# Differences between *in vivo* and *ex vivo* models drive divergent perturbation outcomes

**DOI:** 10.1101/2025.09.02.673692

**Authors:** Aarathy Ravi Sundar Jose Geetha, Cynthia del Valle, Wolfgang Esser-Skala, David Lara-Astiaso, Nikolaus Fortelny

## Abstract

*Ex vivo* experimental models are extensively used as reductionist models, yet the impact of model choice on molecular states and perturbation responses has not been systematically evaluated. Here, we compared the outcomes of genetic perturbations in hematopoietic stem cells between *in vivo* and *ex vivo* model systems. This revealed strong baseline differences characterized by a reduced interferon response and elevated growth and metabolism signatures in *ex vivo* models, a pattern recapitulated in various orthogonal human and mouse systems. We further found substantial differences between perturbation effects between *in vivo* and *ex vivo* models, with some perturbations showing opposite effects. Finally, we evaluated whether AI-based perturbation prediction models can predict *in vivo* perturbation effects from *ex vivo* perturbation effects, which proved challenging. Together, our findings reveal systematic biases in *ex vivo* models, demonstrate how these biases influence perturbation outcomes, suggest approaches to enhance *ex vivo* systems, and provide a test case for computational prediction of perturbation effects.

## Introduction

Biological research utilizes both *in vivo* and *ex vivo* models to study multiple basic and applied biological problems including development, immune responses, and disease conditions. While *in vivo* models are commonly considered to preserve physiological and cellular complexity, performing precise manipulations in cellular states at scale *in vivo* is still a major technological challenge. Hence, *ex vivo* models play a central role in advancing our understanding of cellular physiology and function across multiple tissue contexts^1–4^. For instance, the vast majority of functional genomics maps, which link genes to functions, have been conducted using immortal cancer cell lines^5–10^ followed by primary cell or organoid *ex vivo* models. While *in vivo* studies remain indispensable for understanding cellular behavior within native tissue environments, *ex vivo* models remain essential tools for mechanistic dissection and target discovery. Additionally, *ex vivo* cell cultures are a crucial step in many personalized therapies. For example, autologous (patient-derived) or allogeneic (donor-derived) cells are cultured *ex vivo* to produce next generation medicines such as CAR-T cells or to test the effects of specific drugs or genetic modifications, ultimately enabling treatments tailored to individual patients^11–18^.

Despite the broad application of *ex vivo* models, recent research has highlighted substantial differences in cellular functions and activities of key molecular players between *ex vivo* and *in vivo* model systems (experimental models)^19–23^. Some of these differences, particularly those related to metabolism and mechanosensing, are expected as cells in traditional *ex vivo* cultures are grown with excess of nutrients, constant gas levels, and on 2D surfaces^24–26^. This scenario is very far from the heterogeneous environment found across organs/tissues under physiological *in vivo* conditions. Thus, *ex vivo* models generally cannot recapitulate the intricate microenvironment with cellular interactions, signaling, and mechanical forces that play important roles *in vivo*^27–29^. Strategies to make *ex vivo* models more comparable to *in vivo* systems include co-culture models, additions of small molecules (for example to modulate growth, differentiation, and apoptosis), 3D models, and dynamic culture platforms (enabling fluid shear stress or nutrient gradient) that better mimic physiological conditions^30–37^. Despite these efforts, the precise molecular differences between cells cultured *ex vivo* compared to cells grown *in vivo* have not been systematically studied, limiting the interpretation of *ex vivo* results and improvement of culture systems. This knowledge gap is particularly important given recent FDA initiatives to shift from animal models to human-based research technologies, aimed at a broad scientific and ethical evolution^38,39^, in line with the 3Rs^40^ principles: Replacement, Reduction, and Refinement of animal use in research. While this initiative promotes the use of advanced human models including organoids, tissue chips, and other *in vitro* systems, currently, systematic evaluation of these models against *in vivo* biology remains limited. Our study addresses this gap by establishing a framework for such comparisons and highlighting opportunities to improve *ex vivo/in vitro* platforms, including restoration of altered signaling in the respective systems.

Here, we investigated *ex vivo* to *in vivo* differences in multiple tissue contexts. First, we complemented our recently published *in vivo* screen^41^ on primary hematopoietic progenitors (HPCs) undergoing differentiation with a comparable *ex vivo* dataset. By analyzing unperturbed cells (harboring non-targeting control guide RNAs; NTCs) in these two experimental models, we identified two major differences between *in vivo* and *ex vivo* conditions: interferon pathway and growth/metabolic pathways. To evaluate how generalizable these observations are, we extended our analysis to eight additional, publicly available datasets of *in vivo* and *ex vivo* experimental models spanning diverse tissues. This revealed that the observed *ex vivo–in vivo* transcriptional shifts, particularly in interferon and growth/metabolism associated genes, are not restricted to hematopoiesis but are a general property of *ex vivo* culture. Next, we found that the baseline differences (in unperturbed cells) also lead to differences in the outcome of genetic perturbations between *ex vivo* and *in vivo* conditions. For example, loss of Brd9, a subunit of the ncBAF complex, produces opposing outcomes of lineage differentiation between *in vivo* and *ex vivo* conditions. Analysis of splenic immune cells and glioblastoma models revealed the same baseline biases in interferon and metabolic pathways, again leading to divergent knockout effects between conditions. Finally, we benchmark recent AI tools at predicting *in vivo* KO effects from *ex vivo* KO and NTC cells and *in vivo* NTCs, which demonstrated limitations of current models and potential to develop more accurate models incorporating context-specific constraints to predictive frameworks.

## Results

### *Ex vivo* hematopoietic models exhibit reduced interferon response and increased growth/metabolic expression signatures

To address differences between *ex vivo* and *in vivo* systems during stem cell differentiation we complemented our previous mouse *in vivo* Perturb-seq study in hematopoiesis^41^ with an *ex vivo* Perturb-seq experiment in which we perturbed the same 40 genetic perturbations in the same cell context, the multipotent HPC (Lin-, ckit+, Sca1+), grown under *ex vivo* culture conditions that stimulate hematopoietic differentiation (**Fig. 1A**).

**Figure 1.**
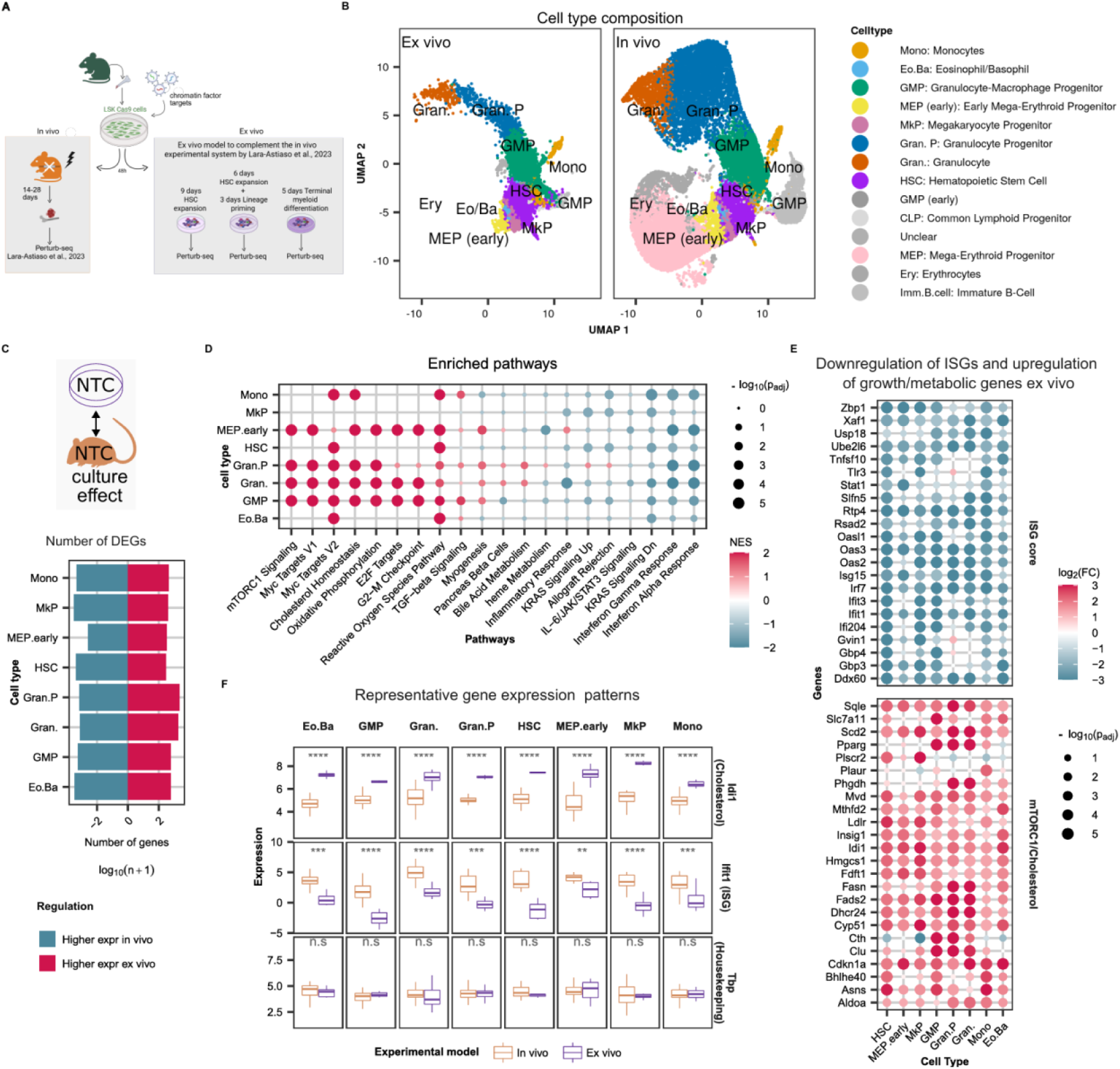
E*x vivo* hematopoietic models exhibit reduced interferon response and increased growth/metabolic expression signatures **A,** Experimental framework. **B,** UMAP of hematopoietic cells differentiated *ex vivo* or *in vivo*. **C,** Number of differentially expressed genes. Results were filtered based on an absolute log2 fold change greater than 1 and an adjusted p-value lower than 0.05. **D-F**, Culture effect on gene expression per cell type. **D,** Gene set enrichment analysis of differentially expressed genes. **E**, Selected significant genes with culture effects. **F,** Gene expression of representative genes *Idi1* (Cholesterol), *Ifit1* (ISG core) and *Tbp* (housekeeping).

Using computational projection of single-cell transcriptomes, we mapped the *ex vivo* generated cellular states onto the *in vivo* hematopoietic system, establishing equivalent cellular states between the two systems (**Fig. 1B** and **Extended Data fig. 1A-B).** The *ex vivo* system recapitulated myeloid differentiation, producing cell types similar to hematopoietic stem cells (HSC), myeloid progenitors (GMPs), granulocytes and their progenitors (Gran and Gran. P), and monocytes (Mono). Additionally, *ex vivo* culture conditions supported early mega-erythroid differentiation, yielding less abundant populations of mega-erythroid progenitors (MEP) and megakaryocytic progenitors (MkP). However, advanced erythroid differentiation and the mature lymphoid branch were largely absent *ex vivo*, likely due to the lack of necessary instructing cytokines and cellular stroma. Then, to assess the “culture effect” (the difference between *in vivo* and *ex vivo* models) in cell types retained prominently in both experimental models, we compared the expression patterns within each cell type between *ex vivo* and *in vivo*. This analysis revealed systematic and consistent differences across all cell types, with a similar magnitude (log fold changes) and number of differentially expressed genes (**Fig. 1C, Extended Data fig. 1C-D, Sup. Table 1**).

Gene set enrichment analysis identified the major pathways and mechanisms impacted by this culture effect and the results were similar for most cell types (**Fig. 1D**). Specifically, across diverse cell types, *ex vivo* conditions consistently showed increased expression of genes from pathways related to growth, such as mTORC1 signaling, DNA replication, and ribosomal machinery as well as other metabolic pathways such as cholesterol homeostasis and oxidative phosphorylation (**Fig. 1D–F, Extended Data fig. 2A, Sup. Table 2**). This enhanced growth and metabolism is expected in nutrient-rich *ex vivo* systems^42^. The upregulation of replication and translation-associated genes, consistent with elevated *Myc* expression (particularly in granulocytes, **Extended Data fig. 2A**), may be driven by the GM-CSF and IL-3 cytokines present in the myeloid media, known activators of *Myc*-dependent growth programs. In strong contrast, *ex vivo* conditions led to a strong downregulation of interferon responses and adhesion programs across all cell types (**Fig. 1D-F**). This is likely due to the absence of stromal cells and extracellular matrix, which provide basal (tonic) interferon levels *in vivo*^43–45^, and the molecular cues that orchestrate cell adhesion (**Extended Data fig. 2A**). Finally, to rule out the possibility that elevated interferon-stimulated genes (ISGs) *in vivo* were due to an inflammatory environment produced by bone marrow pre-conditioning, we examined ISG expression in two independent *in vivo* datasets^46,47^ obtained from animals under steady-state physiological conditions (no pre-conditioning). In both control datasets, ISG expression was similar to the *in vivo* Perturb-seq data and higher than the *ex vivo* perturb-seq data (**Extended Data fig. 2B**). This confirms that the decreased ISG expression observed *ex vivo* is a genuine effect of the *ex vivo* culture conditions.

Altogether, our analysis reveals a experimental model-dependent regulation of metabolic, signaling (interferon) and adhesion mechanisms and highlighting processes that may confound the interpretation of perturbation screens performed *ex vivo*.

### Metabolic programs and interferon responses are dysregulated across multiple *ex vivo* models

To systematically assess the conservation of *ex vivo* culture model-specific effects across tissues, we curated eight publicly available datasets containing matched *in vivo* and *ex vivo* samples and then performed differential gene expression. Correlating gene-level log fold changes, we found clear positive correlations across nearly all comparisons, which demonstrates a conserved transcriptional response to *ex vivo* culture across diverse tissue types in both human and mouse systems (**Extended Data fig. 3A, Sup. Table 3-7**). Moreover, pathway enrichment analyses further revealed that differences affected similar biological programs, particularly, diminished interferon signaling and enhanced metabolic and growth-related pathways in *ex vivo* models (**Extended Data fig. 3B, Sup. Table 8**).

In mouse models, the downregulation of interferon-stimulated genes and upregulation of mTORC1 and metabolic pathways was observed in *ex vivo* cultures used for the expansion of long-term hematopoietic stem cells (**Fig. 2A**) (LT-HSCs)^48,49^, as well as in intestinal organoids derived from primary intestinal tissue^50^ (**Fig. 2B**). Similarly, murine splenic macrophages and T cells exhibited downregulation of interferon-stimulated genes after only 20 hours of *ex vivo* culture^51^ (**Fig. 2C, D**). However, the induction of cholesterol and mTORC1-related pathways was less pronounced in this setting, likely reflecting the limited duration of culture and incomplete metabolic adaptation.

**Figure 2.**
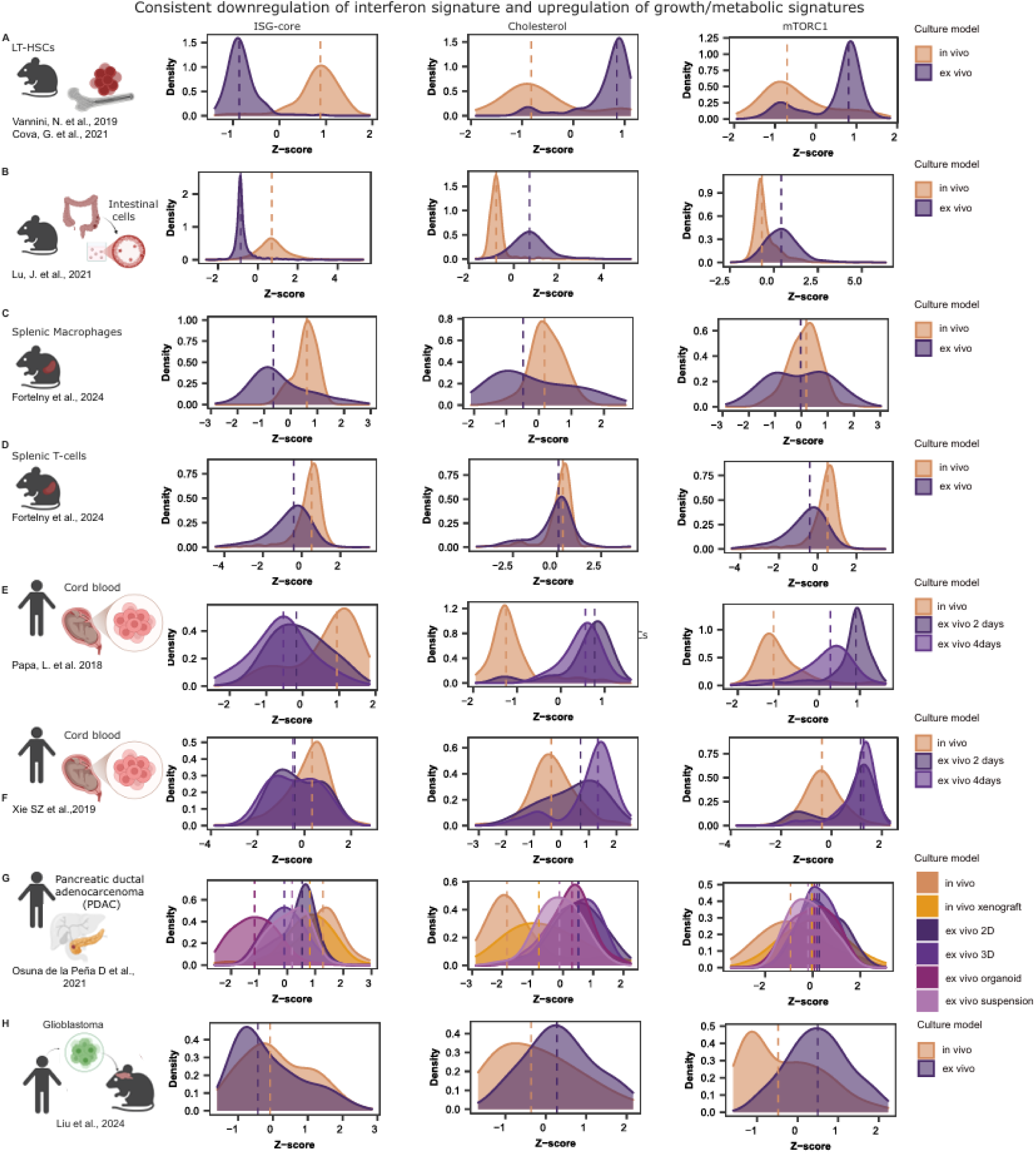
Metabolic programs and interferon responses are dysregulated across multiple *ex vivo* models Consistent culture effects on ISGs and Cholesterol genes in multiple tissue contexts. Genes shown are the top downregulated ISGs and the top upregulated cholesterol-and mTORC1-associated genes identified in the Perturb-seq dataset^41^. Expression values are shown as gene-wise z-scores across all samples in each dataset. Distributions are shown across genes. The vertical lines represent the median z-scores within each experimental model across genes. **A**, Mouse long-term hematopoietic stem cells (LT-HSCs): Transcriptional differences of selected genes between freshly isolated primary long-term hematopoietic stem cells obtained from dataset generated by Vannini and colleagues^48^ versus those cultured *ex vivo* for 2 days, obtained from dataset generated by Cova and colleagues^49^. **B**,Mouse intestinal cells: Comparison of gene expression between epithelial cells directly isolated from *in vivo* intestinal crypts and those cultured *ex vivo* as intestinal organoids (dataset from Lu and colleagues^50^. **C-D,** Mouse splenic macrophages and T-cells: Comparison of gene expression between directly isolated primary splenic macrophages and T-cells to those cultured *ex vivo* for 20 hours ^51^, shows consistent downregulation of ISG core genes, however no evident changes in mTORC1 and cholesterol genes between the experimental models. **E-F,** Human cord blood cells: Comparison of freshly isolated CD34⁺ cells versus those maintained in *ex vivo* culture for 2 days or 4 days **E,** Papa and colleagues^53,54^ and **F,** Xie and colleagues^52^. **G,** Human pancreatic ductal adenocarcinoma (PDAC): Comparison of gene expression between primary PDAC tumour samples (*in vivo*), primary patient-derived xenografts grown in nude mice (*in vivo* xenograft), monocultures of cells derived from the xenografts, prepared as 2D monolayers(*ex vivo* 2D), matrigel-embedded organoids(*ex vivo* organoid), 3D cultures in self-assembling peptide amphiphile (PA) hydrogels(*ex vivo* 3D), monocultures in suspension(*ex vivo* suspension) (dataset obtained from Osuna de la Peña D and colleagues^55^), shows a general downregulation of selected ISGs and upregulation of cholesterol and mTORC1 genes across all *ex vivo* models when compared to *in vivo* models. **H,** Glioblastoma: GL261 cells were grown in *ex vivo* cultures or transplanted intracranially in C57BL/6 mice (*in vivo)*^56^.

In human datasets, we observed a comparable trend. Analyses of cord blood hematopoietic progenitor cells (HPCs) from two independent studies^52–54^showed consistent transcriptional shifts when comparing freshly isolated cells to those cultured *ex vivo* for 2-4 days (**Fig. 2E, F**). A similar pattern was also evident in primary human pancreatic ductal adenocarcinoma (PDAC) samples compared to their corresponding *ex vivo* models, including both 2D cultures and more physiologically relevant organoid systems^55^ (**Fig. 2G**). Xenografts were included as an *in vivo* model from this public dataset but are interpreted as a tumor growth model rather than a physiologically normal tissue state and considered within a spectrum of increasing complexity (2D, organoid, xenograft, primary tumor). Finally, we identified analogous changes in a murine glioblastoma model^56^ where comparison of *in vivo* intracranial grafts derived from GL261 cells with *ex vivo* cultured counterparts revealed reduced interferon responses alongside enhanced metabolic signalling (**Fig. 2H**). While pathway-level trends were consistent with those observed in other systems, the specific interferon-stimulated genes driving these signatures differed in the glioblastoma model (**Fig. 2, Extended Data fig. 4**).

Altogether, our analysis across human and mouse experimental systems from various tissues reinforce the conclusion that the observed transcriptional differences generalize across various tissues, more and less differentiated cells, healthy and cancer cells, and species. These findings underscore the importance of understanding the impact of *ex vivo* culture. This is especially critical given the scientific and regulatory emphasis on transitioning from animal models to human *in vitro* models^38,39,57^. The prominence of this consistently altered gene expression patterns in organoid cultures also suggests that three-dimensional architecture alone is insufficient to maintain *in vivo*-like, physiological ISG expression, and appears to require specific extrinsic signals.

### Genetic perturbations cause distinct effects between *in vivo* and *ex vivo* models

To examine if the differences between *ex vivo* and *in vivo* systems alter the phenotypic outcomes of genetic perturbations, we next compared the KO effects (log₂ fold changes from differential expression analysis comparing KOs to NTCs) for each genetic perturbation between the *ex vivo* and *in vivo* models within each cell type (**Fig. 3A–B**). In the Perturb-seq dataset of hematopoiesis, while some perturbations (37 perturbations) showed similar effects (greater than 0.5) between *in vivo* and *ex vivo* conditions the vast majority (126 perturbations) showed low to no correlation (below 0.5) (**Fig. 3B, Extended Data fig. 5A**). In particular, perturbations of *Brd9, Phf10*, and *Setdb2* showed opposite effects (negative correlation) between *ex vivo* and *in vivo* conditions across most cell types. These similarities and differences in KO responses were not explained by differences in baseline expression of the targeted regulators between *ex vivo* and *in vivo* conditions (**Fig. 3C, Extended Data fig. 5B**), indicating that differential perturbation phenotypes arise from mechanisms other than simple expression differences of the regulators themselves.

**Figure 3.**
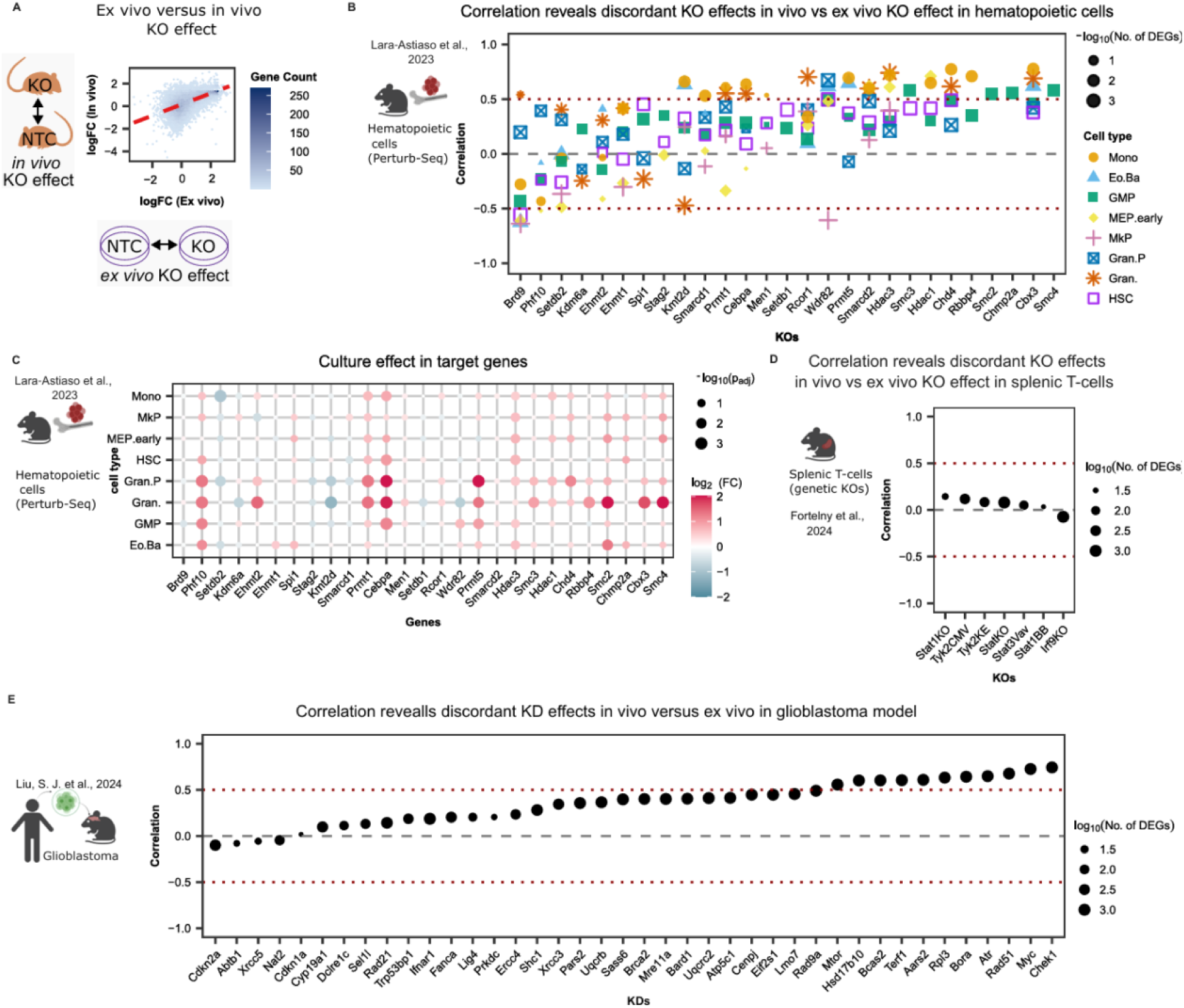
Genetic perturbations cause distinct effects between *in vivo* and *ex vivo* models. A, Pearson correlation in the example of the *Wdr82*KO effect in eosinophils/basophils obtained from *in vivo* versus *ex vivo* model from the perturb-seq dataset of hematopoietic cells^41^.**B, D, E** Pearson correlation of genetic perturbation effects between *in vivo* and *ex vivo* grown cells for all KOs and cell types. The size reflects the number of differentially expressed genes among the maximum observed between *in vivo* and *ex vivo* genetic perturbation effects. The genetic perturbation targets are arranged in x axis in increasing order of Pearson correlation between genetic perturbation effects in *in vivo* and *ex vivo* models (mean across all cell types was used for ordering). **B,** Perturb-seq in hematopoietic cells^41^. **C,** Differential expression statistics of the selected regulators (targeted for KO) at baseline (unperturbed) from perturb-seq dataset of hematopoietic cells^41^. **D,** *In vivo* and *ex vivo* genetic perturbation effects in the genetic KOs of JAK-STAT pathway components in splenic T-cells^51^. **E** *In vivo* and *ex vivo* genetic perturbation effects in the Perturb-seq datasets with knock downs (KDs) in Glioblastoma under no radiotherapy conditions^56^.

To validate the observed differences in responses to genetic perturbations in hematopoietic cells between *in vivo* and *ex vivo* systems in additional tissue contexts, we extended our analysis to two additional datasets where we had already observed consistent culture effects (**Fig. 2D, G**): one dataset of genetic KOs of JAK-STAT pathway components in splenic T-cells^51^ (**Fig. 3D**) and one dataset of CRISPRi-based Perturb-seq knockdown (KD) in a glioblastoma model where we compared perturbations between cultured vs *in vivo* grafted glioblastoma cells (GL261)^56^ (**Fig. 3E**). In both cases, we observed discordant perturbation effects across multiple perturbations when comparing *ex vivo* and *in vivo* models, validating our finding that baseline differences (culture effects) modulate perturbation effects.

### Culture effects anticipate differences in perturbation patterns between *ex vivo* and *in vivo* models

A possible explanation for the variable effects of some perturbations is that targets with strong differences in perturbation effects (*ex vivo* versus *in vivo)* (**Fig. 3B, D, E**) may be functionally related to pathways with strong baseline differences (culture effects). To test this hypothesis, we analyzed the enrichment of *in vivo* perturbation targets among genes with significant culture-effects. Genetic perturbations with more discordant *ex vivo–in vivo* effects showed greater overlap of their target genes with culture-effect genes, resulting in stronger enrichment (**Extended Data fig. 6 A-C, Sup. Table 9-11).** This suggests that, for example, the strong difference in *Brd9*-KO effects (**Fig. 3B**) may be due to its role in interferon signaling^58–60^, which was strongly affected by experimental models in unperturbed cells (culture effects) (**Fig. 1**)

To identify the precise genes and pathways that exhibit differences in genetic perturbation effects between the experimental models, we next calculated interaction effects that statistically test the difference in perturbation outcome between *ex vivo* and *in vivo* models (**Fig. 4A, Sup. Table 12-14**). We performed this interaction analysis across all three perturbation datasets: (1) Perturb-seq in hematopoietic cells^41^, (2) genetic KOs of JAK–STAT pathway components^51^, and (3) Perturb-seq in glioblastoma^56^. This analysis revealed a substantial number of genes showing significant interaction effects for multiple perturbations (**Fig. 4B-D**). For example, all 27 KOs in the Perturb-seq data of hematopoietic cells had interaction effects in more than 10 genes in at least one cell type. As expected, genes with interaction effects were enriched for pathways with strong baseline differences (culture effects) such as interferon signaling, growth, metabolism, and adhesion (**Extended Data fig. 7A, Sup. Table 15**). This indicates that culture effects drive differences in perturbation effects between *ex vivo* and *in vivo*.

**Figure 4.**
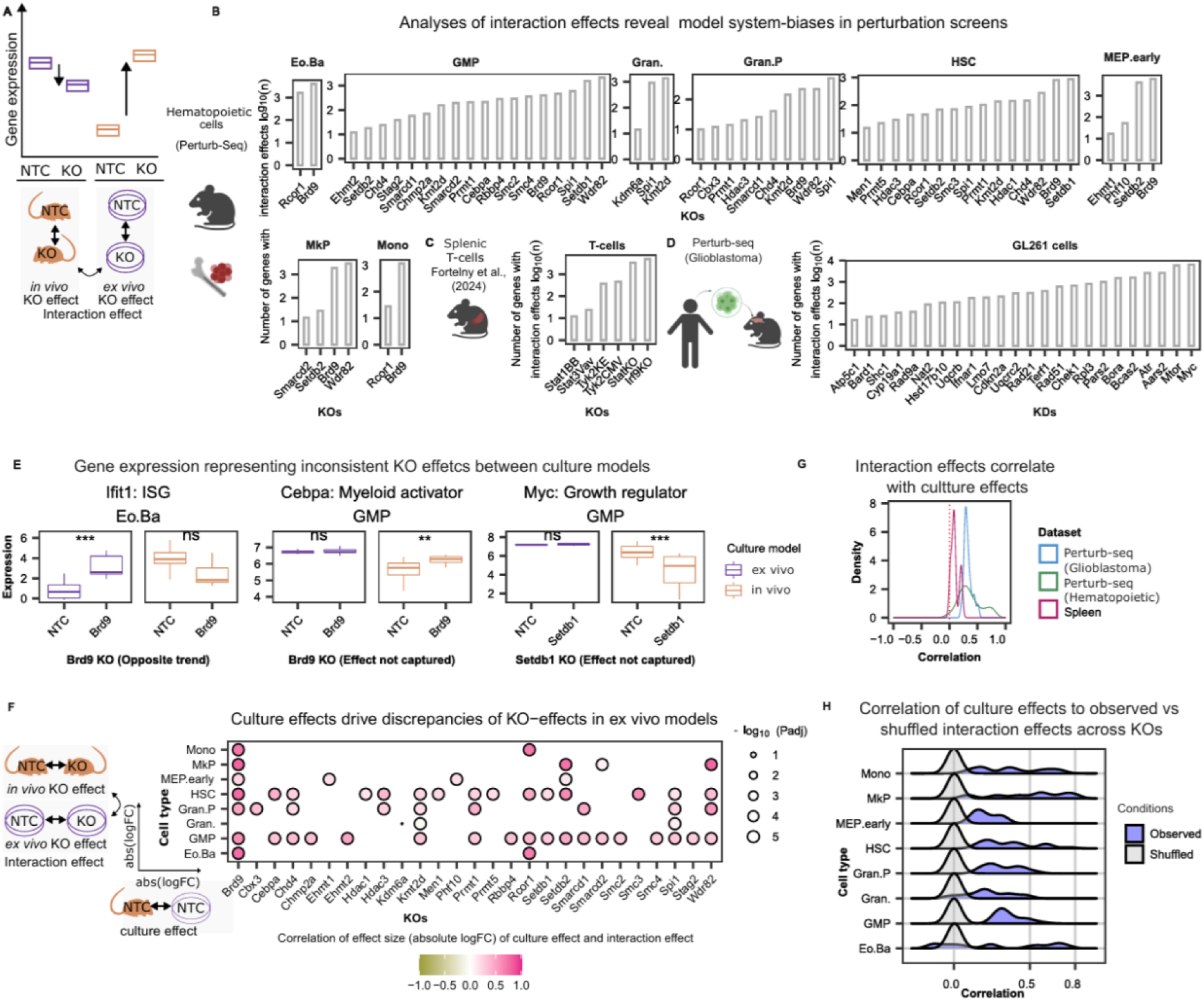
**Culture effects anticipate differences in perturbation patterns between *ex vivo* and *in vivo* models**. Comparison of culture effects, genetic perturbation effects, and interaction effects. Knockouts with at least 10 genes with significant interaction effect are selected here. **A**, Schematic representing of the interaction effect (of experimental model and the genetic perturbation effects) that statistically tests the differences between genetic perturbation effects *ex vivo* versus genetic perturbation effects *in vivo*. **B-D,** Number of genes with significant interaction effects of experimental models and KO effects in **B,** perturb-seq dataset of hematopoiesis^41^, **C,** genetic KOs of Splenic T-cells^51^ and **D,** knockdown (KD) effect in perturb-seq dataset of glioblastoma^56^. genetic perturbations with at least 10 differentially expressed genes are shown. Significant genes were filtered based on an absolute log2 fold change greater than 1 and an adjusted p-value lower than 0.05 **E,** Gene expression representing inconsistent KO effects between *in vivo* and *ex vivo* experimental models in the perturb-seq dataset of hematopoiesis^41^. **F,** Correlation of effect size (absolute logFC) of culture effect and interaction effect in perturb-seq dataset of hematopoiesis^41^.**G,** Pearson correlation of absolute logFC of culture effect versus interaction effect (average across cell types if applicable) across all genetic perturbations within each dataset. **H,** Pearson correlations between the absolute log_2_ fold changes of the culture effects and the absolute log_2_ fold changes for the interaction effects across all genes. Density plots across all KOs within each cell type in perturb-seq dataset of hematopoiesis^41^. The observed correlations are shown in blue. Baseline control correlations obtained after shuffling log2 fold changes are shown in grey

Confirming our results above, *Brd9*-KO, which showed strong differences in KO effects based on our correlation analysis (**Fig. 3B**), also produced most significant interaction effects, especially across top ISGs in multiple cell types (**Fig. 4E, Extended Data fig. 7B**). For example, when comparing gene expression levels of *Ifit1*, a known ISG, we found opposite trends for *Brd9*-KO *in vivo* and *ex vivo* in eosinophil / basophil cells (**Fig. 4E**) that coincide with clear differences between *Ifit1* expression *in vivo* versus *ex vivo* in NTCs. Similarly, the effect of *Brd9* on the regulation of *Cebpa* in GMPs was only prominent *in vivo* but not *ex vivo*. Beyond the identification of pathway regulators, *ex vivo* perturbation approaches can also miss relevant effects. For example, blockade of myeloid differentiation was only revealed upon *Brd9* perturbation *in vivo*^38^ (**Extended Data fig. 7C**). Likewise, *Setdb1* knockout altered gene expression of *Myc* in granulocyte-macrophage progenitors only *in vivo* but not *ex vivo,* which is also reflected in the interaction analysis across multiple *Myc* targets (**Extended Data fig. 8**). Given that *Setdb1*-mediated regulation of *Myc* is being investigated in multiple tumor contexts^61–63^, careful examination of model-specific effects is particularly important. Despite these differences, many KO effects were also consistent across experimental models, for example the expression of *Klk1,* a serine protease or *Atp7b,* a protein involved in copper transport in the knockout of *Cbx3* (**Extended Data fig. 9 A**).

Based on the observation that culture effects and interaction effects influence similar pathways/gene programs, we hypothesized that indeed the culture effects drive differences in perturbation effects between *ex vivo* and *in vivo*. To systematically analyze this across genetic perturbations, we correlated the magnitude of interaction effects to that of culture effects (**Fig. 4F, Extended Data fig. 9B-C**). A significant positive correlation was observed in almost all cases across the three biological systems in question (**Fig. 4F-H, Extended Data fig. 9B-C**), suggesting that the culture effects anticipate interaction effects observed in genetic perturbations.

### *Ex vivo* culture modulates the outcomes of radiotherapy treatment in glioblastoma model

To investigate whether culture conditions also influence responses to non-genetic perturbations, we leveraged the same glioblastoma Perturb-seq dataset previously analyzed for KD effects^56^ where, in addition to genetic perturbations, one of the experimental arms was exposed to radiotherapy (2 Gy × 5 fractions). Analyzing interactions of the culture model and the radiotherapy effects in NTCs (i.e, differences in gene expression outcomes of radiotherapy in *in vivo* vs *ex vivo* models) (**Fig. 5A),** we observed a positive correlation (R = 0.4) of interaction effects and culture effects (*ex vivo* vs *in vivo* at baseline(no radiotherapy)), similar to what we observed above for genetic perturbation. We show that genes with strong culture effects also showed strong differences in the outcomes of both perturbations (genetic and radiotherapy), between *in vivo* and *ex vivo* models (**Fig. 5B, Extended Data fig. 10**).

**Figure 5.**
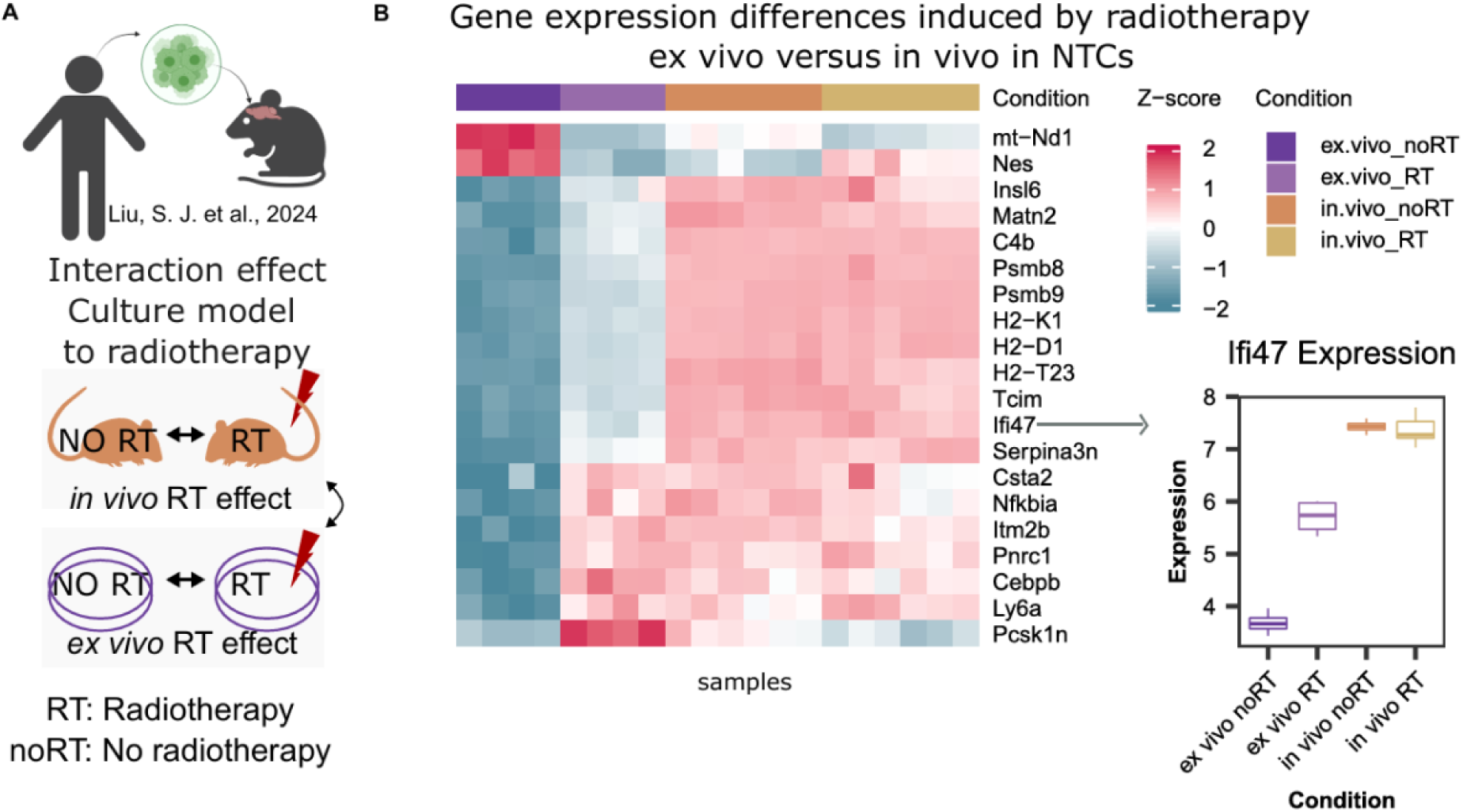
E*x vivo* culture modulates the outcomes of radiotherapy treatment in glioblastoma model. **A-B,** Interaction effect of radiotherapy to experimental model. **B,** Top 20 genes selected based on lowest p-value and strongest absolute log2 fold change respectively, showing interaction effects of radiotherapy to culture model.

This demonstrates that culture-dependent regulatory programs systematically rewire how cells respond to diverse (not only genetic) perturbations, underscoring that perturbation outcomes *ex vivo* should be interpreted by accounting for culture-driven transcriptional programs.

### Integrating *ex vivo* perturbations and paired *in vivo* baseline profiles for predicting *In vivo* perturbation effects using AI models

Artificial Intelligence (AI) holds the potential to unravel intricate gene regulatory relationships and predict responses to novel perturbations. While an ideal predictive model would be trained on large scale, genome-wide *in vivo* perturbation data, generating such data is still a major challenge. Instead, we aimed to determine if *in vivo* perturbation patterns could be predicted by integrating *ex vivo* perturbation data with their *in vivo* counterparts measured under baseline (unperturbed) conditions, which are readily available for multiple tissue types.

To test this hypothesis, we first used linear models^64^ to examine whether relatively simple relationships between *ex vivo* and *in vivo* KO effects exist (**Extended Data fig. 11A**). We first tested models that use the *ex vivo* log₂ fold changes (logFCs) to predict the *in vivo* logFCs across all genes in cross-validation. We then tested more complex models that consider baseline (NTC) differences (culture effects) and interaction terms of *ex vivo* KO effects and culture effects. When trained across all knockouts (thus learning the same coefficient(s) across all KOs), none of these models were able to predict *in vivo* KO effects (top row of **Extended Data fig. 11A**), where correlations of predicted and actual KO effects mirrored the correlations of unmodified *ex vivo* and *in vivo* logFCs (**Fig. 3B**). However, when trained separately on individual knockouts, both models drastically improved correlations, in particular for difficult cases such as Brd9 (bottom row of **Extended Data fig. 11A**). These results show that, in principle, learnable relationships of *ex vivo* and *in vivo* KO effects exist.

While the above approach requires data on *in vivo* KO effects, which are difficult to obtain, we next sought to test options to predict unseen *in vivo* KO effects. We used the recently published and highly performant tools biolord and PeturbNet, which are applicable to predict single-cell perturbations across diverse scenarios^65,66^. We trained both tools to predict *in vivo* KO single-cell transcriptomic profiles from *ex vivo* KO and NTC profiles and *in vivo* NTC profiles (**Fig. 6A**). To assess prediction performance, we then calculated *in vivo* KO effects (KO versus NTCs) from KO profiles predicted by biolord/PerturbNet and compared them to the observed *in vivo* KO effects (log₂FC). As a baseline, we also calculated the correlation between the *ex vivo* KO effects and the observed *in vivo* KO effects. KO effects inferred from the predicted *in vivo* KO expression profiles generated by biolord outperformed *ex vivo* in 34 out of 84 cases while PerturbNet outperformed *ex vivo* in 125 out of 152 cases (**Fig. 6B-C, Extended Data fig. 11 B, C**). Although, PerturbNet consistently outperformed biolord and *ex vivo* in most cases (**Extended Data fig. 11 C)**, the magnitude of improvement was often modest, with performance remaining insufficient to fully recapitulate true *in vivo* responses (19 out of 152 predictions with correlation > 0.8). Particularly for perturbations with divergent *ex vivo* behavior, the prediction remained poor. For instance when examining *Setdb1* and *Brd9* knockouts, which exhibit opposite trends between *in vivo* and *ex vivo* models (**Fig. 4**), we find that both the prediction tools fail to correct for these divergent effects observed *ex vivo* and capture real *in vivo* effect (**Fig. 6D**).

**Figure 6.**
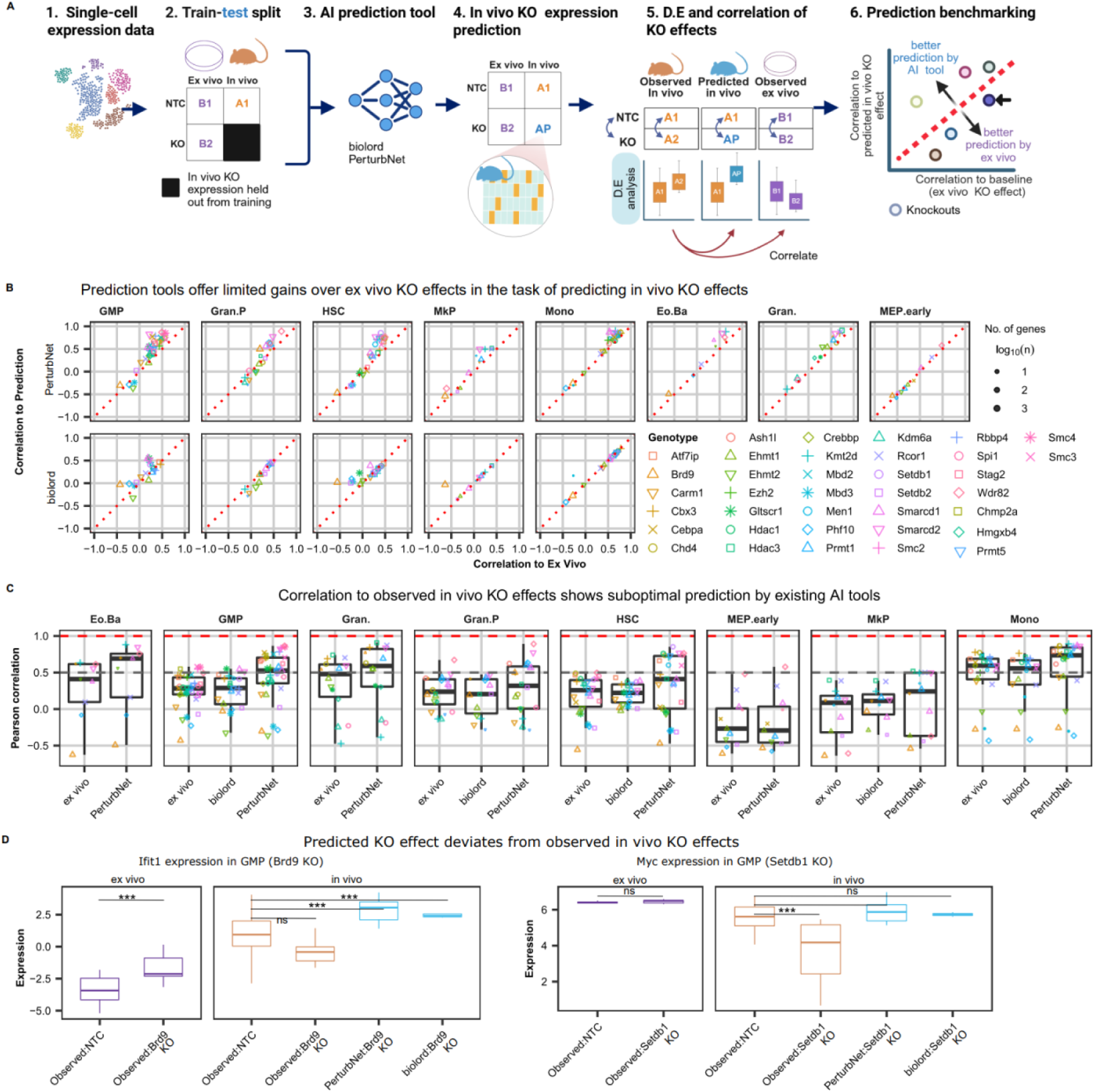
Integrating *ex vivo* perturbations and paired *in vivo* baseline profiles for predicting *In vivo* perturbation effects using AI models **A,** Analysis workflow: **(1)** Single-cell expression data is provided as the input. **(2)** Data is split into training data, which contains (i) *ex vivo* NTCs and (ii) *ex vivo* KOs as well as (iii) *in vivo* NTCs, and test data, which is unseen by the algorithms and contains all *in vivo* KOs, which are thus only used to evaluate the algorithm after training. (**3**) AI prediction tools are trained on training data. **(4)** The *in vivo* KO expression data are predicted by the AI tool. **(5)** Predicted data is used to perform differential gene expression analysis comparing predicted *in vivo* KOs to observed *in vivo* NTCs. Then, to evaluate AI tools, the KO effects (log fold changes across genes) from the predicted *in vivo* KOs are correlated with KO effects from the observed *in vivo* KOs. **(6)** As a baseline for benchmarking, the correlation from (5) is compared to the same correlation using observed *ex vivo* KO effects instead of predicted *in vivo* KO effects. **B,** Pearson correlation of *in vivo* KO effects to predicted/*ex vivo* KO effects in the x-/y-axis. Values above the diagonal indicate better prediction by the respective prediction tool, while below the diagonal shows worse prediction than the baseline. **C,** Pearson correlation of predicted to observed *in vivo* KO effects for the AI tools, compared to baseline (*ex vivo*) performance, i.e. correlation of observed *ex vivo* to *in vivo* KO effect.

In summary, despite the consistent inherent differences between *in vivo* and *ex vivo* experimental models that we identified in bone marrow hematopoietic cells, across multiple cell types, it was challenging to accurately predict *in vivo* KO effects with existing prediction tools. While these models show limitations, improving approaches of incorporating *in vivo*-derived signals and model-specific constraints into predictive frameworks may further enhance performance and help bridge the gap between experimental contexts.

## Discussion

*In vivo* models are increasingly become amendable for precise genetic and environmental manipulations^67–69^. Nevertheless, *ex vivo* models continue to serve a controlled model system for mechanistic dissection, therapeutic target optimization, and conducting detailed functional assays at a scale that is challenging to achieve *in vivo*^70–72^, underscoring the importance of systematic validation of these models.

We report consistent upregulation of genes involved in metabolic and growth-related pathways *ex vivo* which likely stems from the non-physiological conditions of uniformly distributed and excess nutrients common to *ex vivo* models. Perhaps more surprisingly, we observed a consistent downregulation of interferon stimulated genes across multiple biological systems. These *ex vivo*-induced alterations were consistently observed in hematopoietic cells^41,48,49^, cord blood cells^52–54^, pancreatic adenocarcinoma^55^, and intestinal cells^50^ across multiple human and mouse models. Both type I and type II IFN signaling have been shown to influence hematopoietic stem cells (HSC) by modulating self-renewal and differentiation, in part through their interactions with bone marrow stromal cells and niche components^43,44,73–75^. Moreover, constitutive interferon signaling is implicated in priming subsequent responses by maintaining adequate STAT protein levels^76,77^. Thus, the deprivation of IFN signaling *ex vivo* might lead to functional impacts. Furthermore, our observations from the analyzed datasets are consistent with studies in other systems. For example, previous studies have shown that the 4T1 breast cancer cell line exhibits a pronounced downregulation of interferon-stimulated genes when cultured *ex vivo* compared with orthotopic or subcutaneous tumors grown *in vivo*^78^. On the other hand, the reduced ISG signature apparent from gene set enrichment analysis when comparing *ex vivo* models to *in vivo* grown GL261 mouse glioblastoma^56^ resulted from downregulation in a different set of interferon-stimulated genes (ISGs) suggesting that, while the specific genes may differ, reduced interferon signaling may be a general feature of *ex vivo* models. Given the relevance of interferons in modulating tumor microenvironment and cancer therapeutics^79^, excluding their *ex vivo* culture-impact is critical.

Numerous studies have evaluated culture systems to benchmark *ex vivo* conditions against *in vivo* or alternative models^50,80,81^. In contrast, we focused on the molecular differences that arise as a consequence of *ex vivo* culturing to provide a framework for identifying gaps in *ex vivo* models and guiding improvements. For example, based on our results, future work could test whether low-level interferon supplementation or co-culture with interferon-producing cells enhances the physiological relevance of *ex vivo* cultures. Systematic understanding of model-specific effects can also be leveraged to study micro-environment specific signaling. For instance, knowledge about the significance of tumor microenvironment and associated signaling events have led to effective use of *ex vivo* settings as controls, which lack these specific contexts, to specifically identify regulators of tumor micro-environment interactions^19^. Likewise, mapping molecular discrepancies between *ex vivo* and *in vivo* models will inform the development of improved culture systems that more faithfully recapitulate physiological contexts.

Our investigation on the effect of genetic KOs revealed that several KOs induced distinct responses in *ex vivo* cultured cells relative to the *in vivo* model One of the most pronounced differences between the experimental models was the KO of *Brd9,* a gene regulating transcriptional output of ISG^58–60^. Furthermore, our analyses suggests that the differences in basal interferon signaling between the models impact the differences in KO effects. These transcriptional differences are not limited to hematopoietic cells. We validated these findings in splenic T-cells from genetic knockouts of JAK-STAT components^51^ and a perturb-seq dataset on glioblastoma^56^. These datasets complement the Perturb-seq dataset on hematopoietic cells in several ways: The JAK-STAT dataset avoids potential artifacts from viral vectors (used for CRISPR guide RNA delivery) and from inflammatory environment due to bone marrow irradiation. It includes lymphocyte populations (T cells) in splenic cells and thus extends the scope beyond blood derived myeloid cells. Finally, it allowed us to assess interaction effects in KOs of regulators directly involved in interferon signaling response. Perturb-seq dataset on glioblastoma on the other hand, enables the comparison of the same technique in a cancer context. Despite, the complementary characteristics, culture effects in all three datasets were found to be contributing to the discrepancies in perturbation effects between *ex vivo* and *in vivo* settings in the respective systems. For the genetic knockouts of JAK-STAT components^51^ and glioblastoma Perturb-seq^56^ datasets, we included knockouts with n ≥ 2 replicates as available in the original studies. We acknowledge that this limits statistical power for gene-level inference. However, the observed correlation between *ex vivo* and *in vivo* knockout effects, based on consistent expression trends across genes, supports the robustness of the conclusions.

Our findings further suggest that KOs that are functionally linked to pathways that differ between the experimental models (culture effects) are more likely to result in model-specific outcomes (interaction effects). Nevertheless, in some KOs (*Setdb1* in granulocyte macrophage progenitors or *Smc3* in hematopoietic stem cells), we observed many genes with interaction effects or strong correlations between KO effects and culture effects, but a limited overlap between the culture effect and their target genes. This discrepancy may reflect the presence of compensatory mechanisms active *in vivo* specifically in certain regulators^82^.

Given the influence of culture effect in modulating KO effects on transcriptome, we sought to predict *in vivo* KO effects using AI models, using *in vivo* and *ex vivo* NTCs as well as *ex vivo* KOs. Despite the limited success of current models, our analysis points to future directions for improving such models, in particular by incorporating information on the functional proximity of KOs to pathways that are altered by culture effects. Similar to ongoing efforts in the virtual cell challenge^83^, such approaches may enable more accurate *in silico* modeling of *in vivo* perturbation responses. Here, we extend this direction by introducing a complementary benchmark-*in vivo* versus *ex vivo KO transfer*, distinct from the intra-condition perturbation prediction tasks typically targeted by virtual cell models.

Taken together, our study offers valuable insights into the scope and limitations of *ex vivo* models, enabling a more informed and context-appropriate use of these models. It furthermore suggests improvements for *ex vivo* models of hematopoiesis, for example by optimizing nutrient and cytokine supplements. Our analyses were enabled by the availability of Perturb-seq screens that were performed in parallel *ex vivo* and *in vivo* to target the same regulators (chromatin factors) or radiotherapy as a distinct perturbation agent. This limits technical differences (reagents, experimenters, and similar) between the two experimental models as much as possible, so that we expect that the observed differences (culture effects) are driven by differences in the culture model. Nevertheless, technical differences cannot be ruled out completely when comparing different culture model systems. We therefore used additional, unperturbed *in vivo* datasets to support the low expression of interferon response genes *ex vivo*. Furthermore, we validated our key findings in external datasets and different tissue contexts. Technical effects likely play a minor role when analyzing differences in KO effects (interaction effects) as these analyses rely on comparisons of KOs and NTCs (or WT) *within* each model, thus avoiding potential technical effects.

In summary, our framework offers a systematic approach to evaluate any experimental models by assessing cell type composition and gene expression differences, given comparable systems. In the future, we expect that similar approaches will be useful to validate optimized *ex vivo* systems, for example by incorporating cocultures in 3D architectures and/or small molecules. Together with ongoing efforts to develop machine learning prediction tools^83–85^, we expect that advances in *ex vivo* models will strongly improve the prediction of *in vivo* perturbation effects.

## Supplementary Tables

**Supplementary Table 1:** Differential expression statistics of genes with culture effect (*ex vivo* to *in vivo*) in the perturb-seq dataset on hematopoietic cells. logFC - log_2_foldchange; adj.P.Val - adjusted p-value

**Supplementary Table 2:** Gene set enrichment analysis of genes with culture effect in the perturb-seq dataset on hematopoietic cells. padj-adjusted p-value; NES - normalized enrichment score; db - enrichment database; leadingEdge - genes that contribute most to the observed enrichment.

**Supplementary Table 3-7:** Differential expression statistics of genes with culture effects (under no perturbation), analyzed from, publicly available datasets, included in the manuscript.

**Supplementary Table 8:** Gene set enrichment analysis between *in vivo* versus *ex vivo* models in publicly available datasets; NES - normalized enrichment score; leadingEdge - genes that contribute most to the observed enrichment.

**Supplementary Table 9:** Fisher’s exact test statistics when analyzing the overlap between genes with culture effect and regulator (KO) targets (in *in vivo* culture model) in perturb-seq dataset of hematopoiesis. The table is filtered for overlap > 5.

**Supplementary Table 10:** Fisher’s exact test statistics when analyzing the overlap between genes with culture effect and regulator (KO) target (in *in vivo* culture model) in bulk RNA-seq dataset of spleen cells. The table is filtered for overlap > 5

**Supplementary Table 11:** Fisher’s exact test statistics when analyzing the overlap between genes with culture effect and regulator (KO) target (in *in vivo* culture model) in perturb-seq dataset of glioblastoma. The table is filtered for overlap > 5

**Supplementary Table 12:** Differential expression statistics of genes with *ex vivo* KO effects, *in vivo* KO effects and interaction effects in Perturb-seq dataset of hematopoiesis. logFC - log_2_foldchange; adj.P.Val - adjusted p-value interaction, condition-the comparison of interest (interaction/ *ex vivo* KO effect/ *in vivo* KO effect)

**Supplementary Table 13:** Differential expression statistics of genes with culture effect, KO effects or interaction effect in bulk RNA-seq dataset of spleen cells. logFC - log_2_foldchange; adj.P.Val - adjusted p-value, condition-the comparison of interest (interaction/ *ex vivo* KO effect/ *in vivo* KO effect)

**Supplementary Table 14:** Differential expression statistics of genes with culture effect, KO effects or interaction effect of knockouts/radiotherapy to experimental models in perturb-seq dataset of glioblastoma. logFC - log_2_foldchange; adj.P.Val - adjusted p-value; coef – comparison of interest (culture effect: *ex vivo* NTC cells under no radiotherapy versus *ex vivo* NTC cells under no radiotherapy conditions, <genotype>_ex.vivo_noRT: knockout effect of the respective knockout in *ex vivo* model under no radiotherapy, <genotype>_in.vivo_noRT: knockout effect of the respective knockout in *in vivo* model under no radiotherapy, interaction_genotype_<genotype>: interaction effect of genotype and experimental model under no radiotherapy, interaction_RT: Interaction of radiotherapy and experimental model in NTC cells.

**Supplementary Table 15:** Gene set enrichment analysis of genes with interaction effect in Perturb-seq dataset of hematopoietic cells, bulk RNA-seq of spleen cells and perturb-seq of glioblastoma. padj - adjusted p-value; NES - normalized enrichment score; db - enrichment database; leadingEdge - genes that contribute most to the observed enrichment.

**Supplementary Table 16:** Datasets used for comparing gene expression in matched *ex vivo* and *in vivo* models in different tissue contexts.

## Methods

### Data acquisition

Perturb-seq single-cell RNA sequencing (scRNA-seq) data (GSE213511, https://doi.org/10.5281/zenodo.17014352) were obtained from (David Lara et al, 2023)^41^. Experimental procedures are described in detail in the original publications. For *ex vivo* Perturb-seq, data from cells cultured in conditions for HSC expansion, lineage priming and myeloid differentiation as follows: For HPC expansion (9-day, no priming), transduced LSKs were cultured for 9 days in F12 medium supplemented with polyvinyl alcohol (PVA), thrombopoietin (TPO; 100 ng/mL), and stem cell factor (SCF; 10 ng/mL), without lineage priming, followed by Perturb-seq profiling. For HPC expansion with lineage priming (6+3 days), **t**ransduced LSKs were cultured for 6 days in F12 medium supplemented with PVA, TPO (100 ng/mL), and SCF (10 ng/mL), followed by 3 days of lineage priming in F12 medium containing GM-CSF (5 ng/mL), Flt3L (5 ng/mL), IL-6 (5 ng/mL), IL-3 (5 ng/mL), erythropoietin (10 U/mL), TPO (5 ng/mL), and SCF (10 ng/mL). Cells were then subjected to Perturb-seq. For Terminal myeloid differentiation, LSK cells were transduced and differentiated under myeloid-promoting conditions for 5 days in IMDM supplemented with 20% FBS, GM-CSF, IL-3, IL-6, and Flt3L.

Bulk RNA-seq data (GSE204736) of sorted splenic macrophages and T cells were obtained from (Fortelny et. al, 2024)^51^. Experimental procedures are described in detail in the original publication.

### Computational setup

Analyses were performed in R (v4.0.2) if not specified otherwise. Package management was handled using renv (v0.12.0) to ensure reproducibility. Data visualization utilized ggplot2 (v3.3.3) and tidyverse packages (v1.3.0). Initial quality control of single-cell data was conducted using Seurat (v4.0.0) and Monocle 3^86^ (v0.2.3.0), following the workflow described by (David Lara et al, 2023)^41^, with an additional preprocessing step to remove ambient RNA contamination using SoupX^87^ (v1.4.5).

### Cell type assignment and data projection

Cell type annotation for *in vivo* samples were obtained from (David Lara et al, 2023)^41^ where they were based on singleR^88^ (v1.4.1) and CytoTRACE^89^ (v0.3.3).

*Ex vivo* cultured cells from the Perturb-seq dataset and unperturbed *in vivo* cells from Anna Konturek-Ciesla and colleagues^47^ were projected onto the UMAP obtained from *in vivo* grown cells from the Perturb-seq dataset for cell type assignment. We used ProjecTILs^85^ algorithm (v2.0.2) to predict the cell types based on closest cell type with reference to the UMAP coordinates of already annotated *in vivo* cells using a k-nearest neighbour approach with k = 20 neighbours.

For downstream analysis, we used cell types that were prominent in both *ex vivo* and *in vivo* models, with mean confidence of cell type assignment >= 0.7. For external datasets obtained from Izzo and colleagues (GSE124822)^46^, we used the cell type assignments from the original publications

### Comparing interferon/metabolic gene signature across tissues from publicly available datasets

Datasets used to compare interferon signatures and cholesterol/metabolism-associated genes are summarized in **Sup. Table 16.** Top downregulated ISGs and upregulated cholesterol genes and mTORC1 genes in the Perturb-seq dataset^41^ were compared against each dataset. For this top 20 genes were ranked per cell type (from the Perturb-seq dataset) based on increasing p-value and strongest log2 fold change respectively and combined to obtain one list of genes. To enable cross-species comparison, we translated mouse gene symbols to their human counterparts. Ortholog relationships were taken from a BioMart export (Ensemble, July 2021) containing curated human–mouse gene pairs. Cell type assignments for external datasets were obtained from the respective datasets. Normalized counts were either obtained directly from the public databases or generated by reanalyzing the raw data using limma. Z-scores were then calculated for each gene across samples within each dataset.

### Differential expression analysis

For Perturb-seq dataset, pseudobulk samples were generated by summing the counts across cells that belong to the same sample and cell type and possess the same guide RNA. Following this, counts were normalized by calculating normalizing factors using the function calcNormFactors from edgeR^90^ (v3.32.1). Counts were transformed to counts per million (CPM) and log_2_-normalized using limma^91^ (v3.46.0). For comparing global transcriptome simiilarity, voom-normalized pseudobulk expression matrices for *in vivo* and *ex vivo* samples were combined, and pairwise Euclidean distances between samples were computed across genes. Classical multidimensional scaling (MDS) was performed using the cmdscale function, and the first two dimensions were used for visualization. Genes with CPM > 30 in at least one sample were further processed for differential expression analysis using limma. For the analysis of NTC cells, a design matrix was constructed by combining cell type and culture model (*ex vivo*, *in vivo*) into a single factor(group) and fitting a model without intercept (∼0 + group). Normalization and variance modeling were performed with voom (limma), followed by linear modeling with lmFit and empirical Bayes moderation (eBayes). For estimating KO effects, a model which estimates interaction and main effects (∼ experimental model * genotype) for experimental model (*in vivo* and *ex vivo*) and KOs was used. KOs with a minimum of three biological replicates (n ≥ 3) in respective cell type were included for downstream analyses. Genes were considered as significantly differentially expressed if they had an absolute log_2_ fold change (logFC) greater than 1 and a Benjamini-Hochberg adjusted *p*-value less than 0.05. Gene set enrichment was performed using the function fgsea from the fgsea package^92^ (v1.16.0).

For the bulk-sorted data from splenic macrophages and T-cells^51^, raw counts were processed and normalized separately for each cell type, and differential expression analysis was performed with linear mixed models which estimates main effects and interaction effects for experimental model (*in vivo* and *ex vivo*) and KOs and included experiment ID as a random intercept term (1 | experiment_id) to account for batch effects (∼ genotype * experimental model + (1|experiment_id) with other settings as described in the original publication (n≥2). Only T-cells were used for interaction effect analysis as splenic macrophages showed altered marker genes after *ex vivo* cultures.

For Perturb-seq data on glioblastoma^56^, we generated pseudobulk samples according to the annotated treatment conditions. Normalization was performed as described for perturb-seq data for hematopoiesis above, followed by filtering genes based on “filterByExpr” function available from edgeR packageas. A three-way interaction model including experimental model (*in vivo* vs. *ex vivo*), genotype (KDs), and treatment (radiotherapy vs. no radiotherapy) was initially fitted; however only Culture effects, KO effects and radiotherapy treatment effect and not the three-way interactions are included in the final analyses. Normalization and variance modeling were performed as described above. Conditions with a minimum of two biological replicates (n ≥ 2) were included for downstream analyses. Genes were considered significantly differentially expressed if they had an absolute log_2_ fold change (logFC) greater than 1 and a Benjamini-Hochberg adjusted *p*-value less than 0.05. Gene set enrichment was performed using the function fgsea from the fgsea package^92^ (v1.16.0).

### Analysis of interferon and metabolic signatures in additional datasets

For the following datasets, relevant samples enabling *ex vivo* versus *in vivo* comparisons were selected, normalizations were performed as described above for perturb-seq dataset on hematopoietic cells, followed by filtering and differential expression analysis was performed using limma as described above by setting a design matrix constructed by using experimental model (*ex vivo*, *in vivo*) and as the factor.

**1**, Mouse long-term hematopoietic stem cells (LT-HSCs): primary long-term hematopoietic stem cells obtained from dataset generated by Vannini and colleagues^48^ versus those cultured *ex vivo* for 2 days, obtained from dataset generated by Cova and colleagues^49^. **2,** Mouse intestinal cells: Epithelial cells directly isolated from *in vivo* intestinal crypts versus those cultured *ex vivo* as intestinal organoids (dataset from Lu and colleagues^50^). **3, 4** Human cord blood cells: Comparison of freshly isolated CD34⁺ cells versus those maintained in *ex vivo* culture for 2 days or 4 days from Papa and colleagues^53,54^ and, Xie and colleagues^52^. **5,** Human pancreatic ductal adenocarcinoma (PDAC): Comparison primary PDAC tumour samples (*in vivo*), primary patient-derived xenografts grown in nude mice (*in vivo* xenograft), monocultures of cells derived from the xenografts, prepared as 2D monolayers(*ex vivo* 2D), matrigel-embedded organoids(*ex vivo* organoid), 3D cultures in self-assembling peptide amphiphile (PA) hydrogels(*ex vivo* 3D), monocultures in suspension(*ex vivo* suspension) (dataset obtained from Osuna de la Peña D and colleagues^55^), For visualization of culture effect and interaction effects on expression level from perturb-seq datsets on hematopoietic cells and glioblastoma^41,56^, we selected genes belonging to specific pathways. Pathways were selected primarily based on the outcome of enrichment analysis (fgsea) and known associated biological processes. Top significant genes among pathways of interest were filtered, followed by sorting based on increasing adjusted *p-value* and decreasing absolute logFCs respectively. A subset of ISG-core genes as described by (Mostafavi et. al)^93^ were selected with these criteria for representing the culture effects and interaction effects in interferon stimulated genes. Other gene sets for pathways of interest were obtained from Molecular Signatures Database^94^ (MSigDB, 2020 and Reactome Pathway^95^ Database (2022), downloaded using Enrichr^96^.

### Prediction of *in vivo* KO effects

Linear models were trained using the *ex vivo* log₂ fold changes (0 + logFCs) or baseline (NTC) differences (culture effects) and interaction terms of *ex vivo* KO effects and culture effects (ex.vivo_logFC * NTC_logFC) to predict the *in vivo* logFCs across all genes. The full set of genes was randomly partitioned into five equal subsets where 20% of the genes were withheld as a test set, and the remaining 80% were used for model training. The models were trained (fitted) either across all knockouts or within each KO. Model performance was assessed based on Pearson’s correlation coefficient calculated between actual and predicted logFCs for *in vivo* KOs.

The biolord^65^ package (v0.0.3) was used to predict single cell expression profiles in *in vivo* KO samples from *ex vivo* KO and NTC and *in vivo* NTC cells from the Perturb-seq dataset. Genotype, cell type and sample were used as attribute labels while training. Default parameters were used throughout. To assess prediction performance of biolord, Pearson’s correlation coefficient was calculated for predicted versus actual *in vivo* KO effects (log₂FC) obtained after differential expression analysis.

PerturbNet package (v0.0.3b1) was used to predict *in vivo* KO expression profiles. To predict *in vivo* knockout gene expression, we used PerturbNet trained exclusively on *ex vivo* NTC, *ex vivo* KOs and *in vivo* NTC, with all *in vivo* KO samples held out during training. While training the CINN model, we adapted the training factors by including onehot vectors of celltype, experimental model (*ex vivo* or *in vivo*), sample identity and guide in addition to genotypeVae (which was originally used in the PerturbNet-based prediction on genetic perturbations). The model was trained using scVI (v0.7.1) to learn a latent representation of gene expression. All model components were trained without access to *in vivo* KO expression. The *in vivo* KO dataset was used exclusively for evaluation. For the held-out *in vivo* KO cells, predicted gene expression was generated using the trained scVI decoder. Predictions were first obtained as normalized expression profiles at a fixed reference library size. To ensure realistic sequencing depth and enable direct comparison with observed data, predicted expression values were subsequently rescaled to match the observed per-cell library sizes of the corresponding *in vivo* cells.

## Code availability

All analysis scripts are publicly available at https://doi.org/10.5281/zenodo.17011954

## Competing interests

The authors declare no competing interests.

## Acknowledgements

We thank Markus Wiederstein for maintaining the HPC infrastructure and providing technical support. We are grateful to Thomas Decker, Brian Plosky and Rodrigues Nunes da Silva Natalia for their valuable comments and feedback on this work. The data presented in this publication were collected using the EFRE-FFG-funded resesarch infrastructure CELLCOMM [FO999912153]. This research was funded in whole or in part by the Austrian Science Fund (FWF) 10.55776/F100900 [“EpiFlaMe”].

## Author contributions

A.G.: Methodology, Software, Investigation, Resources, Data Curation, Visualization, Writing - Original Draft, Writing - Review & Editing; W.E.S.: Software, Investigation, Writing - Review & Editing; C.V.: Resources; D.L.A.: Conceptualization, Writing - Review & Editing; N.F.: Conceptualization, Methodology, Investigation, Writing - Original Draft, Writing - Review & Editing, Supervision, Project administration.

---

**Extended Data figure 1.**
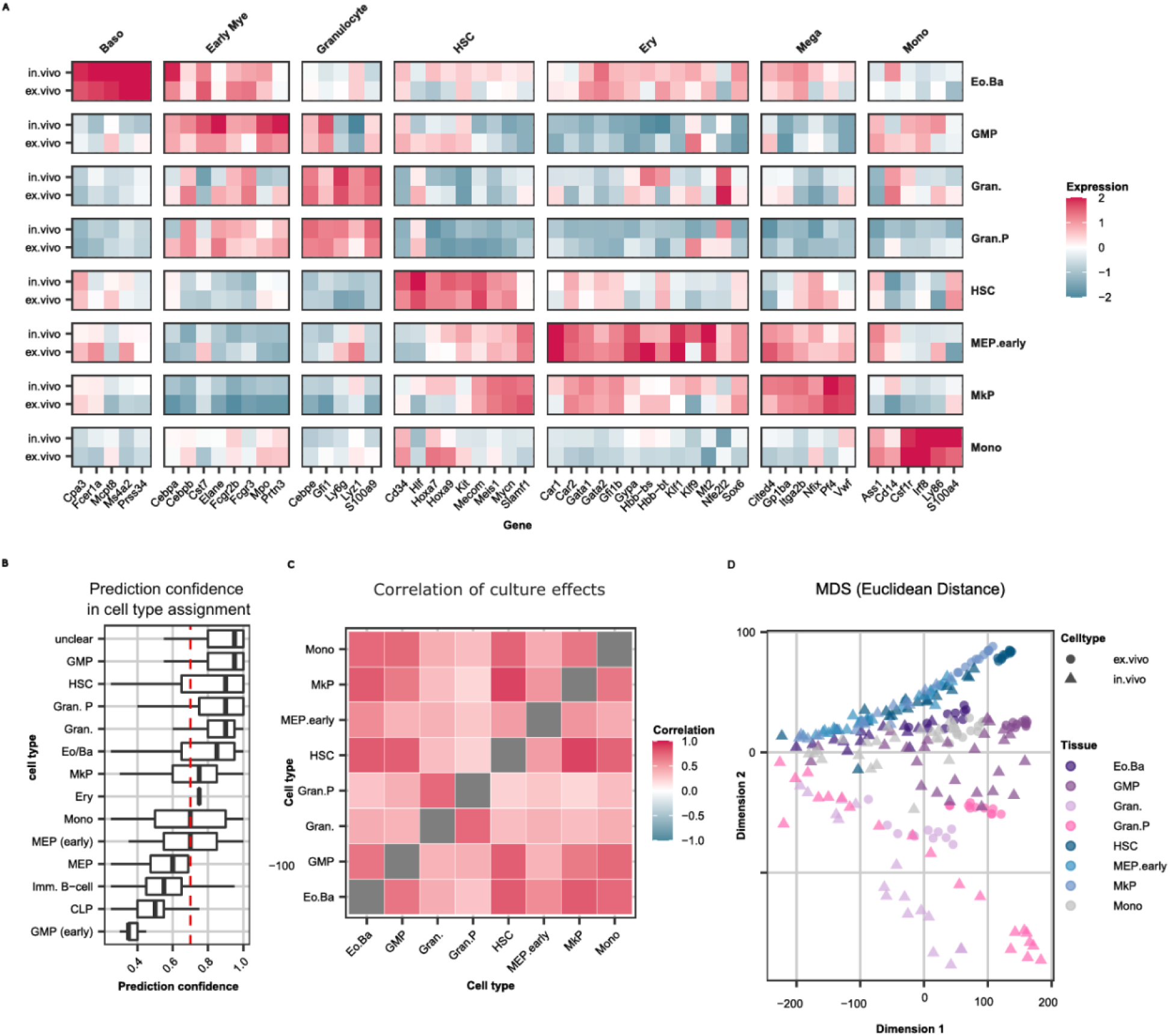
**A,**Normalized log_2_ counts per million (CPM) expression of marker genes of corresponding cell types (marked above the corresponding panels), scaled across cell types. **B,** Prediction confidence of *ex vivo* cell type assignment by projecting *ex vivo* cells on to the UMAP of *in vivo* cells using ProjecTILs^85^. The red line indictae prediction confidence of 0.7**. C,** Correlation of transcriptional culture effects (log_2_FC comparing *ex vivo* versus *in vivo*) between cell types. **D,** MDS plots representing Euclidean distance between samples.

**Extended Data figure 2.**
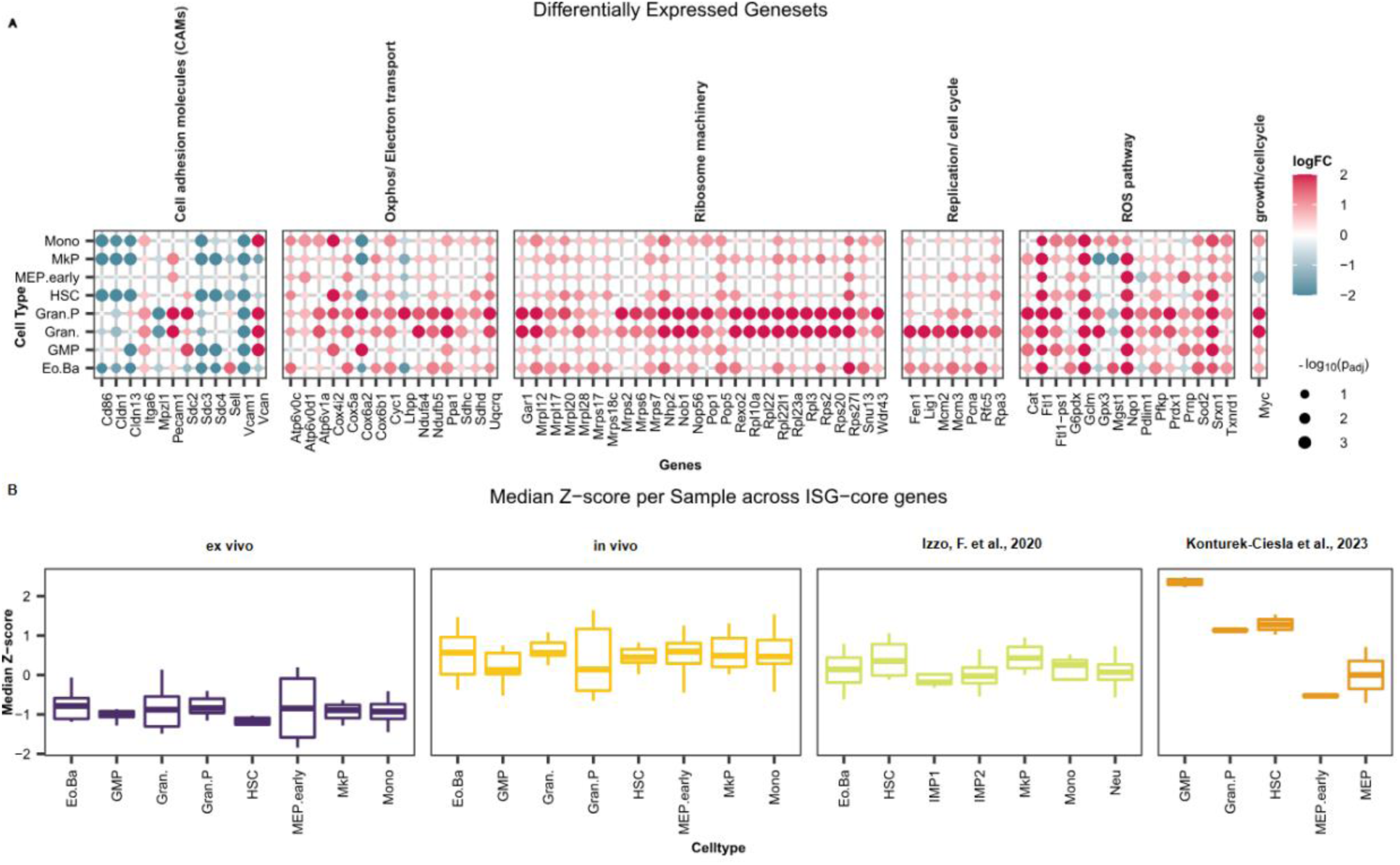
**A,**Differential expression statistics of genes with culture effect. **B,** Aggregated median expression from pseudobulk samples across ISGs represented in Figure 1D, in *ex vivo*, *in vivo* from the Perturb-seq dataset^41^ as well as unperturbed *in vivo* cells from Anna Konturek-Ciesla and colleagues^47^, and from Izzo and colleagues^46^. The cell type assignments were obtained from the original publication in the external dataset 1 (Izzo and colleagues) and transferred from the *in vivo* Perturb-seq data set after projection in the external dataset 2 (Anna Konturek-Ciesla and colleagues) and in the *ex vivo* Perturb-seq dataset, based on a k-nearest neighbour approach with k = 20 neighbours.

**Extended Data figure 3.**
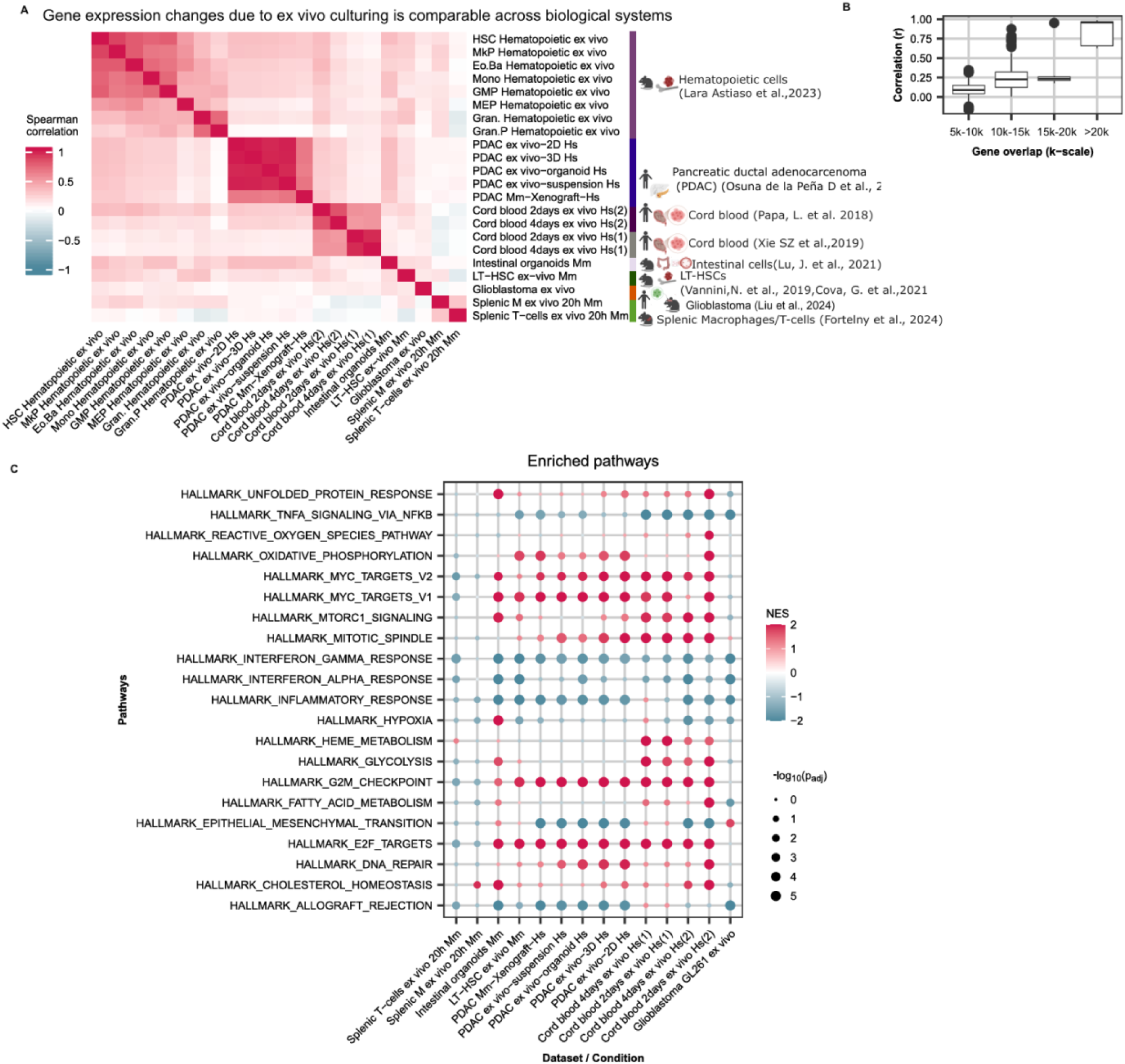
**A,**Correlation of logFC of gene expression comparing cells cultured under *ex vivo* conditions against *in vivo* cells across different biological systems. **B,** Pearson correlation coefficients (r) computed between log₂ fold changes across pair-wise dataset comparisons, stratified by the number of total overlapping genes between the datasets in comparison. **C,** Gene set enrichment analysis of differentially expressed genes between *ex vivo* and *in vivo* models in each dataset representing diverse biological systems. NES: normalized enrichment score

**Extended Data figure 4.**
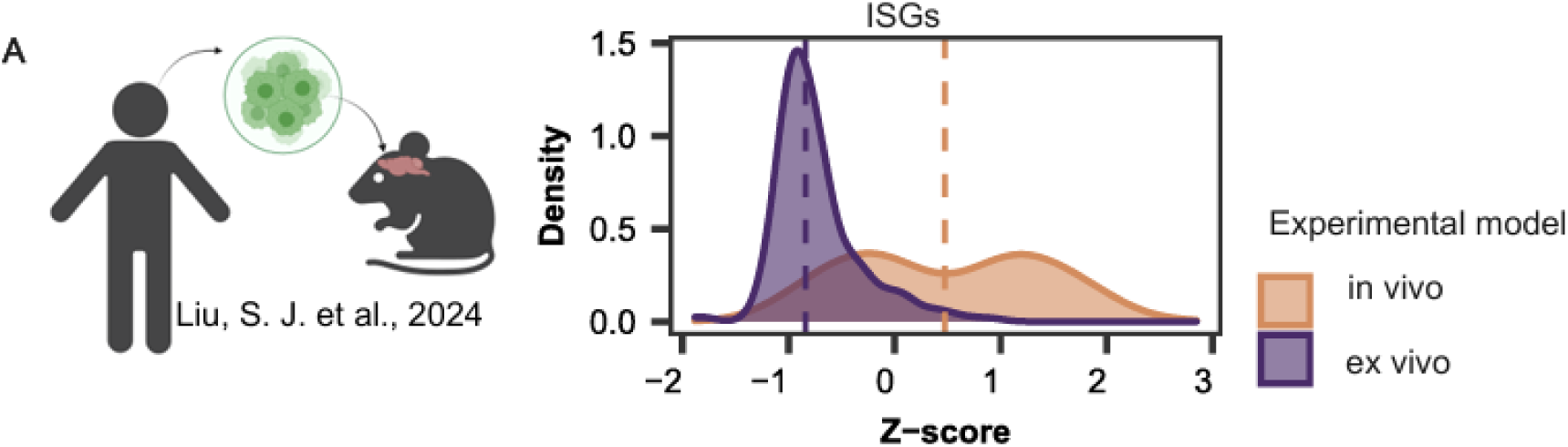
ISGs downregulated in GL261 *ex vivo* cultures of glioblastoma relative to the *in vivo* grown cells. The ISG genes (n = 119) were selected from the enrichment analysis.

**Extended Data fig. 5.**
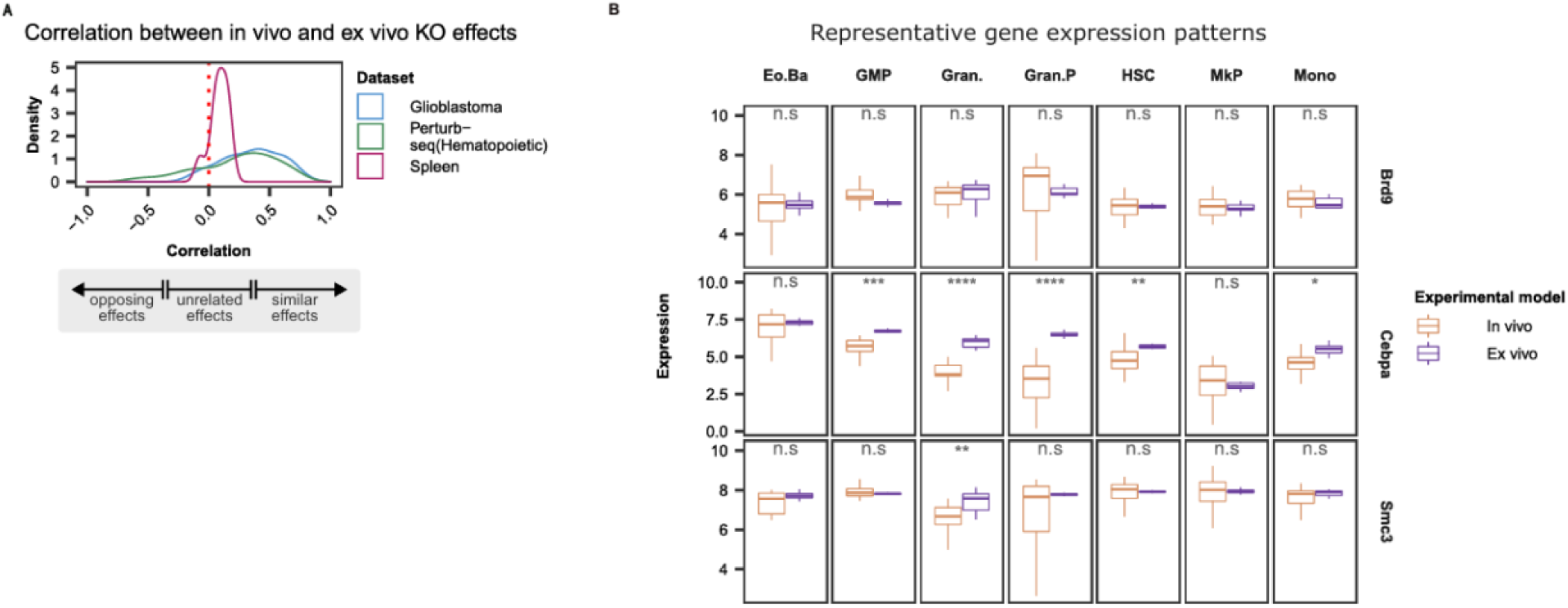
**A,**Pearson correlation of *in vivo* versus *ex vivo* KO effects (average across cell types if applicable) within each dataset. **B,** Baseline expression of selected the regulators in *ex vivo* vs. *in vivo* conditions in perturb-seq dataset on hematopoietic cells^41^.

**Extended Data figure 6.**
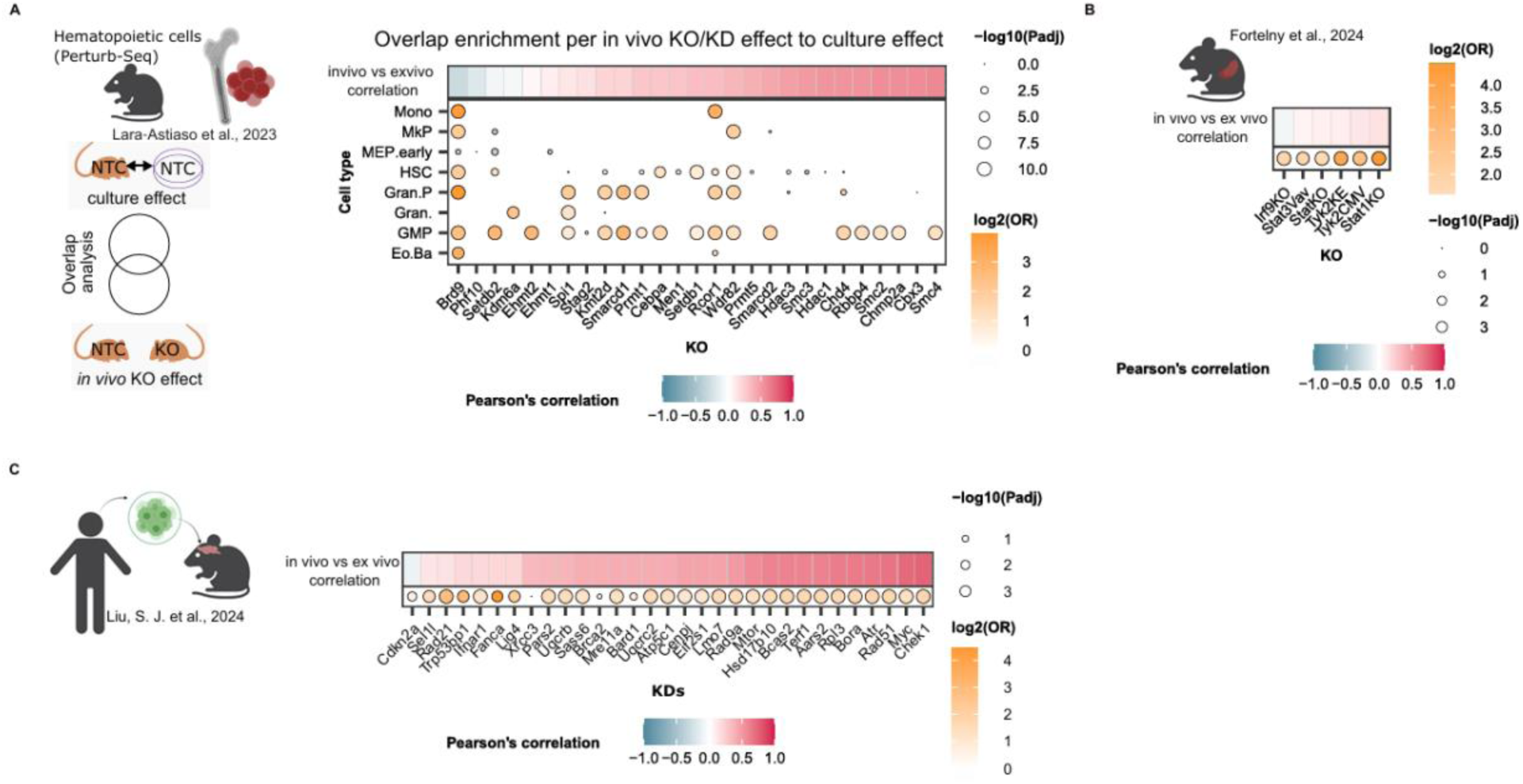
**A-C,**The heatmaps represents correlation of *in vivo* versus *ex vivo* genetic perturbation effects (mean across cell types) in the perturb-seq of hematopoietic cells^41^. The dot plots represent enrichment assessing overlap of target genes of the perturbed regulators (KO/ KDs) in *in vivo* models(actual targets of respective genetic perturbations) and genes contributing to culture effects. **A,** Perturb-seq in hematopoietic cells. **B,** Genetic knockouts of JAK STAT pathway components. **C,** Perturb-seq in Glioblastoma.

**Extended Data figure 7.**
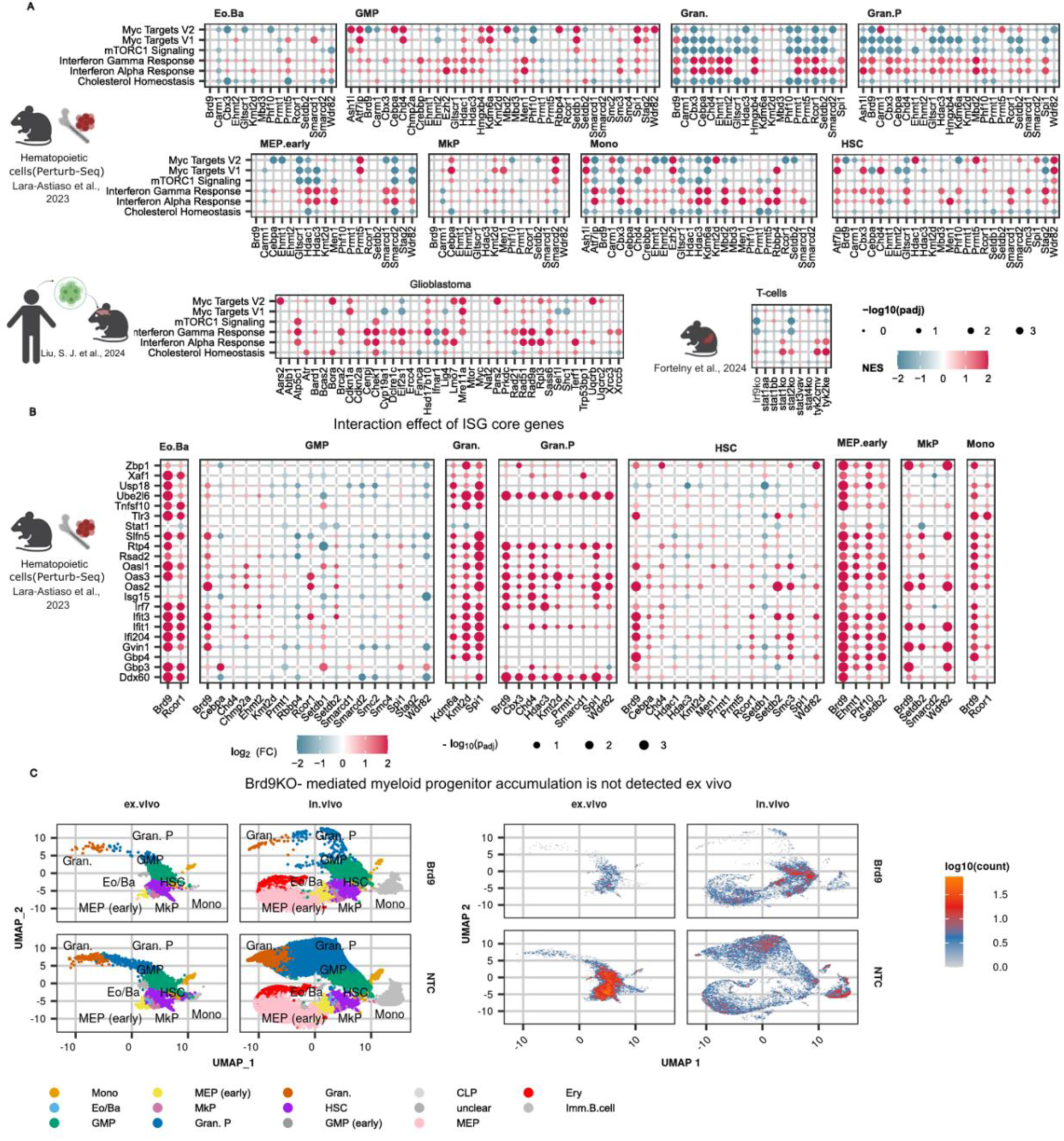
**A,**Gene set enrichment in selected pathways for genes with significant interaction effects in the indicated datasets. **B,** Differential expression statistics of interaction effects of ISGs (y-axis) in respective KOs (x-axis) in the perturb-seq dataset of hematopoietic cells^41^, knockdowns of perturb-seq dataset in glioblastoma model^56^ and genetic knockouts in mouse spenic T-cells^51^. **C,** Cell type distribution in *Brd9*KO and NTC in *in vivo* and *ex vivo* experimental models.

**Extended Data figure 8.**
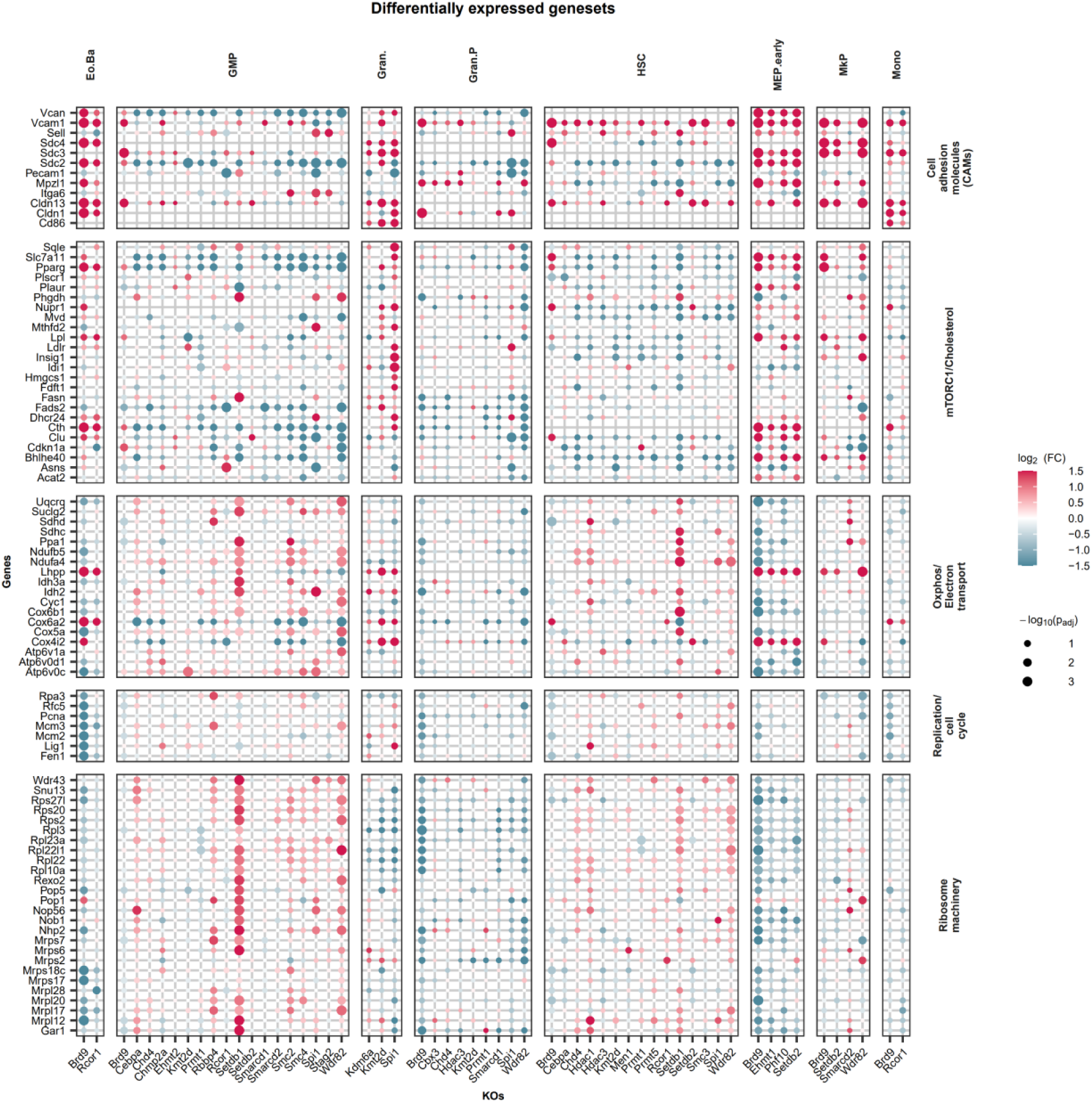
Differential expression statistics of interaction effect analysis. Interaction effects in genes represented in Extended Data fig. 2A, with culture effects.

**Extended Data figure 9.**
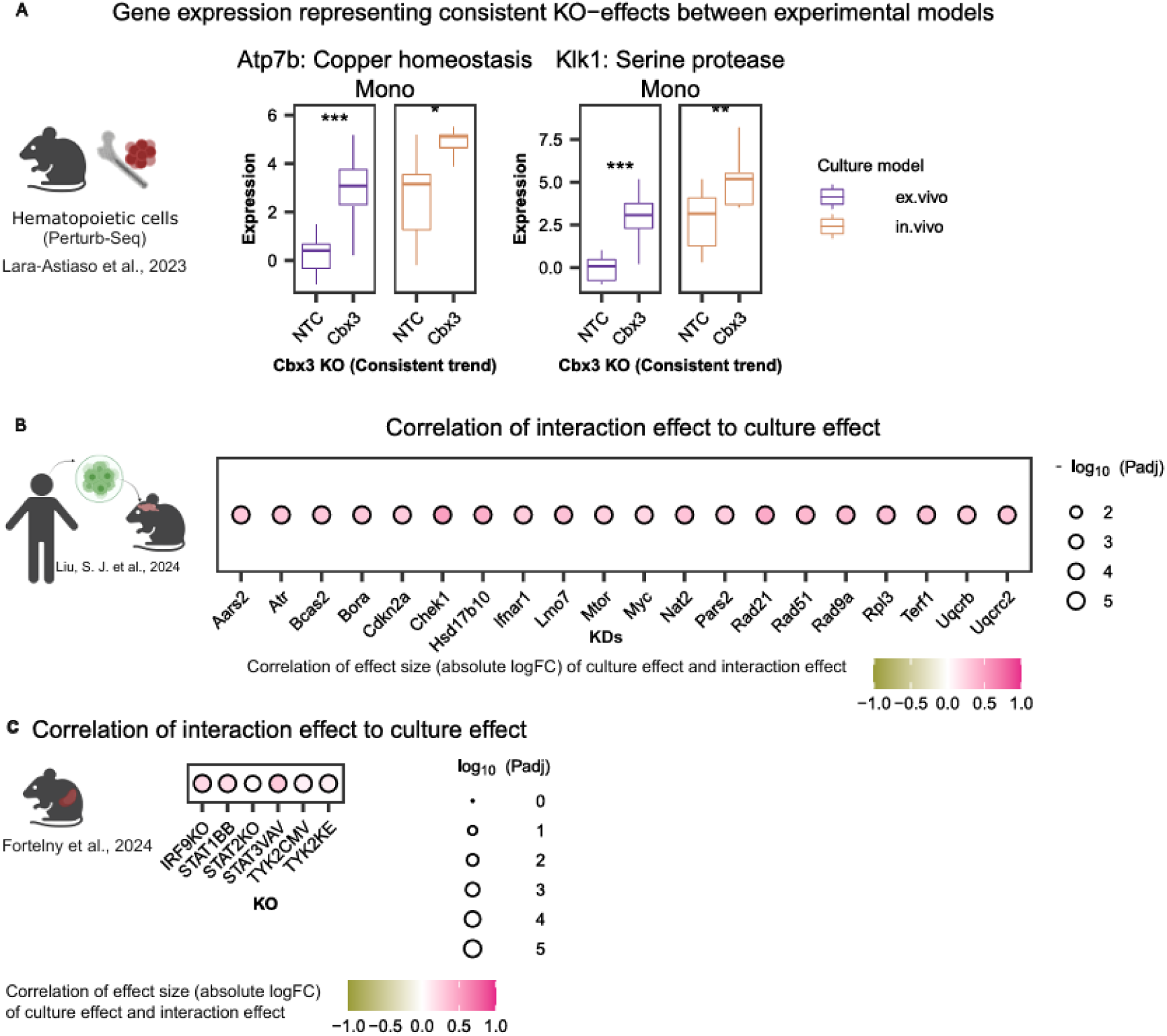
**A,**Gene expression representing consistent perturbation effects between experimental models. **B,** Correlation of effect size (absolute logFC) of culture effect and interaction effect in perturb-seq dataset of glioblastoma(left) and splenic T-cells from genetic knockouts of JAK-STAT components(right).

**Extended Data figure 10.**
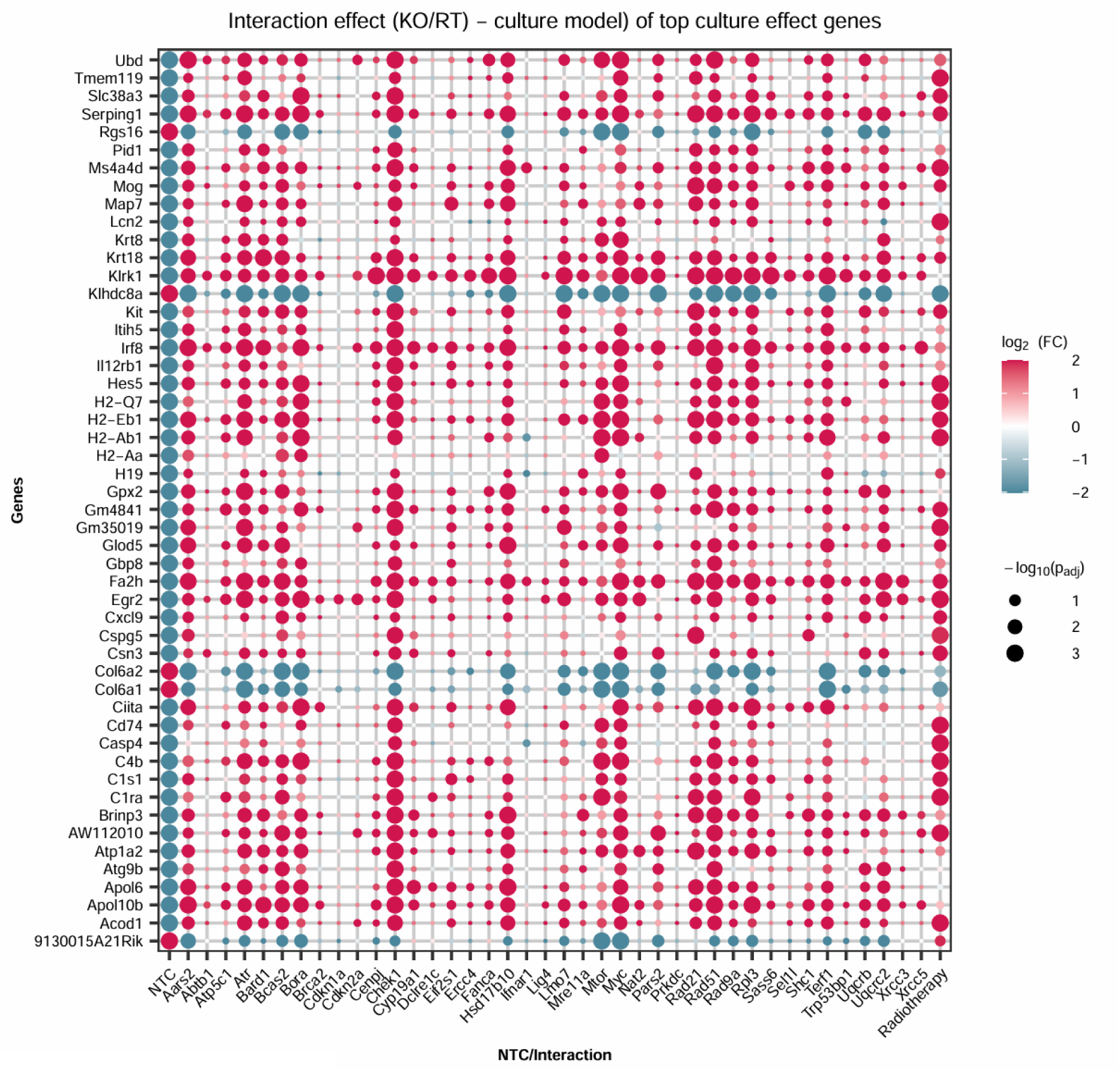
Top 50 significant genes (adjusted p value < 0.05) based on strongest absolute log_2_ fold change for culture effect (NTC) and the log_2_ fold changes of the corresponding genes for interaction effects of KDs and culture model and the interaction effect of radiotherapy and culture effect in NTCs (Ratiotherapy) from the glioblastoma dataset from Liu and colleagues^56^.

**Extended Data figure 11.**
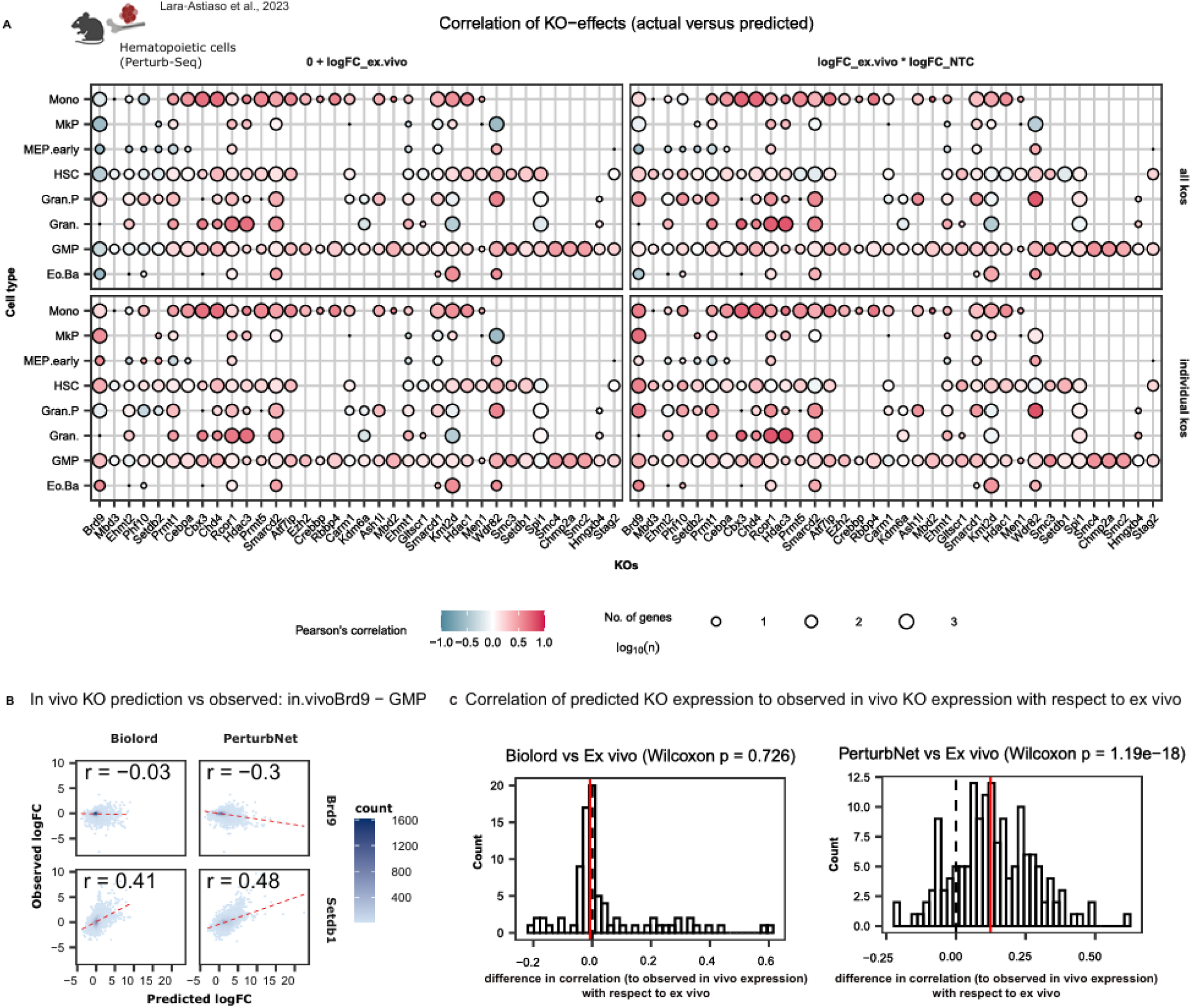
**A,**Pearson’s correlation of actual (*in vivo*) versus predicted log_2_ fold change using the *ex vivo* log₂ fold changes (0 + logFCs) or baseline (NTC) differences (culture effects) and interaction terms of *ex vivo* KO effects and culture effects (*ex.vivo*_logFC * NTC_logFC) to predict the *in vivo* logFCs across all gene when trained across or within knockouts (KO) to predict the gene expression in test set. **B,** Observed versus predicted log_2_ fold change of KO effects of *Brd9* and *Setdb1* in Granulocyte-macrophage progenitor cells (GMP). **C,** Difference in correlation of predicted vs. observed *in vivo* KO expression relative to *ex vivo* KO expression. The red line denotes the median Δ correlation compared to the *ex vivo* baseline.

## Notes

### Competing Interest Statement

The authors have declared no competing interest.

### Summary of Updates

Added/ revised figures and supplementary figures for validation and better clarity, new datasets included

