## Supplementary Table 14 for "Differences between *in vivo* and *ex vivo* models drive divergent perturbation outcomes"

| coef | gene | logFC | adj.P.Val | celltype |
|---|---|---|---|---|
| culture_effect | Ppan | -0.446 | 1.00E-03 | glio |
| Interaction_genotype_Atr | Arsj | -4.305 | 1.00E-03 | glio |
| Interaction_genotype_Mtor | Gm13684 | -3.39 | 1.00E-03 | glio |
| Interaction_genotype_Mtor | Miga2 | -2.568 | 1.00E-03 | glio |
| Interaction_genotype_Mtor | Ewsr1 | 1.507 | 1.00E-03 | glio |
| Interaction_genotype_Myc | Rgs4 | -2.729 | 1.00E-03 | glio |
| Interaction_genotype_Myc | Scaf11 | 1.353 | 1.00E-03 | glio |
| Interaction_genotype_Terf1 | Klrk1 | 3.941 | 1.00E-03 | glio |
| interaction_RT | Pex13 | 0.966 | 1.00E-03 | glio |
| interaction_RT | Rida | 0.987 | 1.00E-03 | glio |
| Aars2_in.vivo_noRT | 1700019D03R | 2.515 | 1.00E-03 | glio |
| Atp5c1_in.vivo_noRT | Gm43672 | 2.668 | 1.00E-03 | glio |
| Atp5c1_in.vivo_noRT | Slc4a9 | 2.136 | 1.00E-03 | glio |
| Atr_in.vivo_noRT | Ppp1r13l | 2.354 | 1.00E-03 | glio |
| Atr_in.vivo_noRT | Bbs9 | 2.106 | 1.00E-03 | glio |
| Atr_in.vivo_noRT | Tagap1 | 1.901 | 1.00E-03 | glio |
| Atr_in.vivo_noRT | Vps33b | 1.825 | 1.00E-03 | glio |
| Bcas2_in.vivo_noRT | Jazf1 | 2.302 | 1.00E-03 | glio |
| Bcas2_in.vivo_noRT | Pfn1 | -0.642 | 1.00E-03 | glio |
| Bora_in.vivo_noRT | Gm10553 | 2.39 | 1.00E-03 | glio |
| Hsd17b10_in.vivo_noRT | Col17a1 | 2.616 | 1.00E-03 | glio |
| Hsd17b10_in.vivo_noRT | Fads3 | 1.537 | 1.00E-03 | glio |
| culture_effect | Mindy3 | 0.723 | 1.00E-04 | glio |
| culture_effect | Ankrd26 | 0.815 | 1.00E-04 | glio |
| Interaction_genotype_Mtor | Snhg4.1 | -3.969 | 1.00E-04 | glio |
| Interaction_genotype_Myc | Fam228a | -4.488 | 1.00E-04 | glio |
| Interaction_genotype_Myc | Gm44699 | -3.96 | 1.00E-04 | glio |
| Interaction_genotype_Myc | Gm13986 | -4.581 | 1.00E-04 | glio |
| Interaction_genotype_Myc | Gm12500 | -4.209 | 1.00E-04 | glio |
| Interaction_genotype_Myc | Gm18280 | -3.259 | 1.00E-04 | glio |
| Interaction_genotype_Myc | Cbx2 | -4.045 | 1.00E-04 | glio |
| interaction_RT | Actn4 | -0.782 | 1.00E-04 | glio |
| Aars2_ex.vivo_noRT | Gm36043 | 3.293 | 1.00E-04 | glio |
| Aars2_ex.vivo_noRT | Tmem52 | 3.293 | 1.00E-04 | glio |
| Aars2_ex.vivo_noRT | Prkcq | 3.293 | 1.00E-04 | glio |
| Atr_in.vivo_noRT | Gm3448 | 2.312 | 1.00E-04 | glio |
| Bcas2_in.vivo_noRT | Wfikkn2 | 2.833 | 1.00E-04 | glio |
| Bcas2_ex.vivo_noRT | Gm12800 | 2.742 | 1.00E-04 | glio |
| Bcas2_ex.vivo_noRT | Trmt61b | 2.742 | 1.00E-04 | glio |
| Bora_in.vivo_noRT | Dhrs13os | 3.1 | 1.00E-04 | glio |
| Bora_in.vivo_noRT | Kif5a | 3.006 | 1.00E-04 | glio |
| Bora_in.vivo_noRT | Sapcd1 | 2.796 | 1.00E-04 | glio |
| Bora_in.vivo_noRT | Adipor1 | -1.272 | 1.00E-04 | glio |
| Chek1_in.vivo_noRT | Abcc10 | 2.852 | 1.00E-04 | glio |
| Chek1_in.vivo_noRT | Zfp974 | 2.912 | 1.00E-04 | glio |
| Interaction_genotype_Mtor | 4930594M22F | -5.318 | 1.00E-05 | glio |
| Interaction_genotype_Mtor | Gm47872 | -5.031 | 1.00E-05 | glio |
| Interaction_genotype_Mtor | Birc6 | 2.366 | 1.00E-05 | glio |
| interaction_RT | Sp140 | 1.525 | 1.00E-05 | glio |
| Aars2_in.vivo_noRT | Plxnb1 | 3.195 | 1.00E-05 | glio |
| Atr_in.vivo_noRT | Moap1 | 2.822 | 1.00E-05 | glio |

| | | | | |
|---|---|---|---|---|
| Atr_in.vivo_noRT | Mutyh | 3.08 | 1.00E-05 | glio |
| Atr_ex.vivo_noRT | H2-Eb2 | 3.164 | 1.00E-05 | glio |
| Chek1_ex.vivo_noRT | Gm47872 | 3.647 | 1.00E-05 | glio |
| Hsd17b10_in.vivo_noRT | Nuak2 | 3.429 | 1.00E-05 | glio |
| Interaction_genotype_Myc | Asphd2 | -5.454 | 1.00E-06 | glio |
| Atr_in.vivo_noRT | Ccdc106 | 3.679 | 1.00E-06 | glio |
| Atr_in.vivo_noRT | Cd93 | 3.5 | 1.00E-06 | glio |
| Atr_in.vivo_noRT | Armc7 | 2.949 | 1.00E-06 | glio |
| Atr_in.vivo_noRT | 8430429K09R | 2.742 | 1.00E-06 | glio |
| Bora_in.vivo_noRT | Trnp1 | 3.495 | 1.00E-06 | glio |
| Atr_in.vivo_noRT | Saxo2 | 3.61 | 1.00E-07 | glio |
| Bcas2_in.vivo_noRT | BC030343 | 3.77 | 1.00E-07 | glio |
| Bcas2_ex.vivo_noRT | Fabp6 | 3.885 | 1.00E-07 | glio |
| Hsd17b10_in.vivo_noRT | Ric3 | 3.356 | 1.00E-07 | glio |
| culture_effect | Pakap.1 | 2.05 | 1.00E-08 | glio |
| Interaction_genotype_Mtor | Serp2 | -6.055 | 1.00E-08 | glio |
| Aars2_in.vivo_noRT | Arhgef19 | 4.148 | 1.00E-08 | glio |
| Bcas2_in.vivo_noRT | 5033403F01Ri | 3.881 | 1.00E-08 | glio |
| Bora_in.vivo_noRT | Sbk3 | 3.768 | 1.00E-08 | glio |
| Chek1_in.vivo_noRT | Vcam1 | 5.057 | 1.00E-08 | glio |
| Chek1_in.vivo_noRT | Gm44096 | 3.773 | 1.00E-08 | glio |
| Aars2_in.vivo_noRT | Hist1h4f | 4.535 | 1.00E-09 | glio |
| Atr_in.vivo_noRT | S100a7a | 4.305 | 1.00E-09 | glio |
| culture_effect | Hoxaas3 | 2.224 | 1.00E-10 | glio |
| Aars2_in.vivo_noRT | Aqp5 | 5.063 | 1.00E-10 | glio |
| Atr_in.vivo_noRT | Gpr37l1 | 4.419 | 1.00E-10 | glio |
| Atr_in.vivo_noRT | Olfr1189 | 3.878 | 1.00E-10 | glio |
| Chek1_in.vivo_noRT | Pard3bos1 | 4.047 | 1.00E-10 | glio |
| culture_effect | Gm50203 | 2.912 | 1.00E-11 | glio |
| Aars2_in.vivo_noRT | Neurl1b | 4.801 | 1.00E-11 | glio |
| Chek1_in.vivo_noRT | E030042O20R | 5.044 | 1.00E-11 | glio |
| Chek1_ex.vivo_noRT | Gm36037 | 4.447 | 1.00E-12 | glio |
| Chek1_ex.vivo_noRT | Slc45a1 | 4.443 | 1.00E-12 | glio |
| Hsd17b10_in.vivo_noRT | Gm13398 | 3.616 | 1.00E-12 | glio |
| Atr_in.vivo_noRT | Olfr279 | 5.004 | 1.00E-13 | glio |
| Chek1_in.vivo_noRT | Ceacam16 | 4.386 | 1.00E-13 | glio |
| Bcas2_in.vivo_noRT | Iqch | 4.543 | 1.00E-14 | glio |
| culture_effect | Irgm2 | -1.539 | 1.00E-15 | glio |
| Bora_in.vivo_noRT | Gm45406 | 4.414 | 1.00E-16 | glio |
| Atr_in.vivo_noRT | Pmp2 | 5.533 | 1.00E-18 | glio |
| Atr_in.vivo_noRT | Tmem74b | 5.269 | 1.00E-19 | glio |
| culture_effect | Svbp | 1.408 | 1.00E-24 | glio |
| culture_effect | Ckmt1 | -1.754 | 1.01E-03 | glio |
| culture_effect | Gcdh | -0.986 | 1.01E-03 | glio |
| culture_effect | Kif1bp | -0.731 | 1.01E-03 | glio |
| culture_effect | Prrg4 | 0.872 | 1.01E-03 | glio |
| culture_effect | Ppp2r2a | -0.439 | 1.01E-03 | glio |
| culture_effect | Znrd2 | 0.42 | 1.01E-03 | glio |
| Interaction_genotype_Aars2 | Gm49694 | -4.102 | 1.01E-03 | glio |
| Interaction_genotype_Atr | 4930588A03R | -2.769 | 1.01E-03 | glio |
| Interaction_genotype_Bora | Gm44752 | -3.911 | 1.01E-03 | glio |
| Interaction_genotype_Cdkn2a | Zscan4-ps2 | 3.615 | 1.01E-03 | glio |

| | | | | |
|---|---|---|---|---|
| Interaction_genotype_Cdkn2a | Ablim2 | 3.368 | 1.01E-03 | glio |
| Interaction_genotype_Mtor | Cxcl1 | -4.099 | 1.01E-03 | glio |
| Interaction_genotype_Mtor | Fes | -3.786 | 1.01E-03 | glio |
| Interaction_genotype_Mtor | Zfp39 | -3.832 | 1.01E-03 | glio |
| Interaction_genotype_Mtor | Slc25a29 | -3.465 | 1.01E-03 | glio |
| Interaction_genotype_Mtor | Bst1 | -3.308 | 1.01E-03 | glio |
| Interaction_genotype_Mtor | 2810408I11Ri | -2.683 | 1.01E-03 | glio |
| Interaction_genotype_Mtor | Emc8 | 1.578 | 1.01E-03 | glio |
| Interaction_genotype_Myc | Hoxa13 | -3.895 | 1.01E-03 | glio |
| Interaction_genotype_Myc | Ank3 | -2.94 | 1.01E-03 | glio |
| Interaction_genotype_Myc | Ldlrad4 | -3.141 | 1.01E-03 | glio |
| Interaction_genotype_Myc | Adnp2 | -2.783 | 1.01E-03 | glio |
| Interaction_genotype_Myc | Exoc6b | -2.79 | 1.01E-03 | glio |
| interaction_RT | Cln3 | 1.378 | 1.01E-03 | glio |
| interaction_RT | Fasn | -0.826 | 1.01E-03 | glio |
| Aars2_in.vivo_noRT | Zfp169 | 2.262 | 1.01E-03 | glio |
| Aars2_in.vivo_noRT | Ddx18 | -1.241 | 1.01E-03 | glio |
| Aars2_ex.vivo_noRT | 9430041J12Ri | 2.499 | 1.01E-03 | glio |
| Atp5c1_in.vivo_noRT | Rhox7a | 2.668 | 1.01E-03 | glio |
| Atr_in.vivo_noRT | Zfp296 | 2.04 | 1.01E-03 | glio |
| Atr_in.vivo_noRT | Aldh4a1 | 2.086 | 1.01E-03 | glio |
| Atr_in.vivo_noRT | Idi1 | -1.841 | 1.01E-03 | glio |
| Bard1_ex.vivo_noRT | Olfr543 | 2.419 | 1.01E-03 | glio |
| Bard1_ex.vivo_noRT | Cfh | 2.415 | 1.01E-03 | glio |
| Bard1_ex.vivo_noRT | Gm28578 | 3.322 | 1.01E-03 | glio |
| Bcas2_in.vivo_noRT | Nop2 | -1.236 | 1.01E-03 | glio |
| Bcas2_ex.vivo_noRT | Apol6 | 2.737 | 1.01E-03 | glio |
| Bora_in.vivo_noRT | Gm14326 | 2.428 | 1.01E-03 | glio |
| Bora_in.vivo_noRT | Hist1h4c | 2.504 | 1.01E-03 | glio |
| Bora_in.vivo_noRT | Fhad1 | 2.211 | 1.01E-03 | glio |
| Bora_in.vivo_noRT | Gm16845 | 2.265 | 1.01E-03 | glio |
| Bora_in.vivo_noRT | Insrr | 2.16 | 1.01E-03 | glio |
| Bora_in.vivo_noRT | Tnfrsf9 | 2.113 | 1.01E-03 | glio |
| Bora_in.vivo_noRT | Bcl2l1 | -1.341 | 1.01E-03 | glio |
| Bora_in.vivo_noRT | Dock9 | -1.443 | 1.01E-03 | glio |
| Bora_in.vivo_noRT | Smarca4 | -1.281 | 1.01E-03 | glio |
| Bora_in.vivo_noRT | Eif3a | -0.555 | 1.01E-03 | glio |
| Bora_ex.vivo_noRT | Gm13449 | 2.414 | 1.01E-03 | glio |
| Bora_ex.vivo_noRT | Gm19276 | 2.188 | 1.01E-03 | glio |
| Bora_ex.vivo_noRT | Sap30bpos | 2.808 | 1.01E-03 | glio |
| Bora_ex.vivo_noRT | Ms4a4d | 3.201 | 1.01E-03 | glio |
| Chek1_in.vivo_noRT | Klhl8 | 2.258 | 1.01E-03 | glio |
| Chek1_in.vivo_noRT | Klhl32 | 2.088 | 1.01E-03 | glio |
| Chek1_ex.vivo_noRT | Krt16 | 3.747 | 1.01E-03 | glio |
| Chek1_ex.vivo_noRT | Abi3bp | 3.072 | 1.01E-03 | glio |
| Eif2s1_in.vivo_noRT | Gm12703 | 1.877 | 1.01E-03 | glio |
| Eif2s1_in.vivo_noRT | Akr1c18 | 2.143 | 1.01E-03 | glio |
| Eif2s1_in.vivo_noRT | 4921504E06Ri | 2.143 | 1.01E-03 | glio |
| Eif2s1_in.vivo_noRT | 4931431C16R | 2.143 | 1.01E-03 | glio |
| Eif2s1_in.vivo_noRT | Maged2 | -0.899 | 1.01E-03 | glio |
| Hsd17b10_in.vivo_noRT | Pdzk1ip1 | 2.626 | 1.01E-03 | glio |
| Hsd17b10_in.vivo_noRT | Ache | 2.57 | 1.01E-03 | glio |

| | | | | |
|---|---|---|---|---|
| culture_effect | Herpud1 | -0.722 | 1.01E-04 | glio |
| Interaction_genotype_Atr | Gm48435 | -3.157 | 1.01E-04 | glio |
| Interaction_genotype_Atr | Zfand5 | 1.435 | 1.01E-04 | glio |
| Interaction_genotype_Chek1 | Larp1b | 2.888 | 1.01E-04 | glio |
| Interaction_genotype_Mtor | Gm5941 | -4.799 | 1.01E-04 | glio |
| Interaction_genotype_Mtor | Cxcl13 | -4.492 | 1.01E-04 | glio |
| Interaction_genotype_Mtor | Iba57 | -4.56 | 1.01E-04 | glio |
| Interaction_genotype_Mtor | Klhl8 | -4.461 | 1.01E-04 | glio |
| Interaction_genotype_Myc | Thada | -4.028 | 1.01E-04 | glio |
| interaction_RT | Cd81 | 1.706 | 1.01E-04 | glio |
| interaction_RT | Cd83 | 3.288 | 1.01E-04 | glio |
| Aars2_in.vivo_noRT | Rad54b | 2.046 | 1.01E-04 | glio |
| Aars2_ex.vivo_noRT | Slc10a1 | 2.899 | 1.01E-04 | glio |
| Atr_in.vivo_noRT | Gba2 | 2.753 | 1.01E-04 | glio |
| Atr_in.vivo_noRT | Zfp974 | 2.879 | 1.01E-04 | glio |
| Atr_in.vivo_noRT | Otof | 2.119 | 1.01E-04 | glio |
| Atr_in.vivo_noRT | Fscn1 | -1.6 | 1.01E-04 | glio |
| Atr_ex.vivo_noRT | Sult5a1 | 3.736 | 1.01E-04 | glio |
| Bcas2_in.vivo_noRT | Hcrtr1 | 2.869 | 1.01E-04 | glio |
| Bcas2_in.vivo_noRT | Mtbp | -1.798 | 1.01E-04 | glio |
| Bcas2_ex.vivo_noRT | Gm5890 | 2.742 | 1.01E-04 | glio |
| Bcas2_ex.vivo_noRT | Zscan4c | 2.742 | 1.01E-04 | glio |
| Bcas2_ex.vivo_noRT | Speer4a | 2.742 | 1.01E-04 | glio |
| Bcas2_ex.vivo_noRT | BC080695 | 2.742 | 1.01E-04 | glio |
| Bora_in.vivo_noRT | Fam71e1 | 2.514 | 1.01E-04 | glio |
| Bora_in.vivo_noRT | Maged1 | -1.259 | 1.01E-04 | glio |
| Bora_ex.vivo_noRT | Gm33973 | 3.201 | 1.01E-04 | glio |
| Chek1_in.vivo_noRT | Ankrd13d | 2.826 | 1.01E-04 | glio |
| Chek1_in.vivo_noRT | Gm45871 | 2.514 | 1.01E-04 | glio |
| Chek1_in.vivo_noRT | Xbp1 | -1.606 | 1.01E-04 | glio |
| Chek1_ex.vivo_noRT | Hsh2d | 3.301 | 1.01E-04 | glio |
| Hsd17b10_in.vivo_noRT | Gm26542 | 2.765 | 1.01E-04 | glio |
| Hsd17b10_in.vivo_noRT | Cldn19 | 2.834 | 1.01E-04 | glio |
| Hsd17b10_in.vivo_noRT | Pcdhga9 | 2.709 | 1.01E-04 | glio |
| culture_effect | Scel | -1.853 | 1.01E-05 | glio |
| Interaction_genotype_Mtor | Tnfrsf13b | -5.601 | 1.01E-05 | glio |
| Interaction_genotype_Mtor | Il17b | -5.823 | 1.01E-05 | glio |
| Aars2_in.vivo_noRT | A830082N09R | 3.245 | 1.01E-05 | glio |
| Aars2_in.vivo_noRT | Rps6ka4 | 2.898 | 1.01E-05 | glio |
| Aars2_ex.vivo_noRT | Klrk1 | 3.293 | 1.01E-05 | glio |
| Atp5c1_in.vivo_noRT | Speer4a | 2.402 | 1.01E-05 | glio |
| Atr_in.vivo_noRT | Gm42372 | 3.534 | 1.01E-05 | glio |
| Atr_in.vivo_noRT | Zfp583 | 3.241 | 1.01E-05 | glio |
| Bcas2_in.vivo_noRT | Mir142hg | 3.214 | 1.01E-05 | glio |
| Bcas2_in.vivo_noRT | Ppp1r18os | 3.395 | 1.01E-05 | glio |
| Bcas2_in.vivo_noRT | Gm41396 | 3.314 | 1.01E-05 | glio |
| Bora_in.vivo_noRT | Mrpl51 | -0.88 | 1.01E-05 | glio |
| Chek1_in.vivo_noRT | B430010I23Ri | 2.957 | 1.01E-05 | glio |
| Chek1_in.vivo_noRT | 1500002C15R | 3.276 | 1.01E-05 | glio |
| Chek1_in.vivo_noRT | D830025C05R | 3.262 | 1.01E-05 | glio |
| Chek1_ex.vivo_noRT | Platr31 | 4.443 | 1.01E-05 | glio |
| Interaction_genotype_Nat2 | Gm13078 | 4.155 | 1.01E-06 | glio |

| | | | | |
|---|---|---|---|---|
| Aars2_in.vivo_noRT | Spon1 | 3.473 | 1.01E-06 | glio |
| Aars2_in.vivo_noRT | B430010I23Ri | 3.24 | 1.01E-06 | glio |
| Aars2_ex.vivo_noRT | Gm13178 | 3.293 | 1.01E-06 | glio |
| Atr_in.vivo_noRT | AA386476 | 3.503 | 1.01E-06 | glio |
| Bora_in.vivo_noRT | Mmp28 | 3.567 | 1.01E-06 | glio |
| Chek1_in.vivo_noRT | Gm30541 | 3.348 | 1.01E-06 | glio |
| Chek1_in.vivo_noRT | Gm26766 | 3.348 | 1.01E-06 | glio |
| Chek1_in.vivo_noRT | Zfp28 | 3.443 | 1.01E-06 | glio |
| Chek1_in.vivo_noRT | Ddrgk1 | -1.563 | 1.01E-06 | glio |
| culture_effect | Zdhhc9 | -1.683 | 1.01E-07 | glio |
| culture_effect | Tubb2b | -0.837 | 1.01E-07 | glio |
| Aars2_in.vivo_noRT | B230319C09R | 4.017 | 1.01E-07 | glio |
| Aars2_in.vivo_noRT | Gm26519 | 4.14 | 1.01E-07 | glio |
| Aars2_in.vivo_noRT | Rala | -1.79 | 1.01E-07 | glio |
| Atr_in.vivo_noRT | Tnfrsf13b | 3.895 | 1.01E-07 | glio |
| Atr_in.vivo_noRT | Gm10602 | 4.126 | 1.01E-07 | glio |
| Atr_in.vivo_noRT | Hist1h2ad | 3.762 | 1.01E-07 | glio |
| Bora_in.vivo_noRT | Gm47972 | 3.642 | 1.01E-07 | glio |
| Bora_in.vivo_noRT | Gm16576 | 3.805 | 1.01E-07 | glio |
| Chek1_ex.vivo_noRT | Pcdhga11 | 3.871 | 1.01E-07 | glio |
| culture_effect | 5033403F01Ri | 2.851 | 1.01E-08 | glio |
| culture_effect | Brix1 | -0.683 | 1.01E-08 | glio |
| Interaction_genotype_Mtor | Selenov | -5.869 | 1.01E-08 | glio |
| Aars2_in.vivo_noRT | Zmynd15 | 3.889 | 1.01E-08 | glio |
| Bcas2_in.vivo_noRT | B020031M17F | 4.807 | 1.01E-08 | glio |
| Bora_in.vivo_noRT | Aldh1l1 | 4.441 | 1.01E-08 | glio |
| Chek1_in.vivo_noRT | Gm34256 | 4.243 | 1.01E-08 | glio |
| Chek1_in.vivo_noRT | Alpl | 3.859 | 1.01E-08 | glio |
| Chek1_in.vivo_noRT | Ipcef1 | 3.23 | 1.01E-08 | glio |
| Chek1_ex.vivo_noRT | Rab17 | 4.948 | 1.01E-08 | glio |
| Chek1_ex.vivo_noRT | Alpl | 4.842 | 1.01E-08 | glio |
| Chek1_ex.vivo_noRT | Ctss | 4.449 | 1.01E-08 | glio |
| Atr_in.vivo_noRT | Gm3604 | 4.251 | 1.01E-09 | glio |
| Atr_in.vivo_noRT | Zfp652os | 4.228 | 1.01E-09 | glio |
| Bora_in.vivo_noRT | Spns3 | 4.279 | 1.01E-09 | glio |
| Chek1_in.vivo_noRT | Ankef1 | 4.304 | 1.01E-09 | glio |
| Chek1_ex.vivo_noRT | Wfdc18 | 6.308 | 1.01E-09 | glio |
| culture_effect | Bcl3 | -3.235 | 1.01E-10 | glio |
| culture_effect | Snhg5 | -1.095 | 1.01E-10 | glio |
| culture_effect | Hspa8 | -0.958 | 1.01E-10 | glio |
| Atr_in.vivo_noRT | A330040F15Ri | 4.754 | 1.01E-10 | glio |
| Bcas2_in.vivo_noRT | Mfap3l | 4.934 | 1.01E-10 | glio |
| Bora_in.vivo_noRT | Cd52 | 4.279 | 1.01E-10 | glio |
| Chek1_in.vivo_noRT | 9630014M24F | 4.265 | 1.01E-10 | glio |
| Chek1_in.vivo_noRT | Gm30938 | 4.398 | 1.01E-10 | glio |
| Chek1_ex.vivo_noRT | Gm19265 | 4.443 | 1.01E-10 | glio |
| culture_effect | Map2 | -1.519 | 1.01E-11 | glio |
| culture_effect | Psip1 | -0.905 | 1.01E-11 | glio |
| Chek1_ex.vivo_noRT | A330076H08R | 4.842 | 1.01E-11 | glio |
| Hsd17b10_in.vivo_noRT | Ush1g | 3.616 | 1.01E-11 | glio |
| Hsd17b10_in.vivo_noRT | Gm31063 | 3.616 | 1.01E-11 | glio |
| Atr_in.vivo_noRT | BC028471 | 5.001 | 1.01E-12 | glio |

| | | | | |
|---|---|---|---|---|
| Chek1_in.vivo_noRT | Ngb | 4.797 | 1.01E-12 | glio |
| Chek1_in.vivo_noRT | 1110046J04Ri | 5.044 | 1.01E-12 | glio |
| Bora_in.vivo_noRT | Snhg4.1 | 4.414 | 1.01E-13 | glio |
| Chek1_in.vivo_noRT | Gm17276 | 4.253 | 1.01E-13 | glio |
| Chek1_ex.vivo_noRT | Slc16a5 | 4.449 | 1.01E-13 | glio |
| Atr_in.vivo_noRT | Ramp3 | 5.267 | 1.01E-14 | glio |
| Atr_ex.vivo_noRT | Gm5286 | 5.228 | 1.01E-14 | glio |
| Chek1_in.vivo_noRT | Tent5c | 4.118 | 1.01E-14 | glio |
| culture_effect | Gas7 | 1.601 | 1.01E-15 | glio |
| Bora_in.vivo_noRT | Gm14540 | 4.414 | 1.01E-16 | glio |
| Atr_in.vivo_noRT | BC147527 | 5.533 | 1.01E-17 | glio |
| Bora_in.vivo_noRT | Gm13288 | 4.936 | 1.01E-17 | glio |
| culture_effect | Sema6a | -3.382 | 1.01E-18 | glio |
| culture_effect | Olfml3 | -2.562 | 1.01E-18 | glio |
| culture_effect | Serinc5 | -4.353 | 1.01E-19 | glio |
| Aars2_in.vivo_noRT | Hist1h2ag | 4.794 | 1.01E-19 | glio |
| Aars2_in.vivo_noRT | Angptl7 | 4.461 | 1.01E-19 | glio |
| Bora_in.vivo_noRT | Clec4d | 4.936 | 1.01E-19 | glio |
| Atr_in.vivo_noRT | Plcl1 | 5.533 | 1.01E-21 | glio |
| Atr_in.vivo_noRT | Gm12703 | 5.267 | 1.01E-25 | glio |
| culture_effect | Nup133 | -0.778 | 1.02E-03 | glio |
| culture_effect | Rpn1 | -0.465 | 1.02E-03 | glio |
| culture_effect | Sf3b2 | -0.385 | 1.02E-03 | glio |
| culture_effect | Selenom | 0.454 | 1.02E-03 | glio |
| culture_effect | Rbm8a | -0.369 | 1.02E-03 | glio |
| Interaction_genotype_Aars2 | Hsbp1 | 1.111 | 1.02E-03 | glio |
| Interaction_genotype_Atr | Luc7l | 1.706 | 1.02E-03 | glio |
| Interaction_genotype_Mtor | Gm15972 | -3.505 | 1.02E-03 | glio |
| Interaction_genotype_Mtor | Fam228a | -3.904 | 1.02E-03 | glio |
| Interaction_genotype_Mtor | Gm26533 | -3.557 | 1.02E-03 | glio |
| Interaction_genotype_Mtor | Gm49708 | -3.51 | 1.02E-03 | glio |
| Interaction_genotype_Myc | Ninj2 | -3.777 | 1.02E-03 | glio |
| Interaction_genotype_Myc | Ttc38 | -3.619 | 1.02E-03 | glio |
| Interaction_genotype_Myc | Gm45184 | -3.679 | 1.02E-03 | glio |
| Interaction_genotype_Myc | Coro2b | -3.104 | 1.02E-03 | glio |
| Interaction_genotype_Myc | Alg9 | -2.579 | 1.02E-03 | glio |
| Interaction_genotype_Myc | Kif13a | -2.923 | 1.02E-03 | glio |
| Interaction_genotype_Myc | Txnl4a | 1.225 | 1.02E-03 | glio |
| Interaction_genotype_Myc | Phf5a | 1.258 | 1.02E-03 | glio |
| Interaction_genotype_Myc | Atp5md | 1.831 | 1.02E-03 | glio |
| Aars2_in.vivo_noRT | Gm31645 | 2.373 | 1.02E-03 | glio |
| Aars2_in.vivo_noRT | Gm19531 | 2.165 | 1.02E-03 | glio |
| Aars2_in.vivo_noRT | Gm7467 | 2.046 | 1.02E-03 | glio |
| Aars2_in.vivo_noRT | Cep128 | 2.107 | 1.02E-03 | glio |
| Atp5c1_in.vivo_noRT | Gm15743 | 2.405 | 1.02E-03 | glio |
| Atp5c1_in.vivo_noRT | Aqp5 | 2.927 | 1.02E-03 | glio |
| Atp5c1_in.vivo_noRT | Syngap1 | 2.508 | 1.02E-03 | glio |
| Atr_in.vivo_noRT | 5430427N15R | 2.26 | 1.02E-03 | glio |
| Atr_in.vivo_noRT | Hk1os | 2.165 | 1.02E-03 | glio |
| Atr_in.vivo_noRT | Krt8 | -2.77 | 1.02E-03 | glio |
| Atr_ex.vivo_noRT | Gm45620 | 2.846 | 1.02E-03 | glio |
| Atr_ex.vivo_noRT | Ift88 | -2.503 | 1.02E-03 | glio |

| | | | | |
|---|---|---|---|---|
| Atr_ex.vivo_noRT | Atf3 | 2.272 | 1.02E-03 | glio |
| Bard1_ex.vivo_noRT | Rhbg | 2.421 | 1.02E-03 | glio |
| Bcas2_in.vivo_noRT | Zfp583 | 2.514 | 1.02E-03 | glio |
| Bcas2_ex.vivo_noRT | 1700012B07R | 3.515 | 1.02E-03 | glio |
| Chek1_in.vivo_noRT | Etaa1os | 2.294 | 1.02E-03 | glio |
| Chek1_in.vivo_noRT | Ccdc8 | 2.329 | 1.02E-03 | glio |
| Chek1_in.vivo_noRT | Klk9 | 2.397 | 1.02E-03 | glio |
| Chek1_in.vivo_noRT | Gm26881 | 2.087 | 1.02E-03 | glio |
| Chek1_in.vivo_noRT | Rilpl1 | 2.082 | 1.02E-03 | glio |
| Chek1_in.vivo_noRT | Coro1c | -1.35 | 1.02E-03 | glio |
| Chek1_in.vivo_noRT | Rpl26 | -0.599 | 1.02E-03 | glio |
| Chek1_ex.vivo_noRT | Gm46447 | 2.707 | 1.02E-03 | glio |
| Chek1_ex.vivo_noRT | Gm50232 | 2.888 | 1.02E-03 | glio |
| Chek1_ex.vivo_noRT | Itga2b | 2.824 | 1.02E-03 | glio |
| Chek1_ex.vivo_noRT | Srebf2 | 1.65 | 1.02E-03 | glio |
| Eif2s1_in.vivo_noRT | Gast | 2.143 | 1.02E-03 | glio |
| Eif2s1_in.vivo_noRT | Gm36172 | 2.141 | 1.02E-03 | glio |
| Hsd17b10_in.vivo_noRT | 2900042K21R | 2.431 | 1.02E-03 | glio |
| Hsd17b10_in.vivo_noRT | Ctsf | 1.948 | 1.02E-03 | glio |
| Hsd17b10_in.vivo_noRT | Zfp157 | -1.826 | 1.02E-03 | glio |
| culture_effect | Mir142hg | 2.083 | 1.02E-04 | glio |
| culture_effect | Stox2 | 0.693 | 1.02E-04 | glio |
| culture_effect | Pold2 | -0.623 | 1.02E-04 | glio |
| Interaction_genotype_Atr | Gm11973 | -3.867 | 1.02E-04 | glio |
| Interaction_genotype_Atr | Tmem60 | 2.058 | 1.02E-04 | glio |
| Interaction_genotype_Mtor | Gm19605 | -4.528 | 1.02E-04 | glio |
| Interaction_genotype_Mtor | Meak7 | -4.451 | 1.02E-04 | glio |
| Interaction_genotype_Mtor | Serf1 | 1.946 | 1.02E-04 | glio |
| Interaction_genotype_Mtor | S100b | 3.113 | 1.02E-04 | glio |
| Interaction_genotype_Mtor | Ndufc2 | 1.932 | 1.02E-04 | glio |
| Interaction_genotype_Myc | E130307A14Ri | -4.095 | 1.02E-04 | glio |
| Interaction_genotype_Myc | Zfp14 | -4.561 | 1.02E-04 | glio |
| Interaction_genotype_Myc | Palb2 | -3.607 | 1.02E-04 | glio |
| Interaction_genotype_Myc | Sucnr1 | -2.729 | 1.02E-04 | glio |
| Interaction_genotype_Myc | Capg | 2.012 | 1.02E-04 | glio |
| Interaction_genotype_Nat2 | Nts | 5.521 | 1.02E-04 | glio |
| Aars2_in.vivo_noRT | Napsa | 2.822 | 1.02E-04 | glio |
| Aars2_in.vivo_noRT | Arf1 | -0.692 | 1.02E-04 | glio |
| Aars2_in.vivo_noRT | Cfl1 | -0.822 | 1.02E-04 | glio |
| Aars2_ex.vivo_noRT | Myt1 | 2.893 | 1.02E-04 | glio |
| Aars2_ex.vivo_noRT | Auts2 | 3.293 | 1.02E-04 | glio |
| Bard1_ex.vivo_noRT | Cxcl5 | 2.814 | 1.02E-04 | glio |
| Bcas2_ex.vivo_noRT | BB287469 | 2.742 | 1.02E-04 | glio |
| Bora_in.vivo_noRT | E230025N22R | 2.589 | 1.02E-04 | glio |
| Bora_in.vivo_noRT | Aldh1a2 | 2.579 | 1.02E-04 | glio |
| Chek1_in.vivo_noRT | Des | 2.793 | 1.02E-04 | glio |
| Eif2s1_in.vivo_noRT | 4930588A03R | 2.143 | 1.02E-04 | glio |
| Eif2s1_in.vivo_noRT | Gm13856 | 2.143 | 1.02E-04 | glio |
| Eif2s1_in.vivo_noRT | Gm21936 | 2.143 | 1.02E-04 | glio |
| culture_effect | Snx9 | 0.87 | 1.02E-05 | glio |
| Interaction_genotype_Mtor | 9430065F17Ri | -5.178 | 1.02E-05 | glio |
| Interaction_genotype_Mtor | Rab37 | -5.112 | 1.02E-05 | glio |

| | | | | |
|---|---|---|---|---|
| Interaction_genotype_Mtor | Ccr5 | -4.861 | 1.02E-05 | glio |
| Interaction_genotype_Mtor | Mtmr11 | -5.388 | 1.02E-05 | glio |
| Interaction_genotype_Mtor | Speer4a | -3.833 | 1.02E-05 | glio |
| Interaction_genotype_Mtor | Capzb | 2.165 | 1.02E-05 | glio |
| Interaction_genotype_Myc | N4bp2os | -5.122 | 1.02E-05 | glio |
| Interaction_genotype_Myc | Sumo2 | 1.828 | 1.02E-05 | glio |
| Aars2_ex.vivo_noRT | Gm12841 | 2.9 | 1.02E-05 | glio |
| Atr_in.vivo_noRT | Scn2a | 3.12 | 1.02E-05 | glio |
| Atr_in.vivo_noRT | 3010001F23Ri | 2.905 | 1.02E-05 | glio |
| Atr_in.vivo_noRT | Gm48855 | 2.763 | 1.02E-05 | glio |
| Bard1_ex.vivo_noRT | Slc14a2 | 2.814 | 1.02E-05 | glio |
| Bora_ex.vivo_noRT | H2-Eb2 | 3.201 | 1.02E-05 | glio |
| Chek1_in.vivo_noRT | A830005F24Ri | 3.127 | 1.02E-05 | glio |
| culture_effect | Limk2 | 1.366 | 1.02E-06 | glio |
| culture_effect | Ywhag | -0.594 | 1.02E-06 | glio |
| Interaction_genotype_Mtor | Hist1h2ai | -6.099 | 1.02E-06 | glio |
| Aars2_in.vivo_noRT | Gm49077 | 3.553 | 1.02E-06 | glio |
| Atr_in.vivo_noRT | 9130008F23Ri | 3.767 | 1.02E-06 | glio |
| Bcas2_in.vivo_noRT | Ltb4r1 | 3.632 | 1.02E-06 | glio |
| Bcas2_in.vivo_noRT | Stk36 | 3.519 | 1.02E-06 | glio |
| Chek1_in.vivo_noRT | Hspb9 | 3.732 | 1.02E-06 | glio |
| Chek1_in.vivo_noRT | Gm29361 | 3.542 | 1.02E-06 | glio |
| culture_effect | Egr3 | -2.227 | 1.02E-07 | glio |
| culture_effect | Lage3 | 0.78 | 1.02E-07 | glio |
| Aars2_in.vivo_noRT | Lamc2 | 3.679 | 1.02E-07 | glio |
| Atr_in.vivo_noRT | Gm34643 | 3.818 | 1.02E-07 | glio |
| Bcas2_in.vivo_noRT | 2210417A02R | 4.016 | 1.02E-07 | glio |
| Bcas2_in.vivo_noRT | Nhsl2 | 3.689 | 1.02E-07 | glio |
| Bora_in.vivo_noRT | Chd5 | 3.691 | 1.02E-07 | glio |
| Chek1_in.vivo_noRT | 6330408M09F | 4.002 | 1.02E-07 | glio |
| Chek1_in.vivo_noRT | Slc16a6 | 3.86 | 1.02E-07 | glio |
| Chek1_in.vivo_noRT | Pde5a | 3.757 | 1.02E-07 | glio |
| culture_effect | Gng10 | 0.691 | 1.02E-08 | glio |
| Interaction_genotype_Mtor | Crhbp | -6.136 | 1.02E-08 | glio |
| Interaction_genotype_Mtor | Glyat | -4.896 | 1.02E-08 | glio |
| interaction_RT | Lcn2 | 11.024 | 1.02E-08 | glio |
| Atr_in.vivo_noRT | C430042M11F | 4.004 | 1.02E-08 | glio |
| Bcas2_in.vivo_noRT | Necab3 | 3.882 | 1.02E-08 | glio |
| Bcas2_in.vivo_noRT | Prox2os | 3.503 | 1.02E-08 | glio |
| Bora_in.vivo_noRT | Wfdc1 | 3.951 | 1.02E-08 | glio |
| Chek1_in.vivo_noRT | Gm48933 | 4.042 | 1.02E-08 | glio |
| Hsd17b10_in.vivo_noRT | Gm36198 | 3.356 | 1.02E-08 | glio |
| culture_effect | Me1 | 1.097 | 1.02E-09 | glio |
| Aars2_in.vivo_noRT | 2900042K21R | 4.142 | 1.02E-09 | glio |
| Aars2_in.vivo_noRT | BC034090 | 3.953 | 1.02E-09 | glio |
| Atr_in.vivo_noRT | Gm43062 | 4.293 | 1.02E-09 | glio |
| Atr_ex.vivo_noRT | Zscan4c | 3.947 | 1.02E-09 | glio |
| Atr_ex.vivo_noRT | Speer4a | 3.947 | 1.02E-09 | glio |
| Bcas2_in.vivo_noRT | Rspo1 | 4.015 | 1.02E-09 | glio |
| Chek1_in.vivo_noRT | 5033403F01Ri | 4.118 | 1.02E-09 | glio |
| Chek1_in.vivo_noRT | Gm14264 | 4.05 | 1.02E-09 | glio |
| Chek1_in.vivo_noRT | Gm46652 | 3.994 | 1.02E-09 | glio |

| | | | | |
|---|---|---|---|---|
| Chek1_in.vivo_noRT | Gm50295 | 4.252 | 1.02E-09 | glio |
| Atr_in.vivo_noRT | Gm48933 | 4.532 | 1.02E-10 | glio |
| Hsd17b10_in.vivo_noRT | Gm18280 | 3.352 | 1.02E-10 | glio |
| Hsd17b10_in.vivo_noRT | Gm4027 | 3.616 | 1.02E-10 | glio |
| Hsd17b10_in.vivo_noRT | AC154200.1 | 3.616 | 1.02E-10 | glio |
| Aars2_in.vivo_noRT | Pcdhgb6 | 4.142 | 1.02E-11 | glio |
| Aars2_in.vivo_noRT | 9930111J21Ri | 4.147 | 1.02E-11 | glio |
| Atr_in.vivo_noRT | Gpr179 | 4.532 | 1.02E-11 | glio |
| Atr_in.vivo_noRT | Gm48610 | 4.751 | 1.02E-11 | glio |
| Atr_in.vivo_noRT | Milr1 | 4.799 | 1.02E-11 | glio |
| Atr_in.vivo_noRT | Ephx1 | 4.635 | 1.02E-11 | glio |
| Bora_in.vivo_noRT | Gm13054 | 4.41 | 1.02E-12 | glio |
| culture_effect | Tmem234 | 0.762 | 1.02E-14 | glio |
| culture_effect | Eif3e | -0.873 | 1.02E-14 | glio |
| Bcas2_in.vivo_noRT | Gm13713 | 4.275 | 1.02E-14 | glio |
| Chek1_in.vivo_noRT | Gm31152 | 5.033 | 1.02E-15 | glio |
| culture_effect | Clcf1 | 3.2 | 1.02E-17 | glio |
| Bcas2_in.vivo_noRT | Tal2 | 4.544 | 1.02E-18 | glio |
| Bcas2_in.vivo_noRT | Prap1 | 4.807 | 1.02E-19 | glio |
| Atr_in.vivo_noRT | 4732463B04R | 5.269 | 1.02E-21 | glio |
| Chek1_in.vivo_noRT | C330002G04R | 5.044 | 1.02E-21 | glio |
| culture_effect | Glod5 | -7.492 | 1.02E-27 | glio |
| culture_effect | Tm4sf1 | 2.201 | 1.02E-28 | glio |
| culture_effect | Ppef1 | -1.925 | 1.03E-03 | glio |
| culture_effect | E530011L22Ri | -1.613 | 1.03E-03 | glio |
| culture_effect | Atxn7 | -0.939 | 1.03E-03 | glio |
| Interaction_genotype_Aars2 | 1700034I23Ri | -2.831 | 1.03E-03 | glio |
| Interaction_genotype_Aars2 | Zfp287 | -3.263 | 1.03E-03 | glio |
| Interaction_genotype_Aars2 | Ppa1 | 1.358 | 1.03E-03 | glio |
| Interaction_genotype_Atr | Mecom | -3.671 | 1.03E-03 | glio |
| Interaction_genotype_Mtor | A830035O19R | -3.741 | 1.03E-03 | glio |
| Interaction_genotype_Mtor | Gm43066 | -3.821 | 1.03E-03 | glio |
| Interaction_genotype_Mtor | Ankrd9 | -2.767 | 1.03E-03 | glio |
| Interaction_genotype_Mtor | Dusp11 | 1.539 | 1.03E-03 | glio |
| Interaction_genotype_Myc | Ltb | -3.612 | 1.03E-03 | glio |
| Interaction_genotype_Myc | Esp24 | -2.729 | 1.03E-03 | glio |
| Interaction_genotype_Myc | Rapgef3 | -4.133 | 1.03E-03 | glio |
| Interaction_genotype_Myc | Wdr5b | -3.528 | 1.03E-03 | glio |
| Interaction_genotype_Myc | Frs3 | -3.685 | 1.03E-03 | glio |
| Interaction_genotype_Myc | Rab18 | 1.412 | 1.03E-03 | glio |
| Interaction_genotype_Myc | Rpl39 | 1.505 | 1.03E-03 | glio |
| Aars2_in.vivo_noRT | Hyal1 | 2.303 | 1.03E-03 | glio |
| Aars2_in.vivo_noRT | Clcn2 | 2.472 | 1.03E-03 | glio |
| Aars2_in.vivo_noRT | Tmem223 | 2.398 | 1.03E-03 | glio |
| Aars2_in.vivo_noRT | BC028528 | 2.395 | 1.03E-03 | glio |
| Aars2_in.vivo_noRT | Gm45902 | 2.328 | 1.03E-03 | glio |
| Aars2_in.vivo_noRT | Nav3 | -0.93 | 1.03E-03 | glio |
| Aars2_in.vivo_noRT | Timm23 | -0.803 | 1.03E-03 | glio |
| Aars2_in.vivo_noRT | Polr1d | -0.644 | 1.03E-03 | glio |
| Aars2_ex.vivo_noRT | Ankrd33b | 3.293 | 1.03E-03 | glio |
| Aars2_ex.vivo_noRT | Gm15941 | 2.897 | 1.03E-03 | glio |
| Atp5c1_in.vivo_noRT | Postn | 2.555 | 1.03E-03 | glio |

| | | | | |
|---|---|---|---|---|
| Atr_in.vivo_noRT | Gm12166 | 2.406 | 1.03E-03 | glio |
| Atr_in.vivo_noRT | Grb7 | 2.356 | 1.03E-03 | glio |
| Atr_in.vivo_noRT | Mtss1 | 2.439 | 1.03E-03 | glio |
| Atr_in.vivo_noRT | Gon4l | -1.302 | 1.03E-03 | glio |
| Atr_in.vivo_noRT | Resf1 | -1.489 | 1.03E-03 | glio |
| Atr_ex.vivo_noRT | 4833407H14R | 3.422 | 1.03E-03 | glio |
| Atr_ex.vivo_noRT | Bco1 | 2.769 | 1.03E-03 | glio |
| Bcas2_in.vivo_noRT | Fam71f2 | 2.463 | 1.03E-03 | glio |
| Bcas2_in.vivo_noRT | Fam76b | -1.543 | 1.03E-03 | glio |
| Bcas2_in.vivo_noRT | Clk3 | -1.562 | 1.03E-03 | glio |
| Bcas2_in.vivo_noRT | Haus1 | -1.583 | 1.03E-03 | glio |
| Bcas2_ex.vivo_noRT | Boc | 2.349 | 1.03E-03 | glio |
| Bora_in.vivo_noRT | Shroom1 | 2.319 | 1.03E-03 | glio |
| Bora_in.vivo_noRT | 2310022A10R | 1.401 | 1.03E-03 | glio |
| Bora_in.vivo_noRT | Huwe1 | -1.025 | 1.03E-03 | glio |
| Bora_in.vivo_noRT | Anp32b | -0.626 | 1.03E-03 | glio |
| Bora_ex.vivo_noRT | Bglap2 | 3.201 | 1.03E-03 | glio |
| Cenpj_in.vivo_noRT | 5330439K02R | 2.816 | 1.03E-03 | glio |
| Cenpj_ex.vivo_noRT | 4930570E03Ri | 2.591 | 1.03E-03 | glio |
| Cenpj_ex.vivo_noRT | Gm5286 | 2.591 | 1.03E-03 | glio |
| Cenpj_ex.vivo_noRT | H2-Eb2 | 2.591 | 1.03E-03 | glio |
| Chek1_in.vivo_noRT | Myo16 | 2.405 | 1.03E-03 | glio |
| Chek1_in.vivo_noRT | Mpv17 | -1.285 | 1.03E-03 | glio |
| Chek1_in.vivo_noRT | Mrpl15 | -0.858 | 1.03E-03 | glio |
| Chek1_in.vivo_noRT | Ywhae | -0.588 | 1.03E-03 | glio |
| Eif2s1_in.vivo_noRT | Cldn5 | 1.877 | 1.03E-03 | glio |
| culture_effect | Gm36888 | -2.301 | 1.03E-04 | glio |
| culture_effect | Stk11ip | -1.614 | 1.03E-04 | glio |
| Interaction_genotype_Atr | Mob3b | -4.163 | 1.03E-04 | glio |
| Interaction_genotype_Bora | Lsm14a | 2.228 | 1.03E-04 | glio |
| Interaction_genotype_Bora | Rpa2 | 2.541 | 1.03E-04 | glio |
| Interaction_genotype_Bora | Asgr1 | -4.971 | 1.03E-04 | glio |
| Interaction_genotype_Cenpj | Klrk1 | 4.688 | 1.03E-04 | glio |
| Interaction_genotype_Chek1 | 4930478M13F | -5.046 | 1.03E-04 | glio |
| Interaction_genotype_Mtor | Iqcn | -4.37 | 1.03E-04 | glio |
| Interaction_genotype_Mtor | Ubash3a | -6.621 | 1.03E-04 | glio |
| Interaction_genotype_Mtor | Gm26981 | -4.527 | 1.03E-04 | glio |
| Interaction_genotype_Mtor | Fhl3 | -4.602 | 1.03E-04 | glio |
| Interaction_genotype_Mtor | Tax1bp1 | 1.897 | 1.03E-04 | glio |
| Interaction_genotype_Mtor | Rpl13a | 1.322 | 1.03E-04 | glio |
| Interaction_genotype_Myc | Gm48772 | -4.022 | 1.03E-04 | glio |
| Interaction_genotype_Myc | Thbs3 | -4.132 | 1.03E-04 | glio |
| Interaction_genotype_Myc | Cd59a | -4.383 | 1.03E-04 | glio |
| Interaction_genotype_Myc | Sgk3 | -4.387 | 1.03E-04 | glio |
| Interaction_genotype_Myc | Nipa1 | -3.619 | 1.03E-04 | glio |
| interaction_RT | Ube2l6 | 0.991 | 1.03E-04 | glio |
| Aars2_in.vivo_noRT | Hist1h4j | 3.207 | 1.03E-04 | glio |
| Aars2_in.vivo_noRT | Igf2r | 2.572 | 1.03E-04 | glio |
| Aars2_in.vivo_noRT | Zfp280b | 2.195 | 1.03E-04 | glio |
| Aars2_in.vivo_noRT | Pa2g4 | -1.177 | 1.03E-04 | glio |
| Atr_in.vivo_noRT | Magi2 | 2.69 | 1.03E-04 | glio |
| Atr_in.vivo_noRT | Foxc2 | 2.798 | 1.03E-04 | glio |

| | | | | |
|---|---|---|---|---|
| Atr_in.vivo_noRT | Gm32031 | 2.409 | 1.03E-04 | glio |
| Bard1_ex.vivo_noRT | Gm47936 | 2.814 | 1.03E-04 | glio |
| Bcas2_in.vivo_noRT | Mfsd7a | 3.201 | 1.03E-04 | glio |
| Bcas2_in.vivo_noRT | Pcp4l1 | 2.849 | 1.03E-04 | glio |
| Bcas2_in.vivo_noRT | Gm43156 | 2.789 | 1.03E-04 | glio |
| Bcas2_in.vivo_noRT | Gm15910 | 2.687 | 1.03E-04 | glio |
| Bcas2_in.vivo_noRT | Nfu1 | -1.393 | 1.03E-04 | glio |
| Bcas2_in.vivo_noRT | Slc38a1 | -1.175 | 1.03E-04 | glio |
| Bcas2_ex.vivo_noRT | H1foo | 2.742 | 1.03E-04 | glio |
| Bora_in.vivo_noRT | Cxcl2 | 4.603 | 1.03E-04 | glio |
| Bora_in.vivo_noRT | Rbm3os | 2.669 | 1.03E-04 | glio |
| Bora_in.vivo_noRT | Borcs8 | -1.446 | 1.03E-04 | glio |
| Bora_ex.vivo_noRT | Gm34776 | 2.808 | 1.03E-04 | glio |
| Chek1_in.vivo_noRT | B230303O12R | 2.794 | 1.03E-04 | glio |
| Chek1_in.vivo_noRT | Rpl28 | -0.681 | 1.03E-04 | glio |
| Chek1_ex.vivo_noRT | Slc9a9 | 3.57 | 1.03E-04 | glio |
| Chek1_ex.vivo_noRT | Dock6 | 3.263 | 1.03E-04 | glio |
| Eif2s1_in.vivo_noRT | Ankrd22 | 2.143 | 1.03E-04 | glio |
| Eif2s1_in.vivo_noRT | Gm47404 | 2.143 | 1.03E-04 | glio |
| Eif2s1_in.vivo_noRT | Gm31717 | 2.143 | 1.03E-04 | glio |
| Eif2s1_in.vivo_noRT | Nos2 | 2.827 | 1.03E-04 | glio |
| Eif2s1_in.vivo_noRT | Gm14742 | 2.143 | 1.03E-04 | glio |
| Eif2s1_in.vivo_noRT | Ctla4 | 2.143 | 1.03E-04 | glio |
| Eif2s1_in.vivo_noRT | Zscan4-ps3 | 2.143 | 1.03E-04 | glio |
| Eif2s1_in.vivo_noRT | A030014E15Ri | 2.143 | 1.03E-04 | glio |
| Eif2s1_in.vivo_noRT | Gm21731 | 2.143 | 1.03E-04 | glio |
| Eif2s1_in.vivo_noRT | Gm11232 | 2.143 | 1.03E-04 | glio |
| Eif2s1_in.vivo_noRT | Cgnl1 | 2.143 | 1.03E-04 | glio |
| Eif2s1_in.vivo_noRT | Tcp11x2 | 2.675 | 1.03E-04 | glio |
| Eif2s1_in.vivo_noRT | Gm26543 | 2.143 | 1.03E-04 | glio |
| Eif2s1_in.vivo_noRT | Dscaml1 | 2.143 | 1.03E-04 | glio |
| Eif2s1_in.vivo_noRT | 2410017I17Ri | 2.143 | 1.03E-04 | glio |
| Eif2s1_in.vivo_noRT | Pspn | 2.143 | 1.03E-04 | glio |
| Eif2s1_in.vivo_noRT | Ephx3 | 2.998 | 1.03E-04 | glio |
| Eif2s1_in.vivo_noRT | Atf5 | 1.062 | 1.03E-04 | glio |
| culture_effect | Shank1 | -2.561 | 1.03E-05 | glio |
| culture_effect | Wdr45 | -1.054 | 1.03E-05 | glio |
| culture_effect | Col11a1 | -0.53 | 1.03E-05 | glio |
| Interaction_genotype_Mtor | Dok2 | -4.549 | 1.03E-05 | glio |
| Interaction_genotype_Myc | Gm49891 | -5.286 | 1.03E-05 | glio |
| Interaction_genotype_Myc | Acoxl | -4.715 | 1.03E-05 | glio |
| Interaction_genotype_Myc | Snx3 | 1.458 | 1.03E-05 | glio |
| Aars2_in.vivo_noRT | Trim14 | 3.383 | 1.03E-05 | glio |
| Aars2_in.vivo_noRT | Cacnb3 | 2.821 | 1.03E-05 | glio |
| Atr_in.vivo_noRT | Ccdc173 | 2.815 | 1.03E-05 | glio |
| Bcas2_in.vivo_noRT | Ablim3 | 3.247 | 1.03E-05 | glio |
| Chek1_ex.vivo_noRT | Gm28221 | 3.733 | 1.03E-05 | glio |
| culture_effect | Fam189a1 | -2.605 | 1.03E-06 | glio |
| culture_effect | Cdh18 | -2.856 | 1.03E-06 | glio |
| culture_effect | Ky | -1.918 | 1.03E-06 | glio |
| culture_effect | Mrps26 | 0.547 | 1.03E-06 | glio |
| Interaction_genotype_Atr | Resf1 | 2.909 | 1.03E-06 | glio |

| | | | | |
|---|---|---|---|---|
| Interaction_genotype_Mtor | Gpr137b | -5.576 | 1.03E-06 | glio |
| Aars2_ex.vivo_noRT | Esp24 | 3.293 | 1.03E-06 | glio |
| Bcas2_in.vivo_noRT | Syngap1 | 3.347 | 1.03E-06 | glio |
| Bcas2_in.vivo_noRT | Plppr3 | 3.421 | 1.03E-06 | glio |
| Bcas2_in.vivo_noRT | Slc17a9 | 3.352 | 1.03E-06 | glio |
| Bora_in.vivo_noRT | Olfr1369-ps1 | 4.195 | 1.03E-06 | glio |
| Chek1_in.vivo_noRT | Rnf208 | 3.651 | 1.03E-06 | glio |
| Chek1_in.vivo_noRT | Lmod1 | 3.612 | 1.03E-06 | glio |
| Chek1_ex.vivo_noRT | Psca | 3.794 | 1.03E-06 | glio |
| Hsd17b10_in.vivo_noRT | Gm13270 | 3.616 | 1.03E-06 | glio |
| culture_effect | D030040B21R | 2.76 | 1.03E-07 | glio |
| Interaction_genotype_Mtor | Ift20 | 2.859 | 1.03E-07 | glio |
| Interaction_genotype_Mtor | Col8a1 | -6.236 | 1.03E-07 | glio |
| Interaction_genotype_Mtor | Gm30692 | -4.492 | 1.03E-07 | glio |
| Aars2_in.vivo_noRT | Zfp972 | 3.882 | 1.03E-07 | glio |
| Atr_in.vivo_noRT | H60c | 3.94 | 1.03E-07 | glio |
| Atr_in.vivo_noRT | 4930405A21R | 3.42 | 1.03E-07 | glio |
| Bcas2_in.vivo_noRT | Sv2a | 3.899 | 1.03E-07 | glio |
| Bora_in.vivo_noRT | Ptp4a1 | 3.242 | 1.03E-07 | glio |
| Bora_in.vivo_noRT | Gm16759 | 3.903 | 1.03E-07 | glio |
| Bora_in.vivo_noRT | Klf4 | -2.875 | 1.03E-07 | glio |
| Chek1_ex.vivo_noRT | Gm49860 | 3.861 | 1.03E-07 | glio |
| Hsd17b10_in.vivo_noRT | Dapk2 | 2.833 | 1.03E-07 | glio |
| culture_effect | Zdhhc14 | -3.007 | 1.03E-08 | glio |
| culture_effect | Cct3 | -0.681 | 1.03E-08 | glio |
| Interaction_genotype_Mtor | D030040B21R | -6.633 | 1.03E-08 | glio |
| Aars2_in.vivo_noRT | Gm10655 | 4.142 | 1.03E-08 | glio |
| Aars2_in.vivo_noRT | Cd93 | 4.084 | 1.03E-08 | glio |
| Atr_ex.vivo_noRT | Dnaaf3 | 4.746 | 1.03E-08 | glio |
| Bcas2_in.vivo_noRT | Spdye4a | 4.149 | 1.03E-08 | glio |
| Bora_in.vivo_noRT | Gm16094 | 3.631 | 1.03E-08 | glio |
| Bora_in.vivo_noRT | Gm20455 | 4.216 | 1.03E-08 | glio |
| Chek1_in.vivo_noRT | Ceacam19 | 4.011 | 1.03E-08 | glio |
| Chek1_in.vivo_noRT | Hba-a2 | 4.132 | 1.03E-08 | glio |
| culture_effect | Atxn1 | 2.208 | 1.03E-09 | glio |
| Atr_ex.vivo_noRT | BB287469 | 3.947 | 1.03E-09 | glio |
| Bcas2_in.vivo_noRT | Has1 | 3.91 | 1.03E-09 | glio |
| Bora_in.vivo_noRT | Mycl | 4.414 | 1.03E-09 | glio |
| Chek1_in.vivo_noRT | Necab3 | 4.118 | 1.03E-09 | glio |
| Bora_in.vivo_noRT | Gm47232 | 4.414 | 1.03E-10 | glio |
| Aars2_in.vivo_noRT | Platr20 | 5.063 | 1.03E-11 | glio |
| Bora_in.vivo_noRT | Serpina3g | 4.263 | 1.03E-11 | glio |
| culture_effect | Bco1 | -3.818 | 1.03E-12 | glio |
| Atr_in.vivo_noRT | Lrfn1 | 4.891 | 1.03E-12 | glio |
| Atr_in.vivo_noRT | 4933406C10R | 4.497 | 1.03E-12 | glio |
| Chek1_in.vivo_noRT | Gm6566 | 4.118 | 1.03E-12 | glio |
| Atr_in.vivo_noRT | 2310067P03R | 5.011 | 1.03E-13 | glio |
| Bora_in.vivo_noRT | Gm16751 | 4.42 | 1.03E-13 | glio |
| Atr_in.vivo_noRT | Ntn1 | 4.632 | 1.03E-14 | glio |
| Atr_in.vivo_noRT | Lce1g | 5.533 | 1.03E-14 | glio |
| Chek1_in.vivo_noRT | Nos2 | 4.784 | 1.03E-14 | glio |
| culture_effect | Tmem119 | -5.614 | 1.03E-15 | glio |

| | | | | |
|---|---|---|---|---|
| Aars2_in.vivo_noRT | A330032B11R | 5.06 | 1.03E-15 | glio |
| Atr_in.vivo_noRT | Gm42918 | 4.891 | 1.03E-15 | glio |
| Bcas2_in.vivo_noRT | 1700017M07F | 4.807 | 1.03E-15 | glio |
| Aars2_in.vivo_noRT | Npy1r | 4.682 | 1.03E-17 | glio |
| Bora_in.vivo_noRT | Edn2 | 4.936 | 1.03E-23 | glio |
| Chek1_in.vivo_noRT | Gm5788 | 5.044 | 1.03E-23 | glio |
| culture_effect | Tent5a | -0.848 | 1.04E-03 | glio |
| culture_effect | Diaph3 | 0.734 | 1.04E-03 | glio |
| culture_effect | Usp9x | -0.497 | 1.04E-03 | glio |
| Interaction_genotype_Aars2 | Gm17399 | -4.29 | 1.04E-03 | glio |
| Interaction_genotype_Aars2 | Yipf4 | 1.299 | 1.04E-03 | glio |
| Interaction_genotype_Aars2 | Eps8 | 1.435 | 1.04E-03 | glio |
| Interaction_genotype_Aars2 | Swi5 | 0.968 | 1.04E-03 | glio |
| Interaction_genotype_Atr | Gm13449 | -3.157 | 1.04E-03 | glio |
| Interaction_genotype_Bcas2 | Akap3 | -4.442 | 1.04E-03 | glio |
| Interaction_genotype_Mtor | Ly6g6c | -3.632 | 1.04E-03 | glio |
| Interaction_genotype_Mtor | Gm42699 | -3.834 | 1.04E-03 | glio |
| Interaction_genotype_Mtor | Lst1 | -3.889 | 1.04E-03 | glio |
| Interaction_genotype_Mtor | Smim41 | -4.267 | 1.04E-03 | glio |
| Interaction_genotype_Mtor | Fxyd4 | -4.192 | 1.04E-03 | glio |
| Interaction_genotype_Mtor | A530017D24R | -3.691 | 1.04E-03 | glio |
| Interaction_genotype_Mtor | Hsd17b10 | 1.812 | 1.04E-03 | glio |
| Interaction_genotype_Mtor | Capns1 | 1.609 | 1.04E-03 | glio |
| Interaction_genotype_Mtor | Mrpl57 | 1.717 | 1.04E-03 | glio |
| Interaction_genotype_Myc | Mfsd4b4 | -3.372 | 1.04E-03 | glio |
| Interaction_genotype_Myc | Btc | -4.482 | 1.04E-03 | glio |
| Interaction_genotype_Myc | Rtl8a | -2.963 | 1.04E-03 | glio |
| Interaction_genotype_Terf1 | Gm11457 | -4.528 | 1.04E-03 | glio |
| Aars2_in.vivo_noRT | Zbtb4 | 2.235 | 1.04E-03 | glio |
| Aars2_in.vivo_noRT | Dzip1l | 1.856 | 1.04E-03 | glio |
| Aars2_in.vivo_noRT | Camk2n2 | 1.945 | 1.04E-03 | glio |
| Aars2_ex.vivo_noRT | Gm45235 | 2.499 | 1.04E-03 | glio |
| Atp5c1_in.vivo_noRT | Gm35696 | 2.408 | 1.04E-03 | glio |
| Atp5c1_in.vivo_noRT | Rftn2 | 2.277 | 1.04E-03 | glio |
| Atp5c1_in.vivo_noRT | Cxcl1 | 2.668 | 1.04E-03 | glio |
| Atr_in.vivo_noRT | Ntn3 | 2.195 | 1.04E-03 | glio |
| Atr_in.vivo_noRT | Tcf20 | -1.477 | 1.04E-03 | glio |
| Atr_in.vivo_noRT | Xpo1 | -1.258 | 1.04E-03 | glio |
| Atr_in.vivo_noRT | Rpl13a | -0.644 | 1.04E-03 | glio |
| Atr_ex.vivo_noRT | Map3k5 | 2.764 | 1.04E-03 | glio |
| Atr_ex.vivo_noRT | Gm16853 | 2.982 | 1.04E-03 | glio |
| Bcas2_in.vivo_noRT | Gm34455 | 3.042 | 1.04E-03 | glio |
| Bcas2_in.vivo_noRT | Pcdhb16 | 2.581 | 1.04E-03 | glio |
| Bcas2_in.vivo_noRT | Tmem53 | 2.547 | 1.04E-03 | glio |
| Bcas2_in.vivo_noRT | Sulf2 | -1.271 | 1.04E-03 | glio |
| Bcas2_in.vivo_noRT | Srsf11 | -0.853 | 1.04E-03 | glio |
| Bcas2_in.vivo_noRT | Anp32b | -0.621 | 1.04E-03 | glio |
| Bora_in.vivo_noRT | Fastkd1 | 2.139 | 1.04E-03 | glio |
| Bora_ex.vivo_noRT | Kcnd1 | 2.802 | 1.04E-03 | glio |
| Bora_ex.vivo_noRT | Ric3 | 2.808 | 1.04E-03 | glio |
| Cenpj_in.vivo_noRT | Ube4bos1 | 2.288 | 1.04E-03 | glio |
| Chek1_in.vivo_noRT | Gpr161 | 2.428 | 1.04E-03 | glio |

| | | | | |
|---|---|---|---|---|
| Chek1_in.vivo_noRT | Ypel4 | 2.209 | 1.04E-03 | glio |
| Chek1_in.vivo_noRT | Depdc7 | -1.666 | 1.04E-03 | glio |
| Hsd17b10_in.vivo_noRT | Gm44096 | 2.346 | 1.04E-03 | glio |
| Hsd17b10_in.vivo_noRT | Scgb1a1 | 2.506 | 1.04E-03 | glio |
| culture_effect | Gm15050 | 1.267 | 1.04E-04 | glio |
| culture_effect | Tgm2 | -1.183 | 1.04E-04 | glio |
| Interaction_genotype_Aars2 | Btf3 | 1.011 | 1.04E-04 | glio |
| Interaction_genotype_Mtor | Gm49123 | -4.857 | 1.04E-04 | glio |
| Interaction_genotype_Mtor | Rpl3 | 1.326 | 1.04E-04 | glio |
| Interaction_genotype_Myc | Cnn1 | -5.411 | 1.04E-04 | glio |
| Interaction_genotype_Myc | Nin | -3.399 | 1.04E-04 | glio |
| interaction_RT | Tiam1 | -1.387 | 1.04E-04 | glio |
| interaction_RT | Gm44386 | 3.607 | 1.04E-04 | glio |
| Aars2_in.vivo_noRT | Ppat | -1.502 | 1.04E-04 | glio |
| Atr_in.vivo_noRT | Pcx | 2.674 | 1.04E-04 | glio |
| Atr_in.vivo_noRT | Npr2 | 3.122 | 1.04E-04 | glio |
| Atr_in.vivo_noRT | Cacnb3 | 2.491 | 1.04E-04 | glio |
| Atr_in.vivo_noRT | Pcdhga6 | 2.558 | 1.04E-04 | glio |
| Atr_in.vivo_noRT | Lipt2 | 2.222 | 1.04E-04 | glio |
| Atr_in.vivo_noRT | Pold2 | -1.543 | 1.04E-04 | glio |
| Atr_in.vivo_noRT | Rpl27 | -0.837 | 1.04E-04 | glio |
| Atr_ex.vivo_noRT | Gm36757 | 3.562 | 1.04E-04 | glio |
| Bard1_ex.vivo_noRT | Gm14540 | 2.818 | 1.04E-04 | glio |
| Bora_in.vivo_noRT | Tc2n | 2.849 | 1.04E-04 | glio |
| Bora_in.vivo_noRT | Gm28417 | 2.616 | 1.04E-04 | glio |
| Bora_in.vivo_noRT | Fam111a | -2.229 | 1.04E-04 | glio |
| Chek1_in.vivo_noRT | Nat8l | 2.453 | 1.04E-04 | glio |
| Chek1_ex.vivo_noRT | 4930483J18Ri | 3.166 | 1.04E-04 | glio |
| Chek1_ex.vivo_noRT | Flnc | 3.444 | 1.04E-04 | glio |
| Chek1_ex.vivo_noRT | Adcy7 | 3.311 | 1.04E-04 | glio |
| Hsd17b10_in.vivo_noRT | Tcam1 | 2.965 | 1.04E-04 | glio |
| culture_effect | Zfp36l2 | -0.837 | 1.04E-05 | glio |
| culture_effect | Parp1 | 0.614 | 1.04E-05 | glio |
| Interaction_genotype_Mtor | Spdye4a | -5.52 | 1.04E-05 | glio |
| Interaction_genotype_Mtor | Crb3 | -4.099 | 1.04E-05 | glio |
| Bcas2_in.vivo_noRT | Gm47061 | 3.502 | 1.04E-05 | glio |
| Bcas2_in.vivo_noRT | Gm16048 | 3.371 | 1.04E-05 | glio |
| Bcas2_in.vivo_noRT | Bc1 | 3.309 | 1.04E-05 | glio |
| Bora_in.vivo_noRT | Zfp661 | 3.186 | 1.04E-05 | glio |
| Bora_in.vivo_noRT | Mia3 | -1.871 | 1.04E-05 | glio |
| Chek1_in.vivo_noRT | 3010001F23Ri | 2.926 | 1.04E-05 | glio |
| culture_effect | Sgsm2 | -1.375 | 1.04E-06 | glio |
| culture_effect | Klhl5 | 1.266 | 1.04E-06 | glio |
| culture_effect | Acaa1a | -1.005 | 1.04E-06 | glio |
| Interaction_genotype_Mtor | Slc45a3 | -5.889 | 1.04E-06 | glio |
| interaction_RT | Zfp560 | 1.391 | 1.04E-06 | glio |
| Aars2_in.vivo_noRT | Icam5 | 3.305 | 1.04E-06 | glio |
| Aars2_in.vivo_noRT | D130062J10Ri | 3.113 | 1.04E-06 | glio |
| Atr_ex.vivo_noRT | Vmn1r88 | 3.962 | 1.04E-06 | glio |
| Bora_in.vivo_noRT | Gm43660 | 3.234 | 1.04E-06 | glio |
| Chek1_in.vivo_noRT | Zfpm1 | 3.527 | 1.04E-06 | glio |
| culture_effect | Tceal9 | 0.839 | 1.04E-07 | glio |

| | | | | |
|---|---|---|---|---|
| Aars2_in.vivo_noRT | 9530018F02Ri | 3.941 | 1.04E-07 | glio |
| Bcas2_in.vivo_noRT | Rab3b | 3.651 | 1.04E-07 | glio |
| Bcas2_in.vivo_noRT | Serpina3i | 3.486 | 1.04E-07 | glio |
| Bora_in.vivo_noRT | C230013L11Ri | 4.024 | 1.04E-07 | glio |
| Bora_in.vivo_noRT | Gm46404 | 3.9 | 1.04E-07 | glio |
| Bora_in.vivo_noRT | Rrad | 3.975 | 1.04E-07 | glio |
| Chek1_ex.vivo_noRT | Rnf208 | 4.86 | 1.04E-07 | glio |
| culture_effect | Slc6a9 | -1.143 | 1.04E-08 | glio |
| Aars2_in.vivo_noRT | Gm31805 | 3.879 | 1.04E-08 | glio |
| Atr_in.vivo_noRT | Ggnbp1 | 3.851 | 1.04E-08 | glio |
| Atr_in.vivo_noRT | Gm30790 | 4.017 | 1.04E-08 | glio |
| Atr_in.vivo_noRT | Alox5ap | 4.428 | 1.04E-08 | glio |
| Atr_in.vivo_noRT | Haglr | 4.29 | 1.04E-08 | glio |
| Bcas2_in.vivo_noRT | Gm14264 | 3.813 | 1.04E-08 | glio |
| Chek1_ex.vivo_noRT | Dcn | 4.449 | 1.04E-08 | glio |
| Hsd17b10_in.vivo_noRT | Tmem74b | 3.352 | 1.04E-08 | glio |
| culture_effect | Gm49694 | 3.284 | 1.04E-09 | glio |
| Aars2_in.vivo_noRT | Fgd3 | 4.591 | 1.04E-09 | glio |
| Atr_in.vivo_noRT | Nppc | 4.534 | 1.04E-09 | glio |
| Aars2_in.vivo_noRT | Gtsf1l | 3.697 | 1.04E-10 | glio |
| Bora_in.vivo_noRT | Gm20633 | 4.145 | 1.04E-10 | glio |
| Chek1_ex.vivo_noRT | Ntn1 | 4.842 | 1.04E-10 | glio |
| Chek1_ex.vivo_noRT | Pmch | 4.842 | 1.04E-10 | glio |
| culture_effect | Gm4890 | 1.944 | 1.04E-12 | glio |
| Interaction_genotype_Mtor | Hbq1b | -9.021 | 1.04E-12 | glio |
| Interaction_genotype_Mtor | Folr1 | -7.669 | 1.04E-12 | glio |
| Atr_in.vivo_noRT | Gipc2 | 5.27 | 1.04E-12 | glio |
| Hsd17b10_in.vivo_noRT | Gm266 | 3.616 | 1.04E-12 | glio |
| culture_effect | Cbx4 | -1.679 | 1.04E-13 | glio |
| Aars2_in.vivo_noRT | Gm43672 | 5.327 | 1.04E-13 | glio |
| Atr_in.vivo_noRT | Gm20275 | 4.179 | 1.04E-13 | glio |
| culture_effect | Smad7 | 1.845 | 1.04E-15 | glio |
| Bcas2_in.vivo_noRT | 1700024I08Ri | 4.547 | 1.04E-15 | glio |
| Bora_in.vivo_noRT | Tmem54 | 4.676 | 1.04E-15 | glio |
| Chek1_in.vivo_noRT | Gm12610 | 4.515 | 1.04E-15 | glio |
| Aars2_in.vivo_noRT | Pnliprp2 | 4.548 | 1.04E-16 | glio |
| Atr_in.vivo_noRT | Fgf8 | 5.017 | 1.04E-16 | glio |
| Bcas2_in.vivo_noRT | Akr1c18 | 4.807 | 1.04E-17 | glio |
| Bcas2_in.vivo_noRT | 4921504E06Ri | 4.807 | 1.04E-17 | glio |
| Bcas2_in.vivo_noRT | Zmynd10 | 5.065 | 1.04E-17 | glio |
| Bcas2_in.vivo_noRT | Gm35082 | 4.807 | 1.04E-18 | glio |
| Bora_in.vivo_noRT | Speer4a | 4.67 | 1.04E-19 | glio |
| Bora_in.vivo_noRT | Ccdc80 | 4.936 | 1.04E-19 | glio |
| culture_effect | Sod3 | 5.275 | 1.04E-20 | glio |
| Bora_in.vivo_noRT | Cd7 | 4.672 | 1.04E-20 | glio |
| Aars2_in.vivo_noRT | Sytl4 | 5.06 | 1.04E-22 | glio |
| Atr_in.vivo_noRT | Ube4bos1 | 5.773 | 1.04E-22 | glio |
| Aars2_in.vivo_noRT | Olfr56 | 5.327 | 1.04E-23 | glio |
| Bora_in.vivo_noRT | Gm14207 | 4.936 | 1.04E-23 | glio |
| Aars2_in.vivo_noRT | Gm30794 | 5.327 | 1.04E-24 | glio |
| Atr_in.vivo_noRT | Sema6d | 5.533 | 1.04E-24 | glio |
| culture_effect | Ttc7 | 1.029 | 1.05E-03 | glio |

| | | | | |
|---|---|---|---|---|
| culture_effect | Miga2 | 0.895 | 1.05E-03 | glio |
| culture_effect | Sft2d2 | -0.671 | 1.05E-03 | glio |
| culture_effect | Vps37a | 0.522 | 1.05E-03 | glio |
| Interaction_genotype_Atr | Gm21936 | -2.765 | 1.05E-03 | glio |
| Interaction_genotype_Atr | Gm13856 | -2.763 | 1.05E-03 | glio |
| Interaction_genotype_Mtor | Mro | -3.869 | 1.05E-03 | glio |
| Interaction_genotype_Mtor | Zfp811 | -3.903 | 1.05E-03 | glio |
| Interaction_genotype_Mtor | Dlx4 | -3.746 | 1.05E-03 | glio |
| Interaction_genotype_Mtor | Gm8013 | -4.041 | 1.05E-03 | glio |
| Interaction_genotype_Mtor | Abat | -4.201 | 1.05E-03 | glio |
| Interaction_genotype_Mtor | Gm45494 | -3.62 | 1.05E-03 | glio |
| Interaction_genotype_Mtor | Dsel | -3.444 | 1.05E-03 | glio |
| Interaction_genotype_Mtor | Vangl2 | -2.92 | 1.05E-03 | glio |
| Interaction_genotype_Mtor | Ikbkg | -2.615 | 1.05E-03 | glio |
| Interaction_genotype_Mtor | Sf3b5 | 1.672 | 1.05E-03 | glio |
| Interaction_genotype_Myc | AF067063 | -2.729 | 1.05E-03 | glio |
| Interaction_genotype_Myc | Ddx19b | -3.426 | 1.05E-03 | glio |
| Interaction_genotype_Myc | Zfp90 | -3.202 | 1.05E-03 | glio |
| Interaction_genotype_Myc | Ipp | -3.045 | 1.05E-03 | glio |
| Interaction_genotype_Myc | Ninl | -3.164 | 1.05E-03 | glio |
| Interaction_genotype_Myc | Mmachc | -2.79 | 1.05E-03 | glio |
| Interaction_genotype_Myc | Mapt | -2.884 | 1.05E-03 | glio |
| interaction_RT | Gch1 | 2.088 | 1.05E-03 | glio |
| interaction_RT | Bysl | -0.89 | 1.05E-03 | glio |
| Atp5c1_in.vivo_noRT | Prickle4 | 2.405 | 1.05E-03 | glio |
| Atr_in.vivo_noRT | Ppef1 | 2.205 | 1.05E-03 | glio |
| Atr_in.vivo_noRT | Myo16 | 2.365 | 1.05E-03 | glio |
| Atr_in.vivo_noRT | Gm14167 | 2.097 | 1.05E-03 | glio |
| Atr_in.vivo_noRT | Zfp971 | 2.138 | 1.05E-03 | glio |
| Atr_in.vivo_noRT | Lzts3 | 2.028 | 1.05E-03 | glio |
| Atr_in.vivo_noRT | Scaf11 | -0.93 | 1.05E-03 | glio |
| Atr_ex.vivo_noRT | Gm15972 | 2.764 | 1.05E-03 | glio |
| Bcas2_in.vivo_noRT | Snph | 2.293 | 1.05E-03 | glio |
| Bcas2_in.vivo_noRT | Gsto2 | 2.423 | 1.05E-03 | glio |
| Bcas2_in.vivo_noRT | Pycard | 2.199 | 1.05E-03 | glio |
| Bcas2_ex.vivo_noRT | Gm17203 | 2.342 | 1.05E-03 | glio |
| Bcas2_ex.vivo_noRT | Otof | 2.742 | 1.05E-03 | glio |
| Bcas2_ex.vivo_noRT | Bok | -2.968 | 1.05E-03 | glio |
| Bora_in.vivo_noRT | Nnt | 2.16 | 1.05E-03 | glio |
| Bora_in.vivo_noRT | Zswim8 | -1.562 | 1.05E-03 | glio |
| Bora_ex.vivo_noRT | Serpinb2 | 2.802 | 1.05E-03 | glio |
| Bora_ex.vivo_noRT | Olfr1189 | 2.802 | 1.05E-03 | glio |
| Cenpj_ex.vivo_noRT | Fam166a | 2.591 | 1.05E-03 | glio |
| Cenpj_ex.vivo_noRT | Gm15894 | 2.591 | 1.05E-03 | glio |
| Chek1_in.vivo_noRT | 6720489N17R | 2.651 | 1.05E-03 | glio |
| Chek1_in.vivo_noRT | Gm38414 | 2.637 | 1.05E-03 | glio |
| Chek1_in.vivo_noRT | Tmem68 | -1.347 | 1.05E-03 | glio |
| Chek1_ex.vivo_noRT | Mt4 | 3.862 | 1.05E-03 | glio |
| Chek1_ex.vivo_noRT | Elobl | 3.106 | 1.05E-03 | glio |
| Chek1_ex.vivo_noRT | Gm44096 | 2.778 | 1.05E-03 | glio |
| Chek1_ex.vivo_noRT | Mettl26 | -1.691 | 1.05E-03 | glio |
| Chek1_ex.vivo_noRT | Fbxl12 | 1.861 | 1.05E-03 | glio |

| | | | | |
|---|---|---|---|---|
| Hsd17b10_in.vivo_noRT | Klhl30 | 2.431 | 1.05E-03 | glio |
| Hsd17b10_in.vivo_noRT | Styk1 | 2.551 | 1.05E-03 | glio |
| Hsd17b10_in.vivo_noRT | Mief1 | -1.658 | 1.05E-03 | glio |
| Hsd17b10_in.vivo_noRT | Chpt1 | -1.055 | 1.05E-03 | glio |
| culture_effect | D830039M14I | 2.286 | 1.05E-04 | glio |
| culture_effect | Focad | -1.268 | 1.05E-04 | glio |
| culture_effect | Nprl2 | -1.113 | 1.05E-04 | glio |
| Interaction_genotype_Atr | Gm14399 | -4.837 | 1.05E-04 | glio |
| Interaction_genotype_Bora | Gm39326 | -4.949 | 1.05E-04 | glio |
| Interaction_genotype_Mtor | 4933400F21Ri | -3.308 | 1.05E-04 | glio |
| Interaction_genotype_Mtor | Angptl7 | -3.233 | 1.05E-04 | glio |
| Interaction_genotype_Mtor | E130208F15Ri | -4.977 | 1.05E-04 | glio |
| Interaction_genotype_Mtor | Gm4724 | -4.144 | 1.05E-04 | glio |
| Interaction_genotype_Myc | G430095P16R | -4.423 | 1.05E-04 | glio |
| Interaction_genotype_Myc | Fbxl2 | -3.965 | 1.05E-04 | glio |
| Interaction_genotype_Myc | Gm8098 | -4.378 | 1.05E-04 | glio |
| Interaction_genotype_Myc | Gimap9 | -4.119 | 1.05E-04 | glio |
| Interaction_genotype_Myc | Gm4419 | -3.612 | 1.05E-04 | glio |
| Interaction_genotype_Myc | Dhx35 | -3.535 | 1.05E-04 | glio |
| Interaction_genotype_Myc | G3bp1 | 1.749 | 1.05E-04 | glio |
| Interaction_genotype_Uqcrc2 | Tesc | 3.313 | 1.05E-04 | glio |
| Interaction_genotype_Uqcrc2 | Rassf6 | -4.688 | 1.05E-04 | glio |
| Aars2_in.vivo_noRT | Gm38973 | 2.731 | 1.05E-04 | glio |
| Aars2_in.vivo_noRT | A730017L22Ri | 2.405 | 1.05E-04 | glio |
| Atr_in.vivo_noRT | Erfe | 2.519 | 1.05E-04 | glio |
| Atr_in.vivo_noRT | Yaf2 | -1.245 | 1.05E-04 | glio |
| Atr_in.vivo_noRT | Mrps7 | -1.232 | 1.05E-04 | glio |
| Atr_ex.vivo_noRT | Kcnq2 | 3.164 | 1.05E-04 | glio |
| Bcas2_in.vivo_noRT | Olfr655 | 3.891 | 1.05E-04 | glio |
| Bora_in.vivo_noRT | Slpi | 3.11 | 1.05E-04 | glio |
| Bora_in.vivo_noRT | Hrk | 2.811 | 1.05E-04 | glio |
| Bora_in.vivo_noRT | Mmp23 | 2.662 | 1.05E-04 | glio |
| Bora_in.vivo_noRT | Shank1 | 2.569 | 1.05E-04 | glio |
| Bora_in.vivo_noRT | Gpt | 2.693 | 1.05E-04 | glio |
| Bora_in.vivo_noRT | Plod2 | -1.968 | 1.05E-04 | glio |
| Bora_in.vivo_noRT | Dek | -1.039 | 1.05E-04 | glio |
| Chek1_in.vivo_noRT | Ccn2 | 3.716 | 1.05E-04 | glio |
| Chek1_in.vivo_noRT | Gm17597 | 3.008 | 1.05E-04 | glio |
| Chek1_in.vivo_noRT | Ccnj | 2.862 | 1.05E-04 | glio |
| Chek1_in.vivo_noRT | Eme2 | 2.63 | 1.05E-04 | glio |
| Chek1_in.vivo_noRT | Hddc3 | 2.661 | 1.05E-04 | glio |
| Chek1_in.vivo_noRT | Tsc1 | 2.19 | 1.05E-04 | glio |
| Chek1_ex.vivo_noRT | Ripply3 | 3.278 | 1.05E-04 | glio |
| Chek1_ex.vivo_noRT | Wdr93 | 3.26 | 1.05E-04 | glio |
| Hsd17b10_in.vivo_noRT | Gm16244 | 2.959 | 1.05E-04 | glio |
| Hsd17b10_in.vivo_noRT | 4933429H19R | 2.704 | 1.05E-04 | glio |
| Hsd17b10_in.vivo_noRT | 2810049E08Ri | 2.579 | 1.05E-04 | glio |
| culture_effect | Cfb | -2.239 | 1.05E-05 | glio |
| culture_effect | Tmem203 | 0.993 | 1.05E-05 | glio |
| culture_effect | Tbk1 | -0.898 | 1.05E-05 | glio |
| culture_effect | Oat | -0.699 | 1.05E-05 | glio |
| culture_effect | Pycr2 | -0.69 | 1.05E-05 | glio |

| | | | | |
|---|---|---|---|---|
| Interaction_genotype_Myc | Gm16222 | -5.766 | 1.05E-05 | glio |
| Interaction_genotype_Nat2 | Ankrd22 | 3.724 | 1.05E-05 | glio |
| Atr_in.vivo_noRT | D830025C05R | 3.228 | 1.05E-05 | glio |
| Atr_in.vivo_noRT | Rpl22 | -0.874 | 1.05E-05 | glio |
| Bard1_ex.vivo_noRT | Gm4340 | 2.814 | 1.05E-05 | glio |
| Bard1_ex.vivo_noRT | Gm16239 | 2.814 | 1.05E-05 | glio |
| Bcas2_in.vivo_noRT | Slc46a1 | 3.291 | 1.05E-05 | glio |
| Bcas2_ex.vivo_noRT | Cyp2c23 | 3.727 | 1.05E-05 | glio |
| Bcas2_ex.vivo_noRT | Gm13191 | 2.742 | 1.05E-05 | glio |
| Bcas2_ex.vivo_noRT | 4933416E03Ri | 2.742 | 1.05E-05 | glio |
| Bora_in.vivo_noRT | Cyp2s1 | 3.363 | 1.05E-05 | glio |
| Chek1_in.vivo_noRT | Gm10244 | 3.223 | 1.05E-05 | glio |
| Hsd17b10_in.vivo_noRT | 4933408J17Ri | 2.827 | 1.05E-05 | glio |
| culture_effect | Mrpl30 | 0.464 | 1.05E-06 | glio |
| Interaction_genotype_Mtor | Caprin1 | 2.361 | 1.05E-06 | glio |
| Interaction_genotype_Mtor | Adgrb3 | -5.573 | 1.05E-06 | glio |
| Interaction_genotype_Mtor | Elobl | -6.223 | 1.05E-06 | glio |
| Interaction_genotype_Mtor | Gjb4 | -5.967 | 1.05E-06 | glio |
| Interaction_genotype_Mtor | Gm42695 | -5.816 | 1.05E-06 | glio |
| Interaction_genotype_Mtor | Gm5788 | -4.099 | 1.05E-06 | glio |
| interaction_RT | Akr1b3 | 1.784 | 1.05E-06 | glio |
| Aars2_ex.vivo_noRT | Gm9758 | 3.293 | 1.05E-06 | glio |
| Atp5c1_in.vivo_noRT | AU019990 | 2.668 | 1.05E-06 | glio |
| Atr_ex.vivo_noRT | Gm15533 | 3.164 | 1.05E-06 | glio |
| Bcas2_in.vivo_noRT | 5031434O11R | 3.928 | 1.05E-06 | glio |
| Chek1_in.vivo_noRT | 4930515G01R | 3.161 | 1.05E-06 | glio |
| Chek1_in.vivo_noRT | 9130008F23Ri | 3.789 | 1.05E-06 | glio |
| Chek1_in.vivo_noRT | Arg2 | 3.403 | 1.05E-06 | glio |
| Chek1_ex.vivo_noRT | Gm44983 | 3.556 | 1.05E-06 | glio |
| Hsd17b10_in.vivo_noRT | Lce1g | 3.616 | 1.05E-06 | glio |
| culture_effect | Mir670hg | 3.034 | 1.05E-07 | glio |
| culture_effect | Eif3h | -0.558 | 1.05E-07 | glio |
| Interaction_genotype_Bcas2 | Folr1 | -5.636 | 1.05E-07 | glio |
| Interaction_genotype_Mtor | Ostc | 2.527 | 1.05E-07 | glio |
| Interaction_genotype_Mtor | Gm45620 | -6.165 | 1.05E-07 | glio |
| Interaction_genotype_Mtor | Prrx2 | -6.397 | 1.05E-07 | glio |
| Interaction_genotype_Mtor | Gtsf1 | -5.262 | 1.05E-07 | glio |
| Interaction_genotype_Mtor | Cldn2 | -4.099 | 1.05E-07 | glio |
| Aars2_in.vivo_noRT | Eng | 4.13 | 1.05E-07 | glio |
| Bora_in.vivo_noRT | Gm26533 | 3.826 | 1.05E-07 | glio |
| Chek1_in.vivo_noRT | Jmjd8 | 3.678 | 1.05E-07 | glio |
| Chek1_in.vivo_noRT | Slc7a3 | 3.777 | 1.05E-07 | glio |
| Chek1_ex.vivo_noRT | Cyp4f40 | 4.449 | 1.05E-07 | glio |
| culture_effect | Pdhb | -0.819 | 1.05E-08 | glio |
| Chek1_in.vivo_noRT | Neurl3 | 4.053 | 1.05E-08 | glio |
| Chek1_ex.vivo_noRT | Slc16a4 | 3.987 | 1.05E-08 | glio |
| culture_effect | Bdnf | 2.154 | 1.05E-09 | glio |
| Aars2_in.vivo_noRT | Npb | 4.159 | 1.05E-09 | glio |
| Bora_in.vivo_noRT | 4933429H19R | 4.024 | 1.05E-09 | glio |
| Bora_in.vivo_noRT | Gm16348 | 4.285 | 1.05E-09 | glio |
| Bora_in.vivo_noRT | Gap43 | 4.777 | 1.05E-09 | glio |
| Chek1_in.vivo_noRT | 3930402G23R | 4.657 | 1.05E-10 | glio |

| | | | | |
|---|---|---|---|---|
| Aars2_in.vivo_noRT | Lyz2 | 4.941 | 1.05E-11 | glio |
| Chek1_in.vivo_noRT | Camk2b | 4.522 | 1.05E-11 | glio |
| culture_effect | Id3 | 1.378 | 1.05E-12 | glio |
| Aars2_in.vivo_noRT | Hbq1b | 5.06 | 1.05E-12 | glio |
| Atr_in.vivo_noRT | Ifi207 | 5.27 | 1.05E-12 | glio |
| Bcas2_in.vivo_noRT | AI463229 | 4.162 | 1.05E-12 | glio |
| Hsd17b10_in.vivo_noRT | Ildr1 | 3.616 | 1.05E-12 | glio |
| Atr_in.vivo_noRT | Gm13822 | 5.008 | 1.05E-13 | glio |
| Bora_in.vivo_noRT | Ntn4 | 4.67 | 1.05E-14 | glio |
| Chek1_in.vivo_noRT | Crybb1 | 4.657 | 1.05E-14 | glio |
| Chek1_in.vivo_noRT | 1700026N04R | 4.784 | 1.05E-14 | glio |
| Hsd17b10_in.vivo_noRT | Stfa3 | 3.616 | 1.05E-14 | glio |
| Hsd17b10_in.vivo_noRT | Olfr787 | 3.616 | 1.05E-14 | glio |
| Atr_in.vivo_noRT | Gm49441 | 4.742 | 1.05E-15 | glio |
| Aars2_in.vivo_noRT | Slc1a2 | 5.063 | 1.05E-17 | glio |
| Bcas2_in.vivo_noRT | 4931431C16R | 4.807 | 1.05E-17 | glio |
| Chek1_in.vivo_noRT | Pdgfb | 4.78 | 1.05E-17 | glio |
| Chek1_in.vivo_noRT | Ccser1 | 5.044 | 1.05E-17 | glio |
| culture_effect | Cope | -1.091 | 1.05E-18 | glio |
| Chek1_in.vivo_noRT | A930019D19R | 5.044 | 1.05E-25 | glio |
| culture_effect | Kit | -5.797 | 1.05E-28 | glio |
| Atr_in.vivo_noRT | Gm5415 | 5.533 | 1.05E-28 | glio |
| culture_effect | Dhh | -0.985 | 1.06E-03 | glio |
| Interaction_genotype_Aars2 | Slc25a29 | -3.702 | 1.06E-03 | glio |
| Interaction_genotype_Aars2 | Camk2b | -3.939 | 1.06E-03 | glio |
| Interaction_genotype_Aars2 | Glyat | -2.831 | 1.06E-03 | glio |
| Interaction_genotype_Aars2 | 4921536K21R | -4.026 | 1.06E-03 | glio |
| Interaction_genotype_Aars2 | Gm43156 | -3.996 | 1.06E-03 | glio |
| Interaction_genotype_Aars2 | Hibadh | 1.913 | 1.06E-03 | glio |
| Interaction_genotype_Aars2 | Tiprl | 1.52 | 1.06E-03 | glio |
| Interaction_genotype_Aars2 | Rps5 | 0.982 | 1.06E-03 | glio |
| Interaction_genotype_Atr | Tut4 | 1.777 | 1.06E-03 | glio |
| Interaction_genotype_Atr | Mtx2 | 1.35 | 1.06E-03 | glio |
| Interaction_genotype_Bcas2 | Eng | -4.769 | 1.06E-03 | glio |
| Interaction_genotype_Bora | Gm47204 | -4.425 | 1.06E-03 | glio |
| Interaction_genotype_Chek1 | Nfkbia | 2.439 | 1.06E-03 | glio |
| Interaction_genotype_Mtor | Sptbn4 | -3.711 | 1.06E-03 | glio |
| Interaction_genotype_Mtor | Nfat5 | 1.877 | 1.06E-03 | glio |
| Interaction_genotype_Mtor | Fam104a | 1.348 | 1.06E-03 | glio |
| Interaction_genotype_Myc | Gm2016 | -2.729 | 1.06E-03 | glio |
| Interaction_genotype_Myc | Plekhg3 | -3.49 | 1.06E-03 | glio |
| Interaction_genotype_Myc | Rnf169 | -2.855 | 1.06E-03 | glio |
| Interaction_genotype_Myc | Nup205 | -2.327 | 1.06E-03 | glio |
| Interaction_genotype_Myc | Eps8 | 1.486 | 1.06E-03 | glio |
| Interaction_genotype_Pars2 | D430001F17R | -4.214 | 1.06E-03 | glio |
| Interaction_genotype_Pars2 | 4930412M03F | -4.338 | 1.06E-03 | glio |
| Interaction_genotype_Pars2 | Gm15890 | -3.831 | 1.06E-03 | glio |
| Interaction_genotype_Pars2 | Dntt | -4.671 | 1.06E-03 | glio |
| Interaction_genotype_Pars2 | Psmd3 | 1.315 | 1.06E-03 | glio |
| Aars2_in.vivo_noRT | Abca13 | 2.595 | 1.06E-03 | glio |
| Aars2_in.vivo_noRT | Mlip | 2.322 | 1.06E-03 | glio |
| Aars2_in.vivo_noRT | Lrp4 | 2.369 | 1.06E-03 | glio |

| | | | | |
|---|---|---|---|---|
| Aars2_in.vivo_noRT | 6430590A07R | 1.811 | 1.06E-03 | glio |
| Aars2_in.vivo_noRT | Tomm70a | -1.104 | 1.06E-03 | glio |
| Aars2_in.vivo_noRT | Tmbim6 | -0.839 | 1.06E-03 | glio |
| Atr_in.vivo_noRT | 4931428F04Ri | 1.992 | 1.06E-03 | glio |
| Atr_in.vivo_noRT | mt-Nd2 | -0.759 | 1.06E-03 | glio |
| Atr_ex.vivo_noRT | C4b | 2.585 | 1.06E-03 | glio |
| Bard1_ex.vivo_noRT | Ush1g | 2.415 | 1.06E-03 | glio |
| Bard1_ex.vivo_noRT | Gm36757 | 3.213 | 1.06E-03 | glio |
| Bard1_ex.vivo_noRT | Pcdh15 | 2.814 | 1.06E-03 | glio |
| Bard1_ex.vivo_noRT | Car10 | 2.814 | 1.06E-03 | glio |
| Bcas2_in.vivo_noRT | Gm42699 | 2.257 | 1.06E-03 | glio |
| Bcas2_in.vivo_noRT | Usp49 | 2.131 | 1.06E-03 | glio |
| Bcas2_ex.vivo_noRT | Hrh2 | 2.742 | 1.06E-03 | glio |
| Bora_in.vivo_noRT | A730063M14F | 2.138 | 1.06E-03 | glio |
| Bora_in.vivo_noRT | Fgfr3 | -2.13 | 1.06E-03 | glio |
| Bora_in.vivo_noRT | Pcmtd2 | -1.713 | 1.06E-03 | glio |
| Bora_in.vivo_noRT | Brd2 | -1.025 | 1.06E-03 | glio |
| Chek1_in.vivo_noRT | 3110001I22Ri | 2.316 | 1.06E-03 | glio |
| Chek1_in.vivo_noRT | Riok3 | -1.407 | 1.06E-03 | glio |
| Chek1_in.vivo_noRT | Ppm1g | -1.091 | 1.06E-03 | glio |
| Chek1_ex.vivo_noRT | Ptp4a1.1 | 2.866 | 1.06E-03 | glio |
| Chek1_ex.vivo_noRT | Nxnl2 | 2.684 | 1.06E-03 | glio |
| Chek1_ex.vivo_noRT | Sft2d1 | -1.768 | 1.06E-03 | glio |
| Chek1_ex.vivo_noRT | Cks2 | -1.284 | 1.06E-03 | glio |
| Eif2s1_in.vivo_noRT | Gm38163 | 2.143 | 1.06E-03 | glio |
| Eif2s1_ex.vivo_noRT | Nmrk2 | 3.373 | 1.06E-03 | glio |
| culture_effect | Gm36371 | 1.742 | 1.06E-04 | glio |
| Interaction_genotype_Aars2 | Gm10300 | -5.012 | 1.06E-04 | glio |
| Interaction_genotype_Aars2 | 2010106G04R | -4.915 | 1.06E-04 | glio |
| Interaction_genotype_Aars2 | Plcl1 | -3.986 | 1.06E-04 | glio |
| Interaction_genotype_Mtor | Gm37584 | -4.657 | 1.06E-04 | glio |
| Interaction_genotype_Mtor | Cyp2c23 | -3.971 | 1.06E-04 | glio |
| Interaction_genotype_Mtor | 2410080I02Ri | -4.383 | 1.06E-04 | glio |
| Interaction_genotype_Mtor | Gm36371 | -4.212 | 1.06E-04 | glio |
| Interaction_genotype_Myc | Olfr552 | -3.128 | 1.06E-04 | glio |
| Interaction_genotype_Myc | Abcg4 | -4.085 | 1.06E-04 | glio |
| Interaction_genotype_Myc | 9430041J12Ri | -3.523 | 1.06E-04 | glio |
| Interaction_genotype_Myc | 2310068J16Ri | -3.658 | 1.06E-04 | glio |
| Interaction_genotype_Myc | Phka2 | -3.776 | 1.06E-04 | glio |
| Interaction_genotype_Nat2 | Tpo | 3.461 | 1.06E-04 | glio |
| Aars2_in.vivo_noRT | Gm29358 | 2.827 | 1.06E-04 | glio |
| Aars2_in.vivo_noRT | Kyat1 | 2.963 | 1.06E-04 | glio |
| Aars2_in.vivo_noRT | Rell2 | 2.849 | 1.06E-04 | glio |
| Aars2_ex.vivo_noRT | Dio1 | 3.293 | 1.06E-04 | glio |
| Atp5c1_in.vivo_noRT | Klk13 | 2.136 | 1.06E-04 | glio |
| Atr_in.vivo_noRT | Ankrd13d | 2.786 | 1.06E-04 | glio |
| Atr_in.vivo_noRT | 2410080I02Ri | 2.62 | 1.06E-04 | glio |
| Atr_in.vivo_noRT | 6330418K02R | 2.258 | 1.06E-04 | glio |
| Atr_in.vivo_noRT | Pcdh17 | -1.958 | 1.06E-04 | glio |
| Bcas2_in.vivo_noRT | Etohd2 | 2.915 | 1.06E-04 | glio |
| Bcas2_in.vivo_noRT | Lrig1 | 2.956 | 1.06E-04 | glio |
| Bcas2_in.vivo_noRT | Naa80 | 2.265 | 1.06E-04 | glio |

| | | | | |
|---|---|---|---|---|
| Bcas2_in.vivo_noRT | Col9a3 | -1.936 | 1.06E-04 | glio |
| Bora_in.vivo_noRT | Gm4129 | 2.767 | 1.06E-04 | glio |
| Bora_ex.vivo_noRT | 4930423D22R | 2.622 | 1.06E-04 | glio |
| Chek1_in.vivo_noRT | Hist2h4 | 2.637 | 1.06E-04 | glio |
| culture_effect | Slc6a19 | -2.711 | 1.06E-05 | glio |
| culture_effect | Bst1 | 1.958 | 1.06E-05 | glio |
| culture_effect | Tiprl | -0.581 | 1.06E-05 | glio |
| Interaction_genotype_Mtor | Itgb6 | -5.285 | 1.06E-05 | glio |
| Interaction_genotype_Mtor | Gm47622 | -3.882 | 1.06E-05 | glio |
| Aars2_in.vivo_noRT | Prkar2b | 3.13 | 1.06E-05 | glio |
| Aars2_in.vivo_noRT | Iba57 | 2.941 | 1.06E-05 | glio |
| Atp5c1_in.vivo_noRT | Tal2 | 2.405 | 1.06E-05 | glio |
| Atr_in.vivo_noRT | C030014I23Ri | 3.083 | 1.06E-05 | glio |
| Atr_in.vivo_noRT | Tsga10ip | 3.044 | 1.06E-05 | glio |
| Atr_ex.vivo_noRT | Vmn2r56 | 3.164 | 1.06E-05 | glio |
| Chek1_ex.vivo_noRT | Krtdap | 3.86 | 1.06E-05 | glio |
| Chek1_ex.vivo_noRT | Acod1 | 4.842 | 1.06E-05 | glio |
| Hsd17b10_in.vivo_noRT | Gm16344 | 2.705 | 1.06E-05 | glio |
| culture_effect | Gm37933 | 2.15 | 1.06E-06 | glio |
| culture_effect | Eif4a2 | -0.712 | 1.06E-06 | glio |
| Interaction_genotype_Aars2 | 4930478M13F | -6.102 | 1.06E-06 | glio |
| Aars2_in.vivo_noRT | Gemin4 | 3.581 | 1.06E-06 | glio |
| Aars2_in.vivo_noRT | Rpl7a | -0.809 | 1.06E-06 | glio |
| Aars2_ex.vivo_noRT | AF067063 | 3.293 | 1.06E-06 | glio |
| Atr_in.vivo_noRT | Gm35339 | 3.474 | 1.06E-06 | glio |
| Atr_in.vivo_noRT | Slc25a29 | 3.099 | 1.06E-06 | glio |
| Atr_in.vivo_noRT | Best1 | 3.074 | 1.06E-06 | glio |
| Bcas2_in.vivo_noRT | 4921523L03Ri | 3.639 | 1.06E-06 | glio |
| Bora_in.vivo_noRT | Snrnp40 | -1.836 | 1.06E-06 | glio |
| Bora_ex.vivo_noRT | Gm266 | 3.201 | 1.06E-06 | glio |
| Bora_ex.vivo_noRT | Ildr1 | 3.201 | 1.06E-06 | glio |
| Chek1_in.vivo_noRT | Gm15535 | 3.727 | 1.06E-06 | glio |
| Chek1_in.vivo_noRT | Ngef | 3.035 | 1.06E-06 | glio |
| Chek1_in.vivo_noRT | Tnni2 | 4.006 | 1.06E-06 | glio |
| Chek1_in.vivo_noRT | Tgfb1i1 | 3.201 | 1.06E-06 | glio |
| Chek1_ex.vivo_noRT | A530013C23R | 3.963 | 1.06E-06 | glio |
| Hsd17b10_in.vivo_noRT | Rab27a | 3.09 | 1.06E-06 | glio |
| culture_effect | 1110006O24R | 2.593 | 1.06E-07 | glio |
| culture_effect | Slc2a8 | -1.727 | 1.06E-07 | glio |
| culture_effect | 1810009A15R | 1.662 | 1.06E-07 | glio |
| culture_effect | Cox7a2 | 0.6 | 1.06E-07 | glio |
| Interaction_genotype_Mtor | Mir670hg | -6.907 | 1.06E-07 | glio |
| Interaction_genotype_Mtor | Gm26671 | -4.492 | 1.06E-07 | glio |
| Interaction_genotype_Mtor | 1700026N04R | -5.761 | 1.06E-07 | glio |
| Interaction_genotype_Nat2 | Gm2016 | 4.734 | 1.06E-07 | glio |
| interaction_RT | 2410006H16R | 1.368 | 1.06E-07 | glio |
| Aars2_in.vivo_noRT | Zfpm1 | 3.81 | 1.06E-07 | glio |
| Atr_ex.vivo_noRT | Prap1 | 3.554 | 1.06E-07 | glio |
| Bcas2_in.vivo_noRT | Gm4221 | 3.765 | 1.06E-07 | glio |
| Bcas2_in.vivo_noRT | Pycrl | -2.227 | 1.06E-07 | glio |
| Bora_in.vivo_noRT | Tmem30c | 3.75 | 1.06E-07 | glio |
| Aars2_in.vivo_noRT | Cyp4f18 | 5.603 | 1.06E-08 | glio |

| | | | | |
|---|---|---|---|---|
| Bcas2_in.vivo_noRT | 4930518J21Ri | 3.882 | 1.06E-08 | glio |
| Bora_in.vivo_noRT | Gm14698 | 4.024 | 1.06E-08 | glio |
| Bora_in.vivo_noRT | Rasl10a | 3.317 | 1.06E-08 | glio |
| Bora_in.vivo_noRT | D430040D24R | 4.019 | 1.06E-08 | glio |
| culture_effect | Btg1 | -0.991 | 1.06E-09 | glio |
| culture_effect | Anapc11 | 0.749 | 1.06E-09 | glio |
| Aars2_in.vivo_noRT | Apol7d | 4.425 | 1.06E-09 | glio |
| Atr_in.vivo_noRT | Dcpp3 | 4.016 | 1.06E-09 | glio |
| Chek1_in.vivo_noRT | Ahrr | 4.243 | 1.06E-09 | glio |
| Chek1_ex.vivo_noRT | Gpr37l1 | 5.249 | 1.06E-09 | glio |
| Aars2_in.vivo_noRT | Ppp1r3g | 4.142 | 1.06E-10 | glio |
| Atr_in.vivo_noRT | Neurl3 | 4.543 | 1.06E-10 | glio |
| Bcas2_in.vivo_noRT | Hoxa6 | 4.292 | 1.06E-10 | glio |
| Aars2_in.vivo_noRT | Gm5039 | 5.327 | 1.06E-11 | glio |
| Atr_in.vivo_noRT | Camkk1 | 4.763 | 1.06E-11 | glio |
| Aars2_in.vivo_noRT | Gm41764 | 4.41 | 1.06E-12 | glio |
| Aars2_in.vivo_noRT | 2310067P03R | 4.804 | 1.06E-12 | glio |
| Atr_in.vivo_noRT | H3f3aos | 4.888 | 1.06E-12 | glio |
| Chek1_ex.vivo_noRT | Cxcl5 | 4.842 | 1.06E-13 | glio |
| Chek1_in.vivo_noRT | Fbxo15 | 4.78 | 1.06E-14 | glio |
| Aars2_in.vivo_noRT | Gm15894 | 4.282 | 1.06E-15 | glio |
| Chek1_in.vivo_noRT | Pik3ap1 | 4.78 | 1.06E-15 | glio |
| Aars2_in.vivo_noRT | Krtcap3 | 4.535 | 1.06E-16 | glio |
| Aars2_in.vivo_noRT | Dpp6 | 4.399 | 1.06E-18 | glio |
| Chek1_in.vivo_noRT | Gtsf1 | 5.044 | 1.06E-18 | glio |
| Atr_in.vivo_noRT | Bco2 | 5.533 | 1.06E-19 | glio |
| Bcas2_in.vivo_noRT | Ank1 | 4.807 | 1.06E-20 | glio |
| Aars2_in.vivo_noRT | 1700019E08Ri | 5.327 | 1.06E-21 | glio |
| Chek1_in.vivo_noRT | Gm15533 | 5.044 | 1.06E-22 | glio |
| Atr_in.vivo_noRT | Gm9758 | 5.273 | 1.06E-23 | glio |
| culture_effect | Macc1 | -1.87 | 1.07E-03 | glio |
| culture_effect | Cacul1 | 0.651 | 1.07E-03 | glio |
| culture_effect | Slc25a51 | 0.584 | 1.07E-03 | glio |
| culture_effect | Clcn5 | 0.568 | 1.07E-03 | glio |
| Interaction_genotype_Aars2 | Gm13986 | -4.253 | 1.07E-03 | glio |
| Interaction_genotype_Aars2 | Htr1b | -3.989 | 1.07E-03 | glio |
| Interaction_genotype_Aars2 | Gm13663 | -3.588 | 1.07E-03 | glio |
| Interaction_genotype_Aars2 | Spcs1 | 1.164 | 1.07E-03 | glio |
| Interaction_genotype_Atr | Gm42695 | -4.086 | 1.07E-03 | glio |
| Interaction_genotype_Bora | 2410080I02Ri | -4.137 | 1.07E-03 | glio |
| Interaction_genotype_Bora | Gm47484 | -4.523 | 1.07E-03 | glio |
| Interaction_genotype_Mtor | Dyrk4 | -4.528 | 1.07E-03 | glio |
| Interaction_genotype_Mtor | Ccdc106 | -4.179 | 1.07E-03 | glio |
| Interaction_genotype_Mtor | Hist1h3b | -4.05 | 1.07E-03 | glio |
| Interaction_genotype_Mtor | Zfp689 | -2.971 | 1.07E-03 | glio |
| Interaction_genotype_Mtor | Adamts7 | -3.272 | 1.07E-03 | glio |
| Interaction_genotype_Myc | Dok2 | -3.179 | 1.07E-03 | glio |
| Interaction_genotype_Myc | Gm39459 | -3.747 | 1.07E-03 | glio |
| Interaction_genotype_Myc | Gm26930 | -3.952 | 1.07E-03 | glio |
| Interaction_genotype_Myc | Gm43707 | -3.324 | 1.07E-03 | glio |
| Interaction_genotype_Myc | A430105J06Ri | -3.416 | 1.07E-03 | glio |
| Interaction_genotype_Myc | Gm48933 | -3.417 | 1.07E-03 | glio |

| | | | | |
|---|---|---|---|---|
| Interaction_genotype_Myc | Gen1 | -3.007 | 1.07E-03 | glio |
| Interaction_genotype_Myc | Gm50340 | -3.199 | 1.07E-03 | glio |
| Interaction_genotype_Myc | Mrpl53 | 1.337 | 1.07E-03 | glio |
| Interaction_genotype_Myc | Dnajc2 | 1.751 | 1.07E-03 | glio |
| Interaction_genotype_Myc | Rps16 | 1.189 | 1.07E-03 | glio |
| Interaction_genotype_Pars2 | 1700013F07Ri | -4.787 | 1.07E-03 | glio |
| Interaction_genotype_Pars2 | Prl3d3 | -3.101 | 1.07E-03 | glio |
| Interaction_genotype_Pars2 | Gm47622 | -3.367 | 1.07E-03 | glio |
| interaction_RT | 4933412E12Ri | 1.524 | 1.07E-03 | glio |
| interaction_RT | Apoc1 | 3.25 | 1.07E-03 | glio |
| interaction_RT | Fh1 | -0.891 | 1.07E-03 | glio |
| interaction_RT | Tnni2 | 3.75 | 1.07E-03 | glio |
| interaction_RT | Jund | 0.746 | 1.07E-03 | glio |
| interaction_RT | Alyref | 0.662 | 1.07E-03 | glio |
| Aars2_in.vivo_noRT | 3300002I08Ri | 2.3 | 1.07E-03 | glio |
| Aars2_in.vivo_noRT | Fhod1 | 1.971 | 1.07E-03 | glio |
| Aars2_in.vivo_noRT | Pex10 | 1.827 | 1.07E-03 | glio |
| Aars2_in.vivo_noRT | Rps5 | -0.607 | 1.07E-03 | glio |
| Aars2_ex.vivo_noRT | Kdm2b | -2.297 | 1.07E-03 | glio |
| Atr_in.vivo_noRT | Klhl32 | 2.048 | 1.07E-03 | glio |
| Atr_in.vivo_noRT | Otud6b | -1.219 | 1.07E-03 | glio |
| Atr_in.vivo_noRT | Ifrd1 | -1.063 | 1.07E-03 | glio |
| Atr_in.vivo_noRT | Gm47283 | -0.864 | 1.07E-03 | glio |
| Atr_in.vivo_noRT | Rpl6 | -0.669 | 1.07E-03 | glio |
| Atr_ex.vivo_noRT | Gm13054 | 2.769 | 1.07E-03 | glio |
| Atr_ex.vivo_noRT | Emc2 | -1.224 | 1.07E-03 | glio |
| Bcas2_in.vivo_noRT | Ankrd42 | 2.424 | 1.07E-03 | glio |
| Bcas2_in.vivo_noRT | Gm4181 | 2.394 | 1.07E-03 | glio |
| Bcas2_in.vivo_noRT | Chrnb1 | 2.237 | 1.07E-03 | glio |
| Bcas2_in.vivo_noRT | Top3a | 1.86 | 1.07E-03 | glio |
| Bora_in.vivo_noRT | Gm9776 | 2.312 | 1.07E-03 | glio |
| Bora_in.vivo_noRT | Rmst | 2.49 | 1.07E-03 | glio |
| Bora_in.vivo_noRT | Tmem214 | -1.483 | 1.07E-03 | glio |
| Bora_in.vivo_noRT | Parp1 | -1.198 | 1.07E-03 | glio |
| Bora_ex.vivo_noRT | Gm21762 | 2.808 | 1.07E-03 | glio |
| Cenpj_in.vivo_noRT | Rgs4 | 2.022 | 1.07E-03 | glio |
| Chek1_in.vivo_noRT | Lipe | 2.35 | 1.07E-03 | glio |
| Chek1_ex.vivo_noRT | Gm40557 | 2.96 | 1.07E-03 | glio |
| Chek1_ex.vivo_noRT | Gh | 3.279 | 1.07E-03 | glio |
| Chek1_ex.vivo_noRT | Ctsa | 1.449 | 1.07E-03 | glio |
| Eif2s1_in.vivo_noRT | Tesc | -1.519 | 1.07E-03 | glio |
| culture_effect | Tmem79 | 1.84 | 1.07E-04 | glio |
| culture_effect | Timm8a1 | 0.583 | 1.07E-04 | glio |
| Interaction_genotype_Aars2 | Hlx | -4.687 | 1.07E-04 | glio |
| Interaction_genotype_Mtor | Bend7 | -4.475 | 1.07E-04 | glio |
| Interaction_genotype_Mtor | Zfp873 | -4.086 | 1.07E-04 | glio |
| Interaction_genotype_Myc | 2510046G10R | -3.544 | 1.07E-04 | glio |
| Interaction_genotype_Myc | Gabarap | 1.412 | 1.07E-04 | glio |
| Aars2_in.vivo_noRT | Gpx1 | -1.096 | 1.07E-04 | glio |
| Aars2_ex.vivo_noRT | Arg2 | 3.676 | 1.07E-04 | glio |
| Atp5c1_in.vivo_noRT | Layn | 2.242 | 1.07E-04 | glio |
| Atr_in.vivo_noRT | Klhl33 | 2.733 | 1.07E-04 | glio |

| | | | | |
|---|---|---|---|---|
| Atr_in.vivo_noRT | Ppargc1b | 2.777 | 1.07E-04 | glio |
| Atr_in.vivo_noRT | Tfam | -1.263 | 1.07E-04 | glio |
| Atr_in.vivo_noRT | Stat3 | -1.34 | 1.07E-04 | glio |
| Atr_in.vivo_noRT | Eif2s3x | -1.204 | 1.07E-04 | glio |
| Atr_in.vivo_noRT | Rps25 | -0.776 | 1.07E-04 | glio |
| Atr_ex.vivo_noRT | 1110017D15R | 3.569 | 1.07E-04 | glio |
| Atr_ex.vivo_noRT | Slc11a1 | 3.722 | 1.07E-04 | glio |
| Bora_in.vivo_noRT | A430018G15R | 2.292 | 1.07E-04 | glio |
| Chek1_in.vivo_noRT | 1110006O24R | 2.842 | 1.07E-04 | glio |
| Chek1_in.vivo_noRT | Gm20337 | 2.826 | 1.07E-04 | glio |
| Chek1_in.vivo_noRT | Supt4a | -0.908 | 1.07E-04 | glio |
| Chek1_ex.vivo_noRT | Gm48357 | 3.036 | 1.07E-04 | glio |
| Chek1_ex.vivo_noRT | Bcl2l14 | 3.738 | 1.07E-04 | glio |
| culture_effect | Ap2a1 | 0.844 | 1.07E-05 | glio |
| culture_effect | Ctbp1 | 0.502 | 1.07E-05 | glio |
| Interaction_genotype_Aars2 | Gm9958 | -5.567 | 1.07E-05 | glio |
| Interaction_genotype_Aars2 | Hist1h2bh | -5.616 | 1.07E-05 | glio |
| Interaction_genotype_Mtor | C130013H08R | -5.741 | 1.07E-05 | glio |
| Interaction_genotype_Mtor | Cldn19 | -5.002 | 1.07E-05 | glio |
| Interaction_genotype_Mtor | Bfsp2 | -5.932 | 1.07E-05 | glio |
| Interaction_genotype_Rad21 | Gpr17 | 5.404 | 1.07E-05 | glio |
| Atr_in.vivo_noRT | Ovca2 | 2.88 | 1.07E-05 | glio |
| Atr_ex.vivo_noRT | Gm30692 | 2.77 | 1.07E-05 | glio |
| Bcas2_in.vivo_noRT | Tcea2 | 3.079 | 1.07E-05 | glio |
| Bora_in.vivo_noRT | Gm34643 | 3.221 | 1.07E-05 | glio |
| Bora_in.vivo_noRT | Supt16 | -1.562 | 1.07E-05 | glio |
| Bora_ex.vivo_noRT | Lmx1a | 3.201 | 1.07E-05 | glio |
| Bora_ex.vivo_noRT | Vmn2r56 | 3.201 | 1.07E-05 | glio |
| Bora_ex.vivo_noRT | Gm26671 | 2.808 | 1.07E-05 | glio |
| Chek1_in.vivo_noRT | Gm10638 | 3.076 | 1.07E-05 | glio |
| Chek1_ex.vivo_noRT | Hpse | 3.877 | 1.07E-05 | glio |
| Chek1_ex.vivo_noRT | Slxl1 | 3.866 | 1.07E-05 | glio |
| Chek1_ex.vivo_noRT | Gm45441 | 3.828 | 1.07E-05 | glio |
| Chek1_ex.vivo_noRT | Flot2 | -2.664 | 1.07E-05 | glio |
| culture_effect | Svip | 1.863 | 1.07E-06 | glio |
| culture_effect | 2810013P06R | 1.504 | 1.07E-06 | glio |
| Interaction_genotype_Atr | Gm49312 | -5.08 | 1.07E-06 | glio |
| Interaction_genotype_Myc | Klk14 | -5.669 | 1.07E-06 | glio |
| Aars2_in.vivo_noRT | Gm37305 | 3.369 | 1.07E-06 | glio |
| Atp5c1_in.vivo_noRT | Gm36793 | 2.668 | 1.07E-06 | glio |
| Atr_in.vivo_noRT | Vgll3 | 3.412 | 1.07E-06 | glio |
| Bcas2_in.vivo_noRT | Gm13283 | 3.444 | 1.07E-06 | glio |
| Bora_in.vivo_noRT | Tmem86a | 3.569 | 1.07E-06 | glio |
| Bora_in.vivo_noRT | Alox5ap | 3.831 | 1.07E-06 | glio |
| Hsd17b10_in.vivo_noRT | Gm43672 | 3.616 | 1.07E-06 | glio |
| Interaction_genotype_Mtor | Stub1 | 2.47 | 1.07E-07 | glio |
| Interaction_genotype_Mtor | Gm44702 | -6.588 | 1.07E-07 | glio |
| Atr_in.vivo_noRT | Gm4673 | 3.735 | 1.07E-07 | glio |
| Atr_ex.vivo_noRT | Gm36026 | 3.962 | 1.07E-07 | glio |
| Bcas2_in.vivo_noRT | Gm26822 | 3.892 | 1.07E-07 | glio |
| Bcas2_in.vivo_noRT | Slc25a35 | 3.697 | 1.07E-07 | glio |
| Bora_in.vivo_noRT | Gm47484 | 3.943 | 1.07E-07 | glio |

| | | | | |
|---|---|---|---|---|
| Chek1_in.vivo_noRT | Lynx1 | 3.74 | 1.07E-07 | glio |
| culture_effect | Ctdnep1 | 0.793 | 1.07E-08 | glio |
| Interaction_genotype_Myc | Man2c1os | -6.863 | 1.07E-08 | glio |
| Aars2_in.vivo_noRT | Gm9484 | 4.168 | 1.07E-08 | glio |
| Aars2_in.vivo_noRT | A530072M11F | 3.998 | 1.07E-08 | glio |
| Atr_in.vivo_noRT | Dct | 4.73 | 1.07E-08 | glio |
| Bcas2_in.vivo_noRT | Milr1 | 4.072 | 1.07E-08 | glio |
| Bcas2_in.vivo_noRT | Gm46652 | 3.758 | 1.07E-08 | glio |
| Bcas2_in.vivo_noRT | Gm47101 | 4.016 | 1.07E-09 | glio |
| Atr_in.vivo_noRT | Rab3b | 4.377 | 1.07E-10 | glio |
| Atr_in.vivo_noRT | Il17b | 4.883 | 1.07E-10 | glio |
| Bcas2_in.vivo_noRT | Col5a2 | 3.495 | 1.07E-10 | glio |
| Bora_in.vivo_noRT | Slc16a7 | 3.825 | 1.07E-10 | glio |
| culture_effect | Hnrnpa2b1 | -0.705 | 1.07E-11 | glio |
| Atr_in.vivo_noRT | Camk1d | 4.732 | 1.07E-11 | glio |
| culture_effect | Akr1b7 | -4.433 | 1.07E-12 | glio |
| Chek1_in.vivo_noRT | Gm43951 | 4.118 | 1.07E-12 | glio |
| Chek1_in.vivo_noRT | Nmrk2 | 4.402 | 1.07E-12 | glio |
| Aars2_in.vivo_noRT | Osm | 4.514 | 1.07E-13 | glio |
| Aars2_in.vivo_noRT | Rhox7a | 5.327 | 1.07E-13 | glio |
| Atr_in.vivo_noRT | Gjb5 | 4.543 | 1.07E-13 | glio |
| Hsd17b10_in.vivo_noRT | Cgnl1 | 3.616 | 1.07E-13 | glio |
| Hsd17b10_in.vivo_noRT | Gm26543 | 3.616 | 1.07E-13 | glio |
| Hsd17b10_in.vivo_noRT | Dscaml1 | 3.616 | 1.07E-13 | glio |
| Hsd17b10_in.vivo_noRT | 2410017I17Ri | 3.616 | 1.07E-13 | glio |
| culture_effect | Slc38a2 | -0.985 | 1.07E-15 | glio |
| Aars2_in.vivo_noRT | Dnah10 | 4.947 | 1.07E-15 | glio |
| Atr_in.vivo_noRT | Gm31152 | 5.011 | 1.07E-15 | glio |
| Bora_in.vivo_noRT | Gm13032 | 4.55 | 1.07E-15 | glio |
| Chek1_in.vivo_noRT | Minar2 | 4.784 | 1.07E-17 | glio |
| Chek1_in.vivo_noRT | H1foo | 4.511 | 1.07E-18 | glio |
| Bcas2_in.vivo_noRT | Stfa3 | 4.807 | 1.07E-24 | glio |
| Bcas2_in.vivo_noRT | Olfr787 | 4.807 | 1.07E-24 | glio |
| culture_effect | Itgbl1 | -1.726 | 1.08E-03 | glio |
| culture_effect | Prox2 | 1.984 | 1.08E-03 | glio |
| culture_effect | Gm15535 | 1.937 | 1.08E-03 | glio |
| culture_effect | Syn3 | 1.599 | 1.08E-03 | glio |
| culture_effect | Gm16537 | 1.333 | 1.08E-03 | glio |
| culture_effect | Trim30d | 0.834 | 1.08E-03 | glio |
| culture_effect | Tiam1 | -0.689 | 1.08E-03 | glio |
| culture_effect | Bcl10 | -0.564 | 1.08E-03 | glio |
| culture_effect | Rab22a | 0.578 | 1.08E-03 | glio |
| culture_effect | Smdt1 | 0.382 | 1.08E-03 | glio |
| Interaction_genotype_Atr | Chic2 | 1.402 | 1.08E-03 | glio |
| Interaction_genotype_Bora | Gm10655 | -4.438 | 1.08E-03 | glio |
| Interaction_genotype_Bora | St8sia1 | -4.452 | 1.08E-03 | glio |
| Interaction_genotype_Bora | Wfdc1 | -3.487 | 1.08E-03 | glio |
| Interaction_genotype_Bora | Gm11627 | -3.664 | 1.08E-03 | glio |
| Interaction_genotype_Bora | 2310067P03R | -4.177 | 1.08E-03 | glio |
| Interaction_genotype_Bora | 1810021B22R | -4.368 | 1.08E-03 | glio |
| Interaction_genotype_Bora | Ccna2 | 2.651 | 1.08E-03 | glio |
| Interaction_genotype_Bora | Hmmr | 2.834 | 1.08E-03 | glio |

| | | | | |
|---|---|---|---|---|
| Interaction_genotype_Mtor | Hpn | -3.809 | 1.08E-03 | glio |
| Interaction_genotype_Mtor | Gorasp1 | -3.408 | 1.08E-03 | glio |
| Interaction_genotype_Mtor | Ttc9 | -2.562 | 1.08E-03 | glio |
| Interaction_genotype_Myc | Scgb1c1 | -3.122 | 1.08E-03 | glio |
| Interaction_genotype_Myc | Strip2 | -2.865 | 1.08E-03 | glio |
| Interaction_genotype_Myc | A530013C23R | -3.491 | 1.08E-03 | glio |
| Interaction_genotype_Myc | Car12 | -3.521 | 1.08E-03 | glio |
| Interaction_genotype_Myc | Kif7 | -3.425 | 1.08E-03 | glio |
| Interaction_genotype_Myc | Fam171a2 | -3.659 | 1.08E-03 | glio |
| Interaction_genotype_Myc | Napepld | -3.213 | 1.08E-03 | glio |
| Interaction_genotype_Myc | Hspa2 | -2.834 | 1.08E-03 | glio |
| Interaction_genotype_Myc | Gm28791 | -3.276 | 1.08E-03 | glio |
| Interaction_genotype_Myc | Hs2st1 | -1.802 | 1.08E-03 | glio |
| Interaction_genotype_Myc | Rps9 | 1.229 | 1.08E-03 | glio |
| Interaction_genotype_Rad21 | Cspg5 | 3.271 | 1.08E-03 | glio |
| interaction_RT | Hist1h3f | -2.153 | 1.08E-03 | glio |
| interaction_RT | Ptn | 3.445 | 1.08E-03 | glio |
| Aars2_in.vivo_noRT | B3gat2 | 2.466 | 1.08E-03 | glio |
| Aars2_in.vivo_noRT | Slc6a20b | 2.319 | 1.08E-03 | glio |
| Aars2_in.vivo_noRT | Gm14418 | 1.833 | 1.08E-03 | glio |
| Aars2_in.vivo_noRT | Rtl8b | 1.872 | 1.08E-03 | glio |
| Aars2_in.vivo_noRT | Bcat1 | -1.663 | 1.08E-03 | glio |
| Aars2_in.vivo_noRT | Arl4a | -1.406 | 1.08E-03 | glio |
| Aars2_in.vivo_noRT | Calr | -0.963 | 1.08E-03 | glio |
| Aars2_ex.vivo_noRT | Pkd1l2 | 2.897 | 1.08E-03 | glio |
| Abtb1_ex.vivo_noRT | Gm29670 | 2.95 | 1.08E-03 | glio |
| Atp5c1_in.vivo_noRT | Snrpn | 2.01 | 1.08E-03 | glio |
| Atp5c1_in.vivo_noRT | Gm32250 | 1.886 | 1.08E-03 | glio |
| Atp5c1_in.vivo_noRT | Gm15080 | 2.734 | 1.08E-03 | glio |
| Atr_in.vivo_noRT | Ypel4 | 2.169 | 1.08E-03 | glio |
| Atr_ex.vivo_noRT | 4930447K03R | 2.376 | 1.08E-03 | glio |
| Atr_ex.vivo_noRT | Nt5e | 2.771 | 1.08E-03 | glio |
| Atr_ex.vivo_noRT | Gm36913 | 3.124 | 1.08E-03 | glio |
| Atr_ex.vivo_noRT | Cited1 | 2.664 | 1.08E-03 | glio |
| Bard1_ex.vivo_noRT | Usp17la | 3.217 | 1.08E-03 | glio |
| Bard1_ex.vivo_noRT | Ntn1 | 2.814 | 1.08E-03 | glio |
| Bard1_ex.vivo_noRT | Pmch | 2.814 | 1.08E-03 | glio |
| Bcas2_in.vivo_noRT | Mmp14 | 2.451 | 1.08E-03 | glio |
| Bcas2_ex.vivo_noRT | Gtsf1l | 2.742 | 1.08E-03 | glio |
| Bora_in.vivo_noRT | Mpz | 2.965 | 1.08E-03 | glio |
| Bora_in.vivo_noRT | Cdkn1c | 2.368 | 1.08E-03 | glio |
| Bora_in.vivo_noRT | A230056P14R | 2.366 | 1.08E-03 | glio |
| Bora_in.vivo_noRT | Fdft1 | -1.82 | 1.08E-03 | glio |
| Bora_in.vivo_noRT | Pdxk | -1.534 | 1.08E-03 | glio |
| Bora_ex.vivo_noRT | Rab27a | 2.806 | 1.08E-03 | glio |
| Bora_ex.vivo_noRT | Tmeff2 | 3.598 | 1.08E-03 | glio |
| Bora_ex.vivo_noRT | Osm | 2.802 | 1.08E-03 | glio |
| Chek1_ex.vivo_noRT | E030042O20R | 3.155 | 1.08E-03 | glio |
| Chek1_ex.vivo_noRT | Gm45620 | 2.784 | 1.08E-03 | glio |
| Chek1_ex.vivo_noRT | Ccdc184 | 3.154 | 1.08E-03 | glio |
| Chek1_ex.vivo_noRT | Gm9967 | 2.777 | 1.08E-03 | glio |
| Hsd17b10_in.vivo_noRT | Flrt1 | 2.641 | 1.08E-03 | glio |

| | | | | |
|---|---|---|---|---|
| Hsd17b10_in.vivo_noRT | Dagla | 2.651 | 1.08E-03 | glio |
| Hsd17b10_in.vivo_noRT | Zfhx2os | 2.567 | 1.08E-03 | glio |
| culture_effect | Noxo1 | -1.949 | 1.08E-04 | glio |
| culture_effect | Ost4 | 0.493 | 1.08E-04 | glio |
| Interaction_genotype_Mtor | Nos3 | -4.627 | 1.08E-04 | glio |
| Interaction_genotype_Mtor | Fam107b | -3.704 | 1.08E-04 | glio |
| Interaction_genotype_Myc | Serinc2 | -4.218 | 1.08E-04 | glio |
| Interaction_genotype_Myc | Nek8 | -4.088 | 1.08E-04 | glio |
| Interaction_genotype_Myc | 9230114K14R | -3.623 | 1.08E-04 | glio |
| Interaction_genotype_Myc | Vmp1 | 1.975 | 1.08E-04 | glio |
| Interaction_genotype_Myc | Spcs1 | 1.626 | 1.08E-04 | glio |
| Interaction_genotype_Myc | Krtcap2 | 1.597 | 1.08E-04 | glio |
| Aars2_in.vivo_noRT | Rtl5 | 2.658 | 1.08E-04 | glio |
| Aars2_ex.vivo_noRT | Gm19265 | 2.893 | 1.08E-04 | glio |
| Atr_in.vivo_noRT | Dennd4b | 2.718 | 1.08E-04 | glio |
| Atr_in.vivo_noRT | B230303O12R | 2.754 | 1.08E-04 | glio |
| Atr_in.vivo_noRT | Hist2h4 | 2.603 | 1.08E-04 | glio |
| Atr_in.vivo_noRT | Kctd7 | 2.531 | 1.08E-04 | glio |
| Atr_in.vivo_noRT | Gm10184 | 2.517 | 1.08E-04 | glio |
| Atr_in.vivo_noRT | Fam214a | 2.175 | 1.08E-04 | glio |
| Atr_ex.vivo_noRT | Insc | 3.164 | 1.08E-04 | glio |
| Atr_ex.vivo_noRT | Icam2 | 3.164 | 1.08E-04 | glio |
| Atr_ex.vivo_noRT | Wnt6 | 4.347 | 1.08E-04 | glio |
| Bcas2_in.vivo_noRT | Gm15706 | 2.388 | 1.08E-04 | glio |
| Bora_in.vivo_noRT | 1600002K03R | 2.873 | 1.08E-04 | glio |
| Bora_in.vivo_noRT | Gm15834 | 2.755 | 1.08E-04 | glio |
| Bora_in.vivo_noRT | Nrap | 2.929 | 1.08E-04 | glio |
| Bora_in.vivo_noRT | Serpinb9 | 2.639 | 1.08E-04 | glio |
| Bora_in.vivo_noRT | Ift46 | -1.791 | 1.08E-04 | glio |
| Bora_in.vivo_noRT | Chst11 | -1.802 | 1.08E-04 | glio |
| Chek1_in.vivo_noRT | 4930455G09R | 3.228 | 1.08E-04 | glio |
| Chek1_in.vivo_noRT | H60b | 2.675 | 1.08E-04 | glio |
| Hsd17b10_in.vivo_noRT | Gm3734 | 3.093 | 1.08E-04 | glio |
| Hsd17b10_in.vivo_noRT | Gm13404 | 2.878 | 1.08E-04 | glio |
| Hsd17b10_in.vivo_noRT | Lyrm1 | 2.848 | 1.08E-04 | glio |
| Hsd17b10_in.vivo_noRT | Taf9 | 2.771 | 1.08E-04 | glio |
| culture_effect | Stamos | 1.793 | 1.08E-05 | glio |
| culture_effect | Acy1 | -1.33 | 1.08E-05 | glio |
| culture_effect | Sem1 | 0.499 | 1.08E-05 | glio |
| Interaction_genotype_Mtor | 2610027K06R | -4.555 | 1.08E-05 | glio |
| Interaction_genotype_Mtor | Gm9967 | -4.944 | 1.08E-05 | glio |
| Interaction_genotype_Mtor | Ube2b | 1.67 | 1.08E-05 | glio |
| Interaction_genotype_Myc | D930019O06R | -5.154 | 1.08E-05 | glio |
| Interaction_genotype_Myc | Pcyox1l | -4.329 | 1.08E-05 | glio |
| Interaction_genotype_Myc | Fbxl12os | -4.831 | 1.08E-05 | glio |
| Interaction_genotype_Myc | Inha | -5.078 | 1.08E-05 | glio |
| Atp5c1_in.vivo_noRT | 1700026N04R | 2.936 | 1.08E-05 | glio |
| Atr_in.vivo_noRT | Trir | -1.105 | 1.08E-05 | glio |
| Atr_in.vivo_noRT | Gabarap | -1.133 | 1.08E-05 | glio |
| Bard1_ex.vivo_noRT | Usp17lc | 3.612 | 1.08E-05 | glio |
| Bard1_ex.vivo_noRT | Hoxc8 | 2.814 | 1.08E-05 | glio |
| Bcas2_in.vivo_noRT | Tpsab1 | 2.857 | 1.08E-05 | glio |

| | | | | |
|---|---|---|---|---|
| Bcas2_in.vivo_noRT | Palm3 | 3.21 | 1.08E-05 | glio |
| Bora_in.vivo_noRT | C920021L13Ri | 3.139 | 1.08E-05 | glio |
| Bora_in.vivo_noRT | Ypel4 | 2.882 | 1.08E-05 | glio |
| Chek1_in.vivo_noRT | Gm14636 | 3.755 | 1.08E-05 | glio |
| Chek1_in.vivo_noRT | Rab17 | 3.049 | 1.08E-05 | glio |
| Chek1_in.vivo_noRT | Slc9a9 | 3.179 | 1.08E-05 | glio |
| Chek1_ex.vivo_noRT | Nrip3 | 3.656 | 1.08E-05 | glio |
| Chek1_ex.vivo_noRT | Gm49086 | 3.279 | 1.08E-05 | glio |
| Chek1_ex.vivo_noRT | Rgs5 | 4.842 | 1.08E-05 | glio |
| Hsd17b10_in.vivo_noRT | Gm14698 | 3.225 | 1.08E-05 | glio |
| culture_effect | Rabep1 | 0.659 | 1.08E-06 | glio |
| Interaction_genotype_Mtor | Aqp5 | -6.629 | 1.08E-06 | glio |
| Interaction_genotype_Mtor | Ripply3 | -5.663 | 1.08E-06 | glio |
| Interaction_genotype_Myc | 8030456M14F | -5.486 | 1.08E-06 | glio |
| interaction_RT | Piezo1 | -1.42 | 1.08E-06 | glio |
| Aars2_ex.vivo_noRT | Gm2016 | 3.293 | 1.08E-06 | glio |
| Atr_in.vivo_noRT | Mmp19 | 3.339 | 1.08E-06 | glio |
| Bcas2_in.vivo_noRT | Ggt5 | 3.174 | 1.08E-06 | glio |
| Chek1_ex.vivo_noRT | Pcdhgb6 | 3.738 | 1.08E-06 | glio |
| culture_effect | Hecw2 | 2.666 | 1.08E-07 | glio |
| Aars2_in.vivo_noRT | Gm15510 | 3.732 | 1.08E-07 | glio |
| Aars2_in.vivo_noRT | Rab40b | 3.59 | 1.08E-07 | glio |
| Atr_in.vivo_noRT | 4833418N02R | 3.832 | 1.08E-07 | glio |
| Bcas2_in.vivo_noRT | Zfp458 | 3.861 | 1.08E-07 | glio |
| Chek1_in.vivo_noRT | 5430400D12R | 3.762 | 1.08E-07 | glio |
| Chek1_ex.vivo_noRT | Gm10324 | 3.48 | 1.08E-07 | glio |
| Hsd17b10_in.vivo_noRT | Gm37214 | 3.877 | 1.08E-07 | glio |
| culture_effect | Esr1 | -3.398 | 1.08E-08 | glio |
| culture_effect | Gar1 | -0.766 | 1.08E-08 | glio |
| Interaction_genotype_Myc | Zeb2os | -6.215 | 1.08E-08 | glio |
| Interaction_genotype_Myc | Rcan2 | -6.877 | 1.08E-08 | glio |
| Bcas2_in.vivo_noRT | Slc23a3 | 3.905 | 1.08E-08 | glio |
| Bora_in.vivo_noRT | Gm8300 | 4.936 | 1.08E-08 | glio |
| Bora_in.vivo_noRT | Inhba | 4.295 | 1.08E-08 | glio |
| Chek1_ex.vivo_noRT | Dreh | 4.449 | 1.08E-08 | glio |
| culture_effect | Gm15706 | 2.206 | 1.08E-09 | glio |
| Bcas2_in.vivo_noRT | Gipc2 | 4.544 | 1.08E-09 | glio |
| Bcas2_in.vivo_noRT | Gm30938 | 4.161 | 1.08E-09 | glio |
| Bcas2_in.vivo_noRT | Gm5039 | 4.807 | 1.08E-09 | glio |
| Chek1_in.vivo_noRT | 4930518J21Ri | 4.118 | 1.08E-09 | glio |
| Chek1_in.vivo_noRT | Mag | 4.004 | 1.08E-09 | glio |
| Chek1_ex.vivo_noRT | Prl7c1 | 3.747 | 1.08E-09 | glio |
| culture_effect | Gm4419 | 2.505 | 1.08E-10 | glio |
| culture_effect | 0610040B10R | 2.23 | 1.08E-10 | glio |
| Bcas2_in.vivo_noRT | Icam2 | 3.941 | 1.08E-10 | glio |
| Bora_in.vivo_noRT | Ipcef1 | 3.647 | 1.08E-10 | glio |
| Bcas2_in.vivo_noRT | Thnsl2 | 3.639 | 1.08E-11 | glio |
| Atr_in.vivo_noRT | Hist2h3c2 | 5.267 | 1.08E-12 | glio |
| Aars2_in.vivo_noRT | Col5a2 | 4.015 | 1.08E-13 | glio |
| Aars2_in.vivo_noRT | Ramp3 | 5.06 | 1.08E-13 | glio |
| Hsd17b10_in.vivo_noRT | Pspn | 3.616 | 1.08E-13 | glio |
| Atr_in.vivo_noRT | Gm43672 | 5.533 | 1.08E-14 | glio |

| | | | | |
|---|---|---|---|---|
| culture_effect | Wtip | 1.342 | 1.08E-15 | glio |
| Chek1_in.vivo_noRT | Psca | 5.044 | 1.08E-15 | glio |
| Aars2_in.vivo_noRT | 2310003N18R | 5.327 | 1.08E-19 | glio |
| culture_effect | Zfp36 | -3.183 | 1.08E-21 | glio |
| culture_effect | Shroom2 | 2.584 | 1.08E-21 | glio |
| Aars2_in.vivo_noRT | Gad1 | 5.06 | 1.08E-23 | glio |
| Bora_in.vivo_noRT | Gm45338 | 4.936 | 1.08E-25 | glio |
| Bora_in.vivo_noRT | Fhl1 | 5.861 | 1.08E-25 | glio |
| Bora_in.vivo_noRT | Scg2 | 4.936 | 1.08E-25 | glio |
| Atr_in.vivo_noRT | Edn2 | 5.533 | 1.08E-28 | glio |
| Atr_in.vivo_noRT | Gm14207 | 5.533 | 1.08E-28 | glio |
| Atr_in.vivo_noRT | Nrip2 | 5.533 | 1.08E-28 | glio |
| culture_effect | Tmem87a | -1.07 | 1.09E-03 | glio |
| culture_effect | Zfp608 | -0.591 | 1.09E-03 | glio |
| Interaction_genotype_Atr | Clns1a | 1.657 | 1.09E-03 | glio |
| Interaction_genotype_Bcas2 | Glt1d1 | -4.168 | 1.09E-03 | glio |
| Interaction_genotype_Bcas2 | Mfsd13a | 3.558 | 1.09E-03 | glio |
| Interaction_genotype_Bcas2 | Mir670hg | -4.653 | 1.09E-03 | glio |
| Interaction_genotype_Bcas2 | Mycl | -4.553 | 1.09E-03 | glio |
| Interaction_genotype_Bcas2 | Srl | -4.466 | 1.09E-03 | glio |
| Interaction_genotype_Bcas2 | Hrct1 | -3.836 | 1.09E-03 | glio |
| Interaction_genotype_Bcas2 | Dscaml1 | -2.978 | 1.09E-03 | glio |
| Interaction_genotype_Bcas2 | Nfix | 2.172 | 1.09E-03 | glio |
| Interaction_genotype_Bcas2 | Numa1 | 1.912 | 1.09E-03 | glio |
| Interaction_genotype_Bora | Bhlhe41 | -5.022 | 1.09E-03 | glio |
| Interaction_genotype_Mtor | Umps | -2.542 | 1.09E-03 | glio |
| Interaction_genotype_Mtor | Ubfd1 | 1.703 | 1.09E-03 | glio |
| Interaction_genotype_Mtor | Sec61a1 | 1.645 | 1.09E-03 | glio |
| Interaction_genotype_Myc | 9530053A07R | -3.709 | 1.09E-03 | glio |
| Interaction_genotype_Myc | Sptbn4 | -3.573 | 1.09E-03 | glio |
| Interaction_genotype_Myc | Glrx | -3.434 | 1.09E-03 | glio |
| Interaction_genotype_Myc | 2610507I01Ri | -3.064 | 1.09E-03 | glio |
| Interaction_genotype_Myc | Grk5 | -3.024 | 1.09E-03 | glio |
| interaction_RT | Cspg4 | -1.549 | 1.09E-03 | glio |
| interaction_RT | Agpat4 | 1.08 | 1.09E-03 | glio |
| interaction_RT | Npepl1 | -0.963 | 1.09E-03 | glio |
| interaction_RT | Ube2w | 0.949 | 1.09E-03 | glio |
| interaction_RT | Utp6 | -0.867 | 1.09E-03 | glio |
| interaction_RT | Tcp1 | -0.604 | 1.09E-03 | glio |
| Aars2_in.vivo_noRT | Gm16230 | 2.309 | 1.09E-03 | glio |
| Aars2_in.vivo_noRT | Il1rapl1 | 2.359 | 1.09E-03 | glio |
| Aars2_in.vivo_noRT | Ccdc14 | 2.164 | 1.09E-03 | glio |
| Aars2_in.vivo_noRT | Cdc25b | 1.676 | 1.09E-03 | glio |
| Aars2_ex.vivo_noRT | H2-M10.1 | 2.706 | 1.09E-03 | glio |
| Atp5c1_in.vivo_noRT | Msl3l2 | 2.554 | 1.09E-03 | glio |
| Atr_in.vivo_noRT | 3110001I22Ri | 2.276 | 1.09E-03 | glio |
| Atr_in.vivo_noRT | Zfp763 | 2.18 | 1.09E-03 | glio |
| Atr_in.vivo_noRT | Glt1d1 | 2.191 | 1.09E-03 | glio |
| Atr_in.vivo_noRT | Gramd4 | 2.307 | 1.09E-03 | glio |
| Atr_in.vivo_noRT | 2210408I21Ri | 2.131 | 1.09E-03 | glio |
| Atr_in.vivo_noRT | Tcerg1 | -1.045 | 1.09E-03 | glio |
| Atr_ex.vivo_noRT | Ttc21b | -2.233 | 1.09E-03 | glio |

| | | | | |
|---|---|---|---|---|
| Bcas2_in.vivo_noRT | Rbp7 | 2.415 | 1.09E-03 | glio |
| Bcas2_in.vivo_noRT | Trp53rkb | 1.821 | 1.09E-03 | glio |
| Bcas2_in.vivo_noRT | Polr3c | 1.413 | 1.09E-03 | glio |
| Bora_in.vivo_noRT | Gm45605 | 2.383 | 1.09E-03 | glio |
| Bora_in.vivo_noRT | Gm10658 | 2.271 | 1.09E-03 | glio |
| Bora_in.vivo_noRT | Pdcd10 | -1.181 | 1.09E-03 | glio |
| Bora_in.vivo_noRT | Rps24 | -0.642 | 1.09E-03 | glio |
| Bora_ex.vivo_noRT | Pcdhgb5 | 2.808 | 1.09E-03 | glio |
| Chek1_in.vivo_noRT | Tab1 | 2.006 | 1.09E-03 | glio |
| Chek1_in.vivo_noRT | Ankrd13b | 1.529 | 1.09E-03 | glio |
| Chek1_ex.vivo_noRT | 4930505N22R | 2.707 | 1.09E-03 | glio |
| Cyp19a1_ex.vivo_noRT | Slc17a6 | 2.159 | 1.09E-03 | glio |
| Eif2s1_in.vivo_noRT | Gm15929 | 2.143 | 1.09E-03 | glio |
| Eif2s1_in.vivo_noRT | Cd7 | 1.879 | 1.09E-03 | glio |
| Eif2s1_in.vivo_noRT | Pitx3 | 2.143 | 1.09E-03 | glio |
| Eif2s1_in.vivo_noRT | Nisch | -0.85 | 1.09E-03 | glio |
| culture_effect | Vasn | -2.199 | 1.09E-04 | glio |
| culture_effect | Gm49708 | 1.593 | 1.09E-04 | glio |
| culture_effect | Mau2 | -0.759 | 1.09E-04 | glio |
| culture_effect | Nop10 | 0.512 | 1.09E-04 | glio |
| Interaction_genotype_Aars2 | Adgrb3 | -4.662 | 1.09E-04 | glio |
| Interaction_genotype_Atr | D630036H23R | -4.583 | 1.09E-04 | glio |
| Interaction_genotype_Atr | Lamtor3 | 2.055 | 1.09E-04 | glio |
| Interaction_genotype_Chek1 | Bex3 | 2.002 | 1.09E-04 | glio |
| Interaction_genotype_Chek1 | Atoh8 | -4.613 | 1.09E-04 | glio |
| Interaction_genotype_Chek1 | Map1lc3b | 1.712 | 1.09E-04 | glio |
| Interaction_genotype_Ifnar1 | Gm5934 | 4.002 | 1.09E-04 | glio |
| Interaction_genotype_Mtor | Olfr1191-ps1 | -4.236 | 1.09E-04 | glio |
| Interaction_genotype_Mtor | Hebp2 | 2.096 | 1.09E-04 | glio |
| Interaction_genotype_Mtor | Rad21 | 2.155 | 1.09E-04 | glio |
| Interaction_genotype_Mtor | Ncor1 | 1.733 | 1.09E-04 | glio |
| Interaction_genotype_Myc | A930033M14R | -3.694 | 1.09E-04 | glio |
| Interaction_genotype_Myc | Aph1b | -3.875 | 1.09E-04 | glio |
| Interaction_genotype_Myc | Col15a1 | -3.717 | 1.09E-04 | glio |
| Aars2_in.vivo_noRT | Myo1h | 2.832 | 1.09E-04 | glio |
| Aars2_in.vivo_noRT | Tsga10 | 2.864 | 1.09E-04 | glio |
| Aars2_in.vivo_noRT | Lrrc20 | 2.494 | 1.09E-04 | glio |
| Aars2_ex.vivo_noRT | Ctsf | 2.9 | 1.09E-04 | glio |
| Atp5c1_in.vivo_noRT | Psca | 2.668 | 1.09E-04 | glio |
| Atp5c1_in.vivo_noRT | Slc16a4 | 2.404 | 1.09E-04 | glio |
| Atr_in.vivo_noRT | Zc3h6 | 2.772 | 1.09E-04 | glio |
| Atr_in.vivo_noRT | Zfp764 | 2.541 | 1.09E-04 | glio |
| Atr_in.vivo_noRT | Rps3 | -0.804 | 1.09E-04 | glio |
| Atr_in.vivo_noRT | Rps19 | -0.764 | 1.09E-04 | glio |
| Bcas2_in.vivo_noRT | Gm9776 | 2.7 | 1.09E-04 | glio |
| Bcas2_ex.vivo_noRT | Zscan4-ps2 | 2.742 | 1.09E-04 | glio |
| Bcas2_ex.vivo_noRT | Gm6566 | 3.147 | 1.09E-04 | glio |
| Bora_in.vivo_noRT | Gm17477 | 2.49 | 1.09E-04 | glio |
| Chek1_in.vivo_noRT | Elf3 | 3.003 | 1.09E-04 | glio |
| Chek1_ex.vivo_noRT | Gm20536 | 3.475 | 1.09E-04 | glio |
| Chek1_ex.vivo_noRT | 1700017M07R | 2.897 | 1.09E-04 | glio |
| culture_effect | Agpat5 | -0.685 | 1.09E-05 | glio |

| | | | | |
|---|---|---|---|---|
| Interaction_genotype_Mtor | BC037032 | -5.615 | 1.09E-05 | glio |
| Interaction_genotype_Myc | K230015D01R | -5.044 | 1.09E-05 | glio |
| Interaction_genotype_Myc | Ubd | 4.872 | 1.09E-05 | glio |
| Interaction_genotype_Myc | Fau | 1.877 | 1.09E-05 | glio |
| Interaction_genotype_Rad51 | Ubd | 5.476 | 1.09E-05 | glio |
| Interaction_genotype_Rad51 | Klrk1 | 4.852 | 1.09E-05 | glio |
| Atr_ex.vivo_noRT | Alpl | 3.947 | 1.09E-05 | glio |
| Chek1_ex.vivo_noRT | A330032B11R | 3.463 | 1.09E-05 | glio |
| Chek1_ex.vivo_noRT | Arid3c | 3.664 | 1.09E-05 | glio |
| Hsd17b10_in.vivo_noRT | Thnsl2 | 2.448 | 1.09E-05 | glio |
| Hsd17b10_in.vivo_noRT | Gm5039 | 3.616 | 1.09E-05 | glio |
| culture_effect | Ccndbp1 | -0.815 | 1.09E-06 | glio |
| Interaction_genotype_Mtor | Nlrx1 | -6.145 | 1.09E-06 | glio |
| Interaction_genotype_Mtor | Gm49172 | -3.839 | 1.09E-06 | glio |
| Aars2_in.vivo_noRT | Lbhd2 | 4.085 | 1.09E-06 | glio |
| Aars2_in.vivo_noRT | 2610507I01Ri | 3.065 | 1.09E-06 | glio |
| Atr_in.vivo_noRT | Gm29361 | 3.508 | 1.09E-06 | glio |
| Atr_in.vivo_noRT | H2-Q10 | 3.365 | 1.09E-06 | glio |
| Bcas2_in.vivo_noRT | Gm10115 | 3.369 | 1.09E-06 | glio |
| Bora_in.vivo_noRT | Gm49494 | 3.591 | 1.09E-06 | glio |
| Bora_in.vivo_noRT | Kcnmb4os2 | 3.499 | 1.09E-06 | glio |
| Chek1_in.vivo_noRT | Lamc2 | 3.396 | 1.09E-06 | glio |
| Chek1_in.vivo_noRT | Ikzf2 | 3.62 | 1.09E-06 | glio |
| Chek1_ex.vivo_noRT | Akr1b7 | 4.443 | 1.09E-06 | glio |
| Hsd17b10_in.vivo_noRT | Rhox7a | 3.616 | 1.09E-06 | glio |
| Interaction_genotype_Mtor | 1110020A21R | -6.45 | 1.09E-07 | glio |
| Aars2_in.vivo_noRT | Gchfr | 4.214 | 1.09E-07 | glio |
| Aars2_in.vivo_noRT | Rfx6 | 3.567 | 1.09E-07 | glio |
| Atr_in.vivo_noRT | Gm13421 | 3.809 | 1.09E-07 | glio |
| Atr_in.vivo_noRT | 9130204L05Ri | 3.939 | 1.09E-07 | glio |
| Bora_in.vivo_noRT | Gm2788 | 3.888 | 1.09E-07 | glio |
| Bora_in.vivo_noRT | Gm48882 | 3.812 | 1.09E-07 | glio |
| Bora_in.vivo_noRT | Zfp473 | 3.512 | 1.09E-07 | glio |
| Bora_in.vivo_noRT | Eps8l2 | 3.834 | 1.09E-07 | glio |
| Chek1_ex.vivo_noRT | Cox7b2 | 4.045 | 1.09E-07 | glio |
| Hsd17b10_in.vivo_noRT | Gm21762 | 3.352 | 1.09E-07 | glio |
| Aars2_in.vivo_noRT | Bfsp2 | 4.464 | 1.09E-08 | glio |
| Aars2_in.vivo_noRT | Syngap1 | 3.867 | 1.09E-08 | glio |
| Bcas2_in.vivo_noRT | Ctsf | 3.139 | 1.09E-08 | glio |
| Bcas2_in.vivo_noRT | Fam228a | 4.019 | 1.09E-08 | glio |
| Bora_in.vivo_noRT | Tnnt3 | 4.302 | 1.09E-08 | glio |
| Interaction_genotype_Mtor | Gm13288 | -6.234 | 1.09E-09 | glio |
| Atr_in.vivo_noRT | Gm50399 | 3.981 | 1.09E-09 | glio |
| Bcas2_in.vivo_noRT | Ifi207 | 4.544 | 1.09E-09 | glio |
| Chek1_ex.vivo_noRT | Gm44646 | 4.256 | 1.09E-09 | glio |
| Atr_in.vivo_noRT | Traf3ip3 | 4.469 | 1.09E-10 | glio |
| Atr_in.vivo_noRT | A430106G13R | 4.742 | 1.09E-10 | glio |
| Bcas2_in.vivo_noRT | 2300002M23F | 3.638 | 1.09E-10 | glio |
| Bcas2_in.vivo_noRT | Dlx4 | 4.281 | 1.09E-10 | glio |
| Bora_in.vivo_noRT | 2610027K06R | 4.28 | 1.09E-10 | glio |
| Atr_in.vivo_noRT | Slc17a9 | 4.588 | 1.09E-11 | glio |
| Atr_in.vivo_noRT | Amdhd1 | 4.49 | 1.09E-11 | glio |

| | | | | |
|---|---|---|---|---|
| Atr_in.vivo_noRT | Spdye4a | 4.876 | 1.09E-11 | glio |
| culture_effect | Hnrnpc | -0.796 | 1.09E-12 | glio |
| Aars2_in.vivo_noRT | Gm13822 | 4.801 | 1.09E-12 | glio |
| Bora_in.vivo_noRT | Noxred1 | 4.733 | 1.09E-12 | glio |
| Aars2_in.vivo_noRT | Vmn1r91 | 4.674 | 1.09E-13 | glio |
| Bora_in.vivo_noRT | Sh3d21 | 4.414 | 1.09E-13 | glio |
| Chek1_in.vivo_noRT | Ttc16 | 5.051 | 1.09E-13 | glio |
| Atr_in.vivo_noRT | Gm50394 | 5.267 | 1.09E-14 | glio |
| Chek1_in.vivo_noRT | Masp1 | 4.522 | 1.09E-14 | glio |
| Chek1_in.vivo_noRT | Ly6d | 4.316 | 1.09E-14 | glio |
| Chek1_in.vivo_noRT | Gm4858 | 5.044 | 1.09E-14 | glio |
| Chek1_in.vivo_noRT | Gm4166 | 4.116 | 1.09E-15 | glio |
| culture_effect | Paics | -1.047 | 1.09E-17 | glio |
| Bcas2_in.vivo_noRT | Gast | 4.807 | 1.09E-17 | glio |
| culture_effect | Ier5 | -2.881 | 1.09E-19 | glio |
| Bora_in.vivo_noRT | Gm30794 | 4.936 | 1.09E-21 | glio |
| culture_effect | Tkt | -1.537 | 1.09E-22 | glio |
| Aars2_in.vivo_noRT | Gm13449 | 5.327 | 1.09E-22 | glio |
| Bora_in.vivo_noRT | Gpc5 | 4.936 | 1.09E-22 | glio |
| culture_effect | Pmp22 | 3.987 | 1.09E-23 | glio |
| Bcas2_in.vivo_noRT | A930019D19R | 4.807 | 1.09E-23 | glio |
| Aars2_in.vivo_noRT | Gm12703 | 5.57 | 1.09E-27 | glio |
| Atr_in.vivo_noRT | Gm5788 | 5.533 | 1.09E-27 | glio |
| culture_effect | Nox1 | -1.822 | 1.10E-03 | glio |
| culture_effect | Mgp | -2.452 | 1.10E-03 | glio |
| culture_effect | Cyld | 0.649 | 1.10E-03 | glio |
| Interaction_genotype_Aars2 | Atoh8 | -3.307 | 1.10E-03 | glio |
| Interaction_genotype_Bora | Zfp873 | -3.831 | 1.10E-03 | glio |
| Interaction_genotype_Mtor | Zhx3 | -4.001 | 1.10E-03 | glio |
| Interaction_genotype_Mtor | Sdr39u1 | -3.139 | 1.10E-03 | glio |
| Interaction_genotype_Mtor | H2-DMb1 | 2.046 | 1.10E-03 | glio |
| Interaction_genotype_Mtor | Ubxn1 | 1.137 | 1.10E-03 | glio |
| Interaction_genotype_Myc | Olfr56 | -2.729 | 1.10E-03 | glio |
| Interaction_genotype_Myc | Pglyrp1 | -3.797 | 1.10E-03 | glio |
| Interaction_genotype_Myc | Nim1k | -3.841 | 1.10E-03 | glio |
| Interaction_genotype_Myc | Cops9 | 1.432 | 1.10E-03 | glio |
| interaction_RT | Pgm5 | 3.18 | 1.10E-03 | glio |
| interaction_RT | Mdc1 | -1.259 | 1.10E-03 | glio |
| interaction_RT | Aven | -1.28 | 1.10E-03 | glio |
| interaction_RT | Elf1 | 1.027 | 1.10E-03 | glio |
| interaction_RT | Nol4l | 1.312 | 1.10E-03 | glio |
| Aars2_in.vivo_noRT | Tmem255a | 2.418 | 1.10E-03 | glio |
| Aars2_ex.vivo_noRT | Esyt3 | 3.104 | 1.10E-03 | glio |
| Atp5c1_in.vivo_noRT | Lrfn1 | 2.548 | 1.10E-03 | glio |
| Atr_in.vivo_noRT | Tspan8 | 2.296 | 1.10E-03 | glio |
| Atr_in.vivo_noRT | Cpeb3 | 2.349 | 1.10E-03 | glio |
| Atr_in.vivo_noRT | Rbm5 | -1.451 | 1.10E-03 | glio |
| Bcas2_in.vivo_noRT | 5730414N17R | 2.537 | 1.10E-03 | glio |
| Bcas2_in.vivo_noRT | Srfbp1 | -1.534 | 1.10E-03 | glio |
| Bcas2_in.vivo_noRT | Hnrnpab | -0.591 | 1.10E-03 | glio |
| Bcas2_ex.vivo_noRT | Masp1 | 2.727 | 1.10E-03 | glio |
| Bcas2_ex.vivo_noRT | AI463229 | 2.742 | 1.10E-03 | glio |

| | | | | |
|---|---|---|---|---|
| Bcas2_ex.vivo_noRT | Nck2 | -2.58 | 1.10E-03 | glio |
| Bora_in.vivo_noRT | Cpne7 | 2.538 | 1.10E-03 | glio |
| Bora_in.vivo_noRT | Pcdhb17 | 2.339 | 1.10E-03 | glio |
| Bora_in.vivo_noRT | Stamos | 2.154 | 1.10E-03 | glio |
| Bora_in.vivo_noRT | 2700081O15R | -1.536 | 1.10E-03 | glio |
| Cenpj_ex.vivo_noRT | Gm49808 | 2.198 | 1.10E-03 | glio |
| Chek1_in.vivo_noRT | Rbakdn | 2.342 | 1.10E-03 | glio |
| Chek1_in.vivo_noRT | Sapcd1 | 2.379 | 1.10E-03 | glio |
| Chek1_ex.vivo_noRT | Foxl2os | 2.856 | 1.10E-03 | glio |
| Chek1_ex.vivo_noRT | Tlr2 | 3.159 | 1.10E-03 | glio |
| Fanca_ex.vivo_noRT | Ubd | 3.315 | 1.10E-03 | glio |
| Fanca_ex.vivo_noRT | Akap14 | 2.919 | 1.10E-03 | glio |
| Hsd17b10_in.vivo_noRT | Gm47484 | 2.623 | 1.10E-03 | glio |
| culture_effect | Smim1 | 1.146 | 1.10E-04 | glio |
| culture_effect | Stmn1 | -0.644 | 1.10E-04 | glio |
| Interaction_genotype_Aars2 | Nxph3 | -4.386 | 1.10E-04 | glio |
| Interaction_genotype_Bcas2 | Rasgrp2 | -5.119 | 1.10E-04 | glio |
| Interaction_genotype_Bcas2 | Daam2 | -5.033 | 1.10E-04 | glio |
| Interaction_genotype_Bcas2 | Jund | 1.667 | 1.10E-04 | glio |
| Interaction_genotype_Bora | Hspa12a | -5.07 | 1.10E-04 | glio |
| Interaction_genotype_Cdkn2a | Gm47510 | 3.894 | 1.10E-04 | glio |
| Interaction_genotype_Cdkn2a | Pnliprp2 | 4.413 | 1.10E-04 | glio |
| Interaction_genotype_Cdkn2a | Gm48617 | 3.898 | 1.10E-04 | glio |
| Interaction_genotype_Fanca | Gpr17 | 5.32 | 1.10E-04 | glio |
| Interaction_genotype_Mtor | Gm26930 | -4.95 | 1.10E-04 | glio |
| Interaction_genotype_Mtor | Tead3 | -4.538 | 1.10E-04 | glio |
| Interaction_genotype_Mtor | 1700007L15Ri | -3.332 | 1.10E-04 | glio |
| Interaction_genotype_Myc | Gm5577 | -4.009 | 1.10E-04 | glio |
| Interaction_genotype_Myc | Pmp2 | -3.851 | 1.10E-04 | glio |
| Interaction_genotype_Myc | Hist1h2bl | -3.923 | 1.10E-04 | glio |
| Interaction_genotype_Myc | Gm32051 | -4.481 | 1.10E-04 | glio |
| Aars2_in.vivo_noRT | Slc25a43 | 2.743 | 1.10E-04 | glio |
| Aars2_in.vivo_noRT | Fam177a | 2.559 | 1.10E-04 | glio |
| Aars2_in.vivo_noRT | Alas1 | 1.969 | 1.10E-04 | glio |
| Aars2_in.vivo_noRT | P4hb | -0.855 | 1.10E-04 | glio |
| Atr_in.vivo_noRT | Rasl12 | 2.486 | 1.10E-04 | glio |
| Atr_in.vivo_noRT | Pcbp3 | 2.325 | 1.10E-04 | glio |
| Atr_in.vivo_noRT | A430005L14Ri | -1.549 | 1.10E-04 | glio |
| Bard1_ex.vivo_noRT | Gm2022 | 2.814 | 1.10E-04 | glio |
| Cenpj_ex.vivo_noRT | Cfb | 3.389 | 1.10E-04 | glio |
| Cenpj_ex.vivo_noRT | Capn13 | 2.591 | 1.10E-04 | glio |
| Cenpj_ex.vivo_noRT | Gm47540 | 2.996 | 1.10E-04 | glio |
| Cenpj_ex.vivo_noRT | Gm13191 | 2.591 | 1.10E-04 | glio |
| Cenpj_ex.vivo_noRT | 4933416E03Ri | 2.591 | 1.10E-04 | glio |
| Chek1_in.vivo_noRT | Gm43773 | 2.623 | 1.10E-04 | glio |
| Chek1_in.vivo_noRT | Pcdhga6 | 2.58 | 1.10E-04 | glio |
| Chek1_in.vivo_noRT | Car7 | 2.719 | 1.10E-04 | glio |
| Chek1_in.vivo_noRT | Rpl35a | -0.666 | 1.10E-04 | glio |
| Chek1_ex.vivo_noRT | Ptges | 3.572 | 1.10E-04 | glio |
| Hsd17b10_in.vivo_noRT | Mro | 2.825 | 1.10E-04 | glio |
| Hsd17b10_in.vivo_noRT | S100a7a | 2.901 | 1.10E-04 | glio |
| culture_effect | Zscan22 | 1.601 | 1.10E-05 | glio |

| | | | | |
|---|---|---|---|---|
| culture_effect | Rpl36a-ps1 | -0.897 | 1.10E-05 | glio |
| culture_effect | Rce1 | -0.728 | 1.10E-05 | glio |
| Interaction_genotype_Mtor | Gm19391 | -4.467 | 1.10E-05 | glio |
| Interaction_genotype_Mtor | Gm13421 | -5.328 | 1.10E-05 | glio |
| Interaction_genotype_Mtor | Tcstv1 | -4.498 | 1.10E-05 | glio |
| Interaction_genotype_Mtor | 4933439K11R | -5.391 | 1.10E-05 | glio |
| Interaction_genotype_Mtor | Gm21319 | -4.099 | 1.10E-05 | glio |
| Interaction_genotype_Mtor | Psma3 | 1.958 | 1.10E-05 | glio |
| Interaction_genotype_Mtor | Ubb | 1.758 | 1.10E-05 | glio |
| interaction_RT | Akr1b8 | 1.383 | 1.10E-05 | glio |
| Aars2_in.vivo_noRT | 6720489N17R | 3.444 | 1.10E-05 | glio |
| Aars2_in.vivo_noRT | Rasl12 | 2.816 | 1.10E-05 | glio |
| Atp5c1_in.vivo_noRT | Ccl3 | 2.408 | 1.10E-05 | glio |
| Atp5c1_in.vivo_noRT | Gm48780 | 2.408 | 1.10E-05 | glio |
| Atp5c1_in.vivo_noRT | Gm15138 | 2.402 | 1.10E-05 | glio |
| Atp5c1_in.vivo_noRT | Pldi | 2.668 | 1.10E-05 | glio |
| Atr_in.vivo_noRT | E430018J23Ri | 2.942 | 1.10E-05 | glio |
| Atr_in.vivo_noRT | Rab15 | 3.024 | 1.10E-05 | glio |
| Atr_ex.vivo_noRT | Gm26671 | 2.771 | 1.10E-05 | glio |
| Bcas2_in.vivo_noRT | Zfhx2os | 3.226 | 1.10E-05 | glio |
| Bora_in.vivo_noRT | Gm10401 | 2.734 | 1.10E-05 | glio |
| Bora_in.vivo_noRT | S100a7a | 3.17 | 1.10E-05 | glio |
| Bora_ex.vivo_noRT | Slc16a5 | 2.808 | 1.10E-05 | glio |
| Chek1_in.vivo_noRT | Nxf2 | 3.23 | 1.10E-05 | glio |
| Chek1_in.vivo_noRT | Mapk15 | 3.059 | 1.10E-05 | glio |
| Chek1_in.vivo_noRT | Tlr2 | 3.146 | 1.10E-05 | glio |
| Chek1_in.vivo_noRT | Nup50 | -1.535 | 1.10E-05 | glio |
| Chek1_ex.vivo_noRT | Gm28592 | 3.679 | 1.10E-05 | glio |
| Chek1_ex.vivo_noRT | Gm30124 | 3.652 | 1.10E-05 | glio |
| Chek1_ex.vivo_noRT | Cfap46 | 3.457 | 1.10E-05 | glio |
| culture_effect | C87436 | 1 | 1.10E-06 | glio |
| Interaction_genotype_Mtor | Boc | -4.228 | 1.10E-06 | glio |
| Interaction_genotype_Mtor | Mmp13 | -4.099 | 1.10E-06 | glio |
| Aars2_in.vivo_noRT | Slc24a1 | 3.465 | 1.10E-06 | glio |
| Bora_in.vivo_noRT | 4930445N18R | 3.492 | 1.10E-06 | glio |
| Bora_in.vivo_noRT | Syne3 | 3.68 | 1.10E-06 | glio |
| Chek1_in.vivo_noRT | Gm15510 | 3.449 | 1.10E-06 | glio |
| Chek1_in.vivo_noRT | Synpr | 3.881 | 1.10E-06 | glio |
| Hsd17b10_in.vivo_noRT | AI854703 | 3.497 | 1.10E-06 | glio |
| Hsd17b10_in.vivo_noRT | Ceacam16 | 2.958 | 1.10E-06 | glio |
| culture_effect | Rmdn3 | -1.297 | 1.10E-07 | glio |
| culture_effect | Rcor2 | -1.187 | 1.10E-07 | glio |
| Interaction_genotype_Mtor | Limch1 | -6.638 | 1.10E-07 | glio |
| Interaction_genotype_Mtor | Tnni3k | -4.886 | 1.10E-07 | glio |
| Interaction_genotype_Mtor | Gm13348 | -5.371 | 1.10E-07 | glio |
| Aars2_in.vivo_noRT | Lmod1 | 3.895 | 1.10E-07 | glio |
| Bora_in.vivo_noRT | A330084C13R | 4.018 | 1.10E-07 | glio |
| Bora_in.vivo_noRT | 1700011M02F | 4.673 | 1.10E-07 | glio |
| Aars2_in.vivo_noRT | Gm49042 | 3.697 | 1.10E-08 | glio |
| Atr_in.vivo_noRT | Vcam1 | 5.017 | 1.10E-08 | glio |
| Atr_in.vivo_noRT | 6430590A07R | 3.001 | 1.10E-08 | glio |
| Chek1_in.vivo_noRT | 4833407H14R | 4.409 | 1.10E-08 | glio |

| | | | | |
|---|---|---|---|---|
| Aars2_in.vivo_noRT | Arg2 | 4.201 | 1.10E-09 | glio |
| Aars2_in.vivo_noRT | Klhl30 | 4.142 | 1.10E-09 | glio |
| Atr_in.vivo_noRT | 1700011D18R | 4.488 | 1.10E-10 | glio |
| Chek1_ex.vivo_noRT | Car10 | 4.842 | 1.10E-10 | glio |
| Aars2_in.vivo_noRT | Ngb | 4.556 | 1.10E-11 | glio |
| Atr_in.vivo_noRT | Mfap3l | 5.142 | 1.10E-11 | glio |
| Aars2_in.vivo_noRT | Abcg4 | 4.536 | 1.10E-12 | glio |
| Aars2_in.vivo_noRT | Gm15579 | 5.327 | 1.10E-12 | glio |
| Aars2_in.vivo_noRT | E130215H24R | 4.536 | 1.10E-12 | glio |
| Chek1_ex.vivo_noRT | Snrpn | 4.842 | 1.10E-12 | glio |
| Chek1_ex.vivo_noRT | Pnliprp2 | 4.842 | 1.10E-12 | glio |
| culture_effect | Maff | -1.972 | 1.10E-13 | glio |
| culture_effect | Afdn | 1.255 | 1.10E-13 | glio |
| Chek1_in.vivo_noRT | Gm41318 | 5.044 | 1.10E-13 | glio |
| Chek1_ex.vivo_noRT | Gm47936 | 4.842 | 1.10E-13 | glio |
| culture_effect | Clic1 | -0.879 | 1.10E-14 | glio |
| Bcas2_in.vivo_noRT | Gm13054 | 4.799 | 1.10E-14 | glio |
| Bora_in.vivo_noRT | Spi1 | 4.936 | 1.10E-14 | glio |
| Chek1_in.vivo_noRT | Acat3 | 4.252 | 1.10E-14 | glio |
| Chek1_in.vivo_noRT | Gm5602 | 4.252 | 1.10E-14 | glio |
| culture_effect | Epb41l3 | 1.81 | 1.10E-15 | glio |
| Bcas2_in.vivo_noRT | Ptprc | 4.543 | 1.10E-16 | glio |
| Chek1_in.vivo_noRT | Gm30042 | 4.784 | 1.10E-16 | glio |
| Chek1_in.vivo_noRT | Gm34939 | 4.78 | 1.10E-16 | glio |
| Bora_in.vivo_noRT | 2010001K21R | 5.46 | 1.10E-17 | glio |
| Aars2_in.vivo_noRT | Trim16 | 5.577 | 1.10E-18 | glio |
| Bcas2_in.vivo_noRT | Gad1 | 4.541 | 1.10E-19 | glio |
| culture_effect | S100a6 | 1.776 | 1.10E-20 | glio |
| Atr_in.vivo_noRT | Gm50321 | 5.533 | 1.10E-20 | glio |
| culture_effect | H2-M3 | -3.157 | 1.10E-24 | glio |
| culture_effect | Gm34590 | 1.793 | 1.11E-03 | glio |
| culture_effect | Clec2d | 1.063 | 1.11E-03 | glio |
| Interaction_genotype_Aars2 | A930001C03R | -3.9 | 1.11E-03 | glio |
| Interaction_genotype_Aars2 | Ttbk1 | -3.962 | 1.11E-03 | glio |
| Interaction_genotype_Bcas2 | Sprr1a | -4.438 | 1.11E-03 | glio |
| Interaction_genotype_Bcas2 | Rapgef3os2 | -3.251 | 1.11E-03 | glio |
| Interaction_genotype_Bcas2 | Gm48435 | -2.852 | 1.11E-03 | glio |
| Interaction_genotype_Bcas2 | 3110082J24Ri | -4.515 | 1.11E-03 | glio |
| Interaction_genotype_Bcas2 | Prl7c1 | -3.16 | 1.11E-03 | glio |
| Interaction_genotype_Mtor | Gm47802 | -3.698 | 1.11E-03 | glio |
| Interaction_genotype_Mtor | Zfp54 | -3.717 | 1.11E-03 | glio |
| Interaction_genotype_Mtor | Mfap5 | -4.016 | 1.11E-03 | glio |
| Interaction_genotype_Myc | Gm48215 | -2.729 | 1.11E-03 | glio |
| Interaction_genotype_Myc | Slc16a6 | -3.864 | 1.11E-03 | glio |
| Interaction_genotype_Myc | Styk1 | -3.7 | 1.11E-03 | glio |
| Interaction_genotype_Myc | Mirg | -3.78 | 1.11E-03 | glio |
| Interaction_genotype_Myc | Plcxd2 | -3.776 | 1.11E-03 | glio |
| Interaction_genotype_Myc | Tmem246 | -3.487 | 1.11E-03 | glio |
| Interaction_genotype_Myc | Ppargc1b | -3.717 | 1.11E-03 | glio |
| Interaction_genotype_Myc | Inava | -3.463 | 1.11E-03 | glio |
| Interaction_genotype_Myc | Zfp647 | -3.154 | 1.11E-03 | glio |
| Interaction_genotype_Myc | Klhl12 | -3.21 | 1.11E-03 | glio |

| | | | | |
|---|---|---|---|---|
| Interaction_genotype_Rad51 | Cebpg | 1.806 | 1.11E-03 | glio |
| Aars2_in.vivo_noRT | 9230112E08Ri | 2.332 | 1.11E-03 | glio |
| Aars2_in.vivo_noRT | Zfr2 | 2.2 | 1.11E-03 | glio |
| Aars2_in.vivo_noRT | Gm43597 | 2.06 | 1.11E-03 | glio |
| Aars2_in.vivo_noRT | Tmco1 | -0.829 | 1.11E-03 | glio |
| Aars2_ex.vivo_noRT | Gm42664 | 2.706 | 1.11E-03 | glio |
| Atp5c1_in.vivo_noRT | Masp1 | 2.146 | 1.11E-03 | glio |
| Atr_in.vivo_noRT | Chn2 | 2.541 | 1.11E-03 | glio |
| Atr_in.vivo_noRT | Wdr78 | 2.501 | 1.11E-03 | glio |
| Atr_in.vivo_noRT | Chdh | 2.245 | 1.11E-03 | glio |
| Atr_in.vivo_noRT | Ddb2 | 1.834 | 1.11E-03 | glio |
| Atr_in.vivo_noRT | Syt1 | 2.21 | 1.11E-03 | glio |
| Bard1_ex.vivo_noRT | AC159140.1 | 2.421 | 1.11E-03 | glio |
| Bcas2_in.vivo_noRT | Gm14273 | 2.317 | 1.11E-03 | glio |
| Bcas2_ex.vivo_noRT | Gm36037 | 2.347 | 1.11E-03 | glio |
| Bora_in.vivo_noRT | Rell2 | 2.458 | 1.11E-03 | glio |
| Bora_in.vivo_noRT | A230087F16Ri | 2.312 | 1.11E-03 | glio |
| Bora_in.vivo_noRT | Fbxl14 | -1.566 | 1.11E-03 | glio |
| Chek1_in.vivo_noRT | Nrap | 2.504 | 1.11E-03 | glio |
| Chek1_in.vivo_noRT | Chn2 | 2.581 | 1.11E-03 | glio |
| Chek1_in.vivo_noRT | Sh3kbp1 | -1.268 | 1.11E-03 | glio |
| Chek1_in.vivo_noRT | Rpl7 | -0.626 | 1.11E-03 | glio |
| Chek1_ex.vivo_noRT | 4921507P07R | 2.867 | 1.11E-03 | glio |
| Eif2s1_in.vivo_noRT | Npy4r | 2.359 | 1.11E-03 | glio |
| Hsd17b10_in.vivo_noRT | Olfr920 | 2.397 | 1.11E-03 | glio |
| culture_effect | Cela1 | -0.75 | 1.11E-04 | glio |
| Interaction_genotype_Bcas2 | Gm13288 | -4.201 | 1.11E-04 | glio |
| Interaction_genotype_Mtor | 1700024B05R | -4.776 | 1.11E-04 | glio |
| Interaction_genotype_Mtor | Gm16244 | -4.66 | 1.11E-04 | glio |
| Interaction_genotype_Mtor | Gm16023 | -5.003 | 1.11E-04 | glio |
| Interaction_genotype_Mtor | Sh3glb1 | 1.831 | 1.11E-04 | glio |
| Interaction_genotype_Myc | Gngt2 | -4.398 | 1.11E-04 | glio |
| Interaction_genotype_Myc | Zgpat | -3.77 | 1.11E-04 | glio |
| interaction_RT | Hagh | 0.892 | 1.11E-04 | glio |
| Aars2_ex.vivo_noRT | Gm20732 | 3.293 | 1.11E-04 | glio |
| Atr_in.vivo_noRT | Gm26620 | 2.454 | 1.11E-04 | glio |
| Atr_ex.vivo_noRT | Gm10827 | 3.688 | 1.11E-04 | glio |
| Bcas2_in.vivo_noRT | Ksr1 | 2.758 | 1.11E-04 | glio |
| Bcas2_in.vivo_noRT | Tmem191c | 2.593 | 1.11E-04 | glio |
| Bora_in.vivo_noRT | Gm20707 | 2.759 | 1.11E-04 | glio |
| Bora_in.vivo_noRT | Slc25a29 | 2.525 | 1.11E-04 | glio |
| Bora_in.vivo_noRT | Naip6 | 2.684 | 1.11E-04 | glio |
| Chek1_in.vivo_noRT | Nudt7 | 2.593 | 1.11E-04 | glio |
| Eif2s1_in.vivo_noRT | Gm5415 | 2.143 | 1.11E-04 | glio |
| Eif2s1_in.vivo_noRT | Edn2 | 2.143 | 1.11E-04 | glio |
| Eif2s1_in.vivo_noRT | Gm14207 | 2.143 | 1.11E-04 | glio |
| Eif2s1_in.vivo_noRT | Gm49698 | 2.143 | 1.11E-04 | glio |
| Eif2s1_in.vivo_noRT | Olfr881 | 2.143 | 1.11E-04 | glio |
| culture_effect | 4930558J18Ri | 2.132 | 1.11E-05 | glio |
| culture_effect | Terf1 | -0.815 | 1.11E-05 | glio |
| culture_effect | Loxl3 | 0.927 | 1.11E-05 | glio |
| culture_effect | Stx5a | -0.638 | 1.11E-05 | glio |

| | | | | |
|---|---|---|---|---|
| Interaction_genotype_Mtor | Gm20707 | -5.104 | 1.11E-05 | glio |
| Interaction_genotype_Mtor | Tomm6os | -5.43 | 1.11E-05 | glio |
| Interaction_genotype_Mtor | 4930505N22R | -4.813 | 1.11E-05 | glio |
| Interaction_genotype_Mtor | Ablim2 | -3.567 | 1.11E-05 | glio |
| Interaction_genotype_Mtor | Wdr93 | -5.025 | 1.11E-05 | glio |
| Interaction_genotype_Myc | Rpl28 | 1.496 | 1.11E-05 | glio |
| Aars2_in.vivo_noRT | Nat8l | 2.736 | 1.11E-05 | glio |
| Atr_in.vivo_noRT | Gm49708 | 2.887 | 1.11E-05 | glio |
| Atr_ex.vivo_noRT | Snrpn | 3.164 | 1.11E-05 | glio |
| Atr_ex.vivo_noRT | Pnliprp2 | 3.164 | 1.11E-05 | glio |
| Bora_in.vivo_noRT | Lrrc75b | 3.28 | 1.11E-05 | glio |
| Bora_in.vivo_noRT | Gstt1 | 3.223 | 1.11E-05 | glio |
| Chek1_in.vivo_noRT | Gm11110 | 3.126 | 1.11E-05 | glio |
| Chek1_in.vivo_noRT | AC241534.1 | 3.184 | 1.11E-05 | glio |
| culture_effect | Rab12 | 0.757 | 1.11E-06 | glio |
| Interaction_genotype_Mtor | Hrct1 | -5.077 | 1.11E-06 | glio |
| Interaction_genotype_Mtor | Gm8016 | -4.099 | 1.11E-06 | glio |
| Aars2_in.vivo_noRT | Tlr6 | 3.492 | 1.11E-06 | glio |
| Aars2_in.vivo_noRT | Vgll2 | 2.991 | 1.11E-06 | glio |
| Aars2_ex.vivo_noRT | Gm14820 | 3.293 | 1.11E-06 | glio |
| Atr_in.vivo_noRT | Cdrt4 | 3.558 | 1.11E-06 | glio |
| Atr_in.vivo_noRT | Ccdc103 | 3.414 | 1.11E-06 | glio |
| Bora_in.vivo_noRT | Gm35066 | 3.444 | 1.11E-06 | glio |
| Bora_in.vivo_noRT | 1700001L05Ri | 3.238 | 1.11E-06 | glio |
| Bora_in.vivo_noRT | D830035M03I | 3.35 | 1.11E-06 | glio |
| culture_effect | 2310030G06R | 2.518 | 1.11E-07 | glio |
| culture_effect | 0610038B21R | 2.182 | 1.11E-07 | glio |
| Interaction_genotype_Mtor | Rab25 | -5.778 | 1.11E-07 | glio |
| Atr_in.vivo_noRT | Grip1 | 3.654 | 1.11E-07 | glio |
| Atr_in.vivo_noRT | Dennd2d | 3.266 | 1.11E-07 | glio |
| Atr_ex.vivo_noRT | Hoxc8 | 3.164 | 1.11E-07 | glio |
| Bora_in.vivo_noRT | Flrt1 | 3.961 | 1.11E-07 | glio |
| Bora_in.vivo_noRT | Rbm11 | 3.538 | 1.11E-07 | glio |
| Chek1_in.vivo_noRT | Arhgef19 | 3.865 | 1.11E-07 | glio |
| Chek1_ex.vivo_noRT | Elfn2 | 4.268 | 1.11E-07 | glio |
| Chek1_ex.vivo_noRT | Dact1 | 4.443 | 1.11E-07 | glio |
| Chek1_ex.vivo_noRT | Mpeg1 | 4.268 | 1.11E-07 | glio |
| Chek1_ex.vivo_noRT | Tmem229b | 4.472 | 1.11E-07 | glio |
| Aars2_in.vivo_noRT | Slc17a9 | 3.872 | 1.11E-08 | glio |
| Aars2_in.vivo_noRT | Guca1b | 4.038 | 1.11E-08 | glio |
| Bcas2_in.vivo_noRT | Hist2h3c2 | 4.541 | 1.11E-09 | glio |
| culture_effect | Vma21 | 0.738 | 1.11E-10 | glio |
| Bora_in.vivo_noRT | Cd83 | 3.936 | 1.11E-10 | glio |
| Chek1_in.vivo_noRT | Gm17146 | 4.126 | 1.11E-10 | glio |
| Aars2_in.vivo_noRT | Gm42984 | 4.676 | 1.11E-11 | glio |
| Aars2_in.vivo_noRT | 4933406C10R | 4.29 | 1.11E-11 | glio |
| Atr_in.vivo_noRT | Ppp1r3g | 4.349 | 1.11E-11 | glio |
| Chek1_ex.vivo_noRT | Gm12589 | 4.443 | 1.11E-11 | glio |
| Atr_in.vivo_noRT | Gm50375 | 5.267 | 1.11E-12 | glio |
| Chek1_in.vivo_noRT | A030005L19Ri | 4.515 | 1.11E-12 | glio |
| culture_effect | Slfn2 | -3.524 | 1.11E-13 | glio |
| culture_effect | P4hb | -0.833 | 1.11E-13 | glio |

| | | | | |
|---|---|---|---|---|
| culture_effect | Gm10399 | 3.699 | 1.11E-14 | glio |
| Bora_in.vivo_noRT | Gm36356 | 4.676 | 1.11E-14 | glio |
| culture_effect | Hes1 | -1.814 | 1.11E-15 | glio |
| Atr_in.vivo_noRT | Cyp4x1 | 5.267 | 1.11E-15 | glio |
| Atr_in.vivo_noRT | 9630014M24F | 5.292 | 1.11E-15 | glio |
| Atr_in.vivo_noRT | Gngt2 | 5.533 | 1.11E-15 | glio |
| Bcas2_in.vivo_noRT | Mecom | 5.336 | 1.11E-15 | glio |
| Bora_in.vivo_noRT | 1700016P04R | 5.194 | 1.11E-15 | glio |
| Chek1_in.vivo_noRT | Gm45235 | 4.784 | 1.11E-17 | glio |
| Aars2_in.vivo_noRT | Dok2 | 5.311 | 1.11E-18 | glio |
| Bora_in.vivo_noRT | Spink6 | 4.936 | 1.11E-18 | glio |
| Bcas2_in.vivo_noRT | Fabp6 | 4.807 | 1.11E-19 | glio |
| Chek1_in.vivo_noRT | Gstt4 | 4.78 | 1.11E-20 | glio |
| culture_effect | Cd47 | -1.682 | 1.11E-22 | glio |
| Atr_in.vivo_noRT | Hoxc8 | 5.533 | 1.11E-29 | glio |
| culture_effect | Cidea | -1.85 | 1.12E-03 | glio |
| culture_effect | Oip5os1 | 0.444 | 1.12E-03 | glio |
| Interaction_genotype_Aars2 | Cd248 | -3.965 | 1.12E-03 | glio |
| Interaction_genotype_Aars2 | 2210406O10R | -4.032 | 1.12E-03 | glio |
| Interaction_genotype_Atr | Olfml2a | -4.008 | 1.12E-03 | glio |
| Interaction_genotype_Atr | Otos | -3.064 | 1.12E-03 | glio |
| Interaction_genotype_Atr | Tmem205 | 1.771 | 1.12E-03 | glio |
| Interaction_genotype_Atr | 5430416N02R | 2.413 | 1.12E-03 | glio |
| Interaction_genotype_Atr | Fos | 2.321 | 1.12E-03 | glio |
| Interaction_genotype_Chek1 | Tax1bp3 | 2.366 | 1.12E-03 | glio |
| Interaction_genotype_Mtor | Lnx2 | -3.569 | 1.12E-03 | glio |
| Interaction_genotype_Mtor | Gstt2 | -3.738 | 1.12E-03 | glio |
| Interaction_genotype_Mtor | Btbd19 | -2.859 | 1.12E-03 | glio |
| Interaction_genotype_Mtor | Tnni1 | -2.749 | 1.12E-03 | glio |
| Interaction_genotype_Myc | Vsig10l | -3.627 | 1.12E-03 | glio |
| Interaction_genotype_Myc | Gm45606 | -3.565 | 1.12E-03 | glio |
| Interaction_genotype_Myc | Inhba | -3.941 | 1.12E-03 | glio |
| Interaction_genotype_Myc | Gtf2ird2 | -3.389 | 1.12E-03 | glio |
| Interaction_genotype_Myc | Kbtbd7 | -3.304 | 1.12E-03 | glio |
| Interaction_genotype_Myc | Ssh2 | -2.967 | 1.12E-03 | glio |
| Interaction_genotype_Myc | Smap2 | -2.171 | 1.12E-03 | glio |
| interaction_RT | Med21 | 0.553 | 1.12E-03 | glio |
| Aars2_in.vivo_noRT | Syde2 | 2.251 | 1.12E-03 | glio |
| Aars2_in.vivo_noRT | Gm5914 | 1.914 | 1.12E-03 | glio |
| Aars2_in.vivo_noRT | Rap2a | -1.533 | 1.12E-03 | glio |
| Aars2_in.vivo_noRT | Atp5j2 | -0.803 | 1.12E-03 | glio |
| Aars2_in.vivo_noRT | Rps25 | -0.64 | 1.12E-03 | glio |
| Aars2_ex.vivo_noRT | Ly6g6c | 2.899 | 1.12E-03 | glio |
| Atr_in.vivo_noRT | Arhgef28 | 2.065 | 1.12E-03 | glio |
| Atr_in.vivo_noRT | Tmem229b | 2.147 | 1.12E-03 | glio |
| Bard1_ex.vivo_noRT | Gm48265 | 2.814 | 1.12E-03 | glio |
| Bcas2_in.vivo_noRT | Ucn2 | 2.313 | 1.12E-03 | glio |
| Bcas2_in.vivo_noRT | Gm10353 | 2.293 | 1.12E-03 | glio |
| Bcas2_ex.vivo_noRT | Slc45a1 | 2.342 | 1.12E-03 | glio |
| Bora_in.vivo_noRT | 9330121J05Ri | 2.364 | 1.12E-03 | glio |
| Bora_ex.vivo_noRT | Bco1 | 2.806 | 1.12E-03 | glio |
| Cenpj_in.vivo_noRT | Esp24 | 2.022 | 1.12E-03 | glio |

| | | | | |
|---|---|---|---|---|
| Cenpj_ex.vivo_noRT | Crb3 | 2.591 | 1.12E-03 | glio |
| Chek1_in.vivo_noRT | Actr10 | -0.987 | 1.12E-03 | glio |
| Chek1_ex.vivo_noRT | Dab2 | 2.732 | 1.12E-03 | glio |
| Eif2s1_in.vivo_noRT | Apobr | 2.922 | 1.12E-03 | glio |
| Hsd17b10_in.vivo_noRT | Apoa1 | 2.583 | 1.12E-03 | glio |
| Hsd17b10_in.vivo_noRT | 4930545E07Ri | 2.583 | 1.12E-03 | glio |
| culture_effect | Thap11 | -0.695 | 1.12E-04 | glio |
| Interaction_genotype_Dclre1c | Gm40538 | 4.349 | 1.12E-04 | glio |
| Interaction_genotype_Mtor | Zfp184 | -4.296 | 1.12E-04 | glio |
| Interaction_genotype_Mtor | Gm20324 | -3.971 | 1.12E-04 | glio |
| Interaction_genotype_Mtor | Dnmt1 | 2.05 | 1.12E-04 | glio |
| Interaction_genotype_Myc | Usp6nl | -3.104 | 1.12E-04 | glio |
| interaction_RT | Pcdh18 | -1.574 | 1.12E-04 | glio |
| Aars2_in.vivo_noRT | Lgr6 | 3.118 | 1.12E-04 | glio |
| Aars2_in.vivo_noRT | Cpne7 | 2.928 | 1.12E-04 | glio |
| Aars2_in.vivo_noRT | Pou5f1 | 2.542 | 1.12E-04 | glio |
| Aars2_in.vivo_noRT | P4ha3 | 2.628 | 1.12E-04 | glio |
| Aars2_ex.vivo_noRT | Vgll2 | 3.293 | 1.12E-04 | glio |
| Aars2_ex.vivo_noRT | Rem2 | 3.293 | 1.12E-04 | glio |
| Atp5c1_in.vivo_noRT | Jph4 | 2.933 | 1.12E-04 | glio |
| Atr_in.vivo_noRT | Gm20597 | 2.952 | 1.12E-04 | glio |
| Atr_in.vivo_noRT | Car7 | 2.685 | 1.12E-04 | glio |
| Atr_in.vivo_noRT | Triap1 | -1.589 | 1.12E-04 | glio |
| Atr_in.vivo_noRT | Rac1 | -1.061 | 1.12E-04 | glio |
| Bard1_ex.vivo_noRT | 4930447F24Ri | 3.432 | 1.12E-04 | glio |
| Bcas2_ex.vivo_noRT | Ccl3 | 2.742 | 1.12E-04 | glio |
| Bcas2_ex.vivo_noRT | AU019990 | 2.742 | 1.12E-04 | glio |
| Bcas2_ex.vivo_noRT | Gm48780 | 2.742 | 1.12E-04 | glio |
| Bcas2_ex.vivo_noRT | Cys1 | 3.131 | 1.12E-04 | glio |
| Bcas2_ex.vivo_noRT | Gm36793 | 2.742 | 1.12E-04 | glio |
| Bcas2_ex.vivo_noRT | Brsk2 | 3.524 | 1.12E-04 | glio |
| Bora_in.vivo_noRT | Dynlt1c | 3.133 | 1.12E-04 | glio |
| Chek1_in.vivo_noRT | Nup210l | 2.917 | 1.12E-04 | glio |
| Chek1_in.vivo_noRT | Fam178b | 2.866 | 1.12E-04 | glio |
| Hsd17b10_in.vivo_noRT | Tspyl3 | 2.533 | 1.12E-04 | glio |
| Hsd17b10_in.vivo_noRT | Vcam1 | 3.621 | 1.12E-04 | glio |
| culture_effect | Pde6h | 1.866 | 1.12E-05 | glio |
| culture_effect | AW011738 | 1.325 | 1.12E-05 | glio |
| culture_effect | Abcd3 | 0.874 | 1.12E-05 | glio |
| culture_effect | Csk | 0.846 | 1.12E-05 | glio |
| culture_effect | Sap30l | 0.602 | 1.12E-05 | glio |
| Interaction_genotype_Mtor | Cyp2u1 | -4.719 | 1.12E-05 | glio |
| Interaction_genotype_Mtor | Gm35066 | -5.086 | 1.12E-05 | glio |
| Interaction_genotype_Mtor | Ccl3 | -3.839 | 1.12E-05 | glio |
| Interaction_genotype_Mtor | Dppa2 | -4.934 | 1.12E-05 | glio |
| Interaction_genotype_Mtor | Speer4e | -4.099 | 1.12E-05 | glio |
| Interaction_genotype_Mtor | Gm48780 | -3.839 | 1.12E-05 | glio |
| Interaction_genotype_Myc | Sfr1 | 1.786 | 1.12E-05 | glio |
| Aars2_in.vivo_noRT | Stac3 | 3.184 | 1.12E-05 | glio |
| Aars2_ex.vivo_noRT | Gm28940 | 3.293 | 1.12E-05 | glio |
| Atr_in.vivo_noRT | D130062J10Ri | 2.805 | 1.12E-05 | glio |
| Bcas2_in.vivo_noRT | Mpeg1 | 2.831 | 1.12E-05 | glio |

| | | | | |
|---|---|---|---|---|
| Bora_in.vivo_noRT | 4930590J08Ri | 3.189 | 1.12E-05 | glio |
| Chek1_in.vivo_noRT | Tdrp | 2.995 | 1.12E-05 | glio |
| Chek1_in.vivo_noRT | E430018J23Ri | 2.964 | 1.12E-05 | glio |
| culture_effect | Mapk12 | -1.202 | 1.12E-06 | glio |
| culture_effect | Atp6v1b2 | -0.792 | 1.12E-06 | glio |
| Interaction_genotype_Aars2 | Ube2e1 | 2.551 | 1.12E-06 | glio |
| interaction_RT | Dbp | 2.03 | 1.12E-06 | glio |
| Atr_in.vivo_noRT | Tspan33 | 3.439 | 1.12E-06 | glio |
| Atr_ex.vivo_noRT | Gpr4 | 3.164 | 1.12E-06 | glio |
| Bcas2_in.vivo_noRT | Gm49891 | 3.61 | 1.12E-06 | glio |
| Chek1_in.vivo_noRT | Dnaaf3 | 3.091 | 1.12E-06 | glio |
| Chek1_in.vivo_noRT | Dmrt2 | 3.634 | 1.12E-06 | glio |
| Chek1_in.vivo_noRT | Gm14327 | 3.538 | 1.12E-06 | glio |
| Chek1_in.vivo_noRT | Cnfn | 3.649 | 1.12E-06 | glio |
| Chek1_in.vivo_noRT | Gm49801 | 3.512 | 1.12E-06 | glio |
| Chek1_in.vivo_noRT | Slc7a14 | 3.273 | 1.12E-06 | glio |
| Chek1_ex.vivo_noRT | Aldh1a2 | 4.049 | 1.12E-06 | glio |
| Interaction_genotype_Mtor | D630036H23R | -6.146 | 1.12E-07 | glio |
| Interaction_genotype_Mtor | Ccdc125 | -5.267 | 1.12E-07 | glio |
| Aars2_in.vivo_noRT | Rnf208 | 3.934 | 1.12E-07 | glio |
| Aars2_in.vivo_noRT | Hsd17b14 | 3.906 | 1.12E-07 | glio |
| Atr_in.vivo_noRT | Mrgprf | 3.737 | 1.12E-07 | glio |
| Bcas2_in.vivo_noRT | Gm8212 | 3.705 | 1.12E-07 | glio |
| Bora_in.vivo_noRT | Gm2012 | 3.772 | 1.12E-07 | glio |
| Bora_in.vivo_noRT | Grip2 | 3.685 | 1.12E-07 | glio |
| Chek1_in.vivo_noRT | Dcpp3 | 3.526 | 1.12E-07 | glio |
| Chek1_in.vivo_noRT | Mrgprf | 3.759 | 1.12E-07 | glio |
| culture_effect | Lbhd2 | 3.343 | 1.12E-08 | glio |
| culture_effect | Got1 | -0.886 | 1.12E-08 | glio |
| Aars2_in.vivo_noRT | 1700124L16Ri | 4.092 | 1.12E-08 | glio |
| Atr_in.vivo_noRT | Gm26879 | 3.526 | 1.12E-08 | glio |
| Atr_in.vivo_noRT | Dnaaf3 | 3.581 | 1.12E-08 | glio |
| Chek1_in.vivo_noRT | Serpina3i | 3.723 | 1.12E-08 | glio |
| Chek1_in.vivo_noRT | Gpr37l1 | 3.93 | 1.12E-08 | glio |
| Aars2_in.vivo_noRT | Hist1h4k | 4.271 | 1.12E-09 | glio |
| Aars2_in.vivo_noRT | Mir670hg | 4.676 | 1.12E-09 | glio |
| Bcas2_in.vivo_noRT | B930094E09Ri | 4.291 | 1.12E-09 | glio |
| Chek1_in.vivo_noRT | Slc23a3 | 4.142 | 1.12E-09 | glio |
| Hsd17b10_in.vivo_noRT | Masp1 | 3.625 | 1.12E-09 | glio |
| culture_effect | Qk | 0.802 | 1.12E-10 | glio |
| Atr_in.vivo_noRT | Nhsl2 | 4.415 | 1.12E-10 | glio |
| Bora_in.vivo_noRT | D830039M14I | 4.67 | 1.12E-10 | glio |
| Hsd17b10_in.vivo_noRT | Gm32250 | 3.365 | 1.12E-10 | glio |
| culture_effect | Bckdk | 1.077 | 1.12E-11 | glio |
| Bcas2_in.vivo_noRT | Zfp810 | 4.807 | 1.12E-12 | glio |
| Bora_in.vivo_noRT | Cfap46 | 4.469 | 1.12E-12 | glio |
| Atr_in.vivo_noRT | Gm15972 | 4.54 | 1.12E-13 | glio |
| Chek1_in.vivo_noRT | Gm44752 | 4.657 | 1.12E-13 | glio |
| Chek1_in.vivo_noRT | Gm27239 | 5.044 | 1.12E-13 | glio |
| Chek1_in.vivo_noRT | 4930412M03F | 5.044 | 1.12E-13 | glio |
| Chek1_in.vivo_noRT | 4930483J18Ri | 4.784 | 1.12E-13 | glio |
| Aars2_in.vivo_noRT | Gm13403 | 5.179 | 1.12E-14 | glio |

| | | | | |
|---|---|---|---|---|
| Atr_in.vivo_noRT | Dppa2 | 5.001 | 1.12E-14 | glio |
| Atr_in.vivo_noRT | Rhox7a | 5.533 | 1.12E-14 | glio |
| Bcas2_in.vivo_noRT | Gm20705 | 4.275 | 1.12E-14 | glio |
| Chek1_in.vivo_noRT | Aloxe3 | 4.515 | 1.12E-14 | glio |
| Chek1_in.vivo_noRT | Gm20767 | 5.044 | 1.12E-14 | glio |
| culture_effect | Mlxipl | -4.097 | 1.12E-15 | glio |
| Bora_in.vivo_noRT | 4933400F21Ri | 4.145 | 1.12E-16 | glio |
| Chek1_ex.vivo_noRT | Hoxc8 | 4.842 | 1.12E-16 | glio |
| Chek1_in.vivo_noRT | Prickle2 | 5.044 | 1.12E-17 | glio |
| Aars2_in.vivo_noRT | Map3k9 | 5.067 | 1.12E-18 | glio |
| Atr_in.vivo_noRT | Nkg7 | 5.273 | 1.12E-18 | glio |
| Bcas2_in.vivo_noRT | Gm15138 | 4.541 | 1.12E-18 | glio |
| Chek1_ex.vivo_noRT | Slc17a6 | 4.842 | 1.12E-19 | glio |
| Chek1_in.vivo_noRT | Slc45a1 | 4.784 | 1.12E-20 | glio |
| Chek1_in.vivo_noRT | Slc19a3 | 5.044 | 1.12E-22 | glio |
| Bora_in.vivo_noRT | Gm49698 | 4.936 | 1.12E-23 | glio |
| Chek1_in.vivo_noRT | Gm19276 | 5.044 | 1.12E-23 | glio |
| culture_effect | Itih5 | -6.179 | 1.12E-24 | glio |
| Aars2_in.vivo_noRT | Ank1 | 5.327 | 1.12E-24 | glio |
| Atr_in.vivo_noRT | Prl3d3 | 5.27 | 1.12E-26 | glio |
| culture_effect | Tpm1 | 1.99 | 1.12E-33 | glio |
| culture_effect | Unkl | -1.284 | 1.13E-03 | glio |
| culture_effect | Eif5a2 | 1.157 | 1.13E-03 | glio |
| culture_effect | Sox2ot | 1.264 | 1.13E-03 | glio |
| culture_effect | Katnal1 | -0.806 | 1.13E-03 | glio |
| culture_effect | Zc3hc1 | -0.629 | 1.13E-03 | glio |
| culture_effect | Chd2 | -0.687 | 1.13E-03 | glio |
| culture_effect | P3h1 | -0.562 | 1.13E-03 | glio |
| culture_effect | Cdkn1a | 0.534 | 1.13E-03 | glio |
| Interaction_genotype_Atr | Pakap | -3.216 | 1.13E-03 | glio |
| Interaction_genotype_Atr | Gm35082 | -3.167 | 1.13E-03 | glio |
| Interaction_genotype_Atr | Agap2 | -3.355 | 1.13E-03 | glio |
| Interaction_genotype_Mtor | Tnnt3 | -4.258 | 1.13E-03 | glio |
| Interaction_genotype_Mtor | Glrx | -3.642 | 1.13E-03 | glio |
| Interaction_genotype_Mtor | Polr2h | 1.797 | 1.13E-03 | glio |
| Interaction_genotype_Mtor | Rab8a | 1.623 | 1.13E-03 | glio |
| Interaction_genotype_Myc | Tagap | -2.469 | 1.13E-03 | glio |
| Interaction_genotype_Myc | 4732463B04R | -2.859 | 1.13E-03 | glio |
| Interaction_genotype_Myc | Stk36 | -3.733 | 1.13E-03 | glio |
| Interaction_genotype_Myc | 9230112E08Ri | -3.736 | 1.13E-03 | glio |
| Interaction_genotype_Myc | Zic2 | -3.075 | 1.13E-03 | glio |
| Interaction_genotype_Myc | Fut10 | -2.951 | 1.13E-03 | glio |
| Interaction_genotype_Myc | Tcaf1 | -2.267 | 1.13E-03 | glio |
| Interaction_genotype_Myc | Vcp | 1.479 | 1.13E-03 | glio |
| Interaction_genotype_Pars2 | Igf2bp1 | -4.322 | 1.13E-03 | glio |
| Interaction_genotype_Pars2 | Olfr552 | -3.197 | 1.13E-03 | glio |
| Interaction_genotype_Pars2 | Nxph4 | -4.072 | 1.13E-03 | glio |
| Interaction_genotype_Pars2 | Zfp446 | -3.954 | 1.13E-03 | glio |
| Interaction_genotype_Pars2 | Gm49312 | -3.521 | 1.13E-03 | glio |
| Interaction_genotype_Pars2 | Cebpa | -4.514 | 1.13E-03 | glio |
| Interaction_genotype_Pars2 | Kctd5 | 2.118 | 1.13E-03 | glio |
| Interaction_genotype_Pars2 | Skida1 | -4.477 | 1.13E-03 | glio |

| | | | | |
|---|---|---|---|---|
| Interaction_genotype_Pars2 | Clec4d | -3.453 | 1.13E-03 | glio |
| Interaction_genotype_Pars2 | Gm14207 | -3.062 | 1.13E-03 | glio |
| Interaction_genotype_Pars2 | Wincr1 | -5.556 | 1.13E-03 | glio |
| Interaction_genotype_Pars2 | Cpne8 | -4.485 | 1.13E-03 | glio |
| Interaction_genotype_Pars2 | Rhox7a | -4.645 | 1.13E-03 | glio |
| Aars2_in.vivo_noRT | Gm15545 | 2.217 | 1.13E-03 | glio |
| Aars2_ex.vivo_noRT | Glod5 | 3.693 | 1.13E-03 | glio |
| Atr_in.vivo_noRT | Rexo5 | 2.235 | 1.13E-03 | glio |
| Atr_in.vivo_noRT | Rassf8 | 1.971 | 1.13E-03 | glio |
| Atr_in.vivo_noRT | Fam117a | 2.083 | 1.13E-03 | glio |
| Atr_in.vivo_noRT | Srr | 1.771 | 1.13E-03 | glio |
| Atr_in.vivo_noRT | Spast | -1.552 | 1.13E-03 | glio |
| Atr_in.vivo_noRT | Mak16 | -1 | 1.13E-03 | glio |
| Atr_ex.vivo_noRT | Gm35082 | 2.367 | 1.13E-03 | glio |
| Bcas2_in.vivo_noRT | Tmem88 | 2.325 | 1.13E-03 | glio |
| Bcas2_in.vivo_noRT | Arhgdib | 2.386 | 1.13E-03 | glio |
| Bcas2_in.vivo_noRT | Grhl2 | 2.368 | 1.13E-03 | glio |
| Bcas2_ex.vivo_noRT | Serping1 | 2.742 | 1.13E-03 | glio |
| Bora_in.vivo_noRT | Gm15133 | 2.33 | 1.13E-03 | glio |
| Bora_in.vivo_noRT | Gm29358 | 2.436 | 1.13E-03 | glio |
| Bora_in.vivo_noRT | Mier1 | -0.971 | 1.13E-03 | glio |
| Chek1_in.vivo_noRT | Gm11713 | 2.101 | 1.13E-03 | glio |
| Chek1_in.vivo_noRT | Ppef1 | 2.227 | 1.13E-03 | glio |
| Chek1_in.vivo_noRT | Carmil1 | 2.229 | 1.13E-03 | glio |
| Chek1_in.vivo_noRT | Fam49a | 1.946 | 1.13E-03 | glio |
| Chek1_ex.vivo_noRT | Vpreb1 | 2.887 | 1.13E-03 | glio |
| Chek1_ex.vivo_noRT | Cdh18 | 2.98 | 1.13E-03 | glio |
| Eif2s1_in.vivo_noRT | Gad1 | 1.877 | 1.13E-03 | glio |
| Hsd17b10_in.vivo_noRT | Catsperd | 2.437 | 1.13E-03 | glio |
| culture_effect | Proscos | 2.089 | 1.13E-04 | glio |
| Interaction_genotype_Mtor | Gm16740 | -4.455 | 1.13E-04 | glio |
| Interaction_genotype_Mtor | Alyref2 | -3.857 | 1.13E-04 | glio |
| Interaction_genotype_Mtor | Vps33a | -3.648 | 1.13E-04 | glio |
| Interaction_genotype_Myc | Gipc3 | -4.517 | 1.13E-04 | glio |
| Interaction_genotype_Myc | Gm43689 | -3.695 | 1.13E-04 | glio |
| Interaction_genotype_Myc | Ipo11 | -3.272 | 1.13E-04 | glio |
| Interaction_genotype_Myc | Ndufa4 | 1.556 | 1.13E-04 | glio |
| Aars2_in.vivo_noRT | Spice1 | 2.331 | 1.13E-04 | glio |
| Aars2_in.vivo_noRT | Kdelr2 | -1.219 | 1.13E-04 | glio |
| Atr_in.vivo_noRT | Zfp595 | 2.903 | 1.13E-04 | glio |
| Atr_in.vivo_noRT | Cd24a | 3.205 | 1.13E-04 | glio |
| Atr_in.vivo_noRT | Tmem128 | -1.126 | 1.13E-04 | glio |
| Atr_ex.vivo_noRT | Ptp4a1 | 3.146 | 1.13E-04 | glio |
| Bcas2_in.vivo_noRT | Rpp21 | -1.57 | 1.13E-04 | glio |
| Bora_in.vivo_noRT | 1700126G02R | 2.803 | 1.13E-04 | glio |
| Chek1_in.vivo_noRT | Usp51 | 2.624 | 1.13E-04 | glio |
| Chek1_in.vivo_noRT | Rchy1 | -1.552 | 1.13E-04 | glio |
| Chek1_ex.vivo_noRT | 1700001L05Ri | 3.287 | 1.13E-04 | glio |
| Eif2s1_in.vivo_noRT | BC061195 | 2.143 | 1.13E-04 | glio |
| Hsd17b10_in.vivo_noRT | Gm24362 | 2.573 | 1.13E-04 | glio |
| culture_effect | Agfg2 | 1.053 | 1.13E-05 | glio |
| culture_effect | Cdc45 | -1.015 | 1.13E-05 | glio |

| | | | | |
|---|---|---|---|---|
| culture_effect | Rnf5 | -0.802 | 1.13E-05 | glio |
| culture_effect | Drg2 | -0.694 | 1.13E-05 | glio |
| Interaction_genotype_Chek1 | Egr2 | 5.557 | 1.13E-05 | glio |
| Interaction_genotype_Mtor | Nat8f1 | -4.896 | 1.13E-05 | glio |
| Aars2_in.vivo_noRT | Ppa1 | -1.276 | 1.13E-05 | glio |
| Aars2_ex.vivo_noRT | Cfb | 3.293 | 1.13E-05 | glio |
| Atr_in.vivo_noRT | Slc9a9 | 3.145 | 1.13E-05 | glio |
| Atr_in.vivo_noRT | Gnl3 | -1.535 | 1.13E-05 | glio |
| Atr_ex.vivo_noRT | Slc16a5 | 2.771 | 1.13E-05 | glio |
| Bcas2_in.vivo_noRT | Msl3l2 | 3.111 | 1.13E-05 | glio |
| Bcas2_ex.vivo_noRT | Gm47540 | 3.147 | 1.13E-05 | glio |
| Bcas2_ex.vivo_noRT | Cldn2 | 2.742 | 1.13E-05 | glio |
| Bora_in.vivo_noRT | Rasl2-9 | 3.499 | 1.13E-05 | glio |
| Bora_in.vivo_noRT | Hist1h2ad | 3.165 | 1.13E-05 | glio |
| Bora_in.vivo_noRT | Gm13421 | 3.212 | 1.13E-05 | glio |
| Bora_in.vivo_noRT | Rit2 | 2.968 | 1.13E-05 | glio |
| Bora_in.vivo_noRT | Slc25a11 | -1.583 | 1.13E-05 | glio |
| Bora_ex.vivo_noRT | Snrpn | 3.201 | 1.13E-05 | glio |
| Chek1_in.vivo_noRT | Gm45833 | 4.196 | 1.13E-05 | glio |
| Cyp19a1_in.vivo_noRT | Serpine2 | 3.318 | 1.13E-05 | glio |
| Hsd17b10_in.vivo_noRT | B230311B06R | 3.353 | 1.13E-05 | glio |
| culture_effect | Appbp2os | 2.646 | 1.13E-06 | glio |
| culture_effect | Tmem59l | 1.932 | 1.13E-06 | glio |
| Interaction_genotype_Mtor | Poln | -5.856 | 1.13E-06 | glio |
| Aars2_ex.vivo_noRT | Gm4166 | 3.293 | 1.13E-06 | glio |
| Bora_in.vivo_noRT | Zfp575 | 3.684 | 1.13E-06 | glio |
| Chek1_ex.vivo_noRT | Gm48742 | 3.467 | 1.13E-06 | glio |
| Chek1_ex.vivo_noRT | Gm9929 | 3.873 | 1.13E-06 | glio |
| culture_effect | Pcdhgb1 | -2.894 | 1.13E-07 | glio |
| Aars2_in.vivo_noRT | Pcdhb18 | 3.576 | 1.13E-07 | glio |
| Aars2_in.vivo_noRT | Apold1 | 4.217 | 1.13E-07 | glio |
| Aars2_in.vivo_noRT | Gm48529 | 3.597 | 1.13E-07 | glio |
| Aars2_in.vivo_noRT | Gadd45gip1 | -1.659 | 1.13E-07 | glio |
| Atr_in.vivo_noRT | Tcam1 | 3.819 | 1.13E-07 | glio |
| Atr_in.vivo_noRT | Cep295nl | 3.655 | 1.13E-07 | glio |
| Bcas2_in.vivo_noRT | Slc7a14 | 3.564 | 1.13E-07 | glio |
| Bcas2_in.vivo_noRT | Rpl3l | 3.389 | 1.13E-07 | glio |
| Chek1_in.vivo_noRT | Hspa12a | 3.743 | 1.13E-07 | glio |
| Atr_in.vivo_noRT | Casz1 | 4.258 | 1.13E-08 | glio |
| Atr_in.vivo_noRT | Tbxa2r | 3.694 | 1.13E-08 | glio |
| Bcas2_in.vivo_noRT | Peli3 | 3.892 | 1.13E-08 | glio |
| Bcas2_in.vivo_noRT | Arl6ip4 | -2.328 | 1.13E-08 | glio |
| Chek1_in.vivo_noRT | 1700011D18R | 3.999 | 1.13E-08 | glio |
| Chek1_in.vivo_noRT | Gm20667 | 4.043 | 1.13E-08 | glio |
| culture_effect | Myo5a | 1.067 | 1.13E-09 | glio |
| Atr_in.vivo_noRT | Tmem179 | 4.375 | 1.13E-09 | glio |
| Bora_in.vivo_noRT | Gm43387 | 4.294 | 1.13E-09 | glio |
| Chek1_in.vivo_noRT | Gm12602 | 4.017 | 1.13E-09 | glio |
| Aars2_in.vivo_noRT | Ptges | 4.671 | 1.13E-10 | glio |
| Atr_in.vivo_noRT | Hba-a2 | 4.621 | 1.13E-10 | glio |
| Bora_in.vivo_noRT | Gm20635 | 4.291 | 1.13E-10 | glio |
| Chek1_in.vivo_noRT | B930094E09Ri | 4.528 | 1.13E-10 | glio |

| | | | | |
|---|---|---|---|---|
| Bcas2_in.vivo_noRT | Gm31518 | 4.807 | 1.13E-11 | glio |
| culture_effect | Igbp1 | -1.19 | 1.13E-12 | glio |
| culture_effect | Pdap1 | 0.793 | 1.13E-12 | glio |
| Aars2_in.vivo_noRT | Cysrt1 | 5.067 | 1.13E-12 | glio |
| Aars2_in.vivo_noRT | Krtdap | 4.936 | 1.13E-12 | glio |
| Chek1_in.vivo_noRT | Accsl | 5.307 | 1.13E-12 | glio |
| Chek1_ex.vivo_noRT | Gm47540 | 4.449 | 1.13E-12 | glio |
| Atr_in.vivo_noRT | Hbq1b | 5.267 | 1.13E-13 | glio |
| Atr_in.vivo_noRT | Lrrc72 | 4.742 | 1.13E-13 | glio |
| Bora_in.vivo_noRT | Itgbl1 | 4.273 | 1.13E-13 | glio |
| Atr_in.vivo_noRT | Stx19 | 4.622 | 1.13E-15 | glio |
| Bcas2_in.vivo_noRT | Gm5934 | 4.544 | 1.13E-17 | glio |
| Bcas2_in.vivo_noRT | Gm5286 | 4.807 | 1.13E-18 | glio |
| Bcas2_in.vivo_noRT | Gm34776 | 4.807 | 1.13E-18 | glio |
| Chek1_in.vivo_noRT | Zscan4d | 5.044 | 1.13E-18 | glio |
| Bcas2_in.vivo_noRT | Gpc5 | 4.807 | 1.13E-21 | glio |
| Aars2_in.vivo_noRT | Gm48215 | 5.327 | 1.13E-23 | glio |
| Aars2_in.vivo_noRT | Gm11583 | 5.315 | 1.13E-23 | glio |
| Bora_in.vivo_noRT | Olfr881 | 4.936 | 1.13E-23 | glio |
| culture_effect | Pdia3 | -1.458 | 1.13E-25 | glio |
| culture_effect | Zfand5 | -1.433 | 1.13E-26 | glio |
| Atr_in.vivo_noRT | Gm12841 | 5.533 | 1.13E-26 | glio |
| culture_effect | Gm32913 | 2.578 | 1.14E-03 | glio |
| culture_effect | 1500009L16Ri | -1.33 | 1.14E-03 | glio |
| culture_effect | Dera | -0.644 | 1.14E-03 | glio |
| culture_effect | Tpm3 | 0.408 | 1.14E-03 | glio |
| Interaction_genotype_Aars2 | Ppil6 | -3.709 | 1.14E-03 | glio |
| Interaction_genotype_Aars2 | Gm10324 | -3.13 | 1.14E-03 | glio |
| Interaction_genotype_Aars2 | 1700113B09R | -4.144 | 1.14E-03 | glio |
| Interaction_genotype_Aars2 | Selenof | 1.216 | 1.14E-03 | glio |
| Interaction_genotype_Atr | Olfm2 | -3.995 | 1.14E-03 | glio |
| Interaction_genotype_Atr | Flacc1 | -3.559 | 1.14E-03 | glio |
| Interaction_genotype_Atr | Mxd4 | 2.04 | 1.14E-03 | glio |
| Interaction_genotype_Atr | Laptm4b | 1.894 | 1.14E-03 | glio |
| Interaction_genotype_Bcas2 | Nrg2 | -4.32 | 1.14E-03 | glio |
| Interaction_genotype_Bcas2 | Col8a1 | -4.197 | 1.14E-03 | glio |
| Interaction_genotype_Bora | Skida1 | -4.301 | 1.14E-03 | glio |
| Interaction_genotype_Bora | 1700017M07F | -3.68 | 1.14E-03 | glio |
| Interaction_genotype_Mtor | Gm12860 | -3.091 | 1.14E-03 | glio |
| Interaction_genotype_Mtor | Hist1h2bb | -3.896 | 1.14E-03 | glio |
| Interaction_genotype_Myc | Gm33051 | -2.873 | 1.14E-03 | glio |
| Interaction_genotype_Myc | Platr20 | -3.935 | 1.14E-03 | glio |
| Interaction_genotype_Myc | 4930426I24Ri | -3.46 | 1.14E-03 | glio |
| Interaction_genotype_Myc | Pdgfra | -3.611 | 1.14E-03 | glio |
| Interaction_genotype_Myc | Hk1os | -3.28 | 1.14E-03 | glio |
| Interaction_genotype_Myc | Tmem33 | 1.646 | 1.14E-03 | glio |
| interaction_RT | Rfc1 | -0.686 | 1.14E-03 | glio |
| Aars2_in.vivo_noRT | Lfng | 2.336 | 1.14E-03 | glio |
| Aars2_in.vivo_noRT | Castor2 | 2.181 | 1.14E-03 | glio |
| Aars2_in.vivo_noRT | Zfp748 | 2.293 | 1.14E-03 | glio |
| Aars2_in.vivo_noRT | Wipf1 | -1.346 | 1.14E-03 | glio |
| Aars2_in.vivo_noRT | Gng10 | -1.004 | 1.14E-03 | glio |

| | | | | |
|---|---|---|---|---|
| Aars2_in.vivo_noRT | Rpl15 | -0.667 | 1.14E-03 | glio |
| Atp5c1_ex.vivo_noRT | Synpo2l | 3.363 | 1.14E-03 | glio |
| Atr_in.vivo_noRT | Nadsyn1 | 2.348 | 1.14E-03 | glio |
| Atr_in.vivo_noRT | Meis3 | 1.943 | 1.14E-03 | glio |
| Atr_in.vivo_noRT | Luc7l | -1.23 | 1.14E-03 | glio |
| Atr_ex.vivo_noRT | Gm48899 | 2.588 | 1.14E-03 | glio |
| Bcas2_in.vivo_noRT | Catsperz | 2.333 | 1.14E-03 | glio |
| Bcas2_in.vivo_noRT | Cd59a | 2.417 | 1.14E-03 | glio |
| Bcas2_in.vivo_noRT | Lman2 | -1.038 | 1.14E-03 | glio |
| Bcas2_ex.vivo_noRT | Mog | 3.54 | 1.14E-03 | glio |
| Bora_in.vivo_noRT | Iapp | 2.773 | 1.14E-03 | glio |
| Bora_in.vivo_noRT | Prrc2c | -0.782 | 1.14E-03 | glio |
| Cenpj_in.vivo_noRT | AF067063 | 2.022 | 1.14E-03 | glio |
| Chek1_in.vivo_noRT | Mtpn | -0.858 | 1.14E-03 | glio |
| Chek1_ex.vivo_noRT | Pax5 | 3.182 | 1.14E-03 | glio |
| Chek1_ex.vivo_noRT | Serpina3m | 3.075 | 1.14E-03 | glio |
| Chek1_ex.vivo_noRT | Tk1 | -2.457 | 1.14E-03 | glio |
| Dclre1c_ex.vivo_noRT | Gm5039 | 3.97 | 1.14E-03 | glio |
| Dclre1c_ex.vivo_noRT | Gm4027 | 2.9 | 1.14E-03 | glio |
| Dclre1c_ex.vivo_noRT | Zscan4c | 2.658 | 1.14E-03 | glio |
| Dclre1c_ex.vivo_noRT | Zscan4d | 2.951 | 1.14E-03 | glio |
| Hsd17b10_in.vivo_noRT | Inha | 2.492 | 1.14E-03 | glio |
| culture_effect | Gm19531 | 1.641 | 1.14E-04 | glio |
| Interaction_genotype_Atr | Pde3a | -4.442 | 1.14E-04 | glio |
| Interaction_genotype_Mtor | Cd83 | -4.079 | 1.14E-04 | glio |
| Interaction_genotype_Mtor | Rell2 | -4.736 | 1.14E-04 | glio |
| Interaction_genotype_Myc | 9530018F02Ri | -4.577 | 1.14E-04 | glio |
| Interaction_genotype_Myc | Nqo1 | -4.442 | 1.14E-04 | glio |
| Interaction_genotype_Myc | Map4k2 | -3.479 | 1.14E-04 | glio |
| interaction_RT | Unc119 | 1.844 | 1.14E-04 | glio |
| Aars2_in.vivo_noRT | Gm32618 | 2.806 | 1.14E-04 | glio |
| Aars2_in.vivo_noRT | Gm29394 | 2.617 | 1.14E-04 | glio |
| Aars2_in.vivo_noRT | Akr1c12 | 2.692 | 1.14E-04 | glio |
| Atr_in.vivo_noRT | H60b | 2.635 | 1.14E-04 | glio |
| Atr_in.vivo_noRT | Tert | 2.547 | 1.14E-04 | glio |
| Atr_in.vivo_noRT | 4732440D04R | 2.473 | 1.14E-04 | glio |
| Atr_ex.vivo_noRT | Col19a1 | 2.771 | 1.14E-04 | glio |
| Bcas2_in.vivo_noRT | Rapgef3 | 3.076 | 1.14E-04 | glio |
| Bcas2_in.vivo_noRT | Neurl1a | 2.999 | 1.14E-04 | glio |
| Bcas2_in.vivo_noRT | Mertk | 2.674 | 1.14E-04 | glio |
| Bcas2_in.vivo_noRT | Gm44190 | 2.537 | 1.14E-04 | glio |
| Bcas2_in.vivo_noRT | Gm9993 | 2.52 | 1.14E-04 | glio |
| Bcas2_in.vivo_noRT | Vps13a | -1.437 | 1.14E-04 | glio |
| Bcas2_ex.vivo_noRT | Spink8 | 2.742 | 1.14E-04 | glio |
| Bcas2_ex.vivo_noRT | Psmb11 | 2.742 | 1.14E-04 | glio |
| Bora_in.vivo_noRT | Glt8d2 | 2.885 | 1.14E-04 | glio |
| Bora_in.vivo_noRT | Cops8 | -1.449 | 1.14E-04 | glio |
| Chek1_in.vivo_noRT | Klhl33 | 2.755 | 1.14E-04 | glio |
| Chek1_in.vivo_noRT | 1110038F14Ri | -1.002 | 1.14E-04 | glio |
| Chek1_ex.vivo_noRT | Bbs1 | 3.4 | 1.14E-04 | glio |
| Chek1_ex.vivo_noRT | Yipf6 | -2.102 | 1.14E-04 | glio |
| Chek1_ex.vivo_noRT | Yif1a | -2.081 | 1.14E-04 | glio |

| | | | | |
|---|---|---|---|---|
| Eif2s1_in.vivo_noRT | Idi2 | 2.143 | 1.14E-04 | glio |
| Eif2s1_in.vivo_noRT | Vmn2r18 | 2.143 | 1.14E-04 | glio |
| Eif2s1_in.vivo_noRT | Gm16178 | 2.143 | 1.14E-04 | glio |
| Eif2s1_in.vivo_noRT | Gm364 | 2.143 | 1.14E-04 | glio |
| Hsd17b10_in.vivo_noRT | Slxl1 | 2.838 | 1.14E-04 | glio |
| Hsd17b10_in.vivo_noRT | 1700012B09R | 3.356 | 1.14E-04 | glio |
| Hsd17b10_in.vivo_noRT | Gm13625 | 2.627 | 1.14E-04 | glio |
| culture_effect | Ypel2 | 1.273 | 1.14E-05 | glio |
| culture_effect | Nfkb2 | -0.871 | 1.14E-05 | glio |
| culture_effect | Tmem176a | -0.977 | 1.14E-05 | glio |
| culture_effect | Nemf | -0.577 | 1.14E-05 | glio |
| culture_effect | Rp9 | 0.53 | 1.14E-05 | glio |
| Interaction_genotype_Mtor | Gm36608 | -5.245 | 1.14E-05 | glio |
| Interaction_genotype_Mtor | Tmem190 | -5.831 | 1.14E-05 | glio |
| Interaction_genotype_Mtor | 1700012B09R | -6.219 | 1.14E-05 | glio |
| Interaction_genotype_Mtor | Ifitm5 | -5.159 | 1.14E-05 | glio |
| Atp5c1_in.vivo_noRT | Spink8 | 2.408 | 1.14E-05 | glio |
| Atr_in.vivo_noRT | Sipa1l2 | 3.091 | 1.14E-05 | glio |
| Atr_in.vivo_noRT | Zfp964 | 3.159 | 1.14E-05 | glio |
| Atr_in.vivo_noRT | Ocm | 2.692 | 1.14E-05 | glio |
| Atr_ex.vivo_noRT | 9430041J12Ri | 3.168 | 1.14E-05 | glio |
| Atr_ex.vivo_noRT | Nppc | 4.274 | 1.14E-05 | glio |
| Bora_in.vivo_noRT | Gm37933 | 2.785 | 1.14E-05 | glio |
| Bora_in.vivo_noRT | Nim1k | 3.261 | 1.14E-05 | glio |
| Bora_in.vivo_noRT | Gm15614 | 3.059 | 1.14E-05 | glio |
| Bora_in.vivo_noRT | Fbn2 | -2.731 | 1.14E-05 | glio |
| Chek1_in.vivo_noRT | Adamts7 | 2.873 | 1.14E-05 | glio |
| Chek1_ex.vivo_noRT | Gbp8 | 3.577 | 1.14E-05 | glio |
| culture_effect | Sema4g | -2.396 | 1.14E-06 | glio |
| culture_effect | Irf3 | -0.806 | 1.14E-06 | glio |
| culture_effect | Erh | 0.844 | 1.14E-06 | glio |
| culture_effect | Aldoa | -0.686 | 1.14E-06 | glio |
| culture_effect | Rps18 | -0.746 | 1.14E-06 | glio |
| Interaction_genotype_Mtor | Bnip3l | 2.763 | 1.14E-06 | glio |
| Interaction_genotype_Mtor | Mcmdc2 | -4.949 | 1.14E-06 | glio |
| Interaction_genotype_Mtor | Slc45a1 | -4.239 | 1.14E-06 | glio |
| Interaction_genotype_Mtor | Chia1 | -4.093 | 1.14E-06 | glio |
| Aars2_in.vivo_noRT | Wif1 | 3.83 | 1.14E-06 | glio |
| Chek1_in.vivo_noRT | 4732471J01Ri | 3.486 | 1.14E-06 | glio |
| Chek1_in.vivo_noRT | Rxra | 3.574 | 1.14E-06 | glio |
| culture_effect | Kmt2c | -0.865 | 1.14E-07 | glio |
| culture_effect | Washc5 | 0.851 | 1.14E-07 | glio |
| culture_effect | Rpl26 | -0.646 | 1.14E-07 | glio |
| interaction_RT | Crem | 1.724 | 1.14E-07 | glio |
| Aars2_ex.vivo_noRT | A330076H08R | 4.069 | 1.14E-07 | glio |
| Atr_in.vivo_noRT | Ctss | 3.162 | 1.14E-07 | glio |
| Bora_in.vivo_noRT | Hsf4 | 3.749 | 1.14E-07 | glio |
| Chek1_in.vivo_noRT | Gm10655 | 3.859 | 1.14E-07 | glio |
| Hsd17b10_in.vivo_noRT | Nos2 | 3.356 | 1.14E-07 | glio |
| culture_effect | Gm5431 | -3.378 | 1.14E-08 | glio |
| culture_effect | Abracl | 0.681 | 1.14E-08 | glio |
| culture_effect | Cd9 | -0.835 | 1.14E-08 | glio |

| | | | | |
|---|---|---|---|---|
| Aars2_in.vivo_noRT | 4932441J04Ri | 4.076 | 1.14E-08 | glio |
| Atr_in.vivo_noRT | 4930515G01R | 3.651 | 1.14E-08 | glio |
| Bcas2_in.vivo_noRT | Gm48610 | 4.025 | 1.14E-08 | glio |
| Bora_in.vivo_noRT | Gm41442 | 3.895 | 1.14E-08 | glio |
| Bora_in.vivo_noRT | Zfp455 | 3.915 | 1.14E-08 | glio |
| Chek1_in.vivo_noRT | Traf3ip3 | 3.98 | 1.14E-08 | glio |
| Fanca_in.vivo_noRT | Tulp2 | -3.495 | 1.14E-08 | glio |
| culture_effect | Car10 | -3.278 | 1.14E-09 | glio |
| Interaction_genotype_Mtor | Eps8l2 | -7.658 | 1.14E-09 | glio |
| Interaction_genotype_Mtor | Gm27239 | -7.203 | 1.14E-09 | glio |
| Aars2_in.vivo_noRT | Ccl8 | 4.801 | 1.14E-09 | glio |
| Aars2_in.vivo_noRT | Gm13610 | 4.17 | 1.14E-09 | glio |
| Atr_in.vivo_noRT | Ggt5 | 3.9 | 1.14E-09 | glio |
| Chek1_in.vivo_noRT | Gm15520 | 3.727 | 1.14E-09 | glio |
| Chek1_ex.vivo_noRT | Arhgef16 | 4.443 | 1.14E-09 | glio |
| interaction_RT | Nfkbia | 2.05 | 1.14E-10 | glio |
| Atr_in.vivo_noRT | Gm17210 | 4.474 | 1.14E-10 | glio |
| Chek1_ex.vivo_noRT | Gm48265 | 4.842 | 1.14E-10 | glio |
| Hsd17b10_in.vivo_noRT | Spink6 | 3.616 | 1.14E-10 | glio |
| culture_effect | Fam114a1 | 1.473 | 1.14E-11 | glio |
| Atr_in.vivo_noRT | Npy4r | 4.166 | 1.14E-11 | glio |
| Atr_in.vivo_noRT | Gm10399 | 4.601 | 1.14E-11 | glio |
| Atr_in.vivo_noRT | Insc | 4.076 | 1.14E-11 | glio |
| Bcas2_in.vivo_noRT | Pla2g2d | 4.544 | 1.14E-12 | glio |
| Bcas2_in.vivo_noRT | Htr1b | 4.807 | 1.14E-12 | glio |
| Atr_in.vivo_noRT | BC048671 | 4.232 | 1.14E-13 | glio |
| Atr_in.vivo_noRT | Gm9999 | 5.096 | 1.14E-14 | glio |
| Atr_in.vivo_noRT | Rnft2 | 5.239 | 1.14E-14 | glio |
| Aars2_in.vivo_noRT | Kpna7 | 5.063 | 1.14E-15 | glio |
| culture_effect | Gatad1 | 1.034 | 1.14E-16 | glio |
| Aars2_in.vivo_noRT | D430001F17R | 5.327 | 1.14E-16 | glio |
| Chek1_in.vivo_noRT | Dpp6 | 4.116 | 1.14E-16 | glio |
| Chek1_in.vivo_noRT | Npr3 | 5.044 | 1.14E-16 | glio |
| culture_effect | Gpnmb | -1.748 | 1.14E-17 | glio |
| Chek1_ex.vivo_noRT | 9430041J12Ri | 5.96 | 1.14E-17 | glio |
| Chek1_ex.vivo_noRT | Stfa3 | 4.842 | 1.14E-17 | glio |
| Chek1_ex.vivo_noRT | Olfr787 | 4.842 | 1.14E-17 | glio |
| culture_effect | Fry | -4.096 | 1.14E-18 | glio |
| Aars2_in.vivo_noRT | BC049987 | 4.544 | 1.14E-19 | glio |
| Atr_in.vivo_noRT | AC163347.1 | 5.27 | 1.14E-20 | glio |
| Chek1_in.vivo_noRT | Gm13449 | 5.044 | 1.14E-20 | glio |
| Aars2_in.vivo_noRT | Gm44899 | 5.327 | 1.14E-21 | glio |
| Aars2_in.vivo_noRT | Olfr543 | 5.067 | 1.14E-21 | glio |
| Aars2_in.vivo_noRT | Rimklb | 5.063 | 1.14E-24 | glio |
| Aars2_in.vivo_noRT | Gm13191 | 5.327 | 1.14E-29 | glio |
| Aars2_in.vivo_noRT | 4933416E03Ri | 5.327 | 1.14E-29 | glio |
| culture_effect | C1ra | -6.421 | 1.14E-37 | glio |
| Interaction_genotype_Atr | A030014E15Ri | -2.763 | 1.15E-03 | glio |
| Interaction_genotype_Bcas2 | Acoxl | -4.051 | 1.15E-03 | glio |
| Interaction_genotype_Bcas2 | Mrpl34 | 1.264 | 1.15E-03 | glio |
| Interaction_genotype_Bora | Lockd | 2.634 | 1.15E-03 | glio |
| Interaction_genotype_Mtor | Platr31 | -4.498 | 1.15E-03 | glio |

| | | | | |
|---|---|---|---|---|
| Interaction_genotype_Mtor | Fpgt | -3.201 | 1.15E-03 | glio |
| Interaction_genotype_Myc | Gm29966 | -2.469 | 1.15E-03 | glio |
| Interaction_genotype_Myc | Gm36266 | -4.095 | 1.15E-03 | glio |
| Interaction_genotype_Myc | Gstt2 | -3.592 | 1.15E-03 | glio |
| Interaction_genotype_Myc | Sh3d19 | -3.446 | 1.15E-03 | glio |
| Interaction_genotype_Myc | 1700030K09R | -2.751 | 1.15E-03 | glio |
| Interaction_genotype_Myc | Ncan | -3.084 | 1.15E-03 | glio |
| Interaction_genotype_Myc | Agfg2 | -2.383 | 1.15E-03 | glio |
| Aars2_in.vivo_noRT | 1700063D05R | 2.414 | 1.15E-03 | glio |
| Aars2_in.vivo_noRT | Gm11772 | 2.328 | 1.15E-03 | glio |
| Aars2_in.vivo_noRT | Tgfbrap1 | 1.48 | 1.15E-03 | glio |
| Aars2_in.vivo_noRT | Ddx24 | -0.841 | 1.15E-03 | glio |
| Aars2_in.vivo_noRT | Psmg4 | -0.76 | 1.15E-03 | glio |
| Aars2_in.vivo_noRT | Fbl | -0.761 | 1.15E-03 | glio |
| Atp5c1_in.vivo_noRT | Aloxe3 | 2.139 | 1.15E-03 | glio |
| Atp5c1_in.vivo_noRT | Ccdc39 | 2.495 | 1.15E-03 | glio |
| Atr_in.vivo_noRT | Utp6 | -1.268 | 1.15E-03 | glio |
| Atr_ex.vivo_noRT | Larp1 | -1.041 | 1.15E-03 | glio |
| Bard1_ex.vivo_noRT | Atp8a2 | 2.825 | 1.15E-03 | glio |
| Bcas2_in.vivo_noRT | Cxcl2 | 3.941 | 1.15E-03 | glio |
| Bcas2_in.vivo_noRT | Fignl2 | 2.541 | 1.15E-03 | glio |
| Bcas2_in.vivo_noRT | Gm12043 | 2.214 | 1.15E-03 | glio |
| Bcas2_in.vivo_noRT | Pigl | 1.867 | 1.15E-03 | glio |
| Bora_in.vivo_noRT | Myo1h | 2.442 | 1.15E-03 | glio |
| Bora_ex.vivo_noRT | Ly6c2 | 3.201 | 1.15E-03 | glio |
| Bora_ex.vivo_noRT | Snhg14 | 3.361 | 1.15E-03 | glio |
| Brca2_in.vivo_noRT | A930003A15R | 2.845 | 1.15E-03 | glio |
| Brca2_ex.vivo_noRT | Bmp15 | 3.221 | 1.15E-03 | glio |
| Cenpj_in.vivo_noRT | Gm2016 | 2.022 | 1.15E-03 | glio |
| Cenpj_in.vivo_noRT | Rapgef3os2 | 2.022 | 1.15E-03 | glio |
| Cenpj_in.vivo_noRT | Gm47622 | 2.022 | 1.15E-03 | glio |
| Chek1_ex.vivo_noRT | Tmem8b | 2.898 | 1.15E-03 | glio |
| Chek1_ex.vivo_noRT | Ifi208 | 3.255 | 1.15E-03 | glio |
| Chek1_ex.vivo_noRT | Nat14 | 2.453 | 1.15E-03 | glio |
| Chek1_ex.vivo_noRT | Cdk1 | -2.026 | 1.15E-03 | glio |
| Eif2s1_in.vivo_noRT | Ntn1 | 2.295 | 1.15E-03 | glio |
| culture_effect | Gm26910 | 1.655 | 1.15E-04 | glio |
| culture_effect | Kif1b | 0.612 | 1.15E-04 | glio |
| culture_effect | Lrrc59 | 0.486 | 1.15E-04 | glio |
| Interaction_genotype_Aars2 | Rab11fip4os1 | -4.932 | 1.15E-04 | glio |
| Interaction_genotype_Aars2 | Gm50334 | -5.244 | 1.15E-04 | glio |
| Interaction_genotype_Cdkn2a | Gjd4 | 3.615 | 1.15E-04 | glio |
| Interaction_genotype_Mtor | Klhl23 | -4.173 | 1.15E-04 | glio |
| Interaction_genotype_Mtor | Hspa5 | 2.422 | 1.15E-04 | glio |
| Interaction_genotype_Myc | 4930599N23R | -3.982 | 1.15E-04 | glio |
| Interaction_genotype_Myc | Gemin4 | -4.462 | 1.15E-04 | glio |
| Aars2_in.vivo_noRT | Hdx | 2.582 | 1.15E-04 | glio |
| Aars2_in.vivo_noRT | Tuba1a | -0.976 | 1.15E-04 | glio |
| Aars2_ex.vivo_noRT | Gm41349 | 3.293 | 1.15E-04 | glio |
| Atp5c1_in.vivo_noRT | H1foo | 2.136 | 1.15E-04 | glio |
| Atr_in.vivo_noRT | Hemk1 | 2.768 | 1.15E-04 | glio |
| Atr_in.vivo_noRT | Tcta | 2.174 | 1.15E-04 | glio |

| | | | | |
|---|---|---|---|---|
| Atr_in.vivo_noRT | Atp6v1c1 | -1.38 | 1.15E-04 | glio |
| Atr_in.vivo_noRT | Magoh | -1.148 | 1.15E-04 | glio |
| Atr_ex.vivo_noRT | 4732463B04R | 2.771 | 1.15E-04 | glio |
| Bcas2_in.vivo_noRT | Zfp960 | 2.912 | 1.15E-04 | glio |
| Bcas2_in.vivo_noRT | Gask1b | 3.012 | 1.15E-04 | glio |
| Bcas2_in.vivo_noRT | Kctd11 | 2.853 | 1.15E-04 | glio |
| Bcas2_ex.vivo_noRT | Vtn | 2.742 | 1.15E-04 | glio |
| Bora_in.vivo_noRT | Trim68 | 2.627 | 1.15E-04 | glio |
| Bora_in.vivo_noRT | Tmem151a | 2.75 | 1.15E-04 | glio |
| Bora_in.vivo_noRT | Aspscr1 | -1.729 | 1.15E-04 | glio |
| Chek1_in.vivo_noRT | Unc5cl | 2.742 | 1.15E-04 | glio |
| Chek1_in.vivo_noRT | Gm20515 | 2.706 | 1.15E-04 | glio |
| Chek1_in.vivo_noRT | Dip2a | 2.627 | 1.15E-04 | glio |
| Chek1_ex.vivo_noRT | Gm44836 | 3.345 | 1.15E-04 | glio |
| culture_effect | Mob3b | 1.657 | 1.15E-05 | glio |
| Interaction_genotype_Bora | Gm30938 | -5.437 | 1.15E-05 | glio |
| Interaction_genotype_Bora | Rassf6 | -4.769 | 1.15E-05 | glio |
| Interaction_genotype_Mtor | Cebpa | -5.285 | 1.15E-05 | glio |
| Interaction_genotype_Mtor | Gm48899 | -4.415 | 1.15E-05 | glio |
| Interaction_genotype_Mtor | Pum2 | 2.237 | 1.15E-05 | glio |
| Interaction_genotype_Myc | H2-Eb1 | 4.811 | 1.15E-05 | glio |
| Interaction_genotype_Myc | Enho | -4.883 | 1.15E-05 | glio |
| Aars2_in.vivo_noRT | Abi3bp | 3.098 | 1.15E-05 | glio |
| Aars2_in.vivo_noRT | Dhtkd1 | 3.364 | 1.15E-05 | glio |
| Atr_in.vivo_noRT | Olfr77 | 2.948 | 1.15E-05 | glio |
| Atr_in.vivo_noRT | Gm28221 | 2.864 | 1.15E-05 | glio |
| Bora_in.vivo_noRT | Ttc38 | 3.069 | 1.15E-05 | glio |
| Bora_ex.vivo_noRT | Rnase4 | 4.046 | 1.15E-05 | glio |
| Chek1_in.vivo_noRT | Phldb2 | 3.309 | 1.15E-05 | glio |
| Chek1_in.vivo_noRT | Lipt2 | 2.505 | 1.15E-05 | glio |
| Chek1_ex.vivo_noRT | Ucn2 | 3.867 | 1.15E-05 | glio |
| Hsd17b10_in.vivo_noRT | Sema3a | 2.938 | 1.15E-05 | glio |
| culture_effect | Srcin1 | 2.193 | 1.15E-06 | glio |
| culture_effect | Pcca | -1.51 | 1.15E-06 | glio |
| Interaction_genotype_Mtor | Tcea2 | -5.634 | 1.15E-06 | glio |
| Bcas2_in.vivo_noRT | Gm10382 | 3.563 | 1.15E-06 | glio |
| Bora_in.vivo_noRT | Gm5089 | 3.898 | 1.15E-06 | glio |
| Bora_ex.vivo_noRT | Dpp6 | 3.201 | 1.15E-06 | glio |
| Hsd17b10_in.vivo_noRT | Cfap46 | 3.149 | 1.15E-06 | glio |
| culture_effect | Nmral1 | -1.461 | 1.15E-07 | glio |
| Interaction_genotype_Myc | Uqcrq | 2.368 | 1.15E-07 | glio |
| Aars2_in.vivo_noRT | Gm49801 | 3.795 | 1.15E-07 | glio |
| Aars2_in.vivo_noRT | 4732471J01Ri | 3.769 | 1.15E-07 | glio |
| Aars2_in.vivo_noRT | Olfr920 | 3.576 | 1.15E-07 | glio |
| Atr_in.vivo_noRT | Gm15056 | 4.099 | 1.15E-07 | glio |
| Atr_in.vivo_noRT | 1700047M11F | 3.344 | 1.15E-07 | glio |
| Atr_ex.vivo_noRT | St18 | 3.962 | 1.15E-07 | glio |
| Bora_ex.vivo_noRT | Gm15908 | 3.201 | 1.15E-07 | glio |
| Bora_ex.vivo_noRT | A930014E10Ri | 3.201 | 1.15E-07 | glio |
| Bora_ex.vivo_noRT | Gm38163 | 3.993 | 1.15E-07 | glio |
| Chek1_in.vivo_noRT | Gm15246 | 3.693 | 1.15E-07 | glio |
| Chek1_in.vivo_noRT | 1810034E14Ri | 3.684 | 1.15E-07 | glio |

| | | | | |
|---|---|---|---|---|
| Chek1_ex.vivo_noRT | Prox2os | 4.049 | 1.15E-07 | glio |
| Chek1_ex.vivo_noRT | Gm36696 | 4.558 | 1.15E-07 | glio |
| Hsd17b10_in.vivo_noRT | 1700026N04R | 3.356 | 1.15E-07 | glio |
| culture_effect | Sertad3 | -1.501 | 1.15E-08 | glio |
| culture_effect | Crlf1 | 1.424 | 1.15E-08 | glio |
| culture_effect | Fech | 1.081 | 1.15E-08 | glio |
| Bcas2_in.vivo_noRT | Camkk1 | 4.037 | 1.15E-08 | glio |
| Chek1_in.vivo_noRT | Tnxb | 3.891 | 1.15E-08 | glio |
| Chek1_in.vivo_noRT | Gm41492 | 3.474 | 1.15E-08 | glio |
| Hsd17b10_in.vivo_noRT | Lrfn1 | 4.019 | 1.15E-08 | glio |
| culture_effect | Gm20513 | -3.031 | 1.15E-09 | glio |
| Aars2_in.vivo_noRT | Gm26822 | 4.412 | 1.15E-09 | glio |
| Bcas2_in.vivo_noRT | 9630014M24F | 4.028 | 1.15E-09 | glio |
| Aars2_in.vivo_noRT | Gm10010 | 4.586 | 1.15E-10 | glio |
| Atr_in.vivo_noRT | Gm15473 | 4.66 | 1.15E-10 | glio |
| Atr_in.vivo_noRT | Gm20667 | 4.533 | 1.15E-10 | glio |
| Atr_in.vivo_noRT | Gm11491 | 4.491 | 1.15E-10 | glio |
| Atr_ex.vivo_noRT | Usp17la | 5.515 | 1.15E-10 | glio |
| Atr_ex.vivo_noRT | BB557941 | 4.76 | 1.15E-10 | glio |
| Atr_ex.vivo_noRT | Aqp9 | 4.725 | 1.15E-10 | glio |
| Bcas2_in.vivo_noRT | Gm49441 | 4.016 | 1.15E-11 | glio |
| Bcas2_in.vivo_noRT | Gm37035 | 3.89 | 1.15E-11 | glio |
| Bora_in.vivo_noRT | Gm39326 | 4.673 | 1.15E-11 | glio |
| Bora_in.vivo_noRT | Dppa2 | 4.404 | 1.15E-11 | glio |
| Aars2_in.vivo_noRT | Tecrl | 5.067 | 1.15E-12 | glio |
| Atr_in.vivo_noRT | Ngb | 4.763 | 1.15E-12 | glio |
| Atr_in.vivo_noRT | Gm20682 | 4.489 | 1.15E-14 | glio |
| Bcas2_in.vivo_noRT | Mcmdc2 | 4.547 | 1.15E-14 | glio |
| Atr_in.vivo_noRT | Acoxl | 5.533 | 1.15E-17 | glio |
| Bora_in.vivo_noRT | Strip2 | 4.672 | 1.15E-17 | glio |
| Bora_in.vivo_noRT | Gm15326 | 4.407 | 1.15E-17 | glio |
| Chek1_in.vivo_noRT | Gm15890 | 5.044 | 1.15E-17 | glio |
| Aars2_in.vivo_noRT | Pldi | 5.327 | 1.15E-20 | glio |
| Aars2_in.vivo_noRT | Npnt | 5.06 | 1.15E-22 | glio |
| Chek1_in.vivo_noRT | Prl3d3 | 4.78 | 1.15E-22 | glio |
| Bora_in.vivo_noRT | Gm4340 | 4.936 | 1.15E-24 | glio |
| Bora_in.vivo_noRT | Gm16239 | 4.936 | 1.15E-24 | glio |
| Atr_in.vivo_noRT | Gm49698 | 5.533 | 1.15E-28 | glio |
| culture_effect | Pycard | -1.874 | 1.16E-03 | glio |
| culture_effect | Nutf2-ps1 | -0.916 | 1.16E-03 | glio |
| culture_effect | Prpsap2 | -0.573 | 1.16E-03 | glio |
| culture_effect | Cdt1 | -0.644 | 1.16E-03 | glio |
| Interaction_genotype_Aars2 | Mrpl53 | 1.354 | 1.16E-03 | glio |
| Interaction_genotype_Atr | Tdrd5 | -3.712 | 1.16E-03 | glio |
| Interaction_genotype_Cdkn2a | AF067063 | 3.615 | 1.16E-03 | glio |
| Interaction_genotype_Mtor | St5 | -3.522 | 1.16E-03 | glio |
| Interaction_genotype_Mtor | Rcan3 | -3.294 | 1.16E-03 | glio |
| Interaction_genotype_Mtor | Nucks1 | 1.774 | 1.16E-03 | glio |
| Interaction_genotype_Myc | D030051J21Ri | -3.58 | 1.16E-03 | glio |
| Interaction_genotype_Myc | Klrk1 | 2.974 | 1.16E-03 | glio |
| Interaction_genotype_Myc | Zfp599 | -3.903 | 1.16E-03 | glio |
| Interaction_genotype_Myc | Relt | -3.76 | 1.16E-03 | glio |

| | | | | |
|---|---|---|---|---|
| Interaction_genotype_Myc | 4931428F04Ri | -3.176 | 1.16E-03 | glio |
| Interaction_genotype_Myc | Rbm45 | -2.945 | 1.16E-03 | glio |
| Interaction_genotype_Myc | A630072M18F | -3.028 | 1.16E-03 | glio |
| Interaction_genotype_Myc | Chsy1 | -2.547 | 1.16E-03 | glio |
| interaction_RT | Pabpn1 | -0.668 | 1.16E-03 | glio |
| Aars2_in.vivo_noRT | Nsun7 | 2.539 | 1.16E-03 | glio |
| Aars2_in.vivo_noRT | 3110045C21R | 2.179 | 1.16E-03 | glio |
| Aars2_in.vivo_noRT | Gm48550 | 2.224 | 1.16E-03 | glio |
| Aars2_in.vivo_noRT | Ppm1l | 1.724 | 1.16E-03 | glio |
| Aars2_in.vivo_noRT | Wbp4 | -1.198 | 1.16E-03 | glio |
| Aars2_ex.vivo_noRT | Txndc2 | 2.505 | 1.16E-03 | glio |
| Atr_in.vivo_noRT | Gm44777 | 2.194 | 1.16E-03 | glio |
| Bard1_ex.vivo_noRT | Gm11131 | 3.217 | 1.16E-03 | glio |
| Bcas2_in.vivo_noRT | Med9os | 2.381 | 1.16E-03 | glio |
| Bcas2_in.vivo_noRT | Rrad | 2.544 | 1.16E-03 | glio |
| Bcas2_in.vivo_noRT | Prdm9 | 2.397 | 1.16E-03 | glio |
| Bcas2_in.vivo_noRT | Sox2ot | 2.202 | 1.16E-03 | glio |
| Bcas2_ex.vivo_noRT | Abra | 2.742 | 1.16E-03 | glio |
| Bcas2_ex.vivo_noRT | Ccl19 | 2.742 | 1.16E-03 | glio |
| Bora_in.vivo_noRT | Zfp599 | 2.503 | 1.16E-03 | glio |
| Bora_ex.vivo_noRT | Map3k5 | 2.802 | 1.16E-03 | glio |
| Cenpj_in.vivo_noRT | Gm11583 | 2.026 | 1.16E-03 | glio |
| Cenpj_in.vivo_noRT | Ephx3 | 2.596 | 1.16E-03 | glio |
| Cenpj_in.vivo_noRT | Gm50138 | 2.275 | 1.16E-03 | glio |
| Chek1_in.vivo_noRT | Cdh24 | 2.567 | 1.16E-03 | glio |
| Chek1_in.vivo_noRT | Ccdc180 | 2.132 | 1.16E-03 | glio |
| Chek1_in.vivo_noRT | Nadsyn1 | 2.382 | 1.16E-03 | glio |
| Chek1_in.vivo_noRT | Ankrd46 | -1.557 | 1.16E-03 | glio |
| Chek1_in.vivo_noRT | Cebpb | -1.395 | 1.16E-03 | glio |
| Chek1_ex.vivo_noRT | Slc46a3 | 3.016 | 1.16E-03 | glio |
| Chek1_ex.vivo_noRT | D030040B21R | 2.673 | 1.16E-03 | glio |
| Chek1_ex.vivo_noRT | Cox10 | -1.928 | 1.16E-03 | glio |
| Chek1_ex.vivo_noRT | Hmgb1 | -0.787 | 1.16E-03 | glio |
| Eif2s1_in.vivo_noRT | Gm10354 | 2.143 | 1.16E-03 | glio |
| Eif2s1_in.vivo_noRT | Ccdc162 | 2.138 | 1.16E-03 | glio |
| Hsd17b10_in.vivo_noRT | 2610016A17R | 2.557 | 1.16E-03 | glio |
| Hsd17b10_in.vivo_noRT | 9030612E09Ri | 2.295 | 1.16E-03 | glio |
| culture_effect | Gm10642 | 2.092 | 1.16E-04 | glio |
| culture_effect | Sec24d | -1.014 | 1.16E-04 | glio |
| Interaction_genotype_Mtor | Col19a1 | -3.575 | 1.16E-04 | glio |
| Interaction_genotype_Mtor | Tmem54 | -3.839 | 1.16E-04 | glio |
| Interaction_genotype_Myc | Ccdc136 | -4.114 | 1.16E-04 | glio |
| interaction_RT | Col11a2 | -1.018 | 1.16E-04 | glio |
| interaction_RT | Zc3hav1 | 0.881 | 1.16E-04 | glio |
| interaction_RT | Atp5g2 | 0.674 | 1.16E-04 | glio |
| interaction_RT | B2m | 0.804 | 1.16E-04 | glio |
| Aars2_in.vivo_noRT | Zfp599 | 2.894 | 1.16E-04 | glio |
| Aars2_ex.vivo_noRT | Abcb5 | 2.499 | 1.16E-04 | glio |
| Atr_in.vivo_noRT | Fam178b | 2.826 | 1.16E-04 | glio |
| Atr_in.vivo_noRT | Bdh2 | 2.707 | 1.16E-04 | glio |
| Atr_in.vivo_noRT | Pacs1 | 2.671 | 1.16E-04 | glio |
| Atr_in.vivo_noRT | Arhgap26 | 2.583 | 1.16E-04 | glio |

| | | | | |
|---|---|---|---|---|
| Atr_in.vivo_noRT | Jrk | 2.542 | 1.16E-04 | glio |
| Atr_in.vivo_noRT | Zfp319 | 1.839 | 1.16E-04 | glio |
| Bcas2_in.vivo_noRT | Hist1h4c | 2.902 | 1.16E-04 | glio |
| Bora_in.vivo_noRT | Gm13830 | 3.006 | 1.16E-04 | glio |
| Bora_in.vivo_noRT | Sept1 | 2.993 | 1.16E-04 | glio |
| Chek1_in.vivo_noRT | Pafah2 | 3.015 | 1.16E-04 | glio |
| Chek1_in.vivo_noRT | 4930594M22F | 2.797 | 1.16E-04 | glio |
| Chek1_in.vivo_noRT | Sqor | 2.472 | 1.16E-04 | glio |
| Chek1_in.vivo_noRT | Nubp2 | -1.576 | 1.16E-04 | glio |
| Interaction_genotype_Mtor | Gm29666 | -5.245 | 1.16E-05 | glio |
| Interaction_genotype_Mtor | 4930592C13R | -5.352 | 1.16E-05 | glio |
| Interaction_genotype_Mtor | Spink8 | -3.839 | 1.16E-05 | glio |
| Interaction_genotype_Mtor | Aoc1 | -5.518 | 1.16E-05 | glio |
| Interaction_genotype_Myc | Fam178b | -5.178 | 1.16E-05 | glio |
| Interaction_genotype_Myc | Dnah10 | -4.485 | 1.16E-05 | glio |
| Atp5c1_in.vivo_noRT | Vtn | 2.408 | 1.16E-05 | glio |
| Atr_ex.vivo_noRT | Gm7102 | 3.963 | 1.16E-05 | glio |
| Bcas2_in.vivo_noRT | Col17a1 | 3.274 | 1.16E-05 | glio |
| Chek1_in.vivo_noRT | Alox8 | 3.787 | 1.16E-05 | glio |
| Chek1_ex.vivo_noRT | Gm35696 | 3.577 | 1.16E-05 | glio |
| culture_effect | Cmas | -0.726 | 1.16E-06 | glio |
| Interaction_genotype_Mtor | Lman1 | 2.419 | 1.16E-06 | glio |
| Bora_ex.vivo_noRT | BC049987 | 3.201 | 1.16E-06 | glio |
| Chek1_in.vivo_noRT | Axdnd1 | 3.732 | 1.16E-06 | glio |
| Chek1_ex.vivo_noRT | B3gnt4 | 3.657 | 1.16E-06 | glio |
| culture_effect | Lars2 | -1.429 | 1.16E-07 | glio |
| culture_effect | Ruvbl2 | -0.792 | 1.16E-07 | glio |
| culture_effect | Ipo7 | -0.629 | 1.16E-07 | glio |
| Interaction_genotype_Mtor | 1810019D21R | -6.206 | 1.16E-07 | glio |
| Aars2_in.vivo_noRT | Hspb9 | 4.015 | 1.16E-07 | glio |
| Aars2_in.vivo_noRT | Gm14327 | 3.821 | 1.16E-07 | glio |
| Atr_in.vivo_noRT | Gm15091 | 3.829 | 1.16E-07 | glio |
| Atr_in.vivo_noRT | Lrrc75b | 3.877 | 1.16E-07 | glio |
| Bora_in.vivo_noRT | Skp2 | -2.389 | 1.16E-07 | glio |
| Chek1_in.vivo_noRT | Gm9484 | 3.885 | 1.16E-07 | glio |
| culture_effect | Lrp11 | -1.366 | 1.16E-08 | glio |
| Aars2_in.vivo_noRT | EU599041 | 4.05 | 1.16E-08 | glio |
| Aars2_in.vivo_noRT | Slurp1 | 5.789 | 1.16E-08 | glio |
| Bcas2_in.vivo_noRT | Gm12602 | 3.78 | 1.16E-08 | glio |
| Hsd17b10_in.vivo_noRT | Nat8f2 | 3.356 | 1.16E-08 | glio |
| Chek1_in.vivo_noRT | Osm | 3.695 | 1.16E-09 | glio |
| Hsd17b10_in.vivo_noRT | Ccser1 | 3.616 | 1.16E-09 | glio |
| culture_effect | Ralgds | -1.92 | 1.16E-10 | glio |
| Chek1_ex.vivo_noRT | Ipcef1 | 4.842 | 1.16E-11 | glio |
| Hsd17b10_in.vivo_noRT | Gm47510 | 3.356 | 1.16E-11 | glio |
| Aars2_in.vivo_noRT | Cd83 | 4.326 | 1.16E-12 | glio |
| Atr_in.vivo_noRT | A530013C23R | 4.615 | 1.16E-12 | glio |
| Atr_in.vivo_noRT | Gm12925 | 5.076 | 1.16E-12 | glio |
| Atr_in.vivo_noRT | Neurl1b | 5.007 | 1.16E-12 | glio |
| Bcas2_in.vivo_noRT | Ccr5 | 4.526 | 1.16E-12 | glio |
| culture_effect | Qars | -1.305 | 1.16E-13 | glio |
| Aars2_in.vivo_noRT | Gm50394 | 5.06 | 1.16E-13 | glio |

| | | | | |
|---|---|---|---|---|
| Atr_in.vivo_noRT | Gm49579 | 4.48 | 1.16E-13 | glio |
| Atr_in.vivo_noRT | Gm20633 | 4.742 | 1.16E-13 | glio |
| Bcas2_in.vivo_noRT | Gm43915 | 4.275 | 1.16E-13 | glio |
| culture_effect | Unc93b1 | -2.348 | 1.16E-14 | glio |
| Atr_in.vivo_noRT | Spn | 5.011 | 1.16E-14 | glio |
| Bora_in.vivo_noRT | 4930502E18Ri | 4.401 | 1.16E-14 | glio |
| Aars2_in.vivo_noRT | AI463229 | 4.682 | 1.16E-15 | glio |
| Atr_ex.vivo_noRT | Speer4e | 5.477 | 1.16E-15 | glio |
| Atr_ex.vivo_noRT | A930014E10Ri | 4.547 | 1.16E-15 | glio |
| Chek1_in.vivo_noRT | 4930570E03Ri | 4.784 | 1.16E-18 | glio |
| Chek1_in.vivo_noRT | 4930447K03R | 4.784 | 1.16E-18 | glio |
| Bora_in.vivo_noRT | Ank1 | 4.936 | 1.16E-21 | glio |
| Atr_in.vivo_noRT | 9430041J12Ri | 5.533 | 1.16E-22 | glio |
| Atr_in.vivo_noRT | Cxcl5 | 5.533 | 1.16E-24 | glio |
| culture_effect | Ly6d | -1.662 | 1.17E-03 | glio |
| culture_effect | Csrp1 | 0.659 | 1.17E-03 | glio |
| culture_effect | Brd3 | 0.533 | 1.17E-03 | glio |
| culture_effect | Ppie | -0.502 | 1.17E-03 | glio |
| Interaction_genotype_Aars2 | Slc4a9 | -3.388 | 1.17E-03 | glio |
| Interaction_genotype_Atr | E030037K01Ri | -3.705 | 1.17E-03 | glio |
| Interaction_genotype_Atr | 2610507I01Ri | -3.44 | 1.17E-03 | glio |
| Interaction_genotype_Atr | Cd164 | 1.487 | 1.17E-03 | glio |
| Interaction_genotype_Atr | 1810058I24Ri | 1.322 | 1.17E-03 | glio |
| Interaction_genotype_Mtor | Acot6 | -3.825 | 1.17E-03 | glio |
| Interaction_genotype_Mtor | Irak3 | -4.076 | 1.17E-03 | glio |
| Interaction_genotype_Mtor | Tmem116 | -4.227 | 1.17E-03 | glio |
| Interaction_genotype_Mtor | Cass4 | -3.9 | 1.17E-03 | glio |
| Interaction_genotype_Mtor | Smarcc1 | 1.868 | 1.17E-03 | glio |
| Interaction_genotype_Myc | B3galt4 | -3.374 | 1.17E-03 | glio |
| Interaction_genotype_Myc | Tcstv1 | -3.128 | 1.17E-03 | glio |
| Interaction_genotype_Myc | Hist1h3g | -3.903 | 1.17E-03 | glio |
| Interaction_genotype_Myc | Lgr6 | -4.056 | 1.17E-03 | glio |
| Interaction_genotype_Myc | Rgs3 | -3.553 | 1.17E-03 | glio |
| interaction_RT | Gm10399 | -2.33 | 1.17E-03 | glio |
| interaction_RT | Tulp2 | 1.404 | 1.17E-03 | glio |
| interaction_RT | Ndel1 | 0.893 | 1.17E-03 | glio |
| interaction_RT | Nup98 | 0.905 | 1.17E-03 | glio |
| interaction_RT | Efhd2 | -1.04 | 1.17E-03 | glio |
| interaction_RT | Nasp | -0.671 | 1.17E-03 | glio |
| Aars2_in.vivo_noRT | Phlda1 | -1.094 | 1.17E-03 | glio |
| Atp5c1_in.vivo_noRT | Cfh | 1.889 | 1.17E-03 | glio |
| Atr_in.vivo_noRT | Zfp597 | 2.127 | 1.17E-03 | glio |
| Atr_ex.vivo_noRT | Igdcc4 | 2.77 | 1.17E-03 | glio |
| Atr_ex.vivo_noRT | Adcy2 | 2.771 | 1.17E-03 | glio |
| Bard1_ex.vivo_noRT | Gm33973 | 2.814 | 1.17E-03 | glio |
| Bcas2_in.vivo_noRT | Nsd3 | -1.256 | 1.17E-03 | glio |
| Bcas2_ex.vivo_noRT | Astn2 | 3.129 | 1.17E-03 | glio |
| Bora_in.vivo_noRT | Nuf2 | -1.761 | 1.17E-03 | glio |
| Bora_in.vivo_noRT | Lamb1 | -1.667 | 1.17E-03 | glio |
| Cenpj_ex.vivo_noRT | Thnsl2 | 2.591 | 1.17E-03 | glio |
| Cenpj_ex.vivo_noRT | Akap14 | 2.591 | 1.17E-03 | glio |
| Chek1_in.vivo_noRT | Slc35d2 | 2.41 | 1.17E-03 | glio |

| | | | | |
|---|---|---|---|---|
| Chek1_in.vivo_noRT | Gm9887 | 2.276 | 1.17E-03 | glio |
| Chek1_in.vivo_noRT | Psmb5 | -0.752 | 1.17E-03 | glio |
| Chek1_ex.vivo_noRT | Klra6 | 3.105 | 1.17E-03 | glio |
| Chek1_ex.vivo_noRT | Gm15234 | 2.874 | 1.17E-03 | glio |
| Chek1_ex.vivo_noRT | Abcg4 | 2.696 | 1.17E-03 | glio |
| Chek1_ex.vivo_noRT | Gm43608 | 2.955 | 1.17E-03 | glio |
| Chek1_ex.vivo_noRT | Tsga10ip | 3.158 | 1.17E-03 | glio |
| Chek1_ex.vivo_noRT | Napb | 2.426 | 1.17E-03 | glio |
| Chek1_ex.vivo_noRT | Pin1 | -0.881 | 1.17E-03 | glio |
| Chek1_ex.vivo_noRT | Gabarap | 0.782 | 1.17E-03 | glio |
| Hsd17b10_in.vivo_noRT | Emid1 | 2.75 | 1.17E-03 | glio |
| Hsd17b10_in.vivo_noRT | C4a | 2.178 | 1.17E-03 | glio |
| culture_effect | Gm10401 | -2.155 | 1.17E-04 | glio |
| Interaction_genotype_Atr | Fgfr1op2 | 1.513 | 1.17E-04 | glio |
| Interaction_genotype_Mtor | Gm15326 | -3.353 | 1.17E-04 | glio |
| Interaction_genotype_Mtor | Parvb | -4.315 | 1.17E-04 | glio |
| Interaction_genotype_Myc | Spon1 | -4.449 | 1.17E-04 | glio |
| Interaction_genotype_Myc | Gm41307 | -3.122 | 1.17E-04 | glio |
| Interaction_genotype_Myc | 9230112J17Ri | -3.71 | 1.17E-04 | glio |
| Interaction_genotype_Myc | Slc38a9 | -3.375 | 1.17E-04 | glio |
| Interaction_genotype_Myc | Pemt | -3.383 | 1.17E-04 | glio |
| Interaction_genotype_Myc | Ncam1 | -2.581 | 1.17E-04 | glio |
| interaction_RT | Grina | 0.697 | 1.17E-04 | glio |
| Aars2_in.vivo_noRT | Spry3 | 2.799 | 1.17E-04 | glio |
| Aars2_in.vivo_noRT | Def6 | 2.624 | 1.17E-04 | glio |
| Aars2_ex.vivo_noRT | Gm5934 | 2.713 | 1.17E-04 | glio |
| Atr_in.vivo_noRT | 4833417C18R | 2.709 | 1.17E-04 | glio |
| Atr_in.vivo_noRT | Gng3 | 2.573 | 1.17E-04 | glio |
| Atr_in.vivo_noRT | Rfesd | 1.968 | 1.17E-04 | glio |
| Atr_in.vivo_noRT | Rfk | 1.327 | 1.17E-04 | glio |
| Atr_ex.vivo_noRT | Clca3a2 | 3.164 | 1.17E-04 | glio |
| Bard1_ex.vivo_noRT | Gm45406 | 2.814 | 1.17E-04 | glio |
| Bcas2_in.vivo_noRT | Gm37986 | 2.47 | 1.17E-04 | glio |
| Bcas2_in.vivo_noRT | Cdc27 | -1.726 | 1.17E-04 | glio |
| Bora_in.vivo_noRT | Rhot1 | -1.658 | 1.17E-04 | glio |
| Bora_in.vivo_noRT | Pigs | -1.856 | 1.17E-04 | glio |
| Bora_ex.vivo_noRT | Kcnq2 | 3.201 | 1.17E-04 | glio |
| Cenpj_ex.vivo_noRT | Cldn2 | 2.591 | 1.17E-04 | glio |
| Chek1_in.vivo_noRT | Gm17259 | 2.593 | 1.17E-04 | glio |
| Chek1_in.vivo_noRT | March10 | 2.687 | 1.17E-04 | glio |
| Chek1_ex.vivo_noRT | Gm10553 | 3.547 | 1.17E-04 | glio |
| Eif2s1_in.vivo_noRT | Ccl9 | 2.143 | 1.17E-04 | glio |
| Interaction_genotype_Mtor | Nrip3 | -5.041 | 1.17E-05 | glio |
| Interaction_genotype_Mtor | Ccno | -5.44 | 1.17E-05 | glio |
| Interaction_genotype_Mtor | Vtn | -3.839 | 1.17E-05 | glio |
| Atp5c1_in.vivo_noRT | Gtsf1 | 2.668 | 1.17E-05 | glio |
| Atr_in.vivo_noRT | Pcdhb16 | 3.307 | 1.17E-05 | glio |
| Atr_in.vivo_noRT | Trim66 | 2.821 | 1.17E-05 | glio |
| Atr_ex.vivo_noRT | Gm40538 | 3.164 | 1.17E-05 | glio |
| Bcas2_in.vivo_noRT | Tbc1d10c | 3.112 | 1.17E-05 | glio |
| Bcas2_in.vivo_noRT | 2610016A17R | 3.221 | 1.17E-05 | glio |
| Bora_in.vivo_noRT | Tph2 | 2.923 | 1.17E-05 | glio |

| | | | | |
|---|---|---|---|---|
| Bora_ex.vivo_noRT | Gm4027 | 3.201 | 1.17E-05 | glio |
| Chek1_in.vivo_noRT | Zfp980 | 3.277 | 1.17E-05 | glio |
| Hsd17b10_in.vivo_noRT | 2310067P03R | 3.093 | 1.17E-05 | glio |
| Hsd17b10_in.vivo_noRT | Olfr279 | 3.087 | 1.17E-05 | glio |
| Hsd17b10_in.vivo_noRT | Syngap1 | 3.101 | 1.17E-05 | glio |
| culture_effect | Camk2n2 | 1.586 | 1.17E-06 | glio |
| culture_effect | Ddx25 | -1.351 | 1.17E-06 | glio |
| culture_effect | Cbx3 | 0.625 | 1.17E-06 | glio |
| Interaction_genotype_Mtor | Gm10734 | -5.809 | 1.17E-06 | glio |
| Interaction_genotype_Mtor | Gm24362 | -5.336 | 1.17E-06 | glio |
| Interaction_genotype_Mtor | Gm5934 | -4.415 | 1.17E-06 | glio |
| Aars2_in.vivo_noRT | Rab17 | 3.332 | 1.17E-06 | glio |
| Atr_in.vivo_noRT | Trim68 | 3.224 | 1.17E-06 | glio |
| Bcas2_in.vivo_noRT | 1700013F07Ri | 3.652 | 1.17E-06 | glio |
| Chek1_in.vivo_noRT | Gm16174 | 3.654 | 1.17E-06 | glio |
| Chek1_in.vivo_noRT | Rpl3l | 3.107 | 1.17E-06 | glio |
| Chek1_ex.vivo_noRT | Amy1 | 4.055 | 1.17E-06 | glio |
| Chek1_ex.vivo_noRT | Nox1 | 3.747 | 1.17E-06 | glio |
| Hsd17b10_in.vivo_noRT | Cxcl1 | 3.616 | 1.17E-06 | glio |
| culture_effect | Ngdn | -0.766 | 1.17E-07 | glio |
| culture_effect | Pgk1 | -0.638 | 1.17E-07 | glio |
| Aars2_in.vivo_noRT | 1700029J07Ri | 3.769 | 1.17E-07 | glio |
| Aars2_in.vivo_noRT | Hrk | 3.738 | 1.17E-07 | glio |
| Bcas2_in.vivo_noRT | Gm26519 | 4.148 | 1.17E-07 | glio |
| Bora_in.vivo_noRT | 9930014A18R | 3.542 | 1.17E-07 | glio |
| Bora_ex.vivo_noRT | Scg2 | 3.201 | 1.17E-07 | glio |
| Chek1_ex.vivo_noRT | Gm48008 | 3.652 | 1.17E-07 | glio |
| Interaction_genotype_Mtor | Adam12 | -5.997 | 1.17E-08 | glio |
| Aars2_in.vivo_noRT | Serinc2 | 3.768 | 1.17E-08 | glio |
| Bcas2_in.vivo_noRT | Zmynd15 | 3.899 | 1.17E-08 | glio |
| Chek1_in.vivo_noRT | Gm45890 | 4.127 | 1.17E-08 | glio |
| Chek1_in.vivo_noRT | Gm11491 | 4.002 | 1.17E-08 | glio |
| Chek1_in.vivo_noRT | Gm28874 | 4.022 | 1.17E-08 | glio |
| Chek1_in.vivo_noRT | 1110020A21R | 4.093 | 1.17E-08 | glio |
| Bcas2_in.vivo_noRT | 4921508D12R | 3.88 | 1.17E-09 | glio |
| Bora_in.vivo_noRT | Igdcc4 | 3.761 | 1.17E-09 | glio |
| Chek1_in.vivo_noRT | Sult5a1 | 4.511 | 1.17E-09 | glio |
| Chek1_in.vivo_noRT | Peli3 | 4.129 | 1.17E-09 | glio |
| Chek1_in.vivo_noRT | H2-M10.1 | 3.589 | 1.17E-09 | glio |
| Chek1_in.vivo_noRT | Mogat1 | 4.522 | 1.17E-09 | glio |
| Chek1_in.vivo_noRT | Gm43258 | 4.159 | 1.17E-09 | glio |
| Chek1_ex.vivo_noRT | BC147527 | 4.842 | 1.17E-09 | glio |
| Hsd17b10_in.vivo_noRT | 4732463B04R | 3.352 | 1.17E-09 | glio |
| Chek1_ex.vivo_noRT | Pcdh15 | 4.842 | 1.17E-10 | glio |
| Aars2_in.vivo_noRT | A530013C23R | 4.409 | 1.17E-11 | glio |
| Atr_in.vivo_noRT | Gm5544 | 4.742 | 1.17E-11 | glio |
| Bora_in.vivo_noRT | Jph4 | 4.67 | 1.17E-11 | glio |
| culture_effect | Prpf19 | -0.968 | 1.17E-12 | glio |
| Aars2_in.vivo_noRT | A930012O16R | 5.067 | 1.17E-12 | glio |
| Bora_in.vivo_noRT | Tcap | 4.557 | 1.17E-12 | glio |
| Atr_in.vivo_noRT | Krtdap | 5.142 | 1.17E-13 | glio |
| Bcas2_in.vivo_noRT | Gm13270 | 5.334 | 1.17E-13 | glio |

| | | | | |
|---|---|---|---|---|
| Bcas2_in.vivo_noRT | Thsd1 | 4.42 | 1.17E-13 | glio |
| Aars2_in.vivo_noRT | Calhm5 | 5.135 | 1.17E-14 | glio |
| Atr_in.vivo_noRT | Gm13288 | 5.533 | 1.17E-21 | glio |
| Atr_in.vivo_noRT | Gm45151 | 5.511 | 1.17E-22 | glio |
| Aars2_in.vivo_noRT | Trim17 | 5.327 | 1.17E-26 | glio |
| Atr_in.vivo_noRT | Olfr881 | 5.533 | 1.17E-28 | glio |
| culture_effect | Cd81 | -4.779 | 1.17E-48 | glio |
| culture_effect | Supt20 | 0.549 | 1.18E-03 | glio |
| culture_effect | Akt1 | 0.483 | 1.18E-03 | glio |
| Interaction_genotype_Atr | Gm47851 | -3.713 | 1.18E-03 | glio |
| Interaction_genotype_Bcas2 | Arf6 | 1.546 | 1.18E-03 | glio |
| Interaction_genotype_Mtor | Gm13610 | -3.875 | 1.18E-03 | glio |
| Interaction_genotype_Mtor | Mok | -4.027 | 1.18E-03 | glio |
| Interaction_genotype_Mtor | Myl4 | -4.012 | 1.18E-03 | glio |
| Interaction_genotype_Mtor | E430018J23Ri | -3.666 | 1.18E-03 | glio |
| Interaction_genotype_Mtor | Gm10184 | -3.71 | 1.18E-03 | glio |
| Interaction_genotype_Mtor | Slc39a9 | -2.454 | 1.18E-03 | glio |
| Interaction_genotype_Mtor | N6amt1 | -1.973 | 1.18E-03 | glio |
| Interaction_genotype_Mtor | Lsm6 | 1.669 | 1.18E-03 | glio |
| Interaction_genotype_Myc | Fabp6 | -2.729 | 1.18E-03 | glio |
| Interaction_genotype_Myc | Olfr1152 | -2.463 | 1.18E-03 | glio |
| Interaction_genotype_Myc | Rasal1 | -4.11 | 1.18E-03 | glio |
| Interaction_genotype_Myc | Zfp936 | -3.452 | 1.18E-03 | glio |
| Interaction_genotype_Myc | 1700086P04R | -3.368 | 1.18E-03 | glio |
| Interaction_genotype_Myc | Zfp605 | -2.99 | 1.18E-03 | glio |
| Interaction_genotype_Myc | Tafa5 | -3.196 | 1.18E-03 | glio |
| Interaction_genotype_Myc | Cox14 | 1.462 | 1.18E-03 | glio |
| Interaction_genotype_Myc | Npm1 | 1.325 | 1.18E-03 | glio |
| interaction_RT | Adgrb3 | -2.948 | 1.18E-03 | glio |
| interaction_RT | Colgalt1 | -0.808 | 1.18E-03 | glio |
| Aars2_in.vivo_noRT | Cacna1a | 1.86 | 1.18E-03 | glio |
| Atr_in.vivo_noRT | Chrnb1 | 2.181 | 1.18E-03 | glio |
| Atr_in.vivo_noRT | Gen1 | 1.874 | 1.18E-03 | glio |
| Atr_in.vivo_noRT | Bysl | -1.297 | 1.18E-03 | glio |
| Atr_ex.vivo_noRT | Cxcr6 | 3.164 | 1.18E-03 | glio |
| Bcas2_in.vivo_noRT | 4930412C18R | 2.471 | 1.18E-03 | glio |
| Bcas2_in.vivo_noRT | Gm41760 | 2.352 | 1.18E-03 | glio |
| Bcas2_in.vivo_noRT | Spink10 | 2.233 | 1.18E-03 | glio |
| Bcas2_in.vivo_noRT | Plekhh2 | 2.151 | 1.18E-03 | glio |
| Bcas2_in.vivo_noRT | Gm5617 | 1.705 | 1.18E-03 | glio |
| Bcas2_in.vivo_noRT | Tmem263 | -1.199 | 1.18E-03 | glio |
| Bora_in.vivo_noRT | Soga1 | 2.446 | 1.18E-03 | glio |
| Bora_in.vivo_noRT | Rab27b | 2.527 | 1.18E-03 | glio |
| Bora_ex.vivo_noRT | Gm15972 | 2.802 | 1.18E-03 | glio |
| Bora_ex.vivo_noRT | 4930447K03R | 2.414 | 1.18E-03 | glio |
| Brca2_in.vivo_noRT | 4930512B01R | 2.296 | 1.18E-03 | glio |
| Cenpj_in.vivo_noRT | Olfr56 | 2.022 | 1.18E-03 | glio |
| Chek1_ex.vivo_noRT | Casp1 | 2.66 | 1.18E-03 | glio |
| Eif2s1_in.vivo_noRT | Gm28940 | 2.154 | 1.18E-03 | glio |
| culture_effect | Hist2h4 | 1.737 | 1.18E-04 | glio |
| Interaction_genotype_Aars2 | Tmem134 | 1.942 | 1.18E-04 | glio |
| Interaction_genotype_Atr | Rtraf | 1.616 | 1.18E-04 | glio |

| | | | | |
|---|---|---|---|---|
| Interaction_genotype_Mtor | Gpr162 | -4.678 | 1.18E-04 | glio |
| Interaction_genotype_Mtor | Gm2762 | -4.333 | 1.18E-04 | glio |
| Interaction_genotype_Myc | Abhd15 | -4.711 | 1.18E-04 | glio |
| Interaction_genotype_Myc | Klk10 | -4.208 | 1.18E-04 | glio |
| Interaction_genotype_Myc | Ndufc2 | 1.725 | 1.18E-04 | glio |
| Interaction_genotype_Myc | Selenow | 1.561 | 1.18E-04 | glio |
| Aars2_in.vivo_noRT | Stard10 | 2.66 | 1.18E-04 | glio |
| Aars2_ex.vivo_noRT | Serpina3g | 3.293 | 1.18E-04 | glio |
| Atp5c1_in.vivo_noRT | Map3k9 | 2.408 | 1.18E-04 | glio |
| Atr_in.vivo_noRT | Cr2 | 2.282 | 1.18E-04 | glio |
| Atr_in.vivo_noRT | Col5a3 | 2.034 | 1.18E-04 | glio |
| Atr_ex.vivo_noRT | Prox2os | 3.154 | 1.18E-04 | glio |
| Atr_ex.vivo_noRT | Mt4 | 4.519 | 1.18E-04 | glio |
| Bcas2_in.vivo_noRT | Rnf144b | 2.75 | 1.18E-04 | glio |
| Bcas2_in.vivo_noRT | BC049715 | 2.583 | 1.18E-04 | glio |
| Bcas2_in.vivo_noRT | Fam136a | -1.232 | 1.18E-04 | glio |
| Bora_in.vivo_noRT | Rab36 | 2.768 | 1.18E-04 | glio |
| Bora_in.vivo_noRT | Slc46a1 | 2.902 | 1.18E-04 | glio |
| Bora_in.vivo_noRT | Hdac10 | 2.395 | 1.18E-04 | glio |
| Chek1_in.vivo_noRT | Gpr146 | 2.79 | 1.18E-04 | glio |
| Chek1_ex.vivo_noRT | Rab15 | 3.47 | 1.18E-04 | glio |
| Chek1_ex.vivo_noRT | Eaf2 | 3.478 | 1.18E-04 | glio |
| Chek1_ex.vivo_noRT | Cks1b | -1.312 | 1.18E-04 | glio |
| Hsd17b10_in.vivo_noRT | Esrp2 | 2.704 | 1.18E-04 | glio |
| culture_effect | Astn2 | -2.627 | 1.18E-05 | glio |
| culture_effect | Rcbtb2 | -0.983 | 1.18E-05 | glio |
| culture_effect | Ciao1 | -0.726 | 1.18E-05 | glio |
| culture_effect | Mrps7 | 0.537 | 1.18E-05 | glio |
| Interaction_genotype_Atr | Klf6 | 2.578 | 1.18E-05 | glio |
| Interaction_genotype_Mtor | Zfp607b | -4.16 | 1.18E-05 | glio |
| Interaction_genotype_Mtor | 4933421A08R | -5.532 | 1.18E-05 | glio |
| Interaction_genotype_Mtor | 7530414M10F | -5.07 | 1.18E-05 | glio |
| Interaction_genotype_Mtor | Olfr171 | -4.099 | 1.18E-05 | glio |
| Interaction_genotype_Mtor | Gapdh | 2.335 | 1.18E-05 | glio |
| Interaction_genotype_Myc | Nrg2 | -4.984 | 1.18E-05 | glio |
| Interaction_genotype_Myc | Abcb5 | -3.523 | 1.18E-05 | glio |
| Interaction_genotype_Myc | Sprr1a | -5.102 | 1.18E-05 | glio |
| Aars2_in.vivo_noRT | Ostc | -1.052 | 1.18E-05 | glio |
| Atr_in.vivo_noRT | B230312C02R | 3.922 | 1.18E-05 | glio |
| Atr_in.vivo_noRT | 4930412C18R | 3.197 | 1.18E-05 | glio |
| Atr_in.vivo_noRT | G430095P16R | 3.079 | 1.18E-05 | glio |
| Atr_in.vivo_noRT | Taco1os | 2.873 | 1.18E-05 | glio |
| culture_effect | Synpr | 2.805 | 1.18E-06 | glio |
| culture_effect | Itgb4 | -1.281 | 1.18E-06 | glio |
| Interaction_genotype_Mtor | Wdr38 | -5.514 | 1.18E-06 | glio |
| interaction_RT | 4930597O21R | 4.499 | 1.18E-06 | glio |
| interaction_RT | Acot5 | 4.351 | 1.18E-06 | glio |
| Aars2_in.vivo_noRT | Ccdc148 | 3.566 | 1.18E-06 | glio |
| Aars2_in.vivo_noRT | Dll1 | 3.602 | 1.18E-06 | glio |
| Aars2_ex.vivo_noRT | Olfr56 | 3.293 | 1.18E-06 | glio |
| Atr_in.vivo_noRT | Gm4221 | 3.429 | 1.18E-06 | glio |
| Atr_in.vivo_noRT | Gatad2a | -1.798 | 1.18E-06 | glio |

| | | | | |
|---|---|---|---|---|
| Bcas2_in.vivo_noRT | 1810041H14R | 3.688 | 1.18E-06 | glio |
| Chek1_in.vivo_noRT | Zfp433 | 3.604 | 1.18E-06 | glio |
| Chek1_in.vivo_noRT | Zfp467 | 3.503 | 1.18E-06 | glio |
| Chek1_in.vivo_noRT | Gm26879 | 3.037 | 1.18E-06 | glio |
| Aars2_in.vivo_noRT | Ikzf2 | 3.903 | 1.18E-07 | glio |
| Aars2_in.vivo_noRT | Gm50012 | 3.726 | 1.18E-07 | glio |
| Chek1_in.vivo_noRT | Gm50399 | 3.491 | 1.18E-07 | glio |
| Chek1_ex.vivo_noRT | Dcpp1 | 4.842 | 1.18E-07 | glio |
| Chek1_in.vivo_noRT | Gm17210 | 3.984 | 1.18E-08 | glio |
| Chek1_in.vivo_noRT | Ccl8 | 4.518 | 1.18E-08 | glio |
| culture_effect | Auts2 | -3.385 | 1.18E-09 | glio |
| Aars2_in.vivo_noRT | Sema3a | 3.889 | 1.18E-09 | glio |
| Aars2_in.vivo_noRT | 1110035H17R | 4.695 | 1.18E-09 | glio |
| Aars2_in.vivo_noRT | Srsf5 | -1.967 | 1.18E-09 | glio |
| Bcas2_in.vivo_noRT | 1110020A21R | 4.375 | 1.18E-09 | glio |
| Bora_in.vivo_noRT | Gm12280 | 4.672 | 1.18E-09 | glio |
| Chek1_in.vivo_noRT | Aqp5 | 4.78 | 1.18E-09 | glio |
| Atr_in.vivo_noRT | Gm28874 | 4.511 | 1.18E-10 | glio |
| Bcas2_in.vivo_noRT | Arhgef16 | 3.771 | 1.18E-10 | glio |
| Chek1_ex.vivo_noRT | Gm18280 | 4.049 | 1.18E-10 | glio |
| Bora_in.vivo_noRT | Spn | 4.414 | 1.18E-11 | glio |
| Chek1_ex.vivo_noRT | Abcb5 | 4.049 | 1.18E-11 | glio |
| Chek1_in.vivo_noRT | Rgs9 | 4.243 | 1.18E-12 | glio |
| Chek1_ex.vivo_noRT | Gm4027 | 4.842 | 1.18E-12 | glio |
| Atr_in.vivo_noRT | Wdr93 | 4.877 | 1.18E-13 | glio |
| Bora_in.vivo_noRT | C130060C02R | 4.672 | 1.18E-13 | glio |
| Chek1_in.vivo_noRT | 1700016P04R | 4.777 | 1.18E-13 | glio |
| culture_effect | Gadd45b | 2.876 | 1.18E-14 | glio |
| Atr_in.vivo_noRT | Usp17la | 5.533 | 1.18E-15 | glio |
| Chek1_in.vivo_noRT | E030037K01Ri | 5.035 | 1.18E-15 | glio |
| Bcas2_in.vivo_noRT | Selenov | 4.807 | 1.18E-16 | glio |
| Bcas2_in.vivo_noRT | A330076H08R | 4.807 | 1.18E-16 | glio |
| Bora_in.vivo_noRT | Gm11899 | 4.414 | 1.18E-16 | glio |
| Aars2_in.vivo_noRT | Gm33051 | 4.674 | 1.18E-17 | glio |
| Bcas2_in.vivo_noRT | Ccl3 | 4.547 | 1.18E-18 | glio |
| culture_effect | Polr3k | -1.468 | 1.18E-19 | glio |
| Aars2_in.vivo_noRT | Scel | 4.409 | 1.18E-20 | glio |
| Atr_in.vivo_noRT | Dpep3 | 5.001 | 1.18E-23 | glio |
| Bora_in.vivo_noRT | 1700003E24Ri | 4.936 | 1.18E-25 | glio |
| culture_effect | D230022J07Ri | 1.785 | 1.19E-03 | glio |
| culture_effect | Gm31597 | 1.098 | 1.19E-03 | glio |
| Interaction_genotype_Atr | A930033H14R | -4.177 | 1.19E-03 | glio |
| Interaction_genotype_Atr | Gm48008 | -3.296 | 1.19E-03 | glio |
| Interaction_genotype_Atr | Gm16835 | -4.009 | 1.19E-03 | glio |
| Interaction_genotype_Atr | Rbm45 | -3.261 | 1.19E-03 | glio |
| Interaction_genotype_Bard1 | Kcnip3 | 4.901 | 1.19E-03 | glio |
| Interaction_genotype_Bard1 | Auts2 | 4.614 | 1.19E-03 | glio |
| Interaction_genotype_Bard1 | Krt18 | 6.9 | 1.19E-03 | glio |
| Interaction_genotype_Bard1 | St18 | 4.204 | 1.19E-03 | glio |
| Interaction_genotype_Bard1 | Art2b | 4.5 | 1.19E-03 | glio |
| Interaction_genotype_Bora | 1810041H14R | -4.553 | 1.19E-03 | glio |
| Interaction_genotype_Mtor | Tfr2 | -3.528 | 1.19E-03 | glio |

| | | | | |
|---|---|---|---|---|
| Interaction_genotype_Mtor | Map3k15 | -3.215 | 1.19E-03 | glio |
| Interaction_genotype_Mtor | Btbd16 | -3.293 | 1.19E-03 | glio |
| Interaction_genotype_Mtor | Nme2 | 1.167 | 1.19E-03 | glio |
| Interaction_genotype_Myc | Dapk2 | -2.744 | 1.19E-03 | glio |
| Interaction_genotype_Myc | Mmaa | -3.181 | 1.19E-03 | glio |
| Interaction_genotype_Myc | Zscan12 | -2.779 | 1.19E-03 | glio |
| Interaction_genotype_Myc | Lsm11 | -2.371 | 1.19E-03 | glio |
| Interaction_genotype_Nat2 | Kmo | 3.926 | 1.19E-03 | glio |
| Interaction_genotype_Uqcrb | Esrp2 | -5.009 | 1.19E-03 | glio |
| Aars2_in.vivo_noRT | Tspan32 | 2.348 | 1.19E-03 | glio |
| Aars2_in.vivo_noRT | Tlr2 | 2.378 | 1.19E-03 | glio |
| Aars2_in.vivo_noRT | Pacc1 | 1.961 | 1.19E-03 | glio |
| Atr_ex.vivo_noRT | Dnmt1 | -1.19 | 1.19E-03 | glio |
| Bcas2_in.vivo_noRT | Nxf2 | 2.464 | 1.19E-03 | glio |
| Bcas2_in.vivo_noRT | Mogat2 | 2.29 | 1.19E-03 | glio |
| Bcas2_in.vivo_noRT | Gm28229 | 2.166 | 1.19E-03 | glio |
| Bcas2_in.vivo_noRT | Mafa | 2.145 | 1.19E-03 | glio |
| Bcas2_in.vivo_noRT | Mrpl33 | -0.641 | 1.19E-03 | glio |
| Bora_in.vivo_noRT | Mb | 2.366 | 1.19E-03 | glio |
| Bora_in.vivo_noRT | Napsa | 2.419 | 1.19E-03 | glio |
| Bora_in.vivo_noRT | Surf1 | -1.335 | 1.19E-03 | glio |
| Bora_in.vivo_noRT | Kdelr1 | -1.009 | 1.19E-03 | glio |
| Bora_ex.vivo_noRT | Gm13054 | 2.806 | 1.19E-03 | glio |
| Bora_ex.vivo_noRT | Col17a1 | 3.345 | 1.19E-03 | glio |
| Brca2_in.vivo_noRT | Zmynd10 | 2.564 | 1.19E-03 | glio |
| Chek1_in.vivo_noRT | Gm37612 | 2.378 | 1.19E-03 | glio |
| Chek1_in.vivo_noRT | Gm12359 | 2.352 | 1.19E-03 | glio |
| Chek1_ex.vivo_noRT | 5930403N24R | 2.974 | 1.19E-03 | glio |
| Chek1_ex.vivo_noRT | Gm49701 | 2.815 | 1.19E-03 | glio |
| Eif2s1_in.vivo_noRT | Cd200r2 | 1.879 | 1.19E-03 | glio |
| culture_effect | C1rl | -1.706 | 1.19E-04 | glio |
| culture_effect | Msh2 | -0.724 | 1.19E-04 | glio |
| culture_effect | Vdac2 | -0.434 | 1.19E-04 | glio |
| Interaction_genotype_Aars2 | Pex10 | -3.889 | 1.19E-04 | glio |
| Interaction_genotype_Aars2 | Ak1 | -4.471 | 1.19E-04 | glio |
| Interaction_genotype_Aars2 | Tmem179 | -4.769 | 1.19E-04 | glio |
| Interaction_genotype_Aars2 | Pygo1 | -4.497 | 1.19E-04 | glio |
| Interaction_genotype_Mtor | Gm15265 | -4.601 | 1.19E-04 | glio |
| Interaction_genotype_Mtor | Akap3 | -4.522 | 1.19E-04 | glio |
| Interaction_genotype_Mtor | Gm39526 | -3.576 | 1.19E-04 | glio |
| Interaction_genotype_Mtor | Cacnb1 | -5.008 | 1.19E-04 | glio |
| Interaction_genotype_Mtor | Slc41a3 | -4.405 | 1.19E-04 | glio |
| Interaction_genotype_Mtor | Gm16685 | -3.84 | 1.19E-04 | glio |
| Interaction_genotype_Mtor | Mrpl42 | 1.74 | 1.19E-04 | glio |
| Interaction_genotype_Mtor | Cdk4 | 1.79 | 1.19E-04 | glio |
| Interaction_genotype_Myc | Stat5b | -3.72 | 1.19E-04 | glio |
| Aars2_in.vivo_noRT | Itgal | 2.855 | 1.19E-04 | glio |
| Aars2_in.vivo_noRT | Slc36a1 | 2.447 | 1.19E-04 | glio |
| Aars2_ex.vivo_noRT | CAAA0106612 | 3.275 | 1.19E-04 | glio |
| Atr_in.vivo_noRT | Gm49123 | 2.874 | 1.19E-04 | glio |
| Bard1_ex.vivo_noRT | H2-Eb2 | 2.814 | 1.19E-04 | glio |
| Bcas2_in.vivo_noRT | 2810414N06R | 2.785 | 1.19E-04 | glio |

| | | | | |
|---|---|---|---|---|
| Bcas2_in.vivo_noRT | Slc47a2 | 2.701 | 1.19E-04 | glio |
| Bcas2_in.vivo_noRT | Plk3 | -1.82 | 1.19E-04 | glio |
| Bora_in.vivo_noRT | Slc22a18 | 2.91 | 1.19E-04 | glio |
| Bora_in.vivo_noRT | Sardh | 2.649 | 1.19E-04 | glio |
| Bora_in.vivo_noRT | Mthfd1l | -1.548 | 1.19E-04 | glio |
| Cenpj_ex.vivo_noRT | Stfa3 | 2.591 | 1.19E-04 | glio |
| Cenpj_ex.vivo_noRT | Olfr787 | 2.591 | 1.19E-04 | glio |
| Chek1_in.vivo_noRT | Btc | 3.484 | 1.19E-04 | glio |
| Chek1_in.vivo_noRT | Gm15063 | 3.06 | 1.19E-04 | glio |
| Chek1_in.vivo_noRT | Gm10501 | 2.901 | 1.19E-04 | glio |
| Chek1_in.vivo_noRT | Gm36445 | 2.754 | 1.19E-04 | glio |
| Chek1_ex.vivo_noRT | Amdhd1 | 3.278 | 1.19E-04 | glio |
| Hsd17b10_in.vivo_noRT | Ankef1 | 2.877 | 1.19E-04 | glio |
| Interaction_genotype_Mtor | Ptgis | -5.187 | 1.19E-05 | glio |
| Interaction_genotype_Mtor | Gm35439 | -4.119 | 1.19E-05 | glio |
| Atr_in.vivo_noRT | Nxf2 | 3.19 | 1.19E-05 | glio |
| Atr_in.vivo_noRT | Tdrp | 2.961 | 1.19E-05 | glio |
| Bora_in.vivo_noRT | Tcte2 | 2.814 | 1.19E-05 | glio |
| Bora_in.vivo_noRT | Hells | -2.332 | 1.19E-05 | glio |
| Bora_ex.vivo_noRT | Gm40538 | 3.201 | 1.19E-05 | glio |
| Chek1_ex.vivo_noRT | Mrps7 | -1.411 | 1.19E-05 | glio |
| Hsd17b10_in.vivo_noRT | Gm13822 | 3.09 | 1.19E-05 | glio |
| Hsd17b10_in.vivo_noRT | Igfbp7 | 2.85 | 1.19E-05 | glio |
| culture_effect | 1700041G16R | 2.338 | 1.19E-06 | glio |
| culture_effect | Arpc3 | -0.628 | 1.19E-06 | glio |
| culture_effect | Btf3 | -0.455 | 1.19E-06 | glio |
| Aars2_in.vivo_noRT | Gm10638 | 3.359 | 1.19E-06 | glio |
| Atr_in.vivo_noRT | Crabp1 | 3.411 | 1.19E-06 | glio |
| Bcas2_in.vivo_noRT | Gm50103 | 3.5 | 1.19E-06 | glio |
| Chek1_in.vivo_noRT | Rab40b | 3.307 | 1.19E-06 | glio |
| culture_effect | Gm34006 | -2.107 | 1.19E-07 | glio |
| culture_effect | 1500015A07R | 2.039 | 1.19E-07 | glio |
| Interaction_genotype_Mtor | Stfa3 | -4.099 | 1.19E-07 | glio |
| Interaction_genotype_Mtor | Olfr787 | -4.099 | 1.19E-07 | glio |
| Atr_in.vivo_noRT | Olfr267 | 3.759 | 1.19E-07 | glio |
| Atr_in.vivo_noRT | Igsf23 | 3.873 | 1.19E-07 | glio |
| Bcas2_in.vivo_noRT | Gm50334 | 4.162 | 1.19E-07 | glio |
| Bora_in.vivo_noRT | Zhx2 | 3.891 | 1.19E-07 | glio |
| Bora_ex.vivo_noRT | 1700003E24Ri | 3.201 | 1.19E-07 | glio |
| Chek1_ex.vivo_noRT | Vwce | 4.056 | 1.19E-07 | glio |
| Atr_in.vivo_noRT | Kcnab3 | 3.999 | 1.19E-08 | glio |
| Atr_in.vivo_noRT | Cyth4 | 3.993 | 1.19E-08 | glio |
| Bcas2_in.vivo_noRT | 4930426I24Ri | 3.926 | 1.19E-08 | glio |
| Chek1_in.vivo_noRT | Hist1h4f | 4.252 | 1.19E-08 | glio |
| Atr_in.vivo_noRT | Jmjd8 | 4.167 | 1.19E-09 | glio |
| Bora_in.vivo_noRT | Vmn1r88 | 3.751 | 1.19E-09 | glio |
| Bora_in.vivo_noRT | Gm13594 | 4.411 | 1.19E-09 | glio |
| Chek1_in.vivo_noRT | Spdye4a | 4.386 | 1.19E-09 | glio |
| Chek1_ex.vivo_noRT | Dph6 | -3.503 | 1.19E-09 | glio |
| culture_effect | Gm5617 | 1.663 | 1.19E-10 | glio |
| Atr_in.vivo_noRT | BC034090 | 4.16 | 1.19E-10 | glio |
| Atr_in.vivo_noRT | Bpifa1 | 4.098 | 1.19E-10 | glio |

| | | | | |
|---|---|---|---|---|
| culture_effect | Tmem63b | -0.946 | 1.19E-11 | glio |
| Bcas2_in.vivo_noRT | 1110046J04Ri | 4.807 | 1.19E-11 | glio |
| Chek1_ex.vivo_noRT | Gm36026 | 4.842 | 1.19E-11 | glio |
| Chek1_in.vivo_noRT | Gm48161 | 4.256 | 1.19E-12 | glio |
| Chek1_in.vivo_noRT | Dcn | 4.653 | 1.19E-13 | glio |
| culture_effect | Guca1a | -2.032 | 1.19E-18 | glio |
| culture_effect | Itm2b | -1.35 | 1.19E-18 | glio |
| culture_effect | Sept7 | -1.161 | 1.19E-18 | glio |
| Atr_in.vivo_noRT | Obscn | 5.533 | 1.19E-18 | glio |
| Chek1_in.vivo_noRT | Ltbp2 | 5.044 | 1.19E-19 | glio |
| Chek1_in.vivo_noRT | Lmx1a | 5.044 | 1.19E-19 | glio |
| culture_effect | Stat3 | -1.449 | 1.19E-21 | glio |
| Chek1_in.vivo_noRT | Gm13288 | 5.555 | 1.19E-21 | glio |
| Atr_in.vivo_noRT | Halr1 | 5.267 | 1.19E-28 | glio |
| Atr_in.vivo_noRT | Capn13 | 5.533 | 1.19E-31 | glio |
| culture_effect | Smim10l1 | 0.488 | 1.20E-03 | glio |
| Interaction_genotype_Aars2 | Rdh10 | -3.137 | 1.20E-03 | glio |
| Interaction_genotype_Atr | Pspn | -2.765 | 1.20E-03 | glio |
| Interaction_genotype_Atr | Apoh | -3.938 | 1.20E-03 | glio |
| Interaction_genotype_Atr | Cep295nl | -4.041 | 1.20E-03 | glio |
| Interaction_genotype_Bcas2 | Dntt | -4.468 | 1.20E-03 | glio |
| Interaction_genotype_Bcas2 | Agbl5 | -3.929 | 1.20E-03 | glio |
| Interaction_genotype_Bcas2 | Gm27253 | -4.238 | 1.20E-03 | glio |
| Interaction_genotype_Bcas2 | D430001F17R | -4.01 | 1.20E-03 | glio |
| Interaction_genotype_Bcas2 | 4930412M03F | -4.142 | 1.20E-03 | glio |
| Interaction_genotype_Bora | Hes5 | 5.409 | 1.20E-03 | glio |
| Interaction_genotype_Mtor | Arvcf | -4.222 | 1.20E-03 | glio |
| Interaction_genotype_Mtor | Hist1h2ad | -3.997 | 1.20E-03 | glio |
| Interaction_genotype_Mtor | Gm4707 | -3.8 | 1.20E-03 | glio |
| Interaction_genotype_Mtor | Gipc3 | -3.933 | 1.20E-03 | glio |
| Interaction_genotype_Mtor | A330074K22R | -3.377 | 1.20E-03 | glio |
| Interaction_genotype_Myc | 4930512B01R | -2.729 | 1.20E-03 | glio |
| Interaction_genotype_Myc | Ltb4r1 | -3.902 | 1.20E-03 | glio |
| Interaction_genotype_Myc | Accsl | -3.914 | 1.20E-03 | glio |
| Interaction_genotype_Myc | 1700013F07Ri | -3.829 | 1.20E-03 | glio |
| Interaction_genotype_Myc | Dis3l | -3.288 | 1.20E-03 | glio |
| Interaction_genotype_Myc | Cdkl2 | -3.237 | 1.20E-03 | glio |
| Interaction_genotype_Myc | Gart | -2.392 | 1.20E-03 | glio |
| Interaction_genotype_Myc | Nek6 | -2.787 | 1.20E-03 | glio |
| Aars2_ex.vivo_noRT | Svopl | 2.9 | 1.20E-03 | glio |
| Aars2_ex.vivo_noRT | Prm1 | 2.893 | 1.20E-03 | glio |
| Atr_in.vivo_noRT | Xrcc3 | 2.246 | 1.20E-03 | glio |
| Atr_in.vivo_noRT | Immt | -1.041 | 1.20E-03 | glio |
| Atr_in.vivo_noRT | 1810058I24Ri | -0.982 | 1.20E-03 | glio |
| Atr_in.vivo_noRT | D8Ertd738e | -0.647 | 1.20E-03 | glio |
| Atr_ex.vivo_noRT | Gm39459 | 3.077 | 1.20E-03 | glio |
| Bcas2_in.vivo_noRT | Aldh3a2 | 1.702 | 1.20E-03 | glio |
| Bora_in.vivo_noRT | Specc1l | -1.641 | 1.20E-03 | glio |
| Bora_in.vivo_noRT | Vps45 | 1.603 | 1.20E-03 | glio |
| Bora_in.vivo_noRT | Set | -0.633 | 1.20E-03 | glio |
| Cenpj_in.vivo_noRT | Gm48215 | 2.022 | 1.20E-03 | glio |
| Cenpj_ex.vivo_noRT | Gm21319 | 2.591 | 1.20E-03 | glio |

| | | | | |
|---|---|---|---|---|
| Chek1_in.vivo_noRT | Tmem117 | 2.416 | 1.20E-03 | glio |
| Chek1_in.vivo_noRT | Tmem219 | -1.516 | 1.20E-03 | glio |
| Chek1_ex.vivo_noRT | 2810408A11R | 2.98 | 1.20E-03 | glio |
| Chek1_ex.vivo_noRT | 4833407H14R | 3.279 | 1.20E-03 | glio |
| Chek1_ex.vivo_noRT | Cd1d1 | 2.754 | 1.20E-03 | glio |
| Eif2s1_ex.vivo_noRT | Ly6d | 3.014 | 1.20E-03 | glio |
| Hsd17b10_in.vivo_noRT | Bfsp2 | 2.771 | 1.20E-03 | glio |
| Hsd17b10_in.vivo_noRT | Pla2g7 | 2.429 | 1.20E-03 | glio |
| Hsd17b10_in.vivo_noRT | Ccdc39 | 2.396 | 1.20E-03 | glio |
| Hsd17b10_in.vivo_noRT | 9130230N09R | 2.118 | 1.20E-03 | glio |
| culture_effect | Ctnnbl1 | -0.742 | 1.20E-04 | glio |
| culture_effect | Osbp | -0.665 | 1.20E-04 | glio |
| Interaction_genotype_Atr | Eps8l2 | -4.756 | 1.20E-04 | glio |
| Interaction_genotype_Bora | Gm50334 | -5.454 | 1.20E-04 | glio |
| Interaction_genotype_Mtor | Arrdc3 | -4.337 | 1.20E-04 | glio |
| Interaction_genotype_Mtor | Bag1 | 2.163 | 1.20E-04 | glio |
| Aars2_in.vivo_noRT | Sh3bgrl2 | 2.718 | 1.20E-04 | glio |
| Aars2_in.vivo_noRT | Zfp296 | 2.369 | 1.20E-04 | glio |
| Aars2_in.vivo_noRT | Zfp704 | 2.355 | 1.20E-04 | glio |
| Aars2_ex.vivo_noRT | Zfp352 | 3.293 | 1.20E-04 | glio |
| Aars2_ex.vivo_noRT | Gm34643 | 3.774 | 1.20E-04 | glio |
| Atr_in.vivo_noRT | Tmprss5 | 2.697 | 1.20E-04 | glio |
| Atr_in.vivo_noRT | Ccdc96 | 2.425 | 1.20E-04 | glio |
| Atr_ex.vivo_noRT | Art2b | 3.164 | 1.20E-04 | glio |
| Bora_in.vivo_noRT | Ckm | 2.662 | 1.20E-04 | glio |
| Chek1_in.vivo_noRT | Mblac1 | 2.898 | 1.20E-04 | glio |
| Chek1_in.vivo_noRT | Cnbd2 | 2.897 | 1.20E-04 | glio |
| Chek1_ex.vivo_noRT | 1700025N23R | 3.091 | 1.20E-04 | glio |
| Chek1_ex.vivo_noRT | Gm8104 | 3.298 | 1.20E-04 | glio |
| Chek1_ex.vivo_noRT | Ifi206 | 3.448 | 1.20E-04 | glio |
| culture_effect | Parp9 | -0.887 | 1.20E-05 | glio |
| culture_effect | Rps10 | -0.623 | 1.20E-05 | glio |
| Interaction_genotype_Mtor | 2210011C24R | -4.68 | 1.20E-05 | glio |
| Interaction_genotype_Mtor | 1700108F19Ri | -4.229 | 1.20E-05 | glio |
| Aars2_in.vivo_noRT | Bpifc | 3.307 | 1.20E-05 | glio |
| Aars2_in.vivo_noRT | 1700028J19Ri | 3.066 | 1.20E-05 | glio |
| Atp5c1_in.vivo_noRT | Zscan4d | 2.668 | 1.20E-05 | glio |
| Atr_in.vivo_noRT | Smim40 | 3.032 | 1.20E-05 | glio |
| Atr_in.vivo_noRT | Gm26910 | 2.844 | 1.20E-05 | glio |
| Atr_in.vivo_noRT | Ube2d2a | -1.226 | 1.20E-05 | glio |
| Atr_ex.vivo_noRT | Prl7c1 | 2.852 | 1.20E-05 | glio |
| Bcas2_in.vivo_noRT | Wdr38 | 3.236 | 1.20E-05 | glio |
| Bora_in.vivo_noRT | Gm42997 | 3.054 | 1.20E-05 | glio |
| Bora_in.vivo_noRT | Gstt2 | 3.151 | 1.20E-05 | glio |
| Chek1_in.vivo_noRT | C430014B12R | 2.995 | 1.20E-05 | glio |
| Chek1_in.vivo_noRT | Smyd2 | -1.583 | 1.20E-05 | glio |
| culture_effect | Gm17501 | 2.385 | 1.20E-06 | glio |
| culture_effect | Plekhh2 | 1.832 | 1.20E-06 | glio |
| culture_effect | Cmc2 | 0.74 | 1.20E-06 | glio |
| culture_effect | Selenok | -0.607 | 1.20E-06 | glio |
| Interaction_genotype_Mtor | 9130230N09R | -5.08 | 1.20E-06 | glio |
| Interaction_genotype_Mtor | 1700015F17Ri | -5.778 | 1.20E-06 | glio |

| | | | | |
|---|---|---|---|---|
| Interaction_genotype_Mtor | Erich6b | -4.099 | 1.20E-06 | glio |
| Atr_in.vivo_noRT | Hsd17b14 | 3.576 | 1.20E-06 | glio |
| Bcas2_in.vivo_noRT | Dchs1 | 3.312 | 1.20E-06 | glio |
| Bora_in.vivo_noRT | 1700025N23R | 3.044 | 1.20E-06 | glio |
| Chek1_in.vivo_noRT | Gm50012 | 3.443 | 1.20E-06 | glio |
| Chek1_in.vivo_noRT | C130013H08R | 3.804 | 1.20E-06 | glio |
| Chek1_in.vivo_noRT | Slc22a5 | 3.124 | 1.20E-06 | glio |
| culture_effect | Gm45336 | -2.807 | 1.20E-07 | glio |
| culture_effect | Glb1 | -1.15 | 1.20E-07 | glio |
| culture_effect | Prmt1 | -0.682 | 1.20E-07 | glio |
| Atr_in.vivo_noRT | Gm46218 | 3.75 | 1.20E-07 | glio |
| Bcas2_in.vivo_noRT | 1700023H06R | 4.016 | 1.20E-07 | glio |
| Chek1_ex.vivo_noRT | Ngef | 4.148 | 1.20E-07 | glio |
| Hsd17b10_in.vivo_noRT | 4930502E18Ri | 3.081 | 1.20E-07 | glio |
| Chek1_in.vivo_noRT | E230001N04R | 3.458 | 1.20E-08 | glio |
| culture_effect | Rps24 | -0.81 | 1.20E-09 | glio |
| Aars2_in.vivo_noRT | Acot6 | 4.081 | 1.20E-09 | glio |
| Aars2_in.vivo_noRT | Cd59b | 4.282 | 1.20E-09 | glio |
| Atr_in.vivo_noRT | Gm43258 | 4.137 | 1.20E-09 | glio |
| Bora_in.vivo_noRT | Flicr | 4.136 | 1.20E-09 | glio |
| Chek1_in.vivo_noRT | Fam228a | 4.256 | 1.20E-09 | glio |
| Atr_in.vivo_noRT | Hipk4 | 4.056 | 1.20E-10 | glio |
| Bcas2_in.vivo_noRT | Fam131a | 4.281 | 1.20E-10 | glio |
| Bcas2_in.vivo_noRT | Elobl | 4.807 | 1.20E-10 | glio |
| Bora_in.vivo_noRT | 4930540M05F | 4.279 | 1.20E-10 | glio |
| Hsd17b10_in.vivo_noRT | B230206H07R | 3.09 | 1.20E-10 | glio |
| Atr_in.vivo_noRT | Sult5a1 | 5.001 | 1.20E-11 | glio |
| Bcas2_in.vivo_noRT | Pcdhgb6 | 4.152 | 1.20E-11 | glio |
| Bora_in.vivo_noRT | Slc22a1 | 3.762 | 1.20E-11 | glio |
| Aars2_in.vivo_noRT | Pcdhga1 | 4.275 | 1.20E-12 | glio |
| Aars2_in.vivo_noRT | Grin3b | 4.804 | 1.20E-12 | glio |
| Bcas2_in.vivo_noRT | Fgf8 | 4.291 | 1.20E-12 | glio |
| Chek1_in.vivo_noRT | Cd248 | 4.777 | 1.20E-12 | glio |
| Bcas2_in.vivo_noRT | Tmem236 | 4.541 | 1.20E-13 | glio |
| Bcas2_in.vivo_noRT | Bcl2l15 | 4.547 | 1.20E-13 | glio |
| Bora_in.vivo_noRT | Olfr90 | 4.672 | 1.20E-14 | glio |
| culture_effect | Myo18a | 1.841 | 1.20E-15 | glio |
| Atr_in.vivo_noRT | Gm47872 | 5.269 | 1.20E-15 | glio |
| Aars2_in.vivo_noRT | Ssh3 | 5.041 | 1.20E-16 | glio |
| Atr_in.vivo_noRT | Pik3ap1 | 5.27 | 1.20E-18 | glio |
| Bcas2_in.vivo_noRT | Gm48780 | 4.547 | 1.20E-18 | glio |
| culture_effect | Yipf1 | -1.61 | 1.20E-20 | glio |
| Aars2_in.vivo_noRT | Tpo | 5.327 | 1.20E-26 | glio |
| culture_effect | Gm50232 | -1.901 | 1.21E-03 | glio |
| culture_effect | Zbtb11os1 | 1.471 | 1.21E-03 | glio |
| culture_effect | Ik | -0.425 | 1.21E-03 | glio |
| Interaction_genotype_Aars2 | Bcl7c | 1.482 | 1.21E-03 | glio |
| Interaction_genotype_Mtor | D630003M21I | -2.821 | 1.21E-03 | glio |
| Interaction_genotype_Myc | Traf1 | -3.682 | 1.21E-03 | glio |
| Interaction_genotype_Myc | Ikzf2 | -3.905 | 1.21E-03 | glio |
| Interaction_genotype_Myc | Gm20234 | -3.89 | 1.21E-03 | glio |
| Interaction_genotype_Myc | 1700063J08Ri | -3.515 | 1.21E-03 | glio |

| | | | | |
|---|---|---|---|---|
| Interaction_genotype_Myc | Anxa9 | -3.364 | 1.21E-03 | glio |
| Interaction_genotype_Myc | Mbd4 | -2.948 | 1.21E-03 | glio |
| Interaction_genotype_Pars2 | Loxl4 | -4.093 | 1.21E-03 | glio |
| Interaction_genotype_Rad21 | Gm20513 | 3.788 | 1.21E-03 | glio |
| interaction_RT | Adk | -0.832 | 1.21E-03 | glio |
| Aars2_in.vivo_noRT | Tlx3 | 2.281 | 1.21E-03 | glio |
| Aars2_in.vivo_noRT | Neurl4 | 1.78 | 1.21E-03 | glio |
| Aars2_in.vivo_noRT | Fbxw11 | -1.491 | 1.21E-03 | glio |
| Aars2_in.vivo_noRT | Tmem43 | -1.477 | 1.21E-03 | glio |
| Aars2_in.vivo_noRT | Tmbim4 | -1.071 | 1.21E-03 | glio |
| Aars2_in.vivo_noRT | Hmgn1 | -0.857 | 1.21E-03 | glio |
| Aars2_in.vivo_noRT | Rpl12 | -0.602 | 1.21E-03 | glio |
| Aars2_ex.vivo_noRT | Grin2d | 2.939 | 1.21E-03 | glio |
| Aars2_ex.vivo_noRT | Gm27241 | 2.312 | 1.21E-03 | glio |
| Atr_in.vivo_noRT | Ccdc180 | 2.092 | 1.21E-03 | glio |
| Atr_in.vivo_noRT | Tpcn2 | 2.302 | 1.21E-03 | glio |
| Atr_in.vivo_noRT | Kctd18 | 1.884 | 1.21E-03 | glio |
| Atr_in.vivo_noRT | Dvl3 | 1.567 | 1.21E-03 | glio |
| Atr_ex.vivo_noRT | Gm43625 | 2.776 | 1.21E-03 | glio |
| Bard1_ex.vivo_noRT | Gm49504 | 2.415 | 1.21E-03 | glio |
| Bcas2_in.vivo_noRT | Slurp1 | 3.513 | 1.21E-03 | glio |
| Bcas2_in.vivo_noRT | Htra1 | 2.637 | 1.21E-03 | glio |
| Bcas2_in.vivo_noRT | Gm42670 | 2.136 | 1.21E-03 | glio |
| Bcas2_in.vivo_noRT | Gm31645 | 2.382 | 1.21E-03 | glio |
| Bcas2_in.vivo_noRT | Mpnd | -1.534 | 1.21E-03 | glio |
| Bora_in.vivo_noRT | Itgal | 2.465 | 1.21E-03 | glio |
| Bora_in.vivo_noRT | Gm32618 | 2.416 | 1.21E-03 | glio |
| Cenpj_ex.vivo_noRT | Gm39526 | 2.591 | 1.21E-03 | glio |
| Cenpj_ex.vivo_noRT | Speer4e | 2.591 | 1.21E-03 | glio |
| Chek1_in.vivo_noRT | Dhtkd1 | 2.566 | 1.21E-03 | glio |
| Chek1_in.vivo_noRT | Gstm7 | 2.355 | 1.21E-03 | glio |
| Chek1_in.vivo_noRT | B230369F24Ri | 2.348 | 1.21E-03 | glio |
| Chek1_in.vivo_noRT | Gm15446 | 2.068 | 1.21E-03 | glio |
| Chek1_in.vivo_noRT | Plec | -1.025 | 1.21E-03 | glio |
| Chek1_ex.vivo_noRT | Taf5 | -1.859 | 1.21E-03 | glio |
| Eif2s1_ex.vivo_noRT | Gm41307 | 2.639 | 1.21E-03 | glio |
| Hsd17b10_in.vivo_noRT | Nek3 | 2.41 | 1.21E-03 | glio |
| Interaction_genotype_Atr | Rapgef3os2 | -3.555 | 1.21E-04 | glio |
| Interaction_genotype_Lmo7 | Klhdc8a | -4.44 | 1.21E-04 | glio |
| Interaction_genotype_Mtor | Acad11 | -3.857 | 1.21E-04 | glio |
| Interaction_genotype_Myc | Ube4bos1 | -3.651 | 1.21E-04 | glio |
| Interaction_genotype_Myc | Neurl2 | -3.793 | 1.21E-04 | glio |
| Interaction_genotype_Nat2 | B020031M17F | 5.625 | 1.21E-04 | glio |
| Interaction_genotype_Nat2 | Gm33466 | 3.605 | 1.21E-04 | glio |
| Interaction_genotype_Rpl3 | 6330415G19R | -5.048 | 1.21E-04 | glio |
| Interaction_genotype_Rpl3 | Gm27239 | -5.023 | 1.21E-04 | glio |
| Aars2_in.vivo_noRT | Gm15614 | 2.682 | 1.21E-04 | glio |
| Atr_in.vivo_noRT | Gm20186 | 2.856 | 1.21E-04 | glio |
| Atr_in.vivo_noRT | Sp1 | -1.527 | 1.21E-04 | glio |
| Atr_in.vivo_noRT | Nol8 | -1.549 | 1.21E-04 | glio |
| Atr_in.vivo_noRT | Ddrgk1 | -1.257 | 1.21E-04 | glio |
| Atr_ex.vivo_noRT | 4933408J17Ri | 3.178 | 1.21E-04 | glio |

| | | | | |
|---|---|---|---|---|
| Bcas2_in.vivo_noRT | Tmem240 | 3.079 | 1.21E-04 | glio |
| Bora_in.vivo_noRT | Plekho1 | -1.614 | 1.21E-04 | glio |
| Bora_ex.vivo_noRT | Insc | 3.201 | 1.21E-04 | glio |
| Bora_ex.vivo_noRT | Gm38477 | 2.802 | 1.21E-04 | glio |
| Cenpj_ex.vivo_noRT | Gm15908 | 2.591 | 1.21E-04 | glio |
| Cenpj_ex.vivo_noRT | A930014E10Ri | 2.591 | 1.21E-04 | glio |
| Cenpj_ex.vivo_noRT | Scg2 | 2.591 | 1.21E-04 | glio |
| Cenpj_ex.vivo_noRT | Slc25a18 | 3.375 | 1.21E-04 | glio |
| Cenpj_ex.vivo_noRT | 1700003E24Ri | 2.591 | 1.21E-04 | glio |
| Cenpj_ex.vivo_noRT | Krt19 | 2.591 | 1.21E-04 | glio |
| Cenpj_ex.vivo_noRT | Gm48155 | 2.591 | 1.21E-04 | glio |
| Cenpj_ex.vivo_noRT | Abra | 3.389 | 1.21E-04 | glio |
| Cenpj_ex.vivo_noRT | Gm34059 | 2.591 | 1.21E-04 | glio |
| Cenpj_ex.vivo_noRT | Halr1 | 2.591 | 1.21E-04 | glio |
| Cenpj_ex.vivo_noRT | Helt | 2.591 | 1.21E-04 | glio |
| Cenpj_ex.vivo_noRT | Gm20625 | 2.591 | 1.21E-04 | glio |
| Cenpj_ex.vivo_noRT | Gm5157 | 2.591 | 1.21E-04 | glio |
| Cenpj_ex.vivo_noRT | Gm44736 | 2.591 | 1.21E-04 | glio |
| Cenpj_ex.vivo_noRT | Fam227b | 2.591 | 1.21E-04 | glio |
| Cenpj_ex.vivo_noRT | Gm20658 | 2.591 | 1.21E-04 | glio |
| Cenpj_ex.vivo_noRT | Olfr810 | 2.591 | 1.21E-04 | glio |
| Cenpj_ex.vivo_noRT | Apoa5 | 2.591 | 1.21E-04 | glio |
| Cenpj_ex.vivo_noRT | Gm49172 | 2.591 | 1.21E-04 | glio |
| Cenpj_ex.vivo_noRT | Gjd4 | 2.591 | 1.21E-04 | glio |
| Chek1_ex.vivo_noRT | Fam13a | 3.379 | 1.21E-04 | glio |
| Eif2s1_in.vivo_noRT | Trim17 | 2.143 | 1.21E-04 | glio |
| Eif2s1_in.vivo_noRT | Tpo | 2.143 | 1.21E-04 | glio |
| Hsd17b10_in.vivo_noRT | Map7 | -2.662 | 1.21E-04 | glio |
| culture_effect | Apbb1ip | -1.708 | 1.21E-05 | glio |
| culture_effect | Carhsp1 | 0.86 | 1.21E-05 | glio |
| culture_effect | mt-Co2 | 0.58 | 1.21E-05 | glio |
| Interaction_genotype_Mtor | Gm14399 | -5.237 | 1.21E-05 | glio |
| Interaction_genotype_Mtor | Eif3m | 2.062 | 1.21E-05 | glio |
| Aars2_in.vivo_noRT | Aarsd1 | -1.697 | 1.21E-05 | glio |
| Atr_in.vivo_noRT | Slamf1 | 3.094 | 1.21E-05 | glio |
| Atr_in.vivo_noRT | Gm50203 | 2.901 | 1.21E-05 | glio |
| Atr_in.vivo_noRT | Rpl23a | -0.891 | 1.21E-05 | glio |
| Bcas2_in.vivo_noRT | Prm1 | 2.797 | 1.21E-05 | glio |
| Bcas2_in.vivo_noRT | Ash2l | -1.413 | 1.21E-05 | glio |
| Bora_in.vivo_noRT | Rasal1 | 3.46 | 1.21E-05 | glio |
| Bora_in.vivo_noRT | Desi2 | -1.725 | 1.21E-05 | glio |
| Chek1_in.vivo_noRT | Vipr2 | 3.101 | 1.21E-05 | glio |
| Chek1_in.vivo_noRT | Hdac11 | 3.159 | 1.21E-05 | glio |
| Chek1_ex.vivo_noRT | Stpg2 | 3.86 | 1.21E-05 | glio |
| Chek1_ex.vivo_noRT | Gm15816 | 3.363 | 1.21E-05 | glio |
| Chek1_ex.vivo_noRT | Ampd3 | 3.857 | 1.21E-05 | glio |
| Interaction_genotype_Mtor | Tac2 | -4.099 | 1.21E-06 | glio |
| Interaction_genotype_Myc | Gm7467 | -4.837 | 1.21E-06 | glio |
| Interaction_genotype_Myc | 4921531C22R | -5.384 | 1.21E-06 | glio |
| Interaction_genotype_Myc | Efhd1 | -5.659 | 1.21E-06 | glio |
| Aars2_in.vivo_noRT | Upk1a | 3.535 | 1.21E-06 | glio |
| Aars2_in.vivo_noRT | Mapk15 | 3.342 | 1.21E-06 | glio |

| | | | | |
|---|---|---|---|---|
| Aars2_ex.vivo_noRT | Gm48215 | 3.293 | 1.21E-06 | glio |
| Bora_in.vivo_noRT | Gm13272 | 3.233 | 1.21E-06 | glio |
| Bora_in.vivo_noRT | Gng13 | 3.343 | 1.21E-06 | glio |
| culture_effect | Rnasek | 0.615 | 1.21E-07 | glio |
| Interaction_genotype_Atr | Matn4 | -6.099 | 1.21E-07 | glio |
| Interaction_genotype_Myc | Alox12 | -6.25 | 1.21E-07 | glio |
| Aars2_in.vivo_noRT | Tpsab1 | 3.377 | 1.21E-07 | glio |
| Aars2_in.vivo_noRT | Gm15535 | 4.01 | 1.21E-07 | glio |
| Aars2_in.vivo_noRT | Zfp467 | 3.786 | 1.21E-07 | glio |
| Aars2_in.vivo_noRT | Mpeg1 | 3.351 | 1.21E-07 | glio |
| Atr_in.vivo_noRT | Cyp2s1 | 3.96 | 1.21E-07 | glio |
| Bcas2_in.vivo_noRT | Gpr162 | 3.759 | 1.21E-07 | glio |
| culture_effect | Sipa1l3 | 1.541 | 1.21E-08 | glio |
| Aars2_in.vivo_noRT | Gm10115 | 3.889 | 1.21E-08 | glio |
| Aars2_in.vivo_noRT | Plppr3 | 3.94 | 1.21E-08 | glio |
| Atr_in.vivo_noRT | Mir155hg | 4.288 | 1.21E-08 | glio |
| Bcas2_in.vivo_noRT | Camk1d | 4.006 | 1.21E-08 | glio |
| culture_effect | Gjb1 | -1.468 | 1.21E-09 | glio |
| Bcas2_in.vivo_noRT | Efhd1 | 4.291 | 1.21E-09 | glio |
| Bora_in.vivo_noRT | Clca3a1 | 3.885 | 1.21E-09 | glio |
| Hsd17b10_in.vivo_noRT | Prickle2 | 3.616 | 1.21E-09 | glio |
| Atr_in.vivo_noRT | Slc25a35 | 4.423 | 1.21E-10 | glio |
| Bora_in.vivo_noRT | Pcdhga1 | 3.885 | 1.21E-10 | glio |
| culture_effect | Sp110 | -2.103 | 1.21E-11 | glio |
| Atr_in.vivo_noRT | Cgn | 5.007 | 1.21E-13 | glio |
| Bcas2_in.vivo_noRT | Gm15520 | 4.554 | 1.21E-13 | glio |
| Bcas2_in.vivo_noRT | Gm45151 | 4.275 | 1.21E-14 | glio |
| Bcas2_in.vivo_noRT | 4930527F14Ri | 4.278 | 1.21E-15 | glio |
| Chek1_in.vivo_noRT | Gm28940 | 4.528 | 1.21E-15 | glio |
| Atr_in.vivo_noRT | Gm11769 | 5.273 | 1.21E-16 | glio |
| culture_effect | Ptpn2 | -1.168 | 1.21E-18 | glio |
| Atr_in.vivo_noRT | 9230112J17Ri | 5.533 | 1.21E-20 | glio |
| Bora_in.vivo_noRT | Gad1 | 4.67 | 1.21E-20 | glio |
| Aars2_in.vivo_noRT | Smtnl1 | 5.327 | 1.21E-22 | glio |
| Aars2_in.vivo_noRT | Speer4a | 5.06 | 1.21E-22 | glio |
| Chek1_in.vivo_noRT | Vmn2r44 | 5.044 | 1.21E-23 | glio |
| Chek1_in.vivo_noRT | Mmp13 | 5.044 | 1.21E-23 | glio |
| Atr_in.vivo_noRT | Gm19276 | 5.533 | 1.21E-27 | glio |
| culture_effect | Apmap | -0.749 | 1.22E-03 | glio |
| Interaction_genotype_Atr | Cldn19 | -3.886 | 1.22E-03 | glio |
| Interaction_genotype_Atr | Vwce | -3.665 | 1.22E-03 | glio |
| Interaction_genotype_Atr | Gm48140 | -2.9 | 1.22E-03 | glio |
| Interaction_genotype_Atr | Gm11627 | -3.251 | 1.22E-03 | glio |
| Interaction_genotype_Lmo7 | Mecom | -4.46 | 1.22E-03 | glio |
| Interaction_genotype_Mtor | Gm11110 | -3.952 | 1.22E-03 | glio |
| Interaction_genotype_Mtor | Gm10457 | -4.607 | 1.22E-03 | glio |
| Interaction_genotype_Mtor | Gm20274 | -3.166 | 1.22E-03 | glio |
| Interaction_genotype_Mtor | Med28 | 1.347 | 1.22E-03 | glio |
| Interaction_genotype_Mtor | Nedd4 | 1.114 | 1.22E-03 | glio |
| Interaction_genotype_Myc | E430018J23Ri | -3.461 | 1.22E-03 | glio |
| Interaction_genotype_Myc | Enc1 | -2.827 | 1.22E-03 | glio |
| Interaction_genotype_Rpl3 | 1700003G18R | -5.085 | 1.22E-03 | glio |

| | | | | |
|---|---|---|---|---|
| interaction_RT | Acyp1 | 0.827 | 1.22E-03 | glio |
| Aars2_in.vivo_noRT | Msantd1 | 2.339 | 1.22E-03 | glio |
| Aars2_in.vivo_noRT | Dcaf12 | -1.186 | 1.22E-03 | glio |
| Aars2_in.vivo_noRT | Mrfap1 | -0.67 | 1.22E-03 | glio |
| Atr_in.vivo_noRT | Dnaic2 | 1.579 | 1.22E-03 | glio |
| Atr_in.vivo_noRT | Oip5os1 | -1.112 | 1.22E-03 | glio |
| Atr_in.vivo_noRT | Lsm3 | -1.235 | 1.22E-03 | glio |
| Atr_in.vivo_noRT | Tpm1 | -1.33 | 1.22E-03 | glio |
| Atr_in.vivo_noRT | Lamp1 | -0.868 | 1.22E-03 | glio |
| Bard1_ex.vivo_noRT | Gm31152 | 2.86 | 1.22E-03 | glio |
| Bcas2_in.vivo_noRT | Slc9a9 | 2.419 | 1.22E-03 | glio |
| Bcas2_in.vivo_noRT | Ccdc78 | 2.267 | 1.22E-03 | glio |
| Bcas2_in.vivo_noRT | Rps24 | -0.634 | 1.22E-03 | glio |
| Bcas2_ex.vivo_noRT | Gm15138 | 2.347 | 1.22E-03 | glio |
| Bcas2_ex.vivo_noRT | Kcnk7 | 2.742 | 1.22E-03 | glio |
| Bcas2_ex.vivo_noRT | Myoz1 | 2.742 | 1.22E-03 | glio |
| Bcas2_ex.vivo_noRT | Usp17lc | 2.742 | 1.22E-03 | glio |
| Bcas2_ex.vivo_noRT | Brinp3 | 2.742 | 1.22E-03 | glio |
| Bora_in.vivo_noRT | Gdap1l1 | 2.583 | 1.22E-03 | glio |
| Bora_in.vivo_noRT | Cavin1 | -1.701 | 1.22E-03 | glio |
| Bora_in.vivo_noRT | Cep192 | -1.535 | 1.22E-03 | glio |
| Bora_in.vivo_noRT | Pura | -1.028 | 1.22E-03 | glio |
| Bora_ex.vivo_noRT | Nt5e | 2.808 | 1.22E-03 | glio |
| Brca2_in.vivo_noRT | Olfr543 | 2.272 | 1.22E-03 | glio |
| Brca2_in.vivo_noRT | Sema6d | 2.288 | 1.22E-03 | glio |
| Chek1_in.vivo_noRT | Wnt10a | 2.347 | 1.22E-03 | glio |
| Chek1_in.vivo_noRT | Atp6v1g2 | 2.166 | 1.22E-03 | glio |
| Chek1_in.vivo_noRT | Myb | 2.327 | 1.22E-03 | glio |
| Chek1_in.vivo_noRT | Serf2 | -0.633 | 1.22E-03 | glio |
| Chek1_ex.vivo_noRT | Efhb | 3.036 | 1.22E-03 | glio |
| Hsd17b10_in.vivo_noRT | Serpina3i | 2.295 | 1.22E-03 | glio |
| Hsd17b10_in.vivo_noRT | Gm2788 | 2.568 | 1.22E-03 | glio |
| Hsd17b10_in.vivo_noRT | Hpse | 2.426 | 1.22E-03 | glio |
| culture_effect | Wdr54 | -1.242 | 1.22E-04 | glio |
| Interaction_genotype_Aars2 | 1700063J08Ri | -4.77 | 1.22E-04 | glio |
| Interaction_genotype_Aars2 | Uck2 | 1.935 | 1.22E-04 | glio |
| Interaction_genotype_Aars2 | Psmg4 | 1.259 | 1.22E-04 | glio |
| Interaction_genotype_Mtor | 5930403N24R | -4.649 | 1.22E-04 | glio |
| Interaction_genotype_Mtor | Gm10353 | -4.526 | 1.22E-04 | glio |
| Interaction_genotype_Mtor | Slc25a18 | -3.833 | 1.22E-04 | glio |
| Interaction_genotype_Myc | Gm10767 | -4.436 | 1.22E-04 | glio |
| Interaction_genotype_Myc | Cul1 | 1.931 | 1.22E-04 | glio |
| Interaction_genotype_Myc | Rrp1 | 1.522 | 1.22E-04 | glio |
| Aars2_in.vivo_noRT | Gm16124 | 3.092 | 1.22E-04 | glio |
| Aars2_in.vivo_noRT | Bbs10 | 2.83 | 1.22E-04 | glio |
| Aars2_in.vivo_noRT | Stpg4 | 2.838 | 1.22E-04 | glio |
| Aars2_in.vivo_noRT | Kctd1 | 2.622 | 1.22E-04 | glio |
| Aars2_in.vivo_noRT | Lsm1 | -1.269 | 1.22E-04 | glio |
| Aars2_ex.vivo_noRT | Slc22a1 | 2.897 | 1.22E-04 | glio |
| Atp5c1_in.vivo_noRT | Psmb11 | 2.145 | 1.22E-04 | glio |
| Atp5c1_in.vivo_noRT | 2010001K21R | 2.668 | 1.22E-04 | glio |
| Atp5c1_in.vivo_noRT | Gm26978 | 2.408 | 1.22E-04 | glio |

| | | | | |
|---|---|---|---|---|
| Atp5c1_in.vivo_noRT | Ppil6 | 2.668 | 1.22E-04 | glio |
| Atr_in.vivo_noRT | 4930506C21R | 2.669 | 1.22E-04 | glio |
| Atr_in.vivo_noRT | St6gal1 | 2.823 | 1.22E-04 | glio |
| Atr_ex.vivo_noRT | Nos2 | 3.164 | 1.22E-04 | glio |
| Atr_ex.vivo_noRT | Efna2 | 4.125 | 1.22E-04 | glio |
| Atr_ex.vivo_noRT | Tmem151a | 3 | 1.22E-04 | glio |
| Bard1_ex.vivo_noRT | Dmrtc1a | 2.814 | 1.22E-04 | glio |
| Bard1_ex.vivo_noRT | Gm11899 | 2.814 | 1.22E-04 | glio |
| Bora_in.vivo_noRT | Gm4189 | 2.734 | 1.22E-04 | glio |
| Bora_in.vivo_noRT | Rwdd3 | 2.886 | 1.22E-04 | glio |
| Chek1_in.vivo_noRT | Gm32816 | 2.984 | 1.22E-04 | glio |
| Chek1_in.vivo_noRT | 1700012D14R | 2.665 | 1.22E-04 | glio |
| Eif2s1_in.vivo_noRT | 1700034I23Ri | 2.143 | 1.22E-04 | glio |
| Hsd17b10_in.vivo_noRT | 4933406C10R | 2.58 | 1.22E-04 | glio |
| Interaction_genotype_Mtor | Fbxl22 | -4.958 | 1.22E-05 | glio |
| Interaction_genotype_Mtor | Ptges | -5.514 | 1.22E-05 | glio |
| Aars2_in.vivo_noRT | Pwp1 | -1.795 | 1.22E-05 | glio |
| Aars2_in.vivo_noRT | Tra2b | -1.623 | 1.22E-05 | glio |
| Aars2_in.vivo_noRT | Nop16 | -1.561 | 1.22E-05 | glio |
| Atr_in.vivo_noRT | 1700020L24Ri | 2.873 | 1.22E-05 | glio |
| Atr_in.vivo_noRT | Mfsd4b4 | 2.808 | 1.22E-05 | glio |
| Atr_ex.vivo_noRT | Htr2b | 3.164 | 1.22E-05 | glio |
| Bcas2_in.vivo_noRT | Hoxa7 | 3.094 | 1.22E-05 | glio |
| Bora_in.vivo_noRT | Prkcq | 2.615 | 1.22E-05 | glio |
| Hsd17b10_in.vivo_noRT | Zic4 | 3.353 | 1.22E-05 | glio |
| culture_effect | Gm16835 | 2.455 | 1.22E-06 | glio |
| culture_effect | Bphl | -1.082 | 1.22E-06 | glio |
| Interaction_genotype_Mtor | Vcp | 2.637 | 1.22E-06 | glio |
| Chek1_ex.vivo_noRT | Klk6 | 3.861 | 1.22E-06 | glio |
| Chek1_ex.vivo_noRT | Epb42 | 4.484 | 1.22E-06 | glio |
| culture_effect | Alad | -0.822 | 1.22E-07 | glio |
| culture_effect | Spcs2 | -0.616 | 1.22E-07 | glio |
| Aars2_in.vivo_noRT | Gm21814 | 3.468 | 1.22E-07 | glio |
| Aars2_in.vivo_noRT | Ucma | 4.328 | 1.22E-07 | glio |
| Atr_in.vivo_noRT | Cbs | 3.807 | 1.22E-07 | glio |
| Atr_ex.vivo_noRT | B020031M17F | 5.622 | 1.22E-07 | glio |
| Bcas2_in.vivo_noRT | Ccdc106 | 3.996 | 1.22E-07 | glio |
| culture_effect | Tbc1d30 | 2.05 | 1.22E-08 | glio |
| Aars2_in.vivo_noRT | Calml4 | 3.766 | 1.22E-08 | glio |
| Atr_in.vivo_noRT | Brsk2 | 3.61 | 1.22E-08 | glio |
| Chek1_in.vivo_noRT | Rab3b | 3.887 | 1.22E-08 | glio |
| Chek1_in.vivo_noRT | Nhsl2 | 3.925 | 1.22E-08 | glio |
| Chek1_ex.vivo_noRT | Ntn5 | 4.859 | 1.22E-08 | glio |
| Hsd17b10_in.vivo_noRT | 4930527F14Ri | 3.087 | 1.22E-08 | glio |
| Chek1_in.vivo_noRT | A830035O19R | 3.994 | 1.22E-09 | glio |
| Hsd17b10_in.vivo_noRT | Car14 | 4.137 | 1.22E-09 | glio |
| Hsd17b10_in.vivo_noRT | Gm15890 | 3.616 | 1.22E-09 | glio |
| Aars2_in.vivo_noRT | Man2c1os | 4.682 | 1.22E-10 | glio |
| Aars2_in.vivo_noRT | Otop1 | 4.142 | 1.22E-10 | glio |
| Atr_ex.vivo_noRT | AC125149.3 | 5.322 | 1.22E-10 | glio |
| Atr_in.vivo_noRT | Mogat1 | 5.011 | 1.22E-11 | glio |
| Bora_in.vivo_noRT | Masp2 | 4.009 | 1.22E-11 | glio |

| | | | | |
|---|---|---|---|---|
| Chek1_ex.vivo_noRT | Vmn2r54 | 4.842 | 1.22E-11 | glio |
| culture_effect | Gm32090 | 3.343 | 1.22E-12 | glio |
| culture_effect | Abhd5 | 1.523 | 1.22E-12 | glio |
| Bcas2_in.vivo_noRT | Hbq1b | 5.07 | 1.22E-12 | glio |
| Bcas2_in.vivo_noRT | Myo5c | 4.807 | 1.22E-12 | glio |
| Chek1_in.vivo_noRT | Skida1 | 5.044 | 1.22E-12 | glio |
| Chek1_in.vivo_noRT | Fbxl2 | 4.398 | 1.22E-12 | glio |
| Chek1_in.vivo_noRT | BC018473 | 5.044 | 1.22E-13 | glio |
| Atr_in.vivo_noRT | Lrrc4b | 4.623 | 1.22E-14 | glio |
| culture_effect | Adamts1 | -1.911 | 1.22E-15 | glio |
| Aars2_in.vivo_noRT | Gm41349 | 5.063 | 1.22E-15 | glio |
| Bora_in.vivo_noRT | 2410141K09R | 4.672 | 1.22E-15 | glio |
| Bcas2_in.vivo_noRT | Kcnk13 | 4.544 | 1.22E-16 | glio |
| culture_effect | Nlrc5 | -2.434 | 1.22E-17 | glio |
| Bora_in.vivo_noRT | Esp15 | 4.936 | 1.22E-17 | glio |
| Bora_in.vivo_noRT | Gm13178 | 4.42 | 1.22E-17 | glio |
| Bora_in.vivo_noRT | Gm10801 | 4.936 | 1.22E-17 | glio |
| Aars2_in.vivo_noRT | Gm49086 | 5.327 | 1.22E-18 | glio |
| Atr_in.vivo_noRT | Gm128 | 5.153 | 1.22E-19 | glio |
| Bora_in.vivo_noRT | Gm21761 | 4.936 | 1.22E-19 | glio |
| Atr_in.vivo_noRT | Gm35439 | 5.154 | 1.22E-20 | glio |
| Bcas2_in.vivo_noRT | Cd36 | 4.807 | 1.22E-20 | glio |
| Bora_in.vivo_noRT | Lncbate10 | 4.936 | 1.22E-22 | glio |
| Aars2_in.vivo_noRT | Gm29670 | 5.315 | 1.22E-23 | glio |
| Atr_in.vivo_noRT | Cldn5 | 5.267 | 1.22E-25 | glio |
| Chek1_in.vivo_noRT | Gm49808 | 5.044 | 1.22E-25 | glio |
| culture_effect | Foxd2 | 1.233 | 1.23E-03 | glio |
| culture_effect | Gstm1 | 1.186 | 1.23E-03 | glio |
| culture_effect | Atpaf1 | -0.869 | 1.23E-03 | glio |
| culture_effect | Tjp2 | 0.736 | 1.23E-03 | glio |
| culture_effect | Plk3 | -0.684 | 1.23E-03 | glio |
| culture_effect | Tpd52 | -0.618 | 1.23E-03 | glio |
| culture_effect | Ext1 | 0.579 | 1.23E-03 | glio |
| Interaction_genotype_Aars2 | 1010001B22R | -3.965 | 1.23E-03 | glio |
| Interaction_genotype_Aars2 | Cerkl | -3.653 | 1.23E-03 | glio |
| Interaction_genotype_Aars2 | Eef1a1 | 1.014 | 1.23E-03 | glio |
| Interaction_genotype_Atr | Olfr1369-ps1 | -4.947 | 1.23E-03 | glio |
| Interaction_genotype_Atr | Stk36 | -4.169 | 1.23E-03 | glio |
| Interaction_genotype_Atr | Ripply3 | -3.933 | 1.23E-03 | glio |
| Interaction_genotype_Atr | Ttbk1 | -3.925 | 1.23E-03 | glio |
| Interaction_genotype_Atr | Slc35e3 | -2.281 | 1.23E-03 | glio |
| Interaction_genotype_Mtor | Magea10 | -3.418 | 1.23E-03 | glio |
| Interaction_genotype_Mtor | Ctrl | -4.127 | 1.23E-03 | glio |
| Interaction_genotype_Mtor | 1810024B03R | -2.76 | 1.23E-03 | glio |
| Interaction_genotype_Myc | 4930553M12F | -2.856 | 1.23E-03 | glio |
| Interaction_genotype_Myc | Mroh7 | -3.595 | 1.23E-03 | glio |
| Interaction_genotype_Myc | Klhl36 | -3.594 | 1.23E-03 | glio |
| Interaction_genotype_Myc | Pnpla7 | -3.443 | 1.23E-03 | glio |
| Interaction_genotype_Myc | Arhgap10 | -2.174 | 1.23E-03 | glio |
| Interaction_genotype_Myc | Atp5g2 | 1.138 | 1.23E-03 | glio |
| interaction_RT | Capn6 | 2.731 | 1.23E-03 | glio |
| Aars2_in.vivo_noRT | Tecr | -1.058 | 1.23E-03 | glio |

| | | | | |
|---|---|---|---|---|
| Aars2_ex.vivo_noRT | Cyp7a1 | 3.093 | 1.23E-03 | glio |
| Aars2_ex.vivo_noRT | Icosl | 2.893 | 1.23E-03 | glio |
| Atr_in.vivo_noRT | Ssc5d | 2.144 | 1.23E-03 | glio |
| Atr_ex.vivo_noRT | Tubb4a | 3.517 | 1.23E-03 | glio |
| Atr_ex.vivo_noRT | Pnpo | 1.74 | 1.23E-03 | glio |
| Bard1_ex.vivo_noRT | Olfr718-ps1 | 2.421 | 1.23E-03 | glio |
| Bcas2_in.vivo_noRT | Zfp534 | 2.465 | 1.23E-03 | glio |
| Bcas2_in.vivo_noRT | Mief2 | 2.164 | 1.23E-03 | glio |
| Bcas2_in.vivo_noRT | Kpnb1 | -1.027 | 1.23E-03 | glio |
| Bora_in.vivo_noRT | Cep83 | -1.408 | 1.23E-03 | glio |
| Bora_in.vivo_noRT | Mettl9 | -1.321 | 1.23E-03 | glio |
| Brca2_in.vivo_noRT | Gm49504 | 2.288 | 1.23E-03 | glio |
| Chek1_in.vivo_noRT | Efr3b | 2.613 | 1.23E-03 | glio |
| Chek1_in.vivo_noRT | Bglap3 | 2.325 | 1.23E-03 | glio |
| Chek1_in.vivo_noRT | Tpbg | 2.302 | 1.23E-03 | glio |
| Chek1_in.vivo_noRT | Esr1 | 2.187 | 1.23E-03 | glio |
| Chek1_in.vivo_noRT | Fst | 2.141 | 1.23E-03 | glio |
| Chek1_in.vivo_noRT | Grhl2 | 2.339 | 1.23E-03 | glio |
| Chek1_in.vivo_noRT | Smap1 | -1.074 | 1.23E-03 | glio |
| Eif2s1_in.vivo_noRT | 1700006H21R | 2.67 | 1.23E-03 | glio |
| Eif2s1_in.vivo_noRT | Pldi | 2.143 | 1.23E-03 | glio |
| Hsd17b10_in.vivo_noRT | 4930515G01R | 2.265 | 1.23E-03 | glio |
| Interaction_genotype_Atr | Afg1l | -4.816 | 1.23E-04 | glio |
| Interaction_genotype_Mtor | AA388235 | -4.721 | 1.23E-04 | glio |
| Interaction_genotype_Mtor | B230312C02R | -5.79 | 1.23E-04 | glio |
| Interaction_genotype_Mtor | Ly6e | 2.682 | 1.23E-04 | glio |
| Interaction_genotype_Myc | Fbxo45 | -3.424 | 1.23E-04 | glio |
| interaction_RT | Bop1 | -1.004 | 1.23E-04 | glio |
| Aars2_in.vivo_noRT | Plcb4 | 2.74 | 1.23E-04 | glio |
| Aars2_in.vivo_noRT | Cd34 | 2.768 | 1.23E-04 | glio |
| Aars2_in.vivo_noRT | 1700010I14Ri | 2.657 | 1.23E-04 | glio |
| Aars2_ex.vivo_noRT | 1700108F19Ri | 2.899 | 1.23E-04 | glio |
| Atr_in.vivo_noRT | Gm7879 | 2.881 | 1.23E-04 | glio |
| Atr_in.vivo_noRT | Gpr146 | 2.75 | 1.23E-04 | glio |
| Atr_in.vivo_noRT | Slc15a3 | 2.514 | 1.23E-04 | glio |
| Atr_in.vivo_noRT | Zfp870 | 2.345 | 1.23E-04 | glio |
| Atr_in.vivo_noRT | Habp4 | 2.171 | 1.23E-04 | glio |
| Atr_ex.vivo_noRT | 1700006H21R | 3.549 | 1.23E-04 | glio |
| Bcas2_in.vivo_noRT | Cyct | -1.625 | 1.23E-04 | glio |
| Bcas2_ex.vivo_noRT | Rgs4 | 2.742 | 1.23E-04 | glio |
| Chek1_in.vivo_noRT | Mzf1 | 2.653 | 1.23E-04 | glio |
| Chek1_in.vivo_noRT | Cd38 | 2.707 | 1.23E-04 | glio |
| Hsd17b10_in.vivo_noRT | Isl1 | 2.813 | 1.23E-04 | glio |
| Hsd17b10_in.vivo_noRT | Ifi214 | 2.825 | 1.23E-04 | glio |
| culture_effect | 1700096K18R | 1.481 | 1.23E-05 | glio |
| Interaction_genotype_Mtor | 4930579C12R | -4.494 | 1.23E-05 | glio |
| Interaction_genotype_Mtor | Gm6803 | -4.099 | 1.23E-05 | glio |
| Interaction_genotype_Myc | Gm37800 | -4.596 | 1.23E-05 | glio |
| Interaction_genotype_Xrcc5 | Slc16a4 | 4.69 | 1.23E-05 | glio |
| Aars2_in.vivo_noRT | Magi2 | 3.019 | 1.23E-05 | glio |
| Aars2_in.vivo_noRT | Cops8 | -1.641 | 1.23E-05 | glio |
| Aars2_ex.vivo_noRT | Dennd2d | 3.676 | 1.23E-05 | glio |

| | | | | |
|---|---|---|---|---|
| Atp5c1_in.vivo_noRT | Gm18280 | 2.404 | 1.23E-05 | glio |
| Atr_in.vivo_noRT | Plac1 | 2.952 | 1.23E-05 | glio |
| Atr_ex.vivo_noRT | Stat4 | 4.035 | 1.23E-05 | glio |
| Bora_in.vivo_noRT | Cbs | 3.21 | 1.23E-05 | glio |
| Bora_in.vivo_noRT | Fes | 3.149 | 1.23E-05 | glio |
| Chek1_in.vivo_noRT | B230317F23Ri | 3.001 | 1.23E-05 | glio |
| Chek1_ex.vivo_noRT | Trim66 | 3.657 | 1.23E-05 | glio |
| Hsd17b10_in.vivo_noRT | A930003A15R | 2.985 | 1.23E-05 | glio |
| culture_effect | Tsga10ip | -3.027 | 1.23E-06 | glio |
| Interaction_genotype_Mtor | Slc39a8 | -6.346 | 1.23E-06 | glio |
| Interaction_genotype_Mtor | Gm12905 | -6.241 | 1.23E-06 | glio |
| Interaction_genotype_Myc | Rpsa | 2.035 | 1.23E-06 | glio |
| Atr_in.vivo_noRT | A930003O13R | 3.734 | 1.23E-06 | glio |
| Atr_in.vivo_noRT | Gstm7 | 3.382 | 1.23E-06 | glio |
| Bora_in.vivo_noRT | Gm20492 | 3.345 | 1.23E-06 | glio |
| Chek1_in.vivo_noRT | Ss18l1 | 2.926 | 1.23E-06 | glio |
| Hsd17b10_in.vivo_noRT | Ngb | 3.377 | 1.23E-06 | glio |
| culture_effect | Gcat | -0.93 | 1.23E-07 | glio |
| culture_effect | Itga6 | 0.771 | 1.23E-07 | glio |
| Aars2_ex.vivo_noRT | Ankrd22 | 3.293 | 1.23E-07 | glio |
| Aars2_ex.vivo_noRT | Gm47404 | 3.293 | 1.23E-07 | glio |
| Aars2_ex.vivo_noRT | Gm31717 | 3.293 | 1.23E-07 | glio |
| Aars2_ex.vivo_noRT | Gm14742 | 3.293 | 1.23E-07 | glio |
| Aars2_ex.vivo_noRT | Scel | 3.293 | 1.23E-07 | glio |
| Aars2_ex.vivo_noRT | Ctla4 | 3.293 | 1.23E-07 | glio |
| Aars2_ex.vivo_noRT | Zscan4-ps3 | 3.293 | 1.23E-07 | glio |
| Aars2_ex.vivo_noRT | Tagap | 3.293 | 1.23E-07 | glio |
| Aars2_ex.vivo_noRT | Gm21731 | 3.293 | 1.23E-07 | glio |
| Aars2_ex.vivo_noRT | Gm11232 | 3.293 | 1.23E-07 | glio |
| Aars2_ex.vivo_noRT | Gm29966 | 3.293 | 1.23E-07 | glio |
| Aars2_ex.vivo_noRT | Olfr1152 | 3.293 | 1.23E-07 | glio |
| Bcas2_in.vivo_noRT | Lrrc46 | 3.677 | 1.23E-07 | glio |
| Bcas2_in.vivo_noRT | Gm41492 | 3.237 | 1.23E-07 | glio |
| Bcas2_in.vivo_noRT | Abhd2 | -2.454 | 1.23E-07 | glio |
| Bora_in.vivo_noRT | Gm50069 | 3.819 | 1.23E-07 | glio |
| Bora_in.vivo_noRT | Ak1 | 3.673 | 1.23E-07 | glio |
| culture_effect | Dlx1 | 2.076 | 1.23E-08 | glio |
| Atr_in.vivo_noRT | Rpl3l | 3.597 | 1.23E-08 | glio |
| Atr_in.vivo_noRT | Tgfb1i1 | 3.69 | 1.23E-08 | glio |
| Bcas2_in.vivo_noRT | Gm48772 | 3.899 | 1.23E-08 | glio |
| Chek1_in.vivo_noRT | Mir670hg | 4.393 | 1.23E-08 | glio |
| Chek1_in.vivo_noRT | Trim67 | 3.475 | 1.23E-08 | glio |
| Chek1_in.vivo_noRT | A430035B10R | 4.119 | 1.23E-08 | glio |
| Chek1_in.vivo_noRT | A230020J21Ri | 4.116 | 1.23E-08 | glio |
| culture_effect | Cd109 | -0.892 | 1.23E-09 | glio |
| Atr_in.vivo_noRT | Stab1 | 4.399 | 1.23E-09 | glio |
| Bcas2_in.vivo_noRT | Accsl | 4.547 | 1.23E-09 | glio |
| Atr_in.vivo_noRT | Gm34256 | 4.733 | 1.23E-10 | glio |
| Chek1_in.vivo_noRT | Gipc2 | 4.78 | 1.23E-10 | glio |
| Chek1_in.vivo_noRT | Efhd1 | 4.528 | 1.23E-10 | glio |
| Chek1_in.vivo_noRT | Tas1r1 | 4.017 | 1.23E-10 | glio |
| Chek1_in.vivo_noRT | Ifi207 | 4.78 | 1.23E-10 | glio |

| | | | | |
|---|---|---|---|---|
| Hsd17b10_in.vivo_noRT | Gm9758 | 3.356 | 1.23E-10 | glio |
| Bora_in.vivo_noRT | Ksr2 | 3.898 | 1.23E-11 | glio |
| Chek1_ex.vivo_noRT | St18 | 4.842 | 1.23E-11 | glio |
| Hsd17b10_in.vivo_noRT | Gpr22 | 3.616 | 1.23E-11 | glio |
| Hsd17b10_in.vivo_noRT | Gm50024 | 3.353 | 1.23E-11 | glio |
| Hsd17b10_in.vivo_noRT | Gm43256 | 4.415 | 1.23E-11 | glio |
| Aars2_in.vivo_noRT | Cgn | 4.801 | 1.23E-12 | glio |
| culture_effect | Myo5b | -1.273 | 1.23E-13 | glio |
| Atr_in.vivo_noRT | Cysrt1 | 5.273 | 1.23E-13 | glio |
| Hsd17b10_in.vivo_noRT | Gm48780 | 3.887 | 1.23E-13 | glio |
| Aars2_in.vivo_noRT | Gngt2 | 5.327 | 1.23E-14 | glio |
| Aars2_in.vivo_noRT | Gm20646 | 5.327 | 1.23E-15 | glio |
| Chek1_in.vivo_noRT | Card9 | 4.78 | 1.23E-15 | glio |
| Aars2_in.vivo_noRT | Thsd1 | 4.94 | 1.23E-16 | glio |
| Atr_in.vivo_noRT | Col19a1 | 4.616 | 1.23E-17 | glio |
| Chek1_in.vivo_noRT | Olfr543 | 4.784 | 1.23E-19 | glio |
| Bcas2_in.vivo_noRT | Gm43701 | 4.807 | 1.23E-20 | glio |
| culture_effect | Apol7d | 1.911 | 1.24E-03 | glio |
| culture_effect | Gm26728 | 1.766 | 1.24E-03 | glio |
| culture_effect | Gm20005 | 0.828 | 1.24E-03 | glio |
| culture_effect | Gpatch11 | 0.696 | 1.24E-03 | glio |
| Interaction_genotype_Aars2 | Abcb5 | -2.828 | 1.24E-03 | glio |
| Interaction_genotype_Aars2 | Gm26674 | -3.816 | 1.24E-03 | glio |
| Interaction_genotype_Aars2 | BC037032 | -4.342 | 1.24E-03 | glio |
| Interaction_genotype_Aars2 | Gpr162 | -4.146 | 1.24E-03 | glio |
| Interaction_genotype_Atr | Phactr4 | 2.457 | 1.24E-03 | glio |
| Interaction_genotype_Bcas2 | Olfm2 | -4.306 | 1.24E-03 | glio |
| Interaction_genotype_Mtor | Nup210l | -4.138 | 1.24E-03 | glio |
| Interaction_genotype_Mtor | Zfp27 | -3.378 | 1.24E-03 | glio |
| Interaction_genotype_Myc | Bpi | -3.301 | 1.24E-03 | glio |
| Interaction_genotype_Myc | Rell2 | -3.731 | 1.24E-03 | glio |
| Interaction_genotype_Myc | Lin52 | -2.764 | 1.24E-03 | glio |
| Interaction_genotype_Pars2 | 1700034I23Ri | -3.066 | 1.24E-03 | glio |
| Interaction_genotype_Pars2 | Idi2 | -3.045 | 1.24E-03 | glio |
| Interaction_genotype_Pars2 | Clcn1 | -3.575 | 1.24E-03 | glio |
| Interaction_genotype_Pars2 | C130060C02R | -3.971 | 1.24E-03 | glio |
| Interaction_genotype_Pars2 | Efhd1 | -4.412 | 1.24E-03 | glio |
| Interaction_genotype_Pars2 | Glyat | -3.066 | 1.24E-03 | glio |
| Interaction_genotype_Pars2 | Gm49086 | -3.831 | 1.24E-03 | glio |
| Interaction_genotype_Pars2 | Cysrt1 | -4.564 | 1.24E-03 | glio |
| Interaction_genotype_Rpl3 | Mecom | -4.17 | 1.24E-03 | glio |
| interaction_RT | Nr4a3 | 2.617 | 1.24E-03 | glio |
| interaction_RT | Sppl2b | -0.932 | 1.24E-03 | glio |
| Aars2_in.vivo_noRT | Gnasas1 | 2.175 | 1.24E-03 | glio |
| Aars2_in.vivo_noRT | Zkscan5 | 1.867 | 1.24E-03 | glio |
| Atp5c1_in.vivo_noRT | Tmem52 | 2.289 | 1.24E-03 | glio |
| Atp5c1_in.vivo_noRT | Rslcan18 | 2.282 | 1.24E-03 | glio |
| Atr_in.vivo_noRT | Col11a1 | -0.799 | 1.24E-03 | glio |
| Bard1_ex.vivo_noRT | 5730507C01R | 2.405 | 1.24E-03 | glio |
| Bard1_ex.vivo_noRT | Gm44899 | 2.492 | 1.24E-03 | glio |
| Bcas2_in.vivo_noRT | Ccdc57 | 2.461 | 1.24E-03 | glio |
| Bcas2_in.vivo_noRT | Gm10138 | 2.185 | 1.24E-03 | glio |

| | | | | |
|---|---|---|---|---|
| Bcas2_in.vivo_noRT | Rpl10a | -0.532 | 1.24E-03 | glio |
| Bcas2_ex.vivo_noRT | Ceacam16 | 2.742 | 1.24E-03 | glio |
| Bcas2_ex.vivo_noRT | Apol10b | 2.742 | 1.24E-03 | glio |
| Bora_in.vivo_noRT | 2810405F17Ri | 2.25 | 1.24E-03 | glio |
| Bora_in.vivo_noRT | Acss2 | 2.098 | 1.24E-03 | glio |
| Bora_in.vivo_noRT | Prdm4 | -1.597 | 1.24E-03 | glio |
| Bora_in.vivo_noRT | Top2b | -1.342 | 1.24E-03 | glio |
| Bora_in.vivo_noRT | Atp1a1 | -1.195 | 1.24E-03 | glio |
| Bora_ex.vivo_noRT | C4b | 2.631 | 1.24E-03 | glio |
| Cenpj_ex.vivo_noRT | Ptprc | 2.591 | 1.24E-03 | glio |
| Chek1_in.vivo_noRT | Prr15l | 2.352 | 1.24E-03 | glio |
| Chek1_in.vivo_noRT | Rwdd3 | 2.461 | 1.24E-03 | glio |
| Chek1_in.vivo_noRT | Itgae | 2.417 | 1.24E-03 | glio |
| Chek1_in.vivo_noRT | Rbm3os | 2.253 | 1.24E-03 | glio |
| Eif2s1_in.vivo_noRT | Gm48140 | 1.88 | 1.24E-03 | glio |
| Eif2s1_in.vivo_noRT | Timp4 | -3.239 | 1.24E-03 | glio |
| Fanca_ex.vivo_noRT | Vmn2r56 | 2.919 | 1.24E-03 | glio |
| Hsd17b10_in.vivo_noRT | 1810062O18R | 2.561 | 1.24E-03 | glio |
| Hsd17b10_in.vivo_noRT | Nrp2 | 0.944 | 1.24E-03 | glio |
| culture_effect | 4933400F21Ri | -1.726 | 1.24E-04 | glio |
| culture_effect | Csnk1d | -0.618 | 1.24E-04 | glio |
| culture_effect | Copb1 | -0.542 | 1.24E-04 | glio |
| Interaction_genotype_Aars2 | Zeb2os | -4.386 | 1.24E-04 | glio |
| Interaction_genotype_Bcas2 | Tanc1 | 2.744 | 1.24E-04 | glio |
| Interaction_genotype_Bora | 1700026J14Ri | -4.525 | 1.24E-04 | glio |
| Interaction_genotype_Mtor | Gm20667 | -4.588 | 1.24E-04 | glio |
| Interaction_genotype_Myc | Tead3 | -4.329 | 1.24E-04 | glio |
| Interaction_genotype_Myc | Necab3 | -4.292 | 1.24E-04 | glio |
| Interaction_genotype_Myc | Gm16348 | -4.256 | 1.24E-04 | glio |
| Atr_in.vivo_noRT | 1700012D14R | 2.631 | 1.24E-04 | glio |
| Atr_ex.vivo_noRT | Gm11131 | 3.566 | 1.24E-04 | glio |
| Bcas2_ex.vivo_noRT | H2-Eb1 | 3.522 | 1.24E-04 | glio |
| Bora_in.vivo_noRT | Shpk | 2.949 | 1.24E-04 | glio |
| Bora_in.vivo_noRT | Bc1 | 2.909 | 1.24E-04 | glio |
| Bora_in.vivo_noRT | Fhdc1 | 2.914 | 1.24E-04 | glio |
| Bora_in.vivo_noRT | Gm48604 | 2.677 | 1.24E-04 | glio |
| Bora_ex.vivo_noRT | Col19a1 | 2.808 | 1.24E-04 | glio |
| Bora_ex.vivo_noRT | Strip2 | 2.802 | 1.24E-04 | glio |
| Bora_ex.vivo_noRT | Icam2 | 3.201 | 1.24E-04 | glio |
| Bora_ex.vivo_noRT | 4732463B04R | 2.808 | 1.24E-04 | glio |
| Bora_ex.vivo_noRT | 2010001K21R | 3.202 | 1.24E-04 | glio |
| Bora_ex.vivo_noRT | Art2b | 3.201 | 1.24E-04 | glio |
| Bora_ex.vivo_noRT | 1110017D15R | 3.6 | 1.24E-04 | glio |
| Chek1_in.vivo_noRT | Krt17 | 4.784 | 1.24E-04 | glio |
| Chek1_in.vivo_noRT | Gm15879 | 3.55 | 1.24E-04 | glio |
| Chek1_in.vivo_noRT | 0610009E02Ri | 2.937 | 1.24E-04 | glio |
| Chek1_ex.vivo_noRT | Pcdhgb4 | 3.072 | 1.24E-04 | glio |
| culture_effect | Hdac9 | 2.389 | 1.24E-05 | glio |
| culture_effect | A730071L15Ri | 2.424 | 1.24E-05 | glio |
| culture_effect | Idh2 | -0.639 | 1.24E-05 | glio |
| Interaction_genotype_Mtor | Fhl2 | -4.492 | 1.24E-05 | glio |
| Interaction_genotype_Mtor | Gm49767 | -5.433 | 1.24E-05 | glio |

| | | | | |
|---|---|---|---|---|
| Interaction_genotype_Myc | Rgl1 | -4.488 | 1.24E-05 | glio |
| Interaction_genotype_Myc | AC163347.1 | -4.048 | 1.24E-05 | glio |
| Aars2_in.vivo_noRT | Gm21154 | 3.284 | 1.24E-05 | glio |
| Aars2_ex.vivo_noRT | Boc | 2.9 | 1.24E-05 | glio |
| Aars2_ex.vivo_noRT | Muc4 | 3.293 | 1.24E-05 | glio |
| Bora_in.vivo_noRT | Mrgprf | 3.14 | 1.24E-05 | glio |
| Bora_ex.vivo_noRT | Spink6 | 3.201 | 1.24E-05 | glio |
| Bora_ex.vivo_noRT | Htr2b | 3.201 | 1.24E-05 | glio |
| Chek1_in.vivo_noRT | C030037D09R | 3.072 | 1.24E-05 | glio |
| Chek1_in.vivo_noRT | Nnt | 2.782 | 1.24E-05 | glio |
| Bora_ex.vivo_noRT | Adam5 | 3.201 | 1.24E-06 | glio |
| Bora_ex.vivo_noRT | 4933400F21Ri | 3.201 | 1.24E-06 | glio |
| Chek1_in.vivo_noRT | Gm48529 | 3.314 | 1.24E-06 | glio |
| culture_effect | Qtrt1 | -1.073 | 1.24E-07 | glio |
| Interaction_genotype_Mtor | Gm45591 | -5.421 | 1.24E-07 | glio |
| Aars2_in.vivo_noRT | Cnfn | 3.932 | 1.24E-07 | glio |
| Aars2_ex.vivo_noRT | 2410017I17Ri | 3.293 | 1.24E-07 | glio |
| Aars2_ex.vivo_noRT | Cgnl1 | 3.293 | 1.24E-07 | glio |
| Aars2_ex.vivo_noRT | Gm26543 | 3.293 | 1.24E-07 | glio |
| Bcas2_in.vivo_noRT | Pax5 | 3.962 | 1.24E-07 | glio |
| Bcas2_in.vivo_noRT | Cd59b | 3.762 | 1.24E-07 | glio |
| Bcas2_in.vivo_noRT | Gm5431 | 3.426 | 1.24E-07 | glio |
| Bora_in.vivo_noRT | Gm13625 | 3.415 | 1.24E-07 | glio |
| Bora_in.vivo_noRT | Ttc41 | 3.924 | 1.24E-07 | glio |
| Bora_in.vivo_noRT | Epb41l4b | -2.204 | 1.24E-07 | glio |
| Hsd17b10_in.vivo_noRT | Lhx1os | 3.09 | 1.24E-07 | glio |
| culture_effect | Timp3 | 1.567 | 1.24E-08 | glio |
| Atr_in.vivo_noRT | Gm5089 | 4.495 | 1.24E-08 | glio |
| Atr_in.vivo_noRT | A430035B10R | 4.097 | 1.24E-08 | glio |
| Bora_in.vivo_noRT | Hecw2 | 4.11 | 1.24E-08 | glio |
| culture_effect | 3110039M20F | 1.193 | 1.24E-10 | glio |
| Aars2_in.vivo_noRT | Spint4 | 4.536 | 1.24E-10 | glio |
| Bora_in.vivo_noRT | Ushbp1 | 4.411 | 1.24E-10 | glio |
| Hsd17b10_in.vivo_noRT | Ablim2 | 3.084 | 1.24E-10 | glio |
| Aars2_in.vivo_noRT | Ptp4a1.1 | 4.53 | 1.24E-11 | glio |
| Aars2_in.vivo_noRT | Gm14968 | 4.947 | 1.24E-11 | glio |
| Chek1_in.vivo_noRT | Amy1 | 4.515 | 1.24E-11 | glio |
| Atr_ex.vivo_noRT | Ctsf | 4.978 | 1.24E-12 | glio |
| Chek1_in.vivo_noRT | Gm34047 | 5.044 | 1.24E-12 | glio |
| Chek1_ex.vivo_noRT | Gm40538 | 4.842 | 1.24E-12 | glio |
| Hsd17b10_in.vivo_noRT | Serpinb9g | 3.616 | 1.24E-12 | glio |
| Aars2_in.vivo_noRT | Cxcl1 | 5.327 | 1.24E-13 | glio |
| Atr_in.vivo_noRT | Iqcn | 5.007 | 1.24E-14 | glio |
| Bora_in.vivo_noRT | Slc25a18 | 4.67 | 1.24E-15 | glio |
| Aars2_in.vivo_noRT | Hcn3 | 5.221 | 1.24E-16 | glio |
| Atr_in.vivo_noRT | Gm17276 | 4.742 | 1.24E-16 | glio |
| Bcas2_in.vivo_noRT | Scel | 3.889 | 1.24E-16 | glio |
| Bora_in.vivo_noRT | Usp17lc | 4.936 | 1.24E-16 | glio |
| Bora_in.vivo_noRT | Gm48008 | 4.672 | 1.24E-16 | glio |
| Chek1_in.vivo_noRT | Gm36198 | 4.784 | 1.24E-16 | glio |
| Bora_in.vivo_noRT | Akap14 | 4.673 | 1.24E-17 | glio |
| Atr_in.vivo_noRT | Gm20711 | 5.533 | 1.24E-20 | glio |

| | | | | |
|---|---|---|---|---|
| Bcas2_in.vivo_noRT | Gm5415 | 4.807 | 1.24E-22 | glio |
| Chek1_in.vivo_noRT | Gpr4 | 5.044 | 1.24E-22 | glio |
| Chek1_in.vivo_noRT | Prl7c1 | 5.044 | 1.24E-22 | glio |
| Chek1_in.vivo_noRT | Gm33466 | 5.044 | 1.24E-22 | glio |
| Bora_in.vivo_noRT | BC061195 | 4.936 | 1.24E-23 | glio |
| Chek1_in.vivo_noRT | Gm8016 | 5.044 | 1.24E-23 | glio |
| Atr_in.vivo_noRT | Gm47936 | 5.533 | 1.24E-24 | glio |
| culture_effect | Hes5 | -8.078 | 1.24E-28 | glio |
| culture_effect | Slc38a3 | -6.025 | 1.24E-28 | glio |
| culture_effect | 4922502N22R | -2.058 | 1.25E-03 | glio |
| culture_effect | Gpat2 | -1.196 | 1.25E-03 | glio |
| culture_effect | Mpv17l2 | 0.539 | 1.25E-03 | glio |
| culture_effect | Ltv1 | -0.574 | 1.25E-03 | glio |
| culture_effect | Dkc1 | -0.452 | 1.25E-03 | glio |
| Interaction_genotype_Aars2 | 8030456M14F | -3.878 | 1.25E-03 | glio |
| Interaction_genotype_Bcas2 | Cabp4 | -4.369 | 1.25E-03 | glio |
| Interaction_genotype_Cdkn2a | Gm40264 | 4.088 | 1.25E-03 | glio |
| Interaction_genotype_Cdkn2a | Slc17a6 | 2.835 | 1.25E-03 | glio |
| Interaction_genotype_Cdkn2a | Gm15138 | 3.505 | 1.25E-03 | glio |
| Interaction_genotype_Cdkn2a | Gad1 | 3.374 | 1.25E-03 | glio |
| Interaction_genotype_Cdkn2a | Scel | 3.227 | 1.25E-03 | glio |
| Interaction_genotype_Cdkn2a | Edn2 | 3.239 | 1.25E-03 | glio |
| Interaction_genotype_Cdkn2a | Halr1 | 3.101 | 1.25E-03 | glio |
| Interaction_genotype_Chek1 | Slc35f1 | -3.35 | 1.25E-03 | glio |
| Interaction_genotype_Chek1 | Otof | 3.751 | 1.25E-03 | glio |
| Interaction_genotype_Chek1 | Smim14 | 1.634 | 1.25E-03 | glio |
| Interaction_genotype_Mtor | Tmem86a | -4.099 | 1.25E-03 | glio |
| Interaction_genotype_Mtor | Zfp960 | -4.112 | 1.25E-03 | glio |
| Interaction_genotype_Mtor | Gm10115 | -3.695 | 1.25E-03 | glio |
| Interaction_genotype_Mtor | Gm20234 | -4.096 | 1.25E-03 | glio |
| Interaction_genotype_Myc | Hist1h4f | -3.884 | 1.25E-03 | glio |
| Interaction_genotype_Myc | Slc13a5 | -3.495 | 1.25E-03 | glio |
| Interaction_genotype_Myc | Ndufb8 | 1.112 | 1.25E-03 | glio |
| Interaction_genotype_Pars2 | Gm10244 | -4.576 | 1.25E-03 | glio |
| Interaction_genotype_Uqcrc2 | Nxph3 | -4.399 | 1.25E-03 | glio |
| Aars2_in.vivo_noRT | Lypd6 | 2.339 | 1.25E-03 | glio |
| Aars2_in.vivo_noRT | B4galnt4 | 2.218 | 1.25E-03 | glio |
| Aars2_in.vivo_noRT | Coro1b | 2.18 | 1.25E-03 | glio |
| Aars2_in.vivo_noRT | Tnfsf12 | 1.911 | 1.25E-03 | glio |
| Aars2_in.vivo_noRT | Cox6b1 | -0.659 | 1.25E-03 | glio |
| Aars2_in.vivo_noRT | Ifitm3 | -0.737 | 1.25E-03 | glio |
| Atp5c1_in.vivo_noRT | Tcp11x2 | 2.146 | 1.25E-03 | glio |
| Atr_in.vivo_noRT | Sncg | 2.501 | 1.25E-03 | glio |
| Atr_in.vivo_noRT | Rpap2 | 1.715 | 1.25E-03 | glio |
| Atr_in.vivo_noRT | Txn2 | -0.873 | 1.25E-03 | glio |
| Atr_ex.vivo_noRT | Pldi | 2.469 | 1.25E-03 | glio |
| Atr_ex.vivo_noRT | Gpx2 | 3.931 | 1.25E-03 | glio |
| Bcas2_in.vivo_noRT | A430105J06Ri | 2.299 | 1.25E-03 | glio |
| Bcas2_in.vivo_noRT | Sall3 | 1.629 | 1.25E-03 | glio |
| Bcas2_ex.vivo_noRT | Gm4841 | 3.54 | 1.25E-03 | glio |
| Bora_in.vivo_noRT | Spry3 | 2.409 | 1.25E-03 | glio |
| Bora_in.vivo_noRT | Stpg4 | 2.448 | 1.25E-03 | glio |

| | | | | |
|---|---|---|---|---|
| Chek1_in.vivo_noRT | Fgl2 | 2.151 | 1.25E-03 | glio |
| Chek1_in.vivo_noRT | Rack1 | -0.645 | 1.25E-03 | glio |
| Chek1_ex.vivo_noRT | Gm44386 | 3.079 | 1.25E-03 | glio |
| Hsd17b10_in.vivo_noRT | Acot6 | 2.371 | 1.25E-03 | glio |
| Hsd17b10_in.vivo_noRT | Glmp | 2.658 | 1.25E-03 | glio |
| Hsd17b10_in.vivo_noRT | Lcor | -1.698 | 1.25E-03 | glio |
| Interaction_genotype_Mtor | Pkd1l2 | -4.23 | 1.25E-04 | glio |
| Interaction_genotype_Mtor | Gm16133 | -4.658 | 1.25E-04 | glio |
| Interaction_genotype_Mtor | Tdrp | -4.526 | 1.25E-04 | glio |
| Interaction_genotype_Mtor | Gpr135 | -4.564 | 1.25E-04 | glio |
| Interaction_genotype_Mtor | Cbx5 | 1.685 | 1.25E-04 | glio |
| Interaction_genotype_Mtor | Atp5b | 1.511 | 1.25E-04 | glio |
| Interaction_genotype_Myc | Baiap2l2 | -4.07 | 1.25E-04 | glio |
| Interaction_genotype_Myc | Gm49860 | -3.711 | 1.25E-04 | glio |
| Interaction_genotype_Myc | Capn13 | -2.729 | 1.25E-04 | glio |
| Interaction_genotype_Myc | Vps25 | -3.915 | 1.25E-04 | glio |
| Interaction_genotype_Myc | Ttl | -3.596 | 1.25E-04 | glio |
| Aars2_ex.vivo_noRT | 2810049E08Ri | 3.293 | 1.25E-04 | glio |
| Atr_in.vivo_noRT | Bod1l | -1.275 | 1.25E-04 | glio |
| Bcas2_in.vivo_noRT | Gm17249 | 2.67 | 1.25E-04 | glio |
| Bcas2_ex.vivo_noRT | Gm32122 | 3.524 | 1.25E-04 | glio |
| Chek1_in.vivo_noRT | Srpr | -1.174 | 1.25E-04 | glio |
| Chek1_ex.vivo_noRT | Trp53inp1 | 3.042 | 1.25E-04 | glio |
| Interaction_genotype_Mtor | 1700019E08Ri | -4.099 | 1.25E-05 | glio |
| Interaction_genotype_Mtor | Necab3 | -5.119 | 1.25E-05 | glio |
| Interaction_genotype_Mtor | BC026762 | -5.006 | 1.25E-05 | glio |
| Interaction_genotype_Myc | Kpna7 | -4.402 | 1.25E-05 | glio |
| Aars2_ex.vivo_noRT | Gm10801 | 3.293 | 1.25E-05 | glio |
| Atr_in.vivo_noRT | Shbg | 3.154 | 1.25E-05 | glio |
| Atr_in.vivo_noRT | Cdiptos | 2.984 | 1.25E-05 | glio |
| Atr_in.vivo_noRT | Zfp534 | 3.192 | 1.25E-05 | glio |
| Bcas2_ex.vivo_noRT | Stfa3 | 2.742 | 1.25E-05 | glio |
| Bcas2_ex.vivo_noRT | Olfr787 | 2.742 | 1.25E-05 | glio |
| Chek1_in.vivo_noRT | Mucl3 | 3.272 | 1.25E-05 | glio |
| Chek1_in.vivo_noRT | Ehhadh | 3.348 | 1.25E-05 | glio |
| Chek1_in.vivo_noRT | Mpv17l | 2.713 | 1.25E-05 | glio |
| Chek1_ex.vivo_noRT | Nr1h4 | 3.736 | 1.25E-05 | glio |
| culture_effect | Tshz1 | -1.183 | 1.25E-06 | glio |
| culture_effect | Tmx3 | -0.771 | 1.25E-06 | glio |
| Aars2_in.vivo_noRT | Gm20522 | 3.521 | 1.25E-06 | glio |
| Aars2_in.vivo_noRT | Myl9 | 3.764 | 1.25E-06 | glio |
| Aars2_in.vivo_noRT | Adgrd1 | 3.17 | 1.25E-06 | glio |
| Aars2_in.vivo_noRT | Scn2a | 3.424 | 1.25E-06 | glio |
| Aars2_ex.vivo_noRT | Gm11583 | 3.293 | 1.25E-06 | glio |
| Atp5c1_in.vivo_noRT | Rgs4 | 2.668 | 1.25E-06 | glio |
| Atr_in.vivo_noRT | Atp8a2 | 2.854 | 1.25E-06 | glio |
| Atr_in.vivo_noRT | 1700113B09R | 3.46 | 1.25E-06 | glio |
| Bcas2_in.vivo_noRT | Gm49042 | 3.178 | 1.25E-06 | glio |
| Bcas2_in.vivo_noRT | Calml4 | 3.246 | 1.25E-06 | glio |
| Bora_in.vivo_noRT | Gm12500 | 3.295 | 1.25E-06 | glio |
| Bora_in.vivo_noRT | Olfml2a | 3.433 | 1.25E-06 | glio |
| Bora_in.vivo_noRT | Etohd2 | 3.577 | 1.25E-06 | glio |

| | | | | |
|---|---|---|---|---|
| Bora_ex.vivo_noRT | Gm12794 | 3.201 | 1.25E-06 | glio |
| Chek1_in.vivo_noRT | Pcdhb18 | 3.293 | 1.25E-06 | glio |
| culture_effect | Gm29994 | -3.028 | 1.25E-07 | glio |
| Interaction_genotype_Mtor | Vdac2 | 2.503 | 1.25E-07 | glio |
| Interaction_genotype_Mtor | Sf3b3 | 2.742 | 1.25E-07 | glio |
| Interaction_genotype_Myc | 1110020A21R | -6.242 | 1.25E-07 | glio |
| Atr_in.vivo_noRT | Arhgap4 | 3.547 | 1.25E-07 | glio |
| Atr_in.vivo_noRT | Cdc42ep4 | 3.443 | 1.25E-07 | glio |
| Bcas2_in.vivo_noRT | 2900042K21R | 3.622 | 1.25E-07 | glio |
| Bora_in.vivo_noRT | Slc46a3 | 3.751 | 1.25E-07 | glio |
| culture_effect | 1700012B07R | -3.602 | 1.25E-08 | glio |
| interaction_RT | Eif3b | -1.097 | 1.25E-08 | glio |
| Atr_in.vivo_noRT | 4921507P07R | 3.769 | 1.25E-08 | glio |
| Bora_in.vivo_noRT | Gm49701 | 3.887 | 1.25E-08 | glio |
| Chek1_in.vivo_noRT | BC030343 | 4.007 | 1.25E-08 | glio |
| culture_effect | Ifi27 | -1.04 | 1.25E-09 | glio |
| culture_effect | Rabac1 | -0.705 | 1.25E-09 | glio |
| interaction_RT | 5430416N02R | 2.092 | 1.25E-09 | glio |
| Aars2_in.vivo_noRT | 2210417A02R | 4.536 | 1.25E-09 | glio |
| Aars2_in.vivo_noRT | Zfp811 | 4.153 | 1.25E-09 | glio |
| Aars2_in.vivo_noRT | Papln | 4.059 | 1.25E-09 | glio |
| Bcas2_in.vivo_noRT | 5930403N24R | 4.281 | 1.25E-09 | glio |
| Hsd17b10_in.vivo_noRT | Mettl7a3 | 4.028 | 1.25E-09 | glio |
| culture_effect | Akap12 | 1.568 | 1.25E-10 | glio |
| Aars2_in.vivo_noRT | Insc | 3.869 | 1.25E-10 | glio |
| Bora_in.vivo_noRT | Iqcn | 4.41 | 1.25E-11 | glio |
| Chek1_ex.vivo_noRT | Aqp9 | 4.842 | 1.25E-11 | glio |
| Chek1_ex.vivo_noRT | Gm15326 | 4.268 | 1.25E-11 | glio |
| Atr_in.vivo_noRT | Accsl | 5.273 | 1.25E-12 | glio |
| Chek1_in.vivo_noRT | Gm21818 | 5.044 | 1.25E-12 | glio |
| Atr_in.vivo_noRT | Tecrl | 5.273 | 1.25E-13 | glio |
| Atr_in.vivo_noRT | Ifitm5 | 5.011 | 1.25E-13 | glio |
| culture_effect | Col22a1 | 2.231 | 1.25E-14 | glio |
| Atr_in.vivo_noRT | Ldhd | 4.616 | 1.25E-14 | glio |
| Bora_in.vivo_noRT | Gm16741 | 4.936 | 1.25E-14 | glio |
| Bcas2_in.vivo_noRT | 4930579C12R | 4.807 | 1.25E-15 | glio |
| Chek1_in.vivo_noRT | Gm2449 | 4.78 | 1.25E-15 | glio |
| Chek1_in.vivo_noRT | Snrpn | 4.385 | 1.25E-15 | glio |
| Bcas2_in.vivo_noRT | 4930512B01R | 4.807 | 1.25E-19 | glio |
| Bora_in.vivo_noRT | Tal2 | 4.673 | 1.25E-19 | glio |
| Bcas2_in.vivo_noRT | Erich6b | 4.807 | 1.25E-21 | glio |
| Bcas2_in.vivo_noRT | Lncbate10 | 4.807 | 1.25E-21 | glio |
| Bcas2_in.vivo_noRT | Edn2 | 4.807 | 1.25E-22 | glio |
| Bcas2_in.vivo_noRT | Gm14207 | 4.807 | 1.25E-22 | glio |
| Bcas2_in.vivo_noRT | Nrip2 | 4.807 | 1.25E-22 | glio |
| Bora_in.vivo_noRT | Tac2 | 4.936 | 1.25E-22 | glio |
| Atr_in.vivo_noRT | Tnni3k | 5.533 | 1.25E-24 | glio |
| Aars2_in.vivo_noRT | Gm34059 | 5.327 | 1.25E-28 | glio |
| culture_effect | Psmb9 | -4.99 | 1.25E-74 | glio |
| Interaction_genotype_Atr | Edn2 | -2.765 | 1.26E-03 | glio |
| Interaction_genotype_Atr | Gm26766 | -3.802 | 1.26E-03 | glio |
| Interaction_genotype_Mtor | Gm13403 | -3.739 | 1.26E-03 | glio |

| | | | | |
|---|---|---|---|---|
| Interaction_genotype_Mtor | Rhbg | -2.897 | 1.26E-03 | glio |
| Interaction_genotype_Mtor | Plpp6 | -3.535 | 1.26E-03 | glio |
| Interaction_genotype_Mtor | Ufsp1 | -2.728 | 1.26E-03 | glio |
| Interaction_genotype_Mtor | Orai2 | -2.827 | 1.26E-03 | glio |
| Interaction_genotype_Mtor | Fbxo38 | -2.415 | 1.26E-03 | glio |
| Interaction_genotype_Mtor | Lzts2 | 1.716 | 1.26E-03 | glio |
| Interaction_genotype_Mtor | Uqcr10 | 1.744 | 1.26E-03 | glio |
| Interaction_genotype_Myc | Gm12089 | -3.779 | 1.26E-03 | glio |
| Interaction_genotype_Myc | Gm28229 | -3.376 | 1.26E-03 | glio |
| Interaction_genotype_Myc | Zfp839 | -3.505 | 1.26E-03 | glio |
| Interaction_genotype_Myc | Tmem151a | -3.654 | 1.26E-03 | glio |
| Interaction_genotype_Myc | Ska1 | -2.435 | 1.26E-03 | glio |
| Interaction_genotype_Nat2 | Gm45151 | 3.59 | 1.26E-03 | glio |
| Interaction_genotype_Pars2 | Car14 | -4.262 | 1.26E-03 | glio |
| Interaction_genotype_Pars2 | Gm10425 | -4.562 | 1.26E-03 | glio |
| Interaction_genotype_Rad51 | H2-Eb1 | 4.093 | 1.26E-03 | glio |
| Interaction_genotype_Rad51 | Asap2 | 2.567 | 1.26E-03 | glio |
| Interaction_genotype_Rad51 | Gm38534 | -4.268 | 1.26E-03 | glio |
| Interaction_genotype_Uqcrc2 | Nxph4 | -4.619 | 1.26E-03 | glio |
| Interaction_genotype_Uqcrc2 | Rcan2 | -4.884 | 1.26E-03 | glio |
| interaction_RT | Nfkbiz | 1.434 | 1.26E-03 | glio |
| interaction_RT | Limk1 | -1.278 | 1.26E-03 | glio |
| interaction_RT | Kifc3 | -0.988 | 1.26E-03 | glio |
| interaction_RT | Nts | 3.55 | 1.26E-03 | glio |
| Aars2_in.vivo_noRT | Kank2 | 2.228 | 1.26E-03 | glio |
| Atr_in.vivo_noRT | Zfp346 | 2.071 | 1.26E-03 | glio |
| Atr_in.vivo_noRT | 4933439C10R | 2.148 | 1.26E-03 | glio |
| Bcas2_in.vivo_noRT | Usp27x | 2.5 | 1.26E-03 | glio |
| Bcas2_in.vivo_noRT | C030014I23Ri | 2.357 | 1.26E-03 | glio |
| Bcas2_in.vivo_noRT | Ankrd28 | -1.674 | 1.26E-03 | glio |
| Bora_in.vivo_noRT | Clec1a | 2.255 | 1.26E-03 | glio |
| Bora_in.vivo_noRT | Slfn3 | 2.127 | 1.26E-03 | glio |
| Bora_in.vivo_noRT | Nipsnap1 | -1.565 | 1.26E-03 | glio |
| Bora_ex.vivo_noRT | Gm48899 | 2.626 | 1.26E-03 | glio |
| Chek1_in.vivo_noRT | Rgl1 | 2.128 | 1.26E-03 | glio |
| Chek1_in.vivo_noRT | Sh3pxd2b | 2.125 | 1.26E-03 | glio |
| Chek1_in.vivo_noRT | Dennd6b | 2.154 | 1.26E-03 | glio |
| Chek1_in.vivo_noRT | Hbp1 | -1.556 | 1.26E-03 | glio |
| Chek1_ex.vivo_noRT | Sox8 | 2.604 | 1.26E-03 | glio |
| Chek1_ex.vivo_noRT | Abtb2 | -2.178 | 1.26E-03 | glio |
| Hsd17b10_in.vivo_noRT | Zhx2 | 2.571 | 1.26E-03 | glio |
| Hsd17b10_in.vivo_noRT | 1700001G11R | 2.115 | 1.26E-03 | glio |
| culture_effect | Nipsnap3b | -0.633 | 1.26E-04 | glio |
| culture_effect | Rlbp1 | -0.568 | 1.26E-04 | glio |
| Interaction_genotype_Mtor | A430057M04F | -4.503 | 1.26E-04 | glio |
| Interaction_genotype_Mtor | Kctd11 | -4.651 | 1.26E-04 | glio |
| Interaction_genotype_Mtor | Tbc1d22b | -4.285 | 1.26E-04 | glio |
| Interaction_genotype_Mtor | Mthfr | -4.001 | 1.26E-04 | glio |
| Interaction_genotype_Mtor | Sec22c | -4.098 | 1.26E-04 | glio |
| Interaction_genotype_Myc | Olfr1369-ps1 | -5.307 | 1.26E-04 | glio |
| Interaction_genotype_Myc | Gm15943 | -3.447 | 1.26E-04 | glio |
| Interaction_genotype_Myc | Fabp3 | -4.807 | 1.26E-04 | glio |

| | | | | |
|---|---|---|---|---|
| Interaction_genotype_Myc | Lynx1 | -4.177 | 1.26E-04 | glio |
| interaction_RT | 2410004B18R | 0.916 | 1.26E-04 | glio |
| interaction_RT | AA467197 | 1.733 | 1.26E-04 | glio |
| interaction_RT | Sparcl1 | 1.401 | 1.26E-04 | glio |
| interaction_RT | Ftsj3 | -0.888 | 1.26E-04 | glio |
| Aars2_in.vivo_noRT | Zscan2 | 2.613 | 1.26E-04 | glio |
| Atr_in.vivo_noRT | Gm49602 | 2.594 | 1.26E-04 | glio |
| Atr_in.vivo_noRT | A330074K22R | 2.457 | 1.26E-04 | glio |
| Atr_in.vivo_noRT | Macc1 | 2.466 | 1.26E-04 | glio |
| Atr_in.vivo_noRT | Gm29170 | 2.623 | 1.26E-04 | glio |
| Atr_in.vivo_noRT | Itga2 | 2.373 | 1.26E-04 | glio |
| Atr_in.vivo_noRT | Fermt2 | -1.649 | 1.26E-04 | glio |
| Bcas2_in.vivo_noRT | Gm16133 | 2.824 | 1.26E-04 | glio |
| Bcas2_in.vivo_noRT | Cep72 | 2.686 | 1.26E-04 | glio |
| Bcas2_ex.vivo_noRT | Gm13178 | 2.742 | 1.26E-04 | glio |
| Bora_in.vivo_noRT | Gm34552 | 2.742 | 1.26E-04 | glio |
| Bora_in.vivo_noRT | Gm26982 | 2.716 | 1.26E-04 | glio |
| Bora_ex.vivo_noRT | Gm41409 | 3.242 | 1.26E-04 | glio |
| Cenpj_in.vivo_noRT | Cys1 | 2.81 | 1.26E-04 | glio |
| Chek1_in.vivo_noRT | Ifit1bl2 | 2.847 | 1.26E-04 | glio |
| Chek1_in.vivo_noRT | Gm42937 | 2.734 | 1.26E-04 | glio |
| Chek1_ex.vivo_noRT | Gm5624 | 3.247 | 1.26E-04 | glio |
| Interaction_genotype_Mtor | Tpt1 | 1.71 | 1.26E-05 | glio |
| Interaction_genotype_Myc | Rnft2 | -4.903 | 1.26E-05 | glio |
| Aars2_in.vivo_noRT | Slc43a1 | 3.125 | 1.26E-05 | glio |
| Aars2_ex.vivo_noRT | Gm17203 | 2.893 | 1.26E-05 | glio |
| Atr_in.vivo_noRT | Kank3 | 2.922 | 1.26E-05 | glio |
| Bcas2_in.vivo_noRT | Gm15672 | 3.084 | 1.26E-05 | glio |
| Bcas2_in.vivo_noRT | Gm44836 | 2.975 | 1.26E-05 | glio |
| Bcas2_in.vivo_noRT | C030037D09R | 3.08 | 1.26E-05 | glio |
| Chek1_in.vivo_noRT | Adgrd1 | 2.887 | 1.26E-05 | glio |
| Chek1_in.vivo_noRT | Fhdc1 | 3.266 | 1.26E-05 | glio |
| Chek1_ex.vivo_noRT | Gm43660 | 3.659 | 1.26E-05 | glio |
| Hsd17b10_in.vivo_noRT | Cgn | 3.09 | 1.26E-05 | glio |
| culture_effect | Myd88 | -1.337 | 1.26E-06 | glio |
| culture_effect | Tle4 | -1.048 | 1.26E-06 | glio |
| Atr_in.vivo_noRT | Vcan | 3.664 | 1.26E-06 | glio |
| Atr_ex.vivo_noRT | Gm26555 | 3.164 | 1.26E-06 | glio |
| Chek1_in.vivo_noRT | Gm34590 | 3.471 | 1.26E-06 | glio |
| Chek1_in.vivo_noRT | Gm19409 | 3.464 | 1.26E-06 | glio |
| culture_effect | Ndufs6 | 0.578 | 1.26E-07 | glio |
| Bcas2_in.vivo_noRT | Npb | 3.64 | 1.26E-07 | glio |
| Bora_in.vivo_noRT | 4933439K11R | 3.751 | 1.26E-07 | glio |
| Bora_in.vivo_noRT | Las1l | -2.099 | 1.26E-07 | glio |
| Chek1_in.vivo_noRT | Gm14273 | 3.581 | 1.26E-07 | glio |
| Interaction_genotype_Aars2 | Klhdc8a | -5.607 | 1.26E-08 | glio |
| Aars2_in.vivo_noRT | Pagr1a | 3.642 | 1.26E-08 | glio |
| Atr_in.vivo_noRT | Hist1h4b | 4.301 | 1.26E-08 | glio |
| Bcas2_in.vivo_noRT | Ifitm1 | 4.541 | 1.26E-08 | glio |
| Bora_in.vivo_noRT | Pom121 | -2.41 | 1.26E-08 | glio |
| culture_effect | Kif3a | -1.265 | 1.26E-09 | glio |
| Atr_in.vivo_noRT | Pde5a | 4.247 | 1.26E-09 | glio |

| | | | | |
|---|---|---|---|---|
| Bcas2_in.vivo_noRT | Fgd3 | 4.599 | 1.26E-09 | glio |
| Bcas2_in.vivo_noRT | Trpm3 | 4.285 | 1.26E-09 | glio |
| Chek1_in.vivo_noRT | Gm48610 | 4.261 | 1.26E-09 | glio |
| Chek1_ex.vivo_noRT | Lncenc1 | 4.845 | 1.26E-09 | glio |
| Aars2_in.vivo_noRT | Zkscan4 | 4.526 | 1.26E-10 | glio |
| Atr_in.vivo_noRT | BC030343 | 4.497 | 1.26E-10 | glio |
| Bora_in.vivo_noRT | Gjb5 | 3.946 | 1.26E-10 | glio |
| Chek1_in.vivo_noRT | Hist2h3c2 | 4.777 | 1.26E-10 | glio |
| Atr_in.vivo_noRT | Lst1 | 4.768 | 1.26E-11 | glio |
| Bcas2_in.vivo_noRT | Gm41409 | 4.275 | 1.26E-11 | glio |
| Bora_in.vivo_noRT | Gm11963 | 4.676 | 1.26E-11 | glio |
| Bcas2_in.vivo_noRT | Necab2 | 4.543 | 1.26E-12 | glio |
| interaction_RT | Spry2 | 2.754 | 1.26E-13 | glio |
| Bcas2_in.vivo_noRT | Nxph3 | 4.807 | 1.26E-13 | glio |
| Bora_in.vivo_noRT | Gm20646 | 4.936 | 1.26E-13 | glio |
| Bora_in.vivo_noRT | Cyp2c23 | 4.414 | 1.26E-13 | glio |
| Atr_in.vivo_noRT | Gm49312 | 4.997 | 1.26E-14 | glio |
| Atr_in.vivo_noRT | Gm42695 | 5.273 | 1.26E-14 | glio |
| Bcas2_in.vivo_noRT | AF067061 | 4.807 | 1.26E-15 | glio |
| Bora_in.vivo_noRT | Cfh | 4.157 | 1.26E-15 | glio |
| Aars2_in.vivo_noRT | Slc4a9 | 4.794 | 1.26E-16 | glio |
| Atr_in.vivo_noRT | Gm47851 | 5.273 | 1.26E-16 | glio |
| Bora_in.vivo_noRT | Gm35439 | 4.557 | 1.26E-16 | glio |
| Atr_in.vivo_noRT | Matn4 | 5.807 | 1.26E-17 | glio |
| Bora_in.vivo_noRT | AC159140.1 | 4.676 | 1.26E-18 | glio |
| Atr_in.vivo_noRT | Gm27241 | 4.877 | 1.26E-20 | glio |
| Bcas2_in.vivo_noRT | 9130019O22R | 4.807 | 1.26E-20 | glio |
| Bora_in.vivo_noRT | Prap1 | 4.936 | 1.26E-20 | glio |
| Chek1_in.vivo_noRT | Smtnl1 | 5.044 | 1.26E-20 | glio |
| Bcas2_in.vivo_noRT | Gm49808 | 4.807 | 1.26E-23 | glio |
| Interaction_genotype_Aars2 | Gm20667 | -4.108 | 1.27E-03 | glio |
| Interaction_genotype_Aars2 | Serf2 | 0.966 | 1.27E-03 | glio |
| Interaction_genotype_Atr | Celsr3 | -3.789 | 1.27E-03 | glio |
| Interaction_genotype_Mtor | Nmrk2 | -3.457 | 1.27E-03 | glio |
| Interaction_genotype_Mtor | Gm12089 | -3.987 | 1.27E-03 | glio |
| Interaction_genotype_Myc | 4930470G03R | -2.856 | 1.27E-03 | glio |
| Interaction_genotype_Myc | 4930579C12R | -3.124 | 1.27E-03 | glio |
| Interaction_genotype_Myc | Gpr22 | -2.729 | 1.27E-03 | glio |
| Interaction_genotype_Myc | 4930518J21Ri | -3.567 | 1.27E-03 | glio |
| Interaction_genotype_Myc | 1010001B22R | -3.487 | 1.27E-03 | glio |
| Interaction_genotype_Myc | Galns | -3.199 | 1.27E-03 | glio |
| Interaction_genotype_Myc | Zfp607a | -3.545 | 1.27E-03 | glio |
| Interaction_genotype_Myc | Mt1 | 2.542 | 1.27E-03 | glio |
| Interaction_genotype_Pars2 | Tnfsf13os | -4.408 | 1.27E-03 | glio |
| Interaction_genotype_Pars2 | Gm10354 | -3.563 | 1.27E-03 | glio |
| Interaction_genotype_Pars2 | Gm16741 | -4.011 | 1.27E-03 | glio |
| Interaction_genotype_Pars2 | Gm5152 | -4.523 | 1.27E-03 | glio |
| interaction_RT | Ezh2 | -0.76 | 1.27E-03 | glio |
| Aars2_in.vivo_noRT | Fyb | 2.28 | 1.27E-03 | glio |
| Aars2_in.vivo_noRT | Ilf3 | -1.364 | 1.27E-03 | glio |
| Aars2_in.vivo_noRT | Zfp105 | 1.729 | 1.27E-03 | glio |
| Aars2_in.vivo_noRT | Dap | -1.378 | 1.27E-03 | glio |

| | | | | |
|---|---|---|---|---|
| Atr_in.vivo_noRT | Atp6v1g2 | 2.126 | 1.27E-03 | glio |
| Atr_in.vivo_noRT | Brd3os | 2.135 | 1.27E-03 | glio |
| Atr_in.vivo_noRT | Crem | -1.555 | 1.27E-03 | glio |
| Atr_in.vivo_noRT | Cnpy3 | -1.525 | 1.27E-03 | glio |
| Atr_ex.vivo_noRT | Dcn | 2.771 | 1.27E-03 | glio |
| Bcas2_in.vivo_noRT | Lztr1 | 1.707 | 1.27E-03 | glio |
| Bora_in.vivo_noRT | Gm38394 | 2.29 | 1.27E-03 | glio |
| Bora_in.vivo_noRT | Cdh18 | 2.198 | 1.27E-03 | glio |
| Cenpj_ex.vivo_noRT | Olfr171 | 2.591 | 1.27E-03 | glio |
| Chek1_in.vivo_noRT | Arrdc2 | 2.535 | 1.27E-03 | glio |
| Chek1_in.vivo_noRT | Arhgap27 | 2.195 | 1.27E-03 | glio |
| Chek1_in.vivo_noRT | Mmp23 | 2.246 | 1.27E-03 | glio |
| Chek1_in.vivo_noRT | 1500015A07R | 2.186 | 1.27E-03 | glio |
| Chek1_in.vivo_noRT | Rhno1 | -1.198 | 1.27E-03 | glio |
| Chek1_in.vivo_noRT | Rps23 | -0.609 | 1.27E-03 | glio |
| Eif2s1_in.vivo_noRT | Trpm3 | 2.682 | 1.27E-03 | glio |
| Interaction_genotype_Mtor | Thsd1 | -3.901 | 1.27E-04 | glio |
| Interaction_genotype_Mtor | Gm15910 | -4.276 | 1.27E-04 | glio |
| Interaction_genotype_Mtor | Axl | 1.965 | 1.27E-04 | glio |
| Interaction_genotype_Myc | Arrdc3 | -4.129 | 1.27E-04 | glio |
| Interaction_genotype_Myc | Hectd2os | -4.196 | 1.27E-04 | glio |
| Interaction_genotype_Myc | Gm45605 | -4.295 | 1.27E-04 | glio |
| Interaction_genotype_Myc | Bloc1s6os | -3.948 | 1.27E-04 | glio |
| Aars2_in.vivo_noRT | Ifi205 | 3.506 | 1.27E-04 | glio |
| Aars2_in.vivo_noRT | Rgs5 | 3.293 | 1.27E-04 | glio |
| Aars2_in.vivo_noRT | Phldb3 | 2.741 | 1.27E-04 | glio |
| Aars2_in.vivo_noRT | Pusl1 | -1.935 | 1.27E-04 | glio |
| Aars2_in.vivo_noRT | Rpl36al | -0.733 | 1.27E-04 | glio |
| Atp5c1_in.vivo_noRT | Gm13178 | 2.152 | 1.27E-04 | glio |
| Atp5c1_in.vivo_noRT | 1700026J14Ri | 2.677 | 1.27E-04 | glio |
| Atr_in.vivo_noRT | 1700019L13Ri | 2.704 | 1.27E-04 | glio |
| Atr_in.vivo_noRT | 0610009E02Ri | 2.897 | 1.27E-04 | glio |
| Atr_in.vivo_noRT | 1700001L19Ri | 2.445 | 1.27E-04 | glio |
| Atr_in.vivo_noRT | E130311K13Ri | 2.241 | 1.27E-04 | glio |
| Atr_ex.vivo_noRT | 9930111J21Ri | 3.164 | 1.27E-04 | glio |
| Bcas2_in.vivo_noRT | Cd79a | 2.816 | 1.27E-04 | glio |
| Bcas2_ex.vivo_noRT | Esp24 | 2.742 | 1.27E-04 | glio |
| Bora_in.vivo_noRT | Gm20628 | 2.988 | 1.27E-04 | glio |
| Bora_in.vivo_noRT | 4930542D17R | 2.656 | 1.27E-04 | glio |
| Chek1_ex.vivo_noRT | 1810013L24Ri | 1.842 | 1.27E-04 | glio |
| Eif2s1_in.vivo_noRT | Clec12b | 2.143 | 1.27E-04 | glio |
| Eif2s1_in.vivo_noRT | Gm14929 | 2.143 | 1.27E-04 | glio |
| Eif2s1_in.vivo_noRT | Vmn1r32 | 2.143 | 1.27E-04 | glio |
| Eif2s1_in.vivo_noRT | Glyat | 2.143 | 1.27E-04 | glio |
| Eif2s1_in.vivo_noRT | Tmem132c | 2.143 | 1.27E-04 | glio |
| Eif2s1_in.vivo_noRT | Strip2 | 2.41 | 1.27E-04 | glio |
| Eif2s1_in.vivo_noRT | Gm44693 | 2.143 | 1.27E-04 | glio |
| Eif2s1_in.vivo_noRT | Gm48742 | 2.554 | 1.27E-04 | glio |
| culture_effect | Rbm42 | -0.665 | 1.27E-05 | glio |
| Interaction_genotype_Aars2 | Mtx1 | 2.425 | 1.27E-05 | glio |
| Interaction_genotype_Myc | 9430065F17Ri | -4.966 | 1.27E-05 | glio |
| Interaction_genotype_Myc | Gm15234 | -4.982 | 1.27E-05 | glio |

| | | | | |
|---|---|---|---|---|
| interaction_RT | Mfap2 | 2.44 | 1.27E-05 | glio |
| Aars2_in.vivo_noRT | Kirrel3 | 3.069 | 1.27E-05 | glio |
| Aars2_in.vivo_noRT | Gm49164 | 3.136 | 1.27E-05 | glio |
| Aars2_in.vivo_noRT | Tbca | -0.82 | 1.27E-05 | glio |
| Atr_in.vivo_noRT | Ccdc57 | 3.187 | 1.27E-05 | glio |
| Bcas2_in.vivo_noRT | Dagla | 3.311 | 1.27E-05 | glio |
| Bcas2_in.vivo_noRT | Gm16759 | 3.241 | 1.27E-05 | glio |
| Bcas2_in.vivo_noRT | Cebpa | 3.146 | 1.27E-05 | glio |
| Bcas2_in.vivo_noRT | Gm9725 | 3.139 | 1.27E-05 | glio |
| Bcas2_in.vivo_noRT | Pabpc1 | -0.693 | 1.27E-05 | glio |
| Bora_in.vivo_noRT | Olfr267 | 3.162 | 1.27E-05 | glio |
| Bora_in.vivo_noRT | Slco2a1 | 2.938 | 1.27E-05 | glio |
| Bora_in.vivo_noRT | 9430060I03Ri | 2.979 | 1.27E-05 | glio |
| Hsd17b10_in.vivo_noRT | Ifitm5 | 3.093 | 1.27E-05 | glio |
| Hsd17b10_in.vivo_noRT | Ccl8 | 3.617 | 1.27E-05 | glio |
| culture_effect | Agpat1 | 0.87 | 1.27E-06 | glio |
| culture_effect | Mmp2 | 1.016 | 1.27E-06 | glio |
| Interaction_genotype_Chek1 | Cfap100 | -5.841 | 1.27E-06 | glio |
| Interaction_genotype_Mtor | Vsig10l | -5.786 | 1.27E-06 | glio |
| Interaction_genotype_Mtor | Luzp2 | -4.371 | 1.27E-06 | glio |
| Aars2_in.vivo_noRT | 1500002C15R | 3.559 | 1.27E-06 | glio |
| Aars2_ex.vivo_noRT | Gm29670 | 3.293 | 1.27E-06 | glio |
| Atp5c1_in.vivo_noRT | Apol7d | 3.755 | 1.27E-06 | glio |
| Atr_in.vivo_noRT | Prpf4 | -1.922 | 1.27E-06 | glio |
| Bcas2_in.vivo_noRT | 4930592C13R | 3.54 | 1.27E-06 | glio |
| Bora_in.vivo_noRT | Krt16 | 4.227 | 1.27E-06 | glio |
| Bora_in.vivo_noRT | 1700063J08Ri | 3.429 | 1.27E-06 | glio |
| Chek1_ex.vivo_noRT | Ifitm1 | 4.842 | 1.27E-06 | glio |
| Hsd17b10_in.vivo_noRT | Gm13054 | 3.09 | 1.27E-06 | glio |
| culture_effect | Cacnb3 | 2.167 | 1.27E-07 | glio |
| Interaction_genotype_Mtor | Gm14326 | -6.413 | 1.27E-07 | glio |
| Interaction_genotype_Mtor | Nid1 | -6.551 | 1.27E-07 | glio |
| Interaction_genotype_Mtor | Slc2a9 | -5.296 | 1.27E-07 | glio |
| Aars2_in.vivo_noRT | Gm19409 | 3.747 | 1.27E-07 | glio |
| Aars2_in.vivo_noRT | Gm34590 | 3.754 | 1.27E-07 | glio |
| Aars2_in.vivo_noRT | Prm1 | 3.316 | 1.27E-07 | glio |
| Atr_in.vivo_noRT | Astn2 | 3.445 | 1.27E-07 | glio |
| Atr_in.vivo_noRT | 1810034E14Ri | 3.65 | 1.27E-07 | glio |
| Atr_in.vivo_noRT | 2310030G06R | 3.701 | 1.27E-07 | glio |
| Chek1_ex.vivo_noRT | Prickle4 | 4.215 | 1.27E-07 | glio |
| Chek1_ex.vivo_noRT | Gbp2b | 4.525 | 1.27E-07 | glio |
| Aars2_in.vivo_noRT | Dchs1 | 3.832 | 1.27E-08 | glio |
| Bcas2_in.vivo_noRT | A830035O19R | 3.758 | 1.27E-08 | glio |
| Hsd17b10_in.vivo_noRT | Pmp2 | 3.616 | 1.27E-08 | glio |
| culture_effect | Prdm1 | -3.608 | 1.27E-09 | glio |
| Aars2_in.vivo_noRT | Ccdc39 | 4.106 | 1.27E-09 | glio |
| Atr_in.vivo_noRT | Itga2b | 4.146 | 1.27E-09 | glio |
| Bcas2_in.vivo_noRT | Gm40557 | 4.284 | 1.27E-09 | glio |
| Chek1_ex.vivo_noRT | Gm16853 | 4.998 | 1.27E-09 | glio |
| Bcas2_in.vivo_noRT | Gm20732 | 4.037 | 1.27E-10 | glio |
| Chek1_in.vivo_noRT | 5930403N24R | 4.518 | 1.27E-10 | glio |
| Chek1_in.vivo_noRT | Wnt16 | 4.253 | 1.27E-10 | glio |

| | | | | |
|---|---|---|---|---|
| Chek1_in.vivo_noRT | Gm43948 | 3.568 | 1.27E-10 | glio |
| Aars2_in.vivo_noRT | Zfp14 | 4.804 | 1.27E-11 | glio |
| Aars2_in.vivo_noRT | Col6a2 | 4.543 | 1.27E-11 | glio |
| Bcas2_in.vivo_noRT | A930003A15R | 4.433 | 1.27E-11 | glio |
| Chek1_in.vivo_noRT | Elobl | 5.044 | 1.27E-11 | glio |
| Hsd17b10_in.vivo_noRT | Gm48617 | 3.352 | 1.27E-11 | glio |
| Aars2_in.vivo_noRT | Wdr93 | 4.671 | 1.27E-12 | glio |
| Aars2_in.vivo_noRT | Ifitm5 | 4.804 | 1.27E-12 | glio |
| Chek1_in.vivo_noRT | B3galt4 | 4.518 | 1.27E-12 | glio |
| Aars2_in.vivo_noRT | 6430562O15R | 4.805 | 1.27E-14 | glio |
| Bcas2_in.vivo_noRT | Slc1a2 | 4.543 | 1.27E-14 | glio |
| Bcas2_in.vivo_noRT | Gm19265 | 4.291 | 1.27E-14 | glio |
| Atr_in.vivo_noRT | Car14 | 5.533 | 1.27E-16 | glio |
| Bcas2_in.vivo_noRT | Gm49223 | 4.802 | 1.27E-16 | glio |
| Chek1_in.vivo_noRT | Tmem74b | 4.78 | 1.27E-16 | glio |
| Atr_in.vivo_noRT | D930028M14I | 5.533 | 1.27E-18 | glio |
| Atr_in.vivo_noRT | Npr3 | 5.533 | 1.27E-19 | glio |
| Bcas2_in.vivo_noRT | Tac2 | 4.807 | 1.27E-21 | glio |
| Atr_in.vivo_noRT | Gm12794 | 5.533 | 1.27E-26 | glio |
| culture_effect | Id1 | 2.647 | 1.27E-36 | glio |
| Interaction_genotype_Mtor | Slc46a3 | -3.949 | 1.28E-03 | glio |
| Interaction_genotype_Mtor | Col20a1 | -3.864 | 1.28E-03 | glio |
| Interaction_genotype_Mtor | Relt | -3.968 | 1.28E-03 | glio |
| Interaction_genotype_Mtor | Gm26549 | -3.225 | 1.28E-03 | glio |
| Interaction_genotype_Mtor | 2500004C02R | -2.923 | 1.28E-03 | glio |
| Interaction_genotype_Mtor | Edil3 | -2.9 | 1.28E-03 | glio |
| Interaction_genotype_Myc | Gm4129 | -3.587 | 1.28E-03 | glio |
| Interaction_genotype_Myc | Kctd6 | -3.395 | 1.28E-03 | glio |
| interaction_RT | Sept11 | -0.767 | 1.28E-03 | glio |
| interaction_RT | Ndufb3 | 0.611 | 1.28E-03 | glio |
| Aars2_in.vivo_noRT | Zfp97 | 2.303 | 1.28E-03 | glio |
| Aars2_in.vivo_noRT | Wdr61 | -1.184 | 1.28E-03 | glio |
| Aars2_in.vivo_noRT | Ppp2ca | -0.709 | 1.28E-03 | glio |
| Aars2_in.vivo_noRT | Dad1 | -0.851 | 1.28E-03 | glio |
| Aars2_in.vivo_noRT | Cct4 | -0.819 | 1.28E-03 | glio |
| Aars2_in.vivo_noRT | Cox5a | -0.671 | 1.28E-03 | glio |
| Atp5c1_in.vivo_noRT | Gm47872 | 2.404 | 1.28E-03 | glio |
| Atr_in.vivo_noRT | Mindy4 | 2.188 | 1.28E-03 | glio |
| Atr_in.vivo_noRT | Pacc1 | 1.943 | 1.28E-03 | glio |
| Atr_in.vivo_noRT | Bzw1 | -1.002 | 1.28E-03 | glio |
| Atr_in.vivo_noRT | Rbm25 | -0.663 | 1.28E-03 | glio |
| Bcas2_in.vivo_noRT | Shbg | 2.428 | 1.28E-03 | glio |
| Bcas2_in.vivo_noRT | Rufy2 | 1.407 | 1.28E-03 | glio |
| Chek1_ex.vivo_noRT | Upb1 | 2.771 | 1.28E-03 | glio |
| Chek1_ex.vivo_noRT | A930012O16R | 2.957 | 1.28E-03 | glio |
| Chek1_ex.vivo_noRT | Tceal1 | 2.766 | 1.28E-03 | glio |
| Eif2s1_in.vivo_noRT | Gtsf1 | 2.143 | 1.28E-03 | glio |
| Hsd17b10_in.vivo_noRT | Sri | -0.928 | 1.28E-03 | glio |
| culture_effect | BC049987 | -1.717 | 1.28E-04 | glio |
| Interaction_genotype_Atr | 5430402O13R | -4.769 | 1.28E-04 | glio |
| Interaction_genotype_Atr | 5330439K02R | -4.776 | 1.28E-04 | glio |
| Interaction_genotype_Mtor | Uhrf1bp1 | -3.43 | 1.28E-04 | glio |

| | | | | |
|---|---|---|---|---|
| Interaction_genotype_Mtor | Pfdn1 | 1.921 | 1.28E-04 | glio |
| Interaction_genotype_Myc | Oxt | -4.34 | 1.28E-04 | glio |
| Interaction_genotype_Myc | Rps13 | 1.369 | 1.28E-04 | glio |
| interaction_RT | Zbp1 | 1.069 | 1.28E-04 | glio |
| Aars2_in.vivo_noRT | Tmc8 | 2.685 | 1.28E-04 | glio |
| Aars2_ex.vivo_noRT | Lncbate10 | 2.505 | 1.28E-04 | glio |
| Aars2_ex.vivo_noRT | Pla2g2f | 3.293 | 1.28E-04 | glio |
| Atp5c1_in.vivo_noRT | Gm41409 | 2.664 | 1.28E-04 | glio |
| Atr_ex.vivo_noRT | Map1lc3b | 1.118 | 1.28E-04 | glio |
| Bora_in.vivo_noRT | Trim14 | 2.993 | 1.28E-04 | glio |
| Bora_in.vivo_noRT | Gm12971 | 2.816 | 1.28E-04 | glio |
| Bora_in.vivo_noRT | Zscan22 | 2.294 | 1.28E-04 | glio |
| Chek1_in.vivo_noRT | Fbxw2 | -1.585 | 1.28E-04 | glio |
| Chek1_in.vivo_noRT | Hnrnpab | -0.722 | 1.28E-04 | glio |
| Hsd17b10_in.vivo_noRT | Fam13a | 2.816 | 1.28E-04 | glio |
| Hsd17b10_in.vivo_noRT | Grm1 | 2.512 | 1.28E-04 | glio |
| Interaction_genotype_Mtor | Gm43728 | -5.397 | 1.28E-05 | glio |
| Interaction_genotype_Myc | Gm10484 | -5.22 | 1.28E-05 | glio |
| Aars2_in.vivo_noRT | Glb1l | 3.075 | 1.28E-05 | glio |
| Atr_in.vivo_noRT | 1700063D05R | 3.158 | 1.28E-05 | glio |
| Atr_in.vivo_noRT | Plscr4 | 2.44 | 1.28E-05 | glio |
| Bcas2_in.vivo_noRT | Prex2 | 3.371 | 1.28E-05 | glio |
| Bcas2_in.vivo_noRT | Fam222a | 3.293 | 1.28E-05 | glio |
| Bora_in.vivo_noRT | 4933406P04R | 3.255 | 1.28E-05 | glio |
| Interaction_genotype_Mtor | Gm48742 | -4.824 | 1.28E-06 | glio |
| Interaction_genotype_Mtor | Npr3 | -5.071 | 1.28E-06 | glio |
| Interaction_genotype_Mtor | 8030423F21Ri | -4.946 | 1.28E-06 | glio |
| Aars2_in.vivo_noRT | Gm11110 | 3.409 | 1.28E-06 | glio |
| Aars2_in.vivo_noRT | Gm10244 | 3.506 | 1.28E-06 | glio |
| Aars2_in.vivo_noRT | C430014B12R | 3.278 | 1.28E-06 | glio |
| Atr_in.vivo_noRT | Sardh | 3.246 | 1.28E-06 | glio |
| Atr_in.vivo_noRT | Arid3c | 3.038 | 1.28E-06 | glio |
| Atr_ex.vivo_noRT | Gm5941 | 4.459 | 1.28E-06 | glio |
| Chek1_in.vivo_noRT | Olfr920 | 3.293 | 1.28E-06 | glio |
| Hsd17b10_in.vivo_noRT | Gm45606 | 3.356 | 1.28E-06 | glio |
| Aars2_in.vivo_noRT | Zfp433 | 3.887 | 1.28E-07 | glio |
| Bcas2_in.vivo_noRT | Edn1 | 4.428 | 1.28E-07 | glio |
| Bcas2_in.vivo_noRT | Olfm2 | 3.9 | 1.28E-07 | glio |
| Bcas2_in.vivo_noRT | Wfdc18 | 4.541 | 1.28E-07 | glio |
| Bcas2_in.vivo_noRT | Pde1a | 3.807 | 1.28E-07 | glio |
| Bcas2_in.vivo_noRT | Sdr16c5 | 3.49 | 1.28E-07 | glio |
| Bcas2_in.vivo_noRT | Ipcef1 | 2.994 | 1.28E-07 | glio |
| Bora_in.vivo_noRT | Shroom3 | 3.657 | 1.28E-07 | glio |
| Chek1_in.vivo_noRT | Sec1 | 3.887 | 1.28E-07 | glio |
| Chek1_in.vivo_noRT | A530072M11F | 3.715 | 1.28E-07 | glio |
| Atr_in.vivo_noRT | Rnase4 | 3.971 | 1.28E-08 | glio |
| Bcas2_in.vivo_noRT | Gm5544 | 4.016 | 1.28E-08 | glio |
| Bora_in.vivo_noRT | 1700012B09R | 4.676 | 1.28E-08 | glio |
| Bora_in.vivo_noRT | Gsx2 | 4.243 | 1.28E-08 | glio |
| Bora_in.vivo_noRT | Prrt3 | 3.522 | 1.28E-08 | glio |
| Aars2_in.vivo_noRT | Gm10230 | 4.446 | 1.28E-09 | glio |
| Bcas2_in.vivo_noRT | Zfp808 | 3.69 | 1.28E-09 | glio |

| | | | | |
|---|---|---|---|---|
| Bcas2_in.vivo_noRT | Pard3bos1 | 3.81 | 1.28E-09 | glio |
| Chek1_in.vivo_noRT | 4933408J17Ri | 3.727 | 1.28E-09 | glio |
| Chek1_in.vivo_noRT | Camkk1 | 4.273 | 1.28E-09 | glio |
| Chek1_in.vivo_noRT | Gm48772 | 4.136 | 1.28E-09 | glio |
| culture_effect | Gm6710 | 1.944 | 1.28E-10 | glio |
| Atr_in.vivo_noRT | 1110020A21R | 4.583 | 1.28E-10 | glio |
| interaction_RT | Itm2b | 1.537 | 1.28E-11 | glio |
| interaction_RT | Pnrc1 | 1.516 | 1.28E-11 | glio |
| interaction_RT | Matn2 | 2.452 | 1.28E-11 | glio |
| Aars2_in.vivo_noRT | Flnc | 4.801 | 1.28E-11 | glio |
| Chek1_in.vivo_noRT | A930003A15R | 4.421 | 1.28E-11 | glio |
| Bora_in.vivo_noRT | Cyp4x1 | 4.67 | 1.28E-12 | glio |
| Atr_in.vivo_noRT | A930012O16R | 5.273 | 1.28E-13 | glio |
| Chek1_in.vivo_noRT | Thap8 | 5.562 | 1.28E-14 | glio |
| Atr_in.vivo_noRT | Gm11457 | 5.533 | 1.28E-16 | glio |
| Bcas2_in.vivo_noRT | Btn1a1 | 4.547 | 1.28E-16 | glio |
| Chek1_in.vivo_noRT | Hrct1 | 5.044 | 1.28E-16 | glio |
| Atr_in.vivo_noRT | Masp1 | 5.011 | 1.28E-17 | glio |
| Chek1_in.vivo_noRT | Olfr56 | 5.044 | 1.28E-21 | glio |
| Aars2_in.vivo_noRT | Gm48764 | 5.327 | 1.28E-25 | glio |
| Aars2_in.vivo_noRT | 1700034I23Ri | 5.327 | 1.28E-26 | glio |
| culture_effect | Serping1 | -7.158 | 1.28E-41 | glio |
| culture_effect | E130102H24R | -1.956 | 1.29E-03 | glio |
| culture_effect | Greb1 | 0.592 | 1.29E-03 | glio |
| culture_effect | Rnf7 | 0.37 | 1.29E-03 | glio |
| Interaction_genotype_Mtor | Pkp3 | -4.193 | 1.29E-03 | glio |
| Interaction_genotype_Mtor | Gstm4 | -3.434 | 1.29E-03 | glio |
| Interaction_genotype_Mtor | Fastkd5 | -3.162 | 1.29E-03 | glio |
| Interaction_genotype_Mtor | 4930599N23R | -3.443 | 1.29E-03 | glio |
| Interaction_genotype_Pars2 | Rab25 | -3.947 | 1.29E-03 | glio |
| Interaction_genotype_Pars2 | D6Ertd527e | -4.377 | 1.29E-03 | glio |
| Interaction_genotype_Rad21 | Ocm | 4.265 | 1.29E-03 | glio |
| Interaction_genotype_Rad21 | Scel | 3.138 | 1.29E-03 | glio |
| interaction_RT | Cspg5 | 1.894 | 1.29E-03 | glio |
| Aars2_in.vivo_noRT | Xylb | 2.278 | 1.29E-03 | glio |
| Aars2_in.vivo_noRT | S100a6 | -1.096 | 1.29E-03 | glio |
| Aars2_in.vivo_noRT | Eef2 | -0.538 | 1.29E-03 | glio |
| Aars2_ex.vivo_noRT | Lrp2bp | 2.774 | 1.29E-03 | glio |
| Aars2_ex.vivo_noRT | Ntrk1 | 3.192 | 1.29E-03 | glio |
| Atr_in.vivo_noRT | Nt5m | 1.245 | 1.29E-03 | glio |
| Bard1_ex.vivo_noRT | Fa2h | 2.814 | 1.29E-03 | glio |
| Bcas2_in.vivo_noRT | Acacb | 2.305 | 1.29E-03 | glio |
| Bcas2_in.vivo_noRT | Nell2 | 2.136 | 1.29E-03 | glio |
| Bcas2_in.vivo_noRT | B3gat2 | 2.474 | 1.29E-03 | glio |
| Bcas2_in.vivo_noRT | Crls1 | -1.263 | 1.29E-03 | glio |
| Bora_in.vivo_noRT | Gata3 | 2.553 | 1.29E-03 | glio |
| Bora_in.vivo_noRT | Zfp600 | 2.327 | 1.29E-03 | glio |
| Cenpj_in.vivo_noRT | Prap1 | 2.022 | 1.29E-03 | glio |
| Cenpj_in.vivo_noRT | Halr1 | 1.756 | 1.29E-03 | glio |
| Chek1_in.vivo_noRT | Nkain4 | 2.179 | 1.29E-03 | glio |
| Chek1_in.vivo_noRT | Tbc1d10a | -1.458 | 1.29E-03 | glio |
| culture_effect | Mllt6 | -1.038 | 1.29E-04 | glio |

| | | | | |
|---|---|---|---|---|
| Interaction_genotype_Chek1 | Brinp3 | 4.346 | 1.29E-04 | glio |
| Interaction_genotype_Mtor | Tacr2 | -4.725 | 1.29E-04 | glio |
| Interaction_genotype_Mtor | Hist1h2ac | -4.432 | 1.29E-04 | glio |
| Interaction_genotype_Mtor | 1600002K03R | -4.646 | 1.29E-04 | glio |
| Interaction_genotype_Mtor | Serinc1 | 1.837 | 1.29E-04 | glio |
| Interaction_genotype_Myc | Gm5089 | -4.958 | 1.29E-04 | glio |
| Interaction_genotype_Myc | Rnf123 | -3.38 | 1.29E-04 | glio |
| Interaction_genotype_Myc | BC049715 | -3.789 | 1.29E-04 | glio |
| Interaction_genotype_Myc | Rps12 | 1.557 | 1.29E-04 | glio |
| interaction_RT | Rundc3a | 2.514 | 1.29E-04 | glio |
| Aars2_in.vivo_noRT | Abhd18 | 2.788 | 1.29E-04 | glio |
| Aars2_in.vivo_noRT | 1600014C10R | 2.105 | 1.29E-04 | glio |
| Atp5c1_in.vivo_noRT | Capn9 | 2.668 | 1.29E-04 | glio |
| Atr_in.vivo_noRT | Cd38 | 2.667 | 1.29E-04 | glio |
| Atr_in.vivo_noRT | Uevld | 2.204 | 1.29E-04 | glio |
| Atr_in.vivo_noRT | Epcam | -2.506 | 1.29E-04 | glio |
| Atr_ex.vivo_noRT | Gm49860 | 2.98 | 1.29E-04 | glio |
| Atr_ex.vivo_noRT | 4930553M12F | 2.77 | 1.29E-04 | glio |
| Bcas2_in.vivo_noRT | Ppp1r26 | 2.898 | 1.29E-04 | glio |
| Bcas2_in.vivo_noRT | Zfp799 | 2.707 | 1.29E-04 | glio |
| Bcas2_ex.vivo_noRT | Gm9758 | 2.742 | 1.29E-04 | glio |
| Bora_in.vivo_noRT | Col17a1 | 2.886 | 1.29E-04 | glio |
| Bora_in.vivo_noRT | Zfp59 | 2.779 | 1.29E-04 | glio |
| Bora_in.vivo_noRT | Clspn | -1.95 | 1.29E-04 | glio |
| Bora_ex.vivo_noRT | Clca3a2 | 3.201 | 1.29E-04 | glio |
| Bora_ex.vivo_noRT | C430002N11R | 2.806 | 1.29E-04 | glio |
| Cenpj_ex.vivo_noRT | A930019D19R | 2.591 | 1.29E-04 | glio |
| Chek1_ex.vivo_noRT | Slc8a3 | 3.586 | 1.29E-04 | glio |
| Hsd17b10_in.vivo_noRT | Milr1 | 2.881 | 1.29E-04 | glio |
| culture_effect | H2-T22 | -0.632 | 1.29E-05 | glio |
| culture_effect | Emc6 | 0.499 | 1.29E-05 | glio |
| Aars2_in.vivo_noRT | Eme2 | 2.913 | 1.29E-05 | glio |
| Aars2_in.vivo_noRT | 1700041G16R | 3.072 | 1.29E-05 | glio |
| Atr_in.vivo_noRT | Fignl2 | 3.268 | 1.29E-05 | glio |
| Atr_in.vivo_noRT | 1700102H20R | 3.009 | 1.29E-05 | glio |
| Chek1_in.vivo_noRT | Gm26930 | 3.354 | 1.29E-05 | glio |
| Chek1_in.vivo_noRT | Vgll2 | 2.708 | 1.29E-05 | glio |
| Chek1_ex.vivo_noRT | Ptgis | 3.754 | 1.29E-05 | glio |
| Chek1_ex.vivo_noRT | Etohd2 | 3.948 | 1.29E-05 | glio |
| Chek1_ex.vivo_noRT | Dock9 | -2.304 | 1.29E-05 | glio |
| Hsd17b10_in.vivo_noRT | Masp2 | 2.689 | 1.29E-05 | glio |
| culture_effect | B020010K11R | -1.751 | 1.29E-06 | glio |
| culture_effect | Jmjd6 | -0.62 | 1.29E-06 | glio |
| Interaction_genotype_Mtor | Gm36495 | -6.107 | 1.29E-06 | glio |
| Interaction_genotype_Mtor | Gm43409 | -6.393 | 1.29E-06 | glio |
| Interaction_genotype_Mtor | Gm34776 | -4.492 | 1.29E-06 | glio |
| Atr_in.vivo_noRT | Zfp458 | 3.521 | 1.29E-06 | glio |
| Bcas2_in.vivo_noRT | Mag | 3.225 | 1.29E-06 | glio |
| Chek1_in.vivo_noRT | Gm10767 | 3.444 | 1.29E-06 | glio |
| culture_effect | Sept1 | 2.6 | 1.29E-07 | glio |
| Aars2_in.vivo_noRT | Gm10767 | 3.727 | 1.29E-07 | glio |
| Bcas2_in.vivo_noRT | E230001N04R | 3.221 | 1.29E-07 | glio |

| | | | | |
|---|---|---|---|---|
| Chek1_in.vivo_noRT | A530083I20Ri | 4.036 | 1.29E-07 | glio |
| Chek1_in.vivo_noRT | 4932441J04Ri | 3.793 | 1.29E-07 | glio |
| Chek1_in.vivo_noRT | Zmynd15 | 3.606 | 1.29E-07 | glio |
| Aars2_in.vivo_noRT | Pcdhgb5 | 3.423 | 1.29E-08 | glio |
| Atr_in.vivo_noRT | Traf1 | 3.929 | 1.29E-08 | glio |
| Bcas2_in.vivo_noRT | Syp | 4.416 | 1.29E-08 | glio |
| Hsd17b10_in.vivo_noRT | Scel | 2.698 | 1.29E-08 | glio |
| Bora_in.vivo_noRT | Rnft2 | 4.133 | 1.29E-09 | glio |
| Chek1_in.vivo_noRT | Svopl | 3.862 | 1.29E-09 | glio |
| culture_effect | Sdc3 | -3.402 | 1.29E-10 | glio |
| Aars2_in.vivo_noRT | Apoh | 4.412 | 1.29E-10 | glio |
| Bcas2_in.vivo_noRT | Camk2b | 4.285 | 1.29E-10 | glio |
| Chek1_in.vivo_noRT | Trpm3 | 4.522 | 1.29E-10 | glio |
| Chek1_in.vivo_noRT | Gm40557 | 4.521 | 1.29E-10 | glio |
| Chek1_in.vivo_noRT | Gm50375 | 4.777 | 1.29E-10 | glio |
| Chek1_ex.vivo_noRT | Prl3d3 | 3.747 | 1.29E-10 | glio |
| Aars2_in.vivo_noRT | Baalc | 4.535 | 1.29E-11 | glio |
| Atr_in.vivo_noRT | Otop1 | 4.349 | 1.29E-11 | glio |
| Chek1_in.vivo_noRT | Prm3 | 4.303 | 1.29E-12 | glio |
| Atr_in.vivo_noRT | Cxcl1 | 5.533 | 1.29E-14 | glio |
| Atr_in.vivo_noRT | D630036H23R | 5.273 | 1.29E-14 | glio |
| Bcas2_in.vivo_noRT | Dapk2 | 4.024 | 1.29E-14 | glio |
| Bora_in.vivo_noRT | Lhx1os | 4.41 | 1.29E-14 | glio |
| Atr_in.vivo_noRT | A930001C03R | 5.39 | 1.29E-15 | glio |
| Bora_in.vivo_noRT | Cers3 | 4.405 | 1.29E-15 | glio |
| Atr_in.vivo_noRT | Ric3 | 5.273 | 1.29E-17 | glio |
| Chek1_in.vivo_noRT | Slc10a1 | 4.784 | 1.29E-17 | glio |
| Chek1_in.vivo_noRT | Gm11973 | 4.777 | 1.29E-17 | glio |
| Chek1_in.vivo_noRT | Angptl7 | 4.178 | 1.29E-17 | glio |
| Aars2_in.vivo_noRT | Gm48242 | 5.327 | 1.29E-18 | glio |
| Aars2_in.vivo_noRT | Gm39526 | 4.804 | 1.29E-18 | glio |
| Atr_in.vivo_noRT | Nat8f2 | 5.273 | 1.29E-19 | glio |
| Atr_in.vivo_noRT | Oit3 | 6.037 | 1.29E-19 | glio |
| Aars2_in.vivo_noRT | 4930570E03Ri | 5.067 | 1.29E-20 | glio |
| Aars2_in.vivo_noRT | 4930447K03R | 5.067 | 1.29E-20 | glio |
| Bcas2_in.vivo_noRT | A330049N07R | 4.541 | 1.29E-20 | glio |
| Bora_in.vivo_noRT | A330049N07R | 4.67 | 1.29E-21 | glio |
| Chek1_in.vivo_noRT | Gm5415 | 5.044 | 1.29E-24 | glio |
| Atr_in.vivo_noRT | BC061195 | 5.533 | 1.29E-28 | glio |
| culture_effect | Spats2l | -2.282 | 1.29E-29 | glio |
| Atr_in.vivo_noRT | Krt19 | 5.533 | 1.29E-30 | glio |
| culture_effect | Apoh | 1.806 | 1.30E-03 | glio |
| culture_effect | Ptcd1 | -1.176 | 1.30E-03 | glio |
| culture_effect | Faf1 | -0.594 | 1.30E-03 | glio |
| Interaction_genotype_Aars2 | B4galnt1 | -3.403 | 1.30E-03 | glio |
| Interaction_genotype_Aars2 | Rab7b | -3.522 | 1.30E-03 | glio |
| Interaction_genotype_Aars2 | Atpif1 | 1.041 | 1.30E-03 | glio |
| Interaction_genotype_Bora | Slpi | -4.677 | 1.30E-03 | glio |
| Interaction_genotype_Bora | Gm26881 | -3.658 | 1.30E-03 | glio |
| Interaction_genotype_Bora | Pcnx3 | 2.193 | 1.30E-03 | glio |
| Interaction_genotype_Mtor | Lncenc1 | -3.59 | 1.30E-03 | glio |
| Interaction_genotype_Mtor | Tmem91 | -4.017 | 1.30E-03 | glio |

| | | | | |
|---|---|---|---|---|
| Interaction_genotype_Mtor | Htra1 | -4.242 | 1.30E-03 | glio |
| Interaction_genotype_Mtor | Rnf38 | -2.585 | 1.30E-03 | glio |
| Interaction_genotype_Mtor | Tasor2 | -1.776 | 1.30E-03 | glio |
| Interaction_genotype_Myc | Retn | -4.717 | 1.30E-03 | glio |
| Interaction_genotype_Myc | Ephx3 | -3.482 | 1.30E-03 | glio |
| Interaction_genotype_Myc | Gm10516 | -3.599 | 1.30E-03 | glio |
| Interaction_genotype_Myc | Sult6b1 | -3.568 | 1.30E-03 | glio |
| Interaction_genotype_Myc | Nupr1l | -4.041 | 1.30E-03 | glio |
| Interaction_genotype_Myc | Rwdd3 | -3.827 | 1.30E-03 | glio |
| Interaction_genotype_Myc | Zfp28 | -3.699 | 1.30E-03 | glio |
| Interaction_genotype_Myc | Tha1 | -3.477 | 1.30E-03 | glio |
| Interaction_genotype_Myc | 4833438C02R | -3.174 | 1.30E-03 | glio |
| Interaction_genotype_Myc | Tgfb2 | -3.057 | 1.30E-03 | glio |
| Interaction_genotype_Myc | Gdi2 | 1.437 | 1.30E-03 | glio |
| Interaction_genotype_Myc | Naca | 1.094 | 1.30E-03 | glio |
| Interaction_genotype_Rpl3 | 4930412M03F | -4.345 | 1.30E-03 | glio |
| Interaction_genotype_Rpl3 | Cpne7 | -4.723 | 1.30E-03 | glio |
| Interaction_genotype_Rpl3 | Prnp | 2.14 | 1.30E-03 | glio |
| interaction_RT | 0610009B22R | 0.778 | 1.30E-03 | glio |
| interaction_RT | Mrpl51 | -0.534 | 1.30E-03 | glio |
| Aars2_in.vivo_noRT | Gm4356 | 2.526 | 1.30E-03 | glio |
| Aars2_in.vivo_noRT | Serpinb1a | 2.91 | 1.30E-03 | glio |
| Aars2_in.vivo_noRT | Dner | 2.127 | 1.30E-03 | glio |
| Aars2_in.vivo_noRT | Rpl19 | -0.624 | 1.30E-03 | glio |
| Atr_in.vivo_noRT | Slx1b | 1.95 | 1.30E-03 | glio |
| Atr_in.vivo_noRT | Slc2a1 | -1.767 | 1.30E-03 | glio |
| Atr_in.vivo_noRT | Celf1 | -1.128 | 1.30E-03 | glio |
| Atr_in.vivo_noRT | Btg1 | -1.284 | 1.30E-03 | glio |
| Atr_in.vivo_noRT | Eif3e | -0.652 | 1.30E-03 | glio |
| Atr_ex.vivo_noRT | Dreh | 2.77 | 1.30E-03 | glio |
| Bcas2_in.vivo_noRT | Sipa1l2 | 2.365 | 1.30E-03 | glio |
| Bora_in.vivo_noRT | Rtl5 | 2.268 | 1.30E-03 | glio |
| Bora_in.vivo_noRT | Hdc | 2.43 | 1.30E-03 | glio |
| Bora_in.vivo_noRT | Hmgcs1 | -1.836 | 1.30E-03 | glio |
| Bora_in.vivo_noRT | Serf1 | -0.971 | 1.30E-03 | glio |
| Bora_in.vivo_noRT | Pcna | -1.11 | 1.30E-03 | glio |
| Cenpj_in.vivo_noRT | Fabp6 | 2.022 | 1.30E-03 | glio |
| Chek1_in.vivo_noRT | Gm12971 | 2.399 | 1.30E-03 | glio |
| Chek1_in.vivo_noRT | Cfap69 | 2.303 | 1.30E-03 | glio |
| Chek1_in.vivo_noRT | Camk2n2 | 1.927 | 1.30E-03 | glio |
| Chek1_in.vivo_noRT | Mt2 | 1.029 | 1.30E-03 | glio |
| Chek1_ex.vivo_noRT | Palm3 | 3.085 | 1.30E-03 | glio |
| Chek1_ex.vivo_noRT | Fendrr | 3.015 | 1.30E-03 | glio |
| Chek1_ex.vivo_noRT | Hist3h2a | 2.795 | 1.30E-03 | glio |
| Eif2s1_in.vivo_noRT | Zscan4d | 2.143 | 1.30E-03 | glio |
| Hsd17b10_in.vivo_noRT | Gm48684 | 2.839 | 1.30E-03 | glio |
| Hsd17b10_in.vivo_noRT | Cks1brt | 2.178 | 1.30E-03 | glio |
| Hsd17b10_in.vivo_noRT | Zfp114 | 2.456 | 1.30E-03 | glio |
| culture_effect | Lcorl | -0.818 | 1.30E-04 | glio |
| culture_effect | Micos10 | 0.377 | 1.30E-04 | glio |
| Interaction_genotype_Aars2 | Matn4 | -4.464 | 1.30E-04 | glio |
| Interaction_genotype_Mtor | Adam33 | -4.617 | 1.30E-04 | glio |

| | | | | |
|---|---|---|---|---|
| Interaction_genotype_Mtor | Tbx15 | -4.073 | 1.30E-04 | glio |
| Interaction_genotype_Myc | Pecam1 | -4.448 | 1.30E-04 | glio |
| Interaction_genotype_Myc | Rnf121 | -2.685 | 1.30E-04 | glio |
| Aars2_in.vivo_noRT | Trpm4 | 2.198 | 1.30E-04 | glio |
| Aars2_ex.vivo_noRT | Chia1 | 2.505 | 1.30E-04 | glio |
| Aars2_ex.vivo_noRT | Gpr17 | 3.293 | 1.30E-04 | glio |
| Atp5c1_in.vivo_noRT | Lrguk | 2.404 | 1.30E-04 | glio |
| Atr_in.vivo_noRT | Nop2 | -1.474 | 1.30E-04 | glio |
| Atr_ex.vivo_noRT | Cidea | 3.164 | 1.30E-04 | glio |
| Atr_ex.vivo_noRT | Gm15326 | 2.589 | 1.30E-04 | glio |
| Bcas2_ex.vivo_noRT | AF067063 | 2.742 | 1.30E-04 | glio |
| Bora_in.vivo_noRT | Gm36210 | 2.735 | 1.30E-04 | glio |
| Bora_in.vivo_noRT | Dhtkd1 | 2.992 | 1.30E-04 | glio |
| Chek1_in.vivo_noRT | Anks6 | 2.57 | 1.30E-04 | glio |
| Eif2s1_in.vivo_noRT | Akap14 | 2.407 | 1.30E-04 | glio |
| Eif2s1_in.vivo_noRT | Gm30692 | 2.143 | 1.30E-04 | glio |
| Hsd17b10_in.vivo_noRT | BC048671 | 2.315 | 1.30E-04 | glio |
| Hsd17b10_in.vivo_noRT | Kcnq2 | 2.453 | 1.30E-04 | glio |
| Hsd17b10_in.vivo_noRT | Coro1c | -1.476 | 1.30E-04 | glio |
| culture_effect | Myc | -0.789 | 1.30E-05 | glio |
| culture_effect | Elob | 0.459 | 1.30E-05 | glio |
| Interaction_genotype_Bora | 4930412M03F | -5.478 | 1.30E-05 | glio |
| Interaction_genotype_Mtor | Eme2 | -4.857 | 1.30E-05 | glio |
| Aars2_in.vivo_noRT | Nts | 3.284 | 1.30E-05 | glio |
| Aars2_in.vivo_noRT | Fndc10 | 3.323 | 1.30E-05 | glio |
| Aars2_in.vivo_noRT | Gm29666 | 3.092 | 1.30E-05 | glio |
| Aars2_in.vivo_noRT | Daam2 | 3.154 | 1.30E-05 | glio |
| Aars2_in.vivo_noRT | Efr3b | 3.41 | 1.30E-05 | glio |
| Atp5c1_in.vivo_noRT | Gm44646 | 2.668 | 1.30E-05 | glio |
| Atr_in.vivo_noRT | Gpatch3 | 2.973 | 1.30E-05 | glio |
| Atr_in.vivo_noRT | Nnt | 2.765 | 1.30E-05 | glio |
| Chek1_in.vivo_noRT | Serpinc1 | 3.227 | 1.30E-05 | glio |
| Hsd17b10_in.vivo_noRT | Gm26674 | 2.971 | 1.30E-05 | glio |
| Hsd17b10_in.vivo_noRT | Gm43625 | 2.838 | 1.30E-05 | glio |
| Hsd17b10_in.vivo_noRT | Ttc16 | 3.093 | 1.30E-05 | glio |
| Hsd17b10_in.vivo_noRT | BC065403 | 3.1 | 1.30E-05 | glio |
| culture_effect | Bcl2l11 | -1.673 | 1.30E-06 | glio |
| Interaction_genotype_Mtor | Olfr655 | -7.935 | 1.30E-06 | glio |
| interaction_RT | Ccl27a | 1.638 | 1.30E-06 | glio |
| interaction_RT | Hist2h2ac | -2.424 | 1.30E-06 | glio |
| interaction_RT | Nop56 | -1.031 | 1.30E-06 | glio |
| interaction_RT | Srsf2 | -0.861 | 1.30E-06 | glio |
| interaction_RT | Gm44867 | 4.308 | 1.30E-06 | glio |
| Aars2_in.vivo_noRT | Fam169a | 3.235 | 1.30E-06 | glio |
| Atr_ex.vivo_noRT | Cd36 | 3.164 | 1.30E-06 | glio |
| Atr_ex.vivo_noRT | Gstt4 | 3.164 | 1.30E-06 | glio |
| Bcas2_in.vivo_noRT | Il2rb | 3.265 | 1.30E-06 | glio |
| Bcas2_in.vivo_noRT | Gm37305 | 3.368 | 1.30E-06 | glio |
| Chek1_ex.vivo_noRT | Ikbke | 4.06 | 1.30E-06 | glio |
| Hsd17b10_in.vivo_noRT | Lyl1 | 2.966 | 1.30E-06 | glio |
| Bcas2_in.vivo_noRT | Zfpm1 | 3.809 | 1.30E-07 | glio |
| Aars2_in.vivo_noRT | A530083I20Ri | 4.319 | 1.30E-08 | glio |

| | | | | |
|---|---|---|---|---|
| Aars2_in.vivo_noRT | Lrrc59 | -1.943 | 1.30E-08 | glio |
| Bcas2_in.vivo_noRT | Gm35986 | 3.759 | 1.30E-08 | glio |
| Bcas2_in.vivo_noRT | Lst1 | 4.041 | 1.30E-08 | glio |
| Chek1_in.vivo_noRT | Slc25a35 | 3.934 | 1.30E-08 | glio |
| Chek1_ex.vivo_noRT | Pcdhga2 | 4.646 | 1.30E-08 | glio |
| Bcas2_in.vivo_noRT | Platr20 | 4.543 | 1.30E-09 | glio |
| Bora_in.vivo_noRT | Aoc1 | 4.676 | 1.30E-10 | glio |
| Bcas2_in.vivo_noRT | Gm45606 | 4.547 | 1.30E-11 | glio |
| Bora_in.vivo_noRT | Olfr1191-ps1 | 4.286 | 1.30E-11 | glio |
| Aars2_in.vivo_noRT | Lrrc72 | 4.536 | 1.30E-12 | glio |
| Atr_in.vivo_noRT | Ptp4a1.1 | 4.736 | 1.30E-12 | glio |
| Bcas2_in.vivo_noRT | Pkd1l2 | 4.543 | 1.30E-12 | glio |
| Bora_in.vivo_noRT | Nrros | 4.818 | 1.30E-12 | glio |
| interaction_RT | H2-T23 | 1.986 | 1.30E-13 | glio |
| Atr_in.vivo_noRT | 1700030L20Ri | 4.877 | 1.30E-13 | glio |
| culture_effect | Cdk14 | 1.877 | 1.30E-14 | glio |
| Aars2_in.vivo_noRT | Usp17la | 5.327 | 1.30E-14 | glio |
| Aars2_in.vivo_noRT | Plk5 | 4.671 | 1.30E-14 | glio |
| Atr_in.vivo_noRT | Arhgef16 | 4.497 | 1.30E-14 | glio |
| Aars2_in.vivo_noRT | Rftn2 | 4.936 | 1.30E-15 | glio |
| Aars2_in.vivo_noRT | Gm4166 | 4.399 | 1.30E-17 | glio |
| culture_effect | Il17rc | -1.857 | 1.30E-18 | glio |
| culture_effect | Socs3 | -2.391 | 1.30E-18 | glio |
| Chek1_in.vivo_noRT | Esm1 | 4.78 | 1.30E-20 | glio |
| culture_effect | Tspan4 | -1.627 | 1.30E-21 | glio |
| culture_effect | Klk6 | -5.25 | 1.30E-22 | glio |
| Bora_in.vivo_noRT | Slc35g3 | 4.936 | 1.30E-22 | glio |
| Aars2_in.vivo_noRT | Cd36 | 5.327 | 1.30E-24 | glio |
| Atr_in.vivo_noRT | Rgs4 | 5.533 | 1.30E-25 | glio |
| Chek1_ex.vivo_noRT | Fam227b | 6.273 | 1.30E-25 | glio |
| culture_effect | Ubash3a | 2.38 | 1.31E-03 | glio |
| culture_effect | Slc25a40 | 1.102 | 1.31E-03 | glio |
| culture_effect | Il17d | 0.971 | 1.31E-03 | glio |
| culture_effect | Rita1 | 0.762 | 1.31E-03 | glio |
| Interaction_genotype_Aars2 | Aqp5 | -4.564 | 1.31E-03 | glio |
| Interaction_genotype_Aars2 | A430057M04F | -4.023 | 1.31E-03 | glio |
| Interaction_genotype_Aars2 | S100a7a | -4.079 | 1.31E-03 | glio |
| Interaction_genotype_Aars2 | 2900042K21R | -3.938 | 1.31E-03 | glio |
| Interaction_genotype_Aars2 | Gm47540 | -2.937 | 1.31E-03 | glio |
| Interaction_genotype_Aars2 | Gm10115 | -3.952 | 1.31E-03 | glio |
| Interaction_genotype_Aars2 | Psma6 | 1.216 | 1.31E-03 | glio |
| Interaction_genotype_Bora | Adamts7 | -3.911 | 1.31E-03 | glio |
| Interaction_genotype_Mtor | Mfsd7a | -4.457 | 1.31E-03 | glio |
| Interaction_genotype_Mtor | Rpl18 | 1.043 | 1.31E-03 | glio |
| Interaction_genotype_Myc | A830009L08Ri | -3.322 | 1.31E-03 | glio |
| Interaction_genotype_Myc | Gm30541 | -3.58 | 1.31E-03 | glio |
| Interaction_genotype_Myc | Spred3 | -3.061 | 1.31E-03 | glio |
| Interaction_genotype_Myc | Vps13d | -3.276 | 1.31E-03 | glio |
| Interaction_genotype_Myc | Grcc10 | 1.194 | 1.31E-03 | glio |
| Interaction_genotype_Rad21 | H2-Eb1 | 4.033 | 1.31E-03 | glio |
| Interaction_genotype_Rad21 | Atp8a2 | 4.112 | 1.31E-03 | glio |
| Aars2_in.vivo_noRT | Tmem231 | 2.225 | 1.31E-03 | glio |

| | | | | |
|---|---|---|---|---|
| Aars2_in.vivo_noRT | Anxa7 | -1.391 | 1.31E-03 | glio |
| Aars2_ex.vivo_noRT | Gm43948 | 2.509 | 1.31E-03 | glio |
| Aars2_ex.vivo_noRT | Tex14 | 2.893 | 1.31E-03 | glio |
| Atp5c1_in.vivo_noRT | Tff2 | 2.408 | 1.31E-03 | glio |
| Atp5c1_in.vivo_noRT | Ucma | 2.982 | 1.31E-03 | glio |
| Atr_in.vivo_noRT | Glrx | 2.091 | 1.31E-03 | glio |
| Atr_in.vivo_noRT | Plekhh2 | 2.096 | 1.31E-03 | glio |
| Bard1_ex.vivo_noRT | Insc | 2.814 | 1.31E-03 | glio |
| Bard1_ex.vivo_noRT | Cox7b2 | 2.815 | 1.31E-03 | glio |
| Bcas2_in.vivo_noRT | Tmem255a | 2.426 | 1.31E-03 | glio |
| Bcas2_in.vivo_noRT | 4930581F22Ri | 1.96 | 1.31E-03 | glio |
| Bcas2_in.vivo_noRT | Gm50340 | 2.102 | 1.31E-03 | glio |
| Bcas2_in.vivo_noRT | S100a3 | 2.368 | 1.31E-03 | glio |
| Bcas2_ex.vivo_noRT | 1700010B08R | 2.742 | 1.31E-03 | glio |
| Bora_in.vivo_noRT | Arid3b | 2.25 | 1.31E-03 | glio |
| Cenpj_ex.vivo_noRT | Npy1r | 2.591 | 1.31E-03 | glio |
| Cenpj_ex.vivo_noRT | Gm6803 | 2.591 | 1.31E-03 | glio |
| Chek1_in.vivo_noRT | Anxa11 | 2.625 | 1.31E-03 | glio |
| Chek1_in.vivo_noRT | 2900089D17R | 2.197 | 1.31E-03 | glio |
| Chek1_in.vivo_noRT | Chst11 | -1.507 | 1.31E-03 | glio |
| Chek1_in.vivo_noRT | Trp53 | -0.766 | 1.31E-03 | glio |
| Chek1_ex.vivo_noRT | Adam4 | 2.755 | 1.31E-03 | glio |
| Chek1_ex.vivo_noRT | Nfe2l3 | 2.919 | 1.31E-03 | glio |
| Hsd17b10_in.vivo_noRT | Cpne9 | 2.579 | 1.31E-03 | glio |
| Hsd17b10_in.vivo_noRT | Ctf1 | 2.019 | 1.31E-03 | glio |
| culture_effect | AA465934 | 1.482 | 1.31E-04 | glio |
| Interaction_genotype_Mtor | Obscn | -4.099 | 1.31E-04 | glio |
| Interaction_genotype_Mtor | Inha | -4.494 | 1.31E-04 | glio |
| Interaction_genotype_Mtor | 9330160F10Ri | -4.098 | 1.31E-04 | glio |
| Interaction_genotype_Mtor | 1700124L16Ri | -4.721 | 1.31E-04 | glio |
| Interaction_genotype_Mtor | Igfbp7 | -4.131 | 1.31E-04 | glio |
| Interaction_genotype_Mtor | Gm15892 | -4.946 | 1.31E-04 | glio |
| Interaction_genotype_Mtor | Myl6b | -4.308 | 1.31E-04 | glio |
| Interaction_genotype_Mtor | Ciart | -4.189 | 1.31E-04 | glio |
| Interaction_genotype_Mtor | Ndufa9 | 1.9 | 1.31E-04 | glio |
| Interaction_genotype_Mtor | Psmc2 | 1.848 | 1.31E-04 | glio |
| Interaction_genotype_Mtor | Srpk2 | 1.946 | 1.31E-04 | glio |
| Interaction_genotype_Mtor | Selenof | 1.898 | 1.31E-04 | glio |
| Interaction_genotype_Myc | Tnfsf13os | -4.339 | 1.31E-04 | glio |
| Interaction_genotype_Myc | Zfp296 | -3.871 | 1.31E-04 | glio |
| interaction_RT | Acot7 | -0.789 | 1.31E-04 | glio |
| Aars2_in.vivo_noRT | Gm904 | 2.764 | 1.31E-04 | glio |
| Aars2_in.vivo_noRT | Gm42941 | 2.534 | 1.31E-04 | glio |
| Aars2_in.vivo_noRT | Tbc1d16 | 2.613 | 1.31E-04 | glio |
| Aars2_ex.vivo_noRT | Gm12610 | 2.893 | 1.31E-04 | glio |
| Atp5c1_in.vivo_noRT | Gm15326 | 2.139 | 1.31E-04 | glio |
| Atp5c1_in.vivo_noRT | Gm37800 | 2.668 | 1.31E-04 | glio |
| Atr_in.vivo_noRT | Fam135b | 3.074 | 1.31E-04 | glio |
| Atr_in.vivo_noRT | H2-DMb2 | 2.566 | 1.31E-04 | glio |
| Atr_in.vivo_noRT | BC049715 | 2.534 | 1.31E-04 | glio |
| Atr_ex.vivo_noRT | Adm | 3.554 | 1.31E-04 | glio |
| Bard1_ex.vivo_noRT | Smtnl1 | 2.814 | 1.31E-04 | glio |

| | | | | |
|---|---|---|---|---|
| Bard1_ex.vivo_noRT | Mpeg1 | 3.412 | 1.31E-04 | glio |
| Bard1_ex.vivo_noRT | Gm49808 | 2.421 | 1.31E-04 | glio |
| Bcas2_in.vivo_noRT | 9130008F23Ri | 3.041 | 1.31E-04 | glio |
| Bora_in.vivo_noRT | Gm36738 | 2.928 | 1.31E-04 | glio |
| Bora_in.vivo_noRT | Pgam5 | -1.641 | 1.31E-04 | glio |
| Chek1_in.vivo_noRT | Eps8l1 | 2.73 | 1.31E-04 | glio |
| Chek1_ex.vivo_noRT | Aldh1l2 | -2.34 | 1.31E-04 | glio |
| culture_effect | Tinagl1 | -0.891 | 1.31E-05 | glio |
| Interaction_genotype_Mtor | Mettl18 | -4.334 | 1.31E-05 | glio |
| Interaction_genotype_Mtor | 4930522L14Ri | -5.27 | 1.31E-05 | glio |
| Interaction_genotype_Mtor | Ccdc184 | -5.634 | 1.31E-05 | glio |
| Interaction_genotype_Mtor | Gm49701 | -5.036 | 1.31E-05 | glio |
| Interaction_genotype_Myc | Clcn1 | -4.036 | 1.31E-05 | glio |
| Aars2_in.vivo_noRT | Zfp619 | 3.124 | 1.31E-05 | glio |
| Atr_in.vivo_noRT | Hmgcs2 | 3.45 | 1.31E-05 | glio |
| Atr_ex.vivo_noRT | Gm41409 | 3.563 | 1.31E-05 | glio |
| Bcas2_in.vivo_noRT | Cacnb3 | 2.807 | 1.31E-05 | glio |
| Chek1_in.vivo_noRT | Usp27x | 3.267 | 1.31E-05 | glio |
| culture_effect | Mrpl34 | 0.554 | 1.31E-06 | glio |
| Interaction_genotype_Mtor | Gm48764 | -4.099 | 1.31E-06 | glio |
| Interaction_genotype_Nat2 | Tmem132c | 4.101 | 1.31E-06 | glio |
| Aars2_in.vivo_noRT | Adamts7 | 3.136 | 1.31E-06 | glio |
| Atr_in.vivo_noRT | Foxa1 | 3.681 | 1.31E-06 | glio |
| Atr_in.vivo_noRT | Gm4129 | 3.364 | 1.31E-06 | glio |
| Atr_ex.vivo_noRT | 9130019O22R | 3.164 | 1.31E-06 | glio |
| Bcas2_in.vivo_noRT | Parvg | 3.495 | 1.31E-06 | glio |
| Hsd17b10_in.vivo_noRT | AI182371 | 2.834 | 1.31E-06 | glio |
| culture_effect | Snrnp70 | -0.66 | 1.31E-07 | glio |
| Aars2_in.vivo_noRT | Gm16174 | 3.937 | 1.31E-07 | glio |
| Aars2_in.vivo_noRT | Gm50255 | 4.419 | 1.31E-07 | glio |
| Aars2_in.vivo_noRT | Fam32a | -1.701 | 1.31E-07 | glio |
| Atp5c1_in.vivo_noRT | Gm13856 | 2.668 | 1.31E-07 | glio |
| Atp5c1_in.vivo_noRT | Gm21936 | 2.668 | 1.31E-07 | glio |
| Bcas2_in.vivo_noRT | Trim67 | 3.238 | 1.31E-07 | glio |
| Bcas2_in.vivo_noRT | Gm31517 | 3.773 | 1.31E-07 | glio |
| Chek1_in.vivo_noRT | Bex3 | -1.823 | 1.31E-07 | glio |
| culture_effect | Relb | -1.269 | 1.31E-08 | glio |
| Bcas2_in.vivo_noRT | Ptges | 4.151 | 1.31E-08 | glio |
| Bcas2_in.vivo_noRT | Guca1b | 4.046 | 1.31E-08 | glio |
| culture_effect | Lhfpl2 | -2.221 | 1.31E-09 | glio |
| Bcas2_in.vivo_noRT | 1110035H17R | 4.702 | 1.31E-09 | glio |
| Bcas2_in.vivo_noRT | Wnt6 | 5.039 | 1.31E-09 | glio |
| Hsd17b10_in.vivo_noRT | Ptprc | 3.352 | 1.31E-09 | glio |
| culture_effect | Daam2 | 3.073 | 1.31E-10 | glio |
| Atr_in.vivo_noRT | Gm8212 | 4.431 | 1.31E-10 | glio |
| Atr_in.vivo_noRT | Gm45890 | 4.617 | 1.31E-10 | glio |
| Atr_in.vivo_noRT | 2900042K21R | 4.348 | 1.31E-10 | glio |
| Bora_in.vivo_noRT | Gm15972 | 3.943 | 1.31E-10 | glio |
| Bora_in.vivo_noRT | Gm35290 | 4.198 | 1.31E-10 | glio |
| culture_effect | Atf5 | -1.353 | 1.31E-11 | glio |
| Aars2_in.vivo_noRT | Gm10425 | 5.063 | 1.31E-12 | glio |
| Atr_in.vivo_noRT | Lyz2 | 5.147 | 1.31E-12 | glio |

| | | | | |
|---|---|---|---|---|
| Atr_in.vivo_noRT | 4833407H14R | 5.436 | 1.31E-12 | glio |
| Atr_in.vivo_noRT | Gm15579 | 5.533 | 1.31E-13 | glio |
| Atr_in.vivo_noRT | Ptprc | 5.269 | 1.31E-21 | glio |
| Atr_in.vivo_noRT | Akr1c18 | 5.533 | 1.31E-22 | glio |
| Chek1_in.vivo_noRT | Edn2 | 5.044 | 1.31E-24 | glio |
| Chek1_in.vivo_noRT | Gm14207 | 5.044 | 1.31E-24 | glio |
| Chek1_in.vivo_noRT | Nrip2 | 5.044 | 1.31E-24 | glio |
| culture_effect | Gbp6 | -3.467 | 1.31E-34 | glio |
| culture_effect | Klhdc8a | 6.896 | 1.31E-43 | glio |
| culture_effect | Kdsr | -0.683 | 1.32E-03 | glio |
| culture_effect | Smarca2 | -0.773 | 1.32E-03 | glio |
| culture_effect | Specc1l | -0.728 | 1.32E-03 | glio |
| culture_effect | Aarsd1 | -0.497 | 1.32E-03 | glio |
| culture_effect | Mrps18a | 0.388 | 1.32E-03 | glio |
| culture_effect | Eif1ad | 0.396 | 1.32E-03 | glio |
| Interaction_genotype_Aars2 | Gm14019 | -3.53 | 1.32E-03 | glio |
| Interaction_genotype_Aars2 | Eif4b | 1.323 | 1.32E-03 | glio |
| Interaction_genotype_Aars2 | Tomm20 | 0.968 | 1.32E-03 | glio |
| Interaction_genotype_Bora | Pygo1 | -3.93 | 1.32E-03 | glio |
| Interaction_genotype_Bora | Gm10734 | -4.238 | 1.32E-03 | glio |
| Interaction_genotype_Mtor | Car11 | -3.277 | 1.32E-03 | glio |
| Interaction_genotype_Myc | Cxcl5 | -2.729 | 1.32E-03 | glio |
| Interaction_genotype_Myc | Serpinb9b | -4.388 | 1.32E-03 | glio |
| Interaction_genotype_Myc | Gm46652 | -3.444 | 1.32E-03 | glio |
| Interaction_genotype_Myc | Fam228b | -3.945 | 1.32E-03 | glio |
| Interaction_genotype_Myc | Gm20522 | -3.54 | 1.32E-03 | glio |
| Interaction_genotype_Myc | Ccar1 | 1.344 | 1.32E-03 | glio |
| Interaction_genotype_Rad21 | H2-DMb2 | 4.659 | 1.32E-03 | glio |
| Interaction_genotype_Rpl3 | Gm49312 | -3.909 | 1.32E-03 | glio |
| Interaction_genotype_Rpl3 | Wfdc1 | -3.826 | 1.32E-03 | glio |
| Interaction_genotype_Terf1 | 5430402O13R | -4.771 | 1.32E-03 | glio |
| interaction_RT | Bik | 1.982 | 1.32E-03 | glio |
| Aars2_in.vivo_noRT | Pla2g2e | 2.518 | 1.32E-03 | glio |
| Aars2_in.vivo_noRT | Med8 | -0.907 | 1.32E-03 | glio |
| Aars2_in.vivo_noRT | Bag1 | -0.86 | 1.32E-03 | glio |
| Atp5c1_in.vivo_noRT | 8030423F21Ri | 2.139 | 1.32E-03 | glio |
| Atp5c1_in.vivo_noRT | Clcn1 | 2.022 | 1.32E-03 | glio |
| Atp5c1_in.vivo_noRT | Cxcl16 | 2.404 | 1.32E-03 | glio |
| Atp5c1_in.vivo_noRT | Slc26a6 | 2.439 | 1.32E-03 | glio |
| Atr_in.vivo_noRT | Gm33782 | 2.039 | 1.32E-03 | glio |
| Atr_in.vivo_noRT | Optn | 2.286 | 1.32E-03 | glio |
| Atr_in.vivo_noRT | Napb | 1.978 | 1.32E-03 | glio |
| Atr_in.vivo_noRT | Gm11808 | -1.164 | 1.32E-03 | glio |
| Atr_ex.vivo_noRT | Gm813 | 3.164 | 1.32E-03 | glio |
| Atr_ex.vivo_noRT | Pcdhb20 | 2.993 | 1.32E-03 | glio |
| Atr_ex.vivo_noRT | Gm2115 | 2.528 | 1.32E-03 | glio |
| Atr_ex.vivo_noRT | Pcyt1b | -1.761 | 1.32E-03 | glio |
| Bard1_ex.vivo_noRT | Prox2os | 2.819 | 1.32E-03 | glio |
| Bcas2_in.vivo_noRT | Syde2 | 2.261 | 1.32E-03 | glio |
| Bcas2_in.vivo_noRT | Nploc4 | -1.351 | 1.32E-03 | glio |
| Bora_in.vivo_noRT | Cd34 | 2.378 | 1.32E-03 | glio |
| Bora_in.vivo_noRT | 0610040B10R | 1.967 | 1.32E-03 | glio |

| | | | | |
|---|---|---|---|---|
| Bora_ex.vivo_noRT | Igdcc4 | 2.808 | 1.32E-03 | glio |
| Bora_ex.vivo_noRT | Adcy2 | 2.808 | 1.32E-03 | glio |
| Bora_ex.vivo_noRT | Cxcr6 | 3.201 | 1.32E-03 | glio |
| Cenpj_ex.vivo_noRT | 1700019E08Ri | 2.591 | 1.32E-03 | glio |
| Chek1_in.vivo_noRT | Ccdc146 | 2.445 | 1.32E-03 | glio |
| Chek1_in.vivo_noRT | Platr3 | 2.453 | 1.32E-03 | glio |
| Chek1_in.vivo_noRT | Arrdc3 | 2.184 | 1.32E-03 | glio |
| Chek1_in.vivo_noRT | Fto | -1.363 | 1.32E-03 | glio |
| Chek1_ex.vivo_noRT | Gm42695 | 2.866 | 1.32E-03 | glio |
| Chek1_ex.vivo_noRT | 9330159F19Ri | 2.68 | 1.32E-03 | glio |
| Hsd17b10_in.vivo_noRT | Cebpa | 2.482 | 1.32E-03 | glio |
| culture_effect | Aaas | -0.797 | 1.32E-04 | glio |
| Interaction_genotype_Aars2 | AC154200.1 | -3.804 | 1.32E-04 | glio |
| Interaction_genotype_Mtor | Rpgr | -4.328 | 1.32E-04 | glio |
| Interaction_genotype_Myc | Fgd3 | -4.739 | 1.32E-04 | glio |
| Interaction_genotype_Myc | C130013H08R | -4.737 | 1.32E-04 | glio |
| Interaction_genotype_Myc | Gm12841 | -3.122 | 1.32E-04 | glio |
| Interaction_genotype_Myc | 1700112D23R | -4.299 | 1.32E-04 | glio |
| Interaction_genotype_Myc | Gm16740 | -4.244 | 1.32E-04 | glio |
| Interaction_genotype_Myc | Ppib | 1.931 | 1.32E-04 | glio |
| interaction_RT | Cdk12 | -0.846 | 1.32E-04 | glio |
| Aars2_in.vivo_noRT | Plekhg5 | 2.596 | 1.32E-04 | glio |
| Atp5c1_in.vivo_noRT | 1700012B09R | 3.46 | 1.32E-04 | glio |
| Atr_in.vivo_noRT | Ccnb1ip1 | 3.002 | 1.32E-04 | glio |
| Atr_in.vivo_noRT | Lhx9 | 2.828 | 1.32E-04 | glio |
| Atr_in.vivo_noRT | A330023F24Ri | 2.755 | 1.32E-04 | glio |
| Atr_in.vivo_noRT | Frs3 | 2.643 | 1.32E-04 | glio |
| Bcas2_ex.vivo_noRT | Gm2016 | 2.742 | 1.32E-04 | glio |
| Bora_in.vivo_noRT | Mill2 | 2.745 | 1.32E-04 | glio |
| Bora_in.vivo_noRT | Tppp | 2.379 | 1.32E-04 | glio |
| Chek1_in.vivo_noRT | C8g | 2.77 | 1.32E-04 | glio |
| Chek1_in.vivo_noRT | Gm19345 | 2.472 | 1.32E-04 | glio |
| culture_effect | Tbx1 | -1.73 | 1.32E-05 | glio |
| Interaction_genotype_Atr | C130060C02R | -4.868 | 1.32E-05 | glio |
| Interaction_genotype_Mtor | Nmi | 2.08 | 1.32E-05 | glio |
| Interaction_genotype_Myc | Cacnb3 | -4.42 | 1.32E-05 | glio |
| Interaction_genotype_Myc | Gm47545 | -4.518 | 1.32E-05 | glio |
| Interaction_genotype_Myc | Ppia | 1.806 | 1.32E-05 | glio |
| Aars2_in.vivo_noRT | Ttc41 | 3.255 | 1.32E-05 | glio |
| Atr_in.vivo_noRT | Cacng7 | 2.939 | 1.32E-05 | glio |
| Atr_ex.vivo_noRT | Gm45151 | 3.164 | 1.32E-05 | glio |
| Atr_ex.vivo_noRT | Gm12860 | 3.164 | 1.32E-05 | glio |
| Bcas2_in.vivo_noRT | Gm26533 | 3.173 | 1.32E-05 | glio |
| Bcas2_in.vivo_noRT | Gm16576 | 3.152 | 1.32E-05 | glio |
| Cenpj_in.vivo_noRT | Osbp2 | 3.818 | 1.32E-05 | glio |
| Cenpj_in.vivo_noRT | Impg2 | 3.077 | 1.32E-05 | glio |
| Cenpj_in.vivo_noRT | Trim17 | 2.548 | 1.32E-05 | glio |
| Cenpj_in.vivo_noRT | Gm14929 | 2.548 | 1.32E-05 | glio |
| Cenpj_in.vivo_noRT | Rtl1 | 3.142 | 1.32E-05 | glio |
| culture_effect | Slco2a1 | -2.933 | 1.32E-06 | glio |
| Interaction_genotype_Myc | Trpm3 | -5.639 | 1.32E-06 | glio |
| Aars2_in.vivo_noRT | Icosl | 3.037 | 1.32E-06 | glio |

| | | | | |
|---|---|---|---|---|
| Atr_in.vivo_noRT | C130013H08R | 3.764 | 1.32E-06 | glio |
| Atr_in.vivo_noRT | Cxcr6 | 3.344 | 1.32E-06 | glio |
| Atr_in.vivo_noRT | Ckm | 3.259 | 1.32E-06 | glio |
| Atr_in.vivo_noRT | Gm26877 | 3.342 | 1.32E-06 | glio |
| Atr_ex.vivo_noRT | Klk13 | 3.164 | 1.32E-06 | glio |
| Chek1_in.vivo_noRT | Gpr21 | 3.178 | 1.32E-06 | glio |
| culture_effect | Atp8a2 | -2.826 | 1.32E-07 | glio |
| culture_effect | Pnp2 | -1.212 | 1.32E-07 | glio |
| Interaction_genotype_Mtor | Vmn2r44 | -4.494 | 1.32E-07 | glio |
| Atr_ex.vivo_noRT | Tmem54 | 3.962 | 1.32E-07 | glio |
| Bcas2_in.vivo_noRT | Klhl30 | 3.622 | 1.32E-07 | glio |
| Bcas2_in.vivo_noRT | Gm13610 | 3.651 | 1.32E-07 | glio |
| Hsd17b10_in.vivo_noRT | Amy1 | 3.619 | 1.32E-07 | glio |
| Hsd17b10_in.vivo_noRT | Gm11457 | 3.616 | 1.32E-07 | glio |
| Bcas2_in.vivo_noRT | Ephx1 | 3.909 | 1.32E-08 | glio |
| Bora_in.vivo_noRT | Gm12840 | 3.678 | 1.32E-08 | glio |
| Chek1_in.vivo_noRT | Apol7d | 4.142 | 1.32E-08 | glio |
| Chek1_ex.vivo_noRT | Gm7247 | 4.045 | 1.32E-08 | glio |
| Hsd17b10_in.vivo_noRT | Ube4bos1 | 3.352 | 1.32E-08 | glio |
| Aars2_in.vivo_noRT | Pde1a | 4.326 | 1.32E-09 | glio |
| Aars2_in.vivo_noRT | Anxa8 | 4.137 | 1.32E-09 | glio |
| Atr_in.vivo_noRT | Lynx1 | 4.229 | 1.32E-09 | glio |
| culture_effect | Fhl2 | 2.711 | 1.32E-10 | glio |
| Atr_in.vivo_noRT | Prob1 | 4.639 | 1.32E-10 | glio |
| Chek1_ex.vivo_noRT | Gm33973 | 4.842 | 1.32E-10 | glio |
| Interaction_genotype_Mtor | Rassf6 | -7.133 | 1.32E-11 | glio |
| Chek1_in.vivo_noRT | 7530414M10F | 4.518 | 1.32E-11 | glio |
| Atr_in.vivo_noRT | AC125149.2 | 4.245 | 1.32E-12 | glio |
| Chek1_in.vivo_noRT | Gm7247 | 4.252 | 1.32E-13 | glio |
| Chek1_ex.vivo_noRT | Gm2022 | 4.842 | 1.32E-13 | glio |
| Hsd17b10_in.vivo_noRT | Gm5415 | 3.616 | 1.32E-13 | glio |
| Hsd17b10_in.vivo_noRT | Edn2 | 3.616 | 1.32E-13 | glio |
| Hsd17b10_in.vivo_noRT | Gm14207 | 3.616 | 1.32E-13 | glio |
| Hsd17b10_in.vivo_noRT | Nrip2 | 3.616 | 1.32E-13 | glio |
| Aars2_in.vivo_noRT | Ms4a6d | 4.539 | 1.32E-14 | glio |
| Chek1_ex.vivo_noRT | Cd36 | 4.842 | 1.32E-14 | glio |
| Chek1_ex.vivo_noRT | Gstt4 | 4.842 | 1.32E-14 | glio |
| Atr_in.vivo_noRT | Aloxe3 | 5.004 | 1.32E-17 | glio |
| Chek1_in.vivo_noRT | BC049987 | 4.261 | 1.32E-17 | glio |
| Atr_in.vivo_noRT | Rassf6 | 5.533 | 1.32E-20 | glio |
| culture_effect | Trib1 | -1.77 | 1.32E-21 | glio |
| Bcas2_in.vivo_noRT | Slc35g3 | 4.807 | 1.32E-21 | glio |
| Atr_in.vivo_noRT | 4931431C16R | 5.533 | 1.32E-22 | glio |
| Bcas2_in.vivo_noRT | Gm49698 | 4.807 | 1.32E-22 | glio |
| Aars2_in.vivo_noRT | Gm43701 | 5.327 | 1.32E-24 | glio |
| Atr_in.vivo_noRT | Vmn2r44 | 5.533 | 1.32E-27 | glio |
| Atr_in.vivo_noRT | Mmp13 | 5.533 | 1.32E-27 | glio |
| culture_effect | Anxa11 | 1.807 | 1.33E-03 | glio |
| culture_effect | Eml1 | 1.387 | 1.33E-03 | glio |
| culture_effect | Snapin | 0.772 | 1.33E-03 | glio |
| culture_effect | Pdcl3 | 0.452 | 1.33E-03 | glio |
| culture_effect | Srsf11 | -0.388 | 1.33E-03 | glio |

| | | | | |
|---|---|---|---|---|
| culture_effect | Psmd7 | -0.406 | 1.33E-03 | glio |
| Interaction_genotype_Aars2 | Lrrc72 | -3.722 | 1.33E-03 | glio |
| Interaction_genotype_Aars2 | 4931431C16R | -3.212 | 1.33E-03 | glio |
| Interaction_genotype_Aars2 | 2010315B03R | -3.217 | 1.33E-03 | glio |
| Interaction_genotype_Aars2 | Tmed7 | 1.742 | 1.33E-03 | glio |
| Interaction_genotype_Aars2 | Tesc | 2.503 | 1.33E-03 | glio |
| Interaction_genotype_Atr | Epcam | 3.625 | 1.33E-03 | glio |
| Interaction_genotype_Mtor | Fam169a | -3.776 | 1.33E-03 | glio |
| Interaction_genotype_Mtor | Prkar2b | -3.94 | 1.33E-03 | glio |
| Interaction_genotype_Mtor | Klk9 | -3.953 | 1.33E-03 | glio |
| Interaction_genotype_Myc | Gm48899 | -3.045 | 1.33E-03 | glio |
| Interaction_genotype_Myc | EU599041 | -3.746 | 1.33E-03 | glio |
| Interaction_genotype_Myc | Gm47936 | -2.729 | 1.33E-03 | glio |
| Interaction_genotype_Myc | Dixdc1 | -3.57 | 1.33E-03 | glio |
| interaction_RT | Serpinb8 | 1.932 | 1.33E-03 | glio |
| interaction_RT | Dnajc8 | -0.698 | 1.33E-03 | glio |
| Aars2_in.vivo_noRT | Ubash3a | 3.375 | 1.33E-03 | glio |
| Aars2_in.vivo_noRT | Gm36696 | 2.187 | 1.33E-03 | glio |
| Aars2_in.vivo_noRT | Cacng7 | 2.223 | 1.33E-03 | glio |
| Aars2_ex.vivo_noRT | Dcpp1 | 3.293 | 1.33E-03 | glio |
| Atp5c1_in.vivo_noRT | Dapk2 | 1.885 | 1.33E-03 | glio |
| Atr_in.vivo_noRT | Fetub | 1.996 | 1.33E-03 | glio |
| Atr_in.vivo_noRT | Zfp626 | 1.788 | 1.33E-03 | glio |
| Atr_in.vivo_noRT | Rae1 | -1.055 | 1.33E-03 | glio |
| Atr_ex.vivo_noRT | 8030423F21Ri | 2.588 | 1.33E-03 | glio |
| Atr_ex.vivo_noRT | Gm30794 | 2.187 | 1.33E-03 | glio |
| Atr_ex.vivo_noRT | Cabp1 | 2.417 | 1.33E-03 | glio |
| Bard1_ex.vivo_noRT | Gm10696 | 3.213 | 1.33E-03 | glio |
| Bcas2_in.vivo_noRT | Hist1h2bf | 2.495 | 1.33E-03 | glio |
| Bcas2_in.vivo_noRT | 4930432K21R | 2.193 | 1.33E-03 | glio |
| Bcas2_in.vivo_noRT | Mlip | 2.32 | 1.33E-03 | glio |
| Bcas2_in.vivo_noRT | Ahi1 | -1.494 | 1.33E-03 | glio |
| Bcas2_in.vivo_noRT | Pgrmc1 | -1.49 | 1.33E-03 | glio |
| Bcas2_in.vivo_noRT | Tomm40 | -0.918 | 1.33E-03 | glio |
| Bcas2_in.vivo_noRT | Cdk4 | -0.685 | 1.33E-03 | glio |
| Bora_in.vivo_noRT | Slc39a8 | 2.573 | 1.33E-03 | glio |
| Bora_in.vivo_noRT | Cul9 | 2.538 | 1.33E-03 | glio |
| Bora_in.vivo_noRT | Efhd2 | -1.626 | 1.33E-03 | glio |
| Cenpj_in.vivo_noRT | 4930512B01R | 2.022 | 1.33E-03 | glio |
| Chek1_in.vivo_noRT | Zan | 2.281 | 1.33E-03 | glio |
| Chek1_in.vivo_noRT | 5330438D12R | 1.918 | 1.33E-03 | glio |
| Chek1_ex.vivo_noRT | Rpl31 | -0.726 | 1.33E-03 | glio |
| Hsd17b10_in.vivo_noRT | Sdr16c5 | 2.299 | 1.33E-03 | glio |
| culture_effect | Tafa5 | 1.251 | 1.33E-04 | glio |
| culture_effect | Lrrfip1 | 0.664 | 1.33E-04 | glio |
| Interaction_genotype_Aars2 | Lbhd2 | -5.512 | 1.33E-04 | glio |
| Interaction_genotype_Aars2 | Ruvbl2 | 2.135 | 1.33E-04 | glio |
| Interaction_genotype_Chek1 | Serf2 | 1.206 | 1.33E-04 | glio |
| Interaction_genotype_Mtor | Gm40040 | -4.664 | 1.33E-04 | glio |
| Interaction_genotype_Mtor | Zc3h12b | -4.301 | 1.33E-04 | glio |
| Interaction_genotype_Mtor | Rassf8 | -3.843 | 1.33E-04 | glio |
| Interaction_genotype_Mtor | Nmrk1 | -4.278 | 1.33E-04 | glio |

| | | | | |
|---|---|---|---|---|
| Interaction_genotype_Mtor | Nop56 | 2.067 | 1.33E-04 | glio |
| Interaction_genotype_Myc | Gm16244 | -4.449 | 1.33E-04 | glio |
| Interaction_genotype_Pars2 | 2010320O07R | -5.02 | 1.33E-04 | glio |
| Aars2_in.vivo_noRT | Itpr2 | -1.767 | 1.33E-04 | glio |
| Aars2_in.vivo_noRT | Prmt1 | -1.084 | 1.33E-04 | glio |
| Aars2_ex.vivo_noRT | Ocm | 3.293 | 1.33E-04 | glio |
| Aars2_ex.vivo_noRT | Zscan4d | 2.899 | 1.33E-04 | glio |
| Bcas2_in.vivo_noRT | Pwwp2a | -1.926 | 1.33E-04 | glio |
| Bora_ex.vivo_noRT | Nos2 | 3.201 | 1.33E-04 | glio |
| Chek1_ex.vivo_noRT | Mterf1b | 3.287 | 1.33E-04 | glio |
| Eif2s1_in.vivo_noRT | Gm26671 | 2.143 | 1.33E-04 | glio |
| Hsd17b10_in.vivo_noRT | Chsy1 | -2.038 | 1.33E-04 | glio |
| culture_effect | Gm16794 | 2.225 | 1.33E-05 | glio |
| culture_effect | 4931406C07R | 0.927 | 1.33E-05 | glio |
| Interaction_genotype_Atr | Ppp4r2 | 2.19 | 1.33E-05 | glio |
| Interaction_genotype_Atr | Loxl4 | -4.991 | 1.33E-05 | glio |
| Interaction_genotype_Myc | Flrt3 | -4.619 | 1.33E-05 | glio |
| Interaction_genotype_Myc | Tlr6 | -4.974 | 1.33E-05 | glio |
| Interaction_genotype_Myc | Tcte2 | -4.347 | 1.33E-05 | glio |
| Interaction_genotype_Myc | Tomm6os | -5.221 | 1.33E-05 | glio |
| Interaction_genotype_Myc | Gm10125 | -5.089 | 1.33E-05 | glio |
| Interaction_genotype_Myc | Armc2 | -4.973 | 1.33E-05 | glio |
| Interaction_genotype_Myc | Eif1 | 1.544 | 1.33E-05 | glio |
| Aars2_in.vivo_noRT | Hddc3 | 2.944 | 1.33E-05 | glio |
| Atp5c1_in.vivo_noRT | Ccdc125 | 2.668 | 1.33E-05 | glio |
| Atr_in.vivo_noRT | Rrad | 3.27 | 1.33E-05 | glio |
| Atr_in.vivo_noRT | Llgl2 | 2.998 | 1.33E-05 | glio |
| Atr_in.vivo_noRT | Gm35315 | 3.032 | 1.33E-05 | glio |
| Bora_in.vivo_noRT | Acox2 | 2.648 | 1.33E-05 | glio |
| Chek1_in.vivo_noRT | Gm20234 | 3.327 | 1.33E-05 | glio |
| Hsd17b10_in.vivo_noRT | Sap30bpos | 2.753 | 1.33E-05 | glio |
| culture_effect | Comp | -3.294 | 1.33E-06 | glio |
| culture_effect | Fam149a | 0.983 | 1.33E-06 | glio |
| culture_effect | Kin | -0.853 | 1.33E-06 | glio |
| Aars2_in.vivo_noRT | AC241534.1 | 3.467 | 1.33E-06 | glio |
| Aars2_in.vivo_noRT | B230317F23Ri | 3.284 | 1.33E-06 | glio |
| Atp5c1_in.vivo_noRT | Esp24 | 2.668 | 1.33E-06 | glio |
| Atr_in.vivo_noRT | Gm15834 | 3.352 | 1.33E-06 | glio |
| Bard1_ex.vivo_noRT | A330049N07R | 3.213 | 1.33E-06 | glio |
| Bcas2_in.vivo_noRT | Slc26a1 | 3.304 | 1.33E-06 | glio |
| Bcas2_in.vivo_noRT | Klhdc8b | 3.511 | 1.33E-06 | glio |
| Bcas2_ex.vivo_noRT | Gm11583 | 3.54 | 1.33E-06 | glio |
| Bora_ex.vivo_noRT | Gm15533 | 3.201 | 1.33E-06 | glio |
| Chek1_ex.vivo_noRT | Traf3ip3 | 4.162 | 1.33E-06 | glio |
| Chek1_ex.vivo_noRT | Slc26a1 | 4.142 | 1.33E-06 | glio |
| Chek1_ex.vivo_noRT | Plac8 | 4.268 | 1.33E-06 | glio |
| Atr_in.vivo_noRT | Cyp2u1 | 3.43 | 1.33E-07 | glio |
| Bcas2_in.vivo_noRT | Gm32051 | 3.831 | 1.33E-07 | glio |
| Chek1_in.vivo_noRT | Ggt5 | 3.411 | 1.33E-07 | glio |
| Chek1_in.vivo_noRT | EU599041 | 3.767 | 1.33E-07 | glio |
| Chek1_ex.vivo_noRT | Esr1 | 4.449 | 1.33E-07 | glio |
| Aars2_in.vivo_noRT | 5330417C22R | 3.932 | 1.33E-08 | glio |

| | | | | |
|---|---|---|---|---|
| Bcas2_in.vivo_noRT | Gm26682 | 3.889 | 1.33E-08 | glio |
| Bcas2_in.vivo_noRT | Gm16222 | 4.547 | 1.33E-08 | glio |
| Bora_in.vivo_noRT | St14 | 3.55 | 1.33E-08 | glio |
| culture_effect | Pcdhb21 | -3.56 | 1.33E-09 | glio |
| Aars2_in.vivo_noRT | S1pr1 | 4.652 | 1.33E-09 | glio |
| Atr_in.vivo_noRT | Cyp1a1 | 5.533 | 1.33E-09 | glio |
| Bcas2_in.vivo_noRT | BC034090 | 3.952 | 1.33E-09 | glio |
| Chek1_in.vivo_noRT | Camk1d | 4.243 | 1.33E-09 | glio |
| Chek1_in.vivo_noRT | Gm13610 | 4.411 | 1.33E-10 | glio |
| Hsd17b10_in.vivo_noRT | 9430041J12Ri | 3.616 | 1.33E-10 | glio |
| Aars2_in.vivo_noRT | Gm44702 | 4.795 | 1.33E-11 | glio |
| Bora_in.vivo_noRT | Gm9967 | 4.549 | 1.33E-11 | glio |
| Bora_in.vivo_noRT | Gm5624 | 4.48 | 1.33E-11 | glio |
| Chek1_in.vivo_noRT | G630030J09Ri | 4.522 | 1.33E-11 | glio |
| Aars2_in.vivo_noRT | Gm20633 | 4.536 | 1.33E-12 | glio |
| Atr_in.vivo_noRT | Platr20 | 5.269 | 1.33E-12 | glio |
| Atr_in.vivo_noRT | Ctsf | 3.865 | 1.33E-12 | glio |
| Atr_in.vivo_noRT | Igdcc4 | 4.358 | 1.33E-12 | glio |
| Atr_in.vivo_noRT | Col6a2 | 4.75 | 1.33E-12 | glio |
| Atr_in.vivo_noRT | Ttc16 | 5.011 | 1.33E-13 | glio |
| Bora_in.vivo_noRT | Fam131a | 4.934 | 1.33E-13 | glio |
| Bora_in.vivo_noRT | Myoz1 | 4.299 | 1.33E-13 | glio |
| Chek1_in.vivo_noRT | C330024C12R | 4.653 | 1.33E-14 | glio |
| Atr_ex.vivo_noRT | Platr31 | 7.303 | 1.33E-15 | glio |
| Atr_ex.vivo_noRT | Tmem132c | 4.867 | 1.33E-15 | glio |
| Bora_in.vivo_noRT | Cys1 | 4.67 | 1.33E-15 | glio |
| Bora_in.vivo_noRT | Mpzl2 | 4.676 | 1.33E-16 | glio |
| Bora_in.vivo_noRT | 4732463B04R | 4.672 | 1.33E-17 | glio |
| culture_effect | Adm | -5.197 | 1.33E-18 | glio |
| Atr_in.vivo_noRT | Gm20646 | 6.071 | 1.33E-19 | glio |
| culture_effect | Plekha2 | -2.969 | 1.33E-20 | glio |
| Bora_in.vivo_noRT | Cd36 | 4.936 | 1.33E-21 | glio |
| Bcas2_in.vivo_noRT | Olfr881 | 4.807 | 1.33E-22 | glio |
| Bora_in.vivo_noRT | Chia1 | 4.936 | 1.33E-22 | glio |
| culture_effect | Fem1b | -0.612 | 1.34E-03 | glio |
| culture_effect | Ccdc12 | -0.522 | 1.34E-03 | glio |
| culture_effect | March6 | -0.58 | 1.34E-03 | glio |
| Interaction_genotype_Aars2 | Gm36198 | -3.329 | 1.34E-03 | glio |
| Interaction_genotype_Aars2 | Zfp316 | -3.409 | 1.34E-03 | glio |
| Interaction_genotype_Aars2 | Gstm5 | 1.89 | 1.34E-03 | glio |
| Interaction_genotype_Aars2 | Smdt1 | 1.162 | 1.34E-03 | glio |
| Interaction_genotype_Atr | Ncbp1 | 1.689 | 1.34E-03 | glio |
| Interaction_genotype_Cdkn2a | Ccl9 | 3.241 | 1.34E-03 | glio |
| Interaction_genotype_Chek1 | Wfdc1 | -3.96 | 1.34E-03 | glio |
| Interaction_genotype_Mtor | Spry3 | -3.999 | 1.34E-03 | glio |
| Interaction_genotype_Mtor | Tha1 | -3.686 | 1.34E-03 | glio |
| Interaction_genotype_Mtor | Ddx11 | -2.957 | 1.34E-03 | glio |
| Interaction_genotype_Mtor | Pxmp2 | -3.091 | 1.34E-03 | glio |
| Interaction_genotype_Mtor | Peg10 | -3.024 | 1.34E-03 | glio |
| Interaction_genotype_Myc | Fam135b | -4.125 | 1.34E-03 | glio |
| Interaction_genotype_Myc | Gm26981 | -3.524 | 1.34E-03 | glio |
| Interaction_genotype_Myc | Gm50295 | -3.461 | 1.34E-03 | glio |

| | | | | |
|---|---|---|---|---|
| Interaction_genotype_Myc | Vwa8 | -3.147 | 1.34E-03 | glio |
| Interaction_genotype_Myc | Tnk2 | -2.679 | 1.34E-03 | glio |
| Interaction_genotype_Myc | Lgi4 | -2.402 | 1.34E-03 | glio |
| Interaction_genotype_Myc | H2afj | 1.183 | 1.34E-03 | glio |
| interaction_RT | Il15ra | 1.507 | 1.34E-03 | glio |
| interaction_RT | Avpi1 | 0.946 | 1.34E-03 | glio |
| Aars2_in.vivo_noRT | Cass4 | 2.318 | 1.34E-03 | glio |
| Aars2_in.vivo_noRT | Atf6 | -1.638 | 1.34E-03 | glio |
| Aars2_in.vivo_noRT | Psmd11 | -0.867 | 1.34E-03 | glio |
| Aars2_in.vivo_noRT | Rpl29 | -0.546 | 1.34E-03 | glio |
| Atp5c1_in.vivo_noRT | Gm43653 | 2.405 | 1.34E-03 | glio |
| Atr_in.vivo_noRT | Trim65 | 2.176 | 1.34E-03 | glio |
| Atr_in.vivo_noRT | Gm20219 | 2.55 | 1.34E-03 | glio |
| Atr_ex.vivo_noRT | Kcnk13 | 2.376 | 1.34E-03 | glio |
| Atr_ex.vivo_noRT | Btn1a1 | 2.38 | 1.34E-03 | glio |
| Atr_ex.vivo_noRT | Gm11769 | 2.776 | 1.34E-03 | glio |
| Bcas2_in.vivo_noRT | Lrrn4cl | 2.533 | 1.34E-03 | glio |
| Bcas2_in.vivo_noRT | Traf7 | -1.252 | 1.34E-03 | glio |
| Bora_in.vivo_noRT | Sh3bgrl2 | 2.327 | 1.34E-03 | glio |
| Bora_in.vivo_noRT | Abhd18 | 2.397 | 1.34E-03 | glio |
| Bora_in.vivo_noRT | 9130023H24R | 2.352 | 1.34E-03 | glio |
| Bora_in.vivo_noRT | Pus1 | -1.425 | 1.34E-03 | glio |
| Bora_in.vivo_noRT | Arid4a | 1.046 | 1.34E-03 | glio |
| Cenpj_ex.vivo_noRT | Gm13856 | 2.198 | 1.34E-03 | glio |
| Chek1_in.vivo_noRT | Gm26877 | 2.315 | 1.34E-03 | glio |
| Chek1_ex.vivo_noRT | Gm42984 | 2.898 | 1.34E-03 | glio |
| Hsd17b10_in.vivo_noRT | Gm36043 | 2.16 | 1.34E-03 | glio |
| culture_effect | Inpp1 | -0.929 | 1.34E-04 | glio |
| culture_effect | Mmp16 | -1.117 | 1.34E-04 | glio |
| culture_effect | Mrpl39 | -0.735 | 1.34E-04 | glio |
| culture_effect | Rpl41 | 0.637 | 1.34E-04 | glio |
| Interaction_genotype_Mtor | Gm15938 | -4.88 | 1.34E-04 | glio |
| Interaction_genotype_Mtor | Chrna1os | -3.546 | 1.34E-04 | glio |
| Interaction_genotype_Myc | Tmem236 | -3.835 | 1.34E-04 | glio |
| Interaction_genotype_Myc | Gm7008 | -4.211 | 1.34E-04 | glio |
| Interaction_genotype_Myc | Smim3 | -3.432 | 1.34E-04 | glio |
| Interaction_genotype_Myc | Fam104a | 1.349 | 1.34E-04 | glio |
| Aars2_in.vivo_noRT | Tnfrsf22 | 2.601 | 1.34E-04 | glio |
| Aars2_in.vivo_noRT | Stk10 | 2.16 | 1.34E-04 | glio |
| Aars2_in.vivo_noRT | Trappc1 | -1.4 | 1.34E-04 | glio |
| Atp5c1_in.vivo_noRT | Gm30042 | 2.408 | 1.34E-04 | glio |
| Atr_in.vivo_noRT | A930024E05Ri | 2.667 | 1.34E-04 | glio |
| Bcas2_in.vivo_noRT | Gm11767 | 2.885 | 1.34E-04 | glio |
| Bcas2_in.vivo_noRT | Gm29394 | 2.626 | 1.34E-04 | glio |
| Bcas2_in.vivo_noRT | Dnajc2 | -1.339 | 1.34E-04 | glio |
| Bcas2_ex.vivo_noRT | Gm14820 | 2.742 | 1.34E-04 | glio |
| Bora_in.vivo_noRT | Myl7 | 2.808 | 1.34E-04 | glio |
| Bora_in.vivo_noRT | Cfap100 | 2.739 | 1.34E-04 | glio |
| Bora_in.vivo_noRT | Mocs2 | -1.495 | 1.34E-04 | glio |
| Bora_in.vivo_noRT | Top1 | -1.122 | 1.34E-04 | glio |
| Bora_in.vivo_noRT | Ccnd2 | -0.961 | 1.34E-04 | glio |
| Chek1_in.vivo_noRT | Ccnb1ip1 | 3.036 | 1.34E-04 | glio |

| | | | | |
|---|---|---|---|---|
| Chek1_in.vivo_noRT | Ppic | -1.491 | 1.34E-04 | glio |
| Chek1_in.vivo_noRT | Igtp | -1.017 | 1.34E-04 | glio |
| Hsd17b10_in.vivo_noRT | Rilp | 3.09 | 1.34E-04 | glio |
| Hsd17b10_in.vivo_noRT | Ccdc114 | 2.698 | 1.34E-04 | glio |
| Hsd17b10_in.vivo_noRT | Gm50295 | 2.824 | 1.34E-04 | glio |
| culture_effect | Gm37986 | -2.475 | 1.34E-05 | glio |
| culture_effect | Noc2l | 0.664 | 1.34E-05 | glio |
| culture_effect | Naa20 | -0.582 | 1.34E-05 | glio |
| culture_effect | Atp5e | 0.56 | 1.34E-05 | glio |
| Interaction_genotype_Mtor | Tagap1 | -4.123 | 1.34E-05 | glio |
| Interaction_genotype_Mtor | Gm9758 | -3.839 | 1.34E-05 | glio |
| Interaction_genotype_Mtor | Lmx1a | -4.099 | 1.34E-05 | glio |
| Interaction_genotype_Nat2 | BC147527 | 4.902 | 1.34E-05 | glio |
| Aars2_ex.vivo_noRT | 2300002M23F | 3.293 | 1.34E-05 | glio |
| Atp5c1_in.vivo_noRT | Fam234b | -2.851 | 1.34E-05 | glio |
| Atr_in.vivo_noRT | Nr1h3 | 2.727 | 1.34E-05 | glio |
| Atr_ex.vivo_noRT | Arl4d | 4.009 | 1.34E-05 | glio |
| Bcas2_in.vivo_noRT | Il20rb | 3.272 | 1.34E-05 | glio |
| Bora_in.vivo_noRT | 5031434O11R | 3.528 | 1.34E-05 | glio |
| Bora_ex.vivo_noRT | Gm5602 | 3.201 | 1.34E-05 | glio |
| Interaction_genotype_Aars2 | Krtcap2 | 1.659 | 1.34E-06 | glio |
| Interaction_genotype_Myc | Bag1 | 2.544 | 1.34E-06 | glio |
| Aars2_in.vivo_noRT | Frmd8os | 3.809 | 1.34E-06 | glio |
| Aars2_in.vivo_noRT | Gm15879 | 4.37 | 1.34E-06 | glio |
| Aars2_in.vivo_noRT | Hdac7 | 3.319 | 1.34E-06 | glio |
| Aars2_in.vivo_noRT | Txn1 | -0.978 | 1.34E-06 | glio |
| Bora_in.vivo_noRT | Casz1 | 3.661 | 1.34E-06 | glio |
| Chek1_in.vivo_noRT | Pate2 | 3.564 | 1.34E-06 | glio |
| culture_effect | Twf1 | 0.75 | 1.34E-07 | glio |
| Interaction_genotype_Mtor | Igf2bp1 | -6.163 | 1.34E-07 | glio |
| interaction_RT | Tmem179b | 1.37 | 1.34E-07 | glio |
| interaction_RT | Cp | 2.81 | 1.34E-07 | glio |
| interaction_RT | Pantr1 | 2.674 | 1.34E-07 | glio |
| Bora_in.vivo_noRT | Dok3 | 3.801 | 1.34E-07 | glio |
| Bora_in.vivo_noRT | Gm10432 | 3.439 | 1.34E-07 | glio |
| Chek1_ex.vivo_noRT | Gm13822 | 4.447 | 1.34E-07 | glio |
| Chek1_ex.vivo_noRT | Zg16 | 4.443 | 1.34E-07 | glio |
| culture_effect | Filip1l | 1.901 | 1.34E-08 | glio |
| Aars2_in.vivo_noRT | Cemip | 3.292 | 1.34E-08 | glio |
| Chek1_in.vivo_noRT | Sv2a | 4.136 | 1.34E-08 | glio |
| Interaction_genotype_Mtor | Otos | -6.322 | 1.34E-09 | glio |
| Interaction_genotype_Mtor | 5430402O13R | -7.644 | 1.34E-09 | glio |
| Interaction_genotype_Mtor | Gm34047 | -7.601 | 1.34E-09 | glio |
| Bcas2_in.vivo_noRT | Gm17146 | 3.89 | 1.34E-09 | glio |
| Bora_in.vivo_noRT | Rims3 | 3.802 | 1.34E-09 | glio |
| Chek1_ex.vivo_noRT | Tcstv1 | 4.443 | 1.34E-09 | glio |
| culture_effect | Cenpv | -1.015 | 1.34E-10 | glio |
| Aars2_in.vivo_noRT | Gm16046 | 4.535 | 1.34E-10 | glio |
| Bcas2_in.vivo_noRT | Kcnk7 | 3.759 | 1.34E-10 | glio |
| Chek1_in.vivo_noRT | Gm34885 | 4.784 | 1.34E-11 | glio |
| culture_effect | Hotairm1 | 1.757 | 1.34E-12 | glio |
| Atr_in.vivo_noRT | Vmn1r88 | 4.348 | 1.34E-12 | glio |

| | | | | |
|---|---|---|---|---|
| Bcas2_in.vivo_noRT | Rbfaos | 3.89 | 1.34E-12 | glio |
| Aars2_in.vivo_noRT | Dppa2 | 4.794 | 1.34E-13 | glio |
| Aars2_in.vivo_noRT | Gm42695 | 5.067 | 1.34E-13 | glio |
| Bcas2_in.vivo_noRT | Gm11627 | 4.807 | 1.34E-13 | glio |
| Bora_in.vivo_noRT | A330032B11R | 4.67 | 1.34E-13 | glio |
| Chek1_in.vivo_noRT | Celsr3 | 4.784 | 1.34E-13 | glio |
| Chek1_ex.vivo_noRT | Gm13078 | 4.842 | 1.34E-15 | glio |
| Chek1_ex.vivo_noRT | Ctsf | 5.57 | 1.34E-15 | glio |
| Bcas2_in.vivo_noRT | Gm38163 | 4.807 | 1.34E-17 | glio |
| Aars2_in.vivo_noRT | A930033M14F | 5.327 | 1.34E-19 | glio |
| culture_effect | Gbp2 | -4.354 | 1.34E-55 | glio |
| culture_effect | Pabpc4l | -2.104 | 1.35E-03 | glio |
| culture_effect | 1700015F17Ri | 1.906 | 1.35E-03 | glio |
| culture_effect | Gm20324 | 1.316 | 1.35E-03 | glio |
| culture_effect | Pcid2 | -0.736 | 1.35E-03 | glio |
| Interaction_genotype_Aars2 | Pde11a | -4.182 | 1.35E-03 | glio |
| Interaction_genotype_Atr | Cited2 | 2.421 | 1.35E-03 | glio |
| Interaction_genotype_Bora | Tdrd5 | -3.876 | 1.35E-03 | glio |
| Interaction_genotype_Mtor | Gm43260 | -3.576 | 1.35E-03 | glio |
| Interaction_genotype_Mtor | B3galt1 | -3.682 | 1.35E-03 | glio |
| Interaction_genotype_Mtor | Plpp7 | -3.54 | 1.35E-03 | glio |
| Interaction_genotype_Mtor | Loxl2 | -4.074 | 1.35E-03 | glio |
| Interaction_genotype_Mtor | Gm11423 | -2.891 | 1.35E-03 | glio |
| Interaction_genotype_Myc | 1700047K16R | -3.746 | 1.35E-03 | glio |
| Interaction_genotype_Myc | Kdsr | -2.357 | 1.35E-03 | glio |
| Interaction_genotype_Myc | Gm14418 | -2.534 | 1.35E-03 | glio |
| Interaction_genotype_Myc | Naa50 | 1.452 | 1.35E-03 | glio |
| Interaction_genotype_Myc | Ifrd1 | 1.539 | 1.35E-03 | glio |
| interaction_RT | Las1l | -0.837 | 1.35E-03 | glio |
| Aars2_in.vivo_noRT | Asnsd1 | -1.222 | 1.35E-03 | glio |
| Aars2_ex.vivo_noRT | Gm50115 | 2.893 | 1.35E-03 | glio |
| Atp5c1_in.vivo_noRT | Gm16751 | 2.152 | 1.35E-03 | glio |
| Atr_in.vivo_noRT | Serpine2 | 2.742 | 1.35E-03 | glio |
| Atr_in.vivo_noRT | Hist1h2bg | 2.392 | 1.35E-03 | glio |
| Bard1_ex.vivo_noRT | Gm12589 | 2.415 | 1.35E-03 | glio |
| Bcas2_in.vivo_noRT | Slamf1 | 2.367 | 1.35E-03 | glio |
| Bcas2_in.vivo_noRT | Cyb561d1 | 2.485 | 1.35E-03 | glio |
| Bcas2_in.vivo_noRT | Gm4841 | 2.137 | 1.35E-03 | glio |
| Bcas2_in.vivo_noRT | Fkbp8 | -0.906 | 1.35E-03 | glio |
| Bcas2_in.vivo_noRT | Atf4 | -0.732 | 1.35E-03 | glio |
| Bcas2_ex.vivo_noRT | Gm45336 | 2.742 | 1.35E-03 | glio |
| Bcas2_ex.vivo_noRT | Gm16344 | 2.742 | 1.35E-03 | glio |
| Cenpj_in.vivo_noRT | Ush1g | 2.022 | 1.35E-03 | glio |
| Cenpj_in.vivo_noRT | Gm31063 | 2.022 | 1.35E-03 | glio |
| Chek1_in.vivo_noRT | Eif1 | -0.591 | 1.35E-03 | glio |
| Chek1_ex.vivo_noRT | Cd79a | 3.043 | 1.35E-03 | glio |
| Chek1_ex.vivo_noRT | 1700109H08R | 3.13 | 1.35E-03 | glio |
| Chek1_ex.vivo_noRT | Rps15a | -0.843 | 1.35E-03 | glio |
| Eif2s1_in.vivo_noRT | Muc4 | 2.144 | 1.35E-03 | glio |
| Eif2s1_in.vivo_noRT | Adam30 | 2.291 | 1.35E-03 | glio |
| Hsd17b10_in.vivo_noRT | Eps8l2 | 2.514 | 1.35E-03 | glio |
| Hsd17b10_in.vivo_noRT | Gm36913 | 2.458 | 1.35E-03 | glio |

| | | | | |
|---|---|---|---|---|
| culture_effect | Slc30a7 | -0.859 | 1.35E-04 | glio |
| culture_effect | Macrod2 | -0.903 | 1.35E-04 | glio |
| culture_effect | Rasa3 | 0.682 | 1.35E-04 | glio |
| Interaction_genotype_Aars2 | Hba-a2 | -4.895 | 1.35E-04 | glio |
| Interaction_genotype_Bcas2 | Ube3a | 1.782 | 1.35E-04 | glio |
| Interaction_genotype_Mtor | Postn | -4.553 | 1.35E-04 | glio |
| Interaction_genotype_Mtor | Mipep | -3.889 | 1.35E-04 | glio |
| Interaction_genotype_Mtor | B630019K06R | -3.5 | 1.35E-04 | glio |
| Interaction_genotype_Mtor | Matr3 | 1.669 | 1.35E-04 | glio |
| Interaction_genotype_Mtor | Rrp1 | 1.649 | 1.35E-04 | glio |
| Interaction_genotype_Mtor | H2-D1 | 1.947 | 1.35E-04 | glio |
| Interaction_genotype_Myc | Gm16133 | -4.448 | 1.35E-04 | glio |
| Aars2_in.vivo_noRT | Rpain | -1.502 | 1.35E-04 | glio |
| Aars2_in.vivo_noRT | Psmd2 | -1.205 | 1.35E-04 | glio |
| Atr_in.vivo_noRT | 2810455O05R | 2.758 | 1.35E-04 | glio |
| Atr_in.vivo_noRT | Ang | 2.758 | 1.35E-04 | glio |
| Atr_in.vivo_noRT | Bco1 | 2.254 | 1.35E-04 | glio |
| Atr_in.vivo_noRT | Ccdc71l | 2.498 | 1.35E-04 | glio |
| Atr_in.vivo_noRT | Oas1g | 2.619 | 1.35E-04 | glio |
| Atr_in.vivo_noRT | C130036L24Ri | 2.568 | 1.35E-04 | glio |
| Atr_in.vivo_noRT | Specc1 | -1.377 | 1.35E-04 | glio |
| Atr_ex.vivo_noRT | 4930470G03R | 2.771 | 1.35E-04 | glio |
| Bora_in.vivo_noRT | Fam171a2 | 2.774 | 1.35E-04 | glio |
| Bora_in.vivo_noRT | Asphd2 | 2.733 | 1.35E-04 | glio |
| Chek1_in.vivo_noRT | Bpifa1 | 2.517 | 1.35E-04 | glio |
| Chek1_ex.vivo_noRT | Ppm1b | -2.211 | 1.35E-04 | glio |
| Hsd17b10_in.vivo_noRT | Gm36757 | 2.754 | 1.35E-04 | glio |
| Hsd17b10_in.vivo_noRT | Rcor1 | -1.597 | 1.35E-04 | glio |
| culture_effect | Deptor | 1.452 | 1.35E-05 | glio |
| culture_effect | Magt1 | -0.648 | 1.35E-05 | glio |
| culture_effect | Prr13 | 0.689 | 1.35E-05 | glio |
| culture_effect | Timm23 | 0.453 | 1.35E-05 | glio |
| Interaction_genotype_Mtor | Car5a | -4.74 | 1.35E-05 | glio |
| Interaction_genotype_Mtor | Alox5ap | -5.848 | 1.35E-05 | glio |
| Atp5c1_in.vivo_noRT | Gm9758 | 2.408 | 1.35E-05 | glio |
| Atp5c1_in.vivo_noRT | Agap2 | 2.668 | 1.35E-05 | glio |
| Atr_in.vivo_noRT | Mr1 | 3.101 | 1.35E-05 | glio |
| Bcas2_in.vivo_noRT | Slc12a5 | 3.025 | 1.35E-05 | glio |
| Bcas2_in.vivo_noRT | Ncln | -1.641 | 1.35E-05 | glio |
| Bora_ex.vivo_noRT | Gm12860 | 3.201 | 1.35E-05 | glio |
| Bora_ex.vivo_noRT | Gm50024 | 2.802 | 1.35E-05 | glio |
| Bora_ex.vivo_noRT | Vmn2r44 | 2.806 | 1.35E-05 | glio |
| Chek1_in.vivo_noRT | Cbr3 | 3.291 | 1.35E-05 | glio |
| Chek1_in.vivo_noRT | Itgb6 | 3.116 | 1.35E-05 | glio |
| Interaction_genotype_Mtor | Gm45235 | -4.632 | 1.35E-06 | glio |
| Bcas2_in.vivo_noRT | 4930558J18Ri | 3.201 | 1.35E-06 | glio |
| culture_effect | Agap1 | -0.997 | 1.35E-07 | glio |
| Interaction_genotype_Mtor | Ubald2 | 3.426 | 1.35E-07 | glio |
| Interaction_genotype_Mtor | Gm29340 | -5.782 | 1.35E-07 | glio |
| Aars2_in.vivo_noRT | Axdnd1 | 4.014 | 1.35E-07 | glio |
| Bora_ex.vivo_noRT | Krt19 | 3.201 | 1.35E-07 | glio |
| Chek1_in.vivo_noRT | Raly | -1.477 | 1.35E-07 | glio |

| | | | | |
|---|---|---|---|---|
| Interaction_genotype_Atr | Folr1 | -5.94 | 1.35E-08 | glio |
| Aars2_in.vivo_noRT | Adam1a | 3.965 | 1.35E-08 | glio |
| Aars2_in.vivo_noRT | Stk36 | 4.038 | 1.35E-08 | glio |
| Aars2_in.vivo_noRT | Zc3hav1 | -2.16 | 1.35E-08 | glio |
| Atr_in.vivo_noRT | 1700011M02F | 5.27 | 1.35E-09 | glio |
| Bora_in.vivo_noRT | Esrp2 | 4.024 | 1.35E-09 | glio |
| Chek1_ex.vivo_noRT | Snhg4.1 | 4.449 | 1.35E-09 | glio |
| Hsd17b10_in.vivo_noRT | Gm50321 | 3.616 | 1.35E-09 | glio |
| culture_effect | Pter | 1.391 | 1.35E-10 | glio |
| Bora_in.vivo_noRT | Fgf1 | 4.678 | 1.35E-10 | glio |
| Bora_in.vivo_noRT | Dnm3 | 4.411 | 1.35E-10 | glio |
| Aars2_in.vivo_noRT | AI854703 | 4.676 | 1.35E-11 | glio |
| Chek1_in.vivo_noRT | CAAA0106612 | 4.252 | 1.35E-11 | glio |
| Aars2_in.vivo_noRT | Ttc16 | 4.804 | 1.35E-12 | glio |
| Aars2_in.vivo_noRT | Gjb5 | 4.336 | 1.35E-12 | glio |
| Chek1_in.vivo_noRT | Gm37864 | 4.264 | 1.35E-12 | glio |
| Chek1_ex.vivo_noRT | Spink6 | 4.842 | 1.35E-12 | glio |
| Chek1_ex.vivo_noRT | Gm15138 | 4.447 | 1.35E-12 | glio |
| Chek1_ex.vivo_noRT | Htr2b | 4.842 | 1.35E-12 | glio |
| culture_effect | Psma1 | -0.913 | 1.35E-13 | glio |
| culture_effect | Gm29340 | 3.638 | 1.35E-14 | glio |
| Bora_in.vivo_noRT | 4930412M03F | 5.46 | 1.35E-15 | glio |
| Bora_in.vivo_noRT | Gm15270 | 4.673 | 1.35E-15 | glio |
| culture_effect | Col12a1 | 3.113 | 1.35E-16 | glio |
| Atr_ex.vivo_noRT | Spink8 | 5.712 | 1.35E-18 | glio |
| Atr_ex.vivo_noRT | Gm33466 | 5.63 | 1.35E-18 | glio |
| Chek1_in.vivo_noRT | Sytl4 | 4.777 | 1.35E-20 | glio |
| Bcas2_in.vivo_noRT | Chia1 | 4.807 | 1.35E-21 | glio |
| Bora_in.vivo_noRT | Gm43701 | 4.936 | 1.35E-21 | glio |
| culture_effect | Baalc | 4.568 | 1.35E-22 | glio |
| culture_effect | Vgll3 | 5.155 | 1.35E-28 | glio |
| Aars2_in.vivo_noRT | Gm48435 | 5.327 | 1.35E-28 | glio |
| culture_effect | Trafd1 | -2.985 | 1.35E-35 | glio |
| culture_effect | Gm43466 | -1.802 | 1.36E-03 | glio |
| culture_effect | Gm20707 | 1.655 | 1.36E-03 | glio |
| culture_effect | Gng7 | 1.457 | 1.36E-03 | glio |
| culture_effect | Cdc20 | -0.782 | 1.36E-03 | glio |
| Interaction_genotype_Aars2 | 1110046J04Ri | -4.046 | 1.36E-03 | glio |
| Interaction_genotype_Aars2 | Tcap | -3.72 | 1.36E-03 | glio |
| Interaction_genotype_Aars2 | Gm30082 | -3.958 | 1.36E-03 | glio |
| Interaction_genotype_Aars2 | Gm10101 | -3.654 | 1.36E-03 | glio |
| Interaction_genotype_Aars2 | Mcmdc2 | -3.398 | 1.36E-03 | glio |
| Interaction_genotype_Aars2 | Lncbate10 | -2.821 | 1.36E-03 | glio |
| Interaction_genotype_Aars2 | Fdx1 | 1.865 | 1.36E-03 | glio |
| Interaction_genotype_Atr | Gm31415 | -3.398 | 1.36E-03 | glio |
| Interaction_genotype_Bcas2 | Gm11627 | -3.584 | 1.36E-03 | glio |
| Interaction_genotype_Chek1 | Dhx40 | 2.296 | 1.36E-03 | glio |
| Interaction_genotype_Chek1 | Ccpg1 | 2.853 | 1.36E-03 | glio |
| Interaction_genotype_Mtor | Tcam1 | -4.071 | 1.36E-03 | glio |
| Interaction_genotype_Mtor | Nek6 | -2.839 | 1.36E-03 | glio |
| Interaction_genotype_Mtor | Mcam | 2.421 | 1.36E-03 | glio |
| Interaction_genotype_Mtor | Rtcb | 1.429 | 1.36E-03 | glio |

| | | | | |
|---|---|---|---|---|
| Interaction_genotype_Myc | 4930542D17R | -3.538 | 1.36E-03 | glio |
| Interaction_genotype_Myc | Zfp575 | -3.815 | 1.36E-03 | glio |
| Interaction_genotype_Myc | Samd15 | -3.969 | 1.36E-03 | glio |
| Interaction_genotype_Myc | Sirt5 | -2.797 | 1.36E-03 | glio |
| Interaction_genotype_Myc | Gm17501 | -3.355 | 1.36E-03 | glio |
| interaction_RT | Wdyhv1 | 1.078 | 1.36E-03 | glio |
| Aars2_in.vivo_noRT | Prkcq | 1.947 | 1.36E-03 | glio |
| Atp5c1_in.vivo_noRT | Dcn | 2.277 | 1.36E-03 | glio |
| Atr_in.vivo_noRT | Sema3d | 2.304 | 1.36E-03 | glio |
| Atr_in.vivo_noRT | A930004D18R | 2.27 | 1.36E-03 | glio |
| Atr_in.vivo_noRT | Kctd12 | 2.198 | 1.36E-03 | glio |
| Atr_in.vivo_noRT | Ccdc136 | 2.186 | 1.36E-03 | glio |
| Atr_in.vivo_noRT | Timm23 | -0.767 | 1.36E-03 | glio |
| Atr_in.vivo_noRT | Lsm4 | -0.784 | 1.36E-03 | glio |
| Bcas2_in.vivo_noRT | 1700063D05R | 2.424 | 1.36E-03 | glio |
| Bcas2_in.vivo_noRT | Rad52 | 1.854 | 1.36E-03 | glio |
| Bcas2_ex.vivo_noRT | Gm48242 | 2.583 | 1.36E-03 | glio |
| Bora_in.vivo_noRT | Mt4 | 2.94 | 1.36E-03 | glio |
| Bora_in.vivo_noRT | Igf2r | 2.181 | 1.36E-03 | glio |
| Bora_in.vivo_noRT | Hist1h2af | 2.142 | 1.36E-03 | glio |
| Bora_in.vivo_noRT | Tmco3 | -1.481 | 1.36E-03 | glio |
| Bora_in.vivo_noRT | Plod1 | -1.429 | 1.36E-03 | glio |
| Bora_ex.vivo_noRT | Gm21818 | 3.201 | 1.36E-03 | glio |
| Bora_ex.vivo_noRT | Gm39459 | 3.117 | 1.36E-03 | glio |
| Bora_ex.vivo_noRT | Ctss | 2.808 | 1.36E-03 | glio |
| Cenpj_in.vivo_noRT | Insc | 2.327 | 1.36E-03 | glio |
| Cenpj_in.vivo_noRT | Chn2 | 2.611 | 1.36E-03 | glio |
| Chek1_ex.vivo_noRT | Hist1h4k | 2.985 | 1.36E-03 | glio |
| Chek1_ex.vivo_noRT | Gm15013 | 2.897 | 1.36E-03 | glio |
| Chek1_ex.vivo_noRT | Gm47232 | 2.851 | 1.36E-03 | glio |
| Eif2s1_in.vivo_noRT | Pcdhga11 | 2.207 | 1.36E-03 | glio |
| Eif2s1_in.vivo_noRT | Gm44646 | 2.143 | 1.36E-03 | glio |
| Eif2s1_in.vivo_noRT | Adam5 | 1.879 | 1.36E-03 | glio |
| Hsd17b10_in.vivo_noRT | Elk4 | -1.59 | 1.36E-03 | glio |
| culture_effect | Cryzl2 | -0.851 | 1.36E-04 | glio |
| culture_effect | B4galt1 | 0.879 | 1.36E-04 | glio |
| Interaction_genotype_Aars2 | Gnas | 1.123 | 1.36E-04 | glio |
| Interaction_genotype_Myc | Cdrt4 | -4.538 | 1.36E-04 | glio |
| Interaction_genotype_Myc | 1700066B19R | -4.067 | 1.36E-04 | glio |
| interaction_RT | Usf1 | 1.154 | 1.36E-04 | glio |
| interaction_RT | Lgals7 | 3.371 | 1.36E-04 | glio |
| Aars2_ex.vivo_noRT | Pla2g2d | 3.293 | 1.36E-04 | glio |
| Atp5c1_in.vivo_noRT | Gm34939 | 2.404 | 1.36E-04 | glio |
| Atr_in.vivo_noRT | Gfap | 2.506 | 1.36E-04 | glio |
| Bcas2_ex.vivo_noRT | Gm4166 | 2.742 | 1.36E-04 | glio |
| Cenpj_ex.vivo_noRT | Slc14a2 | 2.591 | 1.36E-04 | glio |
| Cenpj_ex.vivo_noRT | Gm4340 | 2.591 | 1.36E-04 | glio |
| Cenpj_ex.vivo_noRT | Gm16239 | 2.591 | 1.36E-04 | glio |
| Cenpj_ex.vivo_noRT | 4930527F14Ri | 2.994 | 1.36E-04 | glio |
| Chek1_in.vivo_noRT | Zfp493 | 2.695 | 1.36E-04 | glio |
| Chek1_in.vivo_noRT | Gm15892 | 2.9 | 1.36E-04 | glio |
| Chek1_in.vivo_noRT | Washc5 | -1.602 | 1.36E-04 | glio |

| | | | | |
|---|---|---|---|---|
| culture_effect | Sik2 | -1.185 | 1.36E-05 | glio |
| culture_effect | Epb41l4a | 1.012 | 1.36E-05 | glio |
| culture_effect | Tfrc | 0.774 | 1.36E-05 | glio |
| culture_effect | Lsm2 | 0.58 | 1.36E-05 | glio |
| culture_effect | Acp1 | 0.499 | 1.36E-05 | glio |
| Interaction_genotype_Mtor | Rtn1 | -5.348 | 1.36E-05 | glio |
| Aars2_in.vivo_noRT | Gm43773 | 2.905 | 1.36E-05 | glio |
| Aars2_in.vivo_noRT | Slc25a47 | 3.079 | 1.36E-05 | glio |
| Aars2_ex.vivo_noRT | Esp15 | 3.293 | 1.36E-05 | glio |
| Aars2_ex.vivo_noRT | Gm11884 | 3.693 | 1.36E-05 | glio |
| Atr_in.vivo_noRT | C030037D09R | 3.032 | 1.36E-05 | glio |
| Atr_in.vivo_noRT | Acrbp | 2.604 | 1.36E-05 | glio |
| Atr_in.vivo_noRT | Rps6 | -0.784 | 1.36E-05 | glio |
| Atr_ex.vivo_noRT | Minar2 | 3.164 | 1.36E-05 | glio |
| Atr_ex.vivo_noRT | Gm50024 | 2.764 | 1.36E-05 | glio |
| Bcas2_in.vivo_noRT | Grcc10 | -0.951 | 1.36E-05 | glio |
| Chek1_ex.vivo_noRT | 9930014A18R | 3.651 | 1.36E-05 | glio |
| Chek1_ex.vivo_noRT | Best1 | 3.563 | 1.36E-05 | glio |
| Hsd17b10_in.vivo_noRT | Gm20732 | 2.846 | 1.36E-05 | glio |
| culture_effect | Prrt3 | -2.722 | 1.36E-06 | glio |
| culture_effect | Mcam | -1.098 | 1.36E-06 | glio |
| Bora_in.vivo_noRT | Gm15473 | 3.554 | 1.36E-06 | glio |
| Bora_in.vivo_noRT | Sparc | -1.749 | 1.36E-06 | glio |
| Chek1_in.vivo_noRT | 0610009L18Ri | 3.532 | 1.36E-06 | glio |
| Chek1_ex.vivo_noRT | 4930555A03R | 3.871 | 1.36E-06 | glio |
| Chek1_ex.vivo_noRT | Gm17210 | 4.256 | 1.36E-06 | glio |
| Chek1_ex.vivo_noRT | Kcnip3 | 4.058 | 1.36E-06 | glio |
| culture_effect | Psma6 | -0.581 | 1.36E-07 | glio |
| culture_effect | Rps5 | -0.658 | 1.36E-07 | glio |
| Interaction_genotype_Myc | Gm36356 | -5.238 | 1.36E-07 | glio |
| Atr_in.vivo_noRT | Hyal1 | 3.563 | 1.36E-07 | glio |
| Bcas2_in.vivo_noRT | Lmod1 | 3.893 | 1.36E-07 | glio |
| Bcas2_in.vivo_noRT | Car12 | 3.484 | 1.36E-07 | glio |
| Bcas2_in.vivo_noRT | Gm15486 | 3.342 | 1.36E-07 | glio |
| Bora_in.vivo_noRT | C77080 | 3.583 | 1.36E-07 | glio |
| Hsd17b10_in.vivo_noRT | Gm50147 | 3.616 | 1.36E-07 | glio |
| Atr_in.vivo_noRT | 4930426I24Ri | 3.878 | 1.36E-08 | glio |
| Atr_in.vivo_noRT | 5430400D12R | 4.251 | 1.36E-09 | glio |
| Atr_in.vivo_noRT | Gm15246 | 4.182 | 1.36E-09 | glio |
| Bora_in.vivo_noRT | D6Ertd527e | 4.154 | 1.36E-09 | glio |
| Bora_in.vivo_noRT | Hectd2os | 4.028 | 1.36E-09 | glio |
| Bora_in.vivo_noRT | Sod3 | 4.936 | 1.36E-09 | glio |
| Chek1_ex.vivo_noRT | Rasl10a | 4.449 | 1.36E-09 | glio |
| Atr_in.vivo_noRT | Sema3a | 4.096 | 1.36E-10 | glio |
| Atr_in.vivo_noRT | Baalc | 4.741 | 1.36E-12 | glio |
| Bora_in.vivo_noRT | Gm47872 | 4.672 | 1.36E-12 | glio |
| Hsd17b10_in.vivo_noRT | 4930570E03Ri | 3.887 | 1.36E-12 | glio |
| Hsd17b10_in.vivo_noRT | Gm12841 | 3.616 | 1.36E-12 | glio |
| Hsd17b10_in.vivo_noRT | Gm49698 | 3.616 | 1.36E-13 | glio |
| Hsd17b10_in.vivo_noRT | Olfr881 | 3.616 | 1.36E-13 | glio |
| Aars2_in.vivo_noRT | Cyp4x1 | 5.06 | 1.36E-14 | glio |
| Aars2_in.vivo_noRT | Gm37035 | 4.41 | 1.36E-14 | glio |

| | | | | |
|---|---|---|---|---|
| Chek1_ex.vivo_noRT | 9130019O22R | 4.842 | 1.36E-14 | glio |
| Bora_in.vivo_noRT | Htr2b | 4.413 | 1.36E-15 | glio |
| Atr_in.vivo_noRT | Gm50147 | 5.533 | 1.36E-16 | glio |
| Chek1_in.vivo_noRT | Gm50321 | 5.044 | 1.36E-17 | glio |
| Aars2_in.vivo_noRT | Psmb11 | 4.804 | 1.36E-20 | glio |
| Bcas2_in.vivo_noRT | Gm36037 | 4.807 | 1.36E-20 | glio |
| culture_effect | Tbx2 | 3.12 | 1.36E-21 | glio |
| culture_effect | Scoc | 1.406 | 1.36E-21 | glio |
| culture_effect | Il15ra | -3.086 | 1.36E-22 | glio |
| culture_effect | Klf5 | -2.169 | 1.36E-22 | glio |
| Aars2_in.vivo_noRT | Gm35439 | 5.457 | 1.36E-22 | glio |
| Aars2_in.vivo_noRT | 9130019O22R | 5.327 | 1.36E-24 | glio |
| Chek1_in.vivo_noRT | Slc14a2 | 5.044 | 1.36E-25 | glio |
| Atr_in.vivo_noRT | Gm8016 | 5.533 | 1.36E-27 | glio |
| culture_effect | Rab4a | 0.778 | 1.37E-03 | glio |
| culture_effect | Hprt | -0.504 | 1.37E-03 | glio |
| culture_effect | Mettl9 | 0.503 | 1.37E-03 | glio |
| culture_effect | Exosc5 | -0.503 | 1.37E-03 | glio |
| Interaction_genotype_Aars2 | Upb1 | -3.842 | 1.37E-03 | glio |
| Interaction_genotype_Bora | Cabp4 | -4.303 | 1.37E-03 | glio |
| Interaction_genotype_Bora | Gm35315 | -4.188 | 1.37E-03 | glio |
| Interaction_genotype_Bora | Irf8 | 3.963 | 1.37E-03 | glio |
| Interaction_genotype_Mtor | B430010I23Ri | -3.625 | 1.37E-03 | glio |
| Interaction_genotype_Mtor | Slamf1 | -3.928 | 1.37E-03 | glio |
| Interaction_genotype_Mtor | Cmtm8 | -4.009 | 1.37E-03 | glio |
| Interaction_genotype_Mtor | Apbb1 | -2.775 | 1.37E-03 | glio |
| Interaction_genotype_Mtor | Nop2 | 1.87 | 1.37E-03 | glio |
| Interaction_genotype_Mtor | Dag1 | 1.956 | 1.37E-03 | glio |
| Interaction_genotype_Mtor | Smap1 | 1.601 | 1.37E-03 | glio |
| Interaction_genotype_Mtor | Clic1 | 1.401 | 1.37E-03 | glio |
| Interaction_genotype_Myc | Cnih2 | -3.612 | 1.37E-03 | glio |
| Interaction_genotype_Myc | Phkg1 | -4.953 | 1.37E-03 | glio |
| Interaction_genotype_Myc | Gm26901 | -3.861 | 1.37E-03 | glio |
| Interaction_genotype_Myc | Veph1 | -3.527 | 1.37E-03 | glio |
| Interaction_genotype_Myc | Med18 | -2.885 | 1.37E-03 | glio |
| Interaction_genotype_Myc | Polk | -2.799 | 1.37E-03 | glio |
| Interaction_genotype_Myc | 1600012H06R | -2.767 | 1.37E-03 | glio |
| Interaction_genotype_Rpl3 | Ak1 | -4.206 | 1.37E-03 | glio |
| Aars2_in.vivo_noRT | Taco1os | 2.157 | 1.37E-03 | glio |
| Aars2_in.vivo_noRT | E130102H24R | 2.215 | 1.37E-03 | glio |
| Aars2_in.vivo_noRT | Nub1 | -1.232 | 1.37E-03 | glio |
| Aars2_in.vivo_noRT | Capzb | -1.029 | 1.37E-03 | glio |
| Aars2_ex.vivo_noRT | Prm3 | 2.713 | 1.37E-03 | glio |
| Atp5c1_in.vivo_noRT | Snhg4.1 | 2.145 | 1.37E-03 | glio |
| Atp5c1_in.vivo_noRT | Gm39459 | 2.548 | 1.37E-03 | glio |
| Atp5c1_in.vivo_noRT | Wnt16 | 2.408 | 1.37E-03 | glio |
| Atr_in.vivo_noRT | Fndc5 | 2.295 | 1.37E-03 | glio |
| Bcas2_in.vivo_noRT | Pdap1 | -0.764 | 1.37E-03 | glio |
| Bora_in.vivo_noRT | Hist1h3g | 2.474 | 1.37E-03 | glio |
| Bora_in.vivo_noRT | Dusp15 | 2.087 | 1.37E-03 | glio |
| Bora_in.vivo_noRT | Arhgap26 | 2.215 | 1.37E-03 | glio |
| Bora_ex.vivo_noRT | Pldi | 2.507 | 1.37E-03 | glio |

| | | | | |
|---|---|---|---|---|
| Bora_ex.vivo_noRT | Prl3d3 | 2.106 | 1.37E-03 | glio |
| Bora_ex.vivo_noRT | Sgk2 | 3.206 | 1.37E-03 | glio |
| Bora_ex.vivo_noRT | Pard3bos1 | 2.815 | 1.37E-03 | glio |
| Cenpj_ex.vivo_noRT | Gm21936 | 2.196 | 1.37E-03 | glio |
| Cenpj_ex.vivo_noRT | 4930588A03R | 2.192 | 1.37E-03 | glio |
| Cenpj_ex.vivo_noRT | Lmx1a | 2.591 | 1.37E-03 | glio |
| Cenpj_ex.vivo_noRT | Vmn2r56 | 2.591 | 1.37E-03 | glio |
| Chek1_in.vivo_noRT | Naip6 | 2.258 | 1.37E-03 | glio |
| Chek1_in.vivo_noRT | Gm4419 | 2.033 | 1.37E-03 | glio |
| Chek1_in.vivo_noRT | Sema3d | 2.338 | 1.37E-03 | glio |
| Chek1_in.vivo_noRT | Peg3 | -1.601 | 1.37E-03 | glio |
| Chek1_in.vivo_noRT | Slc29a1 | -1.585 | 1.37E-03 | glio |
| Chek1_ex.vivo_noRT | Rit2 | 2.772 | 1.37E-03 | glio |
| Chek1_ex.vivo_noRT | Lifr | 2.713 | 1.37E-03 | glio |
| Hsd17b10_in.vivo_noRT | Retn | 3.149 | 1.37E-03 | glio |
| Hsd17b10_in.vivo_noRT | A430057M04F | 2.431 | 1.37E-03 | glio |
| culture_effect | Map3k4 | 0.647 | 1.37E-04 | glio |
| Interaction_genotype_Mtor | Fam161b | -4.641 | 1.37E-04 | glio |
| Interaction_genotype_Mtor | Inpp4b | -4.266 | 1.37E-04 | glio |
| Interaction_genotype_Mtor | Mapk3 | 2.386 | 1.37E-04 | glio |
| Interaction_genotype_Myc | Gm6225 | -4.591 | 1.37E-04 | glio |
| Interaction_genotype_Myc | Arhgap31 | -3.579 | 1.37E-04 | glio |
| Aars2_in.vivo_noRT | Amt | 2.944 | 1.37E-04 | glio |
| Aars2_in.vivo_noRT | Oas1g | 2.628 | 1.37E-04 | glio |
| Aars2_ex.vivo_noRT | Cxcr2 | 3.506 | 1.37E-04 | glio |
| Atp5c1_in.vivo_noRT | Zfp352 | 2.668 | 1.37E-04 | glio |
| Atp5c1_in.vivo_noRT | Grip2 | 2.894 | 1.37E-04 | glio |
| Atr_in.vivo_noRT | Ly6g6d | 2.703 | 1.37E-04 | glio |
| Bcas2_in.vivo_noRT | Slc6a19 | 2.638 | 1.37E-04 | glio |
| Bcas2_in.vivo_noRT | Zfp599 | 2.904 | 1.37E-04 | glio |
| Bora_in.vivo_noRT | Bcl2l14 | 2.895 | 1.37E-04 | glio |
| Bora_in.vivo_noRT | Ccdc146 | 2.862 | 1.37E-04 | glio |
| Chek1_in.vivo_noRT | AY074887 | 2.991 | 1.37E-04 | glio |
| Chek1_in.vivo_noRT | Mthfs | 2.368 | 1.37E-04 | glio |
| Chek1_ex.vivo_noRT | Meaf6 | -1.772 | 1.37E-04 | glio |
| culture_effect | 4930556M19F | 1.959 | 1.37E-05 | glio |
| Interaction_genotype_Aars2 | Appbp2os | -5.614 | 1.37E-05 | glio |
| Interaction_genotype_Myc | Fzd1 | -4.708 | 1.37E-05 | glio |
| Interaction_genotype_Myc | Zfp11 | -4.954 | 1.37E-05 | glio |
| Interaction_genotype_Myc | Smdt1 | 1.799 | 1.37E-05 | glio |
| Aars2_in.vivo_noRT | Glmp | 3.324 | 1.37E-05 | glio |
| Aars2_in.vivo_noRT | Zfp418 | 3.271 | 1.37E-05 | glio |
| Aars2_in.vivo_noRT | Gm47591 | 3.101 | 1.37E-05 | glio |
| Aars2_in.vivo_noRT | Tbrg1 | -1.266 | 1.37E-05 | glio |
| Atp5c1_in.vivo_noRT | Plcl1 | 2.668 | 1.37E-05 | glio |
| Atr_in.vivo_noRT | Crybg2 | 3.009 | 1.37E-05 | glio |
| Atr_in.vivo_noRT | Insrr | 2.757 | 1.37E-05 | glio |
| Atr_in.vivo_noRT | Eif3l | -1.252 | 1.37E-05 | glio |
| Atr_ex.vivo_noRT | Vmn2r44 | 2.769 | 1.37E-05 | glio |
| Bcas2_in.vivo_noRT | Gm49135 | 3.124 | 1.37E-05 | glio |
| Bcas2_in.vivo_noRT | Nckipsd | 2.98 | 1.37E-05 | glio |
| Bora_in.vivo_noRT | Lbhd2 | 3.695 | 1.37E-05 | glio |

| | | | | |
|---|---|---|---|---|
| Bora_in.vivo_noRT | Upk3bl | 2.86 | 1.37E-05 | glio |
| Bora_ex.vivo_noRT | Gast | 3.201 | 1.37E-05 | glio |
| Bora_ex.vivo_noRT | Akr1c18 | 3.201 | 1.37E-05 | glio |
| Bora_ex.vivo_noRT | Minar2 | 3.201 | 1.37E-05 | glio |
| Bora_ex.vivo_noRT | Gm45151 | 3.201 | 1.37E-05 | glio |
| Chek1_in.vivo_noRT | Fhad1 | 2.848 | 1.37E-05 | glio |
| culture_effect | Arhgap23 | -1.098 | 1.37E-06 | glio |
| culture_effect | Tubb2a | -0.845 | 1.37E-06 | glio |
| interaction_RT | Tmbim4 | 1.015 | 1.37E-06 | glio |
| Atr_in.vivo_noRT | Fgfr1op2 | -1.395 | 1.37E-06 | glio |
| Atr_ex.vivo_noRT | Sytl4 | 3.164 | 1.37E-06 | glio |
| Bora_in.vivo_noRT | Mir155hg | 3.691 | 1.37E-06 | glio |
| Chek1_in.vivo_noRT | Ccl20 | 5.044 | 1.37E-06 | glio |
| Aars2_in.vivo_noRT | Sept5 | 3.587 | 1.37E-07 | glio |
| Atp5c1_in.vivo_noRT | Gm15929 | 3.19 | 1.37E-07 | glio |
| Atr_in.vivo_noRT | 9530053A07R | 3.719 | 1.37E-07 | glio |
| Atr_in.vivo_noRT | Gm48832 | 3.517 | 1.37E-07 | glio |
| Bcas2_in.vivo_noRT | Gm21814 | 3.478 | 1.37E-07 | glio |
| Bcas2_in.vivo_noRT | Rab36 | 3.702 | 1.37E-07 | glio |
| Bora_in.vivo_noRT | Gm26542 | 3.558 | 1.37E-07 | glio |
| interaction_RT | H2afv | 0.983 | 1.37E-08 | glio |
| Atr_in.vivo_noRT | Myom1 | 4.077 | 1.37E-08 | glio |
| Bcas2_in.vivo_noRT | A530072M11F | 3.996 | 1.37E-08 | glio |
| Bcas2_in.vivo_noRT | Man2c1os | 4.162 | 1.37E-08 | glio |
| Bora_in.vivo_noRT | Kctd21 | 3.899 | 1.37E-08 | glio |
| Chek1_in.vivo_noRT | Gm8212 | 3.942 | 1.37E-08 | glio |
| Chek1_in.vivo_noRT | Cxcr6 | 3.915 | 1.37E-08 | glio |
| Chek1_ex.vivo_noRT | Rtl1 | 4.447 | 1.37E-08 | glio |
| Aars2_in.vivo_noRT | Tomm6os | 4.409 | 1.37E-09 | glio |
| Chek1_in.vivo_noRT | Gm16853 | 4.01 | 1.37E-09 | glio |
| Chek1_in.vivo_noRT | Pcdhgb5 | 3.651 | 1.37E-09 | glio |
| culture_effect | Fam210b | -1.378 | 1.37E-10 | glio |
| culture_effect | Klf6 | -1.207 | 1.37E-10 | glio |
| Atr_in.vivo_noRT | Npb | 4.366 | 1.37E-10 | glio |
| Chek1_in.vivo_noRT | Gm5039 | 5.044 | 1.37E-10 | glio |
| Bcas2_in.vivo_noRT | Nxnl2 | 4.285 | 1.37E-11 | glio |
| Hsd17b10_in.vivo_noRT | Gm12703 | 3.35 | 1.37E-11 | glio |
| Aars2_in.vivo_noRT | Olfr1191-ps1 | 4.676 | 1.37E-13 | glio |
| Atr_in.vivo_noRT | Cd52 | 4.876 | 1.37E-13 | glio |
| culture_effect | Crabp1 | 3.939 | 1.37E-18 | glio |
| Bcas2_in.vivo_noRT | Spink8 | 4.547 | 1.37E-18 | glio |
| Bcas2_in.vivo_noRT | Ush1g | 4.807 | 1.37E-19 | glio |
| culture_effect | B230208H11R | 1.628 | 1.38E-03 | glio |
| Interaction_genotype_Aars2 | Gm16759 | -4.085 | 1.38E-03 | glio |
| Interaction_genotype_Atr | Slc17a6 | -2.369 | 1.38E-03 | glio |
| Interaction_genotype_Atr | Sult6b1 | -3.997 | 1.38E-03 | glio |
| Interaction_genotype_Atr | Trim17 | -2.769 | 1.38E-03 | glio |
| Interaction_genotype_Atr | Ccl9 | -2.762 | 1.38E-03 | glio |
| Interaction_genotype_Atr | D030040B21R | -3.664 | 1.38E-03 | glio |
| Interaction_genotype_Atr | Gm26981 | -3.959 | 1.38E-03 | glio |
| Interaction_genotype_Atr | Tmx4 | 2.021 | 1.38E-03 | glio |
| Interaction_genotype_Atr | Copg1 | 1.826 | 1.38E-03 | glio |

| | | | | |
|---|---|---|---|---|
| Interaction_genotype_Atr | H13 | 1.591 | 1.38E-03 | glio |
| Interaction_genotype_Bora | Selenov | -3.505 | 1.38E-03 | glio |
| Interaction_genotype_Mtor | Dchs1 | -3.79 | 1.38E-03 | glio |
| Interaction_genotype_Mtor | A630072M18R | -3.022 | 1.38E-03 | glio |
| Interaction_genotype_Mtor | Ppib | 1.447 | 1.38E-03 | glio |
| Interaction_genotype_Myc | Gm2022 | -2.729 | 1.38E-03 | glio |
| Interaction_genotype_Myc | Gm42658 | -3.348 | 1.38E-03 | glio |
| Interaction_genotype_Myc | Rit2 | -3.487 | 1.38E-03 | glio |
| Interaction_genotype_Myc | Kank2 | -3.347 | 1.38E-03 | glio |
| Interaction_genotype_Myc | Gm42997 | -3.256 | 1.38E-03 | glio |
| Interaction_genotype_Myc | 2700033N17R | -3.106 | 1.38E-03 | glio |
| Interaction_genotype_Myc | Evc | -3.002 | 1.38E-03 | glio |
| Interaction_genotype_Myc | Rtel1 | -2.58 | 1.38E-03 | glio |
| Interaction_genotype_Myc | Gtf2f2 | -2.203 | 1.38E-03 | glio |
| Interaction_genotype_Myc | Timm17a | 1.276 | 1.38E-03 | glio |
| Interaction_genotype_Pars2 | Olfm2 | -4.062 | 1.38E-03 | glio |
| Interaction_genotype_Pars2 | Abcb5 | -3.062 | 1.38E-03 | glio |
| Interaction_genotype_Pars2 | Gm10324 | -3.365 | 1.38E-03 | glio |
| Interaction_genotype_Pars2 | Fgf2 | -3.636 | 1.38E-03 | glio |
| Interaction_genotype_Rad21 | Gm11627 | -4.046 | 1.38E-03 | glio |
| interaction_RT | Gm13056 | 2.569 | 1.38E-03 | glio |
| interaction_RT | Ap3s1 | 0.846 | 1.38E-03 | glio |
| Aars2_in.vivo_noRT | Helz | -1.235 | 1.38E-03 | glio |
| Atp5c1_in.vivo_noRT | Gm49874 | 2.668 | 1.38E-03 | glio |
| Atp5c1_in.vivo_noRT | Nppc | 2.716 | 1.38E-03 | glio |
| Atr_in.vivo_noRT | Rnf225 | 2.426 | 1.38E-03 | glio |
| Atr_in.vivo_noRT | Meaf6 | -1.265 | 1.38E-03 | glio |
| Atr_ex.vivo_noRT | Slc14a1 | 2.589 | 1.38E-03 | glio |
| Atr_ex.vivo_noRT | C3 | 3.164 | 1.38E-03 | glio |
| Bard1_ex.vivo_noRT | Icam2 | 2.814 | 1.38E-03 | glio |
| Bcas2_in.vivo_noRT | Lmln | 2.272 | 1.38E-03 | glio |
| Bcas2_in.vivo_noRT | Zmynd11 | -1.376 | 1.38E-03 | glio |
| Bora_in.vivo_noRT | Sbk1 | 2.378 | 1.38E-03 | glio |
| Bora_in.vivo_noRT | Phldb3 | 2.35 | 1.38E-03 | glio |
| Bora_in.vivo_noRT | Gm904 | 2.374 | 1.38E-03 | glio |
| Bora_in.vivo_noRT | Zfp668 | 1.927 | 1.38E-03 | glio |
| Bora_in.vivo_noRT | Rab3gap2 | -1.589 | 1.38E-03 | glio |
| Bora_in.vivo_noRT | Usp34 | -0.932 | 1.38E-03 | glio |
| Cenpj_in.vivo_noRT | Gm49172 | 1.762 | 1.38E-03 | glio |
| Chek1_in.vivo_noRT | Myo1b | 2.659 | 1.38E-03 | glio |
| Chek1_in.vivo_noRT | Slc25a42 | 2.361 | 1.38E-03 | glio |
| Chek1_in.vivo_noRT | Gm21154 | 2.486 | 1.38E-03 | glio |
| Chek1_in.vivo_noRT | Hist1h2bg | 2.426 | 1.38E-03 | glio |
| Chek1_in.vivo_noRT | Ift20 | -0.948 | 1.38E-03 | glio |
| Eif2s1_in.vivo_noRT | Ccdc125 | 2.143 | 1.38E-03 | glio |
| Eif2s1_in.vivo_noRT | Nrros | 2.562 | 1.38E-03 | glio |
| Eif2s1_in.vivo_noRT | Fbn2 | -1.478 | 1.38E-03 | glio |
| Hsd17b10_in.vivo_noRT | Gm15941 | 2.197 | 1.38E-03 | glio |
| culture_effect | Slitrk5 | -1.721 | 1.38E-04 | glio |
| culture_effect | Tle6 | -1.406 | 1.38E-04 | glio |
| Interaction_genotype_Bora | Pmp2 | -4.451 | 1.38E-04 | glio |
| Interaction_genotype_Mtor | Rhov | -4.516 | 1.38E-04 | glio |

| | | | | |
|---|---|---|---|---|
| Interaction_genotype_Mtor | Gm12258 | -4.679 | 1.38E-04 | glio |
| Interaction_genotype_Mtor | Mosmo | 1.904 | 1.38E-04 | glio |
| Interaction_genotype_Myc | Gm24362 | -3.966 | 1.38E-04 | glio |
| Interaction_genotype_Myc | Gm16283 | -4.092 | 1.38E-04 | glio |
| Interaction_genotype_Myc | Eng | -4.63 | 1.38E-04 | glio |
| Interaction_genotype_Myc | Tmem192 | 1.951 | 1.38E-04 | glio |
| Interaction_genotype_Myc | Ufm1 | 1.563 | 1.38E-04 | glio |
| Atp5c1_in.vivo_noRT | Gm35439 | 2.289 | 1.38E-04 | glio |
| Atp5c1_in.vivo_noRT | Gm36198 | 2.408 | 1.38E-04 | glio |
| Bard1_ex.vivo_noRT | Ngb | 3.596 | 1.38E-04 | glio |
| Bcas2_in.vivo_noRT | Pcdhga9 | 2.6 | 1.38E-04 | glio |
| Bcas2_ex.vivo_noRT | Smpd5 | 3.328 | 1.38E-04 | glio |
| Bora_in.vivo_noRT | Gstm2 | 2.812 | 1.38E-04 | glio |
| Bora_in.vivo_noRT | B430219N15R | 2.812 | 1.38E-04 | glio |
| Bora_in.vivo_noRT | Cav2 | -1.835 | 1.38E-04 | glio |
| Chek1_in.vivo_noRT | Myl6b | 2.639 | 1.38E-04 | glio |
| Chek1_ex.vivo_noRT | Eepd1 | 3.287 | 1.38E-04 | glio |
| Eif2s1_in.vivo_noRT | Gm13078 | 2.143 | 1.38E-04 | glio |
| culture_effect | Clec1a | -2.619 | 1.38E-05 | glio |
| culture_effect | Fabp7 | -2.248 | 1.38E-05 | glio |
| Interaction_genotype_Aars2 | Col6a2 | -4.092 | 1.38E-05 | glio |
| Interaction_genotype_Aars2 | Gm19276 | -3.839 | 1.38E-05 | glio |
| Interaction_genotype_Mtor | Pafah2 | -5.608 | 1.38E-05 | glio |
| Atr_in.vivo_noRT | Gm43768 | 2.895 | 1.38E-05 | glio |
| Atr_ex.vivo_noRT | Gast | 3.164 | 1.38E-05 | glio |
| Atr_ex.vivo_noRT | B230303O12R | 4.138 | 1.38E-05 | glio |
| Chek1_in.vivo_noRT | Arvcf | 3.271 | 1.38E-05 | glio |
| Chek1_in.vivo_noRT | Clcn2 | 3.216 | 1.38E-05 | glio |
| culture_effect | Slc6a6 | 0.704 | 1.38E-06 | glio |
| Atp5c1_in.vivo_noRT | AF067063 | 2.668 | 1.38E-06 | glio |
| Atr_ex.vivo_noRT | Trmt61b | 3.164 | 1.38E-06 | glio |
| Bora_in.vivo_noRT | Pkhd1 | 3.419 | 1.38E-06 | glio |
| Bora_in.vivo_noRT | Art2b | 2.847 | 1.38E-06 | glio |
| Chek1_in.vivo_noRT | Gm12905 | 3.739 | 1.38E-06 | glio |
| Chek1_ex.vivo_noRT | Fas | 4.715 | 1.38E-06 | glio |
| culture_effect | Ythdf2 | 0.792 | 1.38E-07 | glio |
| Interaction_genotype_Mtor | 9530018F02Ri | -6.735 | 1.38E-07 | glio |
| Interaction_genotype_Mtor | Sprr1a | -6.471 | 1.38E-07 | glio |
| Aars2_in.vivo_noRT | Synpr | 4.164 | 1.38E-07 | glio |
| Aars2_in.vivo_noRT | Hipk4 | 3.34 | 1.38E-07 | glio |
| Atp5c1_in.vivo_noRT | Ankrd22 | 2.668 | 1.38E-07 | glio |
| Atr_in.vivo_noRT | 4933406P04R | 3.852 | 1.38E-07 | glio |
| Atr_in.vivo_noRT | Rxra | 3.832 | 1.38E-07 | glio |
| Bora_ex.vivo_noRT | Gm48155 | 3.201 | 1.38E-07 | glio |
| culture_effect | Zfp622 | -0.819 | 1.38E-08 | glio |
| culture_effect | Rpl14 | -0.668 | 1.38E-08 | glio |
| Bcas2_in.vivo_noRT | Gm16853 | 3.773 | 1.38E-08 | glio |
| Chek1_in.vivo_noRT | Lrrc46 | 3.919 | 1.38E-08 | glio |
| Hsd17b10_in.vivo_noRT | Obscn | 3.616 | 1.38E-08 | glio |
| culture_effect | Gm49864 | 3.025 | 1.38E-09 | glio |
| culture_effect | Dpy19l1 | -1.319 | 1.38E-09 | glio |
| culture_effect | Itgb1 | 0.767 | 1.38E-10 | glio |

| | | | | |
|---|---|---|---|---|
| Bora_in.vivo_noRT | Vmn1r181 | 4.28 | 1.38E-10 | glio |
| Chek1_in.vivo_noRT | 2010320O07R | 4.393 | 1.38E-10 | glio |
| Chek1_in.vivo_noRT | Kcnq2 | 3.88 | 1.38E-10 | glio |
| Bcas2_in.vivo_noRT | 1700026J14Ri | 4.285 | 1.38E-11 | glio |
| Bora_in.vivo_noRT | Gm17173 | 4.549 | 1.38E-11 | glio |
| culture_effect | Eif4a1 | -0.919 | 1.38E-12 | glio |
| Bcas2_in.vivo_noRT | Gm43256 | 4.544 | 1.38E-12 | glio |
| Chek1_in.vivo_noRT | Dntt | 5.044 | 1.38E-12 | glio |
| Aars2_in.vivo_noRT | Spn | 4.805 | 1.38E-13 | glio |
| Bcas2_in.vivo_noRT | Pmp2 | 4.807 | 1.38E-14 | glio |
| Atr_in.vivo_noRT | Stra8 | 5.01 | 1.38E-16 | glio |
| Bora_in.vivo_noRT | Scel | 4.018 | 1.38E-17 | glio |
| Bcas2_in.vivo_noRT | Gm31063 | 4.807 | 1.38E-19 | glio |
| Bora_in.vivo_noRT | Fabp6 | 4.936 | 1.38E-20 | glio |
| Aars2_in.vivo_noRT | Ltbp2 | 5.327 | 1.38E-21 | glio |
| Aars2_in.vivo_noRT | Lmx1a | 5.327 | 1.38E-21 | glio |
| Bora_in.vivo_noRT | 9130019O22R | 4.936 | 1.38E-21 | glio |
| Chek1_in.vivo_noRT | Gm48215 | 5.044 | 1.38E-21 | glio |
| Bora_in.vivo_noRT | Capn13 | 4.936 | 1.38E-26 | glio |
| culture_effect | Adamts12 | 4.23 | 1.38E-29 | glio |
| culture_effect | H2-Ab1 | -10.125 | 1.38E-63 | glio |
| culture_effect | Icam2 | -1.801 | 1.39E-03 | glio |
| culture_effect | Ccr2 | -2.049 | 1.39E-03 | glio |
| culture_effect | 3110040N11R | 0.713 | 1.39E-03 | glio |
| culture_effect | Ppp2r5e | 0.551 | 1.39E-03 | glio |
| Interaction_genotype_Aars2 | Chia1 | -2.821 | 1.39E-03 | glio |
| Interaction_genotype_Atr | Fbxl13 | -3.484 | 1.39E-03 | glio |
| Interaction_genotype_Bora | Ankrd34a | -4.33 | 1.39E-03 | glio |
| Interaction_genotype_Mtor | 4930555A03R | -3.54 | 1.39E-03 | glio |
| Interaction_genotype_Mtor | Frmd4b | -3.968 | 1.39E-03 | glio |
| Interaction_genotype_Mtor | Stk11ip | -3.42 | 1.39E-03 | glio |
| Interaction_genotype_Mtor | Zfp202 | -3.926 | 1.39E-03 | glio |
| Interaction_genotype_Mtor | Palb2 | -3.028 | 1.39E-03 | glio |
| Interaction_genotype_Myc | Zfp808 | -3.176 | 1.39E-03 | glio |
| Interaction_genotype_Myc | 4732471J01Ri | -3.65 | 1.39E-03 | glio |
| Interaction_genotype_Myc | Rabl3 | -3.006 | 1.39E-03 | glio |
| Interaction_genotype_Myc | Plekhf1 | -3.302 | 1.39E-03 | glio |
| Interaction_genotype_Myc | Fadd | -2.452 | 1.39E-03 | glio |
| Interaction_genotype_Myc | 4932438A13R | -2.417 | 1.39E-03 | glio |
| Interaction_genotype_Myc | Ghitm | 1.486 | 1.39E-03 | glio |
| Interaction_genotype_Myc | Ndufc1 | 1.438 | 1.39E-03 | glio |
| Interaction_genotype_Myc | Ube2b | 1.054 | 1.39E-03 | glio |
| interaction_RT | Snhg16 | 1.202 | 1.39E-03 | glio |
| interaction_RT | Nomo1 | -0.746 | 1.39E-03 | glio |
| Aars2_in.vivo_noRT | 4632404H12R | 2.294 | 1.39E-03 | glio |
| Aars2_in.vivo_noRT | Gm9895 | 2.346 | 1.39E-03 | glio |
| Aars2_in.vivo_noRT | B230307C23R | 1.812 | 1.39E-03 | glio |
| Aars2_in.vivo_noRT | Pcgf5 | -1.428 | 1.39E-03 | glio |
| Aars2_ex.vivo_noRT | Bcl2l15 | 2.718 | 1.39E-03 | glio |
| Aars2_ex.vivo_noRT | Sema6d | 2.311 | 1.39E-03 | glio |
| Atp5c1_in.vivo_noRT | Sh3d21 | 2.145 | 1.39E-03 | glio |
| Atr_in.vivo_noRT | Amz1 | 2.205 | 1.39E-03 | glio |

| | | | | |
|---|---|---|---|---|
| Atr_in.vivo_noRT | Gm16537 | 2.037 | 1.39E-03 | glio |
| Atr_in.vivo_noRT | Hacl1 | 2.07 | 1.39E-03 | glio |
| Atr_in.vivo_noRT | Rrs1 | -1.161 | 1.39E-03 | glio |
| Bcas2_in.vivo_noRT | Rab15 | 2.298 | 1.39E-03 | glio |
| Bcas2_in.vivo_noRT | Gja4 | 2.219 | 1.39E-03 | glio |
| Bora_in.vivo_noRT | Stard10 | 2.27 | 1.39E-03 | glio |
| Bora_in.vivo_noRT | Ptpdc1 | 2.152 | 1.39E-03 | glio |
| Bora_in.vivo_noRT | Tmem184c | -1.601 | 1.39E-03 | glio |
| Bora_ex.vivo_noRT | Gm43625 | 2.808 | 1.39E-03 | glio |
| Brca2_in.vivo_noRT | Slc17a6 | 1.763 | 1.39E-03 | glio |
| Cenpj_in.vivo_noRT | Zfp317 | -1.964 | 1.39E-03 | glio |
| Cenpj_ex.vivo_noRT | Snrpn | 2.591 | 1.39E-03 | glio |
| Chek1_in.vivo_noRT | Flrt2 | 2.381 | 1.39E-03 | glio |
| Chek1_in.vivo_noRT | Rbpms2 | 2.331 | 1.39E-03 | glio |
| Chek1_in.vivo_noRT | Supt16 | -1.107 | 1.39E-03 | glio |
| Chek1_in.vivo_noRT | Gbp3 | -0.934 | 1.39E-03 | glio |
| Eif2s1_in.vivo_noRT | Agap2 | 2.143 | 1.39E-03 | glio |
| Hsd17b10_in.vivo_noRT | Sema4a | 2.573 | 1.39E-03 | glio |
| culture_effect | Trim12c | 0.756 | 1.39E-04 | glio |
| culture_effect | Brd8 | -0.618 | 1.39E-04 | glio |
| Interaction_genotype_Mtor | Gm42726 | -4.462 | 1.39E-04 | glio |
| Interaction_genotype_Mtor | 1700001G11R | -3.538 | 1.39E-04 | glio |
| Interaction_genotype_Mtor | Cdkn1b | 2.469 | 1.39E-04 | glio |
| Interaction_genotype_Myc | Amt | -4.691 | 1.39E-04 | glio |
| Interaction_genotype_Myc | Gpr161 | -4.476 | 1.39E-04 | glio |
| Interaction_genotype_Myc | Taf10 | 1.291 | 1.39E-04 | glio |
| Interaction_genotype_Pars2 | Gm27239 | -5.016 | 1.39E-04 | glio |
| Aars2_in.vivo_noRT | Gm13203 | 2.827 | 1.39E-04 | glio |
| Aars2_in.vivo_noRT | Ece2 | 2.558 | 1.39E-04 | glio |
| Aars2_in.vivo_noRT | Fam50a | -1.571 | 1.39E-04 | glio |
| Atr_in.vivo_noRT | Crocc | 2.622 | 1.39E-04 | glio |
| Bard1_ex.vivo_noRT | 4921504E06Ri | 2.899 | 1.39E-04 | glio |
| Bcas2_in.vivo_noRT | Stard10 | 2.67 | 1.39E-04 | glio |
| Bcas2_in.vivo_noRT | Fam210a | -1.815 | 1.39E-04 | glio |
| Bcas2_ex.vivo_noRT | Olfr56 | 2.742 | 1.39E-04 | glio |
| Bora_in.vivo_noRT | Hist1h2ai | 2.84 | 1.39E-04 | glio |
| Bora_ex.vivo_noRT | 9930111J21Ri | 3.201 | 1.39E-04 | glio |
| Bora_ex.vivo_noRT | Gm15326 | 2.626 | 1.39E-04 | glio |
| Cenpj_ex.vivo_noRT | Hoxc8 | 2.591 | 1.39E-04 | glio |
| Chek1_in.vivo_noRT | Gm30074 | 2.676 | 1.39E-04 | glio |
| Chek1_in.vivo_noRT | Ubxn1 | -0.649 | 1.39E-04 | glio |
| Eif2s1_in.vivo_noRT | Gm13964 | 2.143 | 1.39E-04 | glio |
| Hsd17b10_in.vivo_noRT | Ahrr | 2.816 | 1.39E-04 | glio |
| Hsd17b10_in.vivo_noRT | mt-Co1 | -0.779 | 1.39E-04 | glio |
| culture_effect | St3gal6 | -2.566 | 1.39E-05 | glio |
| Interaction_genotype_Mtor | Gm32591 | -5.044 | 1.39E-05 | glio |
| Interaction_genotype_Mtor | Mien1 | 2.171 | 1.39E-05 | glio |
| Aars2_ex.vivo_noRT | Gm36037 | 2.897 | 1.39E-05 | glio |
| Atr_in.vivo_noRT | Qpct | 3.13 | 1.39E-05 | glio |
| Bora_in.vivo_noRT | Itga3 | 3.307 | 1.39E-05 | glio |
| Chek1_in.vivo_noRT | Lrrc75aos2 | 3.147 | 1.39E-05 | glio |
| Chek1_in.vivo_noRT | Fbxl22 | 2.927 | 1.39E-05 | glio |

| | | | | |
|---|---|---|---|---|
| culture_effect | Mrpl36 | 0.728 | 1.39E-06 | glio |
| Interaction_genotype_Aars2 | Rassf6 | -5.068 | 1.39E-06 | glio |
| Aars2_in.vivo_noRT | D830025C05R | 3.531 | 1.39E-06 | glio |
| Atr_in.vivo_noRT | Hrk | 3.408 | 1.39E-06 | glio |
| Atr_in.vivo_noRT | 1810021B22R | 3.464 | 1.39E-06 | glio |
| Bora_in.vivo_noRT | Hist1h4b | 3.704 | 1.39E-06 | glio |
| Bora_in.vivo_noRT | Gm16133 | 3.486 | 1.39E-06 | glio |
| Hsd17b10_in.vivo_noRT | Notch4 | 3.41 | 1.39E-06 | glio |
| Interaction_genotype_Mtor | Gm26812 | -6.256 | 1.39E-07 | glio |
| Interaction_genotype_Mtor | Gpc5 | -4.494 | 1.39E-07 | glio |
| Atr_in.vivo_noRT | Fbln1 | 3.512 | 1.39E-07 | glio |
| Chek1_in.vivo_noRT | Gm31805 | 3.589 | 1.39E-07 | glio |
| culture_effect | Tmem169 | -2.357 | 1.39E-08 | glio |
| culture_effect | March2 | 1.12 | 1.39E-08 | glio |
| culture_effect | Laptm4a | -0.638 | 1.39E-08 | glio |
| culture_effect | Ndufs5 | 0.544 | 1.39E-08 | glio |
| Atr_in.vivo_noRT | Dlx2 | 4.15 | 1.39E-08 | glio |
| Atr_in.vivo_noRT | Gm30541 | 3.838 | 1.39E-08 | glio |
| culture_effect | 2810405F17Ri | 2.64 | 1.39E-09 | glio |
| Chek1_ex.vivo_noRT | Nkg7 | 4.447 | 1.39E-09 | glio |
| Hsd17b10_in.vivo_noRT | Kcnk13 | 3.353 | 1.39E-09 | glio |
| Atr_in.vivo_noRT | A230020J21Ri | 4.606 | 1.39E-10 | glio |
| Chek1_in.vivo_noRT | Plcd4 | 4.133 | 1.39E-10 | glio |
| Chek1_ex.vivo_noRT | Slc22a1 | 4.447 | 1.39E-10 | glio |
| culture_effect | Hexb | -1.398 | 1.39E-12 | glio |
| culture_effect | Nus1 | 0.904 | 1.39E-12 | glio |
| Aars2_in.vivo_noRT | 1700030L20Ri | 4.671 | 1.39E-12 | glio |
| Bora_in.vivo_noRT | Gm44983 | 4.148 | 1.39E-12 | glio |
| Chek1_in.vivo_noRT | Loxl4 | 4.777 | 1.39E-13 | glio |
| Chek1_ex.vivo_noRT | Gm19276 | 4.39 | 1.39E-13 | glio |
| Chek1_ex.vivo_noRT | Klk13 | 4.842 | 1.39E-14 | glio |
| Atr_in.vivo_noRT | Gm10642 | 5.269 | 1.39E-15 | glio |
| Atr_in.vivo_noRT | Tff2 | 5.273 | 1.39E-15 | glio |
| Chek1_ex.vivo_noRT | Gm13964 | 4.842 | 1.39E-15 | glio |
| Aars2_in.vivo_noRT | Prl2c3 | 5.327 | 1.39E-16 | glio |
| Atr_in.vivo_noRT | Gast | 5.533 | 1.39E-22 | glio |
| Aars2_in.vivo_noRT | Meioc | 5.063 | 1.39E-24 | glio |
| Chek1_in.vivo_noRT | Gm49698 | 5.044 | 1.39E-24 | glio |
| culture_effect | Mef2c | 3.742 | 1.39E-38 | glio |
| Interaction_genotype_Aars2 | Galk1 | 1.534 | 1.40E-03 | glio |
| Interaction_genotype_Atr | 4930434B07R | -3.485 | 1.40E-03 | glio |
| Interaction_genotype_Bcas2 | B930094E09Ri | -4.047 | 1.40E-03 | glio |
| Interaction_genotype_Mtor | Mmp19 | -3.875 | 1.40E-03 | glio |
| Interaction_genotype_Mtor | Fcgrt | -4.249 | 1.40E-03 | glio |
| Interaction_genotype_Mtor | Iapp | -4.443 | 1.40E-03 | glio |
| Interaction_genotype_Mtor | Jup | -3.627 | 1.40E-03 | glio |
| Interaction_genotype_Mtor | Lrrc51 | -2.9 | 1.40E-03 | glio |
| Interaction_genotype_Mtor | 9230114K14R | -2.866 | 1.40E-03 | glio |
| Interaction_genotype_Mtor | Lztfl1 | 1.981 | 1.40E-03 | glio |
| Interaction_genotype_Mtor | Pdcd6ip | 1.631 | 1.40E-03 | glio |
| Interaction_genotype_Mtor | Tle5 | 1.394 | 1.40E-03 | glio |
| Interaction_genotype_Mtor | Rpl10a | 0.928 | 1.40E-03 | glio |

| | | | | |
|---|---|---|---|---|
| Interaction_genotype_Myc | Lrrc45 | -3.236 | 1.40E-03 | glio |
| Interaction_genotype_Myc | Gm49708 | -3.306 | 1.40E-03 | glio |
| Interaction_genotype_Myc | Cdkl5 | -2.556 | 1.40E-03 | glio |
| Interaction_genotype_Myc | Dcaf1 | -2.156 | 1.40E-03 | glio |
| Interaction_genotype_Myc | Arhgef6 | -2.59 | 1.40E-03 | glio |
| Interaction_genotype_Myc | Vps52 | -2.112 | 1.40E-03 | glio |
| Interaction_genotype_Rpl3 | 5033417F24Ri | -4.346 | 1.40E-03 | glio |
| Interaction_genotype_Rpl3 | Serping1 | 3.805 | 1.40E-03 | glio |
| interaction_RT | Dock1 | -1.06 | 1.40E-03 | glio |
| interaction_RT | Gspt1 | -0.671 | 1.40E-03 | glio |
| Aars2_in.vivo_noRT | Flrt3 | 2.099 | 1.40E-03 | glio |
| Aars2_in.vivo_noRT | Slc38a9 | 1.69 | 1.40E-03 | glio |
| Aars2_in.vivo_noRT | Timm21 | -1.404 | 1.40E-03 | glio |
| Aars2_in.vivo_noRT | Osr2 | -1.819 | 1.40E-03 | glio |
| Aars2_in.vivo_noRT | Acbd6 | -1.176 | 1.40E-03 | glio |
| Aars2_in.vivo_noRT | Rpl13a | -0.618 | 1.40E-03 | glio |
| Aars2_ex.vivo_noRT | Sema3a | 2.899 | 1.40E-03 | glio |
| Atr_in.vivo_noRT | Sorl1 | 2.364 | 1.40E-03 | glio |
| Atr_in.vivo_noRT | Gm17491 | 1.803 | 1.40E-03 | glio |
| Atr_in.vivo_noRT | Sorbs1 | 1.299 | 1.40E-03 | glio |
| Atr_in.vivo_noRT | Tmem63b | -1.088 | 1.40E-03 | glio |
| Atr_in.vivo_noRT | Sept7 | -0.879 | 1.40E-03 | glio |
| Bcas2_in.vivo_noRT | Qpct | 2.404 | 1.40E-03 | glio |
| Bcas2_in.vivo_noRT | Igsf8 | -1.734 | 1.40E-03 | glio |
| Bcas2_ex.vivo_noRT | Fam189a1 | 2.742 | 1.40E-03 | glio |
| Bora_in.vivo_noRT | Gm29394 | 2.226 | 1.40E-03 | glio |
| Bora_in.vivo_noRT | Eps8l1 | 2.33 | 1.40E-03 | glio |
| Chek1_in.vivo_noRT | Rnf225 | 2.46 | 1.40E-03 | glio |
| Chek1_in.vivo_noRT | Mast3 | -1.665 | 1.40E-03 | glio |
| Chek1_ex.vivo_noRT | Rhpn2 | 2.77 | 1.40E-03 | glio |
| Chek1_ex.vivo_noRT | Ctso | 2.841 | 1.40E-03 | glio |
| Fanca_ex.vivo_noRT | Gm40538 | 2.919 | 1.40E-03 | glio |
| Fanca_ex.vivo_noRT | Ltbp2 | 2.884 | 1.40E-03 | glio |
| Fanca_ex.vivo_noRT | Tulp2 | -2.816 | 1.40E-03 | glio |
| Fanca_ex.vivo_noRT | Minar2 | 2.913 | 1.40E-03 | glio |
| Fanca_ex.vivo_noRT | Gast | 2.913 | 1.40E-03 | glio |
| Fanca_ex.vivo_noRT | Gm26671 | 2.52 | 1.40E-03 | glio |
| Hsd17b10_in.vivo_noRT | Gm48882 | 2.492 | 1.40E-03 | glio |
| culture_effect | Smarcad1 | 0.602 | 1.40E-04 | glio |
| culture_effect | Saraf | -0.587 | 1.40E-04 | glio |
| culture_effect | Desi1 | 0.517 | 1.40E-04 | glio |
| culture_effect | Sarnp | -0.434 | 1.40E-04 | glio |
| Interaction_genotype_Aars2 | Eif3h | 1.15 | 1.40E-04 | glio |
| Interaction_genotype_Mtor | Lsp1 | -5.064 | 1.40E-04 | glio |
| Interaction_genotype_Mtor | Elovl3 | -4.617 | 1.40E-04 | glio |
| Interaction_genotype_Mtor | Zfp112 | -4.433 | 1.40E-04 | glio |
| Aars2_in.vivo_noRT | Psme2 | -1.06 | 1.40E-04 | glio |
| Atr_in.vivo_noRT | Gm15892 | 2.86 | 1.40E-04 | glio |
| Atr_in.vivo_noRT | Csdc2 | 2.086 | 1.40E-04 | glio |
| Bard1_ex.vivo_noRT | Klrk1 | 2.814 | 1.40E-04 | glio |
| Bcas2_in.vivo_noRT | Spry3 | 2.807 | 1.40E-04 | glio |
| Cenpj_in.vivo_noRT | Gm128 | 2.689 | 1.40E-04 | glio |

| | | | | |
|---|---|---|---|---|
| Chek1_in.vivo_noRT | Setmar | 2.728 | 1.40E-04 | glio |
| Chek1_in.vivo_noRT | Car5a | 2.48 | 1.40E-04 | glio |
| Hsd17b10_in.vivo_noRT | Ptp4a1 | 2.448 | 1.40E-04 | glio |
| Interaction_genotype_Mtor | Gm26658 | -5.567 | 1.40E-05 | glio |
| Interaction_genotype_Mtor | Sdcbp2 | -5.514 | 1.40E-05 | glio |
| Interaction_genotype_Mtor | Gm44096 | -4.901 | 1.40E-05 | glio |
| Interaction_genotype_Myc | L3mbtl2 | -3.288 | 1.40E-05 | glio |
| Interaction_genotype_Myc | Fig4 | -3.837 | 1.40E-05 | glio |
| Interaction_genotype_Myc | Dntt | -5.132 | 1.40E-05 | glio |
| Interaction_genotype_Myc | Mrpl30 | 1.496 | 1.40E-05 | glio |
| Aars2_in.vivo_noRT | 1810059H22R | 3.368 | 1.40E-05 | glio |
| Aars2_ex.vivo_noRT | Slc45a1 | 2.893 | 1.40E-05 | glio |
| Aars2_ex.vivo_noRT | Gm47540 | 2.9 | 1.40E-05 | glio |
| Atp5c1_in.vivo_noRT | Npy1r | 2.545 | 1.40E-05 | glio |
| Atr_in.vivo_noRT | Elovl3 | 3.081 | 1.40E-05 | glio |
| Atr_in.vivo_noRT | Rpn1 | -1.524 | 1.40E-05 | glio |
| Bora_in.vivo_noRT | Gm6225 | 3.26 | 1.40E-05 | glio |
| Bora_in.vivo_noRT | Jazf1 | 2.955 | 1.40E-05 | glio |
| Bora_in.vivo_noRT | Mad2l1 | -1.884 | 1.40E-05 | glio |
| Bora_ex.vivo_noRT | Pcdhb21 | 3.976 | 1.40E-05 | glio |
| Bora_ex.vivo_noRT | Gpc5 | 2.806 | 1.40E-05 | glio |
| Chek1_in.vivo_noRT | Rgs16 | 3.834 | 1.40E-05 | glio |
| Chek1_in.vivo_noRT | Zfp11 | 3.077 | 1.40E-05 | glio |
| Chek1_in.vivo_noRT | Hapln4 | 3.276 | 1.40E-05 | glio |
| Hsd17b10_in.vivo_noRT | Gm49701 | 3.094 | 1.40E-05 | glio |
| Interaction_genotype_Mtor | 6330415G19R | -5.3 | 1.40E-06 | glio |
| Interaction_genotype_Mtor | 1700023H06R | -6.174 | 1.40E-06 | glio |
| Interaction_genotype_Mtor | Ttll13 | -5.136 | 1.40E-06 | glio |
| Aars2_ex.vivo_noRT | Fabp6 | 3.293 | 1.40E-06 | glio |
| Bora_in.vivo_noRT | Vps37a | -1.849 | 1.40E-06 | glio |
| Bora_ex.vivo_noRT | Gpr4 | 3.201 | 1.40E-06 | glio |
| Bora_ex.vivo_noRT | Gm33466 | 3.201 | 1.40E-06 | glio |
| Chek1_in.vivo_noRT | Rab11fip1 | 3.75 | 1.40E-06 | glio |
| Chek1_in.vivo_noRT | Cyp4f18 | 4.784 | 1.40E-06 | glio |
| Hsd17b10_in.vivo_noRT | Ptp4a1.1 | 3.35 | 1.40E-06 | glio |
| culture_effect | Gm50334 | 3.052 | 1.40E-07 | glio |
| culture_effect | 5830444B04R | -1.606 | 1.40E-07 | glio |
| Aars2_in.vivo_noRT | Tnni2 | 4.289 | 1.40E-07 | glio |
| Atr_in.vivo_noRT | Gm15952 | 3.768 | 1.40E-07 | glio |
| Atr_in.vivo_noRT | D730045B01R | 3.251 | 1.40E-07 | glio |
| Bard1_ex.vivo_noRT | 1700019E08Ri | 3.987 | 1.40E-07 | glio |
| Bora_in.vivo_noRT | Atoh8 | 3.553 | 1.40E-07 | glio |
| Chek1_in.vivo_noRT | Syngap1 | 3.584 | 1.40E-07 | glio |
| culture_effect | Ddx24 | 0.611 | 1.40E-08 | glio |
| Aars2_in.vivo_noRT | Gm4755 | 3.569 | 1.40E-08 | glio |
| Atr_in.vivo_noRT | Hoxa13 | 4.086 | 1.40E-08 | glio |
| Atr_in.vivo_noRT | Gm26766 | 3.838 | 1.40E-08 | glio |
| Atr_in.vivo_noRT | S1pr1 | 4.349 | 1.40E-08 | glio |
| Aars2_in.vivo_noRT | Catsperd | 4.148 | 1.40E-09 | glio |
| Bora_ex.vivo_noRT | Gm12703 | 3.994 | 1.40E-09 | glio |
| Chek1_in.vivo_noRT | Ephx1 | 4.145 | 1.40E-09 | glio |
| Chek1_ex.vivo_noRT | Gm27241 | 3.862 | 1.40E-09 | glio |

| | | | | |
|---|---|---|---|---|
| Chek1_ex.vivo_noRT | Clca3a1 | 4.842 | 1.40E-09 | glio |
| Aars2_in.vivo_noRT | Sec1 | 4.684 | 1.40E-10 | glio |
| Chek1_in.vivo_noRT | Fgd3 | 4.838 | 1.40E-10 | glio |
| culture_effect | Cysltr2 | -3.939 | 1.40E-11 | glio |
| culture_effect | Ncam1 | 1.498 | 1.40E-11 | glio |
| Hsd17b10_in.vivo_noRT | Cxcl5 | 3.616 | 1.40E-11 | glio |
| Aars2_in.vivo_noRT | Gm15972 | 4.333 | 1.40E-12 | glio |
| Aars2_in.vivo_noRT | BC065403 | 4.811 | 1.40E-12 | glio |
| Bcas2_in.vivo_noRT | Col8a1 | 4.807 | 1.40E-12 | glio |
| Chek1_in.vivo_noRT | A830009L08Ri | 4.521 | 1.40E-12 | glio |
| culture_effect | Rps9 | -0.971 | 1.40E-13 | glio |
| Atr_in.vivo_noRT | BC065403 | 5.017 | 1.40E-13 | glio |
| Chek1_in.vivo_noRT | Rarres1 | 5.044 | 1.40E-13 | glio |
| culture_effect | Cdk5rap3 | -1.372 | 1.40E-14 | glio |
| culture_effect | Lrrk1 | 2.17 | 1.40E-15 | glio |
| culture_effect | Ccnd2 | 0.95 | 1.40E-15 | glio |
| Atr_in.vivo_noRT | Cxcl16 | 5.269 | 1.40E-15 | glio |
| Chek1_in.vivo_noRT | Fam166a | 4.78 | 1.40E-18 | glio |
| Bcas2_in.vivo_noRT | Slc14a2 | 4.807 | 1.40E-23 | glio |
| Aars2_in.vivo_noRT | 4930588A03R | 5.327 | 1.40E-27 | glio |
| culture_effect | Gm20712 | -2.008 | 1.41E-03 | glio |
| culture_effect | Ipo9 | 0.663 | 1.41E-03 | glio |
| culture_effect | Psmc2 | -0.387 | 1.41E-03 | glio |
| Interaction_genotype_Bcas2 | BC065403 | -3.727 | 1.41E-03 | glio |
| Interaction_genotype_Bcas2 | Tkfc | -3.128 | 1.41E-03 | glio |
| Interaction_genotype_Bcas2 | Tlr1 | -4.252 | 1.41E-03 | glio |
| Interaction_genotype_Bcas2 | Gm28731 | -4.541 | 1.41E-03 | glio |
| Interaction_genotype_Bcas2 | Arf2 | 2.393 | 1.41E-03 | glio |
| Interaction_genotype_Bora | 6330415G19R | -3.971 | 1.41E-03 | glio |
| Interaction_genotype_Chek1 | Gm35019 | 3.913 | 1.41E-03 | glio |
| Interaction_genotype_Chek1 | Selenok | 1.467 | 1.41E-03 | glio |
| Interaction_genotype_Mtor | Rgl1 | -3.187 | 1.41E-03 | glio |
| Interaction_genotype_Mtor | Ep400 | 1.815 | 1.41E-03 | glio |
| Interaction_genotype_Myc | Gm16048 | -3.919 | 1.41E-03 | glio |
| Interaction_genotype_Myc | Haglr | -3.923 | 1.41E-03 | glio |
| Interaction_genotype_Myc | Phldb3 | -3.67 | 1.41E-03 | glio |
| Interaction_genotype_Myc | 4930509H03R | -3.637 | 1.41E-03 | glio |
| Interaction_genotype_Myc | Atxn2l | 1.632 | 1.41E-03 | glio |
| Interaction_genotype_Myc | Selenok | 1.349 | 1.41E-03 | glio |
| Interaction_genotype_Rpl3 | Baalc | -3.875 | 1.41E-03 | glio |
| interaction_RT | Cdc42ep5 | 1.141 | 1.41E-03 | glio |
| interaction_RT | Traf4 | 0.94 | 1.41E-03 | glio |
| interaction_RT | Me2 | -0.89 | 1.41E-03 | glio |
| Aars2_in.vivo_noRT | Gm47863 | 2.329 | 1.41E-03 | glio |
| Aars2_in.vivo_noRT | Ndufb4 | -0.928 | 1.41E-03 | glio |
| Aars2_in.vivo_noRT | Ifitm2 | -0.719 | 1.41E-03 | glio |
| Atp5c1_in.vivo_noRT | Upb1 | 2.405 | 1.41E-03 | glio |
| Atr_in.vivo_noRT | Gm10791 | 2.089 | 1.41E-03 | glio |
| Atr_in.vivo_noRT | Serpina3m | 2.214 | 1.41E-03 | glio |
| Atr_in.vivo_noRT | Ppm1f | 1.892 | 1.41E-03 | glio |
| Atr_in.vivo_noRT | Evc | 1.888 | 1.41E-03 | glio |
| Atr_in.vivo_noRT | Brf2 | 1.652 | 1.41E-03 | glio |

| | | | | |
|---|---|---|---|---|
| Atr_in.vivo_noRT | Sgta | -1.107 | 1.41E-03 | glio |
| Atr_in.vivo_noRT | Ckap2 | 1.169 | 1.41E-03 | glio |
| Bcas2_in.vivo_noRT | 1700008J07Ri | 2.125 | 1.41E-03 | glio |
| Bcas2_in.vivo_noRT | Atp6v1g2 | 2.153 | 1.41E-03 | glio |
| Bcas2_in.vivo_noRT | Gdf11 | 1.782 | 1.41E-03 | glio |
| Bcas2_in.vivo_noRT | Ankrd26 | -1.639 | 1.41E-03 | glio |
| Bora_in.vivo_noRT | Nemp2 | 2.513 | 1.41E-03 | glio |
| Bora_in.vivo_noRT | Samd1 | -1.675 | 1.41E-03 | glio |
| Bora_in.vivo_noRT | Lmna | -0.98 | 1.41E-03 | glio |
| Bora_ex.vivo_noRT | Gm34643 | 3.265 | 1.41E-03 | glio |
| Bora_ex.vivo_noRT | Lce1g | 3.203 | 1.41E-03 | glio |
| Bora_ex.vivo_noRT | Gm30794 | 2.225 | 1.41E-03 | glio |
| Bora_ex.vivo_noRT | Slc15a3 | 3.022 | 1.41E-03 | glio |
| Bora_ex.vivo_noRT | H2-Q7 | 2.71 | 1.41E-03 | glio |
| Chek1_ex.vivo_noRT | Slc22a18 | 3.103 | 1.41E-03 | glio |
| Chek1_ex.vivo_noRT | Gm36356 | 2.406 | 1.41E-03 | glio |
| Eif2s1_in.vivo_noRT | Plcl1 | 2.143 | 1.41E-03 | glio |
| Interaction_genotype_Atr | Zic4 | -4.993 | 1.41E-04 | glio |
| Interaction_genotype_Bcas2 | D430040D24R | -5.015 | 1.41E-04 | glio |
| Interaction_genotype_Mtor | Stra8 | -3.971 | 1.41E-04 | glio |
| Interaction_genotype_Mtor | Rbfaos | -3.58 | 1.41E-04 | glio |
| Interaction_genotype_Mtor | Gm17655 | -4.993 | 1.41E-04 | glio |
| Interaction_genotype_Mtor | Spink2 | -4.922 | 1.41E-04 | glio |
| Interaction_genotype_Myc | Slc19a3 | -3.122 | 1.41E-04 | glio |
| Interaction_genotype_Myc | Gm13191 | -2.729 | 1.41E-04 | glio |
| Interaction_genotype_Myc | 4933416E03Ri | -2.729 | 1.41E-04 | glio |
| Interaction_genotype_Myc | Lmln | -4.207 | 1.41E-04 | glio |
| Interaction_genotype_Myc | Swsap1 | -3.85 | 1.41E-04 | glio |
| Interaction_genotype_Myc | Pomgnt2 | -3.397 | 1.41E-04 | glio |
| Interaction_genotype_Myc | Rpl3 | 1.61 | 1.41E-04 | glio |
| Interaction_genotype_Rad21 | BC049987 | 3.788 | 1.41E-04 | glio |
| Interaction_genotype_Rad21 | Mog | 5.811 | 1.41E-04 | glio |
| interaction_RT | Fstl1 | 1.905 | 1.41E-04 | glio |
| interaction_RT | Gpr37l1 | 3.626 | 1.41E-04 | glio |
| Bcas2_in.vivo_noRT | Abhd15 | 2.958 | 1.41E-04 | glio |
| Bcas2_ex.vivo_noRT | Gm48215 | 2.742 | 1.41E-04 | glio |
| Bora_in.vivo_noRT | Matk | 2.518 | 1.41E-04 | glio |
| Chek1_ex.vivo_noRT | A230020J21Ri | 3.474 | 1.41E-04 | glio |
| culture_effect | 4833439L19Ri | 0.717 | 1.41E-05 | glio |
| Interaction_genotype_Mtor | Gm40264 | -4.426 | 1.41E-05 | glio |
| Interaction_genotype_Mtor | Tmed3 | 2.24 | 1.41E-05 | glio |
| Aars2_in.vivo_noRT | A430033K04R | 2.959 | 1.41E-05 | glio |
| Aars2_in.vivo_noRT | Usp51 | 2.907 | 1.41E-05 | glio |
| Aars2_in.vivo_noRT | Des | 3.076 | 1.41E-05 | glio |
| Aars2_in.vivo_noRT | Gm17056 | 3.108 | 1.41E-05 | glio |
| Aars2_in.vivo_noRT | Bola3 | -1.481 | 1.41E-05 | glio |
| Aars2_in.vivo_noRT | Stt3a | -1.529 | 1.41E-05 | glio |
| Aars2_ex.vivo_noRT | Lrguk | 3.293 | 1.41E-05 | glio |
| Aars2_ex.vivo_noRT | Gm20275 | 3.293 | 1.41E-05 | glio |
| Bcas2_in.vivo_noRT | AA388235 | 3.167 | 1.41E-05 | glio |
| Bcas2_in.vivo_noRT | Gm40040 | 3.251 | 1.41E-05 | glio |
| Bcas2_in.vivo_noRT | Moxd1 | -1.123 | 1.41E-05 | glio |

| | | | | |
|---|---|---|---|---|
| Bora_in.vivo_noRT | Glrp1 | 2.667 | 1.41E-05 | glio |
| Chek1_in.vivo_noRT | Lmtk3 | 2.963 | 1.41E-05 | glio |
| Chek1_ex.vivo_noRT | Gabrd | 3.871 | 1.41E-05 | glio |
| Chek1_ex.vivo_noRT | Fam222a | 4.085 | 1.41E-05 | glio |
| interaction_RT | Mt3 | 2.68 | 1.41E-06 | glio |
| Atp5c1_in.vivo_noRT | Gm2016 | 2.668 | 1.41E-06 | glio |
| Atp5c1_in.vivo_noRT | C330002G04R | 2.668 | 1.41E-06 | glio |
| Atp5c1_in.vivo_noRT | Rapgef3os2 | 2.668 | 1.41E-06 | glio |
| Atp5c1_in.vivo_noRT | Gm47622 | 2.668 | 1.41E-06 | glio |
| Atr_in.vivo_noRT | Gm28050 | 3.495 | 1.41E-06 | glio |
| culture_effect | Ifi208 | -3.352 | 1.41E-07 | glio |
| culture_effect | Magi2 | 2.343 | 1.41E-07 | glio |
| Interaction_genotype_Mtor | D830039M14I | -6.527 | 1.41E-07 | glio |
| Interaction_genotype_Mtor | Armc2 | -6.343 | 1.41E-07 | glio |
| Interaction_genotype_Mtor | Gm15908 | -4.099 | 1.41E-07 | glio |
| Interaction_genotype_Mtor | A930014E10Ri | -4.099 | 1.41E-07 | glio |
| Interaction_genotype_Mtor | Gm13449 | -4.886 | 1.41E-07 | glio |
| Atp5c1_in.vivo_noRT | Gm47404 | 2.668 | 1.41E-07 | glio |
| Bcas2_in.vivo_noRT | Osr2 | -2.998 | 1.41E-07 | glio |
| Bora_in.vivo_noRT | Arhgef15 | 3.555 | 1.41E-07 | glio |
| Chek1_in.vivo_noRT | Slc17a9 | 3.589 | 1.41E-07 | glio |
| Chek1_ex.vivo_noRT | Gm17638 | 4.267 | 1.41E-07 | glio |
| Chek1_ex.vivo_noRT | 4933408J17Ri | 4.058 | 1.41E-07 | glio |
| Hsd17b10_in.vivo_noRT | Gm5662 | 3.616 | 1.41E-07 | glio |
| Hsd17b10_in.vivo_noRT | G630030J09Ri | 3.621 | 1.41E-07 | glio |
| Interaction_genotype_Mtor | Cyp1a1 | -8.968 | 1.41E-08 | glio |
| Interaction_genotype_Mtor | Efhd1 | -7.029 | 1.41E-08 | glio |
| Interaction_genotype_Mtor | Kpna7 | -6.194 | 1.41E-08 | glio |
| Interaction_genotype_Mtor | Gm30794 | -5.075 | 1.41E-08 | glio |
| Interaction_genotype_Mtor | Gm41318 | -6.812 | 1.41E-08 | glio |
| Interaction_genotype_Mtor | Gm11973 | -5.596 | 1.41E-08 | glio |
| Interaction_genotype_Mtor | Abcb5 | -4.893 | 1.41E-08 | glio |
| Atr_ex.vivo_noRT | Pdgfb | 3.984 | 1.41E-08 | glio |
| Atr_in.vivo_noRT | Hspa12a | 4.233 | 1.41E-09 | glio |
| Atr_in.vivo_noRT | Gm49042 | 3.904 | 1.41E-09 | glio |
| Chek1_ex.vivo_noRT | Aloxe3 | 4.325 | 1.41E-09 | glio |
| Chek1_ex.vivo_noRT | Slc15a3 | 5.051 | 1.41E-09 | glio |
| Chek1_ex.vivo_noRT | Kcnk13 | 4.055 | 1.41E-09 | glio |
| Hsd17b10_in.vivo_noRT | 9230112J17Ri | 3.616 | 1.41E-09 | glio |
| Chek1_in.vivo_noRT | Oxt | 4.402 | 1.41E-10 | glio |
| Chek1_in.vivo_noRT | Amh | 4.127 | 1.41E-10 | glio |
| Hsd17b10_in.vivo_noRT | Akr1c18 | 3.616 | 1.41E-10 | glio |
| Hsd17b10_in.vivo_noRT | 4921504E06Ri | 3.616 | 1.41E-10 | glio |
| Hsd17b10_in.vivo_noRT | Gm14820 | 3.353 | 1.41E-10 | glio |
| Hsd17b10_in.vivo_noRT | 4931431C16R | 3.616 | 1.41E-10 | glio |
| culture_effect | Ptgds | -4.048 | 1.41E-11 | glio |
| Aars2_in.vivo_noRT | 9530062K07R | 4.947 | 1.41E-11 | glio |
| Bcas2_in.vivo_noRT | Skida1 | 4.807 | 1.41E-11 | glio |
| Chek1_in.vivo_noRT | Scube3 | 4.511 | 1.41E-12 | glio |
| Hsd17b10_in.vivo_noRT | Gm15908 | 3.616 | 1.41E-14 | glio |
| Hsd17b10_in.vivo_noRT | A930014E10Ri | 3.616 | 1.41E-14 | glio |
| Hsd17b10_in.vivo_noRT | Gm45338 | 3.616 | 1.41E-14 | glio |

| | | | | |
|---|---|---|---|---|
| Hsd17b10_in.vivo_noRT | Scg2 | 3.616 | 1.41E-14 | glio |
| Atr_in.vivo_noRT | 4930527F14Ri | 5.004 | 1.41E-20 | glio |
| Bcas2_in.vivo_noRT | Gm12800 | 4.807 | 1.41E-20 | glio |
| Bora_in.vivo_noRT | Vmn2r18 | 4.936 | 1.41E-23 | glio |
| Bora_in.vivo_noRT | Hoxc8 | 4.936 | 1.41E-24 | glio |
| Chek1_in.vivo_noRT | Olfr881 | 5.044 | 1.41E-24 | glio |
| culture_effect | Mrpl11 | 0.401 | 1.42E-03 | glio |
| Interaction_genotype_Aars2 | Apol7d | -4.103 | 1.42E-03 | glio |
| Interaction_genotype_Aars2 | Lamtor2 | 1.247 | 1.42E-03 | glio |
| Interaction_genotype_Mtor | Gm39469 | -3.142 | 1.42E-03 | glio |
| Interaction_genotype_Myc | Gm13449 | -2.728 | 1.42E-03 | glio |
| Interaction_genotype_Myc | Tmem150c | -3.714 | 1.42E-03 | glio |
| Interaction_genotype_Myc | B3glct | -2.676 | 1.42E-03 | glio |
| Interaction_genotype_Xrcc3 | Kctd4 | -4.752 | 1.42E-03 | glio |
| interaction_RT | Hist2h2bb | -1.742 | 1.42E-03 | glio |
| interaction_RT | Speer4e | 2.47 | 1.42E-03 | glio |
| interaction_RT | Med30 | 0.646 | 1.42E-03 | glio |
| interaction_RT | Calr | -0.765 | 1.42E-03 | glio |
| interaction_RT | Rpl31 | -0.575 | 1.42E-03 | glio |
| Aars2_in.vivo_noRT | Zfp324 | 1.96 | 1.42E-03 | glio |
| Atp5c1_in.vivo_noRT | Fbxl22 | 2.382 | 1.42E-03 | glio |
| Atr_in.vivo_noRT | Gpam | 2.112 | 1.42E-03 | glio |
| Atr_in.vivo_noRT | Zfp12 | 1.775 | 1.42E-03 | glio |
| Atr_ex.vivo_noRT | Slc7a4 | 2.966 | 1.42E-03 | glio |
| Atr_ex.vivo_noRT | Atg101 | 1.178 | 1.42E-03 | glio |
| Bard1_ex.vivo_noRT | 1700006H21R | 3.169 | 1.42E-03 | glio |
| Bcas2_in.vivo_noRT | Srcin1 | 2.233 | 1.42E-03 | glio |
| Bcas2_in.vivo_noRT | Zfr2 | 2.187 | 1.42E-03 | glio |
| Bcas2_ex.vivo_noRT | AF067061 | 2.742 | 1.42E-03 | glio |
| Bora_in.vivo_noRT | Cyp51 | -1.576 | 1.42E-03 | glio |
| Bora_ex.vivo_noRT | Dcn | 2.808 | 1.42E-03 | glio |
| Cenpj_in.vivo_noRT | Gpr22 | 2.022 | 1.42E-03 | glio |
| Cenpj_ex.vivo_noRT | Pnliprp2 | 2.591 | 1.42E-03 | glio |
| Chek1_in.vivo_noRT | Gm48604 | 2.26 | 1.42E-03 | glio |
| Chek1_in.vivo_noRT | Tmem230 | -1.347 | 1.42E-03 | glio |
| Chek1_in.vivo_noRT | Snrnp25 | -1.289 | 1.42E-03 | glio |
| Chek1_ex.vivo_noRT | Gm17227 | 2.764 | 1.42E-03 | glio |
| Chek1_ex.vivo_noRT | Ezh2 | -1.205 | 1.42E-03 | glio |
| Hsd17b10_in.vivo_noRT | Kctd4 | 2.344 | 1.42E-03 | glio |
| culture_effect | Foxa3 | -2.162 | 1.42E-04 | glio |
| culture_effect | Gm26658 | 2.118 | 1.42E-04 | glio |
| culture_effect | Rnf14 | 0.585 | 1.42E-04 | glio |
| culture_effect | Lamtor2 | 0.448 | 1.42E-04 | glio |
| culture_effect | Sec11c | -0.556 | 1.42E-04 | glio |
| Interaction_genotype_Mtor | Gm4189 | -4.435 | 1.42E-04 | glio |
| Interaction_genotype_Mtor | Daam2 | -4.061 | 1.42E-04 | glio |
| Interaction_genotype_Myc | Nnt | -3.973 | 1.42E-04 | glio |
| Interaction_genotype_Myc | Tirap | -3.317 | 1.42E-04 | glio |
| Interaction_genotype_Myc | Afap1l2 | -3.195 | 1.42E-04 | glio |
| interaction_RT | Prdx1 | 0.944 | 1.42E-04 | glio |
| Aars2_in.vivo_noRT | Hist1h3g | 2.864 | 1.42E-04 | glio |
| Aars2_in.vivo_noRT | Maats1 | 2.774 | 1.42E-04 | glio |

| | | | | |
|---|---|---|---|---|
| Atp5c1_in.vivo_noRT | Tmem74b | 2.404 | 1.42E-04 | glio |
| Atr_in.vivo_noRT | Mansc1 | 2.86 | 1.42E-04 | glio |
| Atr_ex.vivo_noRT | Obscn | 3.164 | 1.42E-04 | glio |
| Atr_ex.vivo_noRT | H2-Q10 | 3.567 | 1.42E-04 | glio |
| Atr_ex.vivo_noRT | 4931431C16R | 2.783 | 1.42E-04 | glio |
| Bora_in.vivo_noRT | Tsen34 | -1.37 | 1.42E-04 | glio |
| Bora_in.vivo_noRT | Ube2d2a | -0.999 | 1.42E-04 | glio |
| Chek1_in.vivo_noRT | Gm15601 | 2.832 | 1.42E-04 | glio |
| Chek1_in.vivo_noRT | Ehbp1l1 | 2.794 | 1.42E-04 | glio |
| Chek1_in.vivo_noRT | Pdhb | -1.333 | 1.42E-04 | glio |
| Chek1_in.vivo_noRT | Mef2c | 2.045 | 1.42E-04 | glio |
| Chek1_ex.vivo_noRT | Cpsf3 | -2.019 | 1.42E-04 | glio |
| Hsd17b10_in.vivo_noRT | 9530080O11R | 2.699 | 1.42E-04 | glio |
| culture_effect | D930019O06R | 2.279 | 1.42E-05 | glio |
| culture_effect | Colgalt1 | -0.645 | 1.42E-05 | glio |
| Interaction_genotype_Mtor | Olfm2 | -4.922 | 1.42E-05 | glio |
| Interaction_genotype_Mtor | Gm19409 | -5.338 | 1.42E-05 | glio |
| Interaction_genotype_Myc | Lncbate10 | -3.516 | 1.42E-05 | glio |
| Aars2_in.vivo_noRT | Zc3h6 | 3.102 | 1.42E-05 | glio |
| Aars2_in.vivo_noRT | Pcx | 2.982 | 1.42E-05 | glio |
| Atr_in.vivo_noRT | Gm20234 | 3.287 | 1.42E-05 | glio |
| Bcas2_in.vivo_noRT | Hey2 | 2.96 | 1.42E-05 | glio |
| Chek1_in.vivo_noRT | Tmc7 | 3.231 | 1.42E-05 | glio |
| Chek1_in.vivo_noRT | Gm17399 | 3.302 | 1.42E-05 | glio |
| Chek1_ex.vivo_noRT | Tmem30c | 3.877 | 1.42E-05 | glio |
| Chek1_ex.vivo_noRT | Slc22a22 | 3.873 | 1.42E-05 | glio |
| culture_effect | C130040N14R | 2.531 | 1.42E-06 | glio |
| Interaction_genotype_Mtor | Gm8098 | -5.747 | 1.42E-06 | glio |
| Aars2_ex.vivo_noRT | Gm49808 | 2.9 | 1.42E-06 | glio |
| Atr_ex.vivo_noRT | Gm5890 | 3.164 | 1.42E-06 | glio |
| Bora_in.vivo_noRT | Nox1 | 2.954 | 1.42E-06 | glio |
| Chek1_in.vivo_noRT | 1600002K03R | 3.51 | 1.42E-06 | glio |
| Chek1_in.vivo_noRT | Car12 | 3.192 | 1.42E-06 | glio |
| Hsd17b10_in.vivo_noRT | Col6a2 | 3.359 | 1.42E-06 | glio |
| Interaction_genotype_Mtor | Gm15743 | -5.9 | 1.42E-07 | glio |
| Interaction_genotype_Mtor | Slc35g3 | -4.498 | 1.42E-07 | glio |
| Interaction_genotype_Myc | P4hb | 2.287 | 1.42E-07 | glio |
| Atp5c1_in.vivo_noRT | Gm31717 | 2.668 | 1.42E-07 | glio |
| Bcas2_in.vivo_noRT | Gm49801 | 3.794 | 1.42E-07 | glio |
| Bcas2_in.vivo_noRT | Aqp3 | 4.543 | 1.42E-07 | glio |
| Bora_in.vivo_noRT | Appbp2os | 3.873 | 1.42E-07 | glio |
| Aars2_in.vivo_noRT | Tead3 | 3.902 | 1.42E-08 | glio |
| Bard1_ex.vivo_noRT | Fam227b | 3.596 | 1.42E-08 | glio |
| Bard1_ex.vivo_noRT | BC080695 | 3.987 | 1.42E-08 | glio |
| Bard1_ex.vivo_noRT | A930019D19R | 3.612 | 1.42E-08 | glio |
| Bcas2_in.vivo_noRT | Osm | 3.458 | 1.42E-08 | glio |
| Bcas2_in.vivo_noRT | Svopl | 3.626 | 1.42E-08 | glio |
| Bora_in.vivo_noRT | Lncenc1 | 3.63 | 1.42E-08 | glio |
| Chek1_in.vivo_noRT | Sdr16c5 | 3.727 | 1.42E-08 | glio |
| Aars2_in.vivo_noRT | Kctd4 | 4.054 | 1.42E-09 | glio |
| Atr_in.vivo_noRT | Slc16a6 | 4.35 | 1.42E-09 | glio |
| Atr_in.vivo_noRT | Zmynd15 | 4.096 | 1.42E-09 | glio |

| | | | | |
|---|---|---|---|---|
| Bcas2_in.vivo_noRT | Wnt16 | 4.016 | 1.42E-09 | glio |
| Bcas2_in.vivo_noRT | 2010320O07R | 4.157 | 1.42E-09 | glio |
| Bora_in.vivo_noRT | Lrrc24 | 3.936 | 1.42E-09 | glio |
| Chek1_in.vivo_noRT | Guca1b | 4.285 | 1.42E-09 | glio |
| Chek1_in.vivo_noRT | Gm5544 | 4.253 | 1.42E-09 | glio |
| Chek1_in.vivo_noRT | Hoxa6 | 4.011 | 1.42E-09 | glio |
| Chek1_ex.vivo_noRT | Dnaaf3 | 4.842 | 1.42E-09 | glio |
| Hsd17b10_in.vivo_noRT | Btn1a1 | 3.356 | 1.42E-09 | glio |
| Aars2_in.vivo_noRT | A930003A15R | 4.175 | 1.42E-10 | glio |
| Bcas2_in.vivo_noRT | Gm34885 | 4.547 | 1.42E-10 | glio |
| Chek1_ex.vivo_noRT | 1700108F19Ri | 4.449 | 1.42E-10 | glio |
| culture_effect | Plin2 | -1.135 | 1.42E-11 | glio |
| Bora_in.vivo_noRT | D030040B21R | 4.549 | 1.42E-11 | glio |
| Atr_ex.vivo_noRT | Gm13191 | 3.947 | 1.42E-12 | glio |
| Atr_ex.vivo_noRT | 4933416E03Ri | 3.947 | 1.42E-12 | glio |
| Bcas2_in.vivo_noRT | Grin3b | 4.812 | 1.42E-12 | glio |
| Bcas2_in.vivo_noRT | Abcg4 | 4.534 | 1.42E-12 | glio |
| Chek1_in.vivo_noRT | Carns1 | 5.044 | 1.42E-12 | glio |
| Chek1_in.vivo_noRT | Gm48265 | 4.261 | 1.42E-12 | glio |
| Chek1_in.vivo_noRT | Lyl1 | 4.393 | 1.42E-13 | glio |
| Chek1_in.vivo_noRT | Rnf165 | 5.044 | 1.42E-13 | glio |
| culture_effect | Rpl3 | -1.013 | 1.42E-15 | glio |
| Bora_in.vivo_noRT | Gm39929 | 4.479 | 1.42E-15 | glio |
| Bcas2_in.vivo_noRT | 4930553M12F | 4.541 | 1.42E-16 | glio |
| Bora_in.vivo_noRT | 1700108F19Ri | 4.672 | 1.42E-16 | glio |
| culture_effect | Plaat3 | -3.446 | 1.42E-50 | glio |
| culture_effect | Klk13 | -1.467 | 1.43E-03 | glio |
| Interaction_genotype_Atr | Fbxl2 | -3.605 | 1.43E-03 | glio |
| Interaction_genotype_Bcas2 | Rhox7a | -4.441 | 1.43E-03 | glio |
| Interaction_genotype_Chek1 | Ccsap | -3.212 | 1.43E-03 | glio |
| Interaction_genotype_Mtor | Zfp980 | -4.138 | 1.43E-03 | glio |
| Interaction_genotype_Mtor | Gm28050 | -3.734 | 1.43E-03 | glio |
| Interaction_genotype_Mtor | 1110019D14R | -3.613 | 1.43E-03 | glio |
| Interaction_genotype_Mtor | Pam | 1.982 | 1.43E-03 | glio |
| Interaction_genotype_Mtor | Capg | 1.764 | 1.43E-03 | glio |
| Interaction_genotype_Myc | Rimklb | -2.466 | 1.43E-03 | glio |
| Interaction_genotype_Myc | Gm48768 | -3.652 | 1.43E-03 | glio |
| Interaction_genotype_Myc | Dennd2c | -3.625 | 1.43E-03 | glio |
| Interaction_genotype_Myc | Rtkn2 | -2.843 | 1.43E-03 | glio |
| Interaction_genotype_Myc | Gucd1 | -2.865 | 1.43E-03 | glio |
| Interaction_genotype_Myc | Trim23 | -2.964 | 1.43E-03 | glio |
| Interaction_genotype_Myc | Pspc1 | -2.647 | 1.43E-03 | glio |
| Interaction_genotype_Myc | Mrps36 | 1.324 | 1.43E-03 | glio |
| Interaction_genotype_Myc | Ntm | -2.295 | 1.43E-03 | glio |
| Interaction_genotype_Rpl3 | Skida1 | -4.484 | 1.43E-03 | glio |
| Interaction_genotype_Rpl3 | Rbm19 | -3.387 | 1.43E-03 | glio |
| Interaction_genotype_Rpl3 | Klrk1 | 3.632 | 1.43E-03 | glio |
| interaction_RT | Dnttip2 | -0.682 | 1.43E-03 | glio |
| interaction_RT | Atp6v1g1 | 0.556 | 1.43E-03 | glio |
| Aars2_in.vivo_noRT | Gm26910 | 2.128 | 1.43E-03 | glio |
| Aars2_in.vivo_noRT | Tro | 2.239 | 1.43E-03 | glio |
| Aars2_in.vivo_noRT | Gm48855 | 2.053 | 1.43E-03 | glio |

| | | | | |
|---|---|---|---|---|
| Aars2_in.vivo_noRT | Tcaim | 2.151 | 1.43E-03 | glio |
| Aars2_in.vivo_noRT | Mical1 | 2.079 | 1.43E-03 | glio |
| Aars2_in.vivo_noRT | Anp32b | -0.639 | 1.43E-03 | glio |
| Aars2_ex.vivo_noRT | Ampd3 | 3.107 | 1.43E-03 | glio |
| Atr_in.vivo_noRT | Cog1 | 1.731 | 1.43E-03 | glio |
| Bard1_ex.vivo_noRT | Clca3a2 | 2.814 | 1.43E-03 | glio |
| Bcas2_in.vivo_noRT | Mr1 | 2.374 | 1.43E-03 | glio |
| Bcas2_in.vivo_noRT | Pecr | -1.627 | 1.43E-03 | glio |
| Bcas2_in.vivo_noRT | Nt5c2 | 1.471 | 1.43E-03 | glio |
| Bora_in.vivo_noRT | Fam177a | 2.169 | 1.43E-03 | glio |
| Bora_in.vivo_noRT | Zbtb11os1 | 2.247 | 1.43E-03 | glio |
| Bora_in.vivo_noRT | Btbd8 | 1.785 | 1.43E-03 | glio |
| Bora_ex.vivo_noRT | Gm813 | 3.201 | 1.43E-03 | glio |
| Chek1_in.vivo_noRT | A930029G22R | 2.242 | 1.43E-03 | glio |
| Chek1_in.vivo_noRT | Slc46a2 | 2.063 | 1.43E-03 | glio |
| Chek1_in.vivo_noRT | Wdr33 | -1.261 | 1.43E-03 | glio |
| Chek1_in.vivo_noRT | Sec62 | -0.941 | 1.43E-03 | glio |
| culture_effect | Hddc3 | 1.812 | 1.43E-04 | glio |
| culture_effect | Tinf2 | 0.998 | 1.43E-04 | glio |
| culture_effect | Pck2 | -0.787 | 1.43E-04 | glio |
| culture_effect | Ddx17 | -0.56 | 1.43E-04 | glio |
| culture_effect | Atic | -0.583 | 1.43E-04 | glio |
| Interaction_genotype_Aars2 | Fgf7 | -4.569 | 1.43E-04 | glio |
| Interaction_genotype_Mtor | Pax5 | -4.915 | 1.43E-04 | glio |
| Interaction_genotype_Myc | Zfp786 | -4.201 | 1.43E-04 | glio |
| Interaction_genotype_Myc | Zbtb3 | -4.358 | 1.43E-04 | glio |
| interaction_RT | Eva1c | 3.483 | 1.43E-04 | glio |
| Aars2_in.vivo_noRT | Iapp | 3.164 | 1.43E-04 | glio |
| Aars2_in.vivo_noRT | Rps27l | -0.769 | 1.43E-04 | glio |
| Atp5c1_in.vivo_noRT | Spi1 | 2.668 | 1.43E-04 | glio |
| Atr_in.vivo_noRT | Nmnat3 | 2.834 | 1.43E-04 | glio |
| Bcas2_in.vivo_noRT | Gm15938 | 2.878 | 1.43E-04 | glio |
| Bcas2_in.vivo_noRT | Kctd1 | 2.632 | 1.43E-04 | glio |
| Bora_in.vivo_noRT | Plxnb1 | 2.804 | 1.43E-04 | glio |
| Bora_in.vivo_noRT | Ypel5 | -2.063 | 1.43E-04 | glio |
| Chek1_in.vivo_noRT | B230398E01Ri | 2.71 | 1.43E-04 | glio |
| Chek1_in.vivo_noRT | Gm26658 | 3.013 | 1.43E-04 | glio |
| Chek1_in.vivo_noRT | Ddx58 | -1.389 | 1.43E-04 | glio |
| Hsd17b10_in.vivo_noRT | Camkk1 | 2.846 | 1.43E-04 | glio |
| Hsd17b10_in.vivo_noRT | Fam228a | 2.828 | 1.43E-04 | glio |
| Hsd17b10_in.vivo_noRT | Gm47802 | 2.64 | 1.43E-04 | glio |
| Hsd17b10_in.vivo_noRT | 4932441J04Ri | 2.892 | 1.43E-04 | glio |
| culture_effect | Gm17249 | 2.085 | 1.43E-05 | glio |
| Interaction_genotype_Mtor | B230317F23Ri | -4.761 | 1.43E-05 | glio |
| Interaction_genotype_Mtor | Gm30074 | -4.77 | 1.43E-05 | glio |
| Interaction_genotype_Mtor | Zfp772 | -5.348 | 1.43E-05 | glio |
| Interaction_genotype_Mtor | Gm14820 | -3.836 | 1.43E-05 | glio |
| Interaction_genotype_Mtor | Masp1 | -4.374 | 1.43E-05 | glio |
| Interaction_genotype_Myc | Vma21 | 1.844 | 1.43E-05 | glio |
| Aars2_in.vivo_noRT | Gm17259 | 2.876 | 1.43E-05 | glio |
| Atp5c1_in.vivo_noRT | Gm13288 | 2.668 | 1.43E-05 | glio |
| Atp5c1_in.vivo_noRT | Gm33051 | 2.543 | 1.43E-05 | glio |

| | | | | |
|---|---|---|---|---|
| Atp5c1_in.vivo_noRT | BC018473 | 3.196 | 1.43E-05 | glio |
| Bcas2_in.vivo_noRT | Arf6 | -1.434 | 1.43E-05 | glio |
| Bora_in.vivo_noRT | Car10 | 2.593 | 1.43E-05 | glio |
| Chek1_in.vivo_noRT | Hagh | -1.516 | 1.43E-05 | glio |
| Hsd17b10_in.vivo_noRT | Platr31 | 3.616 | 1.43E-05 | glio |
| Interaction_genotype_Mtor | Mmp28 | -5.957 | 1.43E-06 | glio |
| Interaction_genotype_Mtor | Gm15138 | -4.228 | 1.43E-06 | glio |
| Atr_in.vivo_noRT | 4930542D17R | 3.253 | 1.43E-06 | glio |
| Atr_in.vivo_noRT | Tnnc1 | 3.681 | 1.43E-06 | glio |
| Bcas2_in.vivo_noRT | Slc17a7 | 3.375 | 1.43E-06 | glio |
| Hsd17b10_in.vivo_noRT | Efcab9 | 3.35 | 1.43E-06 | glio |
| culture_effect | Upp1 | -2.092 | 1.43E-07 | glio |
| Interaction_genotype_Chek1 | C1ra | 5.562 | 1.43E-07 | glio |
| Atp5c1_in.vivo_noRT | Gm14742 | 2.668 | 1.43E-07 | glio |
| Atr_in.vivo_noRT | Gm39556 | 3.623 | 1.43E-07 | glio |
| Bora_in.vivo_noRT | Fgf7 | 3.572 | 1.43E-07 | glio |
| culture_effect | A030001D20R | 1.499 | 1.43E-08 | glio |
| Interaction_genotype_Mtor | Gm43701 | -5.081 | 1.43E-08 | glio |
| Aars2_in.vivo_noRT | Slc26a1 | 3.824 | 1.43E-08 | glio |
| Aars2_in.vivo_noRT | Tec | 3.65 | 1.43E-08 | glio |
| Atr_in.vivo_noRT | Retn | 5.066 | 1.43E-08 | glio |
| Bcas2_in.vivo_noRT | Gm10010 | 4.066 | 1.43E-08 | glio |
| Hsd17b10_in.vivo_noRT | D930028M14I | 3.616 | 1.43E-08 | glio |
| culture_effect | Pik3ip1 | -1.853 | 1.43E-09 | glio |
| Atr_in.vivo_noRT | Cnnm1 | 4.622 | 1.43E-09 | glio |
| Chek1_in.vivo_noRT | Gm10115 | 4.136 | 1.43E-09 | glio |
| Chek1_in.vivo_noRT | Gm47545 | 3.793 | 1.43E-09 | glio |
| Hsd17b10_in.vivo_noRT | Gm20711 | 3.616 | 1.43E-09 | glio |
| Atr_in.vivo_noRT | Klhl30 | 4.349 | 1.43E-10 | glio |
| Bora_in.vivo_noRT | Nrg2 | 4.414 | 1.43E-10 | glio |
| culture_effect | Zdhhc2 | 1.232 | 1.43E-12 | glio |
| Hsd17b10_in.vivo_noRT | BC061195 | 3.616 | 1.43E-13 | glio |
| Aars2_in.vivo_noRT | AI182371 | 4.544 | 1.43E-15 | glio |
| Atr_in.vivo_noRT | Gm26789 | 4.616 | 1.43E-15 | glio |
| Atr_in.vivo_noRT | E030037K01Ri | 5.002 | 1.43E-15 | glio |
| Aars2_in.vivo_noRT | Ccdc162 | 4.798 | 1.43E-17 | glio |
| Bora_in.vivo_noRT | 9430041J12Ri | 4.936 | 1.43E-18 | glio |
| Aars2_in.vivo_noRT | Pdgfb | 5.063 | 1.43E-19 | glio |
| Bora_in.vivo_noRT | Gm35082 | 4.936 | 1.43E-19 | glio |
| Aars2_in.vivo_noRT | Gtsf1 | 5.327 | 1.43E-20 | glio |
| culture_effect | Kitl | 2.565 | 1.43E-25 | glio |
| Atr_in.vivo_noRT | Idi2 | 5.533 | 1.43E-28 | glio |
| Atr_in.vivo_noRT | Gm48155 | 5.533 | 1.43E-30 | glio |
| culture_effect | Best1 | -1.818 | 1.44E-03 | glio |
| culture_effect | BC024063 | -1.901 | 1.44E-03 | glio |
| culture_effect | Brd3os | 1.295 | 1.44E-03 | glio |
| culture_effect | Sptan1 | 0.511 | 1.44E-03 | glio |
| culture_effect | Sumo3 | 0.38 | 1.44E-03 | glio |
| culture_effect | Lsm1 | 0.412 | 1.44E-03 | glio |
| Interaction_genotype_Aars2 | Sema6d | -3.015 | 1.44E-03 | glio |
| Interaction_genotype_Aars2 | Serping1 | 3.399 | 1.44E-03 | glio |
| Interaction_genotype_Bcas2 | G430095P16R | -4.288 | 1.44E-03 | glio |

| | | | | |
|---|---|---|---|---|
| Interaction_genotype_Mtor | Rab11fip1 | -4.271 | 1.44E-03 | glio |
| Interaction_genotype_Mtor | C330011M18F | -3.738 | 1.44E-03 | glio |
| Interaction_genotype_Mtor | Tmem250-ps | -2.403 | 1.44E-03 | glio |
| Interaction_genotype_Mtor | Slc39a7 | 1.545 | 1.44E-03 | glio |
| Interaction_genotype_Mtor | Ddr1 | 1.651 | 1.44E-03 | glio |
| Interaction_genotype_Mtor | Bbx | 1.766 | 1.44E-03 | glio |
| Interaction_genotype_Myc | Gm40040 | -3.664 | 1.44E-03 | glio |
| Interaction_genotype_Myc | Commd5 | -3.65 | 1.44E-03 | glio |
| Interaction_genotype_Myc | Phactr2 | -2.684 | 1.44E-03 | glio |
| Interaction_genotype_Myc | Pym1 | -2.331 | 1.44E-03 | glio |
| Interaction_genotype_Myc | Phf6 | -2.297 | 1.44E-03 | glio |
| Interaction_genotype_Myc | Snrpg | 1.548 | 1.44E-03 | glio |
| Interaction_genotype_Rad51 | Ntn4 | -3.384 | 1.44E-03 | glio |
| Interaction_genotype_Rad51 | Gm32834 | -4.192 | 1.44E-03 | glio |
| Interaction_genotype_Rpl3 | Gm29340 | -4.119 | 1.44E-03 | glio |
| Interaction_genotype_Rpl3 | Gm10642 | -4.152 | 1.44E-03 | glio |
| interaction_RT | 1110012L19Ri | 0.972 | 1.44E-03 | glio |
| interaction_RT | Dkc1 | -0.733 | 1.44E-03 | glio |
| Aars2_in.vivo_noRT | Zfp94 | 2.188 | 1.44E-03 | glio |
| Aars2_in.vivo_noRT | Tufm | -1.334 | 1.44E-03 | glio |
| Aars2_in.vivo_noRT | Ccdc86 | -1.384 | 1.44E-03 | glio |
| Aars2_in.vivo_noRT | Uqcrq | -0.699 | 1.44E-03 | glio |
| Atp5c1_in.vivo_noRT | Spns3 | 2.538 | 1.44E-03 | glio |
| Atr_in.vivo_noRT | Bach2 | 2.25 | 1.44E-03 | glio |
| Atr_in.vivo_noRT | Ppic | -1.22 | 1.44E-03 | glio |
| Atr_in.vivo_noRT | Rps4x | -0.758 | 1.44E-03 | glio |
| Atr_ex.vivo_noRT | Gm43701 | 2.181 | 1.44E-03 | glio |
| Bcas2_in.vivo_noRT | Cyp2d22 | 2.47 | 1.44E-03 | glio |
| Bcas2_in.vivo_noRT | Dynlt1b | 2.321 | 1.44E-03 | glio |
| Bcas2_in.vivo_noRT | Cadm2 | 2.444 | 1.44E-03 | glio |
| Bcas2_in.vivo_noRT | Smim40 | 2.305 | 1.44E-03 | glio |
| Bcas2_in.vivo_noRT | Psme2b | 2.223 | 1.44E-03 | glio |
| Bora_in.vivo_noRT | 2410022M11F | 2.365 | 1.44E-03 | glio |
| Bora_ex.vivo_noRT | Dreh | 2.808 | 1.44E-03 | glio |
| Chek1_in.vivo_noRT | Gm43329 | 2.17 | 1.44E-03 | glio |
| Chek1_in.vivo_noRT | Acox2 | 1.989 | 1.44E-03 | glio |
| Chek1_in.vivo_noRT | Rps20 | -0.687 | 1.44E-03 | glio |
| Chek1_ex.vivo_noRT | Dnah2os | 2.76 | 1.44E-03 | glio |
| Eif2s1_ex.vivo_noRT | Rtl9 | 3.479 | 1.44E-03 | glio |
| culture_effect | Clca3a2 | -2.121 | 1.44E-04 | glio |
| Interaction_genotype_Chek1 | Baalc | -4.524 | 1.44E-04 | glio |
| Interaction_genotype_Mtor | Jakmip1 | -4.547 | 1.44E-04 | glio |
| Interaction_genotype_Mtor | Gm15080 | -4.924 | 1.44E-04 | glio |
| Interaction_genotype_Mtor | Gm14698 | -4.593 | 1.44E-04 | glio |
| Interaction_genotype_Mtor | Nphp3 | -4.529 | 1.44E-04 | glio |
| Interaction_genotype_Myc | Ramp3 | -4.273 | 1.44E-04 | glio |
| Interaction_genotype_Myc | Ppp1r3f | -4.563 | 1.44E-04 | glio |
| Interaction_genotype_Myc | Smpd2 | -3.627 | 1.44E-04 | glio |
| Interaction_genotype_Myc | Rpgrip1l | -3.408 | 1.44E-04 | glio |
| Interaction_genotype_Myc | Ciao2a | 2.027 | 1.44E-04 | glio |
| Interaction_genotype_Nat2 | Gm5286 | 3.825 | 1.44E-04 | glio |
| Atr_in.vivo_noRT | Kmt2d | -1.641 | 1.44E-04 | glio |

| | | | | |
|---|---|---|---|---|
| Atr_ex.vivo_noRT | Gm35439 | 2.764 | 1.44E-04 | glio |
| Bcas2_in.vivo_noRT | Stpg4 | 2.848 | 1.44E-04 | glio |
| Bora_in.vivo_noRT | Wwp1 | -1.633 | 1.44E-04 | glio |
| Bora_in.vivo_noRT | Cenps | -1.832 | 1.44E-04 | glio |
| Chek1_ex.vivo_noRT | Gm50399 | 3.153 | 1.44E-04 | glio |
| culture_effect | Lrrc73 | 1.405 | 1.44E-05 | glio |
| culture_effect | Cyp7b1 | -1.394 | 1.44E-05 | glio |
| culture_effect | Gm16062 | 1.123 | 1.44E-05 | glio |
| culture_effect | Galk1 | -0.607 | 1.44E-05 | glio |
| Interaction_genotype_Mtor | Gm4027 | -4.099 | 1.44E-05 | glio |
| Interaction_genotype_Myc | Rab3b | -4.865 | 1.44E-05 | glio |
| Aars2_in.vivo_noRT | 4930554I06Ri | 3.267 | 1.44E-05 | glio |
| Aars2_in.vivo_noRT | Cish | 2.94 | 1.44E-05 | glio |
| Atr_ex.vivo_noRT | Gpc5 | 2.769 | 1.44E-05 | glio |
| Bcas2_in.vivo_noRT | Gm15013 | 3.014 | 1.44E-05 | glio |
| Hsd17b10_in.vivo_noRT | Ntn1 | 2.714 | 1.44E-05 | glio |
| Hsd17b10_in.vivo_noRT | Gm47232 | 3.093 | 1.44E-05 | glio |
| culture_effect | Vps35 | 0.579 | 1.44E-06 | glio |
| Interaction_genotype_Mtor | Zhx2 | -6.088 | 1.44E-06 | glio |
| Interaction_genotype_Mtor | Hectd2os | -5.565 | 1.44E-06 | glio |
| Interaction_genotype_Myc | Sod3 | -5.772 | 1.44E-06 | glio |
| Interaction_genotype_Myc | Kcnmb4os2 | -5.479 | 1.44E-06 | glio |
| Atr_in.vivo_noRT | Tc2n | 3.446 | 1.44E-06 | glio |
| Atr_in.vivo_noRT | Rfx2 | 3.133 | 1.44E-06 | glio |
| Atr_ex.vivo_noRT | Mcmdc2 | 3.636 | 1.44E-06 | glio |
| Bcas2_in.vivo_noRT | Tmem202 | 3.365 | 1.44E-06 | glio |
| Bora_in.vivo_noRT | Tnfrsf4 | 3.35 | 1.44E-06 | glio |
| Bora_in.vivo_noRT | Gm16794 | 3.365 | 1.44E-06 | glio |
| Chek1_in.vivo_noRT | Gm21814 | 3.185 | 1.44E-06 | glio |
| Chek1_in.vivo_noRT | Prrx1 | -1.978 | 1.44E-06 | glio |
| culture_effect | Tpi1 | -0.735 | 1.44E-07 | glio |
| Interaction_genotype_Myc | Ftl1 | 1.978 | 1.44E-07 | glio |
| Aars2_in.vivo_noRT | Ppnr | 3.664 | 1.44E-07 | glio |
| Atp5c1_in.vivo_noRT | Ctla4 | 2.668 | 1.44E-07 | glio |
| Atp5c1_in.vivo_noRT | Zscan4-ps3 | 2.668 | 1.44E-07 | glio |
| Atp5c1_in.vivo_noRT | A030014E15Ri | 2.668 | 1.44E-07 | glio |
| Atp5c1_in.vivo_noRT | Gm21731 | 2.668 | 1.44E-07 | glio |
| Bcas2_in.vivo_noRT | Ccno | 3.787 | 1.44E-07 | glio |
| Bcas2_in.vivo_noRT | Zfp811 | 3.633 | 1.44E-07 | glio |
| Hsd17b10_in.vivo_noRT | Gem | 3.616 | 1.44E-07 | glio |
| Hsd17b10_in.vivo_noRT | Ripply3 | 3.616 | 1.44E-07 | glio |
| Hsd17b10_in.vivo_noRT | Rab25 | 3.35 | 1.44E-07 | glio |
| culture_effect | Tnfsf10 | -2.824 | 1.44E-08 | glio |
| Bora_in.vivo_noRT | Esyt3 | 3.877 | 1.44E-08 | glio |
| Bora_in.vivo_noRT | Rhox3g | 3.822 | 1.44E-08 | glio |
| Chek1_in.vivo_noRT | 2210417A02R | 4.253 | 1.44E-08 | glio |
| culture_effect | Txndc12 | 0.826 | 1.44E-09 | glio |
| Atr_in.vivo_noRT | Pcdhgb5 | 3.629 | 1.44E-09 | glio |
| Atr_ex.vivo_noRT | Slc17a6 | 3.164 | 1.44E-09 | glio |
| Atr_ex.vivo_noRT | Hist1h2bc | 4.754 | 1.44E-09 | glio |
| Bcas2_in.vivo_noRT | BC028471 | 4.275 | 1.44E-09 | glio |
| culture_effect | 4930581F22Ri | 2.203 | 1.44E-10 | glio |

| | | | | |
|---|---|---|---|---|
| culture_effect | Hspb11 | -0.883 | 1.44E-10 | glio |
| culture_effect | Eci1 | -0.925 | 1.44E-10 | glio |
| Aars2_in.vivo_noRT | Qrich2 | 4.548 | 1.44E-10 | glio |
| Aars2_in.vivo_noRT | Cpeb1 | 4.681 | 1.44E-10 | glio |
| Atr_in.vivo_noRT | AU022754 | 4.113 | 1.44E-10 | glio |
| Bcas2_in.vivo_noRT | Gm34047 | 4.807 | 1.44E-11 | glio |
| Bcas2_in.vivo_noRT | Podxl | 4.275 | 1.44E-11 | glio |
| Bcas2_in.vivo_noRT | Gm21818 | 4.807 | 1.44E-11 | glio |
| Hsd17b10_in.vivo_noRT | Gm47936 | 3.616 | 1.44E-11 | glio |
| Hsd17b10_in.vivo_noRT | Tnni3k | 3.616 | 1.44E-11 | glio |
| Atr_in.vivo_noRT | Gm10425 | 5.269 | 1.44E-13 | glio |
| Chek1_in.vivo_noRT | Gm39459 | 5.172 | 1.44E-13 | glio |
| Hsd17b10_in.vivo_noRT | 1700003E24Ri | 3.616 | 1.44E-14 | glio |
| Bcas2_in.vivo_noRT | Slc16a4 | 4.543 | 1.44E-15 | glio |
| Bcas2_in.vivo_noRT | Gm15894 | 4.28 | 1.44E-15 | glio |
| Bcas2_in.vivo_noRT | Gm27241 | 4.151 | 1.44E-15 | glio |
| Chek1_in.vivo_noRT | Gm2115 | 5.044 | 1.44E-16 | glio |
| Atr_in.vivo_noRT | Card9 | 5.269 | 1.44E-18 | glio |
| Atr_in.vivo_noRT | Hrct1 | 5.533 | 1.44E-19 | glio |
| Chek1_in.vivo_noRT | Cd7 | 4.78 | 1.44E-21 | glio |
| culture_effect | Sec14l5 | -1.478 | 1.45E-03 | glio |
| culture_effect | Nup214 | 0.629 | 1.45E-03 | glio |
| culture_effect | Ddx27 | 0.454 | 1.45E-03 | glio |
| Interaction_genotype_Bora | Gm47851 | -3.862 | 1.45E-03 | glio |
| Interaction_genotype_Bora | 4930515G01R | -3.782 | 1.45E-03 | glio |
| Interaction_genotype_Bora | Schip1 | 2.128 | 1.45E-03 | glio |
| Interaction_genotype_Mtor | Stx19 | -3.187 | 1.45E-03 | glio |
| Interaction_genotype_Mtor | Fam222a | -4.143 | 1.45E-03 | glio |
| Interaction_genotype_Mtor | Tom1 | -2.537 | 1.45E-03 | glio |
| Interaction_genotype_Myc | BC018473 | -3.526 | 1.45E-03 | glio |
| Interaction_genotype_Myc | Lrrc46 | -3.606 | 1.45E-03 | glio |
| Interaction_genotype_Myc | Acvr1b | -3.661 | 1.45E-03 | glio |
| Interaction_genotype_Myc | Nt5dc3 | -2.575 | 1.45E-03 | glio |
| Interaction_genotype_Myc | Clcf1 | -2.934 | 1.45E-03 | glio |
| Interaction_genotype_Myc | Mrpl52 | 1.516 | 1.45E-03 | glio |
| Interaction_genotype_Myc | Mrpl54 | 1.206 | 1.45E-03 | glio |
| Interaction_genotype_Pars2 | Gm43653 | -4.101 | 1.45E-03 | glio |
| interaction_RT | Arid4a | 0.952 | 1.45E-03 | glio |
| interaction_RT | Srp14 | 0.482 | 1.45E-03 | glio |
| Aars2_in.vivo_noRT | Zfp689 | 1.839 | 1.45E-03 | glio |
| Aars2_in.vivo_noRT | Rap1b | -0.873 | 1.45E-03 | glio |
| Atp5c1_in.vivo_noRT | Cyp2c23 | 2.145 | 1.45E-03 | glio |
| Atr_ex.vivo_noRT | Gm33051 | 2.367 | 1.45E-03 | glio |
| Atr_ex.vivo_noRT | Ccdc114 | 2.863 | 1.45E-03 | glio |
| Bard1_ex.vivo_noRT | 4930527F14Ri | 2.419 | 1.45E-03 | glio |
| Bard1_ex.vivo_noRT | Fhad1 | 3.026 | 1.45E-03 | glio |
| Bard1_ex.vivo_noRT | Art2b | 2.814 | 1.45E-03 | glio |
| Bcas2_in.vivo_noRT | Fcgrt | 2.56 | 1.45E-03 | glio |
| Bcas2_ex.vivo_noRT | Etohd2 | 3.168 | 1.45E-03 | glio |
| Bora_in.vivo_noRT | Tnnc1 | 2.546 | 1.45E-03 | glio |
| Bora_in.vivo_noRT | B4galt4 | 2.374 | 1.45E-03 | glio |
| Bora_in.vivo_noRT | Sgo2a | -1.903 | 1.45E-03 | glio |

| | | | | |
|---|---|---|---|---|
| Bora_in.vivo_noRT | Cog5 | 1.678 | 1.45E-03 | glio |
| Bora_ex.vivo_noRT | Kcnk13 | 2.413 | 1.45E-03 | glio |
| Bora_ex.vivo_noRT | Btn1a1 | 2.417 | 1.45E-03 | glio |
| Cenpj_ex.vivo_noRT | Gm4027 | 2.591 | 1.45E-03 | glio |
| Chek1_in.vivo_noRT | Tnni1 | 1.684 | 1.45E-03 | glio |
| Chek1_in.vivo_noRT | Snx4 | 1.276 | 1.45E-03 | glio |
| Chek1_ex.vivo_noRT | Pdzk1ip1 | 3.042 | 1.45E-03 | glio |
| Chek1_ex.vivo_noRT | Mrgprf | 2.946 | 1.45E-03 | glio |
| Eif2s1_in.vivo_noRT | Gm13288 | 2.143 | 1.45E-03 | glio |
| Eif2s1_in.vivo_noRT | Il17d | -2.123 | 1.45E-03 | glio |
| Hsd17b10_in.vivo_noRT | Nkx2-2os | 2.492 | 1.45E-03 | glio |
| Hsd17b10_in.vivo_noRT | A630052C17R | 2.332 | 1.45E-03 | glio |
| culture_effect | Pros1 | -1.517 | 1.45E-04 | glio |
| Interaction_genotype_Myc | Gabpb2 | -2.765 | 1.45E-04 | glio |
| interaction_RT | Spag7 | -0.886 | 1.45E-04 | glio |
| interaction_RT | Vars | -0.943 | 1.45E-04 | glio |
| Atr_in.vivo_noRT | Ptn | 3.135 | 1.45E-04 | glio |
| Atr_in.vivo_noRT | Gipc3 | 2.764 | 1.45E-04 | glio |
| Atr_ex.vivo_noRT | Rslcan18 | 3.16 | 1.45E-04 | glio |
| Atr_ex.vivo_noRT | Gngt2 | 3.438 | 1.45E-04 | glio |
| Bora_in.vivo_noRT | Cldn10 | 2.729 | 1.45E-04 | glio |
| Chek1_in.vivo_noRT | Gm48768 | 2.792 | 1.45E-04 | glio |
| Chek1_in.vivo_noRT | Arsj | 2.929 | 1.45E-04 | glio |
| Chek1_in.vivo_noRT | Pabpc4l | 2.765 | 1.45E-04 | glio |
| Chek1_in.vivo_noRT | Gipc3 | 2.798 | 1.45E-04 | glio |
| Chek1_in.vivo_noRT | Hnrnpc | -0.888 | 1.45E-04 | glio |
| Hsd17b10_in.vivo_noRT | Gm48610 | 2.834 | 1.45E-04 | glio |
| culture_effect | Nrarp | -2.127 | 1.45E-05 | glio |
| Interaction_genotype_Mtor | Slc46a1 | -5.347 | 1.45E-05 | glio |
| Interaction_genotype_Myc | Trp53bp2 | -3.949 | 1.45E-05 | glio |
| Interaction_genotype_Myc | Serpine2 | -5.872 | 1.45E-05 | glio |
| Aars2_in.vivo_noRT | Spata1 | 3.166 | 1.45E-05 | glio |
| Aars2_in.vivo_noRT | Tert | 2.855 | 1.45E-05 | glio |
| Atr_ex.vivo_noRT | Gm40264 | 3.36 | 1.45E-05 | glio |
| Atr_ex.vivo_noRT | Anxa8 | 3.952 | 1.45E-05 | glio |
| Bcas2_in.vivo_noRT | 1700006J14Ri | 2.897 | 1.45E-05 | glio |
| Bcas2_in.vivo_noRT | Gm49164 | 3.144 | 1.45E-05 | glio |
| Interaction_genotype_Myc | Zscan22 | -4.653 | 1.45E-06 | glio |
| Aars2_in.vivo_noRT | Hdac11 | 3.442 | 1.45E-06 | glio |
| Aars2_in.vivo_noRT | Fbxl22 | 3.21 | 1.45E-06 | glio |
| Aars2_ex.vivo_noRT | 4930512B01R | 3.293 | 1.45E-06 | glio |
| Bora_in.vivo_noRT | S1pr1 | 3.751 | 1.45E-06 | glio |
| Chek1_in.vivo_noRT | Spink2 | 3.847 | 1.45E-06 | glio |
| Chek1_in.vivo_noRT | BC051077 | 3.863 | 1.45E-06 | glio |
| Chek1_in.vivo_noRT | Cebpg | -1.746 | 1.45E-06 | glio |
| Hsd17b10_in.vivo_noRT | Cd248 | 3.35 | 1.45E-06 | glio |
| culture_effect | Esyt1 | -1.025 | 1.45E-07 | glio |
| Aars2_in.vivo_noRT | Pate2 | 3.847 | 1.45E-07 | glio |
| Aars2_in.vivo_noRT | 0610009L18Ri | 3.815 | 1.45E-07 | glio |
| Aars2_ex.vivo_noRT | Ablim2 | 3.293 | 1.45E-07 | glio |
| Aars2_ex.vivo_noRT | Gm5415 | 3.293 | 1.45E-07 | glio |
| Atr_in.vivo_noRT | Gm37170 | 3.69 | 1.45E-07 | glio |

| | | | | |
|---|---|---|---|---|
| Atr_in.vivo_noRT | Mlf2 | -1.553 | 1.45E-07 | glio |
| Hsd17b10_in.vivo_noRT | Gm45620 | 3.616 | 1.45E-07 | glio |
| Hsd17b10_in.vivo_noRT | Gm38534 | 3.616 | 1.45E-07 | glio |
| culture_effect | Socs2 | 1.789 | 1.45E-08 | glio |
| Chek1_ex.vivo_noRT | 2410141K09R | 4.058 | 1.45E-08 | glio |
| Chek1_ex.vivo_noRT | C1ra | 4.08 | 1.45E-08 | glio |
| Bcas2_in.vivo_noRT | Oxt | 4.166 | 1.45E-09 | glio |
| Chek1_ex.vivo_noRT | Gm26978 | 4.256 | 1.45E-09 | glio |
| Chek1_ex.vivo_noRT | Btn1a1 | 4.058 | 1.45E-09 | glio |
| Chek1_ex.vivo_noRT | Zdhhc14 | 4.447 | 1.45E-09 | glio |
| Chek1_in.vivo_noRT | Postn | 4.409 | 1.45E-10 | glio |
| Hsd17b10_in.vivo_noRT | Gast | 3.616 | 1.45E-10 | glio |
| culture_effect | Gvin1 | -4.176 | 1.45E-11 | glio |
| Aars2_in.vivo_noRT | Notch4 | 4.593 | 1.45E-11 | glio |
| Bora_in.vivo_noRT | Osm | 4.111 | 1.45E-11 | glio |
| culture_effect | Rhoc | 0.976 | 1.45E-12 | glio |
| culture_effect | Lsm7 | 0.811 | 1.45E-12 | glio |
| Hsd17b10_in.vivo_noRT | Gm12794 | 3.616 | 1.45E-12 | glio |
| Aars2_in.vivo_noRT | 2300002M23F | 4.158 | 1.45E-13 | glio |
| Atr_in.vivo_noRT | D030068K23R | 5.142 | 1.45E-13 | glio |
| Bcas2_in.vivo_noRT | Slc4a9 | 4.275 | 1.45E-13 | glio |
| Atr_ex.vivo_noRT | Ankrd22 | 4.725 | 1.45E-15 | glio |
| Bcas2_in.vivo_noRT | Vtn | 4.547 | 1.45E-18 | glio |
| Chek1_in.vivo_noRT | Gm44646 | 5.044 | 1.45E-18 | glio |
| culture_effect | Igsf8 | -2.412 | 1.45E-20 | glio |
| Atr_in.vivo_noRT | Gm11583 | 5.008 | 1.45E-21 | glio |
| culture_effect | Wasf3 | 0.799 | 1.46E-03 | glio |
| culture_effect | Mpv17 | -0.508 | 1.46E-03 | glio |
| culture_effect | Grsf1 | -0.424 | 1.46E-03 | glio |
| Interaction_genotype_Aars2 | Tnfrsf23 | -4.346 | 1.46E-03 | glio |
| Interaction_genotype_Aars2 | Ccnb1ip1 | -4.24 | 1.46E-03 | glio |
| Interaction_genotype_Atr | Gm44702 | -4.06 | 1.46E-03 | glio |
| Interaction_genotype_Atr | 4921504E06Ri | -3.184 | 1.46E-03 | glio |
| Interaction_genotype_Atr | Fhl1 | -3.08 | 1.46E-03 | glio |
| Interaction_genotype_Atr | Clec12b | -2.765 | 1.46E-03 | glio |
| Interaction_genotype_Atr | Gm49874 | -3.94 | 1.46E-03 | glio |
| Interaction_genotype_Atr | Tead3 | -3.969 | 1.46E-03 | glio |
| Interaction_genotype_Atr | Pcdh1 | -3.76 | 1.46E-03 | glio |
| Interaction_genotype_Atr | Ier5 | 2.728 | 1.46E-03 | glio |
| Interaction_genotype_Atr | Tmem128 | 1.308 | 1.46E-03 | glio |
| Interaction_genotype_Atr | Tra2a | 1.578 | 1.46E-03 | glio |
| Interaction_genotype_Atr | Mt1 | 1.931 | 1.46E-03 | glio |
| Interaction_genotype_Bora | Itgb6 | -4.238 | 1.46E-03 | glio |
| Interaction_genotype_Cdkn2a | Gm49172 | 3.095 | 1.46E-03 | glio |
| Interaction_genotype_Cdkn2a | Gm7247 | 3.88 | 1.46E-03 | glio |
| Interaction_genotype_Chek1 | Cebpb | 2.257 | 1.46E-03 | glio |
| Interaction_genotype_Mtor | Pdia6 | 1.76 | 1.46E-03 | glio |
| Interaction_genotype_Mtor | Jpt1 | 1.666 | 1.46E-03 | glio |
| Interaction_genotype_Myc | Dmrtc1a | -2.729 | 1.46E-03 | glio |
| Interaction_genotype_Myc | CAAA0106612 | -3.274 | 1.46E-03 | glio |
| Interaction_genotype_Myc | Gm40264 | -3.056 | 1.46E-03 | glio |
| Interaction_genotype_Myc | Gm26688 | -3.552 | 1.46E-03 | glio |

| | | | | |
|---|---|---|---|---|
| Interaction_genotype_Myc | Pnpla6 | -3.209 | 1.46E-03 | glio |
| Interaction_genotype_Myc | Ap4b1 | -2.907 | 1.46E-03 | glio |
| Interaction_genotype_Myc | Zfp574 | -2.734 | 1.46E-03 | glio |
| Interaction_genotype_Myc | Taf8 | -2.655 | 1.46E-03 | glio |
| Interaction_genotype_Myc | Tgif2 | -2.911 | 1.46E-03 | glio |
| Interaction_genotype_Myc | Rorb | 2.377 | 1.46E-03 | glio |
| Interaction_genotype_Rad51 | Gm2115 | -3.887 | 1.46E-03 | glio |
| interaction_RT | Ier3 | 1.088 | 1.46E-03 | glio |
| interaction_RT | Fkbp4 | -0.625 | 1.46E-03 | glio |
| interaction_RT | Calm2 | 0.626 | 1.46E-03 | glio |
| Aars2_in.vivo_noRT | Syngr4 | 2.286 | 1.46E-03 | glio |
| Aars2_in.vivo_noRT | Gm49041 | 2.029 | 1.46E-03 | glio |
| Aars2_in.vivo_noRT | Car13 | 1.658 | 1.46E-03 | glio |
| Aars2_in.vivo_noRT | Map4k2 | 1.725 | 1.46E-03 | glio |
| Aars2_in.vivo_noRT | Scd2 | -1.557 | 1.46E-03 | glio |
| Aars2_in.vivo_noRT | Tomm5 | -0.843 | 1.46E-03 | glio |
| Aars2_ex.vivo_noRT | Spn | 2.899 | 1.46E-03 | glio |
| Aars2_ex.vivo_noRT | Alox15 | 2.897 | 1.46E-03 | glio |
| Aars2_ex.vivo_noRT | Nrn1l | 2.893 | 1.46E-03 | glio |
| Atp5c1_in.vivo_noRT | Rcan2 | 2.668 | 1.46E-03 | glio |
| Atp5c1_in.vivo_noRT | Gm2012 | 2.556 | 1.46E-03 | glio |
| Atr_in.vivo_noRT | Hist1h2br | 2.481 | 1.46E-03 | glio |
| Atr_in.vivo_noRT | Ccdc7b | 2.192 | 1.46E-03 | glio |
| Atr_in.vivo_noRT | CAAA0114733 | 1.977 | 1.46E-03 | glio |
| Atr_in.vivo_noRT | Poll | 1.748 | 1.46E-03 | glio |
| Atr_in.vivo_noRT | Srsf4 | -1.197 | 1.46E-03 | glio |
| Atr_in.vivo_noRT | Usp34 | -0.979 | 1.46E-03 | glio |
| Atr_ex.vivo_noRT | Rtl1 | 2.769 | 1.46E-03 | glio |
| Atr_ex.vivo_noRT | Podxl | 2.77 | 1.46E-03 | glio |
| Atr_ex.vivo_noRT | Cad | -1.844 | 1.46E-03 | glio |
| Bard1_ex.vivo_noRT | Iqch | 2.647 | 1.46E-03 | glio |
| Bcas2_in.vivo_noRT | Vps50 | -1.567 | 1.46E-03 | glio |
| Bcas2_in.vivo_noRT | Cep57 | -1.281 | 1.46E-03 | glio |
| Bcas2_in.vivo_noRT | Mrpl15 | -0.812 | 1.46E-03 | glio |
| Bora_ex.vivo_noRT | 8030423F21Ri | 2.626 | 1.46E-03 | glio |
| Cenpj_in.vivo_noRT | Sema6d | 2.022 | 1.46E-03 | glio |
| Chek1_ex.vivo_noRT | Mlxipl | 2.668 | 1.46E-03 | glio |
| Chek1_ex.vivo_noRT | Sncg | 3.171 | 1.46E-03 | glio |
| Chek1_ex.vivo_noRT | Pdxdc1 | -1.98 | 1.46E-03 | glio |
| Hsd17b10_in.vivo_noRT | Gm15672 | 2.414 | 1.46E-03 | glio |
| culture_effect | Tmem39b | -0.93 | 1.46E-04 | glio |
| culture_effect | Pprc1 | -0.599 | 1.46E-04 | glio |
| Interaction_genotype_Atr | Gm26673 | -4.684 | 1.46E-04 | glio |
| Interaction_genotype_Chek1 | Upf3a | 2.421 | 1.46E-04 | glio |
| Interaction_genotype_Myc | Myo5c | -4.223 | 1.46E-04 | glio |
| Interaction_genotype_Myc | Sh3d21 | -3.704 | 1.46E-04 | glio |
| Interaction_genotype_Myc | Atp6v0e | 1.473 | 1.46E-04 | glio |
| Interaction_genotype_Terf1 | Folr1 | -4.797 | 1.46E-04 | glio |
| interaction_RT | Eif3l | -0.755 | 1.46E-04 | glio |
| interaction_RT | Gm28535 | 3.388 | 1.46E-04 | glio |
| Atr_in.vivo_noRT | Gldc | 2.625 | 1.46E-04 | glio |
| Atr_in.vivo_noRT | Fzd1 | 2.565 | 1.46E-04 | glio |

| | | | | |
|---|---|---|---|---|
| Atr_in.vivo_noRT | Uba3 | -1.428 | 1.46E-04 | glio |
| Atr_in.vivo_noRT | Rps18 | -0.868 | 1.46E-04 | glio |
| Bcas2_in.vivo_noRT | Prob1 | 2.859 | 1.46E-04 | glio |
| Bora_in.vivo_noRT | Sgk2 | 2.72 | 1.46E-04 | glio |
| Bora_ex.vivo_noRT | Cidea | 3.201 | 1.46E-04 | glio |
| Bora_ex.vivo_noRT | Gm49860 | 3.013 | 1.46E-04 | glio |
| Bora_ex.vivo_noRT | 4930470G03R | 2.808 | 1.46E-04 | glio |
| Bora_ex.vivo_noRT | Asap3 | 3.56 | 1.46E-04 | glio |
| Chek1_in.vivo_noRT | Osbp2 | 3.118 | 1.46E-04 | glio |
| Chek1_in.vivo_noRT | Gm14966 | 2.576 | 1.46E-04 | glio |
| culture_effect | Insc | -2.392 | 1.46E-05 | glio |
| culture_effect | Nts | -2.978 | 1.46E-05 | glio |
| culture_effect | Plrg1 | -1.025 | 1.46E-05 | glio |
| Interaction_genotype_Myc | Gm43728 | -5.189 | 1.46E-05 | glio |
| Interaction_genotype_Myc | Gm14327 | -5.105 | 1.46E-05 | glio |
| Atp5c1_in.vivo_noRT | Gm14820 | 2.405 | 1.46E-05 | glio |
| Atr_in.vivo_noRT | N4bp2os | 3.105 | 1.46E-05 | glio |
| Atr_in.vivo_noRT | Gm19744 | 3.088 | 1.46E-05 | glio |
| Bcas2_ex.vivo_noRT | Gm15908 | 2.742 | 1.46E-05 | glio |
| Bcas2_ex.vivo_noRT | A930014E10Ri | 2.742 | 1.46E-05 | glio |
| Hsd17b10_in.vivo_noRT | Cstdc4 | 2.959 | 1.46E-05 | glio |
| Hsd17b10_in.vivo_noRT | AV039307 | 3.225 | 1.46E-05 | glio |
| Hsd17b10_in.vivo_noRT | Rdh12 | 2.971 | 1.46E-05 | glio |
| Hsd17b10_in.vivo_noRT | Gm5152 | 3.229 | 1.46E-05 | glio |
| culture_effect | Hsp90ab1 | -0.677 | 1.46E-06 | glio |
| Interaction_genotype_Mtor | Gm32834 | -5.098 | 1.46E-06 | glio |
| Aars2_in.vivo_noRT | Ndfip1 | -1.271 | 1.46E-06 | glio |
| Chek1_in.vivo_noRT | 2810457G06R | 3.796 | 1.46E-06 | glio |
| Chek1_in.vivo_noRT | Ppnr | 3.381 | 1.46E-06 | glio |
| culture_effect | Slc16a13 | 1.86 | 1.46E-07 | glio |
| Interaction_genotype_Mtor | Rab3b | -6.235 | 1.46E-07 | glio |
| Atp5c1_in.vivo_noRT | Gm11232 | 2.668 | 1.46E-07 | glio |
| Atr_in.vivo_noRT | 1700066B19R | 3.557 | 1.46E-07 | glio |
| Bcas2_in.vivo_noRT | Gm10767 | 3.737 | 1.46E-07 | glio |
| Bcas2_in.vivo_noRT | Gm50012 | 3.724 | 1.46E-07 | glio |
| Chek1_ex.vivo_noRT | Pcdhb18 | 4.464 | 1.46E-07 | glio |
| culture_effect | Wincr1 | 3.742 | 1.46E-08 | glio |
| Aars2_in.vivo_noRT | Dnajc28 | 4.181 | 1.46E-08 | glio |
| Atr_in.vivo_noRT | Olfr1369-ps1 | 4.792 | 1.46E-08 | glio |
| Atr_in.vivo_noRT | Slc7a14 | 3.763 | 1.46E-08 | glio |
| Bora_in.vivo_noRT | Nid1 | 4.132 | 1.46E-08 | glio |
| culture_effect | Lrrc8c | 1.333 | 1.46E-09 | glio |
| Interaction_genotype_Mtor | Rcan2 | -7.707 | 1.46E-09 | glio |
| Interaction_genotype_Mtor | Gm10010 | -7.569 | 1.46E-09 | glio |
| Interaction_genotype_Myc | Asgr1 | -7.126 | 1.46E-09 | glio |
| Interaction_genotype_Myc | Cyp26b1 | -7.416 | 1.46E-09 | glio |
| Chek1_in.vivo_noRT | Nrip3 | 3.999 | 1.46E-09 | glio |
| culture_effect | Dcun1d5 | 0.743 | 1.46E-10 | glio |
| Aars2_in.vivo_noRT | Syp | 4.936 | 1.46E-10 | glio |
| Bora_in.vivo_noRT | Gm49579 | 3.883 | 1.46E-10 | glio |
| Chek1_in.vivo_noRT | Bend7 | 4.265 | 1.46E-10 | glio |
| Chek1_in.vivo_noRT | Pacrg | 4.252 | 1.46E-11 | glio |

| | | | | |
|---|---|---|---|---|
| Aars2_in.vivo_noRT | Cd52 | 4.67 | 1.46E-12 | glio |
| culture_effect | Lztfl1 | -1.326 | 1.46E-13 | glio |
| culture_effect | Snhg12 | -1.247 | 1.46E-13 | glio |
| Aars2_in.vivo_noRT | Gm47872 | 5.063 | 1.46E-14 | glio |
| Chek1_in.vivo_noRT | D030040B21R | 5.186 | 1.46E-14 | glio |
| Atr_in.vivo_noRT | Gm5662 | 5.533 | 1.46E-16 | glio |
| culture_effect | Trib3 | -2.141 | 1.46E-18 | glio |
| Atr_in.vivo_noRT | Gm2449 | 5.27 | 1.46E-18 | glio |
| Bcas2_in.vivo_noRT | Trmt61b | 4.807 | 1.46E-20 | glio |
| Atr_in.vivo_noRT | Gm29670 | 5.011 | 1.46E-21 | glio |
| Atr_in.vivo_noRT | Esp15 | 5.533 | 1.46E-21 | glio |
| Bcas2_in.vivo_noRT | BC061195 | 4.807 | 1.46E-22 | glio |
| Chek1_in.vivo_noRT | Gpc5 | 5.044 | 1.46E-23 | glio |
| culture_effect | Gm47438 | 1.7 | 1.47E-03 | glio |
| culture_effect | Trpv4 | -0.698 | 1.47E-03 | glio |
| culture_effect | Rbm27 | 0.505 | 1.47E-03 | glio |
| Interaction_genotype_Aars2 | Gm44899 | -3.138 | 1.47E-03 | glio |
| Interaction_genotype_Aars2 | Gm10010 | -3.926 | 1.47E-03 | glio |
| Interaction_genotype_Aars2 | Atxn1l | -2.828 | 1.47E-03 | glio |
| Interaction_genotype_Mtor | Mfsd10 | 1.977 | 1.47E-03 | glio |
| Interaction_genotype_Mtor | Arcn1 | 1.732 | 1.47E-03 | glio |
| Interaction_genotype_Myc | Meioc | -2.466 | 1.47E-03 | glio |
| Interaction_genotype_Myc | Slc10a1 | -2.863 | 1.47E-03 | glio |
| Interaction_genotype_Myc | Tmem86b | -3.708 | 1.47E-03 | glio |
| Interaction_genotype_Myc | Bhlhe41 | -4.075 | 1.47E-03 | glio |
| Interaction_genotype_Myc | Cpt1c | -3.315 | 1.47E-03 | glio |
| Interaction_genotype_Myc | 2900089D17R | -3.242 | 1.47E-03 | glio |
| Interaction_genotype_Myc | Uggt2 | -3.086 | 1.47E-03 | glio |
| Interaction_genotype_Myc | Rilpl2 | -2.646 | 1.47E-03 | glio |
| interaction_RT | H3f3a | 0.529 | 1.47E-03 | glio |
| Aars2_in.vivo_noRT | Whrn | 2.291 | 1.47E-03 | glio |
| Aars2_in.vivo_noRT | Kbtbd3 | 1.835 | 1.47E-03 | glio |
| Atr_in.vivo_noRT | 1500004A13R | 2.308 | 1.47E-03 | glio |
| Atr_in.vivo_noRT | Usp5 | -1.359 | 1.47E-03 | glio |
| Bcas2_in.vivo_noRT | Gm32913 | 3.536 | 1.47E-03 | glio |
| Bcas2_in.vivo_noRT | Cdc14a | -1.623 | 1.47E-03 | glio |
| Bcas2_ex.vivo_noRT | Gm45669 | 2.736 | 1.47E-03 | glio |
| Bora_in.vivo_noRT | Kctd1 | 2.231 | 1.47E-03 | glio |
| Bora_in.vivo_noRT | Pou5f1 | 2.151 | 1.47E-03 | glio |
| Bora_in.vivo_noRT | Efna5 | 2.265 | 1.47E-03 | glio |
| Bora_in.vivo_noRT | Pdzd8 | -1.469 | 1.47E-03 | glio |
| Bora_in.vivo_noRT | Dcaf12 | -1.174 | 1.47E-03 | glio |
| Cenpj_ex.vivo_noRT | Gm40538 | 2.591 | 1.47E-03 | glio |
| Chek1_in.vivo_noRT | Myo5a | -1.373 | 1.47E-03 | glio |
| Chek1_ex.vivo_noRT | Zhx2 | 2.988 | 1.47E-03 | glio |
| Dclre1c_ex.vivo_noRT | Myt1 | 2.687 | 1.47E-03 | glio |
| Dclre1c_ex.vivo_noRT | Olfr787 | 2.292 | 1.47E-03 | glio |
| Dclre1c_ex.vivo_noRT | Adam2 | 3.086 | 1.47E-03 | glio |
| Dclre1c_ex.vivo_noRT | Tex14 | 3.292 | 1.47E-03 | glio |
| Hsd17b10_in.vivo_noRT | Ccl20 | 3.616 | 1.47E-03 | glio |
| Hsd17b10_in.vivo_noRT | Gm43608 | 2.442 | 1.47E-03 | glio |
| culture_effect | Klf4 | 0.765 | 1.47E-04 | glio |

| | | | | |
|---|---|---|---|---|
| culture_effect | Tmbim4 | -0.517 | 1.47E-04 | glio |
| Interaction_genotype_Atr | Cdc42ep4 | -4.487 | 1.47E-04 | glio |
| Interaction_genotype_Atr | Lynx1 | -4.596 | 1.47E-04 | glio |
| Interaction_genotype_Mtor | Bc1 | -4.695 | 1.47E-04 | glio |
| Interaction_genotype_Mtor | Ifi27 | 2.654 | 1.47E-04 | glio |
| Interaction_genotype_Mtor | Fus | 1.949 | 1.47E-04 | glio |
| Interaction_genotype_Mtor | Tmed10 | 1.755 | 1.47E-04 | glio |
| Interaction_genotype_Myc | Akap3 | -4.311 | 1.47E-04 | glio |
| Interaction_genotype_Myc | 9130015A21R | -7.112 | 1.47E-04 | glio |
| Interaction_genotype_Myc | Fbxw17 | -2.754 | 1.47E-04 | glio |
| interaction_RT | Ankrd49 | -1.289 | 1.47E-04 | glio |
| interaction_RT | Hsf2 | 1.232 | 1.47E-04 | glio |
| interaction_RT | Rab9 | 0.769 | 1.47E-04 | glio |
| Aars2_in.vivo_noRT | Gm15998 | 2.693 | 1.47E-04 | glio |
| Aars2_in.vivo_noRT | Gm5577 | 2.483 | 1.47E-04 | glio |
| Atr_ex.vivo_noRT | Klf6 | 1.701 | 1.47E-04 | glio |
| Bard1_ex.vivo_noRT | Gm21761 | 2.814 | 1.47E-04 | glio |
| Bcas2_in.vivo_noRT | Usp44 | 2.818 | 1.47E-04 | glio |
| Bcas2_in.vivo_noRT | Gm43017 | 2.693 | 1.47E-04 | glio |
| Bora_in.vivo_noRT | Zbtb8a | 2.875 | 1.47E-04 | glio |
| Bora_in.vivo_noRT | Tuba1b | -1.017 | 1.47E-04 | glio |
| Chek1_in.vivo_noRT | Spcs1 | -0.918 | 1.47E-04 | glio |
| Chek1_ex.vivo_noRT | Grin3b | 3.287 | 1.47E-04 | glio |
| Eif2s1_in.vivo_noRT | Abcb5 | 2.143 | 1.47E-04 | glio |
| Eif2s1_in.vivo_noRT | 4930553M12F | 2.408 | 1.47E-04 | glio |
| Eif2s1_in.vivo_noRT | Btn1a1 | 2.407 | 1.47E-04 | glio |
| Hsd17b10_in.vivo_noRT | Lrrc24 | 2.616 | 1.47E-04 | glio |
| Interaction_genotype_Mtor | Hsf2bp | -4.869 | 1.47E-05 | glio |
| Interaction_genotype_Myc | Gm10354 | -4.024 | 1.47E-05 | glio |
| Atr_in.vivo_noRT | Tmem246 | 2.837 | 1.47E-05 | glio |
| Bcas2_in.vivo_noRT | Pde3a | 3.029 | 1.47E-05 | glio |
| Bcas2_in.vivo_noRT | Pfkfb2 | 3.171 | 1.47E-05 | glio |
| Bora_in.vivo_noRT | 9530053A07R | 3.122 | 1.47E-05 | glio |
| Chek1_in.vivo_noRT | Gm13056 | 3.289 | 1.47E-05 | glio |
| Hsd17b10_in.vivo_noRT | Shisal2a | 3.353 | 1.47E-05 | glio |
| culture_effect | Kif5c | 1.474 | 1.47E-06 | glio |
| Interaction_genotype_Myc | Ak1 | -5.271 | 1.47E-06 | glio |
| Interaction_genotype_Myc | C4b | 4.856 | 1.47E-06 | glio |
| Aars2_in.vivo_noRT | Acsf2 | 2.973 | 1.47E-06 | glio |
| Atr_in.vivo_noRT | 9330179D12R | 3.604 | 1.47E-06 | glio |
| Atr_in.vivo_noRT | Rfx6 | 3.237 | 1.47E-06 | glio |
| Chek1_in.vivo_noRT | Sept5 | 3.304 | 1.47E-06 | glio |
| Chek1_ex.vivo_noRT | Ltb | 4.021 | 1.47E-06 | glio |
| Chek1_ex.vivo_noRT | Has1 | 3.86 | 1.47E-06 | glio |
| culture_effect | Isg20 | -1.018 | 1.47E-07 | glio |
| Interaction_genotype_Myc | 6430590A07R | -4.762 | 1.47E-07 | glio |
| Interaction_genotype_Myc | 2310058D17R | -6.179 | 1.47E-07 | glio |
| Interaction_genotype_Myc | Gm50334 | -6.716 | 1.47E-07 | glio |
| Interaction_genotype_Myc | Appbp2os | -6.31 | 1.47E-07 | glio |
| Aars2_in.vivo_noRT | Hpse | 3.616 | 1.47E-07 | glio |
| Aars2_ex.vivo_noRT | Nrip2 | 3.293 | 1.47E-07 | glio |
| Aars2_ex.vivo_noRT | Gm49698 | 3.293 | 1.47E-07 | glio |

| | | | | |
|---|---|---|---|---|
| Aars2_ex.vivo_noRT | Olfr881 | 3.293 | 1.47E-07 | glio |
| Bora_in.vivo_noRT | Sema4a | 3.893 | 1.47E-07 | glio |
| Chek1_in.vivo_noRT | Plppr3 | 3.657 | 1.47E-07 | glio |
| Chek1_ex.vivo_noRT | Nrros | 4.449 | 1.47E-07 | glio |
| Hsd17b10_in.vivo_noRT | Bmx | 3.349 | 1.47E-07 | glio |
| culture_effect | B9d1 | -1.036 | 1.47E-08 | glio |
| Aars2_in.vivo_noRT | Pbld1 | 4.14 | 1.47E-08 | glio |
| Atr_in.vivo_noRT | 3110082J24Ri | 4.091 | 1.47E-08 | glio |
| Bcas2_in.vivo_noRT | EU599041 | 4.048 | 1.47E-08 | glio |
| Bcas2_in.vivo_noRT | Gm12925 | 4.349 | 1.47E-09 | glio |
| Bcas2_in.vivo_noRT | Lyz2 | 4.421 | 1.47E-09 | glio |
| Chek1_in.vivo_noRT | Lst1 | 4.278 | 1.47E-09 | glio |
| Chek1_in.vivo_noRT | Il2rb | 4.031 | 1.47E-09 | glio |
| Hsd17b10_in.vivo_noRT | Rassf6 | 3.616 | 1.47E-09 | glio |
| Chek1_ex.vivo_noRT | Hfm1 | 4.268 | 1.47E-10 | glio |
| Atr_in.vivo_noRT | Spon1 | 4.733 | 1.47E-11 | glio |
| culture_effect | Tnfaip8l1 | 2.087 | 1.47E-13 | glio |
| Bora_in.vivo_noRT | Kpna7 | 4.672 | 1.47E-13 | glio |
| Aars2_in.vivo_noRT | Gm31152 | 4.805 | 1.47E-14 | glio |
| Atr_in.vivo_noRT | Pecam1 | 5.51 | 1.47E-14 | glio |
| Chek1_ex.vivo_noRT | Sytl4 | 4.842 | 1.47E-14 | glio |
| Chek1_ex.vivo_noRT | Gm12800 | 4.842 | 1.47E-14 | glio |
| culture_effect | BC065403 | 3.876 | 1.47E-15 | glio |
| Bora_in.vivo_noRT | Serpinb2 | 4.936 | 1.47E-15 | glio |
| Atr_in.vivo_noRT | Muc4 | 5.011 | 1.47E-18 | glio |
| Aars2_in.vivo_noRT | Minar2 | 5.067 | 1.47E-19 | glio |
| Atr_in.vivo_noRT | Gm10801 | 5.533 | 1.47E-21 | glio |
| Atr_in.vivo_noRT | Kcnk13 | 5.27 | 1.47E-21 | glio |
| Atr_in.vivo_noRT | Hfm1 | 5.533 | 1.47E-23 | glio |
| Atr_in.vivo_noRT | D630024D03R | 5.533 | 1.47E-23 | glio |
| Aars2_in.vivo_noRT | Gm13856 | 5.327 | 1.47E-27 | glio |
| Atr_in.vivo_noRT | Vmn2r18 | 5.533 | 1.47E-28 | glio |
| culture_effect | Gbp4 | -5.146 | 1.47E-29 | glio |
| culture_effect | Gm43948 | -1.626 | 1.48E-03 | glio |
| culture_effect | Nr2c2 | -0.723 | 1.48E-03 | glio |
| culture_effect | Urm1 | -0.637 | 1.48E-03 | glio |
| culture_effect | Cyfip1 | 0.53 | 1.48E-03 | glio |
| Interaction_genotype_Aars2 | Fa2h | 3.756 | 1.48E-03 | glio |
| Interaction_genotype_Aars2 | Reln | 1.693 | 1.48E-03 | glio |
| Interaction_genotype_Aars2 | Tra2b | 1.649 | 1.48E-03 | glio |
| Interaction_genotype_Aars2 | Ostc | 1.081 | 1.48E-03 | glio |
| Interaction_genotype_Atr | Eme2 | -3.734 | 1.48E-03 | glio |
| Interaction_genotype_Atr | Gm17655 | -4.425 | 1.48E-03 | glio |
| Interaction_genotype_Atr | Gm48170 | -3.167 | 1.48E-03 | glio |
| Interaction_genotype_Atr | 5730507C01R | -3.038 | 1.48E-03 | glio |
| Interaction_genotype_Atr | Vmn1r32 | -2.762 | 1.48E-03 | glio |
| Interaction_genotype_Atr | Pgghg | -3.303 | 1.48E-03 | glio |
| Interaction_genotype_Mtor | Gm32816 | -4.145 | 1.48E-03 | glio |
| Interaction_genotype_Mtor | B3gnt5 | -3.624 | 1.48E-03 | glio |
| Interaction_genotype_Mtor | Anxa11 | -4.034 | 1.48E-03 | glio |
| Interaction_genotype_Mtor | Stag2 | 1.794 | 1.48E-03 | glio |
| Interaction_genotype_Myc | Zc3h3 | -2.888 | 1.48E-03 | glio |

| | | | | |
|---|---|---|---|---|
| Interaction_genotype_Terf1 | Wincr1 | -5.904 | 1.48E-03 | glio |
| interaction_RT | Epor | 2.808 | 1.48E-03 | glio |
| interaction_RT | Epsti1 | 1.173 | 1.48E-03 | glio |
| interaction_RT | Prmt5 | -0.974 | 1.48E-03 | glio |
| interaction_RT | Gadd45gip1 | -0.638 | 1.48E-03 | glio |
| interaction_RT | Atp5md | 0.8 | 1.48E-03 | glio |
| Aars2_ex.vivo_noRT | Prl7c1 | 2.197 | 1.48E-03 | glio |
| Aars2_ex.vivo_noRT | Dusp15 | 2.893 | 1.48E-03 | glio |
| Atp5c1_in.vivo_noRT | Tmem30c | 2.533 | 1.48E-03 | glio |
| Atr_in.vivo_noRT | Ankrd40 | 1.042 | 1.48E-03 | glio |
| Atr_in.vivo_noRT | Sod1 | -0.908 | 1.48E-03 | glio |
| Atr_ex.vivo_noRT | Ubxn10 | 2.764 | 1.48E-03 | glio |
| Bard1_ex.vivo_noRT | Nos2 | 2.814 | 1.48E-03 | glio |
| Bcas2_in.vivo_noRT | Btbd17 | 2.775 | 1.48E-03 | glio |
| Bora_in.vivo_noRT | Dennd3 | 2.416 | 1.48E-03 | glio |
| Cenpj_ex.vivo_noRT | A030014E15Ri | 2.198 | 1.48E-03 | glio |
| Chek1_in.vivo_noRT | Rpl36al | -0.594 | 1.48E-03 | glio |
| Chek1_in.vivo_noRT | H3f3a | -0.55 | 1.48E-03 | glio |
| culture_effect | Fam107b | 1.265 | 1.48E-04 | glio |
| culture_effect | Ppp1r15b | -0.697 | 1.48E-04 | glio |
| Interaction_genotype_Atr | Gm32834 | -4.295 | 1.48E-04 | glio |
| Interaction_genotype_Mtor | Nop16 | 2.032 | 1.48E-04 | glio |
| Interaction_genotype_Myc | 9330121J05Ri | -4.232 | 1.48E-04 | glio |
| Interaction_genotype_Myc | Nsun6 | -3.379 | 1.48E-04 | glio |
| interaction_RT | 8030462N17R | 1.362 | 1.48E-04 | glio |
| Aars2_in.vivo_noRT | Cxcl2 | 4.46 | 1.48E-04 | glio |
| Aars2_in.vivo_noRT | Slc25a53 | 2.89 | 1.48E-04 | glio |
| Aars2_in.vivo_noRT | Fam117a | 2.4 | 1.48E-04 | glio |
| Atp5c1_in.vivo_noRT | Nat8f2 | 2.408 | 1.48E-04 | glio |
| Atr_in.vivo_noRT | Col20a1 | 2.696 | 1.48E-04 | glio |
| Atr_in.vivo_noRT | Hmx3 | 2.458 | 1.48E-04 | glio |
| Atr_in.vivo_noRT | Mpdu1 | -1.511 | 1.48E-04 | glio |
| Atr_ex.vivo_noRT | Tns4 | 3.15 | 1.48E-04 | glio |
| Bcas2_in.vivo_noRT | Gm16740 | 2.712 | 1.48E-04 | glio |
| Bcas2_in.vivo_noRT | Zscan2 | 2.623 | 1.48E-04 | glio |
| Bora_in.vivo_noRT | Gpsm1 | -2.008 | 1.48E-04 | glio |
| Bora_in.vivo_noRT | Mbd2 | -1.613 | 1.48E-04 | glio |
| Chek1_ex.vivo_noRT | Gm26916 | 3.229 | 1.48E-04 | glio |
| Interaction_genotype_Mtor | 5031434O11R | -5.831 | 1.48E-05 | glio |
| Aars2_in.vivo_noRT | Prdx3 | -1.387 | 1.48E-05 | glio |
| Bard1_ex.vivo_noRT | Ankrd22 | 2.814 | 1.48E-05 | glio |
| Bcas2_in.vivo_noRT | Retreg1 | 1.795 | 1.48E-05 | glio |
| Bora_in.vivo_noRT | Coq8a | 2.843 | 1.48E-05 | glio |
| Chek1_in.vivo_noRT | Gadl1 | 2.869 | 1.48E-05 | glio |
| Chek1_in.vivo_noRT | Rabac1 | -1.101 | 1.48E-05 | glio |
| culture_effect | Zfp322a | 1.135 | 1.48E-06 | glio |
| culture_effect | Mrps15 | 0.659 | 1.48E-06 | glio |
| Atr_ex.vivo_noRT | H1foo | 3.164 | 1.48E-06 | glio |
| Bcas2_in.vivo_noRT | Pagr1a | 3.123 | 1.48E-06 | glio |
| Hsd17b10_in.vivo_noRT | Bmp15 | 2.963 | 1.48E-06 | glio |
| culture_effect | Pfn1 | 0.562 | 1.48E-07 | glio |
| Interaction_genotype_Mtor | Gm14968 | -6.306 | 1.48E-07 | glio |

| | | | | |
|---|---|---|---|---|
| Interaction_genotype_Mtor | Scg2 | -4.099 | 1.48E-07 | glio |
| Aars2_in.vivo_noRT | Map3k5 | 3.162 | 1.48E-07 | glio |
| Bora_ex.vivo_noRT | Gm34059 | 3.201 | 1.48E-07 | glio |
| Bora_ex.vivo_noRT | Halr1 | 3.201 | 1.48E-07 | glio |
| culture_effect | Dync1i2 | 0.798 | 1.48E-08 | glio |
| Bcas2_in.vivo_noRT | Nrip3 | 3.763 | 1.48E-08 | glio |
| Bora_in.vivo_noRT | Slc30a5 | -2.525 | 1.48E-08 | glio |
| Chek1_in.vivo_noRT | 2900042K21R | 3.859 | 1.48E-08 | glio |
| Aars2_in.vivo_noRT | Gm26581 | 4.468 | 1.48E-09 | glio |
| Bcas2_in.vivo_noRT | Postn | 4.173 | 1.48E-09 | glio |
| culture_effect | Cr1l | -1.107 | 1.48E-10 | glio |
| Chek1_in.vivo_noRT | BC028471 | 4.511 | 1.48E-10 | glio |
| Hsd17b10_in.vivo_noRT | Cldn5 | 3.35 | 1.48E-11 | glio |
| culture_effect | Coq8a | -4.866 | 1.48E-12 | glio |
| culture_effect | Tuba1a | -1.079 | 1.48E-13 | glio |
| Aars2_in.vivo_noRT | Iqcn | 4.801 | 1.48E-13 | glio |
| Aars2_in.vivo_noRT | Ntn1 | 4.425 | 1.48E-13 | glio |
| Chek1_in.vivo_noRT | 9230112J17Ri | 5.044 | 1.48E-17 | glio |
| Bora_in.vivo_noRT | Ccl3 | 4.676 | 1.48E-19 | glio |
| Atr_in.vivo_noRT | Gm48008 | 5.269 | 1.48E-20 | glio |
| culture_effect | Pnrc1 | -1.471 | 1.48E-21 | glio |
| culture_effect | Coq9 | -0.703 | 1.49E-03 | glio |
| Interaction_genotype_Aars2 | Gm45591 | -3.356 | 1.49E-03 | glio |
| Interaction_genotype_Aars2 | Sept1 | -4.059 | 1.49E-03 | glio |
| Interaction_genotype_Aars2 | Arhgef26 | 2.039 | 1.49E-03 | glio |
| Interaction_genotype_Aars2 | Armc1 | 1.629 | 1.49E-03 | glio |
| Interaction_genotype_Aars2 | Txnl4a | 1.097 | 1.49E-03 | glio |
| Interaction_genotype_Atr | Cdrt4 | -4.179 | 1.49E-03 | glio |
| Interaction_genotype_Atr | Fndc10 | -4.347 | 1.49E-03 | glio |
| Interaction_genotype_Atr | B9d1 | 2.036 | 1.49E-03 | glio |
| Interaction_genotype_Bora | Wwp1 | 2.006 | 1.49E-03 | glio |
| Interaction_genotype_Mtor | Wdr19 | -3.494 | 1.49E-03 | glio |
| Interaction_genotype_Mtor | Man2b1 | -3.154 | 1.49E-03 | glio |
| Interaction_genotype_Mtor | Stxbp4 | -2.912 | 1.49E-03 | glio |
| Interaction_genotype_Myc | Gm48008 | -2.867 | 1.49E-03 | glio |
| Interaction_genotype_Myc | Fgfr1 | -4.224 | 1.49E-03 | glio |
| Interaction_genotype_Myc | Zfp597 | -3.368 | 1.49E-03 | glio |
| Interaction_genotype_Myc | Hhla1 | -3.047 | 1.49E-03 | glio |
| Interaction_genotype_Myc | Adamts20 | -2.883 | 1.49E-03 | glio |
| Interaction_genotype_Myc | Gnb2 | 1.332 | 1.49E-03 | glio |
| interaction_RT | Cox7a1 | 3.14 | 1.49E-03 | glio |
| interaction_RT | Kdelr3 | 1.366 | 1.49E-03 | glio |
| Aars2_in.vivo_noRT | Gm8773 | 2.424 | 1.49E-03 | glio |
| Aars2_in.vivo_noRT | Ovca2 | 2.136 | 1.49E-03 | glio |
| Aars2_in.vivo_noRT | Fabp3 | 2.567 | 1.49E-03 | glio |
| Aars2_in.vivo_noRT | Akna | 2.205 | 1.49E-03 | glio |
| Aars2_in.vivo_noRT | Rpa1 | -1.497 | 1.49E-03 | glio |
| Aars2_ex.vivo_noRT | Dhrs9 | 2.893 | 1.49E-03 | glio |
| Atp5c1_in.vivo_noRT | Ddx25 | -1.843 | 1.49E-03 | glio |
| Atr_in.vivo_noRT | Ell2 | 2.039 | 1.49E-03 | glio |
| Atr_in.vivo_noRT | Ttc12 | 2.158 | 1.49E-03 | glio |
| Atr_in.vivo_noRT | Elk3 | 1.565 | 1.49E-03 | glio |

| | | | | |
|---|---|---|---|---|
| Atr_in.vivo_noRT | Tep1 | 1.701 | 1.49E-03 | glio |
| Atr_in.vivo_noRT | Safb | -1.277 | 1.49E-03 | glio |
| Atr_ex.vivo_noRT | Prrt3 | 2.771 | 1.49E-03 | glio |
| Atr_ex.vivo_noRT | Cacng4 | 2.771 | 1.49E-03 | glio |
| Bcas2_in.vivo_noRT | N4bp2os | 2.379 | 1.49E-03 | glio |
| Bcas2_in.vivo_noRT | Elovl3 | 2.355 | 1.49E-03 | glio |
| Bcas2_in.vivo_noRT | Eif5 | -0.647 | 1.49E-03 | glio |
| Bcas2_ex.vivo_noRT | Adam2 | 2.742 | 1.49E-03 | glio |
| Bcas2_ex.vivo_noRT | Kit | 2.742 | 1.49E-03 | glio |
| Bora_in.vivo_noRT | Nuak2 | 2.533 | 1.49E-03 | glio |
| Bora_in.vivo_noRT | Csad | -1.373 | 1.49E-03 | glio |
| Cenpj_in.vivo_noRT | Cxcl5 | 2.022 | 1.49E-03 | glio |
| Cenpj_in.vivo_noRT | Gm47936 | 2.022 | 1.49E-03 | glio |
| Cenpj_in.vivo_noRT | Tnni3k | 2.022 | 1.49E-03 | glio |
| Cenpj_in.vivo_noRT | Gm14540 | 2.029 | 1.49E-03 | glio |
| Cenpj_in.vivo_noRT | Oit3 | 2.551 | 1.49E-03 | glio |
| Cenpj_in.vivo_noRT | Gm49504 | 2.022 | 1.49E-03 | glio |
| Cenpj_in.vivo_noRT | Sdc3 | 1.777 | 1.49E-03 | glio |
| Chek1_in.vivo_noRT | Gm26901 | 2.457 | 1.49E-03 | glio |
| Chek1_in.vivo_noRT | 1500004A13R | 2.342 | 1.49E-03 | glio |
| Chek1_in.vivo_noRT | Sfi1 | 2.111 | 1.49E-03 | glio |
| culture_effect | Ufl1 | -0.705 | 1.49E-04 | glio |
| Interaction_genotype_Aars2 | 5430402O13R | -4.779 | 1.49E-04 | glio |
| Interaction_genotype_Aars2 | Mettl7b | -4.693 | 1.49E-04 | glio |
| Interaction_genotype_Aars2 | 1110004F10Ri | 1.607 | 1.49E-04 | glio |
| Interaction_genotype_Mtor | Dmrt2 | -4.821 | 1.49E-04 | glio |
| Interaction_genotype_Mtor | Gm15234 | -4.398 | 1.49E-04 | glio |
| Interaction_genotype_Mtor | Tph2 | -3.989 | 1.49E-04 | glio |
| Interaction_genotype_Mtor | Det1 | -3.547 | 1.49E-04 | glio |
| Interaction_genotype_Myc | Medag | -4.193 | 1.49E-04 | glio |
| Interaction_genotype_Myc | Slc6a13 | -4.614 | 1.49E-04 | glio |
| Interaction_genotype_Myc | Clec16a | -3.909 | 1.49E-04 | glio |
| Interaction_genotype_Myc | Ankrd1 | -4.864 | 1.49E-04 | glio |
| Interaction_genotype_Myc | Hist1h2an | -4.195 | 1.49E-04 | glio |
| Interaction_genotype_Myc | Tonsl | -3.576 | 1.49E-04 | glio |
| Interaction_genotype_Myc | Ncs1 | -3.337 | 1.49E-04 | glio |
| Interaction_genotype_Myc | Pdcd5 | 1.453 | 1.49E-04 | glio |
| Interaction_genotype_Myc | Rps26 | 1.583 | 1.49E-04 | glio |
| Interaction_genotype_Rpl3 | Myl6b | -4.982 | 1.49E-04 | glio |
| Interaction_genotype_Rpl3 | Gm36356 | -4.423 | 1.49E-04 | glio |
| Aars2_in.vivo_noRT | Krba1 | 2.71 | 1.49E-04 | glio |
| Aars2_in.vivo_noRT | Uchl3 | -1.429 | 1.49E-04 | glio |
| Atr_ex.vivo_noRT | Gm47622 | 2.589 | 1.49E-04 | glio |
| Atr_ex.vivo_noRT | Vipr2 | 3.582 | 1.49E-04 | glio |
| Bcas2_in.vivo_noRT | Phldb3 | 2.75 | 1.49E-04 | glio |
| Bcas2_in.vivo_noRT | Hp1bp3 | -1.088 | 1.49E-04 | glio |
| Bcas2_in.vivo_noRT | Eny2 | -0.669 | 1.49E-04 | glio |
| Bora_ex.vivo_noRT | Dok3 | 3.659 | 1.49E-04 | glio |
| Chek1_ex.vivo_noRT | Mycl | 3.404 | 1.49E-04 | glio |
| culture_effect | Thnsl2 | -2.103 | 1.49E-05 | glio |
| culture_effect | Atp9a | -1.408 | 1.49E-05 | glio |
| culture_effect | Sema3e | -1.409 | 1.49E-05 | glio |

| | | | | |
|---|---|---|---|---|
| culture_effect | Tram2 | 0.896 | 1.49E-05 | glio |
| culture_effect | Auh | -0.835 | 1.49E-05 | glio |
| Interaction_genotype_Atr | Gstm5 | 2.596 | 1.49E-05 | glio |
| Interaction_genotype_Mtor | C030005K06R | -5.128 | 1.49E-05 | glio |
| interaction_RT | Arhgdia | -0.713 | 1.49E-05 | glio |
| Aars2_in.vivo_noRT | Usp11 | 3.228 | 1.49E-05 | glio |
| Atr_in.vivo_noRT | Meak7 | 3.04 | 1.49E-05 | glio |
| Chek1_ex.vivo_noRT | Rhox3g | 3.696 | 1.49E-05 | glio |
| Hsd17b10_in.vivo_noRT | K230015D01R | 3.35 | 1.49E-05 | glio |
| Interaction_genotype_Mtor | Swi5 | 2.144 | 1.49E-06 | glio |
| Interaction_genotype_Mtor | Gngt2 | -5.768 | 1.49E-06 | glio |
| Aars2_in.vivo_noRT | Gadl1 | 3.152 | 1.49E-06 | glio |
| Atr_in.vivo_noRT | Pde4d | 3.358 | 1.49E-06 | glio |
| Atr_in.vivo_noRT | Map3k8 | 3.595 | 1.49E-06 | glio |
| Bcas2_in.vivo_noRT | Uckl1os | 3.572 | 1.49E-06 | glio |
| Bora_in.vivo_noRT | Pglyrp1 | 3.457 | 1.49E-06 | glio |
| Chek1_in.vivo_noRT | Hacd2 | -2.218 | 1.49E-06 | glio |
| Chek1_ex.vivo_noRT | Ifitm5 | 4.06 | 1.49E-06 | glio |
| Chek1_ex.vivo_noRT | Npm3 | -1.287 | 1.49E-06 | glio |
| Atr_in.vivo_noRT | Gm48684 | 4.229 | 1.49E-07 | glio |
| Chek1_in.vivo_noRT | Tbrg1 | -1.555 | 1.49E-07 | glio |
| Hsd17b10_in.vivo_noRT | Krt80 | 3.35 | 1.49E-07 | glio |
| culture_effect | Gm49066 | 1.565 | 1.49E-08 | glio |
| culture_effect | Arhgef1 | -1.235 | 1.49E-08 | glio |
| culture_effect | Ap1m1 | -1.059 | 1.49E-08 | glio |
| Aars2_in.vivo_noRT | Gpd1 | 4.154 | 1.49E-08 | glio |
| Bcas2_in.vivo_noRT | Pcdhgb5 | 3.431 | 1.49E-08 | glio |
| Bora_in.vivo_noRT | Gm15559 | 4.009 | 1.49E-08 | glio |
| Chek1_ex.vivo_noRT | Podxl | 4.449 | 1.49E-08 | glio |
| Chek1_ex.vivo_noRT | Ubxn10 | 4.443 | 1.49E-08 | glio |
| Chek1_ex.vivo_noRT | Lrp2bp | 4.348 | 1.49E-08 | glio |
| Bora_in.vivo_noRT | Gm8013 | 4.291 | 1.49E-09 | glio |
| Chek1_in.vivo_noRT | Platr20 | 4.78 | 1.49E-10 | glio |
| Chek1_in.vivo_noRT | Mettl7a3 | 4.393 | 1.49E-11 | glio |
| Hsd17b10_in.vivo_noRT | Gm49504 | 3.616 | 1.49E-11 | glio |
| Bcas2_in.vivo_noRT | AI182371 | 4.025 | 1.49E-12 | glio |
| culture_effect | Olfml2b | -4.018 | 1.49E-13 | glio |
| culture_effect | Rdm1 | -1.448 | 1.49E-13 | glio |
| Bcas2_in.vivo_noRT | Gm49086 | 4.807 | 1.49E-15 | glio |
| culture_effect | Txnip | -4.678 | 1.49E-16 | glio |
| Aars2_in.vivo_noRT | BC147527 | 5.327 | 1.49E-16 | glio |
| Atr_in.vivo_noRT | Crybb1 | 5.146 | 1.49E-17 | glio |
| culture_effect | Cd82 | -2.995 | 1.49E-20 | glio |
| culture_effect | Pstpip2 | -4.348 | 1.49E-20 | glio |
| Atr_in.vivo_noRT | Scgb1c1 | 5.533 | 1.49E-20 | glio |
| Chek1_in.vivo_noRT | Npnt | 4.777 | 1.49E-20 | glio |
| Aars2_in.vivo_noRT | Fhl1 | 5.327 | 1.49E-22 | glio |
| Bora_in.vivo_noRT | Gm16178 | 4.936 | 1.49E-23 | glio |
| Bora_in.vivo_noRT | Gm364 | 4.936 | 1.49E-23 | glio |
| Aars2_in.vivo_noRT | Gm36037 | 5.327 | 1.49E-24 | glio |
| culture_effect | Snrpn | -1.593 | 1.50E-03 | glio |
| culture_effect | BC034090 | -1.915 | 1.50E-03 | glio |

| | | | | |
|---|---|---|---|---|
| culture_effect | Gm37305 | 1.653 | 1.50E-03 | glio |
| culture_effect | Gabbr1 | -0.867 | 1.50E-03 | glio |
| culture_effect | Casc4 | 0.552 | 1.50E-03 | glio |
| culture_effect | Malat1 | -0.702 | 1.50E-03 | glio |
| Interaction_genotype_Atr | Asphd2 | -3.695 | 1.50E-03 | glio |
| Interaction_genotype_Bora | Gm36371 | -3.848 | 1.50E-03 | glio |
| Interaction_genotype_Bora | Snrnp40 | 1.763 | 1.50E-03 | glio |
| Interaction_genotype_Mtor | Rab20 | -3.295 | 1.50E-03 | glio |
| Interaction_genotype_Mtor | Mppe1 | -2.8 | 1.50E-03 | glio |
| Interaction_genotype_Myc | Gm17227 | -3.426 | 1.50E-03 | glio |
| Interaction_genotype_Myc | A330040F15Ri | -3.757 | 1.50E-03 | glio |
| Interaction_genotype_Myc | Susd1 | -3.905 | 1.50E-03 | glio |
| Interaction_genotype_Myc | Mycl | -3.633 | 1.50E-03 | glio |
| Interaction_genotype_Myc | Pkdcc | -3.633 | 1.50E-03 | glio |
| Interaction_genotype_Myc | Wfs1 | -3.552 | 1.50E-03 | glio |
| Interaction_genotype_Myc | Ropn1l | -3.506 | 1.50E-03 | glio |
| Interaction_genotype_Myc | Usp12 | -2.197 | 1.50E-03 | glio |
| Interaction_genotype_Rad51 | Serping1 | 3.831 | 1.50E-03 | glio |
| Interaction_genotype_Rad51 | Col6a2 | -3.448 | 1.50E-03 | glio |
| Interaction_genotype_Rad51 | Shld1 | -3.637 | 1.50E-03 | glio |
| Interaction_genotype_Rad51 | D830039M14I | -4.604 | 1.50E-03 | glio |
| interaction_RT | Rrp7a | -0.769 | 1.50E-03 | glio |
| Aars2_in.vivo_noRT | 9330159F19Ri | 2.139 | 1.50E-03 | glio |
| Aars2_in.vivo_noRT | Tmem177 | 1.988 | 1.50E-03 | glio |
| Aars2_in.vivo_noRT | Fam53b | 1.942 | 1.50E-03 | glio |
| Atr_in.vivo_noRT | Ttc30b | 2.425 | 1.50E-03 | glio |
| Atr_in.vivo_noRT | Commd5 | 2.255 | 1.50E-03 | glio |
| Atr_in.vivo_noRT | Gm14325 | 2.289 | 1.50E-03 | glio |
| Atr_in.vivo_noRT | Fxyd7 | 2.069 | 1.50E-03 | glio |
| Atr_in.vivo_noRT | Szt2 | 1.811 | 1.50E-03 | glio |
| Atr_ex.vivo_noRT | Pstpip2 | 2.37 | 1.50E-03 | glio |
| Atr_ex.vivo_noRT | Ormdl3 | 1.369 | 1.50E-03 | glio |
| Bard1_ex.vivo_noRT | 9930111J21Ri | 2.814 | 1.50E-03 | glio |
| Bcas2_in.vivo_noRT | Qars | -1.328 | 1.50E-03 | glio |
| Bcas2_in.vivo_noRT | Plekhj1 | -0.875 | 1.50E-03 | glio |
| Bcas2_in.vivo_noRT | Skap2 | -0.984 | 1.50E-03 | glio |
| Bora_in.vivo_noRT | Irak3 | 2.42 | 1.50E-03 | glio |
| Bora_in.vivo_noRT | Gm14488 | 2.497 | 1.50E-03 | glio |
| Bora_ex.vivo_noRT | Esm1 | 2.226 | 1.50E-03 | glio |
| Cenpj_ex.vivo_noRT | Spink6 | 2.591 | 1.50E-03 | glio |
| Cenpj_ex.vivo_noRT | Htr2b | 2.591 | 1.50E-03 | glio |
| Chek1_in.vivo_noRT | Vdac3 | -0.838 | 1.50E-03 | glio |
| Chek1_ex.vivo_noRT | Usp11 | 3.104 | 1.50E-03 | glio |
| Chek1_ex.vivo_noRT | 1700124L16Ri | 2.963 | 1.50E-03 | glio |
| Eif2s1_in.vivo_noRT | 4930423D22R | 1.883 | 1.50E-03 | glio |
| Eif2s1_in.vivo_noRT | Esp15 | 2.143 | 1.50E-03 | glio |
| Eif2s1_in.vivo_noRT | Boc | 1.879 | 1.50E-03 | glio |
| Eif2s1_in.vivo_noRT | Gm10801 | 2.143 | 1.50E-03 | glio |
| Interaction_genotype_Aars2 | 9130604C24R | -5.121 | 1.50E-04 | glio |
| Interaction_genotype_Atr | Tcim | 2.9 | 1.50E-04 | glio |
| Interaction_genotype_Mtor | Dcn | -4.101 | 1.50E-04 | glio |
| Interaction_genotype_Mtor | Gm50115 | -4.235 | 1.50E-04 | glio |

| | | | | |
|---|---|---|---|---|
| Interaction_genotype_Mtor | Ccdc63 | -4.423 | 1.50E-04 | glio |
| Interaction_genotype_Mtor | D730045A05R | -4.494 | 1.50E-04 | glio |
| Interaction_genotype_Mtor | Micall2 | -4.454 | 1.50E-04 | glio |
| Interaction_genotype_Mtor | 1700120C14R | -3.935 | 1.50E-04 | glio |
| Interaction_genotype_Mtor | 9430060I03Ri | -3.833 | 1.50E-04 | glio |
| Interaction_genotype_Myc | Atp5g3 | 1.446 | 1.50E-04 | glio |
| Interaction_genotype_Nat2 | Gm4340 | 3.282 | 1.50E-04 | glio |
| Interaction_genotype_Nat2 | Gm16239 | 3.282 | 1.50E-04 | glio |
| interaction_RT | Gm9758 | 2.73 | 1.50E-04 | glio |
| Aars2_in.vivo_noRT | Axin2 | 2.757 | 1.50E-04 | glio |
| Aars2_in.vivo_noRT | Ctdp1 | 1.802 | 1.50E-04 | glio |
| Atp5c1_in.vivo_noRT | Gm16741 | 2.668 | 1.50E-04 | glio |
| Atr_in.vivo_noRT | Rorc | 2.872 | 1.50E-04 | glio |
| Atr_in.vivo_noRT | 4933404O12R | 2.628 | 1.50E-04 | glio |
| Atr_in.vivo_noRT | Lca5 | 2.587 | 1.50E-04 | glio |
| Atr_in.vivo_noRT | Tmco1 | -1.012 | 1.50E-04 | glio |
| Chek1_in.vivo_noRT | Ccdc28b | 2.341 | 1.50E-04 | glio |
| Chek1_ex.vivo_noRT | Grhl2 | 3.568 | 1.50E-04 | glio |
| Chek1_ex.vivo_noRT | Prr7 | 3.392 | 1.50E-04 | glio |
| Chek1_ex.vivo_noRT | Casp4 | 3.098 | 1.50E-04 | glio |
| Eif2s1_in.vivo_noRT | Gm5788 | 2.143 | 1.50E-04 | glio |
| culture_effect | Gm11457 | 2.366 | 1.50E-05 | glio |
| culture_effect | Inpp5k | -1.432 | 1.50E-05 | glio |
| culture_effect | Aldh7a1 | -0.886 | 1.50E-05 | glio |
| Interaction_genotype_Myc | Gm36839 | -5 | 1.50E-05 | glio |
| Atr_in.vivo_noRT | Gm2822 | 3.317 | 1.50E-05 | glio |
| Atr_in.vivo_noRT | Gm42701 | 3.017 | 1.50E-05 | glio |
| Atr_in.vivo_noRT | Phex | 2.846 | 1.50E-05 | glio |
| Atr_in.vivo_noRT | Apex1 | -1.634 | 1.50E-05 | glio |
| Atr_in.vivo_noRT | Emd | -1.231 | 1.50E-05 | glio |
| Bard1_ex.vivo_noRT | Gm47404 | 2.814 | 1.50E-05 | glio |
| Bcas2_in.vivo_noRT | Gm50069 | 3.166 | 1.50E-05 | glio |
| Chek1_in.vivo_noRT | 2900041M22F | 3.234 | 1.50E-05 | glio |
| culture_effect | Gtf3a | 0.624 | 1.50E-06 | glio |
| Interaction_genotype_Mtor | G630030J09Ri | -5.472 | 1.50E-06 | glio |
| Aars2_in.vivo_noRT | Qpct | 3.46 | 1.50E-06 | glio |
| Chek1_in.vivo_noRT | 1700055D18R | 3.654 | 1.50E-06 | glio |
| culture_effect | Gna15 | -3.024 | 1.50E-07 | glio |
| Chek1_in.vivo_noRT | Wincr1 | 4.78 | 1.50E-07 | glio |
| Bcas2_in.vivo_noRT | Gm3734 | 4.284 | 1.50E-08 | glio |
| culture_effect | Clstn3 | -2.554 | 1.50E-09 | glio |
| Aars2_in.vivo_noRT | Pax5 | 4.482 | 1.50E-09 | glio |
| Aars2_in.vivo_noRT | Slc26a6 | 4.035 | 1.50E-09 | glio |
| Aars2_in.vivo_noRT | Ccno | 4.306 | 1.50E-09 | glio |
| Chek1_ex.vivo_noRT | Gm16751 | 4.443 | 1.50E-09 | glio |
| culture_effect | Insrr | -3.711 | 1.50E-10 | glio |
| Aars2_in.vivo_noRT | Glis1 | 4.415 | 1.50E-10 | glio |
| Bcas2_in.vivo_noRT | Amy1 | 4.278 | 1.50E-10 | glio |
| Atr_in.vivo_noRT | Gm10010 | 4.792 | 1.50E-11 | glio |
| Bcas2_in.vivo_noRT | Stx19 | 3.895 | 1.50E-11 | glio |
| Bcas2_in.vivo_noRT | Gm14968 | 4.955 | 1.50E-11 | glio |
| Atr_in.vivo_noRT | Zfp14 | 5.01 | 1.50E-12 | glio |

| | | | | |
|---|---|---|---|---|
| Bora_in.vivo_noRT | 6430562O15R | 4.414 | 1.50E-12 | glio |
| Aars2_in.vivo_noRT | Gm42918 | 4.684 | 1.50E-14 | glio |
| Chek1_ex.vivo_noRT | Trmt61b | 4.842 | 1.50E-14 | glio |
| Aars2_in.vivo_noRT | Pcdhga11 | 4.844 | 1.50E-16 | glio |
| Aars2_in.vivo_noRT | Tmem236 | 5.06 | 1.50E-16 | glio |
| Bcas2_in.vivo_noRT | Olfr718-ps1 | 4.543 | 1.50E-17 | glio |
| Bora_in.vivo_noRT | Gm48780 | 4.676 | 1.50E-19 | glio |
| Aars2_in.vivo_noRT | Tal2 | 5.063 | 1.50E-22 | glio |
| Chek1_in.vivo_noRT | Slc17a6 | 5.044 | 1.50E-29 | glio |
| culture_effect | B2m | -1.944 | 1.50E-39 | glio |
| culture_effect | H1foo | -1.467 | 1.51E-03 | glio |
| culture_effect | Mccc1 | -1.01 | 1.51E-03 | glio |
| culture_effect | Chek1 | -0.682 | 1.51E-03 | glio |
| Interaction_genotype_Aars2 | Txnl4b | -3.368 | 1.51E-03 | glio |
| Interaction_genotype_Atr | Gm32250 | -2.96 | 1.51E-03 | glio |
| Interaction_genotype_Atr | Gm16133 | -4.089 | 1.51E-03 | glio |
| Interaction_genotype_Atr | Sox5 | -4.132 | 1.51E-03 | glio |
| Interaction_genotype_Atr | Fam120a | -1.397 | 1.51E-03 | glio |
| Interaction_genotype_Bora | Hecw2 | -4.072 | 1.51E-03 | glio |
| Interaction_genotype_Bora | Sptbn4 | -4.154 | 1.51E-03 | glio |
| Interaction_genotype_Bora | Atoh8 | -3.511 | 1.51E-03 | glio |
| Interaction_genotype_Bora | Gm29083 | -3.859 | 1.51E-03 | glio |
| Interaction_genotype_Mtor | Usp27x | -3.904 | 1.51E-03 | glio |
| Interaction_genotype_Myc | Smtnl1 | -2.729 | 1.51E-03 | glio |
| Interaction_genotype_Myc | Parp16 | -3.413 | 1.51E-03 | glio |
| Interaction_genotype_Myc | Zfp983 | -3.024 | 1.51E-03 | glio |
| Interaction_genotype_Myc | Cpa2 | -3.401 | 1.51E-03 | glio |
| Interaction_genotype_Myc | Zfp12 | -2.857 | 1.51E-03 | glio |
| Interaction_genotype_Myc | Prmt6 | -2.895 | 1.51E-03 | glio |
| Interaction_genotype_Myc | 2010315B03R | -2.856 | 1.51E-03 | glio |
| Interaction_genotype_Myc | B3galt6 | -2.624 | 1.51E-03 | glio |
| Interaction_genotype_Myc | Borcs5 | -2.763 | 1.51E-03 | glio |
| Interaction_genotype_Myc | Arl8b | 1.628 | 1.51E-03 | glio |
| Interaction_genotype_Myc | Ap2s1 | 1.332 | 1.51E-03 | glio |
| interaction_RT | Ddx56 | -1.072 | 1.51E-03 | glio |
| Aars2_in.vivo_noRT | Zfp975 | 2.209 | 1.51E-03 | glio |
| Aars2_in.vivo_noRT | Eri3 | -1.201 | 1.51E-03 | glio |
| Aars2_in.vivo_noRT | Pitpnm1 | 1.641 | 1.51E-03 | glio |
| Aars2_ex.vivo_noRT | Gm20478 | 2.897 | 1.51E-03 | glio |
| Aars2_ex.vivo_noRT | Map3k9 | 2.505 | 1.51E-03 | glio |
| Atp5c1_in.vivo_noRT | Syp | 2.799 | 1.51E-03 | glio |
| Atr_in.vivo_noRT | Creg2 | 2.286 | 1.51E-03 | glio |
| Atr_in.vivo_noRT | Eif3b | -0.969 | 1.51E-03 | glio |
| Bard1_ex.vivo_noRT | Tent5c | 2.421 | 1.51E-03 | glio |
| Bcas2_in.vivo_noRT | Xpo4 | -1.592 | 1.51E-03 | glio |
| Bcas2_in.vivo_noRT | Arih1 | -1.207 | 1.51E-03 | glio |
| Bora_in.vivo_noRT | Cyb561 | 2.462 | 1.51E-03 | glio |
| Bora_in.vivo_noRT | Wdr5b | 2.249 | 1.51E-03 | glio |
| Bora_in.vivo_noRT | Zscan2 | 2.223 | 1.51E-03 | glio |
| Bora_in.vivo_noRT | Maats1 | 2.38 | 1.51E-03 | glio |
| Bora_in.vivo_noRT | Fads2 | -1.584 | 1.51E-03 | glio |
| Bora_in.vivo_noRT | Dusp2 | -2.444 | 1.51E-03 | glio |

| | | | | |
|---|---|---|---|---|
| Bora_ex.vivo_noRT | Slc7a4 | 3.013 | 1.51E-03 | glio |
| Chek1_in.vivo_noRT | 4833408A19R | 2.194 | 1.51E-03 | glio |
| Chek1_in.vivo_noRT | Ptpru | 2.043 | 1.51E-03 | glio |
| Chek1_in.vivo_noRT | Rpsa | -0.628 | 1.51E-03 | glio |
| Chek1_ex.vivo_noRT | 1700030C12R | 2.986 | 1.51E-03 | glio |
| Interaction_genotype_Aars2 | Gm32585 | -4.982 | 1.51E-04 | glio |
| Interaction_genotype_Mtor | Foxl2os | -4.377 | 1.51E-04 | glio |
| Interaction_genotype_Myc | Cldn2 | -2.729 | 1.51E-04 | glio |
| Interaction_genotype_Myc | 2010001K21R | -3.892 | 1.51E-04 | glio |
| interaction_RT | Flna | -0.924 | 1.51E-04 | glio |
| Aars2_in.vivo_noRT | Mpz | 3.356 | 1.51E-04 | glio |
| Aars2_in.vivo_noRT | Ctrl | 2.778 | 1.51E-04 | glio |
| Aars2_in.vivo_noRT | Enho | 2.668 | 1.51E-04 | glio |
| Aars2_in.vivo_noRT | Exosc3 | -1.225 | 1.51E-04 | glio |
| Atr_in.vivo_noRT | Phactr4 | -2.069 | 1.51E-04 | glio |
| Bcas2_in.vivo_noRT | Zfp598 | -1.843 | 1.51E-04 | glio |
| Bcas2_in.vivo_noRT | Hspg2 | -1.759 | 1.51E-04 | glio |
| Bcas2_in.vivo_noRT | Fbxl6 | -1.493 | 1.51E-04 | glio |
| Bora_in.vivo_noRT | Bpifc | 2.916 | 1.51E-04 | glio |
| Bora_in.vivo_noRT | Gm21154 | 2.903 | 1.51E-04 | glio |
| Bora_in.vivo_noRT | Cdc16 | -1.569 | 1.51E-04 | glio |
| Bora_in.vivo_noRT | Paics | -0.866 | 1.51E-04 | glio |
| Chek1_in.vivo_noRT | Nim1k | 2.845 | 1.51E-04 | glio |
| Chek1_in.vivo_noRT | Myl4 | 2.793 | 1.51E-04 | glio |
| Hsd17b10_in.vivo_noRT | Gpd1 | 2.964 | 1.51E-04 | glio |
| Hsd17b10_in.vivo_noRT | 5830454E08Ri | 2.691 | 1.51E-04 | glio |
| Hsd17b10_in.vivo_noRT | Gm20275 | 2.262 | 1.51E-04 | glio |
| Hsd17b10_in.vivo_noRT | Rexo4 | -1.899 | 1.51E-04 | glio |
| culture_effect | Dgcr6 | 0.597 | 1.51E-05 | glio |
| Interaction_genotype_Mtor | Gm10101 | -4.561 | 1.51E-05 | glio |
| Interaction_genotype_Myc | D2hgdh | -4.208 | 1.51E-05 | glio |
| Interaction_genotype_Myc | Notch4 | -4.878 | 1.51E-05 | glio |
| Interaction_genotype_Myc | Gm10324 | -3.826 | 1.51E-05 | glio |
| Aars2_in.vivo_noRT | Cracr2b | 3.202 | 1.51E-05 | glio |
| Aars2_in.vivo_noRT | Bloc1s3 | 2.716 | 1.51E-05 | glio |
| Aars2_in.vivo_noRT | Llph | -1.101 | 1.51E-05 | glio |
| Atp5c1_in.vivo_noRT | Esp15 | 2.668 | 1.51E-05 | glio |
| Atr_in.vivo_noRT | Zfp114 | 3.06 | 1.51E-05 | glio |
| Bard1_ex.vivo_noRT | Gm31717 | 2.814 | 1.51E-05 | glio |
| Bard1_ex.vivo_noRT | Gm14742 | 2.814 | 1.51E-05 | glio |
| Bora_in.vivo_noRT | 1810034E14Ri | 3.053 | 1.51E-05 | glio |
| Bora_in.vivo_noRT | Gm34121 | 2.811 | 1.51E-05 | glio |
| Chek1_in.vivo_noRT | 4930597O21R | 3.316 | 1.51E-05 | glio |
| Chek1_ex.vivo_noRT | Tec | 3.567 | 1.51E-05 | glio |
| culture_effect | Ecscr | -1.495 | 1.51E-06 | glio |
| Atp5c1_in.vivo_noRT | Olfr56 | 2.668 | 1.51E-06 | glio |
| Bcas2_ex.vivo_noRT | Fhl1 | 3.589 | 1.51E-06 | glio |
| Bora_in.vivo_noRT | 1110035H17R | 3.768 | 1.51E-06 | glio |
| Chek1_in.vivo_noRT | Tpsab1 | 3.094 | 1.51E-06 | glio |
| Chek1_in.vivo_noRT | Gm36495 | 3.749 | 1.51E-06 | glio |
| culture_effect | Bbc3 | -1.343 | 1.51E-07 | glio |
| culture_effect | Lima1 | 0.853 | 1.51E-07 | glio |

| | | | | |
|---|---|---|---|---|
| Aars2_ex.vivo_noRT | BC061195 | 3.293 | 1.51E-07 | glio |
| Bcas2_in.vivo_noRT | Acot6 | 3.562 | 1.51E-07 | glio |
| Chek1_in.vivo_noRT | 1700081N11R | 4.19 | 1.51E-07 | glio |
| Atr_in.vivo_noRT | Poln | 4.03 | 1.51E-08 | glio |
| Bcas2_in.vivo_noRT | Gpr179 | 3.806 | 1.51E-08 | glio |
| culture_effect | Ski | 1.324 | 1.51E-09 | glio |
| Bora_in.vivo_noRT | Tnfsf13os | 4.166 | 1.51E-09 | glio |
| Chek1_in.vivo_noRT | 1700040D17R | 4.491 | 1.51E-09 | glio |
| culture_effect | Olfm1 | 2.306 | 1.51E-10 | glio |
| Atr_in.vivo_noRT | Gm13610 | 4.377 | 1.51E-10 | glio |
| Chek1_in.vivo_noRT | Gm26822 | 4.652 | 1.51E-10 | glio |
| culture_effect | Klhl30 | 3.384 | 1.51E-12 | glio |
| Atr_in.vivo_noRT | Mamdc2 | 4.733 | 1.51E-12 | glio |
| Atr_in.vivo_noRT | Gm5039 | 5.533 | 1.51E-12 | glio |
| Atr_in.vivo_noRT | Flnc | 5.008 | 1.51E-12 | glio |
| Chek1_in.vivo_noRT | Tns4 | 4.393 | 1.51E-12 | glio |
| Hsd17b10_in.vivo_noRT | Idi2 | 3.616 | 1.51E-13 | glio |
| Atr_in.vivo_noRT | Kcnk7 | 4.485 | 1.51E-14 | glio |
| Atr_in.vivo_noRT | Gm43653 | 5.27 | 1.51E-15 | glio |
| culture_effect | Jcad | -4.183 | 1.51E-16 | glio |
| Aars2_in.vivo_noRT | Folr1 | 5.327 | 1.51E-18 | glio |
| Bcas2_in.vivo_noRT | Cd200r2 | 4.543 | 1.51E-19 | glio |
| Bora_in.vivo_noRT | Gm36037 | 4.936 | 1.51E-21 | glio |
| culture_effect | Hipk2 | -3.118 | 1.51E-44 | glio |
| culture_effect | Klhdc8b | 1.741 | 1.52E-03 | glio |
| culture_effect | Scly | -0.857 | 1.52E-03 | glio |
| culture_effect | Cyp39a1 | -0.851 | 1.52E-03 | glio |
| culture_effect | 9530068E07Ri | -0.467 | 1.52E-03 | glio |
| culture_effect | Zmat2 | 0.401 | 1.52E-03 | glio |
| Interaction_genotype_Aars2 | Selp | -3.985 | 1.52E-03 | glio |
| Interaction_genotype_Aars2 | Gm13348 | -3.306 | 1.52E-03 | glio |
| Interaction_genotype_Aars2 | Gm13398 | -2.831 | 1.52E-03 | glio |
| Interaction_genotype_Aars2 | Nme4 | 1.886 | 1.52E-03 | glio |
| Interaction_genotype_Atr | Npnt | -2.901 | 1.52E-03 | glio |
| Interaction_genotype_Atr | Ccdc85c | -3.875 | 1.52E-03 | glio |
| Interaction_genotype_Atr | Elk3 | -2.255 | 1.52E-03 | glio |
| Interaction_genotype_Atr | Arf4 | 1.189 | 1.52E-03 | glio |
| Interaction_genotype_Mtor | Gm48265 | -3.316 | 1.52E-03 | glio |
| Interaction_genotype_Mtor | Arhgdib | -3.934 | 1.52E-03 | glio |
| Interaction_genotype_Mtor | Zfp316 | -3.101 | 1.52E-03 | glio |
| Interaction_genotype_Mtor | D830025C05R | -3.615 | 1.52E-03 | glio |
| Interaction_genotype_Mtor | Gm19531 | -3.216 | 1.52E-03 | glio |
| Interaction_genotype_Mtor | Timm10b | 1.534 | 1.52E-03 | glio |
| Interaction_genotype_Myc | Serac1 | -3.455 | 1.52E-03 | glio |
| Interaction_genotype_Myc | 4930579K19R | -3.575 | 1.52E-03 | glio |
| Interaction_genotype_Myc | Sap25 | -3.415 | 1.52E-03 | glio |
| Interaction_genotype_Myc | Nabp1 | -3.439 | 1.52E-03 | glio |
| Interaction_genotype_Myc | Fbxl19 | -3.399 | 1.52E-03 | glio |
| Interaction_genotype_Myc | Ttk | -2.65 | 1.52E-03 | glio |
| Interaction_genotype_Myc | Abhd10 | -2.587 | 1.52E-03 | glio |
| Interaction_genotype_Pars2 | D030040B21R | -4.007 | 1.52E-03 | glio |
| Interaction_genotype_Pars2 | Slc4a9 | -3.622 | 1.52E-03 | glio |

| | | | | |
|---|---|---|---|---|
| Interaction_genotype_Pars2 | Serp2 | -3.006 | 1.52E-03 | glio |
| interaction_RT | Fubp1 | -0.612 | 1.52E-03 | glio |
| Aars2_in.vivo_noRT | Aox1 | 2.1 | 1.52E-03 | glio |
| Aars2_in.vivo_noRT | Gimap9 | 2.106 | 1.52E-03 | glio |
| Aars2_in.vivo_noRT | Zfp90 | 1.928 | 1.52E-03 | glio |
| Atp5c1_in.vivo_noRT | Pde11a | 2.667 | 1.52E-03 | glio |
| Atr_in.vivo_noRT | Gm17484 | 2.147 | 1.52E-03 | glio |
| Atr_in.vivo_noRT | Rac3 | 2.111 | 1.52E-03 | glio |
| Atr_in.vivo_noRT | Shank1 | 2.106 | 1.52E-03 | glio |
| Atr_in.vivo_noRT | 1500015A07R | 2.125 | 1.52E-03 | glio |
| Atr_in.vivo_noRT | Tamm41 | 1.497 | 1.52E-03 | glio |
| Atr_ex.vivo_noRT | Mcm3 | -1.661 | 1.52E-03 | glio |
| Bcas2_in.vivo_noRT | Gm19744 | 2.362 | 1.52E-03 | glio |
| Bcas2_in.vivo_noRT | Bst1 | 2.168 | 1.52E-03 | glio |
| Bcas2_in.vivo_noRT | Xylb | 2.287 | 1.52E-03 | glio |
| Bcas2_in.vivo_noRT | Gm527 | 2.192 | 1.52E-03 | glio |
| Bora_ex.vivo_noRT | Slc14a1 | 2.626 | 1.52E-03 | glio |
| Bora_ex.vivo_noRT | Gm43701 | 2.219 | 1.52E-03 | glio |
| Chek1_in.vivo_noRT | Rac3 | 2.145 | 1.52E-03 | glio |
| Chek1_ex.vivo_noRT | Galnt7 | -2.024 | 1.52E-03 | glio |
| Eif2s1_in.vivo_noRT | Gm26555 | 1.88 | 1.52E-03 | glio |
| Ercc4_ex.vivo_noRT | Rimklb | 2.704 | 1.52E-03 | glio |
| Hsd17b10_in.vivo_noRT | Hlx | 2.36 | 1.52E-03 | glio |
| Hsd17b10_in.vivo_noRT | Gm16348 | 2.432 | 1.52E-03 | glio |
| Hsd17b10_in.vivo_noRT | Slc26a6 | 2.324 | 1.52E-03 | glio |
| Hsd17b10_in.vivo_noRT | Gm43387 | 2.441 | 1.52E-03 | glio |
| Hsd17b10_in.vivo_noRT | Ciao2a | -1.004 | 1.52E-03 | glio |
| culture_effect | Atg16l2 | -1.487 | 1.52E-04 | glio |
| culture_effect | Trim35 | 0.662 | 1.52E-04 | glio |
| culture_effect | Higd1a | 0.549 | 1.52E-04 | glio |
| Interaction_genotype_Bora | Ak1 | -4.709 | 1.52E-04 | glio |
| Interaction_genotype_Bora | Cldn19 | -4.834 | 1.52E-04 | glio |
| Interaction_genotype_Mtor | Pcdhgb6 | -4.019 | 1.52E-04 | glio |
| Interaction_genotype_Mtor | Trim67 | -4.026 | 1.52E-04 | glio |
| Interaction_genotype_Mtor | C230035I16Ri | -4.77 | 1.52E-04 | glio |
| Interaction_genotype_Mtor | Gpi1 | 1.861 | 1.52E-04 | glio |
| Interaction_genotype_Myc | Rbx1 | 1.286 | 1.52E-04 | glio |
| interaction_RT | Pcolce2 | -1.092 | 1.52E-04 | glio |
| Aars2_in.vivo_noRT | Ifnb1 | 5.327 | 1.52E-04 | glio |
| Aars2_in.vivo_noRT | Irak3 | 2.811 | 1.52E-04 | glio |
| Aars2_ex.vivo_noRT | Ccdc125 | 2.893 | 1.52E-04 | glio |
| Aars2_ex.vivo_noRT | BC147527 | 3.293 | 1.52E-04 | glio |
| Atr_in.vivo_noRT | Sgsh | 2.79 | 1.52E-04 | glio |
| Bora_in.vivo_noRT | Ampd3 | 2.654 | 1.52E-04 | glio |
| Bora_in.vivo_noRT | Gm43138 | 2.824 | 1.52E-04 | glio |
| Chek1_in.vivo_noRT | Cldn9 | 2.598 | 1.52E-04 | glio |
| Chek1_ex.vivo_noRT | Gm13288 | 2.707 | 1.52E-04 | glio |
| Hsd17b10_in.vivo_noRT | Gm10010 | 2.875 | 1.52E-04 | glio |
| interaction_RT | Tmem176a | 1.52 | 1.52E-05 | glio |
| Aars2_in.vivo_noRT | Ccnj | 3.145 | 1.52E-05 | glio |
| Aars2_ex.vivo_noRT | Gm34939 | 3.293 | 1.52E-05 | glio |
| Atp5c1_in.vivo_noRT | Gm10801 | 2.668 | 1.52E-05 | glio |

| | | | | |
|---|---|---|---|---|
| Atr_in.vivo_noRT | Hist1h2bf | 3.221 | 1.52E-05 | glio |
| Atr_in.vivo_noRT | Pxt1 | 3.06 | 1.52E-05 | glio |
| Atr_in.vivo_noRT | Fhad1 | 2.808 | 1.52E-05 | glio |
| Bard1_ex.vivo_noRT | Scel | 2.814 | 1.52E-05 | glio |
| Bard1_ex.vivo_noRT | Ctla4 | 2.814 | 1.52E-05 | glio |
| Bard1_ex.vivo_noRT | Tagap | 2.814 | 1.52E-05 | glio |
| Bard1_ex.vivo_noRT | Gm21731 | 2.814 | 1.52E-05 | glio |
| Bard1_ex.vivo_noRT | Gm11232 | 2.814 | 1.52E-05 | glio |
| Bard1_ex.vivo_noRT | Gm29966 | 2.814 | 1.52E-05 | glio |
| Bard1_ex.vivo_noRT | Olfr1152 | 2.814 | 1.52E-05 | glio |
| Bcas2_in.vivo_noRT | Gm43331 | 2.595 | 1.52E-05 | glio |
| Hsd17b10_in.vivo_noRT | Mir670hg | 3.493 | 1.52E-05 | glio |
| interaction_RT | Mlec | -1.056 | 1.52E-06 | glio |
| Aars2_in.vivo_noRT | Zfp781 | 3.225 | 1.52E-06 | glio |
| Bora_in.vivo_noRT | Gm10734 | 3.42 | 1.52E-06 | glio |
| culture_effect | Eepd1 | -2.993 | 1.52E-07 | glio |
| culture_effect | Smarcb1 | -0.702 | 1.52E-07 | glio |
| Interaction_genotype_Aars2 | Ntn4 | -5.181 | 1.52E-07 | glio |
| Interaction_genotype_Aars2 | Folr1 | -5.604 | 1.52E-07 | glio |
| Interaction_genotype_Aars2 | Nxph4 | -6.416 | 1.52E-07 | glio |
| Interaction_genotype_Mtor | Gm49891 | -6.656 | 1.52E-07 | glio |
| Bora_in.vivo_noRT | Gm8104 | 3.517 | 1.52E-07 | glio |
| Chek1_ex.vivo_noRT | Gm47802 | 4.275 | 1.52E-07 | glio |
| Hsd17b10_in.vivo_noRT | Cxcl13 | 3.616 | 1.52E-07 | glio |
| Interaction_genotype_Atr | Col6a2 | -5.531 | 1.52E-08 | glio |
| Bcas2_in.vivo_noRT | 4933408J17Ri | 3.491 | 1.52E-08 | glio |
| Chek1_in.vivo_noRT | Npb | 3.876 | 1.52E-08 | glio |
| Atr_in.vivo_noRT | 1700040D17R | 4.469 | 1.52E-09 | glio |
| Bora_in.vivo_noRT | Gm47802 | 3.961 | 1.52E-09 | glio |
| Bora_in.vivo_noRT | Ptpn22 | 4.673 | 1.52E-09 | glio |
| Atr_in.vivo_noRT | Hist1h4f | 4.741 | 1.52E-10 | glio |
| Aars2_in.vivo_noRT | Igdcc4 | 4.151 | 1.52E-11 | glio |
| Atr_in.vivo_noRT | AI854703 | 4.883 | 1.52E-12 | glio |
| Atr_in.vivo_noRT | 2610027K06R | 4.877 | 1.52E-13 | glio |
| Hsd17b10_in.vivo_noRT | Vmn2r18 | 3.616 | 1.52E-13 | glio |
| Bcas2_in.vivo_noRT | Gm10482 | 4.285 | 1.52E-14 | glio |
| culture_effect | Apod | -4.538 | 1.52E-16 | glio |
| Aars2_in.vivo_noRT | Bcl2l15 | 5.067 | 1.52E-16 | glio |
| Aars2_in.vivo_noRT | Gm50138 | 4.804 | 1.52E-16 | glio |
| Aars2_in.vivo_noRT | Hunk | 5.304 | 1.52E-16 | glio |
| Aars2_in.vivo_noRT | Tent5c | 4.401 | 1.52E-16 | glio |
| Atr_in.vivo_noRT | Gm21762 | 5.269 | 1.52E-17 | glio |
| Chek1_in.vivo_noRT | Gm20711 | 5.044 | 1.52E-17 | glio |
| Chek1_in.vivo_noRT | Ccdc125 | 5.044 | 1.52E-18 | glio |
| Atr_in.vivo_noRT | Cers3 | 5.002 | 1.52E-19 | glio |
| Chek1_in.vivo_noRT | Fhl1 | 5.044 | 1.52E-20 | glio |
| Atr_in.vivo_noRT | Crb3 | 5.533 | 1.52E-23 | glio |
| culture_effect | H2-Q7 | -7.093 | 1.52E-40 | glio |
| culture_effect | Npy1r | -1.579 | 1.53E-03 | glio |
| culture_effect | Gm34256 | 1.918 | 1.53E-03 | glio |
| culture_effect | Airn | -1.008 | 1.53E-03 | glio |
| Interaction_genotype_Atr | C330002G04R | -2.956 | 1.53E-03 | glio |

| | | | | |
|---|---|---|---|---|
| Interaction_genotype_Atr | Gm30692 | -2.763 | 1.53E-03 | glio |
| Interaction_genotype_Bora | Ankrd1 | -4.854 | 1.53E-03 | glio |
| Interaction_genotype_Mtor | Nrn1l | -3.573 | 1.53E-03 | glio |
| Interaction_genotype_Mtor | Fam178b | -3.741 | 1.53E-03 | glio |
| Interaction_genotype_Mtor | B230208H11R | -3.649 | 1.53E-03 | glio |
| Interaction_genotype_Mtor | Gm45051 | -3.094 | 1.53E-03 | glio |
| Interaction_genotype_Mtor | Ikbip | 1.831 | 1.53E-03 | glio |
| Interaction_genotype_Mtor | Ccnd2 | 1.512 | 1.53E-03 | glio |
| Interaction_genotype_Myc | Procr | -4.355 | 1.53E-03 | glio |
| Interaction_genotype_Myc | Hcn2 | -3.782 | 1.53E-03 | glio |
| Interaction_genotype_Myc | Catsper2 | -3.417 | 1.53E-03 | glio |
| Interaction_genotype_Myc | Cenpi | -2.922 | 1.53E-03 | glio |
| Interaction_genotype_Myc | Tmem160 | 1.327 | 1.53E-03 | glio |
| Interaction_genotype_Myc | Cdc42 | 1.26 | 1.53E-03 | glio |
| Interaction_genotype_Rpl3 | C1ra | 3.582 | 1.53E-03 | glio |
| interaction_RT | Vav3 | 1.399 | 1.53E-03 | glio |
| interaction_RT | Junos | 2.736 | 1.53E-03 | glio |
| Aars2_in.vivo_noRT | Nckap1l | 2.259 | 1.53E-03 | glio |
| Aars2_in.vivo_noRT | Rps18 | -0.716 | 1.53E-03 | glio |
| Atr_in.vivo_noRT | Pnpla6 | 1.994 | 1.53E-03 | glio |
| Atr_in.vivo_noRT | Rbm39 | -0.696 | 1.53E-03 | glio |
| Atr_ex.vivo_noRT | Slc23a3 | 2.976 | 1.53E-03 | glio |
| Atr_ex.vivo_noRT | Rbakdn | 3.08 | 1.53E-03 | glio |
| Bcas2_in.vivo_noRT | 1700109H08R | 2.374 | 1.53E-03 | glio |
| Bcas2_ex.vivo_noRT | Dcdc2a | 2.742 | 1.53E-03 | glio |
| Bora_in.vivo_noRT | Hcn2 | 2.483 | 1.53E-03 | glio |
| Bora_in.vivo_noRT | Zfp458 | 2.415 | 1.53E-03 | glio |
| Bora_in.vivo_noRT | Plekhg5 | 2.215 | 1.53E-03 | glio |
| Bora_in.vivo_noRT | Rnpc3 | -1.473 | 1.53E-03 | glio |
| Bora_in.vivo_noRT | Aatf | -1.151 | 1.53E-03 | glio |
| Bora_in.vivo_noRT | Hspa8 | -0.766 | 1.53E-03 | glio |
| Chek1_in.vivo_noRT | Npr2 | 2.632 | 1.53E-03 | glio |
| Chek1_in.vivo_noRT | Mutyh | 2.305 | 1.53E-03 | glio |
| Chek1_in.vivo_noRT | Lrif1 | -1.377 | 1.53E-03 | glio |
| Chek1_in.vivo_noRT | Rps8 | -0.593 | 1.53E-03 | glio |
| Chek1_ex.vivo_noRT | Cadm2 | 3.072 | 1.53E-03 | glio |
| Chek1_ex.vivo_noRT | Brsk1 | 2.467 | 1.53E-03 | glio |
| Chek1_ex.vivo_noRT | Slc25a5 | -1.004 | 1.53E-03 | glio |
| Cyp19a1_ex.vivo_noRT | St18 | 2.933 | 1.53E-03 | glio |
| Eif2s1_in.vivo_noRT | Slc45a1 | 1.883 | 1.53E-03 | glio |
| Hsd17b10_in.vivo_noRT | Gtsf1l | 1.987 | 1.53E-03 | glio |
| Hsd17b10_in.vivo_noRT | Stk24 | -1.407 | 1.53E-03 | glio |
| Interaction_genotype_Chek1 | Resf1 | 2.496 | 1.53E-04 | glio |
| Interaction_genotype_Mtor | Htr2b | -3.576 | 1.53E-04 | glio |
| Interaction_genotype_Mtor | 4933406P04R | -4.545 | 1.53E-04 | glio |
| Interaction_genotype_Mtor | Camk2n2 | -3.631 | 1.53E-04 | glio |
| Interaction_genotype_Myc | Slc1a2 | -3.651 | 1.53E-04 | glio |
| Interaction_genotype_Myc | Rab11fip3 | -3.868 | 1.53E-04 | glio |
| Interaction_genotype_Myc | Snapc5 | -3.674 | 1.53E-04 | glio |
| Interaction_genotype_Myc | Gstz1 | -3.532 | 1.53E-04 | glio |
| Interaction_genotype_Myc | Polr1d | 1.281 | 1.53E-04 | glio |
| Aars2_in.vivo_noRT | Cdc42ep2 | 2.763 | 1.53E-04 | glio |

| | | | | |
|---|---|---|---|---|
| Aars2_in.vivo_noRT | Pcdh1 | 2.699 | 1.53E-04 | glio |
| Atp5c1_in.vivo_noRT | Plk5 | 2.534 | 1.53E-04 | glio |
| Bcas2_in.vivo_noRT | 4930458D05R | 2.999 | 1.53E-04 | glio |
| Bcas2_in.vivo_noRT | Gm15614 | 2.68 | 1.53E-04 | glio |
| Bcas2_in.vivo_noRT | Def6 | 2.609 | 1.53E-04 | glio |
| Bcas2_ex.vivo_noRT | Gm29670 | 2.742 | 1.53E-04 | glio |
| Bora_in.vivo_noRT | Ccdc142os | 2.681 | 1.53E-04 | glio |
| Bora_in.vivo_noRT | Gm17300 | 2.574 | 1.53E-04 | glio |
| Bora_ex.vivo_noRT | Ltbp2 | 2.806 | 1.53E-04 | glio |
| Bora_ex.vivo_noRT | Nmrk2 | 3.201 | 1.53E-04 | glio |
| Eif2s1_in.vivo_noRT | Gm19276 | 2.143 | 1.53E-04 | glio |
| Eif2s1_in.vivo_noRT | Vmn2r44 | 2.143 | 1.53E-04 | glio |
| Eif2s1_in.vivo_noRT | Mmp13 | 2.143 | 1.53E-04 | glio |
| culture_effect | Ndufb11 | -0.447 | 1.53E-05 | glio |
| Interaction_genotype_Mtor | Pik3r3 | -4.765 | 1.53E-05 | glio |
| Interaction_genotype_Myc | 2010110K18R | -4.727 | 1.53E-05 | glio |
| Interaction_genotype_Myc | 4930505N22R | -4.605 | 1.53E-05 | glio |
| Aars2_in.vivo_noRT | Gm20337 | 3.109 | 1.53E-05 | glio |
| Aars2_in.vivo_noRT | Rars | -1.583 | 1.53E-05 | glio |
| Atr_in.vivo_noRT | Cyb561d1 | 3.211 | 1.53E-05 | glio |
| Bcas2_in.vivo_noRT | Nkx2-2os | 3.152 | 1.53E-05 | glio |
| Bora_ex.vivo_noRT | Ms4a15 | 3.201 | 1.53E-05 | glio |
| Chek1_in.vivo_noRT | Sbk2 | 3.078 | 1.53E-05 | glio |
| Interaction_genotype_Mtor | Sult6b1 | -5.727 | 1.53E-06 | glio |
| Interaction_genotype_Myc | Scube3 | -4.347 | 1.53E-06 | glio |
| Interaction_genotype_Myc | Gstm1 | -4.85 | 1.53E-06 | glio |
| Interaction_genotype_Myc | Spx | -5.289 | 1.53E-06 | glio |
| interaction_RT | Sdcbp | 0.964 | 1.53E-06 | glio |
| Atr_in.vivo_noRT | BC051077 | 3.829 | 1.53E-06 | glio |
| Atr_in.vivo_noRT | Pfkfb1 | 3.372 | 1.53E-06 | glio |
| Atr_in.vivo_noRT | 1700126G02R | 3.4 | 1.53E-06 | glio |
| Bcas2_in.vivo_noRT | Gm38979 | 3.633 | 1.53E-06 | glio |
| Bcas2_in.vivo_noRT | Gnaz | 3.509 | 1.53E-06 | glio |
| Bora_in.vivo_noRT | Sptbn4 | 3.388 | 1.53E-06 | glio |
| Atp5c1_in.vivo_noRT | Cgnl1 | 2.668 | 1.53E-07 | glio |
| Atp5c1_in.vivo_noRT | Gm26543 | 2.668 | 1.53E-07 | glio |
| Atp5c1_in.vivo_noRT | Dscaml1 | 2.668 | 1.53E-07 | glio |
| Atp5c1_in.vivo_noRT | 2410017I17Ri | 2.668 | 1.53E-07 | glio |
| Atp5c1_in.vivo_noRT | Pspn | 2.668 | 1.53E-07 | glio |
| Atr_in.vivo_noRT | Lmln | 3.535 | 1.53E-07 | glio |
| Bcas2_in.vivo_noRT | Ccdc39 | 3.587 | 1.53E-07 | glio |
| Bcas2_in.vivo_noRT | Ccdc148 | 3.845 | 1.53E-07 | glio |
| Chek1_ex.vivo_noRT | Prdm1 | 4.449 | 1.53E-07 | glio |
| Hsd17b10_in.vivo_noRT | Vwce | 3.356 | 1.53E-07 | glio |
| Atr_in.vivo_noRT | Hist1h4k | 3.964 | 1.53E-08 | glio |
| Bcas2_in.vivo_noRT | H2-M10.1 | 3.352 | 1.53E-08 | glio |
| Bcas2_in.vivo_noRT | 1700003G18R | 4.547 | 1.53E-08 | glio |
| Aars2_in.vivo_noRT | 1700023H06R | 4.536 | 1.53E-09 | glio |
| Atr_in.vivo_noRT | Syngap1 | 4.073 | 1.53E-09 | glio |
| Atr_in.vivo_noRT | Gm20536 | 4.146 | 1.53E-09 | glio |
| Bora_in.vivo_noRT | Rtn1 | 4.29 | 1.53E-09 | glio |
| Chek1_in.vivo_noRT | B020031M17F | 5.044 | 1.53E-09 | glio |

| | | | | |
|---|---|---|---|---|
| Chek1_in.vivo_noRT | Gpr179 | 4.042 | 1.53E-09 | glio |
| Chek1_ex.vivo_noRT | Irf8 | 4.842 | 1.53E-09 | glio |
| Hsd17b10_in.vivo_noRT | 4930553M12F | 3.35 | 1.53E-09 | glio |
| culture_effect | Cenpb | 1.153 | 1.53E-10 | glio |
| culture_effect | Eid1 | -1.13 | 1.53E-10 | glio |
| Interaction_genotype_Mtor | Krt80 | -7.1 | 1.53E-10 | glio |
| Aars2_in.vivo_noRT | B020031M17F | 5.327 | 1.53E-10 | glio |
| Atr_in.vivo_noRT | Acot6 | 4.288 | 1.53E-10 | glio |
| Atr_in.vivo_noRT | Gm10230 | 4.676 | 1.53E-10 | glio |
| Chek1_ex.vivo_noRT | Gm39556 | 5.611 | 1.53E-10 | glio |
| Aars2_in.vivo_noRT | Olfm2 | 4.929 | 1.53E-11 | glio |
| Atr_in.vivo_noRT | Ptges | 4.877 | 1.53E-11 | glio |
| Bcas2_in.vivo_noRT | Efcab9 | 4.541 | 1.53E-11 | glio |
| Bora_in.vivo_noRT | Pcdhga11 | 3.939 | 1.53E-11 | glio |
| Chek1_in.vivo_noRT | Gm26674 | 4.399 | 1.53E-11 | glio |
| Chek1_in.vivo_noRT | Gm37214 | 4.784 | 1.53E-11 | glio |
| Aars2_in.vivo_noRT | Gm28578 | 5.067 | 1.53E-12 | glio |
| Aars2_in.vivo_noRT | Gm7102 | 5.067 | 1.53E-12 | glio |
| Bcas2_in.vivo_noRT | Gm41318 | 4.807 | 1.53E-12 | glio |
| Bora_in.vivo_noRT | Gm10642 | 4.672 | 1.53E-12 | glio |
| Bora_in.vivo_noRT | Tff2 | 4.676 | 1.53E-12 | glio |
| Hsd17b10_in.vivo_noRT | Olfr171 | 4.137 | 1.53E-13 | glio |
| Hsd17b10_in.vivo_noRT | Gm16178 | 3.616 | 1.53E-13 | glio |
| Hsd17b10_in.vivo_noRT | Gm364 | 3.616 | 1.53E-13 | glio |
| culture_effect | 2900093K20R | 2.406 | 1.53E-14 | glio |
| Chek1_in.vivo_noRT | 2010001K21R | 5.044 | 1.53E-15 | glio |
| Atr_in.vivo_noRT | Gem | 5.533 | 1.53E-16 | glio |
| Chek1_in.vivo_noRT | Fam25c | 4.511 | 1.53E-16 | glio |
| Chek1_in.vivo_noRT | Gm32250 | 4.261 | 1.53E-16 | glio |
| Aars2_in.vivo_noRT | Lrguk | 5.063 | 1.53E-18 | glio |
| Chek1_in.vivo_noRT | Psmb11 | 4.521 | 1.53E-18 | glio |
| Aars2_in.vivo_noRT | Gm45235 | 5.067 | 1.53E-19 | glio |
| culture_effect | Gbp10 | -5.199 | 1.53E-20 | glio |
| Chek1_in.vivo_noRT | Gm30794 | 5.044 | 1.53E-22 | glio |
| Aars2_in.vivo_noRT | Gm21936 | 5.327 | 1.53E-27 | glio |
| culture_effect | Gm45767 | -1.855 | 1.54E-03 | glio |
| culture_effect | Stk32c | -1.741 | 1.54E-03 | glio |
| culture_effect | 4833408A19R | 1.478 | 1.54E-03 | glio |
| culture_effect | Rtl8a | 1.039 | 1.54E-03 | glio |
| culture_effect | Fbxl3 | -0.855 | 1.54E-03 | glio |
| culture_effect | Usp40 | 0.956 | 1.54E-03 | glio |
| culture_effect | Eif2b1 | -0.582 | 1.54E-03 | glio |
| culture_effect | Ipo4 | -0.531 | 1.54E-03 | glio |
| culture_effect | Rps19bp1 | 0.377 | 1.54E-03 | glio |
| Interaction_genotype_Aars2 | Usp30 | -2.919 | 1.54E-03 | glio |
| Interaction_genotype_Atr | Gm29340 | -3.389 | 1.54E-03 | glio |
| Interaction_genotype_Atr | Cnot4 | 1.425 | 1.54E-03 | glio |
| Interaction_genotype_Mtor | Gm11884 | -3.5 | 1.54E-03 | glio |
| Interaction_genotype_Mtor | Slc16a2 | -3.812 | 1.54E-03 | glio |
| Interaction_genotype_Mtor | Akr1c12 | -3.896 | 1.54E-03 | glio |
| Interaction_genotype_Mtor | A930003O13R | -4.134 | 1.54E-03 | glio |
| Interaction_genotype_Mtor | Acox3 | -3.806 | 1.54E-03 | glio |

| | | | | |
|---|---|---|---|---|
| Interaction_genotype_Mtor | Vma21 | 1.484 | 1.54E-03 | glio |
| Interaction_genotype_Mtor | Fkbp4 | 1.565 | 1.54E-03 | glio |
| Interaction_genotype_Mtor | Zfp593 | -1.627 | 1.54E-03 | glio |
| Interaction_genotype_Myc | Gm28578 | -3.714 | 1.54E-03 | glio |
| Interaction_genotype_Myc | B3gat2 | -3.787 | 1.54E-03 | glio |
| Interaction_genotype_Myc | Rtn4ip1 | -3.033 | 1.54E-03 | glio |
| Interaction_genotype_Myc | Xab2 | -2.861 | 1.54E-03 | glio |
| Interaction_genotype_Myc | Eldr | -2.933 | 1.54E-03 | glio |
| Interaction_genotype_Myc | Smad1 | -1.937 | 1.54E-03 | glio |
| Interaction_genotype_Myc | Moxd1 | 1.322 | 1.54E-03 | glio |
| Interaction_genotype_Pars2 | 1700015F17Ri | -4.313 | 1.54E-03 | glio |
| Interaction_genotype_Pars2 | Lncbate10 | -3.056 | 1.54E-03 | glio |
| Aars2_in.vivo_noRT | Plppr2 | 2.311 | 1.54E-03 | glio |
| Aars2_in.vivo_noRT | Ptpre | 1.826 | 1.54E-03 | glio |
| Aars2_ex.vivo_noRT | Gm47872 | 2.897 | 1.54E-03 | glio |
| Atp5c1_in.vivo_noRT | Adam2 | 2.142 | 1.54E-03 | glio |
| Atp5c1_in.vivo_noRT | Gjb5 | 2.208 | 1.54E-03 | glio |
| Atp5c1_in.vivo_noRT | 1700081N11R | 2.866 | 1.54E-03 | glio |
| Atr_in.vivo_noRT | Icam4 | 2.185 | 1.54E-03 | glio |
| Atr_in.vivo_noRT | Arrdc1 | 2.05 | 1.54E-03 | glio |
| Atr_in.vivo_noRT | Ift46 | -1.456 | 1.54E-03 | glio |
| Atr_ex.vivo_noRT | A030005L19Ri | 2.77 | 1.54E-03 | glio |
| Bcas2_in.vivo_noRT | Gm35315 | 2.306 | 1.54E-03 | glio |
| Bcas2_in.vivo_noRT | Upf2 | -1.461 | 1.54E-03 | glio |
| Bcas2_in.vivo_noRT | Twf1 | -1.185 | 1.54E-03 | glio |
| Bcas2_in.vivo_noRT | Cct5 | -0.61 | 1.54E-03 | glio |
| Bora_in.vivo_noRT | Axin2 | 2.367 | 1.54E-03 | glio |
| Bora_in.vivo_noRT | Fhl2 | 2.168 | 1.54E-03 | glio |
| Bora_in.vivo_noRT | Slc36a1 | 2.071 | 1.54E-03 | glio |
| Bora_in.vivo_noRT | Zc4h2 | 1.927 | 1.54E-03 | glio |
| Bora_ex.vivo_noRT | Gm33051 | 2.404 | 1.54E-03 | glio |
| Cenpj_in.vivo_noRT | Gm2022 | 2.022 | 1.54E-03 | glio |
| Cenpj_ex.vivo_noRT | Gm5602 | 2.591 | 1.54E-03 | glio |
| Chek1_in.vivo_noRT | Hykk | 2.279 | 1.54E-03 | glio |
| Chek1_in.vivo_noRT | Cyp4f17 | 2.325 | 1.54E-03 | glio |
| Chek1_in.vivo_noRT | Pex11g | 2.158 | 1.54E-03 | glio |
| Chek1_in.vivo_noRT | Ddx59 | 1.826 | 1.54E-03 | glio |
| Chek1_in.vivo_noRT | Pomgnt1 | -1.619 | 1.54E-03 | glio |
| Chek1_in.vivo_noRT | Tinagl1 | -1.458 | 1.54E-03 | glio |
| Chek1_ex.vivo_noRT | Ppp1r16b | 3.067 | 1.54E-03 | glio |
| Fanca_ex.vivo_noRT | Slc17a6 | 2.124 | 1.54E-03 | glio |
| Fanca_ex.vivo_noRT | Vmn2r44 | 2.524 | 1.54E-03 | glio |
| culture_effect | Dll1 | -2.583 | 1.54E-04 | glio |
| culture_effect | Gm17399 | 2.098 | 1.54E-04 | glio |
| culture_effect | 2900097C17R | -0.686 | 1.54E-04 | glio |
| culture_effect | Gbp7 | -0.532 | 1.54E-04 | glio |
| Interaction_genotype_Aars2 | Gm34590 | -4.762 | 1.54E-04 | glio |
| Interaction_genotype_Aars2 | Fam32a | 1.681 | 1.54E-04 | glio |
| Interaction_genotype_Atr | Myl6b | -4.532 | 1.54E-04 | glio |
| Interaction_genotype_Chek1 | Adam12 | -4.295 | 1.54E-04 | glio |
| Interaction_genotype_Chek1 | H2-Q7 | 4.393 | 1.54E-04 | glio |
| Interaction_genotype_Chek1 | Rasgrp2 | -4.958 | 1.54E-04 | glio |

| | | | | |
|---|---|---|---|---|
| Interaction_genotype_Chek1 | Gm50223 | 4.105 | 1.54E-04 | glio |
| Interaction_genotype_Mtor | Zfp972 | -4.74 | 1.54E-04 | glio |
| Interaction_genotype_Mtor | Gm42670 | -3.949 | 1.54E-04 | glio |
| Interaction_genotype_Mtor | Akap6 | -3.659 | 1.54E-04 | glio |
| Interaction_genotype_Myc | Myl6b | -4.097 | 1.54E-04 | glio |
| Aars2_in.vivo_noRT | 1700123M08F | 2.674 | 1.54E-04 | glio |
| Aars2_in.vivo_noRT | Amd2 | 2.639 | 1.54E-04 | glio |
| Aars2_in.vivo_noRT | Kcnt2 | 2.604 | 1.54E-04 | glio |
| Aars2_in.vivo_noRT | Creb3l1 | 2.656 | 1.54E-04 | glio |
| Atr_in.vivo_noRT | Fanca | 2.288 | 1.54E-04 | glio |
| Bcas2_in.vivo_noRT | A930003O13R | 3.007 | 1.54E-04 | glio |
| Chek1_in.vivo_noRT | Egfl8 | 2.832 | 1.54E-04 | glio |
| Dclre1c_ex.vivo_noRT | Gm34059 | 2.658 | 1.54E-04 | glio |
| Eif2s1_in.vivo_noRT | Gm8016 | 2.143 | 1.54E-04 | glio |
| culture_effect | 4930563E22Ri | 2.185 | 1.54E-05 | glio |
| culture_effect | Eldr | 1.234 | 1.54E-05 | glio |
| Aars2_in.vivo_noRT | 1110006O24R | 3.125 | 1.54E-05 | glio |
| Aars2_in.vivo_noRT | Gm20515 | 2.989 | 1.54E-05 | glio |
| Aars2_in.vivo_noRT | March10 | 2.97 | 1.54E-05 | glio |
| Aars2_ex.vivo_noRT | Gm30042 | 3.293 | 1.54E-05 | glio |
| Atr_in.vivo_noRT | Rpl13 | -0.917 | 1.54E-05 | glio |
| Atr_ex.vivo_noRT | Ms4a15 | 3.164 | 1.54E-05 | glio |
| Bcas2_in.vivo_noRT | Snai1 | 3.289 | 1.54E-05 | glio |
| Bora_ex.vivo_noRT | Slc35g3 | 2.802 | 1.54E-05 | glio |
| culture_effect | Slitrk6 | 1.674 | 1.54E-06 | glio |
| Interaction_genotype_Mtor | Mrps33 | 2.102 | 1.54E-06 | glio |
| Interaction_genotype_Mtor | Clcn1 | -4.613 | 1.54E-06 | glio |
| Interaction_genotype_Mtor | Nwd1 | -4.817 | 1.54E-06 | glio |
| Atr_in.vivo_noRT | Rab36 | 3.365 | 1.54E-06 | glio |
| Atr_in.vivo_noRT | Gm4189 | 3.331 | 1.54E-06 | glio |
| Bcas2_in.vivo_noRT | 2210406O10R | 3.49 | 1.54E-06 | glio |
| Bcas2_in.vivo_noRT | 1700122E12Ri | 3.411 | 1.54E-06 | glio |
| Bora_in.vivo_noRT | Kcnt2 | 3.273 | 1.54E-06 | glio |
| Chek1_in.vivo_noRT | Hpse | 3.333 | 1.54E-06 | glio |
| Chek1_in.vivo_noRT | Anxa8 | 3.318 | 1.54E-06 | glio |
| Chek1_in.vivo_noRT | Mpeg1 | 3.068 | 1.54E-06 | glio |
| Hsd17b10_in.vivo_noRT | Gm42664 | 2.966 | 1.54E-06 | glio |
| Hsd17b10_in.vivo_noRT | 6430562O15R | 3.094 | 1.54E-06 | glio |
| culture_effect | 2810001G20R | 1.199 | 1.54E-07 | glio |
| Interaction_genotype_Mtor | 1700003E24Ri | -4.099 | 1.54E-07 | glio |
| Interaction_genotype_Myc | Gm29340 | -5.717 | 1.54E-07 | glio |
| Interaction_genotype_Myc | Olfml2a | -6.083 | 1.54E-07 | glio |
| Interaction_genotype_Myc | Hlx | -5.925 | 1.54E-07 | glio |
| Atr_in.vivo_noRT | Rex2 | 3.86 | 1.54E-07 | glio |
| Bora_in.vivo_noRT | Gm29340 | 3.62 | 1.54E-07 | glio |
| Interaction_genotype_Mtor | Cnpy2 | 2.605 | 1.54E-08 | glio |
| Interaction_genotype_Mtor | Gm45338 | -4.492 | 1.54E-08 | glio |
| Aars2_in.vivo_noRT | Ltb4r1 | 4.152 | 1.54E-08 | glio |
| Bora_in.vivo_noRT | Gm49694 | 4.21 | 1.54E-08 | glio |
| Chek1_in.vivo_noRT | Gpr162 | 3.995 | 1.54E-08 | glio |
| Chek1_in.vivo_noRT | Vamp8 | 3.22 | 1.54E-08 | glio |
| Chek1_ex.vivo_noRT | Masp2 | 4.449 | 1.54E-09 | glio |

| | | | | |
|---|---|---|---|---|
| Aars2_in.vivo_noRT | 1700061G19R | 4.315 | 1.54E-10 | glio |
| Bora_in.vivo_noRT | Gm15234 | 4.42 | 1.54E-10 | glio |
| Aars2_in.vivo_noRT | Vmn1r88 | 4.142 | 1.54E-11 | glio |
| Aars2_in.vivo_noRT | Gm15941 | 4.444 | 1.54E-12 | glio |
| Bora_in.vivo_noRT | Cxcl16 | 4.672 | 1.54E-12 | glio |
| Bora_in.vivo_noRT | Dnah10 | 4.557 | 1.54E-13 | glio |
| Chek1_in.vivo_noRT | Agrp | 4.261 | 1.54E-13 | glio |
| Chek1_in.vivo_noRT | Rab25 | 4.777 | 1.54E-14 | glio |
| Atr_in.vivo_noRT | Ripply3 | 5.533 | 1.54E-16 | glio |
| Aars2_in.vivo_noRT | Zscan4d | 5.327 | 1.54E-20 | glio |
| culture_effect | Gbp8 | -5.524 | 1.54E-24 | glio |
| culture_effect | Mthfd2l | 2.318 | 1.54E-28 | glio |
| culture_effect | Zfp968 | 1.795 | 1.55E-03 | glio |
| culture_effect | Fam71f1 | 1.168 | 1.55E-03 | glio |
| culture_effect | Tfdp1 | 0.497 | 1.55E-03 | glio |
| culture_effect | Dnajc2 | 0.459 | 1.55E-03 | glio |
| Interaction_genotype_Ifnar1 | Slc17a6 | 2.734 | 1.55E-03 | glio |
| Interaction_genotype_Mtor | Slc17a7 | -3.846 | 1.55E-03 | glio |
| Interaction_genotype_Mtor | Ercc3 | -2.96 | 1.55E-03 | glio |
| Interaction_genotype_Mtor | Mrpl22 | 1.534 | 1.55E-03 | glio |
| Interaction_genotype_Myc | Masp1 | -3.005 | 1.55E-03 | glio |
| Interaction_genotype_Myc | Gm44091 | -3.466 | 1.55E-03 | glio |
| Interaction_genotype_Myc | Ppp1r3e | -3.439 | 1.55E-03 | glio |
| Interaction_genotype_Myc | Mpped2 | -3.613 | 1.55E-03 | glio |
| Interaction_genotype_Myc | Fam214a | -2.907 | 1.55E-03 | glio |
| Interaction_genotype_Myc | Nhej1 | -3.36 | 1.55E-03 | glio |
| Interaction_genotype_Myc | Fancc | -2.638 | 1.55E-03 | glio |
| Interaction_genotype_Myc | Neto1 | 2.018 | 1.55E-03 | glio |
| Interaction_genotype_Myc | Utp11 | 1.354 | 1.55E-03 | glio |
| Interaction_genotype_Pars2 | Gm13404 | -4.412 | 1.55E-03 | glio |
| Interaction_genotype_Pars2 | Chia1 | -3.056 | 1.55E-03 | glio |
| Interaction_genotype_Pars2 | Ppil6 | -3.943 | 1.55E-03 | glio |
| Interaction_genotype_Pars2 | Gm41318 | -4.184 | 1.55E-03 | glio |
| Interaction_genotype_Pars2 | Impg2 | -3.586 | 1.55E-03 | glio |
| interaction_RT | Klhdc8a | -2.313 | 1.55E-03 | glio |
| Aars2_in.vivo_noRT | Fam71f1 | 1.912 | 1.55E-03 | glio |
| Aars2_in.vivo_noRT | Tmem108 | 1.95 | 1.55E-03 | glio |
| Aars2_in.vivo_noRT | Taldo1 | -0.923 | 1.55E-03 | glio |
| Aars2_in.vivo_noRT | Phb2 | -0.716 | 1.55E-03 | glio |
| Aars2_in.vivo_noRT | Ifi27l2a | -1.039 | 1.55E-03 | glio |
| Aars2_ex.vivo_noRT | Tmem30c | 3.127 | 1.55E-03 | glio |
| Atp5c1_in.vivo_noRT | Dok2 | 2.142 | 1.55E-03 | glio |
| Atr_ex.vivo_noRT | Zfp811 | 2.975 | 1.55E-03 | glio |
| Bard1_ex.vivo_noRT | Gm34776 | 2.421 | 1.55E-03 | glio |
| Bard1_ex.vivo_noRT | 4933408J17Ri | 2.812 | 1.55E-03 | glio |
| Bcas2_in.vivo_noRT | 1700102H20R | 2.282 | 1.55E-03 | glio |
| Bcas2_in.vivo_noRT | Limd1 | -1.26 | 1.55E-03 | glio |
| Eif2s1_in.vivo_noRT | Gstt4 | 1.88 | 1.55E-03 | glio |
| Eif2s1_in.vivo_noRT | Slc2a9 | 2.143 | 1.55E-03 | glio |
| Hsd17b10_in.vivo_noRT | Kcnb1 | 2.532 | 1.55E-03 | glio |
| culture_effect | Fgf1 | 2.181 | 1.55E-04 | glio |
| culture_effect | Samd9l | 0.794 | 1.55E-04 | glio |

| | | | | |
|---|---|---|---|---|
| Interaction_genotype_Myc | E130215H24R | -3.986 | 1.55E-04 | glio |
| Interaction_genotype_Myc | Fzd4 | -4.178 | 1.55E-04 | glio |
| Interaction_genotype_Myc | Higd1a | 1.989 | 1.55E-04 | glio |
| Aars2_in.vivo_noRT | Utrn | -1.643 | 1.55E-04 | glio |
| Aars2_in.vivo_noRT | Slc25a5 | -0.987 | 1.55E-04 | glio |
| Atr_in.vivo_noRT | Ebi3 | 3.155 | 1.55E-04 | glio |
| Atr_in.vivo_noRT | Plekho2 | 2.599 | 1.55E-04 | glio |
| Atr_in.vivo_noRT | Hoxaas3 | 2.225 | 1.55E-04 | glio |
| Atr_ex.vivo_noRT | Gm14207 | 2.37 | 1.55E-04 | glio |
| Atr_ex.vivo_noRT | Nat8f4 | 3.174 | 1.55E-04 | glio |
| Bcas2_in.vivo_noRT | Cd34 | 2.766 | 1.55E-04 | glio |
| Bcas2_in.vivo_noRT | Arhgdia | -0.79 | 1.55E-04 | glio |
| Bora_in.vivo_noRT | Nt5dc2 | -2.046 | 1.55E-04 | glio |
| Chek1_in.vivo_noRT | Cnn1 | 3.374 | 1.55E-04 | glio |
| Chek1_in.vivo_noRT | Fam83e | 2.795 | 1.55E-04 | glio |
| Chek1_in.vivo_noRT | Foxl2os | 2.535 | 1.55E-04 | glio |
| Chek1_in.vivo_noRT | Tlcd2 | 3.018 | 1.55E-04 | glio |
| Hsd17b10_in.vivo_noRT | E2f8 | -2.24 | 1.55E-04 | glio |
| culture_effect | Fkbp2 | -0.548 | 1.55E-05 | glio |
| Interaction_genotype_Bora | Gm26533 | -5.624 | 1.55E-05 | glio |
| interaction_RT | Cbx5 | -0.738 | 1.55E-05 | glio |
| Atr_in.vivo_noRT | Krt17 | 5.273 | 1.55E-05 | glio |
| Bard1_ex.vivo_noRT | Cgnl1 | 2.814 | 1.55E-05 | glio |
| Bard1_ex.vivo_noRT | Gm26543 | 2.814 | 1.55E-05 | glio |
| Bcas2_in.vivo_noRT | Tox2 | 3.032 | 1.55E-05 | glio |
| Bora_in.vivo_noRT | Gm20536 | 3.04 | 1.55E-05 | glio |
| Bora_in.vivo_noRT | Gm10353 | 2.955 | 1.55E-05 | glio |
| Chek1_in.vivo_noRT | Gm36608 | 3.113 | 1.55E-05 | glio |
| Chek1_in.vivo_noRT | Nr1h4 | 2.995 | 1.55E-05 | glio |
| Interaction_genotype_Mtor | A730071L15Ri | -5.873 | 1.55E-06 | glio |
| Interaction_genotype_Myc | Cyp3a13 | -5.861 | 1.55E-06 | glio |
| Interaction_genotype_Myc | Apoh | -5.469 | 1.55E-06 | glio |
| Interaction_genotype_Myc | Gm41318 | -5.442 | 1.55E-06 | glio |
| Atp5c1_in.vivo_noRT | Gm48215 | 2.668 | 1.55E-06 | glio |
| Atr_in.vivo_noRT | A930033H14R | 3.509 | 1.55E-06 | glio |
| Interaction_genotype_Mtor | Npnt | -4.63 | 1.55E-07 | glio |
| Aars2_ex.vivo_noRT | Idi2 | 3.293 | 1.55E-07 | glio |
| Aars2_ex.vivo_noRT | Vmn2r18 | 3.293 | 1.55E-07 | glio |
| Aars2_ex.vivo_noRT | Gm16178 | 3.293 | 1.55E-07 | glio |
| Aars2_ex.vivo_noRT | Gm364 | 3.293 | 1.55E-07 | glio |
| Atr_in.vivo_noRT | Gm50218 | 3.94 | 1.55E-07 | glio |
| Hsd17b10_in.vivo_noRT | Gm49223 | 3.093 | 1.55E-07 | glio |
| Aars2_in.vivo_noRT | Limch1 | 4.289 | 1.55E-08 | glio |
| Chek1_ex.vivo_noRT | Prrt3 | 4.449 | 1.55E-08 | glio |
| Chek1_ex.vivo_noRT | Cacng4 | 4.449 | 1.55E-08 | glio |
| culture_effect | Gm36495 | 3.271 | 1.55E-09 | glio |
| Aars2_in.vivo_noRT | Ggt1 | 4.014 | 1.55E-09 | glio |
| Atr_in.vivo_noRT | Arl4d | 4.222 | 1.55E-09 | glio |
| Chek1_in.vivo_noRT | 1700124L16Ri | 4.333 | 1.55E-09 | glio |
| Aars2_in.vivo_noRT | Gm3734 | 4.804 | 1.55E-10 | glio |
| Atr_in.vivo_noRT | Apol7d | 4.632 | 1.55E-10 | glio |
| culture_effect | Sumo1 | 0.707 | 1.55E-11 | glio |

| | | | | |
|---|---|---|---|---|
| Chek1_ex.vivo_noRT | BC048671 | 4.842 | 1.55E-11 | glio |
| Atr_in.vivo_noRT | Gm14968 | 5.154 | 1.55E-12 | glio |
| Bcas2_in.vivo_noRT | Rslcan18 | 4.421 | 1.55E-12 | glio |
| Bcas2_in.vivo_noRT | Gm27239 | 4.807 | 1.55E-12 | glio |
| Bora_in.vivo_noRT | Gm41349 | 4.673 | 1.55E-13 | glio |
| Atr_in.vivo_noRT | Vmn2r54 | 4.372 | 1.55E-14 | glio |
| Bcas2_in.vivo_noRT | Gm15929 | 4.807 | 1.55E-17 | glio |
| Atr_in.vivo_noRT | Dapk2 | 4.75 | 1.55E-19 | glio |
| Atr_in.vivo_noRT | Gm15445 | 4.887 | 1.55E-19 | glio |
| Bora_in.vivo_noRT | Gm5286 | 4.936 | 1.55E-19 | glio |
| Bora_in.vivo_noRT | Gm34776 | 4.936 | 1.55E-19 | glio |
| Aars2_in.vivo_noRT | Fam166a | 5.063 | 1.55E-20 | glio |
| Chek1_in.vivo_noRT | BC061195 | 5.044 | 1.55E-24 | glio |
| culture_effect | H2-D1 | -3.672 | 1.55E-70 | glio |
| culture_effect | Gm26881 | 1.286 | 1.56E-03 | glio |
| culture_effect | Fancl | -0.911 | 1.56E-03 | glio |
| culture_effect | Rbms3 | 0.917 | 1.56E-03 | glio |
| Interaction_genotype_Atr | Gm26671 | -2.762 | 1.56E-03 | glio |
| Interaction_genotype_Atr | Ece2 | -3.818 | 1.56E-03 | glio |
| Interaction_genotype_Mtor | Zfp990 | -3.684 | 1.56E-03 | glio |
| Interaction_genotype_Mtor | Tbxa2r | -3.622 | 1.56E-03 | glio |
| Interaction_genotype_Mtor | Gm49768 | -3.587 | 1.56E-03 | glio |
| Interaction_genotype_Mtor | Ift172 | -3.247 | 1.56E-03 | glio |
| Interaction_genotype_Mtor | Amph | -3.241 | 1.56E-03 | glio |
| Interaction_genotype_Myc | Mcm8 | -2.471 | 1.56E-03 | glio |
| Interaction_genotype_Myc | Stip1 | 1.309 | 1.56E-03 | glio |
| Interaction_genotype_Myc | Rex1bd | 1.327 | 1.56E-03 | glio |
| interaction_RT | Fosl2 | 1.426 | 1.56E-03 | glio |
| interaction_RT | Ola1 | -0.63 | 1.56E-03 | glio |
| Aars2_in.vivo_noRT | Phf20l1 | -1.291 | 1.56E-03 | glio |
| Aars2_ex.vivo_noRT | Magea10 | 2.713 | 1.56E-03 | glio |
| Atr_in.vivo_noRT | Gm38190 | 2.101 | 1.56E-03 | glio |
| Atr_in.vivo_noRT | Kcnk2 | 1.922 | 1.56E-03 | glio |
| Atr_in.vivo_noRT | Hoxd4 | 2.035 | 1.56E-03 | glio |
| Atr_in.vivo_noRT | Scfd1 | -1.274 | 1.56E-03 | glio |
| Bard1_ex.vivo_noRT | Ciita | 2.415 | 1.56E-03 | glio |
| Bcas2_in.vivo_noRT | Cox7a1 | 2.489 | 1.56E-03 | glio |
| Bcas2_in.vivo_noRT | 3010001F23Ri | 2.178 | 1.56E-03 | glio |
| Bcas2_ex.vivo_noRT | 1700024I08Ri | 2.542 | 1.56E-03 | glio |
| Bora_in.vivo_noRT | Mkrn2 | 1.428 | 1.56E-03 | glio |
| Chek1_in.vivo_noRT | Elac1 | 2.181 | 1.56E-03 | glio |
| Chek1_in.vivo_noRT | Slc5a6 | 1.956 | 1.56E-03 | glio |
| Chek1_in.vivo_noRT | Cox8a | -0.587 | 1.56E-03 | glio |
| Chek1_in.vivo_noRT | Rps25 | -0.616 | 1.56E-03 | glio |
| Cyp19a1_ex.vivo_noRT | Gm10801 | 2.959 | 1.56E-03 | glio |
| Dclre1c_ex.vivo_noRT | Gm20767 | 3.344 | 1.56E-03 | glio |
| Hsd17b10_in.vivo_noRT | Tmem30c | 2.43 | 1.56E-03 | glio |
| culture_effect | Kcnq2 | -2.098 | 1.56E-04 | glio |
| culture_effect | Stag1 | 0.591 | 1.56E-04 | glio |
| Interaction_genotype_Atr | Mapk1 | 1.856 | 1.56E-04 | glio |
| Interaction_genotype_Atr | Ier2 | 2.544 | 1.56E-04 | glio |
| Interaction_genotype_Mtor | Spcs1 | 1.84 | 1.56E-04 | glio |

| | | | | |
|---|---|---|---|---|
| Interaction_genotype_Myc | Psat1 | 1.715 | 1.56E-04 | glio |
| interaction_RT | Drosha | -0.864 | 1.56E-04 | glio |
| interaction_RT | H2-Q7 | 2.653 | 1.56E-04 | glio |
| interaction_RT | Msmo1 | 1.103 | 1.56E-04 | glio |
| interaction_RT | Phldb2 | -3.277 | 1.56E-04 | glio |
| Aars2_in.vivo_noRT | Zfp458 | 2.805 | 1.56E-04 | glio |
| Bora_in.vivo_noRT | Stac3 | 2.794 | 1.56E-04 | glio |
| Bora_in.vivo_noRT | Cysltr2 | 2.465 | 1.56E-04 | glio |
| Hsd17b10_in.vivo_noRT | Gm5544 | 2.825 | 1.56E-04 | glio |
| culture_effect | Syt17 | 2.356 | 1.56E-05 | glio |
| culture_effect | Cct2 | -0.434 | 1.56E-05 | glio |
| Interaction_genotype_Mtor | 4930589L23Ri | -4.896 | 1.56E-05 | glio |
| Interaction_genotype_Mtor | Milr1 | -5.316 | 1.56E-05 | glio |
| Interaction_genotype_Myc | Gm49086 | -4.292 | 1.56E-05 | glio |
| Atr_in.vivo_noRT | Lmtk3 | 2.923 | 1.56E-05 | glio |
| Atr_in.vivo_noRT | Prdm6 | 2.76 | 1.56E-05 | glio |
| Atr_ex.vivo_noRT | A530013C23R | 3.655 | 1.56E-05 | glio |
| Bard1_ex.vivo_noRT | 2410017I17Ri | 2.814 | 1.56E-05 | glio |
| Chek1_ex.vivo_noRT | Necab3 | 3.666 | 1.56E-05 | glio |
| Chek1_ex.vivo_noRT | 4933406C10R | 3.464 | 1.56E-05 | glio |
| culture_effect | Mtg1 | -0.97 | 1.56E-06 | glio |
| culture_effect | Trim30a | 0.819 | 1.56E-06 | glio |
| Interaction_genotype_Mtor | Golph3 | 2.223 | 1.56E-06 | glio |
| Interaction_genotype_Mtor | Gm10324 | -4.403 | 1.56E-06 | glio |
| Interaction_genotype_Myc | 1700084E18Ri | -5.763 | 1.56E-06 | glio |
| Interaction_genotype_Myc | Gm50069 | -5.642 | 1.56E-06 | glio |
| Atr_in.vivo_noRT | Stmn4 | 5.017 | 1.56E-06 | glio |
| Atr_in.vivo_noRT | Gm26982 | 3.313 | 1.56E-06 | glio |
| Bcas2_in.vivo_noRT | Cdo1 | 3.519 | 1.56E-06 | glio |
| culture_effect | Mgarp | -3.12 | 1.56E-07 | glio |
| culture_effect | Gm50340 | 1.925 | 1.56E-07 | glio |
| Atr_in.vivo_noRT | Gm6225 | 3.857 | 1.56E-07 | glio |
| Atr_in.vivo_noRT | Sdcbp2 | 3.839 | 1.56E-07 | glio |
| Bora_in.vivo_noRT | A730071L15Ri | 3.876 | 1.56E-07 | glio |
| Atr_in.vivo_noRT | Sv2a | 4.096 | 1.56E-08 | glio |
| Atr_in.vivo_noRT | Lrrc46 | 3.885 | 1.56E-08 | glio |
| Bcas2_in.vivo_noRT | Cpeb1 | 4.162 | 1.56E-08 | glio |
| culture_effect | Denr | 0.691 | 1.56E-09 | glio |
| culture_effect | AI506816 | -0.814 | 1.56E-09 | glio |
| culture_effect | Uqcrq | 0.71 | 1.56E-09 | glio |
| Aars2_in.vivo_noRT | Pla2g7 | 4.14 | 1.56E-09 | glio |
| Atr_in.vivo_noRT | Rps24 | -1.264 | 1.56E-09 | glio |
| Chek1_in.vivo_noRT | Sap30bpos | 3.649 | 1.56E-09 | glio |
| Chek1_in.vivo_noRT | Ptges | 4.388 | 1.56E-09 | glio |
| culture_effect | H2-DMb2 | -3.902 | 1.56E-10 | glio |
| Bora_in.vivo_noRT | Gm28404 | 4.048 | 1.56E-10 | glio |
| Chek1_in.vivo_noRT | Gm12925 | 4.586 | 1.56E-10 | glio |
| Atr_in.vivo_noRT | A930003A15R | 4.381 | 1.56E-11 | glio |
| Atr_in.vivo_noRT | Aqp5 | 5.27 | 1.56E-11 | glio |
| Bcas2_in.vivo_noRT | Cd248 | 4.541 | 1.56E-11 | glio |
| Atr_in.vivo_noRT | Gm44702 | 5.002 | 1.56E-12 | glio |
| Bcas2_in.vivo_noRT | 4930412M03F | 4.807 | 1.56E-12 | glio |

| | | | | |
|---|---|---|---|---|
| Chek1_in.vivo_noRT | 5430402O13R | 5.044 | 1.56E-12 | glio |
| Chek1_in.vivo_noRT | Gm44702 | 5.023 | 1.56E-12 | glio |
| culture_effect | 2510046G10R | 3.13 | 1.56E-14 | glio |
| Bora_in.vivo_noRT | D430001F17R | 4.936 | 1.56E-14 | glio |
| Chek1_in.vivo_noRT | Tcstv3 | 5.044 | 1.56E-14 | glio |
| Chek1_ex.vivo_noRT | Gm13856 | 4.449 | 1.56E-14 | glio |
| Chek1_ex.vivo_noRT | Gm5890 | 4.842 | 1.56E-14 | glio |
| Chek1_ex.vivo_noRT | Zscan4c | 4.842 | 1.56E-14 | glio |
| Chek1_ex.vivo_noRT | Speer4a | 4.842 | 1.56E-14 | glio |
| Chek1_ex.vivo_noRT | BC080695 | 4.842 | 1.56E-14 | glio |
| Chek1_ex.vivo_noRT | 4930588A03R | 4.443 | 1.56E-14 | glio |
| culture_effect | Tmbim1 | -1.461 | 1.56E-15 | glio |
| culture_effect | Prune1 | -1.413 | 1.56E-16 | glio |
| Aars2_in.vivo_noRT | Olfr279 | 5.589 | 1.56E-16 | glio |
| Atr_in.vivo_noRT | Gm44658 | 5.002 | 1.56E-16 | glio |
| Bcas2_in.vivo_noRT | Gm13348 | 4.807 | 1.56E-16 | glio |
| Chek1_in.vivo_noRT | Nat8f2 | 4.784 | 1.56E-16 | glio |
| Bora_in.vivo_noRT | 4930512B01R | 4.936 | 1.56E-20 | glio |
| Aars2_in.vivo_noRT | Gm12800 | 5.327 | 1.56E-24 | glio |
| culture_effect | Gm35019 | -6.745 | 1.56E-36 | glio |
| culture_effect | Hlx | 1.718 | 1.57E-03 | glio |
| culture_effect | Hyou1 | -0.663 | 1.57E-03 | glio |
| culture_effect | Ube3a | 0.4 | 1.57E-03 | glio |
| Interaction_genotype_Aars2 | 1700011D18R | -3.988 | 1.57E-03 | glio |
| Interaction_genotype_Aars2 | Gm28874 | -3.931 | 1.57E-03 | glio |
| Interaction_genotype_Aars2 | D230022J07Ri | -3.977 | 1.57E-03 | glio |
| Interaction_genotype_Aars2 | Hist1h4b | -4.297 | 1.57E-03 | glio |
| Interaction_genotype_Aars2 | Trim30a | 1.944 | 1.57E-03 | glio |
| Interaction_genotype_Aars2 | Rplp2 | 0.945 | 1.57E-03 | glio |
| Interaction_genotype_Atr | Erlec1 | 1.628 | 1.57E-03 | glio |
| Interaction_genotype_Bora | Slc2a9 | -3.456 | 1.57E-03 | glio |
| Interaction_genotype_Bora | Chd4 | 1.334 | 1.57E-03 | glio |
| Interaction_genotype_Mtor | Gm35339 | -4.003 | 1.57E-03 | glio |
| Interaction_genotype_Mtor | Gm43727 | -3.681 | 1.57E-03 | glio |
| Interaction_genotype_Mtor | Igfbp6 | -3.321 | 1.57E-03 | glio |
| Interaction_genotype_Myc | Tex52 | -3.457 | 1.57E-03 | glio |
| Interaction_genotype_Myc | Plekhm3 | -3.706 | 1.57E-03 | glio |
| Interaction_genotype_Myc | Dgka | -3.152 | 1.57E-03 | glio |
| Interaction_genotype_Rpl3 | Slpi | -4.839 | 1.57E-03 | glio |
| Interaction_genotype_Rpl3 | Fa2h | 4.162 | 1.57E-03 | glio |
| interaction_RT | Ifrd2 | -1.026 | 1.57E-03 | glio |
| interaction_RT | Ctsb | 0.852 | 1.57E-03 | glio |
| Aars2_in.vivo_noRT | Zfp937 | 2.396 | 1.57E-03 | glio |
| Aars2_in.vivo_noRT | Kif24 | 2.212 | 1.57E-03 | glio |
| Aars2_ex.vivo_noRT | Sh3d21 | 2.595 | 1.57E-03 | glio |
| Atr_in.vivo_noRT | Gm20300 | 2.275 | 1.57E-03 | glio |
| Atr_in.vivo_noRT | Thbs3 | 2.074 | 1.57E-03 | glio |
| Atr_in.vivo_noRT | Zfp93 | 1.831 | 1.57E-03 | glio |
| Atr_in.vivo_noRT | Il1rap | 1.462 | 1.57E-03 | glio |
| Atr_ex.vivo_noRT | Gm48170 | 2.367 | 1.57E-03 | glio |
| Atr_ex.vivo_noRT | Tal2 | 2.182 | 1.57E-03 | glio |
| Bcas2_in.vivo_noRT | Cdiptos | 2.258 | 1.57E-03 | glio |

| | | | | |
|---|---|---|---|---|
| Bcas2_in.vivo_noRT | Pcdhgb4 | 2.012 | 1.57E-03 | glio |
| Bcas2_in.vivo_noRT | Cacng7 | 2.233 | 1.57E-03 | glio |
| Bcas2_in.vivo_noRT | Ube2i | -0.675 | 1.57E-03 | glio |
| Bora_in.vivo_noRT | Gm9873 | 2.081 | 1.57E-03 | glio |
| Bora_ex.vivo_noRT | Gm26542 | 3.012 | 1.57E-03 | glio |
| Bora_ex.vivo_noRT | Ubxn10 | 2.802 | 1.57E-03 | glio |
| Chek1_in.vivo_noRT | Rab42 | 2.44 | 1.57E-03 | glio |
| Chek1_in.vivo_noRT | A230072E10Ri | 2.277 | 1.57E-03 | glio |
| Chek1_in.vivo_noRT | 2010110K18R | 2.198 | 1.57E-03 | glio |
| Chek1_in.vivo_noRT | Slc25a29 | 2.099 | 1.57E-03 | glio |
| Chek1_in.vivo_noRT | Prdm15 | 1.958 | 1.57E-03 | glio |
| Chek1_in.vivo_noRT | Mfap1b | -1.396 | 1.57E-03 | glio |
| Chek1_in.vivo_noRT | Zc3h15 | -0.815 | 1.57E-03 | glio |
| Chek1_ex.vivo_noRT | Flrt1 | 3.097 | 1.57E-03 | glio |
| Chek1_ex.vivo_noRT | Gm16740 | 2.704 | 1.57E-03 | glio |
| Chek1_ex.vivo_noRT | Ska2 | -1.884 | 1.57E-03 | glio |
| Chek1_ex.vivo_noRT | Eprs | -0.898 | 1.57E-03 | glio |
| Hsd17b10_in.vivo_noRT | Dok3 | 2.481 | 1.57E-03 | glio |
| culture_effect | Cpne8 | 2.194 | 1.57E-04 | glio |
| culture_effect | 2500004C02R | 1.426 | 1.57E-04 | glio |
| culture_effect | Tmem128 | -0.462 | 1.57E-04 | glio |
| Interaction_genotype_Aars2 | Gm20455 | -4.941 | 1.57E-04 | glio |
| Interaction_genotype_Bora | Pip5k1a | 2.231 | 1.57E-04 | glio |
| Interaction_genotype_Mtor | Rnf144b | -4.606 | 1.57E-04 | glio |
| Interaction_genotype_Mtor | Fbxo10 | -4.46 | 1.57E-04 | glio |
| Interaction_genotype_Mtor | Syt6 | -4.081 | 1.57E-04 | glio |
| Interaction_genotype_Myc | Gm36037 | -3.124 | 1.57E-04 | glio |
| Interaction_genotype_Myc | Arhgef28 | -3.87 | 1.57E-04 | glio |
| Interaction_genotype_Myc | 9130230N09R | -3.707 | 1.57E-04 | glio |
| Interaction_genotype_Myc | Zfp335 | -3.318 | 1.57E-04 | glio |
| interaction_RT | Ppef2 | 2.063 | 1.57E-04 | glio |
| interaction_RT | Gm17092 | -1.67 | 1.57E-04 | glio |
| interaction_RT | Katnbl1 | 1.071 | 1.57E-04 | glio |
| interaction_RT | Srrt | -0.904 | 1.57E-04 | glio |
| Aars2_in.vivo_noRT | Btbd6 | 2.444 | 1.57E-04 | glio |
| Aars2_in.vivo_noRT | Cavin2 | 2.806 | 1.57E-04 | glio |
| Atp5c1_in.vivo_noRT | Scube3 | 2.663 | 1.57E-04 | glio |
| Atp5c1_in.vivo_noRT | Tlr1 | 2.93 | 1.57E-04 | glio |
| Atr_ex.vivo_noRT | Slc7a3 | 3.556 | 1.57E-04 | glio |
| Bard1_ex.vivo_noRT | 4930570E03Ri | 2.814 | 1.57E-04 | glio |
| Bard1_ex.vivo_noRT | Gm5286 | 2.814 | 1.57E-04 | glio |
| Bcas2_in.vivo_noRT | C130013H08R | 3.038 | 1.57E-04 | glio |
| Bcas2_in.vivo_noRT | AW046200 | 2.749 | 1.57E-04 | glio |
| Bcas2_in.vivo_noRT | Vmac | 3.013 | 1.57E-04 | glio |
| Bora_in.vivo_noRT | Gm48796 | 2.564 | 1.57E-04 | glio |
| Bora_in.vivo_noRT | Hist1h3d | 2.557 | 1.57E-04 | glio |
| Bora_in.vivo_noRT | Tlx3 | 2.658 | 1.57E-04 | glio |
| Chek1_in.vivo_noRT | Kdelr3 | -2.163 | 1.57E-04 | glio |
| Chek1_in.vivo_noRT | Acaa1a | -1.745 | 1.57E-04 | glio |
| culture_effect | Epor | -2.485 | 1.57E-05 | glio |
| culture_effect | Cspg4 | -1.112 | 1.57E-05 | glio |
| Interaction_genotype_Myc | Rmi1 | -3.726 | 1.57E-05 | glio |

| | | | | |
|---|---|---|---|---|
| interaction_RT | Sesn3 | 1.421 | 1.57E-05 | glio |
| Atr_ex.vivo_noRT | Slc35g3 | 2.764 | 1.57E-05 | glio |
| Bcas2_in.vivo_noRT | Gm47591 | 3.109 | 1.57E-05 | glio |
| Bcas2_in.vivo_noRT | Eme2 | 2.911 | 1.57E-05 | glio |
| Bora_in.vivo_noRT | Pcdhgb4 | 2.665 | 1.57E-05 | glio |
| Chek1_in.vivo_noRT | Tle4 | -2.022 | 1.57E-05 | glio |
| Hsd17b10_in.vivo_noRT | Icam2 | 2.75 | 1.57E-05 | glio |
| Hsd17b10_in.vivo_noRT | Glod5 | -3.341 | 1.57E-05 | glio |
| culture_effect | Poglut2 | -1.048 | 1.57E-06 | glio |
| Interaction_genotype_Myc | A930001C03R | -5.372 | 1.57E-06 | glio |
| Interaction_genotype_Myc | Pitx3 | -4.493 | 1.57E-06 | glio |
| Interaction_genotype_Myc | Rpl32 | 1.979 | 1.57E-06 | glio |
| Bcas2_in.vivo_noRT | Vcam1 | 4.291 | 1.57E-06 | glio |
| Bora_in.vivo_noRT | D430018E03R | 3.21 | 1.57E-06 | glio |
| Bora_ex.vivo_noRT | Gm26555 | 3.201 | 1.57E-06 | glio |
| Chek1_in.vivo_noRT | Cfap100 | 3.376 | 1.57E-06 | glio |
| Interaction_genotype_Mtor | Skp1a | 2.567 | 1.57E-07 | glio |
| Interaction_genotype_Mtor | Atp6v1g1 | 2.437 | 1.57E-07 | glio |
| Interaction_genotype_Mtor | Gm39326 | -6.153 | 1.57E-07 | glio |
| Interaction_genotype_Myc | Poln | -6.19 | 1.57E-07 | glio |
| Bcas2_in.vivo_noRT | Nlrx1 | 3.919 | 1.57E-07 | glio |
| Bcas2_in.vivo_noRT | Synpr | 4.174 | 1.57E-07 | glio |
| Bora_ex.vivo_noRT | Helt | 3.201 | 1.57E-07 | glio |
| Bora_ex.vivo_noRT | Gm20625 | 3.201 | 1.57E-07 | glio |
| Bora_ex.vivo_noRT | Gm5157 | 3.201 | 1.57E-07 | glio |
| Bora_ex.vivo_noRT | Gm44736 | 3.201 | 1.57E-07 | glio |
| Bora_ex.vivo_noRT | Fam227b | 3.201 | 1.57E-07 | glio |
| Bora_ex.vivo_noRT | Gm20658 | 3.201 | 1.57E-07 | glio |
| Bora_ex.vivo_noRT | Olfr810 | 3.201 | 1.57E-07 | glio |
| Bora_ex.vivo_noRT | Apoa5 | 3.201 | 1.57E-07 | glio |
| Bora_ex.vivo_noRT | Gjd4 | 3.201 | 1.57E-07 | glio |
| Bora_ex.vivo_noRT | Gm49172 | 3.201 | 1.57E-07 | glio |
| Chek1_in.vivo_noRT | Pbld1 | 3.857 | 1.57E-07 | glio |
| Chek1_in.vivo_noRT | Gm49042 | 3.414 | 1.57E-07 | glio |
| culture_effect | Ybey | 1.829 | 1.57E-08 | glio |
| Aars2_in.vivo_noRT | Klhdc8b | 4.049 | 1.57E-08 | glio |
| Bora_in.vivo_noRT | C4a | 3.498 | 1.57E-08 | glio |
| Hsd17b10_in.vivo_noRT | 2410141K09R | 3.352 | 1.57E-08 | glio |
| Aars2_in.vivo_noRT | Cnih2 | 4.139 | 1.57E-09 | glio |
| Atr_in.vivo_noRT | Arhgef19 | 4.354 | 1.57E-09 | glio |
| Bcas2_in.vivo_noRT | Tas1r1 | 3.78 | 1.57E-09 | glio |
| Bora_in.vivo_noRT | Sult6b1 | 4.145 | 1.57E-09 | glio |
| Chek1_ex.vivo_noRT | 4930579C12R | 4.447 | 1.57E-09 | glio |
| Chek1_ex.vivo_noRT | Cyp2c23 | 4.447 | 1.57E-09 | glio |
| Chek1_ex.vivo_noRT | Card9 | 4.443 | 1.57E-09 | glio |
| Hsd17b10_in.vivo_noRT | Scgb1c1 | 3.616 | 1.57E-09 | glio |
| culture_effect | Mrps31 | -1.002 | 1.57E-10 | glio |
| Interaction_genotype_Mtor | Gm32090 | -7.216 | 1.57E-10 | glio |
| Aars2_in.vivo_noRT | Gm28424 | 4.315 | 1.57E-10 | glio |
| Chek1_in.vivo_noRT | 5330439K02R | 4.777 | 1.57E-11 | glio |
| Chek1_ex.vivo_noRT | Gm5602 | 4.842 | 1.57E-12 | glio |
| culture_effect | Eif3f | -0.9 | 1.57E-13 | glio |

| | | | | |
|---|---|---|---|---|
| Bora_in.vivo_noRT | Car14 | 4.936 | 1.57E-13 | glio |
| Chek1_ex.vivo_noRT | Gm50024 | 4.443 | 1.57E-13 | glio |
| culture_effect | Misp | -3.857 | 1.57E-14 | glio |
| culture_effect | Rpl7 | -0.949 | 1.57E-14 | glio |
| Bcas2_in.vivo_noRT | Gm48242 | 4.807 | 1.57E-15 | glio |
| Chek1_in.vivo_noRT | Ppil6 | 5.044 | 1.57E-15 | glio |
| Bcas2_in.vivo_noRT | Gm17203 | 4.157 | 1.57E-16 | glio |
| Bcas2_in.vivo_noRT | Aqp9 | 4.807 | 1.57E-16 | glio |
| Chek1_in.vivo_noRT | Pakap | 4.511 | 1.57E-16 | glio |
| Bcas2_in.vivo_noRT | Gm5890 | 4.807 | 1.57E-20 | glio |
| Chek1_in.vivo_noRT | Speer4a | 4.777 | 1.57E-20 | glio |
| Bora_in.vivo_noRT | Gm12800 | 4.936 | 1.57E-21 | glio |
| culture_effect | Gm42418 | -2.704 | 1.57E-22 | glio |
| Chek1_in.vivo_noRT | Gm4340 | 5.044 | 1.57E-25 | glio |
| Chek1_in.vivo_noRT | Gm16239 | 5.044 | 1.57E-25 | glio |
| culture_effect | Pnpla2 | -2.01 | 1.57E-26 | glio |
| culture_effect | Aig1 | -1.315 | 1.57E-26 | glio |
| Atr_in.vivo_noRT | Gm15533 | 5.533 | 1.57E-26 | glio |
| Atr_in.vivo_noRT | Gm16178 | 5.533 | 1.57E-28 | glio |
| Atr_in.vivo_noRT | Gm364 | 5.533 | 1.57E-28 | glio |
| culture_effect | C130013H08R | 1.868 | 1.58E-03 | glio |
| culture_effect | Phactr2 | 0.881 | 1.58E-03 | glio |
| Interaction_genotype_Aars2 | Tsn | 1.023 | 1.58E-03 | glio |
| Interaction_genotype_Atr | Grin3b | -3.911 | 1.58E-03 | glio |
| Interaction_genotype_Atr | Usp39 | 1.944 | 1.58E-03 | glio |
| Interaction_genotype_Bcas2 | Rilp | -4.525 | 1.58E-03 | glio |
| Interaction_genotype_Bcas2 | Lama5 | -4.384 | 1.58E-03 | glio |
| Interaction_genotype_Bcas2 | Abcg4 | -3.939 | 1.58E-03 | glio |
| Interaction_genotype_Bcas2 | Ifitm6 | -4.288 | 1.58E-03 | glio |
| Interaction_genotype_Mtor | Gm21154 | -4.065 | 1.58E-03 | glio |
| Interaction_genotype_Mtor | Fbxl12os | -3.459 | 1.58E-03 | glio |
| Interaction_genotype_Mtor | Gm50340 | -3.106 | 1.58E-03 | glio |
| Interaction_genotype_Mtor | Cnep1r1 | 1.892 | 1.58E-03 | glio |
| Interaction_genotype_Mtor | Cenpf | 3.126 | 1.58E-03 | glio |
| Interaction_genotype_Myc | Rasl2-9 | -4.041 | 1.58E-03 | glio |
| Interaction_genotype_Myc | Tnfrsf10b | -3.439 | 1.58E-03 | glio |
| Interaction_genotype_Myc | Ankle1 | -3.168 | 1.58E-03 | glio |
| interaction_RT | Maff | 1.327 | 1.58E-03 | glio |
| Aars2_in.vivo_noRT | Sdf2 | -0.918 | 1.58E-03 | glio |
| Aars2_in.vivo_noRT | Lsm6 | -1.015 | 1.58E-03 | glio |
| Aars2_ex.vivo_noRT | Gm50138 | 2.534 | 1.58E-03 | glio |
| Atr_in.vivo_noRT | Gm4631 | 2.167 | 1.58E-03 | glio |
| Atr_in.vivo_noRT | Yrdc | -0.954 | 1.58E-03 | glio |
| Bcas2_in.vivo_noRT | Tdrp | 2.235 | 1.58E-03 | glio |
| Bora_in.vivo_noRT | Chaf1a | -1.428 | 1.58E-03 | glio |
| Cenpj_ex.vivo_noRT | Gm12860 | 2.591 | 1.58E-03 | glio |
| Cenpj_ex.vivo_noRT | Dscaml1 | 2.196 | 1.58E-03 | glio |
| Cenpj_ex.vivo_noRT | Pspn | 2.196 | 1.58E-03 | glio |
| Cenpj_ex.vivo_noRT | Akr1c18 | 2.591 | 1.58E-03 | glio |
| Cenpj_ex.vivo_noRT | Minar2 | 2.591 | 1.58E-03 | glio |
| Cenpj_ex.vivo_noRT | Gm45151 | 2.591 | 1.58E-03 | glio |
| Cenpj_ex.vivo_noRT | Gast | 2.591 | 1.58E-03 | glio |

| | | | | |
|---|---|---|---|---|
| Cenpj_ex.vivo_noRT | Gm27241 | 2.408 | 1.58E-03 | glio |
| Cenpj_ex.vivo_noRT | Txndc2 | 2.601 | 1.58E-03 | glio |
| Cenpj_ex.vivo_noRT | Serping1 | 2.591 | 1.58E-03 | glio |
| Chek1_in.vivo_noRT | Tspan2 | 1.952 | 1.58E-03 | glio |
| Chek1_in.vivo_noRT | Ccni | -1.334 | 1.58E-03 | glio |
| Chek1_in.vivo_noRT | Vps4b | 0.986 | 1.58E-03 | glio |
| Chek1_ex.vivo_noRT | C430042M11F | 2.938 | 1.58E-03 | glio |
| Chek1_ex.vivo_noRT | Mroh2b | 2.986 | 1.58E-03 | glio |
| Chek1_ex.vivo_noRT | Sbsn | 2.562 | 1.58E-03 | glio |
| Hsd17b10_in.vivo_noRT | Phf23 | -1.098 | 1.58E-03 | glio |
| culture_effect | Stk10 | 1.159 | 1.58E-04 | glio |
| culture_effect | Scrn1 | 0.68 | 1.58E-04 | glio |
| Interaction_genotype_Mtor | Sv2a | -4.724 | 1.58E-04 | glio |
| Interaction_genotype_Mtor | Itpripl1 | -3.851 | 1.58E-04 | glio |
| Interaction_genotype_Mtor | Lncppara | -4.453 | 1.58E-04 | glio |
| Interaction_genotype_Mtor | Pde8a | -3.417 | 1.58E-04 | glio |
| interaction_RT | Rnpepl1 | -1.265 | 1.58E-04 | glio |
| interaction_RT | Pla2g2e | 3.665 | 1.58E-04 | glio |
| Aars2_in.vivo_noRT | Tmem40 | 2.982 | 1.58E-04 | glio |
| Atr_in.vivo_noRT | Syn3 | 2.68 | 1.58E-04 | glio |
| Atr_in.vivo_noRT | Spata2l | 2.741 | 1.58E-04 | glio |
| Atr_in.vivo_noRT | Ndufa4 | -0.863 | 1.58E-04 | glio |
| Bcas2_in.vivo_noRT | Rimkla | 2.588 | 1.58E-04 | glio |
| Bcas2_in.vivo_noRT | Tnfrsf22 | 2.609 | 1.58E-04 | glio |
| Bora_in.vivo_noRT | Ttc5 | -1.871 | 1.58E-04 | glio |
| Bora_in.vivo_noRT | Ccnt1 | -1.591 | 1.58E-04 | glio |
| Bora_ex.vivo_noRT | Obscn | 3.201 | 1.58E-04 | glio |
| Chek1_in.vivo_noRT | Gm6598 | 2.525 | 1.58E-04 | glio |
| culture_effect | Gm15834 | 2.115 | 1.58E-05 | glio |
| Interaction_genotype_Aars2 | 4930412M03F | -5.253 | 1.58E-05 | glio |
| Interaction_genotype_Atr | Rnf165 | -5.081 | 1.58E-05 | glio |
| Interaction_genotype_Atr | Gm16741 | -4.909 | 1.58E-05 | glio |
| Interaction_genotype_Mtor | Alox8 | -6.293 | 1.58E-05 | glio |
| interaction_RT | Kpnb1 | -0.884 | 1.58E-05 | glio |
| Aars2_in.vivo_noRT | Unc5cl | 3.025 | 1.58E-05 | glio |
| Aars2_in.vivo_noRT | Zfp37 | 2.925 | 1.58E-05 | glio |
| Atr_in.vivo_noRT | Cadm2 | 3.17 | 1.58E-05 | glio |
| Atr_in.vivo_noRT | Srpk2 | -1.405 | 1.58E-05 | glio |
| Atr_in.vivo_noRT | Rpl9 | -0.826 | 1.58E-05 | glio |
| Bcas2_in.vivo_noRT | Zfp990 | 2.906 | 1.58E-05 | glio |
| Bcas2_in.vivo_noRT | Acot5 | 2.923 | 1.58E-05 | glio |
| Bcas2_in.vivo_noRT | Daam2 | 3.152 | 1.58E-05 | glio |
| Bcas2_ex.vivo_noRT | 1700003E24Ri | 2.742 | 1.58E-05 | glio |
| Bora_in.vivo_noRT | 5330426L24Ri | 3.226 | 1.58E-05 | glio |
| Bora_in.vivo_noRT | Gm37170 | 3.093 | 1.58E-05 | glio |
| Hsd17b10_in.vivo_noRT | Lrrc4b | 2.705 | 1.58E-05 | glio |
| culture_effect | Elmsan1 | -1.283 | 1.58E-06 | glio |
| Interaction_genotype_Myc | Ttbk1 | -5.457 | 1.58E-06 | glio |
| Bcas2_in.vivo_noRT | Gm26916 | 3.23 | 1.58E-06 | glio |
| Chek1_in.vivo_noRT | Gm26620 | 3.025 | 1.58E-06 | glio |
| Chek1_in.vivo_noRT | Wdr45b | -1.913 | 1.58E-06 | glio |
| Chek1_ex.vivo_noRT | Atp1a2 | 4.443 | 1.58E-06 | glio |

| | | | | |
|---|---|---|---|---|
| Interaction_genotype_Mtor | Hspa12a | -6.389 | 1.58E-07 | glio |
| Interaction_genotype_Mtor | Mycl | -6.587 | 1.58E-07 | glio |
| Atr_in.vivo_noRT | Rasl2-9 | 4.096 | 1.58E-07 | glio |
| Atr_in.vivo_noRT | Stpg2 | 3.585 | 1.58E-07 | glio |
| Bcas2_in.vivo_noRT | Catsperd | 3.628 | 1.58E-07 | glio |
| Chek1_in.vivo_noRT | Gpd1 | 3.871 | 1.58E-07 | glio |
| Atr_in.vivo_noRT | Arg2 | 3.893 | 1.58E-08 | glio |
| Atr_in.vivo_noRT | Gm11127 | 3.973 | 1.58E-08 | glio |
| Atr_in.vivo_noRT | Gpr21 | 3.668 | 1.58E-08 | glio |
| Atr_ex.vivo_noRT | Capn13 | 3.164 | 1.58E-08 | glio |
| Atr_ex.vivo_noRT | Gm21319 | 3.947 | 1.58E-08 | glio |
| Bcas2_in.vivo_noRT | Spint4 | 4.016 | 1.58E-08 | glio |
| Chek1_ex.vivo_noRT | Prm3 | 4.263 | 1.58E-08 | glio |
| Bora_in.vivo_noRT | Ntn5 | 4.144 | 1.58E-09 | glio |
| Bora_in.vivo_noRT | Stx1b | 4.033 | 1.58E-10 | glio |
| Chek1_in.vivo_noRT | Tspyl3 | 3.961 | 1.58E-10 | glio |
| Atr_in.vivo_noRT | Apoh | 4.618 | 1.58E-11 | glio |
| Bcas2_in.vivo_noRT | Dntt | 4.807 | 1.58E-11 | glio |
| Chek1_ex.vivo_noRT | Lncbate10 | 4.055 | 1.58E-11 | glio |
| culture_effect | Adam10 | 1.32 | 1.58E-12 | glio |
| Atr_in.vivo_noRT | Notch4 | 4.8 | 1.58E-12 | glio |
| Bora_in.vivo_noRT | Prrx2 | 4.936 | 1.58E-12 | glio |
| Chek1_in.vivo_noRT | Matn4 | 4.78 | 1.58E-12 | glio |
| culture_effect | Syne2 | -1.261 | 1.58E-13 | glio |
| Bcas2_in.vivo_noRT | Obscn | 4.807 | 1.58E-14 | glio |
| Chek1_in.vivo_noRT | Abra | 4.518 | 1.58E-14 | glio |
| Chek1_ex.vivo_noRT | Gm21936 | 4.447 | 1.58E-14 | glio |
| Chek1_ex.vivo_noRT | BB287469 | 4.842 | 1.58E-14 | glio |
| Bora_in.vivo_noRT | Gm15445 | 4.29 | 1.58E-15 | glio |
| culture_effect | Uqcr10 | 1.032 | 1.58E-16 | glio |
| Atr_in.vivo_noRT | Gm45620 | 5.533 | 1.58E-16 | glio |
| Atr_in.vivo_noRT | Gm38534 | 5.533 | 1.58E-16 | glio |
| Bcas2_in.vivo_noRT | Pitx3 | 4.807 | 1.58E-17 | glio |
| Chek1_in.vivo_noRT | 5730507C01R | 4.521 | 1.58E-17 | glio |
| Atr_in.vivo_noRT | Gm13032 | 5.147 | 1.58E-19 | glio |
| Bcas2_in.vivo_noRT | Zscan4c | 4.807 | 1.58E-20 | glio |
| Bcas2_in.vivo_noRT | Rimklb | 4.544 | 1.58E-20 | glio |
| Bcas2_in.vivo_noRT | Idi2 | 4.807 | 1.58E-22 | glio |
| culture_effect | Nlrx1 | 1.905 | 1.59E-03 | glio |
| culture_effect | Btbd6 | 1.41 | 1.59E-03 | glio |
| culture_effect | Arl8a | 0.6 | 1.59E-03 | glio |
| culture_effect | Ggh | -0.489 | 1.59E-03 | glio |
| Interaction_genotype_Aars2 | Chrna1os | -3.182 | 1.59E-03 | glio |
| Interaction_genotype_Atr | B430219N15R | -4.042 | 1.59E-03 | glio |
| Interaction_genotype_Atr | Rlbp1 | -1.458 | 1.59E-03 | glio |
| Interaction_genotype_Chek1 | AW112010 | 4.847 | 1.59E-03 | glio |
| Interaction_genotype_Chek1 | Gm27239 | -4.098 | 1.59E-03 | glio |
| Interaction_genotype_Chek1 | Eif4ebp2 | 2.181 | 1.59E-03 | glio |
| Interaction_genotype_Chek1 | Ezr | 1.881 | 1.59E-03 | glio |
| Interaction_genotype_Chek1 | Cd47 | 1.877 | 1.59E-03 | glio |
| Interaction_genotype_Mtor | Pax3 | -3.058 | 1.59E-03 | glio |
| Interaction_genotype_Myc | Aqp3 | -4.428 | 1.59E-03 | glio |

| | | | | |
|---|---|---|---|---|
| Interaction_genotype_Myc | Pde5a | -3.64 | 1.59E-03 | glio |
| Interaction_genotype_Myc | Hcn1 | -3.856 | 1.59E-03 | glio |
| Interaction_genotype_Myc | Vmac | -3.861 | 1.59E-03 | glio |
| Interaction_genotype_Myc | Fam120c | -3.551 | 1.59E-03 | glio |
| Interaction_genotype_Myc | Unc13d | -3.437 | 1.59E-03 | glio |
| Interaction_genotype_Myc | Grb14 | -2.724 | 1.59E-03 | glio |
| Interaction_genotype_Myc | Stx17 | -2.47 | 1.59E-03 | glio |
| Interaction_genotype_Myc | D630045J12Ri | -2.424 | 1.59E-03 | glio |
| Interaction_genotype_Myc | Rpl17 | 1.236 | 1.59E-03 | glio |
| interaction_RT | Uaca | -0.89 | 1.59E-03 | glio |
| Aars2_in.vivo_noRT | Naa25 | -1.311 | 1.59E-03 | glio |
| Aars2_in.vivo_noRT | Ndufb9 | -0.589 | 1.59E-03 | glio |
| Atr_in.vivo_noRT | Spag4 | 1.803 | 1.59E-03 | glio |
| Atr_in.vivo_noRT | Adck2 | 2.271 | 1.59E-03 | glio |
| Bcas2_in.vivo_noRT | Cyb5r1 | -1.634 | 1.59E-03 | glio |
| Bcas2_in.vivo_noRT | Upf3b | -1.537 | 1.59E-03 | glio |
| Bcas2_in.vivo_noRT | Becn1 | -0.79 | 1.59E-03 | glio |
| Bcas2_ex.vivo_noRT | Etl4 | 2.742 | 1.59E-03 | glio |
| Bora_in.vivo_noRT | 9030025P20R | 2.464 | 1.59E-03 | glio |
| Bora_in.vivo_noRT | Gm15998 | 2.303 | 1.59E-03 | glio |
| Bora_in.vivo_noRT | Pcyt1b | -1.471 | 1.59E-03 | glio |
| Bora_in.vivo_noRT | Rab5c | -1.044 | 1.59E-03 | glio |
| Bora_ex.vivo_noRT | Gm40264 | 2.615 | 1.59E-03 | glio |
| Chek1_in.vivo_noRT | Serpina3m | 2.222 | 1.59E-03 | glio |
| Eif2s1_in.vivo_noRT | Pyy | 2.143 | 1.59E-03 | glio |
| Hsd17b10_in.vivo_noRT | Hemk1 | 2.431 | 1.59E-03 | glio |
| Hsd17b10_in.vivo_noRT | C2cd5 | -1.874 | 1.59E-03 | glio |
| culture_effect | Ttbk1 | 1.793 | 1.59E-04 | glio |
| culture_effect | Gnl3 | -0.532 | 1.59E-04 | glio |
| Interaction_genotype_Aars2 | Pyy | -3.914 | 1.59E-04 | glio |
| Interaction_genotype_Atr | Gm17597 | -4.955 | 1.59E-04 | glio |
| Interaction_genotype_Mtor | Hnrnpu | 1.553 | 1.59E-04 | glio |
| interaction_RT | Rabac1 | 0.701 | 1.59E-04 | glio |
| Aars2_in.vivo_noRT | Herpud1 | -1.735 | 1.59E-04 | glio |
| Aars2_in.vivo_noRT | Brd4 | -1.218 | 1.59E-04 | glio |
| Atr_in.vivo_noRT | Snx3 | -0.819 | 1.59E-04 | glio |
| Bcas2_ex.vivo_noRT | Gm45338 | 2.348 | 1.59E-04 | glio |
| Bora_in.vivo_noRT | Fndc10 | 2.932 | 1.59E-04 | glio |
| Bora_in.vivo_noRT | Prelid3a | 2.708 | 1.59E-04 | glio |
| Bora_in.vivo_noRT | Wbp11 | -0.979 | 1.59E-04 | glio |
| Bora_ex.vivo_noRT | Adam4 | 3.309 | 1.59E-04 | glio |
| Chek1_ex.vivo_noRT | Gm44658 | 2.9 | 1.59E-04 | glio |
| Eif2s1_in.vivo_noRT | Gpc5 | 2.143 | 1.59E-04 | glio |
| Hsd17b10_in.vivo_noRT | 1700013F07Ri | 2.988 | 1.59E-04 | glio |
| culture_effect | Gm47061 | 2.492 | 1.59E-05 | glio |
| culture_effect | Cers4 | -1.072 | 1.59E-05 | glio |
| Interaction_genotype_Mtor | Fkbp9 | 2.084 | 1.59E-05 | glio |
| Interaction_genotype_Myc | Stard6 | -5.027 | 1.59E-05 | glio |
| Interaction_genotype_Myc | Arf1 | 1.391 | 1.59E-05 | glio |
| Aars2_in.vivo_noRT | Ankrd13d | 3.094 | 1.59E-05 | glio |
| Atr_in.vivo_noRT | Ston1 | 3.083 | 1.59E-05 | glio |
| Bora_ex.vivo_noRT | Gm10482 | 3.201 | 1.59E-05 | glio |

| | | | | |
|---|---|---|---|---|
| Hsd17b10_in.vivo_noRT | Ccl7 | 3.616 | 1.59E-05 | glio |
| Hsd17b10_in.vivo_noRT | Ushbp1 | 3.09 | 1.59E-05 | glio |
| Interaction_genotype_Myc | Atp5o | 2.245 | 1.59E-06 | glio |
| Atr_in.vivo_noRT | 1600002K03R | 3.47 | 1.59E-06 | glio |
| Bora_in.vivo_noRT | Sgk3 | 3.382 | 1.59E-06 | glio |
| Chek1_in.vivo_noRT | Prkcsh | -1.624 | 1.59E-06 | glio |
| Interaction_genotype_Nat2 | Gm5157 | 4.205 | 1.59E-07 | glio |
| Atr_in.vivo_noRT | Vgll2 | 3.197 | 1.59E-07 | glio |
| Atr_in.vivo_noRT | 2610507I01Ri | 3.273 | 1.59E-07 | glio |
| Bcas2_in.vivo_noRT | Papln | 3.54 | 1.59E-07 | glio |
| Bora_in.vivo_noRT | Dzank1 | 3.761 | 1.59E-07 | glio |
| Bora_in.vivo_noRT | Slc14a1 | 3.051 | 1.59E-07 | glio |
| Chek1_in.vivo_noRT | Adam1a | 3.682 | 1.59E-07 | glio |
| Hsd17b10_in.vivo_noRT | Adam12 | 3.356 | 1.59E-07 | glio |
| culture_effect | Gbp3 | -0.739 | 1.59E-08 | glio |
| Aars2_in.vivo_noRT | Gm26559 | 4.026 | 1.59E-08 | glio |
| interaction_RT | Noct | 2.031 | 1.59E-09 | glio |
| Atr_in.vivo_noRT | A530072M11F | 4.205 | 1.59E-09 | glio |
| Hsd17b10_in.vivo_noRT | Abra | 3.611 | 1.59E-09 | glio |
| Atr_in.vivo_noRT | Papln | 4.266 | 1.59E-10 | glio |
| Bcas2_in.vivo_noRT | 7530414M10F | 4.281 | 1.59E-10 | glio |
| Bcas2_in.vivo_noRT | G630030J09Ri | 4.285 | 1.59E-10 | glio |
| Bora_in.vivo_noRT | Tex52 | 4.286 | 1.59E-10 | glio |
| Atr_in.vivo_noRT | Spint4 | 4.742 | 1.59E-11 | glio |
| culture_effect | Dlk2 | 2.566 | 1.59E-13 | glio |
| Bora_in.vivo_noRT | Dok2 | 4.411 | 1.59E-13 | glio |
| Bora_in.vivo_noRT | Gm11457 | 4.936 | 1.59E-13 | glio |
| Chek1_in.vivo_noRT | Gm50115 | 4.78 | 1.59E-13 | glio |
| Atr_in.vivo_noRT | A330015K06R | 4.876 | 1.59E-14 | glio |
| Bcas2_in.vivo_noRT | Gm36172 | 4.275 | 1.59E-14 | glio |
| Bora_in.vivo_noRT | 9130230L23Ri | 4.676 | 1.59E-14 | glio |
| Aars2_in.vivo_noRT | Car14 | 5.327 | 1.59E-15 | glio |
| Atr_in.vivo_noRT | Pla2g2d | 5.27 | 1.59E-16 | glio |
| Bora_in.vivo_noRT | Gm5934 | 4.673 | 1.59E-18 | glio |
| Bora_in.vivo_noRT | Akr1c18 | 4.936 | 1.59E-18 | glio |
| Bora_in.vivo_noRT | 4921504E06Ri | 4.936 | 1.59E-18 | glio |
| Bcas2_in.vivo_noRT | BC080695 | 4.807 | 1.59E-20 | glio |
| Bora_in.vivo_noRT | Rimklb | 4.673 | 1.59E-21 | glio |
| culture_effect | Ninj1 | 1.592 | 1.59E-22 | glio |
| Bcas2_in.vivo_noRT | Gm4340 | 4.807 | 1.59E-23 | glio |
| Bcas2_in.vivo_noRT | Gm16239 | 4.807 | 1.59E-23 | glio |
| Aars2_in.vivo_noRT | Gm49172 | 5.067 | 1.59E-26 | glio |
| Aars2_in.vivo_noRT | Clec12b | 5.327 | 1.59E-26 | glio |
| culture_effect | Abi1 | 0.559 | 1.60E-03 | glio |
| culture_effect | Cyct | -0.539 | 1.60E-03 | glio |
| Interaction_genotype_Aars2 | 4933406P04R | -4.057 | 1.60E-03 | glio |
| Interaction_genotype_Atr | Gm47622 | -2.944 | 1.60E-03 | glio |
| Interaction_genotype_Atr | Akap3 | -3.963 | 1.60E-03 | glio |
| Interaction_genotype_Atr | Gm20707 | -3.796 | 1.60E-03 | glio |
| Interaction_genotype_Atr | Cdc16 | -1.839 | 1.60E-03 | glio |
| Interaction_genotype_Atr | Mrps25 | 1.489 | 1.60E-03 | glio |
| Interaction_genotype_Atr | Slu7 | 1.665 | 1.60E-03 | glio |

| | | | | |
|---|---|---|---|---|
| Interaction_genotype_Atr | Srpk2 | 1.435 | 1.60E-03 | glio |
| Interaction_genotype_Bcas2 | A430010J10Ri | -5.039 | 1.60E-03 | glio |
| Interaction_genotype_Mtor | Tmem30c | -3.878 | 1.60E-03 | glio |
| Interaction_genotype_Mtor | 1700055D18R | -3.827 | 1.60E-03 | glio |
| Interaction_genotype_Myc | Lmod1 | -3.792 | 1.60E-03 | glio |
| Interaction_genotype_Myc | Gm26982 | -3.56 | 1.60E-03 | glio |
| Interaction_genotype_Myc | Ccdc96 | -3.253 | 1.60E-03 | glio |
| Interaction_genotype_Myc | Tkfc | -2.557 | 1.60E-03 | glio |
| Interaction_genotype_Myc | Xrcc1 | -2.151 | 1.60E-03 | glio |
| Interaction_genotype_Myc | Mef2c | -2.333 | 1.60E-03 | glio |
| Interaction_genotype_Myc | Eif3k | 1.318 | 1.60E-03 | glio |
| Interaction_genotype_Terf1 | Pcdh17 | 2.838 | 1.60E-03 | glio |
| Interaction_genotype_Terf1 | Jkamp | 1.962 | 1.60E-03 | glio |
| Interaction_genotype_Terf1 | Gm43653 | -4.391 | 1.60E-03 | glio |
| Atr_in.vivo_noRT | Prdm15 | 1.924 | 1.60E-03 | glio |
| Atr_ex.vivo_noRT | Gm31063 | 2.24 | 1.60E-03 | glio |
| Atr_ex.vivo_noRT | Gsto2 | 2.955 | 1.60E-03 | glio |
| Atr_ex.vivo_noRT | Gm42047 | -2.622 | 1.60E-03 | glio |
| Bcas2_in.vivo_noRT | Ifi205 | 2.986 | 1.60E-03 | glio |
| Bcas2_in.vivo_noRT | Zfp114 | 2.333 | 1.60E-03 | glio |
| Bcas2_in.vivo_noRT | Pxt1 | 2.334 | 1.60E-03 | glio |
| Bcas2_in.vivo_noRT | Ston1 | 2.357 | 1.60E-03 | glio |
| Bcas2_in.vivo_noRT | Llgl2 | 2.272 | 1.60E-03 | glio |
| Bcas2_in.vivo_noRT | Pdlim1 | -1.666 | 1.60E-03 | glio |
| Bcas2_in.vivo_noRT | Arhgap17 | -1.453 | 1.60E-03 | glio |
| Bcas2_in.vivo_noRT | Dhfr | -1.447 | 1.60E-03 | glio |
| Bora_in.vivo_noRT | 1700120G11R | 2.453 | 1.60E-03 | glio |
| Bora_in.vivo_noRT | 1700021F05Ri | -1.237 | 1.60E-03 | glio |
| Chek1_in.vivo_noRT | Rnf126 | -1.125 | 1.60E-03 | glio |
| Chek1_ex.vivo_noRT | Bpi | 2.59 | 1.60E-03 | glio |
| Chek1_ex.vivo_noRT | Fbxo9 | -1.886 | 1.60E-03 | glio |
| Cyp19a1_ex.vivo_noRT | Ecscr | -2.777 | 1.60E-03 | glio |
| Eif2s1_in.vivo_noRT | Esm1 | 1.879 | 1.60E-03 | glio |
| Hsd17b10_in.vivo_noRT | Amhr2 | 2.74 | 1.60E-03 | glio |
| culture_effect | Gm35066 | 2.005 | 1.60E-04 | glio |
| culture_effect | Pkn2 | 0.484 | 1.60E-04 | glio |
| Interaction_genotype_Chek1 | Tcim | 3.336 | 1.60E-04 | glio |
| Interaction_genotype_Chek1 | Relb | 3.072 | 1.60E-04 | glio |
| Interaction_genotype_Mtor | Has1 | -4.184 | 1.60E-04 | glio |
| Interaction_genotype_Mtor | Zfp273 | -4.499 | 1.60E-04 | glio |
| Interaction_genotype_Mtor | AA386476 | -4.358 | 1.60E-04 | glio |
| Interaction_genotype_Myc | Gm16741 | -3.874 | 1.60E-04 | glio |
| Aars2_in.vivo_noRT | Cyb561 | 2.852 | 1.60E-04 | glio |
| Aars2_in.vivo_noRT | Fam129c | 2.659 | 1.60E-04 | glio |
| Aars2_in.vivo_noRT | Dennd3 | 2.793 | 1.60E-04 | glio |
| Atp5c1_in.vivo_noRT | Ube4bos1 | 2.404 | 1.60E-04 | glio |
| Atp5c1_in.vivo_noRT | D430001F17R | 2.668 | 1.60E-04 | glio |
| Atr_in.vivo_noRT | Ddr1 | -1.382 | 1.60E-04 | glio |
| Bora_ex.vivo_noRT | 4931431C16R | 2.815 | 1.60E-04 | glio |
| Chek1_in.vivo_noRT | Gstt1 | 2.797 | 1.60E-04 | glio |
| Chek1_in.vivo_noRT | Cox4i1 | -0.663 | 1.60E-04 | glio |
| Chek1_ex.vivo_noRT | Slc5a5 | 3.669 | 1.60E-04 | glio |

| | | | | |
|---|---|---|---|---|
| Chek1_ex.vivo_noRT | Galnt12 | -1.985 | 1.60E-04 | glio |
| Hsd17b10_in.vivo_noRT | Pola2 | -1.719 | 1.60E-04 | glio |
| culture_effect | Ldlr | -1.007 | 1.60E-05 | glio |
| culture_effect | Txndc9 | -0.575 | 1.60E-05 | glio |
| Interaction_genotype_Mtor | Sult5a1 | -5.55 | 1.60E-05 | glio |
| Interaction_genotype_Myc | Radil | -4.86 | 1.60E-05 | glio |
| Interaction_genotype_Myc | Tmem40 | -5.365 | 1.60E-05 | glio |
| Interaction_genotype_Myc | Lhx9 | -5.174 | 1.60E-05 | glio |
| Aars2_in.vivo_noRT | Cib2 | 3.445 | 1.60E-05 | glio |
| Aars2_in.vivo_noRT | Gstm6 | 3.26 | 1.60E-05 | glio |
| Aars2_in.vivo_noRT | Anks6 | 2.853 | 1.60E-05 | glio |
| Aars2_in.vivo_noRT | Mzf1 | 2.936 | 1.60E-05 | glio |
| Aars2_in.vivo_noRT | Bpifa1 | 2.8 | 1.60E-05 | glio |
| Aars2_ex.vivo_noRT | Gm15138 | 2.897 | 1.60E-05 | glio |
| Atr_in.vivo_noRT | 9030407P20R | 2.987 | 1.60E-05 | glio |
| Atr_ex.vivo_noRT | Ctss | 3.554 | 1.60E-05 | glio |
| culture_effect | Gm32031 | 1.808 | 1.60E-06 | glio |
| Interaction_genotype_Mtor | 4930558J18Ri | -5.213 | 1.60E-06 | glio |
| Interaction_genotype_Mtor | Nkx2-2os | -5.918 | 1.60E-06 | glio |
| Interaction_genotype_Mtor | Zfp14 | -5.931 | 1.60E-06 | glio |
| Interaction_genotype_Myc | Pyy | -4.609 | 1.60E-06 | glio |
| Atr_ex.vivo_noRT | Zscan4-ps2 | 3.164 | 1.60E-06 | glio |
| Bora_in.vivo_noRT | Rhov | 3.283 | 1.60E-06 | glio |
| Bora_in.vivo_noRT | Tssk6 | 2.974 | 1.60E-06 | glio |
| Bora_ex.vivo_noRT | Npnt | 3.196 | 1.60E-06 | glio |
| Bora_ex.vivo_noRT | Cd36 | 3.201 | 1.60E-06 | glio |
| Bora_ex.vivo_noRT | Gstt4 | 3.201 | 1.60E-06 | glio |
| Chek1_ex.vivo_noRT | AC125149.3 | 4.039 | 1.60E-06 | glio |
| culture_effect | Etl4 | -2.81 | 1.60E-07 | glio |
| culture_effect | Rab26os | 1.702 | 1.60E-07 | glio |
| Interaction_genotype_Mtor | Zfp11 | -6.323 | 1.60E-07 | glio |
| Bora_in.vivo_noRT | Nmb | 3.643 | 1.60E-07 | glio |
| Bora_in.vivo_noRT | Gm45336 | 3.064 | 1.60E-07 | glio |
| Bora_in.vivo_noRT | Otos | 3.407 | 1.60E-07 | glio |
| Chek1_in.vivo_noRT | Icam5 | 3.545 | 1.60E-07 | glio |
| culture_effect | Atf7ip | -0.94 | 1.60E-08 | glio |
| Interaction_genotype_Mtor | P4hb | 2.771 | 1.60E-08 | glio |
| Aars2_in.vivo_noRT | Gm50103 | 4.02 | 1.60E-08 | glio |
| Chek1_in.vivo_noRT | Cd59b | 3.999 | 1.60E-08 | glio |
| Chek1_in.vivo_noRT | Klhl30 | 3.859 | 1.60E-08 | glio |
| culture_effect | Gm9887 | 2.447 | 1.60E-09 | glio |
| Atr_in.vivo_noRT | Pagr1a | 3.849 | 1.60E-09 | glio |
| Bcas2_in.vivo_noRT | Neurl1b | 4.281 | 1.60E-09 | glio |
| Bcas2_in.vivo_noRT | Bend7 | 4.029 | 1.60E-09 | glio |
| Bora_in.vivo_noRT | Prrc2a | -2.25 | 1.60E-09 | glio |
| Aars2_in.vivo_noRT | Mtmr11 | 4.481 | 1.60E-10 | glio |
| Aars2_in.vivo_noRT | Ifitm1 | 5.06 | 1.60E-10 | glio |
| Aars2_in.vivo_noRT | Gm3716 | 4.409 | 1.60E-10 | glio |
| Aars2_in.vivo_noRT | Parvg | 4.551 | 1.60E-10 | glio |
| Atr_in.vivo_noRT | Gpr162 | 4.485 | 1.60E-10 | glio |
| Bcas2_in.vivo_noRT | Gm15579 | 4.807 | 1.60E-10 | glio |
| Aars2_in.vivo_noRT | AC125149.2 | 4.038 | 1.60E-11 | glio |

| | | | | |
|---|---|---|---|---|
| Aars2_in.vivo_noRT | 2610027K06R | 4.671 | 1.60E-12 | glio |
| Bora_in.vivo_noRT | 1700006H21R | 4.936 | 1.60E-12 | glio |
| Bcas2_in.vivo_noRT | Ric3 | 4.547 | 1.60E-13 | glio |
| culture_effect | Bahcc1 | -2.293 | 1.60E-15 | glio |
| Atr_in.vivo_noRT | Gm6566 | 4.608 | 1.60E-15 | glio |
| Chek1_ex.vivo_noRT | Gm16933 | 4.842 | 1.60E-15 | glio |
| Bora_in.vivo_noRT | Tcstv1 | 4.936 | 1.60E-16 | glio |
| Chek1_in.vivo_noRT | Gm14019 | 5.044 | 1.60E-16 | glio |
| Bora_in.vivo_noRT | 4931431C16R | 4.936 | 1.60E-18 | glio |
| culture_effect | Xrcc5 | -1.568 | 1.60E-20 | glio |
| Atr_in.vivo_noRT | Btn1a1 | 5.273 | 1.60E-21 | glio |
| Aars2_in.vivo_noRT | Cd200r2 | 5.063 | 1.60E-23 | glio |
| Aars2_in.vivo_noRT | Prap1 | 5.327 | 1.60E-23 | glio |
| Atr_in.vivo_noRT | Boc | 5.269 | 1.60E-24 | glio |
| Aars2_in.vivo_noRT | Gm14929 | 5.327 | 1.60E-26 | glio |
| culture_effect | H2-Aa | -6.416 | 1.60E-56 | glio |
| Interaction_genotype_Aars2 | 9430065F17Ri | -3.89 | 1.61E-03 | glio |
| Interaction_genotype_Aars2 | Zfp968 | -3.987 | 1.61E-03 | glio |
| Interaction_genotype_Aars2 | 1110038F14Ri | 1.18 | 1.61E-03 | glio |
| Interaction_genotype_Aars2 | Emp3 | 1.18 | 1.61E-03 | glio |
| Interaction_genotype_Atr | Zfp652os | -3.815 | 1.61E-03 | glio |
| Interaction_genotype_Atr | Map7 | 4.1 | 1.61E-03 | glio |
| Interaction_genotype_Atr | Zfp786 | -3.846 | 1.61E-03 | glio |
| Interaction_genotype_Atr | Olfr718-ps1 | -3.009 | 1.61E-03 | glio |
| Interaction_genotype_Atr | Cluap1 | -3.148 | 1.61E-03 | glio |
| Interaction_genotype_Atr | Knstrn | -2.239 | 1.61E-03 | glio |
| Interaction_genotype_Atr | Ccdc174 | 1.862 | 1.61E-03 | glio |
| Interaction_genotype_Atr | Herpud1 | 1.966 | 1.61E-03 | glio |
| Interaction_genotype_Chek1 | Tbrg1 | 1.539 | 1.61E-03 | glio |
| Interaction_genotype_Chek1 | Raly | 1.457 | 1.61E-03 | glio |
| Interaction_genotype_Hsd17b | K230015D01R | -4.939 | 1.61E-03 | glio |
| Interaction_genotype_Hsd17b | Irf8 | 4.473 | 1.61E-03 | glio |
| Interaction_genotype_Mtor | P2ry10b | -3.949 | 1.61E-03 | glio |
| Interaction_genotype_Mtor | Fam241a | -3.919 | 1.61E-03 | glio |
| Interaction_genotype_Mtor | Apol7a | -3.514 | 1.61E-03 | glio |
| Interaction_genotype_Mtor | Lacc1 | -3.12 | 1.61E-03 | glio |
| Interaction_genotype_Mtor | Tbx2 | -2.634 | 1.61E-03 | glio |
| Interaction_genotype_Mtor | Synj2bp | 1.651 | 1.61E-03 | glio |
| Interaction_genotype_Mtor | Mak16 | 1.482 | 1.61E-03 | glio |
| Interaction_genotype_Myc | Gm21761 | -2.729 | 1.61E-03 | glio |
| Interaction_genotype_Myc | Dock6 | -3.467 | 1.61E-03 | glio |
| Interaction_genotype_Rpl3 | Rnf165 | -4.183 | 1.61E-03 | glio |
| Interaction_genotype_Rpl3 | Rgs16 | -5.022 | 1.61E-03 | glio |
| Interaction_genotype_Rpl3 | Gm49864 | -4.287 | 1.61E-03 | glio |
| Interaction_genotype_Rpl3 | Cbl | 2.264 | 1.61E-03 | glio |
| Aars2_in.vivo_noRT | Zfp772 | 2.308 | 1.61E-03 | glio |
| Aars2_in.vivo_noRT | Fbxo48 | 2.306 | 1.61E-03 | glio |
| Aars2_in.vivo_noRT | Arid3a | 1.961 | 1.61E-03 | glio |
| Aars2_in.vivo_noRT | Kidins220 | 1.311 | 1.61E-03 | glio |
| Aars2_in.vivo_noRT | Brix1 | -0.805 | 1.61E-03 | glio |
| Atr_in.vivo_noRT | Procr | 2.757 | 1.61E-03 | glio |
| Atr_in.vivo_noRT | Elac1 | 2.141 | 1.61E-03 | glio |

| | | | | |
|---|---|---|---|---|
| Atr_in.vivo_noRT | Mlxip | 2.033 | 1.61E-03 | glio |
| Atr_in.vivo_noRT | D330050G23R | 1.86 | 1.61E-03 | glio |
| Bard1_in.vivo_noRT | Tal2 | 2.246 | 1.61E-03 | glio |
| Bard1_in.vivo_noRT | Gm15908 | 1.976 | 1.61E-03 | glio |
| Bard1_in.vivo_noRT | Krt8 | -2.005 | 1.61E-03 | glio |
| Bard1_in.vivo_noRT | Agap2 | 2.504 | 1.61E-03 | glio |
| Bcas2_in.vivo_noRT | E130102H24R | 2.225 | 1.61E-03 | glio |
| Bcas2_in.vivo_noRT | Calm3 | -1.055 | 1.61E-03 | glio |
| Bora_ex.vivo_noRT | Cxcl1 | 3.201 | 1.61E-03 | glio |
| Bora_ex.vivo_noRT | Prrt3 | 2.808 | 1.61E-03 | glio |
| Cdkn2a_in.vivo_noRT | Pnliprp2 | -2.453 | 1.61E-03 | glio |
| Cenpj_in.vivo_noRT | Tom1 | -1.86 | 1.61E-03 | glio |
| Chek1_in.vivo_noRT | Ythdf3 | -1.436 | 1.61E-03 | glio |
| Chek1_in.vivo_noRT | Brd8 | -1.261 | 1.61E-03 | glio |
| Chek1_ex.vivo_noRT | Dut | -1.14 | 1.61E-03 | glio |
| Chek1_ex.vivo_noRT | Rpl11 | -0.818 | 1.61E-03 | glio |
| Eif2s1_in.vivo_noRT | Impg2 | 2.143 | 1.61E-03 | glio |
| Hsd17b10_in.vivo_noRT | Dnase1l2 | 2.328 | 1.61E-03 | glio |
| Hsd17b10_in.vivo_noRT | Ggt1 | 2.304 | 1.61E-03 | glio |
| Hsd17b10_in.vivo_noRT | Hsf4 | 2.429 | 1.61E-03 | glio |
| culture_effect | Gm13470 | 1.52 | 1.61E-04 | glio |
| culture_effect | Slc39a7 | -0.485 | 1.61E-04 | glio |
| culture_effect | Tomm22 | 0.371 | 1.61E-04 | glio |
| culture_effect | Calm1 | -0.456 | 1.61E-04 | glio |
| Interaction_genotype_Atr | Gm9484 | -4.898 | 1.61E-04 | glio |
| Interaction_genotype_Mtor | Flrt1 | -4.869 | 1.61E-04 | glio |
| Interaction_genotype_Mtor | Aass | -4.248 | 1.61E-04 | glio |
| Interaction_genotype_Mtor | Thbs3 | -4.215 | 1.61E-04 | glio |
| Interaction_genotype_Mtor | Amz1 | -4.408 | 1.61E-04 | glio |
| Interaction_genotype_Mtor | Wdr83os | 1.81 | 1.61E-04 | glio |
| Interaction_genotype_Pars2 | Gpr179 | -5.097 | 1.61E-04 | glio |
| Aars2_in.vivo_noRT | Ccdc103 | 2.698 | 1.61E-04 | glio |
| Atp5c1_in.vivo_noRT | Gm29670 | 2.146 | 1.61E-04 | glio |
| Atr_in.vivo_noRT | Klhdc1 | 2.621 | 1.61E-04 | glio |
| Atr_in.vivo_noRT | Gstt3 | 2.715 | 1.61E-04 | glio |
| Atr_in.vivo_noRT | Emg1 | -0.951 | 1.61E-04 | glio |
| Atr_ex.vivo_noRT | C330002G04R | 2.577 | 1.61E-04 | glio |
| Bcas2_in.vivo_noRT | BC051077 | 3.103 | 1.61E-04 | glio |
| Bcas2_in.vivo_noRT | Nqo1 | 2.82 | 1.61E-04 | glio |
| Bora_ex.vivo_noRT | Gm14207 | 2.408 | 1.61E-04 | glio |
| Chek1_ex.vivo_noRT | Nppc | 3.64 | 1.61E-04 | glio |
| Hsd17b10_in.vivo_noRT | AC160336.1 | -2.308 | 1.61E-04 | glio |
| Interaction_genotype_Chek1 | Col6a2 | -4.781 | 1.61E-05 | glio |
| Interaction_genotype_Mtor | Gm16283 | -4.85 | 1.61E-05 | glio |
| Interaction_genotype_Mtor | Tm7sf2 | -4.249 | 1.61E-05 | glio |
| Interaction_genotype_Mtor | Adh7 | -5.107 | 1.61E-05 | glio |
| Atr_in.vivo_noRT | Gm19605 | 2.986 | 1.61E-05 | glio |
| Chek1_in.vivo_noRT | Hyal3 | 3.454 | 1.61E-05 | glio |
| culture_effect | Dscc1 | -1.212 | 1.61E-06 | glio |
| interaction_RT | Map1lc3b | 0.962 | 1.61E-06 | glio |
| Aars2_in.vivo_noRT | Itgb6 | 3.399 | 1.61E-06 | glio |
| Atr_in.vivo_noRT | Gm26728 | 3.504 | 1.61E-06 | glio |

| | | | | |
|---|---|---|---|---|
| Bora_in.vivo_noRT | C430042M11F | 3.407 | 1.61E-06 | glio |
| Bora_in.vivo_noRT | Gm30790 | 3.42 | 1.61E-06 | glio |
| Bora_in.vivo_noRT | Spx | 3.21 | 1.61E-06 | glio |
| Bora_ex.vivo_noRT | 9130019O22R | 3.201 | 1.61E-06 | glio |
| culture_effect | Lancl1 | -1.126 | 1.61E-07 | glio |
| culture_effect | Glod4 | -0.832 | 1.61E-07 | glio |
| culture_effect | Rock2 | 0.645 | 1.61E-07 | glio |
| Atr_in.vivo_noRT | Gm31805 | 3.549 | 1.61E-07 | glio |
| Chek1_in.vivo_noRT | 5330417C22R | 3.649 | 1.61E-07 | glio |
| Hsd17b10_in.vivo_noRT | Thap8 | 3.887 | 1.61E-07 | glio |
| culture_effect | Syt1 | -3.342 | 1.61E-08 | glio |
| culture_effect | As3mt | -1.611 | 1.61E-08 | glio |
| Bcas2_in.vivo_noRT | Lmx1b | 4.02 | 1.61E-08 | glio |
| Bora_in.vivo_noRT | Gm26673 | 3.916 | 1.61E-08 | glio |
| Bora_in.vivo_noRT | 1700056N10R | 3.55 | 1.61E-08 | glio |
| culture_effect | Yipf4 | -0.709 | 1.61E-09 | glio |
| Atr_in.vivo_noRT | Gm10655 | 4.348 | 1.61E-09 | glio |
| Bora_in.vivo_noRT | Selenok-ps1 | 3.896 | 1.61E-09 | glio |
| Hsd17b10_in.vivo_noRT | 8030423F21Ri | 3.619 | 1.61E-09 | glio |
| Hsd17b10_in.vivo_noRT | Klk13 | 3.084 | 1.61E-09 | glio |
| culture_effect | Gm7967 | -1.976 | 1.61E-10 | glio |
| Bcas2_in.vivo_noRT | Gm37214 | 4.547 | 1.61E-10 | glio |
| Chek1_in.vivo_noRT | Vsig10l | 4.391 | 1.61E-10 | glio |
| Atr_in.vivo_noRT | Zkscan4 | 4.733 | 1.61E-11 | glio |
| Chek1_ex.vivo_noRT | Tmem54 | 4.842 | 1.61E-11 | glio |
| culture_effect | Tmem88b | -2.923 | 1.61E-12 | glio |
| Aars2_in.vivo_noRT | Gm20275 | 3.972 | 1.61E-12 | glio |
| Chek1_ex.vivo_noRT | Gm12860 | 4.842 | 1.61E-12 | glio |
| Atr_in.vivo_noRT | Gm20635 | 4.888 | 1.61E-13 | glio |
| Bcas2_in.vivo_noRT | Vmn2r56 | 4.025 | 1.61E-13 | glio |
| Aars2_in.vivo_noRT | Gm11457 | 5.327 | 1.61E-15 | glio |
| Bora_in.vivo_noRT | Adgre5 | 4.676 | 1.61E-15 | glio |
| Bora_in.vivo_noRT | Gm48899 | 4.676 | 1.61E-15 | glio |
| Bora_in.vivo_noRT | Klhdc8a | 5.703 | 1.61E-16 | glio |
| Chek1_in.vivo_noRT | Rassf6 | 5.044 | 1.61E-17 | glio |
| Aars2_in.vivo_noRT | 4933400F21Ri | 4.536 | 1.61E-19 | glio |
| Aars2_in.vivo_noRT | Gm48170 | 5.327 | 1.61E-21 | glio |
| Bora_in.vivo_noRT | Trmt61b | 4.936 | 1.61E-21 | glio |
| Aars2_in.vivo_noRT | Trmt61b | 5.327 | 1.61E-24 | glio |
| Aars2_in.vivo_noRT | Helt | 5.327 | 1.61E-28 | glio |
| Aars2_in.vivo_noRT | Gm20625 | 5.327 | 1.61E-28 | glio |
| culture_effect | Pla2g4c | -1.945 | 1.62E-03 | glio |
| culture_effect | Sema4f | 1.573 | 1.62E-03 | glio |
| culture_effect | 6330403L08Ri | 1.511 | 1.62E-03 | glio |
| culture_effect | Uggt2 | -1.121 | 1.62E-03 | glio |
| culture_effect | Scaf11 | 0.372 | 1.62E-03 | glio |
| culture_effect | Ube2b | -0.305 | 1.62E-03 | glio |
| culture_effect | Ranbp1 | 0.376 | 1.62E-03 | glio |
| Interaction_genotype_Aars2 | Gm15816 | -3.513 | 1.62E-03 | glio |
| Interaction_genotype_Aars2 | Gm42658 | -3.67 | 1.62E-03 | glio |
| Interaction_genotype_Aars2 | Ddx18 | 1.589 | 1.62E-03 | glio |
| Interaction_genotype_Atr | 9230112J17Ri | -3.35 | 1.62E-03 | glio |

| | | | | |
|---|---|---|---|---|
| Interaction_genotype_Atr | Gm26579 | -3.726 | 1.62E-03 | glio |
| Interaction_genotype_Mtor | Thnsl2 | -2.931 | 1.62E-03 | glio |
| Interaction_genotype_Mtor | Usp51 | -3.714 | 1.62E-03 | glio |
| Interaction_genotype_Mtor | Galnt2 | 1.909 | 1.62E-03 | glio |
| Interaction_genotype_Mtor | Tomm40 | 1.586 | 1.62E-03 | glio |
| Interaction_genotype_Myc | Gm7879 | -3.726 | 1.62E-03 | glio |
| Interaction_genotype_Myc | Btbd6 | -3.119 | 1.62E-03 | glio |
| Interaction_genotype_Myc | Pex16 | -2.489 | 1.62E-03 | glio |
| Interaction_genotype_Rpl3 | 2010320O07R | -4.117 | 1.62E-03 | glio |
| Atp5c1_in.vivo_noRT | Magea10 | 2.201 | 1.62E-03 | glio |
| Atp5c1_in.vivo_noRT | Cox7b2 | 2.201 | 1.62E-03 | glio |
| Atp5c1_in.vivo_noRT | Gm45606 | 2.408 | 1.62E-03 | glio |
| Atr_in.vivo_noRT | Zswim9 | 1.869 | 1.62E-03 | glio |
| Atr_ex.vivo_noRT | Gm15941 | 2.769 | 1.62E-03 | glio |
| Atr_ex.vivo_noRT | Sh3gl3 | 3.033 | 1.62E-03 | glio |
| Bcas2_in.vivo_noRT | Meak7 | 2.313 | 1.62E-03 | glio |
| Bcas2_in.vivo_noRT | Plac1 | 2.226 | 1.62E-03 | glio |
| Bcas2_in.vivo_noRT | C1ra | -1.713 | 1.62E-03 | glio |
| Bora_ex.vivo_noRT | Rtl1 | 2.806 | 1.62E-03 | glio |
| Bora_ex.vivo_noRT | Pstpip2 | 2.408 | 1.62E-03 | glio |
| Cdkn2a_in.vivo_noRT | Scel | -2.066 | 1.62E-03 | glio |
| Cdkn2a_in.vivo_noRT | Gm44096 | -2.945 | 1.62E-03 | glio |
| Cdkn2a_in.vivo_noRT | Mgp | 2.243 | 1.62E-03 | glio |
| Cdkn2a_in.vivo_noRT | Klk13 | -2.207 | 1.62E-03 | glio |
| Chek1_in.vivo_noRT | Gm26812 | 2.239 | 1.62E-03 | glio |
| Chek1_ex.vivo_noRT | 9530080O11R | 2.764 | 1.62E-03 | glio |
| culture_effect | Ifnz | 1.866 | 1.62E-04 | glio |
| Interaction_genotype_Mtor | Dok3 | -4.701 | 1.62E-04 | glio |
| Interaction_genotype_Mtor | Hipk4 | -4.192 | 1.62E-04 | glio |
| Interaction_genotype_Mtor | Asphd2 | -3.913 | 1.62E-04 | glio |
| Interaction_genotype_Mtor | Ndufs8 | 1.934 | 1.62E-04 | glio |
| Interaction_genotype_Myc | March10 | -4.202 | 1.62E-04 | glio |
| Interaction_genotype_Myc | Cyb5d1 | -4.276 | 1.62E-04 | glio |
| Interaction_genotype_Myc | Gm45902 | -4.284 | 1.62E-04 | glio |
| Interaction_genotype_Myc | Entpd6 | -3.267 | 1.62E-04 | glio |
| interaction_RT | Slc8b1 | 1.876 | 1.62E-04 | glio |
| Aars2_in.vivo_noRT | Tnnc1 | 2.937 | 1.62E-04 | glio |
| Aars2_in.vivo_noRT | S100a10 | -1.141 | 1.62E-04 | glio |
| Aars2_in.vivo_noRT | Eef1g | -0.782 | 1.62E-04 | glio |
| Aars2_in.vivo_noRT | Ptma | -0.764 | 1.62E-04 | glio |
| Aars2_ex.vivo_noRT | Gm50223 | 2.9 | 1.62E-04 | glio |
| Atr_in.vivo_noRT | Fam174b | 2.959 | 1.62E-04 | glio |
| Atr_in.vivo_noRT | Zbtb34 | 2.561 | 1.62E-04 | glio |
| Atr_in.vivo_noRT | Cisd3 | -1.48 | 1.62E-04 | glio |
| Bcas2_in.vivo_noRT | Myom1 | 2.814 | 1.62E-04 | glio |
| Bcas2_in.vivo_noRT | Prrg1 | 2.783 | 1.62E-04 | glio |
| Bcas2_in.vivo_noRT | Adamts4 | 2.408 | 1.62E-04 | glio |
| Bcas2_ex.vivo_noRT | 4930512B01R | 2.742 | 1.62E-04 | glio |
| Bora_ex.vivo_noRT | Vmn1r88 | 3.201 | 1.62E-04 | glio |
| Chek1_ex.vivo_noRT | Gmpr2 | -2.154 | 1.62E-04 | glio |
| Eif2s1_in.vivo_noRT | Erich6b | 2.143 | 1.62E-04 | glio |
| Eif2s1_in.vivo_noRT | Lncbate10 | 2.143 | 1.62E-04 | glio |

| | | | | |
|---|---|---|---|---|
| Eif2s1_in.vivo_noRT | Tac2 | 2.143 | 1.62E-04 | glio |
| Eif2s1_in.vivo_noRT | Slc35g3 | 2.143 | 1.62E-04 | glio |
| Eif2s1_in.vivo_noRT | Chia1 | 2.143 | 1.62E-04 | glio |
| Hsd17b10_in.vivo_noRT | Gm16183 | 2.974 | 1.62E-04 | glio |
| Hsd17b10_in.vivo_noRT | Gltp | -1.396 | 1.62E-04 | glio |
| culture_effect | Crabp2 | 2.097 | 1.62E-05 | glio |
| Interaction_genotype_Atr | Hist1h2bc | 4.593 | 1.62E-05 | glio |
| Interaction_genotype_Mtor | Hist1h3d | -4.26 | 1.62E-05 | glio |
| Interaction_genotype_Myc | Zfp934 | -4.495 | 1.62E-05 | glio |
| Interaction_genotype_Myc | Hcrtr1 | -5.039 | 1.62E-05 | glio |
| Interaction_genotype_Myc | Fgf1 | -5.056 | 1.62E-05 | glio |
| Interaction_genotype_Myc | A330049N07R | -3.405 | 1.62E-05 | glio |
| interaction_RT | Dhrs1 | -1.271 | 1.62E-05 | glio |
| Aars2_in.vivo_noRT | H2-T24 | 2.868 | 1.62E-05 | glio |
| Atp5c1_in.vivo_noRT | Slc2a9 | 2.668 | 1.62E-05 | glio |
| Atr_in.vivo_noRT | Zmynd19 | -1.767 | 1.62E-05 | glio |
| Bard1_ex.vivo_noRT | Gm15326 | 3.037 | 1.62E-05 | glio |
| Bcas2_in.vivo_noRT | Arhgef4 | 3.042 | 1.62E-05 | glio |
| Bcas2_in.vivo_noRT | Krt8 | -4.013 | 1.62E-05 | glio |
| Bora_in.vivo_noRT | Mroh2b | 3.132 | 1.62E-05 | glio |
| Bora_in.vivo_noRT | Gng8 | 2.964 | 1.62E-05 | glio |
| Hsd17b10_in.vivo_noRT | Ap5z1 | -2.13 | 1.62E-05 | glio |
| culture_effect | Pik3r1 | 1.108 | 1.62E-06 | glio |
| culture_effect | Rps3a1 | -0.624 | 1.62E-06 | glio |
| Aars2_in.vivo_noRT | Mucl3 | 3.555 | 1.62E-06 | glio |
| Aars2_in.vivo_noRT | Fen1 | -2.123 | 1.62E-06 | glio |
| Atr_in.vivo_noRT | Nrap | 3.531 | 1.62E-06 | glio |
| Bcas2_in.vivo_noRT | Tec | 3.13 | 1.62E-06 | glio |
| Bora_ex.vivo_noRT | Klk13 | 3.201 | 1.62E-06 | glio |
| Chek1_in.vivo_noRT | Rps24 | -0.948 | 1.62E-06 | glio |
| Chek1_ex.vivo_noRT | Ier5 | 3.013 | 1.62E-06 | glio |
| Atr_in.vivo_noRT | Wnt6 | 4.358 | 1.62E-07 | glio |
| Atr_ex.vivo_noRT | Gm20711 | 3.916 | 1.62E-07 | glio |
| Bcas2_in.vivo_noRT | Gm16174 | 3.935 | 1.62E-07 | glio |
| Chek1_in.vivo_noRT | Ltb4r1 | 3.869 | 1.62E-07 | glio |
| Hsd17b10_in.vivo_noRT | Crybb1 | 3.229 | 1.62E-07 | glio |
| Interaction_genotype_Myc | Gm2115 | -5.765 | 1.62E-08 | glio |
| Aars2_in.vivo_noRT | Gm26916 | 3.75 | 1.62E-08 | glio |
| Aars2_in.vivo_noRT | Gm36210 | 3.916 | 1.62E-08 | glio |
| Atr_in.vivo_noRT | Lamc2 | 3.885 | 1.62E-08 | glio |
| Bcas2_in.vivo_noRT | Zkscan4 | 4.007 | 1.62E-08 | glio |
| Bcas2_in.vivo_noRT | Ankrd1 | 4.807 | 1.62E-08 | glio |
| Bcas2_in.vivo_noRT | Gm47545 | 3.556 | 1.62E-08 | glio |
| Bcas2_in.vivo_noRT | Tomm6os | 4.418 | 1.62E-09 | glio |
| Atr_in.vivo_noRT | Fgd3 | 4.798 | 1.62E-10 | glio |
| Bora_in.vivo_noRT | Ly6g6c | 4.076 | 1.62E-10 | glio |
| Chek1_in.vivo_noRT | Lyz2 | 4.658 | 1.62E-10 | glio |
| Chek1_ex.vivo_noRT | Gm12610 | 4.443 | 1.62E-10 | glio |
| Bora_in.vivo_noRT | Ccdc63 | 4.55 | 1.62E-11 | glio |
| Chek1_in.vivo_noRT | 1700061G19R | 4.556 | 1.62E-11 | glio |
| Chek1_in.vivo_noRT | Itgbl1 | 3.857 | 1.62E-11 | glio |
| Aars2_in.vivo_noRT | Gm16364.1 | 5.327 | 1.62E-12 | glio |

| | | | | |
|---|---|---|---|---|
| Hsd17b10_in.vivo_noRT | Gm15533 | 3.616 | 1.62E-12 | glio |
| Atr_in.vivo_noRT | Gm13594 | 5.545 | 1.62E-14 | glio |
| Atr_in.vivo_noRT | Gm39326 | 5.27 | 1.62E-14 | glio |
| Bora_in.vivo_noRT | B3gnt4 | 4.673 | 1.62E-14 | glio |
| Atr_ex.vivo_noRT | Gm12800 | 5.079 | 1.62E-15 | glio |
| culture_effect | Slc16a3 | -2.967 | 1.62E-16 | glio |
| culture_effect | Fam171a1 | -2.058 | 1.62E-16 | glio |
| Aars2_in.vivo_noRT | Gm40264 | 5.067 | 1.62E-17 | glio |
| Bora_in.vivo_noRT | Pyy | 4.936 | 1.62E-17 | glio |
| Bcas2_in.vivo_noRT | Map3k9 | 5.065 | 1.62E-18 | glio |
| Chek1_in.vivo_noRT | Gm4027 | 5.044 | 1.62E-19 | glio |
| Atr_in.vivo_noRT | Mpzl2 | 5.273 | 1.62E-20 | glio |
| Bcas2_in.vivo_noRT | Vmn2r18 | 4.807 | 1.62E-22 | glio |
| culture_effect | Gm14966 | 1.524 | 1.63E-03 | glio |
| culture_effect | Npc1 | -0.752 | 1.63E-03 | glio |
| culture_effect | Ncbp1 | 0.467 | 1.63E-03 | glio |
| culture_effect | Ganab | -0.403 | 1.63E-03 | glio |
| Interaction_genotype_Aars2 | Gm4181 | -3.968 | 1.63E-03 | glio |
| Interaction_genotype_Aars2 | A430105J06Ri | -3.73 | 1.63E-03 | glio |
| Interaction_genotype_Aars2 | Tspo | 1.206 | 1.63E-03 | glio |
| Interaction_genotype_Atr | Bbc3 | 2.524 | 1.63E-03 | glio |
| Interaction_genotype_Bora | Gm49312 | -3.485 | 1.63E-03 | glio |
| Interaction_genotype_Mtor | Gm5861 | -4.523 | 1.63E-03 | glio |
| Interaction_genotype_Mtor | Ccnj | -3.834 | 1.63E-03 | glio |
| Interaction_genotype_Mtor | Mapre1 | 1.568 | 1.63E-03 | glio |
| Interaction_genotype_Mtor | Pnn | 1.732 | 1.63E-03 | glio |
| Interaction_genotype_Myc | Ttc41 | -3.798 | 1.63E-03 | glio |
| Interaction_genotype_Myc | Prkaa2 | -3.175 | 1.63E-03 | glio |
| Interaction_genotype_Myc | Tmem59l | -3.028 | 1.63E-03 | glio |
| Aars2_in.vivo_noRT | Ccdc13 | 2.313 | 1.63E-03 | glio |
| Aars2_in.vivo_noRT | Eef1akmt3 | 2.11 | 1.63E-03 | glio |
| Aars2_in.vivo_noRT | Trappc2l | -1.263 | 1.63E-03 | glio |
| Aars2_ex.vivo_noRT | Cstdc4 | 2.893 | 1.63E-03 | glio |
| Atr_in.vivo_noRT | 2310001H17R | 1.862 | 1.63E-03 | glio |
| Atr_in.vivo_noRT | Cisd2 | -1.012 | 1.63E-03 | glio |
| Atr_ex.vivo_noRT | Ccn2 | 3.527 | 1.63E-03 | glio |
| Bard1_ex.vivo_noRT | Nmrk2 | 2.814 | 1.63E-03 | glio |
| Bard1_ex.vivo_noRT | Gm48435 | 2.027 | 1.63E-03 | glio |
| Bcas2_in.vivo_noRT | Gpatch3 | 2.246 | 1.63E-03 | glio |
| Bcas2_in.vivo_noRT | Cdt1 | -1.507 | 1.63E-03 | glio |
| Bcas2_in.vivo_noRT | Tpt1 | -0.713 | 1.63E-03 | glio |
| Bora_in.vivo_noRT | Ovgp1 | 2.254 | 1.63E-03 | glio |
| Bora_in.vivo_noRT | Cldn14 | 2.16 | 1.63E-03 | glio |
| Bora_ex.vivo_noRT | Podxl | 2.808 | 1.63E-03 | glio |
| Chek1_in.vivo_noRT | Gm19391 | 2.144 | 1.63E-03 | glio |
| Chek1_in.vivo_noRT | Zfp639 | -1.507 | 1.63E-03 | glio |
| Chek1_in.vivo_noRT | Acbd3 | -1.162 | 1.63E-03 | glio |
| Chek1_ex.vivo_noRT | Usp51 | 2.808 | 1.63E-03 | glio |
| Chek1_ex.vivo_noRT | Gm49801 | 2.949 | 1.63E-03 | glio |
| Eif2s1_in.vivo_noRT | Sytl4 | 1.877 | 1.63E-03 | glio |
| Hsd17b10_in.vivo_noRT | Elovl1 | -0.857 | 1.63E-03 | glio |
| Interaction_genotype_Chek1 | Atg9b | 4.475 | 1.63E-04 | glio |

| | | | | |
|---|---|---|---|---|
| Interaction_genotype_Mtor | Prss22 | -5.607 | 1.63E-04 | glio |
| Interaction_genotype_Mtor | Hcn2 | -4.787 | 1.63E-04 | glio |
| Interaction_genotype_Myc | Wnt16 | -4.117 | 1.63E-04 | glio |
| Interaction_genotype_Myc | Tc2n | -4.399 | 1.63E-04 | glio |
| Interaction_genotype_Myc | Slc25a29 | -3.871 | 1.63E-04 | glio |
| Interaction_genotype_Myc | Stfa3 | -2.729 | 1.63E-04 | glio |
| Interaction_genotype_Myc | A530017D24R | -4.026 | 1.63E-04 | glio |
| Interaction_genotype_Myc | Chchd6 | -3.68 | 1.63E-04 | glio |
| Aars2_in.vivo_noRT | Etaa1os | 2.577 | 1.63E-04 | glio |
| Aars2_in.vivo_noRT | 5330438D12R | 2.201 | 1.63E-04 | glio |
| Aars2_ex.vivo_noRT | 4930502E18Ri | 2.9 | 1.63E-04 | glio |
| Aars2_ex.vivo_noRT | 1700112J16Ri | 2.706 | 1.63E-04 | glio |
| Atp5c1_in.vivo_noRT | Gm11583 | 2.142 | 1.63E-04 | glio |
| Atp5c1_in.vivo_noRT | Nwd1 | 2.404 | 1.63E-04 | glio |
| Atp5c1_in.vivo_noRT | Tmem54 | 2.408 | 1.63E-04 | glio |
| Atp5c1_in.vivo_noRT | Tex52 | 2.791 | 1.63E-04 | glio |
| Atr_in.vivo_noRT | Katnal2 | 2.133 | 1.63E-04 | glio |
| Atr_in.vivo_noRT | Synrg | -1.603 | 1.63E-04 | glio |
| Atr_ex.vivo_noRT | Tmem52 | 3.164 | 1.63E-04 | glio |
| Atr_ex.vivo_noRT | Gm36043 | 3.164 | 1.63E-04 | glio |
| Atr_ex.vivo_noRT | Prkcq | 3.164 | 1.63E-04 | glio |
| Bard1_ex.vivo_noRT | Fam166a | 2.814 | 1.63E-04 | glio |
| Bard1_ex.vivo_noRT | Gm15894 | 2.814 | 1.63E-04 | glio |
| Bard1_ex.vivo_noRT | Pex6 | -2.459 | 1.63E-04 | glio |
| Bora_in.vivo_noRT | Nts | 2.893 | 1.63E-04 | glio |
| Bora_in.vivo_noRT | Ktn1 | -1.474 | 1.63E-04 | glio |
| Bora_in.vivo_noRT | Vim | -0.85 | 1.63E-04 | glio |
| Chek1_ex.vivo_noRT | Cdca8 | -2.156 | 1.63E-04 | glio |
| culture_effect | Pdlim4 | 2.013 | 1.63E-05 | glio |
| culture_effect | Gpr155 | -1.379 | 1.63E-05 | glio |
| culture_effect | Heatr5a | -0.869 | 1.63E-05 | glio |
| Interaction_genotype_Mtor | Hist1h4b | -5.57 | 1.63E-05 | glio |
| Interaction_genotype_Mtor | Setmar | -5.218 | 1.63E-05 | glio |
| Interaction_genotype_Mtor | Zfp808 | -4.545 | 1.63E-05 | glio |
| Interaction_genotype_Myc | Gga3 | -3.72 | 1.63E-05 | glio |
| Interaction_genotype_Myc | Gm39326 | -4.777 | 1.63E-05 | glio |
| Atp5c1_in.vivo_noRT | Gm15127 | 3.17 | 1.63E-05 | glio |
| Bora_in.vivo_noRT | E2f3 | -1.845 | 1.63E-05 | glio |
| Chek1_in.vivo_noRT | Hecw2 | 3.159 | 1.63E-05 | glio |
| Chek1_ex.vivo_noRT | Gm10827 | 4.071 | 1.63E-05 | glio |
| Hsd17b10_in.vivo_noRT | Ldhd | 2.699 | 1.63E-05 | glio |
| Aars2_in.vivo_noRT | Zfp11 | 3.36 | 1.63E-06 | glio |
| Aars2_in.vivo_noRT | Ostf1 | -1.796 | 1.63E-06 | glio |
| Atr_in.vivo_noRT | Gm34552 | 3.339 | 1.63E-06 | glio |
| Hsd17b10_in.vivo_noRT | Ncf1 | 2.963 | 1.63E-06 | glio |
| culture_effect | Gm5914 | 1.682 | 1.63E-07 | glio |
| Interaction_genotype_Mtor | A930012O16R | -6.524 | 1.63E-07 | glio |
| Interaction_genotype_Mtor | Ppil6 | -5.774 | 1.63E-07 | glio |
| Interaction_genotype_Mtor | Olfr552 | -4.498 | 1.63E-07 | glio |
| Bora_in.vivo_noRT | Celf5 | 3.873 | 1.63E-07 | glio |
| Chek1_in.vivo_noRT | Ifit1bl1 | 4.511 | 1.63E-07 | glio |
| Chek1_in.vivo_noRT | Gap43 | 4.127 | 1.63E-07 | glio |

| | | | | |
|---|---|---|---|---|
| Interaction_genotype_Rad21 | Pstpip2 | 5.634 | 1.63E-08 | glio |
| Aars2_in.vivo_noRT | 4921523L03Ri | 4.159 | 1.63E-08 | glio |
| culture_effect | 6430590A07R | 1.893 | 1.63E-09 | glio |
| Atr_in.vivo_noRT | 6330408M09F | 4.492 | 1.63E-09 | glio |
| Chek1_ex.vivo_noRT | Gm33051 | 4.045 | 1.63E-09 | glio |
| Bcas2_in.vivo_noRT | Armc2 | 4.416 | 1.63E-10 | glio |
| Bora_in.vivo_noRT | Gm32591 | 4.414 | 1.63E-10 | glio |
| Bora_in.vivo_noRT | Hsf2bp | 4.144 | 1.63E-10 | glio |
| Atr_in.vivo_noRT | Lmx1b | 4.746 | 1.63E-11 | glio |
| Bcas2_in.vivo_noRT | Carns1 | 4.807 | 1.63E-11 | glio |
| Chek1_ex.vivo_noRT | Clec4d | 4.837 | 1.63E-13 | glio |
| Chek1_ex.vivo_noRT | Vmn2r44 | 4.447 | 1.63E-13 | glio |
| Hsd17b10_in.vivo_noRT | Ccl9 | 3.616 | 1.63E-13 | glio |
| Atr_in.vivo_noRT | Gm9967 | 5.146 | 1.63E-14 | glio |
| culture_effect | A430018G15R | 2.855 | 1.63E-15 | glio |
| culture_effect | Adamtsl5 | 2.349 | 1.63E-15 | glio |
| Bora_in.vivo_noRT | Gm31415 | 4.67 | 1.63E-15 | glio |
| Bora_in.vivo_noRT | B4galnt1 | 4.676 | 1.63E-15 | glio |
| culture_effect | Mgst2 | -3.719 | 1.63E-16 | glio |
| Chek1_in.vivo_noRT | Agap2 | 5.044 | 1.63E-18 | glio |
| Atr_in.vivo_noRT | Gm2115 | 5.533 | 1.63E-19 | glio |
| Chek1_in.vivo_noRT | AC154200.1 | 5.044 | 1.63E-19 | glio |
| Bcas2_in.vivo_noRT | Gm47540 | 4.807 | 1.63E-20 | glio |
| Bcas2_in.vivo_noRT | BB287469 | 4.807 | 1.63E-20 | glio |
| Aars2_in.vivo_noRT | Gm15445 | 5.19 | 1.63E-21 | glio |
| Aars2_in.vivo_noRT | 1700112J16Ri | 5.327 | 1.63E-22 | glio |
| Aars2_in.vivo_noRT | Clec4d | 5.327 | 1.63E-22 | glio |
| Chek1_in.vivo_noRT | Erich6b | 5.044 | 1.63E-23 | glio |
| Chek1_in.vivo_noRT | Lncbate10 | 5.044 | 1.63E-23 | glio |
| culture_effect | Zmiz2 | 0.568 | 1.64E-03 | glio |
| culture_effect | Phf20l1 | 0.495 | 1.64E-03 | glio |
| Interaction_genotype_Mtor | Reck | -3.849 | 1.64E-03 | glio |
| Interaction_genotype_Mtor | 6330403L08Ri | -3.374 | 1.64E-03 | glio |
| Interaction_genotype_Mtor | Zbtb25 | -3.148 | 1.64E-03 | glio |
| Interaction_genotype_Mtor | Bola3 | 1.556 | 1.64E-03 | glio |
| Interaction_genotype_Mtor | Idh3g | 1.44 | 1.64E-03 | glio |
| Interaction_genotype_Myc | Gm47510 | -2.469 | 1.64E-03 | glio |
| Interaction_genotype_Myc | Kif5a | -3.866 | 1.64E-03 | glio |
| Interaction_genotype_Myc | B3gntl1 | -3.477 | 1.64E-03 | glio |
| Interaction_genotype_Myc | Fchsd1 | -3.4 | 1.64E-03 | glio |
| Interaction_genotype_Myc | Gm34256 | -3.631 | 1.64E-03 | glio |
| Interaction_genotype_Myc | Gm14488 | -3.748 | 1.64E-03 | glio |
| Interaction_genotype_Myc | Pdpr | -3.083 | 1.64E-03 | glio |
| Interaction_genotype_Myc | Htt | -1.862 | 1.64E-03 | glio |
| Interaction_genotype_Myc | Anapc1 | -1.839 | 1.64E-03 | glio |
| Aars2_in.vivo_noRT | Rps6kl1 | 2.276 | 1.64E-03 | glio |
| Aars2_in.vivo_noRT | Pdlim2 | 2.049 | 1.64E-03 | glio |
| Aars2_ex.vivo_noRT | Ubd | 2.9 | 1.64E-03 | glio |
| Aars2_ex.vivo_noRT | H2-Eb1 | 2.899 | 1.64E-03 | glio |
| Atr_in.vivo_noRT | Arhgap22 | 2.174 | 1.64E-03 | glio |
| Atr_in.vivo_noRT | Hist1h2bn | 2.166 | 1.64E-03 | glio |
| Atr_in.vivo_noRT | Gm42463 | 2.075 | 1.64E-03 | glio |

| | | | | |
|---|---|---|---|---|
| Bcas2_ex.vivo_noRT | Prap1 | 2.349 | 1.64E-03 | glio |
| Bcas2_ex.vivo_noRT | D730045A05R | 3.129 | 1.64E-03 | glio |
| Bcas2_ex.vivo_noRT | Car14 | 2.957 | 1.64E-03 | glio |
| Bora_in.vivo_noRT | Mtcp1 | 2.24 | 1.64E-03 | glio |
| Bora_in.vivo_noRT | Plpp6 | 2.162 | 1.64E-03 | glio |
| Chek1_in.vivo_noRT | Fgf15 | 2.475 | 1.64E-03 | glio |
| Chek1_in.vivo_noRT | Hpn | 2.225 | 1.64E-03 | glio |
| Chek1_in.vivo_noRT | Gm14418 | 1.787 | 1.64E-03 | glio |
| Chek1_ex.vivo_noRT | Flicr | 2.806 | 1.64E-03 | glio |
| Chek1_ex.vivo_noRT | Zfpm1 | 2.954 | 1.64E-03 | glio |
| Eif2s1_in.vivo_noRT | Selenov | 2.143 | 1.64E-03 | glio |
| Eif2s1_in.vivo_noRT | A330076H08R | 2.143 | 1.64E-03 | glio |
| culture_effect | 9930111J21Ri | -2.114 | 1.64E-04 | glio |
| culture_effect | Tgfbr2 | -1.067 | 1.64E-04 | glio |
| culture_effect | Grhpr | -0.69 | 1.64E-04 | glio |
| culture_effect | Sdf4 | -0.498 | 1.64E-04 | glio |
| Interaction_genotype_Mtor | Kcnd1 | -3.976 | 1.64E-04 | glio |
| Interaction_genotype_Mtor | Usp17la | -4.494 | 1.64E-04 | glio |
| Interaction_genotype_Mtor | St14 | -4.086 | 1.64E-04 | glio |
| Interaction_genotype_Mtor | Gm26581 | -4.54 | 1.64E-04 | glio |
| Interaction_genotype_Mtor | Phtf1os | -4.165 | 1.64E-04 | glio |
| Interaction_genotype_Mtor | Psmb8 | 1.95 | 1.64E-04 | glio |
| Interaction_genotype_Myc | Olfr787 | -2.729 | 1.64E-04 | glio |
| Interaction_genotype_Myc | Gm3604 | -4.331 | 1.64E-04 | glio |
| Interaction_genotype_Myc | Gm47438 | -4.575 | 1.64E-04 | glio |
| Interaction_genotype_Myc | AC132307.1 | -3.929 | 1.64E-04 | glio |
| Interaction_genotype_Myc | Cacna1a | -3.467 | 1.64E-04 | glio |
| Interaction_genotype_Nat2 | AU019990 | 3.605 | 1.64E-04 | glio |
| Aars2_in.vivo_noRT | Arhgef28 | 2.373 | 1.64E-04 | glio |
| Aars2_in.vivo_noRT | Krt8 | -3.673 | 1.64E-04 | glio |
| Aars2_in.vivo_noRT | Tspo | -0.995 | 1.64E-04 | glio |
| Atr_in.vivo_noRT | Gm10419 | 2.881 | 1.64E-04 | glio |
| Atr_in.vivo_noRT | Hr | 2.388 | 1.64E-04 | glio |
| Atr_in.vivo_noRT | Ndufaf2 | -1.343 | 1.64E-04 | glio |
| Bora_in.vivo_noRT | 1810059H22R | 2.978 | 1.64E-04 | glio |
| Bora_in.vivo_noRT | Rab42 | 2.865 | 1.64E-04 | glio |
| Bora_in.vivo_noRT | Psat1 | -0.994 | 1.64E-04 | glio |
| Bora_ex.vivo_noRT | Vipr2 | 3.622 | 1.64E-04 | glio |
| Chek1_in.vivo_noRT | Cntf | 2.675 | 1.64E-04 | glio |
| Chek1_ex.vivo_noRT | Utp11 | -1.14 | 1.64E-04 | glio |
| Hsd17b10_in.vivo_noRT | Aass | 2.615 | 1.64E-04 | glio |
| Interaction_genotype_Mtor | Bbs1 | -5.203 | 1.64E-05 | glio |
| Interaction_genotype_Mtor | Ywhah | 1.974 | 1.64E-05 | glio |
| Interaction_genotype_Myc | Gm11963 | -4.926 | 1.64E-05 | glio |
| Aars2_in.vivo_noRT | 4930594M22F | 3.08 | 1.64E-05 | glio |
| Aars2_in.vivo_noRT | Car5a | 2.763 | 1.64E-05 | glio |
| Aars2_in.vivo_noRT | Zfp974 | 3.182 | 1.64E-05 | glio |
| Atr_in.vivo_noRT | Cd72 | 2.972 | 1.64E-05 | glio |
| Atr_in.vivo_noRT | Jam3 | -1.983 | 1.64E-05 | glio |
| Bcas2_in.vivo_noRT | Map3k5 | 2.643 | 1.64E-05 | glio |
| Bcas2_in.vivo_noRT | 4930554I06Ri | 3.277 | 1.64E-05 | glio |
| Bora_in.vivo_noRT | Arl4d | 3.088 | 1.64E-05 | glio |

| | | | | |
|---|---|---|---|---|
| Chek1_ex.vivo_noRT | Postn | 3.754 | 1.64E-05 | glio |
| culture_effect | Hells | 0.942 | 1.64E-06 | glio |
| culture_effect | Apex1 | -0.715 | 1.64E-06 | glio |
| Aars2_in.vivo_noRT | Serpinc1 | 3.51 | 1.64E-06 | glio |
| Chek1_in.vivo_noRT | Gm5861 | 3.995 | 1.64E-06 | glio |
| Interaction_genotype_Mtor | Gm15127 | -6.213 | 1.64E-07 | glio |
| Atr_in.vivo_noRT | Itga3 | 3.904 | 1.64E-07 | glio |
| Chek1_in.vivo_noRT | Dchs1 | 3.549 | 1.64E-07 | glio |
| Chek1_in.vivo_noRT | Calml4 | 3.483 | 1.64E-07 | glio |
| culture_effect | Sugt1 | -0.897 | 1.64E-08 | glio |
| Interaction_genotype_Mtor | Slc17a6 | -4.099 | 1.64E-08 | glio |
| Aars2_in.vivo_noRT | Gm43330 | 3.427 | 1.64E-08 | glio |
| Aars2_in.vivo_noRT | Gm10382 | 4.083 | 1.64E-08 | glio |
| Atr_in.vivo_noRT | Rab40b | 3.796 | 1.64E-08 | glio |
| Interaction_genotype_Mtor | Gm11627 | -6.578 | 1.64E-09 | glio |
| Interaction_genotype_Mtor | Gm48435 | -4.886 | 1.64E-09 | glio |
| Atr_in.vivo_noRT | Calml4 | 3.973 | 1.64E-09 | glio |
| Bcas2_in.vivo_noRT | Plcd4 | 3.896 | 1.64E-09 | glio |
| Hsd17b10_in.vivo_noRT | Gm49860 | 3.616 | 1.64E-09 | glio |
| Atr_in.vivo_noRT | Ccdc39 | 4.313 | 1.64E-10 | glio |
| Bcas2_in.vivo_noRT | 5330439K02R | 4.541 | 1.64E-10 | glio |
| culture_effect | Fam136a | 0.826 | 1.64E-11 | glio |
| Bora_in.vivo_noRT | Gm43653 | 4.673 | 1.64E-12 | glio |
| Atr_in.vivo_noRT | Stx1b | 4.63 | 1.64E-13 | glio |
| Chek1_ex.vivo_noRT | Gm45406 | 4.842 | 1.64E-13 | glio |
| culture_effect | Ctdspl | -1.746 | 1.64E-14 | glio |
| Atr_in.vivo_noRT | Jph4 | 5.267 | 1.64E-14 | glio |
| Chek1_ex.vivo_noRT | H1foo | 4.842 | 1.64E-14 | glio |
| culture_effect | Bricd5 | -3.408 | 1.64E-15 | glio |
| culture_effect | Prkca | 2.379 | 1.64E-16 | glio |
| Bcas2_in.vivo_noRT | 4930470G03R | 4.541 | 1.64E-16 | glio |
| Bora_in.vivo_noRT | Ptprc | 4.672 | 1.64E-17 | glio |
| Chek1_in.vivo_noRT | Ank1 | 5.044 | 1.64E-22 | glio |
| Atr_in.vivo_noRT | Gm2022 | 5.533 | 1.64E-24 | glio |
| Atr_in.vivo_noRT | Esp24 | 5.533 | 1.64E-25 | glio |
| Atr_in.vivo_noRT | Gpc5 | 5.533 | 1.64E-27 | glio |
| culture_effect | 1810021B22R | 1.698 | 1.65E-03 | glio |
| culture_effect | Zfp94 | 1.433 | 1.65E-03 | glio |
| culture_effect | Kcnk2 | 1.142 | 1.65E-03 | glio |
| culture_effect | Pgap1 | -1.068 | 1.65E-03 | glio |
| culture_effect | Stxbp3 | -0.671 | 1.65E-03 | glio |
| culture_effect | Pet100 | 0.637 | 1.65E-03 | glio |
| Interaction_genotype_Aars2 | Gm31063 | -2.958 | 1.65E-03 | glio |
| Interaction_genotype_Aars2 | Olfr279 | -3.823 | 1.65E-03 | glio |
| Interaction_genotype_Aars2 | Gm44658 | -3.445 | 1.65E-03 | glio |
| Interaction_genotype_Aars2 | Rilp | -4.232 | 1.65E-03 | glio |
| Interaction_genotype_Mtor | B3gat2 | -3.996 | 1.65E-03 | glio |
| Interaction_genotype_Mtor | Plcb4 | -3.712 | 1.65E-03 | glio |
| Interaction_genotype_Mtor | Rsrp1 | 1.831 | 1.65E-03 | glio |
| Interaction_genotype_Mtor | Hnrnpa1 | 1.304 | 1.65E-03 | glio |
| Interaction_genotype_Myc | Gm12280 | -3.944 | 1.65E-03 | glio |
| Interaction_genotype_Myc | Gprin2 | -3.009 | 1.65E-03 | glio |

| | | | | |
|---|---|---|---|---|
| Interaction_genotype_Myc | Ccnj | -3.622 | 1.65E-03 | glio |
| Interaction_genotype_Myc | Wdr7 | -2.949 | 1.65E-03 | glio |
| Interaction_genotype_Myc | Ddx51 | -2.748 | 1.65E-03 | glio |
| Interaction_genotype_Myc | Klhl20 | -2.426 | 1.65E-03 | glio |
| Interaction_genotype_Myc | Prkar1a | 1.231 | 1.65E-03 | glio |
| interaction_RT | Ift80 | 1.413 | 1.65E-03 | glio |
| interaction_RT | Timeless | -1.463 | 1.65E-03 | glio |
| Atr_in.vivo_noRT | Gm50340 | 2.025 | 1.65E-03 | glio |
| Atr_in.vivo_noRT | Myo7a | 1.855 | 1.65E-03 | glio |
| Atr_ex.vivo_noRT | Ftl1 | 0.671 | 1.65E-03 | glio |
| Bora_in.vivo_noRT | Fhod3 | 2.419 | 1.65E-03 | glio |
| Bora_in.vivo_noRT | Enho | 2.278 | 1.65E-03 | glio |
| Bora_in.vivo_noRT | Kctd3 | -1.607 | 1.65E-03 | glio |
| Chek1_in.vivo_noRT | Nek3 | 2.289 | 1.65E-03 | glio |
| Chek1_in.vivo_noRT | Gm50340 | 2.053 | 1.65E-03 | glio |
| Chek1_in.vivo_noRT | Cep78 | 1.87 | 1.65E-03 | glio |
| Chek1_in.vivo_noRT | Gclc | -1.493 | 1.65E-03 | glio |
| Chek1_ex.vivo_noRT | Rdh12 | 2.696 | 1.65E-03 | glio |
| Chek1_ex.vivo_noRT | Vdr | 3.218 | 1.65E-03 | glio |
| Chek1_ex.vivo_noRT | Sh3rf1 | 2.399 | 1.65E-03 | glio |
| Eif2s1_in.vivo_noRT | Npnt | 1.877 | 1.65E-03 | glio |
| Eif2s1_in.vivo_noRT | Gm43915 | 2.141 | 1.65E-03 | glio |
| culture_effect | Cyp27a1 | -2 | 1.65E-04 | glio |
| culture_effect | Rnaseh2b | -0.915 | 1.65E-04 | glio |
| Interaction_genotype_Ifnar1 | Gprin2 | 4.403 | 1.65E-04 | glio |
| Interaction_genotype_Ifnar1 | Gstt4 | 3.784 | 1.65E-04 | glio |
| Interaction_genotype_Mtor | AI854703 | -4.498 | 1.65E-04 | glio |
| Interaction_genotype_Myc | Gm47540 | -3.122 | 1.65E-04 | glio |
| Interaction_genotype_Myc | Cdc42ep4 | -4.054 | 1.65E-04 | glio |
| Interaction_genotype_Myc | Mettl21a | -3.417 | 1.65E-04 | glio |
| interaction_RT | Ablim3 | 3.271 | 1.65E-04 | glio |
| Aars2_in.vivo_noRT | Gm32913 | 4.056 | 1.65E-04 | glio |
| Atr_in.vivo_noRT | Hist4h4 | 2.737 | 1.65E-04 | glio |
| Atr_in.vivo_noRT | Nqo1 | 2.768 | 1.65E-04 | glio |
| Atr_in.vivo_noRT | Slc27a3 | 2.66 | 1.65E-04 | glio |
| Bcas2_in.vivo_noRT | Gata3 | 2.953 | 1.65E-04 | glio |
| Bora_in.vivo_noRT | Ddx49 | -1.5 | 1.65E-04 | glio |
| Chek1_in.vivo_noRT | Ifnz | 2.7 | 1.65E-04 | glio |
| Chek1_ex.vivo_noRT | Kif18a | -2.657 | 1.65E-04 | glio |
| Interaction_genotype_Atr | Gm36356 | -4.495 | 1.65E-05 | glio |
| Interaction_genotype_Mtor | C230037L18Ri | -4.746 | 1.65E-05 | glio |
| Interaction_genotype_Mtor | Tas1r1 | -4.631 | 1.65E-05 | glio |
| Interaction_genotype_Myc | Gm47061 | -5.367 | 1.65E-05 | glio |
| Interaction_genotype_Myc | Atox1 | 2.104 | 1.65E-05 | glio |
| Atr_in.vivo_noRT | Kcna4 | 3.007 | 1.65E-05 | glio |
| Bard1_ex.vivo_noRT | Irf8 | 3.612 | 1.65E-05 | glio |
| Bard1_ex.vivo_noRT | Synpo2l | 3.592 | 1.65E-05 | glio |
| Bard1_ex.vivo_noRT | Ablim2 | 2.814 | 1.65E-05 | glio |
| Bard1_ex.vivo_noRT | Gm5415 | 2.814 | 1.65E-05 | glio |
| Bard1_ex.vivo_noRT | Lrrc4b | 3.592 | 1.65E-05 | glio |
| Bard1_ex.vivo_noRT | Nrip2 | 2.814 | 1.65E-05 | glio |
| Chek1_in.vivo_noRT | 4930524O07R | 3.119 | 1.65E-05 | glio |

| | | | | |
|---|---|---|---|---|
| Hsd17b10_in.vivo_noRT | Gipc2 | 3.353 | 1.65E-05 | glio |
| Hsd17b10_in.vivo_noRT | Ifi207 | 3.353 | 1.65E-05 | glio |
| Aars2_in.vivo_noRT | Gsx2 | 3.588 | 1.65E-06 | glio |
| Chek1_in.vivo_noRT | Prm1 | 3.033 | 1.65E-06 | glio |
| Chek1_in.vivo_noRT | Zfp59 | 3.41 | 1.65E-06 | glio |
| Interaction_genotype_Mtor | D8Ertd738e | 2.211 | 1.65E-07 | glio |
| Aars2_in.vivo_noRT | Tcea2 | 3.599 | 1.65E-07 | glio |
| Atr_in.vivo_noRT | B430010I23Ri | 3.447 | 1.65E-07 | glio |
| Bcas2_in.vivo_noRT | Pelp1 | -2.388 | 1.65E-07 | glio |
| Bora_in.vivo_noRT | Rab11fip4os1 | 3.9 | 1.65E-07 | glio |
| Bora_in.vivo_noRT | Bbof1 | 3.508 | 1.65E-07 | glio |
| Bora_in.vivo_noRT | Slc6a19 | 3.533 | 1.65E-07 | glio |
| Hsd17b10_in.vivo_noRT | Gm43915 | 3.084 | 1.65E-07 | glio |
| culture_effect | Icam1 | -3.074 | 1.65E-08 | glio |
| culture_effect | Fbxl6 | -0.901 | 1.65E-08 | glio |
| Atr_in.vivo_noRT | Ccdc184 | 4.2 | 1.65E-08 | glio |
| Atr_in.vivo_noRT | Aldh1a1 | 4.008 | 1.65E-08 | glio |
| Atr_ex.vivo_noRT | Gm9758 | 3.962 | 1.65E-09 | glio |
| Bcas2_in.vivo_noRT | Vsig10l | 4.154 | 1.65E-09 | glio |
| Hsd17b10_in.vivo_noRT | 4930470G03R | 3.35 | 1.65E-09 | glio |
| Atr_in.vivo_noRT | Zfp811 | 4.359 | 1.65E-10 | glio |
| Chek1_in.vivo_noRT | Gm11884 | 4.045 | 1.65E-10 | glio |
| Chek1_in.vivo_noRT | Neurl1b | 4.518 | 1.65E-10 | glio |
| Chek1_in.vivo_noRT | Lrfn1 | 4.401 | 1.65E-10 | glio |
| Chek1_ex.vivo_noRT | D630024D03F | 4.256 | 1.65E-10 | glio |
| Hsd17b10_in.vivo_noRT | Gm38163 | 3.616 | 1.65E-10 | glio |
| Atr_in.vivo_noRT | Man2c1os | 4.888 | 1.65E-11 | glio |
| Hsd17b10_in.vivo_noRT | Gm2022 | 3.616 | 1.65E-11 | glio |
| culture_effect | Ltbp4 | -1.839 | 1.65E-14 | glio |
| Bora_in.vivo_noRT | Atad3a | -2.983 | 1.65E-14 | glio |
| Bora_in.vivo_noRT | Clcn1 | 4.291 | 1.65E-14 | glio |
| Chek1_in.vivo_noRT | Krt80 | 4.777 | 1.65E-14 | glio |
| interaction_RT | C4b | 5.211 | 1.65E-18 | glio |
| Bcas2_in.vivo_noRT | Gm18280 | 4.543 | 1.65E-18 | glio |
| Atr_in.vivo_noRT | Htr2b | 5.01 | 1.65E-19 | glio |
| Chek1_in.vivo_noRT | 1700112J16Ri | 5.044 | 1.65E-20 | glio |
| Chek1_in.vivo_noRT | Clec4d | 5.044 | 1.65E-20 | glio |
| Aars2_in.vivo_noRT | Gm15138 | 5.06 | 1.65E-22 | glio |
| Chek1_in.vivo_noRT | A330049N07R | 4.777 | 1.65E-22 | glio |
| Interaction_genotype_Aars2 | C230013L11Ri | -4.167 | 1.66E-03 | glio |
| Interaction_genotype_Aars2 | Rims3 | -3.562 | 1.66E-03 | glio |
| Interaction_genotype_Aars2 | Elobl | -4.173 | 1.66E-03 | glio |
| Interaction_genotype_Aars2 | Hist2h3c1 | -3.888 | 1.66E-03 | glio |
| Interaction_genotype_Aars2 | Zdhhc1 | -3.675 | 1.66E-03 | glio |
| Interaction_genotype_Aars2 | E230029C05Ri | -3.528 | 1.66E-03 | glio |
| Interaction_genotype_Aars2 | Gm50240 | -3.108 | 1.66E-03 | glio |
| Interaction_genotype_Atr | Myo5c | -3.863 | 1.66E-03 | glio |
| Interaction_genotype_Atr | Rpgr | -3.76 | 1.66E-03 | glio |
| Interaction_genotype_Bcas2 | Prl3d3 | -2.897 | 1.66E-03 | glio |
| Interaction_genotype_Bcas2 | Ncln | 1.78 | 1.66E-03 | glio |
| Interaction_genotype_Mtor | Hif3a | -3.85 | 1.66E-03 | glio |
| Interaction_genotype_Mtor | Pcsk4 | -3.636 | 1.66E-03 | glio |

| | | | | |
|---|---|---|---|---|
| Interaction_genotype_Mtor | 2310068J16Ri | -2.975 | 1.66E-03 | glio |
| Interaction_genotype_Mtor | Brd1 | 1.537 | 1.66E-03 | glio |
| Interaction_genotype_Mtor | Cbx1 | 1.404 | 1.66E-03 | glio |
| Interaction_genotype_Myc | Adam19 | -3.596 | 1.66E-03 | glio |
| Interaction_genotype_Myc | Gm32857 | -3.225 | 1.66E-03 | glio |
| Interaction_genotype_Myc | 2810002D19R | -2.826 | 1.66E-03 | glio |
| Interaction_genotype_Rpl3 | C330013E15Ri | -4.206 | 1.66E-03 | glio |
| Interaction_genotype_Rpl3 | Gm49874 | -4.459 | 1.66E-03 | glio |
| Interaction_genotype_Rpl3 | Ubr5 | 1.66 | 1.66E-03 | glio |
| interaction_RT | Slfn2 | 2.031 | 1.66E-03 | glio |
| interaction_RT | Tmem38b | 1.247 | 1.66E-03 | glio |
| interaction_RT | Nfkbib | 0.836 | 1.66E-03 | glio |
| Aars2_in.vivo_noRT | Rragd | 2.38 | 1.66E-03 | glio |
| Aars2_in.vivo_noRT | Gm16973 | 2.279 | 1.66E-03 | glio |
| Aars2_in.vivo_noRT | Pnmal2 | 2.206 | 1.66E-03 | glio |
| Aars2_in.vivo_noRT | Xpo1 | -1.215 | 1.66E-03 | glio |
| Aars2_ex.vivo_noRT | BC034090 | 2.9 | 1.66E-03 | glio |
| Atr_in.vivo_noRT | Rhbdl1 | 2.17 | 1.66E-03 | glio |
| Atr_in.vivo_noRT | Cptp | 1.656 | 1.66E-03 | glio |
| Atr_in.vivo_noRT | Mrps9 | -1.206 | 1.66E-03 | glio |
| Atr_in.vivo_noRT | Mycbp2 | -1.285 | 1.66E-03 | glio |
| Atr_in.vivo_noRT | Kcmf1 | -0.987 | 1.66E-03 | glio |
| Atr_in.vivo_noRT | Nasp | -0.888 | 1.66E-03 | glio |
| Atr_in.vivo_noRT | Hspe1 | -0.683 | 1.66E-03 | glio |
| Bora_in.vivo_noRT | Sgms2 | 2.431 | 1.66E-03 | glio |
| Bora_ex.vivo_noRT | Slc23a3 | 3.023 | 1.66E-03 | glio |
| Bora_ex.vivo_noRT | Cacng4 | 2.808 | 1.66E-03 | glio |
| Cenpj_ex.vivo_noRT | Gm38163 | 2.591 | 1.66E-03 | glio |
| Cenpj_ex.vivo_noRT | Edn2 | 2.196 | 1.66E-03 | glio |
| Chek1_ex.vivo_noRT | Cks1brt | 2.571 | 1.66E-03 | glio |
| Eif2s1_in.vivo_noRT | Speer4a | 1.877 | 1.66E-03 | glio |
| Hsd17b10_in.vivo_noRT | Klra6 | 2.506 | 1.66E-03 | glio |
| Hsd17b10_in.vivo_noRT | Rasl2-9 | 2.705 | 1.66E-03 | glio |
| Hsd17b10_in.vivo_noRT | Tmem150c | 2.448 | 1.66E-03 | glio |
| Hsd17b10_in.vivo_noRT | Rnf31 | -1.73 | 1.66E-03 | glio |
| Hsd17b10_in.vivo_noRT | Twf2 | -1.321 | 1.66E-03 | glio |
| Hsd17b10_in.vivo_noRT | Ngdn | -0.984 | 1.66E-03 | glio |
| culture_effect | Gm49463 | 1.36 | 1.66E-04 | glio |
| culture_effect | Gm26797 | 1.325 | 1.66E-04 | glio |
| culture_effect | Aifm2 | -1.014 | 1.66E-04 | glio |
| Interaction_genotype_Mtor | Gm50069 | -4.371 | 1.66E-04 | glio |
| Interaction_genotype_Mtor | Fam114a2 | 1.811 | 1.66E-04 | glio |
| Interaction_genotype_Mtor | Ndufa10 | 1.772 | 1.66E-04 | glio |
| Interaction_genotype_Myc | Casz1 | -4.659 | 1.66E-04 | glio |
| Interaction_genotype_Myc | Necab2 | -3.959 | 1.66E-04 | glio |
| Interaction_genotype_Myc | Gm3734 | -4.67 | 1.66E-04 | glio |
| Interaction_genotype_Myc | Tifa | -3.844 | 1.66E-04 | glio |
| interaction_RT | Tbx2 | -1.954 | 1.66E-04 | glio |
| interaction_RT | Slc26a10 | 1.822 | 1.66E-04 | glio |
| interaction_RT | Sh3pxd2a | -1.496 | 1.66E-04 | glio |
| interaction_RT | H2-T22 | 0.85 | 1.66E-04 | glio |
| interaction_RT | Rpl39 | 0.944 | 1.66E-04 | glio |

| | | | | |
|---|---|---|---|---|
| Aars2_in.vivo_noRT | Hcn2 | 2.873 | 1.66E-04 | glio |
| Aars2_in.vivo_noRT | Nuak2 | 2.923 | 1.66E-04 | glio |
| Aars2_in.vivo_noRT | Mafa | 2.423 | 1.66E-04 | glio |
| Aars2_in.vivo_noRT | Egfl7 | 2.377 | 1.66E-04 | glio |
| Aars2_in.vivo_noRT | Atxn1l | 1.938 | 1.66E-04 | glio |
| Aars2_in.vivo_noRT | Parp12 | -1.487 | 1.66E-04 | glio |
| Aars2_in.vivo_noRT | Eif1a | -1.06 | 1.66E-04 | glio |
| Atr_in.vivo_noRT | Gm15492 | 2.706 | 1.66E-04 | glio |
| Atr_in.vivo_noRT | Nanos3 | 2.405 | 1.66E-04 | glio |
| Atr_in.vivo_noRT | Enah | 2.383 | 1.66E-04 | glio |
| Atr_in.vivo_noRT | Fbxo31 | 1.484 | 1.66E-04 | glio |
| Bora_in.vivo_noRT | Prss22 | 3.338 | 1.66E-04 | glio |
| Bora_in.vivo_noRT | Glmp | 2.933 | 1.66E-04 | glio |
| Bora_in.vivo_noRT | AC125149.3 | 2.531 | 1.66E-04 | glio |
| Chek1_in.vivo_noRT | Gm15492 | 2.74 | 1.66E-04 | glio |
| Chek1_in.vivo_noRT | Eef1a1 | -0.705 | 1.66E-04 | glio |
| Hsd17b10_in.vivo_noRT | Vpreb1 | 2.713 | 1.66E-04 | glio |
| Interaction_genotype_Myc | Net1 | -4.12 | 1.66E-05 | glio |
| Interaction_genotype_Myc | Rnf165 | -4.63 | 1.66E-05 | glio |
| Interaction_genotype_Myc | Gm49701 | -4.828 | 1.66E-05 | glio |
| Aars2_in.vivo_noRT | Gm17597 | 3.291 | 1.66E-05 | glio |
| Aars2_in.vivo_noRT | Micall2 | 3.144 | 1.66E-05 | glio |
| Bard1_ex.vivo_noRT | Gm49698 | 2.814 | 1.66E-05 | glio |
| Bard1_ex.vivo_noRT | Olfr881 | 2.814 | 1.66E-05 | glio |
| Bard1_ex.vivo_noRT | Vmn2r18 | 2.814 | 1.66E-05 | glio |
| Bcas2_in.vivo_noRT | Gm15265 | 3.005 | 1.66E-05 | glio |
| Bcas2_in.vivo_noRT | Glmp | 3.322 | 1.66E-05 | glio |
| Bora_in.vivo_noRT | Gm47689 | 3.071 | 1.66E-05 | glio |
| Chek1_in.vivo_noRT | Shd | 2.956 | 1.66E-05 | glio |
| culture_effect | Cox18 | 0.771 | 1.66E-06 | glio |
| Interaction_genotype_Mtor | Gm266 | -4.099 | 1.66E-06 | glio |
| Interaction_genotype_Myc | 2810405F17Ri | -5.15 | 1.66E-06 | glio |
| Atr_in.vivo_noRT | Slc35d2 | 3.409 | 1.66E-06 | glio |
| Atr_ex.vivo_noRT | Gm45338 | 2.77 | 1.66E-06 | glio |
| Atr_ex.vivo_noRT | Gm41349 | 3.947 | 1.66E-06 | glio |
| Bora_ex.vivo_noRT | Sytl4 | 3.201 | 1.66E-06 | glio |
| Bora_ex.vivo_noRT | Gm12800 | 3.201 | 1.66E-06 | glio |
| Hsd17b10_in.vivo_noRT | Gm31152 | 3.094 | 1.66E-06 | glio |
| Hsd17b10_in.vivo_noRT | Gm17276 | 2.825 | 1.66E-06 | glio |
| Aars2_in.vivo_noRT | Tmeff2 | 4.07 | 1.66E-07 | glio |
| Aars2_in.vivo_noRT | Emilin2 | 3.72 | 1.66E-07 | glio |
| Bcas2_in.vivo_noRT | Hpse | 3.625 | 1.66E-07 | glio |
| Chek1_ex.vivo_noRT | Rslcan18 | 4.055 | 1.66E-07 | glio |
| Bora_in.vivo_noRT | Gm5129 | 4.136 | 1.66E-08 | glio |
| Atr_in.vivo_noRT | Dio1 | 3.697 | 1.66E-09 | glio |
| Chek1_in.vivo_noRT | Man2c1os | 4.399 | 1.66E-09 | glio |
| Chek1_in.vivo_noRT | Gm10010 | 4.303 | 1.66E-09 | glio |
| Aars2_in.vivo_noRT | 2010106G04R | 4.526 | 1.66E-10 | glio |
| Chek1_ex.vivo_noRT | Sbk3 | 5.247 | 1.66E-10 | glio |
| culture_effect | Gm43689 | 2.835 | 1.66E-11 | glio |
| Bcas2_in.vivo_noRT | A030005L19Ri | 4.278 | 1.66E-11 | glio |
| Bora_in.vivo_noRT | 1810019D21R | 4.676 | 1.66E-11 | glio |

| | | | | |
|---|---|---|---|---|
| Chek1_ex.vivo_noRT | Gm47622 | 4.268 | 1.66E-11 | glio |
| Chek1_ex.vivo_noRT | Chia1 | 4.055 | 1.66E-11 | glio |
| Chek1_ex.vivo_noRT | Tent5c | 4.449 | 1.66E-11 | glio |
| Bora_in.vivo_noRT | Ifitm6 | 4.936 | 1.66E-12 | glio |
| Bora_in.vivo_noRT | Gm50147 | 4.936 | 1.66E-13 | glio |
| culture_effect | Cnn3 | 0.974 | 1.66E-14 | glio |
| Aars2_in.vivo_noRT | Gm49441 | 4.536 | 1.66E-14 | glio |
| Aars2_in.vivo_noRT | Acoxl | 5.327 | 1.66E-16 | glio |
| Atr_in.vivo_noRT | Gm49860 | 5.533 | 1.66E-20 | glio |
| Bora_in.vivo_noRT | Luzp2 | 4.934 | 1.66E-20 | glio |
| culture_effect | AV356131 | 4.519 | 1.66E-21 | glio |
| Aars2_in.vivo_noRT | 2410017I17Ri | 5.841 | 1.66E-30 | glio |
| Aars2_in.vivo_noRT | Sucnr1 | 5.327 | 1.66E-30 | glio |
| culture_effect | Col9a1 | -3.024 | 1.66E-37 | glio |
| culture_effect | 6330408M09F | 1.892 | 1.67E-03 | glio |
| culture_effect | Meis1 | -1.115 | 1.67E-03 | glio |
| culture_effect | Znrd1as | 0.758 | 1.67E-03 | glio |
| culture_effect | Rad18 | 0.674 | 1.67E-03 | glio |
| culture_effect | Gid4 | 0.519 | 1.67E-03 | glio |
| culture_effect | Rplp1 | -0.442 | 1.67E-03 | glio |
| Interaction_genotype_Aars2 | Esrp2 | -3.69 | 1.67E-03 | glio |
| Interaction_genotype_Aars2 | Gm43409 | -4.328 | 1.67E-03 | glio |
| Interaction_genotype_Aars2 | Gm15510 | -3.875 | 1.67E-03 | glio |
| Interaction_genotype_Aars2 | Gm13283 | -3.663 | 1.67E-03 | glio |
| Interaction_genotype_Atr | Gm43727 | -3.905 | 1.67E-03 | glio |
| Interaction_genotype_Bcas2 | Cysrt1 | -4.36 | 1.67E-03 | glio |
| Interaction_genotype_Mtor | Zfp114 | -3.684 | 1.67E-03 | glio |
| Interaction_genotype_Mtor | Rrp12 | -3.161 | 1.67E-03 | glio |
| Interaction_genotype_Mtor | Mrps25 | 1.534 | 1.67E-03 | glio |
| Interaction_genotype_Mtor | Gmnn | 1.887 | 1.67E-03 | glio |
| Interaction_genotype_Mtor | Dctn3 | 1.373 | 1.67E-03 | glio |
| Interaction_genotype_Myc | Gm5286 | -2.729 | 1.67E-03 | glio |
| Interaction_genotype_Myc | 1700108F19Ri | -2.859 | 1.67E-03 | glio |
| Interaction_genotype_Myc | Slc25a18 | -2.957 | 1.67E-03 | glio |
| Interaction_genotype_Myc | Prm3 | -3.11 | 1.67E-03 | glio |
| Interaction_genotype_Myc | Gm15672 | -3.5 | 1.67E-03 | glio |
| Interaction_genotype_Myc | Gm12043 | -3.353 | 1.67E-03 | glio |
| Interaction_genotype_Myc | 1600002K03R | -3.635 | 1.67E-03 | glio |
| Interaction_genotype_Myc | Ip6k3 | -3.638 | 1.67E-03 | glio |
| Interaction_genotype_Myc | Ptpdc1 | -3.385 | 1.67E-03 | glio |
| Interaction_genotype_Myc | Hoxa4 | -3.149 | 1.67E-03 | glio |
| Interaction_genotype_Myc | Gtse1 | -2.877 | 1.67E-03 | glio |
| interaction_RT | Ifrd1 | 0.696 | 1.67E-03 | glio |
| Aars2_in.vivo_noRT | Gm12758 | 2.22 | 1.67E-03 | glio |
| Aars2_in.vivo_noRT | 6030442K20R | 2.174 | 1.67E-03 | glio |
| Aars2_in.vivo_noRT | Wdr27 | 1.975 | 1.67E-03 | glio |
| Aars2_in.vivo_noRT | Pdzd7 | 2.282 | 1.67E-03 | glio |
| Aars2_in.vivo_noRT | Gm14139 | 1.66 | 1.67E-03 | glio |
| Aars2_in.vivo_noRT | Dvl3 | 1.525 | 1.67E-03 | glio |
| Aars2_in.vivo_noRT | Ergic3 | -0.964 | 1.67E-03 | glio |
| Aars2_in.vivo_noRT | Tap2 | -0.978 | 1.67E-03 | glio |
| Aars2_ex.vivo_noRT | Impg2 | 2.505 | 1.67E-03 | glio |

| | | | | |
|---|---|---|---|---|
| Aars2_ex.vivo_noRT | Gjb5 | 2.713 | 1.67E-03 | glio |
| Aars2_ex.vivo_noRT | Ccdc80 | 2.324 | 1.67E-03 | glio |
| Atr_in.vivo_noRT | 4931414P19R | 2.155 | 1.67E-03 | glio |
| Atr_in.vivo_noRT | Pfkfb3 | 1.979 | 1.67E-03 | glio |
| Atr_ex.vivo_noRT | Klra6 | 3.13 | 1.67E-03 | glio |
| Atr_ex.vivo_noRT | Mcph1 | -1.783 | 1.67E-03 | glio |
| Atr_ex.vivo_noRT | Hspe1 | -0.796 | 1.67E-03 | glio |
| Bcas2_in.vivo_noRT | Gm29107 | 2.37 | 1.67E-03 | glio |
| Bcas2_ex.vivo_noRT | Bpifa3 | 2.742 | 1.67E-03 | glio |
| Bora_in.vivo_noRT | 1700123M08F | 2.283 | 1.67E-03 | glio |
| Bora_in.vivo_noRT | Gm42941 | 2.143 | 1.67E-03 | glio |
| Bora_in.vivo_noRT | Fzd4 | 2.193 | 1.67E-03 | glio |
| Bora_ex.vivo_noRT | Tal2 | 2.22 | 1.67E-03 | glio |
| Bora_ex.vivo_noRT | Ankrd33b | 3.201 | 1.67E-03 | glio |
| Cenpj_in.vivo_noRT | Bend7 | 2.545 | 1.67E-03 | glio |
| Cenpj_ex.vivo_noRT | B230206H07R | 2.196 | 1.67E-03 | glio |
| Chek1_in.vivo_noRT | Rhbdl1 | 2.204 | 1.67E-03 | glio |
| Chek1_in.vivo_noRT | Gm38190 | 2.123 | 1.67E-03 | glio |
| Chek1_in.vivo_noRT | Pdpk1 | -1.365 | 1.67E-03 | glio |
| Chek1_in.vivo_noRT | Acin1 | -0.981 | 1.67E-03 | glio |
| Chek1_ex.vivo_noRT | Slc36a2 | 2.797 | 1.67E-03 | glio |
| Chek1_ex.vivo_noRT | Tnfrsf4 | 2.868 | 1.67E-03 | glio |
| Chek1_ex.vivo_noRT | Emilin2 | 2.96 | 1.67E-03 | glio |
| Chek1_ex.vivo_noRT | Cxcl2 | 4.787 | 1.67E-03 | glio |
| Chek1_ex.vivo_noRT | Hdac9 | 2.683 | 1.67E-03 | glio |
| Chek1_ex.vivo_noRT | Inpp5b | -2.067 | 1.67E-03 | glio |
| Chek1_ex.vivo_noRT | Eif2s3x | -1.039 | 1.67E-03 | glio |
| Chek1_ex.vivo_noRT | Fubp1 | -0.92 | 1.67E-03 | glio |
| Hsd17b10_in.vivo_noRT | Slc46a3 | 2.431 | 1.67E-03 | glio |
| Hsd17b10_in.vivo_noRT | 4933439K11R | 2.431 | 1.67E-03 | glio |
| Interaction_genotype_Aars2 | Prickle2 | -3.976 | 1.67E-04 | glio |
| Interaction_genotype_Aars2 | 4930540M05F | -4.54 | 1.67E-04 | glio |
| Interaction_genotype_Atr | Hunk | -4.339 | 1.67E-04 | glio |
| Interaction_genotype_Mtor | CAAA0106612 | -4.101 | 1.67E-04 | glio |
| Interaction_genotype_Mtor | Brk1 | 1.532 | 1.67E-04 | glio |
| interaction_RT | Slc25a5 | -0.807 | 1.67E-04 | glio |
| interaction_RT | Ccl20 | 4.933 | 1.67E-04 | glio |
| Aars2_in.vivo_noRT | Epb42 | 2.719 | 1.67E-04 | glio |
| Aars2_ex.vivo_noRT | Clca3a1 | 3.293 | 1.67E-04 | glio |
| Atr_in.vivo_noRT | Gm9828 | 2.619 | 1.67E-04 | glio |
| Atr_ex.vivo_noRT | Auts2 | 3.164 | 1.67E-04 | glio |
| Atr_ex.vivo_noRT | H2-K1 | 1.279 | 1.67E-04 | glio |
| Brca2_ex.vivo_noRT | Fam227b | 2.82 | 1.67E-04 | glio |
| Cenpj_ex.vivo_noRT | Ankrd22 | 2.591 | 1.67E-04 | glio |
| Cenpj_ex.vivo_noRT | Gm47404 | 2.591 | 1.67E-04 | glio |
| Cenpj_ex.vivo_noRT | Gm31717 | 2.591 | 1.67E-04 | glio |
| Cenpj_ex.vivo_noRT | Gm14742 | 2.591 | 1.67E-04 | glio |
| Cenpj_ex.vivo_noRT | Scel | 2.591 | 1.67E-04 | glio |
| Cenpj_ex.vivo_noRT | Ctla4 | 2.591 | 1.67E-04 | glio |
| Cenpj_ex.vivo_noRT | Zscan4-ps3 | 2.591 | 1.67E-04 | glio |
| Cenpj_ex.vivo_noRT | Tagap | 2.591 | 1.67E-04 | glio |
| Cenpj_ex.vivo_noRT | Gm21731 | 2.591 | 1.67E-04 | glio |

| | | | | |
|---|---|---|---|---|
| Cenpj_ex.vivo_noRT | Gm11232 | 2.591 | 1.67E-04 | glio |
| Cenpj_ex.vivo_noRT | Olfr1152 | 2.591 | 1.67E-04 | glio |
| Cenpj_ex.vivo_noRT | Kcnq2 | 3.389 | 1.67E-04 | glio |
| Cenpj_ex.vivo_noRT | Gm14207 | 2.595 | 1.67E-04 | glio |
| Cenpj_ex.vivo_noRT | Cgnl1 | 2.591 | 1.67E-04 | glio |
| Cenpj_ex.vivo_noRT | Gm26543 | 2.591 | 1.67E-04 | glio |
| Cenpj_ex.vivo_noRT | 2410017I17Ri | 2.591 | 1.67E-04 | glio |
| Chek1_in.vivo_noRT | Fndc3a | -1.886 | 1.67E-04 | glio |
| Chek1_ex.vivo_noRT | Rtn2 | 3.044 | 1.67E-04 | glio |
| Hsd17b10_in.vivo_noRT | Gm32834 | 2.699 | 1.67E-04 | glio |
| culture_effect | Arpc1b | -0.526 | 1.67E-05 | glio |
| Interaction_genotype_Mtor | Gm16933 | -3.577 | 1.67E-05 | glio |
| Interaction_genotype_Myc | R3hcc1 | -3.864 | 1.67E-05 | glio |
| Atp5c1_in.vivo_noRT | Pyy | 2.668 | 1.67E-05 | glio |
| Atr_in.vivo_noRT | Htra1 | 3.363 | 1.67E-05 | glio |
| Atr_in.vivo_noRT | Itga7 | 3.058 | 1.67E-05 | glio |
| Bcas2_in.vivo_noRT | Fgf2 | 3.181 | 1.67E-05 | glio |
| Chek1_in.vivo_noRT | Gm43221 | 2.872 | 1.67E-05 | glio |
| Hsd17b10_in.vivo_noRT | Dnm3 | 3.09 | 1.67E-05 | glio |
| Interaction_genotype_Atr | Ormdl3 | 2.618 | 1.67E-06 | glio |
| Interaction_genotype_Myc | Gm3716 | -5.251 | 1.67E-06 | glio |
| Interaction_genotype_Myc | Gm11457 | -5.235 | 1.67E-06 | glio |
| Aars2_ex.vivo_noRT | Hist1h2ag | 3.293 | 1.67E-06 | glio |
| Atr_in.vivo_noRT | Gm48796 | 3.161 | 1.67E-06 | glio |
| Bora_in.vivo_noRT | Gm32122 | 3.108 | 1.67E-06 | glio |
| Bora_ex.vivo_noRT | Trmt61b | 3.201 | 1.67E-06 | glio |
| Bora_ex.vivo_noRT | H2-Ab1 | 3.665 | 1.67E-06 | glio |
| Chek1_in.vivo_noRT | Serpinb9b | 4.046 | 1.67E-06 | glio |
| Chek1_in.vivo_noRT | Gm9958 | 3.474 | 1.67E-06 | glio |
| Chek1_in.vivo_noRT | Ache | 3.471 | 1.67E-06 | glio |
| Interaction_genotype_Mtor | Saxo2 | -6.183 | 1.67E-07 | glio |
| Interaction_genotype_Mtor | Gm10125 | -6.458 | 1.67E-07 | glio |
| Interaction_genotype_Myc | 2210406O10R | -6.153 | 1.67E-07 | glio |
| Interaction_genotype_Myc | Tmem179 | -6.101 | 1.67E-07 | glio |
| Interaction_genotype_Myc | 4930478M13F | -5.989 | 1.67E-07 | glio |
| Interaction_genotype_Myc | 1700026J14Ri | -5.519 | 1.67E-07 | glio |
| Aars2_in.vivo_noRT | Gm12905 | 4.022 | 1.67E-07 | glio |
| Atr_in.vivo_noRT | Rasal1 | 4.057 | 1.67E-07 | glio |
| Bora_in.vivo_noRT | Klra6 | 3.826 | 1.67E-07 | glio |
| Bora_in.vivo_noRT | Nppc | 3.937 | 1.67E-07 | glio |
| Bora_in.vivo_noRT | Gstp3 | 3.636 | 1.67E-07 | glio |
| Cenpj_in.vivo_noRT | Prickle2 | 3.607 | 1.67E-07 | glio |
| Aars2_in.vivo_noRT | Gm49891 | 4.129 | 1.67E-08 | glio |
| Atr_in.vivo_noRT | Mpeg1 | 3.558 | 1.67E-08 | glio |
| Atr_in.vivo_noRT | Tpsab1 | 3.584 | 1.67E-08 | glio |
| Bcas2_ex.vivo_noRT | Scg2 | 3.54 | 1.67E-08 | glio |
| Chek1_in.vivo_noRT | BC034090 | 3.67 | 1.67E-08 | glio |
| Chek1_ex.vivo_noRT | Gm15270 | 4.049 | 1.67E-08 | glio |
| Hsd17b10_in.vivo_noRT | Psca | 3.616 | 1.67E-08 | glio |
| Hsd17b10_in.vivo_noRT | Cys1 | 3.35 | 1.67E-08 | glio |
| Hsd17b10_in.vivo_noRT | Gm15270 | 3.353 | 1.67E-08 | glio |
| culture_effect | Nasp | -0.749 | 1.67E-09 | glio |

| | | | | |
|---|---|---|---|---|
| Atr_in.vivo_noRT | 4930558J18Ri | 3.927 | 1.67E-09 | glio |
| Bora_in.vivo_noRT | Tmem150c | 4.292 | 1.67E-09 | glio |
| Atr_in.vivo_noRT | Cd59b | 4.488 | 1.67E-10 | glio |
| culture_effect | Gm14419 | 2.616 | 1.67E-11 | glio |
| culture_effect | 4930550L24Ri | -0.965 | 1.67E-11 | glio |
| culture_effect | 1110038F14Ri | -0.795 | 1.67E-11 | glio |
| culture_effect | Zcrb1 | 0.681 | 1.67E-11 | glio |
| Chek1_in.vivo_noRT | Gm29083 | 4.511 | 1.67E-12 | glio |
| Hsd17b10_in.vivo_noRT | Slc19a3 | 3.616 | 1.67E-12 | glio |
| Aars2_in.vivo_noRT | Gm10642 | 5.063 | 1.67E-14 | glio |
| Bcas2_in.vivo_noRT | D930028M14I | 4.807 | 1.67E-14 | glio |
| Aars2_in.vivo_noRT | Fgf8 | 4.811 | 1.67E-15 | glio |
| Atr_in.vivo_noRT | Gm43951 | 4.608 | 1.67E-15 | glio |
| Bora_in.vivo_noRT | Gm15816 | 4.936 | 1.67E-15 | glio |
| Chek1_in.vivo_noRT | Hnf1aos1 | 5.044 | 1.67E-16 | glio |
| Atr_in.vivo_noRT | Txndc2 | 4.754 | 1.67E-17 | glio |
| culture_effect | Snhg3 | 2.06 | 1.67E-20 | glio |
| Chek1_in.vivo_noRT | Gad1 | 4.777 | 1.67E-21 | glio |
| culture_effect | Serpini1 | -3.567 | 1.67E-23 | glio |
| Bora_in.vivo_noRT | Ccl9 | 4.936 | 1.67E-23 | glio |
| Chek1_in.vivo_noRT | Tac2 | 5.044 | 1.67E-23 | glio |
| culture_effect | C130026I21Ri | -3.25 | 1.67E-26 | glio |
| Atr_in.vivo_noRT | Slc19a3 | 5.533 | 1.67E-26 | glio |
| culture_effect | Mrpl20 | 0.379 | 1.68E-03 | glio |
| culture_effect | Rpl32 | -0.441 | 1.68E-03 | glio |
| Interaction_genotype_Aars2 | Gtsf1 | -3.197 | 1.68E-03 | glio |
| Interaction_genotype_Atr | 4930563E22Ri | -3.929 | 1.68E-03 | glio |
| Interaction_genotype_Mtor | Gm15446 | -3.379 | 1.68E-03 | glio |
| Interaction_genotype_Mtor | Ppp4c | 1.762 | 1.68E-03 | glio |
| Interaction_genotype_Mtor | Pam16 | 1.556 | 1.68E-03 | glio |
| Interaction_genotype_Myc | Gm26822 | -3.741 | 1.68E-03 | glio |
| Interaction_genotype_Myc | Gm8212 | -3.569 | 1.68E-03 | glio |
| Interaction_genotype_Myc | Gm48617 | -2.466 | 1.68E-03 | glio |
| Interaction_genotype_Myc | Hacl1 | -3.315 | 1.68E-03 | glio |
| Interaction_genotype_Myc | Endov | -3.064 | 1.68E-03 | glio |
| Interaction_genotype_Myc | Slc39a11 | -2.793 | 1.68E-03 | glio |
| Interaction_genotype_Myc | Shc1 | -2.49 | 1.68E-03 | glio |
| Interaction_genotype_Myc | Col27a1 | -2.399 | 1.68E-03 | glio |
| Interaction_genotype_Pars2 | Gm15853 | -4.435 | 1.68E-03 | glio |
| Interaction_genotype_Pars2 | Prickle4 | -4.06 | 1.68E-03 | glio |
| Interaction_genotype_Pars2 | Ccdc39 | -4.234 | 1.68E-03 | glio |
| Interaction_genotype_Pars2 | Gm36198 | -3.564 | 1.68E-03 | glio |
| Interaction_genotype_Pars2 | Gm13398 | -3.066 | 1.68E-03 | glio |
| Interaction_genotype_Pars2 | Matn4 | -3.926 | 1.68E-03 | glio |
| Interaction_genotype_Pars2 | Gm26674 | -4.05 | 1.68E-03 | glio |
| Interaction_genotype_Pars2 | Gm14023 | -4.403 | 1.68E-03 | glio |
| interaction_RT | Chac1 | 1.519 | 1.68E-03 | glio |
| Aars2_in.vivo_noRT | Lrig1 | 2.413 | 1.68E-03 | glio |
| Aars2_in.vivo_noRT | Hrh2 | 1.784 | 1.68E-03 | glio |
| Aars2_in.vivo_noRT | Gm16196 | 2.102 | 1.68E-03 | glio |
| Aars2_in.vivo_noRT | Gm20756 | 2.081 | 1.68E-03 | glio |
| Aars2_in.vivo_noRT | Kbtbd11 | 1.963 | 1.68E-03 | glio |

| | | | | |
|---|---|---|---|---|
| Aars2_in.vivo_noRT | Lgals3bp | -1.117 | 1.68E-03 | glio |
| Aars2_in.vivo_noRT | Cycs | -0.748 | 1.68E-03 | glio |
| Atr_in.vivo_noRT | Rhobtb1 | 2.263 | 1.68E-03 | glio |
| Atr_in.vivo_noRT | Hpn | 2.185 | 1.68E-03 | glio |
| Atr_in.vivo_noRT | Gm48646 | 2.182 | 1.68E-03 | glio |
| Atr_in.vivo_noRT | Ormdl3 | -1.249 | 1.68E-03 | glio |
| Atr_in.vivo_noRT | Cfdp1 | -0.845 | 1.68E-03 | glio |
| Atr_in.vivo_noRT | Gapdh | -0.797 | 1.68E-03 | glio |
| Atr_ex.vivo_noRT | Gm11884 | 2.764 | 1.68E-03 | glio |
| Bard1_ex.vivo_noRT | Pcdhb21 | 3.056 | 1.68E-03 | glio |
| Bcas2_in.vivo_noRT | Hist1h4j | 2.688 | 1.68E-03 | glio |
| Bcas2_in.vivo_noRT | A330009N23R | 2.235 | 1.68E-03 | glio |
| Bcas2_in.vivo_noRT | Ppp1r9b | -1.675 | 1.68E-03 | glio |
| Bcas2_in.vivo_noRT | Orai1 | -1.563 | 1.68E-03 | glio |
| Bcas2_in.vivo_noRT | Dusp19 | -1.51 | 1.68E-03 | glio |
| Bcas2_in.vivo_noRT | Hexa | -1.001 | 1.68E-03 | glio |
| Bora_in.vivo_noRT | Map3k8 | 2.461 | 1.68E-03 | glio |
| Bora_ex.vivo_noRT | Bco2 | 2.633 | 1.68E-03 | glio |
| Cenpj_in.vivo_noRT | Dmrtc1a | 2.022 | 1.68E-03 | glio |
| Cenpj_in.vivo_noRT | Gm12589 | 2.022 | 1.68E-03 | glio |
| Cenpj_ex.vivo_noRT | Cyp7a1 | 3.197 | 1.68E-03 | glio |
| Chek1_in.vivo_noRT | Slc13a5 | 2.166 | 1.68E-03 | glio |
| Chek1_in.vivo_noRT | Pmf1 | -1.304 | 1.68E-03 | glio |
| Eif2s1_in.vivo_noRT | Selenok | -0.622 | 1.68E-03 | glio |
| Hsd17b10_in.vivo_noRT | Acy3 | 2.436 | 1.68E-03 | glio |
| culture_effect | Serpina3i | -2.256 | 1.68E-04 | glio |
| culture_effect | Klhl28 | -0.949 | 1.68E-04 | glio |
| culture_effect | Nans | -0.537 | 1.68E-04 | glio |
| culture_effect | Cox16 | -0.516 | 1.68E-04 | glio |
| Interaction_genotype_Ifnar1 | Speer4a | 3.787 | 1.68E-04 | glio |
| Interaction_genotype_Myc | Gal3st4 | -4.178 | 1.68E-04 | glio |
| Aars2_in.vivo_noRT | Ccdc8 | 2.612 | 1.68E-04 | glio |
| Aars2_in.vivo_noRT | 5830428M24F | 2.673 | 1.68E-04 | glio |
| Aars2_in.vivo_noRT | Crabp1 | 2.695 | 1.68E-04 | glio |
| Aars2_in.vivo_noRT | 3110001I22Ri | 2.606 | 1.68E-04 | glio |
| Aars2_in.vivo_noRT | Mpc1 | -1.271 | 1.68E-04 | glio |
| Atr_in.vivo_noRT | Mocs1 | 2.339 | 1.68E-04 | glio |
| Atr_in.vivo_noRT | Mtdh | -0.742 | 1.68E-04 | glio |
| Atr_ex.vivo_noRT | Hif3a | 3.554 | 1.68E-04 | glio |
| Atr_ex.vivo_noRT | Gm13713 | 2.764 | 1.68E-04 | glio |
| Atr_ex.vivo_noRT | Gm5039 | 3.937 | 1.68E-04 | glio |
| Bora_ex.vivo_noRT | C330002G04R | 2.615 | 1.68E-04 | glio |
| Chek1_in.vivo_noRT | C920021L13Ri | 2.713 | 1.68E-04 | glio |
| Chek1_in.vivo_noRT | Kif7 | 2.512 | 1.68E-04 | glio |
| Chek1_ex.vivo_noRT | Gm1123 | 3.468 | 1.68E-04 | glio |
| Chek1_ex.vivo_noRT | Pcdhb17 | 3.425 | 1.68E-04 | glio |
| Chek1_ex.vivo_noRT | Gm43331 | 2.887 | 1.68E-04 | glio |
| culture_effect | Gm13663 | 1.77 | 1.68E-05 | glio |
| culture_effect | Slc25a30 | -0.867 | 1.68E-05 | glio |
| culture_effect | Zfp292 | -0.631 | 1.68E-05 | glio |
| Interaction_genotype_Aars2 | Baalc | -4.719 | 1.68E-05 | glio |
| Interaction_genotype_Atr | Csnk1d | 2.483 | 1.68E-05 | glio |

| | | | | |
|---|---|---|---|---|
| Aars2_in.vivo_noRT | Derl2 | -1.49 | 1.68E-05 | glio |
| Atr_in.vivo_noRT | Gmnn | -1.74 | 1.68E-05 | glio |
| Bard1_ex.vivo_noRT | BC061195 | 2.814 | 1.68E-05 | glio |
| Bard1_ex.vivo_noRT | Idi2 | 2.814 | 1.68E-05 | glio |
| Bard1_ex.vivo_noRT | Gm364 | 2.814 | 1.68E-05 | glio |
| Bard1_ex.vivo_noRT | Gm14019 | 3.526 | 1.68E-05 | glio |
| Bora_in.vivo_noRT | Arhgap4 | 2.95 | 1.68E-05 | glio |
| Chek1_in.vivo_noRT | Gm4907 | 3.438 | 1.68E-05 | glio |
| Chek1_in.vivo_noRT | Prxl2b | 2.572 | 1.68E-05 | glio |
| Chek1_ex.vivo_noRT | D930028M14I | 3.356 | 1.68E-05 | glio |
| Hsd17b10_in.vivo_noRT | Hist2h3c2 | 3.35 | 1.68E-05 | glio |
| Interaction_genotype_Mtor | Ildr1 | -4.099 | 1.68E-06 | glio |
| Bora_ex.vivo_noRT | Gm5890 | 3.201 | 1.68E-06 | glio |
| Bora_ex.vivo_noRT | Zscan4c | 3.201 | 1.68E-06 | glio |
| Bora_ex.vivo_noRT | Speer4a | 3.201 | 1.68E-06 | glio |
| Bora_ex.vivo_noRT | BC080695 | 3.201 | 1.68E-06 | glio |
| Hsd17b10_in.vivo_noRT | Gm43463 | 2.958 | 1.68E-06 | glio |
| Interaction_genotype_Mtor | Krt19 | -4.099 | 1.68E-07 | glio |
| Interaction_genotype_Mtor | Gm35082 | -4.896 | 1.68E-07 | glio |
| Bora_in.vivo_noRT | Jakmip1 | 3.616 | 1.68E-07 | glio |
| Bcas2_in.vivo_noRT | Amdhd1 | 3.764 | 1.68E-08 | glio |
| Bcas2_in.vivo_noRT | Gm16046 | 4.015 | 1.68E-08 | glio |
| Chek1_in.vivo_noRT | Tomm6os | 4.126 | 1.68E-08 | glio |
| Chek1_in.vivo_noRT | 1700029J07Ri | 4.016 | 1.68E-08 | glio |
| Chek1_ex.vivo_noRT | Gm15929 | 3.866 | 1.68E-08 | glio |
| Chek1_ex.vivo_noRT | Serpinb9 | 4.817 | 1.68E-08 | glio |
| Chek1_ex.vivo_noRT | 5730507C01R | 3.652 | 1.68E-08 | glio |
| Atr_in.vivo_noRT | Guca1b | 4.245 | 1.68E-09 | glio |
| Bcas2_in.vivo_noRT | Amh | 3.89 | 1.68E-09 | glio |
| Bcas2_in.vivo_noRT | Lrfn1 | 4.165 | 1.68E-09 | glio |
| Atr_in.vivo_noRT | Gm26822 | 4.618 | 1.68E-10 | glio |
| Bora_in.vivo_noRT | Cyp4f40 | 4.283 | 1.68E-10 | glio |
| Chek1_in.vivo_noRT | 4930555A03R | 4.306 | 1.68E-11 | glio |
| Chek1_in.vivo_noRT | Gm5941 | 5.044 | 1.68E-12 | glio |
| Chek1_in.vivo_noRT | Gm4724 | 4.388 | 1.68E-12 | glio |
| Bcas2_in.vivo_noRT | Gm50138 | 4.284 | 1.68E-13 | glio |
| Aars2_in.vivo_noRT | Tff2 | 5.067 | 1.68E-14 | glio |
| Aars2_in.vivo_noRT | Cxcl16 | 5.063 | 1.68E-14 | glio |
| Atr_ex.vivo_noRT | Gm47404 | 4.547 | 1.68E-14 | glio |
| Bora_in.vivo_noRT | Cpxm1 | 4.67 | 1.68E-14 | glio |
| Atr_in.vivo_noRT | Slc45a3 | 5.521 | 1.68E-15 | glio |
| Chek1_ex.vivo_noRT | Gm45338 | 4.449 | 1.68E-15 | glio |
| Bora_in.vivo_noRT | Gm48764 | 4.936 | 1.68E-22 | glio |
| culture_effect | 1700023H06R | 1.933 | 1.69E-03 | glio |
| culture_effect | Gm11767 | 1.739 | 1.69E-03 | glio |
| Interaction_genotype_Aars2 | Akr1a1 | 1.037 | 1.69E-03 | glio |
| Interaction_genotype_Bora | H2-Eb1 | 3.764 | 1.69E-03 | glio |
| Interaction_genotype_Bora | Pgam5 | 2.048 | 1.69E-03 | glio |
| interaction_RT | Crip2 | 1.708 | 1.69E-03 | glio |
| interaction_RT | Dpep2 | 2.804 | 1.69E-03 | glio |
| interaction_RT | Plscr2 | -1.354 | 1.69E-03 | glio |
| interaction_RT | Tle5 | 0.563 | 1.69E-03 | glio |

| | | | | |
|---|---|---|---|---|
| Aars2_in.vivo_noRT | Samd14 | 1.918 | 1.69E-03 | glio |
| Atr_in.vivo_noRT | Smpdl3b | -1.605 | 1.69E-03 | glio |
| Atr_in.vivo_noRT | Sccpdh | 1.441 | 1.69E-03 | glio |
| Bard1_ex.vivo_noRT | Prkcq | 2.814 | 1.69E-03 | glio |
| Bcas2_in.vivo_noRT | Ppp2r5c | -1.176 | 1.69E-03 | glio |
| Bcas2_in.vivo_noRT | Anapc11 | -0.905 | 1.69E-03 | glio |
| Bora_in.vivo_noRT | Ccdc103 | 2.308 | 1.69E-03 | glio |
| Bora_in.vivo_noRT | Anapc13 | -1.133 | 1.69E-03 | glio |
| Bora_in.vivo_noRT | Ddx39b | -0.931 | 1.69E-03 | glio |
| Bora_ex.vivo_noRT | Gm48170 | 2.404 | 1.69E-03 | glio |
| Cenpj_ex.vivo_noRT | Ms4a15 | 2.591 | 1.69E-03 | glio |
| Chek1_ex.vivo_noRT | Gm7102 | 2.946 | 1.69E-03 | glio |
| Chek1_ex.vivo_noRT | Notch4 | 2.761 | 1.69E-03 | glio |
| Chek1_ex.vivo_noRT | 5330426L24Ri | 3.029 | 1.69E-03 | glio |
| Chek1_ex.vivo_noRT | 5830444B04R | 2.026 | 1.69E-03 | glio |
| Chek1_ex.vivo_noRT | Dek | -0.956 | 1.69E-03 | glio |
| Hsd17b10_in.vivo_noRT | Lrp11 | -1.671 | 1.69E-03 | glio |
| culture_effect | Gm48027 | -1.757 | 1.69E-04 | glio |
| culture_effect | Ndufc2 | -0.452 | 1.69E-04 | glio |
| Interaction_genotype_Mtor | Eid2b | -3.493 | 1.69E-04 | glio |
| Interaction_genotype_Mtor | Idh3b | 2.208 | 1.69E-04 | glio |
| Interaction_genotype_Myc | Arhgef4 | -4.259 | 1.69E-04 | glio |
| Interaction_genotype_Rad51 | BC065403 | -4.784 | 1.69E-04 | glio |
| interaction_RT | Hivep2 | 1.261 | 1.69E-04 | glio |
| Aars2_in.vivo_noRT | Selenof | -1.011 | 1.69E-04 | glio |
| Aars2_ex.vivo_noRT | Dnaaf3 | 3.293 | 1.69E-04 | glio |
| Atp5c1_in.vivo_noRT | Prl2c3 | 2.668 | 1.69E-04 | glio |
| Atr_in.vivo_noRT | Accs | 2.54 | 1.69E-04 | glio |
| Atr_ex.vivo_noRT | Gm20705 | 2.769 | 1.69E-04 | glio |
| Atr_ex.vivo_noRT | Gm19684 | 3.764 | 1.69E-04 | glio |
| Bcas2_in.vivo_noRT | Auts2 | 2.357 | 1.69E-04 | glio |
| Chek1_in.vivo_noRT | Ppfia3 | 2.6 | 1.69E-04 | glio |
| Eif2s1_ex.vivo_noRT | 9130019O22R | 3.032 | 1.69E-04 | glio |
| Hsd17b10_in.vivo_noRT | Ift20 | -0.885 | 1.69E-04 | glio |
| culture_effect | Rps4x | -0.706 | 1.69E-05 | glio |
| Interaction_genotype_Mtor | Bpi | -4.67 | 1.69E-05 | glio |
| Interaction_genotype_Myc | Gm15127 | -4.844 | 1.69E-05 | glio |
| Bora_in.vivo_noRT | Hhipl2 | 3.377 | 1.69E-05 | glio |
| Bora_in.vivo_noRT | Itga2b | 3.012 | 1.69E-05 | glio |
| Bora_in.vivo_noRT | Psmb5 | -0.996 | 1.69E-05 | glio |
| Chek1_in.vivo_noRT | AC166172.1 | 2.843 | 1.69E-05 | glio |
| Chek1_ex.vivo_noRT | Gm35290 | 3.547 | 1.69E-05 | glio |
| Chek1_ex.vivo_noRT | Tmem74b | 3.079 | 1.69E-05 | glio |
| Hsd17b10_in.vivo_noRT | 4930555A03R | 2.878 | 1.69E-05 | glio |
| Interaction_genotype_Mtor | Gm20633 | -5.344 | 1.69E-06 | glio |
| Interaction_genotype_Mtor | Slc16a4 | -4.691 | 1.69E-06 | glio |
| Atr_in.vivo_noRT | Gm36210 | 3.332 | 1.69E-06 | glio |
| Atr_ex.vivo_noRT | Ccl3 | 3.164 | 1.69E-06 | glio |
| Bcas2_in.vivo_noRT | Gm35867 | 3.354 | 1.69E-06 | glio |
| Bora_in.vivo_noRT | A430035B10R | 3.5 | 1.69E-06 | glio |
| Bora_ex.vivo_noRT | BB287469 | 3.201 | 1.69E-06 | glio |
| Chek1_in.vivo_noRT | Scd3 | 3.656 | 1.69E-06 | glio |

| | | | | |
|---|---|---|---|---|
| Hsd17b10_in.vivo_noRT | Matn4 | 3.352 | 1.69E-06 | glio |
| Interaction_genotype_Nat2 | Gm5662 | 5.734 | 1.69E-07 | glio |
| Interaction_genotype_Nat2 | Gm11232 | 4.295 | 1.69E-07 | glio |
| Interaction_genotype_Nat2 | Gm44693 | 4.373 | 1.69E-07 | glio |
| Aars2_in.vivo_noRT | Rab11fip1 | 4.033 | 1.69E-07 | glio |
| Atr_in.vivo_noRT | Gm49479 | 3.695 | 1.69E-07 | glio |
| Chek1_ex.vivo_noRT | Dok2 | 3.867 | 1.69E-07 | glio |
| Chek1_ex.vivo_noRT | Tns4 | 4.045 | 1.69E-07 | glio |
| Chek1_ex.vivo_noRT | C4a | 4.055 | 1.69E-07 | glio |
| Atr_in.vivo_noRT | Fkbp4 | -1.545 | 1.69E-08 | glio |
| Bcas2_in.vivo_noRT | 5330417C22R | 3.93 | 1.69E-08 | glio |
| Bora_in.vivo_noRT | Gm37584 | 4.025 | 1.69E-08 | glio |
| Chek1_in.vivo_noRT | Pde1a | 4.043 | 1.69E-08 | glio |
| Chek1_ex.vivo_noRT | A030005L19Ri | 4.449 | 1.69E-08 | glio |
| Chek1_in.vivo_noRT | Amdhd1 | 4.001 | 1.69E-09 | glio |
| Aars2_in.vivo_noRT | Gm28731 | 4.941 | 1.69E-11 | glio |
| Aars2_in.vivo_noRT | Gm47232 | 4.804 | 1.69E-12 | glio |
| Aars2_in.vivo_noRT | Gm20635 | 4.681 | 1.69E-12 | glio |
| Bcas2_in.vivo_noRT | BC018473 | 4.807 | 1.69E-12 | glio |
| Bora_in.vivo_noRT | Gm31152 | 4.414 | 1.69E-12 | glio |
| Atr_in.vivo_noRT | Gm35290 | 4.795 | 1.69E-13 | glio |
| Chek1_in.vivo_noRT | Gm7102 | 5.295 | 1.69E-13 | glio |
| Bcas2_in.vivo_noRT | 6430562O15R | 4.803 | 1.69E-14 | glio |
| Atr_in.vivo_noRT | Gm7247 | 4.741 | 1.69E-16 | glio |
| Bcas2_in.vivo_noRT | Gm48764 | 4.807 | 1.69E-21 | glio |
| Aars2_in.vivo_noRT | Ccdc80 | 5.327 | 1.69E-22 | glio |
| Atr_in.vivo_noRT | 4930423D22R | 5.273 | 1.69E-24 | glio |
| Atr_in.vivo_noRT | A330049N07R | 5.267 | 1.69E-26 | glio |
| Aars2_in.vivo_noRT | Gm5157 | 5.327 | 1.69E-28 | glio |
| culture_effect | Twsg1 | -0.755 | 1.70E-03 | glio |
| Interaction_genotype_Atr | Gm24362 | -3.606 | 1.70E-03 | glio |
| Interaction_genotype_Atr | Vmn2r44 | -2.765 | 1.70E-03 | glio |
| Interaction_genotype_Atr | Slc25a35 | -3.866 | 1.70E-03 | glio |
| Interaction_genotype_Atr | Egr1 | 2.936 | 1.70E-03 | glio |
| Interaction_genotype_Atr | Tpt1 | 1.223 | 1.70E-03 | glio |
| Interaction_genotype_Bora | Senp6 | 1.763 | 1.70E-03 | glio |
| Interaction_genotype_Bora | Tram1 | 1.363 | 1.70E-03 | glio |
| Interaction_genotype_Chek1 | Atp6v1f | 1.189 | 1.70E-03 | glio |
| Interaction_genotype_Chek1 | Ubb | 1.094 | 1.70E-03 | glio |
| Interaction_genotype_Mtor | Chadl | 2.589 | 1.70E-03 | glio |
| Interaction_genotype_Myc | Shroom3 | -3.548 | 1.70E-03 | glio |
| Interaction_genotype_Myc | Rac3 | -3.353 | 1.70E-03 | glio |
| Interaction_genotype_Myc | Rpgr | -3.331 | 1.70E-03 | glio |
| Interaction_genotype_Rad51 | Shc3 | -3.586 | 1.70E-03 | glio |
| interaction_RT | Bclaf1 | -0.635 | 1.70E-03 | glio |
| Aars2_in.vivo_noRT | Ttyh3 | 2.042 | 1.70E-03 | glio |
| Aars2_in.vivo_noRT | Traf2 | 1.512 | 1.70E-03 | glio |
| Aars2_in.vivo_noRT | Tmed5 | -1.056 | 1.70E-03 | glio |
| Atp5c1_in.vivo_noRT | Vmn2r54 | 2.037 | 1.70E-03 | glio |
| Atr_in.vivo_noRT | Apoc1 | 2.489 | 1.70E-03 | glio |
| Atr_in.vivo_noRT | Gm43466 | 2.065 | 1.70E-03 | glio |
| Atr_in.vivo_noRT | Scyl1 | 1.463 | 1.70E-03 | glio |

| | | | | |
|---|---|---|---|---|
| Atr_in.vivo_noRT | Mrpl24 | -0.962 | 1.70E-03 | glio |
| Bcas2_in.vivo_noRT | Itga7 | 2.332 | 1.70E-03 | glio |
| Bcas2_in.vivo_noRT | 1700001G11R | 1.996 | 1.70E-03 | glio |
| Bcas2_ex.vivo_noRT | Rangrf | -1.433 | 1.70E-03 | glio |
| Bora_in.vivo_noRT | Ankrd23 | 2.429 | 1.70E-03 | glio |
| Bora_in.vivo_noRT | Dixdc1 | 2.303 | 1.70E-03 | glio |
| Bora_in.vivo_noRT | Stk11 | -1.261 | 1.70E-03 | glio |
| Bora_ex.vivo_noRT | Gm31063 | 2.277 | 1.70E-03 | glio |
| Cenpj_ex.vivo_noRT | A930002C04R | 2.192 | 1.70E-03 | glio |
| Chek1_ex.vivo_noRT | Gm20712 | 2.947 | 1.70E-03 | glio |
| Eif2s1_in.vivo_noRT | 5430402O13R | 2.666 | 1.70E-03 | glio |
| Eif2s1_in.vivo_noRT | Tal2 | 1.88 | 1.70E-03 | glio |
| Hsd17b10_in.vivo_noRT | Gm15903 | 2.304 | 1.70E-03 | glio |
| Hsd17b10_in.vivo_noRT | Arhgef4 | 2.382 | 1.70E-03 | glio |
| Hsd17b10_in.vivo_noRT | Chd5 | 2.371 | 1.70E-03 | glio |
| Hsd17b10_in.vivo_noRT | Zfp981 | 2.282 | 1.70E-03 | glio |
| Hsd17b10_in.vivo_noRT | Misp | -2.115 | 1.70E-03 | glio |
| culture_effect | Prkab2 | -1.263 | 1.70E-04 | glio |
| culture_effect | Nudcd3 | 0.623 | 1.70E-04 | glio |
| Interaction_genotype_Mtor | Hmgn1 | 1.698 | 1.70E-04 | glio |
| Interaction_genotype_Mtor | Metap2 | 1.838 | 1.70E-04 | glio |
| Interaction_genotype_Myc | Spata33 | -4.478 | 1.70E-04 | glio |
| Interaction_genotype_Myc | Gm13421 | -4.331 | 1.70E-04 | glio |
| Interaction_genotype_Myc | Fgd6 | -4.072 | 1.70E-04 | glio |
| Interaction_genotype_Myc | Mtor | -2.813 | 1.70E-04 | glio |
| Aars2_in.vivo_noRT | Eml6 | 2.726 | 1.70E-04 | glio |
| Aars2_in.vivo_noRT | 1700120G11R | 2.843 | 1.70E-04 | glio |
| Aars2_in.vivo_noRT | Hdac1 | -1.073 | 1.70E-04 | glio |
| Aars2_ex.vivo_noRT | Ciita | 2.893 | 1.70E-04 | glio |
| Atr_in.vivo_noRT | Gm5930 | 2.626 | 1.70E-04 | glio |
| Atr_in.vivo_noRT | Pcdhb20 | 2.474 | 1.70E-04 | glio |
| Atr_in.vivo_noRT | Zfp446 | 2.305 | 1.70E-04 | glio |
| Atr_in.vivo_noRT | Med28 | -0.888 | 1.70E-04 | glio |
| Bcas2_in.vivo_noRT | Vcan | 2.938 | 1.70E-04 | glio |
| Bcas2_in.vivo_noRT | Tmem91 | 2.764 | 1.70E-04 | glio |
| Bcas2_in.vivo_noRT | Foxa1 | 2.955 | 1.70E-04 | glio |
| Bcas2_in.vivo_noRT | 4930522L14Ri | 2.789 | 1.70E-04 | glio |
| Bcas2_in.vivo_noRT | Ptcd3 | -1.707 | 1.70E-04 | glio |
| Bora_ex.vivo_noRT | Prkcq | 3.201 | 1.70E-04 | glio |
| Hsd17b10_in.vivo_noRT | Clca3a2 | 2.43 | 1.70E-04 | glio |
| Interaction_genotype_Mtor | Akr1c18 | -4.099 | 1.70E-05 | glio |
| Interaction_genotype_Mtor | 4930434B07R | -4.423 | 1.70E-05 | glio |
| Interaction_genotype_Myc | Gm13398 | -3.526 | 1.70E-05 | glio |
| Aars2_in.vivo_noRT | Zfp60 | 2.308 | 1.70E-05 | glio |
| Aars2_ex.vivo_noRT | Ipcef1 | 3.293 | 1.70E-05 | glio |
| Atp5c1_in.vivo_noRT | A330032B11R | 2.933 | 1.70E-05 | glio |
| Atr_in.vivo_noRT | Gm16845 | 2.862 | 1.70E-05 | glio |
| Atr_in.vivo_noRT | 4732419C18R | 2.945 | 1.70E-05 | glio |
| Bora_in.vivo_noRT | Gm39556 | 3.026 | 1.70E-05 | glio |
| Hsd17b10_in.vivo_noRT | Gm50375 | 3.35 | 1.70E-05 | glio |
| culture_effect | Mrps14 | 0.515 | 1.70E-06 | glio |
| Interaction_genotype_Mtor | Gm30082 | -5.662 | 1.70E-06 | glio |

| | | | | |
|---|---|---|---|---|
| Aars2_in.vivo_noRT | Ehhadh | 3.631 | 1.70E-06 | glio |
| Aars2_in.vivo_noRT | Lrrc75aos2 | 3.43 | 1.70E-06 | glio |
| Aars2_ex.vivo_noRT | Gpr22 | 3.293 | 1.70E-06 | glio |
| Atr_in.vivo_noRT | Hcfc1r1 | 3.735 | 1.70E-06 | glio |
| Atr_ex.vivo_noRT | Gm48780 | 3.164 | 1.70E-06 | glio |
| Atr_ex.vivo_noRT | Fabp4 | 4.44 | 1.70E-06 | glio |
| Interaction_genotype_Mtor | Dntt | -6.502 | 1.70E-07 | glio |
| Aars2_in.vivo_noRT | Ndufs5 | -1.116 | 1.70E-07 | glio |
| Chek1_in.vivo_noRT | 4921523L03Ri | 3.876 | 1.70E-07 | glio |
| Hsd17b10_in.vivo_noRT | Lrp2bp | 3.352 | 1.70E-07 | glio |
| Hsd17b10_in.vivo_noRT | Gm42849 | 3.35 | 1.70E-07 | glio |
| culture_effect | 1700047M11F | -3.268 | 1.70E-08 | glio |
| Bcas2_in.vivo_noRT | Adam1a | 3.963 | 1.70E-08 | glio |
| culture_effect | Pcdh1 | 2.633 | 1.70E-09 | glio |
| Bora_in.vivo_noRT | Ccdc114 | 4.018 | 1.70E-09 | glio |
| Bora_in.vivo_noRT | Gm2762 | 3.897 | 1.70E-09 | glio |
| Bora_in.vivo_noRT | B230311B06R | 4.673 | 1.70E-10 | glio |
| Aars2_in.vivo_noRT | BC048671 | 4.026 | 1.70E-12 | glio |
| Bcas2_in.vivo_noRT | Magea10 | 4.34 | 1.70E-12 | glio |
| Bcas2_in.vivo_noRT | Cox7b2 | 4.34 | 1.70E-12 | glio |
| Bora_in.vivo_noRT | Nxph4 | 4.936 | 1.70E-12 | glio |
| Aars2_in.vivo_noRT | Gm39326 | 5.063 | 1.70E-13 | glio |
| Atr_in.vivo_noRT | Gm47232 | 5.011 | 1.70E-13 | glio |
| Atr_in.vivo_noRT | Gm28578 | 5.273 | 1.70E-13 | glio |
| Atr_in.vivo_noRT | 4930540M05F | 4.876 | 1.70E-13 | glio |
| Aars2_in.vivo_noRT | Gm50147 | 5.327 | 1.70E-15 | glio |
| Aars2_in.vivo_noRT | Gm11769 | 5.067 | 1.70E-15 | glio |
| Bora_in.vivo_noRT | BB557941 | 4.676 | 1.70E-15 | glio |
| Bora_in.vivo_noRT | Gm13713 | 4.404 | 1.70E-15 | glio |
| Aars2_in.vivo_noRT | Pmp2 | 5.327 | 1.70E-17 | glio |
| Atr_in.vivo_noRT | Nos2 | 5.273 | 1.70E-17 | glio |
| Aars2_in.vivo_noRT | Gm32250 | 4.544 | 1.70E-18 | glio |
| Bcas2_in.vivo_noRT | Gm16178 | 4.807 | 1.70E-22 | glio |
| Bcas2_in.vivo_noRT | Gm364 | 4.807 | 1.70E-22 | glio |
| Chek1_in.vivo_noRT | Idi2 | 5.044 | 1.70E-24 | glio |
| Aars2_in.vivo_noRT | Vmn1r32 | 5.327 | 1.70E-26 | glio |
| Aars2_in.vivo_noRT | Glyat | 5.327 | 1.70E-26 | glio |
| Aars2_in.vivo_noRT | Cldn2 | 5.327 | 1.70E-29 | glio |
| culture_effect | Zc2hc1c | -1.709 | 1.71E-03 | glio |
| culture_effect | Ulbp1 | -1.047 | 1.71E-03 | glio |
| culture_effect | Ppa2 | -0.596 | 1.71E-03 | glio |
| culture_effect | Rab3gap1 | -0.517 | 1.71E-03 | glio |
| culture_effect | Vezf1 | -0.542 | 1.71E-03 | glio |
| Interaction_genotype_Aars2 | Gm30262 | -4.087 | 1.71E-03 | glio |
| Interaction_genotype_Atr | 0610040B10R | -3.082 | 1.71E-03 | glio |
| Interaction_genotype_Mtor | Gm527 | -3.696 | 1.71E-03 | glio |
| Interaction_genotype_Mtor | Mlh3 | -2.779 | 1.71E-03 | glio |
| Interaction_genotype_Myc | Paqr8 | -4.042 | 1.71E-03 | glio |
| Interaction_genotype_Myc | Klc2 | -3.313 | 1.71E-03 | glio |
| Interaction_genotype_Myc | Zdhhc18 | -3.14 | 1.71E-03 | glio |
| Interaction_genotype_Myc | BC055324 | -3.129 | 1.71E-03 | glio |
| Interaction_genotype_Myc | Ttll12 | -2.291 | 1.71E-03 | glio |

| | | | | |
|---|---|---|---|---|
| Interaction_genotype_Myc | Rpl8 | 1.168 | 1.71E-03 | glio |
| Interaction_genotype_Rad51 | Gm50334 | -4.839 | 1.71E-03 | glio |
| Interaction_genotype_Rad51 | Igf2bp1 | -4.32 | 1.71E-03 | glio |
| interaction_RT | Clk4 | 0.853 | 1.71E-03 | glio |
| interaction_RT | H2-DMb1 | 0.898 | 1.71E-03 | glio |
| interaction_RT | Hnrnpa1 | -0.624 | 1.71E-03 | glio |
| Aars2_in.vivo_noRT | Dennd2a | 2.253 | 1.71E-03 | glio |
| Aars2_in.vivo_noRT | Cuedc2 | -0.749 | 1.71E-03 | glio |
| Aars2_in.vivo_noRT | Bst2 | -0.741 | 1.71E-03 | glio |
| Aars2_ex.vivo_noRT | B3galt2 | 2.893 | 1.71E-03 | glio |
| Atp5c1_in.vivo_noRT | Tcap | 2.289 | 1.71E-03 | glio |
| Atp5c1_in.vivo_noRT | Fgf8 | 2.152 | 1.71E-03 | glio |
| Atp5c1_in.vivo_noRT | Mogat1 | 2.668 | 1.71E-03 | glio |
| Bard1_ex.vivo_noRT | Vmn1r88 | 2.814 | 1.71E-03 | glio |
| Bard1_ex.vivo_noRT | Rslcan18 | 2.809 | 1.71E-03 | glio |
| Bard1_ex.vivo_noRT | 4732463B04R | 2.421 | 1.71E-03 | glio |
| Bcas2_in.vivo_noRT | Rab34 | -0.818 | 1.71E-03 | glio |
| Bora_in.vivo_noRT | Selenbp1 | 2.498 | 1.71E-03 | glio |
| Bora_in.vivo_noRT | Prx | 2.408 | 1.71E-03 | glio |
| Bora_in.vivo_noRT | Amd2 | 2.249 | 1.71E-03 | glio |
| Chek1_in.vivo_noRT | Snn | 2.308 | 1.71E-03 | glio |
| Chek1_ex.vivo_noRT | Ccdc8 | 2.897 | 1.71E-03 | glio |
| culture_effect | Gpatch8 | -0.545 | 1.71E-04 | glio |
| Interaction_genotype_Atr | Gm40040 | -4.888 | 1.71E-04 | glio |
| Interaction_genotype_Atr | Gm49504 | -3.561 | 1.71E-04 | glio |
| Interaction_genotype_Ifnar1 | Nrip2 | 3.531 | 1.71E-04 | glio |
| Interaction_genotype_Mtor | Pacrg | -4.102 | 1.71E-04 | glio |
| Interaction_genotype_Mtor | Gm45151 | -3.567 | 1.71E-04 | glio |
| Interaction_genotype_Mtor | Plekho2 | -4.368 | 1.71E-04 | glio |
| Interaction_genotype_Mtor | Fblim1 | -4.451 | 1.71E-04 | glio |
| Interaction_genotype_Mtor | Pcdh1 | -4.087 | 1.71E-04 | glio |
| Interaction_genotype_Mtor | Snrpb | 1.563 | 1.71E-04 | glio |
| Aars2_in.vivo_noRT | Gm35339 | 2.73 | 1.71E-04 | glio |
| Aars2_in.vivo_noRT | Afg1l | 2.723 | 1.71E-04 | glio |
| Aars2_in.vivo_noRT | 9030025P20R | 2.854 | 1.71E-04 | glio |
| Aars2_in.vivo_noRT | Intu | 2.544 | 1.71E-04 | glio |
| Aars2_ex.vivo_noRT | Gm48140 | 2.499 | 1.71E-04 | glio |
| Atp5c1_in.vivo_noRT | BC147527 | 2.668 | 1.71E-04 | glio |
| Atp5c1_in.vivo_noRT | Gm29083 | 2.657 | 1.71E-04 | glio |
| Atp5c1_in.vivo_noRT | Lmx1b | 2.944 | 1.71E-04 | glio |
| Atr_in.vivo_noRT | Snx32 | 2.501 | 1.71E-04 | glio |
| Bard1_ex.vivo_noRT | Gm13856 | 2.421 | 1.71E-04 | glio |
| Bcas2_in.vivo_noRT | Iapp | 3.172 | 1.71E-04 | glio |
| Bora_in.vivo_noRT | Slc22a5 | 2.48 | 1.71E-04 | glio |
| Bora_in.vivo_noRT | Tro | 2.632 | 1.71E-04 | glio |
| Bora_in.vivo_noRT | Dpysl2 | -1.728 | 1.71E-04 | glio |
| Bora_in.vivo_noRT | Gpx4 | -0.807 | 1.71E-04 | glio |
| Chek1_ex.vivo_noRT | Tes3-ps | 3.352 | 1.71E-04 | glio |
| Chek1_ex.vivo_noRT | Maats1 | 3.438 | 1.71E-04 | glio |
| Chek1_ex.vivo_noRT | Pabpc1 | -0.725 | 1.71E-04 | glio |
| Eif2s1_in.vivo_noRT | Gm48764 | 2.143 | 1.71E-04 | glio |
| Hsd17b10_in.vivo_noRT | Gm9999 | 2.641 | 1.71E-04 | glio |

| | | | | |
|---|---|---|---|---|
| culture_effect | P2ry10 | -0.914 | 1.71E-05 | glio |
| culture_effect | Usp50 | -1.218 | 1.71E-05 | glio |
| Interaction_genotype_Mtor | Rab40b | -5.054 | 1.71E-05 | glio |
| Interaction_genotype_Myc | Snrpf | 2.031 | 1.71E-05 | glio |
| Interaction_genotype_Rad51 | Sike1 | 2.61 | 1.71E-05 | glio |
| Aars2_in.vivo_noRT | Nup210l | 3.2 | 1.71E-05 | glio |
| Aars2_in.vivo_noRT | Tmem86b | 3.1 | 1.71E-05 | glio |
| Atp5c1_in.vivo_noRT | Impg2 | 2.668 | 1.71E-05 | glio |
| Atr_in.vivo_noRT | Ptprg | 2.605 | 1.71E-05 | glio |
| Atr_ex.vivo_noRT | Hist3h2ba | 3.509 | 1.71E-05 | glio |
| Bard1_ex.vivo_noRT | Gm16178 | 2.814 | 1.71E-05 | glio |
| Bcas2_in.vivo_noRT | 1810059H22R | 3.366 | 1.71E-05 | glio |
| Bora_in.vivo_noRT | Smad4 | -1.754 | 1.71E-05 | glio |
| Chek1_ex.vivo_noRT | 1700061G19R | 3.689 | 1.71E-05 | glio |
| Chek1_ex.vivo_noRT | Gm4755 | 3.467 | 1.71E-05 | glio |
| Hsd17b10_in.vivo_noRT | Nrg2 | 3.093 | 1.71E-05 | glio |
| Hsd17b10_in.vivo_noRT | Npy4r | 2.776 | 1.71E-05 | glio |
| culture_effect | Mmgt2 | 1.241 | 1.71E-06 | glio |
| Interaction_genotype_Mtor | Tomm5 | 2.477 | 1.71E-06 | glio |
| Interaction_genotype_Mtor | Stk36 | -5.898 | 1.71E-06 | glio |
| Interaction_genotype_Mtor | Gm14540 | -4.37 | 1.71E-06 | glio |
| Interaction_genotype_Mtor | F630040K05Ri | -5.173 | 1.71E-06 | glio |
| Atp5c1_in.vivo_noRT | Tagap | 2.408 | 1.71E-06 | glio |
| Atr_ex.vivo_noRT | Lhx1os | 3.552 | 1.71E-06 | glio |
| Atr_ex.vivo_noRT | Gm36793 | 3.164 | 1.71E-06 | glio |
| Chek1_in.vivo_noRT | Spata33 | 3.519 | 1.71E-06 | glio |
| Chek1_in.vivo_noRT | Tcea2 | 3.316 | 1.71E-06 | glio |
| Chek1_ex.vivo_noRT | Amh | 3.871 | 1.71E-06 | glio |
| Hsd17b10_in.vivo_noRT | Synpo2l | 2.971 | 1.71E-06 | glio |
| culture_effect | Mrps24 | 0.531 | 1.71E-07 | glio |
| Interaction_genotype_Chek1 | Ntn4 | -5.249 | 1.71E-07 | glio |
| Atr_ex.vivo_noRT | Gm31717 | 3.164 | 1.71E-07 | glio |
| Bcas2_in.vivo_noRT | Hipk4 | 3.338 | 1.71E-07 | glio |
| Chek1_in.vivo_noRT | E130208F15Ri | 3.991 | 1.71E-07 | glio |
| culture_effect | Metrn | 0.731 | 1.71E-08 | glio |
| culture_effect | Lbh | 0.832 | 1.71E-08 | glio |
| Bcas2_in.vivo_noRT | Kcnj13 | 3.526 | 1.71E-08 | glio |
| Bcas2_in.vivo_noRT | Pbld1 | 4.148 | 1.71E-08 | glio |
| Bora_in.vivo_noRT | Gm31517 | 4.018 | 1.71E-08 | glio |
| culture_effect | Gm30191 | 2.102 | 1.71E-09 | glio |
| Atp5c1_in.vivo_noRT | Glyat | 3.199 | 1.71E-09 | glio |
| Atr_in.vivo_noRT | Gm9484 | 4.375 | 1.71E-09 | glio |
| Atr_in.vivo_noRT | Gm4755 | 3.776 | 1.71E-09 | glio |
| Bcas2_in.vivo_noRT | Platr31 | 4.807 | 1.71E-09 | glio |
| Bora_in.vivo_noRT | Gm11549 | 4.411 | 1.71E-10 | glio |
| Chek1_in.vivo_noRT | H3f3aos | 4.399 | 1.71E-10 | glio |
| culture_effect | P3h2 | 1.972 | 1.71E-11 | glio |
| Chek1_ex.vivo_noRT | Slc25a18 | 4.842 | 1.71E-11 | glio |
| Atr_in.vivo_noRT | Rims3 | 4.399 | 1.71E-12 | glio |
| Bora_in.vivo_noRT | Plk5 | 4.28 | 1.71E-12 | glio |
| Atr_in.vivo_noRT | Gm7102 | 5.273 | 1.71E-13 | glio |
| Atr_in.vivo_noRT | Gm17173 | 5.146 | 1.71E-14 | glio |

| | | | | |
|---|---|---|---|---|
| Bora_in.vivo_noRT | Zmynd10 | 4.676 | 1.71E-15 | glio |
| culture_effect | Cpm | -2.374 | 1.71E-16 | glio |
| Atr_in.vivo_noRT | Gm11627 | 5.533 | 1.71E-17 | glio |
| Bora_in.vivo_noRT | Spink8 | 4.676 | 1.71E-19 | glio |
| Aars2_in.vivo_noRT | Gm15326 | 4.798 | 1.71E-20 | glio |
| Bora_in.vivo_noRT | Ush1g | 4.936 | 1.71E-20 | glio |
| Chek1_in.vivo_noRT | Ccdc80 | 5.044 | 1.71E-20 | glio |
| Chek1_in.vivo_noRT | Gm15908 | 5.044 | 1.71E-26 | glio |
| Chek1_in.vivo_noRT | A930014E10Ri | 5.044 | 1.71E-26 | glio |
| culture_effect | Tmem181a | -0.649 | 1.72E-03 | glio |
| Interaction_genotype_Aars2 | 2210417A02R | -4.098 | 1.72E-03 | glio |
| Interaction_genotype_Aars2 | Zdhhc17 | -2.862 | 1.72E-03 | glio |
| Interaction_genotype_Bcas2 | Clec4d | -3.25 | 1.72E-03 | glio |
| Interaction_genotype_Bcas2 | Gm10425 | -4.359 | 1.72E-03 | glio |
| Interaction_genotype_Bcas2 | Fgd3 | -4.602 | 1.72E-03 | glio |
| Interaction_genotype_Bcas2 | Nsd3 | 1.757 | 1.72E-03 | glio |
| Interaction_genotype_Bora | Gm50147 | -4.001 | 1.72E-03 | glio |
| Interaction_genotype_Mtor | Plekhm3 | -3.915 | 1.72E-03 | glio |
| Interaction_genotype_Mtor | Rab30 | -3.108 | 1.72E-03 | glio |
| Interaction_genotype_Mtor | 1700008J07Ri | -3.026 | 1.72E-03 | glio |
| Interaction_genotype_Mtor | Tuba1a | 1.678 | 1.72E-03 | glio |
| Interaction_genotype_Myc | Gm12703 | -2.463 | 1.72E-03 | glio |
| Interaction_genotype_Myc | Peli3 | -3.483 | 1.72E-03 | glio |
| Interaction_genotype_Myc | Serpinb2 | -3.128 | 1.72E-03 | glio |
| Interaction_genotype_Myc | Slc46a1 | -3.639 | 1.72E-03 | glio |
| Interaction_genotype_Myc | Draxin | -3.548 | 1.72E-03 | glio |
| Interaction_genotype_Myc | Aplf | -2.988 | 1.72E-03 | glio |
| Interaction_genotype_Myc | Nr1h3 | -3.103 | 1.72E-03 | glio |
| Interaction_genotype_Pars2 | Gm11457 | -3.986 | 1.72E-03 | glio |
| Interaction_genotype_Pars2 | 5330438D12R | -3.02 | 1.72E-03 | glio |
| Interaction_genotype_Rad51 | AW112010 | 5.037 | 1.72E-03 | glio |
| interaction_RT | Itgae | 3.011 | 1.72E-03 | glio |
| Aars2_in.vivo_noRT | Ccdc141 | 2.332 | 1.72E-03 | glio |
| Aars2_in.vivo_noRT | D830050J10Ri | 2.113 | 1.72E-03 | glio |
| Aars2_in.vivo_noRT | Spcs2 | -0.776 | 1.72E-03 | glio |
| Aars2_in.vivo_noRT | Rbbp7 | -0.701 | 1.72E-03 | glio |
| Aars2_in.vivo_noRT | Pfn1 | -0.645 | 1.72E-03 | glio |
| Aars2_in.vivo_noRT | Rpl9 | -0.588 | 1.72E-03 | glio |
| Atr_in.vivo_noRT | Gm43560 | 1.92 | 1.72E-03 | glio |
| Atr_in.vivo_noRT | Lmbr1 | 1.718 | 1.72E-03 | glio |
| Bcas2_in.vivo_noRT | Gm50203 | 2.175 | 1.72E-03 | glio |
| Bcas2_in.vivo_noRT | 4732491K20R | 2.093 | 1.72E-03 | glio |
| Bcas2_in.vivo_noRT | Arih2 | -1.463 | 1.72E-03 | glio |
| Bcas2_ex.vivo_noRT | Gm36757 | 3.125 | 1.72E-03 | glio |
| Bora_in.vivo_noRT | Wfs1 | 2.237 | 1.72E-03 | glio |
| Bora_in.vivo_noRT | Akr1e1 | -1.425 | 1.72E-03 | glio |
| Bora_in.vivo_noRT | Sgo1 | -1.827 | 1.72E-03 | glio |
| Chek1_in.vivo_noRT | Ccdc142os | 2.256 | 1.72E-03 | glio |
| Chek1_in.vivo_noRT | Emc2 | -1.045 | 1.72E-03 | glio |
| Eif2s1_in.vivo_noRT | Ccl3 | 1.883 | 1.72E-03 | glio |
| Eif2s1_in.vivo_noRT | Gm48780 | 1.883 | 1.72E-03 | glio |
| Eif2s1_in.vivo_noRT | Gm13348 | 2.143 | 1.72E-03 | glio |

| | | | | |
|---|---|---|---|---|
| Eif2s1_in.vivo_noRT | Aqp9 | 2.143 | 1.72E-03 | glio |
| Eif2s1_in.vivo_noRT | Slc4a9 | 2.141 | 1.72E-03 | glio |
| Eif2s1_in.vivo_noRT | Gm15138 | 1.877 | 1.72E-03 | glio |
| Eif2s1_in.vivo_noRT | Rpp40 | 1.097 | 1.72E-03 | glio |
| Hsd17b10_in.vivo_noRT | Gm33973 | 2.05 | 1.72E-03 | glio |
| culture_effect | Setx | -0.622 | 1.72E-04 | glio |
| Interaction_genotype_Mtor | Cnnm1 | -5.016 | 1.72E-04 | glio |
| Interaction_genotype_Mtor | Zfp352 | -4.099 | 1.72E-04 | glio |
| Interaction_genotype_Mtor | Gm47484 | -4.565 | 1.72E-04 | glio |
| Interaction_genotype_Mtor | Tmeff1 | -3.91 | 1.72E-04 | glio |
| Interaction_genotype_Myc | Gm4189 | -4.25 | 1.72E-04 | glio |
| Interaction_genotype_Myc | Gm13404 | -4.344 | 1.72E-04 | glio |
| Aars2_in.vivo_noRT | Fhod3 | 2.81 | 1.72E-04 | glio |
| Atr_in.vivo_noRT | Hist1h4a | 2.541 | 1.72E-04 | glio |
| Bcas2_in.vivo_noRT | C330013E15Ri | 2.692 | 1.72E-04 | glio |
| Bora_in.vivo_noRT | Zfp418 | 2.881 | 1.72E-04 | glio |
| Chek1_in.vivo_noRT | Ces1a | 2.921 | 1.72E-04 | glio |
| culture_effect | Cox17 | 0.643 | 1.72E-05 | glio |
| Interaction_genotype_Myc | AA386476 | -4.94 | 1.72E-05 | glio |
| Aars2_in.vivo_noRT | Nanos3 | 2.735 | 1.72E-05 | glio |
| Aars2_in.vivo_noRT | Tubb2a | -1.929 | 1.72E-05 | glio |
| Aars2_ex.vivo_noRT | Gm36026 | 3.293 | 1.72E-05 | glio |
| Atr_in.vivo_noRT | Eef1a1 | -0.82 | 1.72E-05 | glio |
| Chek1_in.vivo_noRT | Tes3-ps | 3.009 | 1.72E-05 | glio |
| Chek1_in.vivo_noRT | 2310008N11R | 3.258 | 1.72E-05 | glio |
| Chek1_in.vivo_noRT | Gdnf | 2.186 | 1.72E-05 | glio |
| Chek1_ex.vivo_noRT | Kctd4 | 3.679 | 1.72E-05 | glio |
| culture_effect | Zdhhc15 | -2.271 | 1.72E-06 | glio |
| culture_effect | Gfpt1 | -0.764 | 1.72E-06 | glio |
| culture_effect | Immp1l | 0.656 | 1.72E-06 | glio |
| culture_effect | Rorb | 0.971 | 1.72E-06 | glio |
| Interaction_genotype_Mtor | Hist4h4 | -5.7 | 1.72E-06 | glio |
| Interaction_genotype_Mtor | Lynx1 | -5.544 | 1.72E-06 | glio |
| Interaction_genotype_Mtor | 4921523L03Ri | -5.993 | 1.72E-06 | glio |
| Interaction_genotype_Mtor | Thap8 | -5.785 | 1.72E-06 | glio |
| Aars2_in.vivo_noRT | AC166172.1 | 3.126 | 1.72E-06 | glio |
| Atr_in.vivo_noRT | Kif5a | 3.603 | 1.72E-06 | glio |
| Atr_in.vivo_noRT | AC125149.3 | 3.128 | 1.72E-06 | glio |
| Bcas2_in.vivo_noRT | Gm30262 | 3.515 | 1.72E-06 | glio |
| Bora_ex.vivo_noRT | H1foo | 3.201 | 1.72E-06 | glio |
| culture_effect | Tm7sf3 | -0.807 | 1.72E-07 | glio |
| Bora_in.vivo_noRT | Adgrb3 | 3.63 | 1.72E-07 | glio |
| Aars2_in.vivo_noRT | Tmem202 | 3.885 | 1.72E-08 | glio |
| Aars2_in.vivo_noRT | Slc17a7 | 3.895 | 1.72E-08 | glio |
| Atr_in.vivo_noRT | Pcdhb18 | 3.783 | 1.72E-08 | glio |
| Atr_in.vivo_noRT | Prm1 | 3.523 | 1.72E-08 | glio |
| Bora_in.vivo_noRT | Dact1 | 3.769 | 1.72E-08 | glio |
| Bora_in.vivo_noRT | Zfp551 | 3.902 | 1.72E-08 | glio |
| Chek1_ex.vivo_noRT | Rhov | 4.642 | 1.72E-08 | glio |
| Atr_in.vivo_noRT | Gm10115 | 4.096 | 1.72E-09 | glio |
| Bora_in.vivo_noRT | Cldn19 | 4.154 | 1.72E-09 | glio |
| Bora_in.vivo_noRT | Slc7a4 | 4.046 | 1.72E-09 | glio |

| | | | | |
|---|---|---|---|---|
| culture_effect | Vegfa | -1.546 | 1.72E-10 | glio |
| Aars2_in.vivo_noRT | Lama5 | 4.805 | 1.72E-11 | glio |
| Atr_in.vivo_noRT | Gm16046 | 4.741 | 1.72E-11 | glio |
| Bcas2_in.vivo_noRT | Nmrk2 | 4.165 | 1.72E-11 | glio |
| Chek1_in.vivo_noRT | Gm15579 | 5.044 | 1.72E-11 | glio |
| Chek1_in.vivo_noRT | Pde6b | 4.012 | 1.72E-11 | glio |
| Atr_in.vivo_noRT | Clca3a1 | 4.482 | 1.72E-12 | glio |
| Atr_in.vivo_noRT | Bmx | 4.746 | 1.72E-14 | glio |
| Chek1_in.vivo_noRT | Dcdc2a | 4.784 | 1.72E-15 | glio |
| Chek1_in.vivo_noRT | Tcp11x2 | 5.045 | 1.72E-17 | glio |
| Bora_in.vivo_noRT | Gm31063 | 4.936 | 1.72E-20 | glio |
| culture_effect | 9130023H24R | 1.508 | 1.73E-03 | glio |
| Interaction_genotype_Aars2 | B230319C09R | -4.226 | 1.73E-03 | glio |
| Interaction_genotype_Atr | Tc2n | -4.04 | 1.73E-03 | glio |
| Interaction_genotype_Atr | 6330415G19R | -3.515 | 1.73E-03 | glio |
| Interaction_genotype_Bora | Epb41l2 | 1.795 | 1.73E-03 | glio |
| Interaction_genotype_Mtor | Pla1a | -3.567 | 1.73E-03 | glio |
| Interaction_genotype_Mtor | Traf3ip3 | -3.75 | 1.73E-03 | glio |
| Interaction_genotype_Mtor | Dok7 | -3.78 | 1.73E-03 | glio |
| Interaction_genotype_Mtor | 1700102H20R | -3.591 | 1.73E-03 | glio |
| Interaction_genotype_Mtor | Zfp870 | -3.11 | 1.73E-03 | glio |
| Interaction_genotype_Mtor | Lmbrd1 | 1.919 | 1.73E-03 | glio |
| Interaction_genotype_Myc | Chst12 | -2.91 | 1.73E-03 | glio |
| Interaction_genotype_Myc | Timp1 | 2.416 | 1.73E-03 | glio |
| Interaction_genotype_Nat2 | Speer4a | 3.198 | 1.73E-03 | glio |
| Interaction_genotype_Nat2 | Gm14929 | 3.042 | 1.73E-03 | glio |
| Interaction_genotype_Pars2 | Gm44899 | -3.373 | 1.73E-03 | glio |
| interaction_RT | Cmbl | 1.21 | 1.73E-03 | glio |
| Aars2_in.vivo_noRT | Lsm7 | -0.864 | 1.73E-03 | glio |
| Aars2_ex.vivo_noRT | Pcdhgb1 | 2.718 | 1.73E-03 | glio |
| Atr_in.vivo_noRT | Ezh2 | -1.13 | 1.73E-03 | glio |
| Bcas2_in.vivo_noRT | Susd1 | 2.495 | 1.73E-03 | glio |
| Bcas2_in.vivo_noRT | Moap1 | 2.095 | 1.73E-03 | glio |
| Bcas2_in.vivo_noRT | Impact | -1.573 | 1.73E-03 | glio |
| Bora_in.vivo_noRT | Eml6 | 2.336 | 1.73E-03 | glio |
| Bora_in.vivo_noRT | Gm35339 | 2.339 | 1.73E-03 | glio |
| Bora_in.vivo_noRT | Cfap77 | 2.085 | 1.73E-03 | glio |
| Bora_in.vivo_noRT | Rnf114 | -1.441 | 1.73E-03 | glio |
| Cenpj_in.vivo_noRT | Gm13449 | 2.022 | 1.73E-03 | glio |
| Chek1_in.vivo_noRT | Klf15 | 2.287 | 1.73E-03 | glio |
| Chek1_in.vivo_noRT | Gm50232 | 2.123 | 1.73E-03 | glio |
| Chek1_in.vivo_noRT | 3110021N24R | 1.932 | 1.73E-03 | glio |
| Chek1_in.vivo_noRT | Slc35g1 | 1.775 | 1.73E-03 | glio |
| Chek1_ex.vivo_noRT | Dnah2 | 2.971 | 1.73E-03 | glio |
| Hsd17b10_in.vivo_noRT | Eme2 | 2.265 | 1.73E-03 | glio |
| culture_effect | Gemin8 | -0.785 | 1.73E-04 | glio |
| culture_effect | Tmx1 | -0.518 | 1.73E-04 | glio |
| culture_effect | Cavin3 | 0.543 | 1.73E-04 | glio |
| Interaction_genotype_Mtor | Gm17276 | -3.703 | 1.73E-04 | glio |
| Interaction_genotype_Mtor | H3f3aos | -4.319 | 1.73E-04 | glio |
| Interaction_genotype_Mtor | 1700030K09R | -3.219 | 1.73E-04 | glio |
| Interaction_genotype_Myc | Ccdc9b | -3.229 | 1.73E-04 | glio |

| | | | | |
|---|---|---|---|---|
| Atp5c1_in.vivo_noRT | 2410141K09R | 2.404 | 1.73E-04 | glio |
| Atr_in.vivo_noRT | Tmem171 | 3.379 | 1.73E-04 | glio |
| Atr_in.vivo_noRT | Ramp1 | 3.042 | 1.73E-04 | glio |
| Atr_in.vivo_noRT | Gm10825 | 2.704 | 1.73E-04 | glio |
| Atr_in.vivo_noRT | Gm29093 | 2.617 | 1.73E-04 | glio |
| Atr_in.vivo_noRT | 4930590J08Ri | 2.728 | 1.73E-04 | glio |
| Bora_in.vivo_noRT | Gm43689 | 2.47 | 1.73E-04 | glio |
| Bora_in.vivo_noRT | Hp1bp3 | -1.115 | 1.73E-04 | glio |
| Chek1_in.vivo_noRT | Syt17 | 2.979 | 1.73E-04 | glio |
| Chek1_in.vivo_noRT | Dock11 | 2.501 | 1.73E-04 | glio |
| Interaction_genotype_Mtor | Gm26673 | -5.011 | 1.73E-05 | glio |
| Interaction_genotype_Mtor | Gast | -4.099 | 1.73E-05 | glio |
| Aars2_in.vivo_noRT | Elf3 | 3.286 | 1.73E-05 | glio |
| Aars2_in.vivo_noRT | Gm20186 | 3.186 | 1.73E-05 | glio |
| Bcas2_in.vivo_noRT | Bfsp2 | 3.435 | 1.73E-05 | glio |
| Bcas2_in.vivo_noRT | E230029C05Ri | 2.935 | 1.73E-05 | glio |
| Bora_in.vivo_noRT | Gm26559 | 3.125 | 1.73E-05 | glio |
| Chek1_in.vivo_noRT | Arhgap33 | 3.104 | 1.73E-05 | glio |
| Chek1_in.vivo_noRT | Pcdhb21 | 2.8 | 1.73E-05 | glio |
| Hsd17b10_in.vivo_noRT | Pcdhga11 | 2.619 | 1.73E-05 | glio |
| Interaction_genotype_Mtor | Cers3 | -4.539 | 1.73E-06 | glio |
| Interaction_genotype_Myc | 2010106G04R | -5.611 | 1.73E-06 | glio |
| Aars2_in.vivo_noRT | Gm43221 | 3.155 | 1.73E-06 | glio |
| Atr_in.vivo_noRT | Gm5431 | 3.09 | 1.73E-06 | glio |
| Chek1_in.vivo_noRT | Hipk4 | 3.057 | 1.73E-06 | glio |
| culture_effect | Kazald1 | 1.192 | 1.73E-07 | glio |
| culture_effect | Orc2 | 0.758 | 1.73E-07 | glio |
| Atp5c1_in.vivo_noRT | Gm5415 | 2.668 | 1.73E-07 | glio |
| Atp5c1_in.vivo_noRT | Edn2 | 2.668 | 1.73E-07 | glio |
| Atp5c1_in.vivo_noRT | Gm14207 | 2.668 | 1.73E-07 | glio |
| Atr_in.vivo_noRT | Gm12454 | 3.629 | 1.73E-07 | glio |
| Atr_in.vivo_noRT | Mag | 3.434 | 1.73E-07 | glio |
| Bcas2_in.vivo_noRT | Zfp85os | 3.748 | 1.73E-07 | glio |
| Bcas2_in.vivo_noRT | Zkscan7 | 3.938 | 1.73E-08 | glio |
| Chek1_in.vivo_noRT | 1700023H06R | 4.253 | 1.73E-08 | glio |
| Aars2_in.vivo_noRT | Scgb1a1 | 4.217 | 1.73E-09 | glio |
| Bcas2_in.vivo_noRT | H3f3aos | 4.162 | 1.73E-09 | glio |
| Chek1_in.vivo_noRT | Syp | 4.653 | 1.73E-09 | glio |
| Chek1_ex.vivo_noRT | Pstpip2 | 4.049 | 1.73E-09 | glio |
| culture_effect | Ppp1r14b | 0.7 | 1.73E-10 | glio |
| Aars2_in.vivo_noRT | Gm16222 | 5.067 | 1.73E-10 | glio |
| Bora_in.vivo_noRT | AV039307 | 4.545 | 1.73E-10 | glio |
| Bora_in.vivo_noRT | Gm17638 | 4.144 | 1.73E-10 | glio |
| Chek1_ex.vivo_noRT | Zscan4d | 4.449 | 1.73E-10 | glio |
| Aars2_in.vivo_noRT | Gm30082 | 4.676 | 1.73E-11 | glio |
| Chek1_in.vivo_noRT | Gprin2 | 4.132 | 1.73E-12 | glio |
| Chek1_ex.vivo_noRT | Akr1c18 | 4.842 | 1.73E-12 | glio |
| Chek1_ex.vivo_noRT | Minar2 | 4.842 | 1.73E-12 | glio |
| Chek1_ex.vivo_noRT | Gm45151 | 4.842 | 1.73E-12 | glio |
| Aars2_in.vivo_noRT | Jph4 | 5.06 | 1.73E-13 | glio |
| Chek1_in.vivo_noRT | Muc4 | 4.522 | 1.73E-15 | glio |
| Atr_in.vivo_noRT | Cxcl13 | 5.533 | 1.73E-16 | glio |

| | | | | |
|---|---|---|---|---|
| Bora_in.vivo_noRT | 1700024I08Ri | 4.676 | 1.73E-16 | glio |
| Atr_in.vivo_noRT | Psca | 5.533 | 1.73E-18 | glio |
| Atr_in.vivo_noRT | 1700108F19Ri | 5.269 | 1.73E-20 | glio |
| culture_effect | Maged2 | -1.873 | 1.73E-25 | glio |
| culture_effect | 4933432I03Ri | -1.802 | 1.74E-03 | glio |
| culture_effect | Ceacam1 | -2.001 | 1.74E-03 | glio |
| culture_effect | D830050J10Ri | 1.395 | 1.74E-03 | glio |
| culture_effect | 8430429K09R | 1.094 | 1.74E-03 | glio |
| culture_effect | Dnase2a | 0.931 | 1.74E-03 | glio |
| culture_effect | Ttyh2 | 0.787 | 1.74E-03 | glio |
| Interaction_genotype_Aars2 | Herpud1 | 2.055 | 1.74E-03 | glio |
| Interaction_genotype_Aars2 | Cebpb | 1.861 | 1.74E-03 | glio |
| Interaction_genotype_Aars2 | Ndufa13 | 0.941 | 1.74E-03 | glio |
| Interaction_genotype_Atr | Gpc5 | -2.765 | 1.74E-03 | glio |
| Interaction_genotype_Atr | Ccnj | -4.058 | 1.74E-03 | glio |
| Interaction_genotype_Chek1 | Ciita | 3.508 | 1.74E-03 | glio |
| Interaction_genotype_Chek1 | Gbp10 | 4.063 | 1.74E-03 | glio |
| Interaction_genotype_Mtor | Cks1brt | -3.348 | 1.74E-03 | glio |
| Interaction_genotype_Mtor | Ahrr | -3.691 | 1.74E-03 | glio |
| Interaction_genotype_Mtor | Poll | -2.942 | 1.74E-03 | glio |
| Interaction_genotype_Mtor | Fzd1 | -3.217 | 1.74E-03 | glio |
| Interaction_genotype_Mtor | Gtf2e1 | -2.966 | 1.74E-03 | glio |
| Interaction_genotype_Myc | Kifc5b | -3.017 | 1.74E-03 | glio |
| Interaction_genotype_Myc | 1600014C10R | -2.77 | 1.74E-03 | glio |
| Interaction_genotype_Myc | Ckap2l | -2.809 | 1.74E-03 | glio |
| Aars2_in.vivo_noRT | Adcy9 | 2.027 | 1.74E-03 | glio |
| Aars2_in.vivo_noRT | Swsap1 | 1.965 | 1.74E-03 | glio |
| Aars2_in.vivo_noRT | Nectin3 | 1.562 | 1.74E-03 | glio |
| Aars2_in.vivo_noRT | Ywhag | -0.87 | 1.74E-03 | glio |
| Aars2_ex.vivo_noRT | Amdhd1 | 2.905 | 1.74E-03 | glio |
| Atp5c1_in.vivo_noRT | Efcab9 | 2.402 | 1.74E-03 | glio |
| Atp5c1_in.vivo_noRT | Elobl | 2.668 | 1.74E-03 | glio |
| Atr_in.vivo_noRT | Cep78 | 1.836 | 1.74E-03 | glio |
| Atr_in.vivo_noRT | Nudt21 | -0.843 | 1.74E-03 | glio |
| Atr_ex.vivo_noRT | H2-M10.1 | 2.577 | 1.74E-03 | glio |
| Atr_ex.vivo_noRT | Eef1d | -0.688 | 1.74E-03 | glio |
| Bard1_ex.vivo_noRT | Neurl3 | 3.213 | 1.74E-03 | glio |
| Bard1_ex.vivo_noRT | Gbp10 | 2.814 | 1.74E-03 | glio |
| Bcas2_in.vivo_noRT | Gm10800 | 2.497 | 1.74E-03 | glio |
| Bcas2_in.vivo_noRT | Gm43066 | 2.155 | 1.74E-03 | glio |
| Bcas2_in.vivo_noRT | S100a11 | -1.291 | 1.74E-03 | glio |
| Bcas2_ex.vivo_noRT | Ntn1 | 2.742 | 1.74E-03 | glio |
| Bcas2_ex.vivo_noRT | Pmch | 2.742 | 1.74E-03 | glio |
| Bcas2_ex.vivo_noRT | Olfr543 | 2.347 | 1.74E-03 | glio |
| Bora_in.vivo_noRT | Hist1h2bp | 2.412 | 1.74E-03 | glio |
| Bora_in.vivo_noRT | Fam129c | 2.268 | 1.74E-03 | glio |
| Bora_in.vivo_noRT | Gm43378 | 2.137 | 1.74E-03 | glio |
| Bora_in.vivo_noRT | Creb3l1 | 2.262 | 1.74E-03 | glio |
| Bora_in.vivo_noRT | Ccdc28b | 1.978 | 1.74E-03 | glio |
| Bora_in.vivo_noRT | Ccdc25 | -1.265 | 1.74E-03 | glio |
| Bora_in.vivo_noRT | Dph3 | -0.992 | 1.74E-03 | glio |
| Cenpj_ex.vivo_noRT | Gm20513 | 2.591 | 1.74E-03 | glio |

| | | | | |
|---|---|---|---|---|
| Chek1_in.vivo_noRT | Klhl23 | 2.11 | 1.74E-03 | glio |
| Chek1_in.vivo_noRT | Hdac10 | 1.987 | 1.74E-03 | glio |
| Chek1_in.vivo_noRT | Igsf8 | -1.695 | 1.74E-03 | glio |
| Chek1_in.vivo_noRT | Csnk2a1 | -0.973 | 1.74E-03 | glio |
| Chek1_ex.vivo_noRT | Camkk1 | 2.9 | 1.74E-03 | glio |
| Chek1_ex.vivo_noRT | Klhl24 | 1.814 | 1.74E-03 | glio |
| Hsd17b10_in.vivo_noRT | 6330408M09F | 2.575 | 1.74E-03 | glio |
| Hsd17b10_in.vivo_noRT | 5031434O11R | 2.739 | 1.74E-03 | glio |
| Hsd17b10_in.vivo_noRT | H60c | 2.554 | 1.74E-03 | glio |
| Hsd17b10_in.vivo_noRT | Ssr2 | -0.624 | 1.74E-03 | glio |
| Hsd17b10_in.vivo_noRT | Ppa1 | -0.665 | 1.74E-03 | glio |
| Interaction_genotype_Mtor | Prm3 | -3.937 | 1.74E-04 | glio |
| Interaction_genotype_Mtor | Gm7879 | -4.724 | 1.74E-04 | glio |
| Interaction_genotype_Myc | Gm20646 | -4.103 | 1.74E-04 | glio |
| Interaction_genotype_Myc | AY074887 | -4.529 | 1.74E-04 | glio |
| Interaction_genotype_Myc | Hrct1 | -3.704 | 1.74E-04 | glio |
| Interaction_genotype_Myc | Ube2i | 1.51 | 1.74E-04 | glio |
| Aars2_in.vivo_noRT | Srsf6 | -1.165 | 1.74E-04 | glio |
| Atr_in.vivo_noRT | Rpl19 | -0.734 | 1.74E-04 | glio |
| Bcas2_in.vivo_noRT | Galk1 | -1.245 | 1.74E-04 | glio |
| Bora_in.vivo_noRT | Crtap | -1.584 | 1.74E-04 | glio |
| Eif2s1_in.vivo_noRT | Rab27a | 2.678 | 1.74E-04 | glio |
| culture_effect | Rassf6 | 2.099 | 1.74E-05 | glio |
| Interaction_genotype_Aars2 | Srsf5 | 1.949 | 1.74E-05 | glio |
| Interaction_genotype_Aars2 | Gm29340 | -5.028 | 1.74E-05 | glio |
| Interaction_genotype_Atr | Crem | 2.925 | 1.74E-05 | glio |
| Interaction_genotype_Myc | 4930432K21R | -4.553 | 1.74E-05 | glio |
| Interaction_genotype_Myc | Cpne8 | -5.062 | 1.74E-05 | glio |
| Aars2_in.vivo_noRT | Samd15 | 3.305 | 1.74E-05 | glio |
| Aars2_in.vivo_noRT | Gm42937 | 3.017 | 1.74E-05 | glio |
| Aars2_ex.vivo_noRT | Vmn2r54 | 3.293 | 1.74E-05 | glio |
| Bard1_ex.vivo_noRT | Kcnk13 | 3.199 | 1.74E-05 | glio |
| Bcas2_ex.vivo_noRT | Krt19 | 2.742 | 1.74E-05 | glio |
| Hsd17b10_in.vivo_noRT | Ksr2 | 2.578 | 1.74E-05 | glio |
| Hsd17b10_in.vivo_noRT | Gm15234 | 3.1 | 1.74E-05 | glio |
| culture_effect | Cdc37l1 | 0.697 | 1.74E-06 | glio |
| Interaction_genotype_Mtor | Alox12 | -5.666 | 1.74E-06 | glio |
| Interaction_genotype_Mtor | Gm15445 | -4.438 | 1.74E-06 | glio |
| Interaction_genotype_Mtor | Gm44646 | -4.685 | 1.74E-06 | glio |
| Aars2_in.vivo_noRT | Nr1h4 | 3.278 | 1.74E-06 | glio |
| Atp5c1_in.vivo_noRT | Prap1 | 2.668 | 1.74E-06 | glio |
| Atr_in.vivo_noRT | Dock6 | 3.276 | 1.74E-06 | glio |
| Atr_in.vivo_noRT | Ankrd23 | 3.529 | 1.74E-06 | glio |
| Hsd17b10_in.vivo_noRT | A530013C23R | 3.229 | 1.74E-06 | glio |
| culture_effect | Cyren | 1.409 | 1.74E-07 | glio |
| Interaction_genotype_Mtor | Car14 | -6.092 | 1.74E-07 | glio |
| Interaction_genotype_Mtor | Gm48155 | -4.099 | 1.74E-07 | glio |
| Interaction_genotype_Mtor | Eef2 | 1.57 | 1.74E-07 | glio |
| Atr_in.vivo_noRT | Tead3 | 3.595 | 1.74E-07 | glio |
| Bcas2_in.vivo_noRT | Pla2g7 | 3.62 | 1.74E-07 | glio |
| Bcas2_in.vivo_noRT | Scgb1a1 | 3.697 | 1.74E-07 | glio |
| Atr_in.vivo_noRT | Gm48529 | 3.804 | 1.74E-08 | glio |

| | | | | |
|---|---|---|---|---|
| culture_effect | Ntrk1 | -3.551 | 1.74E-09 | glio |
| Atr_in.vivo_noRT | Gm13283 | 4.179 | 1.74E-09 | glio |
| Atr_in.vivo_noRT | 1700124L16Ri | 4.299 | 1.74E-09 | glio |
| Aars2_in.vivo_noRT | Gm16094 | 4.022 | 1.74E-10 | glio |
| Chek1_ex.vivo_noRT | Gm16094 | 5.056 | 1.74E-10 | glio |
| Hsd17b10_in.vivo_noRT | Npr3 | 4.147 | 1.74E-11 | glio |
| Chek1_in.vivo_noRT | Hunk | 4.511 | 1.74E-12 | glio |
| Bcas2_in.vivo_noRT | Pnliprp2 | 4.028 | 1.74E-13 | glio |
| Bora_in.vivo_noRT | Impg2 | 4.936 | 1.74E-17 | glio |
| Aars2_in.vivo_noRT | Crhbp | 5.327 | 1.74E-18 | glio |
| Chek1_in.vivo_noRT | Plcl1 | 5.044 | 1.74E-18 | glio |
| Bora_in.vivo_noRT | Gm5890 | 4.936 | 1.74E-21 | glio |
| Atr_in.vivo_noRT | Klk13 | 5.001 | 1.74E-22 | glio |
| Chek1_in.vivo_noRT | Slc35g3 | 5.044 | 1.74E-23 | glio |
| Chek1_in.vivo_noRT | Vmn2r18 | 5.044 | 1.74E-24 | glio |
| culture_effect | Ero1l | -2.466 | 1.74E-26 | glio |
| culture_effect | 1810041H14R | 1.883 | 1.75E-03 | glio |
| culture_effect | Dok4 | -1.686 | 1.75E-03 | glio |
| culture_effect | Brip1os | -0.665 | 1.75E-03 | glio |
| culture_effect | Rufy1 | -0.647 | 1.75E-03 | glio |
| culture_effect | Fam98b | 0.565 | 1.75E-03 | glio |
| culture_effect | Rnf11 | 0.539 | 1.75E-03 | glio |
| Interaction_genotype_Aars2 | Zfp810 | -3.816 | 1.75E-03 | glio |
| Interaction_genotype_Aars2 | Slc2a9 | -3.231 | 1.75E-03 | glio |
| Interaction_genotype_Aars2 | Trnau1ap | 1.386 | 1.75E-03 | glio |
| Interaction_genotype_Atr | Anxa11 | -4.314 | 1.75E-03 | glio |
| Interaction_genotype_Atr | Fbxo31 | -2.251 | 1.75E-03 | glio |
| Interaction_genotype_Chek1 | Man2c1os | -4.335 | 1.75E-03 | glio |
| Interaction_genotype_Mtor | Rtel1 | -2.482 | 1.75E-03 | glio |
| Interaction_genotype_Myc | Ntn3 | -3.233 | 1.75E-03 | glio |
| Interaction_genotype_Rpl3 | A730071L15Ri | -4.485 | 1.75E-03 | glio |
| Interaction_genotype_Rpl3 | AV039307 | -4.378 | 1.75E-03 | glio |
| Interaction_genotype_Rpl3 | Gm20492 | -4.166 | 1.75E-03 | glio |
| Interaction_genotype_Rpl3 | Mapk3 | 2.139 | 1.75E-03 | glio |
| Aars2_in.vivo_noRT | Gm26631 | 2.362 | 1.75E-03 | glio |
| Aars2_in.vivo_noRT | Pcdhb14 | 2.293 | 1.75E-03 | glio |
| Aars2_in.vivo_noRT | Me2 | -1.362 | 1.75E-03 | glio |
| Aars2_in.vivo_noRT | Sec11c | -1.041 | 1.75E-03 | glio |
| Aars2_ex.vivo_noRT | Gipc2 | 3.223 | 1.75E-03 | glio |
| Aars2_ex.vivo_noRT | Sept5 | 2.989 | 1.75E-03 | glio |
| Atr_in.vivo_noRT | Drc3 | 2.303 | 1.75E-03 | glio |
| Atr_in.vivo_noRT | Zfp82 | 2.195 | 1.75E-03 | glio |
| Atr_in.vivo_noRT | Tnfsf12 | 1.852 | 1.75E-03 | glio |
| Atr_in.vivo_noRT | Tirap | 1.653 | 1.75E-03 | glio |
| Atr_ex.vivo_noRT | Slc6a13 | 3.107 | 1.75E-03 | glio |
| Bard1_ex.vivo_noRT | Dio1 | 2.814 | 1.75E-03 | glio |
| Bcas2_in.vivo_noRT | Serp2 | 2.07 | 1.75E-03 | glio |
| Bcas2_in.vivo_noRT | Wdr70 | -1.14 | 1.75E-03 | glio |
| Bora_in.vivo_noRT | Crabp1 | 2.305 | 1.75E-03 | glio |
| Bora_in.vivo_noRT | Zfp729b | 1.989 | 1.75E-03 | glio |
| Bora_in.vivo_noRT | Igfbp4 | -2.262 | 1.75E-03 | glio |
| Chek1_in.vivo_noRT | B230312C02R | 2.923 | 1.75E-03 | glio |

| | | | | |
|---|---|---|---|---|
| Eif2s1_in.vivo_noRT | Spink8 | 1.883 | 1.75E-03 | glio |
| culture_effect | Arap2 | -1.201 | 1.75E-04 | glio |
| culture_effect | Sdc4 | 1.171 | 1.75E-04 | glio |
| culture_effect | Nedd4 | -0.362 | 1.75E-04 | glio |
| Interaction_genotype_Mtor | Krtcap3 | -3.5 | 1.75E-04 | glio |
| Interaction_genotype_Mtor | Enah | -4.198 | 1.75E-04 | glio |
| Interaction_genotype_Myc | Ndufa10 | 1.615 | 1.75E-04 | glio |
| Aars2_ex.vivo_noRT | Gm13398 | 2.495 | 1.75E-04 | glio |
| Atp5c1_in.vivo_noRT | Slc25a18 | 2.402 | 1.75E-04 | glio |
| Atr_in.vivo_noRT | Lncppara | 2.859 | 1.75E-04 | glio |
| Atr_in.vivo_noRT | BC065397 | 2.947 | 1.75E-04 | glio |
| Atr_in.vivo_noRT | Ppfia3 | 2.567 | 1.75E-04 | glio |
| Bard1_ex.vivo_noRT | Crb3 | 2.814 | 1.75E-04 | glio |
| Bard1_ex.vivo_noRT | Gm21936 | 2.419 | 1.75E-04 | glio |
| Bard1_ex.vivo_noRT | Thnsl2 | 2.814 | 1.75E-04 | glio |
| Bard1_ex.vivo_noRT | 4930588A03R | 2.415 | 1.75E-04 | glio |
| Bcas2_in.vivo_noRT | Zfp992 | 2.779 | 1.75E-04 | glio |
| Bcas2_in.vivo_noRT | Reck | 2.714 | 1.75E-04 | glio |
| Bcas2_in.vivo_noRT | Maob | 2.594 | 1.75E-04 | glio |
| Bcas2_in.vivo_noRT | Fbln5 | 1.51 | 1.75E-04 | glio |
| Bora_in.vivo_noRT | St18 | 2.097 | 1.75E-04 | glio |
| Chek1_in.vivo_noRT | Batf | 2.673 | 1.75E-04 | glio |
| Chek1_in.vivo_noRT | Osbpl7 | 2.399 | 1.75E-04 | glio |
| Chek1_ex.vivo_noRT | Fbxl2 | 2.961 | 1.75E-04 | glio |
| Eif2s1_in.vivo_noRT | Pygo1 | 2.834 | 1.75E-04 | glio |
| Interaction_genotype_Mtor | 2810414N06R | -5.067 | 1.75E-05 | glio |
| Aars2_in.vivo_noRT | Dock4 | 3.046 | 1.75E-05 | glio |
| Atr_in.vivo_noRT | Ddx11 | 2.181 | 1.75E-05 | glio |
| Atr_ex.vivo_noRT | Gata3 | 4.268 | 1.75E-05 | glio |
| Bcas2_in.vivo_noRT | Zeb2os | 2.894 | 1.75E-05 | glio |
| Bcas2_in.vivo_noRT | Gm17259 | 2.874 | 1.75E-05 | glio |
| Bcas2_in.vivo_noRT | Prune1 | -1.75 | 1.75E-05 | glio |
| Bora_in.vivo_noRT | Hyal1 | 2.972 | 1.75E-05 | glio |
| Chek1_ex.vivo_noRT | Selenov | 3.072 | 1.75E-05 | glio |
| Chek1_ex.vivo_noRT | Tcam1 | 3.957 | 1.75E-05 | glio |
| Chek1_ex.vivo_noRT | Txnip | 4.397 | 1.75E-05 | glio |
| culture_effect | Wls | -0.839 | 1.75E-06 | glio |
| culture_effect | Zfp385b | -0.765 | 1.75E-06 | glio |
| Interaction_genotype_Mtor | Acoxl | -5.293 | 1.75E-06 | glio |
| Interaction_genotype_Myc | Atp5g1 | 1.965 | 1.75E-06 | glio |
| Aars2_in.vivo_noRT | Hmgcs2 | 3.78 | 1.75E-06 | glio |
| Atr_in.vivo_noRT | Glt8d2 | 3.482 | 1.75E-06 | glio |
| Atr_in.vivo_noRT | 1110013H19R | 3.661 | 1.75E-06 | glio |
| Atr_in.vivo_noRT | 8030456M14F | 3.38 | 1.75E-06 | glio |
| Bcas2_in.vivo_noRT | Gm26850 | 3.413 | 1.75E-06 | glio |
| Bcas2_in.vivo_noRT | Gm4755 | 3.05 | 1.75E-06 | glio |
| Bora_in.vivo_noRT | Dlx2 | 3.553 | 1.75E-06 | glio |
| Interaction_genotype_Mtor | Hist2h4 | -5.978 | 1.75E-07 | glio |
| Aars2_in.vivo_noRT | 1700055D18R | 3.937 | 1.75E-07 | glio |
| Aars2_in.vivo_noRT | Gnas | -1.029 | 1.75E-07 | glio |
| Bcas2_in.vivo_noRT | Cnih2 | 3.619 | 1.75E-07 | glio |
| Chek1_in.vivo_noRT | 4930558J18Ri | 3.438 | 1.75E-07 | glio |

| | | | | |
|---|---|---|---|---|
| Aars2_in.vivo_noRT | Amhr2 | 4.45 | 1.75E-08 | glio |
| Aars2_in.vivo_noRT | E130208F15Ri | 4.274 | 1.75E-08 | glio |
| Aars2_in.vivo_noRT | Gm46447 | 3.516 | 1.75E-08 | glio |
| Aars2_in.vivo_noRT | Gm28592 | 3.649 | 1.75E-08 | glio |
| Chek1_in.vivo_noRT | Pax5 | 4.199 | 1.75E-08 | glio |
| Aars2_in.vivo_noRT | Gm50334 | 4.682 | 1.75E-09 | glio |
| Atr_in.vivo_noRT | 4932441J04Ri | 4.283 | 1.75E-09 | glio |
| Chek1_in.vivo_noRT | Stk36 | 4.285 | 1.75E-09 | glio |
| Aars2_in.vivo_noRT | Ltb | 4.267 | 1.75E-10 | glio |
| Atr_ex.vivo_noRT | Slc14a2 | 3.947 | 1.75E-11 | glio |
| culture_effect | Lrrc14 | 1.316 | 1.75E-12 | glio |
| culture_effect | Clip1 | 1.965 | 1.75E-14 | glio |
| culture_effect | Il7 | -4.45 | 1.75E-15 | glio |
| Atr_in.vivo_noRT | Nat8f4 | 5.001 | 1.75E-15 | glio |
| Atr_in.vivo_noRT | 1700026N04R | 5.273 | 1.75E-17 | glio |
| Atr_in.vivo_noRT | Fbxo15 | 5.27 | 1.75E-17 | glio |
| Chek1_ex.vivo_noRT | Trim17 | 5.564 | 1.75E-19 | glio |
| Chek1_ex.vivo_noRT | Meioc | 5.611 | 1.75E-19 | glio |
| culture_effect | Rps6 | -1.106 | 1.75E-21 | glio |
| culture_effect | Psmb11 | -1.457 | 1.76E-03 | glio |
| culture_effect | Ube2t | -0.715 | 1.76E-03 | glio |
| culture_effect | Gnaq | 0.575 | 1.76E-03 | glio |
| Interaction_genotype_Aars2 | Hddc3 | -3.647 | 1.76E-03 | glio |
| Interaction_genotype_Atr | Gm45669 | -3.404 | 1.76E-03 | glio |
| Interaction_genotype_Atr | Slc35g3 | -2.769 | 1.76E-03 | glio |
| Interaction_genotype_Atr | Gm3604 | -3.972 | 1.76E-03 | glio |
| Interaction_genotype_Atr | B230317F23Ri | -3.552 | 1.76E-03 | glio |
| Interaction_genotype_Atr | Nipsnap3b | 1.747 | 1.76E-03 | glio |
| Interaction_genotype_Bcas2 | Fa2h | 3.986 | 1.76E-03 | glio |
| Interaction_genotype_Bora | 6330403L08Ri | -3.912 | 1.76E-03 | glio |
| Interaction_genotype_Bora | Gm44702 | -4.224 | 1.76E-03 | glio |
| Interaction_genotype_Bora | Mtmr2 | 2.2 | 1.76E-03 | glio |
| Interaction_genotype_Chek1 | Gm13283 | -4.135 | 1.76E-03 | glio |
| Interaction_genotype_Chek1 | Cspg5 | 3.462 | 1.76E-03 | glio |
| Interaction_genotype_Chek1 | Rab12 | 2.002 | 1.76E-03 | glio |
| Interaction_genotype_Mtor | Mlip | -3.857 | 1.76E-03 | glio |
| Interaction_genotype_Mtor | Gm5577 | -3.285 | 1.76E-03 | glio |
| Interaction_genotype_Myc | Cldn5 | -2.463 | 1.76E-03 | glio |
| Interaction_genotype_Myc | H60c | -3.798 | 1.76E-03 | glio |
| Interaction_genotype_Myc | Gm11361 | -3.247 | 1.76E-03 | glio |
| Interaction_genotype_Myc | Eme1 | -2.734 | 1.76E-03 | glio |
| Interaction_genotype_Myc | Zfp532 | -2.7 | 1.76E-03 | glio |
| Interaction_genotype_Myc | Hsp90ab1 | 1.284 | 1.76E-03 | glio |
| Aars2_in.vivo_noRT | Cntd1 | 2.252 | 1.76E-03 | glio |
| Aars2_in.vivo_noRT | Sun1 | -1.456 | 1.76E-03 | glio |
| Aars2_in.vivo_noRT | Atpsckmt | -1.335 | 1.76E-03 | glio |
| Atr_in.vivo_noRT | Nhlrc3 | 1.854 | 1.76E-03 | glio |
| Atr_in.vivo_noRT | Ankrd11 | -1.06 | 1.76E-03 | glio |
| Bcas2_in.vivo_noRT | Gm17501 | 2.384 | 1.76E-03 | glio |
| Bcas2_in.vivo_noRT | Gm28221 | 2.138 | 1.76E-03 | glio |
| Bora_in.vivo_noRT | Gm29994 | 2.059 | 1.76E-03 | glio |
| Cenpj_ex.vivo_noRT | Gm43948 | 2.605 | 1.76E-03 | glio |

| | | | | |
|---|---|---|---|---|
| Chek1_in.vivo_noRT | Card14 | 2.332 | 1.76E-03 | glio |
| Chek1_in.vivo_noRT | Naca | -0.535 | 1.76E-03 | glio |
| Hsd17b10_in.vivo_noRT | Mamdc2 | 2.306 | 1.76E-03 | glio |
| Hsd17b10_in.vivo_noRT | Mpeg1 | 2.172 | 1.76E-03 | glio |
| culture_effect | Upp2 | 1.461 | 1.76E-04 | glio |
| culture_effect | Pnma5 | -0.924 | 1.76E-04 | glio |
| culture_effect | Map2k1 | -0.63 | 1.76E-04 | glio |
| culture_effect | Chpt1 | -0.575 | 1.76E-04 | glio |
| Interaction_genotype_Bora | Vps4b | 2.134 | 1.76E-04 | glio |
| Interaction_genotype_Mtor | Des | -4.605 | 1.76E-04 | glio |
| Interaction_genotype_Mtor | Tspan33 | -4.557 | 1.76E-04 | glio |
| Interaction_genotype_Mtor | Tmem79 | -3.984 | 1.76E-04 | glio |
| Aars2_in.vivo_noRT | Atp6v1g2 | 2.456 | 1.76E-04 | glio |
| Aars2_ex.vivo_noRT | Irf8 | 3.293 | 1.76E-04 | glio |
| Atr_in.vivo_noRT | Prdm1 | 2.467 | 1.76E-04 | glio |
| Bcas2_in.vivo_noRT | Cdrt4 | 2.832 | 1.76E-04 | glio |
| Bcas2_in.vivo_noRT | Rhov | 2.621 | 1.76E-04 | glio |
| Bcas2_in.vivo_noRT | Pcdhb19 | 2.515 | 1.76E-04 | glio |
| Bcas2_in.vivo_noRT | Kpna3 | -1.761 | 1.76E-04 | glio |
| Chek1_in.vivo_noRT | Plek2 | 2.593 | 1.76E-04 | glio |
| Hsd17b10_in.vivo_noRT | Spint4 | 2.825 | 1.76E-04 | glio |
| Hsd17b10_in.vivo_noRT | Slc23a3 | 2.714 | 1.76E-04 | glio |
| Hsd17b10_in.vivo_noRT | 9930111J21Ri | 2.436 | 1.76E-04 | glio |
| Interaction_genotype_Mtor | Arhgap22 | -4.891 | 1.76E-05 | glio |
| Interaction_genotype_Mtor | Gask1b | -5.47 | 1.76E-05 | glio |
| interaction_RT | Egr2 | 3.206 | 1.76E-05 | glio |
| interaction_RT | 1110017D15R | 4.044 | 1.76E-05 | glio |
| Aars2_in.vivo_noRT | Ppm1f | 2.492 | 1.76E-05 | glio |
| Aars2_ex.vivo_noRT | Aqp9 | 3.293 | 1.76E-05 | glio |
| Atr_in.vivo_noRT | Gm29107 | 3.096 | 1.76E-05 | glio |
| Bora_in.vivo_noRT | Gm4181 | 3.056 | 1.76E-05 | glio |
| Bora_in.vivo_noRT | Tmem88 | 2.978 | 1.76E-05 | glio |
| Chek1_ex.vivo_noRT | Cd34 | 3.936 | 1.76E-05 | glio |
| Hsd17b10_in.vivo_noRT | Accsl | 3.356 | 1.76E-05 | glio |
| culture_effect | 1110020A21R | 2.578 | 1.76E-06 | glio |
| Interaction_genotype_Chek1 | Apol6 | 5.143 | 1.76E-06 | glio |
| Interaction_genotype_Mtor | B3gnt4 | -5.021 | 1.76E-06 | glio |
| Interaction_genotype_Myc | BC065403 | -4.983 | 1.76E-06 | glio |
| Interaction_genotype_Myc | Gm17259 | -5.08 | 1.76E-06 | glio |
| Interaction_genotype_Myc | Loxl4 | -5.098 | 1.76E-06 | glio |
| interaction_RT | Marcksl1 | 1.19 | 1.76E-06 | glio |
| Aars2_in.vivo_noRT | Gm26930 | 3.637 | 1.76E-06 | glio |
| Aars2_in.vivo_noRT | Gm14636 | 4.038 | 1.76E-06 | glio |
| Aars2_ex.vivo_noRT | Tpo | 3.293 | 1.76E-07 | glio |
| Atp5c1_in.vivo_noRT | Gm49698 | 2.668 | 1.76E-07 | glio |
| Atp5c1_in.vivo_noRT | Olfr881 | 2.668 | 1.76E-07 | glio |
| Bcas2_in.vivo_noRT | Gm49767 | 3.769 | 1.76E-07 | glio |
| Bcas2_in.vivo_noRT | Kctd4 | 3.534 | 1.76E-07 | glio |
| Bcas2_in.vivo_noRT | Wscd1 | -2.33 | 1.76E-07 | glio |
| Chek1_in.vivo_noRT | Icosl | 3.277 | 1.76E-07 | glio |
| culture_effect | Hist4h4 | 2.621 | 1.76E-08 | glio |
| Atr_in.vivo_noRT | Gm15510 | 3.939 | 1.76E-08 | glio |

| | | | | |
|---|---|---|---|---|
| Bcas2_in.vivo_noRT | Qrich2 | 4.029 | 1.76E-08 | glio |
| Atr_in.vivo_noRT | Plppr3 | 4.147 | 1.76E-09 | glio |
| Atr_in.vivo_noRT | EU599041 | 4.256 | 1.76E-09 | glio |
| Bora_in.vivo_noRT | Dhrs9 | 3.874 | 1.76E-09 | glio |
| Atr_in.vivo_noRT | Cyp27b1 | 3.972 | 1.76E-10 | glio |
| Bora_in.vivo_noRT | Gm5152 | 4.549 | 1.76E-10 | glio |
| Chek1_in.vivo_noRT | Gm44672 | 4.279 | 1.76E-10 | glio |
| culture_effect | Ocm | -3.776 | 1.76E-11 | glio |
| Bora_in.vivo_noRT | Gm49874 | 4.936 | 1.76E-11 | glio |
| Chek1_in.vivo_noRT | Gm15127 | 4.784 | 1.76E-12 | glio |
| Hsd17b10_in.vivo_noRT | Gpr4 | 3.616 | 1.76E-12 | glio |
| Hsd17b10_in.vivo_noRT | Prl7c1 | 3.616 | 1.76E-12 | glio |
| Hsd17b10_in.vivo_noRT | Gm33466 | 3.616 | 1.76E-12 | glio |
| Bora_in.vivo_noRT | Gm5662 | 4.936 | 1.76E-13 | glio |
| Chek1_ex.vivo_noRT | Dmrtc1a | 4.842 | 1.76E-13 | glio |
| Chek1_in.vivo_noRT | Col19a1 | 4.127 | 1.76E-14 | glio |
| culture_effect | Hnrnpa1 | -0.952 | 1.76E-15 | glio |
| Aars2_in.vivo_noRT | Gm47851 | 5.067 | 1.76E-15 | glio |
| Atr_in.vivo_noRT | Gm12860 | 4.525 | 1.76E-16 | glio |
| Chek1_ex.vivo_noRT | Gm15908 | 4.842 | 1.76E-17 | glio |
| Chek1_ex.vivo_noRT | A930014E10Ri | 4.842 | 1.76E-17 | glio |
| culture_effect | Arl5a | 1.306 | 1.76E-18 | glio |
| Bora_in.vivo_noRT | Zscan4c | 4.936 | 1.76E-21 | glio |
| Aars2_in.vivo_noRT | Gm5890 | 5.327 | 1.76E-24 | glio |
| Bora_in.vivo_noRT | Krt19 | 4.936 | 1.76E-25 | glio |
| Hsd17b10_in.vivo_noRT | Hsd17b10 | -5.098 | 1.76E-30 | glio |
| Interaction_genotype_Atr | A930012O16R | -4.012 | 1.77E-03 | glio |
| Interaction_genotype_Atr | Gm14305 | -3.261 | 1.77E-03 | glio |
| Interaction_genotype_Mtor | Slc25a43 | -3.82 | 1.77E-03 | glio |
| Interaction_genotype_Mtor | Cfap20 | 1.487 | 1.77E-03 | glio |
| Interaction_genotype_Myc | Hes5 | 4.414 | 1.77E-03 | glio |
| interaction_RT | Naga | 1.557 | 1.77E-03 | glio |
| interaction_RT | Sec24d | 1.344 | 1.77E-03 | glio |
| interaction_RT | Eif2b5 | -1.053 | 1.77E-03 | glio |
| interaction_RT | Prrg1 | 3.155 | 1.77E-03 | glio |
| interaction_RT | Pstk | -0.939 | 1.77E-03 | glio |
| interaction_RT | Pnn | -0.724 | 1.77E-03 | glio |
| Aars2_in.vivo_noRT | Fam189b | 2.319 | 1.77E-03 | glio |
| Atp5c1_in.vivo_noRT | Scd1 | -1.576 | 1.77E-03 | glio |
| Atr_in.vivo_noRT | Pyroxd2 | 2.077 | 1.77E-03 | glio |
| Atr_in.vivo_noRT | Pcdhb22 | 2.112 | 1.77E-03 | glio |
| Atr_in.vivo_noRT | Cerk | 1.908 | 1.77E-03 | glio |
| Atr_in.vivo_noRT | Bckdha | -1.433 | 1.77E-03 | glio |
| Atr_ex.vivo_noRT | Gm45235 | 2.37 | 1.77E-03 | glio |
| Bcas2_in.vivo_noRT | Gm8773 | 2.433 | 1.77E-03 | glio |
| Bcas2_in.vivo_noRT | Zfp345 | 1.572 | 1.77E-03 | glio |
| Bcas2_in.vivo_noRT | Uhrf1 | -1.445 | 1.77E-03 | glio |
| Bora_in.vivo_noRT | Mgat1 | -1.462 | 1.77E-03 | glio |
| Bora_in.vivo_noRT | Nop2 | -1.188 | 1.77E-03 | glio |
| Bora_in.vivo_noRT | Phlda3 | -1.211 | 1.77E-03 | glio |
| Cenpj_in.vivo_noRT | Smtnl1 | 2.022 | 1.77E-03 | glio |
| Cenpj_ex.vivo_noRT | Ccl9 | 2.198 | 1.77E-03 | glio |

| | | | | |
|---|---|---|---|---|
| Chek1_in.vivo_noRT | Senp6 | -1.208 | 1.77E-03 | glio |
| Chek1_in.vivo_noRT | Nans | -1.093 | 1.77E-03 | glio |
| Chek1_ex.vivo_noRT | Itgb7 | 2.763 | 1.77E-03 | glio |
| Eif2s1_in.vivo_noRT | Vtn | 1.883 | 1.77E-03 | glio |
| Hsd17b10_in.vivo_noRT | Cxcr2 | 2.304 | 1.77E-03 | glio |
| Hsd17b10_in.vivo_noRT | Plekha7 | -1.83 | 1.77E-03 | glio |
| culture_effect | Trip6 | 0.654 | 1.77E-04 | glio |
| culture_effect | Ufc1 | -0.448 | 1.77E-04 | glio |
| Interaction_genotype_Aars2 | Rpl4 | 1.105 | 1.77E-04 | glio |
| Interaction_genotype_Mtor | Ttc16 | -4.38 | 1.77E-04 | glio |
| Interaction_genotype_Mtor | Myl9 | -4.654 | 1.77E-04 | glio |
| Interaction_genotype_Myc | Klhl26 | -3.81 | 1.77E-04 | glio |
| Interaction_genotype_Myc | Cavin3 | 1.856 | 1.77E-04 | glio |
| Interaction_genotype_Myc | Pdap1 | 1.523 | 1.77E-04 | glio |
| Aars2_in.vivo_noRT | 1810058I24Ri | -1.153 | 1.77E-04 | glio |
| Aars2_in.vivo_noRT | Srsf11 | -1.013 | 1.77E-04 | glio |
| Aars2_ex.vivo_noRT | Lhx1os | 2.897 | 1.77E-04 | glio |
| Aars2_ex.vivo_noRT | Dtx1 | 3.676 | 1.77E-04 | glio |
| Atp5c1_in.vivo_noRT | Acoxl | 2.668 | 1.77E-04 | glio |
| Atr_in.vivo_noRT | Gm8126 | 2.401 | 1.77E-04 | glio |
| Atr_ex.vivo_noRT | Dio1 | 3.164 | 1.77E-04 | glio |
| Atr_ex.vivo_noRT | Prrx2 | 3.226 | 1.77E-04 | glio |
| Bcas2_in.vivo_noRT | Fgfr1 | 3.17 | 1.77E-04 | glio |
| Bcas2_in.vivo_noRT | AA386476 | 2.777 | 1.77E-04 | glio |
| Bcas2_in.vivo_noRT | Ctrl | 2.788 | 1.77E-04 | glio |
| Bcas2_in.vivo_noRT | Timm8a1 | -1.483 | 1.77E-04 | glio |
| Bora_in.vivo_noRT | Ccn2 | 3.609 | 1.77E-04 | glio |
| Bora_in.vivo_noRT | Agfg1 | -1.369 | 1.77E-04 | glio |
| Eif2s1_in.vivo_noRT | Olfr552 | 2.143 | 1.77E-04 | glio |
| culture_effect | Pgm2 | -0.935 | 1.77E-05 | glio |
| Interaction_genotype_Chek1 | Pnrc1 | 2.032 | 1.77E-05 | glio |
| Interaction_genotype_Mtor | Zic4 | -5.32 | 1.77E-05 | glio |
| Interaction_genotype_Mtor | A530013C23R | -4.861 | 1.77E-05 | glio |
| Interaction_genotype_Myc | Gm46218 | -4.905 | 1.77E-05 | glio |
| Interaction_genotype_Myc | Gm43581 | -4.544 | 1.77E-05 | glio |
| Atp5c1_in.vivo_noRT | Selenov | 2.668 | 1.77E-05 | glio |
| Atp5c1_in.vivo_noRT | A330076H08R | 2.668 | 1.77E-05 | glio |
| Atr_ex.vivo_noRT | Gm20513 | 3.164 | 1.77E-05 | glio |
| Bcas2_in.vivo_noRT | Prdm10 | 2.67 | 1.77E-05 | glio |
| Bcas2_ex.vivo_noRT | Gm48155 | 2.742 | 1.77E-05 | glio |
| Bora_in.vivo_noRT | Gm49479 | 3.098 | 1.77E-05 | glio |
| Bora_ex.vivo_noRT | Gm20513 | 3.201 | 1.77E-05 | glio |
| Bora_ex.vivo_noRT | Olfr552 | 2.802 | 1.77E-05 | glio |
| Chek1_in.vivo_noRT | 4930589L23Ri | 3.165 | 1.77E-05 | glio |
| Chek1_in.vivo_noRT | Tmsb15b2 | 3.323 | 1.77E-05 | glio |
| Chek1_in.vivo_noRT | Myorg | 2.974 | 1.77E-05 | glio |
| Hsd17b10_in.vivo_noRT | Wdr93 | 2.96 | 1.77E-05 | glio |
| Aars2_in.vivo_noRT | Gm37986 | 2.989 | 1.77E-06 | glio |
| Atr_ex.vivo_noRT | Psmb11 | 3.164 | 1.77E-06 | glio |
| Chek1_ex.vivo_noRT | Acot5 | 4.049 | 1.77E-06 | glio |
| Hsd17b10_in.vivo_noRT | Gm15127 | 3.356 | 1.77E-06 | glio |
| Hsd17b10_in.vivo_noRT | Fgf1 | 3.598 | 1.77E-06 | glio |

| | | | | |
|---|---|---|---|---|
| Interaction_genotype_Mtor | Gm32250 | -4.69 | 1.77E-07 | glio |
| Interaction_genotype_Myc | C030037D09R | -5.988 | 1.77E-07 | glio |
| Aars2_in.vivo_noRT | Gm44836 | 3.495 | 1.77E-07 | glio |
| Atr_in.vivo_noRT | Icam5 | 3.511 | 1.77E-07 | glio |
| Chek1_in.vivo_noRT | Gm49891 | 3.846 | 1.77E-07 | glio |
| Atr_in.vivo_noRT | Olfr920 | 3.783 | 1.77E-08 | glio |
| Chek1_in.vivo_noRT | Olfm2 | 4.136 | 1.77E-08 | glio |
| Chek1_in.vivo_noRT | Zfp972 | 4.111 | 1.77E-08 | glio |
| Atr_in.vivo_noRT | Dchs1 | 4.039 | 1.77E-09 | glio |
| Atr_in.vivo_noRT | Fam169a | 3.979 | 1.77E-09 | glio |
| Chek1_in.vivo_noRT | Ifitm1 | 4.777 | 1.77E-09 | glio |
| Chek1_ex.vivo_noRT | Myzap | 4.443 | 1.77E-09 | glio |
| Chek1_ex.vivo_noRT | Kcnq2 | 4.842 | 1.77E-10 | glio |
| Aars2_in.vivo_noRT | 4933429H19R | 4.415 | 1.77E-11 | glio |
| Aars2_in.vivo_noRT | Gm43137 | 4.811 | 1.77E-11 | glio |
| Aars2_in.vivo_noRT | Slc6a13 | 4.947 | 1.77E-11 | glio |
| Bcas2_in.vivo_noRT | 5430402O13R | 4.807 | 1.77E-11 | glio |
| Bcas2_in.vivo_noRT | Sbk3 | 4.416 | 1.77E-11 | glio |
| Atr_in.vivo_noRT | 9530062K07R | 5.154 | 1.77E-12 | glio |
| Bora_in.vivo_noRT | Gm45591 | 4.676 | 1.77E-15 | glio |
| Aars2_in.vivo_noRT | Ccser1 | 5.327 | 1.77E-19 | glio |
| Bora_in.vivo_noRT | BC080695 | 4.936 | 1.77E-21 | glio |
| Aars2_in.vivo_noRT | Fabp6 | 5.327 | 1.77E-23 | glio |
| culture_effect | Tnnt2 | 3.212 | 1.77E-24 | glio |
| culture_effect | Birc3 | -1.735 | 1.78E-03 | glio |
| culture_effect | Trim37 | 0.616 | 1.78E-03 | glio |
| Interaction_genotype_Atr | Cnbp | 1.283 | 1.78E-03 | glio |
| Interaction_genotype_Bcas2 | Ephx1 | -4.066 | 1.78E-03 | glio |
| Interaction_genotype_Bora | Sema6b | -3.955 | 1.78E-03 | glio |
| Interaction_genotype_Mtor | H2-M10.1 | -3.231 | 1.78E-03 | glio |
| Interaction_genotype_Mtor | Rhbdf2 | -3.636 | 1.78E-03 | glio |
| Interaction_genotype_Mtor | Miga1 | -2.441 | 1.78E-03 | glio |
| Interaction_genotype_Mtor | Sdhaf4 | 1.797 | 1.78E-03 | glio |
| Interaction_genotype_Mtor | Sumo3 | 1.408 | 1.78E-03 | glio |
| Interaction_genotype_Mtor | Fubp1 | 1.438 | 1.78E-03 | glio |
| Interaction_genotype_Myc | Slc25a47 | -3.513 | 1.78E-03 | glio |
| Interaction_genotype_Myc | 5730455P16R | -2.441 | 1.78E-03 | glio |
| Interaction_genotype_Myc | Slc25a39 | 1.325 | 1.78E-03 | glio |
| Interaction_genotype_Myc | Tpt1 | 1.269 | 1.78E-03 | glio |
| Interaction_genotype_Pars2 | Gm20635 | -4.029 | 1.78E-03 | glio |
| Interaction_genotype_Pars2 | Ppp1r3g | -3.961 | 1.78E-03 | glio |
| Interaction_genotype_Uqcrc2 | Baalc | -4.016 | 1.78E-03 | glio |
| Interaction_genotype_Uqcrc2 | Gm45620 | -4.513 | 1.78E-03 | glio |
| interaction_RT | Tbc1d30 | -1.883 | 1.78E-03 | glio |
| interaction_RT | Mrps7 | -0.633 | 1.78E-03 | glio |
| Aars2_in.vivo_noRT | Fam120c | 2.261 | 1.78E-03 | glio |
| Atp5c1_in.vivo_noRT | Ank3 | -2.006 | 1.78E-03 | glio |
| Atr_in.vivo_noRT | Get4 | -1.165 | 1.78E-03 | glio |
| Atr_ex.vivo_noRT | Nr1h4 | 2.886 | 1.78E-03 | glio |
| Bard1_ex.vivo_noRT | Chd5 | 3.044 | 1.78E-03 | glio |
| Bcas2_ex.vivo_noRT | Ssh3 | 2.743 | 1.78E-03 | glio |
| Bora_in.vivo_noRT | 6720489N17R | 2.543 | 1.78E-03 | glio |

| | | | | |
|---|---|---|---|---|
| Bora_in.vivo_noRT | 5830428M24F | 2.283 | 1.78E-03 | glio |
| Bora_in.vivo_noRT | Gm44777 | 2.153 | 1.78E-03 | glio |
| Chek1_in.vivo_noRT | 2600006K01R | 2.385 | 1.78E-03 | glio |
| Chek1_in.vivo_noRT | Rpl13 | -0.65 | 1.78E-03 | glio |
| Chek1_ex.vivo_noRT | Ninj2 | 2.887 | 1.78E-03 | glio |
| Chek1_ex.vivo_noRT | Gm16046 | 2.717 | 1.78E-03 | glio |
| Hsd17b10_in.vivo_noRT | D230022J07Ri | 2.459 | 1.78E-03 | glio |
| Hsd17b10_in.vivo_noRT | Ctdp1 | -1.743 | 1.78E-03 | glio |
| Interaction_genotype_Bora | Nxph3 | -4.471 | 1.78E-04 | glio |
| Interaction_genotype_Mtor | Ldhd | -3.892 | 1.78E-04 | glio |
| Interaction_genotype_Mtor | 4930545E07Ri | -4.695 | 1.78E-04 | glio |
| Aars2_in.vivo_noRT | Prx | 2.798 | 1.78E-04 | glio |
| Aars2_in.vivo_noRT | Mapkapk5 | 2.628 | 1.78E-04 | glio |
| Aars2_in.vivo_noRT | Zfp346 | 2.386 | 1.78E-04 | glio |
| Atp5c1_in.vivo_noRT | Gm15270 | 2.405 | 1.78E-04 | glio |
| Atp5c1_in.vivo_noRT | Cys1 | 2.402 | 1.78E-04 | glio |
| Bcas2_in.vivo_noRT | Tnfrsf1b | 2.513 | 1.78E-04 | glio |
| Bora_in.vivo_noRT | Slc43a1 | 2.734 | 1.78E-04 | glio |
| Chek1_ex.vivo_noRT | Pde1a | 3.463 | 1.78E-04 | glio |
| Chek1_ex.vivo_noRT | 2310001H17R | 2.605 | 1.78E-04 | glio |
| Hsd17b10_in.vivo_noRT | 5033403F01Ri | 2.69 | 1.78E-04 | glio |
| culture_effect | Gm4876 | 1.343 | 1.78E-05 | glio |
| culture_effect | Mpdu1 | -0.705 | 1.78E-05 | glio |
| Interaction_genotype_Mtor | Rbm41 | -4.863 | 1.78E-05 | glio |
| Interaction_genotype_Mtor | Prtn3 | -5.36 | 1.78E-05 | glio |
| Interaction_genotype_Myc | Gm49359 | -4.065 | 1.78E-05 | glio |
| Interaction_genotype_Myc | Gm16183 | -5.228 | 1.78E-05 | glio |
| Interaction_genotype_Myc | Rnf7 | 1.759 | 1.78E-05 | glio |
| Aars2_in.vivo_noRT | Gm29170 | 2.924 | 1.78E-05 | glio |
| Bora_in.vivo_noRT | Gm32051 | 3.171 | 1.78E-05 | glio |
| Chek1_in.vivo_noRT | Phf11a | 2.951 | 1.78E-05 | glio |
| Interaction_genotype_Mtor | Hddc3 | -5.267 | 1.78E-06 | glio |
| Interaction_genotype_Mtor | Serpinb9g | -4.099 | 1.78E-06 | glio |
| Interaction_genotype_Mtor | Myo5c | -5.592 | 1.78E-06 | glio |
| Aars2_in.vivo_noRT | Radil | 3.296 | 1.78E-06 | glio |
| Atp5c1_in.vivo_noRT | Gm29966 | 2.408 | 1.78E-06 | glio |
| Atp5c1_in.vivo_noRT | Fabp6 | 2.668 | 1.78E-06 | glio |
| Atr_in.vivo_noRT | Cfap100 | 3.336 | 1.78E-06 | glio |
| Atr_in.vivo_noRT | Asphd2 | 3.33 | 1.78E-06 | glio |
| culture_effect | Dhx58 | -1.35 | 1.78E-07 | glio |
| culture_effect | Mbtd1 | -1.01 | 1.78E-07 | glio |
| Aars2_in.vivo_noRT | Gm9958 | 3.757 | 1.78E-07 | glio |
| Aars2_in.vivo_noRT | Gm36839 | 3.764 | 1.78E-07 | glio |
| Aars2_in.vivo_noRT | Ache | 3.754 | 1.78E-07 | glio |
| Aars2_in.vivo_noRT | Cfap100 | 3.644 | 1.78E-07 | glio |
| Aars2_in.vivo_noRT | Gm19345 | 3.291 | 1.78E-07 | glio |
| Bora_ex.vivo_noRT | A930019D19R | 3.201 | 1.78E-07 | glio |
| Chek1_in.vivo_noRT | Gm10382 | 3.8 | 1.78E-07 | glio |
| Atr_in.vivo_noRT | Car12 | 3.681 | 1.78E-08 | glio |
| Atr_in.vivo_noRT | Zfp972 | 4.089 | 1.78E-08 | glio |
| Atr_in.vivo_noRT | Zfpm1 | 4.017 | 1.78E-08 | glio |
| Atr_in.vivo_noRT | Gm21814 | 3.675 | 1.78E-08 | glio |

| | | | | |
|---|---|---|---|---|
| Chek1_in.vivo_noRT | Zfp811 | 3.87 | 1.78E-08 | glio |
| Chek1_in.vivo_noRT | Acot6 | 3.798 | 1.78E-08 | glio |
| Chek1_ex.vivo_noRT | H2-DMb2 | 4.842 | 1.78E-08 | glio |
| Chek1_ex.vivo_noRT | Gm43951 | 4.049 | 1.78E-08 | glio |
| Aars2_in.vivo_noRT | Gm17227 | 3.944 | 1.78E-09 | glio |
| Atr_in.vivo_noRT | Il2rb | 3.991 | 1.78E-09 | glio |
| culture_effect | Sumo2 | 0.682 | 1.78E-10 | glio |
| Bora_in.vivo_noRT | Gm13403 | 4.279 | 1.78E-10 | glio |
| Chek1_in.vivo_noRT | 2810049E08Ri | 4.007 | 1.78E-10 | glio |
| Chek1_in.vivo_noRT | Slc14a1 | 3.688 | 1.78E-10 | glio |
| Chek1_in.vivo_noRT | Cidea | 4.129 | 1.78E-11 | glio |
| Chek1_ex.vivo_noRT | Stx19 | 4.842 | 1.78E-11 | glio |
| culture_effect | Gpc1 | 2.182 | 1.78E-12 | glio |
| Bcas2_in.vivo_noRT | 4930483J18Ri | 4.547 | 1.78E-12 | glio |
| Aars2_in.vivo_noRT | Gm20682 | 4.282 | 1.78E-13 | glio |
| Atr_in.vivo_noRT | D030040B21R | 5.146 | 1.78E-14 | glio |
| Atr_in.vivo_noRT | 4930447F24Ri | 5.273 | 1.78E-15 | glio |
| Chek1_in.vivo_noRT | Fbxl13 | 4.784 | 1.78E-15 | glio |
| culture_effect | Pcdh15 | -4.509 | 1.78E-17 | glio |
| Bcas2_in.vivo_noRT | Hfm1 | 4.807 | 1.78E-18 | glio |
| Bcas2_in.vivo_noRT | D630024D03F | 4.807 | 1.78E-18 | glio |
| Chek1_in.vivo_noRT | Chia1 | 5.044 | 1.78E-23 | glio |
| Aars2_in.vivo_noRT | Zscan4c | 5.327 | 1.78E-24 | glio |
| Aars2_in.vivo_noRT | Tagap | 5.067 | 1.78E-25 | glio |
| Atr_in.vivo_noRT | Ccl9 | 5.533 | 1.78E-28 | glio |
| culture_effect | Ggt5 | -1.867 | 1.79E-03 | glio |
| culture_effect | Gm16222 | 2.102 | 1.79E-03 | glio |
| culture_effect | 5730409E04Ri | 1.262 | 1.79E-03 | glio |
| culture_effect | Rcan3 | 1.152 | 1.79E-03 | glio |
| culture_effect | Gm11033 | 0.961 | 1.79E-03 | glio |
| culture_effect | Mrps12 | 0.431 | 1.79E-03 | glio |
| Interaction_genotype_Aars2 | D930028M14I | -3.521 | 1.79E-03 | glio |
| Interaction_genotype_Aars2 | 4921504E06Ri | -3.122 | 1.79E-03 | glio |
| Interaction_genotype_Aars2 | B230369F24Ri | -3.916 | 1.79E-03 | glio |
| Interaction_genotype_Aars2 | Fgf2 | -3.717 | 1.79E-03 | glio |
| Interaction_genotype_Aars2 | Wdr38 | -3.649 | 1.79E-03 | glio |
| Interaction_genotype_Atr | Gm11520 | -3.566 | 1.79E-03 | glio |
| Interaction_genotype_Atr | Gpx1 | 1.198 | 1.79E-03 | glio |
| Interaction_genotype_Chek1 | 2610027K06R | -3.939 | 1.79E-03 | glio |
| Interaction_genotype_Myc | Ccdc142os | -3.393 | 1.79E-03 | glio |
| Interaction_genotype_Pars2 | Nxnl2 | -3.905 | 1.79E-03 | glio |
| Interaction_genotype_Rpl3 | Gm26766 | -4.141 | 1.79E-03 | glio |
| interaction_RT | Mospd1 | 1.522 | 1.79E-03 | glio |
| interaction_RT | Vldlr | 1.442 | 1.79E-03 | glio |
| interaction_RT | Arf5 | 0.559 | 1.79E-03 | glio |
| interaction_RT | Rack1 | -0.686 | 1.79E-03 | glio |
| Aars2_in.vivo_noRT | Col28a1 | 2.127 | 1.79E-03 | glio |
| Aars2_in.vivo_noRT | Dnah2 | 2.266 | 1.79E-03 | glio |
| Aars2_in.vivo_noRT | Gm44867 | 2.269 | 1.79E-03 | glio |
| Aars2_in.vivo_noRT | Cox20 | -1.393 | 1.79E-03 | glio |
| Aars2_in.vivo_noRT | Lsm3 | -1.196 | 1.79E-03 | glio |
| Atr_in.vivo_noRT | Gm29488 | 2.167 | 1.79E-03 | glio |

| | | | | |
|---|---|---|---|---|
| Atr_in.vivo_noRT | Apol8 | 1.79 | 1.79E-03 | glio |
| Atr_in.vivo_noRT | Zfp647 | 1.887 | 1.79E-03 | glio |
| Atr_in.vivo_noRT | 9530082P21R | 1.794 | 1.79E-03 | glio |
| Atr_in.vivo_noRT | Qrich1 | -1.262 | 1.79E-03 | glio |
| Atr_in.vivo_noRT | Maged2 | -1.231 | 1.79E-03 | glio |
| Atr_in.vivo_noRT | Hsbp1 | -0.767 | 1.79E-03 | glio |
| Bard1_ex.vivo_noRT | Col19a1 | 2.421 | 1.79E-03 | glio |
| Bard1_ex.vivo_noRT | Gm38477 | 2.415 | 1.79E-03 | glio |
| Bcas2_in.vivo_noRT | Kcna4 | 2.28 | 1.79E-03 | glio |
| Bcas2_ex.vivo_noRT | Gm48265 | 2.742 | 1.79E-03 | glio |
| Bcas2_ex.vivo_noRT | Ush1g | 2.342 | 1.79E-03 | glio |
| Bora_in.vivo_noRT | Gm38414 | 2.529 | 1.79E-03 | glio |
| Bora_in.vivo_noRT | Fubp1 | -0.79 | 1.79E-03 | glio |
| Chek1_in.vivo_noRT | Matk | 2.101 | 1.79E-03 | glio |
| Chek1_in.vivo_noRT | Son | -0.95 | 1.79E-03 | glio |
| Chek1_ex.vivo_noRT | Zfp534 | 3.061 | 1.79E-03 | glio |
| Chek1_ex.vivo_noRT | Pcdhb22 | 2.845 | 1.79E-03 | glio |
| Chek1_ex.vivo_noRT | Tmem151a | 2.594 | 1.79E-03 | glio |
| Eif2s1_in.vivo_noRT | 2310003N18R | 2.143 | 1.79E-03 | glio |
| Hsd17b10_in.vivo_noRT | Grip2 | 2.365 | 1.79E-03 | glio |
| Hsd17b10_in.vivo_noRT | Pla2g4b | 2.319 | 1.79E-03 | glio |
| Hsd17b10_in.vivo_noRT | G3bp2 | -0.677 | 1.79E-03 | glio |
| culture_effect | Vmn1r88 | -2.119 | 1.79E-04 | glio |
| culture_effect | Spink2 | 2.267 | 1.79E-04 | glio |
| culture_effect | Eif4b | -0.469 | 1.79E-04 | glio |
| Aars2_in.vivo_noRT | Zbp1 | -1.603 | 1.79E-04 | glio |
| Atr_in.vivo_noRT | 6430503K07R | 2.474 | 1.79E-04 | glio |
| Atr_in.vivo_noRT | Fyttd1 | -1.27 | 1.79E-04 | glio |
| Atr_in.vivo_noRT | Slc38a2 | -1.007 | 1.79E-04 | glio |
| Atr_ex.vivo_noRT | Hrct1 | 2.975 | 1.79E-04 | glio |
| Bard1_ex.vivo_noRT | Akap14 | 2.814 | 1.79E-04 | glio |
| Bora_in.vivo_noRT | Daam2 | 2.764 | 1.79E-04 | glio |
| Bora_in.vivo_noRT | Tspyl5 | 2.306 | 1.79E-04 | glio |
| Hsd17b10_in.vivo_noRT | Necab3 | 2.691 | 1.79E-04 | glio |
| Hsd17b10_in.vivo_noRT | Pcdhgb6 | 2.431 | 1.79E-04 | glio |
| culture_effect | BC048671 | -2.235 | 1.79E-05 | glio |
| culture_effect | Tox | -2.511 | 1.79E-05 | glio |
| culture_effect | 2210016L21Ri | 0.71 | 1.79E-05 | glio |
| culture_effect | Mrpl15 | -0.498 | 1.79E-05 | glio |
| Interaction_genotype_Aars2 | Gm10642 | -5.06 | 1.79E-05 | glio |
| Interaction_genotype_Myc | Peg10 | -4.284 | 1.79E-05 | glio |
| Interaction_genotype_Myc | Prelid3a | -4.88 | 1.79E-05 | glio |
| Interaction_genotype_Myc | Gm41760 | -4.796 | 1.79E-05 | glio |
| Interaction_genotype_Myc | Rhox7a | -5.105 | 1.79E-05 | glio |
| interaction_RT | Prdx5 | 0.943 | 1.79E-05 | glio |
| Aars2_in.vivo_noRT | Hnrnpa2b1 | -0.804 | 1.79E-05 | glio |
| Atr_ex.vivo_noRT | Ccdc162 | 3.164 | 1.79E-05 | glio |
| Atr_ex.vivo_noRT | Olfr552 | 2.764 | 1.79E-05 | glio |
| Bora_ex.vivo_noRT | Klrk1 | 3.201 | 1.79E-05 | glio |
| Chek1_in.vivo_noRT | Mmp28 | 3.15 | 1.79E-05 | glio |
| Chek1_in.vivo_noRT | Gm28592 | 2.829 | 1.79E-05 | glio |
| Chek1_in.vivo_noRT | Efna4 | 3.086 | 1.79E-05 | glio |

| | | | | |
|---|---|---|---|---|
| culture_effect | Bach1 | -1.356 | 1.79E-06 | glio |
| Interaction_genotype_Mtor | A330032B11R | -5.212 | 1.79E-06 | glio |
| Interaction_genotype_Myc | Eif5a | 2.06 | 1.79E-06 | glio |
| interaction_RT | Cct6a | -0.851 | 1.79E-06 | glio |
| Atr_in.vivo_noRT | Zfp59 | 3.376 | 1.79E-06 | glio |
| Chek1_in.vivo_noRT | Srgn | 3.999 | 1.79E-06 | glio |
| culture_effect | Fam92a | 0.676 | 1.79E-07 | glio |
| culture_effect | Hnrnph1 | -0.65 | 1.79E-07 | glio |
| Atr_in.vivo_noRT | 1700101I11Ri | 3.893 | 1.79E-07 | glio |
| Atr_ex.vivo_noRT | Scel | 3.164 | 1.79E-07 | glio |
| Atr_ex.vivo_noRT | Ctla4 | 3.164 | 1.79E-07 | glio |
| Atr_ex.vivo_noRT | 4930423D22R | 3.382 | 1.79E-07 | glio |
| Atr_ex.vivo_noRT | Zscan4-ps3 | 3.164 | 1.79E-07 | glio |
| Atr_ex.vivo_noRT | Tagap | 3.164 | 1.79E-07 | glio |
| Atr_ex.vivo_noRT | Bbof1 | 4.531 | 1.79E-07 | glio |
| Atr_ex.vivo_noRT | Gm21731 | 3.164 | 1.79E-07 | glio |
| Chek1_in.vivo_noRT | Gm50103 | 3.737 | 1.79E-07 | glio |
| Hsd17b10_in.vivo_noRT | Trim16 | 3.356 | 1.79E-07 | glio |
| culture_effect | Emc2 | -0.783 | 1.79E-08 | glio |
| culture_effect | Nabp2 | 0.732 | 1.79E-08 | glio |
| Bora_in.vivo_noRT | Nt5e | 3.457 | 1.79E-08 | glio |
| Aars2_in.vivo_noRT | Ptp4a1 | 3.632 | 1.79E-09 | glio |
| Aars2_in.vivo_noRT | Ikbke | 4.01 | 1.79E-09 | glio |
| Hsd17b10_in.vivo_noRT | Gm15943 | 3.352 | 1.79E-09 | glio |
| Aars2_in.vivo_noRT | Ly6c2 | 4.424 | 1.79E-10 | glio |
| Bcas2_in.vivo_noRT | Slc14a1 | 3.699 | 1.79E-10 | glio |
| Chek1_in.vivo_noRT | Nrn1l | 4.118 | 1.79E-10 | glio |
| Bora_in.vivo_noRT | Gm14379 | 4.42 | 1.79E-11 | glio |
| culture_effect | Tnfrsf13c | -3.271 | 1.79E-12 | glio |
| Chek1_in.vivo_noRT | Ttll13 | 4.31 | 1.79E-12 | glio |
| Chek1_ex.vivo_noRT | Gast | 4.842 | 1.79E-12 | glio |
| Chek1_in.vivo_noRT | Gm15743 | 4.78 | 1.79E-13 | glio |
| Chek1_in.vivo_noRT | Vwce | 4.784 | 1.79E-14 | glio |
| Hsd17b10_in.vivo_noRT | Krt19 | 3.616 | 1.79E-14 | glio |
| Chek1_in.vivo_noRT | Gm45406 | 4.522 | 1.79E-17 | glio |
| Chek1_in.vivo_noRT | Scgb1c1 | 5.044 | 1.79E-17 | glio |
| Chek1_in.vivo_noRT | Gm14540 | 4.521 | 1.79E-17 | glio |
| Chek1_in.vivo_noRT | 4933400F21Ri | 4.253 | 1.79E-17 | glio |
| Atr_in.vivo_noRT | 4930553M12F | 5.267 | 1.79E-21 | glio |
| Atr_in.vivo_noRT | Gm38163 | 5.533 | 1.79E-22 | glio |
| Atr_in.vivo_noRT | Gm26555 | 5.27 | 1.79E-24 | glio |
| culture_effect | Psme1 | -2.524 | 1.79E-49 | glio |
| culture_effect | Cpq | -2.831 | 1.79E-49 | glio |
| culture_effect | Fam111a | 0.739 | 1.80E-03 | glio |
| culture_effect | Lims1 | 0.558 | 1.80E-03 | glio |
| culture_effect | Tmcc3 | 0.583 | 1.80E-03 | glio |
| culture_effect | Rbck1 | -0.495 | 1.80E-03 | glio |
| culture_effect | Mrpl57 | 0.406 | 1.80E-03 | glio |
| culture_effect | Dnajc21 | 0.395 | 1.80E-03 | glio |
| Interaction_genotype_Bora | Skp2 | 2.113 | 1.80E-03 | glio |
| Interaction_genotype_Bora | Las1l | 1.892 | 1.80E-03 | glio |
| Interaction_genotype_Ifnar1 | Halr1 | 3 | 1.80E-03 | glio |

| | | | | |
|---|---|---|---|---|
| Interaction_genotype_Mtor | Zfp619 | -3.928 | 1.80E-03 | glio |
| Interaction_genotype_Mtor | Gm48427 | -3.625 | 1.80E-03 | glio |
| Interaction_genotype_Mtor | Dffa | -3.035 | 1.80E-03 | glio |
| Interaction_genotype_Mtor | Tcf3 | 1.665 | 1.80E-03 | glio |
| Interaction_genotype_Mtor | Tmem126a | 1.466 | 1.80E-03 | glio |
| Interaction_genotype_Mtor | Uqcc2 | 1.559 | 1.80E-03 | glio |
| Interaction_genotype_Mtor | Ndufs5 | 1.246 | 1.80E-03 | glio |
| Interaction_genotype_Mtor | Ifi27l2a | 2.055 | 1.80E-03 | glio |
| Interaction_genotype_Myc | 6530402F18Ri | -3.401 | 1.80E-03 | glio |
| Interaction_genotype_Myc | Mfsd3 | -2.848 | 1.80E-03 | glio |
| Interaction_genotype_Myc | 2700062C07R | -2.004 | 1.80E-03 | glio |
| Interaction_genotype_Myc | Gas2l1 | -1.768 | 1.80E-03 | glio |
| Interaction_genotype_Myc | Vps35 | 1.295 | 1.80E-03 | glio |
| Interaction_genotype_Rad51 | AV356131 | -3.987 | 1.80E-03 | glio |
| interaction_RT | Tbl3 | -0.911 | 1.80E-03 | glio |
| Aars2_in.vivo_noRT | Med10 | -1.259 | 1.80E-03 | glio |
| Atr_in.vivo_noRT | Lhx6 | 2.117 | 1.80E-03 | glio |
| Atr_in.vivo_noRT | Iba57 | 2.131 | 1.80E-03 | glio |
| Atr_in.vivo_noRT | Napepld | 1.968 | 1.80E-03 | glio |
| Atr_in.vivo_noRT | Klk10 | 2.116 | 1.80E-03 | glio |
| Bard1_ex.vivo_noRT | Strip2 | 2.415 | 1.80E-03 | glio |
| Bcas2_in.vivo_noRT | 1700047M11F | 2.08 | 1.80E-03 | glio |
| Bcas2_ex.vivo_noRT | Car10 | 2.742 | 1.80E-03 | glio |
| Bcas2_ex.vivo_noRT | Pcdh15 | 2.742 | 1.80E-03 | glio |
| Bcas2_ex.vivo_noRT | Gm11033 | 2.028 | 1.80E-03 | glio |
| Bora_in.vivo_noRT | Fhl3 | 2.3 | 1.80E-03 | glio |
| Bora_in.vivo_noRT | Commd3 | -1.166 | 1.80E-03 | glio |
| Chek1_in.vivo_noRT | AI429214 | 2.287 | 1.80E-03 | glio |
| Chek1_in.vivo_noRT | Gm17300 | 2.157 | 1.80E-03 | glio |
| Chek1_in.vivo_noRT | Apba3 | -1.299 | 1.80E-03 | glio |
| Chek1_in.vivo_noRT | Borcs8 | -1.139 | 1.80E-03 | glio |
| Hsd17b10_in.vivo_noRT | Kyat1 | 2.563 | 1.80E-03 | glio |
| culture_effect | Ints9 | -0.954 | 1.80E-04 | glio |
| culture_effect | Ube2d1 | 0.696 | 1.80E-04 | glio |
| Interaction_genotype_Atr | Gal3st4 | -4.607 | 1.80E-04 | glio |
| Interaction_genotype_Atr | Sprr1a | -4.742 | 1.80E-04 | glio |
| Interaction_genotype_Mtor | Rab27a | -3.968 | 1.80E-04 | glio |
| Interaction_genotype_Mtor | C4b | 3.856 | 1.80E-04 | glio |
| interaction_RT | H2-DMa | 1.256 | 1.80E-04 | glio |
| interaction_RT | Copz2 | 0.989 | 1.80E-04 | glio |
| interaction_RT | Itpa | -0.949 | 1.80E-04 | glio |
| interaction_RT | Srp72 | -0.779 | 1.80E-04 | glio |
| interaction_RT | Hbq1b | -3.294 | 1.80E-04 | glio |
| Aars2_in.vivo_noRT | Ankrd23 | 2.819 | 1.80E-04 | glio |
| Atr_in.vivo_noRT | Tmem35b | 2.542 | 1.80E-04 | glio |
| Atr_in.vivo_noRT | Ado | -1.488 | 1.80E-04 | glio |
| Atr_in.vivo_noRT | Snd1 | -1.276 | 1.80E-04 | glio |
| Atr_ex.vivo_noRT | Gm20732 | 3.164 | 1.80E-04 | glio |
| Atr_ex.vivo_noRT | Slc10a1 | 2.77 | 1.80E-04 | glio |
| Atr_ex.vivo_noRT | Vgll2 | 3.164 | 1.80E-04 | glio |
| Atr_ex.vivo_noRT | Rem2 | 3.164 | 1.80E-04 | glio |
| Atr_ex.vivo_noRT | Tmeff2 | 3.941 | 1.80E-04 | glio |

| | | | | |
|---|---|---|---|---|
| Atr_ex.vivo_noRT | Usp51 | 3.336 | 1.80E-04 | glio |
| Atr_ex.vivo_noRT | Lpin1 | 3.096 | 1.80E-04 | glio |
| Bcas2_ex.vivo_noRT | Hist1h2ag | 2.742 | 1.80E-04 | glio |
| Bora_in.vivo_noRT | 4930554I06Ri | 2.876 | 1.80E-04 | glio |
| Bora_ex.vivo_noRT | Thsd1 | 3.012 | 1.80E-04 | glio |
| Cenpj_ex.vivo_noRT | Ablim2 | 2.591 | 1.80E-04 | glio |
| Cenpj_ex.vivo_noRT | Gm5415 | 2.591 | 1.80E-04 | glio |
| Cenpj_ex.vivo_noRT | Nrip2 | 2.591 | 1.80E-04 | glio |
| Cenpj_ex.vivo_noRT | Gm49698 | 2.591 | 1.80E-04 | glio |
| Cenpj_ex.vivo_noRT | Olfr881 | 2.591 | 1.80E-04 | glio |
| Chek1_in.vivo_noRT | Gm46378 | 2.81 | 1.80E-04 | glio |
| Interaction_genotype_Myc | 1700026N04R | -4.391 | 1.80E-05 | glio |
| interaction_RT | Casp1 | 2.744 | 1.80E-05 | glio |
| Aars2_in.vivo_noRT | Gldc | 2.955 | 1.80E-05 | glio |
| Atp5c1_in.vivo_noRT | Olfr543 | 2.408 | 1.80E-05 | glio |
| Atr_in.vivo_noRT | Arhgap33 | 3.07 | 1.80E-05 | glio |
| Bora_in.vivo_noRT | Sumo2 | -0.861 | 1.80E-05 | glio |
| Chek1_ex.vivo_noRT | Wfdc3 | 4.672 | 1.80E-05 | glio |
| culture_effect | Trp53rkb | 1.348 | 1.80E-06 | glio |
| culture_effect | Ubald2 | -0.833 | 1.80E-06 | glio |
| Interaction_genotype_Mtor | 6430562O15R | -5.167 | 1.80E-06 | glio |
| Aars2_in.vivo_noRT | Cbr3 | 3.574 | 1.80E-06 | glio |
| Atr_ex.vivo_noRT | Vtn | 3.164 | 1.80E-06 | glio |
| Bora_in.vivo_noRT | Lsp1 | 3.749 | 1.80E-06 | glio |
| Bora_in.vivo_noRT | Rtn2 | 3.029 | 1.80E-06 | glio |
| Chek1_in.vivo_noRT | Ablim3 | 3.483 | 1.80E-06 | glio |
| Chek1_ex.vivo_noRT | Plcd4 | 3.867 | 1.80E-06 | glio |
| Chek1_ex.vivo_noRT | Wnt16 | 4.235 | 1.80E-07 | glio |
| Hsd17b10_in.vivo_noRT | Zfp810 | 3.616 | 1.80E-07 | glio |
| culture_effect | Gm49874 | 3.177 | 1.80E-08 | glio |
| culture_effect | Ankrd1 | 3.768 | 1.80E-09 | glio |
| culture_effect | 8030456M14F | 2.864 | 1.80E-09 | glio |
| Chek1_ex.vivo_noRT | Gbp10 | 4.842 | 1.80E-09 | glio |
| Atr_in.vivo_noRT | Gpnmb | -3.502 | 1.80E-10 | glio |
| Bora_in.vivo_noRT | Prickle4 | 4.145 | 1.80E-10 | glio |
| Atr_in.vivo_noRT | Gm16094 | 4.228 | 1.80E-11 | glio |
| Bcas2_in.vivo_noRT | Gm26789 | 3.89 | 1.80E-11 | glio |
| Bora_in.vivo_noRT | A330015K06R | 4.279 | 1.80E-11 | glio |
| culture_effect | Ctdsp2 | -1.088 | 1.80E-12 | glio |
| Aars2_in.vivo_noRT | St8sia1 | 5.067 | 1.80E-12 | glio |
| Aars2_in.vivo_noRT | Ldhd | 4.41 | 1.80E-13 | glio |
| Aars2_in.vivo_noRT | Gm9967 | 4.94 | 1.80E-13 | glio |
| Bcas2_in.vivo_noRT | Gm40264 | 4.547 | 1.80E-14 | glio |
| Aars2_in.vivo_noRT | Gm5662 | 5.327 | 1.80E-15 | glio |
| Bcas2_in.vivo_noRT | Folr1 | 4.807 | 1.80E-15 | glio |
| Chek1_in.vivo_noRT | Gm45669 | 4.78 | 1.80E-15 | glio |
| Chek1_ex.vivo_noRT | Gm47510 | 4.842 | 1.80E-15 | glio |
| Chek1_ex.vivo_noRT | Gm5788 | 4.842 | 1.80E-15 | glio |
| Atr_in.vivo_noRT | Cox7b2 | 5.066 | 1.80E-16 | glio |
| Aars2_in.vivo_noRT | Gm12610 | 4.798 | 1.80E-17 | glio |
| Chek1_ex.vivo_noRT | Sucnr1 | 4.842 | 1.80E-18 | glio |
| Aars2_in.vivo_noRT | 5730507C01R | 4.804 | 1.80E-19 | glio |

| | | | | |
|---|---|---|---|---|
| Aars2_in.vivo_noRT | Gm11973 | 5.06 | 1.80E-19 | glio |
| Aars2_in.vivo_noRT | Slc10a1 | 5.067 | 1.80E-19 | glio |
| Atr_in.vivo_noRT | Nwd1 | 5.269 | 1.80E-19 | glio |
| Atr_in.vivo_noRT | Slc2a9 | 5.533 | 1.80E-21 | glio |
| Aars2_in.vivo_noRT | Ccl3 | 5.067 | 1.80E-22 | glio |
| Aars2_in.vivo_noRT | BC080695 | 5.327 | 1.80E-24 | glio |
| culture_effect | Rbakdn | -1.97 | 1.81E-03 | glio |
| culture_effect | Ncmap | -1.863 | 1.81E-03 | glio |
| culture_effect | Zfp277 | -0.871 | 1.81E-03 | glio |
| culture_effect | Pfkl | -0.576 | 1.81E-03 | glio |
| culture_effect | Pak3 | 0.628 | 1.81E-03 | glio |
| Interaction_genotype_Chek1 | Gpx2 | 5.575 | 1.81E-03 | glio |
| Interaction_genotype_Chek1 | 2310030G06R | -4.183 | 1.81E-03 | glio |
| Interaction_genotype_Mtor | Ddah2 | -3.833 | 1.81E-03 | glio |
| Interaction_genotype_Mtor | Mafa | -3.505 | 1.81E-03 | glio |
| Interaction_genotype_Mtor | Slc25a47 | -3.718 | 1.81E-03 | glio |
| Interaction_genotype_Mtor | Prps1l3 | -3.08 | 1.81E-03 | glio |
| Interaction_genotype_Myc | Wif1 | -3.985 | 1.81E-03 | glio |
| Interaction_genotype_Myc | Sstr2 | -3.304 | 1.81E-03 | glio |
| Interaction_genotype_Myc | Nol7 | 1.308 | 1.81E-03 | glio |
| Interaction_genotype_Pars2 | Gm47204 | -4.312 | 1.81E-03 | glio |
| Aars2_in.vivo_noRT | Klf12 | 2.15 | 1.81E-03 | glio |
| Aars2_in.vivo_noRT | 2810429I04Ri | 1.826 | 1.81E-03 | glio |
| Aars2_in.vivo_noRT | Retsat | 1.541 | 1.81E-03 | glio |
| Aars2_in.vivo_noRT | Ndufa4 | -0.684 | 1.81E-03 | glio |
| Aars2_ex.vivo_noRT | Ptgis | 2.981 | 1.81E-03 | glio |
| Atp5c1_in.vivo_noRT | Ptp4a1.1 | 2.393 | 1.81E-03 | glio |
| Atr_in.vivo_noRT | Pif1 | 2.098 | 1.81E-03 | glio |
| Atr_in.vivo_noRT | Zfp729a | 2.052 | 1.81E-03 | glio |
| Atr_in.vivo_noRT | Atp5g1 | -0.65 | 1.81E-03 | glio |
| Atr_ex.vivo_noRT | Svopl | 2.771 | 1.81E-03 | glio |
| Atr_ex.vivo_noRT | 9630013A20R | 2.903 | 1.81E-03 | glio |
| Bard1_ex.vivo_noRT | Gm36043 | 2.814 | 1.81E-03 | glio |
| Bard1_ex.vivo_noRT | Gm49860 | 2.615 | 1.81E-03 | glio |
| Bard1_ex.vivo_noRT | Hes5 | 3.612 | 1.81E-03 | glio |
| Bcas2_in.vivo_noRT | 9030407P20R | 2.26 | 1.81E-03 | glio |
| Bcas2_in.vivo_noRT | Gm19605 | 2.26 | 1.81E-03 | glio |
| Bcas2_in.vivo_noRT | Zfp94 | 2.186 | 1.81E-03 | glio |
| Bcas2_in.vivo_noRT | N4bp2l2 | -1.126 | 1.81E-03 | glio |
| Bcas2_in.vivo_noRT | Eif1a | -0.82 | 1.81E-03 | glio |
| Bora_in.vivo_noRT | Nav1 | -1.662 | 1.81E-03 | glio |
| Bora_in.vivo_noRT | Tprkb | -1.507 | 1.81E-03 | glio |
| Bora_ex.vivo_noRT | Gm15941 | 2.806 | 1.81E-03 | glio |
| Chek1_in.vivo_noRT | Pon3 | 2.293 | 1.81E-03 | glio |
| Chek1_in.vivo_noRT | Gm10282 | 2.267 | 1.81E-03 | glio |
| Chek1_in.vivo_noRT | Art2b | 1.909 | 1.81E-03 | glio |
| Chek1_in.vivo_noRT | Rhobtb1 | 2.285 | 1.81E-03 | glio |
| Chek1_ex.vivo_noRT | Zkscan4 | 2.9 | 1.81E-03 | glio |
| Chek1_ex.vivo_noRT | Msl3l2 | 2.887 | 1.81E-03 | glio |
| Chek1_ex.vivo_noRT | Opalin | 3.861 | 1.81E-03 | glio |
| Eif2s1_in.vivo_noRT | Gm50138 | 2.144 | 1.81E-03 | glio |
| Eif2s1_in.vivo_noRT | Maged1 | -0.724 | 1.81E-03 | glio |

| | | | | |
|---|---|---|---|---|
| Hsd17b10_in.vivo_noRT | Dzank1 | 2.441 | 1.81E-03 | glio |
| Hsd17b10_in.vivo_noRT | A530083I20Ri | 2.608 | 1.81E-03 | glio |
| culture_effect | Ncor2 | 0.788 | 1.81E-04 | glio |
| Interaction_genotype_Atr | Ccnb1ip1 | -5.009 | 1.81E-04 | glio |
| Interaction_genotype_Mtor | Gm10482 | -3.577 | 1.81E-04 | glio |
| Interaction_genotype_Myc | 1700006J14Ri | -4.031 | 1.81E-04 | glio |
| Interaction_genotype_Myc | 4930592C13R | -4.354 | 1.81E-04 | glio |
| Interaction_genotype_Myc | Gm45494 | -4.026 | 1.81E-04 | glio |
| Interaction_genotype_Myc | Gm50399 | -4.046 | 1.81E-04 | glio |
| Interaction_genotype_Myc | Fam110a | -3.608 | 1.81E-04 | glio |
| Interaction_genotype_Myc | Mcph1 | -2.688 | 1.81E-04 | glio |
| Interaction_genotype_Myc | Ddx21 | 1.766 | 1.81E-04 | glio |
| Aars2_ex.vivo_noRT | Gbp10 | 3.293 | 1.81E-04 | glio |
| Bcas2_in.vivo_noRT | Hsd17b14 | 2.849 | 1.81E-04 | glio |
| Chek1_ex.vivo_noRT | Gm35986 | 3.164 | 1.81E-04 | glio |
| Hsd17b10_in.vivo_noRT | Zkscan4 | 2.816 | 1.81E-04 | glio |
| Hsd17b10_in.vivo_noRT | Qrich2 | 2.838 | 1.81E-04 | glio |
| Hsd17b10_in.vivo_noRT | Gm16046 | 2.824 | 1.81E-04 | glio |
| Interaction_genotype_Mtor | Gm14295 | -4.178 | 1.81E-05 | glio |
| Interaction_genotype_Myc | Hspb9 | -5.292 | 1.81E-05 | glio |
| Interaction_genotype_Myc | Wdr83os | 1.909 | 1.81E-05 | glio |
| interaction_RT | Mcm3 | -1.243 | 1.81E-05 | glio |
| Atr_in.vivo_noRT | Gm49768 | 2.78 | 1.81E-05 | glio |
| Bcas2_in.vivo_noRT | 4930417H01R | 3.167 | 1.81E-05 | glio |
| Bcas2_in.vivo_noRT | Reep4 | -2.359 | 1.81E-05 | glio |
| Bora_ex.vivo_noRT | Ly6d | 3.201 | 1.81E-05 | glio |
| Hsd17b10_in.vivo_noRT | 4930478M13F | 3.237 | 1.81E-05 | glio |
| Aars2_in.vivo_noRT | Sbk2 | 3.361 | 1.81E-06 | glio |
| Aars2_in.vivo_noRT | Arvcf | 3.554 | 1.81E-06 | glio |
| Hsd17b10_in.vivo_noRT | Rbfaos | 2.699 | 1.81E-06 | glio |
| culture_effect | Ggta1 | -1.54 | 1.81E-07 | glio |
| culture_effect | Gata6 | -1.263 | 1.81E-07 | glio |
| culture_effect | Sipa1l1 | 0.934 | 1.81E-07 | glio |
| Interaction_genotype_Mtor | C330002G04R | -4.685 | 1.81E-07 | glio |
| Aars2_in.vivo_noRT | Mir142hg | 3.734 | 1.81E-07 | glio |
| Atp5c1_in.vivo_noRT | BC061195 | 2.668 | 1.81E-07 | glio |
| Atr_in.vivo_noRT | Gap43 | 4.087 | 1.81E-07 | glio |
| Interaction_genotype_Myc | 4930589L23Ri | -6.696 | 1.81E-08 | glio |
| Interaction_genotype_Myc | Gm48242 | -5.678 | 1.81E-08 | glio |
| Aars2_in.vivo_noRT | 4930592C13R | 4.06 | 1.81E-08 | glio |
| Aars2_in.vivo_noRT | Vipr2 | 3.92 | 1.81E-08 | glio |
| culture_effect | Ankrd40 | 0.922 | 1.81E-09 | glio |
| Aars2_in.vivo_noRT | Zfp85os | 4.267 | 1.81E-09 | glio |
| Aars2_in.vivo_noRT | Dnase1l2 | 4.038 | 1.81E-09 | glio |
| Atr_in.vivo_noRT | Gm20707 | 4.148 | 1.81E-09 | glio |
| Atr_in.vivo_noRT | Gadl1 | 3.896 | 1.81E-09 | glio |
| Chek1_ex.vivo_noRT | Rpl3l | 4.842 | 1.81E-09 | glio |
| Chek1_ex.vivo_noRT | Gm13684 | 4.449 | 1.81E-09 | glio |
| Atr_in.vivo_noRT | Kctd4 | 4.261 | 1.81E-10 | glio |
| Atr_in.vivo_noRT | Mir670hg | 4.883 | 1.81E-10 | glio |
| Atr_ex.vivo_noRT | Serping1 | 4.746 | 1.81E-10 | glio |
| Atr_ex.vivo_noRT | Gm47510 | 3.947 | 1.81E-10 | glio |

| | | | | |
|---|---|---|---|---|
| Aars2_in.vivo_noRT | Ankrd33b | 4.811 | 1.81E-11 | glio |
| Aars2_in.vivo_noRT | Clca3a1 | 4.275 | 1.81E-11 | glio |
| Aars2_in.vivo_noRT | Apobr | 5.327 | 1.81E-12 | glio |
| culture_effect | Gt(ROSA)26Sor | 1.558 | 1.81E-13 | glio |
| Atr_in.vivo_noRT | Ly6g6c | 4.673 | 1.81E-13 | glio |
| Bcas2_in.vivo_noRT | Krtcap3 | 4.015 | 1.81E-13 | glio |
| Bcas2_in.vivo_noRT | Gm21762 | 4.543 | 1.81E-13 | glio |
| Chek1_in.vivo_noRT | Gm12860 | 4.036 | 1.81E-13 | glio |
| Chek1_in.vivo_noRT | Bmp15 | 4.391 | 1.81E-13 | glio |
| Hsd17b10_in.vivo_noRT | Trim17 | 3.616 | 1.81E-13 | glio |
| Aars2_in.vivo_noRT | Gm43653 | 5.063 | 1.81E-14 | glio |
| Bora_in.vivo_noRT | Vtn | 4.676 | 1.81E-19 | glio |
| culture_effect | Atox1 | 1.259 | 1.81E-22 | glio |
| Chek1_in.vivo_noRT | Gm45338 | 5.044 | 1.81E-26 | glio |
| Aars2_in.vivo_noRT | Gm44736 | 5.327 | 1.81E-28 | glio |
| culture_effect | Amer1 | -0.76 | 1.82E-03 | glio |
| culture_effect | Agpat2 | 0.749 | 1.82E-03 | glio |
| culture_effect | Slc25a17 | -0.466 | 1.82E-03 | glio |
| culture_effect | Vapb | 0.494 | 1.82E-03 | glio |
| Interaction_genotype_Aars2 | Ccdc80 | -3.003 | 1.82E-03 | glio |
| Interaction_genotype_Atr | Tmem179 | -3.69 | 1.82E-03 | glio |
| Interaction_genotype_Atr | H2-Eb1 | 3.619 | 1.82E-03 | glio |
| Interaction_genotype_Atr | Gaa | 1.834 | 1.82E-03 | glio |
| Interaction_genotype_Bcas2 | AC163347.1 | -3.384 | 1.82E-03 | glio |
| Interaction_genotype_Bcas2 | Kbtbd8 | -4.114 | 1.82E-03 | glio |
| Interaction_genotype_Bcas2 | Gm30794 | -3.042 | 1.82E-03 | glio |
| Interaction_genotype_Bcas2 | Gm15743 | -3.877 | 1.82E-03 | glio |
| Interaction_genotype_Bcas2 | Gm49086 | -3.628 | 1.82E-03 | glio |
| Interaction_genotype_Bcas2 | Gm43701 | -3.047 | 1.82E-03 | glio |
| Interaction_genotype_Mtor | Ip6k3 | -3.846 | 1.82E-03 | glio |
| Interaction_genotype_Mtor | Cerk | -3.047 | 1.82E-03 | glio |
| Interaction_genotype_Mtor | Gm17501 | -3.422 | 1.82E-03 | glio |
| Interaction_genotype_Mtor | Nsfl1c | 1.616 | 1.82E-03 | glio |
| Interaction_genotype_Mtor | Cdc123 | 1.453 | 1.82E-03 | glio |
| Interaction_genotype_Myc | Mogat1 | -3.791 | 1.82E-03 | glio |
| Interaction_genotype_Myc | Zmynd15 | -3.424 | 1.82E-03 | glio |
| Interaction_genotype_Myc | 1700066M21F | -3.227 | 1.82E-03 | glio |
| Interaction_genotype_Myc | Pcdhgc3 | -2.824 | 1.82E-03 | glio |
| Interaction_genotype_Myc | Eif2ak3 | -2.688 | 1.82E-03 | glio |
| Interaction_genotype_Myc | Basp1 | -3.667 | 1.82E-03 | glio |
| Interaction_genotype_Nat2 | Prex2 | 4.79 | 1.82E-03 | glio |
| Interaction_genotype_Pars2 | Clk3 | 2.206 | 1.82E-03 | glio |
| Interaction_genotype_Rad51 | Pde3a | -4.129 | 1.82E-03 | glio |
| Interaction_genotype_Rad9a | Pcolce | -3.758 | 1.82E-03 | glio |
| Interaction_genotype_Rpl3 | Rbm4 | 2.244 | 1.82E-03 | glio |
| Aars2_in.vivo_noRT | Cep152 | 1.77 | 1.82E-03 | glio |
| Atr_in.vivo_noRT | Fbxo10 | 2.194 | 1.82E-03 | glio |
| Atr_in.vivo_noRT | Gars | -0.98 | 1.82E-03 | glio |
| Atr_ex.vivo_noRT | Prm1 | 2.764 | 1.82E-03 | glio |
| Bora_in.vivo_noRT | Gm15708 | 2.249 | 1.82E-03 | glio |
| Bora_in.vivo_noRT | Smc5 | -1.567 | 1.82E-03 | glio |
| Cenpj_in.vivo_noRT | Ripply3 | 2.551 | 1.82E-03 | glio |

| | | | | |
|---|---|---|---|---|
| culture_effect | Grcc10 | -0.424 | 1.82E-04 | glio |
| Interaction_genotype_Ifnar1 | Gm16178 | 3.531 | 1.82E-04 | glio |
| Interaction_genotype_Mtor | Rpl7a | 1.146 | 1.82E-04 | glio |
| Interaction_genotype_Myc | D630024D03R | -3.316 | 1.82E-04 | glio |
| Interaction_genotype_Myc | Gm15908 | -2.729 | 1.82E-04 | glio |
| Interaction_genotype_Myc | A930014E10Ri | -2.729 | 1.82E-04 | glio |
| Interaction_genotype_Myc | Plekho2 | -4.16 | 1.82E-04 | glio |
| Interaction_genotype_Myc | Pif1 | -4.065 | 1.82E-04 | glio |
| Interaction_genotype_Myc | Pag1 | -3.878 | 1.82E-04 | glio |
| Interaction_genotype_Myc | Tbc1d4 | -3.643 | 1.82E-04 | glio |
| Interaction_genotype_Myc | Map2k2 | 1.589 | 1.82E-04 | glio |
| interaction_RT | Bpgm | 1.13 | 1.82E-04 | glio |
| Aars2_in.vivo_noRT | Fst | 2.424 | 1.82E-04 | glio |
| Aars2_in.vivo_noRT | Gm16537 | 2.367 | 1.82E-04 | glio |
| Bcas2_in.vivo_noRT | Tnp2 | 2.744 | 1.82E-04 | glio |
| Bcas2_ex.vivo_noRT | Gpr22 | 2.742 | 1.82E-04 | glio |
| Bcas2_ex.vivo_noRT | Napsa | 3.93 | 1.82E-04 | glio |
| Bora_in.vivo_noRT | Rps6ka4 | 2.508 | 1.82E-04 | glio |
| Cenpj_ex.vivo_noRT | BC061195 | 2.591 | 1.82E-04 | glio |
| Hsd17b10_in.vivo_noRT | Gm9970 | 2.971 | 1.82E-04 | glio |
| Hsd17b10_in.vivo_noRT | 4930518J21Ri | 2.691 | 1.82E-04 | glio |
| Hsd17b10_in.vivo_noRT | H2-DMb2 | 2.601 | 1.82E-04 | glio |
| culture_effect | Peg3 | -0.874 | 1.82E-05 | glio |
| Interaction_genotype_Mtor | 0610009L18Ri | -4.739 | 1.82E-05 | glio |
| Interaction_genotype_Mtor | Gm45441 | -5.175 | 1.82E-05 | glio |
| Interaction_genotype_Myc | 4930513N10R | -5.535 | 1.82E-05 | glio |
| Interaction_genotype_Myc | Gm28731 | -5.205 | 1.82E-05 | glio |
| Interaction_genotype_Myc | Rps27a | 1.594 | 1.82E-05 | glio |
| Aars2_in.vivo_noRT | Gm10501 | 3.184 | 1.82E-05 | glio |
| Aars2_in.vivo_noRT | Myl6b | 2.922 | 1.82E-05 | glio |
| Aars2_ex.vivo_noRT | St18 | 3.293 | 1.82E-05 | glio |
| Bcas2_in.vivo_noRT | Aldh1l1 | 3.385 | 1.82E-05 | glio |
| Bcas2_in.vivo_noRT | Usp11 | 3.226 | 1.82E-05 | glio |
| Bcas2_in.vivo_noRT | Anks6 | 2.861 | 1.82E-05 | glio |
| Bora_in.vivo_noRT | Max | -1.657 | 1.82E-05 | glio |
| Bora_ex.vivo_noRT | Ccdc162 | 3.201 | 1.82E-05 | glio |
| Chek1_in.vivo_noRT | Phtf1os | 3.119 | 1.82E-05 | glio |
| Chek1_in.vivo_noRT | Gm48086 | 3.096 | 1.82E-05 | glio |
| Chek1_in.vivo_noRT | 1010001B22R | 3.043 | 1.82E-05 | glio |
| Hsd17b10_in.vivo_noRT | Gm32591 | 3.093 | 1.82E-05 | glio |
| Hsd17b10_in.vivo_noRT | Platr20 | 3.352 | 1.82E-05 | glio |
| culture_effect | Tbc1d15 | -0.727 | 1.82E-06 | glio |
| Interaction_genotype_Myc | Arsj | -5.837 | 1.82E-06 | glio |
| Aars2_in.vivo_noRT | Gm13663 | 3.051 | 1.82E-06 | glio |
| Atr_in.vivo_noRT | C1qtnf12 | 3.685 | 1.82E-06 | glio |
| Atr_in.vivo_noRT | D830050J10Ri | 3.112 | 1.82E-06 | glio |
| Bora_in.vivo_noRT | Kcnab3 | 3.402 | 1.82E-06 | glio |
| Bora_ex.vivo_noRT | Gm20732 | 3.993 | 1.82E-06 | glio |
| Bora_ex.vivo_noRT | Gm45338 | 2.808 | 1.82E-06 | glio |
| Bora_ex.vivo_noRT | Rem2 | 3.994 | 1.82E-06 | glio |
| Bora_ex.vivo_noRT | Zscan4-ps2 | 3.201 | 1.82E-06 | glio |
| Chek1_in.vivo_noRT | Gm41396 | 3.551 | 1.82E-06 | glio |

| | | | | |
|---|---|---|---|---|
| Chek1_ex.vivo_noRT | Gm8013 | 4.256 | 1.82E-06 | glio |
| Interaction_genotype_Mtor | Gm10399 | -5.6 | 1.82E-07 | glio |
| Chek1_in.vivo_noRT | Slc26a1 | 3.541 | 1.82E-07 | glio |
| Hsd17b10_in.vivo_noRT | Htr1b | 3.616 | 1.82E-07 | glio |
| culture_effect | Pole4 | 0.754 | 1.82E-08 | glio |
| Aars2_in.vivo_noRT | Rpl4 | -1.041 | 1.82E-08 | glio |
| Atr_in.vivo_noRT | 4930513N10R | 4.462 | 1.82E-08 | glio |
| Atr_in.vivo_noRT | Gm43689 | 3.577 | 1.82E-08 | glio |
| Bora_in.vivo_noRT | Slc36a2 | 3.817 | 1.82E-08 | glio |
| Chek1_in.vivo_noRT | Gm50334 | 4.399 | 1.82E-08 | glio |
| Chek1_in.vivo_noRT | Spint4 | 4.253 | 1.82E-09 | glio |
| culture_effect | Gm3716 | 3.181 | 1.82E-10 | glio |
| culture_effect | Scrg1 | -2.146 | 1.82E-10 | glio |
| Aars2_in.vivo_noRT | Rax | 4.148 | 1.82E-10 | glio |
| Bora_in.vivo_noRT | Brsk2 | 4.058 | 1.82E-10 | glio |
| Hsd17b10_in.vivo_noRT | Gm15929 | 3.616 | 1.82E-10 | glio |
| Atr_in.vivo_noRT | Alox12 | 5.249 | 1.82E-13 | glio |
| Bcas2_in.vivo_noRT | Gm4858 | 4.807 | 1.82E-13 | glio |
| Atr_in.vivo_noRT | Gm11963 | 5.273 | 1.82E-14 | glio |
| Atr_in.vivo_noRT | Necab2 | 5.269 | 1.82E-16 | glio |
| Bora_in.vivo_noRT | 4930527F14Ri | 4.407 | 1.82E-16 | glio |
| Chek1_in.vivo_noRT | Adam30 | 5.176 | 1.82E-17 | glio |
| Chek1_in.vivo_noRT | Gm20705 | 5.042 | 1.82E-19 | glio |
| culture_effect | Slc26a11 | -1.555 | 1.83E-03 | glio |
| culture_effect | Rhoq | 0.902 | 1.83E-03 | glio |
| culture_effect | Tnip1 | -0.698 | 1.83E-03 | glio |
| Interaction_genotype_Bora | H2-Q7 | 3.632 | 1.83E-03 | glio |
| Interaction_genotype_Mtor | Tmem44 | -3.795 | 1.83E-03 | glio |
| Interaction_genotype_Mtor | Zc3h12c | -3.429 | 1.83E-03 | glio |
| Interaction_genotype_Mtor | Tarsl2 | -2.898 | 1.83E-03 | glio |
| Interaction_genotype_Mtor | Naxe | 1.721 | 1.83E-03 | glio |
| Interaction_genotype_Myc | Gm35986 | -3.359 | 1.83E-03 | glio |
| Interaction_genotype_Myc | Gpd1 | -3.694 | 1.83E-03 | glio |
| Interaction_genotype_Rad21 | Gtsf1l | 3.85 | 1.83E-03 | glio |
| Interaction_genotype_Uqcrb | Mxra7 | 2.775 | 1.83E-03 | glio |
| Interaction_genotype_Uqcrb | Hbq1b | -4.837 | 1.83E-03 | glio |
| interaction_RT | Tpbg | 2.774 | 1.83E-03 | glio |
| interaction_RT | Reep5 | 0.599 | 1.83E-03 | glio |
| Aars2_in.vivo_noRT | Gm5141 | 2.214 | 1.83E-03 | glio |
| Aars2_ex.vivo_noRT | Sbk3 | 2.897 | 1.83E-03 | glio |
| Atp5c1_in.vivo_noRT | Ak1 | 2.456 | 1.83E-03 | glio |
| Atr_in.vivo_noRT | Syn1 | 2.268 | 1.83E-03 | glio |
| Bard1_ex.vivo_noRT | 4930423D22R | 2.235 | 1.83E-03 | glio |
| Bcas2_in.vivo_noRT | Hist1h2bh | 2.342 | 1.83E-03 | glio |
| Bcas2_in.vivo_noRT | Edc4 | -1.472 | 1.83E-03 | glio |
| Bcas2_in.vivo_noRT | Btd | -1.711 | 1.83E-03 | glio |
| Bcas2_in.vivo_noRT | Osgepl1 | 1.381 | 1.83E-03 | glio |
| Bcas2_ex.vivo_noRT | Adm | 3.147 | 1.83E-03 | glio |
| Cenpj_in.vivo_noRT | Fhl1 | 2.022 | 1.83E-03 | glio |
| Chek1_in.vivo_noRT | Vamp3 | -1.286 | 1.83E-03 | glio |
| Chek1_in.vivo_noRT | Rnaseh2c | -1.046 | 1.83E-03 | glio |
| Chek1_ex.vivo_noRT | Map4k4 | -1.268 | 1.83E-03 | glio |

| | | | | |
|---|---|---|---|---|
| Eif2s1_ex.vivo_noRT | Fam25c | 2.798 | 1.83E-03 | glio |
| Eif2s1_ex.vivo_noRT | Sucnr1 | 2.234 | 1.83E-03 | glio |
| Hsd17b10_in.vivo_noRT | Ggt7 | -1.637 | 1.83E-03 | glio |
| culture_effect | Hk1os | 1.571 | 1.83E-04 | glio |
| culture_effect | Nprl3 | 0.814 | 1.83E-04 | glio |
| culture_effect | Rps20 | -0.601 | 1.83E-04 | glio |
| Interaction_genotype_Mtor | Nova2 | -4.195 | 1.83E-04 | glio |
| Interaction_genotype_Myc | Gm26619 | -4.477 | 1.83E-04 | glio |
| Interaction_genotype_Myc | Foxd2 | -3.73 | 1.83E-04 | glio |
| interaction_RT | Mgarp | 3.297 | 1.83E-04 | glio |
| Aars2_in.vivo_noRT | Map3k8 | 2.851 | 1.83E-04 | glio |
| Bard1_ex.vivo_noRT | Nts | 3.987 | 1.83E-04 | glio |
| Bora_in.vivo_noRT | Knstrn | -1.776 | 1.83E-04 | glio |
| Bora_in.vivo_noRT | Aopep | -1.393 | 1.83E-04 | glio |
| Bora_ex.vivo_noRT | Tmem52 | 3.201 | 1.83E-04 | glio |
| Bora_ex.vivo_noRT | Gm36043 | 3.201 | 1.83E-04 | glio |
| Chek1_in.vivo_noRT | Def6 | 2.573 | 1.83E-04 | glio |
| Chek1_ex.vivo_noRT | Ttc16 | 3.247 | 1.83E-04 | glio |
| Chek1_ex.vivo_noRT | Rrp15 | -1.335 | 1.83E-04 | glio |
| Chek1_ex.vivo_noRT | Gtf3a | -1.333 | 1.83E-04 | glio |
| Hsd17b10_in.vivo_noRT | Gm15853 | 2.881 | 1.83E-04 | glio |
| Hsd17b10_in.vivo_noRT | Gm50103 | 2.84 | 1.83E-04 | glio |
| Hsd17b10_in.vivo_noRT | Zfp777 | -1.941 | 1.83E-04 | glio |
| culture_effect | Naglu | -1.262 | 1.83E-05 | glio |
| culture_effect | Retsat | -1.193 | 1.83E-05 | glio |
| culture_effect | Polr1d | 0.406 | 1.83E-05 | glio |
| Interaction_genotype_Mtor | Cyb561d1 | -5.469 | 1.83E-05 | glio |
| Interaction_genotype_Mtor | Gm9844 | -4.943 | 1.83E-05 | glio |
| Interaction_genotype_Mtor | Gm38163 | -4.099 | 1.83E-05 | glio |
| Aars2_in.vivo_noRT | Prss22 | 3.729 | 1.83E-05 | glio |
| Aars2_in.vivo_noRT | Mblac1 | 3.181 | 1.83E-05 | glio |
| Aars2_in.vivo_noRT | Cnbd2 | 3.18 | 1.83E-05 | glio |
| Aars2_in.vivo_noRT | Ang | 3.088 | 1.83E-05 | glio |
| Bcas2_in.vivo_noRT | Gramd2 | 2.969 | 1.83E-05 | glio |
| Bcas2_in.vivo_noRT | Xist | 2.955 | 1.83E-05 | glio |
| Bcas2_in.vivo_noRT | Gstm6 | 3.268 | 1.83E-05 | glio |
| Chek1_in.vivo_noRT | Rab12 | -1.688 | 1.83E-05 | glio |
| culture_effect | Eif3j1 | 0.809 | 1.83E-06 | glio |
| Aars2_in.vivo_noRT | Tmc7 | 3.514 | 1.83E-06 | glio |
| Aars2_in.vivo_noRT | Shd | 3.239 | 1.83E-06 | glio |
| Bcas2_in.vivo_noRT | Rasgrp2 | 3.506 | 1.83E-06 | glio |
| Bora_in.vivo_noRT | Cyth4 | 3.396 | 1.83E-06 | glio |
| Chek1_ex.vivo_noRT | BC018473 | 4.045 | 1.83E-06 | glio |
| Interaction_genotype_Mtor | Gm15929 | -5.075 | 1.83E-07 | glio |
| interaction_RT | Paip2 | 0.855 | 1.83E-07 | glio |
| Aars2_in.vivo_noRT | 2810457G06R | 4.079 | 1.83E-07 | glio |
| Atr_ex.vivo_noRT | Gm11232 | 3.164 | 1.83E-07 | glio |
| Bcas2_in.vivo_noRT | Sema3a | 3.369 | 1.83E-07 | glio |
| Chek1_in.vivo_noRT | Pagr1a | 3.359 | 1.83E-07 | glio |
| Interaction_genotype_Mtor | Gm49312 | -5.918 | 1.83E-08 | glio |
| interaction_RT | Npm3 | -1.05 | 1.83E-08 | glio |
| Hsd17b10_in.vivo_noRT | Gm38477 | 3.094 | 1.83E-08 | glio |

| | | | | |
|---|---|---|---|---|
| Hsd17b10_in.vivo_noRT | Adgre5 | 3.356 | 1.83E-08 | glio |
| Hsd17b10_in.vivo_noRT | Gm48899 | 3.356 | 1.83E-08 | glio |
| culture_effect | Naga | -1.863 | 1.83E-09 | glio |
| Atr_in.vivo_noRT | Tec | 3.857 | 1.83E-09 | glio |
| culture_effect | Stard3nl | 1.218 | 1.83E-10 | glio |
| culture_effect | Eif3l | -0.771 | 1.83E-10 | glio |
| Atr_in.vivo_noRT | 1700061G19R | 4.522 | 1.83E-11 | glio |
| Chek1_in.vivo_noRT | Cstdc4 | 4.387 | 1.83E-11 | glio |
| culture_effect | Peg10 | 2.205 | 1.83E-12 | glio |
| Aars2_in.vivo_noRT | Thap8 | 5.067 | 1.83E-12 | glio |
| Aars2_in.vivo_noRT | Gm35290 | 4.589 | 1.83E-12 | glio |
| Atr_in.vivo_noRT | Col6a3 | 4.497 | 1.83E-13 | glio |
| Hsd17b10_in.vivo_noRT | Tpo | 3.616 | 1.83E-13 | glio |
| Atr_in.vivo_noRT | Rhbg | 3.938 | 1.83E-14 | glio |
| culture_effect | Mog | -5.466 | 1.83E-17 | glio |
| Bora_in.vivo_noRT | Kcnk13 | 4.673 | 1.83E-17 | glio |
| Bcas2_in.vivo_noRT | Crb3 | 4.807 | 1.83E-18 | glio |
| culture_effect | Col18a1 | -2.156 | 1.83E-20 | glio |
| culture_effect | Atp5b | -1.025 | 1.83E-20 | glio |
| Bora_in.vivo_noRT | Gm47540 | 4.936 | 1.83E-21 | glio |
| Bora_in.vivo_noRT | BB287469 | 4.936 | 1.83E-21 | glio |
| Chek1_in.vivo_noRT | Scg2 | 5.044 | 1.83E-26 | glio |
| culture_effect | Efhd2 | 0.631 | 1.84E-03 | glio |
| Interaction_genotype_Myc | Mnt | -2.835 | 1.84E-03 | glio |
| Interaction_genotype_Myc | 8430429K09R | -2.688 | 1.84E-03 | glio |
| Interaction_genotype_Myc | Derl1 | 1.387 | 1.84E-03 | glio |
| Interaction_genotype_Myc | Pkm | 1.503 | 1.84E-03 | glio |
| Interaction_genotype_Pars2 | Gm45591 | -3.591 | 1.84E-03 | glio |
| Interaction_genotype_Pars2 | Gm13348 | -3.541 | 1.84E-03 | glio |
| Interaction_genotype_Pars2 | Gm31063 | -3.192 | 1.84E-03 | glio |
| Interaction_genotype_Pars2 | Upb1 | -4.076 | 1.84E-03 | glio |
| Interaction_genotype_Pars2 | Gm30082 | -4.195 | 1.84E-03 | glio |
| Interaction_genotype_Rpl3 | Gm45589 | -4.29 | 1.84E-03 | glio |
| interaction_RT | Gemin5 | -0.883 | 1.84E-03 | glio |
| interaction_RT | Mgat2 | -0.863 | 1.84E-03 | glio |
| interaction_RT | Xaf1 | 0.711 | 1.84E-03 | glio |
| Aars2_in.vivo_noRT | Stat4 | 2.162 | 1.84E-03 | glio |
| Aars2_in.vivo_noRT | Sobp | 2.123 | 1.84E-03 | glio |
| Atr_in.vivo_noRT | Vax2 | 2.169 | 1.84E-03 | glio |
| Atr_in.vivo_noRT | Zfp382 | 2.065 | 1.84E-03 | glio |
| Atr_in.vivo_noRT | Slc4a1ap | -1.449 | 1.84E-03 | glio |
| Atr_ex.vivo_noRT | Ntrk1 | 3.076 | 1.84E-03 | glio |
| Atr_ex.vivo_noRT | Ccdc28b | 2.644 | 1.84E-03 | glio |
| Bora_in.vivo_noRT | Bbs9 | 2.041 | 1.84E-03 | glio |
| Bora_in.vivo_noRT | Tlr4 | 1.627 | 1.84E-03 | glio |
| Cenpj_ex.vivo_noRT | Ly6d | 2.591 | 1.84E-03 | glio |
| Cenpj_ex.vivo_noRT | Svopl | 2.982 | 1.84E-03 | glio |
| Cenpj_ex.vivo_noRT | Ccdc162 | 2.591 | 1.84E-03 | glio |
| Cenpj_ex.vivo_noRT | Mpzl2 | 2.591 | 1.84E-03 | glio |
| Cenpj_ex.vivo_noRT | Trim17 | 2.192 | 1.84E-03 | glio |
| Cenpj_ex.vivo_noRT | A330049N07R | 2.192 | 1.84E-03 | glio |
| Cenpj_ex.vivo_noRT | Klrk1 | 2.591 | 1.84E-03 | glio |

| | | | | |
|---|---|---|---|---|
| Chek1_in.vivo_noRT | Gm20597 | 2.463 | 1.84E-03 | glio |
| Chek1_in.vivo_noRT | Elof1 | -1.139 | 1.84E-03 | glio |
| Eif2s1_in.vivo_noRT | Gm18280 | 1.879 | 1.84E-03 | glio |
| Eif2s1_in.vivo_noRT | Pcdhb19 | 2.327 | 1.84E-03 | glio |
| Hsd17b10_in.vivo_noRT | Zfp830 | -1.208 | 1.84E-03 | glio |
| culture_effect | Micall2 | 1.944 | 1.84E-04 | glio |
| culture_effect | Mis12 | 0.79 | 1.84E-04 | glio |
| culture_effect | Add3 | 0.821 | 1.84E-04 | glio |
| culture_effect | Ipo5 | -0.608 | 1.84E-04 | glio |
| Interaction_genotype_Atr | Gm10484 | -4.86 | 1.84E-04 | glio |
| Interaction_genotype_Atr | Yars | 2.26 | 1.84E-04 | glio |
| Interaction_genotype_Atr | Ngfr | -2.949 | 1.84E-04 | glio |
| Interaction_genotype_Chek1 | H2-Eb2 | 3.892 | 1.84E-04 | glio |
| Interaction_genotype_Mtor | Gm31805 | -4.178 | 1.84E-04 | glio |
| Interaction_genotype_Mtor | Bri3 | 1.811 | 1.84E-04 | glio |
| Interaction_genotype_Myc | Ankef1 | -4.345 | 1.84E-04 | glio |
| Interaction_genotype_Myc | Cyp4x1 | -4.045 | 1.84E-04 | glio |
| Interaction_genotype_Myc | Gm16835 | -4.164 | 1.84E-04 | glio |
| Aars2_in.vivo_noRT | Hist1h2bp | 2.802 | 1.84E-04 | glio |
| Aars2_in.vivo_noRT | Rbakdn | 2.625 | 1.84E-04 | glio |
| Aars2_in.vivo_noRT | Gbp3 | -1.15 | 1.84E-04 | glio |
| Atr_in.vivo_noRT | Zfp882 | 2.845 | 1.84E-04 | glio |
| Bcas2_in.vivo_noRT | Gm15998 | 2.692 | 1.84E-04 | glio |
| Bcas2_in.vivo_noRT | Rabac1 | -0.915 | 1.84E-04 | glio |
| Bora_in.vivo_noRT | Abcd3 | -1.762 | 1.84E-04 | glio |
| Bora_ex.vivo_noRT | Gm13713 | 2.802 | 1.84E-04 | glio |
| Cenpj_ex.vivo_noRT | Idi2 | 2.591 | 1.84E-04 | glio |
| Hsd17b10_in.vivo_noRT | Peli3 | 2.701 | 1.84E-04 | glio |
| Interaction_genotype_Atr | Fbxo32 | 3.77 | 1.84E-05 | glio |
| Interaction_genotype_Atr | Mkrn1 | 2.272 | 1.84E-05 | glio |
| Interaction_genotype_Atr | 4921523L03Ri | -5.56 | 1.84E-05 | glio |
| Interaction_genotype_Atr | Klk14 | -5.3 | 1.84E-05 | glio |
| Interaction_genotype_Mtor | Tbc1d4 | -4.379 | 1.84E-05 | glio |
| Interaction_genotype_Mtor | Tal1 | -5.103 | 1.84E-05 | glio |
| Interaction_genotype_Mtor | Gm38477 | -3.976 | 1.84E-05 | glio |
| Interaction_genotype_Mtor | Sdhb | 1.902 | 1.84E-05 | glio |
| Interaction_genotype_Myc | Ehd2 | -3.591 | 1.84E-05 | glio |
| Interaction_genotype_Myc | Rab25 | -4.408 | 1.84E-05 | glio |
| interaction_RT | Dhx29 | -1.463 | 1.84E-05 | glio |
| Atr_in.vivo_noRT | Zfp772 | 3.052 | 1.84E-05 | glio |
| Atr_in.vivo_noRT | Abcd4 | 2.65 | 1.84E-05 | glio |
| Atr_ex.vivo_noRT | Mpzl2 | 3.164 | 1.84E-05 | glio |
| Atr_ex.vivo_noRT | Tex52 | 3.742 | 1.84E-05 | glio |
| Bcas2_in.vivo_noRT | Gm10602 | 3.4 | 1.84E-05 | glio |
| Bora_in.vivo_noRT | Gemin4 | 3.191 | 1.84E-05 | glio |
| Hsd17b10_in.vivo_noRT | 1700030L20Ri | 2.96 | 1.84E-05 | glio |
| culture_effect | Map2k4 | 0.859 | 1.84E-06 | glio |
| Interaction_genotype_Mtor | Tmem69 | -4.768 | 1.84E-06 | glio |
| Interaction_genotype_Myc | Cog8 | -4.453 | 1.84E-06 | glio |
| Interaction_genotype_Myc | Mettl18 | -4.761 | 1.84E-06 | glio |
| Interaction_genotype_Myc | A930012O16R | -5.698 | 1.84E-06 | glio |
| Bcas2_in.vivo_noRT | Serpinc1 | 3.519 | 1.84E-06 | glio |

| | | | | |
|---|---|---|---|---|
| Bora_in.vivo_noRT | Tmem116 | 3.758 | 1.84E-06 | glio |
| Aars2_in.vivo_noRT | Msl3l2 | 3.631 | 1.84E-07 | glio |
| Atr_in.vivo_noRT | 4930556M19F | 3.423 | 1.84E-07 | glio |
| Atr_ex.vivo_noRT | Gm29966 | 3.164 | 1.84E-07 | glio |
| Atr_ex.vivo_noRT | Olfr1152 | 3.164 | 1.84E-07 | glio |
| Atr_ex.vivo_noRT | Tex14 | 4.347 | 1.84E-07 | glio |
| Bora_in.vivo_noRT | Drd4 | 3.752 | 1.84E-07 | glio |
| Aars2_in.vivo_noRT | mt-Co1 | -1.282 | 1.84E-08 | glio |
| Atr_in.vivo_noRT | B230319C09R | 4.223 | 1.84E-08 | glio |
| Atr_in.vivo_noRT | Lsp1 | 4.346 | 1.84E-08 | glio |
| Atr_in.vivo_noRT | Rpl18a | -1.049 | 1.84E-08 | glio |
| Hsd17b10_in.vivo_noRT | B4galnt1 | 3.356 | 1.84E-08 | glio |
| culture_effect | Tmem59 | -0.666 | 1.84E-09 | glio |
| Bcas2_in.vivo_noRT | 1700061G19R | 4.325 | 1.84E-10 | glio |
| Hsd17b10_in.vivo_noRT | Pitx3 | 3.616 | 1.84E-10 | glio |
| culture_effect | Dbnl | -1.008 | 1.84E-13 | glio |
| Bora_in.vivo_noRT | Rftn2 | 4.545 | 1.84E-13 | glio |
| Bora_in.vivo_noRT | Gem | 4.936 | 1.84E-13 | glio |
| Chek1_ex.vivo_noRT | Gpc5 | 4.447 | 1.84E-13 | glio |
| Chek1_in.vivo_noRT | Folr1 | 5.822 | 1.84E-21 | glio |
| Chek1_in.vivo_noRT | Gm16178 | 5.044 | 1.84E-24 | glio |
| Chek1_in.vivo_noRT | Gm364 | 5.044 | 1.84E-24 | glio |
| culture_effect | Gm15510 | 1.706 | 1.85E-03 | glio |
| Interaction_genotype_Aars2 | 9330104G04R | -3.815 | 1.85E-03 | glio |
| Interaction_genotype_Aars2 | 1700028J19Ri | -3.831 | 1.85E-03 | glio |
| Interaction_genotype_Aars2 | Kctd11 | -3.989 | 1.85E-03 | glio |
| Interaction_genotype_Aars2 | Lncppara | -4.031 | 1.85E-03 | glio |
| Interaction_genotype_Aars2 | Abcc5 | -3.602 | 1.85E-03 | glio |
| Interaction_genotype_Aars2 | Nop16 | 1.56 | 1.85E-03 | glio |
| Interaction_genotype_Aars2 | Ift20 | 1.328 | 1.85E-03 | glio |
| Interaction_genotype_Aars2 | Ciao2b | 1.135 | 1.85E-03 | glio |
| Interaction_genotype_Atr | E130215H24R | -3.626 | 1.85E-03 | glio |
| Interaction_genotype_Atr | Gm43728 | -4.046 | 1.85E-03 | glio |
| Interaction_genotype_Atr | 3110083C13R | -3.764 | 1.85E-03 | glio |
| Interaction_genotype_Atr | Myo7a | -3.41 | 1.85E-03 | glio |
| Interaction_genotype_Atr | Chchd2 | 0.918 | 1.85E-03 | glio |
| Interaction_genotype_Bora | Gm36356 | -3.447 | 1.85E-03 | glio |
| Interaction_genotype_Cdkn2a | Gm17203 | 3.338 | 1.85E-03 | glio |
| Interaction_genotype_Cdkn2a | Klk13 | 3.368 | 1.85E-03 | glio |
| Interaction_genotype_Mtor | Mogat1 | -4 | 1.85E-03 | glio |
| Interaction_genotype_Myc | Cd7 | -2.466 | 1.85E-03 | glio |
| Interaction_genotype_Myc | Cdo1 | -3.693 | 1.85E-03 | glio |
| Interaction_genotype_Myc | Stpg4 | -3.619 | 1.85E-03 | glio |
| Interaction_genotype_Myc | Nmral1 | -2.746 | 1.85E-03 | glio |
| Interaction_genotype_Myc | Klhl13 | -2.819 | 1.85E-03 | glio |
| Interaction_genotype_Myc | Dtx4 | -2.801 | 1.85E-03 | glio |
| Interaction_genotype_Myc | Zranb2 | 1.303 | 1.85E-03 | glio |
| Interaction_genotype_Pars2 | Gm32585 | -4.419 | 1.85E-03 | glio |
| interaction_RT | Kif21a | 0.92 | 1.85E-03 | glio |
| Aars2_in.vivo_noRT | Rmst | 2.347 | 1.85E-03 | glio |
| Aars2_in.vivo_noRT | Mrpl2 | -0.976 | 1.85E-03 | glio |
| Aars2_ex.vivo_noRT | Nxnl2 | 2.712 | 1.85E-03 | glio |

| | | | | |
|---|---|---|---|---|
| Atr_in.vivo_noRT | Nae1 | -1.256 | 1.85E-03 | glio |
| Atr_in.vivo_noRT | Ckap4 | -1.06 | 1.85E-03 | glio |
| Atr_ex.vivo_noRT | Dcpp1 | 3.164 | 1.85E-03 | glio |
| Bard1_ex.vivo_noRT | C430002N11R | 2.419 | 1.85E-03 | glio |
| Bard1_ex.vivo_noRT | Myt1 | 2.415 | 1.85E-03 | glio |
| Bcas2_in.vivo_noRT | 4930556M19F | 2.16 | 1.85E-03 | glio |
| Bora_in.vivo_noRT | Paox | 2.098 | 1.85E-03 | glio |
| Bora_in.vivo_noRT | Papss1 | -1.321 | 1.85E-03 | glio |
| Brca2_in.vivo_noRT | Meioc | 2.033 | 1.85E-03 | glio |
| Brca2_in.vivo_noRT | Pcdhga1 | 2.537 | 1.85E-03 | glio |
| Cenpj_in.vivo_noRT | Gm45620 | 2.548 | 1.85E-03 | glio |
| Cenpj_in.vivo_noRT | 1700112J16Ri | 2.022 | 1.85E-03 | glio |
| Cenpj_in.vivo_noRT | Clec4d | 2.022 | 1.85E-03 | glio |
| Cenpj_in.vivo_noRT | Ccdc80 | 2.022 | 1.85E-03 | glio |
| Chek1_in.vivo_noRT | Clic1 | -0.668 | 1.85E-03 | glio |
| Chek1_ex.vivo_noRT | Rtl5 | 2.851 | 1.85E-03 | glio |
| Chek1_ex.vivo_noRT | 3110001I22Ri | 2.704 | 1.85E-03 | glio |
| Chek1_ex.vivo_noRT | Srgap1 | 2.473 | 1.85E-03 | glio |
| Eif2s1_in.vivo_noRT | A930033M14F | 2.143 | 1.85E-03 | glio |
| Eif2s1_in.vivo_noRT | Gm35696 | 2.407 | 1.85E-03 | glio |
| Eif2s1_in.vivo_noRT | Gm10638 | 2.491 | 1.85E-03 | glio |
| culture_effect | Ccdc28b | -1.89 | 1.85E-04 | glio |
| culture_effect | Gm6169 | -1.833 | 1.85E-04 | glio |
| culture_effect | Txnl4a | 0.398 | 1.85E-04 | glio |
| Interaction_genotype_Atr | D430001F17R | -4.315 | 1.85E-04 | glio |
| Interaction_genotype_Mtor | Klra6 | -4.726 | 1.85E-04 | glio |
| Aars2_in.vivo_noRT | Unkl | 2.118 | 1.85E-04 | glio |
| Atp5c1_in.vivo_noRT | Adgre5 | 2.408 | 1.85E-04 | glio |
| Atp5c1_in.vivo_noRT | Gm48899 | 2.408 | 1.85E-04 | glio |
| Atr_in.vivo_noRT | Dlg5 | 2.491 | 1.85E-04 | glio |
| Atr_ex.vivo_noRT | 1700034I23Ri | 2.367 | 1.85E-04 | glio |
| Atr_ex.vivo_noRT | Ccl8 | 3.967 | 1.85E-04 | glio |
| Atr_ex.vivo_noRT | Myt1 | 2.764 | 1.85E-04 | glio |
| Atr_ex.vivo_noRT | Ccdc148 | 3.295 | 1.85E-04 | glio |
| Bcas2_in.vivo_noRT | Gm29361 | 2.782 | 1.85E-04 | glio |
| Bora_in.vivo_noRT | Zfp619 | 2.734 | 1.85E-04 | glio |
| Bora_ex.vivo_noRT | Gm20705 | 2.806 | 1.85E-04 | glio |
| Cenpj_ex.vivo_noRT | Gm16178 | 2.591 | 1.85E-04 | glio |
| Cenpj_ex.vivo_noRT | Gm364 | 2.591 | 1.85E-04 | glio |
| Chek1_in.vivo_noRT | Rnasel | 2.433 | 1.85E-04 | glio |
| Chek1_ex.vivo_noRT | Tipin | -1.62 | 1.85E-04 | glio |
| Hsd17b10_in.vivo_noRT | Gm5624 | 2.627 | 1.85E-04 | glio |
| Hsd17b10_in.vivo_noRT | Sec23b | -1.761 | 1.85E-04 | glio |
| culture_effect | Sema4b | -1.176 | 1.85E-05 | glio |
| culture_effect | Ankra2 | 0.903 | 1.85E-05 | glio |
| culture_effect | 5430416N02R | -0.98 | 1.85E-05 | glio |
| culture_effect | Eif2a | -0.65 | 1.85E-05 | glio |
| culture_effect | Ier3ip1 | 0.579 | 1.85E-05 | glio |
| Bcas2_in.vivo_noRT | 4930481B07R | 3.243 | 1.85E-05 | glio |
| Chek1_in.vivo_noRT | Smim13 | -1.945 | 1.85E-05 | glio |
| Interaction_genotype_Mtor | Gm50334 | -6.132 | 1.85E-06 | glio |
| Atr_in.vivo_noRT | Ampd3 | 3.251 | 1.85E-06 | glio |

| | | | | |
|---|---|---|---|---|
| Bcas2_in.vivo_noRT | Retn | 4.34 | 1.85E-06 | glio |
| Bora_in.vivo_noRT | Ripk2 | 1.596 | 1.85E-06 | glio |
| Chek1_in.vivo_noRT | Msl3l2 | 3.348 | 1.85E-06 | glio |
| Chek1_in.vivo_noRT | Gm43331 | 2.832 | 1.85E-06 | glio |
| Hsd17b10_in.vivo_noRT | Zfp786 | 3.352 | 1.85E-06 | glio |
| Atr_in.vivo_noRT | Slc7a7 | 3.585 | 1.85E-07 | glio |
| Atr_in.vivo_noRT | 9530018F02Ri | 4.147 | 1.85E-08 | glio |
| Atr_in.vivo_noRT | Map3k5 | 3.369 | 1.85E-08 | glio |
| Bard1_ex.vivo_noRT | D730045A05R | 5.222 | 1.85E-08 | glio |
| Bcas2_in.vivo_noRT | Apoh | 3.892 | 1.85E-08 | glio |
| Chek1_in.vivo_noRT | Ccdc39 | 3.823 | 1.85E-08 | glio |
| Chek1_in.vivo_noRT | Zfpl1 | -2.116 | 1.85E-08 | glio |
| Atr_ex.vivo_noRT | Esp24 | 3.947 | 1.85E-09 | glio |
| Bcas2_in.vivo_noRT | 9530062K07R | 4.428 | 1.85E-09 | glio |
| Chek1_in.vivo_noRT | Lmx1b | 4.257 | 1.85E-09 | glio |
| Chek1_in.vivo_noRT | Zkscan4 | 4.243 | 1.85E-09 | glio |
| Atr_in.vivo_noRT | Ptp4a1 | 3.839 | 1.85E-10 | glio |
| Atr_in.vivo_noRT | Catsperd | 4.354 | 1.85E-10 | glio |
| Aars2_in.vivo_noRT | Platr31 | 5.327 | 1.85E-11 | glio |
| Atr_in.vivo_noRT | Glis1 | 4.621 | 1.85E-11 | glio |
| Atr_ex.vivo_noRT | Gm4340 | 3.947 | 1.85E-11 | glio |
| Atr_ex.vivo_noRT | Gm16239 | 3.947 | 1.85E-11 | glio |
| Chek1_in.vivo_noRT | Gm13270 | 5.044 | 1.85E-12 | glio |
| Bora_in.vivo_noRT | Ripply3 | 4.936 | 1.85E-13 | glio |
| Atr_in.vivo_noRT | Kcnj13 | 4.762 | 1.85E-14 | glio |
| Bora_in.vivo_noRT | Prl2c3 | 4.936 | 1.85E-14 | glio |
| Bora_in.vivo_noRT | Iqch | 4.672 | 1.85E-15 | glio |
| Chek1_in.vivo_noRT | Gm38393 | 4.784 | 1.85E-15 | glio |
| Aars2_in.vivo_noRT | Flacc1 | 5.327 | 1.85E-18 | glio |
| Atr_in.vivo_noRT | Gm14019 | 5.533 | 1.85E-19 | glio |
| Bora_in.vivo_noRT | Meioc | 4.673 | 1.85E-21 | glio |
| Bora_in.vivo_noRT | Halr1 | 4.67 | 1.85E-23 | glio |
| culture_effect | Tafa4 | -4.3 | 1.85E-24 | glio |
| Bcas2_in.vivo_noRT | Gm15908 | 4.807 | 1.85E-24 | glio |
| Bcas2_in.vivo_noRT | A930014E10Ri | 4.807 | 1.85E-24 | glio |
| Atr_in.vivo_noRT | AF067063 | 5.533 | 1.85E-25 | glio |
| Atr_in.vivo_noRT | Gm13191 | 5.533 | 1.85E-31 | glio |
| Atr_in.vivo_noRT | 4933416E03Ri | 5.533 | 1.85E-31 | glio |
| culture_effect | Nr4a2 | -1.297 | 1.86E-03 | glio |
| culture_effect | Cyb5r4 | 0.533 | 1.86E-03 | glio |
| Interaction_genotype_Aars2 | Copa | 1.443 | 1.86E-03 | glio |
| Interaction_genotype_Atr | Gm15943 | -3.087 | 1.86E-03 | glio |
| Interaction_genotype_Atr | Olfr552 | -2.769 | 1.86E-03 | glio |
| Interaction_genotype_Atr | Ttc21b | -2.843 | 1.86E-03 | glio |
| Interaction_genotype_Bcas2 | Gm14207 | -2.858 | 1.86E-03 | glio |
| Interaction_genotype_Mtor | Myo16 | -3.868 | 1.86E-03 | glio |
| Interaction_genotype_Mtor | Gm16104 | -3.496 | 1.86E-03 | glio |
| Interaction_genotype_Myc | Gm21319 | -2.729 | 1.86E-03 | glio |
| Interaction_genotype_Myc | BC051077 | -4.035 | 1.86E-03 | glio |
| Interaction_genotype_Myc | Gm14305 | -2.829 | 1.86E-03 | glio |
| Interaction_genotype_Myc | Cnot9 | -2.099 | 1.86E-03 | glio |
| Interaction_genotype_Rpl3 | Prox2 | -4.576 | 1.86E-03 | glio |

| | | | | |
|---|---|---|---|---|
| Interaction_genotype_Uqcrc2 | Klhdc8a | -3.435 | 1.86E-03 | glio |
| Aars2_in.vivo_noRT | A230056P14R | 2.232 | 1.86E-03 | glio |
| Aars2_in.vivo_noRT | 2700046G09R | 1.833 | 1.86E-03 | glio |
| Aars2_in.vivo_noRT | Brms1 | -1.19 | 1.86E-03 | glio |
| Aars2_in.vivo_noRT | Sec61g | -0.803 | 1.86E-03 | glio |
| Aars2_in.vivo_noRT | Hsp90aa1 | -0.668 | 1.86E-03 | glio |
| Atp5c1_in.vivo_noRT | Gm17146 | 2.278 | 1.86E-03 | glio |
| Atr_in.vivo_noRT | Eif5 | -0.677 | 1.86E-03 | glio |
| Atr_ex.vivo_noRT | Icosl | 2.764 | 1.86E-03 | glio |
| Atr_ex.vivo_noRT | Bcl3 | 2.78 | 1.86E-03 | glio |
| Bcas2_in.vivo_noRT | Trnt1 | -1.219 | 1.86E-03 | glio |
| Bcas2_in.vivo_noRT | Tapbpl | -1.27 | 1.86E-03 | glio |
| Bcas2_ex.vivo_noRT | Gm33973 | 2.742 | 1.86E-03 | glio |
| Bcas2_ex.vivo_noRT | Rhbg | 2.349 | 1.86E-03 | glio |
| Bora_in.vivo_noRT | Ppp1r13l | 2.281 | 1.86E-03 | glio |
| Bora_in.vivo_noRT | Tst | -1.49 | 1.86E-03 | glio |
| Bora_ex.vivo_noRT | Pkd1l2 | 2.806 | 1.86E-03 | glio |
| Chek1_in.vivo_noRT | Hist1h3d | 2.14 | 1.86E-03 | glio |
| Chek1_in.vivo_noRT | Mmaa | 1.986 | 1.86E-03 | glio |
| Chek1_in.vivo_noRT | Gnb1 | -0.804 | 1.86E-03 | glio |
| Chek1_ex.vivo_noRT | Osbp2 | 3.345 | 1.86E-03 | glio |
| Chek1_ex.vivo_noRT | Hmgcs2 | 3.298 | 1.86E-03 | glio |
| Chek1_ex.vivo_noRT | 1110046J04Ri | 2.831 | 1.86E-03 | glio |
| Chek1_ex.vivo_noRT | Cct6b | 2.763 | 1.86E-03 | glio |
| Chek1_ex.vivo_noRT | A530072M11F | 2.84 | 1.86E-03 | glio |
| Chek1_ex.vivo_noRT | Gm28417 | 2.74 | 1.86E-03 | glio |
| Chek1_ex.vivo_noRT | Acox3 | 2.708 | 1.86E-03 | glio |
| Chek1_ex.vivo_noRT | Naa10 | -0.992 | 1.86E-03 | glio |
| Hsd17b10_in.vivo_noRT | Gm47972 | 2.322 | 1.86E-03 | glio |
| Hsd17b10_in.vivo_noRT | Tnfrsf13b | 2.509 | 1.86E-03 | glio |
| culture_effect | Msra | -1.13 | 1.86E-04 | glio |
| culture_effect | Kdm3a | -0.793 | 1.86E-04 | glio |
| culture_effect | Rnf121 | -0.726 | 1.86E-04 | glio |
| Interaction_genotype_Atr | Camk1d | -4.513 | 1.86E-04 | glio |
| Interaction_genotype_Mtor | Cyp27b1 | -4.082 | 1.86E-04 | glio |
| Interaction_genotype_Mtor | Kctd14 | -4.13 | 1.86E-04 | glio |
| Interaction_genotype_Mtor | Tnfrsf4 | -4.488 | 1.86E-04 | glio |
| Interaction_genotype_Mtor | Eif3j2 | -4.464 | 1.86E-04 | glio |
| Interaction_genotype_Myc | Spata31d1a | -4.32 | 1.86E-04 | glio |
| Interaction_genotype_Myc | Impg2 | -3.517 | 1.86E-04 | glio |
| Interaction_genotype_Myc | Hist1h2bh | -4.283 | 1.86E-04 | glio |
| Interaction_genotype_Myc | Bahd1 | -3.747 | 1.86E-04 | glio |
| Interaction_genotype_Myc | Ampd2 | -2.866 | 1.86E-04 | glio |
| Interaction_genotype_Myc | C1qbp | 1.65 | 1.86E-04 | glio |
| Interaction_genotype_Myc | Pgls | 1.545 | 1.86E-04 | glio |
| Aars2_in.vivo_noRT | Rgl1 | 2.411 | 1.86E-04 | glio |
| Aars2_in.vivo_noRT | Gpn3 | -1.652 | 1.86E-04 | glio |
| Aars2_ex.vivo_noRT | Gm2449 | 2.995 | 1.86E-04 | glio |
| Atp5c1_in.vivo_noRT | B4galnt1 | 2.408 | 1.86E-04 | glio |
| Atr_in.vivo_noRT | Pde6h | 2.459 | 1.86E-04 | glio |
| Atr_in.vivo_noRT | Spa17 | 2.598 | 1.86E-04 | glio |
| Bora_in.vivo_noRT | Sox8 | 2.21 | 1.86E-04 | glio |

| | | | | |
|---|---|---|---|---|
| Chek1_in.vivo_noRT | Gm12258 | 2.837 | 1.86E-04 | glio |
| Chek1_in.vivo_noRT | Agfg1 | -1.364 | 1.86E-04 | glio |
| culture_effect | Slc6a15 | -1.273 | 1.86E-05 | glio |
| culture_effect | Ppl | -1.149 | 1.86E-05 | glio |
| culture_effect | AU040320 | -1.033 | 1.86E-05 | glio |
| Interaction_genotype_Mtor | A930029G22R | -4.528 | 1.86E-05 | glio |
| Interaction_genotype_Mtor | Nyap1 | -5.064 | 1.86E-05 | glio |
| Interaction_genotype_Mtor | Srl | -5.178 | 1.86E-05 | glio |
| Interaction_genotype_Mtor | Hnrnpl | 2.172 | 1.86E-05 | glio |
| Aars2_in.vivo_noRT | Zfp493 | 2.978 | 1.86E-05 | glio |
| Aars2_in.vivo_noRT | Gm14966 | 2.859 | 1.86E-05 | glio |
| Atr_in.vivo_noRT | Gm5141 | 2.958 | 1.86E-05 | glio |
| Bard1_ex.vivo_noRT | Tpo | 2.814 | 1.86E-05 | glio |
| Bcas2_in.vivo_noRT | H2-DMb2 | 2.883 | 1.86E-05 | glio |
| Bora_in.vivo_noRT | Myl9 | 3.373 | 1.86E-05 | glio |
| Bora_in.vivo_noRT | Gm48832 | 2.92 | 1.86E-05 | glio |
| Bora_ex.vivo_noRT | Gm41307 | 2.808 | 1.86E-05 | glio |
| Interaction_genotype_Mtor | Medag | -5.563 | 1.86E-06 | glio |
| Aars2_in.vivo_noRT | Hapln4 | 3.559 | 1.86E-06 | glio |
| Aars2_ex.vivo_noRT | Cxcl5 | 3.293 | 1.86E-06 | glio |
| Atp5c1_in.vivo_noRT | 4930512B01R | 2.668 | 1.86E-06 | glio |
| Chek1_ex.vivo_noRT | Gm15486 | 3.871 | 1.86E-06 | glio |
| Hsd17b10_in.vivo_noRT | 4921536K21R | 3.356 | 1.86E-06 | glio |
| culture_effect | Fhad1 | -3.074 | 1.86E-07 | glio |
| culture_effect | Pdxk | 0.988 | 1.86E-07 | glio |
| culture_effect | Rae1 | -0.701 | 1.86E-07 | glio |
| Aars2_in.vivo_noRT | Gm36495 | 4.032 | 1.86E-07 | glio |
| Atp5c1_in.vivo_noRT | Idi2 | 2.668 | 1.86E-07 | glio |
| Atp5c1_in.vivo_noRT | Vmn2r18 | 2.668 | 1.86E-07 | glio |
| Bcas2_in.vivo_noRT | AU022754 | 3.387 | 1.86E-07 | glio |
| Interaction_genotype_Mtor | Lncbate10 | -4.886 | 1.86E-08 | glio |
| Aars2_in.vivo_noRT | 1700013F07Ri | 4.172 | 1.86E-08 | glio |
| Bcas2_in.vivo_noRT | Cfap58 | 4.807 | 1.86E-08 | glio |
| Bcas2_in.vivo_noRT | Nfix | -2.368 | 1.86E-08 | glio |
| Bora_in.vivo_noRT | Gm26590 | 3.648 | 1.86E-08 | glio |
| Aars2_in.vivo_noRT | Gm15853 | 4.591 | 1.86E-10 | glio |
| Atr_in.vivo_noRT | Gm34590 | 4.498 | 1.86E-10 | glio |
| Chek1_in.vivo_noRT | Gm14968 | 4.664 | 1.86E-10 | glio |
| Bora_in.vivo_noRT | Pla1a | 4.405 | 1.86E-11 | glio |
| Chek1_in.vivo_noRT | 5033417F24Ri | 5.044 | 1.86E-13 | glio |
| Chek1_ex.vivo_noRT | Gm11899 | 4.842 | 1.86E-13 | glio |
| Hsd17b10_in.vivo_noRT | Gm48155 | 3.616 | 1.86E-14 | glio |
| Bora_in.vivo_noRT | Gm38477 | 4.938 | 1.86E-19 | glio |
| Bcas2_in.vivo_noRT | Meioc | 4.544 | 1.86E-20 | glio |
| culture_effect | Ube2l6 | -1.684 | 1.86E-23 | glio |
| Aars2_in.vivo_noRT | Tmem132c | 5.327 | 1.86E-26 | glio |
| Atr_in.vivo_noRT | Erich6b | 5.533 | 1.86E-27 | glio |
| Atr_in.vivo_noRT | Lncbate10 | 5.533 | 1.86E-27 | glio |
| culture_effect | Fos | -3.247 | 1.86E-36 | glio |
| culture_effect | Gstm2 | 1.669 | 1.87E-03 | glio |
| culture_effect | Stx11 | -0.865 | 1.87E-03 | glio |
| culture_effect | Acads | -0.605 | 1.87E-03 | glio |

| | | | | |
|---|---|---|---|---|
| culture_effect | Fbxw2 | 0.512 | 1.87E-03 | glio |
| culture_effect | Nucks1 | 0.411 | 1.87E-03 | glio |
| culture_effect | mt-Cytb | 0.349 | 1.87E-03 | glio |
| Interaction_genotype_Bora | Gm16244 | -4.25 | 1.87E-03 | glio |
| Interaction_genotype_Mtor | Ehbp1l1 | -3.896 | 1.87E-03 | glio |
| Interaction_genotype_Mtor | Dzip1 | -2.904 | 1.87E-03 | glio |
| Interaction_genotype_Mtor | Aup1 | 1.412 | 1.87E-03 | glio |
| Interaction_genotype_Mtor | Eif3f | 1.256 | 1.87E-03 | glio |
| Interaction_genotype_Myc | Tmem223 | -3.613 | 1.87E-03 | glio |
| Interaction_genotype_Myc | Gm10439 | -2.841 | 1.87E-03 | glio |
| Interaction_genotype_Myc | Ptbp2 | -2.508 | 1.87E-03 | glio |
| Interaction_genotype_Myc | Clip1 | -2.327 | 1.87E-03 | glio |
| Interaction_genotype_Myc | Znrd1 | 1.299 | 1.87E-03 | glio |
| interaction_RT | Ralgds | 1.511 | 1.87E-03 | glio |
| interaction_RT | Gm8013 | 3.104 | 1.87E-03 | glio |
| Aars2_in.vivo_noRT | Vmac | 2.491 | 1.87E-03 | glio |
| Aars2_in.vivo_noRT | Gm16536 | 1.762 | 1.87E-03 | glio |
| Aars2_in.vivo_noRT | Epb41 | -1.394 | 1.87E-03 | glio |
| Atr_in.vivo_noRT | Ccdc73 | 1.973 | 1.87E-03 | glio |
| Atr_in.vivo_noRT | Nisch | -1.128 | 1.87E-03 | glio |
| Atr_ex.vivo_noRT | 1700056N10R | 2.688 | 1.87E-03 | glio |
| Atr_ex.vivo_noRT | Speg | 2.955 | 1.87E-03 | glio |
| Atr_ex.vivo_noRT | Col11a1 | -0.888 | 1.87E-03 | glio |
| Bcas2_in.vivo_noRT | Cd72 | 2.246 | 1.87E-03 | glio |
| Bcas2_in.vivo_noRT | Gm43768 | 2.169 | 1.87E-03 | glio |
| Bcas2_in.vivo_noRT | Pdgfa | -1.581 | 1.87E-03 | glio |
| Chek1_in.vivo_noRT | Fam135b | 2.584 | 1.87E-03 | glio |
| Hsd17b10_in.vivo_noRT | Gpm6b | 0.934 | 1.87E-03 | glio |
| culture_effect | 1110065P20R | 0.458 | 1.87E-04 | glio |
| Interaction_genotype_Aars2 | Gm42670 | -4.206 | 1.87E-04 | glio |
| Interaction_genotype_Aars2 | 4930405A21R | -4.333 | 1.87E-04 | glio |
| Interaction_genotype_Aars2 | Trnp1 | -4.511 | 1.87E-04 | glio |
| Interaction_genotype_Aars2 | Crhbp | -4.071 | 1.87E-04 | glio |
| Interaction_genotype_Atr | Gm13421 | -4.76 | 1.87E-04 | glio |
| Interaction_genotype_Myc | Scg2 | -2.729 | 1.87E-04 | glio |
| Interaction_genotype_Myc | Gm36608 | -4.241 | 1.87E-04 | glio |
| Aars2_in.vivo_noRT | Fgl2 | 2.434 | 1.87E-04 | glio |
| Aars2_in.vivo_noRT | Apex1 | -1.402 | 1.87E-04 | glio |
| Aars2_in.vivo_noRT | Ift20 | -1.154 | 1.87E-04 | glio |
| Aars2_in.vivo_noRT | Cstb | -1.49 | 1.87E-04 | glio |
| Aars2_in.vivo_noRT | Lgals1 | -0.948 | 1.87E-04 | glio |
| Atr_in.vivo_noRT | Plcxd2 | 2.687 | 1.87E-04 | glio |
| Atr_in.vivo_noRT | Hist1h2ac | 2.576 | 1.87E-04 | glio |
| Bard1_ex.vivo_noRT | Gm21319 | 2.814 | 1.87E-04 | glio |
| Bora_in.vivo_noRT | Krt17 | 4.676 | 1.87E-04 | glio |
| Bora_in.vivo_noRT | Gm29666 | 2.702 | 1.87E-04 | glio |
| Bora_in.vivo_noRT | Kirrel3 | 2.679 | 1.87E-04 | glio |
| Bora_in.vivo_noRT | Glb1l | 2.685 | 1.87E-04 | glio |
| Bora_ex.vivo_noRT | Auts2 | 3.201 | 1.87E-04 | glio |
| Chek1_in.vivo_noRT | Gm4673 | 2.722 | 1.87E-04 | glio |
| Chek1_ex.vivo_noRT | Lrp1b | 3.483 | 1.87E-04 | glio |
| culture_effect | Kat8 | -0.967 | 1.87E-05 | glio |

| | | | | |
|---|---|---|---|---|
| culture_effect | Commd8 | 0.673 | 1.87E-05 | glio |
| Interaction_genotype_Mtor | Slc25a35 | -4.99 | 1.87E-05 | glio |
| Interaction_genotype_Mtor | Hmg20b | 2.282 | 1.87E-05 | glio |
| Interaction_genotype_Myc | Ifitm6 | -4.952 | 1.87E-05 | glio |
| Interaction_genotype_Pars2 | Gm19276 | -4.073 | 1.87E-05 | glio |
| Aars2_in.vivo_noRT | Paqr8 | 3.43 | 1.87E-05 | glio |
| Aars2_in.vivo_noRT | Ifit1bl2 | 3.13 | 1.87E-05 | glio |
| Aars2_in.vivo_noRT | 4933404O12R | 2.958 | 1.87E-05 | glio |
| Atr_ex.vivo_noRT | Clmn | 3.953 | 1.87E-05 | glio |
| Chek1_in.vivo_noRT | Gm12689 | 3.346 | 1.87E-05 | glio |
| Chek1_ex.vivo_noRT | Smu1 | -1.893 | 1.87E-05 | glio |
| Interaction_genotype_Mtor | Ndufa11 | 2.221 | 1.87E-06 | glio |
| Interaction_genotype_Mtor | Wnt16 | -5.487 | 1.87E-06 | glio |
| Atr_in.vivo_noRT | Abcd2 | 3.694 | 1.87E-06 | glio |
| Atr_in.vivo_noRT | Rps6ka4 | 3.105 | 1.87E-06 | glio |
| Bora_in.vivo_noRT | Gm26519 | 3.75 | 1.87E-06 | glio |
| Bora_in.vivo_noRT | Hoxa13 | 3.489 | 1.87E-06 | glio |
| Bora_in.vivo_noRT | C330013E15Ri | 3.345 | 1.87E-06 | glio |
| Atp5c1_in.vivo_noRT | Gm16178 | 2.668 | 1.87E-07 | glio |
| Atp5c1_in.vivo_noRT | Gm364 | 2.668 | 1.87E-07 | glio |
| Atr_in.vivo_noRT | Hdac7 | 3.553 | 1.87E-07 | glio |
| Bcas2_in.vivo_noRT | Vamp8 | 2.984 | 1.87E-07 | glio |
| Hsd17b10_in.vivo_noRT | C330024C12R | 3.225 | 1.87E-07 | glio |
| Hsd17b10_in.vivo_noRT | Slc4a9 | 3.084 | 1.87E-07 | glio |
| Chek1_ex.vivo_noRT | Zmynd10 | 4.055 | 1.87E-08 | glio |
| Hsd17b10_in.vivo_noRT | Gm31415 | 3.35 | 1.87E-08 | glio |
| Atr_in.vivo_noRT | Gm43330 | 3.633 | 1.87E-09 | glio |
| Chek1_ex.vivo_noRT | Brsk2 | 4.842 | 1.87E-09 | glio |
| Chek1_ex.vivo_noRT | Bmp15 | 4.449 | 1.87E-09 | glio |
| Chek1_ex.vivo_noRT | Slc16a7 | 4.443 | 1.87E-09 | glio |
| Atr_in.vivo_noRT | 1110035H17R | 4.903 | 1.87E-10 | glio |
| Bcas2_in.vivo_noRT | CAAA0106612 | 4.015 | 1.87E-10 | glio |
| Aars2_in.vivo_noRT | Rims3 | 4.193 | 1.87E-11 | glio |
| Atr_in.vivo_noRT | Gm28424 | 4.522 | 1.87E-11 | glio |
| Atr_in.vivo_noRT | Qrich2 | 4.755 | 1.87E-11 | glio |
| Chek1_in.vivo_noRT | Hbq1b | 4.777 | 1.87E-11 | glio |
| culture_effect | Irf2 | -1.184 | 1.87E-12 | glio |
| Bcas2_in.vivo_noRT | 1700016P04R | 4.541 | 1.87E-12 | glio |
| Bora_in.vivo_noRT | 4930447F24Ri | 4.676 | 1.87E-12 | glio |
| Chek1_in.vivo_noRT | BC026762 | 4.664 | 1.87E-12 | glio |
| Atr_in.vivo_noRT | Upb1 | 5.27 | 1.87E-15 | glio |
| Bora_in.vivo_noRT | 1700017M07F | 4.936 | 1.87E-16 | glio |
| Bcas2_in.vivo_noRT | Gpr22 | 4.807 | 1.87E-19 | glio |
| Aars2_in.vivo_noRT | Gm13178 | 4.811 | 1.87E-20 | glio |
| culture_effect | Kif19a | -1.18 | 1.88E-03 | glio |
| culture_effect | Ikbkg | 0.835 | 1.88E-03 | glio |
| culture_effect | 4930453N24R | 0.493 | 1.88E-03 | glio |
| culture_effect | Gm16286 | 0.399 | 1.88E-03 | glio |
| culture_effect | Cpe | 0.54 | 1.88E-03 | glio |
| Interaction_genotype_Aars2 | Ank1 | -2.818 | 1.88E-03 | glio |
| Interaction_genotype_Atr | Gm15853 | -4.02 | 1.88E-03 | glio |
| Interaction_genotype_Atr | Stard3 | 1.994 | 1.88E-03 | glio |

| | | | | |
|---|---|---|---|---|
| Interaction_genotype_Bora | Tra2a | 1.51 | 1.88E-03 | glio |
| Interaction_genotype_Mtor | Gm16344 | -3.188 | 1.88E-03 | glio |
| Interaction_genotype_Mtor | Wfdc12 | -3.975 | 1.88E-03 | glio |
| Interaction_genotype_Mtor | Ythdf2 | 1.825 | 1.88E-03 | glio |
| Interaction_genotype_Myc | Speer4e | -2.729 | 1.88E-03 | glio |
| Interaction_genotype_Myc | Mrgprf | -3.598 | 1.88E-03 | glio |
| Interaction_genotype_Myc | Itga7 | -3.516 | 1.88E-03 | glio |
| Interaction_genotype_Myc | Satb2 | -2.972 | 1.88E-03 | glio |
| Interaction_genotype_Myc | Gm12184 | -3.002 | 1.88E-03 | glio |
| Interaction_genotype_Myc | Hist1h1d | -3.054 | 1.88E-03 | glio |
| Aars2_in.vivo_noRT | Cers2 | -0.805 | 1.88E-03 | glio |
| Atr_ex.vivo_noRT | Fam222a | 3.188 | 1.88E-03 | glio |
| Atr_ex.vivo_noRT | Msrb2 | 2.875 | 1.88E-03 | glio |
| Atr_ex.vivo_noRT | Gm45509 | 3.078 | 1.88E-03 | glio |
| Bcas2_in.vivo_noRT | D130062J10Ri | 2.079 | 1.88E-03 | glio |
| Bcas2_ex.vivo_noRT | AC159140.1 | 2.349 | 1.88E-03 | glio |
| Bora_in.vivo_noRT | Gm5577 | 2.092 | 1.88E-03 | glio |
| Bora_ex.vivo_noRT | 9430041J12Ri | 2.407 | 1.88E-03 | glio |
| Chek1_in.vivo_noRT | Tmem171 | 2.889 | 1.88E-03 | glio |
| Chek1_in.vivo_noRT | Slc25a47 | 2.281 | 1.88E-03 | glio |
| Chek1_in.vivo_noRT | Hoxa7 | 2.267 | 1.88E-03 | glio |
| Chek1_in.vivo_noRT | B3gntl1 | 2.193 | 1.88E-03 | glio |
| Chek1_in.vivo_noRT | Ube2k | -0.805 | 1.88E-03 | glio |
| Chek1_ex.vivo_noRT | Hba-a1 | 3.566 | 1.88E-03 | glio |
| Chek1_ex.vivo_noRT | Timeless | -2.241 | 1.88E-03 | glio |
| Hsd17b10_in.vivo_noRT | Gm12089 | 2.447 | 1.88E-03 | glio |
| Interaction_genotype_Mtor | Dpp6 | -3.172 | 1.88E-04 | glio |
| Interaction_genotype_Mtor | Hspb9 | -4.709 | 1.88E-04 | glio |
| Interaction_genotype_Myc | Gm15853 | -4.368 | 1.88E-04 | glio |
| Interaction_genotype_Myc | Gm38037 | -4.093 | 1.88E-04 | glio |
| Atp5c1_in.vivo_noRT | Luzp2 | 2.142 | 1.88E-04 | glio |
| Atp5c1_in.vivo_noRT | BB557941 | 2.408 | 1.88E-04 | glio |
| Atp5c1_in.vivo_noRT | Zmynd10 | 2.408 | 1.88E-04 | glio |
| Atr_in.vivo_noRT | Rab30 | 2.351 | 1.88E-04 | glio |
| Atr_in.vivo_noRT | Ddx19b | 2.39 | 1.88E-04 | glio |
| Atr_ex.vivo_noRT | Serpina3g | 3.164 | 1.88E-04 | glio |
| Atr_ex.vivo_noRT | Ncf1 | 2.981 | 1.88E-04 | glio |
| Bcas2_in.vivo_noRT | Dpy30 | -1.07 | 1.88E-04 | glio |
| Chek1_ex.vivo_noRT | E030037K01Ri | 2.976 | 1.88E-04 | glio |
| Chek1_ex.vivo_noRT | Crispld2 | 3.079 | 1.88E-04 | glio |
| Aars2_in.vivo_noRT | Gm30074 | 2.959 | 1.88E-05 | glio |
| Aars2_in.vivo_noRT | C8g | 3.053 | 1.88E-05 | glio |
| Aars2_in.vivo_noRT | Zfp595 | 3.21 | 1.88E-05 | glio |
| Aars2_ex.vivo_noRT | Slc6a11 | 3.293 | 1.88E-05 | glio |
| Atr_in.vivo_noRT | Hoxa3 | 2.487 | 1.88E-05 | glio |
| Atr_in.vivo_noRT | Slc38a1 | -1.342 | 1.88E-05 | glio |
| Atr_ex.vivo_noRT | Gm41307 | 2.771 | 1.88E-05 | glio |
| Bcas2_in.vivo_noRT | Gfap | 2.821 | 1.88E-05 | glio |
| Bcas2_ex.vivo_noRT | Gm34059 | 2.742 | 1.88E-05 | glio |
| Bcas2_ex.vivo_noRT | Halr1 | 2.742 | 1.88E-05 | glio |
| Bora_in.vivo_noRT | Gm48357 | 2.621 | 1.88E-05 | glio |
| Bora_ex.vivo_noRT | Mpzl2 | 3.201 | 1.88E-05 | glio |

| | | | | |
|---|---|---|---|---|
| Hsd17b10_in.vivo_noRT | Gm11549 | 3.09 | 1.88E-05 | glio |
| culture_effect | Lynx1 | 2.464 | 1.88E-06 | glio |
| culture_effect | Csnk2b | -0.617 | 1.88E-06 | glio |
| Interaction_genotype_Myc | Cnksr1 | -5.292 | 1.88E-06 | glio |
| Interaction_genotype_Myc | Eny2 | 1.696 | 1.88E-06 | glio |
| Aars2_in.vivo_noRT | BC037704 | 3.278 | 1.88E-06 | glio |
| Atr_in.vivo_noRT | Nat8l | 2.942 | 1.88E-06 | glio |
| Bcas2_in.vivo_noRT | Gsx2 | 3.596 | 1.88E-06 | glio |
| Bcas2_in.vivo_noRT | Gm10101 | 3.238 | 1.88E-06 | glio |
| Interaction_genotype_Mtor | Gm41307 | -4.492 | 1.88E-07 | glio |
| interaction_RT | Nupr1 | 2.247 | 1.88E-07 | glio |
| Aars2_in.vivo_noRT | Spata33 | 3.802 | 1.88E-07 | glio |
| Aars2_in.vivo_noRT | Gstm2 | 3.739 | 1.88E-07 | glio |
| Aars2_in.vivo_noRT | Itgb7 | 3.424 | 1.88E-07 | glio |
| Aars2_ex.vivo_noRT | Rimklb | 3.293 | 1.88E-07 | glio |
| Bcas2_in.vivo_noRT | Slc26a6 | 3.515 | 1.88E-07 | glio |
| Atp5c1_in.vivo_noRT | Sucnr1 | 2.668 | 1.88E-08 | glio |
| Bora_in.vivo_noRT | Smg7 | -2.284 | 1.88E-08 | glio |
| Chek1_in.vivo_noRT | Papln | 3.776 | 1.88E-08 | glio |
| Interaction_genotype_Mtor | Ciao2b | 2.898 | 1.88E-09 | glio |
| Interaction_genotype_Mtor | mt-Cytb | 2.128 | 1.88E-09 | glio |
| Bcas2_in.vivo_noRT | Kcnq2 | 3.643 | 1.88E-09 | glio |
| culture_effect | Scd1 | 1.296 | 1.88E-10 | glio |
| Aars2_in.vivo_noRT | Gm15927 | 4.325 | 1.88E-10 | glio |
| Aars2_in.vivo_noRT | Sh3gl3 | 4.415 | 1.88E-10 | glio |
| Atr_ex.vivo_noRT | Gm48617 | 3.947 | 1.88E-10 | glio |
| Chek1_in.vivo_noRT | Grm1 | 3.94 | 1.88E-10 | glio |
| culture_effect | Asgr1 | 3.462 | 1.88E-11 | glio |
| Chek1_ex.vivo_noRT | Gm34776 | 4.449 | 1.88E-11 | glio |
| Hsd17b10_in.vivo_noRT | Cd7 | 3.352 | 1.88E-11 | glio |
| Chek1_in.vivo_noRT | Zfp786 | 4.78 | 1.88E-12 | glio |
| Bcas2_in.vivo_noRT | Gm20767 | 4.807 | 1.88E-13 | glio |
| Chek1_in.vivo_noRT | Rab27a | 4.518 | 1.88E-13 | glio |
| Chek1_in.vivo_noRT | Txndc2 | 4.264 | 1.88E-14 | glio |
| Chek1_in.vivo_noRT | 8030423F21Ri | 4.515 | 1.88E-14 | glio |
| Bora_in.vivo_noRT | Gm20705 | 4.405 | 1.88E-15 | glio |
| Chek1_in.vivo_noRT | Capn9 | 5.044 | 1.88E-15 | glio |
| Atr_in.vivo_noRT | Rslcan18 | 5.147 | 1.88E-16 | glio |
| Atr_in.vivo_noRT | Pkd1l2 | 5.269 | 1.88E-16 | glio |
| culture_effect | Tgtp1 | -4.656 | 1.88E-17 | glio |
| culture_effect | Idnk | -1.546 | 1.88E-17 | glio |
| Aars2_in.vivo_noRT | Gm30042 | 5.067 | 1.88E-18 | glio |
| Aars2_in.vivo_noRT | Gm34939 | 5.063 | 1.88E-18 | glio |
| Bcas2_in.vivo_noRT | Gm48140 | 4.544 | 1.88E-19 | glio |
| Bcas2_in.vivo_noRT | Ccl9 | 4.807 | 1.88E-22 | glio |
| Aars2_in.vivo_noRT | BB287469 | 5.327 | 1.88E-24 | glio |
| Atr_in.vivo_noRT | Gpr4 | 5.533 | 1.88E-26 | glio |
| Atr_in.vivo_noRT | Prl7c1 | 5.533 | 1.88E-26 | glio |
| Atr_in.vivo_noRT | Gm33466 | 5.533 | 1.88E-26 | glio |
| culture_effect | Sesn2 | -0.717 | 1.89E-03 | glio |
| culture_effect | Sgo1 | 0.73 | 1.89E-03 | glio |
| culture_effect | Atp5o | -0.38 | 1.89E-03 | glio |

| | | | | |
|---|---|---|---|---|
| Interaction_genotype_Aars2 | 1110035H17R | -4.358 | 1.89E-03 | glio |
| Interaction_genotype_Aars2 | D430040D24F | -3.807 | 1.89E-03 | glio |
| Interaction_genotype_Aars2 | Cth | 3.104 | 1.89E-03 | glio |
| Interaction_genotype_Aars2 | Tmx2 | 1.672 | 1.89E-03 | glio |
| Interaction_genotype_Myc | Tgfbr1 | -2.429 | 1.89E-03 | glio |
| Interaction_genotype_Myc | Mcrip1 | 1.293 | 1.89E-03 | glio |
| Interaction_genotype_Pars2 | 6330415G19R | -3.909 | 1.89E-03 | glio |
| Interaction_genotype_Pars2 | Ntn4 | -2.839 | 1.89E-03 | glio |
| Interaction_genotype_Rad21 | Rhbg | 3.423 | 1.89E-03 | glio |
| interaction_RT | Elp1 | -0.91 | 1.89E-03 | glio |
| Aars2_in.vivo_noRT | Cuta | -0.848 | 1.89E-03 | glio |
| Atp5c1_in.vivo_noRT | Matn4 | 2.404 | 1.89E-03 | glio |
| Atr_in.vivo_noRT | Lsm12 | -0.916 | 1.89E-03 | glio |
| Bcas2_in.vivo_noRT | Lgr6 | 2.599 | 1.89E-03 | glio |
| Bcas2_in.vivo_noRT | Trim66 | 2.095 | 1.89E-03 | glio |
| Bcas2_in.vivo_noRT | Asb1 | 1.923 | 1.89E-03 | glio |
| Bora_ex.vivo_noRT | Gm11884 | 2.802 | 1.89E-03 | glio |
| Chek1_in.vivo_noRT | B230206L02Ri | 2.242 | 1.89E-03 | glio |
| Chek1_in.vivo_noRT | 4921531C22R | 2.202 | 1.89E-03 | glio |
| Chek1_in.vivo_noRT | Ttbk1 | 2.246 | 1.89E-03 | glio |
| Chek1_in.vivo_noRT | Xrcc5 | -1.192 | 1.89E-03 | glio |
| Chek1_in.vivo_noRT | Pdcd10 | -1.113 | 1.89E-03 | glio |
| Chek1_in.vivo_noRT | Tbca | -0.549 | 1.89E-03 | glio |
| Chek1_ex.vivo_noRT | Stk32c | 2.719 | 1.89E-03 | glio |
| Hsd17b10_in.vivo_noRT | Rps19bp1 | -0.698 | 1.89E-03 | glio |
| culture_effect | Kiz | -1.018 | 1.89E-04 | glio |
| Interaction_genotype_Mtor | March11 | -4.738 | 1.89E-04 | glio |
| Interaction_genotype_Mtor | 9230112E08Ri | -4.563 | 1.89E-04 | glio |
| Interaction_genotype_Mtor | Mtrr | -3.275 | 1.89E-04 | glio |
| Interaction_genotype_Mtor | Ywhaz | 1.791 | 1.89E-04 | glio |
| Aars2_in.vivo_noRT | Sapcd1 | 2.662 | 1.89E-04 | glio |
| Atp5c1_in.vivo_noRT | Gm31415 | 2.402 | 1.89E-04 | glio |
| Bard1_ex.vivo_noRT | Gm39526 | 2.814 | 1.89E-04 | glio |
| Bcas2_in.vivo_noRT | C030005K06R | 2.708 | 1.89E-04 | glio |
| Bora_in.vivo_noRT | Usp11 | 2.837 | 1.89E-04 | glio |
| Bora_in.vivo_noRT | 1700041G16R | 2.682 | 1.89E-04 | glio |
| Bora_in.vivo_noRT | B4galt3 | -1.542 | 1.89E-04 | glio |
| Chek1_in.vivo_noRT | Siae | 2.428 | 1.89E-04 | glio |
| Chek1_ex.vivo_noRT | Prr15l | 3.463 | 1.89E-04 | glio |
| Aars2_in.vivo_noRT | Fam178b | 3.129 | 1.89E-05 | glio |
| Aars2_in.vivo_noRT | Cnpy2 | -0.97 | 1.89E-05 | glio |
| Chek1_in.vivo_noRT | Kcnb1 | 3.179 | 1.89E-05 | glio |
| Chek1_ex.vivo_noRT | Gm4724 | 3.349 | 1.89E-05 | glio |
| Hsd17b10_in.vivo_noRT | Cd52 | 2.959 | 1.89E-05 | glio |
| culture_effect | Pcdhb19 | -2.831 | 1.89E-06 | glio |
| culture_effect | Tnfaip3 | -2.615 | 1.89E-06 | glio |
| culture_effect | Gm11423 | 1.661 | 1.89E-06 | glio |
| culture_effect | Glce | -1.283 | 1.89E-06 | glio |
| Interaction_genotype_Mtor | Spon1 | -5.819 | 1.89E-06 | glio |
| Interaction_genotype_Myc | Hspa9 | 2.079 | 1.89E-06 | glio |
| Interaction_genotype_Myc | Syt4 | 2.257 | 1.89E-06 | glio |
| Aars2_in.vivo_noRT | Gm36608 | 3.396 | 1.89E-06 | glio |

| | | | | |
|---|---|---|---|---|
| Aars2_in.vivo_noRT | Armc1 | -1.766 | 1.89E-06 | glio |
| Aars2_ex.vivo_noRT | Gm47936 | 3.293 | 1.89E-06 | glio |
| Atr_in.vivo_noRT | Slc46a1 | 3.499 | 1.89E-06 | glio |
| Bcas2_in.vivo_noRT | A630072M18F | 2.847 | 1.89E-06 | glio |
| Chek1_in.vivo_noRT | Ppp1r18os | 3.631 | 1.89E-06 | glio |
| Atr_in.vivo_noRT | Gm37305 | 3.576 | 1.89E-07 | glio |
| culture_effect | Ech1 | -0.858 | 1.89E-08 | glio |
| Hsd17b10_in.vivo_noRT | BB557941 | 3.356 | 1.89E-08 | glio |
| Hsd17b10_in.vivo_noRT | Zmynd10 | 3.356 | 1.89E-08 | glio |
| Aars2_in.vivo_noRT | Hlx | 4.07 | 1.89E-09 | glio |
| Aars2_in.vivo_noRT | Dmrt2 | 4.431 | 1.89E-09 | glio |
| Atr_in.vivo_noRT | Adgrd1 | 3.886 | 1.89E-09 | glio |
| Chek1_in.vivo_noRT | Gm43625 | 3.739 | 1.89E-09 | glio |
| culture_effect | Gm47758 | -3.242 | 1.89E-10 | glio |
| Atr_in.vivo_noRT | Pde1a | 4.533 | 1.89E-10 | glio |
| Atr_in.vivo_noRT | Slc26a6 | 4.241 | 1.89E-10 | glio |
| culture_effect | Tubb4a | -4.464 | 1.89E-11 | glio |
| Bcas2_in.vivo_noRT | Gm44752 | 4.42 | 1.89E-12 | glio |
| Aars2_in.vivo_noRT | Gm17173 | 4.94 | 1.89E-13 | glio |
| Bora_in.vivo_noRT | Gm45620 | 4.936 | 1.89E-13 | glio |
| Bora_in.vivo_noRT | Gm38534 | 4.936 | 1.89E-13 | glio |
| Hsd17b10_in.vivo_noRT | 1700034I23Ri | 3.616 | 1.89E-13 | glio |
| Aars2_in.vivo_noRT | Gem | 5.327 | 1.89E-15 | glio |
| Bcas2_in.vivo_noRT | Gm15943 | 4.543 | 1.89E-16 | glio |
| Aars2_in.vivo_noRT | Prickle2 | 5.327 | 1.89E-19 | glio |
| Chek1_in.vivo_noRT | Tal2 | 4.78 | 1.89E-20 | glio |
| Atr_in.vivo_noRT | Gm45669 | 5.78 | 1.89E-21 | glio |
| Chek1_in.vivo_noRT | Cd36 | 5.044 | 1.89E-22 | glio |
| Aars2_in.vivo_noRT | Gm44693 | 5.327 | 1.89E-26 | glio |
| culture_effect | Gm47484 | 1.853 | 1.90E-03 | glio |
| culture_effect | Asnsd1 | 0.467 | 1.90E-03 | glio |
| culture_effect | Utp14a | -0.465 | 1.90E-03 | glio |
| culture_effect | Nop56 | -0.426 | 1.90E-03 | glio |
| Interaction_genotype_Aars2 | Apoa5 | -2.548 | 1.90E-03 | glio |
| Interaction_genotype_Aars2 | Skap2 | 1.432 | 1.90E-03 | glio |
| Interaction_genotype_Aars2 | Emc4 | 1.343 | 1.90E-03 | glio |
| Interaction_genotype_Bcas2 | Gm50375 | -4.518 | 1.90E-03 | glio |
| Interaction_genotype_Bcas2 | Arsj | -4.375 | 1.90E-03 | glio |
| Interaction_genotype_Bcas2 | Gm16046 | -4.099 | 1.90E-03 | glio |
| Interaction_genotype_Bcas2 | Tnfrsf22 | -3.86 | 1.90E-03 | glio |
| Interaction_genotype_Bcas2 | Rab25 | -3.744 | 1.90E-03 | glio |
| Interaction_genotype_Bcas2 | 4833418N02R | -4.299 | 1.90E-03 | glio |
| Interaction_genotype_Bcas2 | Bnip2 | 1.383 | 1.90E-03 | glio |
| Interaction_genotype_Myc | Gad1 | -2.463 | 1.90E-03 | glio |
| Interaction_genotype_Myc | Gm35439 | -2.749 | 1.90E-03 | glio |
| Interaction_genotype_Myc | Isl1 | -3.508 | 1.90E-03 | glio |
| Interaction_genotype_Myc | Emid1 | -3.866 | 1.90E-03 | glio |
| Interaction_genotype_Myc | Neil2 | -3.25 | 1.90E-03 | glio |
| Interaction_genotype_Myc | Hist1h2ak | -3.088 | 1.90E-03 | glio |
| Interaction_genotype_Myc | Cavin2 | -2.963 | 1.90E-03 | glio |
| Interaction_genotype_Myc | Rpl6 | 1.179 | 1.90E-03 | glio |
| interaction_RT | Cse1l | -0.888 | 1.90E-03 | glio |

| | | | | |
|---|---|---|---|---|
| Atr_in.vivo_noRT | Ttc25 | 2.169 | 1.90E-03 | glio |
| Atr_in.vivo_noRT | Mien1 | -0.964 | 1.90E-03 | glio |
| Atr_in.vivo_noRT | Tcim | -1.461 | 1.90E-03 | glio |
| Bcas2_in.vivo_noRT | Cyb5d1 | 2.271 | 1.90E-03 | glio |
| Bcas2_in.vivo_noRT | Ubox5 | 1.883 | 1.90E-03 | glio |
| Bcas2_ex.vivo_noRT | Ceacam19 | 3.174 | 1.90E-03 | glio |
| Bora_in.vivo_noRT | Grb7 | 2.284 | 1.90E-03 | glio |
| Chek1_in.vivo_noRT | Cracr2b | 2.383 | 1.90E-03 | glio |
| Chek1_in.vivo_noRT | Zfp703 | -1.759 | 1.90E-03 | glio |
| Chek1_ex.vivo_noRT | Gm12454 | 2.887 | 1.90E-03 | glio |
| Chek1_ex.vivo_noRT | 9530062K07R | 2.878 | 1.90E-03 | glio |
| Dclre1c_ex.vivo_noRT | Cxcl5 | 2.666 | 1.90E-03 | glio |
| Interaction_genotype_Aars2 | Rarres1 | -4.597 | 1.90E-04 | glio |
| Interaction_genotype_Atr | Tomm6os | -4.861 | 1.90E-04 | glio |
| Interaction_genotype_Mtor | Pde6b | -3.86 | 1.90E-04 | glio |
| Interaction_genotype_Mtor | Taco1os | -3.982 | 1.90E-04 | glio |
| Interaction_genotype_Mtor | Plekhn1 | -3.895 | 1.90E-04 | glio |
| Interaction_genotype_Myc | Hfm1 | -3.304 | 1.90E-04 | glio |
| Interaction_genotype_Myc | Prickle4 | -3.996 | 1.90E-04 | glio |
| Interaction_genotype_Myc | Hspa12a | -4.224 | 1.90E-04 | glio |
| Interaction_genotype_Myc | Stab1 | -4.46 | 1.90E-04 | glio |
| Interaction_genotype_Myc | Gm13830 | -4.523 | 1.90E-04 | glio |
| Interaction_genotype_Myc | Smad3 | -3.992 | 1.90E-04 | glio |
| Interaction_genotype_Myc | Mrpl33 | 1.39 | 1.90E-04 | glio |
| Interaction_genotype_Myc | mt-Co1 | 1.447 | 1.90E-04 | glio |
| Aars2_ex.vivo_noRT | Hfm1 | 2.718 | 1.90E-04 | glio |
| Atp5c1_in.vivo_noRT | Gm45591 | 2.408 | 1.90E-04 | glio |
| Atr_in.vivo_noRT | Rps15 | -0.682 | 1.90E-04 | glio |
| Atr_ex.vivo_noRT | Gm19265 | 2.764 | 1.90E-04 | glio |
| Atr_ex.vivo_noRT | Gm15559 | 3.539 | 1.90E-04 | glio |
| Bard1_ex.vivo_noRT | Speer4e | 2.814 | 1.90E-04 | glio |
| Hsd17b10_in.vivo_noRT | Gm48772 | 2.708 | 1.90E-04 | glio |
| Hsd17b10_in.vivo_noRT | Ppp1ca | -0.705 | 1.90E-04 | glio |
| Interaction_genotype_Aars2 | Gm30938 | -5.07 | 1.90E-05 | glio |
| Interaction_genotype_Mtor | Ncf1 | -4.427 | 1.90E-05 | glio |
| Interaction_genotype_Myc | 4930447K03R | -3.8 | 1.90E-05 | glio |
| Interaction_genotype_Myc | Sema6d | -3.711 | 1.90E-05 | glio |
| Bora_in.vivo_noRT | Gm13283 | 3.044 | 1.90E-05 | glio |
| Chek1_in.vivo_noRT | Rtl8b | 2.384 | 1.90E-05 | glio |
| Dclre1c_ex.vivo_noRT | BB287469 | 3.224 | 1.90E-05 | glio |
| Interaction_genotype_Mtor | Tmem40 | -6.236 | 1.90E-06 | glio |
| Atp5c1_in.vivo_noRT | Olfr1152 | 2.402 | 1.90E-06 | glio |
| Atp5c1_in.vivo_noRT | Ush1g | 2.668 | 1.90E-06 | glio |
| Atp5c1_in.vivo_noRT | Gm31063 | 2.668 | 1.90E-06 | glio |
| Atr_in.vivo_noRT | Fam171a2 | 3.371 | 1.90E-06 | glio |
| Bora_in.vivo_noRT | Ccdc184 | 3.603 | 1.90E-06 | glio |
| Hsd17b10_in.vivo_noRT | Gm7247 | 2.824 | 1.90E-06 | glio |
| culture_effect | 4632427E13Ri | 0.928 | 1.90E-07 | glio |
| Chek1_in.vivo_noRT | 1700013F07Ri | 3.889 | 1.90E-07 | glio |
| Chek1_ex.vivo_noRT | Catsperd | 4.48 | 1.90E-07 | glio |
| Hsd17b10_in.vivo_noRT | Myo5c | 3.616 | 1.90E-07 | glio |
| Chek1_ex.vivo_noRT | Gm15941 | 4.447 | 1.90E-08 | glio |

| | | | | |
|---|---|---|---|---|
| Atr_in.vivo_noRT | Sec1 | 4.376 | 1.90E-09 | glio |
| Atr_in.vivo_noRT | Gm26559 | 4.26 | 1.90E-09 | glio |
| Bcas2_in.vivo_noRT | Gm44672 | 4.043 | 1.90E-09 | glio |
| Chek1_in.vivo_noRT | Gm16222 | 4.784 | 1.90E-09 | glio |
| Chek1_ex.vivo_noRT | Gm48170 | 4.045 | 1.90E-09 | glio |
| Chek1_ex.vivo_noRT | Gm4858 | 4.842 | 1.90E-09 | glio |
| Chek1_ex.vivo_noRT | Insc | 4.842 | 1.90E-10 | glio |
| Aars2_in.vivo_noRT | Gm11963 | 5.067 | 1.90E-13 | glio |
| Aars2_in.vivo_noRT | Stx19 | 4.415 | 1.90E-14 | glio |
| Atr_in.vivo_noRT | Igfbp7 | 4.768 | 1.90E-14 | glio |
| culture_effect | Wfdc13 | 1.799 | 1.90E-15 | glio |
| Bora_in.vivo_noRT | Gm16685 | 4.801 | 1.90E-16 | glio |
| Chek1_in.vivo_noRT | Gm15326 | 4.515 | 1.90E-18 | glio |
| Interaction_genotype_Aars2 | Tagap1 | -3.143 | 1.91E-03 | glio |
| Interaction_genotype_Aars2 | Bccip | 1.415 | 1.91E-03 | glio |
| Interaction_genotype_Bcas2 | Shisal2a | -4.451 | 1.91E-03 | glio |
| Interaction_genotype_Myc | Zc3h12a | -3.366 | 1.91E-03 | glio |
| Interaction_genotype_Myc | Slc29a2 | -3.471 | 1.91E-03 | glio |
| Interaction_genotype_Myc | Dbndd1 | -3.126 | 1.91E-03 | glio |
| Interaction_genotype_Myc | Map7d2 | -2.975 | 1.91E-03 | glio |
| Interaction_genotype_Myc | B230206L02Ri | -3.142 | 1.91E-03 | glio |
| Interaction_genotype_Myc | Myo18a | -2.057 | 1.91E-03 | glio |
| Interaction_genotype_Myc | U2af2 | 1.599 | 1.91E-03 | glio |
| Aars2_in.vivo_noRT | Dnajc15 | -1.214 | 1.91E-03 | glio |
| Aars2_in.vivo_noRT | Msmo1 | -1.421 | 1.91E-03 | glio |
| Atp5c1_in.vivo_noRT | Asgr1 | 2.54 | 1.91E-03 | glio |
| Atp5c1_in.vivo_noRT | Dpp6 | 1.741 | 1.91E-03 | glio |
| Atp5c1_in.vivo_noRT | Gm15579 | 2.668 | 1.91E-03 | glio |
| Atp5c1_in.vivo_noRT | Gm5861 | 2.93 | 1.91E-03 | glio |
| Atr_in.vivo_noRT | Neurl2 | 1.94 | 1.91E-03 | glio |
| Atr_in.vivo_noRT | Phldb1 | -1.131 | 1.91E-03 | glio |
| Bcas2_in.vivo_noRT | Acad10 | 2.302 | 1.91E-03 | glio |
| Bcas2_in.vivo_noRT | Car13 | 1.645 | 1.91E-03 | glio |
| Bora_in.vivo_noRT | 1700113B09R | 2.326 | 1.91E-03 | glio |
| Cenpj_in.vivo_noRT | Gm21761 | 2.022 | 1.91E-03 | glio |
| Chek1_in.vivo_noRT | Gm12981 | 2.257 | 1.91E-03 | glio |
| Chek1_in.vivo_noRT | Reep5 | -0.871 | 1.91E-03 | glio |
| Chek1_in.vivo_noRT | Ybx3 | -0.715 | 1.91E-03 | glio |
| Chek1_ex.vivo_noRT | Gm24362 | 2.563 | 1.91E-03 | glio |
| Chek1_ex.vivo_noRT | Milr1 | 2.89 | 1.91E-03 | glio |
| Hsd17b10_in.vivo_noRT | Ak1 | 2.353 | 1.91E-03 | glio |
| culture_effect | Alkbh6 | -0.755 | 1.91E-04 | glio |
| Interaction_genotype_Cyp19a | Mmp13 | 3.917 | 1.91E-04 | glio |
| Interaction_genotype_Mtor | Vpreb1 | -4.359 | 1.91E-04 | glio |
| Interaction_genotype_Myc | 1700003E24Ri | -2.729 | 1.91E-04 | glio |
| Interaction_genotype_Myc | Gm9725 | -4.31 | 1.91E-04 | glio |
| Interaction_genotype_Myc | Zfp689 | -3.324 | 1.91E-04 | glio |
| Aars2_in.vivo_noRT | Spata5 | -1.641 | 1.91E-04 | glio |
| Atp5c1_in.vivo_noRT | Nxph3 | 2.668 | 1.91E-04 | glio |
| Atr_in.vivo_noRT | Pmp22 | 2.422 | 1.91E-04 | glio |
| Atr_in.vivo_noRT | Sem1 | -0.926 | 1.91E-04 | glio |
| Bard1_ex.vivo_noRT | Ptprc | 2.814 | 1.91E-04 | glio |

| | | | | |
|---|---|---|---|---|
| culture_effect | Foxn2 | 0.822 | 1.91E-05 | glio |
| Interaction_genotype_Mtor | Snrpc | 1.904 | 1.91E-05 | glio |
| Aars2_in.vivo_noRT | Pcdhb20 | 2.804 | 1.91E-05 | glio |
| Atr_in.vivo_noRT | Gm17655 | 3.341 | 1.91E-05 | glio |
| Bcas2_ex.vivo_noRT | Cfh | 3.141 | 1.91E-05 | glio |
| Bcas2_ex.vivo_noRT | Helt | 2.742 | 1.91E-05 | glio |
| Bcas2_ex.vivo_noRT | Gm20625 | 2.742 | 1.91E-05 | glio |
| Bcas2_ex.vivo_noRT | Gm5157 | 2.742 | 1.91E-05 | glio |
| Bcas2_ex.vivo_noRT | Gm44736 | 2.742 | 1.91E-05 | glio |
| Bcas2_ex.vivo_noRT | Fam227b | 2.742 | 1.91E-05 | glio |
| Bcas2_ex.vivo_noRT | Gm20658 | 2.742 | 1.91E-05 | glio |
| Bcas2_ex.vivo_noRT | Olfr810 | 2.742 | 1.91E-05 | glio |
| Bcas2_ex.vivo_noRT | Gjd4 | 2.742 | 1.91E-05 | glio |
| Bcas2_ex.vivo_noRT | Apoa5 | 2.742 | 1.91E-05 | glio |
| Bcas2_ex.vivo_noRT | Gm49172 | 2.742 | 1.91E-05 | glio |
| Bora_in.vivo_noRT | Spon1 | 3.083 | 1.91E-05 | glio |
| Interaction_genotype_Mtor | Ptprg | -5.072 | 1.91E-06 | glio |
| Interaction_genotype_Mtor | Tcf12 | 2.775 | 1.91E-06 | glio |
| Interaction_genotype_Mtor | Gm10767 | -5.805 | 1.91E-06 | glio |
| Interaction_genotype_Mtor | Prl2c3 | -5.294 | 1.91E-06 | glio |
| Interaction_genotype_Mtor | Gm12794 | -4.099 | 1.91E-06 | glio |
| Bcas2_in.vivo_noRT | Tgfbr3l | 3.503 | 1.91E-06 | glio |
| Bcas2_in.vivo_noRT | Ehhadh | 3.641 | 1.91E-06 | glio |
| culture_effect | Asrgl1 | -0.827 | 1.91E-07 | glio |
| Interaction_genotype_Mtor | Ephx1 | -6.105 | 1.91E-07 | glio |
| Interaction_genotype_Mtor | Gm34059 | -4.099 | 1.91E-07 | glio |
| Bora_in.vivo_noRT | Gm12089 | 3.759 | 1.91E-07 | glio |
| Interaction_genotype_Mtor | Mecom | -6.357 | 1.91E-08 | glio |
| interaction_RT | Irf7 | 1.536 | 1.91E-08 | glio |
| Atr_in.vivo_noRT | Tmem116 | 4.355 | 1.91E-08 | glio |
| culture_effect | Tigd2 | 1.579 | 1.91E-09 | glio |
| Aars2_in.vivo_noRT | Gm49767 | 4.289 | 1.91E-09 | glio |
| Hsd17b10_in.vivo_noRT | H1foo | 3.084 | 1.91E-09 | glio |
| Bcas2_in.vivo_noRT | Gm5941 | 4.807 | 1.91E-11 | glio |
| Chek1_in.vivo_noRT | Adam12 | 4.784 | 1.91E-14 | glio |
| Aars2_in.vivo_noRT | Ripply3 | 5.327 | 1.91E-15 | glio |
| Chek1_ex.vivo_noRT | Gm48617 | 4.842 | 1.91E-15 | glio |
| Chek1_in.vivo_noRT | AC163347.1 | 4.78 | 1.91E-17 | glio |
| culture_effect | Eef1a1 | -1.248 | 1.91E-19 | glio |
| Aars2_in.vivo_noRT | Gm4027 | 5.327 | 1.91E-21 | glio |
| Chek1_in.vivo_noRT | Prap1 | 5.044 | 1.91E-21 | glio |
| Atr_in.vivo_noRT | Tac2 | 5.533 | 1.91E-27 | glio |
| culture_effect | Elk3 | 2.959 | 1.91E-30 | glio |
| culture_effect | Atp8b2 | -0.982 | 1.92E-03 | glio |
| culture_effect | Phkb | 0.755 | 1.92E-03 | glio |
| culture_effect | Psmg1 | -0.743 | 1.92E-03 | glio |
| culture_effect | Pnpla8 | -0.587 | 1.92E-03 | glio |
| culture_effect | Plxnb2 | -0.561 | 1.92E-03 | glio |
| culture_effect | Mrps33 | 0.326 | 1.92E-03 | glio |
| Interaction_genotype_Aars2 | Anks6 | -3.572 | 1.92E-03 | glio |
| Interaction_genotype_Aars2 | Siva1 | 1.253 | 1.92E-03 | glio |
| Interaction_genotype_Atr | Sh3d21 | -3.344 | 1.92E-03 | glio |

| | | | | |
|---|---|---|---|---|
| Interaction_genotype_Atr | Ankef1 | -3.985 | 1.92E-03 | glio |
| Interaction_genotype_Bora | Gm16348 | -4.028 | 1.92E-03 | glio |
| Interaction_genotype_Mtor | S1pr1 | -4.165 | 1.92E-03 | glio |
| Interaction_genotype_Mtor | Card14 | -3.755 | 1.92E-03 | glio |
| Interaction_genotype_Mtor | Polr3g | -2.875 | 1.92E-03 | glio |
| Interaction_genotype_Mtor | Tmx3 | 1.774 | 1.92E-03 | glio |
| Interaction_genotype_Myc | Gm43773 | -3.37 | 1.92E-03 | glio |
| Interaction_genotype_Myc | Trim62 | -2.954 | 1.92E-03 | glio |
| interaction_RT | Pmvk | 1.031 | 1.92E-03 | glio |
| interaction_RT | Elf2 | 0.713 | 1.92E-03 | glio |
| Aars2_in.vivo_noRT | Nxpe3 | 1.877 | 1.92E-03 | glio |
| Atp5c1_in.vivo_noRT | 1700047K16R | 2.538 | 1.92E-03 | glio |
| Atp5c1_in.vivo_noRT | Tcam1 | 2.549 | 1.92E-03 | glio |
| Atp5c1_in.vivo_noRT | Chaf1a | -1.31 | 1.92E-03 | glio |
| Atr_in.vivo_noRT | Acvr1b | 2.212 | 1.92E-03 | glio |
| Atr_in.vivo_noRT | Slc16a2 | 2.108 | 1.92E-03 | glio |
| Atr_in.vivo_noRT | Farsa | -1.218 | 1.92E-03 | glio |
| Atr_ex.vivo_noRT | Tmem236 | 2.59 | 1.92E-03 | glio |
| Bcas2_in.vivo_noRT | Brcc3 | -1.037 | 1.92E-03 | glio |
| Bora_in.vivo_noRT | Adh7 | 2.249 | 1.92E-03 | glio |
| Bora_in.vivo_noRT | Fxyd3 | -2.294 | 1.92E-03 | glio |
| Bora_in.vivo_noRT | Ccnd3 | -0.851 | 1.92E-03 | glio |
| Bora_ex.vivo_noRT | Ubd | 2.808 | 1.92E-03 | glio |
| Cenpj_ex.vivo_noRT | Clec12b | 2.196 | 1.92E-03 | glio |
| Cenpj_ex.vivo_noRT | Vmn1r32 | 2.198 | 1.92E-03 | glio |
| Chek1_in.vivo_noRT | Ptn | 2.646 | 1.92E-03 | glio |
| Chek1_in.vivo_noRT | Gm29488 | 2.188 | 1.92E-03 | glio |
| Chek1_in.vivo_noRT | Mphosph6 | -1.202 | 1.92E-03 | glio |
| Chek1_ex.vivo_noRT | Cfap58 | 3.482 | 1.92E-03 | glio |
| Hsd17b10_in.vivo_noRT | Stab1 | 2.482 | 1.92E-03 | glio |
| Hsd17b10_in.vivo_noRT | Acot5 | 2.263 | 1.92E-03 | glio |
| Hsd17b10_in.vivo_noRT | Cpne7 | 2.528 | 1.92E-03 | glio |
| Hsd17b10_in.vivo_noRT | Gm43773 | 2.246 | 1.92E-03 | glio |
| Interaction_genotype_Aars2 | BC028471 | -4.711 | 1.92E-04 | glio |
| Interaction_genotype_Atr | Gm10447 | -3.643 | 1.92E-04 | glio |
| Interaction_genotype_Mtor | Nim1k | -4.671 | 1.92E-04 | glio |
| Interaction_genotype_Mtor | A330009N23R | -4.367 | 1.92E-04 | glio |
| Interaction_genotype_Mtor | Srp19 | 1.744 | 1.92E-04 | glio |
| Interaction_genotype_Mtor | Arpc2 | 1.627 | 1.92E-04 | glio |
| Interaction_genotype_Mtor | Hspa8 | 1.914 | 1.92E-04 | glio |
| Interaction_genotype_Myc | Dlg5 | -4.025 | 1.92E-04 | glio |
| Interaction_genotype_Myc | Hs1bp3 | -3.613 | 1.92E-04 | glio |
| Interaction_genotype_Rad21 | Prdm1 | 5.149 | 1.92E-04 | glio |
| interaction_RT | Gar1 | -0.824 | 1.92E-04 | glio |
| Aars2_in.vivo_noRT | 1700113B09R | 2.716 | 1.92E-04 | glio |
| Atr_in.vivo_noRT | Zfp28 | 2.662 | 1.92E-04 | glio |
| Atr_in.vivo_noRT | Zkscan7 | 2.68 | 1.92E-04 | glio |
| Bcas2_in.vivo_noRT | Il23a | 2.685 | 1.92E-04 | glio |
| Bcas2_in.vivo_noRT | Gm19684 | 2.848 | 1.92E-04 | glio |
| Bcas2_in.vivo_noRT | Tnnc1 | 2.947 | 1.92E-04 | glio |
| Bcas2_in.vivo_noRT | Eps8 | -1.035 | 1.92E-04 | glio |
| Bora_in.vivo_noRT | Gm4707 | 2.544 | 1.92E-04 | glio |

| | | | | |
|---|---|---|---|---|
| Chek1_in.vivo_noRT | Gm14023 | 2.766 | 1.92E-04 | glio |
| Hsd17b10_in.vivo_noRT | Tnnt3 | 2.981 | 1.92E-04 | glio |
| Hsd17b10_in.vivo_noRT | Astn2 | 2.591 | 1.92E-04 | glio |
| Interaction_genotype_Mtor | Cdkl2 | -4.612 | 1.92E-05 | glio |
| Interaction_genotype_Mtor | Txndc2 | -4.108 | 1.92E-05 | glio |
| Interaction_genotype_Myc | Cct6a | 1.821 | 1.92E-05 | glio |
| Atr_in.vivo_noRT | Ogdhl | 2.978 | 1.92E-05 | glio |
| Bcas2_in.vivo_noRT | Plin1 | 3.094 | 1.92E-05 | glio |
| Bora_in.vivo_noRT | Astn2 | 2.848 | 1.92E-05 | glio |
| culture_effect | Tomm7 | 0.687 | 1.92E-06 | glio |
| culture_effect | Rpl23 | -0.719 | 1.92E-06 | glio |
| Interaction_genotype_Mtor | 1700001L19Ri | -5.295 | 1.92E-06 | glio |
| Aars2_in.vivo_noRT | Gm17399 | 3.585 | 1.92E-06 | glio |
| Atr_in.vivo_noRT | Slc22a18 | 3.507 | 1.92E-06 | glio |
| Bcas2_in.vivo_noRT | Smc2os | 3.411 | 1.92E-06 | glio |
| Bora_ex.vivo_noRT | Ccl3 | 3.201 | 1.92E-06 | glio |
| Bora_ex.vivo_noRT | Gm48780 | 3.201 | 1.92E-06 | glio |
| Bora_ex.vivo_noRT | AU019990 | 3.201 | 1.92E-06 | glio |
| Chek1_in.vivo_noRT | Zfhx2os | 3.463 | 1.92E-06 | glio |
| Chek1_in.vivo_noRT | Tbc1d10c | 3.348 | 1.92E-06 | glio |
| Interaction_genotype_Mtor | Gm44658 | -5.51 | 1.92E-07 | glio |
| Aars2_in.vivo_noRT | 1700006J14Ri | 3.416 | 1.92E-07 | glio |
| Atr_ex.vivo_noRT | Cgnl1 | 3.164 | 1.92E-07 | glio |
| Bora_in.vivo_noRT | 2810408A11R | 3.662 | 1.92E-07 | glio |
| Chek1_ex.vivo_noRT | Gm48832 | 4.447 | 1.92E-07 | glio |
| Chek1_ex.vivo_noRT | Parvg | 4.671 | 1.92E-07 | glio |
| Bora_in.vivo_noRT | Prox2 | 4.21 | 1.92E-08 | glio |
| Bora_in.vivo_noRT | Gm33973 | 3.37 | 1.92E-08 | glio |
| culture_effect | Rps3 | -0.788 | 1.92E-09 | glio |
| Aars2_in.vivo_noRT | Psma4 | -1.69 | 1.92E-09 | glio |
| Bora_in.vivo_noRT | Rem2 | 3.751 | 1.92E-09 | glio |
| Chek1_in.vivo_noRT | Cpeb1 | 4.398 | 1.92E-09 | glio |
| Chek1_ex.vivo_noRT | Sema6d | 3.861 | 1.92E-09 | glio |
| Chek1_in.vivo_noRT | Ccno | 4.547 | 1.92E-10 | glio |
| Chek1_in.vivo_noRT | Gm9929 | 4.265 | 1.92E-11 | glio |
| Aars2_in.vivo_noRT | Gm10300 | 5.067 | 1.92E-12 | glio |
| Aars2_in.vivo_noRT | Stx1b | 4.424 | 1.92E-12 | glio |
| Aars2_in.vivo_noRT | Gm28404 | 4.439 | 1.92E-12 | glio |
| Bcas2_in.vivo_noRT | Rarres1 | 4.807 | 1.92E-12 | glio |
| Chek1_in.vivo_noRT | Lce1g | 5.044 | 1.92E-12 | glio |
| Chek1_ex.vivo_noRT | Ms4a15 | 4.842 | 1.92E-12 | glio |
| Atr_in.vivo_noRT | Tmem54 | 5.273 | 1.92E-19 | glio |
| Bora_in.vivo_noRT | Cox7b2 | 5.778 | 1.92E-20 | glio |
| Chek1_in.vivo_noRT | Gm43701 | 5.044 | 1.92E-22 | glio |
| Chek1_in.vivo_noRT | Hoxc8 | 5.044 | 1.92E-25 | glio |
| culture_effect | Ubd | -8.536 | 1.92E-46 | glio |
| culture_effect | Zfp110 | -0.725 | 1.93E-03 | glio |
| Interaction_genotype_Aars2 | Ifitm2 | 0.984 | 1.93E-03 | glio |
| Interaction_genotype_Mtor | Gm47428 | -3.61 | 1.93E-03 | glio |
| Interaction_genotype_Mtor | Il6 | -4.283 | 1.93E-03 | glio |
| Interaction_genotype_Mtor | Ube2e1 | 1.661 | 1.93E-03 | glio |
| Interaction_genotype_Mtor | Cox7a2 | 1.426 | 1.93E-03 | glio |

| | | | | |
|---|---|---|---|---|
| Interaction_genotype_Myc | Olfr171 | -2.729 | 1.93E-03 | glio |
| Interaction_genotype_Myc | 3930402G23R | -3.555 | 1.93E-03 | glio |
| Interaction_genotype_Myc | Hist1h4n | -3.776 | 1.93E-03 | glio |
| Interaction_genotype_Myc | Zfp975 | -3.416 | 1.93E-03 | glio |
| Interaction_genotype_Myc | 6330403L08Ri | -3.224 | 1.93E-03 | glio |
| Interaction_genotype_Myc | Pccb | -3.317 | 1.93E-03 | glio |
| Interaction_genotype_Myc | Il17ra | -3.062 | 1.93E-03 | glio |
| Interaction_genotype_Myc | Patj | -2.865 | 1.93E-03 | glio |
| Interaction_genotype_Myc | Usp40 | -2.881 | 1.93E-03 | glio |
| Interaction_genotype_Myc | Gramd1a | -2.422 | 1.93E-03 | glio |
| Interaction_genotype_Myc | Plekha8 | -2.254 | 1.93E-03 | glio |
| Interaction_genotype_Myc | Ndst1 | -1.879 | 1.93E-03 | glio |
| Interaction_genotype_Myc | Adk | 1.814 | 1.93E-03 | glio |
| Interaction_genotype_Myc | Vdac2 | 1.312 | 1.93E-03 | glio |
| Interaction_genotype_Myc | Banf1 | 1.274 | 1.93E-03 | glio |
| interaction_RT | Wdr46 | -0.95 | 1.93E-03 | glio |
| Aars2_in.vivo_noRT | Ndufb11 | -0.693 | 1.93E-03 | glio |
| Aars2_ex.vivo_noRT | Gm15890 | 2.506 | 1.93E-03 | glio |
| Atr_ex.vivo_noRT | Gm26590 | 2.783 | 1.93E-03 | glio |
| Bcas2_in.vivo_noRT | Lmtk3 | 2.196 | 1.93E-03 | glio |
| Bcas2_ex.vivo_noRT | Gm49579 | 2.737 | 1.93E-03 | glio |
| Bcas2_ex.vivo_noRT | H2-DMa | 1.392 | 1.93E-03 | glio |
| Bora_ex.vivo_noRT | H2-M10.1 | 2.615 | 1.93E-03 | glio |
| Bora_ex.vivo_noRT | Gm45235 | 2.408 | 1.93E-03 | glio |
| Chek1_in.vivo_noRT | Psmc1 | -1.239 | 1.93E-03 | glio |
| Chek1_in.vivo_noRT | D8Ertd738e | -0.596 | 1.93E-03 | glio |
| Eif2s1_in.vivo_noRT | Gm9758 | 1.883 | 1.93E-03 | glio |
| culture_effect | Rapgefl1 | -2.298 | 1.93E-04 | glio |
| culture_effect | Pea15a | 1.105 | 1.93E-04 | glio |
| Interaction_genotype_Aars2 | Gm5129 | -4.808 | 1.93E-04 | glio |
| Interaction_genotype_Bcas2 | Efhd1 | -4.995 | 1.93E-04 | glio |
| Interaction_genotype_Bcas2 | Gm36356 | -4.189 | 1.93E-04 | glio |
| Interaction_genotype_Bcas2 | Rbpms2 | -4.916 | 1.93E-04 | glio |
| Interaction_genotype_Bcas2 | Gm32090 | -4.387 | 1.93E-04 | glio |
| Interaction_genotype_Bcas2 | Pde3a | -4.639 | 1.93E-04 | glio |
| Interaction_genotype_Bcas2 | 4930524O07R | -4.928 | 1.93E-04 | glio |
| Interaction_genotype_Mtor | Gm26978 | -3.633 | 1.93E-04 | glio |
| Interaction_genotype_Mtor | Dcdc2a | -3.839 | 1.93E-04 | glio |
| Interaction_genotype_Mtor | Rab18 | 1.723 | 1.93E-04 | glio |
| Interaction_genotype_Myc | Ccdc39 | -4.165 | 1.93E-04 | glio |
| Aars2_in.vivo_noRT | Selenbp1 | 2.888 | 1.93E-04 | glio |
| Atr_in.vivo_noRT | Hist2h3b | 2.624 | 1.93E-04 | glio |
| Atr_in.vivo_noRT | Klhdc2 | -1.32 | 1.93E-04 | glio |
| Atr_in.vivo_noRT | Hspa8 | -0.959 | 1.93E-04 | glio |
| Bard1_ex.vivo_noRT | Npy1r | 2.814 | 1.93E-04 | glio |
| Bard1_ex.vivo_noRT | A030014E15Ri | 2.421 | 1.93E-04 | glio |
| Bcas2_in.vivo_noRT | 1700123M08F | 2.672 | 1.93E-04 | glio |
| Bcas2_in.vivo_noRT | Amd2 | 2.637 | 1.93E-04 | glio |
| Bcas2_in.vivo_noRT | Ltbp4 | -1.801 | 1.93E-04 | glio |
| Bcas2_ex.vivo_noRT | Cxcl5 | 2.742 | 1.93E-04 | glio |
| Bora_in.vivo_noRT | Spata1 | 2.776 | 1.93E-04 | glio |
| Bora_in.vivo_noRT | Sec23ip | -1.72 | 1.93E-04 | glio |

| | | | | |
|---|---|---|---|---|
| Chek1_ex.vivo_noRT | 1700026N04R | 2.921 | 1.93E-04 | glio |
| Chek1_ex.vivo_noRT | Bcl6 | 3.47 | 1.93E-04 | glio |
| Chek1_ex.vivo_noRT | Eef1e1 | -1.084 | 1.93E-04 | glio |
| Hsd17b10_in.vivo_noRT | Inhba | 2.975 | 1.93E-04 | glio |
| Hsd17b10_in.vivo_noRT | Rims3 | 2.482 | 1.93E-04 | glio |
| culture_effect | Pag1 | 1.626 | 1.93E-05 | glio |
| culture_effect | Orai1 | -0.974 | 1.93E-05 | glio |
| culture_effect | Wwc1 | 0.948 | 1.93E-05 | glio |
| culture_effect | Mrps9 | -0.679 | 1.93E-05 | glio |
| Interaction_genotype_Mtor | Car12 | -4.891 | 1.93E-05 | glio |
| Aars2_in.vivo_noRT | Rpl11 | -0.926 | 1.93E-05 | glio |
| Atr_in.vivo_noRT | Acad10 | 3.028 | 1.93E-05 | glio |
| Atr_in.vivo_noRT | Myorg | 2.934 | 1.93E-05 | glio |
| Atr_ex.vivo_noRT | Klrk1 | 3.164 | 1.93E-05 | glio |
| Bora_in.vivo_noRT | Gm12454 | 3.032 | 1.93E-05 | glio |
| Bora_in.vivo_noRT | Setd4 | 2.985 | 1.93E-05 | glio |
| Bora_in.vivo_noRT | Impdh2 | -1.032 | 1.93E-05 | glio |
| Chek1_ex.vivo_noRT | Zfp808 | 3.279 | 1.93E-05 | glio |
| Hsd17b10_in.vivo_noRT | 3930402G23R | 3.229 | 1.93E-05 | glio |
| culture_effect | Gm8797 | -1.371 | 1.93E-06 | glio |
| culture_effect | Mtch1 | 0.615 | 1.93E-06 | glio |
| Aars2_in.vivo_noRT | Alox8 | 4.07 | 1.93E-06 | glio |
| Chek1_in.vivo_noRT | Tnc | 3.995 | 1.93E-06 | glio |
| Chek1_in.vivo_noRT | Gm16048 | 3.608 | 1.93E-06 | glio |
| Chek1_in.vivo_noRT | Gm44836 | 3.212 | 1.93E-06 | glio |
| Chek1_ex.vivo_noRT | Ptp4a1 | 3.652 | 1.93E-06 | glio |
| Interaction_genotype_Mtor | Pakap | -4.946 | 1.93E-07 | glio |
| Aars2_in.vivo_noRT | Tbc1d10c | 3.631 | 1.93E-07 | glio |
| Aars2_ex.vivo_noRT | Gm14929 | 3.293 | 1.93E-07 | glio |
| Atp5c1_in.vivo_noRT | Ccl9 | 2.668 | 1.93E-07 | glio |
| Atr_ex.vivo_noRT | Gm26543 | 3.164 | 1.93E-07 | glio |
| Bora_ex.vivo_noRT | Slc14a2 | 3.201 | 1.93E-07 | glio |
| Bora_ex.vivo_noRT | Tff2 | 4.606 | 1.93E-07 | glio |
| Bora_ex.vivo_noRT | Prap1 | 3.601 | 1.93E-07 | glio |
| Bora_ex.vivo_noRT | Gm4340 | 3.201 | 1.93E-07 | glio |
| Bora_ex.vivo_noRT | Gm16239 | 3.201 | 1.93E-07 | glio |
| Hsd17b10_in.vivo_noRT | Odc1 | -1.673 | 1.93E-07 | glio |
| culture_effect | 5033430I15Ri | 2.628 | 1.93E-08 | glio |
| Atr_in.vivo_noRT | 4732471J01Ri | 3.975 | 1.93E-08 | glio |
| Atr_in.vivo_noRT | Lmod1 | 4.102 | 1.93E-08 | glio |
| Bora_in.vivo_noRT | Zfp986 | 3.895 | 1.93E-08 | glio |
| Hsd17b10_in.vivo_noRT | Gm45591 | 3.356 | 1.93E-08 | glio |
| Hsd17b10_in.vivo_noRT | BC049987 | 2.833 | 1.93E-08 | glio |
| culture_effect | Ppp1r16b | -3.63 | 1.93E-09 | glio |
| Aars2_in.vivo_noRT | Styk1 | 4.277 | 1.93E-09 | glio |
| Bcas2_in.vivo_noRT | Zfp14 | 4.284 | 1.93E-09 | glio |
| Bora_in.vivo_noRT | 9530080O11R | 4.019 | 1.93E-09 | glio |
| Chek1_in.vivo_noRT | Gm3734 | 4.521 | 1.93E-09 | glio |
| Atr_in.vivo_noRT | 2210417A02R | 4.742 | 1.93E-10 | glio |
| Atr_in.vivo_noRT | Ccl8 | 5.007 | 1.93E-10 | glio |
| Bora_in.vivo_noRT | Fam13a | 4.66 | 1.93E-11 | glio |
| Bora_in.vivo_noRT | Gm10696 | 4.936 | 1.93E-12 | glio |

| | | | | |
|---|---|---|---|---|
| Bora_in.vivo_noRT | Gm42918 | 4.294 | 1.93E-12 | glio |
| Chek1_ex.vivo_noRT | Zscan4-ps2 | 4.842 | 1.93E-14 | glio |
| Aars2_in.vivo_noRT | Acat3 | 4.535 | 1.93E-16 | glio |
| Chek1_in.vivo_noRT | Ube4bos1 | 4.78 | 1.93E-16 | glio |
| Atr_in.vivo_noRT | Adam30 | 5.154 | 1.93E-17 | glio |
| Bora_in.vivo_noRT | Selenov | 4.936 | 1.93E-17 | glio |
| Bora_in.vivo_noRT | Btn1a1 | 4.676 | 1.93E-17 | glio |
| Bora_in.vivo_noRT | A330076H08R | 4.936 | 1.93E-17 | glio |
| Aars2_in.vivo_noRT | AC154200.1 | 5.327 | 1.93E-21 | glio |
| Bcas2_in.vivo_noRT | Hoxc8 | 4.807 | 1.93E-23 | glio |
| culture_effect | H2-Eb2 | -5.028 | 1.93E-25 | glio |
| Aars2_in.vivo_noRT | Fam227b | 5.327 | 1.93E-28 | glio |
| Aars2_in.vivo_noRT | Gm20658 | 5.327 | 1.93E-28 | glio |
| culture_effect | Trim24 | 0.599 | 1.94E-03 | glio |
| culture_effect | Nup35 | 0.526 | 1.94E-03 | glio |
| Interaction_genotype_Mtor | Gm43660 | -3.579 | 1.94E-03 | glio |
| Interaction_genotype_Mtor | Cd79a | -3.907 | 1.94E-03 | glio |
| Interaction_genotype_Mtor | Arid3b | -3.549 | 1.94E-03 | glio |
| Interaction_genotype_Mtor | Fetub | -3.168 | 1.94E-03 | glio |
| Interaction_genotype_Mtor | Amdhd2 | -2.352 | 1.94E-03 | glio |
| Interaction_genotype_Mtor | Hoxaas3 | -2.705 | 1.94E-03 | glio |
| Interaction_genotype_Mtor | Sys1 | 1.68 | 1.94E-03 | glio |
| Interaction_genotype_Myc | Ercc6l | -2.493 | 1.94E-03 | glio |
| Interaction_genotype_Myc | Ndufb2 | 1.369 | 1.94E-03 | glio |
| Aars2_in.vivo_noRT | Mpped2 | 2.221 | 1.94E-03 | glio |
| Atr_ex.vivo_noRT | Txndc2 | 2.376 | 1.94E-03 | glio |
| Atr_ex.vivo_noRT | Rax | 2.808 | 1.94E-03 | glio |
| Atr_ex.vivo_noRT | Grin2d | 2.794 | 1.94E-03 | glio |
| Atr_ex.vivo_noRT | Cysrt1 | 2.945 | 1.94E-03 | glio |
| Atr_ex.vivo_noRT | AC168977.1 | 3.413 | 1.94E-03 | glio |
| Atr_ex.vivo_noRT | Chl1 | -1.224 | 1.94E-03 | glio |
| Bora_in.vivo_noRT | 9130015A21R | 4.038 | 1.94E-03 | glio |
| Chek1_in.vivo_noRT | Gm45494 | 2.166 | 1.94E-03 | glio |
| Chek1_in.vivo_noRT | Tppp | 1.963 | 1.94E-03 | glio |
| Chek1_ex.vivo_noRT | Dennd3 | 2.974 | 1.94E-03 | glio |
| culture_effect | Ccdc136 | 1.617 | 1.94E-04 | glio |
| culture_effect | Hnrnpf | -0.372 | 1.94E-04 | glio |
| Interaction_genotype_Myc | D430001F17R | -3.886 | 1.94E-04 | glio |
| Interaction_genotype_Myc | Gm50107 | -3.942 | 1.94E-04 | glio |
| Interaction_genotype_Myc | Gm43156 | -4.096 | 1.94E-04 | glio |
| Interaction_genotype_Myc | Ubash3a | -6.044 | 1.94E-04 | glio |
| Aars2_in.vivo_noRT | Alox15 | 2.366 | 1.94E-04 | glio |
| Aars2_in.vivo_noRT | Pfdn1 | -0.951 | 1.94E-04 | glio |
| Aars2_ex.vivo_noRT | Gm17276 | 2.897 | 1.94E-04 | glio |
| Aars2_ex.vivo_noRT | 9130204L05Ri | 3.768 | 1.94E-04 | glio |
| Aars2_ex.vivo_noRT | Gm4858 | 3.293 | 1.94E-04 | glio |
| Aars2_ex.vivo_noRT | Fa2h | 3.293 | 1.94E-04 | glio |
| Atp5c1_in.vivo_noRT | Gm11627 | 2.668 | 1.94E-04 | glio |
| Atp5c1_in.vivo_noRT | Iqch | 2.404 | 1.94E-04 | glio |
| Atr_in.vivo_noRT | E130114P18Ri | 2.781 | 1.94E-04 | glio |
| Atr_in.vivo_noRT | Sap25 | 2.469 | 1.94E-04 | glio |
| Atr_ex.vivo_noRT | Pcdhga9 | 3.371 | 1.94E-04 | glio |

| | | | | |
|---|---|---|---|---|
| Bcas2_in.vivo_noRT | Mkrn3 | 2.55 | 1.94E-04 | glio |
| Bora_in.vivo_noRT | Zbtb24 | 1.955 | 1.94E-04 | glio |
| Chek1_in.vivo_noRT | Hist2h3b | 2.658 | 1.94E-04 | glio |
| culture_effect | Dennd2d | -2.408 | 1.94E-05 | glio |
| culture_effect | Yae1d1 | 0.725 | 1.94E-05 | glio |
| culture_effect | Mtln | 0.609 | 1.94E-05 | glio |
| Interaction_genotype_Mtor | Snai1 | -5.263 | 1.94E-05 | glio |
| Interaction_genotype_Mtor | Jazf1 | -4.896 | 1.94E-05 | glio |
| interaction_RT | Wsb1 | 1.182 | 1.94E-05 | glio |
| Atp5c1_in.vivo_noRT | Gm43915 | 2.657 | 1.94E-05 | glio |
| Atr_in.vivo_noRT | Dusp15 | 2.684 | 1.94E-05 | glio |
| Atr_in.vivo_noRT | Gm10658 | 2.868 | 1.94E-05 | glio |
| Bard1_ex.vivo_noRT | Rimklb | 2.814 | 1.94E-05 | glio |
| Bard1_ex.vivo_noRT | Kcnq2 | 3.612 | 1.94E-05 | glio |
| Bcas2_in.vivo_noRT | 9130604C24R | 3.341 | 1.94E-05 | glio |
| Interaction_genotype_Mtor | Afg1l | -5.754 | 1.94E-06 | glio |
| Aars2_in.vivo_noRT | Txn2 | -1.384 | 1.94E-06 | glio |
| Bcas2_in.vivo_noRT | Dio1 | 2.971 | 1.94E-06 | glio |
| Hsd17b10_in.vivo_noRT | E030037K01Ri | 3.084 | 1.94E-06 | glio |
| culture_effect | Tppp | -2.852 | 1.94E-07 | glio |
| Chek1_ex.vivo_noRT | Crybb1 | 3.873 | 1.94E-07 | glio |
| Atr_in.vivo_noRT | Gm26519 | 4.347 | 1.94E-08 | glio |
| Atr_in.vivo_noRT | Gm49801 | 4.002 | 1.94E-08 | glio |
| Atr_in.vivo_noRT | 5330417C22R | 4.138 | 1.94E-09 | glio |
| Chek1_in.vivo_noRT | Gm16046 | 4.252 | 1.94E-09 | glio |
| Chek1_ex.vivo_noRT | Gm20767 | 4.842 | 1.94E-09 | glio |
| Hsd17b10_in.vivo_noRT | Usp17lc | 3.616 | 1.94E-09 | glio |
| Aars2_in.vivo_noRT | Sbk3 | 4.158 | 1.94E-10 | glio |
| Aars2_in.vivo_noRT | Bglap2 | 4.398 | 1.94E-10 | glio |
| Atr_in.vivo_noRT | Ggt1 | 4.221 | 1.94E-10 | glio |
| Bcas2_in.vivo_noRT | Gm26674 | 4.162 | 1.94E-10 | glio |
| Bora_in.vivo_noRT | Zic4 | 4.673 | 1.94E-10 | glio |
| Chek1_in.vivo_noRT | Gm42984 | 4.393 | 1.94E-10 | glio |
| Chek1_in.vivo_noRT | Rdh12 | 4.398 | 1.94E-11 | glio |
| culture_effect | Mfsd13a | -2.745 | 1.94E-12 | glio |
| Bcas2_in.vivo_noRT | Rnf165 | 4.807 | 1.94E-12 | glio |
| Bora_in.vivo_noRT | Thnsl2 | 3.768 | 1.94E-12 | glio |
| Bora_in.vivo_noRT | Upb1 | 4.673 | 1.94E-12 | glio |
| Chek1_in.vivo_noRT | 4921536K21R | 4.784 | 1.94E-12 | glio |
| Chek1_in.vivo_noRT | Smpd5 | 4.387 | 1.94E-12 | glio |
| Aars2_in.vivo_noRT | Gm5602 | 4.535 | 1.94E-16 | glio |
| Bcas2_in.vivo_noRT | 2310003N18R | 4.807 | 1.94E-16 | glio |
| Chek1_in.vivo_noRT | Gm49860 | 5.044 | 1.94E-17 | glio |
| Atr_in.vivo_noRT | Hnf1aos1 | 5.533 | 1.94E-19 | glio |
| culture_effect | Ubc | -1.656 | 1.94E-25 | glio |
| Bora_in.vivo_noRT | Slc17a6 | 4.936 | 1.94E-28 | glio |
| culture_effect | Timp2 | 2.028 | 1.94E-30 | glio |
| culture_effect | Atp6v1g2 | 1.331 | 1.95E-03 | glio |
| culture_effect | Igsf3 | -1.023 | 1.95E-03 | glio |
| culture_effect | Echdc2 | -0.666 | 1.95E-03 | glio |
| culture_effect | Hook3 | -0.544 | 1.95E-03 | glio |
| Interaction_genotype_Aars2 | St8sia1 | -3.974 | 1.95E-03 | glio |

| | | | | |
|---|---|---|---|---|
| Interaction_genotype_Aars2 | Cox5b | 0.994 | 1.95E-03 | glio |
| Interaction_genotype_Atr | Spata31d1a | -3.96 | 1.95E-03 | glio |
| Interaction_genotype_Mtor | Fam189b | -3.983 | 1.95E-03 | glio |
| Interaction_genotype_Mtor | 1810009A15R | -3.049 | 1.95E-03 | glio |
| Interaction_genotype_Mtor | Cd47 | 1.958 | 1.95E-03 | glio |
| Interaction_genotype_Mtor | Hmgb2 | 3.037 | 1.95E-03 | glio |
| Interaction_genotype_Myc | Rassf7 | -2.808 | 1.95E-03 | glio |
| Interaction_genotype_Myc | Elac2 | -2.209 | 1.95E-03 | glio |
| Interaction_genotype_Myc | H2-DMb1 | 1.796 | 1.95E-03 | glio |
| Aars2_in.vivo_noRT | Gdap1l1 | 2.449 | 1.95E-03 | glio |
| Aars2_in.vivo_noRT | Adamts4 | 1.985 | 1.95E-03 | glio |
| Atr_in.vivo_noRT | Mmaa | 1.946 | 1.95E-03 | glio |
| Bora_in.vivo_noRT | Pxdn | 1.93 | 1.95E-03 | glio |
| Chek1_ex.vivo_noRT | Shisal2a | 2.962 | 1.95E-03 | glio |
| Chek1_ex.vivo_noRT | Foxa3 | 2.865 | 1.95E-03 | glio |
| Hsd17b10_in.vivo_noRT | Cyp4f18 | 3.356 | 1.95E-03 | glio |
| culture_effect | 9130604C24R | 2.153 | 1.95E-04 | glio |
| culture_effect | 2610002M06F | -0.683 | 1.95E-04 | glio |
| culture_effect | Tut4 | -0.55 | 1.95E-04 | glio |
| Interaction_genotype_Myc | Rapgef3os2 | -3.119 | 1.95E-04 | glio |
| Interaction_genotype_Myc | Efcab11 | -3.598 | 1.95E-04 | glio |
| Interaction_genotype_Myc | Cisd1 | 1.595 | 1.95E-04 | glio |
| Bcas2_in.vivo_noRT | Gm15708 | 2.64 | 1.95E-04 | glio |
| Bcas2_ex.vivo_noRT | Gm47936 | 2.742 | 1.95E-04 | glio |
| Bora_ex.vivo_noRT | Dok2 | 3.018 | 1.95E-04 | glio |
| Chek1_in.vivo_noRT | Bod1 | -1.445 | 1.95E-04 | glio |
| Chek1_ex.vivo_noRT | Ctsd | 1.052 | 1.95E-04 | glio |
| Dclre1c_ex.vivo_noRT | Gm5662 | 3.85 | 1.95E-04 | glio |
| Dclre1c_ex.vivo_noRT | Gm21936 | 2.685 | 1.95E-04 | glio |
| Interaction_genotype_Mtor | 5830454E08Ri | -4.464 | 1.95E-05 | glio |
| Interaction_genotype_Mtor | Sema6b | -4.647 | 1.95E-05 | glio |
| Interaction_genotype_Myc | Ephx1 | -4.733 | 1.95E-05 | glio |
| Interaction_genotype_Myc | Ost4 | 1.94 | 1.95E-05 | glio |
| Interaction_genotype_Rad51 | Baalc | -5.193 | 1.95E-05 | glio |
| Atr_in.vivo_noRT | Lbx2 | 3.244 | 1.95E-05 | glio |
| Atr_in.vivo_noRT | Gm44190 | 2.745 | 1.95E-05 | glio |
| Bcas2_in.vivo_noRT | Hoga1 | 2.993 | 1.95E-05 | glio |
| Bcas2_in.vivo_noRT | Tmem86b | 3.108 | 1.95E-05 | glio |
| Chek1_in.vivo_noRT | Zbtb46 | 3.205 | 1.95E-05 | glio |
| Aars2_in.vivo_noRT | 2900041M22F | 3.517 | 1.95E-06 | glio |
| Atr_in.vivo_noRT | Fgfr1 | 3.896 | 1.95E-06 | glio |
| Atr_in.vivo_noRT | Nupr1l | 3.753 | 1.95E-06 | glio |
| Atr_in.vivo_noRT | Sgk2 | 3.317 | 1.95E-06 | glio |
| Atr_in.vivo_noRT | Cct4 | -1.296 | 1.95E-06 | glio |
| Bora_in.vivo_noRT | Eng | 3.74 | 1.95E-06 | glio |
| Bora_in.vivo_noRT | Ggnbp1 | 3.254 | 1.95E-06 | glio |
| Bora_ex.vivo_noRT | Gm36793 | 3.201 | 1.95E-06 | glio |
| Hsd17b10_in.vivo_noRT | Gm49874 | 3.616 | 1.95E-06 | glio |
| Hsd17b10_in.vivo_noRT | Hdac9 | 3.35 | 1.95E-06 | glio |
| Aars2_in.vivo_noRT | Ablim3 | 3.766 | 1.95E-07 | glio |
| Atr_ex.vivo_noRT | Spns3 | 4.708 | 1.95E-07 | glio |
| Atr_ex.vivo_noRT | 2410017I17Ri | 3.164 | 1.95E-07 | glio |

| | | | | |
|---|---|---|---|---|
| Bora_in.vivo_noRT | Gm10125 | 3.735 | 1.95E-07 | glio |
| Bora_in.vivo_noRT | Tmem8b | 3.548 | 1.95E-07 | glio |
| Chek1_in.vivo_noRT | 1810041H14R | 3.925 | 1.95E-07 | glio |
| Chek1_ex.vivo_noRT | D730045A05R | 4.447 | 1.95E-07 | glio |
| Atr_in.vivo_noRT | Gm50012 | 3.933 | 1.95E-08 | glio |
| Atr_in.vivo_noRT | 1700029J07Ri | 3.976 | 1.95E-08 | glio |
| Bcas2_in.vivo_noRT | Dph3 | -1.885 | 1.95E-08 | glio |
| Bora_in.vivo_noRT | D930030I03Ri | 4.23 | 1.95E-08 | glio |
| Bcas2_in.vivo_noRT | Gm42984 | 4.157 | 1.95E-09 | glio |
| Bcas2_in.vivo_noRT | Flnc | 4.281 | 1.95E-09 | glio |
| Bora_in.vivo_noRT | Cpne9 | 4.682 | 1.95E-10 | glio |
| Chek1_ex.vivo_noRT | Ciita | 4.443 | 1.95E-10 | glio |
| Aars2_in.vivo_noRT | Adam33 | 4.684 | 1.95E-11 | glio |
| Bcas2_in.vivo_noRT | Fbxl2 | 4.161 | 1.95E-11 | glio |
| culture_effect | Gpi1 | -0.953 | 1.95E-13 | glio |
| Chek1_in.vivo_noRT | Ptgis | 5.044 | 1.95E-13 | glio |
| Aars2_in.vivo_noRT | E030037K01Ri | 4.795 | 1.95E-14 | glio |
| Atr_in.vivo_noRT | Gm14379 | 5.017 | 1.95E-14 | glio |
| Aars2_in.vivo_noRT | Gm45620 | 5.327 | 1.95E-15 | glio |
| Aars2_in.vivo_noRT | Gm38534 | 5.327 | 1.95E-15 | glio |
| Chek1_in.vivo_noRT | 4930434B07R | 4.78 | 1.95E-15 | glio |
| Chek1_in.vivo_noRT | 1700026J14Ri | 5.045 | 1.95E-15 | glio |
| Aars2_in.vivo_noRT | Gm15890 | 5.327 | 1.95E-19 | glio |
| culture_effect | Skap2 | -1.447 | 1.95E-23 | glio |
| Bora_in.vivo_noRT | Trim17 | 4.936 | 1.95E-23 | glio |
| Aars2_in.vivo_noRT | Ankrd22 | 5.327 | 1.95E-27 | glio |
| culture_effect | Traf5 | -1.639 | 1.96E-03 | glio |
| culture_effect | Exo5 | 0.774 | 1.96E-03 | glio |
| culture_effect | Syf2 | -0.427 | 1.96E-03 | glio |
| Interaction_genotype_Aars2 | 1700024I08Ri | -3.147 | 1.96E-03 | glio |
| Interaction_genotype_Aars2 | Gm26850 | -3.921 | 1.96E-03 | glio |
| Interaction_genotype_Aars2 | Mras | 1.777 | 1.96E-03 | glio |
| Interaction_genotype_Atr | Tmem181a | 2.102 | 1.96E-03 | glio |
| Interaction_genotype_Mtor | Fzd5 | -3.772 | 1.96E-03 | glio |
| Interaction_genotype_Mtor | Gm9803 | -3.43 | 1.96E-03 | glio |
| Interaction_genotype_Mtor | Atp6v1g2 | -3.232 | 1.96E-03 | glio |
| Interaction_genotype_Mtor | Zranb2 | 1.475 | 1.96E-03 | glio |
| Interaction_genotype_Myc | 1700006H21R | -3.516 | 1.96E-03 | glio |
| Interaction_genotype_Myc | Lrrc75aos2 | -3.616 | 1.96E-03 | glio |
| Interaction_genotype_Myc | Gm15879 | -4.509 | 1.96E-03 | glio |
| Interaction_genotype_Myc | Tmem117 | -3.543 | 1.96E-03 | glio |
| Interaction_genotype_Myc | Lypd6 | -3.462 | 1.96E-03 | glio |
| Interaction_genotype_Myc | Ccdc15 | -3.168 | 1.96E-03 | glio |
| Interaction_genotype_Myc | Zfp971 | -3.163 | 1.96E-03 | glio |
| Interaction_genotype_Myc | Etaa1os | -3.201 | 1.96E-03 | glio |
| Interaction_genotype_Myc | Mgst3 | -3.018 | 1.96E-03 | glio |
| Interaction_genotype_Myc | Zfp369 | -3.104 | 1.96E-03 | glio |
| Interaction_genotype_Myc | 2900060B14R | -3.296 | 1.96E-03 | glio |
| Interaction_genotype_Myc | Armc5 | -3.019 | 1.96E-03 | glio |
| Interaction_genotype_Myc | Ptpn9 | -2.299 | 1.96E-03 | glio |
| Interaction_genotype_Myc | Tcim | 2.706 | 1.96E-03 | glio |
| Interaction_genotype_Myc | Odc1 | 2.434 | 1.96E-03 | glio |

| | | | | |
|---|---|---|---|---|
| Aars2_in.vivo_noRT | 9330151L19Ri | 2.007 | 1.96E-03 | glio |
| Aars2_in.vivo_noRT | Gle1 | -1.355 | 1.96E-03 | glio |
| Atr_in.vivo_noRT | Fbxo46 | 1.934 | 1.96E-03 | glio |
| Atr_in.vivo_noRT | Mapk1ip1l | -1.18 | 1.96E-03 | glio |
| Atr_in.vivo_noRT | Gabarapl1 | -1.011 | 1.96E-03 | glio |
| Atr_ex.vivo_noRT | Mxd1 | 2.45 | 1.96E-03 | glio |
| Bcas2_in.vivo_noRT | Rps6kl1 | 2.284 | 1.96E-03 | glio |
| Bora_in.vivo_noRT | Nuak1 | 1.913 | 1.96E-03 | glio |
| Cenpj_ex.vivo_noRT | Apol10b | 2.591 | 1.96E-03 | glio |
| Chek1_ex.vivo_noRT | Gm13203 | 3.061 | 1.96E-03 | glio |
| Chek1_ex.vivo_noRT | Npr2 | 3.344 | 1.96E-03 | glio |
| Chek1_ex.vivo_noRT | Tubb2b | -1.118 | 1.96E-03 | glio |
| Chek1_ex.vivo_noRT | Ftl1 | 0.633 | 1.96E-03 | glio |
| Eif2s1_in.vivo_noRT | Ccser1 | 2.143 | 1.96E-03 | glio |
| Hsd17b10_in.vivo_noRT | Drd4 | 2.432 | 1.96E-03 | glio |
| culture_effect | Anapc7 | -0.907 | 1.96E-04 | glio |
| Interaction_genotype_Atr | Lmbr1 | -3.689 | 1.96E-04 | glio |
| Interaction_genotype_Bcas2 | Nrip1 | 2.6 | 1.96E-04 | glio |
| Interaction_genotype_Mtor | Agrp | -3.712 | 1.96E-04 | glio |
| Interaction_genotype_Mtor | Crybg2 | -4.424 | 1.96E-04 | glio |
| Interaction_genotype_Myc | Recql5 | -3.594 | 1.96E-04 | glio |
| Aars2_in.vivo_noRT | Lap3 | -1.176 | 1.96E-04 | glio |
| Aars2_ex.vivo_noRT | Gm20767 | 3.293 | 1.96E-04 | glio |
| Aars2_ex.vivo_noRT | Brsk2 | 3.293 | 1.96E-04 | glio |
| Atr_in.vivo_noRT | Rnasel | 2.393 | 1.96E-04 | glio |
| Atr_ex.vivo_noRT | Crybb1 | 2.978 | 1.96E-04 | glio |
| Bcas2_in.vivo_noRT | A430010J10Ri | 3.3 | 1.96E-04 | glio |
| Bora_in.vivo_noRT | 4930455G09R | 3.12 | 1.96E-04 | glio |
| Bora_in.vivo_noRT | Gstm6 | 2.869 | 1.96E-04 | glio |
| Bora_in.vivo_noRT | Atp8a2 | 2.257 | 1.96E-04 | glio |
| Bora_in.vivo_noRT | Radil | 2.638 | 1.96E-04 | glio |
| Chek1_in.vivo_noRT | 2510046G10R | 2.414 | 1.96E-04 | glio |
| Chek1_ex.vivo_noRT | Thap8 | 3.259 | 1.96E-04 | glio |
| culture_effect | 4732491K20R | 1.753 | 1.96E-05 | glio |
| culture_effect | Calcrl | -1.404 | 1.96E-05 | glio |
| culture_effect | Lbp | -1.011 | 1.96E-05 | glio |
| Interaction_genotype_Mtor | Ltb | -4.982 | 1.96E-05 | glio |
| Interaction_genotype_Mtor | Hnrnpr | 2.037 | 1.96E-05 | glio |
| Interaction_genotype_Myc | Lama5 | -5.048 | 1.96E-05 | glio |
| Aars2_in.vivo_noRT | Gm6598 | 2.808 | 1.96E-05 | glio |
| Aars2_ex.vivo_noRT | BC048671 | 3.293 | 1.96E-05 | glio |
| Bora_in.vivo_noRT | Ppif | -2.117 | 1.96E-05 | glio |
| Chek1_in.vivo_noRT | C2cd2 | 3.134 | 1.96E-05 | glio |
| Hsd17b10_in.vivo_noRT | Gm20635 | 2.971 | 1.96E-05 | glio |
| Hsd17b10_in.vivo_noRT | 2610027K06R | 2.96 | 1.96E-05 | glio |
| Aars2_in.vivo_noRT | Tagap1 | 2.675 | 1.96E-06 | glio |
| Aars2_in.vivo_noRT | Psmd7 | -1.453 | 1.96E-06 | glio |
| Atr_in.vivo_noRT | Cldn10 | 3.326 | 1.96E-06 | glio |
| Bcas2_in.vivo_noRT | Bbs1 | 3.348 | 1.96E-06 | glio |
| Bora_in.vivo_noRT | Apold1 | 3.826 | 1.96E-06 | glio |
| Hsd17b10_in.vivo_noRT | Pmch | 2.958 | 1.96E-06 | glio |
| Interaction_genotype_Mtor | Fam25c | -4.94 | 1.96E-07 | glio |

| | | | | |
|---|---|---|---|---|
| Interaction_genotype_Myc | Gm43137 | -5.957 | 1.96E-07 | glio |
| Chek1_in.vivo_noRT | 4930592C13R | 3.777 | 1.96E-07 | glio |
| culture_effect | 2610027K06R | 2.803 | 1.96E-08 | glio |
| culture_effect | Dzip1l | 1.765 | 1.96E-08 | glio |
| culture_effect | Lmbrd1 | -1.022 | 1.96E-08 | glio |
| culture_effect | Gna11 | 0.978 | 1.96E-08 | glio |
| Aars2_in.vivo_noRT | 1810041H14R | 4.208 | 1.96E-08 | glio |
| Atr_in.vivo_noRT | Krt16 | 4.824 | 1.96E-08 | glio |
| Chek1_in.vivo_noRT | Catsperd | 3.865 | 1.96E-08 | glio |
| culture_effect | Cyp26b1 | 3.209 | 1.96E-09 | glio |
| culture_effect | Aopep | 0.838 | 1.96E-09 | glio |
| Chek1_ex.vivo_noRT | Gm38393 | 4.449 | 1.96E-09 | glio |
| culture_effect | Pcna | -0.985 | 1.96E-10 | glio |
| Bora_in.vivo_noRT | Vpreb1 | 4.557 | 1.96E-11 | glio |
| Aars2_in.vivo_noRT | D030040B21R | 4.94 | 1.96E-13 | glio |
| Hsd17b10_in.vivo_noRT | Slc17a6 | 3.616 | 1.96E-16 | glio |
| Atr_in.vivo_noRT | C330024C12R | 5.142 | 1.96E-17 | glio |
| Atr_in.vivo_noRT | Gm39929 | 5.076 | 1.96E-19 | glio |
| culture_effect | Csgalnact1 | -3.404 | 1.96E-20 | glio |
| culture_effect | Psme2 | -1.376 | 1.96E-22 | glio |
| Chek1_in.vivo_noRT | 9130019O22R | 5.044 | 1.96E-22 | glio |
| Bcas2_in.vivo_noRT | Gm45338 | 4.807 | 1.96E-24 | glio |
| Bora_in.vivo_noRT | Gm48155 | 4.936 | 1.96E-25 | glio |
| Chek1_in.vivo_noRT | 1700003E24Ri | 5.044 | 1.96E-26 | glio |
| culture_effect | Gm13178 | -1.451 | 1.97E-03 | glio |
| culture_effect | Lamtor1 | 0.406 | 1.97E-03 | glio |
| Interaction_genotype_Aars2 | Gm45890 | -3.985 | 1.97E-03 | glio |
| Interaction_genotype_Aars2 | Gm10602 | -4.294 | 1.97E-03 | glio |
| Interaction_genotype_Atr | Hspa12a | -3.876 | 1.97E-03 | glio |
| Interaction_genotype_Atr | Gas5 | 1.178 | 1.97E-03 | glio |
| Interaction_genotype_Bcas2 | Gm40557 | -4.225 | 1.97E-03 | glio |
| Interaction_genotype_Bcas2 | Gm36495 | -4.263 | 1.97E-03 | glio |
| Interaction_genotype_Bcas2 | 2010001K21R | -3.763 | 1.97E-03 | glio |
| Interaction_genotype_Mtor | Abcc10 | -3.658 | 1.97E-03 | glio |
| Interaction_genotype_Myc | Cd200r2 | -2.466 | 1.97E-03 | glio |
| Interaction_genotype_Myc | Gm6803 | -2.729 | 1.97E-03 | glio |
| Interaction_genotype_Myc | Zfp354c | -3.226 | 1.97E-03 | glio |
| Interaction_genotype_Myc | Snta1 | -2.695 | 1.97E-03 | glio |
| Interaction_genotype_Rad21 | Slc1a1 | 3.967 | 1.97E-03 | glio |
| interaction_RT | Nfe2l2 | 0.961 | 1.97E-03 | glio |
| Aars2_in.vivo_noRT | Nfatc2 | 2.295 | 1.97E-03 | glio |
| Aars2_in.vivo_noRT | Gmpr | 1.746 | 1.97E-03 | glio |
| Aars2_in.vivo_noRT | Iigp1 | -1.836 | 1.97E-03 | glio |
| Aars2_in.vivo_noRT | Rps23 | -0.592 | 1.97E-03 | glio |
| Aars2_ex.vivo_noRT | Gm7247 | 2.495 | 1.97E-03 | glio |
| Aars2_ex.vivo_noRT | A330032B11R | 2.713 | 1.97E-03 | glio |
| Atp5c1_in.vivo_noRT | Gm44658 | 2.136 | 1.97E-03 | glio |
| Atp5c1_in.vivo_noRT | 4921536K21R | 2.408 | 1.97E-03 | glio |
| Atp5c1_in.vivo_noRT | Zfp786 | 2.404 | 1.97E-03 | glio |
| Bard1_ex.vivo_noRT | 4930553M12F | 2.421 | 1.97E-03 | glio |
| Bcas2_in.vivo_noRT | D730045B01R | 2.021 | 1.97E-03 | glio |
| Bcas2_in.vivo_noRT | Hist3h2a | 2.169 | 1.97E-03 | glio |

| | | | | |
|---|---|---|---|---|
| Bcas2_ex.vivo_noRT | 1110038B12R | 0.835 | 1.97E-03 | glio |
| Bora_in.vivo_noRT | Zfp763 | 2.116 | 1.97E-03 | glio |
| Bora_in.vivo_noRT | Ranbp2 | -1.33 | 1.97E-03 | glio |
| Bora_ex.vivo_noRT | Gm42664 | 2.615 | 1.97E-03 | glio |
| Bora_ex.vivo_noRT | Gm43258 | 3.022 | 1.97E-03 | glio |
| Chek1_in.vivo_noRT | Ndufb8 | -0.568 | 1.97E-03 | glio |
| Chek1_ex.vivo_noRT | Rasip1 | 3.063 | 1.97E-03 | glio |
| culture_effect | Rel | -0.936 | 1.97E-04 | glio |
| Interaction_genotype_Mtor | Plcd4 | -4.164 | 1.97E-04 | glio |
| Interaction_genotype_Mtor | Gm10696 | -4.498 | 1.97E-04 | glio |
| Interaction_genotype_Mtor | Tmem240 | -4.724 | 1.97E-04 | glio |
| Interaction_genotype_Mtor | Cmpk1 | 1.911 | 1.97E-04 | glio |
| Interaction_genotype_Mtor | Dazap2 | 1.842 | 1.97E-04 | glio |
| Interaction_genotype_Myc | Zfp661 | -4.326 | 1.97E-04 | glio |
| Interaction_genotype_Myc | Prl2c3 | -3.925 | 1.97E-04 | glio |
| Interaction_genotype_Myc | Klhdc8b | -4.238 | 1.97E-04 | glio |
| Interaction_genotype_Myc | Egfl7 | -3.86 | 1.97E-04 | glio |
| Interaction_genotype_Myc | Gpr173 | -3.538 | 1.97E-04 | glio |
| Interaction_genotype_Myc | Def8 | -3.251 | 1.97E-04 | glio |
| interaction_RT | Hist1h3c | -1.72 | 1.97E-04 | glio |
| Atr_in.vivo_noRT | Dyrk4 | 2.965 | 1.97E-04 | glio |
| Atr_in.vivo_noRT | Sf3b4 | -1.46 | 1.97E-04 | glio |
| Atr_ex.vivo_noRT | Glyat | 2.367 | 1.97E-04 | glio |
| Bcas2_in.vivo_noRT | Adam4 | 2.466 | 1.97E-04 | glio |
| Bora_in.vivo_noRT | Gm17597 | 2.901 | 1.97E-04 | glio |
| Bora_in.vivo_noRT | Itgb7 | 2.519 | 1.97E-04 | glio |
| Bora_in.vivo_noRT | Grcc10 | -0.81 | 1.97E-04 | glio |
| Bora_ex.vivo_noRT | Slc10a1 | 2.808 | 1.97E-04 | glio |
| Bora_ex.vivo_noRT | 1700034I23Ri | 2.404 | 1.97E-04 | glio |
| Chek1_in.vivo_noRT | Gh | 2.988 | 1.97E-04 | glio |
| Eif2s1_in.vivo_noRT | Gm41307 | 2.143 | 1.97E-04 | glio |
| Eif2s1_in.vivo_noRT | Gm13713 | 2.384 | 1.97E-04 | glio |
| culture_effect | Gars | -0.584 | 1.97E-05 | glio |
| Interaction_genotype_Bora | Serping1 | 4.758 | 1.97E-05 | glio |
| Interaction_genotype_Bora | 1810019D21R | -5.427 | 1.97E-05 | glio |
| Interaction_genotype_Myc | Car14 | -4.722 | 1.97E-05 | glio |
| Aars2_in.vivo_noRT | Gm32816 | 3.267 | 1.97E-05 | glio |
| Aars2_in.vivo_noRT | Setmar | 3.011 | 1.97E-05 | glio |
| Atp5c1_in.vivo_noRT | Gm13348 | 2.668 | 1.97E-05 | glio |
| Atp5c1_in.vivo_noRT | Aqp9 | 2.668 | 1.97E-05 | glio |
| Atr_in.vivo_noRT | Fcgrt | 3.286 | 1.97E-05 | glio |
| Atr_in.vivo_noRT | Cox7a1 | 3.215 | 1.97E-05 | glio |
| Atr_in.vivo_noRT | Pmf1 | -1.757 | 1.97E-05 | glio |
| Atr_in.vivo_noRT | Gar1 | -1.412 | 1.97E-05 | glio |
| Atr_ex.vivo_noRT | Ly6c2 | 3.962 | 1.97E-05 | glio |
| Atr_ex.vivo_noRT | Gm35019 | 3.484 | 1.97E-05 | glio |
| Fanca_ex.vivo_noRT | Gm15908 | 2.919 | 1.97E-05 | glio |
| culture_effect | Calu | -0.548 | 1.97E-06 | glio |
| Aars2_in.vivo_noRT | Gm13056 | 3.572 | 1.97E-06 | glio |
| Aars2_in.vivo_noRT | Tes3-ps | 3.292 | 1.97E-06 | glio |
| Atr_in.vivo_noRT | Ccdc78 | 3.177 | 1.97E-06 | glio |
| Bora_in.vivo_noRT | Ucma | 3.938 | 1.97E-06 | glio |

| | | | | |
|---|---|---|---|---|
| Interaction_genotype_Myc | Gm30191 | -5.766 | 1.97E-07 | glio |
| Bora_in.vivo_noRT | Cnnm1 | 4.025 | 1.97E-07 | glio |
| Bora_in.vivo_noRT | Gm43581 | 3.366 | 1.97E-07 | glio |
| Bora_in.vivo_noRT | Armc1 | -2.091 | 1.97E-08 | glio |
| Chek1_ex.vivo_noRT | Gm13032 | 4.443 | 1.97E-10 | glio |
| Bcas2_in.vivo_noRT | Nos2 | 4.547 | 1.97E-13 | glio |
| Bora_in.vivo_noRT | BC147527 | 4.936 | 1.97E-14 | glio |
| Bora_in.vivo_noRT | Dapk2 | 4.153 | 1.97E-15 | glio |
| Aars2_in.vivo_noRT | Klhdc8a | 5.863 | 1.97E-17 | glio |
| Chek1_in.vivo_noRT | Scel | 4.126 | 1.97E-18 | glio |
| Chek1_in.vivo_noRT | Spink6 | 5.044 | 1.97E-19 | glio |
| culture_effect | Egr1 | -3.147 | 1.97E-25 | glio |
| Atr_in.vivo_noRT | Slc17a6 | 5.533 | 1.97E-34 | glio |
| culture_effect | Ocrl | 0.944 | 1.98E-03 | glio |
| culture_effect | Borcs5 | 0.838 | 1.98E-03 | glio |
| culture_effect | Ptpn9 | 0.66 | 1.98E-03 | glio |
| Interaction_genotype_Aars2 | Ksr1 | -3.768 | 1.98E-03 | glio |
| Interaction_genotype_Bcas2 | Gm10354 | -3.36 | 1.98E-03 | glio |
| Interaction_genotype_Mtor | Ccl20 | -5.602 | 1.98E-03 | glio |
| Interaction_genotype_Mtor | Gm10419 | -4.147 | 1.98E-03 | glio |
| Interaction_genotype_Mtor | D830035M03I | -3.586 | 1.98E-03 | glio |
| Interaction_genotype_Myc | Kif5c | -2.748 | 1.98E-03 | glio |
| Aars2_in.vivo_noRT | Gng7 | 2.138 | 1.98E-03 | glio |
| Aars2_ex.vivo_noRT | 2410141K09R | 2.509 | 1.98E-03 | glio |
| Atr_in.vivo_noRT | Gm36988 | 2.214 | 1.98E-03 | glio |
| Atr_in.vivo_noRT | E230016M11F | 2.02 | 1.98E-03 | glio |
| Atr_in.vivo_noRT | Slc9a3r2 | 1.908 | 1.98E-03 | glio |
| Atr_in.vivo_noRT | D330050I16Ri | 1.804 | 1.98E-03 | glio |
| Bard1_ex.vivo_noRT | Vgll2 | 2.814 | 1.98E-03 | glio |
| Bard1_ex.vivo_noRT | Serpina3g | 2.814 | 1.98E-03 | glio |
| Bcas2_in.vivo_noRT | Gm14399 | 2.29 | 1.98E-03 | glio |
| Bcas2_in.vivo_noRT | Mfsd4b4 | 2.082 | 1.98E-03 | glio |
| Bcas2_in.vivo_noRT | Trim6 | 2.354 | 1.98E-03 | glio |
| Bcas2_in.vivo_noRT | Epn2 | -1.178 | 1.98E-03 | glio |
| Bcas2_in.vivo_noRT | Topors | -0.933 | 1.98E-03 | glio |
| Bora_in.vivo_noRT | Chdh | 2.181 | 1.98E-03 | glio |
| Bora_in.vivo_noRT | Slbp | -1.037 | 1.98E-03 | glio |
| Cenpj_in.vivo_noRT | Gm35082 | 2.022 | 1.98E-03 | glio |
| Cenpj_in.vivo_noRT | 9130230L23Ri | 2.292 | 1.98E-03 | glio |
| Cenpj_in.vivo_noRT | Tagap | 1.762 | 1.98E-03 | glio |
| Cenpj_in.vivo_noRT | Pcdhga1 | 2.269 | 1.98E-03 | glio |
| Cenpj_in.vivo_noRT | Gm5286 | 2.022 | 1.98E-03 | glio |
| Cenpj_in.vivo_noRT | Gm34776 | 2.022 | 1.98E-03 | glio |
| Chek1_ex.vivo_noRT | BC003965 | -1.278 | 1.98E-03 | glio |
| Hsd17b10_in.vivo_noRT | Tmem179 | 2.458 | 1.98E-03 | glio |
| Hsd17b10_in.vivo_noRT | Bcl7b | -1.486 | 1.98E-03 | glio |
| culture_effect | Rgl1 | 1.613 | 1.98E-04 | glio |
| Interaction_genotype_Mtor | Slc36a4 | -4.329 | 1.98E-04 | glio |
| Interaction_genotype_Mtor | Taf1 | 2.008 | 1.98E-04 | glio |
| Interaction_genotype_Mtor | Slc25a5 | 1.912 | 1.98E-04 | glio |
| Interaction_genotype_Myc | Spink10 | -3.888 | 1.98E-04 | glio |
| Interaction_genotype_Myc | BC037704 | -3.918 | 1.98E-04 | glio |

| | | | | |
|---|---|---|---|---|
| Interaction_genotype_Rad51 | Klf5 | 3.24 | 1.98E-04 | glio |
| Interaction_genotype_Rad51 | Rcan2 | -5.252 | 1.98E-04 | glio |
| Interaction_genotype_Rad51 | Hbq1b | -4.981 | 1.98E-04 | glio |
| Interaction_genotype_Rad51 | Fa2h | 4.795 | 1.98E-04 | glio |
| Aars2_in.vivo_noRT | Nkain4 | 2.462 | 1.98E-04 | glio |
| Aars2_in.vivo_noRT | Ola1 | -1.132 | 1.98E-04 | glio |
| Aars2_in.vivo_noRT | Rpl5 | -0.731 | 1.98E-04 | glio |
| Atr_in.vivo_noRT | Gh | 2.948 | 1.98E-04 | glio |
| Atr_in.vivo_noRT | Gm41609 | 2.847 | 1.98E-04 | glio |
| Atr_in.vivo_noRT | Slc4a11 | 2.746 | 1.98E-04 | glio |
| Atr_in.vivo_noRT | Slc22a22 | 2.576 | 1.98E-04 | glio |
| Atr_in.vivo_noRT | Nova2 | 2.515 | 1.98E-04 | glio |
| Atr_in.vivo_noRT | A530041M06R | 2.664 | 1.98E-04 | glio |
| Atr_in.vivo_noRT | Gm43813 | 2.463 | 1.98E-04 | glio |
| Atr_in.vivo_noRT | Gm48427 | 2.403 | 1.98E-04 | glio |
| Bcas2_in.vivo_noRT | 5830428M24R | 2.683 | 1.98E-04 | glio |
| Bora_ex.vivo_noRT | Dio1 | 3.201 | 1.98E-04 | glio |
| Chek1_in.vivo_noRT | Gzmm | 2.715 | 1.98E-04 | glio |
| culture_effect | Dram1 | -1.814 | 1.98E-05 | glio |
| Aars2_in.vivo_noRT | Cldn9 | 2.881 | 1.98E-05 | glio |
| Atr_in.vivo_noRT | Gm17501 | 3.11 | 1.98E-05 | glio |
| Bard1_ex.vivo_noRT | Gm14929 | 2.814 | 1.98E-05 | glio |
| Bcas2_in.vivo_noRT | Dyrk3 | -2.325 | 1.98E-05 | glio |
| Bora_in.vivo_noRT | Klhdc8b | 3.122 | 1.98E-05 | glio |
| Bora_in.vivo_noRT | Fam71f2 | 3.115 | 1.98E-05 | glio |
| interaction_RT | N6amt1 | -1.398 | 1.98E-06 | glio |
| interaction_RT | Acsl3 | 1.347 | 1.98E-06 | glio |
| interaction_RT | Cuta | 0.835 | 1.98E-06 | glio |
| Aars2_in.vivo_noRT | Phf11a | 3.234 | 1.98E-06 | glio |
| Atr_in.vivo_noRT | Myl7 | 3.405 | 1.98E-06 | glio |
| Bora_in.vivo_noRT | Gm35551 | 3.12 | 1.98E-06 | glio |
| culture_effect | Lrrc42 | -1.019 | 1.98E-07 | glio |
| culture_effect | Rex1bd | 0.618 | 1.98E-07 | glio |
| Interaction_genotype_Myc | Gm49312 | -5.282 | 1.98E-07 | glio |
| Aars2_ex.vivo_noRT | Meioc | 3.293 | 1.98E-07 | glio |
| Aars2_ex.vivo_noRT | Tmem132c | 3.293 | 1.98E-07 | glio |
| Aars2_ex.vivo_noRT | Gm44693 | 3.293 | 1.98E-07 | glio |
| Aars2_ex.vivo_noRT | Gm47622 | 3.517 | 1.98E-07 | glio |
| Atr_in.vivo_noRT | Gm43066 | 3.391 | 1.98E-07 | glio |
| Atr_in.vivo_noRT | Selenok | -1.544 | 1.98E-07 | glio |
| Bcas2_in.vivo_noRT | Ggt1 | 3.494 | 1.98E-07 | glio |
| Bora_in.vivo_noRT | AV356131 | 3.653 | 1.98E-07 | glio |
| Hsd17b10_in.vivo_noRT | Gm50138 | 3.093 | 1.98E-07 | glio |
| Hsd17b10_in.vivo_noRT | Clcn1 | 2.97 | 1.98E-07 | glio |
| culture_effect | Pou5f1 | -3.326 | 1.98E-08 | glio |
| Interaction_genotype_Mtor | Esrp2 | -6.555 | 1.98E-08 | glio |
| Aars2_in.vivo_noRT | Cabp4 | 4.016 | 1.98E-08 | glio |
| Aars2_in.vivo_noRT | 1700122E12Ri | 3.931 | 1.98E-08 | glio |
| Atr_ex.vivo_noRT | C430002N11R | 3.913 | 1.98E-08 | glio |
| Atr_ex.vivo_noRT | Gm6803 | 3.947 | 1.98E-08 | glio |
| Bora_in.vivo_noRT | 9430037G07R | 4.028 | 1.98E-08 | glio |
| Bora_in.vivo_noRT | Gm36043 | 3.48 | 1.98E-08 | glio |

| | | | | |
|---|---|---|---|---|
| Chek1_in.vivo_noRT | Sema3a | 3.606 | 1.98E-08 | glio |
| Aars2_in.vivo_noRT | Spata31d1a | 4.685 | 1.98E-11 | glio |
| Atr_in.vivo_noRT | Cpeb1 | 4.888 | 1.98E-11 | glio |
| Bcas2_in.vivo_noRT | Gm6566 | 3.882 | 1.98E-11 | glio |
| Chek1_ex.vivo_noRT | C330002G04R | 4.256 | 1.98E-11 | glio |
| Aars2_in.vivo_noRT | A330015K06R | 4.67 | 1.98E-13 | glio |
| Chek1_in.vivo_noRT | Gm42664 | 4.393 | 1.98E-13 | glio |
| Chek1_in.vivo_noRT | Gm35696 | 4.784 | 1.98E-13 | glio |
| Chek1_in.vivo_noRT | Zfp810 | 5.573 | 1.98E-16 | glio |
| Atr_in.vivo_noRT | Abra | 5.008 | 1.98E-17 | glio |
| Chek1_ex.vivo_noRT | Scg2 | 4.842 | 1.98E-17 | glio |
| Atr_in.vivo_noRT | Gm21319 | 5.533 | 1.98E-23 | glio |
| culture_effect | Flot1 | -0.59 | 1.99E-03 | glio |
| culture_effect | Uri1 | 0.507 | 1.99E-03 | glio |
| Interaction_genotype_Aars2 | BC030343 | -3.845 | 1.99E-03 | glio |
| Interaction_genotype_Mtor | 4930412C18R | -3.499 | 1.99E-03 | glio |
| Interaction_genotype_Mtor | 3830403N18R | 1.504 | 1.99E-03 | glio |
| Interaction_genotype_Myc | 1700019E08Ri | -2.729 | 1.99E-03 | glio |
| Interaction_genotype_Myc | 5830428M24F | -3.458 | 1.99E-03 | glio |
| Interaction_genotype_Myc | Gm16364.1 | -3.575 | 1.99E-03 | glio |
| Interaction_genotype_Myc | 4930522L14Ri | -3.468 | 1.99E-03 | glio |
| Interaction_genotype_Myc | Setdb2 | -2.934 | 1.99E-03 | glio |
| Interaction_genotype_Myc | Wdr41 | -2.842 | 1.99E-03 | glio |
| Interaction_genotype_Myc | Adat2 | -2.841 | 1.99E-03 | glio |
| Interaction_genotype_Myc | Urgcp | -2.647 | 1.99E-03 | glio |
| interaction_RT | Sf3b6 | 0.513 | 1.99E-03 | glio |
| Aars2_in.vivo_noRT | Rasip1 | 2.297 | 1.99E-03 | glio |
| Aars2_in.vivo_noRT | Fastkd1 | 2.007 | 1.99E-03 | glio |
| Aars2_in.vivo_noRT | Tmem11 | -1.031 | 1.99E-03 | glio |
| Atr_in.vivo_noRT | Ubox5 | 1.834 | 1.99E-03 | glio |
| Atr_in.vivo_noRT | Sh2b3 | 2.111 | 1.99E-03 | glio |
| Atr_in.vivo_noRT | Fbxo32 | -1.775 | 1.99E-03 | glio |
| Atr_ex.vivo_noRT | Flnb | -1.553 | 1.99E-03 | glio |
| Bcas2_in.vivo_noRT | Mpz | 2.836 | 1.99E-03 | glio |
| Bcas2_in.vivo_noRT | BC025920 | 2.03 | 1.99E-03 | glio |
| Bora_in.vivo_noRT | Cdh24 | 2.459 | 1.99E-03 | glio |
| Bora_in.vivo_noRT | Ppef1 | 2.132 | 1.99E-03 | glio |
| Chek1_in.vivo_noRT | Ebi3 | 2.666 | 1.99E-03 | glio |
| Chek1_in.vivo_noRT | Lrp1b | 2.265 | 1.99E-03 | glio |
| Chek1_in.vivo_noRT | Izumo4 | 2.256 | 1.99E-03 | glio |
| Chek1_in.vivo_noRT | Tceal1 | 2.148 | 1.99E-03 | glio |
| Chek1_in.vivo_noRT | Hdhd5 | 2.021 | 1.99E-03 | glio |
| Chek1_in.vivo_noRT | Zfp113 | 1.907 | 1.99E-03 | glio |
| Chek1_in.vivo_noRT | Fam53b | 1.914 | 1.99E-03 | glio |
| Chek1_in.vivo_noRT | Pak4 | -1.598 | 1.99E-03 | glio |
| Chek1_in.vivo_noRT | Wdr43 | -1.065 | 1.99E-03 | glio |
| Chek1_ex.vivo_noRT | Gm46378 | 2.995 | 1.99E-03 | glio |
| Eif2s1_in.vivo_noRT | Prickle2 | 2.143 | 1.99E-03 | glio |
| Interaction_genotype_Mtor | Fxyd1 | -3.784 | 1.99E-04 | glio |
| Interaction_genotype_Myc | Zfp467 | -4.267 | 1.99E-04 | glio |
| Interaction_genotype_Myc | 2210417A02R | -4.386 | 1.99E-04 | glio |
| interaction_RT | 4930589L23Ri | -2.916 | 1.99E-04 | glio |

| | | | | |
|---|---|---|---|---|
| Aars2_in.vivo_noRT | Mt4 | 3.331 | 1.99E-04 | glio |
| Aars2_in.vivo_noRT | Trim65 | 2.506 | 1.99E-04 | glio |
| Aars2_in.vivo_noRT | Myc | -1.688 | 1.99E-04 | glio |
| Aars2_in.vivo_noRT | Serf2 | -0.753 | 1.99E-04 | glio |
| Aars2_ex.vivo_noRT | Gm43915 | 2.899 | 1.99E-04 | glio |
| Atp5c1_in.vivo_noRT | Hist1h2ag | 2.136 | 1.99E-04 | glio |
| Atp5c1_in.vivo_noRT | Gstp3 | 2.832 | 1.99E-04 | glio |
| Atr_ex.vivo_noRT | 2810049E08Ri | 3.164 | 1.99E-04 | glio |
| Bora_in.vivo_noRT | Paqr8 | 3.04 | 1.99E-04 | glio |
| Bora_in.vivo_noRT | Mettl6 | -1.834 | 1.99E-04 | glio |
| Chek1_in.vivo_noRT | Zfp963 | 2.966 | 1.99E-04 | glio |
| Interaction_genotype_Atr | Adamts12 | -4.094 | 1.99E-05 | glio |
| Interaction_genotype_Atr | Igf2 | -3.648 | 1.99E-05 | glio |
| Interaction_genotype_Myc | Slc4a9 | -4.083 | 1.99E-05 | glio |
| Aars2_in.vivo_noRT | Gm15063 | 3.343 | 1.99E-05 | glio |
| Aars2_in.vivo_noRT | B230398E01Ri | 2.993 | 1.99E-05 | glio |
| Aars2_ex.vivo_noRT | Tmem54 | 3.293 | 1.99E-05 | glio |
| Chek1_in.vivo_noRT | Acvr1b | 2.987 | 1.99E-05 | glio |
| Chek1_in.vivo_noRT | Zfp607b | 2.501 | 1.99E-05 | glio |
| Chek1_ex.vivo_noRT | Mkrn3 | 3.595 | 1.99E-05 | glio |
| Eif2s1_in.vivo_noRT | Slc17a6 | 2.143 | 1.99E-05 | glio |
| Hsd17b10_in.vivo_noRT | Gm9929 | 2.837 | 1.99E-05 | glio |
| Hsd17b10_in.vivo_noRT | 9130204L05Ri | 3.313 | 1.99E-05 | glio |
| Bora_in.vivo_noRT | Rnase4 | 3.374 | 1.99E-06 | glio |
| Bora_in.vivo_noRT | Gm50255 | 4.029 | 1.99E-06 | glio |
| Bora_in.vivo_noRT | Cdk2 | -2.457 | 1.99E-06 | glio |
| Hsd17b10_in.vivo_noRT | Agrp | 2.834 | 1.99E-06 | glio |
| Interaction_genotype_Mtor | Gm18280 | -4.629 | 1.99E-07 | glio |
| Bcas2_in.vivo_noRT | Inha | 3.683 | 1.99E-07 | glio |
| Chek1_in.vivo_noRT | Parvg | 3.732 | 1.99E-07 | glio |
| Chek1_ex.vivo_noRT | B4galnt1 | 3.75 | 1.99E-07 | glio |
| Atr_in.vivo_noRT | Gm14327 | 4.028 | 1.99E-08 | glio |
| Bora_in.vivo_noRT | Gm1123 | 4.029 | 1.99E-08 | glio |
| Chek1_in.vivo_noRT | Olfr1189 | 3.389 | 1.99E-08 | glio |
| Hsd17b10_in.vivo_noRT | Iqch | 3.352 | 1.99E-08 | glio |
| Hsd17b10_in.vivo_noRT | Nat8f4 | 3.611 | 1.99E-08 | glio |
| Hsd17b10_in.vivo_noRT | 2010001K21R | 3.616 | 1.99E-08 | glio |
| Hsd17b10_in.vivo_noRT | Gm27241 | 2.96 | 1.99E-08 | glio |
| Atr_in.vivo_noRT | Slc26a1 | 4.03 | 1.99E-09 | glio |
| Bora_in.vivo_noRT | Mro | 4.145 | 1.99E-09 | glio |
| Chek1_ex.vivo_noRT | Kmo | 4.449 | 1.99E-09 | glio |
| Aars2_in.vivo_noRT | Alox12 | 4.538 | 1.99E-10 | glio |
| Aars2_in.vivo_noRT | Klk14 | 4.795 | 1.99E-11 | glio |
| Atr_in.vivo_noRT | Gm3716 | 4.615 | 1.99E-11 | glio |
| Atr_in.vivo_noRT | Gm30082 | 4.883 | 1.99E-12 | glio |
| Chek1_in.vivo_noRT | Klhdc8a | 5.044 | 1.99E-13 | glio |
| culture_effect | Hagh | -1.117 | 1.99E-14 | glio |
| Atr_in.vivo_noRT | Pla1a | 5.002 | 1.99E-14 | glio |
| Bcas2_in.vivo_noRT | Gm50375 | 5.699 | 1.99E-14 | glio |
| Bora_in.vivo_noRT | Gm35696 | 5.2 | 1.99E-15 | glio |
| Bcas2_in.vivo_noRT | Gm29670 | 4.285 | 1.99E-16 | glio |
| Bcas2_in.vivo_noRT | Gm11583 | 4.281 | 1.99E-16 | glio |

| | | | | |
|---|---|---|---|---|
| Aars2_in.vivo_noRT | Snrpn | 4.668 | 1.99E-17 | glio |
| Aars2_in.vivo_noRT | Obscn | 5.327 | 1.99E-17 | glio |
| culture_effect | Layn | 3.019 | 1.99E-18 | glio |
| culture_effect | Bcas3 | 1.848 | 1.99E-18 | glio |
| Aars2_in.vivo_noRT | AC159140.1 | 5.067 | 1.99E-21 | glio |
| culture_effect | Krt18 | -7.921 | 1.99E-24 | glio |
| Bcas2_in.vivo_noRT | Scg2 | 4.807 | 1.99E-24 | glio |
| culture_effect | Cpeb1 | 1.882 | 2.00E-03 | glio |
| culture_effect | Rbm8a2 | -1.72 | 2.00E-03 | glio |
| culture_effect | Fbxw4 | 0.791 | 2.00E-03 | glio |
| culture_effect | Cpsf3 | -0.594 | 2.00E-03 | glio |
| culture_effect | Iws1 | -0.538 | 2.00E-03 | glio |
| culture_effect | Atp6ap2 | -0.567 | 2.00E-03 | glio |
| Interaction_genotype_Aars2 | Cdkn2a | 0.93 | 2.00E-03 | glio |
| Interaction_genotype_Atr | Gm50069 | -3.863 | 2.00E-03 | glio |
| Interaction_genotype_Atr | Tmx2 | 1.545 | 2.00E-03 | glio |
| Interaction_genotype_Bcas2 | 1700034I23Ri | -2.862 | 2.00E-03 | glio |
| Interaction_genotype_Cdkn2a | Sucnr1 | 2.835 | 2.00E-03 | glio |
| Interaction_genotype_Cdkn2a | Gpc5 | 3.22 | 2.00E-03 | glio |
| Interaction_genotype_Mtor | Nphs1 | -3.563 | 2.00E-03 | glio |
| Interaction_genotype_Mtor | Map3k20 | -3.074 | 2.00E-03 | glio |
| Interaction_genotype_Myc | Gm19744 | -3.579 | 2.00E-03 | glio |
| Interaction_genotype_Myc | Klf1 | -3.618 | 2.00E-03 | glio |
| Interaction_genotype_Pars2 | Gm34047 | -4.172 | 2.00E-03 | glio |
| Interaction_genotype_Rad51 | Gm11457 | -4.152 | 2.00E-03 | glio |
| interaction_RT | Adam12 | -2.567 | 2.00E-03 | glio |
| interaction_RT | Ift27 | 0.943 | 2.00E-03 | glio |
| interaction_RT | Lage3 | -0.762 | 2.00E-03 | glio |
| Atp5c1_in.vivo_noRT | Gm3734 | 2.673 | 2.00E-03 | glio |
| Atr_in.vivo_noRT | Mmp11 | 2.27 | 2.00E-03 | glio |
| Atr_in.vivo_noRT | BC024063 | 2.13 | 2.00E-03 | glio |
| Atr_in.vivo_noRT | Sox6 | -1.248 | 2.00E-03 | glio |
| Atr_in.vivo_noRT | Cpq | -1.088 | 2.00E-03 | glio |
| Atr_in.vivo_noRT | Atad2 | -1.405 | 2.00E-03 | glio |
| Atr_ex.vivo_noRT | Nell2 | 2.783 | 2.00E-03 | glio |
| Bcas2_in.vivo_noRT | Mfsd3 | 1.895 | 2.00E-03 | glio |
| Bcas2_in.vivo_noRT | Tmcc1 | -1.412 | 2.00E-03 | glio |
| Bcas2_in.vivo_noRT | Idh3a | -1.177 | 2.00E-03 | glio |
| Cenpj_in.vivo_noRT | Zfp810 | 2.548 | 2.00E-03 | glio |
| Cenpj_in.vivo_noRT | Gm29966 | 1.762 | 2.00E-03 | glio |
| Cenpj_in.vivo_noRT | Htr1b | 2.548 | 2.00E-03 | glio |
| Chek1_in.vivo_noRT | 1700028J19Ri | 2.246 | 2.00E-03 | glio |
| Chek1_in.vivo_noRT | Rps18 | -0.695 | 2.00E-03 | glio |
| Eif2s1_in.vivo_noRT | Gm15890 | 2.143 | 2.00E-03 | glio |
| Interaction_genotype_Mtor | Gm13054 | -3.968 | 2.00E-04 | glio |
| Interaction_genotype_Mtor | Gm9887 | -3.943 | 2.00E-04 | glio |
| Interaction_genotype_Myc | Gm15520 | -3.76 | 2.00E-04 | glio |
| Interaction_genotype_Myc | Gm32913 | -6.242 | 2.00E-04 | glio |
| Interaction_genotype_Myc | Gm45716 | -3.732 | 2.00E-04 | glio |
| Interaction_genotype_Myc | Slc25a3 | 1.376 | 2.00E-04 | glio |
| Interaction_genotype_Myc | Gapdh | 1.961 | 2.00E-04 | glio |
| Interaction_genotype_Myc | Rplp2 | 1.315 | 2.00E-04 | glio |

| | | | | |
|---|---|---|---|---|
| Atr_in.vivo_noRT | Zik1 | 2.215 | 2.00E-04 | glio |
| Atr_in.vivo_noRT | Tex261 | -1.241 | 2.00E-04 | glio |
| Atr_in.vivo_noRT | Pbrm1 | -1.381 | 2.00E-04 | glio |
| Bard1_ex.vivo_noRT | Olfr171 | 2.814 | 2.00E-04 | glio |
| Bcas2_in.vivo_noRT | 9330179D12R | 2.877 | 2.00E-04 | glio |
| Bcas2_in.vivo_noRT | Asap3 | 2.594 | 2.00E-04 | glio |
| Bora_in.vivo_noRT | Ndufaf6 | -1.355 | 2.00E-04 | glio |
| Bora_ex.vivo_noRT | Myt1 | 2.802 | 2.00E-04 | glio |
| Cenpj_ex.vivo_noRT | Tpo | 2.591 | 2.00E-04 | glio |
| Chek1_in.vivo_noRT | Ccdc9b | 2.403 | 2.00E-04 | glio |
| Eif2s1_in.vivo_noRT | Gm13398 | 2.143 | 2.00E-04 | glio |
| culture_effect | Enho | 2.008 | 2.00E-05 | glio |
| Atr_in.vivo_noRT | Fam161b | 2.986 | 2.00E-05 | glio |
| Atr_in.vivo_noRT | 6330403L08Ri | 2.872 | 2.00E-05 | glio |
| Chek1_in.vivo_noRT | Gm44190 | 2.767 | 2.00E-05 | glio |
| Chek1_ex.vivo_noRT | Gm14379 | 3.501 | 2.00E-05 | glio |
| Chek1_ex.vivo_noRT | Esco2 | -2.953 | 2.00E-05 | glio |
| Cyp19a1_ex.vivo_noRT | Scg2 | 2.959 | 2.00E-05 | glio |
| culture_effect | Chrna1os | 1.827 | 2.00E-06 | glio |
| Interaction_genotype_Mtor | Gtf2h5 | 2.051 | 2.00E-06 | glio |
| Interaction_genotype_Mtor | AI182371 | -4.69 | 2.00E-06 | glio |
| Interaction_genotype_Mtor | Gm20646 | -5.472 | 2.00E-06 | glio |
| Aars2_ex.vivo_noRT | Gm13449 | 3.305 | 2.00E-06 | glio |
| Bora_in.vivo_noRT | Slc1a3 | -2.551 | 2.00E-06 | glio |
| Bora_ex.vivo_noRT | Spink8 | 3.201 | 2.00E-06 | glio |
| Bora_ex.vivo_noRT | Psmb11 | 3.201 | 2.00E-06 | glio |
| Chek1_in.vivo_noRT | Prelid3a | 3.339 | 2.00E-06 | glio |
| Atr_in.vivo_noRT | Icosl | 3.243 | 2.00E-07 | glio |
| Bora_in.vivo_noRT | Bpi | 3.255 | 2.00E-07 | glio |
| Atr_in.vivo_noRT | Rnf208 | 4.141 | 2.00E-08 | glio |
| Bora_in.vivo_noRT | Insm1 | 3.818 | 2.00E-08 | glio |
| Chek1_ex.vivo_noRT | Tubb4a | 5.482 | 2.00E-08 | glio |
| Chek1_ex.vivo_noRT | Pdlim3 | 4.049 | 2.00E-08 | glio |
| Chek1_ex.vivo_noRT | Pkd1l2 | 4.447 | 2.00E-08 | glio |
| culture_effect | 9630013A20R | -3.394 | 2.00E-09 | glio |
| culture_effect | Pgm3 | -1.4 | 2.00E-09 | glio |
| Aars2_in.vivo_noRT | Inha | 4.203 | 2.00E-09 | glio |
| Atr_in.vivo_noRT | Adam1a | 4.172 | 2.00E-09 | glio |
| Chek1_in.vivo_noRT | Ppp1r3g | 3.859 | 2.00E-09 | glio |
| Atr_ex.vivo_noRT | Luzp2 | 4.188 | 2.00E-10 | glio |
| Bora_in.vivo_noRT | Rbms1 | -2.44 | 2.00E-10 | glio |
| Chek1_ex.vivo_noRT | Neurl3 | 5.623 | 2.00E-10 | glio |
| Chek1_in.vivo_noRT | Gm43672 | 5.044 | 2.00E-12 | glio |
| Hsd17b10_in.vivo_noRT | Gm30794 | 3.616 | 2.00E-12 | glio |
| Aars2_in.vivo_noRT | Arhgef16 | 4.29 | 2.00E-13 | glio |
| Chek1_in.vivo_noRT | Gm37800 | 5.044 | 2.00E-15 | glio |
| Chek1_ex.vivo_noRT | Gm12703 | 4.842 | 2.00E-15 | glio |
| Chek1_ex.vivo_noRT | Mmp13 | 4.842 | 2.00E-15 | glio |
| culture_effect | Eif3k | -0.946 | 2.00E-17 | glio |
| culture_effect | Zdhhc20 | -1.588 | 2.00E-18 | glio |
| Chek1_in.vivo_noRT | Gm21761 | 5.044 | 2.00E-20 | glio |
| Aars2_in.vivo_noRT | Gm21761 | 5.327 | 2.00E-22 | glio |

| | | | | |
|---|---|---|---|---|
| Bora_in.vivo_noRT | Tpo | 4.936 | 2.00E-23 | glio |
| Atr_in.vivo_noRT | Cd7 | 5.269 | 2.00E-25 | glio |
| Atr_in.vivo_noRT | Gm2016 | 5.533 | 2.00E-25 | glio |
| Atr_in.vivo_noRT | Slc35g3 | 5.533 | 2.00E-27 | glio |
| culture_effect | Pcdhb14 | -1.994 | 2.01E-03 | glio |
| culture_effect | Bbs2 | -1.472 | 2.01E-03 | glio |
| Interaction_genotype_Atr | Gm20633 | -3.614 | 2.01E-03 | glio |
| Interaction_genotype_Atr | Gm49694 | -3.789 | 2.01E-03 | glio |
| Interaction_genotype_Mtor | 1700006J14Ri | -3.448 | 2.01E-03 | glio |
| Interaction_genotype_Mtor | Tmem117 | -3.748 | 2.01E-03 | glio |
| Interaction_genotype_Mtor | Zc3h6 | -3.442 | 2.01E-03 | glio |
| Interaction_genotype_Mtor | Pmm2 | -2.34 | 2.01E-03 | glio |
| Interaction_genotype_Mtor | S1pr2 | 1.801 | 2.01E-03 | glio |
| Interaction_genotype_Myc | Zfp524 | -2.444 | 2.01E-03 | glio |
| Interaction_genotype_Myc | Cdk14 | -2.216 | 2.01E-03 | glio |
| Interaction_genotype_Rad21 | Fam189a1 | 3.891 | 2.01E-03 | glio |
| Aars2_in.vivo_noRT | Rab27b | 2.385 | 2.01E-03 | glio |
| Aars2_in.vivo_noRT | BC025920 | 1.996 | 2.01E-03 | glio |
| Aars2_in.vivo_noRT | Lpin2 | 1.729 | 2.01E-03 | glio |
| Aars2_in.vivo_noRT | Mcfd2 | -1.329 | 2.01E-03 | glio |
| Aars2_in.vivo_noRT | Mdh1 | -0.859 | 2.01E-03 | glio |
| Aars2_in.vivo_noRT | mt-Nd4l | 1.161 | 2.01E-03 | glio |
| Aars2_ex.vivo_noRT | Mog | 3.293 | 2.01E-03 | glio |
| Atr_in.vivo_noRT | Hao1 | 2.325 | 2.01E-03 | glio |
| Atr_in.vivo_noRT | Oplah | 2.067 | 2.01E-03 | glio |
| Atr_in.vivo_noRT | Bcat1 | -1.563 | 2.01E-03 | glio |
| Atr_ex.vivo_noRT | Mycl | 2.911 | 2.01E-03 | glio |
| Bcas2_in.vivo_noRT | Ccdc159 | 2.384 | 2.01E-03 | glio |
| Bora_in.vivo_noRT | Sptlc2 | -1.505 | 2.01E-03 | glio |
| Bora_ex.vivo_noRT | Prm1 | 2.802 | 2.01E-03 | glio |
| Chek1_in.vivo_noRT | Dhx29 | 1.341 | 2.01E-03 | glio |
| Hsd17b10_in.vivo_noRT | Shroom3 | 2.337 | 2.01E-03 | glio |
| culture_effect | Fip1l1 | 0.51 | 2.01E-04 | glio |
| Interaction_genotype_Mtor | Amh | -4.154 | 2.01E-04 | glio |
| Interaction_genotype_Mtor | Ccdc57 | -4.72 | 2.01E-04 | glio |
| Interaction_genotype_Mtor | Carmil1 | -4.363 | 2.01E-04 | glio |
| Interaction_genotype_Mtor | A430033K04R | -4.241 | 2.01E-04 | glio |
| Interaction_genotype_Mtor | Kdm5a | 1.964 | 2.01E-04 | glio |
| Interaction_genotype_Mtor | Tapbp | 2.004 | 2.01E-04 | glio |
| Interaction_genotype_Myc | B4galnt1 | -3.562 | 2.01E-04 | glio |
| Interaction_genotype_Myc | Nxpe3 | -3.631 | 2.01E-04 | glio |
| Atr_in.vivo_noRT | Sh3bp5 | 2.594 | 2.01E-04 | glio |
| Atr_in.vivo_noRT | Rps17 | -0.776 | 2.01E-04 | glio |
| Bcas2_in.vivo_noRT | Gm26981 | 2.633 | 2.01E-04 | glio |
| Hsd17b10_in.vivo_noRT | Igdcc4 | 2.44 | 2.01E-04 | glio |
| Interaction_genotype_Mtor | Gm26766 | -4.738 | 2.01E-05 | glio |
| Interaction_genotype_Mtor | Gm15535 | -5.385 | 2.01E-05 | glio |
| Interaction_genotype_Mtor | Gm47509 | -5.255 | 2.01E-05 | glio |
| Interaction_genotype_Mtor | Gm13270 | -5.165 | 2.01E-05 | glio |
| Interaction_genotype_Mtor | Nat8f2 | -4.234 | 2.01E-05 | glio |
| Interaction_genotype_Myc | Fbxl14 | -3.28 | 2.01E-05 | glio |
| Aars2_in.vivo_noRT | Zbtb17 | 2.117 | 2.01E-05 | glio |

| | | | | |
|---|---|---|---|---|
| Bard1_ex.vivo_noRT | 4930470G03R | 3.203 | 2.01E-05 | glio |
| Bard1_ex.vivo_noRT | Meioc | 2.814 | 2.01E-05 | glio |
| Bora_in.vivo_noRT | Gm31805 | 2.952 | 2.01E-05 | glio |
| Bora_in.vivo_noRT | Tead3 | 2.998 | 2.01E-05 | glio |
| Chek1_ex.vivo_noRT | 1700016P04R | 3.447 | 2.01E-05 | glio |
| culture_effect | Fbn1 | -2.366 | 2.01E-06 | glio |
| culture_effect | Gm20337 | 2.344 | 2.01E-06 | glio |
| culture_effect | Gm10762 | 1.639 | 2.01E-06 | glio |
| Interaction_genotype_Mtor | 2010320O07R | -5.096 | 2.01E-06 | glio |
| Interaction_genotype_Mtor | Gm15533 | -4.099 | 2.01E-06 | glio |
| Atr_in.vivo_noRT | Dennd5a | -1.958 | 2.01E-06 | glio |
| Bcas2_in.vivo_noRT | Dct | 4.004 | 2.01E-06 | glio |
| Bora_in.vivo_noRT | B230319C09R | 3.626 | 2.01E-06 | glio |
| Bora_in.vivo_noRT | Sv2a | 3.499 | 2.01E-06 | glio |
| Aars2_in.vivo_noRT | Scd3 | 3.939 | 2.01E-07 | glio |
| Aars2_in.vivo_noRT | Acvr1b | 3.555 | 2.01E-07 | glio |
| Bcas2_in.vivo_noRT | Sh2d3c | 3.906 | 2.01E-07 | glio |
| Bcas2_in.vivo_noRT | 1700055D18R | 3.945 | 2.01E-07 | glio |
| Bora_in.vivo_noRT | Tmem44 | 3.678 | 2.01E-07 | glio |
| Hsd17b10_in.vivo_noRT | Ppil6 | 3.616 | 2.01E-08 | glio |
| Aars2_in.vivo_noRT | Ctsf | 3.658 | 2.01E-11 | glio |
| Aars2_in.vivo_noRT | Fabp4 | 4.798 | 2.01E-11 | glio |
| Aars2_in.vivo_noRT | Gm39459 | 4.676 | 2.01E-11 | glio |
| Chek1_in.vivo_noRT | Cysrt1 | 4.784 | 2.01E-11 | glio |
| Chek1_in.vivo_noRT | Rhbg | 3.448 | 2.01E-11 | glio |
| Chek1_ex.vivo_noRT | Gm11884 | 5.564 | 2.01E-12 | glio |
| Atr_in.vivo_noRT | Gm16364.1 | 5.533 | 2.01E-13 | glio |
| Aars2_in.vivo_noRT | Ly6d | 4.599 | 2.01E-16 | glio |
| culture_effect | Cavin2 | 4.108 | 2.01E-20 | glio |
| Atr_in.vivo_noRT | Gm34059 | 5.533 | 2.01E-30 | glio |
| culture_effect | Col6a2 | 6.381 | 2.01E-40 | glio |
| culture_effect | Gm48265 | -1.717 | 2.02E-03 | glio |
| culture_effect | Iqcg | -1.772 | 2.02E-03 | glio |
| culture_effect | Gm6133 | -0.937 | 2.02E-03 | glio |
| culture_effect | Slc35a4 | 0.538 | 2.02E-03 | glio |
| culture_effect | Slc29a1 | -0.609 | 2.02E-03 | glio |
| culture_effect | Atp6v1g1 | 0.336 | 2.02E-03 | glio |
| Interaction_genotype_Aars2 | 5730414N17R | -4.008 | 2.02E-03 | glio |
| Interaction_genotype_Atr | Snhg1 | 1.557 | 2.02E-03 | glio |
| Interaction_genotype_Mtor | Amy1 | -3.565 | 2.02E-03 | glio |
| Interaction_genotype_Mtor | Rfx6 | -3.684 | 2.02E-03 | glio |
| Interaction_genotype_Mtor | Mknk1 | -2.839 | 2.02E-03 | glio |
| Interaction_genotype_Mtor | Lmo7 | -3.015 | 2.02E-03 | glio |
| Interaction_genotype_Myc | Gm47232 | -3.409 | 2.02E-03 | glio |
| Interaction_genotype_Myc | Gm5624 | -3.336 | 2.02E-03 | glio |
| Interaction_genotype_Myc | Abcc10 | -3.446 | 2.02E-03 | glio |
| Interaction_genotype_Myc | Gm26620 | -3.057 | 2.02E-03 | glio |
| Interaction_genotype_Myc | Hddc3 | -3.156 | 2.02E-03 | glio |
| Interaction_genotype_Myc | Sh2b3 | -3.332 | 2.02E-03 | glio |
| Interaction_genotype_Myc | Elf4 | -3.189 | 2.02E-03 | glio |
| Interaction_genotype_Myc | Tnfaip2 | -3.759 | 2.02E-03 | glio |
| Interaction_genotype_Myc | Zfp438 | -2.891 | 2.02E-03 | glio |

| | | | | |
|---|---|---|---|---|
| interaction_RT | Gsk3b | 0.713 | 2.02E-03 | glio |
| Aars2_in.vivo_noRT | Mb | 2.233 | 2.02E-03 | glio |
| Aars2_in.vivo_noRT | Gm12059 | 2.146 | 2.02E-03 | glio |
| Aars2_in.vivo_noRT | Tal1 | 2.208 | 2.02E-03 | glio |
| Aars2_in.vivo_noRT | Tbx15 | 1.947 | 2.02E-03 | glio |
| Aars2_in.vivo_noRT | Armc7 | 1.945 | 2.02E-03 | glio |
| Aars2_in.vivo_noRT | A630089N07R | 1.901 | 2.02E-03 | glio |
| Aars2_in.vivo_noRT | Rpl10 | -0.582 | 2.02E-03 | glio |
| Aars2_ex.vivo_noRT | Gem | 2.9 | 2.02E-03 | glio |
| Aars2_ex.vivo_noRT | Gm15445 | 2.307 | 2.02E-03 | glio |
| Atp5c1_in.vivo_noRT | Llgl2 | 2.367 | 2.02E-03 | glio |
| Bard1_ex.vivo_noRT | Dennd2d | 2.819 | 2.02E-03 | glio |
| Bcas2_in.vivo_noRT | 5930430L01Ri | 2.129 | 2.02E-03 | glio |
| Bcas2_in.vivo_noRT | Adprh | -1.271 | 2.02E-03 | glio |
| Bcas2_ex.vivo_noRT | Gm49504 | 2.342 | 2.02E-03 | glio |
| Chek1_in.vivo_noRT | Eva1c | 2.316 | 2.02E-03 | glio |
| Chek1_in.vivo_noRT | Nsmce2 | -1.017 | 2.02E-03 | glio |
| Chek1_in.vivo_noRT | Dusp6 | -1.297 | 2.02E-03 | glio |
| Hsd17b10_in.vivo_noRT | Slc35f6 | -1.681 | 2.02E-03 | glio |
| culture_effect | Sh3bgrl3 | 0.52 | 2.02E-04 | glio |
| Interaction_genotype_Mtor | Erp27 | -4.56 | 2.02E-04 | glio |
| Interaction_genotype_Mtor | 4930554I06Ri | -4.601 | 2.02E-04 | glio |
| Interaction_genotype_Mtor | Fabp3 | -4.811 | 2.02E-04 | glio |
| Interaction_genotype_Mtor | Gm13056 | -4.071 | 2.02E-04 | glio |
| Interaction_genotype_Myc | Krt19 | -2.729 | 2.02E-04 | glio |
| Interaction_genotype_Myc | Pex26 | -3.754 | 2.02E-04 | glio |
| Interaction_genotype_Myc | Sept6 | -3.329 | 2.02E-04 | glio |
| Interaction_genotype_Myc | 0610009L18Ri | -3.958 | 2.02E-04 | glio |
| interaction_RT | Pttg1 | 1.007 | 2.02E-04 | glio |
| interaction_RT | Tpr | -0.671 | 2.02E-04 | glio |
| Aars2_in.vivo_noRT | Car9 | 2.555 | 2.02E-04 | glio |
| Aars2_in.vivo_noRT | Gpam | 2.442 | 2.02E-04 | glio |
| Bcas2_in.vivo_noRT | Hebp2 | -1.386 | 2.02E-04 | glio |
| Bcas2_ex.vivo_noRT | Gm13449 | 2.753 | 2.02E-04 | glio |
| Bora_in.vivo_noRT | Des | 2.686 | 2.02E-04 | glio |
| Bora_in.vivo_noRT | 4430402I18Ri | 2.374 | 2.02E-04 | glio |
| Chek1_ex.vivo_noRT | Batf2 | 2.824 | 2.02E-04 | glio |
| Eif2s1_in.vivo_noRT | Gm266 | 2.143 | 2.02E-04 | glio |
| Hsd17b10_in.vivo_noRT | 2010106G04R | 2.816 | 2.02E-04 | glio |
| culture_effect | Adipor1 | 0.535 | 2.02E-05 | glio |
| culture_effect | Psmd3 | -0.552 | 2.02E-05 | glio |
| Interaction_genotype_Bora | Ntn4 | -4.123 | 2.02E-05 | glio |
| Interaction_genotype_Bora | Col6a2 | -4.226 | 2.02E-05 | glio |
| Interaction_genotype_Bora | Gm2115 | -4.771 | 2.02E-05 | glio |
| Interaction_genotype_Nat2 | Scd3 | 5.426 | 2.02E-05 | glio |
| Aars2_ex.vivo_noRT | Apol6 | 3.305 | 2.02E-05 | glio |
| Atr_in.vivo_noRT | Alpk2 | 3.817 | 2.02E-05 | glio |
| Atr_in.vivo_noRT | Gm9776 | 2.909 | 2.02E-05 | glio |
| Bard1_ex.vivo_noRT | Tmem132c | 2.814 | 2.02E-05 | glio |
| Bard1_ex.vivo_noRT | Gm44693 | 2.814 | 2.02E-05 | glio |
| Chek1_in.vivo_noRT | Gm43156 | 3.025 | 2.02E-05 | glio |
| Chek1_in.vivo_noRT | Gm16835 | 3.117 | 2.02E-05 | glio |

| | | | | |
|---|---|---|---|---|
| Interaction_genotype_Mtor | Prickle4 | -5.366 | 2.02E-06 | glio |
| Interaction_genotype_Mtor | Tagap | -3.839 | 2.02E-06 | glio |
| Interaction_genotype_Myc | Ran | 2.319 | 2.02E-06 | glio |
| Aars2_in.vivo_noRT | Wnt4 | 3.684 | 2.02E-06 | glio |
| Bcas2_in.vivo_noRT | Gabrd | 3.358 | 2.02E-06 | glio |
| Bora_ex.vivo_noRT | Vtn | 3.201 | 2.02E-06 | glio |
| Interaction_genotype_Mtor | Zfp91 | 2.552 | 2.02E-07 | glio |
| Chek1_ex.vivo_noRT | Igfbp7 | 4.045 | 2.02E-07 | glio |
| Interaction_genotype_Mtor | Mrpl12 | 3.011 | 2.02E-09 | glio |
| Bcas2_in.vivo_noRT | Gm44702 | 4.275 | 2.02E-09 | glio |
| Bcas2_in.vivo_noRT | Gm11884 | 3.808 | 2.02E-09 | glio |
| Chek1_in.vivo_noRT | Apoh | 4.129 | 2.02E-09 | glio |
| Atr_ex.vivo_noRT | Tcstv1 | 4.76 | 2.02E-10 | glio |
| culture_effect | Mocs2 | 1.007 | 2.02E-11 | glio |
| Atr_in.vivo_noRT | Rax | 4.354 | 2.02E-11 | glio |
| Chek1_in.vivo_noRT | Cfb | 3.74 | 2.02E-11 | glio |
| Aars2_in.vivo_noRT | Pcdhga2 | 4.593 | 2.02E-12 | glio |
| Bcas2_in.vivo_noRT | Dcn | 4.416 | 2.02E-12 | glio |
| Chek1_in.vivo_noRT | Gm14002 | 4.522 | 2.02E-12 | glio |
| Bcas2_in.vivo_noRT | 1700026N04R | 4.547 | 2.02E-13 | glio |
| Chek1_in.vivo_noRT | Armc2 | 5.687 | 2.02E-16 | glio |
| Bora_in.vivo_noRT | Gm17203 | 4.286 | 2.02E-17 | glio |
| Aars2_in.vivo_noRT | Gm44646 | 5.327 | 2.02E-20 | glio |
| culture_effect | Epcam | -4.528 | 2.02E-32 | glio |
| culture_effect | Elobl | 1.981 | 2.03E-03 | glio |
| culture_effect | Pate2 | 1.773 | 2.03E-03 | glio |
| Interaction_genotype_Aars2 | Efcab9 | -3.701 | 2.03E-03 | glio |
| Interaction_genotype_Aars2 | Chd4 | 1.284 | 2.03E-03 | glio |
| Interaction_genotype_Atr | Slc1a2 | -3.291 | 2.03E-03 | glio |
| Interaction_genotype_Atr | Gm41307 | -2.762 | 2.03E-03 | glio |
| Interaction_genotype_Atr | Gm15234 | -3.824 | 2.03E-03 | glio |
| Interaction_genotype_Atr | 2210417A02R | -4.028 | 2.03E-03 | glio |
| Interaction_genotype_Bora | Gm26673 | -4.05 | 2.03E-03 | glio |
| Interaction_genotype_Lmo7 | Gm43672 | -4.797 | 2.03E-03 | glio |
| Interaction_genotype_Myc | Bmx | -3.12 | 2.03E-03 | glio |
| Interaction_genotype_Myc | Nptx1 | -3.835 | 2.03E-03 | glio |
| Interaction_genotype_Myc | Lzts3 | -3.11 | 2.03E-03 | glio |
| Interaction_genotype_Myc | Als2 | -3.172 | 2.03E-03 | glio |
| Interaction_genotype_Myc | Rps19bp1 | 1.295 | 2.03E-03 | glio |
| Interaction_genotype_Pars2 | B930094E09Ri | -3.778 | 2.03E-03 | glio |
| Interaction_genotype_Pars2 | Apol7d | -4.328 | 2.03E-03 | glio |
| Interaction_genotype_Pars2 | Plcl1 | -3.424 | 2.03E-03 | glio |
| Interaction_genotype_Pars2 | Ifit1bl1 | -5.055 | 2.03E-03 | glio |
| Interaction_genotype_Pars2 | Gm15816 | -3.748 | 2.03E-03 | glio |
| interaction_RT | Pfkl | -0.969 | 2.03E-03 | glio |
| Aars2_in.vivo_noRT | Hist1h3b | 2.351 | 2.03E-03 | glio |
| Aars2_in.vivo_noRT | Zfp961 | 1.639 | 2.03E-03 | glio |
| Aars2_ex.vivo_noRT | Ercc6 | -2.505 | 2.03E-03 | glio |
| Atp5c1_in.vivo_noRT | Gm27010 | 2.367 | 2.03E-03 | glio |
| Atr_in.vivo_noRT | Fubp1 | -0.827 | 2.03E-03 | glio |
| Bard1_ex.vivo_noRT | Gpr17 | 2.814 | 2.03E-03 | glio |
| Bcas2_in.vivo_noRT | Acy3 | 2.314 | 2.03E-03 | glio |

| | | | | |
|---|---|---|---|---|
| Bcas2_in.vivo_noRT | Gm11131 | 2.189 | 2.03E-03 | glio |
| Bcas2_in.vivo_noRT | Ogdhl | 2.252 | 2.03E-03 | glio |
| Bora_in.vivo_noRT | Gm14167 | 2.024 | 2.03E-03 | glio |
| Bora_in.vivo_noRT | Tmem170 | 1.881 | 2.03E-03 | glio |
| Bora_in.vivo_noRT | Pak1ip1 | -0.999 | 2.03E-03 | glio |
| Cenpj_ex.vivo_noRT | Gm30692 | 2.198 | 2.03E-03 | glio |
| Cenpj_ex.vivo_noRT | Acod1 | 3.737 | 2.03E-03 | glio |
| Chek1_in.vivo_noRT | Radil | 2.222 | 2.03E-03 | glio |
| Chek1_in.vivo_noRT | Ppargc1b | 2.289 | 2.03E-03 | glio |
| Chek1_in.vivo_noRT | Klhl18 | 1.84 | 2.03E-03 | glio |
| Interaction_genotype_Atr | Dntt | -4.772 | 2.03E-04 | glio |
| Interaction_genotype_Atr | Strn3 | 1.98 | 2.03E-04 | glio |
| Interaction_genotype_Bora | Thap8 | -5.006 | 2.03E-04 | glio |
| Interaction_genotype_Mtor | Ccdc162 | -3.57 | 2.03E-04 | glio |
| Interaction_genotype_Mtor | Hist1h4a | -4.139 | 2.03E-04 | glio |
| Interaction_genotype_Mtor | Gm9828 | -3.993 | 2.03E-04 | glio |
| Interaction_genotype_Mtor | Cdk1 | 3.255 | 2.03E-04 | glio |
| Interaction_genotype_Myc | A330032B11R | -3.842 | 2.03E-04 | glio |
| Interaction_genotype_Myc | Pldi | -3.423 | 2.03E-04 | glio |
| Interaction_genotype_Myc | Mfn2 | -2.853 | 2.03E-04 | glio |
| interaction_RT | Ppp1r1b | 2.24 | 2.03E-04 | glio |
| interaction_RT | Casp12 | 1.524 | 2.03E-04 | glio |
| interaction_RT | Dusp14 | 1.188 | 2.03E-04 | glio |
| Aars2_in.vivo_noRT | Slc4a5 | 2.796 | 2.03E-04 | glio |
| Aars2_ex.vivo_noRT | D630024D03F | 2.706 | 2.03E-04 | glio |
| Bora_in.vivo_noRT | Samd15 | 2.914 | 2.03E-04 | glio |
| Bora_in.vivo_noRT | 9330160F10Ri | 2.453 | 2.03E-04 | glio |
| Bora_ex.vivo_noRT | Vgll2 | 3.201 | 2.03E-04 | glio |
| Bora_ex.vivo_noRT | 4930518J21Ri | 3.391 | 2.03E-04 | glio |
| Chek1_in.vivo_noRT | Mrps14 | -0.919 | 2.03E-04 | glio |
| Eif2s1_in.vivo_noRT | Ildr1 | 2.143 | 2.03E-04 | glio |
| interaction_RT | Nrip1 | 1.315 | 2.03E-05 | glio |
| Aars2_in.vivo_noRT | Adam19 | 3.071 | 2.03E-05 | glio |
| Chek1_ex.vivo_noRT | Cldn10 | 3.857 | 2.03E-05 | glio |
| culture_effect | Slc7a6 | -1.059 | 2.03E-06 | glio |
| Interaction_genotype_Mtor | Cyp4x1 | -5.415 | 2.03E-06 | glio |
| Interaction_genotype_Myc | Dpf1 | -4.751 | 2.03E-06 | glio |
| Aars2_in.vivo_noRT | 4930524O07R | 3.402 | 2.03E-06 | glio |
| Aars2_in.vivo_noRT | Vopp1 | 2.989 | 2.03E-06 | glio |
| Chek1_ex.vivo_noRT | Ncf1 | 3.862 | 2.03E-07 | glio |
| Hsd17b10_in.vivo_noRT | C130060C02R | 3.352 | 2.03E-07 | glio |
| Aars2_in.vivo_noRT | Gm10101 | 3.758 | 2.03E-08 | glio |
| Bcas2_in.vivo_noRT | 2010106G04R | 4.007 | 2.03E-08 | glio |
| Hsd17b10_in.vivo_noRT | Gm13032 | 3.23 | 2.03E-08 | glio |
| Hsd17b10_in.vivo_noRT | Gm44983 | 3.355 | 2.03E-08 | glio |
| Bcas2_in.vivo_noRT | Tspyl3 | 3.724 | 2.03E-09 | glio |
| Atr_in.vivo_noRT | Tomm6os | 4.615 | 2.03E-10 | glio |
| Chek1_in.vivo_noRT | Gm24362 | 4.001 | 2.03E-10 | glio |
| Bcas2_in.vivo_noRT | Matn4 | 4.543 | 2.03E-11 | glio |
| Chek1_in.vivo_noRT | Dreh | 4.397 | 2.03E-12 | glio |
| Bcas2_in.vivo_noRT | Fbxo15 | 4.544 | 2.03E-13 | glio |
| culture_effect | Gm14418 | 2.442 | 2.03E-15 | glio |

| | | | | |
|---|---|---|---|---|
| Bora_in.vivo_noRT | Gm45151 | 4.405 | 2.03E-15 | glio |
| Chek1_ex.vivo_noRT | Gm8016 | 4.842 | 2.03E-15 | glio |
| Aars2_in.vivo_noRT | Bco2 | 5.327 | 2.03E-18 | glio |
| Chek1_in.vivo_noRT | Gm13178 | 4.528 | 2.03E-18 | glio |
| Aars2_in.vivo_noRT | 4930512B01R | 5.327 | 2.03E-23 | glio |
| Aars2_in.vivo_noRT | Gm48140 | 5.063 | 2.03E-23 | glio |
| Atr_in.vivo_noRT | Gm10447 | 5.533 | 2.03E-23 | glio |
| culture_effect | C1s1 | -5.749 | 2.03E-24 | glio |
| culture_effect | Elp2 | -0.654 | 2.04E-03 | glio |
| culture_effect | Abhd14a | -0.533 | 2.04E-03 | glio |
| culture_effect | Nr2c2ap | 0.406 | 2.04E-03 | glio |
| Interaction_genotype_Aars2 | Gm9967 | -3.657 | 2.04E-03 | glio |
| Interaction_genotype_Aars2 | Styk1 | -3.908 | 2.04E-03 | glio |
| Interaction_genotype_Aars2 | Wdr5b | -3.702 | 2.04E-03 | glio |
| Interaction_genotype_Aars2 | 4930558J18Ri | -3.429 | 2.04E-03 | glio |
| Interaction_genotype_Aars2 | Mettl18 | -3.289 | 2.04E-03 | glio |
| Interaction_genotype_Aars2 | Pkia | -3.258 | 2.04E-03 | glio |
| Interaction_genotype_Atr | Gm50399 | -3.686 | 2.04E-03 | glio |
| Interaction_genotype_Atr | 1700006J14Ri | -3.672 | 2.04E-03 | glio |
| Interaction_genotype_Bora | Gm34047 | -4.076 | 2.04E-03 | glio |
| Interaction_genotype_Mtor | Shpk | -4.028 | 2.04E-03 | glio |
| Interaction_genotype_Mtor | Stard6 | -3.589 | 2.04E-03 | glio |
| Interaction_genotype_Mtor | Zfp438 | -2.968 | 2.04E-03 | glio |
| Interaction_genotype_Mtor | Rtn4 | 1.484 | 2.04E-03 | glio |
| Interaction_genotype_Myc | Gm35290 | -3.286 | 2.04E-03 | glio |
| Interaction_genotype_Myc | Rslcan18 | -3.131 | 2.04E-03 | glio |
| Interaction_genotype_Myc | Elmod3 | -3.319 | 2.04E-03 | glio |
| Interaction_genotype_Myc | Tmem129 | -2.571 | 2.04E-03 | glio |
| Interaction_genotype_Pars2 | Gm44658 | -3.679 | 2.04E-03 | glio |
| Interaction_genotype_Pars2 | BC065403 | -3.572 | 2.04E-03 | glio |
| interaction_RT | Ptpn18 | 1.017 | 2.04E-03 | glio |
| Aars2_in.vivo_noRT | Usp4 | -1.405 | 2.04E-03 | glio |
| Aars2_in.vivo_noRT | Bod1l | -0.994 | 2.04E-03 | glio |
| Aars2_ex.vivo_noRT | Slc16a4 | 2.437 | 2.04E-03 | glio |
| Atr_in.vivo_noRT | 3300005D01R | 2.384 | 2.04E-03 | glio |
| Atr_in.vivo_noRT | Txndc9 | -1.013 | 2.04E-03 | glio |
| Atr_ex.vivo_noRT | Prm3 | 2.584 | 2.04E-03 | glio |
| Bcas2_in.vivo_noRT | Xpo6 | -1.506 | 2.04E-03 | glio |
| Bora_in.vivo_noRT | Lhfpl4 | 2.038 | 2.04E-03 | glio |
| Bora_in.vivo_noRT | Necap1 | -1.603 | 2.04E-03 | glio |
| Bora_in.vivo_noRT | Mthfd2 | -1.241 | 2.04E-03 | glio |
| Bora_ex.vivo_noRT | Svopl | 2.808 | 2.04E-03 | glio |
| Chek1_in.vivo_noRT | Gm49123 | 2.384 | 2.04E-03 | glio |
| Chek1_in.vivo_noRT | Lrp8 | -1.566 | 2.04E-03 | glio |
| Chek1_in.vivo_noRT | Tmem183a | -1.32 | 2.04E-03 | glio |
| Chek1_in.vivo_noRT | Man1a2 | -1.145 | 2.04E-03 | glio |
| culture_effect | Evi5l | -1.398 | 2.04E-04 | glio |
| culture_effect | Trp53bp1 | -1.092 | 2.04E-04 | glio |
| culture_effect | Rpa1 | 0.743 | 2.04E-04 | glio |
| Interaction_genotype_Aars2 | Pgls | 1.318 | 2.04E-04 | glio |
| Interaction_genotype_Mtor | Gm43463 | -3.834 | 2.04E-04 | glio |
| Interaction_genotype_Mtor | Nrap | -4.648 | 2.04E-04 | glio |

| | | | | |
|---|---|---|---|---|
| Interaction_genotype_Mtor | H3f3a | 1.327 | 2.04E-04 | glio |
| Interaction_genotype_Myc | Gm49479 | -4.143 | 2.04E-04 | glio |
| Interaction_genotype_Myc | Mir142hg | -4.163 | 2.04E-04 | glio |
| Aars2_ex.vivo_noRT | Nat8f2 | 2.897 | 2.04E-04 | glio |
| Bora_in.vivo_noRT | Eva1c | 2.741 | 2.04E-04 | glio |
| Chek1_in.vivo_noRT | N4bp2l1 | 2.748 | 2.04E-04 | glio |
| Chek1_in.vivo_noRT | Zfp74 | 2.062 | 2.04E-04 | glio |
| Interaction_genotype_Myc | Pde8a | -3.867 | 2.04E-05 | glio |
| Atr_in.vivo_noRT | Gm15910 | 2.884 | 2.04E-05 | glio |
| Atr_in.vivo_noRT | Slc1a3 | -2.275 | 2.04E-05 | glio |
| Bora_in.vivo_noRT | Tlr6 | 3.102 | 2.04E-05 | glio |
| Chek1_in.vivo_noRT | Alpk2 | 3.851 | 2.04E-05 | glio |
| Chek1_in.vivo_noRT | Gm13212 | 3.049 | 2.04E-05 | glio |
| Interaction_genotype_Bcas2 | Ntn4 | -4.386 | 2.04E-06 | glio |
| Interaction_genotype_Mtor | Tcap | -5.275 | 2.04E-06 | glio |
| Atr_in.vivo_noRT | Palm3 | 3.406 | 2.04E-06 | glio |
| Chek1_ex.vivo_noRT | Mst1 | 4.049 | 2.04E-06 | glio |
| Aars2_in.vivo_noRT | Hey2 | 3.479 | 2.04E-07 | glio |
| Aars2_in.vivo_noRT | Gm15672 | 3.603 | 2.04E-07 | glio |
| Aars2_in.vivo_noRT | Fam171a2 | 3.674 | 2.04E-07 | glio |
| Atr_in.vivo_noRT | Slc24a1 | 3.671 | 2.04E-07 | glio |
| Atr_in.vivo_noRT | Rab17 | 3.539 | 2.04E-07 | glio |
| Chek1_in.vivo_noRT | Gm47223 | 4.777 | 2.04E-07 | glio |
| Chek1_in.vivo_noRT | Tec | 3.367 | 2.04E-07 | glio |
| culture_effect | Pstk | 1.009 | 2.04E-08 | glio |
| Chek1_in.vivo_noRT | AU022754 | 3.623 | 2.04E-08 | glio |
| Interaction_genotype_Myc | Fis1 | 2.794 | 2.04E-09 | glio |
| Atr_in.vivo_noRT | B230317F23Ri | 4.028 | 2.04E-09 | glio |
| culture_effect | Synm | 2.245 | 2.04E-10 | glio |
| culture_effect | Lrpap1 | -1.185 | 2.04E-10 | glio |
| culture_effect | Atg101 | -0.989 | 2.04E-10 | glio |
| Chek1_in.vivo_noRT | Zfp14 | 4.521 | 2.04E-10 | glio |
| Atr_in.vivo_noRT | Mtmr11 | 4.687 | 2.04E-11 | glio |
| Chek1_in.vivo_noRT | Tecrl | 4.784 | 2.04E-11 | glio |
| Aars2_in.vivo_noRT | Gm13594 | 5.311 | 2.04E-13 | glio |
| Atr_in.vivo_noRT | Vmn1r181 | 4.877 | 2.04E-13 | glio |
| Atr_in.vivo_noRT | St8sia1 | 5.273 | 2.04E-13 | glio |
| Atr_in.vivo_noRT | Hsf2bp | 4.741 | 2.04E-13 | glio |
| Bora_in.vivo_noRT | Cxcl13 | 4.936 | 2.04E-13 | glio |
| Bora_in.vivo_noRT | Gm49223 | 4.414 | 2.04E-14 | glio |
| Bcas2_in.vivo_noRT | Crhbp | 4.807 | 2.04E-15 | glio |
| Atr_in.vivo_noRT | Gstt4 | 5.27 | 2.04E-24 | glio |
| culture_effect | Gm11583 | -1.46 | 2.05E-03 | glio |
| culture_effect | Zfp598 | -0.653 | 2.05E-03 | glio |
| culture_effect | Shmt1 | -0.623 | 2.05E-03 | glio |
| Interaction_genotype_Aars2 | Frg1 | 1.631 | 2.05E-03 | glio |
| Interaction_genotype_Atr | Chd7 | 2.138 | 2.05E-03 | glio |
| Interaction_genotype_Cdkn2a | Olfr1152 | 3.101 | 2.05E-03 | glio |
| Interaction_genotype_Cdkn2a | Gm29966 | 3.095 | 2.05E-03 | glio |
| Interaction_genotype_Chek1 | Ift20 | 1.451 | 2.05E-03 | glio |
| Interaction_genotype_Mtor | Dnmt3b | -3.735 | 2.05E-03 | glio |
| Interaction_genotype_Myc | Gm42918 | -3.059 | 2.05E-03 | glio |

| | | | | |
|---|---|---|---|---|
| Interaction_genotype_Myc | Acsf2 | -3.078 | 2.05E-03 | glio |
| Interaction_genotype_Myc | Tmem136 | -3.041 | 2.05E-03 | glio |
| interaction_RT | Chd1l | -1.726 | 2.05E-03 | glio |
| interaction_RT | Rnf166 | 1.267 | 2.05E-03 | glio |
| interaction_RT | 1810013L24Ri | 1.033 | 2.05E-03 | glio |
| Aars2_in.vivo_noRT | Gm28535 | 2.144 | 2.05E-03 | glio |
| Aars2_in.vivo_noRT | Gm45716 | 2.112 | 2.05E-03 | glio |
| Aars2_in.vivo_noRT | Entpd5 | 2.114 | 2.05E-03 | glio |
| Aars2_in.vivo_noRT | Gm14295 | 1.851 | 2.05E-03 | glio |
| Atp5c1_in.vivo_noRT | Gm36026 | 2.017 | 2.05E-03 | glio |
| Atp5c1_in.vivo_noRT | Gm48742 | 2.018 | 2.05E-03 | glio |
| Bard1_ex.vivo_noRT | Gm41349 | 2.814 | 2.05E-03 | glio |
| Bard1_ex.vivo_noRT | 4931431C16R | 2.434 | 2.05E-03 | glio |
| Bcas2_in.vivo_noRT | Gm16124 | 2.572 | 2.05E-03 | glio |
| Bcas2_in.vivo_noRT | D830050J10Ri | 2.121 | 2.05E-03 | glio |
| Bcas2_in.vivo_noRT | Pigk | -1.377 | 2.05E-03 | glio |
| Bora_in.vivo_noRT | Btbd6 | 2.053 | 2.05E-03 | glio |
| Bora_in.vivo_noRT | Khdrbs2 | 2.028 | 2.05E-03 | glio |
| Bora_in.vivo_noRT | Pgp | -0.914 | 2.05E-03 | glio |
| Chek1_in.vivo_noRT | 4833413G10R | 2.373 | 2.05E-03 | glio |
| Chek1_in.vivo_noRT | Scfd1 | -1.244 | 2.05E-03 | glio |
| Chek1_ex.vivo_noRT | Zfp980 | 3.038 | 2.05E-03 | glio |
| Chek1_ex.vivo_noRT | A530032D15R | 3.015 | 2.05E-03 | glio |
| Eif2s1_in.vivo_noRT | Gm14820 | 1.88 | 2.05E-03 | glio |
| culture_effect | Mphosph10 | -0.49 | 2.05E-04 | glio |
| Interaction_genotype_Aars2 | Lyz2 | -4.858 | 2.05E-04 | glio |
| Interaction_genotype_Atr | Dnah10 | -4.125 | 2.05E-04 | glio |
| Interaction_genotype_Bcas2 | Gm19541 | -4.937 | 2.05E-04 | glio |
| Interaction_genotype_Myc | Slc16a13 | -3.573 | 2.05E-04 | glio |
| interaction_RT | Znhit6 | -0.982 | 2.05E-04 | glio |
| Aars2_in.vivo_noRT | Gm12359 | 2.635 | 2.05E-04 | glio |
| Aars2_in.vivo_noRT | 2900089D17R | 2.48 | 2.05E-04 | glio |
| Aars2_in.vivo_noRT | Slc46a2 | 2.346 | 2.05E-04 | glio |
| Aars2_in.vivo_noRT | Nr2c2ap | -1.249 | 2.05E-04 | glio |
| Atr_in.vivo_noRT | Adamts12 | 2.339 | 2.05E-04 | glio |
| Atr_ex.vivo_noRT | Pla2g2f | 3.164 | 2.05E-04 | glio |
| Bcas2_in.vivo_noRT | Tspan33 | 2.713 | 2.05E-04 | glio |
| Bora_in.vivo_noRT | Btc | 3.377 | 2.05E-04 | glio |
| Bora_in.vivo_noRT | 2410080I02Ri | 2.556 | 2.05E-04 | glio |
| Bora_in.vivo_noRT | Crmp1 | 1.936 | 2.05E-04 | glio |
| Bora_in.vivo_noRT | Rrm2 | -1.925 | 2.05E-04 | glio |
| Bora_in.vivo_noRT | Tpt1 | -0.839 | 2.05E-04 | glio |
| Cenpj_in.vivo_noRT | Sucnr1 | 2.022 | 2.05E-04 | glio |
| Cenpj_in.vivo_noRT | Crb3 | 2.551 | 2.05E-04 | glio |
| Cenpj_in.vivo_noRT | Gm10447 | 2.548 | 2.05E-04 | glio |
| Cenpj_in.vivo_noRT | Ngb | 3.13 | 2.05E-04 | glio |
| Cenpj_in.vivo_noRT | Skida1 | 3.323 | 2.05E-04 | glio |
| Cenpj_ex.vivo_noRT | Gm14929 | 2.591 | 2.05E-04 | glio |
| Cenpj_ex.vivo_noRT | Rimklb | 2.591 | 2.05E-04 | glio |
| Cyp19a1_ex.vivo_noRT | Trmt61b | 2.933 | 2.05E-04 | glio |
| Hsd17b10_in.vivo_noRT | Tead3 | 2.722 | 2.05E-04 | glio |
| Interaction_genotype_Mtor | Fam227b | -3.307 | 2.05E-05 | glio |

| | | | | |
|---|---|---|---|---|
| Interaction_genotype_Myc | Gm10244 | -5.051 | 2.05E-05 | glio |
| interaction_RT | Lrrc59 | -0.855 | 2.05E-05 | glio |
| Atr_in.vivo_noRT | Idh2 | -1.56 | 2.05E-05 | glio |
| Bora_in.vivo_noRT | Srcin1 | 2.895 | 2.05E-05 | glio |
| Chek1_in.vivo_noRT | Stmn4 | 4.528 | 2.05E-05 | glio |
| Chek1_ex.vivo_noRT | Gm9895 | 3.853 | 2.05E-05 | glio |
| Atr_in.vivo_noRT | Bc1 | 3.506 | 2.05E-06 | glio |
| Bora_in.vivo_noRT | Cct6b | 3.18 | 2.05E-06 | glio |
| Chek1_in.vivo_noRT | 2610016A17R | 3.458 | 2.05E-06 | glio |
| Chek1_in.vivo_noRT | Gm15672 | 3.32 | 2.05E-06 | glio |
| Chek1_in.vivo_noRT | Prkar2b | 3.371 | 2.05E-06 | glio |
| culture_effect | 5330438D12R | 1.934 | 2.05E-07 | glio |
| Aars2_in.vivo_noRT | Lyrm1 | 3.614 | 2.05E-07 | glio |
| Atp5c1_in.vivo_noRT | Trim17 | 2.668 | 2.05E-07 | glio |
| Atp5c1_in.vivo_noRT | Tpo | 2.668 | 2.05E-07 | glio |
| Atr_in.vivo_noRT | Gemin4 | 3.788 | 2.05E-07 | glio |
| Atr_in.vivo_noRT | Ankrd42 | 3.66 | 2.05E-07 | glio |
| Atr_in.vivo_noRT | Adamts7 | 3.343 | 2.05E-07 | glio |
| Atr_in.vivo_noRT | Rpl32 | -1.089 | 2.05E-07 | glio |
| Bcas2_ex.vivo_noRT | Chia1 | 3.536 | 2.05E-07 | glio |
| Bcas2_ex.vivo_noRT | Gm48617 | 3.54 | 2.05E-07 | glio |
| Bcas2_ex.vivo_noRT | Angptl7 | 3.54 | 2.05E-07 | glio |
| Bora_in.vivo_noRT | Gm20478 | 3.364 | 2.05E-07 | glio |
| Bora_ex.vivo_noRT | Hoxc8 | 3.201 | 2.05E-07 | glio |
| Bora_ex.vivo_noRT | Ipcef1 | 3.994 | 2.05E-07 | glio |
| culture_effect | Gm27239 | 2.962 | 2.05E-08 | glio |
| culture_effect | Ap3s1 | 0.894 | 2.05E-08 | glio |
| culture_effect | Actr10 | -0.725 | 2.05E-08 | glio |
| culture_effect | Alyref | 0.696 | 2.05E-08 | glio |
| Atr_in.vivo_noRT | Zfp467 | 3.993 | 2.05E-08 | glio |
| Bcas2_in.vivo_noRT | Mtmr11 | 3.961 | 2.05E-08 | glio |
| Bora_in.vivo_noRT | Cks1brt | 3.499 | 2.05E-08 | glio |
| Bcas2_in.vivo_noRT | Ccl7 | 4.807 | 2.05E-09 | glio |
| Chek1_ex.vivo_noRT | Fa2h | 4.842 | 2.05E-09 | glio |
| Chek1_ex.vivo_noRT | Pldi | 4.148 | 2.05E-09 | glio |
| Chek1_ex.vivo_noRT | Icam2 | 4.842 | 2.05E-10 | glio |
| Atr_in.vivo_noRT | Ltb | 4.474 | 2.05E-11 | glio |
| Bcas2_in.vivo_noRT | Gm43951 | 3.881 | 2.05E-11 | glio |
| Bcas2_in.vivo_noRT | A830009L08Ri | 4.284 | 2.05E-11 | glio |
| Bcas2_in.vivo_noRT | Gm48161 | 4.019 | 2.05E-11 | glio |
| Bcas2_in.vivo_noRT | Rgs9 | 4.007 | 2.05E-11 | glio |
| Bora_in.vivo_noRT | Lrrc4b | 4.026 | 2.05E-11 | glio |
| Chek1_in.vivo_noRT | Rhox7a | 5.044 | 2.05E-12 | glio |
| Bora_in.vivo_noRT | Gm11769 | 4.676 | 2.05E-13 | glio |
| Chek1_in.vivo_noRT | Gm36026 | 4.393 | 2.05E-14 | glio |
| Atr_in.vivo_noRT | Gm43256 | 5.27 | 2.05E-16 | glio |
| Aars2_in.vivo_noRT | Gm45406 | 4.805 | 2.05E-19 | glio |
| Atr_in.vivo_noRT | Pyy | 5.533 | 2.05E-21 | glio |
| Chek1_in.vivo_noRT | Ccl9 | 5.044 | 2.05E-24 | glio |
| Chek1_in.vivo_noRT | Halr1 | 4.777 | 2.05E-24 | glio |
| culture_effect | Ldlrad4 | 1.071 | 2.06E-03 | glio |
| culture_effect | Cnot7 | 0.547 | 2.06E-03 | glio |

| | | | | |
|---|---|---|---|---|
| culture_effect | Syt11 | 0.601 | 2.06E-03 | glio |
| Interaction_genotype_Bora | Jun | 1.779 | 2.06E-03 | glio |
| Interaction_genotype_Mtor | Pcdhga1 | -3.225 | 2.06E-03 | glio |
| Interaction_genotype_Mtor | Ifi214 | -3.714 | 2.06E-03 | glio |
| Interaction_genotype_Mtor | Gng13 | -3.697 | 2.06E-03 | glio |
| Interaction_genotype_Mtor | Pla2g7 | -3.646 | 2.06E-03 | glio |
| Interaction_genotype_Mtor | Zfp354a | -3.724 | 2.06E-03 | glio |
| Interaction_genotype_Mtor | Soga1 | -3.797 | 2.06E-03 | glio |
| Interaction_genotype_Mtor | Zik1 | -3.108 | 2.06E-03 | glio |
| Interaction_genotype_Mtor | Ccdc93 | -3.116 | 2.06E-03 | glio |
| Interaction_genotype_Mtor | Gm26670 | -2.991 | 2.06E-03 | glio |
| Interaction_genotype_Mtor | Ppp1r15a | 2.091 | 2.06E-03 | glio |
| Interaction_genotype_Mtor | Eno3 | 2.107 | 2.06E-03 | glio |
| Interaction_genotype_Mtor | Tomm70a | 1.562 | 2.06E-03 | glio |
| Interaction_genotype_Mtor | Mrpl43 | 1.355 | 2.06E-03 | glio |
| Interaction_genotype_Mtor | C1qbp | 1.388 | 2.06E-03 | glio |
| Interaction_genotype_Mtor | Rbm3 | 1.258 | 2.06E-03 | glio |
| Interaction_genotype_Myc | Lmx1a | -2.729 | 2.06E-03 | glio |
| Interaction_genotype_Myc | Gm37214 | -3.565 | 2.06E-03 | glio |
| Interaction_genotype_Myc | Etohd2 | -3.679 | 2.06E-03 | glio |
| Interaction_genotype_Myc | Rex2 | -3.716 | 2.06E-03 | glio |
| Interaction_genotype_Myc | Gm14023 | -3.552 | 2.06E-03 | glio |
| Interaction_genotype_Myc | Aldh1a3 | -3.367 | 2.06E-03 | glio |
| Interaction_genotype_Myc | Ndufb9 | 1.018 | 2.06E-03 | glio |
| Interaction_genotype_Rpl3 | Gm27010 | -4.126 | 2.06E-03 | glio |
| interaction_RT | Nubp1 | -0.955 | 2.06E-03 | glio |
| interaction_RT | Minar2 | 2.446 | 2.06E-03 | glio |
| interaction_RT | Wdr36 | -0.882 | 2.06E-03 | glio |
| interaction_RT | Ubtd1 | 0.994 | 2.06E-03 | glio |
| interaction_RT | Atp5b | -0.551 | 2.06E-03 | glio |
| Aars2_in.vivo_noRT | Ppp1r3f | 2.346 | 2.06E-03 | glio |
| Aars2_in.vivo_noRT | A730063M14F | 1.995 | 2.06E-03 | glio |
| Aars2_in.vivo_noRT | Nnt | 2.019 | 2.06E-03 | glio |
| Aars2_in.vivo_noRT | Plrg1 | -1.551 | 2.06E-03 | glio |
| Aars2_in.vivo_noRT | Mcrs1 | -1.146 | 2.06E-03 | glio |
| Atr_in.vivo_noRT | Uhrf1 | -1.402 | 2.06E-03 | glio |
| Bcas2_in.vivo_noRT | Kyat1 | 2.444 | 2.06E-03 | glio |
| Bcas2_in.vivo_noRT | Fam161b | 2.259 | 2.06E-03 | glio |
| Bcas2_ex.vivo_noRT | Olfr718-ps1 | 2.348 | 2.06E-03 | glio |
| Bcas2_ex.vivo_noRT | Kcnq2 | 2.742 | 2.06E-03 | glio |
| Bora_in.vivo_noRT | Slc35d2 | 2.303 | 2.06E-03 | glio |
| Bora_in.vivo_noRT | Car9 | 2.183 | 2.06E-03 | glio |
| Bora_in.vivo_noRT | Smim4 | -1.64 | 2.06E-03 | glio |
| Bora_in.vivo_noRT | Glrx5 | -0.893 | 2.06E-03 | glio |
| Bora_ex.vivo_noRT | Lrp2bp | 2.699 | 2.06E-03 | glio |
| Cenpj_ex.vivo_noRT | Gm28940 | 2.591 | 2.06E-03 | glio |
| Cenpj_ex.vivo_noRT | Gm26671 | 2.198 | 2.06E-03 | glio |
| Chek1_in.vivo_noRT | Herc3 | 1.666 | 2.06E-03 | glio |
| Chek1_ex.vivo_noRT | Kif23 | -2.195 | 2.06E-03 | glio |
| Chek1_ex.vivo_noRT | Mcm3 | -1.659 | 2.06E-03 | glio |
| Hsd17b10_in.vivo_noRT | Limch1 | 2.578 | 2.06E-03 | glio |
| Interaction_genotype_Atr | B230219D22R | 2.133 | 2.06E-04 | glio |

| | | | | |
|---|---|---|---|---|
| Interaction_genotype_Mtor | Gm14023 | -4.55 | 2.06E-04 | glio |
| Interaction_genotype_Myc | Dynlt1c | -4.754 | 2.06E-04 | glio |
| Interaction_genotype_Myc | Gm48155 | -2.729 | 2.06E-04 | glio |
| Interaction_genotype_Myc | Ppox | -3.191 | 2.06E-04 | glio |
| Interaction_genotype_Myc | Hbegf | 2.395 | 2.06E-04 | glio |
| Aars2_in.vivo_noRT | Gm43707 | 2.318 | 2.06E-04 | glio |
| Atr_in.vivo_noRT | 2500004C02R | 2.161 | 2.06E-04 | glio |
| Atr_in.vivo_noRT | D030028A08R | 2.045 | 2.06E-04 | glio |
| Bcas2_in.vivo_noRT | Vgll3 | 2.685 | 2.06E-04 | glio |
| Chek1_in.vivo_noRT | Syt6 | 2.304 | 2.06E-04 | glio |
| Chek1_in.vivo_noRT | Ube2i | -0.83 | 2.06E-04 | glio |
| Hsd17b10_in.vivo_noRT | Gm14264 | 2.622 | 2.06E-04 | glio |
| Interaction_genotype_Mtor | Gm28875 | -5.306 | 2.06E-05 | glio |
| Aars2_in.vivo_noRT | Fzd1 | 2.868 | 2.06E-05 | glio |
| Aars2_in.vivo_noRT | Csnk1d | -1.702 | 2.06E-05 | glio |
| Aars2_in.vivo_noRT | Csnk2a2 | -1.376 | 2.06E-05 | glio |
| Aars2_ex.vivo_noRT | Slc25a18 | 3.293 | 2.06E-05 | glio |
| Atr_in.vivo_noRT | Dnajc9 | -1.864 | 2.06E-05 | glio |
| Bora_in.vivo_noRT | Slc24a1 | 3.074 | 2.06E-05 | glio |
| Chek1_in.vivo_noRT | Hcrtr1 | 3.105 | 2.06E-05 | glio |
| Chek1_in.vivo_noRT | Gm14326 | 3.065 | 2.06E-05 | glio |
| Chek1_in.vivo_noRT | 2810405F17Ri | 2.887 | 2.06E-05 | glio |
| Aars2_in.vivo_noRT | Fastk | -1.952 | 2.06E-06 | glio |
| Aars2_ex.vivo_noRT | Gm2022 | 3.293 | 2.06E-06 | glio |
| Atr_in.vivo_noRT | Gstm2 | 3.409 | 2.06E-06 | glio |
| Atr_in.vivo_noRT | N4bp2l2 | -1.739 | 2.06E-06 | glio |
| Bcas2_in.vivo_noRT | Speg | 3.43 | 2.06E-06 | glio |
| Bcas2_in.vivo_noRT | Gm28592 | 3.12 | 2.06E-06 | glio |
| Bora_in.vivo_noRT | Trim44 | -1.965 | 2.06E-06 | glio |
| Hsd17b10_in.vivo_noRT | Aqp5 | 3.874 | 2.06E-06 | glio |
| culture_effect | Mrpl53 | 0.614 | 2.06E-07 | glio |
| Aars2_in.vivo_noRT | Gm41396 | 3.834 | 2.06E-07 | glio |
| Atr_in.vivo_noRT | Klhdc8b | 3.719 | 2.06E-07 | glio |
| Chek1_in.vivo_noRT | Slc17a7 | 3.612 | 2.06E-07 | glio |
| Chek1_ex.vivo_noRT | Fbln1 | 4.453 | 2.06E-07 | glio |
| Hsd17b10_in.vivo_noRT | Col8a1 | 3.616 | 2.06E-07 | glio |
| Bcas2_in.vivo_noRT | E130208F15Ri | 4.282 | 2.06E-08 | glio |
| Atr_ex.vivo_noRT | Gm48215 | 3.962 | 2.06E-09 | glio |
| Chek1_in.vivo_noRT | Qrich2 | 4.265 | 2.06E-09 | glio |
| culture_effect | Cadm4 | 1.602 | 2.06E-10 | glio |
| Bora_in.vivo_noRT | Col6a3 | 3.9 | 2.06E-10 | glio |
| Bora_in.vivo_noRT | Tmem190 | 4.936 | 2.06E-10 | glio |
| Chek1_in.vivo_noRT | Hdac9 | 4.777 | 2.06E-12 | glio |
| Chek1_in.vivo_noRT | Gm48742 | 4.393 | 2.06E-14 | glio |
| Atr_in.vivo_noRT | Tmem52 | 5.154 | 2.06E-16 | glio |
| Bora_in.vivo_noRT | Gm4166 | 4.542 | 2.06E-18 | glio |
| Bora_in.vivo_noRT | Gm18280 | 4.672 | 2.06E-19 | glio |
| Atr_in.vivo_noRT | Chia1 | 5.533 | 2.06E-27 | glio |
| culture_effect | Nalcn | -0.793 | 2.07E-03 | glio |
| culture_effect | Tdrkh | -0.554 | 2.07E-03 | glio |
| Interaction_genotype_Aars2 | Ldlr | 2.579 | 2.07E-03 | glio |
| Interaction_genotype_Mtor | 1700113B09R | -3.644 | 2.07E-03 | glio |

| | | | | |
|---|---|---|---|---|
| Interaction_genotype_Mtor | Mtmr9 | -2.888 | 2.07E-03 | glio |
| Interaction_genotype_Mtor | Azin2 | -3.065 | 2.07E-03 | glio |
| Interaction_genotype_Myc | 4930556M19F | -3.066 | 2.07E-03 | glio |
| Interaction_genotype_Myc | Trim45 | -2.972 | 2.07E-03 | glio |
| Aars2_in.vivo_noRT | 2410004B18R | -1.118 | 2.07E-03 | glio |
| Aars2_ex.vivo_noRT | Cxcl13 | 2.9 | 2.07E-03 | glio |
| Aars2_ex.vivo_noRT | Pcdhb19 | 2.897 | 2.07E-03 | glio |
| Atr_in.vivo_noRT | 5830408C22R | 2.031 | 2.07E-03 | glio |
| Atr_in.vivo_noRT | Frg2f1 | 1.906 | 2.07E-03 | glio |
| Bora_in.vivo_noRT | Glt1d1 | 2.118 | 2.07E-03 | glio |
| Bora_in.vivo_noRT | Gab2 | 2.093 | 2.07E-03 | glio |
| Bora_in.vivo_noRT | Axin1 | -1.613 | 2.07E-03 | glio |
| Bora_in.vivo_noRT | Ddx10 | -1.39 | 2.07E-03 | glio |
| Cdkn2a_in.vivo_noRT | Gm15445 | -2.321 | 2.07E-03 | glio |
| Cdkn2a_in.vivo_noRT | Halr1 | -1.941 | 2.07E-03 | glio |
| Chek1_in.vivo_noRT | Gm7879 | 2.391 | 2.07E-03 | glio |
| Chek1_in.vivo_noRT | Gm12940 | 1.918 | 2.07E-03 | glio |
| Chek1_in.vivo_noRT | Zdhhc5 | -1.455 | 2.07E-03 | glio |
| Chek1_ex.vivo_noRT | 1700122E12Ri | 2.821 | 2.07E-03 | glio |
| culture_effect | Fbln1 | -2.275 | 2.07E-04 | glio |
| culture_effect | Fbln5 | -0.863 | 2.07E-04 | glio |
| culture_effect | Atp6v1c1 | 0.507 | 2.07E-04 | glio |
| Interaction_genotype_Chek1 | Hist1h2bc | 4.606 | 2.07E-04 | glio |
| Interaction_genotype_Myc | Tcap | -3.905 | 2.07E-04 | glio |
| Interaction_genotype_Myc | Hunk | -3.83 | 2.07E-04 | glio |
| Interaction_genotype_Myc | Ache | -4.355 | 2.07E-04 | glio |
| Interaction_genotype_Myc | Gm10655 | -4.278 | 2.07E-04 | glio |
| Interaction_genotype_Myc | Nrap | -4.44 | 2.07E-04 | glio |
| Interaction_genotype_Myc | Xkr5 | -3.476 | 2.07E-04 | glio |
| Interaction_genotype_Myc | Alg13 | -2.937 | 2.07E-04 | glio |
| Aars2_in.vivo_noRT | Nrap | 2.787 | 2.07E-04 | glio |
| Aars2_in.vivo_noRT | Tpbg | 2.585 | 2.07E-04 | glio |
| Aars2_in.vivo_noRT | Mmp23 | 2.529 | 2.07E-04 | glio |
| Aars2_in.vivo_noRT | Galk1 | -1.27 | 2.07E-04 | glio |
| Atp5c1_in.vivo_noRT | Abat | 3.074 | 2.07E-04 | glio |
| Bcas2_ex.vivo_noRT | Gm2022 | 2.742 | 2.07E-04 | glio |
| Bora_in.vivo_noRT | Usp38 | -1.777 | 2.07E-04 | glio |
| Chek1_in.vivo_noRT | Camkv | 2.809 | 2.07E-04 | glio |
| Hsd17b10_in.vivo_noRT | Puf60 | -0.903 | 2.07E-04 | glio |
| culture_effect | Tnfrsf1a | -0.704 | 2.07E-05 | glio |
| Interaction_genotype_Aars2 | Gm36356 | -4.532 | 2.07E-05 | glio |
| Interaction_genotype_Aars2 | Pitx3 | -4.312 | 2.07E-05 | glio |
| Interaction_genotype_Aars2 | 1700026J14Ri | -4.824 | 2.07E-05 | glio |
| Interaction_genotype_Mtor | Gipc2 | -5.483 | 2.07E-05 | glio |
| Interaction_genotype_Mtor | Dpep3 | -3.567 | 2.07E-05 | glio |
| Atr_in.vivo_noRT | Fut11 | 2.897 | 2.07E-05 | glio |
| Bora_in.vivo_noRT | Paip1 | -1.772 | 2.07E-05 | glio |
| Chek1_in.vivo_noRT | Card10 | 3.246 | 2.07E-05 | glio |
| Chek1_in.vivo_noRT | Gm15133 | 2.967 | 2.07E-05 | glio |
| Chek1_in.vivo_noRT | Rpl7a | -0.69 | 2.07E-05 | glio |
| Hsd17b10_in.vivo_noRT | Slc45a3 | 3.094 | 2.07E-05 | glio |
| Hsd17b10_in.vivo_noRT | 4930540M05F | 2.958 | 2.07E-05 | glio |

| | | | | |
|---|---|---|---|---|
| culture_effect | Magoh | -0.593 | 2.07E-06 | glio |
| Interaction_genotype_Aars2 | 2310058D17R | -5.997 | 2.07E-06 | glio |
| Interaction_genotype_Mtor | Slpi | -6.248 | 2.07E-06 | glio |
| Interaction_genotype_Mtor | Gm50399 | -5.416 | 2.07E-06 | glio |
| Interaction_genotype_Mtor | Psca | -5.147 | 2.07E-06 | glio |
| Aars2_in.vivo_noRT | 4930597O21R | 3.599 | 2.07E-06 | glio |
| Atr_ex.vivo_noRT | Gm13178 | 3.164 | 2.07E-06 | glio |
| Bcas2_in.vivo_noRT | Prr7 | 3.416 | 2.07E-06 | glio |
| Bcas2_in.vivo_noRT | Gm16283 | 3.283 | 2.07E-06 | glio |
| Bcas2_in.vivo_noRT | Dcpp1 | 3.48 | 2.07E-06 | glio |
| culture_effect | Nav1 | 1.147 | 2.07E-07 | glio |
| Aars2_in.vivo_noRT | Zfhx2os | 3.746 | 2.07E-07 | glio |
| Aars2_in.vivo_noRT | Slc12a5 | 3.545 | 2.07E-07 | glio |
| Aars2_in.vivo_noRT | Myzap | 3.005 | 2.07E-07 | glio |
| Bora_in.vivo_noRT | Gm48682 | 3.625 | 2.07E-07 | glio |
| Bora_in.vivo_noRT | Gm10644 | 3.695 | 2.07E-07 | glio |
| Bora_in.vivo_noRT | 4933433G15R | 3.803 | 2.07E-07 | glio |
| Chek1_in.vivo_noRT | Tmem202 | 3.602 | 2.07E-07 | glio |
| culture_effect | Lamc2 | 2.735 | 2.07E-08 | glio |
| culture_effect | Ccnb1ip1 | 2.85 | 2.07E-08 | glio |
| culture_effect | B230206L02Ri | 2.507 | 2.07E-08 | glio |
| Atr_in.vivo_noRT | Gm26581 | 4.138 | 2.07E-08 | glio |
| Bcas2_in.vivo_noRT | Glis1 | 3.895 | 2.07E-08 | glio |
| Chek1_in.vivo_noRT | Fam169a | 3.731 | 2.07E-08 | glio |
| Aars2_in.vivo_noRT | Mamdc2 | 4.017 | 2.07E-09 | glio |
| Atr_in.vivo_noRT | Stk36 | 4.245 | 2.07E-09 | glio |
| Chek1_ex.vivo_noRT | Gm43463 | 4.449 | 2.07E-09 | glio |
| Hsd17b10_in.vivo_noRT | Gm40538 | 3.353 | 2.07E-09 | glio |
| Atr_ex.vivo_noRT | Gm4858 | 5.228 | 2.07E-10 | glio |
| Bcas2_in.vivo_noRT | Cysrt1 | 4.547 | 2.07E-10 | glio |
| Bora_in.vivo_noRT | Slc45a3 | 4.414 | 2.07E-10 | glio |
| Chek1_in.vivo_noRT | Flnc | 4.518 | 2.07E-10 | glio |
| Chek1_in.vivo_noRT | A930012O16R | 4.784 | 2.07E-11 | glio |
| Hsd17b10_in.vivo_noRT | Gad1 | 3.35 | 2.07E-11 | glio |
| Aars2_in.vivo_noRT | Ly6g6c | 4.467 | 2.07E-12 | glio |
| Chek1_ex.vivo_noRT | Gm38163 | 4.842 | 2.07E-12 | glio |
| Atr_in.vivo_noRT | Thap8 | 5.273 | 2.07E-13 | glio |
| Atr_in.vivo_noRT | Ccdc63 | 5.147 | 2.07E-14 | glio |
| Atr_in.vivo_noRT | Prrx2 | 5.533 | 2.07E-15 | glio |
| Atr_in.vivo_noRT | Gm41409 | 5.002 | 2.07E-15 | glio |
| Bora_in.vivo_noRT | Gm27241 | 4.28 | 2.07E-16 | glio |
| culture_effect | St18 | -4.297 | 2.07E-17 | glio |
| Bora_in.vivo_noRT | Gm38163 | 4.936 | 2.07E-18 | glio |
| Chek1_in.vivo_noRT | Strip2 | 4.78 | 2.07E-18 | glio |
| culture_effect | Celf2 | 1.896 | 2.07E-19 | glio |
| Aars2_in.vivo_noRT | Gm14540 | 4.804 | 2.07E-19 | glio |
| Chek1_in.vivo_noRT | AC159140.1 | 4.784 | 2.07E-19 | glio |
| Atr_in.vivo_noRT | C330002G04R | 5.533 | 2.07E-25 | glio |
| culture_effect | Tuba8 | -1.811 | 2.08E-03 | glio |
| culture_effect | Nme7 | -0.673 | 2.08E-03 | glio |
| culture_effect | Ankrd24 | 0.583 | 2.08E-03 | glio |
| Interaction_genotype_Aars2 | Abcc10 | -3.895 | 2.08E-03 | glio |

| | | | | |
|---|---|---|---|---|
| Interaction_genotype_Aars2 | Npl | -3.439 | 2.08E-03 | glio |
| Interaction_genotype_Atr | Cyp4x1 | -3.685 | 2.08E-03 | glio |
| Interaction_genotype_Bora | Gm16046 | -4.027 | 2.08E-03 | glio |
| Interaction_genotype_Bora | C920021L13Ri | -3.972 | 2.08E-03 | glio |
| Interaction_genotype_Bora | Akap3 | -4.112 | 2.08E-03 | glio |
| Interaction_genotype_Bora | Aldh1l1 | -4.544 | 2.08E-03 | glio |
| Interaction_genotype_Bora | Zfp786 | -3.995 | 2.08E-03 | glio |
| Interaction_genotype_Bora | Arhgef26 | 2.075 | 2.08E-03 | glio |
| Interaction_genotype_Mtor | 9530053A07R | -3.671 | 2.08E-03 | glio |
| Interaction_genotype_Mtor | Gm14403 | -3.225 | 2.08E-03 | glio |
| Interaction_genotype_Mtor | Zfp287 | -2.845 | 2.08E-03 | glio |
| Interaction_genotype_Mtor | Kif3c | -2.818 | 2.08E-03 | glio |
| Interaction_genotype_Myc | Ccn2 | -4.552 | 2.08E-03 | glio |
| Interaction_genotype_Myc | 9330159F19Ri | -3.352 | 2.08E-03 | glio |
| Interaction_genotype_Myc | Rhobtb2 | -2.944 | 2.08E-03 | glio |
| Interaction_genotype_Myc | Mmd | -3.056 | 2.08E-03 | glio |
| Interaction_genotype_Myc | Rcor1 | -2.075 | 2.08E-03 | glio |
| Interaction_genotype_Myc | Dhx32 | -2.084 | 2.08E-03 | glio |
| Interaction_genotype_Myc | Bex4 | 1.832 | 2.08E-03 | glio |
| Aars2_in.vivo_noRT | B3galt2 | 1.978 | 2.08E-03 | glio |
| Aars2_in.vivo_noRT | Paqr7 | 2.117 | 2.08E-03 | glio |
| Atr_in.vivo_noRT | Zfp946 | 2.079 | 2.08E-03 | glio |
| Bora_in.vivo_noRT | Efr3b | 2.505 | 2.08E-03 | glio |
| Bora_in.vivo_noRT | Zfp446 | 1.954 | 2.08E-03 | glio |
| Chek1_in.vivo_noRT | Doc2g | 2.306 | 2.08E-03 | glio |
| Chek1_in.vivo_noRT | 5830448L01Ri | 2.188 | 2.08E-03 | glio |
| Chek1_in.vivo_noRT | Zhx3 | 2.234 | 2.08E-03 | glio |
| Eif2s1_in.vivo_noRT | Gm50321 | 2.143 | 2.08E-03 | glio |
| Eif2s1_in.vivo_noRT | Ppl | -1.837 | 2.08E-03 | glio |
| Hsd17b10_in.vivo_noRT | Gm10125 | 2.415 | 2.08E-03 | glio |
| Hsd17b10_in.vivo_noRT | Gm50218 | 2.554 | 2.08E-03 | glio |
| Interaction_genotype_Atr | Gm14207 | -3.163 | 2.08E-04 | glio |
| Interaction_genotype_Mtor | B230319C09R | -4.706 | 2.08E-04 | glio |
| Interaction_genotype_Mtor | Glg1 | 2.116 | 2.08E-04 | glio |
| Interaction_genotype_Myc | 4930423D22R | -3.049 | 2.08E-04 | glio |
| Interaction_genotype_Myc | A930002C04R | -2.869 | 2.08E-04 | glio |
| Aars2_in.vivo_noRT | Slc35d2 | 2.693 | 2.08E-04 | glio |
| Aars2_in.vivo_noRT | Fbxo24 | 2.608 | 2.08E-04 | glio |
| Atp5c1_in.vivo_noRT | Fbxl2 | 2.544 | 2.08E-04 | glio |
| Atr_in.vivo_noRT | Col6a1 | 2.569 | 2.08E-04 | glio |
| Atr_in.vivo_noRT | Csgalnact2 | 2.118 | 2.08E-04 | glio |
| Atr_in.vivo_noRT | Trp53 | -0.957 | 2.08E-04 | glio |
| Bard1_ex.vivo_noRT | Gm6803 | 2.814 | 2.08E-04 | glio |
| Bora_in.vivo_noRT | 1110006O24R | 2.734 | 2.08E-04 | glio |
| Bora_ex.vivo_noRT | Gm41349 | 3.201 | 2.08E-04 | glio |
| Bora_ex.vivo_noRT | Serpina3g | 3.201 | 2.08E-04 | glio |
| Bora_ex.vivo_noRT | Glyat | 2.404 | 2.08E-04 | glio |
| Chek1_in.vivo_noRT | Gm42372 | 3.044 | 2.08E-04 | glio |
| Chek1_in.vivo_noRT | 9430060I03Ri | 2.562 | 2.08E-04 | glio |
| Hsd17b10_in.vivo_noRT | Vmn1r88 | 2.431 | 2.08E-04 | glio |
| Interaction_genotype_Mtor | Pbld1 | -5.322 | 2.08E-05 | glio |
| Bcas2_in.vivo_noRT | Gm15056 | 3.373 | 2.08E-05 | glio |

| | | | | |
|---|---|---|---|---|
| Chek1_in.vivo_noRT | Pcp4l1 | 3.086 | 2.08E-05 | glio |
| Chek1_in.vivo_noRT | Gm43728 | 3.185 | 2.08E-05 | glio |
| Bcas2_in.vivo_noRT | Tbkbp1 | 3.514 | 2.08E-06 | glio |
| Bora_in.vivo_noRT | Traf1 | 3.332 | 2.08E-06 | glio |
| Interaction_genotype_Mtor | Helt | -4.099 | 2.08E-07 | glio |
| Atp5c1_in.vivo_noRT | 1700034I23Ri | 2.668 | 2.08E-07 | glio |
| Atr_in.vivo_noRT | Tlr6 | 3.699 | 2.08E-07 | glio |
| Bcas2_in.vivo_noRT | Cyp1a1 | 4.807 | 2.08E-07 | glio |
| Bora_in.vivo_noRT | Tmem179 | 3.778 | 2.08E-07 | glio |
| Aars2_in.vivo_noRT | Gm35867 | 3.874 | 2.08E-08 | glio |
| Bora_in.vivo_noRT | Fsbp | 4.016 | 2.08E-08 | glio |
| Chek1_in.vivo_noRT | Gm26581 | 4.16 | 2.08E-08 | glio |
| Atr_in.vivo_noRT | Pla2g7 | 4.346 | 2.08E-10 | glio |
| Bora_in.vivo_noRT | Serpina3i | 4.148 | 2.08E-10 | glio |
| Bora_in.vivo_noRT | BC048671 | 3.635 | 2.08E-10 | glio |
| Chek1_in.vivo_noRT | 9530062K07R | 4.664 | 2.08E-10 | glio |
| culture_effect | Prkcsh | -0.925 | 2.08E-11 | glio |
| Aars2_in.vivo_noRT | Zfp712 | 4.146 | 2.08E-11 | glio |
| Bcas2_in.vivo_noRT | Scube3 | 4.275 | 2.08E-11 | glio |
| Bora_in.vivo_noRT | Rcan2 | 4.936 | 2.08E-11 | glio |
| Aars2_in.vivo_noRT | Ushbp1 | 4.801 | 2.08E-12 | glio |
| Atr_in.vivo_noRT | Lama5 | 5.011 | 2.08E-12 | glio |
| Hsd17b10_in.vivo_noRT | Ank1 | 3.616 | 2.08E-12 | glio |
| Bora_in.vivo_noRT | AC125149.2 | 4.431 | 2.08E-13 | glio |
| Atr_ex.vivo_noRT | BC018473 | 6.529 | 2.08E-16 | glio |
| Chek1_in.vivo_noRT | Kcnd1 | 5.051 | 2.08E-16 | glio |
| culture_effect | Actn4 | 1.075 | 2.08E-17 | glio |
| culture_effect | Cdh6 | 2.117 | 2.08E-18 | glio |
| Chek1_in.vivo_noRT | Gm15138 | 4.777 | 2.08E-20 | glio |
| Chek1_in.vivo_noRT | Rimklb | 4.78 | 2.08E-22 | glio |
| Atr_in.vivo_noRT | Slc45a1 | 5.273 | 2.08E-24 | glio |
| Atr_in.vivo_noRT | Rapgef3os2 | 5.533 | 2.08E-25 | glio |
| culture_effect | C2 | -5.122 | 2.08E-33 | glio |
| culture_effect | Lpcat1 | -0.665 | 2.09E-03 | glio |
| culture_effect | Mecp2 | -0.502 | 2.09E-03 | glio |
| Interaction_genotype_Atr | Gm13283 | -3.387 | 2.09E-03 | glio |
| Interaction_genotype_Atr | Btg2 | 2.597 | 2.09E-03 | glio |
| Interaction_genotype_Lmo7 | A430106G13R | -4.747 | 2.09E-03 | glio |
| Interaction_genotype_Mtor | Mfap3l | -4.047 | 2.09E-03 | glio |
| Interaction_genotype_Mtor | Hpdl | -3.492 | 2.09E-03 | glio |
| Interaction_genotype_Myc | Pla2g4b | -3.413 | 2.09E-03 | glio |
| Interaction_genotype_Myc | Gm6871 | -2.865 | 2.09E-03 | glio |
| Interaction_genotype_Myc | Wdr53 | -2.654 | 2.09E-03 | glio |
| Interaction_genotype_Myc | Zfp746 | -2.797 | 2.09E-03 | glio |
| Interaction_genotype_Pars2 | 4921504E06Ri | -3.357 | 2.09E-03 | glio |
| Interaction_genotype_Rad21 | C3 | 4.697 | 2.09E-03 | glio |
| Interaction_genotype_Rad51 | Lmo7 | -3.818 | 2.09E-03 | glio |
| Interaction_genotype_Rad51 | 2310030G06R | -4.304 | 2.09E-03 | glio |
| Interaction_genotype_Rad51 | Dlst | 2.131 | 2.09E-03 | glio |
| Interaction_genotype_Rad51 | Fxyd3 | 3.697 | 2.09E-03 | glio |
| Interaction_genotype_Rad51 | Ppic | 1.928 | 2.09E-03 | glio |
| Interaction_genotype_Rpl3 | 4930589L23Ri | -4.293 | 2.09E-03 | glio |

| | | | | |
|---|---|---|---|---|
| Interaction_genotype_Rpl3 | AY074887 | -4.606 | 2.09E-03 | glio |
| interaction_RT | Amn1 | 0.997 | 2.09E-03 | glio |
| Aars2_in.vivo_noRT | Mrpl10 | -1.486 | 2.09E-03 | glio |
| Aars2_ex.vivo_noRT | Smpd5 | 2.706 | 2.09E-03 | glio |
| Atr_in.vivo_noRT | 1190007I07Ri | -1.036 | 2.09E-03 | glio |
| Atr_in.vivo_noRT | Cox7c | -0.792 | 2.09E-03 | glio |
| Atr_ex.vivo_noRT | Ccl7 | 3.372 | 2.09E-03 | glio |
| Bcas2_in.vivo_noRT | Gm14326 | 2.299 | 2.09E-03 | glio |
| Bcas2_in.vivo_noRT | Gm26631 | 2.37 | 2.09E-03 | glio |
| Bcas2_in.vivo_noRT | Mettl26 | -1.287 | 2.09E-03 | glio |
| Bcas2_ex.vivo_noRT | A930033M14F | 2.575 | 2.09E-03 | glio |
| Bora_in.vivo_noRT | Zglp1 | 2.46 | 2.09E-03 | glio |
| Bora_in.vivo_noRT | Haus8 | -1.455 | 2.09E-03 | glio |
| Bora_in.vivo_noRT | Lars | -0.844 | 2.09E-03 | glio |
| Chek1_in.vivo_noRT | Gm20186 | 2.367 | 2.09E-03 | glio |
| Chek1_in.vivo_noRT | Zc3h6 | 2.282 | 2.09E-03 | glio |
| Chek1_in.vivo_noRT | Chrna1os | 1.894 | 2.09E-03 | glio |
| Chek1_in.vivo_noRT | Foxc2 | 2.295 | 2.09E-03 | glio |
| Chek1_ex.vivo_noRT | Slc43a1 | 2.93 | 2.09E-03 | glio |
| Chek1_ex.vivo_noRT | Gm42670 | 2.402 | 2.09E-03 | glio |
| Chek1_ex.vivo_noRT | Rgp1 | 2.341 | 2.09E-03 | glio |
| Chek1_ex.vivo_noRT | Samd5 | 2.508 | 2.09E-03 | glio |
| Chek1_ex.vivo_noRT | Ddx39b | -1.117 | 2.09E-03 | glio |
| Hsd17b10_in.vivo_noRT | Csrp1 | -1.602 | 2.09E-03 | glio |
| Interaction_genotype_Mtor | BC147527 | -4.099 | 2.09E-04 | glio |
| Interaction_genotype_Myc | Klhl32 | -3.749 | 2.09E-04 | glio |
| Interaction_genotype_Myc | Hmx3 | -3.885 | 2.09E-04 | glio |
| Interaction_genotype_Myc | Gm10130 | -3.51 | 2.09E-04 | glio |
| Aars2_ex.vivo_noRT | Ank1 | 2.509 | 2.09E-04 | glio |
| Aars2_ex.vivo_noRT | Tpsab1 | 3.293 | 2.09E-04 | glio |
| Aars2_ex.vivo_noRT | Gm29994 | 3.285 | 2.09E-04 | glio |
| Atr_in.vivo_noRT | Cnbp | -1.081 | 2.09E-04 | glio |
| Atr_ex.vivo_noRT | Gpr17 | 3.164 | 2.09E-04 | glio |
| Bora_ex.vivo_noRT | Gm19265 | 2.802 | 2.09E-04 | glio |
| Bora_ex.vivo_noRT | Ctsf | 2.808 | 2.09E-04 | glio |
| Interaction_genotype_Aars2 | Fastk | 2.444 | 2.09E-05 | glio |
| Interaction_genotype_Aars2 | Skida1 | -5.392 | 2.09E-05 | glio |
| Interaction_genotype_Mtor | Adk | 2.58 | 2.09E-05 | glio |
| Interaction_genotype_Myc | Fzd7 | -4.687 | 2.09E-05 | glio |
| Aars2_in.vivo_noRT | Pheta1 | 3.039 | 2.09E-05 | glio |
| Aars2_in.vivo_noRT | Set | -0.93 | 2.09E-05 | glio |
| Aars2_ex.vivo_noRT | Stx19 | 3.293 | 2.09E-05 | glio |
| Atr_in.vivo_noRT | Gm10553 | 2.987 | 2.09E-05 | glio |
| Hsd17b10_in.vivo_noRT | Ms4a15 | 2.447 | 2.09E-05 | glio |
| Hsd17b10_in.vivo_noRT | Nox1 | 2.696 | 2.09E-05 | glio |
| Hsd17b10_in.vivo_noRT | Krt8 | -2.743 | 2.09E-05 | glio |
| culture_effect | Alox12 | 2.586 | 2.09E-06 | glio |
| culture_effect | Tm9sf1 | -0.904 | 2.09E-06 | glio |
| culture_effect | Rictor | -0.727 | 2.09E-06 | glio |
| Interaction_genotype_Myc | Hdac9 | -5.264 | 2.09E-06 | glio |
| Bora_in.vivo_noRT | Tmem91 | 3.417 | 2.09E-06 | glio |
| Interaction_genotype_Mtor | Gm3716 | -5.893 | 2.09E-07 | glio |

| | | | | |
|---|---|---|---|---|
| Atr_ex.vivo_noRT | 4930527F14Ri | 3.566 | 2.09E-07 | glio |
| Bcas2_in.vivo_noRT | Olfr1189 | 3.152 | 2.09E-07 | glio |
| Chek1_ex.vivo_noRT | Slc6a19 | 4.449 | 2.09E-07 | glio |
| culture_effect | Mecom | 2.908 | 2.09E-08 | glio |
| culture_effect | Cct7 | -0.651 | 2.09E-08 | glio |
| Interaction_genotype_Atr | 2010320O07R | -6.536 | 2.09E-08 | glio |
| Bcas2_in.vivo_noRT | Gm15853 | 4.072 | 2.09E-08 | glio |
| Aars2_in.vivo_noRT | Cxcr2 | 4.015 | 2.09E-09 | glio |
| Bcas2_in.vivo_noRT | Nrn1l | 3.882 | 2.09E-09 | glio |
| Atr_in.vivo_noRT | Cnih2 | 4.345 | 2.09E-10 | glio |
| Bcas2_in.vivo_noRT | Celsr3 | 4.547 | 2.09E-12 | glio |
| Atr_in.vivo_noRT | Olfr1191-ps1 | 5.392 | 2.09E-17 | glio |
| Chek1_in.vivo_noRT | Gm11899 | 4.522 | 2.09E-17 | glio |
| Chek1_ex.vivo_noRT | 1700003E24Ri | 4.842 | 2.09E-17 | glio |
| Bora_in.vivo_noRT | Olfr718-ps1 | 4.672 | 2.09E-18 | glio |
| Bcas2_in.vivo_noRT | Olfr552 | 4.807 | 2.09E-21 | glio |
| Aars2_in.vivo_noRT | Gm29966 | 5.067 | 2.09E-25 | glio |
| Aars2_in.vivo_noRT | Olfr810 | 5.327 | 2.09E-28 | glio |
| culture_effect | Rassf8 | 1.131 | 2.10E-03 | glio |
| culture_effect | Nek7 | 0.675 | 2.10E-03 | glio |
| Interaction_genotype_Mtor | Trmu | -2.787 | 2.10E-03 | glio |
| Interaction_genotype_Mtor | Sfxn1 | 1.739 | 2.10E-03 | glio |
| Interaction_genotype_Myc | Mob3c | -3.108 | 2.10E-03 | glio |
| interaction_RT | Tsc22d4 | 0.758 | 2.10E-03 | glio |
| Aars2_in.vivo_noRT | Fut11 | 2.153 | 2.10E-03 | glio |
| Aars2_in.vivo_noRT | Gm813 | 2.156 | 2.10E-03 | glio |
| Aars2_in.vivo_noRT | Tmem14c | -0.823 | 2.10E-03 | glio |
| Atr_in.vivo_noRT | Pald1 | 2.12 | 2.10E-03 | glio |
| Atr_ex.vivo_noRT | Edn1 | 3.021 | 2.10E-03 | glio |
| Bard1_ex.vivo_noRT | Ltbp2 | 2.419 | 2.10E-03 | glio |
| Bcas2_in.vivo_noRT | Fam189b | 2.329 | 2.10E-03 | glio |
| Bcas2_in.vivo_noRT | Jund | -0.896 | 2.10E-03 | glio |
| Bcas2_ex.vivo_noRT | Insc | 2.742 | 2.10E-03 | glio |
| Bora_in.vivo_noRT | Rspo3 | 2.581 | 2.10E-03 | glio |
| Bora_in.vivo_noRT | Srsf7 | -0.755 | 2.10E-03 | glio |
| Cenpj_ex.vivo_noRT | Slc16a5 | 2.198 | 2.10E-03 | glio |
| Chek1_in.vivo_noRT | Gm36371 | 2.135 | 2.10E-03 | glio |
| Chek1_in.vivo_noRT | 9130230N09R | 1.95 | 2.10E-03 | glio |
| Chek1_in.vivo_noRT | Fam13c | 2.42 | 2.10E-03 | glio |
| Chek1_ex.vivo_noRT | Gm44672 | 2.711 | 2.10E-03 | glio |
| Chek1_ex.vivo_noRT | 4930455G09R | 3.359 | 2.10E-03 | glio |
| Chek1_ex.vivo_noRT | Chek1 | -1.936 | 2.10E-03 | glio |
| Chek1_ex.vivo_noRT | Casp12 | 1.921 | 2.10E-03 | glio |
| culture_effect | Serpina3g | -2.131 | 2.10E-04 | glio |
| culture_effect | Mvd | -1.088 | 2.10E-04 | glio |
| culture_effect | Zfp469 | -0.786 | 2.10E-04 | glio |
| Interaction_genotype_Mtor | Lama5 | -4.464 | 2.10E-04 | glio |
| Interaction_genotype_Mtor | Zfp324 | -3.687 | 2.10E-04 | glio |
| Interaction_genotype_Myc | Hoxa6 | -4.048 | 2.10E-04 | glio |
| Interaction_genotype_Myc | Gm20515 | -4.024 | 2.10E-04 | glio |
| Interaction_genotype_Myc | Ephb3 | -3.25 | 2.10E-04 | glio |
| Interaction_genotype_Myc | Birc6 | 1.854 | 2.10E-04 | glio |

| | | | | |
|---|---|---|---|---|
| interaction_RT | Nudcd1 | -0.907 | 2.10E-04 | glio |
| interaction_RT | Sem1 | 0.67 | 2.10E-04 | glio |
| Aars2_in.vivo_noRT | Wnt10a | 2.63 | 2.10E-04 | glio |
| Aars2_in.vivo_noRT | Bglap3 | 2.608 | 2.10E-04 | glio |
| Aars2_in.vivo_noRT | Gstm7 | 2.638 | 2.10E-04 | glio |
| Atp5c1_in.vivo_noRT | Slc1a2 | 2.404 | 2.10E-04 | glio |
| Bard1_ex.vivo_noRT | 8030423F21Ri | 3.037 | 2.10E-04 | glio |
| Bard1_ex.vivo_noRT | Dscaml1 | 2.419 | 2.10E-04 | glio |
| Bcas2_in.vivo_noRT | 5033430I15Ri | 2.628 | 2.10E-04 | glio |
| Bcas2_in.vivo_noRT | C1qtnf12 | 2.959 | 2.10E-04 | glio |
| Bora_in.vivo_noRT | Gm20337 | 2.718 | 2.10E-04 | glio |
| Cenpj_ex.vivo_noRT | Glyat | 2.578 | 2.10E-04 | glio |
| Cenpj_ex.vivo_noRT | Meioc | 2.591 | 2.10E-04 | glio |
| Cenpj_ex.vivo_noRT | Tmem132c | 2.591 | 2.10E-04 | glio |
| Cenpj_ex.vivo_noRT | Gm44693 | 2.591 | 2.10E-04 | glio |
| Chek1_in.vivo_noRT | Cep128 | 2.37 | 2.10E-04 | glio |
| Cyp19a1_ex.vivo_noRT | Magea10 | 3.532 | 2.10E-04 | glio |
| Dclre1c_ex.vivo_noRT | D730045A05R | 3.864 | 2.10E-04 | glio |
| Dclre1c_ex.vivo_noRT | Dscaml1 | 2.691 | 2.10E-04 | glio |
| culture_effect | B230303O12R | -2.733 | 2.10E-05 | glio |
| culture_effect | 4930524O07R | 2.183 | 2.10E-05 | glio |
| culture_effect | Ufsp2 | 0.666 | 2.10E-05 | glio |
| Interaction_genotype_Myc | 2210011C24R | -4.525 | 2.10E-05 | glio |
| Interaction_genotype_Myc | Gm48682 | -4.531 | 2.10E-05 | glio |
| Interaction_genotype_Myc | 3110083C13R | -4.759 | 2.10E-05 | glio |
| Atr_ex.vivo_noRT | Map3k9 | 3.174 | 2.10E-05 | glio |
| Bcas2_ex.vivo_noRT | A930019D19R | 2.742 | 2.10E-05 | glio |
| Bora_in.vivo_noRT | Gm14636 | 3.648 | 2.10E-05 | glio |
| culture_effect | Lpar4 | -1.016 | 2.10E-06 | glio |
| Interaction_genotype_Mtor | Iqch | -4.8 | 2.10E-06 | glio |
| Interaction_genotype_Mtor | Gpr4 | -4.099 | 2.10E-06 | glio |
| Interaction_genotype_Mtor | Gm29966 | -3.839 | 2.10E-06 | glio |
| Interaction_genotype_Mtor | Gm33466 | -4.099 | 2.10E-06 | glio |
| Aars2_in.vivo_noRT | Cyb561d1 | 3.519 | 2.10E-06 | glio |
| Atr_in.vivo_noRT | Col17a1 | 3.483 | 2.10E-06 | glio |
| Bora_in.vivo_noRT | Poln | 3.433 | 2.10E-06 | glio |
| Chek1_in.vivo_noRT | Map3k5 | 2.879 | 2.10E-06 | glio |
| Chek1_ex.vivo_noRT | Kif11 | -3.43 | 2.10E-06 | glio |
| Interaction_genotype_Mtor | Gm20625 | -4.099 | 2.10E-07 | glio |
| Atr_in.vivo_noRT | Gm10638 | 3.566 | 2.10E-07 | glio |
| Hsd17b10_in.vivo_noRT | Grin3b | 3.62 | 2.10E-07 | glio |
| Atr_in.vivo_noRT | Ikzf2 | 4.109 | 2.10E-08 | glio |
| Bcas2_in.vivo_noRT | Ppp1r3g | 3.622 | 2.10E-08 | glio |
| Bora_in.vivo_noRT | Pmvk | -2.707 | 2.10E-08 | glio |
| Chek1_in.vivo_noRT | Kctd4 | 3.771 | 2.10E-08 | glio |
| Bcas2_in.vivo_noRT | Tecrl | 4.547 | 2.10E-10 | glio |
| Bora_in.vivo_noRT | Ephx3 | 4.209 | 2.10E-10 | glio |
| Chek1_ex.vivo_noRT | Clca3a2 | 4.842 | 2.10E-10 | glio |
| Aars2_in.vivo_noRT | Nlrx1 | 4.953 | 2.10E-11 | glio |
| Aars2_in.vivo_noRT | Vmn1r181 | 4.671 | 2.10E-12 | glio |
| culture_effect | Chd7 | -1.438 | 2.10E-14 | glio |
| Chek1_in.vivo_noRT | Nwd1 | 4.78 | 2.10E-16 | glio |

| | | | | |
|---|---|---|---|---|
| Bora_in.vivo_noRT | Pnliprp2 | 4.682 | 2.10E-17 | glio |
| Chek1_in.vivo_noRT | Fabp6 | 5.044 | 2.10E-21 | glio |
| Bora_in.vivo_noRT | Olfr552 | 4.936 | 2.10E-22 | glio |
| culture_effect | Rpl5 | -1.377 | 2.10E-26 | glio |
| culture_effect | Pcif1 | 0.573 | 2.11E-03 | glio |
| culture_effect | Ubac1 | 0.517 | 2.11E-03 | glio |
| Interaction_genotype_Aars2 | Slc26a6 | -3.714 | 2.11E-03 | glio |
| Interaction_genotype_Aars2 | Kcmf1 | 1.324 | 2.11E-03 | glio |
| Interaction_genotype_Mtor | Ndufa7 | 1.171 | 2.11E-03 | glio |
| Interaction_genotype_Myc | Slc25a53 | -3.746 | 2.11E-03 | glio |
| Interaction_genotype_Myc | Ovca2 | -3.299 | 2.11E-03 | glio |
| Interaction_genotype_Myc | Nxf2 | -3.557 | 2.11E-03 | glio |
| Interaction_genotype_Myc | Tab1 | -2.945 | 2.11E-03 | glio |
| Interaction_genotype_Myc | Arhgap9 | -2.851 | 2.11E-03 | glio |
| Interaction_genotype_Myc | Pogz | -2.377 | 2.11E-03 | glio |
| Interaction_genotype_Myc | Haus1 | -2.307 | 2.11E-03 | glio |
| Interaction_genotype_Myc | Fkbp2 | 1.415 | 2.11E-03 | glio |
| Interaction_genotype_Pars2 | Dnm3 | -3.361 | 2.11E-03 | glio |
| interaction_RT | E130102H24R | 2.56 | 2.11E-03 | glio |
| interaction_RT | Cttn | -0.879 | 2.11E-03 | glio |
| interaction_RT | Ppfibp1 | -0.629 | 2.11E-03 | glio |
| Aars2_in.vivo_noRT | Slc9b1 | 2.128 | 2.11E-03 | glio |
| Atp5c1_in.vivo_noRT | Col6a3 | 2.159 | 2.11E-03 | glio |
| Atp5c1_in.vivo_noRT | Surf4 | -0.616 | 2.11E-03 | glio |
| Atr_in.vivo_noRT | Ubr5 | -1.076 | 2.11E-03 | glio |
| Atr_in.vivo_noRT | Tuba1b | -0.882 | 2.11E-03 | glio |
| Atr_ex.vivo_noRT | 1810012K16R | 2.78 | 2.11E-03 | glio |
| Atr_ex.vivo_noRT | Atr | -2.466 | 2.11E-03 | glio |
| Bcas2_in.vivo_noRT | Gm10553 | 2.261 | 2.11E-03 | glio |
| Bcas2_in.vivo_noRT | Zfp658 | 2.26 | 2.11E-03 | glio |
| Bcas2_in.vivo_noRT | Depdc1a | -1.762 | 2.11E-03 | glio |
| Bora_in.vivo_noRT | Tmem117 | 2.309 | 2.11E-03 | glio |
| Bora_in.vivo_noRT | Gm37612 | 2.27 | 2.11E-03 | glio |
| Chek1_in.vivo_noRT | Gm26670 | 2.094 | 2.11E-03 | glio |
| Chek1_in.vivo_noRT | Cysltr2 | 2.04 | 2.11E-03 | glio |
| Chek1_ex.vivo_noRT | Kpna7 | 2.443 | 2.11E-03 | glio |
| Eif2s1_in.vivo_noRT | 9230112J17Ri | 2.143 | 2.11E-03 | glio |
| culture_effect | 9530062K07R | 2.222 | 2.11E-04 | glio |
| culture_effect | Hip1 | 0.949 | 2.11E-04 | glio |
| Interaction_genotype_Atr | Col6a1 | -3.411 | 2.11E-04 | glio |
| Interaction_genotype_Atr | Ptpa | 1.852 | 2.11E-04 | glio |
| Interaction_genotype_Mtor | Gm32051 | -4.445 | 2.11E-04 | glio |
| Interaction_genotype_Myc | 6430562O15R | -3.797 | 2.11E-04 | glio |
| Interaction_genotype_Myc | Hint1 | 1.18 | 2.11E-04 | glio |
| Atr_in.vivo_noRT | Mgst1 | 2.677 | 2.11E-04 | glio |
| Bcas2_in.vivo_noRT | 1110013H19R | 2.935 | 2.11E-04 | glio |
| Bcas2_in.vivo_noRT | Slc41a3 | 2.546 | 2.11E-04 | glio |
| Cenpj_ex.vivo_noRT | Vmn1r88 | 3.375 | 2.11E-04 | glio |
| Chek1_in.vivo_noRT | Fxyd4 | 2.928 | 2.11E-04 | glio |
| Chek1_in.vivo_noRT | Bloc1s6os | 2.524 | 2.11E-04 | glio |
| Chek1_ex.vivo_noRT | Ccdc106 | 3.443 | 2.11E-04 | glio |
| Fanca_ex.vivo_noRT | Gm4027 | 3.279 | 2.11E-04 | glio |

| | | | | |
|---|---|---|---|---|
| Fanca_ex.vivo_noRT | BC049987 | 2.913 | 2.11E-04 | glio |
| Fanca_ex.vivo_noRT | Gm20732 | 3.708 | 2.11E-04 | glio |
| Fanca_ex.vivo_noRT | Ank1 | 2.924 | 2.11E-04 | glio |
| culture_effect | Spc24 | -0.818 | 2.11E-05 | glio |
| Interaction_genotype_Mtor | Gm28229 | -4.746 | 2.11E-05 | glio |
| Interaction_genotype_Mtor | Gm45606 | -4.935 | 2.11E-05 | glio |
| Interaction_genotype_Myc | Zfp873 | -4.308 | 2.11E-05 | glio |
| Interaction_genotype_Myc | Wnt10a | -4.76 | 2.11E-05 | glio |
| Interaction_genotype_Myc | Gm15728 | -5.086 | 2.11E-05 | glio |
| Interaction_genotype_Myc | Rplp1 | 1.647 | 2.11E-05 | glio |
| Atr_in.vivo_noRT | Stat4 | 2.873 | 2.11E-05 | glio |
| Atr_ex.vivo_noRT | Cited4 | 4.126 | 2.11E-05 | glio |
| culture_effect | Trmo | -1.651 | 2.11E-06 | glio |
| Atr_in.vivo_noRT | Gm13830 | 3.603 | 2.11E-06 | glio |
| Atr_in.vivo_noRT | Clns1a | -1.721 | 2.11E-06 | glio |
| Hsd17b10_in.vivo_noRT | Ramp3 | 3.35 | 2.11E-06 | glio |
| Atr_in.vivo_noRT | Dynlt1b | 3.585 | 2.11E-07 | glio |
| Bcas2_in.vivo_noRT | Gm36495 | 4.042 | 2.11E-07 | glio |
| Bora_in.vivo_noRT | Zfp712 | 3.241 | 2.11E-07 | glio |
| Bora_in.vivo_noRT | Rce1 | -2.082 | 2.11E-07 | glio |
| Chek1_in.vivo_noRT | Gm4755 | 3.286 | 2.11E-07 | glio |
| Chek1_in.vivo_noRT | Gm20522 | 3.767 | 2.11E-07 | glio |
| Atr_ex.vivo_noRT | AF067061 | 4.309 | 2.11E-08 | glio |
| Aars2_in.vivo_noRT | 4922502N22R | 4.344 | 2.11E-10 | glio |
| culture_effect | Adam12 | 3.159 | 2.11E-11 | glio |
| Bora_in.vivo_noRT | Zmynd15 | 4.557 | 2.11E-11 | glio |
| Chek1_in.vivo_noRT | Myoz1 | 3.878 | 2.11E-11 | glio |
| Chek1_ex.vivo_noRT | Smtnl1 | 4.842 | 2.11E-13 | glio |
| Aars2_in.vivo_noRT | Ric3 | 5.067 | 2.11E-16 | glio |
| Atr_in.vivo_noRT | Tcp11x2 | 5.011 | 2.11E-17 | glio |
| Bcas2_in.vivo_noRT | Gm10354 | 4.807 | 2.11E-17 | glio |
| culture_effect | Ndufa9 | -1.155 | 2.11E-18 | glio |
| Bora_in.vivo_noRT | Gm48140 | 4.673 | 2.11E-20 | glio |
| Bora_in.vivo_noRT | 1700034I23Ri | 4.936 | 2.11E-23 | glio |
| culture_effect | Fam234a | -0.817 | 2.12E-03 | glio |
| Interaction_genotype_Atr | Foxd2 | -3.392 | 2.12E-03 | glio |
| Interaction_genotype_Bcas2 | Glyat | -2.862 | 2.12E-03 | glio |
| Interaction_genotype_Mtor | Igdcc4 | -3.317 | 2.12E-03 | glio |
| Interaction_genotype_Mtor | Zfp971 | -3.32 | 2.12E-03 | glio |
| Interaction_genotype_Myc | Inpp1 | -2.489 | 2.12E-03 | glio |
| Interaction_genotype_Myc | Edil3 | -2.704 | 2.12E-03 | glio |
| Interaction_genotype_Myc | Timm10b | 1.29 | 2.12E-03 | glio |
| Interaction_genotype_Myc | Rpl13a | 1.113 | 2.12E-03 | glio |
| Interaction_genotype_Pars2 | 1700003G18R | -4.629 | 2.12E-03 | glio |
| Aars2_in.vivo_noRT | Pcdhb21 | 2.059 | 2.12E-03 | glio |
| Aars2_ex.vivo_noRT | Vmn1r181 | 2.897 | 2.12E-03 | glio |
| Aars2_ex.vivo_noRT | Sbk2 | 3.087 | 2.12E-03 | glio |
| Aars2_ex.vivo_noRT | Calhm6 | 2.713 | 2.12E-03 | glio |
| Atp5c1_in.vivo_noRT | Gm16364.1 | 2.668 | 2.12E-03 | glio |
| Atp5c1_in.vivo_noRT | Ramp3 | 2.402 | 2.12E-03 | glio |
| Atp5c1_in.vivo_noRT | D6Ertd527e | 2.413 | 2.12E-03 | glio |
| Atr_in.vivo_noRT | Nagpa | 1.894 | 2.12E-03 | glio |

| | | | | |
|---|---|---|---|---|
| Atr_ex.vivo_noRT | Gm27241 | 2.183 | 2.12E-03 | glio |
| Atr_ex.vivo_noRT | Nap1l1 | -0.745 | 2.12E-03 | glio |
| Bard1_ex.vivo_noRT | Gm35439 | 2.415 | 2.12E-03 | glio |
| Bard1_ex.vivo_noRT | Pacrg | 2.802 | 2.12E-03 | glio |
| Bard1_ex.vivo_noRT | Ocm | 2.814 | 2.12E-03 | glio |
| Bard1_ex.vivo_noRT | Bard1 | -2.572 | 2.12E-03 | glio |
| Bard1_ex.vivo_noRT | Tgm2 | -2.257 | 2.12E-03 | glio |
| Bcas2_in.vivo_noRT | Slc2a6 | 2.419 | 2.12E-03 | glio |
| Bcas2_in.vivo_noRT | Dnah2 | 2.276 | 2.12E-03 | glio |
| Bora_in.vivo_noRT | Puf60 | -0.902 | 2.12E-03 | glio |
| Cenpj_in.vivo_noRT | Olfr1152 | 1.756 | 2.12E-03 | glio |
| Chek1_in.vivo_noRT | Hspbap1 | 1.4 | 2.12E-03 | glio |
| Chek1_in.vivo_noRT | Tpgs1 | 0.964 | 2.12E-03 | glio |
| Chek1_in.vivo_noRT | Gdi2 | -0.87 | 2.12E-03 | glio |
| Chek1_in.vivo_noRT | Sub1 | -0.539 | 2.12E-03 | glio |
| Chek1_ex.vivo_noRT | Prdm2 | 1.489 | 2.12E-03 | glio |
| Eif2s1_in.vivo_noRT | Gm20711 | 2.143 | 2.12E-03 | glio |
| Eif2s1_in.vivo_noRT | Cyp4x1 | 2.408 | 2.12E-03 | glio |
| Hsd17b10_in.vivo_noRT | Myl6b | 2.274 | 2.12E-03 | glio |
| culture_effect | Kcnk7 | -1.983 | 2.12E-04 | glio |
| culture_effect | Mfsd3 | 1.37 | 2.12E-04 | glio |
| Interaction_genotype_Mtor | Cfap58 | -5.459 | 2.12E-04 | glio |
| Interaction_genotype_Mtor | Mrpl2 | 1.783 | 2.12E-04 | glio |
| Aars2_in.vivo_noRT | Adat1 | 2.189 | 2.12E-04 | glio |
| Aars2_ex.vivo_noRT | 2310003N18R | 2.9 | 2.12E-04 | glio |
| Aars2_ex.vivo_noRT | AU022754 | 3.293 | 2.12E-04 | glio |
| Aars2_ex.vivo_noRT | Bcl2l14 | 3.766 | 2.12E-04 | glio |
| Atr_in.vivo_noRT | Ddah2 | 2.658 | 2.12E-04 | glio |
| Bard1_ex.vivo_noRT | Pspn | 2.419 | 2.12E-04 | glio |
| Bcas2_in.vivo_noRT | Abcd2 | 2.968 | 2.12E-04 | glio |
| Bcas2_in.vivo_noRT | Gm43707 | 2.336 | 2.12E-04 | glio |
| Bcas2_in.vivo_noRT | Cnot9 | -1.721 | 2.12E-04 | glio |
| Bcas2_in.vivo_noRT | Arfip2 | -1.663 | 2.12E-04 | glio |
| Bora_in.vivo_noRT | Gm15879 | 3.443 | 2.12E-04 | glio |
| Bora_in.vivo_noRT | Kctd7 | 2.467 | 2.12E-04 | glio |
| Bora_in.vivo_noRT | Rexo1 | -1.602 | 2.12E-04 | glio |
| Chek1_in.vivo_noRT | Egf | 2.852 | 2.12E-04 | glio |
| Chek1_ex.vivo_noRT | Gm37800 | 2.976 | 2.12E-04 | glio |
| Hsd17b10_in.vivo_noRT | Alox12 | 2.828 | 2.12E-04 | glio |
| culture_effect | Slc14a1 | -2.245 | 2.12E-05 | glio |
| culture_effect | Cir1 | -0.716 | 2.12E-05 | glio |
| culture_effect | Rpl30 | -0.556 | 2.12E-05 | glio |
| Interaction_genotype_Mtor | 6430503K07R | -4.709 | 2.12E-05 | glio |
| Interaction_genotype_Mtor | Gm17597 | -5.272 | 2.12E-05 | glio |
| Interaction_genotype_Mtor | Gm29243 | -4.583 | 2.12E-05 | glio |
| Interaction_genotype_Mtor | Myom1 | -5.321 | 2.12E-05 | glio |
| Interaction_genotype_Myc | Cysrt1 | -5.024 | 2.12E-05 | glio |
| Interaction_genotype_Sass6 | Klrk1 | 4.945 | 2.12E-05 | glio |
| Aars2_in.vivo_noRT | Pabpc4l | 3.048 | 2.12E-05 | glio |
| Aars2_in.vivo_noRT | Plvap | 2.954 | 2.12E-05 | glio |
| Atp5c1_in.vivo_noRT | AC159140.1 | 2.408 | 2.12E-05 | glio |
| Bora_in.vivo_noRT | Upk1a | 3.144 | 2.12E-05 | glio |

| | | | | |
|---|---|---|---|---|
| Chek1_in.vivo_noRT | Haglr | 3.263 | 2.12E-05 | glio |
| Chek1_in.vivo_noRT | Mertk | 2.911 | 2.12E-05 | glio |
| Cyp19a1_ex.vivo_noRT | Gm20658 | 2.959 | 2.12E-05 | glio |
| Cyp19a1_ex.vivo_noRT | Olfr56 | 3.336 | 2.12E-05 | glio |
| Cyp19a1_ex.vivo_noRT | 2310003N18R | 3.691 | 2.12E-05 | glio |
| culture_effect | Ptov1 | -0.553 | 2.12E-06 | glio |
| Interaction_genotype_Myc | Ppp1r18os | -5.749 | 2.12E-06 | glio |
| Aars2_in.vivo_noRT | Gm14399 | 3.346 | 2.12E-06 | glio |
| Atr_ex.vivo_noRT | Gm19276 | 2.941 | 2.12E-06 | glio |
| Bcas2_in.vivo_noRT | Gm8098 | 3.304 | 2.12E-06 | glio |
| Bora_ex.vivo_noRT | Gm17276 | 3.598 | 2.12E-06 | glio |
| Chek1_ex.vivo_noRT | Bmp2 | 4.049 | 2.12E-06 | glio |
| Aars2_in.vivo_noRT | Psmc6 | -1.61 | 2.12E-07 | glio |
| Aars2_ex.vivo_noRT | Gm26978 | 3.882 | 2.12E-07 | glio |
| Bcas2_in.vivo_noRT | Spink2 | 4.14 | 2.12E-07 | glio |
| Chek1_in.vivo_noRT | Gm26916 | 3.467 | 2.12E-07 | glio |
| Atr_in.vivo_noRT | Gm46447 | 3.722 | 2.12E-09 | glio |
| Chek1_ex.vivo_noRT | Col6a3 | 4.849 | 2.12E-10 | glio |
| Chek1_ex.vivo_noRT | Gm128 | 4.449 | 2.12E-10 | glio |
| Aars2_in.vivo_noRT | Spns3 | 4.669 | 2.12E-11 | glio |
| Hsd17b10_in.vivo_noRT | Akap14 | 3.884 | 2.12E-12 | glio |
| Bora_in.vivo_noRT | Gm47851 | 4.676 | 2.12E-13 | glio |
| Bcas2_in.vivo_noRT | Npy1r | 4.162 | 2.12E-14 | glio |
| Chek1_in.vivo_noRT | Gm42849 | 4.777 | 2.12E-14 | glio |
| culture_effect | Rgcc | -1.672 | 2.12E-15 | glio |
| Bora_in.vivo_noRT | Mcmdc2 | 4.676 | 2.12E-15 | glio |
| Aars2_in.vivo_noRT | D930028M14I | 5.327 | 2.12E-17 | glio |
| Bora_in.vivo_noRT | 4930553M12F | 4.67 | 2.12E-17 | glio |
| Atr_in.vivo_noRT | Gm47622 | 5.533 | 2.12E-25 | glio |
| culture_effect | Acot1 | -2.453 | 2.12E-29 | glio |
| culture_effect | S100a11 | 2.057 | 2.12E-33 | glio |
| culture_effect | Gm29670 | -1.457 | 2.13E-03 | glio |
| culture_effect | Hdac7 | 1.529 | 2.13E-03 | glio |
| culture_effect | Ccdc180 | 1.292 | 2.13E-03 | glio |
| culture_effect | Ddx60 | -0.758 | 2.13E-03 | glio |
| culture_effect | Spry2 | -0.739 | 2.13E-03 | glio |
| Interaction_genotype_Aars2 | 1810058I24Ri | 1.269 | 2.13E-03 | glio |
| Interaction_genotype_Atr | Myo19 | -3.095 | 2.13E-03 | glio |
| Interaction_genotype_Bora | Slc25a35 | -4.03 | 2.13E-03 | glio |
| Interaction_genotype_Bora | Gm16364.1 | -4.356 | 2.13E-03 | glio |
| Interaction_genotype_Mtor | Slmap | -2.053 | 2.13E-03 | glio |
| Interaction_genotype_Mtor | Usp1 | 1.748 | 2.13E-03 | glio |
| Interaction_genotype_Myc | Chrdl2 | -3.226 | 2.13E-03 | glio |
| Interaction_genotype_Myc | Akr1e1 | -2.094 | 2.13E-03 | glio |
| Aars2_in.vivo_noRT | Stamos | 2.012 | 2.13E-03 | glio |
| Aars2_in.vivo_noRT | Cerkl | 1.946 | 2.13E-03 | glio |
| Aars2_in.vivo_noRT | 9230114K14R | 1.869 | 2.13E-03 | glio |
| Bora_in.vivo_noRT | Gm12359 | 2.244 | 2.13E-03 | glio |
| Bora_in.vivo_noRT | Impdh1 | -1.52 | 2.13E-03 | glio |
| Bora_in.vivo_noRT | Cmc1 | -1.32 | 2.13E-03 | glio |
| Chek1_ex.vivo_noRT | Gm15853 | 2.853 | 2.13E-03 | glio |
| Eif2s1_in.vivo_noRT | Rassf6 | 2.143 | 2.13E-03 | glio |

| | | | | |
|---|---|---|---|---|
| Hsd17b10_in.vivo_noRT | Slc25a43 | 2.346 | 2.13E-03 | glio |
| culture_effect | Btc | 2.412 | 2.13E-04 | glio |
| culture_effect | Nr2f6 | 0.64 | 2.13E-04 | glio |
| Interaction_genotype_Bcas2 | Gm38534 | -4.599 | 2.13E-04 | glio |
| Interaction_genotype_Bcas2 | D030040B21R | -4.587 | 2.13E-04 | glio |
| Interaction_genotype_Bcas2 | Gm43156 | -4.827 | 2.13E-04 | glio |
| Interaction_genotype_Bcas2 | Apoh | -4.805 | 2.13E-04 | glio |
| Interaction_genotype_Bcas2 | 4930589L23Ri | -4.889 | 2.13E-04 | glio |
| Interaction_genotype_Bcas2 | Plcl1 | -4.018 | 2.13E-04 | glio |
| Interaction_genotype_Bcas2 | 1700015F17Ri | -4.906 | 2.13E-04 | glio |
| Interaction_genotype_Mtor | Sgms2 | -4.581 | 2.13E-04 | glio |
| Interaction_genotype_Mtor | 2010110K18R | -4.143 | 2.13E-04 | glio |
| Aars2_in.vivo_noRT | Vma21 | -1.068 | 2.13E-04 | glio |
| Atr_ex.vivo_noRT | Ocm | 3.164 | 2.13E-04 | glio |
| Atr_ex.vivo_noRT | Hist1h2ac | 2.925 | 2.13E-04 | glio |
| Bcas2_in.vivo_noRT | Uchl5 | -1.463 | 2.13E-04 | glio |
| Cenpj_in.vivo_noRT | Capn13 | 2.022 | 2.13E-04 | glio |
| Cenpj_in.vivo_noRT | B3gnt4 | 2.813 | 2.13E-04 | glio |
| Cenpj_in.vivo_noRT | Gm48170 | 2.551 | 2.13E-04 | glio |
| Chek1_in.vivo_noRT | Slc2a13 | 2.631 | 2.13E-04 | glio |
| Cyp19a1_ex.vivo_noRT | Gm15270 | 3.317 | 2.13E-04 | glio |
| culture_effect | Haus8 | 0.822 | 2.13E-05 | glio |
| culture_effect | Creb5 | -0.888 | 2.13E-05 | glio |
| Atp5c1_in.vivo_noRT | 2310003N18R | 2.668 | 2.13E-05 | glio |
| Bora_in.vivo_noRT | Mmp2 | -2.216 | 2.13E-05 | glio |
| Hsd17b10_in.vivo_noRT | 2810410L24Ri | 3.09 | 2.13E-05 | glio |
| Aars2_ex.vivo_noRT | Rpl3l | 4.069 | 2.13E-06 | glio |
| Aars2_ex.vivo_noRT | Gm21936 | 2.897 | 2.13E-06 | glio |
| Aars2_ex.vivo_noRT | 4930588A03R | 2.893 | 2.13E-06 | glio |
| Bcas2_in.vivo_noRT | Gm36608 | 3.404 | 2.13E-06 | glio |
| Chek1_in.vivo_noRT | E030030I06Ril | -2.248 | 2.13E-06 | glio |
| Chek1_ex.vivo_noRT | Gja4 | 4.058 | 2.13E-06 | glio |
| Interaction_genotype_Mtor | Ifitm6 | -6.322 | 2.13E-07 | glio |
| Interaction_genotype_Mtor | 5730507C01R | -4.767 | 2.13E-07 | glio |
| Interaction_genotype_Mtor | Gm5157 | -4.099 | 2.13E-07 | glio |
| Atr_in.vivo_noRT | Mapk15 | 3.549 | 2.13E-07 | glio |
| Atr_in.vivo_noRT | Rbp7 | 3.645 | 2.13E-07 | glio |
| Atr_in.vivo_noRT | Gm19409 | 3.954 | 2.13E-08 | glio |
| Bora_in.vivo_noRT | 4933421A08R | 4.157 | 2.13E-08 | glio |
| Bora_in.vivo_noRT | Gm47428 | 3.76 | 2.13E-08 | glio |
| Bcas2_in.vivo_noRT | A930012O16R | 4.547 | 2.13E-10 | glio |
| Hsd17b10_in.vivo_noRT | Dmrtc1a | 3.616 | 2.13E-11 | glio |
| Hsd17b10_in.vivo_noRT | Gm12589 | 3.616 | 2.13E-11 | glio |
| Atr_in.vivo_noRT | Ushbp1 | 5.008 | 2.13E-13 | glio |
| Chek1_in.vivo_noRT | Mettl7b | 5.044 | 2.13E-13 | glio |
| Bcas2_in.vivo_noRT | Psca | 4.807 | 2.13E-14 | glio |
| Chek1_in.vivo_noRT | Lrp2bp | 4.78 | 2.13E-14 | glio |
| Aars2_in.vivo_noRT | Cxcl13 | 5.327 | 2.13E-15 | glio |
| Aars2_in.vivo_noRT | Nkg7 | 5.067 | 2.13E-17 | glio |
| Atr_in.vivo_noRT | Dcdc2a | 5.273 | 2.13E-18 | glio |
| Bcas2_in.vivo_noRT | Zscan4-ps2 | 4.807 | 2.13E-20 | glio |
| Chek1_in.vivo_noRT | Gm36037 | 5.044 | 2.13E-22 | glio |

| | | | | |
|---|---|---|---|---|
| culture_effect | Cptp | 0.858 | 2.14E-03 | glio |
| culture_effect | Met | -0.879 | 2.14E-03 | glio |
| culture_effect | Ssna1 | 0.393 | 2.14E-03 | glio |
| Interaction_genotype_Aars2 | Gm32591 | -3.764 | 2.14E-03 | glio |
| Interaction_genotype_Aars2 | Taldo1 | 1.266 | 2.14E-03 | glio |
| Interaction_genotype_Atr | 4930522L14Ri | -3.904 | 2.14E-03 | glio |
| Interaction_genotype_Atr | Yae1d1 | 1.843 | 2.14E-03 | glio |
| Interaction_genotype_Mtor | Prob1 | -3.916 | 2.14E-03 | glio |
| Interaction_genotype_Mtor | Armc7 | -3.251 | 2.14E-03 | glio |
| Interaction_genotype_Mtor | Tatdn2 | -3.06 | 2.14E-03 | glio |
| Interaction_genotype_Mtor | 2410022M11F | -3.372 | 2.14E-03 | glio |
| Interaction_genotype_Mtor | Zfp719 | -2.796 | 2.14E-03 | glio |
| Interaction_genotype_Myc | Gm4027 | -2.729 | 2.14E-03 | glio |
| Interaction_genotype_Myc | Gm20633 | -3.185 | 2.14E-03 | glio |
| Interaction_genotype_Myc | Ccdc14 | -3.191 | 2.14E-03 | glio |
| Interaction_genotype_Rad51 | Fbln5 | 2.866 | 2.14E-03 | glio |
| interaction_RT | Zfp87 | 1.749 | 2.14E-03 | glio |
| Atr_in.vivo_noRT | Pola1 | -1.253 | 2.14E-03 | glio |
| Atr_in.vivo_noRT | Eif5b | -1.046 | 2.14E-03 | glio |
| Atr_ex.vivo_noRT | Spn | 2.77 | 2.14E-03 | glio |
| Atr_ex.vivo_noRT | Gm43948 | 2.38 | 2.14E-03 | glio |
| Atr_ex.vivo_noRT | Alox15 | 2.769 | 2.14E-03 | glio |
| Atr_ex.vivo_noRT | Tecrl | 2.865 | 2.14E-03 | glio |
| Bcas2_in.vivo_noRT | Eno1 | -1.148 | 2.14E-03 | glio |
| Bora_ex.vivo_noRT | Dcpp1 | 3.201 | 2.14E-03 | glio |
| Bora_ex.vivo_noRT | Txndc2 | 2.413 | 2.14E-03 | glio |
| Cenpj_ex.vivo_noRT | Magea10 | 2.809 | 2.14E-03 | glio |
| Chek1_in.vivo_noRT | Sap18 | -1.176 | 2.14E-03 | glio |
| Chek1_ex.vivo_noRT | Echs1 | -1.235 | 2.14E-03 | glio |
| Chek1_ex.vivo_noRT | Pdap1 | -0.804 | 2.14E-03 | glio |
| Hsd17b10_in.vivo_noRT | Igsf23 | 2.483 | 2.14E-03 | glio |
| interaction_RT | Per1 | 2.114 | 2.14E-04 | glio |
| Aars2_in.vivo_noRT | Itprid2 | 1.766 | 2.14E-04 | glio |
| Bard1_ex.vivo_noRT | Nt5e | 3.203 | 2.14E-04 | glio |
| Bcas2_in.vivo_noRT | Eml6 | 2.724 | 2.14E-04 | glio |
| Bora_in.vivo_noRT | Nup210l | 2.809 | 2.14E-04 | glio |
| Bora_in.vivo_noRT | Hddc3 | 2.553 | 2.14E-04 | glio |
| Bora_in.vivo_noRT | Car7 | 2.621 | 2.14E-04 | glio |
| Chek1_in.vivo_noRT | March11 | 2.77 | 2.14E-04 | glio |
| Chek1_in.vivo_noRT | Vps35 | -1.149 | 2.14E-04 | glio |
| Interaction_genotype_Mtor | 1700040D17R | -5.398 | 2.14E-05 | glio |
| Aars2_in.vivo_noRT | Gm48768 | 3.075 | 2.14E-05 | glio |
| Aars2_in.vivo_noRT | Gm15601 | 3.115 | 2.14E-05 | glio |
| Aars2_in.vivo_noRT | Trim30a | -1.922 | 2.14E-05 | glio |
| Atr_in.vivo_noRT | Hist1h2af | 2.739 | 2.14E-05 | glio |
| Bcas2_in.vivo_noRT | C77080 | 2.93 | 2.14E-05 | glio |
| Chek1_in.vivo_noRT | 9430065F17Ri | 3.068 | 2.14E-05 | glio |
| Chek1_ex.vivo_noRT | Dppa2 | 3.475 | 2.14E-05 | glio |
| culture_effect | Gpn3 | -0.916 | 2.14E-06 | glio |
| Interaction_genotype_Mtor | Gm50107 | -5.308 | 2.14E-06 | glio |
| Interaction_genotype_Mtor | Psme1 | 2.715 | 2.14E-06 | glio |
| Interaction_genotype_Myc | G630030J09Ri | -5.263 | 2.14E-06 | glio |

| | | | | |
|---|---|---|---|---|
| Atp5c1_in.vivo_noRT | Gpr22 | 2.668 | 2.14E-06 | glio |
| Atr_in.vivo_noRT | Olfr655 | 4.617 | 2.14E-06 | glio |
| Bcas2_in.vivo_noRT | B930059L03Ri | 3.622 | 2.14E-06 | glio |
| Bcas2_ex.vivo_noRT | Slc17a6 | 2.742 | 2.14E-06 | glio |
| culture_effect | Tgfbi | -2.638 | 2.14E-07 | glio |
| Bcas2_in.vivo_noRT | Cyp26b1 | 3.882 | 2.14E-07 | glio |
| Chek1_ex.vivo_noRT | Nat8f4 | 4.055 | 2.14E-07 | glio |
| Interaction_genotype_Mtor | Fgf7 | -6.634 | 2.14E-08 | glio |
| Aars2_in.vivo_noRT | Ifit1bl1 | 4.794 | 2.14E-08 | glio |
| Aars2_in.vivo_noRT | Wnt6 | 4.661 | 2.14E-08 | glio |
| Aars2_in.vivo_noRT | Elfn2 | 3.552 | 2.14E-08 | glio |
| Atr_in.vivo_noRT | Hspb9 | 4.222 | 2.14E-08 | glio |
| Atr_in.vivo_noRT | Gchfr | 4.421 | 2.14E-08 | glio |
| Atr_in.vivo_noRT | Sept5 | 3.794 | 2.14E-08 | glio |
| Bcas2_in.vivo_noRT | Gm43625 | 3.502 | 2.14E-08 | glio |
| Chek1_ex.vivo_noRT | Masp1 | 4.045 | 2.14E-08 | glio |
| culture_effect | Fam104a | 0.6 | 2.14E-09 | glio |
| Bora_in.vivo_noRT | 5830454E08Ri | 4.011 | 2.14E-09 | glio |
| Atr_ex.vivo_noRT | Gm10482 | 4.746 | 2.14E-11 | glio |
| Chek1_in.vivo_noRT | Olfr279 | 4.515 | 2.14E-11 | glio |
| Hsd17b10_in.vivo_noRT | Clec12b | 3.616 | 2.14E-13 | glio |
| Hsd17b10_in.vivo_noRT | Gm14929 | 3.616 | 2.14E-13 | glio |
| Aars2_in.vivo_noRT | Fam25c | 4.794 | 2.14E-18 | glio |
| culture_effect | Snx10 | -1.404 | 2.14E-19 | glio |
| Bcas2_in.vivo_noRT | Sema6d | 4.807 | 2.14E-19 | glio |
| Aars2_in.vivo_noRT | Ccdc125 | 5.327 | 2.14E-20 | glio |
| Aars2_in.vivo_noRT | Spink8 | 5.067 | 2.14E-22 | glio |
| Bcas2_in.vivo_noRT | 1700003E24Ri | 4.807 | 2.14E-24 | glio |
| culture_effect | Hist1h2ag | -1.467 | 2.15E-03 | glio |
| culture_effect | Lyz2 | 1.889 | 2.15E-03 | glio |
| culture_effect | Gm30082 | 1.789 | 2.15E-03 | glio |
| culture_effect | Numbl | 0.81 | 2.15E-03 | glio |
| Interaction_genotype_Mtor | Gm41760 | -3.43 | 2.15E-03 | glio |
| Interaction_genotype_Mtor | Top2a | 2.974 | 2.15E-03 | glio |
| Interaction_genotype_Myc | 5330417C22R | -3.432 | 2.15E-03 | glio |
| Interaction_genotype_Myc | Gm11696 | -3.42 | 2.15E-03 | glio |
| Interaction_genotype_Myc | Icam4 | -3.275 | 2.15E-03 | glio |
| Interaction_genotype_Myc | Brf1 | -2.737 | 2.15E-03 | glio |
| interaction_RT | Nfil3 | 1.483 | 2.15E-03 | glio |
| interaction_RT | Ltb | 2.811 | 2.15E-03 | glio |
| interaction_RT | Mcrip1 | 0.614 | 2.15E-03 | glio |
| Aars2_in.vivo_noRT | Hba-a1 | 2.685 | 2.15E-03 | glio |
| Aars2_in.vivo_noRT | Mtcl1 | 2.144 | 2.15E-03 | glio |
| Aars2_in.vivo_noRT | Pou6f1 | 2.104 | 2.15E-03 | glio |
| Aars2_in.vivo_noRT | Plp2 | -1.099 | 2.15E-03 | glio |
| Aars2_ex.vivo_noRT | Tec | 2.817 | 2.15E-03 | glio |
| Atp5c1_in.vivo_noRT | Kcnd1 | 2.146 | 2.15E-03 | glio |
| Atr_in.vivo_noRT | Dhrs3 | 2.236 | 2.15E-03 | glio |
| Atr_in.vivo_noRT | Stx6 | 1.612 | 2.15E-03 | glio |
| Atr_in.vivo_noRT | Tufm | -1.281 | 2.15E-03 | glio |
| Atr_ex.vivo_noRT | Nrn1l | 2.764 | 2.15E-03 | glio |
| Bcas2_in.vivo_noRT | Myorg | 2.208 | 2.15E-03 | glio |

| | | | | |
|---|---|---|---|---|
| Bcas2_in.vivo_noRT | 2900009J06Ri | 2.186 | 2.15E-03 | glio |
| Bcas2_in.vivo_noRT | Tigd5 | 2.045 | 2.15E-03 | glio |
| Cenpj_ex.vivo_noRT | Muc4 | 2.591 | 2.15E-03 | glio |
| Chek1_in.vivo_noRT | A630052C17R | 2.172 | 2.15E-03 | glio |
| Chek1_in.vivo_noRT | Magi2 | 2.2 | 2.15E-03 | glio |
| Chek1_in.vivo_noRT | Fdft1 | -1.713 | 2.15E-03 | glio |
| Chek1_in.vivo_noRT | Swi5 | -0.595 | 2.15E-03 | glio |
| Chek1_ex.vivo_noRT | Mak16 | -1 | 2.15E-03 | glio |
| Hsd17b10_in.vivo_noRT | Zdhhc20 | -1.148 | 2.15E-03 | glio |
| culture_effect | Etaa1os | 1.723 | 2.15E-04 | glio |
| Interaction_genotype_Myc | Crb3 | -3.272 | 2.15E-04 | glio |
| Aars2_in.vivo_noRT | Amz1 | 2.535 | 2.15E-04 | glio |
| Aars2_ex.vivo_noRT | Tcstv3 | 3.293 | 2.15E-04 | glio |
| Atr_ex.vivo_noRT | Slc22a1 | 2.769 | 2.15E-04 | glio |
| Bora_in.vivo_noRT | Tnfaip2 | 2.729 | 2.15E-04 | glio |
| Bora_in.vivo_noRT | Cpsf3 | -1.618 | 2.15E-04 | glio |
| culture_effect | Gm11713 | -2.38 | 2.15E-05 | glio |
| culture_effect | Gm43303 | -1.955 | 2.15E-05 | glio |
| culture_effect | Tulp2 | -1.195 | 2.15E-05 | glio |
| culture_effect | Ssbp4 | -1.114 | 2.15E-05 | glio |
| Interaction_genotype_Mtor | 1810041H14R | -5.331 | 2.15E-05 | glio |
| Aars2_ex.vivo_noRT | Gm15270 | 3.275 | 2.15E-05 | glio |
| Bcas2_in.vivo_noRT | Dock4 | 3.044 | 2.15E-05 | glio |
| Bcas2_in.vivo_noRT | Gm20186 | 3.174 | 2.15E-05 | glio |
| Chek1_ex.vivo_noRT | Serbp1 | -0.931 | 2.15E-05 | glio |
| culture_effect | Pnp | -0.754 | 2.15E-06 | glio |
| culture_effect | Sox2 | 0.636 | 2.15E-06 | glio |
| Atr_in.vivo_noRT | Shpk | 3.546 | 2.15E-06 | glio |
| Atr_in.vivo_noRT | Plxnb1 | 3.401 | 2.15E-06 | glio |
| Atr_in.vivo_noRT | Itgb7 | 3.116 | 2.15E-06 | glio |
| Bora_in.vivo_noRT | Gm9833 | 3.155 | 2.15E-06 | glio |
| Hsd17b10_in.vivo_noRT | Rcan2 | 3.616 | 2.15E-06 | glio |
| culture_effect | Gm49785 | 2.646 | 2.15E-07 | glio |
| culture_effect | Npr1 | 0.983 | 2.15E-07 | glio |
| interaction_RT | Gpatch4 | -1.256 | 2.15E-07 | glio |
| interaction_RT | Spcs1 | 0.946 | 2.15E-07 | glio |
| Atr_in.vivo_noRT | Ola1 | -1.622 | 2.15E-07 | glio |
| Bora_in.vivo_noRT | 9130604C24R | 4.003 | 2.15E-07 | glio |
| Chek1_ex.vivo_noRT | Npr3 | 3.871 | 2.15E-07 | glio |
| Chek1_ex.vivo_noRT | Pcdhb21 | 4.433 | 2.15E-07 | glio |
| Atr_in.vivo_noRT | Gm10767 | 3.933 | 2.15E-08 | glio |
| Atr_in.vivo_noRT | Mrps25 | -1.878 | 2.15E-08 | glio |
| Chek1_ex.vivo_noRT | Magea10 | 4.263 | 2.15E-08 | glio |
| Aars2_in.vivo_noRT | Rtl9 | 3.885 | 2.15E-09 | glio |
| Atr_in.vivo_noRT | Ccno | 4.513 | 2.15E-10 | glio |
| Aars2_in.vivo_noRT | Hif3a | 4.684 | 2.15E-11 | glio |
| Aars2_in.vivo_noRT | Pecam1 | 4.794 | 2.15E-11 | glio |
| Bcas2_in.vivo_noRT | Lce1g | 4.807 | 2.15E-11 | glio |
| Chek1_in.vivo_noRT | Krtdap | 4.653 | 2.15E-11 | glio |
| culture_effect | Gm4013 | 1.836 | 2.15E-13 | glio |
| Bcas2_in.vivo_noRT | Flacc1 | 4.807 | 2.15E-15 | glio |
| Bora_in.vivo_noRT | Gm19265 | 4.42 | 2.15E-15 | glio |

| | | | | |
|---|---|---|---|---|
| Chek1_ex.vivo_noRT | Angptl7 | 4.842 | 2.15E-15 | glio |
| culture_effect | 9330020H09R | 3.123 | 2.15E-17 | glio |
| Aars2_in.vivo_noRT | Gm36198 | 5.067 | 2.15E-18 | glio |
| Atr_in.vivo_noRT | Magea10 | 5.576 | 2.15E-19 | glio |
| Atr_in.vivo_noRT | Gm13713 | 5.001 | 2.15E-19 | glio |
| culture_effect | Tmbim6 | -1.172 | 2.15E-20 | glio |
| Atr_in.vivo_noRT | 4930470G03R | 5.267 | 2.15E-21 | glio |
| culture_effect | 2610016A17R | 1.73 | 2.16E-03 | glio |
| culture_effect | Abhd16a | -0.561 | 2.16E-03 | glio |
| culture_effect | Tmed1 | -0.563 | 2.16E-03 | glio |
| culture_effect | Iffo2 | 0.623 | 2.16E-03 | glio |
| Interaction_genotype_Atr | Prickle4 | -3.637 | 2.16E-03 | glio |
| Interaction_genotype_Atr | Ifit1bl1 | -4.631 | 2.16E-03 | glio |
| Interaction_genotype_Atr | Sertad1 | 1.983 | 2.16E-03 | glio |
| Interaction_genotype_Mtor | Csnk1e | 1.623 | 2.16E-03 | glio |
| Interaction_genotype_Myc | Nat8f2 | -2.865 | 2.16E-03 | glio |
| Interaction_genotype_Myc | Bcl2l15 | -3.044 | 2.16E-03 | glio |
| Interaction_genotype_Myc | 9030025P20R | -3.653 | 2.16E-03 | glio |
| Interaction_genotype_Myc | Zfp346 | -3.134 | 2.16E-03 | glio |
| Aars2_in.vivo_noRT | Tigd5 | 2.012 | 2.16E-03 | glio |
| Atp5c1_in.vivo_noRT | Gm50394 | 2.402 | 2.16E-03 | glio |
| Atr_in.vivo_noRT | Tra2a | -1.113 | 2.16E-03 | glio |
| Bcas2_ex.vivo_noRT | Icam2 | 2.742 | 2.16E-03 | glio |
| Bora_in.vivo_noRT | Zfp39 | 2.291 | 2.16E-03 | glio |
| Bora_in.vivo_noRT | Gstm7 | 2.248 | 2.16E-03 | glio |
| Bora_in.vivo_noRT | Smarcc2 | -1.324 | 2.16E-03 | glio |
| Bora_in.vivo_noRT | Uchl3 | -1.105 | 2.16E-03 | glio |
| Cenpj_in.vivo_noRT | Hfm1 | 2.022 | 2.16E-03 | glio |
| Cenpj_in.vivo_noRT | D630024D03R | 2.022 | 2.16E-03 | glio |
| Chek1_in.vivo_noRT | Ccr2 | 2.249 | 2.16E-03 | glio |
| Chek1_in.vivo_noRT | Dennd4b | 2.229 | 2.16E-03 | glio |
| Chek1_ex.vivo_noRT | 4833413G10R | 3.072 | 2.16E-03 | glio |
| Chek1_ex.vivo_noRT | Gmds | -1.955 | 2.16E-03 | glio |
| Chek1_ex.vivo_noRT | 5430405H02R | 2.231 | 2.16E-03 | glio |
| Chek1_ex.vivo_noRT | Ran | -0.793 | 2.16E-03 | glio |
| Hsd17b10_in.vivo_noRT | Dct | 2.813 | 2.16E-03 | glio |
| Interaction_genotype_Atr | Gm28424 | -4.283 | 2.16E-04 | glio |
| Interaction_genotype_Mtor | Npy1r | -3.454 | 2.16E-04 | glio |
| Interaction_genotype_Mtor | Hist1h2bl | -3.957 | 2.16E-04 | glio |
| Interaction_genotype_Mtor | A730063M14R | -3.87 | 2.16E-04 | glio |
| Interaction_genotype_Mtor | 1190005I06Ri | -3.808 | 2.16E-04 | glio |
| Interaction_genotype_Mtor | Eif3g | 1.83 | 2.16E-04 | glio |
| Interaction_genotype_Myc | Gm34059 | -2.729 | 2.16E-04 | glio |
| Interaction_genotype_Myc | Cbr3 | -4.399 | 2.16E-04 | glio |
| Interaction_genotype_Myc | Atpif1 | 1.57 | 2.16E-04 | glio |
| Aars2_in.vivo_noRT | Prr15l | 2.635 | 2.16E-04 | glio |
| Atp5c1_in.vivo_noRT | Gm4858 | 2.668 | 2.16E-04 | glio |
| Atp5c1_in.vivo_noRT | Spn | 2.668 | 2.16E-04 | glio |
| Bcas2_in.vivo_noRT | Dhrs13os | 2.971 | 2.16E-04 | glio |
| Bcas2_in.vivo_noRT | Ankrd23 | 2.827 | 2.16E-04 | glio |
| Bora_in.vivo_noRT | Pafah2 | 2.907 | 2.16E-04 | glio |
| Chek1_in.vivo_noRT | Mirg | 2.702 | 2.16E-04 | glio |

| | | | | |
|---|---|---|---|---|
| Chek1_in.vivo_noRT | Tph2 | 2.497 | 2.16E-04 | glio |
| Eif2s1_in.vivo_noRT | Serpinb9g | 2.143 | 2.16E-04 | glio |
| culture_effect | Selenbp1 | -2.974 | 2.16E-05 | glio |
| Aars2_in.vivo_noRT | Sgsh | 3.12 | 2.16E-05 | glio |
| Aars2_in.vivo_noRT | Zfp316 | 2.576 | 2.16E-05 | glio |
| Atr_in.vivo_noRT | A230087F16Ri | 2.909 | 2.16E-05 | glio |
| Bora_in.vivo_noRT | 1500002C15R | 3.169 | 2.16E-05 | glio |
| Bora_in.vivo_noRT | Bpifa3 | 2.522 | 2.16E-05 | glio |
| Chek1_in.vivo_noRT | Ksr1 | 2.995 | 2.16E-05 | glio |
| Interaction_genotype_Mtor | Cyb5d1 | -5.654 | 2.16E-06 | glio |
| Bora_in.vivo_noRT | Abhd15 | 3.611 | 2.16E-06 | glio |
| Bora_in.vivo_noRT | Rhbdf2 | 3.141 | 2.16E-06 | glio |
| Chek1_in.vivo_noRT | Cebpa | 3.382 | 2.16E-06 | glio |
| Chek1_in.vivo_noRT | Gm9725 | 3.376 | 2.16E-06 | glio |
| Chek1_in.vivo_noRT | Slc12a5 | 3.262 | 2.16E-06 | glio |
| Chek1_in.vivo_noRT | Gm36913 | 3.354 | 2.16E-06 | glio |
| Bard1_ex.vivo_noRT | Gpr4 | 3.612 | 2.16E-07 | glio |
| Aars2_in.vivo_noRT | Wincr1 | 5.063 | 2.16E-08 | glio |
| culture_effect | Zfp593 | 0.905 | 2.16E-09 | glio |
| Bcas2_in.vivo_noRT | Gm28731 | 4.421 | 2.16E-09 | glio |
| Chek1_in.vivo_noRT | Dnajc28 | 4.421 | 2.16E-09 | glio |
| Bora_in.vivo_noRT | Gm32090 | 4.144 | 2.16E-10 | glio |
| Chek1_ex.vivo_noRT | Ccdc125 | 4.443 | 2.16E-10 | glio |
| Bcas2_in.vivo_noRT | Prm3 | 4.066 | 2.16E-11 | glio |
| Chek1_in.vivo_noRT | Ncf1 | 4.391 | 2.16E-13 | glio |
| Bcas2_in.vivo_noRT | Gm26978 | 4.547 | 2.16E-15 | glio |
| Bora_in.vivo_noRT | Pmch | 4.802 | 2.16E-15 | glio |
| Aars2_in.vivo_noRT | Gm28940 | 4.811 | 2.16E-17 | glio |
| culture_effect | Ppib | -1.264 | 2.16E-21 | glio |
| Atr_in.vivo_noRT | Gm15929 | 5.533 | 2.16E-22 | glio |
| Aars2_in.vivo_noRT | Gm47404 | 5.327 | 2.16E-27 | glio |
| culture_effect | G730013B05R | -0.708 | 2.17E-03 | glio |
| culture_effect | Arhgap39 | -0.635 | 2.17E-03 | glio |
| culture_effect | Setd5 | -0.562 | 2.17E-03 | glio |
| Interaction_genotype_Atr | Fam228a | -3.883 | 2.17E-03 | glio |
| Interaction_genotype_Atr | Gm30938 | -3.516 | 2.17E-03 | glio |
| Interaction_genotype_Atr | H3f3b | 0.998 | 2.17E-03 | glio |
| Interaction_genotype_Bora | Gm16283 | -3.89 | 2.17E-03 | glio |
| Interaction_genotype_Bora | AV356131 | -3.482 | 2.17E-03 | glio |
| Interaction_genotype_Mtor | A130014A01R | -3.197 | 2.17E-03 | glio |
| Interaction_genotype_Myc | Adam5 | -2.466 | 2.17E-03 | glio |
| Interaction_genotype_Myc | A430035B10R | -3.566 | 2.17E-03 | glio |
| Interaction_genotype_Myc | Eml6 | -3.528 | 2.17E-03 | glio |
| Interaction_genotype_Myc | Entpd5 | -3.295 | 2.17E-03 | glio |
| Interaction_genotype_Myc | Impa2 | -2.499 | 2.17E-03 | glio |
| Interaction_genotype_Myc | Ino80c | -2.011 | 2.17E-03 | glio |
| Interaction_genotype_Myc | Iah1 | 1.619 | 2.17E-03 | glio |
| interaction_RT | Elac2 | -1.049 | 2.17E-03 | glio |
| interaction_RT | Gm10115 | -2.916 | 2.17E-03 | glio |
| interaction_RT | Irf2 | 0.832 | 2.17E-03 | glio |
| Aars2_in.vivo_noRT | Ralgapa1 | -1.446 | 2.17E-03 | glio |
| Aars2_in.vivo_noRT | Eif2ak2 | -1.282 | 2.17E-03 | glio |

| | | | | |
|---|---|---|---|---|
| Aars2_in.vivo_noRT | Hnrnpr | -0.98 | 2.17E-03 | glio |
| Aars2_in.vivo_noRT | Otulin | -0.966 | 2.17E-03 | glio |
| Aars2_in.vivo_noRT | Ywhah | -0.733 | 2.17E-03 | glio |
| Aars2_ex.vivo_noRT | Astn2 | 2.897 | 2.17E-03 | glio |
| Atp5c1_in.vivo_noRT | Bok | -2.239 | 2.17E-03 | glio |
| Atr_in.vivo_noRT | Gm26549 | 1.93 | 2.17E-03 | glio |
| Atr_in.vivo_noRT | Ralgps1 | 1.806 | 2.17E-03 | glio |
| Atr_ex.vivo_noRT | Arhgef1 | -1.939 | 2.17E-03 | glio |
| Bora_in.vivo_noRT | Arrdc2 | 2.428 | 2.17E-03 | glio |
| Bora_in.vivo_noRT | Gm11713 | 1.993 | 2.17E-03 | glio |
| Bora_in.vivo_noRT | Ube2g1 | -1.293 | 2.17E-03 | glio |
| Bora_in.vivo_noRT | Lbh | -1.077 | 2.17E-03 | glio |
| Chek1_in.vivo_noRT | Hemk1 | 2.279 | 2.17E-03 | glio |
| Chek1_ex.vivo_noRT | Arhgap12 | 1.84 | 2.17E-03 | glio |
| Interaction_genotype_Aars2 | Gm50321 | -3.93 | 2.17E-04 | glio |
| Interaction_genotype_Mtor | Insm1 | -4.359 | 2.17E-04 | glio |
| Interaction_genotype_Mtor | Gm11767 | -4.451 | 2.17E-04 | glio |
| Interaction_genotype_Mtor | Galm | -3.924 | 2.17E-04 | glio |
| Interaction_genotype_Mtor | Ndufs2 | 1.811 | 2.17E-04 | glio |
| Interaction_genotype_Myc | 9530062K07R | -4.301 | 2.17E-04 | glio |
| Interaction_genotype_Myc | Tlr1 | -4.114 | 2.17E-04 | glio |
| interaction_RT | Uhrf1 | -1.148 | 2.17E-04 | glio |
| interaction_RT | Mcm5 | -1.115 | 2.17E-04 | glio |
| Atr_in.vivo_noRT | Nup85 | -1.614 | 2.17E-04 | glio |
| Bcas2_in.vivo_noRT | Hist1h2bp | 2.812 | 2.17E-04 | glio |
| Bcas2_in.vivo_noRT | Agap1 | -1.678 | 2.17E-04 | glio |
| Bora_in.vivo_noRT | Cdkn2a | -0.722 | 2.17E-04 | glio |
| Bora_ex.vivo_noRT | Zfp352 | 3.201 | 2.17E-04 | glio |
| Hsd17b10_in.vivo_noRT | Gm20455 | 2.896 | 2.17E-04 | glio |
| Hsd17b10_in.vivo_noRT | Rhbg | 2.021 | 2.17E-04 | glio |
| Hsd17b10_in.vivo_noRT | Ap3b1 | -1.517 | 2.17E-04 | glio |
| culture_effect | Smim19 | 0.799 | 2.17E-05 | glio |
| culture_effect | Acat1 | -0.753 | 2.17E-05 | glio |
| Atr_in.vivo_noRT | Ccdc141 | 3.076 | 2.17E-05 | glio |
| Aars2_in.vivo_noRT | 1010001B22R | 3.326 | 2.17E-06 | glio |
| Aars2_in.vivo_noRT | Tex14 | 2.986 | 2.17E-06 | glio |
| Atp5c1_in.vivo_noRT | A930002C04R | 2.408 | 2.17E-06 | glio |
| Atr_in.vivo_noRT | Prelid3a | 3.305 | 2.17E-06 | glio |
| Bcas2_in.vivo_noRT | Tanc1 | -2.262 | 2.17E-06 | glio |
| Bora_in.vivo_noRT | 9530018F02Ri | 3.55 | 2.17E-06 | glio |
| Bora_in.vivo_noRT | Hdac1 | -1.33 | 2.17E-06 | glio |
| Aars2_in.vivo_noRT | Zfp990 | 3.425 | 2.17E-07 | glio |
| Aars2_in.vivo_noRT | Uck2 | -1.865 | 2.17E-07 | glio |
| Bcas2_in.vivo_noRT | Hlx | 3.551 | 2.17E-07 | glio |
| Bcas2_in.vivo_noRT | Dnase1l2 | 3.519 | 2.17E-07 | glio |
| Bora_in.vivo_noRT | Stab1 | 3.802 | 2.17E-07 | glio |
| Chek1_in.vivo_noRT | Cabp4 | 3.733 | 2.17E-07 | glio |
| Hsd17b10_in.vivo_noRT | CAAA0106612 | 3.355 | 2.17E-07 | glio |
| Hsd17b10_in.vivo_noRT | Gm41318 | 3.616 | 2.17E-07 | glio |
| culture_effect | Ccnl1 | -1.044 | 2.17E-08 | glio |
| Bora_in.vivo_noRT | A330040F15Ri | 4.157 | 2.17E-08 | glio |
| culture_effect | Hspg2 | -1.211 | 2.17E-10 | glio |

| | | | | |
|---|---|---|---|---|
| Aars2_in.vivo_noRT | Gm11696 | 4.46 | 2.17E-10 | glio |
| Aars2_in.vivo_noRT | St14 | 3.941 | 2.17E-10 | glio |
| Aars2_in.vivo_noRT | Gabrd | 4.414 | 2.17E-10 | glio |
| Bora_in.vivo_noRT | 2810410L24Ri | 4.411 | 2.17E-10 | glio |
| Hsd17b10_in.vivo_noRT | Olfr543 | 3.356 | 2.17E-10 | glio |
| culture_effect | A530032D15R | -3.756 | 2.17E-12 | glio |
| culture_effect | Eif3d | -0.807 | 2.17E-12 | glio |
| Atr_in.vivo_noRT | Gm28731 | 5.147 | 2.17E-12 | glio |
| Bcas2_in.vivo_noRT | Loxl4 | 4.541 | 2.17E-12 | glio |
| Atr_in.vivo_noRT | Sema6b | 5.094 | 2.17E-13 | glio |
| Bcas2_in.vivo_noRT | Crybb1 | 4.42 | 2.17E-13 | glio |
| culture_effect | Gstk1 | 1.416 | 2.17E-14 | glio |
| Aars2_in.vivo_noRT | Stra8 | 4.804 | 2.17E-15 | glio |
| Atr_in.vivo_noRT | Speer4e | 5.533 | 2.17E-23 | glio |
| Atr_in.vivo_noRT | Gm48435 | 5.533 | 2.17E-30 | glio |
| culture_effect | Zfp799 | 1.516 | 2.18E-03 | glio |
| culture_effect | Myadm | -0.707 | 2.18E-03 | glio |
| Interaction_genotype_Mtor | Trpm6 | -3.205 | 2.18E-03 | glio |
| Interaction_genotype_Mtor | Cisd1 | 1.509 | 2.18E-03 | glio |
| Interaction_genotype_Myc | Cox7b2 | -3.059 | 2.18E-03 | glio |
| Interaction_genotype_Myc | Gm17203 | -2.478 | 2.18E-03 | glio |
| Interaction_genotype_Myc | Cyth4 | -3.293 | 2.18E-03 | glio |
| Interaction_genotype_Myc | B130055M24F | -3.223 | 2.18E-03 | glio |
| Interaction_genotype_Myc | Mthfs | -3.069 | 2.18E-03 | glio |
| Interaction_genotype_Myc | 9530034E10Ri | -2.843 | 2.18E-03 | glio |
| Interaction_genotype_Myc | Sppl3 | -2.02 | 2.18E-03 | glio |
| Interaction_genotype_Myc | Gabarapl1 | 1.552 | 2.18E-03 | glio |
| interaction_RT | Tgtp1 | 2.397 | 2.18E-03 | glio |
| Aars2_in.vivo_noRT | Acbd7 | 2.294 | 2.18E-03 | glio |
| Aars2_in.vivo_noRT | Cep85 | 2.043 | 2.18E-03 | glio |
| Aars2_in.vivo_noRT | Dctpp1 | -0.948 | 2.18E-03 | glio |
| Aars2_in.vivo_noRT | Rpl36 | -0.729 | 2.18E-03 | glio |
| Atp5c1_in.vivo_noRT | Apobr | 2.668 | 2.18E-03 | glio |
| Atr_in.vivo_noRT | Rnpc3 | -1.4 | 2.18E-03 | glio |
| Atr_in.vivo_noRT | Esf1 | -0.854 | 2.18E-03 | glio |
| Bard1_ex.vivo_noRT | 2810049E08Ri | 2.814 | 2.18E-03 | glio |
| Bard1_ex.vivo_noRT | Pla2g2f | 2.814 | 2.18E-03 | glio |
| Bcas2_in.vivo_noRT | Gm9844 | 2.159 | 2.18E-03 | glio |
| Bcas2_in.vivo_noRT | Rab5if | -0.776 | 2.18E-03 | glio |
| Bora_in.vivo_noRT | Itgae | 2.309 | 2.18E-03 | glio |
| Bora_in.vivo_noRT | Gm26579 | 2.114 | 2.18E-03 | glio |
| Bora_ex.vivo_noRT | Gm45606 | 2.899 | 2.18E-03 | glio |
| Chek1_in.vivo_noRT | Dnajc14 | 1.553 | 2.18E-03 | glio |
| Chek1_ex.vivo_noRT | Zc3h11a | 2.574 | 2.18E-03 | glio |
| Chek1_ex.vivo_noRT | Serinc2 | 2.566 | 2.18E-03 | glio |
| Chek1_ex.vivo_noRT | Rpl9 | -0.711 | 2.18E-03 | glio |
| Hsd17b10_in.vivo_noRT | Olfr1369-ps1 | 2.875 | 2.18E-03 | glio |
| culture_effect | Cmc1 | -0.638 | 2.18E-04 | glio |
| Interaction_genotype_Atr | Gm30794 | -3.346 | 2.18E-04 | glio |
| Interaction_genotype_Atr | Clec4d | -3.554 | 2.18E-04 | glio |
| interaction_RT | Carm1 | -1.226 | 2.18E-04 | glio |
| interaction_RT | Adh5 | -0.895 | 2.18E-04 | glio |

| | | | | |
|---|---|---|---|---|
| Aars2_in.vivo_noRT | Tepsin | 2.198 | 2.18E-04 | glio |
| Atp5c1_in.vivo_noRT | Gm20767 | 2.668 | 2.18E-04 | glio |
| Atr_in.vivo_noRT | Galm | 2.53 | 2.18E-04 | glio |
| Atr_in.vivo_noRT | Pdk2 | 2.66 | 2.18E-04 | glio |
| Atr_in.vivo_noRT | Slc2a13 | 2.597 | 2.18E-04 | glio |
| Atr_in.vivo_noRT | Rsph3a | 2.436 | 2.18E-04 | glio |
| Atr_ex.vivo_noRT | Pla2g2d | 3.164 | 2.18E-04 | glio |
| Bard1_ex.vivo_noRT | Vmn2r56 | 2.814 | 2.18E-04 | glio |
| Bora_in.vivo_noRT | Zc3h6 | 2.698 | 2.18E-04 | glio |
| Chek1_in.vivo_noRT | Gm10131 | 2.61 | 2.18E-04 | glio |
| Chek1_ex.vivo_noRT | Catsperz | 3.34 | 2.18E-04 | glio |
| Hsd17b10_in.vivo_noRT | Cpne8 | 2.881 | 2.18E-04 | glio |
| culture_effect | Neurl2 | 1.524 | 2.18E-05 | glio |
| culture_effect | Crip2 | -1.373 | 2.18E-05 | glio |
| culture_effect | Dedd2 | -1.153 | 2.18E-05 | glio |
| culture_effect | Gnb4 | -0.862 | 2.18E-05 | glio |
| Interaction_genotype_Aars2 | Pmp2 | -4.75 | 2.18E-05 | glio |
| Interaction_genotype_Atr | 4930589L23Ri | -5.172 | 2.18E-05 | glio |
| Interaction_genotype_Atr | Kcnmb4os2 | -5.141 | 2.18E-05 | glio |
| Atr_in.vivo_noRT | Acy3 | 3.04 | 2.18E-05 | glio |
| Atr_ex.vivo_noRT | Cfb | 3.164 | 2.18E-05 | glio |
| Atr_ex.vivo_noRT | Ubd | 3.554 | 2.18E-05 | glio |
| Bcas2_in.vivo_noRT | Gm48684 | 3.503 | 2.18E-05 | glio |
| Bcas2_in.vivo_noRT | Gm28875 | 3.112 | 2.18E-05 | glio |
| Bcas2_in.vivo_noRT | Gm12089 | 3.106 | 2.18E-05 | glio |
| Bora_in.vivo_noRT | Gm37305 | 2.979 | 2.18E-05 | glio |
| Bora_in.vivo_noRT | Gm20522 | 3.13 | 2.18E-05 | glio |
| Chek1_in.vivo_noRT | Igf2r | 2.812 | 2.18E-05 | glio |
| Atr_in.vivo_noRT | Hist1h2ai | 3.437 | 2.18E-06 | glio |
| Bora_in.vivo_noRT | 4930522L14Ri | 3.441 | 2.18E-06 | glio |
| Hsd17b10_in.vivo_noRT | Gm50394 | 3.35 | 2.18E-06 | glio |
| culture_effect | Cemip | -2.969 | 2.18E-08 | glio |
| culture_effect | Nadk2 | -0.988 | 2.18E-08 | glio |
| Chek1_in.vivo_noRT | Pla2g7 | 3.857 | 2.18E-08 | glio |
| Atr_in.vivo_noRT | Csnk1d | -2.404 | 2.18E-09 | glio |
| Bcas2_in.vivo_noRT | Ngb | 4.037 | 2.18E-09 | glio |
| Atr_in.vivo_noRT | Gm17227 | 4.151 | 2.18E-10 | glio |
| Chek1_in.vivo_noRT | Hlx | 4.311 | 2.18E-10 | glio |
| Chek1_in.vivo_noRT | Slc6a11 | 3.612 | 2.18E-10 | glio |
| Atr_in.vivo_noRT | Sbk3 | 4.365 | 2.18E-11 | glio |
| Atr_in.vivo_noRT | 2010106G04R | 4.733 | 2.18E-11 | glio |
| Atr_in.vivo_noRT | St14 | 4.147 | 2.18E-11 | glio |
| Chek1_in.vivo_noRT | 2310067P03R | 4.521 | 2.18E-11 | glio |
| culture_effect | Gm4951 | -1.718 | 2.18E-13 | glio |
| Atr_in.vivo_noRT | Gm10300 | 5.273 | 2.18E-13 | glio |
| culture_effect | Arl4a | -1.368 | 2.18E-14 | glio |
| Chek1_ex.vivo_noRT | Cldn5 | 4.842 | 2.18E-15 | glio |
| Chek1_in.vivo_noRT | Gm13032 | 4.658 | 2.18E-16 | glio |
| culture_effect | Nr4a1 | -1.892 | 2.18E-17 | glio |
| culture_effect | Casp1 | -4.392 | 2.18E-20 | glio |
| culture_effect | Trf | -4.462 | 2.18E-20 | glio |
| culture_effect | Crispld2 | -4.847 | 2.18E-20 | glio |

| | | | | |
|---|---|---|---|---|
| culture_effect | Slc1a1 | -5.161 | 2.18E-21 | glio |
| Atr_in.vivo_noRT | Trim17 | 5.533 | 2.18E-28 | glio |
| culture_effect | Rdh14 | 0.837 | 2.19E-03 | glio |
| culture_effect | Atp5l | 0.361 | 2.19E-03 | glio |
| Interaction_genotype_Bcas2 | Igf2bp1 | -3.952 | 2.19E-03 | glio |
| Interaction_genotype_Bcas2 | Camk2b | -3.97 | 2.19E-03 | glio |
| Interaction_genotype_Bcas2 | Chordc1 | 1.74 | 2.19E-03 | glio |
| Interaction_genotype_Mtor | Ikbke | -3.572 | 2.19E-03 | glio |
| Interaction_genotype_Mtor | Zscan2 | -3.502 | 2.19E-03 | glio |
| Interaction_genotype_Mtor | AI429214 | -3.619 | 2.19E-03 | glio |
| Interaction_genotype_Mtor | Mmaa | -3.152 | 2.19E-03 | glio |
| Interaction_genotype_Myc | Kcnj13 | -3.123 | 2.19E-03 | glio |
| Interaction_genotype_Myc | Dock11 | -3.081 | 2.19E-03 | glio |
| Interaction_genotype_Myc | 4933439C10R | -3.255 | 2.19E-03 | glio |
| Interaction_genotype_Myc | Cfap300 | -2.8 | 2.19E-03 | glio |
| Interaction_genotype_Myc | Hirip3 | -2.429 | 2.19E-03 | glio |
| Aars2_in.vivo_noRT | Ppid | -0.908 | 2.19E-03 | glio |
| Atr_in.vivo_noRT | Mpc2 | -1.12 | 2.19E-03 | glio |
| Atr_in.vivo_noRT | Rpl36 | -0.744 | 2.19E-03 | glio |
| Atr_ex.vivo_noRT | Dhrs9 | 2.764 | 2.19E-03 | glio |
| Atr_ex.vivo_noRT | 7530414M10F | 2.811 | 2.19E-03 | glio |
| Atr_ex.vivo_noRT | Gabrd | 2.99 | 2.19E-03 | glio |
| Atr_ex.vivo_noRT | Tarsl2 | -2.317 | 2.19E-03 | glio |
| Bcas2_in.vivo_noRT | Cdc25a | -1.451 | 2.19E-03 | glio |
| Bora_in.vivo_noRT | Anxa11 | 2.518 | 2.19E-03 | glio |
| Bora_in.vivo_noRT | Fbxo24 | 2.217 | 2.19E-03 | glio |
| Bora_in.vivo_noRT | Mettl18 | 1.92 | 2.19E-03 | glio |
| Bora_in.vivo_noRT | D2hgdh | 1.817 | 2.19E-03 | glio |
| Chek1_in.vivo_noRT | Pcx | 2.184 | 2.19E-03 | glio |
| Chek1_in.vivo_noRT | Pfkp | -1.786 | 2.19E-03 | glio |
| Chek1_ex.vivo_noRT | Gm16973 | 2.81 | 2.19E-03 | glio |
| Chek1_ex.vivo_noRT | Lsm8 | -0.943 | 2.19E-03 | glio |
| Eif2s1_in.vivo_noRT | Scgb1c1 | 2.143 | 2.19E-03 | glio |
| culture_effect | Rem2 | -2.119 | 2.19E-04 | glio |
| Interaction_genotype_Aars2 | Hecw2 | -4.593 | 2.19E-04 | glio |
| Interaction_genotype_Chek1 | Serping1 | 4.119 | 2.19E-04 | glio |
| Interaction_genotype_Mtor | Hemk1 | -4.423 | 2.19E-04 | glio |
| Interaction_genotype_Myc | Syn3 | -4.108 | 2.19E-04 | glio |
| Interaction_genotype_Myc | Armc6 | -3.134 | 2.19E-04 | glio |
| Atp5c1_in.vivo_noRT | 5730507C01R | 2.145 | 2.19E-04 | glio |
| Atr_in.vivo_noRT | Homez | 2.627 | 2.19E-04 | glio |
| Atr_in.vivo_noRT | Cfap74 | 2.332 | 2.19E-04 | glio |
| Atr_in.vivo_noRT | Gm19412 | 1.881 | 2.19E-04 | glio |
| Bcas2_in.vivo_noRT | Fhl3 | 2.689 | 2.19E-04 | glio |
| Chek1_in.vivo_noRT | Mroh2b | 2.716 | 2.19E-04 | glio |
| Chek1_ex.vivo_noRT | Nin | -2.517 | 2.19E-04 | glio |
| Hsd17b10_in.vivo_noRT | mt-Co3 | -0.711 | 2.19E-04 | glio |
| culture_effect | AC125149.3 | -2.537 | 2.19E-05 | glio |
| culture_effect | A530040E14Ri | -2.17 | 2.19E-05 | glio |
| Aars2_in.vivo_noRT | Gstt3 | 3.045 | 2.19E-05 | glio |
| Chek1_ex.vivo_noRT | Ccdc28b | 3.353 | 2.19E-05 | glio |
| Interaction_genotype_Mtor | Olfr1152 | -3.833 | 2.19E-06 | glio |

| | | | | |
|---|---|---|---|---|
| interaction_RT | Rpl5 | -0.94 | 2.19E-06 | glio |
| Bcas2_in.vivo_noRT | Gm42658 | 3.054 | 2.19E-06 | glio |
| Bcas2_in.vivo_noRT | Trim14 | 3.641 | 2.19E-06 | glio |
| Atr_ex.vivo_noRT | Cxcr2 | 4.533 | 2.19E-07 | glio |
| Bora_in.vivo_noRT | Gm3604 | 3.654 | 2.19E-07 | glio |
| Bora_in.vivo_noRT | Gm50107 | 3.421 | 2.19E-07 | glio |
| Bora_in.vivo_noRT | Rtl1 | 3.244 | 2.19E-07 | glio |
| Bora_in.vivo_noRT | 6330415G19R | 3.629 | 2.19E-07 | glio |
| Bora_in.vivo_noRT | 1810012K16R | 3.42 | 2.19E-07 | glio |
| Hsd17b10_in.vivo_noRT | A330032B11R | 3.35 | 2.19E-07 | glio |
| Hsd17b10_in.vivo_noRT | Gm27239 | 3.616 | 2.19E-07 | glio |
| Hsd17b10_in.vivo_noRT | 4930412M03F | 3.616 | 2.19E-07 | glio |
| Aars2_in.vivo_noRT | 2210406O10R | 4.01 | 2.19E-08 | glio |
| Chek1_in.vivo_noRT | Cnih2 | 3.856 | 2.19E-08 | glio |
| Chek1_ex.vivo_noRT | Apol6 | 4.055 | 2.19E-08 | glio |
| culture_effect | 4833438C02R | 2.31 | 2.19E-09 | glio |
| Interaction_genotype_Myc | Swi5 | 2.404 | 2.19E-09 | glio |
| Interaction_genotype_Myc | Matn4 | -6.704 | 2.19E-09 | glio |
| Bora_in.vivo_noRT | D030051J21Ri | 4.033 | 2.19E-09 | glio |
| Atr_in.vivo_noRT | Olfm2 | 4.626 | 2.19E-10 | glio |
| Atr_ex.vivo_noRT | BC080695 | 4.309 | 2.19E-11 | glio |
| Atr_ex.vivo_noRT | Zscan4d | 4.766 | 2.19E-11 | glio |
| Chek1_ex.vivo_noRT | Serping1 | 4.842 | 2.19E-11 | glio |
| Bcas2_in.vivo_noRT | Gm17276 | 4.016 | 2.19E-12 | glio |
| Bora_in.vivo_noRT | E030037K01Ri | 4.405 | 2.19E-12 | glio |
| Chek1_ex.vivo_noRT | Slc35g3 | 4.443 | 2.19E-13 | glio |
| Hsd17b10_in.vivo_noRT | Vmn1r32 | 3.616 | 2.19E-13 | glio |
| Hsd17b10_in.vivo_noRT | Glyat | 3.616 | 2.19E-13 | glio |
| Bora_in.vivo_noRT | Gm43915 | 4.404 | 2.19E-14 | glio |
| Bora_in.vivo_noRT | Acoxl | 4.936 | 2.19E-14 | glio |
| Chek1_ex.vivo_noRT | Ccl3 | 4.842 | 2.19E-14 | glio |
| Chek1_ex.vivo_noRT | Gm48780 | 4.842 | 2.19E-14 | glio |
| Chek1_ex.vivo_noRT | AU019990 | 4.842 | 2.19E-14 | glio |
| Atr_in.vivo_noRT | Ifitm6 | 5.533 | 2.19E-15 | glio |
| Bcas2_in.vivo_noRT | Gm11457 | 5.325 | 2.19E-15 | glio |
| culture_effect | Phactr1 | 1.882 | 2.19E-22 | glio |
| Interaction_genotype_Aars2 | Glis1 | -3.83 | 2.20E-03 | glio |
| Interaction_genotype_Aars2 | Gjb4 | -3.902 | 2.20E-03 | glio |
| Interaction_genotype_Aars2 | 1810041H14R | -4.052 | 2.20E-03 | glio |
| Interaction_genotype_Aars2 | Zfp867 | -3.325 | 2.20E-03 | glio |
| Interaction_genotype_Mtor | Ntn1 | -3.197 | 2.20E-03 | glio |
| Interaction_genotype_Mtor | Gm48796 | -3.595 | 2.20E-03 | glio |
| Interaction_genotype_Mtor | Rps6ka6 | -3.431 | 2.20E-03 | glio |
| Interaction_genotype_Myc | Spink6 | -2.729 | 2.20E-03 | glio |
| Interaction_genotype_Myc | 4930445N18R | -3.5 | 2.20E-03 | glio |
| Interaction_genotype_Myc | Dus3l | -2.717 | 2.20E-03 | glio |
| Interaction_genotype_Myc | Strn | -2.309 | 2.20E-03 | glio |
| interaction_RT | Alkbh6 | 1.008 | 2.20E-03 | glio |
| interaction_RT | Tec | 2.864 | 2.20E-03 | glio |
| interaction_RT | Ripk2 | 1.078 | 2.20E-03 | glio |
| interaction_RT | Srsf3 | -0.603 | 2.20E-03 | glio |
| Aars2_in.vivo_noRT | Tmem191c | 2.071 | 2.20E-03 | glio |

| | | | | |
|---|---|---|---|---|
| Aars2_in.vivo_noRT | 4930578M01F | 1.942 | 2.20E-03 | glio |
| Atp5c1_in.vivo_noRT | Gm42695 | 2.408 | 2.20E-03 | glio |
| Atr_in.vivo_noRT | Gm4707 | 2.096 | 2.20E-03 | glio |
| Bcas2_in.vivo_noRT | Shroom1 | 2.189 | 2.20E-03 | glio |
| Bcas2_in.vivo_noRT | Gm16845 | 2.136 | 2.20E-03 | glio |
| Bcas2_in.vivo_noRT | A230056P14R | 2.242 | 2.20E-03 | glio |
| Bora_in.vivo_noRT | Prr15l | 2.245 | 2.20E-03 | glio |
| Bora_in.vivo_noRT | Tubb4a | 2.312 | 2.20E-03 | glio |
| Chek1_in.vivo_noRT | Trabd | -1.314 | 2.20E-03 | glio |
| Chek1_ex.vivo_noRT | H2afz | -0.83 | 2.20E-03 | glio |
| culture_effect | Hdac2 | 0.476 | 2.20E-04 | glio |
| Interaction_genotype_Aars2 | Gm34256 | -4.886 | 2.20E-04 | glio |
| Interaction_genotype_Atr | 5033417F24Ri | -4.445 | 2.20E-04 | glio |
| Interaction_genotype_Atr | 2210011C24R | -4.021 | 2.20E-04 | glio |
| Interaction_genotype_Atr | Gm35066 | -4.512 | 2.20E-04 | glio |
| Interaction_genotype_Atr | Gm47545 | -4.158 | 2.20E-04 | glio |
| Interaction_genotype_Atr | Gm43701 | -3.352 | 2.20E-04 | glio |
| Interaction_genotype_Atr | Plekhf2 | 2.324 | 2.20E-04 | glio |
| Interaction_genotype_Atr | Sbds | 1.804 | 2.20E-04 | glio |
| Interaction_genotype_Mtor | Gm42849 | -3.912 | 2.20E-04 | glio |
| Interaction_genotype_Mtor | Gm28940 | -3.583 | 2.20E-04 | glio |
| Interaction_genotype_Mtor | Gm26688 | -4.363 | 2.20E-04 | glio |
| Interaction_genotype_Mtor | Hist1h4n | -4.781 | 2.20E-04 | glio |
| Interaction_genotype_Mtor | Commd5 | -4.485 | 2.20E-04 | glio |
| Interaction_genotype_Mtor | Sardh | -3.99 | 2.20E-04 | glio |
| Interaction_genotype_Mtor | Orc6 | 2.085 | 2.20E-04 | glio |
| Interaction_genotype_Mtor | Impdh2 | 1.682 | 2.20E-04 | glio |
| Aars2_in.vivo_noRT | Zfp39 | 2.682 | 2.20E-04 | glio |
| Bcas2_in.vivo_noRT | Gm28050 | 2.769 | 2.20E-04 | glio |
| Bcas2_in.vivo_noRT | Gm4419 | 2.325 | 2.20E-04 | glio |
| Bcas2_in.vivo_noRT | Cmas | -1.287 | 2.20E-04 | glio |
| Bora_in.vivo_noRT | A430033K04R | 2.568 | 2.20E-04 | glio |
| Cenpj_ex.vivo_noRT | Gm40264 | 3.176 | 2.20E-04 | glio |
| Chek1_ex.vivo_noRT | Efna2 | 3.86 | 2.20E-04 | glio |
| Dclre1c_ex.vivo_noRT | Zscan4-ps2 | 2.9 | 2.20E-04 | glio |
| Hsd17b10_in.vivo_noRT | Gm10399 | 2.684 | 2.20E-04 | glio |
| Hsd17b10_in.vivo_noRT | Mtmr11 | 2.77 | 2.20E-04 | glio |
| Interaction_genotype_Mtor | C920021L13Ri | -4.886 | 2.20E-05 | glio |
| Interaction_genotype_Mtor | Axdnd1 | -5.596 | 2.20E-05 | glio |
| Interaction_genotype_Mtor | Serpinb2 | -4.498 | 2.20E-05 | glio |
| Atp5c1_in.vivo_noRT | Gm50138 | 2.667 | 2.20E-05 | glio |
| Atr_in.vivo_noRT | Fbxl12os | 2.892 | 2.20E-05 | glio |
| culture_effect | Clpb | -1.015 | 2.20E-06 | glio |
| culture_effect | Cited4 | -3.317 | 2.20E-07 | glio |
| Interaction_genotype_Mtor | Gm44736 | -4.099 | 2.20E-07 | glio |
| Atr_in.vivo_noRT | C430014B12R | 3.484 | 2.20E-07 | glio |
| Bora_in.vivo_noRT | Gm36757 | 3.543 | 2.20E-07 | glio |
| Atr_in.vivo_noRT | Dmrt2 | 4.124 | 2.20E-08 | glio |
| Bcas2_in.vivo_noRT | Gm3716 | 3.889 | 2.20E-08 | glio |
| Chek1_in.vivo_noRT | Edn1 | 4.664 | 2.20E-08 | glio |
| Atr_in.vivo_noRT | Gm26916 | 3.956 | 2.20E-09 | glio |
| Bora_in.vivo_noRT | A930001C03R | 4.283 | 2.20E-10 | glio |

| | | | | |
|---|---|---|---|---|
| Bora_in.vivo_noRT | Zg16 | 4.283 | 2.20E-10 | glio |
| Hsd17b10_in.vivo_noRT | Gm10354 | 3.616 | 2.20E-10 | glio |
| Chek1_in.vivo_noRT | Ccr5 | 4.244 | 2.20E-11 | glio |
| Atr_in.vivo_noRT | Gm17638 | 4.741 | 2.20E-13 | glio |
| Chek1_in.vivo_noRT | Tdrd5 | 4.658 | 2.20E-13 | glio |
| Aars2_in.vivo_noRT | Prrx2 | 5.327 | 2.20E-14 | glio |
| Aars2_in.vivo_noRT | Pla2g2d | 5.063 | 2.20E-15 | glio |
| Aars2_in.vivo_noRT | Tmem74b | 5.063 | 2.20E-18 | glio |
| culture_effect | Klre1 | -5.287 | 2.20E-21 | glio |
| culture_effect | Cnksr1 | 1.628 | 2.21E-03 | glio |
| culture_effect | Zfp566 | -0.828 | 2.21E-03 | glio |
| Interaction_genotype_Atr | Emid1 | -4.295 | 2.21E-03 | glio |
| Interaction_genotype_Atr | Gm12841 | -2.762 | 2.21E-03 | glio |
| Interaction_genotype_Atr | Gm14023 | -3.981 | 2.21E-03 | glio |
| Interaction_genotype_Atr | Jam3 | 2.264 | 2.21E-03 | glio |
| Interaction_genotype_Atr | Mt2 | 1.878 | 2.21E-03 | glio |
| Interaction_genotype_Bcas2 | 4921536K21R | -4.057 | 2.21E-03 | glio |
| Interaction_genotype_Cdkn2a | Gast | 3.634 | 2.21E-03 | glio |
| Interaction_genotype_Mtor | Lipt2 | -2.742 | 2.21E-03 | glio |
| Interaction_genotype_Myc | Zscan2 | -3.297 | 2.21E-03 | glio |
| Interaction_genotype_Myc | Gm49085 | -3.222 | 2.21E-03 | glio |
| Interaction_genotype_Myc | 1700001L19Ri | -3.13 | 2.21E-03 | glio |
| Interaction_genotype_Myc | F2r | -3.16 | 2.21E-03 | glio |
| Interaction_genotype_Myc | Traip | -2.373 | 2.21E-03 | glio |
| Interaction_genotype_Myc | Psmc3 | 1.424 | 2.21E-03 | glio |
| interaction_RT | Grn | 0.765 | 2.21E-03 | glio |
| Aars2_in.vivo_noRT | Cul9 | 2.396 | 2.21E-03 | glio |
| Aars2_in.vivo_noRT | Veph1 | 2.192 | 2.21E-03 | glio |
| Aars2_in.vivo_noRT | Sphk1 | 2.24 | 2.21E-03 | glio |
| Aars2_in.vivo_noRT | Gipc1 | -1.438 | 2.21E-03 | glio |
| Aars2_in.vivo_noRT | B4galt6 | -1.811 | 2.21E-03 | glio |
| Aars2_in.vivo_noRT | Ufc1 | -0.8 | 2.21E-03 | glio |
| Aars2_ex.vivo_noRT | Zmynd10 | 2.505 | 2.21E-03 | glio |
| Atp5c1_in.vivo_noRT | D630036H23F | 2.408 | 2.21E-03 | glio |
| Atr_ex.vivo_noRT | Hnrnpu | -0.721 | 2.21E-03 | glio |
| Bcas2_in.vivo_noRT | Cpne7 | 2.408 | 2.21E-03 | glio |
| Bcas2_in.vivo_noRT | Sufu | 1.471 | 2.21E-03 | glio |
| Bcas2_in.vivo_noRT | Eif1 | -0.555 | 2.21E-03 | glio |
| Bora_in.vivo_noRT | Bglap3 | 2.218 | 2.21E-03 | glio |
| Bora_in.vivo_noRT | Xrcc3 | 2.173 | 2.21E-03 | glio |
| Cenpj_in.vivo_noRT | Tinagl1 | -1.039 | 2.21E-03 | glio |
| culture_effect | Shisa7 | -1.753 | 2.21E-04 | glio |
| Interaction_genotype_Mtor | Gm14488 | -4.688 | 2.21E-04 | glio |
| Interaction_genotype_Myc | Gm15535 | -4.434 | 2.21E-04 | glio |
| Interaction_genotype_Myc | Gm20075 | -4.04 | 2.21E-04 | glio |
| Interaction_genotype_Myc | Akap7 | -3.346 | 2.21E-04 | glio |
| Aars2_in.vivo_noRT | Shank1 | 2.436 | 2.21E-04 | glio |
| Aars2_in.vivo_noRT | Rac3 | 2.441 | 2.21E-04 | glio |
| Aars2_in.vivo_noRT | Natd1 | 2.078 | 2.21E-04 | glio |
| Atp5c1_in.vivo_noRT | F630040K05Ri | 2.545 | 2.21E-04 | glio |
| Bora_in.vivo_noRT | Gm15063 | 2.952 | 2.21E-04 | glio |
| Bora_in.vivo_noRT | Zfp595 | 2.829 | 2.21E-04 | glio |

| | | | | |
|---|---|---|---|---|
| Bora_ex.vivo_noRT | 2810049E08Ri | 3.201 | 2.21E-04 | glio |
| Chek1_in.vivo_noRT | Comp | 2.821 | 2.21E-04 | glio |
| Chek1_ex.vivo_noRT | Zfp992 | 3.475 | 2.21E-04 | glio |
| Chek1_ex.vivo_noRT | Gm30790 | 3.353 | 2.21E-04 | glio |
| Hsd17b10_in.vivo_noRT | Gsx2 | 2.928 | 2.21E-04 | glio |
| Interaction_genotype_Mtor | Fam177a | -4.622 | 2.21E-05 | glio |
| Interaction_genotype_Mtor | Il20rb | -5.422 | 2.21E-05 | glio |
| Interaction_genotype_Mtor | Actr10 | 2.123 | 2.21E-05 | glio |
| Interaction_genotype_Mtor | Ghitm | 2.329 | 2.21E-05 | glio |
| Interaction_genotype_Nat2 | Gm12800 | 3.908 | 2.21E-05 | glio |
| interaction_RT | Dusp2 | 2.537 | 2.21E-05 | glio |
| interaction_RT | Mettl1 | -1.208 | 2.21E-05 | glio |
| Aars2_in.vivo_noRT | Ssu72 | -1.296 | 2.21E-05 | glio |
| Bcas2_in.vivo_noRT | Gm43727 | 2.98 | 2.21E-05 | glio |
| Bora_in.vivo_noRT | Dennd2d | 2.669 | 2.21E-05 | glio |
| Bora_in.vivo_noRT | Wnt10a | 3.007 | 2.21E-05 | glio |
| Chek1_ex.vivo_noRT | Rps27l | -0.839 | 2.21E-05 | glio |
| culture_effect | Gm19345 | -2.719 | 2.21E-06 | glio |
| culture_effect | Akr1b10 | -1.133 | 2.21E-06 | glio |
| Interaction_genotype_Atr | Rab14 | 1.798 | 2.21E-06 | glio |
| Interaction_genotype_Mtor | Vbp1 | 2.452 | 2.21E-06 | glio |
| Atr_in.vivo_noRT | Prkar2b | 3.337 | 2.21E-06 | glio |
| Atr_in.vivo_noRT | Gstm5 | -2.042 | 2.21E-06 | glio |
| Atr_in.vivo_noRT | Npm1 | -0.937 | 2.21E-06 | glio |
| Atr_ex.vivo_noRT | Gm14820 | 3.164 | 2.21E-06 | glio |
| Bcas2_in.vivo_noRT | Alox8 | 4.078 | 2.21E-06 | glio |
| Bcas2_in.vivo_noRT | Gm46447 | 2.996 | 2.21E-06 | glio |
| Bora_ex.vivo_noRT | Rgs4 | 3.201 | 2.21E-06 | glio |
| Chek1_in.vivo_noRT | Hey2 | 3.196 | 2.21E-06 | glio |
| culture_effect | Nuak1 | 1.743 | 2.21E-07 | glio |
| culture_effect | Rps23 | -0.625 | 2.21E-07 | glio |
| Interaction_genotype_Atr | Capza2 | 1.8 | 2.21E-07 | glio |
| Interaction_genotype_Atr | Alox12 | -6.394 | 2.21E-07 | glio |
| Interaction_genotype_Mtor | Pdgfb | -4.999 | 2.21E-07 | glio |
| Bora_in.vivo_noRT | Tnfrsf13b | 3.831 | 2.21E-07 | glio |
| Interaction_genotype_Mtor | mt-Atp6 | 2.441 | 2.21E-08 | glio |
| Aars2_in.vivo_noRT | Gnaz | 4.028 | 2.21E-08 | glio |
| Chek1_ex.vivo_noRT | Ly6g6c | 4.449 | 2.21E-08 | glio |
| Bcas2_in.vivo_noRT | 2810049E08Ri | 3.77 | 2.21E-09 | glio |
| Chek1_ex.vivo_noRT | Gem | 5.25 | 2.21E-10 | glio |
| Atr_in.vivo_noRT | Syp | 5.142 | 2.21E-11 | glio |
| Bora_in.vivo_noRT | Gm27253 | 4.67 | 2.21E-11 | glio |
| Atr_in.vivo_noRT | Ankrd33b | 5.017 | 2.21E-12 | glio |
| Chek1_in.vivo_noRT | Pdlim3 | 4.264 | 2.21E-13 | glio |
| Chek1_in.vivo_noRT | D730045A05R | 5.044 | 2.21E-13 | glio |
| Chek1_in.vivo_noRT | Tmem54 | 4.784 | 2.21E-16 | glio |
| Cdkn2a_ex.vivo_noRT | Cdkn2a | -1.416 | 2.21E-17 | glio |
| Aars2_in.vivo_noRT | Pakap | 4.794 | 2.21E-18 | glio |
| Bcas2_in.vivo_noRT | Gm16933 | 4.285 | 2.21E-18 | glio |
| Atr_in.vivo_noRT | Pitx3 | 5.533 | 2.21E-22 | glio |
| culture_effect | Gm16023 | 1.923 | 2.22E-03 | glio |
| culture_effect | Slain2 | 0.535 | 2.22E-03 | glio |

| | | | | |
|---|---|---|---|---|
| Interaction_genotype_Aars2 | Rftn2 | -3.414 | 2.22E-03 | glio |
| Interaction_genotype_Aars2 | AW046200 | -3.86 | 2.22E-03 | glio |
| Interaction_genotype_Aars2 | Gm6598 | -3.508 | 2.22E-03 | glio |
| Interaction_genotype_Aars2 | Cop1 | 1.805 | 2.22E-03 | glio |
| Interaction_genotype_Atr | Klf9 | 2.144 | 2.22E-03 | glio |
| Interaction_genotype_Bora | Msi2 | 1.762 | 2.22E-03 | glio |
| Interaction_genotype_Chek1 | Gm49874 | -4.313 | 2.22E-03 | glio |
| Interaction_genotype_Mtor | Gm12689 | -4.068 | 2.22E-03 | glio |
| Interaction_genotype_Mtor | Ywhae | 1.159 | 2.22E-03 | glio |
| Interaction_genotype_Mtor | Rpl13 | 1.114 | 2.22E-03 | glio |
| Interaction_genotype_Myc | Wnt4 | -3.849 | 2.22E-03 | glio |
| Interaction_genotype_Myc | Rps6kl1 | -3.505 | 2.22E-03 | glio |
| Interaction_genotype_Pars2 | Nemp1 | 2.39 | 2.22E-03 | glio |
| Aars2_in.vivo_noRT | P2ry2 | 2.135 | 2.22E-03 | glio |
| Aars2_in.vivo_noRT | Nap1l1 | -0.626 | 2.22E-03 | glio |
| Atr_in.vivo_noRT | Samm50 | -0.975 | 2.22E-03 | glio |
| Bora_in.vivo_noRT | Tpbg | 2.194 | 2.22E-03 | glio |
| Bora_in.vivo_noRT | Mapre3 | 1.962 | 2.22E-03 | glio |
| Bora_in.vivo_noRT | Flii | -1.536 | 2.22E-03 | glio |
| Bora_in.vivo_noRT | Nedd8 | -0.65 | 2.22E-03 | glio |
| Bora_ex.vivo_noRT | Tmem236 | 2.621 | 2.22E-03 | glio |
| Bora_ex.vivo_noRT | Krtdap | 3.011 | 2.22E-03 | glio |
| Chek1_ex.vivo_noRT | Gm36913 | 2.859 | 2.22E-03 | glio |
| Chek1_ex.vivo_noRT | Matk | 2.689 | 2.22E-03 | glio |
| Chek1_ex.vivo_noRT | Pafah1b3 | -1.8 | 2.22E-03 | glio |
| Chek1_ex.vivo_noRT | Heatr3 | -1.865 | 2.22E-03 | glio |
| Chek1_ex.vivo_noRT | Plk2 | 2.025 | 2.22E-03 | glio |
| Hsd17b10_in.vivo_noRT | Gm41492 | 2.046 | 2.22E-03 | glio |
| Interaction_genotype_Aars2 | Sfr1 | 1.279 | 2.22E-04 | glio |
| Interaction_genotype_Mtor | Gm41442 | -4.124 | 2.22E-04 | glio |
| Interaction_genotype_Myc | Gm7102 | -4.366 | 2.22E-04 | glio |
| Interaction_genotype_Myc | Hist2h3c2 | -4.402 | 2.22E-04 | glio |
| Interaction_genotype_Myc | Lamtor1 | 1.669 | 2.22E-04 | glio |
| Interaction_genotype_Myc | Emg1 | 1.571 | 2.22E-04 | glio |
| Atr_in.vivo_noRT | Gm14418 | 2.041 | 2.22E-04 | glio |
| Atr_ex.vivo_noRT | Pcdhga1 | 2.993 | 2.22E-04 | glio |
| Bard1_ex.vivo_noRT | Lmx1a | 2.814 | 2.22E-04 | glio |
| Bcas2_in.vivo_noRT | Cpsf6 | -1.325 | 2.22E-04 | glio |
| Chek1_ex.vivo_noRT | Mb | 3.468 | 2.22E-04 | glio |
| Interaction_genotype_Mtor | Ggnbp2 | 1.948 | 2.22E-05 | glio |
| Interaction_genotype_Myc | Itga2b | -4.771 | 2.22E-05 | glio |
| Interaction_genotype_Myc | 4930405A21R | -4.492 | 2.22E-05 | glio |
| Interaction_genotype_Myc | C330011M18F | -4.933 | 2.22E-05 | glio |
| Interaction_genotype_Myc | Rer1 | 1.685 | 2.22E-05 | glio |
| Atr_in.vivo_noRT | Fam189b | 3.063 | 2.22E-05 | glio |
| Chek1_in.vivo_noRT | Etohd2 | 3.152 | 2.22E-05 | glio |
| Chek1_ex.vivo_noRT | Gm15520 | 3.264 | 2.22E-05 | glio |
| Hsd17b10_in.vivo_noRT | Vmn1r181 | 2.96 | 2.22E-05 | glio |
| Hsd17b10_in.vivo_noRT | 1700081N11R | 3.535 | 2.22E-05 | glio |
| culture_effect | Apeh | -1.164 | 2.22E-06 | glio |
| Aars2_in.vivo_noRT | Gm45833 | 4.479 | 2.22E-06 | glio |
| Atr_in.vivo_noRT | A830082N09R | 3.452 | 2.22E-06 | glio |

| | | | | |
|---|---|---|---|---|
| Bcas2_in.vivo_noRT | Mroh7 | 3.283 | 2.22E-06 | glio |
| Interaction_genotype_Mtor | Wac | 2.503 | 2.22E-07 | glio |
| Interaction_genotype_Mtor | K230015D01R | -6.278 | 2.22E-07 | glio |
| Interaction_genotype_Mtor | Gm16222 | -7.135 | 2.22E-07 | glio |
| Aars2_in.vivo_noRT | Gm36913 | 3.637 | 2.22E-07 | glio |
| Aars2_ex.vivo_noRT | mt-Nd4 | 1.002 | 2.22E-07 | glio |
| Atr_ex.vivo_noRT | Ablim2 | 3.164 | 2.22E-07 | glio |
| Atr_ex.vivo_noRT | Gm5415 | 3.164 | 2.22E-07 | glio |
| Bcas2_in.vivo_noRT | Ikbke | 3.491 | 2.22E-07 | glio |
| Atr_in.vivo_noRT | Gm15728 | 4.084 | 2.22E-08 | glio |
| culture_effect | Faim2 | -3.006 | 2.22E-09 | glio |
| Bora_in.vivo_noRT | Gm9999 | 3.961 | 2.22E-09 | glio |
| Bora_in.vivo_noRT | 1700047K16R | 4.279 | 2.22E-09 | glio |
| Chek1_in.vivo_noRT | Gm13822 | 4.518 | 2.22E-11 | glio |
| culture_effect | Bmp2 | -4.071 | 2.22E-12 | glio |
| Chek1_in.vivo_noRT | Gm128 | 4.126 | 2.22E-13 | glio |
| culture_effect | Pdia6 | -1.073 | 2.22E-14 | glio |
| Aars2_in.vivo_noRT | 1700006H21R | 5.327 | 2.22E-14 | glio |
| Aars2_in.vivo_noRT | Upb1 | 5.063 | 2.22E-14 | glio |
| Hsd17b10_in.vivo_noRT | Gm34059 | 3.616 | 2.22E-14 | glio |
| Atr_in.vivo_noRT | 4930505N22R | 5.273 | 2.22E-16 | glio |
| Bcas2_in.vivo_noRT | Trim17 | 4.807 | 2.22E-22 | glio |
| Chek1_in.vivo_noRT | Gm12800 | 5.044 | 2.22E-22 | glio |
| culture_effect | Apol10b | -7.168 | 2.22E-41 | glio |
| Interaction_genotype_Aars2 | Stx1b | -3.493 | 2.23E-03 | glio |
| Interaction_genotype_Aars2 | Hdgfl2 | 1.879 | 2.23E-03 | glio |
| Interaction_genotype_Atr | Rhox7a | -3.962 | 2.23E-03 | glio |
| Interaction_genotype_Atr | 4930554I06Ri | -4.027 | 2.23E-03 | glio |
| Interaction_genotype_Atr | Gsn | 2.057 | 2.23E-03 | glio |
| Interaction_genotype_Atr | Arl2bp | 1.387 | 2.23E-03 | glio |
| Interaction_genotype_Ifnar1 | Gm49172 | 2.994 | 2.23E-03 | glio |
| Interaction_genotype_Mtor | Rgs3 | -3.588 | 2.23E-03 | glio |
| Interaction_genotype_Mtor | Tnrc6c | 1.809 | 2.23E-03 | glio |
| Interaction_genotype_Mtor | Srp14 | 1.134 | 2.23E-03 | glio |
| Interaction_genotype_Mtor | Pcbp2 | 1.335 | 2.23E-03 | glio |
| Interaction_genotype_Myc | Parvb | -3.189 | 2.23E-03 | glio |
| Interaction_genotype_Myc | Harbi1 | -2.816 | 2.23E-03 | glio |
| Interaction_genotype_Myc | Cep19 | -2.719 | 2.23E-03 | glio |
| Interaction_genotype_Myc | Simc1 | -2.781 | 2.23E-03 | glio |
| Interaction_genotype_Myc | Msrb3 | -2.817 | 2.23E-03 | glio |
| interaction_RT | E230025N22R | 2.801 | 2.23E-03 | glio |
| Aars2_in.vivo_noRT | Usp35 | 2.2 | 2.23E-03 | glio |
| Aars2_in.vivo_noRT | Rmnd5b | 1.433 | 2.23E-03 | glio |
| Aars2_in.vivo_noRT | Nup205 | 1.238 | 2.23E-03 | glio |
| Aars2_in.vivo_noRT | Med28 | -0.711 | 2.23E-03 | glio |
| Aars2_ex.vivo_noRT | Ikbke | 2.887 | 2.23E-03 | glio |
| Atr_in.vivo_noRT | Zfp52 | 1.834 | 2.23E-03 | glio |
| Atr_ex.vivo_noRT | Gm49801 | 2.937 | 2.23E-03 | glio |
| Atr_ex.vivo_noRT | 1700008J07Ri | -2.594 | 2.23E-03 | glio |
| Bora_ex.vivo_noRT | Tex14 | 2.802 | 2.23E-03 | glio |
| Bora_ex.vivo_noRT | 9630014M24F | 2.827 | 2.23E-03 | glio |
| Brca2_ex.vivo_noRT | Prox2os | 3.191 | 2.23E-03 | glio |

| | | | | |
|---|---|---|---|---|
| Brca2_ex.vivo_noRT | Gm15972 | 3.215 | 2.23E-03 | glio |
| Chek1_in.vivo_noRT | Faim2 | 1.781 | 2.23E-03 | glio |
| Chek1_in.vivo_noRT | Minpp1 | -1.26 | 2.23E-03 | glio |
| Chek1_in.vivo_noRT | Rpl13a | -0.589 | 2.23E-03 | glio |
| Chek1_ex.vivo_noRT | Il23a | 2.856 | 2.23E-03 | glio |
| Cyp19a1_ex.vivo_noRT | Gm43951 | 2.942 | 2.23E-03 | glio |
| Cyp19a1_ex.vivo_noRT | Agrp | 2.941 | 2.23E-03 | glio |
| culture_effect | Unc119 | -1.165 | 2.23E-04 | glio |
| culture_effect | Ttc27 | -1.114 | 2.23E-04 | glio |
| culture_effect | Fbn2 | -0.888 | 2.23E-04 | glio |
| Interaction_genotype_Atr | Gm9967 | -4.385 | 2.23E-04 | glio |
| Interaction_genotype_Myc | Ube2k | 1.486 | 2.23E-04 | glio |
| Aars2_ex.vivo_noRT | Itgbl1 | 2.897 | 2.23E-04 | glio |
| Atr_in.vivo_noRT | Fgf21 | 2.767 | 2.23E-04 | glio |
| Atr_in.vivo_noRT | Eml1 | 2.367 | 2.23E-04 | glio |
| Atr_in.vivo_noRT | Ddx5 | -0.681 | 2.23E-04 | glio |
| Atr_ex.vivo_noRT | Gm37986 | 3.164 | 2.23E-04 | glio |
| Bcas2_in.vivo_noRT | Sapcd1 | 2.672 | 2.23E-04 | glio |
| Bora_ex.vivo_noRT | A830035O19R | 3.302 | 2.23E-04 | glio |
| Chek1_in.vivo_noRT | Tex19.2 | 2.58 | 2.23E-04 | glio |
| Chek1_in.vivo_noRT | Frmd4b | 2.66 | 2.23E-04 | glio |
| Chek1_in.vivo_noRT | Slx1b | 2.236 | 2.23E-04 | glio |
| Hsd17b10_in.vivo_noRT | Prtn3 | 2.834 | 2.23E-04 | glio |
| Hsd17b10_in.vivo_noRT | Tbxa2r | 2.557 | 2.23E-04 | glio |
| Hsd17b10_in.vivo_noRT | Gm20707 | 2.722 | 2.23E-04 | glio |
| Interaction_genotype_Mtor | AC241534.1 | -4.892 | 2.23E-05 | glio |
| Interaction_genotype_Mtor | Ubr4 | 2.216 | 2.23E-05 | glio |
| Interaction_genotype_Myc | Tmem79 | -4.462 | 2.23E-05 | glio |
| Interaction_genotype_Myc | Gm32090 | -4.135 | 2.23E-05 | glio |
| Interaction_genotype_Myc | Gm10425 | -5.023 | 2.23E-05 | glio |
| interaction_RT | Hist1h1b | -2.203 | 2.23E-05 | glio |
| Atr_in.vivo_noRT | Gm14326 | 3.025 | 2.23E-05 | glio |
| Atr_in.vivo_noRT | Pcdhb17 | 2.936 | 2.23E-05 | glio |
| Atr_in.vivo_noRT | Gsn | -1.933 | 2.23E-05 | glio |
| Bcas2_in.vivo_noRT | 4930455G09R | 3.521 | 2.23E-05 | glio |
| culture_effect | Pigs | -1.013 | 2.23E-06 | glio |
| Interaction_genotype_Aars2 | 4930589L23Ri | -6.001 | 2.23E-06 | glio |
| Interaction_genotype_Aars2 | Hbq1b | -5.78 | 2.23E-06 | glio |
| Interaction_genotype_Myc | Crhbp | -4.767 | 2.23E-06 | glio |
| Aars2_in.vivo_noRT | Rps6 | -0.844 | 2.23E-06 | glio |
| Atr_in.vivo_noRT | Gm26812 | 3.203 | 2.23E-06 | glio |
| Bcas2_in.vivo_noRT | Cnksr1 | 3.238 | 2.23E-06 | glio |
| Chek1_ex.vivo_noRT | Zfp455 | 4.079 | 2.23E-06 | glio |
| culture_effect | Tgfb2 | 2.017 | 2.23E-07 | glio |
| Aars2_in.vivo_noRT | 2610016A17R | 3.741 | 2.23E-07 | glio |
| Aars2_in.vivo_noRT | Acot5 | 3.443 | 2.23E-07 | glio |
| Atp5c1_in.vivo_noRT | Clec12b | 2.668 | 2.23E-07 | glio |
| Atp5c1_in.vivo_noRT | Gm14929 | 2.668 | 2.23E-07 | glio |
| Bard1_ex.vivo_noRT | Gm26555 | 3.612 | 2.23E-07 | glio |
| Bard1_ex.vivo_noRT | 9130019O22R | 3.612 | 2.23E-07 | glio |
| culture_effect | Cgref1 | 1.483 | 2.23E-08 | glio |
| interaction_RT | Nrp2 | -1.355 | 2.23E-08 | glio |

| | | | | |
|---|---|---|---|---|
| Bcas2_in.vivo_noRT | Gm16183 | 4.165 | 2.23E-08 | glio |
| Chek1_in.vivo_noRT | Slc26a6 | 3.752 | 2.23E-08 | glio |
| Interaction_genotype_Mtor | Ntn4 | -5.321 | 2.23E-09 | glio |
| Bcas2_in.vivo_noRT | Slc6a13 | 4.428 | 2.23E-09 | glio |
| Bora_in.vivo_noRT | Asgr1 | 4.278 | 2.23E-09 | glio |
| Bora_in.vivo_noRT | Slxl1 | 4.158 | 2.23E-09 | glio |
| Chek1_in.vivo_noRT | 1700003G18R | 4.784 | 2.23E-09 | glio |
| Chek1_ex.vivo_noRT | Tpsab1 | 4.842 | 2.23E-09 | glio |
| Aars2_in.vivo_noRT | Gm16183 | 4.684 | 2.23E-10 | glio |
| Bcas2_in.vivo_noRT | Gm15127 | 4.547 | 2.23E-11 | glio |
| Bcas2_in.vivo_noRT | Gm43672 | 4.807 | 2.23E-11 | glio |
| Chek1_in.vivo_noRT | Gm43463 | 4.386 | 2.23E-13 | glio |
| Chek1_in.vivo_noRT | Igf2bp1 | 5.044 | 2.23E-13 | glio |
| Bora_in.vivo_noRT | Gm14002 | 4.938 | 2.23E-14 | glio |
| culture_effect | Rpl9-ps6 | -1.199 | 2.23E-15 | glio |
| Atr_in.vivo_noRT | Fbxl13 | 5.273 | 2.23E-18 | glio |
| culture_effect | Ptpn18 | -1.952 | 2.23E-20 | glio |
| Atr_in.vivo_noRT | 1700024I08Ri | 5.273 | 2.23E-20 | glio |
| Atr_in.vivo_noRT | C430002N11R | 5.533 | 2.23E-23 | glio |
| Atr_in.vivo_noRT | Rimklb | 5.27 | 2.23E-26 | glio |
| culture_effect | Rufy3 | -0.834 | 2.24E-03 | glio |
| Interaction_genotype_Atr | Sucnr1 | -2.369 | 2.24E-03 | glio |
| Interaction_genotype_Atr | 1700112J16Ri | -2.956 | 2.24E-03 | glio |
| Interaction_genotype_Atr | Gm26581 | -3.955 | 2.24E-03 | glio |
| Interaction_genotype_Atr | Myl9 | -4.038 | 2.24E-03 | glio |
| Interaction_genotype_Atr | Zfhx2os | -3.79 | 2.24E-03 | glio |
| Interaction_genotype_Atr | Cpm | 2.655 | 2.24E-03 | glio |
| Interaction_genotype_Atr | Supt4a | 1.007 | 2.24E-03 | glio |
| Interaction_genotype_Bcas2 | Klhl30 | -3.682 | 2.24E-03 | glio |
| Interaction_genotype_Bcas2 | Cyct | 1.896 | 2.24E-03 | glio |
| Interaction_genotype_Mtor | Gm43258 | -3.675 | 2.24E-03 | glio |
| Interaction_genotype_Mtor | Ovgp1 | -3.6 | 2.24E-03 | glio |
| Interaction_genotype_Mtor | Dis3l | -3.325 | 2.24E-03 | glio |
| Interaction_genotype_Mtor | Crip1 | 2.389 | 2.24E-03 | glio |
| Interaction_genotype_Myc | A230056P14R | -3.41 | 2.24E-03 | glio |
| interaction_RT | Sdhaf4 | 0.785 | 2.24E-03 | glio |
| interaction_RT | Leprotl1 | 0.636 | 2.24E-03 | glio |
| Aars2_in.vivo_noRT | Eid3 | 1.904 | 2.24E-03 | glio |
| Atr_in.vivo_noRT | B3gnt5 | 1.987 | 2.24E-03 | glio |
| Atr_in.vivo_noRT | D430020J02Ri | 1.896 | 2.24E-03 | glio |
| Bcas2_in.vivo_noRT | Fhad1 | 2.081 | 2.24E-03 | glio |
| Bcas2_in.vivo_noRT | Klf7 | -1.582 | 2.24E-03 | glio |
| Bcas2_in.vivo_noRT | Marcksl1 | -1.228 | 2.24E-03 | glio |
| Bcas2_ex.vivo_noRT | Clca3a2 | 2.742 | 2.24E-03 | glio |
| Bcas2_ex.vivo_noRT | Art2b | 2.742 | 2.24E-03 | glio |
| Bora_in.vivo_noRT | Mafa | 2.029 | 2.24E-03 | glio |
| Bora_in.vivo_noRT | Klk7 | 3.065 | 2.24E-03 | glio |
| Bora_in.vivo_noRT | Raly | -0.791 | 2.24E-03 | glio |
| Chek1_in.vivo_noRT | Hspe1 | -0.632 | 2.24E-03 | glio |
| Chek1_ex.vivo_noRT | Guca1b | 2.821 | 2.24E-03 | glio |
| Chek1_ex.vivo_noRT | Msrb2 | 2.789 | 2.24E-03 | glio |
| culture_effect | Gtf2h1 | 0.574 | 2.24E-04 | glio |

| | | | | |
|---|---|---|---|---|
| Interaction_genotype_Mtor | 4930553M12F | -3.433 | 2.24E-04 | glio |
| Interaction_genotype_Mtor | Fndc10 | -4.774 | 2.24E-04 | glio |
| Interaction_genotype_Myc | Med21 | 1.408 | 2.24E-04 | glio |
| Aars2_in.vivo_noRT | Fhdc1 | 2.78 | 2.24E-04 | glio |
| Aars2_ex.vivo_noRT | Gm32122 | 3.293 | 2.24E-04 | glio |
| Atr_in.vivo_noRT | Comp | 2.781 | 2.24E-04 | glio |
| Atr_in.vivo_noRT | A130014A01R | 2.335 | 2.24E-04 | glio |
| Atr_in.vivo_noRT | Bst2 | -0.901 | 2.24E-04 | glio |
| Bora_in.vivo_noRT | Pcbp3 | 2.261 | 2.24E-04 | glio |
| Chek1_ex.vivo_noRT | Tmem86a | 3.475 | 2.24E-04 | glio |
| Eif2s1_in.vivo_noRT | Gm12841 | 2.143 | 2.24E-04 | glio |
| culture_effect | Casp6 | -1.066 | 2.24E-05 | glio |
| culture_effect | Hspa13 | -0.738 | 2.24E-05 | glio |
| Interaction_genotype_Mtor | Cabp4 | -5.082 | 2.24E-05 | glio |
| Interaction_genotype_Mtor | Myl7 | -5.257 | 2.24E-05 | glio |
| interaction_RT | Tlnrd1 | -1.353 | 2.24E-05 | glio |
| Bcas2_in.vivo_noRT | Rac1 | -1.075 | 2.24E-05 | glio |
| Bora_in.vivo_noRT | Icam5 | 2.914 | 2.24E-05 | glio |
| Chek1_ex.vivo_noRT | Tnnt3 | 4.049 | 2.24E-05 | glio |
| Hsd17b10_in.vivo_noRT | Ccr5 | 2.817 | 2.24E-05 | glio |
| culture_effect | Msl3 | 0.702 | 2.24E-06 | glio |
| Aars2_in.vivo_noRT | Gm48086 | 3.379 | 2.24E-06 | glio |
| Aars2_in.vivo_noRT | Ppp1cc | -1.709 | 2.24E-06 | glio |
| Atp5c1_in.vivo_noRT | Sema6d | 2.668 | 2.24E-06 | glio |
| Bcas2_in.vivo_noRT | 4922502N22R | 3.288 | 2.24E-06 | glio |
| Bora_in.vivo_noRT | Zfp972 | 3.492 | 2.24E-06 | glio |
| Hsd17b10_in.vivo_noRT | Fgf2 | 3.53 | 2.24E-06 | glio |
| Interaction_genotype_Mtor | 4833419F23Ri | -6.355 | 2.24E-07 | glio |
| Interaction_genotype_Mtor | Fhl1 | -4.809 | 2.24E-07 | glio |
| Interaction_genotype_Mtor | Gm20658 | -4.099 | 2.24E-07 | glio |
| Aars2_in.vivo_noRT | Gm9725 | 3.659 | 2.24E-07 | glio |
| Aars2_in.vivo_noRT | Gm15013 | 3.534 | 2.24E-07 | glio |
| Aars2_in.vivo_noRT | Ppp1r18os | 3.914 | 2.24E-07 | glio |
| Bora_in.vivo_noRT | Zfp652os | 3.631 | 2.24E-07 | glio |
| Bora_in.vivo_noRT | Zfp273 | 3.693 | 2.24E-07 | glio |
| Atr_in.vivo_noRT | Cnfn | 4.138 | 2.24E-08 | glio |
| Atr_in.vivo_noRT | Apold1 | 4.423 | 2.24E-08 | glio |
| Atr_in.vivo_noRT | Gm15535 | 4.216 | 2.24E-08 | glio |
| Hsd17b10_in.vivo_noRT | Capn9 | 3.616 | 2.24E-08 | glio |
| Interaction_genotype_Myc | Mecom | -6.572 | 2.24E-09 | glio |
| Atr_in.vivo_noRT | A530083I20Ri | 4.525 | 2.24E-09 | glio |
| culture_effect | Plod1 | -1.156 | 2.24E-10 | glio |
| Atr_ex.vivo_noRT | Usp17lc | 5.228 | 2.24E-12 | glio |
| Atr_ex.vivo_noRT | Gm13964 | 4.309 | 2.24E-12 | glio |
| Chek1_in.vivo_noRT | S100a14 | 4.243 | 2.24E-12 | glio |
| Chek1_in.vivo_noRT | Hcn3 | 4.401 | 2.24E-12 | glio |
| Hsd17b10_in.vivo_noRT | Cd36 | 3.616 | 2.24E-12 | glio |
| Hsd17b10_in.vivo_noRT | Strip2 | 3.873 | 2.24E-12 | glio |
| Hsd17b10_in.vivo_noRT | Gm43701 | 3.616 | 2.24E-12 | glio |
| Bora_in.vivo_noRT | Adam2 | 4.935 | 2.24E-16 | glio |
| Atr_in.vivo_noRT | Gm36026 | 4.883 | 2.24E-17 | glio |
| Atr_in.vivo_noRT | 4930502E18Ri | 4.998 | 2.24E-18 | glio |

| | | | | |
|---|---|---|---|---|
| Chek1_in.vivo_noRT | Ccl3 | 4.784 | 2.24E-20 | glio |
| Atr_in.vivo_noRT | Impg2 | 5.533 | 2.24E-21 | glio |
| Atr_in.vivo_noRT | Tpo | 5.533 | 2.24E-28 | glio |
| culture_effect | Cd74 | -11.379 | 2.24E-52 | glio |
| culture_effect | Nxt2 | 1.075 | 2.25E-03 | glio |
| culture_effect | Pdzd4 | 1.08 | 2.25E-03 | glio |
| culture_effect | Ilrun | -0.608 | 2.25E-03 | glio |
| culture_effect | Pim3 | -0.516 | 2.25E-03 | glio |
| Interaction_genotype_Atr | Rab3b | -3.722 | 2.25E-03 | glio |
| Interaction_genotype_Atr | Sec1 | -3.946 | 2.25E-03 | glio |
| Interaction_genotype_Bcas2 | Gm30938 | -3.826 | 2.25E-03 | glio |
| Interaction_genotype_Bcas2 | Fgd6 | -3.926 | 2.25E-03 | glio |
| Interaction_genotype_Bora | Atf6 | 2.526 | 2.25E-03 | glio |
| Interaction_genotype_Chek1 | Sec24c | 2.798 | 2.25E-03 | glio |
| Interaction_genotype_Chek1 | Tiam1 | -2.839 | 2.25E-03 | glio |
| Interaction_genotype_Chek1 | Myo5a | 2.094 | 2.25E-03 | glio |
| Interaction_genotype_Chek1 | Hmox1 | 3.469 | 2.25E-03 | glio |
| Interaction_genotype_Mtor | Col15a1 | -2.986 | 2.25E-03 | glio |
| Interaction_genotype_Mtor | Stxbp5 | -2.7 | 2.25E-03 | glio |
| Interaction_genotype_Myc | Nyap2 | -3.494 | 2.25E-03 | glio |
| Interaction_genotype_Myc | Ldlrap1 | -3.324 | 2.25E-03 | glio |
| Interaction_genotype_Myc | Rpusd3 | -2.906 | 2.25E-03 | glio |
| Interaction_genotype_Myc | Chst3 | -2.772 | 2.25E-03 | glio |
| Interaction_genotype_Myc | Zfp951 | -2.693 | 2.25E-03 | glio |
| Interaction_genotype_Myc | Aagab | -3.095 | 2.25E-03 | glio |
| Interaction_genotype_Myc | Spc25 | -2.618 | 2.25E-03 | glio |
| interaction_RT | Tnfaip2 | 2.109 | 2.25E-03 | glio |
| interaction_RT | Dhrs13 | -1.127 | 2.25E-03 | glio |
| interaction_RT | Pold2 | -0.833 | 2.25E-03 | glio |
| Aars2_in.vivo_noRT | Nck2 | 1.701 | 2.25E-03 | glio |
| Aars2_ex.vivo_noRT | Cyp4f40 | 2.9 | 2.25E-03 | glio |
| Atr_in.vivo_noRT | Slc5a5 | 2.28 | 2.25E-03 | glio |
| Atr_in.vivo_noRT | Man2b2 | 2.087 | 2.25E-03 | glio |
| Atr_in.vivo_noRT | Zscan29 | 1.855 | 2.25E-03 | glio |
| Bcas2_in.vivo_noRT | Gm17491 | 1.756 | 2.25E-03 | glio |
| Bcas2_in.vivo_noRT | Adra1b | -1.669 | 2.25E-03 | glio |
| Bcas2_in.vivo_noRT | Zfp106 | -1.071 | 2.25E-03 | glio |
| Bcas2_in.vivo_noRT | Eif5a | -0.582 | 2.25E-03 | glio |
| Bora_in.vivo_noRT | Cd2bp2 | -1.441 | 2.25E-03 | glio |
| Bora_in.vivo_noRT | Cop1 | -1.282 | 2.25E-03 | glio |
| Bora_in.vivo_noRT | Sf3a2 | -1.322 | 2.25E-03 | glio |
| Cenpj_in.vivo_noRT | A930002C04R | 1.762 | 2.25E-03 | glio |
| Chek1_ex.vivo_noRT | Mill2 | 2.746 | 2.25E-03 | glio |
| Chek1_ex.vivo_noRT | Cep83 | -1.565 | 2.25E-03 | glio |
| Eif2s1_in.vivo_noRT | Gm47872 | 2.403 | 2.25E-03 | glio |
| culture_effect | Snx17 | 0.468 | 2.25E-04 | glio |
| Interaction_genotype_Mtor | Bpifc | -4.818 | 2.25E-04 | glio |
| Interaction_genotype_Mtor | Zfp986 | -4.429 | 2.25E-04 | glio |
| Interaction_genotype_Mtor | Camkk1 | -4.478 | 2.25E-04 | glio |
| Interaction_genotype_Mtor | Shroom1 | -4.317 | 2.25E-04 | glio |
| Interaction_genotype_Mtor | Gm15706 | -3.455 | 2.25E-04 | glio |
| Interaction_genotype_Myc | Gm9967 | -3.944 | 2.25E-04 | glio |

| | | | | |
|---|---|---|---|---|
| Interaction_genotype_Myc | Mill2 | -4.206 | 2.25E-04 | glio |
| Interaction_genotype_Myc | Sema4f | -4.071 | 2.25E-04 | glio |
| Interaction_genotype_Myc | Zfp994 | -3.259 | 2.25E-04 | glio |
| Interaction_genotype_Myc | Rab4a | -2.924 | 2.25E-04 | glio |
| Interaction_genotype_Myc | Strap | 1.579 | 2.25E-04 | glio |
| interaction_RT | C2 | 2.055 | 2.25E-04 | glio |
| interaction_RT | Ube2t | 1.299 | 2.25E-04 | glio |
| Atp5c1_in.vivo_noRT | Gm45406 | 2.146 | 2.25E-04 | glio |
| Atr_in.vivo_noRT | Pkp3 | 2.734 | 2.25E-04 | glio |
| Bcas2_in.vivo_noRT | Slpi | 2.981 | 2.25E-04 | glio |
| Bcas2_in.vivo_noRT | Npl | 2.511 | 2.25E-04 | glio |
| Bora_in.vivo_noRT | Nat8l | 2.345 | 2.25E-04 | glio |
| Bora_in.vivo_noRT | 8430429K09R | 2.143 | 2.25E-04 | glio |
| Chek1_in.vivo_noRT | Slc4a3 | 2.444 | 2.25E-04 | glio |
| culture_effect | Gins1 | -1.256 | 2.25E-05 | glio |
| Interaction_genotype_Myc | Vps54 | -3.396 | 2.25E-05 | glio |
| Interaction_genotype_Myc | Rmi2 | -4.137 | 2.25E-05 | glio |
| Interaction_genotype_Myc | Gm26766 | -4.59 | 2.25E-05 | glio |
| Interaction_genotype_Myc | Gm9922 | -4.578 | 2.25E-05 | glio |
| Interaction_genotype_Myc | Camkk2 | -3.748 | 2.25E-05 | glio |
| Interaction_genotype_Myc | Gm44899 | -3.834 | 2.25E-05 | glio |
| Interaction_genotype_Myc | Gm31063 | -3.653 | 2.25E-05 | glio |
| Interaction_genotype_Myc | Eif4g2 | 1.571 | 2.25E-05 | glio |
| Interaction_genotype_Myc | Eif2s2 | 1.599 | 2.25E-05 | glio |
| Interaction_genotype_Myc | Rps23 | 1.652 | 2.25E-05 | glio |
| Aars2_in.vivo_noRT | Prrc1 | -1.839 | 2.25E-05 | glio |
| Aars2_in.vivo_noRT | Srsf9 | -1.266 | 2.25E-05 | glio |
| Aars2_ex.vivo_noRT | Gm43951 | 3.276 | 2.25E-05 | glio |
| Atr_in.vivo_noRT | Zfp37 | 2.865 | 2.25E-05 | glio |
| Atr_ex.vivo_noRT | Gm12841 | 2.771 | 2.25E-05 | glio |
| Bcas2_in.vivo_noRT | Gm6598 | 2.816 | 2.25E-05 | glio |
| Hsd17b10_in.vivo_noRT | Gm45441 | 3.088 | 2.25E-05 | glio |
| Interaction_genotype_Myc | Gm9958 | -5.474 | 2.25E-06 | glio |
| Atr_ex.vivo_noRT | Gm2016 | 3.164 | 2.25E-06 | glio |
| Bora_in.vivo_noRT | Tnni2 | 3.898 | 2.25E-06 | glio |
| Interaction_genotype_Bora | Dnm3 | -6.173 | 2.25E-07 | glio |
| Interaction_genotype_Myc | Tomm20 | 2.088 | 2.25E-07 | glio |
| Interaction_genotype_Myc | Rgs16 | -6.602 | 2.25E-07 | glio |
| Interaction_genotype_Myc | Pfn1 | 2.172 | 2.25E-07 | glio |
| Interaction_genotype_Myc | Lbhd2 | -7.007 | 2.25E-07 | glio |
| Aars2_in.vivo_noRT | Slc22a18 | 3.81 | 2.25E-07 | glio |
| Aars2_ex.vivo_noRT | C430002N11R | 3.697 | 2.25E-07 | glio |
| Bcas2_in.vivo_noRT | Gm15903 | 3.495 | 2.25E-07 | glio |
| Chek1_in.vivo_noRT | Dio1 | 3.208 | 2.25E-07 | glio |
| Chek1_in.vivo_noRT | Nlrx1 | 4.156 | 2.25E-08 | glio |
| Chek1_ex.vivo_noRT | A830009L08Ri | 4.368 | 2.25E-08 | glio |
| culture_effect | 1810058I24Ri | 0.71 | 2.25E-09 | glio |
| interaction_RT | Atp6v1f | 1.019 | 2.25E-09 | glio |
| interaction_RT | Dok1 | -1.817 | 2.25E-09 | glio |
| Aars2_in.vivo_noRT | Lrrn3 | 3.871 | 2.25E-09 | glio |
| Bora_in.vivo_noRT | 2310058D17R | 4.29 | 2.25E-09 | glio |
| Hsd17b10_in.vivo_noRT | Psmb11 | 3.093 | 2.25E-09 | glio |

| | | | | |
|---|---|---|---|---|
| culture_effect | Asphd2 | 2.956 | 2.25E-10 | glio |
| Chek1_in.vivo_noRT | Platr31 | 5.044 | 2.25E-10 | glio |
| Atr_in.vivo_noRT | Gm3734 | 5.01 | 2.25E-11 | glio |
| Bora_in.vivo_noRT | Akap3 | 4.676 | 2.25E-11 | glio |
| Aars2_in.vivo_noRT | Hsf2bp | 4.535 | 2.25E-12 | glio |
| Atr_in.vivo_noRT | Gm20442 | 4.525 | 2.25E-13 | glio |
| Atr_in.vivo_noRT | Apobr | 5.533 | 2.25E-13 | glio |
| Atr_in.vivo_noRT | Gm48742 | 4.883 | 2.25E-17 | glio |
| culture_effect | Pfkp | -0.78 | 2.26E-03 | glio |
| culture_effect | Slc7a5 | -0.873 | 2.26E-03 | glio |
| culture_effect | Ascc3 | 0.5 | 2.26E-03 | glio |
| culture_effect | Sigmar1 | -0.509 | 2.26E-03 | glio |
| Interaction_genotype_Atr | Gm2788 | -4.044 | 2.26E-03 | glio |
| Interaction_genotype_Atr | Gm9958 | -3.732 | 2.26E-03 | glio |
| Interaction_genotype_Atr | Hsbp1 | 1.033 | 2.26E-03 | glio |
| Interaction_genotype_Bcas2 | Plekho2 | -4.026 | 2.26E-03 | glio |
| Interaction_genotype_Bcas2 | 5330439K02R | -4.096 | 2.26E-03 | glio |
| Interaction_genotype_Bcas2 | Gm10324 | -3.162 | 2.26E-03 | glio |
| Interaction_genotype_Mtor | Gm16754 | -3.786 | 2.26E-03 | glio |
| Interaction_genotype_Mtor | Ms4a15 | -2.929 | 2.26E-03 | glio |
| Interaction_genotype_Mtor | Slc25a53 | -3.956 | 2.26E-03 | glio |
| Interaction_genotype_Mtor | AW046200 | -3.603 | 2.26E-03 | glio |
| Interaction_genotype_Myc | Gm20442 | -3.088 | 2.26E-03 | glio |
| Interaction_genotype_Myc | 9930014A18R | -3.326 | 2.26E-03 | glio |
| Interaction_genotype_Myc | Mocs3 | -3.107 | 2.26E-03 | glio |
| Interaction_genotype_Myc | Rpl27 | 1.213 | 2.26E-03 | glio |
| interaction_RT | Pole | -1.318 | 2.26E-03 | glio |
| Aars2_in.vivo_noRT | Nemp2 | 2.38 | 2.26E-03 | glio |
| Aars2_in.vivo_noRT | Rpl14 | -0.583 | 2.26E-03 | glio |
| Atr_in.vivo_noRT | AI597479 | 1.692 | 2.26E-03 | glio |
| Atr_in.vivo_noRT | Hspa9 | -0.772 | 2.26E-03 | glio |
| Bcas2_in.vivo_noRT | Ube2cbp | 1.979 | 2.26E-03 | glio |
| Bcas2_in.vivo_noRT | Saysd1 | -1.501 | 2.26E-03 | glio |
| Bcas2_in.vivo_noRT | Nabp2 | -1.006 | 2.26E-03 | glio |
| Bcas2_ex.vivo_noRT | Gm12589 | 2.342 | 2.26E-03 | glio |
| Cenpj_in.vivo_noRT | Gm21319 | 2.022 | 2.26E-03 | glio |
| Cenpj_in.vivo_noRT | Sbk3 | 2.438 | 2.26E-03 | glio |
| Chek1_in.vivo_noRT | Prpf40b | 1.717 | 2.26E-03 | glio |
| Chek1_ex.vivo_noRT | Klhl36 | 2.683 | 2.26E-03 | glio |
| Hsd17b10_in.vivo_noRT | AY036118 | 1.117 | 2.26E-03 | glio |
| Interaction_genotype_Mtor | Itgae | -4.641 | 2.26E-04 | glio |
| Interaction_genotype_Mtor | Ece2 | -4.217 | 2.26E-04 | glio |
| Interaction_genotype_Mtor | Pex10 | -3.534 | 2.26E-04 | glio |
| Interaction_genotype_Myc | Helt | -2.729 | 2.26E-04 | glio |
| Interaction_genotype_Myc | Cdca2 | -3.318 | 2.26E-04 | glio |
| Atp5c1_in.vivo_noRT | Gm14540 | 2.145 | 2.26E-04 | glio |
| Atr_in.vivo_noRT | Zfp316 | 2.246 | 2.26E-04 | glio |
| Atr_in.vivo_noRT | Scpep1 | -1.487 | 2.26E-04 | glio |
| Bcas2_in.vivo_noRT | Cela1 | -1.737 | 2.26E-04 | glio |
| Bora_in.vivo_noRT | Zfp764 | 2.467 | 2.26E-04 | glio |
| Cenpj_in.vivo_noRT | Gm13191 | 2.022 | 2.26E-04 | glio |
| Cenpj_in.vivo_noRT | 4933416E03Ri | 2.022 | 2.26E-04 | glio |

| | | | | |
|---|---|---|---|---|
| Chek1_in.vivo_noRT | Dll1 | 2.809 | 2.26E-04 | glio |
| Interaction_genotype_Myc | 4933429H19R | -4.418 | 2.26E-05 | glio |
| Aars2_in.vivo_noRT | Myl4 | 3.076 | 2.26E-05 | glio |
| Aars2_in.vivo_noRT | AY074887 | 3.274 | 2.26E-05 | glio |
| Aars2_in.vivo_noRT | Gal3st4 | 2.968 | 2.26E-05 | glio |
| Aars2_in.vivo_noRT | Rorc | 3.202 | 2.26E-05 | glio |
| Aars2_in.vivo_noRT | Tars | -1.245 | 2.26E-05 | glio |
| Aars2_in.vivo_noRT | Vdac1 | -1.264 | 2.26E-05 | glio |
| Bora_in.vivo_noRT | Gm10244 | 3.115 | 2.26E-05 | glio |
| Bora_ex.vivo_noRT | Gm28940 | 3.201 | 2.26E-05 | glio |
| Chek1_in.vivo_noRT | Tmco1 | -1.095 | 2.26E-05 | glio |
| culture_effect | 2310058D17R | 2.515 | 2.26E-06 | glio |
| Interaction_genotype_Mtor | Spink10 | -5.263 | 2.26E-06 | glio |
| Interaction_genotype_Mtor | Zfp599 | -6.073 | 2.26E-06 | glio |
| Interaction_genotype_Mtor | Ikzf2 | -6.075 | 2.26E-06 | glio |
| Bora_in.vivo_noRT | Hspb9 | 3.625 | 2.26E-06 | glio |
| Bora_in.vivo_noRT | Epb41l2 | -1.914 | 2.26E-06 | glio |
| Chek1_in.vivo_noRT | Gm16759 | 3.478 | 2.26E-06 | glio |
| Chek1_in.vivo_noRT | A830082N09R | 3.474 | 2.26E-06 | glio |
| Chek1_ex.vivo_noRT | Napsa | 4.449 | 2.26E-06 | glio |
| Interaction_genotype_Mtor | Gm12841 | -4.492 | 2.26E-07 | glio |
| Aars2_in.vivo_noRT | Prelid3a | 3.609 | 2.26E-07 | glio |
| Atr_in.vivo_noRT | Col28a1 | 3.387 | 2.26E-07 | glio |
| Bcas2_in.vivo_noRT | Gm10484 | 3.806 | 2.26E-07 | glio |
| Bora_in.vivo_noRT | Etl4 | 3.06 | 2.26E-07 | glio |
| Chek1_ex.vivo_noRT | Fam25c | 3.469 | 2.26E-07 | glio |
| culture_effect | Stk19 | -0.875 | 2.26E-08 | glio |
| culture_effect | Fus | -0.745 | 2.26E-08 | glio |
| Bora_in.vivo_noRT | C230037L18Ri | 3.768 | 2.26E-08 | glio |
| culture_effect | Rbm28 | 0.851 | 2.26E-09 | glio |
| Chek1_in.vivo_noRT | Otop1 | 3.859 | 2.26E-09 | glio |
| Bcas2_in.vivo_noRT | Gm16364.1 | 4.807 | 2.26E-10 | glio |
| Atr_in.vivo_noRT | Ly6c2 | 4.63 | 2.26E-11 | glio |
| Bcas2_in.vivo_noRT | Gm30082 | 4.674 | 2.26E-11 | glio |
| culture_effect | Wdr38 | 3.613 | 2.26E-13 | glio |
| Aars2_in.vivo_noRT | Gm14379 | 4.811 | 2.26E-13 | glio |
| Aars2_in.vivo_noRT | Ccdc63 | 4.941 | 2.26E-13 | glio |
| Aars2_in.vivo_noRT | Gm17276 | 4.536 | 2.26E-15 | glio |
| Atr_in.vivo_noRT | Rgs9 | 4.733 | 2.26E-15 | glio |
| Atr_in.vivo_noRT | Nxph4 | 5.533 | 2.26E-15 | glio |
| culture_effect | Efna1 | -3.716 | 2.26E-17 | glio |
| Chek1_in.vivo_noRT | Akap14 | 4.78 | 2.26E-18 | glio |
| culture_effect | Cstb | -1.985 | 2.26E-19 | glio |
| Aars2_in.vivo_noRT | Gm16933 | 4.805 | 2.26E-22 | glio |
| Aars2_in.vivo_noRT | Ush1g | 5.327 | 2.26E-23 | glio |
| Chek1_in.vivo_noRT | Gm48764 | 5.044 | 2.26E-23 | glio |
| culture_effect | Pink1 | -0.668 | 2.27E-03 | glio |
| culture_effect | Mfap1b | -0.585 | 2.27E-03 | glio |
| culture_effect | Ubfd1 | -0.433 | 2.27E-03 | glio |
| culture_effect | Dcaf13 | -0.478 | 2.27E-03 | glio |
| Interaction_genotype_Aars2 | Farsb | 1.677 | 2.27E-03 | glio |
| Interaction_genotype_Mtor | Plppr3 | -3.693 | 2.27E-03 | glio |

| | | | | |
|---|---|---|---|---|
| Interaction_genotype_Mtor | Wrap53 | -2.662 | 2.27E-03 | glio |
| Interaction_genotype_Mtor | Chd7 | 1.993 | 2.27E-03 | glio |
| Interaction_genotype_Mtor | Prkcsh | 1.59 | 2.27E-03 | glio |
| Interaction_genotype_Myc | Gm49785 | -3.19 | 2.27E-03 | glio |
| Interaction_genotype_Myc | Rps8 | 1.055 | 2.27E-03 | glio |
| Aars2_in.vivo_noRT | Fgd6 | 2.082 | 2.27E-03 | glio |
| Aars2_in.vivo_noRT | Cyb5r4 | -1.24 | 2.27E-03 | glio |
| Atr_in.vivo_noRT | Kank4os | 1.703 | 2.27E-03 | glio |
| Atr_in.vivo_noRT | Nup50 | -1.061 | 2.27E-03 | glio |
| Atr_in.vivo_noRT | Myof | 1.395 | 2.27E-03 | glio |
| Bcas2_in.vivo_noRT | Cdkn1c | 2.238 | 2.27E-03 | glio |
| Bcas2_in.vivo_noRT | Gm45605 | 2.254 | 2.27E-03 | glio |
| Bcas2_in.vivo_noRT | Rell2 | 2.329 | 2.27E-03 | glio |
| Brca2_in.vivo_noRT | Dnajc16 | -2.614 | 2.27E-03 | glio |
| Chek1_in.vivo_noRT | Lhx9 | 2.339 | 2.27E-03 | glio |
| Chek1_ex.vivo_noRT | Tuba1b | -0.978 | 2.27E-03 | glio |
| Hsd17b10_in.vivo_noRT | Slc16a6 | 2.432 | 2.27E-03 | glio |
| culture_effect | Dctn2 | -0.491 | 2.27E-04 | glio |
| Interaction_genotype_Aars2 | Gm27253 | -4.55 | 2.27E-04 | glio |
| Interaction_genotype_Aars2 | 1700017M07F | -3.979 | 2.27E-04 | glio |
| Interaction_genotype_Aars2 | Gadd45gip1 | 1.598 | 2.27E-04 | glio |
| Interaction_genotype_Aars2 | Uqcrq | 1.191 | 2.27E-04 | glio |
| Interaction_genotype_Atr | Gm9970 | -4.815 | 2.27E-04 | glio |
| Interaction_genotype_Bcas2 | Gm49479 | -4.874 | 2.27E-04 | glio |
| Interaction_genotype_Bcas2 | Rnft2 | -4.769 | 2.27E-04 | glio |
| Interaction_genotype_Mtor | 8030456M14F | -3.9 | 2.27E-04 | glio |
| Interaction_genotype_Mtor | Tuba1c | 2.755 | 2.27E-04 | glio |
| interaction_RT | Sft2d1 | 1.099 | 2.27E-04 | glio |
| interaction_RT | Nop2 | -0.918 | 2.27E-04 | glio |
| interaction_RT | Junb | 1.38 | 2.27E-04 | glio |
| Aars2_in.vivo_noRT | Ccdc7b | 2.522 | 2.27E-04 | glio |
| Aars2_in.vivo_noRT | Rpl8 | -0.735 | 2.27E-04 | glio |
| Aars2_in.vivo_noRT | Eef1a1 | -0.694 | 2.27E-04 | glio |
| Atr_in.vivo_noRT | Msh5 | 2.766 | 2.27E-04 | glio |
| Bcas2_in.vivo_noRT | 1810021B22R | 2.738 | 2.27E-04 | glio |
| Bcas2_in.vivo_noRT | Mmp19 | 2.613 | 2.27E-04 | glio |
| Bcas2_ex.vivo_noRT | Gm45406 | 2.742 | 2.27E-04 | glio |
| Bora_in.vivo_noRT | 4930594M22F | 2.689 | 2.27E-04 | glio |
| Bora_in.vivo_noRT | Gm10501 | 2.794 | 2.27E-04 | glio |
| Bora_ex.vivo_noRT | Pla2g2f | 3.201 | 2.27E-04 | glio |
| Chek1_in.vivo_noRT | Jazf1 | 2.539 | 2.27E-04 | glio |
| Chek1_in.vivo_noRT | Gm38115 | 2.631 | 2.27E-04 | glio |
| Chek1_in.vivo_noRT | Rpn2 | -0.943 | 2.27E-04 | glio |
| Chek1_in.vivo_noRT | Eif5 | -0.791 | 2.27E-04 | glio |
| Chek1_ex.vivo_noRT | Shroom3 | 3.287 | 2.27E-04 | glio |
| Dclre1c_ex.vivo_noRT | Ankrd22 | 2.658 | 2.27E-04 | glio |
| culture_effect | Mpp6 | 0.601 | 2.27E-05 | glio |
| Interaction_genotype_Mtor | Col11a1 | 1.857 | 2.27E-05 | glio |
| Atr_in.vivo_noRT | Phkg1 | 4.104 | 2.27E-05 | glio |
| Atr_in.vivo_noRT | Cdkn1c | 2.965 | 2.27E-05 | glio |
| Atr_in.vivo_noRT | Gm15133 | 2.927 | 2.27E-05 | glio |
| Atr_in.vivo_noRT | 2810405F17Ri | 2.847 | 2.27E-05 | glio |

| | | | | |
|---|---|---|---|---|
| Atr_in.vivo_noRT | Ube2d3 | -0.865 | 2.27E-05 | glio |
| Bcas2_in.vivo_noRT | Cldn9 | 2.889 | 2.27E-05 | glio |
| Bcas2_in.vivo_noRT | 0610038B21R | 2.767 | 2.27E-05 | glio |
| Bcas2_ex.vivo_noRT | Slc14a2 | 2.742 | 2.27E-05 | glio |
| Bora_in.vivo_noRT | Slc9a9 | 3.081 | 2.27E-05 | glio |
| Chek1_in.vivo_noRT | Gm31517 | 3.067 | 2.27E-05 | glio |
| Chek1_in.vivo_noRT | Tex14 | 2.703 | 2.27E-05 | glio |
| Chek1_in.vivo_noRT | Slc47a2 | 2.938 | 2.27E-05 | glio |
| Aars2_in.vivo_noRT | 4930589L23Ri | 3.448 | 2.27E-06 | glio |
| Aars2_in.vivo_noRT | Phtf1os | 3.402 | 2.27E-06 | glio |
| Atr_in.vivo_noRT | Hoxa7 | 3.294 | 2.27E-06 | glio |
| Bora_in.vivo_noRT | Hist1h4k | 3.367 | 2.27E-06 | glio |
| Chek1_in.vivo_noRT | Gm16576 | 3.389 | 2.27E-06 | glio |
| Hsd17b10_in.vivo_noRT | Gm41409 | 3.084 | 2.27E-06 | glio |
| Interaction_genotype_Bcas2 | Klhdc8a | -5.063 | 2.27E-07 | glio |
| Aars2_in.vivo_noRT | Gm16048 | 3.891 | 2.27E-07 | glio |
| Aars2_in.vivo_noRT | Bcl2l14 | 3.822 | 2.27E-07 | glio |
| Atp5c1_in.vivo_noRT | Vmn1r32 | 2.668 | 2.27E-07 | glio |
| Atr_in.vivo_noRT | Zfp433 | 4.094 | 2.27E-08 | glio |
| Chek1_in.vivo_noRT | Scgb1a1 | 3.934 | 2.27E-08 | glio |
| Hsd17b10_in.vivo_noRT | Tcstv1 | 3.616 | 2.27E-09 | glio |
| Aars2_in.vivo_noRT | 1110017D15R | 4.326 | 2.27E-10 | glio |
| culture_effect | Tusc3 | -1.025 | 2.27E-11 | glio |
| Hsd17b10_in.vivo_noRT | 9130019O22R | 3.616 | 2.27E-12 | glio |
| Aars2_in.vivo_noRT | Bmx | 4.539 | 2.27E-13 | glio |
| Chek1_ex.vivo_noRT | Gm36793 | 4.842 | 2.27E-14 | glio |
| Chek1_ex.vivo_noRT | A030014E15Ri | 4.449 | 2.27E-14 | glio |
| Hsd17b10_in.vivo_noRT | Gm48435 | 3.616 | 2.27E-14 | glio |
| Aars2_in.vivo_noRT | Nxph3 | 5.327 | 2.27E-16 | glio |
| Chek1_in.vivo_noRT | Cfh | 4.264 | 2.27E-16 | glio |
| Aars2_in.vivo_noRT | Pik3ap1 | 5.063 | 2.27E-17 | glio |
| Bcas2_in.vivo_noRT | Gm21319 | 4.807 | 2.27E-18 | glio |
| Aars2_in.vivo_noRT | Vtn | 5.067 | 2.27E-22 | glio |
| Bcas2_in.vivo_noRT | Tpo | 4.807 | 2.27E-22 | glio |
| Aars2_in.vivo_noRT | Gm31063 | 5.327 | 2.27E-23 | glio |
| Aars2_in.vivo_noRT | Gm31717 | 5.327 | 2.27E-27 | glio |
| culture_effect | Rab26 | 1.365 | 2.28E-03 | glio |
| Interaction_genotype_Aars2 | Gm44096 | -3.628 | 2.28E-03 | glio |
| Interaction_genotype_Bcas2 | Cd248 | -3.996 | 2.28E-03 | glio |
| Interaction_genotype_Mtor | Pawr | -3.305 | 2.28E-03 | glio |
| Interaction_genotype_Mtor | Cfap69 | -3.703 | 2.28E-03 | glio |
| Interaction_genotype_Mtor | 4930513N10R | -3.916 | 2.28E-03 | glio |
| Interaction_genotype_Mtor | Suox | -3.335 | 2.28E-03 | glio |
| Interaction_genotype_Mtor | Eif3a | 1.071 | 2.28E-03 | glio |
| Interaction_genotype_Myc | Gng3 | -3.324 | 2.28E-03 | glio |
| Interaction_genotype_Myc | Ago4 | -3.324 | 2.28E-03 | glio |
| interaction_RT | Akirin2 | 0.538 | 2.28E-03 | glio |
| Aars2_in.vivo_noRT | Rsrc2 | -1.095 | 2.28E-03 | glio |
| Atr_in.vivo_noRT | Ide | -1.247 | 2.28E-03 | glio |
| Bcas2_in.vivo_noRT | Rundc1 | 1.838 | 2.28E-03 | glio |
| Bora_in.vivo_noRT | Esr1 | 2.08 | 2.28E-03 | glio |
| Bora_in.vivo_noRT | Gdpd5 | 2.208 | 2.28E-03 | glio |

| | | | | |
|---|---|---|---|---|
| Cenpj_in.vivo_noRT | Speer4e | 2.022 | 2.28E-03 | glio |
| Cenpj_in.vivo_noRT | 1700047K16R | 2.663 | 2.28E-03 | glio |
| Cenpj_in.vivo_noRT | C430002N11R | 2.022 | 2.28E-03 | glio |
| Cenpj_ex.vivo_noRT | Mfap3l | 3.404 | 2.28E-03 | glio |
| Chek1_in.vivo_noRT | Relb | -1.469 | 2.28E-03 | glio |
| Chek1_in.vivo_noRT | Taf5 | -1.378 | 2.28E-03 | glio |
| Chek1_ex.vivo_noRT | 2310008N11R | 2.966 | 2.28E-03 | glio |
| Chek1_ex.vivo_noRT | Resf1 | 1.405 | 2.28E-03 | glio |
| Hsd17b10_in.vivo_noRT | Nmb | 2.322 | 2.28E-03 | glio |
| Hsd17b10_in.vivo_noRT | Trim67 | 2.047 | 2.28E-03 | glio |
| Hsd17b10_in.vivo_noRT | Gm9484 | 2.458 | 2.28E-03 | glio |
| Hsd17b10_in.vivo_noRT | Zfp661 | 2.392 | 2.28E-03 | glio |
| culture_effect | Ccdc117 | -0.867 | 2.28E-04 | glio |
| Interaction_genotype_Mtor | Rwdd1 | 1.441 | 2.28E-04 | glio |
| Interaction_genotype_Myc | Gm20625 | -2.729 | 2.28E-04 | glio |
| Interaction_genotype_Myc | Uckl1os | -4.429 | 2.28E-04 | glio |
| Interaction_genotype_Myc | Kcnmb4 | -3.98 | 2.28E-04 | glio |
| Aars2_in.vivo_noRT | Dnah2os | 2.377 | 2.28E-04 | glio |
| Atr_in.vivo_noRT | Rab11fip5 | 2.693 | 2.28E-04 | glio |
| Atr_in.vivo_noRT | Pgk1 | -0.825 | 2.28E-04 | glio |
| Chek1_in.vivo_noRT | Smim41 | 2.849 | 2.28E-04 | glio |
| Chek1_in.vivo_noRT | Gm32051 | 2.755 | 2.28E-04 | glio |
| Chek1_in.vivo_noRT | Rnf182 | 2.348 | 2.28E-04 | glio |
| Chek1_in.vivo_noRT | Vdac2 | -0.887 | 2.28E-04 | glio |
| Chek1_in.vivo_noRT | Tpt1 | -0.837 | 2.28E-04 | glio |
| Chek1_ex.vivo_noRT | Gm44867 | 3.458 | 2.28E-04 | glio |
| culture_effect | Gm27253 | 2.381 | 2.28E-05 | glio |
| culture_effect | Mettl17 | -1.62 | 2.28E-05 | glio |
| Interaction_genotype_Aars2 | Carns1 | -5.387 | 2.28E-05 | glio |
| Interaction_genotype_Mtor | D030051J21Ri | -4.95 | 2.28E-05 | glio |
| Interaction_genotype_Mtor | Rbm25 | 1.661 | 2.28E-05 | glio |
| Atr_in.vivo_noRT | Gm12059 | 2.862 | 2.28E-05 | glio |
| Atr_in.vivo_noRT | Zbtb8os | -1.582 | 2.28E-05 | glio |
| Bora_ex.vivo_noRT | Cfb | 3.201 | 2.28E-05 | glio |
| Hsd17b10_in.vivo_noRT | Sod3 | 3.616 | 2.28E-05 | glio |
| Aars2_in.vivo_noRT | Mmp28 | 3.433 | 2.28E-06 | glio |
| Aars2_ex.vivo_noRT | Gm45406 | 3.293 | 2.28E-06 | glio |
| Atp5c1_in.vivo_noRT | Cxcl5 | 2.668 | 2.28E-06 | glio |
| Atp5c1_in.vivo_noRT | Hif3a | 3.558 | 2.28E-06 | glio |
| Atp5c1_in.vivo_noRT | Kcnn4 | -3.921 | 2.28E-06 | glio |
| Atr_in.vivo_noRT | Gm36738 | 3.525 | 2.28E-06 | glio |
| Bcas2_in.vivo_noRT | Dynll1 | -0.938 | 2.28E-06 | glio |
| Bora_ex.vivo_noRT | Ocm | 3.994 | 2.28E-06 | glio |
| Aars2_in.vivo_noRT | Gm36738 | 3.855 | 2.28E-07 | glio |
| Aars2_ex.vivo_noRT | Gm13078 | 3.293 | 2.28E-07 | glio |
| Atr_in.vivo_noRT | Slfn3 | 3.245 | 2.28E-07 | glio |
| Bcas2_in.vivo_noRT | Emp1 | -2.141 | 2.28E-07 | glio |
| Chek1_in.vivo_noRT | 1700122E12Ri | 3.648 | 2.28E-07 | glio |
| Chek1_in.vivo_noRT | Sipa1l2 | 3.629 | 2.28E-07 | glio |
| Bora_in.vivo_noRT | Gm16754 | 3.956 | 2.28E-08 | glio |
| Bora_in.vivo_noRT | Slc41a3 | 3.72 | 2.28E-08 | glio |
| Atr_in.vivo_noRT | Ikbke | 4.217 | 2.28E-10 | glio |

| | | | | |
|---|---|---|---|---|
| Aars2_in.vivo_noRT | Flicr | 4.526 | 2.28E-11 | glio |
| Atr_in.vivo_noRT | Gm15927 | 4.532 | 2.28E-11 | glio |
| Bcas2_in.vivo_noRT | Gm37864 | 4.027 | 2.28E-11 | glio |
| Bcas2_in.vivo_noRT | Rhox7a | 4.807 | 2.28E-11 | glio |
| Bora_in.vivo_noRT | Bmx | 4.149 | 2.28E-11 | glio |
| Bora_in.vivo_noRT | 2300002M23F | 3.768 | 2.28E-11 | glio |
| Aars2_in.vivo_noRT | Col6a3 | 4.29 | 2.28E-12 | glio |
| Atr_in.vivo_noRT | C230037L18Ri | 4.902 | 2.28E-13 | glio |
| culture_effect | Lgals7 | -4.827 | 2.28E-14 | glio |
| culture_effect | Eef1b2 | -0.947 | 2.28E-14 | glio |
| Atr_in.vivo_noRT | Lyl1 | 4.883 | 2.28E-16 | glio |
| Bcas2_in.vivo_noRT | Gm9758 | 4.547 | 2.28E-18 | glio |
| Atr_in.vivo_noRT | 2410141K09R | 5.269 | 2.28E-19 | glio |
| Chek1_in.vivo_noRT | Gm48780 | 4.784 | 2.28E-20 | glio |
| culture_effect | Gm36757 | -1.929 | 2.29E-03 | glio |
| culture_effect | 1110046J04Ri | 1.877 | 2.29E-03 | glio |
| culture_effect | Zcwpw1 | 0.954 | 2.29E-03 | glio |
| Interaction_genotype_Atr | Slc19a3 | -2.763 | 2.29E-03 | glio |
| Interaction_genotype_Atr | Hoxa6 | -3.689 | 2.29E-03 | glio |
| Interaction_genotype_Cdkn2a | Capn13 | 2.835 | 2.29E-03 | glio |
| Interaction_genotype_Mtor | Gm20732 | -3.329 | 2.29E-03 | glio |
| Interaction_genotype_Mtor | Gm16094 | -3.38 | 2.29E-03 | glio |
| Interaction_genotype_Mtor | Gm15614 | -3.683 | 2.29E-03 | glio |
| Interaction_genotype_Mtor | Gm47438 | -3.745 | 2.29E-03 | glio |
| Interaction_genotype_Mtor | Inafm1 | -3.07 | 2.29E-03 | glio |
| Interaction_genotype_Mtor | Platr4 | -2.846 | 2.29E-03 | glio |
| Interaction_genotype_Myc | Kifc1 | -2.655 | 2.29E-03 | glio |
| Interaction_genotype_Rpl3 | Tslp | -4.279 | 2.29E-03 | glio |
| interaction_RT | Fxyd1 | 1.672 | 2.29E-03 | glio |
| interaction_RT | Gadd45g | -1.59 | 2.29E-03 | glio |
| Aars2_in.vivo_noRT | Nutf2 | 2.171 | 2.29E-03 | glio |
| Aars2_in.vivo_noRT | Gm38394 | 2.148 | 2.29E-03 | glio |
| Aars2_in.vivo_noRT | C030034I22Ri | 2.017 | 2.29E-03 | glio |
| Atr_in.vivo_noRT | Mrps31 | -1.138 | 2.29E-03 | glio |
| Atr_in.vivo_noRT | Top1 | -0.919 | 2.29E-03 | glio |
| Atr_ex.vivo_noRT | Lrrc72 | 2.641 | 2.29E-03 | glio |
| Bora_in.vivo_noRT | Myo1b | 2.552 | 2.29E-03 | glio |
| Bora_in.vivo_noRT | Platr3 | 2.346 | 2.29E-03 | glio |
| Bora_ex.vivo_noRT | Gm50115 | 2.802 | 2.29E-03 | glio |
| Chek1_in.vivo_noRT | Snapc1 | -1.367 | 2.29E-03 | glio |
| Chek1_ex.vivo_noRT | Gm47484 | 2.837 | 2.29E-03 | glio |
| Hsd17b10_in.vivo_noRT | C77080 | 2.27 | 2.29E-03 | glio |
| culture_effect | Gm15261 | -0.741 | 2.29E-04 | glio |
| culture_effect | Ube2q1 | 0.576 | 2.29E-04 | glio |
| culture_effect | Samm50 | -0.492 | 2.29E-04 | glio |
| Interaction_genotype_Atr | Serf1 | 1.623 | 2.29E-04 | glio |
| Interaction_genotype_Bora | Matn4 | -4.55 | 2.29E-04 | glio |
| Interaction_genotype_Bora | Nid1 | -4.98 | 2.29E-04 | glio |
| Interaction_genotype_Bora | Gm7008 | -4.801 | 2.29E-04 | glio |
| Aars2_in.vivo_noRT | Gm43329 | 2.453 | 2.29E-04 | glio |
| Atr_in.vivo_noRT | Pcdhga12 | 2.537 | 2.29E-04 | glio |
| Atr_in.vivo_noRT | Tha1 | 2.446 | 2.29E-04 | glio |

| | | | | |
|---|---|---|---|---|
| Bcas2_in.vivo_noRT | Selenbp1 | 2.898 | 2.29E-04 | glio |
| Bora_in.vivo_noRT | Mblac1 | 2.79 | 2.29E-04 | glio |
| Bora_in.vivo_noRT | Unc5cl | 2.634 | 2.29E-04 | glio |
| Chek1_in.vivo_noRT | Snx13 | 2.139 | 2.29E-04 | glio |
| Eif2s1_in.vivo_noRT | Gm12794 | 2.143 | 2.29E-04 | glio |
| culture_effect | Rhpn2 | -2.236 | 2.29E-05 | glio |
| Interaction_genotype_Aars2 | Gm43672 | -5.415 | 2.29E-05 | glio |
| Interaction_genotype_Myc | Zfp551 | -4.779 | 2.29E-05 | glio |
| interaction_RT | 1500009L16Ri | 2.712 | 2.29E-05 | glio |
| Aars2_in.vivo_noRT | Cntf | 2.958 | 2.29E-05 | glio |
| Bcas2_in.vivo_noRT | Gm44091 | 2.888 | 2.29E-05 | glio |
| Bcas2_in.vivo_noRT | Trabd | -1.798 | 2.29E-05 | glio |
| culture_effect | Clk4 | -0.781 | 2.29E-06 | glio |
| Interaction_genotype_Aars2 | Kpna7 | -5.291 | 2.29E-06 | glio |
| Atr_in.vivo_noRT | 1700024B05R | 3.329 | 2.29E-06 | glio |
| Bora_in.vivo_noRT | 4930426I24Ri | 3.281 | 2.29E-06 | glio |
| Bora_ex.vivo_noRT | Gm13178 | 3.201 | 2.29E-06 | glio |
| Chek1_in.vivo_noRT | Gm26533 | 3.41 | 2.29E-06 | glio |
| culture_effect | Pcbp3 | -2.771 | 2.29E-07 | glio |
| culture_effect | Gm9993 | 1.974 | 2.29E-07 | glio |
| Chek1_ex.vivo_noRT | Itih5 | 4.642 | 2.29E-07 | glio |
| Hsd17b10_in.vivo_noRT | Kpna7 | 3.352 | 2.29E-07 | glio |
| culture_effect | Gls | -0.818 | 2.29E-08 | glio |
| Bora_in.vivo_noRT | 1700030C12R | 4.011 | 2.29E-08 | glio |
| Hsd17b10_in.vivo_noRT | Slc1a2 | 3.352 | 2.29E-08 | glio |
| culture_effect | 6330415G19R | 3.255 | 2.29E-11 | glio |
| Atr_in.vivo_noRT | Slc6a13 | 5.154 | 2.29E-12 | glio |
| Bcas2_in.vivo_noRT | Nkg7 | 4.547 | 2.29E-14 | glio |
| culture_effect | Calhm6 | -4.401 | 2.29E-16 | glio |
| Bora_in.vivo_noRT | 4930579C12R | 4.936 | 2.29E-16 | glio |
| Aars2_in.vivo_noRT | Slc16a5 | 5.063 | 2.29E-24 | glio |
| culture_effect | Actr8 | -0.718 | 2.30E-03 | glio |
| culture_effect | Mex3c | -0.678 | 2.30E-03 | glio |
| culture_effect | Phpt1 | -0.696 | 2.30E-03 | glio |
| culture_effect | Dstn | 0.377 | 2.30E-03 | glio |
| Interaction_genotype_Atr | Gm11696 | -3.846 | 2.30E-03 | glio |
| Interaction_genotype_Atr | Gm37170 | -3.767 | 2.30E-03 | glio |
| Interaction_genotype_Chek1 | Klre1 | 4.151 | 2.30E-03 | glio |
| Interaction_genotype_Chek1 | Mrpl57 | 1.664 | 2.30E-03 | glio |
| Interaction_genotype_Chek1 | 2410006H16R | 1.531 | 2.30E-03 | glio |
| Interaction_genotype_Myc | Gm36988 | -3.505 | 2.30E-03 | glio |
| Interaction_genotype_Myc | Pigv | -3.595 | 2.30E-03 | glio |
| Interaction_genotype_Myc | Slc36a4 | -3.324 | 2.30E-03 | glio |
| Interaction_genotype_Myc | Gm20033 | -2.91 | 2.30E-03 | glio |
| Interaction_genotype_Myc | Slc26a10 | -2.69 | 2.30E-03 | glio |
| Interaction_genotype_Myc | Trp53cor1 | -2.528 | 2.30E-03 | glio |
| Interaction_genotype_Rpl3 | Gm30938 | -4.029 | 2.30E-03 | glio |
| interaction_RT | Pcdh10 | 2.059 | 2.30E-03 | glio |
| interaction_RT | Tmem60 | 0.75 | 2.30E-03 | glio |
| Aars2_in.vivo_noRT | Arhgap9 | 1.851 | 2.30E-03 | glio |
| Aars2_in.vivo_noRT | Cnbp | -0.867 | 2.30E-03 | glio |
| Atp5c1_in.vivo_noRT | Adgrb3 | 2.413 | 2.30E-03 | glio |

| | | | | |
|---|---|---|---|---|
| Bcas2_in.vivo_noRT | D3Ertd751e | 1.834 | 2.30E-03 | glio |
| Bora_ex.vivo_noRT | Thg1l | -2.212 | 2.30E-03 | glio |
| Cenpj_ex.vivo_noRT | 2300002M23F | 2.591 | 2.30E-03 | glio |
| Chek1_in.vivo_noRT | Cep76 | 1.586 | 2.30E-03 | glio |
| Chek1_in.vivo_noRT | Cct5 | -0.611 | 2.30E-03 | glio |
| Eif2s1_in.vivo_noRT | Gm43653 | 2.407 | 2.30E-03 | glio |
| Eif2s1_in.vivo_noRT | Olfr543 | 1.883 | 2.30E-03 | glio |
| Eif2s1_in.vivo_noRT | Gm10642 | 2.403 | 2.30E-03 | glio |
| Eif2s1_in.vivo_noRT | Clcn1 | 2.021 | 2.30E-03 | glio |
| Eif2s1_in.vivo_noRT | Cxcl16 | 2.403 | 2.30E-03 | glio |
| Eif2s1_in.vivo_noRT | Spry4 | -2.124 | 2.30E-03 | glio |
| Hsd17b10_in.vivo_noRT | Gm10644 | 2.375 | 2.30E-03 | glio |
| Hsd17b10_in.vivo_noRT | S100a4 | -1.612 | 2.30E-03 | glio |
| Hsd17b10_in.vivo_noRT | Adk | -0.891 | 2.30E-03 | glio |
| Interaction_genotype_Atr | 1700034I23Ri | -3.167 | 2.30E-04 | glio |
| Interaction_genotype_Atr | Arpc5l | 1.567 | 2.30E-04 | glio |
| Interaction_genotype_Mtor | Tusc3 | 2.053 | 2.30E-04 | glio |
| Interaction_genotype_Myc | Hdac10 | -3.709 | 2.30E-04 | glio |
| Interaction_genotype_Myc | Rps5 | 1.276 | 2.30E-04 | glio |
| Aars2_in.vivo_noRT | Cdh24 | 2.85 | 2.30E-04 | glio |
| Aars2_in.vivo_noRT | C1qtnf1 | 2.585 | 2.30E-04 | glio |
| Aars2_in.vivo_noRT | A830005F24Ri | 2.632 | 2.30E-04 | glio |
| Aars2_in.vivo_noRT | Oaz1 | -0.702 | 2.30E-04 | glio |
| Aars2_ex.vivo_noRT | Agrp | 2.897 | 2.30E-04 | glio |
| Atp5c1_in.vivo_noRT | Gm40264 | 2.408 | 2.30E-04 | glio |
| Bard1_ex.vivo_noRT | Edn2 | 2.419 | 2.30E-04 | glio |
| Bard1_ex.vivo_noRT | Snrpn | 2.814 | 2.30E-04 | glio |
| Bard1_ex.vivo_noRT | Pnliprp2 | 2.814 | 2.30E-04 | glio |
| Bard1_ex.vivo_noRT | B230206H07R | 2.419 | 2.30E-04 | glio |
| Bard1_ex.vivo_noRT | Tns4 | 3.19 | 2.30E-04 | glio |
| Bcas2_in.vivo_noRT | Fchsd1 | 2.563 | 2.30E-04 | glio |
| Bora_in.vivo_noRT | Tmem86b | 2.709 | 2.30E-04 | glio |
| Bora_in.vivo_noRT | Plekhg6 | 2.73 | 2.30E-04 | glio |
| Bora_in.vivo_noRT | Mrpl41 | -1.338 | 2.30E-04 | glio |
| Cenpj_ex.vivo_noRT | Gm13078 | 2.591 | 2.30E-04 | glio |
| Cenpj_ex.vivo_noRT | Gm5934 | 2.809 | 2.30E-04 | glio |
| Cenpj_ex.vivo_noRT | Gm13964 | 2.591 | 2.30E-04 | glio |
| Chek1_ex.vivo_noRT | Babam2 | -1.797 | 2.30E-04 | glio |
| Interaction_genotype_Myc | Col22a1 | -3.709 | 2.30E-05 | glio |
| Interaction_genotype_Myc | Fam122b | -3.795 | 2.30E-05 | glio |
| Interaction_genotype_Myc | Med14 | -3.602 | 2.30E-05 | glio |
| Interaction_genotype_Myc | Snai1 | -5.051 | 2.30E-05 | glio |
| Interaction_genotype_Myc | Gm43653 | -4.563 | 2.30E-05 | glio |
| Interaction_genotype_Myc | Gm36198 | -4.024 | 2.30E-05 | glio |
| Interaction_genotype_Myc | Mien1 | 1.866 | 2.30E-05 | glio |
| Interaction_genotype_Myc | Ndufa2 | 1.813 | 2.30E-05 | glio |
| Atr_in.vivo_noRT | Syngr2 | -1.69 | 2.30E-05 | glio |
| Bcas2_in.vivo_noRT | H60c | 3.213 | 2.30E-05 | glio |
| Bora_in.vivo_noRT | B430010I23Ri | 2.85 | 2.30E-05 | glio |
| Bora_in.vivo_noRT | Ubxn10 | 2.559 | 2.30E-05 | glio |
| Chek1_in.vivo_noRT | Nop9 | -1.915 | 2.30E-05 | glio |
| Interaction_genotype_Atr | 1700008J07Ri | -5.204 | 2.30E-06 | glio |

| | | | | |
|---|---|---|---|---|
| Interaction_genotype_Myc | Fpgs | -3.926 | 2.30E-06 | glio |
| Interaction_genotype_Myc | Ccdc180 | -4.956 | 2.30E-06 | glio |
| Interaction_genotype_Myc | Tmem234 | 1.711 | 2.30E-06 | glio |
| Interaction_genotype_Myc | Fth1 | 2.751 | 2.30E-06 | glio |
| Atr_in.vivo_noRT | Nudt7 | 3.097 | 2.30E-06 | glio |
| Bcas2_in.vivo_noRT | 4930524O07R | 3.41 | 2.30E-06 | glio |
| Bora_in.vivo_noRT | Cacng5 | 3.086 | 2.30E-06 | glio |
| Hsd17b10_in.vivo_noRT | Gm42695 | 3.356 | 2.30E-06 | glio |
| interaction_RT | Ndufa1 | 1.072 | 2.30E-07 | glio |
| Aars2_in.vivo_noRT | Gm5861 | 4.278 | 2.30E-07 | glio |
| Bcas2_in.vivo_noRT | Rfc5 | -2.441 | 2.30E-07 | glio |
| Chek1_ex.vivo_noRT | Usp17la | 4.447 | 2.30E-07 | glio |
| Chek1_ex.vivo_noRT | Gm35551 | 4.256 | 2.30E-07 | glio |
| Hsd17b10_in.vivo_noRT | BC018473 | 3.616 | 2.30E-07 | glio |
| culture_effect | Lcn2 | -8.278 | 2.30E-08 | glio |
| Interaction_genotype_Mtor | Trim16 | -6.213 | 2.30E-08 | glio |
| Aars2_in.vivo_noRT | Cdo1 | 4.039 | 2.30E-08 | glio |
| Bora_in.vivo_noRT | Medag | 3.769 | 2.30E-08 | glio |
| Chek1_in.vivo_noRT | Wfdc18 | 4.777 | 2.30E-08 | glio |
| Aars2_in.vivo_noRT | A430057M04F | 4.141 | 2.30E-09 | glio |
| Bcas2_in.vivo_noRT | AI854703 | 4.157 | 2.30E-09 | glio |
| Chek1_in.vivo_noRT | Glis1 | 4.132 | 2.30E-09 | glio |
| Atr_in.vivo_noRT | Slc14a1 | 3.648 | 2.30E-10 | glio |
| Atr_in.vivo_noRT | Pax5 | 4.689 | 2.30E-10 | glio |
| Bcas2_in.vivo_noRT | Cstdc4 | 4.15 | 2.30E-10 | glio |
| Interaction_genotype_Mtor | Gm19276 | -5.904 | 2.30E-11 | glio |
| Aars2_in.vivo_noRT | Ccl7 | 5.327 | 2.30E-11 | glio |
| Aars2_in.vivo_noRT | Lrrc24 | 4.326 | 2.30E-11 | glio |
| Chek1_in.vivo_noRT | Gm10425 | 4.78 | 2.30E-11 | glio |
| Aars2_in.vivo_noRT | D830039M14I | 5.06 | 2.30E-12 | glio |
| Aars2_in.vivo_noRT | Dnm3 | 4.801 | 2.30E-12 | glio |
| Atr_in.vivo_noRT | Grip2 | 4.82 | 2.30E-12 | glio |
| Bora_in.vivo_noRT | 2810049E08Ri | 4.424 | 2.30E-12 | glio |
| Bora_in.vivo_noRT | Sprr1a | 4.936 | 2.30E-12 | glio |
| Aars2_in.vivo_noRT | Kcnk7 | 4.278 | 2.30E-13 | glio |
| Hsd17b10_in.vivo_noRT | Tmem132c | 3.616 | 2.30E-13 | glio |
| culture_effect | Aldoc | -4.755 | 2.30E-15 | glio |
| Aars2_in.vivo_noRT | Gm49579 | 4.81 | 2.30E-15 | glio |
| culture_effect | Erp44 | -1.237 | 2.30E-17 | glio |
| Bcas2_in.vivo_noRT | Gm10447 | 4.807 | 2.30E-18 | glio |
| Bcas2_in.vivo_noRT | Halr1 | 4.541 | 2.30E-22 | glio |
| Chek1_in.vivo_noRT | Trmt61b | 5.044 | 2.30E-22 | glio |
| culture_effect | Sulf2 | -2.506 | 2.30E-36 | glio |
| Interaction_genotype_Atr | Ado | 1.724 | 2.31E-03 | glio |
| Interaction_genotype_Bora | Sprr1a | -4.107 | 2.31E-03 | glio |
| Interaction_genotype_Bora | Nav2 | 1.973 | 2.31E-03 | glio |
| Interaction_genotype_Mtor | AC125149.2 | -3.206 | 2.31E-03 | glio |
| Interaction_genotype_Mtor | Guca1b | -3.679 | 2.31E-03 | glio |
| Interaction_genotype_Mtor | Mdm1 | -3.215 | 2.31E-03 | glio |
| Interaction_genotype_Mtor | 5930430L01Ri | -2.981 | 2.31E-03 | glio |
| Interaction_genotype_Mtor | Rhno1 | 1.604 | 2.31E-03 | glio |
| Interaction_genotype_Mtor | Taf10 | 1.173 | 2.31E-03 | glio |

| | | | | |
|---|---|---|---|---|
| Interaction_genotype_Myc | Akr1c18 | -2.729 | 2.31E-03 | glio |
| Interaction_genotype_Myc | Gm4356 | -3.775 | 2.31E-03 | glio |
| Interaction_genotype_Myc | Rad9b | -3.197 | 2.31E-03 | glio |
| Interaction_genotype_Myc | Borcs8 | 1.474 | 2.31E-03 | glio |
| Interaction_genotype_Myc | Itga1 | -2.19 | 2.31E-03 | glio |
| Interaction_genotype_Myc | Ndufa5 | 1.337 | 2.31E-03 | glio |
| interaction_RT | Bcorl1 | 1.443 | 2.31E-03 | glio |
| interaction_RT | Tsc22d3 | 1.228 | 2.31E-03 | glio |
| interaction_RT | Fmc1 | 0.674 | 2.31E-03 | glio |
| Aars2_in.vivo_noRT | Psd | 1.982 | 2.31E-03 | glio |
| Aars2_in.vivo_noRT | Ctf1 | 1.883 | 2.31E-03 | glio |
| Aars2_in.vivo_noRT | 5033406O09R | -1.417 | 2.31E-03 | glio |
| Aars2_in.vivo_noRT | Syt4 | -0.741 | 2.31E-03 | glio |
| Aars2_ex.vivo_noRT | Pdlim3 | 2.499 | 2.31E-03 | glio |
| Atp5c1_in.vivo_noRT | Gm13404 | 2.461 | 2.31E-03 | glio |
| Atr_in.vivo_noRT | Zfp934 | 1.972 | 2.31E-03 | glio |
| Bcas2_in.vivo_noRT | Gdap1l1 | 2.459 | 2.31E-03 | glio |
| Bora_in.vivo_noRT | A830005F24Ri | 2.241 | 2.31E-03 | glio |
| Bora_in.vivo_noRT | Fst | 2.034 | 2.31E-03 | glio |
| Bora_in.vivo_noRT | Anapc1 | -1.25 | 2.31E-03 | glio |
| Chek1_in.vivo_noRT | Bdh2 | 2.218 | 2.31E-03 | glio |
| Chek1_in.vivo_noRT | 4833417C18R | 2.219 | 2.31E-03 | glio |
| Chek1_in.vivo_noRT | Edc3 | 1.637 | 2.31E-03 | glio |
| Chek1_in.vivo_noRT | Hdac1 | -0.81 | 2.31E-03 | glio |
| Eif2s1_in.vivo_noRT | Dcn | 2.279 | 2.31E-03 | glio |
| Fanca_ex.vivo_noRT | A330076H08R | 2.913 | 2.31E-03 | glio |
| Fanca_ex.vivo_noRT | Ipcef1 | 2.919 | 2.31E-03 | glio |
| Fanca_ex.vivo_noRT | 4930502E18Ri | 2.886 | 2.31E-03 | glio |
| Hsd17b10_in.vivo_noRT | Krt16 | 2.907 | 2.31E-03 | glio |
| Hsd17b10_in.vivo_noRT | 4933406P04R | 2.466 | 2.31E-03 | glio |
| culture_effect | Sgk2 | -2.363 | 2.31E-04 | glio |
| culture_effect | Prrx2 | 2.156 | 2.31E-04 | glio |
| culture_effect | Zbed5 | -1.784 | 2.31E-04 | glio |
| culture_effect | 4921531C22R | 1.707 | 2.31E-04 | glio |
| Interaction_genotype_Mtor | C2cd2 | -4.571 | 2.31E-04 | glio |
| Interaction_genotype_Myc | Gm5157 | -2.729 | 2.31E-04 | glio |
| Interaction_genotype_Myc | Acad12 | -4.106 | 2.31E-04 | glio |
| Interaction_genotype_Myc | Pcgf3 | -2.833 | 2.31E-04 | glio |
| Interaction_genotype_Rad21 | Gm813 | 5.391 | 2.31E-04 | glio |
| interaction_RT | Slc35a4 | -1.047 | 2.31E-04 | glio |
| Atr_in.vivo_noRT | 6430548M08F | 2.627 | 2.31E-04 | glio |
| Bcas2_in.vivo_noRT | Pck2 | -1.84 | 2.31E-04 | glio |
| Bora_in.vivo_noRT | Gm20515 | 2.599 | 2.31E-04 | glio |
| Bora_in.vivo_noRT | Cdk13 | -1.509 | 2.31E-04 | glio |
| Chek1_in.vivo_noRT | Gm26621 | 2.641 | 2.31E-04 | glio |
| Chek1_in.vivo_noRT | Saxo2 | 2.583 | 2.31E-04 | glio |
| Chek1_ex.vivo_noRT | Gm13270 | 3.312 | 2.31E-04 | glio |
| Interaction_genotype_Mtor | Gpc1 | -3.636 | 2.31E-05 | glio |
| Interaction_genotype_Mtor | Tubb2a | 2.827 | 2.31E-05 | glio |
| Interaction_genotype_Myc | Gm45338 | -3.122 | 2.31E-05 | glio |
| Aars2_in.vivo_noRT | Ccn2 | 3.999 | 2.31E-05 | glio |
| Aars2_in.vivo_noRT | 2510046G10R | 2.697 | 2.31E-05 | glio |

| | | | | |
|---|---|---|---|---|
| Atp5c1_in.vivo_noRT | Gm5934 | 2.405 | 2.31E-05 | glio |
| Bard1_ex.vivo_noRT | Gm20705 | 3.217 | 2.31E-05 | glio |
| Bard1_ex.vivo_noRT | Gm13078 | 2.814 | 2.31E-05 | glio |
| Bcas2_in.vivo_noRT | Gm30074 | 2.957 | 2.31E-05 | glio |
| Bcas2_ex.vivo_noRT | Gm4340 | 2.742 | 2.31E-05 | glio |
| Bcas2_ex.vivo_noRT | Gm16239 | 2.742 | 2.31E-05 | glio |
| Chek1_ex.vivo_noRT | Dchs1 | 3.657 | 2.31E-05 | glio |
| Chek1_ex.vivo_noRT | Jakmip2 | 3.652 | 2.31E-05 | glio |
| Atp5c1_in.vivo_noRT | Gm47936 | 2.668 | 2.31E-06 | glio |
| Atp5c1_in.vivo_noRT | Tnni3k | 2.668 | 2.31E-06 | glio |
| Atr_in.vivo_noRT | Bcl2l14 | 3.492 | 2.31E-06 | glio |
| Chek1_ex.vivo_noRT | Gm35019 | 3.649 | 2.31E-06 | glio |
| culture_effect | Pim1 | -1.85 | 2.31E-07 | glio |
| Interaction_genotype_Atr | Jund | 2.381 | 2.31E-07 | glio |
| Interaction_genotype_Atr | 6430590A07R | -4.76 | 2.31E-07 | glio |
| Interaction_genotype_Mtor | Gm20711 | -5.29 | 2.31E-07 | glio |
| Interaction_genotype_Mtor | Olfr810 | -4.099 | 2.31E-07 | glio |
| Atr_in.vivo_noRT | Fzd7 | 3.436 | 2.31E-07 | glio |
| Bcas2_in.vivo_noRT | Adam33 | 3.65 | 2.31E-07 | glio |
| Bcas2_in.vivo_noRT | Lyrm1 | 3.623 | 2.31E-07 | glio |
| Chek1_ex.vivo_noRT | Slc2a9 | 3.645 | 2.31E-07 | glio |
| Atr_in.vivo_noRT | Anxa8 | 3.807 | 2.31E-08 | glio |
| Bcas2_in.vivo_noRT | Vipr2 | 3.908 | 2.31E-08 | glio |
| Chek1_in.vivo_noRT | Zfp85os | 3.984 | 2.31E-08 | glio |
| Chek1_ex.vivo_noRT | Ep400 | -2.572 | 2.31E-08 | glio |
| Chek1_in.vivo_noRT | Limch1 | 4.535 | 2.31E-09 | glio |
| Atr_in.vivo_noRT | Gm20513 | 3.437 | 2.31E-10 | glio |
| Hsd17b10_in.vivo_noRT | Gm44693 | 3.616 | 2.31E-13 | glio |
| Aars2_in.vivo_noRT | Ifitm6 | 5.327 | 2.31E-14 | glio |
| Atr_in.vivo_noRT | Ms4a15 | 4.364 | 2.31E-15 | glio |
| Atr_in.vivo_noRT | Gm48161 | 4.745 | 2.31E-15 | glio |
| Aars2_in.vivo_noRT | Agap2 | 5.327 | 2.31E-20 | glio |
| Atr_in.vivo_noRT | Gad1 | 5.267 | 2.31E-25 | glio |
| culture_effect | Snx29 | -1.327 | 2.32E-03 | glio |
| Interaction_genotype_Atr | Gm43672 | -3.807 | 2.32E-03 | glio |
| Interaction_genotype_Atr | P4ha3 | -3.271 | 2.32E-03 | glio |
| Interaction_genotype_Mtor | Inava | -3.488 | 2.32E-03 | glio |
| Interaction_genotype_Myc | 4930471E19Ri | -3.556 | 2.32E-03 | glio |
| Interaction_genotype_Myc | Bambi | -3.325 | 2.32E-03 | glio |
| Interaction_genotype_Myc | Acot13 | 1.628 | 2.32E-03 | glio |
| Interaction_genotype_Myc | Nhp2 | 1.381 | 2.32E-03 | glio |
| Interaction_genotype_Pars2 | Bco2 | -3.628 | 2.32E-03 | glio |
| Interaction_genotype_Pars2 | 4930478M13F | -3.656 | 2.32E-03 | glio |
| Aars2_in.vivo_noRT | Aars2 | 2.092 | 2.32E-03 | glio |
| Aars2_ex.vivo_noRT | Nts | 3.293 | 2.32E-03 | glio |
| Atr_in.vivo_noRT | Trim46 | 2.008 | 2.32E-03 | glio |
| Atr_ex.vivo_noRT | Magea10 | 2.584 | 2.32E-03 | glio |
| Atr_ex.vivo_noRT | Pkd2 | -2.35 | 2.32E-03 | glio |
| Bcas2_in.vivo_noRT | Gm29358 | 2.307 | 2.32E-03 | glio |
| Bora_ex.vivo_noRT | Gm27241 | 2.22 | 2.32E-03 | glio |
| Cenpj_ex.vivo_noRT | Esp15 | 2.591 | 2.32E-03 | glio |
| Cenpj_ex.vivo_noRT | Gm10801 | 2.591 | 2.32E-03 | glio |

| | | | | |
|---|---|---|---|---|
| Chek1_in.vivo_noRT | Ramp1 | 2.553 | 2.32E-03 | glio |
| Chek1_in.vivo_noRT | Mansc1 | 2.371 | 2.32E-03 | glio |
| Chek1_in.vivo_noRT | Ift74 | -1.366 | 2.32E-03 | glio |
| Chek1_ex.vivo_noRT | Cspg5 | 2.227 | 2.32E-03 | glio |
| Hsd17b10_in.vivo_noRT | Arhgef19 | 2.437 | 2.32E-03 | glio |
| Hsd17b10_in.vivo_noRT | Gm16023 | 2.619 | 2.32E-03 | glio |
| Hsd17b10_in.vivo_noRT | Kmo | 1.974 | 2.32E-03 | glio |
| Hsd17b10_in.vivo_noRT | Pcdh10 | -2.099 | 2.32E-03 | glio |
| culture_effect | Lrp1b | -2.353 | 2.32E-04 | glio |
| Interaction_genotype_Myc | Plpp7 | -3.872 | 2.32E-04 | glio |
| Interaction_genotype_Rad21 | Irf8 | 4.146 | 2.32E-04 | glio |
| Interaction_genotype_Rad21 | Kit | 4.312 | 2.32E-04 | glio |
| Interaction_genotype_Rad21 | Art2b | 4.578 | 2.32E-04 | glio |
| Aars2_in.vivo_noRT | Gm26877 | 2.598 | 2.32E-04 | glio |
| Aars2_in.vivo_noRT | Rpl26 | -0.677 | 2.32E-04 | glio |
| Aars2_ex.vivo_noRT | Pla1a | 3.293 | 2.32E-04 | glio |
| Aars2_ex.vivo_noRT | Cys1 | 2.9 | 2.32E-04 | glio |
| Atr_in.vivo_noRT | Plxna2 | 2.816 | 2.32E-04 | glio |
| Atr_ex.vivo_noRT | Gm12610 | 2.764 | 2.32E-04 | glio |
| Bcas2_ex.vivo_noRT | Dmrtc1a | 2.742 | 2.32E-04 | glio |
| Bcas2_ex.vivo_noRT | Gm19541 | -3.567 | 2.32E-04 | glio |
| Bora_in.vivo_noRT | Bdh2 | 2.634 | 2.32E-04 | glio |
| Bora_in.vivo_noRT | Trim37 | -1.76 | 2.32E-04 | glio |
| Chek1_in.vivo_noRT | Slc36a4 | 2.57 | 2.32E-04 | glio |
| Chek1_in.vivo_noRT | Klc4 | 2.162 | 2.32E-04 | glio |
| Chek1_ex.vivo_noRT | Crhbp | 2.805 | 2.32E-04 | glio |
| Chek1_ex.vivo_noRT | Gnl2 | -1.652 | 2.32E-04 | glio |
| culture_effect | Ankrd37 | -1.47 | 2.32E-05 | glio |
| culture_effect | Ttc5 | -0.991 | 2.32E-05 | glio |
| culture_effect | Bex3 | 0.548 | 2.32E-05 | glio |
| Interaction_genotype_Aars2 | Tomm22 | 1.39 | 2.32E-05 | glio |
| Aars2_in.vivo_noRT | Nim1k | 3.128 | 2.32E-05 | glio |
| Aars2_in.vivo_noRT | Scfd1 | -1.725 | 2.32E-05 | glio |
| Atr_in.vivo_noRT | Kctd11 | 3.062 | 2.32E-05 | glio |
| Atr_ex.vivo_noRT | Ccl20 | 5.394 | 2.32E-05 | glio |
| Chek1_in.vivo_noRT | Rnf144b | 2.986 | 2.32E-05 | glio |
| Chek1_in.vivo_noRT | Prph | -2.152 | 2.32E-05 | glio |
| Interaction_genotype_Mtor | Fgd6 | -5.442 | 2.32E-06 | glio |
| Interaction_genotype_Mtor | Gm32585 | -5.885 | 2.32E-06 | glio |
| Interaction_genotype_Mtor | Skida1 | -5.608 | 2.32E-06 | glio |
| Interaction_genotype_Mtor | Ccnd3 | 2.357 | 2.32E-06 | glio |
| Interaction_genotype_Myc | Phkb | -3.754 | 2.32E-06 | glio |
| Aars2_in.vivo_noRT | 2310008N11R | 3.541 | 2.32E-06 | glio |
| Atr_in.vivo_noRT | 4933408B17R | 3.316 | 2.32E-06 | glio |
| Bora_in.vivo_noRT | Rnf208 | 3.544 | 2.32E-06 | glio |
| Chek1_in.vivo_noRT | Gm15013 | 3.251 | 2.32E-06 | glio |
| Hsd17b10_in.vivo_noRT | Nxnl2 | 3.094 | 2.32E-06 | glio |
| Hsd17b10_in.vivo_noRT | D630036H23F | 3.356 | 2.32E-06 | glio |
| Hsd17b10_in.vivo_noRT | 1700026J14Ri | 3.094 | 2.32E-06 | glio |
| Interaction_genotype_Mtor | Rhox7a | -6.475 | 2.32E-07 | glio |
| Interaction_genotype_Myc | Olfm2 | -6.122 | 2.32E-07 | glio |
| Aars2_in.vivo_noRT | Wdr38 | 3.756 | 2.32E-07 | glio |

| | | | | |
|---|---|---|---|---|
| Atp5c1_in.vivo_noRT | Tmem132c | 2.668 | 2.32E-07 | glio |
| Atp5c1_in.vivo_noRT | Gm44693 | 2.668 | 2.32E-07 | glio |
| Bcas2_in.vivo_noRT | Alox12 | 4.018 | 2.32E-08 | glio |
| Bcas2_ex.vivo_noRT | Gm26555 | 3.915 | 2.32E-08 | glio |
| Hsd17b10_in.vivo_noRT | Gm37800 | 3.616 | 2.32E-08 | glio |
| Hsd17b10_in.vivo_noRT | 4933400F21Ri | 2.825 | 2.32E-08 | glio |
| culture_effect | Bri3 | 0.714 | 2.32E-09 | glio |
| Interaction_genotype_Mtor | Col6a2 | -5.666 | 2.32E-09 | glio |
| Interaction_genotype_Mtor | AC154200.1 | -5.869 | 2.32E-09 | glio |
| culture_effect | Nfe2l3 | -3.575 | 2.32E-10 | glio |
| Aars2_in.vivo_noRT | Hist2h3c1 | 4.415 | 2.32E-10 | glio |
| Bcas2_in.vivo_noRT | Gm16094 | 4.02 | 2.32E-10 | glio |
| Aars2_ex.vivo_noRT | mt-Nd2 | 1.512 | 2.32E-11 | glio |
| Atr_in.vivo_noRT | Adam4 | 4.245 | 2.32E-11 | glio |
| culture_effect | Itm2c | -1.188 | 2.32E-12 | glio |
| Atr_in.vivo_noRT | Spata31d1a | 4.892 | 2.32E-12 | glio |
| Atr_in.vivo_noRT | Mfsd10 | -2.984 | 2.32E-12 | glio |
| Atr_in.vivo_noRT | Prickle4 | 4.742 | 2.32E-13 | glio |
| Atr_in.vivo_noRT | Iqcf1 | 4.88 | 2.32E-13 | glio |
| culture_effect | Gbp9 | -2.141 | 2.32E-15 | glio |
| culture_effect | Sh3pxd2a | 1.861 | 2.32E-15 | glio |
| Bora_in.vivo_noRT | AF067061 | 4.936 | 2.32E-16 | glio |
| culture_effect | Shf | -3.102 | 2.32E-37 | glio |
| culture_effect | Macf1 | 0.461 | 2.33E-03 | glio |
| Interaction_genotype_Atr | Npr3 | -3.341 | 2.33E-03 | glio |
| Interaction_genotype_Atr | Aldh1l1 | -4.251 | 2.33E-03 | glio |
| Interaction_genotype_Atr | Spx | -3.588 | 2.33E-03 | glio |
| Interaction_genotype_Atr | Phtf1os | -3.7 | 2.33E-03 | glio |
| Interaction_genotype_Mtor | Icam2 | -3.233 | 2.33E-03 | glio |
| Interaction_genotype_Mtor | Serpinc1 | -3.806 | 2.33E-03 | glio |
| Interaction_genotype_Mtor | H2-Q7 | 3.22 | 2.33E-03 | glio |
| Interaction_genotype_Mtor | Luc7l | 1.727 | 2.33E-03 | glio |
| Interaction_genotype_Mtor | Cdca3 | 2.16 | 2.33E-03 | glio |
| Interaction_genotype_Myc | Gm44672 | -3.307 | 2.33E-03 | glio |
| Interaction_genotype_Myc | E130311K13Ri | -2.962 | 2.33E-03 | glio |
| interaction_RT | Jup | 2.041 | 2.33E-03 | glio |
| Aars2_in.vivo_noRT | Ranbp6 | 1.9 | 2.33E-03 | glio |
| Aars2_in.vivo_noRT | Phf21a | 1.351 | 2.33E-03 | glio |
| Atp5c1_in.vivo_noRT | Rab27a | 2.142 | 2.33E-03 | glio |
| Atp5c1_in.vivo_noRT | Ly6d | 1.941 | 2.33E-03 | glio |
| Atr_in.vivo_noRT | Cldn34c1 | 2.054 | 2.33E-03 | glio |
| Atr_in.vivo_noRT | Uhrf1bp1 | 1.658 | 2.33E-03 | glio |
| Atr_in.vivo_noRT | Abcb10 | 1.625 | 2.33E-03 | glio |
| Atr_in.vivo_noRT | Psmb4 | -0.734 | 2.33E-03 | glio |
| Bcas2_in.vivo_noRT | Pcdhb17 | 2.21 | 2.33E-03 | glio |
| Bcas2_in.vivo_noRT | Fah | 1.819 | 2.33E-03 | glio |
| Bora_in.vivo_noRT | Cfap69 | 2.199 | 2.33E-03 | glio |
| Cenpj_in.vivo_noRT | Abi3bp | 2.499 | 2.33E-03 | glio |
| Cenpj_in.vivo_noRT | Cenpj | -2.187 | 2.33E-03 | glio |
| Cenpj_ex.vivo_noRT | Gm30124 | 2.983 | 2.33E-03 | glio |
| Chek1_in.vivo_noRT | Cpt1b | 2.162 | 2.33E-03 | glio |
| Chek1_ex.vivo_noRT | Gm9908 | 2.959 | 2.33E-03 | glio |

| | | | | |
|---|---|---|---|---|
| Chek1_ex.vivo_noRT | Dcaf15 | -1.963 | 2.33E-03 | glio |
| Hsd17b10_in.vivo_noRT | Gm29340 | 2.3 | 2.33E-03 | glio |
| Interaction_genotype_Atr | Egr2 | 4.434 | 2.33E-04 | glio |
| Interaction_genotype_Mtor | Whrn | -4.402 | 2.33E-04 | glio |
| Interaction_genotype_Mtor | Olfml2a | -4.165 | 2.33E-04 | glio |
| Interaction_genotype_Mtor | Cacybp | 1.724 | 2.33E-04 | glio |
| Interaction_genotype_Myc | Stac3 | -4.351 | 2.33E-04 | glio |
| Interaction_genotype_Myc | Abcb10 | -3.23 | 2.33E-04 | glio |
| Aars2_in.vivo_noRT | Rwdd3 | 2.744 | 2.33E-04 | glio |
| Aars2_in.vivo_noRT | Wdr78 | 2.808 | 2.33E-04 | glio |
| Aars2_in.vivo_noRT | 1700008J07Ri | 2.381 | 2.33E-04 | glio |
| Atr_in.vivo_noRT | 9130015A21R | 4.635 | 2.33E-04 | glio |
| Atr_in.vivo_noRT | Micu1 | 1.544 | 2.33E-04 | glio |
| Bard1_in.vivo_noRT | Gm36793 | 2.504 | 2.33E-04 | glio |
| Bcas2_in.vivo_noRT | Acox3 | 2.58 | 2.33E-04 | glio |
| Chek1_in.vivo_noRT | Zfand5 | -0.928 | 2.33E-04 | glio |
| Hsd17b10_in.vivo_noRT | Cyp1a1 | 3.616 | 2.33E-04 | glio |
| Hsd17b10_in.vivo_noRT | Gm12602 | 2.589 | 2.33E-04 | glio |
| Interaction_genotype_Myc | Parp8 | -3.679 | 2.33E-05 | glio |
| Interaction_genotype_Myc | A330084C13R | -5.026 | 2.33E-05 | glio |
| Interaction_genotype_Myc | Rpl41 | 1.819 | 2.33E-05 | glio |
| interaction_RT | C130026I21Ri | 1.785 | 2.33E-05 | glio |
| Aars2_ex.vivo_noRT | Otof | 3.293 | 2.33E-05 | glio |
| Atr_in.vivo_noRT | Arid3b | 2.851 | 2.33E-05 | glio |
| Bcas2_in.vivo_noRT | Itpr2 | -1.947 | 2.33E-05 | glio |
| Bora_ex.vivo_noRT | A030005L19Ri | 3.601 | 2.33E-05 | glio |
| Chek1_in.vivo_noRT | 4930513N10R | 3.435 | 2.33E-05 | glio |
| Hsd17b10_in.vivo_noRT | Iqcf1 | 2.963 | 2.33E-05 | glio |
| Interaction_genotype_Mtor | Gm15056 | -6.288 | 2.33E-06 | glio |
| Bard1_in.vivo_noRT | Bard1 | -3.474 | 2.33E-06 | glio |
| Bcas2_in.vivo_noRT | Gm17399 | 3.583 | 2.33E-06 | glio |
| Bora_in.vivo_noRT | Kctd14 | 3.012 | 2.33E-06 | glio |
| Bora_ex.vivo_noRT | Esp24 | 3.201 | 2.33E-06 | glio |
| Hsd17b10_in.vivo_noRT | Plk5 | 2.96 | 2.33E-06 | glio |
| Aars2_in.vivo_noRT | Hnrnpa1 | -1.147 | 2.33E-07 | glio |
| Aars2_in.vivo_noRT | Btf3 | -0.855 | 2.33E-07 | glio |
| Aars2_ex.vivo_noRT | Gm13964 | 3.293 | 2.33E-07 | glio |
| Atr_in.vivo_noRT | Upk1a | 3.741 | 2.33E-07 | glio |
| Atr_in.vivo_noRT | Hhipl2 | 3.974 | 2.33E-07 | glio |
| Atr_in.vivo_noRT | Gm11110 | 3.615 | 2.33E-07 | glio |
| Chek1_ex.vivo_noRT | 1110017D15R | 4.449 | 2.33E-07 | glio |
| Chek1_ex.vivo_noRT | Pacsin1 | 4.269 | 2.33E-07 | glio |
| Chek1_in.vivo_noRT | Ggt1 | 3.731 | 2.33E-08 | glio |
| culture_effect | Ndufa2 | 0.743 | 2.33E-10 | glio |
| Aars2_in.vivo_noRT | Nyap2 | 4.415 | 2.33E-10 | glio |
| Atr_in.vivo_noRT | Dnase1l2 | 4.245 | 2.33E-10 | glio |
| Bora_in.vivo_noRT | Gm20442 | 3.928 | 2.33E-10 | glio |
| Chek1_in.vivo_noRT | AI854703 | 4.393 | 2.33E-10 | glio |
| Chek1_in.vivo_noRT | Ptp4a1.1 | 4.247 | 2.33E-10 | glio |
| Bora_in.vivo_noRT | Gm37035 | 4.019 | 2.33E-12 | glio |
| Chek1_in.vivo_noRT | Cxcl1 | 5.044 | 2.33E-12 | glio |
| Atr_in.vivo_noRT | Gm43948 | 4.058 | 2.33E-13 | glio |

| | | | | |
|---|---|---|---|---|
| Bcas2_in.vivo_noRT | Ccdc162 | 4.278 | 2.33E-14 | glio |
| Bcas2_in.vivo_noRT | Bco2 | 4.807 | 2.33E-15 | glio |
| Bcas2_in.vivo_noRT | A930033M14F | 4.807 | 2.33E-16 | glio |
| Atr_in.vivo_noRT | Slc25a18 | 5.267 | 2.33E-19 | glio |
| culture_effect | Wfdc1 | 5.137 | 2.33E-27 | glio |
| culture_effect | Ksr1 | 1.599 | 2.34E-03 | glio |
| culture_effect | Mob3a | -0.902 | 2.34E-03 | glio |
| culture_effect | Exosc3 | -0.401 | 2.34E-03 | glio |
| Interaction_genotype_Aars2 | Ifnz | -3.701 | 2.34E-03 | glio |
| Interaction_genotype_Aars2 | Icam4 | -3.623 | 2.34E-03 | glio |
| Interaction_genotype_Aars2 | Klc3 | -3.327 | 2.34E-03 | glio |
| Interaction_genotype_Aars2 | Itpr2 | 2.169 | 2.34E-03 | glio |
| Interaction_genotype_Aars2 | Ndufb9 | 0.856 | 2.34E-03 | glio |
| Interaction_genotype_Atr | Tcap | -3.546 | 2.34E-03 | glio |
| Interaction_genotype_Myc | Gzmm | -3.525 | 2.34E-03 | glio |
| Interaction_genotype_Myc | Akna | -3.378 | 2.34E-03 | glio |
| Interaction_genotype_Myc | Fndc3c1 | -3.282 | 2.34E-03 | glio |
| Interaction_genotype_Myc | Cep135 | -2.739 | 2.34E-03 | glio |
| Aars2_in.vivo_noRT | Ppih | -1.202 | 2.34E-03 | glio |
| Atr_in.vivo_noRT | Tbc1d24 | 2.067 | 2.34E-03 | glio |
| Atr_in.vivo_noRT | Ciao1 | -1.188 | 2.34E-03 | glio |
| Bcas2_in.vivo_noRT | 9330121J05Ri | 2.235 | 2.34E-03 | glio |
| Bcas2_ex.vivo_noRT | Nos2 | 2.742 | 2.34E-03 | glio |
| Bora_in.vivo_noRT | Fgl2 | 2.043 | 2.34E-03 | glio |
| Bora_in.vivo_noRT | Arhgap27 | 2.087 | 2.34E-03 | glio |
| Cenpj_in.vivo_noRT | Pakap | 2.019 | 2.34E-03 | glio |
| Cenpj_in.vivo_noRT | Olfr171 | 2.022 | 2.34E-03 | glio |
| Cenpj_in.vivo_noRT | Prl3d3 | 1.759 | 2.34E-03 | glio |
| Chek1_ex.vivo_noRT | Gstt1 | 2.961 | 2.34E-03 | glio |
| Eif2s1_in.vivo_noRT | Usp17lc | 2.143 | 2.34E-03 | glio |
| Eif2s1_in.vivo_noRT | 4930447F24Ri | 2.411 | 2.34E-03 | glio |
| Hsd17b10_in.vivo_noRT | Tmem44 | 2.358 | 2.34E-03 | glio |
| Hsd17b10_in.vivo_noRT | Gstp3 | 2.316 | 2.34E-03 | glio |
| Hsd17b10_in.vivo_noRT | E230001N04R | 2.03 | 2.34E-03 | glio |
| Interaction_genotype_Mtor | Foxa1 | -4.881 | 2.34E-04 | glio |
| Interaction_genotype_Mtor | Rpl23a | 1.311 | 2.34E-04 | glio |
| Interaction_genotype_Myc | Prap1 | -3.122 | 2.34E-04 | glio |
| Interaction_genotype_Myc | Gm44736 | -2.729 | 2.34E-04 | glio |
| Interaction_genotype_Myc | 9330160F10Ri | -3.771 | 2.34E-04 | glio |
| Interaction_genotype_Myc | Vav2 | -4.105 | 2.34E-04 | glio |
| Interaction_genotype_Myc | Sem1 | 1.424 | 2.34E-04 | glio |
| Aars2_in.vivo_noRT | Lemd3 | 1.67 | 2.34E-04 | glio |
| Atr_in.vivo_noRT | Naca | -0.651 | 2.34E-04 | glio |
| Bard1_ex.vivo_noRT | A930002C04R | 2.415 | 2.34E-04 | glio |
| Bard1_ex.vivo_noRT | Gm4027 | 2.814 | 2.34E-04 | glio |
| Bard1_ex.vivo_noRT | Dcn | 3.219 | 2.34E-04 | glio |
| Bcas2_in.vivo_noRT | Larp4 | -0.92 | 2.34E-04 | glio |
| Chek1_in.vivo_noRT | Gm47689 | 2.655 | 2.34E-04 | glio |
| Chek1_in.vivo_noRT | Snph | 2.53 | 2.34E-04 | glio |
| Chek1_in.vivo_noRT | Gm7467 | 2.293 | 2.34E-04 | glio |
| Chek1_in.vivo_noRT | Chmp5 | -0.922 | 2.34E-04 | glio |
| Hsd17b10_in.vivo_noRT | Gm46652 | 2.567 | 2.34E-04 | glio |

| | | | | |
|---|---|---|---|---|
| culture_effect | Slc37a3 | -0.992 | 2.34E-05 | glio |
| Interaction_genotype_Myc | AV356131 | -4.028 | 2.34E-05 | glio |
| Interaction_genotype_Myc | Ptges3 | 1.586 | 2.34E-05 | glio |
| interaction_RT | Dek | -0.861 | 2.34E-05 | glio |
| Aars2_in.vivo_noRT | Ifnz | 2.983 | 2.34E-05 | glio |
| Aars2_in.vivo_noRT | Plek2 | 2.876 | 2.34E-05 | glio |
| Aars2_in.vivo_noRT | Ccdc9b | 2.686 | 2.34E-05 | glio |
| Aars2_in.vivo_noRT | Klhdc1 | 2.929 | 2.34E-05 | glio |
| Atr_in.vivo_noRT | Gm45605 | 2.98 | 2.34E-05 | glio |
| Atr_in.vivo_noRT | Rpl35a | -0.753 | 2.34E-05 | glio |
| Bcas2_in.vivo_noRT | 4933433G15R | 3.141 | 2.34E-05 | glio |
| Bora_in.vivo_noRT | Gm26910 | 2.783 | 2.34E-05 | glio |
| Bora_in.vivo_noRT | Ppef2 | 2.479 | 2.34E-05 | glio |
| Chek1_in.vivo_noRT | Gm17249 | 2.906 | 2.34E-05 | glio |
| Hsd17b10_in.vivo_noRT | Gm47428 | 2.967 | 2.34E-05 | glio |
| culture_effect | Wfdc3 | -3.326 | 2.34E-06 | glio |
| culture_effect | Brcc3 | -0.649 | 2.34E-06 | glio |
| Interaction_genotype_Mtor | 1700081N11R | -6.342 | 2.34E-06 | glio |
| Interaction_genotype_Mtor | Pde11a | -5.885 | 2.34E-06 | glio |
| Interaction_genotype_Mtor | Gm15672 | -5.658 | 2.34E-06 | glio |
| Atr_in.vivo_noRT | Ccdc146 | 3.445 | 2.34E-06 | glio |
| Bcas2_in.vivo_noRT | Ntf5 | 3.512 | 2.34E-06 | glio |
| Bcas2_in.vivo_noRT | Gm43330 | 2.907 | 2.34E-06 | glio |
| Chek1_in.vivo_noRT | Gm49135 | 3.361 | 2.34E-06 | glio |
| Chek1_in.vivo_noRT | Dagla | 3.547 | 2.34E-06 | glio |
| culture_effect | 2610507I01Ri | 2.154 | 2.34E-07 | glio |
| Aars2_in.vivo_noRT | Pde3a | 3.549 | 2.34E-07 | glio |
| Bcas2_in.vivo_noRT | A430057M04F | 3.622 | 2.34E-07 | glio |
| Bcas2_ex.vivo_noRT | Ltbp2 | 3.928 | 2.34E-07 | glio |
| Aars2_in.vivo_noRT | Gm26850 | 3.932 | 2.34E-08 | glio |
| Atr_in.vivo_noRT | Ppnr | 3.871 | 2.34E-08 | glio |
| Atr_in.vivo_noRT | Mill2 | 3.879 | 2.34E-08 | glio |
| Bora_in.vivo_noRT | A430106G13R | 4.145 | 2.34E-08 | glio |
| Bora_in.vivo_noRT | Zfp960 | 4.099 | 2.34E-08 | glio |
| culture_effect | Man2c1os | 3.199 | 2.34E-09 | glio |
| Bora_in.vivo_noRT | Gm47204 | 4.291 | 2.34E-09 | glio |
| Aars2_in.vivo_noRT | Prtn3 | 4.544 | 2.34E-10 | glio |
| Atr_in.vivo_noRT | Gm15903 | 4.221 | 2.34E-10 | glio |
| Hsd17b10_in.vivo_noRT | Ntn4 | 3.881 | 2.34E-10 | glio |
| Bora_in.vivo_noRT | Ankef1 | 4.721 | 2.34E-11 | glio |
| Chek1_in.vivo_noRT | Ssh3 | 4.243 | 2.34E-12 | glio |
| Atr_in.vivo_noRT | Nxnl2 | 5.011 | 2.34E-15 | glio |
| Chek1_in.vivo_noRT | Zfp352 | 5.044 | 2.34E-15 | glio |
| Aars2_in.vivo_noRT | Tcap | 5.457 | 2.34E-17 | glio |
| Atr_in.vivo_noRT | Gm38393 | 5.273 | 2.34E-18 | glio |
| Bcas2_in.vivo_noRT | Cxcl5 | 4.807 | 2.34E-19 | glio |
| Atr_in.vivo_noRT | Usp17lc | 5.533 | 2.34E-20 | glio |
| Chek1_in.vivo_noRT | Gm35082 | 5.044 | 2.34E-20 | glio |
| culture_effect | Atf6 | -0.718 | 2.35E-03 | glio |
| Interaction_genotype_Atr | Platr4 | -3.061 | 2.35E-03 | glio |
| Interaction_genotype_Atr | Zscan22 | -3.079 | 2.35E-03 | glio |
| Interaction_genotype_Atr | Strn | -2.494 | 2.35E-03 | glio |

| | | | | |
|---|---|---|---|---|
| Interaction_genotype_Cyp19a | Gm32122 | 4.781 | 2.35E-03 | glio |
| Interaction_genotype_Mtor | A630052C17R | -3.422 | 2.35E-03 | glio |
| Interaction_genotype_Mtor | Clec16a | -3.185 | 2.35E-03 | glio |
| Interaction_genotype_Myc | Ccdc114 | -3.279 | 2.35E-03 | glio |
| Interaction_genotype_Myc | 9130008F23Ri | -3.825 | 2.35E-03 | glio |
| Interaction_genotype_Myc | Sugp2 | -2.768 | 2.35E-03 | glio |
| Interaction_genotype_Myc | Fnip2 | -2.439 | 2.35E-03 | glio |
| Interaction_genotype_Pars2 | Cyp1a1 | -5.19 | 2.35E-03 | glio |
| interaction_RT | 4933432I03Ri | 2.56 | 2.35E-03 | glio |
| Aars2_in.vivo_noRT | Ccdc62 | 2.043 | 2.35E-03 | glio |
| Aars2_in.vivo_noRT | Rad9b | 2.055 | 2.35E-03 | glio |
| Aars2_in.vivo_noRT | Gxylt2 | 1.922 | 2.35E-03 | glio |
| Aars2_in.vivo_noRT | Acp6 | -1.447 | 2.35E-03 | glio |
| Atp5c1_ex.vivo_noRT | Ocm | 3.395 | 2.35E-03 | glio |
| Atp5c1_ex.vivo_noRT | Gm48617 | 2.61 | 2.35E-03 | glio |
| Atr_in.vivo_noRT | Syk | 1.944 | 2.35E-03 | glio |
| Atr_in.vivo_noRT | Gnb1l | 1.933 | 2.35E-03 | glio |
| Atr_in.vivo_noRT | B3galt6 | 1.575 | 2.35E-03 | glio |
| Atr_ex.vivo_noRT | Sertad1 | 1.517 | 2.35E-03 | glio |
| Bcas2_in.vivo_noRT | Myo1h | 2.312 | 2.35E-03 | glio |
| Bcas2_in.vivo_noRT | Gm10658 | 2.142 | 2.35E-03 | glio |
| Bcas2_in.vivo_noRT | Gm13261 | 2.07 | 2.35E-03 | glio |
| Bcas2_in.vivo_noRT | Socs1 | -1.617 | 2.35E-03 | glio |
| Bcas2_in.vivo_noRT | Smg1 | -1.27 | 2.35E-03 | glio |
| Bora_in.vivo_noRT | Reep4 | -1.699 | 2.35E-03 | glio |
| Bora_in.vivo_noRT | Bag3 | -1.42 | 2.35E-03 | glio |
| Chek1_in.vivo_noRT | Fam174b | 2.47 | 2.35E-03 | glio |
| Chek1_in.vivo_noRT | 2410080I02Ri | 2.131 | 2.35E-03 | glio |
| Chek1_in.vivo_noRT | Psat1 | -0.808 | 2.35E-03 | glio |
| Chek1_ex.vivo_noRT | Lims2 | 3.084 | 2.35E-03 | glio |
| Eif2s1_in.vivo_noRT | Cryl1 | -1.558 | 2.35E-03 | glio |
| Eif2s1_ex.vivo_noRT | Capn13 | 2.234 | 2.35E-03 | glio |
| Fanca_ex.vivo_noRT | Fsd2 | 3.488 | 2.35E-03 | glio |
| Hsd17b10_in.vivo_noRT | Sec1 | 2.459 | 2.35E-03 | glio |
| Hsd17b10_in.vivo_noRT | Gm46218 | 2.38 | 2.35E-03 | glio |
| culture_effect | Tnp2 | -2.411 | 2.35E-04 | glio |
| culture_effect | Tfcp2l1 | -1.582 | 2.35E-04 | glio |
| culture_effect | Fkbp3 | -0.435 | 2.35E-04 | glio |
| Interaction_genotype_Mtor | Gm37305 | -4.122 | 2.35E-04 | glio |
| Interaction_genotype_Myc | Car13 | -2.716 | 2.35E-04 | glio |
| Atr_in.vivo_noRT | Gm43696 | 2.672 | 2.35E-04 | glio |
| Atr_ex.vivo_noRT | Selenok-ps1 | 3.183 | 2.35E-04 | glio |
| Bcas2_in.vivo_noRT | Tmem79 | 2.523 | 2.35E-04 | glio |
| Bcas2_in.vivo_noRT | Adcy7 | 2.511 | 2.35E-04 | glio |
| Bcas2_in.vivo_noRT | Ccdc136 | 2.522 | 2.35E-04 | glio |
| Bora_in.vivo_noRT | March10 | 2.579 | 2.35E-04 | glio |
| Interaction_genotype_Mtor | Gm2012 | -5.199 | 2.35E-05 | glio |
| Interaction_genotype_Mtor | Gm13032 | -4.112 | 2.35E-05 | glio |
| Interaction_genotype_Myc | Zfp715 | -4.16 | 2.35E-05 | glio |
| Interaction_genotype_Myc | Irf8 | 4.568 | 2.35E-05 | glio |
| Interaction_genotype_Myc | Pik3ap1 | -4.162 | 2.35E-05 | glio |
| Interaction_genotype_Myc | Ank1 | -3.513 | 2.35E-05 | glio |

| | | | | |
|---|---|---|---|---|
| Atr_in.vivo_noRT | Gm34455 | 3.768 | 2.35E-05 | glio |
| Bard1_ex.vivo_noRT | Gm13964 | 2.814 | 2.35E-05 | glio |
| culture_effect | Gm13261 | -2.644 | 2.35E-06 | glio |
| culture_effect | Neto2 | 0.962 | 2.35E-06 | glio |
| culture_effect | Cav1 | 0.955 | 2.35E-06 | glio |
| culture_effect | Uqcc3 | 0.604 | 2.35E-06 | glio |
| interaction_RT | Gm15261 | -1.621 | 2.35E-06 | glio |
| interaction_RT | Plagl1 | 1.651 | 2.35E-06 | glio |
| Atr_in.vivo_noRT | Gm20628 | 3.585 | 2.35E-06 | glio |
| Bora_in.vivo_noRT | Lmod1 | 3.505 | 2.35E-06 | glio |
| Bora_in.vivo_noRT | Gm15535 | 3.619 | 2.35E-06 | glio |
| Hsd17b10_in.vivo_noRT | Pdlim3 | 2.837 | 2.35E-06 | glio |
| Interaction_genotype_Mtor | Camk2b | -6.004 | 2.35E-07 | glio |
| Interaction_genotype_Mtor | Lrrc72 | -5.787 | 2.35E-07 | glio |
| Interaction_genotype_Myc | Otos | -4.964 | 2.35E-07 | glio |
| culture_effect | Pkp4 | 1.031 | 2.35E-08 | glio |
| Atr_in.vivo_noRT | Hpse | 3.822 | 2.35E-08 | glio |
| Atr_in.vivo_noRT | Rpl5 | -1.167 | 2.35E-08 | glio |
| Bcas2_in.vivo_noRT | Otop1 | 3.623 | 2.35E-08 | glio |
| Chek1_in.vivo_noRT | Mtmr11 | 4.198 | 2.35E-09 | glio |
| Bcas2_in.vivo_noRT | Krtdap | 4.416 | 2.35E-10 | glio |
| interaction_RT | Csta2 | 2.553 | 2.35E-11 | glio |
| Hsd17b10_in.vivo_noRT | Gm13449 | 3.616 | 2.35E-11 | glio |
| Atr_in.vivo_noRT | Gm39459 | 4.883 | 2.35E-12 | glio |
| Hsd17b10_in.vivo_noRT | Tagap | 3.356 | 2.35E-12 | glio |
| Aars2_in.vivo_noRT | Icam2 | 4.46 | 2.35E-13 | glio |
| Atr_in.vivo_noRT | Ifi214 | 5.28 | 2.35E-14 | glio |
| Bora_in.vivo_noRT | Gpr22 | 4.936 | 2.35E-20 | glio |
| culture_effect | Ust | -0.916 | 2.36E-03 | glio |
| Interaction_genotype_Atr | F630040K05Ri | -3.444 | 2.36E-03 | glio |
| Interaction_genotype_Atr | Gm15910 | -3.598 | 2.36E-03 | glio |
| Interaction_genotype_Atr | Tcea2 | -3.646 | 2.36E-03 | glio |
| Interaction_genotype_Atr | A230020J21Ri | -3.991 | 2.36E-03 | glio |
| Interaction_genotype_Atr | Gm39929 | -3.075 | 2.36E-03 | glio |
| Interaction_genotype_Atr | Kcnk4 | -3.873 | 2.36E-03 | glio |
| Interaction_genotype_Atr | Hddc3 | -3.534 | 2.36E-03 | glio |
| Interaction_genotype_Atr | Plpp6 | -3.572 | 2.36E-03 | glio |
| Interaction_genotype_Atr | Gm48772 | -3.425 | 2.36E-03 | glio |
| Interaction_genotype_Atr | Tafa5 | -3.272 | 2.36E-03 | glio |
| Interaction_genotype_Atr | Whamm | 2.196 | 2.36E-03 | glio |
| Interaction_genotype_Atr | Pet2 | 1.462 | 2.36E-03 | glio |
| Interaction_genotype_Atr | Atg101 | 1.454 | 2.36E-03 | glio |
| Interaction_genotype_Atr | Med28 | 1.015 | 2.36E-03 | glio |
| Interaction_genotype_Bcas2 | Tmem190 | -4.59 | 2.36E-03 | glio |
| Interaction_genotype_Bcas2 | Npl | -3.731 | 2.36E-03 | glio |
| Interaction_genotype_Bcas2 | Pmp2 | -3.608 | 2.36E-03 | glio |
| Interaction_genotype_Chek1 | 1110038B12R | 1.31 | 2.36E-03 | glio |
| Interaction_genotype_Mtor | Krt17 | -6.428 | 2.36E-03 | glio |
| Interaction_genotype_Mtor | Syk | -3.263 | 2.36E-03 | glio |
| Interaction_genotype_Mtor | Hyal1 | -3.39 | 2.36E-03 | glio |
| Interaction_genotype_Mtor | Tgfb3 | -2.996 | 2.36E-03 | glio |
| Interaction_genotype_Myc | Gm26555 | -2.466 | 2.36E-03 | glio |

| | | | | |
|---|---|---|---|---|
| Interaction_genotype_Myc | Nadsyn1 | -3.563 | 2.36E-03 | glio |
| Interaction_genotype_Myc | Fetub | -2.915 | 2.36E-03 | glio |
| Interaction_genotype_Myc | Mamld1 | -2.582 | 2.36E-03 | glio |
| Interaction_genotype_Rad51 | Sppl2a | 1.783 | 2.36E-03 | glio |
| Aars2_in.vivo_noRT | 9130023H24R | 2.21 | 2.36E-03 | glio |
| Aars2_in.vivo_noRT | Aatk | 2.263 | 2.36E-03 | glio |
| Aars2_in.vivo_noRT | Eci2 | -1.25 | 2.36E-03 | glio |
| Aars2_ex.vivo_noRT | Ldhd | 2.582 | 2.36E-03 | glio |
| Atp5c1_in.vivo_noRT | Gm39326 | 2.405 | 2.36E-03 | glio |
| Atr_in.vivo_noRT | Zfp981 | 2.123 | 2.36E-03 | glio |
| Atr_in.vivo_noRT | Rps20 | -0.665 | 2.36E-03 | glio |
| Bard1_ex.vivo_noRT | Pla2g2d | 2.814 | 2.36E-03 | glio |
| Bcas2_in.vivo_noRT | Tmem246 | 2.091 | 2.36E-03 | glio |
| Bcas2_in.vivo_noRT | Mrfap1 | -0.586 | 2.36E-03 | glio |
| Bora_in.vivo_noRT | Sema3d | 2.24 | 2.36E-03 | glio |
| Bora_ex.vivo_noRT | Alox15 | 2.806 | 2.36E-03 | glio |
| Cenpj_ex.vivo_noRT | Pcdhgb1 | 2.814 | 2.36E-03 | glio |
| Cenpj_ex.vivo_noRT | Lrguk | 2.591 | 2.36E-03 | glio |
| Cenpj_ex.vivo_noRT | Vmn2r44 | 2.196 | 2.36E-03 | glio |
| Cenpj_ex.vivo_noRT | Gm20275 | 2.591 | 2.36E-03 | glio |
| Cenpj_ex.vivo_noRT | Gm50024 | 2.192 | 2.36E-03 | glio |
| Chek1_ex.vivo_noRT | AC133103.1 | 3.032 | 2.36E-03 | glio |
| Cyp19a1_ex.vivo_noRT | Gm36037 | 2.538 | 2.36E-03 | glio |
| Cyp19a1_ex.vivo_noRT | Sucnr1 | 2.159 | 2.36E-03 | glio |
| Dclre1c_in.vivo_noRT | Fn1 | 1.642 | 2.36E-03 | glio |
| Hsd17b10_in.vivo_noRT | 2810408A11R | 2.342 | 2.36E-03 | glio |
| Hsd17b10_in.vivo_noRT | Gm10655 | 2.431 | 2.36E-03 | glio |
| Hsd17b10_in.vivo_noRT | Tmco1 | -0.622 | 2.36E-03 | glio |
| culture_effect | Cd1d1 | -1.935 | 2.36E-04 | glio |
| culture_effect | Kdm7a | -0.911 | 2.36E-04 | glio |
| Interaction_genotype_Aars2 | Oas1g | -4.43 | 2.36E-04 | glio |
| Interaction_genotype_Aars2 | Jph4 | -4.57 | 2.36E-04 | glio |
| Interaction_genotype_Aars2 | Ostf1 | 1.924 | 2.36E-04 | glio |
| Interaction_genotype_Atr | Gm26682 | -4.488 | 2.36E-04 | glio |
| Interaction_genotype_Bcas2 | A430106G13R | -4.88 | 2.36E-04 | glio |
| Interaction_genotype_Bcas2 | Gm10399 | -4.028 | 2.36E-04 | glio |
| Interaction_genotype_Bcas2 | Pygo1 | -4.529 | 2.36E-04 | glio |
| Interaction_genotype_Myc | C030005K06R | -4.13 | 2.36E-04 | glio |
| Interaction_genotype_Myc | Gm20337 | -4.057 | 2.36E-04 | glio |
| Interaction_genotype_Myc | Rpl26 | 1.268 | 2.36E-04 | glio |
| interaction_RT | Pa2g4 | -0.77 | 2.36E-04 | glio |
| Bcas2_in.vivo_noRT | Cct6b | 2.518 | 2.36E-04 | glio |
| Bcas2_ex.vivo_noRT | Gm11899 | 2.742 | 2.36E-04 | glio |
| Bora_ex.vivo_noRT | Gm5934 | 2.622 | 2.36E-04 | glio |
| Bora_ex.vivo_noRT | Gpr17 | 3.201 | 2.36E-04 | glio |
| Chek1_ex.vivo_noRT | Ccdc7b | 3.345 | 2.36E-04 | glio |
| culture_effect | Gm36043 | -2.391 | 2.36E-05 | glio |
| culture_effect | Mrps10 | 0.507 | 2.36E-05 | glio |
| culture_effect | Mrpl42 | 0.452 | 2.36E-05 | glio |
| culture_effect | Edf1 | 0.425 | 2.36E-05 | glio |
| Atr_in.vivo_noRT | Susd1 | 3.221 | 2.36E-05 | glio |
| Bcas2_in.vivo_noRT | Gm10863 | 2.725 | 2.36E-05 | glio |

| | | | | |
|---|---|---|---|---|
| Bcas2_in.vivo_noRT | Ddx19b | 2.702 | 2.36E-05 | glio |
| Hsd17b10_in.vivo_noRT | Tex52 | 2.966 | 2.36E-05 | glio |
| Interaction_genotype_Mtor | Cd36 | -4.099 | 2.36E-06 | glio |
| Aars2_in.vivo_noRT | Cop1 | -1.944 | 2.36E-06 | glio |
| Aars2_ex.vivo_noRT | Dmrtc1a | 3.293 | 2.36E-06 | glio |
| Atp5c1_in.vivo_noRT | Gm49504 | 2.668 | 2.36E-06 | glio |
| Atr_ex.vivo_noRT | Gm4166 | 3.164 | 2.36E-06 | glio |
| Bora_in.vivo_noRT | Mterf1b | 3.169 | 2.36E-06 | glio |
| Chek1_ex.vivo_noRT | Glrp1 | 3.652 | 2.36E-06 | glio |
| Interaction_genotype_Mtor | Apoa5 | -4.099 | 2.36E-07 | glio |
| Interaction_genotype_Myc | Plekhh2 | -5.502 | 2.36E-07 | glio |
| Bcas2_in.vivo_noRT | Gm17227 | 3.424 | 2.36E-07 | glio |
| Hsd17b10_in.vivo_noRT | Gm41349 | 3.353 | 2.36E-07 | glio |
| culture_effect | Habp4 | 1.77 | 2.36E-08 | glio |
| culture_effect | Mllt11 | 1.051 | 2.36E-08 | glio |
| Chek1_ex.vivo_noRT | Ms4a4d | 4.842 | 2.36E-08 | glio |
| culture_effect | 1810044D09R | 1.566 | 2.36E-09 | glio |
| Bora_in.vivo_noRT | Isl1 | 4.133 | 2.36E-09 | glio |
| Chek1_ex.vivo_noRT | Gng13 | 5.082 | 2.36E-09 | glio |
| Chek1_ex.vivo_noRT | Gm48161 | 4.449 | 2.36E-09 | glio |
| Atr_in.vivo_noRT | Gm45336 | 3.661 | 2.36E-10 | glio |
| Bcas2_in.vivo_noRT | Gm10425 | 4.543 | 2.36E-10 | glio |
| Chek1_in.vivo_noRT | Col6a2 | 4.26 | 2.36E-10 | glio |
| Aars2_in.vivo_noRT | Esrp2 | 4.415 | 2.36E-11 | glio |
| Bcas2_in.vivo_noRT | Zfp786 | 4.543 | 2.36E-11 | glio |
| Chek1_ex.vivo_noRT | Gm10482 | 4.842 | 2.36E-12 | glio |
| Atr_in.vivo_noRT | Dnm3 | 5.008 | 2.36E-13 | glio |
| Aars2_in.vivo_noRT | Nat8f4 | 4.794 | 2.36E-14 | glio |
| Atr_in.vivo_noRT | 1700026J14Ri | 5.011 | 2.36E-15 | glio |
| Bora_in.vivo_noRT | Slc1a2 | 4.672 | 2.36E-15 | glio |
| culture_effect | Wsb1 | -1.472 | 2.36E-16 | glio |
| Bcas2_in.vivo_noRT | 1700034I23Ri | 4.807 | 2.36E-22 | glio |
| Aars2_in.vivo_noRT | Gm30692 | 5.327 | 2.36E-26 | glio |
| culture_effect | Sdr16c5 | -1.856 | 2.37E-03 | glio |
| culture_effect | Pawr | -1.729 | 2.37E-03 | glio |
| culture_effect | 1500002C15R | 1.715 | 2.37E-03 | glio |
| culture_effect | Evl | 0.692 | 2.37E-03 | glio |
| culture_effect | Uba1 | -0.453 | 2.37E-03 | glio |
| Interaction_genotype_Aars2 | Gm20711 | -3.225 | 2.37E-03 | glio |
| Interaction_genotype_Atr | Impg2 | -3.157 | 2.37E-03 | glio |
| Interaction_genotype_Atr | C030005K06R | -3.776 | 2.37E-03 | glio |
| Interaction_genotype_Bard1 | Apol10b | 3.788 | 2.37E-03 | glio |
| Interaction_genotype_Bora | B630019K06R | -3.291 | 2.37E-03 | glio |
| Interaction_genotype_Myc | Guca1b | -3.474 | 2.37E-03 | glio |
| Interaction_genotype_Rpl3 | Nxph3 | -3.856 | 2.37E-03 | glio |
| Interaction_genotype_Rpl3 | Akirin2 | 1.165 | 2.37E-03 | glio |
| Aars2_in.vivo_noRT | Spef1 | 2.202 | 2.37E-03 | glio |
| Aars2_in.vivo_noRT | Gstm5 | -1.297 | 2.37E-03 | glio |
| Aars2_in.vivo_noRT | Rplp1 | -0.614 | 2.37E-03 | glio |
| Atr_in.vivo_noRT | Fam83a | 2.472 | 2.37E-03 | glio |
| Atr_in.vivo_noRT | Tbc1d30 | 1.961 | 2.37E-03 | glio |
| Bcas2_in.vivo_noRT | Cavin1 | -1.609 | 2.37E-03 | glio |

| | | | | |
|---|---|---|---|---|
| Bcas2_in.vivo_noRT | Emg1 | -0.669 | 2.37E-03 | glio |
| Bora_in.vivo_noRT | Cstf3 | -1.539 | 2.37E-03 | glio |
| Bora_in.vivo_noRT | Gfpt2 | -1.445 | 2.37E-03 | glio |
| Bora_in.vivo_noRT | Pes1 | -1.22 | 2.37E-03 | glio |
| Bora_ex.vivo_noRT | Sema3a | 2.808 | 2.37E-03 | glio |
| Bora_ex.vivo_noRT | Gm43948 | 2.417 | 2.37E-03 | glio |
| Cenpj_ex.vivo_noRT | Brinp3 | 2.591 | 2.37E-03 | glio |
| Chek1_in.vivo_noRT | Tmprss5 | 2.208 | 2.37E-03 | glio |
| Chek1_in.vivo_noRT | AC132307.1 | 2.008 | 2.37E-03 | glio |
| Chek1_in.vivo_noRT | Pdlim2 | 2.002 | 2.37E-03 | glio |
| Chek1_in.vivo_noRT | Zc3h11a | 2.008 | 2.37E-03 | glio |
| Chek1_in.vivo_noRT | Sra1 | -1.043 | 2.37E-03 | glio |
| Chek1_ex.vivo_noRT | Jmjd8 | 2.779 | 2.37E-03 | glio |
| Chek1_ex.vivo_noRT | Tas1r1 | 2.491 | 2.37E-03 | glio |
| Chek1_ex.vivo_noRT | Ltbp1 | 2.299 | 2.37E-03 | glio |
| Chek1_ex.vivo_noRT | Gdpgp1 | 2.289 | 2.37E-03 | glio |
| Hsd17b10_in.vivo_noRT | Gm13421 | 2.412 | 2.37E-03 | glio |
| Hsd17b10_in.vivo_noRT | Cep72 | 2.276 | 2.37E-03 | glio |
| Interaction_genotype_Mtor | Gm128 | -3.574 | 2.37E-04 | glio |
| Interaction_genotype_Mtor | Rab11b | 1.825 | 2.37E-04 | glio |
| Interaction_genotype_Mtor | Mrps18b | 1.805 | 2.37E-04 | glio |
| Interaction_genotype_Myc | Fam227b | -2.729 | 2.37E-04 | glio |
| Interaction_genotype_Myc | Psca | -3.777 | 2.37E-04 | glio |
| Interaction_genotype_Myc | Gm20658 | -2.729 | 2.37E-04 | glio |
| Interaction_genotype_Myc | Fam13a | -4.193 | 2.37E-04 | glio |
| Interaction_genotype_Myc | Kcnk4 | -4.233 | 2.37E-04 | glio |
| Interaction_genotype_Myc | Gm26682 | -4.052 | 2.37E-04 | glio |
| Interaction_genotype_Myc | Aox1 | -3.82 | 2.37E-04 | glio |
| Interaction_genotype_Myc | Prodh | -3.704 | 2.37E-04 | glio |
| Interaction_genotype_Myc | Kank1 | -2.766 | 2.37E-04 | glio |
| Interaction_genotype_Pars2 | Thap8 | -5.267 | 2.37E-04 | glio |
| Aars2_in.vivo_noRT | Zfp729a | 2.394 | 2.37E-04 | glio |
| Aars2_in.vivo_noRT | Rpl28 | -0.653 | 2.37E-04 | glio |
| Atr_in.vivo_noRT | Fndc7 | 2.78 | 2.37E-04 | glio |
| Atr_ex.vivo_noRT | Dtx1 | 3.569 | 2.37E-04 | glio |
| Bcas2_in.vivo_noRT | Tc2n | 2.719 | 2.37E-04 | glio |
| Bora_ex.vivo_noRT | Slc22a1 | 2.806 | 2.37E-04 | glio |
| Bora_ex.vivo_noRT | 1700108F19Ri | 2.808 | 2.37E-04 | glio |
| culture_effect | Zfp91 | 0.501 | 2.37E-05 | glio |
| Interaction_genotype_Mtor | 1500002C15R | -5.163 | 2.37E-05 | glio |
| Interaction_genotype_Mtor | Gm10638 | -4.673 | 2.37E-05 | glio |
| Interaction_genotype_Myc | Npm3 | 1.935 | 2.37E-05 | glio |
| Atr_ex.vivo_noRT | Pkd1l2 | 3.566 | 2.37E-05 | glio |
| Bora_in.vivo_noRT | Ccnh | -1.789 | 2.37E-05 | glio |
| Interaction_genotype_Mtor | Gm48933 | -5.463 | 2.37E-06 | glio |
| Interaction_genotype_Myc | Gm50321 | -4.625 | 2.37E-06 | glio |
| interaction_RT | Serf2 | 0.792 | 2.37E-06 | glio |
| Aars2_in.vivo_noRT | Mertk | 3.194 | 2.37E-06 | glio |
| Atp5c1_in.vivo_noRT | Prl3d3 | 2.405 | 2.37E-06 | glio |
| Bcas2_in.vivo_noRT | Olfr1369-ps1 | 4.066 | 2.37E-06 | glio |
| Bcas2_in.vivo_noRT | 1700084E18Ri | 3.565 | 2.37E-06 | glio |
| Bora_ex.vivo_noRT | Gm9758 | 3.201 | 2.37E-06 | glio |

| | | | | |
|---|---|---|---|---|
| Bora_in.vivo_noRT | 1700040D17R | 3.872 | 2.37E-07 | glio |
| Bcas2_in.vivo_noRT | Ly6c2 | 3.904 | 2.37E-08 | glio |
| Bora_in.vivo_noRT | Gm30124 | 3.666 | 2.37E-08 | glio |
| culture_effect | Psmc6 | -0.785 | 2.37E-10 | glio |
| Aars2_in.vivo_noRT | Wfdc1 | 4.342 | 2.37E-10 | glio |
| Bora_in.vivo_noRT | Gm20682 | 3.892 | 2.37E-11 | glio |
| Chek1_ex.vivo_noRT | Otof | 4.842 | 2.37E-11 | glio |
| Chek1_in.vivo_noRT | Ramp3 | 4.777 | 2.37E-12 | glio |
| Hsd17b10_in.vivo_noRT | Gm36037 | 3.616 | 2.37E-12 | glio |
| Chek1_in.vivo_noRT | Gm13403 | 4.909 | 2.37E-13 | glio |
| Chek1_in.vivo_noRT | Gm35439 | 4.664 | 2.37E-17 | glio |
| Aars2_in.vivo_noRT | Gm50321 | 5.327 | 2.37E-19 | glio |
| culture_effect | Hexa | -1.398 | 2.37E-20 | glio |
| Aars2_in.vivo_noRT | Spink6 | 5.327 | 2.37E-21 | glio |
| Chek1_in.vivo_noRT | Cd200r2 | 4.78 | 2.37E-21 | glio |
| Chek1_in.vivo_noRT | 4930512B01R | 5.044 | 2.37E-21 | glio |
| Atr_in.vivo_noRT | Gm30794 | 5.533 | 2.37E-26 | glio |
| Aars2_in.vivo_noRT | Gm14742 | 5.327 | 2.37E-27 | glio |
| Bora_in.vivo_noRT | Gjd4 | 5.46 | 2.37E-28 | glio |
| culture_effect | Slc4a5 | -2.032 | 2.38E-03 | glio |
| culture_effect | Gm28874 | 1.739 | 2.38E-03 | glio |
| culture_effect | Epb41l5 | -1.068 | 2.38E-03 | glio |
| Interaction_genotype_Aars2 | Kdm4a | 1.976 | 2.38E-03 | glio |
| Interaction_genotype_Atr | Fam13a | -3.833 | 2.38E-03 | glio |
| Interaction_genotype_Mtor | 9130204L05Ri | -3.984 | 2.38E-03 | glio |
| Interaction_genotype_Mtor | Ubd | 3.503 | 2.38E-03 | glio |
| Interaction_genotype_Mtor | Lrrc73 | -2.738 | 2.38E-03 | glio |
| Interaction_genotype_Pars2 | Rilp | -4.469 | 2.38E-03 | glio |
| Interaction_genotype_Terf1 | Man2c1os | -4.573 | 2.38E-03 | glio |
| Interaction_genotype_Terf1 | Gm43672 | -4.608 | 2.38E-03 | glio |
| Interaction_genotype_Terf1 | Carns1 | -4.58 | 2.38E-03 | glio |
| Interaction_genotype_Terf1 | Gm11627 | -3.952 | 2.38E-03 | glio |
| Interaction_genotype_Terf1 | H2-Ab1 | 3.571 | 2.38E-03 | glio |
| Interaction_genotype_Terf1 | K230015D01R | -4.576 | 2.38E-03 | glio |
| interaction_RT | Ube2g1 | 0.881 | 2.38E-03 | glio |
| interaction_RT | Tcerg1 | -0.641 | 2.38E-03 | glio |
| Aars2_in.vivo_noRT | Arid3b | 2.107 | 2.38E-03 | glio |
| Aars2_in.vivo_noRT | Dnase2a | 1.684 | 2.38E-03 | glio |
| Aars2_in.vivo_noRT | Rps3a1 | -0.615 | 2.38E-03 | glio |
| Aars2_ex.vivo_noRT | Masp1 | 2.495 | 2.38E-03 | glio |
| Aars2_ex.vivo_noRT | Esr1 | 2.899 | 2.38E-03 | glio |
| Atr_in.vivo_noRT | Ktn1 | -1.172 | 2.38E-03 | glio |
| Atr_in.vivo_noRT | Mcl1 | -1.136 | 2.38E-03 | glio |
| Atr_ex.vivo_noRT | Cstdc4 | 2.764 | 2.38E-03 | glio |
| Bard1_ex.vivo_noRT | Plac8 | 3.037 | 2.38E-03 | glio |
| Bcas2_in.vivo_noRT | Gm15133 | 2.201 | 2.38E-03 | glio |
| Bcas2_in.vivo_noRT | 6330403L08Ri | 2.146 | 2.38E-03 | glio |
| Bcas2_in.vivo_noRT | Chst11 | -1.433 | 2.38E-03 | glio |
| Bcas2_ex.vivo_noRT | Vpreb1 | 2.954 | 2.38E-03 | glio |
| Bora_in.vivo_noRT | Rgl1 | 2.021 | 2.38E-03 | glio |
| Bora_in.vivo_noRT | Nkain4 | 2.072 | 2.38E-03 | glio |
| Bora_in.vivo_noRT | Mindy4 | 2.114 | 2.38E-03 | glio |

| | | | | |
|---|---|---|---|---|
| Bora_in.vivo_noRT | Rbm38 | -1.475 | 2.38E-03 | glio |
| Chek1_in.vivo_noRT | Nmnat3 | 2.344 | 2.38E-03 | glio |
| Chek1_in.vivo_noRT | Rorc | 2.382 | 2.38E-03 | glio |
| Chek1_in.vivo_noRT | Pacs1 | 2.182 | 2.38E-03 | glio |
| Chek1_in.vivo_noRT | 2010109A12R | 2.165 | 2.38E-03 | glio |
| Chek1_in.vivo_noRT | Snx3 | -0.598 | 2.38E-03 | glio |
| Chek1_ex.vivo_noRT | Fbn1 | 2.576 | 2.38E-03 | glio |
| Hsd17b10_in.vivo_noRT | Elovl7 | -1.111 | 2.38E-03 | glio |
| Interaction_genotype_Mtor | BC065397 | -4.914 | 2.38E-04 | glio |
| Interaction_genotype_Mtor | Commd3 | 2.011 | 2.38E-04 | glio |
| Atr_in.vivo_noRT | Rbm45 | 2.039 | 2.38E-04 | glio |
| Atr_ex.vivo_noRT | Lce1g | 3.517 | 2.38E-04 | glio |
| Chek1_in.vivo_noRT | Rpl10-ps3 | -1.289 | 2.38E-04 | glio |
| Chek1_in.vivo_noRT | Zcrb1 | -0.806 | 2.38E-04 | glio |
| Chek1_in.vivo_noRT | Rps5 | -0.676 | 2.38E-04 | glio |
| Dclre1c_ex.vivo_noRT | Zscan4-ps3 | 2.658 | 2.38E-04 | glio |
| Hsd17b10_in.vivo_noRT | Apoh | 2.701 | 2.38E-04 | glio |
| culture_effect | Cldn9 | -2.583 | 2.38E-05 | glio |
| culture_effect | Trim66 | -2.461 | 2.38E-05 | glio |
| culture_effect | Azin2 | 1.694 | 2.38E-05 | glio |
| culture_effect | Tasor | -0.977 | 2.38E-05 | glio |
| Interaction_genotype_Mtor | Tslp | -4.684 | 2.38E-05 | glio |
| Interaction_genotype_Nat2 | Gm2022 | 3.978 | 2.38E-05 | glio |
| interaction_RT | Resf1 | 1.177 | 2.38E-05 | glio |
| Aars2_in.vivo_noRT | Sub1 | -0.774 | 2.38E-05 | glio |
| Bcas2_in.vivo_noRT | Tnfrsf13b | 3.169 | 2.38E-05 | glio |
| Bora_in.vivo_noRT | Gm10638 | 2.969 | 2.38E-05 | glio |
| Interaction_genotype_Mtor | Nrg2 | -5.561 | 2.38E-06 | glio |
| Aars2_in.vivo_noRT | Cdkn1c | 3.295 | 2.38E-06 | glio |
| Bora_in.vivo_noRT | Synpr | 3.773 | 2.38E-06 | glio |
| Bora_in.vivo_noRT | Gm26581 | 3.541 | 2.38E-06 | glio |
| Chek1_in.vivo_noRT | C330011M18F | 3.389 | 2.38E-06 | glio |
| Chek1_in.vivo_noRT | Slc43a1 | 3.371 | 2.38E-06 | glio |
| Chek1_in.vivo_noRT | Pkdcc | 3.392 | 2.38E-06 | glio |
| Chek1_ex.vivo_noRT | Bmx | 3.664 | 2.38E-06 | glio |
| culture_effect | Wrap73 | -1.319 | 2.38E-07 | glio |
| Bora_in.vivo_noRT | 4833418N02R | 3.767 | 2.38E-07 | glio |
| Chek1_in.vivo_noRT | 2210406O10R | 3.727 | 2.38E-07 | glio |
| Chek1_in.vivo_noRT | Gm37986 | 3.23 | 2.38E-07 | glio |
| Atr_in.vivo_noRT | Gm16174 | 4.143 | 2.38E-08 | glio |
| Bcas2_in.vivo_noRT | Gm28424 | 3.796 | 2.38E-08 | glio |
| Bora_in.vivo_noRT | Pcdhga2 | 3.666 | 2.38E-08 | glio |
| Chek1_in.vivo_noRT | Gm49767 | 4.006 | 2.38E-08 | glio |
| Chek1_in.vivo_noRT | 1700006J14Ri | 3.663 | 2.38E-08 | glio |
| Chek1_in.vivo_noRT | Hist1h2af | 3.559 | 2.38E-08 | glio |
| Atr_in.vivo_noRT | Pbld1 | 4.346 | 2.38E-09 | glio |
| Chek1_in.vivo_noRT | 2010106G04R | 4.243 | 2.38E-09 | glio |
| Chek1_ex.vivo_noRT | Scgb1a1 | 5.212 | 2.38E-09 | glio |
| culture_effect | Smox | -1.275 | 2.38E-11 | glio |
| Atr_in.vivo_noRT | Sh3gl3 | 4.621 | 2.38E-11 | glio |
| Hsd17b10_in.vivo_noRT | Hnf1aos1 | 4.143 | 2.38E-11 | glio |
| Aars2_in.vivo_noRT | Gm26789 | 4.409 | 2.38E-14 | glio |

| | | | | |
|---|---|---|---|---|
| Aars2_in.vivo_noRT | Nxph4 | 5.327 | 2.38E-14 | glio |
| Bora_in.vivo_noRT | Col5a2 | 4.149 | 2.38E-14 | glio |
| Aars2_in.vivo_noRT | Masp1 | 4.805 | 2.38E-16 | glio |
| Chek1_ex.vivo_noRT | Ankrd22 | 4.842 | 2.38E-16 | glio |
| culture_effect | Olfr889 | 2.376 | 2.38E-17 | glio |
| Aars2_in.vivo_noRT | Gm35082 | 5.327 | 2.38E-22 | glio |
| Aars2_in.vivo_noRT | Olfr1152 | 5.06 | 2.38E-25 | glio |
| culture_effect | Bcat1 | -0.625 | 2.39E-03 | glio |
| Interaction_genotype_Aars2 | Slc45a3 | -3.824 | 2.39E-03 | glio |
| Interaction_genotype_Aars2 | Gm11767 | -3.909 | 2.39E-03 | glio |
| Interaction_genotype_Aars2 | Zbtb11os1 | -3.64 | 2.39E-03 | glio |
| Interaction_genotype_Aars2 | Gng10 | 1.258 | 2.39E-03 | glio |
| Interaction_genotype_Atr | Mertk | -3.699 | 2.39E-03 | glio |
| Interaction_genotype_Atr | Gm39326 | -3.628 | 2.39E-03 | glio |
| Interaction_genotype_Atr | 1110020A21R | -3.711 | 2.39E-03 | glio |
| Interaction_genotype_Atr | Ppargc1b | -3.858 | 2.39E-03 | glio |
| Interaction_genotype_Atr | Gm3716 | -3.391 | 2.39E-03 | glio |
| Interaction_genotype_Atr | Dis3l2 | -2.916 | 2.39E-03 | glio |
| Interaction_genotype_Atr | Tax1bp3 | 1.8 | 2.39E-03 | glio |
| Interaction_genotype_Atr | Rsrp1 | 1.557 | 2.39E-03 | glio |
| Interaction_genotype_Bora | Gm11973 | -3.232 | 2.39E-03 | glio |
| Interaction_genotype_Mtor | Plaa | 1.562 | 2.39E-03 | glio |
| Interaction_genotype_Myc | Pde11a | -3.474 | 2.39E-03 | glio |
| Interaction_genotype_Myc | Nckap1l | -3.445 | 2.39E-03 | glio |
| Interaction_genotype_Myc | 6030443J06Ri | -3.35 | 2.39E-03 | glio |
| interaction_RT | Mars | -0.885 | 2.39E-03 | glio |
| Aars2_in.vivo_noRT | Cd1d1 | 1.948 | 2.39E-03 | glio |
| Aars2_in.vivo_noRT | Cped1 | 2.038 | 2.39E-03 | glio |
| Aars2_in.vivo_noRT | Ppie | -1.155 | 2.39E-03 | glio |
| Aars2_in.vivo_noRT | C1qbp | -0.709 | 2.39E-03 | glio |
| Atr_in.vivo_noRT | Gm38037 | 2.118 | 2.39E-03 | glio |
| Atr_in.vivo_noRT | Dip2a | 2.09 | 2.39E-03 | glio |
| Atr_in.vivo_noRT | Ulk2 | 1.7 | 2.39E-03 | glio |
| Atr_in.vivo_noRT | Txnl1 | -0.977 | 2.39E-03 | glio |
| Atr_ex.vivo_noRT | Sema6d | 2.182 | 2.39E-03 | glio |
| Atr_ex.vivo_noRT | Naa10 | -0.957 | 2.39E-03 | glio |
| Atr_ex.vivo_noRT | Eef1g | -0.763 | 2.39E-03 | glio |
| Bcas2_in.vivo_noRT | Soga1 | 2.316 | 2.39E-03 | glio |
| Bcas2_in.vivo_noRT | Nfatc3 | -1.555 | 2.39E-03 | glio |
| Bora_in.vivo_noRT | C1qtnf1 | 2.195 | 2.39E-03 | glio |
| Bora_in.vivo_noRT | Zan | 2.173 | 2.39E-03 | glio |
| Bora_ex.vivo_noRT | Prm3 | 2.622 | 2.39E-03 | glio |
| Chek1_in.vivo_noRT | 2210011C24R | 2.153 | 2.39E-03 | glio |
| Chek1_in.vivo_noRT | A930024E05Ri | 2.198 | 2.39E-03 | glio |
| Hsd17b10_in.vivo_noRT | Adgrb3 | 2.31 | 2.39E-03 | glio |
| culture_effect | B130034C11R | 1.356 | 2.39E-04 | glio |
| culture_effect | N4bp1 | -0.536 | 2.39E-04 | glio |
| Interaction_genotype_Mtor | Commd1 | 1.78 | 2.39E-04 | glio |
| Interaction_genotype_Myc | Ush1g | -3.128 | 2.39E-04 | glio |
| Interaction_genotype_Myc | Kif24 | -4.085 | 2.39E-04 | glio |
| Interaction_genotype_Myc | Tmem37 | -4.084 | 2.39E-04 | glio |
| Interaction_genotype_Myc | Mpz | -5.003 | 2.39E-04 | glio |

| | | | | |
|---|---|---|---|---|
| interaction_RT | Lclat1 | -1.316 | 2.39E-04 | glio |
| interaction_RT | Hist1h1a | -1.662 | 2.39E-04 | glio |
| Aars2_in.vivo_noRT | A930029G22R | 2.525 | 2.39E-04 | glio |
| Aars2_in.vivo_noRT | Gm48604 | 2.543 | 2.39E-04 | glio |
| Aars2_in.vivo_noRT | Snhg14 | 2.661 | 2.39E-04 | glio |
| Aars2_in.vivo_noRT | Eif4h | -1.229 | 2.39E-04 | glio |
| Aars2_ex.vivo_noRT | Gm16685 | 2.893 | 2.39E-04 | glio |
| Aars2_ex.vivo_noRT | Insrr | 3.293 | 2.39E-04 | glio |
| Aars2_ex.vivo_noRT | Ccser1 | 2.899 | 2.39E-04 | glio |
| Aars2_ex.vivo_noRT | Vamp8 | 2.893 | 2.39E-04 | glio |
| Atr_in.vivo_noRT | Gm4117 | 2.242 | 2.39E-04 | glio |
| Bcas2_in.vivo_noRT | Dynlt1c | 3.004 | 2.39E-04 | glio |
| Bora_in.vivo_noRT | Eftud2 | -1.397 | 2.39E-04 | glio |
| Cyp19a1_ex.vivo_noRT | Zmynd10 | 3.296 | 2.39E-04 | glio |
| Eif2s1_in.vivo_noRT | Gm15533 | 2.143 | 2.39E-04 | glio |
| culture_effect | Rnf128 | -2.525 | 2.39E-05 | glio |
| Interaction_genotype_Atr | A330084C13R | -5.454 | 2.39E-05 | glio |
| Interaction_genotype_Mtor | Pygo1 | -4.609 | 2.39E-05 | glio |
| Interaction_genotype_Mtor | Ephx3 | -4.851 | 2.39E-05 | glio |
| Atr_in.vivo_noRT | Gm10800 | 3.223 | 2.39E-05 | glio |
| Atr_ex.vivo_noRT | Muc4 | 3.164 | 2.39E-05 | glio |
| Bcas2_in.vivo_noRT | C130040N14R | 3.16 | 2.39E-05 | glio |
| Bcas2_in.vivo_noRT | Tmem35b | 2.879 | 2.39E-05 | glio |
| Chek1_in.vivo_noRT | 2810414N06R | 3.022 | 2.39E-05 | glio |
| Hsd17b10_in.vivo_noRT | Abcg4 | 2.825 | 2.39E-05 | glio |
| culture_effect | 2210406O10R | 2.489 | 2.39E-06 | glio |
| Interaction_genotype_Mtor | 9130019O22R | -4.099 | 2.39E-06 | glio |
| Interaction_genotype_Mtor | Sox10 | 2.365 | 2.39E-06 | glio |
| Interaction_genotype_Mtor | Olfr543 | -4.234 | 2.39E-06 | glio |
| Interaction_genotype_Mtor | Hsp90ab1 | 1.873 | 2.39E-06 | glio |
| interaction_RT | Letm1 | -1.114 | 2.39E-06 | glio |
| interaction_RT | Trappc6a | 1.272 | 2.39E-06 | glio |
| Atr_in.vivo_noRT | Rack1 | -0.973 | 2.39E-06 | glio |
| Atr_ex.vivo_noRT | Agrp | 3.552 | 2.39E-06 | glio |
| Bora_ex.vivo_noRT | AF067063 | 3.201 | 2.39E-06 | glio |
| Chek1_in.vivo_noRT | Pde3a | 3.266 | 2.39E-06 | glio |
| Chek1_ex.vivo_noRT | Lrrc24 | 3.857 | 2.39E-06 | glio |
| Hsd17b10_in.vivo_noRT | Gm42918 | 2.974 | 2.39E-06 | glio |
| Hsd17b10_in.vivo_noRT | BC026762 | 3.237 | 2.39E-06 | glio |
| Aars2_in.vivo_noRT | Serpinb9b | 4.329 | 2.39E-07 | glio |
| Aars2_in.vivo_noRT | Pla2g4b | 3.499 | 2.39E-07 | glio |
| Aars2_in.vivo_noRT | Gba2 | 3.605 | 2.39E-07 | glio |
| Bora_in.vivo_noRT | Fsd2 | 3.553 | 2.39E-07 | glio |
| Chek1_in.vivo_noRT | Gm16023 | 4.047 | 2.39E-07 | glio |
| Aars2_in.vivo_noRT | 1110004F10Ri | -1.717 | 2.39E-08 | glio |
| Bora_in.vivo_noRT | Cenpq | -2.067 | 2.39E-08 | glio |
| Chek1_ex.vivo_noRT | Gjb5 | 4.263 | 2.39E-08 | glio |
| Bora_in.vivo_noRT | Gm13404 | 4.198 | 2.39E-09 | glio |
| Bora_in.vivo_noRT | Cdk11b | -2.095 | 2.39E-09 | glio |
| Bora_in.vivo_noRT | Sox11 | 4.29 | 2.39E-10 | glio |
| Chek1_in.vivo_noRT | A530013C23R | 4.126 | 2.39E-10 | glio |
| Chek1_ex.vivo_noRT | Art2b | 4.842 | 2.39E-10 | glio |

| | | | | |
|---|---|---|---|---|
| Aars2_in.vivo_noRT | Hectd2os | 4.419 | 2.39E-11 | glio |
| culture_effect | S1pr2 | -1.143 | 2.39E-12 | glio |
| Aars2_in.vivo_noRT | Iqcf1 | 4.674 | 2.39E-12 | glio |
| Bcas2_in.vivo_noRT | Gm20682 | 4.28 | 2.39E-13 | glio |
| Bcas2_in.vivo_noRT | Gm33051 | 4.154 | 2.39E-14 | glio |
| Chek1_ex.vivo_noRT | Spink8 | 4.842 | 2.39E-14 | glio |
| culture_effect | Ppfia2 | -4.047 | 2.39E-15 | glio |
| Aars2_in.vivo_noRT | Npr3 | 5.327 | 2.39E-18 | glio |
| Chek1_in.vivo_noRT | Esp15 | 5.044 | 2.39E-18 | glio |
| culture_effect | Acadl | -1.722 | 2.39E-22 | glio |
| Atr_in.vivo_noRT | 4930588A03R | 5.533 | 2.39E-29 | glio |
| culture_effect | Slc8b1 | -0.972 | 2.40E-03 | glio |
| culture_effect | Bcorl1 | -0.882 | 2.40E-03 | glio |
| Interaction_genotype_Aars2 | Gm42849 | -3.432 | 2.40E-03 | glio |
| Interaction_genotype_Aars2 | Mtcp1 | -3.72 | 2.40E-03 | glio |
| Interaction_genotype_Aars2 | 9330160F10Ri | -3.454 | 2.40E-03 | glio |
| Interaction_genotype_Aars2 | Echs1 | 1.471 | 2.40E-03 | glio |
| Interaction_genotype_Atr | 6430562O15R | -3.437 | 2.40E-03 | glio |
| Interaction_genotype_Atr | Mpc2 | 1.473 | 2.40E-03 | glio |
| Interaction_genotype_Bcas2 | Abcb5 | -2.859 | 2.40E-03 | glio |
| Interaction_genotype_Mtor | Ccdc78 | -3.602 | 2.40E-03 | glio |
| Interaction_genotype_Mtor | Gng3 | -3.532 | 2.40E-03 | glio |
| Interaction_genotype_Mtor | Psmb5 | 1.422 | 2.40E-03 | glio |
| Interaction_genotype_Myc | Gstt4 | -2.466 | 2.40E-03 | glio |
| Interaction_genotype_Myc | Dzank1 | -3.545 | 2.40E-03 | glio |
| Interaction_genotype_Myc | Egfl8 | -3.617 | 2.40E-03 | glio |
| Interaction_genotype_Rad51 | Gm4419 | -3.662 | 2.40E-03 | glio |
| interaction_RT | Rpl10 | -0.633 | 2.40E-03 | glio |
| Aars2_in.vivo_noRT | Slfn3 | 1.985 | 2.40E-03 | glio |
| Aars2_in.vivo_noRT | Zfp54 | 2.039 | 2.40E-03 | glio |
| Aars2_in.vivo_noRT | Xndc1 | 1.677 | 2.40E-03 | glio |
| Aars2_in.vivo_noRT | 1110038F14Ri | -0.797 | 2.40E-03 | glio |
| Atr_in.vivo_noRT | Tm6sf1 | 2.4 | 2.40E-03 | glio |
| Atr_in.vivo_noRT | Uhrf1bp1l | -1.391 | 2.40E-03 | glio |
| Atr_in.vivo_noRT | Smdt1 | -0.758 | 2.40E-03 | glio |
| Bcas2_in.vivo_noRT | 2310022A10R | -1.533 | 2.40E-03 | glio |
| Bcas2_ex.vivo_noRT | 9930111J21Ri | 2.742 | 2.40E-03 | glio |
| Bcas2_ex.vivo_noRT | Cidea | 2.742 | 2.40E-03 | glio |
| Bora_in.vivo_noRT | 2900089D17R | 2.09 | 2.40E-03 | glio |
| Bora_in.vivo_noRT | Cd24a | 2.622 | 2.40E-03 | glio |
| Bora_ex.vivo_noRT | Gm17173 | 2.889 | 2.40E-03 | glio |
| Cdkn2a_in.vivo_noRT | Pcp4l1 | -3.105 | 2.40E-03 | glio |
| Cdkn2a_in.vivo_noRT | Gm49172 | -1.935 | 2.40E-03 | glio |
| Chek1_ex.vivo_noRT | Rfx2 | 2.76 | 2.40E-03 | glio |
| Eif2s1_ex.vivo_noRT | Fam131a | 3.415 | 2.40E-03 | glio |
| Hsd17b10_in.vivo_noRT | Zfp273 | 2.37 | 2.40E-03 | glio |
| Hsd17b10_in.vivo_noRT | Gh | 2.592 | 2.40E-03 | glio |
| Hsd17b10_in.vivo_noRT | Rhox5 | -2.82 | 2.40E-03 | glio |
| Hsd17b10_in.vivo_noRT | Ptbp3 | -0.693 | 2.40E-03 | glio |
| culture_effect | Rbm5 | -0.692 | 2.40E-04 | glio |
| Interaction_genotype_Aars2 | 2310067P03R | -4.476 | 2.40E-04 | glio |
| Interaction_genotype_Mtor | D930030I03Ri | -4.396 | 2.40E-04 | glio |

| | | | | |
|---|---|---|---|---|
| Interaction_genotype_Mtor | Tbl1x | 1.708 | 2.40E-04 | glio |
| Interaction_genotype_Myc | Olfr810 | -2.729 | 2.40E-04 | glio |
| Interaction_genotype_Myc | Gm20597 | -4.505 | 2.40E-04 | glio |
| Interaction_genotype_Myc | Grik3 | -3.832 | 2.40E-04 | glio |
| Interaction_genotype_Myc | Rcor2 | -3.02 | 2.40E-04 | glio |
| Interaction_genotype_Myc | Rab7b | -3.644 | 2.40E-04 | glio |
| Interaction_genotype_Myc | Prdx6 | 1.872 | 2.40E-04 | glio |
| Aars2_in.vivo_noRT | Lmo4 | -1.498 | 2.40E-04 | glio |
| Atr_in.vivo_noRT | Zbtb4 | 2.469 | 2.40E-04 | glio |
| Bcas2_in.vivo_noRT | Pcdhgb1 | 2.273 | 2.40E-04 | glio |
| Bcas2_in.vivo_noRT | Mis12 | -1.81 | 2.40E-04 | glio |
| Bcas2_in.vivo_noRT | Fbl | -0.796 | 2.40E-04 | glio |
| Bora_in.vivo_noRT | Gm17259 | 2.485 | 2.40E-04 | glio |
| Chek1_in.vivo_noRT | Mok | 2.683 | 2.40E-04 | glio |
| Chek1_in.vivo_noRT | Cul9 | 2.855 | 2.40E-04 | glio |
| Chek1_ex.vivo_noRT | Rsl1d1 | -1.147 | 2.40E-04 | glio |
| Chek1_ex.vivo_noRT | Npm1 | -0.884 | 2.40E-04 | glio |
| culture_effect | Aldh1a2 | -2.498 | 2.40E-05 | glio |
| Interaction_genotype_Aars2 | Ndufs5 | 1.285 | 2.40E-05 | glio |
| interaction_RT | Lyar | -0.875 | 2.40E-05 | glio |
| Atp5c1_in.vivo_noRT | Pkd1l2 | 2.935 | 2.40E-05 | glio |
| Atr_ex.vivo_noRT | Gm39556 | 3.947 | 2.40E-05 | glio |
| Bcas2_in.vivo_noRT | 9130204L05Ri | 3.213 | 2.40E-05 | glio |
| Bora_in.vivo_noRT | AC241534.1 | 3.076 | 2.40E-05 | glio |
| Bora_in.vivo_noRT | Rab17 | 2.941 | 2.40E-05 | glio |
| Chek1_in.vivo_noRT | D930019O06R | 3.147 | 2.40E-05 | glio |
| Chek1_ex.vivo_noRT | Ccl8 | 4.263 | 2.40E-05 | glio |
| Hsd17b10_in.vivo_noRT | E130215H24R | 2.825 | 2.40E-05 | glio |
| culture_effect | Snhg4 | 1.005 | 2.40E-06 | glio |
| Interaction_genotype_Myc | Gm42670 | -4.902 | 2.40E-06 | glio |
| Interaction_genotype_Myc | Pabpc1 | 1.708 | 2.40E-06 | glio |
| Aars2_in.vivo_noRT | Hyal3 | 3.737 | 2.40E-06 | glio |
| Atr_in.vivo_noRT | Vdr | 3.557 | 2.40E-06 | glio |
| Bora_in.vivo_noRT | Zfpm1 | 3.42 | 2.40E-06 | glio |
| Interaction_genotype_Chek1 | Irf8 | 6.061 | 2.40E-07 | glio |
| Atr_in.vivo_noRT | Nckipsd | 3.465 | 2.40E-07 | glio |
| Atr_in.vivo_noRT | Adh7 | 3.523 | 2.40E-07 | glio |
| Bora_in.vivo_noRT | Adcy2 | 3.183 | 2.40E-07 | glio |
| Chek1_in.vivo_noRT | Uckl1os | 3.809 | 2.40E-07 | glio |
| Chek1_ex.vivo_noRT | Pakap | 3.463 | 2.40E-07 | glio |
| culture_effect | Inka1 | 1.23 | 2.40E-08 | glio |
| Aars2_ex.vivo_noRT | Sucnr1 | 3.293 | 2.40E-08 | glio |
| Bard1_ex.vivo_noRT | Zscan4-ps3 | 3.612 | 2.40E-08 | glio |
| Atr_in.vivo_noRT | Dnajc28 | 4.387 | 2.40E-09 | glio |
| Atr_in.vivo_noRT | Gm42658 | 3.781 | 2.40E-09 | glio |
| Bora_in.vivo_noRT | Gm32585 | 4.411 | 2.40E-09 | glio |
| Chek1_ex.vivo_noRT | Tcstv3 | 4.842 | 2.40E-09 | glio |
| culture_effect | Smpdl3b | -1.557 | 2.40E-10 | glio |
| Atr_in.vivo_noRT | Scgb1a1 | 4.423 | 2.40E-10 | glio |
| Atr_in.vivo_noRT | 1700023H06R | 4.742 | 2.40E-10 | glio |
| culture_effect | 4933421O10R | 2.05 | 2.40E-12 | glio |
| culture_effect | Rab3ip | 1.111 | 2.40E-12 | glio |

| | | | | |
|---|---|---|---|---|
| Chek1_in.vivo_noRT | Snhg4.1 | 4.521 | 2.40E-14 | glio |
| Chek1_in.vivo_noRT | Trim16 | 4.784 | 2.40E-14 | glio |
| Chek1_ex.vivo_noRT | Psmb11 | 4.842 | 2.40E-14 | glio |
| Hsd17b10_in.vivo_noRT | Helt | 3.616 | 2.40E-14 | glio |
| Hsd17b10_in.vivo_noRT | Gm20625 | 3.616 | 2.40E-14 | glio |
| Chek1_ex.vivo_noRT | Dpep3 | 4.842 | 2.40E-15 | glio |
| Chek1_ex.vivo_noRT | Erich6b | 4.842 | 2.40E-15 | glio |
| Bcas2_in.vivo_noRT | Gm40538 | 4.544 | 2.40E-16 | glio |
| Atr_in.vivo_noRT | 1700034I23Ri | 5.533 | 2.40E-28 | glio |
| culture_effect | Gm10655 | 1.769 | 2.41E-03 | glio |
| culture_effect | Gatd3a | -0.723 | 2.41E-03 | glio |
| Interaction_genotype_Atr | Gm36037 | -2.765 | 2.41E-03 | glio |
| Interaction_genotype_Atr | Radil | -3.702 | 2.41E-03 | glio |
| Interaction_genotype_Atr | Gm3448 | -2.692 | 2.41E-03 | glio |
| Interaction_genotype_Mtor | Gm10244 | -3.682 | 2.41E-03 | glio |
| Interaction_genotype_Mtor | Optn | -3.746 | 2.41E-03 | glio |
| Interaction_genotype_Mtor | Hr | -3.175 | 2.41E-03 | glio |
| Interaction_genotype_Mtor | Lefty1 | -2.935 | 2.41E-03 | glio |
| Interaction_genotype_Mtor | Fam71f1 | -2.887 | 2.41E-03 | glio |
| Interaction_genotype_Myc | Gm38163 | -2.729 | 2.41E-03 | glio |
| Interaction_genotype_Myc | Wdr35 | -2.75 | 2.41E-03 | glio |
| interaction_RT | Pop1 | -1.095 | 2.41E-03 | glio |
| interaction_RT | Alkbh5 | 0.681 | 2.41E-03 | glio |
| Aars2_in.vivo_noRT | Lurap1 | 2.148 | 2.41E-03 | glio |
| Atp5c1_in.vivo_noRT | Gm11963 | 2.408 | 2.41E-03 | glio |
| Atp5c1_in.vivo_noRT | Rilp | 2.664 | 2.41E-03 | glio |
| Atr_in.vivo_noRT | Hook3 | -1.28 | 2.41E-03 | glio |
| Atr_ex.vivo_noRT | BC034090 | 2.771 | 2.41E-03 | glio |
| Atr_ex.vivo_noRT | 1700047M11F | 2.771 | 2.41E-03 | glio |
| Bcas2_in.vivo_noRT | Fam241a | 2.288 | 2.41E-03 | glio |
| Bcas2_ex.vivo_noRT | 4930527F14Ri | 2.347 | 2.41E-03 | glio |
| Bora_in.vivo_noRT | Ubtd1 | -1.455 | 2.41E-03 | glio |
| Cenpj_in.vivo_noRT | Gm6803 | 2.022 | 2.41E-03 | glio |
| Cenpj_in.vivo_noRT | A930003A15R | 2.415 | 2.41E-03 | glio |
| Cenpj_in.vivo_noRT | Nos2 | 2.292 | 2.41E-03 | glio |
| Chek1_in.vivo_noRT | Gm36266 | 2.467 | 2.41E-03 | glio |
| Chek1_in.vivo_noRT | A330023F24Ri | 2.265 | 2.41E-03 | glio |
| Chek1_ex.vivo_noRT | Gm26822 | 2.916 | 2.41E-03 | glio |
| Chek1_ex.vivo_noRT | Syt1 | 2.89 | 2.41E-03 | glio |
| Chek1_ex.vivo_noRT | Gm29243 | 2.434 | 2.41E-03 | glio |
| Hsd17b10_in.vivo_noRT | 9930014A18R | 2.222 | 2.41E-03 | glio |
| Hsd17b10_in.vivo_noRT | Cnmd | -2.027 | 2.41E-03 | glio |
| Hsd17b10_in.vivo_noRT | Pdap1 | -0.617 | 2.41E-03 | glio |
| culture_effect | Npw | -2.038 | 2.41E-04 | glio |
| Interaction_genotype_Atr | Tarsl2 | -3.653 | 2.41E-04 | glio |
| Interaction_genotype_Atr | AC163347.1 | -3.688 | 2.41E-04 | glio |
| Interaction_genotype_Atr | Glyat | -3.167 | 2.41E-04 | glio |
| Interaction_genotype_Mtor | Gm43625 | -3.98 | 2.41E-04 | glio |
| Interaction_genotype_Mtor | Muc4 | -3.577 | 2.41E-04 | glio |
| Interaction_genotype_Mtor | Ppp1r26 | -4.63 | 2.41E-04 | glio |
| Interaction_genotype_Mtor | Hist2h3b | -4.369 | 2.41E-04 | glio |
| Interaction_genotype_Myc | Gm15473 | -4.373 | 2.41E-04 | glio |

| | | | | |
|---|---|---|---|---|
| interaction_RT | Rpa1 | -1.193 | 2.41E-04 | glio |
| interaction_RT | Nphs1 | -3.148 | 2.41E-04 | glio |
| Aars2_ex.vivo_noRT | Gm6566 | 2.899 | 2.41E-04 | glio |
| Atp5c1_in.vivo_noRT | Tcstv3 | 2.668 | 2.41E-04 | glio |
| Atr_in.vivo_noRT | Slc36a4 | 2.53 | 2.41E-04 | glio |
| Bcas2_in.vivo_noRT | Lmna | -1.129 | 2.41E-04 | glio |
| Bora_in.vivo_noRT | Wbp1 | -1.828 | 2.41E-04 | glio |
| Cenpj_ex.vivo_noRT | Gm16933 | 2.591 | 2.41E-04 | glio |
| Eif2s1_in.vivo_noRT | Slc19a3 | 2.143 | 2.41E-04 | glio |
| Hsd17b10_in.vivo_noRT | Gm49312 | 2.542 | 2.41E-04 | glio |
| Atr_in.vivo_noRT | Osgin1 | 2.723 | 2.41E-05 | glio |
| Bcas2_in.vivo_noRT | Med29 | -1.573 | 2.41E-05 | glio |
| Hsd17b10_in.vivo_noRT | Lyz2 | 3.23 | 2.41E-05 | glio |
| Hsd17b10_in.vivo_noRT | Gm20682 | 2.572 | 2.41E-05 | glio |
| culture_effect | Itpkc | -1.959 | 2.41E-06 | glio |
| culture_effect | 2310039H08R | 0.943 | 2.41E-06 | glio |
| culture_effect | Trim8 | 0.716 | 2.41E-06 | glio |
| culture_effect | Psmd6 | -0.71 | 2.41E-06 | glio |
| Interaction_genotype_Myc | 4833408A19R | -5.142 | 2.41E-06 | glio |
| Atr_in.vivo_noRT | Stac3 | 3.391 | 2.41E-06 | glio |
| Bcas2_in.vivo_noRT | Cemip | 2.772 | 2.41E-06 | glio |
| Chek1_in.vivo_noRT | Zfp990 | 3.142 | 2.41E-06 | glio |
| Hsd17b10_in.vivo_noRT | Cyp7a1 | 3.356 | 2.41E-06 | glio |
| Bora_in.vivo_noRT | Ccl19 | 2.985 | 2.41E-07 | glio |
| Chek1_ex.vivo_noRT | Enpp3 | 4.443 | 2.41E-07 | glio |
| Interaction_genotype_Mtor | Plcl1 | -6.051 | 2.41E-09 | glio |
| Aars2_in.vivo_noRT | Gm43660 | 4.138 | 2.41E-10 | glio |
| Bcas2_in.vivo_noRT | Rdh12 | 4.162 | 2.41E-10 | glio |
| Chek1_in.vivo_noRT | Baalc | 4.252 | 2.41E-10 | glio |
| Bcas2_in.vivo_noRT | Gm29083 | 4.275 | 2.41E-11 | glio |
| Bcas2_in.vivo_noRT | Gm48265 | 4.024 | 2.41E-11 | glio |
| Bora_in.vivo_noRT | Igfbp7 | 4.171 | 2.41E-11 | glio |
| Chek1_ex.vivo_noRT | Gtsf1l | 4.842 | 2.41E-11 | glio |
| Aars2_in.vivo_noRT | Gm17638 | 4.535 | 2.41E-12 | glio |
| Bora_in.vivo_noRT | AI463229 | 4.824 | 2.41E-16 | glio |
| Chek1_in.vivo_noRT | Gm48008 | 4.78 | 2.41E-17 | glio |
| Bora_in.vivo_noRT | Gm33051 | 4.807 | 2.41E-18 | glio |
| Bora_in.vivo_noRT | Gm15929 | 4.936 | 2.41E-18 | glio |
| Chek1_in.vivo_noRT | Gm10801 | 5.044 | 2.41E-18 | glio |
| Bora_in.vivo_noRT | Zscan4-ps2 | 4.936 | 2.41E-21 | glio |
| culture_effect | Junb | -2.597 | 2.41E-25 | glio |
| culture_effect | 4930513N10R | 1.871 | 2.42E-03 | glio |
| Interaction_genotype_Atr | Zfp46 | -3.319 | 2.42E-03 | glio |
| Interaction_genotype_Atr | Mlf2 | 1.216 | 2.42E-03 | glio |
| Interaction_genotype_Mtor | Rps6ka4 | -3.519 | 2.42E-03 | glio |
| Interaction_genotype_Mtor | Gzmm | -3.734 | 2.42E-03 | glio |
| Interaction_genotype_Mtor | Tnnc1 | -4.051 | 2.42E-03 | glio |
| Interaction_genotype_Mtor | Rexo5 | -3.557 | 2.42E-03 | glio |
| Interaction_genotype_Mtor | Gm17092 | -2.536 | 2.42E-03 | glio |
| Interaction_genotype_Mtor | Drap1 | 1.39 | 2.42E-03 | glio |
| Interaction_genotype_Myc | Cys1 | -2.856 | 2.42E-03 | glio |
| Interaction_genotype_Myc | Jmjd8 | -3.427 | 2.42E-03 | glio |

| | | | | |
|---|---|---|---|---|
| Interaction_genotype_Myc | Pfkfb2 | -3.502 | 2.42E-03 | glio |
| Aars2_in.vivo_noRT | Gm47644 | 2.114 | 2.42E-03 | glio |
| Aars2_in.vivo_noRT | Ropn1l | 2.127 | 2.42E-03 | glio |
| Aars2_in.vivo_noRT | Ndufa2 | -0.721 | 2.42E-03 | glio |
| Bora_in.vivo_noRT | Slc25a42 | 2.253 | 2.42E-03 | glio |
| Bora_in.vivo_noRT | Ndrg4 | 2.282 | 2.42E-03 | glio |
| Bora_ex.vivo_noRT | Ptpn22 | 3.281 | 2.42E-03 | glio |
| Cenpj_in.vivo_noRT | 1700019E08Ri | 2.022 | 2.42E-03 | glio |
| Chek1_in.vivo_noRT | Firre | 2.034 | 2.42E-03 | glio |
| Chek1_in.vivo_noRT | Slc12a2 | -1.613 | 2.42E-03 | glio |
| Chek1_ex.vivo_noRT | D6Ertd527e | 2.728 | 2.42E-03 | glio |
| Chek1_ex.vivo_noRT | Trim16 | 2.469 | 2.42E-03 | glio |
| Eif2s1_ex.vivo_noRT | Gm13191 | 2.234 | 2.42E-03 | glio |
| Eif2s1_ex.vivo_noRT | 4933416E03Ri | 2.234 | 2.42E-03 | glio |
| Eif2s1_ex.vivo_noRT | Cldn2 | 2.234 | 2.42E-03 | glio |
| Eif2s1_ex.vivo_noRT | Stfa3 | 2.234 | 2.42E-03 | glio |
| Eif2s1_ex.vivo_noRT | Olfr787 | 2.234 | 2.42E-03 | glio |
| Eif2s1_ex.vivo_noRT | Esr1 | 3.421 | 2.42E-03 | glio |
| Eif2s1_ex.vivo_noRT | Gm15908 | 2.234 | 2.42E-03 | glio |
| Eif2s1_ex.vivo_noRT | A930014E10Ri | 2.234 | 2.42E-03 | glio |
| Eif2s1_ex.vivo_noRT | Scg2 | 2.234 | 2.42E-03 | glio |
| Eif2s1_ex.vivo_noRT | 1700003E24Ri | 2.234 | 2.42E-03 | glio |
| Eif2s1_ex.vivo_noRT | Hint2 | 1.008 | 2.42E-03 | glio |
| Eif2s1_ex.vivo_noRT | Plcd4 | 3.214 | 2.42E-03 | glio |
| Eif2s1_ex.vivo_noRT | Krt19 | 2.234 | 2.42E-03 | glio |
| Eif2s1_ex.vivo_noRT | Gm48155 | 2.234 | 2.42E-03 | glio |
| Eif2s1_ex.vivo_noRT | Plk5 | 3.015 | 2.42E-03 | glio |
| Eif2s1_ex.vivo_noRT | Gm34059 | 2.234 | 2.42E-03 | glio |
| Eif2s1_ex.vivo_noRT | Halr1 | 2.234 | 2.42E-03 | glio |
| Eif2s1_ex.vivo_noRT | Prex2 | 3.814 | 2.42E-03 | glio |
| Eif2s1_ex.vivo_noRT | Irf8 | 3.014 | 2.42E-03 | glio |
| Eif2s1_ex.vivo_noRT | Helt | 2.234 | 2.42E-03 | glio |
| Eif2s1_ex.vivo_noRT | Gm20625 | 2.234 | 2.42E-03 | glio |
| Eif2s1_ex.vivo_noRT | Gm5157 | 2.234 | 2.42E-03 | glio |
| Eif2s1_ex.vivo_noRT | Gm44736 | 2.234 | 2.42E-03 | glio |
| Eif2s1_ex.vivo_noRT | Fam227b | 2.234 | 2.42E-03 | glio |
| Eif2s1_ex.vivo_noRT | Gm20658 | 2.234 | 2.42E-03 | glio |
| Eif2s1_ex.vivo_noRT | Ift88 | -2.653 | 2.42E-03 | glio |
| Eif2s1_ex.vivo_noRT | Olfr810 | 2.234 | 2.42E-03 | glio |
| Eif2s1_ex.vivo_noRT | Insc | 3.014 | 2.42E-03 | glio |
| Eif2s1_ex.vivo_noRT | Apoa5 | 2.234 | 2.42E-03 | glio |
| Eif2s1_ex.vivo_noRT | Gm49172 | 2.234 | 2.42E-03 | glio |
| Eif2s1_ex.vivo_noRT | C1s1 | 3.125 | 2.42E-03 | glio |
| Eif2s1_ex.vivo_noRT | Gjd4 | 2.234 | 2.42E-03 | glio |
| Hsd17b10_in.vivo_noRT | AV356131 | 2.332 | 2.42E-03 | glio |
| Hsd17b10_in.vivo_noRT | Jakmip1 | 2.295 | 2.42E-03 | glio |
| Hsd17b10_in.vivo_noRT | Rtl9 | 2.175 | 2.42E-03 | glio |
| Hsd17b10_in.vivo_noRT | Cyb5rl | 1.855 | 2.42E-03 | glio |
| Interaction_genotype_Mtor | Mkrn3 | -4.269 | 2.42E-04 | glio |
| Interaction_genotype_Mtor | Gm26621 | -4.36 | 2.42E-04 | glio |
| Interaction_genotype_Myc | 2810454H06R | -3.837 | 2.42E-04 | glio |
| interaction_RT | Atp6ap2 | 1.069 | 2.42E-04 | glio |

| | | | | |
|---|---|---|---|---|
| interaction_RT | Atp6v1d | 0.773 | 2.42E-04 | glio |
| Atr_in.vivo_noRT | B230216N24R | 2.859 | 2.42E-04 | glio |
| Atr_in.vivo_noRT | Dok7 | 2.505 | 2.42E-04 | glio |
| Atr_in.vivo_noRT | Ube2q1 | -1.488 | 2.42E-04 | glio |
| Bcas2_in.vivo_noRT | Hrk | 2.682 | 2.42E-04 | glio |
| Bcas2_in.vivo_noRT | Gm12359 | 2.645 | 2.42E-04 | glio |
| Bcas2_in.vivo_noRT | 1700113B09R | 2.714 | 2.42E-04 | glio |
| Bcas2_ex.vivo_noRT | Idh2 | -1.651 | 2.42E-04 | glio |
| Bora_in.vivo_noRT | Ifit1bl2 | 2.739 | 2.42E-04 | glio |
| Bora_ex.vivo_noRT | Slc38a3 | 3.196 | 2.42E-04 | glio |
| Chek1_in.vivo_noRT | Mrpl18 | -1.272 | 2.42E-04 | glio |
| Chek1_ex.vivo_noRT | Rab25 | 2.898 | 2.42E-04 | glio |
| Eif2s1_in.vivo_noRT | Gm20513 | 2.292 | 2.42E-04 | glio |
| Eif2s1_in.vivo_noRT | Gpr4 | 2.143 | 2.42E-04 | glio |
| Eif2s1_in.vivo_noRT | Prl7c1 | 2.143 | 2.42E-04 | glio |
| Eif2s1_in.vivo_noRT | Gm33466 | 2.143 | 2.42E-04 | glio |
| culture_effect | Snrnp25 | 0.653 | 2.42E-05 | glio |
| culture_effect | Armc1 | 0.599 | 2.42E-05 | glio |
| Interaction_genotype_Myc | Zfp61 | -3.934 | 2.42E-05 | glio |
| Interaction_genotype_Myc | Kif18b | -4.005 | 2.42E-05 | glio |
| Aars2_in.vivo_noRT | Gm26658 | 3.296 | 2.42E-05 | glio |
| Bcas2_in.vivo_noRT | Eid2b | 2.502 | 2.42E-05 | glio |
| Bora_in.vivo_noRT | Phldb2 | 3.202 | 2.42E-05 | glio |
| Bora_in.vivo_noRT | Gm11110 | 3.018 | 2.42E-05 | glio |
| Bora_in.vivo_noRT | Mag | 2.837 | 2.42E-05 | glio |
| Bora_in.vivo_noRT | Pcdhb16 | 3.234 | 2.42E-05 | glio |
| Chek1_in.vivo_noRT | Zfp799 | 2.946 | 2.42E-05 | glio |
| Interaction_genotype_Mtor | Spx | -5.323 | 2.42E-06 | glio |
| Aars2_in.vivo_noRT | Gm43156 | 3.308 | 2.42E-06 | glio |
| Bcas2_in.vivo_noRT | Gm13986 | 3.548 | 2.42E-06 | glio |
| Chek1_in.vivo_noRT | Kirrel3 | 3.31 | 2.42E-06 | glio |
| Hsd17b10_in.vivo_noRT | Podxl | 3.084 | 2.42E-06 | glio |
| Hsd17b10_in.vivo_noRT | Sult6b1 | 3.352 | 2.42E-06 | glio |
| culture_effect | Sod1 | -0.635 | 2.42E-07 | glio |
| Aars2_in.vivo_noRT | Gm49135 | 3.644 | 2.42E-07 | glio |
| Bora_in.vivo_noRT | Gm43062 | 3.682 | 2.42E-07 | glio |
| Chek1_in.vivo_noRT | Tnfrsf23 | 4.125 | 2.42E-07 | glio |
| culture_effect | Ptpn13 | -1.328 | 2.42E-08 | glio |
| culture_effect | Camsap2 | -1.191 | 2.42E-08 | glio |
| culture_effect | Tecr | -0.791 | 2.42E-08 | glio |
| Chek1_in.vivo_noRT | Emilin2 | 3.967 | 2.42E-08 | glio |
| interaction_RT | Rpl4 | -1.004 | 2.42E-09 | glio |
| Bcas2_in.vivo_noRT | Grm1 | 3.703 | 2.42E-09 | glio |
| Aars2_in.vivo_noRT | Mycl | 4.805 | 2.42E-11 | glio |
| Bora_in.vivo_noRT | 4833419F23Ri | 4.936 | 2.42E-12 | glio |
| Hsd17b10_in.vivo_noRT | Gm12800 | 3.616 | 2.42E-12 | glio |
| Bcas2_in.vivo_noRT | Pik3ap1 | 4.544 | 2.42E-14 | glio |
| Hsd17b10_in.vivo_noRT | Gm5157 | 3.616 | 2.42E-14 | glio |
| Bora_in.vivo_noRT | Slc16a4 | 4.672 | 2.42E-16 | glio |
| Atr_in.vivo_noRT | 8030423F21Ri | 5.004 | 2.42E-17 | glio |
| Bcas2_in.vivo_noRT | Speer4e | 4.807 | 2.42E-18 | glio |
| Chek1_in.vivo_noRT | 4732463B04R | 4.78 | 2.42E-18 | glio |

| | | | | |
|---|---|---|---|---|
| culture_effect | Nhsl1 | -1.726 | 2.42E-20 | glio |
| Aars2_in.vivo_noRT | Strip2 | 5.063 | 2.42E-20 | glio |
| Chek1_in.vivo_noRT | Trim17 | 5.044 | 2.42E-24 | glio |
| Chek1_in.vivo_noRT | Clec12b | 5.822 | 2.42E-29 | glio |
| culture_effect | Fam234b | -1.099 | 2.43E-03 | glio |
| culture_effect | Fahd1 | 0.822 | 2.43E-03 | glio |
| Interaction_genotype_Mtor | Gm1123 | -3.774 | 2.43E-03 | glio |
| Interaction_genotype_Mtor | Gm15520 | -3.176 | 2.43E-03 | glio |
| Interaction_genotype_Mtor | R3hdm2 | 1.63 | 2.43E-03 | glio |
| Interaction_genotype_Mtor | Qk | 1.464 | 2.43E-03 | glio |
| Interaction_genotype_Myc | Wfikkn2 | -3.512 | 2.43E-03 | glio |
| Interaction_genotype_Myc | Sgk1 | -2.476 | 2.43E-03 | glio |
| Interaction_genotype_Rpl3 | Gm14636 | -4.887 | 2.43E-03 | glio |
| Aars2_in.vivo_noRT | A930001M01F | 2.166 | 2.43E-03 | glio |
| Aars2_in.vivo_noRT | Gm49085 | 2.048 | 2.43E-03 | glio |
| Aars2_in.vivo_noRT | Maff | -1.69 | 2.43E-03 | glio |
| Aars2_in.vivo_noRT | Igtp | -0.837 | 2.43E-03 | glio |
| Aars2_in.vivo_noRT | Cox8a | -0.572 | 2.43E-03 | glio |
| Aars2_in.vivo_noRT | Rps27a | -0.598 | 2.43E-03 | glio |
| Atp5c1_in.vivo_noRT | Gm16685 | 2.01 | 2.43E-03 | glio |
| Atr_in.vivo_noRT | Chrac1 | -1.149 | 2.43E-03 | glio |
| Bora_in.vivo_noRT | Flrt2 | 2.274 | 2.43E-03 | glio |
| Bora_in.vivo_noRT | Stip1 | -0.73 | 2.43E-03 | glio |
| Cenpj_ex.vivo_noRT | Gpc5 | 2.196 | 2.43E-03 | glio |
| culture_effect | Entr1 | 0.726 | 2.43E-04 | glio |
| Interaction_genotype_Atr | Gm49701 | -4.468 | 2.43E-04 | glio |
| Interaction_genotype_Mtor | H2-Eb1 | 4.298 | 2.43E-04 | glio |
| Interaction_genotype_Myc | Tbc1d13 | -3.506 | 2.43E-04 | glio |
| interaction_RT | Tecr | 0.84 | 2.43E-04 | glio |
| Aars2_in.vivo_noRT | Ccdc136 | 2.489 | 2.43E-04 | glio |
| Bcas2_in.vivo_noRT | Rhbdf2 | 2.487 | 2.43E-04 | glio |
| Bcas2_in.vivo_noRT | Tpbg | 2.594 | 2.43E-04 | glio |
| Bcas2_in.vivo_noRT | 2900089D17R | 2.488 | 2.43E-04 | glio |
| Chek1_ex.vivo_noRT | Isl1 | 3.26 | 2.43E-04 | glio |
| Interaction_genotype_Mtor | Gm10501 | -5.233 | 2.43E-05 | glio |
| Interaction_genotype_Mtor | Plekha6 | -4.403 | 2.43E-05 | glio |
| Interaction_genotype_Mtor | Rgs16 | -5.07 | 2.43E-05 | glio |
| Aars2_in.vivo_noRT | C920021L13Ri | 2.996 | 2.43E-05 | glio |
| Aars2_in.vivo_noRT | Gm46218 | 3.04 | 2.43E-05 | glio |
| Aars2_in.vivo_noRT | Smdt1 | -1.081 | 2.43E-05 | glio |
| Bcas2_ex.vivo_noRT | Hoxc8 | 2.742 | 2.43E-05 | glio |
| Bora_in.vivo_noRT | Smim40 | 2.967 | 2.43E-05 | glio |
| Bora_ex.vivo_noRT | Muc4 | 3.201 | 2.43E-05 | glio |
| Hsd17b10_in.vivo_noRT | Cyp4f40 | 2.963 | 2.43E-05 | glio |
| culture_effect | Gm26533 | 2.431 | 2.43E-06 | glio |
| Interaction_genotype_Mtor | Gm45833 | -7.403 | 2.43E-06 | glio |
| Interaction_genotype_Mtor | 4931431C16R | -4.485 | 2.43E-06 | glio |
| Interaction_genotype_Myc | Hba-a2 | -5.59 | 2.43E-06 | glio |
| Aars2_ex.vivo_noRT | Gm11899 | 3.293 | 2.43E-06 | glio |
| culture_effect | Gm43221 | -3.106 | 2.43E-07 | glio |
| Atr_in.vivo_noRT | Fbxl22 | 3.417 | 2.43E-07 | glio |
| Atr_in.vivo_noRT | Bglap2 | 4.605 | 2.43E-11 | glio |

| | | | | |
|---|---|---|---|---|
| Bcas2_in.vivo_noRT | 4921536K21R | 4.547 | 2.43E-11 | glio |
| Bcas2_in.vivo_noRT | F630040K05Ri | 4.156 | 2.43E-11 | glio |
| Atr_in.vivo_noRT | Klk14 | 5.002 | 2.43E-12 | glio |
| Atr_in.vivo_noRT | Tex52 | 4.883 | 2.43E-13 | glio |
| Bcas2_in.vivo_noRT | Masp1 | 4.285 | 2.43E-13 | glio |
| Chek1_in.vivo_noRT | Ntn4 | 4.777 | 2.43E-15 | glio |
| Aars2_in.vivo_noRT | Gm11627 | 5.327 | 2.43E-16 | glio |
| Atr_in.vivo_noRT | Gm44752 | 5.146 | 2.43E-16 | glio |
| Aars2_in.vivo_noRT | Slc17a6 | 5.327 | 2.43E-32 | glio |
| culture_effect | Loxl4 | 1.687 | 2.44E-03 | glio |
| culture_effect | Gm4117 | -1.411 | 2.44E-03 | glio |
| Interaction_genotype_Atr | Lyrm1 | -3.608 | 2.44E-03 | glio |
| Interaction_genotype_Bcas2 | Nid1 | -4.137 | 2.44E-03 | glio |
| Interaction_genotype_Bcas2 | Dph3 | 1.463 | 2.44E-03 | glio |
| Interaction_genotype_Lmo7 | Pmp2 | -4.132 | 2.44E-03 | glio |
| Interaction_genotype_Mtor | Fam228b | -3.98 | 2.44E-03 | glio |
| Interaction_genotype_Myc | Glis1 | -3.359 | 2.44E-03 | glio |
| Interaction_genotype_Myc | Prss22 | -4.237 | 2.44E-03 | glio |
| Interaction_genotype_Myc | Gm15492 | -3.444 | 2.44E-03 | glio |
| Interaction_genotype_Myc | Gm38246 | -3.38 | 2.44E-03 | glio |
| Interaction_genotype_Myc | Rabgap1l | -2.948 | 2.44E-03 | glio |
| Interaction_genotype_Myc | 3110009E18Ri | -2.355 | 2.44E-03 | glio |
| Interaction_genotype_Myc | Me1 | -1.876 | 2.44E-03 | glio |
| Interaction_genotype_Myc | Hnrnpf | 1.09 | 2.44E-03 | glio |
| Interaction_genotype_Myc | Ptma | 1.242 | 2.44E-03 | glio |
| Interaction_genotype_Pars2 | Gm33782 | -3.755 | 2.44E-03 | glio |
| Interaction_genotype_Pars2 | Gm34885 | -4.181 | 2.44E-03 | glio |
| Interaction_genotype_Pars2 | 4930540M05F | -3.984 | 2.44E-03 | glio |
| Interaction_genotype_Pars2 | 1700126G02R | -4.223 | 2.44E-03 | glio |
| Interaction_genotype_Pars2 | Olfr279 | -4.044 | 2.44E-03 | glio |
| Interaction_genotype_Pars2 | Gm45338 | -2.662 | 2.44E-03 | glio |
| Interaction_genotype_Pars2 | Slc16a4 | -3.385 | 2.44E-03 | glio |
| Interaction_genotype_Pars2 | 5033430I15Ri | -3.471 | 2.44E-03 | glio |
| Interaction_genotype_Rpl3 | Bhlhe41 | -4.944 | 2.44E-03 | glio |
| Interaction_genotype_Rpl3 | Nfkbiz | 3.1 | 2.44E-03 | glio |
| Interaction_genotype_Rpl3 | Tonsl | -3.526 | 2.44E-03 | glio |
| Interaction_genotype_Rpl3 | 8030456M14F | -4.05 | 2.44E-03 | glio |
| Interaction_genotype_Rpl3 | Tmem43 | 2.454 | 2.44E-03 | glio |
| Interaction_genotype_Rpl3 | Myo5a | 1.965 | 2.44E-03 | glio |
| interaction_RT | Platr3 | 3.114 | 2.44E-03 | glio |
| interaction_RT | Ruvbl1 | -0.709 | 2.44E-03 | glio |
| Aars2_in.vivo_noRT | Selenos | -0.884 | 2.44E-03 | glio |
| Atp5c1_in.vivo_noRT | Tmeff2 | 2.703 | 2.44E-03 | glio |
| Atr_in.vivo_noRT | Pigv | 2.246 | 2.44E-03 | glio |
| Atr_in.vivo_noRT | Gm43445 | 2.099 | 2.44E-03 | glio |
| Atr_in.vivo_noRT | Gm12976 | 2.484 | 2.44E-03 | glio |
| Atr_in.vivo_noRT | Bsg | -0.635 | 2.44E-03 | glio |
| Bard1_ex.vivo_noRT | Gm13713 | 2.415 | 2.44E-03 | glio |
| Bcas2_in.vivo_noRT | Itgal | 2.336 | 2.44E-03 | glio |
| Bcas2_in.vivo_noRT | Fbxl12os | 2.166 | 2.44E-03 | glio |
| Bcas2_in.vivo_noRT | Gm45716 | 2.12 | 2.44E-03 | glio |
| Bcas2_in.vivo_noRT | Dkc1 | -0.968 | 2.44E-03 | glio |

| | | | | |
|---|---|---|---|---|
| Bcas2_in.vivo_noRT | Nasp | -0.782 | 2.44E-03 | glio |
| Bcas2_ex.vivo_noRT | Cd72 | 3.145 | 2.44E-03 | glio |
| Bora_ex.vivo_noRT | Spn | 2.808 | 2.44E-03 | glio |
| Cenpj_ex.vivo_noRT | Gm34939 | 2.591 | 2.44E-03 | glio |
| Chek1_in.vivo_noRT | 2810455O05R | 2.268 | 2.44E-03 | glio |
| Chek1_in.vivo_noRT | 9330160F10Ri | 2.037 | 2.44E-03 | glio |
| Chek1_in.vivo_noRT | Rpl23 | -0.641 | 2.44E-03 | glio |
| culture_effect | Thoc5 | -0.767 | 2.44E-04 | glio |
| culture_effect | Paf1 | -0.752 | 2.44E-04 | glio |
| Interaction_genotype_Chek1 | Ms4a4d | 4.94 | 2.44E-04 | glio |
| Interaction_genotype_Mtor | 2810410L24Ri | -4.382 | 2.44E-04 | glio |
| Interaction_genotype_Mtor | 9030612E09Ri | -4.145 | 2.44E-04 | glio |
| Interaction_genotype_Mtor | Rbpms2 | -4.142 | 2.44E-04 | glio |
| Interaction_genotype_Mtor | E130317F20Ri | -3.015 | 2.44E-04 | glio |
| Interaction_genotype_Myc | Apoa5 | -2.729 | 2.44E-04 | glio |
| Interaction_genotype_Myc | Gm48742 | -3.454 | 2.44E-04 | glio |
| Interaction_genotype_Myc | A230020J21Ri | -4.35 | 2.44E-04 | glio |
| Atr_in.vivo_noRT | Erbb3 | -1.777 | 2.44E-04 | glio |
| Bcas2_in.vivo_noRT | Slc8a3 | 2.609 | 2.44E-04 | glio |
| Bora_in.vivo_noRT | Dock4 | 2.655 | 2.44E-04 | glio |
| Chek1_in.vivo_noRT | Wasl | -1.284 | 2.44E-04 | glio |
| Hsd17b10_in.vivo_noRT | Wfdc18 | 3.35 | 2.44E-04 | glio |
| culture_effect | Cdc14a | -0.936 | 2.44E-05 | glio |
| Interaction_genotype_Myc | Gm46620 | -3.586 | 2.44E-05 | glio |
| Interaction_genotype_Myc | Ppil6 | -4.404 | 2.44E-05 | glio |
| Interaction_genotype_Myc | Calu | 1.718 | 2.44E-05 | glio |
| Interaction_genotype_Myc | Rps20 | 1.632 | 2.44E-05 | glio |
| interaction_RT | Ube2h | 1.213 | 2.44E-05 | glio |
| Aars2_in.vivo_noRT | Btc | 3.767 | 2.44E-05 | glio |
| Aars2_ex.vivo_noRT | Gtsf1l | 3.293 | 2.44E-05 | glio |
| Atr_in.vivo_noRT | Alad | -1.606 | 2.44E-05 | glio |
| Atr_in.vivo_noRT | Tbl1x | -1.292 | 2.44E-05 | glio |
| Atr_ex.vivo_noRT | Pacrg | 3.515 | 2.44E-05 | glio |
| Chek1_in.vivo_noRT | D030056L22R | -1.857 | 2.44E-05 | glio |
| Chek1_ex.vivo_noRT | Hcn3 | 3.34 | 2.44E-05 | glio |
| Interaction_genotype_Mtor | Ntf5 | -5.971 | 2.44E-06 | glio |
| Interaction_genotype_Mtor | Gm17146 | -5.254 | 2.44E-06 | glio |
| Interaction_genotype_Mtor | H2afj | 2.159 | 2.44E-06 | glio |
| Atr_in.vivo_noRT | Erg28 | -1.799 | 2.44E-06 | glio |
| Atr_ex.vivo_noRT | Olfr56 | 3.164 | 2.44E-06 | glio |
| Bcas2_in.vivo_noRT | 9330104G04R | 3.327 | 2.44E-06 | glio |
| Bora_in.vivo_noRT | Gm30541 | 3.241 | 2.44E-06 | glio |
| Bora_in.vivo_noRT | 4921507P07R | 3.172 | 2.44E-06 | glio |
| Chek1_in.vivo_noRT | 1810062O18R | 3.458 | 2.44E-06 | glio |
| Chek1_in.vivo_noRT | Acot5 | 3.16 | 2.44E-06 | glio |
| Aars2_in.vivo_noRT | Gm26533 | 3.693 | 2.44E-07 | glio |
| Aars2_in.vivo_noRT | Zeb2os | 3.413 | 2.44E-07 | glio |
| Bcas2_in.vivo_noRT | Bhlhe41 | 4.284 | 2.44E-07 | glio |
| Bcas2_in.vivo_noRT | Elfn2 | 3.562 | 2.44E-08 | glio |
| Bora_in.vivo_noRT | Fhl4 | 3.686 | 2.44E-08 | glio |
| Aars2_in.vivo_noRT | Slc14a1 | 3.441 | 2.44E-09 | glio |
| Atr_in.vivo_noRT | Gpd1 | 4.361 | 2.44E-09 | glio |

| | | | | |
|---|---|---|---|---|
| Bora_in.vivo_noRT | Pde11a | 4.407 | 2.44E-09 | glio |
| culture_effect | Rai14 | 1.455 | 2.44E-13 | glio |
| Chek1_in.vivo_noRT | Gm16751 | 4.528 | 2.44E-14 | glio |
| Chek1_ex.vivo_noRT | Tac2 | 4.842 | 2.44E-15 | glio |
| Atr_in.vivo_noRT | Gm20705 | 5.002 | 2.44E-19 | glio |
| culture_effect | H2-DMb1 | -3.107 | 2.44E-50 | glio |
| culture_effect | Skiv2l | -0.878 | 2.45E-03 | glio |
| Interaction_genotype_Atr | Pik3ap1 | -3.259 | 2.45E-03 | glio |
| Interaction_genotype_Mtor | Wdr27 | -2.963 | 2.45E-03 | glio |
| Interaction_genotype_Mtor | Ptms | 1.611 | 2.45E-03 | glio |
| Interaction_genotype_Myc | BB557941 | -2.869 | 2.45E-03 | glio |
| Interaction_genotype_Myc | Gpam | -3.164 | 2.45E-03 | glio |
| Interaction_genotype_Myc | Trim13 | -3.146 | 2.45E-03 | glio |
| Interaction_genotype_Myc | Eif4a1 | 1.408 | 2.45E-03 | glio |
| Aars2_in.vivo_noRT | Zbtb11os1 | 2.114 | 2.45E-03 | glio |
| Aars2_in.vivo_noRT | Rilpl1 | 1.921 | 2.45E-03 | glio |
| Atp5c1_in.vivo_noRT | Lrrn3 | 2.259 | 2.45E-03 | glio |
| Atr_in.vivo_noRT | Pou3f1 | 2.174 | 2.45E-03 | glio |
| Atr_in.vivo_noRT | 2010109A12R | 2.125 | 2.45E-03 | glio |
| Atr_in.vivo_noRT | Mob3b | 1.89 | 2.45E-03 | glio |
| Atr_in.vivo_noRT | Ube2cbp | 1.929 | 2.45E-03 | glio |
| Atr_in.vivo_noRT | Pfkl | -1.325 | 2.45E-03 | glio |
| Bard1_ex.vivo_noRT | Gtsf1 | 2.449 | 2.45E-03 | glio |
| Bard1_ex.vivo_noRT | Ccdc189 | 2.763 | 2.45E-03 | glio |
| Bcas2_in.vivo_noRT | Ubxn10 | 1.901 | 2.45E-03 | glio |
| Bcas2_in.vivo_noRT | Pcdhb21 | 2.066 | 2.45E-03 | glio |
| Bcas2_in.vivo_noRT | Tmem11 | -0.97 | 2.45E-03 | glio |
| Bcas2_in.vivo_noRT | Eif2s1 | -0.708 | 2.45E-03 | glio |
| Bora_in.vivo_noRT | Rnf225 | 2.353 | 2.45E-03 | glio |
| Bora_in.vivo_noRT | Atp5j | -0.505 | 2.45E-03 | glio |
| Cenpj_in.vivo_noRT | Gm44899 | 2.022 | 2.45E-03 | glio |
| Chek1_in.vivo_noRT | Ang | 2.268 | 2.45E-03 | glio |
| Chek1_in.vivo_noRT | 1700019L13Ri | 2.214 | 2.45E-03 | glio |
| Chek1_in.vivo_noRT | Lysmd4 | 2.031 | 2.45E-03 | glio |
| Chek1_in.vivo_noRT | Got1 | -1.133 | 2.45E-03 | glio |
| Chek1_ex.vivo_noRT | Rbm17 | -0.825 | 2.45E-03 | glio |
| Interaction_genotype_Atr | 5033403F01Ri | -4.199 | 2.45E-04 | glio |
| Interaction_genotype_Atr | Gm15127 | -4.484 | 2.45E-04 | glio |
| Interaction_genotype_Mtor | Zfp433 | -4.706 | 2.45E-04 | glio |
| Interaction_genotype_Mtor | 9030025P20R | -4.651 | 2.45E-04 | glio |
| Interaction_genotype_Mtor | 9130015A21R | -6.696 | 2.45E-04 | glio |
| Interaction_genotype_Myc | Gm20635 | -3.892 | 2.45E-04 | glio |
| Interaction_genotype_Myc | Gstm2 | -4.167 | 2.45E-04 | glio |
| Interaction_genotype_Myc | Vipas39 | -2.782 | 2.45E-04 | glio |
| interaction_RT | Sept4 | 3.776 | 2.45E-04 | glio |
| Aars2_in.vivo_noRT | Gm11520 | 2.397 | 2.45E-04 | glio |
| Atr_in.vivo_noRT | Psmd2 | -1.155 | 2.45E-04 | glio |
| Atr_ex.vivo_noRT | Plac1 | 3.383 | 2.45E-04 | glio |
| Bard1_ex.vivo_noRT | Spink6 | 2.814 | 2.45E-04 | glio |
| Bard1_ex.vivo_noRT | Htr2b | 2.814 | 2.45E-04 | glio |
| Bard1_ex.vivo_noRT | Serping1 | 2.814 | 2.45E-04 | glio |
| Bcas2_in.vivo_noRT | 1600002K03R | 2.744 | 2.45E-04 | glio |

| | | | | |
|---|---|---|---|---|
| Bcas2_in.vivo_noRT | Nrap | 2.797 | 2.45E-04 | glio |
| Bcas2_ex.vivo_noRT | Smtnl1 | 2.742 | 2.45E-04 | glio |
| Bora_ex.vivo_noRT | Abcb5 | 2.407 | 2.45E-04 | glio |
| Cenpj_in.vivo_noRT | Cldn2 | 2.022 | 2.45E-04 | glio |
| Interaction_genotype_Mtor | Gm26737 | -7.426 | 2.45E-05 | glio |
| Interaction_genotype_Mtor | Cys1 | -4.226 | 2.45E-05 | glio |
| Interaction_genotype_Myc | Arhgap19 | -4.095 | 2.45E-05 | glio |
| Bard1_ex.vivo_noRT | Cidea | 3.612 | 2.45E-05 | glio |
| Bard1_ex.vivo_noRT | Gm16933 | 2.814 | 2.45E-05 | glio |
| Bora_in.vivo_noRT | Mapk15 | 2.952 | 2.45E-05 | glio |
| Bora_in.vivo_noRT | Fkbp2 | -1.201 | 2.45E-05 | glio |
| culture_effect | Gm48855 | -2.645 | 2.45E-06 | glio |
| culture_effect | Fabp3 | 2.298 | 2.45E-06 | glio |
| Aars2_in.vivo_noRT | Auts2 | 2.876 | 2.45E-06 | glio |
| Atp5c1_in.vivo_noRT | Gm11899 | 2.677 | 2.45E-06 | glio |
| Bora_in.vivo_noRT | Gm26766 | 3.241 | 2.45E-06 | glio |
| Bora_ex.vivo_noRT | Gm2016 | 3.201 | 2.45E-06 | glio |
| Chek1_in.vivo_noRT | AA388235 | 3.404 | 2.45E-06 | glio |
| Chek1_in.vivo_noRT | A430033K04R | 3.205 | 2.45E-06 | glio |
| culture_effect | Igf1 | 1.764 | 2.45E-07 | glio |
| culture_effect | Dxo | -1.205 | 2.45E-07 | glio |
| Atr_in.vivo_noRT | Gm10244 | 3.712 | 2.45E-07 | glio |
| Atr_ex.vivo_noRT | BC061195 | 3.164 | 2.45E-07 | glio |
| Bora_in.vivo_noRT | 6330408M09F | 3.895 | 2.45E-07 | glio |
| Chek1_in.vivo_noRT | Cdo1 | 3.756 | 2.45E-07 | glio |
| Chek1_in.vivo_noRT | Gm35867 | 3.591 | 2.45E-07 | glio |
| culture_effect | Kars | -0.731 | 2.45E-08 | glio |
| interaction_RT | Tmem119 | 5.131 | 2.45E-08 | glio |
| Chek1_ex.vivo_noRT | Olfr90 | 4.449 | 2.45E-09 | glio |
| culture_effect | Mtpn | 0.679 | 2.45E-10 | glio |
| Chek1_in.vivo_noRT | Grin3b | 4.521 | 2.45E-11 | glio |
| Aars2_in.vivo_noRT | Gm44658 | 4.795 | 2.45E-15 | glio |
| Aars2_in.vivo_noRT | Gm21762 | 5.063 | 2.45E-16 | glio |
| Bora_in.vivo_noRT | Pitx3 | 4.936 | 2.45E-18 | glio |
| culture_effect | Sec1 | 1.786 | 2.46E-03 | glio |
| culture_effect | Dop1b | -1.706 | 2.46E-03 | glio |
| culture_effect | Lamc1 | 0.735 | 2.46E-03 | glio |
| culture_effect | Ppp1r9b | -0.727 | 2.46E-03 | glio |
| culture_effect | Bhlhe22 | 0.709 | 2.46E-03 | glio |
| culture_effect | Trmt6 | 0.483 | 2.46E-03 | glio |
| Interaction_genotype_Aars2 | Lmx1b | -3.911 | 2.46E-03 | glio |
| Interaction_genotype_Atr | B4galnt1 | -3.202 | 2.46E-03 | glio |
| Interaction_genotype_Atr | Pcdh17 | 2.508 | 2.46E-03 | glio |
| Interaction_genotype_Atr | Higd1a | 1.561 | 2.46E-03 | glio |
| Interaction_genotype_Bcas2 | Padi2 | -4.226 | 2.46E-03 | glio |
| Interaction_genotype_Bcas2 | Elk3 | -2.249 | 2.46E-03 | glio |
| Interaction_genotype_Mtor | Pnmal2 | -3.73 | 2.46E-03 | glio |
| Interaction_genotype_Mtor | Serpinb8 | -3.008 | 2.46E-03 | glio |
| Interaction_genotype_Mtor | Zfp706 | 1.241 | 2.46E-03 | glio |
| Interaction_genotype_Myc | Ifi205 | -4.443 | 2.46E-03 | glio |
| Interaction_genotype_Myc | Pknox2 | -3.572 | 2.46E-03 | glio |
| Interaction_genotype_Myc | Cldn11 | -3.298 | 2.46E-03 | glio |

| | | | | |
|---|---|---|---|---|
| Interaction_genotype_Myc | Ttf2 | -2.483 | 2.46E-03 | glio |
| Interaction_genotype_Myc | Dstn | 1.254 | 2.46E-03 | glio |
| Aars2_in.vivo_noRT | Platr16 | 2.339 | 2.46E-03 | glio |
| Bcas2_in.vivo_noRT | Mt4 | 2.811 | 2.46E-03 | glio |
| Bcas2_in.vivo_noRT | Rnh1 | -0.955 | 2.46E-03 | glio |
| Bora_in.vivo_noRT | Rbpms2 | 2.224 | 2.46E-03 | glio |
| Bora_in.vivo_noRT | Sfi1 | 2.028 | 2.46E-03 | glio |
| Bora_in.vivo_noRT | Trf | -2.502 | 2.46E-03 | glio |
| Chek1_in.vivo_noRT | Pheta1 | 2.246 | 2.46E-03 | glio |
| Chek1_in.vivo_noRT | Nfyb | -1.281 | 2.46E-03 | glio |
| Chek1_ex.vivo_noRT | Gsto2 | 2.851 | 2.46E-03 | glio |
| Interaction_genotype_Bora | Pomk | -4.304 | 2.46E-04 | glio |
| Interaction_genotype_Mtor | Mblac1 | -4.583 | 2.46E-04 | glio |
| Interaction_genotype_Myc | Gm16794 | -3.933 | 2.46E-04 | glio |
| Interaction_genotype_Myc | Ggact | -3.625 | 2.46E-04 | glio |
| Aars2_in.vivo_noRT | Zglp1 | 2.851 | 2.46E-04 | glio |
| Aars2_in.vivo_noRT | 4833408A19R | 2.477 | 2.46E-04 | glio |
| Atr_in.vivo_noRT | Mok | 2.643 | 2.46E-04 | glio |
| Atr_in.vivo_noRT | Rab7b | 2.516 | 2.46E-04 | glio |
| Atr_in.vivo_noRT | Mturn | 2.408 | 2.46E-04 | glio |
| Atr_in.vivo_noRT | Galnt4 | 2.467 | 2.46E-04 | glio |
| Atr_in.vivo_noRT | Trappc4 | -1.184 | 2.46E-04 | glio |
| Bcas2_in.vivo_noRT | Gclc | -1.729 | 2.46E-04 | glio |
| Bora_ex.vivo_noRT | Pla2g2d | 3.201 | 2.46E-04 | glio |
| Cenpj_ex.vivo_noRT | Gm47510 | 2.591 | 2.46E-04 | glio |
| Cenpj_ex.vivo_noRT | Gm5788 | 2.591 | 2.46E-04 | glio |
| Cenpj_ex.vivo_noRT | Gm48617 | 2.591 | 2.46E-04 | glio |
| Cenpj_ex.vivo_noRT | Gm12703 | 2.591 | 2.46E-04 | glio |
| Cenpj_ex.vivo_noRT | Gm128 | 2.996 | 2.46E-04 | glio |
| Cenpj_ex.vivo_noRT | Mmp13 | 2.591 | 2.46E-04 | glio |
| Cenpj_ex.vivo_noRT | Angptl7 | 2.591 | 2.46E-04 | glio |
| Cenpj_ex.vivo_noRT | Gm8016 | 2.591 | 2.46E-04 | glio |
| Chek1_in.vivo_noRT | 2810021J22Ri | 2.724 | 2.46E-04 | glio |
| Chek1_in.vivo_noRT | Pcdhga12 | 2.559 | 2.46E-04 | glio |
| Chek1_in.vivo_noRT | Rtcb | -0.826 | 2.46E-04 | glio |
| Chek1_ex.vivo_noRT | Tubb6 | -1.987 | 2.46E-04 | glio |
| culture_effect | Avpi1 | 0.771 | 2.46E-05 | glio |
| Interaction_genotype_Mtor | Otx1 | -4.57 | 2.46E-05 | glio |
| Interaction_genotype_Mtor | Apold1 | -5.801 | 2.46E-05 | glio |
| Aars2_in.vivo_noRT | Ssr2 | -1.12 | 2.46E-05 | glio |
| Atr_in.vivo_noRT | Fam177a | 2.766 | 2.46E-05 | glio |
| Atr_ex.vivo_noRT | Slc19a3 | 2.77 | 2.46E-05 | glio |
| Chek1_in.vivo_noRT | Mfsd7a | 3.437 | 2.46E-05 | glio |
| Chek1_in.vivo_noRT | 9130221H12R | 2.735 | 2.46E-05 | glio |
| Chek1_ex.vivo_noRT | Gm8212 | 3.7 | 2.46E-05 | glio |
| Hsd17b10_in.vivo_noRT | Arhgef16 | 2.58 | 2.46E-05 | glio |
| Hsd17b10_in.vivo_noRT | Ikzf2 | 3.255 | 2.46E-05 | glio |
| culture_effect | Cacng5 | -2.785 | 2.46E-06 | glio |
| Interaction_genotype_Mtor | Platr20 | -6.1 | 2.46E-06 | glio |
| Interaction_genotype_Myc | Gm49864 | -5.294 | 2.46E-06 | glio |
| Interaction_genotype_Myc | 5430402O13R | -5.485 | 2.46E-06 | glio |
| Interaction_genotype_Myc | Mettl7b | -5.388 | 2.46E-06 | glio |

| | | | | |
|---|---|---|---|---|
| Aars2_in.vivo_noRT | Rgs16 | 4.117 | 2.46E-06 | glio |
| Aars2_in.vivo_noRT | Wfikkn2 | 3.352 | 2.46E-06 | glio |
| Interaction_genotype_Mtor | H2-K1 | 3.519 | 2.46E-07 | glio |
| Interaction_genotype_Mtor | Gjd4 | -4.099 | 2.46E-07 | glio |
| Bcas2_in.vivo_noRT | Mamdc2 | 3.497 | 2.46E-07 | glio |
| Interaction_genotype_Mtor | Chchd1 | 2.688 | 2.46E-08 | glio |
| Interaction_genotype_Mtor | Gm10300 | -7.077 | 2.46E-08 | glio |
| Interaction_genotype_Mtor | Gm10447 | -5.373 | 2.46E-08 | glio |
| Interaction_genotype_Mtor | Clec4d | -5.284 | 2.46E-08 | glio |
| Aars2_in.vivo_noRT | Gm16283 | 3.803 | 2.46E-08 | glio |
| culture_effect | Atg4d | 1.154 | 2.46E-09 | glio |
| Aars2_in.vivo_noRT | Edn1 | 4.947 | 2.46E-09 | glio |
| Chek1_in.vivo_noRT | Gm3716 | 4.126 | 2.46E-09 | glio |
| Chek1_ex.vivo_noRT | AU022754 | 4.842 | 2.46E-09 | glio |
| Chek1_in.vivo_noRT | Gm28731 | 4.658 | 2.46E-10 | glio |
| Chek1_in.vivo_noRT | 4933406C10R | 4.007 | 2.46E-10 | glio |
| Atr_in.vivo_noRT | C4a | 4.095 | 2.46E-11 | glio |
| Bora_in.vivo_noRT | Tlr1 | 4.676 | 2.46E-11 | glio |
| culture_effect | Gm26519 | 3.753 | 2.46E-12 | glio |
| culture_effect | Gm19426 | 2.631 | 2.46E-12 | glio |
| Hsd17b10_in.vivo_noRT | Trmt61b | 3.616 | 2.46E-12 | glio |
| Atr_in.vivo_noRT | Podxl | 5.002 | 2.46E-15 | glio |
| Aars2_in.vivo_noRT | Aloxe3 | 4.798 | 2.46E-16 | glio |
| Atr_in.vivo_noRT | Gm49172 | 5.273 | 2.46E-28 | glio |
| culture_effect | Wdr27 | 1.245 | 2.47E-03 | glio |
| Interaction_genotype_Aars2 | Gm50394 | -3.613 | 2.47E-03 | glio |
| Interaction_genotype_Aars2 | Neurl1a | -4.114 | 2.47E-03 | glio |
| Interaction_genotype_Atr | D630024D03F | -2.956 | 2.47E-03 | glio |
| Interaction_genotype_Bcas2 | Gm26674 | -3.847 | 2.47E-03 | glio |
| Interaction_genotype_Bora | Gpt | -4.073 | 2.47E-03 | glio |
| Interaction_genotype_Mtor | E230025N22R | -3.499 | 2.47E-03 | glio |
| Interaction_genotype_Mtor | Wars2 | -3.19 | 2.47E-03 | glio |
| Interaction_genotype_Myc | Adgre5 | -2.863 | 2.47E-03 | glio |
| Interaction_genotype_Myc | Rps6 | 1.013 | 2.47E-03 | glio |
| Interaction_genotype_Uqcrc2 | Kctd2 | 2.366 | 2.47E-03 | glio |
| Interaction_genotype_Uqcrc2 | Rpp25l | 1.283 | 2.47E-03 | glio |
| Aars2_in.vivo_noRT | Nav1 | 1.413 | 2.47E-03 | glio |
| Atp5c1_in.vivo_noRT | Vmn2r56 | 1.886 | 2.47E-03 | glio |
| Atr_in.vivo_noRT | Padi2 | 2.257 | 2.47E-03 | glio |
| Atr_in.vivo_noRT | Zfp790 | 2.004 | 2.47E-03 | glio |
| Atr_in.vivo_noRT | Nmi | -0.838 | 2.47E-03 | glio |
| Bcas2_in.vivo_noRT | Zfp287 | 1.756 | 2.47E-03 | glio |
| Bcas2_in.vivo_noRT | Gm48427 | 2.028 | 2.47E-03 | glio |
| Bcas2_in.vivo_noRT | Poglut2 | -1.497 | 2.47E-03 | glio |
| Bora_in.vivo_noRT | Faap20 | -1.291 | 2.47E-03 | glio |
| Bora_ex.vivo_noRT | Dhrs9 | 2.802 | 2.47E-03 | glio |
| Cenpj_ex.vivo_noRT | Gm30042 | 2.591 | 2.47E-03 | glio |
| Chek1_in.vivo_noRT | 4930506C21R | 2.18 | 2.47E-03 | glio |
| Chek1_in.vivo_noRT | Zfp52 | 1.841 | 2.47E-03 | glio |
| Chek1_in.vivo_noRT | Nr6a1 | 1.919 | 2.47E-03 | glio |
| Eif2s1_in.vivo_noRT | Gm44672 | 2.427 | 2.47E-03 | glio |
| Hsd17b10_in.vivo_noRT | Dnajc28 | 2.47 | 2.47E-03 | glio |

| | | | | |
|---|---|---|---|---|
| Hsd17b10_in.vivo_noRT | Fgf7 | 2.252 | 2.47E-03 | glio |
| Interaction_genotype_Aars2 | Gm39326 | -4.471 | 2.47E-04 | glio |
| Interaction_genotype_Myc | 4833419F23Ri | -4.196 | 2.47E-04 | glio |
| Interaction_genotype_Myc | Kctd1 | -3.95 | 2.47E-04 | glio |
| Atr_in.vivo_noRT | Papola | -1.201 | 2.47E-04 | glio |
| Bcas2_in.vivo_noRT | Gm4129 | 2.638 | 2.47E-04 | glio |
| Bcas2_ex.vivo_noRT | 4930447K03R | 2.753 | 2.47E-04 | glio |
| Chek1_in.vivo_noRT | Msh5 | 2.787 | 2.47E-04 | glio |
| Chek1_ex.vivo_noRT | Gm29083 | 2.955 | 2.47E-04 | glio |
| Hsd17b10_in.vivo_noRT | Zfp964 | 2.781 | 2.47E-04 | glio |
| Interaction_genotype_Mtor | Slc47a2 | -4.916 | 2.47E-05 | glio |
| Interaction_genotype_Mtor | Hmgn3 | 2.306 | 2.47E-05 | glio |
| Aars2_in.vivo_noRT | Mansc1 | 3.164 | 2.47E-05 | glio |
| Aars2_in.vivo_noRT | Hist1h4a | 2.844 | 2.47E-05 | glio |
| Atr_in.vivo_noRT | Ccdc159 | 3.11 | 2.47E-05 | glio |
| Chek1_in.vivo_noRT | Hist1h4c | 3.135 | 2.47E-05 | glio |
| Chek1_in.vivo_noRT | Stat3 | -1.465 | 2.47E-05 | glio |
| Interaction_genotype_Mtor | Gm12800 | -4.099 | 2.47E-06 | glio |
| Atp5c1_in.vivo_noRT | Gm2022 | 2.668 | 2.47E-06 | glio |
| Atr_in.vivo_noRT | Eme2 | 3.119 | 2.47E-06 | glio |
| Atr_in.vivo_noRT | Nyap1 | 3.285 | 2.47E-06 | glio |
| Bcas2_in.vivo_noRT | Zfp968 | 3.43 | 2.47E-06 | glio |
| culture_effect | Gm19412 | 1.397 | 2.47E-07 | glio |
| Atr_in.vivo_noRT | 1500002C15R | 3.766 | 2.47E-07 | glio |
| culture_effect | Arid4a | -1.014 | 2.47E-08 | glio |
| Atr_in.vivo_noRT | Gm50103 | 4.226 | 2.47E-09 | glio |
| Bcas2_in.vivo_noRT | Lama5 | 4.285 | 2.47E-09 | glio |
| culture_effect | Myo10 | 1.655 | 2.47E-10 | glio |
| Aars2_in.vivo_noRT | Gm12840 | 4.068 | 2.47E-10 | glio |
| Aars2_in.vivo_noRT | Rilp | 4.801 | 2.47E-10 | glio |
| Aars2_in.vivo_noRT | C4a | 3.888 | 2.47E-10 | glio |
| Aars2_in.vivo_noRT | Gm8098 | 4.338 | 2.47E-10 | glio |
| Atr_in.vivo_noRT | Hlx | 4.277 | 2.47E-10 | glio |
| Bcas2_in.vivo_noRT | Apobr | 4.807 | 2.47E-10 | glio |
| Aars2_in.vivo_noRT | Rnft2 | 4.523 | 2.47E-11 | glio |
| Chek1_in.vivo_noRT | Gm16364.1 | 5.044 | 2.47E-11 | glio |
| Hsd17b10_in.vivo_noRT | Smtnl1 | 3.616 | 2.47E-11 | glio |
| Hsd17b10_in.vivo_noRT | Cd200r2 | 3.352 | 2.47E-11 | glio |
| Aars2_in.vivo_noRT | Nrg2 | 4.804 | 2.47E-12 | glio |
| Atr_in.vivo_noRT | Fabp4 | 5.004 | 2.47E-12 | glio |
| Bcas2_in.vivo_noRT | Gm50115 | 4.544 | 2.47E-12 | glio |
| Bora_in.vivo_noRT | Rbfaos | 4.019 | 2.47E-13 | glio |
| Aars2_in.vivo_noRT | Gm30938 | 5.19 | 2.47E-14 | glio |
| Chek1_in.vivo_noRT | Oit3 | 5.044 | 2.47E-14 | glio |
| culture_effect | Oxct1 | 1.263 | 2.47E-17 | glio |
| Aars2_in.vivo_noRT | Gm11899 | 4.805 | 2.47E-19 | glio |
| Bcas2_in.vivo_noRT | Gm47936 | 4.807 | 2.47E-19 | glio |
| Aars2_in.vivo_noRT | Plcl1 | 5.327 | 2.47E-20 | glio |
| Atr_in.vivo_noRT | Scel | 4.615 | 2.47E-22 | glio |
| culture_effect | Gadl1 | -1.823 | 2.48E-03 | glio |
| culture_effect | Coro2b | 1.083 | 2.48E-03 | glio |
| culture_effect | Retreg1 | 0.682 | 2.48E-03 | glio |

| | | | | |
|---|---|---|---|---|
| Interaction_genotype_Aars2 | Fen1 | 1.995 | 2.48E-03 | glio |
| Interaction_genotype_Atr | Capn13 | -2.369 | 2.48E-03 | glio |
| Interaction_genotype_Atr | A530072M11F | -3.844 | 2.48E-03 | glio |
| Interaction_genotype_Atr | Snai1 | -3.905 | 2.48E-03 | glio |
| Interaction_genotype_Atr | 1500002C15R | -3.874 | 2.48E-03 | glio |
| Interaction_genotype_Atr | Erg28 | 1.614 | 2.48E-03 | glio |
| Interaction_genotype_Bora | Nrg2 | -3.989 | 2.48E-03 | glio |
| Interaction_genotype_Bora | 4930513N10R | -4.541 | 2.48E-03 | glio |
| Interaction_genotype_Bora | Col8a1 | -3.872 | 2.48E-03 | glio |
| Interaction_genotype_Bora | Gm10125 | -4.094 | 2.48E-03 | glio |
| Interaction_genotype_Bora | Gm47061 | -4.37 | 2.48E-03 | glio |
| Interaction_genotype_Bora | Gm43728 | -4.195 | 2.48E-03 | glio |
| Interaction_genotype_Bora | Gm10399 | -3.442 | 2.48E-03 | glio |
| Interaction_genotype_Bora | Dntt | -4.138 | 2.48E-03 | glio |
| Interaction_genotype_Bora | Mad2l1 | 1.968 | 2.48E-03 | glio |
| Interaction_genotype_Bora | Chst11 | 2.011 | 2.48E-03 | glio |
| Interaction_genotype_Myc | Sytl4 | -2.463 | 2.48E-03 | glio |
| Interaction_genotype_Myc | Ifi207 | -3.639 | 2.48E-03 | glio |
| Interaction_genotype_Myc | Zer1 | -2.82 | 2.48E-03 | glio |
| Interaction_genotype_Myc | Arl4c | -2.966 | 2.48E-03 | glio |
| Interaction_genotype_Myc | Fastkd2 | -2.497 | 2.48E-03 | glio |
| Interaction_genotype_Myc | Eif2b5 | -2.122 | 2.48E-03 | glio |
| Interaction_genotype_Myc | Rpl7 | 1.131 | 2.48E-03 | glio |
| Interaction_genotype_Pars2 | Map3k7 | 1.809 | 2.48E-03 | glio |
| interaction_RT | Lpcat4 | 1.671 | 2.48E-03 | glio |
| interaction_RT | Selenbp1 | 3.394 | 2.48E-03 | glio |
| interaction_RT | Ints5 | -0.947 | 2.48E-03 | glio |
| Aars2_in.vivo_noRT | Pacsin1 | 1.981 | 2.48E-03 | glio |
| Aars2_in.vivo_noRT | Ptprg | 1.889 | 2.48E-03 | glio |
| Aars2_in.vivo_noRT | Cep85l | 1.977 | 2.48E-03 | glio |
| Aars2_in.vivo_noRT | Slc1a3 | -1.498 | 2.48E-03 | glio |
| Atp5c1_in.vivo_noRT | Pnliprp2 | 1.889 | 2.48E-03 | glio |
| Atp5c1_in.vivo_noRT | AI463229 | 2.023 | 2.48E-03 | glio |
| Atp5c1_in.vivo_noRT | Uqcc1 | -1.504 | 2.48E-03 | glio |
| Atr_in.vivo_noRT | Usf3 | 1.502 | 2.48E-03 | glio |
| Atr_in.vivo_noRT | Chmp3 | -1.075 | 2.48E-03 | glio |
| Bcas2_in.vivo_noRT | Rad50 | -1.344 | 2.48E-03 | glio |
| Bora_in.vivo_noRT | A930004D18R | 2.197 | 2.48E-03 | glio |
| Bora_in.vivo_noRT | Elovl6 | -1.517 | 2.48E-03 | glio |
| Bora_in.vivo_noRT | Pbrm1 | -1.084 | 2.48E-03 | glio |
| Chek1_ex.vivo_noRT | Zfp59 | 2.831 | 2.48E-03 | glio |
| Chek1_ex.vivo_noRT | C8g | 2.678 | 2.48E-03 | glio |
| Eif2s1_in.vivo_noRT | Tcstv1 | 2.143 | 2.48E-03 | glio |
| Eif2s1_in.vivo_noRT | Klra6 | 2.615 | 2.48E-03 | glio |
| culture_effect | Ctbs | -0.883 | 2.48E-04 | glio |
| culture_effect | Cdk17 | 0.95 | 2.48E-04 | glio |
| culture_effect | Zfp266 | 0.68 | 2.48E-04 | glio |
| Interaction_genotype_Bora | Pmvk | 2.753 | 2.48E-04 | glio |
| Interaction_genotype_Mtor | Plk5 | -3.836 | 2.48E-04 | glio |
| Interaction_genotype_Mtor | 9330179D12R | -4.7 | 2.48E-04 | glio |
| Interaction_genotype_Mtor | Nanp | -2.939 | 2.48E-04 | glio |
| Interaction_genotype_Myc | Gjd4 | -2.729 | 2.48E-04 | glio |

| | | | | |
|---|---|---|---|---|
| Interaction_genotype_Myc | Lmo7 | -3.591 | 2.48E-04 | glio |
| Interaction_genotype_Myc | Zswim9 | -3.499 | 2.48E-04 | glio |
| interaction_RT | St3gal5 | -1.096 | 2.48E-04 | glio |
| interaction_RT | Chst11 | -1.081 | 2.48E-04 | glio |
| interaction_RT | Wasl | 0.814 | 2.48E-04 | glio |
| Aars2_in.vivo_noRT | Herc6 | -1.344 | 2.48E-04 | glio |
| Bard1_ex.vivo_noRT | Ccl9 | 2.421 | 2.48E-04 | glio |
| Bora_ex.vivo_noRT | Gm12610 | 2.802 | 2.48E-04 | glio |
| Cenpj_in.vivo_noRT | Opalin | 3.371 | 2.48E-04 | glio |
| Cenpj_in.vivo_noRT | Stfa3 | 2.022 | 2.48E-04 | glio |
| Cenpj_in.vivo_noRT | Olfr787 | 2.022 | 2.48E-04 | glio |
| Chek1_in.vivo_noRT | Lcat | 2.71 | 2.48E-04 | glio |
| Chek1_in.vivo_noRT | Lgals4 | 2.408 | 2.48E-04 | glio |
| culture_effect | Nell2 | -2.427 | 2.48E-05 | glio |
| culture_effect | Arhgef19 | 2.31 | 2.48E-05 | glio |
| Atr_ex.vivo_noRT | Arg2 | 3.915 | 2.48E-05 | glio |
| Bora_in.vivo_noRT | 4930412C18R | 3.124 | 2.48E-05 | glio |
| Bora_in.vivo_noRT | A630072M18F | 2.551 | 2.48E-05 | glio |
| Chek1_ex.vivo_noRT | Pagr1a | 3.464 | 2.48E-05 | glio |
| culture_effect | Ctnnbip1 | 0.753 | 2.48E-06 | glio |
| Interaction_genotype_Mtor | Vgll3 | -4.41 | 2.48E-06 | glio |
| Interaction_genotype_Mtor | Pcp4l1 | -5.642 | 2.48E-06 | glio |
| Interaction_genotype_Mtor | Itm2b | 2.876 | 2.48E-06 | glio |
| Interaction_genotype_Mtor | Trmt61b | -4.099 | 2.48E-06 | glio |
| Interaction_genotype_Myc | Ppm1k | -4.26 | 2.48E-06 | glio |
| Bora_in.vivo_noRT | Ikzf2 | 3.512 | 2.48E-06 | glio |
| Bora_ex.vivo_noRT | Gm4166 | 3.201 | 2.48E-06 | glio |
| Interaction_genotype_Mtor | Ppp1r3g | -5.792 | 2.48E-07 | glio |
| Atr_in.vivo_noRT | E130307A14Ri | 3.294 | 2.48E-07 | glio |
| Bcas2_in.vivo_noRT | Cxcr2 | 3.495 | 2.48E-07 | glio |
| Bora_in.vivo_noRT | Flywch1 | -2.072 | 2.48E-07 | glio |
| Hsd17b10_in.vivo_noRT | Pacrg | 3.346 | 2.48E-07 | glio |
| Aars2_in.vivo_noRT | BC049762 | 3.993 | 2.48E-08 | glio |
| Chek1_ex.vivo_noRT | Icosl | 4.443 | 2.48E-08 | glio |
| Chek1_ex.vivo_noRT | Prm1 | 4.443 | 2.48E-08 | glio |
| Interaction_genotype_Atr | Adam12 | -6.463 | 2.48E-09 | glio |
| Bcas2_in.vivo_noRT | Ptp4a1.1 | 4.01 | 2.48E-09 | glio |
| Bcas2_in.vivo_noRT | BC026762 | 4.428 | 2.48E-11 | glio |
| Bora_in.vivo_noRT | Kcnk4 | 4.676 | 2.48E-11 | glio |
| Atr_in.vivo_noRT | Spns3 | 4.876 | 2.48E-12 | glio |
| Atr_in.vivo_noRT | Esrp2 | 4.621 | 2.48E-12 | glio |
| Aars2_in.vivo_noRT | Igfbp7 | 4.561 | 2.48E-13 | glio |
| culture_effect | Ifi35 | -1.031 | 2.48E-14 | glio |
| Chek1_ex.vivo_noRT | Vtn | 4.842 | 2.48E-14 | glio |
| Hsd17b10_in.vivo_noRT | Gm44736 | 3.616 | 2.48E-14 | glio |
| Hsd17b10_in.vivo_noRT | Fam227b | 3.616 | 2.48E-14 | glio |
| Hsd17b10_in.vivo_noRT | Gm20658 | 3.616 | 2.48E-14 | glio |
| Chek1_in.vivo_noRT | 4930502E18Ri | 4.508 | 2.48E-15 | glio |
| Atr_in.vivo_noRT | Zfp810 | 5.533 | 2.48E-16 | glio |
| Atr_in.vivo_noRT | 4930434B07R | 5.27 | 2.48E-18 | glio |
| Bcas2_in.vivo_noRT | C430002N11R | 4.807 | 2.48E-18 | glio |
| Bcas2_in.vivo_noRT | Tnni3k | 4.807 | 2.48E-19 | glio |

| | | | | |
|---|---|---|---|---|
| Chek1_in.vivo_noRT | Gm5890 | 5.044 | 2.48E-22 | glio |
| Chek1_in.vivo_noRT | Meioc | 4.78 | 2.48E-22 | glio |
| Chek1_in.vivo_noRT | Gm10447 | 5.555 | 2.48E-23 | glio |
| Chek1_in.vivo_noRT | Tpo | 5.044 | 2.48E-24 | glio |
| culture_effect | Parp4 | -0.936 | 2.49E-03 | glio |
| culture_effect | Nfkbil1 | 0.665 | 2.49E-03 | glio |
| culture_effect | Uqcrc1 | -0.475 | 2.49E-03 | glio |
| Interaction_genotype_Atr | Gm47540 | -2.762 | 2.49E-03 | glio |
| Interaction_genotype_Atr | A430005L14Ri | 1.718 | 2.49E-03 | glio |
| Interaction_genotype_Myc | Zbtb22 | -2.861 | 2.49E-03 | glio |
| Interaction_genotype_Myc | G3bp2 | 1.26 | 2.49E-03 | glio |
| interaction_RT | Immt | -0.633 | 2.49E-03 | glio |
| Aars2_in.vivo_noRT | Gm19426 | 2.006 | 2.49E-03 | glio |
| Aars2_in.vivo_noRT | Acot7 | -0.963 | 2.49E-03 | glio |
| Atr_ex.vivo_noRT | B3galt2 | 2.764 | 2.49E-03 | glio |
| Atr_ex.vivo_noRT | Gm15879 | 3.462 | 2.49E-03 | glio |
| Bcas2_in.vivo_noRT | Amt | 2.424 | 2.49E-03 | glio |
| Bcas2_in.vivo_noRT | Fbxo31 | -1.409 | 2.49E-03 | glio |
| Bora_ex.vivo_noRT | Dusp15 | 2.802 | 2.49E-03 | glio |
| Cenpj_in.vivo_noRT | Gm16244 | 2.663 | 2.49E-03 | glio |
| Cenpj_ex.vivo_noRT | A330076H08R | 2.591 | 2.49E-03 | glio |
| Cenpj_ex.vivo_noRT | Ipcef1 | 2.591 | 2.49E-03 | glio |
| Chek1_ex.vivo_noRT | Rab19 | 2.902 | 2.49E-03 | glio |
| Chek1_ex.vivo_noRT | Gm14966 | 2.453 | 2.49E-03 | glio |
| Hsd17b10_in.vivo_noRT | Zfp473 | 2.192 | 2.49E-03 | glio |
| Hsd17b10_in.vivo_noRT | Cbs | 2.41 | 2.49E-03 | glio |
| culture_effect | Trip10 | -1.028 | 2.49E-04 | glio |
| Interaction_genotype_Aars2 | Gm10655 | -4.737 | 2.49E-04 | glio |
| Interaction_genotype_Aars2 | 2610016A17R | -4.699 | 2.49E-04 | glio |
| Interaction_genotype_Aars2 | Gm41764 | -4.1 | 2.49E-04 | glio |
| Interaction_genotype_Aars2 | 5033417F24Ri | -4.455 | 2.49E-04 | glio |
| Interaction_genotype_Aars2 | Emc10 | 1.28 | 2.49E-04 | glio |
| Interaction_genotype_Atr | Clcn1 | -3.676 | 2.49E-04 | glio |
| Interaction_genotype_Mtor | Gm26879 | -3.988 | 2.49E-04 | glio |
| Interaction_genotype_Mtor | Mzf1 | -4.13 | 2.49E-04 | glio |
| Interaction_genotype_Mtor | AA465934 | -3.771 | 2.49E-04 | glio |
| Interaction_genotype_Mtor | Oas1c | -3.41 | 2.49E-04 | glio |
| Interaction_genotype_Myc | Acrbp | -3.64 | 2.49E-04 | glio |
| Interaction_genotype_Myc | Zfp408 | -3.747 | 2.49E-04 | glio |
| Interaction_genotype_Myc | Lsm8 | 1.636 | 2.49E-04 | glio |
| Aars2_in.vivo_noRT | Slc25a42 | 2.644 | 2.49E-04 | glio |
| Aars2_in.vivo_noRT | Ccdc146 | 2.728 | 2.49E-04 | glio |
| Aars2_in.vivo_noRT | Paip2 | -0.814 | 2.49E-04 | glio |
| Aars2_ex.vivo_noRT | Npnt | 2.495 | 2.49E-04 | glio |
| Atr_in.vivo_noRT | Tmem186 | 2.471 | 2.49E-04 | glio |
| Bora_in.vivo_noRT | Mzf1 | 2.546 | 2.49E-04 | glio |
| Bora_in.vivo_noRT | Gm42937 | 2.626 | 2.49E-04 | glio |
| Bora_in.vivo_noRT | Elp2 | -1.746 | 2.49E-04 | glio |
| Chek1_in.vivo_noRT | Tnfsf13b | 2.169 | 2.49E-04 | glio |
| Interaction_genotype_Myc | Vac14 | -3.275 | 2.49E-05 | glio |
| interaction_RT | Eif4g2 | -0.673 | 2.49E-05 | glio |
| Aars2_in.vivo_noRT | Rit2 | 2.834 | 2.49E-05 | glio |

| | | | | |
|---|---|---|---|---|
| Atp5c1_in.vivo_noRT | Ccser1 | 2.668 | 2.49E-05 | glio |
| Atr_in.vivo_noRT | Pou5f1 | 2.748 | 2.49E-05 | glio |
| Bcas2_in.vivo_noRT | Taf9 | 2.917 | 2.49E-05 | glio |
| Bora_in.vivo_noRT | Zfp534 | 3.128 | 2.49E-05 | glio |
| Bora_ex.vivo_noRT | Slc19a3 | 2.808 | 2.49E-05 | glio |
| Chek1_in.vivo_noRT | Fzd1 | 2.854 | 2.49E-05 | glio |
| Chek1_in.vivo_noRT | Tsga10 | 3.112 | 2.49E-05 | glio |
| Interaction_genotype_Mtor | Inhba | -6.099 | 2.49E-06 | glio |
| Aars2_in.vivo_noRT | Gm4907 | 3.721 | 2.49E-06 | glio |
| Aars2_in.vivo_noRT | Gm13212 | 3.332 | 2.49E-06 | glio |
| Atr_in.vivo_noRT | Zbtb8a | 3.472 | 2.49E-06 | glio |
| Bcas2_in.vivo_noRT | Gm27010 | 3.206 | 2.49E-06 | glio |
| Bora_ex.vivo_noRT | Gm14820 | 3.201 | 2.49E-06 | glio |
| Chek1_ex.vivo_noRT | Atg9b | 3.659 | 2.49E-06 | glio |
| culture_effect | AU020206 | -1.088 | 2.49E-07 | glio |
| Atp5c1_in.vivo_noRT | Gm30692 | 2.668 | 2.49E-07 | glio |
| Cenpj_ex.vivo_noRT | Gm17203 | 3.774 | 2.49E-07 | glio |
| Chek1_ex.vivo_noRT | Spint4 | 4.553 | 2.49E-07 | glio |
| Hsd17b10_in.vivo_noRT | Rarres1 | 3.616 | 2.49E-07 | glio |
| culture_effect | Cdc5l | -0.714 | 2.49E-08 | glio |
| Aars2_in.vivo_noRT | Bbs1 | 3.867 | 2.49E-08 | glio |
| Atr_in.vivo_noRT | Tkt | -2.072 | 2.49E-08 | glio |
| Atr_ex.vivo_noRT | Cldn2 | 3.164 | 2.49E-08 | glio |
| Aars2_in.vivo_noRT | Gm45336 | 3.454 | 2.49E-09 | glio |
| Bcas2_in.vivo_noRT | Col6a2 | 4.023 | 2.49E-09 | glio |
| Bora_in.vivo_noRT | Ifi214 | 4.145 | 2.49E-09 | glio |
| Bora_in.vivo_noRT | Ctsf | 3.268 | 2.49E-09 | glio |
| culture_effect | Myg1 | -1.122 | 2.49E-10 | glio |
| Bcas2_in.vivo_noRT | Olfr279 | 4.278 | 2.49E-10 | glio |
| culture_effect | Pld3 | -1.317 | 2.49E-12 | glio |
| Aars2_in.vivo_noRT | Tex52 | 4.676 | 2.49E-12 | glio |
| Atr_in.vivo_noRT | Rem2 | 4.349 | 2.49E-12 | glio |
| Atr_in.vivo_noRT | Nmrk2 | 4.891 | 2.49E-15 | glio |
| culture_effect | Abcg2 | 1.899 | 2.49E-18 | glio |
| Atr_in.vivo_noRT | H1foo | 5.001 | 2.49E-22 | glio |
| culture_effect | Gm50223 | -5.241 | 2.49E-26 | glio |
| culture_effect | Catsperz | -1.907 | 2.50E-03 | glio |
| culture_effect | Trpv2 | -1.13 | 2.50E-03 | glio |
| culture_effect | Grk2 | 0.649 | 2.50E-03 | glio |
| Interaction_genotype_Aars2 | Dmrt2 | -4.062 | 2.50E-03 | glio |
| Interaction_genotype_Atr | Trafd1 | 2.071 | 2.50E-03 | glio |
| Interaction_genotype_Hsd17b | Gpx2 | 6.137 | 2.50E-03 | glio |
| Interaction_genotype_Mtor | Arih1 | 1.695 | 2.50E-03 | glio |
| Interaction_genotype_Myc | Gm15246 | -3.439 | 2.50E-03 | glio |
| Interaction_genotype_Myc | Arxes1 | -3.212 | 2.50E-03 | glio |
| Interaction_genotype_Myc | 5930430L01Ri | -2.892 | 2.50E-03 | glio |
| Interaction_genotype_Myc | Sorbs2 | -2.744 | 2.50E-03 | glio |
| Aars2_in.vivo_noRT | Gm14488 | 2.354 | 2.50E-03 | glio |
| Aars2_in.vivo_noRT | 4930509H03R | 2.18 | 2.50E-03 | glio |
| Aars2_in.vivo_noRT | Efna5 | 2.132 | 2.50E-03 | glio |
| Aars2_in.vivo_noRT | Zfp668 | 1.816 | 2.50E-03 | glio |
| Bcas2_in.vivo_noRT | Tsga10 | 2.326 | 2.50E-03 | glio |

| | | | | |
|---|---|---|---|---|
| Bora_in.vivo_noRT | Zfp449 | 1.908 | 2.50E-03 | glio |
| Cenpj_in.vivo_noRT | A330049N07R | 1.756 | 2.50E-03 | glio |
| Cenpj_in.vivo_noRT | Gnaz | 2.649 | 2.50E-03 | glio |
| Cenpj_in.vivo_noRT | Ltbp2 | 2.022 | 2.50E-03 | glio |
| Cenpj_in.vivo_noRT | Lmx1a | 2.022 | 2.50E-03 | glio |
| Chek1_in.vivo_noRT | Surf4 | -0.828 | 2.50E-03 | glio |
| Chek1_in.vivo_noRT | Rpl11 | -0.64 | 2.50E-03 | glio |
| Chek1_ex.vivo_noRT | Myg1 | -1.655 | 2.50E-03 | glio |
| Chek1_ex.vivo_noRT | Rps11 | -0.668 | 2.50E-03 | glio |
| Cyp19a1_ex.vivo_noRT | Ceacam16 | 2.959 | 2.50E-03 | glio |
| Eif2s1_ex.vivo_noRT | Col19a1 | 2.639 | 2.50E-03 | glio |
| Eif2s1_ex.vivo_noRT | A930019D19R | 2.234 | 2.50E-03 | glio |
| Ercc4_ex.vivo_noRT | Gm13032 | 3.082 | 2.50E-03 | glio |
| Hsd17b10_in.vivo_noRT | Fam189a1 | 1.946 | 2.50E-03 | glio |
| Interaction_genotype_Mtor | Cd59b | -4.433 | 2.50E-04 | glio |
| Interaction_genotype_Mtor | Anapc16 | 1.768 | 2.50E-04 | glio |
| Interaction_genotype_Myc | Snhg1 | 2.577 | 2.50E-04 | glio |
| interaction_RT | H2-Ab1 | 2.547 | 2.50E-04 | glio |
| Aars2_ex.vivo_noRT | Cacng5 | 3.293 | 2.50E-04 | glio |
| Aars2_ex.vivo_noRT | Serpina3i | 3.293 | 2.50E-04 | glio |
| Aars2_ex.vivo_noRT | Adgre5 | 2.899 | 2.50E-04 | glio |
| Atp5c1_in.vivo_noRT | Nkg7 | 2.408 | 2.50E-04 | glio |
| Atr_in.vivo_noRT | Gm7467 | 2.252 | 2.50E-04 | glio |
| Bcas2_in.vivo_noRT | Nkain4 | 2.46 | 2.50E-04 | glio |
| Chek1_in.vivo_noRT | Dnaaf4 | 2.862 | 2.50E-04 | glio |
| culture_effect | Ints10 | -1.014 | 2.50E-05 | glio |
| culture_effect | Cd3eap | 0.583 | 2.50E-05 | glio |
| Interaction_genotype_Atr | Nid1 | -5.32 | 2.50E-05 | glio |
| Interaction_genotype_Mtor | Supt4a | 1.808 | 2.50E-05 | glio |
| Interaction_genotype_Mtor | Hint1 | 1.58 | 2.50E-05 | glio |
| Atr_in.vivo_noRT | Gm26737 | 4.43 | 2.50E-05 | glio |
| Atr_in.vivo_noRT | App | -1.351 | 2.50E-05 | glio |
| Bora_in.vivo_noRT | D130062J10Ri | 2.733 | 2.50E-05 | glio |
| Bora_ex.vivo_noRT | Prickle2 | 3.212 | 2.50E-05 | glio |
| Bora_ex.vivo_noRT | Gm39556 | 3.993 | 2.50E-05 | glio |
| Chek1_in.vivo_noRT | Efna5 | 2.909 | 2.50E-05 | glio |
| culture_effect | Dgka | -1.492 | 2.50E-06 | glio |
| culture_effect | Fgfrl1 | -0.622 | 2.50E-06 | glio |
| Aars2_in.vivo_noRT | Tmsb15b2 | 3.606 | 2.50E-06 | glio |
| Atr_in.vivo_noRT | Cpt1b | 3.189 | 2.50E-06 | glio |
| Bora_in.vivo_noRT | Ccl20 | 4.936 | 2.50E-06 | glio |
| Bora_in.vivo_noRT | Arg2 | 3.296 | 2.50E-06 | glio |
| Aars2_ex.vivo_noRT | Gm16933 | 3.293 | 2.50E-07 | glio |
| Atr_in.vivo_noRT | AC241534.1 | 3.674 | 2.50E-07 | glio |
| Atr_in.vivo_noRT | Luc7l2 | -1.416 | 2.50E-07 | glio |
| Chek1_in.vivo_noRT | Cemip | 3.009 | 2.50E-07 | glio |
| culture_effect | Dda1 | 0.699 | 2.50E-08 | glio |
| interaction_RT | Mgst1 | 3.421 | 2.50E-08 | glio |
| Aars2_in.vivo_noRT | Stard6 | 3.956 | 2.50E-08 | glio |
| Bcas2_in.vivo_noRT | Tslp | 4.162 | 2.50E-08 | glio |
| Bcas2_in.vivo_noRT | Sh3gl3 | 3.895 | 2.50E-08 | glio |
| Bcas2_in.vivo_noRT | Cabp4 | 4.014 | 2.50E-08 | glio |

| | | | | |
|---|---|---|---|---|
| Bcas2_in.vivo_noRT | Gm24362 | 3.764 | 2.50E-09 | glio |
| Bcas2_in.vivo_noRT | Styk1 | 4.264 | 2.50E-09 | glio |
| Chek1_in.vivo_noRT | Npy4r | 3.677 | 2.50E-09 | glio |
| Bcas2_in.vivo_noRT | Hunk | 4.275 | 2.50E-11 | glio |
| Chek1_in.vivo_noRT | Cgn | 4.518 | 2.50E-11 | glio |
| Bcas2_in.vivo_noRT | 5033417F24Ri | 4.807 | 2.50E-12 | glio |
| Bora_in.vivo_noRT | Postn | 4.835 | 2.50E-12 | glio |
| Hsd17b10_in.vivo_noRT | Gm29966 | 3.356 | 2.50E-12 | glio |
| Abtb1_in.vivo_noRT | Abtb1 | -4.957 | 2.50E-13 | glio |
| Atr_in.vivo_noRT | Pygo1 | 4.75 | 2.50E-13 | glio |
| Chek1_in.vivo_noRT | Synpo2l | 4.398 | 2.50E-13 | glio |
| Chek1_ex.vivo_noRT | 2310003N18R | 5.25 | 2.50E-13 | glio |
| Bora_in.vivo_noRT | Tmem236 | 4.67 | 2.50E-14 | glio |
| Aars2_in.vivo_noRT | Ceacam16 | 4.669 | 2.50E-15 | glio |
| Aars2_in.vivo_noRT | 4930540M05F | 5.46 | 2.50E-16 | glio |
| Atr_in.vivo_noRT | Gm4858 | 5.533 | 2.50E-17 | glio |
| Bora_in.vivo_noRT | 4930470G03R | 4.67 | 2.50E-17 | glio |
| Bora_in.vivo_noRT | Gm16933 | 4.414 | 2.50E-19 | glio |
| Bora_in.vivo_noRT | Hfm1 | 4.936 | 2.50E-19 | glio |
| Bora_in.vivo_noRT | D630024D03F | 4.936 | 2.50E-19 | glio |
| culture_effect | Cct4 | -1.205 | 2.50E-21 | glio |
| Atr_in.vivo_noRT | Gm45591 | 5.783 | 2.50E-22 | glio |
| Chek1_in.vivo_noRT | Zscan4c | 5.044 | 2.50E-22 | glio |
| Aars2_in.vivo_noRT | Pitx3 | 5.841 | 2.50E-24 | glio |
| culture_effect | Srcap | -0.632 | 2.51E-03 | glio |
| culture_effect | Snrpg | 0.419 | 2.51E-03 | glio |
| Interaction_genotype_Atr | Ifitm6 | -3.795 | 2.51E-03 | glio |
| Interaction_genotype_Bora | Gm15853 | -4.169 | 2.51E-03 | glio |
| Interaction_genotype_Mtor | Zfp964 | -3.804 | 2.51E-03 | glio |
| Interaction_genotype_Mtor | Ankrd13d | -3.695 | 2.51E-03 | glio |
| Interaction_genotype_Mtor | Gm15492 | -3.65 | 2.51E-03 | glio |
| Interaction_genotype_Mtor | Sco1 | -3.127 | 2.51E-03 | glio |
| Interaction_genotype_Mtor | Gga3 | -2.536 | 2.51E-03 | glio |
| Interaction_genotype_Mtor | Anxa5 | 1.807 | 2.51E-03 | glio |
| Interaction_genotype_Myc | Slc22a5 | -3.192 | 2.51E-03 | glio |
| Interaction_genotype_Myc | Sirt7 | -2.249 | 2.51E-03 | glio |
| Interaction_genotype_Myc | Fra10ac1 | -1.811 | 2.51E-03 | glio |
| interaction_RT | Smad7 | -1.165 | 2.51E-03 | glio |
| interaction_RT | Prph | 1.079 | 2.51E-03 | glio |
| Aars2_in.vivo_noRT | Gbp10 | 1.853 | 2.51E-03 | glio |
| Aars2_in.vivo_noRT | Top1 | -0.912 | 2.51E-03 | glio |
| Atr_in.vivo_noRT | Ogt | -1.109 | 2.51E-03 | glio |
| Atr_in.vivo_noRT | Rbm4b | 1.105 | 2.51E-03 | glio |
| Atr_ex.vivo_noRT | Gm17501 | 2.491 | 2.51E-03 | glio |
| Bcas2_in.vivo_noRT | Btf3 | -0.469 | 2.51E-03 | glio |
| Bora_in.vivo_noRT | AI837181 | 1.253 | 2.51E-03 | glio |
| Cenpj_ex.vivo_noRT | Gm7247 | 2.592 | 2.51E-03 | glio |
| Cenpj_ex.vivo_noRT | St18 | 2.591 | 2.51E-03 | glio |
| Chek1_in.vivo_noRT | Phex | 2.076 | 2.51E-03 | glio |
| Chek1_ex.vivo_noRT | Chd5 | 2.761 | 2.51E-03 | glio |
| Chek1_ex.vivo_noRT | Gm5544 | 2.756 | 2.51E-03 | glio |
| Chek1_ex.vivo_noRT | Zfp637 | -1.354 | 2.51E-03 | glio |

| | | | | |
|---|---|---|---|---|
| Hsd17b10_in.vivo_noRT | Gm3604 | 2.334 | 2.51E-03 | glio |
| culture_effect | Sfxn3 | 0.624 | 2.51E-04 | glio |
| Interaction_genotype_Aars2 | Gm26581 | -4.776 | 2.51E-04 | glio |
| Interaction_genotype_Bora | Slc30a5 | 2.603 | 2.51E-04 | glio |
| Interaction_genotype_Bora | D830039M14I | -4.956 | 2.51E-04 | glio |
| interaction_RT | Cluh | -1.304 | 2.51E-04 | glio |
| interaction_RT | Map6 | -1.017 | 2.51E-04 | glio |
| interaction_RT | Mcm6 | -0.996 | 2.51E-04 | glio |
| interaction_RT | Nap1l1 | -0.649 | 2.51E-04 | glio |
| Aars2_in.vivo_noRT | Arrdc2 | 2.818 | 2.51E-04 | glio |
| Aars2_in.vivo_noRT | Sf3b6 | -0.805 | 2.51E-04 | glio |
| Atr_in.vivo_noRT | Adap1 | 2.938 | 2.51E-04 | glio |
| Atr_ex.vivo_noRT | Gm128 | 2.771 | 2.51E-04 | glio |
| Bcas2_in.vivo_noRT | E230025N22R | 2.46 | 2.51E-04 | glio |
| Cenpj_ex.vivo_noRT | Cldn5 | 2.591 | 2.51E-04 | glio |
| Chek1_ex.vivo_noRT | Efr3b | 3.745 | 2.51E-04 | glio |
| Hsd17b10_in.vivo_noRT | Glis1 | 2.704 | 2.51E-04 | glio |
| culture_effect | Gm28221 | -2.494 | 2.51E-05 | glio |
| culture_effect | Fam174a | 0.768 | 2.51E-05 | glio |
| Interaction_genotype_Mtor | Gm10602 | -5.566 | 2.51E-05 | glio |
| Interaction_genotype_Mtor | Tmem236 | -4.413 | 2.51E-05 | glio |
| Interaction_genotype_Mtor | Gm39459 | -5.117 | 2.51E-05 | glio |
| Interaction_genotype_Myc | Yod1 | -4.022 | 2.51E-05 | glio |
| Atp5c1_in.vivo_noRT | Fhl4 | 2.999 | 2.51E-05 | glio |
| Bcas2_in.vivo_noRT | Cyp2s1 | 3.234 | 2.51E-05 | glio |
| Chek1_in.vivo_noRT | Gm10800 | 3.245 | 2.51E-05 | glio |
| Interaction_genotype_Atr | Gm27239 | -5.468 | 2.51E-06 | glio |
| Aars2_in.vivo_noRT | C2cd2 | 3.417 | 2.51E-06 | glio |
| Aars2_in.vivo_noRT | Siva1 | -1.349 | 2.51E-06 | glio |
| Bora_in.vivo_noRT | Lrrc46 | 3.288 | 2.51E-06 | glio |
| Interaction_genotype_Mtor | Slc19a3 | -4.492 | 2.51E-07 | glio |
| Aars2_in.vivo_noRT | Gm16759 | 3.761 | 2.51E-07 | glio |
| Aars2_in.vivo_noRT | C330011M18F | 3.672 | 2.51E-07 | glio |
| Bcas2_in.vivo_noRT | Pdzk1ip1 | 3.817 | 2.51E-07 | glio |
| Hsd17b10_in.vivo_noRT | Rnf165 | 3.616 | 2.51E-07 | glio |
| Hsd17b10_in.vivo_noRT | Aloxe3 | 3.087 | 2.51E-07 | glio |
| culture_effect | Cldn14 | -3.304 | 2.51E-08 | glio |
| culture_effect | 2410006H16R | -0.932 | 2.51E-08 | glio |
| Atr_in.vivo_noRT | Mapk1 | -1.903 | 2.51E-08 | glio |
| Hsd17b10_in.vivo_noRT | 1700017M07F | 3.616 | 2.51E-09 | glio |
| Atr_in.vivo_noRT | Gm33973 | 3.967 | 2.51E-11 | glio |
| Atr_in.vivo_noRT | Flicr | 4.733 | 2.51E-12 | glio |
| Atr_in.vivo_noRT | Hectd2os | 4.625 | 2.51E-12 | glio |
| Bora_in.vivo_noRT | Bcl2l15 | 4.676 | 2.51E-14 | glio |
| Aars2_in.vivo_noRT | Necab2 | 5.063 | 2.51E-15 | glio |
| Aars2_in.vivo_noRT | Gm38477 | 5.341 | 2.51E-22 | glio |
| Atr_in.vivo_noRT | Gm13856 | 5.533 | 2.51E-29 | glio |
| Interaction_genotype_Aars2 | Olfr1369-ps1 | -4.612 | 2.52E-03 | glio |
| Interaction_genotype_Aars2 | Wnt10a | -3.677 | 2.52E-03 | glio |
| Interaction_genotype_Atr | Gm43608 | -3.851 | 2.52E-03 | glio |
| Interaction_genotype_Atr | Slc22a17 | -2.904 | 2.52E-03 | glio |
| Interaction_genotype_Atr | Cog1 | -2.988 | 2.52E-03 | glio |

| | | | | |
|---|---|---|---|---|
| Interaction_genotype_Atr | Ubb | 0.908 | 2.52E-03 | glio |
| Interaction_genotype_Mtor | Cped1 | -3.502 | 2.52E-03 | glio |
| Interaction_genotype_Mtor | Snx17 | 1.471 | 2.52E-03 | glio |
| Interaction_genotype_Myc | Gm16576 | -3.515 | 2.52E-03 | glio |
| Interaction_genotype_Myc | Syt14 | -2.722 | 2.52E-03 | glio |
| interaction_RT | Icam1 | 2.407 | 2.52E-03 | glio |
| interaction_RT | Hes7 | 2.539 | 2.52E-03 | glio |
| interaction_RT | Nol11 | -0.577 | 2.52E-03 | glio |
| Atr_in.vivo_noRT | Oas2 | 2.008 | 2.52E-03 | glio |
| Atr_ex.vivo_noRT | Gm20442 | 2.594 | 2.52E-03 | glio |
| Atr_ex.vivo_noRT | H2-Eb1 | 2.77 | 2.52E-03 | glio |
| Bcas2_in.vivo_noRT | Gm38973 | 2.211 | 2.52E-03 | glio |
| Bcas2_in.vivo_noRT | Ccdc189 | 2.204 | 2.52E-03 | glio |
| Bcas2_in.vivo_noRT | Rif1 | -1.052 | 2.52E-03 | glio |
| Bcas2_in.vivo_noRT | Ndufb10 | -0.71 | 2.52E-03 | glio |
| Bora_in.vivo_noRT | Bach2 | 2.186 | 2.52E-03 | glio |
| Bora_in.vivo_noRT | Entr1 | -1.447 | 2.52E-03 | glio |
| Bora_in.vivo_noRT | Slc29a1 | -1.499 | 2.52E-03 | glio |
| Bora_in.vivo_noRT | Plekhj1 | -0.843 | 2.52E-03 | glio |
| Chek1_in.vivo_noRT | Gm10419 | 2.391 | 2.52E-03 | glio |
| Chek1_in.vivo_noRT | Gm47509 | 2.246 | 2.52E-03 | glio |
| Chek1_ex.vivo_noRT | Tmem91 | 2.889 | 2.52E-03 | glio |
| Chek1_ex.vivo_noRT | Olfr1369-ps1 | 3.275 | 2.52E-03 | glio |
| Eif2s1_in.vivo_noRT | AC159140.1 | 1.883 | 2.52E-03 | glio |
| Hsd17b10_in.vivo_noRT | Sdcbp2 | 2.448 | 2.52E-03 | glio |
| Interaction_genotype_Atr | Klk10 | -4.386 | 2.52E-04 | glio |
| Interaction_genotype_Myc | Frmd8os | -4.668 | 2.52E-04 | glio |
| Interaction_genotype_Myc | Srm | 2.08 | 2.52E-04 | glio |
| Interaction_genotype_Myc | Cd63 | 1.337 | 2.52E-04 | glio |
| Aars2_in.vivo_noRT | Platr3 | 2.736 | 2.52E-04 | glio |
| Aars2_in.vivo_noRT | Got1 | -1.415 | 2.52E-04 | glio |
| Atr_in.vivo_noRT | Atp11a | 2.503 | 2.52E-04 | glio |
| Bcas2_in.vivo_noRT | Gm13830 | 2.876 | 2.52E-04 | glio |
| Bcas2_in.vivo_noRT | Aldh1a2 | 2.45 | 2.52E-04 | glio |
| Bcas2_in.vivo_noRT | Tssk6 | 2.321 | 2.52E-04 | glio |
| Bcas2_in.vivo_noRT | Adap1 | 2.99 | 2.52E-04 | glio |
| Bora_in.vivo_noRT | Ugdh | -1.213 | 2.52E-04 | glio |
| Chek1_in.vivo_noRT | Gm26737 | 3.94 | 2.52E-04 | glio |
| Chek1_in.vivo_noRT | 9130015A21R | 4.675 | 2.52E-04 | glio |
| Chek1_in.vivo_noRT | Gm12043 | 2.45 | 2.52E-04 | glio |
| Chek1_in.vivo_noRT | Etv1 | -1.695 | 2.52E-04 | glio |
| Chek1_in.vivo_noRT | Mpg | -1.514 | 2.52E-04 | glio |
| Cyp19a1_ex.vivo_noRT | Gm32122 | 3.735 | 2.52E-04 | glio |
| Hsd17b10_in.vivo_noRT | Morc4 | -1.757 | 2.52E-04 | glio |
| Interaction_genotype_Mtor | Atp6v0b | 1.799 | 2.52E-05 | glio |
| Interaction_genotype_Myc | Olfr655 | -6.566 | 2.52E-05 | glio |
| Interaction_genotype_Myc | Zfp85os | -4.93 | 2.52E-05 | glio |
| Aars2_in.vivo_noRT | 4930563E22Ri | 3.079 | 2.52E-05 | glio |
| Aars2_in.vivo_noRT | Gstt2 | 3.017 | 2.52E-05 | glio |
| Aars2_in.vivo_noRT | Gm5930 | 2.934 | 2.52E-05 | glio |
| Aars2_in.vivo_noRT | Slc22a22 | 2.906 | 2.52E-05 | glio |
| Bcas2_in.vivo_noRT | Klhdc1 | 2.938 | 2.52E-05 | glio |

| | | | | |
|---|---|---|---|---|
| culture_effect | B230317F23Ri | 2.251 | 2.52E-06 | glio |
| culture_effect | Zfp281 | -1.048 | 2.52E-06 | glio |
| Interaction_genotype_Mtor | Arhgef4 | -5.628 | 2.52E-06 | glio |
| Interaction_genotype_Mtor | Rgs9 | -4.881 | 2.52E-06 | glio |
| Interaction_genotype_Myc | 1110046J04Ri | -5.541 | 2.52E-06 | glio |
| Atr_in.vivo_noRT | Sap30 | -1.905 | 2.52E-06 | glio |
| Chek1_in.vivo_noRT | Pla2g4b | 3.216 | 2.52E-06 | glio |
| culture_effect | D330050I16Ri | 1.641 | 2.52E-07 | glio |
| Aars2_in.vivo_noRT | Tox2 | 3.552 | 2.52E-07 | glio |
| Chek1_ex.vivo_noRT | CAAA0106612 | 4.049 | 2.52E-07 | glio |
| Chek1_ex.vivo_noRT | Mog | 4.842 | 2.52E-07 | glio |
| Chek1_ex.vivo_noRT | Svopl | 4.449 | 2.52E-08 | glio |
| Aars2_in.vivo_noRT | Cebpa | 4.202 | 2.52E-09 | glio |
| Atr_in.vivo_noRT | Ltb4r1 | 4.359 | 2.52E-09 | glio |
| Bcas2_in.vivo_noRT | Gm43137 | 4.291 | 2.52E-09 | glio |
| Bcas2_in.vivo_noRT | 2310067P03R | 4.284 | 2.52E-10 | glio |
| Chek1_in.vivo_noRT | 9930111J21Ri | 3.864 | 2.52E-10 | glio |
| Chek1_in.vivo_noRT | Pcdhgb6 | 3.859 | 2.52E-10 | glio |
| Atr_in.vivo_noRT | Gm15853 | 4.798 | 2.52E-11 | glio |
| Bora_in.vivo_noRT | Icam2 | 4.07 | 2.52E-11 | glio |
| Atr_in.vivo_noRT | Hif3a | 4.891 | 2.52E-12 | glio |
| Aars2_in.vivo_noRT | Vmn2r54 | 4.165 | 2.52E-13 | glio |
| Chek1_in.vivo_noRT | Prl2c2 | 5.044 | 2.52E-14 | glio |
| Bcas2_in.vivo_noRT | Gm38477 | 4.285 | 2.52E-15 | glio |
| Chek1_ex.vivo_noRT | Gm26555 | 5.643 | 2.52E-18 | glio |
| culture_effect | Smoc1 | -2.798 | 2.52E-31 | glio |
| culture_effect | Gm37170 | 1.635 | 2.53E-03 | glio |
| culture_effect | Hist1h2bc | -1.31 | 2.53E-03 | glio |
| culture_effect | Parp14 | -0.775 | 2.53E-03 | glio |
| Interaction_genotype_Aars2 | Gm12925 | -3.823 | 2.53E-03 | glio |
| Interaction_genotype_Aars2 | Gm12758 | -3.725 | 2.53E-03 | glio |
| Interaction_genotype_Aars2 | Kdelr2 | 1.325 | 2.53E-03 | glio |
| Interaction_genotype_Bora | Gm15234 | -3.988 | 2.53E-03 | glio |
| Interaction_genotype_Chek1 | Zyx | 2.202 | 2.53E-03 | glio |
| Interaction_genotype_Mtor | Reep6 | -2.571 | 2.53E-03 | glio |
| Interaction_genotype_Mtor | Rpl14 | 1.089 | 2.53E-03 | glio |
| Interaction_genotype_Myc | Speer4a | -2.463 | 2.53E-03 | glio |
| Interaction_genotype_Myc | Zfp738 | -2.803 | 2.53E-03 | glio |
| interaction_RT | Slc2a13 | 2.821 | 2.53E-03 | glio |
| Aars2_in.vivo_noRT | Vps8 | 1.956 | 2.53E-03 | glio |
| Atp5c1_in.vivo_noRT | Ikzf2 | 2.554 | 2.53E-03 | glio |
| Atp5c1_in.vivo_noRT | Krt8 | -1.666 | 2.53E-03 | glio |
| Atr_in.vivo_noRT | Nat14 | 1.907 | 2.53E-03 | glio |
| Atr_in.vivo_noRT | D3Ertd751e | 1.785 | 2.53E-03 | glio |
| Atr_in.vivo_noRT | Ppp1cb | -1.083 | 2.53E-03 | glio |
| Atr_in.vivo_noRT | Sys1 | -1.103 | 2.53E-03 | glio |
| Atr_ex.vivo_noRT | Gjb5 | 2.584 | 2.53E-03 | glio |
| Bcas2_ex.vivo_noRT | Tent5c | 2.349 | 2.53E-03 | glio |
| Bora_in.vivo_noRT | Npr2 | 2.525 | 2.53E-03 | glio |
| Hsd17b10_in.vivo_noRT | Atoh8 | 2.233 | 2.53E-03 | glio |
| Hsd17b10_in.vivo_noRT | Arhgef15 | 2.235 | 2.53E-03 | glio |
| Interaction_genotype_Mtor | Ubox5 | -3.646 | 2.53E-04 | glio |

| | | | | |
|---|---|---|---|---|
| Interaction_genotype_Myc | Mblac1 | -4.375 | 2.53E-04 | glio |
| Atr_in.vivo_noRT | Gm20712 | 2.564 | 2.53E-04 | glio |
| Atr_ex.vivo_noRT | Gm13032 | 2.764 | 2.53E-04 | glio |
| Atr_ex.vivo_noRT | Npm1 | -0.902 | 2.53E-04 | glio |
| Interaction_genotype_Mtor | Gm9922 | -4.614 | 2.53E-05 | glio |
| Interaction_genotype_Mtor | Gm10131 | -4.938 | 2.53E-05 | glio |
| Interaction_genotype_Mtor | Calm3 | 2.26 | 2.53E-05 | glio |
| interaction_RT | Leprot | 1.058 | 2.53E-05 | glio |
| Atp5c1_in.vivo_noRT | Olfr718-ps1 | 2.404 | 2.53E-05 | glio |
| Atp5c1_in.vivo_noRT | Prickle2 | 2.668 | 2.53E-05 | glio |
| Chek1_ex.vivo_noRT | Gadl1 | 3.556 | 2.53E-05 | glio |
| Hsd17b10_in.vivo_noRT | Gm13403 | 2.958 | 2.53E-05 | glio |
| Interaction_genotype_Mtor | Cpxm1 | -4.937 | 2.53E-06 | glio |
| Interaction_genotype_Mtor | Map3k9 | -4.627 | 2.53E-06 | glio |
| Interaction_genotype_Mtor | Gm5890 | -4.099 | 2.53E-06 | glio |
| Bora_in.vivo_noRT | Dmrt2 | 3.526 | 2.53E-06 | glio |
| Bora_in.vivo_noRT | Cnfn | 3.541 | 2.53E-06 | glio |
| Interaction_genotype_Mtor | Tmem165 | 2.443 | 2.53E-07 | glio |
| Interaction_genotype_Mtor | Zfp551 | -6.149 | 2.53E-07 | glio |
| Aars2_in.vivo_noRT | Srgn | 4.282 | 2.53E-07 | glio |
| Aars2_in.vivo_noRT | Tmx2 | -1.916 | 2.53E-07 | glio |
| Atr_in.vivo_noRT | Vipr2 | 3.59 | 2.53E-07 | glio |
| Atr_ex.vivo_noRT | Idi2 | 3.164 | 2.53E-07 | glio |
| Bora_in.vivo_noRT | Slc16a6 | 3.753 | 2.53E-07 | glio |
| culture_effect | Tmem9b | 1.035 | 2.53E-08 | glio |
| Chek1_ex.vivo_noRT | Gm44899 | 3.738 | 2.53E-08 | glio |
| Hsd17b10_in.vivo_noRT | Zfp352 | 3.616 | 2.53E-08 | glio |
| culture_effect | Tex261 | 0.755 | 2.53E-09 | glio |
| Atr_in.vivo_noRT | Tmem202 | 4.091 | 2.53E-09 | glio |
| Bcas2_in.vivo_noRT | Psmd9 | -2.676 | 2.53E-09 | glio |
| Chek1_ex.vivo_noRT | Synpo2l | 4.447 | 2.53E-09 | glio |
| culture_effect | Anxa4 | -1.116 | 2.53E-10 | glio |
| culture_effect | Psmb2 | -0.757 | 2.53E-10 | glio |
| Bora_in.vivo_noRT | Calhm5 | 4.208 | 2.53E-10 | glio |
| Chek1_ex.vivo_noRT | Vmn1r181 | 5.247 | 2.53E-10 | glio |
| Chek1_ex.vivo_noRT | Nos2 | 4.842 | 2.53E-10 | glio |
| Hsd17b10_in.vivo_noRT | Gm12610 | 3.608 | 2.53E-10 | glio |
| Bora_in.vivo_noRT | 1700015F17Ri | 4.676 | 2.53E-11 | glio |
| Aars2_in.vivo_noRT | Prickle4 | 4.535 | 2.53E-12 | glio |
| Bcas2_in.vivo_noRT | Aloxe3 | 4.278 | 2.53E-13 | glio |
| Aars2_in.vivo_noRT | Crybb1 | 4.94 | 2.53E-16 | glio |
| Chek1_ex.vivo_noRT | Gm47404 | 4.842 | 2.53E-16 | glio |
| Chek1_in.vivo_noRT | BC080695 | 5.044 | 2.53E-22 | glio |
| Aars2_in.vivo_noRT | Stfa3 | 5.327 | 2.53E-29 | glio |
| Aars2_in.vivo_noRT | Olfr787 | 5.327 | 2.53E-29 | glio |
| culture_effect | Gas8 | -0.668 | 2.54E-03 | glio |
| culture_effect | Acadvl | -0.636 | 2.54E-03 | glio |
| culture_effect | Parva | 0.609 | 2.54E-03 | glio |
| Interaction_genotype_Aars2 | Zfp873 | -3.344 | 2.54E-03 | glio |
| Interaction_genotype_Aars2 | Gm26519 | -3.789 | 2.54E-03 | glio |
| Interaction_genotype_Aars2 | Pcolce2 | 1.972 | 2.54E-03 | glio |
| Interaction_genotype_Atr | Gm10101 | -3.378 | 2.54E-03 | glio |

| | | | | |
|---|---|---|---|---|
| Interaction_genotype_Atr | Zfp280c | -2.641 | 2.54E-03 | glio |
| Interaction_genotype_Cdkn2a | Gm13191 | 2.835 | 2.54E-03 | glio |
| Interaction_genotype_Cdkn2a | 4933416E03Ri | 2.835 | 2.54E-03 | glio |
| Interaction_genotype_Chek1 | Tmem179b | 2.01 | 2.54E-03 | glio |
| Interaction_genotype_Mtor | 4921507P07R | -3.517 | 2.54E-03 | glio |
| Interaction_genotype_Mtor | Stradb | -2.894 | 2.54E-03 | glio |
| Interaction_genotype_Mtor | Msrb3 | -2.824 | 2.54E-03 | glio |
| Interaction_genotype_Mtor | Mthfsd | -2.157 | 2.54E-03 | glio |
| Interaction_genotype_Mtor | Gnpda2 | 1.831 | 2.54E-03 | glio |
| Interaction_genotype_Myc | Mir17hg | -3.007 | 2.54E-03 | glio |
| Interaction_genotype_Myc | Col12a1 | -2.592 | 2.54E-03 | glio |
| Interaction_genotype_Uqcrc2 | Gm16741 | -4.194 | 2.54E-03 | glio |
| Interaction_genotype_Uqcrc2 | Gm44702 | -4.663 | 2.54E-03 | glio |
| interaction_RT | Ccpg1os | 1.507 | 2.54E-03 | glio |
| interaction_RT | Peg10 | -1.601 | 2.54E-03 | glio |
| interaction_RT | Baz2b | 0.99 | 2.54E-03 | glio |
| interaction_RT | Slc39a1 | 0.837 | 2.54E-03 | glio |
| interaction_RT | Klf6 | 0.911 | 2.54E-03 | glio |
| interaction_RT | Set | -0.53 | 2.54E-03 | glio |
| Aars2_ex.vivo_noRT | Gm13822 | 2.897 | 2.54E-03 | glio |
| Bard1_ex.vivo_noRT | Gm47622 | 2.239 | 2.54E-03 | glio |
| Bard1_ex.vivo_noRT | BC147527 | 2.814 | 2.54E-03 | glio |
| Bcas2_in.vivo_noRT | Bbs10 | 2.31 | 2.54E-03 | glio |
| Bcas2_in.vivo_noRT | Itch | -1.222 | 2.54E-03 | glio |
| Bcas2_in.vivo_noRT | Rars | -0.925 | 2.54E-03 | glio |
| Bora_in.vivo_noRT | B230208H11R | 2.268 | 2.54E-03 | glio |
| Bora_in.vivo_noRT | Tlr2 | 2.261 | 2.54E-03 | glio |
| Bora_ex.vivo_noRT | Gm20478 | 2.806 | 2.54E-03 | glio |
| Chek1_in.vivo_noRT | Klhdc3 | -1.411 | 2.54E-03 | glio |
| culture_effect | Slc39a6 | 0.575 | 2.54E-04 | glio |
| culture_effect | Isg15 | 0.79 | 2.54E-04 | glio |
| Interaction_genotype_Aars2 | Otos | -3.578 | 2.54E-04 | glio |
| Interaction_genotype_Aars2 | K230015D01R | -4.604 | 2.54E-04 | glio |
| Interaction_genotype_Mtor | Tmpo | 2.131 | 2.54E-04 | glio |
| Interaction_genotype_Myc | Ncf1 | -3.6 | 2.54E-04 | glio |
| Interaction_genotype_Myc | Spag1 | -4.004 | 2.54E-04 | glio |
| interaction_RT | Mbtd1 | 1.154 | 2.54E-04 | glio |
| Aars2_in.vivo_noRT | Folr2 | 2.519 | 2.54E-04 | glio |
| Aars2_in.vivo_noRT | Klhl18 | 2.126 | 2.54E-04 | glio |
| Aars2_in.vivo_noRT | Ywhae | -0.688 | 2.54E-04 | glio |
| Atr_in.vivo_noRT | Mmp14 | 2.659 | 2.54E-04 | glio |
| Atr_in.vivo_noRT | Gas5 | -0.881 | 2.54E-04 | glio |
| Bcas2_in.vivo_noRT | Fam71e1 | 2.385 | 2.54E-04 | glio |
| Bcas2_in.vivo_noRT | 6720489N17R | 2.942 | 2.54E-04 | glio |
| Bcas2_in.vivo_noRT | S100a10 | -1.019 | 2.54E-04 | glio |
| Bora_ex.vivo_noRT | Pcdhga1 | 3.025 | 2.54E-04 | glio |
| Chek1_in.vivo_noRT | Gm14167 | 2.349 | 2.54E-04 | glio |
| Eif2s1_in.vivo_noRT | Gm30794 | 2.143 | 2.54E-04 | glio |
| Hsd17b10_in.vivo_noRT | Gpr179 | 2.615 | 2.54E-04 | glio |
| culture_effect | Gm9856 | 1.342 | 2.54E-05 | glio |
| culture_effect | Morrbid | 1.019 | 2.54E-05 | glio |
| culture_effect | Tspan14 | -0.789 | 2.54E-05 | glio |

| | | | | |
|---|---|---|---|---|
| Interaction_genotype_Aars2 | Gm26766 | -5.049 | 2.54E-05 | glio |
| Interaction_genotype_Mtor | Cpne9 | -5.374 | 2.54E-05 | glio |
| Interaction_genotype_Myc | Bdh2 | -4.706 | 2.54E-05 | glio |
| Aars2_in.vivo_noRT | Pde6h | 2.762 | 2.54E-05 | glio |
| Atr_in.vivo_noRT | Hist1h4c | 3.101 | 2.54E-05 | glio |
| Atr_in.vivo_noRT | Rtl5 | 2.865 | 2.54E-05 | glio |
| Atr_in.vivo_noRT | Napsa | 3.029 | 2.54E-05 | glio |
| Hsd17b10_in.vivo_noRT | Slc22a1 | 2.442 | 2.54E-05 | glio |
| Hsd17b10_in.vivo_noRT | Gm26590 | 2.855 | 2.54E-05 | glio |
| Interaction_genotype_Mtor | Zscan4c | -4.099 | 2.54E-06 | glio |
| Aars2_ex.vivo_noRT | A030014E15Ri | 2.899 | 2.54E-06 | glio |
| Atr_in.vivo_noRT | Zfp994 | 2.542 | 2.54E-06 | glio |
| Chek1_ex.vivo_noRT | Slc11a1 | 4.256 | 2.54E-06 | glio |
| Hsd17b10_in.vivo_noRT | Hectd2os | 3.239 | 2.54E-06 | glio |
| culture_effect | BC037704 | 2.458 | 2.54E-07 | glio |
| culture_effect | Ppp2r1a | -0.763 | 2.54E-07 | glio |
| Aars2_in.vivo_noRT | E230029C05Ri | 3.455 | 2.54E-07 | glio |
| Interaction_genotype_Mtor | Gm13398 | -4.896 | 2.54E-08 | glio |
| interaction_RT | mt-Nd2 | -1.13 | 2.54E-08 | glio |
| Atr_in.vivo_noRT | Axdnd1 | 4.221 | 2.54E-08 | glio |
| culture_effect | Nod1 | -2.735 | 2.54E-09 | glio |
| Bcas2_in.vivo_noRT | Baalc | 4.015 | 2.54E-09 | glio |
| Chek1_in.vivo_noRT | Gm15853 | 4.308 | 2.54E-09 | glio |
| Aars2_in.vivo_noRT | Gm47802 | 4.351 | 2.54E-11 | glio |
| culture_effect | Gale | -2.393 | 2.54E-12 | glio |
| Chek1_in.vivo_noRT | Gm50394 | 4.777 | 2.54E-12 | glio |
| culture_effect | Fam71e1 | 3.031 | 2.54E-13 | glio |
| Bcas2_in.vivo_noRT | Tcstv3 | 4.807 | 2.54E-13 | glio |
| Bcas2_in.vivo_noRT | Lrguk | 4.543 | 2.54E-15 | glio |
| Chek1_in.vivo_noRT | Cers3 | 4.512 | 2.54E-16 | glio |
| Atr_in.vivo_noRT | Cys1 | 5.267 | 2.54E-19 | glio |
| Atr_in.vivo_noRT | Selenov | 5.533 | 2.54E-21 | glio |
| Atr_in.vivo_noRT | Angptl7 | 4.667 | 2.54E-21 | glio |
| Aars2_in.vivo_noRT | Zscan4-ps2 | 5.327 | 2.54E-24 | glio |
| Aars2_in.vivo_noRT | Gm41307 | 5.327 | 2.54E-25 | glio |
| culture_effect | Jrk | 1.3 | 2.55E-03 | glio |
| culture_effect | Sppl3 | -0.582 | 2.55E-03 | glio |
| Interaction_genotype_Atr | Hfm1 | -2.944 | 2.55E-03 | glio |
| Interaction_genotype_Atr | Cstb | 1.826 | 2.55E-03 | glio |
| Interaction_genotype_Bcas2 | Gm20667 | -4.14 | 2.55E-03 | glio |
| Interaction_genotype_Bcas2 | Psenen | 1.078 | 2.55E-03 | glio |
| Interaction_genotype_Myc | 4930527F14Ri | -2.596 | 2.55E-03 | glio |
| Interaction_genotype_Myc | Card10 | -3.307 | 2.55E-03 | glio |
| Interaction_genotype_Myc | Svbp | 1.323 | 2.55E-03 | glio |
| Aars2_in.vivo_noRT | Iqcg | 2.236 | 2.55E-03 | glio |
| Aars2_in.vivo_noRT | 1700017B05R | 1.581 | 2.55E-03 | glio |
| Aars2_ex.vivo_noRT | Zg16 | 2.893 | 2.55E-03 | glio |
| Aars2_ex.vivo_noRT | Tnni2 | 3.517 | 2.55E-03 | glio |
| Atr_in.vivo_noRT | Zfp600 | 2.157 | 2.55E-03 | glio |
| Bcas2_in.vivo_noRT | Sema4f | 2.214 | 2.55E-03 | glio |
| Bcas2_in.vivo_noRT | Gm49768 | 2.054 | 2.55E-03 | glio |
| Bcas2_ex.vivo_noRT | Nmrk2 | 2.742 | 2.55E-03 | glio |

| | | | | |
|---|---|---|---|---|
| Bora_in.vivo_noRT | Gm26901 | 2.35 | 2.55E-03 | glio |
| Bora_in.vivo_noRT | Fetub | 1.923 | 2.55E-03 | glio |
| Bora_in.vivo_noRT | Tesk2 | 2.11 | 2.55E-03 | glio |
| Bora_in.vivo_noRT | Arv1 | 1.677 | 2.55E-03 | glio |
| Bora_in.vivo_noRT | Pard6a | -1.429 | 2.55E-03 | glio |
| Cenpj_in.vivo_noRT | Rimklb | 1.759 | 2.55E-03 | glio |
| Cenpj_ex.vivo_noRT | Slc35g3 | 2.192 | 2.55E-03 | glio |
| Chek1_in.vivo_noRT | 5530601H04R | -1.499 | 2.55E-03 | glio |
| Hsd17b10_in.vivo_noRT | Lrrn3 | 2.16 | 2.55E-03 | glio |
| Hsd17b10_in.vivo_noRT | Pcmt1 | -0.601 | 2.55E-03 | glio |
| culture_effect | Ube3c | 0.568 | 2.55E-04 | glio |
| Interaction_genotype_Mtor | Gm4858 | -4.099 | 2.55E-04 | glio |
| Interaction_genotype_Mtor | Vcan | -4.74 | 2.55E-04 | glio |
| Interaction_genotype_Myc | Kcnk13 | -3.254 | 2.55E-04 | glio |
| Interaction_genotype_Myc | A530072M11F | -4.204 | 2.55E-04 | glio |
| Interaction_genotype_Myc | Gm17484 | -3.967 | 2.55E-04 | glio |
| Interaction_genotype_Myc | Rbm19 | -3.124 | 2.55E-04 | glio |
| Aars2_in.vivo_noRT | Flrt2 | 2.664 | 2.55E-04 | glio |
| Aars2_in.vivo_noRT | Cdc42ep4 | 2.444 | 2.55E-04 | glio |
| Aars2_in.vivo_noRT | Rnf225 | 2.756 | 2.55E-04 | glio |
| Aars2_ex.vivo_noRT | A930003A15R | 3.293 | 2.55E-04 | glio |
| Atr_in.vivo_noRT | Hcn1 | 2.791 | 2.55E-04 | glio |
| Atr_ex.vivo_noRT | Lncbate10 | 2.376 | 2.55E-04 | glio |
| Atr_ex.vivo_noRT | Il12rb1 | 3.159 | 2.55E-04 | glio |
| Bcas2_in.vivo_noRT | Cxcr6 | 2.618 | 2.55E-04 | glio |
| Bora_in.vivo_noRT | AY074887 | 2.884 | 2.55E-04 | glio |
| Chek1_in.vivo_noRT | Gm42670 | 2.373 | 2.55E-04 | glio |
| Hsd17b10_in.vivo_noRT | A830035O19R | 2.567 | 2.55E-04 | glio |
| Interaction_genotype_Mtor | C230014O12R | -4.51 | 2.55E-05 | glio |
| interaction_RT | Lrrc75aos2 | -4.026 | 2.55E-05 | glio |
| Aars2_in.vivo_noRT | Hist1h4n | 3.25 | 2.55E-05 | glio |
| Aars2_ex.vivo_noRT | AI463229 | 3.293 | 2.55E-05 | glio |
| Atp5c1_in.vivo_noRT | Gm15890 | 2.668 | 2.55E-05 | glio |
| Atr_in.vivo_noRT | Tex19.1 | -1.62 | 2.55E-05 | glio |
| Atr_in.vivo_noRT | Jagn1 | -1.622 | 2.55E-05 | glio |
| Atr_ex.vivo_noRT | 2300002M23F | 3.164 | 2.55E-05 | glio |
| Bora_in.vivo_noRT | Zdhhc14 | 2.469 | 2.55E-05 | glio |
| Chek1_ex.vivo_noRT | Pkhd1 | 3.754 | 2.55E-05 | glio |
| Hsd17b10_in.vivo_noRT | Gm14968 | 3.237 | 2.55E-05 | glio |
| Interaction_genotype_Mtor | BC080695 | -4.099 | 2.55E-06 | glio |
| Bcas2_in.vivo_noRT | Enpp3 | 3.24 | 2.55E-06 | glio |
| Bcas2_in.vivo_noRT | Gm26688 | 3.369 | 2.55E-06 | glio |
| Chek1_in.vivo_noRT | Tox2 | 3.269 | 2.55E-06 | glio |
| Chek1_ex.vivo_noRT | Lsm2 | -1.585 | 2.55E-06 | glio |
| Dclre1c_ex.vivo_noRT | Gm21319 | 3.739 | 2.55E-06 | glio |
| Dclre1c_ex.vivo_noRT | Gm2016 | 3.535 | 2.55E-06 | glio |
| Atr_in.vivo_noRT | Lbhd2 | 4.292 | 2.55E-07 | glio |
| Atr_ex.vivo_noRT | Vmn2r18 | 3.164 | 2.55E-07 | glio |
| Aars2_in.vivo_noRT | Gm38979 | 4.152 | 2.55E-08 | glio |
| Aars2_in.vivo_noRT | Cnksr1 | 3.758 | 2.55E-08 | glio |
| Aars2_in.vivo_noRT | H60b | 3.729 | 2.55E-08 | glio |
| culture_effect | Rhox5 | -3.537 | 2.55E-09 | glio |

| | | | | |
|---|---|---|---|---|
| Atr_in.vivo_noRT | Slc17a7 | 4.101 | 2.55E-09 | glio |
| Bcas2_in.vivo_noRT | Ankrd33b | 4.291 | 2.55E-09 | glio |
| Chek1_in.vivo_noRT | Gm28424 | 4.032 | 2.55E-09 | glio |
| culture_effect | Tex19.1 | -0.991 | 2.55E-10 | glio |
| Aars2_in.vivo_noRT | Lncenc1 | 4.021 | 2.55E-10 | glio |
| Aars2_in.vivo_noRT | Tslp | 4.681 | 2.55E-10 | glio |
| Atr_in.vivo_noRT | Ifitm1 | 5.267 | 2.55E-11 | glio |
| Bcas2_in.vivo_noRT | Cxcl1 | 4.807 | 2.55E-11 | glio |
| Bora_in.vivo_noRT | Fgf8 | 4.42 | 2.55E-13 | glio |
| culture_effect | Acan | -3.214 | 2.55E-14 | glio |
| Atr_in.vivo_noRT | Gm15943 | 5.269 | 2.55E-21 | glio |
| culture_effect | Cdh2 | 2.347 | 2.55E-29 | glio |
| culture_effect | Gm48362 | -1.349 | 2.56E-03 | glio |
| culture_effect | Rnf149 | 0.924 | 2.56E-03 | glio |
| culture_effect | Atp7a | 0.836 | 2.56E-03 | glio |
| culture_effect | 2810408I11Ri | 0.756 | 2.56E-03 | glio |
| Interaction_genotype_Atr | H2-Ab1 | 3.131 | 2.56E-03 | glio |
| Interaction_genotype_Bcas2 | Impg2 | -3.371 | 2.56E-03 | glio |
| Interaction_genotype_Myc | Lce1g | -3.52 | 2.56E-03 | glio |
| Interaction_genotype_Myc | C1qtnf12 | -3.718 | 2.56E-03 | glio |
| Interaction_genotype_Myc | Zswim3 | -3.197 | 2.56E-03 | glio |
| Interaction_genotype_Myc | Zfp449 | -2.942 | 2.56E-03 | glio |
| Interaction_genotype_Myc | Zdhhc24 | -2.983 | 2.56E-03 | glio |
| Interaction_genotype_Myc | Lactb | -2.117 | 2.56E-03 | glio |
| Interaction_genotype_Myc | Rpl23a | 1.147 | 2.56E-03 | glio |
| Interaction_genotype_Pars2 | Gm14002 | -3.778 | 2.56E-03 | glio |
| Interaction_genotype_Rad9a | Klhdc8a | -3.651 | 2.56E-03 | glio |
| interaction_RT | Fam161a | 2.077 | 2.56E-03 | glio |
| interaction_RT | Trp53rkb | 1.397 | 2.56E-03 | glio |
| interaction_RT | Mzt2 | 0.973 | 2.56E-03 | glio |
| interaction_RT | Myl12a | 0.578 | 2.56E-03 | glio |
| Atr_in.vivo_noRT | Acad12 | 2.108 | 2.56E-03 | glio |
| Atr_in.vivo_noRT | Clec1a | 2.085 | 2.56E-03 | glio |
| Atr_in.vivo_noRT | Prss56 | 2.271 | 2.56E-03 | glio |
| Atr_in.vivo_noRT | E2f2 | 1.775 | 2.56E-03 | glio |
| Atr_in.vivo_noRT | Gna11 | -1.326 | 2.56E-03 | glio |
| Atr_in.vivo_noRT | Nsmce4a | -1.088 | 2.56E-03 | glio |
| Bard1_ex.vivo_noRT | Slc10a1 | 2.421 | 2.56E-03 | glio |
| Bcas2_in.vivo_noRT | Slc25a43 | 2.224 | 2.56E-03 | glio |
| Bora_in.vivo_noRT | Ccdc7b | 2.128 | 2.56E-03 | glio |
| Bora_in.vivo_noRT | Map3k15 | 1.813 | 2.56E-03 | glio |
| Bora_in.vivo_noRT | Cav1 | -1.68 | 2.56E-03 | glio |
| Bora_in.vivo_noRT | Zfp512 | 1.284 | 2.56E-03 | glio |
| Chek1_ex.vivo_noRT | B230319C09R | 2.912 | 2.56E-03 | glio |
| culture_effect | Ensa | -0.643 | 2.56E-04 | glio |
| culture_effect | Suds3 | 0.491 | 2.56E-04 | glio |
| Interaction_genotype_Myc | Gm9908 | -4.465 | 2.56E-04 | glio |
| Interaction_genotype_Myc | Tlcd2 | -4.649 | 2.56E-04 | glio |
| Aars2_in.vivo_noRT | A430018G15R | 2.149 | 2.56E-04 | glio |
| Aars2_in.vivo_noRT | Col9a1 | -1.718 | 2.56E-04 | glio |
| Atr_in.vivo_noRT | Wnk2 | 2.618 | 2.56E-04 | glio |
| Bcas2_in.vivo_noRT | Glt8d2 | 2.756 | 2.56E-04 | glio |

| | | | | |
|---|---|---|---|---|
| Cenpj_ex.vivo_noRT | Dpep3 | 2.591 | 2.56E-04 | glio |
| Cenpj_ex.vivo_noRT | Erich6b | 2.591 | 2.56E-04 | glio |
| Cenpj_ex.vivo_noRT | Tac2 | 2.591 | 2.56E-04 | glio |
| Interaction_genotype_Myc | Xylt2 | -4.338 | 2.56E-05 | glio |
| Aars2_ex.vivo_noRT | Prap1 | 2.9 | 2.56E-05 | glio |
| Atr_in.vivo_noRT | Gm38973 | 2.937 | 2.56E-05 | glio |
| Bard1_ex.vivo_noRT | Gm47510 | 2.814 | 2.56E-05 | glio |
| Bard1_ex.vivo_noRT | Gm5788 | 2.814 | 2.56E-05 | glio |
| Hsd17b10_in.vivo_noRT | Lrrc72 | 2.825 | 2.56E-05 | glio |
| culture_effect | Sapcd1 | -3.013 | 2.56E-06 | glio |
| culture_effect | Tmem11 | -0.635 | 2.56E-06 | glio |
| Aars2_in.vivo_noRT | Eif3j2 | 3.248 | 2.56E-06 | glio |
| Bcas2_ex.vivo_noRT | Edn2 | 3.145 | 2.56E-06 | glio |
| Chek1_ex.vivo_noRT | Gm37035 | 3.363 | 2.56E-06 | glio |
| Interaction_genotype_Myc | Timm13 | 2.11 | 2.56E-07 | glio |
| Bora_in.vivo_noRT | Babam1 | -2.098 | 2.56E-07 | glio |
| Atr_in.vivo_noRT | Ucma | 4.535 | 2.56E-08 | glio |
| Atr_in.vivo_noRT | Pate2 | 4.054 | 2.56E-08 | glio |
| Bora_in.vivo_noRT | Kmo | 3.295 | 2.56E-08 | glio |
| Hsd17b10_in.vivo_noRT | Gm13178 | 3.1 | 2.56E-09 | glio |
| culture_effect | A930029G22R | 2.747 | 2.56E-10 | glio |
| Aars2_in.vivo_noRT | Gm41442 | 4.285 | 2.56E-10 | glio |
| Bcas2_in.vivo_noRT | Gm9929 | 4.028 | 2.56E-10 | glio |
| Chek1_in.vivo_noRT | Slc6a13 | 4.664 | 2.56E-10 | glio |
| Chek1_ex.vivo_noRT | Gm50223 | 4.449 | 2.56E-10 | glio |
| Hsd17b10_in.vivo_noRT | Gm5890 | 3.616 | 2.56E-12 | glio |
| Atr_in.vivo_noRT | Nrg2 | 5.011 | 2.56E-13 | glio |
| Hsd17b10_in.vivo_noRT | Olfr810 | 3.616 | 2.56E-14 | glio |
| Aars2_in.vivo_noRT | Gm5934 | 5.063 | 2.56E-21 | glio |
| culture_effect | Sertad4 | -1.028 | 2.57E-03 | glio |
| Interaction_genotype_Mtor | Gm9776 | -3.406 | 2.57E-03 | glio |
| Interaction_genotype_Mtor | Zcchc2 | -2.972 | 2.57E-03 | glio |
| Interaction_genotype_Myc | Smim41 | -3.689 | 2.57E-03 | glio |
| Interaction_genotype_Myc | Mertk | -3.263 | 2.57E-03 | glio |
| Interaction_genotype_Myc | 4921523L03Ri | -3.461 | 2.57E-03 | glio |
| Interaction_genotype_Myc | Qpctl | -3.062 | 2.57E-03 | glio |
| Interaction_genotype_Myc | Ccnyl1 | -2.761 | 2.57E-03 | glio |
| Interaction_genotype_Myc | Zbtb33 | -2.42 | 2.57E-03 | glio |
| Interaction_genotype_Myc | Ank2 | -2.141 | 2.57E-03 | glio |
| interaction_RT | Slc52a2 | -1.097 | 2.57E-03 | glio |
| interaction_RT | Xylt1 | -1.023 | 2.57E-03 | glio |
| Aars2_in.vivo_noRT | Zfp354a | 2.081 | 2.57E-03 | glio |
| Aars2_in.vivo_noRT | Cntrob | 1.784 | 2.57E-03 | glio |
| Aars2_in.vivo_noRT | Npm1 | -0.584 | 2.57E-03 | glio |
| Atr_in.vivo_noRT | Cavin2 | 2.271 | 2.57E-03 | glio |
| Atr_in.vivo_noRT | Spats1 | 2.173 | 2.57E-03 | glio |
| Atr_in.vivo_noRT | Pax9 | 2.039 | 2.57E-03 | glio |
| Atr_in.vivo_noRT | Pdxdc1 | -1.461 | 2.57E-03 | glio |
| Atr_in.vivo_noRT | Wasl | -1.046 | 2.57E-03 | glio |
| Bard1_ex.vivo_noRT | Fsd2 | 3.026 | 2.57E-03 | glio |
| Bard1_ex.vivo_noRT | Gm50223 | 2.421 | 2.57E-03 | glio |
| Bcas2_in.vivo_noRT | Cnih4 | -0.681 | 2.57E-03 | glio |

| | | | | |
|---|---|---|---|---|
| Bora_in.vivo_noRT | A930029G22R | 2.135 | 2.57E-03 | glio |
| Bora_in.vivo_noRT | Aida | -1.122 | 2.57E-03 | glio |
| Bora_in.vivo_noRT | Csrp2 | -1.643 | 2.57E-03 | glio |
| Bora_in.vivo_noRT | Gatm | -1.685 | 2.57E-03 | glio |
| Chek1_in.vivo_noRT | Sgsh | 2.301 | 2.57E-03 | glio |
| Chek1_in.vivo_noRT | 4732440D04R | 2.004 | 2.57E-03 | glio |
| Chek1_in.vivo_noRT | Usp5 | -1.301 | 2.57E-03 | glio |
| Chek1_in.vivo_noRT | Cdk5rap3 | -1.253 | 2.57E-03 | glio |
| Chek1_in.vivo_noRT | Tm9sf2 | -1.151 | 2.57E-03 | glio |
| Chek1_in.vivo_noRT | Mrpl28 | -0.836 | 2.57E-03 | glio |
| Chek1_in.vivo_noRT | Rpl3 | -0.59 | 2.57E-03 | glio |
| Eif2s1_in.vivo_noRT | 1700017M07F | 2.143 | 2.57E-03 | glio |
| culture_effect | Gm50069 | 1.978 | 2.57E-04 | glio |
| culture_effect | Scn1b | -2.038 | 2.57E-04 | glio |
| culture_effect | Mcm7 | -0.545 | 2.57E-04 | glio |
| Interaction_genotype_Mtor | Gm20767 | -4.099 | 2.57E-04 | glio |
| Interaction_genotype_Mtor | Anapc13 | 2.022 | 2.57E-04 | glio |
| Aars2_in.vivo_noRT | Mfsd3 | 2.173 | 2.57E-04 | glio |
| Aars2_in.vivo_noRT | Chchd2 | -0.7 | 2.57E-04 | glio |
| Atp5c1_in.vivo_noRT | Gm17203 | 2.018 | 2.57E-04 | glio |
| Atp5c1_in.vivo_noRT | Pik3ap1 | 2.405 | 2.57E-04 | glio |
| Atp5c1_in.vivo_noRT | Gm13032 | 2.282 | 2.57E-04 | glio |
| Atp5c1_in.vivo_noRT | Hcn3 | 2.548 | 2.57E-04 | glio |
| Atr_in.vivo_noRT | Dnajb13 | 2.454 | 2.57E-04 | glio |
| Atr_ex.vivo_noRT | Oit3 | 3.184 | 2.57E-04 | glio |
| Bora_ex.vivo_noRT | Zscan4d | 2.808 | 2.57E-04 | glio |
| Bora_ex.vivo_noRT | Gm37986 | 3.197 | 2.57E-04 | glio |
| Chek1_in.vivo_noRT | Fam71f2 | 2.699 | 2.57E-04 | glio |
| Chek1_ex.vivo_noRT | Krt18 | 4.443 | 2.57E-04 | glio |
| Eif2s1_in.vivo_noRT | Ank1 | 2.143 | 2.57E-04 | glio |
| Interaction_genotype_Mtor | BB557941 | -4.239 | 2.57E-05 | glio |
| Interaction_genotype_Mtor | Agbl3 | -3.81 | 2.57E-05 | glio |
| Interaction_genotype_Mtor | Ammecr1 | 2.62 | 2.57E-05 | glio |
| Interaction_genotype_Mtor | G3bp1 | 2.037 | 2.57E-05 | glio |
| Aars2_in.vivo_noRT | Pgls | -1.046 | 2.57E-05 | glio |
| Bcas2_in.vivo_noRT | Myzap | 2.486 | 2.57E-05 | glio |
| Hsd17b10_in.vivo_noRT | Gm35290 | 2.878 | 2.57E-05 | glio |
| Hsd17b10_in.vivo_noRT | Gm9922 | 3.007 | 2.57E-05 | glio |
| Interaction_genotype_Mtor | Gm9484 | -5.835 | 2.57E-06 | glio |
| Interaction_genotype_Myc | D630036H23F | -5.207 | 2.57E-06 | glio |
| Aars2_in.vivo_noRT | Gm16835 | 3.4 | 2.57E-06 | glio |
| Aars2_in.vivo_noRT | Ksr1 | 3.278 | 2.57E-06 | glio |
| Bora_in.vivo_noRT | Axdnd1 | 3.624 | 2.57E-06 | glio |
| Interaction_genotype_Myc | Anks6 | -5.586 | 2.57E-07 | glio |
| culture_effect | Cyb5r3 | 0.757 | 2.57E-08 | glio |
| Chek1_ex.vivo_noRT | Bglap2 | 4.842 | 2.57E-08 | glio |
| Bcas2_in.vivo_noRT | Gm13822 | 4.281 | 2.57E-10 | glio |
| Bcas2_in.vivo_noRT | Hdac9 | 4.541 | 2.57E-11 | glio |
| Bcas2_in.vivo_noRT | Kcnd1 | 4.285 | 2.57E-12 | glio |
| Bora_in.vivo_noRT | Nat8f4 | 4.404 | 2.57E-12 | glio |
| Hsd17b10_in.vivo_noRT | Zscan4c | 3.616 | 2.57E-12 | glio |
| Atr_in.vivo_noRT | 1810019D21R | 5.273 | 2.57E-14 | glio |

| | | | | |
|---|---|---|---|---|
| Chek1_in.vivo_noRT | Gm36356 | 4.784 | 2.57E-15 | glio |
| Bora_in.vivo_noRT | Crb3 | 4.936 | 2.57E-19 | glio |
| Chek1_in.vivo_noRT | Gm5286 | 5.044 | 2.57E-20 | glio |
| Chek1_in.vivo_noRT | Gm34776 | 5.044 | 2.57E-20 | glio |
| Atr_in.vivo_noRT | Ank1 | 5.533 | 2.57E-26 | glio |
| Atr_in.vivo_noRT | Helt | 5.533 | 2.57E-30 | glio |
| Atr_in.vivo_noRT | Gm20625 | 5.533 | 2.57E-30 | glio |
| culture_effect | Ehd2 | 0.795 | 2.58E-03 | glio |
| Interaction_genotype_Aars2 | Esm1 | -2.746 | 2.58E-03 | glio |
| Interaction_genotype_Atr | Ifi207 | -4.075 | 2.58E-03 | glio |
| Interaction_genotype_Atr | Pldi | -3.064 | 2.58E-03 | glio |
| Interaction_genotype_Bora | 2010320O07R | -3.705 | 2.58E-03 | glio |
| Interaction_genotype_Mtor | 5330426L24Ri | -3.842 | 2.58E-03 | glio |
| Interaction_genotype_Myc | Gm28404 | -3.109 | 2.58E-03 | glio |
| Interaction_genotype_Myc | Ulk3 | -3.413 | 2.58E-03 | glio |
| Interaction_genotype_Myc | Aldh5a1 | -3.132 | 2.58E-03 | glio |
| Interaction_genotype_Myc | Itpkb | -2.851 | 2.58E-03 | glio |
| Interaction_genotype_Myc | Galk2 | -2.351 | 2.58E-03 | glio |
| Interaction_genotype_Myc | Bex1 | 1.803 | 2.58E-03 | glio |
| interaction_RT | Abcf2 | -0.801 | 2.58E-03 | glio |
| interaction_RT | Pold4 | 1.095 | 2.58E-03 | glio |
| Aars2_in.vivo_noRT | Acss1 | 1.978 | 2.58E-03 | glio |
| Aars2_in.vivo_noRT | Gm49774 | 2.09 | 2.58E-03 | glio |
| Aars2_in.vivo_noRT | Eif3b | -0.902 | 2.58E-03 | glio |
| Aars2_ex.vivo_noRT | Cers3 | 2.321 | 2.58E-03 | glio |
| Atr_in.vivo_noRT | Fah | 1.769 | 2.58E-03 | glio |
| Atr_in.vivo_noRT | Hspa4l | -1.24 | 2.58E-03 | glio |
| Atr_ex.vivo_noRT | Rab15 | 2.937 | 2.58E-03 | glio |
| Bora_in.vivo_noRT | Tgfbi | 1.999 | 2.58E-03 | glio |
| Cenpj_ex.vivo_noRT | Vmn2r54 | 2.591 | 2.58E-03 | glio |
| Chek1_in.vivo_noRT | Erfe | 2.03 | 2.58E-03 | glio |
| Chek1_in.vivo_noRT | Tmod3 | -1.321 | 2.58E-03 | glio |
| Chek1_in.vivo_noRT | Ubr2 | -1.218 | 2.58E-03 | glio |
| Ercc4_ex.vivo_noRT | BC049987 | 2.68 | 2.58E-03 | glio |
| Ercc4_ex.vivo_noRT | Magea10 | 3.279 | 2.58E-03 | glio |
| culture_effect | Cln3 | -0.943 | 2.58E-04 | glio |
| culture_effect | Prkce | 0.951 | 2.58E-04 | glio |
| Interaction_genotype_Bcas2 | Baalc | -4.02 | 2.58E-04 | glio |
| Interaction_genotype_Bcas2 | AC154200.1 | -3.836 | 2.58E-04 | glio |
| Interaction_genotype_Mtor | Oaz1 | 1.413 | 2.58E-04 | glio |
| Interaction_genotype_Myc | Syp | -4.565 | 2.58E-04 | glio |
| interaction_RT | mt-Cytb | 0.643 | 2.58E-04 | glio |
| Aars2_ex.vivo_noRT | Tff2 | 3.293 | 2.58E-04 | glio |
| Aars2_ex.vivo_noRT | Fgf8 | 2.983 | 2.58E-04 | glio |
| Bora_in.vivo_noRT | C8g | 2.662 | 2.58E-04 | glio |
| Bora_in.vivo_noRT | Bmpr2 | -1.689 | 2.58E-04 | glio |
| Bora_in.vivo_noRT | Sox2 | -1.179 | 2.58E-04 | glio |
| Chek1_in.vivo_noRT | Gm10353 | 2.529 | 2.58E-04 | glio |
| Hsd17b10_in.vivo_noRT | Rem2 | 2.431 | 2.58E-04 | glio |
| Hsd17b10_in.vivo_noRT | Gm35986 | 2.568 | 2.58E-04 | glio |
| culture_effect | Strn | 1.035 | 2.58E-05 | glio |
| culture_effect | Marf1 | 0.938 | 2.58E-05 | glio |

| | | | | |
|---|---|---|---|---|
| Interaction_genotype_Mtor | Zfpm1 | -5.24 | 2.58E-05 | glio |
| Interaction_genotype_Mtor | Adgre5 | -4.233 | 2.58E-05 | glio |
| Interaction_genotype_Myc | Camk2b | -4.634 | 2.58E-05 | glio |
| Atr_ex.vivo_noRT | Esp15 | 3.164 | 2.58E-05 | glio |
| Cenpj_ex.vivo_noRT | Slc17a6 | 2.591 | 2.58E-05 | glio |
| Chek1_in.vivo_noRT | Gask1b | 3.249 | 2.58E-05 | glio |
| Hsd17b10_in.vivo_noRT | Gm20633 | 2.825 | 2.58E-05 | glio |
| Hsd17b10_in.vivo_noRT | BC030343 | 3.107 | 2.58E-05 | glio |
| Interaction_genotype_Mtor | BB287469 | -4.099 | 2.58E-06 | glio |
| Aars2_ex.vivo_noRT | Smtnl1 | 3.293 | 2.58E-06 | glio |
| Bcas2_in.vivo_noRT | Gm49864 | 3.375 | 2.58E-06 | glio |
| Bcas2_in.vivo_noRT | Sema6b | 3.331 | 2.58E-06 | glio |
| Bora_in.vivo_noRT | Gm14327 | 3.431 | 2.58E-06 | glio |
| Bora_in.vivo_noRT | Gm15510 | 3.341 | 2.58E-06 | glio |
| Bora_in.vivo_noRT | Lamc2 | 3.288 | 2.58E-06 | glio |
| Hsd17b10_in.vivo_noRT | Gm39326 | 3.353 | 2.58E-06 | glio |
| culture_effect | Abcb8 | -1.293 | 2.58E-07 | glio |
| Interaction_genotype_Mtor | Cdc26 | 2.402 | 2.58E-07 | glio |
| Atr_in.vivo_noRT | Zfp781 | 3.431 | 2.58E-07 | glio |
| Atr_ex.vivo_noRT | Tent5c | 3.554 | 2.58E-07 | glio |
| Chek1_in.vivo_noRT | Gm10101 | 3.475 | 2.58E-07 | glio |
| Chek1_in.vivo_noRT | Zfp781 | 3.453 | 2.58E-07 | glio |
| Aars2_in.vivo_noRT | Gm30262 | 4.035 | 2.58E-08 | glio |
| Bora_in.vivo_noRT | Mfap3l | 4.545 | 2.58E-09 | glio |
| Chek1_in.vivo_noRT | Notch4 | 4.31 | 2.58E-10 | glio |
| Chek1_ex.vivo_noRT | Thsd1 | 4.63 | 2.58E-10 | glio |
| Chek1_ex.vivo_noRT | 4930502E18Ri | 4.449 | 2.58E-10 | glio |
| Chek1_in.vivo_noRT | Ifitm5 | 4.521 | 2.58E-11 | glio |
| Chek1_in.vivo_noRT | D030068K23R | 4.653 | 2.58E-11 | glio |
| Hsd17b10_in.vivo_noRT | BC080695 | 3.616 | 2.58E-12 | glio |
| Chek1_in.vivo_noRT | Sh3d21 | 4.521 | 2.58E-14 | glio |
| Aars2_in.vivo_noRT | 9230112J17Ri | 5.327 | 2.58E-19 | glio |
| Chek1_in.vivo_noRT | 9430041J12Ri | 5.044 | 2.58E-19 | glio |
| Bcas2_in.vivo_noRT | AU019990 | 4.807 | 2.58E-20 | glio |
| culture_effect | mt-Nd4l | 1.817 | 2.58E-22 | glio |
| Interaction_genotype_Bora | Hrct1 | -3.505 | 2.59E-03 | glio |
| Interaction_genotype_Mtor | Orc1 | -2.934 | 2.59E-03 | glio |
| Interaction_genotype_Myc | Psmb1 | 1.199 | 2.59E-03 | glio |
| Interaction_genotype_Nat2 | D730045A05R | 4.169 | 2.59E-03 | glio |
| Aars2_in.vivo_noRT | Dpf1 | 1.896 | 2.59E-03 | glio |
| Aars2_in.vivo_noRT | 1700088E04Ri | 1.838 | 2.59E-03 | glio |
| Aars2_in.vivo_noRT | Bod1 | -1.165 | 2.59E-03 | glio |
| Aars2_in.vivo_noRT | Nus1 | -1.012 | 2.59E-03 | glio |
| Aars2_in.vivo_noRT | Naa20 | -0.969 | 2.59E-03 | glio |
| Aars2_in.vivo_noRT | Rbm3 | -0.594 | 2.59E-03 | glio |
| Bcas2_in.vivo_noRT | Slc39a8 | 2.444 | 2.59E-03 | glio |
| Bora_in.vivo_noRT | Pmpca | -1.254 | 2.59E-03 | glio |
| Cenpj_ex.vivo_noRT | Gm36026 | 2.591 | 2.59E-03 | glio |
| Cenpj_ex.vivo_noRT | Kit | 2.591 | 2.59E-03 | glio |
| Chek1_in.vivo_noRT | Ly6g6d | 2.213 | 2.59E-03 | glio |
| Chek1_in.vivo_noRT | Zfp764 | 2.051 | 2.59E-03 | glio |
| Chek1_in.vivo_noRT | Pdcd11 | -1.25 | 2.59E-03 | glio |

| | | | | |
|---|---|---|---|---|
| Chek1_in.vivo_noRT | Slc25a3 | -0.515 | 2.59E-03 | glio |
| Chek1_ex.vivo_noRT | Ptpn22 | 3.071 | 2.59E-03 | glio |
| Chek1_ex.vivo_noRT | Dzank1 | 2.851 | 2.59E-03 | glio |
| Chek1_ex.vivo_noRT | Ube2c | -2.025 | 2.59E-03 | glio |
| Hsd17b10_in.vivo_noRT | Gm48682 | 2.304 | 2.59E-03 | glio |
| Hsd17b10_in.vivo_noRT | Myoz1 | 1.921 | 2.59E-03 | glio |
| Hsd17b10_in.vivo_noRT | Brip1os | -1.449 | 2.59E-03 | glio |
| culture_effect | Tmem267 | 1.24 | 2.59E-04 | glio |
| culture_effect | Ric8a | -0.753 | 2.59E-04 | glio |
| Interaction_genotype_Bora | Flywch1 | 2.218 | 2.59E-04 | glio |
| Interaction_genotype_Mtor | A430105J06Ri | -3.959 | 2.59E-04 | glio |
| Interaction_genotype_Mtor | Sbds | 1.979 | 2.59E-04 | glio |
| Interaction_genotype_Myc | Sardh | -3.843 | 2.59E-04 | glio |
| Interaction_genotype_Myc | Smug1 | -3.018 | 2.59E-04 | glio |
| Bard1_ex.vivo_noRT | Gm5602 | 2.814 | 2.59E-04 | glio |
| Bard1_ex.vivo_noRT | Gm20513 | 2.814 | 2.59E-04 | glio |
| Bcas2_in.vivo_noRT | Gm20707 | 2.629 | 2.59E-04 | glio |
| Chek1_in.vivo_noRT | Gm4181 | 2.631 | 2.59E-04 | glio |
| Chek1_in.vivo_noRT | Gm43597 | 2.302 | 2.59E-04 | glio |
| Chek1_ex.vivo_noRT | Gpr153 | 3.025 | 2.59E-04 | glio |
| Hsd17b10_in.vivo_noRT | Aqp3 | 3.352 | 2.59E-04 | glio |
| Hsd17b10_in.vivo_noRT | Gm16853 | 2.583 | 2.59E-04 | glio |
| culture_effect | 2700038G22R | 0.994 | 2.59E-05 | glio |
| Interaction_genotype_Mtor | Trnp1 | -4.615 | 2.59E-05 | glio |
| Interaction_genotype_Myc | Loxl2 | -5.269 | 2.59E-05 | glio |
| Interaction_genotype_Myc | Gm40557 | -4.889 | 2.59E-05 | glio |
| Aars2_in.vivo_noRT | Reep5 | -1.241 | 2.59E-05 | glio |
| Bcas2_ex.vivo_noRT | Cemip | 3.54 | 2.59E-05 | glio |
| Bora_in.vivo_noRT | Ehhadh | 3.241 | 2.59E-05 | glio |
| Bora_in.vivo_noRT | Slamf1 | 3.02 | 2.59E-05 | glio |
| Bora_in.vivo_noRT | Rbm10 | -2.02 | 2.59E-05 | glio |
| Chek1_in.vivo_noRT | Gm16133 | 3.061 | 2.59E-05 | glio |
| Hsd17b10_in.vivo_noRT | B930094E09Ri | 3.1 | 2.59E-05 | glio |
| Bcas2_in.vivo_noRT | Pih1d1 | -1.952 | 2.59E-06 | glio |
| Chek1_in.vivo_noRT | Pfkfb2 | 3.407 | 2.59E-06 | glio |
| Chek1_ex.vivo_noRT | Dock4 | 4.158 | 2.59E-06 | glio |
| Hsd17b10_in.vivo_noRT | E030042O20R | 3.616 | 2.59E-06 | glio |
| culture_effect | Lman2l | -1.262 | 2.59E-07 | glio |
| Interaction_genotype_Myc | Nxph3 | -5.463 | 2.59E-07 | glio |
| Aars2_in.vivo_noRT | Gm47061 | 4.021 | 2.59E-07 | glio |
| Atp5c1_in.vivo_noRT | Gm26671 | 2.668 | 2.59E-07 | glio |
| Bora_in.vivo_noRT | Pde5a | 3.65 | 2.59E-07 | glio |
| Chek1_ex.vivo_noRT | Mcmdc2 | 3.733 | 2.59E-07 | glio |
| Hsd17b10_in.vivo_noRT | Gm15326 | 3.087 | 2.59E-09 | glio |
| Hsd17b10_in.vivo_noRT | Pdgfb | 3.353 | 2.59E-09 | glio |
| Hsd17b10_in.vivo_noRT | Minar2 | 3.356 | 2.59E-09 | glio |
| Aars2_in.vivo_noRT | Ninj2 | 4.419 | 2.59E-10 | glio |
| Atr_in.vivo_noRT | Zfp85os | 4.474 | 2.59E-10 | glio |
| Atr_in.vivo_noRT | Rtl9 | 4.092 | 2.59E-10 | glio |
| Bcas2_in.vivo_noRT | Gm4724 | 4.151 | 2.59E-11 | glio |
| Atr_in.vivo_noRT | Cyp4f40 | 4.88 | 2.59E-13 | glio |
| culture_effect | Pcdh10 | -3.509 | 2.59E-14 | glio |

| | | | | |
|---|---|---|---|---|
| Aars2_in.vivo_noRT | Pkd1l2 | 5.063 | 2.59E-15 | glio |
| Bora_in.vivo_noRT | Gm10482 | 4.414 | 2.59E-15 | glio |
| Chek1_ex.vivo_noRT | Gm31717 | 4.842 | 2.59E-16 | glio |
| Chek1_in.vivo_noRT | Spink8 | 4.784 | 2.59E-20 | glio |
| Bora_in.vivo_noRT | 1700019E08Ri | 5.46 | 2.59E-22 | glio |
| culture_effect | Pcdh17 | -2.883 | 2.59E-29 | glio |
| culture_effect | Matn2 | -3.71 | 2.59E-40 | glio |
| culture_effect | Phc2 | 0.449 | 2.60E-03 | glio |
| Interaction_genotype_Aars2 | Rpe | 1.976 | 2.60E-03 | glio |
| Interaction_genotype_Atr | Psca | -3.417 | 2.60E-03 | glio |
| Interaction_genotype_Atr | Tigd5 | -3.462 | 2.60E-03 | glio |
| Interaction_genotype_Bora | Gm14327 | -4.11 | 2.60E-03 | glio |
| Interaction_genotype_Lmo7 | Arhgef19 | -4.56 | 2.60E-03 | glio |
| Interaction_genotype_Lmo7 | Gm19276 | -3.221 | 2.60E-03 | glio |
| Interaction_genotype_Mtor | Fam174b | -4.15 | 2.60E-03 | glio |
| Interaction_genotype_Myc | Gm26549 | -2.83 | 2.60E-03 | glio |
| Interaction_genotype_Myc | Chd1l | -2.787 | 2.60E-03 | glio |
| Interaction_genotype_Myc | Dcaf5 | -2.484 | 2.60E-03 | glio |
| Interaction_genotype_Rpl3 | Nxpe3 | -3.61 | 2.60E-03 | glio |
| Interaction_genotype_Rpl3 | Padi2 | -4.347 | 2.60E-03 | glio |
| Interaction_genotype_Rpl3 | Ski | 2.443 | 2.60E-03 | glio |
| interaction_RT | Slc26a11 | 2.27 | 2.60E-03 | glio |
| interaction_RT | Alg6 | -0.808 | 2.60E-03 | glio |
| interaction_RT | Rab18 | 0.61 | 2.60E-03 | glio |
| Aars2_in.vivo_noRT | Gm48427 | 1.989 | 2.60E-03 | glio |
| Aars2_in.vivo_noRT | Irf9 | -1.487 | 2.60E-03 | glio |
| Aars2_ex.vivo_noRT | Prex2 | 3.293 | 2.60E-03 | glio |
| Atp5c1_in.vivo_noRT | 1810019D21R | 2.408 | 2.60E-03 | glio |
| Atp5c1_in.vivo_noRT | Neurl3 | 2.457 | 2.60E-03 | glio |
| Atr_in.vivo_noRT | Trp53inp2 | 1.938 | 2.60E-03 | glio |
| Atr_in.vivo_noRT | Ndufa8 | -0.765 | 2.60E-03 | glio |
| Bcas2_in.vivo_noRT | Acbd7 | 2.302 | 2.60E-03 | glio |
| Bora_in.vivo_noRT | Polr3h | -0.959 | 2.60E-03 | glio |
| Chek1_in.vivo_noRT | Kctd7 | 2.041 | 2.60E-03 | glio |
| Chek1_in.vivo_noRT | Alyref2 | 1.839 | 2.60E-03 | glio |
| Chek1_in.vivo_noRT | Specc1 | -1.068 | 2.60E-03 | glio |
| Chek1_ex.vivo_noRT | Gm16279 | 2.939 | 2.60E-03 | glio |
| Hsd17b10_in.vivo_noRT | 1810041H14R | 2.497 | 2.60E-03 | glio |
| Hsd17b10_in.vivo_noRT | 1700010B08R | 1.929 | 2.60E-03 | glio |
| culture_effect | Anapc2 | -0.707 | 2.60E-04 | glio |
| culture_effect | Ctsh | -0.648 | 2.60E-04 | glio |
| Interaction_genotype_Aars2 | Selenov | -3.804 | 2.60E-04 | glio |
| Interaction_genotype_Myc | A930019D19R | -2.729 | 2.60E-04 | glio |
| Interaction_genotype_Myc | Ttc8 | -3.856 | 2.60E-04 | glio |
| Atr_in.vivo_noRT | Sept1 | 2.807 | 2.60E-04 | glio |
| Atr_ex.vivo_noRT | Chia1 | 2.376 | 2.60E-04 | glio |
| Atr_ex.vivo_noRT | Prl2c3 | 3.113 | 2.60E-04 | glio |
| Bard1_ex.vivo_noRT | Gm12860 | 2.814 | 2.60E-04 | glio |
| Bcas2_in.vivo_noRT | 1700126G02R | 2.674 | 2.60E-04 | glio |
| Chek1_ex.vivo_noRT | Med1 | -1.598 | 2.60E-04 | glio |
| Interaction_genotype_Myc | Rxra | -4.868 | 2.60E-05 | glio |
| Interaction_genotype_Nat2 | Gm14742 | 3.605 | 2.60E-05 | glio |

| | | | | |
|---|---|---|---|---|
| Aars2_in.vivo_noRT | Wfs1 | 2.895 | 2.60E-05 | glio |
| Atr_ex.vivo_noRT | Bcl2l15 | 3.387 | 2.60E-05 | glio |
| Atr_ex.vivo_noRT | Olfml2b | 3.582 | 2.60E-05 | glio |
| Atr_ex.vivo_noRT | S100a7a | 3.462 | 2.60E-05 | glio |
| Bcas2_in.vivo_noRT | Rasl2-9 | 3.369 | 2.60E-05 | glio |
| culture_effect | Fam122a | 1.018 | 2.60E-06 | glio |
| Interaction_genotype_Mtor | Slc4a9 | -4.661 | 2.60E-06 | glio |
| Aars2_in.vivo_noRT | Zbtb46 | 3.488 | 2.60E-06 | glio |
| Bcas2_in.vivo_noRT | Zfp455 | 3.253 | 2.60E-06 | glio |
| Interaction_genotype_Myc | Ccnb1ip1 | -6.513 | 2.60E-07 | glio |
| Interaction_genotype_Myc | mt-Nd1 | 2.428 | 2.60E-07 | glio |
| Aars2_in.vivo_noRT | Psmd6 | -1.999 | 2.60E-07 | glio |
| Atr_ex.vivo_noRT | Gm364 | 3.164 | 2.60E-07 | glio |
| Chek1_ex.vivo_noRT | Olfr1191-ps1 | 4.055 | 2.60E-07 | glio |
| culture_effect | Emc8 | 0.673 | 2.60E-08 | glio |
| Interaction_genotype_Mtor | Dnm3 | -5.853 | 2.60E-08 | glio |
| Bcas2_in.vivo_noRT | Bglap2 | 3.878 | 2.60E-08 | glio |
| Bora_in.vivo_noRT | Gm34256 | 4.136 | 2.60E-08 | glio |
| Chek1_ex.vivo_noRT | Gm32122 | 4.842 | 2.60E-09 | glio |
| Bora_in.vivo_noRT | Arhgef16 | 3.9 | 2.60E-11 | glio |
| Atr_in.vivo_noRT | Gm47802 | 4.558 | 2.60E-12 | glio |
| Atr_in.vivo_noRT | Chd5 | 4.798 | 2.60E-12 | glio |
| Aars2_in.vivo_noRT | 1810019D21R | 5.067 | 2.60E-13 | glio |
| Atr_in.vivo_noRT | Gm20732 | 4.763 | 2.60E-14 | glio |
| Chek1_in.vivo_noRT | 2410141K09R | 4.78 | 2.60E-16 | glio |
| Atr_in.vivo_noRT | Gm20767 | 5.533 | 2.60E-17 | glio |
| Bora_in.vivo_noRT | Gm13348 | 4.936 | 2.60E-17 | glio |
| Chek1_in.vivo_noRT | Usp17lc | 5.044 | 2.60E-17 | glio |
| Chek1_in.vivo_noRT | Mpzl2 | 4.784 | 2.60E-17 | glio |
| culture_effect | B4galt6 | -2.541 | 2.60E-18 | glio |
| Atr_in.vivo_noRT | Lhx1os | 5.007 | 2.60E-18 | glio |
| Atr_in.vivo_noRT | 2010001K21R | 5.533 | 2.60E-18 | glio |
| Aars2_in.vivo_noRT | Gm18280 | 5.063 | 2.60E-22 | glio |
| Aars2_in.vivo_noRT | Gm5286 | 5.327 | 2.60E-22 | glio |
| Aars2_in.vivo_noRT | Gm34776 | 5.327 | 2.60E-22 | glio |
| Chek1_in.vivo_noRT | Gm47540 | 5.044 | 2.60E-22 | glio |
| Chek1_in.vivo_noRT | BB287469 | 5.044 | 2.60E-22 | glio |
| Bora_in.vivo_noRT | C330002G04R | 5.46 | 2.60E-24 | glio |
| culture_effect | Dll3 | -1.979 | 2.61E-03 | glio |
| culture_effect | Mex3a | -0.822 | 2.61E-03 | glio |
| culture_effect | Emp1 | -0.488 | 2.61E-03 | glio |
| Interaction_genotype_Aars2 | Gm45338 | -2.427 | 2.61E-03 | glio |
| Interaction_genotype_Aars2 | Gm16222 | -4.271 | 2.61E-03 | glio |
| Interaction_genotype_Atr | E030042O20R | -4.056 | 2.61E-03 | glio |
| Interaction_genotype_Atr | Mucl3 | -4.001 | 2.61E-03 | glio |
| Interaction_genotype_Atr | Gm34590 | -3.691 | 2.61E-03 | glio |
| Interaction_genotype_Atr | Dcaf12 | 1.508 | 2.61E-03 | glio |
| Interaction_genotype_Bora | Gm50321 | -3.363 | 2.61E-03 | glio |
| Interaction_genotype_Bora | Armc2 | -3.979 | 2.61E-03 | glio |
| Interaction_genotype_Bora | Acoxl | -3.721 | 2.61E-03 | glio |
| Interaction_genotype_Bora | Otos | -3.117 | 2.61E-03 | glio |
| Interaction_genotype_Bora | Utp20 | 2.097 | 2.61E-03 | glio |

| | | | | |
|---|---|---|---|---|
| Interaction_genotype_Mtor | C77080 | -3.595 | 2.61E-03 | glio |
| Interaction_genotype_Mtor | AI506816 | 1.383 | 2.61E-03 | glio |
| Interaction_genotype_Mtor | mt-Nd4 | 1.112 | 2.61E-03 | glio |
| Interaction_genotype_Myc | P4htm | -3.43 | 2.61E-03 | glio |
| Interaction_genotype_Myc | Hao1 | -3.603 | 2.61E-03 | glio |
| Interaction_genotype_Myc | 6030442K20R | -3.249 | 2.61E-03 | glio |
| Interaction_genotype_Myc | Krt8 | 5.904 | 2.61E-03 | glio |
| Interaction_genotype_Myc | 4930523C07R | -3.489 | 2.61E-03 | glio |
| Interaction_genotype_Myc | Tfb1m | -2.818 | 2.61E-03 | glio |
| Interaction_genotype_Rad51 | A430010J10Ri | -5.134 | 2.61E-03 | glio |
| Interaction_genotype_Rad51 | Matn4 | -4.034 | 2.61E-03 | glio |
| Interaction_genotype_Rad51 | Rasgrp2 | -4.286 | 2.61E-03 | glio |
| Aars2_in.vivo_noRT | Tifa | 1.923 | 2.61E-03 | glio |
| Atp5c1_ex.vivo_noRT | Ccdc125 | 2.996 | 2.61E-03 | glio |
| Atr_in.vivo_noRT | 4932422M17F | 1.97 | 2.61E-03 | glio |
| Atr_ex.vivo_noRT | Sdr16c5 | 2.769 | 2.61E-03 | glio |
| Atr_ex.vivo_noRT | Pcdhgb1 | 2.589 | 2.61E-03 | glio |
| Atr_ex.vivo_noRT | Gm8104 | 2.792 | 2.61E-03 | glio |
| Bcas2_in.vivo_noRT | Tmco1 | -0.728 | 2.61E-03 | glio |
| Bora_in.vivo_noRT | Gm4419 | 1.925 | 2.61E-03 | glio |
| Bora_in.vivo_noRT | 1700008J07Ri | 2.009 | 2.61E-03 | glio |
| Bora_in.vivo_noRT | Exoc6 | 1.838 | 2.61E-03 | glio |
| Bora_in.vivo_noRT | Fxr1 | -0.859 | 2.61E-03 | glio |
| Bora_ex.vivo_noRT | Tmem30c | 3.029 | 2.61E-03 | glio |
| culture_effect | Ggt1 | -2.247 | 2.61E-04 | glio |
| Interaction_genotype_Aars2 | 5033403F01Ri | -4.241 | 2.61E-04 | glio |
| Interaction_genotype_Aars2 | Gm15535 | -4.905 | 2.61E-04 | glio |
| Interaction_genotype_Aars2 | Gm37584 | -4.687 | 2.61E-04 | glio |
| Interaction_genotype_Aars2 | Apoa1 | -4.734 | 2.61E-04 | glio |
| Interaction_genotype_Aars2 | Gm31518 | -4.659 | 2.61E-04 | glio |
| Interaction_genotype_Myc | 1700016P04R | -3.858 | 2.61E-04 | glio |
| Interaction_genotype_Myc | Gm43608 | -4.211 | 2.61E-04 | glio |
| Interaction_genotype_Myc | Gm2449 | -3.563 | 2.61E-04 | glio |
| Interaction_genotype_Myc | Btn1a1 | -3.253 | 2.61E-04 | glio |
| Aars2_in.vivo_noRT | 2010110K18R | 2.481 | 2.61E-04 | glio |
| Aars2_ex.vivo_noRT | Col5a2 | 2.706 | 2.61E-04 | glio |
| Aars2_ex.vivo_noRT | Mterf1b | 3.315 | 2.61E-04 | glio |
| Atr_ex.vivo_noRT | Clca3a1 | 3.164 | 2.61E-04 | glio |
| Bard1_ex.vivo_noRT | Gm45151 | 2.814 | 2.61E-04 | glio |
| Bora_in.vivo_noRT | Ly6g6d | 2.639 | 2.61E-04 | glio |
| Bora_in.vivo_noRT | Spry1 | -1.653 | 2.61E-04 | glio |
| Chek1_in.vivo_noRT | Zfp583 | 2.751 | 2.61E-04 | glio |
| Chek1_in.vivo_noRT | Ucn2 | 2.55 | 2.61E-04 | glio |
| Chek1_in.vivo_noRT | Slc35f1 | 2.081 | 2.61E-04 | glio |
| Hsd17b10_in.vivo_noRT | Gm3716 | 2.698 | 2.61E-04 | glio |
| culture_effect | Nme6 | 0.663 | 2.61E-05 | glio |
| culture_effect | Timm50 | -0.653 | 2.61E-05 | glio |
| Interaction_genotype_Chek1 | Egr1 | 4.092 | 2.61E-05 | glio |
| Bcas2_in.vivo_noRT | 2510046G10R | 2.706 | 2.61E-05 | glio |
| Bcas2_in.vivo_noRT | Eef1akmt1 | -1.732 | 2.61E-05 | glio |
| Bora_in.vivo_noRT | Hdac11 | 3.052 | 2.61E-05 | glio |
| culture_effect | Gm32122 | -2.763 | 2.61E-06 | glio |

| | | | | |
|---|---|---|---|---|
| culture_effect | Per1 | -1.69 | 2.61E-06 | glio |
| Aars2_in.vivo_noRT | Pcp4l1 | 3.369 | 2.61E-06 | glio |
| Aars2_in.vivo_noRT | Hcrtr1 | 3.388 | 2.61E-06 | glio |
| Aars2_in.vivo_noRT | Slc47a2 | 3.221 | 2.61E-06 | glio |
| Atr_in.vivo_noRT | Hddc3 | 3.15 | 2.61E-06 | glio |
| Hsd17b10_in.vivo_noRT | A030005L19Ri | 3.087 | 2.61E-06 | glio |
| Hsd17b10_in.vivo_noRT | Jph4 | 3.35 | 2.61E-06 | glio |
| Interaction_genotype_Mtor | 9530062K07R | -6.463 | 2.61E-07 | glio |
| Bcas2_in.vivo_noRT | BC037032 | 3.889 | 2.61E-07 | glio |
| Bora_in.vivo_noRT | 4930405A21R | 3.347 | 2.61E-07 | glio |
| Bora_in.vivo_noRT | Gm15941 | 3.517 | 2.61E-08 | glio |
| Chek1_in.vivo_noRT | Ankrd1 | 5.044 | 2.61E-09 | glio |
| Chek1_in.vivo_noRT | Mrpl57 | -2.041 | 2.61E-09 | glio |
| Atr_in.vivo_noRT | Eng | 4.874 | 2.61E-10 | glio |
| culture_effect | Ei24 | 1.162 | 2.61E-12 | glio |
| Hsd17b10_in.vivo_noRT | Gm47540 | 3.616 | 2.61E-12 | glio |
| Hsd17b10_in.vivo_noRT | BB287469 | 3.616 | 2.61E-12 | glio |
| Bora_in.vivo_noRT | Pla2g2d | 4.673 | 2.61E-13 | glio |
| Hsd17b10_in.vivo_noRT | Gm30692 | 3.616 | 2.61E-13 | glio |
| Atr_in.vivo_noRT | Gm10696 | 5.533 | 2.61E-15 | glio |
| Atr_in.vivo_noRT | Gm15270 | 5.27 | 2.61E-19 | glio |
| Atr_in.vivo_noRT | Sytl4 | 5.267 | 2.61E-24 | glio |
| Atr_in.vivo_noRT | Gm21936 | 5.533 | 2.61E-29 | glio |
| culture_effect | Laptm4b | -2.565 | 2.61E-37 | glio |
| culture_effect | Ciita | -6.425 | 2.61E-37 | glio |
| culture_effect | Kank4os | 0.924 | 2.62E-03 | glio |
| Interaction_genotype_Atr | Cbs | -3.923 | 2.62E-03 | glio |
| Interaction_genotype_Mtor | Mindy4 | -3.539 | 2.62E-03 | glio |
| Interaction_genotype_Myc | Ccl3 | -2.469 | 2.62E-03 | glio |
| Interaction_genotype_Myc | Gm48780 | -2.469 | 2.62E-03 | glio |
| Interaction_genotype_Myc | Ston1 | -3.501 | 2.62E-03 | glio |
| Interaction_genotype_Myc | Cep83os | -3.237 | 2.62E-03 | glio |
| Interaction_genotype_Myc | Gm9844 | -3.211 | 2.62E-03 | glio |
| Interaction_genotype_Myc | Vangl2 | -2.644 | 2.62E-03 | glio |
| Interaction_genotype_Myc | Vrk2 | -2.637 | 2.62E-03 | glio |
| Interaction_genotype_Myc | Trub1 | -2.528 | 2.62E-03 | glio |
| Interaction_genotype_Rad51 | Zfp428 | 2.086 | 2.62E-03 | glio |
| interaction_RT | Slc31a1 | 0.781 | 2.62E-03 | glio |
| Aars2_in.vivo_noRT | Kbtbd7 | 1.905 | 2.62E-03 | glio |
| Aars2_in.vivo_noRT | Rexo2 | -1.069 | 2.62E-03 | glio |
| Aars2_ex.vivo_noRT | Gm15943 | 2.311 | 2.62E-03 | glio |
| Aars2_ex.vivo_noRT | Nrros | 2.899 | 2.62E-03 | glio |
| Aars2_ex.vivo_noRT | H2-T24 | 2.897 | 2.62E-03 | glio |
| Atr_in.vivo_noRT | Atrx | -0.898 | 2.62E-03 | glio |
| Bcas2_in.vivo_noRT | Ensa | -1.295 | 2.62E-03 | glio |
| Bcas2_in.vivo_noRT | Dhx9 | -1.057 | 2.62E-03 | glio |
| Bcas2_ex.vivo_noRT | Obscn | 2.742 | 2.62E-03 | glio |
| Bcas2_ex.vivo_noRT | Gm34776 | 2.348 | 2.62E-03 | glio |
| Bcas2_ex.vivo_noRT | Irf8 | 2.742 | 2.62E-03 | glio |
| Chek1_in.vivo_noRT | Znhit3 | -1.204 | 2.62E-03 | glio |
| Chek1_ex.vivo_noRT | Scn1b | 2.916 | 2.62E-03 | glio |
| Chek1_ex.vivo_noRT | Angptl6 | 1.777 | 2.62E-03 | glio |

| | | | | |
|---|---|---|---|---|
| Eif2s1_ex.vivo_noRT | Slc14a2 | 2.234 | 2.62E-03 | glio |
| Eif2s1_ex.vivo_noRT | Gm4340 | 2.234 | 2.62E-03 | glio |
| Eif2s1_ex.vivo_noRT | Gm16239 | 2.234 | 2.62E-03 | glio |
| Eif2s1_ex.vivo_noRT | Strip2 | 2.615 | 2.62E-03 | glio |
| culture_effect | Thap2 | 0.631 | 2.62E-04 | glio |
| Interaction_genotype_Mtor | Npy4r | -3.922 | 2.62E-04 | glio |
| Interaction_genotype_Mtor | Gm49579 | -3.834 | 2.62E-04 | glio |
| Interaction_genotype_Mtor | B430219N15R | -4.468 | 2.62E-04 | glio |
| Interaction_genotype_Mtor | Glt1d1 | -4.087 | 2.62E-04 | glio |
| Interaction_genotype_Mtor | Kcnk2 | -3.652 | 2.62E-04 | glio |
| Interaction_genotype_Mtor | Sept1 | -4.32 | 2.62E-04 | glio |
| Interaction_genotype_Myc | Fanca | -3.551 | 2.62E-04 | glio |
| Interaction_genotype_Myc | Zfp948 | -3.082 | 2.62E-04 | glio |
| Interaction_genotype_Myc | Itga4 | -3.191 | 2.62E-04 | glio |
| Aars2_in.vivo_noRT | Gm19391 | 2.427 | 2.62E-04 | glio |
| Aars2_in.vivo_noRT | 4931414P19R | 2.485 | 2.62E-04 | glio |
| Atr_in.vivo_noRT | Polr3h | -1.195 | 2.62E-04 | glio |
| Atr_ex.vivo_noRT | Ccdc125 | 2.764 | 2.62E-04 | glio |
| Hsd17b10_in.vivo_noRT | Gm11696 | 2.75 | 2.62E-04 | glio |
| Interaction_genotype_Bora | Smg7 | 2.514 | 2.62E-05 | glio |
| Interaction_genotype_Bora | BC005624 | 2.477 | 2.62E-05 | glio |
| Interaction_genotype_Bora | Armc1 | 2.213 | 2.62E-05 | glio |
| Interaction_genotype_Bora | Adam12 | -4.668 | 2.62E-05 | glio |
| Interaction_genotype_Bora | 5330439K02R | -5.472 | 2.62E-05 | glio |
| Interaction_genotype_Mtor | 4930518J21Ri | -4.937 | 2.62E-05 | glio |
| Interaction_genotype_Myc | Psmb3 | 2.287 | 2.62E-05 | glio |
| Aars2_in.vivo_noRT | Spata45 | 2.89 | 2.62E-05 | glio |
| Bcas2_in.vivo_noRT | Hist1h4a | 2.856 | 2.62E-05 | glio |
| Chek1_in.vivo_noRT | Rapgef3 | 3.312 | 2.62E-05 | glio |
| Interaction_genotype_Mtor | Ccdc114 | -5.437 | 2.62E-06 | glio |
| Interaction_genotype_Nat2 | AF067061 | 4.851 | 2.62E-06 | glio |
| Atr_in.vivo_noRT | Kirrel3 | 3.276 | 2.62E-06 | glio |
| Atr_in.vivo_noRT | Gm43773 | 3.112 | 2.62E-06 | glio |
| Interaction_genotype_Mtor | Gm43137 | -5.915 | 2.62E-07 | glio |
| Interaction_genotype_Mtor | Gm48008 | -5.025 | 2.62E-07 | glio |
| interaction_RT | Rin1 | -2.146 | 2.62E-07 | glio |
| Bora_in.vivo_noRT | Igsf23 | 3.809 | 2.62E-07 | glio |
| Chek1_ex.vivo_noRT | Gm10354 | 3.547 | 2.62E-07 | glio |
| culture_effect | Gm26670 | 2.342 | 2.62E-08 | glio |
| Aars2_in.vivo_noRT | Smc2os | 3.93 | 2.62E-08 | glio |
| Chek1_in.vivo_noRT | Inha | 3.92 | 2.62E-08 | glio |
| culture_effect | Bmp2k | 1.473 | 2.62E-09 | glio |
| culture_effect | Foxd2os | 1.174 | 2.62E-09 | glio |
| culture_effect | Cul4b | -0.921 | 2.62E-09 | glio |
| culture_effect | Bvht | 1.587 | 2.62E-10 | glio |
| culture_effect | Pop1 | 1.359 | 2.62E-10 | glio |
| Bcas2_in.vivo_noRT | Ptgis | 4.807 | 2.62E-12 | glio |
| Atr_in.vivo_noRT | Dcn | 5.142 | 2.62E-16 | glio |
| Chek1_in.vivo_noRT | Gm5934 | 4.78 | 2.62E-19 | glio |
| Aars2_in.vivo_noRT | Ctla4 | 5.327 | 2.62E-27 | glio |
| Aars2_in.vivo_noRT | Gjd4 | 5.327 | 2.62E-28 | glio |
| culture_effect | Tap2 | -4.113 | 2.62E-73 | glio |

| | | | | |
|---|---|---|---|---|
| culture_effect | A930001C03R | 1.708 | 2.63E-03 | glio |
| culture_effect | Tlr2 | -1.909 | 2.63E-03 | glio |
| culture_effect | Zfyve21 | -0.986 | 2.63E-03 | glio |
| culture_effect | Tmem121 | 0.708 | 2.63E-03 | glio |
| culture_effect | Tjap1 | 0.561 | 2.63E-03 | glio |
| Interaction_genotype_Atr | Gm19276 | -2.592 | 2.63E-03 | glio |
| Interaction_genotype_Atr | 1810019D21R | -3.52 | 2.63E-03 | glio |
| Interaction_genotype_Atr | Gm26812 | -3.575 | 2.63E-03 | glio |
| Interaction_genotype_Cdkn2a | Meioc | 3.098 | 2.63E-03 | glio |
| Interaction_genotype_Cdkn2a | Cldn2 | 2.835 | 2.63E-03 | glio |
| Interaction_genotype_Mtor | Napepld | -3.169 | 2.63E-03 | glio |
| Interaction_genotype_Myc | Gm50322 | -3.734 | 2.63E-03 | glio |
| Interaction_genotype_Myc | Elovl4 | -2.416 | 2.63E-03 | glio |
| Aars2_in.vivo_noRT | Rnf32 | 2.07 | 2.63E-03 | glio |
| Aars2_in.vivo_noRT | Ecpas | -1.182 | 2.63E-03 | glio |
| Aars2_in.vivo_noRT | U2surp | -0.935 | 2.63E-03 | glio |
| Aars2_in.vivo_noRT | Cdc42se1 | -1.029 | 2.63E-03 | glio |
| Aars2_in.vivo_noRT | Vdac3 | -0.816 | 2.63E-03 | glio |
| Atr_in.vivo_noRT | Zfp287 | 1.708 | 2.63E-03 | glio |
| Bcas2_in.vivo_noRT | Gm20075 | 2.108 | 2.63E-03 | glio |
| Bcas2_in.vivo_noRT | Elk3 | 1.515 | 2.63E-03 | glio |
| Bora_in.vivo_noRT | B130055M24F | 1.999 | 2.63E-03 | glio |
| Bora_ex.vivo_noRT | Sema6d | 2.22 | 2.63E-03 | glio |
| Cenpj_in.vivo_noRT | Meioc | 1.759 | 2.63E-03 | glio |
| Cenpj_in.vivo_noRT | Gm4027 | 2.022 | 2.63E-03 | glio |
| Cenpj_in.vivo_noRT | AC154200.1 | 2.022 | 2.63E-03 | glio |
| Cenpj_in.vivo_noRT | Cfap61 | -2.092 | 2.63E-03 | glio |
| Cenpj_ex.vivo_noRT | Aqp9 | 2.591 | 2.63E-03 | glio |
| Chek1_in.vivo_noRT | Cacng4 | 1.904 | 2.63E-03 | glio |
| Chek1_in.vivo_noRT | Atp13a1 | -1.422 | 2.63E-03 | glio |
| Chek1_ex.vivo_noRT | Gm12689 | 3.009 | 2.63E-03 | glio |
| Chek1_ex.vivo_noRT | Triqk | 2.716 | 2.63E-03 | glio |
| Chek1_ex.vivo_noRT | Isoc2b | 2.159 | 2.63E-03 | glio |
| Eif2s1_in.vivo_noRT | Pde8a | 1.967 | 2.63E-03 | glio |
| Interaction_genotype_Aars2 | 1700023H06R | -4.901 | 2.63E-04 | glio |
| Interaction_genotype_Mtor | Ggnbp1 | -4.175 | 2.63E-04 | glio |
| Interaction_genotype_Myc | Rbbp7 | 1.571 | 2.63E-04 | glio |
| Interaction_genotype_Myc | Aldoa | 2.032 | 2.63E-04 | glio |
| Interaction_genotype_Rpl3 | Klhdc8a | -3.894 | 2.63E-04 | glio |
| Interaction_genotype_Rpl3 | Gm28424 | -4.79 | 2.63E-04 | glio |
| interaction_RT | C1galt1 | -1.203 | 2.63E-04 | glio |
| interaction_RT | Gabarapl1 | 0.818 | 2.63E-04 | glio |
| interaction_RT | Nol7 | -0.692 | 2.63E-04 | glio |
| Aars2_in.vivo_noRT | Ppil1 | -1.399 | 2.63E-04 | glio |
| Aars2_ex.vivo_noRT | BB557941 | 2.893 | 2.63E-04 | glio |
| Atr_in.vivo_noRT | F8 | 2.395 | 2.63E-04 | glio |
| Atr_in.vivo_noRT | Srrm1 | -1.247 | 2.63E-04 | glio |
| Bcas2_in.vivo_noRT | Ubr5 | -1.245 | 2.63E-04 | glio |
| Interaction_genotype_Mtor | Gm20337 | -4.857 | 2.63E-05 | glio |
| Interaction_genotype_Mtor | Rspo3 | -5.779 | 2.63E-05 | glio |
| Interaction_genotype_Mtor | Atp6v1f | 1.897 | 2.63E-05 | glio |
| Interaction_genotype_Myc | Ecsit | -3.149 | 2.63E-05 | glio |

| | | | | |
|---|---|---|---|---|
| Aars2_in.vivo_noRT | Osbp2 | 3.401 | 2.63E-05 | glio |
| Aars2_in.vivo_noRT | Ttc38 | 2.927 | 2.63E-05 | glio |
| Aars2_in.vivo_noRT | Mrpl3 | -1.672 | 2.63E-05 | glio |
| Atr_in.vivo_noRT | Zfp799 | 2.906 | 2.63E-05 | glio |
| Atr_in.vivo_noRT | Plekhg5 | 2.826 | 2.63E-05 | glio |
| Atr_ex.vivo_noRT | Boc | 2.771 | 2.63E-05 | glio |
| Chek1_ex.vivo_noRT | Macc1 | 3.511 | 2.63E-05 | glio |
| Eif2s1_in.vivo_noRT | Cd14 | 1.903 | 2.63E-05 | glio |
| Hsd17b10_in.vivo_noRT | Zfp433 | 3.239 | 2.63E-05 | glio |
| Interaction_genotype_Mtor | Zeb2os | -5.02 | 2.63E-06 | glio |
| interaction_RT | Gem | 4.054 | 2.63E-06 | glio |
| Bora_in.vivo_noRT | Cyp4f18 | 4.676 | 2.63E-06 | glio |
| Interaction_genotype_Myc | Fxyd7 | -5.454 | 2.63E-07 | glio |
| Interaction_genotype_Myc | Carns1 | -6.082 | 2.63E-07 | glio |
| Aars2_in.vivo_noRT | AA388235 | 3.687 | 2.63E-07 | glio |
| Atr_in.vivo_noRT | Serbp1 | -1.061 | 2.63E-07 | glio |
| Atr_in.vivo_noRT | Tcea2 | 3.805 | 2.63E-08 | glio |
| Bcas2_in.vivo_noRT | Prtn3 | 4.024 | 2.63E-08 | glio |
| Bora_in.vivo_noRT | Fam189a1 | 3.266 | 2.63E-08 | glio |
| Atr_in.vivo_noRT | Gm10382 | 4.29 | 2.63E-09 | glio |
| Bcas2_in.vivo_noRT | Notch4 | 4.074 | 2.63E-09 | glio |
| Bcas2_in.vivo_noRT | Map2 | -3.001 | 2.63E-09 | glio |
| culture_effect | Lclat1 | 1.397 | 2.63E-10 | glio |
| culture_effect | Sapcd2 | 1.379 | 2.63E-11 | glio |
| Aars2_in.vivo_noRT | Gm43409 | 5.063 | 2.63E-11 | glio |
| Chek1_in.vivo_noRT | Abcg4 | 4.253 | 2.63E-11 | glio |
| culture_effect | Ltbp1 | -2.611 | 2.63E-13 | glio |
| Atr_in.vivo_noRT | D830039M14I | 5.267 | 2.63E-13 | glio |
| culture_effect | Ubb | -1.039 | 2.63E-18 | glio |
| Chek1_ex.vivo_noRT | Gm30692 | 5.57 | 2.63E-19 | glio |
| culture_effect | Hspa5 | -1.639 | 2.63E-25 | glio |
| Atr_in.vivo_noRT | Olfr56 | 5.533 | 2.63E-25 | glio |
| Interaction_genotype_Aars2 | Proscos | -3.881 | 2.64E-03 | glio |
| Interaction_genotype_Atr | Gm5129 | -3.825 | 2.64E-03 | glio |
| Interaction_genotype_Bcas2 | 2900042K21R | -3.969 | 2.64E-03 | glio |
| Interaction_genotype_Bcas2 | 5430402O13R | -4.028 | 2.64E-03 | glio |
| Interaction_genotype_Bcas2 | Gm36198 | -3.36 | 2.64E-03 | glio |
| Interaction_genotype_Bcas2 | Lncppara | -4.035 | 2.64E-03 | glio |
| Interaction_genotype_Bcas2 | Hoxa6 | -3.902 | 2.64E-03 | glio |
| Interaction_genotype_Bcas2 | Rab2b | 2.173 | 2.64E-03 | glio |
| Interaction_genotype_Bcas2 | Nfu1 | 1.586 | 2.64E-03 | glio |
| Interaction_genotype_Hsd17b | Serping1 | 3.971 | 2.64E-03 | glio |
| Interaction_genotype_Mtor | Papln | -3.519 | 2.64E-03 | glio |
| Interaction_genotype_Myc | Ddx20 | -2.39 | 2.64E-03 | glio |
| Interaction_genotype_Rad51 | C4b | 3.434 | 2.64E-03 | glio |
| Interaction_genotype_Rad51 | Brinp3 | 3.699 | 2.64E-03 | glio |
| Interaction_genotype_Rad51 | 1110038F14Ri | 1.242 | 2.64E-03 | glio |
| Interaction_genotype_Rpl3 | Gm10101 | -3.901 | 2.64E-03 | glio |
| Aars2_ex.vivo_noRT | Krtcap3 | 2.307 | 2.64E-03 | glio |
| Atr_in.vivo_noRT | 1700120C14R | 2.045 | 2.64E-03 | glio |
| Atr_in.vivo_noRT | Idua | 1.925 | 2.64E-03 | glio |
| Atr_ex.vivo_noRT | Serbp1 | -0.735 | 2.64E-03 | glio |

| | | | | |
|---|---|---|---|---|
| Bcas2_in.vivo_noRT | 2810405F17Ri | 2.121 | 2.64E-03 | glio |
| Bcas2_in.vivo_noRT | Grpel1 | -0.781 | 2.64E-03 | glio |
| Bora_in.vivo_noRT | Gm43329 | 2.062 | 2.64E-03 | glio |
| Bora_in.vivo_noRT | Gm10791 | 2.016 | 2.64E-03 | glio |
| Bora_in.vivo_noRT | Pclaf | -1.794 | 2.64E-03 | glio |
| Bora_in.vivo_noRT | Nudcd2 | -0.678 | 2.64E-03 | glio |
| Chek1_in.vivo_noRT | Adam19 | 2.251 | 2.64E-03 | glio |
| Chek1_in.vivo_noRT | Cacnb3 | 2.002 | 2.64E-03 | glio |
| Chek1_in.vivo_noRT | Rabepk | -1.319 | 2.64E-03 | glio |
| Chek1_in.vivo_noRT | Spag7 | -1.022 | 2.64E-03 | glio |
| Chek1_in.vivo_noRT | Tubb5 | -0.966 | 2.64E-03 | glio |
| Hsd17b10_in.vivo_noRT | 6330415G19R | 2.309 | 2.64E-03 | glio |
| Interaction_genotype_Mtor | C430042M11F | -4.475 | 2.64E-04 | glio |
| Interaction_genotype_Mtor | Shroom3 | -4.375 | 2.64E-04 | glio |
| Interaction_genotype_Mtor | Selenok | 1.812 | 2.64E-04 | glio |
| Interaction_genotype_Myc | Agtpbp1 | -3.272 | 2.64E-04 | glio |
| Atr_in.vivo_noRT | Ripor2 | 2.557 | 2.64E-04 | glio |
| Atr_in.vivo_noRT | Ecm1 | 2.455 | 2.64E-04 | glio |
| Atr_in.vivo_noRT | Sec11a | -0.934 | 2.64E-04 | glio |
| Bard1_ex.vivo_noRT | Akr1c18 | 2.814 | 2.64E-04 | glio |
| Bard1_ex.vivo_noRT | Minar2 | 2.814 | 2.64E-04 | glio |
| Bard1_ex.vivo_noRT | Trim17 | 2.415 | 2.64E-04 | glio |
| Bard1_ex.vivo_noRT | B230311B06R | 3.634 | 2.64E-04 | glio |
| Bcas2_in.vivo_noRT | Slc22a18 | 2.781 | 2.64E-04 | glio |
| Bora_in.vivo_noRT | D730003I15Ri | 2.001 | 2.64E-04 | glio |
| Chek1_in.vivo_noRT | Ankrd42 | 2.661 | 2.64E-04 | glio |
| Chek1_in.vivo_noRT | Tmem88 | 2.561 | 2.64E-04 | glio |
| Chek1_in.vivo_noRT | Ripor2 | 2.591 | 2.64E-04 | glio |
| Eif2s1_in.vivo_noRT | Cd36 | 2.143 | 2.64E-04 | glio |
| Eif2s1_in.vivo_noRT | Gm43701 | 2.143 | 2.64E-04 | glio |
| Eif2s1_in.vivo_noRT | 9130019O22R | 2.143 | 2.64E-04 | glio |
| Eif2s1_in.vivo_noRT | Tmem190 | 3.307 | 2.64E-04 | glio |
| Hsd17b10_in.vivo_noRT | Ly6c2 | 2.713 | 2.64E-04 | glio |
| Interaction_genotype_Atr | AC154200.1 | -4.14 | 2.64E-05 | glio |
| Bcas2_in.vivo_noRT | Fam83e | 3.087 | 2.64E-05 | glio |
| culture_effect | Phip | -0.76 | 2.64E-06 | glio |
| Interaction_genotype_Myc | Gm44190 | -4.835 | 2.64E-06 | glio |
| Interaction_genotype_Myc | Gm41442 | -5.275 | 2.64E-06 | glio |
| Interaction_genotype_Myc | 1700017M07F | -4.675 | 2.64E-06 | glio |
| Atr_in.vivo_noRT | Ifi205 | 4.222 | 2.64E-06 | glio |
| Bora_in.vivo_noRT | Gm16174 | 3.546 | 2.64E-06 | glio |
| Bora_in.vivo_noRT | 4930515G01R | 3.054 | 2.64E-06 | glio |
| Bora_ex.vivo_noRT | Olfr56 | 3.201 | 2.64E-06 | glio |
| Bora_ex.vivo_noRT | Bcl2l15 | 3.787 | 2.64E-06 | glio |
| Chek1_in.vivo_noRT | Gm50069 | 3.403 | 2.64E-06 | glio |
| Interaction_genotype_Mtor | A930002C04R | -4.239 | 2.64E-07 | glio |
| Bcas2_in.vivo_noRT | Apoa1 | 3.773 | 2.64E-07 | glio |
| Chek1_in.vivo_noRT | Gm26850 | 3.649 | 2.64E-07 | glio |
| Interaction_genotype_Chek1 | C4b | 5.703 | 2.64E-08 | glio |
| Interaction_genotype_Chek1 | Apol10b | 6.032 | 2.64E-08 | glio |
| Interaction_genotype_Chek1 | Klhdc8a | -6.257 | 2.64E-08 | glio |
| Atr_ex.vivo_noRT | Gm4027 | 3.947 | 2.64E-08 | glio |

| | | | | |
|---|---|---|---|---|
| Bora_in.vivo_noRT | 1700010B08R | 3.249 | 2.64E-08 | glio |
| Bora_in.vivo_noRT | Selp | 3.892 | 2.64E-08 | glio |
| Chek1_in.vivo_noRT | Dnase1l2 | 3.755 | 2.64E-08 | glio |
| culture_effect | Psph | -0.869 | 2.64E-09 | glio |
| Aars2_in.vivo_noRT | Sh2d3c | 4.425 | 2.64E-09 | glio |
| Aars2_in.vivo_noRT | Zfp473 | 3.902 | 2.64E-09 | glio |
| Chek1_ex.vivo_noRT | AC163347.1 | 4.06 | 2.64E-09 | glio |
| Hsd17b10_in.vivo_noRT | Gm45235 | 3.356 | 2.64E-09 | glio |
| Aars2_in.vivo_noRT | D430040D24R | 4.41 | 2.64E-10 | glio |
| Aars2_in.vivo_noRT | Gm14698 | 4.415 | 2.64E-10 | glio |
| Atr_in.vivo_noRT | B020031M17R | 5.533 | 2.64E-11 | glio |
| Chek1_in.vivo_noRT | Gm28578 | 4.784 | 2.64E-11 | glio |
| Chek1_ex.vivo_noRT | AI463229 | 4.842 | 2.64E-11 | glio |
| Bcas2_in.vivo_noRT | Adam30 | 4.428 | 2.64E-13 | glio |
| Chek1_in.vivo_noRT | Slc25a18 | 4.777 | 2.64E-16 | glio |
| Chek1_ex.vivo_noRT | Gm14742 | 4.842 | 2.64E-16 | glio |
| Bora_in.vivo_noRT | Aqp9 | 4.936 | 2.64E-17 | glio |
| Bora_in.vivo_noRT | Tcp11x2 | 5.197 | 2.64E-18 | glio |
| Aars2_in.vivo_noRT | Tcp11x2 | 5.341 | 2.64E-19 | glio |
| Aars2_in.vivo_noRT | Gm20711 | 5.327 | 2.64E-19 | glio |
| Bcas2_in.vivo_noRT | Gm49504 | 4.807 | 2.64E-19 | glio |
| Chek1_in.vivo_noRT | 1700034I23Ri | 5.044 | 2.64E-24 | glio |
| culture_effect | Gm45406 | -1.457 | 2.65E-03 | glio |
| culture_effect | H2-Q10 | -1.924 | 2.65E-03 | glio |
| culture_effect | Dipk2a | -1.187 | 2.65E-03 | glio |
| culture_effect | Smtn | 0.786 | 2.65E-03 | glio |
| culture_effect | Lsm12 | 0.36 | 2.65E-03 | glio |
| Interaction_genotype_Atr | Gm10399 | -3.136 | 2.65E-03 | glio |
| Interaction_genotype_Atr | Scamp3 | 1.576 | 2.65E-03 | glio |
| Interaction_genotype_Bcas2 | Lncbate10 | -2.852 | 2.65E-03 | glio |
| Interaction_genotype_Mtor | 6030442K20R | -3.454 | 2.65E-03 | glio |
| Interaction_genotype_Mtor | Zswim3 | -3.406 | 2.65E-03 | glio |
| Interaction_genotype_Myc | Lsm12 | 1.268 | 2.65E-03 | glio |
| Interaction_genotype_Uqcrc2 | Gm2115 | -3.906 | 2.65E-03 | glio |
| interaction_RT | D430040D24R | -2.79 | 2.65E-03 | glio |
| interaction_RT | Slc7a11 | 1.674 | 2.65E-03 | glio |
| interaction_RT | Depdc7 | 1.195 | 2.65E-03 | glio |
| interaction_RT | Nkd2 | -1.25 | 2.65E-03 | glio |
| Aars2_in.vivo_noRT | Sinhcaf | -1.11 | 2.65E-03 | glio |
| Aars2_in.vivo_noRT | Tuba1b | -0.841 | 2.65E-03 | glio |
| Bard1_ex.vivo_noRT | Ctsf | 2.421 | 2.65E-03 | glio |
| Bard1_ex.vivo_noRT | Gm19265 | 2.415 | 2.65E-03 | glio |
| Bcas2_in.vivo_noRT | Spink4 | 2.062 | 2.65E-03 | glio |
| Bora_in.vivo_noRT | Foxl2 | 1.831 | 2.65E-03 | glio |
| Bora_in.vivo_noRT | Eif4a3 | -0.881 | 2.65E-03 | glio |
| Chek1_in.vivo_noRT | Tert | 2.058 | 2.65E-03 | glio |
| Chek1_in.vivo_noRT | Pknox2 | 2.221 | 2.65E-03 | glio |
| Chek1_in.vivo_noRT | Sec24b | -1.448 | 2.65E-03 | glio |
| Chek1_ex.vivo_noRT | Rps27rt | -1.573 | 2.65E-03 | glio |
| Cyp19a1_ex.vivo_noRT | Kcnk7 | 2.959 | 2.65E-03 | glio |
| Hsd17b10_in.vivo_noRT | Zfp652os | 2.311 | 2.65E-03 | glio |
| Hsd17b10_in.vivo_noRT | Slc7a3 | 2.349 | 2.65E-03 | glio |

| | | | | |
|---|---|---|---|---|
| culture_effect | Cyth4 | 1.949 | 2.65E-04 | glio |
| Interaction_genotype_Mtor | Doc2g | -4.496 | 2.65E-04 | glio |
| Interaction_genotype_Mtor | 4930432K21R | -3.972 | 2.65E-04 | glio |
| Atp5c1_in.vivo_noRT | Pde6b | 2.416 | 2.65E-04 | glio |
| Bard1_ex.vivo_noRT | Gast | 2.814 | 2.65E-04 | glio |
| Bard1_ex.vivo_noRT | C4b | 2.872 | 2.65E-04 | glio |
| Bora_in.vivo_noRT | Gm26658 | 2.905 | 2.65E-04 | glio |
| Bora_in.vivo_noRT | A330074K22R | 2.384 | 2.65E-04 | glio |
| Hsd17b10_in.vivo_noRT | Stard6 | 2.772 | 2.65E-04 | glio |
| culture_effect | Ms4a15 | -2.104 | 2.65E-05 | glio |
| culture_effect | 9930111J21Ri | -2.114 | 2.65E-05 | glio |
| culture_effect | A630072M18F | 1.568 | 2.65E-05 | glio |
| Interaction_genotype_Myc | Gm10300 | -4.919 | 2.65E-05 | glio |
| Interaction_genotype_Myc | Gm14019 | -4.226 | 2.65E-05 | glio |
| Interaction_genotype_Myc | Rps24 | 1.649 | 2.65E-05 | glio |
| Atr_in.vivo_noRT | Sema4f | 2.94 | 2.65E-05 | glio |
| Bard1_ex.vivo_noRT | Gm12703 | 2.814 | 2.65E-05 | glio |
| Bard1_ex.vivo_noRT | Mmp13 | 2.814 | 2.65E-05 | glio |
| Bard1_ex.vivo_noRT | Obscn | 3.612 | 2.65E-05 | glio |
| Bard1_ex.vivo_noRT | Gm8016 | 2.814 | 2.65E-05 | glio |
| Bora_in.vivo_noRT | Rgs16 | 3.727 | 2.65E-05 | glio |
| Bora_in.vivo_noRT | Gm26930 | 3.247 | 2.65E-05 | glio |
| Bora_in.vivo_noRT | Nav2 | -1.891 | 2.65E-05 | glio |
| Aars2_in.vivo_noRT | Gm17249 | 3.189 | 2.65E-06 | glio |
| Atr_in.vivo_noRT | Glb1l | 3.282 | 2.65E-06 | glio |
| Bora_in.vivo_noRT | Tgfb1i1 | 3.093 | 2.65E-06 | glio |
| Aars2_ex.vivo_noRT | Gm47510 | 3.293 | 2.65E-07 | glio |
| Aars2_ex.vivo_noRT | Gm5788 | 3.293 | 2.65E-07 | glio |
| Atr_in.vivo_noRT | Gm11131 | 3.453 | 2.65E-07 | glio |
| Bora_in.vivo_noRT | Slc7a3 | 3.67 | 2.65E-07 | glio |
| Chek1_in.vivo_noRT | Ikbke | 3.727 | 2.65E-08 | glio |
| culture_effect | Nucb2 | -2.057 | 2.65E-11 | glio |
| Chek1_in.vivo_noRT | E130215H24R | 4.253 | 2.65E-11 | glio |
| Aars2_in.vivo_noRT | Cyp4f40 | 4.674 | 2.65E-12 | glio |
| Hsd17b10_in.vivo_noRT | Olfr1152 | 3.35 | 2.65E-12 | glio |
| Bora_in.vivo_noRT | 5330439K02R | 5.203 | 2.65E-13 | glio |
| Aars2_in.vivo_noRT | Gm6566 | 4.401 | 2.65E-14 | glio |
| Hsd17b10_in.vivo_noRT | Apoa5 | 3.616 | 2.65E-14 | glio |
| Atr_in.vivo_noRT | Armc2 | 5.652 | 2.65E-16 | glio |
| Aars2_in.vivo_noRT | Cfh | 4.547 | 2.65E-18 | glio |
| Aars2_in.vivo_noRT | Akap14 | 5.063 | 2.65E-20 | glio |
| Bora_in.vivo_noRT | Gm14929 | 4.936 | 2.65E-23 | glio |
| Bora_in.vivo_noRT | Gm13191 | 4.936 | 2.65E-26 | glio |
| Bora_in.vivo_noRT | 4933416E03Ri | 4.936 | 2.65E-26 | glio |
| Interaction_genotype_Atr | Lrp2bp | -3.401 | 2.66E-03 | glio |
| Interaction_genotype_Mtor | Rps7 | 1.033 | 2.66E-03 | glio |
| Interaction_genotype_Mtor | Rps16 | 1.035 | 2.66E-03 | glio |
| Interaction_genotype_Myc | Zfp458 | -3.508 | 2.66E-03 | glio |
| Interaction_genotype_Myc | Mcoln1 | -2.629 | 2.66E-03 | glio |
| Interaction_genotype_Rpl3 | Calhm5 | -3.955 | 2.66E-03 | glio |
| Interaction_genotype_Rpl3 | Col8a1 | -4.055 | 2.66E-03 | glio |
| Interaction_genotype_Rpl3 | Gm19276 | -2.931 | 2.66E-03 | glio |

| | | | | |
|---|---|---|---|---|
| Interaction_genotype_Rpl3 | Nat8l | -2.996 | 2.66E-03 | glio |
| interaction_RT | Wdr70 | 0.698 | 2.66E-03 | glio |
| Aars2_in.vivo_noRT | Gnpnat1 | -1.176 | 2.66E-03 | glio |
| Aars2_ex.vivo_noRT | Prdm1 | 2.899 | 2.66E-03 | glio |
| Atr_in.vivo_noRT | St5 | 1.896 | 2.66E-03 | glio |
| Atr_in.vivo_noRT | Itprid2 | 1.466 | 2.66E-03 | glio |
| Bcas2_in.vivo_noRT | Gm14443 | 2.074 | 2.66E-03 | glio |
| Bcas2_in.vivo_noRT | Zcrb1 | -0.629 | 2.66E-03 | glio |
| Bora_in.vivo_noRT | Det1 | 1.667 | 2.66E-03 | glio |
| Chek1_in.vivo_noRT | Pars2 | 2.068 | 2.66E-03 | glio |
| Chek1_in.vivo_noRT | Eif4b | -0.826 | 2.66E-03 | glio |
| Chek1_ex.vivo_noRT | Ldlr | -2.158 | 2.66E-03 | glio |
| Eif2s1_in.vivo_noRT | Tdrd5 | 2.284 | 2.66E-03 | glio |
| Eif2s1_in.vivo_noRT | Gm5934 | 1.88 | 2.66E-03 | glio |
| Eif2s1_in.vivo_noRT | Rtn1 | 2.551 | 2.66E-03 | glio |
| Eif2s1_in.vivo_noRT | 4930579C12R | 2.143 | 2.66E-03 | glio |
| Eif2s1_in.vivo_noRT | AF067061 | 2.143 | 2.66E-03 | glio |
| Eif2s1_in.vivo_noRT | Magea10 | 2.207 | 2.66E-03 | glio |
| Hsd17b10_in.vivo_noRT | Ltb4r1 | 2.441 | 2.66E-03 | glio |
| culture_effect | Elfn2 | -2.134 | 2.66E-04 | glio |
| culture_effect | Glrp1 | -2.014 | 2.66E-04 | glio |
| Interaction_genotype_Mtor | Gm17477 | -4.039 | 2.66E-04 | glio |
| Interaction_genotype_Mtor | Pon3 | -4.496 | 2.66E-04 | glio |
| Interaction_genotype_Mtor | Polr2f | 1.585 | 2.66E-04 | glio |
| Interaction_genotype_Myc | Cxcl16 | -3.952 | 2.66E-04 | glio |
| Interaction_genotype_Myc | Gm11491 | -4.2 | 2.66E-04 | glio |
| Interaction_genotype_Myc | Wdr78 | -4.484 | 2.66E-04 | glio |
| Interaction_genotype_Myc | Uqcr10 | 1.574 | 2.66E-04 | glio |
| Atr_in.vivo_noRT | Bmerb1 | 2.761 | 2.66E-04 | glio |
| Bcas2_in.vivo_noRT | Fhdc1 | 2.788 | 2.66E-04 | glio |
| Bora_in.vivo_noRT | Osbp2 | 3.011 | 2.66E-04 | glio |
| Bora_in.vivo_noRT | H2-T24 | 2.478 | 2.66E-04 | glio |
| Bora_in.vivo_noRT | Bud23 | -1.514 | 2.66E-04 | glio |
| Chek1_ex.vivo_noRT | Rspo1 | 3.076 | 2.66E-04 | glio |
| Chek1_ex.vivo_noRT | Gm46652 | 3.078 | 2.66E-04 | glio |
| Eif2s1_in.vivo_noRT | Gm36037 | 2.143 | 2.66E-04 | glio |
| Eif2s1_in.vivo_noRT | BC026762 | 2.818 | 2.66E-04 | glio |
| Hsd17b10_in.vivo_noRT | Gm49694 | 2.89 | 2.66E-04 | glio |
| Interaction_genotype_Mtor | Tcte2 | -4.189 | 2.66E-05 | glio |
| Atr_in.vivo_noRT | Gm29394 | 2.823 | 2.66E-05 | glio |
| Atr_in.vivo_noRT | Gm20721 | 2.743 | 2.66E-05 | glio |
| Atr_in.vivo_noRT | Efnb1 | 2.656 | 2.66E-05 | glio |
| Bcas2_in.vivo_noRT | Ccnb1ip1 | 3.317 | 2.66E-05 | glio |
| culture_effect | 2700046G09R | 1.59 | 2.66E-06 | glio |
| Bcas2_in.vivo_noRT | Alox5ap | 3.701 | 2.66E-06 | glio |
| Bcas2_in.vivo_noRT | D930030I03Ri | 3.568 | 2.66E-06 | glio |
| Bora_in.vivo_noRT | 4732471J01Ri | 3.378 | 2.66E-06 | glio |
| Interaction_genotype_Mtor | A930019D19R | -4.099 | 2.66E-07 | glio |
| Aars2_in.vivo_noRT | Gm15265 | 3.525 | 2.66E-07 | glio |
| Bcas2_in.vivo_noRT | Cyp27b1 | 3.246 | 2.66E-07 | glio |
| Hsd17b10_in.vivo_noRT | Angptl7 | 2.75 | 2.66E-08 | glio |
| Hsd17b10_in.vivo_noRT | Pldi | 3.616 | 2.66E-10 | glio |

| | | | | |
|---|---|---|---|---|
| culture_effect | Osgep | -1.25 | 2.66E-12 | glio |
| culture_effect | Psmb1 | -0.788 | 2.66E-12 | glio |
| Chek1_in.vivo_noRT | Mecom | 5.044 | 2.66E-14 | glio |
| Aars2_in.vivo_noRT | Gm34885 | 5.603 | 2.66E-15 | glio |
| Atr_in.vivo_noRT | Ppil6 | 5.533 | 2.66E-18 | glio |
| Bcas2_in.vivo_noRT | Gm34939 | 5.061 | 2.66E-18 | glio |
| Chek1_in.vivo_noRT | Gm31063 | 5.044 | 2.66E-21 | glio |
| Interaction_genotype_Atr | 2700081O15R | 2.066 | 2.67E-03 | glio |
| Interaction_genotype_Mtor | Rfx5 | -2.647 | 2.67E-03 | glio |
| Interaction_genotype_Mtor | Slc35b1 | 1.674 | 2.67E-03 | glio |
| Interaction_genotype_Myc | Spink8 | -2.469 | 2.67E-03 | glio |
| Interaction_genotype_Myc | Ppnr | -3.428 | 2.67E-03 | glio |
| Interaction_genotype_Myc | A930003O13R | -3.697 | 2.67E-03 | glio |
| Interaction_genotype_Myc | Aars2 | -3.276 | 2.67E-03 | glio |
| Interaction_genotype_Myc | Gm32618 | -3.36 | 2.67E-03 | glio |
| Interaction_genotype_Myc | Gm12905 | -3.475 | 2.67E-03 | glio |
| Interaction_genotype_Myc | Polh | -2.592 | 2.67E-03 | glio |
| Interaction_genotype_Pars2 | Rassf6 | -3.355 | 2.67E-03 | glio |
| Interaction_genotype_Rad51 | Noc2l | 1.892 | 2.67E-03 | glio |
| Interaction_genotype_Uqcrc2 | D230022J07Ri | -4.536 | 2.67E-03 | glio |
| Aars2_in.vivo_noRT | Grk4 | 2.11 | 2.67E-03 | glio |
| Aars2_in.vivo_noRT | B4galt7 | 1.742 | 2.67E-03 | glio |
| Aars2_in.vivo_noRT | Tbc1d20 | -1.378 | 2.67E-03 | glio |
| Aars2_in.vivo_noRT | Arpc5 | -0.787 | 2.67E-03 | glio |
| Aars2_ex.vivo_noRT | Ms4a6d | 2.505 | 2.67E-03 | glio |
| Atp5c1_in.vivo_noRT | Slc7a4 | 2.316 | 2.67E-03 | glio |
| Atp5c1_in.vivo_noRT | Tinagl1 | -1.008 | 2.67E-03 | glio |
| Atr_in.vivo_noRT | Gm50240 | 1.699 | 2.67E-03 | glio |
| Atr_ex.vivo_noRT | Impg2 | 2.376 | 2.67E-03 | glio |
| Atr_ex.vivo_noRT | Grm1 | 2.586 | 2.67E-03 | glio |
| Atr_ex.vivo_noRT | Gm35696 | 2.696 | 2.67E-03 | glio |
| Atr_ex.vivo_noRT | Mog | 3.164 | 2.67E-03 | glio |
| Atr_ex.vivo_noRT | Zdhhc13 | -2.074 | 2.67E-03 | glio |
| Bora_in.vivo_noRT | Ell2 | 1.975 | 2.67E-03 | glio |
| Bora_in.vivo_noRT | Mrps5 | -1.254 | 2.67E-03 | glio |
| Cenpj_ex.vivo_noRT | Pcdhb19 | 2.994 | 2.67E-03 | glio |
| Chek1_in.vivo_noRT | Dpp7 | 1.818 | 2.67E-03 | glio |
| Chek1_in.vivo_noRT | Crppa | 1.846 | 2.67E-03 | glio |
| Chek1_in.vivo_noRT | Bcap31 | -0.853 | 2.67E-03 | glio |
| Chek1_ex.vivo_noRT | Nasp | -1.018 | 2.67E-03 | glio |
| Eif2s1_ex.vivo_noRT | Hoxc8 | 2.234 | 2.67E-03 | glio |
| culture_effect | Gm26581 | 2.07 | 2.67E-04 | glio |
| culture_effect | Ubtd1 | -0.732 | 2.67E-04 | glio |
| Interaction_genotype_Mtor | E030042O20R | -4.624 | 2.67E-04 | glio |
| Interaction_genotype_Myc | Inca1 | -3.815 | 2.67E-04 | glio |
| Interaction_genotype_Myc | A630001G21R | -3.304 | 2.67E-04 | glio |
| Interaction_genotype_Myc | Srsf7 | 1.664 | 2.67E-04 | glio |
| Aars2_in.vivo_noRT | Tmem97 | -1.421 | 2.67E-04 | glio |
| Atr_in.vivo_noRT | Rbm26 | -1.579 | 2.67E-04 | glio |
| Chek1_in.vivo_noRT | Mcm2 | -1.756 | 2.67E-04 | glio |
| Eif2s1_in.vivo_noRT | Gm12800 | 2.143 | 2.67E-04 | glio |
| Interaction_genotype_Myc | Synm | -4.315 | 2.67E-05 | glio |

| | | | | |
|---|---|---|---|---|
| Interaction_genotype_Myc | Zfp870 | -4.138 | 2.67E-05 | glio |
| Interaction_genotype_Myc | 2410080I02Ri | -4.598 | 2.67E-05 | glio |
| Interaction_genotype_Myc | 4930412C18R | -4.878 | 2.67E-05 | glio |
| Interaction_genotype_Myc | 1700015F17Ri | -4.781 | 2.67E-05 | glio |
| Interaction_genotype_Myc | Vgll3 | -3.826 | 2.67E-05 | glio |
| Interaction_genotype_Myc | Brk1 | 1.529 | 2.67E-05 | glio |
| Aars2_in.vivo_noRT | Atp5b | -0.78 | 2.67E-05 | glio |
| Atr_ex.vivo_noRT | Scd3 | 4.033 | 2.67E-05 | glio |
| Bora_ex.vivo_noRT | 2300002M23F | 3.201 | 2.67E-05 | glio |
| Interaction_genotype_Myc | Gm34047 | -5.436 | 2.67E-06 | glio |
| Aars2_in.vivo_noRT | 9430065F17Ri | 3.351 | 2.67E-06 | glio |
| Bora_in.vivo_noRT | Zfp433 | 3.497 | 2.67E-06 | glio |
| culture_effect | Gm38534 | 2.742 | 2.67E-07 | glio |
| culture_effect | Hint2 | -0.813 | 2.67E-07 | glio |
| culture_effect | Cfap20 | -0.623 | 2.67E-07 | glio |
| Interaction_genotype_Myc | Magi2 | -5.755 | 2.67E-07 | glio |
| Interaction_genotype_Myc | D030040B21R | -5.628 | 2.67E-07 | glio |
| Interaction_genotype_Myc | Gm43672 | -6.11 | 2.67E-07 | glio |
| Bcas2_in.vivo_noRT | 4930545E07Ri | 3.773 | 2.67E-07 | glio |
| Bora_in.vivo_noRT | Tcam1 | 3.747 | 2.67E-07 | glio |
| Chek1_in.vivo_noRT | Gm46447 | 3.233 | 2.67E-07 | glio |
| Bcas2_in.vivo_noRT | Gm11696 | 3.941 | 2.67E-08 | glio |
| Interaction_genotype_Mtor | Mtor | -5.051 | 2.67E-09 | glio |
| Bcas2_in.vivo_noRT | Gm28578 | 4.547 | 2.67E-10 | glio |
| Bcas2_in.vivo_noRT | Gm7102 | 4.547 | 2.67E-10 | glio |
| Chek1_in.vivo_noRT | Lama5 | 4.522 | 2.67E-10 | glio |
| Chek1_in.vivo_noRT | Gm13054 | 4.518 | 2.67E-13 | glio |
| culture_effect | Casp12 | -2.143 | 2.67E-15 | glio |
| Bora_in.vivo_noRT | 4930588A03R | 4.936 | 2.67E-24 | glio |
| Aars2_in.vivo_noRT | Zscan4-ps3 | 5.327 | 2.67E-27 | glio |
| culture_effect | H2-Q4 | -3.672 | 2.67E-34 | glio |
| culture_effect | Slc16a7 | -1.646 | 2.68E-03 | glio |
| culture_effect | Gm41318 | 1.778 | 2.68E-03 | glio |
| culture_effect | A930033H14R | 1.657 | 2.68E-03 | glio |
| Interaction_genotype_Aars2 | Tal2 | -2.752 | 2.68E-03 | glio |
| Interaction_genotype_Bcas2 | Gm32585 | -4.214 | 2.68E-03 | glio |
| Interaction_genotype_Bcas2 | Gm20455 | -4.19 | 2.68E-03 | glio |
| Interaction_genotype_Cdkn2a | Spi1 | 4.023 | 2.68E-03 | glio |
| Interaction_genotype_Cdkn2a | Stfa3 | 2.835 | 2.68E-03 | glio |
| Interaction_genotype_Cdkn2a | Olfr787 | 2.835 | 2.68E-03 | glio |
| Interaction_genotype_Cdkn2a | Gm13032 | 3.621 | 2.68E-03 | glio |
| Interaction_genotype_Mtor | Hhipl2 | -4.045 | 2.68E-03 | glio |
| Interaction_genotype_Mtor | 1810030O07R | -2.294 | 2.68E-03 | glio |
| Interaction_genotype_Myc | Gm13032 | -2.743 | 2.68E-03 | glio |
| Interaction_genotype_Myc | Amd2 | -3.373 | 2.68E-03 | glio |
| Interaction_genotype_Myc | A130051J06Ri | -3.315 | 2.68E-03 | glio |
| Interaction_genotype_Myc | Arhgef10 | -2.886 | 2.68E-03 | glio |
| Interaction_genotype_Myc | Lrrcc1 | -2.697 | 2.68E-03 | glio |
| Interaction_genotype_Myc | Pcyt2 | -2.384 | 2.68E-03 | glio |
| Interaction_genotype_Myc | Ier5l | -2.599 | 2.68E-03 | glio |
| Interaction_genotype_Myc | Eef1a1 | 1.021 | 2.68E-03 | glio |
| Interaction_genotype_Rad21 | Dio1 | 4.056 | 2.68E-03 | glio |

| | | | | |
|---|---|---|---|---|
| Aars2_in.vivo_noRT | Fam220a.1 | 2.079 | 2.68E-03 | glio |
| Aars2_in.vivo_noRT | Cep97 | 1.625 | 2.68E-03 | glio |
| Atr_in.vivo_noRT | Pars2 | 2.034 | 2.68E-03 | glio |
| Atr_ex.vivo_noRT | Ier2 | 1.585 | 2.68E-03 | glio |
| Bcas2_in.vivo_noRT | Efs | -1.677 | 2.68E-03 | glio |
| Bcas2_in.vivo_noRT | Naa35 | -1.354 | 2.68E-03 | glio |
| Cenpj_in.vivo_noRT | Vwce | 2.292 | 2.68E-03 | glio |
| Cenpj_in.vivo_noRT | Rab25 | 2.282 | 2.68E-03 | glio |
| Cenpj_in.vivo_noRT | Gm13713 | 2.019 | 2.68E-03 | glio |
| Cenpj_in.vivo_noRT | Sparcl1 | -1.484 | 2.68E-03 | glio |
| Chek1_in.vivo_noRT | Dyrk4 | 2.476 | 2.68E-03 | glio |
| Chek1_in.vivo_noRT | Lncppara | 2.37 | 2.68E-03 | glio |
| Eif2s1_in.vivo_noRT | E030042O20R | 2.674 | 2.68E-03 | glio |
| Hsd17b10_in.vivo_noRT | Pbld1 | 2.429 | 2.68E-03 | glio |
| Hsd17b10_in.vivo_noRT | 4921523L03Ri | 2.448 | 2.68E-03 | glio |
| Hsd17b10_in.vivo_noRT | 1700124L16Ri | 2.382 | 2.68E-03 | glio |
| Hsd17b10_in.vivo_noRT | Olfr267 | 2.363 | 2.68E-03 | glio |
| Hsd17b10_in.vivo_noRT | Arf2 | -1.585 | 2.68E-03 | glio |
| Interaction_genotype_Atr | Lama5 | -4.688 | 2.68E-04 | glio |
| Interaction_genotype_Mtor | Itga2b | -4.187 | 2.68E-04 | glio |
| Interaction_genotype_Mtor | Gm11273 | -3.733 | 2.68E-04 | glio |
| Interaction_genotype_Mtor | Gm14167 | -3.7 | 2.68E-04 | glio |
| Interaction_genotype_Myc | Ly6e | 2.298 | 2.68E-04 | glio |
| Aars2_in.vivo_noRT | Sbno1 | -1.316 | 2.68E-04 | glio |
| Atp5c1_in.vivo_noRT | Card9 | 2.404 | 2.68E-04 | glio |
| Atp5c1_in.vivo_noRT | Gm2449 | 2.405 | 2.68E-04 | glio |
| Atp5c1_in.vivo_noRT | Prm3 | 2.455 | 2.68E-04 | glio |
| Atp5c1_in.vivo_noRT | Amdhd1 | 2.687 | 2.68E-04 | glio |
| Atr_in.vivo_noRT | Plekhh1 | 2.655 | 2.68E-04 | glio |
| Atr_in.vivo_noRT | Mus81 | 2.009 | 2.68E-04 | glio |
| Bcas2_in.vivo_noRT | Naip6 | 2.551 | 2.68E-04 | glio |
| Bcas2_ex.vivo_noRT | Gm49808 | 2.349 | 2.68E-04 | glio |
| Bora_in.vivo_noRT | Cnn1 | 3.266 | 2.68E-04 | glio |
| Bora_in.vivo_noRT | Tead2 | -1.822 | 2.68E-04 | glio |
| Chek1_in.vivo_noRT | Hmgcs2 | 2.96 | 2.68E-04 | glio |
| Chek1_in.vivo_noRT | Rbp7 | 2.651 | 2.68E-04 | glio |
| Chek1_in.vivo_noRT | Catsperz | 2.569 | 2.68E-04 | glio |
| Chek1_in.vivo_noRT | Zfp169 | 2.503 | 2.68E-04 | glio |
| Chek1_in.vivo_noRT | C230014O12R | 2.3 | 2.68E-04 | glio |
| Chek1_in.vivo_noRT | Ssbp1 | -0.835 | 2.68E-04 | glio |
| Chek1_ex.vivo_noRT | Mrpl20 | -1.054 | 2.68E-04 | glio |
| Eif2s1_in.vivo_noRT | Trmt61b | 2.143 | 2.68E-04 | glio |
| Hsd17b10_in.vivo_noRT | Il17b | 2.966 | 2.68E-04 | glio |
| culture_effect | Csdc2 | 1.259 | 2.68E-05 | glio |
| Interaction_genotype_Mtor | Fam117b | -3.187 | 2.68E-05 | glio |
| Interaction_genotype_Mtor | Ddx50 | 2.272 | 2.68E-05 | glio |
| Aars2_in.vivo_noRT | Gm43813 | 2.771 | 2.68E-05 | glio |
| Atr_ex.vivo_noRT | Lrguk | 3.164 | 2.68E-05 | glio |
| Atr_ex.vivo_noRT | Gm17203 | 2.764 | 2.68E-05 | glio |
| Atr_ex.vivo_noRT | Gm20275 | 3.164 | 2.68E-05 | glio |
| Atr_ex.vivo_noRT | Ly6g6c | 3.554 | 2.68E-05 | glio |
| Bcas2_in.vivo_noRT | Ap3m1 | -1.896 | 2.68E-05 | glio |

| | | | | |
|---|---|---|---|---|
| Bcas2_in.vivo_noRT | Erf | -1.769 | 2.68E-05 | glio |
| Bora_in.vivo_noRT | Ccna2 | -2.394 | 2.68E-05 | glio |
| Bora_ex.vivo_noRT | Boc | 2.808 | 2.68E-05 | glio |
| Bora_in.vivo_noRT | 1700029J07Ri | 3.379 | 2.68E-06 | glio |
| Chek1_in.vivo_noRT | Nkx2-2os | 3.388 | 2.68E-06 | glio |
| Atr_in.vivo_noRT | Gm37986 | 3.196 | 2.68E-07 | glio |
| Atr_in.vivo_noRT | 5730414N17R | 3.801 | 2.68E-07 | glio |
| Chek1_in.vivo_noRT | Gm43330 | 3.144 | 2.68E-07 | glio |
| Chek1_in.vivo_noRT | Mettl18 | 3.055 | 2.68E-07 | glio |
| Chek1_ex.vivo_noRT | Cldn14 | 4.453 | 2.68E-07 | glio |
| Atr_in.vivo_noRT | Gm19684 | 4.084 | 2.68E-08 | glio |
| Atr_in.vivo_noRT | Limch1 | 4.495 | 2.68E-09 | glio |
| Atr_in.vivo_noRT | Cxcr2 | 4.221 | 2.68E-10 | glio |
| Bcas2_in.vivo_noRT | Pde6b | 3.775 | 2.68E-10 | glio |
| Aars2_in.vivo_noRT | D6Ertd527e | 4.544 | 2.68E-11 | glio |
| Bcas2_in.vivo_noRT | Klhdc8a | 4.807 | 2.68E-12 | glio |
| Atr_in.vivo_noRT | Dmrtc1a | 5.533 | 2.68E-24 | glio |
| Atr_in.vivo_noRT | Gm12589 | 5.533 | 2.68E-24 | glio |
| culture_effect | Tent5c | -1.467 | 2.69E-03 | glio |
| Interaction_genotype_Aars2 | Sdhd | 1.447 | 2.69E-03 | glio |
| Interaction_genotype_Bcas2 | Nxnl2 | -3.701 | 2.69E-03 | glio |
| Interaction_genotype_Chek1 | 4930589L23Ri | -4.168 | 2.69E-03 | glio |
| Interaction_genotype_Chek1 | BC065403 | -3.833 | 2.69E-03 | glio |
| Interaction_genotype_Myc | Vtn | -2.469 | 2.69E-03 | glio |
| Interaction_genotype_Myc | Rhox3g | -3.304 | 2.69E-03 | glio |
| Interaction_genotype_Myc | F630040K05Ri | -3.015 | 2.69E-03 | glio |
| Interaction_genotype_Myc | Csgalnact2 | -2.808 | 2.69E-03 | glio |
| Interaction_genotype_Myc | Hist1h3f | -3.437 | 2.69E-03 | glio |
| Interaction_genotype_Myc | Upp2 | -2.921 | 2.69E-03 | glio |
| Aars2_in.vivo_noRT | Fendrr | 2.156 | 2.69E-03 | glio |
| Aars2_in.vivo_noRT | Gab3 | 2.152 | 2.69E-03 | glio |
| Aars2_in.vivo_noRT | Cox18 | -1.238 | 2.69E-03 | glio |
| Atr_in.vivo_noRT | Gm20075 | 2.056 | 2.69E-03 | glio |
| Atr_in.vivo_noRT | Apol7a | 1.884 | 2.69E-03 | glio |
| Bcas2_in.vivo_noRT | B630019K06R | 1.816 | 2.69E-03 | glio |
| Bora_in.vivo_noRT | Slc46a2 | 1.955 | 2.69E-03 | glio |
| Cenpj_in.vivo_noRT | Spink6 | 2.022 | 2.69E-03 | glio |
| Cenpj_in.vivo_noRT | 2810410L24Ri | 2.551 | 2.69E-03 | glio |
| Chek1_ex.vivo_noRT | Ntf5 | 2.855 | 2.69E-03 | glio |
| Chek1_ex.vivo_noRT | Hist1h4b | 2.919 | 2.69E-03 | glio |
| Chek1_ex.vivo_noRT | Ebna1bp2 | -0.86 | 2.69E-03 | glio |
| Hsd17b10_in.vivo_noRT | Wars | -1.399 | 2.69E-03 | glio |
| Interaction_genotype_Atr | Krt80 | -3.952 | 2.69E-04 | glio |
| Interaction_genotype_Atr | Gm49086 | -3.932 | 2.69E-04 | glio |
| Interaction_genotype_Chek1 | Ddit3 | 2.319 | 2.69E-04 | glio |
| Interaction_genotype_Mtor | Plxna2 | -4.802 | 2.69E-04 | glio |
| Interaction_genotype_Mtor | Gm26877 | -4.361 | 2.69E-04 | glio |
| Interaction_genotype_Mtor | Mpp2 | -4.237 | 2.69E-04 | glio |
| Interaction_genotype_Mtor | 0610038B21R | -3.636 | 2.69E-04 | glio |
| Atr_in.vivo_noRT | Tmem82 | 2.772 | 2.69E-04 | glio |
| Atr_in.vivo_noRT | Anxa4 | -1.524 | 2.69E-04 | glio |
| Bcas2_ex.vivo_noRT | Tkfc | -2.584 | 2.69E-04 | glio |

| | | | | |
|---|---|---|---|---|
| Bora_ex.vivo_noRT | Lncbate10 | 2.414 | 2.69E-04 | glio |
| culture_effect | Akap6 | 1.465 | 2.69E-05 | glio |
| culture_effect | Ern1 | -0.942 | 2.69E-05 | glio |
| culture_effect | Ndel1 | -0.707 | 2.69E-05 | glio |
| interaction_RT | Nat8l | -2.471 | 2.69E-05 | glio |
| Aars2_ex.vivo_noRT | 1700047M11F | 3.699 | 2.69E-05 | glio |
| Aars2_ex.vivo_noRT | Hrh2 | 3.293 | 2.69E-05 | glio |
| Atr_in.vivo_noRT | Cela1 | -1.951 | 2.69E-05 | glio |
| Atr_in.vivo_noRT | Gatad2b | -1.559 | 2.69E-05 | glio |
| Bcas2_in.vivo_noRT | Lrrc75b | 3.15 | 2.69E-05 | glio |
| Bcas2_in.vivo_noRT | 4833418N02R | 3.105 | 2.69E-05 | glio |
| Bora_in.vivo_noRT | C430014B12R | 2.887 | 2.69E-05 | glio |
| Chek1_ex.vivo_noRT | 9030612E09Ri | 3.475 | 2.69E-05 | glio |
| Hsd17b10_in.vivo_noRT | 9530062K07R | 3.237 | 2.69E-05 | glio |
| Hsd17b10_in.vivo_noRT | Kcnk7 | 2.568 | 2.69E-05 | glio |
| Hsd17b10_in.vivo_noRT | Cidea | 2.701 | 2.69E-05 | glio |
| Interaction_genotype_Myc | Gm44752 | -4.905 | 2.69E-06 | glio |
| Atr_in.vivo_noRT | A430010J10Ri | 4.027 | 2.69E-06 | glio |
| Atr_in.vivo_noRT | 1700041G16R | 3.279 | 2.69E-06 | glio |
| Bcas2_in.vivo_noRT | Gm10516 | 3.243 | 2.69E-06 | glio |
| Bcas2_in.vivo_noRT | Gm7008 | 3.341 | 2.69E-06 | glio |
| Bcas2_in.vivo_noRT | Trnp1 | 3.365 | 2.69E-06 | glio |
| Bcas2_in.vivo_noRT | Hyal3 | 3.747 | 2.69E-06 | glio |
| Bora_ex.vivo_noRT | Gm48215 | 3.201 | 2.69E-06 | glio |
| Bora_in.vivo_noRT | Gm45441 | 3.877 | 2.69E-08 | glio |
| Bcas2_in.vivo_noRT | A530013C23R | 3.889 | 2.69E-09 | glio |
| Bora_in.vivo_noRT | Prox2os | 3.632 | 2.69E-09 | glio |
| Chek1_in.vivo_noRT | Ly6c2 | 4.141 | 2.69E-09 | glio |
| Aars2_in.vivo_noRT | Rem2 | 4.142 | 2.69E-11 | glio |
| culture_effect | Hmcn1 | -3.756 | 2.69E-12 | glio |
| Bcas2_in.vivo_noRT | Rilp | 5.325 | 2.69E-12 | glio |
| culture_effect | Pls3 | 1.73 | 2.69E-13 | glio |
| Bcas2_in.vivo_noRT | Npr3 | 4.807 | 2.69E-15 | glio |
| Bora_in.vivo_noRT | Gm34059 | 4.936 | 2.69E-25 | glio |
| Atr_in.vivo_noRT | Gm5157 | 5.533 | 2.69E-30 | glio |
| culture_effect | Tox2 | -1.922 | 2.70E-03 | glio |
| Interaction_genotype_Aars2 | Hcn3 | -3.43 | 2.70E-03 | glio |
| Interaction_genotype_Aars2 | Atxn2l | 1.564 | 2.70E-03 | glio |
| Interaction_genotype_Bcas2 | 6430562O15R | -3.651 | 2.70E-03 | glio |
| Interaction_genotype_Bcas2 | Styk1 | -4.073 | 2.70E-03 | glio |
| Interaction_genotype_Bora | A430106G13R | -4.008 | 2.70E-03 | glio |
| Interaction_genotype_Mtor | Zfp781 | -3.493 | 2.70E-03 | glio |
| Interaction_genotype_Mtor | Sipa1 | -2.917 | 2.70E-03 | glio |
| Interaction_genotype_Mtor | Gm10863 | -3.096 | 2.70E-03 | glio |
| Interaction_genotype_Mtor | Map4k4 | 1.657 | 2.70E-03 | glio |
| Interaction_genotype_Myc | Gm16023 | -3.631 | 2.70E-03 | glio |
| Interaction_genotype_Myc | Haspin | -2.753 | 2.70E-03 | glio |
| Interaction_genotype_Myc | Arhgef9 | -2.389 | 2.70E-03 | glio |
| Interaction_genotype_Myc | Rita1 | -2.145 | 2.70E-03 | glio |
| interaction_RT | Iapp | 2.987 | 2.70E-03 | glio |
| Aars2_in.vivo_noRT | Egln3 | 2.118 | 2.70E-03 | glio |
| Aars2_in.vivo_noRT | Bcl2l1 | -1.212 | 2.70E-03 | glio |

| | | | | |
|---|---|---|---|---|
| Aars2_in.vivo_noRT | Higd1a | -1.124 | 2.70E-03 | glio |
| Atr_in.vivo_noRT | Slc25a28 | -0.978 | 2.70E-03 | glio |
| Bard1_ex.vivo_noRT | C330002G04R | 2.227 | 2.70E-03 | glio |
| Bard1_ex.vivo_noRT | Clca3a1 | 2.814 | 2.70E-03 | glio |
| Bard1_ex.vivo_noRT | Rpl3l | 2.814 | 2.70E-03 | glio |
| Bcas2_in.vivo_noRT | Gm38394 | 2.158 | 2.70E-03 | glio |
| Bcas2_in.vivo_noRT | Uqcrb | -0.704 | 2.70E-03 | glio |
| Bora_in.vivo_noRT | Folr2 | 2.128 | 2.70E-03 | glio |
| Bora_in.vivo_noRT | Atp1b3 | -0.987 | 2.70E-03 | glio |
| Bora_in.vivo_noRT | mt-Nd5 | -0.916 | 2.70E-03 | glio |
| Cenpj_in.vivo_noRT | Gm19265 | 2.032 | 2.70E-03 | glio |
| Chek1_in.vivo_noRT | Gm49602 | 2.104 | 2.70E-03 | glio |
| Chek1_in.vivo_noRT | Tmem104 | 1.941 | 2.70E-03 | glio |
| Chek1_in.vivo_noRT | Nek6 | 1.78 | 2.70E-03 | glio |
| Chek1_in.vivo_noRT | Mat2b | -0.955 | 2.70E-03 | glio |
| Chek1_in.vivo_noRT | Rpl14 | -0.566 | 2.70E-03 | glio |
| Chek1_ex.vivo_noRT | Gm49135 | 2.826 | 2.70E-03 | glio |
| Chek1_ex.vivo_noRT | Stx1b | 2.48 | 2.70E-03 | glio |
| Chek1_ex.vivo_noRT | Klf10 | 2.044 | 2.70E-03 | glio |
| Eif2s1_in.vivo_noRT | Baalc | 2.412 | 2.70E-03 | glio |
| culture_effect | Klf16 | -1.065 | 2.70E-04 | glio |
| Interaction_genotype_Atr | Ube4bos1 | -3.795 | 2.70E-04 | glio |
| Interaction_genotype_Bcas2 | Hba-a2 | -4.926 | 2.70E-04 | glio |
| Interaction_genotype_Bcas2 | Alox12 | -4.803 | 2.70E-04 | glio |
| Interaction_genotype_Bcas2 | Pycrl | 2.219 | 2.70E-04 | glio |
| Interaction_genotype_Bcas2 | 4930505N22R | -4.458 | 2.70E-04 | glio |
| Interaction_genotype_Bcas2 | G630030J09Ri | -4.599 | 2.70E-04 | glio |
| Interaction_genotype_Bcas2 | Hdac9 | -4.59 | 2.70E-04 | glio |
| Interaction_genotype_Bcas2 | 1110046J04Ri | -4.877 | 2.70E-04 | glio |
| Interaction_genotype_Mtor | Tgfb1i1 | -4.152 | 2.70E-04 | glio |
| Interaction_genotype_Myc | Gm49504 | -3.128 | 2.70E-04 | glio |
| Interaction_genotype_Myc | Gm11549 | -3.983 | 2.70E-04 | glio |
| Interaction_genotype_Myc | Mcm9 | -3.517 | 2.70E-04 | glio |
| interaction_RT | Cd74 | 3.141 | 2.70E-04 | glio |
| Aars2_in.vivo_noRT | B230208H11R | 2.659 | 2.70E-04 | glio |
| Aars2_in.vivo_noRT | Hykk | 2.562 | 2.70E-04 | glio |
| Atr_in.vivo_noRT | Gm19531 | 2.372 | 2.70E-04 | glio |
| Atr_in.vivo_noRT | Pitpnm2 | 2.331 | 2.70E-04 | glio |
| Atr_in.vivo_noRT | Naf1 | -1.382 | 2.70E-04 | glio |
| Bard1_ex.vivo_noRT | Ms4a15 | 2.814 | 2.70E-04 | glio |
| Bard1_ex.vivo_noRT | Clec12b | 2.419 | 2.70E-04 | glio |
| Bard1_ex.vivo_noRT | Vmn1r32 | 2.421 | 2.70E-04 | glio |
| Bard1_ex.vivo_noRT | Podxl | 3.219 | 2.70E-04 | glio |
| Bcas2_in.vivo_noRT | Pfkfb1 | 2.645 | 2.70E-04 | glio |
| Bcas2_in.vivo_noRT | Ktn1 | -1.387 | 2.70E-04 | glio |
| culture_effect | Ormdl3 | -0.717 | 2.70E-05 | glio |
| Interaction_genotype_Mtor | Rbbp4 | 1.976 | 2.70E-05 | glio |
| Aars2_in.vivo_noRT | Zik1 | 2.487 | 2.70E-05 | glio |
| Atr_in.vivo_noRT | Def6 | 2.832 | 2.70E-05 | glio |
| Atr_in.vivo_noRT | Tmem59l | 2.663 | 2.70E-05 | glio |
| Bard1_ex.vivo_noRT | Angptl7 | 2.814 | 2.70E-05 | glio |
| Bard1_ex.vivo_noRT | Cldn5 | 2.814 | 2.70E-05 | glio |

| | | | | |
|---|---|---|---|---|
| Bcas2_in.vivo_noRT | Gm32618 | 3.064 | 2.70E-05 | glio |
| Bora_in.vivo_noRT | Gm35315 | 2.968 | 2.70E-05 | glio |
| Bora_in.vivo_noRT | Usb1 | -2.005 | 2.70E-05 | glio |
| Bora_ex.vivo_noRT | Esp15 | 3.201 | 2.70E-05 | glio |
| Bora_ex.vivo_noRT | Gm10801 | 3.201 | 2.70E-05 | glio |
| Hsd17b10_in.vivo_noRT | Efhd1 | 3.1 | 2.70E-05 | glio |
| Hsd17b10_in.vivo_noRT | Gm12925 | 3.158 | 2.70E-05 | glio |
| culture_effect | Fhl4 | 2.313 | 2.70E-06 | glio |
| Chek1_in.vivo_noRT | Arhgef4 | 3.278 | 2.70E-06 | glio |
| Chek1_in.vivo_noRT | Zeb2os | 3.13 | 2.70E-06 | glio |
| Chek1_in.vivo_noRT | Ttc41 | 3.502 | 2.70E-06 | glio |
| Hsd17b10_in.vivo_noRT | Gm11963 | 3.356 | 2.70E-06 | glio |
| culture_effect | Idh3b | -0.836 | 2.70E-07 | glio |
| Interaction_genotype_Mtor | Cysrt1 | -6.394 | 2.70E-07 | glio |
| Aars2_in.vivo_noRT | 1810062O18R | 3.741 | 2.70E-07 | glio |
| Bora_in.vivo_noRT | Jmjd8 | 3.57 | 2.70E-07 | glio |
| Aars2_in.vivo_noRT | Mroh7 | 3.803 | 2.70E-08 | glio |
| Chek1_in.vivo_noRT | Gm15903 | 3.732 | 2.70E-08 | glio |
| Chek1_in.vivo_noRT | Gm50255 | 4.66 | 2.70E-08 | glio |
| Chek1_in.vivo_noRT | Mir142hg | 3.98 | 2.70E-08 | glio |
| Aars2_in.vivo_noRT | Gm33973 | 3.76 | 2.70E-10 | glio |
| Atr_in.vivo_noRT | Lrrn3 | 4.077 | 2.70E-10 | glio |
| Atr_in.vivo_noRT | Gm12840 | 4.275 | 2.70E-11 | glio |
| Atr_in.vivo_noRT | Nt5e | 4.055 | 2.70E-11 | glio |
| Atr_in.vivo_noRT | Myoz1 | 3.838 | 2.70E-11 | glio |
| Atr_ex.vivo_noRT | BC147527 | 5.388 | 2.70E-11 | glio |
| Atr_ex.vivo_noRT | Abcb5 | 4.073 | 2.70E-11 | glio |
| Aars2_in.vivo_noRT | Gm15234 | 4.811 | 2.70E-12 | glio |
| Bora_in.vivo_noRT | Gm36172 | 4.404 | 2.70E-15 | glio |
| Atr_in.vivo_noRT | Icam2 | 5.176 | 2.70E-17 | glio |
| Atr_in.vivo_noRT | Rab25 | 5.267 | 2.70E-17 | glio |
| Aars2_in.vivo_noRT | Gm26671 | 5.327 | 2.70E-26 | glio |
| Atr_in.vivo_noRT | Gm48764 | 5.533 | 2.70E-27 | glio |
| Atr_in.vivo_noRT | Cldn2 | 5.533 | 2.70E-31 | glio |
| culture_effect | Loxl2 | 1.606 | 2.71E-03 | glio |
| culture_effect | Actr6 | -0.535 | 2.71E-03 | glio |
| Interaction_genotype_Aars2 | Micall2 | -3.779 | 2.71E-03 | glio |
| Interaction_genotype_Atr | Cxcl16 | -3.592 | 2.71E-03 | glio |
| Interaction_genotype_Atr | Gm16759 | -3.801 | 2.71E-03 | glio |
| Interaction_genotype_Atr | Tmem59l | -3.218 | 2.71E-03 | glio |
| Interaction_genotype_Atr | Mms19 | -2.441 | 2.71E-03 | glio |
| Interaction_genotype_Mtor | Gm16230 | -3.635 | 2.71E-03 | glio |
| Interaction_genotype_Myc | Lrfn1 | -3.374 | 2.71E-03 | glio |
| Interaction_genotype_Myc | Ciita | 2.761 | 2.71E-03 | glio |
| Interaction_genotype_Myc | Zfp287 | -2.637 | 2.71E-03 | glio |
| Interaction_genotype_Myc | Atp10a | -2.748 | 2.71E-03 | glio |
| Interaction_genotype_Myc | Zfp113 | -2.799 | 2.71E-03 | glio |
| Interaction_genotype_Myc | Dcps | -2.209 | 2.71E-03 | glio |
| Interaction_genotype_Myc | Chl1 | 1.459 | 2.71E-03 | glio |
| interaction_RT | L1cam | -0.795 | 2.71E-03 | glio |
| interaction_RT | Crip1 | 0.984 | 2.71E-03 | glio |
| interaction_RT | Atad2 | -0.86 | 2.71E-03 | glio |

| | | | | |
|---|---|---|---|---|
| Aars2_in.vivo_noRT | Slc9a5 | 2.115 | 2.71E-03 | glio |
| Aars2_in.vivo_noRT | Zfp229 | 2.016 | 2.71E-03 | glio |
| Aars2_in.vivo_noRT | Slc25a39 | -0.913 | 2.71E-03 | glio |
| Bard1_ex.vivo_noRT | Dnaaf3 | 2.814 | 2.71E-03 | glio |
| Bcas2_ex.vivo_noRT | C4a | 2.737 | 2.71E-03 | glio |
| Bora_in.vivo_noRT | Gm14325 | 2.215 | 2.71E-03 | glio |
| Bora_in.vivo_noRT | Pigo | 1.953 | 2.71E-03 | glio |
| Bora_in.vivo_noRT | Vwa8 | 1.873 | 2.71E-03 | glio |
| Chek1_in.vivo_noRT | Pcsk4 | 2.195 | 2.71E-03 | glio |
| Chek1_in.vivo_noRT | Acad12 | 2.13 | 2.71E-03 | glio |
| Chek1_in.vivo_noRT | Zfp287 | 1.73 | 2.71E-03 | glio |
| Chek1_ex.vivo_noRT | Hyal3 | 3.089 | 2.71E-03 | glio |
| culture_effect | Mtx1 | -0.558 | 2.71E-04 | glio |
| Interaction_genotype_Bora | Mecom | -4.416 | 2.71E-04 | glio |
| Interaction_genotype_Bora | Plekha3 | 2.314 | 2.71E-04 | glio |
| Interaction_genotype_Bora | Sparc | 1.938 | 2.71E-04 | glio |
| Interaction_genotype_Mtor | Osgin1 | -4.082 | 2.71E-04 | glio |
| Interaction_genotype_Mtor | Ccdc146 | -4.484 | 2.71E-04 | glio |
| Interaction_genotype_Mtor | Tmem104 | -3.833 | 2.71E-04 | glio |
| Interaction_genotype_Nat2 | Gm12794 | 3.461 | 2.71E-04 | glio |
| Aars2_in.vivo_noRT | Mrto4 | -1.023 | 2.71E-04 | glio |
| Atr_in.vivo_noRT | Rnf157 | 2.494 | 2.71E-04 | glio |
| Atr_in.vivo_noRT | C230038L03Ri | 2.364 | 2.71E-04 | glio |
| Atr_in.vivo_noRT | Sf3b3 | -1.174 | 2.71E-04 | glio |
| Bora_ex.vivo_noRT | BC147527 | 3.201 | 2.71E-04 | glio |
| Bora_ex.vivo_noRT | Fa2h | 3.201 | 2.71E-04 | glio |
| Brca2_ex.vivo_noRT | Scel | 2.82 | 2.71E-04 | glio |
| Eif2s1_in.vivo_noRT | Gm5890 | 2.143 | 2.71E-04 | glio |
| Eif2s1_in.vivo_noRT | Zscan4c | 2.143 | 2.71E-04 | glio |
| Eif2s1_in.vivo_noRT | BC080695 | 2.143 | 2.71E-04 | glio |
| Eif2s1_in.vivo_noRT | Gm47540 | 2.143 | 2.71E-04 | glio |
| Eif2s1_in.vivo_noRT | BB287469 | 2.143 | 2.71E-04 | glio |
| culture_effect | Mrfap1 | 0.416 | 2.71E-05 | glio |
| culture_effect | Ndufa13 | -0.45 | 2.71E-05 | glio |
| Interaction_genotype_Myc | Fam177a | -4.41 | 2.71E-05 | glio |
| Atp5c1_in.vivo_noRT | Gm50321 | 2.668 | 2.71E-05 | glio |
| Atr_in.vivo_noRT | Mettl18 | 2.522 | 2.71E-05 | glio |
| culture_effect | Stk40 | -1.048 | 2.71E-06 | glio |
| culture_effect | Smo | 1.053 | 2.71E-06 | glio |
| interaction_RT | Col3a1 | 4.082 | 2.71E-06 | glio |
| Bora_in.vivo_noRT | Pcnx3 | -2.004 | 2.71E-06 | glio |
| culture_effect | Vps51 | 1.524 | 2.71E-07 | glio |
| culture_effect | Ttc1 | 0.671 | 2.71E-07 | glio |
| culture_effect | Rab9 | -0.644 | 2.71E-07 | glio |
| Aars2_in.vivo_noRT | Arhgef4 | 3.561 | 2.71E-07 | glio |
| Aars2_in.vivo_noRT | Fam222a | 3.812 | 2.71E-07 | glio |
| Bora_in.vivo_noRT | Lynx1 | 3.632 | 2.71E-07 | glio |
| Atr_in.vivo_noRT | Gm44836 | 3.701 | 2.71E-08 | glio |
| culture_effect | Rnf114 | -1.21 | 2.71E-09 | glio |
| Interaction_genotype_Myc | Klhdc8a | -5.344 | 2.71E-09 | glio |
| Interaction_genotype_Myc | Gm10399 | -6.567 | 2.71E-09 | glio |
| Aars2_in.vivo_noRT | Gm47972 | 4.032 | 2.71E-09 | glio |

| | | | | |
|---|---|---|---|---|
| Atr_in.vivo_noRT | Elfn2 | 3.759 | 2.71E-09 | glio |
| Chek1_in.vivo_noRT | Gtsf1l | 3.414 | 2.71E-09 | glio |
| Bora_in.vivo_noRT | Has1 | 4.039 | 2.71E-10 | glio |
| Atr_in.vivo_noRT | Cnksr1 | 4.474 | 2.71E-11 | glio |
| Atr_in.vivo_noRT | Lncenc1 | 4.227 | 2.71E-11 | glio |
| Chek1_in.vivo_noRT | Gm13684 | 3.942 | 2.71E-11 | glio |
| Chek1_ex.vivo_noRT | Bcl2l15 | 5.036 | 2.71E-11 | glio |
| Aars2_in.vivo_noRT | Pygo1 | 4.543 | 2.71E-12 | glio |
| culture_effect | F830016B08Ri | -3.354 | 2.71E-14 | glio |
| Aars2_in.vivo_noRT | Magea10 | 4.859 | 2.71E-15 | glio |
| Aars2_in.vivo_noRT | Rslcan18 | 4.941 | 2.71E-15 | glio |
| Aars2_in.vivo_noRT | Cox7b2 | 4.859 | 2.71E-15 | glio |
| Aars2_in.vivo_noRT | Nos2 | 5.067 | 2.71E-16 | glio |
| Chek1_in.vivo_noRT | Htr2b | 4.521 | 2.71E-16 | glio |
| Aars2_in.vivo_noRT | Card9 | 5.063 | 2.71E-17 | glio |
| Aars2_in.vivo_noRT | Hrct1 | 5.327 | 2.71E-18 | glio |
| Bora_in.vivo_noRT | Sema6d | 4.936 | 2.71E-20 | glio |
| culture_effect | Gm11899 | -1.457 | 2.72E-03 | glio |
| culture_effect | Psenen | -0.335 | 2.72E-03 | glio |
| Interaction_genotype_Aars2 | Oit3 | -3.595 | 2.72E-03 | glio |
| Interaction_genotype_Bcas2 | Upb1 | -3.873 | 2.72E-03 | glio |
| Interaction_genotype_Mtor | Cldn11 | -3.507 | 2.72E-03 | glio |
| Interaction_genotype_Mtor | Dock11 | -3.092 | 2.72E-03 | glio |
| Interaction_genotype_Mtor | Synpr | -3.534 | 2.72E-03 | glio |
| Interaction_genotype_Mtor | Fam160b2 | -2.888 | 2.72E-03 | glio |
| Interaction_genotype_Mtor | Naa15 | 1.629 | 2.72E-03 | glio |
| Interaction_genotype_Myc | Zfp781 | -3.284 | 2.72E-03 | glio |
| Interaction_genotype_Myc | Gm4181 | -3.36 | 2.72E-03 | glio |
| Interaction_genotype_Myc | Nsfl1c | -1.646 | 2.72E-03 | glio |
| Aars2_in.vivo_noRT | Rbm25 | -0.629 | 2.72E-03 | glio |
| Atr_in.vivo_noRT | Tmem35a | 2.46 | 2.72E-03 | glio |
| Atr_in.vivo_noRT | 6230400D17R | 2.063 | 2.72E-03 | glio |
| Atr_in.vivo_noRT | Ncald | -1.128 | 2.72E-03 | glio |
| Bcas2_in.vivo_noRT | P4htm | 2.206 | 2.72E-03 | glio |
| Bcas2_in.vivo_noRT | Ndufaf1 | 1.277 | 2.72E-03 | glio |
| Bcas2_in.vivo_noRT | Gtpbp4 | -0.797 | 2.72E-03 | glio |
| Bora_in.vivo_noRT | B230312C02R | 2.816 | 2.72E-03 | glio |
| Bora_in.vivo_noRT | Hykk | 2.171 | 2.72E-03 | glio |
| Bora_in.vivo_noRT | 4833408A19R | 2.086 | 2.72E-03 | glio |
| Bora_in.vivo_noRT | Procr | 2.684 | 2.72E-03 | glio |
| Bora_in.vivo_noRT | Irs4 | 2.35 | 2.72E-03 | glio |
| Bora_in.vivo_noRT | Kctd10 | -1.391 | 2.72E-03 | glio |
| Bora_in.vivo_noRT | Inka1 | -1.73 | 2.72E-03 | glio |
| Bora_in.vivo_noRT | 5033406O09R | 1.247 | 2.72E-03 | glio |
| Bora_in.vivo_noRT | Emd | -0.785 | 2.72E-03 | glio |
| Bora_ex.vivo_noRT | Map3k9 | 2.414 | 2.72E-03 | glio |
| Bora_ex.vivo_noRT | Cstdc4 | 2.802 | 2.72E-03 | glio |
| Bora_ex.vivo_noRT | Gipc2 | 3.139 | 2.72E-03 | glio |
| Bora_ex.vivo_noRT | Nrip3 | 2.8 | 2.72E-03 | glio |
| Bora_ex.vivo_noRT | Gm28592 | 2.831 | 2.72E-03 | glio |
| Bora_ex.vivo_noRT | H2-Eb1 | 2.808 | 2.72E-03 | glio |
| Cdkn2a_in.vivo_noRT | Luzp2 | -2.2 | 2.72E-03 | glio |

| | | | | |
|---|---|---|---|---|
| Hsd17b10_in.vivo_noRT | Pde5a | 2.33 | 2.72E-03 | glio |
| Hsd17b10_in.vivo_noRT | Olfr655 | 3.231 | 2.72E-03 | glio |
| Hsd17b10_in.vivo_noRT | Gm43017 | 2.274 | 2.72E-03 | glio |
| culture_effect | Nisch | -0.536 | 2.72E-04 | glio |
| Interaction_genotype_Atr | Phip | 2.083 | 2.72E-04 | glio |
| Interaction_genotype_Myc | Slc14a2 | -2.729 | 2.72E-04 | glio |
| interaction_RT | Slc26a2 | -0.892 | 2.72E-04 | glio |
| Aars2_in.vivo_noRT | Dnmt3a | 1.569 | 2.72E-04 | glio |
| Atr_in.vivo_noRT | Zbtb3 | 2.593 | 2.72E-04 | glio |
| Atr_in.vivo_noRT | 3110021N24R | 2.187 | 2.72E-04 | glio |
| Bcas2_in.vivo_noRT | Shpk | 2.82 | 2.72E-04 | glio |
| Bcas2_in.vivo_noRT | Nop56 | -1.091 | 2.72E-04 | glio |
| Bora_in.vivo_noRT | Gm15601 | 2.724 | 2.72E-04 | glio |
| Bora_in.vivo_noRT | Sec61b | -0.731 | 2.72E-04 | glio |
| Cenpj_in.vivo_noRT | Gm15908 | 2.022 | 2.72E-04 | glio |
| Cenpj_in.vivo_noRT | A930014E10Ri | 2.022 | 2.72E-04 | glio |
| Cenpj_in.vivo_noRT | Gm45338 | 2.022 | 2.72E-04 | glio |
| Cenpj_in.vivo_noRT | Scg2 | 2.022 | 2.72E-04 | glio |
| Cenpj_in.vivo_noRT | 1700003E24Ri | 2.022 | 2.72E-04 | glio |
| Chek1_in.vivo_noRT | Phkg1 | 3.614 | 2.72E-04 | glio |
| Interaction_genotype_Myc | Fhl3 | -4.817 | 2.72E-05 | glio |
| Interaction_genotype_Myc | Proscos | -4.964 | 2.72E-05 | glio |
| Interaction_genotype_Myc | Gm13056 | -4.821 | 2.72E-05 | glio |
| Interaction_genotype_Myc | Trnp1 | -4.61 | 2.72E-05 | glio |
| Interaction_genotype_Myc | Htr1b | -4.684 | 2.72E-05 | glio |
| Interaction_genotype_Myc | 4921536K21R | -4.721 | 2.72E-05 | glio |
| Interaction_genotype_Myc | Ccdc80 | -3.698 | 2.72E-05 | glio |
| Interaction_genotype_Myc | Gm13348 | -4.002 | 2.72E-05 | glio |
| Interaction_genotype_Myc | Pgp | 1.992 | 2.72E-05 | glio |
| Interaction_genotype_Myc | Atp5h | 1.659 | 2.72E-05 | glio |
| Interaction_genotype_Nat2 | Gm8300 | 6.068 | 2.72E-05 | glio |
| Aars2_in.vivo_noRT | Elf4 | 2.526 | 2.72E-05 | glio |
| Atr_in.vivo_noRT | Slc25a43 | 2.95 | 2.72E-05 | glio |
| Bora_in.vivo_noRT | Arhgef26 | -1.987 | 2.72E-05 | glio |
| Chek1_in.vivo_noRT | Auts2 | 2.593 | 2.72E-05 | glio |
| Hsd17b10_in.vivo_noRT | Olfml3 | -2.259 | 2.72E-05 | glio |
| culture_effect | Nckap5l | -1.442 | 2.72E-06 | glio |
| Atr_in.vivo_noRT | Gm49164 | 3.343 | 2.72E-06 | glio |
| Atr_in.vivo_noRT | Usp51 | 3.114 | 2.72E-06 | glio |
| Atr_in.vivo_noRT | Gm19345 | 2.961 | 2.72E-06 | glio |
| Atr_in.vivo_noRT | Cox7b | -1.114 | 2.72E-06 | glio |
| Bcas2_in.vivo_noRT | Kras | -1.815 | 2.72E-06 | glio |
| Chek1_in.vivo_noRT | Gm15265 | 3.242 | 2.72E-06 | glio |
| Interaction_genotype_Mtor | Rims3 | -5.627 | 2.72E-07 | glio |
| Interaction_genotype_Mtor | Agap2 | -5.084 | 2.72E-07 | glio |
| Interaction_genotype_Myc | Cldn19 | -5.828 | 2.72E-07 | glio |
| Interaction_genotype_Myc | Rassf6 | -4.974 | 2.72E-07 | glio |
| Aars2_in.vivo_noRT | Dagla | 3.83 | 2.72E-07 | glio |
| Aars2_ex.vivo_noRT | Gm48617 | 3.293 | 2.72E-07 | glio |
| Bora_in.vivo_noRT | 5430400D12R | 3.654 | 2.72E-07 | glio |
| Bora_in.vivo_noRT | Arhgef19 | 3.757 | 2.72E-07 | glio |
| Bora_in.vivo_noRT | Gm15091 | 3.757 | 2.72E-07 | glio |

| | | | | |
|---|---|---|---|---|
| Chek1_in.vivo_noRT | Gm30262 | 3.752 | 2.72E-07 | glio |
| Bora_in.vivo_noRT | Hba-a2 | 4.024 | 2.72E-08 | glio |
| Bora_in.vivo_noRT | Gm9922 | 3.801 | 2.72E-08 | glio |
| Hsd17b10_in.vivo_noRT | Fam25c | 3.084 | 2.72E-08 | glio |
| Hsd17b10_in.vivo_noRT | Clca3a1 | 3.627 | 2.72E-08 | glio |
| Aars2_in.vivo_noRT | Wfdc18 | 5.06 | 2.72E-09 | glio |
| Bora_in.vivo_noRT | Hspa12a | 4.681 | 2.72E-11 | glio |
| Chek1_in.vivo_noRT | Apobr | 5.044 | 2.72E-11 | glio |
| Aars2_in.vivo_noRT | 1700003G18R | 5.577 | 2.72E-12 | glio |
| Aars2_in.vivo_noRT | Gm20442 | 4.318 | 2.72E-12 | glio |
| Bora_ex.vivo_noRT | Pnliprp2 | 5.155 | 2.72E-12 | glio |
| Atr_in.vivo_noRT | Grin2d | 4.755 | 2.72E-13 | glio |
| Bcas2_in.vivo_noRT | C330024C12R | 4.416 | 2.72E-13 | glio |
| Chek1_in.vivo_noRT | Spi1 | 5.044 | 2.72E-15 | glio |
| Bcas2_in.vivo_noRT | Gm31717 | 5.325 | 2.72E-26 | glio |
| culture_effect | Igdcc4 | -1.717 | 2.73E-03 | glio |
| culture_effect | Plekha5 | 0.54 | 2.73E-03 | glio |
| Interaction_genotype_Atr | B3gnt4 | -3.292 | 2.73E-03 | glio |
| Interaction_genotype_Atr | 1700016P04R | -3.499 | 2.73E-03 | glio |
| Interaction_genotype_Mtor | Ston1 | -3.71 | 2.73E-03 | glio |
| Interaction_genotype_Myc | Dnajb14 | -1.92 | 2.73E-03 | glio |
| Interaction_genotype_Myc | Csrp1 | -1.953 | 2.73E-03 | glio |
| interaction_RT | H2-Eb1 | 2.546 | 2.73E-03 | glio |
| Atp5c1_in.vivo_noRT | BC026762 | 2.289 | 2.73E-03 | glio |
| Atr_in.vivo_noRT | A630072M18F | 1.845 | 2.73E-03 | glio |
| Atr_in.vivo_noRT | Rhpn2 | 1.928 | 2.73E-03 | glio |
| Atr_in.vivo_noRT | Jmjd1c | -1.065 | 2.73E-03 | glio |
| Bard1_ex.vivo_noRT | Ang | 3.159 | 2.73E-03 | glio |
| Bcas2_in.vivo_noRT | Hist1h3g | 2.344 | 2.73E-03 | glio |
| Bcas2_in.vivo_noRT | Abhd18 | 2.268 | 2.73E-03 | glio |
| Cenpj_ex.vivo_noRT | Slc6a11 | 2.591 | 2.73E-03 | glio |
| Cenpj_ex.vivo_noRT | Olfr552 | 2.192 | 2.73E-03 | glio |
| Chek1_in.vivo_noRT | Col20a1 | 2.207 | 2.73E-03 | glio |
| Chek1_in.vivo_noRT | Dzip1l | 1.726 | 2.73E-03 | glio |
| culture_effect | Gm13986 | 2.084 | 2.73E-04 | glio |
| culture_effect | Syt4 | 0.454 | 2.73E-04 | glio |
| Interaction_genotype_Mtor | 1700025N23R | -3.959 | 2.73E-04 | glio |
| Interaction_genotype_Myc | Oxa1l | -2.556 | 2.73E-04 | glio |
| Interaction_genotype_Myc | Rpl36 | 1.768 | 2.73E-04 | glio |
| interaction_RT | Ky | 2.241 | 2.73E-04 | glio |
| Atp5c1_in.vivo_noRT | 4930527F14Ri | 2.139 | 2.73E-04 | glio |
| Atr_in.vivo_noRT | Gmfb | -1.507 | 2.73E-04 | glio |
| Atr_in.vivo_noRT | Vegfb | -1.389 | 2.73E-04 | glio |
| Bora_in.vivo_noRT | Arsj | 2.822 | 2.73E-04 | glio |
| Bora_in.vivo_noRT | Gm48362 | 2.296 | 2.73E-04 | glio |
| Cenpj_ex.vivo_noRT | Cd7 | 2.591 | 2.73E-04 | glio |
| Cenpj_ex.vivo_noRT | Gm48764 | 2.591 | 2.73E-04 | glio |
| Chek1_in.vivo_noRT | Gm10138 | 2.422 | 2.73E-04 | glio |
| Hsd17b10_in.vivo_noRT | Nrip3 | 2.572 | 2.73E-04 | glio |
| culture_effect | G6pdx | 0.626 | 2.73E-05 | glio |
| culture_effect | BC031181 | 0.49 | 2.73E-05 | glio |
| Interaction_genotype_Mtor | Rab11fip4os1 | -4.816 | 2.73E-05 | glio |

| | | | | |
|---|---|---|---|---|
| Interaction_genotype_Mtor | Gm49874 | -4.776 | 2.73E-05 | glio |
| Bora_in.vivo_noRT | Cdk1 | -2.322 | 2.73E-05 | glio |
| Bora_ex.vivo_noRT | Gm17203 | 2.802 | 2.73E-05 | glio |
| Chek1_in.vivo_noRT | Kyat1 | 3.21 | 2.73E-05 | glio |
| culture_effect | Aldh9a1 | -0.986 | 2.73E-06 | glio |
| Interaction_genotype_Aars2 | 2010320O07R | -5.538 | 2.73E-06 | glio |
| Atr_ex.vivo_noRT | Gm11583 | 3.164 | 2.73E-06 | glio |
| Chek1_in.vivo_noRT | E230029C05Ri | 3.172 | 2.73E-06 | glio |
| Chek1_in.vivo_noRT | H2-T24 | 3.115 | 2.73E-06 | glio |
| Atr_in.vivo_noRT | Hdac11 | 3.649 | 2.73E-07 | glio |
| Atr_in.vivo_noRT | Smim41 | 3.843 | 2.73E-07 | glio |
| Bcas2_in.vivo_noRT | Gm36913 | 3.635 | 2.73E-07 | glio |
| Chek1_ex.vivo_noRT | Pcdhga1 | 3.873 | 2.73E-07 | glio |
| Bcas2_in.vivo_noRT | Gm15927 | 3.806 | 2.73E-08 | glio |
| Bcas2_in.vivo_noRT | Ltb | 3.747 | 2.73E-08 | glio |
| Bora_in.vivo_noRT | 4833407H14R | 4.302 | 2.73E-08 | glio |
| Bora_in.vivo_noRT | Gm32834 | 4.019 | 2.73E-09 | glio |
| culture_effect | 9130015A21R | 5.675 | 2.73E-12 | glio |
| Aars2_in.vivo_noRT | Aoc1 | 5.067 | 2.73E-12 | glio |
| Atr_in.vivo_noRT | Platr31 | 5.533 | 2.73E-12 | glio |
| Bcas2_in.vivo_noRT | Gm15743 | 4.544 | 2.73E-12 | glio |
| Atr_in.vivo_noRT | Kcnd1 | 5.011 | 2.73E-16 | glio |
| Bcas2_in.vivo_noRT | Acat3 | 4.533 | 2.73E-16 | glio |
| Bora_in.vivo_noRT | Gm11583 | 4.411 | 2.73E-17 | glio |
| Bora_in.vivo_noRT | Gm29670 | 4.414 | 2.73E-17 | glio |
| Bcas2_in.vivo_noRT | Gm36793 | 4.807 | 2.73E-20 | glio |
| culture_effect | Gm13684 | -1.643 | 2.74E-03 | glio |
| culture_effect | A130010J15Ri | 0.775 | 2.74E-03 | glio |
| Interaction_genotype_Aars2 | 2410017I17Ri | -2.548 | 2.74E-03 | glio |
| Interaction_genotype_Aars2 | Ctf1 | -3.225 | 2.74E-03 | glio |
| Interaction_genotype_Aars2 | Rab5c | 1.466 | 2.74E-03 | glio |
| Interaction_genotype_Aars2 | Rpl10a | 0.818 | 2.74E-03 | glio |
| Interaction_genotype_Bora | Tspan3 | 1.4 | 2.74E-03 | glio |
| Interaction_genotype_Mtor | Ccl8 | -4.152 | 2.74E-03 | glio |
| Interaction_genotype_Mtor | Gm4181 | -3.566 | 2.74E-03 | glio |
| Interaction_genotype_Mtor | Kctd1 | -3.302 | 2.74E-03 | glio |
| Interaction_genotype_Mtor | Polm | -3.145 | 2.74E-03 | glio |
| Interaction_genotype_Mtor | Zfp626 | -2.765 | 2.74E-03 | glio |
| Interaction_genotype_Mtor | Txnl1 | 1.668 | 2.74E-03 | glio |
| Interaction_genotype_Myc | Nhsl2 | -3.375 | 2.74E-03 | glio |
| Interaction_genotype_Myc | Mob3b | -2.781 | 2.74E-03 | glio |
| interaction_RT | Dtx3 | 1.14 | 2.74E-03 | glio |
| Aars2_in.vivo_noRT | Gm10260 | 2.346 | 2.74E-03 | glio |
| Aars2_in.vivo_noRT | Sik1 | 2.172 | 2.74E-03 | glio |
| Aars2_in.vivo_noRT | Ilf2 | -0.998 | 2.74E-03 | glio |
| Atr_in.vivo_noRT | BC052040 | 1.718 | 2.74E-03 | glio |
| Atr_in.vivo_noRT | Ago2 | -1.21 | 2.74E-03 | glio |
| Bcas2_in.vivo_noRT | Psd | 1.99 | 2.74E-03 | glio |
| Bora_in.vivo_noRT | Exosc7 | -0.942 | 2.74E-03 | glio |
| Bora_in.vivo_noRT | Ppp2ca | -0.637 | 2.74E-03 | glio |
| Bora_ex.vivo_noRT | Ankrd42 | 2.922 | 2.74E-03 | glio |
| Chek1_in.vivo_noRT | Spata2l | 2.252 | 2.74E-03 | glio |

| | | | | |
|---|---|---|---|---|
| Chek1_in.vivo_noRT | Rasl12 | 1.996 | 2.74E-03 | glio |
| Chek1_in.vivo_noRT | Rnf207 | 2.026 | 2.74E-03 | glio |
| Chek1_in.vivo_noRT | Slc36a3os | 1.759 | 2.74E-03 | glio |
| culture_effect | Poln | 1.983 | 2.74E-04 | glio |
| Interaction_genotype_Aars2 | 5430400D12R | -4.583 | 2.74E-04 | glio |
| Interaction_genotype_Mtor | Itga2 | -3.597 | 2.74E-04 | glio |
| Interaction_genotype_Mtor | Cdc5l | 1.745 | 2.74E-04 | glio |
| Interaction_genotype_Mtor | Slc26a2 | 2.233 | 2.74E-04 | glio |
| Interaction_genotype_Myc | Rep15 | -3.79 | 2.74E-04 | glio |
| Aars2_in.vivo_noRT | Rspo3 | 2.972 | 2.74E-04 | glio |
| Aars2_in.vivo_noRT | Anxa11 | 2.908 | 2.74E-04 | glio |
| Aars2_in.vivo_noRT | Mst1 | 2.427 | 2.74E-04 | glio |
| Aars2_in.vivo_noRT | Neil2 | 2.32 | 2.74E-04 | glio |
| Aars2_in.vivo_noRT | Nme2 | -0.664 | 2.74E-04 | glio |
| Aars2_ex.vivo_noRT | Scgb1c1 | 2.9 | 2.74E-04 | glio |
| Cenpj_ex.vivo_noRT | Ccdc125 | 2.989 | 2.74E-04 | glio |
| Chek1_in.vivo_noRT | Fbln1 | 2.513 | 2.74E-04 | glio |
| Interaction_genotype_Aars2 | Cyp26b1 | -5.401 | 2.74E-05 | glio |
| Interaction_genotype_Mtor | D430040D24R | -4.853 | 2.74E-05 | glio |
| Bora_in.vivo_noRT | Serpinc1 | 3.119 | 2.74E-05 | glio |
| Bora_ex.vivo_noRT | Gm20275 | 3.201 | 2.74E-05 | glio |
| Bora_ex.vivo_noRT | Ly6g6c | 3.6 | 2.74E-05 | glio |
| Chek1_in.vivo_noRT | Slc2a6 | 3.186 | 2.74E-05 | glio |
| Hsd17b10_in.vivo_noRT | Gm43581 | 2.826 | 2.74E-05 | glio |
| culture_effect | Eif2d | -0.91 | 2.74E-06 | glio |
| Interaction_genotype_Mtor | 1700016P04R | -5.228 | 2.74E-06 | glio |
| Interaction_genotype_Myc | 5033430I15Ri | -5.138 | 2.74E-06 | glio |
| Interaction_genotype_Myc | A430010J10Ri | -6.486 | 2.74E-06 | glio |
| Aars2_in.vivo_noRT | Pcdhga9 | 3.12 | 2.74E-06 | glio |
| Atr_in.vivo_noRT | Bpifc | 3.513 | 2.74E-06 | glio |
| Atr_in.vivo_noRT | Gm17259 | 3.082 | 2.74E-06 | glio |
| Bora_in.vivo_noRT | Zfp467 | 3.396 | 2.74E-06 | glio |
| Aars2_in.vivo_noRT | Ccl20 | 5.327 | 2.74E-07 | glio |
| Atp5c1_in.vivo_noRT | Gm13078 | 2.668 | 2.74E-07 | glio |
| Bcas2_in.vivo_noRT | Pla2g4b | 3.507 | 2.74E-07 | glio |
| Bora_in.vivo_noRT | Cep295nl | 3.582 | 2.74E-07 | glio |
| Chek1_in.vivo_noRT | Gm42658 | 3.291 | 2.74E-07 | glio |
| Bcas2_in.vivo_noRT | Gm8300 | 4.807 | 2.74E-08 | glio |
| Bora_in.vivo_noRT | Gm48933 | 3.934 | 2.74E-08 | glio |
| Chek1_in.vivo_noRT | Gm17227 | 3.661 | 2.74E-08 | glio |
| Aars2_in.vivo_noRT | Gm10484 | 4.325 | 2.74E-09 | glio |
| Atr_in.vivo_noRT | Gm49891 | 4.336 | 2.74E-09 | glio |
| Aars2_in.vivo_noRT | Ankrd1 | 5.327 | 2.74E-10 | glio |
| Chek1_in.vivo_noRT | Ccl7 | 5.044 | 2.74E-10 | glio |
| Chek1_in.vivo_noRT | Gm43137 | 4.528 | 2.74E-10 | glio |
| Chek1_ex.vivo_noRT | 9930111J21Ri | 4.842 | 2.74E-10 | glio |
| Aars2_in.vivo_noRT | Gm10696 | 5.327 | 2.74E-14 | glio |
| Aars2_in.vivo_noRT | B930094E09Ri | 5.347 | 2.74E-14 | glio |
| Aars2_in.vivo_noRT | Gm41409 | 4.795 | 2.74E-14 | glio |
| Aars2_in.vivo_noRT | Gm43951 | 4.401 | 2.74E-14 | glio |
| Chek1_ex.vivo_noRT | Scel | 4.842 | 2.74E-16 | glio |
| Chek1_ex.vivo_noRT | Ctla4 | 4.842 | 2.74E-16 | glio |

| | | | | |
|---|---|---|---|---|
| Aars2_in.vivo_noRT | Gm2449 | 5.063 | 2.74E-17 | glio |
| Atr_in.vivo_noRT | Olfr718-ps1 | 5.779 | 2.74E-26 | glio |
| Atr_in.vivo_noRT | Meioc | 5.27 | 2.74E-26 | glio |
| culture_effect | Gm36026 | -1.585 | 2.75E-03 | glio |
| culture_effect | A530083I20Ri | 1.853 | 2.75E-03 | glio |
| culture_effect | Oxld1 | 0.991 | 2.75E-03 | glio |
| culture_effect | Fam160b1 | -0.62 | 2.75E-03 | glio |
| Interaction_genotype_Aars2 | Gm10801 | -3.092 | 2.75E-03 | glio |
| Interaction_genotype_Aars2 | Psme1 | 1.252 | 2.75E-03 | glio |
| Interaction_genotype_Atr | Irs1 | -3.434 | 2.75E-03 | glio |
| Interaction_genotype_Atr | Kcnmb4 | -3.418 | 2.75E-03 | glio |
| Interaction_genotype_Mtor | Smpd5 | -3.236 | 2.75E-03 | glio |
| Interaction_genotype_Mtor | Nox1 | -3.212 | 2.75E-03 | glio |
| Interaction_genotype_Mtor | Olfr77 | -3.581 | 2.75E-03 | glio |
| Interaction_genotype_Mtor | Spred3 | -3.023 | 2.75E-03 | glio |
| Interaction_genotype_Mtor | Brpf3 | -3.157 | 2.75E-03 | glio |
| Interaction_genotype_Mtor | Sgsm3 | -2.477 | 2.75E-03 | glio |
| Interaction_genotype_Mtor | Col12a1 | -2.761 | 2.75E-03 | glio |
| Interaction_genotype_Myc | Itga3 | -3.663 | 2.75E-03 | glio |
| Interaction_genotype_Myc | Capza2 | 1.062 | 2.75E-03 | glio |
| Interaction_genotype_Myc | Eif1a | 1.23 | 2.75E-03 | glio |
| Aars2_in.vivo_noRT | Ino80dos | 2.237 | 2.75E-03 | glio |
| Aars2_in.vivo_noRT | Hpdl | 1.98 | 2.75E-03 | glio |
| Aars2_in.vivo_noRT | Tsc22d1 | -1.368 | 2.75E-03 | glio |
| Aars2_in.vivo_noRT | Lamtor5 | -0.975 | 2.75E-03 | glio |
| Atp5c1_in.vivo_noRT | Cytl1 | 2.008 | 2.75E-03 | glio |
| Atr_in.vivo_noRT | Eps8l1 | 2.16 | 2.75E-03 | glio |
| Atr_in.vivo_noRT | Trappc3 | -1.101 | 2.75E-03 | glio |
| Bcas2_in.vivo_noRT | Rtl5 | 2.138 | 2.75E-03 | glio |
| Bcas2_in.vivo_noRT | Nt5dc2 | -1.642 | 2.75E-03 | glio |
| Chek1_in.vivo_noRT | Nqo1 | 2.279 | 2.75E-03 | glio |
| Chek1_in.vivo_noRT | 4932422M17F | 1.992 | 2.75E-03 | glio |
| Chek1_ex.vivo_noRT | Tmem220 | 2.707 | 2.75E-03 | glio |
| Hsd17b10_in.vivo_noRT | 5430400D12R | 2.334 | 2.75E-03 | glio |
| Hsd17b10_in.vivo_noRT | Hax1 | -0.772 | 2.75E-03 | glio |
| Interaction_genotype_Myc | Glb1l | -4.144 | 2.75E-04 | glio |
| Interaction_genotype_Myc | B230312C02R | -5.213 | 2.75E-04 | glio |
| interaction_RT | Nckap5l | 1.802 | 2.75E-04 | glio |
| interaction_RT | Anp32b | -0.657 | 2.75E-04 | glio |
| Atr_in.vivo_noRT | Rplp1 | -0.743 | 2.75E-04 | glio |
| Bcas2_ex.vivo_noRT | Gm21761 | 2.742 | 2.75E-04 | glio |
| Bora_in.vivo_noRT | Ehbp1l1 | 2.686 | 2.75E-04 | glio |
| Bora_in.vivo_noRT | Setmar | 2.621 | 2.75E-04 | glio |
| Bora_in.vivo_noRT | Ccdc71l | 2.422 | 2.75E-04 | glio |
| Cenpj_ex.vivo_noRT | Gad1 | 2.591 | 2.75E-04 | glio |
| Chek1_in.vivo_noRT | Pcdhb16 | 2.817 | 2.75E-04 | glio |
| Chek1_in.vivo_noRT | Arhgdib | 2.623 | 2.75E-04 | glio |
| Chek1_in.vivo_noRT | Mogat2 | 2.526 | 2.75E-04 | glio |
| Chek1_in.vivo_noRT | Nell2 | 2.373 | 2.75E-04 | glio |
| Chek1_in.vivo_noRT | Setd3 | -1.502 | 2.75E-04 | glio |
| Chek1_in.vivo_noRT | Tor3a | -1.772 | 2.75E-04 | glio |
| Chek1_ex.vivo_noRT | Gzmm | 3.29 | 2.75E-04 | glio |

| | | | | |
|---|---|---|---|---|
| Hsd17b10_in.vivo_noRT | Sh3gl3 | 2.704 | 2.75E-04 | glio |
| culture_effect | Nceh1 | 0.842 | 2.75E-05 | glio |
| Interaction_genotype_Mtor | Ipo7 | 1.899 | 2.75E-05 | glio |
| Aars2_in.vivo_noRT | 1700086P04R | 2.918 | 2.75E-05 | glio |
| Aars2_in.vivo_noRT | Bloc1s6os | 2.807 | 2.75E-05 | glio |
| Atr_in.vivo_noRT | Rell2 | 3.055 | 2.75E-05 | glio |
| Bcas2_in.vivo_noRT | Igsf23 | 3.147 | 2.75E-05 | glio |
| Bcas2_in.vivo_noRT | S100a7a | 3.041 | 2.75E-05 | glio |
| Bcas2_in.vivo_noRT | Snapc1 | -1.855 | 2.75E-05 | glio |
| Bcas2_in.vivo_noRT | Numa1 | -1.649 | 2.75E-05 | glio |
| Bora_in.vivo_noRT | B230317F23Ri | 2.894 | 2.75E-05 | glio |
| interaction_RT | Zfand6 | 0.974 | 2.75E-06 | glio |
| Aars2_in.vivo_noRT | Gm29994 | 2.963 | 2.75E-06 | glio |
| Aars2_ex.vivo_noRT | Spi1 | 3.899 | 2.75E-06 | glio |
| Atr_in.vivo_noRT | Hectd2 | 2.89 | 2.75E-06 | glio |
| Bora_in.vivo_noRT | Slc7a14 | 3.166 | 2.75E-06 | glio |
| Bora_in.vivo_noRT | Ngef | 2.928 | 2.75E-06 | glio |
| Aars2_in.vivo_noRT | Tnc | 4.278 | 2.75E-07 | glio |
| Bard1_ex.vivo_noRT | Zscan4-ps2 | 3.612 | 2.75E-07 | glio |
| Chek1_in.vivo_noRT | Cacnb1 | 3.918 | 2.75E-07 | glio |
| Chek1_in.vivo_noRT | Stard6 | 3.673 | 2.75E-07 | glio |
| Atr_in.vivo_noRT | Synpr | 4.37 | 2.75E-08 | glio |
| Hsd17b10_in.vivo_noRT | Spi1 | 3.616 | 2.75E-08 | glio |
| Chek1_ex.vivo_noRT | Lrrc4b | 4.447 | 2.75E-09 | glio |
| Atr_in.vivo_noRT | Gm49767 | 4.495 | 2.75E-10 | glio |
| Atr_in.vivo_noRT | Bpifa3 | 3.657 | 2.75E-10 | glio |
| Bora_in.vivo_noRT | 4930478M13F | 4.557 | 2.75E-10 | glio |
| Chek1_in.vivo_noRT | Gm41764 | 4.127 | 2.75E-11 | glio |
| Bcas2_in.vivo_noRT | Lyl1 | 4.157 | 2.75E-12 | glio |
| Atr_in.vivo_noRT | Zfp808 | 4.416 | 2.75E-13 | glio |
| Bcas2_in.vivo_noRT | Pla1a | 4.805 | 2.75E-13 | glio |
| Atr_ex.vivo_noRT | AU019990 | 5.288 | 2.75E-16 | glio |
| Atr_ex.vivo_noRT | Gm8016 | 5.079 | 2.75E-16 | glio |
| Chek1_in.vivo_noRT | Vtn | 4.784 | 2.75E-20 | glio |
| Atr_in.vivo_noRT | Agrp | 5.543 | 2.75E-21 | glio |
| Aars2_in.vivo_noRT | A030014E15Ri | 5.327 | 2.75E-27 | glio |
| Aars2_in.vivo_noRT | Gm21731 | 5.327 | 2.75E-27 | glio |
| culture_effect | Prcp | -0.575 | 2.76E-03 | glio |
| Interaction_genotype_Aars2 | A930033H14R | -3.849 | 2.76E-03 | glio |
| Interaction_genotype_Atr | Ggps1 | 1.732 | 2.76E-03 | glio |
| Interaction_genotype_Atr | Nsfl1c | 1.484 | 2.76E-03 | glio |
| Interaction_genotype_Atr | Serf2 | 0.897 | 2.76E-03 | glio |
| Interaction_genotype_Mtor | Gramd4 | -3.69 | 2.76E-03 | glio |
| Interaction_genotype_Mtor | Tcaim | -3.253 | 2.76E-03 | glio |
| Interaction_genotype_Myc | Npr3 | -2.912 | 2.76E-03 | glio |
| Interaction_genotype_Myc | Xylb | -3.372 | 2.76E-03 | glio |
| Interaction_genotype_Myc | Cntd1 | -3.426 | 2.76E-03 | glio |
| Interaction_genotype_Myc | Gm49539 | -3.389 | 2.76E-03 | glio |
| Interaction_genotype_Myc | Nyap1 | -3.271 | 2.76E-03 | glio |
| Interaction_genotype_Myc | Foxo3 | -2.412 | 2.76E-03 | glio |
| Interaction_genotype_Myc | Socs7 | -1.866 | 2.76E-03 | glio |
| interaction_RT | Frat2 | 1.473 | 2.76E-03 | glio |

| | | | | |
|---|---|---|---|---|
| interaction_RT | Tenm4 | -1.356 | 2.76E-03 | glio |
| interaction_RT | Cox5b | 0.54 | 2.76E-03 | glio |
| Atr_in.vivo_noRT | Lsm8 | -0.887 | 2.76E-03 | glio |
| Atr_ex.vivo_noRT | Wdr93 | 2.726 | 2.76E-03 | glio |
| Bard1_ex.vivo_noRT | B4galnt1 | 2.52 | 2.76E-03 | glio |
| Bcas2_in.vivo_noRT | Plcb4 | 2.221 | 2.76E-03 | glio |
| Bora_in.vivo_noRT | Fgf15 | 2.368 | 2.76E-03 | glio |
| Bora_in.vivo_noRT | St3gal5 | -1.415 | 2.76E-03 | glio |
| culture_effect | Socs5 | 0.842 | 2.76E-04 | glio |
| Interaction_genotype_Mtor | Tmed8 | -3.43 | 2.76E-04 | glio |
| Interaction_genotype_Mtor | Rpl18a | 1.11 | 2.76E-04 | glio |
| Aars2_in.vivo_noRT | Gm43138 | 2.691 | 2.76E-04 | glio |
| Atr_in.vivo_noRT | Zfp169 | 2.469 | 2.76E-04 | glio |
| Bcas2_in.vivo_noRT | Gm20628 | 2.859 | 2.76E-04 | glio |
| Bcas2_in.vivo_noRT | Gm26877 | 2.606 | 2.76E-04 | glio |
| Bora_in.vivo_noRT | Gm30074 | 2.568 | 2.76E-04 | glio |
| Bora_ex.vivo_noRT | Gm13032 | 2.802 | 2.76E-04 | glio |
| Bora_in.vivo_noRT | Gm20234 | 3.223 | 2.76E-05 | glio |
| Chek1_ex.vivo_noRT | Cct8 | -1.162 | 2.76E-05 | glio |
| Hsd17b10_in.vivo_noRT | A930001C03R | 2.963 | 2.76E-05 | glio |
| Hsd17b10_in.vivo_noRT | Zg16 | 2.963 | 2.76E-05 | glio |
| Hsd17b10_in.vivo_noRT | Sec23a | -2.142 | 2.76E-05 | glio |
| Interaction_genotype_Mtor | Pkdcc | -5.791 | 2.76E-06 | glio |
| Aars2_in.vivo_noRT | Cep72 | 3.206 | 2.76E-06 | glio |
| Aars2_in.vivo_noRT | Foxl2os | 3.355 | 2.76E-07 | glio |
| Bcas2_in.vivo_noRT | BC049762 | 3.714 | 2.76E-07 | glio |
| Hsd17b10_in.vivo_noRT | Gm11769 | 3.356 | 2.76E-07 | glio |
| culture_effect | Ebp | -0.774 | 2.76E-08 | glio |
| Aars2_in.vivo_noRT | Rasgrp2 | 4.026 | 2.76E-08 | glio |
| Bcas2_in.vivo_noRT | Gm9970 | 4.162 | 2.76E-08 | glio |
| Hsd17b10_in.vivo_noRT | Pakap | 3.084 | 2.76E-08 | glio |
| Chek1_ex.vivo_noRT | Gm38477 | 4.443 | 2.76E-11 | glio |
| Bora_in.vivo_noRT | Zfp810 | 4.936 | 2.76E-13 | glio |
| Chek1_ex.vivo_noRT | Zscan4-ps3 | 4.842 | 2.76E-16 | glio |
| Chek1_ex.vivo_noRT | Tagap | 4.842 | 2.76E-16 | glio |
| Aars2_in.vivo_noRT | Nat8f2 | 5.067 | 2.76E-18 | glio |
| culture_effect | Timm13 | 0.327 | 2.77E-03 | glio |
| Interaction_genotype_Bcas2 | Sema6d | -3.047 | 2.77E-03 | glio |
| Interaction_genotype_Bcas2 | Gm15473 | -4.237 | 2.77E-03 | glio |
| Interaction_genotype_Bcas2 | Gm48742 | -3.322 | 2.77E-03 | glio |
| Interaction_genotype_Bcas2 | Gm35315 | -3.986 | 2.77E-03 | glio |
| Interaction_genotype_Bora | Cd151 | -1.675 | 2.77E-03 | glio |
| Interaction_genotype_Mtor | Kcnk7 | -3.051 | 2.77E-03 | glio |
| Interaction_genotype_Mtor | Parvg | -3.759 | 2.77E-03 | glio |
| Interaction_genotype_Mtor | Mff | 1.698 | 2.77E-03 | glio |
| Interaction_genotype_Myc | Amdhd1 | -3.25 | 2.77E-03 | glio |
| Interaction_genotype_Myc | Vmn2r44 | -2.336 | 2.77E-03 | glio |
| Interaction_genotype_Myc | Speg | -3.43 | 2.77E-03 | glio |
| Interaction_genotype_Myc | Mmp14 | -3.43 | 2.77E-03 | glio |
| Interaction_genotype_Myc | Cabp1 | -3.01 | 2.77E-03 | glio |
| Interaction_genotype_Myc | 4930532G15R | -2.967 | 2.77E-03 | glio |
| Interaction_genotype_Myc | Dele1 | -2.477 | 2.77E-03 | glio |

| | | | | |
|---|---|---|---|---|
| Interaction_genotype_Myc | Melk | -2.295 | 2.77E-03 | glio |
| interaction_RT | Atp5mpl | 0.676 | 2.77E-03 | glio |
| Aars2_in.vivo_noRT | Mtcp1 | 2.107 | 2.77E-03 | glio |
| Aars2_ex.vivo_noRT | Sema3e | -2.566 | 2.77E-03 | glio |
| Atp5c1_in.vivo_noRT | Fam13a | 2.396 | 2.77E-03 | glio |
| Atp5c1_in.vivo_noRT | Ifi214 | 2.399 | 2.77E-03 | glio |
| Atr_in.vivo_noRT | AC168977.2 | 2.252 | 2.77E-03 | glio |
| Atr_in.vivo_noRT | Rnf207 | 1.992 | 2.77E-03 | glio |
| Bcas2_in.vivo_noRT | Gm13203 | 2.308 | 2.77E-03 | glio |
| Bcas2_in.vivo_noRT | Phf5a | -0.672 | 2.77E-03 | glio |
| Bora_in.vivo_noRT | A230072E10Ri | 2.17 | 2.77E-03 | glio |
| Bora_in.vivo_noRT | Srfbp1 | -1.41 | 2.77E-03 | glio |
| Bora_ex.vivo_noRT | BC034090 | 2.808 | 2.77E-03 | glio |
| Bora_ex.vivo_noRT | Ptgis | 2.905 | 2.77E-03 | glio |
| Chek1_in.vivo_noRT | Arhgap26 | 2.08 | 2.77E-03 | glio |
| Eif2s1_in.vivo_noRT | Gm49086 | 2.143 | 2.77E-03 | glio |
| Eif2s1_in.vivo_noRT | 4921508D12R | 2.277 | 2.77E-03 | glio |
| culture_effect | Nars | 0.457 | 2.77E-04 | glio |
| Interaction_genotype_Mtor | Otop1 | -4.085 | 2.77E-04 | glio |
| Interaction_genotype_Mtor | Casz1 | -4.693 | 2.77E-04 | glio |
| Interaction_genotype_Mtor | Slc25a3 | 1.206 | 2.77E-04 | glio |
| Interaction_genotype_Myc | Atp6v0c | 1.237 | 2.77E-04 | glio |
| interaction_RT | Dennd2a | 3.216 | 2.77E-04 | glio |
| Aars2_in.vivo_noRT | Klhl23 | 2.393 | 2.77E-04 | glio |
| Aars2_in.vivo_noRT | Gm50232 | 2.406 | 2.77E-04 | glio |
| Aars2_in.vivo_noRT | Faim2 | 2.064 | 2.77E-04 | glio |
| Aars2_in.vivo_noRT | Ccdc120 | 2.395 | 2.77E-04 | glio |
| Atp5c1_in.vivo_noRT | Vmn1r91 | 2.546 | 2.77E-04 | glio |
| Atr_ex.vivo_noRT | Cirbp | 1.973 | 2.77E-04 | glio |
| Bcas2_in.vivo_noRT | Gm17477 | 2.361 | 2.77E-04 | glio |
| Bcas2_in.vivo_noRT | 5031425E22Ri | 1.731 | 2.77E-04 | glio |
| Bora_in.vivo_noRT | Myl6b | 2.532 | 2.77E-04 | glio |
| Chek1_ex.vivo_noRT | Gm12185 | 3.346 | 2.77E-04 | glio |
| culture_effect | Fgd1 | 0.946 | 2.77E-05 | glio |
| Aars2_in.vivo_noRT | Glt1d1 | 2.775 | 2.77E-05 | glio |
| Aars2_in.vivo_noRT | Osbpl6 | 2.094 | 2.77E-05 | glio |
| Aars2_ex.vivo_noRT | Ccl19 | 3.293 | 2.77E-05 | glio |
| Atp5c1_ex.vivo_noRT | 4930512B01R | 3.395 | 2.77E-05 | glio |
| Bard1_ex.vivo_noRT | Dpep3 | 2.814 | 2.77E-05 | glio |
| Bard1_ex.vivo_noRT | Erich6b | 2.814 | 2.77E-05 | glio |
| Bard1_ex.vivo_noRT | Tac2 | 2.814 | 2.77E-05 | glio |
| Bard1_ex.vivo_noRT | Tmem52 | 3.612 | 2.77E-05 | glio |
| Bcas2_in.vivo_noRT | Nop9 | -1.904 | 2.77E-05 | glio |
| culture_effect | Morf4l1 | 0.469 | 2.77E-06 | glio |
| Interaction_genotype_Rad21 | Prkcq | 5.6 | 2.77E-06 | glio |
| Aars2_in.vivo_noRT | Rnf144b | 3.269 | 2.77E-06 | glio |
| Bora_in.vivo_noRT | Tm2d3 | -1.893 | 2.77E-06 | glio |
| Chek1_ex.vivo_noRT | Npy4r | 3.652 | 2.77E-06 | glio |
| Eif2s1_in.vivo_noRT | Id3 | -1.847 | 2.77E-06 | glio |
| culture_effect | Gm29170 | 2.165 | 2.77E-07 | glio |
| Hsd17b10_in.vivo_noRT | Gm40264 | 3.356 | 2.77E-08 | glio |
| Interaction_genotype_Mtor | Nxph4 | -7.688 | 2.77E-10 | glio |

| | | | | |
|---|---|---|---|---|
| Bcas2_in.vivo_noRT | D030068K23R | 4.416 | 2.77E-10 | glio |
| Bora_in.vivo_noRT | Ceacam19 | 4.428 | 2.77E-10 | glio |
| Aars2_in.vivo_noRT | Arhgef15 | 4.455 | 2.77E-11 | glio |
| Chek1_in.vivo_noRT | Wdr93 | 4.388 | 2.77E-11 | glio |
| Atr_in.vivo_noRT | Gm13403 | 4.876 | 2.77E-13 | glio |
| Hsd17b10_in.vivo_noRT | Gjd4 | 3.616 | 2.77E-14 | glio |
| Aars2_in.vivo_noRT | Gm43256 | 5.063 | 2.77E-15 | glio |
| Bora_in.vivo_noRT | Gm49086 | 4.936 | 2.77E-16 | glio |
| Chek1_ex.vivo_noRT | Gm21731 | 4.842 | 2.77E-16 | glio |
| Chek1_in.vivo_noRT | 1700108F19Ri | 4.78 | 2.77E-17 | glio |
| Chek1_ex.vivo_noRT | Krt19 | 4.842 | 2.77E-17 | glio |
| Aars2_in.vivo_noRT | Gm17203 | 4.676 | 2.77E-20 | glio |
| culture_effect | Klrk1 | -6.638 | 2.77E-39 | glio |
| culture_effect | Zc3h15 | 0.348 | 2.78E-03 | glio |
| Interaction_genotype_Atr | 4930447K03R | -2.897 | 2.78E-03 | glio |
| Interaction_genotype_Atr | Triap1 | 1.776 | 2.78E-03 | glio |
| Interaction_genotype_Mtor | 5330438D12R | -2.575 | 2.78E-03 | glio |
| Interaction_genotype_Mtor | Uba2 | 1.554 | 2.78E-03 | glio |
| Interaction_genotype_Pars2 | 4930417H01R | -4.293 | 2.78E-03 | glio |
| Interaction_genotype_Rad51 | Zbtb37 | -3.571 | 2.78E-03 | glio |
| Interaction_genotype_Rad51 | Arf6 | 1.502 | 2.78E-03 | glio |
| interaction_RT | 1700003F12Ri | 1.197 | 2.78E-03 | glio |
| Aars2_in.vivo_noRT | Psd3 | 1.868 | 2.78E-03 | glio |
| Aars2_in.vivo_noRT | Rpl6 | -0.6 | 2.78E-03 | glio |
| Atr_in.vivo_noRT | B4galt4 | 2.204 | 2.78E-03 | glio |
| Atr_in.vivo_noRT | Trim62 | 1.818 | 2.78E-03 | glio |
| Atr_in.vivo_noRT | Gm10827 | 2.793 | 2.78E-03 | glio |
| Atr_in.vivo_noRT | Mff | -1.176 | 2.78E-03 | glio |
| Bcas2_in.vivo_noRT | Sh3bgrl2 | 2.198 | 2.78E-03 | glio |
| Bora_in.vivo_noRT | Tob1 | -1.375 | 2.78E-03 | glio |
| Bora_in.vivo_noRT | Scand1 | -0.987 | 2.78E-03 | glio |
| Chek1_ex.vivo_noRT | B430219N15R | 2.76 | 2.78E-03 | glio |
| Cyp19a1_ex.vivo_noRT | Capn13 | 2.159 | 2.78E-03 | glio |
| Cyp19a1_ex.vivo_noRT | Gm45336 | 2.959 | 2.78E-03 | glio |
| Eif2s1_in.vivo_noRT | Gm48242 | 2.143 | 2.78E-03 | glio |
| Eif2s1_in.vivo_noRT | Olfr718-ps1 | 1.879 | 2.78E-03 | glio |
| Eif2s1_in.vivo_noRT | Serpinc1 | 2.564 | 2.78E-03 | glio |
| culture_effect | Tlr6 | 1.934 | 2.78E-04 | glio |
| Interaction_genotype_Mtor | A930001C03R | -3.995 | 2.78E-04 | glio |
| Interaction_genotype_Mtor | 9330121J05Ri | -4.202 | 2.78E-04 | glio |
| Interaction_genotype_Mtor | Atrip | -3.322 | 2.78E-04 | glio |
| Interaction_genotype_Mtor | D130020L05R | -3.304 | 2.78E-04 | glio |
| Interaction_genotype_Myc | Cd59b | -4.224 | 2.78E-04 | glio |
| Interaction_genotype_Myc | Gm15927 | -3.962 | 2.78E-04 | glio |
| Interaction_genotype_Myc | Arhgap27 | -3.894 | 2.78E-04 | glio |
| Interaction_genotype_Myc | Gm5129 | -4.182 | 2.78E-04 | glio |
| Interaction_genotype_Myc | Sub1 | 1.24 | 2.78E-04 | glio |
| Aars2_in.vivo_noRT | Rpl7 | -0.706 | 2.78E-04 | glio |
| Atr_in.vivo_noRT | Car13 | 1.865 | 2.78E-04 | glio |
| Bcas2_in.vivo_noRT | Trim68 | 2.498 | 2.78E-04 | glio |
| Bcas2_in.vivo_noRT | Rwdd3 | 2.752 | 2.78E-04 | glio |
| Chek1_ex.vivo_noRT | Gm26879 | 2.946 | 2.78E-04 | glio |

| | | | | |
|---|---|---|---|---|
| culture_effect | Gramd2 | 2.08 | 2.78E-05 | glio |
| Interaction_genotype_Bcas2 | Rcan2 | -5.668 | 2.78E-05 | glio |
| Interaction_genotype_Mtor | Pls1 | -5.023 | 2.78E-05 | glio |
| Interaction_genotype_Myc | Shisal2a | -5.115 | 2.78E-05 | glio |
| interaction_RT | C3 | 4.218 | 2.78E-05 | glio |
| Aars2_in.vivo_noRT | Tph2 | 2.78 | 2.78E-05 | glio |
| Aars2_in.vivo_noRT | Pde8a | 2.584 | 2.78E-05 | glio |
| Chek1_in.vivo_noRT | Tox4 | -1.734 | 2.78E-05 | glio |
| Hsd17b10_in.vivo_noRT | 5930403N24R | 3.09 | 2.78E-05 | glio |
| Hsd17b10_in.vivo_noRT | Trpm3 | 3.094 | 2.78E-05 | glio |
| Hsd17b10_in.vivo_noRT | Gm40557 | 3.093 | 2.78E-05 | glio |
| culture_effect | B4galt5 | -1.64 | 2.78E-06 | glio |
| Interaction_genotype_Mtor | Zfp968 | -5.668 | 2.78E-06 | glio |
| Interaction_genotype_Mtor | Gm44672 | -5.466 | 2.78E-06 | glio |
| Aars2_in.vivo_noRT | Gm43728 | 3.468 | 2.78E-06 | glio |
| Atr_in.vivo_noRT | Plekha4 | 3.179 | 2.78E-06 | glio |
| Atr_ex.vivo_noRT | Gm29670 | 3.164 | 2.78E-06 | glio |
| culture_effect | Slc15a2 | -2.346 | 2.78E-07 | glio |
| Interaction_genotype_Myc | Atp6v0b | 1.883 | 2.78E-07 | glio |
| Interaction_genotype_Myc | Atp5b | 2.159 | 2.78E-07 | glio |
| Bcas2_in.vivo_noRT | C230013L11Ri | 3.895 | 2.78E-07 | glio |
| Chek1_in.vivo_noRT | BC049762 | 3.703 | 2.78E-07 | glio |
| interaction_RT | Cct2 | -0.893 | 2.78E-08 | glio |
| Aars2_in.vivo_noRT | Tnfrsf23 | 4.408 | 2.78E-08 | glio |
| Atp5c1_in.vivo_noRT | A330049N07R | 2.93 | 2.78E-08 | glio |
| Chek1_in.vivo_noRT | Alox12 | 4.255 | 2.78E-09 | glio |
| Chek1_ex.vivo_noRT | Gm37864 | 4.449 | 2.78E-09 | glio |
| Bora_in.vivo_noRT | Shisal2a | 4.673 | 2.78E-10 | glio |
| Bora_in.vivo_noRT | Rspo1 | 4.144 | 2.78E-10 | glio |
| Atr_in.vivo_noRT | Gm16222 | 5.273 | 2.78E-11 | glio |
| Atr_in.vivo_noRT | 1110017D15R | 4.533 | 2.78E-11 | glio |
| Hsd17b10_in.vivo_noRT | Fhl1 | 3.616 | 2.78E-11 | glio |
| Bora_in.vivo_noRT | Slc4a9 | 4.404 | 2.78E-14 | glio |
| Chek1_in.vivo_noRT | Lhx1os | 4.518 | 2.78E-15 | glio |
| Chek1_in.vivo_noRT | Olfr90 | 4.78 | 2.78E-15 | glio |
| Atr_in.vivo_noRT | Myo5c | 5.533 | 2.78E-16 | glio |
| Atr_in.vivo_noRT | Clcn1 | 4.888 | 2.78E-18 | glio |
| Atr_in.vivo_noRT | Luzp2 | 5.008 | 2.78E-21 | glio |
| culture_effect | Tssc4 | -4.564 | 2.78E-23 | glio |
| culture_effect | Inf2 | -0.759 | 2.79E-03 | glio |
| culture_effect | Phf13 | -0.698 | 2.79E-03 | glio |
| Interaction_genotype_Aars2 | Gm49123 | -3.954 | 2.79E-03 | glio |
| Interaction_genotype_Aars2 | Snu13 | 0.875 | 2.79E-03 | glio |
| Interaction_genotype_Atr | Ankrd34a | -3.793 | 2.79E-03 | glio |
| Interaction_genotype_Mtor | Gm15998 | -3.617 | 2.79E-03 | glio |
| Interaction_genotype_Myc | Gm38477 | -2.607 | 2.79E-03 | glio |
| Aars2_ex.vivo_noRT | Gm13625 | 2.673 | 2.79E-03 | glio |
| Atp5c1_in.vivo_noRT | 6430562O15R | 2.146 | 2.79E-03 | glio |
| Atr_in.vivo_noRT | Ptpdc1 | 1.999 | 2.79E-03 | glio |
| Bora_in.vivo_noRT | Gm26812 | 2.14 | 2.79E-03 | glio |
| Bora_ex.vivo_noRT | Dmac2 | -2.405 | 2.79E-03 | glio |
| culture_effect | Rflna | -2.235 | 2.79E-04 | glio |

| | | | | |
|---|---|---|---|---|
| culture_effect | Tmem40 | 1.995 | 2.79E-04 | glio |
| culture_effect | C030037D09R | 1.782 | 2.79E-04 | glio |
| culture_effect | Golga1 | 0.977 | 2.79E-04 | glio |
| culture_effect | Mak16 | 0.438 | 2.79E-04 | glio |
| Interaction_genotype_Aars2 | Tdrd5 | -4.175 | 2.79E-04 | glio |
| Interaction_genotype_Mtor | Tcstv3 | -4.099 | 2.79E-04 | glio |
| Interaction_genotype_Mtor | Mrpl51 | 1.575 | 2.79E-04 | glio |
| Interaction_genotype_Myc | Gm4340 | -2.729 | 2.79E-04 | glio |
| Aars2_in.vivo_noRT | Tppp | 2.246 | 2.79E-04 | glio |
| Atp5c1_in.vivo_noRT | Gm15894 | 2.145 | 2.79E-04 | glio |
| Atr_ex.vivo_noRT | 4930502E18Ri | 2.771 | 2.79E-04 | glio |
| Atr_ex.vivo_noRT | Gm50223 | 2.771 | 2.79E-04 | glio |
| Atr_ex.vivo_noRT | Ciita | 2.764 | 2.79E-04 | glio |
| Bora_in.vivo_noRT | Gm48768 | 2.685 | 2.79E-04 | glio |
| Bora_in.vivo_noRT | Gm19345 | 2.364 | 2.79E-04 | glio |
| Bora_in.vivo_noRT | Bpifa1 | 2.41 | 2.79E-04 | glio |
| Bora_in.vivo_noRT | Egfl8 | 2.737 | 2.79E-04 | glio |
| Bora_ex.vivo_noRT | Gm50223 | 2.808 | 2.79E-04 | glio |
| Chek1_in.vivo_noRT | Cep164 | -1.632 | 2.79E-04 | glio |
| culture_effect | Bad | 0.58 | 2.79E-05 | glio |
| Interaction_genotype_Mtor | Sec11a | 1.966 | 2.79E-05 | glio |
| Interaction_genotype_Myc | Gm45591 | -4.052 | 2.79E-05 | glio |
| Aars2_in.vivo_noRT | Gm46378 | 3.093 | 2.79E-05 | glio |
| Aars2_in.vivo_noRT | 9430060I03Ri | 2.845 | 2.79E-05 | glio |
| Aars2_in.vivo_noRT | 4930590J08Ri | 3.036 | 2.79E-05 | glio |
| Aars2_ex.vivo_noRT | Cfh | 2.893 | 2.79E-05 | glio |
| Aars2_ex.vivo_noRT | Olfr543 | 2.897 | 2.79E-05 | glio |
| Aars2_ex.vivo_noRT | Serping1 | 3.293 | 2.79E-05 | glio |
| Atp5c1_in.vivo_noRT | 9230112J17Ri | 2.668 | 2.79E-05 | glio |
| Atr_ex.vivo_noRT | Gm30042 | 3.164 | 2.79E-05 | glio |
| Atr_ex.vivo_noRT | Gm13610 | 3.752 | 2.79E-05 | glio |
| Bcas2_in.vivo_noRT | Gm13421 | 3.083 | 2.79E-05 | glio |
| Bcas2_in.vivo_noRT | Gm26658 | 3.304 | 2.79E-05 | glio |
| Chek1_in.vivo_noRT | Ppp1r26 | 3.135 | 2.79E-05 | glio |
| Hsd17b10_in.vivo_noRT | Gm43409 | 3.353 | 2.79E-05 | glio |
| Hsd17b10_in.vivo_noRT | Nrros | 2.974 | 2.79E-05 | glio |
| Atr_in.vivo_noRT | Gm9873 | 2.945 | 2.79E-06 | glio |
| Hsd17b10_in.vivo_noRT | B3galt4 | 3.09 | 2.79E-06 | glio |
| Interaction_genotype_Myc | Ywhag | 2.297 | 2.79E-07 | glio |
| Aars2_in.vivo_noRT | Apol9a | -2.101 | 2.79E-07 | glio |
| Aars2_ex.vivo_noRT | Mmp13 | 3.293 | 2.79E-07 | glio |
| Bard1_ex.vivo_noRT | AU019990 | 3.612 | 2.79E-07 | glio |
| Bcas2_in.vivo_noRT | Tex14 | 3.244 | 2.79E-07 | glio |
| interaction_RT | Psenen | 0.943 | 2.79E-08 | glio |
| Chek1_in.vivo_noRT | Cyp27b1 | 3.483 | 2.79E-08 | glio |
| Aars2_in.vivo_noRT | Cyp26b1 | 4.401 | 2.79E-09 | glio |
| Chek1_in.vivo_noRT | Insc | 3.586 | 2.79E-09 | glio |
| Chek1_in.vivo_noRT | Ankrd33b | 4.528 | 2.79E-10 | glio |
| culture_effect | Col27a1 | 1.858 | 2.79E-13 | glio |
| Bcas2_in.vivo_noRT | Ly6d | 4.079 | 2.79E-13 | glio |
| Aars2_in.vivo_noRT | 1700026N04R | 5.067 | 2.79E-16 | glio |
| culture_effect | Nfkbia | -1.815 | 2.79E-17 | glio |

| | | | | |
|---|---|---|---|---|
| Bora_in.vivo_noRT | Vmn1r32 | 4.936 | 2.79E-23 | glio |
| Bora_in.vivo_noRT | Glyat | 4.936 | 2.79E-23 | glio |
| Bora_in.vivo_noRT | Gm13856 | 4.936 | 2.79E-24 | glio |
| culture_effect | Hist2h3c2 | 1.892 | 2.80E-03 | glio |
| culture_effect | Vcl | -0.776 | 2.80E-03 | glio |
| Interaction_genotype_Aars2 | Gm41318 | -3.613 | 2.80E-03 | glio |
| Interaction_genotype_Atr | 4930505N22R | -3.447 | 2.80E-03 | glio |
| Interaction_genotype_Atr | Rpl29 | 0.829 | 2.80E-03 | glio |
| Interaction_genotype_Bora | D430001F17R | -3.68 | 2.80E-03 | glio |
| Interaction_genotype_Bora | 2210417A02R | -4.183 | 2.80E-03 | glio |
| Interaction_genotype_Mtor | Myb | -3.74 | 2.80E-03 | glio |
| Interaction_genotype_Mtor | Gm42067 | -2.525 | 2.80E-03 | glio |
| Interaction_genotype_Mtor | Npm1 | 1.06 | 2.80E-03 | glio |
| Interaction_genotype_Myc | Tspyl3 | -3.025 | 2.80E-03 | glio |
| Interaction_genotype_Myc | Serpinb6a | -3.435 | 2.80E-03 | glio |
| Interaction_genotype_Myc | B3galt1 | -3.216 | 2.80E-03 | glio |
| Interaction_genotype_Myc | Abca2 | -2.839 | 2.80E-03 | glio |
| Interaction_genotype_Myc | Ndufs6 | 1.148 | 2.80E-03 | glio |
| Interaction_genotype_Pars2 | Esm1 | -2.98 | 2.80E-03 | glio |
| interaction_RT | Tcp11x2 | -2.484 | 2.80E-03 | glio |
| Aars2_in.vivo_noRT | Echs1 | -1.031 | 2.80E-03 | glio |
| Aars2_in.vivo_noRT | Psme1 | -0.781 | 2.80E-03 | glio |
| Aars2_ex.vivo_noRT | Gm49441 | 2.505 | 2.80E-03 | glio |
| Atr_in.vivo_noRT | Cited4 | 2.17 | 2.80E-03 | glio |
| Atr_in.vivo_noRT | Ckb | -0.807 | 2.80E-03 | glio |
| Bcas2_in.vivo_noRT | Sbk1 | 2.249 | 2.80E-03 | glio |
| Bcas2_in.vivo_noRT | Prox1 | 1.999 | 2.80E-03 | glio |
| Cenpj_in.vivo_noRT | Slc16a5 | 1.758 | 2.80E-03 | glio |
| Cenpj_in.vivo_noRT | 9430041J12Ri | 2.022 | 2.80E-03 | glio |
| Cenpj_in.vivo_noRT | Lrp2bp | 2.287 | 2.80E-03 | glio |
| Cenpj_ex.vivo_noRT | BC048671 | 2.591 | 2.80E-03 | glio |
| Cenpj_ex.vivo_noRT | Gm41307 | 2.198 | 2.80E-03 | glio |
| Chek1_in.vivo_noRT | Zfp882 | 2.355 | 2.80E-03 | glio |
| Chek1_in.vivo_noRT | E130307A14Ri | 2.013 | 2.80E-03 | glio |
| Chek1_ex.vivo_noRT | Cyp4f18 | 3.929 | 2.80E-03 | glio |
| Chek1_ex.vivo_noRT | Zfp712 | 2.46 | 2.80E-03 | glio |
| Hsd17b10_in.vivo_noRT | Hmgcs2 | 2.595 | 2.80E-03 | glio |
| culture_effect | Sardh | 1.764 | 2.80E-04 | glio |
| culture_effect | Arhgap22 | 1.588 | 2.80E-04 | glio |
| culture_effect | Nme1 | 0.403 | 2.80E-04 | glio |
| Interaction_genotype_Mtor | Zfp420 | -4.62 | 2.80E-04 | glio |
| Interaction_genotype_Myc | Gm16239 | -2.729 | 2.80E-04 | glio |
| Interaction_genotype_Myc | Ifit1bl2 | -4.203 | 2.80E-04 | glio |
| interaction_RT | Eif3g | -0.73 | 2.80E-04 | glio |
| Bora_in.vivo_noRT | Adam19 | 2.677 | 2.80E-04 | glio |
| Bora_ex.vivo_noRT | Fabp4 | 3.574 | 2.80E-04 | glio |
| Bora_ex.vivo_noRT | Gm128 | 2.808 | 2.80E-04 | glio |
| Chek1_in.vivo_noRT | Cep112it | 2.6 | 2.80E-04 | glio |
| culture_effect | Carnmt1 | 0.503 | 2.80E-05 | glio |
| Interaction_genotype_Mtor | Igsf23 | -5.241 | 2.80E-05 | glio |
| Aars2_in.vivo_noRT | Atp2a2 | -1.689 | 2.80E-05 | glio |
| Atp5c1_in.vivo_noRT | Gm20711 | 2.668 | 2.80E-05 | glio |

| | | | | |
|---|---|---|---|---|
| Atr_in.vivo_noRT | Rps5 | -0.79 | 2.80E-05 | glio |
| Chek1_in.vivo_noRT | Pola2 | -1.89 | 2.80E-05 | glio |
| culture_effect | Soat1 | 0.696 | 2.80E-06 | glio |
| Interaction_genotype_Mtor | AC159140.1 | -4.232 | 2.80E-06 | glio |
| Interaction_genotype_Mtor | Zscan4-ps2 | -4.099 | 2.80E-06 | glio |
| Interaction_genotype_Mtor | Arl1 | 2.599 | 2.80E-06 | glio |
| Interaction_genotype_Myc | Gm10010 | -5.411 | 2.80E-06 | glio |
| Atr_in.vivo_noRT | Daam2 | 3.361 | 2.80E-06 | glio |
| Bcas2_in.vivo_noRT | Gm5089 | 3.769 | 2.80E-06 | glio |
| Bcas2_in.vivo_noRT | Gm4907 | 3.73 | 2.80E-06 | glio |
| Aars2_in.vivo_noRT | Il20rb | 3.792 | 2.80E-07 | glio |
| Aars2_in.vivo_noRT | Pfkfb2 | 3.69 | 2.80E-07 | glio |
| Aars2_ex.vivo_noRT | Gm8016 | 3.293 | 2.80E-07 | glio |
| Atp5c1_in.vivo_noRT | Gm13964 | 2.668 | 2.80E-07 | glio |
| Atr_in.vivo_noRT | AC166172.1 | 3.333 | 2.80E-07 | glio |
| Bcas2_in.vivo_noRT | Gm43608 | 3.633 | 2.80E-07 | glio |
| Bora_in.vivo_noRT | Gm10655 | 3.751 | 2.80E-07 | glio |
| Bora_in.vivo_noRT | Gm43258 | 3.54 | 2.80E-07 | glio |
| Bora_in.vivo_noRT | Neurl3 | 3.946 | 2.80E-08 | glio |
| Bcas2_in.vivo_noRT | Gm43409 | 4.544 | 2.80E-09 | glio |
| Bcas2_in.vivo_noRT | Klk14 | 4.275 | 2.80E-09 | glio |
| Atr_in.vivo_noRT | Inha | 4.409 | 2.80E-10 | glio |
| Bora_in.vivo_noRT | K230015D01R | 4.67 | 2.80E-10 | glio |
| Atr_in.vivo_noRT | Gm11696 | 4.667 | 2.80E-11 | glio |
| Chek1_in.vivo_noRT | BC065403 | 4.528 | 2.80E-11 | glio |
| culture_effect | Gemin7 | 1.006 | 2.80E-12 | glio |
| Bcas2_in.vivo_noRT | Gm7247 | 4.015 | 2.80E-12 | glio |
| Hsd17b10_in.vivo_noRT | Gm26671 | 3.616 | 2.80E-13 | glio |
| Bora_in.vivo_noRT | Vmn2r56 | 4.154 | 2.80E-14 | glio |
| Aars2_in.vivo_noRT | Fbxo15 | 5.063 | 2.80E-16 | glio |
| Aars2_in.vivo_noRT | AC163347.1 | 5.063 | 2.80E-19 | glio |
| culture_effect | 1700055D18R | 1.782 | 2.81E-03 | glio |
| culture_effect | Tnfsf9 | -1.33 | 2.81E-03 | glio |
| culture_effect | Slc25a22 | -0.87 | 2.81E-03 | glio |
| culture_effect | Lancl2 | 0.797 | 2.81E-03 | glio |
| culture_effect | Efl1 | -0.719 | 2.81E-03 | glio |
| culture_effect | Ifi213 | -0.681 | 2.81E-03 | glio |
| culture_effect | Snhg8 | 0.498 | 2.81E-03 | glio |
| Interaction_genotype_Aars2 | Gm9887 | -3.525 | 2.81E-03 | glio |
| Interaction_genotype_Atr | Cldn2 | -2.369 | 2.81E-03 | glio |
| Interaction_genotype_Atr | Fancl | -2.971 | 2.81E-03 | glio |
| Interaction_genotype_Mtor | Smim40 | -3.68 | 2.81E-03 | glio |
| Interaction_genotype_Mtor | A130051J06Ri | -3.527 | 2.81E-03 | glio |
| Interaction_genotype_Mtor | Rps10 | 1.089 | 2.81E-03 | glio |
| Interaction_genotype_Pars2 | Kpna7 | -3.569 | 2.81E-03 | glio |
| Atr_in.vivo_noRT | Snrpg | -0.799 | 2.81E-03 | glio |
| Bcas2_in.vivo_noRT | 9130023H24R | 2.218 | 2.81E-03 | glio |
| Interaction_genotype_Aars2 | Bhlhe41 | -5.321 | 2.81E-04 | glio |
| Interaction_genotype_Aars2 | Lmo4 | 1.999 | 2.81E-04 | glio |
| Interaction_genotype_Myc | Cbs | -4.283 | 2.81E-04 | glio |
| Interaction_genotype_Myc | C1d | 1.77 | 2.81E-04 | glio |
| Interaction_genotype_Myc | Serf2 | 1.251 | 2.81E-04 | glio |

| | | | | |
|---|---|---|---|---|
| Aars2_ex.vivo_noRT | Acat3 | 2.717 | 2.81E-04 | glio |
| Aars2_ex.vivo_noRT | Cysltr2 | 3.293 | 2.81E-04 | glio |
| Bora_in.vivo_noRT | B230398E01Ri | 2.603 | 2.81E-04 | glio |
| Interaction_genotype_Mtor | Fanca | -4.314 | 2.81E-05 | glio |
| Atp5c1_in.vivo_noRT | B230206H07R | 2.142 | 2.81E-05 | glio |
| Atr_in.vivo_noRT | Myo1h | 3.039 | 2.81E-05 | glio |
| Atr_in.vivo_noRT | Hpdl | 2.69 | 2.81E-05 | glio |
| Bcas2_in.vivo_noRT | Rasal1 | 3.331 | 2.81E-05 | glio |
| Bcas2_in.vivo_noRT | Hist1h2ad | 3.036 | 2.81E-05 | glio |
| Bcas2_in.vivo_noRT | Cntf | 2.956 | 2.81E-05 | glio |
| Bora_in.vivo_noRT | Cbr3 | 3.184 | 2.81E-05 | glio |
| Bora_in.vivo_noRT | Fam169a | 2.845 | 2.81E-05 | glio |
| Bora_ex.vivo_noRT | Lrguk | 3.201 | 2.81E-05 | glio |
| Hsd17b10_in.vivo_noRT | Fhl4 | 2.892 | 2.81E-05 | glio |
| Interaction_genotype_Myc | Zfp94 | -5.111 | 2.81E-06 | glio |
| Atr_in.vivo_noRT | Zfp619 | 3.331 | 2.81E-06 | glio |
| Bcas2_in.vivo_noRT | Kctd21 | 3.247 | 2.81E-06 | glio |
| Bcas2_in.vivo_noRT | Syne3 | 3.551 | 2.81E-06 | glio |
| Bcas2_in.vivo_noRT | Tmem86a | 3.44 | 2.81E-06 | glio |
| Bcas2_in.vivo_noRT | Gm13212 | 3.34 | 2.81E-06 | glio |
| Bora_in.vivo_noRT | Gm50012 | 3.336 | 2.81E-06 | glio |
| culture_effect | Tsc22d1 | 0.889 | 2.81E-07 | glio |
| Hsd17b10_in.vivo_noRT | Gm47851 | 3.356 | 2.81E-07 | glio |
| culture_effect | Nfkbie | -2.161 | 2.81E-08 | glio |
| Atr_in.vivo_noRT | Emilin2 | 3.927 | 2.81E-08 | glio |
| Bcas2_in.vivo_noRT | Zfp11 | 3.881 | 2.81E-08 | glio |
| Chek1_ex.vivo_noRT | Pcdhgb1 | 4.268 | 2.81E-08 | glio |
| culture_effect | Kcnmb4os2 | 2.982 | 2.81E-09 | glio |
| Aars2_in.vivo_noRT | Gm13625 | 3.806 | 2.81E-09 | glio |
| Aars2_in.vivo_noRT | Gm20513 | 3.23 | 2.81E-09 | glio |
| Atr_in.vivo_noRT | Parvg | 4.221 | 2.81E-09 | glio |
| Hsd17b10_in.vivo_noRT | 4930579C12R | 3.616 | 2.81E-09 | glio |
| Bcas2_in.vivo_noRT | Itgbl1 | 3.621 | 2.81E-10 | glio |
| Chek1_ex.vivo_noRT | Col5a2 | 4.256 | 2.81E-10 | glio |
| Bcas2_in.vivo_noRT | Selenok-ps1 | 4.287 | 2.81E-11 | glio |
| Bora_in.vivo_noRT | Camkk1 | 4.699 | 2.81E-11 | glio |
| Chek1_ex.vivo_noRT | Strip2 | 4.443 | 2.81E-11 | glio |
| Atr_in.vivo_noRT | Gm13212 | 4.851 | 2.81E-12 | glio |
| Atr_in.vivo_noRT | Gm15234 | 5.017 | 2.81E-13 | glio |
| Aars2_in.vivo_noRT | Psca | 5.327 | 2.81E-17 | glio |
| Aars2_in.vivo_noRT | Rassf6 | 5.327 | 2.81E-19 | glio |
| Chek1_in.vivo_noRT | Olfr552 | 5.044 | 2.81E-23 | glio |
| culture_effect | Aip | -0.541 | 2.82E-03 | glio |
| Interaction_genotype_Aars2 | Zfp14 | -3.866 | 2.82E-03 | glio |
| Interaction_genotype_Atr | AC166172.1 | -3.372 | 2.82E-03 | glio |
| Interaction_genotype_Bcas2 | Gm44899 | -3.17 | 2.82E-03 | glio |
| Interaction_genotype_Bcas2 | Krt80 | -3.485 | 2.82E-03 | glio |
| Interaction_genotype_Bcas2 | Klhl21 | 1.836 | 2.82E-03 | glio |
| Interaction_genotype_Bora | Spry2 | 2.688 | 2.82E-03 | glio |
| Interaction_genotype_Mtor | Hoga1 | -3.533 | 2.82E-03 | glio |
| Interaction_genotype_Mtor | Gdf11 | -2.57 | 2.82E-03 | glio |
| Interaction_genotype_Myc | Aoc2 | -3.078 | 2.82E-03 | glio |

| | | | | |
|---|---|---|---|---|
| Interaction_genotype_Myc | Ttc30a1 | -3.385 | 2.82E-03 | glio |
| Interaction_genotype_Myc | Synj2 | -2.827 | 2.82E-03 | glio |
| Interaction_genotype_Myc | A030001D20R | -2.497 | 2.82E-03 | glio |
| interaction_RT | Ocm | 2.7 | 2.82E-03 | glio |
| interaction_RT | Ipcef1 | 2.492 | 2.82E-03 | glio |
| interaction_RT | Osbpl9 | 0.953 | 2.82E-03 | glio |
| interaction_RT | Dusp19 | 0.967 | 2.82E-03 | glio |
| Aars2_in.vivo_noRT | Gm9873 | 1.947 | 2.82E-03 | glio |
| Aars2_in.vivo_noRT | Platr4 | 1.736 | 2.82E-03 | glio |
| Aars2_in.vivo_noRT | Pum2 | -1.043 | 2.82E-03 | glio |
| Aars2_in.vivo_noRT | Prdx1 | -0.867 | 2.82E-03 | glio |
| Atr_in.vivo_noRT | Inpp4b | 1.972 | 2.82E-03 | glio |
| Atr_in.vivo_noRT | Fez2 | 1.711 | 2.82E-03 | glio |
| Bcas2_in.vivo_noRT | Gm904 | 2.245 | 2.82E-03 | glio |
| Bcas2_in.vivo_noRT | Nhs | 1.81 | 2.82E-03 | glio |
| Bora_ex.vivo_noRT | 1700047M11F | 2.808 | 2.82E-03 | glio |
| Cenpj_in.vivo_noRT | Ccl4 | 4.584 | 2.82E-03 | glio |
| Cenpj_ex.vivo_noRT | Tmem54 | 2.591 | 2.82E-03 | glio |
| Chek1_ex.vivo_noRT | Gm50431 | 2.432 | 2.82E-03 | glio |
| Chek1_ex.vivo_noRT | Dkc1 | -1.113 | 2.82E-03 | glio |
| Hsd17b10_in.vivo_noRT | Gm8104 | 2.197 | 2.82E-03 | glio |
| culture_effect | Trmt12 | 0.83 | 2.82E-04 | glio |
| culture_effect | Arl16 | 0.748 | 2.82E-04 | glio |
| culture_effect | Tanc2 | 0.686 | 2.82E-04 | glio |
| Interaction_genotype_Atr | B930094E09Ri | -4.262 | 2.82E-04 | glio |
| Interaction_genotype_Mtor | Car13 | -2.794 | 2.82E-04 | glio |
| interaction_RT | Stard3 | 1.159 | 2.82E-04 | glio |
| Aars2_in.vivo_noRT | Hist1h2bg | 2.695 | 2.82E-04 | glio |
| Aars2_in.vivo_noRT | Zfp148 | -1.541 | 2.82E-04 | glio |
| Aars2_in.vivo_noRT | Id3 | -1.807 | 2.82E-04 | glio |
| Atr_in.vivo_noRT | C3 | 2.531 | 2.82E-04 | glio |
| Atr_in.vivo_noRT | Gdf11 | 1.971 | 2.82E-04 | glio |
| Bcas2_in.vivo_noRT | Pafah1b1 | -0.819 | 2.82E-04 | glio |
| Bora_in.vivo_noRT | Pabpc4l | 2.658 | 2.82E-04 | glio |
| Chek1_in.vivo_noRT | Med9os | 2.618 | 2.82E-04 | glio |
| Chek1_in.vivo_noRT | AW146154 | 2.594 | 2.82E-04 | glio |
| Interaction_genotype_Aars2 | Rala | 1.924 | 2.82E-05 | glio |
| Interaction_genotype_Aars2 | 5830454E08Ri | -4.875 | 2.82E-05 | glio |
| Interaction_genotype_Aars2 | AV039307 | -5.286 | 2.82E-05 | glio |
| Interaction_genotype_Aars2 | Atp5o | 1.568 | 2.82E-05 | glio |
| Bcas2_in.vivo_noRT | Tcam1 | 3.093 | 2.82E-05 | glio |
| interaction_RT | Slc31a2 | 2.083 | 2.82E-06 | glio |
| Chek1_ex.vivo_noRT | Cldn9 | 4.045 | 2.82E-06 | glio |
| Interaction_genotype_Mtor | Hlx | -5.959 | 2.82E-07 | glio |
| Interaction_genotype_Mtor | Gm20635 | -5.864 | 2.82E-07 | glio |
| Interaction_genotype_Mtor | D930028M14F | -5.586 | 2.82E-07 | glio |
| Interaction_genotype_Mtor | 4921536K21R | -6.091 | 2.82E-07 | glio |
| Aars2_in.vivo_noRT | Cope | -1.485 | 2.82E-07 | glio |
| Atr_in.vivo_noRT | Tut4 | -2.053 | 2.82E-08 | glio |
| Aars2_in.vivo_noRT | Hhipl2 | 4.558 | 2.82E-09 | glio |
| Bcas2_in.vivo_noRT | Fabp4 | 4.278 | 2.82E-09 | glio |
| Hsd17b10_in.vivo_noRT | AF067061 | 3.616 | 2.82E-09 | glio |

| | | | | |
|---|---|---|---|---|
| Aars2_in.vivo_noRT | Gm49701 | 4.277 | 2.82E-10 | glio |
| Aars2_in.vivo_noRT | Cpne8 | 4.592 | 2.82E-10 | glio |
| Hsd17b10_in.vivo_noRT | Gm48140 | 3.353 | 2.82E-11 | glio |
| culture_effect | Gas5 | -0.988 | 2.82E-12 | glio |
| Bcas2_in.vivo_noRT | Mettl7b | 4.807 | 2.82E-12 | glio |
| Chek1_in.vivo_noRT | Gngt2 | 5.044 | 2.82E-13 | glio |
| Bcas2_in.vivo_noRT | Card9 | 4.543 | 2.82E-14 | glio |
| Bcas2_in.vivo_noRT | Gm12610 | 4.278 | 2.82E-14 | glio |
| Bora_in.vivo_noRT | Pmp2 | 4.936 | 2.82E-15 | glio |
| Aars2_in.vivo_noRT | Lrrc4b | 4.93 | 2.82E-16 | glio |
| Aars2_in.vivo_noRT | Gm13288 | 5.327 | 2.82E-20 | glio |
| culture_effect | 9330104G04R | 1.646 | 2.83E-03 | glio |
| Interaction_genotype_Aars2 | Cerk | -3.192 | 2.83E-03 | glio |
| Interaction_genotype_Mtor | Xlr4c | -3.716 | 2.83E-03 | glio |
| Interaction_genotype_Mtor | Slc27a3 | -3.669 | 2.83E-03 | glio |
| Interaction_genotype_Mtor | Gm26631 | -3.759 | 2.83E-03 | glio |
| Interaction_genotype_Mtor | 4833438C02R | -2.957 | 2.83E-03 | glio |
| Interaction_genotype_Myc | Flnc | -3.37 | 2.83E-03 | glio |
| Interaction_genotype_Myc | 1810058I24Ri | 1.209 | 2.83E-03 | glio |
| Atp5c1_in.vivo_noRT | Akap3 | 2.408 | 2.83E-03 | glio |
| Atp5c1_in.vivo_noRT | Esrp2 | 2.287 | 2.83E-03 | glio |
| Atp5c1_in.vivo_noRT | 1700013F07Ri | 2.56 | 2.83E-03 | glio |
| Atr_in.vivo_noRT | Fbxo2 | -1.39 | 2.83E-03 | glio |
| Atr_ex.vivo_noRT | Ccdc80 | 2.195 | 2.83E-03 | glio |
| Bcas2_ex.vivo_noRT | Fam161a | 2.468 | 2.83E-03 | glio |
| Cenpj_in.vivo_noRT | Akr1c18 | 2.022 | 2.83E-03 | glio |
| Cenpj_in.vivo_noRT | 4921504E06Ri | 2.022 | 2.83E-03 | glio |
| Cenpj_in.vivo_noRT | 4931431C16R | 2.022 | 2.83E-03 | glio |
| Cenpj_in.vivo_noRT | Gm10482 | 2.026 | 2.83E-03 | glio |
| Chek1_in.vivo_noRT | Hist4h4 | 2.247 | 2.83E-03 | glio |
| Chek1_ex.vivo_noRT | Memo1 | -1.365 | 2.83E-03 | glio |
| culture_effect | AY074887 | 2.019 | 2.83E-04 | glio |
| culture_effect | Ptbp1 | -0.624 | 2.83E-04 | glio |
| Interaction_genotype_Atr | Luc7l2 | 1.375 | 2.83E-04 | glio |
| Interaction_genotype_Bcas2 | Mettl7b | -4.724 | 2.83E-04 | glio |
| Interaction_genotype_Myc | Slc16a5 | -2.859 | 2.83E-04 | glio |
| Interaction_genotype_Myc | Gm5152 | -4.195 | 2.83E-04 | glio |
| Aars2_in.vivo_noRT | Cbx2 | 2.324 | 2.83E-04 | glio |
| Aars2_in.vivo_noRT | Ap2b1 | -1.428 | 2.83E-04 | glio |
| Aars2_in.vivo_noRT | Vim | -0.833 | 2.83E-04 | glio |
| Atr_in.vivo_noRT | Wdr83 | 1.485 | 2.83E-04 | glio |
| Bard1_ex.vivo_noRT | Cd72 | 3.592 | 2.83E-04 | glio |
| Bcas2_in.vivo_noRT | Art2b | 2.2 | 2.83E-04 | glio |
| Bcas2_in.vivo_noRT | Kmt2d | -1.59 | 2.83E-04 | glio |
| Bcas2_in.vivo_noRT | Atf5 | -1.747 | 2.83E-04 | glio |
| Cenpj_ex.vivo_noRT | Pla2g2d | 3.375 | 2.83E-04 | glio |
| Cenpj_ex.vivo_noRT | Gm13398 | 2.592 | 2.83E-04 | glio |
| Chek1_in.vivo_noRT | Gm41760 | 2.589 | 2.83E-04 | glio |
| Hsd17b10_in.vivo_noRT | Edn1 | 3.237 | 2.83E-04 | glio |
| culture_effect | 2810402E24Ri | 1.122 | 2.83E-05 | glio |
| Interaction_genotype_Mtor | A230072C01R | -4.59 | 2.83E-05 | glio |
| Interaction_genotype_Myc | Serp2 | -3.809 | 2.83E-05 | glio |

| | | | | |
|---|---|---|---|---|
| Interaction_genotype_Myc | Cd248 | -4.66 | 2.83E-05 | glio |
| Interaction_genotype_Myc | Gm45890 | -4.965 | 2.83E-05 | glio |
| Interaction_genotype_Myc | Sema4a | -5.042 | 2.83E-05 | glio |
| Interaction_genotype_Myc | Ccnd2 | 1.781 | 2.83E-05 | glio |
| Interaction_genotype_Myc | Sec61g | 2.04 | 2.83E-05 | glio |
| interaction_RT | Utp20 | -1.277 | 2.83E-05 | glio |
| Aars2_in.vivo_noRT | Ces1a | 3.204 | 2.83E-05 | glio |
| Aars2_ex.vivo_noRT | Abra | 3.293 | 2.83E-05 | glio |
| Bcas2_in.vivo_noRT | Gm43062 | 3.029 | 2.83E-05 | glio |
| Chek1_in.vivo_noRT | Tctn1 | 3.004 | 2.83E-05 | glio |
| Hsd17b10_in.vivo_noRT | Sox11 | 2.97 | 2.83E-05 | glio |
| Interaction_genotype_Mtor | Btbd6 | -5.041 | 2.83E-06 | glio |
| Bcas2_in.vivo_noRT | Gm49494 | 3.462 | 2.83E-06 | glio |
| Bcas2_in.vivo_noRT | C2cd2 | 3.425 | 2.83E-06 | glio |
| Chek1_ex.vivo_noRT | H2-Q7 | 3.56 | 2.83E-06 | glio |
| culture_effect | Phtf1os | 2.568 | 2.83E-07 | glio |
| Aars2_in.vivo_noRT | Xist | 3.474 | 2.83E-07 | glio |
| Aars2_in.vivo_noRT | Arf5 | -1.263 | 2.83E-07 | glio |
| culture_effect | 1700001G11R | 2.211 | 2.83E-08 | glio |
| Aars2_in.vivo_noRT | Gm27010 | 3.726 | 2.83E-08 | glio |
| Atr_ex.vivo_noRT | Gm5602 | 3.962 | 2.83E-08 | glio |
| Bora_in.vivo_noRT | Rab37 | 4.018 | 2.83E-09 | glio |
| Chek1_in.vivo_noRT | Sh3gl3 | 4.132 | 2.83E-09 | glio |
| Bora_in.vivo_noRT | Il17b | 4.809 | 2.83E-10 | glio |
| Bcas2_in.vivo_noRT | Gm14820 | 4.544 | 2.83E-18 | glio |
| culture_effect | Hmga1b | -1.13 | 2.84E-03 | glio |
| Interaction_genotype_Atr | Prl7c1 | -2.681 | 2.84E-03 | glio |
| Interaction_genotype_Atr | Gm15927 | -3.596 | 2.84E-03 | glio |
| Interaction_genotype_Atr | Cldn34c1 | -3.608 | 2.84E-03 | glio |
| Interaction_genotype_Atr | AV356131 | -2.982 | 2.84E-03 | glio |
| Interaction_genotype_Atr | Jun | 1.632 | 2.84E-03 | glio |
| Interaction_genotype_Bora | Gask1b | -4.266 | 2.84E-03 | glio |
| Interaction_genotype_Bora | Ppef2 | -3.079 | 2.84E-03 | glio |
| Interaction_genotype_Bora | Mier1 | 1.298 | 2.84E-03 | glio |
| Interaction_genotype_Mtor | Cd52 | -3.442 | 2.84E-03 | glio |
| Interaction_genotype_Mtor | Acaca | -2.384 | 2.84E-03 | glio |
| Interaction_genotype_Myc | 1700123M08F | -3.287 | 2.84E-03 | glio |
| Interaction_genotype_Myc | Cnot1 | 1.546 | 2.84E-03 | glio |
| Interaction_genotype_Myc | Tm2d2 | 1.498 | 2.84E-03 | glio |
| Interaction_genotype_Myc | Puf60 | 1.394 | 2.84E-03 | glio |
| interaction_RT | Klhl28 | 1.232 | 2.84E-03 | glio |
| interaction_RT | Sec61g | 0.669 | 2.84E-03 | glio |
| Aars2_in.vivo_noRT | Plpp6 | 2.029 | 2.84E-03 | glio |
| Aars2_in.vivo_noRT | Snw1 | -0.84 | 2.84E-03 | glio |
| Aars2_ex.vivo_noRT | Gm16094 | 2.706 | 2.84E-03 | glio |
| Aars2_ex.vivo_noRT | mt-Nd3 | 1.055 | 2.84E-03 | glio |
| Atr_in.vivo_noRT | Tmem104 | 1.901 | 2.84E-03 | glio |
| Atr_in.vivo_noRT | Dcp1a | 1.634 | 2.84E-03 | glio |
| Atr_in.vivo_noRT | Klk6 | 1.725 | 2.84E-03 | glio |
| Atr_in.vivo_noRT | Utrn | -1.29 | 2.84E-03 | glio |
| Bcas2_in.vivo_noRT | Fgd6 | 2.08 | 2.84E-03 | glio |
| Bcas2_in.vivo_noRT | Nelfcd | -1.526 | 2.84E-03 | glio |

| | | | | |
|---|---|---|---|---|
| Bora_in.vivo_noRT | 2010110K18R | 2.09 | 2.84E-03 | glio |
| Bora_in.vivo_noRT | Tmem17 | 1.868 | 2.84E-03 | glio |
| Bora_in.vivo_noRT | Zfp36l2 | -1.397 | 2.84E-03 | glio |
| Bora_in.vivo_noRT | Sat1 | -1.377 | 2.84E-03 | glio |
| Bora_ex.vivo_noRT | Prl7c1 | 2.106 | 2.84E-03 | glio |
| Chek1_in.vivo_noRT | Hcfc1r1 | 2.453 | 2.84E-03 | glio |
| Chek1_in.vivo_noRT | Gstt3 | 2.225 | 2.84E-03 | glio |
| Chek1_ex.vivo_noRT | Gtf2h4 | -1.854 | 2.84E-03 | glio |
| culture_effect | Mfhas1 | 1.4 | 2.84E-04 | glio |
| Interaction_genotype_Myc | Ralgapa2 | -3.099 | 2.84E-04 | glio |
| interaction_RT | Pola1 | -0.971 | 2.84E-04 | glio |
| Aars2_in.vivo_noRT | Ncam1 | 1.615 | 2.84E-04 | glio |
| Aars2_ex.vivo_noRT | Cd52 | 3.293 | 2.84E-04 | glio |
| Atp5c1_in.vivo_noRT | Mecom | 2.668 | 2.84E-04 | glio |
| Chek1_in.vivo_noRT | Loxl2 | 2.873 | 2.84E-04 | glio |
| culture_effect | Folr2 | -2.572 | 2.84E-05 | glio |
| Interaction_genotype_Mtor | Hoxa13 | -5.265 | 2.84E-05 | glio |
| Interaction_genotype_Mtor | Ubn2 | 2.429 | 2.84E-05 | glio |
| Interaction_genotype_Myc | Tmem17 | -4.176 | 2.84E-05 | glio |
| Aars2_in.vivo_noRT | Eny2 | -0.799 | 2.84E-05 | glio |
| Atr_in.vivo_noRT | Stard10 | 2.867 | 2.84E-05 | glio |
| Atr_in.vivo_noRT | Tpi1 | -1.217 | 2.84E-05 | glio |
| Atr_in.vivo_noRT | Dnaja1 | -0.88 | 2.84E-05 | glio |
| Bcas2_in.vivo_noRT | Gm15091 | 3.103 | 2.84E-05 | glio |
| Bcas2_in.vivo_noRT | Nim1k | 3.126 | 2.84E-05 | glio |
| Chek1_in.vivo_noRT | Tmem240 | 3.315 | 2.84E-05 | glio |
| culture_effect | Cox7c | 0.574 | 2.84E-06 | glio |
| Interaction_genotype_Mtor | Fgd3 | -6.108 | 2.84E-06 | glio |
| Interaction_genotype_Myc | Selenov | -4.5 | 2.84E-06 | glio |
| Aars2_in.vivo_noRT | Tmf1 | -2.03 | 2.84E-06 | glio |
| Aars2_in.vivo_noRT | Gm50069 | 3.686 | 2.84E-07 | glio |
| Atr_in.vivo_noRT | Erlec1 | -1.8 | 2.84E-07 | glio |
| Bora_in.vivo_noRT | Gm9484 | 3.778 | 2.84E-07 | glio |
| culture_effect | Rrp8 | 0.957 | 2.84E-08 | glio |
| culture_effect | Tmem256 | 0.873 | 2.84E-08 | glio |
| Aars2_in.vivo_noRT | Pld6 | 4.07 | 2.84E-08 | glio |
| Atr_in.vivo_noRT | 1700006J14Ri | 3.623 | 2.84E-08 | glio |
| Bcas2_in.vivo_noRT | Npy4r | 3.44 | 2.84E-08 | glio |
| Interaction_genotype_Mtor | Ube2d3 | 2.342 | 2.84E-09 | glio |
| Interaction_genotype_Mtor | Carns1 | -7.452 | 2.84E-09 | glio |
| Aars2_in.vivo_noRT | Gm9908 | 4.41 | 2.84E-09 | glio |
| Aars2_in.vivo_noRT | Gjb4 | 4.67 | 2.84E-11 | glio |
| Bora_in.vivo_noRT | Htr1b | 4.936 | 2.84E-13 | glio |
| Bcas2_in.vivo_noRT | Gm2449 | 4.544 | 2.84E-14 | glio |
| culture_effect | Pafah1b3 | -1.914 | 2.84E-21 | glio |
| culture_effect | Dnm3 | 4.664 | 2.84E-22 | glio |
| Atr_in.vivo_noRT | Olfr171 | 5.533 | 2.84E-23 | glio |
| culture_effect | Rabggta | -0.741 | 2.85E-03 | glio |
| Interaction_genotype_Mtor | Lsm8 | 1.473 | 2.85E-03 | glio |
| Interaction_genotype_Myc | 1700010I14Ri | -3.325 | 2.85E-03 | glio |
| Interaction_genotype_Myc | 2210408F21Ri | -2.543 | 2.85E-03 | glio |
| Interaction_genotype_Rpl3 | Gm50203 | -3.797 | 2.85E-03 | glio |

| | | | | |
|---|---|---|---|---|
| Interaction_genotype_Rpl3 | Gm48882 | -4.229 | 2.85E-03 | glio |
| Interaction_genotype_Rpl3 | Armc2 | -4.135 | 2.85E-03 | glio |
| Interaction_genotype_Rpl3 | 5031434O11R | -4.699 | 2.85E-03 | glio |
| Interaction_genotype_Rpl3 | Efcab11 | -3.554 | 2.85E-03 | glio |
| Interaction_genotype_Rpl3 | Map3k3 | -3.053 | 2.85E-03 | glio |
| Interaction_genotype_Rpl3 | Mdp1 | 1.869 | 2.85E-03 | glio |
| Interaction_genotype_Rpl3 | Emp1 | 1.881 | 2.85E-03 | glio |
| interaction_RT | Nme6 | -0.778 | 2.85E-03 | glio |
| Aars2_ex.vivo_noRT | Jakmip2 | 2.878 | 2.85E-03 | glio |
| Bcas2_in.vivo_noRT | Elavl2 | 2.026 | 2.85E-03 | glio |
| Bcas2_in.vivo_noRT | Pla2g6 | -1.46 | 2.85E-03 | glio |
| Bcas2_in.vivo_noRT | Rbm25 | -0.574 | 2.85E-03 | glio |
| Bora_in.vivo_noRT | Ptpru | 1.935 | 2.85E-03 | glio |
| Bora_in.vivo_noRT | Dnajc21 | -0.842 | 2.85E-03 | glio |
| Cenpj_in.vivo_noRT | Gast | 2.022 | 2.85E-03 | glio |
| Cenpj_in.vivo_noRT | Shfl | 1.571 | 2.85E-03 | glio |
| Chek1_in.vivo_noRT | H2-DMb2 | 2.076 | 2.85E-03 | glio |
| Chek1_in.vivo_noRT | Pnkp | -1.508 | 2.85E-03 | glio |
| Hsd17b10_in.vivo_noRT | mt-Nd6 | -1.685 | 2.85E-03 | glio |
| Interaction_genotype_Atr | BC065403 | -3.911 | 2.85E-04 | glio |
| Interaction_genotype_Atr | N4bp2l2 | 1.863 | 2.85E-04 | glio |
| Interaction_genotype_Mtor | Slc36a2 | -4.233 | 2.85E-04 | glio |
| Interaction_genotype_Myc | Gm29361 | -4.261 | 2.85E-04 | glio |
| Interaction_genotype_Myc | Eid2b | -3.343 | 2.85E-04 | glio |
| Aars2_in.vivo_noRT | Tafa2 | 2.329 | 2.85E-04 | glio |
| Atr_in.vivo_noRT | Cd59a | 2.614 | 2.85E-04 | glio |
| Atr_in.vivo_noRT | Sub1 | -0.67 | 2.85E-04 | glio |
| Bcas2_in.vivo_noRT | Gm4189 | 2.605 | 2.85E-04 | glio |
| Bcas2_in.vivo_noRT | Sardh | 2.52 | 2.85E-04 | glio |
| Bcas2_in.vivo_noRT | Ckm | 2.533 | 2.85E-04 | glio |
| Bcas2_in.vivo_noRT | Rrm2 | -1.885 | 2.85E-04 | glio |
| Bora_in.vivo_noRT | Hist1h4n | 2.86 | 2.85E-04 | glio |
| Chek1_ex.vivo_noRT | 2810410L24Ri | 3.241 | 2.85E-04 | glio |
| Hsd17b10_in.vivo_noRT | AC125149.2 | 2.328 | 2.85E-04 | glio |
| Interaction_genotype_Myc | Hr | -4.305 | 2.85E-05 | glio |
| Interaction_genotype_Myc | Cstb | 3.297 | 2.85E-05 | glio |
| Aars2_in.vivo_noRT | Rack1 | -0.846 | 2.85E-05 | glio |
| Aars2_ex.vivo_noRT | Gm48435 | 2.505 | 2.85E-05 | glio |
| Atp5c1_in.vivo_noRT | Rassf6 | 2.668 | 2.85E-05 | glio |
| Atr_in.vivo_noRT | LTO1 | 1.696 | 2.85E-05 | glio |
| Atr_ex.vivo_noRT | Gm34939 | 3.164 | 2.85E-05 | glio |
| Chek1_in.vivo_noRT | Zscan2 | 2.854 | 2.85E-05 | glio |
| culture_effect | 2610316D01R | -2.406 | 2.85E-06 | glio |
| culture_effect | Zscan18 | -1.702 | 2.85E-06 | glio |
| Interaction_genotype_Mtor | Gm16741 | -5.054 | 2.85E-06 | glio |
| Interaction_genotype_Myc | Npl | -4.98 | 2.85E-06 | glio |
| Interaction_genotype_Myc | Nat8f1 | -5.111 | 2.85E-06 | glio |
| Interaction_genotype_Rad51 | Irf8 | 5.92 | 2.85E-06 | glio |
| Aars2_in.vivo_noRT | Eif3h | -0.923 | 2.85E-06 | glio |
| Bora_ex.vivo_noRT | Ccser1 | 3.601 | 2.85E-06 | glio |
| Bcas2_in.vivo_noRT | C330011M18F | 3.681 | 2.85E-07 | glio |
| Atr_in.vivo_noRT | Gm50255 | 4.626 | 2.85E-08 | glio |

| | | | | |
|---|---|---|---|---|
| Atr_in.vivo_noRT | Spink4 | 3.581 | 2.85E-08 | glio |
| Atr_in.vivo_noRT | 5830448L01Ri | 3.731 | 2.85E-08 | glio |
| culture_effect | Snhg1 | -1.162 | 2.85E-09 | glio |
| Chek1_in.vivo_noRT | Gm16183 | 4.401 | 2.85E-09 | glio |
| culture_effect | Fgf2 | 3.449 | 2.85E-12 | glio |
| culture_effect | Col16a1 | -1.339 | 2.85E-14 | glio |
| Aars2_in.vivo_noRT | Col19a1 | 4.41 | 2.85E-16 | glio |
| Chek1_in.vivo_noRT | Cys1 | 4.777 | 2.85E-16 | glio |
| culture_effect | Smad5 | -0.72 | 2.86E-03 | glio |
| culture_effect | Hipk1 | -0.533 | 2.86E-03 | glio |
| Interaction_genotype_Atr | Gm11549 | -3.605 | 2.86E-03 | glio |
| Interaction_genotype_Atr | Nemf | 1.371 | 2.86E-03 | glio |
| Interaction_genotype_Mtor | Bcap31 | 1.613 | 2.86E-03 | glio |
| Interaction_genotype_Rpl3 | Cd2ap | 1.831 | 2.86E-03 | glio |
| Aars2_in.vivo_noRT | Rusc2 | 1.968 | 2.86E-03 | glio |
| Atr_in.vivo_noRT | Mtx3 | 1.941 | 2.86E-03 | glio |
| Bora_in.vivo_noRT | Phf20l1 | -1.227 | 2.86E-03 | glio |
| Eif2s1_ex.vivo_noRT | Ciita | 2.615 | 2.86E-03 | glio |
| culture_effect | Rnasel | 1.548 | 2.86E-04 | glio |
| culture_effect | Txn1 | -0.423 | 2.86E-04 | glio |
| interaction_RT | Ddx21 | -0.733 | 2.86E-04 | glio |
| Aars2_in.vivo_noRT | Zfp287 | 2.038 | 2.86E-04 | glio |
| Aars2_in.vivo_noRT | Zscan29 | 2.172 | 2.86E-04 | glio |
| Aars2_in.vivo_noRT | Rps6kb1 | -1.309 | 2.86E-04 | glio |
| Atr_in.vivo_noRT | Mfsd6 | 1.724 | 2.86E-04 | glio |
| Atr_in.vivo_noRT | Gm16286 | -1.186 | 2.86E-04 | glio |
| Atr_in.vivo_noRT | Hmgn1 | -1.016 | 2.86E-04 | glio |
| Bcas2_in.vivo_noRT | Gm11520 | 2.406 | 2.86E-04 | glio |
| Chek1_ex.vivo_noRT | Dpp7 | 2.458 | 2.86E-04 | glio |
| Cyp19a1_ex.vivo_noRT | Gm29670 | 2.933 | 2.86E-04 | glio |
| culture_effect | Atp11b | -0.805 | 2.86E-05 | glio |
| Interaction_genotype_Mtor | Zfp652os | -4.769 | 2.86E-05 | glio |
| Interaction_genotype_Myc | Gm20324 | -4.191 | 2.86E-05 | glio |
| Interaction_genotype_Myc | 4930581F22Ri | -3.911 | 2.86E-05 | glio |
| Interaction_genotype_Myc | Crot | -3.646 | 2.86E-05 | glio |
| Aars2_in.vivo_noRT | Fblim1 | 2.862 | 2.86E-05 | glio |
| Atr_in.vivo_noRT | Btbd17 | 3.502 | 2.86E-05 | glio |
| Atr_in.vivo_noRT | Lrrc28 | 2.543 | 2.86E-05 | glio |
| Atr_in.vivo_noRT | Prpf40a | -1.229 | 2.86E-05 | glio |
| Bora_in.vivo_noRT | Pias2 | -1.957 | 2.86E-05 | glio |
| Fanca_ex.vivo_noRT | Fam227b | 2.919 | 2.86E-05 | glio |
| culture_effect | Cyp4v3 | -2.276 | 2.86E-06 | glio |
| culture_effect | Gkap1 | 0.743 | 2.86E-06 | glio |
| culture_effect | Anapc15 | 0.705 | 2.86E-06 | glio |
| Aars2_in.vivo_noRT | Card10 | 3.529 | 2.86E-06 | glio |
| Aars2_in.vivo_noRT | Gm31517 | 3.35 | 2.86E-06 | glio |
| Aars2_ex.vivo_noRT | Dscaml1 | 2.897 | 2.86E-06 | glio |
| Bora_in.vivo_noRT | Dnaaf3 | 2.984 | 2.86E-06 | glio |
| Chek1_in.vivo_noRT | Fndc10 | 3.563 | 2.86E-06 | glio |
| Interaction_genotype_Mtor | Gm10425 | -6.392 | 2.86E-07 | glio |
| Aars2_in.vivo_noRT | Zbtb8a | 3.78 | 2.86E-07 | glio |
| Bcas2_in.vivo_noRT | Gm46404 | 3.771 | 2.86E-07 | glio |

| | | | | |
|---|---|---|---|---|
| culture_effect | Pdlim2 | 2.03 | 2.86E-08 | glio |
| culture_effect | Dusp3 | 1.264 | 2.86E-08 | glio |
| culture_effect | D730045B01R | 2.399 | 2.86E-09 | glio |
| Atr_in.vivo_noRT | Gm10101 | 3.964 | 2.86E-09 | glio |
| Bcas2_in.vivo_noRT | Cgn | 4.281 | 2.86E-10 | glio |
| Bora_in.vivo_noRT | Dlx4 | 4.411 | 2.86E-11 | glio |
| Chek1_ex.vivo_noRT | Col19a1 | 4.449 | 2.86E-11 | glio |
| Aars2_in.vivo_noRT | Gm32591 | 4.804 | 2.86E-12 | glio |
| Atr_in.vivo_noRT | 4930483J18Ri | 5.273 | 2.86E-16 | glio |
| Bora_in.vivo_noRT | Gm15943 | 4.672 | 2.86E-17 | glio |
| Aars2_in.vivo_noRT | 4732463B04R | 5.063 | 2.86E-20 | glio |
| Chek1_in.vivo_noRT | Krt19 | 5.044 | 2.86E-26 | glio |
| culture_effect | Stpg2 | -1.901 | 2.87E-03 | glio |
| culture_effect | Apoa1 | 1.765 | 2.87E-03 | glio |
| culture_effect | Gm32051 | 1.606 | 2.87E-03 | glio |
| culture_effect | Gm20515 | 1.514 | 2.87E-03 | glio |
| culture_effect | Cenpj | -1.033 | 2.87E-03 | glio |
| culture_effect | Dgke | 0.882 | 2.87E-03 | glio |
| culture_effect | Ankrd13a | 0.712 | 2.87E-03 | glio |
| culture_effect | Cwf19l2 | -0.561 | 2.87E-03 | glio |
| culture_effect | Cisd3 | -0.487 | 2.87E-03 | glio |
| culture_effect | Atad1 | -0.48 | 2.87E-03 | glio |
| Interaction_genotype_Aars2 | Gm15929 | -3.01 | 2.87E-03 | glio |
| Interaction_genotype_Aars2 | Fam228a | -3.793 | 2.87E-03 | glio |
| Interaction_genotype_Aars2 | Gm38534 | -3.41 | 2.87E-03 | glio |
| Interaction_genotype_Aars2 | Gm20522 | -3.74 | 2.87E-03 | glio |
| Interaction_genotype_Aars2 | 2610507I01Ri | -3.142 | 2.87E-03 | glio |
| Interaction_genotype_Aars2 | 1700008J07Ri | -3.239 | 2.87E-03 | glio |
| Interaction_genotype_Bora | Spx | -3.752 | 2.87E-03 | glio |
| Interaction_genotype_Bora | Tm2d3 | 1.789 | 2.87E-03 | glio |
| Interaction_genotype_Mtor | 2300002M23F | -2.93 | 2.87E-03 | glio |
| Interaction_genotype_Mtor | Hao1 | -3.811 | 2.87E-03 | glio |
| Interaction_genotype_Myc | Gm9758 | -2.469 | 2.87E-03 | glio |
| Interaction_genotype_Myc | Vmn1r91 | -3.057 | 2.87E-03 | glio |
| Interaction_genotype_Myc | Mical1 | -3.134 | 2.87E-03 | glio |
| Interaction_genotype_Myc | Zfp954 | -2.695 | 2.87E-03 | glio |
| Interaction_genotype_Myc | Gm14295 | -2.64 | 2.87E-03 | glio |
| Interaction_genotype_Myc | Olfm1 | -2.83 | 2.87E-03 | glio |
| Interaction_genotype_Myc | Tbrg1 | 1.286 | 2.87E-03 | glio |
| Interaction_genotype_Rad51 | Rufy1 | 2.417 | 2.87E-03 | glio |
| interaction_RT | Sertad3 | 1.271 | 2.87E-03 | glio |
| interaction_RT | Rrm1 | -0.851 | 2.87E-03 | glio |
| Aars2_in.vivo_noRT | 4930471E19Ri | 2.206 | 2.87E-03 | glio |
| Aars2_in.vivo_noRT | Fzd4 | 2.059 | 2.87E-03 | glio |
| Aars2_in.vivo_noRT | Chmp3 | -1.059 | 2.87E-03 | glio |
| Aars2_ex.vivo_noRT | Gm11973 | 2.306 | 2.87E-03 | glio |
| Atp5c1_in.vivo_noRT | Gm27253 | 2.402 | 2.87E-03 | glio |
| Atr_in.vivo_noRT | B630019K06R | 1.765 | 2.87E-03 | glio |
| Atr_in.vivo_noRT | Aldh1a3 | 2.046 | 2.87E-03 | glio |
| Atr_in.vivo_noRT | Timp3 | -2.375 | 2.87E-03 | glio |
| Bcas2_in.vivo_noRT | Baz2b | 1.138 | 2.87E-03 | glio |
| Bcas2_ex.vivo_noRT | Tmem52 | 2.742 | 2.87E-03 | glio |

| | | | | |
|---|---|---|---|---|
| Bora_in.vivo_noRT | Cdc42ep4 | 2.054 | 2.87E-03 | glio |
| Bora_ex.vivo_noRT | Gjb5 | 2.622 | 2.87E-03 | glio |
| Chek1_in.vivo_noRT | Cep85 | 2.011 | 2.87E-03 | glio |
| Chek1_in.vivo_noRT | Ddost | -0.67 | 2.87E-03 | glio |
| Chek1_ex.vivo_noRT | Fzd4 | 2.581 | 2.87E-03 | glio |
| Chek1_ex.vivo_noRT | Sumo2 | -0.679 | 2.87E-03 | glio |
| Eif2s1_in.vivo_noRT | Folr1 | 2.143 | 2.87E-03 | glio |
| Hsd17b10_in.vivo_noRT | Lynx1 | 2.312 | 2.87E-03 | glio |
| Hsd17b10_in.vivo_noRT | Gm49891 | 2.419 | 2.87E-03 | glio |
| culture_effect | Ppif | -0.824 | 2.87E-04 | glio |
| culture_effect | Lyrm2 | 0.694 | 2.87E-04 | glio |
| culture_effect | Pex13 | -0.662 | 2.87E-04 | glio |
| culture_effect | Zyx | 0.656 | 2.87E-04 | glio |
| culture_effect | Mrpl54 | 0.396 | 2.87E-04 | glio |
| Interaction_genotype_Mtor | Abcd2 | -4.896 | 2.87E-04 | glio |
| Interaction_genotype_Mtor | S100a13 | 1.914 | 2.87E-04 | glio |
| Interaction_genotype_Mtor | Kdelr1 | 1.753 | 2.87E-04 | glio |
| Interaction_genotype_Myc | Mucl3 | -4.361 | 2.87E-04 | glio |
| Interaction_genotype_Rad51 | Sod3 | -5.477 | 2.87E-04 | glio |
| Bora_ex.vivo_noRT | Ccdc125 | 2.802 | 2.87E-04 | glio |
| Cenpj_in.vivo_noRT | Krt19 | 2.022 | 2.87E-04 | glio |
| Cenpj_in.vivo_noRT | Gm44646 | 2.551 | 2.87E-04 | glio |
| Cenpj_in.vivo_noRT | Gm48155 | 2.022 | 2.87E-04 | glio |
| Chek1_in.vivo_noRT | 5730414N17R | 2.774 | 2.87E-04 | glio |
| culture_effect | Gpr137 | 1.121 | 2.87E-05 | glio |
| Interaction_genotype_Mtor | Gm17227 | -4.795 | 2.87E-05 | glio |
| Interaction_genotype_Myc | D830035M03I | -4.713 | 2.87E-05 | glio |
| Bcas2_in.vivo_noRT | Gap43 | 3.361 | 2.87E-05 | glio |
| Bora_in.vivo_noRT | Arvcf | 3.164 | 2.87E-05 | glio |
| Bcas2_in.vivo_noRT | Krt16 | 4.098 | 2.87E-06 | glio |
| Bcas2_in.vivo_noRT | Zfp575 | 3.555 | 2.87E-06 | glio |
| Bcas2_in.vivo_noRT | Iws1 | -1.922 | 2.87E-06 | glio |
| Aars2_in.vivo_noRT | Nkx2-2os | 3.671 | 2.87E-07 | glio |
| Aars2_in.vivo_noRT | Gramd2 | 3.489 | 2.87E-07 | glio |
| Aars2_in.vivo_noRT | Gm20628 | 3.893 | 2.87E-07 | glio |
| Aars2_in.vivo_noRT | Abcc10 | 3.672 | 2.87E-07 | glio |
| Aars2_ex.vivo_noRT | Angptl7 | 3.293 | 2.87E-07 | glio |
| Atr_in.vivo_noRT | Gm43221 | 3.362 | 2.87E-07 | glio |
| Bora_ex.vivo_noRT | Bora | -3.6 | 2.87E-07 | glio |
| Aars2_in.vivo_noRT | 4930579K19R | 4.088 | 2.87E-08 | glio |
| Atp5c1_in.vivo_noRT | Capn13 | 2.668 | 2.87E-08 | glio |
| Bora_in.vivo_noRT | Gm44096 | 3.666 | 2.87E-08 | glio |
| Chek1_in.vivo_noRT | Tmeff2 | 4.316 | 2.87E-08 | glio |
| Aars2_in.vivo_noRT | Chd5 | 4.081 | 2.87E-09 | glio |
| Hsd17b10_in.vivo_noRT | Slc10a1 | 3.356 | 2.87E-09 | glio |
| Chek1_ex.vivo_noRT | Ccl19 | 4.842 | 2.87E-11 | glio |
| Bcas2_in.vivo_noRT | Rab25 | 4.541 | 2.87E-13 | glio |
| Bora_in.vivo_noRT | Nxph3 | 4.936 | 2.87E-14 | glio |
| Chek1_ex.vivo_noRT | Gm11232 | 4.842 | 2.87E-16 | glio |
| Bora_in.vivo_noRT | Thsd1 | 5.073 | 2.87E-17 | glio |
| Chek1_in.vivo_noRT | Akr1c18 | 5.044 | 2.87E-19 | glio |
| Chek1_in.vivo_noRT | 4921504E06Ri | 5.044 | 2.87E-19 | glio |

| | | | | |
|---|---|---|---|---|
| Bcas2_in.vivo_noRT | Gm15326 | 4.796 | 2.87E-20 | glio |
| culture_effect | Rpl7a | -1.101 | 2.87E-24 | glio |
| culture_effect | Gm47644 | -1.806 | 2.88E-03 | glio |
| culture_effect | Hes7 | -1.651 | 2.88E-03 | glio |
| culture_effect | H2-Ke6 | -0.754 | 2.88E-03 | glio |
| culture_effect | Abcd1 | -0.523 | 2.88E-03 | glio |
| Interaction_genotype_Aars2 | Apol6 | 3.15 | 2.88E-03 | glio |
| Interaction_genotype_Aars2 | Ubash3a | -5.349 | 2.88E-03 | glio |
| Interaction_genotype_Aars2 | Gm3716 | -3.399 | 2.88E-03 | glio |
| Interaction_genotype_Aars2 | H2-Eb1 | 3.374 | 2.88E-03 | glio |
| Interaction_genotype_Atr | Tmem74b | -3.072 | 2.88E-03 | glio |
| Interaction_genotype_Cdkn2a | Gm15908 | 2.835 | 2.88E-03 | glio |
| Interaction_genotype_Cdkn2a | A930014E10Ri | 2.835 | 2.88E-03 | glio |
| Interaction_genotype_Mtor | Lipe | -3.569 | 2.88E-03 | glio |
| Interaction_genotype_Myc | Gpx2 | 4.36 | 2.88E-03 | glio |
| Interaction_genotype_Myc | Cdc42ep2 | -3.433 | 2.88E-03 | glio |
| Interaction_genotype_Myc | Aunip | -2.739 | 2.88E-03 | glio |
| Interaction_genotype_Myc | Helq | -2.415 | 2.88E-03 | glio |
| Interaction_genotype_Myc | Clasrp | -2.68 | 2.88E-03 | glio |
| Interaction_genotype_Myc | Rpl15 | 1.151 | 2.88E-03 | glio |
| interaction_RT | Gm9856 | -1.548 | 2.88E-03 | glio |
| Aars2_in.vivo_noRT | Loxl2 | 2.377 | 2.88E-03 | glio |
| Aars2_in.vivo_noRT | Zfp112 | 2.107 | 2.88E-03 | glio |
| Aars2_in.vivo_noRT | Zfp438 | 1.794 | 2.88E-03 | glio |
| Aars2_in.vivo_noRT | Zfp319 | 1.469 | 2.88E-03 | glio |
| Aars2_in.vivo_noRT | Srpk2 | -0.928 | 2.88E-03 | glio |
| Aars2_in.vivo_noRT | Gabarap | -0.687 | 2.88E-03 | glio |
| Aars2_in.vivo_noRT | Ssr4 | -0.663 | 2.88E-03 | glio |
| Atp5c1_in.vivo_noRT | Gm31152 | 2.146 | 2.88E-03 | glio |
| Atr_in.vivo_noRT | Galnt3 | 2.196 | 2.88E-03 | glio |
| Atr_in.vivo_noRT | Pcdh9 | 2.154 | 2.88E-03 | glio |
| Atr_in.vivo_noRT | Gm42869 | 2.074 | 2.88E-03 | glio |
| Atr_in.vivo_noRT | Zfp329 | -1.445 | 2.88E-03 | glio |
| Atr_in.vivo_noRT | Dcun1d1 | -1.316 | 2.88E-03 | glio |
| Atr_in.vivo_noRT | Bvht | 1.553 | 2.88E-03 | glio |
| Atr_in.vivo_noRT | Cbx8 | 1.225 | 2.88E-03 | glio |
| Atr_in.vivo_noRT | Utp11 | -0.817 | 2.88E-03 | glio |
| Atr_in.vivo_noRT | Cct5 | -0.619 | 2.88E-03 | glio |
| Atr_ex.vivo_noRT | Fam131a | 2.764 | 2.88E-03 | glio |
| Bcas2_in.vivo_noRT | Lurap1 | 2.156 | 2.88E-03 | glio |
| Bcas2_in.vivo_noRT | Ndufa6 | -0.739 | 2.88E-03 | glio |
| Cenpj_ex.vivo_noRT | Fam131a | 2.976 | 2.88E-03 | glio |
| Chek1_in.vivo_noRT | Slc15a3 | 2.024 | 2.88E-03 | glio |
| Chek1_ex.vivo_noRT | Lars | -0.97 | 2.88E-03 | glio |
| Interaction_genotype_Mtor | Podxl | -3.961 | 2.88E-04 | glio |
| Interaction_genotype_Mtor | Gm10230 | -4.649 | 2.88E-04 | glio |
| Interaction_genotype_Mtor | Col6a3 | -3.856 | 2.88E-04 | glio |
| Interaction_genotype_Mtor | Nek3 | -4.469 | 2.88E-04 | glio |
| Interaction_genotype_Mtor | Tpk1 | -4.063 | 2.88E-04 | glio |
| Interaction_genotype_Mtor | Plekhg5 | -4.137 | 2.88E-04 | glio |
| Interaction_genotype_Myc | Fbxo24 | -4.091 | 2.88E-04 | glio |
| Interaction_genotype_Myc | Mzf1 | -3.919 | 2.88E-04 | glio |

| | | | | |
|---|---|---|---|---|
| Interaction_genotype_Myc | Ticrr | -3.291 | 2.88E-04 | glio |
| Interaction_genotype_Nat2 | Gm21936 | 3.209 | 2.88E-04 | glio |
| interaction_RT | Stx11 | 1.54 | 2.88E-04 | glio |
| Atr_in.vivo_noRT | Dnttip1 | -1.38 | 2.88E-04 | glio |
| Atr_in.vivo_noRT | Tle5 | -1.042 | 2.88E-04 | glio |
| Hsd17b10_in.vivo_noRT | Cop1 | -1.483 | 2.88E-04 | glio |
| Interaction_genotype_Mtor | G430095P16R | -5 | 2.88E-05 | glio |
| Interaction_genotype_Mtor | Kcnmb4os2 | -4.58 | 2.88E-05 | glio |
| Interaction_genotype_Myc | Krt80 | -4.146 | 2.88E-05 | glio |
| Atr_in.vivo_noRT | Kctd1 | 2.828 | 2.88E-05 | glio |
| Chek1_in.vivo_noRT | Cenpp | -2.121 | 2.88E-05 | glio |
| culture_effect | Prkcd | -1.789 | 2.88E-06 | glio |
| Interaction_genotype_Myc | Gm27010 | -5.211 | 2.88E-06 | glio |
| Interaction_genotype_Myc | Rheb | 1.957 | 2.88E-06 | glio |
| Interaction_genotype_Myc | Selenos | 2.21 | 2.88E-06 | glio |
| Atr_in.vivo_noRT | Ccr2 | 3.276 | 2.88E-06 | glio |
| Bora_in.vivo_noRT | Rab40b | 3.199 | 2.88E-06 | glio |
| Interaction_genotype_Mtor | Slc14a2 | -4.099 | 2.88E-07 | glio |
| Aars2_ex.vivo_noRT | Cldn5 | 3.293 | 2.88E-07 | glio |
| Bcas2_in.vivo_noRT | Gm16348 | 3.623 | 2.88E-07 | glio |
| Aars2_in.vivo_noRT | Cacnb1 | 4.201 | 2.88E-08 | glio |
| Atr_in.vivo_noRT | Rtraf | -1.692 | 2.88E-08 | glio |
| Bora_in.vivo_noRT | 9030612E09Ri | 3.615 | 2.88E-08 | glio |
| Bora_in.vivo_noRT | Ccl8 | 4.41 | 2.88E-08 | glio |
| Chek1_ex.vivo_noRT | Gm45235 | 4.049 | 2.88E-09 | glio |
| Aars2_in.vivo_noRT | Gm9970 | 4.682 | 2.88E-10 | glio |
| Aars2_in.vivo_noRT | 1700056N10R | 3.93 | 2.88E-10 | glio |
| Bcas2_in.vivo_noRT | Gm14002 | 4.285 | 2.88E-11 | glio |
| Bcas2_in.vivo_noRT | Ttll13 | 4.074 | 2.88E-11 | glio |
| Bcas2_in.vivo_noRT | Rnft2 | 4.533 | 2.88E-11 | glio |
| Chek1_ex.vivo_noRT | Dscaml1 | 4.447 | 2.88E-14 | glio |
| Chek1_in.vivo_noRT | Gm39929 | 4.586 | 2.88E-16 | glio |
| Bcas2_in.vivo_noRT | Pldi | 4.807 | 2.88E-17 | glio |
| Bcas2_in.vivo_noRT | Clec12b | 4.807 | 2.88E-22 | glio |
| Bcas2_in.vivo_noRT | Gm14929 | 4.807 | 2.88E-22 | glio |
| culture_effect | Gabarapl1 | -1.511 | 2.88E-24 | glio |
| Atr_in.vivo_noRT | Npnt | 5.267 | 2.88E-24 | glio |
| Aars2_in.vivo_noRT | Gm13398 | 5.327 | 2.88E-25 | glio |
| Atr_in.vivo_noRT | Gm44736 | 5.533 | 2.88E-30 | glio |
| culture_effect | Zdhhc4 | 0.694 | 2.89E-03 | glio |
| culture_effect | Fdx2 | 0.355 | 2.89E-03 | glio |
| Interaction_genotype_Atr | Mcl1 | 1.51 | 2.89E-03 | glio |
| Interaction_genotype_Atr | Ccnd3 | 1.278 | 2.89E-03 | glio |
| Interaction_genotype_Bora | Incenp | 2.3 | 2.89E-03 | glio |
| Interaction_genotype_Chek1 | Cst3 | 1.384 | 2.89E-03 | glio |
| Interaction_genotype_Mtor | Gm10432 | -3.399 | 2.89E-03 | glio |
| Interaction_genotype_Mtor | Ecscr | -3.178 | 2.89E-03 | glio |
| Interaction_genotype_Myc | Nkg7 | -2.865 | 2.89E-03 | glio |
| Interaction_genotype_Myc | Cntf | -3.302 | 2.89E-03 | glio |
| Interaction_genotype_Myc | Gdpd5 | -3.413 | 2.89E-03 | glio |
| Interaction_genotype_Myc | Recql4 | -2.693 | 2.89E-03 | glio |
| Interaction_genotype_Myc | Gm4285 | -2.673 | 2.89E-03 | glio |

| | | | | |
|---|---|---|---|---|
| Interaction_genotype_Pars2 | 1700023H06R | -4.35 | 2.89E-03 | glio |
| Interaction_genotype_Pars2 | Tal2 | -2.987 | 2.89E-03 | glio |
| Interaction_genotype_Pars2 | Cers3 | -3.238 | 2.89E-03 | glio |
| Interaction_genotype_Terf1 | 1810019D21R | -4.303 | 2.89E-03 | glio |
| interaction_RT | Mrpl3 | -0.804 | 2.89E-03 | glio |
| Aars2_in.vivo_noRT | Tctn3 | 2.251 | 2.89E-03 | glio |
| Aars2_in.vivo_noRT | Osgin1 | 1.979 | 2.89E-03 | glio |
| Aars2_in.vivo_noRT | Gm4890 | 1.608 | 2.89E-03 | glio |
| Aars2_in.vivo_noRT | Phf13 | -1.49 | 2.89E-03 | glio |
| Aars2_ex.vivo_noRT | Gm14379 | 2.729 | 2.89E-03 | glio |
| Atr_in.vivo_noRT | Ckmt1 | 1.986 | 2.89E-03 | glio |
| Atr_in.vivo_noRT | Tmem263 | -1.111 | 2.89E-03 | glio |
| Atr_ex.vivo_noRT | Eepd1 | 2.78 | 2.89E-03 | glio |
| Atr_ex.vivo_noRT | H2-Ab1 | 2.451 | 2.89E-03 | glio |
| Bcas2_in.vivo_noRT | Nuak2 | 2.404 | 2.89E-03 | glio |
| Bcas2_in.vivo_noRT | C130036L24Ri | 2.091 | 2.89E-03 | glio |
| Bcas2_in.vivo_noRT | Selenoi | -1.24 | 2.89E-03 | glio |
| Bora_in.vivo_noRT | Rhobtb1 | 2.199 | 2.89E-03 | glio |
| Bora_in.vivo_noRT | Ncaph | -1.659 | 2.89E-03 | glio |
| Chek1_in.vivo_noRT | Syn3 | 2.19 | 2.89E-03 | glio |
| Chek1_in.vivo_noRT | Gldc | 2.136 | 2.89E-03 | glio |
| Chek1_ex.vivo_noRT | Gm49602 | 2.68 | 2.89E-03 | glio |
| culture_effect | Rps8 | -0.479 | 2.89E-04 | glio |
| Interaction_genotype_Atr | Car14 | -4.363 | 2.89E-04 | glio |
| Interaction_genotype_Mtor | D930019O06R | -4.178 | 2.89E-04 | glio |
| Interaction_genotype_Myc | Hoxc8 | -2.729 | 2.89E-04 | glio |
| Interaction_genotype_Myc | Ankrd34a | -4.158 | 2.89E-04 | glio |
| interaction_RT | Sc5d | 1.207 | 2.89E-04 | glio |
| Aars2_in.vivo_noRT | Cnot6 | -1.453 | 2.89E-04 | glio |
| Aars2_ex.vivo_noRT | Bpifa1 | 3.293 | 2.89E-04 | glio |
| Atr_in.vivo_noRT | Prdm9 | 2.594 | 2.89E-04 | glio |
| Atr_in.vivo_noRT | Dus2 | 1.756 | 2.89E-04 | glio |
| Bcas2_in.vivo_noRT | Gm36738 | 2.799 | 2.89E-04 | glio |
| Bora_in.vivo_noRT | Ect2 | -2.028 | 2.89E-04 | glio |
| culture_effect | 2410022M11R | 2.039 | 2.89E-05 | glio |
| culture_effect | Usp46 | 0.866 | 2.89E-05 | glio |
| Atr_in.vivo_noRT | P4htm | 2.932 | 2.89E-05 | glio |
| Bora_in.vivo_noRT | Wnt4 | 3.294 | 2.89E-05 | glio |
| Bora_in.vivo_noRT | Adgrd1 | 2.779 | 2.89E-05 | glio |
| Chek1_in.vivo_noRT | Gm43017 | 2.929 | 2.89E-05 | glio |
| culture_effect | Atp5g1 | 0.51 | 2.89E-06 | glio |
| Interaction_genotype_Myc | Gm20707 | -5.319 | 2.89E-06 | glio |
| Atp5c1_in.vivo_noRT | Dmrtc1a | 2.668 | 2.89E-06 | glio |
| Atp5c1_in.vivo_noRT | Gm12589 | 2.668 | 2.89E-06 | glio |
| Atp5c1_in.vivo_noRT | Rimklb | 2.405 | 2.89E-06 | glio |
| Atr_in.vivo_noRT | Maz | -1.728 | 2.89E-06 | glio |
| Bora_in.vivo_noRT | Gm26877 | 3.266 | 2.89E-06 | glio |
| Bora_ex.vivo_noRT | Ankrd22 | 3.201 | 2.89E-07 | glio |
| Bora_ex.vivo_noRT | Gm47404 | 3.201 | 2.89E-07 | glio |
| Bora_ex.vivo_noRT | Gm31717 | 3.201 | 2.89E-07 | glio |
| Bora_ex.vivo_noRT | Gm14742 | 3.201 | 2.89E-07 | glio |
| Bora_ex.vivo_noRT | Olfr718-ps1 | 3.601 | 2.89E-07 | glio |

| | | | | |
|---|---|---|---|---|
| Bora_ex.vivo_noRT | Scel | 3.201 | 2.89E-07 | glio |
| Bora_ex.vivo_noRT | Ctla4 | 3.201 | 2.89E-07 | glio |
| Bora_ex.vivo_noRT | Zscan4-ps3 | 3.201 | 2.89E-07 | glio |
| Bora_ex.vivo_noRT | Tagap | 3.201 | 2.89E-07 | glio |
| Bora_ex.vivo_noRT | Gm21731 | 3.201 | 2.89E-07 | glio |
| Bora_ex.vivo_noRT | Olfr1152 | 3.201 | 2.89E-07 | glio |
| Bora_ex.vivo_noRT | Gm11232 | 3.201 | 2.89E-07 | glio |
| Chek1_in.vivo_noRT | Bbs1 | 3.584 | 2.89E-07 | glio |
| culture_effect | Ptpn21 | 1.097 | 2.89E-09 | glio |
| Chek1_in.vivo_noRT | Ltb | 3.984 | 2.89E-09 | glio |
| Interaction_genotype_Mtor | Wfdc1 | -7.001 | 2.89E-10 | glio |
| Chek1_in.vivo_noRT | Gm30082 | 4.393 | 2.89E-10 | glio |
| culture_effect | Scand1 | 0.99 | 2.89E-14 | glio |
| Chek1_in.vivo_noRT | Slc2a9 | 5.044 | 2.89E-18 | glio |
| Chek1_in.vivo_noRT | 4931431C16R | 5.044 | 2.89E-19 | glio |
| Atr_in.vivo_noRT | Gm48215 | 5.533 | 2.89E-25 | glio |
| culture_effect | Hist2h3c1 | 1.719 | 2.90E-03 | glio |
| culture_effect | Pcyt1a | 0.792 | 2.90E-03 | glio |
| culture_effect | Zfp105 | 0.835 | 2.90E-03 | glio |
| culture_effect | Tut1 | -0.735 | 2.90E-03 | glio |
| Interaction_genotype_Aars2 | Car12 | -3.579 | 2.90E-03 | glio |
| Interaction_genotype_Aars2 | Sar1a | 1.383 | 2.90E-03 | glio |
| Interaction_genotype_Bcas2 | Nmb | -4.032 | 2.90E-03 | glio |
| Interaction_genotype_Bcas2 | Gm13398 | -2.862 | 2.90E-03 | glio |
| Interaction_genotype_Bcas2 | Gm45591 | -3.388 | 2.90E-03 | glio |
| Interaction_genotype_Bora | Gm11696 | -4.01 | 2.90E-03 | glio |
| Interaction_genotype_Mtor | Csde1 | 1.442 | 2.90E-03 | glio |
| Interaction_genotype_Nat2 | 5033403F01Ri | -3.97 | 2.90E-03 | glio |
| interaction_RT | Mthfsl | 1.508 | 2.90E-03 | glio |
| interaction_RT | Klhl7 | 1.16 | 2.90E-03 | glio |
| interaction_RT | Igfbp6 | 1.562 | 2.90E-03 | glio |
| Aars2_in.vivo_noRT | Pcdh15 | 1.752 | 2.90E-03 | glio |
| Atr_in.vivo_noRT | Mfap3 | 1.118 | 2.90E-03 | glio |
| Cenpj_in.vivo_noRT | Gm47510 | 1.762 | 2.90E-03 | glio |
| Chek1_in.vivo_noRT | Ptges3 | -0.57 | 2.90E-03 | glio |
| culture_effect | Gm16348 | 1.987 | 2.90E-04 | glio |
| Interaction_genotype_Mtor | Gm10484 | -4.394 | 2.90E-04 | glio |
| Interaction_genotype_Mtor | Zfp65 | -3.831 | 2.90E-04 | glio |
| Interaction_genotype_Mtor | Hectd2 | -3.41 | 2.90E-04 | glio |
| Interaction_genotype_Myc | Prr7 | -4.219 | 2.90E-04 | glio |
| Interaction_genotype_Myc | Hras | 1.526 | 2.90E-04 | glio |
| interaction_RT | Etv1 | 1.086 | 2.90E-04 | glio |
| Aars2_ex.vivo_noRT | Tcp11x2 | 2.9 | 2.90E-04 | glio |
| Aars2_ex.vivo_noRT | Gm18280 | 2.499 | 2.90E-04 | glio |
| Atp5c1_in.vivo_noRT | Dcdc2a | 2.408 | 2.90E-04 | glio |
| Bard1_ex.vivo_noRT | Gm30692 | 2.421 | 2.90E-04 | glio |
| Bard1_ex.vivo_noRT | Gm10482 | 2.814 | 2.90E-04 | glio |
| Bora_in.vivo_noRT | Pomc | 2.201 | 2.90E-04 | glio |
| Eif2s1_in.vivo_noRT | Zscan4-ps2 | 2.143 | 2.90E-04 | glio |
| Hsd17b10_in.vivo_noRT | Bglap2 | 2.687 | 2.90E-04 | glio |
| culture_effect | Ralbp1 | 0.518 | 2.90E-05 | glio |
| Interaction_genotype_Bcas2 | Gm10010 | -5.535 | 2.90E-05 | glio |

| | | | | |
|---|---|---|---|---|
| Interaction_genotype_Bcas2 | Gm19276 | -3.87 | 2.90E-05 | glio |
| Interaction_genotype_Mtor | Lhx6 | -4.94 | 2.90E-05 | glio |
| Interaction_genotype_Mtor | 2810457G06R | -5.56 | 2.90E-05 | glio |
| Interaction_genotype_Myc | Gm26674 | -4.511 | 2.90E-05 | glio |
| Interaction_genotype_Rad51 | Apol10b | 4.861 | 2.90E-05 | glio |
| Aars2_ex.vivo_noRT | Ush1g | 2.893 | 2.90E-05 | glio |
| Atr_in.vivo_noRT | Slc2a6 | 3.146 | 2.90E-05 | glio |
| Atr_in.vivo_noRT | 9130023H24R | 2.954 | 2.90E-05 | glio |
| Bora_in.vivo_noRT | Gm10419 | 3.21 | 2.90E-05 | glio |
| Aars2_ex.vivo_noRT | Pspn | 2.897 | 2.90E-06 | glio |
| Bora_ex.vivo_noRT | Gm11583 | 3.201 | 2.90E-06 | glio |
| Chek1_ex.vivo_noRT | Rtn1 | 4.127 | 2.90E-06 | glio |
| culture_effect | Bex2 | -0.918 | 2.90E-07 | glio |
| Chek1_in.vivo_noRT | 4930579K19R | 3.805 | 2.90E-07 | glio |
| Chek1_in.vivo_noRT | Pld6 | 3.787 | 2.90E-07 | glio |
| Chek1_ex.vivo_noRT | Dennd2d | 4.049 | 2.90E-07 | glio |
| Atr_ex.vivo_noRT | Ccser1 | 4.153 | 2.90E-08 | glio |
| Chek1_ex.vivo_noRT | Pcdhb20 | 4.674 | 2.90E-08 | glio |
| Atr_in.vivo_noRT | 4930592C13R | 4.267 | 2.90E-09 | glio |
| Aars2_in.vivo_noRT | 1700081N11R | 4.987 | 2.90E-10 | glio |
| Aars2_in.vivo_noRT | Uckl1os | 4.606 | 2.90E-10 | glio |
| Hsd17b10_in.vivo_noRT | AC159140.1 | 3.356 | 2.90E-10 | glio |
| Chek1_ex.vivo_noRT | Hrh2 | 4.842 | 2.90E-11 | glio |
| Chek1_ex.vivo_noRT | 4732463B04R | 4.449 | 2.90E-11 | glio |
| Chek1_in.vivo_noRT | Gm42695 | 4.784 | 2.90E-12 | glio |
| Chek1_in.vivo_noRT | Slc22a1 | 3.869 | 2.90E-12 | glio |
| Chek1_in.vivo_noRT | Gm15270 | 4.78 | 2.90E-16 | glio |
| culture_effect | Eef2 | -0.962 | 2.90E-20 | glio |
| Bora_in.vivo_noRT | Gm21936 | 4.936 | 2.90E-24 | glio |
| culture_effect | Napa | -0.429 | 2.91E-03 | glio |
| Interaction_genotype_Aars2 | Vsig10l | -3.721 | 2.91E-03 | glio |
| Interaction_genotype_Atr | Lmln | -3.602 | 2.91E-03 | glio |
| Interaction_genotype_Bora | Ccnh | 1.945 | 2.91E-03 | glio |
| Interaction_genotype_Chek1 | Gm38534 | -3.878 | 2.91E-03 | glio |
| Interaction_genotype_Chek1 | Haus8 | 2.252 | 2.91E-03 | glio |
| Interaction_genotype_Lmo7 | Folr1 | -3.81 | 2.91E-03 | glio |
| Interaction_genotype_Lmo7 | Nid1 | -4.661 | 2.91E-03 | glio |
| Interaction_genotype_Mtor | Klrg2 | -3.451 | 2.91E-03 | glio |
| interaction_RT | Trp53inp1 | 1.929 | 2.91E-03 | glio |
| interaction_RT | Dram2 | 1.416 | 2.91E-03 | glio |
| interaction_RT | Cnot3 | -0.691 | 2.91E-03 | glio |
| interaction_RT | Zfp637 | -0.726 | 2.91E-03 | glio |
| interaction_RT | Ndufb11 | 0.508 | 2.91E-03 | glio |
| Aars2_in.vivo_noRT | Fkbp11 | -1.229 | 2.91E-03 | glio |
| Aars2_in.vivo_noRT | Eef1b2 | -0.59 | 2.91E-03 | glio |
| Atr_in.vivo_noRT | Eif4e3 | 1.677 | 2.91E-03 | glio |
| Atr_in.vivo_noRT | Tlr3 | 1.349 | 2.91E-03 | glio |
| Bcas2_in.vivo_noRT | Tvp23a | 1.854 | 2.91E-03 | glio |
| Bora_in.vivo_noRT | Gm11520 | 2.006 | 2.91E-03 | glio |
| Cenpj_ex.vivo_noRT | Stx19 | 2.591 | 2.91E-03 | glio |
| Hsd17b10_in.vivo_noRT | Hspa12a | 2.316 | 2.91E-03 | glio |
| Hsd17b10_in.vivo_noRT | A930005H10R | -1.593 | 2.91E-03 | glio |

| | | | | |
|---|---|---|---|---|
| culture_effect | Kank2 | 1.633 | 2.91E-04 | glio |
| culture_effect | Ttc39a | -1.29 | 2.91E-04 | glio |
| Interaction_genotype_Mtor | Ptpn22 | -4.815 | 2.91E-04 | glio |
| Interaction_genotype_Myc | E030042O20R | -4.416 | 2.91E-04 | glio |
| Interaction_genotype_Myc | Slc7a14 | -3.963 | 2.91E-04 | glio |
| Interaction_genotype_Myc | Cdc42bpg | -3.836 | 2.91E-04 | glio |
| Interaction_genotype_Myc | Meak7 | -4.051 | 2.91E-04 | glio |
| Interaction_genotype_Myc | Tubg2 | -3.182 | 2.91E-04 | glio |
| Interaction_genotype_Myc | Serbp1 | 1.449 | 2.91E-04 | glio |
| Aars2_in.vivo_noRT | Gm26901 | 2.74 | 2.91E-04 | glio |
| Aars2_ex.vivo_noRT | Rasl12 | 3.293 | 2.91E-04 | glio |
| Atr_in.vivo_noRT | Anapc5 | -1.483 | 2.91E-04 | glio |
| Bora_in.vivo_noRT | Myl4 | 2.686 | 2.91E-04 | glio |
| Bora_in.vivo_noRT | L1cam | -1.476 | 2.91E-04 | glio |
| Chek1_in.vivo_noRT | Rps7 | -0.732 | 2.91E-04 | glio |
| Hsd17b10_in.vivo_noRT | Amdhd1 | 2.573 | 2.91E-04 | glio |
| culture_effect | Rgs12 | -1.569 | 2.91E-05 | glio |
| Interaction_genotype_Aars2 | AV356131 | -4.736 | 2.91E-05 | glio |
| Interaction_genotype_Aars2 | Gm49874 | -5.346 | 2.91E-05 | glio |
| Interaction_genotype_Aars2 | Gm44752 | -4.72 | 2.91E-05 | glio |
| Interaction_genotype_Myc | Ddx24 | 1.668 | 2.91E-05 | glio |
| Atr_in.vivo_noRT | Gm32618 | 3.013 | 2.91E-05 | glio |
| Atr_in.vivo_noRT | Mrpl50 | -1.355 | 2.91E-05 | glio |
| Atr_in.vivo_noRT | Rpl7a | -0.707 | 2.91E-05 | glio |
| interaction_RT | Serf1 | 0.975 | 2.91E-06 | glio |
| Bcas2_in.vivo_noRT | Kcnmb4os2 | 3.369 | 2.91E-06 | glio |
| Chek1_ex.vivo_noRT | Chaf1a | -2.548 | 2.91E-06 | glio |
| culture_effect | Gpr146 | -3.324 | 2.91E-07 | glio |
| Interaction_genotype_Mtor | Asgr1 | -5.93 | 2.91E-07 | glio |
| Interaction_genotype_Mtor | Gm5152 | -6.354 | 2.91E-07 | glio |
| Bcas2_in.vivo_noRT | Rtl9 | 3.366 | 2.91E-07 | glio |
| Bora_ex.vivo_noRT | Gm29966 | 3.201 | 2.91E-07 | glio |
| Chek1_in.vivo_noRT | Smc2os | 3.647 | 2.91E-07 | glio |
| Aars2_in.vivo_noRT | Prr7 | 3.936 | 2.91E-08 | glio |
| culture_effect | Psap | -0.78 | 2.91E-10 | glio |
| Bora_in.vivo_noRT | Gm47101 | 4.145 | 2.91E-10 | glio |
| Chek1_ex.vivo_noRT | Cidea | 4.842 | 2.91E-10 | glio |
| Hsd17b10_in.vivo_noRT | 1700112J16Ri | 3.616 | 2.91E-11 | glio |
| Hsd17b10_in.vivo_noRT | Clec4d | 3.616 | 2.91E-11 | glio |
| Aars2_in.vivo_noRT | Grin2d | 4.548 | 2.91E-12 | glio |
| Atr_in.vivo_noRT | Gm32090 | 4.741 | 2.91E-13 | glio |
| Chek1_ex.vivo_noRT | Gm29966 | 4.842 | 2.91E-16 | glio |
| Chek1_ex.vivo_noRT | Olfr1152 | 4.842 | 2.91E-16 | glio |
| Bora_in.vivo_noRT | Gm48435 | 4.936 | 2.91E-25 | glio |
| Interaction_genotype_Aars2 | Stk36 | -3.833 | 2.92E-03 | glio |
| Interaction_genotype_Aars2 | Stx4a | 1.929 | 2.92E-03 | glio |
| Interaction_genotype_Atr | Slc9a5 | -3.677 | 2.92E-03 | glio |
| Interaction_genotype_Mtor | Abca16 | -3.454 | 2.92E-03 | glio |
| Interaction_genotype_Mtor | Atxn1 | -2.739 | 2.92E-03 | glio |
| Interaction_genotype_Mtor | Gpr137 | -2.381 | 2.92E-03 | glio |
| Interaction_genotype_Mtor | Smad4 | 1.711 | 2.92E-03 | glio |
| Interaction_genotype_Mtor | Srsf6 | 1.462 | 2.92E-03 | glio |

| | | | | |
|---|---|---|---|---|
| Aars2_in.vivo_noRT | Cep83os | 2.014 | 2.92E-03 | glio |
| Aars2_in.vivo_noRT | Rsl1 | 2.079 | 2.92E-03 | glio |
| Aars2_in.vivo_noRT | Rnasek | -0.82 | 2.92E-03 | glio |
| Atp5c1_in.vivo_noRT | Kcnk4 | 2.408 | 2.92E-03 | glio |
| Atr_in.vivo_noRT | Gpr17 | 1.82 | 2.92E-03 | glio |
| Atr_ex.vivo_noRT | Cask | -1.849 | 2.92E-03 | glio |
| Bcas2_in.vivo_noRT | Igf2r | 2.052 | 2.92E-03 | glio |
| Bcas2_in.vivo_noRT | Ndufa12 | -0.563 | 2.92E-03 | glio |
| Bora_in.vivo_noRT | Zfp27 | 2.005 | 2.92E-03 | glio |
| Bora_in.vivo_noRT | Maco1 | -1.289 | 2.92E-03 | glio |
| Chek1_in.vivo_noRT | Gm13470 | 1.972 | 2.92E-03 | glio |
| Chek1_in.vivo_noRT | Lifr | 2.052 | 2.92E-03 | glio |
| Chek1_in.vivo_noRT | Rhot1 | -1.296 | 2.92E-03 | glio |
| Chek1_in.vivo_noRT | Mecr | -1.227 | 2.92E-03 | glio |
| Chek1_in.vivo_noRT | Rad23a | -1.246 | 2.92E-03 | glio |
| Chek1_in.vivo_noRT | Nedd8 | -0.643 | 2.92E-03 | glio |
| Eif2s1_in.vivo_noRT | Gm4876 | -2.096 | 2.92E-03 | glio |
| Hsd17b10_in.vivo_noRT | Gmeb2 | -1.81 | 2.92E-03 | glio |
| culture_effect | Vsig10 | -1.529 | 2.92E-04 | glio |
| Interaction_genotype_Mtor | Arl6ip1 | 2.746 | 2.92E-04 | glio |
| interaction_RT | Usp50 | 1.631 | 2.92E-04 | glio |
| interaction_RT | Gstm5 | 0.994 | 2.92E-04 | glio |
| interaction_RT | Irak1 | -0.741 | 2.92E-04 | glio |
| Atr_in.vivo_noRT | Ccdc14 | 2.371 | 2.92E-04 | glio |
| Bcas2_in.vivo_noRT | Gm34552 | 2.613 | 2.92E-04 | glio |
| Chek1_ex.vivo_noRT | Gm2762 | 3.036 | 2.92E-04 | glio |
| Hsd17b10_in.vivo_noRT | Serpina3g | 2.42 | 2.92E-04 | glio |
| culture_effect | 4931406P16R | 0.674 | 2.92E-05 | glio |
| Interaction_genotype_Mtor | Gm49785 | -4.791 | 2.92E-05 | glio |
| Interaction_genotype_Mtor | 9430037G07R | -5.077 | 2.92E-05 | glio |
| Interaction_genotype_Mtor | Gm13713 | -3.966 | 2.92E-05 | glio |
| Aars2_in.vivo_noRT | Gm14023 | 3.049 | 2.92E-05 | glio |
| Aars2_in.vivo_noRT | Gzmm | 2.998 | 2.92E-05 | glio |
| Aars2_in.vivo_noRT | Unc50 | -1.544 | 2.92E-05 | glio |
| Aars2_in.vivo_noRT | Txnl4a | -1.035 | 2.92E-05 | glio |
| Atp5c1_in.vivo_noRT | Ablim2 | 2.136 | 2.92E-05 | glio |
| Atr_in.vivo_noRT | Soga1 | 3.043 | 2.92E-05 | glio |
| Bora_in.vivo_noRT | Hapln4 | 3.169 | 2.92E-05 | glio |
| Chek1_ex.vivo_noRT | Npb | 3.652 | 2.92E-05 | glio |
| Hsd17b10_in.vivo_noRT | Pde1a | 3.143 | 2.92E-05 | glio |
| Atr_in.vivo_noRT | Gm47591 | 3.308 | 2.92E-06 | glio |
| Bora_in.vivo_noRT | Tmeff2 | 3.679 | 2.92E-06 | glio |
| Bora_in.vivo_noRT | Gm34590 | 3.363 | 2.92E-06 | glio |
| Hsd17b10_in.vivo_noRT | A830009L08Ri | 3.093 | 2.92E-06 | glio |
| interaction_RT | Eef1g | -0.973 | 2.92E-07 | glio |
| Aars2_in.vivo_noRT | Gm40040 | 3.77 | 2.92E-07 | glio |
| Bcas2_in.vivo_noRT | Gm47484 | 3.814 | 2.92E-07 | glio |
| Aars2_in.vivo_noRT | Tgfbr3l | 4.023 | 2.92E-08 | glio |
| Atr_in.vivo_noRT | Tnni2 | 4.495 | 2.92E-08 | glio |
| Atr_in.vivo_noRT | Styk1 | 3.947 | 2.92E-08 | glio |
| Atr_ex.vivo_noRT | Stfa3 | 3.164 | 2.92E-08 | glio |
| Atr_ex.vivo_noRT | Olfr787 | 3.164 | 2.92E-08 | glio |

| | | | | |
|---|---|---|---|---|
| Atr_in.vivo_noRT | 5330426L24Ri | 4.327 | 2.92E-09 | glio |
| culture_effect | Tmem179 | 3.226 | 2.92E-10 | glio |
| Bora_in.vivo_noRT | Gm17210 | 4.409 | 2.92E-10 | glio |
| Bora_in.vivo_noRT | E030042O20R | 4.936 | 2.92E-11 | glio |
| culture_effect | Tulp4 | 0.806 | 2.92E-13 | glio |
| Bcas2_in.vivo_noRT | Vmn2r54 | 4.175 | 2.92E-13 | glio |
| Bora_in.vivo_noRT | AI182371 | 4.154 | 2.92E-13 | glio |
| Bora_in.vivo_noRT | Necab2 | 4.672 | 2.92E-13 | glio |
| culture_effect | Prdx6 | -1.082 | 2.92E-14 | glio |
| Atr_in.vivo_noRT | Itgbl1 | 4.347 | 2.92E-14 | glio |
| Atr_in.vivo_noRT | Fbxl2 | 4.887 | 2.92E-15 | glio |
| Atr_in.vivo_noRT | Gm48265 | 4.75 | 2.92E-15 | glio |
| Bora_in.vivo_noRT | Gm9758 | 4.676 | 2.92E-19 | glio |
| culture_effect | Bcan | -3.152 | 2.92E-20 | glio |
| culture_effect | Rcor1 | -0.538 | 2.93E-03 | glio |
| culture_effect | Prmt2 | 0.561 | 2.93E-03 | glio |
| culture_effect | Srp14 | 0.283 | 2.93E-03 | glio |
| Interaction_genotype_Atr | Gm15579 | -4.011 | 2.93E-03 | glio |
| Interaction_genotype_Atr | 1700126G02R | -3.795 | 2.93E-03 | glio |
| Interaction_genotype_Mtor | Zfp322a | -2.19 | 2.93E-03 | glio |
| Interaction_genotype_Myc | Zfp316 | -2.794 | 2.93E-03 | glio |
| Interaction_genotype_Myc | Pgm2l1 | -2.881 | 2.93E-03 | glio |
| Interaction_genotype_Uqcrb | Gm41318 | -4.735 | 2.93E-03 | glio |
| Aars2_in.vivo_noRT | Zbtb42 | 2.077 | 2.93E-03 | glio |
| Atp5c1_in.vivo_noRT | 1700015F17Ri | 2.408 | 2.93E-03 | glio |
| Atr_in.vivo_noRT | A930041C12R | 2.114 | 2.93E-03 | glio |
| Atr_in.vivo_noRT | Dnaja4 | 2.136 | 2.93E-03 | glio |
| Atr_in.vivo_noRT | Pgm5 | 2.3 | 2.93E-03 | glio |
| Bcas2_in.vivo_noRT | 2410022M11F | 2.236 | 2.93E-03 | glio |
| Bcas2_in.vivo_noRT | Crip1 | -1.479 | 2.93E-03 | glio |
| Bcas2_ex.vivo_noRT | Gm36043 | 2.742 | 2.93E-03 | glio |
| Bcas2_ex.vivo_noRT | Prkcq | 2.742 | 2.93E-03 | glio |
| Bora_in.vivo_noRT | Tmem171 | 2.782 | 2.93E-03 | glio |
| Bora_in.vivo_noRT | Nyap1 | 2.184 | 2.93E-03 | glio |
| Bora_in.vivo_noRT | Rcn3 | -1.322 | 2.93E-03 | glio |
| Chek1_in.vivo_noRT | Gm41609 | 2.357 | 2.93E-03 | glio |
| Chek1_in.vivo_noRT | 4933404O12R | 2.139 | 2.93E-03 | glio |
| Chek1_in.vivo_noRT | Crocc | 2.124 | 2.93E-03 | glio |
| Chek1_in.vivo_noRT | Dubr | 1.741 | 2.93E-03 | glio |
| Hsd17b10_in.vivo_noRT | Tmem8b | 2.228 | 2.93E-03 | glio |
| Hsd17b10_in.vivo_noRT | Dock11 | 2.125 | 2.93E-03 | glio |
| Interaction_genotype_Mtor | Lrrc24 | -4.084 | 2.93E-04 | glio |
| Interaction_genotype_Mtor | Gm49801 | -4.468 | 2.93E-04 | glio |
| Interaction_genotype_Mtor | Zfp983 | -3.657 | 2.93E-04 | glio |
| Interaction_genotype_Mtor | Zfp276 | -3.52 | 2.93E-04 | glio |
| Interaction_genotype_Myc | Tmod1 | -3.612 | 2.93E-04 | glio |
| Interaction_genotype_Myc | Ist1 | -2.704 | 2.93E-04 | glio |
| Interaction_genotype_Myc | Ncl | 1.753 | 2.93E-04 | glio |
| interaction_RT | Psmd2 | -0.738 | 2.93E-04 | glio |
| interaction_RT | Cox8a | 0.636 | 2.93E-04 | glio |
| Aars2_ex.vivo_noRT | Gm50375 | 3.623 | 2.93E-04 | glio |
| Atp5c1_in.vivo_noRT | Fbxl13 | 2.408 | 2.93E-04 | glio |

| | | | | |
|---|---|---|---|---|
| Atr_in.vivo_noRT | Rps16 | -0.741 | 2.93E-04 | glio |
| Bcas2_in.vivo_noRT | Gm38190 | 2.419 | 2.93E-04 | glio |
| Bcas2_ex.vivo_noRT | 4930570E03Ri | 2.742 | 2.93E-04 | glio |
| Bcas2_ex.vivo_noRT | Gm5286 | 2.742 | 2.93E-04 | glio |
| Bora_in.vivo_noRT | Jagn1 | -1.398 | 2.93E-04 | glio |
| Bora_ex.vivo_noRT | Clca3a1 | 3.201 | 2.93E-04 | glio |
| Bora_ex.vivo_noRT | Irf8 | 3.201 | 2.93E-04 | glio |
| Eif2s1_in.vivo_noRT | Gm13178 | 2.154 | 2.93E-04 | glio |
| culture_effect | Mb | -2.654 | 2.93E-05 | glio |
| culture_effect | Nek6 | 1.31 | 2.93E-05 | glio |
| culture_effect | Rbm17 | 0.446 | 2.93E-05 | glio |
| Interaction_genotype_Myc | Helb | -3.63 | 2.93E-05 | glio |
| Interaction_genotype_Myc | Gtsf1 | -3.892 | 2.93E-05 | glio |
| Interaction_genotype_Myc | B3gnt4 | -4.194 | 2.93E-05 | glio |
| Interaction_genotype_Myc | Rps19 | 1.56 | 2.93E-05 | glio |
| Aars2_in.vivo_noRT | Odc1 | -1.785 | 2.93E-05 | glio |
| Atr_in.vivo_noRT | Rarres2 | 2.856 | 2.93E-05 | glio |
| culture_effect | Bex4 | -0.759 | 2.93E-06 | glio |
| culture_effect | Rps14 | -0.641 | 2.93E-06 | glio |
| Interaction_genotype_Mtor | Gm35867 | -5.46 | 2.93E-06 | glio |
| Interaction_genotype_Mtor | Med30 | 2.336 | 2.93E-06 | glio |
| Interaction_genotype_Mtor | Spata31d1a | -5.689 | 2.93E-06 | glio |
| Interaction_genotype_Mtor | Gm41409 | -5.111 | 2.93E-06 | glio |
| Bcas2_in.vivo_noRT | 4930445N18R | 3.363 | 2.93E-06 | glio |
| Bora_in.vivo_noRT | Gm19409 | 3.357 | 2.93E-06 | glio |
| Chek1_in.vivo_noRT | Gm15080 | 3.514 | 2.93E-06 | glio |
| Interaction_genotype_Mtor | 4930426I24Ri | -5.987 | 2.93E-07 | glio |
| Bcas2_in.vivo_noRT | A330084C13R | 3.889 | 2.93E-07 | glio |
| Chek1_in.vivo_noRT | BC037704 | 3.525 | 2.93E-07 | glio |
| Hsd17b10_in.vivo_noRT | 5033417F24Ri | 3.616 | 2.93E-07 | glio |
| interaction_RT | Itm2c | 1.438 | 2.93E-08 | glio |
| Bcas2_in.vivo_noRT | Cpne8 | 4.072 | 2.93E-08 | glio |
| Chek1_ex.vivo_noRT | Ly6c2 | 4.842 | 2.93E-08 | glio |
| culture_effect | Dnajc22 | -1.368 | 2.93E-09 | glio |
| Aars2_in.vivo_noRT | Gm43608 | 4.153 | 2.93E-09 | glio |
| Bcas2_in.vivo_noRT | Clca3a2 | 3.621 | 2.93E-09 | glio |
| Bora_in.vivo_noRT | Gm49312 | 3.862 | 2.93E-09 | glio |
| Chek1_in.vivo_noRT | Bglap2 | 4.115 | 2.93E-09 | glio |
| Hsd17b10_in.vivo_noRT | Gm11973 | 3.35 | 2.93E-09 | glio |
| Atr_in.vivo_noRT | Gm50334 | 4.888 | 2.93E-10 | glio |
| Chek1_in.vivo_noRT | Ccl19 | 3.616 | 2.93E-10 | glio |
| Atr_in.vivo_noRT | Wfdc1 | 4.549 | 2.93E-11 | glio |
| Bcas2_in.vivo_noRT | Ramp3 | 4.541 | 2.93E-11 | glio |
| culture_effect | Slc25a3 | -0.751 | 2.93E-13 | glio |
| Bcas2_in.vivo_noRT | Tns4 | 4.934 | 2.93E-15 | glio |
| Bora_in.vivo_noRT | Gm48242 | 4.936 | 2.93E-16 | glio |
| Bcas2_in.vivo_noRT | Slc16a5 | 4.543 | 2.93E-20 | glio |
| culture_effect | Dyrk3 | -2.718 | 2.93E-21 | glio |
| Aars2_in.vivo_noRT | Gm11232 | 5.327 | 2.93E-27 | glio |
| Atr_in.vivo_noRT | Gm14820 | 5.78 | 2.93E-27 | glio |
| Atr_in.vivo_noRT | Esm1 | 5.807 | 2.93E-28 | glio |
| culture_effect | Sav1 | 0.563 | 2.94E-03 | glio |

| | | | | |
|---|---|---|---|---|
| Interaction_genotype_Aars2 | Hspa12b | -4.093 | 2.94E-03 | glio |
| Interaction_genotype_Aars2 | Gtf2h5 | 1.008 | 2.94E-03 | glio |
| Interaction_genotype_Cdkn2a | Scg2 | 2.835 | 2.94E-03 | glio |
| Interaction_genotype_Mtor | CAAA0114733 | -3.245 | 2.94E-03 | glio |
| Interaction_genotype_Mtor | Ddx27 | 1.613 | 2.94E-03 | glio |
| Interaction_genotype_Mtor | Rbm8a | 1.377 | 2.94E-03 | glio |
| Interaction_genotype_Myc | Gm46447 | -3.054 | 2.94E-03 | glio |
| Interaction_genotype_Myc | Mfsd7a | -3.879 | 2.94E-03 | glio |
| Interaction_genotype_Myc | 3300002I08Ri | -3.34 | 2.94E-03 | glio |
| Interaction_genotype_Myc | Zfp866 | -2.965 | 2.94E-03 | glio |
| Interaction_genotype_Myc | Tuba4a | -2.96 | 2.94E-03 | glio |
| Interaction_genotype_Myc | Plcd3 | -2.847 | 2.94E-03 | glio |
| Interaction_genotype_Myc | Shld3 | -2.622 | 2.94E-03 | glio |
| Interaction_genotype_Myc | AC160336.1 | -2.72 | 2.94E-03 | glio |
| Interaction_genotype_Myc | Eya4 | -2.397 | 2.94E-03 | glio |
| interaction_RT | Gm15972 | 2.59 | 2.94E-03 | glio |
| interaction_RT | Rabggtb | 0.68 | 2.94E-03 | glio |
| interaction_RT | Hnrnpm | -0.63 | 2.94E-03 | glio |
| Aars2_in.vivo_noRT | Fam214a | 1.713 | 2.94E-03 | glio |
| Aars2_in.vivo_noRT | Lifr | 2.028 | 2.94E-03 | glio |
| Aars2_in.vivo_noRT | Pank4 | 1.868 | 2.94E-03 | glio |
| Aars2_in.vivo_noRT | Spop | -0.974 | 2.94E-03 | glio |
| Aars2_in.vivo_noRT | Cwc15 | -0.838 | 2.94E-03 | glio |
| Aars2_in.vivo_noRT | Rps10 | -0.634 | 2.94E-03 | glio |
| Atp5c1_in.vivo_noRT | Cacnb1 | 2.605 | 2.94E-03 | glio |
| Atr_in.vivo_noRT | Slu7 | -1.15 | 2.94E-03 | glio |
| Atr_ex.vivo_noRT | Gm36696 | 2.863 | 2.94E-03 | glio |
| Bcas2_in.vivo_noRT | Platr16 | 2.347 | 2.94E-03 | glio |
| Bcas2_in.vivo_noRT | Ptprg | 1.896 | 2.94E-03 | glio |
| Bcas2_in.vivo_noRT | Crot | 1.588 | 2.94E-03 | glio |
| Bcas2_in.vivo_noRT | Sf3a2 | -1.292 | 2.94E-03 | glio |
| Bcas2_ex.vivo_noRT | Gm48435 | 1.954 | 2.94E-03 | glio |
| Bora_ex.vivo_noRT | Mog | 3.201 | 2.94E-03 | glio |
| Cenpj_in.vivo_noRT | Ntn1 | 2.18 | 2.94E-03 | glio |
| Cenpj_in.vivo_noRT | Gm38163 | 2.022 | 2.94E-03 | glio |
| Cenpj_in.vivo_noRT | 1700006H21R | 2.548 | 2.94E-03 | glio |
| culture_effect | D430040D24R | 2.015 | 2.94E-04 | glio |
| Interaction_genotype_Mtor | Tbc1d24 | -4.25 | 2.94E-04 | glio |
| Interaction_genotype_Myc | Smtn | -3.253 | 2.94E-04 | glio |
| interaction_RT | Ly6c1 | 2.819 | 2.94E-04 | glio |
| Aars2_ex.vivo_noRT | Fst | 3.293 | 2.94E-04 | glio |
| Atr_in.vivo_noRT | C230035I16Ri | 2.655 | 2.94E-04 | glio |
| Bora_in.vivo_noRT | Zc3h14 | -1.342 | 2.94E-04 | glio |
| Bora_ex.vivo_noRT | Lck | -2.729 | 2.94E-04 | glio |
| Chek1_ex.vivo_noRT | E230025N22R | 3.096 | 2.94E-04 | glio |
| Interaction_genotype_Atr | Synm | -4.618 | 2.94E-05 | glio |
| Interaction_genotype_Myc | Pde3a | -4.345 | 2.94E-05 | glio |
| Interaction_genotype_Myc | Micall2 | -4.819 | 2.94E-05 | glio |
| Interaction_genotype_Myc | Slc25a10 | -3.788 | 2.94E-05 | glio |
| Interaction_genotype_Myc | Myo1h | -4.891 | 2.94E-05 | glio |
| Interaction_genotype_Myc | Hnrnpa2b1 | 1.652 | 2.94E-05 | glio |
| Aars2_in.vivo_noRT | Syt17 | 3.262 | 2.94E-05 | glio |

| | | | | |
|---|---|---|---|---|
| Aars2_ex.vivo_noRT | Kcnk7 | 3.293 | 2.94E-05 | glio |
| Aars2_ex.vivo_noRT | Myoz1 | 3.293 | 2.94E-05 | glio |
| Atr_in.vivo_noRT | Xrcc5 | -1.599 | 2.94E-05 | glio |
| Atr_ex.vivo_noRT | Gm36037 | 2.769 | 2.94E-05 | glio |
| Bora_in.vivo_noRT | Gm17399 | 3.195 | 2.94E-05 | glio |
| Chek1_in.vivo_noRT | Gm16740 | 2.949 | 2.94E-05 | glio |
| Chek1_ex.vivo_noRT | Ccnl2 | -1.995 | 2.94E-05 | glio |
| Hsd17b10_in.vivo_noRT | Gm28731 | 3.23 | 2.94E-05 | glio |
| Aars2_in.vivo_noRT | Etohd2 | 3.435 | 2.94E-06 | glio |
| Aars2_in.vivo_noRT | 2810414N06R | 3.305 | 2.94E-06 | glio |
| Bora_in.vivo_noRT | Polr3k | -1.743 | 2.94E-06 | glio |
| culture_effect | Tcf12 | -0.739 | 2.94E-07 | glio |
| Interaction_genotype_Mtor | Cyp26b1 | -6.289 | 2.94E-07 | glio |
| Aars2_in.vivo_noRT | Prex2 | 3.89 | 2.94E-07 | glio |
| Atr_in.vivo_noRT | Frmd8os | 4.016 | 2.94E-07 | glio |
| Bora_in.vivo_noRT | Gm15246 | 3.585 | 2.94E-07 | glio |
| Bora_ex.vivo_noRT | Gm26543 | 3.201 | 2.94E-07 | glio |
| culture_effect | Gm44190 | 2.326 | 2.94E-08 | glio |
| culture_effect | Mamld1 | 1.697 | 2.94E-08 | glio |
| culture_effect | Lgals3 | -1.011 | 2.94E-08 | glio |
| Aars2_in.vivo_noRT | Speg | 3.949 | 2.94E-08 | glio |
| Bcas2_in.vivo_noRT | Hist2h3c1 | 3.895 | 2.94E-08 | glio |
| Chek1_ex.vivo_noRT | Gm26542 | 4.66 | 2.94E-08 | glio |
| Aars2_in.vivo_noRT | Bpifa3 | 3.449 | 2.94E-09 | glio |
| Bora_in.vivo_noRT | Gm50295 | 4.144 | 2.94E-09 | glio |
| Bcas2_in.vivo_noRT | Ifitm5 | 4.284 | 2.94E-10 | glio |
| Bcas2_in.vivo_noRT | D730045A05R | 4.807 | 2.94E-12 | glio |
| culture_effect | Zdhhc3 | 1.006 | 2.94E-13 | glio |
| Atr_in.vivo_noRT | Krt80 | 5.267 | 2.94E-17 | glio |
| culture_effect | Myt1 | -4.545 | 2.94E-21 | glio |
| Bora_in.vivo_noRT | AU019990 | 4.936 | 2.94E-21 | glio |
| culture_effect | Reln | -1.727 | 2.94E-25 | glio |
| culture_effect | Cops9 | 1.941 | 2.94E-42 | glio |
| culture_effect | Fosb | -5.134 | 2.94E-42 | glio |
| culture_effect | Ubtd2 | 0.855 | 2.95E-03 | glio |
| culture_effect | 2410004B18R | -0.45 | 2.95E-03 | glio |
| culture_effect | Zfp770 | 0.505 | 2.95E-03 | glio |
| culture_effect | Nudt4 | 0.388 | 2.95E-03 | glio |
| culture_effect | Mt2 | -0.648 | 2.95E-03 | glio |
| culture_effect | Ndfip1 | -0.367 | 2.95E-03 | glio |
| Interaction_genotype_Bcas2 | Gm13348 | -3.337 | 2.95E-03 | glio |
| Interaction_genotype_Bcas2 | Dapp1 | -3.506 | 2.95E-03 | glio |
| Interaction_genotype_Cdkn2a | 1700003E24Ri | 2.835 | 2.95E-03 | glio |
| Interaction_genotype_Myc | Lhfpl3 | -3.244 | 2.95E-03 | glio |
| interaction_RT | Rplp1 | -0.665 | 2.95E-03 | glio |
| Aars2_in.vivo_noRT | Gm20721 | 1.999 | 2.95E-03 | glio |
| Aars2_in.vivo_noRT | Sqor | 1.945 | 2.95E-03 | glio |
| Atr_ex.vivo_noRT | Tspyl3 | 2.568 | 2.95E-03 | glio |
| Bcas2_in.vivo_noRT | Nup50 | -1.037 | 2.95E-03 | glio |
| Bcas2_in.vivo_noRT | Tcim | -1.255 | 2.95E-03 | glio |
| Bora_in.vivo_noRT | Pls1 | 2.15 | 2.95E-03 | glio |
| Bora_in.vivo_noRT | Best1 | 1.974 | 2.95E-03 | glio |

| | | | | |
|---|---|---|---|---|
| Bora_in.vivo_noRT | Dennd2c | 2.164 | 2.95E-03 | glio |
| Bora_in.vivo_noRT | Slc50a1 | -0.957 | 2.95E-03 | glio |
| Chek1_in.vivo_noRT | 4930590J08Ri | 2.238 | 2.95E-03 | glio |
| Chek1_ex.vivo_noRT | E130102H24R | 2.753 | 2.95E-03 | glio |
| Eif2s1_in.vivo_noRT | Crhbp | 2.143 | 2.95E-03 | glio |
| Hsd17b10_in.vivo_noRT | Gm16279 | 2.406 | 2.95E-03 | glio |
| Hsd17b10_in.vivo_noRT | Tubb2b | -0.763 | 2.95E-03 | glio |
| Interaction_genotype_Mtor | Gm13625 | -3.991 | 2.95E-04 | glio |
| Interaction_genotype_Mtor | Ttc30a1 | -4.39 | 2.95E-04 | glio |
| Interaction_genotype_Mtor | Zfp710 | -3.397 | 2.95E-04 | glio |
| Aars2_in.vivo_noRT | Pls1 | 2.541 | 2.95E-04 | glio |
| Aars2_in.vivo_noRT | Irs4 | 2.753 | 2.95E-04 | glio |
| Bcas2_in.vivo_noRT | Tspo | -0.866 | 2.95E-04 | glio |
| Bora_ex.vivo_noRT | Dnaaf3 | 3.201 | 2.95E-04 | glio |
| culture_effect | Ino80c | 0.796 | 2.95E-05 | glio |
| culture_effect | Ubqln2 | -0.723 | 2.95E-05 | glio |
| culture_effect | Srek1ip1 | 0.745 | 2.95E-05 | glio |
| culture_effect | Ndufs4 | -0.46 | 2.95E-05 | glio |
| Aars2_in.vivo_noRT | Gm12258 | 3.12 | 2.95E-05 | glio |
| Aars2_in.vivo_noRT | 4930404N11R | 2.726 | 2.95E-05 | glio |
| Aars2_in.vivo_noRT | Gpr141 | 3.353 | 2.95E-05 | glio |
| Aars2_ex.vivo_noRT | Usp17lc | 3.293 | 2.95E-05 | glio |
| Atp5c1_in.vivo_noRT | Scgb1c1 | 2.668 | 2.95E-05 | glio |
| Atr_in.vivo_noRT | Gm15614 | 2.888 | 2.95E-05 | glio |
| Atr_in.vivo_noRT | 1700010I14Ri | 2.863 | 2.95E-05 | glio |
| Bcas2_in.vivo_noRT | Hist1h4n | 3.258 | 2.95E-05 | glio |
| Chek1_in.vivo_noRT | Rimkla | 2.825 | 2.95E-05 | glio |
| Bora_ex.vivo_noRT | Gm29670 | 3.201 | 2.95E-06 | glio |
| Hsd17b10_in.vivo_noRT | Elobl | 3.616 | 2.95E-06 | glio |
| Chek1_in.vivo_noRT | Gm16283 | 3.52 | 2.95E-07 | glio |
| Chek1_ex.vivo_noRT | Ube4bos1 | 3.657 | 2.95E-07 | glio |
| Chek1_ex.vivo_noRT | Adm | 4.449 | 2.95E-07 | glio |
| Bcas2_in.vivo_noRT | 4933406C10R | 3.771 | 2.95E-09 | glio |
| Chek1_ex.vivo_noRT | Pla1a | 4.842 | 2.95E-09 | glio |
| Chek1_ex.vivo_noRT | Insrr | 4.842 | 2.95E-09 | glio |
| culture_effect | Btg2 | -1.823 | 2.95E-12 | glio |
| Atr_in.vivo_noRT | Cemip | 4.036 | 2.95E-12 | glio |
| Bora_in.vivo_noRT | 9930111J21Ri | 4.281 | 2.95E-12 | glio |
| Chek1_in.vivo_noRT | D630036H23F | 4.784 | 2.95E-12 | glio |
| culture_effect | Dancr | 1.654 | 2.95E-14 | glio |
| Aars2_in.vivo_noRT | Tmem52 | 4.947 | 2.95E-15 | glio |
| Chek1_ex.vivo_noRT | C430002N11R | 5.568 | 2.95E-16 | glio |
| culture_effect | Gm43653 | 1.694 | 2.96E-03 | glio |
| culture_effect | Raet1d | -1.015 | 2.96E-03 | glio |
| culture_effect | Ifi204 | -0.642 | 2.96E-03 | glio |
| culture_effect | Atp6v1d | 0.395 | 2.96E-03 | glio |
| Interaction_genotype_Aars2 | B930059L03Ri | -3.828 | 2.96E-03 | glio |
| Interaction_genotype_Aars2 | Plpp7 | -3.401 | 2.96E-03 | glio |
| Interaction_genotype_Aars2 | Yipf5 | 1.682 | 2.96E-03 | glio |
| Interaction_genotype_Bora | Vsig10l | -3.946 | 2.96E-03 | glio |
| Interaction_genotype_Bora | Notch4 | -3.884 | 2.96E-03 | glio |
| Interaction_genotype_Bora | Micall2 | -3.976 | 2.96E-03 | glio |

| | | | | |
|---|---|---|---|---|
| Interaction_genotype_Mtor | 4930458D05R | -3.999 | 2.96E-03 | glio |
| Interaction_genotype_Myc | Plce1 | -3.095 | 2.96E-03 | glio |
| Aars2_in.vivo_noRT | Gm49539 | 2.172 | 2.96E-03 | glio |
| Aars2_in.vivo_noRT | Tmem68 | -1.217 | 2.96E-03 | glio |
| Atp5c1_in.vivo_noRT | Gm5602 | 1.877 | 2.96E-03 | glio |
| Atp5c1_in.vivo_noRT | Acat3 | 1.876 | 2.96E-03 | glio |
| Atp5c1_in.vivo_noRT | Pmel | -2.102 | 2.96E-03 | glio |
| Atr_in.vivo_noRT | Fam120a | 0.893 | 2.96E-03 | glio |
| Atr_in.vivo_noRT | Rpl36a | -0.691 | 2.96E-03 | glio |
| Bcas2_in.vivo_noRT | 9130015A21R | 3.909 | 2.96E-03 | glio |
| Bcas2_in.vivo_noRT | Hist1h2af | 2.012 | 2.96E-03 | glio |
| Bcas2_in.vivo_noRT | Gpx7 | -1.089 | 2.96E-03 | glio |
| Bcas2_in.vivo_noRT | Ube2d2a | -0.755 | 2.96E-03 | glio |
| Bcas2_in.vivo_noRT | H2afz | -0.652 | 2.96E-03 | glio |
| Cenpj_ex.vivo_noRT | Masp1 | 2.592 | 2.96E-03 | glio |
| Chek1_ex.vivo_noRT | 4933432I03Ri | 2.756 | 2.96E-03 | glio |
| Chek1_ex.vivo_noRT | Phyh | 2.219 | 2.96E-03 | glio |
| Hsd17b10_in.vivo_noRT | Slc7a4 | 2.202 | 2.96E-03 | glio |
| Bard1_ex.vivo_noRT | Gm26671 | 2.421 | 2.96E-04 | glio |
| Bcas2_in.vivo_noRT | Zfp59 | 2.65 | 2.96E-04 | glio |
| Bcas2_ex.vivo_noRT | Gm15894 | 2.742 | 2.96E-04 | glio |
| Chek1_in.vivo_noRT | Opalin | 3.329 | 2.96E-04 | glio |
| Hsd17b10_in.vivo_noRT | Npc1 | -1.942 | 2.96E-04 | glio |
| Interaction_genotype_Mtor | Bmx | -4.49 | 2.96E-05 | glio |
| Interaction_genotype_Mtor | Esp15 | -4.099 | 2.96E-05 | glio |
| Atr_in.vivo_noRT | Sh3bgrl2 | 2.924 | 2.96E-05 | glio |
| Bora_in.vivo_noRT | Elovl3 | 3.009 | 2.96E-05 | glio |
| Hsd17b10_in.vivo_noRT | Slc6a13 | 3.237 | 2.96E-05 | glio |
| culture_effect | Ube2n | 0.48 | 2.96E-06 | glio |
| Interaction_genotype_Mtor | Slc24a1 | -5.715 | 2.96E-06 | glio |
| Interaction_genotype_Mtor | AU019990 | -4.099 | 2.96E-06 | glio |
| Bora_in.vivo_noRT | Sbf1 | -1.972 | 2.96E-06 | glio |
| culture_effect | Pcdhb20 | -3.025 | 2.96E-07 | glio |
| culture_effect | Gm7467 | 1.969 | 2.96E-07 | glio |
| culture_effect | Tmem134 | 0.707 | 2.96E-07 | glio |
| Interaction_genotype_Mtor | Gm48170 | -4.896 | 2.96E-07 | glio |
| Atr_in.vivo_noRT | Gm14443 | 3.338 | 2.96E-07 | glio |
| Atr_ex.vivo_noRT | Gm16344 | 3.947 | 2.96E-07 | glio |
| Bcas2_in.vivo_noRT | Chd5 | 3.562 | 2.96E-07 | glio |
| Bcas2_in.vivo_noRT | Gm47972 | 3.513 | 2.96E-07 | glio |
| Bora_in.vivo_noRT | Gm45833 | 4.856 | 2.96E-07 | glio |
| Bora_in.vivo_noRT | Dcpp3 | 3.419 | 2.96E-07 | glio |
| culture_effect | Acox2 | -3.008 | 2.96E-08 | glio |
| Bcas2_in.vivo_noRT | Nyap2 | 3.895 | 2.96E-08 | glio |
| Aars2_in.vivo_noRT | 9930014A18R | 3.932 | 2.96E-09 | glio |
| Chek1_in.vivo_noRT | Rilp | 4.518 | 2.96E-09 | glio |
| Aars2_in.vivo_noRT | Shroom3 | 4.562 | 2.96E-11 | glio |
| Atr_in.vivo_noRT | Hist2h3c1 | 4.622 | 2.96E-11 | glio |
| Bora_in.vivo_noRT | Kcnk7 | 3.888 | 2.96E-11 | glio |
| Chek1_in.vivo_noRT | Slc16a7 | 3.932 | 2.96E-11 | glio |
| Hsd17b10_in.vivo_noRT | Ccdc80 | 3.616 | 2.96E-11 | glio |
| Bcas2_in.vivo_noRT | Igf2bp1 | 4.807 | 2.96E-12 | glio |

| | | | | |
|---|---|---|---|---|
| Bora_in.vivo_noRT | Gm41409 | 4.405 | 2.96E-12 | glio |
| Chek1_in.vivo_noRT | D6Ertd527e | 4.772 | 2.96E-12 | glio |
| culture_effect | Prelid1 | 0.778 | 2.96E-13 | glio |
| culture_effect | Odc1 | -1.437 | 2.96E-13 | glio |
| Bcas2_in.vivo_noRT | Gm5602 | 4.016 | 2.96E-13 | glio |
| culture_effect | Prdx4 | -1.085 | 2.96E-14 | glio |
| Aars2_in.vivo_noRT | Zfp810 | 5.327 | 2.96E-15 | glio |
| Bcas2_in.vivo_noRT | Ceacam16 | 4.679 | 2.96E-15 | glio |
| Bcas2_in.vivo_noRT | Stra8 | 4.802 | 2.96E-15 | glio |
| Chek1_ex.vivo_noRT | Gm48155 | 4.842 | 2.96E-17 | glio |
| culture_effect | Ly6e | -2.996 | 2.96E-47 | glio |
| culture_effect | Dnm1 | -1.566 | 2.97E-03 | glio |
| culture_effect | Emc3 | -0.444 | 2.97E-03 | glio |
| culture_effect | Ddx39 | -0.402 | 2.97E-03 | glio |
| Interaction_genotype_Bora | Cep295nl | -3.93 | 2.97E-03 | glio |
| Interaction_genotype_Chek1 | Gm30938 | -3.903 | 2.97E-03 | glio |
| Interaction_genotype_Chek1 | Map1lc3a | 1.338 | 2.97E-03 | glio |
| Interaction_genotype_Mtor | Smc2os | -3.453 | 2.97E-03 | glio |
| Interaction_genotype_Mtor | Tex15 | -3.42 | 2.97E-03 | glio |
| Interaction_genotype_Mtor | Lrrc49 | -2.948 | 2.97E-03 | glio |
| Interaction_genotype_Mtor | Nexn | -3.164 | 2.97E-03 | glio |
| Interaction_genotype_Mtor | Sag | -3.499 | 2.97E-03 | glio |
| Interaction_genotype_Mtor | Ppard | -2.659 | 2.97E-03 | glio |
| Interaction_genotype_Mtor | Tgfb2 | -2.94 | 2.97E-03 | glio |
| Aars2_in.vivo_noRT | Arhgap10 | -1.494 | 2.97E-03 | glio |
| Aars2_in.vivo_noRT | Zmynd19 | -1.236 | 2.97E-03 | glio |
| Atr_ex.vivo_noRT | Cxcl13 | 2.771 | 2.97E-03 | glio |
| Atr_ex.vivo_noRT | Pcdhb19 | 2.769 | 2.97E-03 | glio |
| Bard1_ex.vivo_noRT | Slc22a1 | 2.419 | 2.97E-03 | glio |
| Bard1_ex.vivo_noRT | Tpsab1 | 2.814 | 2.97E-03 | glio |
| Bcas2_ex.vivo_noRT | Auts2 | 2.742 | 2.97E-03 | glio |
| Bcas2_ex.vivo_noRT | Col19a1 | 2.349 | 2.97E-03 | glio |
| Bcas2_ex.vivo_noRT | Cdh10 | -2.469 | 2.97E-03 | glio |
| Bora_in.vivo_noRT | Snn | 2.201 | 2.97E-03 | glio |
| Bora_in.vivo_noRT | Gm19391 | 2.037 | 2.97E-03 | glio |
| Bora_ex.vivo_noRT | Pcdhgb1 | 2.626 | 2.97E-03 | glio |
| Cenpj_in.vivo_noRT | Gm50024 | 1.759 | 2.97E-03 | glio |
| Chek1_in.vivo_noRT | B4galt4 | 2.225 | 2.97E-03 | glio |
| Chek1_ex.vivo_noRT | Slc12a5 | 2.711 | 2.97E-03 | glio |
| Chek1_ex.vivo_noRT | Dnaaf4 | 3.006 | 2.97E-03 | glio |
| Hsd17b10_in.vivo_noRT | EU599041 | 2.339 | 2.97E-03 | glio |
| Hsd17b10_in.vivo_noRT | Bbof1 | 2.188 | 2.97E-03 | glio |
| Hsd17b10_in.vivo_noRT | Cyp27b1 | 2.055 | 2.97E-03 | glio |
| culture_effect | Gm35551 | -2.164 | 2.97E-04 | glio |
| culture_effect | Pbdc1 | -0.426 | 2.97E-04 | glio |
| Interaction_genotype_Atr | Gm14968 | -4.387 | 2.97E-04 | glio |
| Interaction_genotype_Atr | Gm10425 | -4.663 | 2.97E-04 | glio |
| Interaction_genotype_Myc | Gm12840 | -3.845 | 2.97E-04 | glio |
| Interaction_genotype_Myc | Serping1 | 3.494 | 2.97E-04 | glio |
| Interaction_genotype_Myc | Gm9873 | -3.693 | 2.97E-04 | glio |
| Interaction_genotype_Rad21 | Fst | 4.868 | 2.97E-04 | glio |
| Interaction_genotype_Rad21 | Gm47936 | 3.784 | 2.97E-04 | glio |

| | | | | |
|---|---|---|---|---|
| interaction_RT | Meiob | 2.07 | 2.97E-04 | glio |
| interaction_RT | Wbp11 | -0.628 | 2.97E-04 | glio |
| Aars2_in.vivo_noRT | Ndufaf2 | -1.278 | 2.97E-04 | glio |
| Atp5c1_in.vivo_noRT | Gm38393 | 2.408 | 2.97E-04 | glio |
| Bora_in.vivo_noRT | Fam83e | 2.687 | 2.97E-04 | glio |
| Bora_in.vivo_noRT | Pdik1l | 1.844 | 2.97E-04 | glio |
| Chek1_in.vivo_noRT | A430105J06Ri | 2.535 | 2.97E-04 | glio |
| Chek1_in.vivo_noRT | Castor2 | 2.428 | 2.97E-04 | glio |
| Chek1_ex.vivo_noRT | Moap1 | 3.008 | 2.97E-04 | glio |
| culture_effect | Dzip3 | 0.932 | 2.97E-05 | glio |
| culture_effect | Prnp | -0.713 | 2.97E-05 | glio |
| Interaction_genotype_Atr | Pygo1 | -4.833 | 2.97E-05 | glio |
| interaction_RT | Mpp6 | -0.968 | 2.97E-05 | glio |
| Aars2_in.vivo_noRT | Rxra | 3.088 | 2.97E-05 | glio |
| Aars2_in.vivo_noRT | Ptn | 3.438 | 2.97E-05 | glio |
| Bora_in.vivo_noRT | Tmc7 | 3.124 | 2.97E-05 | glio |
| Bora_in.vivo_noRT | Lrrc75aos2 | 3.04 | 2.97E-05 | glio |
| Bora_in.vivo_noRT | Aldh2 | -1.855 | 2.97E-05 | glio |
| Aars2_in.vivo_noRT | Haglr | 3.546 | 2.97E-06 | glio |
| Atr_in.vivo_noRT | Nts | 3.49 | 2.97E-06 | glio |
| Bcas2_in.vivo_noRT | 9430065F17Ri | 3.361 | 2.97E-06 | glio |
| Chek1_in.vivo_noRT | Xist | 3.191 | 2.97E-06 | glio |
| Chek1_ex.vivo_noRT | Hist1h2bf | 4.406 | 2.97E-06 | glio |
| Aars2_in.vivo_noRT | Vcp | -1.603 | 2.97E-07 | glio |
| Bcas2_in.vivo_noRT | A230087F16Ri | 3.496 | 2.97E-07 | glio |
| Chek1_ex.vivo_noRT | Mapk15 | 4.452 | 2.97E-07 | glio |
| Hsd17b10_in.vivo_noRT | Gm4166 | 2.689 | 2.97E-07 | glio |
| culture_effect | Acsl4 | 1.243 | 2.97E-08 | glio |
| Chek1_in.vivo_noRT | Gm15927 | 4.042 | 2.97E-09 | glio |
| Bcas2_in.vivo_noRT | Smpd5 | 4.15 | 2.97E-11 | glio |
| Chek1_in.vivo_noRT | Usp17la | 5.044 | 2.97E-13 | glio |
| Atr_in.vivo_noRT | Gm10125 | 5.385 | 2.97E-14 | glio |
| Chek1_ex.vivo_noRT | Pspn | 4.447 | 2.97E-14 | glio |
| Chek1_ex.vivo_noRT | Rgs4 | 4.842 | 2.97E-14 | glio |
| Bcas2_in.vivo_noRT | Ccser1 | 4.807 | 2.97E-16 | glio |
| Chek1_in.vivo_noRT | Vmn2r56 | 4.791 | 2.97E-18 | glio |
| culture_effect | Tapbp | -2.291 | 2.97E-43 | glio |
| Interaction_genotype_Aars2 | Inhba | -4.034 | 2.98E-03 | glio |
| Interaction_genotype_Aars2 | Tnni3k | -2.821 | 2.98E-03 | glio |
| Interaction_genotype_Aars2 | Aloxe3 | -3.143 | 2.98E-03 | glio |
| Interaction_genotype_Aars2 | Gm15445 | -2.883 | 2.98E-03 | glio |
| Interaction_genotype_Aars2 | Zfp629 | -3.328 | 2.98E-03 | glio |
| Interaction_genotype_Aars2 | 0610009L18Ri | -3.554 | 2.98E-03 | glio |
| Interaction_genotype_Aars2 | Trmu | -2.686 | 2.98E-03 | glio |
| Interaction_genotype_Aars2 | Ptrh2 | 1.484 | 2.98E-03 | glio |
| Interaction_genotype_Aars2 | Msmo1 | 1.884 | 2.98E-03 | glio |
| Interaction_genotype_Aars2 | Tmbim4 | 1.357 | 2.98E-03 | glio |
| Interaction_genotype_Atr | Stfa3 | -2.369 | 2.98E-03 | glio |
| Interaction_genotype_Atr | Olfr787 | -2.369 | 2.98E-03 | glio |
| Interaction_genotype_Atr | 4833418N02R | -3.805 | 2.98E-03 | glio |
| Interaction_genotype_Atr | Wls | 1.798 | 2.98E-03 | glio |
| Interaction_genotype_Atr | Tle5 | 1.145 | 2.98E-03 | glio |

| | | | | |
|---|---|---|---|---|
| Interaction_genotype_Mtor | Unc13d | -3.472 | 2.98E-03 | glio |
| Interaction_genotype_Mtor | Cbr3 | -3.575 | 2.98E-03 | glio |
| Interaction_genotype_Mtor | Rusc2 | -3.286 | 2.98E-03 | glio |
| Interaction_genotype_Mtor | Asb16 | -3.284 | 2.98E-03 | glio |
| Interaction_genotype_Mtor | 1600014C10R | -2.639 | 2.98E-03 | glio |
| Interaction_genotype_Mtor | Ldha | 1.93 | 2.98E-03 | glio |
| Interaction_genotype_Mtor | Anxa2 | 1.751 | 2.98E-03 | glio |
| Interaction_genotype_Mtor | Ftl1 | 1.167 | 2.98E-03 | glio |
| Interaction_genotype_Myc | Gm14820 | -2.466 | 2.98E-03 | glio |
| Interaction_genotype_Myc | Card9 | -2.865 | 2.98E-03 | glio |
| Interaction_genotype_Myc | Acad10 | -3.426 | 2.98E-03 | glio |
| Interaction_genotype_Myc | Gm10825 | -3.385 | 2.98E-03 | glio |
| Interaction_genotype_Myc | Fndc7 | -3.503 | 2.98E-03 | glio |
| Interaction_genotype_Myc | Pfdn2 | 1.043 | 2.98E-03 | glio |
| Aars2_in.vivo_noRT | 1110019D14R | 2.067 | 2.98E-03 | glio |
| Aars2_in.vivo_noRT | Zfp36l2 | -1.374 | 2.98E-03 | glio |
| Aars2_in.vivo_noRT | Rnh1 | -0.986 | 2.98E-03 | glio |
| Atp5c1_in.vivo_noRT | Tmem190 | 2.668 | 2.98E-03 | glio |
| Bcas2_in.vivo_noRT | Cyb561 | 2.332 | 2.98E-03 | glio |
| Bcas2_in.vivo_noRT | Irak3 | 2.291 | 2.98E-03 | glio |
| Bcas2_in.vivo_noRT | Mmd2 | 1.763 | 2.98E-03 | glio |
| Bcas2_ex.vivo_noRT | Gm38477 | 2.342 | 2.98E-03 | glio |
| Bcas2_ex.vivo_noRT | 4732463B04R | 2.349 | 2.98E-03 | glio |
| Bora_ex.vivo_noRT | Impg2 | 2.413 | 2.98E-03 | glio |
| Eif2s1_in.vivo_noRT | Flacc1 | 2.143 | 2.98E-03 | glio |
| Hsd17b10_in.vivo_noRT | Lmf2 | -1.186 | 2.98E-03 | glio |
| culture_effect | Ppp6c | -0.567 | 2.98E-04 | glio |
| Interaction_genotype_Aars2 | Gm13270 | -4.685 | 2.98E-04 | glio |
| Interaction_genotype_Aars2 | Psmb2 | 1.322 | 2.98E-04 | glio |
| Interaction_genotype_Aars2 | Ndufa7 | 1.07 | 2.98E-04 | glio |
| interaction_RT | Zdhhc15 | 2.545 | 2.98E-04 | glio |
| Atp5c1_in.vivo_noRT | Gm45669 | 2.405 | 2.98E-04 | glio |
| Atr_ex.vivo_noRT | Rpl3l | 3.164 | 2.98E-04 | glio |
| Bcas2_ex.vivo_noRT | Sap30bpos | 3.131 | 2.98E-04 | glio |
| Bora_in.vivo_noRT | Gm14966 | 2.469 | 2.98E-04 | glio |
| Chek1_ex.vivo_noRT | Pcdhga6 | 3.113 | 2.98E-04 | glio |
| Hsd17b10_in.vivo_noRT | Prox2 | 2.89 | 2.98E-04 | glio |
| culture_effect | Snhg14 | -2.689 | 2.98E-05 | glio |
| Interaction_genotype_Mtor | AC166172.1 | -4.488 | 2.98E-05 | glio |
| Interaction_genotype_Mtor | Gm10801 | -4.099 | 2.98E-05 | glio |
| Atr_in.vivo_noRT | Gm42941 | 2.74 | 2.98E-05 | glio |
| Atr_ex.vivo_noRT | Slc45a1 | 2.764 | 2.98E-05 | glio |
| Bard1_ex.vivo_noRT | Cd7 | 2.814 | 2.98E-05 | glio |
| Bard1_ex.vivo_noRT | Gm48764 | 2.814 | 2.98E-05 | glio |
| Bard1_ex.vivo_noRT | 1700108F19Ri | 3.219 | 2.98E-05 | glio |
| culture_effect | Ptrh2 | -0.669 | 2.98E-06 | glio |
| Aars2_in.vivo_noRT | Rarres2 | 3.186 | 2.98E-06 | glio |
| Atr_in.vivo_noRT | Des | 3.283 | 2.98E-06 | glio |
| Atr_in.vivo_noRT | Gm21154 | 3.48 | 2.98E-06 | glio |
| Atr_in.vivo_noRT | Card14 | 3.331 | 2.98E-06 | glio |
| Bora_in.vivo_noRT | Gm48529 | 3.207 | 2.98E-06 | glio |
| Bora_in.vivo_noRT | Rpl3l | 3 | 2.98E-06 | glio |

| | | | | |
|---|---|---|---|---|
| culture_effect | Scfd1 | -0.836 | 2.98E-07 | glio |
| culture_effect | Plp1 | -1.203 | 2.98E-07 | glio |
| Interaction_genotype_Myc | Zfhx2os | -6.023 | 2.98E-07 | glio |
| Interaction_genotype_Myc | Tnfrsf23 | -6.619 | 2.98E-07 | glio |
| interaction_RT | Gbp9 | 1.957 | 2.98E-07 | glio |
| Bora_ex.vivo_noRT | Cgnl1 | 3.201 | 2.98E-07 | glio |
| Chek1_ex.vivo_noRT | Fam20b | -2.63 | 2.98E-07 | glio |
| culture_effect | Calhm2 | 1.844 | 2.98E-08 | glio |
| Chek1_ex.vivo_noRT | Ms4a6d | 4.055 | 2.98E-08 | glio |
| Hsd17b10_in.vivo_noRT | Gm16741 | 3.616 | 2.98E-08 | glio |
| Atr_in.vivo_noRT | Nyap2 | 4.621 | 2.98E-11 | glio |
| Chek1_ex.vivo_noRT | C330024C12R | 5.218 | 2.98E-12 | glio |
| Bcas2_in.vivo_noRT | Tent5c | 3.881 | 2.98E-13 | glio |
| Bora_in.vivo_noRT | Bglap2 | 5.066 | 2.98E-13 | glio |
| Chek1_ex.vivo_noRT | Gm21761 | 4.842 | 2.98E-13 | glio |
| Chek1_ex.vivo_noRT | Olfr552 | 4.443 | 2.98E-13 | glio |
| Aars2_in.vivo_noRT | 4930505N22R | 5.067 | 2.98E-15 | glio |
| Bcas2_in.vivo_noRT | Hrct1 | 4.807 | 2.98E-15 | glio |
| Bora_in.vivo_noRT | Cxcl5 | 4.936 | 2.98E-20 | glio |
| Atr_in.vivo_noRT | Cd36 | 5.533 | 2.98E-26 | glio |
| Atr_in.vivo_noRT | Clec12b | 5.533 | 2.98E-28 | glio |
| Atr_in.vivo_noRT | Gm14929 | 5.533 | 2.98E-28 | glio |
| Interaction_genotype_Atr | 8030456M14F | -3.481 | 2.99E-03 | glio |
| Interaction_genotype_Mtor | Cidea | -3.184 | 2.99E-03 | glio |
| Interaction_genotype_Mtor | Cep76 | -2.513 | 2.99E-03 | glio |
| Interaction_genotype_Mtor | Ogt | 1.616 | 2.99E-03 | glio |
| Interaction_genotype_Myc | Rimkla | -3.237 | 2.99E-03 | glio |
| Interaction_genotype_Myc | Cdh24 | -3.668 | 2.99E-03 | glio |
| Interaction_genotype_Myc | Smc2os | -3.306 | 2.99E-03 | glio |
| Interaction_genotype_Myc | Usp27x | -3.445 | 2.99E-03 | glio |
| Interaction_genotype_Rad21 | Dpp6 | 3.148 | 2.99E-03 | glio |
| Interaction_genotype_Rad21 | Tpsab1 | 4.17 | 2.99E-03 | glio |
| Interaction_genotype_Rad21 | Mag | 4.32 | 2.99E-03 | glio |
| Aars2_in.vivo_noRT | Acads | -1.37 | 2.99E-03 | glio |
| Atr_in.vivo_noRT | Gm45871 | 1.943 | 2.99E-03 | glio |
| Atr_in.vivo_noRT | 1700052K11R | 1.727 | 2.99E-03 | glio |
| Atr_in.vivo_noRT | Lmf2 | -1.217 | 2.99E-03 | glio |
| Atr_ex.vivo_noRT | Mylip | 2.134 | 2.99E-03 | glio |
| Atr_ex.vivo_noRT | Mt2 | 1.42 | 2.99E-03 | glio |
| Bcas2_in.vivo_noRT | Hcn2 | 2.354 | 2.99E-03 | glio |
| Bcas2_in.vivo_noRT | 1700010I14Ri | 2.137 | 2.99E-03 | glio |
| Bcas2_in.vivo_noRT | Tmem218 | 1.684 | 2.99E-03 | glio |
| Bcas2_ex.vivo_noRT | Strip2 | 2.342 | 2.99E-03 | glio |
| Bora_in.vivo_noRT | Drc3 | 2.239 | 2.99E-03 | glio |
| Chek1_ex.vivo_noRT | Chdh | 2.771 | 2.99E-03 | glio |
| Chek1_ex.vivo_noRT | Cep85l | 2.379 | 2.99E-03 | glio |
| Hsd17b10_in.vivo_noRT | Guca1b | 2.328 | 2.99E-03 | glio |
| culture_effect | Fam76a | 0.758 | 2.99E-04 | glio |
| culture_effect | Sec13 | -0.544 | 2.99E-04 | glio |
| Interaction_genotype_Atr | Gm14418 | -3.23 | 2.99E-04 | glio |
| Interaction_genotype_Atr | Rab25 | -4.048 | 2.99E-04 | glio |
| Interaction_genotype_Mtor | Cdkn1c | -4.339 | 2.99E-04 | glio |

| | | | | |
|---|---|---|---|---|
| Interaction_genotype_Mtor | Gm48646 | -4.335 | 2.99E-04 | glio |
| interaction_RT | Gm10687 | 2.885 | 2.99E-04 | glio |
| interaction_RT | Tuba1b | -0.87 | 2.99E-04 | glio |
| Aars2_in.vivo_noRT | Tigd2 | 1.901 | 2.99E-04 | glio |
| Atr_ex.vivo_noRT | 1700112J16Ri | 2.577 | 2.99E-04 | glio |
| Bcas2_in.vivo_noRT | Platr3 | 2.744 | 2.99E-04 | glio |
| Cenpj_in.vivo_noRT | Gm34059 | 2.022 | 2.99E-04 | glio |
| Cenpj_in.vivo_noRT | Gm48435 | 2.022 | 2.99E-04 | glio |
| Cenpj_in.vivo_noRT | Helt | 2.022 | 2.99E-04 | glio |
| Cenpj_in.vivo_noRT | Gm20625 | 2.022 | 2.99E-04 | glio |
| Cenpj_in.vivo_noRT | Gm5157 | 2.022 | 2.99E-04 | glio |
| Cenpj_in.vivo_noRT | Gm44736 | 2.022 | 2.99E-04 | glio |
| Cenpj_in.vivo_noRT | Fam227b | 2.022 | 2.99E-04 | glio |
| Cenpj_in.vivo_noRT | Gm20658 | 2.022 | 2.99E-04 | glio |
| Cenpj_in.vivo_noRT | Olfr810 | 2.022 | 2.99E-04 | glio |
| Cenpj_ex.vivo_noRT | Cd200r2 | 2.591 | 2.99E-04 | glio |
| Chek1_in.vivo_noRT | C230035I16Ri | 2.689 | 2.99E-04 | glio |
| Chek1_in.vivo_noRT | Elf2 | -1.236 | 2.99E-04 | glio |
| culture_effect | Dpp6 | -1.862 | 2.99E-05 | glio |
| Interaction_genotype_Myc | Tph2 | -4.357 | 2.99E-05 | glio |
| Interaction_genotype_Myc | Gm15579 | -5.169 | 2.99E-05 | glio |
| Interaction_genotype_Myc | 4921504E06Ri | -3.818 | 2.99E-05 | glio |
| interaction_RT | Smo | -1.547 | 2.99E-05 | glio |
| interaction_RT | Hirip3 | -1.439 | 2.99E-05 | glio |
| Aars2_in.vivo_noRT | Gpr135 | 2.912 | 2.99E-05 | glio |
| Atr_in.vivo_noRT | Rpl11 | -0.942 | 2.99E-05 | glio |
| Bcas2_in.vivo_noRT | Lbhd2 | 3.565 | 2.99E-05 | glio |
| Chek1_ex.vivo_noRT | B230303O12R | 3.861 | 2.99E-05 | glio |
| culture_effect | Ufd1 | -0.828 | 2.99E-06 | glio |
| culture_effect | Coa5 | 0.846 | 2.99E-06 | glio |
| Atr_in.vivo_noRT | Ttc41 | 3.462 | 2.99E-06 | glio |
| Atr_in.vivo_noRT | Car5a | 2.969 | 2.99E-06 | glio |
| Chek1_in.vivo_noRT | Gramd2 | 3.206 | 2.99E-06 | glio |
| interaction_RT | Kdm7a | 1.951 | 2.99E-07 | glio |
| Atr_in.vivo_noRT | Gm28592 | 3.319 | 2.99E-07 | glio |
| Atr_in.vivo_noRT | Zfp11 | 3.566 | 2.99E-07 | glio |
| Atr_in.vivo_noRT | Itgb6 | 3.605 | 2.99E-07 | glio |
| Atr_in.vivo_noRT | Pcbp4 | -1.749 | 2.99E-07 | glio |
| Atr_in.vivo_noRT | Ccnd3 | -1.586 | 2.99E-07 | glio |
| Chek1_in.vivo_noRT | Gabrd | 3.594 | 2.99E-07 | glio |
| Bora_in.vivo_noRT | Tbxa2r | 3.621 | 2.99E-08 | glio |
| Bcas2_in.vivo_noRT | Pecam1 | 4.275 | 2.99E-09 | glio |
| Aars2_in.vivo_noRT | Gm20455 | 4.606 | 2.99E-10 | glio |
| Chek1_ex.vivo_noRT | Fhl1 | 4.132 | 2.99E-10 | glio |
| Hsd17b10_in.vivo_noRT | Gm16933 | 3.094 | 2.99E-10 | glio |
| culture_effect | Cacng4 | -3.681 | 2.99E-11 | glio |
| Bcas2_in.vivo_noRT | Gm47802 | 4.359 | 2.99E-11 | glio |
| Chek1_in.vivo_noRT | Fam189a1 | 3.885 | 2.99E-11 | glio |
| Atr_in.vivo_noRT | Gm32591 | 5.011 | 2.99E-13 | glio |
| Chek1_in.vivo_noRT | Cfap46 | 4.576 | 2.99E-13 | glio |
| Bcas2_in.vivo_noRT | Gm30042 | 4.547 | 2.99E-15 | glio |
| Bcas2_in.vivo_noRT | Olfr171 | 4.807 | 2.99E-18 | glio |

| | | | | |
|---|---|---|---|---|
| Atr_in.vivo_noRT | Gm19265 | 5.017 | 2.99E-19 | glio |
| Atr_in.vivo_noRT | Dpp6 | 4.606 | 2.99E-20 | glio |
| culture_effect | Ms4a4d | -6.076 | 2.99E-24 | glio |
| culture_effect | Myo7a | -1.183 | 3.00E-03 | glio |
| culture_effect | Ddx28 | -0.715 | 3.00E-03 | glio |
| culture_effect | Nfia | 0.485 | 3.00E-03 | glio |
| Interaction_genotype_Atr | Ftl1 | 0.823 | 3.00E-03 | glio |
| Interaction_genotype_Mtor | Zfp976 | -3.179 | 3.00E-03 | glio |
| Interaction_genotype_Mtor | Pdlim2 | -2.896 | 3.00E-03 | glio |
| Interaction_genotype_Mtor | Rfesd | -2.444 | 3.00E-03 | glio |
| Interaction_genotype_Mtor | Psme3 | 1.444 | 3.00E-03 | glio |
| Interaction_genotype_Mtor | Usp34 | 1.398 | 3.00E-03 | glio |
| Interaction_genotype_Myc | Gm26789 | -2.793 | 3.00E-03 | glio |
| Interaction_genotype_Myc | Gm47689 | -3.335 | 3.00E-03 | glio |
| Interaction_genotype_Myc | Mon1b | -2.984 | 3.00E-03 | glio |
| Interaction_genotype_Myc | Opa1 | -2.321 | 3.00E-03 | glio |
| Aars2_in.vivo_noRT | Fam151b | 2.134 | 3.00E-03 | glio |
| Atr_in.vivo_noRT | Sqor | 1.935 | 3.00E-03 | glio |
| Atr_in.vivo_noRT | Rock2 | -0.935 | 3.00E-03 | glio |
| Atr_in.vivo_noRT | Phf5a | -0.731 | 3.00E-03 | glio |
| Atr_in.vivo_noRT | Brix1 | -0.75 | 3.00E-03 | glio |
| Bcas2_in.vivo_noRT | Rarres2 | 2.13 | 3.00E-03 | glio |
| Bcas2_in.vivo_noRT | Slfn3 | 1.983 | 3.00E-03 | glio |
| Bora_in.vivo_noRT | Mkrn1 | 0.749 | 3.00E-03 | glio |
| Bora_ex.vivo_noRT | Pomk | -2.573 | 3.00E-03 | glio |
| Hsd17b10_in.vivo_noRT | Hist2h4 | 2.164 | 3.00E-03 | glio |
| Interaction_genotype_Bcas2 | Pyy | -3.945 | 3.00E-04 | glio |
| Interaction_genotype_Mtor | A030005L19Ri | -3.964 | 3.00E-04 | glio |
| Interaction_genotype_Mtor | Pkn2 | 1.757 | 3.00E-04 | glio |
| Interaction_genotype_Myc | Gm50024 | -2.865 | 3.00E-04 | glio |
| Interaction_genotype_Myc | Mov10 | -2.625 | 3.00E-04 | glio |
| interaction_RT | Cyb5r3 | -0.796 | 3.00E-04 | glio |
| Aars2_in.vivo_noRT | Myo1b | 2.942 | 3.00E-04 | glio |
| Bcas2_ex.vivo_noRT | Fam166a | 2.742 | 3.00E-04 | glio |
| Bora_in.vivo_noRT | Tlcd2 | 2.905 | 3.00E-04 | glio |
| Chek1_ex.vivo_noRT | 9630013A20R | 3.163 | 3.00E-04 | glio |
| Eif2s1_in.vivo_noRT | Lrguk | 2.407 | 3.00E-04 | glio |
| Eif2s1_in.vivo_noRT | AU019990 | 2.143 | 3.00E-04 | glio |
| Hsd17b10_in.vivo_noRT | Hist2h3c1 | 2.704 | 3.00E-04 | glio |
| Hsd17b10_in.vivo_noRT | Nid1 | 2.812 | 3.00E-04 | glio |
| Hsd17b10_in.vivo_noRT | Nyap2 | 2.704 | 3.00E-04 | glio |
| Interaction_genotype_Mtor | 4930542D17R | -4.908 | 3.00E-05 | glio |
| Atr_in.vivo_noRT | Spry3 | 3.006 | 3.00E-05 | glio |
| Bcas2_in.vivo_noRT | Lncppara | 3.196 | 3.00E-05 | glio |
| Bora_ex.vivo_noRT | Gm36037 | 2.806 | 3.00E-05 | glio |
| Bora_ex.vivo_noRT | Gm34939 | 3.201 | 3.00E-05 | glio |
| Bora_ex.vivo_noRT | Icosl | 3.595 | 3.00E-05 | glio |
| Interaction_genotype_Mtor | 3110082J24Ri | -5.756 | 3.00E-06 | glio |
| Bcas2_in.vivo_noRT | Gm35066 | 3.315 | 3.00E-06 | glio |
| Hsd17b10_in.vivo_noRT | Gm15894 | 2.571 | 3.00E-06 | glio |
| Hsd17b10_in.vivo_noRT | Scube3 | 3.084 | 3.00E-06 | glio |
| Interaction_genotype_Mtor | Gm4340 | -4.099 | 3.00E-07 | glio |

| | | | | |
|---|---|---|---|---|
| Aars2_in.vivo_noRT | Dock11 | 3.321 | 3.00E-07 | glio |
| Aars2_ex.vivo_noRT | Dpep3 | 3.293 | 3.00E-07 | glio |
| Aars2_ex.vivo_noRT | Erich6b | 3.293 | 3.00E-07 | glio |
| Aars2_ex.vivo_noRT | Tac2 | 3.293 | 3.00E-07 | glio |
| Atr_in.vivo_noRT | Zfp980 | 3.767 | 3.00E-07 | glio |
| Bcas2_in.vivo_noRT | Gm43387 | 3.632 | 3.00E-07 | glio |
| Bcas2_in.vivo_noRT | Flrt1 | 3.832 | 3.00E-07 | glio |
| Bcas2_in.vivo_noRT | Gm2788 | 3.759 | 3.00E-07 | glio |
| Bora_in.vivo_noRT | Col9a1 | -2.581 | 3.00E-07 | glio |
| culture_effect | Zbtb1 | -0.986 | 3.00E-08 | glio |
| Aars2_in.vivo_noRT | Zfp980 | 4.075 | 3.00E-08 | glio |
| Chek1_in.vivo_noRT | Mamdc2 | 3.734 | 3.00E-08 | glio |
| Aars2_in.vivo_noRT | Gm16348 | 4.142 | 3.00E-09 | glio |
| Atr_ex.vivo_noRT | Fabp6 | 3.947 | 3.00E-09 | glio |
| Chek1_ex.vivo_noRT | 9130230L23Ri | 4.449 | 3.00E-09 | glio |
| Hsd17b10_in.vivo_noRT | Gtsf1 | 3.616 | 3.00E-10 | glio |
| Aars2_in.vivo_noRT | Ak1 | 4.573 | 3.00E-11 | glio |
| Bcas2_in.vivo_noRT | Gm35696 | 4.547 | 3.00E-12 | glio |
| culture_effect | AI413582 | 1.156 | 3.00E-13 | glio |
| Bora_in.vivo_noRT | Pkd1l2 | 4.672 | 3.00E-13 | glio |
| Atr_in.vivo_noRT | A030005L19Ri | 5.004 | 3.00E-15 | glio |
| Chek1_in.vivo_noRT | Ptprc | 4.78 | 3.00E-18 | glio |
| Chek1_in.vivo_noRT | Gm48140 | 4.78 | 3.00E-21 | glio |
| culture_effect | Ankrd54 | -0.688 | 3.01E-03 | glio |
| culture_effect | Ddx21 | 0.378 | 3.01E-03 | glio |
| Interaction_genotype_Atr | Mro | -3.724 | 3.01E-03 | glio |
| Interaction_genotype_Atr | Abhd18 | -3.762 | 3.01E-03 | glio |
| Interaction_genotype_Atr | Lamp1 | 1.097 | 3.01E-03 | glio |
| Interaction_genotype_Bora | Synpr | -4.313 | 3.01E-03 | glio |
| Interaction_genotype_Bora | Gm19276 | -2.748 | 3.01E-03 | glio |
| Interaction_genotype_Mtor | Slc36a1 | -3.103 | 3.01E-03 | glio |
| Interaction_genotype_Myc | Zfp473 | -3.153 | 3.01E-03 | glio |
| Interaction_genotype_Myc | Rac1 | 1.172 | 3.01E-03 | glio |
| Interaction_genotype_Rpl3 | Atf4 | 1.129 | 3.01E-03 | glio |
| Aars2_in.vivo_noRT | Cul5 | -1.142 | 3.01E-03 | glio |
| Bcas2_in.vivo_noRT | Mta2 | -1.376 | 3.01E-03 | glio |
| Bora_in.vivo_noRT | Hist1h2bn | 2.093 | 3.01E-03 | glio |
| Bora_in.vivo_noRT | Rest | -1.201 | 3.01E-03 | glio |
| Cenpj_in.vivo_noRT | Gm48617 | 1.758 | 3.01E-03 | glio |
| Cenpj_in.vivo_noRT | Gm15929 | 2.022 | 3.01E-03 | glio |
| Chek1_in.vivo_noRT | Gm32031 | 1.911 | 3.01E-03 | glio |
| Chek1_ex.vivo_noRT | Ovgp1 | 2.646 | 3.01E-03 | glio |
| Interaction_genotype_Mtor | Nos2 | -3.839 | 3.01E-04 | glio |
| Interaction_genotype_Mtor | E030037K01Ri | -3.849 | 3.01E-04 | glio |
| Interaction_genotype_Mtor | 1700063J08Ri | -4.092 | 3.01E-04 | glio |
| Interaction_genotype_Mtor | Tnfrsf22 | -3.811 | 3.01E-04 | glio |
| Interaction_genotype_Myc | Gm12589 | -3.128 | 3.01E-04 | glio |
| Interaction_genotype_Myc | Fem1b | -2.428 | 3.01E-04 | glio |
| Aars2_in.vivo_noRT | Plekhb1 | -1.293 | 3.01E-04 | glio |
| Aars2_in.vivo_noRT | mt-Cytb | 0.552 | 3.01E-04 | glio |
| Bcas2_in.vivo_noRT | 4930542D17R | 2.527 | 3.01E-04 | glio |
| Bcas2_in.vivo_noRT | Cdc42ep4 | 2.452 | 3.01E-04 | glio |

| | | | | |
|---|---|---|---|---|
| Cenpj_in.vivo_noRT | Apoa5 | 2.022 | 3.01E-04 | glio |
| Cenpj_in.vivo_noRT | Gjd4 | 2.022 | 3.01E-04 | glio |
| Cenpj_in.vivo_noRT | Man2c1os | 3.208 | 3.01E-04 | glio |
| Cenpj_in.vivo_noRT | Capn9 | 2.798 | 3.01E-04 | glio |
| Cenpj_in.vivo_noRT | A930019D19R | 2.022 | 3.01E-04 | glio |
| Cenpj_in.vivo_noRT | A030005L19Ri | 2.795 | 3.01E-04 | glio |
| Cenpj_in.vivo_noRT | Ly6c1 | 2.486 | 3.01E-04 | glio |
| Bora_ex.vivo_noRT | Slc45a1 | 2.802 | 3.01E-05 | glio |
| culture_effect | 1700086P04R | 2.204 | 3.01E-06 | glio |
| Interaction_genotype_Mtor | Nasp | 2.398 | 3.01E-06 | glio |
| Interaction_genotype_Mtor | Gm36793 | -4.099 | 3.01E-06 | glio |
| interaction_RT | Slc6a8 | -1.534 | 3.01E-06 | glio |
| Atr_in.vivo_noRT | Gm49085 | 3.046 | 3.01E-06 | glio |
| Atr_ex.vivo_noRT | Prl3d3 | 2.866 | 3.01E-06 | glio |
| Interaction_genotype_Mtor | Anp32b | 2.432 | 3.01E-07 | glio |
| Interaction_genotype_Mtor | Gm16239 | -4.099 | 3.01E-07 | glio |
| Interaction_genotype_Mtor | Gm36037 | -4.494 | 3.01E-07 | glio |
| Aars2_in.vivo_noRT | Rbm11 | 3.395 | 3.01E-07 | glio |
| Hsd17b10_in.vivo_noRT | Ptgis | 3.616 | 3.01E-07 | glio |
| Bora_in.vivo_noRT | 5033403F01Ri | 4.01 | 3.01E-09 | glio |
| Chek1_ex.vivo_noRT | Stra8 | 4.447 | 3.01E-09 | glio |
| Aars2_in.vivo_noRT | Hadha | -2.543 | 3.01E-10 | glio |
| Atr_in.vivo_noRT | Nlrx1 | 4.645 | 3.01E-10 | glio |
| Bora_in.vivo_noRT | 3930402G23R | 4.549 | 3.01E-10 | glio |
| Chek1_in.vivo_noRT | 1700030L20Ri | 4.388 | 3.01E-11 | glio |
| culture_effect | Paip2 | -0.775 | 3.01E-13 | glio |
| Bora_in.vivo_noRT | Gm15894 | 3.891 | 3.01E-13 | glio |
| Bcas2_in.vivo_noRT | Ppil6 | 4.807 | 3.01E-14 | glio |
| Chek1_in.vivo_noRT | C130060C02R | 4.78 | 3.01E-14 | glio |
| Chek1_in.vivo_noRT | Cyp2c23 | 4.521 | 3.01E-14 | glio |
| culture_effect | Prkcq | -4.314 | 3.01E-15 | glio |
| Aars2_in.vivo_noRT | Gm7247 | 4.535 | 3.01E-15 | glio |
| Atr_in.vivo_noRT | 1700016P04R | 5.267 | 3.01E-16 | glio |
| Bora_in.vivo_noRT | Slc16a5 | 4.672 | 3.01E-21 | glio |
| Atr_in.vivo_noRT | Gm30042 | 5.811 | 3.01E-23 | glio |
| culture_effect | Mbp | 2.989 | 3.01E-42 | glio |
| culture_effect | Psmb8 | -3.435 | 3.01E-68 | glio |
| culture_effect | Rnf170 | 0.724 | 3.02E-03 | glio |
| culture_effect | Ing4 | -0.614 | 3.02E-03 | glio |
| culture_effect | Taf10 | 0.287 | 3.02E-03 | glio |
| Interaction_genotype_Atr | 4933429H19R | -3.219 | 3.02E-03 | glio |
| Interaction_genotype_Atr | Nol8 | 1.671 | 3.02E-03 | glio |
| Interaction_genotype_Mtor | Svopl | -3.311 | 3.02E-03 | glio |
| Interaction_genotype_Mtor | Pax9 | -3.468 | 3.02E-03 | glio |
| Interaction_genotype_Mtor | Gm10825 | -3.596 | 3.02E-03 | glio |
| Interaction_genotype_Mtor | Kif7 | -3.268 | 3.02E-03 | glio |
| Interaction_genotype_Mtor | Exog | -3.028 | 3.02E-03 | glio |
| Interaction_genotype_Mtor | Exosc5 | 1.723 | 3.02E-03 | glio |
| Interaction_genotype_Myc | Zfp324 | -2.772 | 3.02E-03 | glio |
| Interaction_genotype_Pars2 | Rcan2 | -3.992 | 3.02E-03 | glio |
| Interaction_genotype_Pars2 | Gm11627 | -3.282 | 3.02E-03 | glio |
| Interaction_genotype_Rad51 | Atg9b | 3.839 | 3.02E-03 | glio |

| | | | | |
|---|---|---|---|---|
| Interaction_genotype_Rad51 | Mecom | -3.902 | 3.02E-03 | glio |
| Aars2_in.vivo_noRT | Klhl36 | 2.084 | 3.02E-03 | glio |
| Atr_in.vivo_noRT | Ess2 | 1.71 | 3.02E-03 | glio |
| Atr_in.vivo_noRT | E2f3 | -1.271 | 3.02E-03 | glio |
| Atr_ex.vivo_noRT | Vmn1r181 | 2.769 | 3.02E-03 | glio |
| Bora_in.vivo_noRT | Klf15 | 2.179 | 3.02E-03 | glio |
| Bora_in.vivo_noRT | Zfp82 | 2.131 | 3.02E-03 | glio |
| Bora_in.vivo_noRT | Hacd1 | -1.2 | 3.02E-03 | glio |
| Cenpj_in.vivo_noRT | Pitx3 | 2.022 | 3.02E-03 | glio |
| Chek1_in.vivo_noRT | Gm10825 | 2.215 | 3.02E-03 | glio |
| Chek1_in.vivo_noRT | 2410006H16R | -0.825 | 3.02E-03 | glio |
| Chek1_ex.vivo_noRT | Tsen34 | -1.155 | 3.02E-03 | glio |
| Chek1_ex.vivo_noRT | Chchd1 | -0.842 | 3.02E-03 | glio |
| Interaction_genotype_Bora | Egr2 | 4.651 | 3.02E-04 | glio |
| Interaction_genotype_Mtor | Gm43707 | -3.901 | 3.02E-04 | glio |
| Interaction_genotype_Myc | Gm41409 | -3.742 | 3.02E-04 | glio |
| Interaction_genotype_Myc | BC049762 | -4.105 | 3.02E-04 | glio |
| Atr_in.vivo_noRT | Gm31645 | 2.579 | 3.02E-04 | glio |
| Atr_in.vivo_noRT | Ptprs | -1.563 | 3.02E-04 | glio |
| Atr_in.vivo_noRT | Rbbp7 | -0.876 | 3.02E-04 | glio |
| Bard1_ex.vivo_noRT | Slc16a5 | 2.421 | 3.02E-04 | glio |
| Bcas2_in.vivo_noRT | Gm36210 | 2.606 | 3.02E-04 | glio |
| Bora_in.vivo_noRT | Srgap1 | 2.432 | 3.02E-04 | glio |
| Chek1_in.vivo_noRT | 4930412C18R | 2.708 | 3.02E-04 | glio |
| Chek1_in.vivo_noRT | Mpp2 | 2.594 | 3.02E-04 | glio |
| Chek1_in.vivo_noRT | Nup153 | -1.589 | 3.02E-04 | glio |
| Chek1_in.vivo_noRT | Uqcrq | -0.764 | 3.02E-04 | glio |
| Chek1_ex.vivo_noRT | Scd3 | 3.541 | 3.02E-04 | glio |
| Interaction_genotype_Atr | Gm35019 | 4.663 | 3.02E-05 | glio |
| Interaction_genotype_Atr | Gm20635 | -4.927 | 3.02E-05 | glio |
| Interaction_genotype_Mtor | Gm4129 | -4.96 | 3.02E-05 | glio |
| Interaction_genotype_Mtor | Cox7b2 | -4.429 | 3.02E-05 | glio |
| Interaction_genotype_Mtor | Mob3b | -4 | 3.02E-05 | glio |
| Interaction_genotype_Myc | Gm9833 | -4.5 | 3.02E-05 | glio |
| Interaction_genotype_Nat2 | Gm21762 | 4.582 | 3.02E-05 | glio |
| Aars2_in.vivo_noRT | Psmb2 | -1.033 | 3.02E-05 | glio |
| Atp5c1_in.vivo_noRT | 9530062K07R | 3.336 | 3.02E-05 | glio |
| Atr_in.vivo_noRT | Cpne7 | 3.135 | 3.02E-05 | glio |
| Bcas2_in.vivo_noRT | Cbs | 3.08 | 3.02E-05 | glio |
| Bcas2_in.vivo_noRT | A930001M01F | 2.926 | 3.02E-05 | glio |
| Bora_ex.vivo_noRT | Gm30042 | 3.201 | 3.02E-05 | glio |
| culture_effect | Ints8 | -0.86 | 3.02E-06 | glio |
| Interaction_genotype_Myc | Nln | -3.762 | 3.02E-06 | glio |
| Interaction_genotype_Myc | Gm20455 | -5.636 | 3.02E-06 | glio |
| Bora_in.vivo_noRT | Gm12905 | 3.631 | 3.02E-06 | glio |
| Bora_in.vivo_noRT | Pcdhb18 | 3.186 | 3.02E-06 | glio |
| Bora_in.vivo_noRT | Pate2 | 3.457 | 3.02E-06 | glio |
| Bora_in.vivo_noRT | Tmem40 | 3.651 | 3.02E-06 | glio |
| Chek1_in.vivo_noRT | Fgf2 | 3.418 | 3.02E-06 | glio |
| Interaction_genotype_Mtor | 4933429H19R | -5.644 | 3.02E-07 | glio |
| Atp5c1_in.vivo_noRT | Abcb5 | 2.668 | 3.02E-07 | glio |
| Atp5c1_in.vivo_noRT | Gm10324 | 2.933 | 3.02E-07 | glio |

| | | | | |
|---|---|---|---|---|
| culture_effect | Nfil3 | -1.643 | 3.02E-08 | glio |
| Bora_in.vivo_noRT | Gm20667 | 3.936 | 3.02E-08 | glio |
| Chek1_ex.vivo_noRT | Tex14 | 4.443 | 3.02E-08 | glio |
| culture_effect | Wbp11 | 0.67 | 3.02E-10 | glio |
| Aars2_in.vivo_noRT | Aass | 4.325 | 3.02E-11 | glio |
| Aars2_in.vivo_noRT | Gm11549 | 4.801 | 3.02E-12 | glio |
| Atr_in.vivo_noRT | D6Ertd527e | 4.751 | 3.02E-12 | glio |
| Atr_in.vivo_noRT | Bmp15 | 4.88 | 3.02E-16 | glio |
| Chek1_in.vivo_noRT | Gm15445 | 4.398 | 3.02E-16 | glio |
| Chek1_in.vivo_noRT | Gast | 5.044 | 3.02E-19 | glio |
| Bcas2_in.vivo_noRT | Vmn1r32 | 4.807 | 3.02E-22 | glio |
| Bcas2_in.vivo_noRT | Glyat | 4.807 | 3.02E-22 | glio |
| culture_effect | Trp53cor1 | 0.878 | 3.03E-03 | glio |
| Interaction_genotype_Atr | Sema4a | -3.884 | 3.03E-03 | glio |
| Interaction_genotype_Atr | Clcf1 | -2.882 | 3.03E-03 | glio |
| Interaction_genotype_Mtor | Glmp | -3.977 | 3.03E-03 | glio |
| Interaction_genotype_Mtor | Ak4 | -2.87 | 3.03E-03 | glio |
| Interaction_genotype_Myc | Zscan18 | -2.764 | 3.03E-03 | glio |
| Interaction_genotype_Myc | Cers2 | 1.249 | 3.03E-03 | glio |
| Interaction_genotype_Rpl3 | Wscd1 | 2.251 | 3.03E-03 | glio |
| interaction_RT | Cdc45 | -1.22 | 3.03E-03 | glio |
| interaction_RT | Gm16062 | 1.295 | 3.03E-03 | glio |
| Aars2_in.vivo_noRT | Gm15708 | 2.116 | 3.03E-03 | glio |
| Aars2_in.vivo_noRT | Tnnt2 | 1.833 | 3.03E-03 | glio |
| Atr_in.vivo_noRT | Pacsin2 | -1.311 | 3.03E-03 | glio |
| Atr_in.vivo_noRT | Top2a | -1.732 | 3.03E-03 | glio |
| Bcas2_in.vivo_noRT | Dusp15 | 1.958 | 3.03E-03 | glio |
| Bora_in.vivo_noRT | 2600006K01R | 2.278 | 3.03E-03 | glio |
| Bora_in.vivo_noRT | Slc13a5 | 2.059 | 3.03E-03 | glio |
| Bora_in.vivo_noRT | Car11 | 1.94 | 3.03E-03 | glio |
| Bora_in.vivo_noRT | Uri1 | -1.238 | 3.03E-03 | glio |
| Bora_in.vivo_noRT | Hebp1 | 1.506 | 3.03E-03 | glio |
| Chek1_in.vivo_noRT | Tbp | -1.424 | 3.03E-03 | glio |
| Chek1_in.vivo_noRT | Commd6 | -1.159 | 3.03E-03 | glio |
| Chek1_ex.vivo_noRT | Hmgb2 | -1.521 | 3.03E-03 | glio |
| Interaction_genotype_Mtor | Ushbp1 | -4.284 | 3.03E-04 | glio |
| Interaction_genotype_Mtor | 4930405A21R | -3.908 | 3.03E-04 | glio |
| Interaction_genotype_Myc | Ifnz | -3.946 | 3.03E-04 | glio |
| Interaction_genotype_Myc | Mettl22 | -2.896 | 3.03E-04 | glio |
| Interaction_genotype_Myc | Nampt | 1.782 | 3.03E-04 | glio |
| Aars2_in.vivo_noRT | Ctsb | -1.532 | 3.03E-04 | glio |
| Aars2_ex.vivo_noRT | Gm5431 | 3.293 | 3.03E-04 | glio |
| Atr_ex.vivo_noRT | Gm34047 | 3.103 | 3.03E-04 | glio |
| Bcas2_in.vivo_noRT | Bcl2l14 | 2.766 | 3.03E-04 | glio |
| Chek1_ex.vivo_noRT | Asap3 | 3.248 | 3.03E-04 | glio |
| Eif2s1_in.vivo_noRT | Gm36793 | 2.143 | 3.03E-04 | glio |
| Hsd17b10_in.vivo_noRT | Gm43948 | 2.141 | 3.03E-04 | glio |
| culture_effect | Gm20522 | 2.196 | 3.03E-05 | glio |
| culture_effect | Cyp3a13 | 2.197 | 3.03E-05 | glio |
| culture_effect | 1700001O22R | -0.993 | 3.03E-05 | glio |
| Interaction_genotype_Mtor | Prap1 | -3.699 | 3.03E-05 | glio |
| Interaction_genotype_Mtor | Rslcan18 | -4.5 | 3.03E-05 | glio |

| | | | | |
|---|---|---|---|---|
| Interaction_genotype_Mtor | Cfh | -3.719 | 3.03E-05 | glio |
| Interaction_genotype_Myc | Gm42851 | -4.108 | 3.03E-05 | glio |
| Interaction_genotype_Myc | C130040N14R | -4.8 | 3.03E-05 | glio |
| Interaction_genotype_Myc | Nwd1 | -3.99 | 3.03E-05 | glio |
| Interaction_genotype_Myc | Lrrc72 | -4.417 | 3.03E-05 | glio |
| Interaction_genotype_Myc | Nxnl2 | -4.366 | 3.03E-05 | glio |
| Atp5c1_in.vivo_noRT | Gm49860 | 2.668 | 3.03E-05 | glio |
| Bcas2_in.vivo_noRT | Gm50218 | 3.213 | 3.03E-05 | glio |
| Bcas2_in.vivo_noRT | Lgals3 | -1.271 | 3.03E-05 | glio |
| Bora_in.vivo_noRT | Aplp2 | -1.459 | 3.03E-05 | glio |
| Interaction_genotype_Myc | Sh3pxd2b | -4.96 | 3.03E-06 | glio |
| Atr_ex.vivo_noRT | 4930512B01R | 3.164 | 3.03E-06 | glio |
| Bora_in.vivo_noRT | Rab11fip1 | 3.642 | 3.03E-06 | glio |
| Bora_in.vivo_noRT | Dmxl1 | -2.016 | 3.03E-06 | glio |
| culture_effect | Gm11131 | -3.158 | 3.03E-07 | glio |
| Interaction_genotype_Mtor | Pde3a | -5.69 | 3.03E-07 | glio |
| Atr_in.vivo_noRT | Phldb2 | 3.799 | 3.03E-07 | glio |
| Bora_in.vivo_noRT | A530083I20Ri | 3.928 | 3.03E-07 | glio |
| Chek1_ex.vivo_noRT | 4922502N22R | 4.447 | 3.03E-07 | glio |
| interaction_RT | 1110038B12R | 1.169 | 3.03E-08 | glio |
| Aars2_in.vivo_noRT | Gm47223 | 5.06 | 3.03E-08 | glio |
| Aars2_in.vivo_noRT | 1700025N23R | 3.435 | 3.03E-08 | glio |
| Atr_in.vivo_noRT | Msl3l2 | 3.838 | 3.03E-08 | glio |
| Bora_in.vivo_noRT | Fgd3 | 4.201 | 3.03E-08 | glio |
| Chek1_in.vivo_noRT | Cxcr2 | 3.732 | 3.03E-08 | glio |
| Atr_in.vivo_noRT | 1700122E12Ri | 4.137 | 3.03E-09 | glio |
| Bora_in.vivo_noRT | Gm16244 | 4.279 | 3.03E-09 | glio |
| Chek1_in.vivo_noRT | Cfap58 | 5.044 | 3.03E-09 | glio |
| culture_effect | Gm13283 | 2.999 | 3.03E-10 | glio |
| Interaction_genotype_Mtor | Rbbp7 | 3.091 | 3.03E-10 | glio |
| Atr_in.vivo_noRT | 1700010B08R | 3.847 | 3.03E-11 | glio |
| interaction_RT | Chic2 | 1.359 | 3.03E-12 | glio |
| Atr_in.vivo_noRT | Mycl | 5.011 | 3.03E-12 | glio |
| Chek1_ex.vivo_noRT | Cd7 | 4.842 | 3.03E-15 | glio |
| Chek1_ex.vivo_noRT | Gm48764 | 4.842 | 3.03E-15 | glio |
| Chek1_ex.vivo_noRT | 4930527F14Ri | 5.247 | 3.03E-15 | glio |
| culture_effect | Itch | -0.49 | 3.04E-03 | glio |
| Interaction_genotype_Aars2 | Rdh12 | -3.535 | 3.04E-03 | glio |
| Interaction_genotype_Atr | Zfp790 | -3.523 | 3.04E-03 | glio |
| Interaction_genotype_Mtor | 4930506C21R | -3.589 | 3.04E-03 | glio |
| Interaction_genotype_Mtor | Tm6sf1 | -4.092 | 3.04E-03 | glio |
| Interaction_genotype_Mtor | Fam49a | -2.958 | 3.04E-03 | glio |
| Interaction_genotype_Mtor | Ccdc12 | -1.673 | 3.04E-03 | glio |
| Interaction_genotype_Myc | Myo1b | -3.878 | 3.04E-03 | glio |
| interaction_RT | Magt1 | -0.744 | 3.04E-03 | glio |
| Aars2_in.vivo_noRT | Dpep2 | 1.999 | 3.04E-03 | glio |
| Aars2_in.vivo_noRT | Tmem167 | -0.71 | 3.04E-03 | glio |
| Atr_in.vivo_noRT | Cpa2 | 2.014 | 3.04E-03 | glio |
| Bcas2_ex.vivo_noRT | Cxcl16 | 2.837 | 3.04E-03 | glio |
| Bora_in.vivo_noRT | Card14 | 2.224 | 3.04E-03 | glio |
| Chek1_in.vivo_noRT | Slc27a3 | 2.17 | 3.04E-03 | glio |
| Chek1_in.vivo_noRT | Zfp386 | -1.544 | 3.04E-03 | glio |

| | | | | |
|---|---|---|---|---|
| Chek1_in.vivo_noRT | Cers4 | -1.65 | 3.04E-03 | glio |
| Chek1_in.vivo_noRT | Shmt1 | -1.435 | 3.04E-03 | glio |
| Chek1_in.vivo_noRT | Rpl10 | -0.556 | 3.04E-03 | glio |
| Eif2s1_in.vivo_noRT | Bco2 | 2.143 | 3.04E-03 | glio |
| Eif2s1_in.vivo_noRT | Tle3 | 1.409 | 3.04E-03 | glio |
| Eif2s1_ex.vivo_noRT | Ankrd22 | 2.234 | 3.04E-03 | glio |
| Eif2s1_ex.vivo_noRT | Tmem52 | 3.014 | 3.04E-03 | glio |
| Eif2s1_ex.vivo_noRT | Gm47404 | 2.234 | 3.04E-03 | glio |
| Eif2s1_ex.vivo_noRT | Gm31717 | 2.234 | 3.04E-03 | glio |
| Eif2s1_ex.vivo_noRT | Gm14742 | 2.234 | 3.04E-03 | glio |
| Eif2s1_ex.vivo_noRT | Scel | 2.234 | 3.04E-03 | glio |
| Eif2s1_ex.vivo_noRT | Ctla4 | 2.234 | 3.04E-03 | glio |
| Eif2s1_ex.vivo_noRT | Zscan4-ps3 | 2.234 | 3.04E-03 | glio |
| Eif2s1_ex.vivo_noRT | Tagap | 2.234 | 3.04E-03 | glio |
| Eif2s1_ex.vivo_noRT | Gm21731 | 2.234 | 3.04E-03 | glio |
| Eif2s1_ex.vivo_noRT | Gm11232 | 2.234 | 3.04E-03 | glio |
| Eif2s1_ex.vivo_noRT | Gm29966 | 2.234 | 3.04E-03 | glio |
| Eif2s1_ex.vivo_noRT | Olfr1152 | 2.234 | 3.04E-03 | glio |
| Interaction_genotype_Atr | Fgf2 | -4.352 | 3.04E-04 | glio |
| Interaction_genotype_Mtor | B3galt4 | -3.952 | 3.04E-04 | glio |
| Interaction_genotype_Mtor | Zcrb1 | 1.529 | 3.04E-04 | glio |
| Interaction_genotype_Myc | Ncaph | -3.153 | 3.04E-04 | glio |
| Interaction_genotype_Pars2 | Folr1 | -4.255 | 3.04E-04 | glio |
| Aars2_ex.vivo_noRT | Gm5662 | 3.293 | 3.04E-04 | glio |
| Bcas2_in.vivo_noRT | Myl7 | 2.679 | 3.04E-04 | glio |
| Cenpj_in.vivo_noRT | Gm49808 | 2.022 | 3.04E-04 | glio |
| Chek1_in.vivo_noRT | Taf1 | -1.296 | 3.04E-04 | glio |
| Hsd17b10_in.vivo_noRT | 4833407H14R | 2.982 | 3.04E-04 | glio |
| Hsd17b10_in.vivo_noRT | Tmem119 | -3.541 | 3.04E-04 | glio |
| culture_effect | Nol3 | -1.967 | 3.04E-05 | glio |
| Interaction_genotype_Mtor | Slc16a5 | -3.436 | 3.04E-05 | glio |
| interaction_RT | Sbds | 0.962 | 3.04E-05 | glio |
| Aars2_in.vivo_noRT | Jazf1 | 2.822 | 3.04E-05 | glio |
| Bard1_ex.vivo_noRT | Gad1 | 2.814 | 3.04E-05 | glio |
| Bard1_ex.vivo_noRT | Auts2 | 3.612 | 3.04E-05 | glio |
| Bora_in.vivo_noRT | Gm13056 | 3.181 | 3.04E-05 | glio |
| Chek1_in.vivo_noRT | Cd34 | 3.009 | 3.04E-05 | glio |
| culture_effect | AI480526 | 2.087 | 3.04E-06 | glio |
| culture_effect | Dctn3 | -0.524 | 3.04E-06 | glio |
| interaction_RT | Gpnmb | 1.453 | 3.04E-06 | glio |
| Aars2_ex.vivo_noRT | Gm21761 | 3.293 | 3.04E-06 | glio |
| Interaction_genotype_Mtor | Cox7a2l | 2.436 | 3.04E-07 | glio |
| Interaction_genotype_Myc | Gpx4 | 2.121 | 3.04E-07 | glio |
| Bcas2_in.vivo_noRT | Eps8l2 | 3.705 | 3.04E-07 | glio |
| Bora_ex.vivo_noRT | 2410017I17Ri | 3.201 | 3.04E-07 | glio |
| Chek1_in.vivo_noRT | Tgfbr3l | 3.74 | 3.04E-07 | glio |
| Hsd17b10_in.vivo_noRT | Tcp11x2 | 3.094 | 3.04E-07 | glio |
| Hsd17b10_in.vivo_noRT | Klhdc8a | 3.616 | 3.04E-07 | glio |
| Bcas2_in.vivo_noRT | Gm20455 | 4.087 | 3.04E-08 | glio |
| Bora_in.vivo_noRT | Hist1h4f | 4.144 | 3.04E-08 | glio |
| Bora_in.vivo_noRT | 1700011D18R | 3.891 | 3.04E-08 | glio |
| Chek1_in.vivo_noRT | Adam33 | 3.887 | 3.04E-08 | glio |

| | | | | |
|---|---|---|---|---|
| Chek1_ex.vivo_noRT | Gm48140 | 4.049 | 3.04E-11 | glio |
| culture_effect | Actn1 | 1.413 | 3.04E-12 | glio |
| Chek1_in.vivo_noRT | Vmn1r91 | 4.391 | 3.04E-12 | glio |
| culture_effect | Ergic1 | 1.031 | 3.04E-13 | glio |
| interaction_RT | Ifi27l2a | 1.979 | 3.04E-14 | glio |
| culture_effect | Nckap1 | 1.173 | 3.04E-15 | glio |
| Bora_in.vivo_noRT | Tmem132c | 4.936 | 3.04E-23 | glio |
| Atr_in.vivo_noRT | Gm43701 | 5.533 | 3.04E-26 | glio |
| culture_effect | Bcas1 | -2.991 | 3.04E-29 | glio |
| culture_effect | Ifi47 | -3.745 | 3.04E-48 | glio |
| culture_effect | Acvr2b | -1.194 | 3.05E-03 | glio |
| Interaction_genotype_Aars2 | Gm14399 | -3.701 | 3.05E-03 | glio |
| Interaction_genotype_Atr | Trim16 | -3.311 | 3.05E-03 | glio |
| Interaction_genotype_Bcas2 | Gm15890 | -3.348 | 3.05E-03 | glio |
| Interaction_genotype_Chek1 | Gna12 | -2.515 | 3.05E-03 | glio |
| Interaction_genotype_Mtor | Pcdhb18 | -3.527 | 3.05E-03 | glio |
| Interaction_genotype_Mtor | Sdf4 | 1.463 | 3.05E-03 | glio |
| Interaction_genotype_Mtor | Eif3k | 1.135 | 3.05E-03 | glio |
| Aars2_in.vivo_noRT | Gm12166 | 2.2 | 3.05E-03 | glio |
| Aars2_in.vivo_noRT | Klk9 | 2.157 | 3.05E-03 | glio |
| Aars2_in.vivo_noRT | Ppp1r13l | 2.147 | 3.05E-03 | glio |
| Aars2_in.vivo_noRT | Gm34397 | 1.977 | 3.05E-03 | glio |
| Aars2_in.vivo_noRT | Psenen | -0.714 | 3.05E-03 | glio |
| Atr_in.vivo_noRT | Slc22a21 | 2.159 | 3.05E-03 | glio |
| Atr_in.vivo_noRT | Rbm8a2 | 2.092 | 3.05E-03 | glio |
| Atr_in.vivo_noRT | Sirt7 | -1.394 | 3.05E-03 | glio |
| Bard1_ex.vivo_noRT | AU022754 | 2.814 | 3.05E-03 | glio |
| Bcas2_in.vivo_noRT | Jpt1 | -0.696 | 3.05E-03 | glio |
| Bora_in.vivo_noRT | Fam135b | 2.477 | 3.05E-03 | glio |
| Bora_in.vivo_noRT | Rpl15 | -0.598 | 3.05E-03 | glio |
| Chek1_in.vivo_noRT | E130114P18Ri | 2.291 | 3.05E-03 | glio |
| Chek1_in.vivo_noRT | Btbd16 | 1.821 | 3.05E-03 | glio |
| Chek1_in.vivo_noRT | Arhgap1 | -1.313 | 3.05E-03 | glio |
| Chek1_ex.vivo_noRT | Noc4l | -1.878 | 3.05E-03 | glio |
| Hsd17b10_in.vivo_noRT | Zfp958 | -1.605 | 3.05E-03 | glio |
| culture_effect | Gm4070 | -2.277 | 3.05E-04 | glio |
| culture_effect | Dync1h1 | -0.485 | 3.05E-04 | glio |
| Interaction_genotype_Bora | D630036H23R | -4.575 | 3.05E-04 | glio |
| Interaction_genotype_Bora | Gchfr | -5.239 | 3.05E-04 | glio |
| Interaction_genotype_Bora | 4930478M13F | -4.626 | 3.05E-04 | glio |
| Interaction_genotype_Bora | Gm14002 | -4.409 | 3.05E-04 | glio |
| Interaction_genotype_Bora | Ypel5 | 2.838 | 3.05E-04 | glio |
| Interaction_genotype_Mtor | Slc2a6 | -4.501 | 3.05E-04 | glio |
| Interaction_genotype_Mtor | mt-Nd1 | 1.635 | 3.05E-04 | glio |
| Interaction_genotype_Myc | Pcsk4 | -4.041 | 3.05E-04 | glio |
| Interaction_genotype_Myc | Dhx37 | -3.518 | 3.05E-04 | glio |
| Interaction_genotype_Myc | Taco1 | -3.496 | 3.05E-04 | glio |
| Aars2_in.vivo_noRT | Ccdc142os | 2.539 | 3.05E-04 | glio |
| Atp5c1_in.vivo_noRT | 4930434B07R | 2.405 | 3.05E-04 | glio |
| Atp5c1_in.vivo_noRT | Gm37864 | 2.416 | 3.05E-04 | glio |
| Atr_in.vivo_noRT | Usp39 | -1.571 | 3.05E-04 | glio |
| Atr_in.vivo_noRT | Mtfr1l | 1.378 | 3.05E-04 | glio |

| | | | | |
|---|---|---|---|---|
| Bcas2_in.vivo_noRT | Mill2 | 2.616 | 3.05E-04 | glio |
| Bora_in.vivo_noRT | Wif1 | 2.93 | 3.05E-04 | glio |
| Bora_in.vivo_noRT | Klhdc1 | 2.557 | 3.05E-04 | glio |
| Bora_ex.vivo_noRT | 4930502E18Ri | 2.808 | 3.05E-04 | glio |
| Chek1_in.vivo_noRT | Fignl2 | 2.778 | 3.05E-04 | glio |
| Chek1_in.vivo_noRT | Zfp964 | 2.669 | 3.05E-04 | glio |
| Interaction_genotype_Mtor | Gm34121 | -4.332 | 3.05E-05 | glio |
| Interaction_genotype_Mtor | Gemin4 | -5.044 | 3.05E-05 | glio |
| Interaction_genotype_Mtor | BC018473 | -4.896 | 3.05E-05 | glio |
| Interaction_genotype_Mtor | Glrx5 | 2.048 | 3.05E-05 | glio |
| Interaction_genotype_Myc | A430106G13R | -4.753 | 3.05E-05 | glio |
| Aars2_in.vivo_noRT | Gm10131 | 2.893 | 3.05E-05 | glio |
| Aars2_in.vivo_noRT | Dok7 | 2.835 | 3.05E-05 | glio |
| Atr_ex.vivo_noRT | Gm43409 | 3.915 | 3.05E-05 | glio |
| Bcas2_in.vivo_noRT | Camta2 | 2.059 | 3.05E-05 | glio |
| culture_effect | Dnpep | -0.968 | 3.05E-06 | glio |
| culture_effect | Serinc3 | -0.881 | 3.05E-06 | glio |
| culture_effect | Ppa1 | -0.591 | 3.05E-06 | glio |
| Interaction_genotype_Myc | Trmt2a | -3.952 | 3.05E-06 | glio |
| Interaction_genotype_Myc | Gm28874 | -5.403 | 3.05E-06 | glio |
| Atp5c1_in.vivo_noRT | Gm13449 | 2.668 | 3.05E-06 | glio |
| Bora_in.vivo_noRT | Gm26879 | 2.929 | 3.05E-06 | glio |
| Interaction_genotype_Myc | Dnm3 | -5.188 | 3.05E-07 | glio |
| Interaction_genotype_Myc | Cyp1a1 | -7.598 | 3.05E-07 | glio |
| Bcas2_in.vivo_noRT | Gm48882 | 3.683 | 3.05E-07 | glio |
| Bcas2_in.vivo_noRT | Lrrn3 | 3.351 | 3.05E-07 | glio |
| Aars2_in.vivo_noRT | Enpp3 | 3.76 | 3.05E-08 | glio |
| Bora_in.vivo_noRT | Mir670hg | 4.286 | 3.05E-08 | glio |
| Bora_in.vivo_noRT | Gm45890 | 4.019 | 3.05E-08 | glio |
| Bora_in.vivo_noRT | Gpr37l1 | 3.822 | 3.05E-08 | glio |
| Chek1_in.vivo_noRT | A430057M04F | 3.858 | 3.05E-08 | glio |
| culture_effect | Ccdc85b | 1.109 | 3.05E-09 | glio |
| Atr_in.vivo_noRT | E130208F15Ri | 4.48 | 3.05E-09 | glio |
| Bora_in.vivo_noRT | Necab3 | 4.011 | 3.05E-09 | glio |
| Bora_in.vivo_noRT | Aqp5 | 4.673 | 3.05E-09 | glio |
| Bora_in.vivo_noRT | Gm14264 | 3.942 | 3.05E-09 | glio |
| Bora_in.vivo_noRT | Ahrr | 4.136 | 3.05E-09 | glio |
| Aars2_in.vivo_noRT | Tnfsf13os | 4.556 | 3.05E-11 | glio |
| Bcas2_in.vivo_noRT | Krt80 | 4.541 | 3.05E-13 | glio |
| Aars2_in.vivo_noRT | Htr1b | 5.327 | 3.05E-15 | glio |
| Bcas2_in.vivo_noRT | Gm15890 | 5.325 | 3.05E-19 | glio |
| Bcas2_in.vivo_noRT | Adam5 | 4.543 | 3.05E-19 | glio |
| Atr_in.vivo_noRT | Fam227b | 5.533 | 3.05E-30 | glio |
| Atr_in.vivo_noRT | Gm20658 | 5.533 | 3.05E-30 | glio |
| culture_effect | Rpl35 | 0.61 | 3.06E-03 | glio |
| Interaction_genotype_Atr | Prl2c2 | -3.548 | 3.06E-03 | glio |
| Interaction_genotype_Atr | Daam2 | -3.465 | 3.06E-03 | glio |
| Interaction_genotype_Bcas2 | Gm13270 | -4.071 | 3.06E-03 | glio |
| Interaction_genotype_Bcas2 | Arl6ip4 | 1.785 | 3.06E-03 | glio |
| Interaction_genotype_Bcas2 | Fbxl6 | 1.741 | 3.06E-03 | glio |
| Interaction_genotype_Mtor | Dleu7 | -3.576 | 3.06E-03 | glio |
| Interaction_genotype_Mtor | Ergic3 | 1.463 | 3.06E-03 | glio |

| | | | | |
|---|---|---|---|---|
| Interaction_genotype_Mtor | Med21 | 1.274 | 3.06E-03 | glio |
| Interaction_genotype_Mtor | AI662270 | 1.383 | 3.06E-03 | glio |
| Interaction_genotype_Myc | Rcan3 | -2.861 | 3.06E-03 | glio |
| Interaction_genotype_Myc | Psmb6 | 1.33 | 3.06E-03 | glio |
| Interaction_genotype_Rad9a | Gm27239 | -4.528 | 3.06E-03 | glio |
| interaction_RT | Pcdh17 | 1.154 | 3.06E-03 | glio |
| interaction_RT | Klhl21 | -0.832 | 3.06E-03 | glio |
| interaction_RT | Ogfr | -0.656 | 3.06E-03 | glio |
| Aars2_in.vivo_noRT | Npff | 2.003 | 3.06E-03 | glio |
| Aars2_in.vivo_noRT | Mocs3 | 1.935 | 3.06E-03 | glio |
| Aars2_in.vivo_noRT | Pfn2 | -1.471 | 3.06E-03 | glio |
| Aars2_in.vivo_noRT | Trafd1 | -1.331 | 3.06E-03 | glio |
| Aars2_in.vivo_noRT | Clpp | -0.894 | 3.06E-03 | glio |
| Aars2_ex.vivo_noRT | Ppfia2 | 2.499 | 3.06E-03 | glio |
| Atr_in.vivo_noRT | Aldh3b1 | 2.129 | 3.06E-03 | glio |
| Atr_in.vivo_noRT | Gm13470 | 1.932 | 3.06E-03 | glio |
| Atr_in.vivo_noRT | Ap3m2 | 1.663 | 3.06E-03 | glio |
| Atr_in.vivo_noRT | Tox4 | -1.231 | 3.06E-03 | glio |
| Bard1_ex.vivo_noRT | Gm4858 | 2.814 | 3.06E-03 | glio |
| Bcas2_in.vivo_noRT | Tifa | 1.933 | 3.06E-03 | glio |
| Bcas2_in.vivo_noRT | Tmsb10 | -1.058 | 3.06E-03 | glio |
| Bora_in.vivo_noRT | Zfp974 | 2.282 | 3.06E-03 | glio |
| Bora_in.vivo_noRT | Mst1 | 2.036 | 3.06E-03 | glio |
| Bora_in.vivo_noRT | Hpn | 2.112 | 3.06E-03 | glio |
| Bora_in.vivo_noRT | Eaf1 | -1.559 | 3.06E-03 | glio |
| Bora_in.vivo_noRT | Eloa | 1.533 | 3.06E-03 | glio |
| Chek1_in.vivo_noRT | Gm14409 | 2.07 | 3.06E-03 | glio |
| Chek1_in.vivo_noRT | Lman1 | -0.768 | 3.06E-03 | glio |
| Chek1_in.vivo_noRT | Rpl32 | -0.6 | 3.06E-03 | glio |
| Chek1_ex.vivo_noRT | Urb2 | -2.041 | 3.06E-03 | glio |
| Chek1_ex.vivo_noRT | Pxdc1 | -1.801 | 3.06E-03 | glio |
| Eif2s1_ex.vivo_noRT | Cgnl1 | 2.234 | 3.06E-03 | glio |
| Eif2s1_ex.vivo_noRT | Gm26543 | 2.234 | 3.06E-03 | glio |
| Eif2s1_ex.vivo_noRT | 2410017I17Ri | 2.234 | 3.06E-03 | glio |
| Hsd17b10_in.vivo_noRT | Arhgdia | -0.531 | 3.06E-03 | glio |
| Interaction_genotype_Hsd17b | Brinp3 | 4.639 | 3.06E-04 | glio |
| Aars2_in.vivo_noRT | 9130230N09R | 2.233 | 3.06E-04 | glio |
| Aars2_in.vivo_noRT | Parp1 | -1.315 | 3.06E-04 | glio |
| Atr_in.vivo_noRT | Gm47438 | 2.79 | 3.06E-04 | glio |
| Atr_in.vivo_noRT | Utp14a | -1.332 | 3.06E-04 | glio |
| Bora_in.vivo_noRT | Trim71 | 2.577 | 3.06E-04 | glio |
| Bora_in.vivo_noRT | Cldn9 | 2.491 | 3.06E-04 | glio |
| Bora_in.vivo_noRT | Twf2 | -1.508 | 3.06E-04 | glio |
| Chek1_in.vivo_noRT | Gm16230 | 2.556 | 3.06E-04 | glio |
| Chek1_ex.vivo_noRT | Dct | 3.857 | 3.06E-04 | glio |
| Hsd17b10_in.vivo_noRT | Zfp689 | 2.126 | 3.06E-04 | glio |
| culture_effect | Rpl27a | -0.565 | 3.06E-05 | glio |
| Interaction_genotype_Mtor | Gm20705 | -3.963 | 3.06E-05 | glio |
| Interaction_genotype_Myc | Ppp1r3g | -4.422 | 3.06E-05 | glio |
| Aars2_in.vivo_noRT | Tex19.2 | 2.863 | 3.06E-05 | glio |
| Aars2_in.vivo_noRT | Lgals4 | 2.691 | 3.06E-05 | glio |
| Aars2_in.vivo_noRT | Ano8 | 2.45 | 3.06E-05 | glio |

| | | | | |
|---|---|---|---|---|
| Bcas2_in.vivo_noRT | Gm42997 | 2.925 | 3.06E-05 | glio |
| culture_effect | Mrpl55 | 0.747 | 3.06E-06 | glio |
| culture_effect | Phactr4 | -0.987 | 3.06E-06 | glio |
| Interaction_genotype_Atr | Gm26533 | -5.735 | 3.06E-06 | glio |
| Aars2_in.vivo_noRT | Acox2 | 2.808 | 3.06E-06 | glio |
| interaction_RT | Plxnd1 | -2.706 | 3.06E-07 | glio |
| Bora_in.vivo_noRT | Pawr | 3.17 | 3.06E-07 | glio |
| Chek1_in.vivo_noRT | Lyrm1 | 3.854 | 3.06E-08 | glio |
| culture_effect | Ctss | -3.45 | 3.06E-10 | glio |
| Bora_in.vivo_noRT | 4921508D12R | 4.009 | 3.06E-10 | glio |
| Chek1_in.vivo_noRT | St8sia1 | 4.784 | 3.06E-11 | glio |
| Bora_in.vivo_noRT | Vsig10l | 4.807 | 3.06E-12 | glio |
| Aars2_in.vivo_noRT | Npy4r | 4.496 | 3.06E-13 | glio |
| Bora_in.vivo_noRT | Gm11627 | 4.936 | 3.06E-14 | glio |
| culture_effect | Rab3d | -1.376 | 3.07E-03 | glio |
| Interaction_genotype_Cdkn2a | Krt19 | 2.835 | 3.07E-03 | glio |
| Interaction_genotype_Cdkn2a | Cd7 | 3.099 | 3.07E-03 | glio |
| Interaction_genotype_Mtor | Scn2a | -3.378 | 3.07E-03 | glio |
| Interaction_genotype_Myc | Tmem8b | -3.286 | 3.07E-03 | glio |
| Interaction_genotype_Myc | Zfp811 | -3.326 | 3.07E-03 | glio |
| Interaction_genotype_Myc | AU041133 | -2.873 | 3.07E-03 | glio |
| Interaction_genotype_Terf1 | C230037L18Ri | -4.196 | 3.07E-03 | glio |
| Interaction_genotype_Terf1 | Pmp2 | -3.943 | 3.07E-03 | glio |
| Aars2_in.vivo_noRT | Ccl4 | 4.483 | 3.07E-03 | glio |
| Aars2_in.vivo_noRT | Pakap.1 | 1.873 | 3.07E-03 | glio |
| Atp5c1_in.vivo_noRT | Gm4166 | 1.741 | 3.07E-03 | glio |
| Atp5c1_in.vivo_noRT | Ceacam16 | 2.01 | 3.07E-03 | glio |
| Atr_ex.vivo_noRT | Nts | 3.164 | 3.07E-03 | glio |
| Atr_ex.vivo_noRT | Olig2 | -2.069 | 3.07E-03 | glio |
| Bora_in.vivo_noRT | Gm20597 | 2.355 | 3.07E-03 | glio |
| Chek1_in.vivo_noRT | Macc1 | 1.977 | 3.07E-03 | glio |
| Chek1_in.vivo_noRT | Aptx | -1.189 | 3.07E-03 | glio |
| culture_effect | Ebna1bp2 | -0.438 | 3.07E-04 | glio |
| Interaction_genotype_Mtor | Ccdc159 | -4.552 | 3.07E-04 | glio |
| Interaction_genotype_Myc | Gm10501 | -4.235 | 3.07E-04 | glio |
| Interaction_genotype_Pars2 | AC154200.1 | -4.039 | 3.07E-04 | glio |
| Interaction_genotype_Pars2 | Gm12925 | -5.191 | 3.07E-04 | glio |
| Interaction_genotype_Rad51 | Gm27239 | -4.748 | 3.07E-04 | glio |
| Atp5c1_in.vivo_noRT | Gm20646 | 2.668 | 3.07E-04 | glio |
| Atr_in.vivo_noRT | 2610203C22R | 2.68 | 3.07E-04 | glio |
| Chek1_in.vivo_noRT | Acacb | 2.542 | 3.07E-04 | glio |
| Interaction_genotype_Chek1 | C1s1 | 5.414 | 3.07E-05 | glio |
| Interaction_genotype_Chek1 | Folr1 | -4.55 | 3.07E-05 | glio |
| Interaction_genotype_Mtor | Clint1 | 1.878 | 3.07E-05 | glio |
| Aars2_in.vivo_noRT | Gm42699 | 2.776 | 3.07E-05 | glio |
| Aars2_in.vivo_noRT | Gm17018 | -1.716 | 3.07E-05 | glio |
| Atr_in.vivo_noRT | Hist1h4j | 3.414 | 3.07E-05 | glio |
| Chek1_ex.vivo_noRT | Gm37214 | 3.747 | 3.07E-05 | glio |
| culture_effect | Rdh5 | 1.692 | 3.07E-06 | glio |
| Interaction_genotype_Nat2 | Gm20767 | 5.183 | 3.07E-06 | glio |
| Atp5c1_in.vivo_noRT | Meioc | 2.405 | 3.07E-06 | glio |
| Bora_in.vivo_noRT | Olfr920 | 3.186 | 3.07E-06 | glio |

| | | | | |
|---|---|---|---|---|
| Bard1_ex.vivo_noRT | Gm48780 | 3.596 | 3.07E-07 | glio |
| culture_effect | Lta4h | -1.107 | 3.07E-08 | glio |
| Bcas2_in.vivo_noRT | Gm16023 | 4.338 | 3.07E-08 | glio |
| Bcas2_in.vivo_noRT | Wnt10a | 3.829 | 3.07E-08 | glio |
| Bora_in.vivo_noRT | Grip1 | 3.84 | 3.07E-08 | glio |
| Atp5c1_in.vivo_noRT | A930003A15R | 4.111 | 3.07E-09 | glio |
| Atp5c1_in.vivo_noRT | Erich6b | 3.19 | 3.07E-09 | glio |
| Atp5c1_in.vivo_noRT | Slc17a6 | 2.668 | 3.07E-09 | glio |
| Bcas2_in.vivo_noRT | St8sia1 | 4.547 | 3.07E-10 | glio |
| Bcas2_in.vivo_noRT | Lncenc1 | 4.028 | 3.07E-10 | glio |
| Chek1_in.vivo_noRT | Klk14 | 4.512 | 3.07E-10 | glio |
| Atr_in.vivo_noRT | B430219N15R | 4.697 | 3.07E-11 | glio |
| Chek1_in.vivo_noRT | Lrrc72 | 4.253 | 3.07E-11 | glio |
| Bora_in.vivo_noRT | Gm31518 | 4.936 | 3.07E-12 | glio |
| Bcas2_ex.vivo_noRT | Bcas2 | -3.029 | 3.07E-14 | glio |
| Atr_in.vivo_noRT | Gprin2 | 4.622 | 3.07E-15 | glio |
| culture_effect | Idh3g | -1.01 | 3.07E-17 | glio |
| Aars2_in.vivo_noRT | Gm2115 | 5.327 | 3.07E-18 | glio |
| Bcas2_in.vivo_noRT | Gm2022 | 4.807 | 3.07E-19 | glio |
| Atr_in.vivo_noRT | BC049987 | 4.75 | 3.07E-21 | glio |
| Bora_in.vivo_noRT | Gm44693 | 4.936 | 3.07E-23 | glio |
| Atr_in.vivo_noRT | Speer4a | 5.267 | 3.07E-24 | glio |
| Bcas2_in.vivo_noRT | Krt19 | 4.807 | 3.07E-24 | glio |
| culture_effect | Gm14124 | 0.845 | 3.08E-03 | glio |
| culture_effect | Pcm1 | 0.493 | 3.08E-03 | glio |
| Interaction_genotype_Aars2 | Ripply3 | -3.598 | 3.08E-03 | glio |
| Interaction_genotype_Bcas2 | Gm45620 | -3.781 | 3.08E-03 | glio |
| Interaction_genotype_Bcas2 | B230307C23R | -3.167 | 3.08E-03 | glio |
| Interaction_genotype_Bcas2 | Abhd2 | 2.093 | 3.08E-03 | glio |
| Interaction_genotype_Bora | Man2c1os | -3.915 | 3.08E-03 | glio |
| Interaction_genotype_Mtor | Gm43330 | -3.18 | 3.08E-03 | glio |
| Interaction_genotype_Mtor | Slc22a18 | -3.821 | 3.08E-03 | glio |
| Interaction_genotype_Mtor | Sfxn5 | -3.012 | 3.08E-03 | glio |
| Interaction_genotype_Mtor | Elmod3 | -3.265 | 3.08E-03 | glio |
| Interaction_genotype_Myc | Esp15 | -2.729 | 3.08E-03 | glio |
| Interaction_genotype_Myc | 1600014C23R | -3.443 | 3.08E-03 | glio |
| Interaction_genotype_Myc | Mamdc2 | -3.26 | 3.08E-03 | glio |
| Interaction_genotype_Myc | Gm47863 | -3.306 | 3.08E-03 | glio |
| Interaction_genotype_Myc | Myo9a | -2.319 | 3.08E-03 | glio |
| Interaction_genotype_Myc | Spsb3 | -2.145 | 3.08E-03 | glio |
| Interaction_genotype_Myc | Zfp64 | -2.309 | 3.08E-03 | glio |
| Interaction_genotype_Myc | Layn | -2.46 | 3.08E-03 | glio |
| Interaction_genotype_Myc | Slc35f1 | -2.213 | 3.08E-03 | glio |
| Interaction_genotype_Myc | Ets1 | -1.828 | 3.08E-03 | glio |
| Interaction_genotype_Myc | Prdx2 | 1.21 | 3.08E-03 | glio |
| Interaction_genotype_Terf1 | Gm36356 | -3.728 | 3.08E-03 | glio |
| Interaction_genotype_Terf1 | C230013L11Ri | -4.695 | 3.08E-03 | glio |
| Interaction_genotype_Terf1 | Thap8 | -4.498 | 3.08E-03 | glio |
| Interaction_genotype_Terf1 | Klhl23 | -4.043 | 3.08E-03 | glio |
| Interaction_genotype_Terf1 | Gm27010 | -4.226 | 3.08E-03 | glio |
| Aars2_in.vivo_noRT | Chn2 | 2.335 | 3.08E-03 | glio |
| Aars2_ex.vivo_noRT | Slc6a19 | 2.9 | 3.08E-03 | glio |

| | | | | |
|---|---|---|---|---|
| Atr_in.vivo_noRT | Ppef2 | 1.781 | 3.08E-03 | glio |
| Bard1_ex.vivo_noRT | Gm20767 | 2.814 | 3.08E-03 | glio |
| Bcas2_in.vivo_noRT | Fam177a | 2.039 | 3.08E-03 | glio |
| Bcas2_in.vivo_noRT | Cdc42ep2 | 2.247 | 3.08E-03 | glio |
| Bcas2_ex.vivo_noRT | C430002N11R | 2.347 | 3.08E-03 | glio |
| Bcas2_ex.vivo_noRT | Dio1 | 2.742 | 3.08E-03 | glio |
| Bora_in.vivo_noRT | Mtch2 | -0.799 | 3.08E-03 | glio |
| Brca2_ex.vivo_noRT | Ccl9 | 2.432 | 3.08E-03 | glio |
| Chek1_in.vivo_noRT | Plekho2 | 2.109 | 3.08E-03 | glio |
| Cyp19a1_ex.vivo_noRT | Gm13191 | 2.159 | 3.08E-03 | glio |
| Cyp19a1_ex.vivo_noRT | 4933416E03Ri | 2.159 | 3.08E-03 | glio |
| culture_effect | Ifit1bl1 | 2.472 | 3.08E-04 | glio |
| Interaction_genotype_Aars2 | 4930515G01R | -4.075 | 3.08E-04 | glio |
| Interaction_genotype_Aars2 | Gm41396 | -4.649 | 3.08E-04 | glio |
| Interaction_genotype_Mtor | Gm50138 | -3.549 | 3.08E-04 | glio |
| Interaction_genotype_Mtor | 1110017D15R | -4.292 | 3.08E-04 | glio |
| Interaction_genotype_Myc | Gm27241 | -3.054 | 3.08E-04 | glio |
| Interaction_genotype_Myc | Hist2h3c1 | -3.975 | 3.08E-04 | glio |
| Interaction_genotype_Myc | Gm4221 | -4.075 | 3.08E-04 | glio |
| Interaction_genotype_Myc | A930033H14R | -4.166 | 3.08E-04 | glio |
| Interaction_genotype_Myc | Cox6c | 1.404 | 3.08E-04 | glio |
| Aars2_in.vivo_noRT | Zscan22 | 2.154 | 3.08E-04 | glio |
| Aars2_in.vivo_noRT | Alkbh1 | -1.541 | 3.08E-04 | glio |
| Atr_in.vivo_noRT | 3300002I08Ri | 2.507 | 3.08E-04 | glio |
| Bora_in.vivo_noRT | Rad1 | -1.623 | 3.08E-04 | glio |
| Brca2_in.vivo_noRT | Gm364 | 2.286 | 3.08E-04 | glio |
| Cenpj_in.vivo_noRT | Slc14a2 | 2.022 | 3.08E-04 | glio |
| Chek1_ex.vivo_noRT | Mycbp | -1.849 | 3.08E-04 | glio |
| culture_effect | Smyd4 | 1.201 | 3.08E-05 | glio |
| Interaction_genotype_Myc | Gm14264 | -4.557 | 3.08E-05 | glio |
| Interaction_genotype_Myc | Arhgdia | 1.59 | 3.08E-05 | glio |
| Aars2_in.vivo_noRT | N4bp2l1 | 3.031 | 3.08E-05 | glio |
| Aars2_in.vivo_noRT | Gm42670 | 2.656 | 3.08E-05 | glio |
| Atr_in.vivo_noRT | Plcb4 | 2.947 | 3.08E-05 | glio |
| Atr_ex.vivo_noRT | Gm47540 | 2.771 | 3.08E-05 | glio |
| culture_effect | Hsd17b10 | -0.683 | 3.08E-06 | glio |
| Interaction_genotype_Myc | Gm14968 | -5.357 | 3.08E-06 | glio |
| Atr_in.vivo_noRT | Eva1c | 3.343 | 3.08E-06 | glio |
| Bora_ex.vivo_noRT | Fabp6 | 3.201 | 3.08E-06 | glio |
| Chek1_ex.vivo_noRT | Dlx4 | 3.877 | 3.08E-06 | glio |
| Chek1_ex.vivo_noRT | Phf11a | 4.055 | 3.08E-06 | glio |
| Chek1_ex.vivo_noRT | Tnp2 | 4.256 | 3.08E-06 | glio |
| Hsd17b10_in.vivo_noRT | Gm45336 | 2.806 | 3.08E-06 | glio |
| Hsd17b10_in.vivo_noRT | mt-Co2 | -0.858 | 3.08E-06 | glio |
| culture_effect | Tcte2 | 2.267 | 3.08E-07 | glio |
| Interaction_genotype_Mtor | Aloxe3 | -5.208 | 3.08E-07 | glio |
| Chek1_in.vivo_noRT | Gm8098 | 3.541 | 3.08E-07 | glio |
| Chek1_in.vivo_noRT | Elfn2 | 3.269 | 3.08E-07 | glio |
| Atr_in.vivo_noRT | Gm9958 | 3.963 | 3.08E-08 | glio |
| Atr_in.vivo_noRT | Ache | 3.96 | 3.08E-08 | glio |
| Bora_in.vivo_noRT | Traf3ip3 | 3.872 | 3.08E-08 | glio |
| culture_effect | Dtd1 | -0.922 | 3.08E-09 | glio |

| | | | | |
|---|---|---|---|---|
| Bcas2_in.vivo_noRT | Ttc16 | 4.284 | 3.08E-10 | glio |
| Bora_in.vivo_noRT | Gm30938 | 4.29 | 3.08E-10 | glio |
| Aars2_in.vivo_noRT | Ccdc114 | 4.409 | 3.08E-11 | glio |
| Bcas2_in.vivo_noRT | Dreh | 4.161 | 3.08E-11 | glio |
| Aars2_in.vivo_noRT | Nxnl2 | 4.805 | 3.08E-14 | glio |
| Chek1_ex.vivo_noRT | Cgnl1 | 4.842 | 3.08E-16 | glio |
| Interaction_genotype_Aars2 | Ankrd1 | -4.181 | 3.09E-03 | glio |
| Interaction_genotype_Aars2 | D330041H03R | -3.116 | 3.09E-03 | glio |
| Interaction_genotype_Atr | Paip2 | 0.93 | 3.09E-03 | glio |
| Interaction_genotype_Cdkn2a | Gm48155 | 2.835 | 3.09E-03 | glio |
| Interaction_genotype_Mtor | Krt16 | -4.485 | 3.09E-03 | glio |
| Interaction_genotype_Mtor | Dynlt1c | -3.928 | 3.09E-03 | glio |
| Interaction_genotype_Mtor | BC039966 | -2.63 | 3.09E-03 | glio |
| Interaction_genotype_Mtor | Col9a3 | 2.414 | 3.09E-03 | glio |
| Interaction_genotype_Myc | Gm10801 | -2.729 | 3.09E-03 | glio |
| Interaction_genotype_Myc | Prim2 | -2.4 | 3.09E-03 | glio |
| Interaction_genotype_Myc | Clec2d | -2.713 | 3.09E-03 | glio |
| Interaction_genotype_Pars2 | Pmp2 | -3.569 | 3.09E-03 | glio |
| Interaction_genotype_Pars2 | Cgn | -4.004 | 3.09E-03 | glio |
| Interaction_genotype_Rad21 | C1ra | 3.262 | 3.09E-03 | glio |
| interaction_RT | Rhod | 1.109 | 3.09E-03 | glio |
| interaction_RT | Frg1 | 0.715 | 3.09E-03 | glio |
| Aars2_in.vivo_noRT | Pla2g4c | 2.102 | 3.09E-03 | glio |
| Atr_in.vivo_noRT | Dusp11 | -0.898 | 3.09E-03 | glio |
| Atr_ex.vivo_noRT | Pkhd1 | 2.889 | 3.09E-03 | glio |
| Bcas2_in.vivo_noRT | Psip1 | -0.846 | 3.09E-03 | glio |
| Bora_in.vivo_noRT | Ptn | 2.538 | 3.09E-03 | glio |
| Cenpj_in.vivo_noRT | Gm12703 | 1.756 | 3.09E-03 | glio |
| Chek1_in.vivo_noRT | Klhdc1 | 2.131 | 3.09E-03 | glio |
| Chek1_ex.vivo_noRT | Gm47851 | 2.456 | 3.09E-03 | glio |
| Cyp19a1_ex.vivo_noRT | Ush1g | 2.559 | 3.09E-03 | glio |
| Cyp19a1_ex.vivo_noRT | Cldn2 | 2.159 | 3.09E-03 | glio |
| Cyp19a1_ex.vivo_noRT | Adam2 | 2.933 | 3.09E-03 | glio |
| Cyp19a1_ex.vivo_noRT | Myt1 | 2.559 | 3.09E-03 | glio |
| Cyp19a1_ex.vivo_noRT | Gm48265 | 2.959 | 3.09E-03 | glio |
| Cyp19a1_ex.vivo_noRT | Irf8 | 2.959 | 3.09E-03 | glio |
| Cyp19a1_ex.vivo_noRT | Stfa3 | 2.159 | 3.09E-03 | glio |
| Cyp19a1_ex.vivo_noRT | Olfr787 | 2.159 | 3.09E-03 | glio |
| Hsd17b10_in.vivo_noRT | Grip1 | 2.264 | 3.09E-03 | glio |
| Interaction_genotype_Aars2 | Ppp1r18os | -4.715 | 3.09E-04 | glio |
| Aars2_in.vivo_noRT | Rab42 | 2.723 | 3.09E-04 | glio |
| Aars2_in.vivo_noRT | Hist1h3d | 2.423 | 3.09E-04 | glio |
| Bcas2_in.vivo_noRT | Fam171a2 | 2.645 | 3.09E-04 | glio |
| Bcas2_in.vivo_noRT | 4833408A19R | 2.475 | 3.09E-04 | glio |
| Bcas2_in.vivo_noRT | Tlnrd1 | -1.753 | 3.09E-04 | glio |
| Cenpj_in.vivo_noRT | Gm4340 | 2.022 | 3.09E-04 | glio |
| Cenpj_in.vivo_noRT | Gm16239 | 2.022 | 3.09E-04 | glio |
| Cenpj_in.vivo_noRT | Adgra3 | -1.078 | 3.09E-04 | glio |
| Chek1_ex.vivo_noRT | Lrrc75b | 3.397 | 3.09E-04 | glio |
| Chek1_ex.vivo_noRT | Mrpl47 | -2.074 | 3.09E-04 | glio |
| Chek1_ex.vivo_noRT | Mrps31 | -1.692 | 3.09E-04 | glio |
| Chek1_ex.vivo_noRT | Csf1 | 2.36 | 3.09E-04 | glio |

| | | | | |
|---|---|---|---|---|
| Hsd17b10_in.vivo_noRT | Syne3 | 2.891 | 3.09E-04 | glio |
| Interaction_genotype_Mtor | 4930570E03Ri | -3.839 | 3.09E-05 | glio |
| Interaction_genotype_Myc | Pygo1 | -4.404 | 3.09E-05 | glio |
| Atr_in.vivo_noRT | 5330438D12R | 2.408 | 3.09E-05 | glio |
| Bcas2_ex.vivo_noRT | Pcdhb19 | 3.928 | 3.09E-05 | glio |
| Bcas2_ex.vivo_noRT | Ankrd22 | 2.742 | 3.09E-05 | glio |
| Bcas2_ex.vivo_noRT | Gm31717 | 2.742 | 3.09E-05 | glio |
| Bcas2_ex.vivo_noRT | Gm47404 | 2.742 | 3.09E-05 | glio |
| Bcas2_ex.vivo_noRT | Gm14742 | 2.742 | 3.09E-05 | glio |
| Bcas2_ex.vivo_noRT | Scel | 2.742 | 3.09E-05 | glio |
| Bcas2_ex.vivo_noRT | Gm44646 | 3.298 | 3.09E-05 | glio |
| Bcas2_ex.vivo_noRT | Ctla4 | 2.742 | 3.09E-05 | glio |
| Bcas2_ex.vivo_noRT | Zscan4-ps3 | 2.742 | 3.09E-05 | glio |
| Bcas2_ex.vivo_noRT | Tagap | 2.742 | 3.09E-05 | glio |
| Bcas2_ex.vivo_noRT | Gm21731 | 2.742 | 3.09E-05 | glio |
| Bcas2_ex.vivo_noRT | Gm11232 | 2.742 | 3.09E-05 | glio |
| Bcas2_ex.vivo_noRT | Gm29966 | 2.742 | 3.09E-05 | glio |
| Bcas2_ex.vivo_noRT | Olfr1152 | 2.742 | 3.09E-05 | glio |
| Chek1_in.vivo_noRT | Gm19325 | 3.08 | 3.09E-05 | glio |
| Hsd17b10_in.vivo_noRT | Gm30541 | 2.965 | 3.09E-05 | glio |
| Interaction_genotype_Bora | K230015D01R | -6.236 | 3.09E-06 | glio |
| Bard1_ex.vivo_noRT | Gm5662 | 4.395 | 3.09E-06 | glio |
| Bard1_ex.vivo_noRT | Sucnr1 | 2.814 | 3.09E-06 | glio |
| Bard1_ex.vivo_noRT | Gm40538 | 3.612 | 3.09E-06 | glio |
| Bcas2_in.vivo_noRT | D830035M03I | 3.221 | 3.09E-06 | glio |
| Bora_in.vivo_noRT | 0610009L18Ri | 3.425 | 3.09E-06 | glio |
| culture_effect | Kcnj10 | -3.1 | 3.09E-07 | glio |
| culture_effect | Fn1 | -1.861 | 3.09E-07 | glio |
| Aars2_in.vivo_noRT | Gm36445 | 3.574 | 3.09E-07 | glio |
| Atr_in.vivo_noRT | Nr1h4 | 3.484 | 3.09E-07 | glio |
| Atr_in.vivo_noRT | 1190005I06Ri | 3.28 | 3.09E-07 | glio |
| Bcas2_in.vivo_noRT | BC004004 | -1.925 | 3.09E-07 | glio |
| Bora_in.vivo_noRT | Sec1 | 3.779 | 3.09E-07 | glio |
| Chek1_in.vivo_noRT | Abat | 3.868 | 3.09E-07 | glio |
| Atr_in.vivo_noRT | Gm36839 | 3.971 | 3.09E-08 | glio |
| Bora_in.vivo_noRT | 1110020A21R | 3.986 | 3.09E-08 | glio |
| Chek1_ex.vivo_noRT | Luzp2 | 4.045 | 3.09E-10 | glio |
| Atr_in.vivo_noRT | 4921523L03Ri | 4.869 | 3.09E-11 | glio |
| Bora_in.vivo_noRT | Gm20732 | 4.166 | 3.09E-11 | glio |
| Bora_in.vivo_noRT | Sult5a1 | 4.937 | 3.09E-11 | glio |
| Bora_in.vivo_noRT | Gm10399 | 4.537 | 3.09E-11 | glio |
| Atr_in.vivo_noRT | Aass | 4.532 | 3.09E-12 | glio |
| Atr_in.vivo_noRT | Prox2os | 4.229 | 3.09E-12 | glio |
| Hsd17b10_in.vivo_noRT | Zscan4-ps2 | 3.616 | 3.09E-12 | glio |
| Atr_in.vivo_noRT | Ephx3 | 4.806 | 3.09E-13 | glio |
| Hsd17b10_in.vivo_noRT | Gm13078 | 3.616 | 3.09E-13 | glio |
| culture_effect | Nrp1 | 2.318 | 3.09E-15 | glio |
| culture_effect | Matk | -1.723 | 3.10E-03 | glio |
| culture_effect | Gm38190 | -1.684 | 3.10E-03 | glio |
| culture_effect | Mrpl14 | 0.384 | 3.10E-03 | glio |
| Interaction_genotype_Aars2 | Eif3i | 0.874 | 3.10E-03 | glio |
| Interaction_genotype_Mtor | Oscp1 | -3.579 | 3.10E-03 | glio |

| | | | | |
|---|---|---|---|---|
| Interaction_genotype_Mtor | Fam118b | -2.502 | 3.10E-03 | glio |
| Interaction_genotype_Mtor | Tmem242 | -2.229 | 3.10E-03 | glio |
| Interaction_genotype_Mtor | Rps27a | 0.991 | 3.10E-03 | glio |
| interaction_RT | 4933433G15R | -2.448 | 3.10E-03 | glio |
| Aars2_in.vivo_noRT | Anxa2 | -0.839 | 3.10E-03 | glio |
| Atr_in.vivo_noRT | Klhl9 | -1.41 | 3.10E-03 | glio |
| Atr_in.vivo_noRT | Gbp7 | -1.02 | 3.10E-03 | glio |
| Atr_ex.vivo_noRT | Gm7247 | 2.367 | 3.10E-03 | glio |
| Atr_ex.vivo_noRT | 2410141K09R | 2.38 | 3.10E-03 | glio |
| Atr_ex.vivo_noRT | Astn2 | 2.769 | 3.10E-03 | glio |
| Bcas2_in.vivo_noRT | Axin2 | 2.237 | 3.10E-03 | glio |
| Chek1_in.vivo_noRT | Gfap | 2.016 | 3.10E-03 | glio |
| Chek1_ex.vivo_noRT | Mro | 2.696 | 3.10E-03 | glio |
| Eif2s1_in.vivo_noRT | Stra8 | 2.147 | 3.10E-03 | glio |
| culture_effect | Dtymk | 0.468 | 3.10E-04 | glio |
| Interaction_genotype_Myc | Prl2c2 | -3.908 | 3.10E-04 | glio |
| Interaction_genotype_Myc | Apol10b | 3.504 | 3.10E-04 | glio |
| Interaction_genotype_Myc | Fgf7 | -3.853 | 3.10E-04 | glio |
| Interaction_genotype_Myc | Hinfp | -2.939 | 3.10E-04 | glio |
| Atp5c1_in.vivo_noRT | Cct6b | 2.6 | 3.10E-04 | glio |
| Atr_in.vivo_noRT | Calm2 | -0.941 | 3.10E-04 | glio |
| Bcas2_in.vivo_noRT | A830082N09R | 2.726 | 3.10E-04 | glio |
| Bcas2_in.vivo_noRT | Cfap100 | 2.61 | 3.10E-04 | glio |
| Chek1_in.vivo_noRT | Rrad | 2.781 | 3.10E-04 | glio |
| Chek1_in.vivo_noRT | C030014I23Ri | 2.593 | 3.10E-04 | glio |
| Chek1_in.vivo_noRT | 0610040B10R | 2.176 | 3.10E-04 | glio |
| Hsd17b10_in.vivo_noRT | Ninj2 | 2.709 | 3.10E-04 | glio |
| Interaction_genotype_Myc | Rbmx | -3.553 | 3.10E-05 | glio |
| Aars2_in.vivo_noRT | Cct3 | -1.075 | 3.10E-05 | glio |
| Atr_in.vivo_noRT | Zfp599 | 3.101 | 3.10E-05 | glio |
| Bcas2_in.vivo_noRT | 4933406P04R | 3.126 | 3.10E-05 | glio |
| Bcas2_in.vivo_noRT | Mrgprf | 3.011 | 3.10E-05 | glio |
| Bcas2_ex.vivo_noRT | Cgnl1 | 2.742 | 3.10E-05 | glio |
| Bcas2_ex.vivo_noRT | Slc16a7 | 3.485 | 3.10E-05 | glio |
| Bcas2_ex.vivo_noRT | Gm26543 | 2.742 | 3.10E-05 | glio |
| Bora_ex.vivo_noRT | A330076H08R | 3.201 | 3.10E-05 | glio |
| Bora_ex.vivo_noRT | Gm47540 | 2.808 | 3.10E-05 | glio |
| Chek1_in.vivo_noRT | Usp44 | 3.055 | 3.10E-05 | glio |
| Chek1_in.vivo_noRT | Proscos | 3.148 | 3.10E-05 | glio |
| Chek1_in.vivo_noRT | Pde6h | 2.748 | 3.10E-05 | glio |
| Interaction_genotype_Bcas2 | Cyp26b1 | -6.209 | 3.10E-06 | glio |
| Bora_in.vivo_noRT | Spink2 | 3.739 | 3.10E-06 | glio |
| Chek1_in.vivo_noRT | Uggt1 | -1.955 | 3.10E-06 | glio |
| Chek1_ex.vivo_noRT | Ceacam19 | 4.132 | 3.10E-06 | glio |
| Chek1_ex.vivo_noRT | Sapcd1 | 4.256 | 3.10E-06 | glio |
| interaction_RT | Cd14 | 3.689 | 3.10E-07 | glio |
| Aars2_in.vivo_noRT | Phldb2 | 4.106 | 3.10E-08 | glio |
| Chek1_in.vivo_noRT | Gm36839 | 3.992 | 3.10E-08 | glio |
| Atr_in.vivo_noRT | Gm35867 | 4.08 | 3.10E-09 | glio |
| Atr_ex.vivo_noRT | Dscaml1 | 3.566 | 3.10E-09 | glio |
| Bcas2_in.vivo_noRT | Spata31d1a | 4.166 | 3.10E-09 | glio |
| Bora_in.vivo_noRT | Gm46652 | 3.887 | 3.10E-09 | glio |

| | | | | |
|---|---|---|---|---|
| Bcas2_in.vivo_noRT | Thap8 | 4.547 | 3.10E-10 | glio |
| Bcas2_in.vivo_noRT | Gm50394 | 4.541 | 3.10E-11 | glio |
| Bcas2_in.vivo_noRT | Gprin2 | 3.895 | 3.10E-11 | glio |
| Chek1_ex.vivo_noRT | Gm13398 | 4.045 | 3.10E-11 | glio |
| Bora_in.vivo_noRT | Stx19 | 4.025 | 3.10E-12 | glio |
| Chek1_in.vivo_noRT | Pmch | 4.386 | 3.10E-13 | glio |
| Chek1_ex.vivo_noRT | Gm26543 | 4.842 | 3.10E-16 | glio |
| Bora_in.vivo_noRT | Gm36793 | 4.936 | 3.10E-21 | glio |
| Aars2_in.vivo_noRT | Gm266 | 5.327 | 3.10E-25 | glio |
| culture_effect | Pdk2 | -1.876 | 3.11E-03 | glio |
| culture_effect | Mgat2 | 0.496 | 3.11E-03 | glio |
| culture_effect | Brd7 | 0.485 | 3.11E-03 | glio |
| Interaction_genotype_Bora | Gm37800 | -3.602 | 3.11E-03 | glio |
| Interaction_genotype_Bora | Mia3 | 1.912 | 3.11E-03 | glio |
| Interaction_genotype_Mtor | Pcdhb16 | -3.856 | 3.11E-03 | glio |
| Interaction_genotype_Mtor | Hist1h4j | -3.973 | 3.11E-03 | glio |
| Interaction_genotype_Mtor | Hist1h2be | -3.13 | 3.11E-03 | glio |
| Interaction_genotype_Myc | 1810012K16R | -3.179 | 3.11E-03 | glio |
| Interaction_genotype_Myc | Sestd1 | -2.721 | 3.11E-03 | glio |
| Interaction_genotype_Myc | Parg | -2.368 | 3.11E-03 | glio |
| Interaction_genotype_Myc | Mxra8 | -2.525 | 3.11E-03 | glio |
| Interaction_genotype_Myc | 1700120C14R | -2.858 | 3.11E-03 | glio |
| Interaction_genotype_Myc | Surf4 | 1.214 | 3.11E-03 | glio |
| Interaction_genotype_Pars2 | Gjb4 | -4.137 | 3.11E-03 | glio |
| Interaction_genotype_Pars2 | Gm16046 | -3.903 | 3.11E-03 | glio |
| interaction_RT | Dcpp1 | 3.165 | 3.11E-03 | glio |
| Aars2_in.vivo_noRT | G6pc3 | -1.233 | 3.11E-03 | glio |
| Bcas2_in.vivo_noRT | Vps37c | 1.62 | 3.11E-03 | glio |
| Bcas2_in.vivo_noRT | Stmn1 | -1.065 | 3.11E-03 | glio |
| Chek1_in.vivo_noRT | A330074K22R | 1.967 | 3.11E-03 | glio |
| Chek1_in.vivo_noRT | Gm15706 | 1.871 | 3.11E-03 | glio |
| Chek1_in.vivo_noRT | Eif3f | -0.671 | 3.11E-03 | glio |
| Ercc4_in.vivo_noRT | 1700019E08Ri | 2.433 | 3.11E-03 | glio |
| Ercc4_in.vivo_noRT | Tcp11x2 | 2.57 | 3.11E-03 | glio |
| Hsd17b10_in.vivo_noRT | Ccn2 | 3.061 | 3.11E-03 | glio |
| Hsd17b10_in.vivo_noRT | Bloc1s6os | 2.159 | 3.11E-03 | glio |
| Interaction_genotype_Atr | Oser1 | 2.134 | 3.11E-04 | glio |
| Interaction_genotype_Myc | Zfp933 | -3.429 | 3.11E-04 | glio |
| Interaction_genotype_Uqcrc2 | Fa2h | 5.032 | 3.11E-04 | glio |
| Aars2_ex.vivo_noRT | Arhgef16 | 2.893 | 3.11E-04 | glio |
| Aars2_ex.vivo_noRT | Tnfrsf1b | 3.293 | 3.11E-04 | glio |
| Atr_in.vivo_noRT | Tmem223 | 2.604 | 3.11E-04 | glio |
| Atr_in.vivo_noRT | Mpp2 | 2.554 | 3.11E-04 | glio |
| Atr_in.vivo_noRT | Castor2 | 2.388 | 3.11E-04 | glio |
| Atr_in.vivo_noRT | Acin1 | -1.172 | 3.11E-04 | glio |
| Bcas2_in.vivo_noRT | Hist1h2ai | 2.711 | 3.11E-04 | glio |
| Bcas2_in.vivo_noRT | B430219N15R | 2.683 | 3.11E-04 | glio |
| Bora_in.vivo_noRT | Stk24 | -1.552 | 3.11E-04 | glio |
| Bora_in.vivo_noRT | Ndufs6 | -0.904 | 3.11E-04 | glio |
| Chek1_ex.vivo_noRT | Zan | 3.279 | 3.11E-04 | glio |
| Chek1_ex.vivo_noRT | Pdha1 | -1.094 | 3.11E-04 | glio |
| Eif2s1_in.vivo_noRT | Dab2 | -2.868 | 3.11E-04 | glio |

| | | | | |
|---|---|---|---|---|
| Interaction_genotype_Mtor | Kdm8 | -4.667 | 3.11E-05 | glio |
| Atr_in.vivo_noRT | Arpc5l | -1.302 | 3.11E-05 | glio |
| Bora_in.vivo_noRT | 4930597O21R | 3.209 | 3.11E-05 | glio |
| Bora_in.vivo_noRT | Vgll2 | 2.6 | 3.11E-05 | glio |
| Chek1_in.vivo_noRT | Gm15938 | 3.114 | 3.11E-05 | glio |
| Interaction_genotype_Mtor | D730045B01R | -4.43 | 3.11E-06 | glio |
| Atr_in.vivo_noRT | Abcc10 | 3.342 | 3.11E-06 | glio |
| Atr_in.vivo_noRT | Thoc2 | -1.828 | 3.11E-06 | glio |
| Bora_in.vivo_noRT | Rfx6 | 3.173 | 3.11E-06 | glio |
| Atp5c1_in.vivo_noRT | A930033M14F | 3.196 | 3.11E-07 | glio |
| Bcas2_in.vivo_noRT | Srgn | 4.28 | 3.11E-07 | glio |
| Bora_in.vivo_noRT | Wincr1 | 4.672 | 3.11E-07 | glio |
| culture_effect | Dtx1 | -3.51 | 3.11E-08 | glio |
| Atr_in.vivo_noRT | Mir142hg | 3.94 | 3.11E-08 | glio |
| Aars2_in.vivo_noRT | Myoz1 | 3.631 | 3.11E-10 | glio |
| Chek1_in.vivo_noRT | Fabp4 | 4.515 | 3.11E-10 | glio |
| Chek1_ex.vivo_noRT | Lhx1os | 4.447 | 3.11E-10 | glio |
| Atr_in.vivo_noRT | Gm41442 | 4.492 | 3.11E-11 | glio |
| Atr_in.vivo_noRT | Fam189a1 | 3.863 | 3.11E-11 | glio |
| Aars2_in.vivo_noRT | 1700026J14Ri | 4.805 | 3.11E-14 | glio |
| Bcas2_in.vivo_noRT | Gm41307 | 4.807 | 3.11E-21 | glio |
| Atr_in.vivo_noRT | Gm10354 | 5.533 | 3.11E-22 | glio |
| Atr_in.vivo_noRT | Tagap | 5.273 | 3.11E-27 | glio |
| culture_effect | Nudt7 | 1.365 | 3.12E-03 | glio |
| culture_effect | Fgd4 | 0.75 | 3.12E-03 | glio |
| Interaction_genotype_Aars2 | Plp2 | 1.515 | 3.12E-03 | glio |
| Interaction_genotype_Atr | 1700017M07F | -3.143 | 3.12E-03 | glio |
| Interaction_genotype_Atr | Tmem140 | 3.004 | 3.12E-03 | glio |
| Interaction_genotype_Atr | Rdm1 | 1.851 | 3.12E-03 | glio |
| Interaction_genotype_Atr | Map2k1 | 1.622 | 3.12E-03 | glio |
| Interaction_genotype_Bora | Abhd15 | -4.24 | 3.12E-03 | glio |
| Interaction_genotype_Bora | 5033417F24Ri | -3.794 | 3.12E-03 | glio |
| Interaction_genotype_Chek1 | Nxph3 | -3.73 | 3.12E-03 | glio |
| Interaction_genotype_Mtor | Osm | -3.149 | 3.12E-03 | glio |
| Interaction_genotype_Mtor | Ccar1 | 1.345 | 3.12E-03 | glio |
| Interaction_genotype_Myc | Cox11 | -2.127 | 3.12E-03 | glio |
| Interaction_genotype_Rpl3 | Gm12602 | -3.9 | 3.12E-03 | glio |
| interaction_RT | R3hcc1 | -1.41 | 3.12E-03 | glio |
| interaction_RT | Bex3 | 0.633 | 3.12E-03 | glio |
| Aars2_in.vivo_noRT | Grb7 | 2.15 | 3.12E-03 | glio |
| Aars2_in.vivo_noRT | Dbn1 | 2.18 | 3.12E-03 | glio |
| Aars2_in.vivo_noRT | Clic1 | -0.645 | 3.12E-03 | glio |
| Atr_in.vivo_noRT | Map3k20 | 1.821 | 3.12E-03 | glio |
| Atr_ex.vivo_noRT | Slc22a17 | -2.174 | 3.12E-03 | glio |
| Bcas2_in.vivo_noRT | 9030025P20R | 2.335 | 3.12E-03 | glio |
| Bcas2_in.vivo_noRT | Wrb | 1.581 | 3.12E-03 | glio |
| Bcas2_ex.vivo_noRT | Gm20732 | 2.742 | 3.12E-03 | glio |
| Bora_in.vivo_noRT | AI429214 | 2.179 | 3.12E-03 | glio |
| Bora_in.vivo_noRT | Syn1 | 2.204 | 3.12E-03 | glio |
| Bora_in.vivo_noRT | Eid2b | 1.822 | 3.12E-03 | glio |
| Bora_ex.vivo_noRT | Ccdc80 | 2.232 | 3.12E-03 | glio |
| Cenpj_ex.vivo_noRT | Gm12841 | 2.198 | 3.12E-03 | glio |

| | | | | |
|---|---|---|---|---|
| Cenpj_ex.vivo_noRT | Hrh2 | 2.591 | 3.12E-03 | glio |
| Cenpj_ex.vivo_noRT | Otof | 2.591 | 3.12E-03 | glio |
| Chek1_in.vivo_noRT | Zfand4 | 2.238 | 3.12E-03 | glio |
| Chek1_in.vivo_noRT | Inafm1 | 1.874 | 3.12E-03 | glio |
| Chek1_ex.vivo_noRT | Dbndd1 | 2.303 | 3.12E-03 | glio |
| culture_effect | Rmdn1 | -0.96 | 3.12E-04 | glio |
| culture_effect | Mterf3 | -0.813 | 3.12E-04 | glio |
| Interaction_genotype_Myc | D6Ertd527e | -4.061 | 3.12E-04 | glio |
| Atr_in.vivo_noRT | Dnajc3 | -1.708 | 3.12E-04 | glio |
| Bora_in.vivo_noRT | Ramp1 | 2.978 | 3.12E-04 | glio |
| Eif2s1_in.vivo_noRT | Gm34939 | 2.407 | 3.12E-04 | glio |
| Hsd17b10_in.vivo_noRT | 1810059H22R | 2.948 | 3.12E-04 | glio |
| Hsd17b10_in.vivo_noRT | Krt18 | -3.248 | 3.12E-04 | glio |
| culture_effect | Pou3f2 | -1.047 | 3.12E-05 | glio |
| Interaction_genotype_Mtor | Appbp2os | -4.872 | 3.12E-05 | glio |
| Interaction_genotype_Mtor | Slc16a6 | -5.234 | 3.12E-05 | glio |
| Interaction_genotype_Mtor | Gm19265 | -3.982 | 3.12E-05 | glio |
| Aars2_ex.vivo_noRT | Ceacam16 | 3.293 | 3.12E-05 | glio |
| Aars2_ex.vivo_noRT | Rhbg | 2.9 | 3.12E-05 | glio |
| Atp5c1_in.vivo_noRT | Gm50012 | 3.145 | 3.12E-05 | glio |
| Atr_in.vivo_noRT | Wdr5b | 2.846 | 3.12E-05 | glio |
| Atr_in.vivo_noRT | Cd34 | 2.975 | 3.12E-05 | glio |
| Atr_in.vivo_noRT | Rps8 | -0.791 | 3.12E-05 | glio |
| Bcas2_in.vivo_noRT | Prrt3 | 2.627 | 3.12E-05 | glio |
| Bcas2_in.vivo_noRT | Gm10401 | 2.604 | 3.12E-05 | glio |
| Bcas2_ex.vivo_noRT | 2410017I17Ri | 2.742 | 3.12E-05 | glio |
| Bora_in.vivo_noRT | Naf1 | -1.782 | 3.12E-06 | glio |
| Bora_in.vivo_noRT | 1700124L16Ri | 3.702 | 3.12E-07 | glio |
| Chek1_ex.vivo_noRT | Gm42918 | 3.871 | 3.12E-07 | glio |
| Bcas2_in.vivo_noRT | Inhba | 4.166 | 3.12E-08 | glio |
| Chek1_in.vivo_noRT | Gm11696 | 4.177 | 3.12E-09 | glio |
| Hsd17b10_in.vivo_noRT | Zscan4d | 3.616 | 3.12E-10 | glio |
| Atr_ex.vivo_noRT | Gm49698 | 5.288 | 3.12E-18 | glio |
| Bora_in.vivo_noRT | Gm41307 | 4.936 | 3.12E-22 | glio |
| Bora_in.vivo_noRT | Tagap | 4.676 | 3.12E-22 | glio |
| culture_effect | Serpinb6b | -1.997 | 3.13E-03 | glio |
| culture_effect | Gm15743 | 1.659 | 3.13E-03 | glio |
| culture_effect | Ewsr1 | -0.357 | 3.13E-03 | glio |
| Interaction_genotype_Bcas2 | Hspa12b | -4.368 | 3.13E-03 | glio |
| Interaction_genotype_Bora | Rapgef3os2 | -2.92 | 3.13E-03 | glio |
| Interaction_genotype_Bora | Tiam1 | 2.304 | 3.13E-03 | glio |
| Interaction_genotype_Lmo7 | Gm47204 | -4.631 | 3.13E-03 | glio |
| Interaction_genotype_Mtor | Zbtb45 | -2.647 | 3.13E-03 | glio |
| Interaction_genotype_Mtor | Wnk1 | 1.584 | 3.13E-03 | glio |
| Interaction_genotype_Myc | Gm50218 | -3.642 | 3.13E-03 | glio |
| Interaction_genotype_Myc | 1700037C18R | -3.214 | 3.13E-03 | glio |
| Interaction_genotype_Myc | 3110045C21R | -3.035 | 3.13E-03 | glio |
| Interaction_genotype_Myc | Zkscan6 | -2.749 | 3.13E-03 | glio |
| Interaction_genotype_Myc | Neat1 | 1.903 | 3.13E-03 | glio |
| Interaction_genotype_Rad21 | Brinp3 | 3.53 | 3.13E-03 | glio |
| Interaction_genotype_Sel1l | Pde1a | 4.688 | 3.13E-03 | glio |
| Aars2_in.vivo_noRT | D630003M21I | 1.861 | 3.13E-03 | glio |

| | | | | |
|---|---|---|---|---|
| Aars2_in.vivo_noRT | Rasl11b | 1.911 | 3.13E-03 | glio |
| Aars2_in.vivo_noRT | Dmd | 1.738 | 3.13E-03 | glio |
| Aars2_in.vivo_noRT | 5830444B04R | 1.443 | 3.13E-03 | glio |
| Aars2_ex.vivo_noRT | Gm45669 | 2.505 | 3.13E-03 | glio |
| Atr_in.vivo_noRT | Phf11b | 2.354 | 3.13E-03 | glio |
| Bcas2_in.vivo_noRT | 1700120G11R | 2.323 | 3.13E-03 | glio |
| Bcas2_in.vivo_noRT | Dapp1 | 1.857 | 3.13E-03 | glio |
| Bcas2_in.vivo_noRT | Proser3 | 1.894 | 3.13E-03 | glio |
| Bcas2_in.vivo_noRT | Rwdd1 | -0.56 | 3.13E-03 | glio |
| Bcas2_ex.vivo_noRT | Vgll2 | 2.742 | 3.13E-03 | glio |
| Bcas2_ex.vivo_noRT | Rem2 | 2.742 | 3.13E-03 | glio |
| Chek1_in.vivo_noRT | Nadk2 | -1.258 | 3.13E-03 | glio |
| Chek1_ex.vivo_noRT | Anp32b | -0.695 | 3.13E-03 | glio |
| Eif2s1_in.vivo_noRT | Npr3 | 2.143 | 3.13E-03 | glio |
| Eif2s1_in.vivo_noRT | Gm15127 | 2.414 | 3.13E-03 | glio |
| culture_effect | Slc31a2 | -1.074 | 3.13E-04 | glio |
| Aars2_in.vivo_noRT | Nkain2 | -1.807 | 3.13E-04 | glio |
| Atr_in.vivo_noRT | Slc6a20b | 2.525 | 3.13E-04 | glio |
| Atr_in.vivo_noRT | Gm15545 | 2.423 | 3.13E-04 | glio |
| Atr_in.vivo_noRT | Cntnap1 | 2.354 | 3.13E-04 | glio |
| Bcas2_in.vivo_noRT | Asphd2 | 2.604 | 3.13E-04 | glio |
| Bcas2_in.vivo_noRT | Rfx6 | 2.511 | 3.13E-04 | glio |
| Bcas2_in.vivo_noRT | Cdk11b | -1.24 | 3.13E-04 | glio |
| Bora_ex.vivo_noRT | 1700112J16Ri | 2.615 | 3.13E-04 | glio |
| Chek1_in.vivo_noRT | Gm47438 | 2.824 | 3.13E-04 | glio |
| culture_effect | Gm43672 | 2.446 | 3.13E-05 | glio |
| culture_effect | Grtp1 | -1.413 | 3.13E-05 | glio |
| culture_effect | Atl3 | 0.728 | 3.13E-05 | glio |
| Interaction_genotype_Myc | Gm26559 | -4.846 | 3.13E-05 | glio |
| Interaction_genotype_Myc | Gm17399 | -4.973 | 3.13E-05 | glio |
| Interaction_genotype_Myc | Lrp2bp | -4.303 | 3.13E-05 | glio |
| Aars2_in.vivo_noRT | Rab7b | 2.846 | 3.13E-05 | glio |
| Aars2_in.vivo_noRT | Exosc1 | -1.459 | 3.13E-05 | glio |
| Atr_in.vivo_noRT | Itgal | 3.062 | 3.13E-05 | glio |
| Atr_in.vivo_noRT | Lhfpl4 | 2.639 | 3.13E-05 | glio |
| Bora_in.vivo_noRT | 2900041M22F | 3.126 | 3.13E-05 | glio |
| Bora_in.vivo_noRT | Fbxl22 | 2.82 | 3.13E-05 | glio |
| Chek1_ex.vivo_noRT | Necab2 | 3.349 | 3.13E-05 | glio |
| Chek1_ex.vivo_noRT | Qrich2 | 3.746 | 3.13E-05 | glio |
| culture_effect | Camkk2 | 1.251 | 3.13E-06 | glio |
| Interaction_genotype_Myc | Rarres1 | -5.293 | 3.13E-06 | glio |
| Aars2_in.vivo_noRT | P4htm | 3.262 | 3.13E-06 | glio |
| Atp5c1_in.vivo_noRT | Smtnl1 | 2.668 | 3.13E-06 | glio |
| Bora_in.vivo_noRT | 2810457G06R | 3.689 | 3.13E-06 | glio |
| Aars2_in.vivo_noRT | Adk | -1.995 | 3.13E-07 | glio |
| Chek1_ex.vivo_noRT | Gm11131 | 4.447 | 3.13E-07 | glio |
| Aars2_in.vivo_noRT | Aqp3 | 5.063 | 3.13E-09 | glio |
| Atr_in.vivo_noRT | Cabp4 | 4.223 | 3.13E-09 | glio |
| Chek1_ex.vivo_noRT | Gm17146 | 4.691 | 3.13E-09 | glio |
| culture_effect | Ddx23 | 0.95 | 3.13E-10 | glio |
| Chek1_in.vivo_noRT | Gm20633 | 4.253 | 3.13E-11 | glio |
| Chek1_in.vivo_noRT | Styk1 | 4.722 | 3.13E-11 | glio |

| | | | | |
|---|---|---|---|---|
| Aars2_in.vivo_noRT | Gm32090 | 4.535 | 3.13E-12 | glio |
| Chek1_in.vivo_noRT | Ksr2 | 4.006 | 3.13E-12 | glio |
| Chek1_ex.vivo_noRT | Gm20513 | 4.842 | 3.13E-12 | glio |
| Bora_in.vivo_noRT | Vmn1r91 | 4.807 | 3.13E-14 | glio |
| Bora_in.vivo_noRT | Krtcap3 | 4.144 | 3.13E-14 | glio |
| Chek1_in.vivo_noRT | Gm16741 | 5.044 | 3.13E-15 | glio |
| Atr_in.vivo_noRT | 9130019O22R | 5.533 | 3.13E-26 | glio |
| Bcas2_in.vivo_noRT | Slc17a6 | 4.807 | 3.13E-27 | glio |
| Bcas2_in.vivo_noRT | Gjd4 | 5.325 | 3.13E-27 | glio |
| Interaction_genotype_Hsd17b | Ms4a4d | 4.703 | 3.14E-03 | glio |
| Interaction_genotype_Hsd17b | H2-Eb2 | 3.655 | 3.14E-03 | glio |
| Interaction_genotype_Mtor | Bmp15 | -3.047 | 3.14E-03 | glio |
| Interaction_genotype_Mtor | Gm15903 | -3.456 | 3.14E-03 | glio |
| Interaction_genotype_Mtor | C030014I23Ri | -3.332 | 3.14E-03 | glio |
| Interaction_genotype_Mtor | Basp1 | -3.911 | 3.14E-03 | glio |
| Interaction_genotype_Mtor | Gm6710 | -2.593 | 3.14E-03 | glio |
| Interaction_genotype_Mtor | Casp7 | -2.447 | 3.14E-03 | glio |
| Interaction_genotype_Mtor | Cavin2 | -2.992 | 3.14E-03 | glio |
| Interaction_genotype_Mtor | Rfc1 | 1.492 | 3.14E-03 | glio |
| Interaction_genotype_Myc | D030028A08R | -2.628 | 3.14E-03 | glio |
| Interaction_genotype_Myc | Zfp626 | -2.56 | 3.14E-03 | glio |
| interaction_RT | Map3k15 | -1.889 | 3.14E-03 | glio |
| interaction_RT | Cytl1 | 3.289 | 3.14E-03 | glio |
| interaction_RT | Guca1a | 1.085 | 3.14E-03 | glio |
| interaction_RT | Cnp | 1.151 | 3.14E-03 | glio |
| interaction_RT | Mcm2 | -0.982 | 3.14E-03 | glio |
| interaction_RT | Cox6a2 | 1.518 | 3.14E-03 | glio |
| Aars2_in.vivo_noRT | Gm44777 | 2.02 | 3.14E-03 | glio |
| Aars2_ex.vivo_noRT | Elfn2 | 2.718 | 3.14E-03 | glio |
| Aars2_ex.vivo_noRT | Mpeg1 | 2.718 | 3.14E-03 | glio |
| Aars2_ex.vivo_noRT | Gm4841 | 3.293 | 3.14E-03 | glio |
| Aars2_ex.vivo_noRT | Abcb7 | -1.787 | 3.14E-03 | glio |
| Atp5c1_in.vivo_noRT | Lst1 | 2.424 | 3.14E-03 | glio |
| Atr_in.vivo_noRT | Pcbp2 | -0.694 | 3.14E-03 | glio |
| Atr_ex.vivo_noRT | Lrp1b | 2.976 | 3.14E-03 | glio |
| Atr_ex.vivo_noRT | Gm38973 | 2.768 | 3.14E-03 | glio |
| Atr_ex.vivo_noRT | Ybx3 | -0.848 | 3.14E-03 | glio |
| Bard1_ex.vivo_noRT | Brsk2 | 2.814 | 3.14E-03 | glio |
| Bcas2_in.vivo_noRT | Acot2 | -1.744 | 3.14E-03 | glio |
| Bora_in.vivo_noRT | Pon3 | 2.186 | 3.14E-03 | glio |
| Bora_in.vivo_noRT | Fbxo10 | 2.13 | 3.14E-03 | glio |
| Bora_ex.vivo_noRT | Postn | 2.905 | 3.14E-03 | glio |
| Chek1_in.vivo_noRT | 1190005I06Ri | 1.999 | 3.14E-03 | glio |
| Hsd17b10_in.vivo_noRT | Gm10382 | 2.372 | 3.14E-03 | glio |
| Interaction_genotype_Mtor | Flrt2 | -4.395 | 3.14E-04 | glio |
| Interaction_genotype_Mtor | Cdc42ep2 | -4.436 | 3.14E-04 | glio |
| Interaction_genotype_Myc | 1700012B09R | -4.849 | 3.14E-04 | glio |
| Interaction_genotype_Myc | 4833418N02R | -4.174 | 3.14E-04 | glio |
| Interaction_genotype_Myc | Zfp219 | -2.737 | 3.14E-04 | glio |
| Aars2_in.vivo_noRT | Snn | 2.591 | 3.14E-04 | glio |
| Aars2_in.vivo_noRT | Klf15 | 2.57 | 3.14E-04 | glio |
| Aars2_in.vivo_noRT | Nelfa | -1.432 | 3.14E-04 | glio |

| | | | | |
|---|---|---|---|---|
| Atr_in.vivo_noRT | Slc22a15 | 2.372 | 3.14E-04 | glio |
| Atr_in.vivo_noRT | Akap17b | 2.302 | 3.14E-04 | glio |
| Bora_in.vivo_noRT | Tmem199 | -1.428 | 3.14E-04 | glio |
| Chek1_in.vivo_noRT | Tmem97 | -1.387 | 3.14E-04 | glio |
| culture_effect | Skida1 | 2.424 | 3.14E-05 | glio |
| Interaction_genotype_Bora | Epb41l4b | 2.457 | 3.14E-05 | glio |
| Interaction_genotype_Mtor | Rab5a | 2.027 | 3.14E-05 | glio |
| Interaction_genotype_Sass6 | Itga4 | -2.98 | 3.14E-05 | glio |
| interaction_RT | Txnip | 3.1 | 3.14E-05 | glio |
| Aars2_in.vivo_noRT | Gm28229 | 2.685 | 3.14E-05 | glio |
| Aars2_ex.vivo_noRT | AC159140.1 | 2.9 | 3.14E-05 | glio |
| Aars2_ex.vivo_noRT | 5730507C01R | 2.901 | 3.14E-05 | glio |
| Chek1_ex.vivo_noRT | Mmp19 | 3.673 | 3.14E-05 | glio |
| culture_effect | Nup43 | -0.708 | 3.14E-06 | glio |
| Aars2_in.vivo_noRT | Fhl2 | 3.068 | 3.14E-06 | glio |
| Atr_in.vivo_noRT | March10 | 3.176 | 3.14E-06 | glio |
| Atr_in.vivo_noRT | H2-T24 | 3.075 | 3.14E-06 | glio |
| Chek1_ex.vivo_noRT | Rab27b | 4.479 | 3.14E-06 | glio |
| Chek1_in.vivo_noRT | Cnksr1 | 3.475 | 3.14E-07 | glio |
| Atr_in.vivo_noRT | Hey2 | 3.686 | 3.14E-08 | glio |
| Bcas2_in.vivo_noRT | Tnnt3 | 4.172 | 3.14E-08 | glio |
| Bcas2_in.vivo_noRT | Stard6 | 3.954 | 3.14E-08 | glio |
| Bora_in.vivo_noRT | Gm28874 | 3.914 | 3.14E-08 | glio |
| Bora_in.vivo_noRT | Gm11491 | 3.894 | 3.14E-08 | glio |
| Bora_in.vivo_noRT | Rpa2 | -2.438 | 3.14E-08 | glio |
| Chek1_in.vivo_noRT | Cyp26b1 | 4.118 | 3.14E-08 | glio |
| Hsd17b10_in.vivo_noRT | Gm49086 | 3.616 | 3.14E-09 | glio |
| Aars2_in.vivo_noRT | Rhox3g | 4.212 | 3.14E-10 | glio |
| Aars2_in.vivo_noRT | Sult6b1 | 4.536 | 3.14E-11 | glio |
| Chek1_in.vivo_noRT | Cd52 | 4.387 | 3.14E-11 | glio |
| Bcas2_in.vivo_noRT | Gm20442 | 4.328 | 3.14E-12 | glio |
| culture_effect | Epn2 | -1.178 | 3.14E-13 | glio |
| Bcas2_in.vivo_noRT | Nat8f4 | 4.792 | 3.14E-14 | glio |
| Atr_ex.vivo_noRT | Rgs4 | 5.288 | 3.14E-16 | glio |
| Atr_ex.vivo_noRT | Gm44693 | 4.985 | 3.14E-16 | glio |
| Atr_ex.vivo_noRT | Gm44736 | 4.725 | 3.14E-16 | glio |
| Chek1_in.vivo_noRT | Gm48155 | 5.044 | 3.14E-26 | glio |
| culture_effect | Fgf7 | 1.6 | 3.15E-03 | glio |
| culture_effect | Sspn | 1.013 | 3.15E-03 | glio |
| Interaction_genotype_Bcas2 | Gm15816 | -3.544 | 3.15E-03 | glio |
| Interaction_genotype_Bcas2 | Gm31063 | -2.989 | 3.15E-03 | glio |
| Interaction_genotype_Bcas2 | Pcp4l1 | -3.971 | 3.15E-03 | glio |
| Interaction_genotype_Bcas2 | Ak1 | -3.643 | 3.15E-03 | glio |
| Interaction_genotype_Bcas2 | Tph2 | -3.285 | 3.15E-03 | glio |
| Interaction_genotype_Bora | Lama5 | -4.054 | 3.15E-03 | glio |
| Interaction_genotype_Bora | Eps8l2 | -3.891 | 3.15E-03 | glio |
| Interaction_genotype_Bora | Tob2 | 1.864 | 3.15E-03 | glio |
| Interaction_genotype_Mtor | Vmn2r54 | -2.938 | 3.15E-03 | glio |
| Interaction_genotype_Mtor | Rapgef3 | -3.917 | 3.15E-03 | glio |
| Interaction_genotype_Myc | Pld6 | -3.551 | 3.15E-03 | glio |
| Atr_ex.vivo_noRT | Gm11627 | 2.282 | 3.15E-03 | glio |
| Bard1_ex.vivo_noRT | Gm14207 | 2.021 | 3.15E-03 | glio |

| | | | | |
|---|---|---|---|---|
| Bcas2_in.vivo_noRT | Wdr5b | 2.12 | 3.15E-03 | glio |
| Bcas2_in.vivo_noRT | Snu13 | -0.528 | 3.15E-03 | glio |
| Bora_in.vivo_noRT | Gm50232 | 2.015 | 3.15E-03 | glio |
| Bora_in.vivo_noRT | Ctdspl2 | -1.407 | 3.15E-03 | glio |
| Chek1_ex.vivo_noRT | Naga | 2.014 | 3.15E-03 | glio |
| Hsd17b10_in.vivo_noRT | Gm10432 | 2.119 | 3.15E-03 | glio |
| culture_effect | Gm5129 | 2.103 | 3.15E-04 | glio |
| culture_effect | Ahcyl2 | -0.701 | 3.15E-04 | glio |
| culture_effect | Gnpda2 | -0.618 | 3.15E-04 | glio |
| culture_effect | Parl | 0.51 | 3.15E-04 | glio |
| Interaction_genotype_Bcas2 | Pitx3 | -3.829 | 3.15E-04 | glio |
| Interaction_genotype_Mtor | Syn3 | -3.942 | 3.15E-04 | glio |
| Interaction_genotype_Mtor | Fpgs | -2.989 | 3.15E-04 | glio |
| Interaction_genotype_Myc | Gm45235 | -3.263 | 3.15E-04 | glio |
| Interaction_genotype_Myc | 1700126G02R | -4.154 | 3.15E-04 | glio |
| Interaction_genotype_Myc | Senp8 | -3.492 | 3.15E-04 | glio |
| Interaction_genotype_Myc | Dtwd1 | -3.461 | 3.15E-04 | glio |
| Atr_in.vivo_noRT | Ctdnep1 | -1.316 | 3.15E-04 | glio |
| Atr_in.vivo_noRT | Rnps1 | -1.018 | 3.15E-04 | glio |
| Atr_in.vivo_noRT | Cbx3 | -1.179 | 3.15E-04 | glio |
| Brca2_ex.vivo_noRT | Gm16178 | 2.82 | 3.15E-04 | glio |
| Chek1_in.vivo_noRT | Sec24c | -1.837 | 3.15E-04 | glio |
| Chek1_ex.vivo_noRT | 2010001K21R | 2.89 | 3.15E-04 | glio |
| Interaction_genotype_Aars2 | Gm28424 | -4.899 | 3.15E-05 | glio |
| Interaction_genotype_Aars2 | Tars | 1.705 | 3.15E-05 | glio |
| Interaction_genotype_Myc | Rpl19 | 1.533 | 3.15E-05 | glio |
| Aars2_in.vivo_noRT | Cnn1 | 3.657 | 3.15E-05 | glio |
| Atr_in.vivo_noRT | Tnfrsf22 | 2.808 | 3.15E-05 | glio |
| Chek1_in.vivo_noRT | Prob1 | 3.096 | 3.15E-05 | glio |
| Chek1_in.vivo_noRT | Cnbp | -1.253 | 3.15E-05 | glio |
| Interaction_genotype_Mtor | Tex2 | -5.093 | 3.15E-06 | glio |
| Atr_in.vivo_noRT | Zfp418 | 3.478 | 3.15E-06 | glio |
| Hsd17b10_in.vivo_noRT | 1810019D21R | 3.356 | 3.15E-06 | glio |
| Atp5c1_in.vivo_noRT | Gm5788 | 2.668 | 3.15E-07 | glio |
| Hsd17b10_in.vivo_noRT | Dnah10 | 3.237 | 3.15E-07 | glio |
| Hsd17b10_in.vivo_noRT | Mettl7b | 3.616 | 3.15E-07 | glio |
| Hsd17b10_in.vivo_noRT | Pla2g2d | 3.353 | 3.15E-07 | glio |
| Bcas2_in.vivo_noRT | 1110017D15R | 3.807 | 3.15E-08 | glio |
| Bora_in.vivo_noRT | Tnxb | 3.783 | 3.15E-08 | glio |
| Aars2_in.vivo_noRT | Gm10432 | 3.83 | 3.15E-09 | glio |
| culture_effect | Ndufv1 | -0.987 | 3.15E-10 | glio |
| Atr_in.vivo_noRT | 0610009L18Ri | 4.531 | 3.15E-10 | glio |
| culture_effect | Tuba1c | -1.25 | 3.15E-11 | glio |
| Atr_in.vivo_noRT | Prtn3 | 4.751 | 3.15E-11 | glio |
| Chek1_ex.vivo_noRT | Ly6d | 4.842 | 3.15E-12 | glio |
| culture_effect | Arhgef26 | 1.459 | 3.15E-15 | glio |
| Chek1_in.vivo_noRT | Gm18280 | 4.78 | 3.15E-20 | glio |
| culture_effect | Xrcc1 | -0.538 | 3.16E-03 | glio |
| culture_effect | Rela | -0.545 | 3.16E-03 | glio |
| culture_effect | Ergic2 | 0.393 | 3.16E-03 | glio |
| Interaction_genotype_Aars2 | Flacc1 | -3.224 | 3.16E-03 | glio |
| Interaction_genotype_Aars2 | Lnpk | -1.816 | 3.16E-03 | glio |

| | | | | |
|---|---|---|---|---|
| Interaction_genotype_Chek1 | Gm43137 | -4.024 | 3.16E-03 | glio |
| Interaction_genotype_Myc | Gm17146 | -3.084 | 3.16E-03 | glio |
| Interaction_genotype_Myc | Mro | -3.295 | 3.16E-03 | glio |
| Interaction_genotype_Myc | Cass4 | -3.327 | 3.16E-03 | glio |
| Interaction_genotype_Myc | Fyb | -3.252 | 3.16E-03 | glio |
| Interaction_genotype_Myc | Tmem94 | -3.098 | 3.16E-03 | glio |
| interaction_RT | Klf11 | 1.905 | 3.16E-03 | glio |
| interaction_RT | AC168977.1 | 2.635 | 3.16E-03 | glio |
| interaction_RT | 2310001H17R | 1.608 | 3.16E-03 | glio |
| interaction_RT | Gstt1 | 2.977 | 3.16E-03 | glio |
| interaction_RT | Cbx8 | 1.056 | 3.16E-03 | glio |
| interaction_RT | Coa7 | -0.98 | 3.16E-03 | glio |
| interaction_RT | Atg4d | -0.968 | 3.16E-03 | glio |
| interaction_RT | Dhfr | -0.903 | 3.16E-03 | glio |
| interaction_RT | Vkorc1 | 0.728 | 3.16E-03 | glio |
| interaction_RT | Psmd7 | -0.607 | 3.16E-03 | glio |
| Aars2_in.vivo_noRT | Myo16 | 2.158 | 3.16E-03 | glio |
| Aars2_ex.vivo_noRT | Gm36757 | 2.893 | 3.16E-03 | glio |
| Bard1_ex.vivo_noRT | Zscan4d | 2.421 | 3.16E-03 | glio |
| Bcas2_in.vivo_noRT | Gm2415 | 2.249 | 3.16E-03 | glio |
| Bcas2_in.vivo_noRT | Agbl5 | 1.865 | 3.16E-03 | glio |
| Bcas2_ex.vivo_noRT | Gvin1 | 3.129 | 3.16E-03 | glio |
| Bora_in.vivo_noRT | Pkdcc | 2.224 | 3.16E-03 | glio |
| Bora_in.vivo_noRT | 1810037I17Ri | -0.744 | 3.16E-03 | glio |
| Chek1_in.vivo_noRT | Rpl36a | -0.662 | 3.16E-03 | glio |
| Chek1_ex.vivo_noRT | Zbtb8a | 2.938 | 3.16E-03 | glio |
| Hsd17b10_in.vivo_noRT | Fsd2 | 2.233 | 3.16E-03 | glio |
| Interaction_genotype_Aars2 | Psmd7 | 1.542 | 3.16E-04 | glio |
| Interaction_genotype_Rpl3 | Gm26533 | -5.014 | 3.16E-04 | glio |
| interaction_RT | Id1 | -1.118 | 3.16E-04 | glio |
| interaction_RT | Ifitm2 | 0.596 | 3.16E-04 | glio |
| Aars2_in.vivo_noRT | Gtf2h5 | -0.883 | 3.16E-04 | glio |
| Atr_in.vivo_noRT | Gm16230 | 2.516 | 3.16E-04 | glio |
| Atr_in.vivo_noRT | Lrp4 | 2.576 | 3.16E-04 | glio |
| Atr_in.vivo_noRT | Syde2 | 2.457 | 3.16E-04 | glio |
| Bora_in.vivo_noRT | Syt17 | 2.871 | 3.16E-04 | glio |
| Hsd17b10_in.vivo_noRT | Gm5129 | 2.816 | 3.16E-04 | glio |
| Aars2_in.vivo_noRT | Galm | 2.837 | 3.16E-05 | glio |
| Atr_in.vivo_noRT | Gm43378 | 2.734 | 3.16E-05 | glio |
| Atr_ex.vivo_noRT | Ipcef1 | 3.164 | 3.16E-05 | glio |
| Interaction_genotype_Myc | Nat8l | -3.803 | 3.16E-06 | glio |
| Bcas2_ex.vivo_noRT | Lhx1os | 3.767 | 3.16E-06 | glio |
| Chek1_in.vivo_noRT | Myzap | 2.722 | 3.16E-06 | glio |
| Atr_ex.vivo_noRT | Rimklb | 3.164 | 3.16E-07 | glio |
| Bcas2_in.vivo_noRT | Gm2012 | 3.643 | 3.16E-07 | glio |
| Bora_in.vivo_noRT | Gm50399 | 3.384 | 3.16E-07 | glio |
| Aars2_in.vivo_noRT | Dcpp1 | 4 | 3.16E-08 | glio |
| Chek1_ex.vivo_noRT | Pcbp3 | 4.271 | 3.16E-08 | glio |
| Aars2_in.vivo_noRT | Pdzk1ip1 | 4.337 | 3.16E-09 | glio |
| Atr_in.vivo_noRT | 1700081N11R | 4.68 | 3.16E-09 | glio |
| Atr_in.vivo_noRT | A430057M04F | 4.348 | 3.16E-10 | glio |
| Bora_in.vivo_noRT | 9630014M24F | 4.158 | 3.16E-10 | glio |

| | | | | |
|---|---|---|---|---|
| Atr_in.vivo_noRT | Gm16183 | 4.891 | 3.16E-11 | glio |
| Bcas2_in.vivo_noRT | Gm39526 | 4.284 | 3.16E-15 | glio |
| Bora_in.vivo_noRT | Gm47936 | 4.936 | 3.16E-20 | glio |
| culture_effect | Gm15910 | 1.439 | 3.17E-03 | glio |
| culture_effect | Ska1 | -0.617 | 3.17E-03 | glio |
| culture_effect | Polr2k | 0.345 | 3.17E-03 | glio |
| Interaction_genotype_Atr | Cyp2s1 | -3.923 | 3.17E-03 | glio |
| Interaction_genotype_Mtor | B230216N24R | -3.905 | 3.17E-03 | glio |
| Interaction_genotype_Mtor | Fastkd1 | -3.076 | 3.17E-03 | glio |
| Interaction_genotype_Myc | Zfp960 | -3.535 | 3.17E-03 | glio |
| interaction_RT | Plekhf2 | 0.976 | 3.17E-03 | glio |
| interaction_RT | Trmt1 | -0.798 | 3.17E-03 | glio |
| interaction_RT | 2810004N23R | -0.666 | 3.17E-03 | glio |
| Aars2_in.vivo_noRT | 8430429K09R | 1.733 | 3.17E-03 | glio |
| Aars2_in.vivo_noRT | Vps29 | -0.988 | 3.17E-03 | glio |
| Aars2_ex.vivo_noRT | Gm15929 | 2.316 | 3.17E-03 | glio |
| Aars2_ex.vivo_noRT | Trim67 | 2.573 | 3.17E-03 | glio |
| Aars2_ex.vivo_noRT | Gm20712 | 2.976 | 3.17E-03 | glio |
| Aars2_ex.vivo_noRT | Klc3 | -2.512 | 3.17E-03 | glio |
| Atp5c1_in.vivo_noRT | E030037K01Ri | 2.136 | 3.17E-03 | glio |
| Atr_in.vivo_noRT | Tex15 | 2.068 | 3.17E-03 | glio |
| Atr_in.vivo_noRT | Gbp10 | 1.807 | 3.17E-03 | glio |
| Atr_in.vivo_noRT | 1700088E04Ri | 1.799 | 3.17E-03 | glio |
| Atr_in.vivo_noRT | Cep55 | 1.443 | 3.17E-03 | glio |
| Atr_in.vivo_noRT | Psmd4 | -0.726 | 3.17E-03 | glio |
| Bcas2_in.vivo_noRT | Pou5f1 | 2.022 | 3.17E-03 | glio |
| Bora_in.vivo_noRT | Gm10282 | 2.16 | 3.17E-03 | glio |
| Bora_in.vivo_noRT | Klhl23 | 2.002 | 3.17E-03 | glio |
| Chek1_in.vivo_noRT | Tex15 | 2.102 | 3.17E-03 | glio |
| Chek1_ex.vivo_noRT | Gm50147 | 2.577 | 3.17E-03 | glio |
| Dclre1c_ex.vivo_noRT | Gm5286 | 2.658 | 3.17E-03 | glio |
| Interaction_genotype_Mtor | Prss56 | -4.673 | 3.17E-04 | glio |
| Aars2_in.vivo_noRT | Slc12a6 | 1.974 | 3.17E-04 | glio |
| Aars2_in.vivo_noRT | Lamtor2 | -1.029 | 3.17E-04 | glio |
| Aars2_in.vivo_noRT | Ndufa12 | -0.746 | 3.17E-04 | glio |
| Atr_in.vivo_noRT | 0610040B10R | 2.148 | 3.17E-04 | glio |
| Atr_in.vivo_noRT | Nup98 | -1.506 | 3.17E-04 | glio |
| Bora_in.vivo_noRT | 4930563E22Ri | 2.688 | 3.17E-04 | glio |
| Bora_in.vivo_noRT | Gal3st4 | 2.577 | 3.17E-04 | glio |
| Bora_in.vivo_noRT | Gm46218 | 2.649 | 3.17E-04 | glio |
| Bora_in.vivo_noRT | Foxl2os | 2.428 | 3.17E-04 | glio |
| Cenpj_in.vivo_noRT | Hoxc8 | 2.022 | 3.17E-04 | glio |
| Chek1_in.vivo_noRT | Stk32c | 2.375 | 3.17E-04 | glio |
| Interaction_genotype_Mtor | Oxt | -4.917 | 3.17E-05 | glio |
| Interaction_genotype_Mtor | Cnih2 | -4.982 | 3.17E-05 | glio |
| Interaction_genotype_Mtor | Akirin2 | 1.742 | 3.17E-05 | glio |
| Interaction_genotype_Myc | 1700113B09R | -4.839 | 3.17E-05 | glio |
| Aars2_in.vivo_noRT | Gm9732 | -1.72 | 3.17E-05 | glio |
| Aars2_in.vivo_noRT | Bccip | -1.381 | 3.17E-05 | glio |
| Aars2_ex.vivo_noRT | Apol10b | 3.293 | 3.17E-05 | glio |
| Atr_in.vivo_noRT | Atg13 | 1.741 | 3.17E-05 | glio |
| Chek1_in.vivo_noRT | Xlr4c | 3.009 | 3.17E-05 | glio |

| | | | | |
|---|---|---|---|---|
| Hsd17b10_in.vivo_noRT | Gm12280 | 3.352 | 3.17E-05 | glio |
| Hsd17b10_in.vivo_noRT | Ephx3 | 2.889 | 3.17E-05 | glio |
| Hsd17b10_in.vivo_noRT | Gldc | 2.909 | 3.17E-05 | glio |
| Interaction_genotype_Mtor | Hecw2 | -5.505 | 3.17E-06 | glio |
| Bora_in.vivo_noRT | A930033H14R | 3.445 | 3.17E-06 | glio |
| Chek1_ex.vivo_noRT | Stat4 | 4.132 | 3.17E-06 | glio |
| Fanca_ex.vivo_noRT | Ceacam16 | 4.152 | 3.17E-06 | glio |
| Fanca_ex.vivo_noRT | Ankrd33b | 4.931 | 3.17E-06 | glio |
| Chek1_in.vivo_noRT | Prr7 | 3.653 | 3.17E-07 | glio |
| Bcas2_in.vivo_noRT | Gm39459 | 4.157 | 3.17E-09 | glio |
| Chek1_ex.vivo_noRT | Gm48899 | 4.267 | 3.17E-09 | glio |
| Bcas2_in.vivo_noRT | BC065403 | 4.291 | 3.17E-10 | glio |
| Bora_in.vivo_noRT | Zfp808 | 3.819 | 3.17E-10 | glio |
| Aars2_in.vivo_noRT | Ntn5 | 4.535 | 3.17E-11 | glio |
| Atr_in.vivo_noRT | Kmo | 3.892 | 3.17E-11 | glio |
| Chek1_in.vivo_noRT | Cd83 | 4.043 | 3.17E-11 | glio |
| Atr_in.vivo_noRT | Gm11549 | 5.008 | 3.17E-13 | glio |
| Atr_in.vivo_noRT | Aoc1 | 5.273 | 3.17E-13 | glio |
| Aars2_in.vivo_noRT | Scgb1c1 | 5.327 | 3.17E-19 | glio |
| Bora_in.vivo_noRT | Tnni3k | 4.936 | 3.17E-20 | glio |
| Aars2_in.vivo_noRT | 9430041J12Ri | 5.327 | 3.17E-21 | glio |
| Aars2_in.vivo_noRT | AU019990 | 5.327 | 3.17E-24 | glio |
| culture_effect | Nedd8 | 0.325 | 3.18E-03 | glio |
| Interaction_genotype_Atr | Gm45890 | -3.807 | 3.18E-03 | glio |
| Interaction_genotype_Atr | 2010106G04R | -3.669 | 3.18E-03 | glio |
| Interaction_genotype_Atr | Pcsk4 | -3.58 | 3.18E-03 | glio |
| Interaction_genotype_Atr | Map1b | 1.476 | 3.18E-03 | glio |
| Interaction_genotype_Atr | Ythdf2 | 1.523 | 3.18E-03 | glio |
| Interaction_genotype_Myc | Gm26658 | -3.461 | 3.18E-03 | glio |
| Interaction_genotype_Myc | Ubiad1 | -2.712 | 3.18E-03 | glio |
| Aars2_in.vivo_noRT | Adh7 | 2.116 | 3.18E-03 | glio |
| Aars2_ex.vivo_noRT | Usp17la | 2.897 | 3.18E-03 | glio |
| Atr_in.vivo_noRT | 1810062G17R | 2.057 | 3.18E-03 | glio |
| Atr_in.vivo_noRT | Gm15706 | 1.837 | 3.18E-03 | glio |
| Atr_in.vivo_noRT | Inafm1 | 1.84 | 3.18E-03 | glio |
| Atr_in.vivo_noRT | Snhg10 | 1.876 | 3.18E-03 | glio |
| Atr_in.vivo_noRT | Gak | -1.316 | 3.18E-03 | glio |
| Atr_in.vivo_noRT | Gkap1 | 1.008 | 3.18E-03 | glio |
| Bcas2_in.vivo_noRT | Gab3 | 2.162 | 3.18E-03 | glio |
| Bcas2_in.vivo_noRT | Syce2 | -1.291 | 3.18E-03 | glio |
| Bcas2_ex.vivo_noRT | Gm41349 | 2.742 | 3.18E-03 | glio |
| Bcas2_ex.vivo_noRT | 4930423D22R | 2.163 | 3.18E-03 | glio |
| Cenpj_in.vivo_noRT | Cldn5 | 1.756 | 3.18E-03 | glio |
| Cenpj_in.vivo_noRT | Gm10354 | 2.022 | 3.18E-03 | glio |
| Cenpj_in.vivo_noRT | Itga4 | 1.608 | 3.18E-03 | glio |
| culture_effect | Gm14273 | -2.239 | 3.18E-04 | glio |
| culture_effect | Dpysl3 | 1.034 | 3.18E-04 | glio |
| Interaction_genotype_Bora | Kpna7 | -4.2 | 3.18E-04 | glio |
| Interaction_genotype_Bora | Rhno1 | 1.93 | 3.18E-04 | glio |
| interaction_RT | Gdf15 | 1.624 | 3.18E-04 | glio |
| interaction_RT | Tcf4 | 0.783 | 3.18E-04 | glio |
| interaction_RT | Emc7 | 0.609 | 3.18E-04 | glio |

| | | | | |
|---|---|---|---|---|
| interaction_RT | Hspa8 | -0.883 | 3.18E-04 | glio |
| Aars2_in.vivo_noRT | Zfp873 | 2.256 | 3.18E-04 | glio |
| Aars2_in.vivo_noRT | Sp1 | -1.439 | 3.18E-04 | glio |
| Atr_ex.vivo_noRT | Brsk2 | 3.164 | 3.18E-04 | glio |
| Atr_ex.vivo_noRT | Gm50138 | 2.778 | 3.18E-04 | glio |
| Atr_ex.vivo_noRT | Fa2h | 3.164 | 3.18E-04 | glio |
| Bora_in.vivo_noRT | Ces1a | 2.814 | 3.18E-04 | glio |
| Bora_in.vivo_noRT | Rbm6 | -1.681 | 3.18E-04 | glio |
| Chek1_in.vivo_noRT | Zfp534 | 2.702 | 3.18E-04 | glio |
| Chek1_ex.vivo_noRT | Unc5cl | 3.298 | 3.18E-04 | glio |
| Chek1_ex.vivo_noRT | Gm46404 | 3.367 | 3.18E-04 | glio |
| Chek1_ex.vivo_noRT | Cd3g | 3.86 | 3.18E-05 | glio |
| culture_effect | Ppp3ca | -0.591 | 3.18E-06 | glio |
| Bora_in.vivo_noRT | Pfkfb1 | 3.307 | 3.18E-06 | glio |
| culture_effect | Ets2 | -2.356 | 3.18E-07 | glio |
| Interaction_genotype_Myc | Folr1 | -5.504 | 3.18E-08 | glio |
| Chek1_in.vivo_noRT | Gm9908 | 4.127 | 3.18E-08 | glio |
| Chek1_ex.vivo_noRT | Ldhd | 4.132 | 3.18E-08 | glio |
| culture_effect | Letm1 | 0.875 | 3.18E-09 | glio |
| Bora_in.vivo_noRT | Mogat1 | 4.414 | 3.18E-09 | glio |
| Bora_in.vivo_noRT | 4930518J21Ri | 4.011 | 3.18E-09 | glio |
| Chek1_in.vivo_noRT | Gnaz | 4.269 | 3.18E-09 | glio |
| culture_effect | Ccpg1 | -1.621 | 3.18E-10 | glio |
| Atr_in.vivo_noRT | Cfb | 4.229 | 3.18E-14 | glio |
| culture_effect | Tubb4b | -1.316 | 3.18E-15 | glio |
| Bcas2_in.vivo_noRT | Prickle2 | 4.807 | 3.18E-16 | glio |
| culture_effect | Schip1 | 1.521 | 3.18E-17 | glio |
| Aars2_in.vivo_noRT | Muc4 | 4.805 | 3.18E-17 | glio |
| Atr_in.vivo_noRT | Ntn4 | 5.267 | 3.18E-18 | glio |
| Bcas2_in.vivo_noRT | Capn13 | 4.807 | 3.18E-25 | glio |
| Atr_in.vivo_noRT | Vmn1r32 | 5.533 | 3.18E-28 | glio |
| Atr_in.vivo_noRT | Glyat | 5.533 | 3.18E-28 | glio |
| Interaction_genotype_Atr | Gm15908 | -2.369 | 3.19E-03 | glio |
| Interaction_genotype_Bcas2 | Zfp968 | -4.013 | 3.19E-03 | glio |
| Interaction_genotype_Bora | Fes | -3.91 | 3.19E-03 | glio |
| Interaction_genotype_Bora | Gm15127 | -3.849 | 3.19E-03 | glio |
| Interaction_genotype_Bora | Gm49701 | -3.834 | 3.19E-03 | glio |
| Interaction_genotype_Bora | Dnah10 | -3.491 | 3.19E-03 | glio |
| Interaction_genotype_Chek1 | Tmem140 | 3.598 | 3.19E-03 | glio |
| Interaction_genotype_Mtor | Gm7324 | -3.394 | 3.19E-03 | glio |
| Interaction_genotype_Myc | Gm5941 | -3.43 | 3.19E-03 | glio |
| Interaction_genotype_Myc | Zfp763 | -3.058 | 3.19E-03 | glio |
| Interaction_genotype_Myc | Ocel1 | -2.711 | 3.19E-03 | glio |
| Interaction_genotype_Myc | Srsf11 | 1.238 | 3.19E-03 | glio |
| Interaction_genotype_Uqcrc2 | Tomm6os | -4.681 | 3.19E-03 | glio |
| Interaction_genotype_Uqcrc2 | Gm48242 | -3.805 | 3.19E-03 | glio |
| Aars2_ex.vivo_noRT | 1110017D15R | 2.9 | 3.19E-03 | glio |
| Bard1_ex.vivo_noRT | Insrr | 2.814 | 3.19E-03 | glio |
| Bora_in.vivo_noRT | Ebi3 | 2.558 | 3.19E-03 | glio |
| Bora_ex.vivo_noRT | Sdr16c5 | 2.806 | 3.19E-03 | glio |
| Cenpj_ex.vivo_noRT | Gtsf1l | 2.591 | 3.19E-03 | glio |
| Chek1_in.vivo_noRT | Slc4a11 | 2.256 | 3.19E-03 | glio |

| | | | | |
|---|---|---|---|---|
| Chek1_in.vivo_noRT | Gm5930 | 2.136 | 3.19E-03 | glio |
| Chek1_in.vivo_noRT | A730017L22Ri | 1.892 | 3.19E-03 | glio |
| Chek1_ex.vivo_noRT | Plin1 | 2.755 | 3.19E-03 | glio |
| Chek1_ex.vivo_noRT | Gm16023 | 2.9 | 3.19E-03 | glio |
| Chek1_ex.vivo_noRT | Zfp773 | 2.418 | 3.19E-03 | glio |
| Chek1_ex.vivo_noRT | Cdc6 | -2.375 | 3.19E-03 | glio |
| Eif2s1_in.vivo_noRT | Pla2g2f | 2.198 | 3.19E-03 | glio |
| Hsd17b10_in.vivo_noRT | Gm38979 | 2.442 | 3.19E-03 | glio |
| Hsd17b10_in.vivo_noRT | Fxyd3 | -2.024 | 3.19E-03 | glio |
| Hsd17b10_in.vivo_noRT | Spats2l | -0.999 | 3.19E-03 | glio |
| Interaction_genotype_Mtor | Gm4166 | -3.172 | 3.19E-04 | glio |
| Interaction_genotype_Mtor | C1qtnf12 | -4.716 | 3.19E-04 | glio |
| Interaction_genotype_Mtor | Pou3f1 | -4.361 | 3.19E-04 | glio |
| Interaction_genotype_Mtor | Gm19426 | -3.335 | 3.19E-04 | glio |
| Interaction_genotype_Myc | Fa2h | 3.851 | 3.19E-04 | glio |
| Interaction_genotype_Myc | Wdr27 | -3.498 | 3.19E-04 | glio |
| Interaction_genotype_Myc | Sec11a | 1.567 | 3.19E-04 | glio |
| Interaction_genotype_Myc | Ube2m | 1.341 | 3.19E-04 | glio |
| Aars2_in.vivo_noRT | Hpn | 2.488 | 3.19E-04 | glio |
| Atr_in.vivo_noRT | Hhex | 2.267 | 3.19E-04 | glio |
| Hsd17b10_in.vivo_noRT | D430040D24R | 2.699 | 3.19E-04 | glio |
| culture_effect | Aspa | -2.14 | 3.19E-05 | glio |
| Interaction_genotype_Mtor | Dlx2 | -5.173 | 3.19E-05 | glio |
| interaction_RT | Ubl5 | 0.754 | 3.19E-05 | glio |
| Atr_in.vivo_noRT | Bbs10 | 3.037 | 3.19E-05 | glio |
| Bard1_ex.vivo_noRT | Gm12610 | 3.213 | 3.19E-05 | glio |
| Chek1_ex.vivo_noRT | Tbxa2r | 3.48 | 3.19E-05 | glio |
| Hsd17b10_in.vivo_noRT | Neurl1b | 3.09 | 3.19E-05 | glio |
| Interaction_genotype_Mtor | Gm48772 | -5.056 | 3.19E-06 | glio |
| Chek1_ex.vivo_noRT | Sult5a1 | 4.289 | 3.19E-06 | glio |
| Eif2s1_in.vivo_noRT | Nrip2 | 2.67 | 3.19E-06 | glio |
| Interaction_genotype_Mtor | Hoxc8 | -4.099 | 3.19E-07 | glio |
| Atr_in.vivo_noRT | Dnaaf4 | 3.861 | 3.19E-07 | glio |
| Bora_in.vivo_noRT | 4932441J04Ri | 3.686 | 3.19E-07 | glio |
| culture_effect | Dimt1 | -0.989 | 3.19E-08 | glio |
| culture_effect | Ppp1r2 | 0.853 | 3.19E-08 | glio |
| Atr_in.vivo_noRT | Tbc1d10c | 3.838 | 3.19E-08 | glio |
| Atr_in.vivo_noRT | Gm12905 | 4.228 | 3.19E-08 | glio |
| Bcas2_in.vivo_noRT | Rax | 3.628 | 3.19E-08 | glio |
| Aars2_in.vivo_noRT | Gm43387 | 4.151 | 3.19E-09 | glio |
| Chek1_ex.vivo_noRT | Ccdc162 | 4.842 | 3.19E-12 | glio |
| Chek1_in.vivo_noRT | Tcap | 4.664 | 3.19E-13 | glio |
| Bora_in.vivo_noRT | Ms4a6d | 4.673 | 3.19E-15 | glio |
| Chek1_ex.vivo_noRT | 2410017I17Ri | 4.842 | 3.19E-16 | glio |
| culture_effect | Gadd45g | 3.089 | 3.19E-18 | glio |
| Atr_in.vivo_noRT | Hist1h2ag | 5.001 | 3.19E-21 | glio |
| Bcas2_in.vivo_noRT | Bcas2 | -3.594 | 3.19E-21 | glio |
| culture_effect | Gpx2 | -8.024 | 3.19E-27 | glio |
| Chek1_in.vivo_noRT | Capn13 | 5.044 | 3.19E-27 | glio |
| culture_effect | Cnp | -3.527 | 3.19E-39 | glio |
| culture_effect | Gm26737 | 2.392 | 3.20E-03 | glio |
| culture_effect | Nat8f1 | 1.447 | 3.20E-03 | glio |

| | | | | |
|---|---|---|---|---|
| culture_effect | Pacs2 | 0.691 | 3.20E-03 | glio |
| culture_effect | Stt3a | -0.414 | 3.20E-03 | glio |
| culture_effect | Smarca5 | 0.345 | 3.20E-03 | glio |
| Interaction_genotype_Atr | Nwd1 | -3.087 | 3.20E-03 | glio |
| Interaction_genotype_Atr | Chd5 | -3.715 | 3.20E-03 | glio |
| Interaction_genotype_Atr | Arhgef19 | -3.671 | 3.20E-03 | glio |
| Interaction_genotype_Atr | Asgr1 | -3.324 | 3.20E-03 | glio |
| Interaction_genotype_Atr | Ero1l | 2.193 | 3.20E-03 | glio |
| Interaction_genotype_Chek1 | Dtl | -2.956 | 3.20E-03 | glio |
| Interaction_genotype_Chek1 | Hes5 | 5.107 | 3.20E-03 | glio |
| Interaction_genotype_Chek1 | Sgpp1 | -3.199 | 3.20E-03 | glio |
| Interaction_genotype_Mtor | Slc6a20b | -3.659 | 3.20E-03 | glio |
| Interaction_genotype_Myc | Gm39929 | -2.646 | 3.20E-03 | glio |
| Interaction_genotype_Myc | 1700122E12Ri | -3.335 | 3.20E-03 | glio |
| Interaction_genotype_Myc | Arhgef17 | -2.888 | 3.20E-03 | glio |
| Interaction_genotype_Myc | Apaf1 | -2.411 | 3.20E-03 | glio |
| Interaction_genotype_Myc | Ypel2 | -2.576 | 3.20E-03 | glio |
| Interaction_genotype_Myc | Gnb1 | 1.129 | 3.20E-03 | glio |
| interaction_RT | Mrpl18 | -0.669 | 3.20E-03 | glio |
| Aars2_in.vivo_noRT | Ythdf1 | -1.198 | 3.20E-03 | glio |
| Atp5c1_in.vivo_noRT | Txndc2 | 1.889 | 3.20E-03 | glio |
| Atr_in.vivo_noRT | Xkr8 | 2.006 | 3.20E-03 | glio |
| Atr_in.vivo_noRT | 9130221H12R | 1.954 | 3.20E-03 | glio |
| Atr_in.vivo_noRT | Prickle3 | 1.912 | 3.20E-03 | glio |
| Atr_in.vivo_noRT | Btbd16 | 1.789 | 3.20E-03 | glio |
| Bcas2_in.vivo_noRT | Acsf2 | 1.924 | 3.20E-03 | glio |
| Bora_in.vivo_noRT | Gm17056 | 2.203 | 3.20E-03 | glio |
| Bora_in.vivo_noRT | Pnp | -1.077 | 3.20E-03 | glio |
| Cenpj_in.vivo_noRT | Lrrc4b | 2.166 | 3.20E-03 | glio |
| Chek1_ex.vivo_noRT | Dhcr7 | -2.053 | 3.20E-03 | glio |
| culture_effect | 2010109A12R | 1.594 | 3.20E-04 | glio |
| culture_effect | Nudt18 | 1.083 | 3.20E-04 | glio |
| culture_effect | Akr1b8 | -0.738 | 3.20E-04 | glio |
| culture_effect | Atp6v0b | 0.364 | 3.20E-04 | glio |
| Interaction_genotype_Myc | 1700101I11Ri | -4.389 | 3.20E-04 | glio |
| Interaction_genotype_Myc | Flrt2 | -4.187 | 3.20E-04 | glio |
| Interaction_genotype_Myc | Gdap1l1 | -4.473 | 3.20E-04 | glio |
| Interaction_genotype_Myc | Pigo | -3.684 | 3.20E-04 | glio |
| Interaction_genotype_Myc | Abhd18 | -4.125 | 3.20E-04 | glio |
| Interaction_genotype_Myc | Sipa1l2 | -3.939 | 3.20E-04 | glio |
| Interaction_genotype_Myc | Wdr11 | -3.274 | 3.20E-04 | glio |
| Interaction_genotype_Myc | Pomp | 1.452 | 3.20E-04 | glio |
| Atr_in.vivo_noRT | Clcn2 | 2.679 | 3.20E-04 | glio |
| Atr_in.vivo_noRT | Gm48550 | 2.432 | 3.20E-04 | glio |
| Atr_in.vivo_noRT | B9d1 | -1.572 | 3.20E-04 | glio |
| Atr_ex.vivo_noRT | Pola1 | -1.721 | 3.20E-04 | glio |
| Bcas2_in.vivo_noRT | Flrt2 | 2.663 | 3.20E-04 | glio |
| Bcas2_in.vivo_noRT | B230208H11R | 2.667 | 3.20E-04 | glio |
| Bcas2_in.vivo_noRT | Cd81 | -1.575 | 3.20E-04 | glio |
| Bora_ex.vivo_noRT | Gbp10 | 3.201 | 3.20E-04 | glio |
| Cenpj_ex.vivo_noRT | Gm266 | 2.591 | 3.20E-04 | glio |
| Cenpj_ex.vivo_noRT | Ildr1 | 2.591 | 3.20E-04 | glio |

| | | | | |
|---|---|---|---|---|
| Chek1_in.vivo_noRT | Prkcq | 2.188 | 3.20E-04 | glio |
| culture_effect | Gm14167 | 1.655 | 3.20E-05 | glio |
| culture_effect | Scamp2 | -0.674 | 3.20E-05 | glio |
| culture_effect | Wwc2 | 0.752 | 3.20E-05 | glio |
| culture_effect | Higd2a | 0.599 | 3.20E-05 | glio |
| culture_effect | Rpl8 | -0.565 | 3.20E-05 | glio |
| Interaction_genotype_Mtor | 2310008N11R | -5.369 | 3.20E-05 | glio |
| Interaction_genotype_Mtor | Snrpd1 | 2.118 | 3.20E-05 | glio |
| Interaction_genotype_Myc | Polm | -4.211 | 3.20E-05 | glio |
| Interaction_genotype_Rad51 | Ciita | 4.639 | 3.20E-05 | glio |
| Atr_ex.vivo_noRT | Irf8 | 3.947 | 3.20E-06 | glio |
| Bcas2_in.vivo_noRT | 1700001L05Ri | 3.109 | 3.20E-06 | glio |
| Bcas2_in.vivo_noRT | Card10 | 3.539 | 3.20E-06 | glio |
| Chek1_ex.vivo_noRT | Ckm | 4.045 | 3.20E-06 | glio |
| Chek1_ex.vivo_noRT | Dnajb13 | 4.058 | 3.20E-06 | glio |
| Chek1_in.vivo_noRT | Mroh7 | 3.52 | 3.20E-07 | glio |
| Aars2_in.vivo_noRT | Zfp455 | 3.773 | 3.20E-08 | glio |
| Aars2_in.vivo_noRT | Abat | 4.151 | 3.20E-08 | glio |
| Bcas2_in.vivo_noRT | Ninj2 | 3.9 | 3.20E-08 | glio |
| Bcas2_in.vivo_noRT | Wfdc1 | 3.822 | 3.20E-08 | glio |
| Bcas2_in.vivo_noRT | Tnfrsf23 | 4.418 | 3.20E-08 | glio |
| Aars2_in.vivo_noRT | Tmem30c | 4.14 | 3.20E-09 | glio |
| Aars2_in.vivo_noRT | Apoa1 | 4.293 | 3.20E-09 | glio |
| Aars2_in.vivo_noRT | Ruvbl2 | -2.256 | 3.20E-09 | glio |
| Chek1_in.vivo_noRT | Prtn3 | 4.261 | 3.20E-09 | glio |
| Aars2_in.vivo_noRT | Gm36043 | 3.871 | 3.20E-10 | glio |
| Atr_in.vivo_noRT | Gm36043 | 4.077 | 3.20E-11 | glio |
| Chek1_in.vivo_noRT | Masp2 | 4.116 | 3.20E-12 | glio |
| Aars2_in.vivo_noRT | mt-Nd2 | 1.328 | 3.20E-13 | glio |
| Bora_in.vivo_noRT | Gm43256 | 4.673 | 3.20E-13 | glio |
| culture_effect | Thbs1 | 3.35 | 3.20E-17 | glio |
| Atr_in.vivo_noRT | Tcstv1 | 5.533 | 3.20E-20 | glio |
| Atp5c1_in.vivo_noRT | Atp5c1 | -3.915 | 3.20E-38 | glio |
| Interaction_genotype_Aars2 | Gngt2 | -3.703 | 3.21E-03 | glio |
| Interaction_genotype_Aars2 | Rrp15 | 1.368 | 3.21E-03 | glio |
| Interaction_genotype_Aars2 | Lamtor5 | 1.296 | 3.21E-03 | glio |
| Interaction_genotype_Aars2 | Psmb1 | 0.972 | 3.21E-03 | glio |
| Interaction_genotype_Aars2 | Bsg | 0.908 | 3.21E-03 | glio |
| Interaction_genotype_Mtor | Gdpd5 | -3.621 | 3.21E-03 | glio |
| Interaction_genotype_Myc | Ptn | -3.93 | 3.21E-03 | glio |
| Interaction_genotype_Myc | Bckdhb | -3.029 | 3.21E-03 | glio |
| Interaction_genotype_Myc | Fech | -1.86 | 3.21E-03 | glio |
| Interaction_genotype_Myc | Sept7 | 1.408 | 3.21E-03 | glio |
| Interaction_genotype_Myc | Psmg4 | 1.05 | 3.21E-03 | glio |
| Interaction_genotype_Rad51 | Ap3m1 | 2.259 | 3.21E-03 | glio |
| Atr_in.vivo_noRT | Cerkl | 1.865 | 3.21E-03 | glio |
| Atr_in.vivo_noRT | Senp8 | 1.816 | 3.21E-03 | glio |
| Atr_in.vivo_noRT | Tmem234 | -0.692 | 3.21E-03 | glio |
| Bcas2_in.vivo_noRT | Chd4 | -0.762 | 3.21E-03 | glio |
| Bora_in.vivo_noRT | Adck5 | -1.545 | 3.21E-03 | glio |
| Chek1_in.vivo_noRT | Rplp1 | -0.593 | 3.21E-03 | glio |
| Chek1_ex.vivo_noRT | Gm10230 | 2.897 | 3.21E-03 | glio |

| | | | | |
|---|---|---|---|---|
| Chek1_ex.vivo_noRT | Mctp2 | 2.409 | 3.21E-03 | glio |
| Eif2s1_in.vivo_noRT | Hrct1 | 2.143 | 3.21E-03 | glio |
| Eif2s1_ex.vivo_noRT | Nrip2 | 2.234 | 3.21E-03 | glio |
| Eif2s1_ex.vivo_noRT | Ablim2 | 2.234 | 3.21E-03 | glio |
| Eif2s1_ex.vivo_noRT | Gm5415 | 2.234 | 3.21E-03 | glio |
| Hsd17b10_in.vivo_noRT | Stk36 | 2.328 | 3.21E-03 | glio |
| culture_effect | Hk1 | 0.838 | 3.21E-04 | glio |
| culture_effect | Hat1 | -0.647 | 3.21E-04 | glio |
| Interaction_genotype_Mtor | 9130221H12R | -4.124 | 3.21E-04 | glio |
| Interaction_genotype_Mtor | Kbtbd7 | -3.881 | 3.21E-04 | glio |
| Interaction_genotype_Mtor | Gm17399 | -4.325 | 3.21E-04 | glio |
| Interaction_genotype_Mtor | Ccdc180 | -3.762 | 3.21E-04 | glio |
| Interaction_genotype_Myc | Cdiptos | -3.997 | 3.21E-04 | glio |
| Atr_in.vivo_noRT | Nek8 | 2.302 | 3.21E-04 | glio |
| Bora_in.vivo_noRT | Ifnz | 2.593 | 3.21E-04 | glio |
| Bora_in.vivo_noRT | Prdm10 | 2.28 | 3.21E-04 | glio |
| Bora_in.vivo_noRT | Greb1 | -1.713 | 3.21E-04 | glio |
| Chek1_in.vivo_noRT | Gja4 | 2.456 | 3.21E-04 | glio |
| Chek1_ex.vivo_noRT | Noxred1 | 3.072 | 3.21E-04 | glio |
| culture_effect | Lypla2 | -1.052 | 3.21E-05 | glio |
| Interaction_genotype_Myc | Slc2a9 | -3.926 | 3.21E-05 | glio |
| Interaction_genotype_Myc | Laptm4a | 1.672 | 3.21E-05 | glio |
| interaction_RT | Srp68 | -1.04 | 3.21E-05 | glio |
| Atr_in.vivo_noRT | Stpg4 | 3.045 | 3.21E-05 | glio |
| Atr_in.vivo_noRT | Cep135 | 2.289 | 3.21E-05 | glio |
| Bora_in.vivo_noRT | Hyal3 | 3.346 | 3.21E-05 | glio |
| Bora_in.vivo_noRT | Zfp781 | 2.834 | 3.21E-05 | glio |
| Chek1_in.vivo_noRT | Olfr655 | 4.127 | 3.21E-05 | glio |
| Chek1_in.vivo_noRT | Pcdhb19 | 2.752 | 3.21E-05 | glio |
| Interaction_genotype_Mtor | 2900089D17R | -5.162 | 3.21E-06 | glio |
| Interaction_genotype_Mtor | Eng | -5.999 | 3.21E-06 | glio |
| Interaction_genotype_Mtor | Tlr1 | -5.483 | 3.21E-06 | glio |
| Atr_in.vivo_noRT | Glmp | 3.53 | 3.21E-06 | glio |
| Bora_ex.vivo_noRT | 4930512B01R | 3.201 | 3.21E-06 | glio |
| interaction_RT | Metrn | -1.059 | 3.21E-07 | glio |
| Atr_in.vivo_noRT | Shd | 3.446 | 3.21E-07 | glio |
| Bora_in.vivo_noRT | A530072M11F | 3.608 | 3.21E-07 | glio |
| Bora_in.vivo_noRT | Guca1b | 3.648 | 3.21E-07 | glio |
| culture_effect | Batf | -3.5 | 3.21E-08 | glio |
| Chek1_in.vivo_noRT | Gm10484 | 4.042 | 3.21E-08 | glio |
| Hsd17b10_in.vivo_noRT | Nkg7 | 3.356 | 3.21E-08 | glio |
| culture_effect | Dusp15 | -3.384 | 3.21E-09 | glio |
| culture_effect | Gm9828 | 2.489 | 3.21E-09 | glio |
| Chek1_in.vivo_noRT | Rax | 3.865 | 3.21E-09 | glio |
| Chek1_ex.vivo_noRT | Pou5f1 | 5.249 | 3.21E-10 | glio |
| culture_effect | Rps7 | -0.896 | 3.21E-11 | glio |
| Aars2_in.vivo_noRT | Hsf4 | 4.65 | 3.21E-11 | glio |
| Atr_in.vivo_noRT | Ipcef1 | 3.72 | 3.21E-11 | glio |
| culture_effect | Atcay | -3.305 | 3.21E-19 | glio |
| Bora_in.vivo_noRT | Gm21319 | 4.936 | 3.21E-19 | glio |
| Atr_in.vivo_noRT | Gm13449 | 5.533 | 3.21E-24 | glio |
| Aars2_in.vivo_noRT | Ildr1 | 5.327 | 3.21E-25 | glio |

| | | | | |
|---|---|---|---|---|
| culture_effect | Hpcal1 | -1.584 | 3.22E-03 | glio |
| culture_effect | Oasl1 | -1.566 | 3.22E-03 | glio |
| Interaction_genotype_Aars2 | Hist1h4f | -3.984 | 3.22E-03 | glio |
| Interaction_genotype_Aars2 | A330009N23R | -3.631 | 3.22E-03 | glio |
| Interaction_genotype_Mtor | Cib2 | -4.072 | 3.22E-03 | glio |
| Interaction_genotype_Mtor | Ube2i | 1.306 | 3.22E-03 | glio |
| Interaction_genotype_Myc | Gm38393 | -2.863 | 3.22E-03 | glio |
| Interaction_genotype_Myc | Foxo4 | -2.761 | 3.22E-03 | glio |
| Interaction_genotype_Myc | Ubl7 | -2.451 | 3.22E-03 | glio |
| Interaction_genotype_Terf1 | Gm30938 | -4.142 | 3.22E-03 | glio |
| interaction_RT | Galnt2 | -0.84 | 3.22E-03 | glio |
| Aars2_in.vivo_noRT | 5430427N15R | 2.054 | 3.22E-03 | glio |
| Aars2_in.vivo_noRT | Aip | -1.293 | 3.22E-03 | glio |
| Atr_in.vivo_noRT | Fam102a | 1.538 | 3.22E-03 | glio |
| Bora_in.vivo_noRT | 4933408B17R | 2.189 | 3.22E-03 | glio |
| Bora_in.vivo_noRT | Lsm2 | -0.945 | 3.22E-03 | glio |
| Brca2_in.vivo_noRT | Sucnr1 | 1.763 | 3.22E-03 | glio |
| Cenpj_in.vivo_noRT | Gm12610 | 2.022 | 3.22E-03 | glio |
| Chek1_in.vivo_noRT | Plcxd2 | 2.198 | 3.22E-03 | glio |
| Chek1_in.vivo_noRT | Zfp870 | 1.869 | 3.22E-03 | glio |
| Chek1_in.vivo_noRT | Zfp959 | 1.745 | 3.22E-03 | glio |
| Chek1_ex.vivo_noRT | Tacr2 | 2.784 | 3.22E-03 | glio |
| Chek1_ex.vivo_noRT | Gtf2h2 | -1.787 | 3.22E-03 | glio |
| Chek1_ex.vivo_noRT | Nfkbia | 1.47 | 3.22E-03 | glio |
| Eif2s1_in.vivo_noRT | Zfp786 | 2.41 | 3.22E-03 | glio |
| Hsd17b10_in.vivo_noRT | Jmjd8 | 2.25 | 3.22E-03 | glio |
| Hsd17b10_in.vivo_noRT | A530072M11F | 2.288 | 3.22E-03 | glio |
| Hsd17b10_in.vivo_noRT | 1110038B12R | 0.601 | 3.22E-03 | glio |
| culture_effect | 3110082I17Ri | 0.9 | 3.22E-04 | glio |
| culture_effect | Ruvbl1 | -0.504 | 3.22E-04 | glio |
| Interaction_genotype_Mtor | Gm49135 | -4.433 | 3.22E-04 | glio |
| Interaction_genotype_Rpl3 | Fxyd3 | 4.265 | 3.22E-04 | glio |
| Interaction_genotype_Rpl3 | Otos | -4.273 | 3.22E-04 | glio |
| Interaction_genotype_Rpl3 | Rassf6 | -4.16 | 3.22E-04 | glio |
| Interaction_genotype_Rpl3 | Slf2 | 2.511 | 3.22E-04 | glio |
| Interaction_genotype_Rpl3 | Klhl30 | -4.646 | 3.22E-04 | glio |
| Interaction_genotype_Rpl3 | Psmc3ip | 2.718 | 3.22E-04 | glio |
| Interaction_genotype_Rpl3 | 1810019D21R | -4.818 | 3.22E-04 | glio |
| Interaction_genotype_Rpl3 | Gm10399 | -4.584 | 3.22E-04 | glio |
| Interaction_genotype_Rpl3 | D630036H23F | -4.758 | 3.22E-04 | glio |
| Atp5c1_in.vivo_noRT | B230311B06R | 2.936 | 3.22E-04 | glio |
| Atr_in.vivo_noRT | Slc8a3 | 2.525 | 3.22E-04 | glio |
| Atr_in.vivo_noRT | Fyb | 2.51 | 3.22E-04 | glio |
| Atr_in.vivo_noRT | Naa50 | -1.211 | 3.22E-04 | glio |
| Bard1_ex.vivo_noRT | Ly6d | 2.814 | 3.22E-04 | glio |
| Bard1_ex.vivo_noRT | Fabp4 | 3.608 | 3.22E-04 | glio |
| Bcas2_in.vivo_noRT | Plxnb1 | 2.675 | 3.22E-04 | glio |
| Bora_in.vivo_noRT | Cntf | 2.567 | 3.22E-04 | glio |
| Bora_in.vivo_noRT | Nat8f1 | 2.471 | 3.22E-04 | glio |
| Chek1_in.vivo_noRT | Ccdc57 | 2.697 | 3.22E-04 | glio |
| Chek1_in.vivo_noRT | Pcdhgb4 | 2.248 | 3.22E-04 | glio |
| Chek1_in.vivo_noRT | Cs | -1.157 | 3.22E-04 | glio |

| | | | | |
|---|---|---|---|---|
| Hsd17b10_in.vivo_noRT | 1700061G19R | 2.605 | 3.22E-04 | glio |
| Interaction_genotype_Mtor | Rps9 | 1.392 | 3.22E-05 | glio |
| interaction_RT | Cct3 | -0.804 | 3.22E-05 | glio |
| Aars2_in.vivo_noRT | Mirg | 2.985 | 3.22E-05 | glio |
| Atr_ex.vivo_noRT | Cd83 | 3.355 | 3.22E-05 | glio |
| Chek1_in.vivo_noRT | Ccdc106 | 3.189 | 3.22E-05 | glio |
| Chek1_in.vivo_noRT | AW046200 | 2.986 | 3.22E-05 | glio |
| Hsd17b10_in.vivo_noRT | Hsf2bp | 2.824 | 3.22E-05 | glio |
| Hsd17b10_in.vivo_noRT | Gm30938 | 2.97 | 3.22E-05 | glio |
| culture_effect | Hbp1 | -0.984 | 3.22E-06 | glio |
| culture_effect | Srp9 | 0.498 | 3.22E-06 | glio |
| Interaction_genotype_Atr | 1700026J14Ri | -5.159 | 3.22E-06 | glio |
| Interaction_genotype_Mtor | Gm43581 | -5.122 | 3.22E-06 | glio |
| Interaction_genotype_Mtor | Rimklb | -3.836 | 3.22E-06 | glio |
| Aars2_in.vivo_noRT | D930019O06R | 3.43 | 3.22E-06 | glio |
| Atr_in.vivo_noRT | Ngfr | 2.639 | 3.22E-06 | glio |
| Aars2_in.vivo_noRT | mt-Nd4 | 0.838 | 3.22E-07 | glio |
| Hsd17b10_in.vivo_noRT | D730045A05R | 3.616 | 3.22E-07 | glio |
| culture_effect | Zfp462 | 1.064 | 3.22E-08 | glio |
| Chek1_ex.vivo_noRT | Gm50115 | 4.443 | 3.22E-08 | glio |
| culture_effect | Eif3g | -0.729 | 3.22E-09 | glio |
| Aars2_in.vivo_noRT | Grip2 | 4.076 | 3.22E-09 | glio |
| Chek1_in.vivo_noRT | Tslp | 4.398 | 3.22E-09 | glio |
| Aars2_in.vivo_noRT | Ephx3 | 4.599 | 3.22E-12 | glio |
| Atr_in.vivo_noRT | Serpina3g | 4.337 | 3.22E-12 | glio |
| Bora_in.vivo_noRT | Stra8 | 4.413 | 3.22E-13 | glio |
| Hsd17b10_in.vivo_noRT | Gm13964 | 3.616 | 3.22E-13 | glio |
| Chek1_ex.vivo_noRT | Gad1 | 4.842 | 3.22E-15 | glio |
| Atr_in.vivo_noRT | Pdlim3 | 4.754 | 3.22E-16 | glio |
| Chek1_in.vivo_noRT | Pyy | 5.044 | 3.22E-18 | glio |
| culture_effect | Col19a1 | -1.459 | 3.23E-03 | glio |
| culture_effect | Asl | -1.211 | 3.23E-03 | glio |
| culture_effect | Rpp40 | -0.686 | 3.23E-03 | glio |
| culture_effect | Trappc2 | 0.508 | 3.23E-03 | glio |
| Interaction_genotype_Aars2 | Gm29083 | -3.381 | 3.23E-03 | glio |
| Interaction_genotype_Aars2 | Spon1 | -3.754 | 3.23E-03 | glio |
| Interaction_genotype_Aars2 | Arhgef19 | -3.701 | 3.23E-03 | glio |
| Interaction_genotype_Aars2 | Gm17655 | -4.089 | 3.23E-03 | glio |
| Interaction_genotype_Aars2 | Atp13a3 | -1.902 | 3.23E-03 | glio |
| Interaction_genotype_Aars2 | Srsf11 | 1.135 | 3.23E-03 | glio |
| Interaction_genotype_Aars2 | Rps11 | 0.846 | 3.23E-03 | glio |
| Interaction_genotype_Bcas2 | Rims3 | -3.593 | 3.23E-03 | glio |
| Interaction_genotype_Bcas2 | Gtsf1 | -3.228 | 3.23E-03 | glio |
| Interaction_genotype_Bcas2 | Gpr179 | -3.832 | 3.23E-03 | glio |
| Interaction_genotype_Bcas2 | Hist2h3c1 | -3.92 | 3.23E-03 | glio |
| Interaction_genotype_Mtor | Pld6 | -3.76 | 3.23E-03 | glio |
| Interaction_genotype_Mtor | Pros1 | -3.155 | 3.23E-03 | glio |
| Interaction_genotype_Mtor | Cct2 | 1.141 | 3.23E-03 | glio |
| Interaction_genotype_Nat2 | Hist1h2ag | 3.219 | 3.23E-03 | glio |
| interaction_RT | Sfpq | -0.712 | 3.23E-03 | glio |
| Aars2_in.vivo_noRT | Klc2 | 1.931 | 3.23E-03 | glio |
| Aars2_in.vivo_noRT | Rbfox2 | -1.181 | 3.23E-03 | glio |

| | | | | |
|---|---|---|---|---|
| Aars2_in.vivo_noRT | Dynlrb1 | -0.771 | 3.23E-03 | glio |
| Bcas2_ex.vivo_noRT | 4930553M12F | 2.348 | 3.23E-03 | glio |
| Bcas2_ex.vivo_noRT | Serpina3g | 2.742 | 3.23E-03 | glio |
| Bora_in.vivo_noRT | 1700030K09R | 1.62 | 3.23E-03 | glio |
| Chek1_in.vivo_noRT | Adap1 | 2.449 | 3.23E-03 | glio |
| Eif2s1_ex.vivo_noRT | Gm49698 | 2.234 | 3.23E-03 | glio |
| Eif2s1_ex.vivo_noRT | Olfr881 | 2.234 | 3.23E-03 | glio |
| Eif2s1_ex.vivo_noRT | Gm19265 | 2.615 | 3.23E-03 | glio |
| Eif2s1_ex.vivo_noRT | BC061195 | 2.234 | 3.23E-03 | glio |
| Eif2s1_ex.vivo_noRT | Idi2 | 2.234 | 3.23E-03 | glio |
| Eif2s1_ex.vivo_noRT | Vmn2r18 | 2.234 | 3.23E-03 | glio |
| Eif2s1_ex.vivo_noRT | Gm16178 | 2.234 | 3.23E-03 | glio |
| Eif2s1_ex.vivo_noRT | Gm364 | 2.234 | 3.23E-03 | glio |
| Hsd17b10_in.vivo_noRT | Gm15246 | 2.265 | 3.23E-03 | glio |
| Hsd17b10_in.vivo_noRT | Clstn3 | -1.93 | 3.23E-03 | glio |
| Interaction_genotype_Mtor | Col5a2 | -3.374 | 3.23E-04 | glio |
| Interaction_genotype_Myc | Slc2a6 | -4.289 | 3.23E-04 | glio |
| Interaction_genotype_Myc | Phykpl | -3.164 | 3.23E-04 | glio |
| Aars2_in.vivo_noRT | Ttc30b | 2.733 | 3.23E-04 | glio |
| Aars2_in.vivo_noRT | 2310010J17Ri | 2.006 | 3.23E-04 | glio |
| Aars2_ex.vivo_noRT | Napsa | 3.676 | 3.23E-04 | glio |
| Bard1_ex.vivo_noRT | Ccdc162 | 2.814 | 3.23E-04 | glio |
| Bard1_ex.vivo_noRT | Mpzl2 | 2.814 | 3.23E-04 | glio |
| Bora_in.vivo_noRT | Tiparp | -1.93 | 3.23E-04 | glio |
| Chek1_in.vivo_noRT | Cops2 | -1.242 | 3.23E-04 | glio |
| Interaction_genotype_Mtor | Lcat | -5.179 | 3.23E-05 | glio |
| Interaction_genotype_Mtor | Ybx3 | 2.073 | 3.23E-05 | glio |
| Aars2_in.vivo_noRT | Gm12043 | 2.733 | 3.23E-05 | glio |
| Aars2_ex.vivo_noRT | 1700010B08R | 3.293 | 3.23E-05 | glio |
| Aars2_ex.vivo_noRT | Brinp3 | 3.293 | 3.23E-05 | glio |
| Atr_ex.vivo_noRT | Vmn2r54 | 3.164 | 3.23E-05 | glio |
| Atr_ex.vivo_noRT | Egr2 | 3.661 | 3.23E-05 | glio |
| Bora_in.vivo_noRT | Pxt1 | 2.987 | 3.23E-05 | glio |
| culture_effect | Pin4 | 0.599 | 3.23E-06 | glio |
| Atr_in.vivo_noRT | Fgf15 | 3.502 | 3.23E-06 | glio |
| Atr_ex.vivo_noRT | Gbp10 | 3.947 | 3.23E-06 | glio |
| culture_effect | Klk14 | 2.793 | 3.23E-07 | glio |
| Aars2_in.vivo_noRT | Hoga1 | 3.512 | 3.23E-07 | glio |
| Atr_in.vivo_noRT | Mucl3 | 3.762 | 3.23E-07 | glio |
| Atr_in.vivo_noRT | Serpinc1 | 3.716 | 3.23E-07 | glio |
| Aars2_in.vivo_noRT | 9330104G04R | 3.847 | 3.23E-08 | glio |
| Aars2_in.vivo_noRT | Hecw2 | 3.978 | 3.23E-08 | glio |
| Atr_in.vivo_noRT | Tmeff2 | 4.276 | 3.23E-08 | glio |
| Bora_in.vivo_noRT | A230020J21Ri | 4.009 | 3.23E-08 | glio |
| Aars2_in.vivo_noRT | Cfap58 | 5.327 | 3.23E-10 | glio |
| Bcas2_in.vivo_noRT | Gm10300 | 4.547 | 3.23E-10 | glio |
| culture_effect | Bank1 | -3.398 | 3.23E-12 | glio |
| Bcas2_in.vivo_noRT | Agrp | 4.024 | 3.23E-12 | glio |
| Bora_in.vivo_noRT | Nxnl2 | 4.414 | 3.23E-12 | glio |
| Bora_in.vivo_noRT | 1110046J04Ri | 4.936 | 3.23E-12 | glio |
| culture_effect | D630003M21I | 2.539 | 3.23E-13 | glio |
| Bora_in.vivo_noRT | Gm50138 | 4.414 | 3.23E-14 | glio |

| | | | | |
|---|---|---|---|---|
| culture_effect | Slc25a28 | -1.091 | 3.23E-15 | glio |
| Aars2_in.vivo_noRT | Adam30 | 4.947 | 3.23E-16 | glio |
| culture_effect | Mapk3 | -1.545 | 3.23E-17 | glio |
| Aars2_in.vivo_noRT | Gm13078 | 5.327 | 3.23E-26 | glio |
| culture_effect | Tnfsf13b | -1.247 | 3.24E-03 | glio |
| culture_effect | Mmd | 1.053 | 3.24E-03 | glio |
| culture_effect | Rhebl1 | -0.697 | 3.24E-03 | glio |
| culture_effect | Asf1b | -0.638 | 3.24E-03 | glio |
| culture_effect | Zeb1 | -0.568 | 3.24E-03 | glio |
| culture_effect | Rps6kb1 | -0.428 | 3.24E-03 | glio |
| Interaction_genotype_Atr | Ccnl1 | 1.74 | 3.24E-03 | glio |
| Interaction_genotype_Bcas2 | Esrp2 | -3.723 | 3.24E-03 | glio |
| Interaction_genotype_Bora | Carns1 | -3.927 | 3.24E-03 | glio |
| Interaction_genotype_Cdkn2a | Gm34059 | 2.835 | 3.24E-03 | glio |
| Interaction_genotype_Mtor | Rae1 | 1.496 | 3.24E-03 | glio |
| Interaction_genotype_Myc | Arhgef15 | -3.238 | 3.24E-03 | glio |
| Interaction_genotype_Myc | Rtn1 | -3.367 | 3.24E-03 | glio |
| Interaction_genotype_Myc | Hist1h2ad | -3.42 | 3.24E-03 | glio |
| Interaction_genotype_Myc | 4930578M01F | -2.788 | 3.24E-03 | glio |
| Interaction_genotype_Pars2 | Lyrm4 | 2.056 | 3.24E-03 | glio |
| interaction_RT | Myoz1 | 2.505 | 3.24E-03 | glio |
| interaction_RT | Polr2l | 1.315 | 3.24E-03 | glio |
| Atr_ex.vivo_noRT | Slc16a4 | 2.308 | 3.24E-03 | glio |
| Atr_ex.vivo_noRT | Pou5f1 | 2.77 | 3.24E-03 | glio |
| Bcas2_in.vivo_noRT | Dpf1 | 1.894 | 3.24E-03 | glio |
| Bora_in.vivo_noRT | Siva1 | -0.756 | 3.24E-03 | glio |
| Chek1_in.vivo_noRT | Ccdc71l | 2.001 | 3.24E-03 | glio |
| Chek1_in.vivo_noRT | B4galt7 | 1.728 | 3.24E-03 | glio |
| Chek1_in.vivo_noRT | Ide | -1.219 | 3.24E-03 | glio |
| Chek1_in.vivo_noRT | Timm23 | -0.674 | 3.24E-03 | glio |
| Chek1_ex.vivo_noRT | Cnnm1 | 3.014 | 3.24E-03 | glio |
| Chek1_ex.vivo_noRT | Trip13 | -1.956 | 3.24E-03 | glio |
| Eif2s1_in.vivo_noRT | Egr1 | -1.286 | 3.24E-03 | glio |
| Hsd17b10_in.vivo_noRT | 4930453N24R | -1.109 | 3.24E-03 | glio |
| culture_effect | 2810468N07R | -1.718 | 3.24E-04 | glio |
| culture_effect | Rbmx | 0.93 | 3.24E-04 | glio |
| culture_effect | Hspa4l | -0.592 | 3.24E-04 | glio |
| Interaction_genotype_Mtor | Ninj2 | -4.354 | 3.24E-04 | glio |
| Interaction_genotype_Mtor | Pde1a | -4.478 | 3.24E-04 | glio |
| Interaction_genotype_Mtor | 6330408M09F | -4.406 | 3.24E-04 | glio |
| Interaction_genotype_Mtor | 2210408F21Ri | -3.192 | 3.24E-04 | glio |
| Interaction_genotype_Myc | Samd4 | -3.233 | 3.24E-04 | glio |
| Aars2_in.vivo_noRT | Atg5 | -1.303 | 3.24E-04 | glio |
| Hsd17b10_in.vivo_noRT | Gm28424 | 2.605 | 3.24E-04 | glio |
| Hsd17b10_in.vivo_noRT | Gabrd | 2.688 | 3.24E-04 | glio |
| Interaction_genotype_Mtor | Gm42997 | -4.541 | 3.24E-05 | glio |
| Interaction_genotype_Mtor | Gm26789 | -4.162 | 3.24E-05 | glio |
| Interaction_genotype_Mtor | Fam166a | -3.835 | 3.24E-05 | glio |
| interaction_RT | Ifi206 | 2.908 | 3.24E-05 | glio |
| Atr_in.vivo_noRT | Lgr6 | 3.325 | 3.24E-05 | glio |
| Chek1_in.vivo_noRT | Serac1 | 2.815 | 3.24E-05 | glio |
| Chek1_in.vivo_noRT | Tnfrsf1b | 2.75 | 3.24E-05 | glio |

| | | | | |
|---|---|---|---|---|
| Hsd17b10_in.vivo_noRT | Ccno | 3.123 | 3.24E-05 | glio |
| Interaction_genotype_Mtor | Gm9725 | -5.68 | 3.24E-06 | glio |
| Chek1_ex.vivo_noRT | Fbxl13 | 3.468 | 3.24E-06 | glio |
| Interaction_genotype_Mtor | A330049N07R | -4.232 | 3.24E-07 | glio |
| Atp5c1_in.vivo_noRT | Gm19276 | 2.668 | 3.24E-07 | glio |
| Atr_ex.vivo_noRT | Gm38477 | 3.562 | 3.24E-07 | glio |
| Bcas2_in.vivo_noRT | Arvcf | 3.813 | 3.24E-07 | glio |
| Hsd17b10_in.vivo_noRT | Igf2bp1 | 3.616 | 3.24E-07 | glio |
| Hsd17b10_in.vivo_noRT | 1600014C23R | 3.758 | 3.24E-07 | glio |
| Bcas2_in.vivo_noRT | Trnau1ap | -1.913 | 3.24E-08 | glio |
| Aars2_in.vivo_noRT | 4930545E07Ri | 4.293 | 3.24E-09 | glio |
| Chek1_ex.vivo_noRT | Sbk2 | 5.212 | 3.24E-09 | glio |
| Chek1_ex.vivo_noRT | A930003A15R | 4.842 | 3.24E-09 | glio |
| Hsd17b10_in.vivo_noRT | Gm48242 | 3.616 | 3.24E-09 | glio |
| Chek1_in.vivo_noRT | Gm10300 | 4.784 | 3.24E-11 | glio |
| Atr_in.vivo_noRT | Has1 | 4.636 | 3.24E-13 | glio |
| Aars2_in.vivo_noRT | Podxl | 4.795 | 3.24E-14 | glio |
| Bcas2_in.vivo_noRT | Gm28940 | 4.291 | 3.24E-14 | glio |
| Chek1_ex.vivo_noRT | Gm13178 | 4.842 | 3.24E-14 | glio |
| culture_effect | Pid1 | -5.307 | 3.24E-22 | glio |
| culture_effect | Sbk2 | -1.918 | 3.25E-03 | glio |
| culture_effect | Zfp979 | -1.309 | 3.25E-03 | glio |
| Interaction_genotype_Atr | Kcnk13 | -2.894 | 3.25E-03 | glio |
| Interaction_genotype_Atr | Ush1g | -2.769 | 3.25E-03 | glio |
| Interaction_genotype_Bora | Prl7c1 | -2.83 | 3.25E-03 | glio |
| Interaction_genotype_Mtor | Gfap | -3.446 | 3.25E-03 | glio |
| Interaction_genotype_Mtor | Fah | -2.946 | 3.25E-03 | glio |
| Interaction_genotype_Mtor | Klhdc8b | -3.329 | 3.25E-03 | glio |
| Interaction_genotype_Myc | A830035O19R | -3.164 | 3.25E-03 | glio |
| Interaction_genotype_Myc | 2410006H16R | 1.527 | 3.25E-03 | glio |
| Aars2_in.vivo_noRT | Klhl8 | 2.017 | 3.25E-03 | glio |
| Aars2_in.vivo_noRT | Fdx1 | -1.258 | 3.25E-03 | glio |
| Atr_in.vivo_noRT | A730017L22Ri | 1.858 | 3.25E-03 | glio |
| Atr_in.vivo_noRT | Unc13d | 1.978 | 3.25E-03 | glio |
| Atr_in.vivo_noRT | Smarcad1 | -1.197 | 3.25E-03 | glio |
| Atr_in.vivo_noRT | Zfand5 | -0.738 | 3.25E-03 | glio |
| Bard1_ex.vivo_noRT | Gm32122 | 2.814 | 3.25E-03 | glio |
| Bcas2_in.vivo_noRT | Fhod3 | 2.29 | 3.25E-03 | glio |
| Chek1_in.vivo_noRT | Gm29093 | 2.127 | 3.25E-03 | glio |
| Chek1_in.vivo_noRT | Scaf11 | -0.774 | 3.25E-03 | glio |
| Chek1_ex.vivo_noRT | Smim40 | 2.76 | 3.25E-03 | glio |
| Cyp19a1_ex.vivo_noRT | Olfml2b | 2.947 | 3.25E-03 | glio |
| culture_effect | Mlycd | -1.111 | 3.25E-04 | glio |
| Interaction_genotype_Mtor | Nop10 | 2.012 | 3.25E-04 | glio |
| Interaction_genotype_Myc | Rab42 | -4.24 | 3.25E-04 | glio |
| Aars2_in.vivo_noRT | Tmem50a | -1.216 | 3.25E-04 | glio |
| Atr_in.vivo_noRT | Il1rapl1 | 2.566 | 3.25E-04 | glio |
| Atr_in.vivo_noRT | 9230112E08Ri | 2.538 | 3.25E-04 | glio |
| Atr_in.vivo_noRT | Ggact | 2.209 | 3.25E-04 | glio |
| Atr_in.vivo_noRT | Aup1 | -1.065 | 3.25E-04 | glio |
| Atr_ex.vivo_noRT | Iqcn | 3.15 | 3.25E-04 | glio |
| Bcas2_in.vivo_noRT | Zbtb8a | 2.746 | 3.25E-04 | glio |

| | | | | |
|---|---|---|---|---|
| Bora_in.vivo_noRT | Marc2 | -1.391 | 3.25E-04 | glio |
| culture_effect | Gm48832 | -2.555 | 3.25E-05 | glio |
| culture_effect | Acsl3 | -0.749 | 3.25E-05 | glio |
| Interaction_genotype_Nat2 | Gm5039 | 5.43 | 3.25E-05 | glio |
| Aars2_in.vivo_noRT | Gpr161 | 2.99 | 3.25E-05 | glio |
| Aars2_in.vivo_noRT | Psmb1 | -0.902 | 3.25E-05 | glio |
| Aars2_ex.vivo_noRT | Gm16344 | 3.293 | 3.25E-05 | glio |
| culture_effect | Mst1 | -3.041 | 3.25E-07 | glio |
| Interaction_genotype_Mtor | Calhm5 | -5.768 | 3.25E-07 | glio |
| Interaction_genotype_Mtor | Upb1 | -5.907 | 3.25E-07 | glio |
| Aars2_in.vivo_noRT | Txnl1 | -1.784 | 3.25E-07 | glio |
| Bcas2_in.vivo_noRT | Grip2 | 3.556 | 3.25E-07 | glio |
| Bcas2_in.vivo_noRT | Zhx2 | 3.762 | 3.25E-07 | glio |
| Bcas2_in.vivo_noRT | Hsf4 | 3.62 | 3.25E-07 | glio |
| Chek1_in.vivo_noRT | 4922502N22R | 3.525 | 3.25E-07 | glio |
| culture_effect | Mb21d2 | -2.2 | 3.25E-09 | glio |
| Atr_in.vivo_noRT | Amhr2 | 4.657 | 3.25E-09 | glio |
| Bora_in.vivo_noRT | 1700026J14Ri | 4.414 | 3.25E-12 | glio |
| Atr_in.vivo_noRT | Sprr1a | 5.533 | 3.25E-15 | glio |
| Bora_in.vivo_noRT | Obscn | 4.936 | 3.25E-15 | glio |
| Atr_in.vivo_noRT | Snhg4.1 | 5.011 | 3.25E-17 | glio |
| Bora_in.vivo_noRT | 2310003N18R | 4.936 | 3.25E-17 | glio |
| culture_effect | Parp11 | -0.485 | 3.26E-03 | glio |
| Interaction_genotype_Aars2 | Gm47851 | -3.361 | 3.26E-03 | glio |
| Interaction_genotype_Aars2 | A830009L08Ri | -3.416 | 3.26E-03 | glio |
| Interaction_genotype_Aars2 | Cfl1 | 0.924 | 3.26E-03 | glio |
| Interaction_genotype_Aars2 | Hint1 | 0.775 | 3.26E-03 | glio |
| Interaction_genotype_Atr | Scg2 | -2.369 | 3.26E-03 | glio |
| Interaction_genotype_Atr | Pold1 | -2.122 | 3.26E-03 | glio |
| Interaction_genotype_Mtor | Gm50218 | -3.851 | 3.26E-03 | glio |
| Interaction_genotype_Mtor | Gm26620 | -3.022 | 3.26E-03 | glio |
| Interaction_genotype_Myc | 1700112J16Ri | -2.52 | 3.26E-03 | glio |
| interaction_RT | Nbeal1 | 0.97 | 3.26E-03 | glio |
| Aars2_in.vivo_noRT | Tmem260 | 1.753 | 3.26E-03 | glio |
| Aars2_in.vivo_noRT | Pdcd5 | -0.697 | 3.26E-03 | glio |
| Atr_in.vivo_noRT | Pnpla7 | 1.951 | 3.26E-03 | glio |
| Atr_ex.vivo_noRT | Fat1 | -1.486 | 3.26E-03 | glio |
| Bcas2_in.vivo_noRT | Sgms2 | 2.302 | 3.26E-03 | glio |
| Bcas2_in.vivo_noRT | Map3k8 | 2.332 | 3.26E-03 | glio |
| Bcas2_in.vivo_noRT | Oscp1 | 2.165 | 3.26E-03 | glio |
| Bcas2_in.vivo_noRT | Platr14 | 1.893 | 3.26E-03 | glio |
| Bcas2_in.vivo_noRT | Fxn | -1.35 | 3.26E-03 | glio |
| Bora_in.vivo_noRT | Ccnb1ip1 | 2.405 | 3.26E-03 | glio |
| Bora_in.vivo_noRT | Slc25a47 | 2.174 | 3.26E-03 | glio |
| culture_effect | Srrd | 0.728 | 3.26E-04 | glio |
| culture_effect | Dtnbp1 | 0.669 | 3.26E-04 | glio |
| Interaction_genotype_Aars2 | Gm17018 | 2.097 | 3.26E-04 | glio |
| Interaction_genotype_Mtor | Gm50375 | -4.601 | 3.26E-04 | glio |
| Interaction_genotype_Myc | Chst14 | -3.229 | 3.26E-04 | glio |
| Aars2_in.vivo_noRT | B3gntl1 | 2.476 | 3.26E-04 | glio |
| Atp5c1_in.vivo_noRT | Gm36356 | 2.408 | 3.26E-04 | glio |
| Atr_in.vivo_noRT | Csnk1e | -1.31 | 3.26E-04 | glio |

| | | | | |
|---|---|---|---|---|
| Atr_in.vivo_noRT | 2810004N23R | -1.268 | 3.26E-04 | glio |
| Bora_in.vivo_noRT | Bag1 | -0.908 | 3.26E-04 | glio |
| culture_effect | Mtm1 | -1.221 | 3.26E-05 | glio |
| culture_effect | Prrg2 | 1.024 | 3.26E-05 | glio |
| Aars2_in.vivo_noRT | Camkv | 3.092 | 3.26E-05 | glio |
| Atp5c1_in.vivo_noRT | Cpeb1 | 3.309 | 3.26E-05 | glio |
| Bora_in.vivo_noRT | Mtmr2 | -1.983 | 3.26E-05 | glio |
| Bora_in.vivo_noRT | Cryl1 | -2.379 | 3.26E-05 | glio |
| Chek1_ex.vivo_noRT | Gpr21 | 3.488 | 3.26E-05 | glio |
| Chek1_ex.vivo_noRT | Zfr2 | 3.591 | 3.26E-05 | glio |
| Interaction_genotype_Mtor | Gm50295 | -5.432 | 3.26E-06 | glio |
| Interaction_genotype_Myc | Gm48435 | -3.516 | 3.26E-06 | glio |
| Bora_in.vivo_noRT | Gpr21 | 3.071 | 3.26E-06 | glio |
| Chek1_ex.vivo_noRT | Bco2 | 3.483 | 3.26E-06 | glio |
| culture_effect | Grina | 0.584 | 3.26E-07 | glio |
| Atr_in.vivo_noRT | 1700055D18R | 4.143 | 3.26E-08 | glio |
| Atr_in.vivo_noRT | Rab11fip1 | 4.239 | 3.26E-08 | glio |
| interaction_RT | Insl6 | 1.743 | 3.26E-10 | glio |
| Atp5c1_in.vivo_noRT | Gm49808 | 3.196 | 3.26E-10 | glio |
| Atr_in.vivo_noRT | Adam33 | 4.376 | 3.26E-10 | glio |
| Atr_in.vivo_noRT | Zfp473 | 4.109 | 3.26E-10 | glio |
| Chek1_in.vivo_noRT | Spata31d1a | 4.402 | 3.26E-10 | glio |
| Hsd17b10_in.vivo_noRT | Gm21761 | 3.616 | 3.26E-11 | glio |
| Atr_in.vivo_noRT | Vwce | 5.273 | 3.26E-17 | glio |
| culture_effect | Gfpt2 | -2.025 | 3.26E-19 | glio |
| Aars2_in.vivo_noRT | Gpr22 | 5.327 | 3.26E-23 | glio |
| culture_effect | Slc35f1 | 3.079 | 3.26E-24 | glio |
| culture_effect | Leo1 | -0.528 | 3.27E-03 | glio |
| culture_effect | Sec61g | 0.411 | 3.27E-03 | glio |
| Interaction_genotype_Aars2 | Pde8a | -2.81 | 3.27E-03 | glio |
| Interaction_genotype_Atr | Yrdc | 1.211 | 3.27E-03 | glio |
| Interaction_genotype_Cdkn2a | Myl7 | -4.028 | 3.27E-03 | glio |
| Interaction_genotype_Cdkn2a | Gm20625 | 2.835 | 3.27E-03 | glio |
| Interaction_genotype_Cdkn2a | Gm5157 | 2.835 | 3.27E-03 | glio |
| Interaction_genotype_Cdkn2a | Gm44736 | 2.835 | 3.27E-03 | glio |
| Interaction_genotype_Cdkn2a | Fam227b | 2.835 | 3.27E-03 | glio |
| Interaction_genotype_Cdkn2a | Gm20658 | 2.835 | 3.27E-03 | glio |
| Interaction_genotype_Cdkn2a | Olfr810 | 2.835 | 3.27E-03 | glio |
| Interaction_genotype_Cdkn2a | 4933400F21Ri | 3.1 | 3.27E-03 | glio |
| Interaction_genotype_Cdkn2a | BC049987 | 3.09 | 3.27E-03 | glio |
| Interaction_genotype_Myc | Strada | -2.421 | 3.27E-03 | glio |
| Interaction_genotype_Rpl3 | Gm49785 | -4.094 | 3.27E-03 | glio |
| Interaction_genotype_Rpl3 | D930030I03Ri | -4.336 | 3.27E-03 | glio |
| Aars2_in.vivo_noRT | Cltb | -1.1 | 3.27E-03 | glio |
| Atr_in.vivo_noRT | Slc4a5 | 2.249 | 3.27E-03 | glio |
| Atr_ex.vivo_noRT | Gm13270 | 2.826 | 3.27E-03 | glio |
| Atr_ex.vivo_noRT | Tnnc1 | 2.991 | 3.27E-03 | glio |
| Bcas2_in.vivo_noRT | Fhl2 | 2.039 | 3.27E-03 | glio |
| Bcas2_in.vivo_noRT | Gm10260 | 2.354 | 3.27E-03 | glio |
| Bora_in.vivo_noRT | Hoxa7 | 2.16 | 3.27E-03 | glio |
| Bora_in.vivo_noRT | Pyroxd2 | 2.004 | 3.27E-03 | glio |
| Cenpj_in.vivo_noRT | Gm48265 | 2.169 | 3.27E-03 | glio |

| | | | | |
|---|---|---|---|---|
| Cenpj_in.vivo_noRT | 9530080O11R | 2.405 | 3.27E-03 | glio |
| Cenpj_in.vivo_noRT | Gsx2 | 2.632 | 3.27E-03 | glio |
| Cenpj_in.vivo_noRT | Fbn2 | -1.352 | 3.27E-03 | glio |
| Cenpj_ex.vivo_noRT | AI463229 | 2.591 | 3.27E-03 | glio |
| Cenpj_ex.vivo_noRT | Ccl19 | 2.591 | 3.27E-03 | glio |
| Chek1_ex.vivo_noRT | Syp | 2.941 | 3.27E-03 | glio |
| Eif2s1_in.vivo_noRT | Gm2115 | 2.143 | 3.27E-03 | glio |
| Fanca_ex.vivo_noRT | Fam131a | 3.309 | 3.27E-03 | glio |
| Fanca_ex.vivo_noRT | Sucnr1 | 2.124 | 3.27E-03 | glio |
| Fanca_ex.vivo_noRT | AI463229 | 2.913 | 3.27E-03 | glio |
| Hsd17b10_in.vivo_noRT | Erfe | 2.075 | 3.27E-03 | glio |
| culture_effect | Taf1b | -1.113 | 3.27E-04 | glio |
| Interaction_genotype_Mtor | Cyp2s1 | -4.496 | 3.27E-04 | glio |
| Interaction_genotype_Mtor | Hist1h2an | -4.162 | 3.27E-04 | glio |
| Interaction_genotype_Mtor | Bdh2 | -4.032 | 3.27E-04 | glio |
| Interaction_genotype_Myc | B630019K06R | -3.242 | 3.27E-04 | glio |
| Aars2_in.vivo_noRT | Tmem30a | -1.239 | 3.27E-04 | glio |
| Atp5c1_in.vivo_noRT | Gm38477 | 2.146 | 3.27E-04 | glio |
| Atr_in.vivo_noRT | Zfp748 | 2.499 | 3.27E-04 | glio |
| Chek1_in.vivo_noRT | Gm2822 | 2.827 | 3.27E-04 | glio |
| Chek1_in.vivo_noRT | Shbg | 2.665 | 3.27E-04 | glio |
| Chek1_in.vivo_noRT | Lmln | 2.508 | 3.27E-04 | glio |
| Chek1_in.vivo_noRT | Lfng | 2.583 | 3.27E-04 | glio |
| Chek1_in.vivo_noRT | Thbs3 | 2.363 | 3.27E-04 | glio |
| Chek1_ex.vivo_noRT | Ybx3 | -0.921 | 3.27E-04 | glio |
| culture_effect | Tpm2 | -0.798 | 3.27E-05 | glio |
| culture_effect | Cmtm4 | 0.67 | 3.27E-05 | glio |
| Interaction_genotype_Aars2 | Gm34885 | -5.279 | 3.27E-05 | glio |
| Interaction_genotype_Mtor | Tnfsf13os | -4.916 | 3.27E-05 | glio |
| Chek1_in.vivo_noRT | Rpusd3 | 2.461 | 3.27E-05 | glio |
| Chek1_in.vivo_noRT | Mrps23 | -1.305 | 3.27E-05 | glio |
| Interaction_genotype_Mtor | Gm26682 | -5.425 | 3.27E-06 | glio |
| Aars2_in.vivo_noRT | Yipf5 | -1.818 | 3.27E-06 | glio |
| Chek1_in.vivo_noRT | Emid1 | 3.651 | 3.27E-06 | glio |
| Chek1_in.vivo_noRT | Hoga1 | 3.229 | 3.27E-06 | glio |
| Interaction_genotype_Mtor | 1700017M07F | -5.251 | 3.27E-07 | glio |
| Interaction_genotype_Mtor | Gm47540 | -4.492 | 3.27E-07 | glio |
| Atr_ex.vivo_noRT | Strip2 | 3.562 | 3.27E-07 | glio |
| Atr_in.vivo_noRT | Zfp990 | 3.632 | 3.27E-08 | glio |
| Bcas2_in.vivo_noRT | D430040D24F | 3.89 | 3.27E-08 | glio |
| Bora_in.vivo_noRT | Slc23a3 | 4.035 | 3.27E-09 | glio |
| Chek1_ex.vivo_noRT | Cacng5 | 4.842 | 3.27E-09 | glio |
| Chek1_ex.vivo_noRT | Nmrk2 | 4.842 | 3.27E-10 | glio |
| Aars2_in.vivo_noRT | Cyp7a1 | 4.536 | 3.27E-11 | glio |
| Chek1_ex.vivo_noRT | 4930447K03R | 4.824 | 3.27E-13 | glio |
| Atr_in.vivo_noRT | Adgre5 | 5.273 | 3.27E-19 | glio |
| Atr_in.vivo_noRT | Gm48899 | 5.273 | 3.27E-19 | glio |
| Bcas2_in.vivo_noRT | Tmem132c | 4.807 | 3.27E-22 | glio |
| culture_effect | Rassf1 | -0.61 | 3.28E-03 | glio |
| culture_effect | Cadm1 | 0.626 | 3.28E-03 | glio |
| Interaction_genotype_Mtor | Rtl9 | -3.368 | 3.28E-03 | glio |
| Interaction_genotype_Mtor | Tes3-ps | -3.563 | 3.28E-03 | glio |

| | | | | |
|---|---|---|---|---|
| Interaction_genotype_Mtor | 5730414N17R | -3.427 | 3.28E-03 | glio |
| Interaction_genotype_Myc | Pax3 | -2.727 | 3.28E-03 | glio |
| Interaction_genotype_Myc | Hist1h2be | -2.915 | 3.28E-03 | glio |
| Interaction_genotype_Myc | Psmc5 | 1.243 | 3.28E-03 | glio |
| Interaction_genotype_Rad51 | 4930478M13F | -4.111 | 3.28E-03 | glio |
| Interaction_genotype_Rpl3 | Phc2 | 1.692 | 3.28E-03 | glio |
| interaction_RT | Gad1 | 2.071 | 3.28E-03 | glio |
| interaction_RT | 2810049E08Ri | 2.715 | 3.28E-03 | glio |
| Aars2_ex.vivo_noRT | Enpp3 | 2.893 | 3.28E-03 | glio |
| Atp5c1_in.vivo_noRT | Lyl1 | 2.018 | 3.28E-03 | glio |
| Atr_in.vivo_noRT | Nipal3 | 2.074 | 3.28E-03 | glio |
| Atr_in.vivo_noRT | Oscp1 | 2.114 | 3.28E-03 | glio |
| Atr_in.vivo_noRT | Zfp213 | 1.698 | 3.28E-03 | glio |
| Atr_ex.vivo_noRT | Ankrd54 | 1.624 | 3.28E-03 | glio |
| Bcas2_ex.vivo_noRT | Zfp352 | 2.742 | 3.28E-03 | glio |
| Bora_ex.vivo_noRT | Fam131a | 2.802 | 3.28E-03 | glio |
| Brca2_in.vivo_noRT | 6430562O15R | 2.553 | 3.28E-03 | glio |
| culture_effect | Opalin | -2.981 | 3.28E-04 | glio |
| Interaction_genotype_Mtor | 4921508D12R | -3.965 | 3.28E-04 | glio |
| Interaction_genotype_Mtor | Traf3ip1 | -3.037 | 3.28E-04 | glio |
| interaction_RT | Fez1 | 2.098 | 3.28E-04 | glio |
| interaction_RT | Dynlrb1 | 0.678 | 3.28E-04 | glio |
| interaction_RT | Cxcr2 | 3.34 | 3.28E-04 | glio |
| Aars2_in.vivo_noRT | Gm12940 | 2.185 | 3.28E-04 | glio |
| Atr_in.vivo_noRT | Prkcq | 2.154 | 3.28E-04 | glio |
| Atr_in.vivo_noRT | Stk32c | 2.341 | 3.28E-04 | glio |
| Atr_ex.vivo_noRT | Tpsab1 | 3.164 | 3.28E-04 | glio |
| Atr_ex.vivo_noRT | Atp1a2 | 3.562 | 3.28E-04 | glio |
| Atr_ex.vivo_noRT | Hnrnpa1 | -0.927 | 3.28E-04 | glio |
| Bcas2_in.vivo_noRT | Anxa11 | 2.916 | 3.28E-04 | glio |
| Bora_in.vivo_noRT | Mrfap1 | -0.709 | 3.28E-04 | glio |
| Chek1_ex.vivo_noRT | Nucks1 | -1.141 | 3.28E-04 | glio |
| Interaction_genotype_Mtor | Pecam1 | -5.029 | 3.28E-05 | glio |
| Atr_in.vivo_noRT | Gm5577 | 2.689 | 3.28E-05 | glio |
| Atr_ex.vivo_noRT | Gm2449 | 3.239 | 3.28E-05 | glio |
| Chek1_in.vivo_noRT | Gm15486 | 2.672 | 3.28E-05 | glio |
| Chek1_in.vivo_noRT | A130051J06Ri | 2.82 | 3.28E-05 | glio |
| Interaction_genotype_Mtor | Spata33 | -5.847 | 3.28E-06 | glio |
| interaction_RT | Gm13889 | 4.17 | 3.28E-06 | glio |
| interaction_RT | Ndufa7 | 0.741 | 3.28E-06 | glio |
| Aars2_ex.vivo_noRT | Edn2 | 2.897 | 3.28E-06 | glio |
| Bora_in.vivo_noRT | Gm36495 | 3.642 | 3.28E-06 | glio |
| Hsd17b10_in.vivo_noRT | Gm29083 | 3.084 | 3.28E-06 | glio |
| culture_effect | 1110008P14R | 0.792 | 3.28E-07 | glio |
| Interaction_genotype_Myc | Sema6b | -5.744 | 3.28E-07 | glio |
| Atr_in.vivo_noRT | Anks6 | 3.597 | 3.28E-08 | glio |
| Bcas2_in.vivo_noRT | 4930579K19R | 4.098 | 3.28E-08 | glio |
| Chek1_ex.vivo_noRT | Gm20442 | 4.244 | 3.28E-08 | glio |
| Bcas2_in.vivo_noRT | 9930111J21Ri | 3.627 | 3.28E-09 | glio |
| Bcas2_in.vivo_noRT | Mycl | 4.285 | 3.28E-09 | glio |
| culture_effect | Hrh2 | -4.478 | 3.28E-18 | glio |
| Atr_in.vivo_noRT | Gm31415 | 5.267 | 3.28E-19 | glio |

| | | | | |
|---|---|---|---|---|
| Bora_in.vivo_noRT | Gm10447 | 4.936 | 3.28E-19 | glio |
| culture_effect | Rack1 | -1.503 | 3.28E-27 | glio |
| Interaction_genotype_Atr | A930002C04R | -2.509 | 3.29E-03 | glio |
| Interaction_genotype_Atr | Btn1a1 | -2.894 | 3.29E-03 | glio |
| Interaction_genotype_Atr | 1700003E24Ri | -2.369 | 3.29E-03 | glio |
| Interaction_genotype_Atr | Gm17056 | -3.764 | 3.29E-03 | glio |
| Interaction_genotype_Atr | Gm48610 | -3.655 | 3.29E-03 | glio |
| Interaction_genotype_Atr | A930004D18R | -3.688 | 3.29E-03 | glio |
| Interaction_genotype_Atr | Rad1 | 1.901 | 3.29E-03 | glio |
| Interaction_genotype_Mtor | Map1lc3b | 1.557 | 3.29E-03 | glio |
| Interaction_genotype_Myc | Magee1 | -2.763 | 3.29E-03 | glio |
| Interaction_genotype_Myc | Qrsl1 | -2.421 | 3.29E-03 | glio |
| Aars2_in.vivo_noRT | Mcm4 | -1.321 | 3.29E-03 | glio |
| Atr_in.vivo_noRT | St13 | -0.749 | 3.29E-03 | glio |
| Atr_ex.vivo_noRT | Dact1 | 2.764 | 3.29E-03 | glio |
| Bcas2_in.vivo_noRT | Afmid | 1.763 | 3.29E-03 | glio |
| Bora_in.vivo_noRT | Socs7 | -1.262 | 3.29E-03 | glio |
| Hsd17b10_in.vivo_noRT | Olfr1189 | 1.961 | 3.29E-03 | glio |
| culture_effect | Utp4 | -0.574 | 3.29E-04 | glio |
| Interaction_genotype_Mtor | Tcp11l1 | -3.057 | 3.29E-04 | glio |
| Aars2_in.vivo_noRT | AI429214 | 2.57 | 3.29E-04 | glio |
| Aars2_in.vivo_noRT | Gm10282 | 2.55 | 3.29E-04 | glio |
| Aars2_in.vivo_noRT | Mob3b | 2.22 | 3.29E-04 | glio |
| Atr_in.vivo_noRT | Ccar1 | -1.025 | 3.29E-04 | glio |
| Atr_in.vivo_noRT | Set | -0.778 | 3.29E-04 | glio |
| Bcas2_in.vivo_noRT | Prss22 | 3.209 | 3.29E-04 | glio |
| Bora_ex.vivo_noRT | Ciita | 2.802 | 3.29E-04 | glio |
| Hsd17b10_in.vivo_noRT | Zfp968 | 2.766 | 3.29E-04 | glio |
| Interaction_genotype_Mtor | Rbm3os | -4.777 | 3.29E-05 | glio |
| Interaction_genotype_Mtor | BC037704 | -4.502 | 3.29E-05 | glio |
| Interaction_genotype_Mtor | 5033430I15Ri | -4.455 | 3.29E-05 | glio |
| Aars2_in.vivo_noRT | Mroh2b | 2.999 | 3.29E-05 | glio |
| Atr_ex.vivo_noRT | Pold2 | -1.98 | 3.29E-05 | glio |
| Bcas2_in.vivo_noRT | Tcte2 | 2.685 | 3.29E-05 | glio |
| Chek1_ex.vivo_noRT | Lgals7 | 4.247 | 3.29E-05 | glio |
| Hsd17b10_in.vivo_noRT | Noxred1 | 2.888 | 3.29E-05 | glio |
| Interaction_genotype_Mtor | Olfr718-ps1 | -4.229 | 3.29E-06 | glio |
| Aars2_in.vivo_noRT | Rimkla | 3.108 | 3.29E-06 | glio |
| Aars2_ex.vivo_noRT | 4930570E03Ri | 3.293 | 3.29E-06 | glio |
| Aars2_ex.vivo_noRT | Gm5286 | 3.293 | 3.29E-06 | glio |
| Aars2_ex.vivo_noRT | B230206H07R | 2.897 | 3.29E-06 | glio |
| Atr_ex.vivo_noRT | Gm47872 | 3.952 | 3.29E-06 | glio |
| culture_effect | Plekha3 | 0.873 | 3.29E-07 | glio |
| Atp5c1_in.vivo_noRT | Vmn2r44 | 2.668 | 3.29E-07 | glio |
| Atp5c1_in.vivo_noRT | Mmp13 | 2.668 | 3.29E-07 | glio |
| culture_effect | Pip4p2 | 0.805 | 3.29E-08 | glio |
| Aars2_in.vivo_noRT | Tbkbp1 | 4.033 | 3.29E-08 | glio |
| Aars2_in.vivo_noRT | Srl | 3.908 | 3.29E-08 | glio |
| Aars2_in.vivo_noRT | Gm10516 | 3.763 | 3.29E-08 | glio |
| culture_effect | Gng5 | 0.649 | 3.29E-09 | glio |
| Chek1_ex.vivo_noRT | Serpina3i | 4.842 | 3.29E-09 | glio |
| Aars2_in.vivo_noRT | Esyt3 | 4.267 | 3.29E-10 | glio |

| | | | | |
|---|---|---|---|---|
| Chek1_ex.vivo_noRT | Abra | 4.842 | 3.29E-11 | glio |
| Atr_in.vivo_noRT | Ccdc114 | 4.615 | 3.29E-12 | glio |
| culture_effect | Mest | 4.645 | 3.29E-42 | glio |
| Interaction_genotype_Cdkn2a | Gm6566 | 3.62 | 3.30E-03 | glio |
| Interaction_genotype_Chek1 | Csf1 | 2.977 | 3.30E-03 | glio |
| Interaction_genotype_Mtor | Synpo2l | -3.057 | 3.30E-03 | glio |
| Interaction_genotype_Mtor | Gm16973 | -3.666 | 3.30E-03 | glio |
| Interaction_genotype_Mtor | AW146154 | -3.554 | 3.30E-03 | glio |
| Interaction_genotype_Mtor | Hagh | 1.542 | 3.30E-03 | glio |
| Interaction_genotype_Myc | A330076H08R | -2.729 | 3.30E-03 | glio |
| Interaction_genotype_Myc | Gm41492 | -2.93 | 3.30E-03 | glio |
| Interaction_genotype_Myc | Ogdhl | -3.273 | 3.30E-03 | glio |
| interaction_RT | Fam210b | 0.983 | 3.30E-03 | glio |
| Aars2_in.vivo_noRT | 4930532G15R | 1.931 | 3.30E-03 | glio |
| Aars2_ex.vivo_noRT | Grip2 | 2.907 | 3.30E-03 | glio |
| Atr_in.vivo_noRT | Ppp1r8 | -1.386 | 3.30E-03 | glio |
| Bora_in.vivo_noRT | B230206L02Ri | 2.134 | 3.30E-03 | glio |
| Bora_in.vivo_noRT | Acvr1b | 2.148 | 3.30E-03 | glio |
| Bora_in.vivo_noRT | Lhx6 | 2.044 | 3.30E-03 | glio |
| Bora_ex.vivo_noRT | Cldn10 | 3.008 | 3.30E-03 | glio |
| Cenpj_ex.vivo_noRT | Slc19a3 | 2.198 | 3.30E-03 | glio |
| Chek1_ex.vivo_noRT | Ccdc114 | 2.586 | 3.30E-03 | glio |
| culture_effect | Trappc1 | -0.531 | 3.30E-04 | glio |
| Interaction_genotype_Atr | Zfp551 | -4.42 | 3.30E-04 | glio |
| Interaction_genotype_Mtor | Ankrd23 | -4.564 | 3.30E-04 | glio |
| Interaction_genotype_Myc | Cpxm1 | -3.568 | 3.30E-04 | glio |
| Interaction_genotype_Myc | Cep85 | -3.898 | 3.30E-04 | glio |
| Aars2_in.vivo_noRT | 4921531C22R | 2.485 | 3.30E-04 | glio |
| Atp5c1_in.vivo_noRT | Car14 | 2.668 | 3.30E-04 | glio |
| Bcas2_in.vivo_noRT | Bpifc | 2.787 | 3.30E-04 | glio |
| Bcas2_in.vivo_noRT | Dock6 | 2.55 | 3.30E-04 | glio |
| Interaction_genotype_Myc | Gm19325 | -4.899 | 3.30E-05 | glio |
| Interaction_genotype_Myc | 5033417F24Ri | -4.554 | 3.30E-05 | glio |
| Bcas2_in.vivo_noRT | Itga3 | 3.178 | 3.30E-05 | glio |
| Bora_in.vivo_noRT | Sbk2 | 2.97 | 3.30E-05 | glio |
| Chek1_in.vivo_noRT | Btbd6 | 2.684 | 3.30E-05 | glio |
| Eif2s1_in.vivo_noRT | Gpr22 | 2.67 | 3.30E-05 | glio |
| Eif2s1_in.vivo_noRT | Sema6d | 2.67 | 3.30E-05 | glio |
| Hsd17b10_in.vivo_noRT | Gm41764 | 2.699 | 3.30E-05 | glio |
| Hsd17b10_in.vivo_noRT | 2300002M23F | 2.447 | 3.30E-05 | glio |
| culture_effect | Ube2f | 0.721 | 3.30E-06 | glio |
| Interaction_genotype_Mtor | Lyrm1 | -5.34 | 3.30E-06 | glio |
| Interaction_genotype_Mtor | Arpc5 | 2.096 | 3.30E-06 | glio |
| Aars2_in.vivo_noRT | Zfp960 | 3.432 | 3.30E-06 | glio |
| Aars2_in.vivo_noRT | Gm15486 | 2.955 | 3.30E-06 | glio |
| Chek1_in.vivo_noRT | Rbm11 | 3.112 | 3.30E-06 | glio |
| Chek1_ex.vivo_noRT | Pard3bos1 | 3.664 | 3.30E-06 | glio |
| culture_effect | Scd2 | 0.906 | 3.30E-07 | glio |
| Interaction_genotype_Mtor | Rragd | -6.492 | 3.30E-07 | glio |
| Aars2_in.vivo_noRT | Cd38 | 3.506 | 3.30E-07 | glio |
| Atp5c1_in.vivo_noRT | Gm8016 | 2.668 | 3.30E-07 | glio |
| Bora_in.vivo_noRT | EU599041 | 3.659 | 3.30E-07 | glio |

| | | | | |
|---|---|---|---|---|
| Bora_in.vivo_noRT | Cyp2u1 | 3.357 | 3.30E-07 | glio |
| Chek1_in.vivo_noRT | Dcpp1 | 3.717 | 3.30E-07 | glio |
| culture_effect | Msn | 0.752 | 3.30E-08 | glio |
| Bcas2_in.vivo_noRT | Insc | 3.35 | 3.30E-08 | glio |
| Chek1_ex.vivo_noRT | Agap2 | 3.857 | 3.30E-08 | glio |
| Aars2_in.vivo_noRT | Pla2g2f | 3.684 | 3.30E-09 | glio |
| Chek1_in.vivo_noRT | Gm47232 | 4.521 | 3.30E-11 | glio |
| Bora_in.vivo_noRT | Gm45606 | 4.676 | 3.30E-12 | glio |
| Bcas2_in.vivo_noRT | Gm2115 | 4.807 | 3.30E-15 | glio |
| Aars2_in.vivo_noRT | C330024C12R | 4.936 | 3.30E-16 | glio |
| Bora_in.vivo_noRT | Gm10354 | 4.936 | 3.30E-18 | glio |
| culture_effect | mt-Nd3 | 1.583 | 3.30E-22 | glio |
| Bcas2_in.vivo_noRT | Gm44693 | 4.807 | 3.30E-22 | glio |
| culture_effect | Tesk2 | 1.284 | 3.31E-03 | glio |
| culture_effect | Mycbp2 | -0.49 | 3.31E-03 | glio |
| Interaction_genotype_Aars2 | Sult6b1 | -3.662 | 3.31E-03 | glio |
| Interaction_genotype_Atr | 1700029J07Ri | -3.67 | 3.31E-03 | glio |
| Interaction_genotype_Bcas2 | Elobl | -4.199 | 3.31E-03 | glio |
| Interaction_genotype_Bcas2 | Dock8 | -3.125 | 3.31E-03 | glio |
| Interaction_genotype_Bora | Gm15890 | -3.298 | 3.31E-03 | glio |
| Interaction_genotype_Bora | Borcs8 | 1.592 | 3.31E-03 | glio |
| Interaction_genotype_Mtor | Ctsf | -2.824 | 3.31E-03 | glio |
| Interaction_genotype_Mtor | Gm10644 | -3.626 | 3.31E-03 | glio |
| Interaction_genotype_Mtor | Gls2 | -3.515 | 3.31E-03 | glio |
| Interaction_genotype_Mtor | Gm32618 | -3.425 | 3.31E-03 | glio |
| Interaction_genotype_Mtor | Alyref | 1.568 | 3.31E-03 | glio |
| Interaction_genotype_Myc | Dlx4 | -3.169 | 3.31E-03 | glio |
| Interaction_genotype_Myc | Mcc | -2.827 | 3.31E-03 | glio |
| Interaction_genotype_Myc | Cyb5rl | -2.677 | 3.31E-03 | glio |
| Aars2_in.vivo_noRT | S100a4 | -1.884 | 3.31E-03 | glio |
| Atr_ex.vivo_noRT | Pgk1 | -0.796 | 3.31E-03 | glio |
| Bcas2_in.vivo_noRT | Tmem214 | -1.342 | 3.31E-03 | glio |
| Bcas2_in.vivo_noRT | Itgb1bp1 | -1.161 | 3.31E-03 | glio |
| Bora_in.vivo_noRT | Ttbk1 | 2.139 | 3.31E-03 | glio |
| Bora_in.vivo_noRT | Cnpy4 | -1.35 | 3.31E-03 | glio |
| Chek1_in.vivo_noRT | Zkscan7 | 2.19 | 3.31E-03 | glio |
| Eif2s1_in.vivo_noRT | Gm10324 | 1.877 | 3.31E-03 | glio |
| Eif2s1_in.vivo_noRT | Gm36026 | 2.023 | 3.31E-03 | glio |
| culture_effect | Pbx3 | -0.736 | 3.31E-04 | glio |
| Interaction_genotype_Mtor | Gm5039 | -4.887 | 3.31E-04 | glio |
| Interaction_genotype_Mtor | Cd38 | -4.236 | 3.31E-04 | glio |
| Aars2_in.vivo_noRT | Preb | -1.462 | 3.31E-04 | glio |
| Atp5c1_in.vivo_noRT | Gm11457 | 2.668 | 3.31E-04 | glio |
| Bora_in.vivo_noRT | Hmmr | -2.249 | 3.31E-04 | glio |
| Chek1_in.vivo_noRT | Uba1 | -1.308 | 3.31E-04 | glio |
| culture_effect | Myl9 | 2.368 | 3.31E-05 | glio |
| Interaction_genotype_Myc | 2310022A10R | -3.379 | 3.31E-05 | glio |
| Interaction_genotype_Myc | Macrod2 | -3.564 | 3.31E-05 | glio |
| Interaction_genotype_Myc | 2900052L18Ri | -3.836 | 3.31E-05 | glio |
| Interaction_genotype_Myc | 9430037G07R | -4.869 | 3.31E-05 | glio |
| Aars2_in.vivo_noRT | Nbeal2 | 3.037 | 3.31E-05 | glio |
| Aars2_in.vivo_noRT | 2310030G06R | 2.957 | 3.31E-05 | glio |

| | | | | |
|---|---|---|---|---|
| Aars2_in.vivo_noRT | Srebf2 | -2.143 | 3.31E-05 | glio |
| Aars2_in.vivo_noRT | Psma7 | -0.882 | 3.31E-05 | glio |
| culture_effect | Gm16046 | 2.519 | 3.31E-06 | glio |
| culture_effect | Nudt8 | 1.032 | 3.31E-06 | glio |
| Aars2_in.vivo_noRT | Rab5c | -1.626 | 3.31E-06 | glio |
| Bcas2_in.vivo_noRT | Fsbp | 3.363 | 3.31E-06 | glio |
| Chek1_ex.vivo_noRT | Gm43625 | 3.657 | 3.31E-06 | glio |
| Interaction_genotype_Mtor | Gm36172 | -4.934 | 3.31E-07 | glio |
| culture_effect | Stat4 | -3.388 | 3.31E-08 | glio |
| interaction_RT | Sat1 | 1.801 | 3.31E-08 | glio |
| interaction_RT | Ncl | -1.074 | 3.31E-08 | glio |
| Atr_in.vivo_noRT | Slc12a5 | 3.752 | 3.31E-08 | glio |
| Aars2_ex.vivo_noRT | Slc17a6 | 3.293 | 3.31E-09 | glio |
| Chek1_in.vivo_noRT | Pecam1 | 4.511 | 3.31E-10 | glio |
| Aars2_in.vivo_noRT | Gm13404 | 5.103 | 3.31E-13 | glio |
| Aars2_in.vivo_noRT | Myo5c | 5.327 | 3.31E-15 | glio |
| Bcas2_in.vivo_noRT | Gm44658 | 4.793 | 3.31E-15 | glio |
| Chek1_in.vivo_noRT | 4930527F14Ri | 4.515 | 3.31E-17 | glio |
| Atr_in.vivo_noRT | Capn9 | 5.533 | 3.31E-18 | glio |
| Atp5c1_ex.vivo_noRT | Atp5c1 | -2.956 | 3.31E-23 | glio |
| Atr_in.vivo_noRT | Olfr810 | 5.533 | 3.31E-30 | glio |
| culture_effect | Ankrd34a | 1.661 | 3.32E-03 | glio |
| culture_effect | Oga | -0.427 | 3.32E-03 | glio |
| Interaction_genotype_Aars2 | Gm10767 | -3.74 | 3.32E-03 | glio |
| Interaction_genotype_Aars2 | Psma7 | 0.954 | 3.32E-03 | glio |
| Interaction_genotype_Atr | 1700084E18Ri | -3.651 | 3.32E-03 | glio |
| Interaction_genotype_Atr | Poglut3 | -3.058 | 3.32E-03 | glio |
| Interaction_genotype_Bora | Cysrt1 | -4.03 | 3.32E-03 | glio |
| Interaction_genotype_Chek1 | Smim13 | 2.377 | 3.32E-03 | glio |
| Interaction_genotype_Mtor | Rnf208 | -3.808 | 3.32E-03 | glio |
| Interaction_genotype_Mtor | Lmtk3 | -3.352 | 3.32E-03 | glio |
| Interaction_genotype_Mtor | Brf1 | -2.632 | 3.32E-03 | glio |
| Interaction_genotype_Mtor | Synm | -2.832 | 3.32E-03 | glio |
| Interaction_genotype_Pars2 | Alox12 | -3.922 | 3.32E-03 | glio |
| Interaction_genotype_Rad21 | 1700025N23R | 3.943 | 3.32E-03 | glio |
| Aars2_in.vivo_noRT | Fam13c | 2.296 | 3.32E-03 | glio |
| Atp5c1_in.vivo_noRT | D030051J21Ri | 2.293 | 3.32E-03 | glio |
| Atr_in.vivo_noRT | Lrrc73 | 1.706 | 3.32E-03 | glio |
| Atr_in.vivo_noRT | Wls | -1.264 | 3.32E-03 | glio |
| Atr_ex.vivo_noRT | Zfp568 | 1.915 | 3.32E-03 | glio |
| Bard1_ex.vivo_noRT | Tcstv3 | 2.814 | 3.32E-03 | glio |
| Cenpj_in.vivo_noRT | Cd7 | 1.758 | 3.32E-03 | glio |
| Chek1_in.vivo_noRT | Bend6 | 2.113 | 3.32E-03 | glio |
| Chek1_in.vivo_noRT | Pcbp3 | 1.835 | 3.32E-03 | glio |
| Chek1_in.vivo_noRT | Nek7 | -1.571 | 3.32E-03 | glio |
| Chek1_ex.vivo_noRT | Ppp2r3c | -1.545 | 3.32E-03 | glio |
| Eif2s1_in.vivo_noRT | Gm10644 | 2.485 | 3.32E-03 | glio |
| culture_effect | Phf11a | -2.239 | 3.32E-04 | glio |
| culture_effect | Sf3b3 | -0.472 | 3.32E-04 | glio |
| Interaction_genotype_Aars2 | Poldip2 | 1.781 | 3.32E-04 | glio |
| Interaction_genotype_Mtor | A830009L08Ri | -3.896 | 3.32E-04 | glio |
| Interaction_genotype_Myc | Tomm40l | -3.222 | 3.32E-04 | glio |

| | | | | |
|---|---|---|---|---|
| interaction_RT | Isg20 | 1.083 | 3.32E-04 | glio |
| Aars2_in.vivo_noRT | Card14 | 2.615 | 3.32E-04 | glio |
| Aars2_ex.vivo_noRT | Snhg4.1 | 2.9 | 3.32E-04 | glio |
| Bard1_ex.vivo_noRT | Gm13610 | 3.4 | 3.32E-04 | glio |
| Bcas2_in.vivo_noRT | Dner | 2.386 | 3.32E-04 | glio |
| Bora_ex.vivo_noRT | Lhx1os | 2.806 | 3.32E-04 | glio |
| Bora_ex.vivo_noRT | Brsk2 | 3.201 | 3.32E-04 | glio |
| Bora_ex.vivo_noRT | Rpl3l | 3.201 | 3.32E-04 | glio |
| Chek1_in.vivo_noRT | Slc7a7 | 2.558 | 3.32E-04 | glio |
| Hsd17b10_in.vivo_noRT | Srl | 2.724 | 3.32E-04 | glio |
| Interaction_genotype_Myc | Lamp1 | 1.665 | 3.32E-05 | glio |
| Atr_in.vivo_noRT | Abhd18 | 2.994 | 3.32E-05 | glio |
| Atr_ex.vivo_noRT | Slc6a11 | 3.164 | 3.32E-05 | glio |
| Atr_ex.vivo_noRT | Gm15138 | 2.769 | 3.32E-05 | glio |
| Atr_ex.vivo_noRT | Sema3a | 3.568 | 3.32E-05 | glio |
| Atr_ex.vivo_noRT | Map3k8 | 3.973 | 3.32E-05 | glio |
| Bcas2_in.vivo_noRT | Rex2 | 3.133 | 3.32E-05 | glio |
| Bora_in.vivo_noRT | Gm36608 | 3.006 | 3.32E-05 | glio |
| Bora_ex.vivo_noRT | Gm15138 | 2.806 | 3.32E-05 | glio |
| Aars2_in.vivo_noRT | Zfp799 | 3.209 | 3.32E-06 | glio |
| Atr_in.vivo_noRT | Gm20337 | 3.315 | 3.32E-06 | glio |
| Bora_in.vivo_noRT | Pde4d | 3.285 | 3.32E-06 | glio |
| Bcas2_in.vivo_noRT | Zfp473 | 3.383 | 3.32E-07 | glio |
| Bora_in.vivo_noRT | Gm50218 | 3.875 | 3.32E-07 | glio |
| Atr_in.vivo_noRT | Tpt1 | -1.311 | 3.32E-08 | glio |
| Bcas2_in.vivo_noRT | Hif3a | 4.165 | 3.32E-09 | glio |
| Bora_in.vivo_noRT | Spdye4a | 4.279 | 3.32E-09 | glio |
| Bora_in.vivo_noRT | Pard3bos1 | 3.939 | 3.32E-10 | glio |
| Bora_in.vivo_noRT | B930094E09Ri | 4.42 | 3.32E-10 | glio |
| Chek1_ex.vivo_noRT | Dapk2 | 4.045 | 3.32E-10 | glio |
| Aars2_in.vivo_noRT | Gm20732 | 4.556 | 3.32E-13 | glio |
| culture_effect | Dclk1 | 1.573 | 3.32E-14 | glio |
| Atr_in.vivo_noRT | B4galnt1 | 5.273 | 3.32E-19 | glio |
| Chek1_in.vivo_noRT | Gm14929 | 5.044 | 3.32E-24 | glio |
| culture_effect | Klk7 | -2.614 | 3.33E-03 | glio |
| culture_effect | 4921523L03Ri | 1.752 | 3.33E-03 | glio |
| culture_effect | Zbtb11 | -0.71 | 3.33E-03 | glio |
| culture_effect | Mmp15 | -0.828 | 3.33E-03 | glio |
| culture_effect | Rrbp1 | 0.347 | 3.33E-03 | glio |
| culture_effect | Cox6c | 0.364 | 3.33E-03 | glio |
| Interaction_genotype_Atr | Ube2q1 | 1.617 | 3.33E-03 | glio |
| Interaction_genotype_Bora | Kctd1 | -3.745 | 3.33E-03 | glio |
| Interaction_genotype_Mtor | Gm43331 | -3.049 | 3.33E-03 | glio |
| Interaction_genotype_Myc | Hist1h2bp | -3.531 | 3.33E-03 | glio |
| Interaction_genotype_Myc | Tmem62 | -3.047 | 3.33E-03 | glio |
| interaction_RT | Mex3d | 1.032 | 3.33E-03 | glio |
| Aars2_in.vivo_noRT | Slc16a13 | 1.836 | 3.33E-03 | glio |
| Aars2_in.vivo_noRT | Eif2d | -1.342 | 3.33E-03 | glio |
| Atp5c1_in.vivo_noRT | Nat8f4 | 2.136 | 3.33E-03 | glio |
| Atr_in.vivo_noRT | Klhl1 | 2.002 | 3.33E-03 | glio |
| Atr_in.vivo_noRT | Fbxl8 | 1.645 | 3.33E-03 | glio |
| Atr_ex.vivo_noRT | 1700017M07F | 2.391 | 3.33E-03 | glio |

| | | | | |
|---|---|---|---|---|
| Bcas2_in.vivo_noRT | Dnajc30 | -1.463 | 3.33E-03 | glio |
| Bcas2_in.vivo_noRT | Syngr2 | -1.125 | 3.33E-03 | glio |
| Bora_in.vivo_noRT | B3gntl1 | 2.086 | 3.33E-03 | glio |
| Bora_in.vivo_noRT | Mns1 | -1.392 | 3.33E-03 | glio |
| Bora_ex.vivo_noRT | AC168977.1 | 3.355 | 3.33E-03 | glio |
| Cenpj_in.vivo_noRT | Pldi | 2.022 | 3.33E-03 | glio |
| Chek1_in.vivo_noRT | Sav1 | -1.38 | 3.33E-03 | glio |
| Eif2s1_in.vivo_noRT | Gm14019 | 2.143 | 3.33E-03 | glio |
| Eif2s1_ex.vivo_noRT | Glyat | 2.235 | 3.33E-03 | glio |
| culture_effect | Rab11fip2 | 0.95 | 3.33E-04 | glio |
| culture_effect | Bag3 | -0.684 | 3.33E-04 | glio |
| Interaction_genotype_Atr | Gm4221 | -4.504 | 3.33E-04 | glio |
| Interaction_genotype_Mtor | Fam71f2 | -4.505 | 3.33E-04 | glio |
| Interaction_genotype_Mtor | Pate2 | -4.286 | 3.33E-04 | glio |
| Interaction_genotype_Mtor | Rpl5 | 1.326 | 3.33E-04 | glio |
| Interaction_genotype_Myc | Slc16a4 | -3.321 | 3.33E-04 | glio |
| Aars2_in.vivo_noRT | Fgf15 | 2.758 | 3.33E-04 | glio |
| Aars2_in.vivo_noRT | Gm45494 | 2.449 | 3.33E-04 | glio |
| Aars2_ex.vivo_noRT | Rasl10a | 2.9 | 3.33E-04 | glio |
| Atr_in.vivo_noRT | Stradb | 2.108 | 3.33E-04 | glio |
| Atr_ex.vivo_noRT | Gm17276 | 2.769 | 3.33E-04 | glio |
| Bcas2_in.vivo_noRT | Ccn2 | 3.48 | 3.33E-04 | glio |
| Bcas2_in.vivo_noRT | Cldn10 | 2.6 | 3.33E-04 | glio |
| Bcas2_ex.vivo_noRT | Crb3 | 2.742 | 3.33E-04 | glio |
| Bcas2_ex.vivo_noRT | Gm17210 | 3.576 | 3.33E-04 | glio |
| Chek1_in.vivo_noRT | Srcin1 | 2.47 | 3.33E-04 | glio |
| Eif2s1_in.vivo_noRT | Rgs4 | 2.143 | 3.33E-04 | glio |
| Aars2_ex.vivo_noRT | Gm45336 | 3.293 | 3.33E-05 | glio |
| Aars2_ex.vivo_noRT | Fam131a | 3.693 | 3.33E-05 | glio |
| Bora_ex.vivo_noRT | Magea10 | 3.414 | 3.33E-05 | glio |
| Bora_ex.vivo_noRT | Gm36026 | 3.201 | 3.33E-05 | glio |
| Chek1_in.vivo_noRT | Maob | 2.83 | 3.33E-05 | glio |
| Interaction_genotype_Mtor | Gm11963 | -5.503 | 3.33E-06 | glio |
| Interaction_genotype_Myc | Cuta | 1.853 | 3.33E-06 | glio |
| Aars2_in.vivo_noRT | Gm16133 | 3.344 | 3.33E-06 | glio |
| Aars2_in.vivo_noRT | Gm11713 | 2.898 | 3.33E-06 | glio |
| Atr_in.vivo_noRT | 0610038B21R | 2.987 | 3.33E-06 | glio |
| Bcas2_in.vivo_noRT | Haglr | 3.555 | 3.33E-06 | glio |
| Bcas2_ex.vivo_noRT | Rpl22l1 | 1.074 | 3.33E-06 | glio |
| culture_effect | Slc22a4 | 1.163 | 3.33E-07 | glio |
| Bcas2_in.vivo_noRT | 1810062O18R | 3.739 | 3.33E-07 | glio |
| Bora_ex.vivo_noRT | Myoz1 | 3.994 | 3.33E-07 | glio |
| Bora_ex.vivo_noRT | Gm5415 | 3.201 | 3.33E-07 | glio |
| Atr_in.vivo_noRT | Spata33 | 4.009 | 3.33E-08 | glio |
| Chek1_ex.vivo_noRT | Gm49441 | 4.055 | 3.33E-08 | glio |
| Aars2_in.vivo_noRT | Spink2 | 4.666 | 3.33E-09 | glio |
| Bcas2_in.vivo_noRT | Spns3 | 4.149 | 3.33E-09 | glio |
| Atr_in.vivo_noRT | Gm13625 | 4.013 | 3.33E-10 | glio |
| Atr_in.vivo_noRT | 1700003G18R | 5.273 | 3.33E-11 | glio |
| Hsd17b10_in.vivo_noRT | A930019D19R | 3.616 | 3.33E-14 | glio |
| Bora_in.vivo_noRT | Map3k9 | 4.676 | 3.33E-16 | glio |
| culture_effect | Osr2 | -5.009 | 3.33E-35 | glio |

| | | | | |
|---|---|---|---|---|
| Interaction_genotype_Bora | Cyp3a13 | -4.074 | 3.34E-03 | glio |
| Interaction_genotype_Bora | Pspn | -2.654 | 3.34E-03 | glio |
| Interaction_genotype_Myc | Gm10184 | -3.134 | 3.34E-03 | glio |
| Interaction_genotype_Myc | Stx16 | -1.981 | 3.34E-03 | glio |
| Interaction_genotype_Rad21 | Clca3a1 | 4.056 | 3.34E-03 | glio |
| Interaction_genotype_Rad21 | Mgll | 4.543 | 3.34E-03 | glio |
| interaction_RT | B230311B06R | 2.848 | 3.34E-03 | glio |
| interaction_RT | Nr1d1 | 1.387 | 3.34E-03 | glio |
| interaction_RT | Foxm1 | -1.004 | 3.34E-03 | glio |
| interaction_RT | Ncln | -0.747 | 3.34E-03 | glio |
| interaction_RT | Tmem205 | 0.734 | 3.34E-03 | glio |
| interaction_RT | Rps8 | -0.622 | 3.34E-03 | glio |
| Aars2_in.vivo_noRT | Lipe | 2.104 | 3.34E-03 | glio |
| Aars2_in.vivo_noRT | Tmem59l | 1.919 | 3.34E-03 | glio |
| Aars2_in.vivo_noRT | Plxna1 | 1.642 | 3.34E-03 | glio |
| Aars2_in.vivo_noRT | Cops5 | -0.971 | 3.34E-03 | glio |
| Aars2_in.vivo_noRT | Pdia6 | -0.879 | 3.34E-03 | glio |
| Bcas2_in.vivo_noRT | M6pr | -0.915 | 3.34E-03 | glio |
| Bora_in.vivo_noRT | Gm12981 | 2.149 | 3.34E-03 | glio |
| Bora_in.vivo_noRT | 4921531C22R | 2.095 | 3.34E-03 | glio |
| Bora_in.vivo_noRT | Ccdc73 | 1.909 | 3.34E-03 | glio |
| Bora_in.vivo_noRT | Nhlrc3 | 1.779 | 3.34E-03 | glio |
| Bora_ex.vivo_noRT | Gem | 2.808 | 3.34E-03 | glio |
| Brca2_ex.vivo_noRT | Trim17 | 2.42 | 3.34E-03 | glio |
| Chek1_in.vivo_noRT | B230216N24R | 2.37 | 3.34E-03 | glio |
| Cyp19a1_ex.vivo_noRT | Gm15908 | 2.159 | 3.34E-03 | glio |
| Cyp19a1_ex.vivo_noRT | A930014E10Ri | 2.159 | 3.34E-03 | glio |
| culture_effect | Nt5e | -2.02 | 3.34E-04 | glio |
| Interaction_genotype_Atr | Cdkn1a | 2.031 | 3.34E-04 | glio |
| Interaction_genotype_Mtor | Tafa2 | -4.096 | 3.34E-04 | glio |
| Interaction_genotype_Myc | Abat | -4.416 | 3.34E-04 | glio |
| Interaction_genotype_Myc | Slfn3 | -3.641 | 3.34E-04 | glio |
| Interaction_genotype_Myc | Gm17435 | -3.901 | 3.34E-04 | glio |
| Interaction_genotype_Myc | Cers6 | -2.367 | 3.34E-04 | glio |
| Interaction_genotype_Myc | Ranbp1 | 1.549 | 3.34E-04 | glio |
| Aars2_in.vivo_noRT | Npr2 | 2.915 | 3.34E-04 | glio |
| Aars2_in.vivo_noRT | Pon3 | 2.576 | 3.34E-04 | glio |
| Aars2_ex.vivo_noRT | Tcstv1 | 2.893 | 3.34E-04 | glio |
| Atr_in.vivo_noRT | Rps27a | -0.713 | 3.34E-04 | glio |
| Cenpj_ex.vivo_noRT | Dpp6 | 2.591 | 3.34E-04 | glio |
| Cenpj_ex.vivo_noRT | Serpinb9g | 2.591 | 3.34E-04 | glio |
| Cenpj_ex.vivo_noRT | BC049987 | 2.591 | 3.34E-04 | glio |
| Chek1_ex.vivo_noRT | Gm48933 | 2.987 | 3.34E-04 | glio |
| Interaction_genotype_Bcas2 | Lbhd2 | -6.343 | 3.34E-05 | glio |
| Interaction_genotype_Bora | Folr1 | -4.513 | 3.34E-05 | glio |
| Interaction_genotype_Mtor | Nudcd2 | 1.782 | 3.34E-05 | glio |
| Aars2_in.vivo_noRT | Gm26621 | 2.924 | 3.34E-05 | glio |
| Atp5c1_in.vivo_noRT | Usp17lc | 2.668 | 3.34E-05 | glio |
| Atr_in.vivo_noRT | Sbk1 | 2.975 | 3.34E-05 | glio |
| Atr_in.vivo_noRT | Gm904 | 2.971 | 3.34E-05 | glio |
| Bcas2_in.vivo_noRT | Pabpn1 | -1.131 | 3.34E-05 | glio |
| Bcas2_in.vivo_noRT | Hnrnpf | -0.803 | 3.34E-05 | glio |

| | | | | |
|---|---|---|---|---|
| Bora_in.vivo_noRT | Gm4907 | 3.33 | 3.34E-05 | glio |
| culture_effect | Adh5 | -0.681 | 3.34E-06 | glio |
| culture_effect | Rpl12 | -0.559 | 3.34E-06 | glio |
| Aars2_in.vivo_noRT | Cd79a | 3.335 | 3.34E-06 | glio |
| Aars2_in.vivo_noRT | Neurl1a | 3.528 | 3.34E-06 | glio |
| Atr_in.vivo_noRT | 4930554I06Ri | 3.473 | 3.34E-06 | glio |
| Aars2_in.vivo_noRT | C77080 | 3.449 | 3.34E-07 | glio |
| Aars2_ex.vivo_noRT | Cd7 | 3.293 | 3.34E-07 | glio |
| Aars2_ex.vivo_noRT | Gm48764 | 3.293 | 3.34E-07 | glio |
| Chek1_ex.vivo_noRT | Fhad1 | 4.256 | 3.34E-07 | glio |
| Aars2_in.vivo_noRT | Nox1 | 3.344 | 3.34E-08 | glio |
| Atr_in.vivo_noRT | Gm15672 | 3.81 | 3.34E-08 | glio |
| Chek1_ex.vivo_noRT | Tnni2 | 5.619 | 3.34E-08 | glio |
| Hsd17b10_in.vivo_noRT | Gm5934 | 3.353 | 3.34E-10 | glio |
| Atr_in.vivo_noRT | Ninj2 | 4.626 | 3.34E-11 | glio |
| culture_effect | Mrpl9 | -0.494 | 3.35E-03 | glio |
| Interaction_genotype_Bcas2 | Synpr | -4.247 | 3.35E-03 | glio |
| Interaction_genotype_Bcas2 | Slc2a9 | -3.262 | 3.35E-03 | glio |
| Interaction_genotype_Bcas2 | Gdap1l1 | -4.339 | 3.35E-03 | glio |
| Interaction_genotype_Bcas2 | Wdr70 | 1.507 | 3.35E-03 | glio |
| Interaction_genotype_Bora | Tmem74b | -3.235 | 3.35E-03 | glio |
| Interaction_genotype_Bora | Gm10767 | -3.966 | 3.35E-03 | glio |
| Interaction_genotype_Chek1 | Gm10399 | -3.745 | 3.35E-03 | glio |
| Interaction_genotype_Mtor | Acrbp | -3.057 | 3.35E-03 | glio |
| Interaction_genotype_Mtor | Ccdc69 | -3.386 | 3.35E-03 | glio |
| Aars2_in.vivo_noRT | Gm10863 | 1.932 | 3.35E-03 | glio |
| Aars2_in.vivo_noRT | Efnb1 | 1.912 | 3.35E-03 | glio |
| Aars2_in.vivo_noRT | Rc3h1 | -1.307 | 3.35E-03 | glio |
| Aars2_in.vivo_noRT | Chd8 | -1.25 | 3.35E-03 | glio |
| Aars2_in.vivo_noRT | Slc35a2 | 1.077 | 3.35E-03 | glio |
| Atr_in.vivo_noRT | Atp1b2 | 2.172 | 3.35E-03 | glio |
| Atr_in.vivo_noRT | Ermard | 1.789 | 3.35E-03 | glio |
| Atr_in.vivo_noRT | Mbip | 1.464 | 3.35E-03 | glio |
| Atr_in.vivo_noRT | Glud1 | -1.255 | 3.35E-03 | glio |
| Bcas2_ex.vivo_noRT | 9530080O11R | 2.831 | 3.35E-03 | glio |
| Bora_in.vivo_noRT | Ppargc1b | 2.194 | 3.35E-03 | glio |
| Cdkn2a_in.vivo_noRT | Clcn1 | -2.32 | 3.35E-03 | glio |
| Cdkn2a_in.vivo_noRT | Olfr1152 | -1.941 | 3.35E-03 | glio |
| Cdkn2a_in.vivo_noRT | Rbp2 | 1.708 | 3.35E-03 | glio |
| Chek1_in.vivo_noRT | Plxna2 | 2.326 | 3.35E-03 | glio |
| Chek1_ex.vivo_noRT | 2500004C02R | 2.051 | 3.35E-03 | glio |
| Interaction_genotype_Bcas2 | Adamts4 | -4.132 | 3.35E-04 | glio |
| Interaction_genotype_Bcas2 | 4930558J18Ri | -4.334 | 3.35E-04 | glio |
| Interaction_genotype_Myc | Ankrd22 | -2.729 | 3.35E-04 | glio |
| Interaction_genotype_Myc | Rsad1 | -3.68 | 3.35E-04 | glio |
| Atr_in.vivo_noRT | Tmem255a | 2.625 | 3.35E-04 | glio |
| Atr_in.vivo_noRT | Lfng | 2.543 | 3.35E-04 | glio |
| Atr_ex.vivo_noRT | Gm48140 | 2.37 | 3.35E-04 | glio |
| Bcas2_in.vivo_noRT | Chrna1os | 2.187 | 3.35E-04 | glio |
| Bcas2_in.vivo_noRT | Smim13 | -1.652 | 3.35E-04 | glio |
| Chek1_ex.vivo_noRT | Gngt2 | 3.173 | 3.35E-04 | glio |
| Chek1_ex.vivo_noRT | Nap1l1 | -0.823 | 3.35E-04 | glio |

| | | | | |
|---|---|---|---|---|
| culture_effect | Runx2 | 1.589 | 3.35E-05 | glio |
| Atr_in.vivo_noRT | Rnaseh2a | -1.867 | 3.35E-05 | glio |
| Chek1_ex.vivo_noRT | Nfatc2 | 3.856 | 3.35E-05 | glio |
| culture_effect | Phf11d | -1.697 | 3.35E-06 | glio |
| Interaction_genotype_Mtor | Esm1 | -4.018 | 3.35E-06 | glio |
| Atr_in.vivo_noRT | Ccnj | 3.352 | 3.35E-06 | glio |
| Chek1_ex.vivo_noRT | Gnaz | 4.256 | 3.35E-06 | glio |
| culture_effect | Aimp2 | -0.849 | 3.35E-07 | glio |
| culture_effect | Ythdc1 | 0.666 | 3.35E-07 | glio |
| Aars2_in.vivo_noRT | Fgf2 | 3.701 | 3.35E-07 | glio |
| Bora_ex.vivo_noRT | Nrip2 | 3.201 | 3.35E-07 | glio |
| Bora_ex.vivo_noRT | Ablim2 | 3.201 | 3.35E-07 | glio |
| culture_effect | Arhgef7 | 1.063 | 3.35E-08 | glio |
| Interaction_genotype_Mtor | 1110046J04Ri | -6.911 | 3.35E-08 | glio |
| Bora_in.vivo_noRT | Nhsl2 | 3.818 | 3.35E-08 | glio |
| Bora_in.vivo_noRT | BC030343 | 3.9 | 3.35E-08 | glio |
| Bora_in.vivo_noRT | Rab3b | 3.78 | 3.35E-08 | glio |
| Chek1_in.vivo_noRT | Gm16094 | 3.739 | 3.35E-09 | glio |
| Aars2_in.vivo_noRT | Sprr1a | 5.327 | 3.35E-14 | glio |
| Bora_in.vivo_noRT | Ric3 | 4.676 | 3.35E-14 | glio |
| culture_effect | Lhfp | 2.142 | 3.35E-15 | glio |
| Bcas2_in.vivo_noRT | Gm36198 | 4.547 | 3.35E-15 | glio |
| culture_effect | Srsf5 | -0.368 | 3.36E-03 | glio |
| Interaction_genotype_Aars2 | Gm26981 | -3.624 | 3.36E-03 | glio |
| Interaction_genotype_Aars2 | Nupr1l | -4.141 | 3.36E-03 | glio |
| Interaction_genotype_Bcas2 | Syce2 | 1.856 | 3.36E-03 | glio |
| interaction_RT | Serpina3m | 2.802 | 3.36E-03 | glio |
| interaction_RT | Lefty1 | 1.407 | 3.36E-03 | glio |
| interaction_RT | Gnl2 | -0.76 | 3.36E-03 | glio |
| interaction_RT | Sumo1 | 0.502 | 3.36E-03 | glio |
| Aars2_in.vivo_noRT | Hk1os | 1.958 | 3.36E-03 | glio |
| Atr_in.vivo_noRT | Rprd1a | -1.332 | 3.36E-03 | glio |
| Atr_ex.vivo_noRT | Gm47232 | 2.755 | 3.36E-03 | glio |
| Bcas2_in.vivo_noRT | Prx | 2.278 | 3.36E-03 | glio |
| Bcas2_in.vivo_noRT | Ovgp1 | 2.125 | 3.36E-03 | glio |
| Bora_ex.vivo_noRT | Gm13610 | 2.881 | 3.36E-03 | glio |
| Chek1_in.vivo_noRT | Syt11 | -1.432 | 3.36E-03 | glio |
| Eif2s1_in.vivo_noRT | Hnf1aos1 | 2.143 | 3.36E-03 | glio |
| Eif2s1_in.vivo_noRT | 4930570E03Ri | 1.883 | 3.36E-03 | glio |
| Eif2s1_in.vivo_noRT | 4930447K03R | 1.883 | 3.36E-03 | glio |
| Eif2s1_ex.vivo_noRT | Clca3a1 | 3.032 | 3.36E-03 | glio |
| Eif2s1_ex.vivo_noRT | Tpo | 2.234 | 3.36E-03 | glio |
| culture_effect | Dlg3 | -1.67 | 3.36E-04 | glio |
| culture_effect | 2310022A10R | -0.828 | 3.36E-04 | glio |
| Interaction_genotype_Bora | Gm9970 | -4.978 | 3.36E-04 | glio |
| Interaction_genotype_Bora | Gm13288 | -3.87 | 3.36E-04 | glio |
| Aars2_in.vivo_noRT | 2310068J16Ri | 2.141 | 3.36E-04 | glio |
| Atp5c1_in.vivo_noRT | Gm50147 | 2.668 | 3.36E-04 | glio |
| Atr_in.vivo_noRT | B3gat2 | 2.673 | 3.36E-04 | glio |
| Bcas2_in.vivo_noRT | Sgk2 | 2.59 | 3.36E-04 | glio |
| Bcas2_in.vivo_noRT | Nrip1 | -1.76 | 3.36E-04 | glio |
| Chek1_in.vivo_noRT | Gm34455 | 3.278 | 3.36E-04 | glio |

| | | | | |
|---|---|---|---|---|
| Chek1_in.vivo_noRT | G430095P16R | 2.59 | 3.36E-04 | glio |
| Chek1_in.vivo_noRT | Rab15 | 2.535 | 3.36E-04 | glio |
| Chek1_in.vivo_noRT | Tax1bp3 | -1.608 | 3.36E-04 | glio |
| culture_effect | Ociad1 | -0.56 | 3.36E-05 | glio |
| Interaction_genotype_Bcas2 | Wfdc1 | -4.702 | 3.36E-05 | glio |
| Interaction_genotype_Ifnar1 | Gm40538 | 4.415 | 3.36E-05 | glio |
| Interaction_genotype_Mtor | Phex | -4.584 | 3.36E-05 | glio |
| Interaction_genotype_Mtor | A330076H08R | -4.099 | 3.36E-05 | glio |
| Interaction_genotype_Myc | Gm14326 | -4.67 | 3.36E-05 | glio |
| Atr_in.vivo_noRT | Rps23 | -0.808 | 3.36E-05 | glio |
| Bcas2_in.vivo_noRT | Gm6225 | 3.131 | 3.36E-05 | glio |
| Chek1_ex.vivo_noRT | Gm20646 | 3.469 | 3.36E-05 | glio |
| Aars2_ex.vivo_noRT | A930002C04R | 2.893 | 3.36E-06 | glio |
| Aars2_ex.vivo_noRT | Klk6 | 3.889 | 3.36E-06 | glio |
| Atp5c1_in.vivo_noRT | Fhl1 | 2.668 | 3.36E-06 | glio |
| Atr_in.vivo_noRT | 1810059H22R | 3.575 | 3.36E-06 | glio |
| Atr_in.vivo_noRT | 1110006O24R | 3.331 | 3.36E-06 | glio |
| Bcas2_in.vivo_noRT | Gng13 | 3.214 | 3.36E-06 | glio |
| Hsd17b10_in.vivo_noRT | Hunk | 3.084 | 3.36E-06 | glio |
| Atr_in.vivo_noRT | Gsx2 | 3.795 | 3.36E-07 | glio |
| Atr_in.vivo_noRT | Sbk2 | 3.567 | 3.36E-07 | glio |
| Aars2_in.vivo_noRT | Rasl10a | 3.184 | 3.36E-08 | glio |
| Bcas2_in.vivo_noRT | 1700012B09R | 4.547 | 3.36E-08 | glio |
| Bora_in.vivo_noRT | 3110082J24Ri | 4.027 | 3.36E-08 | glio |
| Chek1_in.vivo_noRT | Rtl9 | 3.602 | 3.36E-08 | glio |
| Hsd17b10_in.vivo_noRT | Pik3ap1 | 3.353 | 3.36E-08 | glio |
| Atr_in.vivo_noRT | 1810041H14R | 4.415 | 3.36E-09 | glio |
| Chek1_ex.vivo_noRT | Ccdc80 | 3.873 | 3.36E-09 | glio |
| Aars2_in.vivo_noRT | Gm8013 | 4.682 | 3.36E-11 | glio |
| Chek1_ex.vivo_noRT | Myoz1 | 4.842 | 3.36E-11 | glio |
| culture_effect | Rnh1 | -1.056 | 3.36E-13 | glio |
| Bora_in.vivo_noRT | Folr1 | 4.936 | 3.36E-16 | glio |
| Atr_in.vivo_noRT | Gm16751 | 5.017 | 3.36E-17 | glio |
| Chek1_in.vivo_noRT | Kcnk13 | 4.78 | 3.36E-18 | glio |
| Bcas2_in.vivo_noRT | Gm44646 | 5.325 | 3.36E-20 | glio |
| culture_effect | Jam3 | -2.206 | 3.36E-24 | glio |
| culture_effect | Map7 | -6.021 | 3.36E-24 | glio |
| Bcas2_in.vivo_noRT | Gm48155 | 4.807 | 3.36E-24 | glio |
| Atr_in.vivo_noRT | Ankrd22 | 5.533 | 3.36E-29 | glio |
| culture_effect | Zfp568 | -0.905 | 3.37E-03 | glio |
| culture_effect | Tubb3 | -1.153 | 3.37E-03 | glio |
| culture_effect | Rb1cc1 | -0.5 | 3.37E-03 | glio |
| culture_effect | Ptp4a2 | 0.343 | 3.37E-03 | glio |
| Interaction_genotype_Aars2 | Glod5 | 4.363 | 3.37E-03 | glio |
| Interaction_genotype_Aars2 | Cyp1a1 | -4.541 | 3.37E-03 | glio |
| Interaction_genotype_Mtor | Pitrm1 | -1.872 | 3.37E-03 | glio |
| Interaction_genotype_Myc | Zfp623 | -2.437 | 3.37E-03 | glio |
| Interaction_genotype_Myc | Prpf40a | 1.281 | 3.37E-03 | glio |
| interaction_RT | Bms1 | -0.832 | 3.37E-03 | glio |
| interaction_RT | Ybx1 | -0.528 | 3.37E-03 | glio |
| Aars2_in.vivo_noRT | Tspan8 | 2.09 | 3.37E-03 | glio |
| Atp5c1_in.vivo_noRT | Gm34885 | 2.408 | 3.37E-03 | glio |

| | | | | |
|---|---|---|---|---|
| Atr_in.vivo_noRT | Gpd2 | -1.57 | 3.37E-03 | glio |
| Atr_in.vivo_noRT | Lats1 | 1.091 | 3.37E-03 | glio |
| Atr_ex.vivo_noRT | Resf1 | 1.42 | 3.37E-03 | glio |
| Bcas2_ex.vivo_noRT | 2810049E08Ri | 2.742 | 3.37E-03 | glio |
| Bcas2_ex.vivo_noRT | Ocm | 2.742 | 3.37E-03 | glio |
| Bcas2_ex.vivo_noRT | Ftsj3 | -1.153 | 3.37E-03 | glio |
| Bora_in.vivo_noRT | Ccdc191 | 1.92 | 3.37E-03 | glio |
| culture_effect | Dhrs9 | -2.122 | 3.37E-04 | glio |
| culture_effect | Gemin4 | 1.965 | 3.37E-04 | glio |
| culture_effect | 9330151L19Ri | 1.492 | 3.37E-04 | glio |
| culture_effect | St3gal4 | 0.97 | 3.37E-04 | glio |
| culture_effect | Mad2l1bp | -0.772 | 3.37E-04 | glio |
| Interaction_genotype_Myc | Otud5 | -2.05 | 3.37E-04 | glio |
| Interaction_genotype_Pars2 | Pyy | -4.148 | 3.37E-04 | glio |
| Interaction_genotype_Pars2 | Pitx3 | -4.032 | 3.37E-04 | glio |
| Interaction_genotype_Pars2 | 5430402O13R | -5.027 | 3.37E-04 | glio |
| Interaction_genotype_Pars2 | 1110046J04Ri | -5.08 | 3.37E-04 | glio |
| Interaction_genotype_Pars2 | Prickle2 | -4.211 | 3.37E-04 | glio |
| Aars2_in.vivo_noRT | Ppargc1b | 2.597 | 3.37E-04 | glio |
| Atp5c1_in.vivo_noRT | Olfr90 | 2.404 | 3.37E-04 | glio |
| Bcas2_in.vivo_noRT | Grwd1 | -1.75 | 3.37E-04 | glio |
| Chek1_in.vivo_noRT | Slamf1 | 2.604 | 3.37E-04 | glio |
| Hsd17b10_in.vivo_noRT | S1pr1 | 2.963 | 3.37E-04 | glio |
| Interaction_genotype_Myc | Gm13283 | -4.46 | 3.37E-05 | glio |
| Interaction_genotype_Myc | 2900042K21R | -4.633 | 3.37E-05 | glio |
| Aars2_in.vivo_noRT | Ubox5 | 2.418 | 3.37E-05 | glio |
| Atr_in.vivo_noRT | Irs1 | 2.685 | 3.37E-05 | glio |
| Chek1_in.vivo_noRT | Bbs10 | 3.058 | 3.37E-05 | glio |
| Interaction_genotype_Mtor | Rgs4 | -4.099 | 3.37E-06 | glio |
| Bcas2_in.vivo_noRT | Olfml2a | 3.304 | 3.37E-06 | glio |
| Bcas2_in.vivo_noRT | Gm43728 | 3.466 | 3.37E-06 | glio |
| Bora_in.vivo_noRT | Maea | -1.773 | 3.37E-06 | glio |
| Interaction_genotype_Myc | Gm15834 | -5.779 | 3.37E-07 | glio |
| Interaction_genotype_Myc | B930059L03Ri | -6.208 | 3.37E-07 | glio |
| Atr_in.vivo_noRT | BC037704 | 3.485 | 3.37E-07 | glio |
| Atr_in.vivo_noRT | Igf2r | 3.316 | 3.37E-07 | glio |
| Atr_ex.vivo_noRT | Meioc | 3.164 | 3.37E-07 | glio |
| Bcas2_in.vivo_noRT | Ccl20 | 5.325 | 3.37E-07 | glio |
| Bora_ex.vivo_noRT | Serping1 | 3.994 | 3.37E-07 | glio |
| Chek1_ex.vivo_noRT | Slc38a3 | 4.045 | 3.37E-07 | glio |
| Hsd17b10_in.vivo_noRT | Necab2 | 3.352 | 3.37E-07 | glio |
| culture_effect | Pold4 | 1.333 | 3.37E-09 | glio |
| Chek1_in.vivo_noRT | Sbk3 | 3.875 | 3.37E-09 | glio |
| Aars2_in.vivo_noRT | Inhba | 4.685 | 3.37E-10 | glio |
| Aars2_in.vivo_noRT | Gm20492 | 4.271 | 3.37E-10 | glio |
| Bcas2_in.vivo_noRT | Wdr93 | 4.151 | 3.37E-10 | glio |
| Bcas2_in.vivo_noRT | Hcn3 | 4.165 | 3.37E-11 | glio |
| Bcas2_in.vivo_noRT | 1700011M02F | 5.784 | 3.37E-11 | glio |
| Aars2_in.vivo_noRT | Vmn2r56 | 5.081 | 3.37E-20 | glio |
| Atr_in.vivo_noRT | Slc16a4 | 5.269 | 3.37E-20 | glio |
| Bora_in.vivo_noRT | Gm49504 | 4.936 | 3.37E-20 | glio |
| culture_effect | Ergic3 | -1.401 | 3.37E-23 | glio |

| | | | | |
|---|---|---|---|---|
| Atr_in.vivo_noRT | Olfr543 | 5.273 | 3.37E-23 | glio |
| culture_effect | Gm20732 | -1.705 | 3.38E-03 | glio |
| culture_effect | Tspan2 | -1.277 | 3.38E-03 | glio |
| culture_effect | Ccm2 | 0.521 | 3.38E-03 | glio |
| Interaction_genotype_Aars2 | Cyb561d1 | -3.918 | 3.38E-03 | glio |
| Interaction_genotype_Aars2 | Klhdc8b | -3.668 | 3.38E-03 | glio |
| Interaction_genotype_Hsd17b | Ntn4 | -3.382 | 3.38E-03 | glio |
| Interaction_genotype_Hsd17b | Gm27239 | -4.335 | 3.38E-03 | glio |
| Interaction_genotype_Mtor | Gm15107 | -3.308 | 3.38E-03 | glio |
| Interaction_genotype_Mtor | 4930581F22Ri | -2.562 | 3.38E-03 | glio |
| Interaction_genotype_Myc | 6330562C20R | -2.648 | 3.38E-03 | glio |
| Aars2_in.vivo_noRT | Ntn3 | 1.989 | 3.38E-03 | glio |
| Aars2_ex.vivo_noRT | Gng13 | 2.878 | 3.38E-03 | glio |
| Aars2_ex.vivo_noRT | Adm | 2.9 | 3.38E-03 | glio |
| Atr_in.vivo_noRT | Mob1b | -1.202 | 3.38E-03 | glio |
| Atr_in.vivo_noRT | Nktr | -1.071 | 3.38E-03 | glio |
| Atr_in.vivo_noRT | Mrpl15 | -0.785 | 3.38E-03 | glio |
| Atr_ex.vivo_noRT | Gm15445 | 2.178 | 3.38E-03 | glio |
| Atr_ex.vivo_noRT | Tmem229b | 2.776 | 3.38E-03 | glio |
| Atr_ex.vivo_noRT | Tifa | 2.293 | 3.38E-03 | glio |
| Bora_in.vivo_noRT | Flt1 | -1.72 | 3.38E-03 | glio |
| Bora_in.vivo_noRT | Ckb | -0.729 | 3.38E-03 | glio |
| Chek1_in.vivo_noRT | Tpm4 | -0.783 | 3.38E-03 | glio |
| Chek1_ex.vivo_noRT | Prtn3 | 2.799 | 3.38E-03 | glio |
| Chek1_ex.vivo_noRT | Eif2b1 | -1.607 | 3.38E-03 | glio |
| culture_effect | Sun1 | -0.784 | 3.38E-04 | glio |
| Interaction_genotype_Mtor | Adam19 | -4.351 | 3.38E-04 | glio |
| Interaction_genotype_Mtor | Polr2c | 1.636 | 3.38E-04 | glio |
| Interaction_genotype_Mtor | Dut | 2.188 | 3.38E-04 | glio |
| Interaction_genotype_Mtor | Nop58 | 1.746 | 3.38E-04 | glio |
| Interaction_genotype_Mtor | Calm2 | 1.72 | 3.38E-04 | glio |
| Aars2_ex.vivo_noRT | Nkg7 | 2.897 | 3.38E-04 | glio |
| Atr_in.vivo_noRT | Gm11772 | 2.534 | 3.38E-04 | glio |
| Bora_in.vivo_noRT | Lncppara | 2.786 | 3.38E-04 | glio |
| culture_effect | Gm2115 | 2.101 | 3.38E-05 | glio |
| culture_effect | Ctps2 | -1.17 | 3.38E-05 | glio |
| culture_effect | Mrpl18 | 0.568 | 3.38E-05 | glio |
| Interaction_genotype_Atr | Armc2 | -5.123 | 3.38E-05 | glio |
| Interaction_genotype_Atr | Gm10324 | -3.97 | 3.38E-05 | glio |
| Interaction_genotype_Myc | Tbx2 | -3.46 | 3.38E-05 | glio |
| Interaction_genotype_Myc | Srrm2 | 1.479 | 3.38E-05 | glio |
| Interaction_genotype_Myc | Atf4 | 1.779 | 3.38E-05 | glio |
| Interaction_genotype_Myc | Rplp0 | 1.573 | 3.38E-05 | glio |
| Aars2_in.vivo_noRT | Nell2 | 2.656 | 3.38E-05 | glio |
| Aars2_in.vivo_noRT | Pcdhga12 | 2.845 | 3.38E-05 | glio |
| Atr_in.vivo_noRT | Gm11713 | 2.59 | 3.38E-05 | glio |
| Atr_in.vivo_noRT | Gab2 | 2.703 | 3.38E-05 | glio |
| Bard1_ex.vivo_noRT | Cd200r2 | 2.814 | 3.38E-05 | glio |
| Bard1_ex.vivo_noRT | Rem2 | 3.612 | 3.38E-05 | glio |
| Bora_in.vivo_noRT | Htra1 | 3.29 | 3.38E-05 | glio |
| Bora_ex.vivo_noRT | Vmn2r54 | 3.201 | 3.38E-05 | glio |
| Chek1_ex.vivo_noRT | Gm41409 | 3.298 | 3.38E-05 | glio |

| | | | | |
|---|---|---|---|---|
| Chek1_ex.vivo_noRT | 6430562O15R | 3.253 | 3.38E-05 | glio |
| Chek1_ex.vivo_noRT | Gm14273 | 3.657 | 3.38E-05 | glio |
| Chek1_ex.vivo_noRT | Cytl1 | 4.142 | 3.38E-05 | glio |
| Interaction_genotype_Mtor | Meioc | -3.836 | 3.38E-06 | glio |
| Interaction_genotype_Myc | 4930412M03F | -5.16 | 3.38E-06 | glio |
| Bcas2_ex.vivo_noRT | Kmo | 3.93 | 3.38E-06 | glio |
| Chek1_in.vivo_noRT | Dock4 | 3.292 | 3.38E-06 | glio |
| Hsd17b10_in.vivo_noRT | Spdye4a | 3.485 | 3.38E-06 | glio |
| Interaction_genotype_Atr | Mfsd10 | 3.15 | 3.38E-07 | glio |
| Atr_in.vivo_noRT | Tex14 | 3.192 | 3.38E-07 | glio |
| Chek1_ex.vivo_noRT | Gstp3 | 4.447 | 3.38E-07 | glio |
| interaction_RT | Ube2b | 0.817 | 3.38E-08 | glio |
| Atr_in.vivo_noRT | Lyrm1 | 3.82 | 3.38E-08 | glio |
| Atr_in.vivo_noRT | 2210011C24R | 3.696 | 3.38E-08 | glio |
| Hsd17b10_in.vivo_noRT | D430001F17R | 3.616 | 3.38E-08 | glio |
| Bcas2_in.vivo_noRT | Slc6a11 | 3.375 | 3.38E-09 | glio |
| Bora_in.vivo_noRT | A530013C23R | 4.551 | 3.38E-12 | glio |
| Chek1_in.vivo_noRT | Dppa2 | 4.511 | 3.38E-12 | glio |
| Atr_in.vivo_noRT | Gm42664 | 4.883 | 3.38E-16 | glio |
| culture_effect | Mif | -0.403 | 3.39E-03 | glio |
| Interaction_genotype_Aars2 | Pdgfb | -2.934 | 3.39E-03 | glio |
| Interaction_genotype_Aars2 | Nlrx1 | -3.809 | 3.39E-03 | glio |
| Interaction_genotype_Aars2 | Scn2a | -3.634 | 3.39E-03 | glio |
| Interaction_genotype_Aars2 | Stub1 | 1.128 | 3.39E-03 | glio |
| Interaction_genotype_Aars2 | Oaz1 | 0.866 | 3.39E-03 | glio |
| Interaction_genotype_Atr | Ccdc78 | -3.626 | 3.39E-03 | glio |
| Interaction_genotype_Atr | Plpp7 | -3.385 | 3.39E-03 | glio |
| Interaction_genotype_Atr | Emc3 | 1.46 | 3.39E-03 | glio |
| Interaction_genotype_Atr | Atp6v1f | 0.997 | 3.39E-03 | glio |
| Interaction_genotype_Bcas2 | Fen1 | 1.874 | 3.39E-03 | glio |
| Interaction_genotype_Myc | Krtdap | -3.321 | 3.39E-03 | glio |
| Interaction_genotype_Myc | Gm13610 | -3.298 | 3.39E-03 | glio |
| Interaction_genotype_Myc | Tmem150a | -2.619 | 3.39E-03 | glio |
| interaction_RT | Uba1 | -0.725 | 3.39E-03 | glio |
| Aars2_in.vivo_noRT | Serpine2 | 2.536 | 3.39E-03 | glio |
| Aars2_in.vivo_noRT | Ppef1 | 1.999 | 3.39E-03 | glio |
| Aars2_in.vivo_noRT | Adck1 | 1.652 | 3.39E-03 | glio |
| Bard1_ex.vivo_noRT | Gm128 | 2.421 | 3.39E-03 | glio |
| Bcas2_ex.vivo_noRT | Msrb2 | 2.844 | 3.39E-03 | glio |
| Bora_in.vivo_noRT | Dnpep | -1.404 | 3.39E-03 | glio |
| Cenpj_in.vivo_noRT | Gm47851 | 2.289 | 3.39E-03 | glio |
| Chek1_in.vivo_noRT | Fgf21 | 2.277 | 3.39E-03 | glio |
| Chek1_in.vivo_noRT | 9130024F11Ri | 2.052 | 3.39E-03 | glio |
| Interaction_genotype_Mtor | Bcas2 | 1.817 | 3.39E-04 | glio |
| Interaction_genotype_Mtor | Rrbp1 | 1.724 | 3.39E-04 | glio |
| Bora_in.vivo_noRT | Abhd2 | -1.668 | 3.39E-04 | glio |
| Chek1_in.vivo_noRT | Cip2a | -1.887 | 3.39E-04 | glio |
| Chek1_ex.vivo_noRT | C1qtnf1 | 3.218 | 3.39E-04 | glio |
| Interaction_genotype_Myc | Lamc2 | -4.449 | 3.39E-05 | glio |
| Interaction_genotype_Myc | Cdc25b | -3.877 | 3.39E-05 | glio |
| Atr_ex.vivo_noRT | Gm50115 | 3.548 | 3.39E-05 | glio |
| Bora_in.vivo_noRT | Gm11131 | 2.851 | 3.39E-05 | glio |

| | | | | |
|---|---|---|---|---|
| Chek1_ex.vivo_noRT | Elf3 | 4.048 | 3.39E-05 | glio |
| Atr_in.vivo_noRT | Mkrn1 | -1.735 | 3.39E-06 | glio |
| Bcas2_in.vivo_noRT | Casz1 | 3.532 | 3.39E-06 | glio |
| Bcas2_in.vivo_noRT | Gm20492 | 3.215 | 3.39E-06 | glio |
| culture_effect | Gm39469 | 1.912 | 3.39E-07 | glio |
| culture_effect | Sympk | -0.86 | 3.39E-07 | glio |
| culture_effect | Gdf15 | -1.471 | 3.39E-07 | glio |
| Interaction_genotype_Myc | 2500004C02R | -4.844 | 3.39E-07 | glio |
| Interaction_genotype_Myc | Gm13288 | -4.865 | 3.39E-07 | glio |
| Interaction_genotype_Mtor | Ndufb5 | 2.788 | 3.39E-08 | glio |
| Chek1_ex.vivo_noRT | Plk5 | 4.449 | 3.39E-09 | glio |
| Atr_in.vivo_noRT | Gjb4 | 4.876 | 3.39E-12 | glio |
| Bora_in.vivo_noRT | Podxl | 4.405 | 3.39E-12 | glio |
| Atr_in.vivo_noRT | Gm6803 | 5.533 | 3.39E-23 | glio |
| Aars2_in.vivo_noRT | Gm36793 | 5.327 | 3.39E-24 | glio |
| culture_effect | BC028471 | 1.742 | 3.40E-03 | glio |
| Interaction_genotype_Bcas2 | 4921504E06Ri | -3.154 | 3.40E-03 | glio |
| Interaction_genotype_Bcas2 | Slc9a5 | -3.89 | 3.40E-03 | glio |
| Interaction_genotype_Mtor | Map3k8 | -3.909 | 3.40E-03 | glio |
| Interaction_genotype_Mtor | Akr1a1 | 1.243 | 3.40E-03 | glio |
| Interaction_genotype_Myc | Slurp1 | -4.868 | 3.40E-03 | glio |
| Interaction_genotype_Myc | Dgcr8 | -2.498 | 3.40E-03 | glio |
| Interaction_genotype_Myc | Tatdn1 | 1.404 | 3.40E-03 | glio |
| Interaction_genotype_Rpl3 | Dhx37 | -3.514 | 3.40E-03 | glio |
| interaction_RT | Ankrd40 | -0.747 | 3.40E-03 | glio |
| Aars2_in.vivo_noRT | Gm26881 | 1.859 | 3.40E-03 | glio |
| Aars2_ex.vivo_noRT | Cldn14 | 2.902 | 3.40E-03 | glio |
| Bcas2_in.vivo_noRT | Zfp112 | 2.116 | 3.40E-03 | glio |
| Bcas2_in.vivo_noRT | Slc9a5 | 2.114 | 3.40E-03 | glio |
| Bcas2_in.vivo_noRT | Pcdh15 | 1.762 | 3.40E-03 | glio |
| Cdkn2a_in.vivo_noRT | Gm29966 | -1.935 | 3.40E-03 | glio |
| Chek1_in.vivo_noRT | Prdx4 | -0.834 | 3.40E-03 | glio |
| Chek1_ex.vivo_noRT | Nop9 | -1.728 | 3.40E-03 | glio |
| Chek1_ex.vivo_noRT | Rpa2 | -1.507 | 3.40E-03 | glio |
| Interaction_genotype_Atr | Lncbate10 | -3.157 | 3.40E-04 | glio |
| Interaction_genotype_Atr | Vdac2 | 1.34 | 3.40E-04 | glio |
| Interaction_genotype_Bora | Olfm2 | -4.755 | 3.40E-04 | glio |
| Aars2_in.vivo_noRT | B230206L02Ri | 2.525 | 3.40E-04 | glio |
| Atr_in.vivo_noRT | Rab10 | -1.04 | 3.40E-04 | glio |
| Bcas2_in.vivo_noRT | Mmaa | 2.262 | 3.40E-04 | glio |
| Bora_in.vivo_noRT | Gm46378 | 2.703 | 3.40E-04 | glio |
| Hsd17b10_in.vivo_noRT | Gm15927 | 2.615 | 3.40E-04 | glio |
| Interaction_genotype_Mtor | Gm20442 | -4.458 | 3.40E-05 | glio |
| Interaction_genotype_Myc | Tmco1 | 1.632 | 3.40E-05 | glio |
| interaction_RT | H2-M3 | 1.808 | 3.40E-05 | glio |
| interaction_RT | Pold1 | -1.392 | 3.40E-05 | glio |
| interaction_RT | Hs2st1 | -1.037 | 3.40E-05 | glio |
| interaction_RT | Pum3 | -0.877 | 3.40E-05 | glio |
| interaction_RT | Gm44190 | -2.727 | 3.40E-05 | glio |
| Aars2_ex.vivo_noRT | Gm50147 | 3.544 | 3.40E-05 | glio |
| Bora_ex.vivo_noRT | Aqp9 | 3.201 | 3.40E-05 | glio |
| Aars2_ex.vivo_noRT | Fam166a | 3.293 | 3.40E-06 | glio |

| | | | | |
|---|---|---|---|---|
| Atr_in.vivo_noRT | Usp11 | 3.434 | 3.40E-06 | glio |
| Bcas2_in.vivo_noRT | Hrh2 | 2.577 | 3.40E-06 | glio |
| Bora_in.vivo_noRT | Gm5861 | 3.888 | 3.40E-06 | glio |
| Chek1_ex.vivo_noRT | 1700006H21R | 4.055 | 3.40E-06 | glio |
| Interaction_genotype_Myc | Gm10101 | -5.508 | 3.40E-07 | glio |
| Interaction_genotype_Myc | 4930515G01R | -5.323 | 3.40E-07 | glio |
| Interaction_genotype_Myc | Cox8a | 1.883 | 3.40E-07 | glio |
| Interaction_genotype_Myc | mt-Co2 | 1.877 | 3.40E-07 | glio |
| Bora_in.vivo_noRT | Sdcbp2 | 3.774 | 3.40E-07 | glio |
| Atr_in.vivo_noRT | Acot5 | 3.649 | 3.40E-08 | glio |
| culture_effect | Atf3 | -1.787 | 3.40E-09 | glio |
| Chek1_in.vivo_noRT | Hist2h3c1 | 4.132 | 3.40E-09 | glio |
| Aars2_in.vivo_noRT | Cks1brt | 3.889 | 3.40E-10 | glio |
| Bora_in.vivo_noRT | Gipc2 | 4.673 | 3.40E-10 | glio |
| Atr_in.vivo_noRT | D430040D24R | 4.617 | 3.40E-11 | glio |
| Atr_in.vivo_noRT | Gm14698 | 4.621 | 3.40E-11 | glio |
| Bora_in.vivo_noRT | Camk2b | 4.414 | 3.40E-11 | glio |
| Chek1_in.vivo_noRT | Pcdhga1 | 3.992 | 3.40E-11 | glio |
| Aars2_in.vivo_noRT | Rgs9 | 4.526 | 3.40E-14 | glio |
| Chek1_ex.vivo_noRT | Esp24 | 4.842 | 3.40E-14 | glio |
| Chek1_in.vivo_noRT | Tcstv1 | 5.044 | 3.40E-17 | glio |
| culture_effect | AI463229 | -1.579 | 3.41E-03 | glio |
| culture_effect | Naa25 | -0.512 | 3.41E-03 | glio |
| culture_effect | Rnf6 | 0.488 | 3.41E-03 | glio |
| Interaction_genotype_Aars2 | C030034I22Ri | -3.358 | 3.41E-03 | glio |
| Interaction_genotype_Cdkn2a | A930019D19R | 2.835 | 3.41E-03 | glio |
| interaction_RT | Rrp15 | -0.617 | 3.41E-03 | glio |
| Aars2_in.vivo_noRT | Nadsyn1 | 2.141 | 3.41E-03 | glio |
| Aars2_ex.vivo_noRT | Pde6b | 2.499 | 3.41E-03 | glio |
| Atr_in.vivo_noRT | Mgll | 2.05 | 3.41E-03 | glio |
| Atr_in.vivo_noRT | Tm2d2 | -1.118 | 3.41E-03 | glio |
| Atr_ex.vivo_noRT | Zmynd10 | 2.376 | 3.41E-03 | glio |
| Bcas2_in.vivo_noRT | Gm38414 | 2.4 | 3.41E-03 | glio |
| Bcas2_in.vivo_noRT | Otud3 | -1.452 | 3.41E-03 | glio |
| Bcas2_in.vivo_noRT | Sept7 | -0.68 | 3.41E-03 | glio |
| Bora_in.vivo_noRT | Eif2s1 | -0.688 | 3.41E-03 | glio |
| Chek1_in.vivo_noRT | Gm42726 | 2.051 | 3.41E-03 | glio |
| Chek1_in.vivo_noRT | Cyb5d2 | 1.564 | 3.41E-03 | glio |
| Chek1_in.vivo_noRT | Mrpl42 | -0.681 | 3.41E-03 | glio |
| Chek1_ex.vivo_noRT | Hnrnpa3 | -0.901 | 3.41E-03 | glio |
| Dclre1c_in.vivo_noRT | Comp | 2.58 | 3.41E-03 | glio |
| Dclre1c_in.vivo_noRT | Edil3 | -2.616 | 3.41E-03 | glio |
| Hsd17b10_in.vivo_noRT | 4930592C13R | 2.349 | 3.41E-03 | glio |
| culture_effect | Mt3 | -1.373 | 3.41E-04 | glio |
| culture_effect | Slfn9 | 0.695 | 3.41E-04 | glio |
| culture_effect | Ndufb6 | 0.406 | 3.41E-04 | glio |
| Interaction_genotype_Aars2 | A730071L15Ri | -4.616 | 3.41E-04 | glio |
| Interaction_genotype_Ifnar1 | Gm10482 | 4.043 | 3.41E-04 | glio |
| Interaction_genotype_Mtor | Gm36913 | -4.393 | 3.41E-04 | glio |
| Interaction_genotype_Myc | Gm47404 | -2.729 | 3.41E-04 | glio |
| Interaction_genotype_Myc | Gm45871 | -3.686 | 3.41E-04 | glio |
| Interaction_genotype_Myc | Pdik1l | -2.969 | 3.41E-04 | glio |

| | | | | |
|---|---|---|---|---|
| Aars2_in.vivo_noRT | Slc22a5 | 2.347 | 3.41E-04 | glio |
| Atr_ex.vivo_noRT | Gm13398 | 2.367 | 3.41E-04 | glio |
| Bora_ex.vivo_noRT | Gm4858 | 3.201 | 3.41E-04 | glio |
| Bora_ex.vivo_noRT | Tmsb15b2 | 3.685 | 3.41E-04 | glio |
| Chek1_in.vivo_noRT | Htra1 | 2.874 | 3.41E-04 | glio |
| Chek1_in.vivo_noRT | Tmem255a | 2.658 | 3.41E-04 | glio |
| Chek1_in.vivo_noRT | Gnaq | -1.575 | 3.41E-04 | glio |
| Interaction_genotype_Ifnar1 | Tagap | 3.781 | 3.41E-05 | glio |
| interaction_RT | 3830403N18R | 0.82 | 3.41E-05 | glio |
| Bcas2_ex.vivo_noRT | Ablim2 | 2.742 | 3.41E-05 | glio |
| Bcas2_ex.vivo_noRT | Gm5415 | 2.742 | 3.41E-05 | glio |
| Bcas2_ex.vivo_noRT | Cox7b2 | 3.55 | 3.41E-05 | glio |
| Bora_ex.vivo_noRT | St18 | 3.201 | 3.41E-05 | glio |
| Chek1_in.vivo_noRT | Prrg1 | 3.019 | 3.41E-05 | glio |
| Hsd17b10_in.vivo_noRT | Pygo1 | 2.832 | 3.41E-05 | glio |
| Interaction_genotype_Mtor | Gm12840 | -5.214 | 3.41E-06 | glio |
| Interaction_genotype_Rpl3 | Ntn4 | -5.039 | 3.41E-06 | glio |
| Aars2_ex.vivo_noRT | Gm15894 | 3.293 | 3.41E-06 | glio |
| Atr_ex.vivo_noRT | Gm49808 | 2.771 | 3.41E-06 | glio |
| Chek1_in.vivo_noRT | Plin1 | 3.33 | 3.41E-06 | glio |
| Chek1_in.vivo_noRT | Gstm6 | 3.506 | 3.41E-06 | glio |
| culture_effect | B630019K06R | 1.781 | 3.41E-07 | glio |
| culture_effect | Pik3cb | 0.987 | 3.41E-07 | glio |
| Aars2_in.vivo_noRT | Snai1 | 3.809 | 3.41E-07 | glio |
| Bcas2_in.vivo_noRT | Ptp4a1 | 3.112 | 3.41E-07 | glio |
| Interaction_genotype_Mtor | Pmp2 | -6.022 | 3.41E-08 | glio |
| Interaction_genotype_Mtor | 1700026J14Ri | -6.096 | 3.41E-08 | glio |
| Interaction_genotype_Mtor | Gm47851 | -6.226 | 3.41E-08 | glio |
| Atr_ex.vivo_noRT | Gm38163 | 3.962 | 3.41E-08 | glio |
| Chek1_in.vivo_noRT | Ptp4a1 | 3.349 | 3.41E-08 | glio |
| culture_effect | 3110021N24R | 2.019 | 3.41E-09 | glio |
| Bora_in.vivo_noRT | Milr1 | 4.202 | 3.41E-09 | glio |
| Bora_in.vivo_noRT | Gm12602 | 3.909 | 3.41E-09 | glio |
| Chek1_in.vivo_noRT | Nyap2 | 4.132 | 3.41E-09 | glio |
| Atr_in.vivo_noRT | Ablim3 | 4.51 | 3.41E-10 | glio |
| Chek1_in.vivo_noRT | 2610027K06R | 4.388 | 3.41E-11 | glio |
| Atr_in.vivo_noRT | Noxred1 | 4.805 | 3.41E-13 | glio |
| Bora_in.vivo_noRT | 4930505N22R | 4.676 | 3.41E-13 | glio |
| culture_effect | Dnajc19 | 1.045 | 3.41E-17 | glio |
| Chek1_in.vivo_noRT | 1700024I08Ri | 4.784 | 3.41E-17 | glio |
| Atr_in.vivo_noRT | Olfr552 | 5.533 | 3.41E-27 | glio |
| culture_effect | Gm39526 | -1.457 | 3.42E-03 | glio |
| culture_effect | 4930532G15R | 1.254 | 3.42E-03 | glio |
| culture_effect | Gmip | -1.103 | 3.42E-03 | glio |
| culture_effect | Lemd3 | -0.775 | 3.42E-03 | glio |
| culture_effect | Zfr | 0.347 | 3.42E-03 | glio |
| Interaction_genotype_Aars2 | Alox12 | -3.6 | 3.42E-03 | glio |
| Interaction_genotype_Aars2 | D330050G23R | -3.042 | 3.42E-03 | glio |
| Interaction_genotype_Bcas2 | D930028M14I | -3.552 | 3.42E-03 | glio |
| Interaction_genotype_Bcas2 | Gm16023 | -4.307 | 3.42E-03 | glio |
| Interaction_genotype_Hsd17b | Ciita | 3.742 | 3.42E-03 | glio |
| Interaction_genotype_Mtor | Palm3 | -3.73 | 3.42E-03 | glio |

| | | | | |
|---|---|---|---|---|
| Interaction_genotype_Mtor | 0610040B10R | -2.573 | 3.42E-03 | glio |
| Interaction_genotype_Mtor | Cdc40 | 1.609 | 3.42E-03 | glio |
| Interaction_genotype_Myc | C8g | -3.24 | 3.42E-03 | glio |
| Interaction_genotype_Myc | Zfp39 | -3.303 | 3.42E-03 | glio |
| Interaction_genotype_Myc | Zbtb5 | -2.419 | 3.42E-03 | glio |
| Interaction_genotype_Myc | Hoxaas3 | -2.465 | 3.42E-03 | glio |
| Interaction_genotype_Myc | Cotl1 | -2.154 | 3.42E-03 | glio |
| Interaction_genotype_Myc | Ndufa11 | 1.134 | 3.42E-03 | glio |
| Aars2_in.vivo_noRT | Pgap3 | 2.125 | 3.42E-03 | glio |
| Aars2_in.vivo_noRT | Rpl10a | -0.495 | 3.42E-03 | glio |
| Aars2_ex.vivo_noRT | Gpr37l1 | 2.899 | 3.42E-03 | glio |
| Atp5c1_in.vivo_noRT | Nxnl2 | 2.146 | 3.42E-03 | glio |
| Bard1_ex.vivo_noRT | Gm5934 | 2.235 | 3.42E-03 | glio |
| Bcas2_in.vivo_noRT | Ccdc103 | 2.178 | 3.42E-03 | glio |
| Bcas2_in.vivo_noRT | Gpx4 | -0.621 | 3.42E-03 | glio |
| Bora_in.vivo_noRT | Rmdn2 | 1.925 | 3.42E-03 | glio |
| Brca2_ex.vivo_noRT | Gm43948 | 2.831 | 3.42E-03 | glio |
| Brca2_ex.vivo_noRT | Gm10482 | 2.82 | 3.42E-03 | glio |
| Brca2_ex.vivo_noRT | Gm41349 | 3.195 | 3.42E-03 | glio |
| Chek1_ex.vivo_noRT | Tnfsf10 | 2.664 | 3.42E-03 | glio |
| Eif2s1_ex.vivo_noRT | Brsk2 | 3.032 | 3.42E-03 | glio |
| Eif2s1_ex.vivo_noRT | Rimklb | 2.234 | 3.42E-03 | glio |
| Eif2s1_ex.vivo_noRT | Rpl3l | 3.032 | 3.42E-03 | glio |
| Eif2s1_ex.vivo_noRT | Gm14929 | 2.234 | 3.42E-03 | glio |
| Eif2s1_ex.vivo_noRT | Gm13032 | 2.633 | 3.42E-03 | glio |
| Hsd17b10_in.vivo_noRT | 4930581F22Ri | 1.852 | 3.42E-03 | glio |
| Interaction_genotype_Mtor | Rps3a1 | 1.277 | 3.42E-04 | glio |
| Interaction_genotype_Myc | C230037L18Ri | -3.748 | 3.42E-04 | glio |
| Interaction_genotype_Myc | Plagl2 | -2.883 | 3.42E-04 | glio |
| Interaction_genotype_Myc | Pfdn1 | 1.501 | 3.42E-04 | glio |
| Aars2_in.vivo_noRT | Ttbk1 | 2.529 | 3.42E-04 | glio |
| Aars2_in.vivo_noRT | Mettl5os | 2.387 | 3.42E-04 | glio |
| Atp5c1_in.vivo_noRT | Gm5662 | 2.668 | 3.42E-04 | glio |
| Atr_in.vivo_noRT | Psme2b | 2.428 | 3.42E-04 | glio |
| Atr_ex.vivo_noRT | Hfm1 | 2.589 | 3.42E-04 | glio |
| Atr_ex.vivo_noRT | Gm43915 | 2.77 | 3.42E-04 | glio |
| Atr_ex.vivo_noRT | AU022754 | 3.164 | 3.42E-04 | glio |
| Atr_ex.vivo_noRT | Gm15890 | 2.773 | 3.42E-04 | glio |
| Bcas2_in.vivo_noRT | Luc7l3 | -0.935 | 3.42E-04 | glio |
| Bcas2_ex.vivo_noRT | Gm13856 | 2.348 | 3.42E-04 | glio |
| Bcas2_ex.vivo_noRT | Akap14 | 2.742 | 3.42E-04 | glio |
| Bora_in.vivo_noRT | Def6 | 2.481 | 3.42E-04 | glio |
| Bora_in.vivo_noRT | Gm11273 | 2.152 | 3.42E-04 | glio |
| Bora_in.vivo_noRT | Ttc19 | -1.472 | 3.42E-04 | glio |
| Chek1_in.vivo_noRT | Tpd52l2 | -1.193 | 3.42E-04 | glio |
| Chek1_ex.vivo_noRT | Gm4070 | 3.43 | 3.42E-04 | glio |
| Chek1_ex.vivo_noRT | Tmem82 | 3.372 | 3.42E-04 | glio |
| Interaction_genotype_Mtor | Ctsz | 1.979 | 3.42E-05 | glio |
| Atr_ex.vivo_noRT | Ankrd33b | 3.962 | 3.42E-05 | glio |
| Atr_ex.vivo_noRT | Calml4 | 3.628 | 3.42E-05 | glio |
| Bcas2_in.vivo_noRT | Gm12258 | 3.128 | 3.42E-05 | glio |
| Bcas2_ex.vivo_noRT | Nrip2 | 2.742 | 3.42E-05 | glio |

| | | | | |
|---|---|---|---|---|
| Bora_in.vivo_noRT | Nr1h4 | 2.887 | 3.42E-05 | glio |
| Aars2_in.vivo_noRT | Pcdhb19 | 3.035 | 3.42E-06 | glio |
| Aars2_in.vivo_noRT | Sema4f | 3.244 | 3.42E-06 | glio |
| Atp5c1_in.vivo_noRT | Mcmdc2 | 2.936 | 3.42E-06 | glio |
| Atp5c1_in.vivo_noRT | 1700112J16Ri | 2.668 | 3.42E-06 | glio |
| Atp5c1_in.vivo_noRT | Clec4d | 2.668 | 3.42E-06 | glio |
| Atr_ex.vivo_noRT | Hist1h2ag | 3.164 | 3.42E-06 | glio |
| Bcas2_in.vivo_noRT | 1700063J08Ri | 3.3 | 3.42E-06 | glio |
| Bora_in.vivo_noRT | Serpinb9b | 3.939 | 3.42E-06 | glio |
| Bora_in.vivo_noRT | Alox15 | 2.924 | 3.42E-06 | glio |
| Chek1_in.vivo_noRT | Cyp2d22 | 3.767 | 3.42E-07 | glio |
| Hsd17b10_in.vivo_noRT | Pkd1l2 | 3.352 | 3.42E-07 | glio |
| culture_effect | Zeb2os | 2.551 | 3.42E-08 | glio |
| culture_effect | Slc3a2 | -0.75 | 3.42E-08 | glio |
| Interaction_genotype_Mtor | Trir | 2.538 | 3.42E-08 | glio |
| Interaction_genotype_Mtor | 4930412M03F | -6.526 | 3.42E-08 | glio |
| Interaction_genotype_Mtor | Hba-a2 | -6.96 | 3.42E-08 | glio |
| Interaction_genotype_Mtor | Gm49808 | -4.492 | 3.42E-08 | glio |
| Atr_in.vivo_noRT | A430033K04R | 3.957 | 3.42E-09 | glio |
| culture_effect | Rps13 | -0.743 | 3.42E-10 | glio |
| Aars2_in.vivo_noRT | Tnnt3 | 4.692 | 3.42E-10 | glio |
| Aars2_in.vivo_noRT | 1700010B08R | 3.64 | 3.42E-10 | glio |
| Hsd17b10_in.vivo_noRT | Thsd1 | 4.01 | 3.42E-11 | glio |
| Aars2_in.vivo_noRT | Gm49874 | 5.327 | 3.42E-13 | glio |
| Chek1_in.vivo_noRT | Gm44983 | 4.256 | 3.42E-13 | glio |
| culture_effect | Anxa8 | -1.867 | 3.43E-03 | glio |
| culture_effect | Arhgdia | 0.305 | 3.43E-03 | glio |
| Interaction_genotype_Aars2 | Gm49808 | -2.427 | 3.43E-03 | glio |
| Interaction_genotype_Bora | Babam1 | 1.785 | 3.43E-03 | glio |
| Interaction_genotype_Chek1 | D730045B01R | -3.535 | 3.43E-03 | glio |
| Interaction_genotype_Chek1 | Ubxn1 | 0.868 | 3.43E-03 | glio |
| Interaction_genotype_Mtor | Gm48086 | -3.652 | 3.43E-03 | glio |
| Interaction_genotype_Mtor | Gm45902 | -3.458 | 3.43E-03 | glio |
| Interaction_genotype_Mtor | Immp2l | -3.035 | 3.43E-03 | glio |
| Interaction_genotype_Mtor | Kcnmb4 | -3.021 | 3.43E-03 | glio |
| Interaction_genotype_Myc | Rab11fip1 | -3.693 | 3.43E-03 | glio |
| Interaction_genotype_Myc | Inpp4b | -3.058 | 3.43E-03 | glio |
| Interaction_genotype_Myc | Gm45397 | -2.856 | 3.43E-03 | glio |
| Interaction_genotype_Myc | Alas1 | -2.564 | 3.43E-03 | glio |
| Interaction_genotype_Pars2 | Tnni3k | -3.056 | 3.43E-03 | glio |
| Interaction_genotype_Pars2 | Gm15929 | -3.245 | 3.43E-03 | glio |
| Aars2_in.vivo_noRT | Bbs9 | 1.899 | 3.43E-03 | glio |
| Aars2_in.vivo_noRT | Usp25 | -1.269 | 3.43E-03 | glio |
| Atp5c1_in.vivo_noRT | Gm9967 | 2.281 | 3.43E-03 | glio |
| Bcas2_in.vivo_noRT | Gmnn | -1.121 | 3.43E-03 | glio |
| Bcas2_ex.vivo_noRT | Pla2g2f | 2.742 | 3.43E-03 | glio |
| Bora_ex.vivo_noRT | Cxcl13 | 2.808 | 3.43E-03 | glio |
| Bora_ex.vivo_noRT | Pcdhb19 | 2.806 | 3.43E-03 | glio |
| Chek1_in.vivo_noRT | A530041M06F | 2.174 | 3.43E-03 | glio |
| Chek1_in.vivo_noRT | Sh3glb1 | -0.818 | 3.43E-03 | glio |
| Chek1_ex.vivo_noRT | Hars | -1.3 | 3.43E-03 | glio |
| Eif2s1_ex.vivo_noRT | Tmem132c | 2.234 | 3.43E-03 | glio |

| | | | | |
|---|---|---|---|---|
| Eif2s1_ex.vivo_noRT | Gm44693 | 2.234 | 3.43E-03 | glio |
| Eif2s1_ex.vivo_noRT | Gpr146 | 3.421 | 3.43E-03 | glio |
| culture_effect | Rnf125 | -1.791 | 3.43E-04 | glio |
| culture_effect | Atp5k | 0.709 | 3.43E-04 | glio |
| Interaction_genotype_Mtor | Gm45184 | -4.253 | 3.43E-04 | glio |
| Interaction_genotype_Myc | Prkca | -2.923 | 3.43E-04 | glio |
| Interaction_genotype_Myc | Tmem167 | 1.337 | 3.43E-04 | glio |
| Interaction_genotype_Nat2 | Gm15894 | 3.722 | 3.43E-04 | glio |
| Aars2_in.vivo_noRT | Ankrd10 | -1.515 | 3.43E-04 | glio |
| Bora_ex.vivo_noRT | Gm20767 | 3.201 | 3.43E-04 | glio |
| Chek1_ex.vivo_noRT | Pla2g4b | 3.131 | 3.43E-04 | glio |
| culture_effect | Mindy1 | -0.727 | 3.43E-05 | glio |
| Interaction_genotype_Mtor | Gm13470 | -4.22 | 3.43E-05 | glio |
| Interaction_genotype_Mtor | Kcnj13 | -4.493 | 3.43E-05 | glio |
| Interaction_genotype_Mtor | Rab7b | -4.253 | 3.43E-05 | glio |
| Aars2_in.vivo_noRT | Gm47689 | 2.938 | 3.43E-05 | glio |
| Atr_in.vivo_noRT | Zfp988 | 2.901 | 3.43E-05 | glio |
| Atr_in.vivo_noRT | Rps7 | -0.861 | 3.43E-05 | glio |
| Bcas2_ex.vivo_noRT | Gm49698 | 2.742 | 3.43E-05 | glio |
| Bcas2_ex.vivo_noRT | Olfr881 | 2.742 | 3.43E-05 | glio |
| Chek1_in.vivo_noRT | Rhov | 2.858 | 3.43E-05 | glio |
| Chek1_ex.vivo_noRT | Aqp3 | 4.452 | 3.43E-05 | glio |
| culture_effect | Gal3st1 | 1.949 | 3.43E-06 | glio |
| interaction_RT | Loxl3 | -1.629 | 3.43E-06 | glio |
| interaction_RT | Gfap | 3.808 | 3.43E-06 | glio |
| Atr_in.vivo_noRT | Mapkbp1 | 2.808 | 3.43E-06 | glio |
| Bora_in.vivo_noRT | Ppnr | 3.274 | 3.43E-06 | glio |
| Interaction_genotype_Mtor | Rftn2 | -5.479 | 3.43E-07 | glio |
| Aars2_in.vivo_noRT | Gm15080 | 3.797 | 3.43E-07 | glio |
| Bora_in.vivo_noRT | 1700081N11R | 4.083 | 3.43E-07 | glio |
| Interaction_genotype_Mtor | BC065403 | -6.103 | 3.43E-08 | glio |
| Aars2_in.vivo_noRT | 1700001L05Ri | 3.629 | 3.43E-08 | glio |
| Bora_in.vivo_noRT | Apol7d | 4.035 | 3.43E-08 | glio |
| Bora_in.vivo_noRT | Fam228a | 4.149 | 3.43E-09 | glio |
| Bora_in.vivo_noRT | Peli3 | 4.021 | 3.43E-09 | glio |
| Interaction_genotype_Mtor | 4930478M13F | -7.782 | 3.43E-10 | glio |
| interaction_RT | Hnrnpd | -1.303 | 3.43E-10 | glio |
| Bora_in.vivo_noRT | Ifi207 | 4.673 | 3.43E-10 | glio |
| Chek1_in.vivo_noRT | 4933429H19R | 4.132 | 3.43E-10 | glio |
| Aars2_in.vivo_noRT | Gm12280 | 5.063 | 3.43E-11 | glio |
| Aars2_in.vivo_noRT | Serpina3g | 4.13 | 3.43E-11 | glio |
| Atr_in.vivo_noRT | Gm49701 | 4.484 | 3.43E-11 | glio |
| Bcas2_in.vivo_noRT | Bmp15 | 4.154 | 3.43E-12 | glio |
| Chek1_ex.vivo_noRT | Mpzl2 | 4.842 | 3.43E-12 | glio |
| Bcas2_in.vivo_noRT | Tmem74b | 4.543 | 3.43E-15 | glio |
| Bora_in.vivo_noRT | Gm40264 | 4.676 | 3.43E-15 | glio |
| Bcas2_in.vivo_noRT | Luzp2 | 4.281 | 3.43E-16 | glio |
| Atr_in.vivo_noRT | Gm16685 | 5.384 | 3.43E-20 | glio |
| Bcas2_in.vivo_noRT | Gm13398 | 4.807 | 3.43E-21 | glio |
| Bcas2_in.vivo_noRT | Tagap | 4.547 | 3.43E-21 | glio |
| culture_effect | Elp4 | -0.686 | 3.44E-03 | glio |
| Interaction_genotype_Aars2 | Cdkn2c | -2.612 | 3.44E-03 | glio |

| | | | | |
|---|---|---|---|---|
| Interaction_genotype_Bcas2 | Cpeb1 | -4.084 | 3.44E-03 | glio |
| Interaction_genotype_Myc | Spry3 | -3.422 | 3.44E-03 | glio |
| Interaction_genotype_Myc | Gm17056 | -3.329 | 3.44E-03 | glio |
| Interaction_genotype_Rpl3 | Mgst2 | 3.69 | 3.44E-03 | glio |
| Interaction_genotype_Rpl3 | Gm49694 | -4.169 | 3.44E-03 | glio |
| Interaction_genotype_Rpl3 | Fh1 | 1.879 | 3.44E-03 | glio |
| Interaction_genotype_Rpl3 | Cav1 | 2.384 | 3.44E-03 | glio |
| interaction_RT | Pcdh15 | 2.485 | 3.44E-03 | glio |
| Aars2_in.vivo_noRT | Ptdss2 | 1.544 | 3.44E-03 | glio |
| Aars2_ex.vivo_noRT | Gm17638 | 2.717 | 3.44E-03 | glio |
| Atr_in.vivo_noRT | 1700109H08R | 2.162 | 3.44E-03 | glio |
| Atr_in.vivo_noRT | Eepd1 | 1.92 | 3.44E-03 | glio |
| Atr_in.vivo_noRT | Eif4a1 | -0.703 | 3.44E-03 | glio |
| Bcas2_in.vivo_noRT | Dixdc1 | 2.174 | 3.44E-03 | glio |
| Bcas2_in.vivo_noRT | Pcdh1 | 2.172 | 3.44E-03 | glio |
| Bcas2_in.vivo_noRT | Krba1 | 2.175 | 3.44E-03 | glio |
| Bcas2_in.vivo_noRT | Gm49797 | 1.668 | 3.44E-03 | glio |
| Bora_in.vivo_noRT | Gm49123 | 2.276 | 3.44E-03 | glio |
| Bora_in.vivo_noRT | Lrp1b | 2.158 | 3.44E-03 | glio |
| Bora_in.vivo_noRT | Sumf2 | -1.434 | 3.44E-03 | glio |
| Bora_in.vivo_noRT | Cdc42se2 | -1.344 | 3.44E-03 | glio |
| Bora_ex.vivo_noRT | Parvg | 3.021 | 3.44E-03 | glio |
| Cenpj_in.vivo_noRT | Gad1 | 1.756 | 3.44E-03 | glio |
| Cenpj_ex.vivo_noRT | Kcnk7 | 2.591 | 3.44E-03 | glio |
| Cenpj_ex.vivo_noRT | Myoz1 | 2.591 | 3.44E-03 | glio |
| Cyp19a1_ex.vivo_noRT | Gm48435 | 2.171 | 3.44E-03 | glio |
| Cyp19a1_ex.vivo_noRT | 1700003E24Ri | 2.159 | 3.44E-03 | glio |
| culture_effect | Pcdhga9 | -2.176 | 3.44E-04 | glio |
| culture_effect | Dmap1 | -0.738 | 3.44E-04 | glio |
| Interaction_genotype_Mtor | Lhx9 | -4.416 | 3.44E-04 | glio |
| Interaction_genotype_Myc | Gm44646 | -3.316 | 3.44E-04 | glio |
| Interaction_genotype_Myc | Gm31717 | -2.729 | 3.44E-04 | glio |
| Interaction_genotype_Myc | Gm12925 | -4.068 | 3.44E-04 | glio |
| Interaction_genotype_Myc | Cep72 | -3.981 | 3.44E-04 | glio |
| Interaction_genotype_Myc | Cd38 | -4.028 | 3.44E-04 | glio |
| Interaction_genotype_Myc | Hsd17b14 | -4.245 | 3.44E-04 | glio |
| Interaction_genotype_Myc | A930037H05R | -3.823 | 3.44E-04 | glio |
| Interaction_genotype_Myc | Canx | 1.46 | 3.44E-04 | glio |
| Aars2_in.vivo_noRT | Hoxa7 | 2.55 | 3.44E-04 | glio |
| Atr_ex.vivo_noRT | Gm10354 | 2.667 | 3.44E-04 | glio |
| Bcas2_in.vivo_noRT | Fndc10 | 2.803 | 3.44E-04 | glio |
| Eif2s1_in.vivo_noRT | Gm45589 | 2.625 | 3.44E-04 | glio |
| culture_effect | Col5a3 | 1.153 | 3.44E-05 | glio |
| Aars2_ex.vivo_noRT | Gm49504 | 2.893 | 3.44E-05 | glio |
| Bard1_ex.vivo_noRT | Dpp6 | 2.814 | 3.44E-05 | glio |
| Bcas2_in.vivo_noRT | Sdcbp2 | 3.112 | 3.44E-05 | glio |
| Bcas2_in.vivo_noRT | Grip1 | 2.928 | 3.44E-05 | glio |
| Bora_in.vivo_noRT | Dpf1 | 2.564 | 3.44E-05 | glio |
| Bora_in.vivo_noRT | Aif1l | -1.309 | 3.44E-05 | glio |
| Interaction_genotype_Mtor | Gm14327 | -5.681 | 3.44E-06 | glio |
| Bcas2_in.vivo_noRT | Mir155hg | 3.562 | 3.44E-06 | glio |
| Bcas2_in.vivo_noRT | Gm13272 | 3.104 | 3.44E-06 | glio |

| | | | | |
|---|---|---|---|---|
| Bora_ex.vivo_noRT | Gm49698 | 3.201 | 3.44E-07 | glio |
| Bora_ex.vivo_noRT | Olfr881 | 3.201 | 3.44E-07 | glio |
| Aars2_in.vivo_noRT | Gm16576 | 4.181 | 3.44E-09 | glio |
| Atr_in.vivo_noRT | Glrp1 | 3.768 | 3.44E-10 | glio |
| Bora_in.vivo_noRT | D930028M14I | 4.936 | 3.44E-15 | glio |
| Chek1_in.vivo_noRT | Serpinb2 | 5.044 | 3.44E-16 | glio |
| Atr_in.vivo_noRT | Gm36356 | 5.273 | 3.44E-18 | glio |
| Atr_in.vivo_noRT | Sucnr1 | 5.533 | 3.44E-32 | glio |
| Interaction_genotype_Aars2 | Bst1 | -3.255 | 3.45E-03 | glio |
| Interaction_genotype_Aars2 | Prdx3 | 1.318 | 3.45E-03 | glio |
| Interaction_genotype_Mtor | Gm17056 | -3.54 | 3.45E-03 | glio |
| Interaction_genotype_Mtor | Srrm1 | 1.499 | 3.45E-03 | glio |
| Interaction_genotype_Myc | Wdfy2 | -2.905 | 3.45E-03 | glio |
| Interaction_genotype_Myc | Gm26797 | -2.669 | 3.45E-03 | glio |
| Interaction_genotype_Myc | Pnkp | -2.302 | 3.45E-03 | glio |
| Interaction_genotype_Myc | Tra2a | 1.478 | 3.45E-03 | glio |
| Aars2_in.vivo_noRT | Idh3a | -1.117 | 3.45E-03 | glio |
| Aars2_in.vivo_noRT | Cox7a2 | -0.64 | 3.45E-03 | glio |
| Atr_in.vivo_noRT | Plcd3 | 1.743 | 3.45E-03 | glio |
| Bcas2_in.vivo_noRT | Ndufa8 | -0.666 | 3.45E-03 | glio |
| Bcas2_ex.vivo_noRT | Akr1b7 | 3.141 | 3.45E-03 | glio |
| Cenpj_in.vivo_noRT | Gtsf1 | 2.022 | 3.45E-03 | glio |
| Chek1_in.vivo_noRT | 4930599N23R | 1.958 | 3.45E-03 | glio |
| Chek1_in.vivo_noRT | 2700033N17R | 1.884 | 3.45E-03 | glio |
| Chek1_ex.vivo_noRT | Zfp619 | 2.811 | 3.45E-03 | glio |
| Eif2s1_in.vivo_noRT | Rnft2 | 2.393 | 3.45E-03 | glio |
| Hsd17b10_in.vivo_noRT | Zfp971 | 2.04 | 3.45E-03 | glio |
| Interaction_genotype_Mtor | Spint4 | -4.398 | 3.45E-04 | glio |
| Interaction_genotype_Mtor | Myo1h | -4.307 | 3.45E-04 | glio |
| Interaction_genotype_Mtor | Ube2cbp | -3.814 | 3.45E-04 | glio |
| Interaction_genotype_Mtor | 2810013P06R | -3.025 | 3.45E-04 | glio |
| Interaction_genotype_Myc | Rgs9 | -3.511 | 3.45E-04 | glio |
| Interaction_genotype_Myc | Gm13625 | -3.782 | 3.45E-04 | glio |
| Interaction_genotype_Myc | Gm14742 | -2.729 | 3.45E-04 | glio |
| Interaction_genotype_Myc | Zfp53 | -3.457 | 3.45E-04 | glio |
| Interaction_genotype_Myc | Apbb1 | -3.202 | 3.45E-04 | glio |
| Aars2_ex.vivo_noRT | Gm16751 | 2.893 | 3.45E-04 | glio |
| Aars2_ex.vivo_noRT | Fhl1 | 2.582 | 3.45E-04 | glio |
| Aars2_ex.vivo_noRT | Ap4b1 | -2.77 | 3.45E-04 | glio |
| Atr_in.vivo_noRT | Gm42595 | 2.434 | 3.45E-04 | glio |
| Bora_in.vivo_noRT | Nanos3 | 2.332 | 3.45E-04 | glio |
| Bora_in.vivo_noRT | Fas | 2.716 | 3.45E-04 | glio |
| Bora_in.vivo_noRT | Ints2 | -1.754 | 3.45E-04 | glio |
| Bora_in.vivo_noRT | Tgs1 | -1.224 | 3.45E-04 | glio |
| Hsd17b10_in.vivo_noRT | 4933421A08R | 2.837 | 3.45E-04 | glio |
| culture_effect | Cdk5rap2 | 0.811 | 3.45E-05 | glio |
| Interaction_genotype_Myc | Ppp2r1b | -3.277 | 3.45E-05 | glio |
| Interaction_genotype_Myc | Esm1 | -3.441 | 3.45E-05 | glio |
| Interaction_genotype_Myc | Gm44658 | -4.14 | 3.45E-05 | glio |
| Atr_ex.vivo_noRT | Gpr22 | 3.164 | 3.45E-06 | glio |
| Atr_ex.vivo_noRT | Tnni3k | 3.16 | 3.45E-06 | glio |
| Chek1_in.vivo_noRT | Serpinb6a | 3.355 | 3.45E-06 | glio |

| | | | | |
|---|---|---|---|---|
| culture_effect | Rasl10a | -2.684 | 3.45E-07 | glio |
| Aars2_in.vivo_noRT | Gm44091 | 3.408 | 3.45E-07 | glio |
| Aars2_ex.vivo_noRT | Gad1 | 3.293 | 3.45E-07 | glio |
| Bora_ex.vivo_noRT | Brinp3 | 3.994 | 3.45E-07 | glio |
| culture_effect | Dgkq | -1.807 | 3.45E-11 | glio |
| Aars2_in.vivo_noRT | Rtn1 | 4.68 | 3.45E-11 | glio |
| Chek1_in.vivo_noRT | Gm13713 | 4.511 | 3.45E-16 | glio |
| Atr_in.vivo_noRT | Gm37864 | 5.263 | 3.45E-18 | glio |
| Bora_in.vivo_noRT | Speer4e | 4.936 | 3.45E-19 | glio |
| Bora_in.vivo_noRT | Helt | 4.936 | 3.45E-25 | glio |
| culture_effect | Tecrl | 1.791 | 3.46E-03 | glio |
| Interaction_genotype_Aars2 | Pakap | -2.881 | 3.46E-03 | glio |
| Interaction_genotype_Aars2 | Ftl1 | 0.815 | 3.46E-03 | glio |
| Interaction_genotype_Bcas2 | A930033H14R | -4.013 | 3.46E-03 | glio |
| Interaction_genotype_Bora | 4930505N22R | -3.61 | 3.46E-03 | glio |
| Interaction_genotype_Fanca | St18 | 4.175 | 3.46E-03 | glio |
| Interaction_genotype_Mtor | Cyp7a1 | -3.492 | 3.46E-03 | glio |
| Interaction_genotype_Mtor | 1700061G19R | -3.449 | 3.46E-03 | glio |
| Interaction_genotype_Mtor | Pttg1 | 2.052 | 3.46E-03 | glio |
| Interaction_genotype_Mtor | Hnrnpa0 | 1.613 | 3.46E-03 | glio |
| Interaction_genotype_Myc | Aqp9 | -2.729 | 3.46E-03 | glio |
| Interaction_genotype_Myc | Olfr90 | -2.859 | 3.46E-03 | glio |
| Interaction_genotype_Myc | Trim68 | -3.185 | 3.46E-03 | glio |
| Interaction_genotype_Myc | Gm2415 | -3.375 | 3.46E-03 | glio |
| Interaction_genotype_Myc | Dusp4 | -3.097 | 3.46E-03 | glio |
| Interaction_genotype_Myc | Cct5 | 1.212 | 3.46E-03 | glio |
| interaction_RT | C920021L13Ri | 2.277 | 3.46E-03 | glio |
| interaction_RT | Snrnp48 | 0.737 | 3.46E-03 | glio |
| Aars2_in.vivo_noRT | Cript | -1.005 | 3.46E-03 | glio |
| Aars2_in.vivo_noRT | Selenok | -0.748 | 3.46E-03 | glio |
| Bard1_ex.vivo_noRT | Fst | 2.814 | 3.46E-03 | glio |
| Bcas2_in.vivo_noRT | Gm35339 | 2.21 | 3.46E-03 | glio |
| Bcas2_in.vivo_noRT | Eml4 | -1.454 | 3.46E-03 | glio |
| Bora_in.vivo_noRT | Izumo4 | 2.149 | 3.46E-03 | glio |
| Bora_in.vivo_noRT | 4833413G10R | 2.265 | 3.46E-03 | glio |
| Bora_in.vivo_noRT | Ankrd13d | 2.189 | 3.46E-03 | glio |
| Bora_in.vivo_noRT | Gm45494 | 2.058 | 3.46E-03 | glio |
| Eif2s1_in.vivo_noRT | Csn3 | -2.711 | 3.46E-03 | glio |
| culture_effect | Qpct | -2.362 | 3.46E-04 | glio |
| culture_effect | Nr1h3 | 1.394 | 3.46E-04 | glio |
| culture_effect | Rasl11a | -1.417 | 3.46E-04 | glio |
| Interaction_genotype_Myc | Gm13594 | -4.154 | 3.46E-04 | glio |
| Interaction_genotype_Myc | Apbb3 | -3.586 | 3.46E-04 | glio |
| interaction_RT | Cxcl5 | 2.634 | 3.46E-04 | glio |
| Aars2_ex.vivo_noRT | Dcpp3 | 3.293 | 3.46E-04 | glio |
| Atp5c1_in.vivo_noRT | Gem | 2.668 | 3.46E-04 | glio |
| Atr_in.vivo_noRT | Vash2 | 2.569 | 3.46E-04 | glio |
| Atr_in.vivo_noRT | Pdcd2 | -1.386 | 3.46E-04 | glio |
| Atr_ex.vivo_noRT | Fgfrl1 | -1.158 | 3.46E-04 | glio |
| Bora_in.vivo_noRT | Gm29093 | 2.544 | 3.46E-04 | glio |
| Cenpj_ex.vivo_noRT | Adam5 | 2.591 | 3.46E-04 | glio |
| Cenpj_ex.vivo_noRT | 4933400F21Ri | 2.591 | 3.46E-04 | glio |

| | | | | |
|---|---|---|---|---|
| Cenpj_ex.vivo_noRT | Gm43915 | 2.995 | 3.46E-04 | glio |
| Cenpj_ex.vivo_noRT | Gm12794 | 2.591 | 3.46E-04 | glio |
| Hsd17b10_in.vivo_noRT | Nfkb2 | -1.683 | 3.46E-04 | glio |
| culture_effect | Pfdn1 | 0.486 | 3.46E-05 | glio |
| Interaction_genotype_Mtor | 4933433G15R | -4.919 | 3.46E-05 | glio |
| Aars2_in.vivo_noRT | Egf | 3.135 | 3.46E-05 | glio |
| Aars2_in.vivo_noRT | Spink10 | 2.752 | 3.46E-05 | glio |
| Aars2_in.vivo_noRT | Ercc5 | 1.993 | 3.46E-05 | glio |
| Aars2_in.vivo_noRT | Eif3i | -0.792 | 3.46E-05 | glio |
| Atr_ex.vivo_noRT | Cxcl1 | 3.947 | 3.46E-05 | glio |
| Bcas2_in.vivo_noRT | Enho | 2.926 | 3.46E-05 | glio |
| Chek1_ex.vivo_noRT | Lrwd1 | -2.561 | 3.46E-05 | glio |
| culture_effect | Cetn3 | -0.686 | 3.46E-06 | glio |
| Interaction_genotype_Atr | Pmp2 | -5.085 | 3.46E-06 | glio |
| Interaction_genotype_Mtor | Gm26619 | -5.824 | 3.46E-06 | glio |
| Aars2_in.vivo_noRT | Gm43017 | 3.212 | 3.46E-06 | glio |
| Atp5c1_in.vivo_noRT | Ccdc80 | 2.668 | 3.46E-06 | glio |
| Atr_in.vivo_noRT | Isoc2a | 2.929 | 3.46E-06 | glio |
| Bora_ex.vivo_noRT | Tnni3k | 3.206 | 3.46E-06 | glio |
| Bora_ex.vivo_noRT | Gm49808 | 2.808 | 3.46E-06 | glio |
| Hsd17b10_in.vivo_noRT | D030051J21Ri | 3.244 | 3.46E-06 | glio |
| culture_effect | Asxl3 | 1.275 | 3.46E-08 | glio |
| culture_effect | Gnb1 | 0.602 | 3.46E-08 | glio |
| Atr_in.vivo_noRT | Cks1brt | 4.096 | 3.46E-11 | glio |
| Aars2_in.vivo_noRT | AV039307 | 4.936 | 3.46E-12 | glio |
| Atr_in.vivo_noRT | Ccl7 | 5.533 | 3.46E-12 | glio |
| Chek1_in.vivo_noRT | Spn | 4.522 | 3.46E-12 | glio |
| Aars2_in.vivo_noRT | Nmrk2 | 4.685 | 3.46E-14 | glio |
| Aars2_in.vivo_noRT | Gm48161 | 4.539 | 3.46E-14 | glio |
| Chek1_in.vivo_noRT | A330032B11R | 4.777 | 3.46E-14 | glio |
| Chek1_in.vivo_noRT | Gm20646 | 5.044 | 3.46E-14 | glio |
| Atr_in.vivo_noRT | 4833419F23Ri | 5.533 | 3.46E-15 | glio |
| culture_effect | Pde8a | 2.967 | 3.46E-16 | glio |
| Atr_in.vivo_noRT | Gm40538 | 5.27 | 3.46E-21 | glio |
| culture_effect | Mrps6 | 1.589 | 3.46E-23 | glio |
| Bora_in.vivo_noRT | Gm49172 | 4.676 | 3.46E-23 | glio |
| Bora_in.vivo_noRT | Gm20625 | 4.936 | 3.46E-25 | glio |
| culture_effect | Col9a3 | -3.141 | 3.46E-35 | glio |
| culture_effect | Afg1l | 1.512 | 3.47E-03 | glio |
| culture_effect | Atp5d | -0.353 | 3.47E-03 | glio |
| Interaction_genotype_Atr | Krt19 | -2.369 | 3.47E-03 | glio |
| Interaction_genotype_Atr | Bckdha | 2.125 | 3.47E-03 | glio |
| Interaction_genotype_Bcas2 | Bco2 | -3.425 | 3.47E-03 | glio |
| Interaction_genotype_Bcas2 | Ank1 | -2.849 | 3.47E-03 | glio |
| Interaction_genotype_Bcas2 | Olfr279 | -3.841 | 3.47E-03 | glio |
| Interaction_genotype_Bcas2 | Gjd4 | -2.583 | 3.47E-03 | glio |
| Interaction_genotype_Bora | Gm16759 | -3.958 | 3.47E-03 | glio |
| Interaction_genotype_Myc | Zfp677 | -3.064 | 3.47E-03 | glio |
| Interaction_genotype_Myc | Prorsd1 | -2.416 | 3.47E-03 | glio |
| Interaction_genotype_Myc | Zbtb12 | -2.341 | 3.47E-03 | glio |
| Interaction_genotype_Myc | Slc25a13 | -2.094 | 3.47E-03 | glio |
| Interaction_genotype_Myc | Rps11 | 0.956 | 3.47E-03 | glio |

| | | | | |
|---|---|---|---|---|
| Aars2_in.vivo_noRT | Chdh | 2.039 | 3.47E-03 | glio |
| Atp5c1_in.vivo_noRT | Gm17173 | 2.281 | 3.47E-03 | glio |
| Atr_in.vivo_noRT | Sag | 2.155 | 3.47E-03 | glio |
| Bard1_ex.vivo_noRT | Cacng5 | 2.814 | 3.47E-03 | glio |
| Bcas2_in.vivo_noRT | Sbds | -0.971 | 3.47E-03 | glio |
| Bcas2_ex.vivo_noRT | Gpr17 | 2.742 | 3.47E-03 | glio |
| Bora_in.vivo_noRT | Gba2 | 2.156 | 3.47E-03 | glio |
| Bora_in.vivo_noRT | Gm45589 | 2.124 | 3.47E-03 | glio |
| Bora_in.vivo_noRT | Txnrd1 | -0.717 | 3.47E-03 | glio |
| Cenpj_in.vivo_noRT | Muc4 | 2.029 | 3.47E-03 | glio |
| Cenpj_ex.vivo_noRT | Boc | 2.198 | 3.47E-03 | glio |
| Chek1_in.vivo_noRT | Map3k20 | 1.829 | 3.47E-03 | glio |
| Chek1_in.vivo_noRT | Mycbp2 | -1.187 | 3.47E-03 | glio |
| Chek1_ex.vivo_noRT | Ifi207 | 2.874 | 3.47E-03 | glio |
| Chek1_ex.vivo_noRT | Rflna | 2.932 | 3.47E-03 | glio |
| Hsd17b10_in.vivo_noRT | Gm5089 | 2.578 | 3.47E-03 | glio |
| Hsd17b10_in.vivo_noRT | Gm43258 | 2.22 | 3.47E-03 | glio |
| Hsd17b10_in.vivo_noRT | Abat | 2.441 | 3.47E-03 | glio |
| Hsd17b10_in.vivo_noRT | Uckl1os | 2.382 | 3.47E-03 | glio |
| Hsd17b10_in.vivo_noRT | Smg9 | -1.623 | 3.47E-03 | glio |
| Hsd17b10_in.vivo_noRT | Cp | 1.456 | 3.47E-03 | glio |
| culture_effect | Mga | -0.701 | 3.47E-04 | glio |
| culture_effect | Mtfr1l | 0.67 | 3.47E-04 | glio |
| culture_effect | Vti1b | 0.498 | 3.47E-04 | glio |
| Interaction_genotype_Atr | Chia1 | -3.157 | 3.47E-04 | glio |
| Interaction_genotype_Mtor | Moap1 | -4.022 | 3.47E-04 | glio |
| Interaction_genotype_Mtor | Gps1 | 1.888 | 3.47E-04 | glio |
| Interaction_genotype_Mtor | Mier1 | 1.63 | 3.47E-04 | glio |
| Interaction_genotype_Myc | Tep1 | -3.22 | 3.47E-04 | glio |
| Aars2_in.vivo_noRT | Phf12 | -1.5 | 3.47E-04 | glio |
| Atp5c1_in.vivo_noRT | Ripply3 | 2.668 | 3.47E-04 | glio |
| Bcas2_ex.vivo_noRT | Gm21319 | 2.742 | 3.47E-04 | glio |
| Bcas2_ex.vivo_noRT | Gm21936 | 2.347 | 3.47E-04 | glio |
| Bcas2_ex.vivo_noRT | Prex1 | -2.955 | 3.47E-04 | glio |
| Chek1_in.vivo_noRT | Chek1 | -1.739 | 3.47E-04 | glio |
| Chek1_in.vivo_noRT | Ipo8 | -1.666 | 3.47E-04 | glio |
| Eif2s1_in.vivo_noRT | Esp24 | 2.143 | 3.47E-04 | glio |
| culture_effect | Dipk1b | 1.039 | 3.47E-05 | glio |
| Interaction_genotype_Mtor | Zfhx2os | -4.827 | 3.47E-05 | glio |
| Interaction_genotype_Mtor | Slfn3 | -4.449 | 3.47E-05 | glio |
| Interaction_genotype_Myc | Dhx29 | -3.207 | 3.47E-05 | glio |
| Atr_in.vivo_noRT | Phip | -1.695 | 3.47E-05 | glio |
| Bard1_ex.vivo_noRT | Kctd4 | 3.819 | 3.47E-05 | glio |
| Chek1_in.vivo_noRT | Myom1 | 3.05 | 3.47E-05 | glio |
| Chek1_in.vivo_noRT | C330013E15Ri | 2.928 | 3.47E-05 | glio |
| Chek1_ex.vivo_noRT | 1600014C23R | 3.749 | 3.47E-05 | glio |
| Chek1_ex.vivo_noRT | Pyy | 2.963 | 3.47E-05 | glio |
| Chek1_ex.vivo_noRT | Slc26a11 | 3.371 | 3.47E-05 | glio |
| Hsd17b10_in.vivo_noRT | Calhm5 | 2.888 | 3.47E-05 | glio |
| Hsd17b10_in.vivo_noRT | Grin2d | 2.837 | 3.47E-05 | glio |
| Interaction_genotype_Mtor | Psma6 | 2.12 | 3.47E-06 | glio |
| Aars2_in.vivo_noRT | Tnfrsf1b | 3.033 | 3.47E-06 | glio |

| | | | | |
|---|---|---|---|---|
| Hsd17b10_in.vivo_noRT | Nmrk2 | 2.974 | 3.47E-06 | glio |
| Chek1_ex.vivo_noRT | Gm10696 | 4.443 | 3.47E-07 | glio |
| Aars2_in.vivo_noRT | Gm2012 | 4.163 | 3.47E-09 | glio |
| Chek1_ex.vivo_noRT | AC125149.2 | 4.447 | 3.47E-09 | glio |
| Bcas2_in.vivo_noRT | E130215H24R | 4.016 | 3.47E-10 | glio |
| Bcas2_in.vivo_noRT | Gm42695 | 4.547 | 3.47E-11 | glio |
| Bora_in.vivo_noRT | Npy1r | 4.291 | 3.47E-15 | glio |
| Chek1_in.vivo_noRT | Clcn1 | 4.398 | 3.47E-15 | glio |
| Aars2_in.vivo_noRT | Gm4858 | 5.327 | 3.47E-16 | glio |
| Chek1_in.vivo_noRT | Impg2 | 5.044 | 3.47E-18 | glio |
| Chek1_ex.vivo_noRT | Gm49172 | 5.643 | 3.47E-21 | glio |
| culture_effect | Ceacam16 | -1.592 | 3.48E-03 | glio |
| culture_effect | Aldh1a3 | 1.114 | 3.48E-03 | glio |
| Interaction_genotype_Atr | Dlx2 | -3.82 | 3.48E-03 | glio |
| Interaction_genotype_Mtor | Sox11 | -3.453 | 3.48E-03 | glio |
| Interaction_genotype_Mtor | Msl3l2 | -3.566 | 3.48E-03 | glio |
| Interaction_genotype_Myc | Ptprg | -2.923 | 3.48E-03 | glio |
| Interaction_genotype_Myc | Fah | -2.735 | 3.48E-03 | glio |
| Interaction_genotype_Myc | Nexn | -2.955 | 3.48E-03 | glio |
| Interaction_genotype_Pars2 | Gm10300 | -4.097 | 3.48E-03 | glio |
| Interaction_genotype_Pars2 | Gm49808 | -2.661 | 3.48E-03 | glio |
| Interaction_genotype_Pars2 | Tbcb | 1.066 | 3.48E-03 | glio |
| Interaction_genotype_Pars2 | Psmb2 | 0.93 | 3.48E-03 | glio |
| Atr_in.vivo_noRT | 4930599N23R | 1.924 | 3.48E-03 | glio |
| Atr_in.vivo_noRT | Vps13a | -1.089 | 3.48E-03 | glio |
| Bora_in.vivo_noRT | Gm7879 | 2.284 | 3.48E-03 | glio |
| Bora_in.vivo_noRT | 1700028J19Ri | 2.139 | 3.48E-03 | glio |
| Bora_in.vivo_noRT | Fam13c | 2.318 | 3.48E-03 | glio |
| Bora_in.vivo_noRT | Tmc6 | -1.384 | 3.48E-03 | glio |
| Chek1_in.vivo_noRT | Arf4 | -0.747 | 3.48E-03 | glio |
| Aars2_in.vivo_noRT | 2600006K01R | 2.668 | 3.48E-04 | glio |
| Atp5c1_in.vivo_noRT | Gm45620 | 2.668 | 3.48E-04 | glio |
| Atp5c1_in.vivo_noRT | Gm38534 | 2.668 | 3.48E-04 | glio |
| Bcas2_ex.vivo_noRT | 4930588A03R | 2.342 | 3.48E-04 | glio |
| Bcas2_ex.vivo_noRT | Gm39526 | 2.742 | 3.48E-04 | glio |
| Bora_in.vivo_noRT | Gm12258 | 2.73 | 3.48E-04 | glio |
| Hsd17b10_in.vivo_noRT | A330040F15Ri | 2.837 | 3.48E-04 | glio |
| Interaction_genotype_Mtor | Kcnab3 | -5.077 | 3.48E-05 | glio |
| Interaction_genotype_Mtor | 1010001B22R | -4.619 | 3.48E-05 | glio |
| Interaction_genotype_Mtor | Ifit1bl1 | -5.921 | 3.48E-05 | glio |
| Interaction_genotype_Mtor | A430018G15R | -3.556 | 3.48E-05 | glio |
| Aars2_in.vivo_noRT | Cyp3a13 | 3.135 | 3.48E-05 | glio |
| Aars2_in.vivo_noRT | Gm10138 | 2.705 | 3.48E-05 | glio |
| Aars2_in.vivo_noRT | Akr1a1 | -0.951 | 3.48E-05 | glio |
| Aars2_ex.vivo_noRT | Eif4e3 | 3.295 | 3.48E-05 | glio |
| Atr_in.vivo_noRT | Btbd6 | 2.65 | 3.48E-05 | glio |
| Bard1_ex.vivo_noRT | Gm13032 | 3.213 | 3.48E-05 | glio |
| Bcas2_in.vivo_noRT | Cep295nl | 2.929 | 3.48E-05 | glio |
| Bora_in.vivo_noRT | Stmn4 | 4.42 | 3.48E-05 | glio |
| Chek1_ex.vivo_noRT | Grip2 | 3.658 | 3.48E-05 | glio |
| Chek1_ex.vivo_noRT | Nme6 | -1.889 | 3.48E-05 | glio |
| Bcas2_in.vivo_noRT | Gm12500 | 3.166 | 3.48E-06 | glio |

| | | | | |
|---|---|---|---|---|
| Hsd17b10_in.vivo_noRT | Gm15579 | 3.616 | 3.48E-06 | glio |
| Aars2_in.vivo_noRT | Gm48882 | 4.203 | 3.48E-09 | glio |
| Chek1_in.vivo_noRT | Gm43409 | 4.78 | 3.48E-10 | glio |
| Atr_in.vivo_noRT | Tnfsf13os | 4.763 | 3.48E-12 | glio |
| Bora_in.vivo_noRT | Rslcan18 | 4.55 | 3.48E-13 | glio |
| Aars2_in.vivo_noRT | Txndc2 | 4.547 | 3.48E-16 | glio |
| Chek1_in.vivo_noRT | Adgre5 | 4.784 | 3.48E-16 | glio |
| Chek1_in.vivo_noRT | Gm48899 | 4.784 | 3.48E-16 | glio |
| Aars2_in.vivo_noRT | Gm14019 | 5.327 | 3.48E-18 | glio |
| Aars2_in.vivo_noRT | Ube4bos1 | 5.063 | 3.48E-18 | glio |
| Bcas2_in.vivo_noRT | Gm6803 | 4.807 | 3.48E-18 | glio |
| Aars2_in.vivo_noRT | Esp15 | 5.327 | 3.48E-20 | glio |
| Chek1_in.vivo_noRT | Gm16933 | 4.522 | 3.48E-20 | glio |
| Bora_in.vivo_noRT | Gm13398 | 4.936 | 3.48E-22 | glio |
| Atr_in.vivo_noRT | Gm36037 | 5.533 | 3.48E-26 | glio |
| culture_effect | Olig1 | -2.754 | 3.48E-31 | glio |
| culture_effect | Junos | -1.67 | 3.49E-03 | glio |
| culture_effect | Scml2 | -1.521 | 3.49E-03 | glio |
| Interaction_genotype_Mtor | Rassf7 | -2.681 | 3.49E-03 | glio |
| interaction_RT | Lin37 | 0.839 | 3.49E-03 | glio |
| interaction_RT | Mrpl19 | -0.837 | 3.49E-03 | glio |
| interaction_RT | Baz1b | -0.731 | 3.49E-03 | glio |
| Aars2_in.vivo_noRT | Large1 | 1.53 | 3.49E-03 | glio |
| Atr_in.vivo_noRT | Psd3 | 1.821 | 3.49E-03 | glio |
| Atr_in.vivo_noRT | Entr1 | -1.374 | 3.49E-03 | glio |
| Atr_in.vivo_noRT | Far1 | -1.284 | 3.49E-03 | glio |
| Bcas2_in.vivo_noRT | Dhrs7 | -1.32 | 3.49E-03 | glio |
| Bora_in.vivo_noRT | Bag5 | -1.265 | 3.49E-03 | glio |
| Bora_in.vivo_noRT | Birc5 | -1.368 | 3.49E-03 | glio |
| Bora_ex.vivo_noRT | Gm7247 | 2.404 | 3.49E-03 | glio |
| Bora_ex.vivo_noRT | Vmn1r181 | 2.806 | 3.49E-03 | glio |
| Bora_ex.vivo_noRT | 2410141K09R | 2.417 | 3.49E-03 | glio |
| Cdkn2a_in.vivo_noRT | Slc17a6 | -1.675 | 3.49E-03 | glio |
| Cenpj_in.vivo_noRT | Zscan4d | 2.022 | 3.49E-03 | glio |
| Chek1_ex.vivo_noRT | Pcdhb14 | 2.868 | 3.49E-03 | glio |
| Eif2s1_in.vivo_noRT | Cldn19 | 2.422 | 3.49E-03 | glio |
| Hsd17b10_in.vivo_noRT | Phldb2 | 2.413 | 3.49E-03 | glio |
| Hsd17b10_in.vivo_noRT | Gm26930 | 2.458 | 3.49E-03 | glio |
| culture_effect | Fancf | 1.206 | 3.49E-04 | glio |
| Interaction_genotype_Bora | Gm48682 | -4.452 | 3.49E-04 | glio |
| Interaction_genotype_Bora | Cpne8 | -4.864 | 3.49E-04 | glio |
| Interaction_genotype_Bora | Ccnt1 | 2.144 | 3.49E-04 | glio |
| Interaction_genotype_Myc | Cyp2s1 | -4.292 | 3.49E-04 | glio |
| Interaction_genotype_Myc | Napb | -3.543 | 3.49E-04 | glio |
| Interaction_genotype_Myc | Slc36a3 | -3.939 | 3.49E-04 | glio |
| Interaction_genotype_Myc | Lrrc28 | -3.409 | 3.49E-04 | glio |
| Interaction_genotype_Myc | Ankrd39 | -2.898 | 3.49E-04 | glio |
| Interaction_genotype_Myc | Hsd17b4 | -2.699 | 3.49E-04 | glio |
| Interaction_genotype_Myc | Psenen | 1.317 | 3.49E-04 | glio |
| Interaction_genotype_Myc | Atp5e | 1.624 | 3.49E-04 | glio |
| Interaction_genotype_Rad51 | Yeats4 | 2.006 | 3.49E-04 | glio |
| Interaction_genotype_Rad51 | Pmm2 | -3.466 | 3.49E-04 | glio |

| | | | | |
|---|---|---|---|---|
| Interaction_genotype_Rad51 | Olfm2 | -5.033 | 3.49E-04 | glio |
| interaction_RT | Selenom | 0.78 | 3.49E-04 | glio |
| Atr_ex.vivo_noRT | Gm32122 | 3.164 | 3.49E-04 | glio |
| Bcas2_ex.vivo_noRT | Speer4e | 2.742 | 3.49E-04 | glio |
| Bora_in.vivo_noRT | Plek2 | 2.485 | 3.49E-04 | glio |
| Bora_in.vivo_noRT | Plcxd2 | 2.623 | 3.49E-04 | glio |
| Bora_in.vivo_noRT | Pbk | -1.961 | 3.49E-04 | glio |
| Chek1_in.vivo_noRT | Anapc15 | -1.396 | 3.49E-04 | glio |
| Chek1_in.vivo_noRT | Eef1b2 | -0.699 | 3.49E-04 | glio |
| Chek1_ex.vivo_noRT | Gm10644 | 3.282 | 3.49E-04 | glio |
| culture_effect | Gm20492 | 2.105 | 3.49E-05 | glio |
| Atr_in.vivo_noRT | Lrp1 | 2.891 | 3.49E-05 | glio |
| Bora_in.vivo_noRT | Car12 | 3.084 | 3.49E-06 | glio |
| Bora_in.vivo_noRT | Gm26728 | 3.431 | 3.49E-06 | glio |
| Chek1_in.vivo_noRT | C77080 | 3.166 | 3.49E-06 | glio |
| Chek1_in.vivo_noRT | Micall2 | 3.384 | 3.49E-06 | glio |
| culture_effect | Slc8a3 | -3.19 | 3.49E-07 | glio |
| Interaction_genotype_Mtor | Mucl3 | -6.531 | 3.49E-07 | glio |
| Atp5c1_in.vivo_noRT | Gpc5 | 2.668 | 3.49E-07 | glio |
| Bora_ex.vivo_noRT | BC061195 | 3.201 | 3.49E-07 | glio |
| Aars2_in.vivo_noRT | Sema6b | 3.851 | 3.49E-08 | glio |
| Atr_in.vivo_noRT | 2210406O10R | 4.216 | 3.49E-09 | glio |
| Aars2_in.vivo_noRT | Fam189a1 | 3.656 | 3.49E-10 | glio |
| Aars2_in.vivo_noRT | Kmo | 3.685 | 3.49E-10 | glio |
| Aars2_in.vivo_noRT | Zfp808 | 4.21 | 3.49E-12 | glio |
| Hsd17b10_in.vivo_noRT | AU019990 | 3.616 | 3.49E-12 | glio |
| culture_effect | Mprip | 1.148 | 3.49E-13 | glio |
| Bcas2_in.vivo_noRT | Tmem52 | 4.957 | 3.49E-15 | glio |
| culture_effect | Gls2 | -1.519 | 3.50E-03 | glio |
| culture_effect | Tlcd1 | 0.643 | 3.50E-03 | glio |
| Interaction_genotype_Atr | Rhobtb3 | 1.915 | 3.50E-03 | glio |
| Interaction_genotype_Mtor | Cyb5d2 | -2.466 | 3.50E-03 | glio |
| Interaction_genotype_Mtor | Nr6a1 | -2.937 | 3.50E-03 | glio |
| Interaction_genotype_Mtor | Tln1 | 1.538 | 3.50E-03 | glio |
| Interaction_genotype_Mtor | Sar1a | 1.553 | 3.50E-03 | glio |
| Interaction_genotype_Mtor | Polr1d | 1.112 | 3.50E-03 | glio |
| Interaction_genotype_Myc | 9630014M24F | -3.121 | 3.50E-03 | glio |
| Interaction_genotype_Myc | Tsga10 | -3.374 | 3.50E-03 | glio |
| Interaction_genotype_Uqcrb | Mfsd13a | 3.694 | 3.50E-03 | glio |
| Interaction_genotype_Uqcrb | Pmp2 | -4.21 | 3.50E-03 | glio |
| Interaction_genotype_Uqcrb | AV039307 | -4.746 | 3.50E-03 | glio |
| interaction_RT | Aoc1 | 2.832 | 3.50E-03 | glio |
| Aars2_in.vivo_noRT | Zfp971 | 1.932 | 3.50E-03 | glio |
| Aars2_in.vivo_noRT | Atp8a1 | 1.416 | 3.50E-03 | glio |
| Aars2_in.vivo_noRT | Rheb | -0.771 | 3.50E-03 | glio |
| Atp5c1_in.vivo_noRT | Gm37214 | 2.408 | 3.50E-03 | glio |
| Atp5c1_in.vivo_noRT | D030040B21R | 2.281 | 3.50E-03 | glio |
| Atp5c1_in.vivo_noRT | Bmp15 | 2.015 | 3.50E-03 | glio |
| Atr_in.vivo_noRT | 9130024F11Ri | 2.012 | 3.50E-03 | glio |
| Atr_in.vivo_noRT | Csnk2b | -0.913 | 3.50E-03 | glio |
| Bora_in.vivo_noRT | 0610009E02Ri | 2.3 | 3.50E-03 | glio |
| Bora_in.vivo_noRT | Mia2 | -1.24 | 3.50E-03 | glio |

| | | | | |
|---|---|---|---|---|
| Bora_in.vivo_noRT | Xlr | -1.248 | 3.50E-03 | glio |
| Bora_ex.vivo_noRT | Kremen1 | -2.434 | 3.50E-03 | glio |
| Chek1_in.vivo_noRT | Fndc7 | 2.29 | 3.50E-03 | glio |
| Chek1_in.vivo_noRT | Hist1h4a | 2.051 | 3.50E-03 | glio |
| Chek1_in.vivo_noRT | Nipal3 | 2.096 | 3.50E-03 | glio |
| Chek1_in.vivo_noRT | Six5 | 1.877 | 3.50E-03 | glio |
| Chek1_ex.vivo_noRT | Cgref1 | -2.13 | 3.50E-03 | glio |
| Hsd17b10_in.vivo_noRT | Hyal3 | 2.557 | 3.50E-03 | glio |
| Hsd17b10_in.vivo_noRT | Gm20536 | 2.241 | 3.50E-03 | glio |
| Hsd17b10_in.vivo_noRT | Kdelr3 | -1.76 | 3.50E-03 | glio |
| Interaction_genotype_Aars2 | Gm12500 | -4.309 | 3.50E-04 | glio |
| Interaction_genotype_Aars2 | H3f3aos | -4.449 | 3.50E-04 | glio |
| Interaction_genotype_Atr | Ldlrad3 | -4.033 | 3.50E-04 | glio |
| Interaction_genotype_Atr | Sap30 | 2.056 | 3.50E-04 | glio |
| Interaction_genotype_Bora | Eme2 | -4.422 | 3.50E-04 | glio |
| Interaction_genotype_Mtor | Gm48161 | -3.704 | 3.50E-04 | glio |
| Interaction_genotype_Mtor | Acbd7 | -4.549 | 3.50E-04 | glio |
| Interaction_genotype_Mtor | Bbs9 | -3.922 | 3.50E-04 | glio |
| Interaction_genotype_Mtor | Nanos3 | -3.503 | 3.50E-04 | glio |
| Interaction_genotype_Mtor | Helz | 1.982 | 3.50E-04 | glio |
| Interaction_genotype_Mtor | Psma1 | 1.766 | 3.50E-04 | glio |
| Interaction_genotype_Mtor | Sf3b2 | 1.66 | 3.50E-04 | glio |
| Interaction_genotype_Myc | Ctla4 | -2.729 | 3.50E-04 | glio |
| interaction_RT | Hist1h2ab | -2.93 | 3.50E-04 | glio |
| Aars2_in.vivo_noRT | Pkdcc | 2.615 | 3.50E-04 | glio |
| Aars2_in.vivo_noRT | Gm12981 | 2.54 | 3.50E-04 | glio |
| Aars2_in.vivo_noRT | Gm26670 | 2.377 | 3.50E-04 | glio |
| Aars2_in.vivo_noRT | Lnpk | 1.348 | 3.50E-04 | glio |
| Atr_in.vivo_noRT | Abca13 | 2.802 | 3.50E-04 | glio |
| Atr_in.vivo_noRT | Dner | 2.334 | 3.50E-04 | glio |
| Bora_ex.vivo_noRT | Gm48140 | 2.408 | 3.50E-04 | glio |
| Chek1_in.vivo_noRT | 1700001G11R | 2.233 | 3.50E-04 | glio |
| Chek1_in.vivo_noRT | Gng5 | -0.745 | 3.50E-04 | glio |
| Interaction_genotype_Mtor | Ramp3 | -4.85 | 3.50E-05 | glio |
| Interaction_genotype_Mtor | B4galnt1 | -4.139 | 3.50E-05 | glio |
| Atr_in.vivo_noRT | Sik1 | 2.916 | 3.50E-05 | glio |
| Bcas2_ex.vivo_noRT | BC061195 | 2.742 | 3.50E-05 | glio |
| Chek1_ex.vivo_noRT | Clcn1 | 2.89 | 3.50E-05 | glio |
| Chek1_ex.vivo_noRT | Tmem190 | 3.968 | 3.50E-05 | glio |
| interaction_RT | Smarca4 | -1.117 | 3.50E-06 | glio |
| interaction_RT | Ubald2 | 1.262 | 3.50E-06 | glio |
| interaction_RT | Nbdy | 0.993 | 3.50E-06 | glio |
| interaction_RT | Aig1 | 0.843 | 3.50E-06 | glio |
| Atr_in.vivo_noRT | Phf11a | 3.441 | 3.50E-07 | glio |
| Aars2_in.vivo_noRT | Kctd21 | 3.766 | 3.50E-08 | glio |
| Atp5c1_in.vivo_noRT | Gm13191 | 2.668 | 3.50E-08 | glio |
| Atp5c1_in.vivo_noRT | 4933416E03Ri | 2.668 | 3.50E-08 | glio |
| Bcas2_in.vivo_noRT | Gm12280 | 4.543 | 3.50E-09 | glio |
| Bora_in.vivo_noRT | Hist2h3c2 | 4.67 | 3.50E-10 | glio |
| Chek1_in.vivo_noRT | Spns3 | 4.386 | 3.50E-10 | glio |
| Chek1_ex.vivo_noRT | Obscn | 4.842 | 3.50E-10 | glio |
| Aars2_in.vivo_noRT | Prox2os | 4.023 | 3.50E-11 | glio |

| | | | | |
|---|---|---|---|---|
| Atr_in.vivo_noRT | Gm12500 | 4.429 | 3.50E-11 | glio |
| Bora_in.vivo_noRT | Col8a1 | 4.936 | 3.50E-13 | glio |
| Aars2_in.vivo_noRT | 2010320O07R | 5.19 | 3.50E-14 | glio |
| Atr_in.vivo_noRT | Dlx4 | 5.008 | 3.50E-14 | glio |
| culture_effect | Hist1h3d | 3.272 | 3.50E-15 | glio |
| Chek1_in.vivo_noRT | Gm16344 | 4.656 | 3.50E-15 | glio |
| Chek1_in.vivo_noRT | Olfr718-ps1 | 4.78 | 3.50E-19 | glio |
| Chek1_in.vivo_noRT | Zscan4-ps2 | 5.044 | 3.50E-22 | glio |
| Aars2_in.vivo_noRT | Gm13964 | 5.327 | 3.50E-26 | glio |
| culture_effect | Fmc1 | 0.4 | 3.51E-03 | glio |
| Interaction_genotype_Atr | Gm5544 | -3.682 | 3.51E-03 | glio |
| Interaction_genotype_Mtor | Ctf1 | -2.75 | 3.51E-03 | glio |
| Interaction_genotype_Mtor | Dzip1l | -2.479 | 3.51E-03 | glio |
| Interaction_genotype_Myc | Nat8f4 | -2.984 | 3.51E-03 | glio |
| Interaction_genotype_Myc | Gm47802 | -3.121 | 3.51E-03 | glio |
| Aars2_in.vivo_noRT | Calhm2 | 1.738 | 3.51E-03 | glio |
| Bard1_ex.vivo_noRT | Gm17210 | 3.01 | 3.51E-03 | glio |
| Bard1_ex.vivo_noRT | Pla1a | 2.814 | 3.51E-03 | glio |
| Bora_in.vivo_noRT | Plk1 | -1.746 | 3.51E-03 | glio |
| Interaction_genotype_Mtor | Gm13830 | -4.486 | 3.51E-04 | glio |
| Interaction_genotype_Myc | Zscan4-ps3 | -2.729 | 3.51E-04 | glio |
| Interaction_genotype_Myc | Gm49494 | -3.913 | 3.51E-04 | glio |
| Interaction_genotype_Myc | Tfeb | -3.3 | 3.51E-04 | glio |
| Interaction_genotype_Myc | Mrps6 | 1.954 | 3.51E-04 | glio |
| Interaction_genotype_Pars2 | Ifitm6 | -5.021 | 3.51E-04 | glio |
| Interaction_genotype_Pars2 | Gm20455 | -5.176 | 3.51E-04 | glio |
| Interaction_genotype_Pars2 | Gm32090 | -4.049 | 3.51E-04 | glio |
| Interaction_genotype_Pars2 | Crhbp | -4.306 | 3.51E-04 | glio |
| Interaction_genotype_Pars2 | Trpm3 | -4.921 | 3.51E-04 | glio |
| Interaction_genotype_Pars2 | Scube3 | -3.67 | 3.51E-04 | glio |
| Interaction_genotype_Pars2 | Gm50321 | -4.165 | 3.51E-04 | glio |
| Interaction_genotype_Pars2 | Gm44752 | -4.445 | 3.51E-04 | glio |
| interaction_RT | Prpf31 | -0.835 | 3.51E-04 | glio |
| Aars2_in.vivo_noRT | Vax2 | 2.472 | 3.51E-04 | glio |
| Bcas2_in.vivo_noRT | Stac3 | 2.665 | 3.51E-04 | glio |
| Bcas2_in.vivo_noRT | Pls1 | 2.549 | 3.51E-04 | glio |
| Chek1_in.vivo_noRT | Hdac7 | 2.526 | 3.51E-04 | glio |
| Chek1_in.vivo_noRT | 9230112E08Ri | 2.56 | 3.51E-04 | glio |
| Interaction_genotype_Atr | Gm17259 | -4.711 | 3.51E-05 | glio |
| Interaction_genotype_Mtor | Gm17173 | -4.816 | 3.51E-05 | glio |
| Interaction_genotype_Mtor | Necab2 | -4.536 | 3.51E-05 | glio |
| Aars2_in.vivo_noRT | Gucd1 | 2.238 | 3.51E-05 | glio |
| Aars2_ex.vivo_noRT | Fam189a1 | 3.293 | 3.51E-05 | glio |
| Aars2_ex.vivo_noRT | Cemip | 3.293 | 3.51E-05 | glio |
| Aars2_ex.vivo_noRT | Olfr718-ps1 | 2.899 | 3.51E-05 | glio |
| Aars2_ex.vivo_noRT | Kit | 3.293 | 3.51E-05 | glio |
| Aars2_ex.vivo_noRT | Sdr16c5 | 3.674 | 3.51E-05 | glio |
| Aars2_ex.vivo_noRT | Agap2 | 3.107 | 3.51E-05 | glio |
| Aars2_ex.vivo_noRT | AF067061 | 3.293 | 3.51E-05 | glio |
| Bora_in.vivo_noRT | 4930524O07R | 3.012 | 3.51E-05 | glio |
| Hsd17b10_in.vivo_noRT | Zfp14 | 3.093 | 3.51E-05 | glio |
| Hsd17b10_in.vivo_noRT | Nt5e | 2.658 | 3.51E-05 | glio |

| | | | | |
|---|---|---|---|---|
| Bora_in.vivo_noRT | Sept5 | 3.197 | 3.51E-06 | glio |
| Chek1_in.vivo_noRT | Plekhg6 | 3.367 | 3.51E-06 | glio |
| culture_effect | Cbx8 | -1.123 | 3.51E-07 | glio |
| culture_effect | Lyar | 0.657 | 3.51E-07 | glio |
| interaction_RT | Psme2 | 1.073 | 3.51E-07 | glio |
| Bora_in.vivo_noRT | Ifit1bl1 | 4.404 | 3.51E-07 | glio |
| culture_effect | Tssk6 | 2.434 | 3.51E-09 | glio |
| culture_effect | Wwp1 | 1.079 | 3.51E-09 | glio |
| Chek1_in.vivo_noRT | 1110017D15R | 4.043 | 3.51E-09 | glio |
| Chek1_ex.vivo_noRT | Tff2 | 4.842 | 3.51E-09 | glio |
| Bcas2_in.vivo_noRT | Gm14698 | 4.413 | 3.51E-10 | glio |
| Chek1_in.vivo_noRT | Hif3a | 4.401 | 3.51E-10 | glio |
| Bcas2_in.vivo_noRT | Pdgfb | 4.544 | 3.51E-16 | glio |
| Bcas2_in.vivo_noRT | Gtsf1 | 4.807 | 3.51E-17 | glio |
| Atr_in.vivo_noRT | BB557941 | 5.273 | 3.51E-19 | glio |
| Bora_in.vivo_noRT | Adam5 | 4.672 | 3.51E-20 | glio |
| Chek1_in.vivo_noRT | Vmn1r32 | 5.044 | 3.51E-24 | glio |
| Chek1_in.vivo_noRT | Glyat | 5.044 | 3.51E-24 | glio |
| Interaction_genotype_Aars2 | Fam25c | -2.875 | 3.52E-03 | glio |
| Interaction_genotype_Aars2 | Ech1 | 1.545 | 3.52E-03 | glio |
| Interaction_genotype_Atr | Alox5ap | -4.119 | 3.52E-03 | glio |
| Interaction_genotype_Bora | 2610027K06R | -3.52 | 3.52E-03 | glio |
| Interaction_genotype_Lig4 | Scgb1c1 | 4.032 | 3.52E-03 | glio |
| Interaction_genotype_Mtor | Plcb1 | -3.175 | 3.52E-03 | glio |
| Interaction_genotype_Mtor | Ankrd17 | 1.67 | 3.52E-03 | glio |
| Interaction_genotype_Myc | Slc44a2 | -3.259 | 3.52E-03 | glio |
| Interaction_genotype_Myc | Sdhb | 1.139 | 3.52E-03 | glio |
| interaction_RT | Ngfr | -1.342 | 3.52E-03 | glio |
| Aars2_in.vivo_noRT | Gm45767 | 1.988 | 3.52E-03 | glio |
| Aars2_in.vivo_noRT | Tcta | 1.689 | 3.52E-03 | glio |
| Atr_in.vivo_noRT | Hoxa1 | 2.036 | 3.52E-03 | glio |
| Atr_in.vivo_noRT | Pold3 | -1.254 | 3.52E-03 | glio |
| Bcas2_in.vivo_noRT | Gm20721 | 2.007 | 3.52E-03 | glio |
| Bora_in.vivo_noRT | Gm20186 | 2.259 | 3.52E-03 | glio |
| Bora_in.vivo_noRT | Hao1 | 2.252 | 3.52E-03 | glio |
| Bora_in.vivo_noRT | Matr3 | -0.816 | 3.52E-03 | glio |
| Chek1_ex.vivo_noRT | Ift172 | 2.165 | 3.52E-03 | glio |
| Interaction_genotype_Myc | Nbea | -3.118 | 3.52E-04 | glio |
| Atr_in.vivo_noRT | Tlx3 | 2.488 | 3.52E-04 | glio |
| Hsd17b10_in.vivo_noRT | Tacr2 | 2.715 | 3.52E-04 | glio |
| Atp5c1_in.vivo_noRT | Icam2 | 2.746 | 3.52E-05 | glio |
| Atr_in.vivo_noRT | 2410022M11F | 2.962 | 3.52E-05 | glio |
| Atr_in.vivo_noRT | Cdk7 | -1.596 | 3.52E-05 | glio |
| Bcas2_in.vivo_noRT | Lgals4 | 2.698 | 3.52E-05 | glio |
| culture_effect | Maged1 | -0.659 | 3.52E-06 | glio |
| Interaction_genotype_Mtor | Chmp5 | 2.073 | 3.52E-06 | glio |
| Atr_in.vivo_noRT | Mt4 | 4.075 | 3.52E-06 | glio |
| Atr_in.vivo_noRT | Gm36445 | 3.244 | 3.52E-06 | glio |
| Bora_in.vivo_noRT | Gm21814 | 3.077 | 3.52E-06 | glio |
| Chek1_ex.vivo_noRT | Tppp | 3.679 | 3.52E-06 | glio |
| Aars2_in.vivo_noRT | Prrt3 | 3.146 | 3.52E-07 | glio |
| Bora_in.vivo_noRT | Gm41492 | 3.366 | 3.52E-08 | glio |

| | | | | |
|---|---|---|---|---|
| Chek1_in.vivo_noRT | Gm27010 | 3.966 | 3.52E-09 | glio |
| Chek1_ex.vivo_noRT | Cd248 | 5.021 | 3.52E-10 | glio |
| Aars2_in.vivo_noRT | Cldn19 | 4.544 | 3.52E-11 | glio |
| culture_effect | Cdh10 | -2.017 | 3.52E-12 | glio |
| Aars2_in.vivo_noRT | Gm5152 | 4.94 | 3.52E-12 | glio |
| Bcas2_in.vivo_noRT | Adam12 | 4.547 | 3.52E-13 | glio |
| Chek1_in.vivo_noRT | Gm31415 | 4.777 | 3.52E-16 | glio |
| Aars2_in.vivo_noRT | Gm49860 | 5.327 | 3.52E-19 | glio |
| Atr_in.vivo_noRT | Cd200r2 | 5.269 | 3.52E-25 | glio |
| Atr_in.vivo_noRT | Tmem132c | 5.533 | 3.52E-28 | glio |
| culture_effect | Gemin5 | -0.501 | 3.53E-03 | glio |
| Interaction_genotype_Aars2 | Gm48008 | -2.96 | 3.53E-03 | glio |
| Interaction_genotype_Aars2 | Gm35082 | -2.831 | 3.53E-03 | glio |
| Interaction_genotype_Aars2 | Gm32834 | -3.198 | 3.53E-03 | glio |
| Interaction_genotype_Mtor | Sap30bpos | -3.098 | 3.53E-03 | glio |
| Interaction_genotype_Mtor | Hist1h2bp | -3.74 | 3.53E-03 | glio |
| Interaction_genotype_Mtor | Mrps18c | 1.48 | 3.53E-03 | glio |
| Interaction_genotype_Myc | Slc46a3 | -3.371 | 3.53E-03 | glio |
| Interaction_genotype_Myc | Pskh1 | -3.041 | 3.53E-03 | glio |
| Interaction_genotype_Pars2 | Gm4724 | -3.63 | 3.53E-03 | glio |
| Interaction_genotype_Pars2 | Gm15743 | -3.693 | 3.53E-03 | glio |
| Interaction_genotype_Pars2 | Gm14968 | -3.815 | 3.53E-03 | glio |
| Interaction_genotype_Rad51 | Adgrb3 | -4.013 | 3.53E-03 | glio |
| Interaction_genotype_Rad51 | 5033417F24Ri | -4.072 | 3.53E-03 | glio |
| Interaction_genotype_Rpl3 | Abhd18 | -4.121 | 3.53E-03 | glio |
| interaction_RT | Isy1 | -0.805 | 3.53E-03 | glio |
| interaction_RT | Phb2 | -0.547 | 3.53E-03 | glio |
| Aars2_in.vivo_noRT | Zfp763 | 1.974 | 3.53E-03 | glio |
| Aars2_in.vivo_noRT | Ypel4 | 1.963 | 3.53E-03 | glio |
| Aars2_ex.vivo_noRT | 4922502N22R | 2.897 | 3.53E-03 | glio |
| Bcas2_in.vivo_noRT | A130010J15Ri | 1.489 | 3.53E-03 | glio |
| Bora_in.vivo_noRT | Fam178b | 2.229 | 3.53E-03 | glio |
| Chek1_ex.vivo_noRT | Samm50 | -1.143 | 3.53E-03 | glio |
| Interaction_genotype_Mtor | Mtfp1 | -3.211 | 3.53E-04 | glio |
| Interaction_genotype_Myc | Gm26581 | -4.15 | 3.53E-04 | glio |
| interaction_RT | Rlbp1 | -0.858 | 3.53E-04 | glio |
| Aars2_in.vivo_noRT | Mras | -1.551 | 3.53E-04 | glio |
| Aars2_in.vivo_noRT | Alyref | -1.037 | 3.53E-04 | glio |
| Aars2_ex.vivo_noRT | Masp2 | 2.9 | 3.53E-04 | glio |
| Aars2_ex.vivo_noRT | Lyl1 | 2.884 | 3.53E-04 | glio |
| Aars2_in.vivo_noRT | Gm10353 | 2.812 | 3.53E-05 | glio |
| Atp5c1_in.vivo_noRT | 4930570E03Ri | 2.408 | 3.53E-05 | glio |
| Atp5c1_in.vivo_noRT | 4930447K03R | 2.408 | 3.53E-05 | glio |
| Atr_in.vivo_noRT | Ovgp1 | 2.851 | 3.53E-05 | glio |
| Atr_in.vivo_noRT | Dmpk | 2.793 | 3.53E-05 | glio |
| Atr_in.vivo_noRT | Snta1 | 2.301 | 3.53E-05 | glio |
| Atr_ex.vivo_noRT | BC048671 | 3.164 | 3.53E-05 | glio |
| Atr_ex.vivo_noRT | Gm43951 | 3.168 | 3.53E-05 | glio |
| Bora_in.vivo_noRT | Rpl7l1 | -1.273 | 3.53E-05 | glio |
| Bora_ex.vivo_noRT | C3 | 3.993 | 3.53E-05 | glio |
| Bora_ex.vivo_noRT | Krtcap3 | 3.008 | 3.53E-05 | glio |
| Chek1_ex.vivo_noRT | Folr2 | 3.672 | 3.53E-05 | glio |

| | | | | |
|---|---|---|---|---|
| culture_effect | Gtf2b | -0.826 | 3.53E-06 | glio |
| Aars2_in.vivo_noRT | Pcdhgb1 | 2.793 | 3.53E-06 | glio |
| Bcas2_in.vivo_noRT | Zfp551 | 3.24 | 3.53E-06 | glio |
| Atr_in.vivo_noRT | Ehhadh | 3.838 | 3.53E-07 | glio |
| Atr_in.vivo_noRT | Chd7 | -2.297 | 3.53E-07 | glio |
| Bcas2_in.vivo_noRT | 9930014A18R | 3.413 | 3.53E-07 | glio |
| Bcas2_in.vivo_noRT | Slc46a3 | 3.622 | 3.53E-07 | glio |
| Bora_in.vivo_noRT | Slc25a35 | 3.826 | 3.53E-08 | glio |
| Aars2_in.vivo_noRT | Gm46404 | 4.29 | 3.53E-09 | glio |
| Hsd17b10_in.vivo_noRT | Folr1 | 3.616 | 3.53E-09 | glio |
| interaction_RT | Ifi47 | 2.124 | 3.53E-11 | glio |
| Aars2_in.vivo_noRT | Noxred1 | 4.599 | 3.53E-12 | glio |
| Chek1_ex.vivo_noRT | 4930570E03Ri | 4.842 | 3.53E-13 | glio |
| Chek1_ex.vivo_noRT | Gm5286 | 4.842 | 3.53E-13 | glio |
| Chek1_ex.vivo_noRT | Gm41307 | 4.449 | 3.53E-13 | glio |
| Aars2_in.vivo_noRT | 4833419F23Ri | 5.327 | 3.53E-14 | glio |
| Atr_in.vivo_noRT | Celsr3 | 5.273 | 3.53E-16 | glio |
| Chek1_in.vivo_noRT | B4galnt1 | 4.784 | 3.53E-16 | glio |
| culture_effect | Bcl2 | 0.824 | 3.54E-03 | glio |
| culture_effect | Slc35b3 | -0.629 | 3.54E-03 | glio |
| Interaction_genotype_Aars2 | Abcg4 | -3.39 | 3.54E-03 | glio |
| Interaction_genotype_Aars2 | Slc6a13 | -3.919 | 3.54E-03 | glio |
| Interaction_genotype_Aars2 | Gm16133 | -3.753 | 3.54E-03 | glio |
| Interaction_genotype_Aars2 | Isg20 | 1.93 | 3.54E-03 | glio |
| Interaction_genotype_Atr | Cpxm1 | -3.208 | 3.54E-03 | glio |
| Interaction_genotype_Atr | Zfp536 | -2.74 | 3.54E-03 | glio |
| Interaction_genotype_Atr | Sapcd2 | -2.03 | 3.54E-03 | glio |
| Interaction_genotype_Ifnar1 | Gm5602 | 3.534 | 3.54E-03 | glio |
| Interaction_genotype_Ifnar1 | Sucnr1 | 2.734 | 3.54E-03 | glio |
| Interaction_genotype_Ifnar1 | Olfr1152 | 3 | 3.54E-03 | glio |
| Interaction_genotype_Mtor | Cyb5rl | -2.746 | 3.54E-03 | glio |
| Interaction_genotype_Mtor | Map2k4 | -1.736 | 3.54E-03 | glio |
| Interaction_genotype_Myc | Gm15080 | -3.554 | 3.54E-03 | glio |
| Interaction_genotype_Myc | Pqlc3 | -3.228 | 3.54E-03 | glio |
| Interaction_genotype_Myc | Tecpr2 | -2.849 | 3.54E-03 | glio |
| Interaction_genotype_Myc | C230014O12R | -2.886 | 3.54E-03 | glio |
| Interaction_genotype_Pars2 | Gm15927 | -3.881 | 3.54E-03 | glio |
| Interaction_genotype_Uqcrb | Folr1 | -3.9 | 3.54E-03 | glio |
| Interaction_genotype_Uqcrb | Gm44752 | -4.2 | 3.54E-03 | glio |
| Interaction_genotype_Uqcrb | 4930412M03F | -4.449 | 3.54E-03 | glio |
| interaction_RT | Ormdl3 | 0.811 | 3.54E-03 | glio |
| Atr_in.vivo_noRT | Zfp362 | 1.808 | 3.54E-03 | glio |
| Bcas2_in.vivo_noRT | Crabp1 | 2.175 | 3.54E-03 | glio |
| Bcas2_in.vivo_noRT | Fam129c | 2.139 | 3.54E-03 | glio |
| Bcas2_in.vivo_noRT | Gm42941 | 2.014 | 3.54E-03 | glio |
| Bora_ex.vivo_noRT | Smpd5 | 2.615 | 3.54E-03 | glio |
| Bora_ex.vivo_noRT | Calhm6 | 2.622 | 3.54E-03 | glio |
| Chek1_in.vivo_noRT | Tor1a | -1.199 | 3.54E-03 | glio |
| Chek1_in.vivo_noRT | Exosc7 | -0.914 | 3.54E-03 | glio |
| Chek1_ex.vivo_noRT | Pdpr | 2.225 | 3.54E-03 | glio |
| Cyp19a1_ex.vivo_noRT | 4930452G13R | -2.818 | 3.54E-03 | glio |
| Eif2s1_in.vivo_noRT | Fam166a | 1.879 | 3.54E-03 | glio |

| | | | | |
|---|---|---|---|---|
| culture_effect | Ppp1r10 | 0.595 | 3.54E-04 | glio |
| Interaction_genotype_Myc | Ccdc85c | -3.964 | 3.54E-04 | glio |
| Interaction_genotype_Myc | Ap5m1 | -2.957 | 3.54E-04 | glio |
| Atr_in.vivo_noRT | Trdmt1 | 2.206 | 3.54E-04 | glio |
| Atr_ex.vivo_noRT | Gpr146 | 3.554 | 3.54E-04 | glio |
| Bora_in.vivo_noRT | Adgra3 | -1.456 | 3.54E-04 | glio |
| Chek1_in.vivo_noRT | Smim40 | 2.542 | 3.54E-04 | glio |
| Chek1_ex.vivo_noRT | Accsl | 3.402 | 3.54E-04 | glio |
| Eif2s1_in.vivo_noRT | AF067063 | 2.143 | 3.54E-04 | glio |
| culture_effect | 4933433G15R | 2.207 | 3.54E-05 | glio |
| culture_effect | Cdr2l | -0.786 | 3.54E-05 | glio |
| Interaction_genotype_Mtor | Bbs10 | -5.113 | 3.54E-05 | glio |
| Aars2_in.vivo_noRT | Gm16279 | 3.071 | 3.54E-05 | glio |
| Aars2_in.vivo_noRT | Gm32051 | 3.038 | 3.54E-05 | glio |
| Aars2_ex.vivo_noRT | Necab3 | 3.649 | 3.54E-05 | glio |
| Atr_in.vivo_noRT | Gm16124 | 3.299 | 3.54E-05 | glio |
| Atr_in.vivo_noRT | Ppid | -1.247 | 3.54E-05 | glio |
| Bcas2_in.vivo_noRT | 1700101I11Ri | 3.167 | 3.54E-05 | glio |
| Bora_in.vivo_noRT | 2310008N11R | 3.15 | 3.54E-05 | glio |
| Bora_in.vivo_noRT | Cd72 | 2.898 | 3.54E-05 | glio |
| culture_effect | Gm44386 | -2.851 | 3.54E-06 | glio |
| Atp5c1_in.vivo_noRT | Slc16a5 | 2.404 | 3.54E-06 | glio |
| Chek1_ex.vivo_noRT | Gm9999 | 3.866 | 3.54E-06 | glio |
| Hsd17b10_in.vivo_noRT | Gm14002 | 3.094 | 3.54E-06 | glio |
| Chek1_in.vivo_noRT | Selenok | -1.384 | 3.54E-07 | glio |
| Chek1_in.vivo_noRT | Lrrn3 | 3.588 | 3.54E-08 | glio |
| Chek1_ex.vivo_noRT | 4921504E06Ri | 3.754 | 3.54E-08 | glio |
| Hsd17b10_in.vivo_noRT | B3gnt4 | 3.874 | 3.54E-10 | glio |
| culture_effect | Sept4 | -4.363 | 3.54E-11 | glio |
| Bcas2_in.vivo_noRT | D630036H23R | 4.547 | 3.54E-11 | glio |
| Bcas2_in.vivo_noRT | Tdrd5 | 4.421 | 3.54E-12 | glio |
| culture_effect | Enpp2 | -4.275 | 3.54E-13 | glio |
| Chek1_ex.vivo_noRT | Gm9758 | 4.842 | 3.54E-14 | glio |
| Atr_in.vivo_noRT | Adam12 | 5.273 | 3.54E-17 | glio |
| Bcas2_in.vivo_noRT | Gm10324 | 4.541 | 3.54E-17 | glio |
| Bora_in.vivo_noRT | C430002N11R | 4.936 | 3.54E-19 | glio |
| culture_effect | Gbp5 | -3.503 | 3.54E-25 | glio |
| culture_effect | Trp53inp1 | -1.271 | 3.55E-03 | glio |
| Interaction_genotype_Aars2 | Pheta1 | -3.743 | 3.55E-03 | glio |
| Interaction_genotype_Atr | Aqp5 | -4.116 | 3.55E-03 | glio |
| Interaction_genotype_Atr | Fsbp | -3.778 | 3.55E-03 | glio |
| Interaction_genotype_Atr | Gm14403 | -3.261 | 3.55E-03 | glio |
| Interaction_genotype_Myc | Tnfrsf13b | -3.443 | 3.55E-03 | glio |
| Interaction_genotype_Myc | Tex2 | -2.928 | 3.55E-03 | glio |
| Interaction_genotype_Myc | Bcr | -2.516 | 3.55E-03 | glio |
| Interaction_genotype_Rad21 | Cemip | 3.726 | 3.55E-03 | glio |
| Interaction_genotype_Rad21 | Ubxn10 | 3.949 | 3.55E-03 | glio |
| interaction_RT | Mboat1 | -1.211 | 3.55E-03 | glio |
| Aars2_in.vivo_noRT | Cd24a | 2.488 | 3.55E-03 | glio |
| Aars2_in.vivo_noRT | Spag7 | -1.019 | 3.55E-03 | glio |
| Atr_ex.vivo_noRT | Zg16 | 2.764 | 3.55E-03 | glio |
| Atr_ex.vivo_noRT | Pdlim3 | 2.37 | 3.55E-03 | glio |

| | | | | |
|---|---|---|---|---|
| Atr_ex.vivo_noRT | Ldhd | 2.453 | 3.55E-03 | glio |
| Bard1_ex.vivo_noRT | 4930502E18Ri | 2.421 | 3.55E-03 | glio |
| Bcas2_in.vivo_noRT | Fam151b | 2.144 | 3.55E-03 | glio |
| Bcas2_in.vivo_noRT | Plpp6 | 2.027 | 3.55E-03 | glio |
| Bcas2_in.vivo_noRT | Cldn22 | 1.884 | 3.55E-03 | glio |
| Bcas2_in.vivo_noRT | Rapgef6 | -1.241 | 3.55E-03 | glio |
| Bcas2_in.vivo_noRT | Fkbp11 | -1.151 | 3.55E-03 | glio |
| Bcas2_in.vivo_noRT | Asns | -1.058 | 3.55E-03 | glio |
| Chek1_in.vivo_noRT | Mfap5 | 2.218 | 3.55E-03 | glio |
| Chek1_in.vivo_noRT | Myt1 | 1.563 | 3.55E-03 | glio |
| Chek1_in.vivo_noRT | Rnf115 | -1.306 | 3.55E-03 | glio |
| Chek1_in.vivo_noRT | Mrpl2 | -0.883 | 3.55E-03 | glio |
| Chek1_in.vivo_noRT | Psmd8 | -0.839 | 3.55E-03 | glio |
| Chek1_ex.vivo_noRT | Abcd2 | 3.006 | 3.55E-03 | glio |
| Chek1_ex.vivo_noRT | Cttn | -1.527 | 3.55E-03 | glio |
| Hsd17b10_in.vivo_noRT | Tmem126b | -1.547 | 3.55E-03 | glio |
| Interaction_genotype_Myc | Gm21731 | -2.729 | 3.55E-04 | glio |
| Interaction_genotype_Myc | Zfp112 | -3.992 | 3.55E-04 | glio |
| Interaction_genotype_Myc | A930004D18R | -4.048 | 3.55E-04 | glio |
| Interaction_genotype_Myc | Arhgap1 | -2.505 | 3.55E-04 | glio |
| Interaction_genotype_Myc | Bsg | 1.381 | 3.55E-04 | glio |
| Aars2_in.vivo_noRT | Dab2ip | 2.242 | 3.55E-04 | glio |
| Aars2_ex.vivo_noRT | Zdhhc14 | 2.897 | 3.55E-04 | glio |
| Bcas2_in.vivo_noRT | Prkar2b | 2.611 | 3.55E-04 | glio |
| Bcas2_in.vivo_noRT | Ampd3 | 2.525 | 3.55E-04 | glio |
| Chek1_in.vivo_noRT | Dynlt1b | 2.558 | 3.55E-04 | glio |
| Chek1_in.vivo_noRT | Tmem151a | 2.568 | 3.55E-04 | glio |
| Chek1_in.vivo_noRT | Sft2d2 | -1.745 | 3.55E-04 | glio |
| Chek1_in.vivo_noRT | Gnas | -0.657 | 3.55E-04 | glio |
| Interaction_genotype_Myc | Lmnb1 | -3.289 | 3.55E-05 | glio |
| Interaction_genotype_Myc | 1700024I08Ri | -3.843 | 3.55E-05 | glio |
| Chek1_in.vivo_noRT | Adam4 | 2.702 | 3.55E-05 | glio |
| Aars2_in.vivo_noRT | Tctn1 | 3.287 | 3.55E-06 | glio |
| Atr_in.vivo_noRT | 4930594M22F | 3.286 | 3.55E-06 | glio |
| Chek1_in.vivo_noRT | Ankrd34a | 3.396 | 3.55E-06 | glio |
| Chek1_ex.vivo_noRT | Fgl2 | 3.867 | 3.55E-06 | glio |
| Chek1_ex.vivo_noRT | Glcci1 | 3.231 | 3.55E-06 | glio |
| Bora_in.vivo_noRT | 1700066B19R | 3.484 | 3.55E-07 | glio |
| Aars2_in.vivo_noRT | Eps8l2 | 4.225 | 3.55E-09 | glio |
| Aars2_in.vivo_noRT | Gm26590 | 4.038 | 3.55E-10 | glio |
| Atr_in.vivo_noRT | Ccl19 | 3.582 | 3.55E-10 | glio |
| Aars2_in.vivo_noRT | 9530080O11R | 4.409 | 3.55E-11 | glio |
| Atr_in.vivo_noRT | 1700013F07Ri | 4.888 | 3.55E-11 | glio |
| Chek1_in.vivo_noRT | Gm20635 | 4.398 | 3.55E-11 | glio |
| interaction_RT | Psmb9 | 2.14 | 3.55E-13 | glio |
| Atr_in.vivo_noRT | Zmynd10 | 5.273 | 3.55E-19 | glio |
| culture_effect | Vps37d | -1.383 | 3.56E-03 | glio |
| culture_effect | 9330175E14Ri | -1.083 | 3.56E-03 | glio |
| culture_effect | Tspan7 | 0.622 | 3.56E-03 | glio |
| culture_effect | Plekhb1 | -0.427 | 3.56E-03 | glio |
| Interaction_genotype_Aars2 | Oxt | -3.645 | 3.56E-03 | glio |
| Interaction_genotype_Ifnar1 | Gm29966 | 2.994 | 3.56E-03 | glio |

| | | | | |
|---|---|---|---|---|
| Interaction_genotype_Myc | Prkar2b | -3.363 | 3.56E-03 | glio |
| Interaction_genotype_Myc | 5730409E04Ri | -2.864 | 3.56E-03 | glio |
| Aars2_in.vivo_noRT | Gab2 | 1.959 | 3.56E-03 | glio |
| Atp5c1_in.vivo_noRT | Gm42664 | 2.018 | 3.56E-03 | glio |
| Atr_in.vivo_noRT | Six5 | 1.843 | 3.56E-03 | glio |
| Bcas2_in.vivo_noRT | Ptpn14 | -1.462 | 3.56E-03 | glio |
| Bcas2_ex.vivo_noRT | Rcor2 | -2.014 | 3.56E-03 | glio |
| Bora_in.vivo_noRT | Doc2g | 2.199 | 3.56E-03 | glio |
| Bora_in.vivo_noRT | Neurl2 | 1.863 | 3.56E-03 | glio |
| Chek1_in.vivo_noRT | Gm43328 | 2.196 | 3.56E-03 | glio |
| Chek1_in.vivo_noRT | Hist1h2ac | 2.087 | 3.56E-03 | glio |
| Chek1_in.vivo_noRT | Senp8 | 1.824 | 3.56E-03 | glio |
| Chek1_in.vivo_noRT | Scaf4 | -1.339 | 3.56E-03 | glio |
| Chek1_in.vivo_noRT | Tpm3-rs7 | -1.137 | 3.56E-03 | glio |
| Chek1_in.vivo_noRT | Ndufs8 | -0.931 | 3.56E-03 | glio |
| Chek1_in.vivo_noRT | Rps19 | -0.565 | 3.56E-03 | glio |
| Chek1_ex.vivo_noRT | Shpk | 2.903 | 3.56E-03 | glio |
| Hsd17b10_in.vivo_noRT | Fzd3 | -1.729 | 3.56E-03 | glio |
| culture_effect | Pcdhgb4 | -1.959 | 3.56E-04 | glio |
| culture_effect | Fam229b | 0.918 | 3.56E-04 | glio |
| culture_effect | Alkbh8 | 0.794 | 3.56E-04 | glio |
| culture_effect | Arl2 | -0.524 | 3.56E-04 | glio |
| Interaction_genotype_Aars2 | Sptbn4 | -4.453 | 3.56E-04 | glio |
| Interaction_genotype_Aars2 | C230037L18Ri | -4.226 | 3.56E-04 | glio |
| Interaction_genotype_Aars2 | Cebpa | -4.55 | 3.56E-04 | glio |
| Interaction_genotype_Aars2 | Spink2 | -4.973 | 3.56E-04 | glio |
| Interaction_genotype_Aars2 | Igf2bp1 | -4.326 | 3.56E-04 | glio |
| Interaction_genotype_Aars2 | Capn9 | -4.098 | 3.56E-04 | glio |
| Interaction_genotype_Mtor | Gm9999 | -4.1 | 3.56E-04 | glio |
| Interaction_genotype_Mtor | Rsph3b | -4.163 | 3.56E-04 | glio |
| Interaction_genotype_Mtor | Glrx3 | 1.847 | 3.56E-04 | glio |
| Interaction_genotype_Myc | 4930417H01R | -4.224 | 3.56E-04 | glio |
| Aars2_in.vivo_noRT | Smim3 | 1.886 | 3.56E-04 | glio |
| Aars2_ex.vivo_noRT | 4930579C12R | 2.897 | 3.56E-04 | glio |
| Aars2_ex.vivo_noRT | Cyp2c23 | 2.897 | 3.56E-04 | glio |
| Aars2_ex.vivo_noRT | Mag | 3.293 | 3.56E-04 | glio |
| Aars2_ex.vivo_noRT | Gm43221 | 3.293 | 3.56E-04 | glio |
| Atr_in.vivo_noRT | Tspan32 | 2.555 | 3.56E-04 | glio |
| Atr_in.vivo_noRT | Rps12 | -0.829 | 3.56E-04 | glio |
| Atr_ex.vivo_noRT | Fam25c | 2.588 | 3.56E-04 | glio |
| Bcas2_in.vivo_noRT | Nts | 2.764 | 3.56E-04 | glio |
| Bcas2_in.vivo_noRT | Nyap1 | 2.584 | 3.56E-04 | glio |
| Bcas2_in.vivo_noRT | Abca13 | 2.85 | 3.56E-04 | glio |
| Chek1_in.vivo_noRT | Hist1h2bf | 2.731 | 3.56E-04 | glio |
| Chek1_in.vivo_noRT | 1110065P20R | -1.112 | 3.56E-04 | glio |
| Hsd17b10_in.vivo_noRT | Svopl | 2.435 | 3.56E-04 | glio |
| Hsd17b10_in.vivo_noRT | Hist1h4b | 2.911 | 3.56E-04 | glio |
| culture_effect | Tmx2 | -0.606 | 3.56E-05 | glio |
| Interaction_genotype_Mtor | Gm20628 | -5.331 | 3.56E-05 | glio |
| Interaction_genotype_Mtor | Gm7247 | -4.104 | 3.56E-05 | glio |
| Interaction_genotype_Mtor | Gm50203 | -4.142 | 3.56E-05 | glio |
| Interaction_genotype_Mtor | Gm19412 | -3.223 | 3.56E-05 | glio |

| | | | | |
|---|---|---|---|---|
| Interaction_genotype_Mtor | Mtfr1l | -2.617 | 3.56E-05 | glio |
| Interaction_genotype_Myc | Osbpl7 | -4.309 | 3.56E-05 | glio |
| Interaction_genotype_Myc | Myo19 | -3.835 | 3.56E-05 | glio |
| Interaction_genotype_Myc | C330013E15Ri | -4.492 | 3.56E-05 | glio |
| Interaction_genotype_Myc | Gm20667 | -4.804 | 3.56E-05 | glio |
| Interaction_genotype_Myc | Akirin2 | 1.633 | 3.56E-05 | glio |
| Aars2_in.vivo_noRT | Usp14 | -1.575 | 3.56E-05 | glio |
| Atr_in.vivo_noRT | Gm45767 | 2.732 | 3.56E-05 | glio |
| Bard1_ex.vivo_noRT | Gm266 | 2.814 | 3.56E-05 | glio |
| Bcas2_ex.vivo_noRT | Idi2 | 2.742 | 3.56E-05 | glio |
| Bcas2_ex.vivo_noRT | Vmn2r18 | 2.742 | 3.56E-05 | glio |
| Bcas2_ex.vivo_noRT | Gm16178 | 2.742 | 3.56E-05 | glio |
| Bcas2_ex.vivo_noRT | Gm364 | 2.742 | 3.56E-05 | glio |
| Chek1_in.vivo_noRT | Tmem91 | 3.001 | 3.56E-05 | glio |
| Hsd17b10_in.vivo_noRT | Flnc | 3.09 | 3.56E-05 | glio |
| Interaction_genotype_Mtor | Atoh8 | -4.411 | 3.56E-06 | glio |
| Interaction_genotype_Mtor | C230013L11Ri | -5.87 | 3.56E-06 | glio |
| Interaction_genotype_Mtor | Esp24 | -4.099 | 3.56E-06 | glio |
| interaction_RT | Hspd1 | -0.916 | 3.56E-06 | glio |
| Chek1_ex.vivo_noRT | Peli3 | 3.942 | 3.56E-06 | glio |
| Bcas2_in.vivo_noRT | Ak1 | 3.544 | 3.56E-07 | glio |
| Bcas2_in.vivo_noRT | 4933439K11R | 3.622 | 3.56E-07 | glio |
| Bora_in.vivo_noRT | Stpg2 | 3.521 | 3.56E-07 | glio |
| culture_effect | Gm10101 | 2.639 | 3.56E-08 | glio |
| Interaction_genotype_Atr | Scube3 | -5.759 | 3.56E-08 | glio |
| Bcas2_in.vivo_noRT | Pld6 | 4.068 | 3.56E-08 | glio |
| Bcas2_in.vivo_noRT | Gm43660 | 3.634 | 3.56E-08 | glio |
| Atr_ex.vivo_noRT | Esm1 | 3.77 | 3.56E-09 | glio |
| Aars2_in.vivo_noRT | Trnp1 | 4.422 | 3.56E-10 | glio |
| Chek1_in.vivo_noRT | Igdcc4 | 3.868 | 3.56E-10 | glio |
| culture_effect | Cox6a1 | -0.843 | 3.56E-12 | glio |
| Aars2_in.vivo_noRT | Gm12860 | 4.319 | 3.56E-15 | glio |
| Bcas2_in.vivo_noRT | Minar2 | 4.547 | 3.56E-16 | glio |
| culture_effect | Neto1 | -1.578 | 3.56E-17 | glio |
| Atr_in.vivo_noRT | Sh3d21 | 5.011 | 3.56E-17 | glio |
| Aars2_in.vivo_noRT | Olfr718-ps1 | 5.063 | 3.56E-21 | glio |
| Aars2_in.vivo_noRT | Akr1c18 | 5.327 | 3.56E-21 | glio |
| Aars2_in.vivo_noRT | 4921504E06Ri | 5.327 | 3.56E-21 | glio |
| culture_effect | Nrp2 | 2.058 | 3.56E-34 | glio |
| culture_effect | Gm47204 | 1.755 | 3.57E-03 | glio |
| culture_effect | Zfp384 | -0.719 | 3.57E-03 | glio |
| Interaction_genotype_Atr | Gm14325 | -3.713 | 3.57E-03 | glio |
| Interaction_genotype_Atr | Pim3 | 1.647 | 3.57E-03 | glio |
| Interaction_genotype_Bora | Gm5152 | -3.99 | 3.57E-03 | glio |
| Interaction_genotype_Bora | C330011M18F | -3.939 | 3.57E-03 | glio |
| Interaction_genotype_Cdkn2a | Slc14a2 | 2.835 | 3.57E-03 | glio |
| Interaction_genotype_Chek1 | Gm10101 | -3.775 | 3.57E-03 | glio |
| Interaction_genotype_Ifnar1 | Gbp3 | -1.004 | 3.57E-03 | glio |
| Interaction_genotype_Mtor | 2900041M22F | -3.762 | 3.57E-03 | glio |
| Interaction_genotype_Mtor | Frs3 | -3.471 | 3.57E-03 | glio |
| Interaction_genotype_Mtor | Tdrd7 | -2.188 | 3.57E-03 | glio |
| Interaction_genotype_Mtor | Rps17 | 1.12 | 3.57E-03 | glio |

| | | | | |
|---|---|---|---|---|
| Interaction_genotype_Myc | Gm9856 | -2.644 | 3.57E-03 | glio |
| Aars2_in.vivo_noRT | B230369F24Ri | 2.108 | 3.57E-03 | glio |
| Aars2_in.vivo_noRT | Gm14167 | 1.89 | 3.57E-03 | glio |
| Aars2_in.vivo_noRT | Brd3os | 1.956 | 3.57E-03 | glio |
| Atp5c1_in.vivo_noRT | Podxl | 2.136 | 3.57E-03 | glio |
| Atp5c1_in.vivo_noRT | 5330439K02R | 2.402 | 3.57E-03 | glio |
| Atp5c1_in.vivo_noRT | Gm5039 | 2.668 | 3.57E-03 | glio |
| Atr_in.vivo_noRT | Zfp607a | 2.027 | 3.57E-03 | glio |
| Bcas2_in.vivo_noRT | Ece2 | 2.039 | 3.57E-03 | glio |
| Bora_ex.vivo_noRT | Astn2 | 2.806 | 3.57E-03 | glio |
| Fanca_ex.vivo_noRT | Fam189a1 | 2.913 | 3.57E-03 | glio |
| Fanca_ex.vivo_noRT | Kcnk7 | 2.913 | 3.57E-03 | glio |
| culture_effect | Pdcd2l | -0.792 | 3.57E-04 | glio |
| Interaction_genotype_Chek1 | Supt4a | 1.343 | 3.57E-04 | glio |
| Interaction_genotype_Mtor | Txndc17 | 1.623 | 3.57E-04 | glio |
| Interaction_genotype_Myc | Gm28050 | -4.09 | 3.57E-04 | glio |
| Interaction_genotype_Rpl3 | Gchfr | -5.422 | 3.57E-04 | glio |
| Interaction_genotype_Rpl3 | Nxph4 | -4.729 | 3.57E-04 | glio |
| Aars2_ex.vivo_noRT | Card9 | 2.893 | 3.57E-04 | glio |
| Atp5c1_in.vivo_noRT | Cxcl13 | 2.668 | 3.57E-04 | glio |
| Atr_in.vivo_noRT | Dleu7 | 2.471 | 3.57E-04 | glio |
| Bora_ex.vivo_noRT | Gm13398 | 2.404 | 3.57E-04 | glio |
| Eif2s1_in.vivo_noRT | Gm2016 | 2.143 | 3.57E-04 | glio |
| Eif2s1_in.vivo_noRT | Halr1 | 1.877 | 3.57E-04 | glio |
| Eif2s1_in.vivo_noRT | C330002G04R | 2.143 | 3.57E-04 | glio |
| Eif2s1_in.vivo_noRT | Rapgef3os2 | 2.143 | 3.57E-04 | glio |
| Eif2s1_in.vivo_noRT | Gm47622 | 2.143 | 3.57E-04 | glio |
| Bard1_ex.vivo_noRT | Ildr1 | 2.814 | 3.57E-05 | glio |
| Bard1_ex.vivo_noRT | Gm20732 | 3.596 | 3.57E-05 | glio |
| Bcas2_in.vivo_noRT | S1pr1 | 3.622 | 3.57E-06 | glio |
| Chek1_in.vivo_noRT | Bfsp2 | 3.672 | 3.57E-06 | glio |
| Hsd17b10_in.vivo_noRT | Ms4a6d | 2.828 | 3.57E-06 | glio |
| culture_effect | Entpd3 | -2.642 | 3.57E-07 | glio |
| Chek1_ex.vivo_noRT | Sept4 | 4.982 | 3.57E-07 | glio |
| Hsd17b10_in.vivo_noRT | Rftn2 | 3.225 | 3.57E-07 | glio |
| Interaction_genotype_Mtor | Efcab9 | -6.542 | 3.57E-08 | glio |
| Aars2_in.vivo_noRT | BC037032 | 4.409 | 3.57E-09 | glio |
| Atr_in.vivo_noRT | Gnaz | 4.235 | 3.57E-09 | glio |
| interaction_RT | Gabarap | 1.064 | 3.57E-10 | glio |
| Aars2_in.vivo_noRT | Gm5129 | 5.063 | 3.57E-12 | glio |
| Atr_in.vivo_noRT | Sult6b1 | 4.742 | 3.57E-12 | glio |
| Hsd17b10_in.vivo_noRT | A930002C04R | 3.356 | 3.57E-12 | glio |
| Atr_in.vivo_noRT | B3galt4 | 5.008 | 3.57E-15 | glio |
| Atr_in.vivo_noRT | Gm45606 | 5.273 | 3.57E-15 | glio |
| Aars2_in.vivo_noRT | Abra | 4.801 | 3.57E-16 | glio |
| Atr_in.vivo_noRT | Gm37800 | 5.533 | 3.57E-18 | glio |
| Bora_in.vivo_noRT | Gm29966 | 4.676 | 3.57E-22 | glio |
| culture_effect | Nmi | -1.459 | 3.57E-28 | glio |
| Atr_in.vivo_noRT | Gm44693 | 5.533 | 3.57E-28 | glio |
| Interaction_genotype_Aars2 | Cenpv | 1.515 | 3.58E-03 | glio |
| Interaction_genotype_Mtor | Pfdn4 | 1.361 | 3.58E-03 | glio |
| Interaction_genotype_Mtor | Eif5a | 1.336 | 3.58E-03 | glio |

| | | | | |
|---|---|---|---|---|
| Interaction_genotype_Myc | Pax5 | -3.546 | 3.58E-03 | glio |
| Interaction_genotype_Myc | Zfp984 | -2.47 | 3.58E-03 | glio |
| Interaction_genotype_Rad51 | Gchfr | -4.624 | 3.58E-03 | glio |
| Interaction_genotype_Rad51 | Gm50223 | 3.455 | 3.58E-03 | glio |
| Aars2_ex.vivo_noRT | Pagr1a | 2.691 | 3.58E-03 | glio |
| Atr_in.vivo_noRT | Ccl2 | 5.142 | 3.58E-03 | glio |
| Atr_in.vivo_noRT | Cep126 | 1.846 | 3.58E-03 | glio |
| Atr_in.vivo_noRT | Ncl | -0.675 | 3.58E-03 | glio |
| Bcas2_in.vivo_noRT | Pusl1 | -1.517 | 3.58E-03 | glio |
| Bora_in.vivo_noRT | Gar1 | -0.878 | 3.58E-03 | glio |
| Chek1_in.vivo_noRT | Ap2a2 | -1.194 | 3.58E-03 | glio |
| Interaction_genotype_Aars2 | Fhl2 | -3.981 | 3.58E-04 | glio |
| Interaction_genotype_Aars2 | Tbca | 0.981 | 3.58E-04 | glio |
| Interaction_genotype_Mtor | Gm26559 | -4.262 | 3.58E-04 | glio |
| Interaction_genotype_Mtor | Mtx3 | -3.851 | 3.58E-04 | glio |
| Interaction_genotype_Mtor | Ufc1 | 1.735 | 3.58E-04 | glio |
| Interaction_genotype_Myc | Gm11232 | -2.729 | 3.58E-04 | glio |
| Interaction_genotype_Myc | Cacnb1 | -4.431 | 3.58E-04 | glio |
| Aars2_ex.vivo_noRT | Serpina3m | 3.48 | 3.58E-04 | glio |
| Bard1_ex.vivo_noRT | Gm50024 | 2.415 | 3.58E-04 | glio |
| Bard1_ex.vivo_noRT | Vmn2r44 | 2.419 | 3.58E-04 | glio |
| Bcas2_in.vivo_noRT | Cmtm4 | -1.474 | 3.58E-04 | glio |
| Bora_in.vivo_noRT | Rxra | 2.698 | 3.58E-04 | glio |
| Bora_in.vivo_noRT | Cdk16 | -1.357 | 3.58E-04 | glio |
| Chek1_in.vivo_noRT | Wdr62 | 2.174 | 3.58E-04 | glio |
| Hsd17b10_in.vivo_noRT | Wfdc1 | 2.631 | 3.58E-04 | glio |
| culture_effect | Gtf2h5 | 0.428 | 3.58E-05 | glio |
| Interaction_genotype_Mtor | Cyth4 | -4.787 | 3.58E-05 | glio |
| Interaction_genotype_Mtor | EU599041 | -5.115 | 3.58E-05 | glio |
| Interaction_genotype_Mtor | Mcrip1 | 1.941 | 3.58E-05 | glio |
| Interaction_genotype_Mtor | Rpl4 | 1.431 | 3.58E-05 | glio |
| Interaction_genotype_Myc | Rps17 | 1.827 | 3.58E-05 | glio |
| Aars2_in.vivo_noRT | Fxyd4 | 3.211 | 3.58E-05 | glio |
| Bcas2_in.vivo_noRT | 2310034G01R | 2.447 | 3.58E-05 | glio |
| culture_effect | Atp5j | 0.406 | 3.58E-06 | glio |
| Aars2_in.vivo_noRT | Gm16740 | 3.232 | 3.58E-06 | glio |
| Bora_in.vivo_noRT | Micu2 | -1.989 | 3.58E-06 | glio |
| Chek1_ex.vivo_noRT | Anxa8 | 4.049 | 3.58E-06 | glio |
| Chek1_ex.vivo_noRT | Vsig10l | 3.933 | 3.58E-06 | glio |
| culture_effect | Pacsin1 | -2.907 | 3.58E-07 | glio |
| culture_effect | Glrx5 | 0.612 | 3.58E-07 | glio |
| interaction_RT | Nr4a1 | 1.769 | 3.58E-07 | glio |
| interaction_RT | Tmem59 | 0.879 | 3.58E-07 | glio |
| Atr_in.vivo_noRT | Tes3-ps | 3.499 | 3.58E-07 | glio |
| Bora_in.vivo_noRT | Limch1 | 3.898 | 3.58E-07 | glio |
| culture_effect | Folr1 | 2.636 | 3.58E-08 | glio |
| Chek1_ex.vivo_noRT | Rgs9 | 4.06 | 3.58E-08 | glio |
| Hsd17b10_in.vivo_noRT | Cers3 | 3.084 | 3.58E-08 | glio |
| Bora_in.vivo_noRT | Gm48610 | 4.154 | 3.58E-09 | glio |
| Bora_in.vivo_noRT | Gm50375 | 4.67 | 3.58E-10 | glio |
| Bora_in.vivo_noRT | Gm17146 | 4.019 | 3.58E-10 | glio |
| Aars2_in.vivo_noRT | 1600014C23R | 4.947 | 3.58E-12 | glio |

| | | | | |
|---|---|---|---|---|
| Aars2_in.vivo_noRT | Gm20767 | 5.327 | 3.58E-16 | glio |
| Chek1_in.vivo_noRT | Btn1a1 | 4.784 | 3.58E-18 | glio |
| culture_effect | Rps27rt | -1.549 | 3.58E-19 | glio |
| Interaction_genotype_Aars2 | Hunk | -3.337 | 3.59E-03 | glio |
| Interaction_genotype_Cdkn2a | Gm4340 | 2.835 | 3.59E-03 | glio |
| Interaction_genotype_Cdkn2a | Gm16239 | 2.835 | 3.59E-03 | glio |
| Interaction_genotype_Mtor | Mrps21 | 1.456 | 3.59E-03 | glio |
| Interaction_genotype_Myc | Gm13713 | -2.596 | 3.59E-03 | glio |
| Interaction_genotype_Myc | Grin2d | -3.103 | 3.59E-03 | glio |
| Interaction_genotype_Myc | Mtmr1 | -2.374 | 3.59E-03 | glio |
| Interaction_genotype_Myc | Bcl9l | -2.204 | 3.59E-03 | glio |
| Interaction_genotype_Myc | Ubap2 | 1.395 | 3.59E-03 | glio |
| Interaction_genotype_Myc | Tinagl1 | 2.242 | 3.59E-03 | glio |
| Aars2_in.vivo_noRT | Hspa13 | -1.239 | 3.59E-03 | glio |
| Bcas2_in.vivo_noRT | Zfp40 | 1.987 | 3.59E-03 | glio |
| Bcas2_ex.vivo_noRT | Pla2g2d | 2.742 | 3.59E-03 | glio |
| Cenpj_ex.vivo_noRT | Gm9929 | 2.794 | 3.59E-03 | glio |
| Chek1_in.vivo_noRT | Ddah2 | 2.169 | 3.59E-03 | glio |
| Chek1_in.vivo_noRT | Tmem35b | 2.052 | 3.59E-03 | glio |
| Cyp19a1_ex.vivo_noRT | Krt19 | 2.159 | 3.59E-03 | glio |
| Eif2s1_ex.vivo_noRT | Calml4 | 3.062 | 3.59E-03 | glio |
| Interaction_genotype_Atr | Gm40557 | -4.529 | 3.59E-04 | glio |
| Interaction_genotype_Mtor | Cct5 | 1.578 | 3.59E-04 | glio |
| Interaction_genotype_Myc | Klhl18 | -3.431 | 3.59E-04 | glio |
| Atr_in.vivo_noRT | Kank2 | 2.435 | 3.59E-04 | glio |
| Bcas2_in.vivo_noRT | Carf | 2.559 | 3.59E-04 | glio |
| Bora_in.vivo_noRT | Isoc2a | 2.327 | 3.59E-04 | glio |
| Bora_in.vivo_noRT | Fam172a | -1.708 | 3.59E-04 | glio |
| Bora_in.vivo_noRT | Dbi | -0.954 | 3.59E-04 | glio |
| Bora_ex.vivo_noRT | Tpsab1 | 3.201 | 3.59E-04 | glio |
| Hsd17b10_in.vivo_noRT | 9330104G04R | 2.667 | 3.59E-04 | glio |
| culture_effect | Lzts2 | -0.59 | 3.59E-05 | glio |
| Interaction_genotype_Aars2 | Gm16283 | -4.981 | 3.59E-05 | glio |
| Interaction_genotype_Aars2 | Rcan2 | -5.258 | 3.59E-05 | glio |
| Interaction_genotype_Myc | Gm49808 | -3.122 | 3.59E-05 | glio |
| Aars2_in.vivo_noRT | Gm14273 | 2.837 | 3.59E-05 | glio |
| Atr_in.vivo_noRT | Pcbp1 | -1.158 | 3.59E-05 | glio |
| Bard1_ex.vivo_noRT | Ccdc125 | 3.213 | 3.59E-05 | glio |
| Bora_ex.vivo_noRT | Slc6a11 | 3.201 | 3.59E-05 | glio |
| Bora_ex.vivo_noRT | Oit3 | 3.593 | 3.59E-05 | glio |
| Chek1_in.vivo_noRT | Rpl4 | -0.715 | 3.59E-05 | glio |
| culture_effect | Stub1 | 0.524 | 3.59E-06 | glio |
| Interaction_genotype_Mtor | Arsj | -5.804 | 3.59E-06 | glio |
| Aars2_in.vivo_noRT | Art2b | 2.729 | 3.59E-06 | glio |
| Atr_in.vivo_noRT | Cbx1 | -1.306 | 3.59E-06 | glio |
| Aars2_in.vivo_noRT | Tmem134 | -1.874 | 3.59E-07 | glio |
| Bora_in.vivo_noRT | Ggt5 | 3.303 | 3.59E-07 | glio |
| Chek1_ex.vivo_noRT | Tbc1d10c | 4.421 | 3.59E-07 | glio |
| culture_effect | Nr4a3 | -2.825 | 3.59E-08 | glio |
| Atr_in.vivo_noRT | Gm15013 | 3.741 | 3.59E-08 | glio |
| Chek1_in.vivo_noRT | B430219N15R | 3.947 | 3.59E-08 | glio |
| Chek1_ex.vivo_noRT | Sema3a | 4.449 | 3.59E-08 | glio |

| | | | | |
|---|---|---|---|---|
| Atr_in.vivo_noRT | Gm16283 | 4.009 | 3.59E-09 | glio |
| Atr_in.vivo_noRT | Rasl10a | 3.391 | 3.59E-09 | glio |
| Chek1_ex.vivo_noRT | S100a14 | 4.443 | 3.59E-09 | glio |
| Aars2_in.vivo_noRT | Gm16023 | 4.866 | 3.59E-10 | glio |
| Atr_in.vivo_noRT | Gm47972 | 4.239 | 3.59E-10 | glio |
| Bora_in.vivo_noRT | Efhd1 | 4.42 | 3.59E-10 | glio |
| culture_effect | Hook1 | -1.226 | 3.59E-11 | glio |
| Chek1_in.vivo_noRT | Gm39326 | 4.78 | 3.59E-12 | glio |
| Hsd17b10_in.vivo_noRT | Gm36793 | 3.616 | 3.59E-12 | glio |
| Atr_in.vivo_noRT | Pard3bos1 | 4.536 | 3.59E-13 | glio |
| Bcas2_in.vivo_noRT | Capn9 | 4.807 | 3.59E-14 | glio |
| Aars2_in.vivo_noRT | 4931431C16R | 5.327 | 3.59E-21 | glio |
| Aars2_in.vivo_noRT | Cgnl1 | 5.327 | 3.59E-27 | glio |
| culture_effect | Phf6 | 0.647 | 3.60E-03 | glio |
| culture_effect | Nrip1 | -0.576 | 3.60E-03 | glio |
| Interaction_genotype_Mtor | Slc7a3 | -3.603 | 3.60E-03 | glio |
| Interaction_genotype_Mtor | Mtcl1 | -3.491 | 3.60E-03 | glio |
| Interaction_genotype_Mtor | Urb1 | -2.541 | 3.60E-03 | glio |
| Interaction_genotype_Mtor | Brcc3 | 1.473 | 3.60E-03 | glio |
| Interaction_genotype_Mtor | Cnot8 | 1.546 | 3.60E-03 | glio |
| Interaction_genotype_Mtor | Oga | 1.524 | 3.60E-03 | glio |
| Interaction_genotype_Myc | 9130230L23Ri | -2.863 | 3.60E-03 | glio |
| Interaction_genotype_Myc | Gm38414 | -3.565 | 3.60E-03 | glio |
| Interaction_genotype_Myc | Lnx2 | -2.986 | 3.60E-03 | glio |
| Interaction_genotype_Pars2 | Gm40557 | -4.024 | 3.60E-03 | glio |
| Interaction_genotype_Pars2 | Gm10734 | -3.974 | 3.60E-03 | glio |
| Aars2_ex.vivo_noRT | Npr2 | 3.373 | 3.60E-03 | glio |
| Atp5c1_in.vivo_noRT | Ccdc63 | 2.282 | 3.60E-03 | glio |
| Atr_in.vivo_noRT | Ctns | 1.876 | 3.60E-03 | glio |
| Atr_in.vivo_noRT | Eri1 | -1.301 | 3.60E-03 | glio |
| Bcas2_in.vivo_noRT | Cyb5b | -1.135 | 3.60E-03 | glio |
| Bora_ex.vivo_noRT | Nts | 3.201 | 3.60E-03 | glio |
| Cenpj_in.vivo_noRT | Cd200r2 | 1.758 | 3.60E-03 | glio |
| Chek1_ex.vivo_noRT | Zfand1 | -1.818 | 3.60E-03 | glio |
| Interaction_genotype_Mtor | Gm37864 | -3.713 | 3.60E-04 | glio |
| Interaction_genotype_Mtor | Hist1h4k | -4.386 | 3.60E-04 | glio |
| Interaction_genotype_Mtor | Nsd2 | 2.139 | 3.60E-04 | glio |
| Interaction_genotype_Myc | D730045B01R | -3.331 | 3.60E-04 | glio |
| Interaction_genotype_Myc | Lars | 1.638 | 3.60E-04 | glio |
| Interaction_genotype_Pars2 | 1700017M07F | -4.214 | 3.60E-04 | glio |
| Bard1_ex.vivo_noRT | Napsa | 3.594 | 3.60E-04 | glio |
| Bcas2_in.vivo_noRT | Aebp1 | -1.827 | 3.60E-04 | glio |
| Bcas2_ex.vivo_noRT | Ptprc | 2.742 | 3.60E-04 | glio |
| Bora_ex.vivo_noRT | Egr2 | 3.277 | 3.60E-04 | glio |
| Cenpj_ex.vivo_noRT | Gm15533 | 2.591 | 3.60E-04 | glio |
| Chek1_in.vivo_noRT | Qpct | 2.641 | 3.60E-04 | glio |
| Chek1_in.vivo_noRT | Flrt3 | 2.34 | 3.60E-04 | glio |
| Hsd17b10_in.vivo_noRT | Tmem121 | -1.934 | 3.60E-04 | glio |
| Interaction_genotype_Atr | Zfp788 | -3.629 | 3.60E-05 | glio |
| Interaction_genotype_Mtor | Gm35986 | -4.729 | 3.60E-05 | glio |
| Interaction_genotype_Mtor | Gabarapl1 | 2.167 | 3.60E-05 | glio |
| Bora_in.vivo_noRT | Tmsb15b2 | 3.216 | 3.60E-05 | glio |

| | | | | |
|---|---|---|---|---|
| Chek1_in.vivo_noRT | A930003O13R | 3.244 | 3.60E-05 | glio |
| culture_effect | Gm10010 | 2.536 | 3.60E-06 | glio |
| Interaction_genotype_Mtor | Hist1h2bh | -5.649 | 3.60E-06 | glio |
| Interaction_genotype_Nat2 | Gm21818 | 5.534 | 3.60E-06 | glio |
| Hsd17b10_in.vivo_noRT | Gm128 | 2.698 | 3.60E-06 | glio |
| Hsd17b10_in.vivo_noRT | Htr2b | 3.093 | 3.60E-08 | glio |
| Chek1_in.vivo_noRT | Wfdc1 | 4.059 | 3.60E-09 | glio |
| Chek1_in.vivo_noRT | Gm9970 | 4.399 | 3.60E-09 | glio |
| culture_effect | Bpifa3 | -3.349 | 3.60E-10 | glio |
| Hsd17b10_in.vivo_noRT | Gm35082 | 3.616 | 3.60E-11 | glio |
| Bcas2_in.vivo_noRT | Tcp11x2 | 4.285 | 3.60E-13 | glio |
| Aars2_in.vivo_noRT | Gm48008 | 5.063 | 3.60E-19 | glio |
| culture_effect | Tvp23a | -1.211 | 3.61E-03 | glio |
| culture_effect | Cox15 | -0.713 | 3.61E-03 | glio |
| Interaction_genotype_Bora | Loxl4 | -3.564 | 3.61E-03 | glio |
| Interaction_genotype_Bora | 5033403F01Ri | -3.567 | 3.61E-03 | glio |
| Interaction_genotype_Mtor | Tbc1d10c | -3.557 | 3.61E-03 | glio |
| Interaction_genotype_Mtor | Gm12971 | -3.672 | 3.61E-03 | glio |
| Interaction_genotype_Mtor | Ing1 | 1.642 | 3.61E-03 | glio |
| Interaction_genotype_Myc | Ccdc51 | -2.692 | 3.61E-03 | glio |
| Interaction_genotype_Myc | Thap3 | -1.954 | 3.61E-03 | glio |
| Interaction_genotype_Pars2 | Hcn3 | -3.652 | 3.61E-03 | glio |
| interaction_RT | Hist1h2bl | -1.925 | 3.61E-03 | glio |
| interaction_RT | Wwtr1 | 0.938 | 3.61E-03 | glio |
| Aars2_in.vivo_noRT | Ddx11 | 1.563 | 3.61E-03 | glio |
| Aars2_in.vivo_noRT | Rab2a | -0.756 | 3.61E-03 | glio |
| Aars2_in.vivo_noRT | Rpl27 | -0.618 | 3.61E-03 | glio |
| Atr_in.vivo_noRT | Erp27 | 2.133 | 3.61E-03 | glio |
| Atr_in.vivo_noRT | Mfap5 | 2.178 | 3.61E-03 | glio |
| Atr_in.vivo_noRT | Cldn14 | 1.973 | 3.61E-03 | glio |
| Atr_in.vivo_noRT | Gm20045 | 1.898 | 3.61E-03 | glio |
| Atr_in.vivo_noRT | Trit1 | 1.522 | 3.61E-03 | glio |
| Atr_ex.vivo_noRT | S100a3 | 2.283 | 3.61E-03 | glio |
| Bcas2_in.vivo_noRT | Steap1 | -1.581 | 3.61E-03 | glio |
| Chek1_ex.vivo_noRT | Tmem44 | 2.728 | 3.61E-03 | glio |
| Chek1_ex.vivo_noRT | Hey2 | 2.613 | 3.61E-03 | glio |
| Eif2s1_in.vivo_noRT | Gm42695 | 2.411 | 3.61E-03 | glio |
| Hsd17b10_in.vivo_noRT | Otos | 2.087 | 3.61E-03 | glio |
| Interaction_genotype_Aars2 | Cldn19 | -4.333 | 3.61E-04 | glio |
| Interaction_genotype_Mtor | Slxl1 | -4.297 | 3.61E-04 | glio |
| Interaction_genotype_Mtor | Gm6566 | -3.567 | 3.61E-04 | glio |
| Aars2_in.vivo_noRT | Gm45589 | 2.515 | 3.61E-04 | glio |
| Aars2_in.vivo_noRT | Sox8 | 2.104 | 3.61E-04 | glio |
| Bora_in.vivo_noRT | Ppp2r2b | -2.238 | 3.61E-04 | glio |
| Chek1_in.vivo_noRT | Cyb561d1 | 2.722 | 3.61E-04 | glio |
| Hsd17b10_in.vivo_noRT | 1700011M02F | 3.353 | 3.61E-04 | glio |
| Interaction_genotype_Myc | Proser3 | -4.17 | 3.61E-05 | glio |
| Interaction_genotype_Pars2 | Gm27253 | -5.584 | 3.61E-05 | glio |
| Atr_in.vivo_noRT | Ppp1r3f | 3.062 | 3.61E-05 | glio |
| Atr_in.vivo_noRT | Mafa | 2.631 | 3.61E-05 | glio |
| Bora_in.vivo_noRT | Rhbdf1 | -2.04 | 3.61E-05 | glio |
| Chek1_in.vivo_noRT | Rnf114 | -1.869 | 3.61E-05 | glio |

| | | | | |
|---|---|---|---|---|
| Hsd17b10_in.vivo_noRT | Pde6b | 2.584 | 3.61E-05 | glio |
| culture_effect | Blzf1 | -0.94 | 3.61E-06 | glio |
| Atr_in.vivo_noRT | Rpl21 | -0.866 | 3.61E-06 | glio |
| Bcas2_in.vivo_noRT | Bnip2 | -1.419 | 3.61E-06 | glio |
| Chek1_ex.vivo_noRT | Gm11769 | 3.672 | 3.61E-06 | glio |
| interaction_RT | Tmem128 | 0.963 | 3.61E-07 | glio |
| Bcas2_in.vivo_noRT | Cpne9 | 3.77 | 3.61E-07 | glio |
| Bora_in.vivo_noRT | Syngap1 | 3.476 | 3.61E-07 | glio |
| Chek1_in.vivo_noRT | Srl | 3.625 | 3.61E-07 | glio |
| Bcas2_in.vivo_noRT | Cacnb1 | 4.2 | 3.61E-08 | glio |
| Bcas2_in.vivo_noRT | Gtsf1l | 3.178 | 3.61E-08 | glio |
| Bcas2_in.vivo_noRT | Gm41442 | 3.765 | 3.61E-08 | glio |
| culture_effect | Cers6 | 1.254 | 3.61E-09 | glio |
| Bora_in.vivo_noRT | Gm5039 | 4.936 | 3.61E-10 | glio |
| Hsd17b10_in.vivo_noRT | Gm44646 | 3.616 | 3.61E-10 | glio |
| culture_effect | Atp5a1 | -0.731 | 3.61E-11 | glio |
| Atr_in.vivo_noRT | Ntn5 | 4.741 | 3.61E-12 | glio |
| Aars2_in.vivo_noRT | Gm27253 | 5.06 | 3.61E-13 | glio |
| Bora_in.vivo_noRT | Gm44658 | 4.405 | 3.61E-13 | glio |
| Chek1_in.vivo_noRT | Cyp4x1 | 4.777 | 3.61E-13 | glio |
| Chek1_in.vivo_noRT | Gm27241 | 4.388 | 3.61E-17 | glio |
| Atr_in.vivo_noRT | Gm13348 | 5.533 | 3.61E-21 | glio |
| Atr_in.vivo_noRT | Smtnl1 | 5.533 | 3.61E-24 | glio |
| Bora_in.vivo_noRT | Ankrd22 | 4.936 | 3.61E-24 | glio |
| Atr_in.vivo_noRT | Apoa5 | 5.533 | 3.61E-30 | glio |
| culture_effect | Fam178b | 1.514 | 3.62E-03 | glio |
| culture_effect | Tgoln1 | -0.493 | 3.62E-03 | glio |
| culture_effect | Rraga | -0.466 | 3.62E-03 | glio |
| Interaction_genotype_Atr | Cpne7 | -3.775 | 3.62E-03 | glio |
| Interaction_genotype_Mtor | Magee1 | -2.971 | 3.62E-03 | glio |
| Interaction_genotype_Myc | Eml1 | -2.847 | 3.62E-03 | glio |
| Interaction_genotype_Myc | Fbxl8 | -2.63 | 3.62E-03 | glio |
| Interaction_genotype_Myc | Pced1a | -2.505 | 3.62E-03 | glio |
| Interaction_genotype_Myc | Stil | -2.628 | 3.62E-03 | glio |
| interaction_RT | Gtf2h1 | -0.748 | 3.62E-03 | glio |
| interaction_RT | Amd1 | -0.701 | 3.62E-03 | glio |
| interaction_RT | Gps1 | -0.653 | 3.62E-03 | glio |
| Aars2_in.vivo_noRT | Dlgap1 | 1.873 | 3.62E-03 | glio |
| Atr_in.vivo_noRT | 2700033N17R | 1.844 | 3.62E-03 | glio |
| Bard1_ex.vivo_noRT | Serpina3i | 2.814 | 3.62E-03 | glio |
| Bcas2_in.vivo_noRT | Krt18 | -2.578 | 3.62E-03 | glio |
| Bcas2_in.vivo_noRT | Csnk2a1 | -0.852 | 3.62E-03 | glio |
| Chek1_ex.vivo_noRT | Ntrk1 | 2.811 | 3.62E-03 | glio |
| Chek1_ex.vivo_noRT | Trmt10a | -2.245 | 3.62E-03 | glio |
| Interaction_genotype_Aars2 | Txnl1 | 1.73 | 3.62E-04 | glio |
| Interaction_genotype_Aars2 | Ndfip1 | 1.368 | 3.62E-04 | glio |
| Interaction_genotype_Myc | Mir155hg | -4.41 | 3.62E-04 | glio |
| Interaction_genotype_Myc | Hist2h2ac | -3.921 | 3.62E-04 | glio |
| interaction_RT | 2900076A07R | 1.607 | 3.62E-04 | glio |
| interaction_RT | Gm17210 | 3.453 | 3.62E-04 | glio |
| Aars2_in.vivo_noRT | A630072M18F | 2.175 | 3.62E-04 | glio |
| Bcas2_in.vivo_noRT | Pak4 | -1.834 | 3.62E-04 | glio |

| | | | | |
|---|---|---|---|---|
| Bora_in.vivo_noRT | Gm14023 | 2.658 | 3.62E-04 | glio |
| Hsd17b10_in.vivo_noRT | Gm15879 | 3.433 | 3.62E-04 | glio |
| culture_effect | Gm26766 | 2.08 | 3.62E-05 | glio |
| Aars2_in.vivo_noRT | Tmem171 | 3.709 | 3.62E-05 | glio |
| Aars2_in.vivo_noRT | Ube2a | -1.663 | 3.62E-05 | glio |
| Chek1_in.vivo_noRT | 4930522L14Ri | 3.025 | 3.62E-05 | glio |
| Interaction_genotype_Mtor | Tal2 | -4.025 | 3.62E-06 | glio |
| Aars2_in.vivo_noRT | Gm43378 | 3.038 | 3.62E-06 | glio |
| Bora_in.vivo_noRT | 8030456M14F | 3.316 | 3.62E-06 | glio |
| Aars2_in.vivo_noRT | Gm26688 | 3.888 | 3.62E-08 | glio |
| Atr_in.vivo_noRT | 2810457G06R | 4.286 | 3.62E-08 | glio |
| Aars2_in.vivo_noRT | Gm26542 | 3.949 | 3.62E-09 | glio |
| Dclre1c_in.vivo_noRT | Dclre1c | -4.457 | 3.62E-09 | glio |
| Bora_in.vivo_noRT | Vmn2r54 | 3.775 | 3.62E-11 | glio |
| Chek1_ex.vivo_noRT | AF067063 | 4.842 | 3.62E-14 | glio |
| Chek1_ex.vivo_noRT | Gm34059 | 4.842 | 3.62E-17 | glio |
| Chek1_ex.vivo_noRT | Halr1 | 4.842 | 3.62E-17 | glio |
| Aars2_in.vivo_noRT | Ptprc | 5.063 | 3.62E-20 | glio |
| Bcas2_in.vivo_noRT | Gm266 | 4.807 | 3.62E-21 | glio |
| culture_effect | Gm31805 | 1.52 | 3.63E-03 | glio |
| culture_effect | Mboat1 | 0.706 | 3.63E-03 | glio |
| culture_effect | Cetn4 | 0.67 | 3.63E-03 | glio |
| culture_effect | Rbm15 | 0.629 | 3.63E-03 | glio |
| Interaction_genotype_Aars2 | Tnfsf13os | -3.644 | 3.63E-03 | glio |
| Interaction_genotype_Atr | 1700026N04R | -3.233 | 3.63E-03 | glio |
| Interaction_genotype_Cdkn2a | Gstt4 | 3.098 | 3.63E-03 | glio |
| Interaction_genotype_Cdkn2a | Hoxc8 | 2.835 | 3.63E-03 | glio |
| Interaction_genotype_Rad51 | Gm39326 | -4.066 | 3.63E-03 | glio |
| interaction_RT | Prune2 | 1.854 | 3.63E-03 | glio |
| Aars2_in.vivo_noRT | Uevld | 1.73 | 3.63E-03 | glio |
| Atr_in.vivo_noRT | Foxred2 | 1.554 | 3.63E-03 | glio |
| Atr_in.vivo_noRT | Uchl1 | -1.379 | 3.63E-03 | glio |
| Atr_ex.vivo_noRT | Apol6 | 2.376 | 3.63E-03 | glio |
| Bora_in.vivo_noRT | B230303O12R | 2.157 | 3.63E-03 | glio |
| Bora_in.vivo_noRT | Gm14419 | 1.885 | 3.63E-03 | glio |
| Bora_in.vivo_noRT | 4933434E20Ri | -1.216 | 3.63E-03 | glio |
| Bora_in.vivo_noRT | Slc19a1 | 1.288 | 3.63E-03 | glio |
| Bora_ex.vivo_noRT | Slc16a4 | 2.345 | 3.63E-03 | glio |
| Cenpj_ex.vivo_noRT | Usp17lc | 2.591 | 3.63E-03 | glio |
| Chek1_in.vivo_noRT | Draxin | 2.11 | 3.63E-03 | glio |
| Cyp19a1_ex.vivo_noRT | Gm48155 | 2.159 | 3.63E-03 | glio |
| Cyp19a1_ex.vivo_noRT | Gm50223 | 2.54 | 3.63E-03 | glio |
| Cyp19a1_ex.vivo_noRT | Gm34059 | 2.159 | 3.63E-03 | glio |
| Cyp19a1_ex.vivo_noRT | Dlx4 | 3.146 | 3.63E-03 | glio |
| Cyp19a1_ex.vivo_noRT | Halr1 | 2.159 | 3.63E-03 | glio |
| Interaction_genotype_Mtor | Ezh2 | 2.02 | 3.63E-04 | glio |
| Interaction_genotype_Myc | Ids | -3.005 | 3.63E-04 | glio |
| Interaction_genotype_Rpl3 | BC065403 | -4.513 | 3.63E-04 | glio |
| Atp5c1_in.vivo_noRT | Gm39929 | 2.21 | 3.63E-04 | glio |
| Bora_in.vivo_noRT | Tulp3 | -1.362 | 3.63E-04 | glio |
| Chek1_in.vivo_noRT | Mr1 | 2.611 | 3.63E-04 | glio |
| culture_effect | Lrrfip2 | 0.694 | 3.63E-05 | glio |

| | | | | |
|---|---|---|---|---|
| Interaction_genotype_Atr | Grasp | 3.862 | 3.63E-05 | glio |
| Aars2_in.vivo_noRT | Ucn2 | 2.833 | 3.63E-05 | glio |
| Atr_in.vivo_noRT | 4930458D05R | 3.196 | 3.63E-05 | glio |
| Bcas2_in.vivo_noRT | 9530053A07R | 2.993 | 3.63E-05 | glio |
| Bcas2_in.vivo_noRT | Ell2 | 2.627 | 3.63E-05 | glio |
| Bcas2_in.vivo_noRT | Palm | -2.031 | 3.63E-05 | glio |
| Chek1_ex.vivo_noRT | Wnt6 | 4.443 | 3.63E-05 | glio |
| Interaction_genotype_Bcas2 | Kpna7 | -5.322 | 3.63E-06 | glio |
| Interaction_genotype_Myc | 2610027K06R | -4.883 | 3.63E-06 | glio |
| Interaction_genotype_Myc | Trim16 | -4.843 | 3.63E-06 | glio |
| Atp5c1_in.vivo_noRT | Gm21761 | 2.668 | 3.63E-06 | glio |
| Bora_in.vivo_noRT | Hpse | 3.225 | 3.63E-06 | glio |
| Aars2_in.vivo_noRT | Plin1 | 3.613 | 3.63E-07 | glio |
| Atp5c1_in.vivo_noRT | Lncbate10 | 2.668 | 3.63E-07 | glio |
| Atr_in.vivo_noRT | Auts2 | 3.083 | 3.63E-07 | glio |
| Hsd17b10_in.vivo_noRT | Gm16751 | 3.1 | 3.63E-07 | glio |
| Atr_in.vivo_noRT | Zfhx2os | 3.952 | 3.63E-08 | glio |
| culture_effect | Matn4 | 3.041 | 3.63E-09 | glio |
| Atr_in.vivo_noRT | Gm26850 | 4.139 | 3.63E-09 | glio |
| Chek1_in.vivo_noRT | Vmn1r88 | 3.859 | 3.63E-10 | glio |
| Aars2_in.vivo_noRT | Has1 | 4.429 | 3.63E-12 | glio |
| Aars2_in.vivo_noRT | A930001C03R | 4.674 | 3.63E-12 | glio |
| Aars2_in.vivo_noRT | Zg16 | 4.674 | 3.63E-12 | glio |
| Atr_in.vivo_noRT | Gm43409 | 5.27 | 3.63E-12 | glio |
| Atr_in.vivo_noRT | Peli3 | 5.128 | 3.63E-14 | glio |
| Atr_in.vivo_noRT | Gm27253 | 5.267 | 3.63E-14 | glio |
| Hsd17b10_in.vivo_noRT | Gm49808 | 3.616 | 3.63E-14 | glio |
| Bcas2_in.vivo_noRT | Gm14019 | 4.807 | 3.63E-15 | glio |
| Aars2_in.vivo_noRT | Gm13032 | 4.941 | 3.63E-18 | glio |
| Aars2_in.vivo_noRT | Hnf1aos1 | 5.327 | 3.63E-18 | glio |
| Atr_in.vivo_noRT | 1700019E08Ri | 5.533 | 3.63E-23 | glio |
| Bora_in.vivo_noRT | Gm5157 | 4.936 | 3.63E-25 | glio |
| culture_effect | Atp1b2 | -1.766 | 3.64E-03 | glio |
| culture_effect | Nupr1 | -0.878 | 3.64E-03 | glio |
| Interaction_genotype_Aars2 | Gm48768 | -3.746 | 3.64E-03 | glio |
| Interaction_genotype_Aars2 | Zfp983 | -3.137 | 3.64E-03 | glio |
| Interaction_genotype_Mtor | Neurl1b | -3.578 | 3.64E-03 | glio |
| Interaction_genotype_Mtor | Mboat2 | -2.592 | 3.64E-03 | glio |
| Interaction_genotype_Mtor | Rplp2 | 0.96 | 3.64E-03 | glio |
| Interaction_genotype_Myc | Gm19265 | -2.613 | 3.64E-03 | glio |
| Interaction_genotype_Myc | Pkn3 | -2.836 | 3.64E-03 | glio |
| interaction_RT | Yipf3 | 0.701 | 3.64E-03 | glio |
| interaction_RT | Med31 | 0.751 | 3.64E-03 | glio |
| Aars2_in.vivo_noRT | Gm20219 | 2.343 | 3.64E-03 | glio |
| Atr_in.vivo_noRT | Pax3 | 1.769 | 3.64E-03 | glio |
| Atr_in.vivo_noRT | Atp5b | -0.545 | 3.64E-03 | glio |
| Atr_ex.vivo_noRT | Nrros | 2.77 | 3.64E-03 | glio |
| Atr_ex.vivo_noRT | H2-T24 | 2.769 | 3.64E-03 | glio |
| Atr_ex.vivo_noRT | Gm8212 | 2.803 | 3.64E-03 | glio |
| Atr_ex.vivo_noRT | Mb | 2.961 | 3.64E-03 | glio |
| Bcas2_in.vivo_noRT | Kcnt2 | 2.084 | 3.64E-03 | glio |
| Bcas2_in.vivo_noRT | Shox2 | 1.473 | 3.64E-03 | glio |

| | | | | |
|---|---|---|---|---|
| Bora_in.vivo_noRT | E230016M11R | 1.947 | 3.64E-03 | glio |
| Chek1_ex.vivo_noRT | Selp | 2.647 | 3.64E-03 | glio |
| Chek1_ex.vivo_noRT | Gdnf | 1.736 | 3.64E-03 | glio |
| culture_effect | Tmem240 | 2.066 | 3.64E-04 | glio |
| Interaction_genotype_Mtor | Hexa | 1.954 | 3.64E-04 | glio |
| Interaction_genotype_Myc | Gm34776 | -3.122 | 3.64E-04 | glio |
| Interaction_genotype_Myc | Il1rapl1 | -4.125 | 3.64E-04 | glio |
| Aars2_in.vivo_noRT | AC132307.1 | 2.298 | 3.64E-04 | glio |
| Atr_ex.vivo_noRT | 2010001K21R | 2.962 | 3.64E-04 | glio |
| Bora_in.vivo_noRT | Tmem35b | 2.469 | 3.64E-04 | glio |
| Bora_in.vivo_noRT | Cdca7l | -1.631 | 3.64E-04 | glio |
| Bora_in.vivo_noRT | Rae1 | -1.173 | 3.64E-04 | glio |
| Interaction_genotype_Mtor | A330084C13R | -4.958 | 3.64E-05 | glio |
| Interaction_genotype_Mtor | Aqp9 | -4.099 | 3.64E-05 | glio |
| Interaction_genotype_Mtor | Gm50012 | -5.077 | 3.64E-05 | glio |
| Interaction_genotype_Myc | Nanos3 | -4.129 | 3.64E-05 | glio |
| Interaction_genotype_Myc | S100a7a | -4.774 | 3.64E-05 | glio |
| Interaction_genotype_Myc | Gm30082 | -4.658 | 3.64E-05 | glio |
| Interaction_genotype_Myc | Polr3gl | -3.617 | 3.64E-05 | glio |
| Interaction_genotype_Myc | St6galnac4 | -3.208 | 3.64E-05 | glio |
| Interaction_genotype_Myc | Tal2 | -3.448 | 3.64E-05 | glio |
| Atp5c1_in.vivo_noRT | Tcstv1 | 2.668 | 3.64E-05 | glio |
| Atr_in.vivo_noRT | Pde8a | 2.537 | 3.64E-05 | glio |
| Hsd17b10_in.vivo_noRT | Ptpn22 | 3.353 | 3.64E-05 | glio |
| Interaction_genotype_Myc | Atp5j | 1.444 | 3.64E-06 | glio |
| Aars2_in.vivo_noRT | Serac1 | 3.098 | 3.64E-06 | glio |
| Bora_ex.vivo_noRT | Hist1h2ag | 3.201 | 3.64E-06 | glio |
| Atp5c1_in.vivo_noRT | Tac2 | 2.668 | 3.64E-07 | glio |
| Bora_in.vivo_noRT | Slc17a9 | 3.481 | 3.64E-07 | glio |
| Bora_ex.vivo_noRT | Vmn2r18 | 3.201 | 3.64E-07 | glio |
| Hsd17b10_in.vivo_noRT | 4930505N22R | 3.356 | 3.64E-07 | glio |
| Chek1_in.vivo_noRT | Pdzk1ip1 | 4.054 | 3.64E-08 | glio |
| Chek1_in.vivo_noRT | Wdr38 | 4.002 | 3.64E-08 | glio |
| Aars2_in.vivo_noRT | Dact1 | 4.159 | 3.64E-10 | glio |
| Bcas2_in.vivo_noRT | 1700030L20Ri | 4.151 | 3.64E-10 | glio |
| Bora_in.vivo_noRT | Gm8212 | 4.358 | 3.64E-10 | glio |
| Bora_in.vivo_noRT | Elobl | 4.936 | 3.64E-11 | glio |
| Aars2_in.vivo_noRT | Gm43948 | 3.851 | 3.64E-12 | glio |
| Atr_in.vivo_noRT | 4921508D12R | 4.606 | 3.64E-13 | glio |
| Chek1_in.vivo_noRT | Dapk2 | 4.261 | 3.64E-16 | glio |
| culture_effect | Nacc2 | 1.781 | 3.64E-18 | glio |
| Atr_in.vivo_noRT | Gm12800 | 5.533 | 3.64E-26 | glio |
| culture_effect | Atxn1l | 0.825 | 3.65E-03 | glio |
| Interaction_genotype_Aars2 | Cdc25b | -2.616 | 3.65E-03 | glio |
| Interaction_genotype_Atr | A330049N07R | -2.503 | 3.65E-03 | glio |
| Atr_in.vivo_noRT | Gm38115 | 2.093 | 3.65E-03 | glio |
| Bcas2_ex.vivo_noRT | Atp11c | -2.279 | 3.65E-03 | glio |
| Bora_in.vivo_noRT | Pdlim1 | -1.537 | 3.65E-03 | glio |
| Chek1_in.vivo_noRT | Hcn1 | 2.301 | 3.65E-03 | glio |
| Chek1_in.vivo_noRT | Sept1 | 2.318 | 3.65E-03 | glio |
| Chek1_ex.vivo_noRT | Cyb561a3 | 2.344 | 3.65E-03 | glio |
| culture_effect | Pafah1b2 | 0.454 | 3.65E-04 | glio |

| | | | | |
|---|---|---|---|---|
| Interaction_genotype_Myc | AI182371 | -3.32 | 3.65E-04 | glio |
| Interaction_genotype_Myc | Tcea2 | -3.841 | 3.65E-04 | glio |
| Bcas2_in.vivo_noRT | Cib2 | 2.925 | 3.65E-04 | glio |
| Bcas2_in.vivo_noRT | Prelid3a | 2.579 | 3.65E-04 | glio |
| Bora_in.vivo_noRT | Spdl1 | -1.714 | 3.65E-04 | glio |
| Chek1_ex.vivo_noRT | Cgn | 3.111 | 3.65E-04 | glio |
| Eif2s1_in.vivo_noRT | Gm11583 | 2.145 | 3.65E-04 | glio |
| Eif2s1_in.vivo_noRT | Ube4bos1 | 2.403 | 3.65E-04 | glio |
| Hsd17b10_in.vivo_noRT | 1110017D15R | 2.616 | 3.65E-04 | glio |
| culture_effect | Tcirg1 | -1.095 | 3.65E-05 | glio |
| Interaction_genotype_Mtor | Card9 | -4.235 | 3.65E-05 | glio |
| Interaction_genotype_Mtor | Klk13 | -3.567 | 3.65E-05 | glio |
| Interaction_genotype_Mtor | Akap14 | -3.836 | 3.65E-05 | glio |
| Atr_in.vivo_noRT | Psmc3 | -1.389 | 3.65E-05 | glio |
| Bcas2_in.vivo_noRT | Eprs | -1.002 | 3.65E-05 | glio |
| Brca2_in.vivo_noRT | Pakap | 2.813 | 3.65E-05 | glio |
| Brca2_in.vivo_noRT | Gm48140 | 2.561 | 3.65E-05 | glio |
| culture_effect | 4933429H19R | 2.339 | 3.65E-06 | glio |
| Bcas2_in.vivo_noRT | 1700025N23R | 2.915 | 3.65E-06 | glio |
| Bora_in.vivo_noRT | Dock6 | 3.212 | 3.65E-06 | glio |
| Hsd17b10_in.vivo_noRT | Gm9967 | 3.229 | 3.65E-06 | glio |
| culture_effect | 1810019D21R | 2.757 | 3.65E-07 | glio |
| Aars2_in.vivo_noRT | Serpinb6b | 3.685 | 3.65E-07 | glio |
| Atr_in.vivo_noRT | Tmc7 | 3.721 | 3.65E-07 | glio |
| Hsd17b10_in.vivo_noRT | Snhg4.1 | 3.093 | 3.65E-07 | glio |
| culture_effect | Ccl19 | -2.886 | 3.65E-08 | glio |
| Bora_in.vivo_noRT | 2210417A02R | 4.145 | 3.65E-08 | glio |
| culture_effect | 2210408F21Ri | 1.859 | 3.65E-09 | glio |
| Aars2_in.vivo_noRT | Tomm22 | -1.382 | 3.65E-09 | glio |
| Bcas2_in.vivo_noRT | Gm9908 | 4.408 | 3.65E-09 | glio |
| Hsd17b10_in.vivo_noRT | 1700016P04R | 3.877 | 3.65E-09 | glio |
| Atr_in.vivo_noRT | Tslp | 4.888 | 3.65E-11 | glio |
| Chek1_in.vivo_noRT | Jph4 | 4.777 | 3.65E-12 | glio |
| Aars2_in.vivo_noRT | Agrp | 4.544 | 3.65E-15 | glio |
| Atr_in.vivo_noRT | Loxl4 | 5.267 | 3.65E-16 | glio |
| Bora_in.vivo_noRT | Gm14820 | 4.673 | 3.65E-19 | glio |
| Atr_in.vivo_noRT | Gm38477 | 5.011 | 3.65E-20 | glio |
| Aars2_in.vivo_noRT | B4galnt1 | 5.603 | 3.65E-21 | glio |
| Aars2_in.vivo_noRT | Adam5 | 5.063 | 3.65E-23 | glio |
| Bcas2_in.vivo_noRT | 4930588A03R | 4.807 | 3.65E-23 | glio |
| Aars2_in.vivo_noRT | Gm26543 | 5.327 | 3.65E-27 | glio |
| Atr_in.vivo_noRT | Gm29966 | 5.273 | 3.65E-27 | glio |
| Bora_in.vivo_noRT | Gm26543 | 5.46 | 3.65E-27 | glio |
| Bora_in.vivo_noRT | Pspn | 5.46 | 3.65E-27 | glio |
| culture_effect | Svopl | -1.723 | 3.66E-03 | glio |
| Interaction_genotype_Aars2 | Miip | -2.667 | 3.66E-03 | glio |
| Interaction_genotype_Atr | Nrap | -3.82 | 3.66E-03 | glio |
| Interaction_genotype_Atr | C4b | 3.001 | 3.66E-03 | glio |
| Interaction_genotype_Atr | Tox4 | 1.734 | 3.66E-03 | glio |
| Interaction_genotype_Mtor | 4930556M19F | -2.915 | 3.66E-03 | glio |
| Interaction_genotype_Mtor | Timm17a | 1.32 | 3.66E-03 | glio |
| Interaction_genotype_Myc | 8030423F21Ri | -2.776 | 3.66E-03 | glio |

| | | | | |
|---|---|---|---|---|
| Interaction_genotype_Myc | Gm42937 | -3.308 | 3.66E-03 | glio |
| Interaction_genotype_Myc | Tsacc | -2.351 | 3.66E-03 | glio |
| Interaction_genotype_Myc | Lrp8 | -2.294 | 3.66E-03 | glio |
| Interaction_genotype_Myc | Nr2c2ap | 1.352 | 3.66E-03 | glio |
| Interaction_genotype_Myc | Rab10 | 1.13 | 3.66E-03 | glio |
| Interaction_genotype_Rpl3 | Pik3c2a | 2.162 | 3.66E-03 | glio |
| Aars2_ex.vivo_noRT | Gm11131 | 2.897 | 3.66E-03 | glio |
| Atp5c1_in.vivo_noRT | Ncf1 | 2.015 | 3.66E-03 | glio |
| Atr_in.vivo_noRT | Fbxo36 | 1.676 | 3.66E-03 | glio |
| Atr_in.vivo_noRT | Hars | -1.115 | 3.66E-03 | glio |
| Atr_ex.vivo_noRT | Dhodh | -2.182 | 3.66E-03 | glio |
| Atr_ex.vivo_noRT | Fbxo32 | 1.994 | 3.66E-03 | glio |
| Bard1_ex.vivo_noRT | 1700034I23Ri | 2.017 | 3.66E-03 | glio |
| Bcas2_in.vivo_noRT | Tuba8 | 2.102 | 3.66E-03 | glio |
| Chek1_in.vivo_noRT | Rpl23a | -0.572 | 3.66E-03 | glio |
| Chek1_ex.vivo_noRT | Gm42658 | 2.471 | 3.66E-03 | glio |
| Interaction_genotype_Bora | Efemp2 | 1.985 | 3.66E-04 | glio |
| Interaction_genotype_Mtor | Fyb | -4.203 | 3.66E-04 | glio |
| Interaction_genotype_Myc | Il17b | -4.454 | 3.66E-04 | glio |
| Interaction_genotype_Myc | Wdpcp | -3.877 | 3.66E-04 | glio |
| Interaction_genotype_Myc | 2810429I04Ri | -3.345 | 3.66E-04 | glio |
| Interaction_genotype_Myc | Sec61a2 | -2.862 | 3.66E-04 | glio |
| Aars2_in.vivo_noRT | Notch2 | 1.904 | 3.66E-04 | glio |
| Aars2_in.vivo_noRT | Ttc1 | -1.228 | 3.66E-04 | glio |
| Atp5c1_in.vivo_noRT | B3gnt4 | 2.405 | 3.66E-04 | glio |
| Atr_in.vivo_noRT | Nemf | -1.168 | 3.66E-04 | glio |
| Bora_ex.vivo_noRT | Hfm1 | 2.626 | 3.66E-04 | glio |
| Chek1_in.vivo_noRT | Lrrn4cl | 2.77 | 3.66E-04 | glio |
| Bora_in.vivo_noRT | A630052C17R | 2.848 | 3.66E-05 | glio |
| Chek1_ex.vivo_noRT | Qpct | 3.867 | 3.66E-05 | glio |
| Hsd17b10_in.vivo_noRT | Gm44702 | 3.084 | 3.66E-05 | glio |
| Interaction_genotype_Aars2 | Gm48242 | -4.982 | 3.66E-06 | glio |
| Interaction_genotype_Mtor | Bloc1s6os | -4.816 | 3.66E-06 | glio |
| Interaction_genotype_Mtor | AF067063 | -4.099 | 3.66E-06 | glio |
| Aars2_in.vivo_noRT | 4930513N10R | 3.718 | 3.66E-06 | glio |
| Chek1_in.vivo_noRT | D230022J07Ri | 3.355 | 3.66E-06 | glio |
| Aars2_in.vivo_noRT | 4930417H01R | 3.687 | 3.66E-07 | glio |
| Aars2_in.vivo_noRT | Pnp | -2.139 | 3.66E-07 | glio |
| Atr_ex.vivo_noRT | Upb1 | 4.158 | 3.66E-07 | glio |
| Bora_ex.vivo_noRT | Gm16178 | 3.201 | 3.66E-07 | glio |
| Bora_ex.vivo_noRT | Gm364 | 3.201 | 3.66E-07 | glio |
| Chek1_ex.vivo_noRT | Alox15 | 4.447 | 3.66E-08 | glio |
| Hsd17b10_in.vivo_noRT | Card9 | 3.352 | 3.66E-08 | glio |
| Atr_in.vivo_noRT | Zg16 | 4.88 | 3.66E-13 | glio |
| Chek1_ex.vivo_noRT | Gm6566 | 5.249 | 3.66E-13 | glio |
| culture_effect | Tmed10 | -0.967 | 3.66E-16 | glio |
| Atr_in.vivo_noRT | Gm10482 | 5.011 | 3.66E-19 | glio |
| Atr_in.vivo_noRT | Aqp9 | 5.533 | 3.66E-21 | glio |
| Bora_in.vivo_noRT | Gm15138 | 5.194 | 3.66E-23 | glio |
| Bora_in.vivo_noRT | Gm30692 | 4.936 | 3.66E-23 | glio |
| Aars2_in.vivo_noRT | A930019D19R | 5.327 | 3.66E-28 | glio |
| culture_effect | 4932422M17F | 1.22 | 3.67E-03 | glio |

| | | | | |
|---|---|---|---|---|
| Interaction_genotype_Myc | Sgms2 | -3.356 | 3.67E-03 | glio |
| Interaction_genotype_Myc | Snhg6 | 1.366 | 3.67E-03 | glio |
| Aars2_in.vivo_noRT | U2af1 | -0.782 | 3.67E-03 | glio |
| Atp5c1_in.vivo_noRT | A030005L19Ri | 2.139 | 3.67E-03 | glio |
| Atp5c1_in.vivo_noRT | Rab37 | 2.281 | 3.67E-03 | glio |
| Atp5c1_in.vivo_noRT | Gpnmb | 0.794 | 3.67E-03 | glio |
| Atr_in.vivo_noRT | Taf3 | -1.099 | 3.67E-03 | glio |
| Bcas2_in.vivo_noRT | Draxin | 2.122 | 3.67E-03 | glio |
| Bcas2_in.vivo_noRT | Zfyve1 | 1.692 | 3.67E-03 | glio |
| Bora_in.vivo_noRT | 5830448L01Ri | 2.081 | 3.67E-03 | glio |
| Chek1_in.vivo_noRT | Sap25 | 1.999 | 3.67E-03 | glio |
| Chek1_in.vivo_noRT | Slu7 | -1.131 | 3.67E-03 | glio |
| Chek1_ex.vivo_noRT | Foxk1 | 1.945 | 3.67E-03 | glio |
| culture_effect | Tmem250-ps | 0.906 | 3.67E-04 | glio |
| culture_effect | Wnk1 | 0.504 | 3.67E-04 | glio |
| Interaction_genotype_Bcas2 | 1110020A21R | -4.675 | 3.67E-04 | glio |
| Interaction_genotype_Bcas2 | Tmem179 | -4.645 | 3.67E-04 | glio |
| Interaction_genotype_Bcas2 | BC028471 | -4.742 | 3.67E-04 | glio |
| Interaction_genotype_Chek1 | Lgals7 | 5.603 | 3.67E-04 | glio |
| Interaction_genotype_Mtor | Gm15013 | -4.252 | 3.67E-04 | glio |
| Interaction_genotype_Myc | Cgnl1 | -2.729 | 3.67E-04 | glio |
| Interaction_genotype_Myc | 1110035H17R | -4.517 | 3.67E-04 | glio |
| Interaction_genotype_Myc | Afg1l | -4.01 | 3.67E-04 | glio |
| Interaction_genotype_Myc | Gatb | -3.496 | 3.67E-04 | glio |
| Interaction_genotype_Myc | Tnfrsf19 | -4.221 | 3.67E-04 | glio |
| Interaction_genotype_Myc | Cdkn2c | -3.184 | 3.67E-04 | glio |
| Atr_in.vivo_noRT | Msantd1 | 2.546 | 3.67E-04 | glio |
| Atr_in.vivo_noRT | Zfp867 | 2.102 | 3.67E-04 | glio |
| Atr_ex.vivo_noRT | D630024D03R | 2.577 | 3.67E-04 | glio |
| Chek1_in.vivo_noRT | Parvb | 2.439 | 3.67E-04 | glio |
| Interaction_genotype_Myc | Ddb2 | -3.838 | 3.67E-05 | glio |
| interaction_RT | S100a13 | 0.909 | 3.67E-05 | glio |
| Aars2_in.vivo_noRT | Gng8 | 2.822 | 3.67E-05 | glio |
| Atr_in.vivo_noRT | Gm15998 | 2.9 | 3.67E-05 | glio |
| Bcas2_in.vivo_noRT | Upk3bl | 2.731 | 3.67E-05 | glio |
| Bora_in.vivo_noRT | Shd | 2.849 | 3.67E-05 | glio |
| Aars2_in.vivo_noRT | Nova2 | 3.099 | 3.67E-06 | glio |
| Bora_in.vivo_noRT | Srgn | 3.892 | 3.67E-06 | glio |
| Atp5c1_in.vivo_noRT | Slc35g3 | 2.668 | 3.67E-07 | glio |
| Atr_in.vivo_noRT | Cbr3 | 3.781 | 3.67E-07 | glio |
| Atr_in.vivo_noRT | Gm26930 | 3.844 | 3.67E-07 | glio |
| Atr_in.vivo_noRT | Arvcf | 3.761 | 3.67E-07 | glio |
| Chek1_ex.vivo_noRT | Gm2449 | 3.746 | 3.67E-07 | glio |
| Hsd17b10_in.vivo_noRT | Gngt2 | 3.616 | 3.67E-07 | glio |
| Interaction_genotype_Atr | Klhdc8a | -5.172 | 3.67E-08 | glio |
| Hsd17b10_in.vivo_noRT | Gm2449 | 3.353 | 3.67E-08 | glio |
| Hsd17b10_in.vivo_noRT | AC163347.1 | 3.353 | 3.67E-09 | glio |
| culture_effect | Unc5c | 1.688 | 3.67E-10 | glio |
| Chek1_in.vivo_noRT | Mycl | 4.522 | 3.67E-10 | glio |
| Chek1_ex.vivo_noRT | Gm30794 | 3.866 | 3.67E-10 | glio |
| culture_effect | Serf2 | 0.714 | 3.67E-11 | glio |
| Chek1_ex.vivo_noRT | Usp17lc | 4.842 | 3.67E-11 | glio |

| | | | | |
|---|---|---|---|---|
| Bcas2_in.vivo_noRT | Gngt2 | 4.807 | 3.67E-12 | glio |
| Bcas2_in.vivo_noRT | 1700019E08Ri | 4.807 | 3.67E-18 | glio |
| Interaction_genotype_Aars2 | Mier3 | -2.477 | 3.68E-03 | glio |
| Interaction_genotype_Bcas2 | BC030343 | -3.893 | 3.68E-03 | glio |
| Interaction_genotype_Bcas2 | B4galt3 | 1.836 | 3.68E-03 | glio |
| Interaction_genotype_Bcas2 | Cdc27 | 1.947 | 3.68E-03 | glio |
| Interaction_genotype_Bora | B930094E09Ri | -3.649 | 3.68E-03 | glio |
| Interaction_genotype_Bora | Fanca | -3.353 | 3.68E-03 | glio |
| Interaction_genotype_Mtor | Ablim3 | -3.506 | 3.68E-03 | glio |
| Interaction_genotype_Mtor | Sybu | -3.246 | 3.68E-03 | glio |
| Interaction_genotype_Mtor | Pcx | -3.299 | 3.68E-03 | glio |
| Interaction_genotype_Mtor | Dgke | -2.396 | 3.68E-03 | glio |
| Interaction_genotype_Pars2 | Trim16 | -3.585 | 3.68E-03 | glio |
| Interaction_genotype_Pars2 | Gm41764 | -3.538 | 3.68E-03 | glio |
| Interaction_genotype_Rad51 | Bcl2 | -3.279 | 3.68E-03 | glio |
| Interaction_genotype_Rad51 | Synpr | -4.591 | 3.68E-03 | glio |
| Interaction_genotype_Rad51 | Gm43672 | -4.232 | 3.68E-03 | glio |
| interaction_RT | Hist1h2be | -1.68 | 3.68E-03 | glio |
| interaction_RT | Gm11808 | 0.697 | 3.68E-03 | glio |
| Aars2_in.vivo_noRT | Tpcn2 | 2.095 | 3.68E-03 | glio |
| Aars2_in.vivo_noRT | Tfr2 | 1.898 | 3.68E-03 | glio |
| Aars2_in.vivo_noRT | BC024978 | 1.786 | 3.68E-03 | glio |
| Aars2_ex.vivo_noRT | Rapgef3os2 | 2.107 | 3.68E-03 | glio |
| Aars2_ex.vivo_noRT | Tubb4a | 3.301 | 3.68E-03 | glio |
| Atp5c1_in.vivo_noRT | Synpo2l | 2.023 | 3.68E-03 | glio |
| Atp5c1_in.vivo_noRT | Gm32834 | 2.279 | 3.68E-03 | glio |
| Bcas2_in.vivo_noRT | Gm43378 | 2.008 | 3.68E-03 | glio |
| Bcas2_in.vivo_noRT | Dnajc3 | -1.394 | 3.68E-03 | glio |
| Bcas2_in.vivo_noRT | Pnp | -1.061 | 3.68E-03 | glio |
| Bcas2_in.vivo_noRT | Snrpd1 | -0.651 | 3.68E-03 | glio |
| Bora_in.vivo_noRT | Klhl33 | 2.136 | 3.68E-03 | glio |
| Chek1_in.vivo_noRT | Sh3bp5 | 2.104 | 3.68E-03 | glio |
| culture_effect | Chdh | -2.151 | 3.68E-04 | glio |
| Interaction_genotype_Aars2 | Rbm48 | -3.058 | 3.68E-04 | glio |
| Interaction_genotype_Aars2 | Bola3 | 1.628 | 3.68E-04 | glio |
| Interaction_genotype_Aars2 | Banf1 | 1.351 | 3.68E-04 | glio |
| Interaction_genotype_Myc | 1700041G16R | -3.816 | 3.68E-04 | glio |
| Interaction_genotype_Pars2 | Prox2 | -5.187 | 3.68E-04 | glio |
| Aars2_in.vivo_noRT | Izumo4 | 2.539 | 3.68E-04 | glio |
| Aars2_in.vivo_noRT | Lrp1b | 2.548 | 3.68E-04 | glio |
| Aars2_in.vivo_noRT | Polr2e | -0.992 | 3.68E-04 | glio |
| Cenpj_in.vivo_noRT | 4930588A03R | 2.022 | 3.68E-04 | glio |
| Cenpj_in.vivo_noRT | Gm13856 | 2.022 | 3.68E-04 | glio |
| Cenpj_in.vivo_noRT | Gm21936 | 2.022 | 3.68E-04 | glio |
| Eif2s1_in.vivo_noRT | Olfr56 | 2.143 | 3.68E-04 | glio |
| Hsd17b10_in.vivo_noRT | A430106G13R | 2.825 | 3.68E-04 | glio |
| culture_effect | Carns1 | 2.419 | 3.68E-05 | glio |
| culture_effect | Hoxd3os1 | 1.108 | 3.68E-05 | glio |
| Interaction_genotype_Myc | Gas2 | -3.845 | 3.68E-05 | glio |
| Bard1_ex.vivo_noRT | 1700112J16Ri | 3.01 | 3.68E-05 | glio |
| Chek1_in.vivo_noRT | Mkrn3 | 2.787 | 3.68E-05 | glio |
| Chek1_ex.vivo_noRT | Snph | 3.532 | 3.68E-05 | glio |

| | | | | |
|---|---|---|---|---|
| culture_effect | 1190007I07Ri | -0.632 | 3.68E-06 | glio |
| interaction_RT | Gbp4 | 2.627 | 3.68E-06 | glio |
| Atr_in.vivo_noRT | Gm8773 | 3.422 | 3.68E-06 | glio |
| Chek1_in.vivo_noRT | Gm44091 | 3.125 | 3.68E-06 | glio |
| Chek1_in.vivo_noRT | Nat8f1 | 3.108 | 3.68E-06 | glio |
| culture_effect | Ppp2r2b | -1.391 | 3.68E-07 | glio |
| Atp5c1_in.vivo_noRT | Chia1 | 2.668 | 3.68E-07 | glio |
| Atr_in.vivo_noRT | Rad1 | -2.268 | 3.68E-07 | glio |
| Chek1_in.vivo_noRT | B930059L03Ri | 3.859 | 3.68E-07 | glio |
| Chek1_in.vivo_noRT | Enpp3 | 3.477 | 3.68E-07 | glio |
| Chek1_in.vivo_noRT | 9330104G04R | 3.564 | 3.68E-07 | glio |
| Chek1_ex.vivo_noRT | 4921508D12R | 4.049 | 3.68E-07 | glio |
| Bora_in.vivo_noRT | Wdr26 | -2.15 | 3.68E-09 | glio |
| Hsd17b10_in.vivo_noRT | Col19a1 | 3.23 | 3.68E-09 | glio |
| culture_effect | Sephs2 | 1.318 | 3.68E-10 | glio |
| Atr_in.vivo_noRT | Rilp | 5.008 | 3.68E-11 | glio |
| culture_effect | Gm19391 | 2.867 | 3.68E-12 | glio |
| Chek1_in.vivo_noRT | Iqcn | 4.518 | 3.68E-12 | glio |
| culture_effect | BC049762 | 1.608 | 3.69E-03 | glio |
| Interaction_genotype_Aars2 | Prox2 | -3.819 | 3.69E-03 | glio |
| Interaction_genotype_Mtor | Hmgb3 | 1.712 | 3.69E-03 | glio |
| Interaction_genotype_Myc | Gm20705 | -2.593 | 3.69E-03 | glio |
| Interaction_genotype_Myc | Iqcf1 | -3.165 | 3.69E-03 | glio |
| Interaction_genotype_Myc | 4833407H14R | -3.658 | 3.69E-03 | glio |
| Interaction_genotype_Myc | Gm37612 | -3.35 | 3.69E-03 | glio |
| Interaction_genotype_Myc | Gm10791 | -3.06 | 3.69E-03 | glio |
| Interaction_genotype_Myc | Cacng7 | -3.103 | 3.69E-03 | glio |
| Interaction_genotype_Myc | Prdm6 | -2.975 | 3.69E-03 | glio |
| Interaction_genotype_Myc | Fsd1 | -2.927 | 3.69E-03 | glio |
| Interaction_genotype_Myc | Bdh1 | -2.529 | 3.69E-03 | glio |
| Interaction_genotype_Myc | Kbtbd4 | -2.249 | 3.69E-03 | glio |
| Aars2_in.vivo_noRT | Tmem229b | 1.941 | 3.69E-03 | glio |
| Aars2_in.vivo_noRT | Ndufb3 | -0.797 | 3.69E-03 | glio |
| Atr_ex.vivo_noRT | Flrt1 | 3.001 | 3.69E-03 | glio |
| Atr_ex.vivo_noRT | Gm13261 | 2.768 | 3.69E-03 | glio |
| Bcas2_ex.vivo_noRT | 1700047K16R | 3.023 | 3.69E-03 | glio |
| Bora_in.vivo_noRT | Nap1l1 | -0.562 | 3.69E-03 | glio |
| Bora_ex.vivo_noRT | Cyp4f40 | 2.808 | 3.69E-03 | glio |
| Bora_ex.vivo_noRT | Itpr1 | -2.189 | 3.69E-03 | glio |
| Cyp19a1_ex.vivo_noRT | Helt | 2.159 | 3.69E-03 | glio |
| Cyp19a1_ex.vivo_noRT | Gm20625 | 2.159 | 3.69E-03 | glio |
| Cyp19a1_ex.vivo_noRT | Gm5157 | 2.159 | 3.69E-03 | glio |
| Cyp19a1_ex.vivo_noRT | Gm44736 | 2.159 | 3.69E-03 | glio |
| Cyp19a1_ex.vivo_noRT | Fam227b | 2.159 | 3.69E-03 | glio |
| Cyp19a1_ex.vivo_noRT | Olfr810 | 2.159 | 3.69E-03 | glio |
| Cyp19a1_ex.vivo_noRT | Apoa5 | 2.159 | 3.69E-03 | glio |
| Cyp19a1_ex.vivo_noRT | Gm49172 | 2.159 | 3.69E-03 | glio |
| Cyp19a1_ex.vivo_noRT | Rslcan18 | 2.947 | 3.69E-03 | glio |
| Cyp19a1_ex.vivo_noRT | Gjd4 | 2.159 | 3.69E-03 | glio |
| Cyp19a1_ex.vivo_noRT | Vmn1r88 | 2.959 | 3.69E-03 | glio |
| Hsd17b10_in.vivo_noRT | Gm50255 | 2.709 | 3.69E-03 | glio |
| culture_effect | Oplah | -1.956 | 3.69E-04 | glio |

| | | | | |
|---|---|---|---|---|
| culture_effect | Epb41l4b | 0.576 | 3.69E-04 | glio |
| Interaction_genotype_Myc | Gm26543 | -2.729 | 3.69E-04 | glio |
| interaction_RT | Mterf1a | 2.009 | 3.69E-04 | glio |
| interaction_RT | Gm47758 | 2.656 | 3.69E-04 | glio |
| Atr_in.vivo_noRT | Gm19967 | 2.306 | 3.69E-04 | glio |
| Bard1_ex.vivo_noRT | Ifitm1 | 3.987 | 3.69E-04 | glio |
| Bcas2_ex.vivo_noRT | Olfr171 | 2.742 | 3.69E-04 | glio |
| Bora_in.vivo_noRT | Gm42372 | 2.937 | 3.69E-04 | glio |
| Hsd17b10_in.vivo_noRT | Sema6b | 2.672 | 3.69E-04 | glio |
| Hsd17b10_in.vivo_noRT | Tsn | -0.629 | 3.69E-04 | glio |
| Interaction_genotype_Mtor | Lmln | -4.785 | 3.69E-05 | glio |
| Aars2_in.vivo_noRT | Pcdhgb4 | 2.531 | 3.69E-05 | glio |
| Atr_in.vivo_noRT | Gm13203 | 3.034 | 3.69E-05 | glio |
| Atr_in.vivo_noRT | 4833438C02R | 2.706 | 3.69E-05 | glio |
| Atr_in.vivo_noRT | Ppp4c | -1.565 | 3.69E-05 | glio |
| Atr_ex.vivo_noRT | Gm20478 | 3.566 | 3.69E-05 | glio |
| culture_effect | N4bp3 | -1.432 | 3.69E-06 | glio |
| culture_effect | Zrsr2 | 0.549 | 3.69E-06 | glio |
| Atr_in.vivo_noRT | Gm42937 | 3.223 | 3.69E-06 | glio |
| Atr_in.vivo_noRT | Myl6b | 3.129 | 3.69E-06 | glio |
| Bcas2_in.vivo_noRT | Pkhd1 | 3.29 | 3.69E-06 | glio |
| Bora_ex.vivo_noRT | Gpr22 | 3.201 | 3.69E-06 | glio |
| Interaction_genotype_Mtor | Klhl30 | -5.526 | 3.69E-07 | glio |
| culture_effect | Gm45051 | 2.12 | 3.69E-08 | glio |
| Aars2_in.vivo_noRT | Tacr2 | 3.894 | 3.69E-08 | glio |
| Aars2_in.vivo_noRT | Gm49864 | 3.894 | 3.69E-08 | glio |
| Aars2_in.vivo_noRT | Atoh8 | 3.944 | 3.69E-09 | glio |
| Chek1_in.vivo_noRT | Cpne8 | 4.309 | 3.69E-09 | glio |
| Bora_in.vivo_noRT | 5930403N24R | 4.41 | 3.69E-10 | glio |
| Bcas2_in.vivo_noRT | S100a14 | 4.007 | 3.69E-11 | glio |
| Bcas2_in.vivo_noRT | Gm45235 | 4.547 | 3.69E-16 | glio |
| Chek1_in.vivo_noRT | BB557941 | 4.784 | 3.69E-16 | glio |
| Chek1_in.vivo_noRT | Gpr22 | 5.044 | 3.69E-21 | glio |
| Bora_in.vivo_noRT | Gm266 | 4.936 | 3.69E-22 | glio |
| culture_effect | Rab27b | -1.989 | 3.70E-03 | glio |
| Interaction_genotype_Atr | Fgf1 | -3.664 | 3.70E-03 | glio |
| Interaction_genotype_Bora | 1700003G18R | -4.29 | 3.70E-03 | glio |
| Interaction_genotype_Myc | Pdlim3 | -2.744 | 3.70E-03 | glio |
| Interaction_genotype_Myc | Sox12 | -3.16 | 3.70E-03 | glio |
| Interaction_genotype_Myc | Gm16537 | -2.811 | 3.70E-03 | glio |
| Interaction_genotype_Myc | Ppp1ca | 1.145 | 3.70E-03 | glio |
| Interaction_genotype_Pars2 | Mtif3 | 2.359 | 3.70E-03 | glio |
| Aars2_in.vivo_noRT | Xrcc3 | 2.039 | 3.70E-03 | glio |
| Aars2_in.vivo_noRT | Gm26579 | 1.981 | 3.70E-03 | glio |
| Aars2_in.vivo_noRT | Jade3 | 1.968 | 3.70E-03 | glio |
| Aars2_in.vivo_noRT | Hnrnpu | -0.61 | 3.70E-03 | glio |
| Atr_in.vivo_noRT | Arhgef17 | 1.888 | 3.70E-03 | glio |
| Atr_in.vivo_noRT | Zfp697 | 1.886 | 3.70E-03 | glio |
| Atr_in.vivo_noRT | Ift140 | 1.907 | 3.70E-03 | glio |
| Bcas2_in.vivo_noRT | Pigt | -1.186 | 3.70E-03 | glio |
| Bcas2_ex.vivo_noRT | 4931431C16R | 2.345 | 3.70E-03 | glio |
| Bora_in.vivo_noRT | Mettl5os | 1.997 | 3.70E-03 | glio |

| | | | | |
|---|---|---|---|---|
| Bora_in.vivo_noRT | Smchd1 | -1.037 | 3.70E-03 | glio |
| Bora_ex.vivo_noRT | Gm42695 | 2.81 | 3.70E-03 | glio |
| Cdkn2a_in.vivo_noRT | Prl3d3 | -1.938 | 3.70E-03 | glio |
| Cdkn2a_in.vivo_noRT | A330049N07R | -1.941 | 3.70E-03 | glio |
| Cdkn2a_in.vivo_noRT | Otos | -2.676 | 3.70E-03 | glio |
| Chek1_in.vivo_noRT | Slc22a22 | 2.086 | 3.70E-03 | glio |
| Chek1_ex.vivo_noRT | Fgd3 | 2.867 | 3.70E-03 | glio |
| Interaction_genotype_Atr | Atp6v1c1 | 1.716 | 3.70E-04 | glio |
| Interaction_genotype_Myc | Zic4 | -4.186 | 3.70E-04 | glio |
| Interaction_genotype_Myc | Cnksr3 | -3.1 | 3.70E-04 | glio |
| Interaction_genotype_Myc | Zfp26 | -2.768 | 3.70E-04 | glio |
| Interaction_genotype_Myc | Klhl5 | -3.004 | 3.70E-04 | glio |
| Interaction_genotype_Myc | Rock2 | 1.532 | 3.70E-04 | glio |
| Atr_in.vivo_noRT | Zfp292 | -1.3 | 3.70E-04 | glio |
| Atr_ex.vivo_noRT | Pla1a | 3.164 | 3.70E-04 | glio |
| Atr_ex.vivo_noRT | Insrr | 3.164 | 3.70E-04 | glio |
| Bcas2_in.vivo_noRT | Ttc41 | 2.736 | 3.70E-04 | glio |
| Brca2_in.vivo_noRT | Btn1a1 | 2.56 | 3.70E-04 | glio |
| Brca2_in.vivo_noRT | Slc35g3 | 2.296 | 3.70E-04 | glio |
| Brca2_in.vivo_noRT | C430002N11R | 2.546 | 3.70E-04 | glio |
| Chek1_ex.vivo_noRT | Xist | 3.155 | 3.70E-04 | glio |
| Chek1_ex.vivo_noRT | Sik1 | 3.255 | 3.70E-04 | glio |
| Hsd17b10_in.vivo_noRT | Trmo | -2.199 | 3.70E-04 | glio |
| culture_effect | Pdlim7 | 1.033 | 3.70E-05 | glio |
| Interaction_genotype_Mtor | Nepro | -3.901 | 3.70E-05 | glio |
| Interaction_genotype_Mtor | 1700112D23R | -4.876 | 3.70E-05 | glio |
| Interaction_genotype_Mtor | Mcm7 | 2.263 | 3.70E-05 | glio |
| Atr_in.vivo_noRT | Mtcp1 | 2.823 | 3.70E-05 | glio |
| Atr_in.vivo_noRT | Cops4 | -1.594 | 3.70E-05 | glio |
| Bcas2_in.vivo_noRT | Mrpl34 | -1.096 | 3.70E-05 | glio |
| Chek1_ex.vivo_noRT | Ifi214 | 3.645 | 3.70E-05 | glio |
| Chek1_ex.vivo_noRT | Slc17a7 | 3.664 | 3.70E-05 | glio |
| Hsd17b10_in.vivo_noRT | Gm32090 | 2.824 | 3.70E-05 | glio |
| culture_effect | Orai2 | 1.477 | 3.70E-06 | glio |
| Interaction_genotype_Myc | Tssk6 | -4.514 | 3.70E-06 | glio |
| Interaction_genotype_Myc | Gm34885 | -5.438 | 3.70E-06 | glio |
| Interaction_genotype_Myc | Efcab9 | -5.173 | 3.70E-06 | glio |
| Atr_in.vivo_noRT | Micall2 | 3.35 | 3.70E-06 | glio |
| Atr_ex.vivo_noRT | Gm45606 | 4.012 | 3.70E-06 | glio |
| Bora_in.vivo_noRT | Scd3 | 3.548 | 3.70E-06 | glio |
| Chek1_in.vivo_noRT | Aldh1l1 | 3.621 | 3.70E-06 | glio |
| culture_effect | Rexo2 | 0.686 | 3.70E-07 | glio |
| Atp5c1_in.vivo_noRT | Abra | 3.194 | 3.70E-07 | glio |
| Chek1_ex.vivo_noRT | H2-Q10 | 4.464 | 3.70E-07 | glio |
| Atr_in.vivo_noRT | Gm20522 | 4.237 | 3.70E-09 | glio |
| Aars2_in.vivo_noRT | B230311B06R | 5.063 | 3.70E-12 | glio |
| Aars2_in.vivo_noRT | Gm37584 | 4.925 | 3.70E-12 | glio |
| Aars2_in.vivo_noRT | Akap3 | 5.067 | 3.70E-13 | glio |
| Bcas2_in.vivo_noRT | Snrpn | 4.149 | 3.70E-14 | glio |
| Aars2_in.vivo_noRT | Lyl1 | 4.676 | 3.70E-15 | glio |
| Aars2_in.vivo_noRT | Dcn | 4.936 | 3.70E-15 | glio |
| Bora_in.vivo_noRT | Gm26978 | 4.676 | 3.70E-16 | glio |

| | | | | |
|---|---|---|---|---|
| Aars2_in.vivo_noRT | Gm47510 | 5.067 | 3.70E-24 | glio |
| culture_effect | Halr1 | -1.201 | 3.71E-03 | glio |
| culture_effect | Gm26549 | 1.18 | 3.71E-03 | glio |
| culture_effect | Ro60 | 0.499 | 3.71E-03 | glio |
| Interaction_genotype_Aars2 | Nid1 | -3.667 | 3.71E-03 | glio |
| Interaction_genotype_Aars2 | 2810414N06R | -3.606 | 3.71E-03 | glio |
| Interaction_genotype_Atr | Gm43689 | -2.873 | 3.71E-03 | glio |
| Interaction_genotype_Atr | Bod1l | 1.339 | 3.71E-03 | glio |
| Interaction_genotype_Bora | Gm48435 | -2.522 | 3.71E-03 | glio |
| Interaction_genotype_Chek1 | Med6 | -2.334 | 3.71E-03 | glio |
| Interaction_genotype_Myc | 0610038B21R | -2.726 | 3.71E-03 | glio |
| Interaction_genotype_Myc | Skp1a | 1.213 | 3.71E-03 | glio |
| Aars2_in.vivo_noRT | Timm13 | -0.637 | 3.71E-03 | glio |
| Atp5c1_in.vivo_noRT | Parvg | 2.407 | 3.71E-03 | glio |
| Atr_in.vivo_noRT | Draxin | 2.07 | 3.71E-03 | glio |
| Atr_in.vivo_noRT | Ankrd17 | -1.046 | 3.71E-03 | glio |
| Atr_in.vivo_noRT | Prdx2 | -0.641 | 3.71E-03 | glio |
| Cenpj_in.vivo_noRT | Ccdc125 | 2.022 | 3.71E-03 | glio |
| Chek1_in.vivo_noRT | Pcdhb20 | 1.985 | 3.71E-03 | glio |
| Chek1_ex.vivo_noRT | Gm32591 | 2.59 | 3.71E-03 | glio |
| Hsd17b10_in.vivo_noRT | Arrdc2 | 2.421 | 3.71E-03 | glio |
| Interaction_genotype_Mtor | Recql4 | -3.455 | 3.71E-04 | glio |
| Atr_in.vivo_noRT | Myo5b | -1.478 | 3.71E-04 | glio |
| Atr_ex.vivo_noRT | Gm10432 | 3.15 | 3.71E-04 | glio |
| Bcas2_in.vivo_noRT | B3gnt5 | 2.324 | 3.71E-04 | glio |
| Bcas2_in.vivo_noRT | Col9a2 | -1.051 | 3.71E-04 | glio |
| Bora_in.vivo_noRT | Dyrk4 | 2.893 | 3.71E-04 | glio |
| culture_effect | Rpf2 | -0.577 | 3.71E-05 | glio |
| Interaction_genotype_Mtor | Ltb4r1 | -5.271 | 3.71E-05 | glio |
| Interaction_genotype_Mtor | Pik3ap1 | -4.196 | 3.71E-05 | glio |
| Atr_ex.vivo_noRT | Slc25a18 | 3.164 | 3.71E-05 | glio |
| Atr_ex.vivo_noRT | Gm17173 | 3.642 | 3.71E-05 | glio |
| Atr_ex.vivo_noRT | Gm15270 | 3.153 | 3.71E-05 | glio |
| Atr_ex.vivo_noRT | 4833419F23Ri | 3.665 | 3.71E-05 | glio |
| Bora_in.vivo_noRT | 4930589L23Ri | 3.058 | 3.71E-05 | glio |
| Interaction_genotype_Mtor | Stab1 | -5.829 | 3.71E-06 | glio |
| Interaction_genotype_Mtor | Gm3734 | -6.039 | 3.71E-06 | glio |
| Aars2_in.vivo_noRT | Adam4 | 2.985 | 3.71E-06 | glio |
| Aars2_ex.vivo_noRT | Ccl9 | 2.9 | 3.71E-06 | glio |
| Hsd17b10_in.vivo_noRT | Akap3 | 3.356 | 3.71E-06 | glio |
| Aars2_in.vivo_noRT | Plekhg6 | 3.657 | 3.71E-07 | glio |
| culture_effect | Jak2 | -1.247 | 3.71E-08 | glio |
| Bora_in.vivo_noRT | E230001N04R | 3.35 | 3.71E-08 | glio |
| Aars2_in.vivo_noRT | Gm2788 | 4.278 | 3.71E-09 | glio |
| Bora_in.vivo_noRT | Gm15520 | 3.62 | 3.71E-09 | glio |
| Bora_in.vivo_noRT | A830035O19R | 3.887 | 3.71E-09 | glio |
| culture_effect | Rpl17 | -0.825 | 3.71E-10 | glio |
| culture_effect | Npm1 | -0.981 | 3.71E-14 | glio |
| Bora_in.vivo_noRT | Rhox7a | 5.46 | 3.71E-14 | glio |
| Chek1_in.vivo_noRT | F630040K05Ri | 4.904 | 3.71E-15 | glio |
| Chek1_in.vivo_noRT | 9130230L23Ri | 4.784 | 3.71E-15 | glio |
| Atr_in.vivo_noRT | Gm41318 | 5.533 | 3.71E-16 | glio |

| | | | | |
|---|---|---|---|---|
| Bora_in.vivo_noRT | Gm40538 | 4.673 | 3.71E-17 | glio |
| culture_effect | Arhgef16 | -1.571 | 3.72E-03 | glio |
| culture_effect | Tmem106a | -1.279 | 3.72E-03 | glio |
| culture_effect | Chp1 | 0.523 | 3.72E-03 | glio |
| Interaction_genotype_Atr | Ptp4a1.1 | -3.549 | 3.72E-03 | glio |
| Interaction_genotype_Atr | Lst1 | -3.684 | 3.72E-03 | glio |
| Interaction_genotype_Mtor | Pomc | -2.669 | 3.72E-03 | glio |
| Interaction_genotype_Myc | Eif3f | 1.258 | 3.72E-03 | glio |
| Interaction_genotype_Rpl3 | Zic4 | -4.276 | 3.72E-03 | glio |
| Atp5c1_in.vivo_noRT | Hbq1b | 2.402 | 3.72E-03 | glio |
| Atp5c1_in.vivo_noRT | Gm43463 | 2.01 | 3.72E-03 | glio |
| Atr_in.vivo_noRT | Pigx | -1.205 | 3.72E-03 | glio |
| Atr_in.vivo_noRT | Rpl39 | -0.761 | 3.72E-03 | glio |
| Bcas2_in.vivo_noRT | Wdr78 | 2.304 | 3.72E-03 | glio |
| Bcas2_ex.vivo_noRT | Gm15326 | 2.167 | 3.72E-03 | glio |
| Bora_in.vivo_noRT | Tceal1 | 2.034 | 3.72E-03 | glio |
| Eif2s1_ex.vivo_noRT | Gm13078 | 2.234 | 3.72E-03 | glio |
| culture_effect | Ampd3 | -2.232 | 3.72E-04 | glio |
| culture_effect | Wasf2 | 0.523 | 3.72E-04 | glio |
| Interaction_genotype_Bora | Max | 2.02 | 3.72E-04 | glio |
| Interaction_genotype_Mtor | Adcy7 | -4.135 | 3.72E-04 | glio |
| Interaction_genotype_Mtor | Fsbp | -4.347 | 3.72E-04 | glio |
| Interaction_genotype_Myc | 2410017I17Ri | -2.729 | 3.72E-04 | glio |
| Interaction_genotype_Myc | Rptor | -2.488 | 3.72E-04 | glio |
| Aars2_in.vivo_noRT | 5830448L01Ri | 2.471 | 3.72E-04 | glio |
| Aars2_ex.vivo_noRT | Ggt1 | 3.293 | 3.72E-04 | glio |
| Bard1_ex.vivo_noRT | Gpc5 | 2.419 | 3.72E-04 | glio |
| Cenpj_ex.vivo_noRT | Gpr4 | 2.591 | 3.72E-04 | glio |
| Cenpj_ex.vivo_noRT | Gm33466 | 2.591 | 3.72E-04 | glio |
| Chek1_ex.vivo_noRT | Platr16 | 3.551 | 3.72E-04 | glio |
| Interaction_genotype_Mtor | Accsl | -5.287 | 3.72E-05 | glio |
| Interaction_genotype_Myc | Zbtb24 | -3.625 | 3.72E-05 | glio |
| Interaction_genotype_Myc | Atp13a2 | -3.219 | 3.72E-05 | glio |
| interaction_RT | Churc1 | 0.909 | 3.72E-05 | glio |
| Aars2_in.vivo_noRT | Tmem88 | 2.844 | 3.72E-05 | glio |
| Atr_in.vivo_noRT | Kcnt2 | 2.81 | 3.72E-05 | glio |
| Bcas2_in.vivo_noRT | Adk | -1.509 | 3.72E-05 | glio |
| Chek1_in.vivo_noRT | Zfp992 | 3.016 | 3.72E-05 | glio |
| Hsd17b10_in.vivo_noRT | Sh2d3c | 3.236 | 3.72E-05 | glio |
| Interaction_genotype_Myc | D17H6S53E | -4.523 | 3.72E-06 | glio |
| Aars2_in.vivo_noRT | Gask1b | 3.532 | 3.72E-06 | glio |
| Atr_in.vivo_noRT | Cdhr4 | 2.978 | 3.72E-06 | glio |
| Bora_in.vivo_noRT | Styk1 | 3.35 | 3.72E-06 | glio |
| Bora_in.vivo_noRT | Gm5431 | 3.025 | 3.72E-06 | glio |
| Hsd17b10_in.vivo_noRT | Gm27253 | 3.35 | 3.72E-06 | glio |
| Hsd17b10_in.vivo_noRT | Gm17173 | 3.229 | 3.72E-06 | glio |
| interaction_RT | Parp1 | -1.128 | 3.72E-07 | glio |
| Aars2_in.vivo_noRT | Gm26619 | 3.731 | 3.72E-07 | glio |
| Chek1_ex.vivo_noRT | Ggt5 | 4.142 | 3.72E-07 | glio |
| Chek1_ex.vivo_noRT | Gm37986 | 4.045 | 3.72E-07 | glio |
| Chek1_ex.vivo_noRT | Grip1 | 4.384 | 3.72E-07 | glio |
| Atr_in.vivo_noRT | Gm36913 | 3.844 | 3.72E-08 | glio |

| | | | | |
|---|---|---|---|---|
| Bcas2_in.vivo_noRT | Top2b | -2.287 | 3.72E-08 | glio |
| interaction_RT | Cst3 | 1.18 | 3.72E-09 | glio |
| Bcas2_in.vivo_noRT | 4933429H19R | 3.895 | 3.72E-09 | glio |
| Bcas2_in.vivo_noRT | Gm47232 | 4.284 | 3.72E-10 | glio |
| Hsd17b10_in.vivo_noRT | Ccdc125 | 3.616 | 3.72E-10 | glio |
| Atp5c1_in.vivo_noRT | Nrip2 | 3.719 | 3.72E-13 | glio |
| Bora_in.vivo_noRT | Tmem52 | 4.557 | 3.72E-13 | glio |
| Bora_in.vivo_noRT | Magea10 | 4.469 | 3.72E-13 | glio |
| Hsd17b10_in.vivo_noRT | Abcb5 | 3.616 | 3.72E-13 | glio |
| culture_effect | Myrf | -4.243 | 3.72E-16 | glio |
| Chek1_in.vivo_noRT | Zmynd10 | 4.784 | 3.72E-16 | glio |
| Bcas2_in.vivo_noRT | Ildr1 | 4.807 | 3.72E-21 | glio |
| culture_effect | Krt8 | -10.11 | 3.72E-28 | glio |
| culture_effect | Col9a2 | -1.954 | 3.72E-31 | glio |
| Interaction_genotype_Bora | Pdcd10 | 1.52 | 3.73E-03 | glio |
| Interaction_genotype_Mtor | Cpne7 | -3.384 | 3.73E-03 | glio |
| Interaction_genotype_Mtor | Ubtd2 | -2.363 | 3.73E-03 | glio |
| Interaction_genotype_Myc | Cnnm1 | -3.646 | 3.73E-03 | glio |
| Interaction_genotype_Myc | Mlf2 | 1.145 | 3.73E-03 | glio |
| Aars2_in.vivo_noRT | Klhl32 | 1.841 | 3.73E-03 | glio |
| Aars2_in.vivo_noRT | Ddx39 | -0.898 | 3.73E-03 | glio |
| Bora_in.vivo_noRT | Hemk1 | 2.171 | 3.73E-03 | glio |
| Bora_in.vivo_noRT | Ccr2 | 2.142 | 3.73E-03 | glio |
| Bora_in.vivo_noRT | Chek1 | -1.431 | 3.73E-03 | glio |
| Bora_in.vivo_noRT | Taldo1 | -0.807 | 3.73E-03 | glio |
| Chek1_in.vivo_noRT | Pxylp1 | 1.977 | 3.73E-03 | glio |
| Hsd17b10_in.vivo_noRT | Alox5ap | 2.511 | 3.73E-03 | glio |
| culture_effect | Mbd3 | 0.429 | 3.73E-04 | glio |
| Interaction_genotype_Mtor | Gm5662 | -4.099 | 3.73E-04 | glio |
| Interaction_genotype_Mtor | Myl12b | 1.903 | 3.73E-04 | glio |
| Atr_in.vivo_noRT | Fam71f1 | 2.12 | 3.73E-04 | glio |
| Bora_ex.vivo_noRT | Gm43915 | 2.808 | 3.73E-04 | glio |
| Chek1_in.vivo_noRT | Olfr77 | 2.458 | 3.73E-04 | glio |
| Chek1_in.vivo_noRT | Bclaf3 | -1.499 | 3.73E-04 | glio |
| Chek1_in.vivo_noRT | Slc1a3 | -1.824 | 3.73E-04 | glio |
| Hsd17b10_in.vivo_noRT | Gstm6 | 2.839 | 3.73E-04 | glio |
| Bard1_ex.vivo_noRT | Serpinb9g | 2.814 | 3.73E-05 | glio |
| Bard1_ex.vivo_noRT | BC049987 | 2.814 | 3.73E-05 | glio |
| Bora_in.vivo_noRT | Tes3-ps | 2.902 | 3.73E-05 | glio |
| Bora_ex.vivo_noRT | BC048671 | 3.201 | 3.73E-05 | glio |
| Chek1_in.vivo_noRT | Gm15728 | 3.091 | 3.73E-05 | glio |
| Interaction_genotype_Mtor | Gm2016 | -4.099 | 3.73E-06 | glio |
| Aars2_in.vivo_noRT | Atp5o | -1.195 | 3.73E-06 | glio |
| Chek1_in.vivo_noRT | Gm43727 | 3.217 | 3.73E-06 | glio |
| Hsd17b10_in.vivo_noRT | Fbxl2 | 2.97 | 3.73E-06 | glio |
| culture_effect | Frg1 | -0.752 | 3.73E-07 | glio |
| Bcas2_in.vivo_noRT | Rbm11 | 3.394 | 3.73E-07 | glio |
| Aars2_in.vivo_noRT | Zfp968 | 3.949 | 3.73E-08 | glio |
| Aars2_ex.vivo_noRT | Capn13 | 3.293 | 3.73E-08 | glio |
| Atr_in.vivo_noRT | Zeb2os | 3.62 | 3.73E-08 | glio |
| Aars2_in.vivo_noRT | Ccl19 | 3.375 | 3.73E-09 | glio |
| Atr_in.vivo_noRT | Cdo1 | 4.246 | 3.73E-09 | glio |

| | | | | |
|---|---|---|---|---|
| Bcas2_in.vivo_noRT | Cfb | 3.503 | 3.73E-10 | glio |
| Chek1_in.vivo_noRT | Otos | 4.044 | 3.73E-10 | glio |
| Chek1_ex.vivo_noRT | Vmn1r88 | 4.842 | 3.73E-10 | glio |
| interaction_RT | Tcim | 2.431 | 3.73E-11 | glio |
| Atr_in.vivo_noRT | Cyp7a1 | 4.742 | 3.73E-12 | glio |
| Bcas2_in.vivo_noRT | Gm42664 | 4.157 | 3.73E-12 | glio |
| Atr_in.vivo_noRT | Akap3 | 5.273 | 3.73E-14 | glio |
| Chek1_in.vivo_noRT | Gm16685 | 4.385 | 3.73E-14 | glio |
| Bcas2_in.vivo_noRT | Zscan4d | 4.807 | 3.73E-17 | glio |
| culture_effect | Gm28535 | -1.742 | 3.74E-03 | glio |
| Interaction_genotype_Aars2 | Fbxl2 | -3.269 | 3.74E-03 | glio |
| Interaction_genotype_Aars2 | Gm13056 | -3.561 | 3.74E-03 | glio |
| Interaction_genotype_Bora | Gm20033 | -3.261 | 3.74E-03 | glio |
| Interaction_genotype_Mtor | Dynlt1b | -3.516 | 3.74E-03 | glio |
| Interaction_genotype_Myc | Zfp959 | -2.746 | 3.74E-03 | glio |
| Interaction_genotype_Myc | Rrnad1 | -2.52 | 3.74E-03 | glio |
| Interaction_genotype_Myc | Mfap3 | -1.858 | 3.74E-03 | glio |
| Interaction_genotype_Myc | Ywhah | 1.157 | 3.74E-03 | glio |
| Aars2_in.vivo_noRT | Chp1 | -1.3 | 3.74E-03 | glio |
| Aars2_in.vivo_noRT | Mrps11 | -0.99 | 3.74E-03 | glio |
| Atr_in.vivo_noRT | Fblim1 | 2.056 | 3.74E-03 | glio |
| Atr_in.vivo_noRT | Sbds | -1.017 | 3.74E-03 | glio |
| Cenpj_in.vivo_noRT | Gm48140 | 1.759 | 3.74E-03 | glio |
| Eif2s1_ex.vivo_noRT | Gm13964 | 2.234 | 3.74E-03 | glio |
| Hsd17b10_in.vivo_noRT | Chek1 | -1.463 | 3.74E-03 | glio |
| culture_effect | Tm9sf4 | -0.55 | 3.74E-04 | glio |
| culture_effect | Ddx58 | 0.501 | 3.74E-04 | glio |
| Interaction_genotype_Mtor | Gm46652 | -4.021 | 3.74E-04 | glio |
| Interaction_genotype_Myc | Pip5k1c | -2.852 | 3.74E-04 | glio |
| Interaction_genotype_Myc | Ncapg | -2.928 | 3.74E-04 | glio |
| Atp5c1_in.vivo_noRT | Cpxm1 | 2.402 | 3.74E-04 | glio |
| Atp5c1_in.vivo_noRT | Pcdhga1 | 2.391 | 3.74E-04 | glio |
| Bcas2_in.vivo_noRT | Rabl3 | 2.11 | 3.74E-04 | glio |
| Bcas2_in.vivo_noRT | Atp5h | -0.649 | 3.74E-04 | glio |
| Bora_in.vivo_noRT | Tmpo | -1.253 | 3.74E-04 | glio |
| Chek1_ex.vivo_noRT | Gpr162 | 3.223 | 3.74E-04 | glio |
| Eif2s1_in.vivo_noRT | Gm48215 | 2.143 | 3.74E-04 | glio |
| Interaction_genotype_Atr | Gabarapl1 | 1.911 | 3.74E-05 | glio |
| Interaction_genotype_Bcas2 | Gm10642 | -5.092 | 3.74E-05 | glio |
| Chek1_in.vivo_noRT | Ctrl | 3.025 | 3.74E-05 | glio |
| Hsd17b10_in.vivo_noRT | Ly6g6c | 2.756 | 3.74E-05 | glio |
| Hsd17b10_in.vivo_noRT | Oxt | 2.975 | 3.74E-05 | glio |
| Hsd17b10_in.vivo_noRT | Postn | 2.982 | 3.74E-05 | glio |
| culture_effect | Dcp2 | 0.982 | 3.74E-06 | glio |
| Interaction_genotype_Mtor | 3930402G23R | -5.716 | 3.74E-06 | glio |
| Atr_in.vivo_noRT | Gstm6 | 3.466 | 3.74E-06 | glio |
| Atr_in.vivo_noRT | Foxl2os | 3.025 | 3.74E-06 | glio |
| culture_effect | Zfp106 | 0.692 | 3.74E-07 | glio |
| interaction_RT | Rras | 1.221 | 3.74E-07 | glio |
| Aars2_in.vivo_noRT | Gm43727 | 3.5 | 3.74E-07 | glio |
| Aars2_in.vivo_noRT | Serpinb6a | 3.638 | 3.74E-07 | glio |
| culture_effect | Gm16196 | 2.137 | 3.74E-08 | glio |

| | | | | |
|---|---|---|---|---|
| Atr_in.vivo_noRT | Gm41396 | 4.041 | 3.74E-08 | glio |
| Hsd17b10_in.vivo_noRT | Prl2c3 | 3.616 | 3.74E-08 | glio |
| Aars2_in.vivo_noRT | Otos | 3.798 | 3.74E-09 | glio |
| Atr_in.vivo_noRT | 1700025N23R | 3.641 | 3.74E-09 | glio |
| Interaction_genotype_Ifnar1 | Psmb8 | -2.323 | 3.74E-11 | glio |
| Atr_in.vivo_noRT | Rhox3g | 4.419 | 3.74E-11 | glio |
| culture_effect | Cebpb | -1.348 | 3.74E-13 | glio |
| interaction_RT | Psmb8 | 1.729 | 3.74E-14 | glio |
| Chek1_ex.vivo_noRT | Edn2 | 4.447 | 3.74E-14 | glio |
| Atr_in.vivo_noRT | Tns4 | 4.883 | 3.74E-15 | glio |
| Atr_in.vivo_noRT | Ncf1 | 4.88 | 3.74E-16 | glio |
| Atr_in.vivo_noRT | Olfr90 | 5.269 | 3.74E-18 | glio |
| culture_effect | Hadhb | -0.501 | 3.75E-03 | glio |
| culture_effect | Tdg | 0.494 | 3.75E-03 | glio |
| Interaction_genotype_Aars2 | Gm3734 | -3.974 | 3.75E-03 | glio |
| Interaction_genotype_Aars2 | N4bp2os | -3.65 | 3.75E-03 | glio |
| Interaction_genotype_Aars2 | Lgals1 | 1.143 | 3.75E-03 | glio |
| Interaction_genotype_Atr | Rgs9 | -3.152 | 3.75E-03 | glio |
| Interaction_genotype_Atr | Rarres1 | -3.514 | 3.75E-03 | glio |
| Interaction_genotype_Atr | Ovca2 | -3.477 | 3.75E-03 | glio |
| Interaction_genotype_Atr | Crebrf | 1.739 | 3.75E-03 | glio |
| Interaction_genotype_Atr | Sdhaf4 | 1.599 | 3.75E-03 | glio |
| Interaction_genotype_Atr | Cops4 | 1.557 | 3.75E-03 | glio |
| Interaction_genotype_Atr | Cdk4 | 1.086 | 3.75E-03 | glio |
| Interaction_genotype_Mtor | 1110013H19R | -3.816 | 3.75E-03 | glio |
| Interaction_genotype_Mtor | Zfpl1 | 1.568 | 3.75E-03 | glio |
| Interaction_genotype_Myc | 2810408A11R | -3.317 | 3.75E-03 | glio |
| Interaction_genotype_Myc | Hist1h2bm | -2.781 | 3.75E-03 | glio |
| Atp5c1_in.vivo_noRT | Tecrl | 2.408 | 3.75E-03 | glio |
| Atp5c1_in.vivo_noRT | B3galt4 | 2.142 | 3.75E-03 | glio |
| Atr_in.vivo_noRT | Mxd1 | 1.847 | 3.75E-03 | glio |
| Bora_in.vivo_noRT | Dennd4b | 2.121 | 3.75E-03 | glio |
| Bora_in.vivo_noRT | Timp2 | -1.355 | 3.75E-03 | glio |
| Bora_in.vivo_noRT | H2-Ab1 | -1.073 | 3.75E-03 | glio |
| Bora_ex.vivo_noRT | Gm15270 | 2.407 | 3.75E-03 | glio |
| Bora_ex.vivo_noRT | Pou5f1 | 2.808 | 3.75E-03 | glio |
| Cenpj_in.vivo_noRT | Agap2 | 2.022 | 3.75E-03 | glio |
| Cenpj_ex.vivo_noRT | Gm36037 | 2.196 | 3.75E-03 | glio |
| Cenpj_ex.vivo_noRT | Ceacam16 | 2.591 | 3.75E-03 | glio |
| Chek1_in.vivo_noRT | 0610038B21R | 1.96 | 3.75E-03 | glio |
| Chek1_in.vivo_noRT | Zfp420 | 2.237 | 3.75E-03 | glio |
| Chek1_in.vivo_noRT | C130036L24Ri | 2.029 | 3.75E-03 | glio |
| Chek1_in.vivo_noRT | Colgalt1 | -1.03 | 3.75E-03 | glio |
| Chek1_ex.vivo_noRT | Zfp418 | 2.88 | 3.75E-03 | glio |
| Chek1_ex.vivo_noRT | Tent5d | 2.668 | 3.75E-03 | glio |
| Hsd17b10_in.vivo_noRT | Plp1 | -1.449 | 3.75E-03 | glio |
| culture_effect | Gm20478 | -2.112 | 3.75E-04 | glio |
| Interaction_genotype_Mtor | Gm14966 | -3.811 | 3.75E-04 | glio |
| Interaction_genotype_Myc | Syne3 | -4.267 | 3.75E-04 | glio |
| Interaction_genotype_Myc | Ube2cbp | -3.603 | 3.75E-04 | glio |
| Aars2_in.vivo_noRT | Wsb1 | -1.527 | 3.75E-04 | glio |
| Bard1_ex.vivo_noRT | Gm28940 | 2.814 | 3.75E-04 | glio |

| | | | | |
|---|---|---|---|---|
| Bcas2_in.vivo_noRT | Zfp418 | 2.751 | 3.75E-04 | glio |
| Bcas2_in.vivo_noRT | Gm48796 | 2.435 | 3.75E-04 | glio |
| Chek1_ex.vivo_noRT | Arl4d | 3.287 | 3.75E-04 | glio |
| Chek1_ex.vivo_noRT | Wdr37 | 1.91 | 3.75E-04 | glio |
| Interaction_genotype_Bcas2 | Gm11457 | -5.102 | 3.75E-05 | glio |
| Atr_in.vivo_noRT | Enho | 2.875 | 3.75E-05 | glio |
| Atr_in.vivo_noRT | Slc35f1 | 2.317 | 3.75E-05 | glio |
| Atr_ex.vivo_noRT | Stx19 | 3.164 | 3.75E-05 | glio |
| Atr_ex.vivo_noRT | Dusp2 | 3.367 | 3.75E-05 | glio |
| Bora_in.vivo_noRT | Gm49768 | 2.716 | 3.75E-05 | glio |
| Interaction_genotype_Bora | Hbq1b | -5.864 | 3.75E-06 | glio |
| Interaction_genotype_Mtor | 4930579K19R | -5.679 | 3.75E-06 | glio |
| Interaction_genotype_Mtor | Gm9970 | -5.756 | 3.75E-06 | glio |
| Aars2_in.vivo_noRT | Ovgp1 | 3.181 | 3.75E-06 | glio |
| Atp5c1_in.vivo_noRT | Gm128 | 2.797 | 3.75E-06 | glio |
| Atr_in.vivo_noRT | Tmem86b | 3.306 | 3.75E-06 | glio |
| Atr_ex.vivo_noRT | Cxcl5 | 3.164 | 3.75E-06 | glio |
| Interaction_genotype_Mtor | Cyp3a13 | -6.438 | 3.75E-07 | glio |
| Interaction_genotype_Mtor | 9430041J12Ri | -4.893 | 3.75E-07 | glio |
| Bcas2_in.vivo_noRT | Tmem150c | 3.639 | 3.75E-07 | glio |
| Aars2_in.vivo_noRT | Gm7008 | 3.861 | 3.75E-08 | glio |
| Bcas2_in.vivo_noRT | Srl | 3.918 | 3.75E-08 | glio |
| Chek1_in.vivo_noRT | Gm43608 | 3.87 | 3.75E-08 | glio |
| Chek1_in.vivo_noRT | Gm47972 | 3.749 | 3.75E-08 | glio |
| Chek1_in.vivo_noRT | Ninj2 | 4.136 | 3.75E-09 | glio |
| Aars2_in.vivo_noRT | Fgf1 | 4.536 | 3.75E-10 | glio |
| Atr_in.vivo_noRT | Pla2g2f | 3.89 | 3.75E-10 | glio |
| culture_effect | Plod3 | -1.185 | 3.75E-15 | glio |
| Aars2_in.vivo_noRT | Gast | 5.327 | 3.75E-21 | glio |
| Atr_in.vivo_noRT | Gm47404 | 5.533 | 3.75E-29 | glio |
| culture_effect | Gm34885 | 1.774 | 3.76E-03 | glio |
| culture_effect | Lsm3 | 0.433 | 3.76E-03 | glio |
| Interaction_genotype_Aars2 | Mycl | -3.745 | 3.76E-03 | glio |
| Interaction_genotype_Aars2 | Adamts7 | -3.294 | 3.76E-03 | glio |
| Interaction_genotype_Aars2 | 2610027K06R | -3.217 | 3.76E-03 | glio |
| Interaction_genotype_Aars2 | Pde3a | -3.22 | 3.76E-03 | glio |
| Interaction_genotype_Aars2 | Cacnb3 | -3.107 | 3.76E-03 | glio |
| Interaction_genotype_Aars2 | St3gal5 | 1.828 | 3.76E-03 | glio |
| Interaction_genotype_Atr | Helt | -2.369 | 3.76E-03 | glio |
| Interaction_genotype_Atr | Iqch | -3.071 | 3.76E-03 | glio |
| Interaction_genotype_Atr | Phldb2 | -3.496 | 3.76E-03 | glio |
| Interaction_genotype_Bcas2 | Eef1a1 | 0.925 | 3.76E-03 | glio |
| Interaction_genotype_Bora | Gm32834 | -3.489 | 3.76E-03 | glio |
| Interaction_genotype_Mtor | Drd4 | -3.62 | 3.76E-03 | glio |
| Interaction_genotype_Mtor | Gm46378 | -3.72 | 3.76E-03 | glio |
| Interaction_genotype_Mtor | Nr2f1 | -2.283 | 3.76E-03 | glio |
| Interaction_genotype_Myc | E130114P18Ri | -3.415 | 3.76E-03 | glio |
| Interaction_genotype_Myc | Gm14325 | -3.277 | 3.76E-03 | glio |
| Interaction_genotype_Myc | Eif1ad | 1.261 | 3.76E-03 | glio |
| Aars2_in.vivo_noRT | Lhfpl4 | 1.895 | 3.76E-03 | glio |
| Atp5c1_in.vivo_noRT | A930012O16R | 2.408 | 3.76E-03 | glio |
| Atp5c1_in.vivo_noRT | Gm20455 | 2.478 | 3.76E-03 | glio |

| | | | | |
|---|---|---|---|---|
| Atr_in.vivo_noRT | Gm19541 | 2.097 | 3.76E-03 | glio |
| Atr_in.vivo_noRT | Gm7160 | 1.963 | 3.76E-03 | glio |
| Bcas2_in.vivo_noRT | 4930550L24Ri | -0.925 | 3.76E-03 | glio |
| Bora_ex.vivo_noRT | Gm15445 | 2.216 | 3.76E-03 | glio |
| Hsd17b10_in.vivo_noRT | Adam1a | 2.255 | 3.76E-03 | glio |
| culture_effect | Galk2 | 0.925 | 3.76E-04 | glio |
| Interaction_genotype_Bcas2 | Gm34885 | -4.774 | 3.76E-04 | glio |
| Interaction_genotype_Bcas2 | Gm44752 | -4.241 | 3.76E-04 | glio |
| Interaction_genotype_Mtor | Gm42658 | -3.926 | 3.76E-04 | glio |
| Interaction_genotype_Mtor | Snhg10 | -3.413 | 3.76E-04 | glio |
| Interaction_genotype_Mtor | Smarca5 | 1.695 | 3.76E-04 | glio |
| Interaction_genotype_Mtor | Rpl19 | 1.212 | 3.76E-04 | glio |
| Interaction_genotype_Myc | Gm5934 | -3.045 | 3.76E-04 | glio |
| Interaction_genotype_Myc | Spi1 | -3.701 | 3.76E-04 | glio |
| Interaction_genotype_Myc | Chchd2 | 1.285 | 3.76E-04 | glio |
| interaction_RT | Polr2k | 0.654 | 3.76E-04 | glio |
| Aars2_ex.vivo_noRT | Gm13684 | 2.899 | 3.76E-04 | glio |
| Aars2_ex.vivo_noRT | Myzap | 2.893 | 3.76E-04 | glio |
| Aars2_ex.vivo_noRT | Spink4 | 3.293 | 3.76E-04 | glio |
| Bcas2_in.vivo_noRT | Rbbp9 | 1.989 | 3.76E-04 | glio |
| Bcas2_in.vivo_noRT | Ywhag | -0.913 | 3.76E-04 | glio |
| Bora_in.vivo_noRT | Gzmm | 2.608 | 3.76E-04 | glio |
| Bora_in.vivo_noRT | Spata45 | 2.499 | 3.76E-04 | glio |
| Bora_in.vivo_noRT | Dpep2 | 2.391 | 3.76E-04 | glio |
| Chek1_ex.vivo_noRT | Tiam1 | -2.13 | 3.76E-04 | glio |
| culture_effect | Nectin2 | -1.193 | 3.76E-05 | glio |
| culture_effect | Rfc4 | -0.788 | 3.76E-05 | glio |
| culture_effect | Ghitm | -0.569 | 3.76E-05 | glio |
| interaction_RT | Mbp | -1.278 | 3.76E-05 | glio |
| Aars2_in.vivo_noRT | Gm42372 | 3.327 | 3.76E-05 | glio |
| Aars2_in.vivo_noRT | Smim41 | 3.132 | 3.76E-05 | glio |
| Aars2_in.vivo_noRT | Fbln1 | 2.796 | 3.76E-05 | glio |
| Aars2_ex.vivo_noRT | Adam2 | 3.293 | 3.76E-05 | glio |
| Atr_ex.vivo_noRT | Mgll | 3.916 | 3.76E-05 | glio |
| Bora_in.vivo_noRT | Gm43221 | 2.764 | 3.76E-05 | glio |
| Chek1_in.vivo_noRT | Cd93 | 3.01 | 3.76E-05 | glio |
| Chek1_ex.vivo_noRT | Igsf23 | 3.65 | 3.76E-05 | glio |
| Chek1_ex.vivo_noRT | Map1lc3a | 1.176 | 3.76E-05 | glio |
| Atr_in.vivo_noRT | Dock4 | 3.252 | 3.76E-06 | glio |
| culture_effect | B930059L03Ri | 2.79 | 3.76E-07 | glio |
| culture_effect | Fez1 | -1.943 | 3.76E-07 | glio |
| Interaction_genotype_Mtor | Lamc2 | -5.816 | 3.76E-07 | glio |
| Aars2_in.vivo_noRT | Gm13272 | 3.623 | 3.76E-08 | glio |
| Chek1_ex.vivo_noRT | Cxcr6 | 4.842 | 3.76E-08 | glio |
| culture_effect | Oas1c | 1.956 | 3.76E-09 | glio |
| Atr_in.vivo_noRT | Bbs1 | 4.074 | 3.76E-09 | glio |
| Bora_in.vivo_noRT | B020031M17F | 4.936 | 3.76E-09 | glio |
| Bora_in.vivo_noRT | Gm48772 | 4.029 | 3.76E-09 | glio |
| Bora_in.vivo_noRT | Trpm3 | 4.414 | 3.76E-10 | glio |
| Bora_in.vivo_noRT | Gm40557 | 4.414 | 3.76E-10 | glio |
| Hsd17b10_in.vivo_noRT | Gm5286 | 3.616 | 3.76E-11 | glio |
| Hsd17b10_in.vivo_noRT | Adam5 | 3.352 | 3.76E-11 | glio |

| | | | | |
|---|---|---|---|---|
| Hsd17b10_in.vivo_noRT | Gm34776 | 3.616 | 3.76E-11 | glio |
| Chek1_in.vivo_noRT | Adam2 | 4.518 | 3.76E-14 | glio |
| Atr_in.vivo_noRT | Rab27a | 5.007 | 3.76E-16 | glio |
| Aars2_in.vivo_noRT | Cers3 | 4.795 | 3.76E-18 | glio |
| Chek1_in.vivo_noRT | Gm38163 | 5.044 | 3.76E-19 | glio |
| Interaction_genotype_Aars2 | Hectd2os | -3.5 | 3.77E-03 | glio |
| Interaction_genotype_Atr | Gm20625 | -2.369 | 3.77E-03 | glio |
| Interaction_genotype_Atr | AC132307.1 | -3.308 | 3.77E-03 | glio |
| Interaction_genotype_Bcas2 | Gm15535 | -4.154 | 3.77E-03 | glio |
| Interaction_genotype_Mtor | Eml6 | -3.559 | 3.77E-03 | glio |
| Interaction_genotype_Mtor | Gm48768 | -3.489 | 3.77E-03 | glio |
| Interaction_genotype_Mtor | 4732440D04R | -2.875 | 3.77E-03 | glio |
| Interaction_genotype_Pars2 | Gm13449 | -3.056 | 3.77E-03 | glio |
| interaction_RT | Efr3b | 3.046 | 3.77E-03 | glio |
| Atr_in.vivo_noRT | Chchd5 | -1.172 | 3.77E-03 | glio |
| Atr_ex.vivo_noRT | Gm10800 | 2.959 | 3.77E-03 | glio |
| Bcas2_in.vivo_noRT | Adh7 | 2.125 | 3.77E-03 | glio |
| Bora_in.vivo_noRT | Magi2 | 2.093 | 3.77E-03 | glio |
| Bora_in.vivo_noRT | Arhgap12 | 1.443 | 3.77E-03 | glio |
| Interaction_genotype_Bora | Rcan2 | -4.733 | 3.77E-04 | glio |
| Interaction_genotype_Bora | Klk14 | -4.67 | 3.77E-04 | glio |
| Interaction_genotype_Mtor | Gprin2 | -3.585 | 3.77E-04 | glio |
| Interaction_genotype_Mtor | D430018E03R | -3.995 | 3.77E-04 | glio |
| Interaction_genotype_Mtor | Reep5 | 1.696 | 3.77E-04 | glio |
| Interaction_genotype_Mtor | Uqcr11 | 2.017 | 3.77E-04 | glio |
| Bora_in.vivo_noRT | Zfp963 | 2.853 | 3.77E-04 | glio |
| Chek1_in.vivo_noRT | 4732491K20R | 2.33 | 3.77E-04 | glio |
| Hsd17b10_in.vivo_noRT | Icosl | 2.377 | 3.77E-04 | glio |
| culture_effect | Gm50240 | 1.316 | 3.77E-05 | glio |
| culture_effect | Pimreg | 0.968 | 3.77E-05 | glio |
| Interaction_genotype_Atr | Hdac9 | -4.895 | 3.77E-05 | glio |
| Interaction_genotype_Atr | Prickle2 | -4.312 | 3.77E-05 | glio |
| Interaction_genotype_Mtor | Gm15890 | -4.077 | 3.77E-05 | glio |
| Interaction_genotype_Mtor | E230029C05Ri | -4.194 | 3.77E-05 | glio |
| Bcas2_in.vivo_noRT | 5330426L24Ri | 3.097 | 3.77E-05 | glio |
| Bcas2_in.vivo_noRT | Gm10131 | 2.891 | 3.77E-05 | glio |
| Chek1_ex.vivo_noRT | Olfr77 | 3.575 | 3.77E-05 | glio |
| Hsd17b10_in.vivo_noRT | 2010320O07R | 2.966 | 3.77E-05 | glio |
| culture_effect | Col28a1 | -2.756 | 3.77E-06 | glio |
| Interaction_genotype_Mtor | Cpne8 | -5.643 | 3.77E-06 | glio |
| Aars2_in.vivo_noRT | Wdr5b | 3.154 | 3.77E-06 | glio |
| Atr_in.vivo_noRT | 4833413G10R | 3.4 | 3.77E-06 | glio |
| Bora_in.vivo_noRT | Gm36839 | 3.374 | 3.77E-06 | glio |
| Chek1_in.vivo_noRT | Gm10602 | 3.636 | 3.77E-06 | glio |
| Bora_in.vivo_noRT | Pbld1 | 3.749 | 3.77E-07 | glio |
| Atr_in.vivo_noRT | Spink2 | 4.336 | 3.77E-08 | glio |
| Atr_ex.vivo_noRT | Gm15908 | 3.164 | 3.77E-08 | glio |
| Chek1_in.vivo_noRT | Apoa1 | 4.01 | 3.77E-08 | glio |
| Chek1_in.vivo_noRT | Gm15972 | 4.05 | 3.77E-11 | glio |
| Chek1_in.vivo_noRT | 4930540M05F | 4.386 | 3.77E-11 | glio |
| Chek1_in.vivo_noRT | Sh2d3c | 4.921 | 3.77E-11 | glio |
| Aars2_in.vivo_noRT | Slc45a3 | 4.805 | 3.77E-12 | glio |

| | | | | |
|---|---|---|---|---|
| Chek1_ex.vivo_noRT | Gm2016 | 4.842 | 3.77E-14 | glio |
| Bcas2_in.vivo_noRT | Hnf1aos1 | 4.807 | 3.77E-15 | glio |
| Atr_in.vivo_noRT | Thnsl2 | 4.365 | 3.77E-16 | glio |
| Atr_in.vivo_noRT | Gm28404 | 5.155 | 3.77E-16 | glio |
| Atr_in.vivo_noRT | Gm36172 | 5.001 | 3.77E-19 | glio |
| Aars2_in.vivo_noRT | Dscaml1 | 5.327 | 3.77E-27 | glio |
| culture_effect | Arhgef11 | -0.911 | 3.78E-03 | glio |
| Interaction_genotype_Aars2 | Gm45833 | -4.705 | 3.78E-03 | glio |
| Interaction_genotype_Aars2 | Arf1 | 0.764 | 3.78E-03 | glio |
| Interaction_genotype_Atr | Gm45235 | -2.903 | 3.78E-03 | glio |
| Interaction_genotype_Chek1 | Kdelr3 | 3.167 | 3.78E-03 | glio |
| Interaction_genotype_Chek1 | Nubp2 | 2.04 | 3.78E-03 | glio |
| Interaction_genotype_Chek1 | Ddrgk1 | 1.583 | 3.78E-03 | glio |
| Interaction_genotype_Mtor | Psmd3 | 1.46 | 3.78E-03 | glio |
| Interaction_genotype_Myc | Maoa | -2.619 | 3.78E-03 | glio |
| interaction_RT | Pum1 | 0.681 | 3.78E-03 | glio |
| interaction_RT | Clic1 | 0.531 | 3.78E-03 | glio |
| Aars2_in.vivo_noRT | Nipa1 | 1.625 | 3.78E-03 | glio |
| Aars2_in.vivo_noRT | Card19 | -1.266 | 3.78E-03 | glio |
| Aars2_in.vivo_noRT | Irgm1 | -0.855 | 3.78E-03 | glio |
| Atr_in.vivo_noRT | Eaf2 | 2.147 | 3.78E-03 | glio |
| Atr_in.vivo_noRT | Gm9774 | 2.092 | 3.78E-03 | glio |
| Bcas2_in.vivo_noRT | Gramd4 | 2.115 | 3.78E-03 | glio |
| Bora_in.vivo_noRT | Olr1 | 2.337 | 3.78E-03 | glio |
| Bora_ex.vivo_noRT | Traf3ip3 | 2.856 | 3.78E-03 | glio |
| Hsd17b10_in.vivo_noRT | 4930579K19R | 2.378 | 3.78E-03 | glio |
| Hsd17b10_in.vivo_noRT | Gm20522 | 2.331 | 3.78E-03 | glio |
| Interaction_genotype_Myc | Gm37584 | -4.083 | 3.78E-04 | glio |
| Interaction_genotype_Myc | 4930481B07R | -4.223 | 3.78E-04 | glio |
| Interaction_genotype_Myc | Ect2 | -3.131 | 3.78E-04 | glio |
| Aars2_in.vivo_noRT | Dusp6 | -1.559 | 3.78E-04 | glio |
| Atr_in.vivo_noRT | Hrh2 | 1.99 | 3.78E-04 | glio |
| Bard1_ex.vivo_noRT | Cfb | 2.814 | 3.78E-04 | glio |
| Bcas2_in.vivo_noRT | Nsun7 | 2.797 | 3.78E-04 | glio |
| Bora_ex.vivo_noRT | Tcstv3 | 3.201 | 3.78E-04 | glio |
| Chek1_in.vivo_noRT | Elovl3 | 2.592 | 3.78E-04 | glio |
| Chek1_in.vivo_noRT | Ptrh2 | -1.252 | 3.78E-04 | glio |
| Chek1_ex.vivo_noRT | Slc7a3 | 3.278 | 3.78E-04 | glio |
| culture_effect | Rnf182 | 1.495 | 3.78E-05 | glio |
| Interaction_genotype_Myc | Phex | -4.373 | 3.78E-05 | glio |
| Interaction_genotype_Myc | Ciao2b | 1.725 | 3.78E-05 | glio |
| Interaction_genotype_Rpl3 | Dnm3 | -5.301 | 3.78E-05 | glio |
| Interaction_genotype_Rpl3 | D030040B21R | -5.344 | 3.78E-05 | glio |
| Interaction_genotype_Rpl3 | Gm14968 | -5.708 | 3.78E-05 | glio |
| Interaction_genotype_Rpl3 | Col6a2 | -4.339 | 3.78E-05 | glio |
| Interaction_genotype_Rpl3 | Ciita | 4.637 | 3.78E-05 | glio |
| Aars2_in.vivo_noRT | Krt17 | 5.067 | 3.78E-05 | glio |
| Atr_in.vivo_noRT | Fam151b | 2.878 | 3.78E-05 | glio |
| Atr_in.vivo_noRT | Rps3a1 | -0.852 | 3.78E-05 | glio |
| Chek1_in.vivo_noRT | Wac | -1.279 | 3.78E-05 | glio |
| Interaction_genotype_Mtor | Ilf2 | 2.467 | 3.78E-06 | glio |
| Atr_in.vivo_noRT | Gm6598 | 3.015 | 3.78E-06 | glio |

| | | | | |
|---|---|---|---|---|
| Chek1_ex.vivo_noRT | Gm5941 | 4.142 | 3.78E-06 | glio |
| culture_effect | Zfas1 | -0.907 | 3.78E-07 | glio |
| Interaction_genotype_Mtor | 4833418N02R | -6.333 | 3.78E-07 | glio |
| interaction_RT | Tacc1 | -1.837 | 3.78E-07 | glio |
| interaction_RT | Ppp3ca | 1 | 3.78E-07 | glio |
| Bora_in.vivo_noRT | Stk36 | 3.648 | 3.78E-07 | glio |
| Hsd17b10_in.vivo_noRT | Usp17la | 3.616 | 3.78E-07 | glio |
| Atr_in.vivo_noRT | Scd3 | 4.145 | 3.78E-08 | glio |
| Bora_in.vivo_noRT | Trim67 | 3.367 | 3.78E-08 | glio |
| culture_effect | Zfp773 | -2.551 | 3.78E-09 | glio |
| culture_effect | Gm12216 | -2.432 | 3.78E-09 | glio |
| culture_effect | Cox7a2l | -0.712 | 3.78E-09 | glio |
| Bcas2_in.vivo_noRT | Gm13594 | 4.281 | 3.78E-09 | glio |
| Bcas2_in.vivo_noRT | Tmem30c | 4.148 | 3.78E-09 | glio |
| Bora_in.vivo_noRT | Efemp2 | -2.259 | 3.78E-09 | glio |
| Hsd17b10_in.vivo_noRT | Crhbp | 3.616 | 3.78E-09 | glio |
| Atr_in.vivo_noRT | 9930014A18R | 4.139 | 3.78E-10 | glio |
| Bcas2_in.vivo_noRT | D830039M14I | 4.541 | 3.78E-10 | glio |
| Bcas2_in.vivo_noRT | Cd52 | 4.15 | 3.78E-10 | glio |
| culture_effect | Snhg10 | 2.304 | 3.78E-11 | glio |
| Bora_in.vivo_noRT | Gm26789 | 4.019 | 3.78E-12 | glio |
| Chek1_ex.vivo_noRT | Gm48435 | 4.055 | 3.78E-13 | glio |
| Bcas2_in.vivo_noRT | Dcdc2a | 4.547 | 3.78E-14 | glio |
| Aars2_in.vivo_noRT | 4930483J18Ri | 5.067 | 3.78E-15 | glio |
| Atr_in.vivo_noRT | Gm44672 | 5.273 | 3.78E-15 | glio |
| Atr_in.vivo_noRT | Gm27239 | 5.533 | 3.78E-16 | glio |
| culture_effect | Gm1673 | 1.897 | 3.78E-32 | glio |
| culture_effect | Nadk | 0.517 | 3.79E-03 | glio |
| culture_effect | Zmynd19 | 0.481 | 3.79E-03 | glio |
| culture_effect | Cnot1 | -0.439 | 3.79E-03 | glio |
| Interaction_genotype_Aars2 | 4930588A03R | -2.433 | 3.79E-03 | glio |
| Interaction_genotype_Aars2 | 4930434B07R | -3.15 | 3.79E-03 | glio |
| Interaction_genotype_Aars2 | Hist1h3i | -3.376 | 3.79E-03 | glio |
| Interaction_genotype_Aars2 | Rpf2 | 1.41 | 3.79E-03 | glio |
| Interaction_genotype_Atr | Frg1 | 1.504 | 3.79E-03 | glio |
| Interaction_genotype_Bcas2 | Wincr1 | -4.801 | 3.79E-03 | glio |
| Interaction_genotype_Myc | Mlst8 | -2.487 | 3.79E-03 | glio |
| Interaction_genotype_Myc | Gm10138 | -2.722 | 3.79E-03 | glio |
| Interaction_genotype_Myc | Trim12c | -1.972 | 3.79E-03 | glio |
| Interaction_genotype_Myc | Btf3 | 0.918 | 3.79E-03 | glio |
| Interaction_genotype_Rad21 | Gm11884 | 3.967 | 3.79E-03 | glio |
| Aars2_in.vivo_noRT | Fdps | -1.383 | 3.79E-03 | glio |
| Atr_in.vivo_noRT | S100a8 | 3.275 | 3.79E-03 | glio |
| Atr_in.vivo_noRT | Zfp420 | 2.197 | 3.79E-03 | glio |
| Atr_in.vivo_noRT | Gstm4 | 1.854 | 3.79E-03 | glio |
| Atr_in.vivo_noRT | Oxld1 | 1.647 | 3.79E-03 | glio |
| Bcas2_in.vivo_noRT | Gm9828 | 2.109 | 3.79E-03 | glio |
| Bcas2_ex.vivo_noRT | Nuak1 | 2.024 | 3.79E-03 | glio |
| Bora_in.vivo_noRT | Gm26670 | 1.986 | 3.79E-03 | glio |
| Bora_in.vivo_noRT | Zfp873 | 1.875 | 3.79E-03 | glio |
| Bora_in.vivo_noRT | Ppp1r16b | 2.047 | 3.79E-03 | glio |
| Bora_in.vivo_noRT | Cyld | -1.408 | 3.79E-03 | glio |

| | | | | |
|---|---|---|---|---|
| Bora_in.vivo_noRT | Tm9sf1 | -1.331 | 3.79E-03 | glio |
| Bora_ex.vivo_noRT | Dact1 | 2.802 | 3.79E-03 | glio |
| Cenpj_in.vivo_noRT | Plcl1 | 2.022 | 3.79E-03 | glio |
| Chek1_in.vivo_noRT | Homez | 2.137 | 3.79E-03 | glio |
| Eif2s1_in.vivo_noRT | Msln | 1.976 | 3.79E-03 | glio |
| Hsd17b10_in.vivo_noRT | B930059L03Ri | 2.431 | 3.79E-03 | glio |
| Interaction_genotype_Atr | Gm10300 | -4.55 | 3.79E-04 | glio |
| Interaction_genotype_Atr | Gm20646 | -4.28 | 3.79E-04 | glio |
| Interaction_genotype_Myc | Ube2s | 1.511 | 3.79E-04 | glio |
| Aars2_in.vivo_noRT | Myt1 | 1.853 | 3.79E-04 | glio |
| Atr_in.vivo_noRT | Nsun7 | 2.745 | 3.79E-04 | glio |
| Atr_in.vivo_noRT | Xylb | 2.484 | 3.79E-04 | glio |
| Atr_in.vivo_noRT | Lypd6 | 2.546 | 3.79E-04 | glio |
| Atr_in.vivo_noRT | Mon1a | 2.03 | 3.79E-04 | glio |
| Atr_ex.vivo_noRT | Acbd7 | 3.233 | 3.79E-04 | glio |
| Bcas2_ex.vivo_noRT | Npy1r | 2.742 | 3.79E-04 | glio |
| Bcas2_ex.vivo_noRT | H2-Eb2 | 2.742 | 3.79E-04 | glio |
| Bcas2_ex.vivo_noRT | Gm6803 | 2.742 | 3.79E-04 | glio |
| Bcas2_ex.vivo_noRT | A030014E15Ri | 2.348 | 3.79E-04 | glio |
| Bcas2_ex.vivo_noRT | Gm48170 | 2.743 | 3.79E-04 | glio |
| Hsd17b10_in.vivo_noRT | Fgd3 | 2.881 | 3.79E-04 | glio |
| Hsd17b10_in.vivo_noRT | Gm15559 | 2.689 | 3.79E-04 | glio |
| Hsd17b10_in.vivo_noRT | Ltb | 2.556 | 3.79E-04 | glio |
| Hsd17b10_in.vivo_noRT | Elac2 | -1.681 | 3.79E-04 | glio |
| culture_effect | 3830406C13R | 0.916 | 3.79E-05 | glio |
| culture_effect | Mpst | -0.793 | 3.79E-05 | glio |
| culture_effect | Ufm1 | 0.491 | 3.79E-05 | glio |
| Atr_in.vivo_noRT | AI413582 | -1.674 | 3.79E-05 | glio |
| Bard1_ex.vivo_noRT | Ank1 | 2.828 | 3.79E-05 | glio |
| Chek1_in.vivo_noRT | AA386476 | 3.014 | 3.79E-05 | glio |
| interaction_RT | Hist1h2bc | 2.752 | 3.79E-06 | glio |
| Aars2_in.vivo_noRT | Ppp1r26 | 3.418 | 3.79E-06 | glio |
| Bora_in.vivo_noRT | Gm9958 | 3.366 | 3.79E-06 | glio |
| Interaction_genotype_Mtor | Gm12925 | -6.229 | 3.79E-07 | glio |
| Atr_in.vivo_noRT | Hapln4 | 3.766 | 3.79E-07 | glio |
| Bora_in.vivo_noRT | Gpd1 | 3.764 | 3.79E-07 | glio |
| Bora_in.vivo_noRT | Plekha3 | -2.228 | 3.79E-07 | glio |
| Hsd17b10_in.vivo_noRT | Sh3d21 | 3.093 | 3.79E-07 | glio |
| Atr_in.vivo_noRT | Gm9725 | 3.865 | 3.79E-08 | glio |
| Interaction_genotype_Myc | Gm27239 | -6.626 | 3.79E-09 | glio |
| Bora_in.vivo_noRT | Camk1d | 4.135 | 3.79E-09 | glio |
| Chek1_ex.vivo_noRT | Acat3 | 4.267 | 3.79E-10 | glio |
| culture_effect | Dusp14 | -1.346 | 3.79E-12 | glio |
| Chek1_in.vivo_noRT | Kpna7 | 4.78 | 3.79E-14 | glio |
| Atr_in.vivo_noRT | Slc22a1 | 4.359 | 3.79E-15 | glio |
| Chek1_in.vivo_noRT | B3gnt4 | 4.78 | 3.79E-15 | glio |
| Atr_in.vivo_noRT | 4930412M03F | 5.533 | 3.79E-16 | glio |
| Atr_in.vivo_noRT | Tcstv3 | 5.533 | 3.79E-17 | glio |
| culture_effect | Sertad1 | -1.996 | 3.79E-21 | glio |
| Aars2_in.vivo_noRT | Sema6d | 5.327 | 3.79E-23 | glio |
| Aars2_in.vivo_noRT | B230206H07R | 4.801 | 3.79E-23 | glio |
| Bcas2_in.vivo_noRT | Gm13856 | 4.807 | 3.79E-23 | glio |

| | | | | |
|---|---|---|---|---|
| culture_effect | Snhg20 | 2.071 | 3.79E-24 | glio |
| Atr_in.vivo_noRT | Trmt61b | 5.533 | 3.79E-26 | glio |
| Interaction_genotype_Atr | Gm5157 | -2.369 | 3.80E-03 | glio |
| Interaction_genotype_Bcas2 | 1700023H06R | -4.15 | 3.80E-03 | glio |
| Interaction_genotype_Myc | Kank3 | -3.176 | 3.80E-03 | glio |
| Aars2_in.vivo_noRT | Clock | -1.193 | 3.80E-03 | glio |
| Atp5c1_in.vivo_noRT | A830009L08Ri | 2.145 | 3.80E-03 | glio |
| Atp5c1_in.vivo_noRT | Vamp8 | 1.907 | 3.80E-03 | glio |
| Bard1_ex.vivo_noRT | A930003A15R | 2.814 | 3.80E-03 | glio |
| Bcas2_in.vivo_noRT | Cdh24 | 2.33 | 3.80E-03 | glio |
| Bora_in.vivo_noRT | B230307C23R | 1.676 | 3.80E-03 | glio |
| Bora_in.vivo_noRT | Bcl7c | -0.865 | 3.80E-03 | glio |
| Bora_in.vivo_noRT | Tomm5 | -0.701 | 3.80E-03 | glio |
| Cenpj_ex.vivo_noRT | Slc45a1 | 2.192 | 3.80E-03 | glio |
| Chek1_ex.vivo_noRT | 4921536K21R | 2.591 | 3.80E-03 | glio |
| Chek1_ex.vivo_noRT | Ncaph | -2.031 | 3.80E-03 | glio |
| Chek1_ex.vivo_noRT | Lrr1 | -2.026 | 3.80E-03 | glio |
| culture_effect | Adamts6 | -1.591 | 3.80E-04 | glio |
| Interaction_genotype_Atr | Gm48242 | -3.736 | 3.80E-04 | glio |
| Interaction_genotype_Mtor | Dreh | -3.846 | 3.80E-04 | glio |
| Aars2_in.vivo_noRT | Gm20597 | 2.746 | 3.80E-04 | glio |
| Aars2_ex.vivo_noRT | Gm31415 | 2.797 | 3.80E-04 | glio |
| Aars2_ex.vivo_noRT | Bmp15 | 2.899 | 3.80E-04 | glio |
| Atr_ex.vivo_noRT | Itgbl1 | 2.769 | 3.80E-04 | glio |
| Bora_ex.vivo_noRT | Nat8f2 | 2.806 | 3.80E-04 | glio |
| Cenpj_in.vivo_noRT | Ankrd22 | 2.022 | 3.80E-04 | glio |
| Cenpj_in.vivo_noRT | Gm47404 | 2.022 | 3.80E-04 | glio |
| Cenpj_in.vivo_noRT | Gm31717 | 2.022 | 3.80E-04 | glio |
| Cenpj_in.vivo_noRT | Gm14742 | 2.022 | 3.80E-04 | glio |
| Cenpj_in.vivo_noRT | Ctla4 | 2.022 | 3.80E-04 | glio |
| Cenpj_in.vivo_noRT | Zscan4-ps3 | 2.022 | 3.80E-04 | glio |
| Cenpj_in.vivo_noRT | A030014E15Ri | 2.022 | 3.80E-04 | glio |
| Cenpj_in.vivo_noRT | Gm21731 | 2.022 | 3.80E-04 | glio |
| Cenpj_in.vivo_noRT | Rslcan18 | 2.691 | 3.80E-04 | glio |
| Cenpj_in.vivo_noRT | Gm11232 | 2.022 | 3.80E-04 | glio |
| Cenpj_in.vivo_noRT | Cgnl1 | 2.022 | 3.80E-04 | glio |
| Cenpj_in.vivo_noRT | Gm26543 | 2.022 | 3.80E-04 | glio |
| Cenpj_in.vivo_noRT | Dscaml1 | 2.022 | 3.80E-04 | glio |
| Cenpj_in.vivo_noRT | 2410017I17Ri | 2.022 | 3.80E-04 | glio |
| Cenpj_in.vivo_noRT | Pspn | 2.022 | 3.80E-04 | glio |
| Cenpj_in.vivo_noRT | Tes | -1.516 | 3.80E-04 | glio |
| Chek1_ex.vivo_noRT | Gm10244 | 3.191 | 3.80E-04 | glio |
| Interaction_genotype_Aars2 | Gm7008 | -5.1 | 3.80E-05 | glio |
| Interaction_genotype_Aars2 | Kars | 2.063 | 3.80E-05 | glio |
| Interaction_genotype_Aars2 | Trim16 | -4.657 | 3.80E-05 | glio |
| Interaction_genotype_Aars2 | Arf5 | 1.443 | 3.80E-05 | glio |
| Atp5c1_in.vivo_noRT | Fam166a | 2.404 | 3.80E-05 | glio |
| Bora_ex.vivo_noRT | Tmem54 | 3.201 | 3.80E-05 | glio |
| Chek1_in.vivo_noRT | Gm49539 | 2.95 | 3.80E-05 | glio |
| Aars2_in.vivo_noRT | Maob | 3.113 | 3.80E-06 | glio |
| Chek1_in.vivo_noRT | Zfp455 | 3.49 | 3.80E-07 | glio |
| Atr_in.vivo_noRT | Pde3a | 3.755 | 3.80E-08 | glio |

| | | | | |
|---|---|---|---|---|
| Atr_in.vivo_noRT | Pla2g4b | 3.705 | 3.80E-08 | glio |
| Chek1_ex.vivo_noRT | Esyt3 | 4.677 | 3.80E-08 | glio |
| Chek1_ex.vivo_noRT | Pmp2 | 4.075 | 3.80E-08 | glio |
| culture_effect | Vim | -0.883 | 3.80E-09 | glio |
| Aars2_in.vivo_noRT | Gm47484 | 4.333 | 3.80E-09 | glio |
| culture_effect | Pecr | -1.644 | 3.80E-11 | glio |
| Bora_in.vivo_noRT | Lrrc72 | 4.67 | 3.80E-13 | glio |
| Bcas2_in.vivo_noRT | Gm37800 | 4.807 | 3.80E-14 | glio |
| culture_effect | Nes | 1.304 | 3.80E-15 | glio |
| culture_effect | Csn3 | -5.754 | 3.80E-17 | glio |
| Bora_in.vivo_noRT | Ildr1 | 4.936 | 3.80E-22 | glio |
| Chek1_in.vivo_noRT | 4930588A03R | 5.044 | 3.80E-25 | glio |
| culture_effect | Insl6 | -2.496 | 3.80E-35 | glio |
| culture_effect | BC030343 | 1.676 | 3.81E-03 | glio |
| Interaction_genotype_Atr | Ddx11 | -2.582 | 3.81E-03 | glio |
| Interaction_genotype_Chek1 | Acod1 | 5.134 | 3.81E-03 | glio |
| Interaction_genotype_Mtor | Tbc1d32 | -2.422 | 3.81E-03 | glio |
| Interaction_genotype_Mtor | Selenoh | 1.675 | 3.81E-03 | glio |
| Interaction_genotype_Mtor | Cst6 | 2.345 | 3.81E-03 | glio |
| Interaction_genotype_Myc | Ctps2 | -2.494 | 3.81E-03 | glio |
| Interaction_genotype_Myc | Pak1ip1 | 1.334 | 3.81E-03 | glio |
| Interaction_genotype_Terf1 | D830039M14I | -4.448 | 3.81E-03 | glio |
| Interaction_genotype_Terf1 | Gm19276 | -3.032 | 3.81E-03 | glio |
| Interaction_genotype_Uqcrc2 | Gm11457 | -4.171 | 3.81E-03 | glio |
| Interaction_genotype_Uqcrc2 | Pygo1 | -4.118 | 3.81E-03 | glio |
| Interaction_genotype_Uqcrc2 | Krt80 | -3.872 | 3.81E-03 | glio |
| Interaction_genotype_Uqcrc2 | Gm11627 | -3.775 | 3.81E-03 | glio |
| Interaction_genotype_Uqcrc2 | Gm44658 | -3.852 | 3.81E-03 | glio |
| Interaction_genotype_Uqcrc2 | 1110046J04Ri | -4.467 | 3.81E-03 | glio |
| Interaction_genotype_Uqcrc2 | Egr2 | 4.182 | 3.81E-03 | glio |
| interaction_RT | Il13ra1 | 1.447 | 3.81E-03 | glio |
| interaction_RT | Car9 | 2.917 | 3.81E-03 | glio |
| interaction_RT | Angptl2 | -1.36 | 3.81E-03 | glio |
| interaction_RT | Kdm5b | 1.241 | 3.81E-03 | glio |
| interaction_RT | Wbp1 | 1.03 | 3.81E-03 | glio |
| interaction_RT | Atp5a1 | -0.538 | 3.81E-03 | glio |
| Aars2_in.vivo_noRT | Nlk | -1.428 | 3.81E-03 | glio |
| Atr_in.vivo_noRT | 2900093K20R | 1.72 | 3.81E-03 | glio |
| Atr_in.vivo_noRT | Fip1l1 | -1.017 | 3.81E-03 | glio |
| Bcas2_in.vivo_noRT | Hjurp | -1.245 | 3.81E-03 | glio |
| Bora_in.vivo_noRT | Lhx9 | 2.231 | 3.81E-03 | glio |
| Chek1_in.vivo_noRT | Sf1 | -0.979 | 3.81E-03 | glio |
| Eif2s1_in.vivo_noRT | Gm39326 | 2.411 | 3.81E-03 | glio |
| Hsd17b10_in.vivo_noRT | Slc24a1 | 2.281 | 3.81E-03 | glio |
| culture_effect | Cpne7 | 1.89 | 3.81E-04 | glio |
| Interaction_genotype_Atr | Igf2bp1 | -4.256 | 3.81E-04 | glio |
| interaction_RT | E130317F20Ri | -1.64 | 3.81E-04 | glio |
| Aars2_in.vivo_noRT | Ldlrad3 | 2.321 | 3.81E-04 | glio |
| Aars2_in.vivo_noRT | Mrps23 | -1.122 | 3.81E-04 | glio |
| Bcas2_ex.vivo_noRT | 1700019E08Ri | 2.742 | 3.81E-04 | glio |
| Chek1_in.vivo_noRT | Gm17655 | 2.852 | 3.81E-04 | glio |
| Chek1_in.vivo_noRT | Cadm2 | 2.68 | 3.81E-04 | glio |

| | | | | |
|---|---|---|---|---|
| Chek1_in.vivo_noRT | N4bp2os | 2.616 | 3.81E-04 | glio |
| Chek1_in.vivo_noRT | 1700102H20R | 2.519 | 3.81E-04 | glio |
| Chek1_in.vivo_noRT | Cdiptos | 2.495 | 3.81E-04 | glio |
| Chek1_in.vivo_noRT | Tmod2 | 1.542 | 3.81E-04 | glio |
| Hsd17b10_in.vivo_noRT | Itgb7 | 2.489 | 3.81E-04 | glio |
| Atr_ex.vivo_noRT | Dusp15 | 3.548 | 3.81E-05 | glio |
| Bcas2_in.vivo_noRT | N4bp2l1 | 3.029 | 3.81E-05 | glio |
| Bcas2_in.vivo_noRT | Nbeal2 | 3.047 | 3.81E-05 | glio |
| Interaction_genotype_Mtor | Gm43672 | -5.526 | 3.81E-06 | glio |
| Interaction_genotype_Myc | Tnni3k | -4.059 | 3.81E-06 | glio |
| Bora_in.vivo_noRT | Ache | 3.363 | 3.81E-06 | glio |
| Bora_in.vivo_noRT | Vps4b | -1.778 | 3.81E-06 | glio |
| Brca2_ex.vivo_noRT | Brca2 | -4.074 | 3.81E-06 | glio |
| Aars2_in.vivo_noRT | Gm37933 | 3.176 | 3.81E-07 | glio |
| Bora_in.vivo_noRT | Plac8 | 3.173 | 3.81E-07 | glio |
| interaction_RT | Tap2 | 1.411 | 3.81E-08 | glio |
| Bora_in.vivo_noRT | Gm35986 | 3.888 | 3.81E-09 | glio |
| Chek1_ex.vivo_noRT | Gm17276 | 4.447 | 3.81E-10 | glio |
| Aars2_in.vivo_noRT | 1810012K16R | 4.32 | 3.81E-11 | glio |
| Chek1_ex.vivo_noRT | Fam166a | 4.842 | 3.81E-13 | glio |
| culture_effect | Tnik | -1.396 | 3.81E-14 | glio |
| Aars2_in.vivo_noRT | Ablim2 | 4.794 | 3.81E-23 | glio |
| culture_effect | Rtcb | -1.406 | 3.81E-27 | glio |
| culture_effect | Chsy1 | 0.73 | 3.82E-03 | glio |
| culture_effect | Mrps11 | 0.39 | 3.82E-03 | glio |
| Interaction_genotype_Bora | Cyp20a1 | 2.191 | 3.82E-03 | glio |
| Interaction_genotype_Mtor | Snx32 | -3.131 | 3.82E-03 | glio |
| Interaction_genotype_Myc | Gm13403 | -3.161 | 3.82E-03 | glio |
| Interaction_genotype_Myc | Ucma | -3.72 | 3.82E-03 | glio |
| Interaction_genotype_Myc | Srgap1 | -3.057 | 3.82E-03 | glio |
| Interaction_genotype_Myc | Lingo3 | -2.235 | 3.82E-03 | glio |
| interaction_RT | Ifi213 | 1.05 | 3.82E-03 | glio |
| Bcas2_in.vivo_noRT | Ccdc8 | 2.093 | 3.82E-03 | glio |
| Bcas2_in.vivo_noRT | Gm29994 | 1.929 | 3.82E-03 | glio |
| Hsd17b10_in.vivo_noRT | Plppr3 | 2.23 | 3.82E-03 | glio |
| Interaction_genotype_Atr | Fa2h | 4.211 | 3.82E-04 | glio |
| Interaction_genotype_Mtor | Rhobtb2 | -3.524 | 3.82E-04 | glio |
| Interaction_genotype_Myc | Tmem71 | -3.332 | 3.82E-04 | glio |
| Aars2_in.vivo_noRT | A630052C17R | 2.455 | 3.82E-04 | glio |
| Aars2_in.vivo_noRT | Gm15726 | 2.33 | 3.82E-04 | glio |
| Atr_in.vivo_noRT | Zfp97 | 2.509 | 3.82E-04 | glio |
| Atr_in.vivo_noRT | Gm49041 | 2.236 | 3.82E-04 | glio |
| Atr_in.vivo_noRT | Frat1 | 2.441 | 3.82E-04 | glio |
| Atr_in.vivo_noRT | Slc18b1 | 1.975 | 3.82E-04 | glio |
| Bora_in.vivo_noRT | Fxyd4 | 2.821 | 3.82E-04 | glio |
| Bora_in.vivo_noRT | 1700086P04R | 2.527 | 3.82E-04 | glio |
| Bora_in.vivo_noRT | Med17 | -1.498 | 3.82E-04 | glio |
| Bora_ex.vivo_noRT | AU022754 | 3.201 | 3.82E-04 | glio |
| Chek1_in.vivo_noRT | Gm35315 | 2.542 | 3.82E-04 | glio |
| Interaction_genotype_Atr | Slc35f1 | -3.506 | 3.82E-05 | glio |
| Interaction_genotype_Mtor | Apol7d | -4.991 | 3.82E-05 | glio |
| Interaction_genotype_Mtor | Dock6 | -4.833 | 3.82E-05 | glio |

| | | | | |
|---|---|---|---|---|
| Interaction_genotype_Mtor | Selp | -4.89 | 3.82E-05 | glio |
| interaction_RT | Fus | -0.889 | 3.82E-05 | glio |
| Aars2_in.vivo_noRT | 2810021J22Ri | 3.007 | 3.82E-05 | glio |
| Aars2_in.vivo_noRT | Lcat | 2.993 | 3.82E-05 | glio |
| Aars2_in.vivo_noRT | Mogat2 | 2.809 | 3.82E-05 | glio |
| Aars2_in.vivo_noRT | Prkar1b | 2.986 | 3.82E-05 | glio |
| Atr_in.vivo_noRT | Amt | 3.15 | 3.82E-05 | glio |
| Bard1_ex.vivo_noRT | Adam5 | 2.814 | 3.82E-05 | glio |
| Bcas2_in.vivo_noRT | Ywhaz | -0.914 | 3.82E-05 | glio |
| Chek1_in.vivo_noRT | Tnp2 | 2.981 | 3.82E-05 | glio |
| Interaction_genotype_Mtor | 1700003G18R | -6.106 | 3.82E-06 | glio |
| Atr_in.vivo_noRT | Gm42726 | 3.078 | 3.82E-06 | glio |
| Atr_in.vivo_noRT | Bbs5 | 2.576 | 3.82E-06 | glio |
| Atr_ex.vivo_noRT | Gm47936 | 3.164 | 3.82E-06 | glio |
| Atr_ex.vivo_noRT | Tcstv3 | 4.309 | 3.82E-07 | glio |
| Atr_in.vivo_noRT | Cebpa | 3.872 | 3.82E-08 | glio |
| culture_effect | Scara5 | 2.159 | 3.82E-09 | glio |
| Atr_in.vivo_noRT | Gm49874 | 5.533 | 3.82E-14 | glio |
| Bcas2_in.vivo_noRT | Gm50321 | 4.807 | 3.82E-16 | glio |
| Bora_in.vivo_noRT | Crhbp | 4.936 | 3.82E-16 | glio |
| Aars2_in.vivo_noRT | Nwd1 | 5.063 | 3.82E-18 | glio |
| culture_effect | Gm28050 | 1.593 | 3.83E-03 | glio |
| Interaction_genotype_Aars2 | Klk6 | 3.323 | 3.83E-03 | glio |
| Interaction_genotype_Aars2 | Gm16740 | -3.544 | 3.83E-03 | glio |
| Interaction_genotype_Aars2 | AY074887 | -3.854 | 3.83E-03 | glio |
| Interaction_genotype_Aars2 | Slc50a1 | 1.303 | 3.83E-03 | glio |
| Interaction_genotype_Atr | Gm12589 | -2.769 | 3.83E-03 | glio |
| Interaction_genotype_Atr | Slc16a4 | -2.962 | 3.83E-03 | glio |
| Interaction_genotype_Atr | Cd248 | -3.502 | 3.83E-03 | glio |
| Interaction_genotype_Atr | Dmrt2 | -3.884 | 3.83E-03 | glio |
| Interaction_genotype_Atr | Gm32585 | -3.735 | 3.83E-03 | glio |
| Interaction_genotype_Atr | D830039M14I | -3.634 | 3.83E-03 | glio |
| Interaction_genotype_Atr | Npl | -3.244 | 3.83E-03 | glio |
| Interaction_genotype_Atr | Mfsd6 | -2.329 | 3.83E-03 | glio |
| Interaction_genotype_Atr | Ano6 | 1.755 | 3.83E-03 | glio |
| Interaction_genotype_Atr | Snrnp27 | 1.281 | 3.83E-03 | glio |
| Interaction_genotype_Fanca | Klrk1 | 3.823 | 3.83E-03 | glio |
| Interaction_genotype_Mtor | Vill | -3.056 | 3.83E-03 | glio |
| Interaction_genotype_Myc | Pars2 | -3.065 | 3.83E-03 | glio |
| Interaction_genotype_Myc | Cdk2ap1 | 1.632 | 3.83E-03 | glio |
| Interaction_genotype_Myc | Nbdy | 1.331 | 3.83E-03 | glio |
| Aars2_in.vivo_noRT | Ppp1r10 | -1.221 | 3.83E-03 | glio |
| Aars2_in.vivo_noRT | Fkbp8 | -0.848 | 3.83E-03 | glio |
| Aars2_in.vivo_noRT | Tagln2 | -1.091 | 3.83E-03 | glio |
| Atr_in.vivo_noRT | Mrpl57 | -0.938 | 3.83E-03 | glio |
| Atr_ex.vivo_noRT | Ppp1r15a | 1.366 | 3.83E-03 | glio |
| Bard1_ex.vivo_noRT | Tff2 | 2.814 | 3.83E-03 | glio |
| Bard1_ex.vivo_noRT | Glyat | 2.017 | 3.83E-03 | glio |
| Bcas2_in.vivo_noRT | Ppp1r13l | 2.146 | 3.83E-03 | glio |
| Bora_in.vivo_noRT | Pcx | 2.077 | 3.83E-03 | glio |
| Bora_ex.vivo_noRT | Gm43951 | 2.408 | 3.83E-03 | glio |
| Chek1_in.vivo_noRT | Prdm1 | 1.977 | 3.83E-03 | glio |

| | | | | |
|---|---|---|---|---|
| Chek1_ex.vivo_noRT | Elac2 | -1.68 | 3.83E-03 | glio |
| Chek1_ex.vivo_noRT | Topbp1 | -1.406 | 3.83E-03 | glio |
| Chek1_ex.vivo_noRT | Nop56 | -0.98 | 3.83E-03 | glio |
| culture_effect | Yipf6 | -0.646 | 3.83E-04 | glio |
| Interaction_genotype_Atr | Gm12925 | -4.509 | 3.83E-04 | glio |
| Interaction_genotype_Mtor | Tctn1 | -4.386 | 3.83E-04 | glio |
| Interaction_genotype_Mtor | Vapa | 1.749 | 3.83E-04 | glio |
| Interaction_genotype_Myc | Fubp1 | 1.519 | 3.83E-04 | glio |
| Bcas2_in.vivo_noRT | B3gntl1 | 2.486 | 3.83E-04 | glio |
| Bora_in.vivo_noRT | N4bp2l1 | 2.64 | 3.83E-04 | glio |
| Chek1_ex.vivo_noRT | Zfp810 | 3.061 | 3.83E-04 | glio |
| Aars2_in.vivo_noRT | Catsperz | 2.852 | 3.83E-05 | glio |
| Atr_in.vivo_noRT | 1700123M08F | 2.88 | 3.83E-05 | glio |
| Bard1_in.vivo_noRT | Krt18 | -4.485 | 3.83E-05 | glio |
| Chek1_in.vivo_noRT | Dcun1d1 | -1.814 | 3.83E-05 | glio |
| Chek1_ex.vivo_noRT | Cyp27b1 | 3.298 | 3.83E-05 | glio |
| culture_effect | Gm41396 | 2.479 | 3.83E-06 | glio |
| culture_effect | Ncln | -0.702 | 3.83E-06 | glio |
| Atp5c1_in.vivo_noRT | Gm35082 | 2.668 | 3.83E-06 | glio |
| Bora_in.vivo_noRT | C1qtnf12 | 3.612 | 3.83E-06 | glio |
| Chek1_ex.vivo_noRT | Serpinb6b | 4.263 | 3.83E-06 | glio |
| culture_effect | Ramp2 | -1.768 | 3.83E-09 | glio |
| Aars2_in.vivo_noRT | Etl4 | 3.451 | 3.83E-09 | glio |
| Bcas2_in.vivo_noRT | Sod3 | 4.807 | 3.83E-09 | glio |
| Chek1_in.vivo_noRT | Gm14698 | 4.132 | 3.83E-09 | glio |
| Chek1_in.vivo_noRT | D430040D24F | 4.127 | 3.83E-09 | glio |
| Chek1_in.vivo_noRT | Rasgrp2 | 4.273 | 3.83E-09 | glio |
| Chek1_ex.vivo_noRT | Txndc2 | 4.055 | 3.83E-09 | glio |
| Chek1_in.vivo_noRT | Clca3a1 | 3.992 | 3.83E-10 | glio |
| Chek1_in.vivo_noRT | D830039M14I | 4.777 | 3.83E-11 | glio |
| culture_effect | Col3a1 | -3.858 | 3.83E-12 | glio |
| Bora_in.vivo_noRT | Gm41318 | 4.936 | 3.83E-13 | glio |
| Chek1_in.vivo_noRT | Dok2 | 4.518 | 3.83E-14 | glio |
| Chek1_in.vivo_noRT | Tmem132c | 5.044 | 3.83E-24 | glio |
| culture_effect | Spink10 | 1.391 | 3.84E-03 | glio |
| culture_effect | Smyd3 | -0.873 | 3.84E-03 | glio |
| Interaction_genotype_Aars2 | Hcn2 | -3.882 | 3.84E-03 | glio |
| Interaction_genotype_Atr | Lmx1b | -3.733 | 3.84E-03 | glio |
| Interaction_genotype_Bora | A730071L15Ri | -3.932 | 3.84E-03 | glio |
| Interaction_genotype_Bora | Catsper2 | -3.717 | 3.84E-03 | glio |
| Interaction_genotype_Mtor | Gm32031 | -2.737 | 3.84E-03 | glio |
| Interaction_genotype_Mtor | Tmem63b | 1.53 | 3.84E-03 | glio |
| Interaction_genotype_Myc | Gm11423 | -2.52 | 3.84E-03 | glio |
| Interaction_genotype_Myc | Trmt10b | -2.196 | 3.84E-03 | glio |
| Interaction_genotype_Myc | Mrto4 | 1.357 | 3.84E-03 | glio |
| Interaction_genotype_Myc | Psmd8 | 1.292 | 3.84E-03 | glio |
| Interaction_genotype_Myc | Zfr | 1.171 | 3.84E-03 | glio |
| Interaction_genotype_Pars2 | Socs1 | 2.295 | 3.84E-03 | glio |
| Interaction_genotype_Rpl3 | Eme2 | -3.815 | 3.84E-03 | glio |
| Atr_in.vivo_noRT | Hsh2d | 1.871 | 3.84E-03 | glio |
| Atr_in.vivo_noRT | Cyp27a1 | 1.91 | 3.84E-03 | glio |
| Atr_in.vivo_noRT | Atrnl1 | 1.851 | 3.84E-03 | glio |

| | | | | |
|---|---|---|---|---|
| Atr_in.vivo_noRT | Dhfr | -1.314 | 3.84E-03 | glio |
| Atr_in.vivo_noRT | Cdc16 | 0.999 | 3.84E-03 | glio |
| Atr_ex.vivo_noRT | Gm37214 | 2.866 | 3.84E-03 | glio |
| Atr_ex.vivo_noRT | Pomc | 1.978 | 3.84E-03 | glio |
| Bora_in.vivo_noRT | Fam174b | 2.362 | 3.84E-03 | glio |
| Bora_in.vivo_noRT | Gm15892 | 2.263 | 3.84E-03 | glio |
| Bora_in.vivo_noRT | Ctr9 | 1.027 | 3.84E-03 | glio |
| Chek1_ex.vivo_noRT | 1700081N11R | 2.901 | 3.84E-03 | glio |
| culture_effect | Castor2 | 1.553 | 3.84E-04 | glio |
| culture_effect | 2510009E07Ri | -1.339 | 3.84E-04 | glio |
| Interaction_genotype_Mtor | Ppargc1b | -4.292 | 3.84E-04 | glio |
| Interaction_genotype_Myc | Flicr | -3.966 | 3.84E-04 | glio |
| Interaction_genotype_Myc | Dpysl3 | -3.173 | 3.84E-04 | glio |
| Interaction_genotype_Myc | Cfl1 | 1.381 | 3.84E-04 | glio |
| Aars2_in.vivo_noRT | Hao1 | 2.655 | 3.84E-04 | glio |
| Aars2_in.vivo_noRT | Rpp25l | -1.102 | 3.84E-04 | glio |
| Bora_in.vivo_noRT | Camkv | 2.701 | 3.84E-04 | glio |
| Chek1_in.vivo_noRT | Serp2 | 2.306 | 3.84E-04 | glio |
| Chek1_in.vivo_noRT | Tspan32 | 2.577 | 3.84E-04 | glio |
| Chek1_ex.vivo_noRT | Gm49767 | 3.271 | 3.84E-04 | glio |
| Chek1_ex.vivo_noRT | Spc25 | -2.413 | 3.84E-04 | glio |
| Interaction_genotype_Myc | Gm35315 | -4.65 | 3.84E-05 | glio |
| Interaction_genotype_Myc | Rims3 | -4.257 | 3.84E-05 | glio |
| Interaction_genotype_Myc | H2afz | 1.934 | 3.84E-05 | glio |
| Interaction_genotype_Myc | mt-Nd2 | 1.741 | 3.84E-05 | glio |
| Atr_in.vivo_noRT | Hist1h3g | 3.071 | 3.84E-05 | glio |
| Atr_in.vivo_noRT | Coro2b | 2.443 | 3.84E-05 | glio |
| Bora_in.vivo_noRT | Phtf1os | 3.012 | 3.84E-05 | glio |
| Bora_in.vivo_noRT | AC166172.1 | 2.736 | 3.84E-05 | glio |
| Bora_in.vivo_noRT | Gm17655 | 3.268 | 3.84E-05 | glio |
| Bora_in.vivo_noRT | Spata33 | 3.412 | 3.84E-06 | glio |
| culture_effect | Tcta | 1.656 | 3.84E-07 | glio |
| culture_effect | Fkbp11 | -0.911 | 3.84E-08 | glio |
| Atr_in.vivo_noRT | Uckl1os | 4.299 | 3.84E-09 | glio |
| Atr_in.vivo_noRT | Gm10432 | 4.036 | 3.84E-10 | glio |
| Atr_in.vivo_noRT | Etl4 | 3.657 | 3.84E-10 | glio |
| Atr_in.vivo_noRT | Gm16576 | 4.416 | 3.84E-10 | glio |
| interaction_RT | Serpina3n | 2.328 | 3.84E-11 | glio |
| Hsd17b10_in.vivo_noRT | Slc14a2 | 3.616 | 3.84E-14 | glio |
| culture_effect | Rabggtb | -1.145 | 3.84E-16 | glio |
| Bcas2_in.vivo_noRT | Hist1h2ag | 4.275 | 3.84E-16 | glio |
| Interaction_genotype_Aars2 | Hdac9 | -3.403 | 3.85E-03 | glio |
| Interaction_genotype_Aars2 | Serinc2 | -3.387 | 3.85E-03 | glio |
| Interaction_genotype_Aars2 | Ngdn | 1.48 | 3.85E-03 | glio |
| Interaction_genotype_Atr | Fam227b | -2.369 | 3.85E-03 | glio |
| Interaction_genotype_Atr | Gm20658 | -2.369 | 3.85E-03 | glio |
| Interaction_genotype_Atr | 4930445N18R | -3.689 | 3.85E-03 | glio |
| Interaction_genotype_Atr | 2410004B18R | 1.429 | 3.85E-03 | glio |
| Interaction_genotype_Mtor | Gm42941 | -3.34 | 3.85E-03 | glio |
| Interaction_genotype_Mtor | 2810429I04Ri | -2.765 | 3.85E-03 | glio |
| Interaction_genotype_Mtor | Uck2 | -1.469 | 3.85E-03 | glio |
| Interaction_genotype_Myc | Gstm6 | -3.379 | 3.85E-03 | glio |

| | | | | |
|---|---|---|---|---|
| Interaction_genotype_Pars2 | 4930588A03R | -2.668 | 3.85E-03 | glio |
| Interaction_genotype_Pars2 | Noxred1 | -3.841 | 3.85E-03 | glio |
| Interaction_genotype_Pars2 | Cyb5b | 1.459 | 3.85E-03 | glio |
| interaction_RT | Hadh | -0.76 | 3.85E-03 | glio |
| interaction_RT | Ostf1 | 0.689 | 3.85E-03 | glio |
| Aars2_in.vivo_noRT | Tdrd3 | 1.818 | 3.85E-03 | glio |
| Aars2_in.vivo_noRT | Hspd1 | -0.676 | 3.85E-03 | glio |
| Atr_in.vivo_noRT | Gm45184 | 2.077 | 3.85E-03 | glio |
| Atr_in.vivo_noRT | Pfn2 | -1.431 | 3.85E-03 | glio |
| Atr_in.vivo_noRT | Lrrc42 | -1.336 | 3.85E-03 | glio |
| Bcas2_in.vivo_noRT | Spryd7 | -1.392 | 3.85E-03 | glio |
| Bora_ex.vivo_noRT | Zmynd10 | 2.414 | 3.85E-03 | glio |
| Chek1_in.vivo_noRT | Bmerb1 | 2.271 | 3.85E-03 | glio |
| Chek1_in.vivo_noRT | Tmem82 | 2.282 | 3.85E-03 | glio |
| Chek1_in.vivo_noRT | Dlg5 | 2.002 | 3.85E-03 | glio |
| Hsd17b10_in.vivo_noRT | Cabp4 | 2.306 | 3.85E-03 | glio |
| Hsd17b10_in.vivo_noRT | Svip | 2.004 | 3.85E-03 | glio |
| Interaction_genotype_Mtor | Traf1 | -4.26 | 3.85E-04 | glio |
| Interaction_genotype_Mtor | C530044C16R | -3.712 | 3.85E-04 | glio |
| Interaction_genotype_Mtor | Dubr | -3.19 | 3.85E-04 | glio |
| Interaction_genotype_Mtor | Tnnt2 | -3.203 | 3.85E-04 | glio |
| Aars2_ex.vivo_noRT | Ngb | 3.293 | 3.85E-04 | glio |
| Aars2_ex.vivo_noRT | Slc16a7 | 2.893 | 3.85E-04 | glio |
| Atr_in.vivo_noRT | Adamts6 | 2.153 | 3.85E-04 | glio |
| Bora_in.vivo_noRT | Slc4a11 | 2.672 | 3.85E-04 | glio |
| Hsd17b10_in.vivo_noRT | Lsp1 | 2.96 | 3.85E-04 | glio |
| culture_effect | Gm4285 | 1.33 | 3.85E-05 | glio |
| Aars2_in.vivo_noRT | Fam199x | 2.325 | 3.85E-05 | glio |
| Atp5c1_in.vivo_noRT | 1700017M07F | 2.668 | 3.85E-05 | glio |
| culture_effect | Rwdd2a | -2.173 | 3.85E-06 | glio |
| Interaction_genotype_Mtor | Gm11549 | -5.353 | 3.85E-06 | glio |
| Atr_in.vivo_noRT | Cyp2d22 | 3.734 | 3.85E-07 | glio |
| Bora_in.vivo_noRT | Gm10115 | 3.499 | 3.85E-07 | glio |
| Chek1_ex.vivo_noRT | Nts | 4.842 | 3.85E-07 | glio |
| Interaction_genotype_Mtor | Gm29083 | -6.222 | 3.85E-08 | glio |
| Interaction_genotype_Mtor | Capn9 | -6.163 | 3.85E-08 | glio |
| Interaction_genotype_Mtor | Tmem74b | -5.599 | 3.85E-08 | glio |
| Aars2_in.vivo_noRT | D830035M03I | 3.741 | 3.85E-08 | glio |
| Chek1_in.vivo_noRT | Chd5 | 3.798 | 3.85E-08 | glio |
| Chek1_in.vivo_noRT | Gm16348 | 3.859 | 3.85E-08 | glio |
| Aars2_in.vivo_noRT | Gm15559 | 4.399 | 3.85E-10 | glio |
| Bcas2_in.vivo_noRT | Gm41764 | 3.89 | 3.85E-10 | glio |
| Bcas2_in.vivo_noRT | Usp17la | 4.807 | 3.85E-12 | glio |
| Bora_in.vivo_noRT | Gm21762 | 4.672 | 3.85E-14 | glio |
| Chek1_in.vivo_noRT | Dnah10 | 4.664 | 3.85E-14 | glio |
| Bcas2_in.vivo_noRT | Gm29966 | 4.547 | 3.85E-21 | glio |
| Atr_in.vivo_noRT | Tal2 | 5.27 | 3.85E-24 | glio |
| culture_effect | Prph | -2.524 | 3.85E-27 | glio |
| Interaction_genotype_Aars2 | Arrdc3 | -3.434 | 3.86E-03 | glio |
| Interaction_genotype_Aars2 | Rps27l | 0.828 | 3.86E-03 | glio |
| Interaction_genotype_Bcas2 | Rab11fip4os1 | -3.889 | 3.86E-03 | glio |
| Interaction_genotype_Bcas2 | Gramd2 | -3.659 | 3.86E-03 | glio |

| | | | | |
|---|---|---|---|---|
| Interaction_genotype_Bcas2 | Dscc1 | 2.595 | 3.86E-03 | glio |
| Interaction_genotype_Bcas2 | Fis1 | 1.189 | 3.86E-03 | glio |
| Interaction_genotype_Mtor | Gpatch2l | -2.894 | 3.86E-03 | glio |
| Interaction_genotype_Mtor | Trmt112 | 1.22 | 3.86E-03 | glio |
| interaction_RT | Map1b | -0.659 | 3.86E-03 | glio |
| Aars2_in.vivo_noRT | Psmb7 | -0.753 | 3.86E-03 | glio |
| Aars2_ex.vivo_noRT | Gm10696 | 2.893 | 3.86E-03 | glio |
| Atp5c1_in.vivo_noRT | Cdo1 | 2.432 | 3.86E-03 | glio |
| Atr_in.vivo_noRT | Pxylp1 | 1.937 | 3.86E-03 | glio |
| Atr_in.vivo_noRT | Bloc1s3 | 1.865 | 3.86E-03 | glio |
| Atr_in.vivo_noRT | Chchd10 | -2.036 | 3.86E-03 | glio |
| Atr_ex.vivo_noRT | Cpeb1 | 2.743 | 3.86E-03 | glio |
| Atr_ex.vivo_noRT | Gch1 | 2.268 | 3.86E-03 | glio |
| Bcas2_ex.vivo_noRT | BC147527 | 2.742 | 3.86E-03 | glio |
| Bora_in.vivo_noRT | Trim46 | 1.946 | 3.86E-03 | glio |
| Bora_in.vivo_noRT | Pprc1 | -1.124 | 3.86E-03 | glio |
| Chek1_in.vivo_noRT | Col6a1 | 2.079 | 3.86E-03 | glio |
| Chek1_in.vivo_noRT | Rragc | -1.262 | 3.86E-03 | glio |
| Chek1_in.vivo_noRT | Col9a3 | -1.419 | 3.86E-03 | glio |
| Chek1_in.vivo_noRT | Rps11 | -0.518 | 3.86E-03 | glio |
| Chek1_ex.vivo_noRT | B230398E01Ri | 2.726 | 3.86E-03 | glio |
| Cyp19a1_ex.vivo_noRT | A930019D19R | 2.159 | 3.86E-03 | glio |
| Dclre1c_ex.vivo_noRT | Gm31717 | 2.292 | 3.86E-03 | glio |
| Eif2s1_in.vivo_noRT | Jph4 | 2.408 | 3.86E-03 | glio |
| culture_effect | Bhlhe41 | 2.352 | 3.86E-04 | glio |
| culture_effect | Car6 | -0.762 | 3.86E-04 | glio |
| culture_effect | Moxd1 | -0.498 | 3.86E-04 | glio |
| Interaction_genotype_Mtor | Rbm11 | -4.121 | 3.86E-04 | glio |
| Interaction_genotype_Mtor | Tmem208 | 1.7 | 3.86E-04 | glio |
| Interaction_genotype_Mtor | Ywhaq | 1.65 | 3.86E-04 | glio |
| Interaction_genotype_Mtor | Rps3 | 1.272 | 3.86E-04 | glio |
| Aars2_ex.vivo_noRT | Slco2a1 | 3.293 | 3.86E-04 | glio |
| Chek1_in.vivo_noRT | Tfam | -1.148 | 3.86E-04 | glio |
| Chek1_ex.vivo_noRT | Gap43 | 3.652 | 3.86E-04 | glio |
| Interaction_genotype_Mtor | Hist2h3c1 | -4.799 | 3.86E-05 | glio |
| Bora_in.vivo_noRT | Gm48086 | 2.988 | 3.86E-05 | glio |
| Chek1_in.vivo_noRT | Sowahc | -2.084 | 3.86E-05 | glio |
| culture_effect | Slc15a3 | -2.983 | 3.86E-07 | glio |
| culture_effect | Mttp | 1.496 | 3.86E-07 | glio |
| Atr_in.vivo_noRT | Zfp983 | 3.049 | 3.86E-07 | glio |
| Bcas2_in.vivo_noRT | Appbp2os | 3.744 | 3.86E-07 | glio |
| Chek1_in.vivo_noRT | Ntf5 | 3.749 | 3.86E-07 | glio |
| Aars2_in.vivo_noRT | B930059L03Ri | 4.142 | 3.86E-08 | glio |
| Atr_in.vivo_noRT | BC049762 | 4.2 | 3.86E-09 | glio |
| Atr_in.vivo_noRT | 9530080O11R | 4.616 | 3.86E-12 | glio |
| Chek1_in.vivo_noRT | Gm47872 | 4.78 | 3.86E-13 | glio |
| Chek1_ex.vivo_noRT | Gm15894 | 4.842 | 3.86E-13 | glio |
| Chek1_ex.vivo_noRT | B230206H07R | 4.447 | 3.86E-14 | glio |
| Atr_in.vivo_noRT | Gm43463 | 4.876 | 3.86E-16 | glio |
| Chek1_in.vivo_noRT | Selenov | 5.044 | 3.86E-18 | glio |
| Atr_in.vivo_noRT | Iqch | 5.269 | 3.86E-19 | glio |
| culture_effect | Fgfr3 | 2.398 | 3.86E-21 | glio |

| | | | | |
|---|---|---|---|---|
| culture_effect | Id2 | 2.681 | 3.86E-28 | glio |
| Atr_in.vivo_noRT | Stfa3 | 5.533 | 3.86E-31 | glio |
| Atr_in.vivo_noRT | Olfr787 | 5.533 | 3.86E-31 | glio |
| culture_effect | Sall3 | 0.758 | 3.87E-03 | glio |
| Interaction_genotype_Aars2 | Dync2li1 | 2.373 | 3.87E-03 | glio |
| Interaction_genotype_Atr | Lrrc75aos2 | -3.766 | 3.87E-03 | glio |
| Interaction_genotype_Atr | Zik1 | -3.065 | 3.87E-03 | glio |
| Interaction_genotype_Mtor | Amhr2 | -3.994 | 3.87E-03 | glio |
| Interaction_genotype_Mtor | Cdc42ep4 | -3.229 | 3.87E-03 | glio |
| Interaction_genotype_Myc | Sema3d | -3.366 | 3.87E-03 | glio |
| Interaction_genotype_Myc | Gm12258 | -3.328 | 3.87E-03 | glio |
| Interaction_genotype_Myc | Tubgcp3 | -2.426 | 3.87E-03 | glio |
| Interaction_genotype_Rad51 | Gm10642 | -3.878 | 3.87E-03 | glio |
| Interaction_genotype_Rad51 | Gm50394 | -4.009 | 3.87E-03 | glio |
| Aars2_in.vivo_noRT | Ndrg4 | 2.149 | 3.87E-03 | glio |
| Aars2_in.vivo_noRT | Tyk2 | 1.742 | 3.87E-03 | glio |
| Aars2_ex.vivo_noRT | Hnf1aos1 | 2.495 | 3.87E-03 | glio |
| Atr_in.vivo_noRT | Vps45 | 1.437 | 3.87E-03 | glio |
| Bard1_ex.vivo_noRT | Gm10324 | 2.234 | 3.87E-03 | glio |
| Bard1_ex.vivo_noRT | Bpifa1 | 2.814 | 3.87E-03 | glio |
| Bcas2_ex.vivo_noRT | Fsd2 | 2.954 | 3.87E-03 | glio |
| Bora_in.vivo_noRT | Mansc1 | 2.263 | 3.87E-03 | glio |
| Cenpj_ex.vivo_noRT | 1700010B08R | 2.591 | 3.87E-03 | glio |
| Eif2s1_ex.vivo_noRT | Gm16933 | 2.234 | 3.87E-03 | glio |
| Hsd17b10_in.vivo_noRT | Cdo1 | 2.328 | 3.87E-03 | glio |
| culture_effect | Zbed4 | 0.837 | 3.87E-04 | glio |
| Interaction_genotype_Myc | Piwil4 | -3.402 | 3.87E-04 | glio |
| Atr_in.vivo_noRT | Serinc2 | 2.425 | 3.87E-04 | glio |
| Atr_in.vivo_noRT | Rpl30 | -0.667 | 3.87E-04 | glio |
| Atr_ex.vivo_noRT | Cfap58 | 4.011 | 3.87E-04 | glio |
| Bcas2_in.vivo_noRT | Abi3bp | 2.578 | 3.87E-04 | glio |
| Bcas2_in.vivo_noRT | Gm10282 | 2.56 | 3.87E-04 | glio |
| Chek1_in.vivo_noRT | Gm19744 | 2.598 | 3.87E-04 | glio |
| Interaction_genotype_Aars2 | Cnpy2 | 1.34 | 3.87E-05 | glio |
| Bcas2_in.vivo_noRT | Hhipl2 | 3.248 | 3.87E-05 | glio |
| Chek1_in.vivo_noRT | Cdrt4 | 3.069 | 3.87E-05 | glio |
| Interaction_genotype_Atr | A430106G13R | -5.798 | 3.87E-06 | glio |
| Interaction_genotype_Atr | Skida1 | -5.728 | 3.87E-06 | glio |
| Interaction_genotype_Mtor | Prl2c2 | -5.277 | 3.87E-06 | glio |
| interaction_RT | Laptm4b | 1.392 | 3.87E-07 | glio |
| Atr_ex.vivo_noRT | Gipc2 | 4.656 | 3.87E-07 | glio |
| culture_effect | Mast2 | 0.912 | 3.87E-08 | glio |
| Chek1_ex.vivo_noRT | Nrn1l | 4.443 | 3.87E-08 | glio |
| Hsd17b10_in.vivo_noRT | BC147527 | 3.616 | 3.87E-08 | glio |
| Chek1_in.vivo_noRT | Gm20455 | 4.323 | 3.87E-09 | glio |
| Chek1_ex.vivo_noRT | Gm43915 | 4.449 | 3.87E-10 | glio |
| Bora_in.vivo_noRT | Skida1 | 4.936 | 3.87E-12 | glio |
| Chek1_ex.vivo_noRT | Prap1 | 4.449 | 3.87E-12 | glio |
| culture_effect | Ubxn10 | -3.971 | 3.87E-13 | glio |
| Atr_in.vivo_noRT | AV039307 | 5.142 | 3.87E-13 | glio |
| Chek1_in.vivo_noRT | Gm45591 | 4.784 | 3.87E-16 | glio |
| Chek1_in.vivo_noRT | Gm15816 | 5.044 | 3.87E-16 | glio |

| | | | | |
|---|---|---|---|---|
| Bcas2_in.vivo_noRT | Klk13 | 4.275 | 3.87E-17 | glio |
| Chek1_in.vivo_noRT | A330076H08R | 5.044 | 3.87E-18 | glio |
| Atr_in.vivo_noRT | 1700017M07F | 5.533 | 3.87E-20 | glio |
| Chek1_in.vivo_noRT | Gm44693 | 5.044 | 3.87E-24 | glio |
| culture_effect | Sft2d3 | 0.92 | 3.88E-03 | glio |
| Interaction_genotype_Aars2 | Gm21936 | -2.429 | 3.88E-03 | glio |
| Interaction_genotype_Bcas2 | Rac1 | 1.093 | 3.88E-03 | glio |
| Interaction_genotype_Mtor | Irs4 | -3.802 | 3.88E-03 | glio |
| Interaction_genotype_Mtor | Zfp988 | -3.607 | 3.88E-03 | glio |
| Interaction_genotype_Mtor | Gm33782 | -3.103 | 3.88E-03 | glio |
| Interaction_genotype_Mtor | Pigo | -3.101 | 3.88E-03 | glio |
| Interaction_genotype_Myc | Cisd2 | 1.264 | 3.88E-03 | glio |
| Aars2_in.vivo_noRT | Clta | -0.707 | 3.88E-03 | glio |
| Atp5c1_in.vivo_noRT | Alox12 | 2.41 | 3.88E-03 | glio |
| Atr_in.vivo_noRT | Smg7 | -1.174 | 3.88E-03 | glio |
| Atr_in.vivo_noRT | Dut | -1.007 | 3.88E-03 | glio |
| Bard1_ex.vivo_noRT | Aldh1a2 | 2.818 | 3.88E-03 | glio |
| Bcas2_in.vivo_noRT | Etaa1os | 2.058 | 3.88E-03 | glio |
| Bcas2_in.vivo_noRT | Slc1a5 | -1.504 | 3.88E-03 | glio |
| Bora_in.vivo_noRT | Setdb2 | 1.75 | 3.88E-03 | glio |
| Chek1_in.vivo_noRT | Psd3 | 1.829 | 3.88E-03 | glio |
| Chek1_in.vivo_noRT | Uba5 | -1.258 | 3.88E-03 | glio |
| Chek1_in.vivo_noRT | Ppan | -0.939 | 3.88E-03 | glio |
| Chek1_ex.vivo_noRT | Nudt18 | -2.133 | 3.88E-03 | glio |
| Chek1_ex.vivo_noRT | Tcim | 1.767 | 3.88E-03 | glio |
| Dclre1c_ex.vivo_noRT | Ctla4 | 2.292 | 3.88E-03 | glio |
| Fanca_ex.vivo_noRT | Capn13 | 2.124 | 3.88E-03 | glio |
| Interaction_genotype_Mtor | Gm11769 | -3.85 | 3.88E-04 | glio |
| Interaction_genotype_Mtor | Psd | -3.788 | 3.88E-04 | glio |
| Interaction_genotype_Myc | Pcdh9 | -4.106 | 3.88E-04 | glio |
| Aars2_in.vivo_noRT | Lzts2 | -1.268 | 3.88E-04 | glio |
| Atr_ex.vivo_noRT | Cacng5 | 3.164 | 3.88E-04 | glio |
| Interaction_genotype_Mtor | Hist1h4f | -5.256 | 3.88E-05 | glio |
| Chek1_in.vivo_noRT | Vcan | 3.174 | 3.88E-05 | glio |
| Chek1_in.vivo_noRT | 4930458D05R | 3.217 | 3.88E-05 | glio |
| interaction_RT | Irgm1 | 1.058 | 3.88E-06 | glio |
| Aars2_in.vivo_noRT | Sbk1 | 3.305 | 3.88E-06 | glio |
| Atr_in.vivo_noRT | Plekhg6 | 3.327 | 3.88E-06 | glio |
| culture_effect | 1700008J07Ri | 2.161 | 3.88E-08 | glio |
| Bcas2_in.vivo_noRT | Fgf1 | 4.016 | 3.88E-08 | glio |
| Chek1_ex.vivo_noRT | Spn | 4.449 | 3.88E-08 | glio |
| Aars2_in.vivo_noRT | C230013L11Ri | 4.415 | 3.88E-09 | glio |
| Aars2_in.vivo_noRT | Slc46a3 | 4.142 | 3.88E-09 | glio |
| Aars2_in.vivo_noRT | Fgf7 | 3.963 | 3.88E-09 | glio |
| Atr_in.vivo_noRT | Gm8098 | 4.03 | 3.88E-09 | glio |
| Hsd17b10_in.vivo_noRT | Agap2 | 3.616 | 3.88E-10 | glio |
| Aars2_in.vivo_noRT | A030005L19Ri | 4.798 | 3.88E-14 | glio |
| culture_effect | Slc35b2 | 1.114 | 3.88E-16 | glio |
| Interaction_genotype_Bcas2 | AW112010 | 4.287 | 3.89E-03 | glio |
| Interaction_genotype_Mtor | Cdc14b | -2.852 | 3.89E-03 | glio |
| interaction_RT | Ttc33 | 0.672 | 3.89E-03 | glio |
| interaction_RT | Mrpl30 | 0.451 | 3.89E-03 | glio |

| | | | | |
|---|---|---|---|---|
| Aars2_in.vivo_noRT | Ccdc78 | 2.009 | 3.89E-03 | glio |
| Aars2_in.vivo_noRT | Pelp1 | -1.332 | 3.89E-03 | glio |
| Atr_in.vivo_noRT | Ttc30a1 | 2.099 | 3.89E-03 | glio |
| Atr_in.vivo_noRT | Myt1 | 1.523 | 3.89E-03 | glio |
| Bora_in.vivo_noRT | Sncg | 2.297 | 3.89E-03 | glio |
| Cenpj_in.vivo_noRT | Gm13288 | 2.022 | 3.89E-03 | glio |
| Chek1_in.vivo_noRT | Bloc1s5 | 1.541 | 3.89E-03 | glio |
| Chek1_ex.vivo_noRT | Glmp | 2.961 | 3.89E-03 | glio |
| Chek1_ex.vivo_noRT | Spata2l | 2.849 | 3.89E-03 | glio |
| culture_effect | Dclk2 | -1.105 | 3.89E-04 | glio |
| culture_effect | Vav3 | -1.011 | 3.89E-04 | glio |
| Interaction_genotype_Bora | Gm16741 | -4.275 | 3.89E-04 | glio |
| Interaction_genotype_Mtor | Notch4 | -4.051 | 3.89E-04 | glio |
| Interaction_genotype_Mtor | Abhd15 | -4.497 | 3.89E-04 | glio |
| Interaction_genotype_Mtor | Homez | -4.381 | 3.89E-04 | glio |
| Interaction_genotype_Mtor | Plcxd2 | -4.354 | 3.89E-04 | glio |
| Interaction_genotype_Mtor | 4933404O12R | -4.034 | 3.89E-04 | glio |
| Interaction_genotype_Mtor | Dgkz | -3.014 | 3.89E-04 | glio |
| Interaction_genotype_Myc | A830082N09R | -4.208 | 3.89E-04 | glio |
| Interaction_genotype_Myc | Zfp202 | -4.141 | 3.89E-04 | glio |
| Interaction_genotype_Myc | Hectd4 | -2.775 | 3.89E-04 | glio |
| Interaction_genotype_Myc | Klhl42 | -3.019 | 3.89E-04 | glio |
| Interaction_genotype_Myc | Rbm8a | 1.477 | 3.89E-04 | glio |
| Aars2_ex.vivo_noRT | Gm38393 | 2.899 | 3.89E-04 | glio |
| Atr_in.vivo_noRT | A330009N23R | 2.444 | 3.89E-04 | glio |
| Atr_in.vivo_noRT | Ranbp1 | -0.924 | 3.89E-04 | glio |
| Atr_ex.vivo_noRT | Serpina3i | 3.164 | 3.89E-04 | glio |
| Chek1_in.vivo_noRT | Plac1 | 2.462 | 3.89E-04 | glio |
| Atr_in.vivo_noRT | Axin2 | 2.964 | 3.89E-05 | glio |
| Bcas2_ex.vivo_noRT | Tpo | 2.742 | 3.89E-05 | glio |
| Hsd17b10_in.vivo_noRT | Zfp811 | 2.969 | 3.89E-05 | glio |
| Interaction_genotype_Mtor | Gm17249 | -5.166 | 3.89E-06 | glio |
| Interaction_genotype_Mtor | Ache | -5.724 | 3.89E-06 | glio |
| Interaction_genotype_Mtor | Kcnk4 | -5.602 | 3.89E-06 | glio |
| Interaction_genotype_Mtor | H13 | 2.369 | 3.89E-06 | glio |
| Interaction_genotype_Mtor | Sept7 | 2.491 | 3.89E-06 | glio |
| Aars2_in.vivo_noRT | Mfsd7a | 3.72 | 3.89E-06 | glio |
| Atr_in.vivo_noRT | Zfp493 | 3.184 | 3.89E-06 | glio |
| Bcas2_in.vivo_noRT | D930019O06R | 3.428 | 3.89E-06 | glio |
| Bora_in.vivo_noRT | Mpeg1 | 2.961 | 3.89E-06 | glio |
| Bcas2_in.vivo_noRT | Gm13625 | 3.286 | 3.89E-07 | glio |
| Bora_in.vivo_noRT | Ltb4r1 | 3.762 | 3.89E-07 | glio |
| Chek1_in.vivo_noRT | Gm47591 | 3.597 | 3.89E-07 | glio |
| Chek1_ex.vivo_noRT | Bbof1 | 4.267 | 3.89E-07 | glio |
| Aars2_in.vivo_noRT | Gm26673 | 4.306 | 3.89E-10 | glio |
| Aars2_in.vivo_noRT | Dlx4 | 4.801 | 3.89E-13 | glio |
| Bcas2_in.vivo_noRT | Gm42849 | 4.541 | 3.89E-13 | glio |
| Bcas2_in.vivo_noRT | Oit3 | 4.807 | 3.89E-13 | glio |
| Bcas2_in.vivo_noRT | Lrp2bp | 4.543 | 3.89E-13 | glio |
| Bora_in.vivo_noRT | Gm20275 | 4.106 | 3.89E-13 | glio |
| Atr_in.vivo_noRT | Fam228a | 5.283 | 3.89E-14 | glio |
| Aars2_in.vivo_noRT | Col8a1 | 5.327 | 3.89E-15 | glio |

| | | | | |
|---|---|---|---|---|
| Chek1_in.vivo_noRT | D430001F17R | 5.044 | 3.89E-15 | glio |
| Chek1_in.vivo_noRT | Tagap | 4.784 | 3.89E-23 | glio |
| Bora_in.vivo_noRT | Gm44736 | 4.936 | 3.89E-25 | glio |
| Interaction_genotype_Atr | Olfr810 | -2.369 | 3.90E-03 | glio |
| Interaction_genotype_Bora | Gm15579 | -4.175 | 3.90E-03 | glio |
| Interaction_genotype_Bora | Zfp85os | -3.935 | 3.90E-03 | glio |
| Interaction_genotype_Bora | Bysl | 1.809 | 3.90E-03 | glio |
| Interaction_genotype_Bora | Smad4 | 1.754 | 3.90E-03 | glio |
| Interaction_genotype_Bora | Hp1bp3 | 1.249 | 3.90E-03 | glio |
| Interaction_genotype_Mtor | Sept5 | -3.449 | 3.90E-03 | glio |
| Interaction_genotype_Mtor | Rab1a | 1.391 | 3.90E-03 | glio |
| Aars2_in.vivo_noRT | Seh1l | -0.995 | 3.90E-03 | glio |
| Aars2_ex.vivo_noRT | Gm19684 | 3.111 | 3.90E-03 | glio |
| Atr_in.vivo_noRT | Ripk3 | 2.339 | 3.90E-03 | glio |
| Atr_in.vivo_noRT | Hlcs | 1.97 | 3.90E-03 | glio |
| Atr_in.vivo_noRT | Aoc2 | 1.831 | 3.90E-03 | glio |
| Atr_in.vivo_noRT | Gm29243 | 1.775 | 3.90E-03 | glio |
| Atr_in.vivo_noRT | Slc5a8 | 1.566 | 3.90E-03 | glio |
| Atr_in.vivo_noRT | Ctnnbip1 | -1.173 | 3.90E-03 | glio |
| Atr_in.vivo_noRT | Tpd52l2 | -0.966 | 3.90E-03 | glio |
| Bcas2_in.vivo_noRT | Rspo3 | 2.452 | 3.90E-03 | glio |
| Bcas2_in.vivo_noRT | Gtf3c3 | 1.499 | 3.90E-03 | glio |
| Bora_in.vivo_noRT | Trim12c | -1.385 | 3.90E-03 | glio |
| Bora_ex.vivo_noRT | Prr15l | 2.984 | 3.90E-03 | glio |
| Cenpj_in.vivo_noRT | 5430402O13R | 2.548 | 3.90E-03 | glio |
| Cenpj_ex.vivo_noRT | Cemip | 2.591 | 3.90E-03 | glio |
| Chek1_in.vivo_noRT | Arhgef17 | 1.91 | 3.90E-03 | glio |
| Chek1_ex.vivo_noRT | Ginm1 | 1.188 | 3.90E-03 | glio |
| Hsd17b10_in.vivo_noRT | Gnaz | 2.318 | 3.90E-03 | glio |
| Hsd17b10_in.vivo_noRT | Pld6 | 2.36 | 3.90E-03 | glio |
| Hsd17b10_in.vivo_noRT | Tmem82 | 2.384 | 3.90E-03 | glio |
| Interaction_genotype_Lmo7 | Hbq1b | -5.111 | 3.90E-04 | glio |
| Interaction_genotype_Lmo7 | Ntn4 | -3.973 | 3.90E-04 | glio |
| Interaction_genotype_Myc | B430010I23Ri | -3.84 | 3.90E-04 | glio |
| Interaction_genotype_Myc | Gm15601 | -4.101 | 3.90E-04 | glio |
| Interaction_genotype_Myc | Trp53 | 1.414 | 3.90E-04 | glio |
| Aars2_in.vivo_noRT | Sumo1 | -0.886 | 3.90E-04 | glio |
| Aars2_ex.vivo_noRT | Kmo | 2.9 | 3.90E-04 | glio |
| Bora_in.vivo_noRT | Egf | 2.744 | 3.90E-04 | glio |
| Eif2s1_in.vivo_noRT | Gm49172 | 1.883 | 3.90E-04 | glio |
| culture_effect | Gm12905 | 2.368 | 3.90E-05 | glio |
| Interaction_genotype_Mtor | Gm35290 | -4.656 | 3.90E-05 | glio |
| Atr_in.vivo_noRT | Gata3 | 3.15 | 3.90E-05 | glio |
| Bcas2_in.vivo_noRT | Drap1 | -1.045 | 3.90E-05 | glio |
| culture_effect | A430010J10Ri | 2.822 | 3.90E-06 | glio |
| culture_effect | Lats2 | 1.081 | 3.90E-06 | glio |
| Atp5c1_in.vivo_noRT | Nmrk2 | 3.073 | 3.90E-06 | glio |
| Atp5c1_in.vivo_noRT | Gm5286 | 2.668 | 3.90E-06 | glio |
| Atp5c1_in.vivo_noRT | Gm34776 | 2.668 | 3.90E-06 | glio |
| Atp5c1_in.vivo_noRT | Gm47510 | 2.408 | 3.90E-06 | glio |
| Chek1_in.vivo_noRT | 9130604C24R | 3.578 | 3.90E-06 | glio |
| Chek1_in.vivo_noRT | Atp6v1d | -1.573 | 3.90E-06 | glio |

| | | | | |
|---|---|---|---|---|
| Atr_in.vivo_noRT | Gm14399 | 3.553 | 3.90E-07 | glio |
| culture_effect | Ptpn11 | 0.896 | 3.90E-08 | glio |
| Hsd17b10_in.vivo_noRT | Flacc1 | 3.616 | 3.90E-09 | glio |
| Interaction_genotype_Bora | Klhdc8a | -6.246 | 3.90E-10 | glio |
| Bora_in.vivo_noRT | Accsl | 4.676 | 3.90E-10 | glio |
| Hsd17b10_in.vivo_noRT | Snrpn | 3.479 | 3.90E-10 | glio |
| Atr_in.vivo_noRT | H2-M10.1 | 4.078 | 3.90E-12 | glio |
| Atr_in.vivo_noRT | Calhm5 | 4.805 | 3.90E-13 | glio |
| Bora_in.vivo_noRT | Gm27239 | 4.936 | 3.90E-13 | glio |
| Bcas2_in.vivo_noRT | Fbxl13 | 4.547 | 3.90E-14 | glio |
| Atr_in.vivo_noRT | F630040K05Ri | 4.883 | 3.90E-15 | glio |
| Bcas2_in.vivo_noRT | Gm44899 | 4.807 | 3.90E-18 | glio |
| Bcas2_in.vivo_noRT | Gm21936 | 4.807 | 3.90E-23 | glio |
| culture_effect | Cryab | 4.627 | 3.90E-47 | glio |
| Interaction_genotype_Aars2 | Sec61b | 0.942 | 3.91E-03 | glio |
| Interaction_genotype_Bcas2 | D030068K23R | -3.931 | 3.91E-03 | glio |
| Interaction_genotype_Bcas2 | Ifit1bl1 | -4.69 | 3.91E-03 | glio |
| Interaction_genotype_Bcas2 | Gm35867 | -3.789 | 3.91E-03 | glio |
| Interaction_genotype_Bcas2 | St8sia1 | -3.984 | 3.91E-03 | glio |
| Interaction_genotype_Bcas2 | Gm28424 | -3.562 | 3.91E-03 | glio |
| Interaction_genotype_Bcas2 | Rgs16 | -3.894 | 3.91E-03 | glio |
| Interaction_genotype_Bcas2 | Pgrmc1 | 1.946 | 3.91E-03 | glio |
| Interaction_genotype_Bora | Ptcd3 | 1.989 | 3.91E-03 | glio |
| Interaction_genotype_Mtor | Gm26916 | -3.319 | 3.91E-03 | glio |
| Interaction_genotype_Mtor | Tmem246 | -3.271 | 3.91E-03 | glio |
| Interaction_genotype_Mtor | Mpv17l | -3.079 | 3.91E-03 | glio |
| Interaction_genotype_Mtor | Sort1 | -2.152 | 3.91E-03 | glio |
| Interaction_genotype_Myc | Ablim2 | -2.197 | 3.91E-03 | glio |
| Interaction_genotype_Myc | 2810414N06R | -3.145 | 3.91E-03 | glio |
| Interaction_genotype_Myc | Yeats4 | 1.392 | 3.91E-03 | glio |
| Interaction_genotype_Uqcrc2 | Gm34047 | -4.372 | 3.91E-03 | glio |
| Aars2_in.vivo_noRT | Gm26692 | 1.996 | 3.91E-03 | glio |
| Aars2_in.vivo_noRT | Tinagl1 | -1.433 | 3.91E-03 | glio |
| Atp5c1_in.vivo_noRT | Gm10425 | 2.404 | 3.91E-03 | glio |
| Atr_in.vivo_noRT | Tmem37 | 2.013 | 3.91E-03 | glio |
| Atr_in.vivo_noRT | Abl1 | 1.611 | 3.91E-03 | glio |
| Atr_in.vivo_noRT | Ccnd2 | -0.779 | 3.91E-03 | glio |
| Bcas2_in.vivo_noRT | Gm14403 | 1.885 | 3.91E-03 | glio |
| Bcas2_in.vivo_noRT | Alg8 | -1.423 | 3.91E-03 | glio |
| Bcas2_in.vivo_noRT | Eci1 | -0.986 | 3.91E-03 | glio |
| Bcas2_in.vivo_noRT | Cdk1 | -1.435 | 3.91E-03 | glio |
| Bcas2_ex.vivo_noRT | Gm13713 | 2.342 | 3.91E-03 | glio |
| Bcas2_ex.vivo_noRT | Gm20705 | 2.347 | 3.91E-03 | glio |
| Bora_in.vivo_noRT | Zfp451 | -1.14 | 3.91E-03 | glio |
| Bora_in.vivo_noRT | Vdac2 | -0.671 | 3.91E-03 | glio |
| Chek1_in.vivo_noRT | 6430548M08F | 2.138 | 3.91E-03 | glio |
| Chek1_in.vivo_noRT | Snap23 | -1.349 | 3.91E-03 | glio |
| Eif2s1_in.vivo_noRT | Serpinb2 | 2.143 | 3.91E-03 | glio |
| Eif2s1_in.vivo_noRT | Arhgef19 | 2.521 | 3.91E-03 | glio |
| Interaction_genotype_Aars2 | Gm49494 | -4.416 | 3.91E-04 | glio |
| Interaction_genotype_Aars2 | Klhl30 | -4.048 | 3.91E-04 | glio |
| Interaction_genotype_Atr | Calm2 | 1.316 | 3.91E-04 | glio |

| | | | | |
|---|---|---|---|---|
| Aars2_in.vivo_noRT | Eva1c | 2.599 | 3.91E-04 | glio |
| Atr_in.vivo_noRT | Kif5c | 2.006 | 3.91E-04 | glio |
| Bard1_ex.vivo_noRT | Muc4 | 2.814 | 3.91E-04 | glio |
| Bora_in.vivo_noRT | Gm26737 | 3.833 | 3.91E-04 | glio |
| Chek1_ex.vivo_noRT | D430001F17R | 2.897 | 3.91E-04 | glio |
| Hsd17b10_in.vivo_noRT | Gm37584 | 2.704 | 3.91E-04 | glio |
| Interaction_genotype_Myc | Spata45 | -4.563 | 3.91E-05 | glio |
| Interaction_genotype_Myc | Nckipsd | -4.526 | 3.91E-05 | glio |
| Atr_in.vivo_noRT | Fermt3 | 2.723 | 3.91E-05 | glio |
| Bora_ex.vivo_noRT | Nrn1l | 3.594 | 3.91E-05 | glio |
| Bora_ex.vivo_noRT | Slc25a18 | 3.201 | 3.91E-05 | glio |
| Chek1_in.vivo_noRT | Pcdhgb1 | 2.51 | 3.91E-05 | glio |
| Ercc4_in.vivo_noRT | Cnfn | 3.502 | 3.91E-05 | glio |
| Interaction_genotype_Mtor | Adam30 | -4.824 | 3.91E-06 | glio |
| Aars2_ex.vivo_noRT | Crb3 | 3.293 | 3.91E-06 | glio |
| Atr_in.vivo_noRT | Gm30074 | 3.165 | 3.91E-06 | glio |
| Bcas2_in.vivo_noRT | Fibp | -2 | 3.91E-06 | glio |
| Bora_in.vivo_noRT | Mir142hg | 3.343 | 3.91E-06 | glio |
| Bora_in.vivo_noRT | Abcd2 | 3.621 | 3.91E-06 | glio |
| Bora_in.vivo_noRT | Plac1 | 3.138 | 3.91E-06 | glio |
| Hsd17b10_in.vivo_noRT | Tlr1 | 3.356 | 3.91E-06 | glio |
| culture_effect | Ppef2 | 1.693 | 3.91E-07 | glio |
| culture_effect | Dnajc30 | 1.079 | 3.91E-07 | glio |
| culture_effect | Efemp2 | 0.739 | 3.91E-07 | glio |
| Aars2_in.vivo_noRT | Nat8f1 | 3.376 | 3.91E-07 | glio |
| Atr_in.vivo_noRT | 4930524O07R | 3.609 | 3.91E-07 | glio |
| culture_effect | Slc11a1 | -3.462 | 3.91E-08 | glio |
| Interaction_genotype_Mtor | Rabac1 | 2.836 | 3.91E-09 | glio |
| Interaction_genotype_Mtor | Pitx3 | -5.863 | 3.91E-09 | glio |
| Aars2_in.vivo_noRT | Ipcef1 | 3.513 | 3.91E-10 | glio |
| Hsd17b10_in.vivo_noRT | Olfr718-ps1 | 3.352 | 3.91E-10 | glio |
| Atr_in.vivo_noRT | Fam131a | 5.007 | 3.91E-14 | glio |
| Aars2_in.vivo_noRT | Dcdc2a | 5.067 | 3.91E-17 | glio |
| Bora_in.vivo_noRT | Gm12610 | 4.931 | 3.91E-18 | glio |
| culture_effect | Rgs16 | 5.456 | 3.91E-21 | glio |
| Aars2_in.vivo_noRT | Gm9758 | 5.067 | 3.91E-22 | glio |
| Atr_in.vivo_noRT | Gm44899 | 5.533 | 3.91E-23 | glio |
| culture_effect | Rab6a | -0.381 | 3.92E-03 | glio |
| Interaction_genotype_Atr | Cfap46 | -3.288 | 3.92E-03 | glio |
| Interaction_genotype_Atr | D8Ertd738e | 0.819 | 3.92E-03 | glio |
| Interaction_genotype_Bora | Gm10447 | -3.009 | 3.92E-03 | glio |
| Interaction_genotype_Mtor | Fhod1 | -2.881 | 3.92E-03 | glio |
| Interaction_genotype_Mtor | D17H6S53E | -2.615 | 3.92E-03 | glio |
| Aars2_in.vivo_noRT | Lman1 | -0.776 | 3.92E-03 | glio |
| Aars2_in.vivo_noRT | Atpif1 | -0.625 | 3.92E-03 | glio |
| Atr_in.vivo_noRT | Fam228b | 2.227 | 3.92E-03 | glio |
| Atr_in.vivo_noRT | Pnkp | 1.302 | 3.92E-03 | glio |
| Bcas2_in.vivo_noRT | Efr3b | 2.376 | 3.92E-03 | glio |
| Chek1_in.vivo_noRT | Nova2 | 2.026 | 3.92E-03 | glio |
| Chek1_in.vivo_noRT | Nanos3 | 1.915 | 3.92E-03 | glio |
| Interaction_genotype_Aars2 | A530083I20Ri | -4.821 | 3.92E-04 | glio |
| Interaction_genotype_Myc | Nuf2 | -2.996 | 3.92E-04 | glio |

| | | | | |
|---|---|---|---|---|
| Interaction_genotype_Myc | Mrps12 | 1.684 | 3.92E-04 | glio |
| Aars2_in.vivo_noRT | Syk | 2.247 | 3.92E-04 | glio |
| Aars2_in.vivo_noRT | Tle5 | -0.995 | 3.92E-04 | glio |
| Atr_ex.vivo_noRT | Stmn4 | 4.339 | 3.92E-04 | glio |
| Bora_in.vivo_noRT | Cyp3a13 | 2.744 | 3.92E-04 | glio |
| Bora_ex.vivo_noRT | Gm32122 | 3.201 | 3.92E-04 | glio |
| Chek1_in.vivo_noRT | Cd81 | -1.732 | 3.92E-04 | glio |
| Chek1_ex.vivo_noRT | Plek2 | 3.163 | 3.92E-04 | glio |
| Chek1_ex.vivo_noRT | Crebl2 | 2.37 | 3.92E-04 | glio |
| Chek1_ex.vivo_noRT | Egr1 | 2.374 | 3.92E-04 | glio |
| Hsd17b10_in.vivo_noRT | Gm34256 | 2.816 | 3.92E-04 | glio |
| Interaction_genotype_Mtor | Gng8 | -4.666 | 3.92E-05 | glio |
| Interaction_genotype_Mtor | Chd5 | -4.935 | 3.92E-05 | glio |
| Aars2_in.vivo_noRT | Plekhh1 | 2.985 | 3.92E-05 | glio |
| Bcas2_in.vivo_noRT | Gm37170 | 2.964 | 3.92E-05 | glio |
| Bcas2_in.vivo_noRT | Arpc1b | -0.839 | 3.92E-05 | glio |
| Chek1_in.vivo_noRT | C030005K06R | 2.944 | 3.92E-05 | glio |
| Hsd17b10_in.vivo_noRT | Gm28404 | 2.728 | 3.92E-05 | glio |
| culture_effect | 2010320M18F | 0.923 | 3.92E-06 | glio |
| Interaction_genotype_Mtor | 1700084E18Ri | -5.6 | 3.92E-06 | glio |
| Aars2_in.vivo_noRT | Rapgef3 | 3.595 | 3.92E-06 | glio |
| Aars2_in.vivo_noRT | Gm11767 | 3.405 | 3.92E-06 | glio |
| Atr_in.vivo_noRT | Nup210l | 3.406 | 3.92E-06 | glio |
| Bcas2_in.vivo_noRT | Gm16794 | 3.236 | 3.92E-06 | glio |
| Bcas2_in.vivo_noRT | Tnfrsf4 | 3.221 | 3.92E-06 | glio |
| Hsd17b10_in.vivo_noRT | Kcnk4 | 3.356 | 3.92E-06 | glio |
| culture_effect | Hopx | -3.579 | 3.92E-10 | glio |
| Aars2_in.vivo_noRT | Rhbg | 3.731 | 3.92E-13 | glio |
| Atr_in.vivo_noRT | BC037032 | 5.407 | 3.92E-13 | glio |
| Atr_in.vivo_noRT | Grin3b | 5.52 | 3.92E-16 | glio |
| culture_effect | Bckdha | -1.871 | 3.92E-17 | glio |
| Atr_in.vivo_noRT | Psmb11 | 5.01 | 3.92E-22 | glio |
| Chek1_in.vivo_noRT | Gm49172 | 4.784 | 3.92E-24 | glio |
| Interaction_genotype_Atr | Apoa5 | -2.369 | 3.93E-03 | glio |
| Interaction_genotype_Atr | Kpna7 | -3.085 | 3.93E-03 | glio |
| Interaction_genotype_Atr | Gm45833 | -4.645 | 3.93E-03 | glio |
| Interaction_genotype_Atr | Rpl18a | 0.889 | 3.93E-03 | glio |
| Interaction_genotype_Lmo7 | Tdrd5 | -4.086 | 3.93E-03 | glio |
| Interaction_genotype_Mtor | Zfp651 | -2.782 | 3.93E-03 | glio |
| Interaction_genotype_Myc | C230035I16Ri | -3.401 | 3.93E-03 | glio |
| Interaction_genotype_Myc | Narf | -2.446 | 3.93E-03 | glio |
| Interaction_genotype_Myc | Tm2d1 | 1.501 | 3.93E-03 | glio |
| Interaction_genotype_Pars2 | Flacc1 | -3.458 | 3.93E-03 | glio |
| Atp5c1_in.vivo_noRT | Gm48772 | 2.291 | 3.93E-03 | glio |
| Atr_in.vivo_noRT | Parvb | 1.985 | 3.93E-03 | glio |
| Atr_in.vivo_noRT | Btf3 | -0.479 | 3.93E-03 | glio |
| Bard1_ex.vivo_noRT | Tnfrsf1b | 2.814 | 3.93E-03 | glio |
| Bcas2_in.vivo_noRT | Dhtkd1 | 2.33 | 3.93E-03 | glio |
| Bcas2_in.vivo_noRT | Kbtbd8 | 2.039 | 3.93E-03 | glio |
| Bcas2_in.vivo_noRT | Tiam2 | 1.887 | 3.93E-03 | glio |
| Bcas2_in.vivo_noRT | Flot2 | -1.444 | 3.93E-03 | glio |
| Bcas2_in.vivo_noRT | Atf3 | -2.111 | 3.93E-03 | glio |

| | | | | |
|---|---|---|---|---|
| Bcas2_in.vivo_noRT | Vmp1 | 0.726 | 3.93E-03 | glio |
| Bora_in.vivo_noRT | Gm36266 | 2.36 | 3.93E-03 | glio |
| Bora_in.vivo_noRT | 9130230N09R | 1.843 | 3.93E-03 | glio |
| Bora_in.vivo_noRT | B4galt7 | 1.699 | 3.93E-03 | glio |
| Bora_in.vivo_noRT | Abt1 | -1.181 | 3.93E-03 | glio |
| Bora_in.vivo_noRT | D8Ertd738e | -0.537 | 3.93E-03 | glio |
| Brca2_in.vivo_noRT | Capn13 | 1.763 | 3.93E-03 | glio |
| Chek1_in.vivo_noRT | Tmem37 | 2.047 | 3.93E-03 | glio |
| Chek1_in.vivo_noRT | Mrpl21 | -0.822 | 3.93E-03 | glio |
| Chek1_ex.vivo_noRT | Spdye4a | 2.764 | 3.93E-03 | glio |
| Chek1_ex.vivo_noRT | Apoh | 2.54 | 3.93E-03 | glio |
| Eif2s1_in.vivo_noRT | Gm11963 | 2.411 | 3.93E-03 | glio |
| Eif2s1_in.vivo_noRT | 4732463B04R | 1.879 | 3.93E-03 | glio |
| Hsd17b10_in.vivo_noRT | Bpgm | -0.95 | 3.93E-03 | glio |
| Interaction_genotype_Atr | 4930426I24Ri | -4.267 | 3.93E-04 | glio |
| Interaction_genotype_Atr | Gm13398 | -3.167 | 3.93E-04 | glio |
| Interaction_genotype_Mtor | Gm26590 | -3.989 | 3.93E-04 | glio |
| Interaction_genotype_Mtor | Alpk2 | -5.454 | 3.93E-04 | glio |
| Interaction_genotype_Mtor | Ankrd27 | -2.98 | 3.93E-04 | glio |
| Interaction_genotype_Myc | Gm42984 | -4.024 | 3.93E-04 | glio |
| Interaction_genotype_Myc | Phgdh | 1.99 | 3.93E-04 | glio |
| Interaction_genotype_Myc | Rpl13 | 1.388 | 3.93E-04 | glio |
| Aars2_in.vivo_noRT | Fam133b | -1.441 | 3.93E-04 | glio |
| Atr_in.vivo_noRT | Zfp959 | 2.019 | 3.93E-04 | glio |
| Atr_in.vivo_noRT | Golph3l | 1.815 | 3.93E-04 | glio |
| Atr_ex.vivo_noRT | Cys1 | 2.771 | 3.93E-04 | glio |
| Atr_ex.vivo_noRT | Gm16685 | 2.764 | 3.93E-04 | glio |
| Atr_ex.vivo_noRT | Bend7 | 3.182 | 3.93E-04 | glio |
| Bora_in.vivo_noRT | Slc25a39 | -1.072 | 3.93E-04 | glio |
| Bora_ex.vivo_noRT | D630024D03F | 2.615 | 3.93E-04 | glio |
| culture_effect | Angptl7 | -1.801 | 3.93E-05 | glio |
| culture_effect | Slc9b1 | -2.435 | 3.93E-05 | glio |
| Atr_ex.vivo_noRT | Masp1 | 3.164 | 3.93E-05 | glio |
| Atr_ex.vivo_noRT | Stoml2 | -1.163 | 3.93E-05 | glio |
| Bard1_ex.vivo_noRT | 4933400F21Ri | 2.814 | 3.93E-05 | glio |
| Bora_in.vivo_noRT | 1010001B22R | 2.936 | 3.93E-05 | glio |
| Bora_in.vivo_noRT | Kcnb1 | 3.072 | 3.93E-05 | glio |
| Chek1_in.vivo_noRT | Foxa1 | 3.191 | 3.93E-05 | glio |
| culture_effect | Polr2e | -0.579 | 3.93E-06 | glio |
| Atr_in.vivo_noRT | Gm17597 | 3.498 | 3.93E-06 | glio |
| Atr_in.vivo_noRT | 2900041M22F | 3.723 | 3.93E-07 | glio |
| Chek1_in.vivo_noRT | Sema6b | 3.568 | 3.93E-07 | glio |
| Chek1_in.vivo_noRT | Gm10516 | 3.48 | 3.93E-07 | glio |
| Chek1_ex.vivo_noRT | Gm36172 | 3.475 | 3.93E-07 | glio |
| Bora_in.vivo_noRT | Gm11127 | 3.9 | 3.93E-08 | glio |
| Chek1_in.vivo_noRT | BC037032 | 4.126 | 3.93E-08 | glio |
| Aars2_in.vivo_noRT | 4933439K11R | 4.142 | 3.93E-09 | glio |
| Aars2_in.vivo_noRT | Gm8104 | 3.907 | 3.93E-09 | glio |
| Atr_in.vivo_noRT | Stard6 | 4.162 | 3.93E-09 | glio |
| Atr_in.vivo_noRT | 9130230N09R | 3.531 | 3.93E-09 | glio |
| Chek1_in.vivo_noRT | Gm38979 | 4.393 | 3.93E-09 | glio |
| Atr_in.vivo_noRT | 2610016A17R | 4.485 | 3.93E-10 | glio |

| | | | | |
|---|---|---|---|---|
| Cyp19a1_ex.vivo_noRT | Mmp13 | 4.171 | 3.93E-10 | glio |
| Atr_in.vivo_noRT | Cpne8 | 4.799 | 3.93E-11 | glio |
| Bora_in.vivo_noRT | Efcab9 | 4.67 | 3.93E-12 | glio |
| Aars2_in.vivo_noRT | Gm27241 | 4.671 | 3.93E-19 | glio |
| Aars2_in.vivo_noRT | Mpzl2 | 5.067 | 3.93E-19 | glio |
| culture_effect | 2700033N17R | 1.161 | 3.94E-03 | glio |
| culture_effect | Pigk | -0.563 | 3.94E-03 | glio |
| Interaction_genotype_Aars2 | Ramp3 | -3.578 | 3.94E-03 | glio |
| Interaction_genotype_Aars2 | Hnrnpa2b1 | 0.841 | 3.94E-03 | glio |
| Interaction_genotype_Atr | Dennd5a | 1.666 | 3.94E-03 | glio |
| Interaction_genotype_Atr | Slc25a28 | 1.342 | 3.94E-03 | glio |
| Interaction_genotype_Mtor | Crybg3 | -3.601 | 3.94E-03 | glio |
| Interaction_genotype_Mtor | Prrg4 | -2.746 | 3.94E-03 | glio |
| Interaction_genotype_Myc | Tlx3 | -3.246 | 3.94E-03 | glio |
| Interaction_genotype_Myc | Arl1 | 1.304 | 3.94E-03 | glio |
| Interaction_genotype_Rpl3 | Efhd1 | -4.056 | 3.94E-03 | glio |
| Interaction_genotype_Rpl3 | Exosc2 | 2.017 | 3.94E-03 | glio |
| Interaction_genotype_Rpl3 | Dnajc15 | 1.79 | 3.94E-03 | glio |
| Bcas2_in.vivo_noRT | Klhl21 | -1.289 | 3.94E-03 | glio |
| Bora_in.vivo_noRT | Gpr146 | 2.153 | 3.94E-03 | glio |
| Cenpj_ex.vivo_noRT | Gm45336 | 2.591 | 3.94E-03 | glio |
| Chek1_in.vivo_noRT | Zfp65 | 1.937 | 3.94E-03 | glio |
| culture_effect | Rab17 | -2.235 | 3.94E-04 | glio |
| culture_effect | Pitpnc1 | -0.92 | 3.94E-04 | glio |
| culture_effect | Nhp2 | 0.434 | 3.94E-04 | glio |
| Interaction_genotype_Myc | Gm5415 | -2.729 | 3.94E-04 | glio |
| Interaction_genotype_Myc | Gm5544 | -4.03 | 3.94E-04 | glio |
| Interaction_genotype_Myc | Uevld | -3.364 | 3.94E-04 | glio |
| Interaction_genotype_Myc | Llph | 1.319 | 3.94E-04 | glio |
| Interaction_genotype_Myc | Rps18 | 1.58 | 3.94E-04 | glio |
| Aars2_in.vivo_noRT | Apoc1 | 2.797 | 3.94E-04 | glio |
| Aars2_in.vivo_noRT | Setd5 | -1.557 | 3.94E-04 | glio |
| Aars2_ex.vivo_noRT | Ccl8 | 3.847 | 3.94E-04 | glio |
| Atp5c1_in.vivo_noRT | Shisal2a | 2.936 | 3.94E-04 | glio |
| Atr_ex.vivo_noRT | A930003A15R | 3.164 | 3.94E-04 | glio |
| Bcas2_ex.vivo_noRT | Ctss | 3.147 | 3.94E-04 | glio |
| Bora_in.vivo_noRT | Slc22a22 | 2.503 | 3.94E-04 | glio |
| Chek1_in.vivo_noRT | Llgl2 | 2.509 | 3.94E-04 | glio |
| Hsd17b10_in.vivo_noRT | Gm50334 | 2.971 | 3.94E-04 | glio |
| culture_effect | Them4 | 1.291 | 3.94E-05 | glio |
| culture_effect | Pithd1 | -1.053 | 3.94E-05 | glio |
| culture_effect | Ccser2 | 1.001 | 3.94E-05 | glio |
| Interaction_genotype_Atr | Slc38a2 | 1.572 | 3.94E-05 | glio |
| Interaction_genotype_Myc | Trappc9 | -3.289 | 3.94E-05 | glio |
| Aars2_ex.vivo_noRT | Dcdc2a | 3.293 | 3.94E-05 | glio |
| Bard1_ex.vivo_noRT | Gm12794 | 2.814 | 3.94E-05 | glio |
| Chek1_ex.vivo_noRT | Apobr | 3.857 | 3.94E-05 | glio |
| Chek1_ex.vivo_noRT | Vwa5a | 2.228 | 3.94E-05 | glio |
| Hsd17b10_in.vivo_noRT | B020031M17F | 3.616 | 3.94E-05 | glio |
| Bcas2_in.vivo_noRT | Pglyrp1 | 3.328 | 3.94E-06 | glio |
| Bora_in.vivo_noRT | Tnc | 3.887 | 3.94E-06 | glio |
| culture_effect | Pttg1 | -0.861 | 3.94E-07 | glio |

| | | | | |
|---|---|---|---|---|
| Bcas2_in.vivo_noRT | Litaf | -2.036 | 3.94E-07 | glio |
| Chek1_in.vivo_noRT | Zfp968 | 3.666 | 3.94E-07 | glio |
| Bcas2_in.vivo_noRT | Gm49701 | 3.758 | 3.94E-08 | glio |
| Bora_in.vivo_noRT | Wnt16 | 4.145 | 3.94E-10 | glio |
| culture_effect | B930036N10R | 2.069 | 3.94E-11 | glio |
| Aars2_in.vivo_noRT | Gm9999 | 4.352 | 3.94E-11 | glio |
| Atr_in.vivo_noRT | Gm26590 | 4.245 | 3.94E-11 | glio |
| Chek1_ex.vivo_noRT | 4930553M12F | 4.449 | 3.94E-11 | glio |
| Chek1_in.vivo_noRT | Cpxm1 | 4.777 | 3.94E-15 | glio |
| Chek1_in.vivo_noRT | 4930553M12F | 4.777 | 3.94E-18 | glio |
| Bora_in.vivo_noRT | 1700112J16Ri | 5.46 | 3.94E-23 | glio |
| Atr_in.vivo_noRT | Gm31717 | 5.533 | 3.94E-29 | glio |
| culture_effect | Fut8 | 0.776 | 3.95E-03 | glio |
| culture_effect | Nudt3 | 0.374 | 3.95E-03 | glio |
| Interaction_genotype_Bcas2 | Gm37170 | -3.837 | 3.95E-03 | glio |
| Interaction_genotype_Mtor | Gm47223 | -4.63 | 3.95E-03 | glio |
| Interaction_genotype_Mtor | Speg | -3.518 | 3.95E-03 | glio |
| Interaction_genotype_Mtor | Zfp82 | -3.401 | 3.95E-03 | glio |
| Interaction_genotype_Mtor | Dnaja4 | -3.456 | 3.95E-03 | glio |
| Interaction_genotype_Mtor | N4bp2os | -3.289 | 3.95E-03 | glio |
| Interaction_genotype_Mtor | Ccdc73 | -2.912 | 3.95E-03 | glio |
| Interaction_genotype_Mtor | Elavl2 | -3.03 | 3.95E-03 | glio |
| Interaction_genotype_Myc | Tdrd3 | -2.849 | 3.95E-03 | glio |
| Interaction_genotype_Myc | Zkscan8 | -2.721 | 3.95E-03 | glio |
| Interaction_genotype_Myc | Gm3448 | -2.355 | 3.95E-03 | glio |
| Aars2_in.vivo_noRT | Procr | 2.55 | 3.95E-03 | glio |
| Aars2_ex.vivo_noRT | Gm49579 | 2.505 | 3.95E-03 | glio |
| Aars2_ex.vivo_noRT | 4833413G10R | 3.1 | 3.95E-03 | glio |
| Atr_in.vivo_noRT | Pole3 | -0.901 | 3.95E-03 | glio |
| Bcas2_in.vivo_noRT | Snx3 | -0.555 | 3.95E-03 | glio |
| Bora_in.vivo_noRT | Etf1 | -0.769 | 3.95E-03 | glio |
| Brca2_ex.vivo_noRT | Vmn2r44 | 2.425 | 3.95E-03 | glio |
| Cdkn2a_in.vivo_noRT | Tcp11x2 | 1.973 | 3.95E-03 | glio |
| Cdkn2a_in.vivo_noRT | Meioc | -1.938 | 3.95E-03 | glio |
| Cdkn2a_in.vivo_noRT | Gm26916 | -2.724 | 3.95E-03 | glio |
| Cdkn2a_in.vivo_noRT | Slc16a5 | -1.938 | 3.95E-03 | glio |
| Cdkn2a_in.vivo_noRT | Gpx2 | 1.296 | 3.95E-03 | glio |
| Cenpj_ex.vivo_noRT | Gm16344 | 2.591 | 3.95E-03 | glio |
| Chek1_in.vivo_noRT | Phpt1 | -1.463 | 3.95E-03 | glio |
| Chek1_in.vivo_noRT | Bptf | -0.786 | 3.95E-03 | glio |
| Eif2s1_ex.vivo_noRT | Gm47510 | 2.234 | 3.95E-03 | glio |
| Eif2s1_ex.vivo_noRT | Gm5788 | 2.234 | 3.95E-03 | glio |
| Eif2s1_ex.vivo_noRT | Gm48617 | 2.234 | 3.95E-03 | glio |
| Eif2s1_ex.vivo_noRT | Il12rb1 | 2.822 | 3.95E-03 | glio |
| Eif2s1_ex.vivo_noRT | Gm12703 | 2.234 | 3.95E-03 | glio |
| Eif2s1_ex.vivo_noRT | Mmp13 | 2.234 | 3.95E-03 | glio |
| Eif2s1_ex.vivo_noRT | Gm8016 | 2.234 | 3.95E-03 | glio |
| Eif2s1_ex.vivo_noRT | Gm17276 | 2.637 | 3.95E-03 | glio |
| Hsd17b10_in.vivo_noRT | Ipcef1 | 1.803 | 3.95E-03 | glio |
| culture_effect | Gm43156 | 1.827 | 3.95E-04 | glio |
| culture_effect | Btg3 | 0.435 | 3.95E-04 | glio |
| Interaction_genotype_Aars2 | 5330439K02R | -4.462 | 3.95E-04 | glio |

| | | | | |
|---|---|---|---|---|
| Interaction_genotype_Mtor | Dlg5 | -3.99 | 3.95E-04 | glio |
| Interaction_genotype_Rad51 | Sardh | -4.582 | 3.95E-04 | glio |
| Aars2_in.vivo_noRT | Cacng4 | 2.187 | 3.95E-04 | glio |
| Aars2_in.vivo_noRT | Ntmt1 | -1.465 | 3.95E-04 | glio |
| Aars2_in.vivo_noRT | Psmd13 | -1.26 | 3.95E-04 | glio |
| Aars2_in.vivo_noRT | Impdh2 | -0.881 | 3.95E-04 | glio |
| Atp5c1_in.vivo_noRT | Zfp810 | 2.668 | 3.95E-04 | glio |
| Bcas2_in.vivo_noRT | Klf15 | 2.568 | 3.95E-04 | glio |
| Hsd17b10_in.vivo_noRT | Prox2os | 2.312 | 3.95E-04 | glio |
| culture_effect | Nucb1 | -0.602 | 3.95E-05 | glio |
| Interaction_genotype_Mtor | Ptprc | -3.835 | 3.95E-05 | glio |
| Interaction_genotype_Rad21 | Sema3a | 5.21 | 3.95E-05 | glio |
| Atr_in.vivo_noRT | Ctrl | 2.985 | 3.95E-05 | glio |
| Chek1_ex.vivo_noRT | Cenpp | -2.628 | 3.95E-05 | glio |
| Chek1_ex.vivo_noRT | Ptdss1 | -2.622 | 3.95E-05 | glio |
| culture_effect | Dab2 | -1.92 | 3.95E-06 | glio |
| culture_effect | Ptbp3 | -0.573 | 3.95E-06 | glio |
| Interaction_genotype_Atr | Cited1 | 4.795 | 3.95E-06 | glio |
| Interaction_genotype_Mtor | Kif24 | -5.455 | 3.95E-06 | glio |
| Interaction_genotype_Mtor | 4930445N18R | -5.665 | 3.95E-06 | glio |
| Interaction_genotype_Mtor | Zscan4d | -4.492 | 3.95E-06 | glio |
| Bcas2_in.vivo_noRT | Tctn1 | 3.297 | 3.95E-06 | glio |
| culture_effect | Ryk | 0.732 | 3.95E-07 | glio |
| Chek1_ex.vivo_noRT | Gm48086 | 4.461 | 3.95E-07 | glio |
| Hsd17b10_in.vivo_noRT | Vmn2r56 | 2.834 | 3.95E-07 | glio |
| culture_effect | Gm49494 | 2.864 | 3.95E-08 | glio |
| culture_effect | S100a3 | 2.4 | 3.95E-09 | glio |
| Atr_in.vivo_noRT | Nox1 | 3.551 | 3.95E-09 | glio |
| Bora_in.vivo_noRT | Gm34047 | 4.936 | 3.95E-12 | glio |
| Atr_in.vivo_noRT | Gm5152 | 5.146 | 3.95E-13 | glio |
| Bcas2_in.vivo_noRT | Prl2c2 | 4.807 | 3.95E-13 | glio |
| Bora_in.vivo_noRT | Ntn1 | 4.558 | 3.95E-14 | glio |
| Chek1_ex.vivo_noRT | Gm14820 | 4.842 | 3.95E-14 | glio |
| Chek1_ex.vivo_noRT | A930002C04R | 4.443 | 3.95E-14 | glio |
| culture_effect | Gpr17 | -4.648 | 3.95E-17 | glio |
| culture_effect | Ncald | -1.434 | 3.95E-17 | glio |
| Bora_in.vivo_noRT | A930033M14F | 4.936 | 3.95E-17 | glio |
| Chek1_in.vivo_noRT | Gm13856 | 5.044 | 3.95E-25 | glio |
| culture_effect | Rbms2 | 0.5 | 3.96E-03 | glio |
| Interaction_genotype_Aars2 | Cnksr1 | -3.42 | 3.96E-03 | glio |
| Interaction_genotype_Aars2 | Gm4419 | -2.855 | 3.96E-03 | glio |
| Interaction_genotype_Atr | Pdzd4 | -2.969 | 3.96E-03 | glio |
| Interaction_genotype_Bcas2 | Hist2h3c2 | -4.094 | 3.96E-03 | glio |
| Interaction_genotype_Bcas2 | Slc26a6 | -3.746 | 3.96E-03 | glio |
| Interaction_genotype_Bcas2 | Man2c1os | -3.797 | 3.96E-03 | glio |
| Interaction_genotype_Bcas2 | Gm27239 | -3.56 | 3.96E-03 | glio |
| Interaction_genotype_Mtor | Xrcc5 | 1.704 | 3.96E-03 | glio |
| Interaction_genotype_Myc | Syt17 | -3.307 | 3.96E-03 | glio |
| Interaction_genotype_Myc | Tsc1 | -2.785 | 3.96E-03 | glio |
| Interaction_genotype_Myc | Snx11 | -2.235 | 3.96E-03 | glio |
| Interaction_genotype_Myc | Pabpn1 | 1.289 | 3.96E-03 | glio |
| Interaction_genotype_Pars2 | Gm21936 | -2.664 | 3.96E-03 | glio |

| | | | | |
|---|---|---|---|---|
| Interaction_genotype_Pars2 | Sag | -3.915 | 3.96E-03 | glio |
| Interaction_genotype_Rad21 | Prss22 | -4.721 | 3.96E-03 | glio |
| Interaction_genotype_Rad51 | Arhgef10l | -3.325 | 3.96E-03 | glio |
| Interaction_genotype_Rpl3 | Gm48772 | -3.8 | 3.96E-03 | glio |
| interaction_RT | Scn1b | 2.369 | 3.96E-03 | glio |
| Aars2_in.vivo_noRT | Ssc5d | 1.938 | 3.96E-03 | glio |
| Aars2_in.vivo_noRT | Lin37 | -1.2 | 3.96E-03 | glio |
| Atr_in.vivo_noRT | Phf19 | 1.913 | 3.96E-03 | glio |
| Atr_ex.vivo_noRT | Map3k15 | -2.33 | 3.96E-03 | glio |
| Bora_in.vivo_noRT | Rorc | 2.275 | 3.96E-03 | glio |
| Bora_in.vivo_noRT | Nsd2 | -1.205 | 3.96E-03 | glio |
| Cenpj_in.vivo_noRT | Ephx1 | 2.43 | 3.96E-03 | glio |
| Chek1_ex.vivo_noRT | Gm12840 | 2.469 | 3.96E-03 | glio |
| Eif2s1_ex.vivo_noRT | Angptl7 | 2.234 | 3.96E-03 | glio |
| Eif2s1_ex.vivo_noRT | Cldn5 | 2.234 | 3.96E-03 | glio |
| Interaction_genotype_Bora | BC065403 | -4.224 | 3.96E-04 | glio |
| Interaction_genotype_Bora | Efhd1 | -4.665 | 3.96E-04 | glio |
| Interaction_genotype_Bora | Gm15743 | -4.329 | 3.96E-04 | glio |
| Interaction_genotype_Rpl3 | Irf8 | 4.482 | 3.96E-04 | glio |
| Atr_in.vivo_noRT | E130102H24R | 2.422 | 3.96E-04 | glio |
| Bard1_ex.vivo_noRT | Gm20633 | 3.109 | 3.96E-04 | glio |
| Bcas2_in.vivo_noRT | Gm45494 | 2.457 | 3.96E-04 | glio |
| Chek1_ex.vivo_noRT | Atp5j | -0.66 | 3.96E-04 | glio |
| Eif2s1_in.vivo_noRT | Gm15270 | 2.407 | 3.96E-04 | glio |
| Eif2s1_ex.vivo_noRT | Gm47936 | 3.032 | 3.96E-04 | glio |
| Eif2s1_ex.vivo_noRT | Gm49808 | 2.639 | 3.96E-04 | glio |
| culture_effect | Ckap2 | -0.795 | 3.96E-05 | glio |
| culture_effect | Rhno1 | 0.587 | 3.96E-05 | glio |
| Interaction_genotype_Mtor | Zfand4 | -5.151 | 3.96E-05 | glio |
| Interaction_genotype_Mtor | H1foo | -3.567 | 3.96E-05 | glio |
| Interaction_genotype_Mtor | Manf | 2.336 | 3.96E-05 | glio |
| Atr_in.vivo_noRT | Vmac | 3.207 | 3.96E-05 | glio |
| Chek1_ex.vivo_noRT | Lgals4 | 3.396 | 3.96E-05 | glio |
| Interaction_genotype_Mtor | Olfr56 | -4.099 | 3.96E-06 | glio |
| Interaction_genotype_Mtor | Zbtb38 | 2.545 | 3.96E-06 | glio |
| Chek1_in.vivo_noRT | Gm12089 | 3.343 | 3.96E-06 | glio |
| Chek1_in.vivo_noRT | Gm28875 | 3.349 | 3.96E-06 | glio |
| Chek1_in.vivo_noRT | Rgs5 | 3.912 | 3.96E-06 | glio |
| Chek1_ex.vivo_noRT | Gm36198 | 3.287 | 3.96E-06 | glio |
| Hsd17b10_in.vivo_noRT | 1700015F17Ri | 3.356 | 3.96E-06 | glio |
| Atp5c1_in.vivo_noRT | Gm48764 | 2.668 | 3.96E-07 | glio |
| Atr_in.vivo_noRT | Gm17399 | 3.792 | 3.96E-07 | glio |
| Chek1_in.vivo_noRT | Gm26688 | 3.605 | 3.96E-07 | glio |
| Atr_in.vivo_noRT | Mzf1 | 3.68 | 3.96E-08 | glio |
| Chek1_in.vivo_noRT | Zfp473 | 3.619 | 3.96E-08 | glio |
| Bora_in.vivo_noRT | A030005L19Ri | 4.407 | 3.96E-12 | glio |
| Bora_in.vivo_noRT | Nmrk2 | 4.294 | 3.96E-12 | glio |
| Bora_in.vivo_noRT | Gm21818 | 4.936 | 3.96E-12 | glio |
| Aars2_in.vivo_noRT | Prm3 | 4.586 | 3.96E-14 | glio |
| Bcas2_in.vivo_noRT | Gm4166 | 3.88 | 3.96E-14 | glio |
| Bcas2_in.vivo_noRT | Gm45669 | 4.544 | 3.96E-14 | glio |
| Hsd17b10_in.vivo_noRT | Sucnr1 | 3.616 | 3.96E-15 | glio |

| | | | | |
|---|---|---|---|---|
| Atr_in.vivo_noRT | Gm42849 | 5.267 | 3.96E-17 | glio |
| Aars2_in.vivo_noRT | Gm48780 | 5.581 | 3.96E-26 | glio |
| Interaction_genotype_Aars2 | Slc25a39 | 1.168 | 3.97E-03 | glio |
| Interaction_genotype_Mtor | Xxylt1 | -3.077 | 3.97E-03 | glio |
| Interaction_genotype_Myc | Flrt1 | -3.499 | 3.97E-03 | glio |
| Interaction_genotype_Myc | Hcfc1r1 | -3.61 | 3.97E-03 | glio |
| Interaction_genotype_Myc | 1810021B22R | -3.176 | 3.97E-03 | glio |
| Interaction_genotype_Myc | Abcc5 | -2.965 | 3.97E-03 | glio |
| Interaction_genotype_Myc | Iqcc | -2.403 | 3.97E-03 | glio |
| Interaction_genotype_Myc | Hyal2 | -2.391 | 3.97E-03 | glio |
| Interaction_genotype_Myc | Ranbp9 | -1.977 | 3.97E-03 | glio |
| interaction_RT | Mgst2 | 1.9 | 3.97E-03 | glio |
| interaction_RT | Gm45397 | -1.741 | 3.97E-03 | glio |
| Aars2_in.vivo_noRT | Hist1h2br | 2.275 | 3.97E-03 | glio |
| Atp5c1_in.vivo_noRT | Adk | -0.763 | 3.97E-03 | glio |
| Atr_in.vivo_noRT | Chtf18 | 1.622 | 3.97E-03 | glio |
| Bard1_ex.vivo_noRT | Gm1123 | 3.021 | 3.97E-03 | glio |
| Bcas2_in.vivo_noRT | Zglp1 | 2.331 | 3.97E-03 | glio |
| Bcas2_in.vivo_noRT | Gde1 | -1.23 | 3.97E-03 | glio |
| Bcas2_ex.vivo_noRT | Egr2 | 2.83 | 3.97E-03 | glio |
| Bcas2_ex.vivo_noRT | Gm10300 | 2.809 | 3.97E-03 | glio |
| Bcas2_ex.vivo_noRT | Mef2b | 2.652 | 3.97E-03 | glio |
| Bora_in.vivo_noRT | Rcbtb2 | -1.411 | 3.97E-03 | glio |
| Bora_in.vivo_noRT | Tns2 | 1.657 | 3.97E-03 | glio |
| Bora_in.vivo_noRT | Bpnt1 | -1.174 | 3.97E-03 | glio |
| Bora_ex.vivo_noRT | Ldhd | 2.491 | 3.97E-03 | glio |
| Chek1_ex.vivo_noRT | Nxph4 | 2.486 | 3.97E-03 | glio |
| Eif2s1_ex.vivo_noRT | Lhx1os | 2.619 | 3.97E-03 | glio |
| Hsd17b10_in.vivo_noRT | Gm50107 | 2.101 | 3.97E-03 | glio |
| Interaction_genotype_Atr | Sema6d | -3.351 | 3.97E-04 | glio |
| Aars2_in.vivo_noRT | Klhl33 | 2.527 | 3.97E-04 | glio |
| Aars2_in.vivo_noRT | B230303O12R | 2.547 | 3.97E-04 | glio |
| Aars2_in.vivo_noRT | Ptges3 | -0.735 | 3.97E-04 | glio |
| Aars2_ex.vivo_noRT | Gm43463 | 2.9 | 3.97E-04 | glio |
| Atr_ex.vivo_noRT | Ank1 | 2.38 | 3.97E-04 | glio |
| Bard1_ex.vivo_noRT | Slc35g3 | 2.415 | 3.97E-04 | glio |
| Bcas2_in.vivo_noRT | Aox1 | 2.359 | 3.97E-04 | glio |
| Bora_in.vivo_noRT | 4933412E12Ri | 1.804 | 3.97E-04 | glio |
| Bora_ex.vivo_noRT | 2310003N18R | 2.808 | 3.97E-04 | glio |
| Cenpj_in.vivo_noRT | Gm5415 | 2.022 | 3.97E-04 | glio |
| Cenpj_in.vivo_noRT | Edn2 | 2.022 | 3.97E-04 | glio |
| Cenpj_in.vivo_noRT | Gm14207 | 2.022 | 3.97E-04 | glio |
| Cenpj_in.vivo_noRT | Nrip2 | 2.022 | 3.97E-04 | glio |
| Cenpj_in.vivo_noRT | Gm3734 | 3.207 | 3.97E-04 | glio |
| Cenpj_in.vivo_noRT | Gm49698 | 2.022 | 3.97E-04 | glio |
| Cenpj_in.vivo_noRT | Olfr881 | 2.022 | 3.97E-04 | glio |
| Chek1_in.vivo_noRT | Gpatch3 | 2.483 | 3.97E-04 | glio |
| Chek1_in.vivo_noRT | Aox1 | 2.347 | 3.97E-04 | glio |
| Interaction_genotype_Mtor | Cavin3 | 2.309 | 3.97E-05 | glio |
| Interaction_genotype_Myc | Tmem190 | -5.254 | 3.97E-05 | glio |
| Aars2_in.vivo_noRT | Fam71f2 | 2.982 | 3.97E-05 | glio |
| Aars2_in.vivo_noRT | Polr2f | -0.856 | 3.97E-05 | glio |

| | | | | |
|---|---|---|---|---|
| Bora_in.vivo_noRT | Lbx2 | 3.171 | 3.97E-05 | glio |
| Chek1_ex.vivo_noRT | D130062J10Ri | 3.39 | 3.97E-05 | glio |
| culture_effect | Dock5 | 1.609 | 3.97E-06 | glio |
| Aars2_in.vivo_noRT | Gm4419 | 2.825 | 3.97E-06 | glio |
| Bcas2_in.vivo_noRT | Gstt3 | 3.302 | 3.97E-06 | glio |
| Hsd17b10_in.vivo_noRT | Gm49441 | 2.825 | 3.97E-06 | glio |
| Aars2_in.vivo_noRT | Ankrd34a | 3.679 | 3.97E-07 | glio |
| Chek1_in.vivo_noRT | Wnt4 | 3.931 | 3.97E-07 | glio |
| culture_effect | Cd302 | -1.007 | 3.97E-09 | glio |
| Bora_in.vivo_noRT | H2-M10.1 | 3.481 | 3.97E-09 | glio |
| Bcas2_in.vivo_noRT | Lrrc72 | 4.016 | 3.97E-10 | glio |
| Aars2_in.vivo_noRT | 5830454E08Ri | 4.401 | 3.97E-11 | glio |
| culture_effect | Klf11 | -2.973 | 3.97E-12 | glio |
| Chek1_in.vivo_noRT | Gm11963 | 4.784 | 3.97E-12 | glio |
| Atr_in.vivo_noRT | Rspo1 | 4.741 | 3.97E-13 | glio |
| Bora_in.vivo_noRT | A930012O16R | 5.2 | 3.97E-13 | glio |
| culture_effect | Polr1c | -1.281 | 3.97E-17 | glio |
| culture_effect | Elovl7 | -1.539 | 3.97E-18 | glio |
| Bora_in.vivo_noRT | Gm2022 | 4.936 | 3.97E-20 | glio |
| Aars2_in.vivo_noRT | Pspn | 5.327 | 3.97E-27 | glio |
| culture_effect | 9530018F02Ri | 1.702 | 3.98E-03 | glio |
| Interaction_genotype_Aars2 | Gm17259 | -3.243 | 3.98E-03 | glio |
| Interaction_genotype_Bora | Sema4a | -4.047 | 3.98E-03 | glio |
| Interaction_genotype_Bora | Clec12b | -2.654 | 3.98E-03 | glio |
| Interaction_genotype_Mtor | Veph1 | -3.306 | 3.98E-03 | glio |
| Interaction_genotype_Myc | Hist1h2br | -3.582 | 3.98E-03 | glio |
| Interaction_genotype_Myc | Rpa1 | -1.979 | 3.98E-03 | glio |
| Interaction_genotype_Pars2 | Pakap | -3.115 | 3.98E-03 | glio |
| Interaction_genotype_Pars2 | Rdh12 | -3.77 | 3.98E-03 | glio |
| Interaction_genotype_Pars2 | Gm35082 | -3.066 | 3.98E-03 | glio |
| Interaction_genotype_Pars2 | Stk36 | -4.068 | 3.98E-03 | glio |
| Interaction_genotype_Pars2 | Col6a2 | -2.795 | 3.98E-03 | glio |
| Interaction_genotype_Pars2 | Gm45716 | -2.963 | 3.98E-03 | glio |
| Interaction_genotype_Rad51 | K230015D01R | -4.192 | 3.98E-03 | glio |
| Interaction_genotype_Rpl3 | Gm49891 | -4.214 | 3.98E-03 | glio |
| Interaction_genotype_Rpl3 | Gm11549 | -3.964 | 3.98E-03 | glio |
| Interaction_genotype_Rpl3 | Ppme1 | 2.052 | 3.98E-03 | glio |
| Aars2_in.vivo_noRT | Itpa | -1.123 | 3.98E-03 | glio |
| Atp5c1_in.vivo_noRT | Sbk1 | 2.331 | 3.98E-03 | glio |
| Atr_in.vivo_noRT | Slc43a2 | 2.167 | 3.98E-03 | glio |
| Atr_in.vivo_noRT | Trmt6 | -1.169 | 3.98E-03 | glio |
| Atr_in.vivo_noRT | Spr | -1.117 | 3.98E-03 | glio |
| Atr_ex.vivo_noRT | Ccr2 | 2.903 | 3.98E-03 | glio |
| Bcas2_in.vivo_noRT | Ntn3 | 1.998 | 3.98E-03 | glio |
| Bcas2_in.vivo_noRT | Efnb1 | 1.92 | 3.98E-03 | glio |
| Bcas2_in.vivo_noRT | Fen1 | -1.268 | 3.98E-03 | glio |
| Bcas2_in.vivo_noRT | Ppil3 | 0.895 | 3.98E-03 | glio |
| Bora_in.vivo_noRT | 4833417C18R | 2.112 | 3.98E-03 | glio |
| Bora_in.vivo_noRT | Fam3c | -1.217 | 3.98E-03 | glio |
| Bora_ex.vivo_noRT | Prex2 | 3.201 | 3.98E-03 | glio |
| Chek1_ex.vivo_noRT | 4932441J04Ri | 2.756 | 3.98E-03 | glio |
| Chek1_ex.vivo_noRT | 1700030L20Ri | 2.511 | 3.98E-03 | glio |

| | | | | |
|---|---|---|---|---|
| Eif2s1_ex.vivo_noRT | Dpep3 | 2.234 | 3.98E-03 | glio |
| Eif2s1_ex.vivo_noRT | Erich6b | 2.234 | 3.98E-03 | glio |
| Eif2s1_ex.vivo_noRT | Tac2 | 2.234 | 3.98E-03 | glio |
| Hsd17b10_in.vivo_noRT | Slc41a3 | 2.128 | 3.98E-03 | glio |
| Interaction_genotype_Myc | Nrip2 | -2.729 | 3.98E-04 | glio |
| Interaction_genotype_Myc | Fsbp | -4.138 | 3.98E-04 | glio |
| Interaction_genotype_Myc | 3110021N24R | -3.293 | 3.98E-04 | glio |
| Interaction_genotype_Myc | Slc38a2 | 1.464 | 3.98E-04 | glio |
| Aars2_in.vivo_noRT | Farsb | -1.373 | 3.98E-04 | glio |
| Atp5c1_in.vivo_noRT | Htr1b | 2.668 | 3.98E-04 | glio |
| Atp5c1_in.vivo_noRT | A430106G13R | 2.924 | 3.98E-04 | glio |
| Atr_in.vivo_noRT | Apoe | 2.227 | 3.98E-04 | glio |
| Atr_ex.vivo_noRT | Tff2 | 3.164 | 3.98E-04 | glio |
| Chek1_in.vivo_noRT | Crybg2 | 2.519 | 3.98E-04 | glio |
| Chek1_ex.vivo_noRT | 1700120G11R | 3.463 | 3.98E-04 | glio |
| Chek1_ex.vivo_noRT | Gemin5 | -1.824 | 3.98E-04 | glio |
| Interaction_genotype_Mtor | Josd2 | 2.182 | 3.98E-05 | glio |
| Interaction_genotype_Mtor | Psmb2 | 2.087 | 3.98E-05 | glio |
| Aars2_in.vivo_noRT | Ybx3 | -1.022 | 3.98E-05 | glio |
| Atr_in.vivo_noRT | Gm4419 | 2.522 | 3.98E-05 | glio |
| Bcas2_ex.vivo_noRT | 4930470G03R | 3.131 | 3.98E-05 | glio |
| Bora_in.vivo_noRT | Srsf3 | -0.954 | 3.98E-05 | glio |
| Bora_ex.vivo_noRT | Stx19 | 3.201 | 3.98E-05 | glio |
| Hsd17b10_in.vivo_noRT | Vmn2r54 | 2.455 | 3.98E-05 | glio |
| Atr_in.vivo_noRT | Cldn9 | 3.088 | 3.98E-06 | glio |
| Bora_in.vivo_noRT | Acsl3 | -1.949 | 3.98E-06 | glio |
| Aars2_in.vivo_noRT | Car10 | 2.983 | 3.98E-07 | glio |
| Bcas2_in.vivo_noRT | Sema4a | 3.764 | 3.98E-07 | glio |
| Bora_in.vivo_noRT | Adam1a | 3.575 | 3.98E-07 | glio |
| culture_effect | Psma7 | -0.665 | 3.98E-09 | glio |
| Atr_in.vivo_noRT | Mroh7 | 4.009 | 3.98E-09 | glio |
| Atr_in.vivo_noRT | Gabrd | 4.084 | 3.98E-09 | glio |
| Interaction_genotype_Mtor | Gm41764 | -6.965 | 3.98E-10 | glio |
| culture_effect | Cd274 | -2.771 | 3.98E-14 | glio |
| Aars2_in.vivo_noRT | 1700016P04R | 5.06 | 3.98E-15 | glio |
| Bora_in.vivo_noRT | Gm47404 | 4.936 | 3.98E-24 | glio |
| Bora_in.vivo_noRT | Cldn2 | 4.936 | 3.98E-26 | glio |
| culture_effect | Gpx7 | -0.436 | 3.99E-03 | glio |
| Interaction_genotype_Atr | Gjd4 | -2.369 | 3.99E-03 | glio |
| Interaction_genotype_Atr | Gm27241 | -2.694 | 3.99E-03 | glio |
| Interaction_genotype_Atr | Gm50103 | -3.528 | 3.99E-03 | glio |
| Interaction_genotype_Atr | Uevld | -2.934 | 3.99E-03 | glio |
| Interaction_genotype_Chek1 | Zeb2os | -3.686 | 3.99E-03 | glio |
| Interaction_genotype_Myc | Gm43260 | -2.998 | 3.99E-03 | glio |
| Interaction_genotype_Myc | Vmn1r139 | -2.92 | 3.99E-03 | glio |
| Interaction_genotype_Myc | Id3 | 2.023 | 3.99E-03 | glio |
| Interaction_genotype_Rpl3 | Limch1 | -4.073 | 3.99E-03 | glio |
| Aars2_in.vivo_noRT | Sema3d | 2.098 | 3.99E-03 | glio |
| Aars2_in.vivo_noRT | Wbp2 | -1.089 | 3.99E-03 | glio |
| Aars2_in.vivo_noRT | Tmem128 | -0.805 | 3.99E-03 | glio |
| Aars2_ex.vivo_noRT | Ssh3 | 2.495 | 3.99E-03 | glio |
| Aars2_ex.vivo_noRT | Ngef | 2.598 | 3.99E-03 | glio |

| | | | | |
|---|---|---|---|---|
| Atr_in.vivo_noRT | Zfp202 | 2.121 | 3.99E-03 | glio |
| Bcas2_in.vivo_noRT | Gm5577 | 1.963 | 3.99E-03 | glio |
| Bcas2_ex.vivo_noRT | Gata3 | 3.257 | 3.99E-03 | glio |
| Bora_in.vivo_noRT | Chrna1os | 1.787 | 3.99E-03 | glio |
| Ercc4_ex.vivo_noRT | Arhgef16 | 3.082 | 3.99E-03 | glio |
| Hsd17b10_in.vivo_noRT | 1810012K16R | 2.1 | 3.99E-03 | glio |
| culture_effect | Scarb2 | -1.032 | 3.99E-04 | glio |
| culture_effect | mt-Atp6 | 0.497 | 3.99E-04 | glio |
| Interaction_genotype_Mtor | Stac3 | -4.385 | 3.99E-04 | glio |
| Interaction_genotype_Myc | Cers3 | -3.169 | 3.99E-04 | glio |
| Interaction_genotype_Myc | 9530080O11R | -3.89 | 3.99E-04 | glio |
| Interaction_genotype_Myc | Lsp1 | -4.486 | 3.99E-04 | glio |
| Interaction_genotype_Myc | Rab20 | -3.515 | 3.99E-04 | glio |
| Aars2_in.vivo_noRT | Emc10 | -0.879 | 3.99E-04 | glio |
| Atr_in.vivo_noRT | Pced1a | 1.907 | 3.99E-04 | glio |
| Chek1_in.vivo_noRT | Gimap9 | 2.353 | 3.99E-04 | glio |
| Chek1_ex.vivo_noRT | Snhg14 | 3.417 | 3.99E-04 | glio |
| culture_effect | Nsmce1 | -0.625 | 3.99E-05 | glio |
| Aars2_in.vivo_noRT | Ankrd42 | 2.944 | 3.99E-05 | glio |
| Aars2_ex.vivo_noRT | Gm12589 | 2.893 | 3.99E-05 | glio |
| Interaction_genotype_Mtor | Gm48682 | -4.966 | 3.99E-06 | glio |
| Interaction_genotype_Mtor | Stpg4 | -5.777 | 3.99E-06 | glio |
| Interaction_genotype_Mtor | 2310058D17R | -6.756 | 3.99E-08 | glio |
| Interaction_genotype_Mtor | Gm14002 | -6.249 | 3.99E-08 | glio |
| Atr_in.vivo_noRT | E230029C05Ri | 3.662 | 3.99E-08 | glio |
| Hsd17b10_in.vivo_noRT | Plcl1 | 3.616 | 3.99E-10 | glio |
| Aars2_in.vivo_noRT | Nrros | 4.684 | 3.99E-12 | glio |
| Aars2_in.vivo_noRT | Fbxl2 | 4.681 | 3.99E-14 | glio |
| Atr_in.vivo_noRT | Pde6b | 4.501 | 3.99E-14 | glio |
| culture_effect | Erap1 | -2.936 | 3.99E-26 | glio |
| culture_effect | Bbof1 | -1.788 | 4.00E-03 | glio |
| culture_effect | Gstt2 | 1.512 | 4.00E-03 | glio |
| culture_effect | Gdnf | 0.824 | 4.00E-03 | glio |
| culture_effect | Crbn | -0.716 | 4.00E-03 | glio |
| Interaction_genotype_Atr | Gm2115 | -3.005 | 4.00E-03 | glio |
| Interaction_genotype_Atr | Gm15133 | -3.577 | 4.00E-03 | glio |
| Interaction_genotype_Bcas2 | Iws1 | 1.837 | 4.00E-03 | glio |
| Interaction_genotype_Bora | Zscan22 | -3.12 | 4.00E-03 | glio |
| Interaction_genotype_Mtor | Rax | -3.281 | 4.00E-03 | glio |
| Interaction_genotype_Mtor | Zfp763 | -3.196 | 4.00E-03 | glio |
| Interaction_genotype_Myc | Wdr59 | -2.749 | 4.00E-03 | glio |
| Interaction_genotype_Myc | Gpatch2l | -2.67 | 4.00E-03 | glio |
| Interaction_genotype_Myc | Kcnc3 | -2.607 | 4.00E-03 | glio |
| Aars2_in.vivo_noRT | Cfap69 | 2.056 | 4.00E-03 | glio |
| Aars2_in.vivo_noRT | Mindy4 | 1.981 | 4.00E-03 | glio |
| Aars2_in.vivo_noRT | Ccdc180 | 1.885 | 4.00E-03 | glio |
| Aars2_in.vivo_noRT | Cyb5rl | 1.718 | 4.00E-03 | glio |
| Aars2_in.vivo_noRT | Flt3l | 1.778 | 4.00E-03 | glio |
| Aars2_in.vivo_noRT | Leng1 | -1.247 | 4.00E-03 | glio |
| Aars2_in.vivo_noRT | Klhl12 | 1.715 | 4.00E-03 | glio |
| Bcas2_in.vivo_noRT | Rbakdn | 2.106 | 4.00E-03 | glio |
| Bora_ex.vivo_noRT | Pdlim3 | 2.407 | 4.00E-03 | glio |

| | | | | |
|---|---|---|---|---|
| Cenpj_ex.vivo_noRT | D730045A05R | 2.994 | 4.00E-03 | glio |
| Chek1_in.vivo_noRT | Chst12 | 1.718 | 4.00E-03 | glio |
| Hsd17b10_in.vivo_noRT | Tcte2 | 2.025 | 4.00E-03 | glio |
| Hsd17b10_in.vivo_noRT | Samd9l | 1.12 | 4.00E-03 | glio |
| Hsd17b10_in.vivo_noRT | Tmed3 | -0.699 | 4.00E-03 | glio |
| culture_effect | Sap30 | -0.573 | 4.00E-04 | glio |
| culture_effect | Hmgb3 | -0.551 | 4.00E-04 | glio |
| Interaction_genotype_Bcas2 | Tecrl | -4.791 | 4.00E-04 | glio |
| Interaction_genotype_Mtor | Slc22a21 | -4.35 | 4.00E-04 | glio |
| Interaction_genotype_Mtor | Zfp729a | -3.92 | 4.00E-04 | glio |
| Interaction_genotype_Mtor | Sord | -3.858 | 4.00E-04 | glio |
| Interaction_genotype_Mtor | Dus4l | -3.284 | 4.00E-04 | glio |
| Interaction_genotype_Myc | AA388235 | -4.144 | 4.00E-04 | glio |
| Interaction_genotype_Myc | Zfp518b | -3.306 | 4.00E-04 | glio |
| Interaction_genotype_Myc | Acbd4 | -2.973 | 4.00E-04 | glio |
| Bcas2_in.vivo_noRT | Btc | 3.248 | 4.00E-04 | glio |
| Bcas2_ex.vivo_noRT | Dscaml1 | 2.347 | 4.00E-04 | glio |
| Bora_in.vivo_noRT | March11 | 2.663 | 4.00E-04 | glio |
| Cenpj_ex.vivo_noRT | Gm26555 | 2.591 | 4.00E-04 | glio |
| Interaction_genotype_Atr | Esrp2 | -4.825 | 4.00E-05 | glio |
| Interaction_genotype_Mtor | Cdrt4 | -5.116 | 4.00E-05 | glio |
| Interaction_genotype_Mtor | Myorg | -4.899 | 4.00E-05 | glio |
| Interaction_genotype_Mtor | Gm12610 | -3.97 | 4.00E-05 | glio |
| Atr_in.vivo_noRT | Fam129c | 2.865 | 4.00E-05 | glio |
| Cyp19a1_ex.vivo_noRT | Scel | 2.933 | 4.00E-05 | glio |
| interaction_RT | Gm48619 | 1.57 | 4.00E-06 | glio |
| Atr_in.vivo_noRT | Wbp11 | -1.267 | 4.00E-06 | glio |
| Aars2_in.vivo_noRT | D230022J07Ri | 3.638 | 4.00E-07 | glio |
| Bora_in.vivo_noRT | E130208F15Ri | 3.883 | 4.00E-07 | glio |
| Chek1_ex.vivo_noRT | Arg2 | 4.449 | 4.00E-07 | glio |
| Atp5c1_in.vivo_noRT | Cldn2 | 2.668 | 4.00E-08 | glio |
| Chek1_in.vivo_noRT | Gm41442 | 4.002 | 4.00E-09 | glio |
| Bora_in.vivo_noRT | Amy1 | 4.407 | 4.00E-11 | glio |
| Aars2_in.vivo_noRT | 2810410L24Ri | 4.801 | 4.00E-12 | glio |
| Bora_in.vivo_noRT | Cd248 | 4.67 | 4.00E-12 | glio |
| Chek1_in.vivo_noRT | Pcdhga11 | 4.047 | 4.00E-12 | glio |
| Atr_in.vivo_noRT | Scube3 | 5.001 | 4.00E-15 | glio |
| Atr_in.vivo_noRT | A830009L08Ri | 5.011 | 4.00E-15 | glio |
| Atr_in.vivo_noRT | Lrp2bp | 5.269 | 4.00E-17 | glio |
| Aars2_in.vivo_noRT | 4930527F14Ri | 4.798 | 4.00E-19 | glio |
| Bcas2_in.vivo_noRT | Gm30692 | 4.807 | 4.00E-22 | glio |
| Chek1_in.vivo_noRT | Slc16a5 | 4.78 | 4.00E-22 | glio |
| culture_effect | Gm45151 | -1.466 | 4.01E-03 | glio |
| Interaction_genotype_Atr | Pkd2 | -2.919 | 4.01E-03 | glio |
| Interaction_genotype_Mtor | Hsd17b7 | -3.096 | 4.01E-03 | glio |
| Interaction_genotype_Myc | Zbtb25 | -2.689 | 4.01E-03 | glio |
| Interaction_genotype_Myc | Wwox | -2.403 | 4.01E-03 | glio |
| Interaction_genotype_Myc | Becn1 | 1.194 | 4.01E-03 | glio |
| Atp5c1_in.vivo_noRT | Pmch | 2.01 | 4.01E-03 | glio |
| Atp5c1_in.vivo_noRT | Gm28578 | 2.408 | 4.01E-03 | glio |
| Atp5c1_in.vivo_noRT | Gm7102 | 2.408 | 4.01E-03 | glio |
| Atp5c1_ex.vivo_noRT | Gpr4 | 2.597 | 4.01E-03 | glio |

| | | | | |
|---|---|---|---|---|
| Atr_in.vivo_noRT | Fancf | 1.701 | 4.01E-03 | glio |
| Atr_in.vivo_noRT | Gjc3 | -0.727 | 4.01E-03 | glio |
| Bora_in.vivo_noRT | Zfp981 | 2.059 | 4.01E-03 | glio |
| Cenpj_in.vivo_noRT | Esp15 | 2.022 | 4.01E-03 | glio |
| Cenpj_in.vivo_noRT | Gm10801 | 2.022 | 4.01E-03 | glio |
| Cenpj_in.vivo_noRT | Krt18 | -2.821 | 4.01E-03 | glio |
| Chek1_in.vivo_noRT | Dgcr6 | -1.021 | 4.01E-03 | glio |
| Chek1_ex.vivo_noRT | Parp2 | -1.821 | 4.01E-03 | glio |
| Interaction_genotype_Atr | Prpf4 | 1.979 | 4.01E-04 | glio |
| Interaction_genotype_Myc | Gm49698 | -2.729 | 4.01E-04 | glio |
| Interaction_genotype_Myc | Gpr162 | -4.1 | 4.01E-04 | glio |
| Interaction_genotype_Myc | 9130604C24R | -4.232 | 4.01E-04 | glio |
| Interaction_genotype_Myc | Tbc1d24 | -3.899 | 4.01E-04 | glio |
| Interaction_genotype_Myc | Gm36495 | -4.031 | 4.01E-04 | glio |
| Interaction_genotype_Myc | Rpl12 | 1.243 | 4.01E-04 | glio |
| interaction_RT | Heca | 1.415 | 4.01E-04 | glio |
| interaction_RT | 2900041M22F | 3.381 | 4.01E-04 | glio |
| interaction_RT | Rpl7l1 | -0.707 | 4.01E-04 | glio |
| interaction_RT | Purb | -0.587 | 4.01E-04 | glio |
| interaction_RT | Serbp1 | -0.626 | 4.01E-04 | glio |
| Aars2_in.vivo_noRT | Dennd4b | 2.512 | 4.01E-04 | glio |
| Atr_in.vivo_noRT | Aprt | -0.861 | 4.01E-04 | glio |
| Bcas2_in.vivo_noRT | AC125149.3 | 2.402 | 4.01E-04 | glio |
| Bcas2_in.vivo_noRT | Napg | -1.651 | 4.01E-04 | glio |
| Bcas2_ex.vivo_noRT | Lmx1a | 2.742 | 4.01E-04 | glio |
| Bcas2_ex.vivo_noRT | Vmn2r56 | 2.742 | 4.01E-04 | glio |
| Bora_in.vivo_noRT | Gm15952 | 2.633 | 4.01E-04 | glio |
| Bora_in.vivo_noRT | Id3 | -1.728 | 4.01E-04 | glio |
| Chek1_in.vivo_noRT | 1700047M11F | 2.317 | 4.01E-04 | glio |
| Chek1_ex.vivo_noRT | Cct3 | -1.001 | 4.01E-04 | glio |
| Eif2s1_in.vivo_noRT | Prap1 | 2.143 | 4.01E-04 | glio |
| Atp5c1_in.vivo_noRT | Gm44752 | 2.809 | 4.01E-05 | glio |
| Atr_in.vivo_noRT | Rps15a | -0.89 | 4.01E-05 | glio |
| Bcas2_in.vivo_noRT | Timm23 | -0.946 | 4.01E-05 | glio |
| Hsd17b10_in.vivo_noRT | Lama5 | 3.094 | 4.01E-05 | glio |
| Hsd17b10_in.vivo_noRT | Gm43137 | 3.1 | 4.01E-05 | glio |
| Hsd17b10_in.vivo_noRT | Parvg | 3.084 | 4.01E-05 | glio |
| Hsd17b10_in.vivo_noRT | Sidt2 | -1.898 | 4.01E-05 | glio |
| Interaction_genotype_Mtor | Celf5 | -5.853 | 4.01E-06 | glio |
| Aars2_in.vivo_noRT | Rhov | 3.141 | 4.01E-06 | glio |
| Bora_in.vivo_noRT | Tcea2 | 3.208 | 4.01E-06 | glio |
| Bora_in.vivo_noRT | Cyp20a1 | -2.144 | 4.01E-06 | glio |
| Bora_ex.vivo_noRT | Cxcl5 | 3.201 | 4.01E-06 | glio |
| Chek1_in.vivo_noRT | Samd15 | 3.545 | 4.01E-06 | glio |
| Chek1_ex.vivo_noRT | Tox2 | 4.055 | 4.01E-06 | glio |
| Hsd17b10_in.vivo_noRT | Ptges | 3.487 | 4.01E-06 | glio |
| culture_effect | Lxn | 1.603 | 4.01E-07 | glio |
| Atr_in.vivo_noRT | Hecw2 | 3.648 | 4.01E-07 | glio |
| Atr_in.vivo_noRT | Shroom1 | 3.425 | 4.01E-07 | glio |
| Chek1_in.vivo_noRT | Dct | 4.24 | 4.01E-07 | glio |
| Chek1_in.vivo_noRT | Tacr2 | 3.611 | 4.01E-07 | glio |
| Chek1_in.vivo_noRT | Gm49864 | 3.611 | 4.01E-07 | glio |

| | | | | |
|---|---|---|---|---|
| Chek1_in.vivo_noRT | 1700084E18Ri | 3.802 | 4.01E-07 | glio |
| Chek1_in.vivo_noRT | A430018G15R | 2.926 | 4.01E-07 | glio |
| Interaction_genotype_Rad21 | Serping1 | 5.539 | 4.01E-08 | glio |
| Atr_in.vivo_noRT | Slc43a1 | 3.869 | 4.01E-08 | glio |
| Aars2_in.vivo_noRT | Flrt1 | 4.352 | 4.01E-09 | glio |
| culture_effect | Kdm6b | -1.204 | 4.01E-10 | glio |
| Chek1_in.vivo_noRT | Ushbp1 | 4.518 | 4.01E-11 | glio |
| Aars2_in.vivo_noRT | Sox11 | 4.68 | 4.01E-12 | glio |
| Aars2_in.vivo_noRT | Gm37864 | 4.547 | 4.01E-14 | glio |
| Chek1_in.vivo_noRT | Gm41349 | 4.78 | 4.01E-14 | glio |
| culture_effect | Jmjd1c | -1.633 | 4.01E-24 | glio |
| culture_effect | Bglap2 | -1.863 | 4.02E-03 | glio |
| culture_effect | Gm41611 | 1.235 | 4.02E-03 | glio |
| culture_effect | Ccdc9 | -0.67 | 4.02E-03 | glio |
| culture_effect | Otud3 | 0.605 | 4.02E-03 | glio |
| Interaction_genotype_Aars2 | Osr2 | 3.06 | 4.02E-03 | glio |
| Interaction_genotype_Mtor | Apobec1 | -3.532 | 4.02E-03 | glio |
| Interaction_genotype_Myc | Wdr62 | -2.8 | 4.02E-03 | glio |
| Interaction_genotype_Myc | 1810062G17R | -3.079 | 4.02E-03 | glio |
| Interaction_genotype_Myc | Ercc4 | -2.944 | 4.02E-03 | glio |
| Interaction_genotype_Myc | Tlr4 | -2.316 | 4.02E-03 | glio |
| Interaction_genotype_Myc | Cryzl2 | -2.202 | 4.02E-03 | glio |
| Interaction_genotype_Myc | Ndufa7 | 0.92 | 4.02E-03 | glio |
| Interaction_genotype_Rpl3 | Tmem216 | -2.818 | 4.02E-03 | glio |
| Aars2_in.vivo_noRT | Zbtb45 | 1.57 | 4.02E-03 | glio |
| Atr_in.vivo_noRT | Prdm10 | 1.832 | 4.02E-03 | glio |
| Atr_in.vivo_noRT | Glis3 | -1.316 | 4.02E-03 | glio |
| Bard1_ex.vivo_noRT | Cd52 | 2.814 | 4.02E-03 | glio |
| Bard1_ex.vivo_noRT | Cysltr2 | 2.814 | 4.02E-03 | glio |
| Cenpj_in.vivo_noRT | Rnf128 | -2.686 | 4.02E-03 | glio |
| Chek1_in.vivo_noRT | Gpr135 | 2.093 | 4.02E-03 | glio |
| Chek1_in.vivo_noRT | Gm19541 | 2.119 | 4.02E-03 | glio |
| Chek1_in.vivo_noRT | Hgs | -1.215 | 4.02E-03 | glio |
| Chek1_in.vivo_noRT | Pole4 | -0.995 | 4.02E-03 | glio |
| Eif2s1_in.vivo_noRT | Gm15816 | 2.143 | 4.02E-03 | glio |
| Eif2s1_ex.vivo_noRT | Ptgis | 3.101 | 4.02E-03 | glio |
| Hsd17b10_in.vivo_noRT | 5330417C22R | 2.221 | 4.02E-03 | glio |
| Hsd17b10_in.vivo_noRT | Zmynd15 | 2.179 | 4.02E-03 | glio |
| Hsd17b10_in.vivo_noRT | 2210406O10R | 2.299 | 4.02E-03 | glio |
| Hsd17b10_in.vivo_noRT | Arvcf | 2.375 | 4.02E-03 | glio |
| Hsd17b10_in.vivo_noRT | Cald1 | 0.939 | 4.02E-03 | glio |
| culture_effect | Pla2g2e | -2.446 | 4.02E-04 | glio |
| culture_effect | Srxn1 | 0.918 | 4.02E-04 | glio |
| Interaction_genotype_Mtor | Gm10516 | -4.176 | 4.02E-04 | glio |
| Interaction_genotype_Myc | Olfr881 | -2.729 | 4.02E-04 | glio |
| Interaction_genotype_Myc | Cgn | -3.935 | 4.02E-04 | glio |
| Interaction_genotype_Myc | Cbx3 | 1.7 | 4.02E-04 | glio |
| Aars2_in.vivo_noRT | 9330160F10Ri | 2.32 | 4.02E-04 | glio |
| Aars2_in.vivo_noRT | Dnajc7 | -1.191 | 4.02E-04 | glio |
| Interaction_genotype_Bcas2 | BC004004 | 2.232 | 4.02E-05 | glio |
| Aars2_in.vivo_noRT | Setd4 | 2.851 | 4.02E-05 | glio |
| Chek1_in.vivo_noRT | Asap3 | 2.825 | 4.02E-05 | glio |

| | | | | |
|---|---|---|---|---|
| Chek1_ex.vivo_noRT | Shd | 3.563 | 4.02E-05 | glio |
| Interaction_genotype_Mtor | Slc7a14 | -5.332 | 4.02E-06 | glio |
| Interaction_genotype_Mtor | Gm15270 | -4.63 | 4.02E-06 | glio |
| Interaction_genotype_Myc | Gm26533 | -5.299 | 4.02E-06 | glio |
| Chek1_in.vivo_noRT | AI480526 | 3.155 | 4.02E-06 | glio |
| culture_effect | Kif13a | 1.463 | 4.02E-07 | glio |
| culture_effect | Ctsl | -0.964 | 4.02E-07 | glio |
| culture_effect | Rpa3 | -0.969 | 4.02E-08 | glio |
| Bcas2_in.vivo_noRT | Abat | 4.15 | 4.02E-08 | glio |
| Atr_in.vivo_noRT | Tssk6 | 3.585 | 4.02E-09 | glio |
| Bora_ex.vivo_noRT | Gm12841 | 3.968 | 4.02E-09 | glio |
| Interaction_genotype_Mtor | Gm11457 | -7.4 | 4.02E-10 | glio |
| Bcas2_in.vivo_noRT | Gm10399 | 5.155 | 4.02E-14 | glio |
| Chek1_in.vivo_noRT | Iqch | 4.78 | 4.02E-16 | glio |
| Chek1_in.vivo_noRT | 1700017M07F | 5.044 | 4.02E-17 | glio |
| Atr_in.vivo_noRT | Htr1b | 6.071 | 4.02E-19 | glio |
| culture_effect | Gm48226 | 1.165 | 4.03E-03 | glio |
| Interaction_genotype_Aars2 | Cript | 1.328 | 4.03E-03 | glio |
| Interaction_genotype_Myc | Gem | -3.122 | 4.03E-03 | glio |
| Interaction_genotype_Myc | Gm15265 | -3.231 | 4.03E-03 | glio |
| Interaction_genotype_Myc | Gm38979 | -3.469 | 4.03E-03 | glio |
| Interaction_genotype_Myc | Zfp446 | -2.84 | 4.03E-03 | glio |
| Interaction_genotype_Myc | Plxnb1 | -3.147 | 4.03E-03 | glio |
| Interaction_genotype_Myc | Cblb | -3.071 | 4.03E-03 | glio |
| Interaction_genotype_Myc | Rttn | -2.599 | 4.03E-03 | glio |
| Interaction_genotype_Myc | Pex19 | -2.068 | 4.03E-03 | glio |
| Interaction_genotype_Myc | Znfx1 | -1.998 | 4.03E-03 | glio |
| Interaction_genotype_Myc | Pbdc1 | 1.222 | 4.03E-03 | glio |
| Interaction_genotype_Rpl3 | C130060C02R | -3.625 | 4.03E-03 | glio |
| Aars2_in.vivo_noRT | Rap1gap | -1.453 | 4.03E-03 | glio |
| Aars2_in.vivo_noRT | Kifc1 | 1.495 | 4.03E-03 | glio |
| Aars2_in.vivo_noRT | Mef2c | 1.657 | 4.03E-03 | glio |
| Aars2_in.vivo_noRT | Slc35b1 | -1.054 | 4.03E-03 | glio |
| Aars2_in.vivo_noRT | Mrpl27 | -0.866 | 4.03E-03 | glio |
| Aars2_in.vivo_noRT | Rps26 | -0.594 | 4.03E-03 | glio |
| Atr_in.vivo_noRT | Nexn | 1.895 | 4.03E-03 | glio |
| Bcas2_in.vivo_noRT | Gm26881 | 1.867 | 4.03E-03 | glio |
| Bcas2_in.vivo_noRT | Relch | -1.518 | 4.03E-03 | glio |
| Bcas2_in.vivo_noRT | Virma | -1.121 | 4.03E-03 | glio |
| Bcas2_in.vivo_noRT | Zfp24 | -1.105 | 4.03E-03 | glio |
| Bcas2_in.vivo_noRT | Krtcap2 | -0.605 | 4.03E-03 | glio |
| Bcas2_ex.vivo_noRT | Dnaaf3 | 2.742 | 4.03E-03 | glio |
| Bora_in.vivo_noRT | Smap2 | -1.312 | 4.03E-03 | glio |
| Bora_ex.vivo_noRT | Apol6 | 2.414 | 4.03E-03 | glio |
| Interaction_genotype_Bcas2 | Phldb2 | -4.757 | 4.03E-04 | glio |
| Interaction_genotype_Myc | Dlx2 | -4.168 | 4.03E-04 | glio |
| Interaction_genotype_Myc | Gpr135 | -3.987 | 4.03E-04 | glio |
| Interaction_genotype_Myc | Zfp868 | -3.536 | 4.03E-04 | glio |
| Interaction_genotype_Myc | Rpl27a | 1.324 | 4.03E-04 | glio |
| Aars2_in.vivo_noRT | Doc2g | 2.589 | 4.03E-04 | glio |
| Aars2_in.vivo_noRT | Lysmd4 | 2.314 | 4.03E-04 | glio |
| Atp5c1_in.vivo_noRT | K230015D01R | 2.93 | 4.03E-04 | glio |

| | | | | |
|---|---|---|---|---|
| Atr_in.vivo_noRT | Bok | 2.175 | 4.03E-04 | glio |
| Bcas2_in.vivo_noRT | Slc43a1 | 2.605 | 4.03E-04 | glio |
| Bora_in.vivo_noRT | 2510046G10R | 2.306 | 4.03E-04 | glio |
| Cenpj_in.vivo_noRT | BC061195 | 2.022 | 4.03E-04 | glio |
| culture_effect | Tcp1 | -0.457 | 4.03E-05 | glio |
| interaction_RT | Hbp1 | 1.347 | 4.03E-05 | glio |
| interaction_RT | Tomm40 | -0.84 | 4.03E-05 | glio |
| Aars2_in.vivo_noRT | Rbp7 | 2.934 | 4.03E-05 | glio |
| Bora_in.vivo_noRT | Zbtb46 | 3.097 | 4.03E-05 | glio |
| Chek1_in.vivo_noRT | Hsd17b14 | 3.086 | 4.03E-05 | glio |
| Chek1_in.vivo_noRT | Gm10791 | 2.66 | 4.03E-05 | glio |
| Hsd17b10_in.vivo_noRT | Ankrd33b | 3.1 | 4.03E-05 | glio |
| culture_effect | Rfc3 | -0.793 | 4.03E-06 | glio |
| Interaction_genotype_Mtor | Gm48215 | -4.099 | 4.03E-06 | glio |
| Interaction_genotype_Myc | Cfap20 | 2.154 | 4.03E-06 | glio |
| Interaction_genotype_Myc | Tecrl | -5.455 | 4.03E-06 | glio |
| Aars2_ex.vivo_noRT | A330049N07R | 2.893 | 4.03E-06 | glio |
| Aars2_ex.vivo_noRT | Thnsl2 | 3.293 | 4.03E-06 | glio |
| Atr_in.vivo_noRT | Cib2 | 3.651 | 4.03E-06 | glio |
| Bcas2_in.vivo_noRT | Serac1 | 3.108 | 4.03E-06 | glio |
| Chek1_in.vivo_noRT | Zfp595 | 3.448 | 4.03E-06 | glio |
| Hsd17b10_in.vivo_noRT | Ccdc63 | 3.23 | 4.03E-06 | glio |
| Aars2_in.vivo_noRT | Taf9 | 3.436 | 4.03E-07 | glio |
| Chek1_ex.vivo_noRT | Cd83 | 3.862 | 4.03E-07 | glio |
| Chek1_ex.vivo_noRT | Dhrs9 | 4.443 | 4.03E-08 | glio |
| Atr_in.vivo_noRT | Gm27010 | 3.932 | 4.03E-09 | glio |
| Bora_in.vivo_noRT | Gm5544 | 4.145 | 4.03E-09 | glio |
| Atr_in.vivo_noRT | 1600014C23R | 5.154 | 4.03E-13 | glio |
| Aars2_in.vivo_noRT | Gm45606 | 5.067 | 4.03E-14 | glio |
| Bora_in.vivo_noRT | Flacc1 | 4.936 | 4.03E-16 | glio |
| Atr_in.vivo_noRT | Vmn2r56 | 4.751 | 4.03E-18 | glio |
| Atr_in.vivo_noRT | Gjd4 | 5.533 | 4.03E-30 | glio |
| culture_effect | Htr2b | -1.457 | 4.04E-03 | glio |
| culture_effect | Sult2b1 | -1.163 | 4.04E-03 | glio |
| Interaction_genotype_Atr | Cox14 | 1.272 | 4.04E-03 | glio |
| Interaction_genotype_Bcas2 | Ccdc146 | -4.036 | 4.04E-03 | glio |
| Interaction_genotype_Mtor | Rcc1l | -2.741 | 4.04E-03 | glio |
| Atp5c1_in.vivo_noRT | Gm4841 | -2.403 | 4.04E-03 | glio |
| Atp5c1_ex.vivo_noRT | Gm15270 | 2.974 | 4.04E-03 | glio |
| Atr_in.vivo_noRT | Siae | 1.89 | 4.04E-03 | glio |
| Atr_in.vivo_noRT | Nsl1 | 1.681 | 4.04E-03 | glio |
| Bcas2_in.vivo_noRT | Iscu | -1.302 | 4.04E-03 | glio |
| Bcas2_in.vivo_noRT | Pcbp1 | -0.696 | 4.04E-03 | glio |
| Bora_in.vivo_noRT | Tra2a | -0.964 | 4.04E-03 | glio |
| Bora_in.vivo_noRT | Ppig | -0.823 | 4.04E-03 | glio |
| Cenpj_in.vivo_noRT | Adam5 | 1.758 | 4.04E-03 | glio |
| Cyp19a1_ex.vivo_noRT | Slc14a2 | 2.159 | 4.04E-03 | glio |
| Cyp19a1_ex.vivo_noRT | Gm4340 | 2.159 | 4.04E-03 | glio |
| Cyp19a1_ex.vivo_noRT | Gm16239 | 2.159 | 4.04E-03 | glio |
| Cyp19a1_ex.vivo_noRT | Pacrg | 2.941 | 4.04E-03 | glio |
| Hsd17b10_in.vivo_noRT | Gm30262 | 2.324 | 4.04E-03 | glio |
| Interaction_genotype_Mtor | Gm10138 | -3.505 | 4.04E-04 | glio |

| | | | | |
|---|---|---|---|---|
| Aars2_in.vivo_noRT | St18 | 1.964 | 4.04E-04 | glio |
| Aars2_in.vivo_noRT | Fzd5 | 2.432 | 4.04E-04 | glio |
| Aars2_in.vivo_noRT | Fkbp1a | -0.76 | 4.04E-04 | glio |
| Atr_in.vivo_noRT | Klhdc10 | -1.61 | 4.04E-04 | glio |
| Atr_in.vivo_noRT | Ano6 | -1.526 | 4.04E-04 | glio |
| Atr_ex.vivo_noRT | Taf1d | 1.142 | 4.04E-04 | glio |
| Bard1_ex.vivo_noRT | Gm45606 | 3.3 | 4.04E-04 | glio |
| Bard1_ex.vivo_noRT | Il7 | 3.19 | 4.04E-04 | glio |
| Bora_in.vivo_noRT | Nbeal2 | 2.647 | 4.04E-04 | glio |
| culture_effect | Gm20559 | 1.172 | 4.04E-05 | glio |
| Aars2_in.vivo_noRT | 1700001G11R | 2.516 | 4.04E-05 | glio |
| Bcas2_in.vivo_noRT | Glrp1 | 2.538 | 4.04E-05 | glio |
| Bcas2_ex.vivo_noRT | Rimklb | 2.742 | 4.04E-05 | glio |
| Chek1_in.vivo_noRT | Gm26981 | 2.869 | 4.04E-05 | glio |
| Chek1_in.vivo_noRT | Alox15 | 2.606 | 4.04E-05 | glio |
| Interaction_genotype_Mtor | 2410141K09R | -4.619 | 4.04E-06 | glio |
| Interaction_genotype_Myc | Tslp | -5.361 | 4.04E-06 | glio |
| Interaction_genotype_Myc | Lyz2 | -5.553 | 4.04E-06 | glio |
| Interaction_genotype_Myc | Prox2 | -5.648 | 4.04E-06 | glio |
| Interaction_genotype_Myc | 2310067P03R | -5.171 | 4.04E-06 | glio |
| Interaction_genotype_Myc | BC030343 | -5.34 | 4.04E-06 | glio |
| Interaction_genotype_Myc | Nap1l1 | 2.013 | 4.04E-06 | glio |
| Aars2_in.vivo_noRT | Soga1 | 3.35 | 4.04E-06 | glio |
| Bcas2_in.vivo_noRT | Gm10734 | 3.291 | 4.04E-06 | glio |
| Aars2_in.vivo_noRT | Trim71 | 3.504 | 4.04E-07 | glio |
| Chek1_ex.vivo_noRT | Spns3 | 4.443 | 4.04E-07 | glio |
| Bora_ex.vivo_noRT | Trim17 | 3.594 | 4.04E-08 | glio |
| Bora_in.vivo_noRT | Svopl | 3.755 | 4.04E-09 | glio |
| Aars2_in.vivo_noRT | Mkrn3 | 4.114 | 4.04E-10 | glio |
| Chek1_in.vivo_noRT | Flicr | 4.243 | 4.04E-10 | glio |
| Chek1_in.vivo_noRT | AC125149.2 | 3.755 | 4.04E-10 | glio |
| Atr_in.vivo_noRT | Esyt3 | 4.474 | 4.04E-11 | glio |
| Bcas2_in.vivo_noRT | Gm38393 | 4.547 | 4.04E-14 | glio |
| Bcas2_in.vivo_noRT | Rgs4 | 4.807 | 4.04E-20 | glio |
| culture_effect | Maob | -1.781 | 4.05E-03 | glio |
| culture_effect | Gm20635 | 1.623 | 4.05E-03 | glio |
| Interaction_genotype_Bcas2 | Efcab9 | -3.71 | 4.05E-03 | glio |
| Interaction_genotype_Myc | Erich3 | -2.74 | 4.05E-03 | glio |
| Interaction_genotype_Myc | Fdxr | -2.334 | 4.05E-03 | glio |
| Interaction_genotype_Myc | D330050I16Ri | -2.391 | 4.05E-03 | glio |
| Interaction_genotype_Myc | Hdac1 | 1.238 | 4.05E-03 | glio |
| interaction_RT | Gata3 | 2.966 | 4.05E-03 | glio |
| Aars2_in.vivo_noRT | A930004D18R | 2.064 | 4.05E-03 | glio |
| Aars2_in.vivo_noRT | Pld1 | 2.093 | 4.05E-03 | glio |
| Aars2_in.vivo_noRT | Tmem170 | 1.747 | 4.05E-03 | glio |
| Aars2_in.vivo_noRT | Eif3k | -0.575 | 4.05E-03 | glio |
| Abtb1_in.vivo_noRT | Shfl | 1.542 | 4.05E-03 | glio |
| Atr_ex.vivo_noRT | Ms4a6d | 2.376 | 4.05E-03 | glio |
| Atr_ex.vivo_noRT | Procr | 3.001 | 4.05E-03 | glio |
| Bora_ex.vivo_noRT | Esr1 | 2.808 | 4.05E-03 | glio |
| Bora_ex.vivo_noRT | Soga3 | -2.245 | 4.05E-03 | glio |
| Cenpj_ex.vivo_noRT | Gm15138 | 2.196 | 4.05E-03 | glio |

| | | | | |
|---|---|---|---|---|
| Chek1_ex.vivo_noRT | Gm26688 | 2.617 | 4.05E-03 | glio |
| Chek1_ex.vivo_noRT | Bckdha | 1.565 | 4.05E-03 | glio |
| Eif2s1_in.vivo_noRT | Slxl1 | 2.413 | 4.05E-03 | glio |
| Hsd17b10_in.vivo_noRT | Rgl1 | 2.014 | 4.05E-03 | glio |
| Interaction_genotype_Aars2 | Tmf1 | 2.201 | 4.05E-04 | glio |
| Interaction_genotype_Chek1 | Wdr45b | 2.321 | 4.05E-04 | glio |
| Interaction_genotype_Mtor | Vmn2r56 | -3.317 | 4.05E-04 | glio |
| Interaction_genotype_Myc | 2310030G06R | -3.792 | 4.05E-04 | glio |
| Atr_ex.vivo_noRT | Vamp8 | 2.764 | 4.05E-04 | glio |
| Bora_in.vivo_noRT | Pigl | 2.006 | 4.05E-04 | glio |
| Bora_ex.vivo_noRT | Ccdc63 | 3.283 | 4.05E-04 | glio |
| Interaction_genotype_Mtor | Zmynd15 | -4.797 | 4.05E-05 | glio |
| Interaction_genotype_Myc | Gm32585 | -4.881 | 4.05E-05 | glio |
| Atr_in.vivo_noRT | Irak3 | 3.018 | 4.05E-05 | glio |
| Bcas2_in.vivo_noRT | Arl4d | 2.959 | 4.05E-05 | glio |
| Chek1_in.vivo_noRT | Slc41a3 | 2.783 | 4.05E-05 | glio |
| Interaction_genotype_Mtor | Gm7102 | -5.735 | 4.05E-06 | glio |
| Interaction_genotype_Mtor | Gm14019 | -4.803 | 4.05E-06 | glio |
| interaction_RT | Selenop | 1.82 | 4.05E-06 | glio |
| Aars2_ex.vivo_noRT | Akap14 | 3.293 | 4.05E-06 | glio |
| Bora_in.vivo_noRT | Gm41396 | 3.444 | 4.05E-06 | glio |
| Bora_in.vivo_noRT | Ablim3 | 3.376 | 4.05E-06 | glio |
| Bora_in.vivo_noRT | 5330417C22R | 3.541 | 4.05E-07 | glio |
| Interaction_genotype_Mtor | 2310067P03R | -6.541 | 4.05E-08 | glio |
| Bcas2_in.vivo_noRT | Gm20633 | 4.016 | 4.05E-10 | glio |
| Bcas2_in.vivo_noRT | Gm16344 | 3.896 | 4.05E-11 | glio |
| Bora_in.vivo_noRT | Gm34885 | 4.676 | 4.05E-11 | glio |
| Bcas2_in.vivo_noRT | 4930505N22R | 5.065 | 4.05E-15 | glio |
| Bora_in.vivo_noRT | Gm49441 | 4.67 | 4.05E-15 | glio |
| Bora_in.vivo_noRT | Ccdc162 | 4.407 | 4.05E-15 | glio |
| culture_effect | Sri | -1.66 | 4.05E-22 | glio |
| culture_effect | Gca | -0.747 | 4.06E-03 | glio |
| Interaction_genotype_Atr | Kcnj13 | -3.273 | 4.06E-03 | glio |
| Interaction_genotype_Atr | Zfp972 | -3.803 | 4.06E-03 | glio |
| Interaction_genotype_Atr | Rasgrp2 | -3.328 | 4.06E-03 | glio |
| Interaction_genotype_Bora | Gm10300 | -3.921 | 4.06E-03 | glio |
| Interaction_genotype_Bora | Myo5c | -3.752 | 4.06E-03 | glio |
| Interaction_genotype_Bora | Sgo2a | 2.479 | 4.06E-03 | glio |
| Interaction_genotype_Bora | Trim44 | 1.753 | 4.06E-03 | glio |
| Interaction_genotype_Chek1 | Zranb1 | 2.197 | 4.06E-03 | glio |
| Interaction_genotype_Mtor | Pdlim4 | -3.567 | 4.06E-03 | glio |
| Interaction_genotype_Mtor | Pml | -2.288 | 4.06E-03 | glio |
| Interaction_genotype_Mtor | Nfic | 1.696 | 4.06E-03 | glio |
| Interaction_genotype_Mtor | Prelid1 | 1.237 | 4.06E-03 | glio |
| Interaction_genotype_Myc | Ric3 | -2.863 | 4.06E-03 | glio |
| Interaction_genotype_Myc | Gorasp1 | -2.85 | 4.06E-03 | glio |
| Interaction_genotype_Myc | Mus81 | -2.633 | 4.06E-03 | glio |
| Interaction_genotype_Myc | Dynlt1a | -2.672 | 4.06E-03 | glio |
| Interaction_genotype_Myc | Ccdc77 | -2.139 | 4.06E-03 | glio |
| Interaction_genotype_Pars2 | Ripply3 | -3.832 | 4.06E-03 | glio |
| Interaction_genotype_Pars2 | Gm28424 | -3.615 | 4.06E-03 | glio |
| Interaction_genotype_Pars2 | Vwce | -3.564 | 4.06E-03 | glio |

| | | | | |
|---|---|---|---|---|
| Interaction_genotype_Pars2 | 4930483J18Ri | -3.685 | 4.06E-03 | glio |
| Interaction_genotype_Pars2 | Gm48008 | -3.195 | 4.06E-03 | glio |
| Aars2_in.vivo_noRT | Phf10 | -1.298 | 4.06E-03 | glio |
| Atp5c1_in.vivo_noRT | Prtn3 | 2.413 | 4.06E-03 | glio |
| Atr_ex.vivo_noRT | Gm16094 | 2.577 | 4.06E-03 | glio |
| Atr_ex.vivo_noRT | Gm48832 | 2.769 | 4.06E-03 | glio |
| Bard1_ex.vivo_noRT | Rasl12 | 2.814 | 4.06E-03 | glio |
| Bcas2_in.vivo_noRT | Chek1 | -1.424 | 4.06E-03 | glio |
| Bcas2_in.vivo_noRT | Galnt1 | -1.252 | 4.06E-03 | glio |
| Bcas2_in.vivo_noRT | Hnrnpu | -0.568 | 4.06E-03 | glio |
| Bora_in.vivo_noRT | Ttc37 | -1.39 | 4.06E-03 | glio |
| Bora_ex.vivo_noRT | Gm13822 | 2.806 | 4.06E-03 | glio |
| Cenpj_ex.vivo_noRT | Fam189a1 | 2.591 | 4.06E-03 | glio |
| Chek1_in.vivo_noRT | Mmp14 | 2.17 | 4.06E-03 | glio |
| Chek1_in.vivo_noRT | Adamts1 | -1.59 | 4.06E-03 | glio |
| Eif2s1_ex.vivo_noRT | Pla1a | 3.014 | 4.06E-03 | glio |
| Interaction_genotype_Myc | Tesk2 | -3.781 | 4.06E-04 | glio |
| Interaction_genotype_Myc | Ier3ip1 | 1.724 | 4.06E-04 | glio |
| Interaction_genotype_Nat2 | Gm4858 | 4.373 | 4.06E-04 | glio |
| Interaction_genotype_Pars2 | Selenov | -4.039 | 4.06E-04 | glio |
| Aars2_in.vivo_noRT | Ccr2 | 2.532 | 4.06E-04 | glio |
| Atp5c1_in.vivo_noRT | Myo5c | 2.668 | 4.06E-04 | glio |
| Bora_in.vivo_noRT | Pgls | -0.834 | 4.06E-04 | glio |
| Cenpj_ex.vivo_noRT | Cd36 | 2.591 | 4.06E-04 | glio |
| Cenpj_ex.vivo_noRT | Gstt4 | 2.591 | 4.06E-04 | glio |
| Chek1_in.vivo_noRT | Fcgrt | 2.797 | 4.06E-04 | glio |
| Chek1_in.vivo_noRT | Gm50203 | 2.411 | 4.06E-04 | glio |
| Chek1_in.vivo_noRT | Gm28221 | 2.374 | 4.06E-04 | glio |
| Chek1_in.vivo_noRT | Gm43066 | 2.392 | 4.06E-04 | glio |
| Chek1_ex.vivo_noRT | Dck | -2.013 | 4.06E-04 | glio |
| culture_effect | Erf | -0.704 | 4.06E-05 | glio |
| Interaction_genotype_Myc | 1700102H20R | -4.597 | 4.06E-05 | glio |
| Interaction_genotype_Myc | Tnnt2 | -3.483 | 4.06E-05 | glio |
| Aars2_in.vivo_noRT | Sdhd | -1.388 | 4.06E-05 | glio |
| Atr_in.vivo_noRT | Slc25a53 | 3.096 | 4.06E-05 | glio |
| Atr_in.vivo_noRT | Platr16 | 3.083 | 4.06E-05 | glio |
| Bcas2_in.vivo_noRT | Gm49077 | 3.047 | 4.06E-05 | glio |
| culture_effect | Ctxn1 | 1.025 | 4.06E-06 | glio |
| Aars2_in.vivo_noRT | Xlr4c | 3.292 | 4.06E-06 | glio |
| Aars2_ex.vivo_noRT | Cd200r2 | 3.293 | 4.06E-07 | glio |
| Atr_in.vivo_noRT | 1010001B22R | 3.533 | 4.06E-07 | glio |
| Bcas2_in.vivo_noRT | Gm26542 | 3.429 | 4.06E-07 | glio |
| Bora_in.vivo_noRT | 4921523L03Ri | 3.769 | 4.06E-07 | glio |
| Atr_in.vivo_noRT | Gm50322 | 4.147 | 4.06E-08 | glio |
| Bora_in.vivo_noRT | Sdr16c5 | 3.619 | 4.06E-08 | glio |
| Chek1_in.vivo_noRT | Gm43387 | 3.868 | 4.06E-08 | glio |
| interaction_RT | Yipf4 | 1.136 | 4.06E-10 | glio |
| Bora_in.vivo_noRT | Tas1r1 | 3.909 | 4.06E-10 | glio |
| Chek1_in.vivo_noRT | Gm35290 | 4.306 | 4.06E-11 | glio |
| culture_effect | Pcolce | -2.133 | 4.06E-12 | glio |
| Atr_in.vivo_noRT | Cldn19 | 4.751 | 4.06E-12 | glio |
| Bcas2_in.vivo_noRT | Ncf1 | 4.154 | 4.06E-12 | glio |

| | | | | |
|---|---|---|---|---|
| Chek1_in.vivo_noRT | Gm9967 | 4.657 | 4.06E-12 | glio |
| Aars2_in.vivo_noRT | Postn | 5.202 | 4.06E-14 | glio |
| culture_effect | Dnajc25 | 0.474 | 4.07E-03 | glio |
| culture_effect | Gatad2a | 0.412 | 4.07E-03 | glio |
| Interaction_genotype_Aars2 | Myo5c | -3.527 | 4.07E-03 | glio |
| Interaction_genotype_Aars2 | Dync1h1 | 1.362 | 4.07E-03 | glio |
| Interaction_genotype_Atr | Gm38979 | -3.898 | 4.07E-03 | glio |
| Interaction_genotype_Bora | Cdca5 | 2.392 | 4.07E-03 | glio |
| Interaction_genotype_Myc | Glod5 | 3.828 | 4.07E-03 | glio |
| interaction_RT | Bcas3 | -0.989 | 4.07E-03 | glio |
| Aars2_in.vivo_noRT | Ago4 | 1.944 | 4.07E-03 | glio |
| Aars2_in.vivo_noRT | 9330020H09R | 1.949 | 4.07E-03 | glio |
| Atr_in.vivo_noRT | Zadh2 | 1.595 | 4.07E-03 | glio |
| Bcas2_in.vivo_noRT | Slc35d2 | 2.174 | 4.07E-03 | glio |
| Bcas2_ex.vivo_noRT | Clca3a1 | 2.742 | 4.07E-03 | glio |
| Bcas2_ex.vivo_noRT | Oit3 | 2.762 | 4.07E-03 | glio |
| Bora_in.vivo_noRT | Cpt1b | 2.055 | 4.07E-03 | glio |
| Hsd17b10_in.vivo_noRT | 1700066B19R | 2.171 | 4.07E-03 | glio |
| Interaction_genotype_Atr | Adam19 | -4.534 | 4.07E-04 | glio |
| Interaction_genotype_Atr | Gm20522 | -4.407 | 4.07E-04 | glio |
| Interaction_genotype_Atr | Hecw2 | -4.367 | 4.07E-04 | glio |
| Interaction_genotype_Atr | Trnp1 | -4.245 | 4.07E-04 | glio |
| Interaction_genotype_Atr | Pmp22 | -3.515 | 4.07E-04 | glio |
| Interaction_genotype_Cenpj | Man2c1os | -5.396 | 4.07E-04 | glio |
| Interaction_genotype_Mtor | Stmn4 | -6.259 | 4.07E-04 | glio |
| Interaction_genotype_Mtor | Cdon | -3.766 | 4.07E-04 | glio |
| Interaction_genotype_Mtor | 2810405F17Ri | -3.525 | 4.07E-04 | glio |
| Aars2_in.vivo_noRT | Aurkaip1 | -0.934 | 4.07E-04 | glio |
| Atr_ex.vivo_noRT | Gm6566 | 2.77 | 4.07E-04 | glio |
| Bcas2_in.vivo_noRT | 4632404H12R | 2.552 | 4.07E-04 | glio |
| Bora_in.vivo_noRT | Immp1l | -1.288 | 4.07E-04 | glio |
| Bora_ex.vivo_noRT | Itgbl1 | 2.806 | 4.07E-04 | glio |
| Cenpj_in.vivo_noRT | Idi2 | 2.022 | 4.07E-04 | glio |
| Cenpj_in.vivo_noRT | Vmn2r18 | 2.022 | 4.07E-04 | glio |
| Cenpj_in.vivo_noRT | Gm16178 | 2.022 | 4.07E-04 | glio |
| Cenpj_in.vivo_noRT | Gm364 | 2.022 | 4.07E-04 | glio |
| Chek1_in.vivo_noRT | Zfp114 | 2.57 | 4.07E-04 | glio |
| culture_effect | A430105J06Ri | 1.938 | 4.07E-05 | glio |
| Interaction_genotype_Mtor | Gm26982 | -4.93 | 4.07E-05 | glio |
| Interaction_genotype_Mtor | Gm15834 | -4.625 | 4.07E-05 | glio |
| Interaction_genotype_Mtor | 0610009E02Ri | -5.315 | 4.07E-05 | glio |
| Interaction_genotype_Mtor | Pmp22 | -3.729 | 4.07E-05 | glio |
| Interaction_genotype_Myc | Zfp949 | -4.123 | 4.07E-05 | glio |
| Interaction_genotype_Myc | Gpc1 | -3.525 | 4.07E-05 | glio |
| Interaction_genotype_Myc | Antxr1 | -4.208 | 4.07E-05 | glio |
| Interaction_genotype_Myc | Gm34590 | -4.668 | 4.07E-05 | glio |
| Interaction_genotype_Myc | Trmu | -3.546 | 4.07E-05 | glio |
| Atr_in.vivo_noRT | Fst | 2.631 | 4.07E-05 | glio |
| Atr_in.vivo_noRT | 4930471E19Ri | 2.95 | 4.07E-05 | glio |
| Atr_in.vivo_noRT | Cyb5rl | 2.372 | 4.07E-05 | glio |
| Bcas2_in.vivo_noRT | Gm45833 | 3.96 | 4.07E-05 | glio |
| Bora_in.vivo_noRT | Acad10 | 2.954 | 4.07E-05 | glio |

| | | | | |
|---|---|---|---|---|
| Hsd17b10_in.vivo_noRT | Stx1b | 2.713 | 4.07E-05 | glio |
| Interaction_genotype_Nat2 | Gm21319 | 4.409 | 4.07E-06 | glio |
| Aars2_ex.vivo_noRT | Trim17 | 2.893 | 4.07E-06 | glio |
| Interaction_genotype_Rad21 | Klrk1 | 5.164 | 4.07E-07 | glio |
| Atr_in.vivo_noRT | Gm13056 | 3.778 | 4.07E-07 | glio |
| Chek1_in.vivo_noRT | Gm13986 | 3.785 | 4.07E-07 | glio |
| Chek1_in.vivo_noRT | Gm45605 | 3.551 | 4.07E-07 | glio |
| culture_effect | Klhl21 | 0.998 | 4.07E-09 | glio |
| Atr_in.vivo_noRT | 4922502N22R | 4.014 | 4.07E-09 | glio |
| Bora_in.vivo_noRT | Ephx1 | 4.038 | 4.07E-09 | glio |
| Aars2_in.vivo_noRT | 4921508D12R | 4.399 | 4.07E-12 | glio |
| Atr_in.vivo_noRT | Nrros | 4.891 | 4.07E-13 | glio |
| culture_effect | Cytl1 | -4.934 | 4.07E-14 | glio |
| culture_effect | Csta2 | -2.233 | 4.07E-17 | glio |
| Aars2_in.vivo_noRT | Fbxl13 | 5.067 | 4.07E-17 | glio |
| Aars2_in.vivo_noRT | Tmem54 | 5.067 | 4.07E-18 | glio |
| Aars2_in.vivo_noRT | Gm50024 | 5.063 | 4.07E-24 | glio |
| Bora_in.vivo_noRT | Gm5415 | 5.46 | 4.07E-27 | glio |
| Bora_in.vivo_noRT | Nrip2 | 5.46 | 4.07E-27 | glio |
| culture_effect | Dand5 | 0.766 | 4.08E-03 | glio |
| Interaction_genotype_Aars2 | Ccnj | -3.722 | 4.08E-03 | glio |
| Interaction_genotype_Bora | C330002G04R | -2.845 | 4.08E-03 | glio |
| Interaction_genotype_Bora | Gm26877 | -3.848 | 4.08E-03 | glio |
| Interaction_genotype_Mtor | Neurl1a | -3.689 | 4.08E-03 | glio |
| Interaction_genotype_Mtor | Efna5 | -3.284 | 4.08E-03 | glio |
| Interaction_genotype_Mtor | Pemt | -2.56 | 4.08E-03 | glio |
| Interaction_genotype_Mtor | Rdh5 | -2.954 | 4.08E-03 | glio |
| Interaction_genotype_Mtor | 2900093K20R | -2.476 | 4.08E-03 | glio |
| Interaction_genotype_Mtor | Polr3e | -2.149 | 4.08E-03 | glio |
| Interaction_genotype_Myc | Gpr19 | -2.141 | 4.08E-03 | glio |
| Interaction_genotype_Myc | Ndufs7 | 1.175 | 4.08E-03 | glio |
| Interaction_genotype_Pars2 | Nim1k | -4.162 | 4.08E-03 | glio |
| Interaction_genotype_Uqcrc2 | Spink2 | -4.856 | 4.08E-03 | glio |
| Interaction_genotype_Uqcrc2 | Slc25a35 | -4.242 | 4.08E-03 | glio |
| Interaction_genotype_Uqcrc2 | Apobr | -4.746 | 4.08E-03 | glio |
| Interaction_genotype_Uqcrc2 | Pdrg1 | 1.44 | 4.08E-03 | glio |
| interaction_RT | Klhl24 | 1.169 | 4.08E-03 | glio |
| Aars2_in.vivo_noRT | Ankrd52 | 1.499 | 4.08E-03 | glio |
| Aars2_in.vivo_noRT | Tmem109 | -1.145 | 4.08E-03 | glio |
| Atr_in.vivo_noRT | Tnpo1 | -1.137 | 4.08E-03 | glio |
| Atr_in.vivo_noRT | Snx21 | 1.488 | 4.08E-03 | glio |
| Atr_in.vivo_noRT | Cd9 | -0.754 | 4.08E-03 | glio |
| Bcas2_in.vivo_noRT | Gm39469 | 1.911 | 4.08E-03 | glio |
| Bora_in.vivo_noRT | Dip2a | 2.026 | 4.08E-03 | glio |
| Hsd17b10_in.vivo_noRT | Espl1 | -1.648 | 4.08E-03 | glio |
| Hsd17b10_in.vivo_noRT | Usp14 | -0.923 | 4.08E-03 | glio |
| Interaction_genotype_Mtor | Mmachc | -3.121 | 4.08E-04 | glio |
| Interaction_genotype_Myc | BC061195 | -2.729 | 4.08E-04 | glio |
| Interaction_genotype_Myc | Soga1 | -4.156 | 4.08E-04 | glio |
| interaction_RT | Tmsb15b2 | 3.596 | 4.08E-04 | glio |
| Atr_in.vivo_noRT | Cass4 | 2.525 | 4.08E-04 | glio |
| Bcas2_in.vivo_noRT | Gm15879 | 3.314 | 4.08E-04 | glio |

| | | | | |
|---|---|---|---|---|
| Bcas2_in.vivo_noRT | Cdca7 | -1.559 | 4.08E-04 | glio |
| Cenpj_ex.vivo_noRT | 9130019O22R | 2.591 | 4.08E-04 | glio |
| Cenpj_ex.vivo_noRT | Klk13 | 2.591 | 4.08E-04 | glio |
| Cyp19a1_ex.vivo_noRT | Smtnl1 | 2.959 | 4.08E-04 | glio |
| Fanca_ex.vivo_noRT | Acat3 | 3.133 | 4.08E-04 | glio |
| Hsd17b10_in.vivo_noRT | Cd82 | -1.897 | 4.08E-04 | glio |
| culture_effect | Depdc1b | -1.485 | 4.08E-05 | glio |
| Bcas2_in.vivo_noRT | Gm34121 | 2.681 | 4.08E-05 | glio |
| Bora_in.vivo_noRT | Gm28592 | 2.722 | 4.08E-05 | glio |
| Hsd17b10_in.vivo_noRT | Vsig10l | 2.963 | 4.08E-05 | glio |
| culture_effect | Adsl | -0.746 | 4.08E-06 | glio |
| culture_effect | Mcrip1 | -0.562 | 4.08E-06 | glio |
| Interaction_genotype_Mtor | Tnxb | -5.448 | 4.08E-06 | glio |
| Aars2_in.vivo_noRT | AW046200 | 3.269 | 4.08E-06 | glio |
| Atr_in.vivo_noRT | Cyb5d1 | 3.507 | 4.08E-07 | glio |
| Chek1_in.vivo_noRT | Gm43660 | 3.341 | 4.08E-07 | glio |
| Chek1_in.vivo_noRT | Pigk | -2.203 | 4.08E-07 | glio |
| Hsd17b10_in.vivo_noRT | Pnliprp2 | 2.837 | 4.08E-07 | glio |
| Hsd17b10_in.vivo_noRT | 4930483J18Ri | 3.356 | 4.08E-07 | glio |
| Atr_in.vivo_noRT | Smc2os | 4.137 | 4.08E-09 | glio |
| Bcas2_in.vivo_noRT | Flicr | 4.007 | 4.08E-09 | glio |
| Bora_in.vivo_noRT | Gm26682 | 4.018 | 4.08E-09 | glio |
| Aars2_in.vivo_noRT | Slc36a2 | 4.208 | 4.08E-10 | glio |
| Bcas2_in.vivo_noRT | 2610027K06R | 4.151 | 4.08E-10 | glio |
| Bcas2_in.vivo_noRT | Ms4a15 | 3.637 | 4.08E-11 | glio |
| Chek1_ex.vivo_noRT | Ubd | 5.25 | 4.08E-11 | glio |
| Atr_in.vivo_noRT | Sox11 | 4.887 | 4.08E-13 | glio |
| culture_effect | 4930478M13F | 3.91 | 4.08E-14 | glio |
| Aars2_in.vivo_noRT | Htr2b | 4.804 | 4.08E-18 | glio |
| Bora_in.vivo_noRT | Gm26671 | 4.936 | 4.08E-23 | glio |
| Chek1_in.vivo_noRT | Gm21936 | 5.044 | 4.08E-25 | glio |
| culture_effect | Dock11 | 1.373 | 4.09E-03 | glio |
| culture_effect | Slc25a23 | 0.809 | 4.09E-03 | glio |
| Interaction_genotype_Aars2 | Sulf2 | 1.81 | 4.09E-03 | glio |
| Interaction_genotype_Aars2 | Mrps11 | 1.32 | 4.09E-03 | glio |
| Interaction_genotype_Atr | Slc16a5 | -2.499 | 4.09E-03 | glio |
| Interaction_genotype_Bora | 1700028J19Ri | -3.795 | 4.09E-03 | glio |
| Interaction_genotype_Mtor | Zbtb3 | -3.405 | 4.09E-03 | glio |
| Interaction_genotype_Mtor | Gm26579 | -3.18 | 4.09E-03 | glio |
| Interaction_genotype_Mtor | Stn1 | -2.796 | 4.09E-03 | glio |
| Interaction_genotype_Mtor | Stat3 | 1.587 | 4.09E-03 | glio |
| Interaction_genotype_Myc | Zgrf1 | -2.442 | 4.09E-03 | glio |
| Interaction_genotype_Uqcrc2 | Trpm3 | -4.315 | 4.09E-03 | glio |
| Aars2_in.vivo_noRT | Cct2 | -0.566 | 4.09E-03 | glio |
| Aars2_in.vivo_noRT | Rpl22 | -0.554 | 4.09E-03 | glio |
| Aars2_ex.vivo_noRT | Ifitm1 | 3.293 | 4.09E-03 | glio |
| Bard1_ex.vivo_noRT | Gm17276 | 2.419 | 4.09E-03 | glio |
| Bora_in.vivo_noRT | Tmprss5 | 2.1 | 4.09E-03 | glio |
| Bora_in.vivo_noRT | Camta1 | -1.355 | 4.09E-03 | glio |
| Bora_in.vivo_noRT | Tmx1 | -0.974 | 4.09E-03 | glio |
| Eif2s1_in.vivo_noRT | Ptprc | 1.879 | 4.09E-03 | glio |
| Eif2s1_in.vivo_noRT | Gm26789 | 2.005 | 4.09E-03 | glio |

| | | | | |
|---|---|---|---|---|
| culture_effect | Artn | 1.047 | 4.09E-04 | glio |
| Interaction_genotype_Atr | Gm15520 | -3.937 | 4.09E-04 | glio |
| Interaction_genotype_Bcas2 | Gm34256 | -4.918 | 4.09E-04 | glio |
| interaction_RT | Steap1 | 1.341 | 4.09E-04 | glio |
| interaction_RT | Grb2 | 0.724 | 4.09E-04 | glio |
| Bcas2_in.vivo_noRT | Slc25a47 | 2.574 | 4.09E-04 | glio |
| Bora_in.vivo_noRT | Mirg | 2.594 | 4.09E-04 | glio |
| Bora_in.vivo_noRT | Cbx5 | -0.771 | 4.09E-04 | glio |
| Chek1_in.vivo_noRT | Pxt1 | 2.571 | 4.09E-04 | glio |
| Eif2s1_in.vivo_noRT | Fabp6 | 2.143 | 4.09E-04 | glio |
| culture_effect | 4732440D04R | 1.645 | 4.09E-05 | glio |
| culture_effect | Pkmyt1 | -1.238 | 4.09E-05 | glio |
| culture_effect | Chst11 | 0.731 | 4.09E-05 | glio |
| Atr_in.vivo_noRT | Dixdc1 | 2.9 | 4.09E-05 | glio |
| Bora_in.vivo_noRT | Spink4 | 2.724 | 4.09E-05 | glio |
| Interaction_genotype_Myc | Adamts7 | -4.905 | 4.09E-06 | glio |
| Atp5c1_in.vivo_noRT | Gm50024 | 2.405 | 4.09E-06 | glio |
| Bora_ex.vivo_noRT | Gm47936 | 3.201 | 4.09E-06 | glio |
| culture_effect | Uchl1 | -0.968 | 4.09E-07 | glio |
| Bora_in.vivo_noRT | 1700101I11Ri | 3.82 | 4.09E-07 | glio |
| culture_effect | Ppdpf | 0.784 | 4.09E-08 | glio |
| Atr_ex.vivo_noRT | Scg2 | 3.164 | 4.09E-08 | glio |
| Bora_in.vivo_noRT | 2900042K21R | 3.751 | 4.09E-08 | glio |
| Bora_in.vivo_noRT | Gpr162 | 3.888 | 4.09E-08 | glio |
| Bora_in.vivo_noRT | Tpsab1 | 3.519 | 4.09E-08 | glio |
| Bora_in.vivo_noRT | Gm43331 | 3.248 | 4.09E-08 | glio |
| Chek1_in.vivo_noRT | Tmem30c | 3.857 | 4.09E-08 | glio |
| Chek1_in.vivo_noRT | Car10 | 3.23 | 4.09E-08 | glio |
| culture_effect | Maf | 1.826 | 4.09E-09 | glio |
| Bora_in.vivo_noRT | Gm16853 | 3.903 | 4.09E-09 | glio |
| Chek1_ex.vivo_noRT | Ceacam16 | 4.842 | 4.09E-11 | glio |
| Atr_in.vivo_noRT | Gm8013 | 4.888 | 4.09E-12 | glio |
| culture_effect | Tmem106c | 1.281 | 4.09E-13 | glio |
| Aars2_in.vivo_noRT | Gm47204 | 5.191 | 4.09E-13 | glio |
| Chek1_ex.vivo_noRT | Gm4166 | 4.842 | 4.09E-14 | glio |
| culture_effect | Lyrm9 | 1.714 | 4.09E-15 | glio |
| Aars2_in.vivo_noRT | Kcnd1 | 4.805 | 4.09E-15 | glio |
| Atr_in.vivo_noRT | Npy1r | 4.888 | 4.09E-19 | glio |
| culture_effect | Mcts1 | -0.439 | 4.10E-03 | glio |
| Interaction_genotype_Aars2 | Gm14419 | -2.859 | 4.10E-03 | glio |
| Interaction_genotype_Aars2 | Atp6v0c | 0.848 | 4.10E-03 | glio |
| Interaction_genotype_Chek1 | Invs | -3.064 | 4.10E-03 | glio |
| Interaction_genotype_Mtor | Gtf2ird2 | -3.173 | 4.10E-03 | glio |
| Interaction_genotype_Mtor | Srd5a1 | -2.566 | 4.10E-03 | glio |
| Interaction_genotype_Mtor | Tm7sf3 | 1.62 | 4.10E-03 | glio |
| Interaction_genotype_Myc | Ppp1r11 | 1.157 | 4.10E-03 | glio |
| interaction_RT | Gm42726 | -2.653 | 4.10E-03 | glio |
| interaction_RT | Cbx4 | 1.04 | 4.10E-03 | glio |
| Aars2_in.vivo_noRT | Zfp607b | 1.738 | 4.10E-03 | glio |
| Aars2_in.vivo_noRT | Tle3 | 1.537 | 4.10E-03 | glio |
| Aars2_in.vivo_noRT | Exoc1 | 1.265 | 4.10E-03 | glio |
| Bard1_ex.vivo_noRT | Bmp2 | 2.803 | 4.10E-03 | glio |

| | | | | |
|---|---|---|---|---|
| Bcas2_in.vivo_noRT | Pura | -0.899 | 4.10E-03 | glio |
| Bora_in.vivo_noRT | A330023F24Ri | 2.158 | 4.10E-03 | glio |
| Bora_in.vivo_noRT | Gm50223 | 1.545 | 4.10E-03 | glio |
| Bora_ex.vivo_noRT | Zg16 | 2.802 | 4.10E-03 | glio |
| Chek1_ex.vivo_noRT | Gm16174 | 2.875 | 4.10E-03 | glio |
| Eif2s1_ex.vivo_noRT | Cd7 | 2.234 | 4.10E-03 | glio |
| Eif2s1_ex.vivo_noRT | Gm48764 | 2.234 | 4.10E-03 | glio |
| Interaction_genotype_Aars2 | Gm48610 | -4.464 | 4.10E-04 | glio |
| Interaction_genotype_Atr | Gm36198 | -3.665 | 4.10E-04 | glio |
| Interaction_genotype_Mtor | Gm17259 | -3.556 | 4.10E-04 | glio |
| Atp5c1_in.vivo_noRT | Fam25c | 2.136 | 4.10E-04 | glio |
| Atr_in.vivo_noRT | Vav2 | 2.497 | 4.10E-04 | glio |
| Atr_in.vivo_noRT | Alyref2 | 2.097 | 4.10E-04 | glio |
| Atr_in.vivo_noRT | Rpl27a | -0.731 | 4.10E-04 | glio |
| Bcas2_in.vivo_noRT | Gm26670 | 2.386 | 4.10E-04 | glio |
| Chek1_in.vivo_noRT | Meak7 | 2.55 | 4.10E-04 | glio |
| culture_effect | 4921524J17Ri | 0.617 | 4.10E-05 | glio |
| culture_effect | Selenof | 0.481 | 4.10E-05 | glio |
| Interaction_genotype_Mtor | Cnn1 | -5.988 | 4.10E-05 | glio |
| Interaction_genotype_Mtor | Zfp810 | -4.72 | 4.10E-05 | glio |
| Interaction_genotype_Myc | Gm26621 | -4.694 | 4.10E-05 | glio |
| culture_effect | Basp1 | 2.085 | 4.10E-06 | glio |
| Interaction_genotype_Atr | Junb | 3.37 | 4.10E-06 | glio |
| Interaction_genotype_Atr | Sod3 | -6.023 | 4.10E-06 | glio |
| Interaction_genotype_Myc | Jph4 | -5.265 | 4.10E-06 | glio |
| Atr_in.vivo_noRT | C8g | 3.259 | 4.10E-06 | glio |
| Hsd17b10_in.vivo_noRT | Man2c1os | 3.498 | 4.10E-06 | glio |
| Atr_in.vivo_noRT | Usp27x | 3.73 | 4.10E-07 | glio |
| Chek1_in.vivo_noRT | Retn | 4.576 | 4.10E-07 | glio |
| Hsd17b10_in.vivo_noRT | Cyp2c23 | 3.093 | 4.10E-07 | glio |
| Aars2_ex.vivo_noRT | Gm13191 | 3.293 | 4.10E-08 | glio |
| Aars2_ex.vivo_noRT | 4933416E03Ri | 3.293 | 4.10E-08 | glio |
| Chek1_ex.vivo_noRT | Gm32250 | 3.469 | 4.10E-08 | glio |
| culture_effect | Gnai2 | 0.798 | 4.10E-09 | glio |
| Hsd17b10_in.vivo_noRT | Gm29670 | 3.094 | 4.10E-09 | glio |
| Hsd17b10_in.vivo_noRT | Bco2 | 3.616 | 4.10E-09 | glio |
| culture_effect | Gm42067 | 2.181 | 4.10E-14 | glio |
| Chek1_in.vivo_noRT | Gm45151 | 4.512 | 4.10E-16 | glio |
| Aars2_in.vivo_noRT | Kcnk13 | 5.063 | 4.10E-20 | glio |
| culture_effect | Atp1a2 | -7.624 | 4.10E-33 | glio |
| culture_effect | St8sia1 | 1.782 | 4.11E-03 | glio |
| Interaction_genotype_Atr | AI182371 | -2.96 | 4.11E-03 | glio |
| Interaction_genotype_Atr | A930019D19R | -2.369 | 4.11E-03 | glio |
| Interaction_genotype_Atr | 1700066B19R | -3.348 | 4.11E-03 | glio |
| Interaction_genotype_Bora | Hells | 2.186 | 4.11E-03 | glio |
| Interaction_genotype_Mtor | Atg4c | -2.658 | 4.11E-03 | glio |
| Interaction_genotype_Mtor | Oasl2 | 1.719 | 4.11E-03 | glio |
| Interaction_genotype_Mtor | Grina | 1.322 | 4.11E-03 | glio |
| Interaction_genotype_Myc | Cxcl13 | -3.122 | 4.11E-03 | glio |
| Interaction_genotype_Myc | Zfp972 | -3.374 | 4.11E-03 | glio |
| Interaction_genotype_Myc | Sema4d | -2.824 | 4.11E-03 | glio |
| Interaction_genotype_Myc | Ccnf | -2.458 | 4.11E-03 | glio |

| | | | | |
|---|---|---|---|---|
| interaction_RT | Itfg2 | -1.063 | 4.11E-03 | glio |
| interaction_RT | Msrb1 | 0.898 | 4.11E-03 | glio |
| interaction_RT | Rrp1b | -0.838 | 4.11E-03 | glio |
| Atp5c1_in.vivo_noRT | Platr31 | 2.668 | 4.11E-03 | glio |
| Atr_in.vivo_noRT | Ufc1 | -0.772 | 4.11E-03 | glio |
| Atr_ex.vivo_noRT | Cers3 | 2.192 | 4.11E-03 | glio |
| Atr_ex.vivo_noRT | Gm4841 | 3.164 | 4.11E-03 | glio |
| Bard1_ex.vivo_noRT | AC125149.3 | 2.809 | 4.11E-03 | glio |
| Bcas2_ex.vivo_noRT | Myt1 | 2.342 | 4.11E-03 | glio |
| Bcas2_ex.vivo_noRT | Emg1 | -0.762 | 4.11E-03 | glio |
| Brca2_in.vivo_noRT | Boc | 2.024 | 4.11E-03 | glio |
| Brca2_in.vivo_noRT | Gm13191 | 1.763 | 4.11E-03 | glio |
| Brca2_in.vivo_noRT | 4933416E03Ri | 1.763 | 4.11E-03 | glio |
| Cenpj_ex.vivo_noRT | AF067061 | 2.591 | 4.11E-03 | glio |
| Chek1_ex.vivo_noRT | Urah | 2.316 | 4.11E-03 | glio |
| culture_effect | Gm49479 | 1.874 | 4.11E-04 | glio |
| culture_effect | Sms | 0.538 | 4.11E-04 | glio |
| Interaction_genotype_Mtor | Plxnb1 | -4.092 | 4.11E-04 | glio |
| Aars2_ex.vivo_noRT | Sox11 | 3.293 | 4.11E-04 | glio |
| Atp5c1_in.vivo_noRT | Bcl2l15 | 2.408 | 4.11E-04 | glio |
| Atr_in.vivo_noRT | Phtf1 | 1.731 | 4.11E-04 | glio |
| Bcas2_ex.vivo_noRT | Gm14929 | 2.742 | 4.11E-05 | glio |
| Bora_in.vivo_noRT | C2cd2 | 3.027 | 4.11E-05 | glio |
| Bora_in.vivo_noRT | Casc4 | -1.745 | 4.11E-05 | glio |
| Chek1_in.vivo_noRT | Gm4117 | 2.481 | 4.11E-05 | glio |
| Chek1_in.vivo_noRT | Uba3 | -1.551 | 4.11E-05 | glio |
| Bcas2_in.vivo_noRT | Sptbn4 | 3.259 | 4.11E-06 | glio |
| Chek1_ex.vivo_noRT | Gm20682 | 3.362 | 4.11E-06 | glio |
| Chek1_in.vivo_noRT | Kctd21 | 3.483 | 4.11E-07 | glio |
| Hsd17b10_in.vivo_noRT | Gm4340 | 3.616 | 4.11E-14 | glio |
| Hsd17b10_in.vivo_noRT | Gm16239 | 3.616 | 4.11E-14 | glio |
| Atr_in.vivo_noRT | Ttll13 | 4.8 | 4.11E-15 | glio |
| culture_effect | Ddost | -1.084 | 4.11E-20 | glio |
| Atr_in.vivo_noRT | Gm14742 | 5.533 | 4.11E-29 | glio |
| culture_effect | Fbxl12os | 1.413 | 4.12E-03 | glio |
| Interaction_genotype_Bcas2 | Gjb4 | -3.934 | 4.12E-03 | glio |
| Interaction_genotype_Bcas2 | 1810041H14R | -4.084 | 4.12E-03 | glio |
| Interaction_genotype_Myc | Fbxo46 | -2.865 | 4.12E-03 | glio |
| Interaction_genotype_Myc | Rragb | -2.63 | 4.12E-03 | glio |
| Interaction_genotype_Myc | Cmtm6 | -2.689 | 4.12E-03 | glio |
| Interaction_genotype_Rad51 | Egr2 | 3.998 | 4.12E-03 | glio |
| Aars2_in.vivo_noRT | Paqr6 | 2.052 | 4.12E-03 | glio |
| Aars2_in.vivo_noRT | Ccdc12 | -1.127 | 4.12E-03 | glio |
| Atp5c1_in.vivo_noRT | Hunk | 2.136 | 4.12E-03 | glio |
| Atr_ex.vivo_noRT | Catsperd | 2.784 | 4.12E-03 | glio |
| Bcas2_in.vivo_noRT | Zfp971 | 1.942 | 4.12E-03 | glio |
| Cenpj_in.vivo_noRT | Mycl | 2.56 | 4.12E-03 | glio |
| Cenpj_in.vivo_noRT | Slc2a9 | 2.022 | 4.12E-03 | glio |
| Cenpj_in.vivo_noRT | Gm43672 | 2.548 | 4.12E-03 | glio |
| Chek1_in.vivo_noRT | 1700112D23R | 2.08 | 4.12E-03 | glio |
| Chek1_in.vivo_noRT | Bco1 | 1.765 | 4.12E-03 | glio |
| Chek1_in.vivo_noRT | Gm45184 | 2.099 | 4.12E-03 | glio |

| | | | | |
|---|---|---|---|---|
| Chek1_ex.vivo_noRT | Tecrl | 2.658 | 4.12E-03 | glio |
| Hsd17b10_in.vivo_noRT | Usp27x | 2.354 | 4.12E-03 | glio |
| Interaction_genotype_Mtor | Rhox3g | -4.131 | 4.12E-04 | glio |
| Interaction_genotype_Mtor | S100a14 | -3.698 | 4.12E-04 | glio |
| Interaction_genotype_Mtor | Abra | -3.573 | 4.12E-04 | glio |
| Interaction_genotype_Mtor | Pnliprp2 | -3.32 | 4.12E-04 | glio |
| Aars2_in.vivo_noRT | Gm49123 | 2.667 | 4.12E-04 | glio |
| Aars2_in.vivo_noRT | 4833413G10R | 2.656 | 4.12E-04 | glio |
| Atr_ex.vivo_noRT | Bpifa1 | 3.164 | 4.12E-04 | glio |
| Bcas2_ex.vivo_noRT | Pspn | 2.347 | 4.12E-04 | glio |
| interaction_RT | 2610037D02R | -2.133 | 4.12E-05 | glio |
| Aars2_in.vivo_noRT | Cep112it | 2.883 | 4.12E-05 | glio |
| Atr_in.vivo_noRT | Ccdc8 | 2.819 | 4.12E-05 | glio |
| Atr_in.vivo_noRT | Ppp1r3e | 2.72 | 4.12E-05 | glio |
| Atr_ex.vivo_noRT | Otof | 3.164 | 4.12E-05 | glio |
| Bora_in.vivo_noRT | Catsper2 | 2.854 | 4.12E-05 | glio |
| culture_effect | BC039966 | 1.449 | 4.12E-06 | glio |
| Aars2_in.vivo_noRT | Gm19325 | 3.363 | 4.12E-06 | glio |
| Atr_in.vivo_noRT | Dock11 | 2.991 | 4.12E-06 | glio |
| Bora_in.vivo_noRT | Ppp1r18os | 3.524 | 4.12E-06 | glio |
| culture_effect | Mbnl1 | 0.765 | 4.12E-08 | glio |
| Chek1_ex.vivo_noRT | Smpd5 | 4.256 | 4.12E-08 | glio |
| Aars2_in.vivo_noRT | Tlr1 | 5.067 | 4.12E-13 | glio |
| Bora_in.vivo_noRT | Ceacam16 | 4.279 | 4.12E-13 | glio |
| Bcas2_in.vivo_noRT | 9230112J17Ri | 4.807 | 4.12E-16 | glio |
| culture_effect | Gjc3 | -1.151 | 4.12E-19 | glio |
| Bcas2_in.vivo_noRT | Dscaml1 | 5.325 | 4.12E-26 | glio |
| culture_effect | Mettl25 | -0.904 | 4.13E-03 | glio |
| culture_effect | Secisbp2l | 0.594 | 4.13E-03 | glio |
| culture_effect | Lockd | 0.644 | 4.13E-03 | glio |
| Interaction_genotype_Aars2 | 4930447F24Ri | -3.435 | 4.13E-03 | glio |
| Interaction_genotype_Aars2 | Gm15672 | -3.594 | 4.13E-03 | glio |
| Interaction_genotype_Aars2 | Krt18 | 4.952 | 4.13E-03 | glio |
| Interaction_genotype_Aars2 | Arsj | -3.697 | 4.13E-03 | glio |
| Interaction_genotype_Aars2 | Eef2 | 0.77 | 4.13E-03 | glio |
| Interaction_genotype_Atr | 4933421A08R | -3.797 | 4.13E-03 | glio |
| Interaction_genotype_Bcas2 | Stx1b | -3.524 | 4.13E-03 | glio |
| Interaction_genotype_Bcas2 | Rmst | -4.03 | 4.13E-03 | glio |
| Interaction_genotype_Mtor | Tmbim4 | 1.544 | 4.13E-03 | glio |
| Interaction_genotype_Myc | Ptp4a1.1 | -3.113 | 4.13E-03 | glio |
| Interaction_genotype_Myc | Dock3 | -2.806 | 4.13E-03 | glio |
| Interaction_genotype_Pars2 | Gm38534 | -3.596 | 4.13E-03 | glio |
| Interaction_genotype_Pars2 | Celsr3 | -3.688 | 4.13E-03 | glio |
| Interaction_genotype_Pars2 | Cldn19 | -3.768 | 4.13E-03 | glio |
| Interaction_genotype_Pars2 | Gm43672 | -3.938 | 4.13E-03 | glio |
| Interaction_genotype_Pars2 | Gm50138 | -3.304 | 4.13E-03 | glio |
| Interaction_genotype_Pars2 | A030014E15Ri | -2.662 | 4.13E-03 | glio |
| Interaction_genotype_Pars2 | D630036H23R | -3.695 | 4.13E-03 | glio |
| Interaction_genotype_Rad51 | Rnf165 | -3.915 | 4.13E-03 | glio |
| Interaction_genotype_Rad51 | Fam71f1 | -3.486 | 4.13E-03 | glio |
| Interaction_genotype_Rad51 | Taok1 | 1.826 | 4.13E-03 | glio |
| interaction_RT | Lztfl1 | 0.829 | 4.13E-03 | glio |

| | | | | |
|---|---|---|---|---|
| interaction_RT | Gcn1 | -0.748 | 4.13E-03 | glio |
| interaction_RT | Dars | -0.736 | 4.13E-03 | glio |
| Aars2_in.vivo_noRT | E530011L22Ri | 1.799 | 4.13E-03 | glio |
| Aars2_ex.vivo_noRT | Spns3 | 2.893 | 4.13E-03 | glio |
| Aars2_ex.vivo_noRT | mt-Nd4l | 1.197 | 4.13E-03 | glio |
| Atr_in.vivo_noRT | Tmem260 | 1.707 | 4.13E-03 | glio |
| Atr_in.vivo_noRT | Cluap1 | 1.594 | 4.13E-03 | glio |
| Bcas2_in.vivo_noRT | Arrdc2 | 2.299 | 4.13E-03 | glio |
| Bcas2_in.vivo_noRT | Ube2j2 | -0.837 | 4.13E-03 | glio |
| Bcas2_ex.vivo_noRT | Gm47622 | 2.167 | 4.13E-03 | glio |
| Bora_in.vivo_noRT | Pacs1 | 2.074 | 4.13E-03 | glio |
| Bora_in.vivo_noRT | Cdc6 | -1.827 | 4.13E-03 | glio |
| Bora_in.vivo_noRT | Tob2 | -1.257 | 4.13E-03 | glio |
| Bora_in.vivo_noRT | Moxd1 | -0.729 | 4.13E-03 | glio |
| Chek1_in.vivo_noRT | Galm | 2.04 | 4.13E-03 | glio |
| Chek1_in.vivo_noRT | Rps15a | -0.608 | 4.13E-03 | glio |
| Hsd17b10_in.vivo_noRT | Slc1a1 | -1.946 | 4.13E-03 | glio |
| culture_effect | Pop7 | 0.552 | 4.13E-04 | glio |
| Interaction_genotype_Aars2 | Wincr1 | -5.534 | 4.13E-04 | glio |
| Interaction_genotype_Aars2 | Irf8 | 4.177 | 4.13E-04 | glio |
| Interaction_genotype_Bcas2 | 1700017M07F | -4.01 | 4.13E-04 | glio |
| Interaction_genotype_Myc | Rpl29 | 1.155 | 4.13E-04 | glio |
| Aars2_in.vivo_noRT | 2410080I02Ri | 2.414 | 4.13E-04 | glio |
| Atp5c1_in.vivo_noRT | Pakap | 2.136 | 4.13E-04 | glio |
| Atr_in.vivo_noRT | Angpt2 | 2.294 | 4.13E-04 | glio |
| Cenpj_ex.vivo_noRT | Sytl4 | 2.591 | 4.13E-04 | glio |
| Cenpj_ex.vivo_noRT | Gm12800 | 2.591 | 4.13E-04 | glio |
| Cenpj_ex.vivo_noRT | Trmt61b | 2.591 | 4.13E-04 | glio |
| Cenpj_ex.vivo_noRT | Gm5890 | 2.591 | 4.13E-04 | glio |
| Cenpj_ex.vivo_noRT | Zscan4c | 2.591 | 4.13E-04 | glio |
| Cenpj_ex.vivo_noRT | Speer4a | 2.591 | 4.13E-04 | glio |
| Cenpj_ex.vivo_noRT | BC080695 | 2.591 | 4.13E-04 | glio |
| Interaction_genotype_Mtor | Dnajc16 | -4.337 | 4.13E-05 | glio |
| Interaction_genotype_Mtor | Isl1 | -4.885 | 4.13E-05 | glio |
| Interaction_genotype_Myc | Gm12940 | -4.119 | 4.13E-05 | glio |
| Interaction_genotype_Myc | Esrp2 | -4.396 | 4.13E-05 | glio |
| interaction_RT | Grcc10 | 0.729 | 4.13E-05 | glio |
| Atp5c1_in.vivo_noRT | 4930579C12R | 2.668 | 4.13E-05 | glio |
| Atr_in.vivo_noRT | Gm2415 | 2.975 | 4.13E-05 | glio |
| Chek1_in.vivo_noRT | Glt1d1 | 2.729 | 4.13E-05 | glio |
| Chek1_ex.vivo_noRT | B3galt4 | 3.672 | 4.13E-06 | glio |
| Chek1_ex.vivo_noRT | Polr1d | -0.946 | 4.13E-06 | glio |
| Chek1_in.vivo_noRT | Gm7008 | 3.578 | 4.13E-07 | glio |
| interaction_RT | Rps15 | -1.026 | 4.13E-08 | glio |
| Atr_in.vivo_noRT | Gm49135 | 3.851 | 4.13E-08 | glio |
| Hsd17b10_in.vivo_noRT | Gm11583 | 3.09 | 4.13E-09 | glio |
| culture_effect | Lmf1 | -1.676 | 4.13E-10 | glio |
| Bora_in.vivo_noRT | 2010320O07R | 4.286 | 4.13E-10 | glio |
| Bora_in.vivo_noRT | Platr20 | 4.672 | 4.13E-10 | glio |
| Bora_in.vivo_noRT | Gm6566 | 4.011 | 4.13E-12 | glio |
| Hsd17b10_in.vivo_noRT | Gm5788 | 3.616 | 4.13E-13 | glio |
| Bcas2_in.vivo_noRT | Nat8f2 | 4.547 | 4.13E-15 | glio |

| | | | | |
|---|---|---|---|---|
| Aars2_in.vivo_noRT | Gm45669 | 5.063 | 4.13E-17 | glio |
| Bora_in.vivo_noRT | Gm31717 | 4.936 | 4.13E-24 | glio |
| Bora_in.vivo_noRT | Cd200r2 | 5.197 | 4.13E-24 | glio |
| culture_effect | Cspg5 | -5.598 | 4.13E-39 | glio |
| culture_effect | Garnl3 | -1.22 | 4.14E-03 | glio |
| Interaction_genotype_Aars2 | Cpeb1 | -3.717 | 4.14E-03 | glio |
| Interaction_genotype_Atr | Zfp575 | -3.824 | 4.14E-03 | glio |
| Interaction_genotype_Bora | 3110082J24Ri | -3.925 | 4.14E-03 | glio |
| Interaction_genotype_Bora | Adam19 | -3.923 | 4.14E-03 | glio |
| Interaction_genotype_Bora | Rgs16 | -3.952 | 4.14E-03 | glio |
| Interaction_genotype_Mtor | Nupl2 | -2.563 | 4.14E-03 | glio |
| Interaction_genotype_Mtor | Rnmt | 1.537 | 4.14E-03 | glio |
| Interaction_genotype_Myc | Slc41a2 | -2.553 | 4.14E-03 | glio |
| Interaction_genotype_Myc | Cog2 | -2.624 | 4.14E-03 | glio |
| Interaction_genotype_Myc | Cad | -2.15 | 4.14E-03 | glio |
| Interaction_genotype_Myc | Paip1 | -1.669 | 4.14E-03 | glio |
| Interaction_genotype_Myc | Plxnd1 | -2.31 | 4.14E-03 | glio |
| Interaction_genotype_Myc | Josd2 | 1.405 | 4.14E-03 | glio |
| Interaction_genotype_Nat2 | Olfr787 | 2.678 | 4.14E-03 | glio |
| Interaction_genotype_Nat2 | Fst | 3.974 | 4.14E-03 | glio |
| interaction_RT | Pcnp | 0.685 | 4.14E-03 | glio |
| interaction_RT | Ndufb1-ps | 0.848 | 4.14E-03 | glio |
| Aars2_in.vivo_noRT | Lrrc28 | 1.799 | 4.14E-03 | glio |
| Atr_in.vivo_noRT | Tmem53 | 2.21 | 4.14E-03 | glio |
| Bora_in.vivo_noRT | 2810455O05R | 2.161 | 4.14E-03 | glio |
| Brca2_ex.vivo_noRT | Slc17a6 | 2.03 | 4.14E-03 | glio |
| Chek1_in.vivo_noRT | Med28 | -0.632 | 4.14E-03 | glio |
| Hsd17b10_in.vivo_noRT | Ucma | 2.617 | 4.14E-03 | glio |
| Hsd17b10_in.vivo_noRT | Emb | -1.602 | 4.14E-03 | glio |
| Interaction_genotype_Aars2 | Gm11457 | -4.158 | 4.14E-04 | glio |
| Interaction_genotype_Mtor | Arhgap33 | -4.418 | 4.14E-04 | glio |
| interaction_RT | Neurl3 | 3.401 | 4.14E-04 | glio |
| Atr_in.vivo_noRT | Zfp94 | 2.394 | 4.14E-04 | glio |
| Atr_ex.vivo_noRT | Acox2 | 2.98 | 4.14E-04 | glio |
| Cenpj_in.vivo_noRT | Ccl9 | 2.022 | 4.14E-04 | glio |
| Cenpj_ex.vivo_noRT | BB287469 | 2.591 | 4.14E-04 | glio |
| Chek1_in.vivo_noRT | Ston1 | 2.594 | 4.14E-04 | glio |
| Chek1_ex.vivo_noRT | Gm21814 | 3.033 | 4.14E-04 | glio |
| culture_effect | 2300002M23F | -2.103 | 4.14E-05 | glio |
| Aars2_ex.vivo_noRT | Etl4 | 3.293 | 4.14E-05 | glio |
| Aars2_ex.vivo_noRT | Dact1 | 3.693 | 4.14E-05 | glio |
| Atp5c1_in.vivo_noRT | AF067061 | 2.668 | 4.14E-05 | glio |
| Bard1_ex.vivo_noRT | Gm15533 | 2.814 | 4.14E-05 | glio |
| Bora_in.vivo_noRT | Myorg | 2.861 | 4.14E-05 | glio |
| Bora_in.vivo_noRT | Ccdc62 | 2.711 | 4.14E-05 | glio |
| Chek1_in.vivo_noRT | Nhlrc3 | 2.387 | 4.14E-05 | glio |
| Atr_in.vivo_noRT | Elf3 | 3.493 | 4.14E-06 | glio |
| Chek1_in.vivo_noRT | Taf9 | 3.153 | 4.14E-06 | glio |
| Interaction_genotype_Myc | 5330439K02R | -5.933 | 4.14E-07 | glio |
| Atr_in.vivo_noRT | Tox2 | 3.758 | 4.14E-08 | glio |
| Atr_in.vivo_noRT | Wdr38 | 3.962 | 4.14E-08 | glio |
| Hsd17b10_in.vivo_noRT | Acoxl | 3.616 | 4.14E-08 | glio |

| | | | | |
|---|---|---|---|---|
| culture_effect | Eef1d | -0.613 | 4.14E-09 | glio |
| Aars2_in.vivo_noRT | A330084C13R | 4.409 | 4.14E-09 | glio |
| Bora_in.vivo_noRT | 4933408J17Ri | 3.62 | 4.14E-09 | glio |
| Bora_in.vivo_noRT | Lst1 | 4.171 | 4.14E-09 | glio |
| Chek1_in.vivo_noRT | Inhba | 4.402 | 4.14E-09 | glio |
| Hsd17b10_in.vivo_noRT | Gm48008 | 3.352 | 4.14E-09 | glio |
| Atr_in.vivo_noRT | Sh2d3c | 4.632 | 4.14E-10 | glio |
| Atr_in.vivo_noRT | Gm43608 | 4.359 | 4.14E-10 | glio |
| Atr_in.vivo_noRT | Gm10484 | 4.532 | 4.14E-10 | glio |
| Atr_ex.vivo_noRT | Cfh | 4.347 | 4.14E-11 | glio |
| culture_effect | Wdr83os | 0.347 | 4.15E-03 | glio |
| Interaction_genotype_Abtb1 | Trmt61b | 3.412 | 4.15E-03 | glio |
| Interaction_genotype_Mtor | Nabp1 | -3.282 | 4.15E-03 | glio |
| Interaction_genotype_Myc | Nptxr | -2.5 | 4.15E-03 | glio |
| Interaction_genotype_Rad9a | Hdac9 | -4.488 | 4.15E-03 | glio |
| interaction_RT | Heatr1 | -1.023 | 4.15E-03 | glio |
| Aars2_ex.vivo_noRT | Speg | 2.843 | 4.15E-03 | glio |
| Atr_in.vivo_noRT | Exog | 1.717 | 4.15E-03 | glio |
| Atr_in.vivo_noRT | Eif5a | -0.591 | 4.15E-03 | glio |
| Bcas2_in.vivo_noRT | Rxylt1 | -0.933 | 4.15E-03 | glio |
| Bora_in.vivo_noRT | Ang | 2.161 | 4.15E-03 | glio |
| Cdkn2a_ex.vivo_noRT | Tnfrsf1b | 3.369 | 4.15E-03 | glio |
| Chek1_in.vivo_noRT | Gm43813 | 1.974 | 4.15E-03 | glio |
| Chek1_in.vivo_noRT | Gm11423 | 1.737 | 4.15E-03 | glio |
| Cyp19a1_ex.vivo_noRT | Hoxc8 | 2.159 | 4.15E-03 | glio |
| culture_effect | 1700026J14Ri | 1.855 | 4.15E-04 | glio |
| Interaction_genotype_Myc | Idi2 | -2.729 | 4.15E-04 | glio |
| Interaction_genotype_Myc | Luzp2 | -3.001 | 4.15E-04 | glio |
| Interaction_genotype_Myc | C430002N11R | -3.124 | 4.15E-04 | glio |
| Interaction_genotype_Myc | Upk3bl | -3.741 | 4.15E-04 | glio |
| Interaction_genotype_Myc | Cdon | -3.558 | 4.15E-04 | glio |
| Interaction_genotype_Pars2 | 2310067P03R | -4.71 | 4.15E-04 | glio |
| Atr_in.vivo_noRT | 9230114K14R | 2.114 | 4.15E-04 | glio |
| Interaction_genotype_Atr | Atoh8 | -4.294 | 4.15E-05 | glio |
| Interaction_genotype_Mtor | Gm5624 | -4.705 | 4.15E-05 | glio |
| Interaction_genotype_Mtor | Cltb | 2.392 | 4.15E-05 | glio |
| Interaction_genotype_Mtor | Rpl15 | 1.524 | 4.15E-05 | glio |
| Interaction_genotype_Myc | Wdr38 | -4.405 | 4.15E-05 | glio |
| interaction_RT | Gpr17 | 3.596 | 4.15E-05 | glio |
| Aars2_in.vivo_noRT | Gm41760 | 2.872 | 4.15E-05 | glio |
| Chek1_in.vivo_noRT | Gata3 | 3.172 | 4.15E-05 | glio |
| Chek1_in.vivo_noRT | Slc4a5 | 3.073 | 4.15E-05 | glio |
| Chek1_ex.vivo_noRT | 4930417H01R | 3.656 | 4.15E-05 | glio |
| Aars2_ex.vivo_noRT | Gm21319 | 3.293 | 4.15E-06 | glio |
| Atr_in.vivo_noRT | Nat8f1 | 3.068 | 4.15E-06 | glio |
| Atr_in.vivo_noRT | Inca1 | 2.931 | 4.15E-06 | glio |
| Atr_in.vivo_noRT | Zfp689 | 2.583 | 4.15E-06 | glio |
| Aars2_in.vivo_noRT | 4930481B07R | 3.763 | 4.15E-07 | glio |
| Atr_in.vivo_noRT | Lrrn4cl | 3.797 | 4.15E-07 | glio |
| Bcas2_in.vivo_noRT | Gm10432 | 3.31 | 4.15E-07 | glio |
| Bcas2_in.vivo_noRT | Atoh8 | 3.424 | 4.15E-07 | glio |
| Bora_in.vivo_noRT | Npb | 3.769 | 4.15E-08 | glio |

| | | | | |
|---|---|---|---|---|
| Aars2_in.vivo_noRT | Bpi | 3.645 | 4.15E-09 | glio |
| culture_effect | Phldb2 | 3.177 | 4.15E-10 | glio |
| Aars2_in.vivo_noRT | Mro | 4.535 | 4.15E-11 | glio |
| Aars2_in.vivo_noRT | D030051J21Ri | 4.424 | 4.15E-11 | glio |
| Aars2_in.vivo_noRT | Rspo1 | 4.535 | 4.15E-12 | glio |
| Aars2_in.vivo_noRT | Pard3bos1 | 4.33 | 4.15E-12 | glio |
| interaction_RT | H2-K1 | 2.034 | 4.15E-14 | glio |
| Bcas2_in.vivo_noRT | Gm49172 | 4.547 | 4.15E-22 | glio |
| culture_effect | Psma5 | -1.493 | 4.15E-24 | glio |
| Atr_in.vivo_noRT | Gm5890 | 5.533 | 4.15E-26 | glio |
| culture_effect | Coro1b | -1.601 | 4.16E-03 | glio |
| Interaction_genotype_Bora | Zfp764 | -3.527 | 4.16E-03 | glio |
| Interaction_genotype_Chek1 | Gnl2 | -2.038 | 4.16E-03 | glio |
| Interaction_genotype_Chek1 | Srebf2 | 2.204 | 4.16E-03 | glio |
| Interaction_genotype_Pars2 | Gngt2 | -3.937 | 4.16E-03 | glio |
| Interaction_genotype_Rad51 | 2010320O07R | -3.858 | 4.16E-03 | glio |
| Interaction_genotype_Rpl3 | Hexim1 | 2.191 | 4.16E-03 | glio |
| Interaction_genotype_Uqcrc2 | Gm49086 | -3.739 | 4.16E-03 | glio |
| Interaction_genotype_Uqcrc2 | BC028471 | -4.331 | 4.16E-03 | glio |
| Interaction_genotype_Uqcrc2 | G630030J09Ri | -4.073 | 4.16E-03 | glio |
| Interaction_genotype_Uqcrc2 | Rarres1 | -4.218 | 4.16E-03 | glio |
| Interaction_genotype_Uqcrc2 | Rab25 | -3.855 | 4.16E-03 | glio |
| Interaction_genotype_Uqcrc2 | Bphl | 2.245 | 4.16E-03 | glio |
| Aars2_in.vivo_noRT | Maml1 | 1.551 | 4.16E-03 | glio |
| Atp5c1_in.vivo_noRT | Gm14002 | 2.146 | 4.16E-03 | glio |
| Bard1_ex.vivo_noRT | Insm1 | 2.855 | 4.16E-03 | glio |
| Bcas2_in.vivo_noRT | Tmem117 | 2.179 | 4.16E-03 | glio |
| Bcas2_in.vivo_noRT | Glt1d1 | 1.994 | 4.16E-03 | glio |
| Chek1_in.vivo_noRT | Prkar1b | 2.167 | 4.16E-03 | glio |
| Chek1_in.vivo_noRT | Gm38246 | 2.086 | 4.16E-03 | glio |
| Chek1_in.vivo_noRT | Atxn1 | 1.841 | 4.16E-03 | glio |
| Eif2s1_ex.vivo_noRT | Gad1 | 2.234 | 4.16E-03 | glio |
| culture_effect | Plekho1 | 0.558 | 4.16E-04 | glio |
| Interaction_genotype_Mtor | Spn | -3.97 | 4.16E-04 | glio |
| Interaction_genotype_Mtor | Gm50255 | -5.144 | 4.16E-04 | glio |
| Interaction_genotype_Mtor | Akap17b | -3.885 | 4.16E-04 | glio |
| Interaction_genotype_Mtor | S100a3 | -3.989 | 4.16E-04 | glio |
| Aars2_in.vivo_noRT | Hist1h1d | 2.091 | 4.16E-04 | glio |
| Atp5c1_in.vivo_noRT | Grin3b | 2.676 | 4.16E-04 | glio |
| Atr_in.vivo_noRT | Kcnmb4 | 2.517 | 4.16E-04 | glio |
| Atr_in.vivo_noRT | Syngr1 | -1.786 | 4.16E-04 | glio |
| Bcas2_in.vivo_noRT | Zfp619 | 2.605 | 4.16E-04 | glio |
| Bcas2_ex.vivo_noRT | Snrpn | 2.742 | 4.16E-04 | glio |
| Bcas2_ex.vivo_noRT | Pnliprp2 | 2.742 | 4.16E-04 | glio |
| Bora_in.vivo_noRT | Gm16279 | 2.681 | 4.16E-04 | glio |
| Bora_in.vivo_noRT | Dll1 | 2.701 | 4.16E-04 | glio |
| Bora_ex.vivo_noRT | Insrr | 3.201 | 4.16E-04 | glio |
| Interaction_genotype_Myc | Smarcd3 | -4.129 | 4.16E-05 | glio |
| Interaction_genotype_Myc | Itgb6 | -4.708 | 4.16E-05 | glio |
| Aars2_in.vivo_noRT | Vapa | -1.252 | 4.16E-05 | glio |
| Atr_in.vivo_noRT | Rgl1 | 2.618 | 4.16E-05 | glio |
| Bcas2_in.vivo_noRT | Itpka | 2.863 | 4.16E-05 | glio |

| | | | | |
|---|---|---|---|---|
| Bora_in.vivo_noRT | Mlf2 | -1.047 | 4.16E-05 | glio |
| Hsd17b10_in.vivo_noRT | Rspo1 | 2.824 | 4.16E-05 | glio |
| Interaction_genotype_Mtor | Ywhag | 2.251 | 4.16E-06 | glio |
| Interaction_genotype_Myc | Spink2 | -5.931 | 4.16E-06 | glio |
| Interaction_genotype_Myc | D8Ertd738e | 1.558 | 4.16E-06 | glio |
| Aars2_in.vivo_noRT | Tcte2 | 3.204 | 4.16E-07 | glio |
| Bora_in.vivo_noRT | Gm47223 | 4.67 | 4.16E-07 | glio |
| interaction_RT | Zdhhc2 | -1.582 | 4.16E-08 | glio |
| Aars2_in.vivo_noRT | 1700084E18Ri | 4.085 | 4.16E-08 | glio |
| Atr_in.vivo_noRT | Gm20515 | 3.733 | 4.16E-08 | glio |
| Chek1_in.vivo_noRT | Speg | 4.19 | 4.16E-09 | glio |
| Aars2_in.vivo_noRT | Kcnk4 | 5.067 | 4.16E-13 | glio |
| Aars2_in.vivo_noRT | Rcan2 | 5.327 | 4.16E-13 | glio |
| Bora_in.vivo_noRT | 4930483J18Ri | 4.676 | 4.16E-13 | glio |
| Atr_in.vivo_noRT | CAAA0106612 | 4.741 | 4.16E-14 | glio |
| Aars2_in.vivo_noRT | 2010001K21R | 5.327 | 4.16E-17 | glio |
| culture_effect | Tcp11l2 | -3.35 | 4.16E-22 | glio |
| Atr_in.vivo_noRT | Olfr1152 | 5.267 | 4.16E-27 | glio |
| Interaction_genotype_Aars2 | Gm48140 | -2.564 | 4.17E-03 | glio |
| Interaction_genotype_Aars2 | Zfp786 | -3.495 | 4.17E-03 | glio |
| Interaction_genotype_Aars2 | Gm13830 | -3.84 | 4.17E-03 | glio |
| Interaction_genotype_Mtor | Ago4 | -3.358 | 4.17E-03 | glio |
| Interaction_genotype_Mtor | Setdb2 | -2.895 | 4.17E-03 | glio |
| Interaction_genotype_Myc | Gm21762 | -2.859 | 4.17E-03 | glio |
| Interaction_genotype_Myc | Timeless | -2.545 | 4.17E-03 | glio |
| Interaction_genotype_Myc | Sra1 | 1.376 | 4.17E-03 | glio |
| interaction_RT | BC049715 | -1.732 | 4.17E-03 | glio |
| interaction_RT | Bet1 | 0.599 | 4.17E-03 | glio |
| Aars2_in.vivo_noRT | Mettl18 | 1.778 | 4.17E-03 | glio |
| Atr_in.vivo_noRT | B4galt7 | 1.662 | 4.17E-03 | glio |
| Bard1_ex.vivo_noRT | Gm43915 | 2.421 | 4.17E-03 | glio |
| Bcas2_in.vivo_noRT | Zkscan3 | -1.267 | 4.17E-03 | glio |
| Bcas2_ex.vivo_noRT | Lrp2bp | 2.62 | 4.17E-03 | glio |
| Bcas2_ex.vivo_noRT | Eef1a1 | 0.692 | 4.17E-03 | glio |
| Bora_in.vivo_noRT | 2210011C24R | 2.046 | 4.17E-03 | glio |
| Chek1_in.vivo_noRT | Smc2 | -1.272 | 4.17E-03 | glio |
| Eif2s1_in.vivo_noRT | Kcnk13 | 1.88 | 4.17E-03 | glio |
| Hsd17b10_in.vivo_noRT | Psmd2 | -0.743 | 4.17E-03 | glio |
| Interaction_genotype_Mtor | Atp5g3 | 1.537 | 4.17E-04 | glio |
| Interaction_genotype_Myc | Vmn2r18 | -2.729 | 4.17E-04 | glio |
| Interaction_genotype_Myc | Igsf9b | -3.519 | 4.17E-04 | glio |
| Interaction_genotype_Myc | Nt5dc1 | -3.519 | 4.17E-04 | glio |
| Aars2_in.vivo_noRT | Fam135b | 2.867 | 4.17E-04 | glio |
| Aars2_in.vivo_noRT | Hemk1 | 2.562 | 4.17E-04 | glio |
| Aars2_ex.vivo_noRT | Alpl | 3.293 | 4.17E-04 | glio |
| Atr_in.vivo_noRT | 4632404H12R | 2.5 | 4.17E-04 | glio |
| Bcas2_in.vivo_noRT | Polr2f | -0.668 | 4.17E-04 | glio |
| Bora_in.vivo_noRT | Smim41 | 2.742 | 4.17E-04 | glio |
| Bora_in.vivo_noRT | Bloc1s6os | 2.417 | 4.17E-04 | glio |
| Cenpj_ex.vivo_noRT | H1foo | 2.591 | 4.17E-04 | glio |
| Chek1_in.vivo_noRT | Ifnb1 | 5.044 | 4.17E-04 | glio |
| Chek1_in.vivo_noRT | Lbx2 | 2.754 | 4.17E-04 | glio |

| | | | | |
|---|---|---|---|---|
| Chek1_in.vivo_noRT | Mapre1 | -1.163 | 4.17E-04 | glio |
| culture_effect | Rwdd4a | 0.536 | 4.17E-05 | glio |
| Bcas2_ex.vivo_noRT | Gm35439 | 3.125 | 4.17E-05 | glio |
| Bcas2_ex.vivo_noRT | Meioc | 2.742 | 4.17E-05 | glio |
| Bcas2_ex.vivo_noRT | Tmem132c | 2.742 | 4.17E-05 | glio |
| Bcas2_ex.vivo_noRT | Gm44693 | 2.742 | 4.17E-05 | glio |
| Aars2_in.vivo_noRT | Tssk6 | 2.841 | 4.17E-06 | glio |
| Hsd17b10_in.vivo_noRT | Rab37 | 3.229 | 4.17E-06 | glio |
| culture_effect | Gm10138 | 2.178 | 4.17E-07 | glio |
| culture_effect | Klhl7 | -1.188 | 4.17E-07 | glio |
| Interaction_genotype_Mtor | Gm13283 | -5.547 | 4.17E-07 | glio |
| Interaction_genotype_Myc | Wincr1 | -7.406 | 4.17E-07 | glio |
| Atr_in.vivo_noRT | Faim2 | 2.808 | 4.17E-07 | glio |
| Chek1_ex.vivo_noRT | Calhm6 | 4.263 | 4.17E-08 | glio |
| culture_effect | Gm14636 | 3.436 | 4.17E-09 | glio |
| Aars2_in.vivo_noRT | Nid1 | 4.522 | 4.17E-10 | glio |
| Aars2_in.vivo_noRT | Ntf5 | 4.568 | 4.17E-10 | glio |
| Chek1_in.vivo_noRT | Rims3 | 3.91 | 4.17E-10 | glio |
| Chek1_in.vivo_noRT | Gm49579 | 3.991 | 4.17E-11 | glio |
| culture_effect | Cd3g | -4.928 | 4.17E-16 | glio |
| Aars2_in.vivo_noRT | Cxcl5 | 5.327 | 4.17E-23 | glio |
| Bora_in.vivo_noRT | Fam227b | 4.936 | 4.17E-25 | glio |
| Bora_in.vivo_noRT | Gm20658 | 4.936 | 4.17E-25 | glio |
| culture_effect | Mpp2 | 1.399 | 4.18E-03 | glio |
| culture_effect | Sema4d | 1.115 | 4.18E-03 | glio |
| culture_effect | Cul2 | -0.469 | 4.18E-03 | glio |
| Interaction_genotype_Mtor | Ppp1r3f | -3.556 | 4.18E-03 | glio |
| Interaction_genotype_Mtor | Vmac | -3.653 | 4.18E-03 | glio |
| Interaction_genotype_Mtor | Rps8 | 0.916 | 4.18E-03 | glio |
| Interaction_genotype_Myc | Slc25a12 | -2.045 | 4.18E-03 | glio |
| Interaction_genotype_Myc | Cnih4 | 1.171 | 4.18E-03 | glio |
| Aars2_in.vivo_noRT | Edil3 | 1.727 | 4.18E-03 | glio |
| Atp5c1_in.vivo_noRT | St8sia1 | 2.408 | 4.18E-03 | glio |
| Atp5c1_in.vivo_noRT | AV356131 | 2.328 | 4.18E-03 | glio |
| Atr_in.vivo_noRT | Rnd3 | 1.626 | 4.18E-03 | glio |
| Bcas2_in.vivo_noRT | Aoc2 | 1.855 | 4.18E-03 | glio |
| Chek1_in.vivo_noRT | Haus8 | -1.357 | 4.18E-03 | glio |
| Interaction_genotype_Atr | Ube2d2a | 1.356 | 4.18E-04 | glio |
| Interaction_genotype_Mtor | Dagla | -4.586 | 4.18E-04 | glio |
| Interaction_genotype_Mtor | Golgb1 | 1.931 | 4.18E-04 | glio |
| Atp5c1_in.vivo_noRT | Tmem236 | 2.402 | 4.18E-04 | glio |
| Atr_in.vivo_noRT | Aox1 | 2.307 | 4.18E-04 | glio |
| Atr_in.vivo_noRT | Fam53b | 2.127 | 4.18E-04 | glio |
| Hsd17b10_in.vivo_noRT | Kctd5 | -1.571 | 4.18E-04 | glio |
| culture_effect | Pin1 | 0.468 | 4.18E-05 | glio |
| Interaction_genotype_Myc | Zfp114 | -4.691 | 4.18E-05 | glio |
| Aars2_in.vivo_noRT | Zfp583 | 3.034 | 4.18E-05 | glio |
| Atr_in.vivo_noRT | Lamtor3 | -1.602 | 4.18E-05 | glio |
| Atr_in.vivo_noRT | Fundc2 | -1.042 | 4.18E-05 | glio |
| Interaction_genotype_Myc | Tdrd5 | -4.87 | 4.18E-06 | glio |
| Atr_in.vivo_noRT | Gm10501 | 3.391 | 4.18E-06 | glio |
| Atr_in.vivo_noRT | Samd15 | 3.511 | 4.18E-06 | glio |

| | | | | |
|---|---|---|---|---|
| Atr_in.vivo_noRT | Mertk | 3.4 | 4.18E-07 | glio |
| Bora_in.vivo_noRT | Gm49042 | 3.307 | 4.18E-07 | glio |
| Aars2_in.vivo_noRT | Gm13986 | 4.068 | 4.18E-08 | glio |
| Aars2_in.vivo_noRT | Gm12500 | 3.685 | 4.18E-08 | glio |
| Chek1_in.vivo_noRT | Tnnt3 | 4.409 | 4.18E-09 | glio |
| Atr_in.vivo_noRT | Gm9970 | 4.888 | 4.18E-11 | glio |
| Atr_in.vivo_noRT | Tlr1 | 5.273 | 4.18E-14 | glio |
| Atr_in.vivo_noRT | BC018473 | 5.533 | 4.18E-16 | glio |
| culture_effect | Gm35867 | 1.588 | 4.19E-03 | glio |
| culture_effect | Zfp697 | 1.18 | 4.19E-03 | glio |
| Interaction_genotype_Atr | Prrc2c | 1.133 | 4.19E-03 | glio |
| Interaction_genotype_Bcas2 | Paqr8 | -4.422 | 4.19E-03 | glio |
| Interaction_genotype_Bcas2 | Zhx2 | -4.054 | 4.19E-03 | glio |
| Interaction_genotype_Bcas2 | Gm44096 | -3.66 | 4.19E-03 | glio |
| Interaction_genotype_Bcas2 | 2310067P03R | -3.709 | 4.19E-03 | glio |
| Interaction_genotype_Bcas2 | Gm20711 | -3.256 | 4.19E-03 | glio |
| Interaction_genotype_Bcas2 | Camk1d | -3.686 | 4.19E-03 | glio |
| Interaction_genotype_Bcas2 | Lrp4 | -3.95 | 4.19E-03 | glio |
| Interaction_genotype_Bcas2 | Loxl4 | -3.514 | 4.19E-03 | glio |
| Interaction_genotype_Bcas2 | C130060C02R | -3.421 | 4.19E-03 | glio |
| Interaction_genotype_Bcas2 | Gm5129 | -3.929 | 4.19E-03 | glio |
| Interaction_genotype_Bcas2 | Serp2 | -3.038 | 4.19E-03 | glio |
| Interaction_genotype_Bcas2 | H2-Eb1 | 3.371 | 4.19E-03 | glio |
| Interaction_genotype_Bcas2 | Terf1 | 1.855 | 4.19E-03 | glio |
| Interaction_genotype_Bcas2 | Prune1 | 1.798 | 4.19E-03 | glio |
| Interaction_genotype_Bcas2 | Cdc5l | 1.264 | 4.19E-03 | glio |
| Interaction_genotype_Bcas2 | Kras | 1.602 | 4.19E-03 | glio |
| Interaction_genotype_Cdkn2a | Ankrd22 | 2.835 | 4.19E-03 | glio |
| Interaction_genotype_Cdkn2a | Ccl3 | 3.095 | 4.19E-03 | glio |
| Interaction_genotype_Cdkn2a | Htr2b | 3.358 | 4.19E-03 | glio |
| Interaction_genotype_Cdkn2a | Gm47404 | 2.835 | 4.19E-03 | glio |
| Interaction_genotype_Cdkn2a | Gm31717 | 2.835 | 4.19E-03 | glio |
| Interaction_genotype_Cdkn2a | Spink8 | 3.095 | 4.19E-03 | glio |
| Interaction_genotype_Cdkn2a | Gm14742 | 2.835 | 4.19E-03 | glio |
| Interaction_genotype_Cdkn2a | Vtn | 3.095 | 4.19E-03 | glio |
| Interaction_genotype_Cenpj | Gm49694 | -4.951 | 4.19E-03 | glio |
| Interaction_genotype_Mtor | Gm35315 | -3.274 | 4.19E-03 | glio |
| Interaction_genotype_Mtor | Lypd6 | -3.358 | 4.19E-03 | glio |
| Aars2_in.vivo_noRT | Kctd12 | 1.992 | 4.19E-03 | glio |
| Aars2_in.vivo_noRT | Rnf157 | 2.01 | 4.19E-03 | glio |
| Aars2_in.vivo_noRT | Rassf7 | 1.712 | 4.19E-03 | glio |
| Aars2_in.vivo_noRT | Tbc1d32 | 1.55 | 4.19E-03 | glio |
| Aars2_in.vivo_noRT | Cubn | 1.616 | 4.19E-03 | glio |
| Aars2_in.vivo_noRT | Pelo | -1.372 | 4.19E-03 | glio |
| Aars2_in.vivo_noRT | Tns2 | 1.622 | 4.19E-03 | glio |
| Aars2_in.vivo_noRT | Rtraf | -0.744 | 4.19E-03 | glio |
| Atp5c1_in.vivo_noRT | Thap8 | 2.408 | 4.19E-03 | glio |
| Atr_ex.vivo_noRT | Gm15943 | 2.182 | 4.19E-03 | glio |
| Bcas2_in.vivo_noRT | Gm14167 | 1.9 | 4.19E-03 | glio |
| Bcas2_ex.vivo_noRT | Gm47101 | 2.78 | 4.19E-03 | glio |
| Bora_in.vivo_noRT | Pheta1 | 2.139 | 4.19E-03 | glio |
| Chek1_ex.vivo_noRT | Gm2381 | 2.502 | 4.19E-03 | glio |

| | | | | |
|---|---|---|---|---|
| culture_effect | Ccp110 | 0.912 | 4.19E-04 | glio |
| culture_effect | S100b | 0.679 | 4.19E-04 | glio |
| culture_effect | Rpl18 | -0.431 | 4.19E-04 | glio |
| Interaction_genotype_Atr | Rcan2 | -4.37 | 4.19E-04 | glio |
| Interaction_genotype_Mtor | Dixdc1 | -4.152 | 4.19E-04 | glio |
| Interaction_genotype_Mtor | Chtf8 | -3.771 | 4.19E-04 | glio |
| Interaction_genotype_Myc | Rps6ka6 | -3.716 | 4.19E-04 | glio |
| Aars2_in.vivo_noRT | Cpt1b | 2.445 | 4.19E-04 | glio |
| Aars2_in.vivo_noRT | Pcdhga6 | 2.351 | 4.19E-04 | glio |
| Atr_in.vivo_noRT | Gimap9 | 2.313 | 4.19E-04 | glio |
| Bora_in.vivo_noRT | Camk2n2 | 2.092 | 4.19E-04 | glio |
| Chek1_ex.vivo_noRT | Gm29093 | 3.226 | 4.19E-04 | glio |
| Chek1_ex.vivo_noRT | Sec14l5 | 2.903 | 4.19E-04 | glio |
| culture_effect | Rubcn | -0.837 | 4.19E-05 | glio |
| culture_effect | Dpysl2 | 0.734 | 4.19E-05 | glio |
| Bcas2_in.vivo_noRT | Afg1l | 2.981 | 4.19E-05 | glio |
| Atp5c1_in.vivo_noRT | Gm48617 | 2.404 | 4.19E-06 | glio |
| Atr_in.vivo_noRT | Mblac1 | 3.387 | 4.19E-06 | glio |
| Chek1_ex.vivo_noRT | Drd4 | 4.146 | 4.19E-06 | glio |
| culture_effect | Ccrl2 | -3.18 | 4.19E-07 | glio |
| culture_effect | 2610306M01F | 1.306 | 4.19E-07 | glio |
| Aars2_in.vivo_noRT | Emid1 | 3.934 | 4.19E-07 | glio |
| Bcas2_in.vivo_noRT | Arhgef15 | 3.426 | 4.19E-07 | glio |
| Bcas2_in.vivo_noRT | Serpinb6b | 3.693 | 4.19E-07 | glio |
| Chek1_in.vivo_noRT | Trnp1 | 3.602 | 4.19E-07 | glio |
| Chek1_in.vivo_noRT | Adgra3 | -2.189 | 4.19E-07 | glio |
| Dclre1c_ex.vivo_noRT | Gm40538 | 4.552 | 4.19E-10 | glio |
| culture_effect | Gbp2b | -3.956 | 4.19E-12 | glio |
| Atr_in.vivo_noRT | Kcnq2 | 4.37 | 4.19E-13 | glio |
| Aars2_in.vivo_noRT | 1700108F19Ri | 5.063 | 4.19E-19 | glio |
| Interaction_genotype_Aars2 | A030014E15Ri | -2.427 | 4.20E-03 | glio |
| Interaction_genotype_Aars2 | Cpne9 | -3.846 | 4.20E-03 | glio |
| Interaction_genotype_Aars2 | Gm3604 | -3.636 | 4.20E-03 | glio |
| Interaction_genotype_Cdkn2a | Zscan4-ps3 | 2.835 | 4.20E-03 | glio |
| Interaction_genotype_Cdkn2a | Gm21731 | 2.835 | 4.20E-03 | glio |
| Interaction_genotype_Cdkn2a | Gm11232 | 2.835 | 4.20E-03 | glio |
| Interaction_genotype_Cdkn2a | Snhg4.1 | 3.577 | 4.20E-03 | glio |
| Interaction_genotype_Mtor | Tmlhe | -3.141 | 4.20E-03 | glio |
| Interaction_genotype_Mtor | Rpl8 | 0.994 | 4.20E-03 | glio |
| Interaction_genotype_Myc | Gm12610 | -2.6 | 4.20E-03 | glio |
| Interaction_genotype_Rad51 | Asphd2 | -3.95 | 4.20E-03 | glio |
| Interaction_genotype_Rad51 | Gm12185 | 4.255 | 4.20E-03 | glio |
| interaction_RT | Ube2d2a | 0.521 | 4.20E-03 | glio |
| Bcas2_in.vivo_noRT | Gm37612 | 2.141 | 4.20E-03 | glio |
| Bcas2_ex.vivo_noRT | Gm10447 | 2.266 | 4.20E-03 | glio |
| Bora_ex.vivo_noRT | H2-T24 | 2.806 | 4.20E-03 | glio |
| Bora_ex.vivo_noRT | Olfr920 | 2.793 | 4.20E-03 | glio |
| Bora_ex.vivo_noRT | Cd3g | 3.011 | 4.20E-03 | glio |
| Cenpj_in.vivo_noRT | Pyy | 2.022 | 4.20E-03 | glio |
| Chek1_ex.vivo_noRT | Trpm3 | 2.59 | 4.20E-03 | glio |
| Hsd17b10_in.vivo_noRT | Selenok-ps1 | 2.052 | 4.20E-03 | glio |
| Interaction_genotype_Mtor | Gm30124 | -4.02 | 4.20E-04 | glio |

| | | | | |
|---|---|---|---|---|
| Interaction_genotype_Mtor | Slc23a3 | -4.168 | 4.20E-04 | glio |
| Interaction_genotype_Myc | Arl10 | -3.423 | 4.20E-04 | glio |
| Atr_in.vivo_noRT | Recql4 | 1.916 | 4.20E-04 | glio |
| Bora_in.vivo_noRT | Gm49077 | 2.648 | 4.20E-04 | glio |
| Bora_in.vivo_noRT | Eno2 | 2.282 | 4.20E-04 | glio |
| Chek1_in.vivo_noRT | 1700020L24Ri | 2.383 | 4.20E-04 | glio |
| Hsd17b10_in.vivo_noRT | Hist1h4f | 2.824 | 4.20E-04 | glio |
| Hsd17b10_in.vivo_noRT | 9430037G07R | 2.708 | 4.20E-04 | glio |
| Hsd17b10_in.vivo_noRT | Gm1123 | 2.709 | 4.20E-04 | glio |
| Interaction_genotype_Bora | Rce1 | 2.483 | 4.20E-05 | glio |
| Interaction_genotype_Chek1 | Kit | 4.661 | 4.20E-05 | glio |
| Bcas2_in.vivo_noRT | Gng8 | 2.832 | 4.20E-05 | glio |
| Chek1_in.vivo_noRT | Fhl3 | 2.937 | 4.20E-05 | glio |
| Chek1_ex.vivo_noRT | Gm41609 | 4.018 | 4.20E-05 | glio |
| Hsd17b10_in.vivo_noRT | Mir142hg | 3.074 | 4.20E-05 | glio |
| Bcas2_in.vivo_noRT | Ncaph2 | -1.561 | 4.20E-06 | glio |
| Chek1_in.vivo_noRT | Nyap1 | 3.235 | 4.20E-06 | glio |
| Hsd17b10_in.vivo_noRT | Tns4 | 2.966 | 4.20E-06 | glio |
| Atr_in.vivo_noRT | Gm36608 | 4.14 | 4.20E-09 | glio |
| Atr_ex.vivo_noRT | Gm20767 | 5.159 | 4.20E-10 | glio |
| Atr_in.vivo_noRT | Gm47101 | 4.742 | 4.20E-13 | glio |
| culture_effect | Naca | -0.875 | 4.20E-16 | glio |
| culture_effect | Mettl7b | 1.724 | 4.21E-03 | glio |
| culture_effect | C130060C02R | 1.564 | 4.21E-03 | glio |
| culture_effect | Gm41442 | 1.611 | 4.21E-03 | glio |
| culture_effect | Med31 | -0.46 | 4.21E-03 | glio |
| Interaction_genotype_Aars2 | 3110082J24Ri | -3.685 | 4.21E-03 | glio |
| Interaction_genotype_Aars2 | Ppia | 0.909 | 4.21E-03 | glio |
| Interaction_genotype_Atr | Aoc1 | -3.789 | 4.21E-03 | glio |
| Interaction_genotype_Atr | Klhdc1 | -3.493 | 4.21E-03 | glio |
| Interaction_genotype_Mtor | Gm49774 | -3.124 | 4.21E-03 | glio |
| Interaction_genotype_Myc | Gm14403 | -2.825 | 4.21E-03 | glio |
| Interaction_genotype_Myc | Wrap73 | -2.345 | 4.21E-03 | glio |
| Interaction_genotype_Myc | Usp34 | 1.187 | 4.21E-03 | glio |
| Aars2_in.vivo_noRT | Scara5 | 1.878 | 4.21E-03 | glio |
| Aars2_in.vivo_noRT | Myl6 | -0.575 | 4.21E-03 | glio |
| Aars2_in.vivo_noRT | Rplp0 | -0.579 | 4.21E-03 | glio |
| Aars2_ex.vivo_noRT | Hif3a | 2.899 | 4.21E-03 | glio |
| Atp5c1_ex.vivo_noRT | Gm48780 | 2.61 | 4.21E-03 | glio |
| Atr_in.vivo_noRT | Trpt1 | 1.609 | 4.21E-03 | glio |
| Bard1_ex.vivo_noRT | Gm5431 | 2.814 | 4.21E-03 | glio |
| Bard1_ex.vivo_noRT | Ppp1r16b | 2.994 | 4.21E-03 | glio |
| Bcas2_in.vivo_noRT | Stard3 | -1.272 | 4.21E-03 | glio |
| Bcas2_in.vivo_noRT | Acat1 | -1.258 | 4.21E-03 | glio |
| Bcas2_ex.vivo_noRT | Mogat1 | 3.114 | 4.21E-03 | glio |
| Bora_in.vivo_noRT | Ldlrad3 | 1.931 | 4.21E-03 | glio |
| Bora_in.vivo_noRT | Dtl | -1.526 | 4.21E-03 | glio |
| Chek1_ex.vivo_noRT | Gm42699 | 2.557 | 4.21E-03 | glio |
| Hsd17b10_in.vivo_noRT | Rasgrp2 | 2.315 | 4.21E-03 | glio |
| culture_effect | Sh3gl1 | -0.648 | 4.21E-04 | glio |
| culture_effect | Nono | -0.426 | 4.21E-04 | glio |
| Interaction_genotype_Cdkn2a | Ly6d | 4.208 | 4.21E-04 | glio |

| | | | | |
|---|---|---|---|---|
| Interaction_genotype_Mtor | Gm43948 | -3.408 | 4.21E-04 | glio |
| Interaction_genotype_Mtor | Prkar1b | -4.281 | 4.21E-04 | glio |
| Interaction_genotype_Myc | Gm16178 | -2.729 | 4.21E-04 | glio |
| Aars2_in.vivo_noRT | Car7 | 2.479 | 4.21E-04 | glio |
| Atr_in.vivo_noRT | Tro | 2.446 | 4.21E-04 | glio |
| Atr_in.vivo_noRT | Clec16a | 2.288 | 4.21E-04 | glio |
| Atr_in.vivo_noRT | Tecr | -1.224 | 4.21E-04 | glio |
| Atr_ex.vivo_noRT | Adgre5 | 2.77 | 4.21E-04 | glio |
| Atr_ex.vivo_noRT | Adam30 | 2.857 | 4.21E-04 | glio |
| Bcas2_in.vivo_noRT | Serpina3n | -1.571 | 4.21E-04 | glio |
| Bora_in.vivo_noRT | Gm10131 | 2.503 | 4.21E-04 | glio |
| Chek1_in.vivo_noRT | Gm42701 | 2.527 | 4.21E-04 | glio |
| Eif2s1_in.vivo_noRT | 4930512B01R | 2.143 | 4.21E-04 | glio |
| culture_effect | Pcdhgc3 | -1.383 | 4.21E-05 | glio |
| Interaction_genotype_Mtor | Gm38393 | -4.233 | 4.21E-05 | glio |
| Aars2_in.vivo_noRT | Med9os | 2.901 | 4.21E-05 | glio |
| Atr_in.vivo_noRT | Car9 | 2.784 | 4.21E-05 | glio |
| Bcas2_in.vivo_noRT | Gm26559 | 2.996 | 4.21E-05 | glio |
| Bcas2_in.vivo_noRT | Car10 | 2.464 | 4.21E-05 | glio |
| Chek1_in.vivo_noRT | 5033430I15Ri | 2.864 | 4.21E-05 | glio |
| Interaction_genotype_Mtor | Ak1 | -4.92 | 4.21E-06 | glio |
| Atr_in.vivo_noRT | Cnbd2 | 3.387 | 4.21E-06 | glio |
| Cenpj_ex.vivo_noRT | Gm10482 | 3.763 | 4.21E-06 | glio |
| culture_effect | Edn1 | 3.17 | 4.21E-07 | glio |
| culture_effect | Hmox1 | 1.581 | 4.21E-07 | glio |
| Atr_in.vivo_noRT | Gm14966 | 3.603 | 4.21E-08 | glio |
| Chek1_ex.vivo_noRT | Gm21818 | 4.842 | 4.21E-08 | glio |
| Chek1_ex.vivo_noRT | Gm20478 | 4.447 | 4.21E-08 | glio |
| Atr_in.vivo_noRT | Gm30262 | 4.241 | 4.21E-09 | glio |
| Bcas2_in.vivo_noRT | Gm20635 | 4.162 | 4.21E-10 | glio |
| Chek1_in.vivo_noRT | Rtl1 | 3.881 | 4.21E-10 | glio |
| culture_effect | Mrpl24 | -0.886 | 4.21E-12 | glio |
| Atr_in.vivo_noRT | Rtn1 | 4.887 | 4.21E-12 | glio |
| Bcas2_in.vivo_noRT | Gm20711 | 4.807 | 4.21E-16 | glio |
| culture_effect | Gm49312 | 3.804 | 4.21E-17 | glio |
| Atr_in.vivo_noRT | Zscan4c | 5.533 | 4.21E-26 | glio |
| culture_effect | 1700094D03R | -1.191 | 4.22E-03 | glio |
| culture_effect | Ggnbp2 | -0.355 | 4.22E-03 | glio |
| culture_effect | Fgfr1op2 | 0.326 | 4.22E-03 | glio |
| Interaction_genotype_Atr | Ppil6 | -3.246 | 4.22E-03 | glio |
| Interaction_genotype_Ifnar1 | Capn13 | 2.734 | 4.22E-03 | glio |
| Interaction_genotype_Mtor | Chmp2b | 1.631 | 4.22E-03 | glio |
| Interaction_genotype_Mtor | Cox4i1 | 1.038 | 4.22E-03 | glio |
| Interaction_genotype_Myc | Elovl3 | -3.245 | 4.22E-03 | glio |
| Interaction_genotype_Myc | Foxj2 | -2.439 | 4.22E-03 | glio |
| Interaction_genotype_Myc | Met | -2.539 | 4.22E-03 | glio |
| Interaction_genotype_Myc | Rps10 | 1.142 | 4.22E-03 | glio |
| interaction_RT | Hnrnpr | -0.606 | 4.22E-03 | glio |
| interaction_RT | Chchd10 | 1.301 | 4.22E-03 | glio |
| interaction_RT | Hnrnpc | -0.533 | 4.22E-03 | glio |
| Aars2_in.vivo_noRT | Neu1 | -1.349 | 4.22E-03 | glio |
| Aars2_in.vivo_noRT | Uqcr10 | -0.722 | 4.22E-03 | glio |

| | | | | |
|---|---|---|---|---|
| Atr_in.vivo_noRT | Snx5 | -0.879 | 4.22E-03 | glio |
| Atr_ex.vivo_noRT | Gm49441 | 2.376 | 4.22E-03 | glio |
| Bcas2_ex.vivo_noRT | Ctsf | 2.349 | 4.22E-03 | glio |
| Bcas2_ex.vivo_noRT | Gm19265 | 2.342 | 4.22E-03 | glio |
| Bora_in.vivo_noRT | 1700019L13Ri | 2.107 | 4.22E-03 | glio |
| Chek1_in.vivo_noRT | Pmp22 | 1.932 | 4.22E-03 | glio |
| Chek1_in.vivo_noRT | Zfp710 | 1.483 | 4.22E-03 | glio |
| culture_effect | B9d2 | -0.858 | 4.22E-04 | glio |
| Interaction_genotype_Aars2 | Thap8 | -4.506 | 4.22E-04 | glio |
| Interaction_genotype_Aars2 | Gm44702 | -4.523 | 4.22E-04 | glio |
| Interaction_genotype_Mtor | Gm21814 | -4.05 | 4.22E-04 | glio |
| Interaction_genotype_Mtor | Gm41349 | -3.836 | 4.22E-04 | glio |
| Interaction_genotype_Mtor | Mndal | 2.379 | 4.22E-04 | glio |
| Interaction_genotype_Myc | Gm364 | -2.729 | 4.22E-04 | glio |
| Interaction_genotype_Myc | Padi2 | -4.094 | 4.22E-04 | glio |
| Interaction_genotype_Rad51 | Gm36356 | -4.149 | 4.22E-04 | glio |
| Aars2_ex.vivo_noRT | Gm48161 | 2.9 | 4.22E-04 | glio |
| Atr_in.vivo_noRT | Bex2 | -1.55 | 4.22E-04 | glio |
| Bora_ex.vivo_noRT | Ank1 | 2.417 | 4.22E-04 | glio |
| Hsd17b10_in.vivo_noRT | Gm41442 | 2.575 | 4.22E-04 | glio |
| Interaction_genotype_Myc | D930028M14I | -4.216 | 4.22E-05 | glio |
| Aars2_in.vivo_noRT | A430105J06Ri | 2.818 | 4.22E-05 | glio |
| Atr_in.vivo_noRT | Slc39a8 | 3.17 | 4.22E-05 | glio |
| Bora_in.vivo_noRT | Card10 | 3.139 | 4.22E-05 | glio |
| Chek1_in.vivo_noRT | Tmf1 | -1.79 | 4.22E-05 | glio |
| Chek1_ex.vivo_noRT | Dtl | -2.569 | 4.22E-05 | glio |
| culture_effect | Fas | -3.072 | 4.22E-06 | glio |
| Aars2_ex.vivo_noRT | Gm39526 | 3.293 | 4.22E-06 | glio |
| Atr_in.vivo_noRT | Setmar | 3.218 | 4.22E-06 | glio |
| Atr_in.vivo_noRT | Fhl2 | 3.302 | 4.22E-07 | glio |
| Bora_in.vivo_noRT | Dchs1 | 3.442 | 4.22E-07 | glio |
| Aars2_in.vivo_noRT | Slc6a19 | 3.672 | 4.22E-08 | glio |
| Hsd17b10_in.vivo_noRT | Gm13713 | 3.084 | 4.22E-08 | glio |
| culture_effect | Uap1l1 | 1.19 | 4.22E-09 | glio |
| Bora_in.vivo_noRT | Oxt | 4.295 | 4.22E-10 | glio |
| Bcas2_in.vivo_noRT | Gm43463 | 4.149 | 4.22E-12 | glio |
| Atr_in.vivo_noRT | Kcnk4 | 5.273 | 4.22E-14 | glio |
| Bora_in.vivo_noRT | Gm4858 | 4.936 | 4.22E-14 | glio |
| Atr_in.vivo_noRT | Gm4724 | 4.877 | 4.22E-15 | glio |
| Chek1_in.vivo_noRT | Efcab9 | 5.289 | 4.22E-15 | glio |
| Aars2_in.vivo_noRT | Rab25 | 5.06 | 4.22E-16 | glio |
| Bcas2_in.vivo_noRT | Slc10a1 | 4.547 | 4.22E-16 | glio |
| Chek1_ex.vivo_noRT | Ablim2 | 4.842 | 4.22E-16 | glio |
| Chek1_ex.vivo_noRT | Gm5415 | 4.842 | 4.22E-16 | glio |
| culture_effect | 1700109H08R | -1.824 | 4.23E-03 | glio |
| Interaction_genotype_Mtor | Cfb | -2.795 | 4.23E-03 | glio |
| Interaction_genotype_Mtor | Arglu1 | 1.503 | 4.23E-03 | glio |
| Interaction_genotype_Myc | Acsf3 | -2.867 | 4.23E-03 | glio |
| Interaction_genotype_Myc | Nsl1 | -2.646 | 4.23E-03 | glio |
| Interaction_genotype_Myc | BC030867 | -2.491 | 4.23E-03 | glio |
| Interaction_genotype_Myc | Ehmt2 | -1.977 | 4.23E-03 | glio |
| Interaction_genotype_Uqcrc2 | Mecom | -3.915 | 4.23E-03 | glio |

| | | | | |
|---|---|---|---|---|
| Interaction_genotype_Uqcrc2 | Gm49312 | -3.467 | 4.23E-03 | glio |
| Interaction_genotype_Uqcrc2 | Pyy | -3.534 | 4.23E-03 | glio |
| Interaction_genotype_Uqcrc2 | Hbq1b | -4.011 | 4.23E-03 | glio |
| Interaction_genotype_Uqcrc2 | Gm20492 | -4.042 | 4.23E-03 | glio |
| Interaction_genotype_Uqcrc2 | Crhbp | -3.692 | 4.23E-03 | glio |
| Interaction_genotype_Uqcrc2 | Gm27253 | -4.172 | 4.23E-03 | glio |
| Interaction_genotype_Uqcrc2 | Adgrb3 | -4.034 | 4.23E-03 | glio |
| Interaction_genotype_Uqcrc2 | Pitx3 | -3.418 | 4.23E-03 | glio |
| interaction_RT | Ndfip2 | 0.84 | 4.23E-03 | glio |
| Aars2_in.vivo_noRT | Rab20 | 1.804 | 4.23E-03 | glio |
| Aars2_in.vivo_noRT | Dclre1a | 1.491 | 4.23E-03 | glio |
| Aars2_in.vivo_noRT | Ech1 | -1.074 | 4.23E-03 | glio |
| Atr_in.vivo_noRT | Tulp4 | -0.808 | 4.23E-03 | glio |
| Atr_ex.vivo_noRT | Krtcap3 | 2.178 | 4.23E-03 | glio |
| Atr_ex.vivo_noRT | Klc4 | 2.126 | 4.23E-03 | glio |
| Bcas2_in.vivo_noRT | Myo1b | 2.422 | 4.23E-03 | glio |
| Bcas2_in.vivo_noRT | Serpine2 | 2.534 | 4.23E-03 | glio |
| Bcas2_in.vivo_noRT | 3110001I22Ri | 2.067 | 4.23E-03 | glio |
| Bcas2_in.vivo_noRT | Gab2 | 1.967 | 4.23E-03 | glio |
| Bcas2_in.vivo_noRT | Sema7a | 1.974 | 4.23E-03 | glio |
| Bcas2_in.vivo_noRT | Ninj1 | -1.229 | 4.23E-03 | glio |
| Bcas2_in.vivo_noRT | Capza1 | -1.068 | 4.23E-03 | glio |
| Bcas2_in.vivo_noRT | Naa50 | -0.906 | 4.23E-03 | glio |
| Chek1_in.vivo_noRT | Plekhh1 | 2.166 | 4.23E-03 | glio |
| Chek1_in.vivo_noRT | Wnk2 | 2.128 | 4.23E-03 | glio |
| Chek1_in.vivo_noRT | Sec61b | -0.591 | 4.23E-03 | glio |
| culture_effect | Gm50295 | 1.983 | 4.23E-04 | glio |
| Interaction_genotype_Mtor | Hoxa4 | -3.802 | 4.23E-04 | glio |
| Interaction_genotype_Myc | E230016M11F | -3.629 | 4.23E-04 | glio |
| Interaction_genotype_Myc | Tiam2 | -3.561 | 4.23E-04 | glio |
| Interaction_genotype_Myc | Chic2 | 1.578 | 4.23E-04 | glio |
| Interaction_genotype_Myc | Nars | 1.579 | 4.23E-04 | glio |
| interaction_RT | Rcc2 | -0.718 | 4.23E-04 | glio |
| Aars2_in.vivo_noRT | Gm7879 | 2.674 | 4.23E-04 | glio |
| Aars2_in.vivo_noRT | Hist2h4 | 2.397 | 4.23E-04 | glio |
| Aars2_in.vivo_noRT | Cct8 | -0.854 | 4.23E-04 | glio |
| Aars2_in.vivo_noRT | Psma2 | -0.856 | 4.23E-04 | glio |
| Atp5c1_in.vivo_noRT | Col8a1 | 2.668 | 4.23E-04 | glio |
| Atr_in.vivo_noRT | Eif4b | -1.046 | 4.23E-04 | glio |
| Bora_in.vivo_noRT | Msi2 | -1.431 | 4.23E-04 | glio |
| Bora_in.vivo_noRT | Use1 | -1.123 | 4.23E-04 | glio |
| Chek1_in.vivo_noRT | D130062J10Ri | 2.316 | 4.23E-04 | glio |
| Interaction_genotype_Myc | Ino80 | -3.33 | 4.23E-05 | glio |
| Interaction_genotype_Myc | Hmgn1 | 1.629 | 4.23E-05 | glio |
| Aars2_in.vivo_noRT | Dnaaf4 | 3.145 | 4.23E-05 | glio |
| Bcas2_in.vivo_noRT | Myl9 | 3.244 | 4.23E-05 | glio |
| Bora_ex.vivo_noRT | 4930483J18Ri | 3.478 | 4.23E-05 | glio |
| Bora_ex.vivo_noRT | Arid3c | 3.608 | 4.23E-05 | glio |
| Chek1_in.vivo_noRT | Gpam | 2.683 | 4.23E-05 | glio |
| Chek1_in.vivo_noRT | Zranb1 | -1.784 | 4.23E-05 | glio |
| culture_effect | Myef2 | 0.753 | 4.23E-06 | glio |
| Aars2_in.vivo_noRT | Snd1 | -1.585 | 4.23E-06 | glio |

| | | | | |
|---|---|---|---|---|
| Atr_in.vivo_noRT | B230398E01Ri | 3.2 | 4.23E-06 | glio |
| Atr_in.vivo_noRT | Rps14 | -0.971 | 4.23E-06 | glio |
| Hsd17b10_in.vivo_noRT | Gm16364.1 | 3.616 | 4.23E-06 | glio |
| Interaction_genotype_Mtor | Gm45890 | -6.335 | 4.23E-07 | glio |
| Bcas2_in.vivo_noRT | A730071L15Ri | 3.747 | 4.23E-07 | glio |
| Bcas2_in.vivo_noRT | Fgf7 | 3.443 | 4.23E-07 | glio |
| Bcas2_in.vivo_noRT | Gm15080 | 3.796 | 4.23E-07 | glio |
| Interaction_genotype_Mtor | Ank1 | -4.883 | 4.23E-08 | glio |
| Bora_in.vivo_noRT | Cd59b | 3.891 | 4.23E-08 | glio |
| Chek1_in.vivo_noRT | C230013L11Ri | 4.132 | 4.23E-08 | glio |
| Atr_in.vivo_noRT | Gm16348 | 4.349 | 4.23E-10 | glio |
| Chek1_in.vivo_noRT | Ergic2 | -2.221 | 4.23E-10 | glio |
| Atr_in.vivo_noRT | Gm20455 | 4.813 | 4.23E-11 | glio |
| Bcas2_in.vivo_noRT | Gm11973 | 4.541 | 4.23E-16 | glio |
| culture_effect | Strn3 | 0.433 | 4.24E-03 | glio |
| culture_effect | Mafg | -0.39 | 4.24E-03 | glio |
| Interaction_genotype_Bcas2 | Ankrd1 | -4.366 | 4.24E-03 | glio |
| Interaction_genotype_Bcas2 | Gm14968 | -3.498 | 4.24E-03 | glio |
| Interaction_genotype_Bora | Gm49086 | -3.298 | 4.24E-03 | glio |
| Interaction_genotype_Bora | Tlr6 | -3.811 | 4.24E-03 | glio |
| Interaction_genotype_Bora | Commd4 | 1.814 | 4.24E-03 | glio |
| Interaction_genotype_Myc | Pex11g | -3.067 | 4.24E-03 | glio |
| Aars2_in.vivo_noRT | Zbtb25 | 1.691 | 4.24E-03 | glio |
| Atp5c1_in.vivo_noRT | Gm10300 | 2.408 | 4.24E-03 | glio |
| Atr_in.vivo_noRT | Gm38246 | 2.046 | 4.24E-03 | glio |
| Atr_in.vivo_noRT | Adamts10 | 1.858 | 4.24E-03 | glio |
| Bora_in.vivo_noRT | BC065397 | 2.345 | 4.24E-03 | glio |
| Chek1_in.vivo_noRT | Gsto2 | 2.132 | 4.24E-03 | glio |
| Chek1_in.vivo_noRT | Tmem53 | 2.244 | 4.24E-03 | glio |
| Chek1_in.vivo_noRT | Klrg2 | 2.002 | 4.24E-03 | glio |
| Chek1_ex.vivo_noRT | 4732419C18R | 2.419 | 4.24E-03 | glio |
| Fanca_ex.vivo_noRT | Gm13191 | 2.124 | 4.24E-03 | glio |
| Fanca_ex.vivo_noRT | 4933416E03Ri | 2.124 | 4.24E-03 | glio |
| Fanca_ex.vivo_noRT | Etl4 | 2.919 | 4.24E-03 | glio |
| culture_effect | Anp32a | -0.459 | 4.24E-04 | glio |
| Interaction_genotype_Myc | Lrrn3 | -3.762 | 4.24E-04 | glio |
| Bcas2_in.vivo_noRT | Gm17597 | 2.772 | 4.24E-04 | glio |
| Bcas2_in.vivo_noRT | Gm29666 | 2.573 | 4.24E-04 | glio |
| Bcas2_in.vivo_noRT | Gm43689 | 2.341 | 4.24E-04 | glio |
| Bora_in.vivo_noRT | Galc | 2.097 | 4.24E-04 | glio |
| culture_effect | Gm15894 | -1.98 | 4.24E-05 | glio |
| culture_effect | Adcy2 | -2.295 | 4.24E-05 | glio |
| Interaction_genotype_Chek1 | Plagl1 | 3.463 | 4.24E-05 | glio |
| Atr_in.vivo_noRT | Fgl2 | 2.64 | 4.24E-05 | glio |
| Bcas2_in.vivo_noRT | Itga2b | 2.882 | 4.24E-05 | glio |
| Bora_in.vivo_noRT | Wfikkn2 | 2.962 | 4.24E-05 | glio |
| Bora_in.vivo_noRT | Gm16835 | 3.009 | 4.24E-05 | glio |
| Bora_in.vivo_noRT | Rasl12 | 2.656 | 4.24E-05 | glio |
| culture_effect | Zfp428 | 0.781 | 4.24E-06 | glio |
| Aars2_in.vivo_noRT | Gm15938 | 3.397 | 4.24E-06 | glio |
| Aars2_ex.vivo_noRT | Speer4e | 3.293 | 4.24E-06 | glio |
| Bora_in.vivo_noRT | Gm16048 | 3.5 | 4.24E-06 | glio |

| | | | | |
|---|---|---|---|---|
| Bcas2_in.vivo_noRT | Serpinb6a | 3.648 | 4.24E-07 | glio |
| Bora_in.vivo_noRT | Gm49891 | 3.739 | 4.24E-07 | glio |
| culture_effect | Usp54 | -2.43 | 4.24E-08 | glio |
| culture_effect | Nsun3 | 2.324 | 4.24E-12 | glio |
| Chek1_in.vivo_noRT | Gm17173 | 4.657 | 4.24E-12 | glio |
| Aars2_in.vivo_noRT | 1700015F17Ri | 5.067 | 4.24E-13 | glio |
| Bora_in.vivo_noRT | BC018473 | 4.936 | 4.24E-13 | glio |
| culture_effect | D730003I15Ri | 2.499 | 4.24E-18 | glio |
| culture_effect | Bola3 | 1.188 | 4.24E-20 | glio |
| Chek1_in.vivo_noRT | Hfm1 | 5.044 | 4.24E-20 | glio |
| Chek1_in.vivo_noRT | D630024D03R | 5.044 | 4.24E-20 | glio |
| Bora_in.vivo_noRT | Gm14742 | 4.936 | 4.24E-24 | glio |
| Atr_in.vivo_noRT | Prap1 | 5.533 | 4.24E-25 | glio |
| culture_effect | Hist1h3b | 1.392 | 4.25E-03 | glio |
| culture_effect | Cpd | -0.492 | 4.25E-03 | glio |
| culture_effect | Sars | -0.37 | 4.25E-03 | glio |
| Interaction_genotype_Atr | Cpne8 | -3.625 | 4.25E-03 | glio |
| Interaction_genotype_Atr | Atpif1 | 0.942 | 4.25E-03 | glio |
| Interaction_genotype_Mtor | Usp11 | -3.714 | 4.25E-03 | glio |
| Interaction_genotype_Mtor | Rpl17 | 0.995 | 4.25E-03 | glio |
| Interaction_genotype_Myc | Rpp40 | -2.189 | 4.25E-03 | glio |
| interaction_RT | Gm19412 | 1.232 | 4.25E-03 | glio |
| interaction_RT | Ubxn8 | -1.14 | 4.25E-03 | glio |
| interaction_RT | Acot1 | 0.952 | 4.25E-03 | glio |
| Aars2_in.vivo_noRT | 1500004A13R | 2.101 | 4.25E-03 | glio |
| Atp5c1_in.vivo_noRT | AI182371 | 1.886 | 4.25E-03 | glio |
| Atp5c1_in.vivo_noRT | Gm42918 | 2.026 | 4.25E-03 | glio |
| Bard1_ex.vivo_noRT | Abcb5 | 2.02 | 4.25E-03 | glio |
| Bcas2_in.vivo_noRT | Zfp39 | 2.162 | 4.25E-03 | glio |
| Bcas2_in.vivo_noRT | Trp53bp2 | 1.564 | 4.25E-03 | glio |
| Bcas2_in.vivo_noRT | Stag1 | -1.161 | 4.25E-03 | glio |
| Bora_in.vivo_noRT | H60b | 2.038 | 4.25E-03 | glio |
| Chek1_in.vivo_noRT | Isoc2a | 1.901 | 4.25E-03 | glio |
| Hsd17b10_in.vivo_noRT | Shbg | 2.282 | 4.25E-03 | glio |
| Hsd17b10_in.vivo_noRT | Cep68 | -1.72 | 4.25E-03 | glio |
| Hsd17b10_in.vivo_noRT | Mef2b | 2.032 | 4.25E-03 | glio |
| culture_effect | Cul5 | 0.53 | 4.25E-04 | glio |
| Interaction_genotype_Bora | Trpm3 | -4.645 | 4.25E-04 | glio |
| Interaction_genotype_Bora | Cenpq | 1.972 | 4.25E-04 | glio |
| Interaction_genotype_Mtor | Emilin2 | -4.374 | 4.25E-04 | glio |
| Interaction_genotype_Mtor | Slurp1 | -6.238 | 4.25E-04 | glio |
| Interaction_genotype_Mtor | Gm36988 | -4.332 | 4.25E-04 | glio |
| Interaction_genotype_Myc | Mettl2 | -2.916 | 4.25E-04 | glio |
| Aars2_ex.vivo_noRT | Luzp2 | 2.495 | 4.25E-04 | glio |
| Bcas2_in.vivo_noRT | Dscc1 | -1.999 | 4.25E-04 | glio |
| Bora_in.vivo_noRT | Frmd4b | 2.552 | 4.25E-04 | glio |
| Bora_in.vivo_noRT | Rab26 | 2.368 | 4.25E-04 | glio |
| Chek1_in.vivo_noRT | Nptx1 | 2.78 | 4.25E-04 | glio |
| Chek1_in.vivo_noRT | Hist1h2br | 2.76 | 4.25E-04 | glio |
| Chek1_ex.vivo_noRT | 2900041M22R | 3.377 | 4.25E-04 | glio |
| Chek1_ex.vivo_noRT | Gm5861 | 3.757 | 4.25E-04 | glio |
| Interaction_genotype_Mtor | Acat3 | -3.883 | 4.25E-05 | glio |

| | | | | |
|---|---|---|---|---|
| Atr_in.vivo_noRT | Alox15 | 2.572 | 4.25E-05 | glio |
| Bcas2_in.vivo_noRT | Gm39556 | 2.897 | 4.25E-05 | glio |
| Hsd17b10_in.vivo_noRT | Gm47101 | 2.825 | 4.25E-05 | glio |
| culture_effect | Platr14 | -2.227 | 4.25E-06 | glio |
| Aars2_ex.vivo_noRT | Clec12b | 2.897 | 4.25E-06 | glio |
| Atr_in.vivo_noRT | Gm14636 | 4.245 | 4.25E-07 | glio |
| Bcas2_in.vivo_noRT | Gm37933 | 3.185 | 4.25E-07 | glio |
| Bora_in.vivo_noRT | Cdca7 | -2.21 | 4.25E-07 | glio |
| culture_effect | 5033406O09R | 1.061 | 4.25E-08 | glio |
| Aars2_in.vivo_noRT | Gng13 | 3.734 | 4.25E-08 | glio |
| Atr_in.vivo_noRT | Ppp1r18os | 4.121 | 4.25E-08 | glio |
| Atr_in.vivo_noRT | Gm26533 | 3.899 | 4.25E-08 | glio |
| Bora_in.vivo_noRT | Prm1 | 3.459 | 4.25E-08 | glio |
| Bcas2_in.vivo_noRT | Gjb4 | 4.15 | 4.25E-09 | glio |
| culture_effect | H13 | -0.845 | 4.25E-10 | glio |
| Bcas2_in.vivo_noRT | Gm13684 | 3.705 | 4.25E-10 | glio |
| Bora_in.vivo_noRT | 7530414M10F | 4.41 | 4.25E-11 | glio |
| Bora_in.vivo_noRT | G630030J09Ri | 4.414 | 4.25E-11 | glio |
| culture_effect | Spry1 | -1.368 | 4.25E-13 | glio |
| Bora_in.vivo_noRT | Gm44752 | 4.549 | 4.25E-13 | glio |
| Atr_in.vivo_noRT | Gm50115 | 5.27 | 4.25E-16 | glio |
| Aars2_in.vivo_noRT | Ppil6 | 5.327 | 4.25E-17 | glio |
| Aars2_in.vivo_noRT | Gm38393 | 5.067 | 4.25E-17 | glio |
| Atr_in.vivo_noRT | Cyp2c23 | 5.011 | 4.25E-17 | glio |
| Atr_in.vivo_noRT | Zfp352 | 5.533 | 4.25E-18 | glio |
| culture_effect | Ccdc9b | 3.701 | 4.25E-20 | glio |
| Aars2_in.vivo_noRT | Serpinb9g | 5.327 | 4.25E-25 | glio |
| Aars2_in.vivo_noRT | A930002C04R | 5.067 | 4.25E-25 | glio |
| Chek1_in.vivo_noRT | Gm34059 | 5.044 | 4.25E-26 | glio |
| Interaction_genotype_Aars2 | Gm43597 | -3.247 | 4.26E-03 | glio |
| Interaction_genotype_Aars2 | Gchfr | -3.876 | 4.26E-03 | glio |
| Interaction_genotype_Aars2 | Fbxo10 | -3.556 | 4.26E-03 | glio |
| Interaction_genotype_Aars2 | Mtch1 | 1.277 | 4.26E-03 | glio |
| Interaction_genotype_Mtor | Tent5d | -3.478 | 4.26E-03 | glio |
| Interaction_genotype_Mtor | Ywhab | 1.333 | 4.26E-03 | glio |
| Interaction_genotype_Myc | Eme2 | -2.937 | 4.26E-03 | glio |
| Interaction_genotype_Myc | Fam83d | -2.851 | 4.26E-03 | glio |
| Interaction_genotype_Rpl3 | Poln | -4.044 | 4.26E-03 | glio |
| Interaction_genotype_Terf1 | Calhm5 | -4.056 | 4.26E-03 | glio |
| Interaction_genotype_Terf1 | Col8a1 | -4.156 | 4.26E-03 | glio |
| Interaction_genotype_Uqcrb | Skida1 | -4.59 | 4.26E-03 | glio |
| Interaction_genotype_Uqcrc2 | Taldo1 | 1.129 | 4.26E-03 | glio |
| Bcas2_in.vivo_noRT | Itgae | 2.18 | 4.26E-03 | glio |
| Bcas2_in.vivo_noRT | Arhgap23 | -1.387 | 4.26E-03 | glio |
| Bcas2_in.vivo_noRT | Aldh1l2 | -1.436 | 4.26E-03 | glio |
| Bcas2_in.vivo_noRT | Anapc10 | -1.084 | 4.26E-03 | glio |
| Bora_in.vivo_noRT | Hist2h4 | 2.006 | 4.26E-03 | glio |
| Bora_in.vivo_noRT | Gm5617 | 1.564 | 4.26E-03 | glio |
| Chek1_in.vivo_noRT | Fancb | 1.683 | 4.26E-03 | glio |
| Hsd17b10_in.vivo_noRT | Gm10115 | 2.178 | 4.26E-03 | glio |
| Interaction_genotype_Mtor | Acacb | -4.128 | 4.26E-04 | glio |
| interaction_RT | Gbp10 | 2.748 | 4.26E-04 | glio |

| | | | | |
|---|---|---|---|---|
| interaction_RT | Ms4a4d | 3.123 | 4.26E-04 | glio |
| interaction_RT | Hsbp1 | 0.595 | 4.26E-04 | glio |
| Bcas2_in.vivo_noRT | Glb1l | 2.556 | 4.26E-04 | glio |
| Bcas2_in.vivo_noRT | Kirrel3 | 2.549 | 4.26E-04 | glio |
| Bora_ex.vivo_noRT | Agrp | 2.806 | 4.26E-04 | glio |
| Chek1_in.vivo_noRT | Eif3i | -0.637 | 4.26E-04 | glio |
| Eif2s1_in.vivo_noRT | Ush1g | 2.143 | 4.26E-04 | glio |
| Eif2s1_in.vivo_noRT | Gm31063 | 2.143 | 4.26E-04 | glio |
| Eif2s1_in.vivo_noRT | Aldoc | -3.12 | 4.26E-04 | glio |
| culture_effect | R74862 | -1.912 | 4.26E-05 | glio |
| Interaction_genotype_Mtor | March10 | -4.779 | 4.26E-05 | glio |
| Aars2_in.vivo_noRT | 4930432K21R | 2.699 | 4.26E-05 | glio |
| Aars2_in.vivo_noRT | Dync1h1 | -1.37 | 4.26E-05 | glio |
| Atr_in.vivo_noRT | 5830428M24F | 2.88 | 4.26E-05 | glio |
| Chek1_in.vivo_noRT | Vgll3 | 2.922 | 4.26E-05 | glio |
| Aars2_in.vivo_noRT | Prob1 | 3.379 | 4.26E-06 | glio |
| Aars2_ex.vivo_noRT | Vmn1r32 | 2.9 | 4.26E-06 | glio |
| Atp5c1_in.vivo_noRT | Man2c1os | 3.604 | 4.26E-06 | glio |
| Atr_in.vivo_noRT | Gaa | -1.839 | 4.26E-06 | glio |
| culture_effect | Nop58 | -0.604 | 4.26E-07 | glio |
| Atr_in.vivo_noRT | Vdac2 | -1.334 | 4.26E-07 | glio |
| Bcas2_in.vivo_noRT | Gm26619 | 3.739 | 4.26E-07 | glio |
| Chek1_ex.vivo_noRT | Gm45669 | 4.055 | 4.26E-08 | glio |
| Chek1_ex.vivo_noRT | Gm15091 | 4.992 | 4.26E-08 | glio |
| Chek1_ex.vivo_noRT | Ppfia2 | 4.049 | 4.26E-08 | glio |
| Bora_in.vivo_noRT | Hoxa6 | 3.904 | 4.26E-09 | glio |
| Bcas2_in.vivo_noRT | Gm39326 | 4.544 | 4.26E-11 | glio |
| Aars2_in.vivo_noRT | Gm48265 | 4.544 | 4.26E-14 | glio |
| Bcas2_in.vivo_noRT | 4930434B07R | 4.544 | 4.26E-14 | glio |
| culture_effect | Serp2 | 3.586 | 4.26E-19 | glio |
| Chek1_in.vivo_noRT | AU019990 | 5.044 | 4.26E-22 | glio |
| Chek1_in.vivo_noRT | Gm41307 | 5.044 | 4.26E-23 | glio |
| Atr_in.vivo_noRT | BC080695 | 5.533 | 4.26E-26 | glio |
| Interaction_genotype_Atr | Gm50024 | -2.506 | 4.27E-03 | glio |
| Interaction_genotype_Mtor | Srsf1 | 1.452 | 4.27E-03 | glio |
| Interaction_genotype_Myc | Rnf225 | -3.463 | 4.27E-03 | glio |
| Interaction_genotype_Myc | Bend6 | -3.146 | 4.27E-03 | glio |
| Interaction_genotype_Myc | AI480526 | -2.879 | 4.27E-03 | glio |
| Interaction_genotype_Myc | Ptpn4 | -2.433 | 4.27E-03 | glio |
| Interaction_genotype_Myc | Tmtc3 | -2.206 | 4.27E-03 | glio |
| Interaction_genotype_Myc | Wdr24 | -2.275 | 4.27E-03 | glio |
| Interaction_genotype_Myc | Slc35c1 | -2.455 | 4.27E-03 | glio |
| Interaction_genotype_Myc | Ssbp3 | -2.301 | 4.27E-03 | glio |
| Interaction_genotype_Rpl3 | Rasgrp2 | -3.93 | 4.27E-03 | glio |
| Interaction_genotype_Rpl3 | Srp68 | 1.684 | 4.27E-03 | glio |
| Interaction_genotype_Rpl3 | Khdrbs1 | 1.262 | 4.27E-03 | glio |
| Aars2_in.vivo_noRT | Dhx9 | -1.031 | 4.27E-03 | glio |
| Aars2_in.vivo_noRT | Atp5c1 | -0.688 | 4.27E-03 | glio |
| Atr_ex.vivo_noRT | Slc6a19 | 2.771 | 4.27E-03 | glio |
| Atr_ex.vivo_noRT | Capn9 | 2.483 | 4.27E-03 | glio |
| Atr_ex.vivo_noRT | Kctd12 | 2.59 | 4.27E-03 | glio |
| Bcas2_in.vivo_noRT | Snhg14 | 2.157 | 4.27E-03 | glio |

| | | | | |
|---|---|---|---|---|
| Bora_in.vivo_noRT | Padi2 | 2.184 | 4.27E-03 | glio |
| Bora_in.vivo_noRT | Kif15 | -1.516 | 4.27E-03 | glio |
| Bora_in.vivo_noRT | Araf | -0.938 | 4.27E-03 | glio |
| Chek1_in.vivo_noRT | Pak2 | -1.112 | 4.27E-03 | glio |
| Chek1_in.vivo_noRT | Dnmt1 | -0.96 | 4.27E-03 | glio |
| Chek1_in.vivo_noRT | Rps27a | -0.564 | 4.27E-03 | glio |
| Bard1_ex.vivo_noRT | 2300002M23R | 2.814 | 4.27E-04 | glio |
| Bard1_ex.vivo_noRT | Sema3a | 3.219 | 4.27E-04 | glio |
| Bcas2_ex.vivo_noRT | Gm4027 | 2.742 | 4.27E-04 | glio |
| Bora_in.vivo_noRT | 2310030G06R | 2.566 | 4.27E-04 | glio |
| Bora_in.vivo_noRT | Rbbp7 | -0.754 | 4.27E-04 | glio |
| Bora_in.vivo_noRT | Uqcc2 | -0.783 | 4.27E-04 | glio |
| culture_effect | Itprid1 | -2.071 | 4.27E-05 | glio |
| Interaction_genotype_Bora | Rbms1 | 2.209 | 4.27E-05 | glio |
| Interaction_genotype_Myc | Gm15952 | -4.7 | 4.27E-05 | glio |
| culture_effect | Smim12 | 0.802 | 4.27E-06 | glio |
| Atr_in.vivo_noRT | 2510046G10R | 2.903 | 4.27E-06 | glio |
| Bard1_ex.vivo_noRT | Capn13 | 2.814 | 4.27E-06 | glio |
| Bard1_ex.vivo_noRT | Gm12841 | 3.219 | 4.27E-06 | glio |
| Bcas2_in.vivo_noRT | Sgk3 | 3.253 | 4.27E-06 | glio |
| Hsd17b10_in.vivo_noRT | F630040K05Ri | 2.965 | 4.27E-06 | glio |
| culture_effect | Tnfrsf23 | 2.956 | 4.27E-07 | glio |
| culture_effect | Mettl5 | -0.907 | 4.27E-07 | glio |
| culture_effect | Ndufaf8 | 0.576 | 4.27E-07 | glio |
| Interaction_genotype_Atr | K230015D01R | -6.87 | 4.27E-08 | glio |
| Interaction_genotype_Mtor | 5033417F24Ri | -6.526 | 4.27E-08 | glio |
| Interaction_genotype_Mtor | BC028471 | -6.776 | 4.27E-08 | glio |
| Interaction_genotype_Mtor | AC163347.1 | -5.418 | 4.27E-08 | glio |
| Atr_in.vivo_noRT | Gm16048 | 4.097 | 4.27E-08 | glio |
| Atr_ex.vivo_noRT | 1700003E24Ri | 3.164 | 4.27E-08 | glio |
| Chek1_in.vivo_noRT | Gm48882 | 3.92 | 4.27E-08 | glio |
| Chek1_in.vivo_noRT | Grip2 | 3.793 | 4.27E-08 | glio |
| Aars2_in.vivo_noRT | Tmem150c | 4.159 | 4.27E-09 | glio |
| Hsd17b10_in.vivo_noRT | Mpzl2 | 3.356 | 4.27E-09 | glio |
| Aars2_in.vivo_noRT | Ms4a15 | 4.157 | 4.27E-14 | glio |
| culture_effect | Tmem30a | -0.4 | 4.28E-03 | glio |
| Interaction_genotype_Aars2 | 4930592C13R | -3.671 | 4.28E-03 | glio |
| Interaction_genotype_Aars2 | Asnsd1 | 1.482 | 4.28E-03 | glio |
| Interaction_genotype_Aars2 | Tmem126a | 1.111 | 4.28E-03 | glio |
| Interaction_genotype_Atr | Gm15473 | -3.74 | 4.28E-03 | glio |
| Interaction_genotype_Bcas2 | Gm42849 | -3.463 | 4.28E-03 | glio |
| Interaction_genotype_Bora | Gm12925 | -3.865 | 4.28E-03 | glio |
| Interaction_genotype_Bora | Fhl3 | -3.823 | 4.28E-03 | glio |
| Interaction_genotype_Bora | Tslp | -3.752 | 4.28E-03 | glio |
| Interaction_genotype_Bora | Pcmtd2 | 2.581 | 4.28E-03 | glio |
| Interaction_genotype_Bora | Cenps | 2.06 | 4.28E-03 | glio |
| Interaction_genotype_Mtor | Magi2 | -2.919 | 4.28E-03 | glio |
| Interaction_genotype_Myc | Bend7 | -3.105 | 4.28E-03 | glio |
| Interaction_genotype_Rad51 | Ifi44 | 1.73 | 4.28E-03 | glio |
| Aars2_in.vivo_noRT | Zbtb20 | 0.973 | 4.28E-03 | glio |
| Aars2_ex.vivo_noRT | Gm20646 | 2.696 | 4.28E-03 | glio |
| Atp5c1_in.vivo_noRT | Mir142hg | 2.375 | 4.28E-03 | glio |

| | | | | |
|---|---|---|---|---|
| Atr_in.vivo_noRT | Larp1b | -1.289 | 4.28E-03 | glio |
| Bora_in.vivo_noRT | 4930506C21R | 2.072 | 4.28E-03 | glio |
| Bora_in.vivo_noRT | Gm47509 | 2.138 | 4.28E-03 | glio |
| Chek1_in.vivo_noRT | Tubd1 | 1.792 | 4.28E-03 | glio |
| Chek1_ex.vivo_noRT | Cercam | 2.458 | 4.28E-03 | glio |
| Chek1_ex.vivo_noRT | Dusp2 | 2.694 | 4.28E-03 | glio |
| Hsd17b10_in.vivo_noRT | Cops5 | -0.775 | 4.28E-03 | glio |
| culture_effect | Mapk8 | -0.884 | 4.28E-04 | glio |
| culture_effect | Syce2 | -0.628 | 4.28E-04 | glio |
| Interaction_genotype_Mtor | Olfr920 | -4.16 | 4.28E-04 | glio |
| Interaction_genotype_Mtor | Ifi35 | 1.851 | 4.28E-04 | glio |
| Interaction_genotype_Mtor | Ssr4 | 1.638 | 4.28E-04 | glio |
| Interaction_genotype_Myc | Akap17b | -3.675 | 4.28E-04 | glio |
| Bcas2_in.vivo_noRT | Spata1 | 2.647 | 4.28E-04 | glio |
| Bora_in.vivo_noRT | Comp | 2.708 | 4.28E-04 | glio |
| Bora_ex.vivo_noRT | Serpina3i | 3.201 | 4.28E-04 | glio |
| Cenpj_in.vivo_noRT | Strip2 | 2.288 | 4.28E-04 | glio |
| Cenpj_in.vivo_noRT | Tpo | 2.022 | 4.28E-04 | glio |
| Chek1_in.vivo_noRT | D730045B01R | 2.257 | 4.28E-04 | glio |
| Chek1_in.vivo_noRT | Arl1 | -1.07 | 4.28E-04 | glio |
| culture_effect | Ints6 | -0.813 | 4.28E-05 | glio |
| Interaction_genotype_Mtor | Gm5089 | -5.536 | 4.28E-05 | glio |
| Interaction_genotype_Mtor | Pde5a | -5.01 | 4.28E-05 | glio |
| Interaction_genotype_Mtor | Slc6a13 | -5.191 | 4.28E-05 | glio |
| Interaction_genotype_Mtor | Psmb11 | -3.576 | 4.28E-05 | glio |
| Interaction_genotype_Mtor | Nfix | 2.475 | 4.28E-05 | glio |
| Interaction_genotype_Mtor | Srsf7 | 2.037 | 4.28E-05 | glio |
| Interaction_genotype_Myc | Tnxb | -4.621 | 4.28E-05 | glio |
| Chek1_in.vivo_noRT | Npl | 2.748 | 4.28E-05 | glio |
| Chek1_ex.vivo_noRT | Maf1 | -1.399 | 4.28E-05 | glio |
| Interaction_genotype_Mtor | Gm47510 | -3.839 | 4.28E-06 | glio |
| Aars2_in.vivo_noRT | C330013E15Ri | 3.211 | 4.28E-06 | glio |
| Aars2_in.vivo_noRT | Slc2a6 | 3.453 | 4.28E-06 | glio |
| Bcas2_in.vivo_noRT | Gm30790 | 3.291 | 4.28E-06 | glio |
| Bcas2_in.vivo_noRT | C430042M11F | 3.278 | 4.28E-06 | glio |
| Chek1_in.vivo_noRT | Gm15056 | 3.609 | 4.28E-06 | glio |
| Hsd17b10_in.vivo_noRT | E130208F15Ri | 3.626 | 4.28E-06 | glio |
| culture_effect | Psmd14 | -0.699 | 4.28E-07 | glio |
| Interaction_genotype_Myc | Dctn3 | 2.211 | 4.28E-07 | glio |
| culture_effect | Psmc4 | -0.749 | 4.28E-08 | glio |
| interaction_RT | Cystm1 | 1.155 | 4.28E-08 | glio |
| Chek1_in.vivo_noRT | St14 | 3.658 | 4.28E-09 | glio |
| Chek1_in.vivo_noRT | Gm13594 | 4.518 | 4.28E-10 | glio |
| Bora_in.vivo_noRT | Gm43951 | 4.01 | 4.28E-12 | glio |
| culture_effect | Syne1 | -1.422 | 4.28E-13 | glio |
| Interaction_genotype_Bcas2 | Gm31717 | -2.583 | 4.29E-03 | glio |
| Interaction_genotype_Bcas2 | 4930478M13F | -3.472 | 4.29E-03 | glio |
| Interaction_genotype_Myc | Usp17lc | -2.729 | 4.29E-03 | glio |
| Interaction_genotype_Myc | Cul5 | -1.545 | 4.29E-03 | glio |
| Aars2_in.vivo_noRT | Bach2 | 2.044 | 4.29E-03 | glio |
| Atr_in.vivo_noRT | Arpc1b | -0.646 | 4.29E-03 | glio |
| Bcas2_in.vivo_noRT | Trim24 | 1.115 | 4.29E-03 | glio |

| | | | | |
|---|---|---|---|---|
| Hsd17b10_in.vivo_noRT | Prrt3 | 1.963 | 4.29E-03 | glio |
| Interaction_genotype_Mtor | Zfp607a | -4.125 | 4.29E-04 | glio |
| Aars2_ex.vivo_noRT | Olfr90 | 2.9 | 4.29E-04 | glio |
| Atr_in.vivo_noRT | Jpx | 1.781 | 4.29E-04 | glio |
| Atr_in.vivo_noRT | Acot13 | -1.458 | 4.29E-04 | glio |
| Bora_in.vivo_noRT | 6430548M08F | 2.563 | 4.29E-04 | glio |
| culture_effect | Gm36738 | 2.183 | 4.29E-05 | glio |
| Interaction_genotype_Mtor | Pdlim3 | -4.114 | 4.29E-05 | glio |
| Interaction_genotype_Myc | Eif1ax | 1.777 | 4.29E-05 | glio |
| Atp5c1_in.vivo_noRT | 5430402O13R | 3.196 | 4.29E-05 | glio |
| Atp5c1_in.vivo_noRT | Strip2 | 2.404 | 4.29E-05 | glio |
| Bcas2_in.vivo_noRT | Fam71f1 | 2.431 | 4.29E-05 | glio |
| Chek1_ex.vivo_noRT | Gm11713 | 3.345 | 4.29E-05 | glio |
| Chek1_ex.vivo_noRT | Cited4 | 3.862 | 4.29E-05 | glio |
| culture_effect | Poc1a | -1.551 | 4.29E-06 | glio |
| Aars2_in.vivo_noRT | Tmem240 | 3.598 | 4.29E-06 | glio |
| Aars2_in.vivo_noRT | Snu13 | -0.909 | 4.29E-06 | glio |
| Bora_in.vivo_noRT | Msl3l2 | 3.241 | 4.29E-06 | glio |
| Chek1_in.vivo_noRT | Msn | -1.668 | 4.29E-06 | glio |
| Interaction_genotype_Mtor | 4930423D22R | -4.418 | 4.29E-07 | glio |
| interaction_RT | Gng11 | 1.428 | 4.29E-07 | glio |
| interaction_RT | Gng12 | 1.105 | 4.29E-07 | glio |
| interaction_RT | Anxa1 | 1.205 | 4.29E-07 | glio |
| Atp5c1_in.vivo_noRT | Olfr552 | 2.668 | 4.29E-07 | glio |
| Bora_in.vivo_noRT | Calml4 | 3.376 | 4.29E-07 | glio |
| Bcas2_in.vivo_noRT | Nid1 | 4.003 | 4.29E-08 | glio |
| Bcas2_in.vivo_noRT | Tacr2 | 3.902 | 4.29E-08 | glio |
| Chek1_in.vivo_noRT | Eps8l2 | 3.942 | 4.29E-08 | glio |
| Hsd17b10_in.vivo_noRT | Gm47204 | 4.022 | 4.29E-08 | glio |
| culture_effect | Bpifa1 | -3.461 | 4.29E-09 | glio |
| Atr_ex.vivo_noRT | Gm10696 | 5.415 | 4.29E-10 | glio |
| Atr_in.vivo_noRT | 2810410L24Ri | 5.008 | 4.29E-13 | glio |
| Bora_in.vivo_noRT | Ttc16 | 4.937 | 4.29E-13 | glio |
| Aars2_in.vivo_noRT | Slc2a9 | 5.327 | 4.29E-20 | glio |
| Chek1_in.vivo_noRT | Sema6d | 5.044 | 4.29E-21 | glio |
| culture_effect | St6galnac6 | 0.905 | 4.30E-03 | glio |
| culture_effect | Sgms1 | 0.848 | 4.30E-03 | glio |
| Interaction_genotype_Bora | Rab25 | -3.414 | 4.30E-03 | glio |
| Interaction_genotype_Rpl3 | Gm16741 | -3.665 | 4.30E-03 | glio |
| Interaction_genotype_Rpl3 | Hnrnpc | 1.077 | 4.30E-03 | glio |
| interaction_RT | Gnptg | 1.113 | 4.30E-03 | glio |
| interaction_RT | Mcm10 | -1.166 | 4.30E-03 | glio |
| interaction_RT | Tmsb4x | 1.931 | 4.30E-03 | glio |
| Atp5c1_in.vivo_noRT | Cyb561 | 2.415 | 4.30E-03 | glio |
| Atr_in.vivo_noRT | Cad | -1.339 | 4.30E-03 | glio |
| Bora_in.vivo_noRT | Gipc3 | 2.167 | 4.30E-03 | glio |
| Cenpj_in.vivo_noRT | Gm50115 | 2.288 | 4.30E-03 | glio |
| Cenpj_in.vivo_noRT | Zfhx2os | 2.515 | 4.30E-03 | glio |
| Cenpj_ex.vivo_noRT | Adam2 | 2.591 | 4.30E-03 | glio |
| Chek1_in.vivo_noRT | Egr2 | -1.8 | 4.30E-03 | glio |
| Chek1_ex.vivo_noRT | Lmx1b | 2.693 | 4.30E-03 | glio |
| Hsd17b10_in.vivo_noRT | Slc17a9 | 2.161 | 4.30E-03 | glio |

| | | | | |
|---|---|---|---|---|
| culture_effect | Copb2 | -0.486 | 4.30E-04 | glio |
| Interaction_genotype_Aars2 | Lynx1 | -4.299 | 4.30E-04 | glio |
| Interaction_genotype_Aars2 | Ndufb8 | 1.005 | 4.30E-04 | glio |
| Interaction_genotype_Myc | 1700124L16Ri | -4.143 | 4.30E-04 | glio |
| Interaction_genotype_Myc | Gm9484 | -4.091 | 4.30E-04 | glio |
| Interaction_genotype_Myc | Rhbdl1 | -3.914 | 4.30E-04 | glio |
| Interaction_genotype_Myc | 4933404O12R | -3.831 | 4.30E-04 | glio |
| Interaction_genotype_Myc | Pomt1 | -3.356 | 4.30E-04 | glio |
| Interaction_genotype_Myc | Atxn1 | -3.315 | 4.30E-04 | glio |
| Interaction_genotype_Myc | Nme2 | 1.262 | 4.30E-04 | glio |
| interaction_RT | Ebp | 0.797 | 4.30E-04 | glio |
| Aars2_in.vivo_noRT | Ccnb1ip1 | 2.795 | 4.30E-04 | glio |
| Aars2_in.vivo_noRT | 2210011C24R | 2.436 | 4.30E-04 | glio |
| Aars2_in.vivo_noRT | Bdh2 | 2.501 | 4.30E-04 | glio |
| Aars2_in.vivo_noRT | Erfe | 2.313 | 4.30E-04 | glio |
| Aars2_in.vivo_noRT | Gtf2b | -1.527 | 4.30E-04 | glio |
| Atr_ex.vivo_noRT | Pyy | 2.667 | 4.30E-04 | glio |
| Bora_in.vivo_noRT | 1700112D23R | 2.505 | 4.30E-04 | glio |
| Bora_ex.vivo_noRT | Cys1 | 2.808 | 4.30E-04 | glio |
| Chek1_in.vivo_noRT | Cox7a1 | 2.725 | 4.30E-04 | glio |
| Chek1_in.vivo_noRT | Trim66 | 2.332 | 4.30E-04 | glio |
| Eif2s1_in.vivo_noRT | Gm45591 | 2.407 | 4.30E-04 | glio |
| culture_effect | Gm14296 | 1.112 | 4.30E-05 | glio |
| Interaction_genotype_Myc | Rpl14 | 1.647 | 4.30E-05 | glio |
| Bcas2_in.vivo_noRT | Arhgap4 | 2.821 | 4.30E-05 | glio |
| Bcas2_in.vivo_noRT | Fbln1 | 2.806 | 4.30E-05 | glio |
| Bora_in.vivo_noRT | Haglr | 3.155 | 4.30E-05 | glio |
| Bora_in.vivo_noRT | Gm43728 | 3.078 | 4.30E-05 | glio |
| Bora_in.vivo_noRT | Gm44190 | 2.672 | 4.30E-05 | glio |
| Chek1_in.vivo_noRT | Gm19684 | 3.085 | 4.30E-05 | glio |
| Chek1_in.vivo_noRT | Pdk2 | 2.943 | 4.30E-05 | glio |
| Chek1_in.vivo_noRT | Tmem186 | 2.757 | 4.30E-05 | glio |
| culture_effect | Gm17597 | 2.434 | 4.30E-06 | glio |
| Hsd17b10_in.vivo_noRT | Gm12860 | 2.608 | 4.30E-06 | glio |
| Atr_in.vivo_noRT | Gm48086 | 3.585 | 4.30E-07 | glio |
| Bora_in.vivo_noRT | Gm10382 | 3.693 | 4.30E-07 | glio |
| Chek1_in.vivo_noRT | Bhlhe41 | 4.521 | 4.30E-08 | glio |
| Chek1_in.vivo_noRT | Cyp1a1 | 5.044 | 4.30E-08 | glio |
| Chek1_in.vivo_noRT | Gm47484 | 4.05 | 4.30E-08 | glio |
| Chek1_in.vivo_noRT | 9930014A18R | 3.649 | 4.30E-08 | glio |
| Atr_in.vivo_noRT | Gm43660 | 3.831 | 4.30E-09 | glio |
| culture_effect | Itm2a | -2.036 | 4.30E-10 | glio |
| culture_effect | Adrb1 | 1.394 | 4.30E-11 | glio |
| Aars2_in.vivo_noRT | Zic4 | 5.063 | 4.30E-12 | glio |
| Bora_in.vivo_noRT | Gm17276 | 4.145 | 4.30E-13 | glio |
| Aars2_in.vivo_noRT | Gm36026 | 4.676 | 4.30E-16 | glio |
| Chek1_ex.vivo_noRT | Helt | 4.842 | 4.30E-17 | glio |
| Chek1_ex.vivo_noRT | Gm20625 | 4.842 | 4.30E-17 | glio |
| Atr_in.vivo_noRT | Gm15138 | 5.267 | 4.30E-24 | glio |
| culture_effect | Gm49172 | -1.194 | 4.31E-03 | glio |
| Interaction_genotype_Bcas2 | Gm21887 | 1.988 | 4.31E-03 | glio |
| Bora_in.vivo_noRT | Prorsd1 | 1.445 | 4.31E-03 | glio |

| | | | | |
|---|---|---|---|---|
| Hsd17b10_in.vivo_noRT | 1700084E18Ri | 2.374 | 4.31E-03 | glio |
| Hsd17b10_in.vivo_noRT | Tgfbr3l | 2.312 | 4.31E-03 | glio |
| Hsd17b10_in.vivo_noRT | Lrrn4cl | 2.387 | 4.31E-03 | glio |
| culture_effect | Otulin | 0.435 | 4.31E-04 | glio |
| Interaction_genotype_Atr | Gm44899 | -3.474 | 4.31E-04 | glio |
| Interaction_genotype_Mtor | 4930597O21R | -4.638 | 4.31E-04 | glio |
| Interaction_genotype_Mtor | Itgb1 | 1.672 | 4.31E-04 | glio |
| Interaction_genotype_Myc | BC037032 | -4.242 | 4.31E-04 | glio |
| Interaction_genotype_Myc | D930030I03Ri | -4.101 | 4.31E-04 | glio |
| Atp5c1_in.vivo_noRT | BC065403 | 2.683 | 4.31E-04 | glio |
| Atr_in.vivo_noRT | Gm4356 | 2.732 | 4.31E-04 | glio |
| Atr_in.vivo_noRT | Gm9895 | 2.553 | 4.31E-04 | glio |
| Atr_in.vivo_noRT | Zdhhc15 | 2.124 | 4.31E-04 | glio |
| Bard1_ex.vivo_noRT | Esp15 | 2.814 | 4.31E-04 | glio |
| Bora_in.vivo_noRT | Prkg2 | -2.058 | 4.31E-04 | glio |
| Cenpj_in.vivo_noRT | 1700034I23Ri | 2.022 | 4.31E-04 | glio |
| culture_effect | Mpz | 2.505 | 4.31E-05 | glio |
| Interaction_genotype_Myc | AC241534.1 | -4.672 | 4.31E-05 | glio |
| Aars2_in.vivo_noRT | Lyst | 2.258 | 4.31E-05 | glio |
| Chek1_in.vivo_noRT | Tspan33 | 2.949 | 4.31E-05 | glio |
| culture_effect | Ahnak2 | 1.174 | 4.31E-06 | glio |
| Aars2_in.vivo_noRT | Proscos | 3.431 | 4.31E-06 | glio |
| Chek1_in.vivo_noRT | Wrb | 2.265 | 4.31E-06 | glio |
| Interaction_genotype_Mtor | Ankrd22 | -4.099 | 4.31E-07 | glio |
| Bora_in.vivo_noRT | Atad2 | -2.427 | 4.31E-07 | glio |
| Hsd17b10_in.vivo_noRT | Dcdc2a | 3.356 | 4.31E-08 | glio |
| Aars2_in.vivo_noRT | Bbof1 | 3.898 | 4.31E-09 | glio |
| Interaction_genotype_Mtor | Arf1 | 2.536 | 4.31E-10 | glio |
| Hsd17b10_in.vivo_noRT | Gm13288 | 3.616 | 4.31E-10 | glio |
| Atr_in.vivo_noRT | Dact1 | 4.366 | 4.31E-11 | glio |
| culture_effect | Prdx2 | -0.804 | 4.31E-12 | glio |
| Aars2_in.vivo_noRT | Fam131a | 4.801 | 4.31E-13 | glio |
| Atr_in.vivo_noRT | Camk2b | 5.011 | 4.31E-14 | glio |
| Atr_in.vivo_noRT | 1700015F17Ri | 5.273 | 4.31E-14 | glio |
| Aars2_in.vivo_noRT | Gm48742 | 4.676 | 4.31E-16 | glio |
| Aars2_in.vivo_noRT | 8030423F21Ri | 4.798 | 4.31E-16 | glio |
| Atr_in.vivo_noRT | Gm4166 | 4.606 | 4.31E-19 | glio |
| culture_effect | Mfap2 | -1.069 | 4.32E-03 | glio |
| Interaction_genotype_Aars2 | Phldb2 | -3.398 | 4.32E-03 | glio |
| Interaction_genotype_Atr | Mcph1 | -2.142 | 4.32E-03 | glio |
| Interaction_genotype_Atr | Tmf1 | 1.703 | 4.32E-03 | glio |
| Interaction_genotype_Bcas2 | Ggnbp1 | -3.726 | 4.32E-03 | glio |
| Interaction_genotype_Mtor | Gm26692 | -3.208 | 4.32E-03 | glio |
| Interaction_genotype_Myc | 4930570E03Ri | -2.469 | 4.32E-03 | glio |
| Interaction_genotype_Myc | Tdrp | -3.157 | 4.32E-03 | glio |
| Interaction_genotype_Pars2 | Masp1 | -3.32 | 4.32E-03 | glio |
| Interaction_genotype_Pars2 | Gm48140 | -2.799 | 4.32E-03 | glio |
| Interaction_genotype_Pars2 | Pspn | -2.664 | 4.32E-03 | glio |
| interaction_RT | Flot1 | 0.875 | 4.32E-03 | glio |
| Aars2_in.vivo_noRT | Mtpn | -0.755 | 4.32E-03 | glio |
| Aars2_ex.vivo_noRT | Moap1 | 2.612 | 4.32E-03 | glio |
| Atp5c1_in.vivo_noRT | Mtmr11 | 2.344 | 4.32E-03 | glio |

| | | | | |
|---|---|---|---|---|
| Atp5c1_in.vivo_noRT | Ccl7 | 2.668 | 4.32E-03 | glio |
| Atp5c1_in.vivo_noRT | Gm49694 | 2.473 | 4.32E-03 | glio |
| Atp5c1_in.vivo_noRT | Gm3604 | 2.33 | 4.32E-03 | glio |
| Atp5c1_in.vivo_noRT | Tgfbr3l | 2.415 | 4.32E-03 | glio |
| Atp5c1_in.vivo_noRT | BC048671 | 1.898 | 4.32E-03 | glio |
| Atp5c1_in.vivo_noRT | 1700086P04R | 2.256 | 4.32E-03 | glio |
| Atp5c1_in.vivo_noRT | Lgals7 | -2.485 | 4.32E-03 | glio |
| Atr_in.vivo_noRT | Klc2 | 1.872 | 4.32E-03 | glio |
| Bora_in.vivo_noRT | Sgsh | 2.193 | 4.32E-03 | glio |
| Bora_in.vivo_noRT | Jrk | 1.968 | 4.32E-03 | glio |
| Bora_in.vivo_noRT | Gm12940 | 1.795 | 4.32E-03 | glio |
| Chek1_in.vivo_noRT | Gna12 | 1.165 | 4.32E-03 | glio |
| Chek1_ex.vivo_noRT | Dagla | 2.859 | 4.32E-03 | glio |
| Interaction_genotype_Mtor | Txn1 | 1.522 | 4.32E-04 | glio |
| Interaction_genotype_Myc | 1500004A13R | -3.994 | 4.32E-04 | glio |
| Aars2_in.vivo_noRT | 0610009E02Ri | 2.69 | 4.32E-04 | glio |
| Aars2_in.vivo_noRT | 4833417C18R | 2.502 | 4.32E-04 | glio |
| Bcas2_in.vivo_noRT | 1700041G16R | 2.553 | 4.32E-04 | glio |
| Bora_in.vivo_noRT | Tex19.2 | 2.472 | 4.32E-04 | glio |
| Bora_in.vivo_noRT | Gopc | -1.7 | 4.32E-04 | glio |
| Bora_in.vivo_noRT | Taf5 | -1.58 | 4.32E-04 | glio |
| culture_effect | Cyth2 | -0.58 | 4.32E-05 | glio |
| Interaction_genotype_Mtor | Slc36a3 | -4.815 | 4.32E-05 | glio |
| Aars2_in.vivo_noRT | Ppp1r9a | 2.343 | 4.32E-05 | glio |
| Aars2_in.vivo_noRT | Paxx | 2.098 | 4.32E-05 | glio |
| Atr_ex.vivo_noRT | Gtsf1l | 3.164 | 4.32E-05 | glio |
| Bard1_ex.vivo_noRT | 1700047K16R | 3.896 | 4.32E-05 | glio |
| Bard1_ex.vivo_noRT | Gm33466 | 2.814 | 4.32E-05 | glio |
| Bcas2_in.vivo_noRT | Gm49479 | 2.969 | 4.32E-05 | glio |
| Bora_in.vivo_noRT | Gm43156 | 2.918 | 4.32E-05 | glio |
| Bora_in.vivo_noRT | Hcrtr1 | 2.998 | 4.32E-05 | glio |
| Bora_in.vivo_noRT | 6330403L08Ri | 2.799 | 4.32E-05 | glio |
| Bora_in.vivo_noRT | Gm813 | 2.825 | 4.32E-05 | glio |
| Hsd17b10_in.vivo_noRT | 9630014M24F | 2.837 | 4.32E-05 | glio |
| Interaction_genotype_Myc | Hspd1 | 2.357 | 4.32E-06 | glio |
| Atr_in.vivo_noRT | Ccdc9b | 2.893 | 4.32E-06 | glio |
| Chek1_in.vivo_noRT | Trim71 | 3.208 | 4.32E-06 | glio |
| Interaction_genotype_Mtor | Atp5f1 | 2.368 | 4.32E-07 | glio |
| Aars2_in.vivo_noRT | Glrp1 | 3.057 | 4.32E-07 | glio |
| Chek1_ex.vivo_noRT | Hif3a | 4.449 | 4.32E-07 | glio |
| Chek1_in.vivo_noRT | Gm2788 | 3.995 | 4.32E-08 | glio |
| Chek1_in.vivo_noRT | Gm2012 | 3.88 | 4.32E-08 | glio |
| Bora_in.vivo_noRT | Plppr3 | 4.083 | 4.32E-09 | glio |
| Bora_in.vivo_noRT | BC028471 | 4.404 | 4.32E-10 | glio |
| Bora_in.vivo_noRT | Ptges | 4.804 | 4.32E-11 | glio |
| Chek1_in.vivo_noRT | Thnsl2 | 3.876 | 4.32E-13 | glio |
| Bora_in.vivo_noRT | 1700026N04R | 4.676 | 4.32E-14 | glio |
| Chek1_ex.vivo_noRT | Capn13 | 4.842 | 4.32E-18 | glio |
| culture_effect | Pmch | -1.592 | 4.33E-03 | glio |
| culture_effect | Gm16741 | 1.605 | 4.33E-03 | glio |
| Interaction_genotype_Mtor | Ddx24 | 1.278 | 4.33E-03 | glio |
| Interaction_genotype_Myc | Gm35867 | -3.048 | 4.33E-03 | glio |

| | | | | |
|---|---|---|---|---|
| Interaction_genotype_Myc | L2hgdh | -2.389 | 4.33E-03 | glio |
| Interaction_genotype_Rpl3 | Gm50069 | -4.038 | 4.33E-03 | glio |
| Interaction_genotype_Rpl3 | Wincr1 | -5.004 | 4.33E-03 | glio |
| Aars2_in.vivo_noRT | Arrdc3 | 1.937 | 4.33E-03 | glio |
| Aars2_in.vivo_noRT | Jmjd6 | -0.883 | 4.33E-03 | glio |
| Aars2_ex.vivo_noRT | Gm35551 | 2.706 | 4.33E-03 | glio |
| Bcas2_ex.vivo_noRT | Rpl3l | 2.742 | 4.33E-03 | glio |
| Bora_in.vivo_noRT | Oas2 | 1.944 | 4.33E-03 | glio |
| Bora_in.vivo_noRT | Asna1 | -0.878 | 4.33E-03 | glio |
| Chek1_ex.vivo_noRT | Exosc8 | -1.199 | 4.33E-03 | glio |
| Hsd17b10_in.vivo_noRT | Gm20478 | 2.044 | 4.33E-03 | glio |
| culture_effect | 1700056E22Ri | 1.245 | 4.33E-04 | glio |
| culture_effect | Fndc3b | 0.601 | 4.33E-04 | glio |
| culture_effect | Notch1 | 0.668 | 4.33E-04 | glio |
| Interaction_genotype_Atr | Gm14019 | -3.866 | 4.33E-04 | glio |
| Interaction_genotype_Bcas2 | Gm32591 | -4.595 | 4.33E-04 | glio |
| Interaction_genotype_Bcas2 | Jph4 | -4.601 | 4.33E-04 | glio |
| Interaction_genotype_Mtor | Hist1h3g | -4.48 | 4.33E-04 | glio |
| Interaction_genotype_Mtor | 4921531C22R | -3.851 | 4.33E-04 | glio |
| Interaction_genotype_Myc | 5031434O11R | -4.465 | 4.33E-04 | glio |
| Aars2_ex.vivo_noRT | Tnni3k | 2.505 | 4.33E-04 | glio |
| Aars2_ex.vivo_noRT | C330024C12R | 2.9 | 4.33E-04 | glio |
| Atr_in.vivo_noRT | Gm47863 | 2.535 | 4.33E-04 | glio |
| Atr_in.vivo_noRT | Ercc6l2 | 1.678 | 4.33E-04 | glio |
| Bcas2_in.vivo_noRT | Radil | 2.515 | 4.33E-04 | glio |
| Bora_ex.vivo_noRT | Gm16685 | 2.802 | 4.33E-04 | glio |
| Bora_ex.vivo_noRT | Cacng5 | 3.201 | 4.33E-04 | glio |
| Chek1_ex.vivo_noRT | Fth1 | 1.128 | 4.33E-04 | glio |
| Hsd17b10_in.vivo_noRT | Ppp1r3g | 2.431 | 4.33E-04 | glio |
| Hsd17b10_in.vivo_noRT | Gm10516 | 2.579 | 4.33E-04 | glio |
| culture_effect | 9430065F17Ri | 2.098 | 4.33E-05 | glio |
| culture_effect | Hmgn3 | 0.56 | 4.33E-05 | glio |
| Interaction_genotype_Mtor | Anks6 | -4.46 | 4.33E-05 | glio |
| Interaction_genotype_Mtor | Ifnz | -4.703 | 4.33E-05 | glio |
| Interaction_genotype_Mtor | Gm28578 | -5.083 | 4.33E-05 | glio |
| Aars2_in.vivo_noRT | Fndc7 | 3.083 | 4.33E-05 | glio |
| Aars2_ex.vivo_noRT | Bpifa3 | 3.293 | 4.33E-05 | glio |
| Chek1_ex.vivo_noRT | Pycard | 3.445 | 4.33E-05 | glio |
| Interaction_genotype_Myc | Rbm3 | 2.007 | 4.33E-06 | glio |
| Atr_in.vivo_noRT | 4930597O21R | 3.806 | 4.33E-07 | glio |
| Atr_in.vivo_noRT | Chchd3 | -1.455 | 4.33E-07 | glio |
| Hsd17b10_in.vivo_noRT | Krtcap3 | 2.824 | 4.33E-07 | glio |
| Aars2_in.vivo_noRT | Kcnmb4os2 | 3.889 | 4.33E-08 | glio |
| Aars2_in.vivo_noRT | 4930445N18R | 3.882 | 4.33E-08 | glio |
| culture_effect | Kcnip3 | -3.416 | 4.33E-09 | glio |
| Bcas2_in.vivo_noRT | Shroom3 | 4.057 | 4.33E-09 | glio |
| Bora_in.vivo_noRT | Gm10230 | 4.586 | 4.33E-10 | glio |
| Bcas2_in.vivo_noRT | Jph4 | 4.541 | 4.33E-11 | glio |
| Atr_in.vivo_noRT | 5830454E08Ri | 4.608 | 4.33E-12 | glio |
| Bcas2_in.vivo_noRT | Zfp352 | 4.807 | 4.33E-14 | glio |
| Bora_in.vivo_noRT | Gm20767 | 4.936 | 4.33E-14 | glio |
| Bora_in.vivo_noRT | Fbxo15 | 4.673 | 4.33E-14 | glio |

| | | | | |
|---|---|---|---|---|
| culture_effect | Myof | 1.681 | 4.33E-15 | glio |
| Aars2_in.vivo_noRT | Gm41318 | 5.327 | 4.33E-15 | glio |
| Atr_in.vivo_noRT | Gm16344 | 4.622 | 4.33E-15 | glio |
| Bora_in.vivo_noRT | Olfr171 | 4.936 | 4.33E-19 | glio |
| Aars2_in.vivo_noRT | Gm49808 | 5.327 | 4.33E-28 | glio |
| culture_effect | Zdhhc12 | -0.565 | 4.34E-03 | glio |
| culture_effect | Smim7 | -0.403 | 4.34E-03 | glio |
| Interaction_genotype_Aars2 | Dscaml1 | -2.429 | 4.34E-03 | glio |
| Interaction_genotype_Chek1 | Adm | 4.061 | 4.34E-03 | glio |
| Interaction_genotype_Mtor | Gm13212 | -3.507 | 4.34E-03 | glio |
| Interaction_genotype_Mtor | Ercc4 | -3.018 | 4.34E-03 | glio |
| Atr_in.vivo_noRT | Arap1 | 2.067 | 4.34E-03 | glio |
| Atr_in.vivo_noRT | Pim2 | 1.626 | 4.34E-03 | glio |
| Cenpj_in.vivo_noRT | Selenov | 2.022 | 4.34E-03 | glio |
| Cenpj_in.vivo_noRT | A330076H08R | 2.022 | 4.34E-03 | glio |
| Cenpj_in.vivo_noRT | 4930423D22R | 1.762 | 4.34E-03 | glio |
| Cenpj_in.vivo_noRT | Gm10399 | 2.397 | 4.34E-03 | glio |
| Chek1_in.vivo_noRT | Mark2 | -1.22 | 4.34E-03 | glio |
| Chek1_ex.vivo_noRT | Cbx1 | -0.834 | 4.34E-03 | glio |
| Fanca_ex.vivo_noRT | Mlst8 | -2.316 | 4.34E-03 | glio |
| Hsd17b10_in.vivo_noRT | Rnf145 | -1.398 | 4.34E-03 | glio |
| culture_effect | Zfp358 | 1.009 | 4.34E-04 | glio |
| culture_effect | Dnttip2 | -0.459 | 4.34E-04 | glio |
| Interaction_genotype_Mtor | D230022J07Ri | -4.074 | 4.34E-04 | glio |
| interaction_RT | Arf4 | 0.687 | 4.34E-04 | glio |
| Aars2_in.vivo_noRT | Dnajb5 | 2.219 | 4.34E-04 | glio |
| Atr_in.vivo_noRT | Npdc1 | 2.03 | 4.34E-04 | glio |
| Atr_in.vivo_noRT | Pllp | -2.052 | 4.34E-04 | glio |
| Bcas2_in.vivo_noRT | Samd15 | 2.785 | 4.34E-04 | glio |
| Bcas2_ex.vivo_noRT | Gm40538 | 2.742 | 4.34E-04 | glio |
| Chek1_in.vivo_noRT | Gm26549 | 2.219 | 4.34E-04 | glio |
| Atr_in.vivo_noRT | Iapp | 3.371 | 4.34E-05 | glio |
| Atr_in.vivo_noRT | Erp29 | -1.294 | 4.34E-05 | glio |
| Bora_in.vivo_noRT | Gm13212 | 2.941 | 4.34E-05 | glio |
| Bora_in.vivo_noRT | Celf1 | -1.424 | 4.34E-05 | glio |
| Bora_ex.vivo_noRT | Masp1 | 3.196 | 4.34E-05 | glio |
| culture_effect | Atxn2l | -0.666 | 4.34E-06 | glio |
| Bora_in.vivo_noRT | Zfhx2os | 3.355 | 4.34E-06 | glio |
| culture_effect | Gpat3 | -1.6 | 4.34E-07 | glio |
| culture_effect | Erlec1 | -0.71 | 4.34E-07 | glio |
| Interaction_genotype_Mtor | Gm15943 | -4.817 | 4.34E-07 | glio |
| Atr_in.vivo_noRT | Gm15265 | 3.731 | 4.34E-08 | glio |
| Atr_ex.vivo_noRT | Ric3 | 4.331 | 4.34E-08 | glio |
| culture_effect | Kifc3 | 1.152 | 4.34E-10 | glio |
| Bcas2_in.vivo_noRT | Aoc1 | 4.547 | 4.34E-10 | glio |
| Bcas2_in.vivo_noRT | Trim16 | 4.547 | 4.34E-13 | glio |
| Atr_in.vivo_noRT | Efcab9 | 5.267 | 4.34E-15 | glio |
| culture_effect | Ifit1 | 2.229 | 4.34E-18 | glio |
| Interaction_genotype_Aars2 | Zmynd15 | -3.524 | 4.35E-03 | glio |
| Interaction_genotype_Aars2 | Gm20492 | -3.309 | 4.35E-03 | glio |
| Interaction_genotype_Atr | 2900042K21R | -3.476 | 4.35E-03 | glio |
| Interaction_genotype_Myc | Sgtb | -3.125 | 4.35E-03 | glio |

| | | | | |
|---|---|---|---|---|
| Interaction_genotype_Myc | Isca1 | -2.36 | 4.35E-03 | glio |
| Aars2_in.vivo_noRT | 4931428F04Ri | 1.762 | 4.35E-03 | glio |
| Aars2_in.vivo_noRT | Rab1b | -1.087 | 4.35E-03 | glio |
| Aars2_in.vivo_noRT | Hmgb1 | -0.6 | 4.35E-03 | glio |
| Bcas2_in.vivo_noRT | Btbd6 | 1.924 | 4.35E-03 | glio |
| Bora_in.vivo_noRT | Srsf2 | -0.667 | 4.35E-03 | glio |
| Fanca_ex.vivo_noRT | Cldn2 | 2.124 | 4.35E-03 | glio |
| culture_effect | Rfx1 | -1.188 | 4.35E-04 | glio |
| culture_effect | Rmc1 | -0.951 | 4.35E-04 | glio |
| Interaction_genotype_Mtor | 3010001F23Ri | -3.588 | 4.35E-04 | glio |
| Interaction_genotype_Mtor | Actb | 1.527 | 4.35E-04 | glio |
| Aars2_in.vivo_noRT | Tmcc2 | 1.888 | 4.35E-04 | glio |
| Aars2_ex.vivo_noRT | Synpo2l | 2.897 | 4.35E-04 | glio |
| Aars2_ex.vivo_noRT | Dapk2 | 2.495 | 4.35E-04 | glio |
| Atr_in.vivo_noRT | Pdlim2 | 2.257 | 4.35E-04 | glio |
| Bcas2_ex.vivo_noRT | Nat8f4 | 3.128 | 4.35E-04 | glio |
| Chek1_ex.vivo_noRT | Dpy30 | -1.085 | 4.35E-04 | glio |
| Interaction_genotype_Mtor | Atp5g2 | 1.656 | 4.35E-05 | glio |
| Interaction_genotype_Myc | Clcn7 | -3.773 | 4.35E-05 | glio |
| Interaction_genotype_Myc | 4930588A03R | -3.128 | 4.35E-05 | glio |
| Interaction_genotype_Aars2 | 1700003G18R | -6.503 | 4.35E-06 | glio |
| Aars2_in.vivo_noRT | Ldlr | -2.515 | 4.35E-06 | glio |
| Bcas2_ex.vivo_noRT | Thnsl2 | 3.524 | 4.35E-06 | glio |
| Bora_in.vivo_noRT | Gm47061 | 3.631 | 4.35E-06 | glio |
| Bora_in.vivo_noRT | Hipk4 | 2.95 | 4.35E-06 | glio |
| Interaction_genotype_Mtor | Eif3h | 1.976 | 4.35E-07 | glio |
| Bora_in.vivo_noRT | 2610507I01Ri | 3.183 | 4.35E-07 | glio |
| Chek1_in.vivo_noRT | Wfikkn2 | 3.593 | 4.35E-07 | glio |
| Atr_in.vivo_noRT | C330011M18I | 3.878 | 4.35E-08 | glio |
| Atr_in.vivo_noRT | Abi3bp | 3.808 | 4.35E-08 | glio |
| Bora_in.vivo_noRT | 1700023H06R | 4.145 | 4.35E-08 | glio |
| Chek1_ex.vivo_noRT | Cd52 | 4.842 | 4.35E-09 | glio |
| Atr_in.vivo_noRT | Cyp26b1 | 4.608 | 4.35E-10 | glio |
| Aars2_in.vivo_noRT | Cpne9 | 4.826 | 4.35E-11 | glio |
| Chek1_ex.vivo_noRT | 4930470G03R | 4.449 | 4.35E-11 | glio |
| Hsd17b10_in.vivo_noRT | Gm19276 | 3.616 | 4.35E-13 | glio |
| Chek1_in.vivo_noRT | Car14 | 5.044 | 4.35E-14 | glio |
| Bcas2_in.vivo_noRT | Olfr1152 | 4.541 | 4.35E-21 | glio |
| culture_effect | Itprid2 | 0.715 | 4.36E-03 | glio |
| culture_effect | Src | -0.718 | 4.36E-03 | glio |
| Interaction_genotype_Aars2 | Pspn | -2.429 | 4.36E-03 | glio |
| Interaction_genotype_Aars2 | Gm36172 | -2.869 | 4.36E-03 | glio |
| Interaction_genotype_Aars2 | Mea1 | 1.489 | 4.36E-03 | glio |
| Interaction_genotype_Atr | Icam5 | -3.496 | 4.36E-03 | glio |
| Interaction_genotype_Atr | Carns1 | -3.515 | 4.36E-03 | glio |
| Interaction_genotype_Atr | Abcb10 | -2.665 | 4.36E-03 | glio |
| Interaction_genotype_Atr | Sat1 | 1.748 | 4.36E-03 | glio |
| Interaction_genotype_Mtor | Ankrd33b | -3.583 | 4.36E-03 | glio |
| Interaction_genotype_Mtor | Gm42679 | -3.376 | 4.36E-03 | glio |
| Interaction_genotype_Mtor | Entpd5 | -3.258 | 4.36E-03 | glio |
| Interaction_genotype_Mtor | Zfp207 | 1.49 | 4.36E-03 | glio |
| Interaction_genotype_Myc | P4ha2 | -2.502 | 4.36E-03 | glio |

| | | | | |
|---|---|---|---|---|
| Interaction_genotype_Myc | Trir | 1.021 | 4.36E-03 | glio |
| Interaction_genotype_Pars2 | Zic4 | -4.095 | 4.36E-03 | glio |
| Aars2_in.vivo_noRT | Gm14325 | 2.082 | 4.36E-03 | glio |
| Aars2_in.vivo_noRT | Dnaaf2 | -1.23 | 4.36E-03 | glio |
| Atr_in.vivo_noRT | Klrg2 | 1.962 | 4.36E-03 | glio |
| Atr_in.vivo_noRT | Atxn1 | 1.801 | 4.36E-03 | glio |
| Atr_in.vivo_noRT | Atxn7l2 | 1.708 | 4.36E-03 | glio |
| Atr_in.vivo_noRT | Tmx2 | -1.027 | 4.36E-03 | glio |
| Bcas2_in.vivo_noRT | Pbrm1 | -0.988 | 4.36E-03 | glio |
| Bcas2_ex.vivo_noRT | C330002G04R | 2.155 | 4.36E-03 | glio |
| Brca2_in.vivo_noRT | Cldn2 | 1.763 | 4.36E-03 | glio |
| Chek1_in.vivo_noRT | Rnd3 | 1.636 | 4.36E-03 | glio |
| Chek1_ex.vivo_noRT | Zfp143 | -1.971 | 4.36E-03 | glio |
| Chek1_ex.vivo_noRT | Trim28 | -1.132 | 4.36E-03 | glio |
| Interaction_genotype_Aars2 | Gm35066 | -4.312 | 4.36E-04 | glio |
| Interaction_genotype_Mtor | Bub3 | 2.003 | 4.36E-04 | glio |
| Atr_ex.vivo_noRT | Cd52 | 3.164 | 4.36E-04 | glio |
| Bard1_ex.vivo_noRT | Olfr552 | 2.415 | 4.36E-04 | glio |
| Bora_in.vivo_noRT | Phkg1 | 3.507 | 4.36E-04 | glio |
| Bora_ex.vivo_noRT | Gm17210 | 3.407 | 4.36E-04 | glio |
| Eif2s1_in.vivo_noRT | Slc1a3 | -1.197 | 4.36E-04 | glio |
| culture_effect | Dusp22 | 0.932 | 4.36E-05 | glio |
| culture_effect | Tmem248 | 0.56 | 4.36E-05 | glio |
| Atr_in.vivo_noRT | Ncor1 | -1.144 | 4.36E-05 | glio |
| Hsd17b10_in.vivo_noRT | Fabp4 | 3.087 | 4.36E-05 | glio |
| Hsd17b10_in.vivo_noRT | Klk14 | 3.084 | 4.36E-05 | glio |
| Hsd17b10_in.vivo_noRT | Cd83 | 2.616 | 4.36E-05 | glio |
| Hsd17b10_in.vivo_noRT | Cnih2 | 2.959 | 4.36E-05 | glio |
| Chek1_ex.vivo_noRT | Gm48242 | 3.324 | 4.36E-06 | glio |
| culture_effect | Socs1 | -1.276 | 4.36E-08 | glio |
| Interaction_genotype_Mtor | Dnase1l2 | -6.541 | 4.36E-08 | glio |
| Interaction_genotype_Mtor | Sema6d | -5.08 | 4.36E-08 | glio |
| Atr_in.vivo_noRT | Gm38979 | 4.359 | 4.36E-09 | glio |
| Chek1_in.vivo_noRT | Rnft2 | 4.24 | 4.36E-10 | glio |
| Chek1_in.vivo_noRT | Ak1 | 4.304 | 4.36E-10 | glio |
| Chek1_ex.vivo_noRT | Nat8f2 | 4.447 | 4.36E-10 | glio |
| Bcas2_in.vivo_noRT | Gm12860 | 3.799 | 4.36E-12 | glio |
| Bora_in.vivo_noRT | Dntt | 4.936 | 4.36E-12 | glio |
| Chek1_in.vivo_noRT | Crb3 | 5.044 | 4.36E-20 | glio |
| Aars2_in.vivo_noRT | Gm48617 | 5.063 | 4.36E-24 | glio |
| culture_effect | Zfp827 | -1.579 | 4.37E-03 | glio |
| culture_effect | Tmem39a | -0.606 | 4.37E-03 | glio |
| culture_effect | Wbp1 | 0.616 | 4.37E-03 | glio |
| Interaction_genotype_Aars2 | Gm48170 | -2.831 | 4.37E-03 | glio |
| Interaction_genotype_Aars2 | A930033M14F | -2.999 | 4.37E-03 | glio |
| Interaction_genotype_Aars2 | Stab1 | -3.764 | 4.37E-03 | glio |
| Interaction_genotype_Aars2 | Gm24362 | -3.271 | 4.37E-03 | glio |
| Interaction_genotype_Aars2 | March10 | -3.507 | 4.37E-03 | glio |
| Interaction_genotype_Aars2 | Arhgef4 | -3.563 | 4.37E-03 | glio |
| Interaction_genotype_Aars2 | Ciita | 2.942 | 4.37E-03 | glio |
| Interaction_genotype_Aars2 | Gm35019 | 3.277 | 4.37E-03 | glio |
| Interaction_genotype_Aars2 | Gm11520 | -3.252 | 4.37E-03 | glio |

| | | | | |
|---|---|---|---|---|
| Interaction_genotype_Aars2 | Mpz | -4.298 | 4.37E-03 | glio |
| Interaction_genotype_Aars2 | Neurl4 | -2.806 | 4.37E-03 | glio |
| Interaction_genotype_Aars2 | Fth1 | 1.303 | 4.37E-03 | glio |
| Interaction_genotype_Atr | Ccdc180 | -3.177 | 4.37E-03 | glio |
| Interaction_genotype_Atr | Gm49708 | -3.176 | 4.37E-03 | glio |
| Interaction_genotype_Bora | Tmem190 | -4.26 | 4.37E-03 | glio |
| Interaction_genotype_Bora | Slx1b | -3.312 | 4.37E-03 | glio |
| Interaction_genotype_Cdkn2a | Gm9758 | 3.095 | 4.37E-03 | glio |
| Interaction_genotype_Cdkn2a | Dapk2 | 3.096 | 4.37E-03 | glio |
| Interaction_genotype_Chek1 | Klhl30 | -3.72 | 4.37E-03 | glio |
| Interaction_genotype_Chek1 | 3010001F23Ri | -3.661 | 4.37E-03 | glio |
| Interaction_genotype_Chek1 | Washc5 | 1.935 | 4.37E-03 | glio |
| Interaction_genotype_Mtor | Vegfb | 1.602 | 4.37E-03 | glio |
| Interaction_genotype_Myc | Acy3 | -3.326 | 4.37E-03 | glio |
| Interaction_genotype_Myc | A630052C17R | -3.032 | 4.37E-03 | glio |
| Interaction_genotype_Myc | Pacc1 | -2.759 | 4.37E-03 | glio |
| Interaction_genotype_Myc | Aldh4a1 | -2.86 | 4.37E-03 | glio |
| Interaction_genotype_Pars2 | Rpia | 2.275 | 4.37E-03 | glio |
| Interaction_genotype_Terf1 | A430018G15R | -3.031 | 4.37E-03 | glio |
| interaction_RT | Fam160b1 | 0.983 | 4.37E-03 | glio |
| Aars2_in.vivo_noRT | Arhgap11a | 1.21 | 4.37E-03 | glio |
| Atr_in.vivo_noRT | Pld2 | 1.881 | 4.37E-03 | glio |
| Cdkn2a_in.vivo_noRT | Gm48617 | -1.938 | 4.37E-03 | glio |
| Cdkn2a_in.vivo_noRT | Gm47510 | -1.935 | 4.37E-03 | glio |
| Cdkn2a_in.vivo_noRT | Htr2b | -2.198 | 4.37E-03 | glio |
| Cenpj_in.vivo_noRT | Boc | 1.758 | 4.37E-03 | glio |
| Cenpj_in.vivo_noRT | Ly6a | 1.425 | 4.37E-03 | glio |
| Eif2s1_in.vivo_noRT | Mtm1 | -1.797 | 4.37E-03 | glio |
| Hsd17b10_in.vivo_noRT | Tmc7 | 2.331 | 4.37E-03 | glio |
| Interaction_genotype_Myc | Vax2 | -3.85 | 4.37E-04 | glio |
| Interaction_genotype_Myc | Lfng | -3.875 | 4.37E-04 | glio |
| Aars2_in.vivo_noRT | Pacs1 | 2.465 | 4.37E-04 | glio |
| Atp5c1_in.vivo_noRT | Gm41318 | 2.668 | 4.37E-04 | glio |
| Bcas2_in.vivo_noRT | Abcc10 | 2.616 | 4.37E-04 | glio |
| Bcas2_in.vivo_noRT | Rps6ka4 | 2.378 | 4.37E-04 | glio |
| Chek1_in.vivo_noRT | Eef1akmt3 | 2.357 | 4.37E-04 | glio |
| Chek1_ex.vivo_noRT | Nkain4 | 3.066 | 4.37E-04 | glio |
| Aars2_in.vivo_noRT | Gja4 | 2.739 | 4.37E-05 | glio |
| Atr_in.vivo_noRT | Cyb561 | 3.059 | 4.37E-05 | glio |
| Atr_in.vivo_noRT | Esr1 | 2.677 | 4.37E-05 | glio |
| Atr_ex.vivo_noRT | Smpd5 | 3.375 | 4.37E-05 | glio |
| Bcas2_in.vivo_noRT | Gemin4 | 3.062 | 4.37E-05 | glio |
| Bcas2_in.vivo_noRT | Tmem40 | 3.24 | 4.37E-05 | glio |
| culture_effect | Gm49207 | -2.47 | 4.37E-06 | glio |
| Chek1_in.vivo_noRT | 4933433G15R | 3.378 | 4.37E-06 | glio |
| Chek1_in.vivo_noRT | Gal3st4 | 3.214 | 4.37E-06 | glio |
| Hsd17b10_in.vivo_noRT | Lst1 | 3.382 | 4.37E-06 | glio |
| culture_effect | Galm | 2.205 | 4.37E-07 | glio |
| Interaction_genotype_Mtor | Gm49494 | -6.713 | 4.37E-08 | glio |
| Bora_in.vivo_noRT | Klhl30 | 3.751 | 4.37E-08 | glio |
| culture_effect | Fbl | -0.687 | 4.37E-10 | glio |
| Bcas2_in.vivo_noRT | B230311B06R | 5.073 | 4.37E-12 | glio |

| | | | | |
|---|---|---|---|---|
| Chek1_ex.vivo_noRT | Nrip2 | 4.842 | 4.37E-16 | glio |
| culture_effect | Slc25a39 | 1.058 | 4.37E-18 | glio |
| Interaction_genotype_Atr | Mob1b | 1.626 | 4.38E-03 | glio |
| Interaction_genotype_Mtor | Slc35d2 | -3.558 | 4.38E-03 | glio |
| Interaction_genotype_Mtor | Sod2 | 1.495 | 4.38E-03 | glio |
| Interaction_genotype_Myc | Magea10 | -2.841 | 4.38E-03 | glio |
| Interaction_genotype_Pars2 | Rab3b | -3.784 | 4.38E-03 | glio |
| Aars2_in.vivo_noRT | Prpf8 | -1.007 | 4.38E-03 | glio |
| Atr_in.vivo_noRT | Asb11 | 1.875 | 4.38E-03 | glio |
| Atr_ex.vivo_noRT | Gm13404 | 2.755 | 4.38E-03 | glio |
| Bcas2_in.vivo_noRT | Poll | 1.606 | 4.38E-03 | glio |
| Bora_in.vivo_noRT | Faim2 | 1.674 | 4.38E-03 | glio |
| Bora_in.vivo_noRT | Asap2 | -1.455 | 4.38E-03 | glio |
| Chek1_in.vivo_noRT | Taldo1 | -0.814 | 4.38E-03 | glio |
| Chek1_ex.vivo_noRT | Myl4 | 2.699 | 4.38E-03 | glio |
| Interaction_genotype_Myc | Zfp729a | -3.734 | 4.38E-04 | glio |
| Interaction_genotype_Rpl3 | D730045B01R | -4.192 | 4.38E-04 | glio |
| Bcas2_in.vivo_noRT | Ppargc1b | 2.582 | 4.38E-04 | glio |
| Chek1_in.vivo_noRT | Itga7 | 2.568 | 4.38E-04 | glio |
| Chek1_in.vivo_noRT | 4930556M19F | 2.396 | 4.38E-04 | glio |
| Chek1_ex.vivo_noRT | 4930518J21Ri | 3.079 | 4.38E-04 | glio |
| culture_effect | Eea1 | 0.644 | 4.38E-05 | glio |
| Atp5c1_in.vivo_noRT | Gm49086 | 2.668 | 4.38E-05 | glio |
| Atp5c1_in.vivo_noRT | Akap14 | 2.405 | 4.38E-05 | glio |
| Bcas2_in.vivo_noRT | 2810021J22Ri | 3.017 | 4.38E-05 | glio |
| Bora_in.vivo_noRT | Pcp4l1 | 2.978 | 4.38E-05 | glio |
| Chek1_ex.vivo_noRT | Mgarp | 3.707 | 4.38E-05 | glio |
| culture_effect | Npl | 2.111 | 4.38E-06 | glio |
| Interaction_genotype_Myc | C230013L11Ri | -5.662 | 4.38E-06 | glio |
| Interaction_genotype_Myc | Cpeb1 | -5.546 | 4.38E-06 | glio |
| Aars2_ex.vivo_noRT | Ptprc | 3.293 | 4.38E-06 | glio |
| Atp5c1_in.vivo_noRT | Gm12703 | 2.402 | 4.38E-06 | glio |
| Bcas2_in.vivo_noRT | Lsp1 | 3.62 | 4.38E-06 | glio |
| Chek1_in.vivo_noRT | Kpnb1 | -1.493 | 4.38E-06 | glio |
| Aars2_in.vivo_noRT | Gm28875 | 3.632 | 4.38E-07 | glio |
| Atr_in.vivo_noRT | Wnt4 | 3.891 | 4.38E-07 | glio |
| Atr_in.vivo_noRT | Myl9 | 4.508 | 4.38E-09 | glio |
| Chek1_in.vivo_noRT | Gm49701 | 3.994 | 4.38E-09 | glio |
| Chek1_ex.vivo_noRT | Gm43701 | 3.86 | 4.38E-10 | glio |
| culture_effect | Stx3 | -1.398 | 4.38E-12 | glio |
| Aars2_in.vivo_noRT | Gm47101 | 4.535 | 4.38E-12 | glio |
| Hsd17b10_in.vivo_noRT | Prl3d3 | 3.353 | 4.38E-12 | glio |
| Chek1_in.vivo_noRT | Gm10642 | 4.78 | 4.38E-13 | glio |
| Chek1_in.vivo_noRT | Tff2 | 4.784 | 4.38E-13 | glio |
| Atr_in.vivo_noRT | 4930570E03Ri | 5.273 | 4.38E-22 | glio |
| Atr_in.vivo_noRT | 4930447K03R | 5.273 | 4.38E-22 | glio |
| culture_effect | Gstp3 | -1.839 | 4.39E-03 | glio |
| culture_effect | 5430427N15R | -1.705 | 4.39E-03 | glio |
| culture_effect | Tmco4 | -1.086 | 4.39E-03 | glio |
| culture_effect | Abtb2 | -0.71 | 4.39E-03 | glio |
| culture_effect | Prim1 | -0.569 | 4.39E-03 | glio |
| culture_effect | Hnrnpab | 0.289 | 4.39E-03 | glio |

| | | | | |
|---|---|---|---|---|
| Interaction_genotype_Aars2 | Fam50a | 1.654 | 4.39E-03 | glio |
| Interaction_genotype_Aars2 | Rps6 | 0.838 | 4.39E-03 | glio |
| Interaction_genotype_Atr | Gm10354 | -2.866 | 4.39E-03 | glio |
| Interaction_genotype_Atr | Nudt7 | -3.251 | 4.39E-03 | glio |
| Interaction_genotype_Atr | Chit1 | 4.121 | 4.39E-03 | glio |
| Interaction_genotype_Atr | Col5a3 | -2.434 | 4.39E-03 | glio |
| Interaction_genotype_Atr | Rpl15 | 0.972 | 4.39E-03 | glio |
| Interaction_genotype_Bora | Greb1 | 1.936 | 4.39E-03 | glio |
| Interaction_genotype_Mtor | Tmem150c | -3.499 | 4.39E-03 | glio |
| Interaction_genotype_Mtor | A230072E10Ri | -3.411 | 4.39E-03 | glio |
| Interaction_genotype_Mtor | Pitpnm2 | -2.995 | 4.39E-03 | glio |
| Interaction_genotype_Myc | Siah1b | -2.239 | 4.39E-03 | glio |
| Interaction_genotype_Rad21 | Cxcr6 | 4.409 | 4.39E-03 | glio |
| interaction_RT | Arhgef16 | 2.437 | 4.39E-03 | glio |
| interaction_RT | Pold3 | -0.859 | 4.39E-03 | glio |
| Bard1_ex.vivo_noRT | 2310003N18R | 2.421 | 4.39E-03 | glio |
| Bard1_ex.vivo_noRT | Slco2a1 | 2.814 | 4.39E-03 | glio |
| Bcas2_in.vivo_noRT | Prr15l | 2.116 | 4.39E-03 | glio |
| Bcas2_in.vivo_noRT | Gm45767 | 1.986 | 4.39E-03 | glio |
| Bcas2_in.vivo_noRT | Ywhaq | -0.603 | 4.39E-03 | glio |
| Bora_in.vivo_noRT | Pcdhga6 | 1.961 | 4.39E-03 | glio |
| Bora_in.vivo_noRT | Atg3 | -1.298 | 4.39E-03 | glio |
| Bora_in.vivo_noRT | Htra2 | -1.334 | 4.39E-03 | glio |
| Bora_in.vivo_noRT | Ppil1 | -1.101 | 4.39E-03 | glio |
| Bora_ex.vivo_noRT | Tmem86a | 2.995 | 4.39E-03 | glio |
| Cenpj_in.vivo_noRT | Maged2 | -0.791 | 4.39E-03 | glio |
| Cenpj_ex.vivo_noRT | Adm | 2.982 | 4.39E-03 | glio |
| Chek1_ex.vivo_noRT | Llgl2 | 2.618 | 4.39E-03 | glio |
| Interaction_genotype_Chek1 | Gm4841 | 5.401 | 4.39E-04 | glio |
| Interaction_genotype_Myc | Lmx1b | -4.092 | 4.39E-04 | glio |
| Aars2_in.vivo_noRT | Kctd7 | 2.324 | 4.39E-04 | glio |
| Aars2_in.vivo_noRT | Pcmt1 | -0.885 | 4.39E-04 | glio |
| Atp5c1_in.vivo_noRT | Gm27239 | 2.668 | 4.39E-04 | glio |
| Atp5c1_in.vivo_noRT | 4930412M03R | 2.668 | 4.39E-04 | glio |
| Atr_in.vivo_noRT | Plvap | 2.482 | 4.39E-04 | glio |
| Atr_in.vivo_noRT | Synm | 2.247 | 4.39E-04 | glio |
| Bard1_ex.vivo_noRT | Gm20275 | 2.814 | 4.39E-04 | glio |
| Cenpj_ex.vivo_noRT | Zscan4-ps2 | 2.591 | 4.39E-04 | glio |
| Chek1_in.vivo_noRT | Gm29107 | 2.607 | 4.39E-04 | glio |
| Cyp19a1_ex.vivo_noRT | Dmrtc1a | 2.933 | 4.39E-04 | glio |
| Aars2_in.vivo_noRT | Ebf4 | 2.95 | 4.39E-05 | glio |
| Atp5c1_in.vivo_noRT | Gm16933 | 2.146 | 4.39E-05 | glio |
| Bora_in.vivo_noRT | Gm15910 | 2.81 | 4.39E-05 | glio |
| Bora_in.vivo_noRT | Commd4 | -1.72 | 4.39E-05 | glio |
| Cenpj_in.vivo_noRT | Gm20705 | 2.789 | 4.39E-05 | glio |
| Interaction_genotype_Mtor | Gramd2 | -5.16 | 4.39E-06 | glio |
| Chek1_ex.vivo_noRT | Gm43256 | 3.672 | 4.39E-06 | glio |
| Hsd17b10_in.vivo_noRT | Slc6a11 | 2.715 | 4.39E-06 | glio |
| culture_effect | Tnni1 | 1.345 | 4.39E-07 | glio |
| Aars2_in.vivo_noRT | Rrp15 | -1.668 | 4.39E-07 | glio |
| Bora_in.vivo_noRT | Gm50103 | 3.629 | 4.39E-07 | glio |
| culture_effect | 1700120C14R | 2.361 | 4.39E-09 | glio |

| | | | | |
|---|---|---|---|---|
| culture_effect | Ndufc1 | 0.745 | 4.39E-09 | glio |
| Aars2_in.vivo_noRT | Dok3 | 4.191 | 4.39E-09 | glio |
| Bora_in.vivo_noRT | Nrip3 | 3.892 | 4.39E-09 | glio |
| culture_effect | C3 | -3.937 | 4.39E-10 | glio |
| Chek1_ex.vivo_noRT | Esm1 | 3.867 | 4.39E-10 | glio |
| Chek1_in.vivo_noRT | Vmn1r181 | 4.388 | 4.39E-11 | glio |
| Atr_in.vivo_noRT | B230311B06R | 5.27 | 4.39E-13 | glio |
| Bora_in.vivo_noRT | Nkg7 | 4.676 | 4.39E-15 | glio |
| Chek1_ex.vivo_noRT | Gtsf1 | 5.455 | 4.39E-15 | glio |
| Chek1_ex.vivo_noRT | Cd200r2 | 4.842 | 4.39E-15 | glio |
| Chek1_in.vivo_noRT | Gm19265 | 4.528 | 4.39E-16 | glio |
| Interaction_genotype_Chek1 | Kansl1 | 1.91 | 4.40E-03 | glio |
| Interaction_genotype_Ifnar1 | 1110017D15R | 4.153 | 4.40E-03 | glio |
| Interaction_genotype_Ifnar1 | Gm13191 | 2.734 | 4.40E-03 | glio |
| Interaction_genotype_Ifnar1 | 4933416E03Ri | 2.734 | 4.40E-03 | glio |
| Interaction_genotype_Ifnar1 | Tap1 | -1.497 | 4.40E-03 | glio |
| Interaction_genotype_Mtor | Nbeal2 | -3.63 | 4.40E-03 | glio |
| Interaction_genotype_Mtor | Ccl2 | -8.224 | 4.40E-03 | glio |
| Interaction_genotype_Myc | Ms4a6d | -2.729 | 4.40E-03 | glio |
| interaction_RT | Gm15417 | 1.372 | 4.40E-03 | glio |
| Aars2_in.vivo_noRT | Eif4a1 | -0.675 | 4.40E-03 | glio |
| Atr_in.vivo_noRT | Gm11423 | 1.697 | 4.40E-03 | glio |
| Atr_in.vivo_noRT | Pecr | -1.43 | 4.40E-03 | glio |
| Bard1_ex.vivo_noRT | Spink4 | 2.814 | 4.40E-03 | glio |
| Bcas2_in.vivo_noRT | Fbxo24 | 2.088 | 4.40E-03 | glio |
| Bcas2_in.vivo_noRT | Lhfpl4 | 1.905 | 4.40E-03 | glio |
| Bcas2_in.vivo_noRT | Acad11 | 1.743 | 4.40E-03 | glio |
| Bcas2_ex.vivo_noRT | Gbp10 | 2.742 | 4.40E-03 | glio |
| Chek1_ex.vivo_noRT | Bcap29 | -1.22 | 4.40E-03 | glio |
| Chek1_ex.vivo_noRT | Bex1 | -1.442 | 4.40E-03 | glio |
| Chek1_ex.vivo_noRT | Sdhb | -0.816 | 4.40E-03 | glio |
| Hsd17b10_in.vivo_noRT | Tbkbp1 | 2.323 | 4.40E-03 | glio |
| Interaction_genotype_Ifnar1 | Psmb9 | -2.128 | 4.40E-04 | glio |
| Interaction_genotype_Mtor | Spink6 | -3.307 | 4.40E-04 | glio |
| Interaction_genotype_Mtor | Gba2 | -4.331 | 4.40E-04 | glio |
| Interaction_genotype_Mtor | Pcbp1 | 1.679 | 4.40E-04 | glio |
| Aars2_in.vivo_noRT | Zfp764 | 2.334 | 4.40E-04 | glio |
| Aars2_ex.vivo_noRT | Lrrc4b | 2.897 | 4.40E-04 | glio |
| Aars2_ex.vivo_noRT | Gm44646 | 2.706 | 4.40E-04 | glio |
| Atr_in.vivo_noRT | 2700046G09R | 2.04 | 4.40E-04 | glio |
| Atr_ex.vivo_noRT | A330032B11R | 2.957 | 4.40E-04 | glio |
| Bcas2_in.vivo_noRT | Elf3 | 2.767 | 4.40E-04 | glio |
| Bora_in.vivo_noRT | Gm26621 | 2.534 | 4.40E-04 | glio |
| Bora_in.vivo_noRT | Plxna2 | 2.743 | 4.40E-04 | glio |
| Bora_in.vivo_noRT | Jpx | 1.795 | 4.40E-04 | glio |
| Chek1_in.vivo_noRT | Atpif1 | -0.761 | 4.40E-04 | glio |
| Chek1_ex.vivo_noRT | Gm39459 | 3.174 | 4.40E-04 | glio |
| culture_effect | Phf5a | 0.435 | 4.40E-05 | glio |
| culture_effect | Rpl13 | -0.587 | 4.40E-05 | glio |
| Bora_in.vivo_noRT | Slc11a1 | 2.867 | 4.40E-05 | glio |
| Hsd17b10_in.vivo_noRT | C230013L11Ri | 3.235 | 4.40E-05 | glio |
| culture_effect | Pxmp2 | 1.503 | 4.40E-06 | glio |

| | | | | |
|---|---|---|---|---|
| Interaction_genotype_Mtor | Trpm3 | -5.424 | 4.40E-06 | glio |
| Bard1_ex.vivo_noRT | Gm38163 | 3.596 | 4.40E-06 | glio |
| Bard1_ex.vivo_noRT | Apol10b | 3.596 | 4.40E-06 | glio |
| Chek1_in.vivo_noRT | Gm10401 | 2.841 | 4.40E-06 | glio |
| Chek1_ex.vivo_noRT | Rarres2 | 4.049 | 4.40E-06 | glio |
| Chek1_ex.vivo_noRT | Comp | 4.449 | 4.40E-06 | glio |
| Bcas2_in.vivo_noRT | Celf5 | 3.744 | 4.40E-07 | glio |
| Chek1_in.vivo_noRT | 1700001L05Ri | 3.346 | 4.40E-07 | glio |
| Hsd17b10_in.vivo_noRT | Fbxl13 | 3.356 | 4.40E-08 | glio |
| Chek1_in.vivo_noRT | Gsx2 | 4.358 | 4.40E-09 | glio |
| culture_effect | Echs1 | -0.931 | 4.40E-11 | glio |
| culture_effect | Mfsd10 | -1.258 | 4.40E-13 | glio |
| Atr_in.vivo_noRT | Tas1r1 | 4.506 | 4.40E-13 | glio |
| Chek1_in.vivo_noRT | Gm11457 | 5.044 | 4.40E-14 | glio |
| Bora_in.vivo_noRT | Lrguk | 4.672 | 4.40E-16 | glio |
| Bora_in.vivo_noRT | Bco2 | 4.936 | 4.40E-16 | glio |
| Chek1_in.vivo_noRT | Rapgef3os2 | 5.555 | 4.40E-25 | glio |
| Chek1_in.vivo_noRT | Gm47622 | 5.555 | 4.40E-25 | glio |
| Interaction_genotype_Aars2 | Fhl1 | -2.744 | 4.41E-03 | glio |
| Interaction_genotype_Aars2 | Zfp712 | -3.235 | 4.41E-03 | glio |
| Interaction_genotype_Aars2 | Cdkn1c | -3.603 | 4.41E-03 | glio |
| Interaction_genotype_Aars2 | Lsm1 | 1.282 | 4.41E-03 | glio |
| Interaction_genotype_Atr | Gm34776 | -2.763 | 4.41E-03 | glio |
| Interaction_genotype_Bcas2 | Gm45338 | -2.458 | 4.41E-03 | glio |
| Interaction_genotype_Bcas2 | Zfp652os | -3.694 | 4.41E-03 | glio |
| Interaction_genotype_Bcas2 | Gm16794 | -3.643 | 4.41E-03 | glio |
| Interaction_genotype_Bora | 4930481B07R | -4.018 | 4.41E-03 | glio |
| Interaction_genotype_Mtor | Gsto2 | -3.566 | 4.41E-03 | glio |
| Interaction_genotype_Mtor | Fchsd1 | -3.236 | 4.41E-03 | glio |
| Interaction_genotype_Myc | Pard3bos1 | -2.911 | 4.41E-03 | glio |
| Interaction_genotype_Myc | E230001N04R | -2.831 | 4.41E-03 | glio |
| Interaction_genotype_Myc | Ankrd44 | -2.327 | 4.41E-03 | glio |
| Interaction_genotype_Pars2 | Gm31415 | -3.297 | 4.41E-03 | glio |
| Interaction_genotype_Pars2 | 5830454E08Ri | -3.427 | 4.41E-03 | glio |
| Interaction_genotype_Pars2 | Gm29340 | -3.367 | 4.41E-03 | glio |
| Aars2_in.vivo_noRT | Commd5 | 2.048 | 4.41E-03 | glio |
| Aars2_in.vivo_noRT | Gm33782 | 1.833 | 4.41E-03 | glio |
| Aars2_in.vivo_noRT | Optn | 2.056 | 4.41E-03 | glio |
| Aars2_in.vivo_noRT | Ddb2 | 1.626 | 4.41E-03 | glio |
| Aars2_in.vivo_noRT | Tarbp1 | 1.708 | 4.41E-03 | glio |
| Aars2_in.vivo_noRT | Abhd17a | -0.919 | 4.41E-03 | glio |
| Aars2_in.vivo_noRT | Ranbp1 | -0.718 | 4.41E-03 | glio |
| Bcas2_in.vivo_noRT | Klhl32 | 1.849 | 4.41E-03 | glio |
| Bora_in.vivo_noRT | Gh | 2.351 | 4.41E-03 | glio |
| Bora_in.vivo_noRT | Lysmd4 | 1.923 | 4.41E-03 | glio |
| Bora_in.vivo_noRT | Prdm5 | -1.446 | 4.41E-03 | glio |
| Bora_in.vivo_noRT | Tmem128 | -0.768 | 4.41E-03 | glio |
| Cenpj_in.vivo_noRT | Gm26555 | 1.759 | 4.41E-03 | glio |
| Cenpj_ex.vivo_noRT | Dcdc2a | 2.591 | 4.41E-03 | glio |
| Interaction_genotype_Aars2 | Pcx | -4.313 | 4.41E-04 | glio |
| Interaction_genotype_Aars2 | Gm45605 | -4.389 | 4.41E-04 | glio |
| Interaction_genotype_Myc | Gm26673 | -3.877 | 4.41E-04 | glio |

| | | | | |
|---|---|---|---|---|
| interaction_RT | Sult2b1 | 2.199 | 4.41E-04 | glio |
| interaction_RT | Stip1 | -0.638 | 4.41E-04 | glio |
| interaction_RT | Cox6b1 | 0.603 | 4.41E-04 | glio |
| Aars2_in.vivo_noRT | Ybx1 | -0.678 | 4.41E-04 | glio |
| Bcas2_ex.vivo_noRT | Spink6 | 2.742 | 4.41E-04 | glio |
| Bcas2_ex.vivo_noRT | Gm40264 | 2.937 | 4.41E-04 | glio |
| Bcas2_ex.vivo_noRT | B230206H07R | 2.347 | 4.41E-04 | glio |
| Bcas2_ex.vivo_noRT | Htr2b | 2.742 | 4.41E-04 | glio |
| Chek1_in.vivo_noRT | Vav2 | 2.519 | 4.41E-04 | glio |
| Chek1_ex.vivo_noRT | Fam169a | 3.074 | 4.41E-04 | glio |
| Fanca_ex.vivo_noRT | Gm5941 | 3.971 | 4.41E-04 | glio |
| Hsd17b10_in.vivo_noRT | Gm47545 | 2.365 | 4.41E-04 | glio |
| Interaction_genotype_Mtor | Ndufb2 | 2.048 | 4.41E-05 | glio |
| Interaction_genotype_Rpl3 | AV356131 | -5.171 | 4.41E-05 | glio |
| Atr_in.vivo_noRT | Gm26692 | 2.74 | 4.41E-05 | glio |
| Atr_ex.vivo_noRT | Hist3h2a | 3.215 | 4.41E-05 | glio |
| culture_effect | Ccdc7b | -2.778 | 4.41E-06 | glio |
| Aars2_in.vivo_noRT | Prkaa2 | 2.789 | 4.41E-06 | glio |
| Bcas2_in.vivo_noRT | Nox1 | 2.824 | 4.41E-06 | glio |
| Interaction_genotype_Mtor | Styk1 | -6.236 | 4.41E-07 | glio |
| Bcas2_in.vivo_noRT | Rab11fip4os1 | 3.771 | 4.41E-07 | glio |
| Bora_in.vivo_noRT | Gm13610 | 3.78 | 4.41E-08 | glio |
| Aars2_in.vivo_noRT | Gm29340 | 4.011 | 4.41E-09 | glio |
| Chek1_in.vivo_noRT | Gm12840 | 3.785 | 4.41E-09 | glio |
| culture_effect | Gm32834 | 3.199 | 4.41E-11 | glio |
| culture_effect | Zfp672 | 1.512 | 4.41E-11 | glio |
| Chek1_in.vivo_noRT | Dnm3 | 4.518 | 4.41E-11 | glio |
| Chek1_in.vivo_noRT | Cxcl16 | 4.78 | 4.41E-13 | glio |
| culture_effect | Nbdy | 1.123 | 4.41E-16 | glio |
| Atr_in.vivo_noRT | Slc6a11 | 4.639 | 4.41E-16 | glio |
| Bcas2_in.vivo_noRT | Abra | 4.809 | 4.41E-16 | glio |
| Bcas2_in.vivo_noRT | Rassf6 | 4.807 | 4.41E-16 | glio |
| Chek1_ex.vivo_noRT | Gm5157 | 4.842 | 4.41E-17 | glio |
| Aars2_in.vivo_noRT | Btn1a1 | 5.067 | 4.41E-20 | glio |
| Atr_in.vivo_noRT | 4933400F21Ri | 4.742 | 4.41E-21 | glio |
| Interaction_genotype_Aars2 | Gm4189 | -3.555 | 4.42E-03 | glio |
| Interaction_genotype_Aars2 | Dynlrb1 | 1.038 | 4.42E-03 | glio |
| Interaction_genotype_Chek1 | Arl1 | 1.429 | 4.42E-03 | glio |
| Interaction_genotype_Ifnar1 | Slc22a1 | 3.513 | 4.42E-03 | glio |
| Interaction_genotype_Mtor | Ebp | 1.495 | 4.42E-03 | glio |
| Interaction_genotype_Myc | Ksr2 | -2.667 | 4.42E-03 | glio |
| Interaction_genotype_Myc | Gm10282 | -3.209 | 4.42E-03 | glio |
| Interaction_genotype_Myc | Ccdc103 | -3.178 | 4.42E-03 | glio |
| Interaction_genotype_Myc | Gm20045 | -2.722 | 4.42E-03 | glio |
| Interaction_genotype_Myc | Oas1g | -2.826 | 4.42E-03 | glio |
| interaction_RT | Stat4 | 2.73 | 4.42E-03 | glio |
| interaction_RT | Foxo3 | 1.26 | 4.42E-03 | glio |
| interaction_RT | 2310009B15R | 0.912 | 4.42E-03 | glio |
| interaction_RT | Espn | -0.601 | 4.42E-03 | glio |
| Atr_in.vivo_noRT | Jakmip2 | 1.892 | 4.42E-03 | glio |
| Bcas2_in.vivo_noRT | Ccdc146 | 2.209 | 4.42E-03 | glio |
| Bcas2_in.vivo_noRT | Bglap3 | 2.088 | 4.42E-03 | glio |

| | | | | |
|---|---|---|---|---|
| Chek1_in.vivo_noRT | Zfp983 | 1.812 | 4.42E-03 | glio |
| culture_effect | Nae1 | -0.58 | 4.42E-04 | glio |
| culture_effect | Plin3 | -0.574 | 4.42E-04 | glio |
| Interaction_genotype_Mtor | Gm38115 | -4.34 | 4.42E-04 | glio |
| Interaction_genotype_Myc | Spsb4 | -3.727 | 4.42E-04 | glio |
| Aars2_in.vivo_noRT | Tsfm | -1.222 | 4.42E-04 | glio |
| Atr_in.vivo_noRT | Syngr4 | 2.492 | 4.42E-04 | glio |
| Atr_ex.vivo_noRT | Cysltr2 | 3.164 | 4.42E-04 | glio |
| Bcas2_in.vivo_noRT | Abca16 | 2.398 | 4.42E-04 | glio |
| culture_effect | Nid1 | 2.31 | 4.42E-05 | glio |
| culture_effect | Vasp | -0.789 | 4.42E-05 | glio |
| Interaction_genotype_Mtor | 9330104G04R | -4.726 | 4.42E-05 | glio |
| Interaction_genotype_Mtor | Kpnb1 | 2.017 | 4.42E-05 | glio |
| Interaction_genotype_Myc | Tmod2 | -2.819 | 4.42E-05 | glio |
| Aars2_in.vivo_noRT | Acacb | 2.825 | 4.42E-05 | glio |
| Bcas2_in.vivo_noRT | Fxyd4 | 3.209 | 4.42E-05 | glio |
| culture_effect | Gm45833 | 3.162 | 4.42E-06 | glio |
| Interaction_genotype_Mtor | Gm47404 | -4.099 | 4.42E-07 | glio |
| Interaction_genotype_Mtor | 1700112J16Ri | -4.685 | 4.42E-07 | glio |
| Atr_in.vivo_noRT | Phtf1os | 3.609 | 4.42E-07 | glio |
| Bcas2_in.vivo_noRT | Gm49694 | 4.081 | 4.42E-08 | glio |
| Chek1_ex.vivo_noRT | Kcnd1 | 4.443 | 4.42E-09 | glio |
| Bora_in.vivo_noRT | Plcd4 | 4.026 | 4.42E-10 | glio |
| Chek1_ex.vivo_noRT | Olfr56 | 4.842 | 4.42E-14 | glio |
| Atr_in.vivo_noRT | Gm47540 | 5.533 | 4.42E-26 | glio |
| Atr_in.vivo_noRT | BB287469 | 5.533 | 4.42E-26 | glio |
| culture_effect | Ntm | 3.419 | 4.42E-35 | glio |
| culture_effect | Apol6 | -6.278 | 4.42E-35 | glio |
| culture_effect | Tap1 | -3.514 | 4.42E-58 | glio |
| culture_effect | H2-T23 | -3.501 | 4.42E-58 | glio |
| Interaction_genotype_Aars2 | Ankef1 | -3.65 | 4.43E-03 | glio |
| Interaction_genotype_Aars2 | Gm36210 | -3.507 | 4.43E-03 | glio |
| Interaction_genotype_Aars2 | Gm43728 | -3.694 | 4.43E-03 | glio |
| Interaction_genotype_Bora | Clec4d | -2.92 | 4.43E-03 | glio |
| Interaction_genotype_Bora | Gm34121 | -3.372 | 4.43E-03 | glio |
| Interaction_genotype_Bora | Styk1 | -3.873 | 4.43E-03 | glio |
| Interaction_genotype_Bora | AC163347.1 | -3.054 | 4.43E-03 | glio |
| Interaction_genotype_Bora | Cox7b2 | -3.374 | 4.43E-03 | glio |
| Interaction_genotype_Bora | Gm37584 | -3.878 | 4.43E-03 | glio |
| Interaction_genotype_Bora | Mettl6 | 2.223 | 4.43E-03 | glio |
| Interaction_genotype_Mtor | Matk | -3.31 | 4.43E-03 | glio |
| Interaction_genotype_Mtor | Zfp583 | -3.591 | 4.43E-03 | glio |
| Interaction_genotype_Myc | Gpkow | -2.401 | 4.43E-03 | glio |
| Interaction_genotype_Myc | Mtfp1 | -2.438 | 4.43E-03 | glio |
| interaction_RT | Rbmx | -1.227 | 4.43E-03 | glio |
| Bcas2_in.vivo_noRT | Intu | 2.009 | 4.43E-03 | glio |
| Bcas2_in.vivo_noRT | Zfp608 | -1.258 | 4.43E-03 | glio |
| Bora_in.vivo_noRT | Cd38 | 2.07 | 4.43E-03 | glio |
| Cenpj_in.vivo_noRT | Slc45a1 | 1.762 | 4.43E-03 | glio |
| Chek1_in.vivo_noRT | Mphosph8 | -1.135 | 4.43E-03 | glio |
| Chek1_ex.vivo_noRT | Ifitm6 | 2.619 | 4.43E-03 | glio |
| culture_effect | Gm15952 | 1.824 | 4.43E-04 | glio |

| | | | | |
|---|---|---|---|---|
| culture_effect | Ssb | -0.391 | 4.43E-04 | glio |
| Interaction_genotype_Aars2 | Pecam1 | -4.549 | 4.43E-04 | glio |
| Interaction_genotype_Chek1 | Slc38a3 | 4.366 | 4.43E-04 | glio |
| Interaction_genotype_Mtor | 1700047K16R | -4.324 | 4.43E-04 | glio |
| Interaction_genotype_Myc | Tpo | -2.729 | 4.43E-04 | glio |
| Interaction_genotype_Myc | 1700012D14R | -3.675 | 4.43E-04 | glio |
| Interaction_genotype_Myc | Lclat1 | -2.408 | 4.43E-04 | glio |
| interaction_RT | Polr1a | -0.934 | 4.43E-04 | glio |
| Aars2_in.vivo_noRT | Gpr146 | 2.544 | 4.43E-04 | glio |
| Aars2_in.vivo_noRT | Tmprss5 | 2.491 | 4.43E-04 | glio |
| Aars2_in.vivo_noRT | Ngfr | 2.074 | 4.43E-04 | glio |
| Atp5c1_in.vivo_noRT | Gm1123 | 2.808 | 4.43E-04 | glio |
| Atr_in.vivo_noRT | Atp5j | -0.628 | 4.43E-04 | glio |
| Bora_in.vivo_noRT | Ppard | 1.838 | 4.43E-04 | glio |
| Bora_ex.vivo_noRT | A930003A15R | 3.201 | 4.43E-04 | glio |
| Bora_ex.vivo_noRT | Vamp8 | 2.802 | 4.43E-04 | glio |
| Cenpj_ex.vivo_noRT | Scgb1c1 | 2.996 | 4.43E-04 | glio |
| Chek1_in.vivo_noRT | Kank3 | 2.418 | 4.43E-04 | glio |
| Chek1_in.vivo_noRT | Rpgr | 2.388 | 4.43E-04 | glio |
| Chek1_ex.vivo_noRT | Slf1 | -2.281 | 4.43E-04 | glio |
| Interaction_genotype_Myc | Elob | 1.538 | 4.43E-05 | glio |
| Atr_in.vivo_noRT | Eml6 | 2.933 | 4.43E-05 | glio |
| Bora_in.vivo_noRT | Gm37986 | 2.599 | 4.43E-05 | glio |
| Bora_in.vivo_noRT | Gm28535 | 2.798 | 4.43E-05 | glio |
| Chek1_in.vivo_noRT | Kctd12 | 2.777 | 4.43E-05 | glio |
| Chek1_ex.vivo_noRT | Hsf4 | 3.672 | 4.43E-05 | glio |
| Bcas2_in.vivo_noRT | D430018E03R | 3.081 | 4.43E-06 | glio |
| Bora_in.vivo_noRT | Schip1 | -2.176 | 4.43E-06 | glio |
| culture_effect | Tpm4 | 0.641 | 4.43E-08 | glio |
| Interaction_genotype_Mtor | Jph4 | -6.635 | 4.43E-08 | glio |
| Hsd17b10_in.vivo_noRT | Gm20705 | 3.084 | 4.43E-08 | glio |
| Atp5c1_in.vivo_noRT | Gm13398 | 3.196 | 4.43E-09 | glio |
| Bcas2_in.vivo_noRT | Ushbp1 | 4.281 | 4.43E-10 | glio |
| Chek1_in.vivo_noRT | Gm12280 | 4.78 | 4.43E-10 | glio |
| culture_effect | Gm13056 | 3.343 | 4.43E-11 | glio |
| Aars2_in.vivo_noRT | Gm27239 | 5.327 | 4.43E-15 | glio |
| culture_effect | Tspan13 | -1.502 | 4.43E-17 | glio |
| Atr_in.vivo_noRT | Serpinb2 | 5.533 | 4.43E-19 | glio |
| culture_effect | Hmga2 | 1.507 | 4.43E-21 | glio |
| Bcas2_in.vivo_noRT | Gm26671 | 4.807 | 4.43E-22 | glio |
| Bora_in.vivo_noRT | Sucnr1 | 4.936 | 4.43E-27 | glio |
| Bora_in.vivo_noRT | Idi2 | 5.46 | 4.43E-27 | glio |
| Interaction_genotype_Aars2 | Aarsd1 | 1.574 | 4.44E-03 | glio |
| Interaction_genotype_Atr | Hoxc8 | -2.369 | 4.44E-03 | glio |
| Interaction_genotype_Chek1 | Gbp2b | 4 | 4.44E-03 | glio |
| Interaction_genotype_Mtor | Dact1 | -3.331 | 4.44E-03 | glio |
| Interaction_genotype_Mtor | Actl6a | 1.518 | 4.44E-03 | glio |
| Interaction_genotype_Myc | 4930594M22F | -3.156 | 4.44E-03 | glio |
| interaction_RT | Gga1 | -0.74 | 4.44E-03 | glio |
| Aars2_in.vivo_noRT | Tex30 | -1.124 | 4.44E-03 | glio |
| Aars2_in.vivo_noRT | Tubb2b | -0.882 | 4.44E-03 | glio |
| Atr_in.vivo_noRT | D130040H23F | 1.93 | 4.44E-03 | glio |

| | | | | |
|---|---|---|---|---|
| Atr_in.vivo_noRT | Tubd1 | 1.759 | 4.44E-03 | glio |
| Atr_ex.vivo_noRT | Elfn2 | 2.589 | 4.44E-03 | glio |
| Atr_ex.vivo_noRT | Grip2 | 2.777 | 4.44E-03 | glio |
| Atr_ex.vivo_noRT | Sema3d | 2.738 | 4.44E-03 | glio |
| Cdkn2a_in.vivo_noRT | Wfdc12 | 1.798 | 4.44E-03 | glio |
| Cenpj_ex.vivo_noRT | Pcdh15 | 2.591 | 4.44E-03 | glio |
| culture_effect | Nebl | -0.878 | 4.44E-04 | glio |
| Interaction_genotype_Myc | Gjc1 | -3.015 | 4.44E-04 | glio |
| Atp5c1_in.vivo_noRT | Ric3 | 2.408 | 4.44E-04 | glio |
| Atp5c1_in.vivo_noRT | Accsl | 2.939 | 4.44E-04 | glio |
| Atp5c1_in.vivo_noRT | Ifitm5 | 2.667 | 4.44E-04 | glio |
| Atr_in.vivo_noRT | Hspa12b | 2.726 | 4.44E-04 | glio |
| Atr_in.vivo_noRT | Klf1 | 2.56 | 4.44E-04 | glio |
| Bora_in.vivo_noRT | Ccdc148 | 2.662 | 4.44E-04 | glio |
| Bora_in.vivo_noRT | Gpr135 | 2.509 | 4.44E-04 | glio |
| Chek1_ex.vivo_noRT | Lpin1 | 3.061 | 4.44E-04 | glio |
| Interaction_genotype_Mtor | Gm14636 | -5.358 | 4.44E-05 | glio |
| Interaction_genotype_Mtor | 2510046G10R | -3.783 | 4.44E-05 | glio |
| Interaction_genotype_Myc | Btbd16 | -4 | 4.44E-05 | glio |
| Interaction_genotype_Myc | Tyw3 | -3.79 | 4.44E-05 | glio |
| Interaction_genotype_Myc | 4933421A08R | -4.955 | 4.44E-05 | glio |
| Atp5c1_in.vivo_noRT | Gm48242 | 2.668 | 4.44E-05 | glio |
| Hsd17b10_in.vivo_noRT | Gm42984 | 2.966 | 4.44E-05 | glio |
| culture_effect | Pepd | -1.069 | 4.44E-06 | glio |
| Chek1_in.vivo_noRT | H60c | 3.45 | 4.44E-06 | glio |
| Chek1_in.vivo_noRT | Paqr8 | 3.677 | 4.44E-06 | glio |
| Hsd17b10_in.vivo_noRT | Camk1d | 3.342 | 4.44E-06 | glio |
| Interaction_genotype_Rad51 | Dnm3 | -6.19 | 4.44E-07 | glio |
| Atr_in.vivo_noRT | Aebp2 | -1.916 | 4.44E-07 | glio |
| Bcas2_in.vivo_noRT | Gm29340 | 3.491 | 4.44E-07 | glio |
| Bcas2_in.vivo_noRT | Gm12840 | 3.549 | 4.44E-08 | glio |
| Bora_in.vivo_noRT | Pde1a | 3.936 | 4.44E-08 | glio |
| Chek1_in.vivo_noRT | Gm13625 | 3.523 | 4.44E-08 | glio |
| culture_effect | Tmem183a | 1.041 | 4.44E-10 | glio |
| culture_effect | Meiob | -2.619 | 4.44E-11 | glio |
| Aars2_in.vivo_noRT | 4930412M03F | 5.327 | 4.44E-15 | glio |
| Atr_in.vivo_noRT | Tmem190 | 6.071 | 4.44E-15 | glio |
| Bora_in.vivo_noRT | Psca | 4.936 | 4.44E-15 | glio |
| Chek1_in.vivo_noRT | Gm15929 | 5.044 | 4.44E-19 | glio |
| Bcas2_in.vivo_noRT | Gm47510 | 4.547 | 4.44E-20 | glio |
| culture_effect | Slc35f5 | -0.823 | 4.45E-03 | glio |
| culture_effect | Mns1 | -0.559 | 4.45E-03 | glio |
| culture_effect | Cggbp1 | 0.385 | 4.45E-03 | glio |
| Interaction_genotype_Aars2 | Tmeff1 | -3.156 | 4.45E-03 | glio |
| Interaction_genotype_Bcas2 | Esm1 | -2.777 | 4.45E-03 | glio |
| Interaction_genotype_Myc | Tns4 | -2.876 | 4.45E-03 | glio |
| Interaction_genotype_Myc | Fam166a | -2.466 | 4.45E-03 | glio |
| Interaction_genotype_Myc | Ccdc71l | -2.944 | 4.45E-03 | glio |
| interaction_RT | Slc14a1 | 2.435 | 4.45E-03 | glio |
| Aars2_in.vivo_noRT | Slc44a2 | 2.017 | 4.45E-03 | glio |
| Aars2_in.vivo_noRT | Arv1 | 1.574 | 4.45E-03 | glio |
| Bora_in.vivo_noRT | Ddx5 | -0.48 | 4.45E-03 | glio |

| | | | | |
|---|---|---|---|---|
| Bora_in.vivo_noRT | H2afz | -0.627 | 4.45E-03 | glio |
| Ercc4_ex.vivo_noRT | Ercc4 | -3.319 | 4.45E-03 | glio |
| Hsd17b10_in.vivo_noRT | 4930513N10R | 2.535 | 4.45E-03 | glio |
| culture_effect | Phex | 1.504 | 4.45E-04 | glio |
| Interaction_genotype_Myc | Adam33 | -4.039 | 4.45E-04 | glio |
| Interaction_genotype_Myc | Fam151b | -3.981 | 4.45E-04 | glio |
| Interaction_genotype_Myc | Nhs | -3.353 | 4.45E-04 | glio |
| Interaction_genotype_Myc | Atp23 | -2.214 | 4.45E-04 | glio |
| Interaction_genotype_Myc | Bdnf | -3.373 | 4.45E-04 | glio |
| interaction_RT | Arl6ip5 | 1.178 | 4.45E-04 | glio |
| Bora_in.vivo_noRT | Carmil1 | 2.388 | 4.45E-04 | glio |
| Bora_ex.vivo_noRT | Gm6566 | 2.808 | 4.45E-04 | glio |
| Chek1_ex.vivo_noRT | Rack1 | -0.917 | 4.45E-04 | glio |
| culture_effect | Hps5 | 0.998 | 4.45E-05 | glio |
| Interaction_genotype_Mtor | 1810059H22R | -5.247 | 4.45E-05 | glio |
| Interaction_genotype_Myc | Arhgef39 | -4.272 | 4.45E-05 | glio |
| Bcas2_in.vivo_noRT | Tmem116 | 3.629 | 4.45E-06 | glio |
| Bcas2_in.vivo_noRT | Dlx2 | 3.424 | 4.45E-06 | glio |
| Bora_in.vivo_noRT | Tbc1d10c | 3.241 | 4.45E-06 | glio |
| Chek1_ex.vivo_noRT | B020031M17F | 4.842 | 4.45E-06 | glio |
| Hsd17b10_in.vivo_noRT | Apobr | 3.616 | 4.45E-06 | glio |
| Interaction_genotype_Myc | Gchfr | -6.502 | 4.45E-07 | glio |
| culture_effect | Snd1 | -0.758 | 4.45E-08 | glio |
| Atr_ex.vivo_noRT | Ly6d | 3.962 | 4.45E-08 | glio |
| Chek1_in.vivo_noRT | Gm29666 | 3.837 | 4.45E-08 | glio |
| Chek1_in.vivo_noRT | Esrp2 | 4.132 | 4.45E-10 | glio |
| Chek1_in.vivo_noRT | Hsf4 | 4.38 | 4.45E-10 | glio |
| Bcas2_in.vivo_noRT | Muc4 | 4.285 | 4.45E-14 | glio |
| Bora_in.vivo_noRT | Crybb1 | 4.549 | 4.45E-14 | glio |
| Chek1_in.vivo_noRT | Col5a2 | 4.255 | 4.45E-15 | glio |
| Chek1_in.vivo_noRT | Gm49223 | 4.521 | 4.45E-15 | glio |
| Aars2_in.vivo_noRT | Gm47936 | 5.327 | 4.45E-23 | glio |
| culture_effect | Phyh | -1.022 | 4.46E-03 | glio |
| Interaction_genotype_Aars2 | Cyp51 | 1.932 | 4.46E-03 | glio |
| Interaction_genotype_Aars2 | Ndufa11 | 1.007 | 4.46E-03 | glio |
| Interaction_genotype_Bora | 4921536K21R | -3.727 | 4.46E-03 | glio |
| Interaction_genotype_Bora | Spata31d1a | -3.858 | 4.46E-03 | glio |
| Interaction_genotype_Terf1 | Pde3a | -3.991 | 4.46E-03 | glio |
| Interaction_genotype_Terf1 | Adamts9 | -4.073 | 4.46E-03 | glio |
| Interaction_genotype_Terf1 | 1700081N11R | -4.798 | 4.46E-03 | glio |
| Atr_in.vivo_noRT | 1700037C18R | 1.944 | 4.46E-03 | glio |
| Atr_in.vivo_noRT | Pomc | 1.738 | 4.46E-03 | glio |
| Atr_in.vivo_noRT | Cct8 | -0.683 | 4.46E-03 | glio |
| Bcas2_in.vivo_noRT | Il15ra | -1.44 | 4.46E-03 | glio |
| Bora_in.vivo_noRT | Ldlr | -1.597 | 4.46E-03 | glio |
| Chek1_in.vivo_noRT | Ccdc88a | -0.963 | 4.46E-03 | glio |
| Interaction_genotype_Myc | Oas1c | -3.23 | 4.46E-04 | glio |
| Interaction_genotype_Rpl3 | Gm2115 | -4.162 | 4.46E-04 | glio |
| interaction_RT | Mybl2 | -1.243 | 4.46E-04 | glio |
| interaction_RT | Fnip1 | 1.109 | 4.46E-04 | glio |
| Atr_ex.vivo_noRT | Napsa | 3.554 | 4.46E-04 | glio |
| Chek1_in.vivo_noRT | Gm43768 | 2.405 | 4.46E-04 | glio |

| | | | | |
|---|---|---|---|---|
| Chek1_in.vivo_noRT | Hist1h2be | 2.157 | 4.46E-04 | glio |
| Chek1_in.vivo_noRT | Bcat1 | -1.792 | 4.46E-04 | glio |
| Chek1_in.vivo_noRT | Ptpn2 | -1.049 | 4.46E-04 | glio |
| Cyp19a1_ex.vivo_noRT | Gm15894 | 2.959 | 4.46E-04 | glio |
| Aars2_ex.vivo_noRT | 4930527F14Ri | 2.897 | 4.46E-05 | glio |
| Atp5c1_in.vivo_noRT | 4732463B04R | 2.404 | 4.46E-05 | glio |
| Atr_in.vivo_noRT | Afg1l | 2.93 | 4.46E-05 | glio |
| Atr_in.vivo_noRT | Fhl3 | 2.897 | 4.46E-05 | glio |
| Atr_in.vivo_noRT | Slc46a2 | 2.552 | 4.46E-05 | glio |
| Atr_in.vivo_noRT | 1700008J07Ri | 2.611 | 4.46E-05 | glio |
| Atr_in.vivo_noRT | Azin2 | 2.602 | 4.46E-05 | glio |
| Atr_in.vivo_noRT | Ero1l | -1.974 | 4.46E-05 | glio |
| Aars2_in.vivo_noRT | Prrg1 | 3.302 | 4.46E-06 | glio |
| Bora_ex.vivo_noRT | Gm2022 | 3.201 | 4.46E-06 | glio |
| Bora_ex.vivo_noRT | Rimklb | 3.201 | 4.46E-07 | glio |
| Bora_in.vivo_noRT | Gm49801 | 3.938 | 4.46E-08 | glio |
| Chek1_in.vivo_noRT | A330084C13R | 4.126 | 4.46E-08 | glio |
| culture_effect | Ddx49 | -0.979 | 4.46E-09 | glio |
| Atr_in.vivo_noRT | Enpp3 | 3.967 | 4.46E-09 | glio |
| Bcas2_in.vivo_noRT | Clca3a1 | 3.756 | 4.46E-09 | glio |
| culture_effect | Car13 | 2.089 | 4.46E-13 | glio |
| Chek1_in.vivo_noRT | Plk5 | 4.388 | 4.46E-13 | glio |
| Bora_in.vivo_noRT | Cyp7a1 | 5.19 | 4.46E-14 | glio |
| Atr_in.vivo_noRT | Cd248 | 5.267 | 4.46E-15 | glio |
| culture_effect | Cript | 0.387 | 4.47E-03 | glio |
| Interaction_genotype_Aars2 | Phex | -3.296 | 4.47E-03 | glio |
| Interaction_genotype_Aars2 | Snrpd2 | 0.869 | 4.47E-03 | glio |
| Interaction_genotype_Atr | Fxyd4 | -3.862 | 4.47E-03 | glio |
| Interaction_genotype_Bora | Gm15601 | -3.902 | 4.47E-03 | glio |
| Interaction_genotype_Bora | Ankef1 | -3.875 | 4.47E-03 | glio |
| Interaction_genotype_Bora | Gatad2a | 1.506 | 4.47E-03 | glio |
| Interaction_genotype_Bora | Cstb | -1.584 | 4.47E-03 | glio |
| Interaction_genotype_Mtor | Bsg | 1.134 | 4.47E-03 | glio |
| Interaction_genotype_Myc | Gm38973 | -3.238 | 4.47E-03 | glio |
| Interaction_genotype_Myc | Gh | -3.616 | 4.47E-03 | glio |
| Interaction_genotype_Myc | Smagp | -3.185 | 4.47E-03 | glio |
| Interaction_genotype_Myc | BC065397 | -3.541 | 4.47E-03 | glio |
| Interaction_genotype_Myc | Ppl | -2.515 | 4.47E-03 | glio |
| Interaction_genotype_Rpl3 | Kdelr3 | 2.798 | 4.47E-03 | glio |
| Atr_in.vivo_noRT | Tnfsf13 | 1.925 | 4.47E-03 | glio |
| Bcas2_in.vivo_noRT | Gm12971 | 2.163 | 4.47E-03 | glio |
| Bcas2_in.vivo_noRT | Rgs10 | -1.415 | 4.47E-03 | glio |
| Chek1_in.vivo_noRT | 1700037C18R | 1.978 | 4.47E-03 | glio |
| Chek1_ex.vivo_noRT | Olfr279 | 2.538 | 4.47E-03 | glio |
| Hsd17b10_in.vivo_noRT | Gm13986 | 2.357 | 4.47E-03 | glio |
| Hsd17b10_in.vivo_noRT | Gm2762 | 2.044 | 4.47E-03 | glio |
| culture_effect | Prr15l | -2.247 | 4.47E-04 | glio |
| Interaction_genotype_Mtor | Grip1 | -4.25 | 4.47E-04 | glio |
| Interaction_genotype_Mtor | Fam151b | -4.189 | 4.47E-04 | glio |
| Interaction_genotype_Myc | Rgs2 | -4.339 | 4.47E-04 | glio |
| Interaction_genotype_Myc | Zfp93 | -3.259 | 4.47E-04 | glio |
| interaction_RT | Atp6v0c | 0.558 | 4.47E-04 | glio |

| | | | | |
|---|---|---|---|---|
| Bora_in.vivo_noRT | Saxo2 | 2.475 | 4.47E-04 | glio |
| Bora_in.vivo_noRT | Tm9sf2 | -1.329 | 4.47E-04 | glio |
| Interaction_genotype_Mtor | Ccdc103 | -4.91 | 4.47E-05 | glio |
| Interaction_genotype_Mtor | Tspyl3 | -4.395 | 4.47E-05 | glio |
| Atp5c1_in.vivo_noRT | Gm13270 | 3.199 | 4.47E-05 | glio |
| Bcas2_in.vivo_noRT | Gm14636 | 3.518 | 4.47E-05 | glio |
| Chek1_in.vivo_noRT | Prrt3 | 2.863 | 4.47E-06 | glio |
| Chek1_in.vivo_noRT | Rit2 | 3.081 | 4.47E-06 | glio |
| Chek1_ex.vivo_noRT | Rpp30 | -2.289 | 4.47E-06 | glio |
| Aars2_in.vivo_noRT | Copz1 | -1.708 | 4.47E-07 | glio |
| Atr_in.vivo_noRT | Gm29994 | 3.165 | 4.47E-07 | glio |
| Chek1_ex.vivo_noRT | Fbxo15 | 3.861 | 4.47E-07 | glio |
| Atp5c1_in.vivo_noRT | Stfa3 | 2.668 | 4.47E-08 | glio |
| Atp5c1_in.vivo_noRT | Olfr787 | 2.668 | 4.47E-08 | glio |
| Chek1_in.vivo_noRT | Flrt1 | 4.069 | 4.47E-08 | glio |
| Hsd17b10_in.vivo_noRT | Gm45669 | 3.353 | 4.47E-08 | glio |
| culture_effect | Rcan2 | 3.466 | 4.47E-10 | glio |
| culture_effect | Serf1 | -0.835 | 4.47E-10 | glio |
| Atr_in.vivo_noRT | Gm9908 | 4.617 | 4.47E-10 | glio |
| Bcas2_in.vivo_noRT | 4930540M05F | 4.149 | 4.47E-10 | glio |
| culture_effect | Kirrel | 1.918 | 4.47E-12 | glio |
| Aars2_in.vivo_noRT | Tnni3k | 5.327 | 4.47E-23 | glio |
| Chek1_in.vivo_noRT | Gm29966 | 4.784 | 4.47E-23 | glio |
| Aars2_in.vivo_noRT | Abcb5 | 5.327 | 4.47E-26 | glio |
| Interaction_genotype_Aars2 | Edn2 | -2.429 | 4.48E-03 | glio |
| Interaction_genotype_Atr | Vgll3 | -2.728 | 4.48E-03 | glio |
| Interaction_genotype_Myc | Pcdhga11 | -2.703 | 4.48E-03 | glio |
| Aars2_in.vivo_noRT | Nek3 | 2.061 | 4.48E-03 | glio |
| Aars2_in.vivo_noRT | 6030443J06Ri | 1.981 | 4.48E-03 | glio |
| Aars2_in.vivo_noRT | Cyp51 | -1.389 | 4.48E-03 | glio |
| Atp5c1_in.vivo_noRT | D830039M14I | 2.402 | 4.48E-03 | glio |
| Atp5c1_in.vivo_noRT | Pcdhb18 | 2.228 | 4.48E-03 | glio |
| Atp5c1_in.vivo_noRT | Kif7 | 2.08 | 4.48E-03 | glio |
| Atr_in.vivo_noRT | Myb | 2.038 | 4.48E-03 | glio |
| Atr_in.vivo_noRT | Sox5 | 2.117 | 4.48E-03 | glio |
| Atr_in.vivo_noRT | Zfp408 | 1.794 | 4.48E-03 | glio |
| Bcas2_in.vivo_noRT | Hmces | -1.473 | 4.48E-03 | glio |
| Bcas2_in.vivo_noRT | Trf | -2.362 | 4.48E-03 | glio |
| Bcas2_in.vivo_noRT | Vars | -1.046 | 4.48E-03 | glio |
| Bcas2_in.vivo_noRT | Tcf25 | -0.755 | 4.48E-03 | glio |
| Bora_in.vivo_noRT | Prelid2 | 2.066 | 4.48E-03 | glio |
| Chek1_ex.vivo_noRT | Oxt | 2.59 | 4.48E-03 | glio |
| Chek1_ex.vivo_noRT | Tmprss5 | 2.717 | 4.48E-03 | glio |
| Chek1_ex.vivo_noRT | Zfp119a | 1.952 | 4.48E-03 | glio |
| Hsd17b10_in.vivo_noRT | Dcpp3 | 2.099 | 4.48E-03 | glio |
| Hsd17b10_in.vivo_noRT | Mfsd7a | 2.541 | 4.48E-03 | glio |
| culture_effect | Ifngr1 | -0.816 | 4.48E-04 | glio |
| Interaction_genotype_Bcas2 | Gm47204 | -4.755 | 4.48E-04 | glio |
| Interaction_genotype_Myc | Grpel1 | 1.454 | 4.48E-04 | glio |
| Atr_in.vivo_noRT | Whrn | 2.498 | 4.48E-04 | glio |
| Atr_ex.vivo_noRT | Rhov | 3.135 | 4.48E-04 | glio |
| Bard1_ex.vivo_noRT | Lrguk | 2.814 | 4.48E-04 | glio |

| | | | | |
|---|---|---|---|---|
| Interaction_genotype_Myc | Zfp273 | -4.714 | 4.48E-05 | glio |
| Interaction_genotype_Myc | Gm32834 | -4.13 | 4.48E-05 | glio |
| Interaction_genotype_Myc | B230317F23Ri | -4.331 | 4.48E-05 | glio |
| Interaction_genotype_Myc | Rpl22 | 1.602 | 4.48E-05 | glio |
| Atr_ex.vivo_noRT | AI463229 | 3.164 | 4.48E-05 | glio |
| Aars2_in.vivo_noRT | Trmu | 2.213 | 4.48E-06 | glio |
| Aars2_in.vivo_noRT | Psmb5 | -1.138 | 4.48E-06 | glio |
| Atr_in.vivo_noRT | Nedd4 | -0.978 | 4.48E-06 | glio |
| culture_effect | Agbl3 | 1.607 | 4.48E-07 | glio |
| culture_effect | Psmd12 | -0.617 | 4.48E-08 | glio |
| Interaction_genotype_Mtor | Ybx1 | 2.829 | 4.48E-08 | glio |
| Interaction_genotype_Mtor | Gm12602 | -6.35 | 4.48E-08 | glio |
| Interaction_genotype_Mtor | Gm20455 | -7.006 | 4.48E-08 | glio |
| Interaction_genotype_Mtor | Gm34885 | -6.808 | 4.48E-08 | glio |
| Interaction_genotype_Mtor | 4930588A03R | -4.498 | 4.48E-08 | glio |
| Hsd17b10_in.vivo_noRT | Gm38393 | 3.356 | 4.48E-08 | glio |
| Hsd17b10_in.vivo_noRT | Gm4724 | 3.487 | 4.48E-08 | glio |
| Bcas2_in.vivo_noRT | Igdcc4 | 3.631 | 4.48E-09 | glio |
| Hsd17b10_in.vivo_noRT | 1700108F19Ri | 3.352 | 4.48E-09 | glio |
| Bora_in.vivo_noRT | Gm12925 | 4.479 | 4.48E-10 | glio |
| Chek1_in.vivo_noRT | Hectd2os | 4.136 | 4.48E-10 | glio |
| Chek1_ex.vivo_noRT | Prkcq | 4.842 | 4.48E-10 | glio |
| Bcas2_in.vivo_noRT | Vmn1r91 | 4.154 | 4.48E-11 | glio |
| Chek1_in.vivo_noRT | Aoc1 | 4.784 | 4.48E-11 | glio |
| Bcas2_in.vivo_noRT | Gm34059 | 4.807 | 4.48E-24 | glio |
| culture_effect | Gm43813 | -1.674 | 4.49E-03 | glio |
| culture_effect | Rragd | 1.458 | 4.49E-03 | glio |
| culture_effect | Mypop | 0.63 | 4.49E-03 | glio |
| Interaction_genotype_Aars2 | Spata31d1a | -3.624 | 4.49E-03 | glio |
| Interaction_genotype_Aars2 | Wnt16 | -3.422 | 4.49E-03 | glio |
| Interaction_genotype_Aars2 | Gm26533 | -3.468 | 4.49E-03 | glio |
| Interaction_genotype_Cdkn2a | Plk5 | 3.625 | 4.49E-03 | glio |
| Interaction_genotype_Myc | Rsph3a | -2.997 | 4.49E-03 | glio |
| Interaction_genotype_Rad51 | Apol7d | -4.218 | 4.49E-03 | glio |
| Interaction_genotype_Rad51 | AV039307 | -4.103 | 4.49E-03 | glio |
| Interaction_genotype_Rad51 | Ifi27l2a | 1.507 | 4.49E-03 | glio |
| Aars2_in.vivo_noRT | Gm1976 | 1.749 | 4.49E-03 | glio |
| Aars2_in.vivo_noRT | Ginm1 | 0.903 | 4.49E-03 | glio |
| Atr_in.vivo_noRT | Gm9887 | 1.987 | 4.49E-03 | glio |
| Atr_in.vivo_noRT | Sema7a | 1.924 | 4.49E-03 | glio |
| Atr_in.vivo_noRT | Mroh1 | 1.465 | 4.49E-03 | glio |
| Atr_in.vivo_noRT | Mif | -0.669 | 4.49E-03 | glio |
| Atr_ex.vivo_noRT | Enpp3 | 2.764 | 4.49E-03 | glio |
| Bard1_ex.vivo_noRT | Vamp8 | 2.415 | 4.49E-03 | glio |
| Bard1_ex.vivo_noRT | Gm43221 | 2.814 | 4.49E-03 | glio |
| Bcas2_ex.vivo_noRT | Gm4858 | 2.742 | 4.49E-03 | glio |
| Bcas2_ex.vivo_noRT | Lims2 | 3.095 | 4.49E-03 | glio |
| Bora_in.vivo_noRT | 1700012D14R | 2.034 | 4.49E-03 | glio |
| Bora_in.vivo_noRT | AC132307.1 | 1.895 | 4.49E-03 | glio |
| Bora_in.vivo_noRT | Mbnl2 | -0.954 | 4.49E-03 | glio |
| Cenpj_in.vivo_noRT | Gstt4 | 1.759 | 4.49E-03 | glio |
| Interaction_genotype_Mtor | Ly6d | -3.371 | 4.49E-04 | glio |

| | | | | |
|---|---|---|---|---|
| Interaction_genotype_Mtor | Gm16279 | -4.516 | 4.49E-04 | glio |
| Interaction_genotype_Mtor | Gpatch3 | -4.131 | 4.49E-04 | glio |
| Interaction_genotype_Mtor | E130114P18Ri | -4.413 | 4.49E-04 | glio |
| Interaction_genotype_Myc | Acacb | -3.918 | 4.49E-04 | glio |
| Interaction_genotype_Myc | Tspan32 | -4.027 | 4.49E-04 | glio |
| Interaction_genotype_Myc | Tti1 | -3.136 | 4.49E-04 | glio |
| Cenpj_ex.vivo_noRT | Ccl3 | 2.591 | 4.49E-04 | glio |
| Cenpj_ex.vivo_noRT | AU019990 | 2.591 | 4.49E-04 | glio |
| Cenpj_ex.vivo_noRT | Gm48780 | 2.591 | 4.49E-04 | glio |
| Chek1_in.vivo_noRT | Mfsd4b4 | 2.319 | 4.49E-04 | glio |
| Chek1_in.vivo_noRT | Klf1 | 2.594 | 4.49E-04 | glio |
| Chek1_in.vivo_noRT | 4632404H12R | 2.522 | 4.49E-04 | glio |
| Interaction_genotype_Myc | Thap8 | -6.001 | 4.49E-07 | glio |
| Atr_in.vivo_noRT | Mmp28 | 3.64 | 4.49E-07 | glio |
| Bora_in.vivo_noRT | Il2rb | 3.394 | 4.49E-07 | glio |
| Chek1_ex.vivo_noRT | Gm20711 | 3.652 | 4.49E-07 | glio |
| Bora_in.vivo_noRT | Syp | 4.545 | 4.49E-09 | glio |
| Chek1_ex.vivo_noRT | Cysltr2 | 4.842 | 4.49E-09 | glio |
| Chek1_ex.vivo_noRT | Bpifa1 | 4.842 | 4.49E-09 | glio |
| Bora_in.vivo_noRT | Carns1 | 4.936 | 4.49E-12 | glio |
| Bora_in.vivo_noRT | Fbxl2 | 4.29 | 4.49E-12 | glio |
| Atr_in.vivo_noRT | Gm17146 | 4.616 | 4.49E-13 | glio |
| Atr_in.vivo_noRT | Synpo2l | 4.888 | 4.49E-16 | glio |
| Aars2_in.vivo_noRT | 4930434B07R | 5.063 | 4.49E-17 | glio |
| Atr_in.vivo_noRT | Fhl1 | 5.533 | 4.49E-24 | glio |
| culture_effect | Rassf4 | 2.524 | 4.49E-25 | glio |
| culture_effect | Tmtc4 | -0.957 | 4.50E-03 | glio |
| culture_effect | Sept11 | 0.416 | 4.50E-03 | glio |
| Interaction_genotype_Aars2 | 9230112J17Ri | -3.015 | 4.50E-03 | glio |
| Interaction_genotype_Atr | Gtf2b | 1.704 | 4.50E-03 | glio |
| Interaction_genotype_Bcas2 | Gm15520 | -3.377 | 4.50E-03 | glio |
| Interaction_genotype_Bcas2 | Psd | -3.437 | 4.50E-03 | glio |
| Interaction_genotype_Bcas2 | BC049762 | -3.804 | 4.50E-03 | glio |
| Interaction_genotype_Bora | Vps37a | 1.667 | 4.50E-03 | glio |
| Interaction_genotype_Pars2 | Edn2 | -2.664 | 4.50E-03 | glio |
| interaction_RT | Ppm1g | -0.627 | 4.50E-03 | glio |
| Aars2_in.vivo_noRT | Gtpbp10 | 1.381 | 4.50E-03 | glio |
| Atr_in.vivo_noRT | Grhl2 | 2.05 | 4.50E-03 | glio |
| Cenpj_in.vivo_noRT | D930030I03Ri | 2.614 | 4.50E-03 | glio |
| Chek1_in.vivo_noRT | Gucd1 | 1.628 | 4.50E-03 | glio |
| Chek1_in.vivo_noRT | Reep4 | -1.59 | 4.50E-03 | glio |
| Eif2s1_in.vivo_noRT | Fam13a | 2.392 | 4.50E-03 | glio |
| culture_effect | Elovl1 | -0.471 | 4.50E-04 | glio |
| Interaction_genotype_Myc | Gm15445 | -3.069 | 4.50E-04 | glio |
| Interaction_genotype_Myc | Ccdc148 | -4.183 | 4.50E-04 | glio |
| Cenpj_in.vivo_noRT | Clec12b | 2.022 | 4.50E-04 | glio |
| Cenpj_in.vivo_noRT | Vmn1r32 | 2.022 | 4.50E-04 | glio |
| Cenpj_in.vivo_noRT | Glyat | 2.022 | 4.50E-04 | glio |
| Chek1_in.vivo_noRT | Map1lc3b | -1.008 | 4.50E-04 | glio |
| Chek1_in.vivo_noRT | Gm4221 | 2.939 | 4.50E-05 | glio |
| Interaction_genotype_Mtor | Gm31717 | -4.099 | 4.50E-07 | glio |
| Bora_in.vivo_noRT | 1700013F07Ri | 3.782 | 4.50E-07 | glio |

| | | | | |
|---|---|---|---|---|
| Interaction_genotype_Mtor | Nxph3 | -6.043 | 4.50E-08 | glio |
| Atr_in.vivo_noRT | Arhgef4 | 3.768 | 4.50E-08 | glio |
| Atr_in.vivo_noRT | Ifit1bl1 | 5.001 | 4.50E-09 | glio |
| Atr_in.vivo_noRT | Rasgrp2 | 4.233 | 4.50E-09 | glio |
| Bcas2_in.vivo_noRT | Ptpn22 | 4.544 | 4.50E-09 | glio |
| Bora_in.vivo_noRT | Bend7 | 4.158 | 4.50E-10 | glio |
| Bora_in.vivo_noRT | Gm26822 | 4.545 | 4.50E-10 | glio |
| Aars2_in.vivo_noRT | Gm5624 | 4.337 | 4.50E-11 | glio |
| Bcas2_in.vivo_noRT | Pdlim3 | 4.028 | 4.50E-12 | glio |
| culture_effect | mt-Nd1 | 1.039 | 4.50E-17 | glio |
| Aars2_in.vivo_noRT | Dapk2 | 4.544 | 4.50E-18 | glio |
| Atr_in.vivo_noRT | Pnliprp2 | 4.755 | 4.50E-18 | glio |
| Atr_in.vivo_noRT | Pldi | 5.533 | 4.50E-22 | glio |
| culture_effect | Anks6 | 1.38 | 4.51E-03 | glio |
| culture_effect | Cct6a | -0.324 | 4.51E-03 | glio |
| Interaction_genotype_Aars2 | Hyal1 | -3.438 | 4.51E-03 | glio |
| Interaction_genotype_Aars2 | Rpl29 | 0.757 | 4.51E-03 | glio |
| Interaction_genotype_Atr | Glt8d2 | -3.67 | 4.51E-03 | glio |
| Interaction_genotype_Atr | Prpf38b | 1.333 | 4.51E-03 | glio |
| Interaction_genotype_Myc | Dok3 | -3.331 | 4.51E-03 | glio |
| Interaction_genotype_Myc | Apold1 | -3.643 | 4.51E-03 | glio |
| Interaction_genotype_Myc | Usp28 | -3 | 4.51E-03 | glio |
| Interaction_genotype_Myc | Gpn2 | -2.565 | 4.51E-03 | glio |
| Interaction_genotype_Myc | Csde1 | 1.203 | 4.51E-03 | glio |
| Interaction_genotype_Uqcrc2 | 2310067P03R | -4.096 | 4.51E-03 | glio |
| Atr_in.vivo_noRT | Carmil1 | 1.94 | 4.51E-03 | glio |
| Atr_in.vivo_noRT | Smad3 | 1.89 | 4.51E-03 | glio |
| Atr_in.vivo_noRT | Gpr153 | 1.923 | 4.51E-03 | glio |
| Atr_in.vivo_noRT | Mbnl2 | -0.978 | 4.51E-03 | glio |
| Bard1_ex.vivo_noRT | Lncbate10 | 2.027 | 4.51E-03 | glio |
| Bard1_ex.vivo_noRT | Chia1 | 2.027 | 4.51E-03 | glio |
| Bcas2_in.vivo_noRT | Gm21154 | 2.25 | 4.51E-03 | glio |
| Bcas2_in.vivo_noRT | Tmem120a | -1.331 | 4.51E-03 | glio |
| Bcas2_ex.vivo_noRT | Gm20767 | 2.742 | 4.51E-03 | glio |
| Bora_in.vivo_noRT | Acad12 | 2.035 | 4.51E-03 | glio |
| Cdkn2a_in.vivo_noRT | Shfl | 1.287 | 4.51E-03 | glio |
| Cenpj_in.vivo_noRT | Gm13348 | 2.022 | 4.51E-03 | glio |
| Cenpj_in.vivo_noRT | Aqp9 | 2.022 | 4.51E-03 | glio |
| Chek1_in.vivo_noRT | Abca13 | 2.312 | 4.51E-03 | glio |
| Chek1_in.vivo_noRT | Zbtb3 | 2.103 | 4.51E-03 | glio |
| Chek1_in.vivo_noRT | Fbxo6 | -1.407 | 4.51E-03 | glio |
| Chek1_in.vivo_noRT | Akap13 | -1.4 | 4.51E-03 | glio |
| Chek1_in.vivo_noRT | Cyfip1 | -1.208 | 4.51E-03 | glio |
| Chek1_in.vivo_noRT | Ttc33 | -1.016 | 4.51E-03 | glio |
| Chek1_ex.vivo_noRT | Ehhadh | 2.897 | 4.51E-03 | glio |
| Eif2s1_ex.vivo_noRT | Cd200r2 | 2.234 | 4.51E-03 | glio |
| culture_effect | 6030458C11R | 0.612 | 4.51E-04 | glio |
| Interaction_genotype_Eif2s1 | Dab2 | 4.702 | 4.51E-04 | glio |
| Interaction_genotype_Myc | Zfp850 | -3.373 | 4.51E-04 | glio |
| Interaction_genotype_Myc | Svip | -3.572 | 4.51E-04 | glio |
| Interaction_genotype_Myc | Kmt5c | -3.088 | 4.51E-04 | glio |
| Interaction_genotype_Myc | Rpap2 | -3.069 | 4.51E-04 | glio |

| | | | | |
|---|---|---|---|---|
| Interaction_genotype_Myc | Plaat3 | 2.361 | 4.51E-04 | glio |
| Interaction_genotype_Myc | Bri3 | 1.434 | 4.51E-04 | glio |
| interaction_RT | Ube2e3 | 0.859 | 4.51E-04 | glio |
| interaction_RT | Taldo1 | -0.78 | 4.51E-04 | glio |
| Bard1_ex.vivo_noRT | Gm41307 | 2.421 | 4.51E-04 | glio |
| Chek1_ex.vivo_noRT | Cdk8 | -1.694 | 4.51E-04 | glio |
| culture_effect | Rab10 | 0.459 | 4.51E-05 | glio |
| Interaction_genotype_Aars2 | Gm47204 | -5.233 | 4.51E-05 | glio |
| Interaction_genotype_Aars2 | Cope | 1.689 | 4.51E-05 | glio |
| Aars2_in.vivo_noRT | Rbm19 | 2.158 | 4.51E-05 | glio |
| Aars2_in.vivo_noRT | Polr3k | -1.537 | 4.51E-05 | glio |
| Atp5c1_in.vivo_noRT | Cyp4x1 | 2.93 | 4.51E-05 | glio |
| Chek1_in.vivo_noRT | Adcy7 | 2.748 | 4.51E-05 | glio |
| Chek1_in.vivo_noRT | Slc39a8 | 3.192 | 4.51E-05 | glio |
| Chek1_ex.vivo_noRT | Slc4a9 | 2.956 | 4.51E-05 | glio |
| Interaction_genotype_Myc | Dnaic2 | -3.893 | 4.51E-06 | glio |
| Atr_ex.vivo_noRT | Gm38393 | 3.915 | 4.51E-07 | glio |
| Atr_ex.vivo_noRT | Synpo2l | 3.941 | 4.51E-07 | glio |
| Chek1_in.vivo_noRT | 1700025N23R | 3.152 | 4.51E-07 | glio |
| Hsd17b10_in.vivo_noRT | Celsr3 | 3.356 | 4.51E-07 | glio |
| Aars2_in.vivo_noRT | Tmem86a | 3.959 | 4.51E-08 | glio |
| Aars2_ex.vivo_noRT | Cldn2 | 3.293 | 4.51E-08 | glio |
| Bcas2_in.vivo_noRT | Gm15559 | 3.88 | 4.51E-08 | glio |
| culture_effect | Mat2a | 0.862 | 4.51E-09 | glio |
| Bora_in.vivo_noRT | Ifitm1 | 4.67 | 4.51E-09 | glio |
| Bora_in.vivo_noRT | Amh | 4.019 | 4.51E-10 | glio |
| Atr_in.vivo_noRT | Osm | 4.184 | 4.51E-12 | glio |
| Bora_in.vivo_noRT | Rgs9 | 4.136 | 4.51E-12 | glio |
| Bora_in.vivo_noRT | B3galt4 | 4.411 | 4.51E-12 | glio |
| Bora_in.vivo_noRT | Gm48161 | 4.148 | 4.51E-12 | glio |
| culture_effect | H19 | -5.847 | 4.51E-13 | glio |
| Hsd17b10_in.vivo_noRT | Vmn2r44 | 3.616 | 4.51E-13 | glio |
| Hsd17b10_in.vivo_noRT | Mmp13 | 3.616 | 4.51E-13 | glio |
| Hsd17b10_in.vivo_noRT | Halr1 | 3.35 | 4.51E-13 | glio |
| Hsd17b10_in.vivo_noRT | Gm8016 | 3.616 | 4.51E-13 | glio |
| Bora_in.vivo_noRT | Olfr810 | 4.936 | 4.51E-25 | glio |
| Atr_in.vivo_noRT | Slc16a5 | 5.269 | 4.51E-26 | glio |
| Atr_in.vivo_noRT | Gm30692 | 5.533 | 4.51E-28 | glio |
| Interaction_genotype_Aars2 | Hspa12a | -3.548 | 4.52E-03 | glio |
| Interaction_genotype_Atr | Serpine2 | -4.171 | 4.52E-03 | glio |
| Interaction_genotype_Atr | Gng7 | -3.342 | 4.52E-03 | glio |
| Interaction_genotype_Atr | Nnt | -3.241 | 4.52E-03 | glio |
| Interaction_genotype_Chek1 | Zfr | 1.275 | 4.52E-03 | glio |
| Interaction_genotype_Myc | Lrfn4 | -2.423 | 4.52E-03 | glio |
| Interaction_genotype_Pars2 | Gpx2 | 5.088 | 4.52E-03 | glio |
| Interaction_genotype_Pars2 | Fbxl2 | -3.504 | 4.52E-03 | glio |
| Interaction_genotype_Pars2 | Edn1 | -4.353 | 4.52E-03 | glio |
| interaction_RT | Litaf | 0.739 | 4.52E-03 | glio |
| Aars2_in.vivo_noRT | Cyp4f17 | 2.079 | 4.52E-03 | glio |
| Aars2_in.vivo_noRT | Arrdc1 | 1.87 | 4.52E-03 | glio |
| Aars2_in.vivo_noRT | Tmem126a | -0.821 | 4.52E-03 | glio |
| Atr_in.vivo_noRT | Dlgap1 | 1.827 | 4.52E-03 | glio |

| | | | | |
|---|---|---|---|---|
| Atr_in.vivo_noRT | Plod2 | -1.42 | 4.52E-03 | glio |
| Bcas2_ex.vivo_noRT | Fa2h | 2.742 | 4.52E-03 | glio |
| Bora_in.vivo_noRT | Mob3b | 1.817 | 4.52E-03 | glio |
| Chek1_in.vivo_noRT | Rab7b | 2.027 | 4.52E-03 | glio |
| Eif2s1_in.vivo_noRT | 4930470G03R | 1.877 | 4.52E-03 | glio |
| Eif2s1_in.vivo_noRT | Pmp2 | 2.143 | 4.52E-03 | glio |
| Eif2s1_in.vivo_noRT | Gm43625 | 2.14 | 4.52E-03 | glio |
| Interaction_genotype_Cdkn2a | Gpr17 | 4.505 | 4.52E-04 | glio |
| Interaction_genotype_Mtor | Gm34552 | -3.823 | 4.52E-04 | glio |
| Interaction_genotype_Myc | Map3k9 | -3.257 | 4.52E-04 | glio |
| Aars2_in.vivo_noRT | B230312C02R | 3.206 | 4.52E-04 | glio |
| Aars2_ex.vivo_noRT | Gm37864 | 2.899 | 4.52E-04 | glio |
| Atr_in.vivo_noRT | Crybg3 | 2.582 | 4.52E-04 | glio |
| Atr_in.vivo_noRT | Pdhb | -1.231 | 4.52E-04 | glio |
| Bcas2_in.vivo_noRT | Slc35a2 | -1.503 | 4.52E-04 | glio |
| Bcas2_ex.vivo_noRT | A930002C04R | 2.342 | 4.52E-04 | glio |
| Bora_in.vivo_noRT | Dnaaf4 | 2.755 | 4.52E-04 | glio |
| Chek1_ex.vivo_noRT | Usp35 | 3.117 | 4.52E-04 | glio |
| culture_effect | Ptpn14 | -0.887 | 4.52E-05 | glio |
| Interaction_genotype_Myc | Gm29666 | -4.668 | 4.52E-05 | glio |
| Bcas2_in.vivo_noRT | Nup153 | -1.79 | 4.52E-05 | glio |
| Bora_in.vivo_noRT | 9430065F17Ri | 2.961 | 4.52E-05 | glio |
| Chek1_in.vivo_noRT | H2-Q10 | 2.876 | 4.52E-05 | glio |
| Chek1_in.vivo_noRT | Fchsd1 | 2.8 | 4.52E-05 | glio |
| Chek1_in.vivo_noRT | Tmem79 | 2.759 | 4.52E-05 | glio |
| Chek1_in.vivo_noRT | Nuak2 | 3.17 | 4.52E-05 | glio |
| Hsd17b10_in.vivo_noRT | Gjb5 | 2.626 | 4.52E-05 | glio |
| Hsd17b10_in.vivo_noRT | Pecam1 | 3.084 | 4.52E-05 | glio |
| culture_effect | Gm33973 | -2.501 | 4.52E-06 | glio |
| culture_effect | Traf4 | -0.827 | 4.52E-06 | glio |
| Interaction_genotype_Mtor | Gm48617 | -3.835 | 4.52E-06 | glio |
| Interaction_genotype_Mtor | Abhd17a | 2.235 | 4.52E-06 | glio |
| Aars2_in.vivo_noRT | Reck | 3.233 | 4.52E-06 | glio |
| Bcas2_in.vivo_noRT | Gchfr | 3.694 | 4.52E-06 | glio |
| Chek1_in.vivo_noRT | Gm49785 | 3.347 | 4.52E-06 | glio |
| Chek1_in.vivo_noRT | C130040N14R | 3.396 | 4.52E-06 | glio |
| Chek1_in.vivo_noRT | Tnfrsf13b | 3.406 | 4.52E-06 | glio |
| Aars2_ex.vivo_noRT | Gm266 | 3.293 | 4.52E-07 | glio |
| Atr_ex.vivo_noRT | Gm16933 | 3.164 | 4.52E-07 | glio |
| Atr_in.vivo_noRT | Gm16759 | 3.967 | 4.52E-08 | glio |
| Atr_in.vivo_noRT | Slpi | 4.245 | 4.52E-08 | glio |
| Bcas2_in.vivo_noRT | Esyt3 | 3.747 | 4.52E-08 | glio |
| Bora_in.vivo_noRT | Olfm2 | 4.029 | 4.52E-08 | glio |
| Bora_in.vivo_noRT | Man2c1os | 4.291 | 4.52E-09 | glio |
| Chek1_in.vivo_noRT | Gjb4 | 4.387 | 4.52E-10 | glio |
| Atr_ex.vivo_noRT | AF067063 | 4.309 | 4.52E-11 | glio |
| culture_effect | Zfp637 | 1.068 | 4.52E-13 | glio |
| Atr_in.vivo_noRT | Gm49223 | 5.011 | 4.52E-18 | glio |
| Bcas2_in.vivo_noRT | Dpep3 | 4.275 | 4.52E-18 | glio |
| Chek1_in.vivo_noRT | Pitx3 | 5.044 | 4.52E-19 | glio |
| culture_effect | H2-DMa | -3.919 | 4.52E-53 | glio |
| culture_effect | Gm44867 | -1.808 | 4.53E-03 | glio |

| | | | | |
|---|---|---|---|---|
| culture_effect | Creb3l1 | -1.759 | 4.53E-03 | glio |
| Interaction_genotype_Chek1 | Mpv17 | 1.872 | 4.53E-03 | glio |
| Interaction_genotype_Mtor | Zfp341 | -3.183 | 4.53E-03 | glio |
| Interaction_genotype_Myc | Gm47101 | -3.043 | 4.53E-03 | glio |
| Interaction_genotype_Pars2 | Zfp810 | -3.783 | 4.53E-03 | glio |
| interaction_RT | Isg15 | 0.979 | 4.53E-03 | glio |
| Aars2_in.vivo_noRT | Gm10791 | 1.883 | 4.53E-03 | glio |
| Aars2_in.vivo_noRT | Fetub | 1.789 | 4.53E-03 | glio |
| Aars2_in.vivo_noRT | Rnf20 | -1.159 | 4.53E-03 | glio |
| Atr_in.vivo_noRT | AU020206 | -1.355 | 4.53E-03 | glio |
| Bcas2_in.vivo_noRT | Lyrm2 | 1.131 | 4.53E-03 | glio |
| Bcas2_ex.vivo_noRT | Gm47972 | 2.84 | 4.53E-03 | glio |
| Brca2_in.vivo_noRT | Stfa3 | 1.763 | 4.53E-03 | glio |
| Brca2_in.vivo_noRT | Olfr787 | 1.763 | 4.53E-03 | glio |
| Chek1_in.vivo_noRT | 2610203C22R | 2.19 | 4.53E-03 | glio |
| Chek1_in.vivo_noRT | Dtx1 | 2.076 | 4.53E-03 | glio |
| Chek1_in.vivo_noRT | Dok7 | 2.015 | 4.53E-03 | glio |
| Cyp19a1_ex.vivo_noRT | A030005L19Ri | 2.943 | 4.53E-03 | glio |
| Eif2s1_ex.vivo_noRT | Gm5431 | 3.032 | 4.53E-03 | glio |
| Hsd17b10_in.vivo_noRT | D930030I03Ri | 2.377 | 4.53E-03 | glio |
| Interaction_genotype_Myc | Gm37035 | -3.292 | 4.53E-04 | glio |
| interaction_RT | Gm35019 | 2.378 | 4.53E-04 | glio |
| Aars2_in.vivo_noRT | Nudt7 | 2.353 | 4.53E-04 | glio |
| Atr_ex.vivo_noRT | Fst | 3.164 | 4.53E-04 | glio |
| Bcas2_in.vivo_noRT | Ccnj | 2.625 | 4.53E-04 | glio |
| Cenpj_ex.vivo_noRT | Gm36793 | 2.591 | 4.53E-04 | glio |
| Chek1_in.vivo_noRT | 9030407P20R | 2.497 | 4.53E-04 | glio |
| Chek1_in.vivo_noRT | Kcna4 | 2.517 | 4.53E-04 | glio |
| Hsd17b10_in.vivo_noRT | Apol6 | -1.984 | 4.53E-04 | glio |
| Interaction_genotype_Myc | Gm13856 | -3.122 | 4.53E-05 | glio |
| Interaction_genotype_Myc | Gm21936 | -3.124 | 4.53E-05 | glio |
| Atr_in.vivo_noRT | Pcdh1 | 2.878 | 4.53E-05 | glio |
| Atr_in.vivo_noRT | Pheta1 | 3.245 | 4.53E-06 | glio |
| Bcas2_in.vivo_noRT | Gm8104 | 3.387 | 4.53E-07 | glio |
| Bcas2_in.vivo_noRT | D230022J07Ri | 3.648 | 4.53E-07 | glio |
| Atr_ex.vivo_noRT | Gm21762 | 4.331 | 4.53E-08 | glio |
| Hsd17b10_in.vivo_noRT | Gm19265 | 3.1 | 4.53E-08 | glio |
| culture_effect | Pxdc1 | 1.173 | 4.53E-09 | glio |
| culture_effect | Zcchc17 | 0.803 | 4.53E-09 | glio |
| culture_effect | Tubb5 | -0.973 | 4.53E-09 | glio |
| Bora_in.vivo_noRT | Gm47545 | 3.685 | 4.53E-09 | glio |
| Bora_in.vivo_noRT | Gpr179 | 3.935 | 4.53E-09 | glio |
| Chek1_ex.vivo_noRT | 1700010B08R | 4.842 | 4.53E-11 | glio |
| Aars2_in.vivo_noRT | B3galt4 | 4.801 | 4.53E-14 | glio |
| Chek1_ex.vivo_noRT | Gm49698 | 4.842 | 4.53E-16 | glio |
| Chek1_in.vivo_noRT | Gm9758 | 4.784 | 4.53E-20 | glio |
| culture_effect | Irgm1 | -1.563 | 4.53E-24 | glio |
| Chek1_in.vivo_noRT | Gm48435 | 5.044 | 4.53E-26 | glio |
| culture_effect | Gm10482 | -1.457 | 4.54E-03 | glio |
| culture_effect | A730063M14F | 1.215 | 4.54E-03 | glio |
| Interaction_genotype_Aars2 | Gm10125 | -3.594 | 4.54E-03 | glio |
| Interaction_genotype_Aars2 | C130040N14R | -3.545 | 4.54E-03 | glio |

| | | | | |
|---|---|---|---|---|
| Interaction_genotype_Aars2 | Gnai2 | 1.389 | 4.54E-03 | glio |
| Interaction_genotype_Atr | Gm16576 | -3.693 | 4.54E-03 | glio |
| Interaction_genotype_Bcas2 | Neurl1a | -4.136 | 4.54E-03 | glio |
| Interaction_genotype_Cdkn2a | Usp17la | 4.133 | 4.54E-03 | glio |
| Interaction_genotype_Chek1 | Tet3 | 2.64 | 4.54E-03 | glio |
| Interaction_genotype_Mtor | Rsph3a | -3.202 | 4.54E-03 | glio |
| Interaction_genotype_Mtor | Zfp600 | -3.203 | 4.54E-03 | glio |
| Interaction_genotype_Mtor | B2m | 1.091 | 4.54E-03 | glio |
| Aars2_in.vivo_noRT | Gm20300 | 2.068 | 4.54E-03 | glio |
| Bcas2_ex.vivo_noRT | Tinagl1 | -1.483 | 4.54E-03 | glio |
| Bora_ex.vivo_noRT | Ms4a6d | 2.414 | 4.54E-03 | glio |
| Cdkn2a_in.vivo_noRT | Cd7 | -1.938 | 4.54E-03 | glio |
| Chek1_in.vivo_noRT | D5Ertd579e | -1.14 | 4.54E-03 | glio |
| Chek1_ex.vivo_noRT | Nlgn2 | 2.222 | 4.54E-03 | glio |
| Chek1_ex.vivo_noRT | Clspn | -1.783 | 4.54E-03 | glio |
| Hsd17b10_in.vivo_noRT | Prmt1 | -0.629 | 4.54E-03 | glio |
| Interaction_genotype_Abtb1 | Prrg1 | 5.281 | 4.54E-04 | glio |
| Interaction_genotype_Abtb1 | Lce1g | 5.048 | 4.54E-04 | glio |
| Interaction_genotype_Abtb1 | Gm29670 | 3.933 | 4.54E-04 | glio |
| Interaction_genotype_Myc | Dmrt2 | -4.243 | 4.54E-04 | glio |
| Interaction_genotype_Myc | Gm16759 | -3.995 | 4.54E-04 | glio |
| Interaction_genotype_Myc | Gnat2 | -3.542 | 4.54E-04 | glio |
| Interaction_genotype_Myc | Sprtn | -3.141 | 4.54E-04 | glio |
| Interaction_genotype_Myc | Son | 1.566 | 4.54E-04 | glio |
| Aars2_in.vivo_noRT | Stub1 | -0.989 | 4.54E-04 | glio |
| Atr_in.vivo_noRT | Nphp3 | 2.459 | 4.54E-04 | glio |
| Atr_in.vivo_noRT | Tmem205 | -1.352 | 4.54E-04 | glio |
| Atr_in.vivo_noRT | Fbn2 | -2.155 | 4.54E-04 | glio |
| Bora_in.vivo_noRT | Nalcn | -1.944 | 4.54E-04 | glio |
| Cenpj_in.vivo_noRT | Tmem132c | 2.022 | 4.54E-04 | glio |
| Cenpj_in.vivo_noRT | Gm44693 | 2.022 | 4.54E-04 | glio |
| Chek1_in.vivo_noRT | Gm19605 | 2.496 | 4.54E-04 | glio |
| Interaction_genotype_Mtor | Amt | -5.268 | 4.54E-05 | glio |
| Interaction_genotype_Mtor | Stx1b | -4.396 | 4.54E-05 | glio |
| Aars2_in.vivo_noRT | Chd4 | -1.239 | 4.54E-05 | glio |
| Atr_in.vivo_noRT | Tmem40 | 3.188 | 4.54E-05 | glio |
| Chek1_in.vivo_noRT | Fgfr1 | 3.406 | 4.54E-05 | glio |
| Chek1_in.vivo_noRT | Cct6b | 2.754 | 4.54E-05 | glio |
| Aars2_in.vivo_noRT | Abhd15 | 3.478 | 4.54E-06 | glio |
| Aars2_ex.vivo_noRT | Olfr171 | 3.293 | 4.54E-06 | glio |
| Atp5c1_in.vivo_noRT | Hfm1 | 2.668 | 4.54E-06 | glio |
| Atp5c1_in.vivo_noRT | D630024D03R | 2.668 | 4.54E-06 | glio |
| Atp5c1_in.vivo_noRT | Cldn5 | 2.402 | 4.54E-06 | glio |
| Atp5c1_in.vivo_noRT | Gm39526 | 2.667 | 4.54E-06 | glio |
| Atp5c1_in.vivo_noRT | Crb3 | 2.668 | 4.54E-06 | glio |
| Bcas2_in.vivo_noRT | Spx | 3.081 | 4.54E-06 | glio |
| Chek1_in.vivo_noRT | Srrt | -1.733 | 4.54E-06 | glio |
| culture_effect | Smu1 | -0.729 | 4.54E-07 | glio |
| culture_effect | Amd1 | 0.729 | 4.54E-07 | glio |
| Aars2_ex.vivo_noRT | Ildr1 | 3.293 | 4.54E-07 | glio |
| Chek1_in.vivo_noRT | D830035M03I | 3.458 | 4.54E-07 | glio |
| Atr_in.vivo_noRT | Xist | 3.681 | 4.54E-08 | glio |

| | | | | |
|---|---|---|---|---|
| Aars2_in.vivo_noRT | Gm35066 | 4.349 | 4.54E-10 | glio |
| Atr_in.vivo_noRT | Tmem30c | 4.347 | 4.54E-10 | glio |
| Bora_in.vivo_noRT | Dcn | 4.545 | 4.54E-13 | glio |
| Atr_in.vivo_noRT | 4933429H19R | 5.159 | 4.54E-15 | glio |
| Atr_in.vivo_noRT | Gm31518 | 5.533 | 4.54E-15 | glio |
| Chek1_ex.vivo_noRT | Olfr881 | 4.842 | 4.54E-16 | glio |
| Bora_in.vivo_noRT | Pldi | 4.936 | 4.54E-18 | glio |
| Chek1_in.vivo_noRT | Gm36793 | 5.044 | 4.54E-22 | glio |
| Atr_in.vivo_noRT | 4921504E06Ri | 6.043 | 4.54E-26 | glio |
| culture_effect | Rpa2 | -0.5 | 4.55E-03 | glio |
| culture_effect | Creld2 | -0.46 | 4.55E-03 | glio |
| Interaction_genotype_Bora | Clcn1 | -3.042 | 4.55E-03 | glio |
| Interaction_genotype_Bora | Gm41318 | -3.656 | 4.55E-03 | glio |
| Interaction_genotype_Bora | Tns2 | -3.133 | 4.55E-03 | glio |
| Interaction_genotype_Bora | Nfix | 1.861 | 4.55E-03 | glio |
| Interaction_genotype_Bora | Copz2 | 1.705 | 4.55E-03 | glio |
| Interaction_genotype_Mtor | Hspa12b | -3.695 | 4.55E-03 | glio |
| Interaction_genotype_Mtor | Fads3 | -2.187 | 4.55E-03 | glio |
| Interaction_genotype_Mtor | Leo1 | -1.725 | 4.55E-03 | glio |
| Interaction_genotype_Mtor | Naca | 0.896 | 4.55E-03 | glio |
| Interaction_genotype_Myc | Pip4k2a | -2.698 | 4.55E-03 | glio |
| Interaction_genotype_Myc | Hist4h4 | -2.916 | 4.55E-03 | glio |
| Interaction_genotype_Myc | Ankrd13a | -2.165 | 4.55E-03 | glio |
| Aars2_in.vivo_noRT | Tsen15 | -1.245 | 4.55E-03 | glio |
| Atr_in.vivo_noRT | Mgat5 | 1.744 | 4.55E-03 | glio |
| Atr_ex.vivo_noRT | Gm45669 | 2.376 | 4.55E-03 | glio |
| Bcas2_in.vivo_noRT | Micos13 | -0.67 | 4.55E-03 | glio |
| Bora_in.vivo_noRT | Cdv3 | -0.678 | 4.55E-03 | glio |
| Cenpj_in.vivo_noRT | Gm49694 | 2.597 | 4.55E-03 | glio |
| Cenpj_ex.vivo_noRT | Etl4 | 2.591 | 4.55E-03 | glio |
| culture_effect | Adarb1 | -0.984 | 4.55E-04 | glio |
| culture_effect | R3hdm4 | 0.531 | 4.55E-04 | glio |
| Interaction_genotype_Aars2 | Gm16244 | -4.549 | 4.55E-04 | glio |
| Interaction_genotype_Mtor | Rasal1 | -4.687 | 4.55E-04 | glio |
| Interaction_genotype_Mtor | Zfp28 | -4.276 | 4.55E-04 | glio |
| Interaction_genotype_Myc | Ankrd16 | -3.245 | 4.55E-04 | glio |
| Interaction_genotype_Myc | Ube2r2 | 1.352 | 4.55E-04 | glio |
| Interaction_genotype_Uqcrc2 | Gm14968 | -5.17 | 4.55E-04 | glio |
| Interaction_genotype_Uqcrc2 | Gm9967 | -4.973 | 4.55E-04 | glio |
| interaction_RT | Tmem11 | 0.765 | 4.55E-04 | glio |
| Aars2_in.vivo_noRT | Cdkn2b | -1.633 | 4.55E-04 | glio |
| Aars2_in.vivo_noRT | Znhit3 | -1.407 | 4.55E-04 | glio |
| Atp5c1_in.vivo_noRT | Htr2b | 2.145 | 4.55E-04 | glio |
| culture_effect | Eps15l1 | -0.901 | 4.55E-05 | glio |
| culture_effect | Tmem214 | -0.776 | 4.55E-05 | glio |
| culture_effect | Atp5h | -0.439 | 4.55E-05 | glio |
| Bora_in.vivo_noRT | 4930513N10R | 3.328 | 4.55E-05 | glio |
| Chek1_in.vivo_noRT | 9330179D12R | 3.114 | 4.55E-05 | glio |
| Chek1_in.vivo_noRT | Fam71e1 | 2.621 | 4.55E-05 | glio |
| Interaction_genotype_Mtor | Nxf2 | -5.722 | 4.55E-06 | glio |
| Aars2_in.vivo_noRT | 9130008F23Ri | 3.561 | 4.55E-06 | glio |
| Atr_in.vivo_noRT | Pafah2 | 3.504 | 4.55E-06 | glio |

| | | | | |
|---|---|---|---|---|
| Bcas2_in.vivo_noRT | Gm50255 | 3.9 | 4.55E-06 | glio |
| Interaction_genotype_Mtor | Mettl7b | -5.965 | 4.55E-07 | glio |
| Interaction_genotype_Mtor | Flacc1 | -5.289 | 4.55E-07 | glio |
| Interaction_genotype_Mtor | Gm14742 | -4.099 | 4.55E-07 | glio |
| Bcas2_in.vivo_noRT | Ankrd34a | 3.687 | 4.55E-07 | glio |
| Interaction_genotype_Myc | 5033403F01Ri | -6.514 | 4.55E-09 | glio |
| Chek1_ex.vivo_noRT | Sap30bpos | 4.449 | 4.55E-09 | glio |
| Bora_in.vivo_noRT | Armc2 | 4.545 | 4.55E-11 | glio |
| Chek1_ex.vivo_noRT | Gm16344 | 4.842 | 4.55E-11 | glio |
| Atr_in.vivo_noRT | Nid1 | 5.233 | 4.55E-13 | glio |
| Aars2_in.vivo_noRT | Krt80 | 5.06 | 4.55E-16 | glio |
| Atr_in.vivo_noRT | Krtcap3 | 4.741 | 4.55E-18 | glio |
| Aars2_in.vivo_noRT | Hfm1 | 5.327 | 4.55E-22 | glio |
| Aars2_in.vivo_noRT | D630024D03F | 5.327 | 4.55E-22 | glio |
| culture_effect | Fes | 1.5 | 4.56E-03 | glio |
| culture_effect | Rbmxl1 | -0.444 | 4.56E-03 | glio |
| Interaction_genotype_Aars2 | Tmem236 | -3.14 | 4.56E-03 | glio |
| Interaction_genotype_Atr | A330040F15Ri | -3.766 | 4.56E-03 | glio |
| Interaction_genotype_Atr | Rilp | -3.785 | 4.56E-03 | glio |
| Interaction_genotype_Bcas2 | 4933433G15R | -3.827 | 4.56E-03 | glio |
| Interaction_genotype_Mtor | Vps29 | 1.499 | 4.56E-03 | glio |
| Interaction_genotype_Myc | Rspo3 | -3.621 | 4.56E-03 | glio |
| Interaction_genotype_Myc | Zfp979 | -2.898 | 4.56E-03 | glio |
| Interaction_genotype_Myc | Notch2 | -2.411 | 4.56E-03 | glio |
| Interaction_genotype_Rad51 | 1700003G18R | -4.544 | 4.56E-03 | glio |
| Interaction_genotype_Rpl3 | Gm13288 | -3.261 | 4.56E-03 | glio |
| interaction_RT | Pelp1 | -0.974 | 4.56E-03 | glio |
| interaction_RT | Tomm7 | 0.668 | 4.56E-03 | glio |
| Aars2_in.vivo_noRT | Srr | 1.569 | 4.56E-03 | glio |
| Cdkn2a_in.vivo_noRT | Mgll | -2.614 | 4.56E-03 | glio |
| Chek1_in.vivo_noRT | Cd59a | 2.124 | 4.56E-03 | glio |
| Chek1_in.vivo_noRT | Tma7 | -0.657 | 4.56E-03 | glio |
| Chek1_ex.vivo_noRT | Mrpl12 | -0.72 | 4.56E-03 | glio |
| Interaction_genotype_Mtor | Zfp575 | -4.398 | 4.56E-04 | glio |
| Interaction_genotype_Mtor | Dennd3 | -4.495 | 4.56E-04 | glio |
| Interaction_genotype_Myc | Zfp382 | -3.742 | 4.56E-04 | glio |
| Interaction_genotype_Myc | Gm4876 | -3.042 | 4.56E-04 | glio |
| Interaction_genotype_Myc | Cltb | 1.757 | 4.56E-04 | glio |
| Interaction_genotype_Myc | Lyar | 1.735 | 4.56E-04 | glio |
| Interaction_genotype_Myc | Lbh | 1.843 | 4.56E-04 | glio |
| Bcas2_in.vivo_noRT | Dnajc22 | -1.718 | 4.56E-04 | glio |
| Bora_in.vivo_noRT | B230216N24R | 2.786 | 4.56E-04 | glio |
| Bora_in.vivo_noRT | Dok7 | 2.44 | 4.56E-04 | glio |
| Bora_in.vivo_noRT | Btbd16 | 2.112 | 4.56E-04 | glio |
| Hsd17b10_in.vivo_noRT | Otop1 | 2.432 | 4.56E-04 | glio |
| Interaction_genotype_Mtor | Mvb12a | 2.171 | 4.56E-05 | glio |
| Interaction_genotype_Myc | Gpr179 | -4.496 | 4.56E-05 | glio |
| culture_effect | Fam126a | 1.25 | 4.56E-06 | glio |
| Interaction_genotype_Mtor | Gm12500 | -5.214 | 4.56E-06 | glio |
| Interaction_genotype_Mtor | Fabp6 | -4.099 | 4.56E-06 | glio |
| Atr_in.vivo_noRT | Copg1 | -1.815 | 4.56E-06 | glio |
| Chek1_ex.vivo_noRT | Eif4e3 | 4.045 | 4.56E-08 | glio |

| | | | | |
|---|---|---|---|---|
| Chek1_ex.vivo_noRT | AW112010 | 5.212 | 4.56E-08 | glio |
| culture_effect | Dnttip1 | -0.898 | 4.56E-09 | glio |
| Atr_in.vivo_noRT | Gm43387 | 4.358 | 4.56E-10 | glio |
| Bora_in.vivo_noRT | Lyz2 | 4.55 | 4.56E-10 | glio |
| culture_effect | Arhgap10 | 1.729 | 4.56E-17 | glio |
| Atr_in.vivo_noRT | Gm15894 | 4.488 | 4.56E-17 | glio |
| Interaction_genotype_Atr | Gm31152 | -3.222 | 4.57E-03 | glio |
| Interaction_genotype_Bcas2 | Tal2 | -2.783 | 4.57E-03 | glio |
| Interaction_genotype_Bcas2 | Zfhx2os | -3.777 | 4.57E-03 | glio |
| Interaction_genotype_Mtor | Kyat3 | -3.295 | 4.57E-03 | glio |
| Interaction_genotype_Mtor | Rbm12b1 | -3.012 | 4.57E-03 | glio |
| Interaction_genotype_Mtor | Gm36738 | -3.22 | 4.57E-03 | glio |
| Interaction_genotype_Myc | Slc25a42 | -3.223 | 4.57E-03 | glio |
| Interaction_genotype_Myc | Cep112 | -2.596 | 4.57E-03 | glio |
| Interaction_genotype_Myc | Hps6 | -2.603 | 4.57E-03 | glio |
| Aars2_ex.vivo_noRT | Mier3 | -1.901 | 4.57E-03 | glio |
| Atr_in.vivo_noRT | Zfp46 | 1.758 | 4.57E-03 | glio |
| Atr_in.vivo_noRT | Parp14 | -1.608 | 4.57E-03 | glio |
| Atr_in.vivo_noRT | Psmd12 | -0.695 | 4.57E-03 | glio |
| Bcas2_ex.vivo_noRT | Slc22a1 | 2.347 | 4.57E-03 | glio |
| Bcas2_ex.vivo_noRT | 1700108F19Ri | 2.348 | 4.57E-03 | glio |
| Bora_in.vivo_noRT | Fzd5 | 2.017 | 4.57E-03 | glio |
| Bora_in.vivo_noRT | 2310068J16Ri | 1.752 | 4.57E-03 | glio |
| Bora_in.vivo_noRT | Edc4 | -1.343 | 4.57E-03 | glio |
| Bora_in.vivo_noRT | Azi2 | -1.23 | 4.57E-03 | glio |
| Chek1_in.vivo_noRT | Ice1 | 1.437 | 4.57E-03 | glio |
| Chek1_ex.vivo_noRT | Tmem150c | 2.69 | 4.57E-03 | glio |
| Chek1_ex.vivo_noRT | Gm44702 | 2.622 | 4.57E-03 | glio |
| Chek1_ex.vivo_noRT | Aoc2 | 2.368 | 4.57E-03 | glio |
| Chek1_ex.vivo_noRT | Hsd17b10 | -1.213 | 4.57E-03 | glio |
| Chek1_ex.vivo_noRT | Eny2 | -0.614 | 4.57E-03 | glio |
| culture_effect | Mapt | 1.068 | 4.57E-04 | glio |
| Interaction_genotype_Myc | Cbr4 | -3.063 | 4.57E-04 | glio |
| Atr_in.vivo_noRT | Eef1akmt3 | 2.317 | 4.57E-04 | glio |
| Atr_in.vivo_noRT | Faf2 | -1.231 | 4.57E-04 | glio |
| Atr_in.vivo_noRT | Psme1 | -0.969 | 4.57E-04 | glio |
| Atr_in.vivo_noRT | Eif1 | -0.672 | 4.57E-04 | glio |
| Chek1_in.vivo_noRT | Btbd17 | 3.012 | 4.57E-04 | glio |
| Chek1_in.vivo_noRT | Bsg | -0.686 | 4.57E-04 | glio |
| culture_effect | S100a10 | 0.6 | 4.57E-05 | glio |
| Interaction_genotype_Mtor | Gm13178 | -3.583 | 4.57E-05 | glio |
| interaction_RT | Npc2 | 0.887 | 4.57E-05 | glio |
| Atr_ex.vivo_noRT | Snhg1 | 1.508 | 4.57E-05 | glio |
| Bcas2_in.vivo_noRT | 4930405A21R | 2.693 | 4.57E-05 | glio |
| Bora_ex.vivo_noRT | Otof | 3.201 | 4.57E-05 | glio |
| Interaction_genotype_Mtor | Zmynd10 | -4.627 | 4.57E-06 | glio |
| Aars2_in.vivo_noRT | Aldh1a1 | 3.298 | 4.57E-06 | glio |
| Atr_in.vivo_noRT | Egfl8 | 3.347 | 4.57E-06 | glio |
| culture_effect | Cep164 | 0.892 | 4.57E-07 | glio |
| Interaction_genotype_Mtor | Shisal2a | -6.485 | 4.57E-07 | glio |
| interaction_RT | Gabarapl2 | 0.966 | 4.57E-07 | glio |
| interaction_RT | Rpl12 | -0.959 | 4.57E-07 | glio |

| | | | | |
|---|---|---|---|---|
| Bora_in.vivo_noRT | 1810041H14R | 3.818 | 4.57E-07 | glio |
| Bora_in.vivo_noRT | Cxcr6 | 3.513 | 4.57E-07 | glio |
| Bora_ex.vivo_noRT | Gm14929 | 3.201 | 4.57E-07 | glio |
| Chek1_in.vivo_noRT | D930030I03Ri | 3.804 | 4.57E-07 | glio |
| Chek1_in.vivo_noRT | Cep72 | 3.446 | 4.57E-07 | glio |
| culture_effect | Napsa | -3.583 | 4.57E-08 | glio |
| culture_effect | Ier5l | 1.555 | 4.57E-08 | glio |
| culture_effect | Dph3 | 0.774 | 4.57E-10 | glio |
| Chek1_in.vivo_noRT | Gm15559 | 4.64 | 4.57E-11 | glio |
| Chek1_ex.vivo_noRT | Apol10b | 4.842 | 4.57E-11 | glio |
| Atr_in.vivo_noRT | Pacrg | 4.742 | 4.57E-14 | glio |
| Atr_in.vivo_noRT | Ctla4 | 5.533 | 4.57E-29 | glio |
| culture_effect | Slfn4 | -1.322 | 4.58E-03 | glio |
| culture_effect | Ap1g2 | -1.29 | 4.58E-03 | glio |
| culture_effect | Tgfb1 | -0.865 | 4.58E-03 | glio |
| culture_effect | Pik3c3 | -0.742 | 4.58E-03 | glio |
| Interaction_genotype_Atr | Tmem202 | -3.564 | 4.58E-03 | glio |
| Interaction_genotype_Atr | D430018E03R | -3.419 | 4.58E-03 | glio |
| Interaction_genotype_Atr | Gm10010 | -3.439 | 4.58E-03 | glio |
| Interaction_genotype_Bcas2 | Top2b | 1.763 | 4.58E-03 | glio |
| Interaction_genotype_Cdkn2a | Gm49698 | 2.835 | 4.58E-03 | glio |
| Interaction_genotype_Cdkn2a | Olfr881 | 2.835 | 4.58E-03 | glio |
| Interaction_genotype_Ifnar1 | Cldn2 | 2.734 | 4.58E-03 | glio |
| Interaction_genotype_Mtor | Zglp1 | -3.808 | 4.58E-03 | glio |
| Interaction_genotype_Mtor | Gm38973 | -3.447 | 4.58E-03 | glio |
| Interaction_genotype_Mtor | Rit2 | -3.271 | 4.58E-03 | glio |
| Interaction_genotype_Mtor | Slitrk5 | -3.312 | 4.58E-03 | glio |
| Interaction_genotype_Mtor | Shroom4 | -2.795 | 4.58E-03 | glio |
| Interaction_genotype_Mtor | Kansl1 | 1.589 | 4.58E-03 | glio |
| Interaction_genotype_Myc | Gm49441 | -2.726 | 4.58E-03 | glio |
| interaction_RT | Eftud2 | -0.696 | 4.58E-03 | glio |
| interaction_RT | Ndufa3 | 0.801 | 4.58E-03 | glio |
| interaction_RT | Aimp1 | -0.552 | 4.58E-03 | glio |
| Aars2_in.vivo_noRT | Hdhd2 | -1.135 | 4.58E-03 | glio |
| Bcas2_in.vivo_noRT | A830005F24Ri | 2.112 | 4.58E-03 | glio |
| Bcas2_in.vivo_noRT | Utp11 | -0.721 | 4.58E-03 | glio |
| Bora_in.vivo_noRT | Hcfc1r1 | 2.345 | 4.58E-03 | glio |
| Brca2_in.vivo_noRT | Gm15908 | 1.763 | 4.58E-03 | glio |
| Brca2_in.vivo_noRT | A930014E10Ri | 1.763 | 4.58E-03 | glio |
| Brca2_in.vivo_noRT | Gm45338 | 1.763 | 4.58E-03 | glio |
| Brca2_in.vivo_noRT | Scg2 | 1.763 | 4.58E-03 | glio |
| Brca2_in.vivo_noRT | 1700003E24Ri | 1.763 | 4.58E-03 | glio |
| Brca2_in.vivo_noRT | Gm14820 | 2.025 | 4.58E-03 | glio |
| Brca2_in.vivo_noRT | Bglap2 | 2.71 | 4.58E-03 | glio |
| Brca2_in.vivo_noRT | Krt19 | 1.763 | 4.58E-03 | glio |
| Brca2_in.vivo_noRT | Gm48155 | 1.763 | 4.58E-03 | glio |
| Brca2_in.vivo_noRT | Gm15890 | 2.286 | 4.58E-03 | glio |
| Brca2_in.vivo_noRT | Gm34059 | 1.763 | 4.58E-03 | glio |
| Brca2_in.vivo_noRT | Gm48435 | 1.763 | 4.58E-03 | glio |
| Brca2_in.vivo_noRT | Helt | 1.763 | 4.58E-03 | glio |
| Brca2_in.vivo_noRT | Cd48 | 1.947 | 4.58E-03 | glio |
| Brca2_in.vivo_noRT | Gm20625 | 1.763 | 4.58E-03 | glio |

| | | | | |
|---|---|---|---|---|
| Brca2_in.vivo_noRT | Gm5157 | 1.763 | 4.58E-03 | glio |
| Brca2_in.vivo_noRT | Scgb1c1 | 2.288 | 4.58E-03 | glio |
| Brca2_in.vivo_noRT | Gm44736 | 1.763 | 4.58E-03 | glio |
| Brca2_in.vivo_noRT | Fam227b | 1.763 | 4.58E-03 | glio |
| Brca2_in.vivo_noRT | Gm20658 | 1.763 | 4.58E-03 | glio |
| Brca2_in.vivo_noRT | Olfr810 | 1.763 | 4.58E-03 | glio |
| Chek1_in.vivo_noRT | Inca1 | 1.904 | 4.58E-03 | glio |
| Hsd17b10_in.vivo_noRT | Dhrs9 | 2.029 | 4.58E-03 | glio |
| Hsd17b10_in.vivo_noRT | 1700122E12Ri | 2.22 | 4.58E-03 | glio |
| Hsd17b10_in.vivo_noRT | 2310008N11R | 2.361 | 4.58E-03 | glio |
| culture_effect | Pabpc1 | -0.337 | 4.58E-04 | glio |
| Interaction_genotype_Myc | Gm14929 | -2.729 | 4.58E-04 | glio |
| Interaction_genotype_Myc | Gm11769 | -3.64 | 4.58E-04 | glio |
| Interaction_genotype_Myc | Spdye4a | -4.15 | 4.58E-04 | glio |
| Interaction_genotype_Myc | Ptpru | -3.482 | 4.58E-04 | glio |
| Aars2_in.vivo_noRT | Lhx9 | 2.622 | 4.58E-04 | glio |
| Aars2_in.vivo_noRT | 4930506C21R | 2.462 | 4.58E-04 | glio |
| Atr_ex.vivo_noRT | Scgb1c1 | 2.771 | 4.58E-04 | glio |
| Atr_ex.vivo_noRT | Spi1 | 2.99 | 4.58E-04 | glio |
| Atr_ex.vivo_noRT | Rasl12 | 3.164 | 4.58E-04 | glio |
| Bcas2_in.vivo_noRT | Des | 2.557 | 4.58E-04 | glio |
| Bora_in.vivo_noRT | 2810021J22Ri | 2.617 | 4.58E-04 | glio |
| Bora_in.vivo_noRT | Ino80dos | 2.614 | 4.58E-04 | glio |
| Bora_in.vivo_noRT | Hk1os | 2.334 | 4.58E-04 | glio |
| Cenpj_ex.vivo_noRT | Spink8 | 2.591 | 4.58E-04 | glio |
| Cenpj_ex.vivo_noRT | Psmb11 | 2.591 | 4.58E-04 | glio |
| culture_effect | Ncstn | -0.725 | 4.58E-05 | glio |
| Interaction_genotype_Atr | Gm41442 | -4.915 | 4.58E-05 | glio |
| Interaction_genotype_Atr | Spon1 | -5.143 | 4.58E-05 | glio |
| Interaction_genotype_Atr | Pde11a | -5.317 | 4.58E-05 | glio |
| Chek1_ex.vivo_noRT | Ccr5 | 3.281 | 4.58E-05 | glio |
| Bcas2_in.vivo_noRT | Ucma | 3.808 | 4.58E-06 | glio |
| Atr_in.vivo_noRT | 9330121J05Ri | 3.498 | 4.58E-07 | glio |
| Bora_in.vivo_noRT | Tox4 | -2.087 | 4.58E-07 | glio |
| Chek1_in.vivo_noRT | Tmem86a | 3.676 | 4.58E-07 | glio |
| Hsd17b10_in.vivo_noRT | Ly6d | 2.889 | 4.58E-07 | glio |
| Aars2_in.vivo_noRT | Psma6 | -1.448 | 4.58E-08 | glio |
| Atr_in.vivo_noRT | Gm10401 | 3.331 | 4.58E-08 | glio |
| Bcas2_in.vivo_noRT | Vmn1r88 | 3.622 | 4.58E-09 | glio |
| Bcas2_in.vivo_noRT | D6Ertd527e | 4.024 | 4.58E-09 | glio |
| Atp5c1_in.vivo_noRT | 4930588A03R | 3.199 | 4.58E-10 | glio |
| Chek1_in.vivo_noRT | Lrrc24 | 4.043 | 4.58E-10 | glio |
| Interaction_genotype_Aars2 | Fhl3 | -3.524 | 4.59E-03 | glio |
| Interaction_genotype_Aars2 | Rheb | 1.033 | 4.59E-03 | glio |
| Interaction_genotype_Mtor | Zfp595 | -3.637 | 4.59E-03 | glio |
| Interaction_genotype_Mtor | Pih1d2 | -3.033 | 4.59E-03 | glio |
| Interaction_genotype_Mtor | 9530077C05R | -2.805 | 4.59E-03 | glio |
| Interaction_genotype_Myc | Tmbim4 | 1.275 | 4.59E-03 | glio |
| Aars2_in.vivo_noRT | Ttc12 | 1.951 | 4.59E-03 | glio |
| Aars2_in.vivo_noRT | Reln | -1.021 | 4.59E-03 | glio |
| Aars2_in.vivo_noRT | Sept7 | -0.716 | 4.59E-03 | glio |
| Bcas2_in.vivo_noRT | Leng9 | 1.981 | 4.59E-03 | glio |

| | | | | |
|---|---|---|---|---|
| Bcas2_ex.vivo_noRT | A330015K06R | 2.622 | 4.59E-03 | glio |
| Bora_in.vivo_noRT | Lrp10 | -1.337 | 4.59E-03 | glio |
| Bora_in.vivo_noRT | Mrpl1 | 0.923 | 4.59E-03 | glio |
| Bora_in.vivo_noRT | Tsn | -0.607 | 4.59E-03 | glio |
| Brca2_in.vivo_noRT | Apoa5 | 1.763 | 4.59E-03 | glio |
| Cenpj_in.vivo_noRT | Esm1 | 1.758 | 4.59E-03 | glio |
| Chek1_ex.vivo_noRT | Yars | -1.532 | 4.59E-03 | glio |
| Dclre1c_ex.vivo_noRT | Ush1g | 2.511 | 4.59E-03 | glio |
| Eif2s1_in.vivo_noRT | Gm15943 | 1.879 | 4.59E-03 | glio |
| Hsd17b10_in.vivo_noRT | Pla2g2f | 1.973 | 4.59E-03 | glio |
| Hsd17b10_in.vivo_noRT | Psma3 | -0.484 | 4.59E-03 | glio |
| Interaction_genotype_Mtor | Eml1 | -3.613 | 4.59E-04 | glio |
| Interaction_genotype_Mtor | Erlec1 | 1.766 | 4.59E-04 | glio |
| Bcas2_in.vivo_noRT | Hs1bp3 | 2.101 | 4.59E-04 | glio |
| Cenpj_in.vivo_noRT | Kcnk13 | 2.288 | 4.59E-04 | glio |
| Cenpj_ex.vivo_noRT | Vtn | 2.591 | 4.59E-04 | glio |
| Cenpj_ex.vivo_noRT | Amy1 | 3.386 | 4.59E-04 | glio |
| Bora_ex.vivo_noRT | Gtsf1l | 3.201 | 4.59E-05 | glio |
| Chek1_in.vivo_noRT | Mmp19 | 2.85 | 4.59E-05 | glio |
| Chek1_ex.vivo_noRT | Tex52 | 3.464 | 4.59E-05 | glio |
| Interaction_genotype_Mtor | Gm49223 | -4.558 | 4.59E-06 | glio |
| Interaction_genotype_Mtor | Efr3a | 2.255 | 4.59E-06 | glio |
| Aars2_in.vivo_noRT | Chrna1os | 2.714 | 4.59E-06 | glio |
| Bcas2_in.vivo_noRT | Xlr4c | 3.3 | 4.59E-06 | glio |
| Chek1_ex.vivo_noRT | Rassf6 | 3.239 | 4.59E-06 | glio |
| Hsd17b10_in.vivo_noRT | Spn | 3.094 | 4.59E-06 | glio |
| Aars2_in.vivo_noRT | Efna2 | 4.055 | 4.59E-07 | glio |
| Bora_in.vivo_noRT | 4930558J18Ri | 3.33 | 4.59E-07 | glio |
| Aars2_in.vivo_noRT | Rtl1 | 3.635 | 4.59E-09 | glio |
| Atr_in.vivo_noRT | Prr7 | 4.142 | 4.59E-09 | glio |
| Atr_in.vivo_noRT | Otos | 4.004 | 4.59E-10 | glio |
| culture_effect | Gm813 | -4.105 | 4.59E-11 | glio |
| Atr_in.vivo_noRT | D030051J21Ri | 4.63 | 4.59E-12 | glio |
| Aars2_in.vivo_noRT | Itgbl1 | 4.14 | 4.59E-13 | glio |
| Bcas2_in.vivo_noRT | 4930570E03Ri | 4.547 | 4.59E-17 | glio |
| Bcas2_in.vivo_noRT | 4930447K03R | 4.547 | 4.59E-17 | glio |
| Chek1_ex.vivo_noRT | Gm44736 | 4.842 | 4.59E-17 | glio |
| Bcas2_in.vivo_noRT | Ltbp2 | 4.807 | 4.59E-18 | glio |
| Bcas2_in.vivo_noRT | Lmx1a | 4.807 | 4.59E-18 | glio |
| culture_effect | Ccdc162 | -1.463 | 4.60E-03 | glio |
| Interaction_genotype_Aars2 | Gal3st4 | -3.472 | 4.60E-03 | glio |
| Interaction_genotype_Aars2 | Gm20707 | -3.448 | 4.60E-03 | glio |
| Interaction_genotype_Myc | Postn | -3.183 | 4.60E-03 | glio |
| Interaction_genotype_Myc | Nars2 | -2.709 | 4.60E-03 | glio |
| Atr_in.vivo_noRT | 1700019D03R | 2.185 | 4.60E-03 | glio |
| Atr_in.vivo_noRT | 6530402F18Ri | 1.939 | 4.60E-03 | glio |
| Bcas2_in.vivo_noRT | Gm26692 | 2.006 | 4.60E-03 | glio |
| Bora_in.vivo_noRT | Mgat4b | -1.233 | 4.60E-03 | glio |
| Bora_ex.vivo_noRT | Cers3 | 2.229 | 4.60E-03 | glio |
| Bora_ex.vivo_noRT | Gm37214 | 2.899 | 4.60E-03 | glio |
| Bora_ex.vivo_noRT | Qpct | 3.018 | 4.60E-03 | glio |
| Chek1_in.vivo_noRT | Rab26 | 1.943 | 4.60E-03 | glio |

| | | | | |
|---|---|---|---|---|
| Eif2s1_in.vivo_noRT | B230206H07R | 1.617 | 4.60E-03 | glio |
| Aars2_in.vivo_noRT | Ngdn | -1.25 | 4.60E-04 | glio |
| Atr_in.vivo_noRT | Nckap1l | 2.465 | 4.60E-04 | glio |
| Atr_in.vivo_noRT | Emc9 | 1.709 | 4.60E-04 | glio |
| Hsd17b10_in.vivo_noRT | 2210417A02R | 2.825 | 4.60E-04 | glio |
| Atr_in.vivo_noRT | Hcn2 | 3.08 | 4.60E-05 | glio |
| Chek1_in.vivo_noRT | Rhbdf2 | 2.724 | 4.60E-05 | glio |
| Interaction_genotype_Mtor | Olfr1369-ps1 | -6.677 | 4.60E-06 | glio |
| Interaction_genotype_Mtor | Gm11491 | -5.57 | 4.60E-06 | glio |
| Interaction_genotype_Myc | Gnas | 1.697 | 4.60E-06 | glio |
| Atr_in.vivo_noRT | Trim71 | 3.174 | 4.60E-06 | glio |
| Chek1_in.vivo_noRT | Kcnmb4os2 | 3.606 | 4.60E-07 | glio |
| culture_effect | C530044C16R | 2.215 | 4.60E-08 | glio |
| Hsd17b10_in.vivo_noRT | Gm45151 | 3.084 | 4.60E-08 | glio |
| interaction_RT | H2-D1 | 1.636 | 4.60E-12 | glio |
| Aars2_in.vivo_noRT | Gm31518 | 5.327 | 4.60E-14 | glio |
| Aars2_in.vivo_noRT | Celsr3 | 5.067 | 4.60E-15 | glio |
| Atr_in.vivo_noRT | Gm48140 | 5.27 | 4.60E-25 | glio |
| culture_effect | Cabp4 | 1.633 | 4.61E-03 | glio |
| culture_effect | Ddx18 | 0.417 | 4.61E-03 | glio |
| Interaction_genotype_Aars2 | Hspb9 | -3.797 | 4.61E-03 | glio |
| Interaction_genotype_Aars2 | Rab30 | -3.082 | 4.61E-03 | glio |
| Interaction_genotype_Atr | Gm44672 | -3.442 | 4.61E-03 | glio |
| Interaction_genotype_Atr | Pdhb | 1.423 | 4.61E-03 | glio |
| Interaction_genotype_Myc | Iqcn | -3.001 | 4.61E-03 | glio |
| Interaction_genotype_Myc | Appl2 | -2.311 | 4.61E-03 | glio |
| Atp5c1_in.vivo_noRT | Aoc1 | 2.408 | 4.61E-03 | glio |
| Atr_in.vivo_noRT | Mcrs1 | -1.055 | 4.61E-03 | glio |
| Atr_ex.vivo_noRT | Folr2 | 2.791 | 4.61E-03 | glio |
| Atr_ex.vivo_noRT | Knstrn | -1.55 | 4.61E-03 | glio |
| Chek1_in.vivo_noRT | Tanc2 | -1.378 | 4.61E-03 | glio |
| Dclre1c_ex.vivo_noRT | Ifnb1 | 6.028 | 4.61E-03 | glio |
| Fanca_ex.vivo_noRT | Stfa3 | 2.124 | 4.61E-03 | glio |
| Fanca_ex.vivo_noRT | Olfr787 | 2.124 | 4.61E-03 | glio |
| Interaction_genotype_Mtor | Gal3st4 | -4.014 | 4.61E-04 | glio |
| Interaction_genotype_Myc | Ltbp2 | -3.124 | 4.61E-04 | glio |
| Aars2_in.vivo_noRT | Zfp870 | 2.165 | 4.61E-04 | glio |
| Aars2_in.vivo_noRT | Clcn7 | 1.942 | 4.61E-04 | glio |
| Atr_ex.vivo_noRT | Col5a2 | 2.577 | 4.61E-04 | glio |
| Bcas2_in.vivo_noRT | Pi4k2b | -1.345 | 4.61E-04 | glio |
| Chek1_in.vivo_noRT | Slurp1 | 3.75 | 4.61E-04 | glio |
| Atr_in.vivo_noRT | Arrdc3 | 2.681 | 4.61E-05 | glio |
| interaction_RT | Jdp2 | 1.65 | 4.61E-07 | glio |
| Chek1_in.vivo_noRT | 4930445N18R | 3.599 | 4.61E-07 | glio |
| Chek1_in.vivo_noRT | Tspan13 | -2.203 | 4.61E-07 | glio |
| Bcas2_in.vivo_noRT | Rhox3g | 3.692 | 4.61E-08 | glio |
| Aars2_in.vivo_noRT | Gm43581 | 3.757 | 4.61E-09 | glio |
| Bora_in.vivo_noRT | Gm10010 | 4.195 | 4.61E-09 | glio |
| Atr_in.vivo_noRT | Edn1 | 5.154 | 4.61E-10 | glio |
| Bora_in.vivo_noRT | Ccdc39 | 4.24 | 4.61E-10 | glio |
| Aars2_in.vivo_noRT | Zhx2 | 4.792 | 4.61E-11 | glio |
| Bcas2_in.vivo_noRT | Dppa2 | 4.275 | 4.61E-11 | glio |

| | | | | |
|---|---|---|---|---|
| Chek1_ex.vivo_noRT | Brinp3 | 4.842 | 4.61E-11 | glio |
| Chek1_ex.vivo_noRT | Gm5934 | 4.263 | 4.61E-11 | glio |
| Chek1_ex.vivo_noRT | Cfh | 4.443 | 4.61E-12 | glio |
| Atr_ex.vivo_noRT | Zfp352 | 5.712 | 4.61E-13 | glio |
| Atr_in.vivo_noRT | Mcmdc2 | 5.273 | 4.61E-19 | glio |
| Interaction_genotype_Aars2 | Cbr3 | -3.722 | 4.62E-03 | glio |
| Interaction_genotype_Cdkn2a | BC061195 | 2.835 | 4.62E-03 | glio |
| Interaction_genotype_Mtor | Timm23 | 1.227 | 4.62E-03 | glio |
| Interaction_genotype_Myc | Dmpk | -3.161 | 4.62E-03 | glio |
| Interaction_genotype_Rad51 | Apol6 | 3.449 | 4.62E-03 | glio |
| Interaction_genotype_Rad51 | Prox2 | -4.302 | 4.62E-03 | glio |
| interaction_RT | Gpr153 | 2.139 | 4.62E-03 | glio |
| interaction_RT | Mapk3 | 0.838 | 4.62E-03 | glio |
| Aars2_in.vivo_noRT | Napb | 1.77 | 4.62E-03 | glio |
| Aars2_ex.vivo_noRT | Cd72 | 2.897 | 4.62E-03 | glio |
| Atr_in.vivo_noRT | Cd68 | 2.187 | 4.62E-03 | glio |
| Atr_in.vivo_noRT | Kirrel | 1.582 | 4.62E-03 | glio |
| Atr_in.vivo_noRT | Slc35g1 | 1.618 | 4.62E-03 | glio |
| Atr_in.vivo_noRT | Nap1l1 | -0.591 | 4.62E-03 | glio |
| Atr_ex.vivo_noRT | Hmgcs2 | 3.18 | 4.62E-03 | glio |
| Bard1_ex.vivo_noRT | Agrp | 2.419 | 4.62E-03 | glio |
| Bcas2_ex.vivo_noRT | Gm42918 | 2.552 | 4.62E-03 | glio |
| Bcas2_ex.vivo_noRT | Hpse | 2.919 | 4.62E-03 | glio |
| Bora_in.vivo_noRT | Zfp882 | 2.248 | 4.62E-03 | glio |
| Bora_in.vivo_noRT | Nqo1 | 2.171 | 4.62E-03 | glio |
| Bora_in.vivo_noRT | Erfe | 1.922 | 4.62E-03 | glio |
| Bora_in.vivo_noRT | Nudt7 | 1.963 | 4.62E-03 | glio |
| Bora_in.vivo_noRT | 4732440D04R | 1.9 | 4.62E-03 | glio |
| Bora_in.vivo_noRT | Tspan9 | -1.591 | 4.62E-03 | glio |
| Bora_in.vivo_noRT | Gtf2f2 | -1.235 | 4.62E-03 | glio |
| Brca2_in.vivo_noRT | Gjd4 | 1.763 | 4.62E-03 | glio |
| culture_effect | Otud7b | -0.606 | 4.62E-04 | glio |
| Interaction_genotype_Aars2 | Atp6v1g2 | -4.037 | 4.62E-04 | glio |
| Interaction_genotype_Bcas2 | Gm41764 | -4.131 | 4.62E-04 | glio |
| Interaction_genotype_Mtor | Calr | 2.206 | 4.62E-04 | glio |
| Interaction_genotype_Myc | Boc | -2.859 | 4.62E-04 | glio |
| Interaction_genotype_Myc | Gm31152 | -3.582 | 4.62E-04 | glio |
| Interaction_genotype_Myc | A430057M04F | -3.923 | 4.62E-04 | glio |
| Interaction_genotype_Myc | Pnn | 1.802 | 4.62E-04 | glio |
| Aars2_in.vivo_noRT | Aprt | -0.856 | 4.62E-04 | glio |
| Atr_in.vivo_noRT | Rpgr | 2.348 | 4.62E-04 | glio |
| Atr_in.vivo_noRT | Syngr3 | 2.154 | 4.62E-04 | glio |
| Atr_in.vivo_noRT | Eef1b2 | -0.706 | 4.62E-04 | glio |
| Atr_ex.vivo_noRT | Gm5431 | 3.164 | 4.62E-04 | glio |
| Bcas2_in.vivo_noRT | Pafah2 | 2.778 | 4.62E-04 | glio |
| Bora_ex.vivo_noRT | Pik3ap1 | 2.84 | 4.62E-04 | glio |
| Bora_ex.vivo_noRT | Adgre5 | 2.808 | 4.62E-04 | glio |
| Chek1_in.vivo_noRT | Gas5 | -0.801 | 4.62E-04 | glio |
| Hsd17b10_in.vivo_noRT | Abhd15 | 2.83 | 4.62E-04 | glio |
| Interaction_genotype_Mtor | Emc10 | 1.772 | 4.62E-05 | glio |
| Interaction_genotype_Myc | Arrb2 | -3.899 | 4.62E-05 | glio |
| Atp5c1_in.vivo_noRT | Folr1 | 2.668 | 4.62E-05 | glio |

| | | | | |
|---|---|---|---|---|
| Atr_in.vivo_noRT | Rbakdn | 2.832 | 4.62E-05 | glio |
| Atr_in.vivo_noRT | Nkain4 | 2.669 | 4.62E-05 | glio |
| Bcas2_in.vivo_noRT | Setd4 | 2.861 | 4.62E-05 | glio |
| Interaction_genotype_Chek1 | Ubd | 5.415 | 4.62E-06 | glio |
| Aars2_in.vivo_noRT | Ccdc106 | 3.472 | 4.62E-06 | glio |
| Aars2_in.vivo_noRT | Tmem91 | 3.284 | 4.62E-06 | glio |
| Chek1_in.vivo_noRT | Gm35066 | 3.551 | 4.62E-07 | glio |
| Atr_in.vivo_noRT | AA388235 | 3.893 | 4.62E-08 | glio |
| Atr_in.vivo_noRT | Unc5cl | 3.769 | 4.62E-08 | glio |
| Aars2_in.vivo_noRT | Insm1 | 4.208 | 4.62E-10 | glio |
| Bcas2_in.vivo_noRT | Cd83 | 3.807 | 4.62E-10 | glio |
| Hsd17b10_in.vivo_noRT | Gm39526 | 3.625 | 4.62E-11 | glio |
| Chek1_in.vivo_noRT | Gm50147 | 5.044 | 4.62E-14 | glio |
| Chek1_in.vivo_noRT | Rftn2 | 4.653 | 4.62E-14 | glio |
| Hsd17b10_in.vivo_noRT | Hoxc8 | 3.616 | 4.62E-14 | glio |
| Atr_in.vivo_noRT | Trim16 | 5.273 | 4.62E-17 | glio |
| Atr_in.vivo_noRT | Rbfaos | 4.616 | 4.62E-17 | glio |
| Bora_in.vivo_noRT | Erich6b | 5.46 | 4.62E-26 | glio |
| Interaction_genotype_Atr | Nsmce2 | 1.286 | 4.63E-03 | glio |
| Interaction_genotype_Bcas2 | Sv2a | -4.012 | 4.63E-03 | glio |
| Interaction_genotype_Mtor | Tgs1 | 1.445 | 4.63E-03 | glio |
| Interaction_genotype_Rpl3 | Gm11520 | -3.644 | 4.63E-03 | glio |
| Atr_in.vivo_noRT | Sh3rf1 | 1.75 | 4.63E-03 | glio |
| Bcas2_in.vivo_noRT | Rbm3os | 2.016 | 4.63E-03 | glio |
| Bcas2_ex.vivo_noRT | Deptor | 2.062 | 4.63E-03 | glio |
| Bora_in.vivo_noRT | Spata2l | 2.144 | 4.63E-03 | glio |
| Brca2_ex.vivo_noRT | Gm10801 | 2.82 | 4.63E-03 | glio |
| culture_effect | Mfsd12 | -1.028 | 4.63E-04 | glio |
| Aars2_in.vivo_noRT | Chchd1 | -0.875 | 4.63E-04 | glio |
| Atp5c1_in.vivo_noRT | Cers3 | 2.136 | 4.63E-04 | glio |
| Atr_in.vivo_noRT | Kif24 | 2.419 | 4.63E-04 | glio |
| Chek1_in.vivo_noRT | Nckipsd | 2.438 | 4.63E-04 | glio |
| Chek1_in.vivo_noRT | Kif24 | 2.453 | 4.63E-04 | glio |
| Chek1_in.vivo_noRT | Cdc14b | 2.032 | 4.63E-04 | glio |
| Chek1_in.vivo_noRT | Zfp239 | 1.177 | 4.63E-04 | glio |
| Atr_in.vivo_noRT | Arhgap27 | 2.684 | 4.63E-05 | glio |
| Atr_in.vivo_noRT | Gm26579 | 2.697 | 4.63E-05 | glio |
| Hsd17b10_in.vivo_noRT | Gm15972 | 2.623 | 4.63E-05 | glio |
| Interaction_genotype_Myc | Sephs1 | -3.556 | 4.63E-06 | glio |
| Interaction_genotype_Myc | Hectd2 | -4.603 | 4.63E-06 | glio |
| Interaction_genotype_Myc | Gm47204 | -5.419 | 4.63E-06 | glio |
| Interaction_genotype_Myc | Gm47851 | -4.856 | 4.63E-06 | glio |
| Aars2_in.vivo_noRT | Myom1 | 3.333 | 4.63E-06 | glio |
| Aars2_in.vivo_noRT | Slc41a3 | 3.066 | 4.63E-06 | glio |
| Interaction_genotype_Mtor | Snx3 | 1.98 | 4.63E-07 | glio |
| Interaction_genotype_Myc | Cycs | 2.255 | 4.63E-07 | glio |
| Bcas2_ex.vivo_noRT | Gpr4 | 3.54 | 4.63E-07 | glio |
| Hsd17b10_in.vivo_noRT | 4930434B07R | 3.353 | 4.63E-08 | glio |
| Chek1_in.vivo_noRT | Gm8300 | 5.044 | 4.63E-09 | glio |
| Bcas2_in.vivo_noRT | Slc16a7 | 3.696 | 4.63E-10 | glio |
| Bcas2_in.vivo_noRT | Rhbg | 3.212 | 4.63E-10 | glio |
| Bora_in.vivo_noRT | Gm37214 | 4.676 | 4.63E-11 | glio |

| | | | | |
|---|---|---|---|---|
| Bora_in.vivo_noRT | CAAA0106612 | 4.144 | 4.63E-11 | glio |
| culture_effect | Sbds | -1.246 | 4.63E-17 | glio |
| Aars2_in.vivo_noRT | Gm39929 | 4.869 | 4.63E-18 | glio |
| culture_effect | Coq3 | -0.621 | 4.64E-03 | glio |
| culture_effect | Hdlbp | 0.352 | 4.64E-03 | glio |
| Interaction_genotype_Aars2 | Nrap | -3.745 | 4.64E-03 | glio |
| Interaction_genotype_Atr | Zfp82 | -3.523 | 4.64E-03 | glio |
| Interaction_genotype_Atr | Zfp101 | -2.709 | 4.64E-03 | glio |
| Interaction_genotype_Atr | Fermt2 | 1.691 | 4.64E-03 | glio |
| Interaction_genotype_Bcas2 | Atf5 | 2.031 | 4.64E-03 | glio |
| Interaction_genotype_Mtor | Tmem167b | -2.21 | 4.64E-03 | glio |
| Interaction_genotype_Rpl3 | Bnip1 | 2.274 | 4.64E-03 | glio |
| Aars2_in.vivo_noRT | Pask | 1.995 | 4.64E-03 | glio |
| Aars2_in.vivo_noRT | Apol9b | -1.214 | 4.64E-03 | glio |
| Atr_in.vivo_noRT | Nme2 | -0.527 | 4.64E-03 | glio |
| Atr_ex.vivo_noRT | Bcl2l11 | 2.055 | 4.64E-03 | glio |
| Bcas2_in.vivo_noRT | Npr2 | 2.395 | 4.64E-03 | glio |
| Bora_in.vivo_noRT | Pcsk4 | 2.088 | 4.64E-03 | glio |
| Bora_in.vivo_noRT | Hint1 | -0.475 | 4.64E-03 | glio |
| Hsd17b10_in.vivo_noRT | Dcpp1 | 2.289 | 4.64E-03 | glio |
| culture_effect | 2900052L18Ri | 1.143 | 4.64E-04 | glio |
| Interaction_genotype_Mtor | Lrrc46 | -4.183 | 4.64E-04 | glio |
| Interaction_genotype_Myc | Slc45a1 | -2.869 | 4.64E-04 | glio |
| Interaction_genotype_Myc | Gm4724 | -3.567 | 4.64E-04 | glio |
| Interaction_genotype_Myc | Efnb1 | -3.624 | 4.64E-04 | glio |
| Interaction_genotype_Myc | Zc3h12c | -3.645 | 4.64E-04 | glio |
| Interaction_genotype_Myc | Vps8 | -3.612 | 4.64E-04 | glio |
| Aars2_in.vivo_noRT | Psmc4 | -1.156 | 4.64E-04 | glio |
| Atr_ex.vivo_noRT | Gm5662 | 3.164 | 4.64E-04 | glio |
| Bcas2_ex.vivo_noRT | Gm5602 | 2.742 | 4.64E-04 | glio |
| Bcas2_ex.vivo_noRT | Gm21818 | 3.54 | 4.64E-04 | glio |
| Bora_in.vivo_noRT | Lcat | 2.603 | 4.64E-04 | glio |
| Bora_in.vivo_noRT | Gm42699 | 2.386 | 4.64E-04 | glio |
| Chek1_in.vivo_noRT | 5930430L01Ri | 2.368 | 4.64E-04 | glio |
| interaction_RT | AW112010 | 3.505 | 4.64E-05 | glio |
| Bora_in.vivo_noRT | Ksr1 | 2.888 | 4.64E-05 | glio |
| Interaction_genotype_Bcas2 | Gm2115 | -5.101 | 4.64E-06 | glio |
| Interaction_genotype_Mtor | Gm12703 | -3.833 | 4.64E-06 | glio |
| Interaction_genotype_Mtor | U2surp | 2.201 | 4.64E-06 | glio |
| Bora_in.vivo_noRT | Gm44836 | 3.104 | 4.64E-06 | glio |
| Chek1_ex.vivo_noRT | Prex2 | 4.842 | 4.64E-07 | glio |
| Hsd17b10_in.vivo_noRT | Adam2 | 3.09 | 4.64E-07 | glio |
| culture_effect | Dync2li1 | -1.403 | 4.64E-08 | glio |
| Aars2_in.vivo_noRT | Selenok-ps1 | 3.763 | 4.64E-09 | glio |
| Aars2_in.vivo_noRT | Ptpn22 | 5.063 | 4.64E-11 | glio |
| culture_effect | Kif13b | -2.23 | 4.64E-13 | glio |
| culture_effect | Ttc26 | -1.234 | 4.65E-03 | glio |
| Interaction_genotype_Atr | Cers3 | -2.81 | 4.65E-03 | glio |
| Interaction_genotype_Mtor | Arhgap9 | -2.749 | 4.65E-03 | glio |
| Interaction_genotype_Myc | Slc25a43 | -3.243 | 4.65E-03 | glio |
| Interaction_genotype_Myc | Trim71 | -3.151 | 4.65E-03 | glio |
| Interaction_genotype_Myc | Agfg1 | -1.388 | 4.65E-03 | glio |

| | | | | |
|---|---|---|---|---|
| Interaction_genotype_Pars2 | Oxt | -3.879 | 4.65E-03 | glio |
| Atp5c1_in.vivo_noRT | Dock6 | 2.241 | 4.65E-03 | glio |
| Atr_in.vivo_noRT | Serpinb1a | 2.58 | 4.65E-03 | glio |
| Atr_in.vivo_noRT | Ppfia2 | 1.627 | 4.65E-03 | glio |
| Bard1_ex.vivo_noRT | Cys1 | 2.421 | 4.65E-03 | glio |
| Bard1_ex.vivo_noRT | Pik3ap1 | 2.459 | 4.65E-03 | glio |
| Bard1_ex.vivo_noRT | Catsperz | 2.893 | 4.65E-03 | glio |
| Bcas2_in.vivo_noRT | Fis1 | -0.803 | 4.65E-03 | glio |
| Bora_in.vivo_noRT | Col20a1 | 2.099 | 4.65E-03 | glio |
| Bora_in.vivo_noRT | Ndufb11 | -0.605 | 4.65E-03 | glio |
| Bora_ex.vivo_noRT | Tnfrsf4 | 2.811 | 4.65E-03 | glio |
| Bora_ex.vivo_noRT | Tm6sf1 | 2.983 | 4.65E-03 | glio |
| Chek1_in.vivo_noRT | Arap1 | 2.089 | 4.65E-03 | glio |
| Chek1_in.vivo_noRT | Dnaic2 | 1.415 | 4.65E-03 | glio |
| Eif2s1_in.vivo_noRT | Obscn | 2.143 | 4.65E-03 | glio |
| Hsd17b10_in.vivo_noRT | Tnni2 | 2.578 | 4.65E-03 | glio |
| Hsd17b10_in.vivo_noRT | Ntf5 | 2.321 | 4.65E-03 | glio |
| Hsd17b10_in.vivo_noRT | Vamp8 | 1.793 | 4.65E-03 | glio |
| culture_effect | Plac1 | -2.153 | 4.65E-04 | glio |
| culture_effect | Calhm5 | 1.895 | 4.65E-04 | glio |
| culture_effect | Pan2 | -0.89 | 4.65E-04 | glio |
| culture_effect | Grn | -0.55 | 4.65E-04 | glio |
| Interaction_genotype_Bora | Pde3a | -4.306 | 4.65E-04 | glio |
| Interaction_genotype_Mtor | Gm16853 | -4.03 | 4.65E-04 | glio |
| Interaction_genotype_Myc | Zfp212 | -3.087 | 4.65E-04 | glio |
| interaction_RT | Anapc15 | -0.872 | 4.65E-04 | glio |
| Aars2_in.vivo_noRT | 1700019L13Ri | 2.497 | 4.65E-04 | glio |
| Aars2_ex.vivo_noRT | Ceacam19 | 3.382 | 4.65E-04 | glio |
| Atr_in.vivo_noRT | Zfp975 | 2.405 | 4.65E-04 | glio |
| Bcas2_in.vivo_noRT | Gm15063 | 2.823 | 4.65E-04 | glio |
| Bcas2_in.vivo_noRT | 1110006O24R | 2.605 | 4.65E-04 | glio |
| Bcas2_in.vivo_noRT | Orc2 | -1.316 | 4.65E-04 | glio |
| Bcas2_ex.vivo_noRT | Gm12860 | 2.742 | 4.65E-04 | glio |
| culture_effect | Vmp1 | -0.631 | 4.65E-05 | glio |
| Aars2_in.vivo_noRT | Pcdhb16 | 3.1 | 4.65E-05 | glio |
| Aars2_in.vivo_noRT | C030014I23Ri | 2.876 | 4.65E-05 | glio |
| Aars2_in.vivo_noRT | Srcin1 | 2.753 | 4.65E-05 | glio |
| Aars2_ex.vivo_noRT | Pou5f1 | 3.676 | 4.65E-05 | glio |
| Bora_in.vivo_noRT | Mertk | 2.803 | 4.65E-05 | glio |
| Bora_ex.vivo_noRT | AI463229 | 3.201 | 4.65E-05 | glio |
| Chek1_in.vivo_noRT | Ifi205 | 3.734 | 4.65E-05 | glio |
| Chek1_in.vivo_noRT | Wnt6 | 3.868 | 4.65E-06 | glio |
| Chek1_in.vivo_noRT | 9130204L05Ri | 3.45 | 4.65E-06 | glio |
| Hsd17b10_in.vivo_noRT | Dppa2 | 3.084 | 4.65E-06 | glio |
| Bora_in.vivo_noRT | Slc26a1 | 3.433 | 4.65E-07 | glio |
| Interaction_genotype_Mtor | Gm13856 | -4.492 | 4.65E-08 | glio |
| Interaction_genotype_Mtor | Gm21936 | -4.494 | 4.65E-08 | glio |
| Bcas2_in.vivo_noRT | St14 | 3.421 | 4.65E-08 | glio |
| Bcas2_in.vivo_noRT | Synpo2l | 4.162 | 4.65E-12 | glio |
| culture_effect | Ddah1 | 1.845 | 4.65E-13 | glio |
| culture_effect | Dhrs7 | -1.356 | 4.65E-13 | glio |
| Chek1_in.vivo_noRT | Gm43915 | 5.023 | 4.65E-18 | glio |

| | | | | |
|---|---|---|---|---|
| Chek1_in.vivo_noRT | 4930470G03R | 4.777 | 4.65E-18 | glio |
| Bora_in.vivo_noRT | Ctla4 | 4.936 | 4.65E-24 | glio |
| culture_effect | Tmed4 | -0.393 | 4.66E-03 | glio |
| culture_effect | Mrps30 | -0.436 | 4.66E-03 | glio |
| Interaction_genotype_Aars2 | Gm49860 | -3.015 | 4.66E-03 | glio |
| Interaction_genotype_Aars2 | Zfp661 | -3.631 | 4.66E-03 | glio |
| Interaction_genotype_Aars2 | Gm19391 | -3.037 | 4.66E-03 | glio |
| Interaction_genotype_Atr | Fam3a | -2.279 | 4.66E-03 | glio |
| Interaction_genotype_Mtor | A930041C12R | -3.443 | 4.66E-03 | glio |
| Interaction_genotype_Mtor | Ccnyl1 | -2.702 | 4.66E-03 | glio |
| Interaction_genotype_Mtor | Cfdp1 | 1.247 | 4.66E-03 | glio |
| Interaction_genotype_Myc | Gm16933 | -2.207 | 4.66E-03 | glio |
| Interaction_genotype_Myc | Mtmr11 | -3.223 | 4.66E-03 | glio |
| Interaction_genotype_Myc | Gm49768 | -3.01 | 4.66E-03 | glio |
| Interaction_genotype_Terf1 | Mecom | -3.901 | 4.66E-03 | glio |
| Interaction_genotype_Terf1 | Zbp1 | 1.801 | 4.66E-03 | glio |
| interaction_RT | Nfia | 0.769 | 4.66E-03 | glio |
| Atr_in.vivo_noRT | Poglut3 | 1.656 | 4.66E-03 | glio |
| Atr_ex.vivo_noRT | Gpr37l1 | 2.77 | 4.66E-03 | glio |
| Atr_ex.vivo_noRT | Pitx3 | 2.183 | 4.66E-03 | glio |
| Atr_ex.vivo_noRT | Cldn14 | 2.772 | 4.66E-03 | glio |
| Bcas2_ex.vivo_noRT | Tpsab1 | 2.742 | 4.66E-03 | glio |
| Bcas2_ex.vivo_noRT | Itpr1 | -2.174 | 4.66E-03 | glio |
| Bora_in.vivo_noRT | Irf5 | 2.412 | 4.66E-03 | glio |
| Bora_in.vivo_noRT | Sec61a2 | -1.482 | 4.66E-03 | glio |
| Bora_in.vivo_noRT | Calu | -0.702 | 4.66E-03 | glio |
| Bora_ex.vivo_noRT | Gm15943 | 2.22 | 4.66E-03 | glio |
| Bora_ex.vivo_noRT | Gm16094 | 2.615 | 4.66E-03 | glio |
| Chek1_in.vivo_noRT | 3110083C13R | 2.04 | 4.66E-03 | glio |
| Chek1_in.vivo_noRT | Fmr1 | -1.19 | 4.66E-03 | glio |
| Interaction_genotype_Aars2 | Lrrc59 | 1.635 | 4.66E-04 | glio |
| Interaction_genotype_Mtor | Fabp4 | -4.358 | 4.66E-04 | glio |
| Interaction_genotype_Myc | Dsel | -3.478 | 4.66E-04 | glio |
| Interaction_genotype_Myc | Hax1 | 1.582 | 4.66E-04 | glio |
| Interaction_genotype_Rad51 | Klf7 | 2.811 | 4.66E-04 | glio |
| interaction_RT | Hnrnpa3 | -0.712 | 4.66E-04 | glio |
| Aars2_in.vivo_noRT | Gm26620 | 2.247 | 4.66E-04 | glio |
| Atr_in.vivo_noRT | Syt4 | -0.84 | 4.66E-04 | glio |
| culture_effect | Ddx39b | -0.523 | 4.66E-05 | glio |
| Interaction_genotype_Myc | Eif3e | 1.68 | 4.66E-05 | glio |
| Aars2_in.vivo_noRT | Dpy19l1 | -1.898 | 4.66E-05 | glio |
| culture_effect | Smarcd1 | 0.787 | 4.66E-06 | glio |
| Interaction_genotype_Nat2 | Gm4027 | 4.443 | 4.66E-06 | glio |
| Bcas2_in.vivo_noRT | Apold1 | 3.697 | 4.66E-06 | glio |
| Bcas2_in.vivo_noRT | Gm19325 | 3.371 | 4.66E-06 | glio |
| Chek1_in.vivo_noRT | Gm37933 | 2.893 | 4.66E-06 | glio |
| Chek1_in.vivo_noRT | Gm49494 | 3.698 | 4.66E-07 | glio |
| Chek1_in.vivo_noRT | Slc6a19 | 3.398 | 4.66E-07 | glio |
| Hsd17b10_in.vivo_noRT | Loxl4 | 3.35 | 4.66E-07 | glio |
| culture_effect | Pdia5 | -1.601 | 4.66E-08 | glio |
| Atr_in.vivo_noRT | Gramd2 | 3.695 | 4.66E-08 | glio |
| Chek1_in.vivo_noRT | Gm47061 | 4.268 | 4.66E-08 | glio |

| | | | | |
|---|---|---|---|---|
| Chek1_in.vivo_noRT | Fam222a | 4.053 | 4.66E-08 | glio |
| culture_effect | Plekha8 | 1.312 | 4.66E-09 | glio |
| Atr_ex.vivo_noRT | D730045A05R | 5.343 | 4.66E-10 | glio |
| Bcas2_in.vivo_noRT | Gm11963 | 4.547 | 4.66E-11 | glio |
| Hsd17b10_in.vivo_noRT | Rgs4 | 3.616 | 4.66E-12 | glio |
| Bora_in.vivo_noRT | Pik3ap1 | 4.673 | 4.66E-15 | glio |
| Chek1_in.vivo_noRT | Prl2c3 | 5.044 | 4.66E-15 | glio |
| Bcas2_in.vivo_noRT | Serpinb9g | 4.807 | 4.66E-21 | glio |
| Atr_in.vivo_noRT | Zscan4-ps3 | 5.533 | 4.66E-29 | glio |
| culture_effect | Zfp664 | 0.526 | 4.67E-03 | glio |
| Interaction_genotype_Mtor | Rnf225 | -3.671 | 4.67E-03 | glio |
| Interaction_genotype_Mtor | 4932422M17F | -2.898 | 4.67E-03 | glio |
| Interaction_genotype_Myc | Tmem240 | -3.34 | 4.67E-03 | glio |
| Interaction_genotype_Myc | Fhl2 | -2.572 | 4.67E-03 | glio |
| Interaction_genotype_Myc | Cyb5r4 | -1.711 | 4.67E-03 | glio |
| Interaction_genotype_Myc | Psma6 | 1.142 | 4.67E-03 | glio |
| interaction_RT | Exosc10 | -0.906 | 4.67E-03 | glio |
| interaction_RT | Pigx | 0.806 | 4.67E-03 | glio |
| Aars2_in.vivo_noRT | Dmxl2 | 1.674 | 4.67E-03 | glio |
| Bcas2_in.vivo_noRT | Gm11713 | 1.864 | 4.67E-03 | glio |
| Chek1_in.vivo_noRT | Ik | -0.874 | 4.67E-03 | glio |
| Chek1_ex.vivo_noRT | Rpl7 | -0.704 | 4.67E-03 | glio |
| Hsd17b10_in.vivo_noRT | Usb1 | -1.435 | 4.67E-03 | glio |
| culture_effect | Traf2 | -0.953 | 4.67E-04 | glio |
| Interaction_genotype_Atr | Rnft2 | -4.255 | 4.67E-04 | glio |
| Interaction_genotype_Atr | Hdac7 | -4.199 | 4.67E-04 | glio |
| Interaction_genotype_Atr | Gm27253 | -4.266 | 4.67E-04 | glio |
| Interaction_genotype_Myc | Adam30 | -3.454 | 4.67E-04 | glio |
| Hsd17b10_in.vivo_noRT | 1700030C12R | 2.691 | 4.67E-04 | glio |
| culture_effect | Gk5 | -1.411 | 4.67E-05 | glio |
| Interaction_genotype_Mtor | Gm6598 | -4.419 | 4.67E-05 | glio |
| Aars2_in.vivo_noRT | Lmln | 2.791 | 4.67E-05 | glio |
| Atr_in.vivo_noRT | Fhod3 | 3.016 | 4.67E-05 | glio |
| Bcas2_in.vivo_noRT | Spon1 | 2.954 | 4.67E-05 | glio |
| Hsd17b10_in.vivo_noRT | Mycl | 3.094 | 4.67E-05 | glio |
| Interaction_genotype_Myc | Gm50394 | -5.098 | 4.67E-06 | glio |
| Atr_in.vivo_noRT | Pabpc4l | 3.255 | 4.67E-06 | glio |
| culture_effect | Nanos3 | 2.05 | 4.67E-07 | glio |
| Atr_in.vivo_noRT | Speg | 4.156 | 4.67E-09 | glio |
| culture_effect | Sfr1 | -0.687 | 4.67E-10 | glio |
| Chek1_in.vivo_noRT | Gm28404 | 4.156 | 4.67E-11 | glio |
| culture_effect | Eif4g2 | 0.691 | 4.67E-12 | glio |
| Chek1_ex.vivo_noRT | Ush1g | 4.443 | 4.67E-12 | glio |
| Chek1_ex.vivo_noRT | Gm48215 | 4.842 | 4.67E-14 | glio |
| Atr_ex.vivo_noRT | Gm48764 | 4.867 | 4.67E-15 | glio |
| Chek1_in.vivo_noRT | Gm11583 | 4.518 | 4.67E-18 | glio |
| Chek1_in.vivo_noRT | Gm29670 | 4.522 | 4.67E-18 | glio |
| Interaction_genotype_Atr | Tslp | -3.269 | 4.68E-03 | glio |
| Interaction_genotype_Atr | Synrg | 1.723 | 4.68E-03 | glio |
| Interaction_genotype_Mtor | Traf5 | -3.394 | 4.68E-03 | glio |
| Interaction_genotype_Mtor | Aff1 | -2.127 | 4.68E-03 | glio |
| Interaction_genotype_Mtor | Bad | 1.485 | 4.68E-03 | glio |

| | | | | |
|---|---|---|---|---|
| Interaction_genotype_Mtor | Slbp | 1.517 | 4.68E-03 | glio |
| Interaction_genotype_Myc | March11 | -3.368 | 4.68E-03 | glio |
| Interaction_genotype_Rpl3 | Cryl1 | 2.629 | 4.68E-03 | glio |
| interaction_RT | Ctnnal1 | -1.338 | 4.68E-03 | glio |
| interaction_RT | Enpp2 | 2.355 | 4.68E-03 | glio |
| Aars2_in.vivo_noRT | Gm17484 | 1.94 | 4.68E-03 | glio |
| Aars2_in.vivo_noRT | Borcs7 | -1.306 | 4.68E-03 | glio |
| Aars2_in.vivo_noRT | Dmwd | -1.166 | 4.68E-03 | glio |
| Atr_in.vivo_noRT | Lyar | -0.835 | 4.68E-03 | glio |
| Bard1_ex.vivo_noRT | Hfm1 | 2.239 | 4.68E-03 | glio |
| Bard1_ex.vivo_noRT | Gm20711 | 2.421 | 4.68E-03 | glio |
| Bcas2_in.vivo_noRT | Actb | -0.623 | 4.68E-03 | glio |
| Cenpj_in.vivo_noRT | Sytl4 | 1.756 | 4.68E-03 | glio |
| Cenpj_in.vivo_noRT | 2310003N18R | 2.022 | 4.68E-03 | glio |
| Hsd17b10_in.vivo_noRT | Mtg2 | -1.585 | 4.68E-03 | glio |
| Interaction_genotype_Mtor | Tex52 | -4.034 | 4.68E-04 | glio |
| Interaction_genotype_Mtor | Serac1 | -4.032 | 4.68E-04 | glio |
| Interaction_genotype_Myc | Hnrnpu | 1.339 | 4.68E-04 | glio |
| Aars2_in.vivo_noRT | Ebi3 | 2.949 | 4.68E-04 | glio |
| Atp5c1_in.vivo_noRT | Rarres1 | 2.668 | 4.68E-04 | glio |
| Atp5c1_in.vivo_noRT | Gm21762 | 2.404 | 4.68E-04 | glio |
| Atr_ex.vivo_noRT | Npnt | 2.367 | 4.68E-04 | glio |
| Bora_in.vivo_noRT | 6430590A07R | 1.942 | 4.68E-04 | glio |
| Bora_in.vivo_noRT | Letm1 | -1.318 | 4.68E-04 | glio |
| Cenpj_in.vivo_noRT | Gm30692 | 2.022 | 4.68E-04 | glio |
| Chek1_in.vivo_noRT | Cd72 | 2.482 | 4.68E-04 | glio |
| Eif2s1_ex.vivo_noRT | C330024C12R | 3.421 | 4.68E-04 | glio |
| Fanca_ex.vivo_noRT | Arhgef16 | 3.309 | 4.68E-04 | glio |
| culture_effect | Mdh2 | -0.532 | 4.68E-05 | glio |
| interaction_RT | Kdm6b | 1.269 | 4.68E-05 | glio |
| Chek1_in.vivo_noRT | Eml6 | 2.955 | 4.68E-05 | glio |
| culture_effect | Rhbg | -2.137 | 4.68E-06 | glio |
| Aars2_ex.vivo_noRT | Gm30692 | 2.899 | 4.68E-06 | glio |
| Atr_in.vivo_noRT | Ehbp1l1 | 3.283 | 4.68E-06 | glio |
| culture_effect | Eef1g | -0.647 | 4.68E-08 | glio |
| Interaction_genotype_Myc | Metap2 | 2.53 | 4.68E-08 | glio |
| Atr_in.vivo_noRT | Rbm11 | 3.602 | 4.68E-08 | glio |
| Bora_in.vivo_noRT | Kcnq2 | 3.773 | 4.68E-10 | glio |
| Bcas2_in.vivo_noRT | Spn | 4.285 | 4.68E-11 | glio |
| Chek1_in.vivo_noRT | Mcmdc2 | 4.784 | 4.68E-16 | glio |
| Atr_in.vivo_noRT | Gm33051 | 4.88 | 4.68E-19 | glio |
| Atr_in.vivo_noRT | Fabp6 | 5.533 | 4.68E-25 | glio |
| Interaction_genotype_Aars2 | Tecrl | -3.583 | 4.69E-03 | glio |
| Interaction_genotype_Bora | Cd248 | -3.666 | 4.69E-03 | glio |
| Interaction_genotype_Bora | D030040B21R | -3.476 | 4.69E-03 | glio |
| Interaction_genotype_Hsd17b | mt-Nd2 | -1.144 | 4.69E-03 | glio |
| Interaction_genotype_Mtor | Gm43608 | -3.384 | 4.69E-03 | glio |
| Interaction_genotype_Myc | Rhov | -3.146 | 4.69E-03 | glio |
| Interaction_genotype_Myc | Alpk2 | -4.085 | 4.69E-03 | glio |
| Interaction_genotype_Myc | Frg2f1 | -2.666 | 4.69E-03 | glio |
| Interaction_genotype_Myc | Zfp426 | -2.289 | 4.69E-03 | glio |
| Interaction_genotype_Myc | Orai2 | -2.387 | 4.69E-03 | glio |

| | | | | |
|---|---|---|---|---|
| Interaction_genotype_Rad51 | Gm9922 | -3.851 | 4.69E-03 | glio |
| interaction_RT | Ctsl | 0.864 | 4.69E-03 | glio |
| Aars2_in.vivo_noRT | Rpap1 | 1.436 | 4.69E-03 | glio |
| Aars2_in.vivo_noRT | Pdcd10 | -1.041 | 4.69E-03 | glio |
| Atr_in.vivo_noRT | Ndrg2 | 2.263 | 4.69E-03 | glio |
| Atr_ex.vivo_noRT | Ppfia2 | 2.37 | 4.69E-03 | glio |
| Bard1_ex.vivo_noRT | Dcpp3 | 2.814 | 4.69E-03 | glio |
| Bora_in.vivo_noRT | Tert | 1.95 | 4.69E-03 | glio |
| Bora_in.vivo_noRT | 1700120C14R | 1.972 | 4.69E-03 | glio |
| Chek1_in.vivo_noRT | Prdm9 | 2.104 | 4.69E-03 | glio |
| Chek1_ex.vivo_noRT | Gm14698 | 2.645 | 4.69E-03 | glio |
| Chek1_ex.vivo_noRT | Cops7b | -1.699 | 4.69E-03 | glio |
| Chek1_ex.vivo_noRT | Vrk1 | -1.535 | 4.69E-03 | glio |
| Eif2s1_in.vivo_noRT | D930028M14I | 2.143 | 4.69E-03 | glio |
| Eif2s1_in.vivo_noRT | Gm5602 | 1.883 | 4.69E-03 | glio |
| Eif2s1_in.vivo_noRT | Zfand2b | -1.504 | 4.69E-03 | glio |
| Eif2s1_in.vivo_noRT | Spcs1 | -0.543 | 4.69E-03 | glio |
| Hsd17b10_in.vivo_noRT | Srp54a | -1.547 | 4.69E-03 | glio |
| Hsd17b10_in.vivo_noRT | Ap2s1 | -0.639 | 4.69E-03 | glio |
| Interaction_genotype_Atr | Gm31063 | -3.293 | 4.69E-04 | glio |
| Interaction_genotype_Myc | Tmem132c | -2.729 | 4.69E-04 | glio |
| Interaction_genotype_Myc | Pdzrn3 | -2.972 | 4.69E-04 | glio |
| Aars2_in.vivo_noRT | Ap1s1 | -1.126 | 4.69E-04 | glio |
| Aars2_ex.vivo_noRT | 9130230L23Ri | 2.9 | 4.69E-04 | glio |
| Atp5c1_in.vivo_noRT | Rnf165 | 2.668 | 4.69E-04 | glio |
| Atr_in.vivo_noRT | Lbhd1 | 1.771 | 4.69E-04 | glio |
| Atr_in.vivo_noRT | Rif1 | -1.265 | 4.69E-04 | glio |
| Bcas2_in.vivo_noRT | Nup210l | 2.68 | 4.69E-04 | glio |
| Bcas2_in.vivo_noRT | Itgb7 | 2.39 | 4.69E-04 | glio |
| culture_effect | Gm39326 | 2.28 | 4.69E-05 | glio |
| Interaction_genotype_Mtor | Tc2n | -4.976 | 4.69E-05 | glio |
| Atr_in.vivo_noRT | 1700120G11R | 3.05 | 4.69E-05 | glio |
| Bard1_ex.vivo_noRT | Lhx1os | 3.201 | 4.69E-05 | glio |
| Chek1_in.vivo_noRT | Aldh1a2 | 2.687 | 4.69E-05 | glio |
| Aars2_in.vivo_noRT | Dixdc1 | 3.23 | 4.69E-06 | glio |
| Bard1_ex.vivo_noRT | Gm13191 | 2.814 | 4.69E-06 | glio |
| Bard1_ex.vivo_noRT | 4933416E03Ri | 2.814 | 4.69E-06 | glio |
| Chek1_ex.vivo_noRT | D030051J21Ri | 3.857 | 4.69E-06 | glio |
| Hsd17b10_in.vivo_noRT | Cpeb1 | 3.492 | 4.69E-06 | glio |
| Interaction_genotype_Mtor | Ctla4 | -4.099 | 4.69E-07 | glio |
| Hsd17b10_in.vivo_noRT | Dok2 | 3.09 | 4.69E-07 | glio |
| culture_effect | Rnf165 | 2.922 | 4.69E-08 | glio |
| culture_effect | Raet1e | -1.161 | 4.69E-08 | glio |
| Chek1_in.vivo_noRT | Lncenc1 | 3.738 | 4.69E-09 | glio |
| Chek1_in.vivo_noRT | Nrg2 | 4.521 | 4.69E-11 | glio |
| Atr_ex.vivo_noRT | Gm14742 | 4.309 | 4.69E-13 | glio |
| Atr_in.vivo_noRT | Rcan2 | 5.533 | 4.69E-14 | glio |
| Bcas2_in.vivo_noRT | Dmrtc1a | 4.807 | 4.69E-19 | glio |
| Bcas2_in.vivo_noRT | Gm12589 | 4.807 | 4.69E-19 | glio |
| Bora_in.vivo_noRT | Gm47510 | 4.676 | 4.69E-21 | glio |
| Aars2_in.vivo_noRT | Crb3 | 5.327 | 4.69E-22 | glio |
| culture_effect | Tet2 | -1.179 | 4.70E-03 | glio |

| | | | | |
|---|---|---|---|---|
| culture_effect | Etv6 | -0.441 | 4.70E-03 | glio |
| culture_effect | Ppp2r2d | 0.429 | 4.70E-03 | glio |
| Interaction_genotype_Rpl3 | 2010106G04R | -4.007 | 4.70E-03 | glio |
| interaction_RT | Stat3 | 0.68 | 4.70E-03 | glio |
| Aars2_in.vivo_noRT | Ly6e | -0.959 | 4.70E-03 | glio |
| Atr_in.vivo_noRT | 1500002F19Ri | 1.953 | 4.70E-03 | glio |
| Bard1_ex.vivo_noRT | Ccser1 | 2.421 | 4.70E-03 | glio |
| Bard1_ex.vivo_noRT | Fgf8 | 2.503 | 4.70E-03 | glio |
| Bard1_ex.vivo_noRT | Slc1a2 | 2.427 | 4.70E-03 | glio |
| Bard1_ex.vivo_noRT | Gm16685 | 2.415 | 4.70E-03 | glio |
| Bard1_ex.vivo_noRT | Mag | 2.814 | 4.70E-03 | glio |
| Bcas2_in.vivo_noRT | 1700019D03R | 2.234 | 4.70E-03 | glio |
| Bora_in.vivo_noRT | 4932422M17F | 1.898 | 4.70E-03 | glio |
| Bora_ex.vivo_noRT | Gm49441 | 2.414 | 4.70E-03 | glio |
| Bora_ex.vivo_noRT | Gm48832 | 2.806 | 4.70E-03 | glio |
| Bora_ex.vivo_noRT | Snx18 | -1.954 | 4.70E-03 | glio |
| Cenpj_in.vivo_noRT | Gm15941 | 2.187 | 4.70E-03 | glio |
| Chek1_in.vivo_noRT | Atp6ap1 | -1.038 | 4.70E-03 | glio |
| Interaction_genotype_Myc | Gm44693 | -2.729 | 4.70E-04 | glio |
| Interaction_genotype_Myc | Pex5 | -2.885 | 4.70E-04 | glio |
| Atp5c1_in.vivo_noRT | Gm27241 | 2.012 | 4.70E-04 | glio |
| Atr_in.vivo_noRT | Adcy9 | 2.234 | 4.70E-04 | glio |
| Bard1_ex.vivo_noRT | Kit | 2.814 | 4.70E-04 | glio |
| Bcas2_in.vivo_noRT | Gm20337 | 2.589 | 4.70E-04 | glio |
| Bcas2_in.vivo_noRT | Zc3h6 | 2.575 | 4.70E-04 | glio |
| Chek1_in.vivo_noRT | Gm17501 | 2.621 | 4.70E-04 | glio |
| culture_effect | 2610318N02R | 0.827 | 4.70E-05 | glio |
| Interaction_genotype_Mtor | P4ha3 | -4.057 | 4.70E-05 | glio |
| Atr_in.vivo_noRT | Nuak2 | 3.13 | 4.70E-05 | glio |
| Atr_in.vivo_noRT | Magohb | -1.657 | 4.70E-05 | glio |
| Chek1_ex.vivo_noRT | Plcl1 | 2.89 | 4.70E-05 | glio |
| culture_effect | Tuft1 | -1.735 | 4.70E-06 | glio |
| Interaction_genotype_Myc | Col6a1 | -3.918 | 4.70E-06 | glio |
| Interaction_genotype_Myc | Lncppara | -5.543 | 4.70E-06 | glio |
| Atr_in.vivo_noRT | Kif7 | 2.982 | 4.70E-06 | glio |
| Bora_in.vivo_noRT | Rex2 | 4.322 | 4.70E-09 | glio |
| Aars2_in.vivo_noRT | Gm47428 | 4.151 | 4.70E-10 | glio |
| Chek1_in.vivo_noRT | Stx1b | 4.141 | 4.70E-11 | glio |
| Hsd17b10_in.vivo_noRT | Nwd1 | 3.879 | 4.70E-11 | glio |
| Chek1_ex.vivo_noRT | Klrk1 | 4.842 | 4.70E-12 | glio |
| Chek1_ex.vivo_noRT | Olfr543 | 4.447 | 4.70E-12 | glio |
| Chek1_in.vivo_noRT | Gm43653 | 4.78 | 4.70E-13 | glio |
| Aars2_in.vivo_noRT | 4930502E18Ri | 4.791 | 4.70E-17 | glio |
| culture_effect | H2afj | 0.948 | 4.70E-18 | glio |
| Chek1_in.vivo_noRT | Gm30692 | 5.044 | 4.70E-24 | glio |
| Interaction_genotype_Atr | Hcn3 | -3.229 | 4.71E-03 | glio |
| Interaction_genotype_Atr | Sorbs2 | -2.999 | 4.71E-03 | glio |
| Interaction_genotype_Chek1 | Ctsa | 1.934 | 4.71E-03 | glio |
| Interaction_genotype_Ifnar1 | Dpep3 | 3.004 | 4.71E-03 | glio |
| Interaction_genotype_Ifnar1 | Stfa3 | 2.734 | 4.71E-03 | glio |
| Interaction_genotype_Ifnar1 | Olfr787 | 2.734 | 4.71E-03 | glio |
| Interaction_genotype_Ifnar1 | Iigp1 | -2.348 | 4.71E-03 | glio |

| | | | | |
|---|---|---|---|---|
| Interaction_genotype_Myc | AF067061 | -2.729 | 4.71E-03 | glio |
| Interaction_genotype_Myc | Slc24a1 | -3.191 | 4.71E-03 | glio |
| Interaction_genotype_Myc | Pald1 | -3.067 | 4.71E-03 | glio |
| Interaction_genotype_Pars2 | Arl6ip6 | 1.476 | 4.71E-03 | glio |
| Interaction_genotype_Rpl3 | 1700015F17Ri | -3.967 | 4.71E-03 | glio |
| Interaction_genotype_Rpl3 | Ppp1r18os | -4.142 | 4.71E-03 | glio |
| Interaction_genotype_Rpl3 | Tppp3 | 2.733 | 4.71E-03 | glio |
| Aars2_in.vivo_noRT | Gm7072 | 1.389 | 4.71E-03 | glio |
| Atp5c1_in.vivo_noRT | D930030I03Ri | 2.476 | 4.71E-03 | glio |
| Atr_in.vivo_noRT | Mfsd8 | 1.762 | 4.71E-03 | glio |
| Bcas2_in.vivo_noRT | B230312C02R | 2.687 | 4.71E-03 | glio |
| Bora_in.vivo_noRT | Ppp1r18 | -1.33 | 4.71E-03 | glio |
| Cenpj_in.vivo_noRT | Angptl7 | 1.685 | 4.71E-03 | glio |
| Cenpj_in.vivo_noRT | Npnt | 1.756 | 4.71E-03 | glio |
| Cenpj_ex.vivo_noRT | Bpifa3 | 2.591 | 4.71E-03 | glio |
| Chek1_ex.vivo_noRT | Eme1 | -2.051 | 4.71E-03 | glio |
| Eif2s1_ex.vivo_noRT | Gm266 | 2.234 | 4.71E-03 | glio |
| Eif2s1_ex.vivo_noRT | Ildr1 | 2.234 | 4.71E-03 | glio |
| Interaction_genotype_Aars2 | Gm26673 | -4.349 | 4.71E-04 | glio |
| Interaction_genotype_Aars2 | Gm49891 | -4.591 | 4.71E-04 | glio |
| Interaction_genotype_Myc | Spice1 | -3.463 | 4.71E-04 | glio |
| Interaction_genotype_Myc | Dpm1 | -1.635 | 4.71E-04 | glio |
| Interaction_genotype_Pars2 | Gm43137 | -4.507 | 4.71E-04 | glio |
| Interaction_genotype_Pars2 | Tdrd5 | -4.409 | 4.71E-04 | glio |
| Interaction_genotype_Pars2 | K230015D01R | -4.731 | 4.71E-04 | glio |
| Interaction_genotype_Pars2 | Gm31518 | -4.893 | 4.71E-04 | glio |
| interaction_RT | Ubtf | -0.823 | 4.71E-04 | glio |
| interaction_RT | Eif4g1 | -0.781 | 4.71E-04 | glio |
| Aars2_in.vivo_noRT | A330023F24Ri | 2.548 | 4.71E-04 | glio |
| Aars2_in.vivo_noRT | 1700012D14R | 2.425 | 4.71E-04 | glio |
| Aars2_in.vivo_noRT | Miip | 1.812 | 4.71E-04 | glio |
| Aars2_ex.vivo_noRT | Stra8 | 2.897 | 4.71E-04 | glio |
| Bora_in.vivo_noRT | Pdrg1 | -1.283 | 4.71E-04 | glio |
| Interaction_genotype_Myc | Srcin1 | -4.273 | 4.71E-05 | glio |
| Interaction_genotype_Myc | Camkv | -4.857 | 4.71E-05 | glio |
| Interaction_genotype_Mtor | 4930512B01R | -4.099 | 4.71E-06 | glio |
| Aars2_in.vivo_noRT | Fam71e1 | 2.904 | 4.71E-06 | glio |
| Atr_ex.vivo_noRT | Gm45406 | 3.164 | 4.71E-06 | glio |
| Bcas2_in.vivo_noRT | Eng | 3.611 | 4.71E-06 | glio |
| culture_effect | Slc5a1 | -2.231 | 4.71E-07 | glio |
| culture_effect | Taok3 | 1.057 | 4.71E-08 | glio |
| Aars2_in.vivo_noRT | Zfp551 | 3.76 | 4.71E-08 | glio |
| Aars2_in.vivo_noRT | Nmb | 4.033 | 4.71E-09 | glio |
| Aars2_in.vivo_noRT | H2-M10.1 | 3.872 | 4.71E-11 | glio |
| Chek1_ex.vivo_noRT | Ltbp2 | 4.447 | 4.71E-11 | glio |
| Aars2_in.vivo_noRT | Gm15908 | 5.327 | 4.71E-29 | glio |
| Aars2_in.vivo_noRT | A930014E10Ri | 5.327 | 4.71E-29 | glio |
| culture_effect | Nsfl1c | -0.405 | 4.72E-03 | glio |
| Interaction_genotype_Aars2 | 9430041J12Ri | -2.828 | 4.72E-03 | glio |
| Interaction_genotype_Aars2 | Ube4bos1 | -2.956 | 4.72E-03 | glio |
| Interaction_genotype_Aars2 | Gm16023 | -3.85 | 4.72E-03 | glio |
| Interaction_genotype_Aars2 | Gm19541 | -3.481 | 4.72E-03 | glio |

| | | | | |
|---|---|---|---|---|
| Interaction_genotype_Aars2 | Apol9a | 1.591 | 4.72E-03 | glio |
| Interaction_genotype_Aars2 | Psmb5 | 1.032 | 4.72E-03 | glio |
| Interaction_genotype_Atr | Gm36495 | -3.383 | 4.72E-03 | glio |
| Interaction_genotype_Cdkn2a | Gm16178 | 2.835 | 4.72E-03 | glio |
| Interaction_genotype_Cdkn2a | Gm364 | 2.835 | 4.72E-03 | glio |
| Interaction_genotype_Mtor | Cptp | -2.446 | 4.72E-03 | glio |
| Interaction_genotype_Mtor | Hspb6 | -2.444 | 4.72E-03 | glio |
| Interaction_genotype_Rad21 | Ciita | 3.199 | 4.72E-03 | glio |
| Interaction_genotype_Rad21 | Ngef | 3.819 | 4.72E-03 | glio |
| Aars2_in.vivo_noRT | Xrcc1 | -1.218 | 4.72E-03 | glio |
| Atr_in.vivo_noRT | Kbtbd2 | -1.271 | 4.72E-03 | glio |
| Atr_in.vivo_noRT | Dlc1 | 1.444 | 4.72E-03 | glio |
| Atr_ex.vivo_noRT | Eif4e3 | 2.367 | 4.72E-03 | glio |
| Atr_ex.vivo_noRT | Hist2h3c2 | 2.779 | 4.72E-03 | glio |
| Atr_ex.vivo_noRT | Car7 | 2.578 | 4.72E-03 | glio |
| Atr_ex.vivo_noRT | Arsk | 2.062 | 4.72E-03 | glio |
| Bcas2_in.vivo_noRT | Vps33b | 1.621 | 4.72E-03 | glio |
| Bora_in.vivo_noRT | Gm49602 | 1.997 | 4.72E-03 | glio |
| Cenpj_ex.vivo_noRT | Ntn1 | 2.591 | 4.72E-03 | glio |
| Chek1_in.vivo_noRT | Acyp2 | -1.537 | 4.72E-03 | glio |
| Hsd17b10_in.vivo_noRT | 1110035H17R | 2.448 | 4.72E-03 | glio |
| Hsd17b10_in.vivo_noRT | Tbc1d16 | 2.066 | 4.72E-03 | glio |
| Interaction_genotype_Aars2 | Prrx2 | -4.347 | 4.72E-04 | glio |
| Interaction_genotype_Aars2 | 1110020A21R | -4.37 | 4.72E-04 | glio |
| Interaction_genotype_Aars2 | Vcp | 1.526 | 4.72E-04 | glio |
| Interaction_genotype_Aars2 | Nme2 | 1.008 | 4.72E-04 | glio |
| Interaction_genotype_Myc | Bbs12 | -3.375 | 4.72E-04 | glio |
| Aars2_in.vivo_noRT | E130307A14Ri | 2.296 | 4.72E-04 | glio |
| Atp5c1_in.vivo_noRT | Nos2 | 2.408 | 4.72E-04 | glio |
| Atp5c1_in.vivo_noRT | Efna2 | 3.119 | 4.72E-04 | glio |
| Bcas2_in.vivo_noRT | 9330160F10Ri | 2.329 | 4.72E-04 | glio |
| Bcas2_ex.vivo_noRT | Ccl9 | 2.349 | 4.72E-04 | glio |
| Bcas2_ex.vivo_noRT | Akr1c18 | 2.742 | 4.72E-04 | glio |
| Bcas2_ex.vivo_noRT | Minar2 | 2.742 | 4.72E-04 | glio |
| Bcas2_ex.vivo_noRT | Gm45151 | 2.742 | 4.72E-04 | glio |
| Bcas2_ex.vivo_noRT | Pabpc4l | 3.573 | 4.72E-04 | glio |
| Hsd17b10_in.vivo_noRT | Gm15520 | 2.3 | 4.72E-04 | glio |
| Aars2_in.vivo_noRT | mt-Nd1 | 0.742 | 4.72E-05 | glio |
| Atr_ex.vivo_noRT | Ptgis | 3.657 | 4.72E-05 | glio |
| Atr_ex.vivo_noRT | Hrh2 | 3.164 | 4.72E-05 | glio |
| Chek1_in.vivo_noRT | Zfp661 | 3.293 | 4.72E-06 | glio |
| Atr_in.vivo_noRT | Wfikkn2 | 3.559 | 4.72E-07 | glio |
| Bcas2_in.vivo_noRT | Anp32a | -1.601 | 4.72E-07 | glio |
| Chek1_in.vivo_noRT | Pcdhga9 | 3.348 | 4.72E-07 | glio |
| Chek1_ex.vivo_noRT | Slc1a2 | 3.657 | 4.72E-07 | glio |
| Aars2_in.vivo_noRT | Slc7a4 | 3.912 | 4.72E-09 | glio |
| Aars2_in.vivo_noRT | Gm49694 | 4.601 | 4.72E-10 | glio |
| Aars2_in.vivo_noRT | Zfp986 | 4.285 | 4.72E-10 | glio |
| Chek1_in.vivo_noRT | Cxcl5 | 5.044 | 4.72E-21 | glio |
| Atr_in.vivo_noRT | Ccl3 | 5.273 | 4.72E-24 | glio |
| culture_effect | Msto1 | 0.654 | 4.73E-03 | glio |
| Interaction_genotype_Aars2 | Tcte2 | -3.028 | 4.73E-03 | glio |

| | | | | |
|---|---|---|---|---|
| Interaction_genotype_Bcas2 | Brd4 | 1.28 | 4.73E-03 | glio |
| Interaction_genotype_Bora | Tmem199 | 1.614 | 4.73E-03 | glio |
| Interaction_genotype_Myc | Zfp937 | -3.333 | 4.73E-03 | glio |
| Interaction_genotype_Pars2 | Olfr655 | -5.219 | 4.73E-03 | glio |
| interaction_RT | Aco2 | -0.536 | 4.73E-03 | glio |
| Aars2_in.vivo_noRT | Uba1 | -1.033 | 4.73E-03 | glio |
| Aars2_ex.vivo_noRT | Prox2os | 2.499 | 4.73E-03 | glio |
| Bcas2_in.vivo_noRT | Mmp23 | 2.009 | 4.73E-03 | glio |
| Bcas2_in.vivo_noRT | Nfrkb | -1.37 | 4.73E-03 | glio |
| Bora_ex.vivo_noRT | D730045A05R | 2.806 | 4.73E-03 | glio |
| Chek1_in.vivo_noRT | Aph1b | 1.778 | 4.73E-03 | glio |
| culture_effect | Mettl3 | -1.045 | 4.73E-04 | glio |
| culture_effect | A930005H10R | 0.762 | 4.73E-04 | glio |
| Interaction_genotype_Atr | Btbd6 | -3.931 | 4.73E-04 | glio |
| Interaction_genotype_Myc | Gm11520 | -3.57 | 4.73E-04 | glio |
| Interaction_genotype_Myc | Gm26670 | -3.449 | 4.73E-04 | glio |
| Atr_in.vivo_noRT | Gm20324 | 2.189 | 4.73E-04 | glio |
| Chek1_in.vivo_noRT | Bzw2 | -1.266 | 4.73E-04 | glio |
| Interaction_genotype_Atr | Nxph4 | -4.809 | 4.73E-05 | glio |
| Atr_ex.vivo_noRT | Fsd2 | 3.722 | 4.73E-05 | glio |
| Bora_in.vivo_noRT | Opalin | 3.689 | 4.73E-05 | glio |
| Chek1_ex.vivo_noRT | Col11a1 | -1.093 | 4.73E-05 | glio |
| Chek1_in.vivo_noRT | Larp1b | -2.047 | 4.73E-06 | glio |
| culture_effect | Gm9958 | 2.599 | 4.73E-07 | glio |
| Bora_ex.vivo_noRT | Meioc | 3.201 | 4.73E-07 | glio |
| Atr_in.vivo_noRT | Zfp455 | 3.98 | 4.73E-09 | glio |
| Atr_in.vivo_noRT | Pld6 | 4.277 | 4.73E-09 | glio |
| Bcas2_in.vivo_noRT | Esrp2 | 3.895 | 4.73E-09 | glio |
| Atr_in.vivo_noRT | Bpi | 3.852 | 4.73E-10 | glio |
| Aars2_in.vivo_noRT | Camk2b | 4.805 | 4.73E-13 | glio |
| Bora_in.vivo_noRT | Rgs4 | 4.936 | 4.73E-21 | glio |
| culture_effect | Hint3 | 0.572 | 4.74E-03 | glio |
| culture_effect | Apba3 | -0.492 | 4.74E-03 | glio |
| culture_effect | Ndufs2 | -0.36 | 4.74E-03 | glio |
| culture_effect | Pfdn6 | -0.336 | 4.74E-03 | glio |
| culture_effect | Atp5c1 | -0.342 | 4.74E-03 | glio |
| Interaction_genotype_Aars2 | Gm32250 | -2.625 | 4.74E-03 | glio |
| Interaction_genotype_Aars2 | Trim17 | -2.433 | 4.74E-03 | glio |
| Interaction_genotype_Aars2 | Egf | -3.793 | 4.74E-03 | glio |
| Interaction_genotype_Aars2 | D930030I03Ri | -3.666 | 4.74E-03 | glio |
| Interaction_genotype_Aars2 | Tex15 | -3.437 | 4.74E-03 | glio |
| Interaction_genotype_Aars2 | Cox8a | 0.813 | 4.74E-03 | glio |
| Interaction_genotype_Atr | Gm28404 | -3.259 | 4.74E-03 | glio |
| Interaction_genotype_Mtor | Ngb | -3.329 | 4.74E-03 | glio |
| Interaction_genotype_Mtor | Exoc3l4 | -2.59 | 4.74E-03 | glio |
| Interaction_genotype_Mtor | Lyn | -2.363 | 4.74E-03 | glio |
| Interaction_genotype_Mtor | Gtf3c6 | 1.51 | 4.74E-03 | glio |
| Interaction_genotype_Myc | Akap14 | -2.466 | 4.74E-03 | glio |
| Interaction_genotype_Myc | Ehbp1l1 | -3.319 | 4.74E-03 | glio |
| Interaction_genotype_Myc | Fmn2 | -2.766 | 4.74E-03 | glio |
| Interaction_genotype_Myc | Fam118a | -2.423 | 4.74E-03 | glio |
| Interaction_genotype_Rad51 | Mog | 4.473 | 4.74E-03 | glio |

| | | | | |
|---|---|---|---|---|
| Interaction_genotype_Uqcrb | Eif4ebp1 | 1.231 | 4.74E-03 | glio |
| Atr_in.vivo_noRT | Gm45051 | 1.862 | 4.74E-03 | glio |
| Atr_in.vivo_noRT | Pomk | 1.778 | 4.74E-03 | glio |
| Atr_in.vivo_noRT | Rpl34 | -0.642 | 4.74E-03 | glio |
| Bcas2_in.vivo_noRT | Esr1 | 1.95 | 4.74E-03 | glio |
| Bcas2_in.vivo_noRT | Tbx15 | 1.829 | 4.74E-03 | glio |
| Bcas2_in.vivo_noRT | Gpd1l | 1.606 | 4.74E-03 | glio |
| Bora_in.vivo_noRT | Cacnb3 | 1.894 | 4.74E-03 | glio |
| Bora_in.vivo_noRT | Gm6871 | 1.658 | 4.74E-03 | glio |
| Bora_in.vivo_noRT | Cyp39a1 | -1.566 | 4.74E-03 | glio |
| Bora_in.vivo_noRT | Rrs1 | -1.001 | 4.74E-03 | glio |
| Bora_in.vivo_noRT | Cks2 | -1.008 | 4.74E-03 | glio |
| Bora_in.vivo_noRT | Hnrnpf | -0.548 | 4.74E-03 | glio |
| Chek1_in.vivo_noRT | Zbtb39 | 1.884 | 4.74E-03 | glio |
| Chek1_in.vivo_noRT | H2-Ke6 | -1.494 | 4.74E-03 | glio |
| Chek1_in.vivo_noRT | Selenon | 1.505 | 4.74E-03 | glio |
| Chek1_in.vivo_noRT | Apex1 | -1.013 | 4.74E-03 | glio |
| Chek1_ex.vivo_noRT | Hoxa6 | 2.491 | 4.74E-03 | glio |
| Eif2s1_in.vivo_noRT | Ablim2 | 1.61 | 4.74E-03 | glio |
| Eif2s1_ex.vivo_noRT | Gm13398 | 2.217 | 4.74E-03 | glio |
| Eif2s1_ex.vivo_noRT | Spi1 | 2.842 | 4.74E-03 | glio |
| Eif2s1_ex.vivo_noRT | Dpp6 | 2.234 | 4.74E-03 | glio |
| Eif2s1_ex.vivo_noRT | Serpinb9g | 2.234 | 4.74E-03 | glio |
| Eif2s1_ex.vivo_noRT | BC049987 | 2.234 | 4.74E-03 | glio |
| Eif2s1_ex.vivo_noRT | Tnfrsf19 | -3.086 | 4.74E-03 | glio |
| Ercc4_in.vivo_noRT | Ercc4 | -2.829 | 4.74E-03 | glio |
| Interaction_genotype_Bcas2 | Trim16 | -4.179 | 4.74E-04 | glio |
| Interaction_genotype_Myc | Sh2b1 | -2.788 | 4.74E-04 | glio |
| Aars2_in.vivo_noRT | Gm15892 | 2.653 | 4.74E-04 | glio |
| Aars2_in.vivo_noRT | Hibadh | -1.454 | 4.74E-04 | glio |
| Atr_ex.vivo_noRT | Gm36172 | 2.594 | 4.74E-04 | glio |
| Atr_ex.vivo_noRT | Tnfrsf1b | 3.164 | 4.74E-04 | glio |
| Chek1_in.vivo_noRT | Cxcl2 | 4.177 | 4.74E-04 | glio |
| Chek1_in.vivo_noRT | Carf | 2.501 | 4.74E-04 | glio |
| Interaction_genotype_Mtor | Gm47232 | -4.775 | 4.74E-05 | glio |
| Interaction_genotype_Myc | Aqp5 | -5.259 | 4.74E-05 | glio |
| Atr_in.vivo_noRT | 9030025P20R | 3.061 | 4.74E-05 | glio |
| Chek1_ex.vivo_noRT | E230001N04R | 3.155 | 4.74E-05 | glio |
| Atr_in.vivo_noRT | Gm47509 | 3.245 | 4.74E-06 | glio |
| Bcas2_in.vivo_noRT | Ifitm2 | -0.999 | 4.74E-06 | glio |
| Chek1_in.vivo_noRT | Gm48684 | 3.739 | 4.74E-06 | glio |
| Interaction_genotype_Mtor | Zscan4-ps3 | -4.099 | 4.74E-07 | glio |
| Atr_in.vivo_noRT | Pcdhga9 | 3.326 | 4.74E-07 | glio |
| Bora_in.vivo_noRT | 4930592C13R | 3.67 | 4.74E-07 | glio |
| culture_effect | Ptpn12 | 1.046 | 4.74E-08 | glio |
| culture_effect | Ndufv3 | 0.748 | 4.74E-08 | glio |
| Aars2_ex.vivo_noRT | Stfa3 | 3.293 | 4.74E-08 | glio |
| Aars2_ex.vivo_noRT | Olfr787 | 3.293 | 4.74E-08 | glio |
| Bora_ex.vivo_noRT | Gm30692 | 3.601 | 4.74E-08 | glio |
| culture_effect | Hebp1 | -1.94 | 4.74E-10 | glio |
| Hsd17b10_in.vivo_noRT | Dpep3 | 3.084 | 4.74E-10 | glio |
| Chek1_in.vivo_noRT | Gm13398 | 5.044 | 4.74E-23 | glio |

| | | | | |
|---|---|---|---|---|
| Bora_in.vivo_noRT | Zscan4-ps3 | 4.936 | 4.74E-24 | glio |
| culture_effect | Pno1 | 0.38 | 4.75E-03 | glio |
| Interaction_genotype_Atr | B4galt4 | -3.665 | 4.75E-03 | glio |
| Interaction_genotype_Bora | Mrpl44 | 1.888 | 4.75E-03 | glio |
| Interaction_genotype_Mtor | Nuak2 | -3.613 | 4.75E-03 | glio |
| interaction_RT | 5830408C22R | 1.984 | 4.75E-03 | glio |
| interaction_RT | Telo2 | -1.128 | 4.75E-03 | glio |
| interaction_RT | Mprip | -0.748 | 4.75E-03 | glio |
| interaction_RT | Tpm3 | -0.583 | 4.75E-03 | glio |
| Aars2_in.vivo_noRT | Ppp1r8 | -1.339 | 4.75E-03 | glio |
| Aars2_in.vivo_noRT | Arpc5l | -0.843 | 4.75E-03 | glio |
| Aars2_ex.vivo_noRT | Tex52 | 2.714 | 4.75E-03 | glio |
| Aars2_ex.vivo_noRT | Car9 | 2.899 | 4.75E-03 | glio |
| Atr_in.vivo_noRT | Taf5 | -1.263 | 4.75E-03 | glio |
| Bora_in.vivo_noRT | Trnt1 | -1.116 | 4.75E-03 | glio |
| Cenpj_in.vivo_noRT | Speer4a | 1.756 | 4.75E-03 | glio |
| Chek1_in.vivo_noRT | Tgfb2 | 1.915 | 4.75E-03 | glio |
| Cyp19a1_ex.vivo_noRT | Ankrd22 | 2.159 | 4.75E-03 | glio |
| Cyp19a1_ex.vivo_noRT | Cxcr2 | 3.148 | 4.75E-03 | glio |
| Cyp19a1_ex.vivo_noRT | Gm47404 | 2.159 | 4.75E-03 | glio |
| Cyp19a1_ex.vivo_noRT | Rpl3l | 2.959 | 4.75E-03 | glio |
| Cyp19a1_ex.vivo_noRT | Gm31717 | 2.159 | 4.75E-03 | glio |
| Cyp19a1_ex.vivo_noRT | Gm14742 | 2.159 | 4.75E-03 | glio |
| Cyp19a1_ex.vivo_noRT | Ctla4 | 2.159 | 4.75E-03 | glio |
| Cyp19a1_ex.vivo_noRT | Zscan4-ps3 | 2.159 | 4.75E-03 | glio |
| Cyp19a1_ex.vivo_noRT | Tagap | 2.159 | 4.75E-03 | glio |
| Cyp19a1_ex.vivo_noRT | Gm21731 | 2.159 | 4.75E-03 | glio |
| Cyp19a1_ex.vivo_noRT | Olfr1152 | 2.159 | 4.75E-03 | glio |
| Cyp19a1_ex.vivo_noRT | Gm11232 | 2.159 | 4.75E-03 | glio |
| Cyp19a1_ex.vivo_noRT | Gm29966 | 2.159 | 4.75E-03 | glio |
| Cyp19a1_ex.vivo_noRT | Cgnl1 | 2.159 | 4.75E-03 | glio |
| Interaction_genotype_Atr | Map3k15 | -3.683 | 4.75E-04 | glio |
| Interaction_genotype_Atr | Ppp1r3g | -4.062 | 4.75E-04 | glio |
| Interaction_genotype_Atr | Acot13 | 1.971 | 4.75E-04 | glio |
| Interaction_genotype_Myc | Icam5 | -3.855 | 4.75E-04 | glio |
| Interaction_genotype_Myc | Anxa11 | -4.305 | 4.75E-04 | glio |
| Interaction_genotype_Myc | Txnrd2 | -3.385 | 4.75E-04 | glio |
| Interaction_genotype_Myc | Xk | -3.742 | 4.75E-04 | glio |
| Interaction_genotype_Myc | Zfp213 | -3.168 | 4.75E-04 | glio |
| Interaction_genotype_Myc | Zswim1 | -2.78 | 4.75E-04 | glio |
| Bcas2_in.vivo_noRT | A430005L14Ri | -1.413 | 4.75E-04 | glio |
| Bcas2_in.vivo_noRT | Alad | -1.332 | 4.75E-04 | glio |
| Bcas2_ex.vivo_noRT | Gast | 2.742 | 4.75E-04 | glio |
| Bora_in.vivo_noRT | Pigk | -1.545 | 4.75E-04 | glio |
| Cenpj_ex.vivo_noRT | Arhgef16 | 2.989 | 4.75E-04 | glio |
| Interaction_genotype_Mtor | Gm36266 | -5.464 | 4.75E-05 | glio |
| Aars2_in.vivo_noRT | Cox4i1 | -0.733 | 4.75E-05 | glio |
| Atp5c1_in.vivo_noRT | Ptprc | 2.404 | 4.75E-05 | glio |
| Atr_in.vivo_noRT | Gatb | 2.365 | 4.75E-05 | glio |
| Eif2s1_in.vivo_noRT | 1700112J16Ri | 2.674 | 4.75E-05 | glio |
| Eif2s1_in.vivo_noRT | Sucnr1 | 2.143 | 4.75E-05 | glio |
| culture_effect | Kpna7 | 2.322 | 4.75E-06 | glio |

| | | | | |
|---|---|---|---|---|
| Interaction_genotype_Mtor | 1700024I08Ri | -4.42 | 4.75E-06 | glio |
| Aars2_in.vivo_noRT | Tesc | -2.666 | 4.75E-06 | glio |
| Aars2_ex.vivo_noRT | Npy1r | 3.293 | 4.75E-06 | glio |
| Aars2_ex.vivo_noRT | Gm6803 | 3.293 | 4.75E-06 | glio |
| Atr_in.vivo_noRT | Gm32816 | 3.474 | 4.75E-06 | glio |
| Bcas2_in.vivo_noRT | Ccdc184 | 3.474 | 4.75E-06 | glio |
| culture_effect | Sdsl | -1.751 | 4.75E-07 | glio |
| culture_effect | Psma3 | -0.53 | 4.75E-07 | glio |
| Atr_in.vivo_noRT | Alox8 | 4.277 | 4.75E-07 | glio |
| Atr_in.vivo_noRT | Slc47a2 | 3.427 | 4.75E-07 | glio |
| Atr_in.vivo_noRT | Gm17249 | 3.396 | 4.75E-07 | glio |
| Bora_ex.vivo_noRT | Tmem132c | 3.201 | 4.75E-07 | glio |
| Bora_ex.vivo_noRT | Gm44693 | 3.201 | 4.75E-07 | glio |
| culture_effect | Ankrd10 | -0.986 | 4.75E-08 | glio |
| Chek1_in.vivo_noRT | Zhx2 | 3.999 | 4.75E-08 | glio |
| Bcas2_in.vivo_noRT | Hectd2os | 3.899 | 4.75E-09 | glio |
| Bora_in.vivo_noRT | Kcnj13 | 3.655 | 4.75E-09 | glio |
| Chek1_ex.vivo_noRT | Fst | 4.842 | 4.75E-09 | glio |
| Chek1_ex.vivo_noRT | 8030423F21Ri | 4.267 | 4.75E-09 | glio |
| culture_effect | Msantd3 | 1.588 | 4.75E-12 | glio |
| Bcas2_in.vivo_noRT | Gm36026 | 4.156 | 4.75E-13 | glio |
| Aars2_in.vivo_noRT | Loxl4 | 5.06 | 4.75E-15 | glio |
| culture_effect | Gm13830 | 1.648 | 4.76E-03 | glio |
| culture_effect | Fbxo31 | -0.607 | 4.76E-03 | glio |
| Interaction_genotype_Aars2 | Ccl9 | -2.427 | 4.76E-03 | glio |
| Interaction_genotype_Aars2 | Gm47232 | -3.503 | 4.76E-03 | glio |
| Interaction_genotype_Aars2 | Gm15234 | -3.487 | 4.76E-03 | glio |
| Interaction_genotype_Aars2 | Zfp11 | -3.482 | 4.76E-03 | glio |
| Interaction_genotype_Aars2 | H3f3a | 0.782 | 4.76E-03 | glio |
| Interaction_genotype_Bora | Aqp5 | -4.265 | 4.76E-03 | glio |
| Interaction_genotype_Chek1 | Cd74 | 4.23 | 4.76E-03 | glio |
| Interaction_genotype_Mtor | 3300005D01R | -3.844 | 4.76E-03 | glio |
| Interaction_genotype_Mtor | Hunk | -2.956 | 4.76E-03 | glio |
| Interaction_genotype_Mtor | Sap18b | -2.784 | 4.76E-03 | glio |
| Interaction_genotype_Myc | Jakmip1 | -3.177 | 4.76E-03 | glio |
| Interaction_genotype_Myc | Fbxo10 | -3.097 | 4.76E-03 | glio |
| Interaction_genotype_Myc | Fam57a | -2.564 | 4.76E-03 | glio |
| interaction_RT | Gtsf1l | 2.396 | 4.76E-03 | glio |
| Bcas2_in.vivo_noRT | Slc25a42 | 2.124 | 4.76E-03 | glio |
| Bcas2_in.vivo_noRT | Tmem170 | 1.745 | 4.76E-03 | glio |
| Bcas2_in.vivo_noRT | Ndor1 | 1.576 | 4.76E-03 | glio |
| Bora_in.vivo_noRT | Hist4h4 | 2.139 | 4.76E-03 | glio |
| Bora_in.vivo_noRT | Incenp | -1.559 | 4.76E-03 | glio |
| Eif2s1_in.vivo_noRT | Gm40538 | 1.88 | 4.76E-03 | glio |
| culture_effect | Gstm5 | -0.601 | 4.76E-04 | glio |
| Interaction_genotype_Bora | Tox4 | 2.263 | 4.76E-04 | glio |
| Interaction_genotype_Mtor | Tlr6 | -3.978 | 4.76E-04 | glio |
| Interaction_genotype_Myc | Gm9887 | -3.487 | 4.76E-04 | glio |
| Atr_in.vivo_noRT | Plppr2 | 2.518 | 4.76E-04 | glio |
| Bora_in.vivo_noRT | Nup88 | -1.306 | 4.76E-04 | glio |
| Hsd17b10_in.vivo_noRT | Gm26673 | 2.596 | 4.76E-04 | glio |
| culture_effect | 4930412M03F | 2.285 | 4.76E-05 | glio |

| | | | | |
|---|---|---|---|---|
| culture_effect | Taf13 | 0.775 | 4.76E-05 | glio |
| Interaction_genotype_Myc | Glt8d2 | -4.813 | 4.76E-05 | glio |
| Atr_in.vivo_noRT | Paqr8 | 3.637 | 4.76E-06 | glio |
| Bora_in.vivo_noRT | Wdr38 | 3.365 | 4.76E-06 | glio |
| Bora_in.vivo_noRT | Lyrm1 | 3.223 | 4.76E-06 | glio |
| Interaction_genotype_Mtor | Loxl4 | -5.559 | 4.76E-07 | glio |
| Bcas2_in.vivo_noRT | Emid1 | 3.944 | 4.76E-07 | glio |
| Bcas2_in.vivo_noRT | Trp53cor1 | 2.521 | 4.76E-07 | glio |
| Chek1_in.vivo_noRT | Gm13272 | 3.34 | 4.76E-07 | glio |
| Atr_in.vivo_noRT | 1700001L05Ri | 3.835 | 4.76E-09 | glio |
| Atr_in.vivo_noRT | Gm26542 | 4.155 | 4.76E-10 | glio |
| Atr_in.vivo_noRT | 2310003N18R | 5.533 | 4.76E-21 | glio |
| Bcas2_in.vivo_noRT | Gm48435 | 4.807 | 4.76E-24 | glio |
| culture_effect | Ly6c2 | -1.837 | 4.77E-03 | glio |
| Interaction_genotype_Mtor | Mirg | -3.385 | 4.77E-03 | glio |
| interaction_RT | Synm | -1.813 | 4.77E-03 | glio |
| interaction_RT | Unc50 | 0.672 | 4.77E-03 | glio |
| interaction_RT | Rps25 | -0.588 | 4.77E-03 | glio |
| Aars2_in.vivo_noRT | Dennd2c | 2.031 | 4.77E-03 | glio |
| Aars2_in.vivo_noRT | Gm26812 | 1.998 | 4.77E-03 | glio |
| Aars2_in.vivo_noRT | Azin2 | 1.858 | 4.77E-03 | glio |
| Aars2_in.vivo_noRT | 4932438A13R | -1.356 | 4.77E-03 | glio |
| Atr_in.vivo_noRT | Osbpl7 | 1.828 | 4.77E-03 | glio |
| Atr_in.vivo_noRT | Usp28 | 1.854 | 4.77E-03 | glio |
| Bora_in.vivo_noRT | Acap3 | 1.575 | 4.77E-03 | glio |
| Chek1_ex.vivo_noRT | Gm42701 | 2.677 | 4.77E-03 | glio |
| Chek1_ex.vivo_noRT | Olr1 | 2.928 | 4.77E-03 | glio |
| Interaction_genotype_Bcas2 | Selenov | -3.836 | 4.77E-04 | glio |
| Interaction_genotype_Mtor | Fgf15 | -4.606 | 4.77E-04 | glio |
| Interaction_genotype_Mtor | Gm4419 | -3.154 | 4.77E-04 | glio |
| Interaction_genotype_Myc | Pafah2 | -4.236 | 4.77E-04 | glio |
| Atr_in.vivo_noRT | Pop5 | -1.024 | 4.77E-04 | glio |
| Bard1_ex.vivo_noRT | Gm30042 | 2.814 | 4.77E-04 | glio |
| Bcas2_in.vivo_noRT | Wbp4 | -1.289 | 4.77E-04 | glio |
| Aars2_ex.vivo_noRT | Ntn1 | 3.293 | 4.77E-05 | glio |
| Aars2_ex.vivo_noRT | Tent5c | 2.9 | 4.77E-05 | glio |
| Aars2_ex.vivo_noRT | Pmch | 3.293 | 4.77E-05 | glio |
| Bcas2_in.vivo_noRT | Gm12454 | 2.902 | 4.77E-05 | glio |
| Bcas2_in.vivo_noRT | Gm48832 | 2.791 | 4.77E-05 | glio |
| Bcas2_in.vivo_noRT | Stpg2 | 2.859 | 4.77E-05 | glio |
| Bora_in.vivo_noRT | Bco1 | 2.424 | 4.77E-05 | glio |
| Chek1_ex.vivo_noRT | Tcap | 3.287 | 4.77E-05 | glio |
| Interaction_genotype_Mtor | Gm48882 | -5.515 | 4.77E-06 | glio |
| Interaction_genotype_Mtor | Gm31152 | -4.952 | 4.77E-06 | glio |
| Bora_in.vivo_noRT | Gm15672 | 3.213 | 4.77E-06 | glio |
| Aars2_in.vivo_noRT | Aldh1l1 | 3.904 | 4.77E-07 | glio |
| Bora_ex.vivo_noRT | Gm47622 | 3.418 | 4.77E-07 | glio |
| Hsd17b10_in.vivo_noRT | Nxph3 | 3.616 | 4.77E-08 | glio |
| culture_effect | D5Ertd579e | 0.912 | 4.77E-09 | glio |
| Chek1_ex.vivo_noRT | Gm45336 | 4.842 | 4.77E-11 | glio |
| Bcas2_in.vivo_noRT | Txndc2 | 4.028 | 4.77E-13 | glio |
| Bora_in.vivo_noRT | Masp1 | 4.414 | 4.77E-14 | glio |

| | | | | |
|---|---|---|---|---|
| Interaction_genotype_Aars2 | Gm36839 | -3.516 | 4.78E-03 | glio |
| Interaction_genotype_Atr | Noxred1 | -3.412 | 4.78E-03 | glio |
| Interaction_genotype_Atr | Prl3d3 | -2.404 | 4.78E-03 | glio |
| Interaction_genotype_Atr | Upk1a | -3.717 | 4.78E-03 | glio |
| Interaction_genotype_Atr | Tspan33 | -3.619 | 4.78E-03 | glio |
| Interaction_genotype_Mtor | Vav3 | -2.618 | 4.78E-03 | glio |
| Interaction_genotype_Mtor | Bcl9 | -1.951 | 4.78E-03 | glio |
| Interaction_genotype_Mtor | Plod2 | 2.188 | 4.78E-03 | glio |
| Interaction_genotype_Mtor | Myc | 1.859 | 4.78E-03 | glio |
| Interaction_genotype_Mtor | Snhg12 | 1.541 | 4.78E-03 | glio |
| Interaction_genotype_Rad21 | Apol6 | 3.31 | 4.78E-03 | glio |
| Aars2_in.vivo_noRT | C230038L03Ri | 1.88 | 4.78E-03 | glio |
| Aars2_in.vivo_noRT | Gbp2b | 1.852 | 4.78E-03 | glio |
| Aars2_in.vivo_noRT | 0610009B22R | -1.079 | 4.78E-03 | glio |
| Aars2_ex.vivo_noRT | Wfdc18 | 3.459 | 4.78E-03 | glio |
| Aars2_ex.vivo_noRT | 1700120G11R | 3.044 | 4.78E-03 | glio |
| Atp5c1_in.vivo_noRT | Tent5c | 1.742 | 4.78E-03 | glio |
| Atp5c1_in.vivo_noRT | Pcdhgb6 | 2.014 | 4.78E-03 | glio |
| Bcas2_in.vivo_noRT | C1qtnf1 | 2.066 | 4.78E-03 | glio |
| Bcas2_in.vivo_noRT | Ppp2cb | -1.192 | 4.78E-03 | glio |
| Bora_in.vivo_noRT | Cdca3 | -1.21 | 4.78E-03 | glio |
| Chek1_in.vivo_noRT | Fktn | 1.202 | 4.78E-03 | glio |
| Chek1_ex.vivo_noRT | Ptbp2 | -1.894 | 4.78E-03 | glio |
| Chek1_ex.vivo_noRT | Relb | 1.604 | 4.78E-03 | glio |
| Cyp19a1_ex.vivo_noRT | 2410017I17Ri | 2.159 | 4.78E-03 | glio |
| Cyp19a1_ex.vivo_noRT | Gm26543 | 2.159 | 4.78E-03 | glio |
| Hsd17b10_in.vivo_noRT | Zfp575 | 2.364 | 4.78E-03 | glio |
| Hsd17b10_in.vivo_noRT | Pag1 | 1.945 | 4.78E-03 | glio |
| Interaction_genotype_Aars2 | Pam16 | 1.613 | 4.78E-04 | glio |
| Interaction_genotype_Atr | Nxnl2 | -4.006 | 4.78E-04 | glio |
| Aars2_in.vivo_noRT | Aqp11 | 2.495 | 4.78E-04 | glio |
| Atr_in.vivo_noRT | Pltp | 2.261 | 4.78E-04 | glio |
| Bard1_ex.vivo_noRT | Brinp3 | 2.814 | 4.78E-04 | glio |
| Chek1_in.vivo_noRT | Blzf1 | -1.575 | 4.78E-04 | glio |
| Interaction_genotype_Mtor | Gm4221 | -4.83 | 4.78E-05 | glio |
| Interaction_genotype_Mtor | Olfr90 | -4.228 | 4.78E-05 | glio |
| interaction_RT | Fos | 1.447 | 4.78E-05 | glio |
| Aars2_in.vivo_noRT | 5730414N17R | 3.057 | 4.78E-05 | glio |
| Bora_in.vivo_noRT | Mfsd7a | 3.33 | 4.78E-05 | glio |
| Chek1_ex.vivo_noRT | Vipr2 | 3.678 | 4.78E-05 | glio |
| Interaction_genotype_Myc | Skida1 | -5.299 | 4.78E-06 | glio |
| Interaction_genotype_Myc | Elobl | -5.645 | 4.78E-06 | glio |
| Interaction_genotype_Myc | Prl7c1 | -3.824 | 4.78E-06 | glio |
| Aars2_in.vivo_noRT | 4930522L14Ri | 3.308 | 4.78E-06 | glio |
| Aars2_in.vivo_noRT | Ptpru | 2.84 | 4.78E-06 | glio |
| Aars2_in.vivo_noRT | Eps8 | -1.435 | 4.78E-06 | glio |
| Atr_in.vivo_noRT | Gm48768 | 3.282 | 4.78E-06 | glio |
| Chek1_in.vivo_noRT | Zfp273 | 3.277 | 4.78E-06 | glio |
| Hsd17b10_in.vivo_noRT | Iqcn | 3.09 | 4.78E-06 | glio |
| culture_effect | Slc24a5 | 0.959 | 4.78E-08 | glio |
| Bora_in.vivo_noRT | BC034090 | 3.563 | 4.78E-08 | glio |
| Atr_in.vivo_noRT | Tgfbr3l | 4.229 | 4.78E-09 | glio |

| | | | | |
|---|---|---|---|---|
| Hsd17b10_in.vivo_noRT | Hrct1 | 3.616 | 4.78E-09 | glio |
| Bora_in.vivo_noRT | Gm43948 | 3.461 | 4.78E-10 | glio |
| Chek1_in.vivo_noRT | Ntn1 | 4.142 | 4.78E-12 | glio |
| culture_effect | Chchd10 | -2.191 | 4.78E-14 | glio |
| Bora_in.vivo_noRT | Ldhd | 4.543 | 4.78E-14 | glio |
| Interaction_genotype_Atr | Cks1brt | -3.204 | 4.79E-03 | glio |
| Interaction_genotype_Atr | Zfp273 | -3.557 | 4.79E-03 | glio |
| Interaction_genotype_Atr | Gm43156 | -3.356 | 4.79E-03 | glio |
| Interaction_genotype_Atr | Zfp13 | -3.04 | 4.79E-03 | glio |
| Interaction_genotype_Myc | Klra6 | -3.356 | 4.79E-03 | glio |
| Interaction_genotype_Myc | Mrtfb | -2.844 | 4.79E-03 | glio |
| Interaction_genotype_Myc | Atg9a | -2.543 | 4.79E-03 | glio |
| interaction_RT | Wnk4 | 1.232 | 4.79E-03 | glio |
| interaction_RT | Rgmb | 0.871 | 4.79E-03 | glio |
| Aars2_in.vivo_noRT | Rab5if | -0.758 | 4.79E-03 | glio |
| Atp5c1_in.vivo_noRT | Gm17276 | 1.877 | 4.79E-03 | glio |
| Atr_in.vivo_noRT | Parp3 | -1.195 | 4.79E-03 | glio |
| Atr_in.vivo_noRT | Ing1 | -1.104 | 4.79E-03 | glio |
| Atr_in.vivo_noRT | Rps27 | -0.828 | 4.79E-03 | glio |
| Bcas2_ex.vivo_noRT | Tcstv3 | 2.742 | 4.79E-03 | glio |
| Bcas2_ex.vivo_noRT | Gm12610 | 2.342 | 4.79E-03 | glio |
| Bora_in.vivo_noRT | Csnk2b | -0.827 | 4.79E-03 | glio |
| Cenpj_in.vivo_noRT | Ltb4r1 | 2.552 | 4.79E-03 | glio |
| Chek1_in.vivo_noRT | Otud6b | -1.066 | 4.79E-03 | glio |
| Chek1_ex.vivo_noRT | Zfp433 | 2.795 | 4.79E-03 | glio |
| Chek1_ex.vivo_noRT | Cfap100 | -2.465 | 4.79E-03 | glio |
| Hsd17b10_in.vivo_noRT | Gchfr | 2.503 | 4.79E-03 | glio |
| Interaction_genotype_Myc | Neurl1a | -4.272 | 4.79E-04 | glio |
| Aars2_in.vivo_noRT | 2810455O05R | 2.551 | 4.79E-04 | glio |
| Atp5c1_in.vivo_noRT | Platr20 | 2.926 | 4.79E-04 | glio |
| Bora_in.vivo_noRT | Gamt | -1.764 | 4.79E-04 | glio |
| Bora_in.vivo_noRT | Fkbp4 | -0.806 | 4.79E-04 | glio |
| Cenpj_in.vivo_noRT | Gm26671 | 2.022 | 4.79E-04 | glio |
| Interaction_genotype_Myc | 1110002L01Ri | -3.595 | 4.79E-05 | glio |
| Bard1_ex.vivo_noRT | Klk13 | 2.814 | 4.79E-05 | glio |
| Bard1_ex.vivo_noRT | Cd36 | 2.814 | 4.79E-05 | glio |
| Bard1_ex.vivo_noRT | Gstt4 | 2.814 | 4.79E-05 | glio |
| Bcas2_in.vivo_noRT | Cep112it | 2.891 | 4.79E-05 | glio |
| Bcas2_in.vivo_noRT | Zbtb3 | 2.91 | 4.79E-05 | glio |
| Bcas2_ex.vivo_noRT | Gm13078 | 2.742 | 4.79E-05 | glio |
| Interaction_genotype_Mtor | Ssr1 | 2.371 | 4.79E-06 | glio |
| Aars2_ex.vivo_noRT | Gm26671 | 2.9 | 4.79E-06 | glio |
| Bcas2_in.vivo_noRT | Kcnab3 | 3.273 | 4.79E-06 | glio |
| Atr_in.vivo_noRT | Ksr1 | 3.485 | 4.79E-07 | glio |
| Bcas2_in.vivo_noRT | Gstp3 | 3.507 | 4.79E-07 | glio |
| culture_effect | Herc6 | -0.938 | 4.79E-10 | glio |
| Bora_in.vivo_noRT | Neurl1b | 4.41 | 4.79E-10 | glio |
| Aars2_in.vivo_noRT | Slxl1 | 4.548 | 4.79E-11 | glio |
| Atr_in.vivo_noRT | Gm5624 | 4.544 | 4.79E-12 | glio |
| Aars2_in.vivo_noRT | Bmp15 | 4.674 | 4.79E-15 | glio |
| Aars2_in.vivo_noRT | Gm38163 | 5.327 | 4.79E-21 | glio |
| Bora_in.vivo_noRT | Gm13078 | 4.936 | 4.79E-23 | glio |

| | | | | |
|---|---|---|---|---|
| culture_effect | Eef1e1 | 0.339 | 4.80E-03 | glio |
| Interaction_genotype_Aars2 | Utrn | 1.711 | 4.80E-03 | glio |
| Interaction_genotype_Aars2 | Tmem11 | 1.31 | 4.80E-03 | glio |
| Interaction_genotype_Atr | Dus2 | -2.432 | 4.80E-03 | glio |
| Interaction_genotype_Bcas2 | Gm15927 | -3.666 | 4.80E-03 | glio |
| Interaction_genotype_Bcas2 | Gm45184 | -3.804 | 4.80E-03 | glio |
| Interaction_genotype_Bcas2 | Ppp4r1 | 1.971 | 4.80E-03 | glio |
| Interaction_genotype_Mtor | Lrrc27 | -2.868 | 4.80E-03 | glio |
| Interaction_genotype_Mtor | Hacd1 | 1.658 | 4.80E-03 | glio |
| Interaction_genotype_Mtor | H2afy | 1.336 | 4.80E-03 | glio |
| Interaction_genotype_Myc | Cul9 | -3.429 | 4.80E-03 | glio |
| Interaction_genotype_Rpl3 | BC024978 | -3.414 | 4.80E-03 | glio |
| Interaction_genotype_Rpl3 | Gm43156 | -3.888 | 4.80E-03 | glio |
| Interaction_genotype_Rpl3 | 4930478M13R | -3.814 | 4.80E-03 | glio |
| Interaction_genotype_Rpl3 | Gm26581 | -4.131 | 4.80E-03 | glio |
| Interaction_genotype_Rpl3 | Apoh | -3.866 | 4.80E-03 | glio |
| interaction_RT | Chmp2b | 0.714 | 4.80E-03 | glio |
| Atr_in.vivo_noRT | Plekhm3 | 2.092 | 4.80E-03 | glio |
| Bcas2_in.vivo_noRT | Lyrm4 | -1.419 | 4.80E-03 | glio |
| Bora_in.vivo_noRT | Gm41609 | 2.25 | 4.80E-03 | glio |
| Bora_in.vivo_noRT | Gstt3 | 2.118 | 4.80E-03 | glio |
| Bora_in.vivo_noRT | Eloc | -0.878 | 4.80E-03 | glio |
| culture_effect | Gm49890 | 1.489 | 4.80E-04 | glio |
| Interaction_genotype_Mtor | Gm5602 | -3.308 | 4.80E-04 | glio |
| Interaction_genotype_Myc | Gm14540 | -3 | 4.80E-04 | glio |
| Interaction_genotype_Myc | Ccdc184 | -4.264 | 4.80E-04 | glio |
| Interaction_genotype_Myc | Entpd2 | -2.914 | 4.80E-04 | glio |
| Atp5c1_in.vivo_noRT | Esyt3 | 2.66 | 4.80E-04 | glio |
| Atr_in.vivo_noRT | Hist1h2bh | 2.54 | 4.80E-04 | glio |
| Atr_in.vivo_noRT | Chchd1 | -0.913 | 4.80E-04 | glio |
| Atr_ex.vivo_noRT | Tsga10ip | 3.436 | 4.80E-04 | glio |
| Bcas2_in.vivo_noRT | Haus3 | -1.47 | 4.80E-04 | glio |
| Bcas2_ex.vivo_noRT | Dct | 3.905 | 4.80E-04 | glio |
| Chek1_ex.vivo_noRT | Top2a | -2.236 | 4.80E-04 | glio |
| Aars2_in.vivo_noRT | Slc9a9 | 2.939 | 4.80E-05 | glio |
| Aars2_in.vivo_noRT | Zfp964 | 2.952 | 4.80E-05 | glio |
| Aars2_in.vivo_noRT | Atp5g2 | -0.792 | 4.80E-05 | glio |
| Bcas2_in.vivo_noRT | 4833438C02R | 2.688 | 4.80E-05 | glio |
| Interaction_genotype_Mtor | Cox6a1 | 2.285 | 4.80E-06 | glio |
| Atr_in.vivo_noRT | 2310008N11R | 3.747 | 4.80E-07 | glio |
| Chek1_ex.vivo_noRT | Pom121 | -2.524 | 4.80E-07 | glio |
| Aars2_in.vivo_noRT | Adcy2 | 3.573 | 4.80E-09 | glio |
| Chek1_ex.vivo_noRT | Gm35439 | 4.443 | 4.80E-11 | glio |
| Aars2_in.vivo_noRT | Gprin2 | 4.415 | 4.80E-14 | glio |
| Aars2_in.vivo_noRT | Gm49504 | 5.327 | 4.80E-23 | glio |
| Atr_in.vivo_noRT | Ltbp2 | 5.533 | 4.80E-23 | glio |
| Atr_in.vivo_noRT | Lmx1a | 5.533 | 4.80E-23 | glio |
| Bcas2_in.vivo_noRT | Ankrd22 | 4.807 | 4.80E-23 | glio |
| culture_effect | Afp | -1.415 | 4.81E-03 | glio |
| Interaction_genotype_Aars2 | C330002G04R | -2.62 | 4.81E-03 | glio |
| Interaction_genotype_Aars2 | AC166172.1 | -3.193 | 4.81E-03 | glio |
| Interaction_genotype_Aars2 | Cstb | 1.744 | 4.81E-03 | glio |

| | | | | |
|---|---|---|---|---|
| Interaction_genotype_Atr | Ano10 | -2.197 | 4.81E-03 | glio |
| Interaction_genotype_Atr | Bpgm | 1.666 | 4.81E-03 | glio |
| Interaction_genotype_Myc | Acat3 | -2.513 | 4.81E-03 | glio |
| Interaction_genotype_Myc | Celf5 | -3.317 | 4.81E-03 | glio |
| Interaction_genotype_Myc | Pgghg | -2.615 | 4.81E-03 | glio |
| Interaction_genotype_Myc | Eif2b3 | -1.912 | 4.81E-03 | glio |
| Interaction_genotype_Myc | Tomm22 | 0.959 | 4.81E-03 | glio |
| Aars2_in.vivo_noRT | Gpr137c | 1.959 | 4.81E-03 | glio |
| Aars2_ex.vivo_noRT | D330041H03R | -2.338 | 4.81E-03 | glio |
| Atp5c1_in.vivo_noRT | Gm11696 | 2.329 | 4.81E-03 | glio |
| Atr_in.vivo_noRT | Mettl15 | 1.526 | 4.81E-03 | glio |
| Bcas2_in.vivo_noRT | Zan | 2.044 | 4.81E-03 | glio |
| Bcas2_ex.vivo_noRT | Zscan4d | 2.348 | 4.81E-03 | glio |
| Cenpj_in.vivo_noRT | A930033M14R | 2.022 | 4.81E-03 | glio |
| Hsd17b10_in.vivo_noRT | Gm26850 | 2.222 | 4.81E-03 | glio |
| Hsd17b10_in.vivo_noRT | Tmem202 | 2.174 | 4.81E-03 | glio |
| Hsd17b10_in.vivo_noRT | Map9 | -1.819 | 4.81E-03 | glio |
| culture_effect | Aatf | -0.513 | 4.81E-04 | glio |
| Interaction_genotype_Myc | Hcn3 | -3.589 | 4.81E-04 | glio |
| Atp5c1_in.vivo_noRT | Fbxo15 | 2.405 | 4.81E-04 | glio |
| Atr_in.vivo_noRT | Cdkn1a | -1.487 | 4.81E-04 | glio |
| Bcas2_in.vivo_noRT | Micall2 | 2.624 | 4.81E-04 | glio |
| culture_effect | Arsj | 2.173 | 4.81E-05 | glio |
| Atr_in.vivo_noRT | Sgms2 | 3.028 | 4.81E-05 | glio |
| Chek1_ex.vivo_noRT | Rad18 | -2.371 | 4.81E-05 | glio |
| Interaction_genotype_Myc | 1700003G18R | -5.899 | 4.81E-06 | glio |
| Interaction_genotype_Myc | 2610016A17R | -5.394 | 4.81E-06 | glio |
| Aars2_in.vivo_noRT | Stmn4 | 4.811 | 4.81E-06 | glio |
| Aars2_ex.vivo_noRT | 1700019E08Ri | 3.293 | 4.81E-06 | glio |
| Atr_in.vivo_noRT | Plek2 | 3.082 | 4.81E-06 | glio |
| Bora_in.vivo_noRT | A830082N09R | 3.378 | 4.81E-06 | glio |
| Chek1_ex.vivo_noRT | Clmn | 4.049 | 4.81E-06 | glio |
| Chek1_ex.vivo_noRT | Emg1 | -1.294 | 4.81E-07 | glio |
| Aars2_in.vivo_noRT | Olfml2a | 3.824 | 4.81E-08 | glio |
| Hsd17b10_in.vivo_noRT | Dreh | 3.501 | 4.81E-08 | glio |
| Aars2_in.vivo_noRT | Bhlhe41 | 4.804 | 4.81E-09 | glio |
| Atr_in.vivo_noRT | Gm16023 | 4.536 | 4.81E-09 | glio |
| Atr_in.vivo_noRT | 4930579K19R | 4.295 | 4.81E-09 | glio |
| Bcas2_in.vivo_noRT | Gm8013 | 4.162 | 4.81E-09 | glio |
| Bora_in.vivo_noRT | Gm16222 | 4.676 | 4.81E-09 | glio |
| Interaction_genotype_Mtor | Tssk6 | -6.676 | 4.81E-10 | glio |
| Interaction_genotype_Mtor | Klk14 | -7.827 | 4.81E-10 | glio |
| Chek1_ex.vivo_noRT | Gm36043 | 4.842 | 4.81E-10 | glio |
| Chek1_ex.vivo_noRT | Tmem52 | 4.842 | 4.81E-10 | glio |
| Bcas2_in.vivo_noRT | Gm43948 | 3.85 | 4.81E-12 | glio |
| culture_effect | Ube2m | 0.754 | 4.81E-13 | glio |
| Chek1_in.vivo_noRT | Prrx2 | 5.044 | 4.81E-13 | glio |
| Bcas2_in.vivo_noRT | Gm50024 | 4.544 | 4.81E-20 | glio |
| Bcas2_in.vivo_noRT | Esp24 | 4.807 | 4.81E-20 | glio |
| Atr_in.vivo_noRT | A030014E15Ri | 5.533 | 4.81E-29 | glio |
| Atr_in.vivo_noRT | Gm21731 | 5.533 | 4.81E-29 | glio |
| culture_effect | Ecpas | -0.454 | 4.82E-03 | glio |

| | | | | |
|---|---|---|---|---|
| Interaction_genotype_Atr | Ppp4c | 1.503 | 4.82E-03 | glio |
| Interaction_genotype_Bcas2 | Col6a1 | -2.76 | 4.82E-03 | glio |
| Interaction_genotype_Chek1 | Dusp2 | 4.084 | 4.82E-03 | glio |
| Interaction_genotype_Chek1 | Peg3 | 2.415 | 4.82E-03 | glio |
| Interaction_genotype_Myc | 2500002B13R | -2.307 | 4.82E-03 | glio |
| Interaction_genotype_Myc | Ndufa13 | 1.03 | 4.82E-03 | glio |
| interaction_RT | Ly6d | 2.3 | 4.82E-03 | glio |
| Aars2_in.vivo_noRT | Fxyd7 | 1.862 | 4.82E-03 | glio |
| Aars2_ex.vivo_noRT | D130062J10Ri | 2.618 | 4.82E-03 | glio |
| Atr_in.vivo_noRT | Itpka | 2.014 | 4.82E-03 | glio |
| Bcas2_ex.vivo_noRT | AU022754 | 2.742 | 4.82E-03 | glio |
| Cenpj_in.vivo_noRT | Tal2 | 1.759 | 4.82E-03 | glio |
| Chek1_in.vivo_noRT | Hsp90b1 | -0.677 | 4.82E-03 | glio |
| culture_effect | Stx19 | -1.846 | 4.82E-04 | glio |
| Interaction_genotype_Mtor | 4930527F14Ri | -3.173 | 4.82E-04 | glio |
| Interaction_genotype_Mtor | Zfp597 | -3.945 | 4.82E-04 | glio |
| Aars2_in.vivo_noRT | Cox5b | -0.773 | 4.82E-04 | glio |
| Bora_in.vivo_noRT | Alpk2 | 3.22 | 4.82E-04 | glio |
| Bora_in.vivo_noRT | Zfp583 | 2.643 | 4.82E-04 | glio |
| Chek1_ex.vivo_noRT | Kctd14 | 2.887 | 4.82E-04 | glio |
| Atp5c1_in.vivo_noRT | Crhbp | 2.668 | 4.82E-05 | glio |
| Atp5c1_in.vivo_noRT | Gm10642 | 2.932 | 4.82E-05 | glio |
| Interaction_genotype_Nat2 | Klrk1 | 4.361 | 4.82E-06 | glio |
| Interaction_genotype_Nat2 | Gm21731 | 3.839 | 4.82E-06 | glio |
| Aars2_in.vivo_noRT | Wdr90 | 2.877 | 4.82E-06 | glio |
| culture_effect | Etfa | -0.73 | 4.82E-07 | glio |
| Interaction_genotype_Mtor | Spata45 | -5.933 | 4.82E-07 | glio |
| Interaction_genotype_Mtor | Gm21731 | -4.099 | 4.82E-07 | glio |
| interaction_RT | Flnb | -1.401 | 4.82E-07 | glio |
| Chek1_in.vivo_noRT | Syne3 | 3.787 | 4.82E-07 | glio |
| Aars2_in.vivo_noRT | Gm20478 | 3.754 | 4.82E-09 | glio |
| Atr_in.vivo_noRT | 5031434O11R | 4.917 | 4.82E-10 | glio |
| Bcas2_in.vivo_noRT | Dnm3 | 4.281 | 4.82E-10 | glio |
| Aars2_in.vivo_noRT | 2410141K09R | 5.063 | 4.82E-18 | glio |
| Aars2_in.vivo_noRT | Usp17lc | 5.327 | 4.82E-19 | glio |
| Atr_in.vivo_noRT | Gm48780 | 5.273 | 4.82E-24 | glio |
| Interaction_genotype_Atr | Gm43256 | -3.277 | 4.83E-03 | glio |
| Interaction_genotype_Ifnar1 | Gm47510 | 2.994 | 4.83E-03 | glio |
| Interaction_genotype_Mtor | Ccn2 | -4.395 | 4.83E-03 | glio |
| Interaction_genotype_Mtor | Trim32 | -2.233 | 4.83E-03 | glio |
| Interaction_genotype_Mtor | Sdhaf2 | -1.459 | 4.83E-03 | glio |
| Interaction_genotype_Mtor | Cox8a | 1.021 | 4.83E-03 | glio |
| Interaction_genotype_Myc | Irgq | -2.218 | 4.83E-03 | glio |
| Interaction_genotype_Uqcrc2 | Sod3 | -4.178 | 4.83E-03 | glio |
| Interaction_genotype_Uqcrc2 | Carns1 | -4.223 | 4.83E-03 | glio |
| Interaction_genotype_Uqcrc2 | 1700017M07F | -3.6 | 4.83E-03 | glio |
| Interaction_genotype_Uqcrc2 | Mkrn1 | 1.339 | 4.83E-03 | glio |
| Atp5c1_in.vivo_noRT | Dppa2 | 2.136 | 4.83E-03 | glio |
| Atr_in.vivo_noRT | Pdia4 | -0.99 | 4.83E-03 | glio |
| Atr_in.vivo_noRT | Polr3k | -1.019 | 4.83E-03 | glio |
| Bcas2_in.vivo_noRT | Usp36 | -1.179 | 4.83E-03 | glio |
| Brca2_in.vivo_noRT | A930019D19R | 1.763 | 4.83E-03 | glio |

| | | | | |
|---|---|---|---|---|
| Eif2s1_in.vivo_noRT | Kcnk4 | 2.407 | 4.83E-03 | glio |
| Hsd17b10_in.vivo_noRT | Apold1 | 2.506 | 4.83E-03 | glio |
| Interaction_genotype_Aars2 | C330013E15Ri | -4.205 | 4.83E-04 | glio |
| Interaction_genotype_Aars2 | Ccdc9b | -3.575 | 4.83E-04 | glio |
| Interaction_genotype_Aars2 | Iah1 | 1.788 | 4.83E-04 | glio |
| Interaction_genotype_Mtor | Cracr2b | -4.502 | 4.83E-04 | glio |
| Aars2_in.vivo_noRT | Prickle3 | 2.242 | 4.83E-04 | glio |
| Atr_in.vivo_noRT | Abracl | -1.082 | 4.83E-04 | glio |
| Bora_in.vivo_noRT | 2810468N07R | 2.301 | 4.83E-04 | glio |
| Chek1_ex.vivo_noRT | Serpinb1a | 4.078 | 4.83E-04 | glio |
| culture_effect | Myh9 | 0.686 | 4.83E-05 | glio |
| culture_effect | Atpif1 | -0.47 | 4.83E-05 | glio |
| Interaction_genotype_Atr | A430033K04R | -4.888 | 4.83E-05 | glio |
| Interaction_genotype_Atr | Gm50321 | -4.266 | 4.83E-05 | glio |
| Interaction_genotype_Atr | Map1lc3b | 1.565 | 4.83E-05 | glio |
| Interaction_genotype_Mtor | Gm45716 | -4.155 | 4.83E-05 | glio |
| Interaction_genotype_Myc | Tapbpl | 2.813 | 4.83E-05 | glio |
| Interaction_genotype_Myc | Anxa1 | 2.268 | 4.83E-05 | glio |
| Chek1_in.vivo_noRT | Wars | -2.012 | 4.83E-05 | glio |
| Atr_ex.vivo_noRT | Gm44646 | 3.375 | 4.83E-06 | glio |
| Bcas2_in.vivo_noRT | Hoxa13 | 3.36 | 4.83E-06 | glio |
| Aars2_in.vivo_noRT | Hist1h2bh | 3.573 | 4.83E-07 | glio |
| Chek1_in.vivo_noRT | Alox5ap | 3.938 | 4.83E-07 | glio |
| Bora_in.vivo_noRT | Zfp811 | 3.762 | 4.83E-08 | glio |
| Chek1_in.vivo_noRT | Gm46404 | 4.531 | 4.83E-10 | glio |
| Hsd17b10_in.vivo_noRT | Esp15 | 3.616 | 4.83E-10 | glio |
| Hsd17b10_in.vivo_noRT | Tmem54 | 3.883 | 4.83E-11 | glio |
| Atr_in.vivo_noRT | Mro | 4.742 | 4.83E-12 | glio |
| Atr_in.vivo_noRT | Gm43915 | 5.001 | 4.83E-18 | glio |
| Chek1_in.vivo_noRT | Ankrd22 | 5.044 | 4.83E-25 | glio |
| culture_effect | Ngfr | 4.024 | 4.83E-35 | glio |
| culture_effect | Chadl | -4.356 | 4.83E-35 | glio |
| culture_effect | Ang | -1.847 | 4.84E-03 | glio |
| culture_effect | Gnpda1 | 0.589 | 4.84E-03 | glio |
| culture_effect | Foxm1 | 0.59 | 4.84E-03 | glio |
| Interaction_genotype_Aars2 | Frmd8os | -3.972 | 4.84E-03 | glio |
| Interaction_genotype_Aars2 | Stard6 | -3.555 | 4.84E-03 | glio |
| Interaction_genotype_Aars2 | Fdps | 1.854 | 4.84E-03 | glio |
| Interaction_genotype_Aars2 | Laptm4a | 0.965 | 4.84E-03 | glio |
| Interaction_genotype_Bora | Hlx | -3.596 | 4.84E-03 | glio |
| Interaction_genotype_Bora | Spry1 | 1.989 | 4.84E-03 | glio |
| Interaction_genotype_Bora | Adipor1 | 1.32 | 4.84E-03 | glio |
| Interaction_genotype_Mtor | Ccdc28b | -3.144 | 4.84E-03 | glio |
| Interaction_genotype_Mtor | Zbtb40 | -2.961 | 4.84E-03 | glio |
| Interaction_genotype_Mtor | Lnpep | 1.666 | 4.84E-03 | glio |
| Interaction_genotype_Mtor | Snx6 | 1.506 | 4.84E-03 | glio |
| Interaction_genotype_Myc | Socs4 | -2.206 | 4.84E-03 | glio |
| Interaction_genotype_Myc | Maf | -2.623 | 4.84E-03 | glio |
| Aars2_in.vivo_noRT | Rhobtb1 | 2.057 | 4.84E-03 | glio |
| Aars2_in.vivo_noRT | Irs1 | 1.919 | 4.84E-03 | glio |
| Atp5c1_in.vivo_noRT | Iqcn | 2.142 | 4.84E-03 | glio |
| Atr_in.vivo_noRT | Gpt | 2.004 | 4.84E-03 | glio |

| | | | | |
|---|---|---|---|---|
| Bcas2_in.vivo_noRT | Arhgap27 | 1.958 | 4.84E-03 | glio |
| Bora_in.vivo_noRT | Amotl1 | -1.352 | 4.84E-03 | glio |
| Bora_ex.vivo_noRT | Slc6a19 | 2.808 | 4.84E-03 | glio |
| Bora_ex.vivo_noRT | 1700016P04R | 2.599 | 4.84E-03 | glio |
| culture_effect | 3110056K07R | 1.007 | 4.84E-04 | glio |
| Interaction_genotype_Mtor | Ccdc96 | -3.83 | 4.84E-04 | glio |
| Bard1_ex.vivo_noRT | A330076H08R | 2.814 | 4.84E-04 | glio |
| Cenpj_ex.vivo_noRT | Rgs4 | 2.591 | 4.84E-04 | glio |
| Chek1_in.vivo_noRT | Cyb5d1 | 2.508 | 4.84E-04 | glio |
| Dclre1c_ex.vivo_noRT | Gm13078 | 2.658 | 4.84E-04 | glio |
| Hsd17b10_in.vivo_noRT | Zfp469 | -1.736 | 4.84E-04 | glio |
| Interaction_genotype_Mtor | Gm49085 | -4.595 | 4.84E-05 | glio |
| Interaction_genotype_Myc | Pidd1 | -3.667 | 4.84E-05 | glio |
| Interaction_genotype_Myc | Slc17a6 | -2.729 | 4.84E-05 | glio |
| Aars2_in.vivo_noRT | Dleu7 | 2.801 | 4.84E-05 | glio |
| Atr_in.vivo_noRT | Slurp1 | 4.239 | 4.84E-05 | glio |
| Interaction_genotype_Mtor | Dapk2 | -4.113 | 4.84E-06 | glio |
| Bcas2_in.vivo_noRT | Cyth4 | 3.267 | 4.84E-06 | glio |
| Bcas2_in.vivo_noRT | Gm32122 | 2.979 | 4.84E-06 | glio |
| Hsd17b10_in.vivo_noRT | Smpd5 | 2.959 | 4.84E-06 | glio |
| Atr_in.vivo_noRT | A430018G15R | 2.893 | 4.84E-07 | glio |
| culture_effect | Lncppara | 2.674 | 4.84E-08 | glio |
| Aars2_in.vivo_noRT | D930030I03Ri | 4.087 | 4.84E-08 | glio |
| Bcas2_in.vivo_noRT | Gm5129 | 4.007 | 4.84E-08 | glio |
| Chek1_in.vivo_noRT | Gm10432 | 3.547 | 4.84E-08 | glio |
| Atr_in.vivo_noRT | Lrrc75aos2 | 4.146 | 4.84E-09 | glio |
| Atr_in.vivo_noRT | 1110046J04Ri | 5.533 | 4.84E-15 | glio |
| culture_effect | Serpine2 | 1.665 | 4.85E-03 | glio |
| culture_effect | Pde12 | -0.685 | 4.85E-03 | glio |
| Interaction_genotype_Aars2 | Necab2 | -3.264 | 4.85E-03 | glio |
| Interaction_genotype_Atr | Slc4a9 | -2.925 | 4.85E-03 | glio |
| Interaction_genotype_Bora | Gm16183 | -3.965 | 4.85E-03 | glio |
| Interaction_genotype_Bora | Prl3d3 | -2.567 | 4.85E-03 | glio |
| Interaction_genotype_Ifnar1 | Ly6d | 3.462 | 4.85E-03 | glio |
| Interaction_genotype_Ifnar1 | Gm15908 | 2.734 | 4.85E-03 | glio |
| Interaction_genotype_Ifnar1 | A930014E10Ri | 2.734 | 4.85E-03 | glio |
| Interaction_genotype_Myc | Abhd5 | -1.987 | 4.85E-03 | glio |
| Interaction_genotype_Pars2 | Slc6a13 | -4.153 | 4.85E-03 | glio |
| Interaction_genotype_Pars2 | Trim17 | -2.668 | 4.85E-03 | glio |
| Interaction_genotype_Rpl3 | Atp1a2 | 4.272 | 4.85E-03 | glio |
| interaction_RT | Ctdsp1 | -0.948 | 4.85E-03 | glio |
| interaction_RT | Nipsnap3b | 0.767 | 4.85E-03 | glio |
| Aars2_in.vivo_noRT | Gm4631 | 1.959 | 4.85E-03 | glio |
| Aars2_in.vivo_noRT | Emp3 | -0.752 | 4.85E-03 | glio |
| Aars2_in.vivo_noRT | Eif5 | -0.616 | 4.85E-03 | glio |
| Aars2_ex.vivo_noRT | Col6a3 | 2.499 | 4.85E-03 | glio |
| Atr_ex.vivo_noRT | Gm17638 | 2.588 | 4.85E-03 | glio |
| Atr_ex.vivo_noRT | Gm37800 | 2.469 | 4.85E-03 | glio |
| Bcas2_in.vivo_noRT | Lrrc28 | 1.809 | 4.85E-03 | glio |
| Bora_in.vivo_noRT | Sinhcaf | -1.031 | 4.85E-03 | glio |
| Bora_ex.vivo_noRT | Ninj2 | 2.831 | 4.85E-03 | glio |
| Hsd17b10_in.vivo_noRT | Shank1 | 2.008 | 4.85E-03 | glio |

| | | | | |
|---|---|---|---|---|
| Hsd17b10_in.vivo_noRT | Tada2a | -1.451 | 4.85E-03 | glio |
| Interaction_genotype_Atr | Gm13348 | -3.642 | 4.85E-04 | glio |
| Interaction_genotype_Atr | Gm20667 | -4.444 | 4.85E-04 | glio |
| Interaction_genotype_Atr | Wfdc1 | -3.377 | 4.85E-04 | glio |
| Interaction_genotype_Atr | Tkt | 1.93 | 4.85E-04 | glio |
| Interaction_genotype_Mtor | Pla2g2d | -3.836 | 4.85E-04 | glio |
| Interaction_genotype_Mtor | AW011738 | -3.019 | 4.85E-04 | glio |
| Atr_in.vivo_noRT | 5930430L01Ri | 2.328 | 4.85E-04 | glio |
| Atr_in.vivo_noRT | Nabp1 | 2.393 | 4.85E-04 | glio |
| Chek1_in.vivo_noRT | Hist1h2bh | 2.574 | 4.85E-04 | glio |
| Chek1_ex.vivo_noRT | Gm49042 | 2.956 | 4.85E-04 | glio |
| Hsd17b10_in.vivo_noRT | Kcnj13 | 2.335 | 4.85E-04 | glio |
| Hsd17b10_in.vivo_noRT | Cd79a | 2.67 | 4.85E-04 | glio |
| Atr_in.vivo_noRT | Prx | 3.005 | 4.85E-05 | glio |
| Atr_ex.vivo_noRT | Ccl19 | 3.164 | 4.85E-05 | glio |
| Atr_ex.vivo_noRT | Gm50147 | 3.454 | 4.85E-05 | glio |
| Bcas2_in.vivo_noRT | Ecpas | -1.598 | 4.85E-05 | glio |
| Chek1_in.vivo_noRT | Gm28050 | 3.006 | 4.85E-05 | glio |
| Chek1_in.vivo_noRT | Gm28417 | 2.724 | 4.85E-05 | glio |
| Hsd17b10_in.vivo_noRT | Col5a2 | 2.304 | 4.85E-05 | glio |
| Hsd17b10_in.vivo_noRT | Papln | 2.87 | 4.85E-05 | glio |
| culture_effect | Camta1 | 0.882 | 4.85E-06 | glio |
| Interaction_genotype_Mtor | Cldn5 | -3.833 | 4.85E-06 | glio |
| Atr_ex.vivo_noRT | Dmrtc1a | 3.164 | 4.85E-06 | glio |
| Chek1_in.vivo_noRT | Mtbp | -2.091 | 4.85E-06 | glio |
| culture_effect | Snx3 | 0.461 | 4.85E-07 | glio |
| Atr_in.vivo_noRT | Prrt3 | 3.353 | 4.85E-08 | glio |
| Atr_ex.vivo_noRT | Krt19 | 3.164 | 4.85E-08 | glio |
| Chek1_in.vivo_noRT | Slc46a3 | 3.859 | 4.85E-08 | glio |
| Atr_in.vivo_noRT | Gm10516 | 3.97 | 4.85E-09 | glio |
| Atr_in.vivo_noRT | Apoa1 | 4.5 | 4.85E-10 | glio |
| Atr_in.vivo_noRT | Pdzk1ip1 | 4.543 | 4.85E-10 | glio |
| Atr_in.vivo_noRT | Atoh8 | 4.15 | 4.85E-10 | glio |
| culture_effect | Cp | -2.447 | 4.85E-11 | glio |
| culture_effect | Fam120a | 0.93 | 4.85E-11 | glio |
| Bora_in.vivo_noRT | Rarres1 | 4.936 | 4.85E-13 | glio |
| Atr_in.vivo_noRT | Gm29083 | 5.001 | 4.85E-15 | glio |
| Chek1_ex.vivo_noRT | BC061195 | 4.842 | 4.85E-16 | glio |
| Aars2_in.vivo_noRT | Pyy | 5.327 | 4.85E-20 | glio |
| Interaction_genotype_Aars2 | Gm28731 | -3.711 | 4.86E-03 | glio |
| Interaction_genotype_Aars2 | Arpc2 | 0.991 | 4.86E-03 | glio |
| Interaction_genotype_Atr | Gm14820 | -2.616 | 4.86E-03 | glio |
| Interaction_genotype_Atr | Selp | -3.523 | 4.86E-03 | glio |
| Interaction_genotype_Atr | Rspo3 | -4.05 | 4.86E-03 | glio |
| Interaction_genotype_Atr | Gm12905 | -3.716 | 4.86E-03 | glio |
| Interaction_genotype_Mtor | Svip | -2.809 | 4.86E-03 | glio |
| Interaction_genotype_Myc | Gm9929 | -2.92 | 4.86E-03 | glio |
| Interaction_genotype_Myc | Foxa1 | -3.512 | 4.86E-03 | glio |
| Interaction_genotype_Myc | Gpr176 | -2.595 | 4.86E-03 | glio |
| Interaction_genotype_Pars2 | Ccl9 | -2.661 | 4.86E-03 | glio |
| Interaction_genotype_Pars2 | Fhl1 | -2.979 | 4.86E-03 | glio |
| Interaction_genotype_Pars2 | Gm7008 | -3.477 | 4.86E-03 | glio |

| | | | | |
|---|---|---|---|---|
| Aars2_in.vivo_noRT | Ell2 | 1.833 | 4.86E-03 | glio |
| Aars2_in.vivo_noRT | Csnk1g2 | -0.955 | 4.86E-03 | glio |
| Aars2_in.vivo_noRT | Nedd8 | -0.633 | 4.86E-03 | glio |
| Atr_ex.vivo_noRT | Rnd1 | 2.186 | 4.86E-03 | glio |
| Bcas2_in.vivo_noRT | Gm26901 | 2.221 | 4.86E-03 | glio |
| Bora_in.vivo_noRT | Cacng4 | 1.797 | 4.86E-03 | glio |
| Bora_in.vivo_noRT | Fkbp1b | 1.914 | 4.86E-03 | glio |
| Bora_in.vivo_noRT | Ythdf2 | -1.128 | 4.86E-03 | glio |
| Chek1_in.vivo_noRT | B3gat2 | 2.183 | 4.86E-03 | glio |
| Chek1_in.vivo_noRT | Zbtb4 | 1.971 | 4.86E-03 | glio |
| Chek1_in.vivo_noRT | Aldh18a1 | -1.392 | 4.86E-03 | glio |
| Chek1_in.vivo_noRT | Aimp1 | -0.693 | 4.86E-03 | glio |
| Chek1_ex.vivo_noRT | Gars | -1.063 | 4.86E-03 | glio |
| Eif2s1_in.vivo_noRT | Tigd2 | 1.301 | 4.86E-03 | glio |
| Eif2s1_in.vivo_noRT | Aif1l | -0.635 | 4.86E-03 | glio |
| culture_effect | Ddx42 | 0.533 | 4.86E-04 | glio |
| Interaction_genotype_Atr | Gm10115 | -4.287 | 4.86E-04 | glio |
| Interaction_genotype_Myc | Gabarapl2 | 1.468 | 4.86E-04 | glio |
| Interaction_genotype_Rad21 | Fa2h | 4.279 | 4.86E-04 | glio |
| Aars2_in.vivo_noRT | Gm49602 | 2.387 | 4.86E-04 | glio |
| Aars2_in.vivo_noRT | Ccdc32 | 1.616 | 4.86E-04 | glio |
| Bora_in.vivo_noRT | Tmem82 | 2.708 | 4.86E-04 | glio |
| Interaction_genotype_Mtor | 2010106G04R | -4.785 | 4.86E-05 | glio |
| Atr_in.vivo_noRT | 2900089D17R | 2.687 | 4.86E-05 | glio |
| Atr_in.vivo_noRT | Art2b | 2.399 | 4.86E-05 | glio |
| Bcas2_ex.vivo_noRT | Gm13964 | 2.742 | 4.86E-05 | glio |
| Interaction_genotype_Myc | Gm11973 | -4.226 | 4.86E-06 | glio |
| Atr_in.vivo_noRT | Gal3st4 | 3.174 | 4.86E-06 | glio |
| Bora_in.vivo_noRT | Prkar2b | 3.263 | 4.86E-06 | glio |
| Interaction_genotype_Mtor | Gm16046 | -5.968 | 4.86E-07 | glio |
| Interaction_genotype_Mtor | Gpr179 | -5.866 | 4.86E-07 | glio |
| Atr_ex.vivo_noRT | Gm5788 | 3.164 | 4.86E-07 | glio |
| culture_effect | Cdk6 | 1.109 | 4.86E-10 | glio |
| Hsd17b10_in.vivo_noRT | Gm10801 | 3.616 | 4.86E-10 | glio |
| Atr_in.vivo_noRT | Gm12280 | 5.269 | 4.86E-12 | glio |
| Chek1_in.vivo_noRT | Isl1 | 4.77 | 4.86E-12 | glio |
| Aars2_in.vivo_noRT | Gm128 | 4.409 | 4.86E-15 | glio |
| Aars2_in.vivo_noRT | BC018473 | 5.327 | 4.86E-15 | glio |
| culture_effect | Eif3i | -0.94 | 4.86E-19 | glio |
| Interaction_genotype_Aars2 | Ccdc39 | -3.47 | 4.87E-03 | glio |
| Interaction_genotype_Mtor | Dgcr8 | -2.397 | 4.87E-03 | glio |
| Interaction_genotype_Mtor | Faf2 | 1.424 | 4.87E-03 | glio |
| Interaction_genotype_Myc | Zbtb46 | -3.343 | 4.87E-03 | glio |
| Interaction_genotype_Myc | Gm12530 | -2.565 | 4.87E-03 | glio |
| Aars2_in.vivo_noRT | Hsp90ab1 | -0.589 | 4.87E-03 | glio |
| Atr_ex.vivo_noRT | 4922502N22R | 2.769 | 4.87E-03 | glio |
| Bcas2_in.vivo_noRT | Rbpms2 | 2.095 | 4.87E-03 | glio |
| Bcas2_in.vivo_noRT | Fst | 1.905 | 4.87E-03 | glio |
| Bcas2_in.vivo_noRT | Slc16a3 | -1.787 | 4.87E-03 | glio |
| Bcas2_in.vivo_noRT | Gmfb | -1.173 | 4.87E-03 | glio |
| Bcas2_in.vivo_noRT | Axl | -0.913 | 4.87E-03 | glio |
| Bcas2_in.vivo_noRT | Lars | -0.775 | 4.87E-03 | glio |

| | | | | |
|---|---|---|---|---|
| Bora_ex.vivo_noRT | Gadl1 | 2.707 | 4.87E-03 | glio |
| Chek1_ex.vivo_noRT | Kbtbd7 | 2.281 | 4.87E-03 | glio |
| Eif2s1_ex.vivo_noRT | Adam5 | 2.234 | 4.87E-03 | glio |
| Eif2s1_ex.vivo_noRT | 4933400F21Ri | 2.234 | 4.87E-03 | glio |
| Eif2s1_ex.vivo_noRT | Gm4841 | 3.373 | 4.87E-03 | glio |
| Eif2s1_ex.vivo_noRT | Gm12794 | 2.234 | 4.87E-03 | glio |
| culture_effect | Hmces | -0.789 | 4.87E-04 | glio |
| Interaction_genotype_Aars2 | Akap3 | -4.411 | 4.87E-04 | glio |
| Interaction_genotype_Mtor | Tmem202 | -4.132 | 4.87E-04 | glio |
| Interaction_genotype_Mtor | Trim65 | -3.86 | 4.87E-04 | glio |
| Chek1_in.vivo_noRT | Susd1 | 2.732 | 4.87E-04 | glio |
| Chek1_in.vivo_noRT | Rabggta | -1.871 | 4.87E-04 | glio |
| Chek1_ex.vivo_noRT | Coq8a | 3.894 | 4.87E-04 | glio |
| Eif2s1_in.vivo_noRT | Cxcl5 | 2.143 | 4.87E-04 | glio |
| Interaction_genotype_Mtor | Grm1 | -4.371 | 4.87E-05 | glio |
| Aars2_in.vivo_noRT | 4930412C18R | 2.991 | 4.87E-05 | glio |
| Aars2_in.vivo_noRT | Serp2 | 2.589 | 4.87E-05 | glio |
| Atp5c1_in.vivo_noRT | Kcnk13 | 2.405 | 4.87E-05 | glio |
| Atr_in.vivo_noRT | Sapcd1 | 2.869 | 4.87E-05 | glio |
| Atr_in.vivo_noRT | Klk9 | 2.873 | 4.87E-05 | glio |
| Atr_in.vivo_noRT | B230369F24Ri | 2.852 | 4.87E-05 | glio |
| Atr_in.vivo_noRT | Phb2 | -0.941 | 4.87E-05 | glio |
| Bard1_ex.vivo_noRT | Sytl4 | 2.814 | 4.87E-05 | glio |
| Bard1_ex.vivo_noRT | Fbxo15 | 3.414 | 4.87E-05 | glio |
| Chek1_in.vivo_noRT | Rbm19 | 2.155 | 4.87E-05 | glio |
| Atr_ex.vivo_noRT | Gm13856 | 2.77 | 4.87E-06 | glio |
| Interaction_genotype_Mtor | Gm31063 | -5.023 | 4.87E-08 | glio |
| Atr_in.vivo_noRT | 1810062O18R | 3.947 | 4.87E-08 | glio |
| culture_effect | Adk | -0.901 | 4.87E-09 | glio |
| Aars2_in.vivo_noRT | Sod3 | 5.327 | 4.87E-11 | glio |
| Bora_in.vivo_noRT | Prm3 | 4.195 | 4.87E-12 | glio |
| Chek1_in.vivo_noRT | 1700006H21R | 5.044 | 4.87E-13 | glio |
| Aars2_in.vivo_noRT | 1110046J04Ri | 5.327 | 4.87E-14 | glio |
| Aars2_in.vivo_noRT | Efcab9 | 5.06 | 4.87E-14 | glio |
| Bcas2_in.vivo_noRT | Fam25c | 4.275 | 4.87E-15 | glio |
| Atr_in.vivo_noRT | Gm15743 | 5.27 | 4.87E-16 | glio |
| culture_effect | Ssrp1 | 0.304 | 4.88E-03 | glio |
| Interaction_genotype_Aars2 | Npnt | -2.565 | 4.88E-03 | glio |
| Interaction_genotype_Mtor | Gm43062 | -3.505 | 4.88E-03 | glio |
| Interaction_genotype_Mtor | Gm48529 | -3.356 | 4.88E-03 | glio |
| Interaction_genotype_Mtor | Fkrp | -2.502 | 4.88E-03 | glio |
| Bard1_ex.vivo_noRT | Ggt1 | 2.814 | 4.88E-03 | glio |
| Bcas2_in.vivo_noRT | Gm43138 | 2.171 | 4.88E-03 | glio |
| Bora_in.vivo_noRT | Fam32a | -0.854 | 4.88E-03 | glio |
| Bora_ex.vivo_noRT | Dnajc28 | 2.975 | 4.88E-03 | glio |
| Chek1_in.vivo_noRT | Tmem223 | 2.115 | 4.88E-03 | glio |
| Chek1_in.vivo_noRT | C3 | 2.042 | 4.88E-03 | glio |
| Chek1_in.vivo_noRT | Uqcrb | -0.677 | 4.88E-03 | glio |
| Chek1_ex.vivo_noRT | Pcdhb16 | 2.843 | 4.88E-03 | glio |
| Hsd17b10_in.vivo_noRT | Rab17 | 2.148 | 4.88E-03 | glio |
| culture_effect | Tfg | -0.579 | 4.88E-04 | glio |
| culture_effect | Rtp4 | 0.491 | 4.88E-04 | glio |

| | | | | |
|---|---|---|---|---|
| culture_effect | Cs | -0.469 | 4.88E-04 | glio |
| Interaction_genotype_Mtor | Gm36026 | -3.448 | 4.88E-04 | glio |
| Interaction_genotype_Mtor | Pabpc1 | 1.11 | 4.88E-04 | glio |
| Interaction_genotype_Myc | Gm14698 | -4.015 | 4.88E-04 | glio |
| Aars2_in.vivo_noRT | Nmnat3 | 2.627 | 4.88E-04 | glio |
| Atr_in.vivo_noRT | Kcnip3 | 2.268 | 4.88E-04 | glio |
| Atr_in.vivo_noRT | Tedc1 | 1.66 | 4.88E-04 | glio |
| Atr_in.vivo_noRT | Rab18 | -1.1 | 4.88E-04 | glio |
| Chek1_in.vivo_noRT | Ocm | 2.203 | 4.88E-04 | glio |
| Chek1_in.vivo_noRT | 6030442K20R | 2.415 | 4.88E-04 | glio |
| Chek1_ex.vivo_noRT | Cyp2d22 | 3.41 | 4.88E-04 | glio |
| Chek1_ex.vivo_noRT | Krt80 | 2.695 | 4.88E-04 | glio |
| Chek1_ex.vivo_noRT | Gm10767 | 3.108 | 4.88E-04 | glio |
| Chek1_ex.vivo_noRT | Gpx2 | 4.049 | 4.88E-04 | glio |
| Interaction_genotype_Myc | Prickle2 | -3.876 | 4.88E-05 | glio |
| Atr_ex.vivo_noRT | Calhm6 | 3.368 | 4.88E-05 | glio |
| Bora_in.vivo_noRT | Tiam1 | -2.046 | 4.88E-05 | glio |
| Chek1_ex.vivo_noRT | Hapln4 | 3.874 | 4.88E-05 | glio |
| culture_effect | Arntl2 | -1.537 | 4.88E-06 | glio |
| Interaction_genotype_Aars2 | Adam12 | -4.951 | 4.88E-06 | glio |
| Interaction_genotype_Mtor | D6Ertd527e | -5.431 | 4.88E-06 | glio |
| Atr_in.vivo_noRT | Gm15601 | 3.321 | 4.88E-06 | glio |
| Bcas2_in.vivo_noRT | Proscos | 3.439 | 4.88E-06 | glio |
| culture_effect | Gpr37 | -1.85 | 4.88E-07 | glio |
| Aars2_in.vivo_noRT | Vcam1 | 4.811 | 4.88E-08 | glio |
| Chek1_ex.vivo_noRT | Rasl12 | 4.842 | 4.88E-09 | glio |
| Aars2_in.vivo_noRT | Gm30124 | 4.056 | 4.88E-10 | glio |
| Bora_in.vivo_noRT | 5430402O13R | 4.936 | 4.88E-12 | glio |
| Bcas2_in.vivo_noRT | Scgb1c1 | 4.807 | 4.88E-16 | glio |
| Aars2_in.vivo_noRT | Gm31415 | 5.575 | 4.88E-21 | glio |
| Bora_in.vivo_noRT | A030014E15Ri | 4.936 | 4.88E-24 | glio |
| Bora_in.vivo_noRT | Gm21731 | 4.936 | 4.88E-24 | glio |
| culture_effect | 5430400D12R | 1.615 | 4.89E-03 | glio |
| culture_effect | Crtc2 | -0.716 | 4.89E-03 | glio |
| Interaction_genotype_Aars2 | Gm14023 | -3.645 | 4.89E-03 | glio |
| Interaction_genotype_Aars2 | Pif1 | -3.37 | 4.89E-03 | glio |
| Interaction_genotype_Aars2 | 1700041G16R | -3.353 | 4.89E-03 | glio |
| Interaction_genotype_Aars2 | Ppp1cc | 1.432 | 4.89E-03 | glio |
| Interaction_genotype_Atr | Gm37035 | -2.932 | 4.89E-03 | glio |
| Interaction_genotype_Atr | Lyz2 | -3.611 | 4.89E-03 | glio |
| Interaction_genotype_Myc | Fignl2 | -3.436 | 4.89E-03 | glio |
| Interaction_genotype_Myc | Pou3f1 | -3.16 | 4.89E-03 | glio |
| Interaction_genotype_Rad51 | Pde8a | -3.308 | 4.89E-03 | glio |
| Aars2_in.vivo_noRT | Klhl3 | 1.881 | 4.89E-03 | glio |
| Aars2_in.vivo_noRT | Tm7sf2 | 1.622 | 4.89E-03 | glio |
| Atp5c1_in.vivo_noRT | Prox2 | 2.464 | 4.89E-03 | glio |
| Atr_in.vivo_noRT | Prkcz | 1.851 | 4.89E-03 | glio |
| Bora_in.vivo_noRT | A930041C12R | 2.05 | 4.89E-03 | glio |
| Bora_in.vivo_noRT | Utp11 | -0.732 | 4.89E-03 | glio |
| Brca2_in.vivo_noRT | Gm49808 | 1.763 | 4.89E-03 | glio |
| Brca2_in.vivo_noRT | Slc14a2 | 1.763 | 4.89E-03 | glio |
| Brca2_in.vivo_noRT | Sox11 | 2.594 | 4.89E-03 | glio |

| | | | | |
|---|---|---|---|---|
| Brca2_in.vivo_noRT | Gm4340 | 1.763 | 4.89E-03 | glio |
| Brca2_in.vivo_noRT | Gm16239 | 1.763 | 4.89E-03 | glio |
| Cenpj_in.vivo_noRT | Ccl3 | 1.762 | 4.89E-03 | glio |
| Cenpj_in.vivo_noRT | Gm48780 | 1.762 | 4.89E-03 | glio |
| Chek1_in.vivo_noRT | Gm31645 | 2.09 | 4.89E-03 | glio |
| Interaction_genotype_Mtor | Wdr78 | -4.518 | 4.89E-04 | glio |
| Interaction_genotype_Myc | Cks1brt | -3.563 | 4.89E-04 | glio |
| Interaction_genotype_Myc | Dad1 | 1.498 | 4.89E-04 | glio |
| Interaction_genotype_Myc | Eef1g | 1.546 | 4.89E-04 | glio |
| Interaction_genotype_Rad51 | Plxnb1 | -4.872 | 4.89E-04 | glio |
| Interaction_genotype_Rad51 | Gm4841 | 5.543 | 4.89E-04 | glio |
| interaction_RT | Gnptab | -1.092 | 4.89E-04 | glio |
| interaction_RT | Tmpo | -0.844 | 4.89E-04 | glio |
| interaction_RT | Mrto4 | -0.696 | 4.89E-04 | glio |
| Atr_in.vivo_noRT | 6030442K20R | 2.381 | 4.89E-04 | glio |
| Atr_ex.vivo_noRT | Vmn1r91 | 2.966 | 4.89E-04 | glio |
| Hsd17b10_in.vivo_noRT | Esyt3 | 2.556 | 4.89E-04 | glio |
| culture_effect | Rfc1 | 0.52 | 4.89E-05 | glio |
| Interaction_genotype_Atr | Elof1 | 2.012 | 4.89E-05 | glio |
| Interaction_genotype_Mtor | Ccdc39 | -4.743 | 4.89E-05 | glio |
| Aars2_in.vivo_noRT | 3010001F23Ri | 2.698 | 4.89E-05 | glio |
| Atp5c1_in.vivo_noRT | Btn1a1 | 2.408 | 4.89E-05 | glio |
| Bcas2_in.vivo_noRT | Gm10230 | 3.412 | 4.89E-06 | glio |
| Bora_in.vivo_noRT | Rpp30 | -1.899 | 4.89E-06 | glio |
| Interaction_genotype_Mtor | Gm11232 | -4.099 | 4.89E-07 | glio |
| Aars2_ex.vivo_noRT | Dpp6 | 3.293 | 4.89E-07 | glio |
| Aars2_ex.vivo_noRT | Serpinb9g | 3.293 | 4.89E-07 | glio |
| Aars2_ex.vivo_noRT | BC049987 | 3.293 | 4.89E-07 | glio |
| Bard1_ex.vivo_noRT | Nat8f2 | 4 | 4.89E-07 | glio |
| Chek1_in.vivo_noRT | Shroom3 | 3.765 | 4.89E-08 | glio |
| Chek1_ex.vivo_noRT | Serpinb2 | 4.443 | 4.89E-09 | glio |
| Chek1_ex.vivo_noRT | Ric3 | 4.449 | 4.89E-09 | glio |
| Chek1_ex.vivo_noRT | Slc14a1 | 4.268 | 4.89E-09 | glio |
| Aars2_in.vivo_noRT | Mtx1 | -2.337 | 4.89E-10 | glio |
| Chek1_ex.vivo_noRT | Gm14540 | 4.049 | 4.89E-10 | glio |
| Chek1_in.vivo_noRT | Ly6g6c | 4.184 | 4.89E-11 | glio |
| Chek1_in.vivo_noRT | Iqcf1 | 4.391 | 4.89E-11 | glio |
| Chek1_ex.vivo_noRT | Gm28940 | 4.842 | 4.89E-12 | glio |
| Chek1_ex.vivo_noRT | Gm20658 | 4.842 | 4.89E-17 | glio |
| Bora_in.vivo_noRT | Apoa5 | 4.936 | 4.89E-25 | glio |
| Aars2_in.vivo_noRT | Slc14a2 | 5.327 | 4.89E-28 | glio |
| Interaction_genotype_Aars2 | Gm50399 | -3.351 | 4.90E-03 | glio |
| Interaction_genotype_Atr | Ccdc191 | -3.205 | 4.90E-03 | glio |
| Interaction_genotype_Bard1 | Irf8 | 3.893 | 4.90E-03 | glio |
| Interaction_genotype_Hsd17b | Col6a2 | -3.374 | 4.90E-03 | glio |
| Interaction_genotype_Mtor | Tor1aip1 | 1.474 | 4.90E-03 | glio |
| Interaction_genotype_Myc | Maml1 | -2.455 | 4.90E-03 | glio |
| Interaction_genotype_Myc | Rpl4 | 1.072 | 4.90E-03 | glio |
| Interaction_genotype_Myc | Rpl9 | 0.951 | 4.90E-03 | glio |
| interaction_RT | Prrc2c | -0.531 | 4.90E-03 | glio |
| Aars2_in.vivo_noRT | Gm38190 | 1.895 | 4.90E-03 | glio |
| Aars2_in.vivo_noRT | Fam104a | -0.651 | 4.90E-03 | glio |

| | | | | |
|---|---|---|---|---|
| Atr_in.vivo_noRT | Map3k14 | 1.833 | 4.90E-03 | glio |
| Atr_in.vivo_noRT | Flt3l | 1.738 | 4.90E-03 | glio |
| Bcas2_in.vivo_noRT | Fgl2 | 1.914 | 4.90E-03 | glio |
| Bcas2_in.vivo_noRT | Cenpc1 | -1.251 | 4.90E-03 | glio |
| Bcas2_ex.vivo_noRT | Ccl7 | 3.298 | 4.90E-03 | glio |
| Bora_in.vivo_noRT | Syn3 | 2.083 | 4.90E-03 | glio |
| Bora_in.vivo_noRT | Tppp3 | -1.817 | 4.90E-03 | glio |
| Bora_in.vivo_noRT | Angptl6 | 1.38 | 4.90E-03 | glio |
| Bora_ex.vivo_noRT | Crispld2 | 2.598 | 4.90E-03 | glio |
| Cenpj_in.vivo_noRT | Kcnk7 | 2.028 | 4.90E-03 | glio |
| Dclre1c_ex.vivo_noRT | Akr1c18 | 2.666 | 4.90E-03 | glio |
| Hsd17b10_in.vivo_noRT | Dchs1 | 2.121 | 4.90E-03 | glio |
| Hsd17b10_in.vivo_noRT | Gm50399 | 2.064 | 4.90E-03 | glio |
| Hsd17b10_in.vivo_noRT | Zfp418 | 2.341 | 4.90E-03 | glio |
| Hsd17b10_in.vivo_noRT | Khsrp | -1.325 | 4.90E-03 | glio |
| Interaction_genotype_Myc | Sv2a | -4.147 | 4.90E-04 | glio |
| Bcas2_ex.vivo_noRT | Slc36a2 | 3.26 | 4.90E-04 | glio |
| Bora_ex.vivo_noRT | BB557941 | 2.802 | 4.90E-04 | glio |
| Bora_ex.vivo_noRT | Slc26a1 | 3.293 | 4.90E-04 | glio |
| Bora_ex.vivo_noRT | Cd52 | 3.201 | 4.90E-04 | glio |
| Bora_ex.vivo_noRT | Amy1 | 3.207 | 4.90E-04 | glio |
| Bora_ex.vivo_noRT | Col5a2 | 2.615 | 4.90E-04 | glio |
| Chek1_ex.vivo_noRT | Gm30541 | 3.065 | 4.90E-04 | glio |
| interaction_RT | Rnf5 | 1.173 | 4.90E-05 | glio |
| Atp5c1_in.vivo_noRT | Krtcap3 | 2.407 | 4.90E-05 | glio |
| Eif2s1_ex.vivo_noRT | A330076H08R | 3.656 | 4.90E-05 | glio |
| Aars2_in.vivo_noRT | Zfp992 | 3.299 | 4.90E-06 | glio |
| Atr_in.vivo_noRT | Cntf | 3.164 | 4.90E-06 | glio |
| Bora_ex.vivo_noRT | Nkg7 | 3.598 | 4.90E-06 | glio |
| Chek1_in.vivo_noRT | Gm34643 | 3.328 | 4.90E-06 | glio |
| Chek1_in.vivo_noRT | Prss22 | 3.975 | 4.90E-06 | glio |
| Interaction_genotype_Myc | Prrx2 | -5.819 | 4.90E-07 | glio |
| Interaction_genotype_Mtor | Pin1 | 2.693 | 4.90E-08 | glio |
| Interaction_genotype_Mtor | Anapc11 | 2.85 | 4.90E-08 | glio |
| interaction_RT | Col11a1 | -1.085 | 4.90E-08 | glio |
| Aars2_in.vivo_noRT | 1700063J08Ri | 3.819 | 4.90E-08 | glio |
| Chek1_in.vivo_noRT | 4933439K11R | 3.859 | 4.90E-08 | glio |
| Aars2_in.vivo_noRT | Jakmip1 | 4.006 | 4.90E-09 | glio |
| Atr_in.vivo_noRT | Wincr1 | 5.269 | 4.90E-09 | glio |
| Chek1_in.vivo_noRT | 4930545E07Ri | 4.521 | 4.90E-10 | glio |
| culture_effect | Samhd1 | -1.032 | 4.90E-11 | glio |
| interaction_RT | Cebpb | 1.814 | 4.90E-11 | glio |
| Bora_in.vivo_noRT | Mettl7a3 | 4.286 | 4.90E-11 | glio |
| Bora_in.vivo_noRT | Gm15579 | 4.936 | 4.90E-11 | glio |
| Chek1_in.vivo_noRT | Gm5662 | 5.044 | 4.90E-14 | glio |
| culture_effect | Gm12185 | -4.88 | 4.90E-17 | glio |
| Aars2_in.vivo_noRT | 4930553M12F | 5.06 | 4.90E-20 | glio |
| Bora_in.vivo_noRT | Serpinb9g | 4.936 | 4.90E-22 | glio |
| Interaction_genotype_Aars2 | Gm7102 | -3.67 | 4.91E-03 | glio |
| Interaction_genotype_Aars2 | Aldh1l1 | -3.916 | 4.91E-03 | glio |
| Interaction_genotype_Aars2 | Gm47484 | -3.645 | 4.91E-03 | glio |
| Interaction_genotype_Aars2 | 9530062K07R | -3.612 | 4.91E-03 | glio |

| | | | | |
|---|---|---|---|---|
| Interaction_genotype_Aars2 | Gars | 1.237 | 4.91E-03 | glio |
| Interaction_genotype_Mtor | Cep78 | -2.767 | 4.91E-03 | glio |
| Interaction_genotype_Myc | Maf1 | 1.138 | 4.91E-03 | glio |
| interaction_RT | Ergic3 | 0.628 | 4.91E-03 | glio |
| interaction_RT | Capns1 | -0.58 | 4.91E-03 | glio |
| Aars2_in.vivo_noRT | Drc3 | 2.096 | 4.91E-03 | glio |
| Aars2_in.vivo_noRT | Arhgap22 | 1.967 | 4.91E-03 | glio |
| Aars2_in.vivo_noRT | Elac1 | 1.935 | 4.91E-03 | glio |
| Aars2_in.vivo_noRT | Atp13a2 | 1.328 | 4.91E-03 | glio |
| Aars2_in.vivo_noRT | Vps72 | -1.207 | 4.91E-03 | glio |
| Atr_in.vivo_noRT | Dnmt3b | 1.996 | 4.91E-03 | glio |
| Atr_in.vivo_noRT | Tgfb2 | 1.875 | 4.91E-03 | glio |
| Bora_in.vivo_noRT | mt-Atp8 | -1.511 | 4.91E-03 | glio |
| Cenpj_ex.vivo_noRT | Mapk15 | 2.983 | 4.91E-03 | glio |
| Chek1_in.vivo_noRT | Gpnmb | -1.043 | 4.91E-03 | glio |
| Chek1_ex.vivo_noRT | Ipo9 | -1.713 | 4.91E-03 | glio |
| Eif2s1_in.vivo_noRT | Psca | 2.143 | 4.91E-03 | glio |
| Interaction_genotype_Mtor | Nupr1l | -4.621 | 4.91E-04 | glio |
| interaction_RT | Ddx27 | -0.799 | 4.91E-04 | glio |
| interaction_RT | Igtp | 0.865 | 4.91E-04 | glio |
| interaction_RT | Ggct | -0.987 | 4.91E-04 | glio |
| interaction_RT | Slc25a18 | 2.846 | 4.91E-04 | glio |
| Aars2_in.vivo_noRT | Gm47509 | 2.529 | 4.91E-04 | glio |
| Aars2_in.vivo_noRT | Cks1b | -1.093 | 4.91E-04 | glio |
| Atp5c1_in.vivo_noRT | 2210417A02R | 2.924 | 4.91E-04 | glio |
| Atr_ex.vivo_noRT | Acat3 | 2.588 | 4.91E-04 | glio |
| Bcas2_in.vivo_noRT | Gm32816 | 2.748 | 4.91E-04 | glio |
| Bcas2_in.vivo_noRT | Sorbs2 | 2.416 | 4.91E-04 | glio |
| Bcas2_in.vivo_noRT | Rnd3 | 1.929 | 4.91E-04 | glio |
| Eif2s1_in.vivo_noRT | Gm47936 | 2.143 | 4.91E-04 | glio |
| Eif2s1_in.vivo_noRT | Tnni3k | 2.143 | 4.91E-04 | glio |
| Hsd17b10_in.vivo_noRT | Bhlhe41 | 3.093 | 4.91E-04 | glio |
| Interaction_genotype_Mtor | Sh3bgrl2 | -4.834 | 4.91E-05 | glio |
| Interaction_genotype_Pars2 | Klhdc8a | -4.005 | 4.91E-05 | glio |
| Aars2_in.vivo_noRT | Nxf2 | 2.984 | 4.91E-05 | glio |
| Atr_in.vivo_noRT | Rbm3os | 2.742 | 4.91E-05 | glio |
| Chek1_in.vivo_noRT | Gm20712 | 2.853 | 4.91E-05 | glio |
| Aars2_in.vivo_noRT | Gm26981 | 3.152 | 4.91E-06 | glio |
| Aars2_ex.vivo_noRT | Slc16a5 | 2.9 | 4.91E-06 | glio |
| Aars2_ex.vivo_noRT | H2-Eb2 | 3.293 | 4.91E-06 | glio |
| Aars2_in.vivo_noRT | Gm10602 | 3.919 | 4.91E-07 | glio |
| Bcas2_in.vivo_noRT | Bbof1 | 3.379 | 4.91E-07 | glio |
| Bcas2_in.vivo_noRT | Otos | 3.278 | 4.91E-07 | glio |
| culture_effect | Lsm6 | 0.681 | 4.91E-08 | glio |
| Chek1_ex.vivo_noRT | St14 | 4.238 | 4.91E-08 | glio |
| Atr_in.vivo_noRT | Gm36495 | 4.776 | 4.91E-10 | glio |
| Bora_in.vivo_noRT | Rnf165 | 4.936 | 4.91E-13 | glio |
| Atr_in.vivo_noRT | Cidea | 4.618 | 4.91E-14 | glio |
| Atr_in.vivo_noRT | S100a14 | 4.733 | 4.91E-15 | glio |
| Aars2_in.vivo_noRT | Slc25a18 | 5.06 | 4.91E-18 | glio |
| Interaction_genotype_Aars2 | Clec12b | -2.429 | 4.92E-03 | glio |
| Interaction_genotype_Aars2 | Plekhg5 | -3.371 | 4.92E-03 | glio |

| | | | | |
|---|---|---|---|---|
| Interaction_genotype_Chek1 | Icam1 | 4.22 | 4.92E-03 | glio |
| Interaction_genotype_Mtor | 9030407P20R | -3.319 | 4.92E-03 | glio |
| Interaction_genotype_Mtor | Pdzd7 | -3.491 | 4.92E-03 | glio |
| Interaction_genotype_Mtor | Inafm2 | -3.004 | 4.92E-03 | glio |
| Interaction_genotype_Myc | Ccdc57 | -3.35 | 4.92E-03 | glio |
| Interaction_genotype_Myc | Eid3 | -2.842 | 4.92E-03 | glio |
| Interaction_genotype_Myc | Fbxo44 | -2.367 | 4.92E-03 | glio |
| Interaction_genotype_Pars2 | Gm36172 | -3.103 | 4.92E-03 | glio |
| Interaction_genotype_Rpl3 | Gm41318 | -3.839 | 4.92E-03 | glio |
| Interaction_genotype_Uqcrc2 | Gm43672 | -4.238 | 4.92E-03 | glio |
| Interaction_genotype_Uqcrc2 | Gm50394 | -4.027 | 4.92E-03 | glio |
| Atr_in.vivo_noRT | Akr1b8 | -1.381 | 4.92E-03 | glio |
| Bcas2_in.vivo_noRT | Recql | -1.346 | 4.92E-03 | glio |
| Bcas2_ex.vivo_noRT | Prickle2 | 2.381 | 4.92E-03 | glio |
| Bora_in.vivo_noRT | Gm15492 | 2.109 | 4.92E-03 | glio |
| Cenpj_ex.vivo_noRT | Gm48265 | 2.591 | 4.92E-03 | glio |
| Cenpj_ex.vivo_noRT | Car10 | 2.591 | 4.92E-03 | glio |
| Interaction_genotype_Aars2 | Tle5 | 1.321 | 4.92E-04 | glio |
| Interaction_genotype_Mtor | 1700066B19R | -3.706 | 4.92E-04 | glio |
| Interaction_genotype_Myc | Cwf19l1 | -2.696 | 4.92E-04 | glio |
| Interaction_genotype_Myc | Ctsl | 2.37 | 4.92E-04 | glio |
| Interaction_genotype_Rad51 | Phldb2 | -4.963 | 4.92E-04 | glio |
| Interaction_genotype_Rad51 | Adam12 | -4.153 | 4.92E-04 | glio |
| Bcas2_in.vivo_noRT | Pfkp | -2.029 | 4.92E-04 | glio |
| Chek1_ex.vivo_noRT | Nyap2 | 3.166 | 4.92E-04 | glio |
| Hsd17b10_in.vivo_noRT | Prmt7 | -1.587 | 4.92E-04 | glio |
| Interaction_genotype_Mtor | Gm40538 | -3.836 | 4.92E-05 | glio |
| Interaction_genotype_Mtor | Ksr1 | -4.679 | 4.92E-05 | glio |
| Interaction_genotype_Mtor | AI480526 | -4.262 | 4.92E-05 | glio |
| Aars2_in.vivo_noRT | Sipa1l2 | 2.884 | 4.92E-05 | glio |
| Atr_in.vivo_noRT | Mpz | 3.562 | 4.92E-05 | glio |
| Bora_ex.vivo_noRT | Hrh2 | 3.201 | 4.92E-05 | glio |
| Bora_ex.vivo_noRT | B3galt2 | 3.595 | 4.92E-05 | glio |
| Bora_in.vivo_noRT | Cebpa | 3.275 | 4.92E-06 | glio |
| Chek1_in.vivo_noRT | Tcte2 | 2.921 | 4.92E-06 | glio |
| Chek1_in.vivo_noRT | Spata45 | 3.136 | 4.92E-06 | glio |
| culture_effect | Snta1 | 1.571 | 4.92E-07 | glio |
| Aars2_in.vivo_noRT | Slc27a3 | 3.498 | 4.92E-07 | glio |
| Atr_in.vivo_noRT | Zfp706 | -1.168 | 4.92E-07 | glio |
| Bora_in.vivo_noRT | Pagr1a | 3.252 | 4.92E-07 | glio |
| Atr_in.vivo_noRT | 4930545E07Ri | 4.5 | 4.92E-10 | glio |
| Atr_in.vivo_noRT | Ak1 | 4.27 | 4.92E-10 | glio |
| Bora_in.vivo_noRT | Lrfn1 | 4.294 | 4.92E-10 | glio |
| Aars2_in.vivo_noRT | 9630014M24F | 4.548 | 4.92E-12 | glio |
| Bcas2_in.vivo_noRT | Cfap46 | 4.34 | 4.92E-12 | glio |
| Bora_in.vivo_noRT | Celsr3 | 4.676 | 4.92E-13 | glio |
| Atr_in.vivo_noRT | Rarres1 | 5.533 | 4.92E-16 | glio |
| Chek1_in.vivo_noRT | Slc16a4 | 4.78 | 4.92E-17 | glio |
| Atr_in.vivo_noRT | 1700112J16Ri | 5.533 | 4.92E-24 | glio |
| Atr_in.vivo_noRT | Clec4d | 5.533 | 4.92E-24 | glio |
| culture_effect | Mta3 | -1.007 | 4.93E-03 | glio |
| Interaction_genotype_Bora | Gm10354 | -3.03 | 4.93E-03 | glio |

| | | | | |
|---|---|---|---|---|
| Interaction_genotype_Myc | Ptprc | -2.466 | 4.93E-03 | glio |
| Interaction_genotype_Myc | Lhx1os | -2.599 | 4.93E-03 | glio |
| Interaction_genotype_Myc | Upk1a | -3.288 | 4.93E-03 | glio |
| Interaction_genotype_Myc | Ssc5d | -3.012 | 4.93E-03 | glio |
| Interaction_genotype_Myc | Zbtb40 | -2.75 | 4.93E-03 | glio |
| Atr_in.vivo_noRT | Ergic3 | -0.853 | 4.93E-03 | glio |
| Chek1_in.vivo_noRT | Pycard | 1.906 | 4.93E-03 | glio |
| Chek1_in.vivo_noRT | Chtop | -0.996 | 4.93E-03 | glio |
| Chek1_in.vivo_noRT | Eif2s2 | -0.531 | 4.93E-03 | glio |
| culture_effect | Senp8 | 1.24 | 4.93E-04 | glio |
| culture_effect | Ggct | -0.61 | 4.93E-04 | glio |
| Interaction_genotype_Atr | Hba-a2 | -4.432 | 4.93E-04 | glio |
| Interaction_genotype_Myc | Dcun1d3 | -3.526 | 4.93E-04 | glio |
| Aars2_in.vivo_noRT | Egfl8 | 2.603 | 4.93E-04 | glio |
| Aars2_in.vivo_noRT | Slc15a3 | 2.307 | 4.93E-04 | glio |
| Aars2_in.vivo_noRT | Mapk1ip1l | -1.333 | 4.93E-04 | glio |
| Atr_in.vivo_noRT | Chic2 | -1.095 | 4.93E-04 | glio |
| Atr_in.vivo_noRT | Rpl8 | -0.701 | 4.93E-04 | glio |
| Aars2_in.vivo_noRT | Slc7a7 | 2.84 | 4.93E-05 | glio |
| Aars2_in.vivo_noRT | Rab15 | 2.818 | 4.93E-05 | glio |
| Atr_in.vivo_noRT | B230219D22R | -1.688 | 4.93E-05 | glio |
| Bcas2_in.vivo_noRT | Tlr6 | 2.973 | 4.93E-05 | glio |
| Bora_in.vivo_noRT | Slc47a2 | 2.83 | 4.93E-05 | glio |
| Hsd17b10_in.vivo_noRT | P4ha3 | 2.764 | 4.93E-05 | glio |
| Aars2_in.vivo_noRT | Il23a | 3.205 | 4.93E-06 | glio |
| Atp5c1_in.vivo_noRT | Gm21319 | 2.668 | 4.93E-06 | glio |
| Atr_in.vivo_noRT | Pycard | 2.933 | 4.93E-06 | glio |
| Bora_in.vivo_noRT | Gm9725 | 3.268 | 4.93E-06 | glio |
| Chek1_in.vivo_noRT | Gm46218 | 3.287 | 4.93E-06 | glio |
| Aars2_in.vivo_noRT | 9130604C24R | 3.861 | 4.93E-07 | glio |
| Atr_ex.vivo_noRT | 4930483J18Ri | 4.062 | 4.93E-07 | glio |
| Atr_ex.vivo_noRT | Pmch | 3.947 | 4.93E-07 | glio |
| Chek1_in.vivo_noRT | Olfr1369-ps1 | 4.303 | 4.93E-07 | glio |
| Chek1_in.vivo_noRT | Gng13 | 3.451 | 4.93E-07 | glio |
| Hsd17b10_in.vivo_noRT | Prrx2 | 3.616 | 4.93E-07 | glio |
| Hsd17b10_in.vivo_noRT | Gm36026 | 2.965 | 4.93E-07 | glio |
| Hsd17b10_in.vivo_noRT | Gm48742 | 2.966 | 4.93E-07 | glio |
| Atr_in.vivo_noRT | Gm29666 | 3.809 | 4.93E-08 | glio |
| Chek1_ex.vivo_noRT | Cstdc4 | 4.443 | 4.93E-08 | glio |
| Bcas2_in.vivo_noRT | Rtn1 | 4.161 | 4.93E-09 | glio |
| Atr_in.vivo_noRT | Arhgef15 | 4.152 | 4.93E-10 | glio |
| Aars2_in.vivo_noRT | Gm42658 | 4.591 | 4.93E-13 | glio |
| Atr_in.vivo_noRT | Gm16933 | 5.011 | 4.93E-24 | glio |
| Interaction_genotype_Atr | B230319C09R | -3.779 | 4.94E-03 | glio |
| Interaction_genotype_Bora | 1700013F07Ri | -3.989 | 4.94E-03 | glio |
| Interaction_genotype_Bora | Aplp2 | 1.44 | 4.94E-03 | glio |
| Interaction_genotype_Mtor | Plp1 | 2.537 | 4.94E-03 | glio |
| Interaction_genotype_Mtor | Acin1 | 1.426 | 4.94E-03 | glio |
| Interaction_genotype_Myc | Bpifc | -3.448 | 4.94E-03 | glio |
| Interaction_genotype_Myc | Tamm41 | -2.248 | 4.94E-03 | glio |
| Aars2_in.vivo_noRT | Hist1h2bn | 1.959 | 4.94E-03 | glio |
| Atr_in.vivo_noRT | Ccdc69 | 2.003 | 4.94E-03 | glio |

| | | | | |
|---|---|---|---|---|
| Bcas2_in.vivo_noRT | Kctd12 | 2.002 | 4.94E-03 | glio |
| Bora_in.vivo_noRT | Rrp36 | -1.169 | 4.94E-03 | glio |
| Bora_ex.vivo_noRT | Fbln1 | 2.804 | 4.94E-03 | glio |
| Bora_ex.vivo_noRT | Gm4841 | 3.201 | 4.94E-03 | glio |
| Cenpj_in.vivo_noRT | Gm15138 | 1.756 | 4.94E-03 | glio |
| Chek1_in.vivo_noRT | A130014A01R | 1.853 | 4.94E-03 | glio |
| Chek1_in.vivo_noRT | Mark4 | 1.438 | 4.94E-03 | glio |
| Chek1_in.vivo_noRT | Arl2bp | -0.849 | 4.94E-03 | glio |
| Interaction_genotype_Aars2 | 2010001K21R | -3.986 | 4.94E-04 | glio |
| Interaction_genotype_Myc | Rnf207 | -3.703 | 4.94E-04 | glio |
| Atr_in.vivo_noRT | Cstb | -1.361 | 4.94E-04 | glio |
| Cenpj_in.vivo_noRT | Gm13078 | 2.022 | 4.94E-04 | glio |
| Eif2s1_in.vivo_noRT | Gm49504 | 2.143 | 4.94E-04 | glio |
| Eif2s1_in.vivo_noRT | Sparcl1 | -1.755 | 4.94E-04 | glio |
| Interaction_genotype_Mtor | Vmn1r91 | -4.427 | 4.94E-05 | glio |
| Chek1_in.vivo_noRT | 1810021B22R | 2.975 | 4.94E-05 | glio |
| Interaction_genotype_Mtor | Znrd1 | 2.147 | 4.94E-06 | glio |
| interaction_RT | Trf | 3.047 | 4.94E-06 | glio |
| Cenpj_ex.vivo_noRT | Gm45338 | 2.982 | 4.94E-06 | glio |
| Chek1_in.vivo_noRT | Cyp2s1 | 3.471 | 4.94E-06 | glio |
| Hsd17b10_in.vivo_noRT | Prm3 | 2.875 | 4.94E-06 | glio |
| Interaction_genotype_Mtor | Sod3 | -5.983 | 4.94E-07 | glio |
| Bcas2_in.vivo_noRT | Adgrb3 | 3.501 | 4.94E-07 | glio |
| Aars2_in.vivo_noRT | Gm32122 | 3.499 | 4.94E-08 | glio |
| Hsd17b10_in.vivo_noRT | Gm11627 | 3.616 | 4.94E-08 | glio |
| Atr_in.vivo_noRT | 9330104G04R | 4.053 | 4.94E-09 | glio |
| Chek1_in.vivo_noRT | Fgf1 | 4.253 | 4.94E-09 | glio |
| Atr_in.vivo_noRT | Zfp712 | 3.838 | 4.94E-10 | glio |
| Chek1_in.vivo_noRT | Gm47802 | 4.068 | 4.94E-10 | glio |
| Bora_in.vivo_noRT | A830009L08Ri | 4.414 | 4.94E-12 | glio |
| culture_effect | S100a4 | 3.091 | 4.94E-39 | glio |
| culture_effect | Snrnp48 | -0.44 | 4.95E-03 | glio |
| Interaction_genotype_Aars2 | Cyp4x1 | -3.35 | 4.95E-03 | glio |
| Interaction_genotype_Aars2 | Gm2788 | -3.709 | 4.95E-03 | glio |
| Interaction_genotype_Atr | Gm14327 | -3.573 | 4.95E-03 | glio |
| Interaction_genotype_Atr | Ric8a | 2 | 4.95E-03 | glio |
| Interaction_genotype_Atr | Fign | 1.9 | 4.95E-03 | glio |
| Interaction_genotype_Atr | Stk19 | 1.45 | 4.95E-03 | glio |
| Interaction_genotype_Ifnar1 | Scg2 | 2.734 | 4.95E-03 | glio |
| Interaction_genotype_Myc | 4930502E18Ri | -2.587 | 4.95E-03 | glio |
| Interaction_genotype_Myc | Vav3 | -2.4 | 4.95E-03 | glio |
| Interaction_genotype_Myc | Dhrs13 | -2.086 | 4.95E-03 | glio |
| Interaction_genotype_Myc | Utp6 | -1.671 | 4.95E-03 | glio |
| Aars2_in.vivo_noRT | Rpusd3 | 1.717 | 4.95E-03 | glio |
| Aars2_in.vivo_noRT | Nsmce1 | -1.047 | 4.95E-03 | glio |
| Atp5c1_in.vivo_noRT | Ttpal | -1.9 | 4.95E-03 | glio |
| Atr_in.vivo_noRT | Tesk2 | 1.942 | 4.95E-03 | glio |
| Atr_in.vivo_noRT | 2310034G01R | 1.72 | 4.95E-03 | glio |
| Atr_ex.vivo_noRT | Gm15929 | 2.187 | 4.95E-03 | glio |
| Atr_ex.vivo_noRT | Gm14273 | 2.762 | 4.95E-03 | glio |
| Atr_ex.vivo_noRT | Ccdc28a | 1.787 | 4.95E-03 | glio |
| Atr_ex.vivo_noRT | Ncl | -0.778 | 4.95E-03 | glio |

| | | | | |
|---|---|---|---|---|
| Bora_in.vivo_noRT | Gm29170 | 2.012 | 4.95E-03 | glio |
| Bora_in.vivo_noRT | Rnf207 | 1.919 | 4.95E-03 | glio |
| Chek1_ex.vivo_noRT | 4930445N18R | 2.633 | 4.95E-03 | glio |
| Chek1_ex.vivo_noRT | Sgpp1 | -2.01 | 4.95E-03 | glio |
| Hsd17b10_in.vivo_noRT | Tdrp | 2.106 | 4.95E-03 | glio |
| Atr_ex.vivo_noRT | Tcp11x2 | 2.771 | 4.95E-04 | glio |
| Atr_ex.vivo_noRT | Kcnip3 | 3.163 | 4.95E-04 | glio |
| Bora_in.vivo_noRT | Nop16 | -1.21 | 4.95E-04 | glio |
| Chek1_in.vivo_noRT | 2900052L18Ri | 2.027 | 4.95E-04 | glio |
| Hsd17b10_in.vivo_noRT | Hba-a2 | 2.704 | 4.95E-04 | glio |
| culture_effect | Zbtb10 | -1.407 | 4.95E-05 | glio |
| Atr_ex.vivo_noRT | Abra | 3.164 | 4.95E-05 | glio |
| Atr_ex.vivo_noRT | Nrip3 | 3.491 | 4.95E-05 | glio |
| Bora_in.vivo_noRT | D930019O06F | 3.04 | 4.95E-05 | glio |
| Chek1_in.vivo_noRT | Gm17477 | 2.598 | 4.95E-05 | glio |
| Atp5c1_in.vivo_noRT | Gm10447 | 2.668 | 4.95E-06 | glio |
| Atr_in.vivo_noRT | Gm15063 | 3.549 | 4.95E-06 | glio |
| Chek1_in.vivo_noRT | Zfp575 | 3.792 | 4.95E-07 | glio |
| Bora_in.vivo_noRT | Acot6 | 3.691 | 4.95E-08 | glio |
| Aars2_in.vivo_noRT | Gm43331 | 3.651 | 4.95E-10 | glio |
| culture_effect | Syvn1 | -1.396 | 4.95E-11 | glio |
| Atr_in.vivo_noRT | Ankrd1 | 5.533 | 4.95E-11 | glio |
| Bcas2_in.vivo_noRT | Gm128 | 3.889 | 4.95E-12 | glio |
| Bcas2_in.vivo_noRT | Tcap | 4.428 | 4.95E-12 | glio |
| Bora_in.vivo_noRT | Aloxe3 | 4.407 | 4.95E-14 | glio |
| Bcas2_in.vivo_noRT | Ccdc125 | 4.807 | 4.95E-17 | glio |
| Atr_in.vivo_noRT | Gm39526 | 5.011 | 4.95E-20 | glio |
| Atr_in.vivo_noRT | 4930579C12R | 5.533 | 4.95E-20 | glio |
| Atr_in.vivo_noRT | Map3k9 | 5.273 | 4.95E-20 | glio |
| culture_effect | Ugt8a | -1 | 4.96E-03 | glio |
| culture_effect | Zfp771 | 0.467 | 4.96E-03 | glio |
| culture_effect | Atp5j2 | 0.353 | 4.96E-03 | glio |
| Interaction_genotype_Bcas2 | Map3k9 | -3.111 | 4.96E-03 | glio |
| Interaction_genotype_Bcas2 | Gm15929 | -3.042 | 4.96E-03 | glio |
| Interaction_genotype_Bcas2 | Dis3l | -3.408 | 4.96E-03 | glio |
| Interaction_genotype_Mtor | Zfp937 | -3.538 | 4.96E-03 | glio |
| Interaction_genotype_Myc | 5930403N24R | -3.172 | 4.96E-03 | glio |
| Interaction_genotype_Myc | Kyat1 | -3.46 | 4.96E-03 | glio |
| Interaction_genotype_Myc | Dennd4b | -3.143 | 4.96E-03 | glio |
| Interaction_genotype_Myc | Celsr2 | -3.03 | 4.96E-03 | glio |
| Interaction_genotype_Myc | Dnhd1 | -2.719 | 4.96E-03 | glio |
| Interaction_genotype_Myc | Ski | -1.896 | 4.96E-03 | glio |
| Interaction_genotype_Myc | Rack1 | 1.146 | 4.96E-03 | glio |
| Interaction_genotype_Myc | Lgals1 | 1.303 | 4.96E-03 | glio |
| Interaction_genotype_Pars2 | Hdac9 | -3.636 | 4.96E-03 | glio |
| Interaction_genotype_Pars2 | 5730507C01R | -2.937 | 4.96E-03 | glio |
| Interaction_genotype_Pars2 | Gm46218 | -3.783 | 4.96E-03 | glio |
| Interaction_genotype_Pars2 | Fam53a | 1.75 | 4.96E-03 | glio |
| interaction_RT | Mat2a | -0.689 | 4.96E-03 | glio |
| Aars2_in.vivo_noRT | Pigo | 1.811 | 4.96E-03 | glio |
| Aars2_in.vivo_noRT | D330050G23F | 1.68 | 4.96E-03 | glio |
| Atr_in.vivo_noRT | 4833439L19Ri | 1.003 | 4.96E-03 | glio |

| | | | | |
|---|---|---|---|---|
| Bora_in.vivo_noRT | Gldc | 2.028 | 4.96E-03 | glio |
| Bora_in.vivo_noRT | Lyar | -0.788 | 4.96E-03 | glio |
| Cenpj_ex.vivo_noRT | Pmch | 2.591 | 4.96E-03 | glio |
| Chek1_in.vivo_noRT | Dnajb13 | 1.964 | 4.96E-03 | glio |
| Chek1_in.vivo_noRT | Dnmt3b | 2.03 | 4.96E-03 | glio |
| culture_effect | Smim3 | 0.965 | 4.96E-04 | glio |
| culture_effect | Nradd | -0.814 | 4.96E-04 | glio |
| culture_effect | Adprh | -0.602 | 4.96E-04 | glio |
| Interaction_genotype_Atr | Sqstm1 | 1.758 | 4.96E-04 | glio |
| Interaction_genotype_Bcas2 | Skida1 | -4.625 | 4.96E-04 | glio |
| Interaction_genotype_Bcas2 | Gm31518 | -4.69 | 4.96E-04 | glio |
| Interaction_genotype_Chek1 | Ginm1 | 2.078 | 4.96E-04 | glio |
| Interaction_genotype_Rpl3 | Gm11627 | -4.221 | 4.96E-04 | glio |
| interaction_RT | Gtf3a | -0.746 | 4.96E-04 | glio |
| interaction_RT | Snx3 | 0.519 | 4.96E-04 | glio |
| Aars2_in.vivo_noRT | 6330418K02R | 2.056 | 4.96E-04 | glio |
| Bcas2_ex.vivo_noRT | A330049N07R | 2.342 | 4.96E-04 | glio |
| Bcas2_ex.vivo_noRT | Gm38163 | 2.742 | 4.96E-04 | glio |
| Chek1_in.vivo_noRT | Acad10 | 2.538 | 4.96E-04 | glio |
| culture_effect | Tcof1 | 0.622 | 4.96E-05 | glio |
| Interaction_genotype_Mtor | Gm15853 | -4.948 | 4.96E-05 | glio |
| Interaction_genotype_Mtor | Gm2449 | -4.14 | 4.96E-05 | glio |
| Interaction_genotype_Myc | Ccdc69 | -4.581 | 4.96E-05 | glio |
| Bora_in.vivo_noRT | Rnf144b | 2.879 | 4.96E-05 | glio |
| Chek1_in.vivo_noRT | Fbxo24 | 2.854 | 4.96E-05 | glio |
| Hsd17b10_in.vivo_noRT | Wnt16 | 2.825 | 4.96E-05 | glio |
| Aars2_in.vivo_noRT | Tnp2 | 3.264 | 4.96E-06 | glio |
| Aars2_in.vivo_noRT | Best1 | 2.901 | 4.96E-06 | glio |
| Bora_ex.vivo_noRT | Gm13856 | 2.808 | 4.96E-06 | glio |
| Bora_ex.vivo_noRT | Gm45406 | 3.201 | 4.96E-06 | glio |
| culture_effect | Mia3 | -0.845 | 4.96E-07 | glio |
| Hsd17b10_in.vivo_noRT | Gm50115 | 3.353 | 4.96E-07 | glio |
| culture_effect | Rps17 | -0.687 | 4.96E-08 | glio |
| Interaction_genotype_Myc | 2010320O07R | -6.101 | 4.96E-08 | glio |
| Interaction_genotype_Myc | Gm19276 | -4.534 | 4.96E-08 | glio |
| Aars2_in.vivo_noRT | Gstp3 | 4.027 | 4.96E-09 | glio |
| Bcas2_in.vivo_noRT | Col19a1 | 3.89 | 4.96E-13 | glio |
| Chek1_in.vivo_noRT | BC147527 | 5.044 | 4.96E-15 | glio |
| Bora_in.vivo_noRT | Klk13 | 4.404 | 4.96E-18 | glio |
| Interaction_genotype_Atr | Plekha4 | -3.371 | 4.97E-03 | glio |
| Interaction_genotype_Myc | Egf | -3.334 | 4.97E-03 | glio |
| Interaction_genotype_Myc | Eno2 | -2.927 | 4.97E-03 | glio |
| interaction_RT | Ier5 | 1.318 | 4.97E-03 | glio |
| Aars2_ex.vivo_noRT | Gm44899 | 2.188 | 4.97E-03 | glio |
| Aars2_ex.vivo_noRT | B3gntl1 | 2.605 | 4.97E-03 | glio |
| Bard1_ex.vivo_noRT | D630024D03R | 2.227 | 4.97E-03 | glio |
| Bora_in.vivo_noRT | H2-DMb2 | 1.969 | 4.97E-03 | glio |
| Bora_in.vivo_noRT | D10Wsu102e | -1.222 | 4.97E-03 | glio |
| Bora_in.vivo_noRT | Rnf4 | -0.823 | 4.97E-03 | glio |
| Cenpj_in.vivo_noRT | Masp1 | 2.026 | 4.97E-03 | glio |
| Cenpj_in.vivo_noRT | Spink8 | 1.762 | 4.97E-03 | glio |
| culture_effect | Smim11 | 0.463 | 4.97E-04 | glio |

| | | | | |
|---|---|---|---|---|
| culture_effect | Tipin | 0.53 | 4.97E-04 | glio |
| culture_effect | Etnk1 | -0.581 | 4.97E-04 | glio |
| Interaction_genotype_Aars2 | Gm17597 | -4.603 | 4.97E-04 | glio |
| Interaction_genotype_Mtor | Gm14418 | -2.852 | 4.97E-04 | glio |
| interaction_RT | Tulp3 | -0.812 | 4.97E-04 | glio |
| Atr_in.vivo_noRT | Fam160a2 | 2.095 | 4.97E-04 | glio |
| Atr_in.vivo_noRT | Rbfa | -1.25 | 4.97E-04 | glio |
| Bcas2_in.vivo_noRT | Gm10501 | 2.665 | 4.97E-04 | glio |
| Bora_in.vivo_noRT | Ankrd42 | 2.553 | 4.97E-04 | glio |
| Bora_in.vivo_noRT | Wnk2 | 2.545 | 4.97E-04 | glio |
| Bora_ex.vivo_noRT | Bpifa1 | 3.201 | 4.97E-04 | glio |
| Bora_ex.vivo_noRT | Napsa | 3.6 | 4.97E-04 | glio |
| Bora_ex.vivo_noRT | Rasl12 | 3.201 | 4.97E-04 | glio |
| Cenpj_ex.vivo_noRT | Gm13178 | 2.591 | 4.97E-04 | glio |
| Chek1_ex.vivo_noRT | Ckmt1 | 3.035 | 4.97E-04 | glio |
| Interaction_genotype_Atr | Gm44752 | -4.546 | 4.97E-05 | glio |
| Interaction_genotype_Mtor | Gm34256 | -4.998 | 4.97E-05 | glio |
| Interaction_genotype_Mtor | Rilp | -5.142 | 4.97E-05 | glio |
| Interaction_genotype_Mtor | Gm16048 | -5.289 | 4.97E-05 | glio |
| Interaction_genotype_Myc | Atp6v1e1 | 1.734 | 4.97E-05 | glio |
| Bcas2_in.vivo_noRT | 3110082J24Ri | 3.365 | 4.97E-06 | glio |
| Bora_in.vivo_noRT | Gm36913 | 3.247 | 4.97E-06 | glio |
| Atr_in.vivo_noRT | C2cd2 | 3.624 | 4.97E-07 | glio |
| culture_effect | Gm28424 | 2.729 | 4.97E-08 | glio |
| culture_effect | Timm10b | 0.646 | 4.97E-08 | glio |
| Interaction_genotype_Mtor | Card10 | -6.5 | 4.97E-08 | glio |
| Interaction_genotype_Mtor | Tecrl | -6.824 | 4.97E-08 | glio |
| Interaction_genotype_Mtor | Gm10354 | -5.394 | 4.97E-08 | glio |
| Atr_in.vivo_noRT | Pfkfb2 | 3.897 | 4.97E-08 | glio |
| Atr_in.vivo_noRT | Dhrs13os | 4.207 | 4.97E-08 | glio |
| Interaction_genotype_Mtor | Rnf165 | -7.955 | 4.97E-11 | glio |
| Aars2_in.vivo_noRT | Gm49312 | 4.253 | 4.97E-11 | glio |
| Atr_in.vivo_noRT | Gm15941 | 4.114 | 4.97E-11 | glio |
| Aars2_in.vivo_noRT | Gm17146 | 4.409 | 4.97E-12 | glio |
| Hsd17b10_in.vivo_noRT | Gpc5 | 3.616 | 4.97E-13 | glio |
| Atr_in.vivo_noRT | Gm15326 | 5.004 | 4.97E-22 | glio |
| culture_effect | Gen1 | 0.927 | 4.98E-03 | glio |
| Interaction_genotype_Atr | Gm31717 | -2.369 | 4.98E-03 | glio |
| Interaction_genotype_Atr | Calhm5 | -3.243 | 4.98E-03 | glio |
| Interaction_genotype_Ifnar1 | 1700003E24Ri | 2.734 | 4.98E-03 | glio |
| Interaction_genotype_Mtor | Chtop | 1.473 | 4.98E-03 | glio |
| Aars2_in.vivo_noRT | Rhbdl1 | 1.963 | 4.98E-03 | glio |
| Aars2_in.vivo_noRT | Thbs3 | 1.868 | 4.98E-03 | glio |
| Atp5c1_in.vivo_noRT | Fxyd3 | -1.694 | 4.98E-03 | glio |
| Atr_in.vivo_noRT | Kif14 | 1.652 | 4.98E-03 | glio |
| Bcas2_in.vivo_noRT | Cfap300 | 1.623 | 4.98E-03 | glio |
| Bora_in.vivo_noRT | E130307A14Ri | 1.905 | 4.98E-03 | glio |
| Bora_in.vivo_noRT | Zfp37 | 2 | 4.98E-03 | glio |
| Bora_in.vivo_noRT | Cnot10 | -1.226 | 4.98E-03 | glio |
| Bora_in.vivo_noRT | Snx30 | 1.364 | 4.98E-03 | glio |
| Bora_in.vivo_noRT | Lsm3 | -1.034 | 4.98E-03 | glio |
| Cenpj_in.vivo_noRT | Ccser1 | 2.022 | 4.98E-03 | glio |

| | | | | |
|---|---|---|---|---|
| Cyp19a1_ex.vivo_noRT | Gm40264 | 2.701 | 4.98E-03 | glio |
| Cyp19a1_ex.vivo_noRT | Ablim2 | 2.159 | 4.98E-03 | glio |
| Cyp19a1_ex.vivo_noRT | Nrip2 | 2.159 | 4.98E-03 | glio |
| Cyp19a1_ex.vivo_noRT | Gm5415 | 2.159 | 4.98E-03 | glio |
| Cyp19a1_ex.vivo_noRT | Insrr | 2.933 | 4.98E-03 | glio |
| Cyp19a1_ex.vivo_noRT | Slxl1 | 3.107 | 4.98E-03 | glio |
| Cyp19a1_ex.vivo_noRT | Gm49698 | 2.159 | 4.98E-03 | glio |
| Cyp19a1_ex.vivo_noRT | Olfr881 | 2.159 | 4.98E-03 | glio |
| culture_effect | Ldlrad3 | 1.526 | 4.98E-04 | glio |
| Interaction_genotype_Myc | Aoc1 | -4.148 | 4.98E-04 | glio |
| Interaction_genotype_Myc | D430018E03R | -3.778 | 4.98E-04 | glio |
| Aars2_in.vivo_noRT | Ly6g6d | 2.496 | 4.98E-04 | glio |
| Bora_in.vivo_noRT | Gm14273 | 2.446 | 4.98E-04 | glio |
| Chek1_in.vivo_noRT | Gm11131 | 2.426 | 4.98E-04 | glio |
| Chek1_in.vivo_noRT | Adamts4 | 2.232 | 4.98E-04 | glio |
| Bard1_ex.vivo_noRT | Gm12800 | 2.814 | 4.98E-05 | glio |
| Chek1_in.vivo_noRT | 1110013H19R | 3.172 | 4.98E-05 | glio |
| Interaction_genotype_Myc | St13 | 1.906 | 4.98E-06 | glio |
| Atr_ex.vivo_noRT | Gm11899 | 3.164 | 4.98E-06 | glio |
| culture_effect | Ndufa5 | 0.626 | 4.98E-07 | glio |
| Atr_in.vivo_noRT | Kcnb1 | 3.669 | 4.98E-07 | glio |
| Chek1_in.vivo_noRT | Gm40040 | 4.011 | 4.98E-08 | glio |
| Interaction_genotype_Mtor | mt-Co3 | 2.804 | 4.98E-09 | glio |
| Chek1_ex.vivo_noRT | Gm5431 | 4.842 | 4.98E-09 | glio |
| Hsd17b10_in.vivo_noRT | 1700024I08Ri | 3.356 | 4.98E-09 | glio |
| Bcas2_in.vivo_noRT | Gm35290 | 4.069 | 4.98E-10 | glio |
| Chek1_in.vivo_noRT | Amhr2 | 4.91 | 4.98E-10 | glio |
| Atr_in.vivo_noRT | Inhba | 4.892 | 4.98E-11 | glio |
| Atr_in.vivo_noRT | Gm26673 | 4.513 | 4.98E-11 | glio |
| Bcas2_in.vivo_noRT | Iqcn | 4.281 | 4.98E-11 | glio |
| Chek1_in.vivo_noRT | Ms4a6d | 4.256 | 4.98E-13 | glio |
| Chek1_in.vivo_noRT | 4930579C12R | 5.044 | 4.98E-17 | glio |
| culture_effect | Adcy9 | -1.551 | 4.99E-03 | glio |
| Interaction_genotype_Atr | Pyy | -2.866 | 4.99E-03 | glio |
| Interaction_genotype_Bcas2 | Zfp14 | -3.897 | 4.99E-03 | glio |
| Interaction_genotype_Cdkn2a | Tpo | 2.835 | 4.99E-03 | glio |
| Interaction_genotype_Mtor | Maats1 | -3.57 | 4.99E-03 | glio |
| Interaction_genotype_Myc | Rhobtb1 | -3.186 | 4.99E-03 | glio |
| Interaction_genotype_Myc | 2310034G01R | -2.335 | 4.99E-03 | glio |
| interaction_RT | Tmem256 | 0.726 | 4.99E-03 | glio |
| Atp5c1_in.vivo_noRT | Ampd3 | 2.216 | 4.99E-03 | glio |
| Atr_in.vivo_noRT | Tti1 | 1.581 | 4.99E-03 | glio |
| Atr_in.vivo_noRT | Alkbh4 | 1.41 | 4.99E-03 | glio |
| Bora_ex.vivo_noRT | Gm36757 | 2.802 | 4.99E-03 | glio |
| Interaction_genotype_Atr | 1110046J04Ri | -4.398 | 4.99E-04 | glio |
| Interaction_genotype_Atr | Pcbp1 | 1.358 | 4.99E-04 | glio |
| Interaction_genotype_Mtor | A330040F15Ri | -4.339 | 4.99E-04 | glio |
| Interaction_genotype_Mtor | H2-Ab1 | 3.754 | 4.99E-04 | glio |
| Interaction_genotype_Myc | Dnmt3a | -2.629 | 4.99E-04 | glio |
| Interaction_genotype_Myc | Lap3 | 1.937 | 4.99E-04 | glio |
| Aars2_in.vivo_noRT | Pcbp3 | 2.118 | 4.99E-04 | glio |
| Aars2_in.vivo_noRT | Rpl31 | -0.683 | 4.99E-04 | glio |

| | | | | |
|---|---|---|---|---|
| Aars2_ex.vivo_noRT | Plk5 | 2.9 | 4.99E-04 | glio |
| Atp5c1_in.vivo_noRT | Gm13713 | 2.136 | 4.99E-04 | glio |
| Atp5c1_in.vivo_noRT | Mettl7a3 | 2.539 | 4.99E-04 | glio |
| Atp5c1_in.vivo_noRT | Gm19265 | 2.152 | 4.99E-04 | glio |
| Atp5c1_in.vivo_noRT | Rhox3g | 2.616 | 4.99E-04 | glio |
| Bora_in.vivo_noRT | Lgals4 | 2.3 | 4.99E-04 | glio |
| Cenpj_in.vivo_noRT | Gm13964 | 2.022 | 4.99E-04 | glio |
| Aars2_ex.vivo_noRT | 4930423D22R | 2.713 | 4.99E-05 | glio |
| Aars2_ex.vivo_noRT | Gm48265 | 3.293 | 4.99E-05 | glio |
| Atr_ex.vivo_noRT | Amdhd1 | 3.543 | 4.99E-05 | glio |
| Chek1_in.vivo_noRT | 1700113B09R | 2.957 | 4.99E-05 | glio |
| Aars2_in.vivo_noRT | C030005K06R | 3.227 | 4.99E-06 | glio |
| Aars2_ex.vivo_noRT | Vmn2r56 | 3.293 | 4.99E-06 | glio |
| Bcas2_in.vivo_noRT | B230319C09R | 3.497 | 4.99E-06 | glio |
| Bcas2_in.vivo_noRT | Gstt1 | 3.339 | 4.99E-06 | glio |
| Bcas2_in.vivo_noRT | Lgals1 | -1.119 | 4.99E-06 | glio |
| Bora_in.vivo_noRT | Glod5 | 2.436 | 4.99E-06 | glio |
| Chek1_in.vivo_noRT | S100a7a | 3.278 | 4.99E-06 | glio |
| Aars2_in.vivo_noRT | Mturn | 3.246 | 4.99E-07 | glio |
| Chek1_in.vivo_noRT | Gm20492 | 3.452 | 4.99E-07 | glio |
| Chek1_ex.vivo_noRT | Rtl9 | 4.132 | 4.99E-07 | glio |
| Hsd17b10_in.vivo_noRT | Rslcan18 | 3.23 | 4.99E-07 | glio |
| Atr_in.vivo_noRT | Gm5861 | 4.485 | 4.99E-08 | glio |
| Bcas2_in.vivo_noRT | Gm26673 | 3.787 | 4.99E-08 | glio |
| Atr_in.vivo_noRT | Gm2012 | 4.369 | 4.99E-10 | glio |
| Atr_in.vivo_noRT | Hsf4 | 4.346 | 4.99E-10 | glio |
| Bora_in.vivo_noRT | Pacrg | 4.145 | 4.99E-11 | glio |
| Bora_in.vivo_noRT | Scube3 | 4.404 | 4.99E-12 | glio |
| Chek1_in.vivo_noRT | AI463229 | 4.399 | 4.99E-14 | glio |
| Atr_in.vivo_noRT | Rnf165 | 5.533 | 4.99E-16 | glio |
| Interaction_genotype_Bora | Gm30794 | -2.711 | 5.00E-03 | glio |
| Interaction_genotype_Bora | Gm20667 | -3.809 | 5.00E-03 | glio |
| Interaction_genotype_Bora | Gmnn | 1.687 | 5.00E-03 | glio |
| Interaction_genotype_Ifnar1 | Cd7 | 2.998 | 5.00E-03 | glio |
| Interaction_genotype_Mtor | Map3k10 | -2.072 | 5.00E-03 | glio |
| Interaction_genotype_Myc | Des | -3.235 | 5.00E-03 | glio |
| Interaction_genotype_Myc | Plekha4 | -3.036 | 5.00E-03 | glio |
| Interaction_genotype_Myc | Flad1 | -2.456 | 5.00E-03 | glio |
| Interaction_genotype_Rpl3 | Gm11423 | -3.197 | 5.00E-03 | glio |
| interaction_RT | Sap30 | 0.748 | 5.00E-03 | glio |
| Atp5c1_in.vivo_noRT | Speg | 2.342 | 5.00E-03 | glio |
| Atr_in.vivo_noRT | Sh3pxd2b | 1.846 | 5.00E-03 | glio |
| Atr_in.vivo_noRT | C1qbp | -0.657 | 5.00E-03 | glio |
| Chek1_in.vivo_noRT | Cenph | -1.351 | 5.00E-03 | glio |
| culture_effect | Smpd2 | -1.062 | 5.00E-04 | glio |
| Interaction_genotype_Mtor | Ndufb11 | 1.405 | 5.00E-04 | glio |
| Interaction_genotype_Myc | A830082K12R | -3.493 | 5.00E-04 | glio |
| interaction_RT | Arsj | -3.034 | 5.00E-04 | glio |
| interaction_RT | Hist1h1e | -1.393 | 5.00E-04 | glio |
| Atr_in.vivo_noRT | Arhgap9 | 2.081 | 5.00E-04 | glio |
| Bcas2_in.vivo_noRT | Mblac1 | 2.661 | 5.00E-04 | glio |
| Bora_in.vivo_noRT | D830025C05R | 2.631 | 5.00E-04 | glio |

| | | | | |
|---|---|---|---|---|
| Bora_in.vivo_noRT | Gm12043 | 2.343 | 5.00E-04 | glio |
| Eif2s1_in.vivo_noRT | Phldb2 | 2.942 | 5.00E-04 | glio |
| Atp5c1_in.vivo_noRT | Bco2 | 2.668 | 5.00E-05 | glio |
| Bard1_ex.vivo_noRT | Trmt61b | 2.814 | 5.00E-05 | glio |
| Bora_in.vivo_noRT | Platr4 | 2.311 | 5.00E-05 | glio |
| Bora_in.vivo_noRT | Rpgrip1l | 2.202 | 5.00E-05 | glio |
| Interaction_genotype_Myc | Tbc1d32 | -4.005 | 5.00E-06 | glio |
| Interaction_genotype_Myc | St8sia1 | -5.446 | 5.00E-06 | glio |
| Aars2_in.vivo_noRT | AA386476 | 3.297 | 5.00E-06 | glio |
| Brca2_in.vivo_noRT | H19 | 2.717 | 5.00E-07 | glio |
| Chek1_in.vivo_noRT | Gm12500 | 3.402 | 5.00E-07 | glio |
| Aars2_in.vivo_noRT | Appbp2os | 4.264 | 5.00E-09 | glio |
| Aars2_in.vivo_noRT | Isl1 | 4.523 | 5.00E-11 | glio |
| Chek1_ex.vivo_noRT | Fam189a1 | 4.842 | 5.00E-11 | glio |
| Chek1_ex.vivo_noRT | Cfb | 4.842 | 5.00E-12 | glio |
| Bcas2_in.vivo_noRT | Spi1 | 4.807 | 5.00E-14 | glio |
| Hsd17b10_in.vivo_noRT | Sema6d | 4.376 | 5.00E-15 | glio |
| Bcas2_in.vivo_noRT | B230206H07R | 4.281 | 5.00E-19 | glio |
| culture_effect | Ccz1 | 0.424 | 5.01E-03 | glio |
| culture_effect | Imp3 | -0.402 | 5.01E-03 | glio |
| culture_effect | Srpr | -0.36 | 5.01E-03 | glio |
| Interaction_genotype_Atr | Slc45a3 | -3.497 | 5.01E-03 | glio |
| Interaction_genotype_Atr | Zfp458 | -3.663 | 5.01E-03 | glio |
| Interaction_genotype_Atr | Txndc17 | 1.063 | 5.01E-03 | glio |
| Interaction_genotype_Bora | Gm43701 | -2.717 | 5.01E-03 | glio |
| Interaction_genotype_Bora | Spink2 | -4.144 | 5.01E-03 | glio |
| Interaction_genotype_Mtor | Setd4 | -3.3 | 5.01E-03 | glio |
| Interaction_genotype_Mtor | Ssc5d | -3.221 | 5.01E-03 | glio |
| Interaction_genotype_Mtor | Gpr161 | -3.451 | 5.01E-03 | glio |
| Interaction_genotype_Mtor | Endod1 | -3.309 | 5.01E-03 | glio |
| Interaction_genotype_Mtor | Gm12758 | -3.225 | 5.01E-03 | glio |
| Interaction_genotype_Mtor | U2af1 | 1.343 | 5.01E-03 | glio |
| Interaction_genotype_Mtor | Nap1l1 | 1.176 | 5.01E-03 | glio |
| Interaction_genotype_Myc | Cfh | -2.349 | 5.01E-03 | glio |
| Interaction_genotype_Myc | Gm35696 | -2.946 | 5.01E-03 | glio |
| Interaction_genotype_Myc | Rab6b | -2.848 | 5.01E-03 | glio |
| Interaction_genotype_Myc | Ccdc73 | -2.726 | 5.01E-03 | glio |
| Interaction_genotype_Myc | Lrrc27 | -2.665 | 5.01E-03 | glio |
| Interaction_genotype_Myc | Hibch | -2.39 | 5.01E-03 | glio |
| Interaction_genotype_Myc | Asb1 | -2.629 | 5.01E-03 | glio |
| interaction_RT | Herpud1 | 0.847 | 5.01E-03 | glio |
| Aars2_in.vivo_noRT | Gm48646 | 1.975 | 5.01E-03 | glio |
| Aars2_in.vivo_noRT | Heca | 1.305 | 5.01E-03 | glio |
| Aars2_in.vivo_noRT | Rer1 | -0.682 | 5.01E-03 | glio |
| Atr_in.vivo_noRT | Sgtb | 1.97 | 5.01E-03 | glio |
| Atr_in.vivo_noRT | Ak3 | 1.328 | 5.01E-03 | glio |
| Atr_in.vivo_noRT | Osbpl8 | 1.29 | 5.01E-03 | glio |
| Bcas2_in.vivo_noRT | Rgl1 | 1.892 | 5.01E-03 | glio |
| Bcas2_in.vivo_noRT | Irs4 | 2.226 | 5.01E-03 | glio |
| Bcas2_in.vivo_noRT | Nol6 | -1.287 | 5.01E-03 | glio |
| Bcas2_in.vivo_noRT | Poldip3 | -1.08 | 5.01E-03 | glio |
| Bcas2_ex.vivo_noRT | Ccdc63 | 2.813 | 5.01E-03 | glio |

| | | | | |
|---|---|---|---|---|
| Bora_in.vivo_noRT | Gm50240 | 1.624 | 5.01E-03 | glio |
| Bora_in.vivo_noRT | Srek1ip1 | -1.306 | 5.01E-03 | glio |
| Cenpj_in.vivo_noRT | Vtn | 1.762 | 5.01E-03 | glio |
| Cyp19a1_ex.vivo_noRT | BC061195 | 2.159 | 5.01E-03 | glio |
| Cyp19a1_ex.vivo_noRT | 1700034I23Ri | 2.161 | 5.01E-03 | glio |
| Cyp19a1_ex.vivo_noRT | Idi2 | 2.159 | 5.01E-03 | glio |
| Cyp19a1_ex.vivo_noRT | Vmn2r18 | 2.159 | 5.01E-03 | glio |
| Cyp19a1_ex.vivo_noRT | Gm16178 | 2.159 | 5.01E-03 | glio |
| Cyp19a1_ex.vivo_noRT | Gm364 | 2.159 | 5.01E-03 | glio |
| Cyp19a1_ex.vivo_noRT | Prag1 | -2.81 | 5.01E-03 | glio |
| Hsd17b10_in.vivo_noRT | Speg | 2.239 | 5.01E-03 | glio |
| Hsd17b10_in.vivo_noRT | Smc2os | 2.22 | 5.01E-03 | glio |
| Hsd17b10_in.vivo_noRT | Rab5c | -0.779 | 5.01E-03 | glio |
| Hsd17b10_in.vivo_noRT | Col9a1 | -0.989 | 5.01E-03 | glio |
| Interaction_genotype_Aars2 | Mrps18a | 1.469 | 5.01E-04 | glio |
| Atr_in.vivo_noRT | Fbxo48 | 2.513 | 5.01E-04 | glio |
| Bora_ex.vivo_noRT | Scgb1c1 | 2.808 | 5.01E-04 | glio |
| Hsd17b10_in.vivo_noRT | 1700023H06R | 2.825 | 5.01E-04 | glio |
| Aars2_in.vivo_noRT | Moap1 | 2.615 | 5.01E-05 | glio |
| Aars2_in.vivo_noRT | Mpp2 | 2.861 | 5.01E-05 | glio |
| Aars2_in.vivo_noRT | Clec16a | 2.618 | 5.01E-05 | glio |
| Aars2_in.vivo_noRT | Snx17 | -1.34 | 5.01E-05 | glio |
| Atr_in.vivo_noRT | Hist1h2bp | 3.009 | 5.01E-05 | glio |
| Bcas2_in.vivo_noRT | Tead3 | 2.868 | 5.01E-05 | glio |
| Chek1_in.vivo_noRT | Sdha | -1.952 | 5.01E-05 | glio |
| Aars2_in.vivo_noRT | 4930458D05R | 3.526 | 5.01E-06 | glio |
| Atr_ex.vivo_noRT | Gm21936 | 2.769 | 5.01E-06 | glio |
| Atr_ex.vivo_noRT | 4930588A03R | 2.764 | 5.01E-06 | glio |
| Aars2_in.vivo_noRT | 4933433G15R | 3.661 | 5.01E-07 | glio |
| Atp5c1_in.vivo_noRT | Gm41307 | 2.668 | 5.01E-07 | glio |
| Atr_in.vivo_noRT | Cep72 | 3.412 | 5.01E-07 | glio |
| Atr_in.vivo_noRT | Rpl26 | -0.946 | 5.01E-07 | glio |
| Atr_in.vivo_noRT | Car10 | 3.19 | 5.01E-08 | glio |
| Atr_in.vivo_noRT | Slc36a2 | 4.414 | 5.01E-11 | glio |
| Aars2_in.vivo_noRT | Cd248 | 5.06 | 5.01E-14 | glio |
| Aars2_in.vivo_noRT | Vwce | 5.067 | 5.01E-16 | glio |
| Atr_in.vivo_noRT | AF067061 | 5.533 | 5.01E-20 | glio |
| Chek1_in.vivo_noRT | Olfr1152 | 4.777 | 5.01E-23 | glio |
| Interaction_genotype_Aars2 | Vmn1r32 | -2.427 | 5.02E-03 | glio |
| Interaction_genotype_Atr | Leng1 | 1.709 | 5.02E-03 | glio |
| Interaction_genotype_Atr | Chkb | 1.562 | 5.02E-03 | glio |
| Interaction_genotype_Mtor | Gm16124 | -3.844 | 5.02E-03 | glio |
| Interaction_genotype_Mtor | 1700056E22Ri | -2.594 | 5.02E-03 | glio |
| Interaction_genotype_Myc | Gm49042 | -2.986 | 5.02E-03 | glio |
| Interaction_genotype_Myc | Zfp729b | -2.793 | 5.02E-03 | glio |
| Interaction_genotype_Myc | Mboat1 | -2.158 | 5.02E-03 | glio |
| Interaction_genotype_Pars2 | Akap3 | -3.859 | 5.02E-03 | glio |
| Atr_in.vivo_noRT | Rwdd4a | -0.944 | 5.02E-03 | glio |
| Atr_ex.vivo_noRT | Cspg4 | -2.141 | 5.02E-03 | glio |
| Bard1_ex.vivo_noRT | BB557941 | 2.415 | 5.02E-03 | glio |
| Bcas2_in.vivo_noRT | Nptn | -1.029 | 5.02E-03 | glio |
| Bcas2_ex.vivo_noRT | Pla1a | 2.742 | 5.02E-03 | glio |

| | | | | |
|---|---|---|---|---|
| Bcas2_ex.vivo_noRT | Insrr | 2.742 | 5.02E-03 | glio |
| Bora_in.vivo_noRT | 4933404O12R | 2.031 | 5.02E-03 | glio |
| Bora_ex.vivo_noRT | Rgs9 | 2.411 | 5.02E-03 | glio |
| Brca2_in.vivo_noRT | Hoxc8 | 1.763 | 5.02E-03 | glio |
| Eif2s1_ex.vivo_noRT | Gm15533 | 2.234 | 5.02E-03 | glio |
| Fanca_in.vivo_noRT | Cytl1 | 2.04 | 5.02E-03 | glio |
| Hsd17b10_in.vivo_noRT | Gm42463 | 1.984 | 5.02E-03 | glio |
| Interaction_genotype_Mtor | Irx2 | -3.39 | 5.02E-04 | glio |
| Atr_in.vivo_noRT | Gm12758 | 2.426 | 5.02E-04 | glio |
| Atr_in.vivo_noRT | Mcm2 | -1.66 | 5.02E-04 | glio |
| Bcas2_in.vivo_noRT | Cnbd2 | 2.661 | 5.02E-04 | glio |
| Bcas2_in.vivo_noRT | Hddc3 | 2.424 | 5.02E-04 | glio |
| Bcas2_in.vivo_noRT | Car7 | 2.487 | 5.02E-04 | glio |
| Bcas2_ex.vivo_noRT | Trim17 | 2.342 | 5.02E-04 | glio |
| culture_effect | Rnf122 | -1.92 | 5.02E-05 | glio |
| Atr_in.vivo_noRT | Mmp23 | 2.735 | 5.02E-05 | glio |
| Bcas2_in.vivo_noRT | Slc24a1 | 2.945 | 5.02E-05 | glio |
| Bcas2_in.vivo_noRT | Astn2 | 2.719 | 5.02E-05 | glio |
| Chek1_in.vivo_noRT | C1qtnf12 | 3.196 | 5.02E-05 | glio |
| Chek1_in.vivo_noRT | Gm50322 | 3.148 | 5.02E-05 | glio |
| culture_effect | Gpr19 | 1.045 | 5.02E-06 | glio |
| interaction_RT | Ramp2 | 2.097 | 5.02E-06 | glio |
| culture_effect | Mag | -3.034 | 5.02E-07 | glio |
| Interaction_genotype_Bcas2 | Mecom | -6.437 | 5.02E-08 | glio |
| Interaction_genotype_Bcas2 | Hbq1b | -6.719 | 5.02E-08 | glio |
| Bora_ex.vivo_noRT | Sucnr1 | 3.201 | 5.02E-08 | glio |
| Atr_in.vivo_noRT | Grasp | -3.514 | 5.02E-09 | glio |
| Bcas2_in.vivo_noRT | Tnfsf13os | 4.036 | 5.02E-09 | glio |
| Bcas2_in.vivo_noRT | Rims3 | 3.673 | 5.02E-09 | glio |
| Chek1_in.vivo_noRT | C4a | 3.605 | 5.02E-09 | glio |
| Interaction_genotype_Myc | Col6a2 | -6.607 | 5.02E-12 | glio |
| Chek1_in.vivo_noRT | Ccdc63 | 4.658 | 5.02E-12 | glio |
| culture_effect | Cdc34 | 0.877 | 5.02E-13 | glio |
| Chek1_in.vivo_noRT | Gm47936 | 5.044 | 5.02E-21 | glio |
| Chek1_in.vivo_noRT | Gm13191 | 5.044 | 5.02E-27 | glio |
| Chek1_in.vivo_noRT | 4933416E03Ri | 5.044 | 5.02E-27 | glio |
| culture_effect | Sfxn4 | -0.929 | 5.03E-03 | glio |
| Interaction_genotype_Mtor | Glrp1 | -3.02 | 5.03E-03 | glio |
| Interaction_genotype_Mtor | N4bp2l1 | -3.558 | 5.03E-03 | glio |
| Interaction_genotype_Mtor | Gstm2 | -3.166 | 5.03E-03 | glio |
| Interaction_genotype_Myc | Tpbg | -3.157 | 5.03E-03 | glio |
| interaction_RT | Pcp4l1 | 2.377 | 5.03E-03 | glio |
| interaction_RT | Unc93b1 | 1.334 | 5.03E-03 | glio |
| interaction_RT | Ramac | 0.692 | 5.03E-03 | glio |
| Aars2_ex.vivo_noRT | Neurl3 | 2.893 | 5.03E-03 | glio |
| Atr_in.vivo_noRT | Mrpl54 | -0.726 | 5.03E-03 | glio |
| Bcas2_in.vivo_noRT | 1500004A13R | 2.111 | 5.03E-03 | glio |
| Bcas2_ex.vivo_noRT | Rbpms2 | -2.821 | 5.03E-03 | glio |
| Bora_in.vivo_noRT | E130114P18Ri | 2.184 | 5.03E-03 | glio |
| Bora_in.vivo_noRT | Ptpmt1 | -0.909 | 5.03E-03 | glio |
| Bora_ex.vivo_noRT | Usp17la | 2.806 | 5.03E-03 | glio |
| Bora_ex.vivo_noRT | Cited4 | 3.013 | 5.03E-03 | glio |

| | | | | |
|---|---|---|---|---|
| Chek1_ex.vivo_noRT | 4930540M05F | 2.469 | 5.03E-03 | glio |
| Atp5c1_in.vivo_noRT | C430042M11F | 2.72 | 5.03E-04 | glio |
| Cenpj_ex.vivo_noRT | Esp24 | 2.591 | 5.03E-04 | glio |
| Hsd17b10_in.vivo_noRT | Pglyrp1 | 2.664 | 5.03E-04 | glio |
| Hsd17b10_in.vivo_noRT | Gpx2 | -2.953 | 5.03E-04 | glio |
| Interaction_genotype_Myc | Kcnab2 | -4.25 | 5.03E-05 | glio |
| Interaction_genotype_Myc | Gm20711 | -3.92 | 5.03E-05 | glio |
| Chek1_in.vivo_noRT | Nupr1l | 3.263 | 5.03E-05 | glio |
| Interaction_genotype_Mtor | 4732419C18R | -5.346 | 5.03E-06 | glio |
| Atp5c1_in.vivo_noRT | Speer4e | 2.668 | 5.03E-06 | glio |
| Bora_in.vivo_noRT | Icosl | 3.17 | 5.03E-07 | glio |
| Aars2_in.vivo_noRT | Bcl7c | -1.768 | 5.03E-08 | glio |
| Atr_in.vivo_noRT | Fam222a | 4.019 | 5.03E-08 | glio |
| Bora_in.vivo_noRT | Gchfr | 4.347 | 5.03E-08 | glio |
| Bora_in.vivo_noRT | Ccno | 3.916 | 5.03E-08 | glio |
| Chek1_in.vivo_noRT | Rhox3g | 3.929 | 5.03E-09 | glio |
| Bora_in.vivo_noRT | Gm26674 | 4.291 | 5.03E-11 | glio |
| Chek1_in.vivo_noRT | Hsf2bp | 4.252 | 5.03E-11 | glio |
| Aars2_in.vivo_noRT | Amdhd1 | 4.793 | 5.03E-13 | glio |
| Bora_in.vivo_noRT | Tent5c | 4.01 | 5.03E-14 | glio |
| Bcas2_in.vivo_noRT | Pakap | 4.275 | 5.03E-15 | glio |
| Bora_in.vivo_noRT | Gm39526 | 4.414 | 5.03E-16 | glio |
| Aars2_in.vivo_noRT | Gm14820 | 5.063 | 5.03E-22 | glio |
| Chek1_in.vivo_noRT | Gm266 | 5.044 | 5.03E-23 | glio |
| culture_effect | Gga2 | -0.728 | 5.04E-03 | glio |
| Interaction_genotype_Myc | Fkbp1a | 0.96 | 5.04E-03 | glio |
| Interaction_genotype_Pars2 | Clec12b | -2.664 | 5.04E-03 | glio |
| Interaction_genotype_Pars2 | Slc17a6 | -2.268 | 5.04E-03 | glio |
| Interaction_genotype_Pars2 | Ccr5 | -3.56 | 5.04E-03 | glio |
| Interaction_genotype_Pars2 | Gm48682 | -3.481 | 5.04E-03 | glio |
| Interaction_genotype_Pars2 | Gm10399 | -3.141 | 5.04E-03 | glio |
| interaction_RT | Oser1 | 0.826 | 5.04E-03 | glio |
| Bcas2_ex.vivo_noRT | Gm13032 | 2.342 | 5.04E-03 | glio |
| Bcas2_ex.vivo_noRT | Gm50223 | 2.349 | 5.04E-03 | glio |
| Chek1_in.vivo_noRT | Mturn | 1.918 | 5.04E-03 | glio |
| Chek1_in.vivo_noRT | Brd3os | 1.915 | 5.04E-03 | glio |
| culture_effect | Pola2 | -0.685 | 5.04E-04 | glio |
| Interaction_genotype_Aars2 | Gm16364.1 | -4.67 | 5.04E-04 | glio |
| Interaction_genotype_Aars2 | Ybx3 | 1.23 | 5.04E-04 | glio |
| Interaction_genotype_Bcas2 | Carns1 | -4.636 | 5.04E-04 | glio |
| Aars2_in.vivo_noRT | Tlcd2 | 2.771 | 5.04E-04 | glio |
| Aars2_in.vivo_noRT | Rps2 | -0.78 | 5.04E-04 | glio |
| Atp5c1_in.vivo_noRT | 5033417F24Ri | 2.668 | 5.04E-04 | glio |
| Bora_in.vivo_noRT | Rbp7 | 2.544 | 5.04E-04 | glio |
| Bora_in.vivo_noRT | Ucn2 | 2.443 | 5.04E-04 | glio |
| Bora_in.vivo_noRT | Dnaaf2 | -1.5 | 5.04E-04 | glio |
| Chek1_in.vivo_noRT | Gm12758 | 2.46 | 5.04E-04 | glio |
| Eif2s1_in.vivo_noRT | Gm10399 | 2.769 | 5.04E-04 | glio |
| Interaction_genotype_Chek1 | Rnf114 | 3.106 | 5.04E-05 | glio |
| Interaction_genotype_Chek1 | Zfand5 | 1.694 | 5.04E-05 | glio |
| Interaction_genotype_Chek1 | Gabarap | 1.406 | 5.04E-05 | glio |
| Interaction_genotype_Mtor | Gm47061 | -5.15 | 5.04E-05 | glio |

| | | | | |
|---|---|---|---|---|
| Interaction_genotype_Mtor | 1700006H21R | -4.886 | 5.04E-05 | glio |
| Interaction_genotype_Mtor | Dync2li1 | -3.534 | 5.04E-05 | glio |
| Interaction_genotype_Myc | Mars2 | -3.441 | 5.04E-05 | glio |
| Interaction_genotype_Myc | Hnf1aos1 | -4.069 | 5.04E-05 | glio |
| Atp5c1_in.vivo_noRT | 4930447F24Ri | 2.939 | 5.04E-05 | glio |
| Atp5c1_in.vivo_noRT | Stra8 | 2.676 | 5.04E-05 | glio |
| Bcas2_in.vivo_noRT | Upk1a | 3.015 | 5.04E-05 | glio |
| Interaction_genotype_Myc | Gm47484 | -5.517 | 5.04E-06 | glio |
| Interaction_genotype_Myc | Cox6b1 | 1.808 | 5.04E-06 | glio |
| interaction_RT | Mybbp1a | -1.106 | 5.04E-07 | glio |
| Hsd17b10_in.vivo_noRT | Ifitm6 | 3.616 | 5.04E-07 | glio |
| Atr_in.vivo_noRT | Gm50069 | 3.892 | 5.04E-08 | glio |
| Chek1_in.vivo_noRT | 1700011M02F | 4.78 | 5.04E-08 | glio |
| Chek1_ex.vivo_noRT | H2-Eb1 | 4.449 | 5.04E-08 | glio |
| culture_effect | Mrps21 | 0.718 | 5.04E-09 | glio |
| Bora_in.vivo_noRT | Amdhd1 | 3.893 | 5.04E-09 | glio |
| Chek1_ex.vivo_noRT | Gm5662 | 4.842 | 5.04E-09 | glio |
| Chek1_in.vivo_noRT | Gm15234 | 4.528 | 5.04E-11 | glio |
| culture_effect | Rars | -1.093 | 5.04E-13 | glio |
| Bcas2_in.vivo_noRT | Snhg4.1 | 4.284 | 5.04E-13 | glio |
| Chek1_in.vivo_noRT | Ifitm6 | 5.044 | 5.04E-13 | glio |
| Aars2_in.vivo_noRT | A830009L08Ri | 4.804 | 5.04E-14 | glio |
| Aars2_in.vivo_noRT | Scube3 | 4.794 | 5.04E-14 | glio |
| Chek1_in.vivo_noRT | AF067061 | 5.044 | 5.04E-17 | glio |
| Bora_in.vivo_noRT | Gm10324 | 4.67 | 5.04E-18 | glio |
| Chek1_in.vivo_noRT | Tnni3k | 5.044 | 5.04E-21 | glio |
| Chek1_in.vivo_noRT | Gm48170 | 5.573 | 5.04E-23 | glio |
| culture_effect | Gm16094 | -1.653 | 5.05E-03 | glio |
| culture_effect | Map3k20 | 1.029 | 5.05E-03 | glio |
| culture_effect | Map3k10 | 0.665 | 5.05E-03 | glio |
| culture_effect | Gm11808 | -0.417 | 5.05E-03 | glio |
| Interaction_genotype_Aars2 | 1700012D14R | -3.275 | 5.05E-03 | glio |
| Interaction_genotype_Bcas2 | Zfp273 | -3.798 | 5.05E-03 | glio |
| Interaction_genotype_Myc | Kif3c | -2.521 | 5.05E-03 | glio |
| Interaction_genotype_Myc | Gxylt2 | -2.396 | 5.05E-03 | glio |
| interaction_RT | 1810037I17Ri | 0.531 | 5.05E-03 | glio |
| Aars2_in.vivo_noRT | Tcf7l1 | 1.729 | 5.05E-03 | glio |
| Aars2_in.vivo_noRT | H2afz | -0.635 | 5.05E-03 | glio |
| Aars2_ex.vivo_noRT | A930033M14F | 2.328 | 5.05E-03 | glio |
| Aars2_ex.vivo_noRT | B230311B06R | 2.935 | 5.05E-03 | glio |
| Atr_ex.vivo_noRT | Pde6b | 2.37 | 5.05E-03 | glio |
| Atr_ex.vivo_noRT | Hrk | 2.647 | 5.05E-03 | glio |
| Bcas2_in.vivo_noRT | Tmem171 | 2.653 | 5.05E-03 | glio |
| Bcas2_in.vivo_noRT | Tlr2 | 2.126 | 5.05E-03 | glio |
| Bcas2_in.vivo_noRT | Cnksr3 | 1.527 | 5.05E-03 | glio |
| Bcas2_in.vivo_noRT | Cdc5l | -0.846 | 5.05E-03 | glio |
| Bora_in.vivo_noRT | Gm10825 | 2.107 | 5.05E-03 | glio |
| Bora_in.vivo_noRT | Lrrfip2 | -1.195 | 5.05E-03 | glio |
| Bora_ex.vivo_noRT | Elfn2 | 2.626 | 5.05E-03 | glio |
| Cenpj_in.vivo_noRT | Gm15890 | 2.022 | 5.05E-03 | glio |
| Cenpj_in.vivo_noRT | Snrpn | 1.89 | 5.05E-03 | glio |
| Chek1_in.vivo_noRT | Psmf1 | -1.29 | 5.05E-03 | glio |

| | | | | |
|---|---|---|---|---|
| Chek1_in.vivo_noRT | H2-K1 | -0.729 | 5.05E-03 | glio |
| Interaction_genotype_Aars2 | Phf10 | 2.126 | 5.05E-04 | glio |
| Interaction_genotype_Mtor | Nmb | -4.129 | 5.05E-04 | glio |
| Interaction_genotype_Mtor | Mettl5os | -3.622 | 5.05E-04 | glio |
| interaction_RT | Bhlhe40 | 2.18 | 5.05E-04 | glio |
| Atr_ex.vivo_noRT | Hist1h2bf | 3.505 | 5.05E-04 | glio |
| Atr_ex.vivo_noRT | Map7 | 3.356 | 5.05E-04 | glio |
| Aars2_ex.vivo_noRT | Gm34776 | 2.899 | 5.05E-05 | glio |
| Atr_ex.vivo_noRT | Apol10b | 3.164 | 5.05E-05 | glio |
| Bard1_ex.vivo_noRT | Gm5890 | 2.814 | 5.05E-05 | glio |
| Bard1_ex.vivo_noRT | Zscan4c | 2.814 | 5.05E-05 | glio |
| Bard1_ex.vivo_noRT | Speer4a | 2.814 | 5.05E-05 | glio |
| Bard1_ex.vivo_noRT | H1foo | 2.814 | 5.05E-05 | glio |
| Bcas2_in.vivo_noRT | D830025C05R | 3.03 | 5.05E-05 | glio |
| Atr_in.vivo_noRT | Rit2 | 3.041 | 5.05E-06 | glio |
| Atr_ex.vivo_noRT | Atp8a2 | 3.959 | 5.05E-07 | glio |
| Atr_in.vivo_noRT | Cacnb1 | 4.408 | 5.05E-09 | glio |
| Atr_in.vivo_noRT | Gm43156 | 4.052 | 5.05E-09 | glio |
| culture_effect | Ak1 | 3.201 | 5.05E-11 | glio |
| Hsd17b10_in.vivo_noRT | Hfm1 | 3.616 | 5.05E-11 | glio |
| Hsd17b10_in.vivo_noRT | D630024D03R | 3.616 | 5.05E-11 | glio |
| Atr_in.vivo_noRT | Spi1 | 5.533 | 5.05E-18 | glio |
| Interaction_genotype_Atr | Ctla4 | -2.369 | 5.06E-03 | glio |
| Interaction_genotype_Atr | Tmem186 | -3.346 | 5.06E-03 | glio |
| Interaction_genotype_Atr | Arl6ip1 | 1.675 | 5.06E-03 | glio |
| Interaction_genotype_Hsd17b | Map7 | 4.649 | 5.06E-03 | glio |
| Interaction_genotype_Mtor | Car7 | -3.347 | 5.06E-03 | glio |
| Interaction_genotype_Mtor | Dlx1 | -2.632 | 5.06E-03 | glio |
| Interaction_genotype_Mtor | Ier3ip1 | 1.564 | 5.06E-03 | glio |
| interaction_RT | Crebrf | 0.847 | 5.06E-03 | glio |
| Aars2_in.vivo_noRT | Aco1 | -1.333 | 5.06E-03 | glio |
| Aars2_in.vivo_noRT | Nras | -1.08 | 5.06E-03 | glio |
| Aars2_in.vivo_noRT | Hnrnpc | -0.688 | 5.06E-03 | glio |
| Atr_in.vivo_noRT | Tlr2 | 2.075 | 5.06E-03 | glio |
| Bora_in.vivo_noRT | Dnaja4 | 2.063 | 5.06E-03 | glio |
| Bora_in.vivo_noRT | Tssc4 | -1.791 | 5.06E-03 | glio |
| Bora_in.vivo_noRT | Ehmt2 | -1.35 | 5.06E-03 | glio |
| Bora_ex.vivo_noRT | Mpeg1 | 2.626 | 5.06E-03 | glio |
| Bora_ex.vivo_noRT | Cfap58 | 3.426 | 5.06E-03 | glio |
| Chek1_in.vivo_noRT | Lrp4 | 2.086 | 5.06E-03 | glio |
| Hsd17b10_in.vivo_noRT | Fam214b | -1.6 | 5.06E-03 | glio |
| Hsd17b10_in.vivo_noRT | Ppp1r9a | 1.697 | 5.06E-03 | glio |
| culture_effect | Zfp324 | 1.294 | 5.06E-04 | glio |
| Interaction_genotype_Mtor | Rpsa | 1.313 | 5.06E-04 | glio |
| Interaction_genotype_Myc | Gm13078 | -2.729 | 5.06E-04 | glio |
| Interaction_genotype_Myc | Upb1 | -3.742 | 5.06E-04 | glio |
| Interaction_genotype_Myc | Enpp1 | -3.627 | 5.06E-04 | glio |
| Interaction_genotype_Myc | Lipt2 | -3.122 | 5.06E-04 | glio |
| Interaction_genotype_Myc | Bcs1l | -2.792 | 5.06E-04 | glio |
| Interaction_genotype_Myc | Rnpepl1 | -2.35 | 5.06E-04 | glio |
| Interaction_genotype_Myc | Sar1a | 1.642 | 5.06E-04 | glio |
| Interaction_genotype_Myc | Calm1 | 1.497 | 5.06E-04 | glio |

| | | | | |
|---|---|---|---|---|
| Atr_in.vivo_noRT | Rps6kl1 | 2.482 | 5.06E-04 | glio |
| Hsd17b10_in.vivo_noRT | Zfp551 | 2.58 | 5.06E-04 | glio |
| Hsd17b10_in.vivo_noRT | Gm49785 | 2.7 | 5.06E-04 | glio |
| culture_effect | Gm15486 | -2.336 | 5.06E-05 | glio |
| culture_effect | 2810454H06R | 1.568 | 5.06E-05 | glio |
| Interaction_genotype_Myc | Gm45833 | -6.03 | 5.06E-05 | glio |
| Interaction_genotype_Myc | Gm9970 | -4.818 | 5.06E-05 | glio |
| Interaction_genotype_Myc | 4930540M05F | -4.445 | 5.06E-05 | glio |
| Interaction_genotype_Myc | Chm | -2.785 | 5.06E-05 | glio |
| Interaction_genotype_Myc | Ssbp1 | 1.68 | 5.06E-05 | glio |
| Aars2_in.vivo_noRT | Vash2 | 2.898 | 5.06E-05 | glio |
| Aars2_in.vivo_noRT | Ddx39b | -1.289 | 5.06E-05 | glio |
| Aars2_ex.vivo_noRT | Car10 | 3.293 | 5.06E-05 | glio |
| Bcas2_in.vivo_noRT | Gm31805 | 2.823 | 5.06E-05 | glio |
| Bcas2_in.vivo_noRT | Mcm3 | -1.835 | 5.06E-05 | glio |
| Bcas2_ex.vivo_noRT | Pcdhga2 | 3.718 | 5.06E-05 | glio |
| Bcas2_ex.vivo_noRT | Gm16933 | 2.742 | 5.06E-05 | glio |
| Bora_in.vivo_noRT | 2810414N06R | 2.914 | 5.06E-05 | glio |
| culture_effect | Slc6a11 | -2.367 | 5.06E-06 | glio |
| culture_effect | Myo1c | 0.875 | 5.06E-06 | glio |
| Atp5c1_in.vivo_noRT | C430002N11R | 2.668 | 5.06E-06 | glio |
| Bcas2_in.vivo_noRT | 1700056N10R | 2.897 | 5.06E-06 | glio |
| Chek1_in.vivo_noRT | Ttc38 | 3.173 | 5.06E-06 | glio |
| Atr_in.vivo_noRT | Gm16835 | 3.606 | 5.06E-07 | glio |
| Atr_in.vivo_noRT | Nkx2-2os | 3.878 | 5.06E-08 | glio |
| Bora_in.vivo_noRT | Vamp8 | 3.113 | 5.06E-08 | glio |
| culture_effect | Tomm6 | 0.684 | 5.06E-09 | glio |
| Aars2_in.vivo_noRT | C230037L18Ri | 4.158 | 5.06E-10 | glio |
| Bcas2_in.vivo_noRT | Gm9967 | 4.42 | 5.06E-11 | glio |
| Bcas2_in.vivo_noRT | Ube4bos1 | 4.543 | 5.06E-15 | glio |
| Bcas2_in.vivo_noRT | Dpp6 | 3.88 | 5.06E-15 | glio |
| culture_effect | Dusp2 | -3.793 | 5.06E-19 | glio |
| culture_effect | Casp4 | -5.422 | 5.06E-27 | glio |
| culture_effect | Il27ra | -1.487 | 5.07E-03 | glio |
| culture_effect | Idua | -1.376 | 5.07E-03 | glio |
| culture_effect | Zfp689 | 1.072 | 5.07E-03 | glio |
| culture_effect | Ash2l | 0.37 | 5.07E-03 | glio |
| Interaction_genotype_Atr | Abat | -3.796 | 5.07E-03 | glio |
| Interaction_genotype_Mtor | Wdr25 | -2.998 | 5.07E-03 | glio |
| Interaction_genotype_Mtor | Tti2 | -2.508 | 5.07E-03 | glio |
| interaction_RT | Gm16897 | 1.323 | 5.07E-03 | glio |
| Atr_in.vivo_noRT | Rsph3b | 1.937 | 5.07E-03 | glio |
| Atr_in.vivo_noRT | Nbr1 | -1.235 | 5.07E-03 | glio |
| Bcas2_in.vivo_noRT | BC028528 | 2.115 | 5.07E-03 | glio |
| Bcas2_in.vivo_noRT | Tdg | -1.204 | 5.07E-03 | glio |
| Bcas2_in.vivo_noRT | Slc25a17 | -1.022 | 5.07E-03 | glio |
| Bora_in.vivo_noRT | Slc15a3 | 1.917 | 5.07E-03 | glio |
| Bora_in.vivo_noRT | Aldh3b1 | 2.065 | 5.07E-03 | glio |
| Chek1_ex.vivo_noRT | Ero1lb | 2.419 | 5.07E-03 | glio |
| Eif2s1_in.vivo_noRT | Minar2 | 1.883 | 5.07E-03 | glio |
| Eif2s1_in.vivo_noRT | Slc6a13 | 2.528 | 5.07E-03 | glio |
| Eif2s1_ex.vivo_noRT | Tmem229b | 3.001 | 5.07E-03 | glio |

| | | | | |
|---|---|---|---|---|
| Eif2s1_ex.vivo_noRT | Gpr4 | 2.234 | 5.07E-03 | glio |
| Eif2s1_ex.vivo_noRT | Gm33466 | 2.234 | 5.07E-03 | glio |
| Eif2s1_ex.vivo_noRT | Msh3 | -2.256 | 5.07E-03 | glio |
| Hsd17b10_in.vivo_noRT | P4hb | -0.508 | 5.07E-03 | glio |
| culture_effect | Glcci1 | -1.348 | 5.07E-04 | glio |
| culture_effect | Mnd1 | -1.326 | 5.07E-04 | glio |
| culture_effect | Srrm1 | 0.459 | 5.07E-04 | glio |
| interaction_RT | Dtx1 | 3.428 | 5.07E-04 | glio |
| Aars2_ex.vivo_noRT | AC125149.2 | 2.897 | 5.07E-04 | glio |
| Atr_in.vivo_noRT | Zfp937 | 2.603 | 5.07E-04 | glio |
| Bcas2_in.vivo_noRT | Hspa12b | 2.753 | 5.07E-04 | glio |
| Bora_in.vivo_noRT | Prkar1b | 2.583 | 5.07E-04 | glio |
| Chek1_ex.vivo_noRT | Inhba | 3.321 | 5.07E-04 | glio |
| Interaction_genotype_Aars2 | Ube2a | 2.281 | 5.07E-05 | glio |
| Interaction_genotype_Aars2 | Gm13288 | -4.169 | 5.07E-05 | glio |
| Interaction_genotype_Mtor | Vcam1 | -6.175 | 5.07E-05 | glio |
| Interaction_genotype_Mtor | Reep3 | 2.036 | 5.07E-05 | glio |
| Chek1_ex.vivo_noRT | Lrfn1 | 3.556 | 5.07E-05 | glio |
| Interaction_genotype_Mtor | B930059L03Ri | -5.628 | 5.07E-06 | glio |
| Interaction_genotype_Myc | Gm29083 | -4.853 | 5.07E-06 | glio |
| Atp5c1_in.vivo_noRT | Cd7 | 2.404 | 5.07E-06 | glio |
| Bora_in.vivo_noRT | Slc12a5 | 3.154 | 5.07E-06 | glio |
| Bora_ex.vivo_noRT | Gm21936 | 2.806 | 5.07E-06 | glio |
| Bora_ex.vivo_noRT | Dmrtc1a | 3.201 | 5.07E-06 | glio |
| Bora_ex.vivo_noRT | 4930588A03R | 2.802 | 5.07E-06 | glio |
| culture_effect | Angptl2 | -1.387 | 5.07E-07 | glio |
| culture_effect | Tmem43 | -1.02 | 5.07E-07 | glio |
| Atr_in.vivo_noRT | Pcp4l1 | 3.575 | 5.07E-07 | glio |
| Atr_ex.vivo_noRT | Gm12703 | 3.164 | 5.07E-07 | glio |
| Atr_ex.vivo_noRT | Mmp13 | 3.164 | 5.07E-07 | glio |
| Bcas2_in.vivo_noRT | Cnnm1 | 3.895 | 5.07E-07 | glio |
| Bcas2_in.vivo_noRT | Drd4 | 3.623 | 5.07E-07 | glio |
| Bora_in.vivo_noRT | Llgl2 | 3.46 | 5.07E-07 | glio |
| Chek1_ex.vivo_noRT | Il12rb1 | 4.055 | 5.07E-07 | glio |
| Aars2_in.vivo_noRT | Fsbp | 3.883 | 5.07E-08 | glio |
| Aars2_in.vivo_noRT | Kcnb1 | 3.976 | 5.07E-08 | glio |
| Atr_in.vivo_noRT | Dagla | 4.037 | 5.07E-08 | glio |
| Hsd17b10_in.vivo_noRT | Gm10482 | 3.094 | 5.07E-08 | glio |
| Interaction_genotype_Mtor | 5330439K02R | -7.303 | 5.07E-09 | glio |
| Interaction_genotype_Mtor | Pyy | -5.979 | 5.07E-09 | glio |
| Chek1_in.vivo_noRT | Esyt3 | 3.984 | 5.07E-09 | glio |
| Bora_in.vivo_noRT | Ggt1 | 4.156 | 5.07E-10 | glio |
| Bora_in.vivo_noRT | H3f3aos | 4.291 | 5.07E-10 | glio |
| Chek1_ex.vivo_noRT | Auts2 | 4.842 | 5.07E-10 | glio |
| Atr_in.vivo_noRT | Tnnt3 | 4.899 | 5.07E-11 | glio |
| Chek1_in.vivo_noRT | Tex52 | 4.393 | 5.07E-11 | glio |
| Bora_in.vivo_noRT | Gm6803 | 4.936 | 5.07E-19 | glio |
| Aars2_in.vivo_noRT | Gm45338 | 5.327 | 5.07E-29 | glio |
| Interaction_genotype_Atr | Zscan4-ps3 | -2.369 | 5.08E-03 | glio |
| Interaction_genotype_Atr | Glod5 | 4.188 | 5.08E-03 | glio |
| Interaction_genotype_Atr | Apol6 | 2.974 | 5.08E-03 | glio |
| Interaction_genotype_Atr | Hcrtr1 | -3.535 | 5.08E-03 | glio |

| | | | | |
|---|---|---|---|---|
| Interaction_genotype_Atr | Rnf181 | 1.534 | 5.08E-03 | glio |
| Interaction_genotype_Bcas2 | 9530053A07R | -3.834 | 5.08E-03 | glio |
| Interaction_genotype_Mtor | B020031M17F | -4.099 | 5.08E-03 | glio |
| Interaction_genotype_Mtor | Hpcal1 | -3.449 | 5.08E-03 | glio |
| Interaction_genotype_Mtor | Gm20939 | -3.119 | 5.08E-03 | glio |
| Interaction_genotype_Mtor | Hhex | -2.806 | 5.08E-03 | glio |
| Interaction_genotype_Mtor | Dolk | -2.398 | 5.08E-03 | glio |
| Interaction_genotype_Mtor | Fkbp2 | 1.454 | 5.08E-03 | glio |
| Interaction_genotype_Myc | Iqch | -2.635 | 5.08E-03 | glio |
| Interaction_genotype_Myc | Gm14443 | -2.888 | 5.08E-03 | glio |
| Interaction_genotype_Myc | Mtr | -2.442 | 5.08E-03 | glio |
| Interaction_genotype_Myc | Mrm2 | -2.178 | 5.08E-03 | glio |
| Interaction_genotype_Myc | Chchd10 | 2.689 | 5.08E-03 | glio |
| Interaction_genotype_Myc | Denr | 1.125 | 5.08E-03 | glio |
| Interaction_genotype_Myc | Larp4 | 1.125 | 5.08E-03 | glio |
| Interaction_genotype_Myc | Brd2 | 1.271 | 5.08E-03 | glio |
| Aars2_in.vivo_noRT | Glipr2 | 2.198 | 5.08E-03 | glio |
| Aars2_in.vivo_noRT | Gprasp2 | 1.86 | 5.08E-03 | glio |
| Aars2_in.vivo_noRT | Stard8 | 1.85 | 5.08E-03 | glio |
| Aars2_in.vivo_noRT | Ciao2b | -0.734 | 5.08E-03 | glio |
| Atr_in.vivo_noRT | 2210016L21Ri | -1.168 | 5.08E-03 | glio |
| Atr_ex.vivo_noRT | Gstp3 | 2.769 | 5.08E-03 | glio |
| Bcas2_ex.vivo_noRT | Gm128 | 2.349 | 5.08E-03 | glio |
| Chek1_in.vivo_noRT | Agpat5 | -1.08 | 5.08E-03 | glio |
| Chek1_in.vivo_noRT | Manf | -0.731 | 5.08E-03 | glio |
| Chek1_ex.vivo_noRT | Pde5a | 2.645 | 5.08E-03 | glio |
| Chek1_ex.vivo_noRT | 1810019D21R | 2.462 | 5.08E-03 | glio |
| Chek1_ex.vivo_noRT | Prpf38a | -1.061 | 5.08E-03 | glio |
| Chek1_ex.vivo_noRT | Dnajc19 | -0.915 | 5.08E-03 | glio |
| Cyp19a1_ex.vivo_noRT | Gm128 | 2.565 | 5.08E-03 | glio |
| Interaction_genotype_Mtor | Ndufv1 | 2.012 | 5.08E-04 | glio |
| Interaction_genotype_Myc | Ranbp17 | -3.876 | 5.08E-04 | glio |
| Interaction_genotype_Myc | Morf4l1 | 1.234 | 5.08E-04 | glio |
| Aars2_in.vivo_noRT | H2-DMb2 | 2.359 | 5.08E-04 | glio |
| Aars2_in.vivo_noRT | Gm42679 | 2.32 | 5.08E-04 | glio |
| Atr_in.vivo_noRT | Tigd5 | 2.246 | 5.08E-04 | glio |
| Cenpj_ex.vivo_noRT | Gm9758 | 2.591 | 5.08E-04 | glio |
| Eif2s1_in.vivo_noRT | Gm2022 | 2.143 | 5.08E-04 | glio |
| Hsd17b10_in.vivo_noRT | Gm20492 | 2.556 | 5.08E-04 | glio |
| Hsd17b10_in.vivo_noRT | Dio1 | 2.307 | 5.08E-04 | glio |
| culture_effect | Yars | -0.705 | 5.08E-05 | glio |
| Bard1_ex.vivo_noRT | BB287469 | 2.814 | 5.08E-05 | glio |
| Bcas2_in.vivo_noRT | Scn2a | 2.924 | 5.08E-05 | glio |
| Interaction_genotype_Myc | Gm14002 | -4.879 | 5.08E-06 | glio |
| Interaction_genotype_Myc | Clint1 | 1.898 | 5.08E-06 | glio |
| Atr_in.vivo_noRT | Myl4 | 3.283 | 5.08E-06 | glio |
| Atr_in.vivo_noRT | Klhl8 | 3.016 | 5.08E-06 | glio |
| Hsd17b10_in.vivo_noRT | Gm34885 | 3.356 | 5.08E-06 | glio |
| Hsd17b10_in.vivo_noRT | Tmem52 | 3.237 | 5.08E-07 | glio |
| Bcas2_in.vivo_noRT | Gm37584 | 3.895 | 5.08E-08 | glio |
| Bora_in.vivo_noRT | Edn1 | 4.557 | 5.08E-08 | glio |
| Chek1_in.vivo_noRT | Gm26542 | 3.666 | 5.08E-08 | glio |

| | | | | |
|---|---|---|---|---|
| Bcas2_in.vivo_noRT | Pcdhga1 | 3.756 | 5.08E-10 | glio |
| culture_effect | Rras2 | 1.344 | 5.08E-12 | glio |
| Bcas2_in.vivo_noRT | Ablim2 | 4.275 | 5.08E-19 | glio |
| Interaction_genotype_Aars2 | 4930445N18R | -3.6 | 5.09E-03 | glio |
| Interaction_genotype_Atr | Gm21731 | -2.369 | 5.09E-03 | glio |
| Interaction_genotype_Atr | 1110035H17R | -3.912 | 5.09E-03 | glio |
| Interaction_genotype_Atr | Cbr3 | -3.662 | 5.09E-03 | glio |
| Interaction_genotype_Bcas2 | Gm50321 | -3.163 | 5.09E-03 | glio |
| Interaction_genotype_Bcas2 | Gm26930 | -4.077 | 5.09E-03 | glio |
| Interaction_genotype_Bcas2 | Tdrd5 | -3.424 | 5.09E-03 | glio |
| Interaction_genotype_Bcas2 | Tnni3k | -2.852 | 5.09E-03 | glio |
| Interaction_genotype_Bcas2 | Vsig10l | -3.752 | 5.09E-03 | glio |
| Interaction_genotype_Bcas2 | Gm32913 | -5.578 | 5.09E-03 | glio |
| Interaction_genotype_Bcas2 | Gm26532 | 1.868 | 5.09E-03 | glio |
| Interaction_genotype_Bcas2 | Rpp21 | 1.643 | 5.09E-03 | glio |
| Interaction_genotype_Mtor | Gm44699 | -2.951 | 5.09E-03 | glio |
| Interaction_genotype_Myc | Gm14966 | -2.827 | 5.09E-03 | glio |
| Atr_in.vivo_noRT | AW146154 | 2.023 | 5.09E-03 | glio |
| Atr_ex.vivo_noRT | Gsx2 | 2.993 | 5.09E-03 | glio |
| Bcas2_ex.vivo_noRT | Aldh1a1 | 3.018 | 5.09E-03 | glio |
| Bora_in.vivo_noRT | Taf1d | -0.898 | 5.09E-03 | glio |
| Hsd17b10_in.vivo_noRT | Gatm | -1.259 | 5.09E-03 | glio |
| Interaction_genotype_Mtor | Gm12043 | -3.93 | 5.09E-04 | glio |
| Interaction_genotype_Myc | Usp24 | -2.722 | 5.09E-04 | glio |
| Aars2_in.vivo_noRT | Gipc3 | 2.558 | 5.09E-04 | glio |
| Aars2_in.vivo_noRT | Rnf207 | 2.295 | 5.09E-04 | glio |
| Atp5c1_in.vivo_noRT | Gm20705 | 2.136 | 5.09E-04 | glio |
| Bcas2_in.vivo_noRT | 4930594M22R | 2.56 | 5.09E-04 | glio |
| Bora_ex.vivo_noRT | Fst | 3.201 | 5.09E-04 | glio |
| Cenpj_ex.vivo_noRT | AF067063 | 2.591 | 5.09E-04 | glio |
| Chek1_in.vivo_noRT | Gm9844 | 2.401 | 5.09E-04 | glio |
| culture_effect | AV039307 | 2.317 | 5.09E-05 | glio |
| Interaction_genotype_Bcas2 | Psmd9 | 2.75 | 5.09E-05 | glio |
| Aars2_in.vivo_noRT | Akap17b | 2.575 | 5.09E-05 | glio |
| Bcas2_in.vivo_noRT | 1500002C15R | 3.04 | 5.09E-05 | glio |
| Bora_in.vivo_noRT | Galk1 | -1.39 | 5.09E-05 | glio |
| Hsd17b10_in.vivo_noRT | Gm13594 | 3.09 | 5.09E-05 | glio |
| Hsd17b10_in.vivo_noRT | Adam33 | 2.99 | 5.09E-05 | glio |
| Chek1_ex.vivo_noRT | Tmem236 | 3.47 | 5.09E-06 | glio |
| Bcas2_in.vivo_noRT | Pla2g2f | 3.164 | 5.09E-07 | glio |
| Aars2_in.vivo_noRT | D430018E03R | 3.601 | 5.09E-08 | glio |
| Chek1_in.vivo_noRT | Prex2 | 4.137 | 5.09E-08 | glio |
| Aars2_in.vivo_noRT | Dhrs9 | 3.74 | 5.09E-09 | glio |
| Bcas2_in.vivo_noRT | Lrrc24 | 3.807 | 5.09E-09 | glio |
| Bora_in.vivo_noRT | Spint4 | 4.145 | 5.09E-09 | glio |
| Atr_in.vivo_noRT | Shroom3 | 4.254 | 5.09E-10 | glio |
| Aars2_in.vivo_noRT | Gm15520 | 4.01 | 5.09E-11 | glio |
| Bcas2_in.vivo_noRT | Tph2 | 4.246 | 5.09E-11 | glio |
| Bora_in.vivo_noRT | Gm37864 | 4.157 | 5.09E-12 | glio |
| Chek1_ex.vivo_noRT | Crb3 | 4.842 | 5.09E-13 | glio |
| Bora_in.vivo_noRT | Ly6d | 4.209 | 5.09E-14 | glio |
| Chek1_ex.vivo_noRT | Ccl9 | 4.449 | 5.09E-14 | glio |

| | | | | |
|---|---|---|---|---|
| Bora_in.vivo_noRT | Gm50024 | 4.673 | 5.09E-21 | glio |
| Interaction_genotype_Aars2 | Radil | -3.377 | 5.10E-03 | glio |
| Interaction_genotype_Aars2 | Ufc1 | 1.04 | 5.10E-03 | glio |
| Interaction_genotype_Atr | Tgfbr3l | -3.702 | 5.10E-03 | glio |
| Interaction_genotype_Atr | Rtn1 | -3.618 | 5.10E-03 | glio |
| Interaction_genotype_Atr | Mzf1 | -3.31 | 5.10E-03 | glio |
| Interaction_genotype_Atr | Cdc42bpg | -3.206 | 5.10E-03 | glio |
| Interaction_genotype_Myc | Gm2762 | -2.963 | 5.10E-03 | glio |
| Interaction_genotype_Myc | Khdrbs1 | 1.219 | 5.10E-03 | glio |
| Interaction_genotype_Pars2 | Vmn1r32 | -2.661 | 5.10E-03 | glio |
| Atp5c1_in.vivo_noRT | Gm7247 | 1.876 | 5.10E-03 | glio |
| Atr_in.vivo_noRT | Fgl1 | 2.062 | 5.10E-03 | glio |
| Atr_in.vivo_noRT | Gm11273 | 1.704 | 5.10E-03 | glio |
| Atr_in.vivo_noRT | Socs2 | 1.705 | 5.10E-03 | glio |
| Atr_in.vivo_noRT | Zfp644 | -1.041 | 5.10E-03 | glio |
| Atr_ex.vivo_noRT | Ifitm1 | 3.164 | 5.10E-03 | glio |
| Bcas2_ex.vivo_noRT | Gm14207 | 1.949 | 5.10E-03 | glio |
| Bora_in.vivo_noRT | Sucla2 | -1.256 | 5.10E-03 | glio |
| Cenpj_in.vivo_noRT | Hif3a | 2.434 | 5.10E-03 | glio |
| Chek1_in.vivo_noRT | Schip1 | -1.353 | 5.10E-03 | glio |
| Eif2s1_in.vivo_noRT | Gm45235 | 1.883 | 5.10E-03 | glio |
| Hsd17b10_in.vivo_noRT | Mir155hg | 2.371 | 5.10E-03 | glio |
| Hsd17b10_in.vivo_noRT | Card10 | 2.35 | 5.10E-03 | glio |
| Interaction_genotype_Mtor | Adam2 | -3.573 | 5.10E-04 | glio |
| Aars2_in.vivo_noRT | Lipt2 | 2.015 | 5.10E-04 | glio |
| Aars2_ex.vivo_noRT | Gm14540 | 2.499 | 5.10E-04 | glio |
| Atr_in.vivo_noRT | Gm9844 | 2.367 | 5.10E-04 | glio |
| Bora_ex.vivo_noRT | Gm5431 | 3.201 | 5.10E-04 | glio |
| culture_effect | Dnajc7 | -0.564 | 5.10E-05 | glio |
| Atp5c1_in.vivo_noRT | Tdrd5 | 2.81 | 5.10E-05 | glio |
| Atr_in.vivo_noRT | Mfsd3 | 2.381 | 5.10E-05 | glio |
| Atr_in.vivo_noRT | Ccnl2 | -1.591 | 5.10E-05 | glio |
| Bora_in.vivo_noRT | Gm38973 | 2.873 | 5.10E-05 | glio |
| Chek1_in.vivo_noRT | Abcd2 | 3.205 | 5.10E-05 | glio |
| culture_effect | Dner | -2.637 | 5.10E-06 | glio |
| Interaction_genotype_Myc | Zfp119a | -4.221 | 5.10E-06 | glio |
| Interaction_genotype_Myc | Gm31518 | -5.354 | 5.10E-06 | glio |
| Atr_in.vivo_noRT | Ybx3 | -1.158 | 5.10E-06 | glio |
| Chek1_in.vivo_noRT | Hist1h2ad | 3.272 | 5.10E-06 | glio |
| culture_effect | Gdf11 | 1.464 | 5.10E-07 | glio |
| Atr_ex.vivo_noRT | Ltbp2 | 3.552 | 5.10E-07 | glio |
| Bora_in.vivo_noRT | Rab15 | 3.728 | 5.10E-08 | glio |
| Hsd17b10_in.vivo_noRT | Gm2115 | 3.616 | 5.10E-09 | glio |
| Atr_in.vivo_noRT | Gm48882 | 4.409 | 5.10E-10 | glio |
| Atr_in.vivo_noRT | Gm8104 | 4.114 | 5.10E-10 | glio |
| Bcas2_in.vivo_noRT | Nrg2 | 4.284 | 5.10E-10 | glio |
| Bcas2_in.vivo_noRT | Vmn1r181 | 4.151 | 5.10E-10 | glio |
| Bora_in.vivo_noRT | Loxl4 | 4.67 | 5.10E-13 | glio |
| Chek1_in.vivo_noRT | Adam5 | 4.78 | 5.10E-21 | glio |
| culture_effect | Gm128 | -1.46 | 5.11E-03 | glio |
| culture_effect | Cog7 | -0.871 | 5.11E-03 | glio |
| Interaction_genotype_Aars2 | Gm20628 | -3.773 | 5.11E-03 | glio |

| | | | | |
|---|---|---|---|---|
| Interaction_genotype_Rad51 | 5330439K02R | -4.056 | 5.11E-03 | glio |
| Interaction_genotype_Rad51 | N6amt1 | 2.019 | 5.11E-03 | glio |
| Aars2_ex.vivo_noRT | Gm49223 | 2.311 | 5.11E-03 | glio |
| Atr_in.vivo_noRT | C530044C16R | 1.934 | 5.11E-03 | glio |
| Atr_in.vivo_noRT | Eef1akmt1 | -1.145 | 5.11E-03 | glio |
| Bcas2_in.vivo_noRT | Gm48604 | 2.024 | 5.11E-03 | glio |
| Bcas2_in.vivo_noRT | Sra1 | -0.934 | 5.11E-03 | glio |
| Chek1_in.vivo_noRT | Hyal1 | 2.02 | 5.11E-03 | glio |
| Eif2s1_in.vivo_noRT | Pdgfb | 1.88 | 5.11E-03 | glio |
| Interaction_genotype_Aars2 | D830039M14I | -4.455 | 5.11E-04 | glio |
| Interaction_genotype_Mtor | Def6 | -4.116 | 5.11E-04 | glio |
| Interaction_genotype_Mtor | Emp3 | 1.891 | 5.11E-04 | glio |
| Interaction_genotype_Myc | Gm43256 | -3.637 | 5.11E-04 | glio |
| Interaction_genotype_Myc | Gm13964 | -2.729 | 5.11E-04 | glio |
| Interaction_genotype_Myc | Gm48684 | -4.569 | 5.11E-04 | glio |
| Atp5c1_in.vivo_noRT | Ptgis | 2.668 | 5.11E-04 | glio |
| Atr_in.vivo_noRT | Ccdc13 | 2.519 | 5.11E-04 | glio |
| Atr_in.vivo_noRT | Rhod | 1.609 | 5.11E-04 | glio |
| Bora_in.vivo_noRT | Gm28229 | 2.295 | 5.11E-04 | glio |
| Bora_in.vivo_noRT | H2afv | -0.74 | 5.11E-04 | glio |
| Chek1_ex.vivo_noRT | Ccdc24 | 3.094 | 5.11E-04 | glio |
| culture_effect | Ubn2 | -0.638 | 5.11E-05 | glio |
| Interaction_genotype_Myc | Reps1 | -3.398 | 5.11E-05 | glio |
| Aars2_in.vivo_noRT | G430095P16R | 2.873 | 5.11E-05 | glio |
| Atr_in.vivo_noRT | Lipe | 2.82 | 5.11E-05 | glio |
| Bora_in.vivo_noRT | Gm17249 | 2.799 | 5.11E-05 | glio |
| Interaction_genotype_Mtor | Copz1 | 2.326 | 5.11E-06 | glio |
| Atr_in.vivo_noRT | Cenpx | -1.865 | 5.11E-06 | glio |
| Interaction_genotype_Mtor | 5033403F01Ri | -5.562 | 5.11E-07 | glio |
| Hsd17b10_in.vivo_noRT | Nxph4 | 3.616 | 5.11E-07 | glio |
| culture_effect | Ptrh1 | 1.641 | 5.11E-08 | glio |
| Aars2_in.vivo_noRT | Syne3 | 4.07 | 5.11E-08 | glio |
| Hsd17b10_in.vivo_noRT | D030040B21R | 3.76 | 5.11E-08 | glio |
| interaction_RT | Smim14 | 1.447 | 5.11E-12 | glio |
| Bcas2_in.vivo_noRT | Gm16751 | 4.291 | 5.11E-13 | glio |
| Chek1_in.vivo_noRT | Gem | 5.044 | 5.11E-14 | glio |
| Aars2_in.vivo_noRT | D630036H23R | 5.577 | 5.11E-16 | glio |
| culture_effect | Ly6a | -3.639 | 5.11E-17 | glio |
| Bcas2_in.vivo_noRT | Gm48617 | 4.543 | 5.11E-20 | glio |
| Atr_in.vivo_noRT | Ccdc80 | 5.533 | 5.11E-24 | glio |
| Interaction_genotype_Aars2 | A430035B10R | -3.659 | 5.12E-03 | glio |
| Interaction_genotype_Atr | Gm11232 | -2.369 | 5.12E-03 | glio |
| Interaction_genotype_Bcas2 | Msmo1 | 1.897 | 5.12E-03 | glio |
| Interaction_genotype_Mtor | Tpbg | -3.366 | 5.12E-03 | glio |
| Interaction_genotype_Mtor | Sf3a3 | 1.333 | 5.12E-03 | glio |
| Interaction_genotype_Myc | Selp | -3.096 | 5.12E-03 | glio |
| Interaction_genotype_Myc | Zfp764 | -2.839 | 5.12E-03 | glio |
| Interaction_genotype_Myc | Gpr89 | -2.135 | 5.12E-03 | glio |
| Interaction_genotype_Myc | Ndufb6 | 1.11 | 5.12E-03 | glio |
| Interaction_genotype_Pars2 | Gm47509 | -3.954 | 5.12E-03 | glio |
| Interaction_genotype_Pars2 | Syngr2 | 1.386 | 5.12E-03 | glio |
| Interaction_genotype_Rad51 | Ubr2 | 1.828 | 5.12E-03 | glio |

| | | | | |
|---|---|---|---|---|
| Interaction_genotype_Rpl3 | 4930426I24Ri | -3.807 | 5.12E-03 | glio |
| Aars2_in.vivo_noRT | 4933408B17R | 2.056 | 5.12E-03 | glio |
| Aars2_in.vivo_noRT | Aasdh | 1.583 | 5.12E-03 | glio |
| Aars2_in.vivo_noRT | Stxbp5 | 1.518 | 5.12E-03 | glio |
| Aars2_in.vivo_noRT | Rnf14 | -1.082 | 5.12E-03 | glio |
| Atp5c1_in.vivo_noRT | Agrp | 1.886 | 5.12E-03 | glio |
| Atp5c1_in.vivo_noRT | Gm20275 | 1.845 | 5.12E-03 | glio |
| Atr_in.vivo_noRT | Pank1 | 1.467 | 5.12E-03 | glio |
| Atr_in.vivo_noRT | Sae1 | -1.04 | 5.12E-03 | glio |
| Bcas2_in.vivo_noRT | Rab42 | 2.203 | 5.12E-03 | glio |
| Bora_in.vivo_noRT | Rbbp4 | -0.742 | 5.12E-03 | glio |
| Chek1_in.vivo_noRT | Wdpcp | 1.956 | 5.12E-03 | glio |
| Eif2s1_in.vivo_noRT | Ptges3l | -1.904 | 5.12E-03 | glio |
| Eif2s1_in.vivo_noRT | Tinagl1 | -0.944 | 5.12E-03 | glio |
| Hsd17b10_in.vivo_noRT | Prr7 | 2.225 | 5.12E-03 | glio |
| Interaction_genotype_Atr | Eml1 | -3.893 | 5.12E-04 | glio |
| Interaction_genotype_Atr | Plcl1 | -3.539 | 5.12E-04 | glio |
| Interaction_genotype_Atr | Gm45397 | -3.834 | 5.12E-04 | glio |
| Interaction_genotype_Atr | Ank1 | -3.153 | 5.12E-04 | glio |
| Interaction_genotype_Atr | Pdcd2 | 1.915 | 5.12E-04 | glio |
| Interaction_genotype_Mtor | Galk1 | 1.895 | 5.12E-04 | glio |
| Interaction_genotype_Myc | BC031181 | 1.407 | 5.12E-04 | glio |
| Interaction_genotype_Rad51 | Nxph3 | -4.38 | 5.12E-04 | glio |
| interaction_RT | Josd2 | 0.749 | 5.12E-04 | glio |
| Atr_in.vivo_noRT | Gm15867 | 2.411 | 5.12E-04 | glio |
| Bard1_ex.vivo_noRT | Ipcef1 | 2.814 | 5.12E-04 | glio |
| Bcas2_in.vivo_noRT | Plekhg6 | 2.601 | 5.12E-04 | glio |
| Bcas2_in.vivo_noRT | Atp8a2 | 2.128 | 5.12E-04 | glio |
| Bcas2_in.vivo_noRT | Dip2a | 2.405 | 5.12E-04 | glio |
| Bcas2_ex.vivo_noRT | Ms4a15 | 2.742 | 5.12E-04 | glio |
| Cenpj_ex.vivo_noRT | Rps6ka4 | 3.286 | 5.12E-04 | glio |
| Cenpj_ex.vivo_noRT | Gm2016 | 2.591 | 5.12E-04 | glio |
| Chek1_in.vivo_noRT | Nxt1 | -1.376 | 5.12E-04 | glio |
| Chek1_in.vivo_noRT | Csnk2a2 | -1.058 | 5.12E-04 | glio |
| Chek1_in.vivo_noRT | Glod5 | 2.098 | 5.12E-04 | glio |
| Chek1_in.vivo_noRT | Eif4ebp1 | -1.058 | 5.12E-04 | glio |
| Chek1_ex.vivo_noRT | Rpl5 | -0.871 | 5.12E-04 | glio |
| culture_effect | Kmo | -2.183 | 5.12E-05 | glio |
| culture_effect | 2310068J16Ri | 1.401 | 5.12E-05 | glio |
| culture_effect | Ccdc28a | -1.174 | 5.12E-05 | glio |
| culture_effect | Ggt7 | -0.988 | 5.12E-05 | glio |
| Interaction_genotype_Atr | 5031434O11R | -5.686 | 5.12E-05 | glio |
| Chek1_in.vivo_noRT | Kif5a | 3.114 | 5.12E-05 | glio |
| Aars2_in.vivo_noRT | Gm49785 | 3.63 | 5.12E-07 | glio |
| Atr_in.vivo_noRT | Hcrtr1 | 3.595 | 5.12E-07 | glio |
| Chek1_ex.vivo_noRT | Pik3ap1 | 3.69 | 5.12E-07 | glio |
| Aars2_ex.vivo_noRT | Slc19a3 | 3.699 | 5.12E-08 | glio |
| Aars2_ex.vivo_noRT | Gm15908 | 3.293 | 5.12E-08 | glio |
| Aars2_ex.vivo_noRT | A930014E10Ri | 3.293 | 5.12E-08 | glio |
| Aars2_in.vivo_noRT | Adgrb3 | 4.021 | 5.12E-09 | glio |
| Bora_in.vivo_noRT | Gm3734 | 4.413 | 5.12E-09 | glio |
| Bora_in.vivo_noRT | Sap30bpos | 3.542 | 5.12E-09 | glio |

| | | | | |
|---|---|---|---|---|
| culture_effect | Fst | -3.837 | 5.12E-11 | glio |
| Atr_in.vivo_noRT | Gm15559 | 4.606 | 5.12E-11 | glio |
| culture_effect | Atxn10 | -1.065 | 5.12E-12 | glio |
| Atr_in.vivo_noRT | Hunk | 5.001 | 5.12E-15 | glio |
| Aars2_in.vivo_noRT | Tcstv3 | 5.327 | 5.12E-16 | glio |
| Bora_in.vivo_noRT | Ccser1 | 4.936 | 5.12E-17 | glio |
| Chek1_ex.vivo_noRT | Olfr810 | 4.842 | 5.12E-17 | glio |
| culture_effect | Lactb | 2.234 | 5.12E-20 | glio |
| Aars2_in.vivo_noRT | Gm12841 | 5.327 | 5.12E-25 | glio |
| Interaction_genotype_Bora | 2610016A17R | -3.771 | 5.13E-03 | glio |
| Interaction_genotype_Bora | Rbm10 | 2.122 | 5.13E-03 | glio |
| Interaction_genotype_Myc | 1810059H22R | -3.325 | 5.13E-03 | glio |
| Interaction_genotype_Myc | Ppp1r16a | -2.323 | 5.13E-03 | glio |
| Interaction_genotype_Myc | Cep290 | -1.99 | 5.13E-03 | glio |
| interaction_RT | Slc27a1 | 1.172 | 5.13E-03 | glio |
| interaction_RT | Ubb | 0.535 | 5.13E-03 | glio |
| Aars2_in.vivo_noRT | Zfp82 | 1.989 | 5.13E-03 | glio |
| Bcas2_in.vivo_noRT | Spryd3 | -1.267 | 5.13E-03 | glio |
| Bora_in.vivo_noRT | Adap1 | 2.341 | 5.13E-03 | glio |
| Bora_in.vivo_noRT | Ndc80 | -1.5 | 5.13E-03 | glio |
| Bora_in.vivo_noRT | Zfp326 | -0.967 | 5.13E-03 | glio |
| Chek1_ex.vivo_noRT | Chrnb1 | 2.531 | 5.13E-03 | glio |
| culture_effect | Sod2 | -0.463 | 5.13E-04 | glio |
| Interaction_genotype_Mtor | Gm12166 | -4.396 | 5.13E-04 | glio |
| Interaction_genotype_Mtor | Atp6v0c | 1.431 | 5.13E-04 | glio |
| Interaction_genotype_Myc | Rpl34 | 1.437 | 5.13E-04 | glio |
| Aars2_in.vivo_noRT | Psmb6 | -0.898 | 5.13E-04 | glio |
| Bcas2_in.vivo_noRT | A430033K04R | 2.439 | 5.13E-04 | glio |
| Chek1_in.vivo_noRT | Hist1h4j | 2.924 | 5.13E-04 | glio |
| Chek1_in.vivo_noRT | Gm14399 | 2.526 | 5.13E-04 | glio |
| Interaction_genotype_Mtor | Lrp2bp | -4.337 | 5.13E-05 | glio |
| Interaction_genotype_Mtor | Otub2 | -4.378 | 5.13E-05 | glio |
| Interaction_genotype_Mtor | Galc | -4 | 5.13E-05 | glio |
| Interaction_genotype_Mtor | Fxyd7 | -4.258 | 5.13E-05 | glio |
| Aars2_in.vivo_noRT | Fignl2 | 3.061 | 5.13E-05 | glio |
| Atp5c1_in.vivo_noRT | 4930553M12F | 2.402 | 5.13E-05 | glio |
| Bora_in.vivo_noRT | Tead4 | -2.057 | 5.13E-05 | glio |
| Chek1_in.vivo_noRT | Tc2n | 2.956 | 5.13E-05 | glio |
| Chek1_in.vivo_noRT | A930033H14R | 3.019 | 5.13E-05 | glio |
| Chek1_in.vivo_noRT | Rpl18a | -0.726 | 5.13E-05 | glio |
| Interaction_genotype_Mtor | Psme2 | 2.586 | 5.13E-06 | glio |
| Aars2_in.vivo_noRT | Gtf3c6 | -1.61 | 5.13E-06 | glio |
| Atr_in.vivo_noRT | 4930455G09R | 3.717 | 5.13E-06 | glio |
| Chek1_in.vivo_noRT | Gm43062 | 3.266 | 5.13E-06 | glio |
| Chek1_in.vivo_noRT | Lrrc75b | 3.387 | 5.13E-06 | glio |
| Atr_in.vivo_noRT | Slc6a19 | 3.364 | 5.13E-07 | glio |
| Hsd17b10_in.vivo_noRT | Crb3 | 3.616 | 5.13E-11 | glio |
| Bora_in.vivo_noRT | Npr3 | 4.936 | 5.13E-16 | glio |
| Chek1_in.vivo_noRT | Ildr1 | 5.044 | 5.13E-23 | glio |
| Atr_in.vivo_noRT | Gm11232 | 5.533 | 5.13E-29 | glio |
| culture_effect | Jun | -2.206 | 5.13E-30 | glio |
| Interaction_genotype_Bcas2 | Aloxe3 | -3.175 | 5.14E-03 | glio |

| | | | | |
|---|---|---|---|---|
| Interaction_genotype_Bcas2 | Gm3716 | -3.425 | 5.14E-03 | glio |
| Interaction_genotype_Lmo7 | Kpna7 | -3.896 | 5.14E-03 | glio |
| Interaction_genotype_Mtor | Fignl2 | -3.645 | 5.14E-03 | glio |
| Interaction_genotype_Mtor | Pacs2 | -2.096 | 5.14E-03 | glio |
| Interaction_genotype_Mtor | Pdxk | -1.911 | 5.14E-03 | glio |
| Interaction_genotype_Myc | D730045A05R | -3.124 | 5.14E-03 | glio |
| Interaction_genotype_Myc | Psd | -2.788 | 5.14E-03 | glio |
| Interaction_genotype_Myc | Pi4kb | -2.52 | 5.14E-03 | glio |
| Interaction_genotype_Myc | Azin2 | -2.692 | 5.14E-03 | glio |
| interaction_RT | Etfbkmt | 1.547 | 5.14E-03 | glio |
| interaction_RT | Lmnb1 | -0.909 | 5.14E-03 | glio |
| interaction_RT | U2surp | -0.576 | 5.14E-03 | glio |
| Aars2_in.vivo_noRT | Pnpla6 | 1.786 | 5.14E-03 | glio |
| Bcas2_in.vivo_noRT | Xrcc1 | -1.224 | 5.14E-03 | glio |
| Bcas2_in.vivo_noRT | Cd44 | -0.953 | 5.14E-03 | glio |
| Bcas2_ex.vivo_noRT | Col9a3 | -2.128 | 5.14E-03 | glio |
| Bora_in.vivo_noRT | Slc27a3 | 2.062 | 5.14E-03 | glio |
| Bora_in.vivo_noRT | Gm26620 | 1.857 | 5.14E-03 | glio |
| Bora_in.vivo_noRT | Tmem104 | 1.827 | 5.14E-03 | glio |
| Bora_in.vivo_noRT | Zfp977 | 1.792 | 5.14E-03 | glio |
| Chek1_ex.vivo_noRT | Pld6 | 2.764 | 5.14E-03 | glio |
| culture_effect | Tlr1 | 2.034 | 5.14E-04 | glio |
| Interaction_genotype_Mtor | Zfp467 | -4.11 | 5.14E-04 | glio |
| Interaction_genotype_Mtor | Gm16759 | -4.06 | 5.14E-04 | glio |
| Interaction_genotype_Myc | Ptges | -4.144 | 5.14E-04 | glio |
| Aars2_in.vivo_noRT | Ccdc96 | 2.218 | 5.14E-04 | glio |
| Atr_in.vivo_noRT | Gpatch8 | -1.248 | 5.14E-04 | glio |
| Atr_ex.vivo_noRT | Traf1 | 3.172 | 5.14E-04 | glio |
| Bcas2_in.vivo_noRT | 4932422M17R | 2.285 | 5.14E-04 | glio |
| Bcas2_ex.vivo_noRT | Gm10482 | 2.742 | 5.14E-04 | glio |
| Bcas2_ex.vivo_noRT | Clec12b | 2.347 | 5.14E-04 | glio |
| Bcas2_ex.vivo_noRT | Vmn1r32 | 2.349 | 5.14E-04 | glio |
| Chek1_in.vivo_noRT | Mrpl32 | -1.003 | 5.14E-04 | glio |
| Interaction_genotype_Mtor | 4930515G01R | -4.194 | 5.14E-05 | glio |
| Bcas2_in.vivo_noRT | B4galt3 | -1.667 | 5.14E-05 | glio |
| Bcas2_in.vivo_noRT | Rad23b | -1.329 | 5.14E-05 | glio |
| Interaction_genotype_Mtor | 1700020L24Ri | -5.101 | 5.14E-06 | glio |
| Bora_in.vivo_noRT | Dagla | 3.44 | 5.14E-06 | glio |
| Interaction_genotype_Mtor | Gm50024 | -4.235 | 5.14E-07 | glio |
| Interaction_genotype_Nat2 | Zscan4d | 4.79 | 5.14E-07 | glio |
| Interaction_genotype_Nat2 | BB287469 | 4.443 | 5.14E-07 | glio |
| Chek1_ex.vivo_noRT | Gm43062 | 4.438 | 5.14E-07 | glio |
| Bora_in.vivo_noRT | BC005624 | -2.191 | 5.14E-08 | glio |
| Atr_in.vivo_noRT | Dcpp1 | 4.206 | 5.14E-09 | glio |
| Aars2_in.vivo_noRT | Gm8300 | 5.327 | 5.14E-10 | glio |
| Chek1_in.vivo_noRT | Tnfsf13os | 4.273 | 5.14E-10 | glio |
| Hsd17b10_in.vivo_noRT | Boc | 3.352 | 5.14E-11 | glio |
| Aars2_in.vivo_noRT | AU022754 | 4.443 | 5.14E-12 | glio |
| Aars2_in.vivo_noRT | Tas1r1 | 4.3 | 5.14E-12 | glio |
| Bcas2_in.vivo_noRT | Boc | 4.543 | 5.14E-19 | glio |
| Chek1_ex.vivo_noRT | Gm14207 | 5.402 | 5.14E-19 | glio |
| Aars2_in.vivo_noRT | Dpep3 | 4.794 | 5.14E-22 | glio |

| | | | | |
|---|---|---|---|---|
| Bora_in.vivo_noRT | Gm13964 | 4.936 | 5.14E-23 | glio |
| Interaction_genotype_Chek1 | Vgll3 | -3.555 | 5.15E-03 | glio |
| Interaction_genotype_Chek1 | Elf2 | 1.67 | 5.15E-03 | glio |
| Interaction_genotype_Mtor | Smu1 | 1.562 | 5.15E-03 | glio |
| Interaction_genotype_Pars2 | Npnt | -2.8 | 5.15E-03 | glio |
| Aars2_in.vivo_noRT | Gm42463 | 1.868 | 5.15E-03 | glio |
| Bcas2_in.vivo_noRT | H2-Eb2 | 1.492 | 5.15E-03 | glio |
| Bora_ex.vivo_noRT | Adm | 2.808 | 5.15E-03 | glio |
| Chek1_in.vivo_noRT | Star | 2.008 | 5.15E-03 | glio |
| Chek1_in.vivo_noRT | Ift172 | 1.694 | 5.15E-03 | glio |
| Chek1_ex.vivo_noRT | Gbp9 | 1.809 | 5.15E-03 | glio |
| Chek1_ex.vivo_noRT | Hnrnpa1 | -0.742 | 5.15E-03 | glio |
| Ercc4_ex.vivo_noRT | Hist1h2ag | 2.68 | 5.15E-03 | glio |
| Hsd17b10_in.vivo_noRT | Haglr | 2.362 | 5.15E-03 | glio |
| Hsd17b10_in.vivo_noRT | Itga7 | 2.203 | 5.15E-03 | glio |
| Hsd17b10_in.vivo_noRT | Anapc2 | -1.262 | 5.15E-03 | glio |
| Hsd17b10_in.vivo_noRT | Calm1 | -0.544 | 5.15E-03 | glio |
| culture_effect | Sap30bpos | -1.936 | 5.15E-04 | glio |
| Interaction_genotype_Mtor | Tmem86b | -4.286 | 5.15E-04 | glio |
| Interaction_genotype_Mtor | 2010109A12R | -3.882 | 5.15E-04 | glio |
| Interaction_genotype_Myc | Deptor | -3.455 | 5.15E-04 | glio |
| Interaction_genotype_Myc | Srp9 | 1.291 | 5.15E-04 | glio |
| Atp5c1_in.vivo_noRT | Klhdc8a | 2.668 | 5.15E-04 | glio |
| Atr_in.vivo_noRT | Tk2 | 2.263 | 5.15E-04 | glio |
| Chek1_in.vivo_noRT | Insrr | 2.267 | 5.15E-04 | glio |
| Interaction_genotype_Myc | Med28 | 1.509 | 5.15E-05 | glio |
| Aars2_in.vivo_noRT | Zfp534 | 2.985 | 5.15E-05 | glio |
| Chek1_ex.vivo_noRT | Gm5039 | 4.055 | 5.15E-05 | glio |
| Chek1_ex.vivo_noRT | Col17a1 | 3.833 | 5.15E-05 | glio |
| Hsd17b10_in.vivo_noRT | Gm46404 | 3.111 | 5.15E-05 | glio |
| interaction_RT | Vmn1r181 | 4.077 | 5.15E-06 | glio |
| Aars2_in.vivo_noRT | Gm15728 | 3.374 | 5.15E-06 | glio |
| Aars2_in.vivo_noRT | 5033430I15Ri | 3.147 | 5.15E-06 | glio |
| Aars2_in.vivo_noRT | Aldh1a2 | 2.97 | 5.15E-06 | glio |
| Atr_in.vivo_noRT | Prss22 | 3.935 | 5.15E-06 | glio |
| culture_effect | Klf10 | -1.41 | 5.15E-07 | glio |
| Aars2_in.vivo_noRT | Fhl4 | 4.076 | 5.15E-10 | glio |
| Atr_in.vivo_noRT | Fgf7 | 4.169 | 5.15E-10 | glio |
| Atr_in.vivo_noRT | Ksr2 | 4.495 | 5.15E-15 | glio |
| Interaction_genotype_Aars2 | Rpl7a | 0.766 | 5.16E-03 | glio |
| Interaction_genotype_Ifnar1 | Krt19 | 2.734 | 5.16E-03 | glio |
| Interaction_genotype_Ifnar1 | Gm48155 | 2.734 | 5.16E-03 | glio |
| Interaction_genotype_Ifnar1 | Thsd1 | 3.569 | 5.16E-03 | glio |
| Interaction_genotype_Ifnar1 | Cd200r2 | 2.998 | 5.16E-03 | glio |
| Interaction_genotype_Ifnar1 | Ccr5 | 3.746 | 5.16E-03 | glio |
| Interaction_genotype_Ifnar1 | 9330175E14Ri | -3.165 | 5.16E-03 | glio |
| Interaction_genotype_Ifnar1 | Helt | 2.734 | 5.16E-03 | glio |
| Interaction_genotype_Ifnar1 | Gm20625 | 2.734 | 5.16E-03 | glio |
| Interaction_genotype_Ifnar1 | Gm5157 | 2.734 | 5.16E-03 | glio |
| Interaction_genotype_Ifnar1 | Gm44736 | 2.734 | 5.16E-03 | glio |
| Interaction_genotype_Ifnar1 | Ankrd33b | 4.209 | 5.16E-03 | glio |
| Interaction_genotype_Ifnar1 | Fam227b | 2.734 | 5.16E-03 | glio |

| | | | | |
|---|---|---|---|---|
| Interaction_genotype_Ifnar1 | Gm20658 | 2.734 | 5.16E-03 | glio |
| Interaction_genotype_Ifnar1 | Gm4951 | -2.188 | 5.16E-03 | glio |
| Interaction_genotype_Mtor | Frk | -3.395 | 5.16E-03 | glio |
| Interaction_genotype_Mtor | Lgr6 | -3.6 | 5.16E-03 | glio |
| Interaction_genotype_Myc | Sh3bgrl2 | -3.091 | 5.16E-03 | glio |
| Interaction_genotype_Myc | Zfp82 | -3.087 | 5.16E-03 | glio |
| Interaction_genotype_Myc | Cmtr2 | -2.491 | 5.16E-03 | glio |
| Interaction_genotype_Myc | Dand5 | -2.227 | 5.16E-03 | glio |
| Interaction_genotype_Myc | Sgf29 | -1.996 | 5.16E-03 | glio |
| Atr_in.vivo_noRT | Actn4 | -0.907 | 5.16E-03 | glio |
| Bora_in.vivo_noRT | Bdp1 | -1.276 | 5.16E-03 | glio |
| Bora_ex.vivo_noRT | Enpp3 | 2.802 | 5.16E-03 | glio |
| Cdkn2a_in.vivo_noRT | Sucnr1 | -1.675 | 5.16E-03 | glio |
| Hsd17b10_in.vivo_noRT | Slc26a1 | 2.113 | 5.16E-03 | glio |
| Interaction_genotype_Nat2 | Gm49808 | 3.081 | 5.16E-04 | glio |
| Atr_ex.vivo_noRT | Arhgef16 | 2.764 | 5.16E-04 | glio |
| Bora_in.vivo_noRT | Gm19531 | 2.308 | 5.16E-04 | glio |
| Hsd17b10_in.vivo_noRT | Osm | 2.267 | 5.16E-04 | glio |
| Interaction_genotype_Mtor | Ksr2 | -4.037 | 5.16E-05 | glio |
| Atr_in.vivo_noRT | Pigo | 2.527 | 5.16E-05 | glio |
| Atr_in.vivo_noRT | Tcof1 | -1.65 | 5.16E-05 | glio |
| Bora_in.vivo_noRT | 2610306M01F | 2.008 | 5.16E-05 | glio |
| Chek1_in.vivo_noRT | Hrk | 2.919 | 5.16E-05 | glio |
| Chek1_in.vivo_noRT | Gm4129 | 2.874 | 5.16E-05 | glio |
| Hsd17b10_in.vivo_noRT | Bend7 | 2.838 | 5.16E-05 | glio |
| Bcas2_in.vivo_noRT | Aldh1a1 | 3.306 | 5.16E-06 | glio |
| Chek1_in.vivo_noRT | Gm13421 | 3.319 | 5.16E-06 | glio |
| Chek1_in.vivo_noRT | Ghitm | -1.484 | 5.16E-06 | glio |
| Chek1_ex.vivo_noRT | Pcdhga9 | 3.877 | 5.16E-06 | glio |
| Chek1_ex.vivo_noRT | Ggps1 | -2.32 | 5.16E-06 | glio |
| Atr_in.vivo_noRT | 9430065F17Ri | 3.558 | 5.16E-07 | glio |
| Bora_in.vivo_noRT | Slc17a7 | 3.504 | 5.16E-07 | glio |
| Interaction_genotype_Mtor | Pnisr | 2.801 | 5.16E-08 | glio |
| Chek1_in.vivo_noRT | Gm45336 | 3.171 | 5.16E-08 | glio |
| culture_effect | Cyp1a1 | 3.935 | 5.16E-09 | glio |
| culture_effect | Wdr89 | -0.963 | 5.16E-09 | glio |
| interaction_RT | Mest | -2.846 | 5.16E-09 | glio |
| Bora_in.vivo_noRT | Lmx1b | 4.149 | 5.16E-09 | glio |
| Chek1_ex.vivo_noRT | Gm21762 | 4.449 | 5.16E-09 | glio |
| Chek1_ex.vivo_noRT | Gm43948 | 4.058 | 5.16E-09 | glio |
| Atr_in.vivo_noRT | Ssh3 | 4.733 | 5.16E-15 | glio |
| Chek1_ex.vivo_noRT | Idi2 | 4.842 | 5.16E-16 | glio |
| culture_effect | Mctp2 | -4.053 | 5.16E-19 | glio |
| Bora_in.vivo_noRT | Gm11232 | 4.936 | 5.16E-24 | glio |
| culture_effect | Gm48882 | 1.642 | 5.17E-03 | glio |
| culture_effect | Stard8 | 1.126 | 5.17E-03 | glio |
| culture_effect | Sept6 | -0.877 | 5.17E-03 | glio |
| Interaction_genotype_Atr | Atf3 | 2.513 | 5.17E-03 | glio |
| Interaction_genotype_Bcas2 | Glod5 | 4.492 | 5.17E-03 | glio |
| Interaction_genotype_Bcas2 | Kirrel3 | -3.754 | 5.17E-03 | glio |
| Interaction_genotype_Bora | Pde11a | -3.891 | 5.17E-03 | glio |
| Interaction_genotype_Bora | Apol10b | 3.189 | 5.17E-03 | glio |

| | | | | |
|---|---|---|---|---|
| Interaction_genotype_Bora | Dmxl1 | 1.876 | 5.17E-03 | glio |
| Interaction_genotype_Bora | Mocs2 | 1.565 | 5.17E-03 | glio |
| Interaction_genotype_Chek1 | Fbxo34 | -2.807 | 5.17E-03 | glio |
| Interaction_genotype_Mtor | Rhobtb1 | -3.391 | 5.17E-03 | glio |
| Interaction_genotype_Myc | 1700056N10R | -2.877 | 5.17E-03 | glio |
| Interaction_genotype_Myc | Pask | -3.154 | 5.17E-03 | glio |
| Interaction_genotype_Myc | Abhd3 | -2.903 | 5.17E-03 | glio |
| Interaction_genotype_Myc | Pus7l | -2.484 | 5.17E-03 | glio |
| Interaction_genotype_Myc | Csk | -1.847 | 5.17E-03 | glio |
| Interaction_genotype_Uqcrb | Rassf6 | -3.734 | 5.17E-03 | glio |
| Aars2_in.vivo_noRT | Syn1 | 2.062 | 5.17E-03 | glio |
| Aars2_in.vivo_noRT | mt-Co3 | -0.582 | 5.17E-03 | glio |
| Atp5c1_in.vivo_noRT | Gm14379 | 2.152 | 5.17E-03 | glio |
| Atr_in.vivo_noRT | Msrb3 | 1.768 | 5.17E-03 | glio |
| Atr_in.vivo_noRT | Lrrc8c | 1.335 | 5.17E-03 | glio |
| Atr_in.vivo_noRT | U2af2 | -1.046 | 5.17E-03 | glio |
| Chek1_ex.vivo_noRT | Spata31d1a | 2.611 | 5.17E-03 | glio |
| Fanca_ex.vivo_noRT | A930014E10Ri | 2.124 | 5.17E-03 | glio |
| Fanca_ex.vivo_noRT | Scg2 | 2.124 | 5.17E-03 | glio |
| Fanca_ex.vivo_noRT | Prox2os | 2.915 | 5.17E-03 | glio |
| Fanca_ex.vivo_noRT | 1700003E24Ri | 2.124 | 5.17E-03 | glio |
| Interaction_genotype_Atr | Gm10642 | -4.013 | 5.17E-04 | glio |
| Interaction_genotype_Myc | D730003I15Ri | -2.781 | 5.17E-04 | glio |
| interaction_RT | 4632427E13Ri | -1.014 | 5.17E-04 | glio |
| Aars2_in.vivo_noRT | Macc1 | 2.26 | 5.17E-04 | glio |
| Aars2_in.vivo_noRT | Cited4 | 2.5 | 5.17E-04 | glio |
| Atr_in.vivo_noRT | Zfp658 | 2.469 | 5.17E-04 | glio |
| Bora_in.vivo_noRT | Catsperz | 2.462 | 5.17E-04 | glio |
| Bora_in.vivo_noRT | Ripor2 | 2.484 | 5.17E-04 | glio |
| culture_effect | Coprs | 0.696 | 5.17E-05 | glio |
| Interaction_genotype_Mtor | 9130230L23Ri | -4.232 | 5.17E-05 | glio |
| Interaction_genotype_Mtor | Uqcrb | 2.054 | 5.17E-05 | glio |
| Atr_ex.vivo_noRT | Kcnk7 | 3.164 | 5.17E-05 | glio |
| Atr_ex.vivo_noRT | Myoz1 | 3.164 | 5.17E-05 | glio |
| Bcas2_in.vivo_noRT | Gstm7 | 2.893 | 5.17E-05 | glio |
| Hsd17b10_in.vivo_noRT | Pcdhga1 | 2.565 | 5.17E-05 | glio |
| Hsd17b10_in.vivo_noRT | BC034090 | 2.77 | 5.17E-05 | glio |
| Aars2_in.vivo_noRT | Rhbdf2 | 3.007 | 5.17E-06 | glio |
| Atr_in.vivo_noRT | Il20rb | 3.998 | 5.17E-08 | glio |
| Atr_in.vivo_noRT | Srl | 4.114 | 5.17E-09 | glio |
| Bora_in.vivo_noRT | Cstb | 1.589 | 5.17E-09 | glio |
| Chek1_ex.vivo_noRT | Gm15445 | 3.857 | 5.17E-09 | glio |
| Aars2_in.vivo_noRT | CAAA0106612 | 4.535 | 5.17E-13 | glio |
| Bcas2_in.vivo_noRT | H1foo | 4.275 | 5.17E-17 | glio |
| Bcas2_in.vivo_noRT | Gm13078 | 4.807 | 5.17E-22 | glio |
| culture_effect | Gm4841 | -6.536 | 5.17E-23 | glio |
| Aars2_in.vivo_noRT | Scg2 | 5.327 | 5.17E-29 | glio |
| culture_effect | Nop16 | -0.4 | 5.18E-03 | glio |
| Interaction_genotype_Ifnar1 | Olfr810 | 2.734 | 5.18E-03 | glio |
| Interaction_genotype_Ifnar1 | Apoa5 | 2.734 | 5.18E-03 | glio |
| Interaction_genotype_Ifnar1 | Lhx1os | 3.394 | 5.18E-03 | glio |
| Interaction_genotype_Mtor | Nt5e | -3.013 | 5.18E-03 | glio |

| | | | | |
|---|---|---|---|---|
| Interaction_genotype_Mtor | Dnaaf4 | -3.731 | 5.18E-03 | glio |
| Interaction_genotype_Mtor | Med8 | 1.283 | 5.18E-03 | glio |
| Interaction_genotype_Mtor | Smim14 | 1.438 | 5.18E-03 | glio |
| Interaction_genotype_Mtor | Vdac1 | 1.412 | 5.18E-03 | glio |
| Interaction_genotype_Myc | Dpep3 | -2.197 | 5.18E-03 | glio |
| Interaction_genotype_Myc | Dnmt3b | -3.158 | 5.18E-03 | glio |
| Interaction_genotype_Myc | Erp27 | -3.194 | 5.18E-03 | glio |
| Interaction_genotype_Myc | Ogfrl1 | -2.383 | 5.18E-03 | glio |
| Interaction_genotype_Myc | Ppp1r37 | -2.288 | 5.18E-03 | glio |
| Interaction_genotype_Rpl3 | Trnp1 | -3.82 | 5.18E-03 | glio |
| Interaction_genotype_Rpl3 | Igf2bp1 | -3.795 | 5.18E-03 | glio |
| Interaction_genotype_Rpl3 | Yaf2 | 1.3 | 5.18E-03 | glio |
| interaction_RT | Col6a1 | -1.726 | 5.18E-03 | glio |
| interaction_RT | Papss2 | -1.557 | 5.18E-03 | glio |
| Atp5c1_in.vivo_noRT | Col19a1 | 1.751 | 5.18E-03 | glio |
| Atr_in.vivo_noRT | Rps6kc1 | 1.753 | 5.18E-03 | glio |
| Atr_in.vivo_noRT | Gapvd1 | -1.053 | 5.18E-03 | glio |
| Atr_in.vivo_noRT | Ndufb9 | -0.538 | 5.18E-03 | glio |
| Bcas2_in.vivo_noRT | Rhob | -1.52 | 5.18E-03 | glio |
| Bcas2_in.vivo_noRT | Sap30bp | -1.047 | 5.18E-03 | glio |
| Bcas2_ex.vivo_noRT | A930003A15R | 2.742 | 5.18E-03 | glio |
| Bcas2_ex.vivo_noRT | Mcmdc2 | 2.414 | 5.18E-03 | glio |
| Bcas2_ex.vivo_noRT | Slc4a5 | 3.119 | 5.18E-03 | glio |
| Bora_ex.vivo_noRT | Gm45669 | 2.413 | 5.18E-03 | glio |
| Bora_ex.vivo_noRT | Gm12454 | 2.831 | 5.18E-03 | glio |
| Bora_ex.vivo_noRT | Ppfia2 | 2.407 | 5.18E-03 | glio |
| Chek1_in.vivo_noRT | Mlip | 2.039 | 5.18E-03 | glio |
| Chek1_ex.vivo_noRT | Gm26674 | 2.416 | 5.18E-03 | glio |
| Hsd17b10_in.vivo_noRT | Tmem116 | 2.438 | 5.18E-03 | glio |
| Hsd17b10_in.vivo_noRT | Pole | -1.584 | 5.18E-03 | glio |
| culture_effect | Olfml2a | 1.876 | 5.18E-04 | glio |
| culture_effect | Ist1 | -0.708 | 5.18E-04 | glio |
| Interaction_genotype_Myc | 4931431C16R | -3.112 | 5.18E-04 | glio |
| Interaction_genotype_Myc | 4933439K11R | -4.021 | 5.18E-04 | glio |
| Aars2_in.vivo_noRT | Fam174b | 2.753 | 5.18E-04 | glio |
| Aars2_in.vivo_noRT | Spcs1 | -0.863 | 5.18E-04 | glio |
| Aars2_ex.vivo_noRT | S100a14 | 2.893 | 5.18E-04 | glio |
| Bora_in.vivo_noRT | Spink10 | 2.362 | 5.18E-04 | glio |
| Bora_in.vivo_noRT | Gm42670 | 2.265 | 5.18E-04 | glio |
| Chek1_in.vivo_noRT | Ogdhl | 2.489 | 5.18E-04 | glio |
| Chek1_in.vivo_noRT | Kansl1 | -1.384 | 5.18E-04 | glio |
| culture_effect | Gm46218 | 2.03 | 5.18E-05 | glio |
| interaction_RT | Rnd2 | 1.645 | 5.18E-05 | glio |
| Aars2_in.vivo_noRT | 1700047M11F | 2.6 | 5.18E-05 | glio |
| Atr_in.vivo_noRT | Tpbg | 2.791 | 5.18E-05 | glio |
| Chek1_in.vivo_noRT | Stk32a | -1.884 | 5.18E-05 | glio |
| Aars2_in.vivo_noRT | Tmem79 | 3.042 | 5.18E-06 | glio |
| Aars2_ex.vivo_noRT | Lmx1a | 3.293 | 5.18E-06 | glio |
| Interaction_genotype_Mtor | Cgnl1 | -4.099 | 5.18E-07 | glio |
| interaction_RT | Gbp2 | 1.733 | 5.18E-07 | glio |
| Atr_in.vivo_noRT | Rnf144b | 3.476 | 5.18E-07 | glio |
| Bcas2_in.vivo_noRT | Slc7a4 | 3.393 | 5.18E-07 | glio |

| | | | | |
|---|---|---|---|---|
| Bora_in.vivo_noRT | Tmem202 | 3.494 | 5.18E-07 | glio |
| Interaction_genotype_Mtor | Gm43689 | -5.27 | 5.18E-08 | glio |
| Atr_in.vivo_noRT | Gm47061 | 4.228 | 5.18E-08 | glio |
| Chek1_ex.vivo_noRT | Pcdhgb5 | 4.449 | 5.18E-09 | glio |
| Bcas2_in.vivo_noRT | Gjb5 | 3.817 | 5.18E-10 | glio |
| Interaction_genotype_Mtor | Nfatc2 | -3.625 | 5.19E-03 | glio |
| Interaction_genotype_Mtor | Gm29394 | -3.284 | 5.19E-03 | glio |
| Interaction_genotype_Mtor | Ankrd11 | 1.507 | 5.19E-03 | glio |
| Interaction_genotype_Myc | Dclk2 | -2.559 | 5.19E-03 | glio |
| interaction_RT | Lcorl | 0.999 | 5.19E-03 | glio |
| interaction_RT | Pdha1 | -0.559 | 5.19E-03 | glio |
| Aars2_in.vivo_noRT | Zfp27 | 1.862 | 5.19E-03 | glio |
| Aars2_in.vivo_noRT | Ppm1m | 1.308 | 5.19E-03 | glio |
| Aars2_in.vivo_noRT | Depdc7 | -1.416 | 5.19E-03 | glio |
| Atr_in.vivo_noRT | Tent5d | 1.938 | 5.19E-03 | glio |
| Bard1_ex.vivo_noRT | BC018473 | 2.799 | 5.19E-03 | glio |
| Bcas2_in.vivo_noRT | A930029G22R | 2.006 | 5.19E-03 | glio |
| Bora_in.vivo_noRT | Zfand4 | 2.131 | 5.19E-03 | glio |
| Cenpj_in.vivo_noRT | Gm18280 | 1.758 | 5.19E-03 | glio |
| Chek1_in.vivo_noRT | Isca2 | -1.217 | 5.19E-03 | glio |
| Chek1_ex.vivo_noRT | Hectd2os | 2.468 | 5.19E-03 | glio |
| Chek1_ex.vivo_noRT | Rpl22l1 | -0.793 | 5.19E-03 | glio |
| culture_effect | Atf7 | -0.618 | 5.19E-04 | glio |
| Interaction_genotype_Mtor | Gm17300 | -4.075 | 5.19E-04 | glio |
| Interaction_genotype_Myc | Gm9776 | -3.772 | 5.19E-04 | glio |
| Interaction_genotype_Myc | Nanp | -3.036 | 5.19E-04 | glio |
| Interaction_genotype_Myc | Trip13 | -2.872 | 5.19E-04 | glio |
| Atp5c1_in.vivo_noRT | Gm45151 | 2.136 | 5.19E-04 | glio |
| culture_effect | 2900076A07R | 1.165 | 5.19E-05 | glio |
| Aars2_in.vivo_noRT | Slamf1 | 2.887 | 5.19E-05 | glio |
| Bora_in.vivo_noRT | Gask1b | 3.141 | 5.19E-05 | glio |
| Bora_in.vivo_noRT | Rapgef3 | 3.205 | 5.19E-05 | glio |
| culture_effect | 6430503K07R | 1.797 | 5.19E-06 | glio |
| Interaction_genotype_Myc | 5430400D12R | -5.278 | 5.19E-06 | glio |
| Aars2_in.vivo_noRT | Fhl3 | 3.227 | 5.19E-06 | glio |
| Chek1_ex.vivo_noRT | Prl2c3 | 3.647 | 5.19E-06 | glio |
| Atr_in.vivo_noRT | Pcdhgb1 | 2.999 | 5.19E-07 | glio |
| Aars2_in.vivo_noRT | Dct | 4.523 | 5.19E-08 | glio |
| Chek1_in.vivo_noRT | Atoh8 | 3.661 | 5.19E-08 | glio |
| Chek1_ex.vivo_noRT | Trim67 | 4.146 | 5.19E-08 | glio |
| Atr_in.vivo_noRT | Kctd21 | 3.973 | 5.19E-09 | glio |
| Bora_in.vivo_noRT | Zkscan4 | 4.136 | 5.19E-09 | glio |
| Hsd17b10_in.vivo_noRT | 4930423D22R | 3.356 | 5.19E-11 | glio |
| Chek1_in.vivo_noRT | Nxph4 | 5.044 | 5.19E-13 | glio |
| Bora_in.vivo_noRT | B230206H07R | 4.411 | 5.19E-20 | glio |
| culture_effect | Os9 | -1.868 | 5.19E-22 | glio |
| Atr_in.vivo_noRT | Gm26671 | 5.533 | 5.19E-28 | glio |
| Atr_in.vivo_noRT | Gm49504 | 6.325 | 5.19E-30 | glio |
| Interaction_genotype_Mtor | Fastkd3 | -2.604 | 5.20E-03 | glio |
| Interaction_genotype_Mtor | Slc35e1 | -2.12 | 5.20E-03 | glio |
| Interaction_genotype_Mtor | Clcf1 | -2.634 | 5.20E-03 | glio |
| Interaction_genotype_Mtor | Mcrip2 | -1.727 | 5.20E-03 | glio |

| | | | | |
|---|---|---|---|---|
| Interaction_genotype_Myc | C330024C12R | -2.731 | 5.20E-03 | glio |
| Interaction_genotype_Myc | Etv2 | -2.821 | 5.20E-03 | glio |
| Interaction_genotype_Myc | Eral1 | -2.544 | 5.20E-03 | glio |
| Interaction_genotype_Myc | Stamos | -2.653 | 5.20E-03 | glio |
| Aars2_in.vivo_noRT | Srsf3 | -0.657 | 5.20E-03 | glio |
| Aars2_ex.vivo_noRT | Fhad1 | 2.706 | 5.20E-03 | glio |
| Atr_in.vivo_noRT | Dlat | -1.096 | 5.20E-03 | glio |
| Bcas2_in.vivo_noRT | Fancl | -1.667 | 5.20E-03 | glio |
| Bcas2_ex.vivo_noRT | Ciita | 2.342 | 5.20E-03 | glio |
| Chek1_ex.vivo_noRT | Mr1 | 2.665 | 5.20E-03 | glio |
| Cyp19a1_ex.vivo_noRT | Tpo | 2.159 | 5.20E-03 | glio |
| Hsd17b10_in.vivo_noRT | 4930589L23Ri | 2.269 | 5.20E-03 | glio |
| Interaction_genotype_Aars2 | D030068K23R | -4.414 | 5.20E-04 | glio |
| Interaction_genotype_Mtor | Gm42918 | -3.636 | 5.20E-04 | glio |
| Interaction_genotype_Mtor | Phf19 | -3.908 | 5.20E-04 | glio |
| Interaction_genotype_Mtor | Armcx1 | -3.391 | 5.20E-04 | glio |
| Interaction_genotype_Mtor | Emc2 | 1.819 | 5.20E-04 | glio |
| Interaction_genotype_Mtor | Utrn | 2.133 | 5.20E-04 | glio |
| Interaction_genotype_Mtor | Eif3e | 1.272 | 5.20E-04 | glio |
| Aars2_in.vivo_noRT | A330074K22R | 2.25 | 5.20E-04 | glio |
| Aars2_in.vivo_noRT | D130020L05R | 2.146 | 5.20E-04 | glio |
| Atr_in.vivo_noRT | Efna4 | 2.515 | 5.20E-04 | glio |
| Bcas2_in.vivo_noRT | Unc5cl | 2.505 | 5.20E-04 | glio |
| Bcas2_in.vivo_noRT | Cnp | -1.726 | 5.20E-04 | glio |
| Cenpj_ex.vivo_noRT | Gm14820 | 2.591 | 5.20E-04 | glio |
| Interaction_genotype_Bora | Pom121 | 2.525 | 5.20E-05 | glio |
| Interaction_genotype_Bora | Prrc2a | 2.16 | 5.20E-05 | glio |
| Interaction_genotype_Bora | Hdac9 | -5.059 | 5.20E-05 | glio |
| Aars2_ex.vivo_noRT | Atp8a2 | 3.305 | 5.20E-05 | glio |
| Aars2_ex.vivo_noRT | Pcdh15 | 3.293 | 5.20E-05 | glio |
| Bcas2_in.vivo_noRT | Gm20522 | 3.001 | 5.20E-05 | glio |
| Bora_in.vivo_noRT | Cep72 | 2.815 | 5.20E-05 | glio |
| Bora_ex.vivo_noRT | Ccl19 | 3.201 | 5.20E-05 | glio |
| Chek1_ex.vivo_noRT | Il2rb | 3.464 | 5.20E-05 | glio |
| Aars2_in.vivo_noRT | Cct6b | 3.037 | 5.20E-06 | glio |
| Aars2_in.vivo_noRT | E230025N22R | 2.98 | 5.20E-06 | glio |
| Bora_in.vivo_noRT | Fam222a | 3.422 | 5.20E-06 | glio |
| Atr_in.vivo_noRT | Cracr2b | 3.664 | 5.20E-07 | glio |
| Chek1_in.vivo_noRT | Olfml2a | 3.541 | 5.20E-07 | glio |
| Bora_in.vivo_noRT | Cpeb1 | 4.291 | 5.20E-09 | glio |
| Aars2_in.vivo_noRT | Medag | 4.159 | 5.20E-10 | glio |
| Atr_in.vivo_noRT | Zic4 | 5.27 | 5.20E-13 | glio |
| Atr_in.vivo_noRT | Amy1 | 5.004 | 5.20E-14 | glio |
| culture_effect | Eif4e3 | -4.531 | 5.20E-19 | glio |
| Atr_in.vivo_noRT | Gm15816 | 5.533 | 5.20E-19 | glio |
| culture_effect | Hist1h2bh | 1.408 | 5.21E-03 | glio |
| Interaction_genotype_Mtor | Gpr37l1 | -3.378 | 5.21E-03 | glio |
| Interaction_genotype_Mtor | AC125149.3 | -3.266 | 5.21E-03 | glio |
| Interaction_genotype_Mtor | Fcsk | -3.111 | 5.21E-03 | glio |
| Interaction_genotype_Rad51 | Gm48682 | -3.813 | 5.21E-03 | glio |
| Interaction_genotype_Rad51 | Pcdh17 | -2.815 | 5.21E-03 | glio |
| Atp5c1_in.vivo_noRT | Zic4 | 2.405 | 5.21E-03 | glio |

| | | | | |
|---|---|---|---|---|
| Bcas2_in.vivo_noRT | Fgf15 | 2.238 | 5.21E-03 | glio |
| Bcas2_in.vivo_noRT | Trim65 | 1.967 | 5.21E-03 | glio |
| Bcas2_in.vivo_noRT | Galk2 | 1.442 | 5.21E-03 | glio |
| Bora_in.vivo_noRT | Phf11b | 2.281 | 5.21E-03 | glio |
| Bora_in.vivo_noRT | Gm13470 | 1.868 | 5.21E-03 | glio |
| Bora_in.vivo_noRT | 4930578M01F | 1.808 | 5.21E-03 | glio |
| Bora_in.vivo_noRT | Fbxo5 | -1.552 | 5.21E-03 | glio |
| Cenpj_in.vivo_noRT | Gm50321 | 2.022 | 5.21E-03 | glio |
| Chek1_in.vivo_noRT | Il1rapl1 | 2.076 | 5.21E-03 | glio |
| Chek1_in.vivo_noRT | Zfp748 | 2.023 | 5.21E-03 | glio |
| Chek1_in.vivo_noRT | Gm45902 | 2.039 | 5.21E-03 | glio |
| Chek1_in.vivo_noRT | Alkbh5 | -0.942 | 5.21E-03 | glio |
| Fanca_ex.vivo_noRT | 4930527F14Ri | 2.524 | 5.21E-03 | glio |
| Interaction_genotype_Mtor | 2810021J22Ri | -4.162 | 5.21E-04 | glio |
| Interaction_genotype_Myc | Tmem202 | -3.924 | 5.21E-04 | glio |
| Interaction_genotype_Myc | Col8a1 | -3.704 | 5.21E-04 | glio |
| Interaction_genotype_Myc | Setd4 | -3.883 | 5.21E-04 | glio |
| Interaction_genotype_Myc | Gm19531 | -3.438 | 5.21E-04 | glio |
| Interaction_genotype_Rpl3 | Hbq1b | -4.624 | 5.21E-04 | glio |
| Interaction_genotype_Rpl3 | Gm48242 | -4.074 | 5.21E-04 | glio |
| Atr_in.vivo_noRT | Paqr7 | 2.351 | 5.21E-04 | glio |
| Bcas2_in.vivo_noRT | Cnn1 | 3.137 | 5.21E-04 | glio |
| Bcas2_in.vivo_noRT | Slc33a1 | -1.856 | 5.21E-04 | glio |
| Bora_in.vivo_noRT | Arhgdib | 2.515 | 5.21E-04 | glio |
| Hsd17b10_in.vivo_noRT | Sbk3 | 2.447 | 5.21E-04 | glio |
| Interaction_genotype_Myc | A030014E15Ri | -3.122 | 5.21E-05 | glio |
| Chek1_in.vivo_noRT | Dhrs13os | 3.208 | 5.21E-05 | glio |
| Hsd17b10_in.vivo_noRT | Has1 | 2.719 | 5.21E-05 | glio |
| Bora_ex.vivo_noRT | Gm11899 | 3.201 | 5.21E-06 | glio |
| Chek1_ex.vivo_noRT | Pid1 | 4.055 | 5.21E-07 | glio |
| Bora_in.vivo_noRT | Papln | 3.669 | 5.21E-08 | glio |
| Atr_in.vivo_noRT | Rtl1 | 3.841 | 5.21E-10 | glio |
| Bora_in.vivo_noRT | Gm14968 | 4.557 | 5.21E-10 | glio |
| culture_effect | Asns | -1.175 | 5.21E-13 | glio |
| Chek1_in.vivo_noRT | Ms4a15 | 4.805 | 5.21E-18 | glio |
| Atr_in.vivo_noRT | Slc1a2 | 5.269 | 5.21E-19 | glio |
| Interaction_genotype_Aars2 | Gm30692 | -2.427 | 5.22E-03 | glio |
| Interaction_genotype_Aars2 | Prickle4 | -3.301 | 5.22E-03 | glio |
| Interaction_genotype_Aars2 | Dnase2a | -2.619 | 5.22E-03 | glio |
| Interaction_genotype_Atr | Mettl7a3 | -3.314 | 5.22E-03 | glio |
| Interaction_genotype_Atr | E230029C05Ri | -3.011 | 5.22E-03 | glio |
| Interaction_genotype_Mtor | Cd302 | 1.674 | 5.22E-03 | glio |
| Interaction_genotype_Myc | 1700011D18R | -3.099 | 5.22E-03 | glio |
| Interaction_genotype_Myc | 2610203C22R | -3.348 | 5.22E-03 | glio |
| Interaction_genotype_Myc | Gm6598 | -2.799 | 5.22E-03 | glio |
| Interaction_genotype_Myc | 9430015G10R | -2.577 | 5.22E-03 | glio |
| Interaction_genotype_Myc | Gal3st1 | -3.055 | 5.22E-03 | glio |
| Interaction_genotype_Myc | Ankrd11 | 1.325 | 5.22E-03 | glio |
| Aars2_in.vivo_noRT | Ube2f | -1.131 | 5.22E-03 | glio |
| Aars2_in.vivo_noRT | Clns1a | -0.979 | 5.22E-03 | glio |
| Aars2_in.vivo_noRT | Ssbp1 | -0.655 | 5.22E-03 | glio |
| Atr_in.vivo_noRT | Gm49741 | 2.164 | 5.22E-03 | glio |

| | | | | |
|---|---|---|---|---|
| Atr_in.vivo_noRT | Zfp341 | 1.837 | 5.22E-03 | glio |
| Atr_in.vivo_noRT | Ubap2 | -0.989 | 5.22E-03 | glio |
| Bard1_ex.vivo_noRT | Scgb1c1 | 2.421 | 5.22E-03 | glio |
| Bcas2_ex.vivo_noRT | Cacng5 | 2.742 | 5.22E-03 | glio |
| Bcas2_ex.vivo_noRT | Serpina3i | 2.742 | 5.22E-03 | glio |
| Bcas2_ex.vivo_noRT | Gm5934 | 2.163 | 5.22E-03 | glio |
| Bcas2_ex.vivo_noRT | Fbxo15 | 2.542 | 5.22E-03 | glio |
| Chek1_in.vivo_noRT | Zfp341 | 1.871 | 5.22E-03 | glio |
| Chek1_ex.vivo_noRT | Ttc26 | 2.237 | 5.22E-03 | glio |
| Chek1_ex.vivo_noRT | Atad1 | -1.352 | 5.22E-03 | glio |
| Hsd17b10_in.vivo_noRT | Gm35867 | 2.163 | 5.22E-03 | glio |
| Hsd17b10_in.vivo_noRT | Cfap100 | 2.179 | 5.22E-03 | glio |
| culture_effect | Mov10 | -0.724 | 5.22E-04 | glio |
| culture_effect | Cse1l | 0.601 | 5.22E-04 | glio |
| culture_effect | Uaca | 0.597 | 5.22E-04 | glio |
| Interaction_genotype_Mtor | Samd15 | -4.547 | 5.22E-04 | glio |
| Bcas2_in.vivo_noRT | Gm32031 | 2.208 | 5.22E-04 | glio |
| Bora_in.vivo_noRT | Ap4s1 | -1.637 | 5.22E-04 | glio |
| Chek1_ex.vivo_noRT | Fhdc1 | 3.384 | 5.22E-04 | glio |
| Hsd17b10_in.vivo_noRT | Rax | 2.437 | 5.22E-04 | glio |
| Interaction_genotype_Mtor | Proscos | -4.927 | 5.22E-05 | glio |
| interaction_RT | Plec | -0.869 | 5.22E-05 | glio |
| Aars2_in.vivo_noRT | Shbg | 2.948 | 5.22E-05 | glio |
| Aars2_in.vivo_noRT | Ccdc57 | 2.98 | 5.22E-05 | glio |
| Atp5c1_in.vivo_noRT | Npr3 | 2.668 | 5.22E-05 | glio |
| Bcas2_in.vivo_noRT | Kank2 | 2.772 | 5.22E-05 | glio |
| Bora_in.vivo_noRT | Tex14 | 2.595 | 5.22E-05 | glio |
| Hsd17b10_in.vivo_noRT | Ankrd1 | 3.616 | 5.22E-05 | glio |
| culture_effect | Rbm43 | -1.102 | 5.22E-06 | glio |
| interaction_RT | Itga6 | -1.079 | 5.22E-06 | glio |
| interaction_RT | Laptm4a | 0.802 | 5.22E-06 | glio |
| Aars2_in.vivo_noRT | Zfp661 | 3.576 | 5.22E-07 | glio |
| Bcas2_in.vivo_noRT | Gstm2 | 3.625 | 5.22E-07 | glio |
| Chek1_in.vivo_noRT | Nox1 | 3.061 | 5.22E-07 | glio |
| Bora_in.vivo_noRT | Wfdc18 | 4.67 | 5.22E-08 | glio |
| culture_effect | Psma2 | -0.705 | 5.22E-09 | glio |
| Aars2_in.vivo_noRT | Dzank1 | 4.151 | 5.22E-09 | glio |
| Bcas2_in.vivo_noRT | Cyp4x1 | 4.541 | 5.22E-12 | glio |
| Chek1_in.vivo_noRT | Gm32834 | 4.65 | 5.22E-12 | glio |
| Aars2_in.vivo_noRT | F630040K05Ri | 4.676 | 5.22E-14 | glio |
| Bcas2_in.vivo_noRT | 8030423F21Ri | 4.808 | 5.22E-16 | glio |
| Chek1_ex.vivo_noRT | Vmn2r18 | 4.842 | 5.22E-16 | glio |
| Aars2_in.vivo_noRT | Capn9 | 5.327 | 5.22E-17 | glio |
| Atr_in.vivo_noRT | 9130230L23Ri | 5.273 | 5.22E-18 | glio |
| culture_effect | Sec23b | -0.566 | 5.23E-03 | glio |
| Interaction_genotype_Chek1 | 1700026J14Ri | -3.514 | 5.23E-03 | glio |
| Interaction_genotype_Chek1 | Nek6 | -3.242 | 5.23E-03 | glio |
| Interaction_genotype_Myc | Poc1a | -2.656 | 5.23E-03 | glio |
| Interaction_genotype_Nat2 | Scel | 2.808 | 5.23E-03 | glio |
| Atr_in.vivo_noRT | Wdpcp | 1.916 | 5.23E-03 | glio |
| Atr_in.vivo_noRT | Star | 1.968 | 5.23E-03 | glio |
| Chek1_in.vivo_noRT | 4930404N11R | 1.907 | 5.23E-03 | glio |

| | | | | |
|---|---|---|---|---|
| Chek1_in.vivo_noRT | Slitrk6 | 1.826 | 5.23E-03 | glio |
| Chek1_in.vivo_noRT | Hspa4 | -0.979 | 5.23E-03 | glio |
| Chek1_in.vivo_noRT | Laptm4a | -0.625 | 5.23E-03 | glio |
| Eif2s1_in.vivo_noRT | Slc10a1 | 1.883 | 5.23E-03 | glio |
| culture_effect | Tmem60 | 0.53 | 5.23E-04 | glio |
| Aars2_in.vivo_noRT | Skp1a | -1.039 | 5.23E-04 | glio |
| Atp5c1_in.vivo_noRT | Cstdc4 | 2.539 | 5.23E-04 | glio |
| Atp5c1_in.vivo_noRT | Mettl7b | 2.668 | 5.23E-04 | glio |
| Atr_ex.vivo_noRT | S100a13 | 1.189 | 5.23E-04 | glio |
| Bard1_ex.vivo_noRT | Vmn2r54 | 2.814 | 5.23E-04 | glio |
| Bora_ex.vivo_noRT | Gm5662 | 3.201 | 5.23E-04 | glio |
| Bora_ex.vivo_noRT | Tcp11x2 | 2.808 | 5.23E-04 | glio |
| Bora_ex.vivo_noRT | Gm8013 | 3.407 | 5.23E-04 | glio |
| Bora_ex.vivo_noRT | Cfap46 | 2.976 | 5.23E-04 | glio |
| Interaction_genotype_Myc | Acvr1 | -3.267 | 5.23E-05 | glio |
| Bcas2_in.vivo_noRT | Dnaaf4 | 3.143 | 5.23E-05 | glio |
| Chek1_in.vivo_noRT | Ifi47 | -1.781 | 5.23E-05 | glio |
| Interaction_genotype_Mtor | Hcn3 | -4.959 | 5.23E-06 | glio |
| Bcas2_in.vivo_noRT | Tnni2 | 3.769 | 5.23E-06 | glio |
| Chek1_ex.vivo_noRT | Diaph3 | -2.948 | 5.23E-06 | glio |
| culture_effect | Tex14 | -2.876 | 5.23E-07 | glio |
| Interaction_genotype_Mtor | Gm26543 | -4.099 | 5.23E-07 | glio |
| Interaction_genotype_Myc | Gm17249 | -5.503 | 5.23E-07 | glio |
| Chek1_ex.vivo_noRT | Cd72 | 4.447 | 5.23E-07 | glio |
| culture_effect | Hsh2d | -3.055 | 5.23E-08 | glio |
| Atr_in.vivo_noRT | Gm13272 | 3.83 | 5.23E-09 | glio |
| Bora_in.vivo_noRT | Tspyl3 | 3.853 | 5.23E-10 | glio |
| Interaction_genotype_Mtor | Gm2115 | -7.135 | 5.23E-11 | glio |
| Chek1_in.vivo_noRT | Gm13348 | 5.044 | 5.23E-18 | glio |
| culture_effect | Mex3d | -0.599 | 5.24E-03 | glio |
| Interaction_genotype_Aars2 | Slurp1 | -5.13 | 5.24E-03 | glio |
| Interaction_genotype_Cdkn2a | Gm14929 | 2.835 | 5.24E-03 | glio |
| Interaction_genotype_Mtor | Piwil4 | -2.818 | 5.24E-03 | glio |
| Interaction_genotype_Mtor | Ppp1r1b | -3.109 | 5.24E-03 | glio |
| Interaction_genotype_Mtor | Zranb3 | -2.522 | 5.24E-03 | glio |
| Interaction_genotype_Myc | Dmac2 | -2.554 | 5.24E-03 | glio |
| Interaction_genotype_Myc | Arpc5l | 1.161 | 5.24E-03 | glio |
| interaction_RT | Pik3r3 | 1.826 | 5.24E-03 | glio |
| interaction_RT | Nsf | -0.926 | 5.24E-03 | glio |
| Atp5c1_in.vivo_noRT | Krtdap | 2.277 | 5.24E-03 | glio |
| Atr_ex.vivo_noRT | Hsf4 | 2.791 | 5.24E-03 | glio |
| Bcas2_in.vivo_noRT | Hyou1 | -1.357 | 5.24E-03 | glio |
| Bcas2_in.vivo_noRT | Polr2e | -0.728 | 5.24E-03 | glio |
| Bora_in.vivo_noRT | Crocc | 2.011 | 5.24E-03 | glio |
| Bora_in.vivo_noRT | Tyms | -1.327 | 5.24E-03 | glio |
| Cdkn2a_in.vivo_noRT | 4933400F21Ri | -1.939 | 5.24E-03 | glio |
| Chek1_in.vivo_noRT | Gm43707 | 1.811 | 5.24E-03 | glio |
| Interaction_genotype_Mtor | Dennd4b | -4.141 | 5.24E-04 | glio |
| Interaction_genotype_Mtor | Trafd1 | 2.47 | 5.24E-04 | glio |
| Interaction_genotype_Myc | Noxred1 | -3.772 | 5.24E-04 | glio |
| Interaction_genotype_Myc | Ccno | -4.07 | 5.24E-04 | glio |
| Aars2_in.vivo_noRT | Pcsk4 | 2.478 | 5.24E-04 | glio |

| | | | | |
|---|---|---|---|---|
| Aars2_in.vivo_noRT | Rbis | -1.315 | 5.24E-04 | glio |
| Atr_ex.vivo_noRT | Dcpp3 | 3.164 | 5.24E-04 | glio |
| Cenpj_in.vivo_noRT | Abcb5 | 2.022 | 5.24E-04 | glio |
| culture_effect | Krt80 | 2.067 | 5.24E-05 | glio |
| Interaction_genotype_Myc | Gm36210 | -4.588 | 5.24E-05 | glio |
| Aars2_in.vivo_noRT | A330009N23R | 2.774 | 5.24E-05 | glio |
| Bora_in.vivo_noRT | Ptcd3 | -1.832 | 5.24E-05 | glio |
| Interaction_genotype_Pars2 | Gm2115 | -5.304 | 5.24E-06 | glio |
| Interaction_genotype_Pars2 | Sod3 | -6.137 | 5.24E-06 | glio |
| Interaction_genotype_Pars2 | Gm48242 | -5.217 | 5.24E-06 | glio |
| Bora_in.vivo_noRT | Abi3bp | 3.231 | 5.24E-06 | glio |
| Aars2_in.vivo_noRT | Zfp273 | 3.56 | 5.24E-07 | glio |
| Aars2_in.vivo_noRT | Slco2a1 | 3.328 | 5.24E-07 | glio |
| Chek1_in.vivo_noRT | Dok3 | 3.908 | 5.24E-08 | glio |
| Chek1_in.vivo_noRT | Arhgef15 | 3.663 | 5.24E-08 | glio |
| Chek1_in.vivo_noRT | Il20rb | 4.02 | 5.24E-08 | glio |
| Chek1_in.vivo_noRT | Tmem86b | 3.877 | 5.24E-08 | glio |
| Atr_in.vivo_noRT | Tnfrsf23 | 4.615 | 5.24E-09 | glio |
| Chek1_in.vivo_noRT | Gm8013 | 4.399 | 5.24E-10 | glio |
| Chek1_in.vivo_noRT | A330015K06R | 4.387 | 5.24E-12 | glio |
| Bcas2_in.vivo_noRT | Gm47404 | 4.807 | 5.24E-23 | glio |
| Chek1_in.vivo_noRT | Helt | 5.044 | 5.24E-26 | glio |
| Chek1_in.vivo_noRT | Gm20625 | 5.044 | 5.24E-26 | glio |
| culture_effect | Chst12 | 0.889 | 5.25E-03 | glio |
| Interaction_genotype_Atr | Cgnl1 | -2.369 | 5.25E-03 | glio |
| Interaction_genotype_Atr | Gm15445 | -2.709 | 5.25E-03 | glio |
| Interaction_genotype_Atr | Gm26543 | -2.369 | 5.25E-03 | glio |
| Interaction_genotype_Atr | 5830454E08Ri | -3.076 | 5.25E-03 | glio |
| Interaction_genotype_Atr | Fip1l1 | 1.309 | 5.25E-03 | glio |
| Interaction_genotype_Mtor | BC048671 | -2.798 | 5.25E-03 | glio |
| Interaction_genotype_Mtor | Dnah2os | -3.24 | 5.25E-03 | glio |
| Interaction_genotype_Mtor | Dhrs3 | -3.327 | 5.25E-03 | glio |
| Interaction_genotype_Mtor | Gm49463 | -2.59 | 5.25E-03 | glio |
| Interaction_genotype_Myc | Mafa | -2.928 | 5.25E-03 | glio |
| Interaction_genotype_Myc | Nmnat3 | -3.218 | 5.25E-03 | glio |
| Interaction_genotype_Myc | Zfp110 | -2.105 | 5.25E-03 | glio |
| Aars2_in.vivo_noRT | Rbm34 | -1.054 | 5.25E-03 | glio |
| Aars2_in.vivo_noRT | Myo1e | 1.198 | 5.25E-03 | glio |
| Atr_in.vivo_noRT | Slc25a20 | 1.483 | 5.25E-03 | glio |
| Bcas2_in.vivo_noRT | Commd5 | 2.056 | 5.25E-03 | glio |
| Chek1_in.vivo_noRT | 1700063D05R | 2.131 | 5.25E-03 | glio |
| Interaction_genotype_Bcas2 | K230015D01R | -4.61 | 5.25E-04 | glio |
| Interaction_genotype_Mtor | Ldlrad3 | -3.572 | 5.25E-04 | glio |
| Interaction_genotype_Mtor | Mrps18a | 1.601 | 5.25E-04 | glio |
| Interaction_genotype_Myc | Gm37305 | -3.733 | 5.25E-04 | glio |
| interaction_RT | Cct8 | -0.666 | 5.25E-04 | glio |
| Atr_in.vivo_noRT | Skp1a | -1.046 | 5.25E-04 | glio |
| Bora_in.vivo_noRT | 5730414N17R | 2.667 | 5.25E-04 | glio |
| Cenpj_ex.vivo_noRT | Gm4166 | 2.591 | 5.25E-04 | glio |
| Interaction_genotype_Myc | Gm49694 | -4.587 | 5.25E-05 | glio |
| Interaction_genotype_Myc | BC028471 | -4.611 | 5.25E-05 | glio |
| interaction_RT | Nacc2 | -1.36 | 5.25E-05 | glio |

| | | | | |
|---|---|---|---|---|
| Aars2_in.vivo_noRT | Rrad | 3.064 | 5.25E-05 | glio |
| Aars2_in.vivo_noRT | Gm20324 | 2.519 | 5.25E-05 | glio |
| Interaction_genotype_Mtor | Cacnb3 | -4.515 | 5.25E-06 | glio |
| Atr_in.vivo_noRT | Fam83e | 3.284 | 5.25E-06 | glio |
| Bard1_ex.vivo_noRT | Cldn2 | 2.814 | 5.25E-06 | glio |
| Bard1_ex.vivo_noRT | Slc45a1 | 3.213 | 5.25E-06 | glio |
| Bcas2_in.vivo_noRT | Rnase4 | 3.245 | 5.25E-06 | glio |
| Chek1_in.vivo_noRT | 4930563E22Ri | 3.325 | 5.25E-06 | glio |
| culture_effect | Arl6ip4 | 0.787 | 5.25E-07 | glio |
| culture_effect | Srp19 | 0.562 | 5.25E-07 | glio |
| Interaction_genotype_Nat2 | Tcstv3 | 5.472 | 5.25E-07 | glio |
| Interaction_genotype_Nat2 | Gm6803 | 4.712 | 5.25E-07 | glio |
| interaction_RT | Sapcd2 | -1.681 | 5.25E-07 | glio |
| interaction_RT | Cd9 | 1.142 | 5.25E-07 | glio |
| Aars2_in.vivo_noRT | Upk3bl | 3.25 | 5.25E-07 | glio |
| Aars2_in.vivo_noRT | Spx | 3.601 | 5.25E-08 | glio |
| Aars2_ex.vivo_noRT | Scg2 | 3.293 | 5.25E-08 | glio |
| Aars2_ex.vivo_noRT | 1700003E24Ri | 3.293 | 5.25E-08 | glio |
| Chek1_ex.vivo_noRT | Rab27a | 4.447 | 5.25E-09 | glio |
| Atr_in.vivo_noRT | Eps8l2 | 4.431 | 5.25E-10 | glio |
| Bora_in.vivo_noRT | Acat3 | 4.144 | 5.25E-14 | glio |
| Chek1_in.vivo_noRT | Slc1a2 | 4.78 | 5.25E-16 | glio |
| Bcas2_in.vivo_noRT | Gm48170 | 4.807 | 5.25E-18 | glio |
| Bcas2_in.vivo_noRT | AF067063 | 4.807 | 5.25E-20 | glio |
| culture_effect | Aldh1a1 | -1.859 | 5.26E-03 | glio |
| Interaction_genotype_Aars2 | Hebp1 | 2.714 | 5.26E-03 | glio |
| Interaction_genotype_Aars2 | Mrpl3 | 1.605 | 5.26E-03 | glio |
| Interaction_genotype_Atr | Slc46a1 | -3.636 | 5.26E-03 | glio |
| Interaction_genotype_Mtor | Gm28809 | -3.047 | 5.26E-03 | glio |
| Atr_in.vivo_noRT | Ftsj3 | -1.025 | 5.26E-03 | glio |
| Bcas2_in.vivo_noRT | Tapbp | -0.79 | 5.26E-03 | glio |
| Bcas2_ex.vivo_noRT | Ascl4 | 2.958 | 5.26E-03 | glio |
| Brca2_in.vivo_noRT | 4930570E03Ri | 2.036 | 5.26E-03 | glio |
| Brca2_in.vivo_noRT | 4930588A03R | 1.763 | 5.26E-03 | glio |
| Brca2_in.vivo_noRT | Rab27b | 2.498 | 5.26E-03 | glio |
| Brca2_in.vivo_noRT | Gm13856 | 1.763 | 5.26E-03 | glio |
| Brca2_in.vivo_noRT | Flacc1 | 2.288 | 5.26E-03 | glio |
| Brca2_in.vivo_noRT | Gm38534 | 2.546 | 5.26E-03 | glio |
| Brca2_in.vivo_noRT | Gm21936 | 1.763 | 5.26E-03 | glio |
| Chek1_in.vivo_noRT | Slc6a20b | 2.036 | 5.26E-03 | glio |
| Chek1_in.vivo_noRT | Rhbdf1 | -1.409 | 5.26E-03 | glio |
| Cyp19a1_ex.vivo_noRT | Gm14929 | 2.159 | 5.26E-03 | glio |
| Cyp19a1_ex.vivo_noRT | Rimklb | 2.159 | 5.26E-03 | glio |
| Cyp19a1_ex.vivo_noRT | Phf11d | -2.738 | 5.26E-03 | glio |
| Eif2s1_in.vivo_noRT | 2010001K21R | 2.143 | 5.26E-03 | glio |
| Hsd17b10_in.vivo_noRT | Cmc2 | -1.026 | 5.26E-03 | glio |
| culture_effect | Mff | 0.524 | 5.26E-04 | glio |
| Interaction_genotype_Aars2 | Snd1 | 1.732 | 5.26E-04 | glio |
| Interaction_genotype_Myc | Dym | -2.549 | 5.26E-04 | glio |
| interaction_RT | Ptgds | 2.921 | 5.26E-04 | glio |
| Aars2_in.vivo_noRT | Kif5c | 1.967 | 5.26E-04 | glio |
| Atp5c1_in.vivo_noRT | Nrg2 | 2.673 | 5.26E-04 | glio |

| | | | | |
|---|---|---|---|---|
| Atr_in.vivo_noRT | Xndc1 | 1.88 | 5.26E-04 | glio |
| Bard1_ex.vivo_noRT | Aqp9 | 2.814 | 5.26E-04 | glio |
| Bora_in.vivo_noRT | Ldhb | -2.099 | 5.26E-05 | glio |
| Chek1_in.vivo_noRT | Gm15834 | 2.863 | 5.26E-05 | glio |
| Bora_in.vivo_noRT | Hey2 | 3.089 | 5.26E-06 | glio |
| Chek1_in.vivo_noRT | Igsf23 | 3.384 | 5.26E-06 | glio |
| Atr_ex.vivo_noRT | Angptl7 | 3.164 | 5.26E-07 | glio |
| Hsd17b10_in.vivo_noRT | Gm15743 | 3.353 | 5.26E-07 | glio |
| Chek1_ex.vivo_noRT | BC034090 | 4.449 | 5.26E-08 | glio |
| Bcas2_in.vivo_noRT | Sult6b1 | 4.016 | 5.26E-09 | glio |
| Chek1_ex.vivo_noRT | Tnfrsf1b | 4.842 | 5.26E-09 | glio |
| Bcas2_in.vivo_noRT | Gm17173 | 4.42 | 5.26E-11 | glio |
| Aars2_in.vivo_noRT | Kcnq2 | 4.163 | 5.26E-12 | glio |
| Bora_in.vivo_noRT | Matn4 | 4.672 | 5.26E-12 | glio |
| Bora_in.vivo_noRT | Gm5941 | 4.936 | 5.26E-12 | glio |
| Hsd17b10_in.vivo_noRT | Esp24 | 3.616 | 5.26E-12 | glio |
| Bora_in.vivo_noRT | Gm30042 | 4.676 | 5.26E-16 | glio |
| Bcas2_in.vivo_noRT | Agap2 | 4.807 | 5.26E-17 | glio |
| Bora_in.vivo_noRT | Dpep3 | 4.404 | 5.26E-19 | glio |
| Aars2_in.vivo_noRT | Impg2 | 5.327 | 5.26E-20 | glio |
| Interaction_genotype_Aars2 | Gm9908 | -3.769 | 5.27E-03 | glio |
| Interaction_genotype_Aars2 | Gm45871 | -3.162 | 5.27E-03 | glio |
| Interaction_genotype_Atr | Arhgef17 | -3.134 | 5.27E-03 | glio |
| Bcas2_in.vivo_noRT | Mrpl44 | -1.232 | 5.27E-03 | glio |
| Bcas2_in.vivo_noRT | Gjc3 | -0.63 | 5.27E-03 | glio |
| Bcas2_ex.vivo_noRT | Gm43330 | 2.559 | 5.27E-03 | glio |
| Bora_in.vivo_noRT | Zfp553 | 1.418 | 5.27E-03 | glio |
| Bora_in.vivo_noRT | Tmcc3 | 1.08 | 5.27E-03 | glio |
| Bora_in.vivo_noRT | Larp4 | -0.703 | 5.27E-03 | glio |
| culture_effect | Dcaf17 | -0.951 | 5.27E-04 | glio |
| culture_effect | Dip2b | 0.544 | 5.27E-04 | glio |
| culture_effect | Reps2 | -0.595 | 5.27E-04 | glio |
| Interaction_genotype_Bora | 4933429H19R | -4.216 | 5.27E-04 | glio |
| Interaction_genotype_Bora | Cyp1a1 | -5.812 | 5.27E-04 | glio |
| Interaction_genotype_Mtor | Plek2 | -4.128 | 5.27E-04 | glio |
| Interaction_genotype_Mtor | Ranbp17 | -4.084 | 5.27E-04 | glio |
| Interaction_genotype_Myc | Traf3ip3 | -3.965 | 5.27E-04 | glio |
| Atp5c1_in.vivo_noRT | Krt16 | 3.343 | 5.27E-04 | glio |
| Atr_in.vivo_noRT | Adamts4 | 2.192 | 5.27E-04 | glio |
| Bard1_ex.vivo_noRT | St18 | 2.814 | 5.27E-04 | glio |
| Chek1_in.vivo_noRT | Bcl7a | 2.529 | 5.27E-04 | glio |
| Hsd17b10_in.vivo_noRT | Gm45890 | 2.699 | 5.27E-04 | glio |
| Aars2_in.vivo_noRT | Smim40 | 2.825 | 5.27E-05 | glio |
| Atr_ex.vivo_noRT | Ceacam16 | 3.164 | 5.27E-05 | glio |
| Bcas2_ex.vivo_noRT | Gm47510 | 2.742 | 5.27E-05 | glio |
| Bcas2_ex.vivo_noRT | Gm5788 | 2.742 | 5.27E-05 | glio |
| culture_effect | Gm20045 | 1.771 | 5.27E-06 | glio |
| Aars2_in.vivo_noRT | Asap3 | 3.094 | 5.27E-06 | glio |
| Bora_ex.vivo_noRT | Card9 | 3.594 | 5.27E-06 | glio |
| Atr_in.vivo_noRT | Ece2 | 3.302 | 5.27E-07 | glio |
| Bora_in.vivo_noRT | Cabp4 | 3.626 | 5.27E-07 | glio |
| Chek1_in.vivo_noRT | Fsbp | 3.6 | 5.27E-07 | glio |

| | | | | |
|---|---|---|---|---|
| Chek1_ex.vivo_noRT | Cxcr2 | 4.256 | 5.27E-07 | glio |
| interaction_RT | Gm43303 | 3.411 | 5.27E-08 | glio |
| culture_effect | Eif2ak2 | 0.971 | 5.27E-10 | glio |
| Atr_in.vivo_noRT | Wfdc18 | 5.267 | 5.27E-10 | glio |
| Atr_in.vivo_noRT | Adcy2 | 3.78 | 5.27E-10 | glio |
| Aars2_in.vivo_noRT | 1700047K16R | 4.669 | 5.27E-11 | glio |
| Hsd17b10_in.vivo_noRT | Erich6b | 3.616 | 5.27E-13 | glio |
| Hsd17b10_in.vivo_noRT | Lncbate10 | 3.616 | 5.27E-13 | glio |
| Bora_in.vivo_noRT | Gm5602 | 4.145 | 5.27E-14 | glio |
| Chek1_in.vivo_noRT | Gm45620 | 5.044 | 5.27E-14 | glio |
| Chek1_in.vivo_noRT | Gm38534 | 5.044 | 5.27E-14 | glio |
| culture_effect | Plxnd1 | 2.665 | 5.27E-17 | glio |
| Bora_in.vivo_noRT | Ablim2 | 4.404 | 5.27E-20 | glio |
| culture_effect | Ift27 | -1.934 | 5.27E-24 | glio |
| culture_effect | Uggt1 | -0.493 | 5.28E-03 | glio |
| Interaction_genotype_Atr | 2410017I17Ri | -2.369 | 5.28E-03 | glio |
| Interaction_genotype_Atr | Adam1a | -3.558 | 5.28E-03 | glio |
| Interaction_genotype_Atr | Gm14296 | -2.522 | 5.28E-03 | glio |
| Interaction_genotype_Myc | Zfp418 | -3.405 | 5.28E-03 | glio |
| Interaction_genotype_Myc | Gm49041 | -2.722 | 5.28E-03 | glio |
| Interaction_genotype_Myc | Mdp1 | -1.699 | 5.28E-03 | glio |
| Interaction_genotype_Rpl3 | Gm43137 | -3.794 | 5.28E-03 | glio |
| Aars2_in.vivo_noRT | Carf | 2.016 | 5.28E-03 | glio |
| Atr_in.vivo_noRT | Gm45902 | 1.999 | 5.28E-03 | glio |
| Atr_in.vivo_noRT | Tnfrsf10b | 1.913 | 5.28E-03 | glio |
| Atr_in.vivo_noRT | Gm15446 | 1.786 | 5.28E-03 | glio |
| Atr_in.vivo_noRT | Ankle1 | 1.787 | 5.28E-03 | glio |
| Eif2s1_in.vivo_noRT | Ppil6 | 2.143 | 5.28E-03 | glio |
| Interaction_genotype_Mtor | Gjb5 | -3.688 | 5.28E-04 | glio |
| Atp5c1_in.vivo_noRT | D730045A05R | 2.668 | 5.28E-04 | glio |
| Bard1_ex.vivo_noRT | Stx19 | 2.814 | 5.28E-04 | glio |
| Cenpj_in.vivo_noRT | Ivns1abp | 1.119 | 5.28E-04 | glio |
| Aars2_in.vivo_noRT | Olfr77 | 2.741 | 5.28E-05 | glio |
| Aars2_in.vivo_noRT | Arid3c | 2.576 | 5.28E-05 | glio |
| Atr_in.vivo_noRT | Gm12359 | 2.841 | 5.28E-05 | glio |
| Atr_in.vivo_noRT | Rpl24 | -0.821 | 5.28E-05 | glio |
| Bora_in.vivo_noRT | Taf11 | -1.668 | 5.28E-05 | glio |
| Chek1_in.vivo_noRT | Tcam1 | 3.33 | 5.28E-06 | glio |
| Chek1_in.vivo_noRT | Gm42997 | 3.161 | 5.28E-06 | glio |
| Chek1_in.vivo_noRT | Gstt2 | 3.257 | 5.28E-06 | glio |
| Aars2_in.vivo_noRT | C130040N14R | 3.679 | 5.28E-07 | glio |
| Chek1_in.vivo_noRT | Gm5089 | 4.006 | 5.28E-07 | glio |
| Atr_ex.vivo_noRT | Gm2022 | 3.947 | 5.28E-09 | glio |
| Aars2_in.vivo_noRT | Asgr1 | 4.668 | 5.28E-11 | glio |
| culture_effect | Plp2 | -1.249 | 5.28E-16 | glio |
| Atr_in.vivo_noRT | Gm44983 | 4.745 | 5.28E-16 | glio |
| Bora_in.vivo_noRT | Gm34939 | 4.672 | 5.28E-16 | glio |
| Bcas2_in.vivo_noRT | 2010001K21R | 5.334 | 5.28E-17 | glio |
| culture_effect | Casp8 | -0.682 | 5.29E-03 | glio |
| culture_effect | Fam102b | -0.655 | 5.29E-03 | glio |
| Interaction_genotype_Aars2 | Gm26671 | -2.427 | 5.29E-03 | glio |
| Interaction_genotype_Aars2 | Psmb3 | 1.142 | 5.29E-03 | glio |

| | | | | |
|---|---|---|---|---|
| Interaction_genotype_Mtor | Dnm1 | -3.467 | 5.29E-03 | glio |
| Interaction_genotype_Mtor | Gm47863 | -3.263 | 5.29E-03 | glio |
| Interaction_genotype_Mtor | B130034C11R | -2.586 | 5.29E-03 | glio |
| Interaction_genotype_Mtor | 2900076A07R | -2.394 | 5.29E-03 | glio |
| Interaction_genotype_Rad51 | H2-DMb1 | 1.819 | 5.29E-03 | glio |
| Interaction_genotype_Rpl3 | A930001C03R | -3.768 | 5.29E-03 | glio |
| Atr_in.vivo_noRT | Pim3 | -1.159 | 5.29E-03 | glio |
| Bcas2_in.vivo_noRT | Nek3 | 2.071 | 5.29E-03 | glio |
| Bora_in.vivo_noRT | Fzd1 | 1.968 | 5.29E-03 | glio |
| Bora_in.vivo_noRT | Plekho2 | 2.002 | 5.29E-03 | glio |
| Cdkn2a_in.vivo_noRT | BC049987 | -1.93 | 5.29E-03 | glio |
| Cenpj_in.vivo_noRT | 9230112J17Ri | 2.022 | 5.29E-03 | glio |
| Chek1_in.vivo_noRT | 3300002I08Ri | 2.017 | 5.29E-03 | glio |
| Chek1_in.vivo_noRT | Cfap74 | 1.835 | 5.29E-03 | glio |
| Chek1_in.vivo_noRT | Yipf6 | -1.227 | 5.29E-03 | glio |
| Chek1_ex.vivo_noRT | Pclaf | -1.817 | 5.29E-03 | glio |
| Hsd17b10_in.vivo_noRT | Serpinb9b | 2.619 | 5.29E-03 | glio |
| Hsd17b10_in.vivo_noRT | Gm26519 | 2.43 | 5.29E-03 | glio |
| Hsd17b10_in.vivo_noRT | Casz1 | 2.341 | 5.29E-03 | glio |
| Interaction_genotype_Atr | Cyp26b1 | -4.342 | 5.29E-04 | glio |
| Interaction_genotype_Atr | Gtsf1 | -3.532 | 5.29E-04 | glio |
| Interaction_genotype_Mtor | Ifit1bl2 | -4.166 | 5.29E-04 | glio |
| Interaction_genotype_Myc | Fermt3 | -3.767 | 5.29E-04 | glio |
| Interaction_genotype_Myc | Fblim1 | -3.871 | 5.29E-04 | glio |
| Atp5c1_in.vivo_noRT | Igf2bp1 | 2.668 | 5.29E-04 | glio |
| Atp5c1_in.vivo_noRT | Gm10482 | 2.146 | 5.29E-04 | glio |
| Atp5c1_in.vivo_noRT | Rab25 | 2.402 | 5.29E-04 | glio |
| Atp5c1_in.vivo_noRT | 1110020A21R | 2.78 | 5.29E-04 | glio |
| Bcas2_in.vivo_noRT | Ddx5 | -0.571 | 5.29E-04 | glio |
| Bora_ex.vivo_noRT | Acat3 | 2.626 | 5.29E-04 | glio |
| Chek1_ex.vivo_noRT | Nadsyn1 | 3.197 | 5.29E-04 | glio |
| Chek1_in.vivo_noRT | Serpinb9 | 2.737 | 5.29E-05 | glio |
| Chek1_ex.vivo_noRT | Gm42849 | 3.178 | 5.29E-05 | glio |
| Chek1_ex.vivo_noRT | Gm43387 | 3.645 | 5.29E-05 | glio |
| Chek1_ex.vivo_noRT | Hrct1 | 3.072 | 5.29E-05 | glio |
| Bora_in.vivo_noRT | Prex2 | 3.5 | 5.29E-06 | glio |
| Bora_in.vivo_noRT | 1700006J14Ri | 3.026 | 5.29E-06 | glio |
| Chek1_in.vivo_noRT | 1700086P04R | 3.164 | 5.29E-06 | glio |
| Dclre1c_ex.vivo_noRT | Gm8300 | 5.406 | 5.29E-06 | glio |
| Aars2_ex.vivo_noRT | Adam5 | 3.293 | 5.29E-07 | glio |
| Aars2_ex.vivo_noRT | 4933400F21Ri | 3.293 | 5.29E-07 | glio |
| Chek1_in.vivo_noRT | 1700063J08Ri | 3.536 | 5.29E-07 | glio |
| Chek1_in.vivo_noRT | Zfp551 | 3.477 | 5.29E-07 | glio |
| Chek1_ex.vivo_noRT | Gm13348 | 3.57 | 5.29E-07 | glio |
| Aars2_in.vivo_noRT | Zfp575 | 4.075 | 5.29E-08 | glio |
| Atr_in.vivo_noRT | Serpinb9b | 4.536 | 5.29E-08 | glio |
| Chek1_in.vivo_noRT | Itgb3bp | -2.152 | 5.29E-08 | glio |
| Chek1_ex.vivo_noRT | Gm13449 | 4.055 | 5.29E-10 | glio |
| culture_effect | Efs | -1.794 | 5.29E-11 | glio |
| Atr_in.vivo_noRT | Fgf1 | 4.742 | 5.29E-11 | glio |
| culture_effect | Cops5 | -1.011 | 5.29E-13 | glio |
| Bcas2_in.vivo_noRT | Gm20646 | 4.807 | 5.29E-13 | glio |

| | | | | |
|---|---|---|---|---|
| Aars2_in.vivo_noRT | Gm13713 | 4.794 | 5.29E-18 | glio |
| Chek1_in.vivo_noRT | Aqp9 | 5.044 | 5.29E-18 | glio |
| Aars2_in.vivo_noRT | 1700024I08Ri | 5.067 | 5.29E-19 | glio |
| culture_effect | Sbk3 | -1.708 | 5.30E-03 | glio |
| culture_effect | Acvr1 | 0.694 | 5.30E-03 | glio |
| culture_effect | Phlda3 | 0.459 | 5.30E-03 | glio |
| Interaction_genotype_Aars2 | 1700012B09R | -4.154 | 5.30E-03 | glio |
| Interaction_genotype_Aars2 | Gm26728 | -3.558 | 5.30E-03 | glio |
| Interaction_genotype_Aars2 | Exoc3 | 1.773 | 5.30E-03 | glio |
| Interaction_genotype_Atr | Dhrs1 | -1.646 | 5.30E-03 | glio |
| Interaction_genotype_Bora | 4933421A08R | -3.961 | 5.30E-03 | glio |
| Interaction_genotype_Mtor | Riox1 | -2.936 | 5.30E-03 | glio |
| Interaction_genotype_Pars2 | Gm30692 | -2.662 | 5.30E-03 | glio |
| Interaction_genotype_Pars2 | 9230112J17Ri | -3.249 | 5.30E-03 | glio |
| Atp5c1_in.vivo_noRT | Sod3 | 2.668 | 5.30E-03 | glio |
| Atr_in.vivo_noRT | Zfp449 | 1.739 | 5.30E-03 | glio |
| Atr_in.vivo_noRT | Oip5 | 1.478 | 5.30E-03 | glio |
| Atr_in.vivo_noRT | Eif4h | -0.945 | 5.30E-03 | glio |
| Bcas2_ex.vivo_noRT | 4930502E18Ri | 2.349 | 5.30E-03 | glio |
| Bcas2_ex.vivo_noRT | Tff2 | 2.742 | 5.30E-03 | glio |
| Bora_in.vivo_noRT | Rccd1 | 1.619 | 5.30E-03 | glio |
| Brca2_in.vivo_noRT | Ankrd22 | 1.763 | 5.30E-03 | glio |
| Brca2_in.vivo_noRT | Gm47404 | 1.763 | 5.30E-03 | glio |
| Brca2_in.vivo_noRT | Gm31717 | 1.763 | 5.30E-03 | glio |
| Brca2_in.vivo_noRT | Fam166a | 2.024 | 5.30E-03 | glio |
| Brca2_in.vivo_noRT | Gm14742 | 1.763 | 5.30E-03 | glio |
| Brca2_in.vivo_noRT | Ctla4 | 1.763 | 5.30E-03 | glio |
| Brca2_in.vivo_noRT | Zscan4-ps3 | 1.763 | 5.30E-03 | glio |
| Brca2_in.vivo_noRT | A030014E15Ri | 1.763 | 5.30E-03 | glio |
| Brca2_in.vivo_noRT | Gm21731 | 1.763 | 5.30E-03 | glio |
| Brca2_in.vivo_noRT | Gm11232 | 1.763 | 5.30E-03 | glio |
| Cyp19a1_ex.vivo_noRT | Fst | 2.933 | 5.30E-03 | glio |
| Fanca_ex.vivo_noRT | Krt19 | 2.124 | 5.30E-03 | glio |
| Fanca_ex.vivo_noRT | Gm48155 | 2.124 | 5.30E-03 | glio |
| Fanca_ex.vivo_noRT | Halr1 | 2.124 | 5.30E-03 | glio |
| Fanca_ex.vivo_noRT | Gm34059 | 2.124 | 5.30E-03 | glio |
| Fanca_ex.vivo_noRT | Gm34776 | 2.52 | 5.30E-03 | glio |
| Fanca_ex.vivo_noRT | Helt | 2.124 | 5.30E-03 | glio |
| Fanca_ex.vivo_noRT | Gm20625 | 2.124 | 5.30E-03 | glio |
| Fanca_ex.vivo_noRT | Gm5157 | 2.124 | 5.30E-03 | glio |
| Fanca_ex.vivo_noRT | Gm44736 | 2.124 | 5.30E-03 | glio |
| Fanca_ex.vivo_noRT | Gm20658 | 2.124 | 5.30E-03 | glio |
| Fanca_ex.vivo_noRT | 4732463B04R | 2.526 | 5.30E-03 | glio |
| Fanca_ex.vivo_noRT | Nmrk2 | 2.913 | 5.30E-03 | glio |
| Fanca_ex.vivo_noRT | Olfr810 | 2.124 | 5.30E-03 | glio |
| Fanca_ex.vivo_noRT | Col19a1 | 2.52 | 5.30E-03 | glio |
| Fanca_ex.vivo_noRT | Apoa5 | 2.124 | 5.30E-03 | glio |
| Fanca_ex.vivo_noRT | Gm49172 | 2.124 | 5.30E-03 | glio |
| Fanca_ex.vivo_noRT | Gjd4 | 2.124 | 5.30E-03 | glio |
| Fanca_ex.vivo_noRT | Gm13054 | 2.884 | 5.30E-03 | glio |
| Interaction_genotype_Myc | Gm15138 | -2.858 | 5.30E-04 | glio |
| Interaction_genotype_Myc | Hyal1 | -3.755 | 5.30E-04 | glio |

| | | | | |
|---|---|---|---|---|
| Interaction_genotype_Rad51 | Gm30938 | -4.554 | 5.30E-04 | glio |
| Bcas2_in.vivo_noRT | Gm20515 | 2.47 | 5.30E-04 | glio |
| Bcas2_in.vivo_noRT | Gm43773 | 2.386 | 5.30E-04 | glio |
| Bcas2_in.vivo_noRT | Tmprss5 | 2.499 | 5.30E-04 | glio |
| Chek1_in.vivo_noRT | Fam161b | 2.496 | 5.30E-04 | glio |
| Atr_in.vivo_noRT | Fbxo24 | 2.814 | 5.30E-05 | glio |
| Atr_in.vivo_noRT | Car11 | 2.537 | 5.30E-05 | glio |
| Chek1_in.vivo_noRT | Gm26728 | 3.014 | 5.30E-05 | glio |
| Hsd17b10_in.vivo_noRT | Cnp | -1.774 | 5.30E-05 | glio |
| culture_effect | Cmc4 | 1.215 | 5.30E-06 | glio |
| Aars2_in.vivo_noRT | Cdrt4 | 3.352 | 5.30E-06 | glio |
| Aars2_ex.vivo_noRT | Gm12794 | 3.293 | 5.30E-07 | glio |
| Atr_in.vivo_noRT | Gm15486 | 3.161 | 5.30E-07 | glio |
| Atr_ex.vivo_noRT | Cldn5 | 3.164 | 5.30E-07 | glio |
| Hsd17b10_in.vivo_noRT | Gm17638 | 3.355 | 5.30E-07 | glio |
| culture_effect | Rpl7l1 | 0.665 | 5.30E-08 | glio |
| Aars2_in.vivo_noRT | Gm2762 | 3.754 | 5.30E-09 | glio |
| Bcas2_in.vivo_noRT | Ntn5 | 4.015 | 5.30E-09 | glio |
| Chek1_ex.vivo_noRT | Ank1 | 4.058 | 5.30E-11 | glio |
| culture_effect | Kdm4a | -0.545 | 5.31E-03 | glio |
| culture_effect | Ppil3 | 0.443 | 5.31E-03 | glio |
| culture_effect | Cst6 | -0.518 | 5.31E-03 | glio |
| Interaction_genotype_Atr | Usp27x | -3.618 | 5.31E-03 | glio |
| Interaction_genotype_Atr | Snhg12 | 1.32 | 5.31E-03 | glio |
| Interaction_genotype_Bora | Sod3 | -3.559 | 5.31E-03 | glio |
| Interaction_genotype_Mtor | Efnb1 | -3.04 | 5.31E-03 | glio |
| Interaction_genotype_Mtor | Gdi1 | 1.483 | 5.31E-03 | glio |
| Interaction_genotype_Mtor | Taok1 | 1.606 | 5.31E-03 | glio |
| Interaction_genotype_Myc | Bccip | 1.29 | 5.31E-03 | glio |
| interaction_RT | Zfp579 | 1.176 | 5.31E-03 | glio |
| Aars2_in.vivo_noRT | Gm29488 | 1.96 | 5.31E-03 | glio |
| Aars2_in.vivo_noRT | Psme2b | 1.939 | 5.31E-03 | glio |
| Atr_in.vivo_noRT | Atp6v1e1 | -0.766 | 5.31E-03 | glio |
| Bcas2_in.vivo_noRT | Dohh | -0.982 | 5.31E-03 | glio |
| Bcas2_in.vivo_noRT | Stat2 | 1.118 | 5.31E-03 | glio |
| Bora_in.vivo_noRT | Junos | 1.974 | 5.31E-03 | glio |
| Bora_in.vivo_noRT | 9630013A20R | 1.856 | 5.31E-03 | glio |
| Cenpj_in.vivo_noRT | Gm20711 | 2.022 | 5.31E-03 | glio |
| Cenpj_in.vivo_noRT | Abra | 2.026 | 5.31E-03 | glio |
| Chek1_in.vivo_noRT | Mga | -1.265 | 5.31E-03 | glio |
| Chek1_in.vivo_noRT | Cd44 | -0.967 | 5.31E-03 | glio |
| Chek1_ex.vivo_noRT | D030068K23R | 2.585 | 5.31E-03 | glio |
| Chek1_ex.vivo_noRT | Prrg1 | 2.707 | 5.31E-03 | glio |
| culture_effect | Usp24 | -0.755 | 5.31E-04 | glio |
| Interaction_genotype_Atr | Zeb2os | -3.899 | 5.31E-04 | glio |
| Interaction_genotype_Mtor | Slc16a13 | -3.313 | 5.31E-04 | glio |
| Bora_in.vivo_noRT | Zbtb3 | 2.52 | 5.31E-04 | glio |
| culture_effect | St13 | 0.454 | 5.31E-05 | glio |
| Aars2_ex.vivo_noRT | Gm33973 | 3.293 | 5.31E-05 | glio |
| Atr_in.vivo_noRT | Cfap69 | 2.8 | 5.31E-05 | glio |
| Aars2_in.vivo_noRT | Cav2 | -2.204 | 5.31E-06 | glio |
| Bora_in.vivo_noRT | Catsperd | 3.757 | 5.31E-08 | glio |

| | | | | |
|---|---|---|---|---|
| Chek1_ex.vivo_noRT | Pde6b | 4.049 | 5.31E-08 | glio |
| Aars2_in.vivo_noRT | Sema4a | 4.283 | 5.31E-09 | glio |
| Atr_ex.vivo_noRT | Nrip2 | 3.962 | 5.31E-11 | glio |
| Chek1_in.vivo_noRT | 1810019D21R | 4.784 | 5.31E-12 | glio |
| Aars2_in.vivo_noRT | Cfb | 4.023 | 5.31E-13 | glio |
| Bcas2_in.vivo_noRT | Gm49860 | 4.807 | 5.31E-16 | glio |
| Chek1_in.vivo_noRT | Gm10482 | 4.522 | 5.31E-16 | glio |
| Aars2_in.vivo_noRT | Olfr552 | 5.841 | 5.31E-29 | glio |
| Aars2_in.vivo_noRT | 1700003E24Ri | 5.327 | 5.31E-29 | glio |
| culture_effect | G630030J09Ri | 1.599 | 5.32E-03 | glio |
| culture_effect | Cebpzos | 0.504 | 5.32E-03 | glio |
| Interaction_genotype_Aars2 | A230020J21Ri | -3.655 | 5.32E-03 | glio |
| Interaction_genotype_Cdkn2a | Tmem132c | 2.835 | 5.32E-03 | glio |
| Interaction_genotype_Cdkn2a | Gm44693 | 2.835 | 5.32E-03 | glio |
| Interaction_genotype_Chek1 | Hrh2 | 3.342 | 5.32E-03 | glio |
| Interaction_genotype_Mtor | Ppp1r3e | -3.224 | 5.32E-03 | glio |
| Interaction_genotype_Myc | Fancb | -2.542 | 5.32E-03 | glio |
| Interaction_genotype_Myc | Sh3bp5l | -2.095 | 5.32E-03 | glio |
| Interaction_genotype_Pars2 | Adam12 | -3.064 | 5.32E-03 | glio |
| Aars2_in.vivo_noRT | Fbxo10 | 1.988 | 5.32E-03 | glio |
| Atr_in.vivo_noRT | Zfp628 | 1.599 | 5.32E-03 | glio |
| Atr_in.vivo_noRT | Eed | -1.135 | 5.32E-03 | glio |
| Atr_in.vivo_noRT | Rfx3 | 1.292 | 5.32E-03 | glio |
| Atr_in.vivo_noRT | Gatm | -1.569 | 5.32E-03 | glio |
| Brca2_in.vivo_noRT | Cgnl1 | 1.763 | 5.32E-03 | glio |
| Brca2_in.vivo_noRT | Gm26543 | 1.763 | 5.32E-03 | glio |
| Brca2_in.vivo_noRT | Dscaml1 | 1.763 | 5.32E-03 | glio |
| Brca2_in.vivo_noRT | 2410017I17Ri | 1.763 | 5.32E-03 | glio |
| Brca2_in.vivo_noRT | Pspn | 1.763 | 5.32E-03 | glio |
| Interaction_genotype_Mtor | A930033H14R | -4.126 | 5.32E-04 | glio |
| Interaction_genotype_Mtor | Marcksl1 | 2.06 | 5.32E-04 | glio |
| Interaction_genotype_Myc | Trim3 | -3.169 | 5.32E-04 | glio |
| interaction_RT | Plac8 | 3.281 | 5.32E-04 | glio |
| Aars2_in.vivo_noRT | Col20a1 | 2.49 | 5.32E-04 | glio |
| Aars2_in.vivo_noRT | Xkr8 | 2.336 | 5.32E-04 | glio |
| Aars2_in.vivo_noRT | Laptm4a | -0.813 | 5.32E-04 | glio |
| Interaction_genotype_Mtor | Itga7 | -4.889 | 5.32E-05 | glio |
| Interaction_genotype_Mtor | Wdpcp | -4.611 | 5.32E-05 | glio |
| Interaction_genotype_Mtor | BC031181 | 1.881 | 5.32E-05 | glio |
| Interaction_genotype_Mtor | Rpl21 | 1.302 | 5.32E-05 | glio |
| Interaction_genotype_Myc | Iqce | -3.883 | 5.32E-05 | glio |
| Atr_ex.vivo_noRT | BC037032 | 3.62 | 5.32E-05 | glio |
| Bora_in.vivo_noRT | Tub | 2.795 | 5.32E-05 | glio |
| Interaction_genotype_Mtor | Gpr22 | -4.099 | 5.32E-06 | glio |
| Bcas2_in.vivo_noRT | Ggnbp1 | 3.125 | 5.32E-06 | glio |
| Chek1_in.vivo_noRT | Gm28229 | 2.932 | 5.32E-06 | glio |
| Interaction_genotype_Myc | Nid1 | -5.974 | 5.32E-07 | glio |
| Aars2_in.vivo_noRT | H60c | 3.733 | 5.32E-07 | glio |
| Aars2_in.vivo_noRT | Ehbp1l1 | 3.591 | 5.32E-07 | glio |
| Atr_in.vivo_noRT | Zbtb46 | 3.694 | 5.32E-07 | glio |
| Atr_in.vivo_noRT | Rpl4 | -0.919 | 5.32E-07 | glio |
| Aars2_in.vivo_noRT | Pkhd1 | 3.809 | 5.32E-08 | glio |

| | | | | |
|---|---|---|---|---|
| Chek1_ex.vivo_noRT | Oit3 | 4.401 | 5.32E-08 | glio |
| Hsd17b10_in.vivo_noRT | Gm36356 | 3.356 | 5.32E-08 | glio |
| Bora_in.vivo_noRT | Chpf | -2.431 | 5.32E-09 | glio |
| Chek1_in.vivo_noRT | 1700012B09R | 4.784 | 5.32E-09 | glio |
| culture_effect | Gm49741 | 2.95 | 5.32E-11 | glio |
| Interaction_genotype_Myc | Hbq1b | -7.651 | 5.32E-11 | glio |
| Atr_in.vivo_noRT | 4930445N18R | 4.626 | 5.32E-11 | glio |
| Chek1_ex.vivo_noRT | Gm12841 | 4.449 | 5.32E-13 | glio |
| Chek1_ex.vivo_noRT | Dcdc2a | 5.611 | 5.32E-14 | glio |
| Aars2_in.vivo_noRT | Gm42664 | 4.676 | 5.32E-15 | glio |
| Atr_in.vivo_noRT | Gm42984 | 5.392 | 5.32E-15 | glio |
| Atr_in.vivo_noRT | B3gnt4 | 5.27 | 5.32E-18 | glio |
| Chek1_in.vivo_noRT | Gm15943 | 4.78 | 5.32E-18 | glio |
| Chek1_in.vivo_noRT | Gm26671 | 5.044 | 5.32E-24 | glio |
| Chek1_in.vivo_noRT | Gm47404 | 5.044 | 5.32E-25 | glio |
| culture_effect | Ntn4 | 4.913 | 5.32E-30 | glio |
| Cdkn1a_ex.vivo_noRT | Cdkn1a | -3.728 | 5.32E-32 | glio |
| culture_effect | Gm10734 | 1.568 | 5.33E-03 | glio |
| culture_effect | Reep6 | 0.886 | 5.33E-03 | glio |
| Interaction_genotype_Atr | Zfp11 | -3.422 | 5.33E-03 | glio |
| Interaction_genotype_Atr | Gm49864 | -3.176 | 5.33E-03 | glio |
| Interaction_genotype_Myc | Ttc12 | -3.051 | 5.33E-03 | glio |
| Interaction_genotype_Myc | Gm49123 | -3.238 | 5.33E-03 | glio |
| Interaction_genotype_Rpl3 | Gm32090 | -3.376 | 5.33E-03 | glio |
| Aars2_in.vivo_noRT | Cs | -0.868 | 5.33E-03 | glio |
| Atp5c1_in.vivo_noRT | Usp33 | -1.703 | 5.33E-03 | glio |
| Atp5c1_ex.vivo_noRT | Gm48215 | 2.61 | 5.33E-03 | glio |
| Atr_in.vivo_noRT | Eif5a2 | 1.717 | 5.33E-03 | glio |
| Atr_in.vivo_noRT | Mmachc | 1.536 | 5.33E-03 | glio |
| Atr_in.vivo_noRT | AI987944 | 1.591 | 5.33E-03 | glio |
| culture_effect | Pdcd2 | -0.556 | 5.33E-04 | glio |
| culture_effect | Tmem251 | 0.545 | 5.33E-04 | glio |
| Interaction_genotype_Mtor | Sgk3 | -4.056 | 5.33E-04 | glio |
| Interaction_genotype_Pars2 | Uchl3 | 1.666 | 5.33E-04 | glio |
| Aars2_in.vivo_noRT | Gm36266 | 2.75 | 5.33E-04 | glio |
| Aars2_in.vivo_noRT | Swi5 | -0.698 | 5.33E-04 | glio |
| Atp5c1_in.vivo_noRT | Acox2 | 2.219 | 5.33E-04 | glio |
| Bcas2_in.vivo_noRT | Ifit1bl2 | 2.61 | 5.33E-04 | glio |
| Bcas2_in.vivo_noRT | Gm36445 | 2.518 | 5.33E-04 | glio |
| Bora_in.vivo_noRT | Cep112it | 2.492 | 5.33E-04 | glio |
| Bora_in.vivo_noRT | Med9os | 2.51 | 5.33E-04 | glio |
| Bora_in.vivo_noRT | Fbln1 | 2.406 | 5.33E-04 | glio |
| Bora_in.vivo_noRT | Mogat2 | 2.419 | 5.33E-04 | glio |
| Bora_in.vivo_noRT | Fggy | 2.247 | 5.33E-04 | glio |
| Bora_ex.vivo_noRT | Tnfrsf1b | 3.201 | 5.33E-04 | glio |
| Chek1_in.vivo_noRT | Gm16845 | 2.373 | 5.33E-04 | glio |
| culture_effect | Pisd | -0.818 | 5.33E-05 | glio |
| culture_effect | Atp6v1a | 0.562 | 5.33E-05 | glio |
| Interaction_genotype_Mtor | 4930502E18Ri | -3.957 | 5.33E-05 | glio |
| Aars2_in.vivo_noRT | D730045B01R | 2.54 | 5.33E-05 | glio |
| Aars2_in.vivo_noRT | Nol12 | -1.276 | 5.33E-05 | glio |
| Atr_in.vivo_noRT | Zfp27 | 2.607 | 5.33E-05 | glio |

| | | | | |
|---|---|---|---|---|
| Bcas2_ex.vivo_noRT | Slc10a1 | 3.131 | 5.33E-05 | glio |
| Interaction_genotype_Aars2 | 4933429H19R | -5.308 | 5.33E-06 | glio |
| Interaction_genotype_Aars2 | Calhm5 | -5.4 | 5.33E-06 | glio |
| Interaction_genotype_Aars2 | Fhl4 | -5.282 | 5.33E-06 | glio |
| Aars2_in.vivo_noRT | Gm29361 | 3.301 | 5.33E-06 | glio |
| Atr_in.vivo_noRT | C920021L13Ri | 3.203 | 5.33E-06 | glio |
| Bora_in.vivo_noRT | Gm49135 | 3.254 | 5.33E-06 | glio |
| Chek1_in.vivo_noRT | Gm15091 | 3.34 | 5.33E-06 | glio |
| Interaction_genotype_Mtor | 2410017I17Ri | -4.099 | 5.33E-07 | glio |
| Atr_in.vivo_noRT | Tmsb15b2 | 3.813 | 5.33E-07 | glio |
| interaction_RT | Wfdc3 | 4.919 | 5.33E-08 | glio |
| Bora_in.vivo_noRT | Gm10767 | 3.861 | 5.33E-08 | glio |
| Chek1_in.vivo_noRT | Cpne9 | 4.007 | 5.33E-08 | glio |
| Bcas2_in.vivo_noRT | C130060C02R | 4.543 | 5.33E-13 | glio |
| Hsd17b10_in.vivo_noRT | Tac2 | 3.616 | 5.33E-13 | glio |
| Chek1_in.vivo_noRT | Rbfaos | 4.127 | 5.33E-14 | glio |
| Aars2_in.vivo_noRT | Cys1 | 5.06 | 5.33E-18 | glio |
| culture_effect | Npdc1 | 0.891 | 5.34E-03 | glio |
| culture_effect | Ikbip | -0.443 | 5.34E-03 | glio |
| culture_effect | Top2a | -0.656 | 5.34E-03 | glio |
| Interaction_genotype_Atr | Acoxl | -3.211 | 5.34E-03 | glio |
| Interaction_genotype_Atr | Arhgef1 | -2.319 | 5.34E-03 | glio |
| Interaction_genotype_Bora | Gm10115 | -3.653 | 5.34E-03 | glio |
| Interaction_genotype_Bora | BC037704 | -3.454 | 5.34E-03 | glio |
| Interaction_genotype_Mtor | Gm6225 | -3.441 | 5.34E-03 | glio |
| Interaction_genotype_Myc | Wdr26 | 1.309 | 5.34E-03 | glio |
| Interaction_genotype_Myc | Dynlrb1 | 1.104 | 5.34E-03 | glio |
| interaction_RT | Tcof1 | -0.713 | 5.34E-03 | glio |
| Aars2_in.vivo_noRT | Fbxl19 | 1.862 | 5.34E-03 | glio |
| Bcas2_in.vivo_noRT | Cyp4f17 | 2.088 | 5.34E-03 | glio |
| Bcas2_in.vivo_noRT | Rfc3 | -1.182 | 5.34E-03 | glio |
| Eif2s1_ex.vivo_noRT | Gm26555 | 2.234 | 5.34E-03 | glio |
| Eif2s1_ex.vivo_noRT | Snhg4.1 | 2.639 | 5.34E-03 | glio |
| Eif2s1_ex.vivo_noRT | Ggt1 | 3.014 | 5.34E-03 | glio |
| Eif2s1_ex.vivo_noRT | Cd36 | 2.234 | 5.34E-03 | glio |
| Eif2s1_ex.vivo_noRT | Gstt4 | 2.234 | 5.34E-03 | glio |
| Interaction_genotype_Aars2 | Gm9922 | -4.118 | 5.34E-04 | glio |
| Interaction_genotype_Atr | Ccdc80 | -3.338 | 5.34E-04 | glio |
| Interaction_genotype_Myc | Fxyd4 | -4.231 | 5.34E-04 | glio |
| Interaction_genotype_Myc | Tbc1d16 | -3.851 | 5.34E-04 | glio |
| Interaction_genotype_Myc | Zscan20 | -3.777 | 5.34E-04 | glio |
| Interaction_genotype_Myc | Bbs4 | -3.642 | 5.34E-04 | glio |
| Interaction_genotype_Myc | Zfp629 | -3.49 | 5.34E-04 | glio |
| Interaction_genotype_Myc | Ccdc93 | -3.33 | 5.34E-04 | glio |
| Aars2_in.vivo_noRT | Otof | 1.911 | 5.34E-04 | glio |
| Aars2_in.vivo_noRT | Nudt21 | -0.953 | 5.34E-04 | glio |
| Atp5c1_in.vivo_noRT | Bmx | 2.402 | 5.34E-04 | glio |
| Atr_in.vivo_noRT | Trim6 | 2.551 | 5.34E-04 | glio |
| Bora_in.vivo_noRT | Zfp748 | 2.452 | 5.34E-04 | glio |
| Bora_in.vivo_noRT | Bvht | 1.777 | 5.34E-04 | glio |
| Chek1_in.vivo_noRT | A430010J10Ri | 3.537 | 5.34E-05 | glio |
| culture_effect | Clp1 | -0.955 | 5.34E-06 | glio |

| | | | | |
|---|---|---|---|---|
| Interaction_genotype_Mtor | Cdiptos | -5.367 | 5.34E-06 | glio |
| Aars2_in.vivo_noRT | Fchsd1 | 3.083 | 5.34E-06 | glio |
| Bora_in.vivo_noRT | Utp20 | -2.248 | 5.34E-07 | glio |
| Interaction_genotype_Mtor | Prdx2 | 2.771 | 5.34E-08 | glio |
| Interaction_genotype_Mtor | 2010001K21R | -6.051 | 5.34E-08 | glio |
| Atr_in.vivo_noRT | Spata1 | 3.91 | 5.34E-08 | glio |
| Chek1_in.vivo_noRT | Fgf7 | 3.68 | 5.34E-08 | glio |
| culture_effect | Trap1 | -1.205 | 5.34E-09 | glio |
| Bora_in.vivo_noRT | Gm11884 | 3.937 | 5.34E-10 | glio |
| Aars2_in.vivo_noRT | Gm32834 | 4.41 | 5.34E-11 | glio |
| Chek1_in.vivo_noRT | Gm49504 | 5.044 | 5.34E-21 | glio |
| Bcas2_in.vivo_noRT | Gm13191 | 4.807 | 5.34E-25 | glio |
| Bcas2_in.vivo_noRT | 4933416E03Ri | 4.807 | 5.34E-25 | glio |
| culture_effect | Atp6v0c | 0.281 | 5.35E-03 | glio |
| Interaction_genotype_Mtor | Rps19 | 0.931 | 5.35E-03 | glio |
| Interaction_genotype_Myc | Mterf2 | -2.543 | 5.35E-03 | glio |
| Aars2_in.vivo_noRT | Ndufb5 | -0.695 | 5.35E-03 | glio |
| Bcas2_in.vivo_noRT | Tpm4 | -0.734 | 5.35E-03 | glio |
| Bora_ex.vivo_noRT | Grip2 | 2.809 | 5.35E-03 | glio |
| Bora_ex.vivo_noRT | Eif4e3 | 2.404 | 5.35E-03 | glio |
| Cenpj_in.vivo_noRT | Rassf6 | 2.022 | 5.35E-03 | glio |
| culture_effect | Fam50a | -0.568 | 5.35E-04 | glio |
| Interaction_genotype_Aars2 | Gm50103 | -4.336 | 5.35E-04 | glio |
| Interaction_genotype_Mtor | Peli3 | -4.061 | 5.35E-04 | glio |
| Interaction_genotype_Myc | Zcrb1 | 1.209 | 5.35E-04 | glio |
| Aars2_in.vivo_noRT | Ramp1 | 2.836 | 5.35E-04 | glio |
| Aars2_in.vivo_noRT | Lyar | -1.084 | 5.35E-04 | glio |
| Atr_in.vivo_noRT | Zmat1 | 2.323 | 5.35E-04 | glio |
| Interaction_genotype_Myc | Gm26579 | -4.374 | 5.35E-05 | glio |
| Interaction_genotype_Myc | Olfr279 | -4.505 | 5.35E-05 | glio |
| Aars2_in.vivo_noRT | Dynlt1b | 2.841 | 5.35E-05 | glio |
| Atr_in.vivo_noRT | Zkscan8 | 2.429 | 5.35E-05 | glio |
| Bcas2_in.vivo_noRT | Acsl5 | -1.52 | 5.35E-05 | glio |
| Bora_ex.vivo_noRT | Abra | 3.201 | 5.35E-05 | glio |
| Hsd17b10_in.vivo_noRT | Gm13684 | 2.514 | 5.35E-05 | glio |
| interaction_RT | Cnot4 | 0.937 | 5.35E-06 | glio |
| Bcas2_in.vivo_noRT | Cyp4f18 | 4.547 | 5.35E-06 | glio |
| culture_effect | Ldhb | -1.211 | 5.35E-08 | glio |
| Aars2_in.vivo_noRT | Gm16794 | 3.756 | 5.35E-08 | glio |
| Hsd17b10_in.vivo_noRT | Gm36172 | 3.084 | 5.35E-08 | glio |
| culture_effect | Ncan | 2.153 | 5.35E-09 | glio |
| Aars2_in.vivo_noRT | 1700012B09R | 5.067 | 5.35E-10 | glio |
| Atr_in.vivo_noRT | Gm46404 | 4.497 | 5.35E-10 | glio |
| Chek1_in.vivo_noRT | Sult6b1 | 4.253 | 5.35E-10 | glio |
| culture_effect | Ifi30 | -1.564 | 5.35E-11 | glio |
| Bcas2_in.vivo_noRT | Frmd8os | 5.1 | 5.35E-11 | glio |
| Chek1_in.vivo_noRT | Gm32591 | 4.521 | 5.35E-11 | glio |
| Chek1_in.vivo_noRT | Gm17638 | 4.252 | 5.35E-11 | glio |
| Chek1_in.vivo_noRT | Cyp4f40 | 4.391 | 5.35E-11 | glio |
| Chek1_in.vivo_noRT | Zfp986 | 4.526 | 5.35E-11 | glio |
| Bcas2_in.vivo_noRT | 4930423D22R | 4.547 | 5.35E-19 | glio |
| Bcas2_in.vivo_noRT | Gm13449 | 4.807 | 5.35E-19 | glio |

| | | | | |
|---|---|---|---|---|
| Atr_in.vivo_noRT | Fam166a | 5.269 | 5.35E-22 | glio |
| Aars2_in.vivo_noRT | Rgs4 | 5.327 | 5.35E-24 | glio |
| Interaction_genotype_Aars2 | Apobr | -3.791 | 5.36E-03 | glio |
| Interaction_genotype_Aars2 | Gm26737 | -5.098 | 5.36E-03 | glio |
| Interaction_genotype_Atr | Sptbn4 | -3.406 | 5.36E-03 | glio |
| Interaction_genotype_Chek1 | Pced1a | -3.042 | 5.36E-03 | glio |
| Interaction_genotype_Chek1 | Eif4e3 | 3.396 | 5.36E-03 | glio |
| Interaction_genotype_Chek1 | Tomm40l | -3.091 | 5.36E-03 | glio |
| Interaction_genotype_Chek1 | P4ha3 | -3.572 | 5.36E-03 | glio |
| Interaction_genotype_Chek1 | H2-Ab1 | 3.187 | 5.36E-03 | glio |
| Interaction_genotype_Chek1 | Tmod2 | -2.295 | 5.36E-03 | glio |
| Interaction_genotype_Mtor | Arhgef18 | -2.05 | 5.36E-03 | glio |
| Interaction_genotype_Myc | Mxd3 | -2.78 | 5.36E-03 | glio |
| Aars2_in.vivo_noRT | Gm43466 | 1.858 | 5.36E-03 | glio |
| Aars2_ex.vivo_noRT | Bbof1 | 2.717 | 5.36E-03 | glio |
| Aars2_ex.vivo_noRT | Nwd1 | 2.311 | 5.36E-03 | glio |
| Atr_in.vivo_noRT | Fkbp9 | -0.934 | 5.36E-03 | glio |
| Atr_in.vivo_noRT | Junb | -1.487 | 5.36E-03 | glio |
| Bora_ex.vivo_noRT | Gpr37l1 | 2.808 | 5.36E-03 | glio |
| Cenpj_ex.vivo_noRT | Prap1 | 2.198 | 5.36E-03 | glio |
| Cenpj_ex.vivo_noRT | Gm33973 | 2.591 | 5.36E-03 | glio |
| Cenpj_ex.vivo_noRT | Insc | 2.591 | 5.36E-03 | glio |
| Chek1_in.vivo_noRT | Vps33b | 1.595 | 5.36E-03 | glio |
| Chek1_in.vivo_noRT | 4933431E20Ri | 1.617 | 5.36E-03 | glio |
| Chek1_in.vivo_noRT | Cdip1 | -1.483 | 5.36E-03 | glio |
| Chek1_in.vivo_noRT | Abhd12 | -1.208 | 5.36E-03 | glio |
| Chek1_in.vivo_noRT | Hras | -0.863 | 5.36E-03 | glio |
| Dclre1c_ex.vivo_noRT | Gm14929 | 2.292 | 5.36E-03 | glio |
| Eif2s1_in.vivo_noRT | Gm11973 | 1.877 | 5.36E-03 | glio |
| Eif2s1_in.vivo_noRT | Lyl1 | 2.016 | 5.36E-03 | glio |
| Interaction_genotype_Mtor | Syngap1 | -4.036 | 5.36E-04 | glio |
| Interaction_genotype_Mtor | BC030343 | -3.964 | 5.36E-04 | glio |
| Interaction_genotype_Mtor | Gm7160 | -3.865 | 5.36E-04 | glio |
| Aars2_in.vivo_noRT | Gm45871 | 2.247 | 5.36E-04 | glio |
| Aars2_in.vivo_noRT | Rpusd1 | 1.781 | 5.36E-04 | glio |
| Aars2_in.vivo_noRT | Fam89a | -1.421 | 5.36E-04 | glio |
| Chek1_in.vivo_noRT | Acy3 | 2.55 | 5.36E-04 | glio |
| culture_effect | Cept1 | -0.903 | 5.36E-05 | glio |
| culture_effect | Tmem126a | 0.469 | 5.36E-05 | glio |
| culture_effect | Gm10076 | 1.539 | 5.36E-05 | glio |
| Interaction_genotype_Atr | Gm44190 | -4.458 | 5.36E-05 | glio |
| Interaction_genotype_Atr | Rxra | -5.116 | 5.36E-05 | glio |
| Interaction_genotype_Atr | BC028471 | -5.046 | 5.36E-05 | glio |
| Atr_in.vivo_noRT | Bglap3 | 2.815 | 5.36E-05 | glio |
| Atr_in.vivo_noRT | Tctn3 | 2.967 | 5.36E-05 | glio |
| Chek1_ex.vivo_noRT | Tnxb | 3.461 | 5.36E-05 | glio |
| Chek1_ex.vivo_noRT | Oas2 | 3.396 | 5.36E-05 | glio |
| culture_effect | Igf2bp1 | 2.534 | 5.36E-06 | glio |
| Interaction_genotype_Myc | Gm20274 | -4.437 | 5.36E-06 | glio |
| Interaction_genotype_Myc | Pate2 | -5.437 | 5.36E-06 | glio |
| Atr_in.vivo_noRT | Adam19 | 3.251 | 5.36E-06 | glio |
| Atp5c1_in.vivo_noRT | Gm266 | 2.668 | 5.36E-07 | glio |

| | | | | |
|---|---|---|---|---|
| Atr_in.vivo_noRT | Hoga1 | 3.719 | 5.36E-08 | glio |
| Atr_in.vivo_noRT | Gm37933 | 3.382 | 5.36E-08 | glio |
| culture_effect | Vgll2 | -3.27 | 5.36E-09 | glio |
| Bcas2_in.vivo_noRT | AC125149.2 | 3.519 | 5.36E-09 | glio |
| Atr_in.vivo_noRT | Zwint | -2.477 | 5.36E-12 | glio |
| Atr_in.vivo_noRT | Dreh | 4.887 | 5.36E-15 | glio |
| Chek1_in.vivo_noRT | Thsd1 | 4.657 | 5.36E-15 | glio |
| Chek1_ex.vivo_noRT | Gm266 | 4.842 | 5.36E-15 | glio |
| culture_effect | Rsrp1 | -1.886 | 5.36E-27 | glio |
| Interaction_genotype_Atr | Dlc1 | -2.078 | 5.37E-03 | glio |
| Interaction_genotype_Myc | Birc2 | -2.332 | 5.37E-03 | glio |
| interaction_RT | Aatf | -0.684 | 5.37E-03 | glio |
| Atr_in.vivo_noRT | Basp1 | 2.313 | 5.37E-03 | glio |
| Bcas2_in.vivo_noRT | Gm20300 | 2.078 | 5.37E-03 | glio |
| Bcas2_in.vivo_noRT | Mrpl50 | -0.844 | 5.37E-03 | glio |
| Bora_in.vivo_noRT | Uqcrfs1 | -0.837 | 5.37E-03 | glio |
| Bora_in.vivo_noRT | Nolc1 | -0.756 | 5.37E-03 | glio |
| Cenpj_in.vivo_noRT | Gm9758 | 1.762 | 5.37E-03 | glio |
| Cenpj_in.vivo_noRT | Tcp11x2 | 2.026 | 5.37E-03 | glio |
| Chek1_ex.vivo_noRT | Tbl1x | -0.947 | 5.37E-03 | glio |
| Eif2s1_ex.vivo_noRT | Klk13 | 2.234 | 5.37E-03 | glio |
| Eif2s1_ex.vivo_noRT | Ngb | 3.014 | 5.37E-03 | glio |
| Eif2s1_ex.vivo_noRT | Sytl4 | 2.234 | 5.37E-03 | glio |
| Eif2s1_ex.vivo_noRT | Gm12800 | 2.234 | 5.37E-03 | glio |
| Eif2s1_ex.vivo_noRT | Trmt61b | 2.234 | 5.37E-03 | glio |
| Eif2s1_ex.vivo_noRT | Gm5890 | 2.234 | 5.37E-03 | glio |
| Eif2s1_ex.vivo_noRT | Cyp2c23 | 2.637 | 5.37E-03 | glio |
| Eif2s1_ex.vivo_noRT | Zscan4c | 2.234 | 5.37E-03 | glio |
| Eif2s1_ex.vivo_noRT | Speer4a | 2.234 | 5.37E-03 | glio |
| Eif2s1_ex.vivo_noRT | BC080695 | 2.234 | 5.37E-03 | glio |
| culture_effect | Adgrl3 | -1.107 | 5.37E-04 | glio |
| Interaction_genotype_Aars2 | Zc3hav1 | 1.836 | 5.37E-04 | glio |
| Interaction_genotype_Bcas2 | Pelp1 | 2.467 | 5.37E-04 | glio |
| Interaction_genotype_Myc | Gdf11 | -2.916 | 5.37E-04 | glio |
| Atr_in.vivo_noRT | Rragd | 2.586 | 5.37E-04 | glio |
| Bard1_ex.vivo_noRT | Gm7247 | 2.799 | 5.37E-04 | glio |
| Bora_in.vivo_noRT | C230035I16Ri | 2.591 | 5.37E-04 | glio |
| Cenpj_in.vivo_noRT | Gm5788 | 2.022 | 5.37E-04 | glio |
| Cenpj_ex.vivo_noRT | Olfr56 | 2.591 | 5.37E-04 | glio |
| Interaction_genotype_Mtor | Ttbk1 | -4.631 | 5.37E-05 | glio |
| Atr_in.vivo_noRT | Ube2r2 | -1.128 | 5.37E-05 | glio |
| Bcas2_ex.vivo_noRT | Gm12703 | 2.742 | 5.37E-05 | glio |
| Bcas2_ex.vivo_noRT | Mmp13 | 2.742 | 5.37E-05 | glio |
| Bcas2_ex.vivo_noRT | Gm8016 | 2.742 | 5.37E-05 | glio |
| Bcas2_ex.vivo_noRT | Vmn1r88 | 3.524 | 5.37E-05 | glio |
| Chek1_in.vivo_noRT | P4ha3 | 2.752 | 5.37E-05 | glio |
| culture_effect | Cox10 | 0.932 | 5.37E-06 | glio |
| interaction_RT | Vegfb | 1.105 | 5.37E-06 | glio |
| interaction_RT | Ndufa11 | 0.804 | 5.37E-06 | glio |
| Interaction_genotype_Atr | Gpnmb | 3.713 | 5.37E-07 | glio |
| Atr_in.vivo_noRT | Hyal3 | 3.943 | 5.37E-07 | glio |
| Bcas2_in.vivo_noRT | Gm10125 | 3.606 | 5.37E-07 | glio |

| | | | | |
|---|---|---|---|---|
| Atr_in.vivo_noRT | Gm40040 | 3.977 | 5.37E-08 | glio |
| Atr_in.vivo_noRT | C77080 | 3.656 | 5.37E-08 | glio |
| Bcas2_in.vivo_noRT | Gm15972 | 3.813 | 5.37E-10 | glio |
| Bora_in.vivo_noRT | Gm44672 | 4.172 | 5.37E-10 | glio |
| Chek1_in.vivo_noRT | Rtn1 | 4.397 | 5.37E-10 | glio |
| Chek1_ex.vivo_noRT | Itgbl1 | 4.447 | 5.37E-10 | glio |
| Bcas2_in.vivo_noRT | Olfr1191-ps1 | 4.157 | 5.37E-11 | glio |
| culture_effect | Vat1 | 1.325 | 5.37E-16 | glio |
| Interaction_genotype_Aars2 | Rpl13a | 0.889 | 5.38E-03 | glio |
| Interaction_genotype_Atr | Ttc16 | -3.443 | 5.38E-03 | glio |
| Interaction_genotype_Atr | Ldlrad4 | -2.872 | 5.38E-03 | glio |
| Interaction_genotype_Mtor | Pex12 | -2.482 | 5.38E-03 | glio |
| Interaction_genotype_Mtor | Arpc1a | 1.539 | 5.38E-03 | glio |
| Interaction_genotype_Rad51 | Cpne7 | -4.208 | 5.38E-03 | glio |
| interaction_RT | Mmd2 | 1.657 | 5.38E-03 | glio |
| Aars2_ex.vivo_noRT | Ksr2 | 2.317 | 5.38E-03 | glio |
| Aars2_ex.vivo_noRT | Mgll | 2.9 | 5.38E-03 | glio |
| Atp5c1_in.vivo_noRT | Gm49312 | 2.122 | 5.38E-03 | glio |
| Bcas2_in.vivo_noRT | Gm10791 | 1.89 | 5.38E-03 | glio |
| Bora_in.vivo_noRT | Xkr8 | 1.942 | 5.38E-03 | glio |
| Bora_in.vivo_noRT | Zfp354a | 1.964 | 5.38E-03 | glio |
| Chek1_in.vivo_noRT | Wdr25 | 1.82 | 5.38E-03 | glio |
| Chek1_ex.vivo_noRT | Lrrc72 | 2.364 | 5.38E-03 | glio |
| Chek1_ex.vivo_noRT | Pla2g12a | -1.19 | 5.38E-03 | glio |
| Cyp19a1_ex.vivo_noRT | Meioc | 2.159 | 5.38E-03 | glio |
| Cyp19a1_ex.vivo_noRT | Tmem132c | 2.159 | 5.38E-03 | glio |
| Cyp19a1_ex.vivo_noRT | Gm44693 | 2.159 | 5.38E-03 | glio |
| Eif2s1_ex.vivo_noRT | BB287469 | 2.234 | 5.38E-03 | glio |
| Interaction_genotype_Myc | 1700001G11R | -3.143 | 5.38E-04 | glio |
| interaction_RT | Zfp706 | 0.57 | 5.38E-04 | glio |
| Atr_ex.vivo_noRT | Smim40 | 3.253 | 5.38E-04 | glio |
| Bcas2_in.vivo_noRT | AY074887 | 2.755 | 5.38E-04 | glio |
| Bora_in.vivo_noRT | Ncl | -0.747 | 5.38E-04 | glio |
| Hsd17b10_in.vivo_noRT | Zfp986 | 2.575 | 5.38E-04 | glio |
| Hsd17b10_in.vivo_noRT | Gng13 | 2.544 | 5.38E-04 | glio |
| culture_effect | 1110032A03R | 0.979 | 5.38E-05 | glio |
| Interaction_genotype_Mtor | Hnrnpab | 1.698 | 5.38E-05 | glio |
| Interaction_genotype_Myc | Pcx | -4.494 | 5.38E-05 | glio |
| Interaction_genotype_Myc | Rab26 | -4.24 | 5.38E-05 | glio |
| Atp5c1_in.vivo_noRT | Ildr1 | 2.668 | 5.38E-07 | glio |
| Atp5c1_in.vivo_noRT | Gm15908 | 2.668 | 5.38E-08 | glio |
| Atp5c1_in.vivo_noRT | A930014E10Ri | 2.668 | 5.38E-08 | glio |
| Atp5c1_in.vivo_noRT | Gm45338 | 2.668 | 5.38E-08 | glio |
| Atp5c1_in.vivo_noRT | Scg2 | 2.668 | 5.38E-08 | glio |
| Bora_in.vivo_noRT | Qrich2 | 4.681 | 5.38E-11 | glio |
| Chek1_ex.vivo_noRT | Olfr1189 | 5.243 | 5.38E-12 | glio |
| Chek1_in.vivo_noRT | Gm11769 | 4.784 | 5.38E-14 | glio |
| culture_effect | Fat1 | -1.402 | 5.38E-15 | glio |
| Atr_in.vivo_noRT | Tdrd5 | 5.147 | 5.38E-16 | glio |
| Bcas2_in.vivo_noRT | Fam166a | 4.543 | 5.38E-17 | glio |
| culture_effect | Irf8 | -8.232 | 5.38E-45 | glio |
| culture_effect | Lgi4 | 0.674 | 5.39E-03 | glio |

| | | | | |
|---|---|---|---|---|
| culture_effect | E2f1 | 0.687 | 5.39E-03 | glio |
| Interaction_genotype_Mtor | B3gntl1 | -3.262 | 5.39E-03 | glio |
| Interaction_genotype_Mtor | Zfp799 | -3.01 | 5.39E-03 | glio |
| Interaction_genotype_Myc | Usp17la | -3.124 | 5.39E-03 | glio |
| Interaction_genotype_Myc | Gm41611 | -2.944 | 5.39E-03 | glio |
| Interaction_genotype_Myc | Tmem119 | 4.08 | 5.39E-03 | glio |
| Interaction_genotype_Myc | Rnf146 | -1.615 | 5.39E-03 | glio |
| Interaction_genotype_Pars2 | Gm26671 | -2.661 | 5.39E-03 | glio |
| Interaction_genotype_Uqcrb | Kpna7 | -3.957 | 5.39E-03 | glio |
| Interaction_genotype_Uqcrb | A530083I20Ri | -4.786 | 5.39E-03 | glio |
| interaction_RT | Plod3 | 0.683 | 5.39E-03 | glio |
| Atr_in.vivo_noRT | Galk1 | -0.958 | 5.39E-03 | glio |
| Atr_ex.vivo_noRT | Zfp808 | 2.384 | 5.39E-03 | glio |
| Atr_ex.vivo_noRT | 1700015F17Ri | 2.604 | 5.39E-03 | glio |
| Chek1_ex.vivo_noRT | Cdca4 | -1.702 | 5.39E-03 | glio |
| Chek1_ex.vivo_noRT | Uqcrc2 | -1.145 | 5.39E-03 | glio |
| Dclre1c_ex.vivo_noRT | Gm128 | 2.693 | 5.39E-03 | glio |
| Hsd17b10_in.vivo_noRT | Hist1h4d | -1.852 | 5.39E-03 | glio |
| culture_effect | Slc12a7 | -1.065 | 5.39E-04 | glio |
| Interaction_genotype_Aars2 | Hrct1 | -3.804 | 5.39E-04 | glio |
| Interaction_genotype_Atr | Yaf2 | 1.498 | 5.39E-04 | glio |
| Aars2_in.vivo_noRT | Gm10419 | 2.674 | 5.39E-04 | glio |
| Atp5c1_in.vivo_noRT | Krt80 | 2.402 | 5.39E-04 | glio |
| Atp5c1_in.vivo_noRT | Vwce | 2.408 | 5.39E-04 | glio |
| Atr_in.vivo_noRT | Bcl7a | 2.489 | 5.39E-04 | glio |
| Atr_in.vivo_noRT | 9330151L19Ri | 2.214 | 5.39E-04 | glio |
| Atr_in.vivo_noRT | Fastkd1 | 2.214 | 5.39E-04 | glio |
| Bora_in.vivo_noRT | Gm41760 | 2.482 | 5.39E-04 | glio |
| Bora_in.vivo_noRT | Fbxw11 | -1.608 | 5.39E-04 | glio |
| Chek1_in.vivo_noRT | 2610507I01Ri | 2.246 | 5.39E-04 | glio |
| Chek1_in.vivo_noRT | Pcdhb14 | 2.539 | 5.39E-04 | glio |
| Atp5c1_in.vivo_noRT | 4930470G03R | 2.402 | 5.39E-05 | glio |
| Atr_in.vivo_noRT | Wnt10a | 2.836 | 5.39E-05 | glio |
| Bcas2_in.vivo_noRT | Gm10244 | 2.986 | 5.39E-05 | glio |
| Interaction_genotype_Mtor | Apoh | -5.254 | 5.39E-06 | glio |
| Atp5c1_in.vivo_noRT | Gad1 | 2.402 | 5.39E-06 | glio |
| Atp5c1_in.vivo_noRT | Olfr171 | 2.668 | 5.39E-06 | glio |
| Atr_in.vivo_noRT | Nim1k | 3.335 | 5.39E-06 | glio |
| Bora_in.vivo_noRT | C330011M18F | 3.281 | 5.39E-06 | glio |
| Interaction_genotype_Mtor | Specc1 | 2.747 | 5.39E-07 | glio |
| Interaction_genotype_Mtor | Gm15579 | -6.539 | 5.39E-07 | glio |
| Interaction_genotype_Mtor | A930033M14F | -5.064 | 5.39E-07 | glio |
| Atr_in.vivo_noRT | Fkbp11 | -2.144 | 5.39E-07 | glio |
| Bora_in.vivo_noRT | Tomm6os | 4.542 | 5.39E-10 | glio |
| Chek1_in.vivo_noRT | Ntn5 | 4.252 | 5.39E-10 | glio |
| Aars2_in.vivo_noRT | Ifi214 | 4.536 | 5.39E-11 | glio |
| Atr_in.vivo_noRT | 4933408J17Ri | 4.217 | 5.39E-12 | glio |
| Bora_in.vivo_noRT | Gm48265 | 4.153 | 5.39E-12 | glio |
| Chek1_in.vivo_noRT | Gm14379 | 4.528 | 5.39E-12 | glio |
| interaction_RT | Ypel3 | 2.078 | 5.39E-13 | glio |
| Chek1_in.vivo_noRT | 4930447F24Ri | 4.784 | 5.39E-13 | glio |
| Aars2_in.vivo_noRT | Adam12 | 5.067 | 5.39E-16 | glio |

| | | | | |
|---|---|---|---|---|
| Chek1_ex.vivo_noRT | Gm16178 | 4.842 | 5.39E-16 | glio |
| Chek1_ex.vivo_noRT | Gm364 | 4.842 | 5.39E-16 | glio |
| Aars2_in.vivo_noRT | Lhx1os | 4.801 | 5.39E-17 | glio |
| Chek1_ex.vivo_noRT | Apoa5 | 4.842 | 5.39E-17 | glio |
| culture_effect | Zfp235 | 0.952 | 5.40E-03 | glio |
| Interaction_genotype_Aars2 | Gm44672 | -3.401 | 5.40E-03 | glio |
| Interaction_genotype_Myc | Olfr1191-ps1 | -2.866 | 5.40E-03 | glio |
| Interaction_genotype_Myc | Gm43915 | -2.59 | 5.40E-03 | glio |
| Interaction_genotype_Myc | Tgfbr3l | -3.266 | 5.40E-03 | glio |
| Interaction_genotype_Myc | Gm20756 | -2.937 | 5.40E-03 | glio |
| Interaction_genotype_Myc | Rap1gds1 | -2.114 | 5.40E-03 | glio |
| Interaction_genotype_Myc | Fktn | -1.862 | 5.40E-03 | glio |
| Interaction_genotype_Myc | Clta | 1.072 | 5.40E-03 | glio |
| interaction_RT | Gpr137b | 2.009 | 5.40E-03 | glio |
| interaction_RT | Nsd3 | 0.73 | 5.40E-03 | glio |
| Aars2_ex.vivo_noRT | 4921504E06Ri | 2.204 | 5.40E-03 | glio |
| Aars2_ex.vivo_noRT | Tns4 | 2.495 | 5.40E-03 | glio |
| Atr_in.vivo_noRT | 2310002F09Ri | 1.844 | 5.40E-03 | glio |
| Atr_in.vivo_noRT | Naa16 | -1.399 | 5.40E-03 | glio |
| Atr_in.vivo_noRT | Qk | -0.907 | 5.40E-03 | glio |
| Atr_in.vivo_noRT | Nsmce2 | -0.924 | 5.40E-03 | glio |
| Atr_in.vivo_noRT | Bcl7c | -0.851 | 5.40E-03 | glio |
| Bcas2_ex.vivo_noRT | Gm1123 | 2.949 | 5.40E-03 | glio |
| Chek1_ex.vivo_noRT | Hmox1 | 2.183 | 5.40E-03 | glio |
| Hsd17b10_in.vivo_noRT | Bpi | 1.935 | 5.40E-03 | glio |
| culture_effect | 4632404H12R | 1.627 | 5.40E-04 | glio |
| culture_effect | Rps27a | -0.491 | 5.40E-04 | glio |
| Interaction_genotype_Bora | Apoh | -4.475 | 5.40E-04 | glio |
| Interaction_genotype_Mtor | Cstb | 3.257 | 5.40E-04 | glio |
| Interaction_genotype_Myc | Sdcbp2 | -4.144 | 5.40E-04 | glio |
| Aars2_in.vivo_noRT | Gfap | 2.299 | 5.40E-04 | glio |
| Aars2_in.vivo_noRT | 1190005I06Ri | 2.282 | 5.40E-04 | glio |
| Atr_in.vivo_noRT | Ubxn10 | 2.098 | 5.40E-04 | glio |
| Bard1_ex.vivo_noRT | Slc19a3 | 2.421 | 5.40E-04 | glio |
| Bard1_ex.vivo_noRT | Hrh2 | 2.814 | 5.40E-04 | glio |
| Chek1_ex.vivo_noRT | Rps6ka4 | 2.994 | 5.40E-04 | glio |
| Interaction_genotype_Myc | Rftn2 | -4.109 | 5.40E-05 | glio |
| Bard1_ex.vivo_noRT | Gm18280 | 2.818 | 5.40E-05 | glio |
| Bard1_ex.vivo_noRT | Gm36793 | 2.814 | 5.40E-05 | glio |
| Hsd17b10_in.vivo_noRT | Gm30082 | 2.966 | 5.40E-05 | glio |
| Interaction_genotype_Myc | Pip4k2b | -4.365 | 5.40E-06 | glio |
| Aars2_in.vivo_noRT | A930003O13R | 3.527 | 5.40E-06 | glio |
| Aars2_in.vivo_noRT | Rtcb | -1.173 | 5.40E-06 | glio |
| culture_effect | Hacd2 | -1.147 | 5.40E-08 | glio |
| Bcas2_in.vivo_noRT | D030040B21R | 4.42 | 5.40E-11 | glio |
| Bcas2_in.vivo_noRT | Sh3d21 | 4.284 | 5.40E-13 | glio |
| Atr_in.vivo_noRT | Lrrc24 | 5.07 | 5.40E-15 | glio |
| Interaction_genotype_Aars2 | Zfp51 | -3.005 | 5.41E-03 | glio |
| Interaction_genotype_Chek1 | Rabac1 | 1.256 | 5.41E-03 | glio |
| Interaction_genotype_Mtor | Ccdc181 | -3.01 | 5.41E-03 | glio |
| Interaction_genotype_Mtor | Trappc12 | -2.162 | 5.41E-03 | glio |
| Interaction_genotype_Mtor | Prdx5 | 1.563 | 5.41E-03 | glio |

| | | | | |
|---|---|---|---|---|
| Interaction_genotype_Myc | 6230400D17R | -2.908 | 5.41E-03 | glio |
| Interaction_genotype_Myc | Nkrf | -2.342 | 5.41E-03 | glio |
| Interaction_genotype_Myc | Ap3m1 | -1.892 | 5.41E-03 | glio |
| interaction_RT | Ang | 2.824 | 5.41E-03 | glio |
| Aars2_in.vivo_noRT | Pcdhb22 | 1.905 | 5.41E-03 | glio |
| Aars2_in.vivo_noRT | Pmvk | -1.357 | 5.41E-03 | glio |
| Aars2_in.vivo_noRT | Spry1 | -1.247 | 5.41E-03 | glio |
| Aars2_in.vivo_noRT | Nampt | -0.97 | 5.41E-03 | glio |
| Aars2_ex.vivo_noRT | C4a | 2.505 | 5.41E-03 | glio |
| Bcas2_in.vivo_noRT | Hykk | 2.042 | 5.41E-03 | glio |
| Bcas2_in.vivo_noRT | Alox15 | 1.846 | 5.41E-03 | glio |
| Bcas2_in.vivo_noRT | Cdkn3 | -1.717 | 5.41E-03 | glio |
| Bora_in.vivo_noRT | Sugct | 1.753 | 5.41E-03 | glio |
| Chek1_in.vivo_noRT | Vash2 | 2.079 | 5.41E-03 | glio |
| Eif2s1_ex.vivo_noRT | H1foo | 2.234 | 5.41E-03 | glio |
| Interaction_genotype_Mtor | Olfr1189 | -3.644 | 5.41E-04 | glio |
| Interaction_genotype_Mtor | Gm3604 | -4.116 | 5.41E-04 | glio |
| Interaction_genotype_Mtor | Msantd1 | -4.304 | 5.41E-04 | glio |
| Interaction_genotype_Myc | Nmb | -3.921 | 5.41E-04 | glio |
| Interaction_genotype_Myc | Ctdp1 | -2.893 | 5.41E-04 | glio |
| interaction_RT | Timm17b | 0.792 | 5.41E-04 | glio |
| Bcas2_in.vivo_noRT | Usp51 | 2.388 | 5.41E-04 | glio |
| Bora_in.vivo_noRT | Mbp | -1.926 | 5.41E-04 | glio |
| interaction_RT | Nus1 | -0.829 | 5.41E-05 | glio |
| Aars2_in.vivo_noRT | Prrc2b | -1.547 | 5.41E-05 | glio |
| Atp5c1_in.vivo_noRT | Hrct1 | 2.668 | 5.41E-05 | glio |
| Chek1_ex.vivo_noRT | Cmss1 | -2.269 | 5.41E-05 | glio |
| Hsd17b10_in.vivo_noRT | Itgbl1 | 2.43 | 5.41E-05 | glio |
| culture_effect | Vamp8 | -2.358 | 5.41E-06 | glio |
| Bora_in.vivo_noRT | Il20rb | 3.401 | 5.41E-06 | glio |
| Hsd17b10_in.vivo_noRT | Gm48161 | 2.828 | 5.41E-06 | glio |
| Hsd17b10_in.vivo_noRT | Lmx1b | 3.356 | 5.41E-06 | glio |
| Aars2_in.vivo_noRT | Ptrh1 | 2.668 | 5.41E-07 | glio |
| Bora_in.vivo_noRT | Tnfrsf23 | 4.018 | 5.41E-07 | glio |
| Bora_in.vivo_noRT | Gm16023 | 3.939 | 5.41E-07 | glio |
| Bora_in.vivo_noRT | Tec | 3.26 | 5.41E-07 | glio |
| Chek1_in.vivo_noRT | Sod3 | 5.044 | 5.41E-10 | glio |
| Hsd17b10_in.vivo_noRT | Slc2a9 | 3.616 | 5.41E-10 | glio |
| Hsd17b10_in.vivo_noRT | Gm26555 | 3.353 | 5.41E-11 | glio |
| Hsd17b10_in.vivo_noRT | Slc25a18 | 3.877 | 5.41E-11 | glio |
| culture_effect | Pde3a | 3.221 | 5.41E-12 | glio |
| Aars2_in.vivo_noRT | Slc25a35 | 4.753 | 5.41E-12 | glio |
| culture_effect | E2f3 | 0.498 | 5.42E-03 | glio |
| Interaction_genotype_Aars2 | Lfng | -3.414 | 5.42E-03 | glio |
| Interaction_genotype_Bcas2 | Ripply3 | -3.629 | 5.42E-03 | glio |
| Interaction_genotype_Mtor | Ggt5 | -3.167 | 5.42E-03 | glio |
| Interaction_genotype_Mtor | Tnfrsf10b | -3.224 | 5.42E-03 | glio |
| Interaction_genotype_Myc | Atg4a | -2.441 | 5.42E-03 | glio |
| Interaction_genotype_Rad9a | Klk6 | 4.195 | 5.42E-03 | glio |
| Interaction_genotype_Rad9a | Nxph3 | -4.161 | 5.42E-03 | glio |
| Interaction_genotype_Uqcrc2 | Lyrm1 | -3.949 | 5.42E-03 | glio |
| Aars2_in.vivo_noRT | Zfand2b | 1.249 | 5.42E-03 | glio |

| | | | | |
|---|---|---|---|---|
| Aars2_in.vivo_noRT | Ppp2r1a | -1.006 | 5.42E-03 | glio |
| Aars2_in.vivo_noRT | Plpp2 | -0.819 | 5.42E-03 | glio |
| Atp5c1_in.vivo_noRT | Eps8l2 | 2.346 | 5.42E-03 | glio |
| Atr_in.vivo_noRT | Ip6k3 | 2.036 | 5.42E-03 | glio |
| Atr_in.vivo_noRT | Smarcc1 | -1.06 | 5.42E-03 | glio |
| Bard1_ex.vivo_noRT | Sox11 | 2.814 | 5.42E-03 | glio |
| Bard1_ex.vivo_noRT | Tcp11x2 | 2.421 | 5.42E-03 | glio |
| Bora_in.vivo_noRT | Gm5930 | 2.029 | 5.42E-03 | glio |
| Bora_in.vivo_noRT | 9130221H12R | 1.89 | 5.42E-03 | glio |
| Bora_in.vivo_noRT | Mrps27 | -1.524 | 5.42E-03 | glio |
| Bora_in.vivo_noRT | Spsb3 | -1.33 | 5.42E-03 | glio |
| Bora_in.vivo_noRT | Ifit1 | -2.154 | 5.42E-03 | glio |
| Chek1_in.vivo_noRT | Ssr4 | -0.609 | 5.42E-03 | glio |
| Hsd17b10_in.vivo_noRT | Rapgef3 | 2.412 | 5.42E-03 | glio |
| Interaction_genotype_Myc | Gm5788 | -2.729 | 5.42E-04 | glio |
| Interaction_genotype_Myc | Hspa12b | -4.278 | 5.42E-04 | glio |
| Interaction_genotype_Myc | Sapcd2 | -2.463 | 5.42E-04 | glio |
| Aars2_in.vivo_noRT | Mpp6 | -1.288 | 5.42E-04 | glio |
| Atr_ex.vivo_noRT | Gm43221 | 3.164 | 5.42E-04 | glio |
| Atr_ex.vivo_noRT | Acot5 | 3.168 | 5.42E-04 | glio |
| Atr_ex.vivo_noRT | Aloxe3 | 2.698 | 5.42E-04 | glio |
| Bcas2_in.vivo_noRT | March10 | 2.45 | 5.42E-04 | glio |
| Chek1_in.vivo_noRT | Ccnc | -1.07 | 5.42E-04 | glio |
| Interaction_genotype_Mtor | Hsd17b14 | -5.069 | 5.42E-05 | glio |
| Aars2_in.vivo_noRT | Tdrp | 2.754 | 5.42E-05 | glio |
| Chek1_in.vivo_noRT | Gm20707 | 2.866 | 5.42E-05 | glio |
| culture_effect | Stard3 | -0.904 | 5.42E-06 | glio |
| Interaction_genotype_Mtor | Strip2 | -4.235 | 5.42E-06 | glio |
| Aars2_in.vivo_noRT | Vgll3 | 3.205 | 5.42E-06 | glio |
| Bora_ex.vivo_noRT | Rasl10a | 4.393 | 5.42E-08 | glio |
| culture_effect | 5930430L01Ri | 2.394 | 5.42E-09 | glio |
| Bcas2_in.vivo_noRT | Cyp7a1 | 4.016 | 5.42E-09 | glio |
| Bora_in.vivo_noRT | Gm16046 | 4.144 | 5.42E-09 | glio |
| Bora_in.vivo_noRT | Adam30 | 4.557 | 5.42E-14 | glio |
| Bora_in.vivo_noRT | Gm48617 | 4.672 | 5.42E-21 | glio |
| culture_effect | Iigp1 | -2.691 | 5.42E-24 | glio |
| Aars2_in.vivo_noRT | Gm5788 | 5.327 | 5.42E-26 | glio |
| Atr_in.vivo_noRT | Gm41307 | 5.533 | 5.42E-27 | glio |
| Interaction_genotype_Aars2 | Gm18280 | -2.564 | 5.43E-03 | glio |
| Interaction_genotype_Bcas2 | Gm49312 | -2.851 | 5.43E-03 | glio |
| Interaction_genotype_Mtor | Npb | -3.329 | 5.43E-03 | glio |
| Interaction_genotype_Mtor | Gm15879 | -4.294 | 5.43E-03 | glio |
| Interaction_genotype_Mtor | Serpinh1 | 1.56 | 5.43E-03 | glio |
| Interaction_genotype_Mtor | Rars | 1.697 | 5.43E-03 | glio |
| Interaction_genotype_Myc | Ccdc63 | -3.054 | 5.43E-03 | glio |
| Interaction_genotype_Myc | Brf2 | -2.334 | 5.43E-03 | glio |
| Interaction_genotype_Myc | Ginm1 | 1.436 | 5.43E-03 | glio |
| Interaction_genotype_Myc | Ppp2ca | 0.995 | 5.43E-03 | glio |
| Aars2_in.vivo_noRT | Zfp12 | 1.573 | 5.43E-03 | glio |
| Atr_in.vivo_noRT | Gm43707 | 1.771 | 5.43E-03 | glio |
| Atr_in.vivo_noRT | Upp2 | 1.797 | 5.43E-03 | glio |
| Bcas2_in.vivo_noRT | Folr2 | 1.999 | 5.43E-03 | glio |

| | | | | |
|---|---|---|---|---|
| Chek1_ex.vivo_noRT | Ngdn | -1.241 | 5.43E-03 | glio |
| culture_effect | Ccnb2 | -0.768 | 5.43E-04 | glio |
| Interaction_genotype_Myc | Gm10642 | -3.552 | 5.43E-04 | glio |
| Interaction_genotype_Myc | Selenom | 1.67 | 5.43E-04 | glio |
| Atp5c1_in.vivo_noRT | Il1rapl1 | 2.55 | 5.43E-04 | glio |
| Atr_in.vivo_noRT | Tmem259 | -1.222 | 5.43E-04 | glio |
| Bcas2_in.vivo_noRT | Osbp2 | 2.882 | 5.43E-04 | glio |
| Bora_in.vivo_noRT | Gm10138 | 2.315 | 5.43E-04 | glio |
| Hsd17b10_in.vivo_noRT | Apol7d | 2.714 | 5.43E-04 | glio |
| Hsd17b10_in.vivo_noRT | 2610044O15R | -1.9 | 5.43E-04 | glio |
| culture_effect | Gm7008 | 2.132 | 5.43E-05 | glio |
| culture_effect | Nfkbib | -0.667 | 5.43E-05 | glio |
| Interaction_genotype_Mtor | Cabp1 | -4.376 | 5.43E-05 | glio |
| Chek1_in.vivo_noRT | Trim68 | 2.734 | 5.43E-05 | glio |
| Chek1_in.vivo_noRT | Gm11520 | 2.637 | 5.43E-05 | glio |
| Chek1_ex.vivo_noRT | Ccsap | -2.581 | 5.43E-05 | glio |
| Chek1_ex.vivo_noRT | Gm15246 | 3.961 | 5.43E-06 | glio |
| culture_effect | Sox4 | 0.854 | 5.43E-07 | glio |
| Interaction_genotype_Atr | Gabarap | 1.764 | 5.43E-07 | glio |
| Chek1_ex.vivo_noRT | Pabpc4l | 4.533 | 5.43E-07 | glio |
| Bora_in.vivo_noRT | Anxa8 | 3.743 | 5.43E-08 | glio |
| Bcas2_in.vivo_noRT | Gm15234 | 4.291 | 5.43E-10 | glio |
| Chek1_ex.vivo_noRT | Cemip | 4.842 | 5.43E-11 | glio |
| culture_effect | Art2b | -4.069 | 5.43E-14 | glio |
| Chek1_ex.vivo_noRT | Gm11583 | 4.842 | 5.43E-14 | glio |
| Atr_in.vivo_noRT | 4930512B01R | 5.533 | 5.43E-25 | glio |
| culture_effect | Olfr1152 | -1.201 | 5.44E-03 | glio |
| culture_effect | Gm28940 | -1.451 | 5.44E-03 | glio |
| culture_effect | Zfp251 | 0.534 | 5.44E-03 | glio |
| culture_effect | Mlkl | -0.461 | 5.44E-03 | glio |
| Interaction_genotype_Atr | 9330179D12R | -3.763 | 5.44E-03 | glio |
| Interaction_genotype_Bora | Gm9833 | -3.506 | 5.44E-03 | glio |
| Interaction_genotype_Bora | Farsa | 1.712 | 5.44E-03 | glio |
| Atr_in.vivo_noRT | Sart1 | 1.052 | 5.44E-03 | glio |
| Bcas2_in.vivo_noRT | Gm43329 | 1.933 | 5.44E-03 | glio |
| Bora_in.vivo_noRT | Gfap | 1.909 | 5.44E-03 | glio |
| Bora_in.vivo_noRT | AV099323 | 1.627 | 5.44E-03 | glio |
| Bora_in.vivo_noRT | Snx21 | 1.464 | 5.44E-03 | glio |
| Bora_in.vivo_noRT | Lig1 | -1.335 | 5.44E-03 | glio |
| Bora_in.vivo_noRT | Nucks1 | -0.802 | 5.44E-03 | glio |
| Bora_in.vivo_noRT | Hspd1 | -0.607 | 5.44E-03 | glio |
| Cenpj_in.vivo_noRT | Scgb1c1 | 2.022 | 5.44E-03 | glio |
| Chek1_in.vivo_noRT | Zfp984 | 1.508 | 5.44E-03 | glio |
| Eif2s1_ex.vivo_noRT | Psca | 2.739 | 5.44E-03 | glio |
| culture_effect | Cd27 | -1.768 | 5.44E-04 | glio |
| culture_effect | Tmem258 | -0.458 | 5.44E-04 | glio |
| Interaction_genotype_Aars2 | Sprr1a | -4.406 | 5.44E-04 | glio |
| Interaction_genotype_Aars2 | Gm10484 | -4.524 | 5.44E-04 | glio |
| Interaction_genotype_Myc | Afap1l1 | -2.884 | 5.44E-04 | glio |
| Aars2_in.vivo_noRT | Dtx4 | 1.885 | 5.44E-04 | glio |
| Aars2_ex.vivo_noRT | Anxa8 | 3.298 | 5.44E-04 | glio |
| Atp5c1_in.vivo_noRT | Gm32591 | 2.673 | 5.44E-04 | glio |

| | | | | |
|---|---|---|---|---|
| Atr_in.vivo_noRT | Sumo2 | -0.756 | 5.44E-04 | glio |
| culture_effect | 5033417F24Ri | 2.285 | 5.44E-05 | glio |
| Interaction_genotype_Mtor | Gm47204 | -4.835 | 5.44E-05 | glio |
| Interaction_genotype_Mtor | Lhx1os | -3.968 | 5.44E-05 | glio |
| Interaction_genotype_Myc | 1700020L24Ri | -4.278 | 5.44E-05 | glio |
| Interaction_genotype_Myc | E230029C05Ri | -4.135 | 5.44E-05 | glio |
| Atr_in.vivo_noRT | Snhg14 | 2.891 | 5.44E-05 | glio |
| Atr_ex.vivo_noRT | Olfr543 | 2.769 | 5.44E-05 | glio |
| Atr_ex.vivo_noRT | Cxcl2 | 5.654 | 5.44E-05 | glio |
| Interaction_genotype_Myc | Capn9 | -4.793 | 5.44E-06 | glio |
| Bora_in.vivo_noRT | 1810062O18R | 3.35 | 5.44E-06 | glio |
| Bora_in.vivo_noRT | Gm15013 | 3.144 | 5.44E-06 | glio |
| Chek1_ex.vivo_noRT | Osm | 4.443 | 5.44E-09 | glio |
| culture_effect | Mlf2 | 0.771 | 5.44E-12 | glio |
| Chek1_ex.vivo_noRT | Thnsl2 | 4.842 | 5.44E-13 | glio |
| Hsd17b10_in.vivo_noRT | Slc35g3 | 3.616 | 5.44E-13 | glio |
| culture_effect | Mvp | -2.103 | 5.44E-26 | glio |
| Chek1_in.vivo_noRT | Gm5157 | 5.044 | 5.44E-26 | glio |
| Interaction_genotype_Aars2 | Slc1a2 | -2.956 | 5.45E-03 | glio |
| Interaction_genotype_Aars2 | Gm15943 | -2.752 | 5.45E-03 | glio |
| Interaction_genotype_Aars2 | Kcnk4 | -3.538 | 5.45E-03 | glio |
| Interaction_genotype_Aars2 | Hoxa6 | -3.353 | 5.45E-03 | glio |
| Interaction_genotype_Aars2 | Rassf7 | -2.826 | 5.45E-03 | glio |
| Interaction_genotype_Aars2 | Wbp4 | 1.394 | 5.45E-03 | glio |
| Interaction_genotype_Aars2 | Vapa | 1.168 | 5.45E-03 | glio |
| Interaction_genotype_Aars2 | Psme2 | 1.157 | 5.45E-03 | glio |
| Interaction_genotype_Aars2 | Eef1b2 | 0.883 | 5.45E-03 | glio |
| Interaction_genotype_Aars2 | mt-Atp6 | 0.872 | 5.45E-03 | glio |
| Atp5c1_in.vivo_noRT | Pla1a | 2.136 | 5.45E-03 | glio |
| Atr_in.vivo_noRT | Cep128 | 1.799 | 5.45E-03 | glio |
| Bard1_ex.vivo_noRT | Alpl | 2.814 | 5.45E-03 | glio |
| Bard1_ex.vivo_noRT | Ceacam19 | 2.902 | 5.45E-03 | glio |
| Bard1_ex.vivo_noRT | Ms4a4d | 2.814 | 5.45E-03 | glio |
| Bora_in.vivo_noRT | Macc1 | 1.869 | 5.45E-03 | glio |
| Chek1_in.vivo_noRT | Cdc42bpg | 1.88 | 5.45E-03 | glio |
| Chek1_in.vivo_noRT | Pex5 | -1.463 | 5.45E-03 | glio |
| Eif2s1_in.vivo_noRT | Nxnl2 | 2.148 | 5.45E-03 | glio |
| Hsd17b10_in.vivo_noRT | 4930524O07R | 2.219 | 5.45E-03 | glio |
| Hsd17b10_in.vivo_noRT | Sugt1 | -0.904 | 5.45E-03 | glio |
| culture_effect | A930003A15R | -2.087 | 5.45E-04 | glio |
| culture_effect | Col8a1 | 1.995 | 5.45E-04 | glio |
| culture_effect | Irgq | 0.916 | 5.45E-04 | glio |
| culture_effect | Exoc7 | -0.739 | 5.45E-04 | glio |
| Interaction_genotype_Myc | Tctn2 | -2.751 | 5.45E-04 | glio |
| Aars2_in.vivo_noRT | Cdkn2a | -0.711 | 5.45E-04 | glio |
| Atr_in.vivo_noRT | Pet2 | -1.218 | 5.45E-04 | glio |
| Atr_ex.vivo_noRT | Snhg4.1 | 2.771 | 5.45E-04 | glio |
| Bard1_ex.vivo_noRT | Slc6a11 | 2.814 | 5.45E-04 | glio |
| Bcas2_in.vivo_noRT | 0610009E02Ri | 2.689 | 5.45E-04 | glio |
| Cenpj_in.vivo_noRT | Gm19276 | 2.022 | 5.45E-04 | glio |
| Cenpj_ex.vivo_noRT | Gm48215 | 2.591 | 5.45E-04 | glio |
| Chek1_in.vivo_noRT | Irf2 | -1.375 | 5.45E-04 | glio |

| | | | | |
|---|---|---|---|---|
| Hsd17b10_in.vivo_noRT | Gm34643 | 2.681 | 5.45E-04 | glio |
| culture_effect | Dus4l | 1.24 | 5.45E-05 | glio |
| Interaction_genotype_Mtor | Adap1 | -5.511 | 5.45E-05 | glio |
| Interaction_genotype_Mtor | Gm7008 | -4.6 | 5.45E-05 | glio |
| Interaction_genotype_Mtor | Olfr267 | -5.002 | 5.45E-05 | glio |
| Interaction_genotype_Mtor | Minar2 | -3.839 | 5.45E-05 | glio |
| Interaction_genotype_Mtor | Mrgprf | -4.968 | 5.45E-05 | glio |
| Bora_in.vivo_noRT | Lrrc51 | 2.246 | 5.45E-05 | glio |
| Interaction_genotype_Aars2 | Krt80 | -5.035 | 5.45E-06 | glio |
| culture_effect | C330013E15Ri | 2.413 | 5.45E-07 | glio |
| Interaction_genotype_Nat2 | Gm5890 | 4.409 | 5.45E-07 | glio |
| Chek1_in.vivo_noRT | Tssk6 | 3.088 | 5.45E-07 | glio |
| Hsd17b10_in.vivo_noRT | Prickle4 | 3.356 | 5.45E-07 | glio |
| Atp5c1_in.vivo_noRT | 1700003E24Ri | 2.668 | 5.45E-08 | glio |
| Aars2_in.vivo_noRT | Pdlim3 | 4.547 | 5.45E-15 | glio |
| Aars2_in.vivo_noRT | Gm15270 | 5.063 | 5.45E-18 | glio |
| culture_effect | Epsti1 | -3.388 | 5.45E-38 | glio |
| culture_effect | Csrp2 | 0.565 | 5.46E-03 | glio |
| Interaction_genotype_Aars2 | Gm26658 | -3.717 | 5.46E-03 | glio |
| Interaction_genotype_Aars2 | BC037704 | -3.218 | 5.46E-03 | glio |
| Interaction_genotype_Atr | Wbp11 | 1.044 | 5.46E-03 | glio |
| Interaction_genotype_Bcas2 | Gm13594 | -3.859 | 5.46E-03 | glio |
| Interaction_genotype_Bcas2 | Gm41318 | -3.605 | 5.46E-03 | glio |
| Interaction_genotype_Myc | Gm40538 | -2.466 | 5.46E-03 | glio |
| Interaction_genotype_Myc | Cdk5rap1 | -2.947 | 5.46E-03 | glio |
| Interaction_genotype_Rad21 | 4933400F21Ri | 3.011 | 5.46E-03 | glio |
| Interaction_genotype_Rad51 | Gmpr2 | 2.278 | 5.46E-03 | glio |
| Aars2_in.vivo_noRT | Lhx6 | 1.911 | 5.46E-03 | glio |
| Aars2_in.vivo_noRT | Clptm1l | -0.995 | 5.46E-03 | glio |
| Chek1_in.vivo_noRT | Gm4356 | 2.243 | 5.46E-03 | glio |
| Chek1_in.vivo_noRT | 1700088E04Ri | 1.731 | 5.46E-03 | glio |
| Eif2s1_in.vivo_noRT | 1700026J14Ri | 2.148 | 5.46E-03 | glio |
| Eif2s1_in.vivo_noRT | Capn9 | 2.143 | 5.46E-03 | glio |
| Hsd17b10_in.vivo_noRT | Gm49494 | 2.271 | 5.46E-03 | glio |
| Hsd17b10_in.vivo_noRT | Eng | 2.42 | 5.46E-03 | glio |
| Interaction_genotype_Bcas2 | Adam12 | -3.848 | 5.46E-04 | glio |
| Interaction_genotype_Mtor | Spcs2 | 1.649 | 5.46E-04 | glio |
| Interaction_genotype_Myc | Gm49767 | -4.063 | 5.46E-04 | glio |
| Atr_in.vivo_noRT | Cntd1 | 2.459 | 5.46E-04 | glio |
| Bcas2_in.vivo_noRT | 4930506C21R | 2.47 | 5.46E-04 | glio |
| Bcas2_in.vivo_noRT | Ghitm | -1.06 | 5.46E-04 | glio |
| Chek1_in.vivo_noRT | Gm10553 | 2.497 | 5.46E-04 | glio |
| Chek1_in.vivo_noRT | Gm10184 | 2.296 | 5.46E-04 | glio |
| Interaction_genotype_Myc | Fbxl22 | -4.382 | 5.46E-05 | glio |
| Interaction_genotype_Myc | Cfap46 | -4.19 | 5.46E-05 | glio |
| Interaction_genotype_Myc | 1700030L20Ri | -4.404 | 5.46E-05 | glio |
| Interaction_genotype_Myc | E130208F15Ri | -5.01 | 5.46E-05 | glio |
| Interaction_genotype_Myc | Smcr8 | -4.009 | 5.46E-05 | glio |
| Interaction_genotype_Myc | Gm15929 | -3.706 | 5.46E-05 | glio |
| Atr_in.vivo_noRT | Gm37612 | 2.867 | 5.46E-05 | glio |
| Bcas2_ex.vivo_noRT | Cldn5 | 2.742 | 5.46E-05 | glio |
| Bcas2_ex.vivo_noRT | Sod3 | 3.524 | 5.46E-05 | glio |

| | | | | |
|---|---|---|---|---|
| Chek1_in.vivo_noRT | C1qtnf1 | 2.832 | 5.46E-05 | glio |
| Hsd17b10_in.vivo_noRT | Gm44672 | 2.852 | 5.46E-05 | glio |
| Interaction_genotype_Mtor | Cd7 | -3.835 | 5.46E-06 | glio |
| Atr_in.vivo_noRT | Rps10 | -0.988 | 5.46E-06 | glio |
| Bcas2_in.vivo_noRT | Poln | 3.304 | 5.46E-06 | glio |
| Bcas2_in.vivo_noRT | Rasl10a | 2.665 | 5.46E-06 | glio |
| Chek1_in.vivo_noRT | Rasl2-9 | 3.606 | 5.46E-06 | glio |
| Chek1_in.vivo_noRT | Slco2a1 | 3.045 | 5.46E-06 | glio |
| Chek1_ex.vivo_noRT | Gm50138 | 3.284 | 5.46E-06 | glio |
| culture_effect | Dnaaf3 | -2.887 | 5.46E-07 | glio |
| Interaction_genotype_Mtor | 1700030L20Ri | -5.774 | 5.46E-07 | glio |
| Chek1_ex.vivo_noRT | Faim2 | 3.652 | 5.46E-07 | glio |
| Atr_in.vivo_noRT | Gm44091 | 3.614 | 5.46E-08 | glio |
| Atr_in.vivo_noRT | Tbkbp1 | 4.24 | 5.46E-09 | glio |
| Chek1_in.vivo_noRT | Nid1 | 4.239 | 5.46E-09 | glio |
| Chek1_in.vivo_noRT | Tbkbp1 | 4.262 | 5.46E-09 | glio |
| Hsd17b10_in.vivo_noRT | Gm14019 | 3.616 | 5.46E-09 | glio |
| Chek1_ex.vivo_noRT | Ildr1 | 4.842 | 5.46E-15 | glio |
| culture_effect | Vamp5 | 1.539 | 5.46E-18 | glio |
| culture_effect | Muc4 | -1.457 | 5.47E-03 | glio |
| Interaction_genotype_Myc | Tnnc1 | -3.474 | 5.47E-03 | glio |
| Interaction_genotype_Myc | C630043F03Ri | -2.489 | 5.47E-03 | glio |
| interaction_RT | S1pr1 | 2.897 | 5.47E-03 | glio |
| interaction_RT | Hint2 | 0.721 | 5.47E-03 | glio |
| Atp5c1_in.vivo_noRT | 4933421A08R | 2.416 | 5.47E-03 | glio |
| Atp5c1_in.vivo_noRT | Slco2a1 | 2.147 | 5.47E-03 | glio |
| Atr_ex.vivo_noRT | Il23a | 2.76 | 5.47E-03 | glio |
| Bard1_ex.vivo_noRT | Gm48140 | 2.021 | 5.47E-03 | glio |
| Bcas2_ex.vivo_noRT | Pawr | 2.568 | 5.47E-03 | glio |
| Bora_in.vivo_noRT | Thtpa | 1.653 | 5.47E-03 | glio |
| Bora_in.vivo_noRT | Ubac2 | -1.22 | 5.47E-03 | glio |
| Chek1_ex.vivo_noRT | Mlycd | 1.937 | 5.47E-03 | glio |
| Interaction_genotype_Bora | Aldh2 | 2.304 | 5.47E-04 | glio |
| Interaction_genotype_Myc | Zfp974 | -4.118 | 5.47E-04 | glio |
| Aars2_ex.vivo_noRT | H2-DMb2 | 3.293 | 5.47E-04 | glio |
| Atr_ex.vivo_noRT | Rasl10a | 2.771 | 5.47E-04 | glio |
| Atr_ex.vivo_noRT | Spink4 | 3.164 | 5.47E-04 | glio |
| Cenpj_in.vivo_noRT | Vmn2r44 | 2.022 | 5.47E-04 | glio |
| Cenpj_in.vivo_noRT | Mmp13 | 2.022 | 5.47E-04 | glio |
| Cenpj_in.vivo_noRT | Gm8016 | 2.022 | 5.47E-04 | glio |
| Hsd17b10_in.vivo_noRT | Rhox3g | 2.501 | 5.47E-04 | glio |
| culture_effect | Gm34121 | 1.863 | 5.47E-05 | glio |
| interaction_RT | Erbb3 | -1.23 | 5.47E-05 | glio |
| Bora_in.vivo_noRT | Cd79a | 2.945 | 5.47E-05 | glio |
| Interaction_genotype_Mtor | Upk3bl | -5.11 | 5.47E-06 | glio |
| interaction_RT | Tmbim6 | 0.863 | 5.47E-06 | glio |
| Aars2_in.vivo_noRT | Eif5a | -0.949 | 5.47E-06 | glio |
| Bcas2_in.vivo_noRT | 9530018F02Ri | 3.421 | 5.47E-06 | glio |
| Bora_in.vivo_noRT | Map3k5 | 2.772 | 5.47E-06 | glio |
| Chek1_ex.vivo_noRT | Gm43330 | 3.862 | 5.47E-07 | glio |
| interaction_RT | Tubb4a | 5.356 | 5.47E-08 | glio |
| culture_effect | Limch1 | 3.8 | 5.47E-13 | glio |

| | | | | |
|---|---|---|---|---|
| Bora_in.vivo_noRT | Card9 | 4.672 | 5.47E-15 | glio |
| culture_effect | Stk39 | 0.62 | 5.48E-03 | glio |
| Interaction_genotype_Atr | Aass | -3.311 | 5.48E-03 | glio |
| Interaction_genotype_Bcas2 | Flacc1 | -3.255 | 5.48E-03 | glio |
| Interaction_genotype_Cdkn2a | Abcb5 | 2.84 | 5.48E-03 | glio |
| Interaction_genotype_Mtor | Pus10 | -2.35 | 5.48E-03 | glio |
| Interaction_genotype_Mtor | Hspe1-rs1 | -1.996 | 5.48E-03 | glio |
| Interaction_genotype_Myc | Hypk | 1.338 | 5.48E-03 | glio |
| Aars2_in.vivo_noRT | Lnx2 | 1.849 | 5.48E-03 | glio |
| Aars2_in.vivo_noRT | Pyroxd2 | 1.871 | 5.48E-03 | glio |
| Aars2_in.vivo_noRT | Ccdc117 | -1.374 | 5.48E-03 | glio |
| Atp5c1_in.vivo_noRT | Pdlim3 | 1.889 | 5.48E-03 | glio |
| Atr_in.vivo_noRT | Eif2s2 | -0.547 | 5.48E-03 | glio |
| Bcas2_in.vivo_noRT | Gm16638 | 1.955 | 5.48E-03 | glio |
| Bcas2_in.vivo_noRT | Cbx4 | -1.319 | 5.48E-03 | glio |
| Bcas2_in.vivo_noRT | Cisd2 | -0.838 | 5.48E-03 | glio |
| Bcas2_in.vivo_noRT | Ipo7 | -0.734 | 5.48E-03 | glio |
| Bcas2_in.vivo_noRT | Zfp706 | -0.534 | 5.48E-03 | glio |
| culture_effect | Polk | 0.996 | 5.48E-04 | glio |
| Interaction_genotype_Mtor | Fam131a | -3.973 | 5.48E-04 | glio |
| Interaction_genotype_Mtor | Gm8212 | -4.146 | 5.48E-04 | glio |
| Interaction_genotype_Myc | Inpp5a | -3.527 | 5.48E-04 | glio |
| Aars2_in.vivo_noRT | Srsf1 | -1.17 | 5.48E-04 | glio |
| Atr_in.vivo_noRT | Hist1h2an | 2.345 | 5.48E-04 | glio |
| Atr_in.vivo_noRT | Mrpl53 | -1.058 | 5.48E-04 | glio |
| Bora_in.vivo_noRT | Scn2a | 2.523 | 5.48E-04 | glio |
| Bora_in.vivo_noRT | Pid1 | 2.022 | 5.48E-04 | glio |
| Interaction_genotype_Myc | Dscaml1 | -3.124 | 5.48E-05 | glio |
| Atp5c1_in.vivo_noRT | Ikbke | 2.933 | 5.48E-05 | glio |
| Atr_in.vivo_noRT | Serpinb9 | 2.703 | 5.48E-05 | glio |
| Atr_in.vivo_noRT | Tubg1 | -1.559 | 5.48E-05 | glio |
| Chek1_in.vivo_noRT | Slpi | 3.218 | 5.48E-05 | glio |
| Aars2_in.vivo_noRT | Gm28417 | 3.007 | 5.48E-06 | glio |
| Bora_in.vivo_noRT | Gm26916 | 3.359 | 5.48E-07 | glio |
| Hsd17b10_in.vivo_noRT | Gm10696 | 3.616 | 5.48E-07 | glio |
| Bcas2_in.vivo_noRT | Prox2 | 4.081 | 5.48E-08 | glio |
| Chek1_ex.vivo_noRT | Gm813 | 4.842 | 5.48E-08 | glio |
| culture_effect | Pnpo | -1.412 | 5.48E-09 | glio |
| Aars2_in.vivo_noRT | 9430037G07R | 4.418 | 5.48E-10 | glio |
| Aars2_in.vivo_noRT | Gm46652 | 4.277 | 5.48E-11 | glio |
| Bora_in.vivo_noRT | Hist1h2ag | 4.404 | 5.48E-17 | glio |
| Bora_in.vivo_noRT | Prickle2 | 4.936 | 5.48E-17 | glio |
| culture_effect | Gm6598 | 1.339 | 5.49E-03 | glio |
| culture_effect | Lgi3 | -1.121 | 5.49E-03 | glio |
| Interaction_genotype_Mtor | 2810049E08Ri | -3.062 | 5.49E-03 | glio |
| Interaction_genotype_Mtor | Gm20275 | -2.745 | 5.49E-03 | glio |
| Interaction_genotype_Mtor | Cntf | -3.264 | 5.49E-03 | glio |
| Interaction_genotype_Mtor | Gnat2 | -2.959 | 5.49E-03 | glio |
| Interaction_genotype_Mtor | Pag1 | -2.815 | 5.49E-03 | glio |
| Interaction_genotype_Mtor | Sipa1l1 | 1.806 | 5.49E-03 | glio |
| Interaction_genotype_Pars2 | Gm45669 | -3.317 | 5.49E-03 | glio |
| interaction_RT | Vdac1 | -0.569 | 5.49E-03 | glio |

| | | | | |
|---|---|---|---|---|
| Aars2_in.vivo_noRT | Prdm15 | 1.718 | 5.49E-03 | glio |
| Atp5c1_in.vivo_noRT | D030068K23R | 2.277 | 5.49E-03 | glio |
| Bora_in.vivo_noRT | Tex15 | 1.995 | 5.49E-03 | glio |
| Chek1_in.vivo_noRT | Fgl1 | 2.084 | 5.49E-03 | glio |
| Chek1_in.vivo_noRT | Mex3c | -1.368 | 5.49E-03 | glio |
| Chek1_ex.vivo_noRT | 4833419F23Ri | 2.587 | 5.49E-03 | glio |
| Hsd17b10_in.vivo_noRT | Il2rb | 2.074 | 5.49E-03 | glio |
| Interaction_genotype_Myc | Limk1 | -2.455 | 5.49E-04 | glio |
| interaction_RT | AY036118 | -1.751 | 5.49E-04 | glio |
| interaction_RT | Acp1 | -0.649 | 5.49E-04 | glio |
| Atp5c1_in.vivo_noRT | Adam12 | 2.408 | 5.49E-04 | glio |
| Atr_in.vivo_noRT | AI480526 | 3.094 | 5.49E-06 | glio |
| culture_effect | D630036H23R | 2.698 | 5.49E-07 | glio |
| culture_effect | Sh3bgrl | 0.823 | 5.49E-07 | glio |
| Aars2_in.vivo_noRT | Arhgap27 | 3.536 | 5.49E-08 | glio |
| Aars2_in.vivo_noRT | Esr1 | 3.515 | 5.49E-08 | glio |
| Aars2_in.vivo_noRT | Tmem8b | 3.939 | 5.49E-09 | glio |
| Hsd17b10_in.vivo_noRT | Gm17203 | 2.966 | 5.49E-09 | glio |
| Bora_in.vivo_noRT | Tns4 | 4.286 | 5.49E-12 | glio |
| Hsd17b10_in.vivo_noRT | Chia1 | 3.616 | 5.49E-13 | glio |
| Chek1_in.vivo_noRT | Gm21319 | 5.044 | 5.49E-20 | glio |
| Atr_in.vivo_noRT | Gm13178 | 5.017 | 5.49E-22 | glio |
| Bcas2_in.vivo_noRT | Helt | 4.807 | 5.49E-24 | glio |
| Bcas2_in.vivo_noRT | Gm20625 | 4.807 | 5.49E-24 | glio |
| culture_effect | Acot7 | 0.356 | 5.50E-03 | glio |
| Interaction_genotype_Atr | Kif7 | -3.225 | 5.50E-03 | glio |
| Interaction_genotype_Myc | Tnfsf12 | -2.385 | 5.50E-03 | glio |
| Interaction_genotype_Myc | Gm4890 | -2.152 | 5.50E-03 | glio |
| Interaction_genotype_Pars2 | Gm41442 | -3.665 | 5.50E-03 | glio |
| Interaction_genotype_Pars2 | Gm49860 | -3.25 | 5.50E-03 | glio |
| Interaction_genotype_Pars2 | Ube4bos1 | -3.19 | 5.50E-03 | glio |
| Interaction_genotype_Pars2 | Glis1 | -3.797 | 5.50E-03 | glio |
| Interaction_genotype_Pars2 | Tmem236 | -3.374 | 5.50E-03 | glio |
| Aars2_in.vivo_noRT | Pbk | -1.524 | 5.50E-03 | glio |
| Aars2_ex.vivo_noRT | Gvin1 | 2.897 | 5.50E-03 | glio |
| Atr_in.vivo_noRT | Dok4 | 2.004 | 5.50E-03 | glio |
| Atr_in.vivo_noRT | Ppm1l | 1.494 | 5.50E-03 | glio |
| Bcas2_in.vivo_noRT | Creld2 | -1.096 | 5.50E-03 | glio |
| Bora_in.vivo_noRT | Gm4673 | 2.085 | 5.50E-03 | glio |
| Bora_ex.vivo_noRT | Gm15929 | 2.225 | 5.50E-03 | glio |
| Bora_ex.vivo_noRT | Slc17a7 | 2.815 | 5.50E-03 | glio |
| Chek1_in.vivo_noRT | Gm11772 | 2.045 | 5.50E-03 | glio |
| Fanca_ex.vivo_noRT | A930019D19R | 2.124 | 5.50E-03 | glio |
| Interaction_genotype_Mtor | Bcl2l15 | -3.622 | 5.50E-04 | glio |
| interaction_RT | Bach2 | 2.992 | 5.50E-04 | glio |
| Chek1_in.vivo_noRT | Kctd11 | 2.572 | 5.50E-04 | glio |
| Chek1_in.vivo_noRT | Shroom1 | 2.426 | 5.50E-04 | glio |
| culture_effect | Tmsb10 | 0.687 | 5.50E-05 | glio |
| Interaction_genotype_Mtor | Son | 2.071 | 5.50E-05 | glio |
| Interaction_genotype_Myc | Cfap100 | -3.9 | 5.50E-05 | glio |
| Aars2_in.vivo_noRT | Gm28221 | 2.657 | 5.50E-05 | glio |
| Aars2_in.vivo_noRT | Ddost | -0.99 | 5.50E-05 | glio |

| | | | | |
|---|---|---|---|---|
| Bcas2_in.vivo_noRT | Rgs16 | 3.598 | 5.50E-05 | glio |
| culture_effect | Sgcz | -1.662 | 5.50E-06 | glio |
| interaction_RT | A330084C13R | -4.045 | 5.50E-06 | glio |
| Bora_ex.vivo_noRT | Smtnl1 | 3.201 | 5.50E-06 | glio |
| culture_effect | 2700081O15R | -1.245 | 5.50E-09 | glio |
| Chek1_ex.vivo_noRT | Gm40264 | 4.256 | 5.50E-09 | glio |
| Atr_in.vivo_noRT | Vamp8 | 3.71 | 5.50E-11 | glio |
| Bcas2_in.vivo_noRT | Masp2 | 3.88 | 5.50E-11 | glio |
| culture_effect | Cdv3 | 0.812 | 5.50E-15 | glio |
| Bcas2_in.vivo_noRT | Nwd1 | 4.543 | 5.50E-15 | glio |
| Atr_in.vivo_noRT | Adam2 | 5.008 | 5.50E-17 | glio |
| Bora_in.vivo_noRT | Pdgfb | 4.673 | 5.50E-17 | glio |
| Bcas2_in.vivo_noRT | Gm12841 | 4.807 | 5.50E-21 | glio |
| Atr_in.vivo_noRT | A930019D19R | 5.533 | 5.50E-30 | glio |
| culture_effect | Fam83d | -0.813 | 5.51E-03 | glio |
| Interaction_genotype_Aars2 | Prl2c3 | -3.229 | 5.51E-03 | glio |
| Interaction_genotype_Aars2 | Guca1b | -3.568 | 5.51E-03 | glio |
| Interaction_genotype_Aars2 | Slc15a4 | -2.676 | 5.51E-03 | glio |
| Interaction_genotype_Myc | Gm31517 | -3.259 | 5.51E-03 | glio |
| Interaction_genotype_Myc | Mtx3 | -2.837 | 5.51E-03 | glio |
| Interaction_genotype_Myc | Abr | -2.868 | 5.51E-03 | glio |
| Interaction_genotype_Myc | Cox6b2 | -2.455 | 5.51E-03 | glio |
| interaction_RT | Gm12185 | 2.379 | 5.51E-03 | glio |
| interaction_RT | Triqk | 2.43 | 5.51E-03 | glio |
| interaction_RT | Gipc2 | 2.746 | 5.51E-03 | glio |
| interaction_RT | Zswim4 | 1.55 | 5.51E-03 | glio |
| interaction_RT | Aen | -0.906 | 5.51E-03 | glio |
| interaction_RT | Orc2 | -0.673 | 5.51E-03 | glio |
| interaction_RT | Nup43 | -0.707 | 5.51E-03 | glio |
| Aars2_in.vivo_noRT | Ss18l2 | -0.744 | 5.51E-03 | glio |
| Bora_in.vivo_noRT | Zkscan7 | 2.083 | 5.51E-03 | glio |
| Bora_in.vivo_noRT | Igf2bp3 | -1.213 | 5.51E-03 | glio |
| Bora_in.vivo_noRT | Rtraf | -0.697 | 5.51E-03 | glio |
| Bora_in.vivo_noRT | Rtcb | -0.613 | 5.51E-03 | glio |
| Brca2_in.vivo_noRT | Gm5415 | 1.763 | 5.51E-03 | glio |
| Brca2_in.vivo_noRT | Edn2 | 1.763 | 5.51E-03 | glio |
| Brca2_in.vivo_noRT | Gm14207 | 1.763 | 5.51E-03 | glio |
| Brca2_in.vivo_noRT | Nrip2 | 1.763 | 5.51E-03 | glio |
| Brca2_in.vivo_noRT | Gm49698 | 1.763 | 5.51E-03 | glio |
| Brca2_in.vivo_noRT | Olfr881 | 1.763 | 5.51E-03 | glio |
| Brca2_in.vivo_noRT | BC061195 | 1.763 | 5.51E-03 | glio |
| Brca2_in.vivo_noRT | Gm43951 | 2.144 | 5.51E-03 | glio |
| Chek1_in.vivo_noRT | Mgarp | 1.955 | 5.51E-03 | glio |
| Chek1_in.vivo_noRT | Hspa13 | -1.191 | 5.51E-03 | glio |
| Chek1_in.vivo_noRT | Smim11 | -0.945 | 5.51E-03 | glio |
| Chek1_in.vivo_noRT | Lars | -0.755 | 5.51E-03 | glio |
| Eif2s1_ex.vivo_noRT | Gm13684 | 2.639 | 5.51E-03 | glio |
| Hsd17b10_in.vivo_noRT | Tmem240 | 2.419 | 5.51E-03 | glio |
| Hsd17b10_in.vivo_noRT | Gm43728 | 2.289 | 5.51E-03 | glio |
| Hsd17b10_in.vivo_noRT | 1500011B03R | -1.023 | 5.51E-03 | glio |
| Interaction_genotype_Myc | Sf3b5 | 1.445 | 5.51E-04 | glio |
| interaction_RT | Dctn3 | 0.622 | 5.51E-04 | glio |

| | | | | |
|---|---|---|---|---|
| Atr_in.vivo_noRT | Pcdhb14 | 2.499 | 5.51E-04 | glio |
| Bcas2_in.vivo_noRT | Pacs1 | 2.463 | 5.51E-04 | glio |
| Chek1_in.vivo_noRT | Ccdc159 | 2.621 | 5.51E-04 | glio |
| culture_effect | Vmn2r54 | -2.096 | 5.51E-05 | glio |
| Interaction_genotype_Mtor | 4933406C10R | -4.441 | 5.51E-05 | glio |
| Interaction_genotype_Mtor | Zfp94 | -4.513 | 5.51E-05 | glio |
| Interaction_genotype_Mtor | Gm26519 | -4.619 | 5.51E-05 | glio |
| Atr_in.vivo_noRT | Gpr137c | 2.704 | 5.51E-05 | glio |
| Bcas2_in.vivo_noRT | Gm16230 | 2.831 | 5.51E-05 | glio |
| culture_effect | Rarres2 | -2.818 | 5.51E-06 | glio |
| Aars2_in.vivo_noRT | Gm14419 | 2.811 | 5.51E-06 | glio |
| Aars2_ex.vivo_noRT | Snrpn | 3.293 | 5.51E-06 | glio |
| Aars2_ex.vivo_noRT | Pnliprp2 | 3.293 | 5.51E-06 | glio |
| Chek1_in.vivo_noRT | Ppp1r2 | -1.846 | 5.51E-06 | glio |
| culture_effect | 0610012G03R | 0.641 | 5.51E-07 | glio |
| Aars2_in.vivo_noRT | S100a7a | 3.561 | 5.51E-07 | glio |
| Hsd17b10_in.vivo_noRT | Gm35696 | 3.356 | 5.51E-07 | glio |
| Aars2_in.vivo_noRT | 4930405A21R | 3.75 | 5.51E-09 | glio |
| Bcas2_in.vivo_noRT | Gm36356 | 4.547 | 5.51E-14 | glio |
| Bora_in.vivo_noRT | Gm2449 | 4.673 | 5.51E-15 | glio |
| culture_effect | 9330160F10Ri | 1.262 | 5.52E-03 | glio |
| culture_effect | Etaa1 | 0.601 | 5.52E-03 | glio |
| Interaction_genotype_Aars2 | Spice1 | -3.027 | 5.52E-03 | glio |
| Interaction_genotype_Atr | Gm5415 | -2.369 | 5.52E-03 | glio |
| Interaction_genotype_Mtor | Zfp874a | -3.2 | 5.52E-03 | glio |
| Interaction_genotype_Mtor | Gm10762 | -2.516 | 5.52E-03 | glio |
| Interaction_genotype_Myc | Krt18 | 4.257 | 5.52E-03 | glio |
| Interaction_genotype_Myc | 1700019L13Ri | -3.041 | 5.52E-03 | glio |
| Interaction_genotype_Myc | Rragd | -3.167 | 5.52E-03 | glio |
| Interaction_genotype_Myc | A230072C01R | -2.796 | 5.52E-03 | glio |
| Interaction_genotype_Myc | Faap100 | -2.261 | 5.52E-03 | glio |
| Interaction_genotype_Myc | Cables2 | -2.499 | 5.52E-03 | glio |
| Interaction_genotype_Myc | Dpy19l4 | -1.981 | 5.52E-03 | glio |
| Interaction_genotype_Pars2 | Elobl | -4.035 | 5.52E-03 | glio |
| Aars2_in.vivo_noRT | Klk10 | 1.908 | 5.52E-03 | glio |
| Aars2_in.vivo_noRT | Arntl | 1.496 | 5.52E-03 | glio |
| Aars2_in.vivo_noRT | Rsrp1 | -1.043 | 5.52E-03 | glio |
| Aars2_in.vivo_noRT | Calm1 | -0.619 | 5.52E-03 | glio |
| Atr_in.vivo_noRT | Pdlim5 | -1.017 | 5.52E-03 | glio |
| Bora_in.vivo_noRT | 1190005I06Ri | 1.891 | 5.52E-03 | glio |
| Bora_in.vivo_noRT | Tcerg1 | -0.868 | 5.52E-03 | glio |
| Bora_ex.vivo_noRT | Piezo2 | -1.625 | 5.52E-03 | glio |
| Hsd17b10_in.vivo_noRT | Cul4a | -1.3 | 5.52E-03 | glio |
| culture_effect | Mmachc | 1.077 | 5.52E-04 | glio |
| Interaction_genotype_Myc | Eri2 | -3.351 | 5.52E-04 | glio |
| interaction_RT | Lgi3 | 2.093 | 5.52E-04 | glio |
| Aars2_in.vivo_noRT | Spata2l | 2.535 | 5.52E-04 | glio |
| Atr_in.vivo_noRT | Rapgefl1 | 2.399 | 5.52E-04 | glio |
| Atr_in.vivo_noRT | Zc3h12a | 2.298 | 5.52E-04 | glio |
| Bcas2_in.vivo_noRT | Ppp2r3d | -1.796 | 5.52E-04 | glio |
| Chek1_in.vivo_noRT | Pde9a | 2.359 | 5.52E-04 | glio |
| Chek1_in.vivo_noRT | Rars | -1.089 | 5.52E-04 | glio |

| | | | | |
|---|---|---|---|---|
| Interaction_genotype_Mtor | Gm41396 | -4.766 | 5.52E-05 | glio |
| Interaction_genotype_Mtor | Usp17lc | -4.099 | 5.52E-05 | glio |
| Atr_in.vivo_noRT | Selenbp1 | 3.095 | 5.52E-05 | glio |
| Bora_ex.vivo_noRT | Sbk3 | 3.599 | 5.52E-05 | glio |
| Aars2_in.vivo_noRT | Car11 | 2.844 | 5.52E-06 | glio |
| Chek1_ex.vivo_noRT | H2-T24 | 5.216 | 5.52E-10 | glio |
| Aars2_in.vivo_noRT | Fam13a | 4.526 | 5.52E-11 | glio |
| Atr_in.vivo_noRT | Gm9833 | 4.29 | 5.52E-11 | glio |
| Aars2_in.vivo_noRT | Gm50115 | 5.063 | 5.52E-15 | glio |
| Aars2_in.vivo_noRT | Rab27a | 4.801 | 5.52E-15 | glio |
| Chek1_in.vivo_noRT | Gm36172 | 4.511 | 5.52E-16 | glio |
| Atr_in.vivo_noRT | Gm32250 | 4.751 | 5.52E-20 | glio |
| culture_effect | Itga1 | 2.523 | 5.52E-24 | glio |
| culture_effect | Ate1 | 0.529 | 5.53E-03 | glio |
| culture_effect | Arl2bp | -0.363 | 5.53E-03 | glio |
| Interaction_genotype_Aars2 | Fam13a | -3.498 | 5.53E-03 | glio |
| Interaction_genotype_Aars2 | Adh7 | -3.426 | 5.53E-03 | glio |
| Interaction_genotype_Mtor | Camkv | -3.337 | 5.53E-03 | glio |
| Interaction_genotype_Mtor | Sh3d19 | -3.161 | 5.53E-03 | glio |
| Interaction_genotype_Mtor | Aldh5a1 | -3.097 | 5.53E-03 | glio |
| Interaction_genotype_Mtor | Gpr137c | -3.159 | 5.53E-03 | glio |
| Interaction_genotype_Mtor | Fzd7 | -3.047 | 5.53E-03 | glio |
| Interaction_genotype_Myc | Itgae | -3.271 | 5.53E-03 | glio |
| Interaction_genotype_Rpl3 | 1700063J08Ri | -3.862 | 5.53E-03 | glio |
| Atr_in.vivo_noRT | Tsn | -0.619 | 5.53E-03 | glio |
| Atr_ex.vivo_noRT | Tbxa2r | 2.585 | 5.53E-03 | glio |
| Bora_in.vivo_noRT | Fgf21 | 2.17 | 5.53E-03 | glio |
| Bora_ex.vivo_noRT | Gm17638 | 2.626 | 5.53E-03 | glio |
| Bora_ex.vivo_noRT | 4922502N22R | 2.806 | 5.53E-03 | glio |
| Brca2_in.vivo_noRT | Idi2 | 1.763 | 5.53E-03 | glio |
| Brca2_in.vivo_noRT | Vmn2r18 | 1.763 | 5.53E-03 | glio |
| Interaction_genotype_Atr | A930001C03R | -4.103 | 5.53E-04 | glio |
| Interaction_genotype_Mtor | Dapp1 | -3.601 | 5.53E-04 | glio |
| Interaction_genotype_Rpl3 | 4933429H19R | -4.399 | 5.53E-04 | glio |
| Atr_in.vivo_noRT | Lrig1 | 2.619 | 5.53E-04 | glio |
| Atr_in.vivo_noRT | Catsper2 | 2.406 | 5.53E-04 | glio |
| Atr_in.vivo_noRT | Rtcb | -0.824 | 5.53E-04 | glio |
| Bard1_ex.vivo_noRT | Gpr146 | 3.563 | 5.53E-04 | glio |
| Bcas2_ex.vivo_noRT | Gm30692 | 2.348 | 5.53E-04 | glio |
| Bora_in.vivo_noRT | Fignl2 | 2.671 | 5.53E-04 | glio |
| Hsd17b10_in.vivo_noRT | Pax5 | 2.771 | 5.53E-04 | glio |
| Hsd17b10_in.vivo_noRT | A230020J21Ri | 2.689 | 5.53E-04 | glio |
| culture_effect | Tph2 | 1.848 | 5.53E-05 | glio |
| Aars2_in.vivo_noRT | Gm20234 | 3.08 | 5.53E-05 | glio |
| Atr_in.vivo_noRT | Gstt1 | 3.287 | 5.53E-06 | glio |
| Bcas2_in.vivo_noRT | Cd93 | 3.303 | 5.53E-06 | glio |
| Bora_in.vivo_noRT | AA388235 | 3.296 | 5.53E-06 | glio |
| Hsd17b10_in.vivo_noRT | Gm26682 | 3.225 | 5.53E-06 | glio |
| Hsd17b10_in.vivo_noRT | Gm37035 | 2.699 | 5.53E-06 | glio |
| Interaction_genotype_Atr | Ubc | 2.457 | 5.53E-07 | glio |
| Interaction_genotype_Atr | 4930478M13F | -6.042 | 5.53E-07 | glio |
| Bcas2_in.vivo_noRT | Gm45336 | 2.935 | 5.53E-07 | glio |

| | | | | |
|---|---|---|---|---|
| Atr_in.vivo_noRT | Srgn | 4.489 | 5.53E-08 | glio |
| Atr_in.vivo_noRT | Doc2g | 3.849 | 5.53E-08 | glio |
| Chek1_ex.vivo_noRT | 1700047M11R | 4.449 | 5.53E-08 | glio |
| Chek1_ex.vivo_noRT | B3galt2 | 4.443 | 5.53E-08 | glio |
| Aars2_in.vivo_noRT | Gm1123 | 4.419 | 5.53E-10 | glio |
| Chek1_in.vivo_noRT | Aqp3 | 5.309 | 5.53E-10 | glio |
| Chek1_in.vivo_noRT | Cyp7a1 | 4.253 | 5.53E-10 | glio |
| Aars2_in.vivo_noRT | Celf5 | 4.8 | 5.53E-11 | glio |
| Aars2_in.vivo_noRT | Rab37 | 4.409 | 5.53E-11 | glio |
| Chek1_ex.vivo_noRT | Akap14 | 4.842 | 5.53E-13 | glio |
| culture_effect | Wnk4 | -2.137 | 5.53E-16 | glio |
| Bora_in.vivo_noRT | Gtsf1 | 4.936 | 5.53E-18 | glio |
| Bcas2_in.vivo_noRT | Gm13964 | 4.807 | 5.53E-22 | glio |
| culture_effect | Cthrc1 | 2.997 | 5.53E-24 | glio |
| Interaction_genotype_Atr | Metrn | -1.078 | 5.54E-03 | glio |
| Interaction_genotype_Bora | Gm20635 | -3.5 | 5.54E-03 | glio |
| Interaction_genotype_Bora | Dnajc9 | 1.809 | 5.54E-03 | glio |
| Interaction_genotype_Chek1 | Spcs1 | 1.154 | 5.54E-03 | glio |
| Interaction_genotype_Mtor | Fermt3 | -3.183 | 5.54E-03 | glio |
| Interaction_genotype_Myc | Galm | -2.733 | 5.54E-03 | glio |
| interaction_RT | LTO1 | -0.954 | 5.54E-03 | glio |
| Aars2_in.vivo_noRT | Il6ra | 1.956 | 5.54E-03 | glio |
| Aars2_in.vivo_noRT | Zfp809 | 1.697 | 5.54E-03 | glio |
| Atp5c1_in.vivo_noRT | Abcc10 | 2.307 | 5.54E-03 | glio |
| Atr_in.vivo_noRT | B230209E15Ri | 2.208 | 5.54E-03 | glio |
| Atr_in.vivo_noRT | Zcchc7 | -1.09 | 5.54E-03 | glio |
| Atr_in.vivo_noRT | Ciapin1 | -1.088 | 5.54E-03 | glio |
| Cenpj_in.vivo_noRT | Gm49860 | 2.022 | 5.54E-03 | glio |
| Chek1_in.vivo_noRT | Gm19531 | 1.882 | 5.54E-03 | glio |
| Chek1_in.vivo_noRT | Ifi204 | -1.483 | 5.54E-03 | glio |
| Eif2s1_in.vivo_noRT | Gm37800 | 2.143 | 5.54E-03 | glio |
| Interaction_genotype_Mtor | Cox14 | 1.837 | 5.54E-04 | glio |
| Atr_in.vivo_noRT | Fam120c | 2.467 | 5.54E-04 | glio |
| Atr_ex.vivo_noRT | Nkg7 | 2.769 | 5.54E-04 | glio |
| Bcas2_in.vivo_noRT | Fkbp1a | -0.678 | 5.54E-04 | glio |
| Bora_in.vivo_noRT | Nell2 | 2.265 | 5.54E-04 | glio |
| Bora_in.vivo_noRT | Hexim2 | 1.604 | 5.54E-04 | glio |
| Bora_in.vivo_noRT | Prrg4 | 1.761 | 5.54E-04 | glio |
| Interaction_genotype_Myc | 4930447F24Ri | -4.416 | 5.54E-05 | glio |
| Interaction_genotype_Myc | Pspn | -3.124 | 5.54E-05 | glio |
| Atr_in.vivo_noRT | Cct2 | -0.793 | 5.54E-05 | glio |
| culture_effect | Capza1 | 0.651 | 5.54E-06 | glio |
| Atr_in.vivo_noRT | Bend6 | 3.112 | 5.54E-06 | glio |
| culture_effect | Hist1h2ap | 2.463 | 5.54E-07 | glio |
| Atr_in.vivo_noRT | Gm12689 | 3.843 | 5.54E-07 | glio |
| Bcas2_ex.vivo_noRT | Sytl4 | 3.54 | 5.54E-07 | glio |
| Bcas2_in.vivo_noRT | Dact1 | 3.64 | 5.54E-08 | glio |
| Bcas2_in.vivo_noRT | C4a | 3.369 | 5.54E-08 | glio |
| Aars2_in.vivo_noRT | Tnfrsf4 | 4.251 | 5.54E-10 | glio |
| Atr_in.vivo_noRT | Abat | 4.895 | 5.54E-11 | glio |
| Hsd17b10_in.vivo_noRT | A330049N07R | 3.35 | 5.54E-12 | glio |
| Atr_in.vivo_noRT | Gm35696 | 5.273 | 5.54E-16 | glio |

| | | | | |
|---|---|---|---|---|
| Aars2_in.vivo_noRT | 4930447F24Ri | 5.581 | 5.54E-17 | glio |
| culture_effect | Grasp | -3.118 | 5.54E-30 | glio |
| culture_effect | Cd34 | -1.818 | 5.55E-03 | glio |
| culture_effect | Arglu1 | 0.387 | 5.55E-03 | glio |
| Interaction_genotype_Atr | Ppp1r26 | -3.693 | 5.55E-03 | glio |
| Interaction_genotype_Bcas2 | Mettl7a3 | -3.537 | 5.55E-03 | glio |
| Interaction_genotype_Bcas2 | Thap8 | -3.755 | 5.55E-03 | glio |
| Interaction_genotype_Bcas2 | Adap1 | -4.255 | 5.55E-03 | glio |
| Aars2_in.vivo_noRT | Ccl2 | 4.936 | 5.55E-03 | glio |
| Aars2_in.vivo_noRT | Mmp11 | 2.064 | 5.55E-03 | glio |
| Aars2_in.vivo_noRT | Ttc25 | 1.962 | 5.55E-03 | glio |
| Aars2_in.vivo_noRT | Ero1lb | 1.829 | 5.55E-03 | glio |
| Aars2_in.vivo_noRT | Wdr24 | 1.417 | 5.55E-03 | glio |
| Aars2_in.vivo_noRT | Cyb5r3 | -0.972 | 5.55E-03 | glio |
| Atp5c1_in.vivo_noRT | Gm15056 | 2.52 | 5.55E-03 | glio |
| Atr_in.vivo_noRT | Mpv17l2 | -1.11 | 5.55E-03 | glio |
| Atr_in.vivo_noRT | Znhit6 | -1.114 | 5.55E-03 | glio |
| Bora_in.vivo_noRT | Arrb1 | 1.51 | 5.55E-03 | glio |
| Brca2_in.vivo_noRT | Gm16178 | 1.763 | 5.55E-03 | glio |
| Eif2s1_in.vivo_noRT | Gm37214 | 2.411 | 5.55E-03 | glio |
| culture_effect | Oas3 | -0.79 | 5.55E-04 | glio |
| Atr_in.vivo_noRT | Zfp982 | 2.354 | 5.55E-04 | glio |
| Atr_in.vivo_noRT | Dlx1 | 2.174 | 5.55E-04 | glio |
| Atr_in.vivo_noRT | Gm4890 | 1.827 | 5.55E-04 | glio |
| Atr_ex.vivo_noRT | Rangrf | -1.482 | 5.55E-04 | glio |
| Bcas2_in.vivo_noRT | E130307A14Ri | 2.306 | 5.55E-04 | glio |
| Bcas2_in.vivo_noRT | Zfp764 | 2.332 | 5.55E-04 | glio |
| Bora_in.vivo_noRT | Gm34455 | 3.171 | 5.55E-04 | glio |
| Bora_in.vivo_noRT | Sart3 | -1.632 | 5.55E-04 | glio |
| Chek1_in.vivo_noRT | Zfp982 | 2.388 | 5.55E-04 | glio |
| interaction_RT | Nudt9 | 1.064 | 5.55E-05 | glio |
| interaction_RT | Hacd1 | 1.008 | 5.55E-05 | glio |
| Aars2_in.vivo_noRT | Usp18 | -2.001 | 5.55E-05 | glio |
| Atp5c1_in.vivo_noRT | Gm15943 | 2.404 | 5.55E-05 | glio |
| Bora_in.vivo_noRT | Olfr655 | 4.02 | 5.55E-05 | glio |
| Bora_in.vivo_noRT | Pcdhga9 | 2.729 | 5.55E-05 | glio |
| Bora_in.vivo_noRT | Sema4f | 2.868 | 5.55E-05 | glio |
| Chek1_ex.vivo_noRT | Gsx2 | 3.873 | 5.55E-05 | glio |
| Interaction_genotype_Mtor | Ankrd34a | -5.534 | 5.55E-06 | glio |
| Interaction_genotype_Mtor | Nxnl2 | -4.943 | 5.55E-06 | glio |
| Atr_in.vivo_noRT | Arl2bp | -1.469 | 5.55E-06 | glio |
| culture_effect | Ptpmt1 | 0.748 | 5.55E-09 | glio |
| Aars2_in.vivo_noRT | Gm50107 | 3.812 | 5.55E-09 | glio |
| Bcas2_in.vivo_noRT | Gm47872 | 4.543 | 5.55E-12 | glio |
| Hsd17b10_in.vivo_noRT | AF067063 | 3.616 | 5.55E-12 | glio |
| Chek1_in.vivo_noRT | Acoxl | 5.044 | 5.55E-15 | glio |
| Bcas2_in.vivo_noRT | Plcl1 | 4.807 | 5.55E-17 | glio |
| Atr_in.vivo_noRT | Pdgfb | 5.27 | 5.55E-21 | glio |
| culture_effect | Nup210 | -0.938 | 5.56E-03 | glio |
| culture_effect | Etv3 | -0.836 | 5.56E-03 | glio |
| Interaction_genotype_Atr | 1700081N11R | -3.844 | 5.56E-03 | glio |
| Interaction_genotype_Atr | AA386476 | -3.42 | 5.56E-03 | glio |

| | | | | |
|---|---|---|---|---|
| Interaction_genotype_Bora | Gm14207 | -2.528 | 5.56E-03 | glio |
| Interaction_genotype_Mtor | Gm15941 | -3.075 | 5.56E-03 | glio |
| Interaction_genotype_Mtor | Calml4 | -3.225 | 5.56E-03 | glio |
| Interaction_genotype_Mtor | Adam1a | -3.334 | 5.56E-03 | glio |
| Interaction_genotype_Mtor | Gm47689 | -3.303 | 5.56E-03 | glio |
| Interaction_genotype_Mtor | 9130023H24R | -3.096 | 5.56E-03 | glio |
| Interaction_genotype_Myc | Slc37a4 | -2.185 | 5.56E-03 | glio |
| Interaction_genotype_Myc | Mrfap1 | 0.953 | 5.56E-03 | glio |
| Aars2_in.vivo_noRT | Rpf1 | -1.06 | 5.56E-03 | glio |
| Aars2_in.vivo_noRT | Prdx5 | -0.923 | 5.56E-03 | glio |
| Atr_ex.vivo_noRT | Pbld1 | 2.719 | 5.56E-03 | glio |
| Bora_in.vivo_noRT | Esco2 | -1.587 | 5.56E-03 | glio |
| Chek1_ex.vivo_noRT | Cul3 | -1.159 | 5.56E-03 | glio |
| Eif2s1_ex.vivo_noRT | Zscan4-ps2 | 2.234 | 5.56E-03 | glio |
| Hsd17b10_in.vivo_noRT | B430010I23Ri | 2.051 | 5.56E-03 | glio |
| interaction_RT | Cldn11 | 2.25 | 5.56E-04 | glio |
| Aars2_ex.vivo_noRT | Phf11a | 3.281 | 5.56E-04 | glio |
| Aars2_ex.vivo_noRT | E230025N22R | 3.124 | 5.56E-04 | glio |
| Atp5c1_in.vivo_noRT | Dpep3 | 2.136 | 5.56E-05 | glio |
| Bcas2_ex.vivo_noRT | 1700026N04R | 3.357 | 5.56E-05 | glio |
| Bcas2_ex.vivo_noRT | Dpep3 | 2.742 | 5.56E-05 | glio |
| Bcas2_ex.vivo_noRT | Erich6b | 2.742 | 5.56E-05 | glio |
| Interaction_genotype_Mtor | 4833408A19R | -4.99 | 5.56E-06 | glio |
| Interaction_genotype_Mtor | Htr1b | -5.262 | 5.56E-06 | glio |
| Atr_in.vivo_noRT | AY074887 | 3.481 | 5.56E-06 | glio |
| Bcas2_in.vivo_noRT | Traf1 | 3.203 | 5.56E-06 | glio |
| Aars2_in.vivo_noRT | Gm34643 | 3.611 | 5.56E-07 | glio |
| Atr_ex.vivo_noRT | Dpep3 | 3.164 | 5.56E-07 | glio |
| Interaction_genotype_Myc | Atoh8 | -5.755 | 5.56E-08 | glio |
| Chek1_in.vivo_noRT | Pla2g2f | 3.401 | 5.56E-08 | glio |
| culture_effect | Ankrd49 | 1.475 | 5.56E-12 | glio |
| Atr_in.vivo_noRT | Spink8 | 5.273 | 5.56E-24 | glio |
| Chek1_in.vivo_noRT | Gm31717 | 5.044 | 5.56E-25 | glio |
| Interaction_genotype_Aars2 | Gm50295 | -3.373 | 5.57E-03 | glio |
| Interaction_genotype_Aars2 | Eif3j2 | -3.451 | 5.57E-03 | glio |
| Interaction_genotype_Myc | Taf4b | -2.54 | 5.57E-03 | glio |
| Interaction_genotype_Myc | Paics | 1.17 | 5.57E-03 | glio |
| Interaction_genotype_Rpl3 | Polr3c | 2.235 | 5.57E-03 | glio |
| interaction_RT | Neb | -0.911 | 5.57E-03 | glio |
| Aars2_in.vivo_noRT | Psmg3 | -1.052 | 5.57E-03 | glio |
| Aars2_in.vivo_noRT | Micos10 | -0.576 | 5.57E-03 | glio |
| Atr_in.vivo_noRT | Celsr2 | 1.91 | 5.57E-03 | glio |
| Atr_in.vivo_noRT | Wdr25 | 1.78 | 5.57E-03 | glio |
| Atr_in.vivo_noRT | Usp6nl | -1.363 | 5.57E-03 | glio |
| Bard1_ex.vivo_noRT | Gm13398 | 2.017 | 5.57E-03 | glio |
| Bcas2_in.vivo_noRT | Ptn | 2.409 | 5.57E-03 | glio |
| Bcas2_in.vivo_noRT | Fam135b | 2.348 | 5.57E-03 | glio |
| Bcas2_in.vivo_noRT | Glod5 | -2.15 | 5.57E-03 | glio |
| Bora_in.vivo_noRT | Vps8 | 1.833 | 5.57E-03 | glio |
| Eif2s1_in.vivo_noRT | Gm41409 | 2.135 | 5.57E-03 | glio |
| culture_effect | Rlf | -0.654 | 5.57E-04 | glio |
| Interaction_genotype_Mtor | Slc22a1 | -3.32 | 5.57E-04 | glio |

| | | | | |
|---|---|---|---|---|
| Interaction_genotype_Mtor | Asf1a | 1.781 | 5.57E-04 | glio |
| Interaction_genotype_Myc | Mmp13 | -2.729 | 5.57E-04 | glio |
| Interaction_genotype_Myc | Aida | -1.695 | 5.57E-04 | glio |
| Aars2_in.vivo_noRT | Tex15 | 2.398 | 5.57E-04 | glio |
| Atr_in.vivo_noRT | Mydgf | -1.136 | 5.57E-04 | glio |
| Chek1_in.vivo_noRT | Rpl12 | -0.634 | 5.57E-04 | glio |
| Chek1_ex.vivo_noRT | Gm16133 | 3.089 | 5.57E-04 | glio |
| Interaction_genotype_Chek1 | Ptdss1 | -3.875 | 5.57E-05 | glio |
| Atp5c1_in.vivo_noRT | Gm2115 | 2.668 | 5.57E-05 | glio |
| Bcas2_ex.vivo_noRT | Tac2 | 2.742 | 5.57E-05 | glio |
| Bora_in.vivo_noRT | Tmem240 | 3.208 | 5.57E-05 | glio |
| Bora_in.vivo_noRT | Zfp799 | 2.832 | 5.57E-05 | glio |
| culture_effect | Adamts10 | 1.761 | 5.57E-06 | glio |
| Interaction_genotype_Myc | mt-Co3 | 1.737 | 5.57E-06 | glio |
| Bcas2_in.vivo_noRT | Hspb9 | 3.495 | 5.57E-06 | glio |
| Bcas2_in.vivo_noRT | Fas | 3.308 | 5.57E-06 | glio |
| Bora_in.vivo_noRT | Pde3a | 3.158 | 5.57E-06 | glio |
| Hsd17b10_in.vivo_noRT | Rgs9 | 2.816 | 5.57E-06 | glio |
| Atr_in.vivo_noRT | Gm4907 | 3.927 | 5.57E-07 | glio |
| Atr_in.vivo_noRT | Gm31517 | 3.557 | 5.57E-07 | glio |
| Atr_in.vivo_noRT | Nptx1 | 3.807 | 5.57E-07 | glio |
| Atr_ex.vivo_noRT | Erich6b | 3.164 | 5.57E-07 | glio |
| Aars2_in.vivo_noRT | Alox5ap | 4.221 | 5.57E-08 | glio |
| Chek1_ex.vivo_noRT | Gm49223 | 3.861 | 5.57E-08 | glio |
| Chek1_in.vivo_noRT | Gm20275 | 3.689 | 5.57E-11 | glio |
| culture_effect | Gm45716 | 2.822 | 5.57E-12 | glio |
| Chek1_in.vivo_noRT | Lrrc4b | 4.133 | 5.57E-12 | glio |
| Chek1_in.vivo_noRT | Gm47851 | 4.784 | 5.57E-14 | glio |
| culture_effect | Irf1 | -2.2 | 5.57E-18 | glio |
| culture_effect | Xpr1 | -0.764 | 5.58E-03 | glio |
| Interaction_genotype_Bora | Slc4a9 | -3.089 | 5.58E-03 | glio |
| Interaction_genotype_Mtor | Ndrg2 | -3.866 | 5.58E-03 | glio |
| Interaction_genotype_Mtor | Tmem60 | 1.554 | 5.58E-03 | glio |
| Interaction_genotype_Myc | Gm47428 | -3.033 | 5.58E-03 | glio |
| Interaction_genotype_Myc | Arid3b | -2.967 | 5.58E-03 | glio |
| Atp5c1_in.vivo_noRT | Gm14698 | 2.287 | 5.58E-03 | glio |
| Atr_in.vivo_noRT | Tmem43 | -1.27 | 5.58E-03 | glio |
| Atr_in.vivo_noRT | Eif3h | -0.55 | 5.58E-03 | glio |
| Brca2_in.vivo_noRT | Ccl9 | 1.763 | 5.58E-03 | glio |
| Brca2_in.vivo_noRT | Ankef1 | 2.606 | 5.58E-03 | glio |
| Interaction_genotype_Mtor | Zfp661 | -4.904 | 5.58E-05 | glio |
| Interaction_genotype_Myc | Gm27253 | -4.46 | 5.58E-05 | glio |
| Interaction_genotype_Myc | Hist1h4b | -4.992 | 5.58E-05 | glio |
| Atr_in.vivo_noRT | Prr15l | 2.842 | 5.58E-05 | glio |
| Bora_in.vivo_noRT | Slc2a6 | 3.081 | 5.58E-05 | glio |
| Bora_in.vivo_noRT | Eme2 | 3.046 | 5.58E-06 | glio |
| Aars2_in.vivo_noRT | Gm15056 | 3.892 | 5.58E-07 | glio |
| Bora_in.vivo_noRT | 1700003G18R | 4.676 | 5.58E-09 | glio |
| Chek1_in.vivo_noRT | Tmem150c | 4.4 | 5.58E-10 | glio |
| Chek1_in.vivo_noRT | Pla1a | 4.512 | 5.58E-12 | glio |
| culture_effect | Atp5md | 1.127 | 5.58E-13 | glio |
| Bora_in.vivo_noRT | C330024C12R | 4.545 | 5.58E-14 | glio |

| | | | | |
|---|---|---|---|---|
| culture_effect | Rpl18a | -1.002 | 5.58E-15 | glio |
| culture_effect | Fbxo32 | -2.449 | 5.58E-16 | glio |
| Bora_in.vivo_noRT | Minar2 | 4.676 | 5.58E-17 | glio |
| Bora_in.vivo_noRT | Gm15890 | 4.936 | 5.58E-17 | glio |
| Atr_in.vivo_noRT | Cpxm1 | 5.267 | 5.58E-18 | glio |
| culture_effect | H2-K1 | -4.295 | 5.58E-72 | glio |
| culture_effect | Trmt1l | -0.586 | 5.59E-03 | glio |
| Interaction_genotype_Atr | Pfkfb1 | -3.531 | 5.59E-03 | glio |
| Interaction_genotype_Atr | Cox6a2 | 2.955 | 5.59E-03 | glio |
| Atr_in.vivo_noRT | Zc2hc1c | 2.012 | 5.59E-03 | glio |
| Atr_in.vivo_noRT | Micos13 | -0.706 | 5.59E-03 | glio |
| Bcas2_in.vivo_noRT | Tmem17 | 1.756 | 5.59E-03 | glio |
| Bora_in.vivo_noRT | A730017L22Ri | 1.794 | 5.59E-03 | glio |
| Bora_in.vivo_noRT | Slc36a3 | 1.91 | 5.59E-03 | glio |
| Bora_in.vivo_noRT | Snap29 | 1.543 | 5.59E-03 | glio |
| Cyp19a1_ex.vivo_noRT | Gm13032 | 2.533 | 5.59E-03 | glio |
| Interaction_genotype_Mtor | Grk4 | -4.008 | 5.59E-04 | glio |
| Interaction_genotype_Mtor | Zfp408 | -3.713 | 5.59E-04 | glio |
| Aars2_in.vivo_noRT | Eif3l | -0.954 | 5.59E-04 | glio |
| Atr_in.vivo_noRT | Dnah2 | 2.473 | 5.59E-04 | glio |
| Atr_in.vivo_noRT | Gm44867 | 2.476 | 5.59E-04 | glio |
| Atr_in.vivo_noRT | Bcorl1 | 1.688 | 5.59E-04 | glio |
| Bard1_ex.vivo_noRT | BC048671 | 2.814 | 5.59E-04 | glio |
| Bard1_ex.vivo_noRT | Eepd1 | 3.214 | 5.59E-04 | glio |
| Chek1_in.vivo_noRT | Slc11a1 | 2.459 | 5.59E-04 | glio |
| culture_effect | Dcpp3 | -2.452 | 5.59E-05 | glio |
| Interaction_genotype_Mtor | Pitpna | 1.856 | 5.59E-05 | glio |
| Interaction_genotype_Myc | 5330438D12R | -3.642 | 5.59E-05 | glio |
| Interaction_genotype_Myc | Gm5914 | -3.762 | 5.59E-05 | glio |
| Interaction_genotype_Myc | Fip1l1 | 1.915 | 5.59E-05 | glio |
| Interaction_genotype_Myc | Ube2d3 | 1.288 | 5.59E-05 | glio |
| Aars2_in.vivo_noRT | Plac1 | 2.745 | 5.59E-05 | glio |
| Aars2_in.vivo_noRT | Gm43066 | 2.675 | 5.59E-05 | glio |
| Bora_ex.vivo_noRT | Kcnk7 | 3.201 | 5.59E-05 | glio |
| Chek1_in.vivo_noRT | Pdxdc1 | -1.951 | 5.59E-05 | glio |
| Chek1_ex.vivo_noRT | Aldh1a1 | 3.738 | 5.59E-05 | glio |
| Chek1_ex.vivo_noRT | Klre1 | 3.469 | 5.59E-05 | glio |
| Atp5c1_in.vivo_noRT | Misp | -3.063 | 5.59E-06 | glio |
| Atr_in.vivo_noRT | Bloc1s6os | 3.014 | 5.59E-06 | glio |
| Hsd17b10_in.vivo_noRT | Ephx1 | 3.239 | 5.59E-06 | glio |
| Hsd17b10_in.vivo_noRT | Gm44752 | 3.229 | 5.59E-07 | glio |
| Hsd17b10_in.vivo_noRT | Acat3 | 2.824 | 5.59E-07 | glio |
| Hsd17b10_in.vivo_noRT | Gm5602 | 2.825 | 5.59E-07 | glio |
| Interaction_genotype_Myc | Paip2 | 2.175 | 5.59E-08 | glio |
| Interaction_genotype_Myc | B930094E09Ri | -6.227 | 5.59E-08 | glio |
| Chek1_ex.vivo_noRT | Gm31517 | 4.791 | 5.59E-08 | glio |
| Atr_in.vivo_noRT | Slc46a3 | 4.348 | 5.59E-10 | glio |
| Atr_in.vivo_noRT | Bbof1 | 4.105 | 5.59E-10 | glio |
| Atr_in.vivo_noRT | Gm43581 | 3.963 | 5.59E-10 | glio |
| Atr_in.vivo_noRT | Ppp4r2 | -2.315 | 5.59E-10 | glio |
| Bcas2_in.vivo_noRT | Idh2 | -2.369 | 5.59E-10 | glio |
| culture_effect | 2210011C24R | 2.817 | 5.59E-11 | glio |

| | | | | |
|---|---|---|---|---|
| Chek1_in.vivo_noRT | Prickle4 | 4.252 | 5.59E-11 | glio |
| Chek1_in.vivo_noRT | BC048671 | 3.743 | 5.59E-11 | glio |
| Chek1_ex.vivo_noRT | AF067061 | 4.842 | 5.59E-11 | glio |
| Atr_in.vivo_noRT | A330084C13R | 5.153 | 5.59E-12 | glio |
| Bora_in.vivo_noRT | Gm7247 | 4.144 | 5.59E-13 | glio |
| Aars2_in.vivo_noRT | Gm37800 | 5.327 | 5.59E-17 | glio |
| Interaction_genotype_Aars2 | Ifitm6 | -3.48 | 5.60E-03 | glio |
| Interaction_genotype_Mtor | Thnsl1 | -2.636 | 5.60E-03 | glio |
| interaction_RT | Nup85 | -0.839 | 5.60E-03 | glio |
| Atr_in.vivo_noRT | Cited1 | -2.131 | 5.60E-03 | glio |
| Bcas2_in.vivo_noRT | 4930550C14R | 1.826 | 5.60E-03 | glio |
| Bcas2_in.vivo_noRT | Sigmar1 | -1.123 | 5.60E-03 | glio |
| Bcas2_in.vivo_noRT | Phb2 | -0.581 | 5.60E-03 | glio |
| Bora_in.vivo_noRT | Msh5 | 2.169 | 5.60E-03 | glio |
| Bora_in.vivo_noRT | Zfp28 | 2.065 | 5.60E-03 | glio |
| Bora_in.vivo_noRT | Gm9828 | 2.017 | 5.60E-03 | glio |
| Bora_in.vivo_noRT | Map3k20 | 1.746 | 5.60E-03 | glio |
| Bora_in.vivo_noRT | Bora | -1.51 | 5.60E-03 | glio |
| Bora_in.vivo_noRT | Ptges2 | -1.362 | 5.60E-03 | glio |
| Bora_in.vivo_noRT | Lsm8 | -0.766 | 5.60E-03 | glio |
| Chek1_in.vivo_noRT | Hspa12b | 2.236 | 5.60E-03 | glio |
| Chek1_in.vivo_noRT | Lmo7 | 1.799 | 5.60E-03 | glio |
| Chek1_in.vivo_noRT | Zfp449 | 1.761 | 5.60E-03 | glio |
| Chek1_in.vivo_noRT | Ylpm1 | -1.129 | 5.60E-03 | glio |
| Chek1_in.vivo_noRT | Ly6e | -0.855 | 5.60E-03 | glio |
| Cyp19a1_ex.vivo_noRT | Pclo | -1.934 | 5.60E-03 | glio |
| Hsd17b10_in.vivo_noRT | Calml4 | 2.055 | 5.60E-03 | glio |
| Hsd17b10_in.vivo_noRT | Bbs1 | 2.157 | 5.60E-03 | glio |
| Hsd17b10_in.vivo_noRT | Tnni1 | -1.647 | 5.60E-03 | glio |
| culture_effect | Mterf1a | 1.258 | 5.60E-04 | glio |
| Interaction_genotype_Bora | Dmac2 | -3.766 | 5.60E-04 | glio |
| Interaction_genotype_Myc | Gm8016 | -2.729 | 5.60E-04 | glio |
| Interaction_genotype_Myc | Cebpa | -3.913 | 5.60E-04 | glio |
| Interaction_genotype_Myc | Jazf1 | -3.695 | 5.60E-04 | glio |
| Interaction_genotype_Myc | Nup37 | 1.898 | 5.60E-04 | glio |
| Aars2_in.vivo_noRT | Klhl1 | 2.332 | 5.60E-04 | glio |
| Bcas2_in.vivo_noRT | Nat8l | 2.216 | 5.60E-04 | glio |
| Bora_in.vivo_noRT | Fen1 | -1.526 | 5.60E-04 | glio |
| Chek1_in.vivo_noRT | Fam120c | 2.501 | 5.60E-04 | glio |
| Aars2_in.vivo_noRT | Cdiptos | 2.778 | 5.60E-05 | glio |
| Bora_ex.vivo_noRT | Usp17lc | 3.201 | 5.60E-05 | glio |
| Bora_ex.vivo_noRT | Apol10b | 3.201 | 5.60E-05 | glio |
| Chek1_in.vivo_noRT | 1700126G02R | 2.911 | 5.60E-05 | glio |
| Interaction_genotype_Myc | A330074K22R | -4.753 | 5.60E-06 | glio |
| Aars2_in.vivo_noRT | Tspan33 | 3.232 | 5.60E-06 | glio |
| Chek1_ex.vivo_noRT | Tex19.2 | 3.957 | 5.60E-06 | glio |
| Hsd17b10_in.vivo_noRT | 5330439K02R | 3.35 | 5.60E-06 | glio |
| Hsd17b10_in.vivo_noRT | Gm37864 | 2.837 | 5.60E-06 | glio |
| Interaction_genotype_Myc | Cox6a1 | 2.216 | 5.60E-07 | glio |
| Bcas2_in.vivo_noRT | AV356131 | 3.523 | 5.60E-07 | glio |
| Hsd17b10_in.vivo_noRT | Hcn3 | 3.495 | 5.60E-08 | glio |
| culture_effect | Jakmip2 | -3.385 | 5.60E-09 | glio |

| | | | | |
|---|---|---|---|---|
| Chek1_in.vivo_noRT | Gm11549 | 4.518 | 5.60E-11 | glio |
| Bora_in.vivo_noRT | Kcnd1 | 4.414 | 5.60E-13 | glio |
| culture_effect | Tagap | -1.194 | 5.61E-03 | glio |
| Interaction_genotype_Bcas2 | Gngt2 | -3.734 | 5.61E-03 | glio |
| Interaction_genotype_Mtor | Mfsd4b4 | -2.973 | 5.61E-03 | glio |
| Interaction_genotype_Myc | Nprl2 | -2.299 | 5.61E-03 | glio |
| Aars2_in.vivo_noRT | Ehd2 | 1.416 | 5.61E-03 | glio |
| Atp5c1_in.vivo_noRT | Tns4 | 2.018 | 5.61E-03 | glio |
| Bora_in.vivo_noRT | Ddr2 | -1.233 | 5.61E-03 | glio |
| Chek1_ex.vivo_noRT | Wnt4 | 2.849 | 5.61E-03 | glio |
| Chek1_ex.vivo_noRT | Rpl29 | -0.578 | 5.61E-03 | glio |
| Interaction_genotype_Atr | Gm14419 | -3.607 | 5.61E-04 | glio |
| Interaction_genotype_Atr | Gm15816 | -3.849 | 5.61E-04 | glio |
| Interaction_genotype_Mtor | Gm50103 | -3.845 | 5.61E-04 | glio |
| Interaction_genotype_Myc | Tspan33 | -3.979 | 5.61E-04 | glio |
| Interaction_genotype_Myc | Katnb1 | -2.697 | 5.61E-04 | glio |
| Bcas2_in.vivo_noRT | 1700012D14R | 2.433 | 5.61E-04 | glio |
| Bcas2_in.vivo_noRT | Gm29170 | 2.423 | 5.61E-04 | glio |
| Bcas2_in.vivo_noRT | Pax3 | 2.106 | 5.61E-04 | glio |
| Cenpj_ex.vivo_noRT | Gm11583 | 2.591 | 5.61E-04 | glio |
| Chek1_in.vivo_noRT | Mta1 | -1.474 | 5.61E-04 | glio |
| Interaction_genotype_Mtor | Castor2 | -4.39 | 5.61E-05 | glio |
| interaction_RT | Arid4b | 0.86 | 5.61E-05 | glio |
| interaction_RT | Fbl | -0.722 | 5.61E-05 | glio |
| Aars2_in.vivo_noRT | Usp27x | 3.02 | 5.61E-05 | glio |
| Chek1_ex.vivo_noRT | Cnfn | 3.862 | 5.61E-05 | glio |
| Interaction_genotype_Mtor | Lrrn3 | -5.132 | 5.61E-06 | glio |
| Atr_in.vivo_noRT | 2810414N06R | 3.511 | 5.61E-07 | glio |
| Atr_ex.vivo_noRT | Tac2 | 3.164 | 5.61E-07 | glio |
| Chek1_in.vivo_noRT | Gm8104 | 3.624 | 5.61E-08 | glio |
| Hsd17b10_in.vivo_noRT | Olfr90 | 3.352 | 5.61E-08 | glio |
| Bora_in.vivo_noRT | Hbq1b | 4.67 | 5.61E-11 | glio |
| Atr_in.vivo_noRT | Mettl7a3 | 4.883 | 5.61E-14 | glio |
| Atr_in.vivo_noRT | Gm26978 | 5.273 | 5.61E-20 | glio |
| Bora_in.vivo_noRT | Gm50321 | 5.46 | 5.61E-20 | glio |
| culture_effect | Adck5 | -0.628 | 5.62E-03 | glio |
| Interaction_genotype_Atr | Zfp960 | -3.713 | 5.62E-03 | glio |
| Interaction_genotype_Bcas2 | Tmem191c | -3.59 | 5.62E-03 | glio |
| Interaction_genotype_Mtor | Rnf126 | 1.483 | 5.62E-03 | glio |
| Interaction_genotype_Mtor | Calu | 1.253 | 5.62E-03 | glio |
| Interaction_genotype_Myc | 9330179D12R | -3.327 | 5.62E-03 | glio |
| Interaction_genotype_Myc | Gjc3 | 1.219 | 5.62E-03 | glio |
| interaction_RT | Ankrd9 | 1.263 | 5.62E-03 | glio |
| Atr_in.vivo_noRT | Mtmr9 | 1.619 | 5.62E-03 | glio |
| Atr_in.vivo_noRT | Asxl1 | -1.369 | 5.62E-03 | glio |
| Atr_in.vivo_noRT | Zfhx3 | -1.119 | 5.62E-03 | glio |
| Bcas2_in.vivo_noRT | Gpam | 1.904 | 5.62E-03 | glio |
| Bora_in.vivo_noRT | Bend6 | 2.006 | 5.62E-03 | glio |
| Bora_in.vivo_noRT | Oscp1 | 2.041 | 5.62E-03 | glio |
| Bora_in.vivo_noRT | 4933432I03Ri | 1.965 | 5.62E-03 | glio |
| Bora_in.vivo_noRT | Tmed4 | -0.937 | 5.62E-03 | glio |
| Bora_ex.vivo_noRT | Gm11131 | 2.806 | 5.62E-03 | glio |

| | | | | |
|---|---|---|---|---|
| Chek1_in.vivo_noRT | Celsr2 | 1.944 | 5.62E-03 | glio |
| Chek1_ex.vivo_noRT | Nup37 | -1.089 | 5.62E-03 | glio |
| Chek1_ex.vivo_noRT | Gtf2a2 | -0.799 | 5.62E-03 | glio |
| Interaction_genotype_Atr | G630030J09Ri | -4.105 | 5.62E-04 | glio |
| Interaction_genotype_Atr | Pip5k1a | 1.9 | 5.62E-04 | glio |
| Interaction_genotype_Mtor | Gm17638 | -3.883 | 5.62E-04 | glio |
| Interaction_genotype_Myc | Pakap.1 | -3.161 | 5.62E-04 | glio |
| Bard1_ex.vivo_noRT | Otof | 2.814 | 5.62E-04 | glio |
| Bcas2_in.vivo_noRT | Gm42937 | 2.497 | 5.62E-04 | glio |
| culture_effect | Arpc2 | 0.433 | 5.62E-05 | glio |
| Interaction_genotype_Mtor | 5330417C22R | -4.798 | 5.62E-05 | glio |
| Interaction_genotype_Mtor | Odc1 | 3.491 | 5.62E-05 | glio |
| Interaction_genotype_Myc | Atp6v1g2 | -4.206 | 5.62E-05 | glio |
| Interaction_genotype_Myc | Selenoh | 2.075 | 5.62E-05 | glio |
| Hsd17b10_in.vivo_noRT | Spata31d1a | 2.975 | 5.62E-05 | glio |
| interaction_RT | Ube2r2 | 0.792 | 5.62E-06 | glio |
| Atr_in.vivo_noRT | Gstt2 | 3.223 | 5.62E-06 | glio |
| Bcas2_in.vivo_noRT | Gm35551 | 2.991 | 5.62E-06 | glio |
| Interaction_genotype_Mtor | 4930447F24Ri | -5.785 | 5.62E-07 | glio |
| culture_effect | Tm9sf2 | -0.878 | 5.62E-08 | glio |
| Aars2_in.vivo_noRT | A730071L15Ri | 4.266 | 5.62E-09 | glio |
| Atr_in.vivo_noRT | Gm2788 | 4.485 | 5.62E-10 | glio |
| Bora_in.vivo_noRT | 4930555A03R | 4.198 | 5.62E-11 | glio |
| Bcas2_in.vivo_noRT | Gm26555 | 4.544 | 5.62E-19 | glio |
| Bcas2_in.vivo_noRT | Gm2016 | 4.807 | 5.62E-20 | glio |
| Bcas2_in.vivo_noRT | Gm12703 | 4.541 | 5.62E-20 | glio |
| culture_effect | Gm15398 | -1.308 | 5.63E-03 | glio |
| culture_effect | Cops7b | 0.609 | 5.63E-03 | glio |
| culture_effect | Zfp148 | 0.491 | 5.63E-03 | glio |
| culture_effect | Trim28 | -0.392 | 5.63E-03 | glio |
| Interaction_genotype_Aars2 | Klk14 | -3.358 | 5.63E-03 | glio |
| Interaction_genotype_Mtor | Ppp2r3a | -2.903 | 5.63E-03 | glio |
| Interaction_genotype_Myc | Ggnbp2 | 1.168 | 5.63E-03 | glio |
| Atr_in.vivo_noRT | Adamts5 | 1.882 | 5.63E-03 | glio |
| Atr_in.vivo_noRT | Cdk20 | 1.684 | 5.63E-03 | glio |
| Eif2s1_in.vivo_noRT | Gm42664 | 2.02 | 5.63E-03 | glio |
| Hsd17b10_in.vivo_noRT | Tmem86a | 2.249 | 5.63E-03 | glio |
| culture_effect | Vmn2r56 | -1.717 | 5.63E-04 | glio |
| Interaction_genotype_Aars2 | Hist1h3d | -3.836 | 5.63E-04 | glio |
| Interaction_genotype_Mtor | Gm32913 | -6.027 | 5.63E-04 | glio |
| Interaction_genotype_Mtor | 1500002F19Ri | -3.975 | 5.63E-04 | glio |
| Interaction_genotype_Mtor | Gdpgp1 | -3.556 | 5.63E-04 | glio |
| Interaction_genotype_Mtor | Tsen34 | 1.808 | 5.63E-04 | glio |
| Interaction_genotype_Myc | Psme2 | 1.628 | 5.63E-04 | glio |
| Atp5c1_in.vivo_noRT | Gngt2 | 2.668 | 5.63E-04 | glio |
| Atr_ex.vivo_noRT | Ggt1 | 3.164 | 5.63E-04 | glio |
| Bora_in.vivo_noRT | Nxf2 | 2.593 | 5.63E-04 | glio |
| culture_effect | Trub2 | 0.951 | 5.63E-05 | glio |
| Interaction_genotype_Mtor | Gm11696 | -4.789 | 5.63E-05 | glio |
| Atr_ex.vivo_noRT | Fgf8 | 3.199 | 5.63E-05 | glio |
| Bcas2_in.vivo_noRT | Phldb2 | 3.073 | 5.63E-05 | glio |
| culture_effect | Akap8l | -0.916 | 5.63E-06 | glio |

| | | | | |
|---|---|---|---|---|
| Interaction_genotype_Mtor | BC049762 | -5.481 | 5.63E-06 | glio |
| Chek1_in.vivo_noRT | Glrp1 | 2.774 | 5.63E-06 | glio |
| Aars2_in.vivo_noRT | Hist1h2ad | 3.555 | 5.63E-07 | glio |
| Aars2_in.vivo_noRT | Gm34121 | 3.201 | 5.63E-07 | glio |
| Bora_in.vivo_noRT | 1700122E12Ri | 3.54 | 5.63E-07 | glio |
| Interaction_genotype_Mtor | E130215H24R | -6.155 | 5.63E-08 | glio |
| Atr_in.vivo_noRT | Gm30124 | 4.263 | 5.63E-11 | glio |
| Aars2_in.vivo_noRT | Pde6b | 4.295 | 5.63E-13 | glio |
| Atr_in.vivo_noRT | Tspyl3 | 4.45 | 5.63E-13 | glio |
| Interaction_genotype_Aars2 | Gm11973 | -2.754 | 5.64E-03 | glio |
| Interaction_genotype_Aars2 | Eno3 | 1.883 | 5.64E-03 | glio |
| Interaction_genotype_Aars2 | Rtcb | 1.071 | 5.64E-03 | glio |
| Interaction_genotype_Atr | Gen1 | -2.744 | 5.64E-03 | glio |
| Interaction_genotype_Atr | Mapk1ip1l | 1.48 | 5.64E-03 | glio |
| Interaction_genotype_Atr | Psmd8 | 1.15 | 5.64E-03 | glio |
| Interaction_genotype_Myc | Ifi214 | -3.136 | 5.64E-03 | glio |
| Interaction_genotype_Myc | Zbtb14 | -2.581 | 5.64E-03 | glio |
| Interaction_genotype_Myc | Nfkbil1 | -1.822 | 5.64E-03 | glio |
| Interaction_genotype_Myc | Polr2j | 1.088 | 5.64E-03 | glio |
| Interaction_genotype_Myc | Cdkn2a | 0.991 | 5.64E-03 | glio |
| Aars2_in.vivo_noRT | Gm36988 | 2.008 | 5.64E-03 | glio |
| Aars2_in.vivo_noRT | 6720427I07Ri | 1.729 | 5.64E-03 | glio |
| Atr_in.vivo_noRT | Topbp1 | -1.206 | 5.64E-03 | glio |
| Bora_in.vivo_noRT | Cenpb | -1.294 | 5.64E-03 | glio |
| Chek1_in.vivo_noRT | Syde2 | 1.968 | 5.64E-03 | glio |
| Chek1_in.vivo_noRT | Fyb | 2.013 | 5.64E-03 | glio |
| Chek1_in.vivo_noRT | Pam16 | -0.881 | 5.64E-03 | glio |
| Hsd17b10_in.vivo_noRT | Gm5861 | 2.568 | 5.64E-03 | glio |
| Hsd17b10_in.vivo_noRT | 4921531C22R | 2.088 | 5.64E-03 | glio |
| Hsd17b10_in.vivo_noRT | Uba1 | -0.883 | 5.64E-03 | glio |
| Hsd17b10_in.vivo_noRT | Snu13 | -0.451 | 5.64E-03 | glio |
| Interaction_genotype_Atr | Zfp114 | -4.331 | 5.64E-04 | glio |
| Interaction_genotype_Atr | Slc2a9 | -3.567 | 5.64E-04 | glio |
| Interaction_genotype_Mtor | Upk1a | -4.286 | 5.64E-04 | glio |
| Interaction_genotype_Mtor | Fam83e | -4.246 | 5.64E-04 | glio |
| Atr_in.vivo_noRT | Homer1 | 1.71 | 5.64E-04 | glio |
| Atr_ex.vivo_noRT | Lmbr1l | 2.079 | 5.64E-04 | glio |
| Bcas2_in.vivo_noRT | Pheta1 | 2.539 | 5.64E-04 | glio |
| Bcas2_ex.vivo_noRT | Gm26671 | 2.349 | 5.64E-04 | glio |
| Bcas2_ex.vivo_noRT | Gm20513 | 2.742 | 5.64E-04 | glio |
| Bcas2_ex.vivo_noRT | Ly6d | 2.742 | 5.64E-04 | glio |
| Bora_in.vivo_noRT | Mutyh | 2.489 | 5.64E-04 | glio |
| Bora_in.vivo_noRT | Rab9 | -1.005 | 5.64E-04 | glio |
| Cenpj_in.vivo_noRT | Gpc5 | 2.022 | 5.64E-04 | glio |
| Atr_in.vivo_noRT | Zan | 2.77 | 5.64E-05 | glio |
| Interaction_genotype_Mtor | Hcrtr1 | -5.447 | 5.64E-06 | glio |
| Interaction_genotype_Myc | Gm10734 | -5.232 | 5.64E-06 | glio |
| Atr_ex.vivo_noRT | Sbk3 | 3.941 | 5.64E-06 | glio |
| Aars2_in.vivo_noRT | 9130204L05Ri | 3.733 | 5.64E-07 | glio |
| Aars2_in.vivo_noRT | Gm43062 | 3.549 | 5.64E-07 | glio |
| Bcas2_in.vivo_noRT | Tmem44 | 3.549 | 5.64E-07 | glio |
| Bcas2_in.vivo_noRT | Efna2 | 4.053 | 5.64E-07 | glio |

| | | | | |
|---|---|---|---|---|
| Bora_ex.vivo_noRT | Gm13078 | 3.201 | 5.64E-07 | glio |
| Interaction_genotype_Mtor | A030014E15Ri | -4.492 | 5.64E-08 | glio |
| Aars2_in.vivo_noRT | Gm16754 | 3.823 | 5.64E-08 | glio |
| Bora_in.vivo_noRT | Dnajc28 | 4.323 | 5.64E-09 | glio |
| Hsd17b10_in.vivo_noRT | Luzp2 | 3.09 | 5.64E-09 | glio |
| Atr_in.vivo_noRT | Selenok-ps1 | 3.969 | 5.64E-10 | glio |
| Bcas2_in.vivo_noRT | Iqcf1 | 4.154 | 5.64E-10 | glio |
| Chek1_in.vivo_noRT | Ccdc114 | 4.126 | 5.64E-10 | glio |
| Aars2_in.vivo_noRT | Tmem190 | 5.327 | 5.64E-12 | glio |
| Aars2_in.vivo_noRT | Amy1 | 4.798 | 5.64E-13 | glio |
| Aars2_in.vivo_noRT | Pacrg | 4.535 | 5.64E-13 | glio |
| Atr_in.vivo_noRT | Gm30938 | 4.887 | 5.64E-13 | glio |
| Chek1_in.vivo_noRT | Upb1 | 4.78 | 5.64E-13 | glio |
| Atr_in.vivo_noRT | 7530414M10F | 5.007 | 5.64E-14 | glio |
| culture_effect | Rbbp7 | -0.875 | 5.64E-15 | glio |
| culture_effect | Zfp395 | -0.913 | 5.65E-03 | glio |
| culture_effect | Prkag1 | -0.493 | 5.65E-03 | glio |
| culture_effect | Rpl24 | -0.335 | 5.65E-03 | glio |
| Interaction_genotype_Aars2 | Cib1 | 1.61 | 5.65E-03 | glio |
| Interaction_genotype_Myc | Gm20300 | -3.101 | 5.65E-03 | glio |
| Interaction_genotype_Myc | Tinf2 | -2.272 | 5.65E-03 | glio |
| Interaction_genotype_Pars2 | Psph | 1.113 | 5.65E-03 | glio |
| Aars2_in.vivo_noRT | Zfp267 | 1.431 | 5.65E-03 | glio |
| Aars2_in.vivo_noRT | Rbm48 | 1.315 | 5.65E-03 | glio |
| Atr_in.vivo_noRT | Pla2g2e | 2.188 | 5.65E-03 | glio |
| Bard1_ex.vivo_noRT | Mfsd2a | -2.055 | 5.65E-03 | glio |
| Bcas2_in.vivo_noRT | Azin2 | 1.866 | 5.65E-03 | glio |
| Bcas2_in.vivo_noRT | St6galnac6 | 1.642 | 5.65E-03 | glio |
| Bcas2_in.vivo_noRT | Ift74 | -1.255 | 5.65E-03 | glio |
| Bcas2_in.vivo_noRT | Nudc | -1.092 | 5.65E-03 | glio |
| Chek1_ex.vivo_noRT | Dmrt2 | 2.719 | 5.65E-03 | glio |
| Eif2s1_in.vivo_noRT | AC163347.1 | 1.88 | 5.65E-03 | glio |
| Eif2s1_in.vivo_noRT | Mcc | -1.997 | 5.65E-03 | glio |
| Eif2s1_in.vivo_noRT | Snhg12 | -0.734 | 5.65E-03 | glio |
| Hsd17b10_in.vivo_noRT | Synpr | 2.453 | 5.65E-03 | glio |
| Hsd17b10_in.vivo_noRT | Eps15l1 | -1.403 | 5.65E-03 | glio |
| Hsd17b10_in.vivo_noRT | Sec23ip | -1.306 | 5.65E-03 | glio |
| culture_effect | Agl | 1.184 | 5.65E-04 | glio |
| Interaction_genotype_Aars2 | Tomm6os | -4.525 | 5.65E-04 | glio |
| Interaction_genotype_Aars2 | Rapgef3os2 | -3.219 | 5.65E-04 | glio |
| Interaction_genotype_Bora | Adgrb3 | -4.37 | 5.65E-04 | glio |
| Interaction_genotype_Bora | A630052C17R | -4.405 | 5.65E-04 | glio |
| Interaction_genotype_Bora | A930012O16R | -4.684 | 5.65E-04 | glio |
| Interaction_genotype_Bora | A930001C03R | -4.378 | 5.65E-04 | glio |
| Interaction_genotype_Bora | Hdac1 | 1.432 | 5.65E-04 | glio |
| Interaction_genotype_Myc | Pa2g4 | 1.72 | 5.65E-04 | glio |
| Aars2_in.vivo_noRT | Syn3 | 2.473 | 5.65E-04 | glio |
| Aars2_in.vivo_noRT | Nqo1 | 2.562 | 5.65E-04 | glio |
| Bard1_ex.vivo_noRT | Tmem54 | 2.814 | 5.65E-04 | glio |
| Bard1_ex.vivo_noRT | Boc | 2.421 | 5.65E-04 | glio |
| Bcas2_ex.vivo_noRT | Ccdc162 | 2.742 | 5.65E-04 | glio |
| Chek1_ex.vivo_noRT | Usp1 | -1.417 | 5.65E-04 | glio |

| | | | | |
|---|---|---|---|---|
| Eif2s1_in.vivo_noRT | Dmrtc1a | 2.143 | 5.65E-04 | glio |
| Eif2s1_in.vivo_noRT | Gm12589 | 2.143 | 5.65E-04 | glio |
| Chek1_in.vivo_noRT | Cdc42ep4 | 2.691 | 5.65E-05 | glio |
| Aars2_in.vivo_noRT | Mmp19 | 3.133 | 5.65E-06 | glio |
| Atr_in.vivo_noRT | Arsj | 3.419 | 5.65E-06 | glio |
| Atr_in.vivo_noRT | Ttc38 | 3.133 | 5.65E-06 | glio |
| Hsd17b10_in.vivo_noRT | Gm14379 | 3.1 | 5.65E-06 | glio |
| interaction_RT | Rheb | 0.884 | 5.65E-07 | glio |
| Chek1_ex.vivo_noRT | Calml4 | 4.163 | 5.65E-07 | glio |
| Atr_in.vivo_noRT | 4933439K11R | 4.349 | 5.65E-10 | glio |
| Aars2_in.vivo_noRT | Gm14264 | 4.333 | 5.65E-11 | glio |
| Atr_in.vivo_noRT | Gm47428 | 4.358 | 5.65E-11 | glio |
| culture_effect | Hectd2 | 2.533 | 5.65E-12 | glio |
| Chek1_in.vivo_noRT | Gm37035 | 4.127 | 5.65E-13 | glio |
| Aars2_in.vivo_noRT | Gm4724 | 4.671 | 5.65E-14 | glio |
| culture_effect | Ifnar2 | -1.547 | 5.65E-16 | glio |
| Atr_in.vivo_noRT | Dok2 | 5.008 | 5.65E-17 | glio |
| Atr_in.vivo_noRT | Gm48170 | 5.533 | 5.65E-23 | glio |
| culture_effect | Gldc | 1.346 | 5.66E-03 | glio |
| Interaction_genotype_Aars2 | Ppm1f | -2.817 | 5.66E-03 | glio |
| Interaction_genotype_Aars2 | Mpc1 | 1.3 | 5.66E-03 | glio |
| Interaction_genotype_Bcas2 | Synm | -2.911 | 5.66E-03 | glio |
| Interaction_genotype_Mtor | Eif1 | 0.961 | 5.66E-03 | glio |
| Aars2_in.vivo_noRT | Zfp827 | 1.922 | 5.66E-03 | glio |
| Aars2_in.vivo_noRT | Arl2 | -0.995 | 5.66E-03 | glio |
| Bcas2_in.vivo_noRT | Zfp867 | 1.712 | 5.66E-03 | glio |
| Bora_in.vivo_noRT | Spire1 | -1.405 | 5.66E-03 | glio |
| Bora_ex.vivo_noRT | Gm11963 | 2.698 | 5.66E-03 | glio |
| Cenpj_ex.vivo_noRT | Spns3 | 2.989 | 5.66E-03 | glio |
| Chek1_in.vivo_noRT | Ssrp1 | -0.682 | 5.66E-03 | glio |
| Eif2s1_in.vivo_noRT | Zfp352 | 2.143 | 5.66E-03 | glio |
| Hsd17b10_in.vivo_noRT | Zfp712 | 1.921 | 5.66E-03 | glio |
| culture_effect | Frat2 | -1.056 | 5.66E-04 | glio |
| Interaction_genotype_Mtor | Mroh7 | -4.057 | 5.66E-04 | glio |
| Interaction_genotype_Mtor | D730003I15Ri | -2.805 | 5.66E-04 | glio |
| Atr_in.vivo_noRT | A730063M14F | 2.202 | 5.66E-04 | glio |
| Atr_in.vivo_noRT | B3galt2 | 2.185 | 5.66E-04 | glio |
| Bora_in.vivo_noRT | Zfp964 | 2.562 | 5.66E-04 | glio |
| Hsd17b10_in.vivo_noRT | 1110020A21R | 2.666 | 5.66E-04 | glio |
| culture_effect | Sp1 | -0.675 | 5.66E-05 | glio |
| Interaction_genotype_Myc | Ksr1 | -4.474 | 5.66E-05 | glio |
| Aars2_in.vivo_noRT | Gm50203 | 2.694 | 5.66E-05 | glio |
| Atr_in.vivo_noRT | Cdc26 | -1.156 | 5.66E-05 | glio |
| Bora_ex.vivo_noRT | Cfh | 2.802 | 5.66E-05 | glio |
| Bora_ex.vivo_noRT | Olfr543 | 2.806 | 5.66E-05 | glio |
| Interaction_genotype_Mtor | Gm26674 | -5.089 | 5.66E-06 | glio |
| Interaction_genotype_Mtor | Gm47101 | -5.181 | 5.66E-06 | glio |
| Interaction_genotype_Mtor | Cxcl5 | -4.099 | 5.66E-06 | glio |
| Interaction_genotype_Mtor | Gad1 | -3.833 | 5.66E-06 | glio |
| Bard1_ex.vivo_noRT | Stfa3 | 2.814 | 5.66E-06 | glio |
| Bard1_ex.vivo_noRT | Olfr787 | 2.814 | 5.66E-06 | glio |
| Bora_in.vivo_noRT | Gmnn | -1.848 | 5.66E-06 | glio |

| | | | | |
|---|---|---|---|---|
| Hsd17b10_in.vivo_noRT | Tnfrsf23 | 3.749 | 5.66E-06 | glio |
| Aars2_ex.vivo_noRT | Gm15533 | 3.293 | 5.66E-07 | glio |
| Interaction_genotype_Myc | Gtf2a2 | 2.323 | 5.66E-08 | glio |
| Interaction_genotype_Myc | Limch1 | -6.675 | 5.66E-08 | glio |
| Interaction_genotype_Myc | Camk1d | -6.291 | 5.66E-08 | glio |
| Atr_in.vivo_noRT | Pcdhga2 | 4.263 | 5.66E-11 | glio |
| Atr_in.vivo_noRT | Gm41492 | 3.963 | 5.66E-11 | glio |
| Aars2_in.vivo_noRT | Rarres1 | 5.327 | 5.66E-15 | glio |
| Atr_in.vivo_noRT | Pmch | 4.876 | 5.66E-16 | glio |
| Aars2_in.vivo_noRT | Ntn4 | 5.06 | 5.66E-17 | glio |
| Atr_in.vivo_noRT | Gtsf1 | 5.533 | 5.66E-22 | glio |
| culture_effect | Arhgef18 | 0.69 | 5.67E-03 | glio |
| Interaction_genotype_Aars2 | Nudt7 | -3.2 | 5.67E-03 | glio |
| Interaction_genotype_Atr | BC061195 | -2.369 | 5.67E-03 | glio |
| Interaction_genotype_Atr | Mamdc2 | -3.409 | 5.67E-03 | glio |
| Interaction_genotype_Atr | Tlr1 | -3.382 | 5.67E-03 | glio |
| Interaction_genotype_Atr | 1500011B03R | 1.548 | 5.67E-03 | glio |
| Interaction_genotype_Mtor | Fdxacb1 | -3.029 | 5.67E-03 | glio |
| Interaction_genotype_Mtor | Nat2 | -2.847 | 5.67E-03 | glio |
| Interaction_genotype_Mtor | Actr3 | 1.385 | 5.67E-03 | glio |
| Interaction_genotype_Myc | Pde9a | -2.918 | 5.67E-03 | glio |
| Interaction_genotype_Myc | Ercc6 | -2.576 | 5.67E-03 | glio |
| Interaction_genotype_Myc | Prepl | -2.264 | 5.67E-03 | glio |
| interaction_RT | Gm4841 | 2.733 | 5.67E-03 | glio |
| Bora_in.vivo_noRT | Fndc7 | 2.183 | 5.67E-03 | glio |
| Bora_in.vivo_noRT | Prickle3 | 1.839 | 5.67E-03 | glio |
| Bora_ex.vivo_noRT | Pde6b | 2.408 | 5.67E-03 | glio |
| Chek1_in.vivo_noRT | Nfib | -0.956 | 5.67E-03 | glio |
| Chek1_ex.vivo_noRT | Gm43813 | 2.512 | 5.67E-03 | glio |
| Eif2s1_in.vivo_noRT | Lrrc24 | 2.191 | 5.67E-03 | glio |
| Eif2s1_in.vivo_noRT | Mob3c | -2.084 | 5.67E-03 | glio |
| culture_effect | Snip1 | -0.748 | 5.67E-04 | glio |
| culture_effect | Bzw2 | -0.51 | 5.67E-04 | glio |
| Interaction_genotype_Atr | Gm16046 | -4.22 | 5.67E-04 | glio |
| Interaction_genotype_Mtor | Acsf2 | -3.658 | 5.67E-04 | glio |
| Interaction_genotype_Mtor | 6530402F18Ri | -3.956 | 5.67E-04 | glio |
| Interaction_genotype_Mtor | Ppp1r12b | -3.785 | 5.67E-04 | glio |
| Interaction_genotype_Mtor | Dnajc21 | 1.771 | 5.67E-04 | glio |
| Interaction_genotype_Mtor | Rack1 | 1.224 | 5.67E-04 | glio |
| Interaction_genotype_Myc | Pbx1 | -3.049 | 5.67E-04 | glio |
| Interaction_genotype_Myc | Rpl18a | 1.149 | 5.67E-04 | glio |
| Bcas2_in.vivo_noRT | Sqor | 2.247 | 5.67E-04 | glio |
| interaction_RT | Hist1h3e | -1.919 | 5.67E-05 | glio |
| Atr_ex.vivo_noRT | Ush1g | 2.764 | 5.67E-05 | glio |
| Hsd17b10_in.vivo_noRT | Cfap58 | 3.616 | 5.67E-05 | glio |
| Hsd17b10_in.vivo_noRT | Gm49579 | 2.563 | 5.67E-05 | glio |
| Aars2_in.vivo_noRT | Vcan | 3.457 | 5.67E-06 | glio |
| Aars2_in.vivo_noRT | Serpinb9 | 3.033 | 5.67E-06 | glio |
| Aars2_in.vivo_noRT | Psmd12 | -1.195 | 5.67E-06 | glio |
| Atp5c1_in.vivo_noRT | Gm6803 | 2.668 | 5.67E-06 | glio |
| Atr_in.vivo_noRT | 4930404N11R | 2.934 | 5.67E-06 | glio |
| Chek1_in.vivo_noRT | Cbs | 3.317 | 5.67E-06 | glio |

| | | | | |
|---|---|---|---|---|
| Hsd17b10_in.vivo_noRT | Ttll13 | 2.883 | 5.67E-06 | glio |
| culture_effect | Stat1 | -0.777 | 5.67E-07 | glio |
| Bcas2_in.vivo_noRT | Tmem8b | 3.419 | 5.67E-07 | glio |
| Bcas2_in.vivo_noRT | Tmem179 | 3.649 | 5.67E-07 | glio |
| Bcas2_in.vivo_noRT | Il17b | 4.157 | 5.67E-08 | glio |
| culture_effect | Slc6a8 | 1.176 | 5.67E-09 | glio |
| culture_effect | Psmb6 | -0.757 | 5.67E-10 | glio |
| Bcas2_in.vivo_noRT | Pmch | 4.149 | 5.67E-12 | glio |
| Bcas2_in.vivo_noRT | Zic4 | 5.062 | 5.67E-12 | glio |
| Chek1_in.vivo_noRT | Cxcl13 | 5.044 | 5.67E-14 | glio |
| Chek1_ex.vivo_noRT | Gm29670 | 4.842 | 5.67E-14 | glio |
| Aars2_in.vivo_noRT | Snhg4.1 | 4.804 | 5.67E-16 | glio |
| Atr_in.vivo_noRT | Gm13054 | 5.007 | 5.67E-16 | glio |
| culture_effect | Pdzd2 | 3.135 | 5.67E-25 | glio |
| culture_effect | Il18bp | -4.498 | 5.67E-27 | glio |
| culture_effect | Gm19684 | -1.912 | 5.68E-03 | glio |
| culture_effect | Gm10863 | 1.195 | 5.68E-03 | glio |
| culture_effect | Cdpf1 | 0.591 | 5.68E-03 | glio |
| culture_effect | Mrpl47 | -0.567 | 5.68E-03 | glio |
| culture_effect | Ndufb5 | -0.33 | 5.68E-03 | glio |
| Interaction_genotype_Atr | Gm14540 | -2.641 | 5.68E-03 | glio |
| Interaction_genotype_Bcas2 | Gm47851 | -3.394 | 5.68E-03 | glio |
| Interaction_genotype_Mtor | Gap43 | -3.844 | 5.68E-03 | glio |
| Interaction_genotype_Mtor | Klf15 | -2.986 | 5.68E-03 | glio |
| Interaction_genotype_Mtor | Pdia3 | 1.38 | 5.68E-03 | glio |
| Interaction_genotype_Myc | Ssh3 | -2.726 | 5.68E-03 | glio |
| Interaction_genotype_Rad21 | Gprin2 | 3.524 | 5.68E-03 | glio |
| Interaction_genotype_Rpl3 | Mir142hg | -3.892 | 5.68E-03 | glio |
| Interaction_genotype_Rpl3 | B930059L03Ri | -4.072 | 5.68E-03 | glio |
| Aars2_in.vivo_noRT | Zfp976 | 1.802 | 5.68E-03 | glio |
| Aars2_in.vivo_noRT | Psma5 | -0.768 | 5.68E-03 | glio |
| Atr_in.vivo_noRT | H2-Eb2 | 1.44 | 5.68E-03 | glio |
| Cenpj_in.vivo_noRT | Gm14820 | 1.759 | 5.68E-03 | glio |
| Cenpj_ex.vivo_noRT | Olfr543 | 2.196 | 5.68E-03 | glio |
| Chek1_ex.vivo_noRT | Hunk | 2.34 | 5.68E-03 | glio |
| Cyp19a1_ex.vivo_noRT | Gjb5 | 2.757 | 5.68E-03 | glio |
| Cyp19a1_ex.vivo_noRT | Pla1a | 2.959 | 5.68E-03 | glio |
| Cyp19a1_ex.vivo_noRT | Gm13078 | 2.159 | 5.68E-03 | glio |
| Cyp19a1_ex.vivo_noRT | Gm13964 | 2.159 | 5.68E-03 | glio |
| Cyp19a1_ex.vivo_noRT | Gm43915 | 2.565 | 5.68E-03 | glio |
| Eif2s1_ex.vivo_noRT | Ccl3 | 2.234 | 5.68E-03 | glio |
| Eif2s1_ex.vivo_noRT | AU019990 | 2.234 | 5.68E-03 | glio |
| Eif2s1_ex.vivo_noRT | Gm48780 | 2.234 | 5.68E-03 | glio |
| Hsd17b10_in.vivo_noRT | Rtl1 | 1.924 | 5.68E-03 | glio |
| Hsd17b10_in.vivo_noRT | Rnf25 | -1.41 | 5.68E-03 | glio |
| culture_effect | Hook2 | -1.142 | 5.68E-04 | glio |
| interaction_RT | Ccndbp1 | 0.938 | 5.68E-04 | glio |
| Atr_in.vivo_noRT | Dgat1 | 1.705 | 5.68E-04 | glio |
| Chek1_in.vivo_noRT | Stamos | 2.258 | 5.68E-04 | glio |
| Chek1_in.vivo_noRT | Ggh | -1.312 | 5.68E-04 | glio |
| Interaction_genotype_Mtor | Ttc41 | -5.168 | 5.68E-05 | glio |
| Interaction_genotype_Mtor | Susd1 | -5.274 | 5.68E-05 | glio |

| | | | | |
|---|---|---|---|---|
| Interaction_genotype_Mtor | Gm29670 | -3.577 | 5.68E-05 | glio |
| Interaction_genotype_Mtor | Cr2 | -4.168 | 5.68E-05 | glio |
| Interaction_genotype_Mtor | Rpn2 | 1.926 | 5.68E-05 | glio |
| Interaction_genotype_Nat2 | Gm13964 | 3.605 | 5.68E-05 | glio |
| Interaction_genotype_Nat2 | AF067063 | 3.825 | 5.68E-05 | glio |
| Aars2_ex.vivo_noRT | Vwce | 3.315 | 5.68E-05 | glio |
| Atr_in.vivo_noRT | Gm11520 | 2.603 | 5.68E-05 | glio |
| Bcas2_in.vivo_noRT | Icam5 | 2.785 | 5.68E-05 | glio |
| Bcas2_in.vivo_noRT | Bpgm | -1.599 | 5.68E-05 | glio |
| Bora_in.vivo_noRT | Slc25a43 | 2.877 | 5.68E-05 | glio |
| Interaction_genotype_Atr | Gm38534 | -5.248 | 5.68E-06 | glio |
| Interaction_genotype_Mtor | Mir142hg | -5.963 | 5.68E-07 | glio |
| Interaction_genotype_Myc | Ybx1 | 2.057 | 5.68E-07 | glio |
| Hsd17b10_in.vivo_noRT | Gm15445 | 2.97 | 5.68E-08 | glio |
| Atr_in.vivo_noRT | Slxl1 | 4.755 | 5.68E-12 | glio |
| Atr_in.vivo_noRT | G630030J09Ri | 5.011 | 5.68E-14 | glio |
| Atr_ex.vivo_noRT | Tpo | 4.725 | 5.68E-15 | glio |
| Bora_in.vivo_noRT | Esp24 | 4.936 | 5.68E-21 | glio |
| Bora_in.vivo_noRT | Stfa3 | 4.936 | 5.68E-26 | glio |
| Bora_in.vivo_noRT | Olfr787 | 4.936 | 5.68E-26 | glio |
| culture_effect | Gm29966 | -1.194 | 5.69E-03 | glio |
| culture_effect | Rpusd1 | 0.771 | 5.69E-03 | glio |
| culture_effect | Baz1a | -0.481 | 5.69E-03 | glio |
| Interaction_genotype_Bcas2 | Celsr3 | -3.485 | 5.69E-03 | glio |
| Interaction_genotype_Bcas2 | Gm12500 | -3.542 | 5.69E-03 | glio |
| Interaction_genotype_Bcas2 | Gm29340 | -3.202 | 5.69E-03 | glio |
| Interaction_genotype_Cdkn2a | Gm30794 | 2.909 | 5.69E-03 | glio |
| Interaction_genotype_Cdkn2a | Gm13078 | 2.835 | 5.69E-03 | glio |
| Interaction_genotype_Mtor | Kcnq2 | -2.935 | 5.69E-03 | glio |
| Interaction_genotype_Mtor | Tmem88 | -3.38 | 5.69E-03 | glio |
| Interaction_genotype_Mtor | Egf | -3.479 | 5.69E-03 | glio |
| Interaction_genotype_Mtor | Gab2 | -3.141 | 5.69E-03 | glio |
| Interaction_genotype_Mtor | Cdv3 | 1.171 | 5.69E-03 | glio |
| Interaction_genotype_Myc | Irf6 | -2.433 | 5.69E-03 | glio |
| Interaction_genotype_Myc | Camta2 | -2.225 | 5.69E-03 | glio |
| Interaction_genotype_Terf1 | Sel1l | 2.055 | 5.69E-03 | glio |
| interaction_RT | Pcdhgc3 | -1.702 | 5.69E-03 | glio |
| interaction_RT | Lancl2 | -1.257 | 5.69E-03 | glio |
| Atr_in.vivo_noRT | Tbccd1 | 1.769 | 5.69E-03 | glio |
| Atr_in.vivo_noRT | Hc | 1.697 | 5.69E-03 | glio |
| Bcas2_in.vivo_noRT | Ebi3 | 2.429 | 5.69E-03 | glio |
| Bcas2_in.vivo_noRT | Shank1 | 1.908 | 5.69E-03 | glio |
| Brca2_in.vivo_noRT | Trim17 | 1.763 | 5.69E-03 | glio |
| Brca2_in.vivo_noRT | Tpo | 1.763 | 5.69E-03 | glio |
| Cenpj_ex.vivo_noRT | Cfh | 2.192 | 5.69E-03 | glio |
| Hsd17b10_in.vivo_noRT | Klf15 | 2.149 | 5.69E-03 | glio |
| Interaction_genotype_Aars2 | Nrg2 | -4.288 | 5.69E-04 | glio |
| Interaction_genotype_Aars2 | A930012O16R | -4.459 | 5.69E-04 | glio |
| interaction_RT | Rrp9 | -0.925 | 5.69E-04 | glio |
| Atr_in.vivo_noRT | Gm26631 | 2.568 | 5.69E-04 | glio |
| Bora_in.vivo_noRT | A430105J06Ri | 2.428 | 5.69E-04 | glio |
| Chek1_in.vivo_noRT | Lgr6 | 2.835 | 5.69E-04 | glio |

| | | | | |
|---|---|---|---|---|
| Interaction_genotype_Mtor | Gm11583 | -3.573 | 5.69E-05 | glio |
| Interaction_genotype_Mtor | Pcdhga11 | -4.073 | 5.69E-05 | glio |
| Aars2_in.vivo_noRT | 1700102H20R | 2.802 | 5.69E-05 | glio |
| Aars2_in.vivo_noRT | Mavs | 2.103 | 5.69E-05 | glio |
| Aars2_in.vivo_noRT | Parp9 | -1.914 | 5.69E-05 | glio |
| Atr_in.vivo_noRT | Naip6 | 2.748 | 5.69E-05 | glio |
| Bora_in.vivo_noRT | Ppp1r26 | 3.028 | 5.69E-05 | glio |
| Chek1_in.vivo_noRT | Pde4d | 2.868 | 5.69E-05 | glio |
| Chek1_in.vivo_noRT | Sardh | 2.757 | 5.69E-05 | glio |
| Chek1_ex.vivo_noRT | Gm34643 | 3.746 | 5.69E-05 | glio |
| Interaction_genotype_Mtor | Stmn1 | 3.063 | 5.69E-06 | glio |
| Bcas2_in.vivo_noRT | Zfp972 | 3.363 | 5.69E-06 | glio |
| Bora_in.vivo_noRT | Gm40040 | 3.38 | 5.69E-06 | glio |
| Atr_in.vivo_noRT | Gm43728 | 3.675 | 5.69E-07 | glio |
| Bora_in.vivo_noRT | Uckl1os | 3.702 | 5.69E-07 | glio |
| Bcas2_in.vivo_noRT | Dok3 | 4.189 | 5.69E-09 | glio |
| Bora_in.vivo_noRT | Nrn1l | 4.011 | 5.69E-10 | glio |
| Atr_in.vivo_noRT | Insm1 | 4.415 | 5.69E-11 | glio |
| Atr_in.vivo_noRT | Col5a2 | 4.222 | 5.69E-15 | glio |
| Aars2_in.vivo_noRT | Gm15929 | 5.327 | 5.69E-21 | glio |
| Bcas2_in.vivo_noRT | Gm5157 | 4.807 | 5.69E-24 | glio |
| culture_effect | Rcl1 | -0.469 | 5.70E-03 | glio |
| Interaction_genotype_Aars2 | Vmn2r44 | -2.429 | 5.70E-03 | glio |
| Interaction_genotype_Aars2 | Dok2 | -2.994 | 5.70E-03 | glio |
| Interaction_genotype_Aars2 | Ifi207 | -3.739 | 5.70E-03 | glio |
| Interaction_genotype_Aars2 | Slc13a5 | -3.336 | 5.70E-03 | glio |
| Interaction_genotype_Atr | Vpreb1 | -3.421 | 5.70E-03 | glio |
| Interaction_genotype_Atr | Mtap | -1.119 | 5.70E-03 | glio |
| Interaction_genotype_Bcas2 | Agap2 | -3.05 | 5.70E-03 | glio |
| Interaction_genotype_Bcas2 | D630036H23F | -3.489 | 5.70E-03 | glio |
| Interaction_genotype_Myc | Zfp433 | -3.337 | 5.70E-03 | glio |
| Interaction_genotype_Myc | Usp33 | -2.325 | 5.70E-03 | glio |
| Interaction_genotype_Pars2 | Lyz2 | -3.923 | 5.70E-03 | glio |
| Interaction_genotype_Pars2 | Hecw2 | -3.472 | 5.70E-03 | glio |
| Interaction_genotype_Uqcrc2 | 4921523L03Ri | -4.341 | 5.70E-03 | glio |
| interaction_RT | Pde10a | 1.442 | 5.70E-03 | glio |
| interaction_RT | Scnm1 | 0.747 | 5.70E-03 | glio |
| interaction_RT | Chchd7 | 0.604 | 5.70E-03 | glio |
| Aars2_in.vivo_noRT | Crcp | -1.166 | 5.70E-03 | glio |
| Aars2_in.vivo_noRT | Mbp | -1.537 | 5.70E-03 | glio |
| Aars2_in.vivo_noRT | Tmem258 | -0.714 | 5.70E-03 | glio |
| Atr_ex.vivo_noRT | Ngef | 2.469 | 5.70E-03 | glio |
| Bard1_in.vivo_noRT | Gm364 | 1.98 | 5.70E-03 | glio |
| Bard1_in.vivo_noRT | 4732463B04R | 2.235 | 5.70E-03 | glio |
| Bora_in.vivo_noRT | A530041M06F | 2.067 | 5.70E-03 | glio |
| culture_effect | Dctn4 | 0.588 | 5.70E-04 | glio |
| Aars2_ex.vivo_noRT | Sap30bpos | 2.899 | 5.70E-04 | glio |
| Bcas2_in.vivo_noRT | Gm42869 | 2.411 | 5.70E-04 | glio |
| Bcas2_ex.vivo_noRT | Slc16a5 | 2.349 | 5.70E-04 | glio |
| Bcas2_ex.vivo_noRT | Epb42 | 3.574 | 5.70E-04 | glio |
| Bora_ex.vivo_noRT | Arhgef16 | 2.802 | 5.70E-04 | glio |
| Atr_ex.vivo_noRT | 1700010B08R | 3.164 | 5.70E-05 | glio |

| | | | | |
|---|---|---|---|---|
| Bard1_ex.vivo_noRT | Adgre5 | 3.219 | 5.70E-05 | glio |
| Bard1_ex.vivo_noRT | Ccl3 | 2.814 | 5.70E-05 | glio |
| interaction_RT | Zswim6 | 1.484 | 5.70E-06 | glio |
| Bora_in.vivo_noRT | Nlrx1 | 4.048 | 5.70E-08 | glio |
| Atr_in.vivo_noRT | Gm26688 | 4.095 | 5.70E-09 | glio |
| Atr_in.vivo_noRT | D830035M03I | 3.947 | 5.70E-09 | glio |
| Chek1_in.vivo_noRT | Nt5e | 3.565 | 5.70E-09 | glio |
| culture_effect | 1700007L15Ri | 2.103 | 5.70E-10 | glio |
| culture_effect | Cited1 | -2.112 | 5.70E-10 | glio |
| Aars2_in.vivo_noRT | Gm41492 | 3.757 | 5.70E-10 | glio |
| culture_effect | 0610009L18Ri | 3.285 | 5.70E-11 | glio |
| Bora_in.vivo_noRT | F630040K05Ri | 4.285 | 5.70E-12 | glio |
| Chek1_ex.vivo_noRT | Srgn | 7.729 | 5.70E-16 | glio |
| Atr_in.vivo_noRT | Minar2 | 5.273 | 5.70E-21 | glio |
| culture_effect | Atrx | -0.375 | 5.71E-03 | glio |
| Interaction_genotype_Chek1 | Dab2 | 3.635 | 5.71E-03 | glio |
| Interaction_genotype_Chek1 | Klk6 | 3.603 | 5.71E-03 | glio |
| Interaction_genotype_Chek1 | Fbxo5 | -2.99 | 5.71E-03 | glio |
| Interaction_genotype_Chek1 | Kdm6b | 2.184 | 5.71E-03 | glio |
| Interaction_genotype_Chek1 | Tbc1d10a | 2.082 | 5.71E-03 | glio |
| Interaction_genotype_Chek1 | Sbds | 1.62 | 5.71E-03 | glio |
| Interaction_genotype_Chek1 | S100a13 | 1.458 | 5.71E-03 | glio |
| Interaction_genotype_Myc | Gm12981 | -3.162 | 5.71E-03 | glio |
| Interaction_genotype_Myc | Apol6 | 2.615 | 5.71E-03 | glio |
| Interaction_genotype_Myc | AI429214 | -3.032 | 5.71E-03 | glio |
| interaction_RT | D930028M14I | 2.389 | 5.71E-03 | glio |
| Bcas2_in.vivo_noRT | Slc46a2 | 1.826 | 5.71E-03 | glio |
| Bcas2_in.vivo_noRT | Rhobtb1 | 2.067 | 5.71E-03 | glio |
| Bcas2_in.vivo_noRT | Plpp7 | 1.895 | 5.71E-03 | glio |
| Bcas2_in.vivo_noRT | Anapc5 | -1.147 | 5.71E-03 | glio |
| Bcas2_in.vivo_noRT | Cox7b | -0.562 | 5.71E-03 | glio |
| Bcas2_ex.vivo_noRT | Cd52 | 2.742 | 5.71E-03 | glio |
| Bcas2_ex.vivo_noRT | Gm5431 | 2.742 | 5.71E-03 | glio |
| Brca2_in.vivo_noRT | 1700034I23Ri | 1.763 | 5.71E-03 | glio |
| Chek1_in.vivo_noRT | Zfp316 | 1.757 | 5.71E-03 | glio |
| Chek1_in.vivo_noRT | Fmc1 | -0.942 | 5.71E-03 | glio |
| Chek1_ex.vivo_noRT | Tdrd5 | 2.316 | 5.71E-03 | glio |
| Chek1_ex.vivo_noRT | Dynlt1b | 2.579 | 5.71E-03 | glio |
| Chek1_ex.vivo_noRT | Rnf123 | 1.869 | 5.71E-03 | glio |
| Chek1_ex.vivo_noRT | Sox2 | -0.994 | 5.71E-03 | glio |
| Eif2s1_in.vivo_noRT | 5330439K02R | 2.4 | 5.71E-03 | glio |
| Eif2s1_in.vivo_noRT | Rasl2-9 | 2.622 | 5.71E-03 | glio |
| Eif2s1_ex.vivo_noRT | Gm36793 | 2.234 | 5.71E-03 | glio |
| Eif2s1_ex.vivo_noRT | Spint4 | 3.099 | 5.71E-03 | glio |
| culture_effect | Gm28187 | -1.383 | 5.71E-04 | glio |
| Interaction_genotype_Atr | Trmt10c | 2.056 | 5.71E-04 | glio |
| Interaction_genotype_Chek1 | Vwa5a | 2.888 | 5.71E-04 | glio |
| Interaction_genotype_Myc | AI854703 | -3.92 | 5.71E-04 | glio |
| Interaction_genotype_Myc | Fam83e | -4.036 | 5.71E-04 | glio |
| Atr_in.vivo_noRT | Platr4 | 1.97 | 5.71E-04 | glio |
| Cenpj_in.vivo_noRT | Erich6b | 2.022 | 5.71E-04 | glio |
| Cenpj_in.vivo_noRT | Lncbate10 | 2.022 | 5.71E-04 | glio |

| | | | | |
|---|---|---|---|---|
| Cenpj_in.vivo_noRT | Tac2 | 2.022 | 5.71E-04 | glio |
| Cenpj_ex.vivo_noRT | Bmp15 | 2.995 | 5.71E-04 | glio |
| Chek1_ex.vivo_noRT | Ramp3 | 3.033 | 5.71E-04 | glio |
| culture_effect | Lipt2 | 1.409 | 5.71E-05 | glio |
| Interaction_genotype_Myc | Edn1 | -5.25 | 5.71E-05 | glio |
| Interaction_genotype_Myc | Mrpl43 | 1.564 | 5.71E-05 | glio |
| Aars2_in.vivo_noRT | Mr1 | 2.894 | 5.71E-05 | glio |
| Aars2_in.vivo_noRT | Trim66 | 2.615 | 5.71E-05 | glio |
| Bard1_ex.vivo_noRT | Gm6566 | 3.219 | 5.71E-05 | glio |
| Hsd17b10_in.vivo_noRT | Slc16a7 | 2.505 | 5.71E-05 | glio |
| Aars2_in.vivo_noRT | C130013H08R | 3.557 | 5.71E-06 | glio |
| culture_effect | Tulp3 | 0.72 | 5.71E-07 | glio |
| Interaction_genotype_Mtor | Lsm4 | 2.702 | 5.71E-08 | glio |
| Interaction_genotype_Mtor | Sucnr1 | -4.099 | 5.71E-08 | glio |
| Atr_in.vivo_noRT | Prex2 | 4.097 | 5.71E-08 | glio |
| Hsd17b10_in.vivo_noRT | Slc45a1 | 3.356 | 5.71E-11 | glio |
| Hsd17b10_in.vivo_noRT | Gstt4 | 3.353 | 5.71E-11 | glio |
| Hsd17b10_in.vivo_noRT | Gm21319 | 3.616 | 5.71E-11 | glio |
| Atr_ex.vivo_noRT | Gm21818 | 6.252 | 5.71E-13 | glio |
| Bcas2_in.vivo_noRT | Gm16741 | 4.807 | 5.71E-14 | glio |
| culture_effect | Ddx5 | -0.777 | 5.71E-16 | glio |
| Bora_in.vivo_noRT | Hrct1 | 4.936 | 5.71E-16 | glio |
| Bcas2_in.vivo_noRT | Olfr543 | 4.547 | 5.71E-18 | glio |
| Bora_in.vivo_noRT | Gm44899 | 4.936 | 5.71E-19 | glio |
| culture_effect | Atp1a3 | -1.183 | 5.72E-03 | glio |
| culture_effect | Klhl42 | 0.756 | 5.72E-03 | glio |
| Interaction_genotype_Atr | Idi2 | -2.369 | 5.72E-03 | glio |
| Interaction_genotype_Atr | Gm16348 | -3.348 | 5.72E-03 | glio |
| Interaction_genotype_Cdkn2a | Gm13964 | 2.835 | 5.72E-03 | glio |
| Interaction_genotype_Mtor | Aldh1l1 | -3.602 | 5.72E-03 | glio |
| Interaction_genotype_Mtor | A830005F24Ri | -3.323 | 5.72E-03 | glio |
| Interaction_genotype_Myc | Plppr2 | -3.219 | 5.72E-03 | glio |
| Interaction_genotype_Myc | Nme1 | 1.085 | 5.72E-03 | glio |
| Interaction_genotype_Pars2 | Wnt16 | -3.656 | 5.72E-03 | glio |
| Interaction_genotype_Pars2 | Cpne9 | -4.073 | 5.72E-03 | glio |
| Interaction_genotype_Uqcrb | C130060C02R | -3.993 | 5.72E-03 | glio |
| interaction_RT | Ppp1r3g | -2.525 | 5.72E-03 | glio |
| interaction_RT | Sp110 | 1.352 | 5.72E-03 | glio |
| interaction_RT | Noc2l | -0.709 | 5.72E-03 | glio |
| interaction_RT | Snhg1 | 0.89 | 5.72E-03 | glio |
| interaction_RT | Elob | 0.481 | 5.72E-03 | glio |
| Aars2_in.vivo_noRT | Eri1 | -1.242 | 5.72E-03 | glio |
| Aars2_in.vivo_noRT | Col11a1 | -0.652 | 5.72E-03 | glio |
| Bcas2_in.vivo_noRT | Ttc12 | 1.949 | 5.72E-03 | glio |
| Chek1_in.vivo_noRT | Msantd1 | 2.056 | 5.72E-03 | glio |
| Chek1_in.vivo_noRT | Ifi203 | -1.507 | 5.72E-03 | glio |
| Interaction_genotype_Aars2 | Prl7c1 | -3.129 | 5.72E-04 | glio |
| Interaction_genotype_Aars2 | Asgr1 | -4.129 | 5.72E-04 | glio |
| Interaction_genotype_Aars2 | Col8a1 | -4.164 | 5.72E-04 | glio |
| Interaction_genotype_Aars2 | Limch1 | -4.364 | 5.72E-04 | glio |
| Interaction_genotype_Aars2 | Gm48435 | -2.821 | 5.72E-04 | glio |
| Interaction_genotype_Aars2 | Pnp | 2.027 | 5.72E-04 | glio |

| | | | | |
|---|---|---|---|---|
| Interaction_genotype_Aars2 | Fkbp1a | 1.04 | 5.72E-04 | glio |
| Interaction_genotype_Mtor | Lrrc4b | -3.584 | 5.72E-04 | glio |
| Interaction_genotype_Myc | Zfp318 | -3.169 | 5.72E-04 | glio |
| Interaction_genotype_Myc | Slc7a6 | -2.605 | 5.72E-04 | glio |
| Interaction_genotype_Myc | Fam181b | -3.527 | 5.72E-04 | glio |
| Interaction_genotype_Myc | Pcnt | -2.403 | 5.72E-04 | glio |
| Interaction_genotype_Myc | Plekhb2 | -2.826 | 5.72E-04 | glio |
| Interaction_genotype_Myc | Trim28 | -1.754 | 5.72E-04 | glio |
| Interaction_genotype_Nat2 | Gm20625 | 3.042 | 5.72E-04 | glio |
| interaction_RT | Syncrip | -0.842 | 5.72E-04 | glio |
| Atp5c1_in.vivo_noRT | Usp17la | 2.668 | 5.72E-04 | glio |
| Atr_in.vivo_noRT | Rabggtb | -1.135 | 5.72E-04 | glio |
| Cenpj_in.vivo_noRT | Slc35g3 | 2.022 | 5.72E-04 | glio |
| Cenpj_in.vivo_noRT | Chia1 | 2.022 | 5.72E-04 | glio |
| Chek1_in.vivo_noRT | Zfp39 | 2.922 | 5.72E-05 | glio |
| Chek1_ex.vivo_noRT | Gm16754 | 3.652 | 5.72E-05 | glio |
| culture_effect | Cybrd1 | -2.355 | 5.72E-06 | glio |
| Aars2_ex.vivo_noRT | Gm4027 | 3.293 | 5.72E-06 | glio |
| culture_effect | Serpina3m | -3.024 | 5.72E-07 | glio |
| Interaction_genotype_Myc | Gm50103 | -5.831 | 5.72E-07 | glio |
| Bora_in.vivo_noRT | Gm4755 | 3.179 | 5.72E-07 | glio |
| Bora_in.vivo_noRT | Rhno1 | -1.863 | 5.72E-07 | glio |
| Bora_in.vivo_noRT | Sema3a | 3.499 | 5.72E-08 | glio |
| culture_effect | Tmed3 | -0.817 | 5.72E-10 | glio |
| Chek1_ex.vivo_noRT | Agrp | 4.447 | 5.72E-10 | glio |
| Aars2_in.vivo_noRT | Ttll13 | 4.593 | 5.72E-14 | glio |
| culture_effect | Rpl10a | -0.967 | 5.72E-19 | glio |
| Aars2_in.vivo_noRT | Gm4340 | 5.327 | 5.72E-28 | glio |
| Aars2_in.vivo_noRT | Gm16239 | 5.327 | 5.72E-28 | glio |
| Interaction_genotype_Aars2 | Rbp7 | -3.52 | 5.73E-03 | glio |
| Interaction_genotype_Atr | Ccnl2 | 1.582 | 5.73E-03 | glio |
| Interaction_genotype_Bora | Gm43672 | -3.713 | 5.73E-03 | glio |
| Interaction_genotype_Myc | Minar2 | -2.469 | 5.73E-03 | glio |
| Interaction_genotype_Myc | Rab30 | -2.63 | 5.73E-03 | glio |
| Interaction_genotype_Myc | Acad9 | -2.253 | 5.73E-03 | glio |
| Interaction_genotype_Myc | Kcmf1 | 1.22 | 5.73E-03 | glio |
| Interaction_genotype_Myc | Eef1b2 | 1.083 | 5.73E-03 | glio |
| Aars2_in.vivo_noRT | Mpg | -1.147 | 5.73E-03 | glio |
| Atr_in.vivo_noRT | Hdgf | -0.678 | 5.73E-03 | glio |
| Bard1_ex.vivo_noRT | Snhg4.1 | 2.421 | 5.73E-03 | glio |
| Bcas2_in.vivo_noRT | 2010110K18R | 1.961 | 5.73E-03 | glio |
| Bora_in.vivo_noRT | Rab11fip5 | 2.111 | 5.73E-03 | glio |
| Bora_in.vivo_noRT | Copz2 | -1.161 | 5.73E-03 | glio |
| Eif2s1_in.vivo_noRT | Gm16933 | 1.621 | 5.73E-03 | glio |
| Eif2s1_in.vivo_noRT | Gm43463 | 2.016 | 5.73E-03 | glio |
| Eif2s1_in.vivo_noRT | A030005L19Ri | 2.142 | 5.73E-03 | glio |
| Eif2s1_ex.vivo_noRT | Spink8 | 2.234 | 5.73E-03 | glio |
| Eif2s1_ex.vivo_noRT | Psmb11 | 2.234 | 5.73E-03 | glio |
| culture_effect | Pvr | -0.699 | 5.73E-04 | glio |
| Interaction_genotype_Myc | Bst1 | -3.36 | 5.73E-04 | glio |
| Interaction_genotype_Myc | Ganc | -3.211 | 5.73E-04 | glio |
| Aars2_in.vivo_noRT | Hist4h4 | 2.53 | 5.73E-04 | glio |

| | | | | |
|---|---|---|---|---|
| Atr_in.vivo_noRT | A230056P14R | 2.438 | 5.73E-04 | glio |
| Bcas2_in.vivo_noRT | C8g | 2.533 | 5.73E-04 | glio |
| Cenpj_in.vivo_noRT | Klrk1 | -2.097 | 5.73E-04 | glio |
| Atr_in.vivo_noRT | Loxl2 | 3.121 | 5.73E-05 | glio |
| Chek1_in.vivo_noRT | Pfkfb1 | 2.882 | 5.73E-05 | glio |
| Eif2s1_ex.vivo_noRT | Meioc | 3.014 | 5.73E-05 | glio |
| Atr_ex.vivo_noRT | Igf2 | -2.823 | 5.73E-06 | glio |
| Bora_in.vivo_noRT | Gnaz | 3.638 | 5.73E-07 | glio |
| Bcas2_in.vivo_noRT | Cldn19 | 4.025 | 5.73E-09 | glio |
| Bora_in.vivo_noRT | Gm42984 | 4.286 | 5.73E-10 | glio |
| Chek1_in.vivo_noRT | 2300002M23F | 3.875 | 5.73E-12 | glio |
| Chek1_in.vivo_noRT | Ldhd | 4.127 | 5.73E-12 | glio |
| Chek1_ex.vivo_noRT | A330049N07R | 4.443 | 5.73E-14 | glio |
| Aars2_in.vivo_noRT | Gm12794 | 5.327 | 5.73E-25 | glio |
| culture_effect | Pygo1 | 1.529 | 5.74E-03 | glio |
| culture_effect | Gm45605 | 1.42 | 5.74E-03 | glio |
| Interaction_genotype_Aars2 | Slc17a6 | -2.034 | 5.74E-03 | glio |
| Interaction_genotype_Aars2 | 4921523L03Ri | -3.587 | 5.74E-03 | glio |
| Interaction_genotype_Aars2 | Rpain | 1.622 | 5.74E-03 | glio |
| Interaction_genotype_Atr | Vmn2r18 | -2.369 | 5.74E-03 | glio |
| Interaction_genotype_Atr | Rnf8 | 1.931 | 5.74E-03 | glio |
| Interaction_genotype_Bora | Cacnb3 | -3.253 | 5.74E-03 | glio |
| Interaction_genotype_Myc | Tnfrsf4 | -3.118 | 5.74E-03 | glio |
| Interaction_genotype_Myc | Gm15706 | -2.404 | 5.74E-03 | glio |
| Interaction_genotype_Myc | Nprl3 | -1.879 | 5.74E-03 | glio |
| Interaction_genotype_Rad51 | Nr4a3 | 4.028 | 5.74E-03 | glio |
| Interaction_genotype_Rad51 | Tulp3 | 1.647 | 5.74E-03 | glio |
| interaction_RT | Auts2 | 2.534 | 5.74E-03 | glio |
| Aars2_in.vivo_noRT | Prkar1a | -0.722 | 5.74E-03 | glio |
| Atr_in.vivo_noRT | Mvb12a | -0.963 | 5.74E-03 | glio |
| Bcas2_in.vivo_noRT | Gm20597 | 2.226 | 5.74E-03 | glio |
| Bcas2_in.vivo_noRT | Tent5d | 1.965 | 5.74E-03 | glio |
| culture_effect | Slc16a2 | -2.022 | 5.74E-04 | glio |
| culture_effect | Pwwp2a | -0.768 | 5.74E-04 | glio |
| culture_effect | Ccnh | -0.604 | 5.74E-04 | glio |
| Interaction_genotype_Chek1 | Sowahc | 3.124 | 5.74E-04 | glio |
| Interaction_genotype_Myc | Eif3j2 | -3.887 | 5.74E-04 | glio |
| Atr_in.vivo_noRT | Slc11a1 | 2.419 | 5.74E-04 | glio |
| Bcas2_in.vivo_noRT | Mzf1 | 2.417 | 5.74E-04 | glio |
| Chek1_in.vivo_noRT | Gm9776 | 2.419 | 5.74E-04 | glio |
| Chek1_in.vivo_noRT | Gm16973 | 2.494 | 5.74E-04 | glio |
| Chek1_ex.vivo_noRT | Gm16230 | 3.069 | 5.74E-04 | glio |
| Hsd17b10_in.vivo_noRT | Axdnd1 | 2.831 | 5.74E-04 | glio |
| Interaction_genotype_Chek1 | Hbq1b | -5.123 | 5.74E-05 | glio |
| Interaction_genotype_Myc | Ubox5 | -3.861 | 5.74E-05 | glio |
| interaction_RT | Chmp5 | 0.71 | 5.74E-05 | glio |
| Chek1_in.vivo_noRT | Glt8d2 | 2.992 | 5.74E-05 | glio |
| Bora_in.vivo_noRT | Usp51 | 3.05 | 5.74E-06 | glio |
| Interaction_genotype_Mtor | Gm20492 | -5.734 | 5.74E-07 | glio |
| Bora_in.vivo_noRT | 2210406O10R | 3.619 | 5.74E-07 | glio |
| Bcas2_in.vivo_noRT | Jakmip1 | 4.016 | 5.74E-09 | glio |
| Bcas2_in.vivo_noRT | Gm32591 | 4.284 | 5.74E-10 | glio |

| | | | | |
|---|---|---|---|---|
| Chek1_ex.vivo_noRT | Tal2 | 3.861 | 5.74E-10 | glio |
| culture_effect | Gm48682 | 3.187 | 5.74E-11 | glio |
| culture_effect | Puf60 | -0.857 | 5.74E-11 | glio |
| Interaction_genotype_Mtor | Gm38534 | -7.776 | 5.74E-11 | glio |
| culture_effect | Kcnmb4 | 2.873 | 5.74E-12 | glio |
| Aars2_in.vivo_noRT | Rnf165 | 5.327 | 5.74E-15 | glio |
| culture_effect | Lap3 | -1.294 | 5.74E-19 | glio |
| Bcas2_in.vivo_noRT | C330002G04R | 4.807 | 5.74E-20 | glio |
| culture_effect | Zfp600 | 1.302 | 5.75E-03 | glio |
| Interaction_genotype_Aars2 | Ttll13 | -3.071 | 5.75E-03 | glio |
| Interaction_genotype_Atr | 6330408M09F | -3.614 | 5.75E-03 | glio |
| Interaction_genotype_Bora | 1700112J16Ri | -2.845 | 5.75E-03 | glio |
| Interaction_genotype_Bora | Ccdc62 | -3.366 | 5.75E-03 | glio |
| Interaction_genotype_Cdkn2a | Gm34643 | 4.112 | 5.75E-03 | glio |
| Interaction_genotype_Mtor | Gm42372 | -3.878 | 5.75E-03 | glio |
| Interaction_genotype_Mtor | Tinf2 | -2.324 | 5.75E-03 | glio |
| Interaction_genotype_Mtor | Canx | 1.21 | 5.75E-03 | glio |
| Interaction_genotype_Myc | Tmem177 | -2.723 | 5.75E-03 | glio |
| Interaction_genotype_Myc | Pak4 | -2.211 | 5.75E-03 | glio |
| Interaction_genotype_Pars2 | Gm18280 | -2.799 | 5.75E-03 | glio |
| Aars2_in.vivo_noRT | Rps16 | -0.559 | 5.75E-03 | glio |
| Atr_in.vivo_noRT | Vamp4 | 0.926 | 5.75E-03 | glio |
| Chek1_in.vivo_noRT | Btg3 | -0.821 | 5.75E-03 | glio |
| Interaction_genotype_Mtor | 1810034E14Ri | -3.954 | 5.75E-04 | glio |
| Interaction_genotype_Myc | Ubtd2 | -2.741 | 5.75E-04 | glio |
| Atp5c1_in.vivo_noRT | Lrp2bp | 2.404 | 5.75E-04 | glio |
| Atr_in.vivo_noRT | Matr3 | -0.993 | 5.75E-04 | glio |
| Bcas2_in.vivo_noRT | 6330418K02R | 2.053 | 5.75E-04 | glio |
| Interaction_genotype_Mtor | Fbxl6 | 2.357 | 5.75E-05 | glio |
| Interaction_genotype_Mtor | Rpl7 | 1.41 | 5.75E-05 | glio |
| Aars2_in.vivo_noRT | Gm2822 | 3.11 | 5.75E-05 | glio |
| Chek1_in.vivo_noRT | Ckm | 2.769 | 5.75E-05 | glio |
| Chek1_in.vivo_noRT | Grasp | -2.347 | 5.75E-05 | glio |
| culture_effect | Canx | -0.512 | 5.75E-06 | glio |
| Aars2_in.vivo_noRT | Zfp28 | 3.208 | 5.75E-06 | glio |
| Chek1_ex.vivo_noRT | Gm13610 | 3.918 | 5.75E-06 | glio |
| Atr_in.vivo_noRT | Rimkla | 3.315 | 5.75E-07 | glio |
| Bcas2_in.vivo_noRT | Gm10644 | 3.566 | 5.75E-07 | glio |
| Bora_ex.vivo_noRT | Gm13964 | 3.201 | 5.75E-07 | glio |
| Chek1_in.vivo_noRT | Pkhd1 | 3.526 | 5.75E-07 | glio |
| Bcas2_in.vivo_noRT | Slc36a2 | 3.688 | 5.75E-08 | glio |
| Chek1_in.vivo_noRT | Snai1 | 4.055 | 5.75E-08 | glio |
| Chek1_ex.vivo_noRT | Bend7 | 4.489 | 5.75E-08 | glio |
| culture_effect | Edil3 | 1.943 | 5.75E-09 | glio |
| Chek1_ex.vivo_noRT | Gjd4 | 4.842 | 5.75E-17 | glio |
| culture_effect | Rpl10-ps3 | -1.584 | 5.75E-24 | glio |
| culture_effect | Egr2 | -6.693 | 5.75E-32 | glio |
| Interaction_genotype_Aars2 | Hsdl1 | -2.568 | 5.76E-03 | glio |
| Interaction_genotype_Atr | 1700040D17R | -3.677 | 5.76E-03 | glio |
| Interaction_genotype_Atr | Mettl6 | 1.969 | 5.76E-03 | glio |
| Interaction_genotype_Bora | Ppp1r3g | -3.428 | 5.76E-03 | glio |
| Interaction_genotype_Bora | Gm20522 | -3.706 | 5.76E-03 | glio |

| | | | | |
|---|---|---|---|---|
| Interaction_genotype_Mtor | Slc26a1 | -3.297 | 5.76E-03 | glio |
| Interaction_genotype_Myc | Gm11772 | -3.125 | 5.76E-03 | glio |
| Interaction_genotype_Myc | Rnf38 | -2.324 | 5.76E-03 | glio |
| Interaction_genotype_Rpl3 | D430040D24R | -3.828 | 5.76E-03 | glio |
| Interaction_genotype_Rpl3 | Gm32585 | -4.073 | 5.76E-03 | glio |
| Interaction_genotype_Rpl3 | Pde11a | -4.074 | 5.76E-03 | glio |
| Interaction_genotype_Rpl3 | Nova2 | -3.702 | 5.76E-03 | glio |
| Interaction_genotype_Rpl3 | Asgr1 | -3.765 | 5.76E-03 | glio |
| Interaction_genotype_Rpl3 | Phtf1os | -3.849 | 5.76E-03 | glio |
| Interaction_genotype_Rpl3 | 1110046J04Ri | -3.938 | 5.76E-03 | glio |
| Interaction_genotype_Rpl3 | Azin2 | -3.487 | 5.76E-03 | glio |
| Interaction_genotype_Rpl3 | Elmsan1 | -2.855 | 5.76E-03 | glio |
| interaction_RT | Tspan8 | 2.604 | 5.76E-03 | glio |
| Atr_in.vivo_noRT | Gm49359 | 1.598 | 5.76E-03 | glio |
| Bard1_ex.vivo_noRT | Arhgef16 | 2.415 | 5.76E-03 | glio |
| Bcas2_ex.vivo_noRT | Bpifa1 | 2.742 | 5.76E-03 | glio |
| Bcas2_ex.vivo_noRT | Cysltr2 | 2.742 | 5.76E-03 | glio |
| Chek1_ex.vivo_noRT | Pmf1 | -1.434 | 5.76E-03 | glio |
| Chek1_ex.vivo_noRT | Drg1 | -0.914 | 5.76E-03 | glio |
| Eif2s1_in.vivo_noRT | B3galt4 | 2.148 | 5.76E-03 | glio |
| Eif2s1_in.vivo_noRT | Spi1 | 2.143 | 5.76E-03 | glio |
| Eif2s1_ex.vivo_noRT | Vtn | 2.234 | 5.76E-03 | glio |
| Fanca_ex.vivo_noRT | Slc14a2 | 2.124 | 5.76E-03 | glio |
| culture_effect | Slpi | 2.007 | 5.76E-04 | glio |
| culture_effect | Wrb | 0.955 | 5.76E-04 | glio |
| Interaction_genotype_Bora | Gm43137 | -4.405 | 5.76E-04 | glio |
| Interaction_genotype_Bora | C4b | 3.802 | 5.76E-04 | glio |
| Interaction_genotype_Bora | Ciita | 3.755 | 5.76E-04 | glio |
| Interaction_genotype_Bora | Lmnb1 | 2.435 | 5.76E-04 | glio |
| interaction_RT | Trim44 | -0.888 | 5.76E-04 | glio |
| Atr_in.vivo_noRT | Frk | 2.451 | 5.76E-04 | glio |
| Bcas2_in.vivo_noRT | Dut | -1.111 | 5.76E-04 | glio |
| Bcas2_ex.vivo_noRT | Mpzl2 | 2.742 | 5.76E-04 | glio |
| Chek1_ex.vivo_noRT | Tgfb1i1 | 2.946 | 5.76E-04 | glio |
| Interaction_genotype_Mtor | Gm16835 | -4.6 | 5.76E-05 | glio |
| Interaction_genotype_Myc | Rilp | -4.931 | 5.76E-05 | glio |
| Interaction_genotype_Myc | Stx1b | -4.188 | 5.76E-05 | glio |
| Aars2_in.vivo_noRT | Acp2 | 1.92 | 5.76E-05 | glio |
| Aars2_in.vivo_noRT | Hsbp1 | -0.972 | 5.76E-05 | glio |
| Aars2_in.vivo_noRT | H3f3a | -0.71 | 5.76E-05 | glio |
| Bcas2_in.vivo_noRT | AC241534.1 | 2.947 | 5.76E-05 | glio |
| Interaction_genotype_Aars2 | Tslp | -5.816 | 5.76E-06 | glio |
| Interaction_genotype_Mtor | Gm47936 | -4.099 | 5.76E-06 | glio |
| Chek1_ex.vivo_noRT | Gprin2 | 3.652 | 5.76E-07 | glio |
| culture_effect | Katna1 | -0.916 | 5.76E-09 | glio |
| Chek1_in.vivo_noRT | Pawr | 3.807 | 5.76E-10 | glio |
| Bcas2_in.vivo_noRT | Rapgef3os2 | 4.807 | 5.76E-20 | glio |
| Chek1_in.vivo_noRT | Gm44736 | 5.044 | 5.76E-26 | glio |
| culture_effect | C4a | -1.585 | 5.77E-03 | glio |
| Interaction_genotype_Atr | 4931431C16R | -2.75 | 5.77E-03 | glio |
| Interaction_genotype_Atr | Gm364 | -2.369 | 5.77E-03 | glio |
| Interaction_genotype_Atr | Il1rap | -2.171 | 5.77E-03 | glio |

| | | | | |
|---|---|---|---|---|
| Interaction_genotype_Chek1 | Nat8l | -3.019 | 5.77E-03 | glio |
| Interaction_genotype_Chek1 | Cbx4 | 2.413 | 5.77E-03 | glio |
| Interaction_genotype_Chek1 | Me2 | -2.3 | 5.77E-03 | glio |
| Interaction_genotype_Chek1 | Prkcsh | 1.664 | 5.77E-03 | glio |
| Interaction_genotype_Ifnar1 | Slc14a2 | 2.734 | 5.77E-03 | glio |
| Interaction_genotype_Ifnar1 | Zbp1 | -1.28 | 5.77E-03 | glio |
| Interaction_genotype_Mtor | Gadl1 | -3.211 | 5.77E-03 | glio |
| Interaction_genotype_Mtor | Cdh24 | -3.635 | 5.77E-03 | glio |
| Interaction_genotype_Mtor | Ucma | -3.686 | 5.77E-03 | glio |
| Interaction_genotype_Mtor | Hectd3 | -2.977 | 5.77E-03 | glio |
| Interaction_genotype_Myc | Gm48960 | -2.906 | 5.77E-03 | glio |
| Interaction_genotype_Myc | Dffa | -2.588 | 5.77E-03 | glio |
| Interaction_genotype_Myc | Adcy6 | -2.35 | 5.77E-03 | glio |
| Interaction_genotype_Myc | Cwc22 | -2.129 | 5.77E-03 | glio |
| Interaction_genotype_Myc | Arid1a | -2.039 | 5.77E-03 | glio |
| Interaction_genotype_Myc | Tmem140 | 2.447 | 5.77E-03 | glio |
| Interaction_genotype_Myc | Sar1b | 1.292 | 5.77E-03 | glio |
| Interaction_genotype_Myc | Sap30l | 1.301 | 5.77E-03 | glio |
| Aars2_in.vivo_noRT | Slc16a2 | 1.902 | 5.77E-03 | glio |
| Aars2_in.vivo_noRT | Iah1 | -1.018 | 5.77E-03 | glio |
| Aars2_ex.vivo_noRT | Clec4d | 2.108 | 5.77E-03 | glio |
| Bcas2_in.vivo_noRT | Rnf11 | -1.223 | 5.77E-03 | glio |
| Bcas2_in.vivo_noRT | Ostf1 | -1 | 5.77E-03 | glio |
| Bora_in.vivo_noRT | AI480526 | 1.987 | 5.77E-03 | glio |
| Bora_in.vivo_noRT | Hdlbp | -0.793 | 5.77E-03 | glio |
| Brca2_in.vivo_noRT | Col5a2 | 1.981 | 5.77E-03 | glio |
| Brca2_in.vivo_noRT | Clec12b | 1.763 | 5.77E-03 | glio |
| Brca2_in.vivo_noRT | Gm14929 | 1.763 | 5.77E-03 | glio |
| Brca2_in.vivo_noRT | Vmn1r32 | 1.763 | 5.77E-03 | glio |
| Brca2_in.vivo_noRT | Glyat | 1.763 | 5.77E-03 | glio |
| Atr_in.vivo_noRT | Ubash3a | 3.582 | 5.77E-04 | glio |
| Bora_in.vivo_noRT | Dnah2os | 2.254 | 5.77E-04 | glio |
| Cenpj_ex.vivo_noRT | Gm29670 | 2.591 | 5.77E-04 | glio |
| Chek1_in.vivo_noRT | Gm32913 | 3.773 | 5.77E-04 | glio |
| Chek1_in.vivo_noRT | Ctss | 2.135 | 5.77E-04 | glio |
| culture_effect | Mpeg1 | -2.335 | 5.77E-05 | glio |
| culture_effect | 4930515G01R | 1.906 | 5.77E-05 | glio |
| culture_effect | Polr2h | -0.582 | 5.77E-05 | glio |
| Chek1_in.vivo_noRT | Gm14419 | 2.522 | 5.77E-05 | glio |
| Atp5c1_in.vivo_noRT | 1700019E08Ri | 2.668 | 5.77E-06 | glio |
| Chek1_in.vivo_noRT | Upk3bl | 2.967 | 5.77E-06 | glio |
| Chek1_ex.vivo_noRT | Gm15559 | 4.045 | 5.77E-06 | glio |
| culture_effect | Tnfrsf1b | -3.229 | 5.77E-08 | glio |
| Chek1_in.vivo_noRT | Ptpn22 | 4.78 | 5.77E-10 | glio |
| Aars2_in.vivo_noRT | Vpreb1 | 4.424 | 5.77E-11 | glio |
| Hsd17b10_in.vivo_noRT | Gm10447 | 3.616 | 5.77E-11 | glio |
| Bora_in.vivo_noRT | Gm13270 | 4.936 | 5.77E-12 | glio |
| Bcas2_in.vivo_noRT | Tmem54 | 4.547 | 5.77E-15 | glio |
| Bora_in.vivo_noRT | Gm28940 | 4.42 | 5.77E-15 | glio |
| Bcas2_in.vivo_noRT | Gm14742 | 4.807 | 5.77E-23 | glio |
| Chek1_in.vivo_noRT | Gm14742 | 5.044 | 5.77E-25 | glio |
| Aars2_in.vivo_noRT | Gm47540 | 5.836 | 5.77E-28 | glio |

| | | | | |
|---|---|---|---|---|
| culture_effect | Glipr2 | -1.814 | 5.78E-03 | glio |
| culture_effect | Gm29243 | -1.251 | 5.78E-03 | glio |
| culture_effect | Hist1h2ab | -1.271 | 5.78E-03 | glio |
| Interaction_genotype_Bcas2 | Gm49808 | -2.458 | 5.78E-03 | glio |
| Interaction_genotype_Bcas2 | Sult6b1 | -3.693 | 5.78E-03 | glio |
| Interaction_genotype_Bora | Upb1 | -3.543 | 5.78E-03 | glio |
| Interaction_genotype_Mtor | Izumo4 | -3.123 | 5.78E-03 | glio |
| Interaction_genotype_Mtor | Bud13 | -2.724 | 5.78E-03 | glio |
| Interaction_genotype_Mtor | P3h2 | -2.314 | 5.78E-03 | glio |
| Interaction_genotype_Myc | Gm43951 | -2.597 | 5.78E-03 | glio |
| Interaction_genotype_Myc | Amph | -2.666 | 5.78E-03 | glio |
| interaction_RT | D630045J12Ri | -1.039 | 5.78E-03 | glio |
| interaction_RT | Sipa1l1 | -0.853 | 5.78E-03 | glio |
| Atr_in.vivo_noRT | Nme1 | -0.6 | 5.78E-03 | glio |
| Bard1_ex.vivo_noRT | H2-DMb2 | 2.814 | 5.78E-03 | glio |
| Chek1_in.vivo_noRT | Upp2 | 1.819 | 5.78E-03 | glio |
| Chek1_in.vivo_noRT | Ypel2 | 1.604 | 5.78E-03 | glio |
| Chek1_in.vivo_noRT | Msi2 | -1.108 | 5.78E-03 | glio |
| Chek1_in.vivo_noRT | Rwdd1 | -0.536 | 5.78E-03 | glio |
| Chek1_ex.vivo_noRT | Cep19 | 1.852 | 5.78E-03 | glio |
| Interaction_genotype_Aars2 | Acoxl | -4.02 | 5.78E-04 | glio |
| Interaction_genotype_Mtor | Lmod1 | -4.37 | 5.78E-04 | glio |
| Interaction_genotype_Mtor | Ppfia3 | -3.996 | 5.78E-04 | glio |
| Interaction_genotype_Myc | Mettl7a3 | -3.674 | 5.78E-04 | glio |
| Interaction_genotype_Myc | Rspo1 | -3.704 | 5.78E-04 | glio |
| Interaction_genotype_Myc | Gm14399 | -3.864 | 5.78E-04 | glio |
| Interaction_genotype_Myc | D3Ertd751e | -3.195 | 5.78E-04 | glio |
| Interaction_genotype_Myc | Tbc1d15 | 1.805 | 5.78E-04 | glio |
| Aars2_in.vivo_noRT | Lncppara | 2.653 | 5.78E-04 | glio |
| Atp5c1_in.vivo_noRT | Gm42849 | 2.402 | 5.78E-04 | glio |
| Atr_in.vivo_noRT | Pdzd4 | 2.045 | 5.78E-04 | glio |
| Bard1_ex.vivo_noRT | Slc25a18 | 2.814 | 5.78E-04 | glio |
| Bcas2_in.vivo_noRT | Nmnat3 | 2.637 | 5.78E-04 | glio |
| Hsd17b10_in.vivo_noRT | Gm26822 | 2.701 | 5.78E-04 | glio |
| culture_effect | Espn | 0.507 | 5.78E-05 | glio |
| culture_effect | Ybx3 | 0.456 | 5.78E-05 | glio |
| Interaction_genotype_Mtor | Ms4a6d | -4.098 | 5.78E-05 | glio |
| Interaction_genotype_Mtor | Zfp932 | -3.359 | 5.78E-05 | glio |
| Aars2_in.vivo_noRT | 1700020L24Ri | 2.666 | 5.78E-05 | glio |
| Aars2_in.vivo_noRT | Msl3 | -1.54 | 5.78E-05 | glio |
| Bora_in.vivo_noRT | Alyref | -1.124 | 5.78E-05 | glio |
| Chek1_in.vivo_noRT | Rab36 | 2.875 | 5.78E-05 | glio |
| Chek1_ex.vivo_noRT | Gm17477 | 3.257 | 5.78E-05 | glio |
| Hsd17b10_in.vivo_noRT | Gm39459 | 2.966 | 5.78E-05 | glio |
| Interaction_genotype_Mtor | Cd248 | -5.238 | 5.78E-06 | glio |
| Interaction_genotype_Mtor | Gm44983 | -4.598 | 5.78E-06 | glio |
| Atr_in.vivo_noRT | Pcdhb19 | 3.242 | 5.78E-07 | glio |
| culture_effect | Aga | -1.119 | 5.78E-09 | glio |
| Chek1_in.vivo_noRT | Gm26673 | 4.023 | 5.78E-09 | glio |
| Hsd17b10_in.vivo_noRT | Gm2016 | 3.616 | 5.78E-12 | glio |
| Bora_in.vivo_noRT | Lyl1 | 4.286 | 5.78E-13 | glio |
| culture_effect | Snx30 | -2.246 | 5.78E-14 | glio |

| | | | | |
|---|---|---|---|---|
| Bora_in.vivo_noRT | Gm45235 | 4.676 | 5.78E-17 | glio |
| culture_effect | Triqk | -1.513 | 5.79E-03 | glio |
| culture_effect | Cyhr1 | 0.36 | 5.79E-03 | glio |
| Interaction_genotype_Aars2 | Csnk1d | 1.604 | 5.79E-03 | glio |
| Interaction_genotype_Aars2 | Cav2 | 1.841 | 5.79E-03 | glio |
| Interaction_genotype_Aars2 | Rars | 1.441 | 5.79E-03 | glio |
| Interaction_genotype_Atr | 1810034E14Ri | -3.375 | 5.79E-03 | glio |
| Interaction_genotype_Bcas2 | Anp32a | 1.316 | 5.79E-03 | glio |
| Interaction_genotype_Mtor | Hist1h2bf | -3.654 | 5.79E-03 | glio |
| Interaction_genotype_Myc | Gm10800 | -3.417 | 5.79E-03 | glio |
| Interaction_genotype_Myc | Rnf32 | -3.043 | 5.79E-03 | glio |
| Interaction_genotype_Myc | Zc3h12b | -2.92 | 5.79E-03 | glio |
| Interaction_genotype_Myc | Ppp1r9a | -2.528 | 5.79E-03 | glio |
| Interaction_genotype_Myc | Dus4l | -2.404 | 5.79E-03 | glio |
| Interaction_genotype_Myc | Hmgcs1 | -2.457 | 5.79E-03 | glio |
| Interaction_genotype_Myc | Rps28 | 1.72 | 5.79E-03 | glio |
| Interaction_genotype_Pars2 | Nrg2 | -3.737 | 5.79E-03 | glio |
| Interaction_genotype_Pars2 | Gm26879 | -3.467 | 5.79E-03 | glio |
| Aars2_ex.vivo_noRT | Gm26789 | 2.312 | 5.79E-03 | glio |
| Atr_in.vivo_noRT | Exosc7 | -0.893 | 5.79E-03 | glio |
| Bcas2_in.vivo_noRT | Snn | 2.071 | 5.79E-03 | glio |
| Bcas2_in.vivo_noRT | Arhgap22 | 1.977 | 5.79E-03 | glio |
| Bcas2_in.vivo_noRT | Cdca3 | -1.184 | 5.79E-03 | glio |
| Bcas2_in.vivo_noRT | Psme2 | -0.686 | 5.79E-03 | glio |
| Bcas2_ex.vivo_noRT | 1700034I23Ri | 1.945 | 5.79E-03 | glio |
| Bora_in.vivo_noRT | Ndst1 | -1.297 | 5.79E-03 | glio |
| Chek1_in.vivo_noRT | Gm15545 | 1.934 | 5.79E-03 | glio |
| Chek1_ex.vivo_noRT | Gm44752 | 2.279 | 5.79E-03 | glio |
| culture_effect | Atp6v0e2 | 0.555 | 5.79E-04 | glio |
| Interaction_genotype_Aars2 | A430106G13R | -4.33 | 5.79E-04 | glio |
| Interaction_genotype_Aars2 | 1810019D21R | -4.148 | 5.79E-04 | glio |
| interaction_RT | Akr1b7 | 3.328 | 5.79E-04 | glio |
| Aars2_in.vivo_noRT | Plekho2 | 2.392 | 5.79E-04 | glio |
| Bora_ex.vivo_noRT | Gm18280 | 2.407 | 5.79E-04 | glio |
| Atp5c1_in.vivo_noRT | Gm14019 | 2.668 | 5.79E-05 | glio |
| Bcas2_in.vivo_noRT | Zfp980 | 3.041 | 5.79E-05 | glio |
| Bcas2_in.vivo_noRT | Lrp4 | 2.891 | 5.79E-05 | glio |
| Bcas2_ex.vivo_noRT | AW112010 | 4.284 | 5.79E-05 | glio |
| Atr_in.vivo_noRT | 4930563E22Ri | 3.285 | 5.79E-06 | glio |
| Atr_in.vivo_noRT | 9430060I03Ri | 3.052 | 5.79E-06 | glio |
| Bcas2_in.vivo_noRT | Slc25a53 | 3.433 | 5.79E-06 | glio |
| Bora_in.vivo_noRT | Zfp990 | 3.035 | 5.79E-06 | glio |
| Chek1_in.vivo_noRT | Olfr267 | 3.269 | 5.79E-06 | glio |
| Chek1_ex.vivo_noRT | Gm15056 | 4.117 | 5.79E-06 | glio |
| Aars2_in.vivo_noRT | 4833418N02R | 3.625 | 5.79E-07 | glio |
| Chek1_ex.vivo_noRT | Adam30 | 3.738 | 5.79E-07 | glio |
| culture_effect | 5031425E22Ri | 1.298 | 5.79E-08 | glio |
| Bora_in.vivo_noRT | Apoh | 4.021 | 5.79E-09 | glio |
| Chek1_ex.vivo_noRT | Dio1 | 4.842 | 5.79E-10 | glio |
| Hsd17b10_in.vivo_noRT | Pyy | 3.616 | 5.79E-10 | glio |
| Aars2_in.vivo_noRT | Gm43258 | 4.467 | 5.79E-11 | glio |
| culture_effect | Steap1 | -1.756 | 5.79E-13 | glio |

| | | | | |
|---|---|---|---|---|
| Atr_in.vivo_noRT | 4930555A03R | 4.795 | 5.79E-14 | glio |
| culture_effect | Itga4 | 2.359 | 5.79E-15 | glio |
| culture_effect | Gpx8 | 1.414 | 5.79E-15 | glio |
| Chek1_in.vivo_noRT | Gm14820 | 4.78 | 5.79E-20 | glio |
| culture_effect | C1galt1c1 | -0.596 | 5.80E-03 | glio |
| culture_effect | Eno3 | -0.541 | 5.80E-03 | glio |
| Interaction_genotype_Atp5c1 | Gm11627 | -4.383 | 5.80E-03 | glio |
| Interaction_genotype_Atp5c1 | Gm42670 | -4.338 | 5.80E-03 | glio |
| Interaction_genotype_Atr | Fbxo15 | -3.088 | 5.80E-03 | glio |
| Interaction_genotype_Atr | Slc45a1 | -2.509 | 5.80E-03 | glio |
| Interaction_genotype_Atr | Boc | -2.499 | 5.80E-03 | glio |
| Interaction_genotype_Atr | Fhl4 | -3.052 | 5.80E-03 | glio |
| Interaction_genotype_Atr | 2210406O10R | -3.326 | 5.80E-03 | glio |
| Interaction_genotype_Atr | Lfng | -3.365 | 5.80E-03 | glio |
| Interaction_genotype_Atr | Gmfb | 1.584 | 5.80E-03 | glio |
| Interaction_genotype_Atr | Sod1 | 1.114 | 5.80E-03 | glio |
| Interaction_genotype_Mtor | Asb11 | -3.187 | 5.80E-03 | glio |
| Interaction_genotype_Mtor | Anln | 2.011 | 5.80E-03 | glio |
| Aars2_in.vivo_noRT | Rbbp9 | 1.596 | 5.80E-03 | glio |
| Atr_in.vivo_noRT | Lmo7 | 1.759 | 5.80E-03 | glio |
| Atr_in.vivo_noRT | Spsb2 | -1.14 | 5.80E-03 | glio |
| Atr_ex.vivo_noRT | Usp11 | 2.864 | 5.80E-03 | glio |
| Brca2_in.vivo_noRT | Tmem132c | 1.763 | 5.80E-03 | glio |
| Brca2_in.vivo_noRT | Gm44693 | 1.763 | 5.80E-03 | glio |
| Eif2s1_in.vivo_noRT | Mpzl2 | 1.883 | 5.80E-03 | glio |
| Eif2s1_in.vivo_noRT | Gm48008 | 1.879 | 5.80E-03 | glio |
| Fanca_ex.vivo_noRT | Gm4340 | 2.124 | 5.80E-03 | glio |
| Fanca_ex.vivo_noRT | Gm16239 | 2.124 | 5.80E-03 | glio |
| Hsd17b10_in.vivo_noRT | Srgn | 2.572 | 5.80E-03 | glio |
| Interaction_genotype_Atr | Smarcd3 | -3.802 | 5.80E-04 | glio |
| Interaction_genotype_Bora | Gm28424 | -4.238 | 5.80E-04 | glio |
| Interaction_genotype_Mtor | Shd | -4.091 | 5.80E-04 | glio |
| Interaction_genotype_Myc | Hrk | -3.962 | 5.80E-04 | glio |
| Atr_in.vivo_noRT | Fabp3 | 2.747 | 5.80E-04 | glio |
| Atr_ex.vivo_noRT | Masp2 | 2.771 | 5.80E-04 | glio |
| Atr_ex.vivo_noRT | Gm16751 | 2.764 | 5.80E-04 | glio |
| Bcas2_in.vivo_noRT | Arsj | 2.693 | 5.80E-04 | glio |
| Bora_in.vivo_noRT | Kcnn4 | -2.715 | 5.80E-04 | glio |
| Chek1_in.vivo_noRT | Gm26631 | 2.602 | 5.80E-04 | glio |
| Aars2_in.vivo_noRT | Gm35315 | 2.825 | 5.80E-05 | glio |
| Aars2_in.vivo_noRT | Gpatch3 | 2.766 | 5.80E-05 | glio |
| Atr_in.vivo_noRT | Tmem117 | 2.906 | 5.80E-05 | glio |
| Atr_in.vivo_noRT | Larp4 | -1.109 | 5.80E-05 | glio |
| Chek1_ex.vivo_noRT | Cxcl16 | 3.356 | 5.80E-05 | glio |
| Aars2_in.vivo_noRT | Gm26737 | 4.76 | 5.80E-06 | glio |
| Bcas2_in.vivo_noRT | Gm15535 | 3.49 | 5.80E-06 | glio |
| Hsd17b10_in.vivo_noRT | Gm48265 | 2.833 | 5.80E-06 | glio |
| Chek1_ex.vivo_noRT | Car9 | 4.449 | 5.80E-07 | glio |
| Bora_in.vivo_noRT | Kctd4 | 3.664 | 5.80E-08 | glio |
| culture_effect | Clmn | -3.926 | 5.80E-11 | glio |
| Atr_in.vivo_noRT | Gm46652 | 4.484 | 5.80E-12 | glio |
| Aars2_in.vivo_noRT | Gm2022 | 5.327 | 5.80E-23 | glio |

| | | | | |
|---|---|---|---|---|
| culture_effect | Pphln1 | 0.46 | 5.81E-03 | glio |
| culture_effect | Nudt9 | -0.461 | 5.81E-03 | glio |
| Interaction_genotype_Aars2 | Gpc5 | -2.429 | 5.81E-03 | glio |
| Interaction_genotype_Aars2 | Tsga10 | -3.582 | 5.81E-03 | glio |
| Interaction_genotype_Myc | Pkd1l2 | -2.861 | 5.81E-03 | glio |
| Interaction_genotype_Rad51 | Tslp | -4.03 | 5.81E-03 | glio |
| Interaction_genotype_Rad51 | Mrps22 | 1.903 | 5.81E-03 | glio |
| Interaction_genotype_Terf1 | Rcan2 | -4.236 | 5.81E-03 | glio |
| Aars2_in.vivo_noRT | Rps15a | -0.588 | 5.81E-03 | glio |
| Atr_in.vivo_noRT | Ndufa3 | -1.269 | 5.81E-03 | glio |
| Atr_ex.vivo_noRT | Slc16a6 | 2.788 | 5.81E-03 | glio |
| Bcas2_in.vivo_noRT | Icam4 | 1.977 | 5.81E-03 | glio |
| Bcas2_in.vivo_noRT | Glipr2 | 2.209 | 5.81E-03 | glio |
| Bcas2_in.vivo_noRT | Atxn3 | -1.409 | 5.81E-03 | glio |
| Bcas2_in.vivo_noRT | Rpl10 | -0.515 | 5.81E-03 | glio |
| Bora_in.vivo_noRT | Ubash3a | 2.985 | 5.81E-03 | glio |
| Bora_in.vivo_noRT | Map1a | 1.59 | 5.81E-03 | glio |
| Bora_in.vivo_noRT | Mrpl45 | -1.062 | 5.81E-03 | glio |
| Bora_in.vivo_noRT | Tnfrsf12a | -1.208 | 5.81E-03 | glio |
| Hsd17b10_in.vivo_noRT | Zfp619 | 2.194 | 5.81E-03 | glio |
| Atr_ex.vivo_noRT | Tec | 3.033 | 5.81E-04 | glio |
| Atr_ex.vivo_noRT | Efr3b | 3.633 | 5.81E-04 | glio |
| Bora_in.vivo_noRT | Ifnb1 | 4.936 | 5.81E-04 | glio |
| Chek1_in.vivo_noRT | Gm10658 | 2.378 | 5.81E-04 | glio |
| Bcas2_in.vivo_noRT | Cstb | -1.442 | 5.81E-05 | glio |
| culture_effect | Arl1 | -0.603 | 5.81E-06 | glio |
| Interaction_genotype_Bcas2 | Scube3 | -4.875 | 5.81E-06 | glio |
| Atr_in.vivo_noRT | Gfpt1 | -1.785 | 5.81E-06 | glio |
| Bora_in.vivo_noRT | Pfkfb2 | 3.3 | 5.81E-06 | glio |
| Bora_in.vivo_noRT | 9530062K07R | 4.557 | 5.81E-10 | glio |
| Bcas2_in.vivo_noRT | Ksr2 | 3.769 | 5.81E-11 | glio |
| Bora_in.vivo_noRT | Gm15127 | 4.676 | 5.81E-12 | glio |
| Chek1_in.vivo_noRT | Kcnj13 | 4.286 | 5.81E-12 | glio |
| Bora_in.vivo_noRT | Gm50115 | 4.673 | 5.81E-13 | glio |
| Interaction_genotype_Aars2 | Kif24 | -3.39 | 5.82E-03 | glio |
| Interaction_genotype_Bcas2 | Pdgfb | -2.965 | 5.82E-03 | glio |
| Interaction_genotype_Ifnar1 | Gm4340 | 2.734 | 5.82E-03 | glio |
| Interaction_genotype_Ifnar1 | Gm16239 | 2.734 | 5.82E-03 | glio |
| Interaction_genotype_Myc | 4930452G13R | -2.854 | 5.82E-03 | glio |
| Interaction_genotype_Pars2 | Vmn2r44 | -2.664 | 5.82E-03 | glio |
| Interaction_genotype_Pars2 | Banp | 2.254 | 5.82E-03 | glio |
| Aars2_in.vivo_noRT | Zcchc14 | 1.559 | 5.82E-03 | glio |
| Aars2_in.vivo_noRT | Slc25a3 | -0.479 | 5.82E-03 | glio |
| Atr_in.vivo_noRT | Hsp90ab1 | -0.584 | 5.82E-03 | glio |
| Brca2_in.vivo_noRT | Tec | 2.329 | 5.82E-03 | glio |
| Chek1_in.vivo_noRT | Kcnmb4 | 2.048 | 5.82E-03 | glio |
| Interaction_genotype_Mtor | Zscan22 | -3.198 | 5.82E-04 | glio |
| Interaction_genotype_Myc | Erich6b | -2.729 | 5.82E-04 | glio |
| Interaction_genotype_Myc | Syngap1 | -3.827 | 5.82E-04 | glio |
| Interaction_genotype_Myc | Zfp772 | -3.978 | 5.82E-04 | glio |
| Interaction_genotype_Myc | Dnajc17 | -3.152 | 5.82E-04 | glio |
| Interaction_genotype_Myc | Fut8 | -2.856 | 5.82E-04 | glio |

| | | | | |
|---|---|---|---|---|
| Interaction_genotype_Myc | Cmc4 | -2.881 | 5.82E-04 | glio |
| Interaction_genotype_Rad21 | Ccl19 | 4.172 | 5.82E-04 | glio |
| Atr_ex.vivo_noRT | Zdhhc14 | 2.769 | 5.82E-04 | glio |
| Chek1_ex.vivo_noRT | Fam71f2 | 3.213 | 5.82E-04 | glio |
| Hsd17b10_in.vivo_noRT | Neurl3 | 2.626 | 5.82E-04 | glio |
| Interaction_genotype_Mtor | Nmnat3 | -5.012 | 5.82E-05 | glio |
| Aars2_in.vivo_noRT | Guk1 | -1.143 | 5.82E-05 | glio |
| Atr_in.vivo_noRT | Jund | -1.376 | 5.82E-05 | glio |
| Chek1_ex.vivo_noRT | Gm19684 | 3.861 | 5.82E-05 | glio |
| Chek1_ex.vivo_noRT | Amhr2 | 4.047 | 5.82E-05 | glio |
| Interaction_genotype_Mtor | Gm40557 | -5.467 | 5.82E-06 | glio |
| Interaction_genotype_Mtor | Gm35696 | -5.105 | 5.82E-06 | glio |
| Aars2_in.vivo_noRT | Foxa1 | 3.474 | 5.82E-06 | glio |
| Aars2_in.vivo_noRT | Gm2415 | 3.283 | 5.82E-06 | glio |
| Bcas2_in.vivo_noRT | Rnf208 | 3.415 | 5.82E-06 | glio |
| Chek1_ex.vivo_noRT | Slc7a4 | 3.861 | 5.82E-06 | glio |
| Interaction_genotype_Mtor | Cct4 | 2.497 | 5.82E-07 | glio |
| Interaction_genotype_Mtor | Matn4 | -5.509 | 5.82E-07 | glio |
| Interaction_genotype_Mtor | Gm34590 | -6.034 | 5.82E-07 | glio |
| Interaction_genotype_Mtor | Rarres1 | -5.87 | 5.82E-07 | glio |
| Interaction_genotype_Mtor | Fbxl13 | -5.214 | 5.82E-07 | glio |
| Chek1_in.vivo_noRT | Aass | 4.042 | 5.82E-10 | glio |
| Aars2_in.vivo_noRT | 4933408J17Ri | 4.01 | 5.82E-11 | glio |
| culture_effect | Cdc123 | -0.344 | 5.83E-03 | glio |
| Interaction_genotype_Atr | 2810457G06R | -3.839 | 5.83E-03 | glio |
| Interaction_genotype_Atr | H3f3aos | -3.366 | 5.83E-03 | glio |
| Interaction_genotype_Atr | Dlx1 | -2.823 | 5.83E-03 | glio |
| Interaction_genotype_Atr | Hspa4l | 1.562 | 5.83E-03 | glio |
| Interaction_genotype_Bcas2 | A930001C03R | -3.535 | 5.83E-03 | glio |
| Interaction_genotype_Bcas2 | Gm26981 | -3.655 | 5.83E-03 | glio |
| Interaction_genotype_Myc | Zfp940 | -3.088 | 5.83E-03 | glio |
| Interaction_genotype_Myc | Ralgps1 | -2.689 | 5.83E-03 | glio |
| Interaction_genotype_Myc | Camk2n2 | -2.445 | 5.83E-03 | glio |
| Interaction_genotype_Pars2 | AW112010 | 4.245 | 5.83E-03 | glio |
| Interaction_genotype_Pars2 | Ddost | 0.874 | 5.83E-03 | glio |
| interaction_RT | Rdm1 | 0.86 | 5.83E-03 | glio |
| Bcas2_in.vivo_noRT | Zfp974 | 2.152 | 5.83E-03 | glio |
| Hsd17b10_in.vivo_noRT | Col6a1 | 2.093 | 5.83E-03 | glio |
| Hsd17b10_in.vivo_noRT | Creld1 | -1.727 | 5.83E-03 | glio |
| Hsd17b10_in.vivo_noRT | Snrpb | -0.465 | 5.83E-03 | glio |
| culture_effect | Map2k2 | 0.4 | 5.83E-04 | glio |
| Aars2_in.vivo_noRT | Tcp1 | -0.81 | 5.83E-04 | glio |
| Bora_in.vivo_noRT | 2610203C22R | 2.607 | 5.83E-04 | glio |
| Chek1_ex.vivo_noRT | Eif4h | -1.316 | 5.83E-04 | glio |
| Interaction_genotype_Myc | Gm17092 | -3.553 | 5.83E-05 | glio |
| Atr_in.vivo_noRT | Zfp39 | 2.888 | 5.83E-05 | glio |
| Atr_in.vivo_noRT | Gm10260 | 3.062 | 5.83E-05 | glio |
| Bora_in.vivo_noRT | Nfix | -1.689 | 5.83E-05 | glio |
| Bora_ex.vivo_noRT | Gata3 | 4.113 | 5.83E-05 | glio |
| Hsd17b10_in.vivo_noRT | Hif3a | 2.974 | 5.83E-05 | glio |
| culture_effect | Stk16 | 0.596 | 5.83E-06 | glio |
| Interaction_genotype_Mtor | 4732463B04R | -4.228 | 5.83E-06 | glio |

| | | | | |
|---|---|---|---|---|
| Bora_in.vivo_noRT | Acot5 | 3.052 | 5.83E-06 | glio |
| Bora_in.vivo_noRT | Tram1 | -1.33 | 5.83E-06 | glio |
| Chek1_in.vivo_noRT | Hist1h4n | 3.497 | 5.83E-06 | glio |
| culture_effect | Nxph3 | 2.594 | 5.83E-07 | glio |
| Chek1_in.vivo_noRT | Gm29340 | 3.728 | 5.83E-08 | glio |
| Bcas2_in.vivo_noRT | Tex52 | 4.157 | 5.83E-10 | glio |
| Bcas2_in.vivo_noRT | Nmb | 4.288 | 5.83E-10 | glio |
| Bora_in.vivo_noRT | Platr31 | 4.936 | 5.83E-10 | glio |
| Chek1_in.vivo_noRT | Upf3a | -2.463 | 5.83E-10 | glio |
| Atr_in.vivo_noRT | Gm26674 | 4.888 | 5.83E-14 | glio |
| Aars2_in.vivo_noRT | Ncf1 | 4.674 | 5.83E-15 | glio |
| Bora_in.vivo_noRT | Gm12841 | 4.936 | 5.83E-22 | glio |
| Interaction_genotype_Atr | Camkk1 | -3.542 | 5.84E-03 | glio |
| Interaction_genotype_Atr | Gm10655 | -3.491 | 5.84E-03 | glio |
| Interaction_genotype_Atr | Xrcc5 | 1.625 | 5.84E-03 | glio |
| Interaction_genotype_Mtor | Ntn5 | -3.298 | 5.84E-03 | glio |
| Interaction_genotype_Myc | Gm43560 | -2.746 | 5.84E-03 | glio |
| Interaction_genotype_Myc | Rpl38 | 1.527 | 5.84E-03 | glio |
| interaction_RT | Stom | 1.634 | 5.84E-03 | glio |
| Atr_in.vivo_noRT | Amigo1 | 1.908 | 5.84E-03 | glio |
| Atr_in.vivo_noRT | Pskh1 | 1.824 | 5.84E-03 | glio |
| Bcas2_in.vivo_noRT | Azin1 | -0.849 | 5.84E-03 | glio |
| Bora_in.vivo_noRT | Hist2h3b | 2.027 | 5.84E-03 | glio |
| Bora_in.vivo_noRT | Gm42726 | 1.943 | 5.84E-03 | glio |
| Bora_in.vivo_noRT | Sun1 | -1.314 | 5.84E-03 | glio |
| Chek1_in.vivo_noRT | Ccdc14 | 1.881 | 5.84E-03 | glio |
| Chek1_in.vivo_noRT | Ppef2 | 1.697 | 5.84E-03 | glio |
| Chek1_in.vivo_noRT | Pcmt1 | -0.67 | 5.84E-03 | glio |
| Chek1_ex.vivo_noRT | Gm10642 | 2.353 | 5.84E-03 | glio |
| Chek1_ex.vivo_noRT | Rnf26 | -1.896 | 5.84E-03 | glio |
| Interaction_genotype_Bora | Vgll3 | -3.827 | 5.84E-04 | glio |
| Interaction_genotype_Mtor | Zfp455 | -4.109 | 5.84E-04 | glio |
| Interaction_genotype_Myc | Tac2 | -2.729 | 5.84E-04 | glio |
| Interaction_genotype_Myc | Gm15270 | -3.26 | 5.84E-04 | glio |
| Aars2_in.vivo_noRT | Gm15492 | 2.5 | 5.84E-04 | glio |
| Atr_in.vivo_noRT | Chtf8 | 2.115 | 5.84E-04 | glio |
| Atr_ex.vivo_noRT | 4930579C12R | 2.769 | 5.84E-04 | glio |
| Atr_ex.vivo_noRT | Cyp2c23 | 2.769 | 5.84E-04 | glio |
| Chek1_in.vivo_noRT | Tmx1 | -1.153 | 5.84E-04 | glio |
| Interaction_genotype_Mtor | Gpd1 | -5.067 | 5.84E-05 | glio |
| Interaction_genotype_Mtor | C1ra | 3.953 | 5.84E-05 | glio |
| Interaction_genotype_Myc | Chd4 | 1.829 | 5.84E-05 | glio |
| Atr_in.vivo_noRT | C1qtnf1 | 2.792 | 5.84E-05 | glio |
| Atr_in.vivo_noRT | Fam3a | 1.873 | 5.84E-05 | glio |
| Atr_ex.vivo_noRT | Gm17210 | 3.749 | 5.84E-05 | glio |
| Bora_in.vivo_noRT | Kyat1 | 3.097 | 5.84E-05 | glio |
| culture_effect | 1190005I06Ri | 1.907 | 5.84E-06 | glio |
| Aars2_in.vivo_noRT | Gm4221 | 3.222 | 5.84E-06 | glio |
| Chek1_ex.vivo_noRT | Neurl1b | 4.045 | 5.84E-06 | glio |
| Chek1_in.vivo_noRT | Cd79a | 3.582 | 5.84E-07 | glio |
| Hsd17b10_in.vivo_noRT | Fgf8 | 3.1 | 5.84E-07 | glio |
| Hsd17b10_in.vivo_noRT | Dcn | 3.225 | 5.84E-07 | glio |

| | | | | |
|---|---|---|---|---|
| culture_effect | Psmb7 | -0.638 | 5.84E-08 | glio |
| Bora_in.vivo_noRT | AU022754 | 3.516 | 5.84E-08 | glio |
| Chek1_in.vivo_noRT | 4930417H01R | 3.933 | 5.84E-08 | glio |
| Atr_in.vivo_noRT | Tacr2 | 4.101 | 5.84E-09 | glio |
| Atr_in.vivo_noRT | Gm49864 | 4.101 | 5.84E-09 | glio |
| Atr_in.vivo_noRT | 1700056N10R | 3.623 | 5.84E-09 | glio |
| Bcas2_in.vivo_noRT | Pawr | 3.819 | 5.84E-10 | glio |
| Bcas2_in.vivo_noRT | Gm49312 | 4.261 | 5.84E-11 | glio |
| Bcas2_in.vivo_noRT | Gm47622 | 4.807 | 5.84E-20 | glio |
| culture_effect | Akr1c13 | -0.921 | 5.85E-03 | glio |
| Interaction_genotype_Aars2 | Slc35g3 | -2.433 | 5.85E-03 | glio |
| Interaction_genotype_Aars2 | Mucl3 | -3.666 | 5.85E-03 | glio |
| Interaction_genotype_Aars2 | Zfhx2os | -3.397 | 5.85E-03 | glio |
| Interaction_genotype_Aars2 | A330084C13R | -3.518 | 5.85E-03 | glio |
| Interaction_genotype_Aars2 | Zscan22 | -2.782 | 5.85E-03 | glio |
| Interaction_genotype_Aars2 | Itprid2 | -2.154 | 5.85E-03 | glio |
| Interaction_genotype_Aars2 | Snx10 | 1.398 | 5.85E-03 | glio |
| Interaction_genotype_Atr | Gm50295 | -3.331 | 5.85E-03 | glio |
| Interaction_genotype_Myc | Sphk1 | -3.207 | 5.85E-03 | glio |
| Interaction_genotype_Myc | Sh3bp2 | -2.954 | 5.85E-03 | glio |
| Interaction_genotype_Myc | Ccdc66 | -2.458 | 5.85E-03 | glio |
| Interaction_genotype_Myc | 2310022B05R | -2.392 | 5.85E-03 | glio |
| Interaction_genotype_Myc | Zfp932 | -2.107 | 5.85E-03 | glio |
| Interaction_genotype_Rpl3 | Gm30082 | -3.849 | 5.85E-03 | glio |
| Interaction_genotype_Rpl3 | Gm9958 | -3.86 | 5.85E-03 | glio |
| Atr_in.vivo_noRT | Crtap | -1.164 | 5.85E-03 | glio |
| Bora_in.vivo_noRT | Tma7 | -0.629 | 5.85E-03 | glio |
| Cenpj_ex.vivo_noRT | Ush1g | 2.192 | 5.85E-03 | glio |
| Chek1_in.vivo_noRT | Lypd6 | 2.056 | 5.85E-03 | glio |
| Chek1_in.vivo_noRT | 4930432K21R | 1.902 | 5.85E-03 | glio |
| Chek1_in.vivo_noRT | Acsl5 | -1.039 | 5.85E-03 | glio |
| Chek1_in.vivo_noRT | Arhgef12 | -0.888 | 5.85E-03 | glio |
| Chek1_in.vivo_noRT | Plekhj1 | -0.765 | 5.85E-03 | glio |
| Chek1_ex.vivo_noRT | Dnmt1 | -1.099 | 5.85E-03 | glio |
| Eif2s1_in.vivo_noRT | Tecrl | 2.411 | 5.85E-03 | glio |
| Aars2_ex.vivo_noRT | Kcnd1 | 2.893 | 5.85E-04 | glio |
| Atr_ex.vivo_noRT | Slco2a1 | 3.164 | 5.85E-04 | glio |
| Bcas2_in.vivo_noRT | 2310001H17R | 2.042 | 5.85E-04 | glio |
| Bora_in.vivo_noRT | Gm2822 | 2.72 | 5.85E-04 | glio |
| Bora_in.vivo_noRT | C030014I23Ri | 2.486 | 5.85E-04 | glio |
| Bora_in.vivo_noRT | Ccdc14 | 2.298 | 5.85E-04 | glio |
| Chek1_ex.vivo_noRT | D830035M03I | 2.996 | 5.85E-04 | glio |
| Chek1_ex.vivo_noRT | Atic | -1.53 | 5.85E-04 | glio |
| culture_effect | Rbpms2 | 1.916 | 5.85E-05 | glio |
| Interaction_genotype_Bora | 5430402O13R | -5.28 | 5.85E-05 | glio |
| Bora_ex.vivo_noRT | Ceacam16 | 3.201 | 5.85E-05 | glio |
| Interaction_genotype_Mtor | Ptp4a1.1 | -5.279 | 5.85E-06 | glio |
| Interaction_genotype_Myc | Slc25a20 | -4.176 | 5.85E-06 | glio |
| Interaction_genotype_Myc | Tnfrsf22 | -4.907 | 5.85E-06 | glio |
| interaction_RT | Myl12b | 0.99 | 5.85E-06 | glio |
| Aars2_ex.vivo_noRT | Gm40538 | 3.293 | 5.85E-06 | glio |
| Atr_in.vivo_noRT | Tomm20 | -0.967 | 5.85E-06 | glio |

| | | | | |
|---|---|---|---|---|
| Atr_in.vivo_noRT | Acox2 | 2.982 | 5.85E-07 | glio |
| Bcas2_in.vivo_noRT | Gm48682 | 3.495 | 5.85E-07 | glio |
| Bcas2_in.vivo_noRT | Stab1 | 3.673 | 5.85E-07 | glio |
| Bcas2_in.vivo_noRT | Gm49785 | 3.638 | 5.85E-07 | glio |
| Bcas2_in.vivo_noRT | Slco2a1 | 3.338 | 5.85E-07 | glio |
| Chek1_in.vivo_noRT | Gm15473 | 3.661 | 5.85E-07 | glio |
| Aars2_in.vivo_noRT | Klra6 | 4.217 | 5.85E-09 | glio |
| Bora_in.vivo_noRT | Zfp14 | 4.413 | 5.85E-10 | glio |
| Bora_in.vivo_noRT | Gm29083 | 4.404 | 5.85E-12 | glio |
| culture_effect | Limk1 | 1.893 | 5.85E-15 | glio |
| Bcas2_in.vivo_noRT | Smtnl1 | 4.807 | 5.85E-19 | glio |
| Interaction_genotype_Mtor | Dr1 | -1.651 | 5.86E-03 | glio |
| Interaction_genotype_Pars2 | Spata31d1a | -3.859 | 5.86E-03 | glio |
| Interaction_genotype_Pars2 | Rgs4 | -2.798 | 5.86E-03 | glio |
| Interaction_genotype_Pars2 | Lrp10 | 1.842 | 5.86E-03 | glio |
| Interaction_genotype_Uqcrc2 | Selenov | -3.425 | 5.86E-03 | glio |
| Chek1_ex.vivo_noRT | Actb | -0.751 | 5.86E-03 | glio |
| Cyp19a1_ex.vivo_noRT | Gm16933 | 2.159 | 5.86E-03 | glio |
| Hsd17b10_in.vivo_noRT | Wdhd1 | -1.882 | 5.86E-03 | glio |
| Hsd17b10_in.vivo_noRT | Hgh1 | -1.325 | 5.86E-03 | glio |
| Interaction_genotype_Myc | Scn2a | -3.829 | 5.86E-04 | glio |
| Interaction_genotype_Myc | Mettl5os | -3.571 | 5.86E-04 | glio |
| Interaction_genotype_Myc | Zfp691 | -3.394 | 5.86E-04 | glio |
| Interaction_genotype_Myc | Eefsec | -2.736 | 5.86E-04 | glio |
| Interaction_genotype_Myc | Fam219b | -2.645 | 5.86E-04 | glio |
| Interaction_genotype_Myc | Rad23a | 1.914 | 5.86E-04 | glio |
| Atp5c1_in.vivo_noRT | Trim16 | 2.408 | 5.86E-04 | glio |
| Bora_in.vivo_noRT | Acacb | 2.434 | 5.86E-04 | glio |
| Cenpj_ex.vivo_noRT | Cenpj | -2.983 | 5.86E-04 | glio |
| Eif2s1_in.vivo_noRT | Gm13449 | 2.143 | 5.86E-04 | glio |
| Bora_in.vivo_noRT | Rarres2 | 2.792 | 5.86E-05 | glio |
| Eif2s1_in.vivo_noRT | Pik3ap1 | 2.934 | 5.86E-05 | glio |
| Interaction_genotype_Atr | Idnk | 2.735 | 5.86E-06 | glio |
| Atr_in.vivo_noRT | Gm42670 | 2.862 | 5.86E-06 | glio |
| Aars2_in.vivo_noRT | Gm48357 | 3.011 | 5.86E-07 | glio |
| Bora_in.vivo_noRT | Cdo1 | 3.649 | 5.86E-07 | glio |
| Atr_in.vivo_noRT | Gm7008 | 4.068 | 5.86E-09 | glio |
| Aars2_in.vivo_noRT | 9030612E09Ri | 4.006 | 5.86E-10 | glio |
| culture_effect | Fads3 | 1.619 | 5.86E-11 | glio |
| culture_effect | Plod2 | -1.329 | 5.86E-11 | glio |
| Bcas2_in.vivo_noRT | Gm32250 | 4.024 | 5.86E-15 | glio |
| Aars2_in.vivo_noRT | 4930470G03R | 5.06 | 5.86E-20 | glio |
| culture_effect | Cfh | -1.315 | 5.87E-03 | glio |
| Interaction_genotype_Bora | E230029C05Ri | -3.335 | 5.87E-03 | glio |
| Interaction_genotype_Mtor | Srgn | -4.036 | 5.87E-03 | glio |
| Interaction_genotype_Mtor | Gbp2 | 2.215 | 5.87E-03 | glio |
| Interaction_genotype_Myc | Crybg2 | -3.059 | 5.87E-03 | glio |
| Interaction_genotype_Myc | Gm3550 | -3.067 | 5.87E-03 | glio |
| Interaction_genotype_Myc | Polr2f | 1.017 | 5.87E-03 | glio |
| Interaction_genotype_Rpl3 | Gm20455 | -4.033 | 5.87E-03 | glio |
| Interaction_genotype_Rpl3 | Add1 | 1.94 | 5.87E-03 | glio |
| Atr_in.vivo_noRT | Galk2 | 1.407 | 5.87E-03 | glio |

| | | | | |
|---|---|---|---|---|
| Atr_in.vivo_noRT | Timm8b | -0.829 | 5.87E-03 | glio |
| Atr_ex.vivo_noRT | Eva1c | 2.713 | 5.87E-03 | glio |
| Atr_ex.vivo_noRT | Mms19 | -1.744 | 5.87E-03 | glio |
| Bard1_ex.vivo_noRT | E030037K01Ri | 2.529 | 5.87E-03 | glio |
| Bcas2_in.vivo_noRT | Card14 | 2.095 | 5.87E-03 | glio |
| Bcas2_ex.vivo_noRT | Fst | 2.742 | 5.87E-03 | glio |
| Bora_in.vivo_noRT | Hcn1 | 2.194 | 5.87E-03 | glio |
| Bora_in.vivo_noRT | Gm43328 | 2.088 | 5.87E-03 | glio |
| Bora_in.vivo_noRT | 4930599N23R | 1.86 | 5.87E-03 | glio |
| Chek1_in.vivo_noRT | Tlx3 | 1.998 | 5.87E-03 | glio |
| culture_effect | Parvb | 1.48 | 5.87E-04 | glio |
| Interaction_genotype_Aars2 | BC052040 | -3.546 | 5.87E-04 | glio |
| Interaction_genotype_Myc | Sult5a1 | -4.18 | 5.87E-04 | glio |
| Interaction_genotype_Myc | Wdr5 | -2.302 | 5.87E-04 | glio |
| Atr_in.vivo_noRT | Gng7 | 2.345 | 5.87E-04 | glio |
| Bcas2_in.vivo_noRT | Abcd4 | 2.2 | 5.87E-04 | glio |
| Bcas2_in.vivo_noRT | Wtap | -1.415 | 5.87E-04 | glio |
| Bora_in.vivo_noRT | Lockd | -2.01 | 5.87E-04 | glio |
| Bora_in.vivo_noRT | Smarcad1 | -1.409 | 5.87E-04 | glio |
| Chek1_in.vivo_noRT | Cst3 | -0.968 | 5.87E-04 | glio |
| Aars2_in.vivo_noRT | Llgl2 | 2.792 | 5.87E-05 | glio |
| Atp5c1_in.vivo_noRT | Hnf1aos1 | 2.668 | 5.87E-05 | glio |
| Atr_ex.vivo_noRT | Gm45336 | 3.164 | 5.87E-05 | glio |
| Bora_ex.vivo_noRT | Ush1g | 2.802 | 5.87E-05 | glio |
| culture_effect | Gja4 | -2.738 | 5.87E-06 | glio |
| Atr_in.vivo_noRT | Tnfrsf1b | 3.239 | 5.87E-07 | glio |
| Bcas2_in.vivo_noRT | Zfp273 | 3.57 | 5.87E-07 | glio |
| Bora_in.vivo_noRT | Pla2g7 | 3.749 | 5.87E-08 | glio |
| Chek1_in.vivo_noRT | Cldn19 | 4.261 | 5.87E-10 | glio |
| Chek1_in.vivo_noRT | AV039307 | 4.653 | 5.87E-11 | glio |
| Hsd17b10_in.vivo_noRT | C330002G04R | 3.616 | 5.87E-12 | glio |
| Hsd17b10_in.vivo_noRT | Rapgef3os2 | 3.616 | 5.87E-12 | glio |
| Eif2s1_in.vivo_noRT | Eif2s1 | -2.176 | 5.87E-27 | glio |
| Interaction_genotype_Aars2 | Ninj1 | 1.532 | 5.88E-03 | glio |
| Interaction_genotype_Mtor | Cep112it | -3.385 | 5.88E-03 | glio |
| Interaction_genotype_Myc | Ktn1 | 1.452 | 5.88E-03 | glio |
| interaction_RT | Plp2 | 0.7 | 5.88E-03 | glio |
| Aars2_in.vivo_noRT | Napepld | 1.761 | 5.88E-03 | glio |
| Aars2_in.vivo_noRT | Ttf2 | 1.368 | 5.88E-03 | glio |
| Atr_in.vivo_noRT | B4galnt4 | 1.911 | 5.88E-03 | glio |
| Atr_in.vivo_noRT | Hectd3 | 1.822 | 5.88E-03 | glio |
| Atr_ex.vivo_noRT | Gm49579 | 2.376 | 5.88E-03 | glio |
| Bcas2_in.vivo_noRT | Pigo | 1.819 | 5.88E-03 | glio |
| Cenpj_in.vivo_noRT | Usp17lc | 2.022 | 5.88E-03 | glio |
| Chek1_in.vivo_noRT | Tbpl1 | -1.083 | 5.88E-03 | glio |
| Chek1_in.vivo_noRT | Eef2 | -0.456 | 5.88E-03 | glio |
| Hsd17b10_in.vivo_noRT | Fam169a | 2.056 | 5.88E-03 | glio |
| culture_effect | AC166172.1 | 1.65 | 5.88E-04 | glio |
| Interaction_genotype_Chek1 | Fth1 | 1.672 | 5.88E-04 | glio |
| Interaction_genotype_Mtor | Rps24 | 1.185 | 5.88E-04 | glio |
| Interaction_genotype_Myc | Timm23 | 1.323 | 5.88E-04 | glio |
| Atr_ex.vivo_noRT | Card9 | 2.764 | 5.88E-04 | glio |

| | | | | |
|---|---|---|---|---|
| Bora_in.vivo_noRT | Ptpn1 | -1.633 | 5.88E-04 | glio |
| Interaction_genotype_Mtor | Gm37214 | -4.935 | 5.88E-05 | glio |
| interaction_RT | Aarsd1 | -1 | 5.88E-05 | glio |
| Aars2_in.vivo_noRT | Kank3 | 2.714 | 5.88E-05 | glio |
| Bcas2_in.vivo_noRT | Gm10638 | 2.84 | 5.88E-05 | glio |
| Bcas2_in.vivo_noRT | G430095P16R | 2.882 | 5.88E-05 | glio |
| Chek1_ex.vivo_noRT | Ccpg1 | 2.279 | 5.88E-05 | glio |
| culture_effect | Bglap3 | -2.86 | 5.88E-06 | glio |
| culture_effect | Znhit2 | 0.807 | 5.88E-06 | glio |
| culture_effect | Ifi203 | -0.992 | 5.88E-06 | glio |
| Atp5c1_in.vivo_noRT | Gm44899 | 2.668 | 5.88E-06 | glio |
| Atp5c1_in.vivo_noRT | Cd200r2 | 2.404 | 5.88E-06 | glio |
| culture_effect | Ilf2 | -0.755 | 5.88E-08 | glio |
| Aars2_in.vivo_noRT | Sptbn4 | 3.778 | 5.88E-08 | glio |
| Bora_in.vivo_noRT | Aqp3 | 4.672 | 5.88E-08 | glio |
| Bora_in.vivo_noRT | Cnih2 | 3.748 | 5.88E-08 | glio |
| Chek1_in.vivo_noRT | Gm49694 | 4.318 | 5.88E-09 | glio |
| Atr_in.vivo_noRT | Gm20478 | 3.961 | 5.88E-10 | glio |
| Atr_in.vivo_noRT | Cfap58 | 5.533 | 5.88E-11 | glio |
| Atr_in.vivo_noRT | E230001N04R | 3.947 | 5.88E-11 | glio |
| Bora_in.vivo_noRT | Tcstv3 | 4.936 | 5.88E-14 | glio |
| Atr_in.vivo_noRT | Nxph3 | 6.325 | 5.88E-22 | glio |
| Interaction_genotype_Aars2 | Sema3e | -3.038 | 5.89E-03 | glio |
| Interaction_genotype_Aars2 | Ppp2ca | 0.857 | 5.89E-03 | glio |
| Interaction_genotype_Mtor | C1qtnf1 | -3.369 | 5.89E-03 | glio |
| Interaction_genotype_Mtor | Gm12981 | -3.37 | 5.89E-03 | glio |
| Interaction_genotype_Mtor | Zfp13 | -2.76 | 5.89E-03 | glio |
| Interaction_genotype_Myc | Gm50255 | -3.774 | 5.89E-03 | glio |
| Interaction_genotype_Myc | Gm36738 | -3.042 | 5.89E-03 | glio |
| Interaction_genotype_Myc | Chrna1os | -2.371 | 5.89E-03 | glio |
| Interaction_genotype_Pars2 | Gpc5 | -2.664 | 5.89E-03 | glio |
| interaction_RT | Sema6a | 1.619 | 5.89E-03 | glio |
| Atp5c1_in.vivo_noRT | Gm4724 | 2.012 | 5.89E-03 | glio |
| Atr_in.vivo_noRT | Rpl37 | -0.744 | 5.89E-03 | glio |
| Bard1_ex.vivo_noRT | Col5a2 | 2.227 | 5.89E-03 | glio |
| Bora_in.vivo_noRT | Sag | 2.081 | 5.89E-03 | glio |
| Bora_ex.vivo_noRT | Gstp3 | 2.806 | 5.89E-03 | glio |
| Chek1_ex.vivo_noRT | Gm16124 | 2.89 | 5.89E-03 | glio |
| Hsd17b10_in.vivo_noRT | Lrp1b | 2.148 | 5.89E-03 | glio |
| Hsd17b10_in.vivo_noRT | Vcl | -1.494 | 5.89E-03 | glio |
| Interaction_genotype_Mtor | Ankef1 | -4.13 | 5.89E-04 | glio |
| Interaction_genotype_Mtor | Ube2s | 1.666 | 5.89E-04 | glio |
| Interaction_genotype_Myc | 1110006O24R | -3.7 | 5.89E-04 | glio |
| Interaction_genotype_Myc | Brpf3 | -3.379 | 5.89E-04 | glio |
| Interaction_genotype_Myc | Kat8 | -2.697 | 5.89E-04 | glio |
| Interaction_genotype_Myc | Polr2k | 1.376 | 5.89E-04 | glio |
| Atr_in.vivo_noRT | Gm16973 | 2.459 | 5.89E-04 | glio |
| Cenpj_ex.vivo_noRT | Fabp6 | 2.591 | 5.89E-04 | glio |
| Chek1_in.vivo_noRT | Cdkn1c | 2.475 | 5.89E-04 | glio |
| Chek1_in.vivo_noRT | Gpn1 | -1.579 | 5.89E-04 | glio |
| Chek1_ex.vivo_noRT | Gm48529 | 3.063 | 5.89E-04 | glio |
| Hsd17b10_in.vivo_noRT | Gm48933 | 2.614 | 5.89E-04 | glio |

| | | | | |
|---|---|---|---|---|
| Hsd17b10_in.vivo_noRT | Tnc | 3.088 | 5.89E-04 | glio |
| Aars2_in.vivo_noRT | C030037D09R | 2.825 | 5.89E-05 | glio |
| Chek1_in.vivo_noRT | Rasal1 | 3.568 | 5.89E-06 | glio |
| Atr_ex.vivo_noRT | Halr1 | 3.164 | 5.89E-08 | glio |
| Bcas2_in.vivo_noRT | Ccdc114 | 3.889 | 5.89E-09 | glio |
| Bcas2_in.vivo_noRT | AV039307 | 4.416 | 5.89E-10 | glio |
| culture_effect | K230015D01R | 3.566 | 5.89E-11 | glio |
| culture_effect | Cyp2j9 | -1.884 | 5.89E-11 | glio |
| Chek1_in.vivo_noRT | B230311B06R | 4.78 | 5.89E-11 | glio |
| Atr_in.vivo_noRT | Isl1 | 4.73 | 5.89E-12 | glio |
| Chek1_ex.vivo_noRT | Gm21319 | 4.842 | 5.89E-13 | glio |
| culture_effect | Itga2 | 1.078 | 5.90E-03 | glio |
| culture_effect | Mtfr1 | 0.509 | 5.90E-03 | glio |
| Interaction_genotype_Aars2 | Cysrt1 | -3.53 | 5.90E-03 | glio |
| Interaction_genotype_Aars2 | Gm26621 | -3.456 | 5.90E-03 | glio |
| Interaction_genotype_Aars2 | 4930426I24Ri | -3.289 | 5.90E-03 | glio |
| Interaction_genotype_Aars2 | Zfp37 | -3.36 | 5.90E-03 | glio |
| Interaction_genotype_Aars2 | 9130015A21R | -5.09 | 5.90E-03 | glio |
| Interaction_genotype_Aars2 | Atf6 | 2.146 | 5.90E-03 | glio |
| Interaction_genotype_Mtor | 5830428M24F | -3.247 | 5.90E-03 | glio |
| Interaction_genotype_Mtor | Ngly1 | -3.118 | 5.90E-03 | glio |
| Interaction_genotype_Mtor | Rps23 | 0.922 | 5.90E-03 | glio |
| Interaction_genotype_Myc | Amy1 | -2.988 | 5.90E-03 | glio |
| Interaction_genotype_Myc | Riox1 | -2.72 | 5.90E-03 | glio |
| Interaction_genotype_Myc | Miip | -2.357 | 5.90E-03 | glio |
| Interaction_genotype_Rad51 | Gm27253 | -3.895 | 5.90E-03 | glio |
| Interaction_genotype_Rad51 | Gm16364.1 | -4.265 | 5.90E-03 | glio |
| Interaction_genotype_Rad51 | Wars | 2.381 | 5.90E-03 | glio |
| Aars2_in.vivo_noRT | Psmd1 | -1.007 | 5.90E-03 | glio |
| Aars2_in.vivo_noRT | Rps14 | -0.569 | 5.90E-03 | glio |
| Atr_in.vivo_noRT | Gm43597 | 1.757 | 5.90E-03 | glio |
| Atr_in.vivo_noRT | Nop56 | -0.842 | 5.90E-03 | glio |
| Atr_ex.vivo_noRT | Gm34643 | 2.85 | 5.90E-03 | glio |
| Chek1_in.vivo_noRT | Fhod1 | 1.708 | 5.90E-03 | glio |
| Chek1_in.vivo_noRT | Psmd7 | -0.705 | 5.90E-03 | glio |
| Interaction_genotype_Atr | Rims3 | -3.898 | 5.90E-04 | glio |
| Interaction_genotype_Atr | Btg1 | 1.91 | 5.90E-04 | glio |
| Interaction_genotype_Mtor | Lurap1 | -4.095 | 5.90E-04 | glio |
| Interaction_genotype_Myc | 2410141K09R | -3.25 | 5.90E-04 | glio |
| Interaction_genotype_Myc | Fam129c | -3.922 | 5.90E-04 | glio |
| Interaction_genotype_Myc | Enox2 | -3.322 | 5.90E-04 | glio |
| Interaction_genotype_Myc | Psd3 | -3.256 | 5.90E-04 | glio |
| Atp5c1_in.vivo_noRT | Thsd1 | 2.281 | 5.90E-04 | glio |
| Atr_in.vivo_noRT | Rfc3 | -1.452 | 5.90E-04 | glio |
| Bcas2_in.vivo_noRT | 8430429K09R | 2.016 | 5.90E-04 | glio |
| Bcas2_in.vivo_noRT | Gm11110 | 2.889 | 5.90E-05 | glio |
| Bora_in.vivo_noRT | Gm11767 | 3.014 | 5.90E-05 | glio |
| Aars2_in.vivo_noRT | Gm13421 | 3.602 | 5.90E-07 | glio |
| culture_effect | Lamp2 | -0.642 | 5.90E-08 | glio |
| culture_effect | Bod1 | 0.852 | 5.90E-09 | glio |
| Chek1_in.vivo_noRT | Pygo1 | 4.26 | 5.90E-11 | glio |
| Chek1_in.vivo_noRT | Gm49874 | 5.044 | 5.90E-12 | glio |

| | | | | |
|---|---|---|---|---|
| Chek1_in.vivo_noRT | Gm10696 | 5.044 | 5.90E-13 | glio |
| Atr_in.vivo_noRT | Smpd5 | 5.414 | 5.90E-18 | glio |
| culture_effect | Slc2a1 | -2.523 | 5.90E-24 | glio |
| culture_effect | Gm48427 | -1.408 | 5.91E-03 | glio |
| culture_effect | Hist1h4a | 1.301 | 5.91E-03 | glio |
| Interaction_genotype_Aars2 | F630040K05Ri | -3.108 | 5.91E-03 | glio |
| Interaction_genotype_Aars2 | Gm11491 | -3.505 | 5.91E-03 | glio |
| Interaction_genotype_Aars2 | Glt1d1 | -3.222 | 5.91E-03 | glio |
| Interaction_genotype_Cdkn2a | Gm16933 | 2.83 | 5.91E-03 | glio |
| Interaction_genotype_Hsd17b | Krt18 | 6.027 | 5.91E-03 | glio |
| Interaction_genotype_Hsd17b | Gm30938 | -4.14 | 5.91E-03 | glio |
| Interaction_genotype_Hsd17b | Nxph3 | -3.967 | 5.91E-03 | glio |
| Interaction_genotype_Mtor | Hpse | -3.353 | 5.91E-03 | glio |
| Interaction_genotype_Mtor | Tmem177 | -2.937 | 5.91E-03 | glio |
| Interaction_genotype_Mtor | Smim8 | 1.572 | 5.91E-03 | glio |
| Interaction_genotype_Myc | Ubl5 | 1.029 | 5.91E-03 | glio |
| Aars2_in.vivo_noRT | BC024063 | 1.924 | 5.91E-03 | glio |
| Aars2_in.vivo_noRT | Itprip | 1.768 | 5.91E-03 | glio |
| Atr_in.vivo_noRT | Rp2 | 1.774 | 5.91E-03 | glio |
| Bard1_ex.vivo_noRT | Rasl10a | 2.421 | 5.91E-03 | glio |
| Bard1_ex.vivo_noRT | Zdhhc14 | 2.419 | 5.91E-03 | glio |
| Bard1_ex.vivo_noRT | 9530080O11R | 2.692 | 5.91E-03 | glio |
| Bcas2_in.vivo_noRT | Rassf4 | -1.624 | 5.91E-03 | glio |
| Cyp19a1_ex.vivo_noRT | Tff2 | 2.959 | 5.91E-03 | glio |
| Cyp19a1_ex.vivo_noRT | Slc26a1 | 3.034 | 5.91E-03 | glio |
| Cyp19a1_ex.vivo_noRT | Gm47510 | 2.159 | 5.91E-03 | glio |
| Cyp19a1_ex.vivo_noRT | Gm5788 | 2.159 | 5.91E-03 | glio |
| Eif2s1_in.vivo_noRT | Gm16741 | 2.143 | 5.91E-03 | glio |
| Hsd17b10_in.vivo_noRT | Tspan9 | -1.609 | 5.91E-03 | glio |
| Interaction_genotype_Myc | Gask1b | -4.106 | 5.91E-04 | glio |
| Interaction_genotype_Myc | Eva1b | -2.63 | 5.91E-04 | glio |
| Aars2_ex.vivo_noRT | Ric3 | 2.899 | 5.91E-04 | glio |
| Bard1_ex.vivo_noRT | Gm17203 | 2.415 | 5.91E-04 | glio |
| Bora_in.vivo_noRT | Ccdc57 | 2.59 | 5.91E-04 | glio |
| culture_effect | Rab34 | -0.468 | 5.91E-05 | glio |
| Aars2_in.vivo_noRT | Elovl3 | 2.875 | 5.91E-05 | glio |
| Aars2_ex.vivo_noRT | Gm38477 | 2.893 | 5.91E-05 | glio |
| Bora_in.vivo_noRT | 1700010I14Ri | 2.799 | 5.91E-05 | glio |
| Chek1_in.vivo_noRT | Folr2 | 2.765 | 5.91E-05 | glio |
| Bora_in.vivo_noRT | Tox2 | 3.161 | 5.91E-06 | glio |
| Bora_in.vivo_noRT | Pla2g4b | 3.108 | 5.91E-06 | glio |
| Bora_in.vivo_noRT | Gm49164 | 3.269 | 5.91E-06 | glio |
| Bora_in.vivo_noRT | Gm43773 | 3.039 | 5.91E-06 | glio |
| Interaction_genotype_Mtor | Gm14379 | -5.724 | 5.91E-07 | glio |
| Interaction_genotype_Mtor | Ube4bos1 | -5.021 | 5.91E-07 | glio |
| Atp5c1_in.vivo_noRT | Serpinb9g | 2.668 | 5.91E-07 | glio |
| Bora_in.vivo_noRT | Gm38979 | 3.762 | 5.91E-07 | glio |
| Chek1_in.vivo_noRT | Casz1 | 3.768 | 5.91E-07 | glio |
| Bcas2_in.vivo_noRT | 9430037G07R | 3.899 | 5.91E-08 | glio |
| Chek1_ex.vivo_noRT | A930033M14F | 3.877 | 5.91E-08 | glio |
| Atr_in.vivo_noRT | Gm47484 | 4.54 | 5.91E-10 | glio |
| Chek1_ex.vivo_noRT | Cys1 | 4.449 | 5.91E-10 | glio |

| | | | | |
|---|---|---|---|---|
| Aars2_in.vivo_noRT | 5033403F01Ri | 4.401 | 5.91E-11 | glio |
| Atr_ex.vivo_noRT | Gm48155 | 3.947 | 5.91E-12 | glio |
| Hsd17b10_in.vivo_noRT | Gm47622 | 3.616 | 5.91E-12 | glio |
| Atr_in.vivo_noRT | Gm16741 | 5.533 | 5.91E-18 | glio |
| Interaction_genotype_Aars2 | Gm26688 | -3.452 | 5.92E-03 | glio |
| Interaction_genotype_Bcas2 | Gm16364.1 | -3.92 | 5.92E-03 | glio |
| Interaction_genotype_Mtor | Ifitm1 | -3.833 | 5.92E-03 | glio |
| Interaction_genotype_Mtor | Mrpl23 | 1.342 | 5.92E-03 | glio |
| Interaction_genotype_Rpl3 | Cdca8 | 2.069 | 5.92E-03 | glio |
| Interaction_genotype_Uqcrc2 | Gm47851 | -3.781 | 5.92E-03 | glio |
| Aars2_in.vivo_noRT | Fam83a | 2.265 | 5.92E-03 | glio |
| Atr_in.vivo_noRT | Dsel | 1.703 | 5.92E-03 | glio |
| Bcas2_in.vivo_noRT | Cd109 | -0.931 | 5.92E-03 | glio |
| Bcas2_ex.vivo_noRT | Rasl12 | 2.742 | 5.92E-03 | glio |
| Brca2_in.vivo_noRT | Gm30692 | 1.763 | 5.92E-03 | glio |
| Cenpj_in.vivo_noRT | Slc26a10 | -1.831 | 5.92E-03 | glio |
| Chek1_ex.vivo_noRT | Nhp2 | -0.737 | 5.92E-03 | glio |
| culture_effect | Coasy | -0.667 | 5.92E-04 | glio |
| Interaction_genotype_Aars2 | Dntt | -4.437 | 5.92E-04 | glio |
| Interaction_genotype_Aars2 | Kdm2b | -3.186 | 5.92E-04 | glio |
| Interaction_genotype_Mtor | Rnase4 | -4.216 | 5.92E-04 | glio |
| Atr_in.vivo_noRT | Rgs5 | 2.963 | 5.92E-04 | glio |
| Bcas2_in.vivo_noRT | Gm15601 | 2.595 | 5.92E-04 | glio |
| Bora_in.vivo_noRT | Cenph | -1.64 | 5.92E-04 | glio |
| Bora_in.vivo_noRT | Top2a | -1.861 | 5.92E-04 | glio |
| Chek1_in.vivo_noRT | Gm15910 | 2.394 | 5.92E-04 | glio |
| Chek1_in.vivo_noRT | 6330403L08Ri | 2.383 | 5.92E-04 | glio |
| culture_effect | Gm4166 | -1.862 | 5.92E-05 | glio |
| culture_effect | BC025920 | 1.551 | 5.92E-05 | glio |
| Bard1_ex.vivo_noRT | Spink8 | 2.814 | 5.92E-05 | glio |
| Bard1_ex.vivo_noRT | Psmb11 | 2.814 | 5.92E-05 | glio |
| Bcas2_in.vivo_noRT | B430010I23Ri | 2.721 | 5.92E-05 | glio |
| Bcas2_in.vivo_noRT | E430018J23Ri | 2.743 | 5.92E-05 | glio |
| Chek1_in.vivo_noRT | Mfsd3 | 2.381 | 5.92E-05 | glio |
| Chek1_ex.vivo_noRT | Gm50321 | 2.946 | 5.92E-05 | glio |
| culture_effect | 5430402O13R | 2.61 | 5.92E-06 | glio |
| Interaction_genotype_Mtor | 9530080O11R | -5.26 | 5.92E-06 | glio |
| Atr_in.vivo_noRT | 1700086P04R | 3.124 | 5.92E-06 | glio |
| Interaction_genotype_Mtor | Gm5415 | -4.099 | 5.92E-07 | glio |
| Chek1_in.vivo_noRT | Tnfrsf4 | 3.458 | 5.92E-07 | glio |
| Aars2_in.vivo_noRT | Gm15473 | 3.944 | 5.92E-08 | glio |
| Aars2_ex.vivo_noRT | Krt19 | 3.293 | 5.92E-08 | glio |
| Chek1_in.vivo_noRT | Sema4a | 4 | 5.92E-08 | glio |
| Bora_in.vivo_noRT | Flnc | 4.411 | 5.92E-10 | glio |
| Chek1_in.vivo_noRT | Rem2 | 3.859 | 5.92E-10 | glio |
| culture_effect | Nxph4 | 3.448 | 5.92E-11 | glio |
| culture_effect | Plekha7 | -2.007 | 5.92E-11 | glio |
| culture_effect | Uqcc2 | 0.741 | 5.92E-11 | glio |
| Aars2_in.vivo_noRT | 4930478M13F | 4.947 | 5.92E-12 | glio |
| Aars2_in.vivo_noRT | Gm44752 | 5.449 | 5.92E-18 | glio |
| Aars2_in.vivo_noRT | Gm20705 | 4.795 | 5.92E-18 | glio |
| Bora_in.vivo_noRT | Zscan4d | 4.936 | 5.92E-18 | glio |

| | | | | |
|---|---|---|---|---|
| Aars2_in.vivo_noRT | Selenov | 5.327 | 5.92E-20 | glio |
| Chek1_in.vivo_noRT | Speer4e | 5.044 | 5.92E-20 | glio |
| Interaction_genotype_Aars2 | Phtf1os | -3.328 | 5.93E-03 | glio |
| Interaction_genotype_Atr | Necab3 | -3.389 | 5.93E-03 | glio |
| Interaction_genotype_Atr | Oas1g | -3.185 | 5.93E-03 | glio |
| Interaction_genotype_Bcas2 | Pakap | -2.912 | 5.93E-03 | glio |
| Interaction_genotype_Bcas2 | Gm50147 | -3.549 | 5.93E-03 | glio |
| Interaction_genotype_Mtor | Gm43329 | -3.203 | 5.93E-03 | glio |
| Interaction_genotype_Myc | Hpdl | -2.914 | 5.93E-03 | glio |
| Atr_in.vivo_noRT | Gm43260 | 1.796 | 5.93E-03 | glio |
| Bard1_ex.vivo_noRT | Trak2 | -1.76 | 5.93E-03 | glio |
| Bcas2_in.vivo_noRT | 2610002M06F | -1.225 | 5.93E-03 | glio |
| Cenpj_ex.vivo_noRT | Il12rb1 | 2.587 | 5.93E-03 | glio |
| Cenpj_ex.vivo_noRT | Irf8 | 2.591 | 5.93E-03 | glio |
| Eif2s1_in.vivo_noRT | Cysrt1 | 2.407 | 5.93E-03 | glio |
| Eif2s1_in.vivo_noRT | 1700108F19Ri | 1.879 | 5.93E-03 | glio |
| Hsd17b10_in.vivo_noRT | Gask1b | 2.342 | 5.93E-03 | glio |
| culture_effect | Lman2 | -0.464 | 5.93E-04 | glio |
| Interaction_genotype_Bcas2 | Gm30082 | -4.508 | 5.93E-04 | glio |
| Interaction_genotype_Bcas2 | Gm44658 | -3.994 | 5.93E-04 | glio |
| Interaction_genotype_Mtor | Gm26822 | -4.319 | 5.93E-04 | glio |
| Interaction_genotype_Mtor | Ero1lb | -3.919 | 5.93E-04 | glio |
| interaction_RT | Ly96 | 1.274 | 5.93E-04 | glio |
| interaction_RT | Tmem243 | 1.145 | 5.93E-04 | glio |
| interaction_RT | Mcam | 1.263 | 5.93E-04 | glio |
| interaction_RT | Hist1h4d | -1.45 | 5.93E-04 | glio |
| Aars2_in.vivo_noRT | Hsh2d | 2.201 | 5.93E-04 | glio |
| Aars2_in.vivo_noRT | Ndufs6 | -0.915 | 5.93E-04 | glio |
| Aars2_in.vivo_noRT | Aldoa | -1 | 5.93E-04 | glio |
| Chek1_in.vivo_noRT | Tep1 | 1.828 | 5.93E-04 | glio |
| Chek1_in.vivo_noRT | Fau | -0.745 | 5.93E-04 | glio |
| Interaction_genotype_Mtor | Cnp | 3.348 | 5.93E-05 | glio |
| Atr_in.vivo_noRT | Zfp873 | 2.486 | 5.93E-05 | glio |
| Bora_in.vivo_noRT | Tctn1 | 2.897 | 5.93E-05 | glio |
| Interaction_genotype_Mtor | 2210406O10R | -5.327 | 5.93E-06 | glio |
| Atr_in.vivo_noRT | Lgals4 | 2.897 | 5.93E-06 | glio |
| culture_effect | Gm48933 | 2.625 | 5.93E-07 | glio |
| culture_effect | Vgf | -2.613 | 5.93E-08 | glio |
| culture_effect | Fh1 | -0.861 | 5.93E-08 | glio |
| culture_effect | E230029C05Ri | 2.861 | 5.93E-10 | glio |
| Bcas2_in.vivo_noRT | Slc22a1 | 3.633 | 5.93E-11 | glio |
| Aars2_in.vivo_noRT | A330076H08R | 5.327 | 5.93E-20 | glio |
| Aars2_in.vivo_noRT | Cldn5 | 5.06 | 5.93E-24 | glio |
| culture_effect | Tnfrsf11b | 0.72 | 5.94E-03 | glio |
| culture_effect | Tra2a | -0.41 | 5.94E-03 | glio |
| Interaction_genotype_Bcas2 | Gm48610 | -3.697 | 5.94E-03 | glio |
| Interaction_genotype_Bora | Gm10324 | -2.831 | 5.94E-03 | glio |
| Interaction_genotype_Bora | Nxnl2 | -3.371 | 5.94E-03 | glio |
| Interaction_genotype_Bora | Gm14019 | -3.231 | 5.94E-03 | glio |
| Interaction_genotype_Bora | Plekha4 | -3.627 | 5.94E-03 | glio |
| Interaction_genotype_Bora | Ift46 | 1.995 | 5.94E-03 | glio |
| Interaction_genotype_Mtor | Adamts9 | -2.98 | 5.94E-03 | glio |

| | | | | |
|---|---|---|---|---|
| Interaction_genotype_Mtor | Ubtd1 | -2.122 | 5.94E-03 | glio |
| Interaction_genotype_Mtor | Ift27 | 1.987 | 5.94E-03 | glio |
| Interaction_genotype_Myc | Osbpl6 | -2.253 | 5.94E-03 | glio |
| interaction_RT | Fam136a | -0.562 | 5.94E-03 | glio |
| Atr_in.vivo_noRT | Eif3d | -0.665 | 5.94E-03 | glio |
| Atr_ex.vivo_noRT | Ssh3 | 2.367 | 5.94E-03 | glio |
| Atr_ex.vivo_noRT | 5930403N24R | 2.714 | 5.94E-03 | glio |
| Bcas2_in.vivo_noRT | 5330438D12R | 1.681 | 5.94E-03 | glio |
| Bcas2_in.vivo_noRT | Rab2b | -1.339 | 5.94E-03 | glio |
| Bora_in.vivo_noRT | Med21 | -0.634 | 5.94E-03 | glio |
| Chek1_ex.vivo_noRT | Ddx18 | -1.079 | 5.94E-03 | glio |
| Cyp19a1_ex.vivo_noRT | Gm12703 | 2.159 | 5.94E-03 | glio |
| Interaction_genotype_Aars2 | Nkain2 | 2.454 | 5.94E-04 | glio |
| Interaction_genotype_Mtor | Lfng | -3.909 | 5.94E-04 | glio |
| Atr_in.vivo_noRT | Stamos | 2.218 | 5.94E-04 | glio |
| Atr_in.vivo_noRT | Eid3 | 2.105 | 5.94E-04 | glio |
| Interaction_genotype_Ifnar1 | Meioc | 3.784 | 5.94E-05 | glio |
| Interaction_genotype_Mtor | Rab36 | -4.766 | 5.94E-05 | glio |
| Atr_in.vivo_noRT | A930029G22R | 2.732 | 5.94E-05 | glio |
| Bcas2_in.vivo_noRT | Zfp964 | 2.95 | 5.94E-05 | glio |
| Atr_ex.vivo_noRT | A030014E15Ri | 2.77 | 5.94E-06 | glio |
| culture_effect | Iah1 | -0.764 | 5.94E-07 | glio |
| Interaction_genotype_Mtor | Olfr279 | -5.874 | 5.94E-07 | glio |
| Atr_in.vivo_noRT | Etohd2 | 3.641 | 5.94E-07 | glio |
| Bora_in.vivo_noRT | Vipr2 | 3.517 | 5.94E-07 | glio |
| Bora_in.vivo_noRT | Gadl1 | 3.286 | 5.94E-07 | glio |
| Bcas2_in.vivo_noRT | Gm1123 | 3.9 | 5.94E-08 | glio |
| Bcas2_in.vivo_noRT | Olfr90 | 4.543 | 5.94E-14 | glio |
| Chek1_in.vivo_noRT | Tmem236 | 4.777 | 5.94E-15 | glio |
| Bora_in.vivo_noRT | Gm36198 | 4.676 | 5.94E-16 | glio |
| culture_effect | Capn6 | -1.5 | 5.95E-03 | glio |
| culture_effect | Itsn1 | 0.59 | 5.95E-03 | glio |
| Interaction_genotype_Atr | Thsd1 | -2.964 | 5.95E-03 | glio |
| Interaction_genotype_Bcas2 | Adam1a | -3.772 | 5.95E-03 | glio |
| Interaction_genotype_Bcas2 | Tnrc6b | 1.484 | 5.95E-03 | glio |
| Interaction_genotype_Chek1 | C130036L24Ri | -3.615 | 5.95E-03 | glio |
| Interaction_genotype_Mtor | Gm34455 | -4.265 | 5.95E-03 | glio |
| Interaction_genotype_Myc | Gm10696 | -3.128 | 5.95E-03 | glio |
| Interaction_genotype_Myc | Scd3 | -3.412 | 5.95E-03 | glio |
| Interaction_genotype_Myc | C2cd2 | -3.199 | 5.95E-03 | glio |
| Interaction_genotype_Myc | Ripor1 | -2.426 | 5.95E-03 | glio |
| Interaction_genotype_Myc | Tnfrsf12a | 1.595 | 5.95E-03 | glio |
| Interaction_genotype_Pars2 | Slc35g3 | -2.668 | 5.95E-03 | glio |
| Aars2_ex.vivo_noRT | Igfbp7 | 2.495 | 5.95E-03 | glio |
| Atr_ex.vivo_noRT | Nasp | -0.915 | 5.95E-03 | glio |
| Bard1_ex.vivo_noRT | Masp2 | 2.421 | 5.95E-03 | glio |
| Bard1_ex.vivo_noRT | Selenbp1 | 3.213 | 5.95E-03 | glio |
| Bard1_ex.vivo_noRT | Tmeff2 | 3.217 | 5.95E-03 | glio |
| Chek1_in.vivo_noRT | Zik1 | 1.711 | 5.95E-03 | glio |
| Chek1_in.vivo_noRT | Leprotl1 | -0.816 | 5.95E-03 | glio |
| Chek1_ex.vivo_noRT | Miga1 | -1.884 | 5.95E-03 | glio |
| Cenpj_ex.vivo_noRT | Ggt1 | 3.389 | 5.95E-04 | glio |

| | | | | |
|---|---|---|---|---|
| Bcas2_ex.vivo_noRT | Cd7 | 2.742 | 5.95E-05 | glio |
| Bcas2_ex.vivo_noRT | Gm48764 | 2.742 | 5.95E-05 | glio |
| Chek1_in.vivo_noRT | Dynlt1c | 3.24 | 5.95E-05 | glio |
| Hsd17b10_in.vivo_noRT | Mfap3l | 3.225 | 5.95E-05 | glio |
| Hsd17b10_in.vivo_noRT | 4921508D12R | 2.689 | 5.95E-05 | glio |
| Chek1_in.vivo_noRT | Gm16794 | 3.473 | 5.95E-07 | glio |
| Chek1_in.vivo_noRT | Rasl10a | 2.901 | 5.95E-07 | glio |
| Hsd17b10_in.vivo_noRT | Gm4858 | 3.616 | 5.95E-08 | glio |
| Bcas2_in.vivo_noRT | Gm49874 | 4.807 | 5.95E-11 | glio |
| Chek1_in.vivo_noRT | Gm5152 | 4.657 | 5.95E-11 | glio |
| Chek1_in.vivo_noRT | Col6a3 | 4.007 | 5.95E-11 | glio |
| Hsd17b10_in.vivo_noRT | Speer4e | 3.616 | 5.95E-11 | glio |
| Atr_in.vivo_noRT | Gm45235 | 5.273 | 5.95E-21 | glio |
| Atr_in.vivo_noRT | A930033M14F | 5.533 | 5.95E-21 | glio |
| Aars2_in.vivo_noRT | Prl3d3 | 5.063 | 5.95E-25 | glio |
| Aars2_in.vivo_noRT | Gm5415 | 5.327 | 5.95E-27 | glio |
| Interaction_genotype_Atr | Gm11963 | -3.394 | 5.96E-03 | glio |
| Interaction_genotype_Bcas2 | Fam25c | -2.906 | 5.96E-03 | glio |
| Interaction_genotype_Bcas2 | Gm35082 | -2.862 | 5.96E-03 | glio |
| Interaction_genotype_Bcas2 | Gm48008 | -2.992 | 5.96E-03 | glio |
| Interaction_genotype_Bcas2 | Wars2 | -3.271 | 5.96E-03 | glio |
| Interaction_genotype_Bcas2 | Cdkn2aip | 2.188 | 5.96E-03 | glio |
| Interaction_genotype_Mtor | 1700011M02F | -4.229 | 5.96E-03 | glio |
| Interaction_genotype_Mtor | Synpo | -3.199 | 5.96E-03 | glio |
| Interaction_genotype_Mtor | Cbr4 | -2.407 | 5.96E-03 | glio |
| Interaction_genotype_Myc | Tmem18 | -2.349 | 5.96E-03 | glio |
| Interaction_genotype_Rpl3 | Appbp2os | -3.928 | 5.96E-03 | glio |
| interaction_RT | Csn3 | 2.462 | 5.96E-03 | glio |
| Aars2_in.vivo_noRT | Rrbp1 | -0.744 | 5.96E-03 | glio |
| Cenpj_ex.vivo_noRT | Car9 | 2.995 | 5.96E-03 | glio |
| Chek1_in.vivo_noRT | Vps13a | -1.04 | 5.96E-03 | glio |
| Fanca_ex.vivo_noRT | Hoxc8 | 2.124 | 5.96E-03 | glio |
| interaction_RT | Mllt3 | 0.952 | 5.96E-04 | glio |
| Bcas2_in.vivo_noRT | Kyat3 | 2.243 | 5.96E-04 | glio |
| Bora_in.vivo_noRT | Slc4a7 | -1.632 | 5.96E-04 | glio |
| Bora_ex.vivo_noRT | Dcpp3 | 3.201 | 5.96E-04 | glio |
| Cenpj_in.vivo_noRT | Gm48764 | 2.022 | 5.96E-04 | glio |
| interaction_RT | Dll3 | 4.207 | 5.96E-05 | glio |
| Bora_ex.vivo_noRT | 4930553M12F | 3.601 | 5.96E-07 | glio |
| Aars2_in.vivo_noRT | Plpp7 | 3.435 | 5.96E-08 | glio |
| Bcas2_in.vivo_noRT | Zfp986 | 3.765 | 5.96E-08 | glio |
| Chek1_in.vivo_noRT | Dact1 | 3.876 | 5.96E-09 | glio |
| Chek1_in.vivo_noRT | Gm33973 | 3.477 | 5.96E-09 | glio |
| Aars2_in.vivo_noRT | E230001N04R | 3.741 | 5.96E-10 | glio |
| Bcas2_in.vivo_noRT | Gm5152 | 4.42 | 5.96E-10 | glio |
| culture_effect | Nop53 | -0.98 | 5.96E-14 | glio |
| culture_effect | Cyb5a | 0.544 | 5.97E-03 | glio |
| Interaction_genotype_Aars2 | Cfap100 | -2.744 | 5.97E-03 | glio |
| Interaction_genotype_Aars2 | Ndufs8 | 1.295 | 5.97E-03 | glio |
| Interaction_genotype_Bcas2 | Pde11a | -3.84 | 5.97E-03 | glio |
| Interaction_genotype_Mtor | Nxpe3 | -2.806 | 5.97E-03 | glio |
| Interaction_genotype_Mtor | Ptov1 | 1.238 | 5.97E-03 | glio |

| | | | | |
|---|---|---|---|---|
| Aars2_in.vivo_noRT | Dhrs3 | 2.03 | 5.97E-03 | glio |
| Aars2_in.vivo_noRT | Zfp946 | 1.883 | 5.97E-03 | glio |
| Aars2_in.vivo_noRT | Casc4 | -1.178 | 5.97E-03 | glio |
| Atr_in.vivo_noRT | Pole4 | -0.978 | 5.97E-03 | glio |
| Bora_in.vivo_noRT | Ddah2 | 2.061 | 5.97E-03 | glio |
| Bora_in.vivo_noRT | Topors | -0.853 | 5.97E-03 | glio |
| Chek1_in.vivo_noRT | Myadm | -1.531 | 5.97E-03 | glio |
| Eif2s1_in.vivo_noRT | Sox11 | 2.276 | 5.97E-03 | glio |
| Interaction_genotype_Bora | Gfpt2 | 2.599 | 5.97E-04 | glio |
| Interaction_genotype_Mtor | Cd59a | -3.907 | 5.97E-04 | glio |
| interaction_RT | Xkr6 | 1.464 | 5.97E-04 | glio |
| interaction_RT | Lrrc8a | 1.201 | 5.97E-04 | glio |
| interaction_RT | Hnrnpu | -0.57 | 5.97E-04 | glio |
| Aars2_ex.vivo_noRT | Serpinb2 | 2.893 | 5.97E-04 | glio |
| Aars2_ex.vivo_noRT | Olfr1189 | 2.893 | 5.97E-04 | glio |
| Chek1_ex.vivo_noRT | Rnase4 | 3.163 | 5.97E-04 | glio |
| culture_effect | Heca | -0.99 | 5.97E-05 | glio |
| culture_effect | 2300009A05R | 0.87 | 5.97E-05 | glio |
| culture_effect | Ccnl2 | -0.643 | 5.97E-05 | glio |
| Interaction_genotype_Atr | 4930412M03F | -4.791 | 5.97E-05 | glio |
| Aars2_in.vivo_noRT | Mfsd4b4 | 2.602 | 5.97E-05 | glio |
| Aars2_in.vivo_noRT | Ocm | 2.486 | 5.97E-05 | glio |
| Aars2_in.vivo_noRT | Copa | -1.339 | 5.97E-05 | glio |
| culture_effect | Tars | -0.546 | 5.97E-06 | glio |
| Interaction_genotype_Myc | 1700023H06R | -5.597 | 5.97E-06 | glio |
| Interaction_genotype_Myc | Mrpl42 | 1.838 | 5.97E-06 | glio |
| Bora_in.vivo_noRT | Gm47591 | 3.244 | 5.97E-06 | glio |
| Chek1_ex.vivo_noRT | Acot6 | 3.871 | 5.97E-06 | glio |
| Hsd17b10_in.vivo_noRT | Pla1a | 3.084 | 5.97E-06 | glio |
| Hsd17b10_in.vivo_noRT | Vmn1r91 | 2.963 | 5.97E-06 | glio |
| Interaction_genotype_Mtor | D630024D03F | -4.685 | 5.97E-07 | glio |
| Aars2_ex.vivo_noRT | Gm48155 | 3.293 | 5.97E-08 | glio |
| Aars2_in.vivo_noRT | Cyp1a1 | 5.327 | 5.97E-09 | glio |
| Bora_in.vivo_noRT | Gm43625 | 3.631 | 5.97E-09 | glio |
| culture_effect | Tgtp2 | -2.741 | 5.97E-13 | glio |
| Atr_in.vivo_noRT | Vtn | 5.273 | 5.97E-24 | glio |
| Aars2_in.vivo_noRT | Edn2 | 5.327 | 5.97E-27 | glio |
| Aars2_in.vivo_noRT | Gm10801 | 6.385 | 5.97E-27 | glio |
| Aars2_in.vivo_noRT | Gm14207 | 5.327 | 5.97E-27 | glio |
| Aars2_in.vivo_noRT | Nrip2 | 5.327 | 5.97E-27 | glio |
| Interaction_genotype_Bora | Hist1h4b | -3.998 | 5.98E-03 | glio |
| Interaction_genotype_Chek1 | Gm50334 | -4.188 | 5.98E-03 | glio |
| Interaction_genotype_Ifnar1 | Hoxc8 | 2.734 | 5.98E-03 | glio |
| Interaction_genotype_Myc | Arntl | -2.441 | 5.98E-03 | glio |
| Interaction_genotype_Rad51 | Hes5 | 4.992 | 5.98E-03 | glio |
| Aars2_in.vivo_noRT | Rfx2 | 1.873 | 5.98E-03 | glio |
| Aars2_in.vivo_noRT | Ppme1 | -1.199 | 5.98E-03 | glio |
| Aars2_in.vivo_noRT | Sugt1 | -1.065 | 5.98E-03 | glio |
| Aars2_in.vivo_noRT | Stoml2 | -0.774 | 5.98E-03 | glio |
| Aars2_in.vivo_noRT | Cct5 | -0.561 | 5.98E-03 | glio |
| Aars2_ex.vivo_noRT | Nat8f4 | 2.505 | 5.98E-03 | glio |
| Atr_in.vivo_noRT | Mink1 | 1.455 | 5.98E-03 | glio |

| | | | | |
|---|---|---|---|---|
| Atr_in.vivo_noRT | Abhd11 | -1.076 | 5.98E-03 | glio |
| Atr_ex.vivo_noRT | Synm | -2.371 | 5.98E-03 | glio |
| Bcas2_in.vivo_noRT | Ccdc142os | 2.019 | 5.98E-03 | glio |
| Bora_in.vivo_noRT | Hist1h4a | 1.944 | 5.98E-03 | glio |
| Bora_in.vivo_noRT | Ugcg | -1.193 | 5.98E-03 | glio |
| Chek1_in.vivo_noRT | BC051226 | 1.415 | 5.98E-03 | glio |
| culture_effect | Pnliprp2 | -1.713 | 5.98E-04 | glio |
| culture_effect | Gm16759 | 1.916 | 5.98E-04 | glio |
| culture_effect | Ipo13 | 0.749 | 5.98E-04 | glio |
| culture_effect | Gclm | 0.614 | 5.98E-04 | glio |
| culture_effect | Ppia | -0.448 | 5.98E-04 | glio |
| Interaction_genotype_Mtor | Gm49539 | -4.145 | 5.98E-04 | glio |
| Interaction_genotype_Myc | Hpn | -3.772 | 5.98E-04 | glio |
| Aars2_in.vivo_noRT | Six5 | 2.151 | 5.98E-04 | glio |
| Aars2_ex.vivo_noRT | Zfp534 | 3.465 | 5.98E-04 | glio |
| Atr_in.vivo_noRT | Pigh | 1.804 | 5.98E-04 | glio |
| Atr_in.vivo_noRT | Psmd11 | -0.907 | 5.98E-04 | glio |
| Bora_in.vivo_noRT | Smarca5 | -0.872 | 5.98E-04 | glio |
| Chek1_ex.vivo_noRT | Shank1 | 3.042 | 5.98E-04 | glio |
| culture_effect | 1700066B19R | 1.987 | 5.98E-05 | glio |
| Interaction_genotype_Myc | Oxld1 | -3.613 | 5.98E-05 | glio |
| Interaction_genotype_Myc | Edn2 | -3.124 | 5.98E-05 | glio |
| interaction_RT | Ppid | -0.853 | 5.98E-05 | glio |
| interaction_RT | Ampd3 | 3.8 | 5.98E-05 | glio |
| Atp5c1_in.vivo_noRT | Gm40538 | 2.405 | 5.98E-05 | glio |
| Atr_in.vivo_noRT | 4833408A19R | 2.683 | 5.98E-05 | glio |
| Atr_in.vivo_noRT | Chrna1os | 2.384 | 5.98E-05 | glio |
| Atr_in.vivo_noRT | Gm41611 | 2.701 | 5.98E-05 | glio |
| Atr_in.vivo_noRT | Nsfl1c | -1.4 | 5.98E-05 | glio |
| Bard1_ex.vivo_noRT | Vtn | 2.814 | 5.98E-05 | glio |
| Bcas2_in.vivo_noRT | Rab17 | 2.812 | 5.98E-05 | glio |
| Chek1_in.vivo_noRT | Gm4189 | 2.842 | 5.98E-05 | glio |
| Bcas2_in.vivo_noRT | Hist1h4k | 3.237 | 5.98E-06 | glio |
| Bora_ex.vivo_noRT | A030014E15Ri | 2.808 | 5.98E-06 | glio |
| Aars2_ex.vivo_noRT | Gpr4 | 3.293 | 5.98E-07 | glio |
| Aars2_ex.vivo_noRT | Gm33466 | 3.293 | 5.98E-07 | glio |
| Aars2_in.vivo_noRT | Gm17477 | 3.395 | 5.98E-08 | glio |
| Aars2_in.vivo_noRT | 2810408A11R | 4.052 | 5.98E-09 | glio |
| Bcas2_in.vivo_noRT | Prrx2 | 4.807 | 5.98E-12 | glio |
| Aars2_in.vivo_noRT | Skida1 | 5.327 | 5.98E-14 | glio |
| culture_effect | Ndufs3 | -1.114 | 5.98E-20 | glio |
| Atr_in.vivo_noRT | 5730507C01R | 5.01 | 5.98E-21 | glio |
| culture_effect | Eif3m | -1.445 | 5.98E-29 | glio |
| Interaction_genotype_Aars2 | Npr3 | -3.006 | 5.99E-03 | glio |
| Interaction_genotype_Aars2 | Gm15520 | -3.065 | 5.99E-03 | glio |
| Interaction_genotype_Aars2 | Ap4b1 | -2.835 | 5.99E-03 | glio |
| Interaction_genotype_Atr | Hist1h1c | 2.245 | 5.99E-03 | glio |
| Interaction_genotype_Myc | Slc12a5 | -3.079 | 5.99E-03 | glio |
| Interaction_genotype_Myc | Tmem186 | -2.918 | 5.99E-03 | glio |
| Atr_in.vivo_noRT | Lars | -0.78 | 5.99E-03 | glio |
| Atr_ex.vivo_noRT | Gm35551 | 2.577 | 5.99E-03 | glio |
| Atr_ex.vivo_noRT | Evi5l | 2.061 | 5.99E-03 | glio |

| | | | | |
|---|---|---|---|---|
| Bcas2_in.vivo_noRT | Etf1 | -0.723 | 5.99E-03 | glio |
| Cenpj_ex.vivo_noRT | Prox2os | 2.595 | 5.99E-03 | glio |
| culture_effect | Gm5602 | -1.726 | 5.99E-04 | glio |
| culture_effect | Mapk6 | 0.675 | 5.99E-04 | glio |
| Interaction_genotype_Myc | Tmem69 | -3.172 | 5.99E-04 | glio |
| interaction_RT | Nacc1 | -0.992 | 5.99E-04 | glio |
| interaction_RT | Tfrc | -1.033 | 5.99E-04 | glio |
| interaction_RT | Esr1 | 3.236 | 5.99E-04 | glio |
| Chek1_in.vivo_noRT | Pcdhb17 | 2.447 | 5.99E-04 | glio |
| Chek1_ex.vivo_noRT | Rhbdf2 | 2.985 | 5.99E-04 | glio |
| Eif2s1_in.vivo_noRT | Smtnl1 | 2.143 | 5.99E-04 | glio |
| Interaction_genotype_Atr | Selenov | -4.14 | 5.99E-05 | glio |
| Aars2_in.vivo_noRT | Crybg2 | 2.802 | 5.99E-05 | glio |
| Aars2_in.vivo_noRT | Ifi203 | -2.26 | 5.99E-05 | glio |
| Atr_in.vivo_noRT | A830005F24Ri | 2.838 | 5.99E-05 | glio |
| Atr_in.vivo_noRT | Gm48604 | 2.75 | 5.99E-05 | glio |
| Atr_ex.vivo_noRT | Brinp3 | 3.164 | 5.99E-05 | glio |
| Bora_in.vivo_noRT | 9130008F23Ri | 3.17 | 5.99E-05 | glio |
| Bora_in.vivo_noRT | Mfsd5 | -1.526 | 5.99E-05 | glio |
| Chek1_in.vivo_noRT | Gm13830 | 3.113 | 5.99E-05 | glio |
| Interaction_genotype_Myc | Crabp1 | -4.849 | 5.99E-06 | glio |
| Atr_in.vivo_noRT | Rpl17 | -0.944 | 5.99E-06 | glio |
| Bcas2_in.vivo_noRT | Gm26581 | 3.412 | 5.99E-06 | glio |
| Chek1_in.vivo_noRT | 4933406P04R | 3.362 | 5.99E-06 | glio |
| culture_effect | Card10 | 2.634 | 5.99E-07 | glio |
| culture_effect | Bax | 0.581 | 5.99E-07 | glio |
| Chek1_in.vivo_noRT | Gm16754 | 3.54 | 5.99E-07 | glio |
| Chek1_ex.vivo_noRT | 1700024I08Ri | 3.469 | 5.99E-07 | glio |
| culture_effect | Slc27a1 | -1.407 | 5.99E-08 | glio |
| Bcas2_in.vivo_noRT | Gm11549 | 4.281 | 5.99E-10 | glio |
| Bcas2_in.vivo_noRT | 1600014C23R | 4.428 | 5.99E-10 | glio |
| Bora_in.vivo_noRT | Lce1g | 4.936 | 5.99E-12 | glio |
| Chek1_in.vivo_noRT | Bcl2l15 | 4.784 | 5.99E-15 | glio |
| culture_effect | Ftl1 | 0.907 | 5.99E-17 | glio |
| Aars2_in.vivo_noRT | Gm21319 | 5.327 | 5.99E-22 | glio |
| Aars2_in.vivo_noRT | Gm19276 | 5.327 | 5.99E-26 | glio |
| Interaction_genotype_Atr | 2410141K09R | -2.89 | 6.00E-03 | glio |
| Interaction_genotype_Atr | Pcx | -3.336 | 6.00E-03 | glio |
| Interaction_genotype_Myc | Slc8b1 | -2.44 | 6.00E-03 | glio |
| Interaction_genotype_Myc | Rnft1 | -1.954 | 6.00E-03 | glio |
| Interaction_genotype_Myc | Lgals3bp | 1.395 | 6.00E-03 | glio |
| Atp5c1_in.vivo_noRT | Dreh | 2.022 | 6.00E-03 | glio |
| Atr_in.vivo_noRT | Trim7 | 1.819 | 6.00E-03 | glio |
| Atr_in.vivo_noRT | Malat1 | -0.951 | 6.00E-03 | glio |
| Atr_ex.vivo_noRT | Maff | 1.779 | 6.00E-03 | glio |
| Bcas2_in.vivo_noRT | Fxyd7 | 1.86 | 6.00E-03 | glio |
| Bcas2_ex.vivo_noRT | Gm5662 | 2.742 | 6.00E-03 | glio |
| Bcas2_ex.vivo_noRT | Gm17276 | 2.347 | 6.00E-03 | glio |
| Bora_in.vivo_noRT | Dapp1 | 1.732 | 6.00E-03 | glio |
| Bora_ex.vivo_noRT | Gm10696 | 2.802 | 6.00E-03 | glio |
| Cdkn2a_in.vivo_noRT | Tff2 | 2.142 | 6.00E-03 | glio |
| Cdkn2a_in.vivo_noRT | Capn13 | -1.675 | 6.00E-03 | glio |

| | | | | |
|---|---|---|---|---|
| Cdkn2a_in.vivo_noRT | Serpine2 | 2.199 | 6.00E-03 | glio |
| Chek1_in.vivo_noRT | 2500004C02R | 1.671 | 6.00E-03 | glio |
| Chek1_in.vivo_noRT | Sec11c | -0.863 | 6.00E-03 | glio |
| Hsd17b10_in.vivo_noRT | Gm49864 | 2.184 | 6.00E-03 | glio |
| culture_effect | Ttc7b | 1.039 | 6.00E-04 | glio |
| culture_effect | Bloc1s1 | 0.832 | 6.00E-04 | glio |
| Interaction_genotype_Myc | Lysmd4 | -3.536 | 6.00E-04 | glio |
| Interaction_genotype_Myc | Nup210 | -3.268 | 6.00E-04 | glio |
| Aars2_in.vivo_noRT | Pcdh9 | 2.462 | 6.00E-04 | glio |
| Atr_in.vivo_noRT | Ptprz1 | -1.731 | 6.00E-04 | glio |
| Bcas2_in.vivo_noRT | Msn | -1.197 | 6.00E-04 | glio |
| Bora_ex.vivo_noRT | Gm15816 | 2.883 | 6.00E-04 | glio |
| Chek1_ex.vivo_noRT | Dhrs13 | -2.051 | 6.00E-04 | glio |
| Aars2_in.vivo_noRT | Hist1h2bf | 3.014 | 6.00E-05 | glio |
| Aars2_in.vivo_noRT | Ptpn2 | -1.248 | 6.00E-05 | glio |
| Aars2_in.vivo_noRT | Mrpl53 | -1.235 | 6.00E-05 | glio |
| Aars2_ex.vivo_noRT | Gm48832 | 3.697 | 6.00E-05 | glio |
| Aars2_ex.vivo_noRT | Iqch | 3.127 | 6.00E-05 | glio |
| Atr_ex.vivo_noRT | Gem | 3.554 | 6.00E-05 | glio |
| Atr_ex.vivo_noRT | Cemip | 3.164 | 6.00E-05 | glio |
| Chek1_in.vivo_noRT | Slc46a1 | 3.01 | 6.00E-05 | glio |
| Hsd17b10_in.vivo_noRT | Ikbke | 2.831 | 6.00E-05 | glio |
| Interaction_genotype_Myc | Gm20492 | -4.98 | 6.00E-06 | glio |
| culture_effect | Mtmr2 | 0.983 | 6.00E-07 | glio |
| Atr_ex.vivo_noRT | Gm5934 | 3.382 | 6.00E-07 | glio |
| culture_effect | Klhl24 | -1.391 | 6.00E-08 | glio |
| Aars2_in.vivo_noRT | Gm10734 | 3.81 | 6.00E-08 | glio |
| Aars2_in.vivo_noRT | Necab3 | 4.401 | 6.00E-11 | glio |
| Bora_in.vivo_noRT | Cstdc4 | 4.279 | 6.00E-11 | glio |
| Bora_in.vivo_noRT | Cysrt1 | 4.676 | 6.00E-11 | glio |
| Atr_in.vivo_noRT | Gm32834 | 4.616 | 6.00E-12 | glio |
| Atr_in.vivo_noRT | Skida1 | 5.533 | 6.00E-15 | glio |
| Atr_in.vivo_noRT | C130060C02R | 5.269 | 6.00E-17 | glio |
| culture_effect | Slc26a1 | -1.737 | 6.01E-03 | glio |
| culture_effect | N4bp2os | 1.458 | 6.01E-03 | glio |
| culture_effect | Znhit1 | 0.392 | 6.01E-03 | glio |
| Interaction_genotype_Bcas2 | Gm50138 | -3.101 | 6.01E-03 | glio |
| Interaction_genotype_Pars2 | Gm2012 | -3.898 | 6.01E-03 | glio |
| Interaction_genotype_Pars2 | Necab2 | -3.498 | 6.01E-03 | glio |
| Interaction_genotype_Terf1 | Kpna7 | -3.696 | 6.01E-03 | glio |
| interaction_RT | Rpl22l1 | 0.536 | 6.01E-03 | glio |
| Aars2_in.vivo_noRT | Hdac10 | 1.74 | 6.01E-03 | glio |
| Aars2_in.vivo_noRT | Cerk | 1.702 | 6.01E-03 | glio |
| Atp5c1_in.vivo_noRT | Dlx4 | 2.142 | 6.01E-03 | glio |
| Bora_in.vivo_noRT | Mtss1 | 2.115 | 6.01E-03 | glio |
| Bora_in.vivo_noRT | Kctd2 | -1.348 | 6.01E-03 | glio |
| Chek1_in.vivo_noRT | Cul1 | -1.014 | 6.01E-03 | glio |
| Chek1_ex.vivo_noRT | Aqp5 | 2.85 | 6.01E-03 | glio |
| Chek1_ex.vivo_noRT | Zfp639 | -1.67 | 6.01E-03 | glio |
| Fanca_ex.vivo_noRT | Fam214a | -2.456 | 6.01E-03 | glio |
| Interaction_genotype_Aars2 | Gm15853 | -4.468 | 6.01E-04 | glio |
| Interaction_genotype_Aars2 | Rnft2 | -4.208 | 6.01E-04 | glio |

| | | | | |
|---|---|---|---|---|
| Interaction_genotype_Myc | Gm10863 | -3.464 | 6.01E-04 | glio |
| Aars2_in.vivo_noRT | Hnrnpk | -0.783 | 6.01E-04 | glio |
| Atr_ex.vivo_noRT | Myzap | 2.764 | 6.01E-04 | glio |
| Atr_ex.vivo_noRT | Mst1 | 3.168 | 6.01E-04 | glio |
| Brca2_in.vivo_noRT | Gm15533 | 2.288 | 6.01E-04 | glio |
| Brca2_in.vivo_noRT | Gpr4 | 2.288 | 6.01E-04 | glio |
| Cenpj_ex.vivo_noRT | 4930512B01R | 2.591 | 6.01E-04 | glio |
| Chek1_in.vivo_noRT | A230087F16Ri | 2.42 | 6.01E-04 | glio |
| Chek1_in.vivo_noRT | Clip1 | 1.772 | 6.01E-04 | glio |
| Aars2_in.vivo_noRT | Rpsa | -0.803 | 6.01E-05 | glio |
| Hsd17b10_in.vivo_noRT | Spns3 | 2.958 | 6.01E-05 | glio |
| Interaction_genotype_Mtor | Gm15816 | -4.786 | 6.01E-06 | glio |
| Aars2_in.vivo_noRT | 2310015A10R | 3.298 | 6.01E-06 | glio |
| Bora_in.vivo_noRT | Nkx2-2os | 3.281 | 6.01E-06 | glio |
| culture_effect | Bloc1s6os | 2.235 | 6.01E-07 | glio |
| culture_effect | Hid1 | -1.807 | 6.01E-07 | glio |
| Interaction_genotype_Mtor | Nrip2 | -4.099 | 6.01E-07 | glio |
| Atr_in.vivo_noRT | Card10 | 3.736 | 6.01E-07 | glio |
| Bora_in.vivo_noRT | Scgb1a1 | 3.826 | 6.01E-08 | glio |
| Bora_in.vivo_noRT | Pax5 | 4.616 | 6.01E-10 | glio |
| Chek1_in.vivo_noRT | 1600014C23R | 4.664 | 6.01E-11 | glio |
| culture_effect | Pcmt1 | -0.912 | 6.01E-14 | glio |
| Bcas2_in.vivo_noRT | Gm44736 | 4.807 | 6.01E-24 | glio |
| culture_effect | Rflnb | -0.86 | 6.02E-03 | glio |
| Interaction_genotype_Atr | 0610038B21R | -2.884 | 6.02E-03 | glio |
| Interaction_genotype_Bora | 1700034I23Ri | -2.532 | 6.02E-03 | glio |
| Interaction_genotype_Myc | Gm42664 | -2.665 | 6.02E-03 | glio |
| Interaction_genotype_Myc | Serpinc1 | -3.226 | 6.02E-03 | glio |
| Interaction_genotype_Myc | Epb41l5 | -2.596 | 6.02E-03 | glio |
| Interaction_genotype_Myc | Wdr34 | -2.572 | 6.02E-03 | glio |
| Interaction_genotype_Pars2 | Sh3d21 | -3.243 | 6.02E-03 | glio |
| Interaction_genotype_Rad51 | C1ra | 3.353 | 6.02E-03 | glio |
| Interaction_genotype_Rpl3 | Gm11457 | -3.628 | 6.02E-03 | glio |
| interaction_RT | C1s1 | 2.295 | 6.02E-03 | glio |
| interaction_RT | 6430590A07R | -1.428 | 6.02E-03 | glio |
| Aars2_ex.vivo_noRT | Tacr2 | 2.813 | 6.02E-03 | glio |
| Brca2_in.vivo_noRT | Gm26671 | 1.763 | 6.02E-03 | glio |
| Interaction_genotype_Mtor | AI463229 | -3.454 | 6.02E-04 | glio |
| Interaction_genotype_Mtor | 1700001L05Ri | -3.957 | 6.02E-04 | glio |
| Interaction_genotype_Mtor | 1810021B22R | -3.987 | 6.02E-04 | glio |
| Interaction_genotype_Mtor | Cd63 | 1.42 | 6.02E-04 | glio |
| Interaction_genotype_Nat2 | Gm10696 | 4.503 | 6.02E-04 | glio |
| Bora_in.vivo_noRT | Mpp2 | 2.48 | 6.02E-04 | glio |
| Atr_in.vivo_noRT | Tppp | 2.452 | 6.02E-05 | glio |
| Chek1_in.vivo_noRT | Slc22a18 | 3.018 | 6.02E-05 | glio |
| Bcas2_in.vivo_noRT | Gm9833 | 3.026 | 6.02E-06 | glio |
| culture_effect | Ly75 | -1.686 | 6.02E-08 | glio |
| Interaction_genotype_Nat2 | Zscan4c | 4.795 | 6.02E-08 | glio |
| Interaction_genotype_Nat2 | Zscan4-ps3 | 4.475 | 6.02E-08 | glio |
| Interaction_genotype_Nat2 | Usp17lc | 5.397 | 6.02E-08 | glio |
| Interaction_genotype_Nat2 | Zscan4-ps2 | 4.755 | 6.02E-08 | glio |
| Bora_in.vivo_noRT | Zfp85os | 3.877 | 6.02E-08 | glio |

| | | | | |
|---|---|---|---|---|
| culture_effect | Pmpcb | -0.779 | 6.02E-09 | glio |
| interaction_RT | Tmem176b | 1.907 | 6.02E-09 | glio |
| Atr_in.vivo_noRT | Fhl4 | 4.283 | 6.02E-11 | glio |
| Atr_in.vivo_noRT | Plcd4 | 4.623 | 6.02E-13 | glio |
| Chek1_ex.vivo_noRT | Clec12b | 4.447 | 6.02E-14 | glio |
| culture_effect | Wnt7b | -3.919 | 6.02E-16 | glio |
| culture_effect | Plek2 | -1.706 | 6.03E-03 | glio |
| culture_effect | Tomm20 | 0.286 | 6.03E-03 | glio |
| Interaction_genotype_Aars2 | Cox5a | 0.858 | 6.03E-03 | glio |
| Interaction_genotype_Atr | Ptprg | -3.073 | 6.03E-03 | glio |
| Interaction_genotype_Bcas2 | Tead3 | -3.665 | 6.03E-03 | glio |
| Interaction_genotype_Bcas2 | Radil | -3.582 | 6.03E-03 | glio |
| Interaction_genotype_Mtor | Sub1 | 1.026 | 6.03E-03 | glio |
| Interaction_genotype_Mtor | Rps20 | 0.997 | 6.03E-03 | glio |
| Interaction_genotype_Myc | Hhipl2 | -3.468 | 6.03E-03 | glio |
| Interaction_genotype_Myc | Ifit1bl1 | -3.579 | 6.03E-03 | glio |
| Interaction_genotype_Myc | Tex9 | -2.469 | 6.03E-03 | glio |
| interaction_RT | Hist3h2ba | 1.852 | 6.03E-03 | glio |
| Atr_in.vivo_noRT | Pik3r3 | 1.829 | 6.03E-03 | glio |
| Atr_in.vivo_noRT | Tsfm | -0.946 | 6.03E-03 | glio |
| Bcas2_ex.vivo_noRT | Glyat | 1.945 | 6.03E-03 | glio |
| Bora_in.vivo_noRT | Hist1h2ac | 1.979 | 6.03E-03 | glio |
| Brca2_in.vivo_noRT | Gm16933 | 1.764 | 6.03E-03 | glio |
| Brca2_in.vivo_noRT | Nlrx1 | 2.7 | 6.03E-03 | glio |
| Cdkn2a_in.vivo_noRT | Gstt4 | -1.938 | 6.03E-03 | glio |
| Cdkn2a_in.vivo_noRT | Slc45a1 | -1.935 | 6.03E-03 | glio |
| Cdkn2a_in.vivo_noRT | Opcml | -1.94 | 6.03E-03 | glio |
| Cdkn2a_in.vivo_noRT | Selp | -2.719 | 6.03E-03 | glio |
| Cdkn2a_in.vivo_noRT | Gm13191 | -1.675 | 6.03E-03 | glio |
| Cdkn2a_in.vivo_noRT | 4933416E03Ri | -1.675 | 6.03E-03 | glio |
| Cenpj_ex.vivo_noRT | Rhbg | 2.198 | 6.03E-03 | glio |
| Chek1_in.vivo_noRT | Dleu7 | 1.981 | 6.03E-03 | glio |
| Cyp19a1_ex.vivo_noRT | Angptl7 | 2.159 | 6.03E-03 | glio |
| Cyp19a1_ex.vivo_noRT | Gm8016 | 2.159 | 6.03E-03 | glio |
| Hsd17b10_in.vivo_noRT | Ggt5 | 1.983 | 6.03E-03 | glio |
| Interaction_genotype_Atr | 1700003G18R | -4.746 | 6.03E-04 | glio |
| Interaction_genotype_Myc | Nf2 | -2.487 | 6.03E-04 | glio |
| Aars2_ex.vivo_noRT | Gm21762 | 2.899 | 6.03E-04 | glio |
| Bora_ex.vivo_noRT | Snhg4.1 | 2.808 | 6.03E-04 | glio |
| Bora_ex.vivo_noRT | AC125149.3 | 3.19 | 6.03E-04 | glio |
| Cyp19a1_ex.vivo_noRT | Olfr171 | 2.959 | 6.03E-04 | glio |
| Cyp19a1_ex.vivo_noRT | Crb3 | 2.933 | 6.03E-04 | glio |
| Hsd17b10_in.vivo_noRT | Zfp972 | 2.703 | 6.03E-04 | glio |
| culture_effect | Pcdhb22 | -2.359 | 6.03E-05 | glio |
| culture_effect | Rxra | 2.073 | 6.03E-05 | glio |
| culture_effect | Gmppb | -0.961 | 6.03E-05 | glio |
| culture_effect | Gsap | 0.898 | 6.03E-05 | glio |
| Interaction_genotype_Aars2 | Psma4 | 1.62 | 6.03E-05 | glio |
| Aars2_in.vivo_noRT | Rbm3os | 3.05 | 6.03E-06 | glio |
| Interaction_genotype_Myc | Oaz1 | 2.108 | 6.03E-08 | glio |
| Chek1_ex.vivo_noRT | Gm20732 | 4.842 | 6.03E-10 | glio |
| Bcas2_in.vivo_noRT | 1810019D21R | 4.547 | 6.03E-11 | glio |

| | | | | |
|---|---|---|---|---|
| Aars2_in.vivo_noRT | Gm42849 | 5.06 | 6.03E-16 | glio |
| Bora_in.vivo_noRT | Boc | 4.672 | 6.03E-20 | glio |
| Bora_in.vivo_noRT | Dmrtc1a | 4.936 | 6.03E-20 | glio |
| Bora_in.vivo_noRT | Gm12589 | 4.936 | 6.03E-20 | glio |
| Bora_in.vivo_noRT | Gm12703 | 4.67 | 6.03E-21 | glio |
| Interaction_genotype_Aars2 | Gm37933 | -2.91 | 6.04E-03 | glio |
| Interaction_genotype_Mtor | Sdr16c5 | -3.177 | 6.04E-03 | glio |
| Interaction_genotype_Myc | Depdc1b | -2.721 | 6.04E-03 | glio |
| Interaction_genotype_Myc | Rbm12b1 | -2.794 | 6.04E-03 | glio |
| Atr_ex.vivo_noRT | D6Ertd527e | 2.632 | 6.04E-03 | glio |
| Bcas2_in.vivo_noRT | Slc25a29 | 1.863 | 6.04E-03 | glio |
| Cenpj_ex.vivo_noRT | Platr31 | 3.363 | 6.04E-03 | glio |
| Cenpj_ex.vivo_noRT | AC159140.1 | 2.198 | 6.04E-03 | glio |
| Chek1_ex.vivo_noRT | Ndufab1 | -0.668 | 6.04E-03 | glio |
| Hsd17b10_in.vivo_noRT | Gm26688 | 2.178 | 6.04E-03 | glio |
| Aars2_in.vivo_noRT | Rps11 | -0.617 | 6.04E-04 | glio |
| Atr_in.vivo_noRT | Rmst | 2.554 | 6.04E-04 | glio |
| Atr_in.vivo_noRT | Gm45716 | 2.318 | 6.04E-04 | glio |
| Bcas2_ex.vivo_noRT | Klrk1 | 2.742 | 6.04E-04 | glio |
| Chek1_ex.vivo_noRT | Capn9 | 2.779 | 6.04E-04 | glio |
| Chek1_ex.vivo_noRT | 9130204L05Ri | 3.363 | 6.04E-04 | glio |
| culture_effect | Gnao1 | -1.678 | 6.04E-05 | glio |
| culture_effect | Cdyl | 0.957 | 6.04E-05 | glio |
| Interaction_genotype_Mtor | Hist1h4c | -4.913 | 6.04E-05 | glio |
| Aars2_in.vivo_noRT | A830082K12R | 2.498 | 6.04E-05 | glio |
| Aars2_in.vivo_noRT | Fam204a | -1.545 | 6.04E-05 | glio |
| Aars2_in.vivo_noRT | Atxn2l | -1.448 | 6.04E-05 | glio |
| Chek1_in.vivo_noRT | Prkaa2 | 2.472 | 6.04E-05 | glio |
| Eif2s1_in.vivo_noRT | Capn13 | 2.143 | 6.04E-05 | glio |
| Interaction_genotype_Mtor | Rxra | -5.382 | 6.04E-06 | glio |
| Interaction_genotype_Myc | Gm41396 | -5.355 | 6.04E-06 | glio |
| Atr_in.vivo_noRT | Rab26 | 2.942 | 6.04E-06 | glio |
| Bcas2_in.vivo_noRT | Gm11127 | 3.246 | 6.04E-06 | glio |
| Chek1_ex.vivo_noRT | Aass | 3.7 | 6.04E-06 | glio |
| Interaction_genotype_Mtor | Gm13272 | -5.348 | 6.04E-07 | glio |
| Atr_in.vivo_noRT | Gm29358 | 3.571 | 6.04E-07 | glio |
| Bcas2_in.vivo_noRT | Gm43581 | 3.237 | 6.04E-07 | glio |
| Atr_in.vivo_noRT | Tcte2 | 3.411 | 6.04E-08 | glio |
| Chek1_ex.vivo_noRT | C3 | 4.842 | 6.04E-08 | glio |
| Hsd17b10_in.vivo_noRT | Gm35439 | 3.237 | 6.04E-09 | glio |
| Hsd17b10_in.vivo_noRT | C430002N11R | 3.616 | 6.04E-11 | glio |
| Bcas2_in.vivo_noRT | B3galt4 | 4.799 | 6.04E-14 | glio |
| Atr_in.vivo_noRT | AC159140.1 | 5.273 | 6.04E-23 | glio |
| culture_effect | Tnpo2 | -0.547 | 6.05E-03 | glio |
| culture_effect | Sqle | -0.564 | 6.05E-03 | glio |
| Interaction_genotype_Mtor | Cyp4f18 | -4.753 | 6.05E-03 | glio |
| Interaction_genotype_Mtor | Cd48 | -3.11 | 6.05E-03 | glio |
| Interaction_genotype_Mtor | Mon1a | -2.772 | 6.05E-03 | glio |
| Interaction_genotype_Mtor | Fosl2 | -2.29 | 6.05E-03 | glio |
| Interaction_genotype_Myc | Gm5039 | -3.517 | 6.05E-03 | glio |
| Interaction_genotype_Myc | Zfp420 | -3.247 | 6.05E-03 | glio |
| Interaction_genotype_Myc | Gm4673 | -3.028 | 6.05E-03 | glio |

| | | | | |
|---|---|---|---|---|
| Interaction_genotype_Myc | Spsb1 | -2.505 | 6.05E-03 | glio |
| Interaction_genotype_Rad51 | Jph4 | -3.908 | 6.05E-03 | glio |
| Aars2_in.vivo_noRT | Ccdc73 | 1.766 | 6.05E-03 | glio |
| Bcas2_in.vivo_noRT | Ptpru | 1.806 | 6.05E-03 | glio |
| Bcas2_in.vivo_noRT | Acox2 | 1.752 | 6.05E-03 | glio |
| Bora_ex.vivo_noRT | Ifitm1 | 3.201 | 6.05E-03 | glio |
| Brca2_in.vivo_noRT | Gm13078 | 1.763 | 6.05E-03 | glio |
| Cdkn2a_in.vivo_noRT | Gm15138 | -1.941 | 6.05E-03 | glio |
| Cdkn2a_in.vivo_noRT | Cyp26b1 | 2.241 | 6.05E-03 | glio |
| Cdkn2a_in.vivo_noRT | Cldn2 | -1.675 | 6.05E-03 | glio |
| Cdkn2a_in.vivo_noRT | Gm39929 | -2.132 | 6.05E-03 | glio |
| Cdkn2a_in.vivo_noRT | Ccl3 | -1.935 | 6.05E-03 | glio |
| Cdkn2a_in.vivo_noRT | Gm18280 | -1.938 | 6.05E-03 | glio |
| Cdkn2a_in.vivo_noRT | Spink8 | -1.935 | 6.05E-03 | glio |
| Cdkn2a_in.vivo_noRT | Stfa3 | -1.675 | 6.05E-03 | glio |
| Cdkn2a_in.vivo_noRT | Olfr787 | -1.675 | 6.05E-03 | glio |
| Cdkn2a_in.vivo_noRT | Vtn | -1.935 | 6.05E-03 | glio |
| Chek1_in.vivo_noRT | 3110045C21R | 1.896 | 6.05E-03 | glio |
| Chek1_in.vivo_noRT | Antxr2 | 1.699 | 6.05E-03 | glio |
| Chek1_ex.vivo_noRT | Aldh3b1 | 2.612 | 6.05E-03 | glio |
| Chek1_ex.vivo_noRT | Plscr4 | 1.964 | 6.05E-03 | glio |
| Eif2s1_ex.vivo_noRT | Rgs4 | 2.234 | 6.05E-03 | glio |
| Eif2s1_ex.vivo_noRT | Gm13178 | 2.234 | 6.05E-03 | glio |
| culture_effect | Slc5a5 | -2.246 | 6.05E-04 | glio |
| culture_effect | Efhd1 | 1.995 | 6.05E-04 | glio |
| Interaction_genotype_Myc | Thsd1 | -3.324 | 6.05E-04 | glio |
| Interaction_genotype_Myc | Rab37 | -3.745 | 6.05E-04 | glio |
| Interaction_genotype_Myc | Gm35066 | -3.719 | 6.05E-04 | glio |
| Interaction_genotype_Rad51 | Fam193a | 2.195 | 6.05E-04 | glio |
| Aars2_in.vivo_noRT | 9130024F11Ri | 2.342 | 6.05E-04 | glio |
| Aars2_in.vivo_noRT | Hmx3 | 2.251 | 6.05E-04 | glio |
| Atr_in.vivo_noRT | Calhm6 | 2.025 | 6.05E-04 | glio |
| Atr_in.vivo_noRT | Hsp90aa1 | -0.742 | 6.05E-04 | glio |
| Bcas2_in.vivo_noRT | Ehbp1l1 | 2.557 | 6.05E-04 | glio |
| Bora_in.vivo_noRT | Shbg | 2.557 | 6.05E-04 | glio |
| Chek1_in.vivo_noRT | Dnah2 | 2.495 | 6.05E-04 | glio |
| culture_effect | Gm50103 | 2.167 | 6.05E-05 | glio |
| Interaction_genotype_Mtor | Layn | -3.668 | 6.05E-05 | glio |
| Interaction_genotype_Myc | Nbas | -3.462 | 6.05E-05 | glio |
| Interaction_genotype_Myc | Rpl24 | 1.511 | 6.05E-05 | glio |
| Atr_ex.vivo_noRT | Rhbg | 2.771 | 6.05E-05 | glio |
| Bcas2_in.vivo_noRT | Dennd2d | 2.54 | 6.05E-05 | glio |
| Chek1_ex.vivo_noRT | Map7 | 3.652 | 6.05E-05 | glio |
| Aars2_ex.vivo_noRT | Gm50024 | 2.893 | 6.05E-06 | glio |
| Aars2_ex.vivo_noRT | Spink6 | 3.293 | 6.05E-06 | glio |
| Aars2_ex.vivo_noRT | Vmn2r44 | 2.897 | 6.05E-06 | glio |
| Aars2_ex.vivo_noRT | Htr2b | 3.293 | 6.05E-06 | glio |
| Atr_in.vivo_noRT | Gm26658 | 3.502 | 6.05E-06 | glio |
| Interaction_genotype_Nat2 | Gm8016 | 4.252 | 6.05E-07 | glio |
| Hsd17b10_in.vivo_noRT | Sprr1a | 3.616 | 6.05E-07 | glio |
| Atr_in.vivo_noRT | Gm29340 | 4.218 | 6.05E-10 | glio |
| Bcas2_in.vivo_noRT | Gm49579 | 3.754 | 6.05E-10 | glio |

| | | | | |
|---|---|---|---|---|
| Atr_in.vivo_noRT | Zfp986 | 4.492 | 6.05E-11 | glio |
| Bcas2_in.vivo_noRT | 1700006H21R | 4.807 | 6.05E-12 | glio |
| Chek1_in.vivo_noRT | Fam227b | 5.044 | 6.05E-26 | glio |
| Chek1_in.vivo_noRT | Gm20658 | 5.044 | 6.05E-26 | glio |
| culture_effect | Ddb1 | -0.359 | 6.06E-03 | glio |
| culture_effect | Lsm8 | 0.342 | 6.06E-03 | glio |
| Interaction_genotype_Myc | Erfe | -2.848 | 6.06E-03 | glio |
| Interaction_genotype_Myc | Sugct | -2.61 | 6.06E-03 | glio |
| Interaction_genotype_Myc | Eef1akmt4 | -2.197 | 6.06E-03 | glio |
| Atr_in.vivo_noRT | Zfr2 | 1.87 | 6.06E-03 | glio |
| Atr_in.vivo_noRT | Gm48226 | 1.881 | 6.06E-03 | glio |
| Atr_in.vivo_noRT | Ndufaf6 | -1 | 6.06E-03 | glio |
| Cenpj_ex.vivo_noRT | Icam2 | 2.591 | 6.06E-03 | glio |
| Chek1_in.vivo_noRT | Zfp97 | 2.02 | 6.06E-03 | glio |
| Chek1_in.vivo_noRT | Ufsp1 | 1.451 | 6.06E-03 | glio |
| Chek1_in.vivo_noRT | Hist1h3i | 1.788 | 6.06E-03 | glio |
| culture_effect | Ppp1r13b | -1.375 | 6.06E-04 | glio |
| culture_effect | Usp1 | 0.509 | 6.06E-04 | glio |
| Interaction_genotype_Aars2 | Armc2 | -4.278 | 6.06E-04 | glio |
| Interaction_genotype_Aars2 | D430001F17R | -3.979 | 6.06E-04 | glio |
| Interaction_genotype_Aars2 | Eri3 | 1.768 | 6.06E-04 | glio |
| Interaction_genotype_Mtor | Gm43773 | -3.948 | 6.06E-04 | glio |
| Interaction_genotype_Mtor | Gm16183 | -4.222 | 6.06E-04 | glio |
| Interaction_genotype_Mtor | Pcdh9 | -4.148 | 6.06E-04 | glio |
| Interaction_genotype_Mtor | Ell2 | -3.737 | 6.06E-04 | glio |
| Interaction_genotype_Mtor | Abhd18 | -4.093 | 6.06E-04 | glio |
| Interaction_genotype_Mtor | Tmem59l | -3.385 | 6.06E-04 | glio |
| Aars2_in.vivo_noRT | Zfp882 | 2.638 | 6.06E-04 | glio |
| Aars2_in.vivo_noRT | Gm10825 | 2.498 | 6.06E-04 | glio |
| Aars2_in.vivo_noRT | Impdh1 | -1.662 | 6.06E-04 | glio |
| Aars2_in.vivo_noRT | Rpf2 | -1.187 | 6.06E-04 | glio |
| Bora_in.vivo_noRT | Sipa1l2 | 2.494 | 6.06E-04 | glio |
| Chek1_in.vivo_noRT | Ubash3a | 3.622 | 6.06E-04 | glio |
| Chek1_in.vivo_noRT | Gm49768 | 2.291 | 6.06E-04 | glio |
| Interaction_genotype_Aars2 | Gm14968 | -5.039 | 6.06E-05 | glio |
| Interaction_genotype_Myc | Gm14379 | -4.355 | 6.06E-05 | glio |
| Aars2_in.vivo_noRT | N4bp2os | 2.899 | 6.06E-05 | glio |
| Bcas2_in.vivo_noRT | Mindy1 | -1.753 | 6.06E-05 | glio |
| Chek1_in.vivo_noRT | Grip1 | 3.165 | 6.06E-06 | glio |
| Interaction_genotype_Myc | C130060C02R | -5.228 | 6.06E-07 | glio |
| Hsd17b10_in.vivo_noRT | Gm20767 | 3.616 | 6.06E-08 | glio |
| Atr_in.vivo_noRT | Zfp968 | 4.156 | 6.06E-09 | glio |
| Hsd17b10_in.vivo_noRT | Impg2 | 3.616 | 6.06E-10 | glio |
| Aars2_in.vivo_noRT | Gm43463 | 4.669 | 6.06E-15 | glio |
| Bcas2_in.vivo_noRT | Gm12794 | 4.807 | 6.06E-21 | glio |
| Interaction_genotype_Aars2 | C330011M18F | -3.461 | 6.07E-03 | glio |
| Interaction_genotype_Atr | Gm26559 | -3.442 | 6.07E-03 | glio |
| Interaction_genotype_Atr | Gm10863 | -3.08 | 6.07E-03 | glio |
| Interaction_genotype_Atr | Dnaic2 | -2.3 | 6.07E-03 | glio |
| Interaction_genotype_Atr | Parp3 | 1.612 | 6.07E-03 | glio |
| Interaction_genotype_Atr | Med9 | -1.485 | 6.07E-03 | glio |
| Interaction_genotype_Atr | Sgta | 1.317 | 6.07E-03 | glio |

| | | | | |
|---|---|---|---|---|
| Interaction_genotype_Atr | Sox4 | 1.637 | 6.07E-03 | glio |
| Interaction_genotype_Atr | H2-D1 | 1.127 | 6.07E-03 | glio |
| Interaction_genotype_Mtor | Kpna3 | -1.871 | 6.07E-03 | glio |
| Atr_in.vivo_noRT | 2610044O15R | 1.364 | 6.07E-03 | glio |
| Atr_in.vivo_noRT | Capza1 | -1.031 | 6.07E-03 | glio |
| Bcas2_in.vivo_noRT | Gm19391 | 1.908 | 6.07E-03 | glio |
| Bcas2_in.vivo_noRT | Rps19bp1 | -0.737 | 6.07E-03 | glio |
| Bora_in.vivo_noRT | Mfap5 | 2.105 | 6.07E-03 | glio |
| Bora_in.vivo_noRT | Ldlrad4 | 1.711 | 6.07E-03 | glio |
| Bora_in.vivo_noRT | Glyr1 | -1.131 | 6.07E-03 | glio |
| Chek1_in.vivo_noRT | 1110012L19Ri | -1.247 | 6.07E-03 | glio |
| Chek1_in.vivo_noRT | Rpl29 | -0.465 | 6.07E-03 | glio |
| Eif2s1_in.vivo_noRT | 1700024I08Ri | 1.883 | 6.07E-03 | glio |
| culture_effect | Pik3cd | -1.549 | 6.07E-04 | glio |
| Interaction_genotype_Hsd17b | Apol6 | 4.375 | 6.07E-04 | glio |
| Interaction_genotype_Hsd17b | Gltp | 2.073 | 6.07E-04 | glio |
| Interaction_genotype_Hsd17b | Glod5 | 6.119 | 6.07E-04 | glio |
| Interaction_genotype_Mtor | 1700030C12R | -4.237 | 6.07E-04 | glio |
| Interaction_genotype_Mtor | Eaf2 | -4.359 | 6.07E-04 | glio |
| Interaction_genotype_Mtor | Grasp | 3.157 | 6.07E-04 | glio |
| Interaction_genotype_Myc | Gm15816 | -3.42 | 6.07E-04 | glio |
| Interaction_genotype_Myc | Ttc25 | -3.82 | 6.07E-04 | glio |
| Interaction_genotype_Myc | Fhod1 | -3.281 | 6.07E-04 | glio |
| Interaction_genotype_Myc | Kif14 | -3.358 | 6.07E-04 | glio |
| Interaction_genotype_Myc | Gpat4 | -2.454 | 6.07E-04 | glio |
| interaction_RT | Siah1a | 1.184 | 6.07E-04 | glio |
| interaction_RT | Btbd17 | 3.664 | 6.07E-04 | glio |
| interaction_RT | Hes5 | 3.922 | 6.07E-04 | glio |
| interaction_RT | Mrps36 | 0.653 | 6.07E-04 | glio |
| interaction_RT | Ccnd1 | -0.877 | 6.07E-04 | glio |
| Interaction_genotype_Aars2 | Txn2 | 1.616 | 6.07E-05 | glio |
| Interaction_genotype_Mtor | Gm28731 | -4.99 | 6.07E-05 | glio |
| Interaction_genotype_Mtor | Trappc6b | 2.105 | 6.07E-05 | glio |
| Interaction_genotype_Myc | Tln2 | -3.717 | 6.07E-05 | glio |
| Bcas2_in.vivo_noRT | Gm26930 | 3.118 | 6.07E-05 | glio |
| Bcas2_in.vivo_noRT | Mapk15 | 2.823 | 6.07E-05 | glio |
| Bcas2_in.vivo_noRT | 9130230N09R | 2.488 | 6.07E-05 | glio |
| Chek1_in.vivo_noRT | Mst1 | 2.673 | 6.07E-05 | glio |
| Chek1_ex.vivo_noRT | Gm19345 | 4.055 | 6.07E-07 | glio |
| interaction_RT | Gbp6 | 1.976 | 6.07E-08 | glio |
| Chek1_ex.vivo_noRT | Vgll2 | 4.842 | 6.07E-10 | glio |
| Bcas2_in.vivo_noRT | Ccdc63 | 4.421 | 6.07E-11 | glio |
| Aars2_in.vivo_noRT | 7530414M10F | 4.801 | 6.07E-13 | glio |
| Aars2_in.vivo_noRT | Kars | -2.503 | 6.07E-13 | glio |
| Bcas2_in.vivo_noRT | A330032B11R | 4.541 | 6.07E-13 | glio |
| Atr_in.vivo_noRT | Hcn3 | 4.891 | 6.07E-15 | glio |
| Aars2_in.vivo_noRT | Lrp2bp | 5.063 | 6.07E-16 | glio |
| Bcas2_in.vivo_noRT | Gm4027 | 4.807 | 6.07E-18 | glio |
| Chek1_in.vivo_noRT | C430002N11R | 5.044 | 6.07E-20 | glio |
| Atr_in.vivo_noRT | Gm21761 | 5.533 | 6.07E-24 | glio |
| culture_effect | Zfp113 | 0.982 | 6.08E-03 | glio |
| culture_effect | Larp1b | -0.532 | 6.08E-03 | glio |

| | | | | |
|---|---|---|---|---|
| Interaction_genotype_Aars2 | Gm17501 | -3.399 | 6.08E-03 | glio |
| Interaction_genotype_Bora | Mettl7b | -3.601 | 6.08E-03 | glio |
| Interaction_genotype_Mtor | P2ry2 | -3.29 | 6.08E-03 | glio |
| Interaction_genotype_Myc | Plppr3 | -3.115 | 6.08E-03 | glio |
| Interaction_genotype_Myc | Myl6 | 0.95 | 6.08E-03 | glio |
| Interaction_genotype_Rpl3 | Gm34047 | -3.85 | 6.08E-03 | glio |
| Interaction_genotype_Rpl3 | Adam12 | -3.264 | 6.08E-03 | glio |
| Aars2_in.vivo_noRT | Slc5a5 | 2.073 | 6.08E-03 | glio |
| Atr_ex.vivo_noRT | Rapgef3os2 | 1.978 | 6.08E-03 | glio |
| Atr_ex.vivo_noRT | Cd72 | 2.769 | 6.08E-03 | glio |
| Bcas2_in.vivo_noRT | Zfp595 | 2.176 | 6.08E-03 | glio |
| Bcas2_in.vivo_noRT | Pkdcc | 2.095 | 6.08E-03 | glio |
| Chek1_in.vivo_noRT | Plekha1 | 1.09 | 6.08E-03 | glio |
| Chek1_in.vivo_noRT | Sfr1 | -0.611 | 6.08E-03 | glio |
| Chek1_in.vivo_noRT | Gabarap | -0.625 | 6.08E-03 | glio |
| Cyp19a1_ex.vivo_noRT | Erich6b | 2.159 | 6.08E-03 | glio |
| Cyp19a1_ex.vivo_noRT | Cldn5 | 2.159 | 6.08E-03 | glio |
| Cyp19a1_ex.vivo_noRT | Dpep3 | 2.159 | 6.08E-03 | glio |
| Interaction_genotype_Aars2 | Gm50107 | -4.036 | 6.08E-04 | glio |
| Interaction_genotype_Mtor | Ces1a | -4.553 | 6.08E-04 | glio |
| Interaction_genotype_Mtor | Gm4673 | -3.866 | 6.08E-04 | glio |
| Interaction_genotype_Mtor | Enho | -3.726 | 6.08E-04 | glio |
| Interaction_genotype_Mtor | Rilpl2 | -2.96 | 6.08E-04 | glio |
| Atp5c1_in.vivo_noRT | Ccdc162 | 2.139 | 6.08E-04 | glio |
| Atr_in.vivo_noRT | Arpc1a | -1.067 | 6.08E-04 | glio |
| Bora_in.vivo_noRT | Usp27x | 2.63 | 6.08E-04 | glio |
| Bora_in.vivo_noRT | Atp5a1 | -0.699 | 6.08E-04 | glio |
| Bora_ex.vivo_noRT | Tcstv1 | 2.802 | 6.08E-04 | glio |
| Chek1_ex.vivo_noRT | Traf1 | 3.085 | 6.08E-04 | glio |
| Bcas2_in.vivo_noRT | Lypd6 | 2.885 | 6.08E-05 | glio |
| Bora_in.vivo_noRT | P4htm | 2.858 | 6.08E-05 | glio |
| Aars2_in.vivo_noRT | Gm19684 | 3.368 | 6.08E-06 | glio |
| Aars2_in.vivo_noRT | Slc25a29 | 2.897 | 6.08E-06 | glio |
| Aars2_in.vivo_noRT | Arhgef26 | -2.12 | 6.08E-06 | glio |
| Chek1_ex.vivo_noRT | Gm15890 | 3.279 | 6.08E-06 | glio |
| Interaction_genotype_Mtor | Syne3 | -6.432 | 6.08E-07 | glio |
| Bora_in.vivo_noRT | Frmd8os | 3.952 | 6.08E-07 | glio |
| Chek1_in.vivo_noRT | Mir155hg | 3.799 | 6.08E-07 | glio |
| Aars2_in.vivo_noRT | Pglyrp1 | 3.847 | 6.08E-08 | glio |
| Atr_in.vivo_noRT | Tnc | 4.484 | 6.08E-08 | glio |
| Bora_in.vivo_noRT | Olfr1189 | 3.281 | 6.08E-08 | glio |
| Chek1_ex.vivo_noRT | Ripor2 | 4.839 | 6.08E-08 | glio |
| Bcas2_in.vivo_noRT | Gm28404 | 3.919 | 6.08E-10 | glio |
| Bcas2_in.vivo_noRT | Cyp4f40 | 4.154 | 6.08E-10 | glio |
| Bora_in.vivo_noRT | Tmem74b | 4.672 | 6.08E-16 | glio |
| Aars2_in.vivo_noRT | Gm36356 | 5.067 | 6.08E-17 | glio |
| Atr_in.vivo_noRT | Zscan4-ps2 | 5.533 | 6.08E-26 | glio |
| culture_effect | Yif1b | -0.36 | 6.09E-03 | glio |
| Interaction_genotype_Aars2 | Gm49701 | -3.333 | 6.09E-03 | glio |
| Interaction_genotype_Aars2 | Zfp458 | -3.602 | 6.09E-03 | glio |
| Interaction_genotype_Aars2 | Psmd8 | 1.089 | 6.09E-03 | glio |
| Interaction_genotype_Aars2 | Ifitm3 | 0.975 | 6.09E-03 | glio |

| | | | | |
|---|---|---|---|---|
| Interaction_genotype_Bcas2 | 2810414N06R | -3.654 | 6.09E-03 | glio |
| Interaction_genotype_Bcas2 | BC049715 | -3.037 | 6.09E-03 | glio |
| Interaction_genotype_Chek1 | Gm49312 | -3.368 | 6.09E-03 | glio |
| Interaction_genotype_Chek1 | Pid1 | 3.642 | 6.09E-03 | glio |
| Interaction_genotype_Hsd17b | Tslp | -4.405 | 6.09E-03 | glio |
| Interaction_genotype_Hsd17b | Fa2h | 3.95 | 6.09E-03 | glio |
| Interaction_genotype_Mtor | Fbxo6 | -1.976 | 6.09E-03 | glio |
| Interaction_genotype_Myc | Neto2 | -2.086 | 6.09E-03 | glio |
| Interaction_genotype_Pars2 | Gm15234 | -3.724 | 6.09E-03 | glio |
| Aars2_in.vivo_noRT | Marc2 | -1.057 | 6.09E-03 | glio |
| Atr_in.vivo_noRT | Dop1b | 1.872 | 6.09E-03 | glio |
| Bcas2_in.vivo_noRT | AI429214 | 2.05 | 6.09E-03 | glio |
| Bcas2_in.vivo_noRT | 4933408B17R | 2.064 | 6.09E-03 | glio |
| Bcas2_in.vivo_noRT | Rbm15 | -1.441 | 6.09E-03 | glio |
| Bcas2_in.vivo_noRT | Bag6 | -1.012 | 6.09E-03 | glio |
| Bcas2_ex.vivo_noRT | Tnfrsf1b | 2.742 | 6.09E-03 | glio |
| Brca2_in.vivo_noRT | Gm13964 | 1.763 | 6.09E-03 | glio |
| Chek1_in.vivo_noRT | Eif3c | -0.693 | 6.09E-03 | glio |
| Chek1_ex.vivo_noRT | Gm43066 | 2.468 | 6.09E-03 | glio |
| Eif2s1_in.vivo_noRT | Gm10425 | 2.407 | 6.09E-03 | glio |
| culture_effect | Nkain4 | -2.022 | 6.09E-04 | glio |
| Interaction_genotype_Atr | Gm45338 | -2.763 | 6.09E-04 | glio |
| Interaction_genotype_Bcas2 | Gm14002 | -4.215 | 6.09E-04 | glio |
| Interaction_genotype_Myc | Gm48764 | -2.729 | 6.09E-04 | glio |
| Chek1_ex.vivo_noRT | Rbakdn | 3.177 | 6.09E-04 | glio |
| culture_effect | AA386476 | 2.072 | 6.09E-05 | glio |
| Aars2_ex.vivo_noRT | Col19a1 | 2.9 | 6.09E-05 | glio |
| Atr_in.vivo_noRT | Itgae | 2.906 | 6.09E-05 | glio |
| Atr_in.vivo_noRT | Hdac10 | 2.456 | 6.09E-05 | glio |
| Bcas2_ex.vivo_noRT | Gad1 | 2.742 | 6.09E-05 | glio |
| Chek1_in.vivo_noRT | 4930542D17R | 2.764 | 6.09E-05 | glio |
| Interaction_genotype_Myc | Gm48882 | -5.306 | 6.09E-06 | glio |
| Aars2_in.vivo_noRT | Dtx1 | 3.149 | 6.09E-06 | glio |
| Atr_in.vivo_noRT | Fes | 3.213 | 6.09E-06 | glio |
| Atr_in.vivo_noRT | Gm28229 | 2.892 | 6.09E-06 | glio |
| Chek1_in.vivo_noRT | Gm50218 | 3.45 | 6.09E-06 | glio |
| Bora_in.vivo_noRT | Gm35867 | 3.483 | 6.09E-07 | glio |
| Bora_ex.vivo_noRT | Gm16933 | 3.201 | 6.09E-07 | glio |
| Atr_in.vivo_noRT | Fgf2 | 3.907 | 6.09E-08 | glio |
| Aars2_ex.vivo_noRT | Gm12703 | 4.092 | 6.09E-10 | glio |
| culture_effect | Dok1 | 1.373 | 6.09E-13 | glio |
| Aars2_in.vivo_noRT | G630030J09Ri | 4.805 | 6.09E-13 | glio |
| culture_effect | P4ha3 | 3.404 | 6.09E-17 | glio |
| culture_effect | Rwdd1 | 0.782 | 6.09E-17 | glio |
| Interaction_genotype_Mtor | Ulk1 | -2.681 | 6.10E-03 | glio |
| Interaction_genotype_Terf1 | Loxl4 | -3.849 | 6.10E-03 | glio |
| interaction_RT | Syce2 | 0.803 | 6.10E-03 | glio |
| interaction_RT | Spsb2 | 0.764 | 6.10E-03 | glio |
| Aars2_in.vivo_noRT | Tceal1 | 1.901 | 6.10E-03 | glio |
| Atr_in.vivo_noRT | Pycr2 | -1.063 | 6.10E-03 | glio |
| Bcas2_in.vivo_noRT | Cracr2b | 2.146 | 6.10E-03 | glio |
| Bcas2_in.vivo_noRT | Pon3 | 2.057 | 6.10E-03 | glio |

| | | | | |
|---|---|---|---|---|
| Bcas2_in.vivo_noRT | Syn1 | 2.071 | 6.10E-03 | glio |
| Bcas2_in.vivo_noRT | Psmb5 | -0.609 | 6.10E-03 | glio |
| Bcas2_ex.vivo_noRT | Gm43915 | 2.348 | 6.10E-03 | glio |
| Bcas2_ex.vivo_noRT | Itgb8 | 1.925 | 6.10E-03 | glio |
| Aars2_in.vivo_noRT | Oscp1 | 2.444 | 6.10E-04 | glio |
| Bcas2_in.vivo_noRT | Ptprn | 2.315 | 6.10E-04 | glio |
| Bora_in.vivo_noRT | Slc6a20b | 2.452 | 6.10E-04 | glio |
| Bora_in.vivo_noRT | Lgalsl | -1.606 | 6.10E-04 | glio |
| Chek1_ex.vivo_noRT | Nhsl2 | 3.079 | 6.10E-04 | glio |
| Eif2s1_in.vivo_noRT | Tagap | 1.883 | 6.10E-04 | glio |
| Interaction_genotype_Myc | Hist1h2ai | -4.729 | 6.10E-05 | glio |
| Interaction_genotype_Myc | Hnrnpk | 1.587 | 6.10E-05 | glio |
| Aars2_in.vivo_noRT | Gm19744 | 2.881 | 6.10E-05 | glio |
| Bcas2_in.vivo_noRT | Mucl3 | 3.036 | 6.10E-05 | glio |
| Bcas2_in.vivo_noRT | Slc7a7 | 2.839 | 6.10E-05 | glio |
| Interaction_genotype_Myc | Gramd2 | -4.948 | 6.10E-06 | glio |
| Bcas2_in.vivo_noRT | Pola1 | -1.862 | 6.10E-06 | glio |
| culture_effect | Pmel | -1.71 | 6.10E-08 | glio |
| culture_effect | Plagl1 | -1.182 | 6.10E-08 | glio |
| Chek1_in.vivo_noRT | Nmb | 3.75 | 6.10E-08 | glio |
| Interaction_genotype_Mtor | Prl3d3 | -4.931 | 6.10E-09 | glio |
| Aars2_in.vivo_noRT | Gm49494 | 4.496 | 6.10E-10 | glio |
| Bora_in.vivo_noRT | Tecrl | 4.676 | 6.10E-11 | glio |
| Bora_in.vivo_noRT | Gm4724 | 4.28 | 6.10E-12 | glio |
| Bora_in.vivo_noRT | Hunk | 4.404 | 6.10E-12 | glio |
| Aars2_in.vivo_noRT | Gm29083 | 4.794 | 6.10E-14 | glio |
| Cdkn1a_in.vivo_noRT | Cdkn1a | -4.087 | 6.10E-47 | glio |
| culture_effect | Zfp334 | -0.92 | 6.11E-03 | glio |
| Interaction_genotype_Aars2 | 6430562O15R | -3.102 | 6.11E-03 | glio |
| Interaction_genotype_Aars2 | Dpf1 | -2.897 | 6.11E-03 | glio |
| Interaction_genotype_Myc | H60b | -3.045 | 6.11E-03 | glio |
| Interaction_genotype_Rpl3 | Epsti1 | 2.56 | 6.11E-03 | glio |
| Atr_in.vivo_noRT | AA465934 | 1.772 | 6.11E-03 | glio |
| Atr_in.vivo_noRT | Nupl2 | 1.546 | 6.11E-03 | glio |
| Bcas2_in.vivo_noRT | Cyb5r3 | -0.931 | 6.11E-03 | glio |
| Chek1_ex.vivo_noRT | Adhfe1 | 2.664 | 6.11E-03 | glio |
| Eif2s1_in.vivo_noRT | Neil2 | -2.18 | 6.11E-03 | glio |
| culture_effect | Phldb1 | -0.505 | 6.11E-04 | glio |
| Interaction_genotype_Bcas2 | Gm29083 | -4.189 | 6.11E-04 | glio |
| Aars2_ex.vivo_noRT | Rab27a | 2.897 | 6.11E-04 | glio |
| Aars2_ex.vivo_noRT | Osm | 2.893 | 6.11E-04 | glio |
| Bcas2_in.vivo_noRT | Zfp493 | 2.458 | 6.11E-04 | glio |
| Chek1_in.vivo_noRT | 9330121J05Ri | 2.471 | 6.11E-04 | glio |
| Bcas2_in.vivo_noRT | Gm28417 | 3.016 | 6.11E-06 | glio |
| Interaction_genotype_Mtor | Gm49698 | -4.099 | 6.11E-07 | glio |
| Aars2_in.vivo_noRT | Lrrc75b | 3.67 | 6.11E-07 | glio |
| Aars2_in.vivo_noRT | Fam83e | 3.592 | 6.11E-07 | glio |
| Aars2_in.vivo_noRT | Saxo2 | 3.402 | 6.11E-07 | glio |
| Atr_in.vivo_noRT | Ptpru | 3.07 | 6.11E-07 | glio |
| Chek1_in.vivo_noRT | Bbof1 | 3.615 | 6.11E-08 | glio |
| culture_effect | Gm21887 | 1.169 | 6.11E-09 | glio |
| Chek1_ex.vivo_noRT | Rem2 | 4.842 | 6.11E-10 | glio |

| | | | | |
|---|---|---|---|---|
| Chek1_ex.vivo_noRT | Gm16685 | 4.443 | 6.11E-10 | glio |
| Chek1_ex.vivo_noRT | Muc4 | 4.842 | 6.11E-12 | glio |
| Chek1_ex.vivo_noRT | Gm39526 | 4.842 | 6.11E-13 | glio |
| Chek1_ex.vivo_noRT | Fabp6 | 4.842 | 6.11E-14 | glio |
| Atr_in.vivo_noRT | Ush1g | 5.533 | 6.11E-25 | glio |
| culture_effect | Fam49b | 0.371 | 6.12E-03 | glio |
| Interaction_genotype_Atr | Srebf2 | 1.84 | 6.12E-03 | glio |
| Interaction_genotype_Bora | Rnf4 | 1.154 | 6.12E-03 | glio |
| Interaction_genotype_Ifnar1 | Ube2l6 | -1.238 | 6.12E-03 | glio |
| Interaction_genotype_Mtor | Cnbd2 | -3.419 | 6.12E-03 | glio |
| Interaction_genotype_Mtor | Phldb1 | 1.457 | 6.12E-03 | glio |
| Interaction_genotype_Mtor | Bod1l | 1.372 | 6.12E-03 | glio |
| Interaction_genotype_Myc | C330018D20R | -2.79 | 6.12E-03 | glio |
| Atp5c1_in.vivo_noRT | Tubb4a | 2.004 | 6.12E-03 | glio |
| Atr_in.vivo_noRT | Map3k6 | 2.036 | 6.12E-03 | glio |
| Atr_in.vivo_noRT | Tmem181a | -1.29 | 6.12E-03 | glio |
| Atr_in.vivo_noRT | Seh1l | -0.945 | 6.12E-03 | glio |
| Atr_in.vivo_noRT | Nav3 | -0.773 | 6.12E-03 | glio |
| Chek1_ex.vivo_noRT | Isoc1 | -1.267 | 6.12E-03 | glio |
| Chek1_ex.vivo_noRT | Gps1 | -1.057 | 6.12E-03 | glio |
| Chek1_ex.vivo_noRT | Ranbp1 | -0.728 | 6.12E-03 | glio |
| culture_effect | Zan | -2.134 | 6.12E-04 | glio |
| Interaction_genotype_Myc | Dynll2 | 1.519 | 6.12E-04 | glio |
| Interaction_genotype_Rad21 | Prune2 | -3.972 | 6.12E-04 | glio |
| Aars2_in.vivo_noRT | Kcmf1 | -1.088 | 6.12E-04 | glio |
| Aars2_in.vivo_noRT | Xrn2 | -0.939 | 6.12E-04 | glio |
| Chek1_ex.vivo_noRT | Gm2115 | 2.607 | 6.12E-04 | glio |
| Interaction_genotype_Aars2 | BC049762 | -5.113 | 6.12E-05 | glio |
| Interaction_genotype_Aars2 | Txn1 | 1.272 | 6.12E-05 | glio |
| Aars2_in.vivo_noRT | 4930556M19F | 2.679 | 6.12E-05 | glio |
| Aars2_ex.vivo_noRT | Strip2 | 2.893 | 6.12E-05 | glio |
| Aars2_ex.vivo_noRT | 4732463B04R | 2.9 | 6.12E-05 | glio |
| Atr_ex.vivo_noRT | AC159140.1 | 2.771 | 6.12E-05 | glio |
| Bora_in.vivo_noRT | Syce2 | -1.759 | 6.12E-05 | glio |
| Hsd17b10_in.vivo_noRT | AU022754 | 2.727 | 6.12E-05 | glio |
| Interaction_genotype_Mtor | Flicr | -5.336 | 6.12E-06 | glio |
| Aars2_in.vivo_noRT | Banf1 | -1.212 | 6.12E-06 | glio |
| Chek1_ex.vivo_noRT | Sh3d21 | 3.345 | 6.12E-06 | glio |
| Interaction_genotype_Mtor | Tnrc6b | 2.901 | 6.12E-07 | glio |
| Interaction_genotype_Mtor | Gm20522 | -6.07 | 6.12E-07 | glio |
| Interaction_genotype_Mtor | Olfr881 | -4.099 | 6.12E-07 | glio |
| Interaction_genotype_Mtor | mt-Co2 | 1.933 | 6.12E-07 | glio |
| culture_effect | Myzap | -2.857 | 6.12E-08 | glio |
| Aars2_in.vivo_noRT | Rab11fip4os1 | 4.29 | 6.12E-09 | glio |
| Aars2_in.vivo_noRT | Prox2 | 4.601 | 6.12E-10 | glio |
| Atr_in.vivo_noRT | Aqp3 | 5.269 | 6.12E-10 | glio |
| Bora_in.vivo_noRT | Gm44702 | 4.405 | 6.12E-10 | glio |
| Aars2_in.vivo_noRT | 2310058D17R | 4.662 | 6.12E-11 | glio |
| Aars2_in.vivo_noRT | Gm34047 | 5.327 | 6.12E-14 | glio |
| Hsd17b10_in.vivo_noRT | Capn13 | 3.616 | 6.12E-15 | glio |
| culture_effect | Crebrf | -1.573 | 6.12E-17 | glio |
| Chek1_in.vivo_noRT | Gm49086 | 5.044 | 6.12E-17 | glio |

| | | | | |
|---|---|---|---|---|
| Atr_in.vivo_noRT | Zscan4d | 5.533 | 6.12E-22 | glio |
| culture_effect | Zfp952 | -1.02 | 6.13E-03 | glio |
| Interaction_genotype_Mtor | Ulbp1 | -2.827 | 6.13E-03 | glio |
| Interaction_genotype_Mtor | Jrk | -2.874 | 6.13E-03 | glio |
| Interaction_genotype_Mtor | Rdx | 1.324 | 6.13E-03 | glio |
| Interaction_genotype_Pars2 | Gm47232 | -3.737 | 6.13E-03 | glio |
| Aars2_in.vivo_noRT | Nhp2 | -0.694 | 6.13E-03 | glio |
| Aars2_ex.vivo_noRT | Cog8 | -2.173 | 6.13E-03 | glio |
| Atp5c1_in.vivo_noRT | Smpd5 | 2.011 | 6.13E-03 | glio |
| Atp5c1_in.vivo_noRT | Gipc2 | 2.405 | 6.13E-03 | glio |
| Atp5c1_in.vivo_noRT | Ifi207 | 2.405 | 6.13E-03 | glio |
| Atp5c1_in.vivo_noRT | Fam131a | 2.142 | 6.13E-03 | glio |
| Atp5c1_in.vivo_noRT | Rem2 | 2.014 | 6.13E-03 | glio |
| Atp5c1_in.vivo_noRT | Cep250 | -1.826 | 6.13E-03 | glio |
| Bcas2_in.vivo_noRT | Slc13a5 | 1.929 | 6.13E-03 | glio |
| Cenpj_in.vivo_noRT | Prkg2 | -1.163 | 6.13E-03 | glio |
| Chek1_in.vivo_noRT | Dhrs11 | 1.896 | 6.13E-03 | glio |
| Chek1_ex.vivo_noRT | Gm15127 | 2.468 | 6.13E-03 | glio |
| Chek1_ex.vivo_noRT | Gm15265 | 2.538 | 6.13E-03 | glio |
| Chek1_ex.vivo_noRT | Slc9b1 | 2.608 | 6.13E-03 | glio |
| culture_effect | Tmem173 | -0.884 | 6.13E-04 | glio |
| culture_effect | Ppil4 | -0.677 | 6.13E-04 | glio |
| Interaction_genotype_Mtor | Vrk2 | -3.22 | 6.13E-04 | glio |
| Interaction_genotype_Mtor | Smc6 | 1.877 | 6.13E-04 | glio |
| Interaction_genotype_Mtor | Pnp | 2.157 | 6.13E-04 | glio |
| Bcas2_in.vivo_noRT | Dennd5a | -1.435 | 6.13E-04 | glio |
| Bora_in.vivo_noRT | Ccar1 | -0.918 | 6.13E-04 | glio |
| Chek1_ex.vivo_noRT | Fbxo34 | -2.176 | 6.13E-04 | glio |
| Aars2_ex.vivo_noRT | Kcnq2 | 3.293 | 6.13E-05 | glio |
| Bcas2_in.vivo_noRT | Gm26910 | 2.652 | 6.13E-05 | glio |
| culture_effect | Abhd17a | 0.568 | 6.13E-06 | glio |
| Interaction_genotype_Mtor | Gm33051 | -4.243 | 6.13E-06 | glio |
| Interaction_genotype_Myc | Gm13270 | -5.38 | 6.13E-06 | glio |
| interaction_RT | Ggh | 1.088 | 6.13E-06 | glio |
| Hsd17b10_in.vivo_noRT | Hbq1b | 3.35 | 6.13E-06 | glio |
| Bcas2_in.vivo_noRT | Gm3604 | 3.525 | 6.13E-07 | glio |
| Cenpj_ex.vivo_noRT | Rem2 | 4.535 | 6.13E-07 | glio |
| Cenpj_ex.vivo_noRT | Gm29966 | 3.389 | 6.13E-07 | glio |
| Aars2_in.vivo_noRT | Sgk3 | 3.772 | 6.13E-08 | glio |
| Bora_in.vivo_noRT | Ppp1r3g | 3.751 | 6.13E-09 | glio |
| Aars2_in.vivo_noRT | Gm44096 | 4.056 | 6.13E-10 | glio |
| Atr_in.vivo_noRT | Gm37584 | 4.622 | 6.13E-11 | glio |
| Hsd17b10_in.vivo_noRT | Rimklb | 3.353 | 6.13E-12 | glio |
| Aars2_in.vivo_noRT | E030042O20R | 5.327 | 6.13E-13 | glio |
| Atr_in.vivo_noRT | Amh | 4.616 | 6.13E-13 | glio |
| Chek1_in.vivo_noRT | Gm49441 | 4.253 | 6.13E-13 | glio |
| Chek1_in.vivo_noRT | Slc4a9 | 4.511 | 6.13E-15 | glio |
| culture_effect | Cd14 | -4.14 | 6.13E-17 | glio |
| Aars2_in.vivo_noRT | Clcn1 | 4.681 | 6.13E-17 | glio |
| Bcas2_in.vivo_noRT | AC154200.1 | 4.807 | 6.13E-18 | glio |
| Aars2_in.vivo_noRT | Gm10447 | 5.327 | 6.13E-22 | glio |
| Interaction_genotype_Bcas2 | Slc6a13 | -3.95 | 6.14E-03 | glio |

| | | | | |
|---|---|---|---|---|
| Interaction_genotype_Bcas2 | C2cd2 | -3.858 | 6.14E-03 | glio |
| Interaction_genotype_Chek1 | Cpsf3 | -2.331 | 6.14E-03 | glio |
| Interaction_genotype_Mtor | Kmo | -2.85 | 6.14E-03 | glio |
| Interaction_genotype_Mtor | Zfp51 | -2.815 | 6.14E-03 | glio |
| Interaction_genotype_Mtor | Tlr4 | -2.348 | 6.14E-03 | glio |
| Interaction_genotype_Rad51 | Scube3 | -3.215 | 6.14E-03 | glio |
| interaction_RT | Pafah1b3 | 0.901 | 6.14E-03 | glio |
| Aars2_in.vivo_noRT | St6gal1 | 2.079 | 6.14E-03 | glio |
| Aars2_ex.vivo_noRT | 9230112J17Ri | 2.312 | 6.14E-03 | glio |
| Atp5c1_in.vivo_noRT | Shfl | 1.509 | 6.14E-03 | glio |
| Atr_in.vivo_noRT | Itpkb | 1.609 | 6.14E-03 | glio |
| Atr_in.vivo_noRT | Bcap29 | -0.985 | 6.14E-03 | glio |
| Atr_in.vivo_noRT | Psma3 | -0.559 | 6.14E-03 | glio |
| Bora_in.vivo_noRT | Eaf2 | 2.083 | 6.14E-03 | glio |
| Chek1_in.vivo_noRT | Atp6v1f | -0.647 | 6.14E-03 | glio |
| Chek1_ex.vivo_noRT | Nol7 | -0.788 | 6.14E-03 | glio |
| Hsd17b10_in.vivo_noRT | Glrx | 1.898 | 6.14E-03 | glio |
| Interaction_genotype_Myc | A330009N23R | -3.79 | 6.14E-04 | glio |
| Interaction_genotype_Myc | Ube2o | -2.365 | 6.14E-04 | glio |
| Atr_in.vivo_noRT | Gm28535 | 2.35 | 6.14E-04 | glio |
| Bcas2_in.vivo_noRT | Gm48768 | 2.556 | 6.14E-04 | glio |
| Chek1_ex.vivo_noRT | Dnah10 | 2.707 | 6.14E-04 | glio |
| Interaction_genotype_Atr | Efcab9 | -4.813 | 6.14E-05 | glio |
| Interaction_genotype_Atr | Htr1b | -4.862 | 6.14E-05 | glio |
| Interaction_genotype_Mtor | Sem1 | 1.971 | 6.14E-05 | glio |
| Bora_ex.vivo_noRT | 1700010B08R | 3.201 | 6.14E-05 | glio |
| Chek1_in.vivo_noRT | Gm26982 | 2.823 | 6.14E-05 | glio |
| Interaction_genotype_Mtor | 1700126G02R | -5.524 | 6.14E-06 | glio |
| Interaction_genotype_Mtor | Gm2022 | -4.099 | 6.14E-06 | glio |
| Chek1_in.vivo_noRT | Cep295nl | 3.166 | 6.14E-06 | glio |
| culture_effect | Gm36696 | -2.99 | 6.14E-07 | glio |
| Aars2_in.vivo_noRT | Cyp2s1 | 3.754 | 6.14E-07 | glio |
| Bcas2_in.vivo_noRT | 4933421A08R | 4.028 | 6.14E-08 | glio |
| Aars2_in.vivo_noRT | Adcy7 | 3.822 | 6.14E-09 | glio |
| Atr_in.vivo_noRT | Trnp1 | 4.092 | 6.14E-09 | glio |
| Bcas2_in.vivo_noRT | Dzank1 | 4.161 | 6.14E-09 | glio |
| Hsd17b10_in.vivo_noRT | Slc16a4 | 3.352 | 6.14E-09 | glio |
| Atr_in.vivo_noRT | Gm34047 | 5.533 | 6.14E-15 | glio |
| Atr_in.vivo_noRT | Gm31063 | 5.533 | 6.14E-25 | glio |
| culture_effect | Nat8l | 5.127 | 6.14E-36 | glio |
| Interaction_genotype_Aars2 | Myl12b | 1.272 | 6.15E-03 | glio |
| Atr_in.vivo_noRT | Entpd2 | 1.677 | 6.15E-03 | glio |
| Bcas2_in.vivo_noRT | Best1 | 1.844 | 6.15E-03 | glio |
| Bcas2_in.vivo_noRT | Sh3kbp1 | -1.053 | 6.15E-03 | glio |
| Brca2_ex.vivo_noRT | Gm17210 | 3.398 | 6.15E-03 | glio |
| Chek1_in.vivo_noRT | Lipt1 | 1.812 | 6.15E-03 | glio |
| Eif2s1_in.vivo_noRT | D430001F17R | 2.143 | 6.15E-03 | glio |
| Ercc4_in.vivo_noRT | Lcn2 | 2.815 | 6.15E-03 | glio |
| Interaction_genotype_Mtor | Tgfbr3l | -4.27 | 6.15E-04 | glio |
| Interaction_genotype_Mtor | Fam160a2 | -3.469 | 6.15E-04 | glio |
| Interaction_genotype_Mtor | Tspan4 | 2.161 | 6.15E-04 | glio |
| Bora_ex.vivo_noRT | Gm43221 | 3.201 | 6.15E-04 | glio |

| | | | | |
|---|---|---|---|---|
| Hsd17b10_in.vivo_noRT | Slc36a2 | 2.497 | 6.15E-04 | glio |
| Interaction_genotype_Mtor | Kif5a | -5.236 | 6.15E-05 | glio |
| Atr_in.vivo_noRT | Ndrg4 | 2.893 | 6.15E-05 | glio |
| Interaction_genotype_Mtor | Gm43256 | -5.007 | 6.15E-06 | glio |
| culture_effect | Grhl2 | -3.143 | 6.15E-07 | glio |
| Interaction_genotype_Mtor | 9230112J17Ri | -5.08 | 6.15E-07 | glio |
| Atr_in.vivo_noRT | Cdc42bpg | 3.153 | 6.15E-07 | glio |
| Aars2_in.vivo_noRT | Retn | 4.859 | 6.15E-08 | glio |
| Aars2_in.vivo_noRT | Gm35551 | 3.511 | 6.15E-08 | glio |
| Bora_in.vivo_noRT | Slc26a6 | 3.644 | 6.15E-08 | glio |
| culture_effect | Rundc3a | -2.781 | 6.15E-10 | glio |
| Aars2_in.vivo_noRT | Gm30790 | 4.347 | 6.15E-10 | glio |
| Aars2_in.vivo_noRT | D030068K23R | 5.45 | 6.15E-15 | glio |
| Bcas2_in.vivo_noRT | Gm13032 | 4.421 | 6.15E-15 | glio |
| Interaction_genotype_Bora | Cebpa | -3.714 | 6.16E-03 | glio |
| Interaction_genotype_Bora | Gm48772 | -3.387 | 6.16E-03 | glio |
| Interaction_genotype_Bora | Taf11 | 1.684 | 6.16E-03 | glio |
| Interaction_genotype_Myc | 1700113A16R | -2.54 | 6.16E-03 | glio |
| Interaction_genotype_Myc | Rhoj | 1.711 | 6.16E-03 | glio |
| Atr_in.vivo_noRT | C730034F03Ri | 1.824 | 6.16E-03 | glio |
| Chek1_in.vivo_noRT | Xylb | 1.995 | 6.16E-03 | glio |
| Chek1_in.vivo_noRT | Tpk1 | 1.849 | 6.16E-03 | glio |
| Chek1_in.vivo_noRT | Cox7a2l | -0.676 | 6.16E-03 | glio |
| Interaction_genotype_Bora | Gm11457 | -4.243 | 6.16E-04 | glio |
| Interaction_genotype_Bora | Maea | 1.905 | 6.16E-04 | glio |
| Interaction_genotype_Lmo7 | 9330104G04R | -5.058 | 6.16E-04 | glio |
| Interaction_genotype_Lmo7 | Gm48242 | -4.365 | 6.16E-04 | glio |
| Interaction_genotype_Lmo7 | Klrk1 | 4.147 | 6.16E-04 | glio |
| Interaction_genotype_Myc | Fbxo15 | -3.448 | 6.16E-04 | glio |
| Interaction_genotype_Myc | Zxdc | -2.559 | 6.16E-04 | glio |
| Interaction_genotype_Myc | Psmd4 | 1.344 | 6.16E-04 | glio |
| Interaction_genotype_Myc | Ybx3 | 1.434 | 6.16E-04 | glio |
| Aars2_in.vivo_noRT | Dyrk4 | 2.759 | 6.16E-04 | glio |
| Atr_ex.vivo_noRT | Fhl1 | 2.453 | 6.16E-04 | glio |
| Bard1_ex.vivo_noRT | Gtsf1l | 2.814 | 6.16E-04 | glio |
| Bcas2_in.vivo_noRT | Setmar | 2.491 | 6.16E-04 | glio |
| Chek1_in.vivo_noRT | Rps16 | -0.687 | 6.16E-04 | glio |
| Hsd17b10_in.vivo_noRT | Gm20667 | 2.616 | 6.16E-04 | glio |
| Interaction_genotype_Atr | Gm35867 | -4.892 | 6.16E-05 | glio |
| Bora_in.vivo_noRT | Gm43017 | 2.822 | 6.16E-05 | glio |
| Bora_in.vivo_noRT | Pcdhgb5 | 3.032 | 6.16E-07 | glio |
| Aars2_in.vivo_noRT | Drd4 | 4.142 | 6.16E-09 | glio |
| Aars2_in.vivo_noRT | AV356131 | 4.043 | 6.16E-09 | glio |
| Bcas2_in.vivo_noRT | Hsf2bp | 4.015 | 6.16E-10 | glio |
| Bcas2_in.vivo_noRT | Stx1b | 3.904 | 6.16E-10 | glio |
| Bcas2_in.vivo_noRT | Fosb | -3.489 | 6.16E-10 | glio |
| Aars2_in.vivo_noRT | Gm21818 | 5.327 | 6.16E-14 | glio |
| Bcas2_in.vivo_noRT | 5730507C01R | 4.284 | 6.16E-16 | glio |
| Chek1_in.vivo_noRT | Gm10354 | 5.044 | 6.16E-19 | glio |
| culture_effect | B230219D22R | 0.467 | 6.17E-03 | glio |
| Interaction_genotype_Atr | Sh2d3c | -3.665 | 6.17E-03 | glio |
| Interaction_genotype_Mtor | Fsd2 | -3.302 | 6.17E-03 | glio |

| | | | | |
|---|---|---|---|---|
| Interaction_genotype_Mtor | Junos | -3.364 | 6.17E-03 | glio |
| Interaction_genotype_Mtor | Pdhb | 1.486 | 6.17E-03 | glio |
| Interaction_genotype_Myc | Pfkfb1 | -3.095 | 6.17E-03 | glio |
| Interaction_genotype_Myc | Gpr137c | -2.901 | 6.17E-03 | glio |
| Interaction_genotype_Myc | Cks1b | 1.38 | 6.17E-03 | glio |
| interaction_RT | 2900009J06Ri | 2.238 | 6.17E-03 | glio |
| interaction_RT | Nsmce2 | 0.605 | 6.17E-03 | glio |
| Aars2_in.vivo_noRT | Pald1 | 1.913 | 6.17E-03 | glio |
| Aars2_in.vivo_noRT | E230016M11F | 1.813 | 6.17E-03 | glio |
| Atr_in.vivo_noRT | Fhod1 | 1.668 | 6.17E-03 | glio |
| Atr_in.vivo_noRT | Elmo2 | 1.395 | 6.17E-03 | glio |
| Atr_ex.vivo_noRT | Tmem74b | 2.198 | 6.17E-03 | glio |
| Bcas2_in.vivo_noRT | Gm17056 | 2.074 | 6.17E-03 | glio |
| Chek1_in.vivo_noRT | Dbnl | -0.893 | 6.17E-03 | glio |
| Fanca_ex.vivo_noRT | Gm41349 | 2.913 | 6.17E-03 | glio |
| culture_effect | Ndfip2 | -0.612 | 6.17E-04 | glio |
| Interaction_genotype_Mtor | Retn | -5.294 | 6.17E-04 | glio |
| Interaction_genotype_Mtor | Upp2 | -3.485 | 6.17E-04 | glio |
| Interaction_genotype_Terf1 | Col6a1 | -3.595 | 6.17E-04 | glio |
| Atr_in.vivo_noRT | Timm50 | -1.327 | 6.17E-04 | glio |
| Atr_ex.vivo_noRT | Gm13684 | 2.77 | 6.17E-04 | glio |
| Chek1_ex.vivo_noRT | Ltb4r1 | 3.264 | 6.17E-04 | glio |
| culture_effect | Fgf21 | -2.709 | 6.17E-05 | glio |
| Interaction_genotype_Chek1 | Gm32834 | -4.858 | 6.17E-05 | glio |
| Interaction_genotype_Myc | Ankrd13b | -3.197 | 6.17E-05 | glio |
| Interaction_genotype_Myc | Mrps16 | 1.634 | 6.17E-05 | glio |
| Aars2_in.vivo_noRT | Ubb | -0.765 | 6.17E-05 | glio |
| Bcas2_in.vivo_noRT | Col11a1 | -0.879 | 6.17E-05 | glio |
| Chek1_in.vivo_noRT | Gm34552 | 2.85 | 6.17E-05 | glio |
| Aars2_in.vivo_noRT | Naip6 | 3.078 | 6.17E-06 | glio |
| culture_effect | Cnpy4 | -1.033 | 6.17E-07 | glio |
| Bora_in.vivo_noRT | Gm36371 | 3.315 | 6.17E-07 | glio |
| culture_effect | AC133103.1 | -3.441 | 6.17E-09 | glio |
| Chek1_ex.vivo_noRT | Vamp8 | 4.443 | 6.17E-10 | glio |
| culture_effect | Arrb1 | -0.865 | 6.18E-03 | glio |
| Interaction_genotype_Aars2 | Glb1l | -3.449 | 6.18E-03 | glio |
| Interaction_genotype_Mtor | Pycard | -3.167 | 6.18E-03 | glio |
| Interaction_genotype_Mtor | Sphk1 | -3.415 | 6.18E-03 | glio |
| Interaction_genotype_Mtor | Adamts12 | -2.336 | 6.18E-03 | glio |
| Aars2_in.vivo_noRT | Chl1 | -0.838 | 6.18E-03 | glio |
| Atr_ex.vivo_noRT | Platr20 | 2.842 | 6.18E-03 | glio |
| Fanca_ex.vivo_noRT | Gm37035 | 2.6 | 6.18E-03 | glio |
| Fanca_ex.vivo_noRT | 2810049E08Ri | 2.919 | 6.18E-03 | glio |
| Interaction_genotype_Myc | 2010109A12R | -3.683 | 6.18E-04 | glio |
| Atp5c1_in.vivo_noRT | Mgst1 | 2.101 | 6.18E-04 | glio |
| Atr_in.vivo_noRT | Zfp57 | 2.366 | 6.18E-04 | glio |
| Atr_in.vivo_noRT | Ifi47 | -1.549 | 6.18E-04 | glio |
| Bora_in.vivo_noRT | Gm15545 | 2.35 | 6.18E-04 | glio |
| Chek1_in.vivo_noRT | Fbxl12os | 2.403 | 6.18E-04 | glio |
| Interaction_genotype_Mtor | Gm29358 | -5.008 | 6.18E-05 | glio |
| Interaction_genotype_Myc | Tbc1d25 | -3.677 | 6.18E-05 | glio |
| Aars2_in.vivo_noRT | Htra1 | 3.157 | 6.18E-05 | glio |

| | | | | |
|---|---|---|---|---|
| Atr_ex.vivo_noRT | Fam189a1 | 3.164 | 6.18E-05 | glio |
| Bora_in.vivo_noRT | Gm49767 | 3.898 | 6.18E-08 | glio |
| Hsd17b10_in.vivo_noRT | Hist1h2ag | 3.084 | 6.18E-09 | glio |
| Atr_ex.vivo_noRT | Nat8f2 | 4.764 | 6.18E-11 | glio |
| Bcas2_in.vivo_noRT | Gm32834 | 4.419 | 6.18E-11 | glio |
| Atr_in.vivo_noRT | Gm14264 | 4.539 | 6.18E-12 | glio |
| Bora_in.vivo_noRT | Zfp786 | 4.672 | 6.18E-12 | glio |
| Bora_in.vivo_noRT | Rab25 | 4.67 | 6.18E-14 | glio |
| Aars2_in.vivo_noRT | Sh3d21 | 4.804 | 6.18E-16 | glio |
| culture_effect | Nmrk2 | -1.576 | 6.19E-03 | glio |
| culture_effect | Rtl9 | -1.665 | 6.19E-03 | glio |
| culture_effect | Mogat2 | -1.701 | 6.19E-03 | glio |
| culture_effect | Snx15 | -0.468 | 6.19E-03 | glio |
| Interaction_genotype_Mtor | Eif1b | 1.52 | 6.19E-03 | glio |
| Interaction_genotype_Myc | Ambra1 | -2.664 | 6.19E-03 | glio |
| Interaction_genotype_Myc | Rpap1 | -2.314 | 6.19E-03 | glio |
| Interaction_genotype_Myc | Mark3 | 1.337 | 6.19E-03 | glio |
| Atp5c1_in.vivo_noRT | Camk2b | 2.146 | 6.19E-03 | glio |
| Atp5c1_in.vivo_noRT | Hist2h3c2 | 2.402 | 6.19E-03 | glio |
| Atp5c1_in.vivo_noRT | Gm44983 | 1.88 | 6.19E-03 | glio |
| Atp5c1_in.vivo_noRT | Tacr2 | 2.283 | 6.19E-03 | glio |
| Bard1_ex.vivo_noRT | Acat3 | 2.239 | 6.19E-03 | glio |
| Bcas2_in.vivo_noRT | Rhbdl1 | 1.961 | 6.19E-03 | glio |
| Bcas2_in.vivo_noRT | Eif4e | -0.842 | 6.19E-03 | glio |
| Bcas2_in.vivo_noRT | Usp34 | -0.798 | 6.19E-03 | glio |
| Cenpj_in.vivo_noRT | Tcstv1 | 2.022 | 6.19E-03 | glio |
| Chek1_ex.vivo_noRT | Gfap | 2.468 | 6.19E-03 | glio |
| culture_effect | Plxnb1 | 1.803 | 6.19E-04 | glio |
| culture_effect | 5730414N17R | 1.839 | 6.19E-04 | glio |
| Interaction_genotype_Myc | Gm47972 | -3.835 | 6.19E-04 | glio |
| Interaction_genotype_Myc | Gm30262 | -3.994 | 6.19E-04 | glio |
| Interaction_genotype_Myc | Snupn | -3.038 | 6.19E-04 | glio |
| Aars2_in.vivo_noRT | Bco1 | 2.048 | 6.19E-04 | glio |
| Aars2_in.vivo_noRT | Rps3 | -0.698 | 6.19E-04 | glio |
| Aars2_ex.vivo_noRT | 4930447K03R | 2.505 | 6.19E-04 | glio |
| Atr_in.vivo_noRT | Hint2 | -1.317 | 6.19E-04 | glio |
| Atr_in.vivo_noRT | Frg1 | -1.239 | 6.19E-04 | glio |
| Bard1_ex.vivo_noRT | Gm5039 | 3.778 | 6.19E-04 | glio |
| Bard1_ex.vivo_noRT | Acox2 | 3.006 | 6.19E-04 | glio |
| Bcas2_in.vivo_noRT | Ptms | -0.969 | 6.19E-04 | glio |
| Bora_in.vivo_noRT | Car13 | 1.79 | 6.19E-04 | glio |
| Bora_in.vivo_noRT | Col9a2 | -1.039 | 6.19E-04 | glio |
| Chek1_in.vivo_noRT | Gm16124 | 2.809 | 6.19E-04 | glio |
| Interaction_genotype_Mtor | Nat8f4 | -4.354 | 6.19E-05 | glio |
| culture_effect | Atp13a1 | -0.915 | 6.19E-06 | glio |
| culture_effect | D8Ertd738e | 0.471 | 6.19E-07 | glio |
| Interaction_genotype_Atr | Nxph3 | -5.518 | 6.19E-07 | glio |
| Bora_in.vivo_noRT | Dio1 | 3.1 | 6.19E-07 | glio |
| Bcas2_in.vivo_noRT | Gm26590 | 3.519 | 6.19E-08 | glio |
| Chek1_ex.vivo_noRT | Map3k5 | 4.443 | 6.19E-09 | glio |
| culture_effect | Car9 | -3.828 | 6.19E-10 | glio |
| Atr_in.vivo_noRT | Dhrs9 | 3.946 | 6.19E-10 | glio |

| | | | | |
|---|---|---|---|---|
| Atr_in.vivo_noRT | Sap30bpos | 4.139 | 6.19E-12 | glio |
| Bcas2_in.vivo_noRT | Tff2 | 4.547 | 6.19E-12 | glio |
| Bcas2_in.vivo_noRT | Gm10642 | 4.543 | 6.19E-12 | glio |
| Chek1_ex.vivo_noRT | Speer4e | 4.842 | 6.19E-13 | glio |
| Atr_in.vivo_noRT | Gm21818 | 5.533 | 6.19E-15 | glio |
| Chek1_in.vivo_noRT | Pnliprp2 | 4.265 | 6.19E-15 | glio |
| Bcas2_in.vivo_noRT | AC163347.1 | 4.544 | 6.19E-16 | glio |
| Atr_in.vivo_noRT | Tent5c | 4.608 | 6.19E-18 | glio |
| culture_effect | Pllp | -3.625 | 6.19E-24 | glio |
| Interaction_genotype_Aars2 | Prr7 | -3.524 | 6.20E-03 | glio |
| Interaction_genotype_Aars2 | Gm15927 | -3.299 | 6.20E-03 | glio |
| Interaction_genotype_Atr | Sf3b4 | 1.49 | 6.20E-03 | glio |
| Interaction_genotype_Bcas2 | Tnfsf13 | -3.462 | 6.20E-03 | glio |
| Interaction_genotype_Myc | Pias3 | -2.503 | 6.20E-03 | glio |
| Interaction_genotype_Myc | Ndufa8 | 1.101 | 6.20E-03 | glio |
| Interaction_genotype_Rad51 | Gm50103 | -3.953 | 6.20E-03 | glio |
| Interaction_genotype_Rad51 | Herc1 | 2.27 | 6.20E-03 | glio |
| interaction_RT | Smim26 | 0.747 | 6.20E-03 | glio |
| Aars2_in.vivo_noRT | Prpf40a | -0.757 | 6.20E-03 | glio |
| Aars2_ex.vivo_noRT | Pcdhga11 | 2.321 | 6.20E-03 | glio |
| Atr_in.vivo_noRT | Zscan20 | 1.851 | 6.20E-03 | glio |
| Bcas2_in.vivo_noRT | Mapk7 | 1.427 | 6.20E-03 | glio |
| Bcas2_in.vivo_noRT | Nmi | -0.718 | 6.20E-03 | glio |
| Bora_in.vivo_noRT | Bmerb1 | 2.164 | 6.20E-03 | glio |
| Bora_in.vivo_noRT | Sh3bp5 | 1.997 | 6.20E-03 | glio |
| Bora_in.vivo_noRT | Pkp3 | 2.123 | 6.20E-03 | glio |
| Bora_in.vivo_noRT | Rit1 | -1.309 | 6.20E-03 | glio |
| Chek1_ex.vivo_noRT | Slc45a3 | 2.53 | 6.20E-03 | glio |
| Interaction_genotype_Aars2 | Gm45620 | -4.093 | 6.20E-04 | glio |
| Interaction_genotype_Mtor | 9130024F11Ri | -3.886 | 6.20E-04 | glio |
| Interaction_genotype_Myc | Gm19605 | -3.772 | 6.20E-04 | glio |
| Interaction_genotype_Myc | Myl9 | -3.93 | 6.20E-04 | glio |
| Interaction_genotype_Rad51 | Il12rb1 | 4.479 | 6.20E-04 | glio |
| Bcas2_in.vivo_noRT | Ikzf4 | 2.733 | 6.20E-05 | glio |
| Chek1_in.vivo_noRT | Krt16 | 4.334 | 6.20E-07 | glio |
| culture_effect | Pak1 | -1.354 | 6.20E-10 | glio |
| Atr_in.vivo_noRT | Slc7a4 | 4.119 | 6.20E-10 | glio |
| Bora_in.vivo_noRT | Grm1 | 3.832 | 6.20E-10 | glio |
| Atr_in.vivo_noRT | Masp2 | 4.606 | 6.20E-15 | glio |
| Atr_in.vivo_noRT | Gm14002 | 5.011 | 6.20E-15 | glio |
| Atr_in.vivo_noRT | Ccdc162 | 5.004 | 6.20E-19 | glio |
| Atr_in.vivo_noRT | Gm13398 | 5.533 | 6.20E-27 | glio |
| Interaction_genotype_Atr | Dynlt1b | -3.485 | 6.21E-03 | glio |
| Interaction_genotype_Bcas2 | Itch | 1.604 | 6.21E-03 | glio |
| Interaction_genotype_Mtor | Ncl | 1.365 | 6.21E-03 | glio |
| Atr_ex.vivo_noRT | Car9 | 2.77 | 6.21E-03 | glio |
| Atr_ex.vivo_noRT | Hoxa5 | 1.615 | 6.21E-03 | glio |
| Bcas2_ex.vivo_noRT | Nat8f2 | 2.347 | 6.21E-03 | glio |
| Chek1_in.vivo_noRT | 1700024B05R | 2.048 | 6.21E-03 | glio |
| Interaction_genotype_Atr | Spry1 | -2.538 | 6.21E-04 | glio |
| Interaction_genotype_Myc | Zcwpw1 | -3.206 | 6.21E-04 | glio |
| Aars2_in.vivo_noRT | Gh | 2.742 | 6.21E-04 | glio |

| | | | | |
|---|---|---|---|---|
| Aars2_ex.vivo_noRT | Pcdhgb5 | 2.899 | 6.21E-04 | glio |
| Atr_in.vivo_noRT | Txnrd1 | -0.907 | 6.21E-04 | glio |
| Bard1_ex.vivo_noRT | Fam131a | 3.213 | 6.21E-04 | glio |
| Bcas2_in.vivo_noRT | H2-T24 | 2.349 | 6.21E-04 | glio |
| Chek1_in.vivo_noRT | Kras | -1.32 | 6.21E-04 | glio |
| Atr_in.vivo_noRT | Cxcl2 | 4.667 | 6.21E-05 | glio |
| Atr_in.vivo_noRT | Gm38414 | 3.126 | 6.21E-05 | glio |
| Bora_in.vivo_noRT | Bbs10 | 2.973 | 6.21E-05 | glio |
| Bora_in.vivo_noRT | Mvb12a | -1.406 | 6.21E-05 | glio |
| Interaction_genotype_Mtor | Cd200r2 | -3.835 | 6.21E-06 | glio |
| Interaction_genotype_Myc | Zcchc8 | -3.622 | 6.21E-06 | glio |
| Interaction_genotype_Myc | Hist1h4a | -4.965 | 6.21E-06 | glio |
| Interaction_genotype_Myc | Mir670hg | -5.543 | 6.21E-06 | glio |
| Atp5c1_in.vivo_noRT | Ltbp2 | 2.668 | 6.21E-06 | glio |
| Atp5c1_in.vivo_noRT | Lmx1a | 2.668 | 6.21E-06 | glio |
| Bora_ex.vivo_noRT | Gm35082 | 3.197 | 6.21E-06 | glio |
| Aars2_in.vivo_noRT | Gm15091 | 3.623 | 6.21E-07 | glio |
| Atr_in.vivo_noRT | Gm45833 | 4.686 | 6.21E-07 | glio |
| Bcas2_in.vivo_noRT | 6330415G19R | 3.5 | 6.21E-07 | glio |
| Chek1_in.vivo_noRT | Hist1h4b | 3.811 | 6.21E-07 | glio |
| Chek1_ex.vivo_noRT | Bco1 | 4.447 | 6.21E-09 | glio |
| Aars2_in.vivo_noRT | Trim67 | 3.758 | 6.21E-10 | glio |
| Bora_in.vivo_noRT | Ngb | 4.166 | 6.21E-10 | glio |
| culture_effect | B930094E09Ri | 3.359 | 6.21E-11 | glio |
| Bora_in.vivo_noRT | Gm43672 | 4.936 | 6.21E-12 | glio |
| Chek1_in.vivo_noRT | Gm20682 | 3.999 | 6.21E-12 | glio |
| Bcas2_in.vivo_noRT | Gstt4 | 4.544 | 6.21E-19 | glio |
| Atr_in.vivo_noRT | Gm49086 | 5.533 | 6.21E-20 | glio |
| Bora_in.vivo_noRT | Cgnl1 | 4.936 | 6.21E-24 | glio |
| culture_effect | Pomgnt1 | 0.611 | 6.22E-03 | glio |
| culture_effect | Atxn2 | -0.435 | 6.22E-03 | glio |
| Interaction_genotype_Atr | Gm44658 | -2.982 | 6.22E-03 | glio |
| Interaction_genotype_Atr | 1700030L20Ri | -3.261 | 6.22E-03 | glio |
| Interaction_genotype_Atr | 4930447F24Ri | -3.272 | 6.22E-03 | glio |
| Interaction_genotype_Cdkn2a | Mmp13 | 2.835 | 6.22E-03 | glio |
| Interaction_genotype_Cdkn2a | Gm8016 | 2.835 | 6.22E-03 | glio |
| Interaction_genotype_Mtor | Zfp213 | -2.5 | 6.22E-03 | glio |
| Interaction_genotype_Myc | Rad54l | -2.356 | 6.22E-03 | glio |
| Interaction_genotype_Myc | Rufy2 | -1.922 | 6.22E-03 | glio |
| Interaction_genotype_Myc | Mkrn1 | 1.468 | 6.22E-03 | glio |
| Atr_in.vivo_noRT | Vwa8 | 1.697 | 6.22E-03 | glio |
| Bora_in.vivo_noRT | Snap23 | -1.287 | 6.22E-03 | glio |
| Chek1_in.vivo_noRT | Igf2bp3 | -1.198 | 6.22E-03 | glio |
| Chek1_ex.vivo_noRT | Dhx57 | 1.94 | 6.22E-03 | glio |
| Cyp19a1_ex.vivo_noRT | Tac2 | 2.159 | 6.22E-03 | glio |
| Eif2s1_ex.vivo_noRT | Esp24 | 2.234 | 6.22E-03 | glio |
| Hsd17b10_in.vivo_noRT | Klhl2 | -1.527 | 6.22E-03 | glio |
| Interaction_genotype_Mtor | Ttc25 | -4.028 | 6.22E-04 | glio |
| Interaction_genotype_Myc | Pla2g7 | -3.862 | 6.22E-04 | glio |
| Interaction_genotype_Myc | A830005F24Ri | -3.907 | 6.22E-04 | glio |
| Atr_in.vivo_noRT | D130020L05R | 2.109 | 6.22E-04 | glio |
| Atr_in.vivo_noRT | Gnas | -0.675 | 6.22E-04 | glio |

| | | | | |
|---|---|---|---|---|
| Atr_in.vivo_noRT | Rbx1 | -0.651 | 6.22E-04 | glio |
| Bcas2_in.vivo_noRT | Adam19 | 2.544 | 6.22E-04 | glio |
| Chek1_ex.vivo_noRT | Jakmip1 | 3.074 | 6.22E-04 | glio |
| Interaction_genotype_Myc | Cerk | -3.892 | 6.22E-05 | glio |
| Interaction_genotype_Myc | Gm48140 | -3.259 | 6.22E-05 | glio |
| Aars2_in.vivo_noRT | Mrps18a | -1.237 | 6.22E-05 | glio |
| Interaction_genotype_Mtor | 1110065P20R | 2.171 | 6.22E-06 | glio |
| Interaction_genotype_Myc | Gm26519 | -5.467 | 6.22E-06 | glio |
| Interaction_genotype_Myc | Rpl36al | 1.704 | 6.22E-06 | glio |
| Atp5c1_in.vivo_noRT | Ntn4 | 2.933 | 6.22E-06 | glio |
| Atp5c1_in.vivo_noRT | Gm48140 | 2.405 | 6.22E-06 | glio |
| Atp5c1_in.vivo_noRT | Itgbl1 | 2.781 | 6.22E-06 | glio |
| Bora_in.vivo_noRT | Arhgef4 | 3.171 | 6.22E-06 | glio |
| Hsd17b10_in.vivo_noRT | Stx19 | 2.704 | 6.22E-06 | glio |
| Interaction_genotype_Myc | A430018G15R | -4.504 | 6.22E-07 | glio |
| Chek1_ex.vivo_noRT | Fgf8 | 3.733 | 6.22E-07 | glio |
| Aars2_ex.vivo_noRT | Gm34059 | 3.293 | 6.22E-08 | glio |
| Aars2_ex.vivo_noRT | Halr1 | 3.293 | 6.22E-08 | glio |
| Aars2_ex.vivo_noRT | Helt | 3.293 | 6.22E-08 | glio |
| Aars2_ex.vivo_noRT | Gm20625 | 3.293 | 6.22E-08 | glio |
| Aars2_ex.vivo_noRT | Gm5157 | 3.293 | 6.22E-08 | glio |
| Aars2_ex.vivo_noRT | Gm44736 | 3.293 | 6.22E-08 | glio |
| Aars2_ex.vivo_noRT | Fam227b | 3.293 | 6.22E-08 | glio |
| Aars2_ex.vivo_noRT | Gm20658 | 3.293 | 6.22E-08 | glio |
| Aars2_ex.vivo_noRT | Apoa5 | 3.293 | 6.22E-08 | glio |
| Aars2_ex.vivo_noRT | Olfr810 | 3.293 | 6.22E-08 | glio |
| Atp5c1_in.vivo_noRT | Krt19 | 2.668 | 6.22E-08 | glio |
| culture_effect | Nfix | -1.071 | 6.22E-10 | glio |
| Bcas2_in.vivo_noRT | Ly6g6c | 3.947 | 6.22E-10 | glio |
| Atr_in.vivo_noRT | Trim67 | 3.964 | 6.22E-11 | glio |
| culture_effect | 1810059H22R | 1.616 | 6.23E-03 | glio |
| Interaction_genotype_Bcas2 | Tnfsf13os | -3.675 | 6.23E-03 | glio |
| Interaction_genotype_Mtor | Pcdhga12 | -3.309 | 6.23E-03 | glio |
| Interaction_genotype_Mtor | Zfp715 | -2.72 | 6.23E-03 | glio |
| Interaction_genotype_Mtor | Cenpa | 2.419 | 6.23E-03 | glio |
| Interaction_genotype_Mtor | Btf3 | 0.863 | 6.23E-03 | glio |
| Interaction_genotype_Myc | Wnk2 | -3.173 | 6.23E-03 | glio |
| Interaction_genotype_Myc | Dctpp1 | 1.348 | 6.23E-03 | glio |
| interaction_RT | Atp1a3 | 1.765 | 6.23E-03 | glio |
| interaction_RT | Cdk18 | -1.554 | 6.23E-03 | glio |
| interaction_RT | Eif4b | -0.58 | 6.23E-03 | glio |
| interaction_RT | Atp5e | 0.579 | 6.23E-03 | glio |
| interaction_RT | Nme1 | -0.505 | 6.23E-03 | glio |
| Aars2_in.vivo_noRT | Ndufc2 | -0.651 | 6.23E-03 | glio |
| Atp5c1_in.vivo_noRT | Gm50375 | 2.402 | 6.23E-03 | glio |
| Atr_in.vivo_noRT | Rmnd5a | -1.301 | 6.23E-03 | glio |
| Atr_in.vivo_noRT | Fuom | -1.184 | 6.23E-03 | glio |
| Bcas2_in.vivo_noRT | Mst1 | 1.907 | 6.23E-03 | glio |
| Bcas2_in.vivo_noRT | Gm48646 | 1.973 | 6.23E-03 | glio |
| Bora_in.vivo_noRT | Cyp27a1 | 1.849 | 6.23E-03 | glio |
| Bora_in.vivo_noRT | AU041133 | 1.729 | 6.23E-03 | glio |
| Bora_ex.vivo_noRT | Hsf4 | 2.823 | 6.23E-03 | glio |

| | | | | |
|---|---|---|---|---|
| Chek1_in.vivo_noRT | Cass4 | 2.035 | 6.23E-03 | glio |
| Chek1_in.vivo_noRT | Ebna1bp2 | -0.641 | 6.23E-03 | glio |
| Chek1_ex.vivo_noRT | Pla2g4c | 2.628 | 6.23E-03 | glio |
| Chek1_ex.vivo_noRT | Cfap69 | 2.527 | 6.23E-03 | glio |
| Chek1_ex.vivo_noRT | Tmem218 | 1.924 | 6.23E-03 | glio |
| Aars2_ex.vivo_noRT | Ms4a4d | 3.293 | 6.23E-04 | glio |
| Bora_in.vivo_noRT | Gja4 | 2.348 | 6.23E-04 | glio |
| Chek1_in.vivo_noRT | Rell2 | 2.566 | 6.23E-04 | glio |
| Interaction_genotype_Mtor | Gm49441 | -4.095 | 6.23E-05 | glio |
| Atr_in.vivo_noRT | Fhdc1 | 2.987 | 6.23E-05 | glio |
| Bcas2_in.vivo_noRT | Bpifa3 | 2.393 | 6.23E-05 | glio |
| Bora_in.vivo_noRT | Gm16740 | 2.841 | 6.23E-05 | glio |
| Interaction_genotype_Mtor | Set | 2.098 | 6.23E-06 | glio |
| Aars2_in.vivo_noRT | Zan | 3.1 | 6.23E-06 | glio |
| Atr_ex.vivo_noRT | Cd7 | 3.164 | 6.23E-07 | glio |
| Atr_in.vivo_noRT | C230013L11Ri | 4.621 | 6.23E-10 | glio |
| Atr_in.vivo_noRT | Tmem150c | 4.366 | 6.23E-10 | glio |
| Hsd17b10_in.vivo_noRT | Esm1 | 3.352 | 6.23E-11 | glio |
| Bcas2_in.vivo_noRT | Cxcl16 | 4.543 | 6.23E-12 | glio |
| Bcas2_in.vivo_noRT | Ifitm6 | 4.807 | 6.23E-12 | glio |
| Chek1_ex.vivo_noRT | Rhbg | 4.449 | 6.23E-12 | glio |
| Bcas2_in.vivo_noRT | Cyp2c23 | 4.284 | 6.23E-13 | glio |
| Aars2_in.vivo_noRT | Gm15743 | 5.063 | 6.23E-15 | glio |
| Bora_in.vivo_noRT | Nos2 | 5.209 | 6.23E-17 | glio |
| Bora_in.vivo_noRT | AF067063 | 4.936 | 6.23E-21 | glio |
| culture_effect | Gpr176 | 0.886 | 6.24E-03 | glio |
| Interaction_genotype_Aars2 | Rack1 | 0.923 | 6.24E-03 | glio |
| Interaction_genotype_Aars2 | Rps3 | 0.902 | 6.24E-03 | glio |
| Interaction_genotype_Atr | Zmynd10 | -2.897 | 6.24E-03 | glio |
| Interaction_genotype_Bcas2 | Zmynd10 | -3.111 | 6.24E-03 | glio |
| Interaction_genotype_Bcas2 | 4930588A03R | -2.464 | 6.24E-03 | glio |
| Interaction_genotype_Bcas2 | Grin3b | -3.625 | 6.24E-03 | glio |
| Interaction_genotype_Bcas2 | Gm41760 | -3.586 | 6.24E-03 | glio |
| Interaction_genotype_Bora | Glyat | -2.532 | 6.24E-03 | glio |
| Interaction_genotype_Bora | Plxnb1 | -3.68 | 6.24E-03 | glio |
| Interaction_genotype_Mtor | Ccdc126 | -2.807 | 6.24E-03 | glio |
| Interaction_genotype_Myc | Gm1123 | -3.197 | 6.24E-03 | glio |
| Interaction_genotype_Myc | Lrrc8b | -2.02 | 6.24E-03 | glio |
| Interaction_genotype_Myc | Spata6 | -1.951 | 6.24E-03 | glio |
| Aars2_in.vivo_noRT | Txnl4b | 1.643 | 6.24E-03 | glio |
| Aars2_in.vivo_noRT | Scaf11 | -0.765 | 6.24E-03 | glio |
| Atr_ex.vivo_noRT | Ccnd1 | -1.011 | 6.24E-03 | glio |
| Bcas2_in.vivo_noRT | Chchd10 | -1.852 | 6.24E-03 | glio |
| Bcas2_in.vivo_noRT | Rpl21 | -0.493 | 6.24E-03 | glio |
| Cenpj_in.vivo_noRT | Gm45606 | 2.291 | 6.24E-03 | glio |
| Chek1_in.vivo_noRT | Kank2 | 1.945 | 6.24E-03 | glio |
| Hsd17b10_in.vivo_noRT | Trnp1 | 2.175 | 6.24E-03 | glio |
| Hsd17b10_in.vivo_noRT | Gm8098 | 2.113 | 6.24E-03 | glio |
| Hsd17b10_in.vivo_noRT | Serp2 | 1.924 | 6.24E-03 | glio |
| Hsd17b10_in.vivo_noRT | Klhl33 | 2.129 | 6.24E-03 | glio |
| Hsd17b10_in.vivo_noRT | 5033406O09R | 1.043 | 6.24E-03 | glio |
| culture_effect | Tfap2a | 0.704 | 6.24E-04 | glio |

| | | | | |
|---|---|---|---|---|
| Interaction_genotype_Atr | Gm9922 | -4.026 | 6.24E-04 | glio |
| Interaction_genotype_Atr | Gpx2 | 5.534 | 6.24E-04 | glio |
| Interaction_genotype_Myc | Akap6 | -3.173 | 6.24E-04 | glio |
| Atr_in.vivo_noRT | Tal1 | 2.415 | 6.24E-04 | glio |
| Bard1_ex.vivo_noRT | Smpd5 | 3.01 | 6.24E-04 | glio |
| Cenpj_in.vivo_noRT | Olfr552 | 2.022 | 6.24E-04 | glio |
| Eif2s1_in.vivo_noRT | Fhl1 | 2.143 | 6.24E-04 | glio |
| Hsd17b10_in.vivo_noRT | Gm14636 | 3.108 | 6.24E-04 | glio |
| Interaction_genotype_Aars2 | Gm43137 | -4.896 | 6.24E-05 | glio |
| Interaction_genotype_Aars2 | BC065403 | -4.54 | 6.24E-05 | glio |
| Interaction_genotype_Myc | Kdelr2 | 1.898 | 6.24E-05 | glio |
| culture_effect | Mlf1 | -1.254 | 6.24E-06 | glio |
| Bcas2_in.vivo_noRT | Gm15728 | 3.372 | 6.24E-06 | glio |
| Chek1_ex.vivo_noRT | Cnih2 | 3.893 | 6.24E-06 | glio |
| Interaction_genotype_Mtor | 1700011D18R | -6.053 | 6.24E-07 | glio |
| Aars2_in.vivo_noRT | Mrgprf | 3.531 | 6.24E-07 | glio |
| Hsd17b10_in.vivo_noRT | 4833419F23Ri | 3.616 | 6.24E-07 | glio |
| culture_effect | Gm48772 | 2.915 | 6.24E-09 | glio |
| Bora_in.vivo_noRT | Ankrd1 | 4.936 | 6.24E-09 | glio |
| Aars2_in.vivo_noRT | Gm12602 | 4.3 | 6.24E-11 | glio |
| Bora_in.vivo_noRT | Cidea | 4.021 | 6.24E-11 | glio |
| Bora_in.vivo_noRT | Pde6b | 3.904 | 6.24E-11 | glio |
| culture_effect | Efnb3 | -1.422 | 6.25E-03 | glio |
| culture_effect | Cenpp | -0.628 | 6.25E-03 | glio |
| Interaction_genotype_Aars2 | 1700086P04R | -3.219 | 6.25E-03 | glio |
| Interaction_genotype_Mtor | Tex19.2 | -3.321 | 6.25E-03 | glio |
| Interaction_genotype_Mtor | Slc35f3 | -2.344 | 6.25E-03 | glio |
| Interaction_genotype_Myc | Ikbke | -2.995 | 6.25E-03 | glio |
| interaction_RT | Ahsa1 | -0.684 | 6.25E-03 | glio |
| Aars2_in.vivo_noRT | Pfkfb3 | 1.749 | 6.25E-03 | glio |
| Aars2_in.vivo_noRT | Cep78 | 1.629 | 6.25E-03 | glio |
| Aars2_in.vivo_noRT | Fancc | 1.426 | 6.25E-03 | glio |
| Brca2_in.vivo_noRT | Psca | 2.288 | 6.25E-03 | glio |
| Dclre1c_ex.vivo_noRT | Adm | 3.06 | 6.25E-03 | glio |
| Dclre1c_ex.vivo_noRT | Gm48617 | 2.296 | 6.25E-03 | glio |
| Dclre1c_ex.vivo_noRT | Gm5788 | 2.292 | 6.25E-03 | glio |
| Interaction_genotype_Myc | Zmynd10 | -3.257 | 6.25E-04 | glio |
| Interaction_genotype_Myc | Gm19409 | -3.968 | 6.25E-04 | glio |
| Interaction_genotype_Myc | Dtnb | -2.746 | 6.25E-04 | glio |
| Bcas2_in.vivo_noRT | Pabpc4l | 2.528 | 6.25E-04 | glio |
| Chek1_in.vivo_noRT | Cpne7 | 2.645 | 6.25E-04 | glio |
| Chek1_in.vivo_noRT | Hspa9 | -0.826 | 6.25E-04 | glio |
| Eif2s1_in.vivo_noRT | Gm29966 | 1.883 | 6.25E-04 | glio |
| culture_effect | Antxr2 | 1.438 | 6.25E-05 | glio |
| Interaction_genotype_Mtor | Gm26728 | -4.846 | 6.25E-05 | glio |
| Interaction_genotype_Myc | 5830454E08Ri | -4.144 | 6.25E-05 | glio |
| interaction_RT | Got2 | -0.87 | 6.25E-05 | glio |
| Aars2_in.vivo_noRT | Med19 | -1.275 | 6.25E-05 | glio |
| Atr_in.vivo_noRT | Gm19391 | 2.634 | 6.25E-05 | glio |
| Bcas2_in.vivo_noRT | Hdac11 | 2.923 | 6.25E-05 | glio |
| Bora_in.vivo_noRT | Proscos | 3.041 | 6.25E-05 | glio |
| Interaction_genotype_Mtor | Apobr | -5.863 | 6.25E-06 | glio |

| | | | | |
|---|---|---|---|---|
| Interaction_genotype_Mtor | Gm49042 | -5.125 | 6.25E-06 | glio |
| Interaction_genotype_Mtor | Fbxo15 | -4.817 | 6.25E-06 | glio |
| Chek1_in.vivo_noRT | Fes | 3.235 | 6.25E-06 | glio |
| culture_effect | Smpdl3a | -2.02 | 6.25E-07 | glio |
| Chek1_in.vivo_noRT | D430018E03R | 3.318 | 6.25E-07 | glio |
| Bcas2_in.vivo_noRT | A330040F15Ri | 4.028 | 6.25E-08 | glio |
| Chek1_in.vivo_noRT | Dzank1 | 3.868 | 6.25E-08 | glio |
| Chek1_in.vivo_noRT | Gm37584 | 4.132 | 6.25E-09 | glio |
| Atr_in.vivo_noRT | Rab37 | 4.615 | 6.25E-12 | glio |
| Atr_in.vivo_noRT | Wnt16 | 4.742 | 6.25E-13 | glio |
| Bora_in.vivo_noRT | Muc4 | 4.947 | 6.25E-18 | glio |
| culture_effect | Gsn | -1.866 | 6.25E-22 | glio |
| culture_effect | Esrp2 | 1.521 | 6.26E-03 | glio |
| Interaction_genotype_Bora | Btbd6 | -3.287 | 6.26E-03 | glio |
| Interaction_genotype_Lmo7 | 1500009L16Ri | 3.466 | 6.26E-03 | glio |
| Interaction_genotype_Mtor | Kars | 1.367 | 6.26E-03 | glio |
| Interaction_genotype_Myc | AW011738 | -2.432 | 6.26E-03 | glio |
| Interaction_genotype_Myc | Ints2 | -2.029 | 6.26E-03 | glio |
| Interaction_genotype_Pars2 | C230037L18Ri | -3.45 | 6.26E-03 | glio |
| Aars2_in.vivo_noRT | AI662270 | -0.777 | 6.26E-03 | glio |
| Atr_in.vivo_noRT | Gm44699 | 1.74 | 6.26E-03 | glio |
| Bcas2_in.vivo_noRT | Fbxo10 | 1.998 | 6.26E-03 | glio |
| Bcas2_in.vivo_noRT | 4931414P19R | 1.946 | 6.26E-03 | glio |
| Bcas2_in.vivo_noRT | Pxmp2 | 1.709 | 6.26E-03 | glio |
| Bcas2_in.vivo_noRT | 2810013P06R | 1.623 | 6.26E-03 | glio |
| Bcas2_in.vivo_noRT | Rhbdf1 | -1.392 | 6.26E-03 | glio |
| Bcas2_in.vivo_noRT | Tex30 | -1.066 | 6.26E-03 | glio |
| Brca2_in.vivo_noRT | Abcb5 | 1.763 | 6.26E-03 | glio |
| Chek1_in.vivo_noRT | 1700007L15Ri | 1.672 | 6.26E-03 | glio |
| Eif2s1_ex.vivo_noRT | Gm9758 | 2.234 | 6.26E-03 | glio |
| Hsd17b10_in.vivo_noRT | Klrk1 | -1.556 | 6.26E-03 | glio |
| Interaction_genotype_Atr | 9130230N09R | -3.652 | 6.26E-04 | glio |
| Interaction_genotype_Atr | Zfp706 | 1.126 | 6.26E-04 | glio |
| Interaction_genotype_Myc | Lrrc49 | -3.177 | 6.26E-04 | glio |
| Atr_ex.vivo_noRT | Bmp15 | 2.77 | 6.26E-04 | glio |
| Bcas2_in.vivo_noRT | Wif1 | 2.801 | 6.26E-04 | glio |
| Bcas2_ex.vivo_noRT | C77080 | 3.31 | 6.26E-04 | glio |
| culture_effect | Abce1 | 0.476 | 6.26E-05 | glio |
| culture_effect | Sh3glb1 | 0.473 | 6.26E-05 | glio |
| Atp5c1_in.vivo_noRT | Ngb | 2.945 | 6.26E-05 | glio |
| Aars2_in.vivo_noRT | Sgms2 | 3.358 | 6.26E-06 | glio |
| Atr_in.vivo_noRT | Gm42699 | 2.983 | 6.26E-06 | glio |
| Bcas2_in.vivo_noRT | Ikzf2 | 3.383 | 6.26E-06 | glio |
| Interaction_genotype_Mtor | Hfm1 | -4.674 | 6.26E-07 | glio |
| Bcas2_in.vivo_noRT | Gm34643 | 3.621 | 6.26E-07 | glio |
| Interaction_genotype_Mtor | C130040N14R | -6.772 | 6.26E-08 | glio |
| Interaction_genotype_Mtor | Dscaml1 | -4.494 | 6.26E-08 | glio |
| culture_effect | Lman1 | -0.858 | 6.26E-12 | glio |
| Bcas2_in.vivo_noRT | Gm13288 | 4.807 | 6.26E-17 | glio |
| culture_effect | Sp140 | -2.019 | 6.26E-18 | glio |
| Atr_in.vivo_noRT | Gm13078 | 5.533 | 6.26E-28 | glio |
| Interaction_genotype_Myc | Zbtb7b | -2.453 | 6.27E-03 | glio |

| | | | | |
|---|---|---|---|---|
| Bcas2_in.vivo_noRT | Fnbp1 | 1.2 | 6.27E-03 | glio |
| Bcas2_in.vivo_noRT | D8Ertd738e | -0.502 | 6.27E-03 | glio |
| Bora_in.vivo_noRT | Rpain | -1.101 | 6.27E-03 | glio |
| Cenpj_ex.vivo_noRT | Clca3a2 | 2.591 | 6.27E-03 | glio |
| Cenpj_ex.vivo_noRT | Art2b | 2.591 | 6.27E-03 | glio |
| Chek1_ex.vivo_noRT | Camk2b | 2.416 | 6.27E-03 | glio |
| Hsd17b10_in.vivo_noRT | 4930558J18Ri | 2.01 | 6.27E-03 | glio |
| Interaction_genotype_Myc | Usf3 | -2.745 | 6.27E-04 | glio |
| Atr_in.vivo_noRT | Epg5 | 2.285 | 6.27E-04 | glio |
| Bora_in.vivo_noRT | Lmln | 2.401 | 6.27E-04 | glio |
| Hsd17b10_in.vivo_noRT | Tomm6os | 2.698 | 6.27E-04 | glio |
| culture_effect | Sptssa | 0.598 | 6.27E-05 | glio |
| culture_effect | Gm9922 | 2.325 | 6.27E-06 | glio |
| Aars2_in.vivo_noRT | Gata3 | 3.453 | 6.27E-06 | glio |
| Aars2_in.vivo_noRT | Igsf23 | 3.667 | 6.27E-07 | glio |
| Bcas2_in.vivo_noRT | 1700040D17R | 3.743 | 6.27E-07 | glio |
| Bcas2_in.vivo_noRT | Gm20478 | 3.235 | 6.27E-07 | glio |
| culture_effect | Vps13b | -1.35 | 6.27E-08 | glio |
| Atr_in.vivo_noRT | Gm43727 | 3.706 | 6.27E-08 | glio |
| Chek1_ex.vivo_noRT | Nwd1 | 3.861 | 6.27E-08 | glio |
| Chek1_in.vivo_noRT | Gm5129 | 4.243 | 6.27E-09 | glio |
| Chek1_ex.vivo_noRT | Rapgef3os2 | 3.657 | 6.27E-09 | glio |
| Chek1_ex.vivo_noRT | BC026762 | 4.735 | 6.27E-09 | glio |
| Bora_in.vivo_noRT | Snrpn | 4.278 | 6.27E-15 | glio |
| Chek1_ex.vivo_noRT | Slc6a11 | 5.643 | 6.27E-15 | glio |
| Bcas2_in.vivo_noRT | Slc45a1 | 4.547 | 6.27E-19 | glio |
| culture_effect | Alg8 | -0.62 | 6.28E-03 | glio |
| culture_effect | Rhod | 0.645 | 6.28E-03 | glio |
| Interaction_genotype_Aars2 | Ptpn2 | 1.261 | 6.28E-03 | glio |
| Interaction_genotype_Mtor | Catsperd | -3.29 | 6.28E-03 | glio |
| Interaction_genotype_Mtor | Batf3 | -3.283 | 6.28E-03 | glio |
| Aars2_in.vivo_noRT | Emc9 | 1.386 | 6.28E-03 | glio |
| Aars2_ex.vivo_noRT | Gm50240 | -2.153 | 6.28E-03 | glio |
| Bard1_ex.vivo_noRT | Gm16751 | 2.415 | 6.28E-03 | glio |
| Bora_in.vivo_noRT | Cul7 | -1.337 | 6.28E-03 | glio |
| Eif2s1_ex.vivo_noRT | AF067063 | 2.234 | 6.28E-03 | glio |
| culture_effect | Shq1 | -0.976 | 6.28E-04 | glio |
| Interaction_genotype_Aars2 | Apol10b | 3.689 | 6.28E-04 | glio |
| Interaction_genotype_Mtor | Syt17 | -4.134 | 6.28E-04 | glio |
| Interaction_genotype_Myc | 9130024F11Ri | -3.681 | 6.28E-04 | glio |
| Interaction_genotype_Myc | Wdr19 | -3.676 | 6.28E-04 | glio |
| Aars2_in.vivo_noRT | Gm42726 | 2.334 | 6.28E-04 | glio |
| Aars2_in.vivo_noRT | Serbp1 | -0.662 | 6.28E-04 | glio |
| Atr_in.vivo_noRT | Prkd2 | 2.225 | 6.28E-04 | glio |
| Bard1_ex.vivo_noRT | Gm36037 | 2.419 | 6.28E-04 | glio |
| Bora_in.vivo_noRT | Slc7a7 | 2.45 | 6.28E-04 | glio |
| Chek1_in.vivo_noRT | Agpat4 | -1.62 | 6.28E-04 | glio |
| Hsd17b10_in.vivo_noRT | Olfm2 | 2.709 | 6.28E-04 | glio |
| Interaction_genotype_Mtor | Nyap2 | -4.863 | 6.28E-05 | glio |
| Interaction_genotype_Myc | Gm28424 | -4.19 | 6.28E-05 | glio |
| Interaction_genotype_Myc | Nudt7 | -4.234 | 6.28E-05 | glio |
| Interaction_genotype_Myc | Slc26a6 | -4.41 | 6.28E-05 | glio |

| | | | | |
|---|---|---|---|---|
| Interaction_genotype_Myc | Atxn1l | -3.438 | 6.28E-05 | glio |
| Hsd17b10_in.vivo_noRT | Gm17146 | 2.699 | 6.28E-05 | glio |
| Hsd17b10_in.vivo_noRT | Cysrt1 | 3.356 | 6.28E-06 | glio |
| Interaction_genotype_Mtor | BC061195 | -4.099 | 6.28E-07 | glio |
| Interaction_genotype_Myc | D830039M14I | -5.95 | 6.28E-07 | glio |
| Atp5c1_in.vivo_noRT | Gm12841 | 2.668 | 6.28E-07 | glio |
| Atr_in.vivo_noRT | Rgs16 | 4.324 | 6.28E-07 | glio |
| Chek1_ex.vivo_noRT | Iqcn | 4.045 | 6.28E-07 | glio |
| Chek1_ex.vivo_noRT | Pawr | 3.871 | 6.28E-07 | glio |
| Atr_in.vivo_noRT | Pcsk4 | 3.738 | 6.28E-08 | glio |
| Chek1_in.vivo_noRT | Gm26590 | 3.755 | 6.28E-09 | glio |
| Chek1_in.vivo_noRT | Grin2d | 4.265 | 6.28E-11 | glio |
| Bora_in.vivo_noRT | Spata31d1a | 4.828 | 6.28E-12 | glio |
| Bora_in.vivo_noRT | BC026762 | 4.557 | 6.28E-12 | glio |
| Chek1_in.vivo_noRT | Fgf8 | 4.528 | 6.28E-14 | glio |
| Bora_in.vivo_noRT | 4930423D22R | 4.676 | 6.28E-20 | glio |
| Bcas2_in.vivo_noRT | Ctla4 | 4.807 | 6.28E-23 | glio |
| Chek1_in.vivo_noRT | Gm13078 | 5.044 | 6.28E-24 | glio |
| culture_effect | Glo1 | 0.419 | 6.29E-03 | glio |
| culture_effect | Fkbp4 | -0.33 | 6.29E-03 | glio |
| Interaction_genotype_Aars2 | Zfp324 | -2.868 | 6.29E-03 | glio |
| Interaction_genotype_Bora | Gm4181 | -3.688 | 6.29E-03 | glio |
| Interaction_genotype_Bora | Mns1 | 1.928 | 6.29E-03 | glio |
| Interaction_genotype_Bora | Supt16 | 1.439 | 6.29E-03 | glio |
| Interaction_genotype_Bora | Bnip3l | 1.515 | 6.29E-03 | glio |
| Interaction_genotype_Mtor | Grin3b | -3.178 | 6.29E-03 | glio |
| Interaction_genotype_Myc | Enah | -2.832 | 6.29E-03 | glio |
| Interaction_genotype_Myc | Gm47283 | 1.211 | 6.29E-03 | glio |
| Interaction_genotype_Rpl3 | Tnfrsf23 | -4.237 | 6.29E-03 | glio |
| Interaction_genotype_Rpl3 | Gm34590 | -3.854 | 6.29E-03 | glio |
| Aars2_in.vivo_noRT | Mageb18 | 1.708 | 6.29E-03 | glio |
| Aars2_in.vivo_noRT | Txnrd1 | -0.712 | 6.29E-03 | glio |
| Atr_in.vivo_noRT | Asb16 | 1.84 | 6.29E-03 | glio |
| Atr_in.vivo_noRT | Ufsp1 | 1.423 | 6.29E-03 | glio |
| Bora_in.vivo_noRT | Pdk2 | 2.058 | 6.29E-03 | glio |
| Bora_in.vivo_noRT | Nol6 | -1.25 | 6.29E-03 | glio |
| Cenpj_in.vivo_noRT | BC034090 | 2.202 | 6.29E-03 | glio |
| Chek1_in.vivo_noRT | Galc | 1.668 | 6.29E-03 | glio |
| Chek1_ex.vivo_noRT | Pla2g7 | 2.523 | 6.29E-03 | glio |
| Interaction_genotype_Mtor | H2afv | 1.392 | 6.29E-04 | glio |
| Interaction_genotype_Myc | Ttc16 | -3.802 | 6.29E-04 | glio |
| Bora_in.vivo_noRT | G430095P16R | 2.482 | 6.29E-04 | glio |
| Brca2_ex.vivo_noRT | Cd200r2 | 2.825 | 6.29E-04 | glio |
| Brca2_ex.vivo_noRT | Serpinb9g | 2.82 | 6.29E-04 | glio |
| Brca2_ex.vivo_noRT | Cd82 | -2.732 | 6.29E-04 | glio |
| Brca2_ex.vivo_noRT | Lhx1os | 3.219 | 6.29E-04 | glio |
| Hsd17b10_in.vivo_noRT | Selp | 2.572 | 6.29E-04 | glio |
| Hsd17b10_in.vivo_noRT | Ceacam19 | 2.584 | 6.29E-04 | glio |
| culture_effect | Epb42 | -2.577 | 6.29E-05 | glio |
| culture_effect | Med9os | 1.968 | 6.29E-05 | glio |
| Interaction_genotype_Mtor | Apoa1 | -4.846 | 6.29E-05 | glio |
| Interaction_genotype_Mtor | Tmem158 | -4.062 | 6.29E-05 | glio |

| | | | | |
|---|---|---|---|---|
| Interaction_genotype_Mtor | Tsg101 | 2.032 | 6.29E-05 | glio |
| Interaction_genotype_Mtor | Cystm1 | 2.1 | 6.29E-05 | glio |
| Interaction_genotype_Myc | B230319C09R | -4.922 | 6.29E-05 | glio |
| Interaction_genotype_Myc | Gm37170 | -4.51 | 6.29E-05 | glio |
| Interaction_genotype_Myc | Zkscan14 | -4.03 | 6.29E-05 | glio |
| Atr_in.vivo_noRT | 6720489N17R | 3.14 | 6.29E-05 | glio |
| Atr_in.vivo_noRT | Elof1 | -1.468 | 6.29E-05 | glio |
| Bcas2_in.vivo_noRT | Pnpla2 | -1.667 | 6.29E-05 | glio |
| Chek1_in.vivo_noRT | 8030456M14F | 2.891 | 6.29E-05 | glio |
| culture_effect | Stxbp2 | -1.304 | 6.29E-06 | glio |
| culture_effect | Sh3kbp1 | 0.671 | 6.29E-06 | glio |
| Interaction_genotype_Atr | Rassf6 | -4.62 | 6.29E-06 | glio |
| Interaction_genotype_Myc | 6330415G19R | -4.893 | 6.29E-06 | glio |
| Interaction_genotype_Myc | Cyp2u1 | -4.749 | 6.29E-06 | glio |
| Interaction_genotype_Myc | Cabp4 | -5.297 | 6.29E-06 | glio |
| Aars2_in.vivo_noRT | Snrpd1 | -1.163 | 6.29E-06 | glio |
| Bora_in.vivo_noRT | Snai1 | 3.418 | 6.29E-06 | glio |
| Interaction_genotype_Mtor | Gm15510 | -5.947 | 6.29E-07 | glio |
| Interaction_genotype_Mtor | Gm28874 | -5.98 | 6.29E-07 | glio |
| Interaction_genotype_Myc | Rabggtb | 2.579 | 6.29E-07 | glio |
| Interaction_genotype_Myc | Odf2 | -4.069 | 6.29E-07 | glio |
| Interaction_genotype_Myc | Psma7 | 2.272 | 6.29E-07 | glio |
| Interaction_genotype_Myc | Gm11627 | -5.014 | 6.29E-07 | glio |
| Chek1_in.vivo_noRT | A730071L15Ri | 3.983 | 6.29E-08 | glio |
| Chek1_ex.vivo_noRT | Ccser1 | 4.449 | 6.29E-10 | glio |
| Chek1_in.vivo_noRT | Gm47510 | 4.784 | 6.29E-22 | glio |
| Interaction_genotype_Bora | G630030J09Ri | -3.477 | 6.30E-03 | glio |
| Interaction_genotype_Myc | Rfx1 | -2.476 | 6.30E-03 | glio |
| Interaction_genotype_Myc | Lrrk1 | -1.893 | 6.30E-03 | glio |
| Aars2_in.vivo_noRT | Polm | 1.774 | 6.30E-03 | glio |
| Atr_in.vivo_noRT | Col9a1 | -1.239 | 6.30E-03 | glio |
| Bora_in.vivo_noRT | Homez | 2.03 | 6.30E-03 | glio |
| Bora_in.vivo_noRT | Dleu2 | -1.489 | 6.30E-03 | glio |
| culture_effect | Slc22a22 | -2.122 | 6.30E-04 | glio |
| culture_effect | Ppp6r3 | -0.613 | 6.30E-04 | glio |
| Interaction_genotype_Atr | Sgk3 | -4.279 | 6.30E-04 | glio |
| Interaction_genotype_Mtor | Paip2 | 1.366 | 6.30E-04 | glio |
| Aars2_in.vivo_noRT | Gm29093 | 2.41 | 6.30E-04 | glio |
| Atp5c1_in.vivo_noRT | BC049987 | 1.885 | 6.30E-04 | glio |
| Atr_in.vivo_noRT | Tnnt1 | 2.228 | 6.30E-04 | glio |
| Atr_in.vivo_noRT | Rpl14 | -0.67 | 6.30E-04 | glio |
| Cenpj_in.vivo_noRT | Ifi207 | 3.057 | 6.30E-04 | glio |
| Interaction_genotype_Atr | Maff | 3.267 | 6.30E-05 | glio |
| Interaction_genotype_Atr | S100a13 | 1.776 | 6.30E-05 | glio |
| Interaction_genotype_Mtor | Sumo2 | 1.835 | 6.30E-05 | glio |
| interaction_RT | Pld3 | 1.176 | 6.30E-05 | glio |
| Aars2_in.vivo_noRT | Nrbp1 | -1.351 | 6.30E-05 | glio |
| Bora_in.vivo_noRT | Chchd3 | -1.097 | 6.30E-05 | glio |
| Bcas2_in.vivo_noRT | Axdnd1 | 3.495 | 6.30E-06 | glio |
| Hsd17b10_in.vivo_noRT | Tslp | 3.498 | 6.30E-06 | glio |
| Hsd17b10_in.vivo_noRT | Slc46a1 | 3.413 | 6.30E-06 | glio |
| Chek1_in.vivo_noRT | Pglyrp1 | 3.564 | 6.30E-07 | glio |

| | | | | |
|---|---|---|---|---|
| Chek1_in.vivo_noRT | Zfp960 | 3.672 | 6.30E-07 | glio |
| Hsd17b10_in.vivo_noRT | 1700006H21R | 4.143 | 6.30E-09 | glio |
| Chek1_ex.vivo_noRT | Dpp6 | 4.842 | 6.30E-15 | glio |
| Bcas2_in.vivo_noRT | Fam227b | 4.807 | 6.30E-24 | glio |
| Bcas2_in.vivo_noRT | Gm20658 | 4.807 | 6.30E-24 | glio |
| culture_effect | Dact1 | -1.703 | 6.31E-03 | glio |
| culture_effect | Uba7 | -1.115 | 6.31E-03 | glio |
| culture_effect | Saysd1 | -0.615 | 6.31E-03 | glio |
| Interaction_genotype_Bcas2 | Vmac | -3.985 | 6.31E-03 | glio |
| Interaction_genotype_Bora | Ccnd2 | 1.032 | 6.31E-03 | glio |
| Interaction_genotype_Mtor | Macc1 | -3.163 | 6.31E-03 | glio |
| Interaction_genotype_Mtor | Smarcd3 | -2.783 | 6.31E-03 | glio |
| Interaction_genotype_Pars2 | Baalc | -2.893 | 6.31E-03 | glio |
| Aars2_in.vivo_noRT | Mga | -1.226 | 6.31E-03 | glio |
| Aars2_ex.vivo_noRT | Gm16576 | 2.783 | 6.31E-03 | glio |
| Atr_in.vivo_noRT | Ccl4 | 4.153 | 6.31E-03 | glio |
| Atr_in.vivo_noRT | Ascl4 | 1.99 | 6.31E-03 | glio |
| Atr_in.vivo_noRT | Gnasas1 | 1.861 | 6.31E-03 | glio |
| Bard1_ex.vivo_noRT | 4930579C12R | 2.419 | 6.31E-03 | glio |
| Bard1_ex.vivo_noRT | Cyp2c23 | 2.419 | 6.31E-03 | glio |
| Bcas2_ex.vivo_noRT | Atic | -1.126 | 6.31E-03 | glio |
| Chek1_in.vivo_noRT | Gm9895 | 2.063 | 6.31E-03 | glio |
| Dclre1c_ex.vivo_noRT | Plac8 | 3.116 | 6.31E-03 | glio |
| Hsd17b10_in.vivo_noRT | Gm10501 | 2.254 | 6.31E-03 | glio |
| culture_effect | Fars2 | -0.704 | 6.31E-04 | glio |
| Interaction_genotype_Mtor | Yipf4 | 1.518 | 6.31E-04 | glio |
| Interaction_genotype_Myc | Tmem116 | -4.257 | 6.31E-04 | glio |
| Interaction_genotype_Myc | Rpl31 | 1.307 | 6.31E-04 | glio |
| Aars2_in.vivo_noRT | Zfp963 | 2.719 | 6.31E-04 | glio |
| Atp5c1_in.vivo_noRT | Slxl1 | 2.664 | 6.31E-04 | glio |
| Bcas2_in.vivo_noRT | B230398E01Ri | 2.474 | 6.31E-04 | glio |
| Chek1_in.vivo_noRT | Atg101 | -1.293 | 6.31E-04 | glio |
| Ercc4_ex.vivo_noRT | Tnni3k | 3.071 | 6.31E-04 | glio |
| Ercc4_ex.vivo_noRT | Gjd4 | 2.704 | 6.31E-04 | glio |
| Interaction_genotype_Atr | Gm45591 | -4.201 | 6.31E-05 | glio |
| Interaction_genotype_Myc | Cpne7 | -4.759 | 6.31E-05 | glio |
| Bora_in.vivo_noRT | Gm15938 | 3.007 | 6.31E-05 | glio |
| Interaction_genotype_Mtor | Ppp1r18os | -5.535 | 6.31E-06 | glio |
| Interaction_genotype_Mtor | 1700041G16R | -5.176 | 6.31E-06 | glio |
| Interaction_genotype_Mtor | Gm11520 | -4.933 | 6.31E-06 | glio |
| Interaction_genotype_Mtor | Gm9833 | -5.077 | 6.31E-06 | glio |
| Interaction_genotype_Mtor | Cgn | -5.305 | 6.31E-06 | glio |
| Interaction_genotype_Mtor | Pdrg1 | 2.5 | 6.31E-06 | glio |
| Chek1_ex.vivo_noRT | Glod5 | 4.443 | 6.31E-06 | glio |
| culture_effect | Arhgap6 | 0.975 | 6.31E-07 | glio |
| Bcas2_in.vivo_noRT | Zfp652os | 3.502 | 6.31E-07 | glio |
| Interaction_genotype_Mtor | Gm31518 | -6.723 | 6.31E-08 | glio |
| culture_effect | Stx7 | 0.95 | 6.31E-09 | glio |
| Bcas2_in.vivo_noRT | Tmem190 | 4.807 | 6.31E-10 | glio |
| Bcas2_in.vivo_noRT | 4930502E18Ri | 4.272 | 6.31E-14 | glio |
| Chek1_in.vivo_noRT | Gm15894 | 3.999 | 6.31E-14 | glio |
| culture_effect | Ccdc62 | 1.224 | 6.32E-03 | glio |

| | | | | |
|---|---|---|---|---|
| Interaction_genotype_Atr | Gm16364.1 | -3.575 | 6.32E-03 | glio |
| Interaction_genotype_Atr | Kank3 | -3.354 | 6.32E-03 | glio |
| Interaction_genotype_Atr | Nat8f1 | -3.169 | 6.32E-03 | glio |
| Interaction_genotype_Mtor | 4930532G15R | -2.762 | 6.32E-03 | glio |
| Interaction_genotype_Myc | Kirrel3 | -3.007 | 6.32E-03 | glio |
| Interaction_genotype_Myc | Slx1b | -2.628 | 6.32E-03 | glio |
| Interaction_genotype_Myc | Abcc4 | -2.506 | 6.32E-03 | glio |
| Interaction_genotype_Nat2 | C4a | 3.709 | 6.32E-03 | glio |
| Interaction_genotype_Pars2 | Armc2 | -3.716 | 6.32E-03 | glio |
| Interaction_genotype_Pars2 | Zmynd15 | -3.662 | 6.32E-03 | glio |
| interaction_RT | 1700029J07Ri | -2.489 | 6.32E-03 | glio |
| Aars2_in.vivo_noRT | Topors | -0.865 | 6.32E-03 | glio |
| Atr_in.vivo_noRT | Tpk1 | 1.809 | 6.32E-03 | glio |
| Atr_in.vivo_noRT | Antxr2 | 1.659 | 6.32E-03 | glio |
| Atr_in.vivo_noRT | Lrrc51 | 1.585 | 6.32E-03 | glio |
| Chek1_in.vivo_noRT | March7 | -1.16 | 6.32E-03 | glio |
| Hsd17b10_in.vivo_noRT | Gm26916 | 2.039 | 6.32E-03 | glio |
| Interaction_genotype_Atr | Prox2 | -4.505 | 6.32E-04 | glio |
| Interaction_genotype_Atr | Bco2 | -3.729 | 6.32E-04 | glio |
| Interaction_genotype_Bora | Plcl1 | -3.687 | 6.32E-04 | glio |
| Atr_in.vivo_noRT | Nfatc2 | 2.502 | 6.32E-04 | glio |
| Atr_in.vivo_noRT | Mb | 2.439 | 6.32E-04 | glio |
| Bcas2_in.vivo_noRT | Rorc | 2.663 | 6.32E-04 | glio |
| Hsd17b10_in.vivo_noRT | Wincr1 | 3.352 | 6.32E-04 | glio |
| culture_effect | Cdk11b | 0.536 | 6.32E-05 | glio |
| Interaction_genotype_Mtor | Pafah1b1 | 1.632 | 6.32E-05 | glio |
| Interaction_genotype_Myc | Nlrx1 | -4.773 | 6.32E-05 | glio |
| Interaction_genotype_Myc | Gm32591 | -4.47 | 6.32E-05 | glio |
| Interaction_genotype_Myc | Gm43409 | -5.023 | 6.32E-05 | glio |
| Aars2_in.vivo_noRT | Cadm2 | 2.963 | 6.32E-05 | glio |
| Aars2_in.vivo_noRT | Gm43768 | 2.688 | 6.32E-05 | glio |
| Bora_in.vivo_noRT | Gm19325 | 2.972 | 6.32E-05 | glio |
| Chek1_in.vivo_noRT | Rfx6 | 2.747 | 6.32E-05 | glio |
| Chek1_ex.vivo_noRT | Mertk | 3.328 | 6.32E-05 | glio |
| Interaction_genotype_Myc | BC024978 | -4.494 | 6.32E-06 | glio |
| Interaction_genotype_Myc | Rpl30 | 1.585 | 6.32E-06 | glio |
| Bcas2_in.vivo_noRT | H2-Q10 | 3.169 | 6.32E-06 | glio |
| Bora_in.vivo_noRT | Gm15265 | 3.134 | 6.32E-06 | glio |
| Bora_in.vivo_noRT | Abcc10 | 3.278 | 6.32E-06 | glio |
| Hsd17b10_in.vivo_noRT | Tecrl | 3.356 | 6.32E-06 | glio |
| culture_effect | Myoz1 | -2.63 | 6.32E-07 | glio |
| Bora_in.vivo_noRT | Cacnb1 | 3.811 | 6.32E-07 | glio |
| Interaction_genotype_Mtor | AV356131 | -5.778 | 6.32E-08 | glio |
| Interaction_genotype_Mtor | Man2c1os | -6.648 | 6.32E-08 | glio |
| Interaction_genotype_Mtor | Pspn | -4.494 | 6.32E-08 | glio |
| Interaction_genotype_Mtor | Gm50147 | -6.365 | 6.32E-08 | glio |
| culture_effect | Ctso | -3.063 | 6.32E-09 | glio |
| Aars2_in.vivo_noRT | Gm48682 | 4.015 | 6.32E-09 | glio |
| Atr_in.vivo_noRT | Gng13 | 3.94 | 6.32E-09 | glio |
| Atr_in.vivo_noRT | Flrt1 | 4.558 | 6.32E-10 | glio |
| Atr_in.vivo_noRT | Zhx2 | 4.488 | 6.32E-10 | glio |
| culture_effect | Vrk3 | -0.507 | 6.33E-03 | glio |

| | | | | |
|---|---|---|---|---|
| Interaction_genotype_Aars2 | Pdzk1ip1 | -3.645 | 6.33E-03 | glio |
| Interaction_genotype_Bard1 | Atp8a2 | 4.055 | 6.33E-03 | glio |
| Interaction_genotype_Bora | Gm36198 | -3.03 | 6.33E-03 | glio |
| Interaction_genotype_Bora | Fgf7 | -3.477 | 6.33E-03 | glio |
| Interaction_genotype_Mtor | Usp44 | -3.367 | 6.33E-03 | glio |
| Interaction_genotype_Mtor | Sgce | 1.697 | 6.33E-03 | glio |
| Interaction_genotype_Myc | Hist1h2bg | -3.202 | 6.33E-03 | glio |
| Interaction_genotype_Myc | Asb16 | -2.825 | 6.33E-03 | glio |
| Aars2_in.vivo_noRT | Irak1bp1 | -1.13 | 6.33E-03 | glio |
| Atr_in.vivo_noRT | Trim13 | 1.761 | 6.33E-03 | glio |
| Chek1_in.vivo_noRT | Asb16 | 1.874 | 6.33E-03 | glio |
| Chek1_ex.vivo_noRT | Pa2g4 | -0.845 | 6.33E-03 | glio |
| Fanca_ex.vivo_noRT | Gpr17 | 2.919 | 6.33E-03 | glio |
| Hsd17b10_in.vivo_noRT | Rbm10 | -1.397 | 6.33E-03 | glio |
| Interaction_genotype_Bora | Krt80 | -3.944 | 6.33E-04 | glio |
| Interaction_genotype_Myc | Cyb561d1 | -4.099 | 6.33E-04 | glio |
| Interaction_genotype_Myc | Stk10 | -3.18 | 6.33E-04 | glio |
| interaction_RT | Rps27l | -0.536 | 6.33E-04 | glio |
| Aars2_in.vivo_noRT | Emc2 | -1.175 | 6.33E-04 | glio |
| Eif2s1_in.vivo_noRT | Clec4d | 2.143 | 6.33E-04 | glio |
| Eif2s1_in.vivo_noRT | Ccdc80 | 2.143 | 6.33E-04 | glio |
| Eif2s1_in.vivo_noRT | Gm48161 | 2.404 | 6.33E-04 | glio |
| culture_effect | Adamts4 | 1.61 | 6.33E-05 | glio |
| Interaction_genotype_Mtor | Fgf2 | -4.266 | 6.33E-05 | glio |
| Aars2_in.vivo_noRT | Meak7 | 2.833 | 6.33E-05 | glio |
| Bora_ex.vivo_noRT | Gm45336 | 3.201 | 6.33E-05 | glio |
| Bora_ex.vivo_noRT | Rhbg | 2.808 | 6.33E-05 | glio |
| Chek1_in.vivo_noRT | Bc1 | 3.016 | 6.33E-05 | glio |
| Chek1_ex.vivo_noRT | Platr3 | 3.829 | 6.33E-05 | glio |
| Chek1_ex.vivo_noRT | Med6 | -2.012 | 6.33E-05 | glio |
| Interaction_genotype_Mtor | Gm20597 | -5.874 | 6.33E-06 | glio |
| Aars2_in.vivo_noRT | BC051077 | 3.623 | 6.33E-06 | glio |
| Atr_in.vivo_noRT | Hist1h4n | 3.457 | 6.33E-06 | glio |
| Bcas2_in.vivo_noRT | Cnfn | 3.412 | 6.33E-06 | glio |
| Chek1_ex.vivo_noRT | Gm47101 | 3.738 | 6.33E-06 | glio |
| Hsd17b10_in.vivo_noRT | Magea10 | 3.149 | 6.33E-07 | glio |
| Hsd17b10_in.vivo_noRT | Cox7b2 | 3.149 | 6.33E-07 | glio |
| Bcas2_in.vivo_noRT | Insm1 | 3.688 | 6.33E-08 | glio |
| culture_effect | Rab11fip4os1 | 3.097 | 6.33E-09 | glio |
| Interaction_genotype_Mtor | Tmem179 | -8.26 | 6.33E-11 | glio |
| Bora_in.vivo_noRT | Rdh12 | 4.291 | 6.33E-11 | glio |
| Chek1_in.vivo_noRT | Ctla4 | 5.044 | 6.33E-25 | glio |
| culture_effect | Gm6566 | -1.467 | 6.34E-03 | glio |
| culture_effect | Amot | -0.901 | 6.34E-03 | glio |
| culture_effect | Caprin1 | 0.338 | 6.34E-03 | glio |
| Interaction_genotype_Aars2 | Gm49785 | -3.314 | 6.34E-03 | glio |
| Interaction_genotype_Mtor | Borcs5 | -2.423 | 6.34E-03 | glio |
| Interaction_genotype_Mtor | Srsf10 | 1.269 | 6.34E-03 | glio |
| Interaction_genotype_Mtor | Larp4 | 1.229 | 6.34E-03 | glio |
| Interaction_genotype_Myc | Fxyd5 | -2.593 | 6.34E-03 | glio |
| Aars2_in.vivo_noRT | Gm45397 | 1.725 | 6.34E-03 | glio |
| Aars2_in.vivo_noRT | Mvb12a | -0.94 | 6.34E-03 | glio |

| | | | | |
|---|---|---|---|---|
| Atr_in.vivo_noRT | 1700007L15Ri | 1.638 | 6.34E-03 | glio |
| Bora_ex.vivo_noRT | Arg2 | 2.808 | 6.34E-03 | glio |
| Chek1_in.vivo_noRT | Rpl39 | -0.696 | 6.34E-03 | glio |
| Eif2s1_ex.vivo_noRT | Gm2016 | 2.234 | 6.34E-03 | glio |
| Hsd17b10_in.vivo_noRT | Gm15473 | 2.233 | 6.34E-03 | glio |
| Hsd17b10_in.vivo_noRT | Gm49042 | 1.987 | 6.34E-03 | glio |
| culture_effect | Ablim2 | -1.467 | 6.34E-04 | glio |
| Interaction_genotype_Atr | Trim14 | -4.704 | 6.34E-04 | glio |
| Interaction_genotype_Myc | Adam1a | -3.918 | 6.34E-04 | glio |
| Aars2_in.vivo_noRT | Ptrh2 | -1.212 | 6.34E-04 | glio |
| Atp5c1_in.vivo_noRT | C130060C02R | 2.404 | 6.34E-04 | glio |
| Interaction_genotype_Mtor | Fam13a | -4.771 | 6.34E-05 | glio |
| Atr_in.vivo_noRT | Folr2 | 2.725 | 6.34E-05 | glio |
| Bora_in.vivo_noRT | Tnfrsf22 | 2.744 | 6.34E-05 | glio |
| Atr_in.vivo_noRT | Jazf1 | 3.029 | 6.34E-06 | glio |
| Chek1_in.vivo_noRT | Gm34121 | 2.918 | 6.34E-06 | glio |
| Bcas2_in.vivo_noRT | Nt5e | 3.328 | 6.34E-08 | glio |
| Aars2_in.vivo_noRT | Gm36757 | 3.933 | 6.34E-09 | glio |
| Aars2_in.vivo_noRT | 4930518J21Ri | 4.401 | 6.34E-11 | glio |
| Atr_in.vivo_noRT | Medag | 4.366 | 6.34E-11 | glio |
| Chek1_ex.vivo_noRT | 4930423D22R | 4.263 | 6.34E-12 | glio |
| Chek1_ex.vivo_noRT | 1700034I23Ri | 4.045 | 6.34E-12 | glio |
| Chek1_ex.vivo_noRT | AC159140.1 | 4.449 | 6.34E-12 | glio |
| Aars2_in.vivo_noRT | Gm16751 | 5.32 | 6.34E-19 | glio |
| Atr_in.vivo_noRT | Cgnl1 | 5.533 | 6.34E-29 | glio |
| culture_effect | Naa10 | -0.351 | 6.35E-03 | glio |
| Interaction_genotype_Mtor | 4732471J01Ri | -3.25 | 6.35E-03 | glio |
| Interaction_genotype_Mtor | Tmod1 | -2.855 | 6.35E-03 | glio |
| Interaction_genotype_Myc | Gpr155 | -2.515 | 6.35E-03 | glio |
| interaction_RT | Dag1 | -0.782 | 6.35E-03 | glio |
| Aars2_in.vivo_noRT | Mmaa | 1.74 | 6.35E-03 | glio |
| Aars2_in.vivo_noRT | Ccdc88c | 1.942 | 6.35E-03 | glio |
| Bora_in.vivo_noRT | Nrp2 | -1.169 | 6.35E-03 | glio |
| Cenpj_ex.vivo_noRT | 4933408J17Ri | 2.605 | 6.35E-03 | glio |
| Hsd17b10_in.vivo_noRT | Ccdc159 | 2.238 | 6.35E-03 | glio |
| Interaction_genotype_Myc | Jrk | -3.553 | 6.35E-04 | glio |
| Bcas2_in.vivo_noRT | Myl4 | 2.557 | 6.35E-04 | glio |
| Bcas2_in.vivo_noRT | Myl6b | 2.403 | 6.35E-04 | glio |
| Bcas2_in.vivo_noRT | Gm36266 | 2.76 | 6.35E-04 | glio |
| Bcas2_in.vivo_noRT | Eif4ebp1 | -1.057 | 6.35E-04 | glio |
| Chek1_ex.vivo_noRT | Mamdc2 | 3.002 | 6.35E-04 | glio |
| Interaction_genotype_Mtor | Amdhd1 | -4.62 | 6.35E-05 | glio |
| Aars2_in.vivo_noRT | Alpk2 | 3.611 | 6.35E-05 | glio |
| Aars2_in.vivo_noRT | Lrrn4cl | 3.053 | 6.35E-05 | glio |
| Aars2_in.vivo_noRT | Marveld1 | 1.954 | 6.35E-05 | glio |
| Aars2_ex.vivo_noRT | Insc | 3.293 | 6.35E-05 | glio |
| Bora_ex.vivo_noRT | Gm16344 | 3.201 | 6.35E-05 | glio |
| Bora_ex.vivo_noRT | Gm48435 | 2.414 | 6.35E-05 | glio |
| Bora_ex.vivo_noRT | AC159140.1 | 2.808 | 6.35E-05 | glio |
| Interaction_genotype_Myc | Hecw2 | -5.176 | 6.35E-06 | glio |
| Bcas2_in.vivo_noRT | Dmrt2 | 3.397 | 6.35E-06 | glio |
| Bora_in.vivo_noRT | Gm15080 | 3.407 | 6.35E-06 | glio |

| | | | | |
|---|---|---|---|---|
| Bora_in.vivo_noRT | Dnajc9 | -1.986 | 6.35E-06 | glio |
| Bcas2_in.vivo_noRT | Bpi | 3.126 | 6.35E-07 | glio |
| culture_effect | Fen1 | -0.964 | 6.35E-08 | glio |
| Atp5c1_in.vivo_noRT | Gm48155 | 2.668 | 6.35E-08 | glio |
| Chek1_ex.vivo_noRT | Pdgfb | 3.679 | 6.35E-08 | glio |
| Aars2_in.vivo_noRT | Wnt16 | 4.536 | 6.35E-12 | glio |
| Bora_in.vivo_noRT | 4921536K21R | 4.676 | 6.35E-12 | glio |
| Aars2_in.vivo_noRT | Mettl7a3 | 4.676 | 6.35E-13 | glio |
| Hsd17b10_in.vivo_noRT | Gm48764 | 3.616 | 6.35E-13 | glio |
| culture_effect | Rap1gap | -2.167 | 6.35E-19 | glio |
| Bcas2_in.vivo_noRT | Zscan4-ps3 | 4.807 | 6.35E-23 | glio |
| culture_effect | Tex19.2 | -1.713 | 6.36E-03 | glio |
| Interaction_genotype_Rpl3 | Mettl7b | -3.784 | 6.36E-03 | glio |
| interaction_RT | Adamts6 | 1.883 | 6.36E-03 | glio |
| interaction_RT | Cfl2 | 0.708 | 6.36E-03 | glio |
| interaction_RT | Ndufb6 | 0.506 | 6.36E-03 | glio |
| Aars2_ex.vivo_noRT | Ccr5 | 2.508 | 6.36E-03 | glio |
| Atr_in.vivo_noRT | Echdc3 | 1.59 | 6.36E-03 | glio |
| Bora_in.vivo_noRT | Gpc6 | -1.523 | 6.36E-03 | glio |
| Cenpj_in.vivo_noRT | Olfr543 | 1.762 | 6.36E-03 | glio |
| Cenpj_in.vivo_noRT | 1700017M07F | 2.022 | 6.36E-03 | glio |
| Cenpj_ex.vivo_noRT | Nos2 | 2.591 | 6.36E-03 | glio |
| Chek1_in.vivo_noRT | Crybg3 | 2.092 | 6.36E-03 | glio |
| culture_effect | Amacr | -0.927 | 6.36E-04 | glio |
| Interaction_genotype_Aars2 | 5930430L01Ri | -3.873 | 6.36E-04 | glio |
| Interaction_genotype_Myc | Dab2ip | -3.539 | 6.36E-04 | glio |
| Atp5c1_in.vivo_noRT | Gm28940 | 2.152 | 6.36E-04 | glio |
| Atr_ex.vivo_noRT | Slc16a7 | 2.764 | 6.36E-04 | glio |
| Bard1_ex.vivo_noRT | Krtcap3 | 2.611 | 6.36E-04 | glio |
| Bcas2_in.vivo_noRT | 1190005I06Ri | 2.291 | 6.36E-04 | glio |
| Chek1_ex.vivo_noRT | Olfr920 | 3.032 | 6.36E-04 | glio |
| Hsd17b10_in.vivo_noRT | C330013E15Ri | 2.552 | 6.36E-04 | glio |
| Interaction_genotype_Myc | Car5a | -4.163 | 6.36E-05 | glio |
| Aars2_in.vivo_noRT | Zfp114 | 2.853 | 6.36E-05 | glio |
| Bora_in.vivo_noRT | Usp44 | 2.947 | 6.36E-05 | glio |
| Bora_in.vivo_noRT | Auts2 | 2.486 | 6.36E-05 | glio |
| Hsd17b10_in.vivo_noRT | Gm20442 | 2.607 | 6.36E-05 | glio |
| Interaction_genotype_Aars2 | 6330415G19R | -5.424 | 6.36E-06 | glio |
| Aars2_in.vivo_noRT | Trim68 | 3.017 | 6.36E-06 | glio |
| Chek1_ex.vivo_noRT | Olfml2b | 3.678 | 6.36E-06 | glio |
| Hsd17b10_in.vivo_noRT | A930012O16R | 3.356 | 6.36E-06 | glio |
| Hsd17b10_in.vivo_noRT | Slc17a7 | 3.247 | 6.36E-06 | glio |
| Aars2_in.vivo_noRT | Casz1 | 4.051 | 6.36E-08 | glio |
| Aars2_in.vivo_noRT | C430042M11F | 3.798 | 6.36E-08 | glio |
| Bcas2_in.vivo_noRT | Aass | 3.806 | 6.36E-09 | glio |
| Chek1_ex.vivo_noRT | Gm15972 | 4.443 | 6.36E-09 | glio |
| Hsd17b10_in.vivo_noRT | Gm49172 | 3.356 | 6.36E-13 | glio |
| Atr_in.vivo_noRT | Gm13684 | 4.431 | 6.36E-14 | glio |
| Bora_in.vivo_noRT | Gm2115 | 4.936 | 6.36E-16 | glio |
| culture_effect | Pdgfa | 1.696 | 6.36E-20 | glio |
| Chek1_in.vivo_noRT | Gm2022 | 5.044 | 6.36E-21 | glio |
| culture_effect | Esyt2 | -0.557 | 6.37E-03 | glio |

| | | | | |
|---|---|---|---|---|
| culture_effect | Trmt10c | -0.5 | 6.37E-03 | glio |
| Interaction_genotype_Bcas2 | Tslp | -3.497 | 6.37E-03 | glio |
| Interaction_genotype_Mtor | Birc3 | -3.299 | 6.37E-03 | glio |
| Interaction_genotype_Mtor | Ldlrap1 | -3.114 | 6.37E-03 | glio |
| Interaction_genotype_Nat2 | Gm44736 | 2.678 | 6.37E-03 | glio |
| Aars2_in.vivo_noRT | Plec | -0.823 | 6.37E-03 | glio |
| Atp5c1_in.vivo_noRT | Hist1h4f | 2.404 | 6.37E-03 | glio |
| Atr_in.vivo_noRT | Oxsm | 2.057 | 6.37E-03 | glio |
| Atr_in.vivo_noRT | Gm4544 | 1.81 | 6.37E-03 | glio |
| Atr_ex.vivo_noRT | Necab2 | 2.454 | 6.37E-03 | glio |
| Bard1_ex.vivo_noRT | Card9 | 2.415 | 6.37E-03 | glio |
| Bcas2_in.vivo_noRT | Car11 | 1.811 | 6.37E-03 | glio |
| Bcas2_in.vivo_noRT | Vax2 | 1.972 | 6.37E-03 | glio |
| Chek1_in.vivo_noRT | Oxsm | 2.097 | 6.37E-03 | glio |
| Chek1_ex.vivo_noRT | Mblac1 | 2.623 | 6.37E-03 | glio |
| Chek1_ex.vivo_noRT | Thoc7 | -1.177 | 6.37E-03 | glio |
| Bard1_ex.vivo_noRT | Gem | 3.219 | 6.37E-04 | glio |
| Interaction_genotype_Mtor | Gm12280 | -5.314 | 6.37E-05 | glio |
| Atr_in.vivo_noRT | Gm14419 | 2.482 | 6.37E-05 | glio |
| Bard1_ex.vivo_noRT | Spn | 3.594 | 6.37E-05 | glio |
| Chek1_in.vivo_noRT | Shpk | 3.056 | 6.37E-05 | glio |
| Chek1_ex.vivo_noRT | A830035O19R | 3.359 | 6.37E-05 | glio |
| culture_effect | Lims2 | -2.782 | 6.37E-06 | glio |
| culture_effect | Twf2 | -0.821 | 6.37E-06 | glio |
| Chek1_ex.vivo_noRT | Prrx2 | 3.778 | 6.37E-06 | glio |
| Aars2_in.vivo_noRT | Gm5089 | 4.289 | 6.37E-08 | glio |
| Chek1_ex.vivo_noRT | Cxcl1 | 4.842 | 6.37E-08 | glio |
| Interaction_genotype_Mtor | Lbhd2 | -8.377 | 6.37E-09 | glio |
| Aars2_in.vivo_noRT | Gm49698 | 5.327 | 6.37E-27 | glio |
| culture_effect | Cenpc1 | -0.524 | 6.38E-03 | glio |
| Interaction_genotype_Aars2 | Ccdc106 | -3.699 | 6.38E-03 | glio |
| Interaction_genotype_Aars2 | Gm48882 | -3.434 | 6.38E-03 | glio |
| Interaction_genotype_Bcas2 | Gm31415 | -3.094 | 6.38E-03 | glio |
| Interaction_genotype_Mtor | 5430427N15R | -3.328 | 6.38E-03 | glio |
| interaction_RT | Mak16 | -0.548 | 6.38E-03 | glio |
| interaction_RT | Fis1 | 0.561 | 6.38E-03 | glio |
| interaction_RT | Impdh2 | -0.556 | 6.38E-03 | glio |
| Atr_in.vivo_noRT | Galc | 1.634 | 6.38E-03 | glio |
| Atr_in.vivo_noRT | Fign | -1.339 | 6.38E-03 | glio |
| Atr_ex.vivo_noRT | Ptpn13 | -1.975 | 6.38E-03 | glio |
| Bora_in.vivo_noRT | Nrf1 | -1.463 | 6.38E-03 | glio |
| Chek1_in.vivo_noRT | Phactr2 | 1.476 | 6.38E-03 | glio |
| Chek1_ex.vivo_noRT | Slc7a7 | 2.586 | 6.38E-03 | glio |
| Chek1_ex.vivo_noRT | Glmn | -1.711 | 6.38E-03 | glio |
| Hsd17b10_in.vivo_noRT | Kcnmb4os2 | 2.179 | 6.38E-03 | glio |
| Hsd17b10_in.vivo_noRT | Gm16283 | 2.092 | 6.38E-03 | glio |
| Hsd17b10_in.vivo_noRT | D730045B01R | 1.892 | 6.38E-03 | glio |
| culture_effect | Mrpl1 | 0.547 | 6.38E-04 | glio |
| Interaction_genotype_Mtor | 1700028J19Ri | -3.964 | 6.38E-04 | glio |
| Interaction_genotype_Mtor | Psmb10 | 1.855 | 6.38E-04 | glio |
| Aars2_in.vivo_noRT | Bend6 | 2.396 | 6.38E-04 | glio |
| Atr_in.vivo_noRT | Gm813 | 2.363 | 6.38E-04 | glio |

| | | | | |
|---|---|---|---|---|
| Atr_in.vivo_noRT | Ldlrad4 | 2.051 | 6.38E-04 | glio |
| Bcas2_in.vivo_noRT | Bcat1 | -1.749 | 6.38E-04 | glio |
| culture_effect | Olfr655 | 2.902 | 6.38E-05 | glio |
| Interaction_genotype_Aars2 | Hist1h4a | -4.779 | 6.38E-05 | glio |
| Interaction_genotype_Myc | Gm26850 | -4.617 | 6.38E-05 | glio |
| Interaction_genotype_Myc | Gm35082 | -3.526 | 6.38E-05 | glio |
| Interaction_genotype_Myc | Micos10 | 1.437 | 6.38E-05 | glio |
| Atr_ex.vivo_noRT | Gm48435 | 2.376 | 6.38E-05 | glio |
| Atr_ex.vivo_noRT | Gm49504 | 2.764 | 6.38E-05 | glio |
| Bcas2_in.vivo_noRT | Stmn4 | 4.291 | 6.38E-05 | glio |
| Hsd17b10_in.vivo_noRT | Zfp990 | 2.777 | 6.38E-05 | glio |
| Interaction_genotype_Mtor | Gm14264 | -5.135 | 6.38E-06 | glio |
| Interaction_genotype_Myc | Fbrsl1 | -3.495 | 6.38E-06 | glio |
| Interaction_genotype_Myc | Gm49874 | -5.256 | 6.38E-06 | glio |
| Atp5c1_in.vivo_noRT | 4933400F21Ri | 2.408 | 6.38E-06 | glio |
| Bcas2_in.vivo_noRT | Tmem107 | -1.95 | 6.38E-06 | glio |
| Bcas2_in.vivo_noRT | Zfp661 | 3.574 | 6.38E-07 | glio |
| Aars2_in.vivo_noRT | Cidea | 4.412 | 6.38E-13 | glio |
| Chek1_in.vivo_noRT | Krtcap3 | 4.252 | 6.38E-15 | glio |
| culture_effect | Tnfrsf9 | -1.19 | 6.39E-03 | glio |
| Interaction_genotype_Atr | D030068K23R | -3.438 | 6.39E-03 | glio |
| Interaction_genotype_Atr | Jph4 | -3.323 | 6.39E-03 | glio |
| Interaction_genotype_Atr | Shisal2a | -3.611 | 6.39E-03 | glio |
| Interaction_genotype_Atr | Zhx2 | -3.575 | 6.39E-03 | glio |
| Interaction_genotype_Atr | Gm32591 | -3.317 | 6.39E-03 | glio |
| Interaction_genotype_Atr | Hmx3 | -3.143 | 6.39E-03 | glio |
| Interaction_genotype_Atr | Gm14326 | -3.305 | 6.39E-03 | glio |
| Interaction_genotype_Atr | Parvb | -3.202 | 6.39E-03 | glio |
| Interaction_genotype_Atr | Chtf18 | -2.676 | 6.39E-03 | glio |
| Interaction_genotype_Atr | Gon4l | 1.521 | 6.39E-03 | glio |
| Interaction_genotype_Cdkn2a | Gm4166 | 2.99 | 6.39E-03 | glio |
| Interaction_genotype_Cdkn2a | Slc25a18 | 3.373 | 6.39E-03 | glio |
| Interaction_genotype_Myc | Tcp11x2 | -2.6 | 6.39E-03 | glio |
| Interaction_genotype_Myc | Rhbdd1 | -2.349 | 6.39E-03 | glio |
| Interaction_genotype_Pars2 | 3930402G23R | -3.881 | 6.39E-03 | glio |
| Atr_ex.vivo_noRT | Agap2 | 2.178 | 6.39E-03 | glio |
| Bcas2_in.vivo_noRT | Iba57 | 1.934 | 6.39E-03 | glio |
| Bcas2_in.vivo_noRT | Cpsf4 | -1.457 | 6.39E-03 | glio |
| Bcas2_in.vivo_noRT | Hspa9 | -0.642 | 6.39E-03 | glio |
| Bora_in.vivo_noRT | E2f4 | -1.275 | 6.39E-03 | glio |
| Chek1_in.vivo_noRT | Gm4544 | 1.844 | 6.39E-03 | glio |
| Chek1_in.vivo_noRT | Tle5 | -0.737 | 6.39E-03 | glio |
| Chek1_in.vivo_noRT | Rpl9 | -0.511 | 6.39E-03 | glio |
| Chek1_ex.vivo_noRT | Glipr2 | 2.884 | 6.39E-03 | glio |
| Chek1_ex.vivo_noRT | Mrps26 | -0.852 | 6.39E-03 | glio |
| culture_effect | Kat6b | -0.777 | 6.39E-04 | glio |
| culture_effect | Guk1 | 0.392 | 6.39E-04 | glio |
| Interaction_genotype_Atr | D930028M14I | -3.856 | 6.39E-04 | glio |
| Interaction_genotype_Atr | Hectd2 | -3.419 | 6.39E-04 | glio |
| Interaction_genotype_Mtor | Gabarap | 1.488 | 6.39E-04 | glio |
| Interaction_genotype_Rad51 | Acaa1a | 2.846 | 6.39E-04 | glio |
| Interaction_genotype_Rad51 | Tnks2 | 2.097 | 6.39E-04 | glio |

| | | | | |
|---|---|---|---|---|
| interaction_RT | Cd302 | 0.925 | 6.39E-04 | glio |
| interaction_RT | Nsun2 | -0.89 | 6.39E-04 | glio |
| interaction_RT | Zfp239 | -0.97 | 6.39E-04 | glio |
| Atr_in.vivo_noRT | Rasip1 | 2.503 | 6.39E-04 | glio |
| Bora_in.vivo_noRT | Cdk19 | -1.443 | 6.39E-04 | glio |
| Bora_ex.vivo_noRT | Fhl1 | 2.491 | 6.39E-04 | glio |
| Bora_ex.vivo_noRT | Mag | 3.201 | 6.39E-04 | glio |
| Bora_ex.vivo_noRT | Spink4 | 3.201 | 6.39E-04 | glio |
| Eif2s1_in.vivo_noRT | B230311B06R | 2.941 | 6.39E-04 | glio |
| culture_effect | D930030I03Ri | 2.276 | 6.39E-05 | glio |
| Aars2_in.vivo_noRT | Pxt1 | 2.854 | 6.39E-05 | glio |
| Atr_in.vivo_noRT | Gm12971 | 2.889 | 6.39E-05 | glio |
| Atr_in.vivo_noRT | Tmem106b | -1.738 | 6.39E-05 | glio |
| Atr_in.vivo_noRT | Psme3 | -1.329 | 6.39E-05 | glio |
| Bcas2_in.vivo_noRT | Gm20234 | 3.09 | 6.39E-05 | glio |
| Bcas2_in.vivo_noRT | Hdac7 | 2.807 | 6.39E-05 | glio |
| Bora_in.vivo_noRT | Hmgn2 | -1.848 | 6.39E-05 | glio |
| Chek1_ex.vivo_noRT | Ang | 3.749 | 6.39E-05 | glio |
| Atp5c1_in.vivo_noRT | Gm48170 | 2.668 | 6.39E-06 | glio |
| Interaction_genotype_Mtor | Hrk | -6.132 | 6.39E-07 | glio |
| culture_effect | Lhfpl3 | -2.111 | 6.39E-08 | glio |
| Aars2_in.vivo_noRT | Pawr | 3.56 | 6.39E-09 | glio |
| Chek1_in.vivo_noRT | Slc36a2 | 3.925 | 6.39E-09 | glio |
| Chek1_in.vivo_noRT | Gm36043 | 3.588 | 6.39E-09 | glio |
| Atr_in.vivo_noRT | 4930589L23Ri | 4.447 | 6.39E-10 | glio |
| Bcas2_in.vivo_noRT | Nxph4 | 4.807 | 6.39E-12 | glio |
| Bora_in.vivo_noRT | 5033417F24Ri | 4.936 | 6.39E-13 | glio |
| Bora_in.vivo_noRT | Ppil6 | 4.936 | 6.39E-15 | glio |
| Chek1_ex.vivo_noRT | Serpinb9g | 4.842 | 6.39E-15 | glio |
| Chek1_ex.vivo_noRT | BC049987 | 4.842 | 6.39E-15 | glio |
| Bcas2_in.vivo_noRT | BC049987 | 4.024 | 6.39E-16 | glio |
| culture_effect | Zdhhc6 | 0.458 | 6.40E-03 | glio |
| Interaction_genotype_Bcas2 | Fbxl13 | -3.18 | 6.40E-03 | glio |
| Interaction_genotype_Lmo7 | Gm3716 | -4.058 | 6.40E-03 | glio |
| Interaction_genotype_Myc | AW112010 | 3.411 | 6.40E-03 | glio |
| Interaction_genotype_Myc | As3mt | -2.362 | 6.40E-03 | glio |
| Aars2_in.vivo_noRT | Kctd9 | 1.38 | 6.40E-03 | glio |
| Bcas2_in.vivo_noRT | Aar2 | 1.288 | 6.40E-03 | glio |
| Chek1_in.vivo_noRT | Gm8773 | 2.141 | 6.40E-03 | glio |
| Chek1_ex.vivo_noRT | 5033417F24Ri | 2.422 | 6.40E-03 | glio |
| culture_effect | Otud6b | 0.503 | 6.40E-04 | glio |
| Interaction_genotype_Chek1 | Gm11627 | -4.075 | 6.40E-04 | glio |
| Interaction_genotype_Chek1 | Atp1a2 | 4.964 | 6.40E-04 | glio |
| Interaction_genotype_Myc | Nxt2 | -3.154 | 6.40E-04 | glio |
| Interaction_genotype_Myc | Vdac3 | 1.44 | 6.40E-04 | glio |
| Aars2_in.vivo_noRT | Ctns | 2.177 | 6.40E-04 | glio |
| Atp5c1_in.vivo_noRT | Gm12610 | 2.139 | 6.40E-04 | glio |
| Atr_in.vivo_noRT | Smc4 | -1.621 | 6.40E-04 | glio |
| Atr_in.vivo_noRT | Srpr | -1.056 | 6.40E-04 | glio |
| Bora_in.vivo_noRT | Hnrnpab | -0.624 | 6.40E-04 | glio |
| Bora_ex.vivo_noRT | Masp2 | 2.808 | 6.40E-04 | glio |
| Bora_ex.vivo_noRT | Gm16751 | 2.802 | 6.40E-04 | glio |

| | | | | |
|---|---|---|---|---|
| Interaction_genotype_Mtor | Cxcl16 | -4.53 | 6.40E-05 | glio |
| Interaction_genotype_Mtor | Gm43915 | -3.96 | 6.40E-05 | glio |
| Interaction_genotype_Mtor | Pabpn1 | 1.98 | 6.40E-05 | glio |
| Atr_ex.vivo_noRT | Spry1 | -2.162 | 6.40E-05 | glio |
| Bora_in.vivo_noRT | Prob1 | 2.989 | 6.40E-05 | glio |
| Bora_in.vivo_noRT | Smc1a | -1.232 | 6.40E-05 | glio |
| Chek1_in.vivo_noRT | Gm36210 | 2.842 | 6.40E-05 | glio |
| Hsd17b10_in.vivo_noRT | Col6a3 | 2.58 | 6.40E-05 | glio |
| Interaction_genotype_Mtor | Lsm1 | 2.258 | 6.40E-06 | glio |
| interaction_RT | Ybx3 | -0.817 | 6.40E-06 | glio |
| Atr_in.vivo_noRT | Plin1 | 3.82 | 6.40E-08 | glio |
| Aars2_in.vivo_noRT | Vamp8 | 3.503 | 6.40E-10 | glio |
| culture_effect | Mettl18 | 1.097 | 6.41E-03 | glio |
| Interaction_genotype_Atr | Gm15138 | -2.499 | 6.41E-03 | glio |
| Interaction_genotype_Bcas2 | Gm21936 | -2.46 | 6.41E-03 | glio |
| Interaction_genotype_Bcas2 | Gm13856 | -2.458 | 6.41E-03 | glio |
| Interaction_genotype_Bcas2 | 4930434B07R | -3.181 | 6.41E-03 | glio |
| Interaction_genotype_Bcas2 | Hectd2os | -3.532 | 6.41E-03 | glio |
| Interaction_genotype_Mtor | Clca3a2 | -2.913 | 6.41E-03 | glio |
| Interaction_genotype_Mtor | Gm10282 | -3.294 | 6.41E-03 | glio |
| Interaction_genotype_Mtor | Psma4 | 1.364 | 6.41E-03 | glio |
| Interaction_genotype_Myc | Gm50115 | -2.865 | 6.41E-03 | glio |
| Interaction_genotype_Myc | Cd83 | -2.71 | 6.41E-03 | glio |
| Interaction_genotype_Myc | Pank1 | -2.279 | 6.41E-03 | glio |
| Interaction_genotype_Myc | Lin9 | -1.841 | 6.41E-03 | glio |
| Interaction_genotype_Myc | Rabif | -1.87 | 6.41E-03 | glio |
| interaction_RT | Cstf2 | -0.861 | 6.41E-03 | glio |
| Aars2_in.vivo_noRT | Coro2b | 1.69 | 6.41E-03 | glio |
| Aars2_in.vivo_noRT | Traf7 | -1.06 | 6.41E-03 | glio |
| Atp5c1_in.vivo_noRT | Vps33a | -1.781 | 6.41E-03 | glio |
| Atr_in.vivo_noRT | Camkk2 | 1.471 | 6.41E-03 | glio |
| Bcas2_in.vivo_noRT | Gm50232 | 1.886 | 6.41E-03 | glio |
| Bcas2_in.vivo_noRT | Sfxn3 | -1.163 | 6.41E-03 | glio |
| Bcas2_in.vivo_noRT | Ssbp1 | -0.597 | 6.41E-03 | glio |
| Bcas2_ex.vivo_noRT | 2310003N18R | 2.349 | 6.41E-03 | glio |
| Bora_in.vivo_noRT | Ripk3 | 2.266 | 6.41E-03 | glio |
| Bora_in.vivo_noRT | Pwp1 | -1.143 | 6.41E-03 | glio |
| Bora_in.vivo_noRT | Nans | -0.937 | 6.41E-03 | glio |
| Chek1_ex.vivo_noRT | Slamf1 | 2.581 | 6.41E-03 | glio |
| Eif2s1_in.vivo_noRT | Prl2c3 | 2.143 | 6.41E-03 | glio |
| Eif2s1_ex.vivo_noRT | Gm14820 | 2.234 | 6.41E-03 | glio |
| Interaction_genotype_Mtor | Arhgef16 | -3.462 | 6.41E-04 | glio |
| Interaction_genotype_Mtor | Radil | -3.907 | 6.41E-04 | glio |
| Interaction_genotype_Mtor | Txndc12 | 1.806 | 6.41E-04 | glio |
| Bcas2_in.vivo_noRT | Gipc3 | 2.556 | 6.41E-04 | glio |
| Chek1_in.vivo_noRT | Myo1h | 2.549 | 6.41E-04 | glio |
| Chek1_in.vivo_noRT | Spink4 | 2.299 | 6.41E-04 | glio |
| Chek1_in.vivo_noRT | Mb | 2.473 | 6.41E-04 | glio |
| Atr_in.vivo_noRT | 2010110K18R | 2.687 | 6.41E-05 | glio |
| Bard1_ex.vivo_noRT | Rgs4 | 2.814 | 6.41E-05 | glio |
| Bcas2_in.vivo_noRT | Esco2 | -2.241 | 6.41E-05 | glio |
| Bora_in.vivo_noRT | Ccdc106 | 3.081 | 6.41E-05 | glio |

| | | | | |
|---|---|---|---|---|
| Bora_in.vivo_noRT | Plcb4 | 2.873 | 6.41E-05 | glio |
| Hsd17b10_in.vivo_noRT | Hey2 | 2.814 | 6.41E-05 | glio |
| Hsd17b10_in.vivo_noRT | Gm36608 | 2.976 | 6.41E-05 | glio |
| Interaction_genotype_Myc | H3f3aos | -5.145 | 6.41E-06 | glio |
| Atr_in.vivo_noRT | 1700112D23R | 3.107 | 6.41E-06 | glio |
| interaction_RT | Pdap1 | -0.871 | 6.41E-07 | glio |
| culture_effect | Xbp1 | -0.909 | 6.41E-08 | glio |
| Aars2_ex.vivo_noRT | Gm49172 | 3.293 | 6.41E-08 | glio |
| Aars2_ex.vivo_noRT | Gjd4 | 3.293 | 6.41E-08 | glio |
| Aars2_in.vivo_noRT | Gm32585 | 4.801 | 6.41E-11 | glio |
| Aars2_in.vivo_noRT | Klk13 | 4.794 | 6.41E-21 | glio |
| Aars2_in.vivo_noRT | Capn13 | 5.327 | 6.41E-30 | glio |
| culture_effect | Pogz | -0.663 | 6.42E-03 | glio |
| Interaction_genotype_Aars2 | Trappc2l | 1.465 | 6.42E-03 | glio |
| Interaction_genotype_Myc | Mpzl2 | -2.469 | 6.42E-03 | glio |
| Interaction_genotype_Myc | Dcaf15 | -2.161 | 6.42E-03 | glio |
| Interaction_genotype_Pars2 | Rnft2 | -3.646 | 6.42E-03 | glio |
| Aars2_in.vivo_noRT | Itga6 | -1.053 | 6.42E-03 | glio |
| Aars2_in.vivo_noRT | Ubc | -0.772 | 6.42E-03 | glio |
| Atr_in.vivo_noRT | Tle2 | 2.062 | 6.42E-03 | glio |
| Bora_in.vivo_noRT | Acbd6 | -0.997 | 6.42E-03 | glio |
| Bora_in.vivo_noRT | Cnp | 0.938 | 6.42E-03 | glio |
| Bora_ex.vivo_noRT | Hnf1aos1 | 2.404 | 6.42E-03 | glio |
| Cenpj_ex.vivo_noRT | Gm49504 | 2.192 | 6.42E-03 | glio |
| Cenpj_ex.vivo_noRT | 4921504E06Ri | 2.3 | 6.42E-03 | glio |
| Cyp19a1_ex.vivo_noRT | Gm6566 | 2.565 | 6.42E-03 | glio |
| Cyp19a1_ex.vivo_noRT | D630024D03F | 2.372 | 6.42E-03 | glio |
| Cyp19a1_ex.vivo_noRT | Cd7 | 2.159 | 6.42E-03 | glio |
| Cyp19a1_ex.vivo_noRT | Gm48764 | 2.159 | 6.42E-03 | glio |
| Interaction_genotype_Myc | Ovgp1 | -3.815 | 6.42E-04 | glio |
| Aars2_in.vivo_noRT | Klhl25 | 2.214 | 6.42E-04 | glio |
| Chek1_in.vivo_noRT | P4ha1 | -1.514 | 6.42E-04 | glio |
| Hsd17b10_in.vivo_noRT | Insm1 | 2.497 | 6.42E-04 | glio |
| Hsd17b10_in.vivo_noRT | Lgals7 | -2.783 | 6.42E-04 | glio |
| Aars2_in.vivo_noRT | Gm42701 | 2.81 | 6.42E-05 | glio |
| Atr_in.vivo_noRT | Rwdd3 | 2.951 | 6.42E-05 | glio |
| Bora_in.vivo_noRT | Rimkla | 2.718 | 6.42E-05 | glio |
| Interaction_genotype_Mtor | Btc | -7.445 | 6.42E-07 | glio |
| Interaction_genotype_Mtor | Tdrd5 | -5.447 | 6.42E-07 | glio |
| Interaction_genotype_Mtor | Idi2 | -4.099 | 6.42E-07 | glio |
| Interaction_genotype_Mtor | Ush1g | -4.498 | 6.42E-07 | glio |
| Aars2_in.vivo_noRT | Grip1 | 3.448 | 6.42E-07 | glio |
| Bora_ex.vivo_noRT | Gm47510 | 3.201 | 6.42E-07 | glio |
| Bora_ex.vivo_noRT | Gm5788 | 3.201 | 6.42E-07 | glio |
| Bora_ex.vivo_noRT | Chia1 | 3.207 | 6.42E-07 | glio |
| Chek1_in.vivo_noRT | Spx | 3.318 | 6.42E-07 | glio |
| Chek1_in.vivo_noRT | Gstp3 | 3.744 | 6.42E-08 | glio |
| Chek1_in.vivo_noRT | Jakmip1 | 3.723 | 6.42E-08 | glio |
| Interaction_genotype_Mtor | Prl7c1 | -5.194 | 6.42E-09 | glio |
| Aars2_in.vivo_noRT | Tmem44 | 4.068 | 6.42E-09 | glio |
| Aars2_in.vivo_noRT | Nppc | 4.328 | 6.42E-09 | glio |
| Aars2_in.vivo_noRT | 1700011M02F | 5.063 | 6.42E-09 | glio |

| | | | | |
|---|---|---|---|---|
| Interaction_genotype_Mtor | Hdac9 | -7.422 | 6.42E-10 | glio |
| Bcas2_in.vivo_noRT | Gm13403 | 4.149 | 6.42E-10 | glio |
| Chek1_ex.vivo_noRT | Gm41349 | 4.842 | 6.42E-10 | glio |
| culture_effect | Upf3a | 0.432 | 6.43E-03 | glio |
| Interaction_genotype_Aars2 | 4930563E22Ri | -3.393 | 6.43E-03 | glio |
| Interaction_genotype_Atr | Jmjd1c | 1.408 | 6.43E-03 | glio |
| Interaction_genotype_Mtor | Ccdc141 | -3.416 | 6.43E-03 | glio |
| Interaction_genotype_Mtor | Phpt1 | -2.045 | 6.43E-03 | glio |
| Interaction_genotype_Myc | Gnasas1 | -2.927 | 6.43E-03 | glio |
| Interaction_genotype_Pars2 | St8sia1 | -3.836 | 6.43E-03 | glio |
| interaction_RT | Rbakdn | 2.66 | 6.43E-03 | glio |
| interaction_RT | Becn1 | 0.533 | 6.43E-03 | glio |
| interaction_RT | Rps6 | -0.519 | 6.43E-03 | glio |
| Atr_in.vivo_noRT | Rtn2 | 1.802 | 6.43E-03 | glio |
| Atr_in.vivo_noRT | Pcnx4 | -1.326 | 6.43E-03 | glio |
| Bora_in.vivo_noRT | Pcdhb20 | 1.877 | 6.43E-03 | glio |
| Bora_in.vivo_noRT | Zswim7 | -1.26 | 6.43E-03 | glio |
| Bora_in.vivo_noRT | Ndufa13 | -0.515 | 6.43E-03 | glio |
| Bora_ex.vivo_noRT | Spns3 | 2.802 | 6.43E-03 | glio |
| Brca2_in.vivo_noRT | Gm5788 | 1.763 | 6.43E-03 | glio |
| Chek1_in.vivo_noRT | Gm47863 | 2.046 | 6.43E-03 | glio |
| Chek1_ex.vivo_noRT | Tox | 2.584 | 6.43E-03 | glio |
| Chek1_ex.vivo_noRT | Optn | 2.53 | 6.43E-03 | glio |
| Chek1_ex.vivo_noRT | Golm1 | -1.861 | 6.43E-03 | glio |
| culture_effect | Ascl4 | -2.129 | 6.43E-04 | glio |
| Interaction_genotype_Mtor | Fgfr1 | -4.801 | 6.43E-04 | glio |
| interaction_RT | Cdc25a | -1.082 | 6.43E-04 | glio |
| Atp5c1_in.vivo_noRT | 2610027K06R | 2.54 | 6.43E-04 | glio |
| culture_effect | Usf1 | -0.767 | 6.43E-05 | glio |
| Interaction_genotype_Mtor | Gm14443 | -4.437 | 6.43E-05 | glio |
| culture_effect | Rin1 | 1.17 | 6.43E-06 | glio |
| interaction_RT | Eef2 | -0.74 | 6.43E-06 | glio |
| Aars2_ex.vivo_noRT | Gpc5 | 2.897 | 6.43E-06 | glio |
| Bora_ex.vivo_noRT | Gm26978 | 3.407 | 6.43E-06 | glio |
| Atr_in.vivo_noRT | Gm43017 | 3.419 | 6.43E-07 | glio |
| Atr_in.vivo_noRT | Serac1 | 3.304 | 6.43E-07 | glio |
| Bora_in.vivo_noRT | Gm26850 | 3.542 | 6.43E-07 | glio |
| culture_effect | Arhgap28 | 1.602 | 6.43E-09 | glio |
| Aars2_in.vivo_noRT | 1700030C12R | 4.401 | 6.43E-10 | glio |
| Chek1_in.vivo_noRT | 9530080O11R | 4.126 | 6.43E-10 | glio |
| Atr_ex.vivo_noRT | Gm16178 | 3.962 | 6.43E-11 | glio |
| Interaction_genotype_Mtor | Klhdc8a | -6.942 | 6.43E-12 | glio |
| Interaction_genotype_Mtor | Gm36356 | -7.396 | 6.43E-12 | glio |
| Chek1_in.vivo_noRT | Zscan4-ps3 | 5.044 | 6.43E-25 | glio |
| Interaction_genotype_Aars2 | Rad54b | -2.614 | 6.44E-03 | glio |
| Interaction_genotype_Myc | Cox7b | 1.079 | 6.44E-03 | glio |
| Interaction_genotype_Myc | Rps14 | 1.033 | 6.44E-03 | glio |
| interaction_RT | Zc3h18 | -0.679 | 6.44E-03 | glio |
| Aars2_in.vivo_noRT | Jup | 1.853 | 6.44E-03 | glio |
| Aars2_in.vivo_noRT | Spred3 | 1.651 | 6.44E-03 | glio |
| Bora_in.vivo_noRT | Khdrbs1 | -0.765 | 6.44E-03 | glio |
| Bora_ex.vivo_noRT | Hif3a | 2.808 | 6.44E-03 | glio |

| | | | | |
|---|---|---|---|---|
| Cdkn2a_in.vivo_noRT | Ksr2 | -2.186 | 6.44E-03 | glio |
| Cdkn2a_in.vivo_noRT | Gm9758 | -1.935 | 6.44E-03 | glio |
| Cdkn2a_in.vivo_noRT | Gm15908 | -1.675 | 6.44E-03 | glio |
| Cdkn2a_in.vivo_noRT | A930014E10Ri | -1.675 | 6.44E-03 | glio |
| Hsd17b10_in.vivo_noRT | Tes3-ps | 2.109 | 6.44E-03 | glio |
| Hsd17b10_in.vivo_noRT | Fbxl22 | 2.031 | 6.44E-03 | glio |
| culture_effect | Gpatch4 | 0.523 | 6.44E-04 | glio |
| Aars2_in.vivo_noRT | Gm41609 | 2.64 | 6.44E-04 | glio |
| Aars2_in.vivo_noRT | Gm20045 | 2.206 | 6.44E-04 | glio |
| Atp5c1_in.vivo_noRT | Ntn1 | 2.294 | 6.44E-04 | glio |
| Atr_in.vivo_noRT | Tmem191c | 2.278 | 6.44E-04 | glio |
| Atr_in.vivo_noRT | 2310068J16Ri | 2.037 | 6.44E-04 | glio |
| Atr_in.vivo_noRT | Noc2l | -1.354 | 6.44E-04 | glio |
| Bora_ex.vivo_noRT | Zdhhc14 | 2.806 | 6.44E-04 | glio |
| Brca2_ex.vivo_noRT | Klk13 | 2.825 | 6.44E-04 | glio |
| Atr_in.vivo_noRT | Rbpms2 | 2.821 | 6.44E-05 | glio |
| Bcas2_in.vivo_noRT | Ift27 | -1.695 | 6.44E-05 | glio |
| Bora_in.vivo_noRT | E230029C05Ri | 3.065 | 6.44E-06 | glio |
| Bora_in.vivo_noRT | Zeb2os | 3.023 | 6.44E-06 | glio |
| Interaction_genotype_Mtor | Vwce | -5.395 | 6.44E-07 | glio |
| Interaction_genotype_Mtor | Vmn2r18 | -4.099 | 6.44E-07 | glio |
| Bcas2_in.vivo_noRT | 6330408M09F | 3.765 | 6.44E-07 | glio |
| Bora_in.vivo_noRT | Lsm14a | -2.08 | 6.44E-08 | glio |
| Atr_in.vivo_noRT | Gm20492 | 3.942 | 6.44E-09 | glio |
| interaction_RT | Ost4 | 1.207 | 6.44E-10 | glio |
| Hsd17b10_in.vivo_noRT | Sytl4 | 3.35 | 6.44E-11 | glio |
| Atr_in.vivo_noRT | Slc4a9 | 5.001 | 6.44E-18 | glio |
| Bcas2_in.vivo_noRT | Cldn5 | 4.541 | 6.44E-20 | glio |
| Interaction_genotype_Atr | Zw10 | -2.575 | 6.45E-03 | glio |
| Interaction_genotype_Bcas2 | Gm26673 | -3.598 | 6.45E-03 | glio |
| Interaction_genotype_Myc | Bach2 | -3.097 | 6.45E-03 | glio |
| Interaction_genotype_Myc | Cenpl | -2.06 | 6.45E-03 | glio |
| Interaction_genotype_Rad51 | Gbp8 | 3.738 | 6.45E-03 | glio |
| Aars2_ex.vivo_noRT | Ggt5 | 2.592 | 6.45E-03 | glio |
| Aars2_ex.vivo_noRT | Akr1b7 | 2.893 | 6.45E-03 | glio |
| Atr_in.vivo_noRT | Zfp516 | -1.218 | 6.45E-03 | glio |
| Bcas2_in.vivo_noRT | Gm49123 | 2.147 | 6.45E-03 | glio |
| Bcas2_in.vivo_noRT | Zbtb22 | 1.664 | 6.45E-03 | glio |
| Bora_in.vivo_noRT | Cdc20 | -1.657 | 6.45E-03 | glio |
| Bora_in.vivo_noRT | Mien1 | 0.648 | 6.45E-03 | glio |
| Cenpj_ex.vivo_noRT | Rslcan18 | 2.601 | 6.45E-03 | glio |
| Eif2s1_ex.vivo_noRT | Gm4166 | 2.234 | 6.45E-03 | glio |
| Hsd17b10_in.vivo_noRT | BC065397 | 2.343 | 6.45E-03 | glio |
| Hsd17b10_in.vivo_noRT | Adam10 | 1.021 | 6.45E-03 | glio |
| Interaction_genotype_Myc | H2-Ab1 | 3.387 | 6.45E-04 | glio |
| Aars2_in.vivo_noRT | Zfand4 | 2.521 | 6.45E-04 | glio |
| Aars2_in.vivo_noRT | Lgals9 | -1.345 | 6.45E-04 | glio |
| Aars2_in.vivo_noRT | Arpc1b | -0.742 | 6.45E-04 | glio |
| Bora_ex.vivo_noRT | BC018473 | 3.196 | 6.45E-04 | glio |
| Bora_ex.vivo_noRT | 4930579C12R | 2.806 | 6.45E-04 | glio |
| Bora_ex.vivo_noRT | Cyp2c23 | 2.806 | 6.45E-04 | glio |
| Hsd17b10_in.vivo_noRT | 1700101I11Ri | 2.756 | 6.45E-04 | glio |

| | | | | |
|---|---|---|---|---|
| Hsd17b10_in.vivo_noRT | Fes | 2.596 | 6.45E-04 | glio |
| Interaction_genotype_Mtor | Rpl10 | 1.285 | 6.45E-05 | glio |
| Interaction_genotype_Mtor | Emid1 | -6.024 | 6.45E-06 | glio |
| Interaction_genotype_Mtor | Scp2 | 2.194 | 6.45E-06 | glio |
| Interaction_genotype_Myc | 1500002C15R | -5.379 | 6.45E-06 | glio |
| interaction_RT | Dbndd2 | 1.5 | 6.45E-06 | glio |
| Bora_ex.vivo_noRT | Gm21761 | 3.201 | 6.45E-06 | glio |
| Chek1_ex.vivo_noRT | Adam2 | 4.842 | 6.45E-11 | glio |
| Chek1_in.vivo_noRT | Bmx | 4.256 | 6.45E-12 | glio |
| Bora_in.vivo_noRT | Gm12794 | 4.936 | 6.45E-22 | glio |
| Aars2_in.vivo_noRT | Olfr881 | 5.327 | 6.45E-27 | glio |
| culture_effect | Fa2h | -7.515 | 6.45E-39 | glio |
| Interaction_genotype_Mtor | Maob | -3.253 | 6.46E-03 | glio |
| Interaction_genotype_Mtor | Mknk2 | -2.221 | 6.46E-03 | glio |
| Interaction_genotype_Mtor | Dcaf10 | -1.838 | 6.46E-03 | glio |
| Interaction_genotype_Mtor | Rc3h2 | 1.58 | 6.46E-03 | glio |
| Interaction_genotype_Mtor | Aldh2 | 1.712 | 6.46E-03 | glio |
| Interaction_genotype_Rpl3 | Ndfip2 | 1.958 | 6.46E-03 | glio |
| Aars2_in.vivo_noRT | Gm4707 | 1.89 | 6.46E-03 | glio |
| Atp5c1_in.vivo_noRT | Gm15559 | 2.272 | 6.46E-03 | glio |
| Atr_ex.vivo_noRT | 5430416N02R | 1.615 | 6.46E-03 | glio |
| Bcas2_in.vivo_noRT | Ttbk1 | 2.01 | 6.46E-03 | glio |
| Bcas2_in.vivo_noRT | B230206L02Ri | 2.005 | 6.46E-03 | glio |
| Bcas2_in.vivo_noRT | Lhx6 | 1.919 | 6.46E-03 | glio |
| Bcas2_in.vivo_noRT | Psmd7 | -0.683 | 6.46E-03 | glio |
| Bora_in.vivo_noRT | Coasy | -1.239 | 6.46E-03 | glio |
| Cdkn2a_in.vivo_noRT | Gm45338 | -1.675 | 6.46E-03 | glio |
| Cdkn2a_in.vivo_noRT | Scg2 | -1.675 | 6.46E-03 | glio |
| Cdkn2a_in.vivo_noRT | 1700003E24Ri | -1.675 | 6.46E-03 | glio |
| Cdkn2a_in.vivo_noRT | Pdlim4 | 1.82 | 6.46E-03 | glio |
| Chek1_in.vivo_noRT | Coa6 | -1.192 | 6.46E-03 | glio |
| Hsd17b10_in.vivo_noRT | Gm7008 | 2.15 | 6.46E-03 | glio |
| Hsd17b10_in.vivo_noRT | 4930445N18R | 2.172 | 6.46E-03 | glio |
| Hsd17b10_in.vivo_noRT | Cep104 | -1.276 | 6.46E-03 | glio |
| culture_effect | Gm10553 | -2.186 | 6.46E-04 | glio |
| culture_effect | BFP | 1.145 | 6.46E-04 | glio |
| Interaction_genotype_Myc | Ifitm5 | -3.79 | 6.46E-04 | glio |
| Interaction_genotype_Myc | Zfp658 | -3.888 | 6.46E-04 | glio |
| Interaction_genotype_Myc | Txn1 | 1.376 | 6.46E-04 | glio |
| Interaction_genotype_Pars2 | Gm49874 | -4.781 | 6.46E-04 | glio |
| Aars2_in.vivo_noRT | 1810062G17R | 2.36 | 6.46E-04 | glio |
| Atr_ex.vivo_noRT | Kmo | 2.771 | 6.46E-04 | glio |
| Bora_in.vivo_noRT | Hist1h2bf | 2.624 | 6.46E-04 | glio |
| culture_effect | mt-Nd6 | -0.938 | 6.46E-05 | glio |
| Atr_ex.vivo_noRT | Cyp4f40 | 3.569 | 6.46E-05 | glio |
| Atr_ex.vivo_noRT | Kit | 3.164 | 6.46E-05 | glio |
| culture_effect | Ep400 | -0.779 | 6.46E-07 | glio |
| culture_effect | Fdps | -0.945 | 6.46E-07 | glio |
| Bora_in.vivo_noRT | Gm30262 | 3.644 | 6.46E-07 | glio |
| Aars2_in.vivo_noRT | Gm10125 | 4.125 | 6.46E-09 | glio |
| Atr_in.vivo_noRT | Ntf5 | 4.238 | 6.46E-09 | glio |
| culture_effect | Tnrc18 | 1.295 | 6.46E-11 | glio |

| | | | | |
|---|---|---|---|---|
| Aars2_in.vivo_noRT | Tcstv1 | 5.327 | 6.46E-19 | glio |
| Atr_in.vivo_noRT | Gm26543 | 5.533 | 6.46E-29 | glio |
| culture_effect | Serpina3n | -3.435 | 6.46E-38 | glio |
| Interaction_genotype_Bora | Calhm5 | -3.403 | 6.47E-03 | glio |
| Interaction_genotype_Bora | Sptlc2 | 2.035 | 6.47E-03 | glio |
| Interaction_genotype_Mtor | Daglb | -2.138 | 6.47E-03 | glio |
| Interaction_genotype_Mtor | Rgcc | 2.156 | 6.47E-03 | glio |
| Interaction_genotype_Mtor | Aig1 | 1.146 | 6.47E-03 | glio |
| interaction_RT | Calu | -0.53 | 6.47E-03 | glio |
| Atr_in.vivo_noRT | Dnah2os | 1.792 | 6.47E-03 | glio |
| Atr_in.vivo_noRT | Ralgps2 | -1.303 | 6.47E-03 | glio |
| Atr_in.vivo_noRT | Uqcrfs1 | -0.871 | 6.47E-03 | glio |
| Atr_ex.vivo_noRT | Fgf21 | 2.97 | 6.47E-03 | glio |
| Bcas2_in.vivo_noRT | Lnx2 | 1.857 | 6.47E-03 | glio |
| Bora_in.vivo_noRT | Psmd8 | -0.771 | 6.47E-03 | glio |
| Cenpj_ex.vivo_noRT | 9930111J21Ri | 2.591 | 6.47E-03 | glio |
| Cenpj_ex.vivo_noRT | Olfr718-ps1 | 2.198 | 6.47E-03 | glio |
| Chek1_ex.vivo_noRT | Fam161b | 2.553 | 6.47E-03 | glio |
| Cyp19a1_ex.vivo_noRT | Gm48140 | 2.165 | 6.47E-03 | glio |
| Cyp19a1_ex.vivo_noRT | Gad1 | 2.159 | 6.47E-03 | glio |
| culture_effect | Cercam | -1.566 | 6.47E-04 | glio |
| culture_effect | Zfand3 | 0.654 | 6.47E-04 | glio |
| Interaction_genotype_Mtor | Rabep1 | 1.778 | 6.47E-04 | glio |
| Interaction_genotype_Rpl3 | Adgrb3 | -4.553 | 6.47E-04 | glio |
| Aars2_ex.vivo_noRT | Gm35082 | 2.495 | 6.47E-04 | glio |
| Atr_in.vivo_noRT | Psd | 2.188 | 6.47E-04 | glio |
| Chek1_in.vivo_noRT | Gm14443 | 2.311 | 6.47E-04 | glio |
| Chek1_in.vivo_noRT | Mcfd2 | -1.472 | 6.47E-04 | glio |
| Eif2s1_in.vivo_noRT | Gm38393 | 2.407 | 6.47E-04 | glio |
| Eif2s1_in.vivo_noRT | Gm21761 | 2.143 | 6.47E-04 | glio |
| Eif2s1_in.vivo_noRT | Olfr1152 | 1.877 | 6.47E-04 | glio |
| Interaction_genotype_Myc | E130317F20Ri | -3.406 | 6.47E-05 | glio |
| Chek1_ex.vivo_noRT | Gm20633 | 3.607 | 6.47E-06 | glio |
| Interaction_genotype_Myc | AC166172.1 | -5.314 | 6.47E-07 | glio |
| Hsd17b10_in.vivo_noRT | Selenov | 3.616 | 6.47E-10 | glio |
| Hsd17b10_in.vivo_noRT | A330076H08R | 3.616 | 6.47E-10 | glio |
| culture_effect | Tpgs2 | 1.465 | 6.47E-12 | glio |
| Aars2_in.vivo_noRT | Plcd4 | 4.416 | 6.47E-12 | glio |
| Bcas2_in.vivo_noRT | A030014E15Ri | 4.807 | 6.47E-23 | glio |
| Bcas2_in.vivo_noRT | Gm21731 | 4.807 | 6.47E-23 | glio |
| Aars2_in.vivo_noRT | Vmn2r44 | 5.327 | 6.47E-26 | glio |
| Aars2_in.vivo_noRT | Mmp13 | 5.327 | 6.47E-26 | glio |
| culture_effect | Mlh3 | 0.755 | 6.48E-03 | glio |
| Interaction_genotype_Bcas2 | Rab11fip5 | -3.79 | 6.48E-03 | glio |
| Interaction_genotype_Bora | Cfap100 | -3.138 | 6.48E-03 | glio |
| Interaction_genotype_Mtor | Rnasel | -2.675 | 6.48E-03 | glio |
| Interaction_genotype_Mtor | Bag5 | -1.653 | 6.48E-03 | glio |
| Interaction_genotype_Mtor | Tcf4 | 1.299 | 6.48E-03 | glio |
| interaction_RT | Fry | 2.05 | 6.48E-03 | glio |
| interaction_RT | Wdcp | 1.06 | 6.48E-03 | glio |
| interaction_RT | Xpo5 | -0.87 | 6.48E-03 | glio |
| interaction_RT | Akirin1 | 0.685 | 6.48E-03 | glio |

| | | | | |
|---|---|---|---|---|
| Aars2_ex.vivo_noRT | Gm49860 | 2.311 | 6.48E-03 | glio |
| Atr_in.vivo_noRT | Atxn10 | -1.021 | 6.48E-03 | glio |
| Bard1_ex.vivo_noRT | Gm10382 | 2.806 | 6.48E-03 | glio |
| Bcas2_in.vivo_noRT | Gm12981 | 2.02 | 6.48E-03 | glio |
| Bcas2_in.vivo_noRT | Klhl23 | 1.873 | 6.48E-03 | glio |
| Bcas2_in.vivo_noRT | Gm17300 | 1.921 | 6.48E-03 | glio |
| Bcas2_in.vivo_noRT | Pold3 | -1.198 | 6.48E-03 | glio |
| Bcas2_in.vivo_noRT | Ube2t | -1.479 | 6.48E-03 | glio |
| Bcas2_in.vivo_noRT | Mrpl1 | -1.124 | 6.48E-03 | glio |
| Bora_ex.vivo_noRT | Ngef | 2.507 | 6.48E-03 | glio |
| Chek1_ex.vivo_noRT | Cep295nl | 2.472 | 6.48E-03 | glio |
| Interaction_genotype_Bora | Hba-a2 | -4.596 | 6.48E-04 | glio |
| Interaction_genotype_Bora | Fgf1 | -4.591 | 6.48E-04 | glio |
| interaction_RT | Scrg1 | 1.88 | 6.48E-04 | glio |
| interaction_RT | Mbd3 | -0.678 | 6.48E-04 | glio |
| Aars2_in.vivo_noRT | E130114P18Ri | 2.574 | 6.48E-04 | glio |
| Aars2_in.vivo_noRT | Lrp1 | 2.429 | 6.48E-04 | glio |
| Bard1_ex.vivo_noRT | Gm47540 | 2.421 | 6.48E-04 | glio |
| Dclre1c_in.vivo_noRT | Magea10 | 2.8 | 6.48E-04 | glio |
| Interaction_genotype_Mtor | Ric3 | -4.233 | 6.48E-05 | glio |
| Interaction_genotype_Myc | Gm50240 | -3.585 | 6.48E-05 | glio |
| interaction_RT | Pabpc1 | -0.615 | 6.48E-05 | glio |
| interaction_RT | Gm47223 | 4.673 | 6.48E-05 | glio |
| Bcas2_in.vivo_noRT | Wnt4 | 3.164 | 6.48E-05 | glio |
| Bcas2_in.vivo_noRT | Cbr3 | 3.054 | 6.48E-05 | glio |
| Interaction_genotype_Myc | Prl3d3 | -3.561 | 6.48E-06 | glio |
| Aars2_in.vivo_noRT | Gm37612 | 3.197 | 6.48E-06 | glio |
| Atr_in.vivo_noRT | Gm12043 | 2.94 | 6.48E-06 | glio |
| Atr_ex.vivo_noRT | Gad1 | 3.164 | 6.48E-07 | glio |
| Bcas2_in.vivo_noRT | Gm48357 | 3.021 | 6.48E-07 | glio |
| Bora_in.vivo_noRT | Gm9929 | 4.158 | 6.48E-11 | glio |
| Chek1_ex.vivo_noRT | Kit | 4.842 | 6.48E-11 | glio |
| culture_effect | Nfkbiz | -2.186 | 6.48E-15 | glio |
| Chek1_in.vivo_noRT | Gm48242 | 5.044 | 6.48E-17 | glio |
| Interaction_genotype_Atr | Gm15952 | -3.33 | 6.49E-03 | glio |
| Interaction_genotype_Atr | Gpatch8 | 1.379 | 6.49E-03 | glio |
| Interaction_genotype_Lmo7 | Gm10399 | -3.599 | 6.49E-03 | glio |
| Interaction_genotype_Mtor | BC034090 | -3.119 | 6.49E-03 | glio |
| Interaction_genotype_Mtor | Cadm2 | -3.513 | 6.49E-03 | glio |
| Interaction_genotype_Mtor | Zc3h12a | -3.151 | 6.49E-03 | glio |
| Interaction_genotype_Mtor | Tcf7l2 | -1.856 | 6.49E-03 | glio |
| Interaction_genotype_Mtor | Ctbp1 | 1.19 | 6.49E-03 | glio |
| Interaction_genotype_Myc | 4931414P19R | -2.947 | 6.49E-03 | glio |
| Interaction_genotype_Myc | Pin1 | 1.123 | 6.49E-03 | glio |
| Interaction_genotype_Rad51 | Tbc1d19 | -2.826 | 6.49E-03 | glio |
| Aars2_in.vivo_noRT | Ttc7b | 1.532 | 6.49E-03 | glio |
| Atr_ex.vivo_noRT | Slc2a13 | 2.715 | 6.49E-03 | glio |
| Atr_ex.vivo_noRT | Gm5089 | 2.882 | 6.49E-03 | glio |
| Bcas2_in.vivo_noRT | 4833413G10R | 2.136 | 6.49E-03 | glio |
| Bora_in.vivo_noRT | Col6a1 | 1.972 | 6.49E-03 | glio |
| Brca2_in.vivo_noRT | Gm19276 | 1.763 | 6.49E-03 | glio |
| Brca2_in.vivo_noRT | Vmn2r44 | 1.763 | 6.49E-03 | glio |

| | | | | |
|---|---|---|---|---|
| Brca2_in.vivo_noRT | Mmp13 | 1.763 | 6.49E-03 | glio |
| Brca2_in.vivo_noRT | Gm8016 | 1.763 | 6.49E-03 | glio |
| Cenpj_in.vivo_noRT | Fth1 | -0.781 | 6.49E-03 | glio |
| Fanca_ex.vivo_noRT | Ankrd22 | 2.124 | 6.49E-03 | glio |
| Fanca_ex.vivo_noRT | Gm47404 | 2.124 | 6.49E-03 | glio |
| Fanca_ex.vivo_noRT | Gm31717 | 2.124 | 6.49E-03 | glio |
| Fanca_ex.vivo_noRT | Gm14742 | 2.124 | 6.49E-03 | glio |
| Fanca_ex.vivo_noRT | Scel | 2.124 | 6.49E-03 | glio |
| Fanca_ex.vivo_noRT | Ctla4 | 2.124 | 6.49E-03 | glio |
| Fanca_ex.vivo_noRT | Zscan4-ps3 | 2.124 | 6.49E-03 | glio |
| Fanca_ex.vivo_noRT | Tagap | 2.124 | 6.49E-03 | glio |
| Fanca_ex.vivo_noRT | Gm21731 | 2.124 | 6.49E-03 | glio |
| Fanca_ex.vivo_noRT | Olfr1152 | 2.124 | 6.49E-03 | glio |
| Fanca_ex.vivo_noRT | Gm11232 | 2.124 | 6.49E-03 | glio |
| Fanca_ex.vivo_noRT | Gm29966 | 2.124 | 6.49E-03 | glio |
| Fanca_ex.vivo_noRT | Cgnl1 | 2.124 | 6.49E-03 | glio |
| Fanca_ex.vivo_noRT | Gm14207 | 2.126 | 6.49E-03 | glio |
| Fanca_ex.vivo_noRT | Gm26543 | 2.124 | 6.49E-03 | glio |
| Fanca_ex.vivo_noRT | 2410017I17Ri | 2.124 | 6.49E-03 | glio |
| Interaction_genotype_Atr | E130208F15Ri | -4.651 | 6.49E-04 | glio |
| Interaction_genotype_Atr | Snhg16 | 2.606 | 6.49E-04 | glio |
| Interaction_genotype_Atr | Aebp2 | 1.811 | 6.49E-04 | glio |
| Interaction_genotype_Mtor | Tubb5 | 2.163 | 6.49E-04 | glio |
| Interaction_genotype_Myc | Ano8 | -3.364 | 6.49E-04 | glio |
| Aars2_in.vivo_noRT | AI480526 | 2.378 | 6.49E-04 | glio |
| Chek1_in.vivo_noRT | Fgd6 | 2.329 | 6.49E-04 | glio |
| Chek1_in.vivo_noRT | Ermard | 2.075 | 6.49E-04 | glio |
| Chek1_ex.vivo_noRT | 1700047K16R | 3.168 | 6.49E-04 | glio |
| culture_effect | Lysmd3 | -0.89 | 6.49E-05 | glio |
| Chek1_in.vivo_noRT | Rex2 | 3.37 | 6.49E-06 | glio |
| Atp5c1_in.vivo_noRT | Gm12794 | 2.668 | 6.49E-07 | glio |
| Chek1_in.vivo_noRT | 1700056N10R | 3.133 | 6.49E-07 | glio |
| Atr_in.vivo_noRT | Gm2762 | 3.961 | 6.49E-10 | glio |
| Aars2_in.vivo_noRT | Gm50295 | 4.535 | 6.49E-11 | glio |
| Aars2_in.vivo_noRT | Pde11a | 4.798 | 6.49E-11 | glio |
| Atr_in.vivo_noRT | Gm9929 | 4.755 | 6.49E-14 | glio |
| Bora_in.vivo_noRT | Abcb5 | 4.936 | 6.49E-23 | glio |
| Chek1_in.vivo_noRT | Olfr810 | 5.044 | 6.49E-26 | glio |
| Interaction_genotype_Atr | 2810410L24Ri | -3.445 | 6.50E-03 | glio |
| Interaction_genotype_Atr | 1810021B22R | -3.399 | 6.50E-03 | glio |
| Interaction_genotype_Atr | Tsc22d1 | 1.58 | 6.50E-03 | glio |
| Interaction_genotype_Ifnar1 | Htr2b | 3.257 | 6.50E-03 | glio |
| Interaction_genotype_Mtor | Zfp534 | -3.539 | 6.50E-03 | glio |
| Interaction_genotype_Myc | Ankrd13d | -3.115 | 6.50E-03 | glio |
| Bcas2_in.vivo_noRT | Gm7879 | 2.155 | 6.50E-03 | glio |
| Bcas2_in.vivo_noRT | Kcnmb4 | 2.035 | 6.50E-03 | glio |
| Chek1_in.vivo_noRT | AA465934 | 1.794 | 6.50E-03 | glio |
| Eif2s1_in.vivo_noRT | BC147527 | 2.143 | 6.50E-03 | glio |
| culture_effect | Phlda2 | -0.923 | 6.50E-04 | glio |
| Interaction_genotype_Ifnar1 | H2-D1 | -1.324 | 6.50E-04 | glio |
| Interaction_genotype_Terf1 | Dnm3 | -4.759 | 6.50E-04 | glio |
| Aars2_ex.vivo_noRT | Bglap2 | 3.293 | 6.50E-04 | glio |

| | | | | |
|---|---|---|---|---|
| Atr_in.vivo_noRT | Rpsa | -0.692 | 6.50E-04 | glio |
| Chek1_ex.vivo_noRT | Ccna2 | -2.284 | 6.50E-04 | glio |
| Interaction_genotype_Mtor | Jmjd8 | -4.796 | 6.50E-05 | glio |
| interaction_RT | Spidr | 1.973 | 6.50E-05 | glio |
| Atr_in.vivo_noRT | Gm32913 | 4.262 | 6.50E-05 | glio |
| Atr_in.vivo_noRT | Zscan22 | 2.356 | 6.50E-05 | glio |
| Aars2_ex.vivo_noRT | Gm5602 | 3.293 | 6.50E-06 | glio |
| Bora_ex.vivo_noRT | Cysltr2 | 3.993 | 6.50E-06 | glio |
| Chek1_ex.vivo_noRT | Sgk2 | 4.055 | 6.50E-06 | glio |
| Chek1_ex.vivo_noRT | Gm16576 | 4.35 | 6.50E-07 | glio |
| culture_effect | D330023K18R | 1.622 | 6.50E-08 | glio |
| Hsd17b10_in.vivo_noRT | 9130230L23Ri | 3.356 | 6.50E-08 | glio |
| Bora_in.vivo_noRT | Slc6a13 | 5.08 | 6.50E-12 | glio |
| Aars2_in.vivo_noRT | Speer4e | 5.327 | 6.50E-22 | glio |
| Interaction_genotype_Aars2 | Sema4a | -3.57 | 6.51E-03 | glio |
| Interaction_genotype_Aars2 | Mbp | 1.904 | 6.51E-03 | glio |
| Interaction_genotype_Cyp19a | Ecscr | -3.5 | 6.51E-03 | glio |
| Interaction_genotype_Cyp19a | Gm45336 | 4.05 | 6.51E-03 | glio |
| Interaction_genotype_Mtor | Hist1h2bg | -3.413 | 6.51E-03 | glio |
| Interaction_genotype_Myc | B930095G15R | -2.912 | 6.51E-03 | glio |
| Interaction_genotype_Myc | Erlec1 | 1.313 | 6.51E-03 | glio |
| Interaction_genotype_Rad21 | Gm40538 | 3.281 | 6.51E-03 | glio |
| interaction_RT | Aup1 | -0.541 | 6.51E-03 | glio |
| Aars2_ex.vivo_noRT | Ephx3 | 2.612 | 6.51E-03 | glio |
| Atr_in.vivo_noRT | 1500011B03R | -1.126 | 6.51E-03 | glio |
| Atr_ex.vivo_noRT | Batf | 2.873 | 6.51E-03 | glio |
| Bcas2_in.vivo_noRT | Ttc25 | 1.972 | 6.51E-03 | glio |
| Bcas2_in.vivo_noRT | 2810002D19R | 1.591 | 6.51E-03 | glio |
| Chek1_in.vivo_noRT | A330009N23R | 1.954 | 6.51E-03 | glio |
| Chek1_in.vivo_noRT | Ints14 | -1.264 | 6.51E-03 | glio |
| Chek1_in.vivo_noRT | Coq2 | -1.132 | 6.51E-03 | glio |
| Chek1_in.vivo_noRT | Tpr | -0.655 | 6.51E-03 | glio |
| Chek1_in.vivo_noRT | Rpl30 | -0.509 | 6.51E-03 | glio |
| culture_effect | Myo1e | 0.692 | 6.51E-04 | glio |
| Interaction_genotype_Bora | Bmpr2 | 2.347 | 6.51E-04 | glio |
| Interaction_genotype_Mtor | Cyc1 | 1.726 | 6.51E-04 | glio |
| Interaction_genotype_Pars2 | Hbq1b | -4.391 | 6.51E-04 | glio |
| Aars2_in.vivo_noRT | Hcfc1r1 | 2.736 | 6.51E-04 | glio |
| Aars2_in.vivo_noRT | Zfpl1 | -1.285 | 6.51E-04 | glio |
| Aars2_in.vivo_noRT | Cyp2d22 | 2.989 | 6.51E-05 | glio |
| Hsd17b10_in.vivo_noRT | Appbp2os | 3.085 | 6.51E-05 | glio |
| Chek1_in.vivo_noRT | Gm32122 | 3.216 | 6.51E-07 | glio |
| culture_effect | Tmem229b | -3.15 | 6.51E-08 | glio |
| Atr_in.vivo_noRT | Snai1 | 4.015 | 6.51E-08 | glio |
| Atr_in.vivo_noRT | Dok3 | 4.398 | 6.51E-10 | glio |
| Bcas2_in.vivo_noRT | Gm17638 | 4.015 | 6.51E-10 | glio |
| Bora_in.vivo_noRT | Gm15743 | 4.673 | 6.51E-13 | glio |
| Bora_in.vivo_noRT | Agrp | 4.154 | 6.51E-13 | glio |
| Interaction_genotype_Atr | Gm50394 | -3.191 | 6.52E-03 | glio |
| Interaction_genotype_Mtor | Clca3a1 | -3.048 | 6.52E-03 | glio |
| Interaction_genotype_Mtor | Sec22b | 1.553 | 6.52E-03 | glio |
| Interaction_genotype_Myc | Naif1 | -2.925 | 6.52E-03 | glio |

| | | | | |
|---|---|---|---|---|
| interaction_RT | Cndp2 | -1.137 | 6.52E-03 | glio |
| Aars2_in.vivo_noRT | Fbxl5 | -1.168 | 6.52E-03 | glio |
| Atr_in.vivo_noRT | G530011O06R | -1.252 | 6.52E-03 | glio |
| Atr_in.vivo_noRT | Gatad1 | -0.888 | 6.52E-03 | glio |
| Bard1_ex.vivo_noRT | Myzap | 2.415 | 6.52E-03 | glio |
| Bora_in.vivo_noRT | Fbxo2 | -1.178 | 6.52E-03 | glio |
| Eif2s1_in.vivo_noRT | Sat1 | -0.994 | 6.52E-03 | glio |
| culture_effect | Mbd2 | 0.567 | 6.52E-04 | glio |
| interaction_RT | Flywch2 | 1.582 | 6.52E-04 | glio |
| interaction_RT | Neto1 | 1.035 | 6.52E-04 | glio |
| Aars2_in.vivo_noRT | Plcxd2 | 2.481 | 6.52E-04 | glio |
| Aars2_in.vivo_noRT | Siae | 2.22 | 6.52E-04 | glio |
| Aars2_in.vivo_noRT | Plekhf1 | 2.188 | 6.52E-04 | glio |
| Atr_in.vivo_noRT | Mtcl1 | 2.351 | 6.52E-04 | glio |
| Chek1_in.vivo_noRT | Vmac | 2.738 | 6.52E-04 | glio |
| Chek1_in.vivo_noRT | Sars | -0.957 | 6.52E-04 | glio |
| Chek1_ex.vivo_noRT | Slc1a1 | 2.976 | 6.52E-04 | glio |
| culture_effect | Thap8 | 2.337 | 6.52E-05 | glio |
| culture_effect | Alkbh7 | -0.832 | 6.52E-05 | glio |
| Interaction_genotype_Mtor | Hyal3 | -5.442 | 6.52E-05 | glio |
| Interaction_genotype_Mtor | 5430400D12R | -4.694 | 6.52E-05 | glio |
| Eif2s1_in.vivo_noRT | Gm13191 | 2.143 | 6.52E-05 | glio |
| Eif2s1_in.vivo_noRT | 4933416E03Ri | 2.143 | 6.52E-05 | glio |
| Eif2s1_in.vivo_noRT | Gm44899 | 2.67 | 6.52E-05 | glio |
| Hsd17b10_in.vivo_noRT | Ifitm1 | 3.35 | 6.52E-05 | glio |
| Hsd17b10_in.vivo_noRT | Gm2012 | 2.979 | 6.52E-05 | glio |
| Bora_in.vivo_noRT | 5330438D12R | 2.87 | 6.52E-07 | glio |
| Chek1_in.vivo_noRT | S1pr1 | 3.859 | 6.52E-07 | glio |
| Chek1_in.vivo_noRT | Gm35986 | 4.519 | 6.52E-12 | glio |
| Aars2_in.vivo_noRT | Gm45151 | 4.795 | 6.52E-18 | glio |
| Interaction_genotype_Bora | Slc25a11 | 1.452 | 6.53E-03 | glio |
| Interaction_genotype_Cdkn2a | Dscaml1 | 2.713 | 6.53E-03 | glio |
| Interaction_genotype_Mtor | Slc39a13 | -1.921 | 6.53E-03 | glio |
| Interaction_genotype_Mtor | Srek1 | 1.506 | 6.53E-03 | glio |
| Interaction_genotype_Myc | 4930458D05R | -3.423 | 6.53E-03 | glio |
| Interaction_genotype_Myc | Eva1c | -3.209 | 6.53E-03 | glio |
| Interaction_genotype_Myc | Pomk | -2.654 | 6.53E-03 | glio |
| Interaction_genotype_Myc | H2-DMa | 1.945 | 6.53E-03 | glio |
| Interaction_genotype_Myc | Cops4 | 1.466 | 6.53E-03 | glio |
| interaction_RT | Zfp746 | -1.362 | 6.53E-03 | glio |
| Atp5c1_in.vivo_noRT | Rbfaos | 1.751 | 6.53E-03 | glio |
| Bora_in.vivo_noRT | 0610038B21R | 1.853 | 6.53E-03 | glio |
| Bora_in.vivo_noRT | Fam228b | 2.154 | 6.53E-03 | glio |
| Bora_in.vivo_noRT | Gm7160 | 1.89 | 6.53E-03 | glio |
| Interaction_genotype_Mtor | Gm20721 | -3.875 | 6.53E-04 | glio |
| Interaction_genotype_Mtor | Mrpl58 | 1.761 | 6.53E-04 | glio |
| Hsd17b10_in.vivo_noRT | Gm28874 | 2.594 | 6.53E-04 | glio |
| culture_effect | Gm48242 | 2.014 | 6.53E-05 | glio |
| culture_effect | Hnrnpk | -0.427 | 6.53E-05 | glio |
| Interaction_genotype_Mtor | Sec1 | -4.867 | 6.53E-05 | glio |
| Interaction_genotype_Mtor | AF067061 | -4.099 | 6.53E-05 | glio |
| Atr_in.vivo_noRT | Klhl23 | 2.6 | 6.53E-05 | glio |

| | | | | |
|---|---|---|---|---|
| Atr_in.vivo_noRT | Clcf1 | 2.626 | 6.53E-05 | glio |
| Bcas2_in.vivo_noRT | Olfr77 | 2.739 | 6.53E-05 | glio |
| Chek1_in.vivo_noRT | Mill2 | 2.852 | 6.53E-05 | glio |
| Chek1_ex.vivo_noRT | Ephx3 | 3.362 | 6.53E-05 | glio |
| culture_effect | Pla2g2f | -2.577 | 6.53E-06 | glio |
| culture_effect | Mrpl17 | 0.53 | 6.53E-06 | glio |
| Interaction_genotype_Mtor | Lrrk2 | -4.015 | 6.53E-06 | glio |
| Interaction_genotype_Mtor | Pgk1 | 2.337 | 6.53E-06 | glio |
| Atr_ex.vivo_noRT | B230311B06R | 4.213 | 6.53E-06 | glio |
| Aars2_in.vivo_noRT | Cep295nl | 3.449 | 6.53E-07 | glio |
| Aars2_in.vivo_noRT | Olfr267 | 3.552 | 6.53E-07 | glio |
| Bora_ex.vivo_noRT | Gm48617 | 3.201 | 6.53E-07 | glio |
| Interaction_genotype_Mtor | Gm49694 | -6.734 | 6.53E-08 | glio |
| Interaction_genotype_Mtor | Gm28424 | -6.179 | 6.53E-08 | glio |
| Interaction_genotype_Mtor | Gm30938 | -6.216 | 6.53E-08 | glio |
| Interaction_genotype_Mtor | St8sia1 | -6.816 | 6.53E-08 | glio |
| Interaction_genotype_Mtor | Gm44899 | -5.203 | 6.53E-08 | glio |
| Atr_ex.vivo_noRT | Helt | 3.164 | 6.53E-08 | glio |
| Atr_in.vivo_noRT | Trim14 | 4.643 | 6.53E-10 | glio |
| Aars2_in.vivo_noRT | Svopl | 4.145 | 6.53E-11 | glio |
| Chek1_ex.vivo_noRT | Slc19a3 | 4.449 | 6.53E-13 | glio |
| culture_effect | Sytl2 | -4.5 | 6.53E-21 | glio |
| culture_effect | Ccdc8 | -1.704 | 6.54E-03 | glio |
| Interaction_genotype_Atr | Rmst | -3.536 | 6.54E-03 | glio |
| Interaction_genotype_Bcas2 | Zmynd11 | 1.67 | 6.54E-03 | glio |
| Interaction_genotype_Myc | Dusp3 | -2.031 | 6.54E-03 | glio |
| Interaction_genotype_Myc | Cox7c | 1.127 | 6.54E-03 | glio |
| Atp5c1_in.vivo_noRT | G630030J09Ri | 2.146 | 6.54E-03 | glio |
| Atp5c1_in.vivo_noRT | Amy1 | 2.139 | 6.54E-03 | glio |
| Atp5c1_in.vivo_noRT | Wfdc1 | 2.211 | 6.54E-03 | glio |
| Atr_ex.vivo_noRT | Selenow | 0.786 | 6.54E-03 | glio |
| Bora_ex.vivo_noRT | Pabpc4l | 2.884 | 6.54E-03 | glio |
| Chek1_in.vivo_noRT | Traf4 | -1.215 | 6.54E-03 | glio |
| Chek1_ex.vivo_noRT | Gm20667 | 2.536 | 6.54E-03 | glio |
| Chek1_ex.vivo_noRT | Kirrel3 | 2.467 | 6.54E-03 | glio |
| Hsd17b10_in.vivo_noRT | 4922502N22R | 2.097 | 6.54E-03 | glio |
| Hsd17b10_in.vivo_noRT | Adcy9 | 1.897 | 6.54E-03 | glio |
| Aars2_in.vivo_noRT | P2ry1 | 2.107 | 6.54E-04 | glio |
| Cenpj_in.vivo_noRT | 6430562O15R | 2.555 | 6.54E-04 | glio |
| Chek1_in.vivo_noRT | Zfp599 | 2.611 | 6.54E-04 | glio |
| Chek1_in.vivo_noRT | H2afz | -0.777 | 6.54E-04 | glio |
| Interaction_genotype_Mtor | Recql5 | -3.895 | 6.54E-05 | glio |
| Atr_in.vivo_noRT | Tex19.2 | 3.069 | 6.54E-06 | glio |
| Atr_in.vivo_noRT | Nell2 | 2.862 | 6.54E-06 | glio |
| Bora_ex.vivo_noRT | Pspn | 2.806 | 6.54E-06 | glio |
| Hsd17b10_in.vivo_noRT | Olfr1191-ps1 | 2.966 | 6.54E-06 | glio |
| Interaction_genotype_Bora | Atad3a | 2.824 | 6.54E-07 | glio |
| Bcas2_in.vivo_noRT | Gm50107 | 3.292 | 6.54E-07 | glio |
| culture_effect | Snx1 | -1.053 | 6.54E-10 | glio |
| Atr_in.vivo_noRT | Nmb | 4.24 | 6.54E-10 | glio |
| Aars2_in.vivo_noRT | Ankef1 | 4.587 | 6.54E-11 | glio |
| culture_effect | 9230114K14R | 2.279 | 6.54E-12 | glio |

| | | | | |
|---|---|---|---|---|
| Atr_in.vivo_noRT | Gm13404 | 4.795 | 6.54E-12 | glio |
| culture_effect | 2310034G01R | 2.631 | 6.54E-16 | glio |
| Interaction_genotype_Atr | Apool | -1.905 | 6.55E-03 | glio |
| Interaction_genotype_Bora | D230022J07Ri | -3.662 | 6.55E-03 | glio |
| Interaction_genotype_Bora | Gdap1l1 | -4.003 | 6.55E-03 | glio |
| Interaction_genotype_Bora | Gm29340 | -3.195 | 6.55E-03 | glio |
| Interaction_genotype_Bora | Sgo1 | 2.276 | 6.55E-03 | glio |
| Interaction_genotype_Myc | Ube3b | -2.012 | 6.55E-03 | glio |
| Interaction_genotype_Myc | Znhit3 | 1.495 | 6.55E-03 | glio |
| Interaction_genotype_Myc | Hnrnpc | 1.121 | 6.55E-03 | glio |
| Interaction_genotype_Myc | Ppp3ca | 1.201 | 6.55E-03 | glio |
| Interaction_genotype_Myc | Romo1 | 1.452 | 6.55E-03 | glio |
| Interaction_genotype_Myc | Atp5j2 | 1.127 | 6.55E-03 | glio |
| interaction_RT | A330076H08R | 2.266 | 6.55E-03 | glio |
| interaction_RT | Csrnp1 | 1.284 | 6.55E-03 | glio |
| Aars2_in.vivo_noRT | Ap1ar | -1.238 | 6.55E-03 | glio |
| Aars2_in.vivo_noRT | Uqcr11 | -0.728 | 6.55E-03 | glio |
| Atr_in.vivo_noRT | Gpr141 | 2.26 | 6.55E-03 | glio |
| Atr_in.vivo_noRT | Lman2 | -0.902 | 6.55E-03 | glio |
| Bard1_ex.vivo_noRT | Bmp15 | 2.421 | 6.55E-03 | glio |
| Chek1_in.vivo_noRT | Cacng7 | 1.94 | 6.55E-03 | glio |
| Chek1_in.vivo_noRT | Tmem9 | -1.332 | 6.55E-03 | glio |
| Hsd17b10_in.vivo_noRT | Zfp275 | -1.349 | 6.55E-03 | glio |
| Hsd17b10_in.vivo_noRT | Gm26532 | 0.938 | 6.55E-03 | glio |
| Interaction_genotype_Mtor | Pde4d | -4.187 | 6.55E-04 | glio |
| Interaction_genotype_Mtor | Ube2c | 3.773 | 6.55E-04 | glio |
| Interaction_genotype_Myc | Ptgis | -3.818 | 6.55E-04 | glio |
| Aars2_in.vivo_noRT | Mnat1 | -1.372 | 6.55E-04 | glio |
| Bcas2_in.vivo_noRT | Cdkn2aip | -1.761 | 6.55E-04 | glio |
| culture_effect | Gm16364.1 | 2.478 | 6.55E-05 | glio |
| Interaction_genotype_Myc | Pakap | -3.576 | 6.55E-05 | glio |
| Chek1_ex.vivo_noRT | Ppil6 | 3.168 | 6.55E-05 | glio |
| Chek1_ex.vivo_noRT | Gm15938 | 3.703 | 6.55E-05 | glio |
| Interaction_genotype_Mtor | Crybb1 | -4.681 | 6.55E-06 | glio |
| Interaction_genotype_Myc | Tmed3 | 2.205 | 6.55E-06 | glio |
| Chek1_ex.vivo_noRT | Cpxm1 | 3.738 | 6.55E-07 | glio |
| Interaction_genotype_Mtor | Prickle2 | -6.041 | 6.55E-09 | glio |
| Atr_ex.vivo_noRT | Smtnl1 | 3.962 | 6.55E-09 | glio |
| culture_effect | Pdia4 | -0.897 | 6.55E-10 | glio |
| culture_effect | Ybx1 | 0.681 | 6.55E-10 | glio |
| Aars2_in.vivo_noRT | Alpl | 4.142 | 6.55E-10 | glio |
| Aars2_in.vivo_noRT | Gm9922 | 4.191 | 6.55E-10 | glio |
| Hsd17b10_in.vivo_noRT | Gm10324 | 3.35 | 6.55E-10 | glio |
| Atr_in.vivo_noRT | AI463229 | 4.888 | 6.55E-17 | glio |
| culture_effect | Fam89a | -1.933 | 6.55E-25 | glio |
| culture_effect | Zfp870 | 1.086 | 6.56E-03 | glio |
| Interaction_genotype_Aars2 | Fbxo24 | -3.396 | 6.56E-03 | glio |
| Interaction_genotype_Aars2 | Vgll3 | -2.648 | 6.56E-03 | glio |
| Interaction_genotype_Aars2 | Csnk2a2 | 1.21 | 6.56E-03 | glio |
| Interaction_genotype_Atr | Maz | 1.442 | 6.56E-03 | glio |
| Interaction_genotype_Bcas2 | Trabd | 1.697 | 6.56E-03 | glio |
| Interaction_genotype_Myc | Aass | -2.875 | 6.56E-03 | glio |

| | | | | |
|---|---|---|---|---|
| Interaction_genotype_Myc | Gm10941 | -2.756 | 6.56E-03 | glio |
| Interaction_genotype_Myc | Dpp7 | -2.477 | 6.56E-03 | glio |
| Aars2_in.vivo_noRT | BC065397 | 2.203 | 6.56E-03 | glio |
| Aars2_in.vivo_noRT | Ttc39a | 1.536 | 6.56E-03 | glio |
| Bcas2_ex.vivo_noRT | Gm8300 | 3.524 | 6.56E-03 | glio |
| Bora_in.vivo_noRT | Tnip1 | 1.175 | 6.56E-03 | glio |
| Chek1_in.vivo_noRT | Dcps | -1.261 | 6.56E-03 | glio |
| Chek1_ex.vivo_noRT | Ppef1 | 2.505 | 6.56E-03 | glio |
| Eif2s1_ex.vivo_noRT | Olfr56 | 2.234 | 6.56E-03 | glio |
| Hsd17b10_in.vivo_noRT | Mroh7 | 2.092 | 6.56E-03 | glio |
| Hsd17b10_in.vivo_noRT | Gm17655 | 2.369 | 6.56E-03 | glio |
| culture_effect | Lfng | 1.622 | 6.56E-04 | glio |
| Interaction_genotype_Myc | Zfp810 | -3.722 | 6.56E-04 | glio |
| Interaction_genotype_Myc | Fastkd1 | -3.441 | 6.56E-04 | glio |
| Atr_in.vivo_noRT | Tsga10 | 2.534 | 6.56E-04 | glio |
| Atr_ex.vivo_noRT | 1700024I08Ri | 2.574 | 6.56E-04 | glio |
| Bcas2_in.vivo_noRT | Syt17 | 2.742 | 6.56E-04 | glio |
| Chek1_in.vivo_noRT | Chchd1 | -0.834 | 6.56E-04 | glio |
| Aars2_in.vivo_noRT | Phkg1 | 3.897 | 6.56E-05 | glio |
| Interaction_genotype_Mtor | Gm10642 | -4.93 | 6.56E-06 | glio |
| Interaction_genotype_Mtor | Gm10382 | -5.654 | 6.56E-06 | glio |
| Interaction_genotype_Mtor | Lyl1 | -4.634 | 6.56E-06 | glio |
| Atr_in.vivo_noRT | Gm46378 | 3.3 | 6.56E-06 | glio |
| Bcas2_in.vivo_noRT | Mterf1b | 3.04 | 6.56E-06 | glio |
| Interaction_genotype_Mtor | Tnfrsf23 | -6.586 | 6.56E-07 | glio |
| Interaction_genotype_Mtor | Gm27253 | -5.829 | 6.56E-07 | glio |
| Interaction_genotype_Mtor | A330015K06R | -5.518 | 6.56E-07 | glio |
| Interaction_genotype_Mtor | Gm16178 | -4.099 | 6.56E-07 | glio |
| Interaction_genotype_Mtor | Gm364 | -4.099 | 6.56E-07 | glio |
| Atr_in.vivo_noRT | D930019O06F | 3.637 | 6.56E-07 | glio |
| Bora_in.vivo_noRT | Gm10101 | 3.367 | 6.56E-07 | glio |
| Chek1_ex.vivo_noRT | Gm29994 | 4.058 | 6.56E-07 | glio |
| Interaction_genotype_Myc | Wfdc1 | -5.485 | 6.56E-08 | glio |
| Interaction_genotype_Myc | Adgrb3 | -6.157 | 6.56E-08 | glio |
| Atr_ex.vivo_noRT | Gm20625 | 3.164 | 6.56E-08 | glio |
| Chek1_in.vivo_noRT | Slc7a4 | 3.629 | 6.56E-08 | glio |
| Atr_in.vivo_noRT | Gm35066 | 4.041 | 6.56E-09 | glio |
| Bora_in.vivo_noRT | Glis1 | 4.024 | 6.56E-09 | glio |
| Bcas2_in.vivo_noRT | Car14 | 4.807 | 6.56E-13 | glio |
| culture_effect | Med9 | 0.457 | 6.57E-03 | glio |
| culture_effect | Cox4i1 | -0.286 | 6.57E-03 | glio |
| Interaction_genotype_Aars2 | Gm41307 | -2.427 | 6.57E-03 | glio |
| Interaction_genotype_Mtor | Mkrn1 | 1.629 | 6.57E-03 | glio |
| Interaction_genotype_Myc | Gm16751 | -2.613 | 6.57E-03 | glio |
| Interaction_genotype_Myc | Gne | -2.392 | 6.57E-03 | glio |
| Aars2_in.vivo_noRT | Dera | -1.231 | 6.57E-03 | glio |
| Aars2_in.vivo_noRT | Cdc123 | -0.8 | 6.57E-03 | glio |
| Bard1_ex.vivo_noRT | Gm13684 | 2.421 | 6.57E-03 | glio |
| Brca2_ex.vivo_noRT | Agrp | 2.8 | 6.57E-03 | glio |
| Chek1_in.vivo_noRT | Dynll2 | -0.87 | 6.57E-03 | glio |
| Chek1_in.vivo_noRT | Bpgm | -1.126 | 6.57E-03 | glio |
| Chek1_ex.vivo_noRT | Hus1 | -1.726 | 6.57E-03 | glio |

| | | | | |
|---|---|---|---|---|
| Chek1_ex.vivo_noRT | Asf1b | -1.72 | 6.57E-03 | glio |
| Chek1_ex.vivo_noRT | Pus7 | -1.699 | 6.57E-03 | glio |
| Chek1_ex.vivo_noRT | Mrpl15 | -0.874 | 6.57E-03 | glio |
| Hsd17b10_in.vivo_noRT | B230319C09R | 2.306 | 6.57E-03 | glio |
| Interaction_genotype_Bora | Scube3 | -3.529 | 6.57E-04 | glio |
| Interaction_genotype_Chek1 | Il12rb1 | 4.34 | 6.57E-04 | glio |
| Interaction_genotype_Myc | 2810006K23R | -2.76 | 6.57E-04 | glio |
| Aars2_ex.vivo_noRT | Bco1 | 2.897 | 6.57E-04 | glio |
| Atr_ex.vivo_noRT | Gm43463 | 2.771 | 6.57E-04 | glio |
| Bcas2_ex.vivo_noRT | Gm28940 | 2.742 | 6.57E-04 | glio |
| Bora_in.vivo_noRT | Cyb561d1 | 2.614 | 6.57E-04 | glio |
| Chek1_in.vivo_noRT | Mybbp1a | -1.198 | 6.57E-04 | glio |
| Aars2_in.vivo_noRT | Hdgfl2 | -1.734 | 6.57E-05 | glio |
| Aars2_in.vivo_noRT | Tfrc | -1.8 | 6.57E-05 | glio |
| Atr_in.vivo_noRT | Mst1 | 2.633 | 6.57E-05 | glio |
| Atr_in.vivo_noRT | Mrps18b | -1.281 | 6.57E-05 | glio |
| Atr_ex.vivo_noRT | Sbk2 | 3.764 | 6.57E-05 | glio |
| Bora_in.vivo_noRT | Xlr4c | 2.902 | 6.57E-05 | glio |
| Chek1_ex.vivo_noRT | Insm1 | 3.463 | 6.57E-05 | glio |
| Interaction_genotype_Myc | Fastkd3 | -4.224 | 6.57E-06 | glio |
| Aars2_ex.vivo_noRT | Gm12860 | 3.293 | 6.57E-06 | glio |
| Bcas2_in.vivo_noRT | Cacng5 | 2.957 | 6.57E-06 | glio |
| Interaction_genotype_Mtor | 4930447K03R | -4.627 | 6.57E-07 | glio |
| Atr_in.vivo_noRT | Gm16133 | 3.551 | 6.57E-07 | glio |
| Chek1_ex.vivo_noRT | Ksr2 | 3.867 | 6.57E-08 | glio |
| Aars2_in.vivo_noRT | 3930402G23R | 4.94 | 6.57E-12 | glio |
| Aars2_in.vivo_noRT | Shisal2a | 5.063 | 6.57E-12 | glio |
| Atr_in.vivo_noRT | Fam13a | 4.733 | 6.57E-12 | glio |
| Bora_in.vivo_noRT | Ttll13 | 4.203 | 6.57E-12 | glio |
| Atr_in.vivo_noRT | Gm49808 | 5.533 | 6.57E-30 | glio |
| culture_effect | Sh3bp4 | -1.004 | 6.58E-03 | glio |
| Interaction_genotype_Atr | Sema4f | -3.295 | 6.58E-03 | glio |
| Interaction_genotype_Bcas2 | Pbld1 | -3.826 | 6.58E-03 | glio |
| Interaction_genotype_Mtor | Zkscan7 | -3.197 | 6.58E-03 | glio |
| Interaction_genotype_Mtor | Psat1 | 1.417 | 6.58E-03 | glio |
| Interaction_genotype_Myc | Igfbp7 | -2.761 | 6.58E-03 | glio |
| Interaction_genotype_Myc | Pax6 | -2.202 | 6.58E-03 | glio |
| Aars2_in.vivo_noRT | Bcap31 | -0.763 | 6.58E-03 | glio |
| Atr_ex.vivo_noRT | Neurl3 | 2.764 | 6.58E-03 | glio |
| Bcas2_in.vivo_noRT | Eva1c | 2.079 | 6.58E-03 | glio |
| Bcas2_in.vivo_noRT | 4921531C22R | 1.966 | 6.58E-03 | glio |
| Bcas2_ex.vivo_noRT | Dcpp3 | 2.742 | 6.58E-03 | glio |
| Bora_in.vivo_noRT | Gm47438 | 2.193 | 6.58E-03 | glio |
| Bora_in.vivo_noRT | Gm45184 | 2.003 | 6.58E-03 | glio |
| Eif2s1_in.vivo_noRT | Slc16a4 | 1.879 | 6.58E-03 | glio |
| Eif2s1_in.vivo_noRT | Col9a2 | -0.651 | 6.58E-03 | glio |
| Interaction_genotype_Myc | Insr | -2.897 | 6.58E-04 | glio |
| Aars2_in.vivo_noRT | 0610010K14R | -1.277 | 6.58E-04 | glio |
| Atr_in.vivo_noRT | Eif3j2 | 2.397 | 6.58E-04 | glio |
| Atr_in.vivo_noRT | Pabpc1 | -0.586 | 6.58E-04 | glio |
| Bora_in.vivo_noRT | Gm11772 | 2.461 | 6.58E-04 | glio |
| Bora_in.vivo_noRT | Ocrl | 1.922 | 6.58E-04 | glio |

| | | | | |
|---|---|---|---|---|
| Chek1_in.vivo_noRT | Soga1 | 2.553 | 6.58E-04 | glio |
| Chek1_in.vivo_noRT | Ccdc62 | 2.29 | 6.58E-04 | glio |
| Chek1_ex.vivo_noRT | Gm49041 | 2.569 | 6.58E-04 | glio |
| Interaction_genotype_Mtor | Gm31415 | -3.967 | 6.58E-05 | glio |
| Interaction_genotype_Mtor | Ippk | -3.668 | 6.58E-05 | glio |
| Aars2_in.vivo_noRT | Gnai2 | -1.456 | 6.58E-05 | glio |
| Chek1_in.vivo_noRT | Col17a1 | 2.993 | 6.58E-05 | glio |
| Chek1_ex.vivo_noRT | Cct2 | -0.88 | 6.58E-05 | glio |
| Interaction_genotype_Myc | Ccdc92 | -3.753 | 6.58E-06 | glio |
| Chek1_in.vivo_noRT | Gm6225 | 3.368 | 6.58E-06 | glio |
| Atr_in.vivo_noRT | Cd79a | 3.542 | 6.58E-07 | glio |
| Bcas2_in.vivo_noRT | 1810012K16R | 3.291 | 6.58E-07 | glio |
| Chek1_in.vivo_noRT | Celf5 | 3.98 | 6.58E-08 | glio |
| Atr_in.vivo_noRT | Vpreb1 | 4.63 | 6.58E-12 | glio |
| Aars2_in.vivo_noRT | Gm26674 | 4.682 | 6.58E-13 | glio |
| Aars2_in.vivo_noRT | Zfp352 | 5.327 | 6.58E-17 | glio |
| culture_effect | Rimklb | -1.198 | 6.59E-03 | glio |
| culture_effect | Ppp1r15a | -0.54 | 6.59E-03 | glio |
| culture_effect | Brd2 | 0.362 | 6.59E-03 | glio |
| Interaction_genotype_Cenpj | Otof | 3.995 | 6.59E-03 | glio |
| Interaction_genotype_Myc | Cd48 | -2.893 | 6.59E-03 | glio |
| Aars2_in.vivo_noRT | Clybl | 1.647 | 6.59E-03 | glio |
| Atp5c1_in.vivo_noRT | 7530414M10F | 2.142 | 6.59E-03 | glio |
| Atr_in.vivo_noRT | Chchd2 | -0.533 | 6.59E-03 | glio |
| Atr_ex.vivo_noRT | Ubald2 | 1.186 | 6.59E-03 | glio |
| Cenpj_ex.vivo_noRT | Cidea | 2.591 | 6.59E-03 | glio |
| culture_effect | Ctsa | -0.609 | 6.59E-04 | glio |
| culture_effect | Oasl2 | 0.528 | 6.59E-04 | glio |
| Interaction_genotype_Mtor | Cdo1 | -4.271 | 6.59E-04 | glio |
| Aars2_in.vivo_noRT | 0610038B21R | 2.243 | 6.59E-04 | glio |
| Aars2_in.vivo_noRT | Ncoa4 | -1.477 | 6.59E-04 | glio |
| Bora_in.vivo_noRT | Lrrn4cl | 2.662 | 6.59E-04 | glio |
| Cenpj_ex.vivo_noRT | Hist1h2ag | 2.591 | 6.59E-04 | glio |
| Chek1_in.vivo_noRT | Gm38973 | 2.448 | 6.59E-04 | glio |
| Hsd17b10_in.vivo_noRT | Ctss | 2.186 | 6.59E-04 | glio |
| Interaction_genotype_Mtor | C430002N11R | -3.701 | 6.59E-05 | glio |
| Aars2_in.vivo_noRT | Ston1 | 2.877 | 6.59E-05 | glio |
| Atr_in.vivo_noRT | Cdh24 | 3.056 | 6.59E-05 | glio |
| Atr_in.vivo_noRT | Slc25a42 | 2.85 | 6.59E-05 | glio |
| Atr_in.vivo_noRT | Gm50232 | 2.612 | 6.59E-05 | glio |
| Atr_in.vivo_noRT | 6030443J06Ri | 2.697 | 6.59E-05 | glio |
| Chek1_ex.vivo_noRT | Hopx | 3.468 | 6.59E-05 | glio |
| Aars2_in.vivo_noRT | Tmed7 | -1.736 | 6.59E-06 | glio |
| Chek1_in.vivo_noRT | Itga3 | 3.414 | 6.59E-06 | glio |
| culture_effect | Tcf4 | -0.658 | 6.59E-07 | glio |
| Atr_in.vivo_noRT | Tph2 | 3.524 | 6.59E-08 | glio |
| Chek1_ex.vivo_noRT | Fbxo5 | -3.545 | 6.59E-08 | glio |
| Bora_in.vivo_noRT | Gm24362 | 3.893 | 6.59E-10 | glio |
| Aars2_in.vivo_noRT | Amh | 4.41 | 6.59E-12 | glio |
| Atr_in.vivo_noRT | Svopl | 4.352 | 6.59E-12 | glio |
| Bcas2_in.vivo_noRT | Pacrg | 4.545 | 6.59E-13 | glio |
| Bcas2_in.vivo_noRT | Kpna7 | 4.543 | 6.59E-13 | glio |

| | | | | |
|---|---|---|---|---|
| Aars2_in.vivo_noRT | Olfr90 | 5.063 | 6.59E-17 | glio |
| Chek1_in.vivo_noRT | A030014E15Ri | 5.044 | 6.59E-25 | glio |
| Chek1_in.vivo_noRT | Gm21731 | 5.044 | 6.59E-25 | glio |
| Interaction_genotype_Atr | BC026762 | -3.277 | 6.60E-03 | glio |
| Interaction_genotype_Mtor | Ribc1 | -2.933 | 6.60E-03 | glio |
| Interaction_genotype_Myc | Adh7 | -2.943 | 6.60E-03 | glio |
| interaction_RT | Tmem41a | 1.232 | 6.60E-03 | glio |
| Aars2_in.vivo_noRT | Pigv | 2.039 | 6.60E-03 | glio |
| Aars2_in.vivo_noRT | Cox6a1 | -0.648 | 6.60E-03 | glio |
| Aars2_ex.vivo_noRT | Olfr1191-ps1 | 2.505 | 6.60E-03 | glio |
| Aars2_ex.vivo_noRT | Pacrg | 2.499 | 6.60E-03 | glio |
| Bcas2_in.vivo_noRT | Acot8 | -1.354 | 6.60E-03 | glio |
| Bora_in.vivo_noRT | Dlg5 | 1.894 | 6.60E-03 | glio |
| Bora_in.vivo_noRT | Kif7 | 1.875 | 6.60E-03 | glio |
| Bora_in.vivo_noRT | Prdm1 | 1.87 | 6.60E-03 | glio |
| Bora_in.vivo_noRT | Mthfs | 1.758 | 6.60E-03 | glio |
| Bora_in.vivo_noRT | Hes5 | -2.208 | 6.60E-03 | glio |
| Bora_in.vivo_noRT | Tubb4b | -0.905 | 6.60E-03 | glio |
| Chek1_in.vivo_noRT | Serinc2 | 1.936 | 6.60E-03 | glio |
| Chek1_in.vivo_noRT | Map3k6 | 2.057 | 6.60E-03 | glio |
| Chek1_in.vivo_noRT | Eid2b | 1.687 | 6.60E-03 | glio |
| culture_effect | Sec61a1 | -0.451 | 6.60E-04 | glio |
| Interaction_genotype_Aars2 | Gm29170 | -3.97 | 6.60E-04 | glio |
| Interaction_genotype_Aars2 | Psph | 1.561 | 6.60E-04 | glio |
| Interaction_genotype_Mtor | H60c | -4.375 | 6.60E-04 | glio |
| Aars2_in.vivo_noRT | Ppfia3 | 2.359 | 6.60E-04 | glio |
| Aars2_ex.vivo_noRT | Map3k5 | 2.893 | 6.60E-04 | glio |
| Bcas2_in.vivo_noRT | Arhgef17 | 2.225 | 6.60E-04 | glio |
| Cenpj_ex.vivo_noRT | Gpr22 | 2.591 | 6.60E-04 | glio |
| Cenpj_ex.vivo_noRT | Tnni3k | 2.588 | 6.60E-04 | glio |
| Hsd17b10_in.vivo_noRT | Dmrt2 | 2.727 | 6.60E-04 | glio |
| Aars2_in.vivo_noRT | Ubxn10 | 2.406 | 6.60E-05 | glio |
| Aars2_in.vivo_noRT | Parn | 1.913 | 6.60E-05 | glio |
| Bora_in.vivo_noRT | Mrps16 | -1.077 | 6.60E-05 | glio |
| culture_effect | C230038L03Ri | -2.524 | 6.60E-06 | glio |
| interaction_RT | Fbxo2 | 1.254 | 6.60E-06 | glio |
| Bora_ex.vivo_noRT | Dscaml1 | 2.806 | 6.60E-06 | glio |
| Interaction_genotype_Aars2 | Mecom | -5.877 | 6.60E-07 | glio |
| Interaction_genotype_Mtor | Nudt7 | -5.606 | 6.60E-07 | glio |
| Interaction_genotype_Myc | Igf2bp1 | -6.198 | 6.60E-08 | glio |
| Bcas2_in.vivo_noRT | Ube3a | -1.768 | 6.60E-08 | glio |
| Bora_in.vivo_noRT | Mtmr11 | 4.09 | 6.60E-09 | glio |
| Aars2_in.vivo_noRT | Brsk2 | 3.918 | 6.60E-10 | glio |
| Aars2_in.vivo_noRT | Tspyl3 | 4.244 | 6.60E-12 | glio |
| Aars2_in.vivo_noRT | K230015D01R | 5.06 | 6.60E-12 | glio |
| Bcas2_in.vivo_noRT | Gm43653 | 4.544 | 6.60E-12 | glio |
| Hsd17b10_in.vivo_noRT | Olfr56 | 3.616 | 6.60E-12 | glio |
| Atr_ex.vivo_noRT | Lmx1a | 4.985 | 6.60E-13 | glio |
| Bora_in.vivo_noRT | 4930570E03Ri | 4.676 | 6.60E-18 | glio |
| Bora_in.vivo_noRT | 4930447K03R | 4.676 | 6.60E-18 | glio |
| culture_effect | Dusp11 | -0.345 | 6.61E-03 | glio |
| culture_effect | Arf6 | 0.364 | 6.61E-03 | glio |

| | | | | |
|---|---|---|---|---|
| Interaction_genotype_Aars2 | Impg2 | -2.822 | 6.61E-03 | glio |
| Interaction_genotype_Aars2 | 1700112J16Ri | -2.62 | 6.61E-03 | glio |
| Interaction_genotype_Hsd17b | Slc35f6 | 2.635 | 6.61E-03 | glio |
| Interaction_genotype_Mtor | Tctn2 | -2.26 | 6.61E-03 | glio |
| Interaction_genotype_Mtor | Pkd1 | -2.411 | 6.61E-03 | glio |
| Interaction_genotype_Myc | Gramd4 | -3.113 | 6.61E-03 | glio |
| interaction_RT | Gm34643 | 2.714 | 6.61E-03 | glio |
| Aars2_in.vivo_noRT | Scaper | 1.523 | 6.61E-03 | glio |
| Atp5c1_in.vivo_noRT | Gm14636 | 2.692 | 6.61E-03 | glio |
| Atr_in.vivo_noRT | Epb41 | -1.22 | 6.61E-03 | glio |
| Atr_in.vivo_noRT | Nipsnap3b | -1.075 | 6.61E-03 | glio |
| Bcas2_in.vivo_noRT | Trim45 | 1.733 | 6.61E-03 | glio |
| Bcas2_in.vivo_noRT | E2f3 | -1.194 | 6.61E-03 | glio |
| Chek1_in.vivo_noRT | Zfp263 | 1.422 | 6.61E-03 | glio |
| culture_effect | Gm16344 | -1.846 | 6.61E-04 | glio |
| culture_effect | Tent2 | 0.668 | 6.61E-04 | glio |
| culture_effect | Pabpc4 | 0.481 | 6.61E-04 | glio |
| Atp5c1_in.vivo_noRT | Prrx2 | 2.668 | 6.61E-04 | glio |
| Bcas2_in.vivo_noRT | Psmg3 | -1.289 | 6.61E-04 | glio |
| Bcas2_ex.vivo_noRT | Cfb | 2.742 | 6.61E-04 | glio |
| Hsd17b10_in.vivo_noRT | Dact1 | 2.449 | 6.61E-04 | glio |
| Interaction_genotype_Mtor | Zfp473 | -4.52 | 6.61E-05 | glio |
| Interaction_genotype_Mtor | Dpf1 | -3.928 | 6.61E-05 | glio |
| Chek1_in.vivo_noRT | Gm20628 | 3.095 | 6.61E-05 | glio |
| Interaction_genotype_Mtor | Lysmd4 | -4.91 | 6.61E-06 | glio |
| Atr_in.vivo_noRT | Gzmm | 3.205 | 6.61E-06 | glio |
| Atr_in.vivo_noRT | Gm10131 | 3.1 | 6.61E-06 | glio |
| Bard1_ex.vivo_noRT | Gm15908 | 2.814 | 6.61E-06 | glio |
| Chek1_in.vivo_noRT | 5031434O11R | 3.635 | 6.61E-06 | glio |
| culture_effect | H2-T24 | -2.998 | 6.61E-07 | glio |
| Atr_in.vivo_noRT | Taf9 | 3.643 | 6.61E-08 | glio |
| Atr_in.vivo_noRT | H13 | -1.864 | 6.61E-08 | glio |
| Chek1_in.vivo_noRT | Adgrb3 | 3.738 | 6.61E-08 | glio |
| Hsd17b10_in.vivo_noRT | Ccdc162 | 3.087 | 6.61E-08 | glio |
| Atr_in.vivo_noRT | Gm32122 | 3.705 | 6.61E-09 | glio |
| Bcas2_in.vivo_noRT | Mro | 4.016 | 6.61E-09 | glio |
| Bora_in.vivo_noRT | 2010106G04R | 4.136 | 6.61E-09 | glio |
| Atr_in.vivo_noRT | 1700047K16R | 4.876 | 6.61E-12 | glio |
| Atr_in.vivo_noRT | 2810408A11R | 4.768 | 6.61E-12 | glio |
| Aars2_in.vivo_noRT | Boc | 5.063 | 6.61E-23 | glio |
| Aars2_in.vivo_noRT | Esp24 | 5.327 | 6.61E-24 | glio |
| Bora_in.vivo_noRT | Dscaml1 | 4.936 | 6.61E-24 | glio |
| Interaction_genotype_Aars2 | Pik3ap1 | -2.924 | 6.62E-03 | glio |
| Interaction_genotype_Atr | Gm11767 | -3.461 | 6.62E-03 | glio |
| Interaction_genotype_Bora | Rrp36 | 1.659 | 6.62E-03 | glio |
| Interaction_genotype_Myc | Nr6a1 | -2.645 | 6.62E-03 | glio |
| Aars2_in.vivo_noRT | Man2b2 | 1.88 | 6.62E-03 | glio |
| Atr_ex.vivo_noRT | Cox7b2 | 2.367 | 6.62E-03 | glio |
| Bcas2_in.vivo_noRT | Ankrd13d | 2.06 | 6.62E-03 | glio |
| Cenpj_in.vivo_noRT | Efcab9 | 2.285 | 6.62E-03 | glio |
| Cenpj_in.vivo_noRT | 4930579C12R | 2.022 | 6.62E-03 | glio |
| Cenpj_in.vivo_noRT | AF067061 | 2.022 | 6.62E-03 | glio |

| | | | | |
|---|---|---|---|---|
| Eif2s1_in.vivo_noRT | Acoxl | 2.143 | 6.62E-03 | glio |
| culture_effect | 1010001B22R | 1.773 | 6.62E-04 | glio |
| culture_effect | Mindy2 | 0.742 | 6.62E-04 | glio |
| interaction_RT | Arl8a | 1.035 | 6.62E-04 | glio |
| Atr_ex.vivo_noRT | 1810026B05R | 1.694 | 6.62E-04 | glio |
| Bcas2_in.vivo_noRT | Ccdc171 | 2.291 | 6.62E-04 | glio |
| Bora_in.vivo_noRT | Mis12 | -1.681 | 6.62E-04 | glio |
| Eif2s1_in.vivo_noRT | Shfl | 1.721 | 6.62E-04 | glio |
| Aars2_ex.vivo_noRT | Icam2 | 3.293 | 6.62E-05 | glio |
| Bard1_ex.vivo_noRT | Gm13178 | 2.814 | 6.62E-05 | glio |
| Aars2_in.vivo_noRT | Pcolce2 | -2.149 | 6.62E-06 | glio |
| Atp5c1_in.vivo_noRT | Rtl1 | 2.988 | 6.62E-06 | glio |
| Bcas2_in.vivo_noRT | Gm30541 | 3.111 | 6.62E-06 | glio |
| Chek1_in.vivo_noRT | Sgk3 | 3.489 | 6.62E-07 | glio |
| culture_effect | Hsp90b1 | -0.91 | 6.62E-11 | glio |
| Bora_in.vivo_noRT | Krtdap | 4.545 | 6.62E-11 | glio |
| Atr_in.vivo_noRT | Asgr1 | 4.875 | 6.62E-12 | glio |
| culture_effect | Nupl2 | 0.885 | 6.63E-03 | glio |
| Interaction_genotype_Atr | Kbtbd2 | 1.752 | 6.63E-03 | glio |
| Interaction_genotype_Bcas2 | Pmp22 | -2.839 | 6.63E-03 | glio |
| Aars2_in.vivo_noRT | Padi2 | 2.051 | 6.63E-03 | glio |
| Atp5c1_in.vivo_noRT | Tesc | -1.329 | 6.63E-03 | glio |
| Atr_in.vivo_noRT | Nphs1 | 1.828 | 6.63E-03 | glio |
| Atr_in.vivo_noRT | Hnrnpr | -0.839 | 6.63E-03 | glio |
| Bcas2_in.vivo_noRT | Lrp1b | 2.029 | 6.63E-03 | glio |
| Bcas2_in.vivo_noRT | Matk | 1.865 | 6.63E-03 | glio |
| Bcas2_in.vivo_noRT | Wasl | -0.935 | 6.63E-03 | glio |
| Chek1_in.vivo_noRT | Rfc4 | -1.294 | 6.63E-03 | glio |
| culture_effect | Ube2v1 | 0.452 | 6.63E-04 | glio |
| culture_effect | Fbxo7 | -0.485 | 6.63E-04 | glio |
| Interaction_genotype_Myc | Psmd12 | 1.432 | 6.63E-04 | glio |
| Interaction_genotype_Myc | Eif3c | 1.563 | 6.63E-04 | glio |
| interaction_RT | Crispld2 | 2.579 | 6.63E-04 | glio |
| Atp5c1_in.vivo_noRT | 1700006H21R | 2.668 | 6.63E-04 | glio |
| Chek1_in.vivo_noRT | Mpz | 3.073 | 6.63E-04 | glio |
| culture_effect | Gm50394 | 2.222 | 6.63E-05 | glio |
| culture_effect | Rint1 | 0.771 | 6.63E-05 | glio |
| Atr_in.vivo_noRT | Paqr6 | 2.768 | 6.63E-05 | glio |
| Atr_in.vivo_noRT | Nek3 | 2.805 | 6.63E-05 | glio |
| Atr_in.vivo_noRT | 2900052L18Ri | 2.265 | 6.63E-05 | glio |
| Bcas2_in.vivo_noRT | 1700020L24Ri | 2.676 | 6.63E-05 | glio |
| Cyp19a1_ex.vivo_noRT | Gm30692 | 2.943 | 6.63E-05 | glio |
| Aars2_in.vivo_noRT | 9330179D12R | 3.397 | 6.63E-06 | glio |
| Aars2_in.vivo_noRT | Gm4129 | 3.157 | 6.63E-06 | glio |
| Bcas2_in.vivo_noRT | Gm14327 | 3.302 | 6.63E-06 | glio |
| Bcas2_in.vivo_noRT | Gm26766 | 3.111 | 6.63E-06 | glio |
| Interaction_genotype_Mtor | Gm15952 | -6.065 | 6.63E-07 | glio |
| Bcas2_in.vivo_noRT | Dhrs9 | 3.22 | 6.63E-07 | glio |
| Bora_ex.vivo_noRT | Gm35439 | 3.595 | 6.63E-07 | glio |
| Bora_ex.vivo_noRT | Mmp13 | 3.201 | 6.63E-07 | glio |
| Atr_in.vivo_noRT | Serpinb6b | 3.891 | 6.63E-08 | glio |
| Chek1_ex.vivo_noRT | Dcpp3 | 4.842 | 6.63E-09 | glio |

| | | | | |
|---|---|---|---|---|
| Aars2_in.vivo_noRT | 4933421A08R | 4.547 | 6.63E-10 | glio |
| Bora_ex.vivo_noRT | Idi2 | 3.993 | 6.63E-10 | glio |
| Bora_in.vivo_noRT | H1foo | 4.404 | 6.63E-18 | glio |
| Bora_in.vivo_noRT | Gm26555 | 4.673 | 6.63E-20 | glio |
| Bora_in.vivo_noRT | 2410017I17Ri | 4.936 | 6.63E-24 | glio |
| culture_effect | Taf11 | 0.456 | 6.64E-03 | glio |
| Interaction_genotype_Aars2 | Cxcl16 | -3.257 | 6.64E-03 | glio |
| Interaction_genotype_Atr | Ankrd42 | -3.522 | 6.64E-03 | glio |
| Interaction_genotype_Mtor | Gm8300 | -4.099 | 6.64E-03 | glio |
| Interaction_genotype_Myc | Cox15 | -2.155 | 6.64E-03 | glio |
| Interaction_genotype_Rad21 | Vgll2 | 3.779 | 6.64E-03 | glio |
| interaction_RT | Klk13 | 1.99 | 6.64E-03 | glio |
| interaction_RT | Plxna3 | 1.213 | 6.64E-03 | glio |
| Atr_in.vivo_noRT | Akna | 1.902 | 6.64E-03 | glio |
| Bcas2_ex.vivo_noRT | Itgbl1 | 2.347 | 6.64E-03 | glio |
| Bora_in.vivo_noRT | Mok | 2.046 | 6.64E-03 | glio |
| Bora_ex.vivo_noRT | Gm49579 | 2.414 | 6.64E-03 | glio |
| Bora_ex.vivo_noRT | Gm43062 | 2.81 | 6.64E-03 | glio |
| Brca2_in.vivo_noRT | Gpc5 | 1.763 | 6.64E-03 | glio |
| Cenpj_in.vivo_noRT | Cmas | -0.698 | 6.64E-03 | glio |
| Chek1_in.vivo_noRT | C1ra | -1.483 | 6.64E-03 | glio |
| Eif2s1_ex.vivo_noRT | Gm48215 | 2.234 | 6.64E-03 | glio |
| Eif2s1_ex.vivo_noRT | AI182371 | 2.441 | 6.64E-03 | glio |
| Hsd17b10_in.vivo_noRT | Gm10101 | 2.047 | 6.64E-03 | glio |
| Hsd17b10_in.vivo_noRT | C130013H08R | 2.378 | 6.64E-03 | glio |
| Aars2_in.vivo_noRT | Kifc5b | 2.012 | 6.64E-04 | glio |
| Bcas2_ex.vivo_noRT | Vmn2r44 | 2.347 | 6.64E-04 | glio |
| Bcas2_ex.vivo_noRT | Gm50024 | 2.342 | 6.64E-04 | glio |
| Interaction_genotype_Myc | Gm15910 | -4.308 | 6.64E-05 | glio |
| Bcas2_in.vivo_noRT | Pnp2 | -2.116 | 6.64E-05 | glio |
| Chek1_ex.vivo_noRT | Kcnj13 | 3.168 | 6.64E-05 | glio |
| Chek1_ex.vivo_noRT | Maob | 3.475 | 6.64E-05 | glio |
| Interaction_genotype_Myc | Phldb2 | -5.257 | 6.64E-06 | glio |
| Interaction_genotype_Myc | Gm15510 | -5.123 | 6.64E-06 | glio |
| Interaction_genotype_Myc | Ccdc59 | 1.788 | 6.64E-06 | glio |
| Interaction_genotype_Mtor | Gm49860 | -5.08 | 6.64E-07 | glio |
| Interaction_genotype_Myc | Hist1h3d | -5.739 | 6.64E-08 | glio |
| Atr_in.vivo_noRT | Gm48242 | 5.533 | 6.64E-20 | glio |
| culture_effect | Fzd1 | 1.29 | 6.65E-03 | glio |
| culture_effect | Rnf31 | -0.73 | 6.65E-03 | glio |
| Interaction_genotype_Aars2 | Magi2 | -3.097 | 6.65E-03 | glio |
| Interaction_genotype_Atr | Prm3 | -3 | 6.65E-03 | glio |
| Interaction_genotype_Atr | Nfkbia | 1.734 | 6.65E-03 | glio |
| Interaction_genotype_Bora | Elobl | -3.859 | 6.65E-03 | glio |
| Interaction_genotype_Bora | Apol6 | 3.085 | 6.65E-03 | glio |
| Interaction_genotype_Mtor | Snn | -3.173 | 6.65E-03 | glio |
| Interaction_genotype_Myc | Mief2 | -2.759 | 6.65E-03 | glio |
| interaction_RT | Sema3b | -1.338 | 6.65E-03 | glio |
| Aars2_in.vivo_noRT | Trip6 | -1.201 | 6.65E-03 | glio |
| Atr_in.vivo_noRT | Chml | 1.934 | 6.65E-03 | glio |
| Atr_in.vivo_noRT | Cnksr3 | 1.465 | 6.65E-03 | glio |
| Bard1_ex.vivo_noRT | Slc16a7 | 2.415 | 6.65E-03 | glio |

| | | | | |
|---|---|---|---|---|
| Bcas2_in.vivo_noRT | Tnfrsf19 | 2.031 | 6.65E-03 | glio |
| Bcas2_in.vivo_noRT | Mindy3 | -1.239 | 6.65E-03 | glio |
| Bcas2_in.vivo_noRT | Cwc25 | -1.231 | 6.65E-03 | glio |
| Bcas2_in.vivo_noRT | Fam120a | -1.006 | 6.65E-03 | glio |
| Bora_in.vivo_noRT | Mmp14 | 2.062 | 6.65E-03 | glio |
| Bora_in.vivo_noRT | Disp1 | 1.699 | 6.65E-03 | glio |
| Bora_ex.vivo_noRT | Rapgef3os2 | 2.016 | 6.65E-03 | glio |
| Bora_ex.vivo_noRT | Gm29093 | 2.769 | 6.65E-03 | glio |
| Cenpj_in.vivo_noRT | Tcap | 2.172 | 6.65E-03 | glio |
| Chek1_in.vivo_noRT | Cntnap1 | 1.85 | 6.65E-03 | glio |
| Hsd17b10_in.vivo_noRT | Spink2 | 2.419 | 6.65E-03 | glio |
| Hsd17b10_in.vivo_noRT | Ccdc184 | 2.283 | 6.65E-03 | glio |
| Hsd17b10_in.vivo_noRT | Shroom1 | 2.061 | 6.65E-03 | glio |
| Hsd17b10_in.vivo_noRT | Plekhj1 | -0.645 | 6.65E-03 | glio |
| Interaction_genotype_Mtor | Pfkfb1 | -4.099 | 6.65E-04 | glio |
| Interaction_genotype_Mtor | Mansc1 | -4.368 | 6.65E-04 | glio |
| Bcas2_in.vivo_noRT | Bpifa1 | 2.28 | 6.65E-04 | glio |
| Bcas2_in.vivo_noRT | Clec2d | 2.017 | 6.65E-04 | glio |
| Bora_ex.vivo_noRT | Slco2a1 | 3.201 | 6.65E-04 | glio |
| Interaction_genotype_Myc | Aloxe3 | -3.839 | 6.65E-05 | glio |
| Interaction_genotype_Myc | Sox10 | 1.77 | 6.65E-05 | glio |
| Aars2_ex.vivo_noRT | Gm15326 | 2.718 | 6.65E-05 | glio |
| Atr_ex.vivo_noRT | Adam2 | 3.164 | 6.65E-05 | glio |
| Bcas2_ex.vivo_noRT | Snhg8 | 1.514 | 6.65E-06 | glio |
| culture_effect | Notch2 | 1.362 | 6.65E-07 | glio |
| Bora_ex.vivo_noRT | Gm8016 | 3.201 | 6.65E-07 | glio |
| Bora_in.vivo_noRT | Bysl | -2.162 | 6.65E-08 | glio |
| Bcas2_in.vivo_noRT | 9530080O11R | 3.89 | 6.65E-09 | glio |
| Aars2_in.vivo_noRT | Slc23a3 | 4.425 | 6.65E-11 | glio |
| Bora_in.vivo_noRT | Krt80 | 4.67 | 6.65E-14 | glio |
| Chek1_in.vivo_noRT | 2310003N18R | 5.044 | 6.65E-18 | glio |
| Interaction_genotype_Aars2 | Cacnb1 | -3.735 | 6.66E-03 | glio |
| Interaction_genotype_Aars2 | Ywhag | 1.066 | 6.66E-03 | glio |
| Interaction_genotype_Atr | Dppa2 | -3.205 | 6.66E-03 | glio |
| Interaction_genotype_Atr | Gm41396 | -3.383 | 6.66E-03 | glio |
| Interaction_genotype_Atr | Eif4a2 | 1.319 | 6.66E-03 | glio |
| Interaction_genotype_Myc | Slc47a2 | -2.928 | 6.66E-03 | glio |
| Interaction_genotype_Myc | Nova2 | -2.824 | 6.66E-03 | glio |
| Interaction_genotype_Myc | Cep41 | -2.215 | 6.66E-03 | glio |
| Interaction_genotype_Rpl3 | Fgf7 | -3.661 | 6.66E-03 | glio |
| Interaction_genotype_Terf1 | Ephx1 | -4.026 | 6.66E-03 | glio |
| Aars2_in.vivo_noRT | St3gal5 | -1.275 | 6.66E-03 | glio |
| Atp5c1_in.vivo_noRT | 1600014C23R | 2.289 | 6.66E-03 | glio |
| Bcas2_ex.vivo_noRT | Spata2l | 2.957 | 6.66E-03 | glio |
| Cenpj_ex.vivo_noRT | 4930432K21R | -2.62 | 6.66E-03 | glio |
| Chek1_in.vivo_noRT | Gm36696 | 1.904 | 6.66E-03 | glio |
| Chek1_ex.vivo_noRT | Fkbp4 | -0.767 | 6.66E-03 | glio |
| culture_effect | Chd3 | 1.054 | 6.66E-04 | glio |
| culture_effect | Eloa | 0.87 | 6.66E-04 | glio |
| Aars2_in.vivo_noRT | Golt1b | -1.32 | 6.66E-04 | glio |
| Atr_in.vivo_noRT | Hist1h3b | 2.558 | 6.66E-04 | glio |
| Bora_in.vivo_noRT | Qpct | 2.533 | 6.66E-04 | glio |

| | | | | |
|---|---|---|---|---|
| Interaction_genotype_Myc | Fam25c | -3.57 | 6.66E-05 | glio |
| Interaction_genotype_Myc | Rhoa | 1.495 | 6.66E-05 | glio |
| Atr_in.vivo_noRT | Matk | 2.591 | 6.66E-05 | glio |
| Atr_in.vivo_noRT | Rpl23 | -0.854 | 6.66E-05 | glio |
| Bora_ex.vivo_noRT | Fam189a1 | 3.201 | 6.66E-05 | glio |
| Bora_ex.vivo_noRT | Cemip | 3.201 | 6.66E-05 | glio |
| Chek1_in.vivo_noRT | Trim14 | 3.1 | 6.66E-05 | glio |
| Interaction_genotype_Myc | Bloc1s5 | -4.059 | 6.66E-06 | glio |
| Bcas2_in.vivo_noRT | Psenen | -1.041 | 6.66E-06 | glio |
| Chek1_ex.vivo_noRT | B230311B06R | 4.119 | 6.66E-06 | glio |
| Bora_in.vivo_noRT | Stard6 | 3.565 | 6.66E-07 | glio |
| Chek1_in.vivo_noRT | Klra6 | 3.934 | 6.66E-08 | glio |
| culture_effect | Psma4 | -0.693 | 6.66E-09 | glio |
| culture_effect | Gm13272 | 2.839 | 6.66E-10 | glio |
| culture_effect | Tm2d1 | 0.936 | 6.66E-10 | glio |
| Aars2_in.vivo_noRT | Serpina3i | 4.006 | 6.66E-10 | glio |
| Chek1_in.vivo_noRT | AI182371 | 4.261 | 6.66E-14 | glio |
| culture_effect | Pde4b | -2.844 | 6.66E-19 | glio |
| Interaction_genotype_Atr | Dnajb4 | 1.744 | 6.67E-03 | glio |
| Interaction_genotype_Bcas2 | Gm49223 | -3.042 | 6.67E-03 | glio |
| Interaction_genotype_Bcas2 | 4930513N10R | -4.089 | 6.67E-03 | glio |
| Interaction_genotype_Myc | Kbtbd3 | -2.558 | 6.67E-03 | glio |
| Aars2_in.vivo_noRT | Tmem35a | 2.253 | 6.67E-03 | glio |
| Aars2_in.vivo_noRT | Sipa1l3 | 1.467 | 6.67E-03 | glio |
| Aars2_ex.vivo_noRT | Tmem205 | 1.08 | 6.67E-03 | glio |
| Atr_in.vivo_noRT | Eid2b | 1.653 | 6.67E-03 | glio |
| Atr_in.vivo_noRT | Cmas | -0.971 | 6.67E-03 | glio |
| Bcas2_in.vivo_noRT | Gm36988 | 2.016 | 6.67E-03 | glio |
| Bora_in.vivo_noRT | Nova2 | 1.918 | 6.67E-03 | glio |
| Bora_in.vivo_noRT | Rgcc | -1.166 | 6.67E-03 | glio |
| Bora_in.vivo_noRT | Psme2 | -0.67 | 6.67E-03 | glio |
| Cdkn2a_in.vivo_noRT | Gm26682 | -2.593 | 6.67E-03 | glio |
| Chek1_in.vivo_noRT | Parp10 | -1.232 | 6.67E-03 | glio |
| Chek1_in.vivo_noRT | Rpl17 | -0.553 | 6.67E-03 | glio |
| culture_effect | Khdc4 | -0.728 | 6.67E-04 | glio |
| Interaction_genotype_Aars2 | Gm16835 | -4.248 | 6.67E-04 | glio |
| Aars2_in.vivo_noRT | Gm4673 | 2.476 | 6.67E-04 | glio |
| Aars2_in.vivo_noRT | Gm36888 | 2.385 | 6.67E-04 | glio |
| Aars2_in.vivo_noRT | Mmachc | 1.85 | 6.67E-04 | glio |
| Aars2_in.vivo_noRT | Tomm6 | -0.965 | 6.67E-04 | glio |
| Aars2_in.vivo_noRT | Snrpe | -0.651 | 6.67E-04 | glio |
| Atp5c1_in.vivo_noRT | Milr1 | 2.713 | 6.67E-04 | glio |
| Eif2s1_in.vivo_noRT | Gm35082 | 2.143 | 6.67E-04 | glio |
| Eif2s1_in.vivo_noRT | Gm36356 | 2.414 | 6.67E-04 | glio |
| interaction_RT | Atf3 | 1.835 | 6.67E-05 | glio |
| interaction_RT | Adgra3 | -1.057 | 6.67E-05 | glio |
| Aars2_in.vivo_noRT | Rplp2 | -0.745 | 6.67E-05 | glio |
| Bora_in.vivo_noRT | Yipf6 | -1.727 | 6.67E-05 | glio |
| Chek1_in.vivo_noRT | St18 | 2.211 | 6.67E-05 | glio |
| Interaction_genotype_Mtor | 2810408A11R | -5.487 | 6.67E-06 | glio |
| Aars2_in.vivo_noRT | Gm28050 | 3.289 | 6.67E-06 | glio |
| Bcas2_in.vivo_noRT | Arg2 | 3.167 | 6.67E-06 | glio |

| | | | | |
|---|---|---|---|---|
| Atr_in.vivo_noRT | 4930417H01R | 3.893 | 6.67E-08 | glio |
| Chek1_in.vivo_noRT | Rab11fip4os1 | 4.007 | 6.67E-08 | glio |
| Chek1_in.vivo_noRT | Appbp2os | 4.511 | 6.67E-10 | glio |
| Chek1_in.vivo_noRT | Ctsf | 3.375 | 6.67E-10 | glio |
| Chek1_in.vivo_noRT | Ephx3 | 4.316 | 6.67E-11 | glio |
| Atr_in.vivo_noRT | 5033403F01Ri | 4.608 | 6.67E-12 | glio |
| Bcas2_in.vivo_noRT | Gm44983 | 4.019 | 6.67E-12 | glio |
| Aars2_in.vivo_noRT | Smpd5 | 4.67 | 6.67E-14 | glio |
| Atr_in.vivo_noRT | BC026762 | 5.154 | 6.67E-15 | glio |
| Aars2_in.vivo_noRT | Gm8016 | 5.327 | 6.67E-26 | glio |
| Atr_in.vivo_noRT | Dscaml1 | 5.533 | 6.67E-29 | glio |
| culture_effect | Gm13889 | -1.587 | 6.68E-03 | glio |
| culture_effect | Cdkal1 | 0.646 | 6.68E-03 | glio |
| culture_effect | Nr1h2 | -0.419 | 6.68E-03 | glio |
| Interaction_genotype_Atr | Slfn3 | -2.936 | 6.68E-03 | glio |
| Interaction_genotype_Mtor | Gm49741 | -2.964 | 6.68E-03 | glio |
| Interaction_genotype_Myc | Gm42869 | -3.023 | 6.68E-03 | glio |
| Aars2_in.vivo_noRT | Tmem131l | 1.559 | 6.68E-03 | glio |
| Aars2_in.vivo_noRT | Rpl17 | -0.554 | 6.68E-03 | glio |
| Aars2_ex.vivo_noRT | E230001N04R | 2.405 | 6.68E-03 | glio |
| Atr_ex.vivo_noRT | Bccip | -0.939 | 6.68E-03 | glio |
| Bcas2_in.vivo_noRT | Fam178b | 2.099 | 6.68E-03 | glio |
| Bcas2_in.vivo_noRT | Izumo4 | 2.02 | 6.68E-03 | glio |
| Bcas2_in.vivo_noRT | Gba2 | 2.027 | 6.68E-03 | glio |
| Bcas2_in.vivo_noRT | Chchd6 | 1.662 | 6.68E-03 | glio |
| Bcas2_in.vivo_noRT | Ppp1r16a | 1.501 | 6.68E-03 | glio |
| Bcas2_in.vivo_noRT | Abhd14a | -1.142 | 6.68E-03 | glio |
| Bcas2_in.vivo_noRT | 9130401M01F | -0.812 | 6.68E-03 | glio |
| Brca2_in.vivo_noRT | Erich6b | 1.763 | 6.68E-03 | glio |
| Brca2_in.vivo_noRT | Lncbate10 | 1.763 | 6.68E-03 | glio |
| Brca2_in.vivo_noRT | Tac2 | 1.763 | 6.68E-03 | glio |
| Cdkn2a_in.vivo_noRT | Krt19 | -1.675 | 6.68E-03 | glio |
| Cdkn2a_in.vivo_noRT | Gm10399 | 2.042 | 6.68E-03 | glio |
| Cenpj_in.vivo_noRT | Zfp455 | 2.297 | 6.68E-03 | glio |
| Interaction_genotype_Aars2 | Gm9970 | -4.477 | 6.68E-04 | glio |
| interaction_RT | Il6 | 2.715 | 6.68E-04 | glio |
| interaction_RT | Socs1 | 1.244 | 6.68E-04 | glio |
| interaction_RT | Neto2 | -1.16 | 6.68E-04 | glio |
| Aars2_in.vivo_noRT | Slc4a11 | 2.539 | 6.68E-04 | glio |
| Aars2_in.vivo_noRT | Zkscan7 | 2.473 | 6.68E-04 | glio |
| Aars2_in.vivo_noRT | Sh3rf1 | 2.08 | 6.68E-04 | glio |
| Bard1_ex.vivo_noRT | AI463229 | 2.814 | 6.68E-04 | glio |
| Bora_in.vivo_noRT | Hdac7 | 2.419 | 6.68E-04 | glio |
| Bora_ex.vivo_noRT | Zfp455 | 3.222 | 6.68E-04 | glio |
| culture_effect | Adgrd1 | -2.381 | 6.68E-05 | glio |
| Interaction_genotype_Myc | Ccl9 | -3.122 | 6.68E-05 | glio |
| Interaction_genotype_Myc | Trim17 | -3.128 | 6.68E-05 | glio |
| Atr_ex.vivo_noRT | Olfr718-ps1 | 2.77 | 6.68E-05 | glio |
| Chek1_in.vivo_noRT | Asphd2 | 2.84 | 6.68E-05 | glio |
| Eif2s1_in.vivo_noRT | Cldn2 | 2.143 | 6.68E-05 | glio |
| Eif2s1_in.vivo_noRT | Stfa3 | 2.143 | 6.68E-05 | glio |
| Eif2s1_in.vivo_noRT | Olfr787 | 2.143 | 6.68E-05 | glio |

| | | | | |
|---|---|---|---|---|
| Eif2s1_in.vivo_noRT | Oit3 | 3.197 | 6.68E-05 | glio |
| Eif2s1_in.vivo_noRT | Gm15908 | 2.143 | 6.68E-05 | glio |
| Eif2s1_in.vivo_noRT | A930014E10Ri | 2.143 | 6.68E-05 | glio |
| Eif2s1_in.vivo_noRT | Gm10010 | 3.399 | 6.68E-05 | glio |
| Eif2s1_in.vivo_noRT | Gm45338 | 2.143 | 6.68E-05 | glio |
| Eif2s1_in.vivo_noRT | Scg2 | 2.143 | 6.68E-05 | glio |
| Eif2s1_in.vivo_noRT | Neb | 1.193 | 6.68E-05 | glio |
| Interaction_genotype_Myc | Snu13 | 1.677 | 6.68E-06 | glio |
| Aars2_in.vivo_noRT | 1810021B22R | 3.258 | 6.68E-06 | glio |
| Aars2_in.vivo_noRT | Tapbp | -1.517 | 6.68E-06 | glio |
| Bard1_ex.vivo_noRT | A930014E10Ri | 2.814 | 6.68E-06 | glio |
| Chek1_ex.vivo_noRT | A330015K06R | 3.567 | 6.68E-06 | glio |
| Interaction_genotype_Myc | Sf3b6 | 2.068 | 6.68E-08 | glio |
| Chek1_in.vivo_noRT | Cks1brt | 3.606 | 6.68E-09 | glio |
| Atr_in.vivo_noRT | 9030612E09Ri | 4.212 | 6.68E-11 | glio |
| Atr_in.vivo_noRT | Ptpn22 | 5.27 | 6.68E-12 | glio |
| Chek1_ex.vivo_noRT | 4930512B01R | 4.842 | 6.68E-14 | glio |
| Bcas2_in.vivo_noRT | Angptl7 | 3.941 | 6.68E-16 | glio |
| Chek1_in.vivo_noRT | Serpinb9g | 5.044 | 6.68E-23 | glio |
| Atr_in.vivo_noRT | 2410017I17Ri | 5.533 | 6.68E-29 | glio |
| Interaction_genotype_Aars2 | Psca | -3.082 | 6.69E-03 | glio |
| Bard1_ex.vivo_noRT | Kmo | 2.421 | 6.69E-03 | glio |
| Bcas2_in.vivo_noRT | Ramp1 | 2.316 | 6.69E-03 | glio |
| Bcas2_in.vivo_noRT | Hist1h3d | 1.904 | 6.69E-03 | glio |
| Bora_ex.vivo_noRT | Ssh3 | 2.404 | 6.69E-03 | glio |
| Bora_ex.vivo_noRT | 1700040D17R | 2.793 | 6.69E-03 | glio |
| Brca2_in.vivo_noRT | Chia1 | 1.763 | 6.69E-03 | glio |
| Brca2_in.vivo_noRT | Zfp661 | 2.495 | 6.69E-03 | glio |
| Hsd17b10_in.vivo_noRT | Shd | 2.06 | 6.69E-03 | glio |
| Hsd17b10_in.vivo_noRT | Gm48357 | 1.832 | 6.69E-03 | glio |
| Interaction_genotype_Myc | Gm47872 | -3.661 | 6.69E-04 | glio |
| Interaction_genotype_Myc | Ptprn | -3.638 | 6.69E-04 | glio |
| Aars2_in.vivo_noRT | Mtmr9 | 1.961 | 6.69E-04 | glio |
| Aars2_ex.vivo_noRT | Gm15972 | 2.893 | 6.69E-04 | glio |
| Atp5c1_in.vivo_noRT | Ifitm6 | 2.668 | 6.69E-04 | glio |
| Atr_in.vivo_noRT | Zfp106 | -1.219 | 6.69E-04 | glio |
| Bcas2_in.vivo_noRT | Ces1a | 2.685 | 6.69E-04 | glio |
| Bora_in.vivo_noRT | Cep57 | -1.365 | 6.69E-04 | glio |
| Fanca_ex.vivo_noRT | Fanca | -3.277 | 6.69E-04 | glio |
| culture_effect | Ndufa6 | -0.488 | 6.69E-05 | glio |
| interaction_RT | Gpr146 | 3.97 | 6.69E-05 | glio |
| Aars2_in.vivo_noRT | Manf | -1.134 | 6.69E-05 | glio |
| Eif2s1_in.vivo_noRT | 1700003E24Ri | 2.143 | 6.69E-05 | glio |
| Bora_in.vivo_noRT | Fgf2 | 3.31 | 6.69E-06 | glio |
| Chek1_in.vivo_noRT | Sdcbp2 | 3.349 | 6.69E-06 | glio |
| Chek1_in.vivo_noRT | 9530053A07R | 3.23 | 6.69E-06 | glio |
| Chek1_in.vivo_noRT | Leprot | -1.768 | 6.69E-06 | glio |
| Interaction_genotype_Mtor | Ctsd | 2.573 | 6.69E-07 | glio |
| Interaction_genotype_Mtor | Gm10655 | -6.802 | 6.69E-08 | glio |
| Chek1_in.vivo_noRT | Serpinb6b | 3.913 | 6.69E-08 | glio |
| Atr_in.vivo_noRT | Gm11884 | 4.535 | 6.69E-13 | glio |
| Bora_in.vivo_noRT | Ptgis | 4.936 | 6.69E-13 | glio |

| | | | | |
|---|---|---|---|---|
| Aars2_in.vivo_noRT | Gm16344 | 4.952 | 6.69E-17 | glio |
| Atr_in.vivo_noRT | Prl2c2 | 5.533 | 6.69E-17 | glio |
| Aars2_in.vivo_noRT | Adgre5 | 5.067 | 6.69E-18 | glio |
| Bora_in.vivo_noRT | Gm2016 | 4.936 | 6.69E-21 | glio |
| Aars2_in.vivo_noRT | C430002N11R | 5.327 | 6.69E-22 | glio |
| culture_effect | Pyroxd1 | -0.739 | 6.70E-03 | glio |
| Interaction_genotype_Bora | Ldhb | 2.295 | 6.70E-03 | glio |
| Interaction_genotype_Hsd17b | Apol10b | 3.651 | 6.70E-03 | glio |
| Interaction_genotype_Mtor | Sarnp | 1.261 | 6.70E-03 | glio |
| Interaction_genotype_Myc | Sspn | -2.474 | 6.70E-03 | glio |
| Interaction_genotype_Nat2 | Cfh | 3.058 | 6.70E-03 | glio |
| Aars2_in.vivo_noRT | Tmem266 | 2.026 | 6.70E-03 | glio |
| Aars2_in.vivo_noRT | Shisa4 | -1.026 | 6.70E-03 | glio |
| Bora_in.vivo_noRT | Fcsk | 1.849 | 6.70E-03 | glio |
| Bora_in.vivo_noRT | Atg7 | 1.746 | 6.70E-03 | glio |
| Hsd17b10_in.vivo_noRT | Dennd2d | 1.881 | 6.70E-03 | glio |
| Hsd17b10_in.vivo_noRT | Uchl3 | -0.822 | 6.70E-03 | glio |
| culture_effect | Alpl | -2.119 | 6.70E-04 | glio |
| culture_effect | Med20 | -0.725 | 6.70E-04 | glio |
| Interaction_genotype_Mtor | Kctd4 | -3.989 | 6.70E-04 | glio |
| Interaction_genotype_Mtor | Gfer | 1.721 | 6.70E-04 | glio |
| Interaction_genotype_Mtor | Cnih4 | 1.61 | 6.70E-04 | glio |
| Atr_in.vivo_noRT | Gdap1l1 | 2.655 | 6.70E-04 | glio |
| Bcas2_in.vivo_noRT | Ppp4r1 | -1.631 | 6.70E-04 | glio |
| Bora_in.vivo_noRT | Dynlt1b | 2.45 | 6.70E-04 | glio |
| culture_effect | Vps28 | -0.577 | 6.70E-05 | glio |
| Aars2_in.vivo_noRT | Vav2 | 2.805 | 6.70E-05 | glio |
| Aars2_in.vivo_noRT | Rrs1 | -1.48 | 6.70E-05 | glio |
| Bora_in.vivo_noRT | AW046200 | 2.878 | 6.70E-05 | glio |
| Hsd17b10_in.vivo_noRT | Gm16222 | 3.356 | 6.70E-05 | glio |
| culture_effect | 3010001F23Ri | 2.014 | 6.70E-06 | glio |
| Atr_in.vivo_noRT | Pcdhgb4 | 2.738 | 6.70E-06 | glio |
| Atr_in.vivo_noRT | Serpinb6a | 3.845 | 6.70E-08 | glio |
| Atr_ex.vivo_noRT | Gm5157 | 3.164 | 6.70E-08 | glio |
| Bora_in.vivo_noRT | Gprin2 | 4.025 | 6.70E-12 | glio |
| Bcas2_in.vivo_noRT | 2410141K09R | 4.543 | 6.70E-15 | glio |
| Aars2_in.vivo_noRT | Gm48899 | 5.067 | 6.70E-18 | glio |
| Interaction_genotype_Aars2 | Gm47545 | -3.046 | 6.71E-03 | glio |
| Interaction_genotype_Atr | Fat1 | -1.788 | 6.71E-03 | glio |
| Interaction_genotype_Bcas2 | Gm49694 | -3.549 | 6.71E-03 | glio |
| Interaction_genotype_Mtor | 4930509H03R | -3.278 | 6.71E-03 | glio |
| interaction_RT | Gm48215 | 2.038 | 6.71E-03 | glio |
| Aars2_in.vivo_noRT | Pafah1b2 | -0.888 | 6.71E-03 | glio |
| Atr_in.vivo_noRT | Spag1 | 1.871 | 6.71E-03 | glio |
| Bcas2_ex.vivo_noRT | Abcb5 | 1.948 | 6.71E-03 | glio |
| Eif2s1_ex.vivo_noRT | Gm11583 | 2.234 | 6.71E-03 | glio |
| Eif2s1_ex.vivo_noRT | Elfn2 | 2.798 | 6.71E-03 | glio |
| Hsd17b10_in.vivo_noRT | Adcy2 | 1.862 | 6.71E-03 | glio |
| culture_effect | Dcps | -0.666 | 6.71E-04 | glio |
| Aars2_in.vivo_noRT | Tmem35b | 2.335 | 6.71E-04 | glio |
| Aars2_in.vivo_noRT | Rab11fip5 | 2.513 | 6.71E-04 | glio |
| Aars2_in.vivo_noRT | Hlcs | 2.3 | 6.71E-04 | glio |

| | | | | |
|---|---|---|---|---|
| Aars2_in.vivo_noRT | Gm10762 | 2.034 | 6.71E-04 | glio |
| Aars2_in.vivo_noRT | Eno3 | -1.632 | 6.71E-04 | glio |
| Aars2_in.vivo_noRT | Rps24 | -0.678 | 6.71E-04 | glio |
| Bcas2_in.vivo_noRT | Rnf125 | 2.247 | 6.71E-04 | glio |
| Bora_in.vivo_noRT | Atp23 | -1.372 | 6.71E-04 | glio |
| Chek1_in.vivo_noRT | Krtcap2 | -0.741 | 6.71E-04 | glio |
| Interaction_genotype_Aars2 | Man2c1os | -5.034 | 6.71E-05 | glio |
| interaction_RT | Adamts12 | -2.279 | 6.71E-05 | glio |
| Atr_in.vivo_noRT | Snrnp27 | -1.287 | 6.71E-05 | glio |
| Hsd17b10_in.vivo_noRT | Nppc | 3.148 | 6.71E-05 | glio |
| Interaction_genotype_Mtor | Gm13594 | -5.53 | 6.71E-06 | glio |
| Interaction_genotype_Myc | AV039307 | -5.192 | 6.71E-06 | glio |
| Aars2_in.vivo_noRT | Gm15834 | 3.146 | 6.71E-06 | glio |
| Aars2_in.vivo_noRT | Cxcr6 | 3.138 | 6.71E-06 | glio |
| culture_effect | Exosc7 | -0.674 | 6.71E-07 | glio |
| Aars2_ex.vivo_noRT | Gm26555 | 3.293 | 6.71E-07 | glio |
| Bcas2_in.vivo_noRT | Zfp712 | 3.112 | 6.71E-07 | glio |
| Aars2_in.vivo_noRT | Mir155hg | 4.082 | 6.71E-08 | glio |
| Bora_in.vivo_noRT | Parvg | 4.157 | 6.71E-09 | glio |
| Chek1_ex.vivo_noRT | Gm13054 | 4.447 | 6.71E-09 | glio |
| Brca2_in.vivo_noRT | Brca2 | -4.486 | 6.71E-12 | glio |
| Atr_in.vivo_noRT | Gm266 | 5.533 | 6.71E-27 | glio |
| culture_effect | Evc | 0.923 | 6.72E-03 | glio |
| culture_effect | Ssc4d | -0.714 | 6.72E-03 | glio |
| Interaction_genotype_Bcas2 | Ramp3 | -3.609 | 6.72E-03 | glio |
| Interaction_genotype_Bcas2 | Zfand3 | 1.793 | 6.72E-03 | glio |
| Interaction_genotype_Bora | Gm34256 | -3.795 | 6.72E-03 | glio |
| Interaction_genotype_Bora | Rab37 | -3.54 | 6.72E-03 | glio |
| Interaction_genotype_Bora | Slc46a1 | -3.799 | 6.72E-03 | glio |
| Interaction_genotype_Bora | Gm36839 | -3.633 | 6.72E-03 | glio |
| Interaction_genotype_Bora | Taco1os | -3.314 | 6.72E-03 | glio |
| Interaction_genotype_Bora | Bag1 | 1.089 | 6.72E-03 | glio |
| Interaction_genotype_Mtor | Mill2 | -3.199 | 6.72E-03 | glio |
| Interaction_genotype_Mtor | Ccdc14 | -2.977 | 6.72E-03 | glio |
| Interaction_genotype_Mtor | Tab1 | -2.724 | 6.72E-03 | glio |
| Interaction_genotype_Mtor | Tmem17 | -2.738 | 6.72E-03 | glio |
| Interaction_genotype_Mtor | Stom | -2.855 | 6.72E-03 | glio |
| Interaction_genotype_Mtor | Inpp5a | -2.81 | 6.72E-03 | glio |
| Interaction_genotype_Mtor | Gm20045 | -2.637 | 6.72E-03 | glio |
| Interaction_genotype_Myc | Hist2h3b | -3.005 | 6.72E-03 | glio |
| Interaction_genotype_Myc | Rmst | -3.109 | 6.72E-03 | glio |
| Interaction_genotype_Myc | Atg4c | -2.342 | 6.72E-03 | glio |
| Atp5c1_in.vivo_noRT | Ms4a6d | 1.88 | 6.72E-03 | glio |
| Atr_in.vivo_noRT | Tmem231 | 1.895 | 6.72E-03 | glio |
| Bcas2_in.vivo_noRT | Foxc2 | 2.064 | 6.72E-03 | glio |
| Bora_in.vivo_noRT | Mrpl52 | -0.822 | 6.72E-03 | glio |
| Chek1_in.vivo_noRT | E130102H24R | 1.932 | 6.72E-03 | glio |
| Chek1_ex.vivo_noRT | Nup210l | 2.684 | 6.72E-03 | glio |
| Chek1_ex.vivo_noRT | Dram1 | 2.145 | 6.72E-03 | glio |
| culture_effect | Bcl2l1 | 0.534 | 6.72E-04 | glio |
| culture_effect | Rpl22 | -0.422 | 6.72E-04 | glio |
| Interaction_genotype_Mtor | 4732491K20R | -3.428 | 6.72E-04 | glio |

| | | | | |
|---|---|---|---|---|
| Interaction_genotype_Myc | B230369F24Ri | -3.816 | 6.72E-04 | glio |
| Interaction_genotype_Myc | Zik1 | -3.264 | 6.72E-04 | glio |
| Chek1_in.vivo_noRT | Sema4f | 2.451 | 6.72E-04 | glio |
| Hsd17b10_in.vivo_noRT | 1700011D18R | 2.571 | 6.72E-04 | glio |
| culture_effect | Angptl4 | -1.742 | 6.72E-05 | glio |
| Interaction_genotype_Myc | D030068K23R | -4.595 | 6.72E-05 | glio |
| Atr_in.vivo_noRT | Slc13a5 | 2.656 | 6.72E-05 | glio |
| Atr_in.vivo_noRT | Mcm5 | -1.872 | 6.72E-05 | glio |
| Hsd17b10_in.vivo_noRT | Cnnm1 | 3.236 | 6.72E-05 | glio |
| Aars2_in.vivo_noRT | Gstt1 | 3.59 | 6.72E-07 | glio |
| Bcas2_in.vivo_noRT | Gm2762 | 3.235 | 6.72E-07 | glio |
| Atr_ex.vivo_noRT | Gm28940 | 3.947 | 6.72E-08 | glio |
| interaction_RT | Il15 | 2.826 | 6.72E-09 | glio |
| Aars2_in.vivo_noRT | A830035O19R | 4.277 | 6.72E-11 | glio |
| Aars2_in.vivo_noRT | Ahrr | 4.526 | 6.72E-11 | glio |
| Atr_in.vivo_noRT | Gm12602 | 4.506 | 6.72E-12 | glio |
| culture_effect | Dlc1 | 1.767 | 6.72E-14 | glio |
| culture_effect | Acod1 | -6.8 | 6.72E-21 | glio |
| culture_effect | Slc25a5 | -1.291 | 6.72E-22 | glio |
| Interaction_genotype_Atr | Gm46404 | -3.609 | 6.73E-03 | glio |
| Interaction_genotype_Atr | Adh5 | -1.316 | 6.73E-03 | glio |
| Interaction_genotype_Mtor | Gm13663 | -2.654 | 6.73E-03 | glio |
| Interaction_genotype_Myc | Cpt1b | -2.967 | 6.73E-03 | glio |
| Interaction_genotype_Myc | Gars | 1.353 | 6.73E-03 | glio |
| Interaction_genotype_Pars2 | Ppp2r5a | 1.917 | 6.73E-03 | glio |
| interaction_RT | Top1mt | -1.212 | 6.73E-03 | glio |
| Atr_in.vivo_noRT | 4930581F22Ri | 1.645 | 6.73E-03 | glio |
| Atr_in.vivo_noRT | Ahcyl2 | -1.195 | 6.73E-03 | glio |
| Bcas2_in.vivo_noRT | 1700028J19Ri | 2.01 | 6.73E-03 | glio |
| Bcas2_ex.vivo_noRT | Gm43221 | 2.742 | 6.73E-03 | glio |
| Chek1_in.vivo_noRT | Zfp937 | 2.113 | 6.73E-03 | glio |
| Chek1_ex.vivo_noRT | Gm2012 | 2.579 | 6.73E-03 | glio |
| Interaction_genotype_Rpl3 | Apol10b | 4.046 | 6.73E-04 | glio |
| Aars2_ex.vivo_noRT | Prl3d3 | 2.197 | 6.73E-04 | glio |
| Atp5c1_in.vivo_noRT | Kpna7 | 2.404 | 6.73E-04 | glio |
| Atr_in.vivo_noRT | Fgd6 | 2.289 | 6.73E-04 | glio |
| Bcas2_in.vivo_noRT | Gm19345 | 2.235 | 6.73E-04 | glio |
| Bora_ex.vivo_noRT | Alpl | 3.201 | 6.73E-04 | glio |
| Hsd17b10_in.vivo_noRT | Git2 | -1.608 | 6.73E-04 | glio |
| Interaction_genotype_Myc | Acap3 | -3.701 | 6.73E-05 | glio |
| Interaction_genotype_Myc | Nop10 | 2.009 | 6.73E-05 | glio |
| Bard1_ex.vivo_noRT | Esp24 | 2.814 | 6.73E-05 | glio |
| Hsd17b10_in.vivo_noRT | Baalc | 2.824 | 6.73E-05 | glio |
| Atp5c1_in.vivo_noRT | Gm4027 | 2.668 | 6.73E-06 | glio |
| Aars2_in.vivo_noRT | Kctd14 | 3.403 | 6.73E-08 | glio |
| Aars2_in.vivo_noRT | Gm10644 | 4.086 | 6.73E-09 | glio |
| Aars2_in.vivo_noRT | Gm12089 | 4.14 | 6.73E-09 | glio |
| Chek1_ex.vivo_noRT | Mag | 4.842 | 6.73E-09 | glio |
| Aars2_in.vivo_noRT | Gm10401 | 4.182 | 6.73E-12 | glio |
| Bora_in.vivo_noRT | Slc10a1 | 4.676 | 6.73E-17 | glio |
| Bora_in.vivo_noRT | Gm11973 | 4.67 | 6.73E-17 | glio |
| Bcas2_in.vivo_noRT | Olfr810 | 4.807 | 6.73E-24 | glio |

| | | | | |
|---|---|---|---|---|
| culture_effect | Meioc | -1.198 | 6.74E-03 | glio |
| Interaction_genotype_Atr | Gm49223 | -2.828 | 6.74E-03 | glio |
| Interaction_genotype_Atr | Pigo | -3.051 | 6.74E-03 | glio |
| Interaction_genotype_Atr | Dgcr8 | -2.555 | 6.74E-03 | glio |
| Interaction_genotype_Atr | Rpl26 | 0.831 | 6.74E-03 | glio |
| Interaction_genotype_Mtor | Ccdc13 | -3.203 | 6.74E-03 | glio |
| Interaction_genotype_Mtor | Pard6g | -2.953 | 6.74E-03 | glio |
| Interaction_genotype_Rpl3 | Gtf2e2 | 1.637 | 6.74E-03 | glio |
| interaction_RT | Upk3bl | 2.252 | 6.74E-03 | glio |
| interaction_RT | A530040E14Ri | 1.954 | 6.74E-03 | glio |
| Atr_in.vivo_noRT | Stat5a | 1.58 | 6.74E-03 | glio |
| Bcas2_in.vivo_noRT | Tlcd2 | 2.252 | 6.74E-03 | glio |
| Bcas2_in.vivo_noRT | Gm45589 | 1.995 | 6.74E-03 | glio |
| Bcas2_ex.vivo_noRT | Agrp | 2.347 | 6.74E-03 | glio |
| Eif2s1_in.vivo_noRT | Dpep3 | 1.61 | 6.74E-03 | glio |
| culture_effect | Arg2 | -2.181 | 6.74E-04 | glio |
| Interaction_genotype_Mtor | Gm45871 | -3.655 | 6.74E-04 | glio |
| Interaction_genotype_Myc | Zscan4d | -3.122 | 6.74E-04 | glio |
| Interaction_genotype_Myc | Vpreb1 | -3.781 | 6.74E-04 | glio |
| Interaction_genotype_Myc | Gm34121 | -3.341 | 6.74E-04 | glio |
| Interaction_genotype_Myc | Fyco1 | -3.14 | 6.74E-04 | glio |
| Interaction_genotype_Myc | Trpt1 | -2.973 | 6.74E-04 | glio |
| Interaction_genotype_Myc | Bet1 | 1.49 | 6.74E-04 | glio |
| Atp5c1_in.vivo_noRT | Nxph4 | 2.668 | 6.74E-04 | glio |
| Cenpj_in.vivo_noRT | Gm41307 | 2.022 | 6.74E-04 | glio |
| Chek1_in.vivo_noRT | Gm32618 | 2.523 | 6.74E-04 | glio |
| Chek1_ex.vivo_noRT | Racgap1 | -2.102 | 6.74E-04 | glio |
| Hsd17b10_in.vivo_noRT | Nlrx1 | 2.728 | 6.74E-04 | glio |
| Interaction_genotype_Atr | Gm9833 | -4.677 | 6.74E-05 | glio |
| Bcas2_in.vivo_noRT | C430014B12R | 2.758 | 6.74E-05 | glio |
| culture_effect | Gm14326 | 2.172 | 6.74E-06 | glio |
| Bcas2_in.vivo_noRT | Zfp433 | 3.367 | 6.74E-06 | glio |
| Atr_ex.vivo_noRT | Gm21761 | 4.867 | 6.74E-13 | glio |
| Chek1_ex.vivo_noRT | Ptprc | 4.842 | 6.74E-13 | glio |
| culture_effect | Tle5 | 0.869 | 6.74E-15 | glio |
| Atr_in.vivo_noRT | Thsd1 | 5.146 | 6.74E-18 | glio |
| culture_effect | Bcl2l14 | -1.871 | 6.75E-03 | glio |
| culture_effect | Gm13270 | 1.716 | 6.75E-03 | glio |
| culture_effect | Gfap | -1.588 | 6.75E-03 | glio |
| culture_effect | Fam192a | -0.39 | 6.75E-03 | glio |
| Interaction_genotype_Atr | Ephx1 | -3.225 | 6.75E-03 | glio |
| Interaction_genotype_Atr | 1700013F07Ri | -3.551 | 6.75E-03 | glio |
| Interaction_genotype_Atr | AV039307 | -3.308 | 6.75E-03 | glio |
| Interaction_genotype_Atr | D930019O06R | -3.375 | 6.75E-03 | glio |
| Interaction_genotype_Atr | Pdrg1 | 1.345 | 6.75E-03 | glio |
| Interaction_genotype_Myc | Snhg4.1 | -2.6 | 6.75E-03 | glio |
| Interaction_genotype_Myc | Klk13 | -2.197 | 6.75E-03 | glio |
| Interaction_genotype_Myc | Zfp986 | -3.06 | 6.75E-03 | glio |
| Interaction_genotype_Myc | Zfp968 | -2.959 | 6.75E-03 | glio |
| Interaction_genotype_Rad51 | Tmem43 | 2.262 | 6.75E-03 | glio |
| Interaction_genotype_Rad51 | Crcp | 1.923 | 6.75E-03 | glio |
| interaction_RT | Pdcd10 | 0.658 | 6.75E-03 | glio |

| | | | | |
|---|---|---|---|---|
| Aars2_in.vivo_noRT | Kit | 1.509 | 6.75E-03 | glio |
| Bora_in.vivo_noRT | Dbn1 | 2.046 | 6.75E-03 | glio |
| Interaction_genotype_Mtor | Prelid3a | -3.928 | 6.75E-04 | glio |
| Interaction_genotype_Mtor | Mief2 | -3.705 | 6.75E-04 | glio |
| Interaction_genotype_Mtor | Stoml2 | 1.598 | 6.75E-04 | glio |
| Atr_in.vivo_noRT | Rab27b | 2.591 | 6.75E-04 | glio |
| Atr_in.vivo_noRT | Ttc9c | 1.099 | 6.75E-04 | glio |
| Bora_in.vivo_noRT | Mr1 | 2.504 | 6.75E-04 | glio |
| Bora_in.vivo_noRT | Icam4 | 2.372 | 6.75E-04 | glio |
| interaction_RT | Klf9 | 1.354 | 6.75E-05 | glio |
| interaction_RT | Ginm1 | 0.993 | 6.75E-05 | glio |
| Aars2_in.vivo_noRT | Insrr | 2.55 | 6.75E-05 | glio |
| Bcas2_in.vivo_noRT | Hapln4 | 3.04 | 6.75E-05 | glio |
| Bora_in.vivo_noRT | Gstm1 | 2.413 | 6.75E-05 | glio |
| Bora_ex.vivo_noRT | AF067061 | 3.201 | 6.75E-05 | glio |
| Chek1_in.vivo_noRT | Myl7 | 2.916 | 6.75E-05 | glio |
| Chek1_in.vivo_noRT | Fam171a2 | 2.881 | 6.75E-05 | glio |
| Interaction_genotype_Myc | Fgf2 | -4.923 | 6.75E-06 | glio |
| Interaction_genotype_Myc | Pik3r3 | -4.798 | 6.75E-06 | glio |
| Interaction_genotype_Myc | Apol7d | -5.329 | 6.75E-06 | glio |
| Bcas2_in.vivo_noRT | 4921507P07R | 3.043 | 6.75E-06 | glio |
| Aars2_in.vivo_noRT | Gm48684 | 4.022 | 6.75E-07 | glio |
| Atr_in.vivo_noRT | Gm16740 | 3.438 | 6.75E-07 | glio |
| Interaction_genotype_Myc | AC154200.1 | -5.042 | 6.75E-08 | glio |
| culture_effect | Fxyd3 | -1.702 | 6.75E-09 | glio |
| culture_effect | Uqcr11 | 0.801 | 6.75E-10 | glio |
| Bora_in.vivo_noRT | Hdac9 | 4.67 | 6.75E-12 | glio |
| Aars2_in.vivo_noRT | Slc22a1 | 4.152 | 6.75E-14 | glio |
| Chek1_ex.vivo_noRT | Tpo | 4.842 | 6.75E-16 | glio |
| Interaction_genotype_Atr | Fam178b | -3.436 | 6.76E-03 | glio |
| Interaction_genotype_Atr | Gnl3 | 1.333 | 6.76E-03 | glio |
| Interaction_genotype_Mtor | Elf3 | -3.653 | 6.76E-03 | glio |
| Interaction_genotype_Mtor | Zfp458 | -3.345 | 6.76E-03 | glio |
| Interaction_genotype_Mtor | Cog8 | -2.533 | 6.76E-03 | glio |
| Interaction_genotype_Mtor | Ivd | -2.636 | 6.76E-03 | glio |
| Aars2_in.vivo_noRT | Zfp981 | 1.916 | 6.76E-03 | glio |
| Aars2_in.vivo_noRT | Ruvbl1 | -0.93 | 6.76E-03 | glio |
| Bard1_ex.vivo_noRT | Gm38393 | 2.421 | 6.76E-03 | glio |
| Bora_in.vivo_noRT | Snx32 | 1.891 | 6.76E-03 | glio |
| Bora_ex.vivo_noRT | Atp8a2 | 2.414 | 6.76E-03 | glio |
| Cdkn2a_in.vivo_noRT | Gm48155 | -1.675 | 6.76E-03 | glio |
| Hsd17b10_in.vivo_noRT | Fsbp | 2.172 | 6.76E-03 | glio |
| Hsd17b10_in.vivo_noRT | Gm16845 | 2.008 | 6.76E-03 | glio |
| culture_effect | Mitd1 | 0.554 | 6.76E-04 | glio |
| interaction_RT | Tnfaip3 | 2.781 | 6.76E-04 | glio |
| interaction_RT | Serpini1 | 1.896 | 6.76E-04 | glio |
| Eif2s1_in.vivo_noRT | Gm5286 | 2.143 | 6.76E-04 | glio |
| Eif2s1_in.vivo_noRT | Gm34776 | 2.143 | 6.76E-04 | glio |
| Atr_ex.vivo_noRT | Pla2g2e | 4.087 | 6.76E-05 | glio |
| Chek1_in.vivo_noRT | Gm36738 | 3.035 | 6.76E-05 | glio |
| Chek1_ex.vivo_noRT | Ddit3 | 1.696 | 6.76E-05 | glio |
| Hsd17b10_in.vivo_noRT | Gm28592 | 2.715 | 6.76E-05 | glio |

| | | | | |
|---|---|---|---|---|
| Interaction_genotype_Mtor | Arhgef19 | -5.39 | 6.76E-06 | glio |
| Interaction_genotype_Mtor | Gm26850 | -5.382 | 6.76E-06 | glio |
| Interaction_genotype_Mtor | Gm42984 | -5.393 | 6.76E-06 | glio |
| Interaction_genotype_Mtor | Snrpn | -4.241 | 6.76E-06 | glio |
| Interaction_genotype_Mtor | 2310003N18R | -4.492 | 6.76E-06 | glio |
| Atr_in.vivo_noRT | Gm14023 | 3.255 | 6.76E-06 | glio |
| Bora_in.vivo_noRT | Mrpl44 | -1.94 | 6.76E-06 | glio |
| culture_effect | Mcts2 | 0.952 | 6.76E-07 | glio |
| Hsd17b10_in.vivo_noRT | Gm16685 | 2.958 | 6.76E-07 | glio |
| culture_effect | Dhx36 | 0.61 | 6.76E-08 | glio |
| Atr_in.vivo_noRT | Jakmip1 | 4.213 | 6.76E-10 | glio |
| culture_effect | Psmb5 | -0.756 | 6.76E-11 | glio |
| Chek1_in.vivo_noRT | Gm32090 | 4.252 | 6.76E-11 | glio |
| Chek1_in.vivo_noRT | Zic4 | 4.78 | 6.76E-11 | glio |
| Atr_in.vivo_noRT | Necab3 | 4.608 | 6.76E-12 | glio |
| culture_effect | Cdkn1b | -1.473 | 6.76E-14 | glio |
| Chek1_in.vivo_noRT | Gm13964 | 5.044 | 6.76E-24 | glio |
| Interaction_genotype_Aars2 | Gm39929 | -2.739 | 6.77E-03 | glio |
| Interaction_genotype_Aars2 | Mir155hg | -3.715 | 6.77E-03 | glio |
| Interaction_genotype_Aars2 | A730017L22Ri | -2.95 | 6.77E-03 | glio |
| Interaction_genotype_Atr | 2810408A11R | -3.467 | 6.77E-03 | glio |
| Interaction_genotype_Bora | Mpz | -4.382 | 6.77E-03 | glio |
| Interaction_genotype_Mtor | Clip1 | -2.14 | 6.77E-03 | glio |
| Interaction_genotype_Myc | Fez2 | -2.368 | 6.77E-03 | glio |
| Interaction_genotype_Myc | Zfp319 | -2.04 | 6.77E-03 | glio |
| Aars2_in.vivo_noRT | Idi1 | -1.525 | 6.77E-03 | glio |
| Atp5c1_in.vivo_noRT | Gm5152 | 2.281 | 6.77E-03 | glio |
| Bcas2_in.vivo_noRT | Doc2g | 2.07 | 6.77E-03 | glio |
| Cdkn2a_in.vivo_noRT | Cfh | -1.928 | 6.77E-03 | glio |
| Chek1_in.vivo_noRT | Syngr4 | 2.003 | 6.77E-03 | glio |
| Chek1_ex.vivo_noRT | Hnrnpr | -0.976 | 6.77E-03 | glio |
| Interaction_genotype_Atr | D430040D24F | -4.174 | 6.77E-04 | glio |
| Interaction_genotype_Atr | 9630014M24F | -4.087 | 6.77E-04 | glio |
| Interaction_genotype_Atr | Cfap100 | -3.612 | 6.77E-04 | glio |
| Interaction_genotype_Atr | Spsb2 | 1.941 | 6.77E-04 | glio |
| Interaction_genotype_Mtor | C8g | -3.998 | 6.77E-04 | glio |
| Interaction_genotype_Mtor | Psmc5 | 1.599 | 6.77E-04 | glio |
| Interaction_genotype_Mtor | Dynll1 | 1.574 | 6.77E-04 | glio |
| Interaction_genotype_Myc | Gm49223 | -3.188 | 6.77E-04 | glio |
| Interaction_genotype_Myc | Nectin3 | -2.824 | 6.77E-04 | glio |
| Interaction_genotype_Myc | Rev1 | -2.639 | 6.77E-04 | glio |
| Aars2_in.vivo_noRT | Zfp988 | 2.431 | 6.77E-04 | glio |
| Aars2_ex.vivo_noRT | Nt5e | 2.9 | 6.77E-04 | glio |
| Bora_in.vivo_noRT | Lin9 | -1.524 | 6.77E-04 | glio |
| Bora_in.vivo_noRT | Ncapd3 | -1.555 | 6.77E-04 | glio |
| Interaction_genotype_Mtor | Gm21762 | -4.229 | 6.77E-05 | glio |
| Atp5c1_in.vivo_noRT | Minar2 | 2.408 | 6.77E-05 | glio |
| Atr_in.vivo_noRT | Idnk | -1.705 | 6.77E-05 | glio |
| Bcas2_in.vivo_noRT | Lpar4 | -2.147 | 6.77E-05 | glio |
| Hsd17b10_in.vivo_noRT | Stra8 | 3.093 | 6.77E-07 | glio |
| Atr_in.vivo_noRT | Gm47545 | 4.282 | 6.77E-12 | glio |
| Atr_in.vivo_noRT | Gm4027 | 5.533 | 6.77E-23 | glio |

| | | | | |
|---|---|---|---|---|
| Interaction_genotype_Bora | Camk2n2 | -2.824 | 6.78E-03 | glio |
| Interaction_genotype_Cdkn2a | Gm48764 | 2.835 | 6.78E-03 | glio |
| Interaction_genotype_Mtor | Gpr141 | -3.922 | 6.78E-03 | glio |
| Interaction_genotype_Mtor | Ogg1 | -2.073 | 6.78E-03 | glio |
| Interaction_genotype_Pars2 | 9130604C24R | -3.938 | 6.78E-03 | glio |
| Atr_in.vivo_noRT | Adhfe1 | 1.964 | 6.78E-03 | glio |
| Atr_in.vivo_noRT | Fkbp1a | -0.577 | 6.78E-03 | glio |
| Bora_in.vivo_noRT | Prnp | -1.142 | 6.78E-03 | glio |
| Chek1_ex.vivo_noRT | Cdca3 | -1.373 | 6.78E-03 | glio |
| culture_effect | 2810049E08Ri | -1.972 | 6.78E-04 | glio |
| Aars2_ex.vivo_noRT | Gm13054 | 2.897 | 6.78E-04 | glio |
| culture_effect | Rhbdf1 | 0.816 | 6.78E-05 | glio |
| Interaction_genotype_Myc | Sos2 | -3.235 | 6.78E-05 | glio |
| Chek1_ex.vivo_noRT | Slc17a9 | 3.48 | 6.78E-05 | glio |
| Aars2_ex.vivo_noRT | Akr1c18 | 3.293 | 6.78E-06 | glio |
| Aars2_ex.vivo_noRT | Minar2 | 3.293 | 6.78E-06 | glio |
| Aars2_ex.vivo_noRT | Gm45151 | 3.293 | 6.78E-06 | glio |
| Atr_ex.vivo_noRT | 4930570E03Ri | 3.164 | 6.78E-06 | glio |
| Bcas2_in.vivo_noRT | Gm15510 | 3.212 | 6.78E-06 | glio |
| Bcas2_in.vivo_noRT | Kctd14 | 2.883 | 6.78E-06 | glio |
| Bora_in.vivo_noRT | Ccnj | 3.287 | 6.78E-06 | glio |
| Interaction_genotype_Mtor | Cox6b1 | 2.354 | 6.78E-07 | glio |
| Atr_in.vivo_noRT | Phldb3 | 3.485 | 6.78E-07 | glio |
| Chek1_in.vivo_noRT | Gm30790 | 3.528 | 6.78E-07 | glio |
| Aars2_in.vivo_noRT | Olfr1369-ps1 | 4.586 | 6.78E-08 | glio |
| culture_effect | Tesc | 1.216 | 6.78E-09 | glio |
| Bcas2_in.vivo_noRT | Prickle4 | 4.016 | 6.78E-10 | glio |
| Bora_in.vivo_noRT | Olfr279 | 4.407 | 6.78E-11 | glio |
| Chek1_in.vivo_noRT | Ripply3 | 5.812 | 6.78E-18 | glio |
| Interaction_genotype_Aars2 | Rnasek | 1.019 | 6.79E-03 | glio |
| Interaction_genotype_Chek1 | Gm47061 | -4.158 | 6.79E-03 | glio |
| Interaction_genotype_Chek1 | C2 | 3.222 | 6.79E-03 | glio |
| Interaction_genotype_Mtor | Atg14 | -2.707 | 6.79E-03 | glio |
| Interaction_genotype_Mtor | Fdps | 2.025 | 6.79E-03 | glio |
| Atp5c1_in.vivo_noRT | Gm37584 | 2.278 | 6.79E-03 | glio |
| Atr_in.vivo_noRT | Tbc1d2b | 1.902 | 6.79E-03 | glio |
| Atr_ex.vivo_noRT | Gm47223 | 3.539 | 6.79E-03 | glio |
| Bcas2_in.vivo_noRT | Tyw3 | 1.569 | 6.79E-03 | glio |
| Bcas2_in.vivo_noRT | Tpr | -0.638 | 6.79E-03 | glio |
| Bora_in.vivo_noRT | Hsh2d | 1.797 | 6.79E-03 | glio |
| Cenpj_ex.vivo_noRT | Fa2h | 2.591 | 6.79E-03 | glio |
| Chek1_in.vivo_noRT | Whrn | 2.008 | 6.79E-03 | glio |
| Chek1_in.vivo_noRT | Dpm3 | -1.198 | 6.79E-03 | glio |
| culture_effect | Trpm3 | 1.975 | 6.79E-04 | glio |
| interaction_RT | Tk1 | -1.37 | 6.79E-04 | glio |
| interaction_RT | Cd72 | 3.295 | 6.79E-04 | glio |
| Bora_in.vivo_noRT | Bst1 | 2.297 | 6.79E-04 | glio |
| Chek1_in.vivo_noRT | Slc25a43 | 2.46 | 6.79E-04 | glio |
| Hsd17b10_in.vivo_noRT | Gm11491 | 2.574 | 6.79E-04 | glio |
| culture_effect | Flt1 | -1.122 | 6.79E-05 | glio |
| culture_effect | C1ql1 | 0.869 | 6.79E-05 | glio |
| Aars2_in.vivo_noRT | Nckipsd | 2.721 | 6.79E-05 | glio |

| | | | | |
|---|---|---|---|---|
| Aars2_in.vivo_noRT | Klf1 | 2.89 | 6.79E-05 | glio |
| Bcas2_in.vivo_noRT | Gpn1 | -1.823 | 6.79E-05 | glio |
| Hsd17b10_in.vivo_noRT | Gjb4 | 2.959 | 6.79E-05 | glio |
| Atr_in.vivo_noRT | Gm36266 | 3.494 | 6.79E-06 | glio |
| Chek1_ex.vivo_noRT | Lce1g | 4.051 | 6.79E-06 | glio |
| Chek1_ex.vivo_noRT | Slc23a3 | 3.871 | 6.79E-06 | glio |
| culture_effect | Sema6b | 2.869 | 6.79E-09 | glio |
| Bora_ex.vivo_noRT | Tpo | 3.994 | 6.79E-10 | glio |
| Chek1_ex.vivo_noRT | Serpina3g | 4.842 | 6.79E-10 | glio |
| Atr_in.vivo_noRT | Grm1 | 4.429 | 6.79E-13 | glio |
| Bora_in.vivo_noRT | A930002C04R | 4.676 | 6.79E-22 | glio |
| culture_effect | Ngb | -1.705 | 6.80E-03 | glio |
| culture_effect | Lrig3 | -0.938 | 6.80E-03 | glio |
| culture_effect | Hinfp | -0.724 | 6.80E-03 | glio |
| Interaction_genotype_Atr | Hoga1 | -3.388 | 6.80E-03 | glio |
| Interaction_genotype_Mtor | Jtb | 1.271 | 6.80E-03 | glio |
| Interaction_genotype_Myc | Obscn | -2.729 | 6.80E-03 | glio |
| Interaction_genotype_Myc | Top3b | -2.492 | 6.80E-03 | glio |
| Interaction_genotype_Myc | Msto1 | -2.058 | 6.80E-03 | glio |
| Aars2_in.vivo_noRT | Sult2b1 | 1.635 | 6.80E-03 | glio |
| Aars2_in.vivo_noRT | Ttc7 | 1.598 | 6.80E-03 | glio |
| Atp5c1_in.vivo_noRT | AV039307 | 2.277 | 6.80E-03 | glio |
| Atr_in.vivo_noRT | Gvin1 | 1.897 | 6.80E-03 | glio |
| Atr_in.vivo_noRT | Ccdc174 | -1.173 | 6.80E-03 | glio |
| Bcas2_ex.vivo_noRT | Mag | 2.742 | 6.80E-03 | glio |
| Bcas2_ex.vivo_noRT | Best1 | 2.618 | 6.80E-03 | glio |
| Bora_in.vivo_noRT | Wars2 | 1.716 | 6.80E-03 | glio |
| Bora_in.vivo_noRT | Cct6a | -0.636 | 6.80E-03 | glio |
| Chek1_in.vivo_noRT | Dner | 1.844 | 6.80E-03 | glio |
| Chek1_in.vivo_noRT | Pskh1 | 1.824 | 6.80E-03 | glio |
| Fanca_ex.vivo_noRT | Ablim2 | 2.124 | 6.80E-03 | glio |
| Fanca_ex.vivo_noRT | Gm5415 | 2.124 | 6.80E-03 | glio |
| Fanca_ex.vivo_noRT | Nrip2 | 2.124 | 6.80E-03 | glio |
| Fanca_ex.vivo_noRT | Brsk2 | 2.913 | 6.80E-03 | glio |
| Fanca_ex.vivo_noRT | Gm49698 | 2.124 | 6.80E-03 | glio |
| Fanca_ex.vivo_noRT | Olfr881 | 2.124 | 6.80E-03 | glio |
| Interaction_genotype_Mtor | Pla2g4b | -3.991 | 6.80E-04 | glio |
| Interaction_genotype_Mtor | Tpm3 | 1.655 | 6.80E-04 | glio |
| Interaction_genotype_Myc | Gtpbp8 | -3.212 | 6.80E-04 | glio |
| Interaction_genotype_Myc | Dzip1l | -2.805 | 6.80E-04 | glio |
| Interaction_genotype_Myc | Imp3 | 1.689 | 6.80E-04 | glio |
| interaction_RT | 4930581F22Ri | -1.779 | 6.80E-04 | glio |
| interaction_RT | Ttc39c | 1.504 | 6.80E-04 | glio |
| interaction_RT | Tnrc6b | 0.87 | 6.80E-04 | glio |
| Chek1_in.vivo_noRT | Itgal | 2.573 | 6.80E-04 | glio |
| Interaction_genotype_Bcas2 | Dnm3 | -4.621 | 6.80E-05 | glio |
| Interaction_genotype_Bcas2 | AV039307 | -5.317 | 6.80E-05 | glio |
| Interaction_genotype_Mtor | Gm2788 | -4.981 | 6.80E-05 | glio |
| Interaction_genotype_Myc | Aig1 | 1.562 | 6.80E-05 | glio |
| Interaction_genotype_Rpl3 | 1700001G11R | -4.564 | 6.80E-05 | glio |
| Atp5c1_in.vivo_noRT | Pdgfb | 2.405 | 6.80E-05 | glio |
| Bcas2_ex.vivo_noRT | Cd200r2 | 2.742 | 6.80E-05 | glio |

| | | | | |
|---|---|---|---|---|
| Bcas2_ex.vivo_noRT | Slc2a13 | 3.664 | 6.80E-05 | glio |
| Interaction_genotype_Mtor | Fgf8 | -4.693 | 6.80E-06 | glio |
| interaction_RT | Hmga2 | -1.11 | 6.80E-06 | glio |
| Chek1_in.vivo_noRT | 2310030G06R | 3.203 | 6.80E-06 | glio |
| Interaction_genotype_Mtor | Slc26a6 | -5.779 | 6.80E-07 | glio |
| Bora_in.vivo_noRT | 4930579K19R | 3.698 | 6.80E-07 | glio |
| Hsd17b10_in.vivo_noRT | Npnt | 3.35 | 6.80E-11 | glio |
| culture_effect | Arf4 | -0.916 | 6.80E-14 | glio |
| culture_effect | Rpl23a | -1.104 | 6.80E-18 | glio |
| Atr_in.vivo_noRT | 1700006H21R | 6.037 | 6.80E-18 | glio |
| Bora_in.vivo_noRT | Ltbp2 | 4.936 | 6.80E-19 | glio |
| Bora_in.vivo_noRT | Lmx1a | 4.936 | 6.80E-19 | glio |
| Bcas2_in.vivo_noRT | Gm11232 | 4.807 | 6.80E-23 | glio |
| Interaction_genotype_Aars2 | Tecr | 1.257 | 6.81E-03 | glio |
| Interaction_genotype_Bora | Gpr179 | -3.502 | 6.81E-03 | glio |
| Interaction_genotype_Bora | Gm41442 | -3.488 | 6.81E-03 | glio |
| Interaction_genotype_Bora | 4833418N02R | -3.709 | 6.81E-03 | glio |
| Interaction_genotype_Bora | Mbd2 | 1.674 | 6.81E-03 | glio |
| Interaction_genotype_Mtor | Prrg1 | -3.417 | 6.81E-03 | glio |
| Interaction_genotype_Mtor | Tmprss5 | -3.388 | 6.81E-03 | glio |
| Interaction_genotype_Mtor | Kctd6 | -2.995 | 6.81E-03 | glio |
| Interaction_genotype_Mtor | Ipo4 | 1.601 | 6.81E-03 | glio |
| Aars2_in.vivo_noRT | Tada2b | 1.271 | 6.81E-03 | glio |
| Atr_in.vivo_noRT | Ap5s1 | 1.395 | 6.81E-03 | glio |
| Bcas2_in.vivo_noRT | Acvr1b | 2.004 | 6.81E-03 | glio |
| Bcas2_in.vivo_noRT | Nudt2 | -1.186 | 6.81E-03 | glio |
| Bcas2_in.vivo_noRT | Rfc2 | -1.174 | 6.81E-03 | glio |
| Bora_in.vivo_noRT | Dnaja2 | -0.766 | 6.81E-03 | glio |
| Chek1_in.vivo_noRT | Rpap2 | 1.488 | 6.81E-03 | glio |
| Chek1_ex.vivo_noRT | Kmt5a | -0.915 | 6.81E-03 | glio |
| Interaction_genotype_Aars2 | Cpne8 | -4.386 | 6.81E-04 | glio |
| Interaction_genotype_Aars2 | Rab3b | -4.17 | 6.81E-04 | glio |
| Interaction_genotype_Atr | Zfp810 | -4.151 | 6.81E-04 | glio |
| Interaction_genotype_Atr | Man2c1os | -4.168 | 6.81E-04 | glio |
| Interaction_genotype_Atr | Mien1 | 1.426 | 6.81E-04 | glio |
| Interaction_genotype_Bora | 1110046J04Ri | -4.547 | 6.81E-04 | glio |
| Aars2_in.vivo_noRT | Hist1h2ac | 2.37 | 6.81E-04 | glio |
| Aars2_in.vivo_noRT | Nipal3 | 2.377 | 6.81E-04 | glio |
| Bcas2_in.vivo_noRT | Gm26737 | 3.704 | 6.81E-04 | glio |
| Chek1_ex.vivo_noRT | Egln3 | 3.032 | 6.81E-04 | glio |
| Interaction_genotype_Mtor | 2310030G06R | -4.417 | 6.81E-05 | glio |
| Interaction_genotype_Mtor | Dmrtc1a | -4.099 | 6.81E-06 | glio |
| Interaction_genotype_Myc | Gm50147 | -4.995 | 6.81E-06 | glio |
| Hsd17b10_in.vivo_noRT | Gm10425 | 3.352 | 6.81E-06 | glio |
| culture_effect | Spink4 | -3.679 | 6.81E-10 | glio |
| Chek1_ex.vivo_noRT | Gm13191 | 4.842 | 6.81E-18 | glio |
| Chek1_ex.vivo_noRT | 4933416E03Ri | 4.842 | 6.81E-18 | glio |
| Atr_in.vivo_noRT | Gm13964 | 5.533 | 6.81E-28 | glio |
| culture_effect | Gtf3c5 | -0.9 | 6.82E-03 | glio |
| culture_effect | Gnai3 | -0.444 | 6.82E-03 | glio |
| culture_effect | Bnip3l | -0.418 | 6.82E-03 | glio |
| culture_effect | Ptma | -0.363 | 6.82E-03 | glio |

| | | | | |
|---|---|---|---|---|
| Interaction_genotype_Aars2 | 4930532G15R | -3.046 | 6.82E-03 | glio |
| Interaction_genotype_Abtb1 | B230303O12R | 4.647 | 6.82E-03 | glio |
| Interaction_genotype_Atr | AW046200 | -3.398 | 6.82E-03 | glio |
| Interaction_genotype_Mtor | Zkscan4 | -3.288 | 6.82E-03 | glio |
| Interaction_genotype_Myc | Suox | -2.752 | 6.82E-03 | glio |
| Aars2_in.vivo_noRT | Pou3f1 | 1.967 | 6.82E-03 | glio |
| Aars2_in.vivo_noRT | Fam241a | 2.03 | 6.82E-03 | glio |
| Aars2_in.vivo_noRT | Psmc2 | -0.719 | 6.82E-03 | glio |
| Atr_ex.vivo_noRT | Mob3b | -2.274 | 6.82E-03 | glio |
| Bcas2_ex.vivo_noRT | Fgf8 | 2.431 | 6.82E-03 | glio |
| Cenpj_in.vivo_noRT | Gm49086 | 2.022 | 6.82E-03 | glio |
| Cenpj_in.vivo_noRT | Platr20 | 2.534 | 6.82E-03 | glio |
| Eif2s1_in.vivo_noRT | Gm14264 | 2.203 | 6.82E-03 | glio |
| Interaction_genotype_Mtor | Tigd5 | -3.722 | 6.82E-04 | glio |
| Interaction_genotype_Mtor | Gm10053 | -3.307 | 6.82E-04 | glio |
| Interaction_genotype_Mtor | Ube2d2a | 1.452 | 6.82E-04 | glio |
| Interaction_genotype_Myc | Axdnd1 | -4.226 | 6.82E-04 | glio |
| Interaction_genotype_Pars2 | Gm11973 | -3.766 | 6.82E-04 | glio |
| Interaction_genotype_Pars2 | Gm50147 | -4.534 | 6.82E-04 | glio |
| Atp5c1_in.vivo_noRT | Gm41349 | 2.405 | 6.82E-04 | glio |
| Bard1_ex.vivo_noRT | Gm15138 | 2.419 | 6.82E-04 | glio |
| Chek1_ex.vivo_noRT | Kcnk4 | 3.079 | 6.82E-04 | glio |
| Chek1_ex.vivo_noRT | Hist2h3c2 | 3.18 | 6.82E-04 | glio |
| Chek1_ex.vivo_noRT | Timm8a1 | -1.421 | 6.82E-04 | glio |
| culture_effect | Polr2a | 0.642 | 6.82E-05 | glio |
| Interaction_genotype_Myc | Zfp652os | -4.383 | 6.82E-05 | glio |
| Atp5c1_in.vivo_noRT | Gm45235 | 2.408 | 6.82E-05 | glio |
| Bard1_ex.vivo_noRT | Gm9758 | 2.814 | 6.82E-05 | glio |
| Bard1_ex.vivo_noRT | AF067063 | 2.814 | 6.82E-05 | glio |
| Bard1_ex.vivo_noRT | Acod1 | 4.419 | 6.82E-05 | glio |
| Hsd17b10_in.vivo_noRT | Plcd4 | 2.705 | 6.82E-05 | glio |
| Aars2_in.vivo_noRT | Icam4 | 3.037 | 6.82E-06 | glio |
| Atr_ex.vivo_noRT | Pspn | 2.769 | 6.82E-06 | glio |
| Bcas2_ex.vivo_noRT | Sucnr1 | 2.742 | 6.82E-06 | glio |
| Atr_in.vivo_noRT | Cdk4 | -1.343 | 6.82E-07 | glio |
| Bora_in.vivo_noRT | Pld6 | 3.68 | 6.82E-07 | glio |
| Bora_in.vivo_noRT | Chd4 | -1.418 | 6.82E-07 | glio |
| Bora_ex.vivo_noRT | Angptl7 | 3.201 | 6.82E-07 | glio |
| Bcas2_in.vivo_noRT | Gm47428 | 3.631 | 6.82E-08 | glio |
| Atr_in.vivo_noRT | B930059L03Ri | 4.349 | 6.82E-09 | glio |
| Chek1_ex.vivo_noRT | 4930434B07R | 4.264 | 6.82E-09 | glio |
| Bora_in.vivo_noRT | AI854703 | 4.286 | 6.82E-10 | glio |
| Bora_in.vivo_noRT | Gm28731 | 4.55 | 6.82E-10 | glio |
| Aars2_in.vivo_noRT | 4930555A03R | 4.589 | 6.82E-13 | glio |
| Bcas2_in.vivo_noRT | Slc25a18 | 4.541 | 6.82E-15 | glio |
| Chek1_in.vivo_noRT | Pmp2 | 5.044 | 6.82E-16 | glio |
| Chek1_in.vivo_noRT | B230206H07R | 4.518 | 6.82E-21 | glio |
| Interaction_genotype_Atr | Capn9 | -3.05 | 6.83E-03 | glio |
| Interaction_genotype_Atr | Gm48933 | -3.172 | 6.83E-03 | glio |
| Interaction_genotype_Atr | A930029G22R | -2.957 | 6.83E-03 | glio |
| Interaction_genotype_Bora | Abcb5 | -2.529 | 6.83E-03 | glio |
| Interaction_genotype_Bora | Gm45591 | -3.057 | 6.83E-03 | glio |

| | | | | |
|---|---|---|---|---|
| Interaction_genotype_Bora | Aass | -3.443 | 6.83E-03 | glio |
| Interaction_genotype_Bora | Wincr1 | -4.459 | 6.83E-03 | glio |
| Interaction_genotype_Myc | Gm44777 | -2.877 | 6.83E-03 | glio |
| Interaction_genotype_Myc | Ring1 | -2.317 | 6.83E-03 | glio |
| Interaction_genotype_Myc | Prkce | -2.111 | 6.83E-03 | glio |
| Interaction_genotype_Myc | Map1lc3b | 1.17 | 6.83E-03 | glio |
| Interaction_genotype_Rpl3 | Gm14419 | -3.252 | 6.83E-03 | glio |
| Interaction_genotype_Rpl3 | Scube3 | -3.019 | 6.83E-03 | glio |
| Aars2_in.vivo_noRT | Lars2 | -1.636 | 6.83E-03 | glio |
| Bora_ex.vivo_noRT | Gm43387 | 2.796 | 6.83E-03 | glio |
| Bora_ex.vivo_noRT | Bcl3 | 2.589 | 6.83E-03 | glio |
| Interaction_genotype_Myc | Wls | 1.915 | 6.83E-04 | glio |
| Aars2_in.vivo_noRT | Prdm1 | 2.26 | 6.83E-04 | glio |
| Bcas2_in.vivo_noRT | Plekho2 | 2.402 | 6.83E-04 | glio |
| Bora_ex.vivo_noRT | Gm13684 | 2.808 | 6.83E-04 | glio |
| Bora_ex.vivo_noRT | Myzap | 2.802 | 6.83E-04 | glio |
| Atr_in.vivo_noRT | Flrt2 | 2.871 | 6.83E-05 | glio |
| Aars2_in.vivo_noRT | Sardh | 3.04 | 6.83E-06 | glio |
| Atr_in.vivo_noRT | Maob | 3.32 | 6.83E-07 | glio |
| Bcas2_in.vivo_noRT | 1700030C12R | 3.882 | 6.83E-08 | glio |
| Chek1_in.vivo_noRT | Map3k9 | 4.784 | 6.83E-17 | glio |
| interaction_RT | Gm26981 | -2.54 | 6.84E-03 | glio |
| Atr_in.vivo_noRT | Cryzl1 | -1.197 | 6.84E-03 | glio |
| Bcas2_in.vivo_noRT | Huwe1 | -0.808 | 6.84E-03 | glio |
| Bora_in.vivo_noRT | Col4a5 | -1.359 | 6.84E-03 | glio |
| Chek1_in.vivo_noRT | Tro | 1.956 | 6.84E-03 | glio |
| Fanca_ex.vivo_noRT | BC061195 | 2.124 | 6.84E-03 | glio |
| Fanca_ex.vivo_noRT | Idi2 | 2.124 | 6.84E-03 | glio |
| Fanca_ex.vivo_noRT | Gm364 | 2.124 | 6.84E-03 | glio |
| Fanca_ex.vivo_noRT | Vmn2r18 | 2.124 | 6.84E-03 | glio |
| Interaction_genotype_Aars2 | Gm16183 | -4.533 | 6.84E-04 | glio |
| Interaction_genotype_Aars2 | Dap | 1.959 | 6.84E-04 | glio |
| Interaction_genotype_Atr | C230013L11Ri | -4.504 | 6.84E-04 | glio |
| Chek1_in.vivo_noRT | Iapp | 2.881 | 6.84E-04 | glio |
| Chek1_in.vivo_noRT | Zfp600 | 2.436 | 6.84E-04 | glio |
| Chek1_in.vivo_noRT | Arf5 | -0.772 | 6.84E-04 | glio |
| Eif2s1_ex.vivo_noRT | Gm39526 | 3.032 | 6.84E-04 | glio |
| Interaction_genotype_Aars2 | Gm42997 | -4.746 | 6.84E-05 | glio |
| Interaction_genotype_Mtor | Phb2 | 1.898 | 6.84E-05 | glio |
| Chek1_ex.vivo_noRT | Hsf2bp | 3.281 | 6.84E-05 | glio |
| Chek1_ex.vivo_noRT | Batf | 3.754 | 6.84E-05 | glio |
| Interaction_genotype_Myc | Gm37933 | -5.025 | 6.84E-07 | glio |
| Aars2_ex.vivo_noRT | Gm13032 | 3.693 | 6.84E-07 | glio |
| Aars2_ex.vivo_noRT | Cd36 | 3.293 | 6.84E-07 | glio |
| Aars2_ex.vivo_noRT | Gstt4 | 3.293 | 6.84E-07 | glio |
| Bora_ex.vivo_noRT | Cldn5 | 3.201 | 6.84E-07 | glio |
| Chek1_in.vivo_noRT | 1110035H17R | 3.876 | 6.84E-07 | glio |
| Aars2_in.vivo_noRT | 6330415G19R | 4.02 | 6.84E-09 | glio |
| Chek1_in.vivo_noRT | Mro | 4.252 | 6.84E-10 | glio |
| culture_effect | Gm31518 | 1.69 | 6.85E-03 | glio |
| Interaction_genotype_Aars2 | Mpp6 | 1.354 | 6.85E-03 | glio |
| Interaction_genotype_Atr | Gm5788 | -2.369 | 6.85E-03 | glio |

| | | | | |
|---|---|---|---|---|
| Interaction_genotype_Atr | Gm37800 | -3.064 | 6.85E-03 | glio |
| Interaction_genotype_Atr | 2310067P03R | -3.229 | 6.85E-03 | glio |
| Interaction_genotype_Atr | Pstk | -1.662 | 6.85E-03 | glio |
| Interaction_genotype_Atr | Tomm20 | 0.837 | 6.85E-03 | glio |
| Interaction_genotype_Mtor | Ptprn | -2.996 | 6.85E-03 | glio |
| Interaction_genotype_Mtor | Mecp2 | 1.548 | 6.85E-03 | glio |
| Interaction_genotype_Mtor | Rps13 | 0.842 | 6.85E-03 | glio |
| Interaction_genotype_Myc | Pcid2 | -2.136 | 6.85E-03 | glio |
| Interaction_genotype_Myc | Nsrp1 | 1.24 | 6.85E-03 | glio |
| interaction_RT | Mb | 2.572 | 6.85E-03 | glio |
| Aars2_in.vivo_noRT | Gm38037 | 1.912 | 6.85E-03 | glio |
| Aars2_in.vivo_noRT | Prss56 | 2.063 | 6.85E-03 | glio |
| Aars2_in.vivo_noRT | Hmgn2 | -1.239 | 6.85E-03 | glio |
| Atr_in.vivo_noRT | A230072C01R | 1.778 | 6.85E-03 | glio |
| Atr_ex.vivo_noRT | Pa2g4 | -0.882 | 6.85E-03 | glio |
| Bcas2_in.vivo_noRT | Mmp11 | 2.062 | 6.85E-03 | glio |
| Bcas2_in.vivo_noRT | Klhdc2 | -0.942 | 6.85E-03 | glio |
| Bora_in.vivo_noRT | Cdh2 | -1.378 | 6.85E-03 | glio |
| Bora_ex.vivo_noRT | Gm35551 | 2.615 | 6.85E-03 | glio |
| Cenpj_in.vivo_noRT | AC159140.1 | 1.762 | 6.85E-03 | glio |
| Chek1_ex.vivo_noRT | Ckap2l | -1.998 | 6.85E-03 | glio |
| Interaction_genotype_Aars2 | Ubd | 4.078 | 6.85E-04 | glio |
| Interaction_genotype_Aars2 | Gm37800 | -3.901 | 6.85E-04 | glio |
| Interaction_genotype_Aars2 | Gm45716 | -3.9 | 6.85E-04 | glio |
| Interaction_genotype_Aars2 | Tmem74b | -3.534 | 6.85E-04 | glio |
| Interaction_genotype_Myc | Hemk1 | -3.849 | 6.85E-04 | glio |
| Interaction_genotype_Myc | Srsf2 | 1.429 | 6.85E-04 | glio |
| Aars2_in.vivo_noRT | Hoxaas3 | 2.023 | 6.85E-04 | glio |
| Atr_in.vivo_noRT | Rcc2 | -1.044 | 6.85E-04 | glio |
| Bcas2_in.vivo_noRT | Trim71 | 2.448 | 6.85E-04 | glio |
| Chek1_ex.vivo_noRT | Gm50375 | 3.282 | 6.85E-04 | glio |
| Interaction_genotype_Mtor | Hist1h2ag | -3.567 | 6.85E-05 | glio |
| Bcas2_in.vivo_noRT | Prkag1 | -1.697 | 6.85E-05 | glio |
| Interaction_genotype_Mtor | A430106G13R | -5.209 | 6.85E-06 | glio |
| Interaction_genotype_Mtor | Plekhj1 | 2.236 | 6.85E-07 | glio |
| Bcas2_in.vivo_noRT | Slc16a6 | 3.623 | 6.85E-07 | glio |
| Bcas2_in.vivo_noRT | Fsd2 | 3.424 | 6.85E-07 | glio |
| Chek1_in.vivo_noRT | Gm29358 | 3.571 | 6.85E-07 | glio |
| Aars2_in.vivo_noRT | Sap30bpos | 3.932 | 6.85E-11 | glio |
| Chek1_in.vivo_noRT | Slc45a3 | 4.522 | 6.85E-11 | glio |
| culture_effect | Romo1 | 1.121 | 6.85E-14 | glio |
| Bora_in.vivo_noRT | Abra | 4.411 | 6.85E-14 | glio |
| Atr_in.vivo_noRT | AC154200.1 | 5.533 | 6.85E-23 | glio |
| culture_effect | Fsd2 | -1.731 | 6.86E-03 | glio |
| culture_effect | Gm40040 | 1.584 | 6.86E-03 | glio |
| culture_effect | Wbp1l | 0.54 | 6.86E-03 | glio |
| Interaction_genotype_Aars2 | 1700016P04R | -3.163 | 6.86E-03 | glio |
| Interaction_genotype_Mtor | Cluap1 | -2.612 | 6.86E-03 | glio |
| Interaction_genotype_Mtor | Mrto4 | 1.365 | 6.86E-03 | glio |
| Aars2_in.vivo_noRT | A930024E05Ri | 1.951 | 6.86E-03 | glio |
| Atr_in.vivo_noRT | BC037039 | 1.644 | 6.86E-03 | glio |
| Atr_in.vivo_noRT | Gch1 | 1.723 | 6.86E-03 | glio |

| | | | | |
|---|---|---|---|---|
| Atr_in.vivo_noRT | Nudt6 | 1.621 | 6.86E-03 | glio |
| Bcas2_in.vivo_noRT | Ephx4 | -1.256 | 6.86E-03 | glio |
| Bora_in.vivo_noRT | Tmem88b | 1.593 | 6.86E-03 | glio |
| Cenpj_in.vivo_noRT | Gm48242 | 2.022 | 6.86E-03 | glio |
| Chek1_in.vivo_noRT | Olig1 | -1.314 | 6.86E-03 | glio |
| Chek1_in.vivo_noRT | Trir | -0.574 | 6.86E-03 | glio |
| Chek1_ex.vivo_noRT | Alcam | 2.205 | 6.86E-03 | glio |
| Eif2s1_in.vivo_noRT | Map3k9 | 1.883 | 6.86E-03 | glio |
| Eif2s1_in.vivo_noRT | Gm16222 | 2.656 | 6.86E-03 | glio |
| Interaction_genotype_Bora | Slc38a3 | 4.147 | 6.86E-04 | glio |
| Interaction_genotype_Bora | Alox12 | -4.463 | 6.86E-04 | glio |
| Interaction_genotype_Bora | Atad2 | 2.314 | 6.86E-04 | glio |
| Aars2_in.vivo_noRT | Hist2h3b | 2.417 | 6.86E-04 | glio |
| Atr_in.vivo_noRT | Ccdc62 | 2.25 | 6.86E-04 | glio |
| Chek1_in.vivo_noRT | Usp35 | 2.446 | 6.86E-04 | glio |
| culture_effect | Cldn19 | 2.164 | 6.86E-05 | glio |
| culture_effect | Zfp652os | 2.115 | 6.86E-05 | glio |
| Bcas2_in.vivo_noRT | B230317F23Ri | 2.765 | 6.86E-05 | glio |
| culture_effect | Tnfrsf22 | 2.032 | 6.86E-06 | glio |
| Bora_in.vivo_noRT | Efna2 | 3.664 | 6.86E-06 | glio |
| Bora_in.vivo_noRT | Emid1 | 3.544 | 6.86E-06 | glio |
| Bora_in.vivo_noRT | Cracr2b | 3.335 | 6.86E-06 | glio |
| culture_effect | Trim7 | -2.989 | 6.86E-08 | glio |
| culture_effect | Itpripl2 | 1.266 | 6.86E-08 | glio |
| Hsd17b10_in.vivo_noRT | Cpxm1 | 3.35 | 6.86E-08 | glio |
| Chek1_in.vivo_noRT | Gm20442 | 4.035 | 6.86E-11 | glio |
| Hsd17b10_in.vivo_noRT | Meioc | 3.353 | 6.86E-12 | glio |
| Hsd17b10_in.vivo_noRT | Gm48215 | 3.616 | 6.86E-12 | glio |
| Bora_in.vivo_noRT | Iqcf1 | 4.816 | 6.86E-13 | glio |
| Atr_in.vivo_noRT | Lrguk | 5.269 | 6.86E-20 | glio |
| Aars2_in.vivo_noRT | Apoa5 | 5.841 | 6.86E-32 | glio |
| culture_effect | Depdc7 | -3.202 | 6.86E-35 | glio |
| culture_effect | Gm34552 | 1.438 | 6.87E-03 | glio |
| Interaction_genotype_Myc | Gm16174 | -3.306 | 6.87E-03 | glio |
| Interaction_genotype_Myc | Hipk4 | -2.822 | 6.87E-03 | glio |
| Interaction_genotype_Myc | Cd68 | -3.347 | 6.87E-03 | glio |
| Interaction_genotype_Pars2 | Gm41307 | -2.661 | 6.87E-03 | glio |
| Interaction_genotype_Terf1 | Cebpg | 1.529 | 6.87E-03 | glio |
| interaction_RT | Aida | -0.656 | 6.87E-03 | glio |
| Aars2_in.vivo_noRT | Xpa | -1.174 | 6.87E-03 | glio |
| Atr_in.vivo_noRT | Ap1s3 | 1.925 | 6.87E-03 | glio |
| Atr_in.vivo_noRT | Atp6v1g1 | -0.672 | 6.87E-03 | glio |
| Chek1_in.vivo_noRT | A230072C01R | 1.812 | 6.87E-03 | glio |
| Chek1_ex.vivo_noRT | Pde4d | 2.579 | 6.87E-03 | glio |
| Fanca_ex.vivo_noRT | Gm16178 | 2.124 | 6.87E-03 | glio |
| Hsd17b10_in.vivo_noRT | 4930594M22F | 2.15 | 6.87E-03 | glio |
| Interaction_genotype_Aars2 | Notch4 | -4.183 | 6.87E-04 | glio |
| Interaction_genotype_Aars2 | Tlr1 | -4.226 | 6.87E-04 | glio |
| Interaction_genotype_Aars2 | Gm11696 | -4.309 | 6.87E-04 | glio |
| Interaction_genotype_Myc | Gm266 | -2.729 | 6.87E-04 | glio |
| Aars2_in.vivo_noRT | Kif7 | 2.266 | 6.87E-04 | glio |
| Bcas2_in.vivo_noRT | 4930563E22Ri | 2.559 | 6.87E-04 | glio |

| | | | | |
|---|---|---|---|---|
| Bcas2_in.vivo_noRT | Brd4 | -1.062 | 6.87E-04 | glio |
| Bora_in.vivo_noRT | Mrps34 | -1.249 | 6.87E-04 | glio |
| Eif2s1_in.vivo_noRT | A930002C04R | 1.883 | 6.87E-04 | glio |
| Eif2s1_in.vivo_noRT | Gm11457 | 2.67 | 6.87E-04 | glio |
| Hsd17b10_in.vivo_noRT | H2-M10.1 | 2.161 | 6.87E-04 | glio |
| Interaction_genotype_Mtor | Mpzl2 | -3.839 | 6.87E-05 | glio |
| Interaction_genotype_Mtor | Inca1 | -4.392 | 6.87E-05 | glio |
| Interaction_genotype_Mtor | A430035B10R | -4.932 | 6.87E-05 | glio |
| Interaction_genotype_Mtor | Uckl1os | -5.006 | 6.87E-05 | glio |
| Interaction_genotype_Mtor | 3110045C21R | -4.405 | 6.87E-05 | glio |
| Interaction_genotype_Mtor | Mfsd3 | -3.879 | 6.87E-05 | glio |
| Interaction_genotype_Myc | Lyrm1 | -4.339 | 6.87E-05 | glio |
| Interaction_genotype_Myc | Gm15545 | -4.322 | 6.87E-05 | glio |
| interaction_RT | Cct5 | -0.82 | 6.87E-06 | glio |
| Aars2_ex.vivo_noRT | Gast | 3.293 | 6.87E-06 | glio |
| Atr_in.vivo_noRT | Tctn1 | 3.494 | 6.87E-07 | glio |
| Bora_in.vivo_noRT | Rasgrp2 | 3.636 | 6.87E-07 | glio |
| Chek1_in.vivo_noRT | Best1 | 3.141 | 6.87E-07 | glio |
| Bora_in.vivo_noRT | Gm10425 | 4.672 | 6.87E-11 | glio |
| culture_effect | Unc80 | -1.978 | 6.87E-12 | glio |
| Bcas2_in.vivo_noRT | Gm45406 | 4.285 | 6.87E-16 | glio |
| culture_effect | 1700029J07Ri | 1.538 | 6.88E-03 | glio |
| culture_effect | Resf1 | -0.495 | 6.88E-03 | glio |
| culture_effect | Polr2m | 0.335 | 6.88E-03 | glio |
| culture_effect | Phb2 | -0.311 | 6.88E-03 | glio |
| Interaction_genotype_Mtor | Zfp52 | -2.758 | 6.88E-03 | glio |
| Interaction_genotype_Mtor | Tmem35b | -2.923 | 6.88E-03 | glio |
| Interaction_genotype_Mtor | Pms1 | -2.706 | 6.88E-03 | glio |
| Interaction_genotype_Mtor | Gm26881 | -2.619 | 6.88E-03 | glio |
| Interaction_genotype_Myc | Slc4a11 | -3.235 | 6.88E-03 | glio |
| Interaction_genotype_Rad51 | Stard10 | -3.8 | 6.88E-03 | glio |
| Interaction_genotype_Rpl3 | Rilp | -4.123 | 6.88E-03 | glio |
| Interaction_genotype_Rpl3 | Hmgxb3 | 2.171 | 6.88E-03 | glio |
| Bcas2_in.vivo_noRT | Zfp950 | -1.289 | 6.88E-03 | glio |
| Bcas2_in.vivo_noRT | Znrd1 | -0.761 | 6.88E-03 | glio |
| Bora_in.vivo_noRT | Pde6h | 1.862 | 6.88E-03 | glio |
| Chek1_in.vivo_noRT | Tmem17 | 1.709 | 6.88E-03 | glio |
| Interaction_genotype_Mtor | Angpt2 | -3.895 | 6.88E-04 | glio |
| Interaction_genotype_Myc | Acaca | -2.795 | 6.88E-04 | glio |
| Interaction_genotype_Xrcc5 | Scel | 3.572 | 6.88E-04 | glio |
| Aars2_in.vivo_noRT | Parvb | 2.315 | 6.88E-04 | glio |
| Aars2_ex.vivo_noRT | Ly6c2 | 3.293 | 6.88E-04 | glio |
| Bcas2_in.vivo_noRT | Map4 | -1.585 | 6.88E-04 | glio |
| Cenpj_in.vivo_noRT | Gm13398 | 2.022 | 6.88E-04 | glio |
| Aars2_in.vivo_noRT | Insr | 1.948 | 6.88E-05 | glio |
| Aars2_in.vivo_noRT | Krtcap2 | -0.923 | 6.88E-05 | glio |
| Atp5c1_in.vivo_noRT | Baalc | 2.928 | 6.88E-05 | glio |
| Atp5c1_in.vivo_noRT | Ninj2 | 3.052 | 6.88E-05 | glio |
| Aars2_in.vivo_noRT | 4732491K20R | 2.874 | 6.88E-06 | glio |
| Bcas2_in.vivo_noRT | Lamc2 | 3.159 | 6.88E-06 | glio |
| Aars2_in.vivo_noRT | Selp | 4.282 | 6.88E-10 | glio |
| Bora_in.vivo_noRT | 2310067P03R | 4.414 | 6.88E-11 | glio |

| | | | | |
|---|---|---|---|---|
| Chek1_in.vivo_noRT | Igfbp7 | 4.278 | 6.88E-12 | glio |
| Bora_in.vivo_noRT | Aass | 4.702 | 6.88E-13 | glio |
| Aars2_in.vivo_noRT | Dntt | 5.327 | 6.88E-14 | glio |
| Aars2_in.vivo_noRT | Matn4 | 5.063 | 6.88E-14 | glio |
| Chek1_in.vivo_noRT | Pla2g2d | 4.78 | 6.88E-14 | glio |
| Atr_in.vivo_noRT | 5033417F24Ri | 5.533 | 6.88E-16 | glio |
| Chek1_in.vivo_noRT | Gm50024 | 4.78 | 6.88E-22 | glio |
| culture_effect | Mfsd2a | -0.873 | 6.89E-03 | glio |
| culture_effect | Snf8 | 0.339 | 6.89E-03 | glio |
| Interaction_genotype_Atr | Gm50375 | -3.663 | 6.89E-03 | glio |
| Interaction_genotype_Atr | Kctd21 | -3.374 | 6.89E-03 | glio |
| Interaction_genotype_Atr | Poln | -3.331 | 6.89E-03 | glio |
| Interaction_genotype_Bora | Ptpn2 | 1.3 | 6.89E-03 | glio |
| Interaction_genotype_Mtor | Ddx25 | -2.521 | 6.89E-03 | glio |
| Interaction_genotype_Mtor | Zbp1 | 1.883 | 6.89E-03 | glio |
| Interaction_genotype_Myc | Gm16230 | -3.057 | 6.89E-03 | glio |
| interaction_RT | Insig1 | 0.999 | 6.89E-03 | glio |
| Aars2_in.vivo_noRT | Zdhhc18 | 1.73 | 6.89E-03 | glio |
| Atr_in.vivo_noRT | Lrrc61 | 1.842 | 6.89E-03 | glio |
| Atr_in.vivo_noRT | Rasa4 | 1.74 | 6.89E-03 | glio |
| Bard1_ex.vivo_noRT | Gm43463 | 2.421 | 6.89E-03 | glio |
| Bcas2_ex.vivo_noRT | Ccser1 | 2.348 | 6.89E-03 | glio |
| Bcas2_ex.vivo_noRT | Ggt1 | 2.742 | 6.89E-03 | glio |
| Bcas2_ex.vivo_noRT | Acod1 | 3.524 | 6.89E-03 | glio |
| Bora_in.vivo_noRT | Plekhh1 | 2.058 | 6.89E-03 | glio |
| Bora_in.vivo_noRT | Ogfod3 | -1.393 | 6.89E-03 | glio |
| Brca2_in.vivo_noRT | Gm48764 | 1.763 | 6.89E-03 | glio |
| Brca2_in.vivo_noRT | Gm48742 | 2.061 | 6.89E-03 | glio |
| Chek1_in.vivo_noRT | Frat1 | 1.937 | 6.89E-03 | glio |
| Chek1_in.vivo_noRT | Rasa4 | 1.774 | 6.89E-03 | glio |
| Chek1_in.vivo_noRT | Rock1 | -0.802 | 6.89E-03 | glio |
| Chek1_in.vivo_noRT | 1110038B12R | -0.693 | 6.89E-03 | glio |
| Chek1_in.vivo_noRT | Rps3 | -0.551 | 6.89E-03 | glio |
| Chek1_ex.vivo_noRT | Fmc1 | -1.086 | 6.89E-03 | glio |
| Eif2s1_ex.vivo_noRT | Gm29670 | 2.234 | 6.89E-03 | glio |
| Hsd17b10_in.vivo_noRT | Ppp1r26 | 2.234 | 6.89E-03 | glio |
| Aars2_in.vivo_noRT | Zfp13 | 2.062 | 6.89E-04 | glio |
| Aars2_in.vivo_noRT | Rpl35a | -0.594 | 6.89E-04 | glio |
| Cenpj_in.vivo_noRT | Mpzl2 | 2.292 | 6.89E-04 | glio |
| Cenpj_in.vivo_noRT | Gm266 | 2.022 | 6.89E-04 | glio |
| Chek1_in.vivo_noRT | Gtf2f1 | -1.49 | 6.89E-04 | glio |
| Chek1_ex.vivo_noRT | Gm43017 | 3.096 | 6.89E-04 | glio |
| Interaction_genotype_Myc | Gm26728 | -4.641 | 6.89E-05 | glio |
| Atr_ex.vivo_noRT | Dcdc2a | 3.164 | 6.89E-05 | glio |
| Atr_ex.vivo_noRT | Gm4724 | 3.252 | 6.89E-05 | glio |
| Interaction_genotype_Myc | Gm30938 | -4.847 | 6.89E-06 | glio |
| Bora_in.vivo_noRT | Inha | 3.812 | 6.89E-08 | glio |
| Bora_in.vivo_noRT | Otop1 | 3.752 | 6.89E-09 | glio |
| Aars2_in.vivo_noRT | Il17b | 4.676 | 6.89E-10 | glio |
| Chek1_in.vivo_noRT | Arhgef16 | 4.007 | 6.89E-12 | glio |
| Bora_in.vivo_noRT | Dpp6 | 4.009 | 6.89E-16 | glio |
| Aars2_in.vivo_noRT | Gm15943 | 5.063 | 6.89E-20 | glio |

| | | | | |
|---|---|---|---|---|
| Atr_in.vivo_noRT | Gm18280 | 5.269 | 6.89E-24 | glio |
| culture_effect | Mapk15 | -1.728 | 6.90E-03 | glio |
| Interaction_genotype_Aars2 | Wdr61 | 1.369 | 6.90E-03 | glio |
| Interaction_genotype_Bcas2 | Itm2a | 2.728 | 6.90E-03 | glio |
| Interaction_genotype_Chek1 | Fgf2 | -3.784 | 6.90E-03 | glio |
| Interaction_genotype_Myc | Gm26631 | -3.179 | 6.90E-03 | glio |
| Interaction_genotype_Myc | Cdyl | -2.044 | 6.90E-03 | glio |
| Interaction_genotype_Rpl3 | Rictor | 1.668 | 6.90E-03 | glio |
| Aars2_in.vivo_noRT | Tbc1d24 | 1.861 | 6.90E-03 | glio |
| Bard1_ex.vivo_noRT | Fgl2 | 2.637 | 6.90E-03 | glio |
| Interaction_genotype_Aars2 | Dnah10 | -3.79 | 6.90E-04 | glio |
| Interaction_genotype_Myc | Ildr1 | -2.729 | 6.90E-04 | glio |
| Interaction_genotype_Myc | Milr1 | -3.947 | 6.90E-04 | glio |
| Interaction_genotype_Myc | Mutyh | -3.916 | 6.90E-04 | glio |
| Interaction_genotype_Myc | Plpp6 | -3.563 | 6.90E-04 | glio |
| interaction_RT | Mast2 | -0.949 | 6.90E-04 | glio |
| Aars2_in.vivo_noRT | A530041M06R | 2.457 | 6.90E-04 | glio |
| Aars2_in.vivo_noRT | Zfp51 | 2.073 | 6.90E-04 | glio |
| Bcas2_in.vivo_noRT | Gm14966 | 2.34 | 6.90E-04 | glio |
| Cenpj_in.vivo_noRT | Ildr1 | 2.022 | 6.90E-04 | glio |
| Interaction_genotype_Atr | Gm32090 | -4.311 | 6.90E-05 | glio |
| Aars2_in.vivo_noRT | 9030407P20R | 2.78 | 6.90E-05 | glio |
| Chek1_in.vivo_noRT | Palm3 | 2.917 | 6.90E-05 | glio |
| Interaction_genotype_Rad21 | Myoz1 | 5.063 | 6.90E-06 | glio |
| Bcas2_in.vivo_noRT | 4732471J01Ri | 3.249 | 6.90E-06 | glio |
| Bcas2_ex.vivo_noRT | 8030423F21Ri | 3.748 | 6.90E-06 | glio |
| culture_effect | Nfat5 | -1.053 | 6.90E-11 | glio |
| culture_effect | Arid4b | -0.952 | 6.90E-13 | glio |
| Chek1_in.vivo_noRT | Shisal2a | 5.31 | 6.90E-13 | glio |
| Aars2_in.vivo_noRT | Pla1a | 5.305 | 6.90E-16 | glio |
| Bcas2_in.vivo_noRT | Gm14540 | 4.284 | 6.90E-16 | glio |
| Bcas2_in.vivo_noRT | Vwce | 5.065 | 6.90E-16 | glio |
| Bcas2_in.vivo_noRT | Usp17lc | 4.807 | 6.90E-16 | glio |
| Chek1_in.vivo_noRT | Ablim2 | 4.511 | 6.90E-21 | glio |
| culture_effect | 4833413G10R | -1.835 | 6.91E-03 | glio |
| Interaction_genotype_Myc | Ccdc159 | -3.186 | 6.91E-03 | glio |
| Interaction_genotype_Rad21 | Slc14a1 | 3.531 | 6.91E-03 | glio |
| Interaction_genotype_Rad51 | Col8a1 | -3.781 | 6.91E-03 | glio |
| Aars2_in.vivo_noRT | Trim46 | 1.806 | 6.91E-03 | glio |
| Aars2_in.vivo_noRT | Gm31597 | 1.622 | 6.91E-03 | glio |
| Aars2_in.vivo_noRT | Pdcd6 | -0.83 | 6.91E-03 | glio |
| Atr_ex.vivo_noRT | Col6a3 | 2.37 | 6.91E-03 | glio |
| Atr_ex.vivo_noRT | Ola1 | -0.961 | 6.91E-03 | glio |
| Bcas2_ex.vivo_noRT | Gm16685 | 2.342 | 6.91E-03 | glio |
| Chek1_in.vivo_noRT | Zfp975 | 1.935 | 6.91E-03 | glio |
| Chek1_in.vivo_noRT | Zfp385b | -1.042 | 6.91E-03 | glio |
| Hsd17b10_in.vivo_noRT | Hspb9 | 2.304 | 6.91E-03 | glio |
| Interaction_genotype_Myc | Sumo1 | 1.245 | 6.91E-04 | glio |
| interaction_RT | H2-Q4 | 1.427 | 6.91E-04 | glio |
| interaction_RT | Cox6b2 | 1.404 | 6.91E-04 | glio |
| interaction_RT | Mki67 | -1.328 | 6.91E-04 | glio |
| Interaction_genotype_Aars2 | Reep5 | 1.614 | 6.91E-05 | glio |

| | | | | |
|---|---|---|---|---|
| Interaction_genotype_Mtor | Gm19541 | -4.649 | 6.91E-05 | glio |
| Bora_ex.vivo_noRT | Kit | 3.201 | 6.91E-05 | glio |
| Chek1_in.vivo_noRT | Dock6 | 2.786 | 6.91E-05 | glio |
| Aars2_ex.vivo_noRT | Slc35g3 | 2.893 | 6.91E-06 | glio |
| Bora_ex.vivo_noRT | 4930570E03Ri | 3.201 | 6.91E-06 | glio |
| Bora_ex.vivo_noRT | Gm5286 | 3.201 | 6.91E-06 | glio |
| interaction_RT | mt-Nd3 | -1.248 | 6.91E-07 | glio |
| culture_effect | Gm42851 | 1.852 | 6.91E-08 | glio |
| Aars2_in.vivo_noRT | Hist1h4b | 4.094 | 6.91E-08 | glio |
| Bcas2_in.vivo_noRT | Abcb5 | 4.807 | 6.91E-22 | glio |
| Atr_in.vivo_noRT | Pspn | 5.533 | 6.91E-29 | glio |
| Interaction_genotype_Chek1 | Sulf2 | 2.093 | 6.92E-03 | glio |
| Interaction_genotype_Chek1 | Igtp | 1.569 | 6.92E-03 | glio |
| Interaction_genotype_Myc | Ttc7b | -2.313 | 6.92E-03 | glio |
| Interaction_genotype_Pars2 | Vsig10l | -3.698 | 6.92E-03 | glio |
| Aars2_in.vivo_noRT | B3gnt5 | 1.781 | 6.92E-03 | glio |
| Aars2_in.vivo_noRT | Snip1 | -1.313 | 6.92E-03 | glio |
| Chek1_in.vivo_noRT | Gnb2 | -0.707 | 6.92E-03 | glio |
| Hsd17b10_in.vivo_noRT | Ccdc106 | 2.288 | 6.92E-03 | glio |
| Hsd17b10_in.vivo_noRT | Actn4 | -0.683 | 6.92E-03 | glio |
| culture_effect | 4930452G13R | -1.353 | 6.92E-04 | glio |
| culture_effect | Capn1 | -0.986 | 6.92E-04 | glio |
| culture_effect | Ttc28 | 0.911 | 6.92E-04 | glio |
| culture_effect | Tsc22d4 | 0.518 | 6.92E-04 | glio |
| Bora_ex.vivo_noRT | Bmp15 | 2.808 | 6.92E-04 | glio |
| culture_effect | Ngef | -2.249 | 6.92E-05 | glio |
| Interaction_genotype_Mtor | Gm16537 | -4.17 | 6.92E-05 | glio |
| Aars2_in.vivo_noRT | Gm19605 | 2.779 | 6.92E-05 | glio |
| Aars2_in.vivo_noRT | Lmtk3 | 2.716 | 6.92E-05 | glio |
| Aars2_in.vivo_noRT | Ppia | -0.778 | 6.92E-05 | glio |
| Aars2_in.vivo_noRT | Tc2n | 3.239 | 6.92E-06 | glio |
| Bora_in.vivo_noRT | Xist | 3.084 | 6.92E-06 | glio |
| Interaction_genotype_Mtor | Morf4l1 | 2.145 | 6.92E-07 | glio |
| Aars2_ex.vivo_noRT | 9130019O22R | 3.293 | 6.92E-07 | glio |
| Bcas2_in.vivo_noRT | Wincr1 | 4.543 | 6.92E-07 | glio |
| Bcas2_in.vivo_noRT | Rtl1 | 3.115 | 6.92E-07 | glio |
| Bora_ex.vivo_noRT | Erich6b | 3.201 | 6.92E-07 | glio |
| Chek1_ex.vivo_noRT | Mgll | 4.449 | 6.92E-07 | glio |
| Chek1_ex.vivo_noRT | C1s1 | 4.226 | 6.92E-07 | glio |
| Atr_in.vivo_noRT | Gm26619 | 3.937 | 6.92E-08 | glio |
| Bcas2_in.vivo_noRT | Gm15473 | 3.952 | 6.92E-08 | glio |
| Chek1_in.vivo_noRT | Gm20513 | 2.947 | 6.92E-08 | glio |
| culture_effect | Tmem50a | -0.794 | 6.92E-09 | glio |
| Atr_in.vivo_noRT | Gstp3 | 4.233 | 6.92E-10 | glio |
| culture_effect | Fstl1 | -0.884 | 6.93E-03 | glio |
| culture_effect | Dynlrb1 | 0.325 | 6.93E-03 | glio |
| Interaction_genotype_Mtor | 3110083C13R | -2.996 | 6.93E-03 | glio |
| Interaction_genotype_Myc | Mindy4 | -2.961 | 6.93E-03 | glio |
| Interaction_genotype_Myc | Cited2 | 1.889 | 6.93E-03 | glio |
| Interaction_genotype_Rad21 | Dusp15 | 3.893 | 6.93E-03 | glio |
| Aars2_in.vivo_noRT | Dip2a | 1.883 | 6.93E-03 | glio |
| Bcas2_ex.vivo_noRT | Vamp8 | 2.342 | 6.93E-03 | glio |

| | | | | |
|---|---|---|---|---|
| Bcas2_ex.vivo_noRT | Spink4 | 2.742 | 6.93E-03 | glio |
| Chek1_in.vivo_noRT | Taco1os | 1.874 | 6.93E-03 | glio |
| Interaction_genotype_Bora | Rhox7a | -4.635 | 6.93E-04 | glio |
| Interaction_genotype_Mtor | Rmdn2 | -3.669 | 6.93E-04 | glio |
| Interaction_genotype_Mtor | mt-Nd5 | 1.936 | 6.93E-04 | glio |
| Interaction_genotype_Myc | Ccdc125 | -3.128 | 6.93E-04 | glio |
| Interaction_genotype_Myc | Elp4 | -2.684 | 6.93E-04 | glio |
| interaction_RT | Ctsf | 2.688 | 6.93E-04 | glio |
| interaction_RT | Psme1 | 0.789 | 6.93E-04 | glio |
| Aars2_in.vivo_noRT | Coq7 | -1.359 | 6.93E-04 | glio |
| Aars2_ex.vivo_noRT | Gm48899 | 2.717 | 6.93E-04 | glio |
| Atr_in.vivo_noRT | Veph1 | 2.398 | 6.93E-04 | glio |
| Interaction_genotype_Myc | 5730414N17R | -4.707 | 6.93E-05 | glio |
| Aars2_in.vivo_noRT | Itga7 | 2.851 | 6.93E-05 | glio |
| Aars2_in.vivo_noRT | B630019K06R | 2.312 | 6.93E-05 | glio |
| Bard1_ex.vivo_noRT | Gm2016 | 2.814 | 6.93E-05 | glio |
| Bcas2_in.vivo_noRT | Tmc7 | 2.995 | 6.93E-05 | glio |
| Interaction_genotype_Mtor | Ppp2ca | 1.833 | 6.93E-06 | glio |
| Bora_in.vivo_noRT | Gramd2 | 3.098 | 6.93E-06 | glio |
| Aars2_in.vivo_noRT | Fes | 3.521 | 6.93E-07 | glio |
| Atp5c1_in.vivo_noRT | Gm15533 | 2.668 | 6.93E-07 | glio |
| Bora_in.vivo_noRT | Gm14443 | 3.261 | 6.93E-07 | glio |
| Aars2_in.vivo_noRT | Ggnbp1 | 3.644 | 6.93E-08 | glio |
| Bora_in.vivo_noRT | Emilin2 | 3.854 | 6.93E-08 | glio |
| Bora_in.vivo_noRT | Gm15853 | 4.201 | 6.93E-09 | glio |
| Aars2_in.vivo_noRT | Gm45441 | 4.267 | 6.93E-10 | glio |
| Atr_in.vivo_noRT | Gm49694 | 4.807 | 6.93E-11 | glio |
| Bcas2_in.vivo_noRT | Gm50147 | 4.807 | 6.93E-13 | glio |
| Atr_in.vivo_noRT | Dnah10 | 5.154 | 6.93E-17 | glio |
| Aars2_in.vivo_noRT | Hoxc8 | 5.327 | 6.93E-28 | glio |
| Interaction_genotype_Bcas2 | Gm48140 | -2.595 | 6.94E-03 | glio |
| Interaction_genotype_Bcas2 | A030014E15Ri | -2.458 | 6.94E-03 | glio |
| Interaction_genotype_Myc | Cyp2c23 | -2.602 | 6.94E-03 | glio |
| Interaction_genotype_Myc | Has1 | -2.814 | 6.94E-03 | glio |
| Aars2_in.vivo_noRT | Nbdy | -0.96 | 6.94E-03 | glio |
| Atr_in.vivo_noRT | Kcnn4 | -2.118 | 6.94E-03 | glio |
| Atr_in.vivo_noRT | Mrpl58 | -0.907 | 6.94E-03 | glio |
| Bcas2_ex.vivo_noRT | Hfm1 | 2.167 | 6.94E-03 | glio |
| Bcas2_ex.vivo_noRT | Slco2a1 | 2.742 | 6.94E-03 | glio |
| Bcas2_ex.vivo_noRT | Lncbate10 | 1.954 | 6.94E-03 | glio |
| Bcas2_ex.vivo_noRT | Gm47223 | 3.697 | 6.94E-03 | glio |
| Bcas2_ex.vivo_noRT | Traf3ip3 | 2.754 | 6.94E-03 | glio |
| Bora_in.vivo_noRT | Ccnyl1 | 1.614 | 6.94E-03 | glio |
| Cenpj_ex.vivo_noRT | Nmrk2 | 2.591 | 6.94E-03 | glio |
| Chek1_in.vivo_noRT | Plppr2 | 2.028 | 6.94E-03 | glio |
| Chek1_ex.vivo_noRT | Gm42463 | 2.389 | 6.94E-03 | glio |
| Hsd17b10_in.vivo_noRT | Rhox2h | -0.876 | 6.94E-03 | glio |
| culture_effect | Rpl19 | -0.458 | 6.94E-04 | glio |
| Interaction_genotype_Atr | Gm34256 | -4.439 | 6.94E-04 | glio |
| Interaction_genotype_Myc | Gm33782 | -3.434 | 6.94E-04 | glio |
| Atr_in.vivo_noRT | Stoml1 | 2.094 | 6.94E-04 | glio |
| Bcas2_in.vivo_noRT | Gm46218 | 2.52 | 6.94E-04 | glio |

| | | | | |
|---|---|---|---|---|
| Bcas2_ex.vivo_noRT | Gpc5 | 2.347 | 6.94E-04 | glio |
| Chek1_in.vivo_noRT | Spry3 | 2.516 | 6.94E-04 | glio |
| culture_effect | Trappc2l | 0.609 | 6.94E-05 | glio |
| Interaction_genotype_Myc | Fam71e1 | -3.714 | 6.94E-05 | glio |
| Interaction_genotype_Myc | Gm16196 | -4.071 | 6.94E-05 | glio |
| Aars2_in.vivo_noRT | 1700024B05R | 2.845 | 6.94E-05 | glio |
| Aars2_in.vivo_noRT | Frmd4a | -1.81 | 6.94E-05 | glio |
| Aars2_ex.vivo_noRT | Clca3a2 | 3.293 | 6.94E-05 | glio |
| Aars2_ex.vivo_noRT | Art2b | 3.293 | 6.94E-05 | glio |
| Bora_in.vivo_noRT | Ece2 | 2.692 | 6.94E-05 | glio |
| Chek1_in.vivo_noRT | Gstm2 | 2.92 | 6.94E-05 | glio |
| Aars2_in.vivo_noRT | Olfr655 | 4.41 | 6.94E-06 | glio |
| Atr_in.vivo_noRT | Osbp2 | 3.608 | 6.94E-06 | glio |
| Chek1_in.vivo_noRT | Gm20536 | 3.147 | 6.94E-06 | glio |
| Bora_in.vivo_noRT | Itgb6 | 3.532 | 6.94E-07 | glio |
| culture_effect | Gm30938 | 2.768 | 6.94E-08 | glio |
| Aars2_in.vivo_noRT | A330040F15Ri | 5.057 | 6.94E-12 | glio |
| Interaction_genotype_Bcas2 | 4933404O12R | -3.523 | 6.95E-03 | glio |
| Interaction_genotype_Bora | Gm36210 | -3.594 | 6.95E-03 | glio |
| Interaction_genotype_Mtor | Gm43387 | -3.328 | 6.95E-03 | glio |
| Interaction_genotype_Mtor | Ctsa | 1.717 | 6.95E-03 | glio |
| Interaction_genotype_Mtor | Lsm14a | 1.495 | 6.95E-03 | glio |
| Interaction_genotype_Myc | Tnks1bp1 | -2.096 | 6.95E-03 | glio |
| Interaction_genotype_Rad21 | Rasl12 | 3.967 | 6.95E-03 | glio |
| interaction_RT | A430106G13R | -2.606 | 6.95E-03 | glio |
| interaction_RT | Ecm1 | 1.921 | 6.95E-03 | glio |
| interaction_RT | Gm30025 | 1.28 | 6.95E-03 | glio |
| Aars2_in.vivo_noRT | 2010109A12R | 1.919 | 6.95E-03 | glio |
| Aars2_in.vivo_noRT | Rell1 | 1.388 | 6.95E-03 | glio |
| Atp5c1_in.vivo_noRT | Olfr1191-ps1 | 2.018 | 6.95E-03 | glio |
| Bcas2_in.vivo_noRT | Tmem168 | 1.414 | 6.95E-03 | glio |
| Bora_in.vivo_noRT | Galm | 1.932 | 6.95E-03 | glio |
| Bora_in.vivo_noRT | Washc5 | 0.923 | 6.95E-03 | glio |
| Chek1_in.vivo_noRT | Ninl | 1.672 | 6.95E-03 | glio |
| Chek1_in.vivo_noRT | Zfp84 | 1.379 | 6.95E-03 | glio |
| Interaction_genotype_Aars2 | Med9os | -4.159 | 6.95E-04 | glio |
| Interaction_genotype_Aars2 | Gm10399 | -3.735 | 6.95E-04 | glio |
| Interaction_genotype_Aars2 | Ssu72 | 1.444 | 6.95E-04 | glio |
| Interaction_genotype_Aars2 | Llph | 1.202 | 6.95E-04 | glio |
| Interaction_genotype_Aars2 | Atp5g1 | 1.047 | 6.95E-04 | glio |
| interaction_RT | 5830444B04R | 1.637 | 6.95E-04 | glio |
| Atr_in.vivo_noRT | Aars2 | 2.299 | 6.95E-04 | glio |
| Chek1_in.vivo_noRT | Nutf2 | 2.418 | 6.95E-04 | glio |
| Bard1_ex.vivo_noRT | Scg2 | 2.814 | 6.95E-06 | glio |
| Bora_in.vivo_noRT | Anks6 | 2.986 | 6.95E-06 | glio |
| Aars2_ex.vivo_noRT | Klk13 | 3.293 | 6.95E-07 | glio |
| Chek1_ex.vivo_noRT | 7530414M10F | 4.114 | 6.95E-07 | glio |
| Bcas2_in.vivo_noRT | Gm36043 | 3.351 | 6.95E-08 | glio |
| Aars2_in.vivo_noRT | Gm3604 | 4.044 | 6.95E-09 | glio |
| Atr_in.vivo_noRT | Kcnmb4os2 | 4.096 | 6.95E-09 | glio |
| Aars2_in.vivo_noRT | Gm47545 | 4.076 | 6.95E-11 | glio |
| Aars2_in.vivo_noRT | Gm35986 | 4.278 | 6.95E-11 | glio |

| | | | | |
|---|---|---|---|---|
| Aars2_in.vivo_noRT | Peli3 | 4.412 | 6.95E-11 | glio |
| Atr_in.vivo_noRT | Gm5129 | 4.733 | 6.95E-11 | glio |
| Bcas2_in.vivo_noRT | Gm41349 | 4.544 | 6.95E-13 | glio |
| Bora_in.vivo_noRT | Rapgef3os2 | 4.936 | 6.95E-21 | glio |
| Bora_in.vivo_noRT | Cldn5 | 4.67 | 6.95E-21 | glio |
| Atr_in.vivo_noRT | Ildr1 | 5.533 | 6.95E-27 | glio |
| culture_effect | Wdr77 | -0.442 | 6.96E-03 | glio |
| culture_effect | Ostc | 0.296 | 6.96E-03 | glio |
| Interaction_genotype_Aars2 | Rmnd5b | -2.099 | 6.96E-03 | glio |
| Interaction_genotype_Atr | Ccdc125 | -2.769 | 6.96E-03 | glio |
| Interaction_genotype_Ifnar1 | Ankrd22 | 2.734 | 6.96E-03 | glio |
| Interaction_genotype_Ifnar1 | Gm34776 | 3.137 | 6.96E-03 | glio |
| Interaction_genotype_Mtor | Blcap | -2.352 | 6.96E-03 | glio |
| Aars2_in.vivo_noRT | Gm26549 | 1.724 | 6.96E-03 | glio |
| Aars2_in.vivo_noRT | Cldn34c1 | 1.847 | 6.96E-03 | glio |
| Atr_in.vivo_noRT | Nudt22 | 1.357 | 6.96E-03 | glio |
| Atr_in.vivo_noRT | Cox18 | 0.947 | 6.96E-03 | glio |
| Bcas2_in.vivo_noRT | Fam174b | 2.233 | 6.96E-03 | glio |
| Bcas2_in.vivo_noRT | B230303O12R | 2.028 | 6.96E-03 | glio |
| Bcas2_in.vivo_noRT | BC024063 | 1.932 | 6.96E-03 | glio |
| Bcas2_in.vivo_noRT | Eral1 | 1.557 | 6.96E-03 | glio |
| Bora_in.vivo_noRT | Acaca | -1.482 | 6.96E-03 | glio |
| Bora_ex.vivo_noRT | Cd72 | 2.806 | 6.96E-03 | glio |
| Chek1_in.vivo_noRT | Zfp385a | 1.56 | 6.96E-03 | glio |
| culture_effect | Slc25a10 | 0.913 | 6.96E-04 | glio |
| Interaction_genotype_Mtor | Gne | -3.188 | 6.96E-04 | glio |
| Interaction_genotype_Mtor | Hdac1 | 1.577 | 6.96E-04 | glio |
| Atr_in.vivo_noRT | Zfp280c | 1.706 | 6.96E-04 | glio |
| Atr_ex.vivo_noRT | Gm48161 | 2.771 | 6.96E-04 | glio |
| Aars2_in.vivo_noRT | Gm17655 | 3.135 | 6.96E-05 | glio |
| Aars2_in.vivo_noRT | Tmem246 | 2.63 | 6.96E-05 | glio |
| Atr_in.vivo_noRT | Dhtkd1 | 3.056 | 6.96E-05 | glio |
| Atr_in.vivo_noRT | Hykk | 2.768 | 6.96E-05 | glio |
| Aars2_in.vivo_noRT | Gm20707 | 3.149 | 6.96E-06 | glio |
| Aars2_in.vivo_noRT | Slc39a8 | 3.473 | 6.96E-06 | glio |
| Hsd17b10_in.vivo_noRT | Gm26789 | 2.699 | 6.96E-06 | glio |
| Interaction_genotype_Mtor | Edn2 | -4.494 | 6.96E-08 | glio |
| Atr_in.vivo_noRT | Dynlt1c | 4.234 | 6.96E-08 | glio |
| culture_effect | Eid2b | 1.999 | 6.96E-09 | glio |
| Atr_in.vivo_noRT | 2810049E08Ri | 4.496 | 6.96E-13 | glio |
| Aars2_in.vivo_noRT | Synpo2l | 4.681 | 6.96E-15 | glio |
| Atr_in.vivo_noRT | Dntt | 5.533 | 6.96E-15 | glio |
| Chek1_in.vivo_noRT | Npy1r | 4.399 | 6.96E-16 | glio |
| Chek1_in.vivo_noRT | Rgs4 | 5.044 | 6.96E-22 | glio |
| Aars2_in.vivo_noRT | Gm15533 | 5.327 | 6.96E-25 | glio |
| culture_effect | Tent5d | -1.555 | 6.97E-03 | glio |
| culture_effect | Gcsh | 0.375 | 6.97E-03 | glio |
| Interaction_genotype_Aars2 | D830025C05R | -3.359 | 6.97E-03 | glio |
| Interaction_genotype_Atr | Acod1 | 4.279 | 6.97E-03 | glio |
| Interaction_genotype_Bora | B930059L03Ri | -3.729 | 6.97E-03 | glio |
| Interaction_genotype_Bora | Ccdc9b | -2.734 | 6.97E-03 | glio |
| Interaction_genotype_Bora | Timp2 | 1.698 | 6.97E-03 | glio |

| | | | | |
|---|---|---|---|---|
| Interaction_genotype_Mtor | Sema3a | -3.055 | 6.97E-03 | glio |
| Interaction_genotype_Mtor | Ovca2 | -3.084 | 6.97E-03 | glio |
| Interaction_genotype_Mtor | Pank4 | -2.95 | 6.97E-03 | glio |
| Interaction_genotype_Mtor | Slc4a8 | -2.787 | 6.97E-03 | glio |
| Interaction_genotype_Myc | Ppp3cc | -2.902 | 6.97E-03 | glio |
| Interaction_genotype_Myc | Pex10 | -2.401 | 6.97E-03 | glio |
| Interaction_genotype_Myc | Bok | -2.743 | 6.97E-03 | glio |
| Interaction_genotype_Myc | Zfp358 | -2.352 | 6.97E-03 | glio |
| Interaction_genotype_Myc | Tada2a | -1.868 | 6.97E-03 | glio |
| Interaction_genotype_Myc | Gga1 | -1.367 | 6.97E-03 | glio |
| Interaction_genotype_Myc | Rps3 | 1.067 | 6.97E-03 | glio |
| interaction_RT | Trafd1 | 0.945 | 6.97E-03 | glio |
| interaction_RT | Naf1 | -0.676 | 6.97E-03 | glio |
| Aars2_in.vivo_noRT | Phex | 1.847 | 6.97E-03 | glio |
| Atr_in.vivo_noRT | Itgb8 | 1.588 | 6.97E-03 | glio |
| Bard1_ex.vivo_noRT | Gm48161 | 2.421 | 6.97E-03 | glio |
| Bcas2_ex.vivo_noRT | Ngb | 2.742 | 6.97E-03 | glio |
| Bcas2_ex.vivo_noRT | Pdzk1ip1 | 2.843 | 6.97E-03 | glio |
| Bcas2_ex.vivo_noRT | Sulf2 | -1.533 | 6.97E-03 | glio |
| Bora_in.vivo_noRT | Gsto2 | 2.025 | 6.97E-03 | glio |
| Chek1_in.vivo_noRT | Slc27a1 | -1.512 | 6.97E-03 | glio |
| Chek1_in.vivo_noRT | Slc44a1 | 1.226 | 6.97E-03 | glio |
| Chek1_in.vivo_noRT | Nudt5 | -0.927 | 6.97E-03 | glio |
| Hsd17b10_in.vivo_noRT | Dlx2 | 2.233 | 6.97E-03 | glio |
| Hsd17b10_in.vivo_noRT | 2810455O05R | 2.154 | 6.97E-03 | glio |
| Interaction_genotype_Aars2 | Lyrm1 | -4.095 | 6.97E-04 | glio |
| Interaction_genotype_Aars2 | Ercc6 | -3.584 | 6.97E-04 | glio |
| Interaction_genotype_Aars2 | Grcc10 | 1.104 | 6.97E-04 | glio |
| Interaction_genotype_Mtor | Cstdc4 | -3.841 | 6.97E-04 | glio |
| Interaction_genotype_Mtor | 1810062O18R | -4.23 | 6.97E-04 | glio |
| Interaction_genotype_Myc | Cst6 | 2.154 | 6.97E-04 | glio |
| interaction_RT | Cd3eap | -0.772 | 6.97E-04 | glio |
| Aars2_in.vivo_noRT | Gm9828 | 2.389 | 6.97E-04 | glio |
| Aars2_in.vivo_noRT | Gtdc1 | 1.961 | 6.97E-04 | glio |
| Atr_in.vivo_noRT | Dhps | -1.133 | 6.97E-04 | glio |
| Atr_ex.vivo_noRT | Ceacam19 | 3.251 | 6.97E-04 | glio |
| Bcas2_in.vivo_noRT | D330041H03F | 1.987 | 6.97E-04 | glio |
| Bora_in.vivo_noRT | Lrrc8a | -1.628 | 6.97E-04 | glio |
| culture_effect | Bcl9l | 0.93 | 6.97E-05 | glio |
| culture_effect | Emp3 | 0.503 | 6.97E-05 | glio |
| Interaction_genotype_Mtor | Aars2 | -4.646 | 6.97E-05 | glio |
| Atr_in.vivo_noRT | Platr3 | 2.943 | 6.97E-05 | glio |
| Bora_in.vivo_noRT | Siah1b | -2.024 | 6.97E-05 | glio |
| culture_effect | Serpinf1 | -1.901 | 6.97E-06 | glio |
| Bcas2_in.vivo_noRT | 1700029J07Ri | 3.25 | 6.97E-06 | glio |
| culture_effect | Msmo1 | -0.906 | 6.97E-07 | glio |
| Bora_in.vivo_noRT | Vcam1 | 4.42 | 6.97E-07 | glio |
| Bora_in.vivo_noRT | Zfp11 | 3.493 | 6.97E-07 | glio |
| Interaction_genotype_Mtor | Dnah10 | -5.855 | 6.97E-08 | glio |
| Bora_in.vivo_noRT | Gm3716 | 4.018 | 6.97E-09 | glio |
| Bcas2_in.vivo_noRT | Gm14379 | 4.291 | 6.97E-11 | glio |
| Bora_in.vivo_noRT | Gm16364.1 | 4.936 | 6.97E-11 | glio |

| | | | | |
|---|---|---|---|---|
| culture_effect | Hivep2 | -1.493 | 6.97E-13 | glio |
| Atr_in.vivo_noRT | 4930478M13F | 5.154 | 6.97E-13 | glio |
| Chek1_in.vivo_noRT | 6430562O15R | 5.051 | 6.97E-16 | glio |
| Chek1_in.vivo_noRT | Gm40538 | 4.78 | 6.97E-18 | glio |
| Bcas2_in.vivo_noRT | Olfr56 | 4.807 | 6.97E-20 | glio |
| Chek1_in.vivo_noRT | Gm11232 | 5.044 | 6.97E-25 | glio |
| culture_effect | Gm49891 | 1.622 | 6.98E-03 | glio |
| culture_effect | Gask1b | 1.597 | 6.98E-03 | glio |
| culture_effect | Zfp580 | 0.763 | 6.98E-03 | glio |
| culture_effect | Rnf168 | 0.618 | 6.98E-03 | glio |
| culture_effect | Serinc1 | -0.356 | 6.98E-03 | glio |
| Interaction_genotype_Bora | Tob1 | 1.806 | 6.98E-03 | glio |
| Interaction_genotype_Bora | Reep5 | 1.082 | 6.98E-03 | glio |
| Interaction_genotype_Myc | Hdac4 | -2.09 | 6.98E-03 | glio |
| Interaction_genotype_Rad21 | Etl4 | 3.571 | 6.98E-03 | glio |
| Atr_in.vivo_noRT | Hnrnpl | -0.912 | 6.98E-03 | glio |
| Bcas2_in.vivo_noRT | Dhrs3 | 2.039 | 6.98E-03 | glio |
| Bora_in.vivo_noRT | Prrc1 | -1.2 | 6.98E-03 | glio |
| Cyp19a1_ex.vivo_noRT | Cd200r2 | 2.159 | 6.98E-03 | glio |
| Interaction_genotype_Mtor | Gldc | -3.815 | 6.98E-04 | glio |
| Interaction_genotype_Myc | Kctd11 | -3.889 | 6.98E-04 | glio |
| Interaction_genotype_Myc | Ddah1 | -2.629 | 6.98E-04 | glio |
| interaction_RT | Fam174a | 1.003 | 6.98E-04 | glio |
| Atr_in.vivo_noRT | Svip | 2.22 | 6.98E-04 | glio |
| Bard1_ex.vivo_noRT | Ccl19 | 2.814 | 6.98E-04 | glio |
| Bcas2_in.vivo_noRT | Klf2 | -2.418 | 6.98E-04 | glio |
| Bora_in.vivo_noRT | Cadm2 | 2.573 | 6.98E-04 | glio |
| culture_effect | Rps16 | -0.553 | 6.98E-05 | glio |
| Interaction_genotype_Myc | Chaf1b | -3.725 | 6.98E-05 | glio |
| Atr_in.vivo_noRT | Ccdc191 | 2.469 | 6.98E-05 | glio |
| Bcas2_in.vivo_noRT | Gm13056 | 3.052 | 6.98E-05 | glio |
| Bora_in.vivo_noRT | Gm16124 | 3.226 | 6.98E-05 | glio |
| Chek1_ex.vivo_noRT | Mrpl19 | -1.995 | 6.98E-05 | glio |
| Aars2_in.vivo_noRT | Ckm | 3.052 | 6.98E-06 | glio |
| Chek1_in.vivo_noRT | Bpi | 3.362 | 6.98E-08 | glio |
| Chek1_ex.vivo_noRT | Gm20705 | 4.447 | 6.98E-11 | glio |
| Atr_in.vivo_noRT | Gm45406 | 5.011 | 6.98E-21 | glio |
| Chek1_in.vivo_noRT | Apoa5 | 5.044 | 6.98E-26 | glio |
| culture_effect | Mpg | 0.451 | 6.99E-03 | glio |
| Interaction_genotype_Aars2 | Kdm8 | -3.139 | 6.99E-03 | glio |
| Interaction_genotype_Aars2 | Parp9 | 1.963 | 6.99E-03 | glio |
| Interaction_genotype_Bora | Gm13348 | -3.007 | 6.99E-03 | glio |
| Interaction_genotype_Terf1 | Mob3b | -3.549 | 6.99E-03 | glio |
| Interaction_genotype_Terf1 | Gm16741 | -3.767 | 6.99E-03 | glio |
| Interaction_genotype_Terf1 | Slc26a6 | -3.967 | 6.99E-03 | glio |
| Interaction_genotype_Terf1 | Gm13288 | -3.362 | 6.99E-03 | glio |
| Interaction_genotype_Terf1 | Card19 | 1.752 | 6.99E-03 | glio |
| Aars2_in.vivo_noRT | Urb1 | 1.42 | 6.99E-03 | glio |
| Aars2_in.vivo_noRT | Pafah1b3 | -1.212 | 6.99E-03 | glio |
| Aars2_in.vivo_noRT | Hebp2 | -1.01 | 6.99E-03 | glio |
| Atp5c1_in.vivo_noRT | Phka1 | -1.674 | 6.99E-03 | glio |
| Bcas2_in.vivo_noRT | Hspa13 | -1.154 | 6.99E-03 | glio |

| | | | | |
|---|---|---|---|---|
| Bcas2_ex.vivo_noRT | Adgre5 | 2.348 | 6.99E-03 | glio |
| Bcas2_ex.vivo_noRT | Acot5 | 2.731 | 6.99E-03 | glio |
| Brca2_in.vivo_noRT | F8 | -2.428 | 6.99E-03 | glio |
| Cenpj_ex.vivo_noRT | Gm26789 | 2.408 | 6.99E-03 | glio |
| Interaction_genotype_Aars2 | Rhox7a | -4.41 | 6.99E-04 | glio |
| Interaction_genotype_Mtor | Ptp4a1 | -3.594 | 6.99E-04 | glio |
| Interaction_genotype_Mtor | Dnajc28 | -4.34 | 6.99E-04 | glio |
| Interaction_genotype_Mtor | Sipa1l2 | -3.829 | 6.99E-04 | glio |
| Interaction_genotype_Myc | Olfr543 | -2.865 | 6.99E-04 | glio |
| Aars2_in.vivo_noRT | Zbtb39 | 2.198 | 6.99E-04 | glio |
| Aars2_in.vivo_noRT | Bsg | -0.694 | 6.99E-04 | glio |
| Bcas2_in.vivo_noRT | Pih1d2 | 2.131 | 6.99E-04 | glio |
| Cenpj_ex.vivo_noRT | Cxcl5 | 2.591 | 6.99E-04 | glio |
| culture_effect | Foxf1 | -1.669 | 6.99E-05 | glio |
| Interaction_genotype_Myc | Smyd5 | -3.843 | 6.99E-05 | glio |
| Interaction_genotype_Myc | Agap2 | -3.715 | 6.99E-05 | glio |
| Interaction_genotype_Myc | 1700056E22Ri | -3.754 | 6.99E-05 | glio |
| Interaction_genotype_Myc | Agbl3 | -3.677 | 6.99E-05 | glio |
| Aars2_in.vivo_noRT | Kcna4 | 2.8 | 6.99E-05 | glio |
| Bora_ex.vivo_noRT | Gm49504 | 2.802 | 6.99E-05 | glio |
| Bora_ex.vivo_noRT | Adam2 | 3.201 | 6.99E-05 | glio |
| Atr_in.vivo_noRT | Ces1a | 3.411 | 6.99E-06 | glio |
| Chek1_in.vivo_noRT | Lbhd2 | 3.802 | 6.99E-06 | glio |
| Atr_in.vivo_noRT | Neurl1a | 3.735 | 6.99E-07 | glio |
| Bcas2_in.vivo_noRT | Gm33973 | 3.24 | 6.99E-08 | glio |
| Chek1_in.vivo_noRT | Gm26619 | 3.959 | 6.99E-08 | glio |
| Bora_in.vivo_noRT | Smpd5 | 4.279 | 6.99E-12 | glio |
| Bcas2_in.vivo_noRT | Lhx1os | 4.281 | 6.99E-14 | glio |
| Bcas2_in.vivo_noRT | D430001F17R | 4.807 | 6.99E-14 | glio |
| Bcas2_in.vivo_noRT | Fhl1 | 4.807 | 6.99E-19 | glio |
| Aars2_in.vivo_noRT | 4930423D22R | 5.067 | 6.99E-23 | glio |
| culture_effect | Gm10432 | -1.634 | 7.00E-03 | glio |
| culture_effect | 4933404O12R | 1.322 | 7.00E-03 | glio |
| Interaction_genotype_Atr | Mmp13 | -2.369 | 7.00E-03 | glio |
| Interaction_genotype_Mtor | Zdhhc23 | -3.331 | 7.00E-03 | glio |
| Interaction_genotype_Myc | H1foo | -2.197 | 7.00E-03 | glio |
| Interaction_genotype_Myc | Insm1 | -2.99 | 7.00E-03 | glio |
| Interaction_genotype_Myc | Zfp59 | -3.107 | 7.00E-03 | glio |
| Interaction_genotype_Myc | Kbtbd8 | -2.939 | 7.00E-03 | glio |
| Interaction_genotype_Myc | Polr3g | -2.358 | 7.00E-03 | glio |
| Interaction_genotype_Myc | Cox10 | -1.931 | 7.00E-03 | glio |
| Interaction_genotype_Myc | Micall1 | -2.118 | 7.00E-03 | glio |
| Interaction_genotype_Myc | Ywhaq | 1.049 | 7.00E-03 | glio |
| Interaction_genotype_Pars2 | Slc16a6 | -3.933 | 7.00E-03 | glio |
| Interaction_genotype_Pars2 | Prl2c3 | -3.464 | 7.00E-03 | glio |
| Aars2_in.vivo_noRT | Cacybp | -0.619 | 7.00E-03 | glio |
| Atr_in.vivo_noRT | Chn1 | 1.624 | 7.00E-03 | glio |
| Bora_in.vivo_noRT | Mrpl55 | -1.155 | 7.00E-03 | glio |
| Hsd17b10_in.vivo_noRT | Gm31517 | 2.161 | 7.00E-03 | glio |
| culture_effect | Atp11c | -0.911 | 7.00E-04 | glio |
| culture_effect | Smim15 | 0.641 | 7.00E-04 | glio |
| Interaction_genotype_Mtor | Tsga10 | -4.18 | 7.00E-04 | glio |

| | | | | |
|---|---|---|---|---|
| Interaction_genotype_Mtor | Maged1 | 1.89 | 7.00E-04 | glio |
| interaction_RT | Khdrbs2 | 2.997 | 7.00E-04 | glio |
| Aars2_in.vivo_noRT | Itga2 | 2.127 | 7.00E-04 | glio |
| Atr_in.vivo_noRT | Gtf2ird2 | 2.169 | 7.00E-04 | glio |
| Atr_in.vivo_noRT | Cul7 | 1.414 | 7.00E-04 | glio |
| Bora_ex.vivo_noRT | Mst1 | 3.199 | 7.00E-04 | glio |
| Chek1_ex.vivo_noRT | Nuf2 | -2.291 | 7.00E-04 | glio |
| culture_effect | Haus7 | -0.862 | 7.00E-05 | glio |
| Interaction_genotype_Atr | Gm47204 | -5.059 | 7.00E-05 | glio |
| Interaction_genotype_Mtor | Gm15246 | -4.809 | 7.00E-05 | glio |
| Interaction_genotype_Myc | Atxn7l1 | -3.691 | 7.00E-05 | glio |
| Interaction_genotype_Myc | Gpr4 | -3.272 | 7.00E-05 | glio |
| Hsd17b10_in.vivo_noRT | Amh | 2.699 | 7.00E-05 | glio |
| Hsd17b10_in.vivo_noRT | Gm17227 | 2.761 | 7.00E-05 | glio |
| Interaction_genotype_Atr | Sema6b | -5.254 | 7.00E-06 | glio |
| Chek1_ex.vivo_noRT | Bglap3 | 4.049 | 7.00E-06 | glio |
| Hsd17b10_in.vivo_noRT | Hoxa6 | 3.105 | 7.00E-06 | glio |
| Aars2_in.vivo_noRT | S100a14 | 4.526 | 7.00E-14 | glio |
| Atr_in.vivo_noRT | Slc16a7 | 4.422 | 7.00E-14 | glio |
| Atr_in.vivo_noRT | A330032B11R | 5.267 | 7.00E-17 | glio |
| culture_effect | Cfap100 | 4.176 | 7.00E-20 | glio |
| culture_effect | Olig2 | -2.715 | 7.00E-29 | glio |
| culture_effect | Osgin2 | -0.563 | 7.01E-03 | glio |
| culture_effect | Rars2 | -0.484 | 7.01E-03 | glio |
| culture_effect | Ivns1abp | 0.551 | 7.01E-03 | glio |
| Interaction_genotype_Ifnar1 | Gm31717 | 2.734 | 7.01E-03 | glio |
| Interaction_genotype_Ifnar1 | Gm14742 | 2.734 | 7.01E-03 | glio |
| Interaction_genotype_Ifnar1 | Psme2 | -0.902 | 7.01E-03 | glio |
| Interaction_genotype_Mtor | Anapc5 | 1.593 | 7.01E-03 | glio |
| Interaction_genotype_Mtor | H3f3b | 1.199 | 7.01E-03 | glio |
| Aars2_in.vivo_noRT | Plekha4 | 1.919 | 7.01E-03 | glio |
| Atr_in.vivo_noRT | Sspn | 1.613 | 7.01E-03 | glio |
| Atr_in.vivo_noRT | Cic | -1.19 | 7.01E-03 | glio |
| Cdkn2a_in.vivo_noRT | Gm34059 | -1.675 | 7.01E-03 | glio |
| Eif2s1_in.vivo_noRT | Gm26978 | 1.883 | 7.01E-03 | glio |
| Eif2s1_in.vivo_noRT | Nxph3 | 2.143 | 7.01E-03 | glio |
| culture_effect | Ankrd39 | 0.809 | 7.01E-04 | glio |
| culture_effect | Ciao2b | -0.392 | 7.01E-04 | glio |
| Bora_in.vivo_noRT | Cyp2d22 | 2.599 | 7.01E-04 | glio |
| Bora_ex.vivo_noRT | Slc16a7 | 2.802 | 7.01E-04 | glio |
| Bora_ex.vivo_noRT | Sox11 | 3.201 | 7.01E-04 | glio |
| culture_effect | AU022754 | -2.355 | 7.01E-05 | glio |
| Interaction_genotype_Atr | Dusp2 | 4.387 | 7.01E-05 | glio |
| Interaction_genotype_Myc | Rasa4 | -4.039 | 7.01E-05 | glio |
| Atr_in.vivo_noRT | Cyp4f17 | 2.823 | 7.01E-05 | glio |
| Chek1_ex.vivo_noRT | 4732471J01Ri | 3.48 | 7.01E-05 | glio |
| Interaction_genotype_Myc | Ap3m2 | -4.208 | 7.01E-06 | glio |
| Interaction_genotype_Myc | A730071L15Ri | -5.299 | 7.01E-06 | glio |
| Aars2_in.vivo_noRT | Slc13a5 | 2.985 | 7.01E-06 | glio |
| Bcas2_in.vivo_noRT | Tmeff2 | 3.55 | 7.01E-06 | glio |
| Interaction_genotype_Mtor | Gm14419 | -5.004 | 7.01E-07 | glio |
| Bcas2_in.vivo_noRT | Atpif1 | -1.06 | 7.01E-07 | glio |

| | | | | |
|---|---|---|---|---|
| Hsd17b10_in.vivo_noRT | Txndc2 | 2.837 | 7.01E-07 | glio |
| Interaction_genotype_Mtor | Ccdc80 | -5.068 | 7.01E-08 | glio |
| Atr_in.vivo_noRT | Eef2 | -0.953 | 7.01E-08 | glio |
| Atr_ex.vivo_noRT | Gm42664 | 4.159 | 7.01E-08 | glio |
| culture_effect | Pxn | 1.284 | 7.01E-10 | glio |
| Bora_in.vivo_noRT | Gm13822 | 4.411 | 7.01E-11 | glio |
| Hsd17b10_in.vivo_noRT | 4930588A03R | 3.616 | 7.01E-14 | glio |
| interaction_RT | Pcsk1n | 3.254 | 7.01E-15 | glio |
| Atr_in.vivo_noRT | Gm14540 | 5.011 | 7.01E-21 | glio |
| Atr_in.vivo_noRT | Gm11973 | 5.267 | 7.01E-21 | glio |
| culture_effect | Rpl10 | -1.245 | 7.01E-22 | glio |
| culture_effect | Naa50 | 0.36 | 7.02E-03 | glio |
| Interaction_genotype_Aars2 | Ndufa6 | 0.969 | 7.02E-03 | glio |
| Interaction_genotype_Bora | Mfsd5 | 1.518 | 7.02E-03 | glio |
| Interaction_genotype_Chek1 | Nupr1 | 2.956 | 7.02E-03 | glio |
| Interaction_genotype_Lmo7 | 2010320O07R | -4.004 | 7.02E-03 | glio |
| Interaction_genotype_Lmo7 | Gm2115 | -3.675 | 7.02E-03 | glio |
| Interaction_genotype_Mtor | Rnh1 | 1.541 | 7.02E-03 | glio |
| Interaction_genotype_Myc | Eogt | -2.582 | 7.02E-03 | glio |
| Aars2_in.vivo_noRT | Nlgn2 | 1.734 | 7.02E-03 | glio |
| Aars2_ex.vivo_noRT | Col28a1 | 2.714 | 7.02E-03 | glio |
| Atp5c1_in.vivo_noRT | Zfp988 | 2.036 | 7.02E-03 | glio |
| Bcas2_in.vivo_noRT | Xpr1 | -1.519 | 7.02E-03 | glio |
| Cenpj_ex.vivo_noRT | Gm12589 | 2.192 | 7.02E-03 | glio |
| Interaction_genotype_Chek1 | Tmf1 | 2.395 | 7.02E-04 | glio |
| Aars2_in.vivo_noRT | Dlg5 | 2.285 | 7.02E-04 | glio |
| Atp5c1_in.vivo_noRT | Gm10696 | 2.668 | 7.02E-04 | glio |
| Atr_in.vivo_noRT | Usp35 | 2.406 | 7.02E-04 | glio |
| Atr_in.vivo_noRT | Yae1d1 | -1.512 | 7.02E-04 | glio |
| Bard1_ex.vivo_noRT | Bpifa3 | 2.814 | 7.02E-04 | glio |
| Bcas2_in.vivo_noRT | Slc27a3 | 2.463 | 7.02E-04 | glio |
| Bcas2_ex.vivo_noRT | Muc4 | 2.742 | 7.02E-04 | glio |
| Bcas2_ex.vivo_noRT | Ccdc80 | 2.555 | 7.02E-04 | glio |
| Bora_in.vivo_noRT | 3010001F23Ri | 2.307 | 7.02E-04 | glio |
| Bora_in.vivo_noRT | E430018J23Ri | 2.345 | 7.02E-04 | glio |
| culture_effect | Cyp2u1 | 1.881 | 7.02E-05 | glio |
| Aars2_ex.vivo_noRT | 4930553M12F | 2.899 | 7.02E-05 | glio |
| Atp5c1_in.vivo_noRT | Cd248 | 2.933 | 7.02E-05 | glio |
| Atr_in.vivo_noRT | Yars | -1.689 | 7.02E-05 | glio |
| Bcas2_in.vivo_noRT | Itgb6 | 2.879 | 7.02E-05 | glio |
| Bcas2_in.vivo_noRT | Adamts7 | 2.636 | 7.02E-05 | glio |
| culture_effect | Srprb | -0.679 | 7.02E-06 | glio |
| Atp5c1_in.vivo_noRT | Slc19a3 | 2.668 | 7.02E-07 | glio |
| Chek1_ex.vivo_noRT | Ankrd33b | 4.842 | 7.02E-08 | glio |
| Chek1_ex.vivo_noRT | Hnf1aos1 | 4.045 | 7.02E-08 | glio |
| Aars2_in.vivo_noRT | Zfp652os | 4.021 | 7.02E-09 | glio |
| Hsd17b10_in.vivo_noRT | Speer4a | 3.35 | 7.02E-11 | glio |
| culture_effect | Tslp | 3.647 | 7.02E-12 | glio |
| Bora_in.vivo_noRT | Gm13449 | 4.936 | 7.02E-20 | glio |
| Bora_in.vivo_noRT | Gm47622 | 4.936 | 7.02E-21 | glio |
| culture_effect | Hspb2 | 3.085 | 7.02E-35 | glio |
| Interaction_genotype_Atr | Gm44983 | -2.868 | 7.03E-03 | glio |

| | | | | |
|---|---|---|---|---|
| Interaction_genotype_Atr | Klhdc10 | 1.812 | 7.03E-03 | glio |
| Interaction_genotype_Ifnar1 | Zscan4-ps3 | 2.734 | 7.03E-03 | glio |
| Interaction_genotype_Ifnar1 | Gm21731 | 2.734 | 7.03E-03 | glio |
| Interaction_genotype_Ifnar1 | Gm11232 | 2.734 | 7.03E-03 | glio |
| Interaction_genotype_Ifnar1 | Gm11583 | 2.999 | 7.03E-03 | glio |
| Interaction_genotype_Mtor | Unc5cl | -3.341 | 7.03E-03 | glio |
| Interaction_genotype_Mtor | Gm16174 | -3.516 | 7.03E-03 | glio |
| Interaction_genotype_Myc | Fbxo25 | -2.266 | 7.03E-03 | glio |
| Aars2_in.vivo_noRT | Nhlrc3 | 1.624 | 7.03E-03 | glio |
| Chek1_in.vivo_noRT | Lrig1 | 2.13 | 7.03E-03 | glio |
| Chek1_in.vivo_noRT | Efr3a | -0.838 | 7.03E-03 | glio |
| Chek1_ex.vivo_noRT | Lrrn3 | 2.353 | 7.03E-03 | glio |
| Eif2s1_ex.vivo_noRT | Fabp6 | 2.234 | 7.03E-03 | glio |
| Interaction_genotype_Mtor | B230206L02Ri | -3.529 | 7.03E-04 | glio |
| Bcas2_ex.vivo_noRT | Phf1 | -2.872 | 7.03E-04 | glio |
| Chek1_in.vivo_noRT | Stpg4 | 2.555 | 7.03E-04 | glio |
| Chek1_in.vivo_noRT | Rtl5 | 2.375 | 7.03E-04 | glio |
| Chek1_in.vivo_noRT | Dusp15 | 2.195 | 7.03E-04 | glio |
| Chek1_in.vivo_noRT | Brca1 | -1.846 | 7.03E-04 | glio |
| Chek1_in.vivo_noRT | Med8 | -0.911 | 7.03E-04 | glio |
| Hsd17b10_in.vivo_noRT | Traf3ip3 | 2.552 | 7.03E-04 | glio |
| Atr_ex.vivo_noRT | Gm26542 | 3.564 | 7.03E-05 | glio |
| Bora_in.vivo_noRT | Prrg1 | 2.912 | 7.03E-05 | glio |
| Bora_in.vivo_noRT | Serac1 | 2.707 | 7.03E-05 | glio |
| Bora_in.vivo_noRT | Mtrex | -1.625 | 7.03E-05 | glio |
| Hsd17b10_in.vivo_noRT | Armc2 | 3.225 | 7.03E-06 | glio |
| Bard1_ex.vivo_noRT | Slc17a6 | 2.814 | 7.03E-07 | glio |
| Bcas2_in.vivo_noRT | Rcan2 | 4.807 | 7.03E-11 | glio |
| Bcas2_in.vivo_noRT | Pcdhga11 | 3.81 | 7.03E-11 | glio |
| Chek1_in.vivo_noRT | Icam2 | 4.177 | 7.03E-12 | glio |
| Chek1_ex.vivo_noRT | 2300002M23F | 4.842 | 7.03E-12 | glio |
| culture_effect | Gm32591 | 1.595 | 7.04E-03 | glio |
| culture_effect | Nptxr | -0.911 | 7.04E-03 | glio |
| Interaction_genotype_Aars2 | Slc7a14 | -3.267 | 7.04E-03 | glio |
| Interaction_genotype_Aars2 | Slc35e2 | -2.197 | 7.04E-03 | glio |
| Interaction_genotype_Atr | Rab40b | -3.325 | 7.04E-03 | glio |
| Interaction_genotype_Rpl3 | Gtf2i | 1.786 | 7.04E-03 | glio |
| interaction_RT | Kars | -0.599 | 7.04E-03 | glio |
| Atr_in.vivo_noRT | Larp1 | -0.794 | 7.04E-03 | glio |
| Atr_ex.vivo_noRT | Fhad1 | 2.577 | 7.04E-03 | glio |
| Bcas2_in.vivo_noRT | Ccdc73 | 1.776 | 7.04E-03 | glio |
| Bora_in.vivo_noRT | Tbcc | -1.274 | 7.04E-03 | glio |
| Chek1_in.vivo_noRT | Morf4l1 | -0.56 | 7.04E-03 | glio |
| Interaction_genotype_Mtor | C630043F03Ri | -3.235 | 7.04E-04 | glio |
| Aars2_in.vivo_noRT | Cr2 | 2.037 | 7.04E-04 | glio |
| Aars2_in.vivo_noRT | Abhd12 | -1.438 | 7.04E-04 | glio |
| Atr_in.vivo_noRT | Gm38394 | 2.354 | 7.04E-04 | glio |
| Bcas2_in.vivo_noRT | Car5a | 2.243 | 7.04E-04 | glio |
| Chek1_in.vivo_noRT | Acbd7 | 2.54 | 7.04E-04 | glio |
| Chek1_ex.vivo_noRT | Gm26682 | 2.927 | 7.04E-04 | glio |
| Chek1_ex.vivo_noRT | Ube2s | -0.836 | 7.04E-04 | glio |
| Eif2s1_in.vivo_noRT | Gem | 2.674 | 7.04E-04 | glio |

| | | | | |
|---|---|---|---|---|
| culture_effect | Cd59a | 1.886 | 7.04E-05 | glio |
| Interaction_genotype_Mtor | Fam71e1 | -3.679 | 7.04E-05 | glio |
| interaction_RT | Casp4 | 2.604 | 7.04E-05 | glio |
| Aars2_in.vivo_noRT | Crybg3 | 2.912 | 7.04E-05 | glio |
| Aars2_in.vivo_noRT | Sar1a | -1.341 | 7.04E-05 | glio |
| Chek1_in.vivo_noRT | Bcl2l14 | 3.003 | 7.04E-05 | glio |
| Chek1_ex.vivo_noRT | Icam1 | 3.705 | 7.04E-05 | glio |
| culture_effect | Tgif1 | -0.821 | 7.04E-06 | glio |
| Interaction_genotype_Mtor | Cfap100 | -4.51 | 7.04E-06 | glio |
| Bcas2_in.vivo_noRT | Tbxa2r | 2.968 | 7.04E-06 | glio |
| Bora_in.vivo_noRT | Smc2os | 3.54 | 7.04E-07 | glio |
| Atp5c1_in.vivo_noRT | Gm34059 | 2.668 | 7.04E-08 | glio |
| Atr_in.vivo_noRT | Cfap46 | 5.066 | 7.04E-16 | glio |
| Interaction_genotype_Mtor | Slc17a9 | -3.215 | 7.05E-03 | glio |
| Interaction_genotype_Mtor | Rps4x | 1.044 | 7.05E-03 | glio |
| Aars2_in.vivo_noRT | Skap2 | -0.867 | 7.05E-03 | glio |
| Atr_in.vivo_noRT | Aqp11 | 1.96 | 7.05E-03 | glio |
| Cdkn2a_in.vivo_noRT | Gm48435 | -1.675 | 7.05E-03 | glio |
| Cenpj_ex.vivo_noRT | Obscn | 2.591 | 7.05E-03 | glio |
| Chek1_in.vivo_noRT | Tent2 | -1.268 | 7.05E-03 | glio |
| Chek1_in.vivo_noRT | Ezr | -1.016 | 7.05E-03 | glio |
| Chek1_ex.vivo_noRT | Gm12602 | 2.334 | 7.05E-03 | glio |
| Hsd17b10_in.vivo_noRT | Gm35066 | 2.124 | 7.05E-03 | glio |
| Hsd17b10_in.vivo_noRT | Gar1 | -0.679 | 7.05E-03 | glio |
| Interaction_genotype_Mtor | Gm13822 | -3.969 | 7.05E-04 | glio |
| Interaction_genotype_Myc | Ppp1r26 | -4.052 | 7.05E-04 | glio |
| Interaction_genotype_Myc | Bax | 1.412 | 7.05E-04 | glio |
| Interaction_genotype_Rpl3 | Fbln5 | 2.952 | 7.05E-04 | glio |
| Atr_in.vivo_noRT | H3f3a | -0.608 | 7.05E-04 | glio |
| Bard1_in.vivo_noRT | Vmn2r18 | 2.228 | 7.05E-04 | glio |
| Bora_in.vivo_noRT | A930003O13R | 3.137 | 7.05E-05 | glio |
| Chek1_in.vivo_noRT | Hist1h2ai | 2.947 | 7.05E-05 | glio |
| Bard1_ex.vivo_noRT | 1700003E24Ri | 2.814 | 7.05E-06 | glio |
| Chek1_ex.vivo_noRT | Adgrd1 | 4.132 | 7.05E-07 | glio |
| Atr_in.vivo_noRT | Gm50107 | 4.018 | 7.05E-10 | glio |
| Chek1_in.vivo_noRT | Rcan2 | 5.044 | 7.05E-12 | glio |
| Chek1_in.vivo_noRT | Nat8f4 | 4.511 | 7.05E-13 | glio |
| Atr_in.vivo_noRT | Slc10a1 | 5.273 | 7.05E-21 | glio |
| Interaction_genotype_Mtor | Bok | -2.949 | 7.06E-03 | glio |
| Interaction_genotype_Myc | Slc27a3 | -3.092 | 7.06E-03 | glio |
| Interaction_genotype_Myc | Ccdc106 | -3.228 | 7.06E-03 | glio |
| interaction_RT | Hmgb3 | 0.693 | 7.06E-03 | glio |
| Atr_in.vivo_noRT | Dcun1d2 | 1.428 | 7.06E-03 | glio |
| Cenpj_in.vivo_noRT | Folr1 | 2.022 | 7.06E-03 | glio |
| Eif2s1_ex.vivo_noRT | Gm44646 | 2.427 | 7.06E-03 | glio |
| Eif2s1_ex.vivo_noRT | Gm26978 | 2.446 | 7.06E-03 | glio |
| Eif2s1_ex.vivo_noRT | Gipc2 | 3.117 | 7.06E-03 | glio |
| Eif2s1_ex.vivo_noRT | 4930512B01R | 2.234 | 7.06E-03 | glio |
| culture_effect | Egln1 | 0.553 | 7.06E-04 | glio |
| Interaction_genotype_Mtor | Gm45605 | -3.888 | 7.06E-04 | glio |
| Interaction_genotype_Myc | Rad51b | -3.479 | 7.06E-04 | glio |
| Interaction_genotype_Rpl3 | Krt80 | -4.127 | 7.06E-04 | glio |

| | | | | |
|---|---|---|---|---|
| Atr_in.vivo_noRT | Dpf1 | 2.103 | 7.06E-04 | glio |
| Bora_in.vivo_noRT | Nsun7 | 2.673 | 7.06E-04 | glio |
| Bora_ex.vivo_noRT | Gm38393 | 2.808 | 7.06E-04 | glio |
| Cenpj_ex.vivo_noRT | Gm47936 | 2.591 | 7.06E-04 | glio |
| Chek1_ex.vivo_noRT | 4930597O21R | 3.344 | 7.06E-04 | glio |
| Eif2s1_in.vivo_noRT | Nr4a1 | -1.167 | 7.06E-04 | glio |
| Bcas2_in.vivo_noRT | 4930597O21R | 3.079 | 7.06E-05 | glio |
| Interaction_genotype_Mtor | 4930470G03R | -4.226 | 7.06E-06 | glio |
| Atr_ex.vivo_noRT | Fam166a | 3.164 | 7.06E-06 | glio |
| Bcas2_in.vivo_noRT | Zfp467 | 3.267 | 7.06E-06 | glio |
| Hsd17b10_in.vivo_noRT | BC028471 | 3.611 | 7.06E-07 | glio |
| Atr_ex.vivo_noRT | Fam227b | 3.164 | 7.06E-08 | glio |
| Atr_ex.vivo_noRT | Gm20658 | 3.164 | 7.06E-08 | glio |
| Bcas2_in.vivo_noRT | 1700047K16R | 4.149 | 7.06E-09 | glio |
| Bcas2_in.vivo_noRT | 2810408A11R | 4.062 | 7.06E-09 | glio |
| Bora_in.vivo_noRT | Ccl7 | 4.936 | 7.06E-10 | glio |
| Bora_in.vivo_noRT | Ptp4a1.1 | 4.139 | 7.06E-10 | glio |
| Chek1_ex.vivo_noRT | Adgre5 | 4.449 | 7.06E-10 | glio |
| Aars2_in.vivo_noRT | Dreh | 4.68 | 7.06E-14 | glio |
| Aars2_in.vivo_noRT | Gm35696 | 5.067 | 7.06E-15 | glio |
| culture_effect | Anapc13 | 0.391 | 7.07E-03 | glio |
| Interaction_genotype_Atr | Ifi213 | 2.083 | 7.07E-03 | glio |
| Interaction_genotype_Myc | Rnf4 | 1.128 | 7.07E-03 | glio |
| interaction_RT | Lamp1 | 0.509 | 7.07E-03 | glio |
| Aars2_in.vivo_noRT | Nup107 | -1.294 | 7.07E-03 | glio |
| Aars2_in.vivo_noRT | Mrps18b | -0.85 | 7.07E-03 | glio |
| Brca2_in.vivo_noRT | Olfr552 | 1.763 | 7.07E-03 | glio |
| Chek1_in.vivo_noRT | Rragd | 2.097 | 7.07E-03 | glio |
| Chek1_in.vivo_noRT | Pole3 | -0.838 | 7.07E-03 | glio |
| Chek1_in.vivo_noRT | Rsl1d1 | -0.731 | 7.07E-03 | glio |
| Chek1_ex.vivo_noRT | Frmd4b | 2.59 | 7.07E-03 | glio |
| Hsd17b10_in.vivo_noRT | Tmeff2 | 2.359 | 7.07E-03 | glio |
| Interaction_genotype_Mtor | Birc5 | 2.75 | 7.07E-04 | glio |
| Interaction_genotype_Mtor | Luc7l3 | 1.66 | 7.07E-04 | glio |
| Aars2_in.vivo_noRT | Zfp983 | 2.095 | 7.07E-04 | glio |
| Aars2_in.vivo_noRT | Chmp6 | -1.491 | 7.07E-04 | glio |
| Aars2_ex.vivo_noRT | Selenbp1 | 3.67 | 7.07E-04 | glio |
| Atp5c1_in.vivo_noRT | Crybb1 | 2.281 | 7.07E-04 | glio |
| Atr_ex.vivo_noRT | Olfr90 | 2.771 | 7.07E-04 | glio |
| Bcas2_in.vivo_noRT | Gal3st4 | 2.448 | 7.07E-04 | glio |
| Bora_in.vivo_noRT | N4bp2os | 2.508 | 7.07E-04 | glio |
| Chek1_ex.vivo_noRT | Plekhh1 | 3.209 | 7.07E-04 | glio |
| Hsd17b10_in.vivo_noRT | Gm17210 | 2.556 | 7.07E-04 | glio |
| Interaction_genotype_Mtor | Bhlhe41 | -5.432 | 7.07E-05 | glio |
| Aars2_in.vivo_noRT | Snrpd2 | -0.832 | 7.07E-05 | glio |
| Atp5c1_in.vivo_noRT | Cfap46 | 2.732 | 7.07E-05 | glio |
| Bard1_ex.vivo_noRT | Gm14820 | 2.814 | 7.07E-05 | glio |
| Bora_in.vivo_noRT | Pcdhb19 | 2.645 | 7.07E-05 | glio |
| Hsd17b10_in.vivo_noRT | Gm8300 | 3.616 | 7.07E-05 | glio |
| Hsd17b10_in.vivo_noRT | A730071L15Ri | 3.083 | 7.07E-05 | glio |
| Hsd17b10_in.vivo_noRT | Ifit1bl1 | 3.605 | 7.07E-05 | glio |
| Interaction_genotype_Atr | 2610027K06R | -6.107 | 7.07E-08 | glio |

| | | | | |
|---|---|---|---|---|
| Aars2_in.vivo_noRT | Fsd2 | 3.944 | 7.07E-09 | glio |
| Atr_in.vivo_noRT | Gm47223 | 5.267 | 7.07E-09 | glio |
| Atr_in.vivo_noRT | Myzap | 3.749 | 7.07E-11 | glio |
| Chek1_in.vivo_noRT | Gm27253 | 4.777 | 7.07E-12 | glio |
| culture_effect | Gm11627 | 3.74 | 7.07E-15 | glio |
| culture_effect | Gm12976 | 1.372 | 7.08E-03 | glio |
| Interaction_genotype_Aars2 | Gm12841 | -2.427 | 7.08E-03 | glio |
| Interaction_genotype_Aars2 | Eng | -3.589 | 7.08E-03 | glio |
| Interaction_genotype_Aars2 | Ndufaf2 | 1.319 | 7.08E-03 | glio |
| Interaction_genotype_Bcas2 | Gm12258 | -3.801 | 7.08E-03 | glio |
| Interaction_genotype_Bora | Zfp810 | -3.517 | 7.08E-03 | glio |
| Interaction_genotype_Mtor | Mapkapk5 | -3.213 | 7.08E-03 | glio |
| Interaction_genotype_Mtor | Disp1 | -2.587 | 7.08E-03 | glio |
| Interaction_genotype_Mtor | Ttc23 | -2.345 | 7.08E-03 | glio |
| Interaction_genotype_Mtor | Gm17435 | -2.813 | 7.08E-03 | glio |
| Interaction_genotype_Myc | C430042M11R | -3.105 | 7.08E-03 | glio |
| Interaction_genotype_Myc | Txnl4b | -2.508 | 7.08E-03 | glio |
| Interaction_genotype_Myc | Ssb | 1.073 | 7.08E-03 | glio |
| Aars2_in.vivo_noRT | Zfp600 | 1.951 | 7.08E-03 | glio |
| Atr_in.vivo_noRT | Acsf3 | 1.678 | 7.08E-03 | glio |
| Bard1_ex.vivo_noRT | Npnt | 2.017 | 7.08E-03 | glio |
| Bcas2_in.vivo_noRT | Klhl33 | 2.007 | 7.08E-03 | glio |
| Chek1_in.vivo_noRT | Zfp94 | 1.905 | 7.08E-03 | glio |
| Hsd17b10_in.vivo_noRT | Gm15535 | 2.299 | 7.08E-03 | glio |
| culture_effect | Gtf2a2 | 0.385 | 7.08E-04 | glio |
| Interaction_genotype_Chek1 | Ep400 | -2.47 | 7.08E-04 | glio |
| Interaction_genotype_Mtor | Sh3gl3 | -4.101 | 7.08E-04 | glio |
| Interaction_genotype_Mtor | 9930014A18R | -3.903 | 7.08E-04 | glio |
| Interaction_genotype_Mtor | Hoxa6 | -3.833 | 7.08E-04 | glio |
| Interaction_genotype_Mtor | Grpel1 | 1.628 | 7.08E-04 | glio |
| Atp5c1_in.vivo_noRT | Muc4 | 2.146 | 7.08E-04 | glio |
| Atp5c1_in.vivo_noRT | Fgd3 | 2.877 | 7.08E-04 | glio |
| Atr_in.vivo_noRT | Pacsin1 | 2.188 | 7.08E-04 | glio |
| Atr_in.vivo_noRT | Ndufb5 | -0.874 | 7.08E-04 | glio |
| Bora_in.vivo_noRT | Cd302 | -1.385 | 7.08E-04 | glio |
| Interaction_genotype_Mtor | Serpinb9b | -5.758 | 7.08E-05 | glio |
| Aars2_in.vivo_noRT | Cd72 | 2.765 | 7.08E-05 | glio |
| Aars2_in.vivo_noRT | Rpgr | 2.652 | 7.08E-05 | glio |
| Atp5c1_in.vivo_noRT | Slc10a1 | 2.408 | 7.08E-05 | glio |
| Bcas2_in.vivo_noRT | Lrrc75aos2 | 2.911 | 7.08E-05 | glio |
| Bcas2_in.vivo_noRT | Rnf122 | 2.468 | 7.08E-05 | glio |
| Bcas2_in.vivo_noRT | Paics | -0.884 | 7.08E-05 | glio |
| Bora_in.vivo_noRT | Myom1 | 2.943 | 7.08E-05 | glio |
| Chek1_in.vivo_noRT | Nat14 | 2.478 | 7.08E-05 | glio |
| Hsd17b10_in.vivo_noRT | Gm8013 | 2.971 | 7.08E-05 | glio |
| Hsd17b10_in.vivo_noRT | Rab11fip4os1 | 3.107 | 7.08E-05 | glio |
| Hsd17b10_in.vivo_noRT | Celf5 | 3.084 | 7.08E-05 | glio |
| culture_effect | Dnajb1 | -0.802 | 7.08E-06 | glio |
| Aars2_in.vivo_noRT | Sfr1 | -1.059 | 7.08E-06 | glio |
| Chek1_in.vivo_noRT | Gng8 | 3.068 | 7.08E-06 | glio |
| Interaction_genotype_Mtor | Gm36210 | -5.958 | 7.08E-07 | glio |
| Bora_in.vivo_noRT | Bbs1 | 3.477 | 7.08E-07 | glio |

| | | | | |
|---|---|---|---|---|
| Bora_ex.vivo_noRT | Dpep3 | 3.201 | 7.08E-07 | glio |
| Chek1_in.vivo_noRT | Gm43581 | 3.474 | 7.08E-08 | glio |
| Bcas2_in.vivo_noRT | Asgr1 | 4.149 | 7.08E-09 | glio |
| Atr_in.vivo_noRT | 1810012K16R | 4.017 | 7.08E-10 | glio |
| Aars2_in.vivo_noRT | Slc6a11 | 3.895 | 7.08E-12 | glio |
| Aars2_in.vivo_noRT | Carns1 | 5.327 | 7.08E-14 | glio |
| Aars2_in.vivo_noRT | Gm19265 | 4.811 | 7.08E-18 | glio |
| Interaction_genotype_Atr | Gldc | -3.231 | 7.09E-03 | glio |
| Interaction_genotype_Atr | Prdm10 | -2.997 | 7.09E-03 | glio |
| Interaction_genotype_Atr | 0610009L18Ri | -2.949 | 7.09E-03 | glio |
| Interaction_genotype_Mtor | Vmn1r88 | -2.914 | 7.09E-03 | glio |
| Interaction_genotype_Mtor | Esyt3 | -3.212 | 7.09E-03 | glio |
| Interaction_genotype_Mtor | Zfp992 | -3.439 | 7.09E-03 | glio |
| Interaction_genotype_Mtor | Zfp57 | -3.153 | 7.09E-03 | glio |
| Interaction_genotype_Mtor | Zfp346 | -2.916 | 7.09E-03 | glio |
| Interaction_genotype_Mtor | Mgst3 | -2.784 | 7.09E-03 | glio |
| Interaction_genotype_Mtor | Vps37c | -2.41 | 7.09E-03 | glio |
| Interaction_genotype_Myc | Pcdhga1 | -2.647 | 7.09E-03 | glio |
| Interaction_genotype_Myc | Xlr4c | -3.139 | 7.09E-03 | glio |
| Interaction_genotype_Myc | Gm10382 | -3.13 | 7.09E-03 | glio |
| Interaction_genotype_Myc | Shroom1 | -2.945 | 7.09E-03 | glio |
| Interaction_genotype_Myc | Nek3 | -3.1 | 7.09E-03 | glio |
| Interaction_genotype_Myc | Slc37a1 | -2.389 | 7.09E-03 | glio |
| Interaction_genotype_Rad51 | Gm20337 | -3.863 | 7.09E-03 | glio |
| Aars2_in.vivo_noRT | Spats1 | 1.966 | 7.09E-03 | glio |
| Aars2_in.vivo_noRT | Gm12976 | 2.239 | 7.09E-03 | glio |
| Aars2_in.vivo_noRT | Zkscan8 | 1.684 | 7.09E-03 | glio |
| Atr_in.vivo_noRT | Gm4070 | 1.978 | 7.09E-03 | glio |
| Atr_in.vivo_noRT | Rnf182 | 1.751 | 7.09E-03 | glio |
| Bcas2_in.vivo_noRT | Hemk1 | 2.042 | 7.09E-03 | glio |
| Bora_in.vivo_noRT | Abca13 | 2.205 | 7.09E-03 | glio |
| Bora_in.vivo_noRT | Myt1 | 1.459 | 7.09E-03 | glio |
| Bora_ex.vivo_noRT | Cox7b2 | 2.404 | 7.09E-03 | glio |
| Chek1_in.vivo_noRT | Nckap1l | 1.976 | 7.09E-03 | glio |
| Chek1_in.vivo_noRT | Slc26a11 | 1.808 | 7.09E-03 | glio |
| Chek1_in.vivo_noRT | Wdr24 | 1.396 | 7.09E-03 | glio |
| Interaction_genotype_Atr | C230037L18Ri | -3.92 | 7.09E-04 | glio |
| Interaction_genotype_Atr | Gm13594 | -4.354 | 7.09E-04 | glio |
| Interaction_genotype_Bcas2 | Tnfrsf23 | -4.927 | 7.09E-04 | glio |
| Interaction_genotype_Myc | 1810062O18R | -4.022 | 7.09E-04 | glio |
| Interaction_genotype_Myc | Vcan | -4.163 | 7.09E-04 | glio |
| Interaction_genotype_Myc | D11Wsu47e | -2.928 | 7.09E-04 | glio |
| Interaction_genotype_Myc | Polr3c | -2.516 | 7.09E-04 | glio |
| Atr_in.vivo_noRT | Pcdhb21 | 2.239 | 7.09E-04 | glio |
| Bora_in.vivo_noRT | Slurp1 | 3.642 | 7.09E-04 | glio |
| Bora_ex.vivo_noRT | Kmo | 2.808 | 7.09E-04 | glio |
| Chek1_in.vivo_noRT | Pcdh15 | 1.993 | 7.09E-04 | glio |
| culture_effect | Ints6l | -1.385 | 7.09E-05 | glio |
| culture_effect | Shisa8 | -1.9 | 7.09E-05 | glio |
| culture_effect | Xlr3b | -0.851 | 7.09E-05 | glio |
| culture_effect | Nup107 | -0.828 | 7.09E-05 | glio |
| Bcas2_in.vivo_noRT | Fam169a | 2.716 | 7.09E-05 | glio |

| | | | | |
|---|---|---|---|---|
| Bcas2_in.vivo_noRT | Gm2822 | 3.108 | 7.09E-05 | glio |
| Bcas2_in.vivo_noRT | Uqcc2 | -0.891 | 7.09E-05 | glio |
| Atr_in.vivo_noRT | Gm12258 | 3.327 | 7.09E-06 | glio |
| Hsd17b10_in.vivo_noRT | Cacnb1 | 3.542 | 7.09E-06 | glio |
| Atr_in.vivo_noRT | Etaa1os | 3.321 | 7.09E-07 | glio |
| Chek1_in.vivo_noRT | Selenok-ps1 | 3.48 | 7.09E-08 | glio |
| culture_effect | Ndst1 | 1.104 | 7.09E-09 | glio |
| culture_effect | Cers2 | -0.827 | 7.09E-13 | glio |
| culture_effect | Rarres1 | 1.629 | 7.10E-03 | glio |
| Interaction_genotype_Aars2 | Lsp1 | -3.791 | 7.10E-03 | glio |
| Interaction_genotype_Aars2 | Smc2os | -3.396 | 7.10E-03 | glio |
| Interaction_genotype_Aars2 | 1700029J07Ri | -3.373 | 7.10E-03 | glio |
| Interaction_genotype_Bcas2 | Apol10b | 3.105 | 7.10E-03 | glio |
| Interaction_genotype_Bora | Gm44899 | -2.839 | 7.10E-03 | glio |
| Interaction_genotype_Bora | Zfp551 | -3.525 | 7.10E-03 | glio |
| Interaction_genotype_Bora | Lypd6 | -3.634 | 7.10E-03 | glio |
| Interaction_genotype_Bora | Kcnmb4os2 | -3.455 | 7.10E-03 | glio |
| Interaction_genotype_Bora | Fbn2 | 2.553 | 7.10E-03 | glio |
| Atr_ex.vivo_noRT | Il2rb | 2.583 | 7.10E-03 | glio |
| Bard1_in.vivo_noRT | Npy4r | 2.459 | 7.10E-03 | glio |
| Bard1_ex.vivo_noRT | Ly6c2 | 2.814 | 7.10E-03 | glio |
| Bard1_ex.vivo_noRT | Gm11973 | 2.193 | 7.10E-03 | glio |
| Bcas2_in.vivo_noRT | Sgta | -0.905 | 7.10E-03 | glio |
| Bora_in.vivo_noRT | Pcdhga12 | 1.94 | 7.10E-03 | glio |
| Bora_in.vivo_noRT | D630045J12Ri | -1.383 | 7.10E-03 | glio |
| Bora_in.vivo_noRT | Msh2 | -1.189 | 7.10E-03 | glio |
| Bora_in.vivo_noRT | Tceal9 | -1.135 | 7.10E-03 | glio |
| Chek1_in.vivo_noRT | Osbpl11 | 1.699 | 7.10E-03 | glio |
| Eif2s1_in.vivo_noRT | Gm11627 | 2.143 | 7.10E-03 | glio |
| Eif2s1_in.vivo_noRT | Fam71e1 | 2.036 | 7.10E-03 | glio |
| Interaction_genotype_Atr | Olfr279 | -4.145 | 7.10E-04 | glio |
| Interaction_genotype_Myc | B230206H07R | -2.599 | 7.10E-04 | glio |
| Atr_in.vivo_noRT | Sphk1 | 2.447 | 7.10E-04 | glio |
| Atr_in.vivo_noRT | Foxg1 | -1.647 | 7.10E-04 | glio |
| Eif2s1_in.vivo_noRT | Adgra3 | -0.972 | 7.10E-04 | glio |
| culture_effect | Ccsap | 1.016 | 7.10E-05 | glio |
| Interaction_genotype_Aars2 | D630036H23R | -4.751 | 7.10E-05 | glio |
| Interaction_genotype_Aars2 | Gm16741 | -4.574 | 7.10E-05 | glio |
| Interaction_genotype_Mtor | Gm5544 | -5.402 | 7.10E-06 | glio |
| Atr_ex.vivo_noRT | Gm15894 | 3.164 | 7.10E-06 | glio |
| Atp5c1_in.vivo_noRT | Gm48435 | 2.668 | 7.10E-08 | glio |
| Chek1_ex.vivo_noRT | Nt5e | 4.449 | 7.10E-09 | glio |
| Chek1_in.vivo_noRT | A930001C03R | 4.391 | 7.10E-11 | glio |
| Chek1_in.vivo_noRT | Zg16 | 4.391 | 7.10E-11 | glio |
| Chek1_in.vivo_noRT | Clca3a2 | 4.369 | 7.10E-13 | glio |
| Bora_in.vivo_noRT | Gm14019 | 4.936 | 7.10E-16 | glio |
| Atr_in.vivo_noRT | Mecom | 5.533 | 7.10E-17 | glio |
| culture_effect | Entpd2 | 2.741 | 7.10E-20 | glio |
| Chek1_in.vivo_noRT | Cldn2 | 5.044 | 7.10E-27 | glio |
| culture_effect | Hist1h4c | 1.465 | 7.11E-03 | glio |
| Interaction_genotype_Aars2 | Cyp2s1 | -3.586 | 7.11E-03 | glio |
| Interaction_genotype_Aars2 | Prdx6 | 1.209 | 7.11E-03 | glio |

| | | | | |
|---|---|---|---|---|
| Aars2_in.vivo_noRT | Dnaja1 | -0.558 | 7.11E-03 | glio |
| Bora_in.vivo_noRT | Gm43813 | 1.866 | 7.11E-03 | glio |
| Bora_in.vivo_noRT | Eif3l | -0.683 | 7.11E-03 | glio |
| Bora_ex.vivo_noRT | Gm16754 | 2.803 | 7.11E-03 | glio |
| Cenpj_in.vivo_noRT | Gm15127 | 2.292 | 7.11E-03 | glio |
| Cenpj_in.vivo_noRT | Gm5152 | 2.405 | 7.11E-03 | glio |
| Cenpj_in.vivo_noRT | Crabp1 | 2.244 | 7.11E-03 | glio |
| Cenpj_ex.vivo_noRT | Prkcq | 2.591 | 7.11E-03 | glio |
| interaction_RT | Higd2a | 0.793 | 7.11E-04 | glio |
| Aars2_in.vivo_noRT | Arfrp1 | -1.27 | 7.11E-04 | glio |
| Interaction_genotype_Mtor | Stard10 | -4.594 | 7.11E-05 | glio |
| Interaction_genotype_Mtor | Neurl2 | -3.867 | 7.11E-05 | glio |
| Interaction_genotype_Mtor | Rras | 2.266 | 7.11E-05 | glio |
| Bora_in.vivo_noRT | Hmgb2 | -1.741 | 7.11E-05 | glio |
| culture_effect | Gm14295 | 1.709 | 7.11E-07 | glio |
| culture_effect | Naa38 | 0.666 | 7.11E-07 | glio |
| Aars2_in.vivo_noRT | Gm9833 | 3.546 | 7.11E-08 | glio |
| Aars2_ex.vivo_noRT | A930019D19R | 3.293 | 7.11E-08 | glio |
| Chek1_ex.vivo_noRT | Zfp352 | 4.842 | 7.11E-10 | glio |
| Aars2_in.vivo_noRT | 4930558J18Ri | 4.257 | 7.11E-11 | glio |
| Aars2_in.vivo_noRT | Kcnj13 | 4.046 | 7.11E-11 | glio |
| Chek1_in.vivo_noRT | Sprr1a | 5.044 | 7.11E-13 | glio |
| Atr_in.vivo_noRT | Gm15127 | 5.273 | 7.11E-15 | glio |
| culture_effect | Gatm | -2.239 | 7.11E-20 | glio |
| Interaction_genotype_Aars2 | Odc1 | 1.689 | 7.12E-03 | glio |
| Interaction_genotype_Bcas2 | Zfp786 | -3.527 | 7.12E-03 | glio |
| Interaction_genotype_Bcas2 | Slc25a35 | -3.561 | 7.12E-03 | glio |
| Interaction_genotype_Bcas2 | 2310058D17R | -3.478 | 7.12E-03 | glio |
| Interaction_genotype_Mtor | Zc4h2 | -2.686 | 7.12E-03 | glio |
| Interaction_genotype_Mtor | Brd3os | -2.803 | 7.12E-03 | glio |
| Interaction_genotype_Mtor | Eef1akmt4 | -2.148 | 7.12E-03 | glio |
| Interaction_genotype_Mtor | Prkca | -2.144 | 7.12E-03 | glio |
| Interaction_genotype_Rpl3 | Hist2h3c1 | -3.779 | 7.12E-03 | glio |
| interaction_RT | Tbc1d14 | -0.934 | 7.12E-03 | glio |
| interaction_RT | Arf6 | 0.576 | 7.12E-03 | glio |
| Aars2_in.vivo_noRT | Pim3 | -1.114 | 7.12E-03 | glio |
| Atp5c1_in.vivo_noRT | Gm49441 | 1.877 | 7.12E-03 | glio |
| Bcas2_in.vivo_noRT | Gm15892 | 2.134 | 7.12E-03 | glio |
| Bora_in.vivo_noRT | Siae | 1.817 | 7.12E-03 | glio |
| Bora_ex.vivo_noRT | Agap2 | 2.216 | 7.12E-03 | glio |
| Chek1_in.vivo_noRT | Rnaseh2a | -1.199 | 7.12E-03 | glio |
| Interaction_genotype_Mtor | Gm43156 | -3.749 | 7.12E-04 | glio |
| Atr_in.vivo_noRT | Nutf2 | 2.378 | 7.12E-04 | glio |
| interaction_RT | Lig1 | -1.215 | 7.12E-05 | glio |
| Aars2_in.vivo_noRT | Gm29107 | 2.89 | 7.12E-05 | glio |
| culture_effect | Rgs5 | -3.505 | 7.12E-06 | glio |
| Atr_in.vivo_noRT | Spink10 | 2.959 | 7.12E-06 | glio |
| Bcas2_in.vivo_noRT | Wipf1 | -1.86 | 7.12E-06 | glio |
| Bora_in.vivo_noRT | Gm26619 | 3.34 | 7.12E-06 | glio |
| Chek1_in.vivo_noRT | 1700101I11Ri | 3.403 | 7.12E-06 | glio |
| Chek1_in.vivo_noRT | Wif1 | 3.567 | 7.12E-06 | glio |
| Chek1_in.vivo_noRT | Nbeal2 | 3.284 | 7.12E-06 | glio |

| | | | | |
|---|---|---|---|---|
| Chek1_ex.vivo_noRT | Otop1 | 3.672 | 7.12E-06 | glio |
| Interaction_genotype_Mtor | Gm16364.1 | -6.352 | 7.12E-07 | glio |
| Interaction_genotype_Mtor | 1700013F07Ri | -6.353 | 7.12E-07 | glio |
| Interaction_genotype_Mtor | Tpo | -4.099 | 7.12E-07 | glio |
| Aars2_ex.vivo_noRT | Sytl4 | 3.293 | 7.12E-07 | glio |
| Aars2_ex.vivo_noRT | Gm12800 | 3.293 | 7.12E-07 | glio |
| Bora_in.vivo_noRT | Abat | 3.761 | 7.12E-07 | glio |
| Bcas2_in.vivo_noRT | Cks1brt | 3.369 | 7.12E-08 | glio |
| Bcas2_in.vivo_noRT | Gm16754 | 3.821 | 7.12E-08 | glio |
| Interaction_genotype_Mtor | Gm14207 | -4.892 | 7.12E-09 | glio |
| Atr_ex.vivo_noRT | Prex2 | 5.622 | 7.12E-09 | glio |
| Bora_in.vivo_noRT | Col6a2 | 4.153 | 7.12E-10 | glio |
| Atr_in.vivo_noRT | E030042O20R | 5.533 | 7.12E-14 | glio |
| Atr_in.vivo_noRT | Cstdc4 | 4.876 | 7.12E-14 | glio |
| Hsd17b10_in.vivo_noRT | Gm13856 | 3.616 | 7.12E-14 | glio |
| Aars2_in.vivo_noRT | Tns4 | 5.213 | 7.12E-17 | glio |
| culture_effect | Ptprz1 | -3.98 | 7.12E-40 | glio |
| culture_effect | Zfp787 | 0.576 | 7.13E-03 | glio |
| Interaction_genotype_Bcas2 | H2-M3 | 2.622 | 7.13E-03 | glio |
| Interaction_genotype_Mtor | 6430590A07R | -2.176 | 7.13E-03 | glio |
| Interaction_genotype_Pars2 | Lcat | -3.878 | 7.13E-03 | glio |
| interaction_RT | Oxct1 | -0.675 | 7.13E-03 | glio |
| Aars2_in.vivo_noRT | Zfp934 | 1.766 | 7.13E-03 | glio |
| Atr_in.vivo_noRT | Pgghg | 1.572 | 7.13E-03 | glio |
| Atr_in.vivo_noRT | Pdgfa | 1.033 | 7.13E-03 | glio |
| Bard1_ex.vivo_noRT | Bglap2 | 2.814 | 7.13E-03 | glio |
| Bcas2_in.vivo_noRT | Lhx9 | 2.102 | 7.13E-03 | glio |
| Bcas2_in.vivo_noRT | Slc5a5 | 2.083 | 7.13E-03 | glio |
| Bcas2_in.vivo_noRT | Tead4 | -1.409 | 7.13E-03 | glio |
| Bcas2_ex.vivo_noRT | 2310040G24R | -2.472 | 7.13E-03 | glio |
| Bora_in.vivo_noRT | Alkbh1 | -1.16 | 7.13E-03 | glio |
| Chek1_ex.vivo_noRT | Srcap | -1.666 | 7.13E-03 | glio |
| Eif2s1_ex.vivo_noRT | Rab27a | 2.637 | 7.13E-03 | glio |
| Eif2s1_ex.vivo_noRT | Gm17638 | 2.798 | 7.13E-03 | glio |
| Eif2s1_ex.vivo_noRT | Map7 | 2.998 | 7.13E-03 | glio |
| culture_effect | Med26 | -0.885 | 7.13E-04 | glio |
| culture_effect | Mrto4 | -0.416 | 7.13E-04 | glio |
| Aars2_in.vivo_noRT | Hspa5 | -0.947 | 7.13E-04 | glio |
| Atr_in.vivo_noRT | Peli1 | -1.517 | 7.13E-04 | glio |
| Bcas2_in.vivo_noRT | Ifnz | 2.464 | 7.13E-04 | glio |
| Bora_in.vivo_noRT | Gm12689 | 2.709 | 7.13E-04 | glio |
| Chek1_ex.vivo_noRT | Mroh7 | 3 | 7.13E-04 | glio |
| Bora_in.vivo_noRT | Gm13203 | 2.97 | 7.13E-05 | glio |
| culture_effect | Dffb | -1.538 | 7.13E-06 | glio |
| Bora_ex.vivo_noRT | Synpo2l | 3.599 | 7.13E-06 | glio |
| culture_effect | Adamts13 | -2.615 | 7.13E-07 | glio |
| Atp5c1_in.vivo_noRT | Halr1 | 2.402 | 7.13E-07 | glio |
| Atp5c1_in.vivo_noRT | Gpr4 | 2.668 | 7.13E-07 | glio |
| Atp5c1_in.vivo_noRT | Prl7c1 | 2.668 | 7.13E-07 | glio |
| Atp5c1_in.vivo_noRT | Gm33466 | 2.668 | 7.13E-07 | glio |
| Atr_in.vivo_noRT | Cyp4f18 | 5.273 | 7.13E-08 | glio |
| Atr_ex.vivo_noRT | Txnip | 5.182 | 7.13E-09 | glio |

| | | | | |
|---|---|---|---|---|
| culture_effect | Snhg9 | 1.186 | 7.13E-10 | glio |
| Atr_in.vivo_noRT | Adgrb3 | 4.227 | 7.13E-10 | glio |
| Bcas2_in.vivo_noRT | Pygo1 | 4.023 | 7.13E-10 | glio |
| Chek1_in.vivo_noRT | 5830454E08Ri | 4.118 | 7.13E-10 | glio |
| culture_effect | Dock9 | -1.259 | 7.13E-12 | glio |
| culture_effect | Adamts14 | -0.975 | 7.14E-03 | glio |
| Interaction_genotype_Bcas2 | Iah1 | 1.392 | 7.14E-03 | glio |
| Interaction_genotype_Bcas2 | AI662270 | 1.043 | 7.14E-03 | glio |
| Interaction_genotype_Myc | Shc3 | -2.416 | 7.14E-03 | glio |
| Interaction_genotype_Myc | Acsl1 | -2.508 | 7.14E-03 | glio |
| interaction_RT | Hsf1 | -0.694 | 7.14E-03 | glio |
| Bora_in.vivo_noRT | Rab30 | 1.778 | 7.14E-03 | glio |
| Brca2_in.vivo_noRT | Spi1 | 2.286 | 7.14E-03 | glio |
| Chek1_in.vivo_noRT | Pum2 | -0.938 | 7.14E-03 | glio |
| Chek1_ex.vivo_noRT | Arhgdib | 2.587 | 7.14E-03 | glio |
| Interaction_genotype_Mtor | Lrp4 | -4.151 | 7.14E-04 | glio |
| Interaction_genotype_Myc | Msh6 | -2.928 | 7.14E-04 | glio |
| Interaction_genotype_Myc | Timm8b | 1.39 | 7.14E-04 | glio |
| Aars2_in.vivo_noRT | Rps13 | -0.612 | 7.14E-04 | glio |
| Atr_in.vivo_noRT | Acbd7 | 2.5 | 7.14E-04 | glio |
| Atr_ex.vivo_noRT | C330024C12R | 2.771 | 7.14E-04 | glio |
| Atr_ex.vivo_noRT | Gpd1 | 3.192 | 7.14E-04 | glio |
| Atr_ex.vivo_noRT | Fkbp4 | -1.032 | 7.14E-04 | glio |
| Bora_in.vivo_noRT | Taco1os | 2.299 | 7.14E-04 | glio |
| Chek1_ex.vivo_noRT | 4930447F24Ri | 2.896 | 7.14E-04 | glio |
| Hsd17b10_in.vivo_noRT | Gm11127 | 2.587 | 7.14E-04 | glio |
| culture_effect | 4930579K19R | 2.231 | 7.14E-05 | glio |
| Interaction_genotype_Mtor | A530072M11F | -4.781 | 7.14E-05 | glio |
| Atr_in.vivo_noRT | Zglp1 | 3.057 | 7.14E-05 | glio |
| Atr_in.vivo_noRT | A230072E10Ri | 2.767 | 7.14E-05 | glio |
| Chek1_in.vivo_noRT | Lrrc75a | 2.529 | 7.14E-05 | glio |
| Chek1_ex.vivo_noRT | Dpep2 | 3.469 | 7.14E-05 | glio |
| Chek1_ex.vivo_noRT | Gm28404 | 3.574 | 7.14E-06 | glio |
| Aars2_in.vivo_noRT | Zdhhc14 | 2.859 | 7.14E-07 | glio |
| Atr_in.vivo_noRT | Rpl28 | -0.901 | 7.14E-07 | glio |
| Hsd17b10_in.vivo_noRT | Gm31518 | 3.616 | 7.14E-07 | glio |
| culture_effect | S100a16 | -1.901 | 7.14E-08 | glio |
| Aars2_in.vivo_noRT | Gm42997 | 3.958 | 7.14E-09 | glio |
| Chek1_in.vivo_noRT | Insm1 | 3.925 | 7.14E-09 | glio |
| Bcas2_in.vivo_noRT | Dnah10 | 4.428 | 7.14E-13 | glio |
| culture_effect | Ncoa4 | -0.549 | 7.15E-03 | glio |
| Interaction_genotype_Bcas2 | Gm20522 | -3.615 | 7.15E-03 | glio |
| Interaction_genotype_Mtor | Glt8d2 | -3.194 | 7.15E-03 | glio |
| Interaction_genotype_Mtor | Mavs | -2.253 | 7.15E-03 | glio |
| Interaction_genotype_Myc | 4921507P07R | -2.94 | 7.15E-03 | glio |
| Interaction_genotype_Myc | Pdzk1ip1 | -3.173 | 7.15E-03 | glio |
| Interaction_genotype_Myc | Zfp217 | -2.851 | 7.15E-03 | glio |
| Interaction_genotype_Myc | Tal1 | -2.941 | 7.15E-03 | glio |
| Interaction_genotype_Myc | Ehbp1 | -2.96 | 7.15E-03 | glio |
| Interaction_genotype_Myc | Kdm1b | -2.508 | 7.15E-03 | glio |
| Interaction_genotype_Myc | Zfp941 | -2.101 | 7.15E-03 | glio |
| Interaction_genotype_Myc | Guk1 | 1.1 | 7.15E-03 | glio |

| | | | | |
|---|---|---|---|---|
| Interaction_genotype_Myc | Hmgb1 | 1.095 | 7.15E-03 | glio |
| Aars2_in.vivo_noRT | Cpe | -1.226 | 7.15E-03 | glio |
| Aars2_in.vivo_noRT | Anapc16 | -0.807 | 7.15E-03 | glio |
| Atr_in.vivo_noRT | Psma4 | -0.662 | 7.15E-03 | glio |
| Eif2s1_in.vivo_noRT | D830039M14I | 2.4 | 7.15E-03 | glio |
| Eif2s1_in.vivo_noRT | Flnb | -0.918 | 7.15E-03 | glio |
| Fanca_ex.vivo_noRT | Insrr | 2.913 | 7.15E-03 | glio |
| Fanca_ex.vivo_noRT | Tpo | 2.124 | 7.15E-03 | glio |
| Fanca_ex.vivo_noRT | Ccdc125 | 2.52 | 7.15E-03 | glio |
| Aars2_in.vivo_noRT | Zranb1 | -1.476 | 7.15E-04 | glio |
| Atr_in.vivo_noRT | Hdc | 2.505 | 7.15E-04 | glio |
| Bcas2_in.vivo_noRT | Txndc5 | -1.352 | 7.15E-04 | glio |
| Chek1_ex.vivo_noRT | 1700056N10R | 2.784 | 7.15E-04 | glio |
| Chek1_ex.vivo_noRT | Gm48796 | 2.974 | 7.15E-04 | glio |
| Interaction_genotype_Mtor | A230020J21Ri | -4.927 | 7.15E-05 | glio |
| Interaction_genotype_Mtor | 4930563E22Ri | -4.654 | 7.15E-05 | glio |
| Bora_in.vivo_noRT | Tnfrsf1b | 2.642 | 7.15E-05 | glio |
| Chek1_ex.vivo_noRT | Shbg | 3.738 | 7.15E-05 | glio |
| culture_effect | Rpl28 | -0.593 | 7.15E-07 | glio |
| Bora_ex.vivo_noRT | Tac2 | 3.201 | 7.15E-07 | glio |
| Interaction_genotype_Mtor | Tmem59 | 2.475 | 7.15E-08 | glio |
| Chek1_in.vivo_noRT | 2310058D17R | 3.865 | 7.15E-08 | glio |
| Chek1_ex.vivo_noRT | Gm43221 | 4.842 | 7.15E-09 | glio |
| Atr_in.vivo_noRT | Pawr | 3.767 | 7.15E-10 | glio |
| Chek1_ex.vivo_noRT | A930019D19R | 4.842 | 7.15E-17 | glio |
| Interaction_genotype_Bcas2 | Pspn | -2.46 | 7.16E-03 | glio |
| Interaction_genotype_Mtor | Slc43a2 | -3.492 | 7.16E-03 | glio |
| Interaction_genotype_Mtor | Flrt3 | -2.819 | 7.16E-03 | glio |
| Aars2_in.vivo_noRT | Ninj1 | -1.163 | 7.16E-03 | glio |
| Aars2_in.vivo_noRT | D8Ertd738e | -0.529 | 7.16E-03 | glio |
| Bcas2_in.vivo_noRT | Ccr2 | 2.013 | 7.16E-03 | glio |
| Bcas2_in.vivo_noRT | 5830448L01Ri | 1.951 | 7.16E-03 | glio |
| Cdkn2a_in.vivo_noRT | Gm20625 | -1.675 | 7.16E-03 | glio |
| Cdkn2a_in.vivo_noRT | Gm5157 | -1.675 | 7.16E-03 | glio |
| Cdkn2a_in.vivo_noRT | Gm44736 | -1.675 | 7.16E-03 | glio |
| Cdkn2a_in.vivo_noRT | Fam227b | -1.675 | 7.16E-03 | glio |
| Cdkn2a_in.vivo_noRT | Gm20658 | -1.675 | 7.16E-03 | glio |
| Cdkn2a_in.vivo_noRT | Gm5934 | -1.938 | 7.16E-03 | glio |
| Cdkn2a_in.vivo_noRT | Olfr810 | -1.675 | 7.16E-03 | glio |
| Cdkn2a_in.vivo_noRT | Hpse | 1.927 | 7.16E-03 | glio |
| Chek1_in.vivo_noRT | Ndufb5 | -0.635 | 7.16E-03 | glio |
| Chek1_ex.vivo_noRT | Iqcg | 2.72 | 7.16E-03 | glio |
| Interaction_genotype_Mtor | Nkg7 | -3.442 | 7.16E-04 | glio |
| Interaction_genotype_Pars2 | Gm48435 | -3.056 | 7.16E-04 | glio |
| Interaction_genotype_Pars2 | Auh | 2.24 | 7.16E-04 | glio |
| Interaction_genotype_Pars2 | Rftn2 | -4.177 | 7.16E-04 | glio |
| Aars2_ex.vivo_noRT | Snhg14 | 3.445 | 7.16E-04 | glio |
| Bcas2_in.vivo_noRT | Nt5c | -1.34 | 7.16E-04 | glio |
| Interaction_genotype_Mtor | Tmem37 | -4.661 | 7.16E-05 | glio |
| Interaction_genotype_Mtor | Rab2a | 2 | 7.16E-05 | glio |
| Interaction_genotype_Myc | Ttc21b | -3.261 | 7.16E-05 | glio |
| Aars2_in.vivo_noRT | Fhad1 | 2.601 | 7.16E-05 | glio |

| | | | | |
|---|---|---|---|---|
| Bard1_ex.vivo_noRT | Gm4166 | 2.814 | 7.16E-05 | glio |
| Aars2_ex.vivo_noRT | D730045A05R | 4.074 | 7.16E-06 | glio |
| Interaction_genotype_Mtor | Gm39929 | -4.804 | 7.16E-07 | glio |
| interaction_RT | Ssrp1 | -0.829 | 7.16E-07 | glio |
| Aars2_ex.vivo_noRT | Trmt61b | 3.293 | 7.16E-07 | glio |
| Bcas2_in.vivo_noRT | Paqr8 | 3.954 | 7.16E-07 | glio |
| interaction_RT | Selenow | 1.214 | 7.16E-11 | glio |
| Bcas2_in.vivo_noRT | Psmb11 | 4.284 | 7.16E-17 | glio |
| Bora_in.vivo_noRT | 9230112J17Ri | 4.936 | 7.16E-17 | glio |
| Bcas2_in.vivo_noRT | A930002C04R | 4.547 | 7.16E-21 | glio |
| Bcas2_in.vivo_noRT | Gm15533 | 4.807 | 7.16E-21 | glio |
| Atr_in.vivo_noRT | Gm15908 | 5.533 | 7.16E-31 | glio |
| Atr_in.vivo_noRT | A930014E10Ri | 5.533 | 7.16E-31 | glio |
| culture_effect | Pik3r3 | 1.134 | 7.17E-03 | glio |
| Interaction_genotype_Mtor | Mrpl54 | 1.21 | 7.17E-03 | glio |
| Interaction_genotype_Myc | Tmco6 | -2.726 | 7.17E-03 | glio |
| Atr_in.vivo_noRT | Gm42851 | 1.701 | 7.17E-03 | glio |
| Atr_ex.vivo_noRT | Gvin1 | 2.769 | 7.17E-03 | glio |
| Bcas2_in.vivo_noRT | Mansc1 | 2.134 | 7.17E-03 | glio |
| Bcas2_in.vivo_noRT | Abcb8 | -1.421 | 7.17E-03 | glio |
| Chek1_ex.vivo_noRT | Cabp4 | 2.488 | 7.17E-03 | glio |
| Chek1_ex.vivo_noRT | Sgo2a | -1.98 | 7.17E-03 | glio |
| Interaction_genotype_Myc | Serpinb9g | -2.729 | 7.17E-04 | glio |
| Interaction_genotype_Myc | 7530414M10F | -3.701 | 7.17E-04 | glio |
| Interaction_genotype_Myc | Gm16638 | -3.702 | 7.17E-04 | glio |
| Interaction_genotype_Myc | Tagap1 | -3.138 | 7.17E-04 | glio |
| Interaction_genotype_Myc | 4632404H12R | -3.707 | 7.17E-04 | glio |
| Interaction_genotype_Myc | Serpinb8 | -3.26 | 7.17E-04 | glio |
| Interaction_genotype_Myc | D130020L05R | -3.189 | 7.17E-04 | glio |
| Interaction_genotype_Myc | Pcdh1 | -3.565 | 7.17E-04 | glio |
| Interaction_genotype_Myc | Cc2d1b | -2.87 | 7.17E-04 | glio |
| Interaction_genotype_Myc | Taok2 | -2.537 | 7.17E-04 | glio |
| Bora_in.vivo_noRT | Olfr77 | 2.351 | 7.17E-04 | glio |
| Bora_in.vivo_noRT | Zfp827 | 2.33 | 7.17E-04 | glio |
| Hsd17b10_in.vivo_noRT | Gm16094 | 2.311 | 7.17E-04 | glio |
| Hsd17b10_in.vivo_noRT | Siva1 | -0.726 | 7.17E-04 | glio |
| Interaction_genotype_Aars2 | Gm46218 | -4.999 | 7.17E-05 | glio |
| Interaction_genotype_Mtor | Scel | -3.181 | 7.17E-05 | glio |
| Interaction_genotype_Nat2 | Usp17la | 4.841 | 7.17E-05 | glio |
| culture_effect | Bcap31 | -0.596 | 7.17E-06 | glio |
| Atr_in.vivo_noRT | N4bp2l1 | 3.237 | 7.17E-06 | glio |
| Chek1_ex.vivo_noRT | Hes5 | 4.842 | 7.17E-06 | glio |
| Bcas2_in.vivo_noRT | C230037L18Ri | 3.638 | 7.17E-08 | glio |
| Atr_in.vivo_noRT | Zfp551 | 3.966 | 7.17E-09 | glio |
| Bcas2_in.vivo_noRT | Slc45a3 | 4.285 | 7.17E-10 | glio |
| Atr_in.vivo_noRT | Gm47204 | 4.888 | 7.17E-12 | glio |
| Bora_in.vivo_noRT | Vwce | 4.676 | 7.17E-14 | glio |
| Chek1_ex.vivo_noRT | Vmn1r32 | 4.449 | 7.17E-14 | glio |
| Aars2_in.vivo_noRT | BC061195 | 5.327 | 7.17E-27 | glio |
| Interaction_genotype_Atr | Lyl1 | -2.904 | 7.18E-03 | glio |
| Interaction_genotype_Mtor | Alkbh4 | -2.343 | 7.18E-03 | glio |
| Interaction_genotype_Myc | Efna5 | -2.895 | 7.18E-03 | glio |

| | | | | |
|---|---|---|---|---|
| Interaction_genotype_Myc | Ddx25 | -2.313 | 7.18E-03 | glio |
| Aars2_in.vivo_noRT | Apobec1 | 1.946 | 7.18E-03 | glio |
| Aars2_in.vivo_noRT | Zc3h12a | 1.835 | 7.18E-03 | glio |
| Atr_in.vivo_noRT | Plekho1 | -1.105 | 7.18E-03 | glio |
| Bcas2_in.vivo_noRT | Rab6b | 1.73 | 7.18E-03 | glio |
| Bora_in.vivo_noRT | Slc18b1 | 1.565 | 7.18E-03 | glio |
| Cenpj_in.vivo_noRT | Gm5934 | 1.759 | 7.18E-03 | glio |
| Cenpj_in.vivo_noRT | Crhbp | 2.022 | 7.18E-03 | glio |
| Chek1_ex.vivo_noRT | C230038L03Ri | 2.471 | 7.18E-03 | glio |
| Interaction_genotype_Myc | Gm31805 | -3.6 | 7.18E-04 | glio |
| Aars2_in.vivo_noRT | Gm39469 | 2.18 | 7.18E-04 | glio |
| Aars2_in.vivo_noRT | Nudt22 | 1.625 | 7.18E-04 | glio |
| Bcas2_in.vivo_noRT | Gnl3 | -1.095 | 7.18E-04 | glio |
| Chek1_in.vivo_noRT | Ccl27a | 1.373 | 7.18E-04 | glio |
| Eif2s1_in.vivo_noRT | Gm39526 | 2.148 | 7.18E-04 | glio |
| Aars2_ex.vivo_noRT | 4933408J17Ri | 3.308 | 7.18E-05 | glio |
| Atr_in.vivo_noRT | Lnx2 | 2.56 | 7.18E-05 | glio |
| Bora_in.vivo_noRT | Aldh1a1 | 2.907 | 7.18E-05 | glio |
| Bora_in.vivo_noRT | Maob | 2.723 | 7.18E-05 | glio |
| Interaction_genotype_Mtor | Pdzk1ip1 | -5.71 | 7.18E-06 | glio |
| Aars2_in.vivo_noRT | Tmem117 | 3.236 | 7.18E-06 | glio |
| Bora_ex.vivo_noRT | Fam166a | 3.201 | 7.18E-06 | glio |
| Chek1_ex.vivo_noRT | Gm11127 | 4.049 | 7.18E-06 | glio |
| Interaction_genotype_Mtor | Gm16348 | -5.86 | 7.18E-07 | glio |
| Aars2_ex.vivo_noRT | Gm5890 | 3.293 | 7.18E-07 | glio |
| Aars2_ex.vivo_noRT | Zscan4c | 3.293 | 7.18E-07 | glio |
| Aars2_ex.vivo_noRT | Gm45338 | 2.899 | 7.18E-07 | glio |
| Aars2_ex.vivo_noRT | Speer4a | 3.293 | 7.18E-07 | glio |
| Aars2_ex.vivo_noRT | BC080695 | 3.293 | 7.18E-07 | glio |
| Atr_in.vivo_noRT | Cish | 3.317 | 7.18E-07 | glio |
| Bcas2_in.vivo_noRT | Pde5a | 3.521 | 7.18E-07 | glio |
| Bora_in.vivo_noRT | Gm46447 | 3.125 | 7.18E-07 | glio |
| culture_effect | Gm8251 | -1.893 | 7.18E-08 | glio |
| Bcas2_in.vivo_noRT | Rem2 | 3.622 | 7.18E-09 | glio |
| culture_effect | Ogfod3 | -2.004 | 7.18E-16 | glio |
| Aars2_in.vivo_noRT | BB557941 | 5.067 | 7.18E-18 | glio |
| culture_effect | Scube3 | 4.439 | 7.18E-23 | glio |
| Interaction_genotype_Bcas2 | Ccr5 | -3.345 | 7.19E-03 | glio |
| Interaction_genotype_Bcas2 | Otos | -2.692 | 7.19E-03 | glio |
| Interaction_genotype_Bora | Ttc41 | -3.863 | 7.19E-03 | glio |
| Interaction_genotype_Mtor | Tm9sf2 | 1.681 | 7.19E-03 | glio |
| Aars2_in.vivo_noRT | Dhx40 | 0.989 | 7.19E-03 | glio |
| Atr_in.vivo_noRT | Naa40 | 1.193 | 7.19E-03 | glio |
| Atr_ex.vivo_noRT | Arvcf | 2.851 | 7.19E-03 | glio |
| Bora_in.vivo_noRT | Pomk | 1.731 | 7.19E-03 | glio |
| Bora_in.vivo_noRT | Lrrc27 | 1.639 | 7.19E-03 | glio |
| Chek1_in.vivo_noRT | Zfp772 | 2.025 | 7.19E-03 | glio |
| Eif2s1_in.vivo_noRT | Apoh | 2.277 | 7.19E-03 | glio |
| Eif2s1_in.vivo_noRT | Gchfr | 2.613 | 7.19E-03 | glio |
| Hsd17b10_in.vivo_noRT | Insrr | 1.902 | 7.19E-03 | glio |
| Interaction_genotype_Myc | Prtn3 | -3.991 | 7.19E-04 | glio |
| Interaction_genotype_Myc | Apobr | -4.119 | 7.19E-04 | glio |

| | | | | |
|---|---|---|---|---|
| Aars2_in.vivo_noRT | Sh3bp5 | 2.387 | 7.19E-04 | glio |
| Aars2_in.vivo_noRT | Ddx27 | -1.248 | 7.19E-04 | glio |
| Atr_in.vivo_noRT | Ybey | 1.972 | 7.19E-04 | glio |
| Bcas2_in.vivo_noRT | Ccdc25 | -1.365 | 7.19E-04 | glio |
| Hsd17b10_in.vivo_noRT | Gm12840 | 2.358 | 7.19E-04 | glio |
| Interaction_genotype_Mtor | E230001N04R | -4.2 | 7.19E-05 | glio |
| Interaction_genotype_Myc | Gm14636 | -5.145 | 7.19E-05 | glio |
| Aars2_in.vivo_noRT | 5930430L01Ri | 2.635 | 7.19E-05 | glio |
| Hsd17b10_in.vivo_noRT | Rtn1 | 2.97 | 7.19E-05 | glio |
| interaction_RT | Eprs | -0.814 | 7.19E-06 | glio |
| interaction_RT | Nolc1 | -0.933 | 7.19E-06 | glio |
| Chek1_in.vivo_noRT | Gm37170 | 3.201 | 7.19E-06 | glio |
| Chek1_in.vivo_noRT | Gm15952 | 3.264 | 7.19E-06 | glio |
| Bora_in.vivo_noRT | Tgfbr3l | 3.632 | 7.19E-07 | glio |
| Interaction_genotype_Mtor | Cers2 | 2.776 | 7.19E-09 | glio |
| Atr_in.vivo_noRT | 4930518J21Ri | 4.608 | 7.19E-12 | glio |
| Bora_in.vivo_noRT | Cxcl1 | 4.936 | 7.19E-12 | glio |
| Bcas2_in.vivo_noRT | Ntn4 | 5.068 | 7.19E-17 | glio |
| culture_effect | Iqgap1 | -0.326 | 7.20E-03 | glio |
| Interaction_genotype_Mtor | Ino80dos | -3.348 | 7.20E-03 | glio |
| Interaction_genotype_Myc | Mdm1 | -2.634 | 7.20E-03 | glio |
| Interaction_genotype_Myc | Pbk | -2.342 | 7.20E-03 | glio |
| interaction_RT | Ypel5 | 0.993 | 7.20E-03 | glio |
| Atr_in.vivo_noRT | Zfp40 | 1.806 | 7.20E-03 | glio |
| Chek1_in.vivo_noRT | Fbxo48 | 2.023 | 7.20E-03 | glio |
| Chek1_in.vivo_noRT | Zfp12 | 1.545 | 7.20E-03 | glio |
| Hsd17b10_in.vivo_noRT | 2810457G06R | 2.369 | 7.20E-03 | glio |
| Hsd17b10_in.vivo_noRT | Ksr1 | 2.099 | 7.20E-03 | glio |
| Interaction_genotype_Mtor | Ceacam16 | -3.441 | 7.20E-04 | glio |
| Interaction_genotype_Myc | A430033K04R | -3.661 | 7.20E-04 | glio |
| Atr_in.vivo_noRT | Hnrnpab | -0.676 | 7.20E-04 | glio |
| Bora_in.vivo_noRT | Gm19744 | 2.491 | 7.20E-04 | glio |
| Interaction_genotype_Mtor | Gm5129 | -4.76 | 7.20E-05 | glio |
| Chek1_in.vivo_noRT | Cldn10 | 2.836 | 7.20E-05 | glio |
| Chek1_in.vivo_noRT | Pls1 | 2.787 | 7.20E-05 | glio |
| Bard1_ex.vivo_noRT | Zfp352 | 3.987 | 7.20E-06 | glio |
| Bard1_ex.vivo_noRT | Bcl2l15 | 3.821 | 7.20E-06 | glio |
| Bard1_ex.vivo_noRT | Krt19 | 2.814 | 7.20E-06 | glio |
| Bard1_ex.vivo_noRT | Gm34939 | 3.596 | 7.20E-06 | glio |
| Bard1_ex.vivo_noRT | Gm36026 | 3.612 | 7.20E-06 | glio |
| Bard1_ex.vivo_noRT | Gm48155 | 2.814 | 7.20E-06 | glio |
| Bora_in.vivo_noRT | BC049762 | 3.59 | 7.20E-07 | glio |
| Chek1_in.vivo_noRT | Noxred1 | 4.316 | 7.20E-11 | glio |
| Bcas2_in.vivo_noRT | Gm10696 | 4.807 | 7.20E-12 | glio |
| Hsd17b10_in.vivo_noRT | Olfr552 | 3.616 | 7.20E-13 | glio |
| Hsd17b10_in.vivo_noRT | Gm21936 | 3.616 | 7.20E-14 | glio |
| Atr_in.vivo_noRT | Carns1 | 5.533 | 7.20E-15 | glio |
| culture_effect | Epha2 | 1.366 | 7.21E-03 | glio |
| Interaction_genotype_Mtor | Galk2 | -2.067 | 7.21E-03 | glio |
| Interaction_genotype_Myc | Inafm1 | -2.51 | 7.21E-03 | glio |
| Interaction_genotype_Myc | Slc3a2 | 1.289 | 7.21E-03 | glio |
| interaction_RT | Slc38a3 | 2.205 | 7.21E-03 | glio |

| | | | | |
|---|---|---|---|---|
| interaction_RT | Slc3a2 | 0.604 | 7.21E-03 | glio |
| Aars2_in.vivo_noRT | Chic2 | -0.83 | 7.21E-03 | glio |
| Aars2_in.vivo_noRT | Gnb2 | -0.684 | 7.21E-03 | glio |
| Atr_in.vivo_noRT | Camk2n2 | 1.617 | 7.21E-03 | glio |
| Atr_in.vivo_noRT | Arl6ip6 | -1.053 | 7.21E-03 | glio |
| Bora_in.vivo_noRT | Tbl3 | -1.127 | 7.21E-03 | glio |
| Chek1_in.vivo_noRT | Plvap | 1.979 | 7.21E-03 | glio |
| Interaction_genotype_Atr | Rab11fip4os1 | -4.23 | 7.21E-04 | glio |
| Interaction_genotype_Bcas2 | Cnp | 2.751 | 7.21E-04 | glio |
| Interaction_genotype_Mtor | Cotl1 | -2.764 | 7.21E-04 | glio |
| Interaction_genotype_Rad21 | Atp1a2 | 4.865 | 7.21E-04 | glio |
| Aars2_in.vivo_noRT | Fth1 | -0.993 | 7.21E-04 | glio |
| Bora_in.vivo_noRT | Fcgrt | 2.689 | 7.21E-04 | glio |
| Bora_in.vivo_noRT | 1700001G11R | 2.125 | 7.21E-04 | glio |
| Bora_in.vivo_noRT | Efcab14 | -1.35 | 7.21E-04 | glio |
| Interaction_genotype_Myc | Fbxo17 | -3.859 | 7.21E-05 | glio |
| Interaction_genotype_Myc | 9330151L19Ri | -4.002 | 7.21E-05 | glio |
| Interaction_genotype_Myc | Clec12b | -3.124 | 7.21E-05 | glio |
| Interaction_genotype_Myc | Exo1 | -3.618 | 7.21E-05 | glio |
| Interaction_genotype_Myc | Eif4ebp1 | 2.14 | 7.21E-05 | glio |
| Chek1_in.vivo_noRT | Plxnb1 | 2.912 | 7.21E-05 | glio |
| Hsd17b10_in.vivo_noRT | 1700003G18R | 3.356 | 7.21E-05 | glio |
| Hsd17b10_in.vivo_noRT | Gm32585 | 3.09 | 7.21E-05 | glio |
| Bora_ex.vivo_noRT | Gm15894 | 3.201 | 7.21E-06 | glio |
| Interaction_genotype_Myc | Gm17597 | -6.098 | 7.21E-07 | glio |
| Aars2_in.vivo_noRT | Cacng5 | 3.476 | 7.21E-08 | glio |
| Atr_in.vivo_noRT | D230022J07Ri | 3.845 | 7.21E-08 | glio |
| Atr_in.vivo_noRT | Gm13986 | 4.275 | 7.21E-09 | glio |
| Atr_in.vivo_noRT | 1700084E18Ri | 4.292 | 7.21E-09 | glio |
| Atr_in.vivo_noRT | D430018E03R | 3.807 | 7.21E-09 | glio |
| Chek1_in.vivo_noRT | 2810410L24Ri | 4.518 | 7.21E-11 | glio |
| Chek1_in.vivo_noRT | Akap3 | 4.784 | 7.21E-12 | glio |
| Chek1_ex.vivo_noRT | Tnni3k | 4.855 | 7.21E-14 | glio |
| culture_effect | Dnlz | 1.224 | 7.21E-15 | glio |
| culture_effect | Sorbs2 | 3.475 | 7.21E-19 | glio |
| culture_effect | Atoh8 | 4.277 | 7.21E-21 | glio |
| Interaction_genotype_Aars2 | 1700122E12Ri | -3.429 | 7.22E-03 | glio |
| Interaction_genotype_Aars2 | 4930417H01R | -3.529 | 7.22E-03 | glio |
| Interaction_genotype_Aars2 | Gm8773 | -3.597 | 7.22E-03 | glio |
| Interaction_genotype_Aars2 | Rasgrp2 | -3.132 | 7.22E-03 | glio |
| Interaction_genotype_Aars2 | 2700097O09R | -2.507 | 7.22E-03 | glio |
| Interaction_genotype_Atr | Rapgef3 | -3.773 | 7.22E-03 | glio |
| Interaction_genotype_Atr | Gm7008 | -3.228 | 7.22E-03 | glio |
| Interaction_genotype_Bora | Gm44658 | -3.146 | 7.22E-03 | glio |
| Interaction_genotype_Bora | Sema6d | -2.716 | 7.22E-03 | glio |
| Interaction_genotype_Bora | Ranbp10 | 2.163 | 7.22E-03 | glio |
| Interaction_genotype_Mtor | Slc4a11 | -3.443 | 7.22E-03 | glio |
| Interaction_genotype_Myc | 3110040N11R | -2.016 | 7.22E-03 | glio |
| Interaction_genotype_Uqcrc2 | Ntn4 | -2.815 | 7.22E-03 | glio |
| interaction_RT | Gm43305 | -1.036 | 7.22E-03 | glio |
| Atp5c1_in.vivo_noRT | B020031M17F | 2.668 | 7.22E-03 | glio |
| Bcas2_in.vivo_noRT | Mrpl51 | -0.513 | 7.22E-03 | glio |

| | | | | |
|---|---|---|---|---|
| Hsd17b10_in.vivo_noRT | Acad12 | 2.021 | 7.22E-03 | glio |
| culture_effect | Ccni | 0.547 | 7.22E-04 | glio |
| Interaction_genotype_Mtor | Gm19744 | -4.156 | 7.22E-04 | glio |
| interaction_RT | Slc5a5 | 3.331 | 7.22E-04 | glio |
| interaction_RT | Abcf1 | -0.647 | 7.22E-04 | glio |
| Aars2_ex.vivo_noRT | Gm30794 | 2.316 | 7.22E-04 | glio |
| Bcas2_in.vivo_noRT | C920021L13Ri | 2.477 | 7.22E-04 | glio |
| Bora_in.vivo_noRT | 1700102H20R | 2.412 | 7.22E-04 | glio |
| Bora_ex.vivo_noRT | Gm43463 | 2.808 | 7.22E-04 | glio |
| Chek1_in.vivo_noRT | P4htm | 2.442 | 7.22E-04 | glio |
| Interaction_genotype_Mtor | Nr1h4 | -4.728 | 7.22E-05 | glio |
| Interaction_genotype_Mtor | Gm15473 | -4.951 | 7.22E-05 | glio |
| Interaction_genotype_Mtor | Wnt10a | -4.542 | 7.22E-05 | glio |
| Interaction_genotype_Mtor | Trim28 | 2.044 | 7.22E-05 | glio |
| Interaction_genotype_Mtor | Smc4 | 2.828 | 7.22E-05 | glio |
| Atr_in.vivo_noRT | Arrdc2 | 3.025 | 7.22E-05 | glio |
| Atr_ex.vivo_noRT | Gm8300 | 4.547 | 7.22E-05 | glio |
| Atr_ex.vivo_noRT | Etl4 | 3.164 | 7.22E-05 | glio |
| Hsd17b10_in.vivo_noRT | Pard3bos1 | 2.619 | 7.22E-05 | glio |
| culture_effect | Samd4 | 1.347 | 7.22E-06 | glio |
| Aars2_in.vivo_noRT | A930033H14R | 3.302 | 7.22E-06 | glio |
| Aars2_in.vivo_noRT | 4933406P04R | 3.645 | 7.22E-07 | glio |
| Bora_in.vivo_noRT | Dnase1l2 | 3.648 | 7.22E-08 | glio |
| culture_effect | Arhgap5 | 0.853 | 7.22E-11 | glio |
| Atr_in.vivo_noRT | Gm44096 | 4.263 | 7.22E-11 | glio |
| Chek1_ex.vivo_noRT | Gm13713 | 4.443 | 7.22E-11 | glio |
| Bcas2_in.vivo_noRT | Apoa5 | 4.807 | 7.22E-24 | glio |
| culture_effect | Acot8 | -0.621 | 7.23E-03 | glio |
| Interaction_genotype_Aars2 | Gm12602 | -3.151 | 7.23E-03 | glio |
| Interaction_genotype_Aars2 | Fastkd1 | -3.011 | 7.23E-03 | glio |
| Interaction_genotype_Aars2 | Med10 | 1.525 | 7.23E-03 | glio |
| Interaction_genotype_Atr | Gpr135 | -3.339 | 7.23E-03 | glio |
| Interaction_genotype_Mtor | Ciapin1 | 1.586 | 7.23E-03 | glio |
| Interaction_genotype_Myc | Ell2 | -2.733 | 7.23E-03 | glio |
| Interaction_genotype_Myc | Zdhhc1 | -2.79 | 7.23E-03 | glio |
| Interaction_genotype_Rpl3 | 3010001F23Ri | -3.461 | 7.23E-03 | glio |
| Aars2_in.vivo_noRT | Pgm2 | -1.31 | 7.23E-03 | glio |
| Bcas2_in.vivo_noRT | Dennd4b | 1.992 | 7.23E-03 | glio |
| Chek1_ex.vivo_noRT | Itgal | 2.709 | 7.23E-03 | glio |
| Chek1_ex.vivo_noRT | Vegfd | 2.315 | 7.23E-03 | glio |
| Interaction_genotype_Myc | Nrip3 | -3.671 | 7.23E-04 | glio |
| Interaction_genotype_Myc | Setmar | -3.848 | 7.23E-04 | glio |
| Bora_in.vivo_noRT | Tdrp | 2.364 | 7.23E-04 | glio |
| Interaction_genotype_Myc | Apoa1 | -4.64 | 7.23E-05 | glio |
| Bora_in.vivo_noRT | C130013H08R | 3.167 | 7.23E-05 | glio |
| Chek1_in.vivo_noRT | Sgk2 | 2.827 | 7.23E-05 | glio |
| Atp5c1_in.vivo_noRT | Spink6 | 2.668 | 7.23E-06 | glio |
| Atr_in.vivo_noRT | Btc | 3.974 | 7.23E-06 | glio |
| culture_effect | Clybl | -2.048 | 7.23E-07 | glio |
| culture_effect | Farsa | -0.79 | 7.23E-07 | glio |
| culture_effect | Slc39a10 | -1.588 | 7.23E-13 | glio |
| Chek1_in.vivo_noRT | Gm50138 | 4.521 | 7.23E-15 | glio |

| | | | | |
|---|---|---|---|---|
| Interaction_genotype_Aars2 | Frk | -3.429 | 7.24E-03 | glio |
| Interaction_genotype_Aars2 | Mars2 | -2.472 | 7.24E-03 | glio |
| Interaction_genotype_Aars2 | Myd88 | -2.594 | 7.24E-03 | glio |
| Interaction_genotype_Atr | Gm44096 | -3.181 | 7.24E-03 | glio |
| Interaction_genotype_Atr | Rassf4 | -2.13 | 7.24E-03 | glio |
| Interaction_genotype_Mtor | Col18a1 | 2.009 | 7.24E-03 | glio |
| Interaction_genotype_Myc | Ice2 | -2.361 | 7.24E-03 | glio |
| Interaction_genotype_Myc | Asap2 | -1.942 | 7.24E-03 | glio |
| Interaction_genotype_Pars2 | Jph4 | -3.635 | 7.24E-03 | glio |
| Aars2_in.vivo_noRT | Acad12 | 1.902 | 7.24E-03 | glio |
| Aars2_in.vivo_noRT | Dync2li1 | -1.401 | 7.24E-03 | glio |
| Atr_in.vivo_noRT | Il27ra | 1.769 | 7.24E-03 | glio |
| Bard1_ex.vivo_noRT | Olfr90 | 2.421 | 7.24E-03 | glio |
| Bcas2_in.vivo_noRT | E230016M11R | 1.821 | 7.24E-03 | glio |
| Bcas2_in.vivo_noRT | Zkscan8 | 1.713 | 7.24E-03 | glio |
| Bcas2_in.vivo_noRT | Rida | -1.28 | 7.24E-03 | glio |
| Bora_in.vivo_noRT | Naa30 | -1.379 | 7.24E-03 | glio |
| Chek1_in.vivo_noRT | Ccdc13 | 2.03 | 7.24E-03 | glio |
| Cyp19a1_ex.vivo_noRT | Ildr1 | 2.159 | 7.24E-03 | glio |
| Hsd17b10_in.vivo_noRT | Plpp2 | -0.656 | 7.24E-03 | glio |
| Interaction_genotype_Myc | Tmem59 | 1.317 | 7.24E-04 | glio |
| Atr_in.vivo_noRT | Tmem143 | 1.967 | 7.24E-04 | glio |
| Interaction_genotype_Atr | Cdkn1b | 2.582 | 7.24E-05 | glio |
| Interaction_genotype_Mtor | Lrfn1 | -4.744 | 7.24E-05 | glio |
| Interaction_genotype_Myc | Plcb4 | -4.537 | 7.24E-05 | glio |
| Interaction_genotype_Myc | Gm41764 | -4 | 7.24E-05 | glio |
| Bora_in.vivo_noRT | 4930458D05R | 3.123 | 7.24E-05 | glio |
| Bora_in.vivo_noRT | Ddx47 | -1.367 | 7.24E-05 | glio |
| Chek1_in.vivo_noRT | Gm43138 | 2.937 | 7.24E-05 | glio |
| Interaction_genotype_Atr | Irf8 | 5.207 | 7.24E-06 | glio |
| Interaction_genotype_Myc | Gm34552 | -5.102 | 7.24E-06 | glio |
| interaction_RT | Kcnq1ot1 | -1.661 | 7.24E-06 | glio |
| Bora_in.vivo_noRT | Serpinb6b | 3.294 | 7.24E-06 | glio |
| Bora_in.vivo_noRT | Gm42658 | 3.184 | 7.24E-07 | glio |
| Bora_in.vivo_noRT | Cemip | 2.902 | 7.24E-07 | glio |
| culture_effect | 1700084E18Ri | 2.888 | 7.24E-08 | glio |
| culture_effect | Trnp1 | 2.897 | 7.24E-09 | glio |
| Bora_in.vivo_noRT | Cfap58 | 4.936 | 7.24E-09 | glio |
| Hsd17b10_in.vivo_noRT | Fbxo15 | 3.874 | 7.24E-10 | glio |
| Atr_in.vivo_noRT | A830035O19R | 4.484 | 7.24E-12 | glio |
| Chek1_ex.vivo_noRT | Olfr171 | 4.842 | 7.24E-13 | glio |
| culture_effect | C130036L24Ri | 2.916 | 7.24E-14 | glio |
| Bcas2_in.vivo_noRT | Esm1 | 4.543 | 7.24E-19 | glio |
| Aars2_in.vivo_noRT | Mcmdc2 | 5.581 | 7.24E-21 | glio |
| culture_effect | Gm41760 | 1.385 | 7.25E-03 | glio |
| Interaction_genotype_Aars2 | B3gnt4 | -2.956 | 7.25E-03 | glio |
| Interaction_genotype_Atr | Erich6b | -2.369 | 7.25E-03 | glio |
| Interaction_genotype_Bcas2 | 3110001I22Ri | -3.585 | 7.25E-03 | glio |
| Interaction_genotype_Bcas2 | Ifi27l2a | 1.255 | 7.25E-03 | glio |
| Interaction_genotype_Myc | Trmo | -2.556 | 7.25E-03 | glio |
| Aars2_in.vivo_noRT | S100a8 | 3.069 | 7.25E-03 | glio |
| Aars2_in.vivo_noRT | Tmem243 | 1.153 | 7.25E-03 | glio |

| | | | | |
|---|---|---|---|---|
| Bcas2_in.vivo_noRT | Prkg2 | -1.55 | 7.25E-03 | glio |
| Bora_in.vivo_noRT | Smyd4 | 1.504 | 7.25E-03 | glio |
| Cenpj_in.vivo_noRT | Flacc1 | 2.022 | 7.25E-03 | glio |
| Chek1_in.vivo_noRT | Nfkbib | -1.087 | 7.25E-03 | glio |
| Interaction_genotype_Hsd17b | Klk6 | 4.676 | 7.25E-04 | glio |
| Interaction_genotype_Myc | Lsm1 | 1.516 | 7.25E-04 | glio |
| Atr_in.vivo_noRT | Tifa | 2.129 | 7.25E-04 | glio |
| Bcas2_in.vivo_noRT | Mmab | 1.789 | 7.25E-04 | glio |
| Atr_in.vivo_noRT | Rpl3 | -0.798 | 7.25E-05 | glio |
| Bcas2_in.vivo_noRT | 2900041M22R | 2.997 | 7.25E-05 | glio |
| Interaction_genotype_Myc | Cdkal1 | -3.397 | 7.25E-06 | glio |
| Interaction_genotype_Myc | Gm42849 | -4.67 | 7.25E-06 | glio |
| Atr_in.vivo_noRT | Ccn2 | 4.206 | 7.25E-06 | glio |
| Bora_in.vivo_noRT | 4930417H01R | 3.296 | 7.25E-06 | glio |
| Interaction_genotype_Myc | 4933433G15R | -5.871 | 7.25E-07 | glio |
| Aars2_ex.vivo_noRT | BB287469 | 3.293 | 7.25E-07 | glio |
| Chek1_in.vivo_noRT | Drd4 | 3.859 | 7.25E-08 | glio |
| culture_effect | Kdelr3 | -1.567 | 7.25E-09 | glio |
| culture_effect | mt-Nd4 | 0.674 | 7.25E-10 | glio |
| Aars2_in.vivo_noRT | Efhd1 | 4.811 | 7.25E-12 | glio |
| culture_effect | Rab7b | 3.146 | 7.25E-14 | glio |
| Aars2_in.vivo_noRT | Zmynd10 | 5.067 | 7.25E-18 | glio |
| culture_effect | Maf1 | -0.326 | 7.26E-03 | glio |
| Interaction_genotype_Atr | Selenow | 0.973 | 7.26E-03 | glio |
| Interaction_genotype_Ifnar1 | 2410017I17Ri | 2.734 | 7.26E-03 | glio |
| Interaction_genotype_Ifnar1 | Gm38477 | 3.127 | 7.26E-03 | glio |
| Interaction_genotype_Ifnar1 | Ablim2 | 2.74 | 7.26E-03 | glio |
| Interaction_genotype_Mtor | Gm10260 | -3.489 | 7.26E-03 | glio |
| Interaction_genotype_Mtor | Aox1 | -2.815 | 7.26E-03 | glio |
| Interaction_genotype_Myc | Dcn | -2.731 | 7.26E-03 | glio |
| Interaction_genotype_Myc | B230208H11R | -2.931 | 7.26E-03 | glio |
| Interaction_genotype_Myc | Mios | -2.378 | 7.26E-03 | glio |
| Interaction_genotype_Rpl3 | Meak7 | -3.77 | 7.26E-03 | glio |
| Aars2_in.vivo_noRT | Sept6 | 1.435 | 7.26E-03 | glio |
| Aars2_in.vivo_noRT | Arhgdia | -0.595 | 7.26E-03 | glio |
| Aars2_ex.vivo_noRT | Atp1a2 | 2.893 | 7.26E-03 | glio |
| Chek1_in.vivo_noRT | Trim6 | 2.061 | 7.26E-03 | glio |
| Interaction_genotype_Aars2 | Gm26682 | -4.152 | 7.26E-04 | glio |
| Interaction_genotype_Myc | Gipc2 | -4.113 | 7.26E-04 | glio |
| Interaction_genotype_Myc | Crybb1 | -3.311 | 7.26E-04 | glio |
| Interaction_genotype_Myc | Saxo2 | -3.647 | 7.26E-04 | glio |
| Interaction_genotype_Myc | S100a3 | -3.744 | 7.26E-04 | glio |
| Interaction_genotype_Myc | Pde12 | -2.726 | 7.26E-04 | glio |
| Interaction_genotype_Myc | Fnbp1 | -2.614 | 7.26E-04 | glio |
| Interaction_genotype_Myc | Mdh2 | 1.678 | 7.26E-04 | glio |
| Interaction_genotype_Myc | Cox7a2 | 1.373 | 7.26E-04 | glio |
| Interaction_genotype_Myc | Pcbp2 | 1.363 | 7.26E-04 | glio |
| Atp5c1_in.vivo_noRT | Vmn1r181 | 2.534 | 7.26E-04 | glio |
| Atr_ex.vivo_noRT | Gm37864 | 2.77 | 7.26E-04 | glio |
| Bcas2_in.vivo_noRT | Gstt2 | 2.497 | 7.26E-04 | glio |
| Bcas2_in.vivo_noRT | Bdp1 | -1.53 | 7.26E-04 | glio |
| Bora_in.vivo_noRT | Cdiptos | 2.387 | 7.26E-04 | glio |

| | | | | |
|---|---|---|---|---|
| Bora_in.vivo_noRT | Etfa | -1.214 | 7.26E-04 | glio |
| Chek1_in.vivo_noRT | Gm44386 | 2.398 | 7.26E-04 | glio |
| Aars2_in.vivo_noRT | Npl | 3.541 | 7.26E-08 | glio |
| Aars2_in.vivo_noRT | Cyp27b1 | 4.28 | 7.26E-12 | glio |
| culture_effect | Il13ra1 | -2.762 | 7.26E-17 | glio |
| culture_effect | Rpl4 | -0.895 | 7.26E-17 | glio |
| culture_effect | Reep3 | 0.368 | 7.27E-03 | glio |
| culture_effect | Dst | -0.384 | 7.27E-03 | glio |
| Interaction_genotype_Atr | Tac2 | -2.369 | 7.27E-03 | glio |
| Interaction_genotype_Atr | Gm2012 | -3.469 | 7.27E-03 | glio |
| Interaction_genotype_Atr | Pcbp4 | 1.27 | 7.27E-03 | glio |
| Interaction_genotype_Mtor | Eif4a3 | 1.396 | 7.27E-03 | glio |
| Interaction_genotype_Mtor | Nucb1 | 1.399 | 7.27E-03 | glio |
| Interaction_genotype_Rpl3 | Esrp2 | -3.582 | 7.27E-03 | glio |
| Aars2_in.vivo_noRT | Rps7 | -0.55 | 7.27E-03 | glio |
| Aars2_ex.vivo_noRT | Gpr146 | 2.899 | 7.27E-03 | glio |
| Atr_in.vivo_noRT | Agpat4 | -1.267 | 7.27E-03 | glio |
| Bora_in.vivo_noRT | Katnbl1 | -1.169 | 7.27E-03 | glio |
| Bora_in.vivo_noRT | Smc2 | -1.137 | 7.27E-03 | glio |
| Bora_ex.vivo_noRT | Car9 | 2.808 | 7.27E-03 | glio |
| Chek1_in.vivo_noRT | Adhfe1 | 1.986 | 7.27E-03 | glio |
| Chek1_in.vivo_noRT | Gvin1 | 1.919 | 7.27E-03 | glio |
| Chek1_in.vivo_noRT | Pbrm1 | -0.946 | 7.27E-03 | glio |
| Chek1_ex.vivo_noRT | Iqgap3 | -2.031 | 7.27E-03 | glio |
| Chek1_ex.vivo_noRT | Mcrip2 | -1.442 | 7.27E-03 | glio |
| culture_effect | Gm10300 | 2.044 | 7.27E-04 | glio |
| Chek1_in.vivo_noRT | Vps25 | 2.053 | 7.27E-04 | glio |
| Interaction_genotype_Mtor | Phf3 | 2.01 | 7.27E-05 | glio |
| Aars2_in.vivo_noRT | Pam16 | -1.319 | 7.27E-05 | glio |
| Chek1_ex.vivo_noRT | C77080 | 3.469 | 7.27E-05 | glio |
| Hsd17b10_in.vivo_noRT | Gm28578 | 3.356 | 7.27E-06 | glio |
| Hsd17b10_in.vivo_noRT | Gm7102 | 3.356 | 7.27E-06 | glio |
| Interaction_genotype_Mtor | Pcmt1 | 2.494 | 7.27E-07 | glio |
| Interaction_genotype_Chek1 | Fa2h | 6.379 | 7.27E-08 | glio |
| Interaction_genotype_Chek1 | Klrk1 | 5.502 | 7.27E-08 | glio |
| Atr_in.vivo_noRT | Ankrd34a | 3.885 | 7.27E-08 | glio |
| Bora_in.vivo_noRT | Hlx | 3.68 | 7.27E-08 | glio |
| Bora_in.vivo_noRT | 1700061G19R | 3.925 | 7.27E-09 | glio |
| Chek1_ex.vivo_noRT | Esp15 | 4.842 | 7.27E-12 | glio |
| Bora_in.vivo_noRT | Mettl7b | 4.936 | 7.27E-13 | glio |
| Bora_in.vivo_noRT | Bmp15 | 4.283 | 7.27E-13 | glio |
| Aars2_in.vivo_noRT | Tdrd5 | 4.941 | 7.27E-15 | glio |
| culture_effect | Fam161a | -1.195 | 7.28E-03 | glio |
| culture_effect | Prcd | 0.773 | 7.28E-03 | glio |
| Interaction_genotype_Aars2 | Apol8 | -2.735 | 7.28E-03 | glio |
| Interaction_genotype_Bcas2 | Gm36172 | -2.9 | 7.28E-03 | glio |
| Interaction_genotype_Bcas2 | Acot2 | 2.461 | 7.28E-03 | glio |
| Interaction_genotype_Bcas2 | Wscd1 | 1.888 | 7.28E-03 | glio |
| Interaction_genotype_Mtor | Ccdc71l | -2.979 | 7.28E-03 | glio |
| Interaction_genotype_Mtor | Pim2 | -2.632 | 7.28E-03 | glio |
| Aars2_in.vivo_noRT | AC168977.2 | 2.045 | 7.28E-03 | glio |
| Aars2_in.vivo_noRT | Eif4e2 | -0.831 | 7.28E-03 | glio |

| | | | | |
|---|---|---|---|---|
| Aars2_in.vivo_noRT | Serf1 | -0.826 | 7.28E-03 | glio |
| Aars2_in.vivo_noRT | Pomp | -0.607 | 7.28E-03 | glio |
| Aars2_ex.vivo_noRT | Slc38a3 | 2.495 | 7.28E-03 | glio |
| Atr_in.vivo_noRT | Tnnt2 | 1.677 | 7.28E-03 | glio |
| Bard1_ex.vivo_noRT | C330024C12R | 2.421 | 7.28E-03 | glio |
| Bard1_ex.vivo_noRT | Pcdhga2 | 2.647 | 7.28E-03 | glio |
| Bcas2_in.vivo_noRT | AW011738 | 1.599 | 7.28E-03 | glio |
| Cdkn2a_in.vivo_noRT | Ly6e | 0.59 | 7.28E-03 | glio |
| culture_effect | 4933406P04R | 1.888 | 7.28E-04 | glio |
| Atr_in.vivo_noRT | Sec61b | -0.76 | 7.28E-04 | glio |
| Bard1_ex.vivo_noRT | Abra | 2.814 | 7.28E-04 | glio |
| Bard1_ex.vivo_noRT | Cyp4f40 | 3.219 | 7.28E-04 | glio |
| Bcas2_ex.vivo_noRT | Magea10 | 2.961 | 7.28E-04 | glio |
| Atp5c1_in.vivo_noRT | Gm11973 | 2.402 | 7.28E-05 | glio |
| Atr_in.vivo_noRT | Ifnb1 | 5.533 | 7.28E-05 | glio |
| Bora_in.vivo_noRT | Amt | 3.086 | 7.28E-05 | glio |
| Bora_in.vivo_noRT | Tmem229b | 2.608 | 7.28E-05 | glio |
| Eif2s1_in.vivo_noRT | Krt19 | 2.143 | 7.28E-05 | glio |
| Atp5c1_in.vivo_noRT | Helt | 2.668 | 7.28E-08 | glio |
| Atp5c1_in.vivo_noRT | Gm20625 | 2.668 | 7.28E-08 | glio |
| Atp5c1_in.vivo_noRT | Gm5157 | 2.668 | 7.28E-08 | glio |
| Atp5c1_in.vivo_noRT | Gm44736 | 2.668 | 7.28E-08 | glio |
| Atp5c1_in.vivo_noRT | Fam227b | 2.668 | 7.28E-08 | glio |
| Atp5c1_in.vivo_noRT | Gm20658 | 2.668 | 7.28E-08 | glio |
| Chek1_in.vivo_noRT | Wdr27 | 3.252 | 7.28E-08 | glio |
| Aars2_in.vivo_noRT | Dio1 | 4.005 | 7.28E-11 | glio |
| Chek1_ex.vivo_noRT | Gm17203 | 4.443 | 7.28E-13 | glio |
| Atr_in.vivo_noRT | Ptgis | 5.533 | 7.28E-16 | glio |
| culture_effect | Col6a1 | 6.433 | 7.28E-46 | glio |
| culture_effect | Srpk2 | -0.358 | 7.29E-03 | glio |
| Interaction_genotype_Aars2 | Ifi203 | 2.174 | 7.29E-03 | glio |
| Interaction_genotype_Atr | Gm15726 | -3.172 | 7.29E-03 | glio |
| Interaction_genotype_Atr | Rnpc3 | 1.806 | 7.29E-03 | glio |
| Interaction_genotype_Bcas2 | C330011M18F | -3.625 | 7.29E-03 | glio |
| Interaction_genotype_Mtor | Ccdc51 | -2.655 | 7.29E-03 | glio |
| Interaction_genotype_Myc | Lrrc75a | -2.747 | 7.29E-03 | glio |
| Interaction_genotype_Myc | Tom1l1 | -2.62 | 7.29E-03 | glio |
| Interaction_genotype_Myc | Armc9 | -2.644 | 7.29E-03 | glio |
| Interaction_genotype_Myc | Zfp955a | -2.607 | 7.29E-03 | glio |
| Interaction_genotype_Myc | Gabpb1 | -1.963 | 7.29E-03 | glio |
| interaction_RT | Srgn | 3.003 | 7.29E-03 | glio |
| interaction_RT | Smpd4 | -1.466 | 7.29E-03 | glio |
| interaction_RT | Cdca7 | -0.809 | 7.29E-03 | glio |
| Aars2_in.vivo_noRT | Rnf187 | -0.782 | 7.29E-03 | glio |
| Atr_ex.vivo_noRT | Set | -0.684 | 7.29E-03 | glio |
| Bcas2_in.vivo_noRT | Ofd1 | -1.252 | 7.29E-03 | glio |
| Bcas2_in.vivo_noRT | Cpm | -1.539 | 7.29E-03 | glio |
| Cdkn2a_in.vivo_noRT | Gjd4 | -1.675 | 7.29E-03 | glio |
| Chek1_in.vivo_noRT | Sh2b3 | 1.89 | 7.29E-03 | glio |
| culture_effect | Hspbap1 | -0.849 | 7.29E-04 | glio |
| culture_effect | Prkaa1 | -0.591 | 7.29E-04 | glio |
| Interaction_genotype_Mtor | Gm4907 | -4.646 | 7.29E-04 | glio |

| | | | | |
|---|---|---|---|---|
| Interaction_genotype_Mtor | Gm28404 | -3.686 | 7.29E-04 | glio |
| Interaction_genotype_Mtor | 4930590J08Ri | -4.283 | 7.29E-04 | glio |
| Interaction_genotype_Mtor | Snta1 | -3.024 | 7.29E-04 | glio |
| Interaction_genotype_Mtor | Nup43 | 1.954 | 7.29E-04 | glio |
| Interaction_genotype_Mtor | Anxa1 | 2.149 | 7.29E-04 | glio |
| Interaction_genotype_Myc | Bco2 | -3.294 | 7.29E-04 | glio |
| Aars2_in.vivo_noRT | Fgf21 | 2.56 | 7.29E-04 | glio |
| Atp5c1_in.vivo_noRT | Adam30 | 2.289 | 7.29E-04 | glio |
| Bcas2_in.vivo_noRT | Gm46378 | 2.574 | 7.29E-04 | glio |
| Bora_in.vivo_noRT | C030037D09R | 2.435 | 7.29E-04 | glio |
| Cenpj_in.vivo_noRT | Serpinb9g | 2.022 | 7.29E-04 | glio |
| Aars2_in.vivo_noRT | Gm12689 | 3.099 | 7.29E-05 | glio |
| Aars2_in.vivo_noRT | Fcgrt | 3.08 | 7.29E-05 | glio |
| Atr_in.vivo_noRT | B230208H11R | 2.865 | 7.29E-05 | glio |
| Hsd17b10_in.vivo_noRT | Pde11a | 3.087 | 7.29E-05 | glio |
| Interaction_genotype_Myc | Klhl23 | -5.299 | 7.29E-07 | glio |
| Interaction_genotype_Myc | mt-Atp6 | 1.827 | 7.29E-07 | glio |
| Aars2_in.vivo_noRT | 1810034E14Ri | 3.444 | 7.29E-07 | glio |
| Bora_in.vivo_noRT | Mucl3 | 3.689 | 7.29E-07 | glio |
| Atr_ex.vivo_noRT | Olfr810 | 3.164 | 7.29E-08 | glio |
| Bcas2_in.vivo_noRT | 2310058D17R | 4.156 | 7.29E-09 | glio |
| Chek1_ex.vivo_noRT | Map3k9 | 4.055 | 7.29E-09 | glio |
| Chek1_in.vivo_noRT | Gm9999 | 4.069 | 7.29E-10 | glio |
| Bora_in.vivo_noRT | Dreh | 4.29 | 7.29E-12 | glio |
| Bcas2_in.vivo_noRT | Gm5662 | 4.807 | 7.29E-13 | glio |
| Chek1_in.vivo_noRT | Gm48617 | 4.78 | 7.29E-22 | glio |
| Interaction_genotype_Aars2 | Gm15890 | -2.82 | 7.30E-03 | glio |
| Interaction_genotype_Aars2 | Crabp1 | -2.848 | 7.30E-03 | glio |
| Interaction_genotype_Bcas2 | Gm13404 | -3.68 | 7.30E-03 | glio |
| Interaction_genotype_Bcas2 | A130010J15Ri | -2.3 | 7.30E-03 | glio |
| Aars2_in.vivo_noRT | Dhx57 | 1.536 | 7.30E-03 | glio |
| Bora_in.vivo_noRT | Pmp22 | 1.825 | 7.30E-03 | glio |
| Chek1_in.vivo_noRT | Gm26910 | 1.845 | 7.30E-03 | glio |
| Chek1_in.vivo_noRT | Rffl | 1.759 | 7.30E-03 | glio |
| Chek1_in.vivo_noRT | Ddx5 | -0.461 | 7.30E-03 | glio |
| Interaction_genotype_Chek1 | Wdr27 | -3.941 | 7.30E-04 | glio |
| Interaction_genotype_Myc | Camkk1 | -3.901 | 7.30E-04 | glio |
| Aars2_in.vivo_noRT | Ddah2 | 2.452 | 7.30E-04 | glio |
| Bora_in.vivo_noRT | Dnaja1 | -0.654 | 7.30E-04 | glio |
| Interaction_genotype_Mtor | Tmem191c | -4.583 | 7.30E-05 | glio |
| Interaction_genotype_Mtor | 9130604C24R | -4.991 | 7.30E-05 | glio |
| Interaction_genotype_Mtor | Sap30l | 2.086 | 7.30E-05 | glio |
| Bora_in.vivo_noRT | Atf6 | -2.029 | 7.30E-05 | glio |
| Chek1_in.vivo_noRT | 4930405A21R | 2.93 | 7.30E-06 | glio |
| Bora_in.vivo_noRT | 1700055D18R | 4.079 | 7.30E-08 | glio |
| Chek1_ex.vivo_noRT | Sdr16c5 | 4.447 | 7.30E-08 | glio |
| Hsd17b10_in.vivo_noRT | Map3k9 | 3.356 | 7.30E-09 | glio |
| Bora_in.vivo_noRT | Baalc | 4.144 | 7.30E-10 | glio |
| Bcas2_in.vivo_noRT | A330015K06R | 4.15 | 7.30E-11 | glio |
| Bora_in.vivo_noRT | Rab27a | 4.41 | 7.30E-13 | glio |
| Atr_in.vivo_noRT | Pcdhga11 | 4.536 | 7.30E-15 | glio |
| culture_effect | Atg9b | -5.354 | 7.30E-26 | glio |

| | | | | |
|---|---|---|---|---|
| culture_effect | Atat1 | -0.658 | 7.31E-03 | glio |
| Interaction_genotype_Aars2 | Zfp955b | -2.686 | 7.31E-03 | glio |
| Interaction_genotype_Atr | Thap8 | -3.328 | 7.31E-03 | glio |
| Interaction_genotype_Atr | Stk35 | -2.564 | 7.31E-03 | glio |
| Interaction_genotype_Atr | Rit1 | 1.753 | 7.31E-03 | glio |
| Interaction_genotype_Bora | Trim37 | 1.908 | 7.31E-03 | glio |
| Interaction_genotype_Chek1 | Gm30074 | -3.619 | 7.31E-03 | glio |
| Interaction_genotype_Chek1 | Mad2l1bp | 2.262 | 7.31E-03 | glio |
| Interaction_genotype_Chek1 | Ggh | 1.733 | 7.31E-03 | glio |
| Interaction_genotype_Myc | Gm13178 | -2.213 | 7.31E-03 | glio |
| Interaction_genotype_Myc | Slc22a18 | -3.243 | 7.31E-03 | glio |
| Interaction_genotype_Myc | Mpv17l | -2.677 | 7.31E-03 | glio |
| Interaction_genotype_Myc | Mmgt1 | -2.182 | 7.31E-03 | glio |
| Interaction_genotype_Myc | Rap2c | -2.16 | 7.31E-03 | glio |
| Interaction_genotype_Rpl3 | 1700112D23R | -3.743 | 7.31E-03 | glio |
| interaction_RT | H13 | 0.609 | 7.31E-03 | glio |
| Atr_in.vivo_noRT | Myh14 | 1.697 | 7.31E-03 | glio |
| Atr_in.vivo_noRT | Zfp948 | 1.382 | 7.31E-03 | glio |
| Bcas2_in.vivo_noRT | Magi2 | 1.963 | 7.31E-03 | glio |
| Bcas2_in.vivo_noRT | Rnf214 | 1.169 | 7.31E-03 | glio |
| Bora_in.vivo_noRT | Gm20712 | 1.967 | 7.31E-03 | glio |
| Chek1_in.vivo_noRT | Helz | -1.044 | 7.31E-03 | glio |
| Interaction_genotype_Mtor | Gm48357 | -3.59 | 7.31E-04 | glio |
| Interaction_genotype_Myc | Emc6 | 1.371 | 7.31E-04 | glio |
| Aars2_in.vivo_noRT | Accs | 2.31 | 7.31E-04 | glio |
| Bcas2_in.vivo_noRT | BC065397 | 2.724 | 7.31E-04 | glio |
| Interaction_genotype_Mtor | Gm37933 | -3.868 | 7.31E-05 | glio |
| Interaction_genotype_Mtor | Gm20682 | -4.534 | 7.31E-06 | glio |
| Atp5c1_in.vivo_noRT | Adam5 | 2.404 | 7.31E-06 | glio |
| Atp5c1_in.vivo_noRT | 9130230L23Ri | 2.936 | 7.31E-06 | glio |
| Atp5c1_in.vivo_noRT | Cysrt1 | 3.455 | 7.31E-06 | glio |
| Atr_in.vivo_noRT | Frmd4b | 3.149 | 7.31E-06 | glio |
| Chek1_ex.vivo_noRT | Mogat2 | 3.892 | 7.31E-06 | glio |
| Bora_in.vivo_noRT | Ikbke | 3.62 | 7.31E-08 | glio |
| Interaction_genotype_Atr | Zwint | 2.789 | 7.31E-09 | glio |
| Aars2_in.vivo_noRT | Tmem179 | 4.168 | 7.31E-09 | glio |
| Bcas2_in.vivo_noRT | Slxl1 | 4.029 | 7.31E-09 | glio |
| Bcas2_in.vivo_noRT | 5830454E08Ri | 3.882 | 7.31E-09 | glio |
| Chek1_in.vivo_noRT | D030051J21Ri | 4.141 | 7.31E-10 | glio |
| culture_effect | Pdzrn3 | 2.358 | 7.31E-14 | glio |
| Interaction_genotype_Atr | Mrpl53 | 1.108 | 7.32E-03 | glio |
| Interaction_genotype_Bora | Tlr1 | -3.543 | 7.32E-03 | glio |
| Interaction_genotype_Mtor | Cd27 | -3.266 | 7.32E-03 | glio |
| Interaction_genotype_Mtor | Ttc8 | -2.954 | 7.32E-03 | glio |
| Aars2_in.vivo_noRT | Tmem237 | -1.324 | 7.32E-03 | glio |
| Atr_in.vivo_noRT | 2810013P06R | 1.566 | 7.32E-03 | glio |
| Bcas2_in.vivo_noRT | Polm | 1.784 | 7.32E-03 | glio |
| Bcas2_in.vivo_noRT | Nagk | 1.467 | 7.32E-03 | glio |
| Bcas2_ex.vivo_noRT | BB557941 | 2.342 | 7.32E-03 | glio |
| Bcas2_ex.vivo_noRT | Sox11 | 2.742 | 7.32E-03 | glio |
| Cenpj_in.vivo_noRT | 1700023H06R | 2.529 | 7.32E-03 | glio |
| Cenpj_in.vivo_noRT | Gm11110 | 2.318 | 7.32E-03 | glio |

| | | | | |
|---|---|---|---|---|
| culture_effect | Rbm19 | 1.044 | 7.32E-04 | glio |
| interaction_RT | Rab11a | 0.682 | 7.32E-04 | glio |
| Aars2_in.vivo_noRT | Isoc2a | 2.184 | 7.32E-04 | glio |
| Aars2_in.vivo_noRT | Zfp446 | 2.097 | 7.32E-04 | glio |
| Aars2_in.vivo_noRT | Rab11fip3 | 2.098 | 7.32E-04 | glio |
| Atr_in.vivo_noRT | Pcdh15 | 1.959 | 7.32E-04 | glio |
| Interaction_genotype_Mtor | Rex2 | -5.085 | 7.32E-05 | glio |
| Interaction_genotype_Mtor | Arhgap27 | -4.471 | 7.32E-05 | glio |
| Interaction_genotype_Mtor | Shc3 | -3.846 | 7.32E-05 | glio |
| Interaction_genotype_Mtor | Snx11 | -3.254 | 7.32E-05 | glio |
| Interaction_genotype_Mtor | Moxd1 | 2.11 | 7.32E-05 | glio |
| interaction_RT | Rpl13a | -0.92 | 7.32E-06 | glio |
| Bcas2_in.vivo_noRT | Slc7a3 | 3.54 | 7.32E-07 | glio |
| Bcas2_in.vivo_noRT | Olfr267 | 3.562 | 7.32E-07 | glio |
| Cenpj_ex.vivo_noRT | Vmn2r18 | 3.389 | 7.32E-07 | glio |
| Chek1_ex.vivo_noRT | Zfp986 | 4.271 | 7.32E-07 | glio |
| Chek1_ex.vivo_noRT | Gm10801 | 4.842 | 7.32E-12 | glio |
| culture_effect | Mob3c | -1.098 | 7.33E-03 | glio |
| culture_effect | Mvb12b | 0.57 | 7.33E-03 | glio |
| Interaction_genotype_Bora | 4930540M05F | -3.45 | 7.33E-03 | glio |
| Interaction_genotype_Mtor | Egfl8 | -3.401 | 7.33E-03 | glio |
| Interaction_genotype_Mtor | Tspyl5 | -2.846 | 7.33E-03 | glio |
| Interaction_genotype_Myc | Ccsap | -2.112 | 7.33E-03 | glio |
| Interaction_genotype_Pars2 | Gm36839 | -3.652 | 7.33E-03 | glio |
| Aars2_in.vivo_noRT | Clec1a | 1.878 | 7.33E-03 | glio |
| Aars2_in.vivo_noRT | Zfp940 | 1.89 | 7.33E-03 | glio |
| culture_effect | Timm17b | -0.483 | 7.33E-04 | glio |
| Interaction_genotype_Myc | Gm44983 | -3.228 | 7.33E-04 | glio |
| Atr_in.vivo_noRT | Brd4 | -1.114 | 7.33E-04 | glio |
| Chek1_in.vivo_noRT | Amt | 2.661 | 7.33E-04 | glio |
| Chek1_in.vivo_noRT | Pnp | -1.21 | 7.33E-04 | glio |
| Interaction_genotype_Myc | Shroom2 | -2.832 | 7.33E-05 | glio |
| Aars2_in.vivo_noRT | Gm34455 | 3.561 | 7.33E-05 | glio |
| Aars2_in.vivo_noRT | Lbx2 | 3.037 | 7.33E-05 | glio |
| Aars2_in.vivo_noRT | Arhgap33 | 2.863 | 7.33E-05 | glio |
| Aars2_in.vivo_noRT | Gm11131 | 2.709 | 7.33E-05 | glio |
| Aars2_ex.vivo_noRT | Nos2 | 3.293 | 7.33E-05 | glio |
| Aars2_ex.vivo_noRT | 4930470G03R | 2.9 | 7.33E-05 | glio |
| Atr_in.vivo_noRT | Polm | 2.49 | 7.33E-05 | glio |
| Atr_in.vivo_noRT | Svbp | -1.412 | 7.33E-05 | glio |
| Eif2s1_in.vivo_noRT | Gm48155 | 2.143 | 7.33E-05 | glio |
| Atr_in.vivo_noRT | 3110083C13R | 3.067 | 7.33E-06 | glio |
| Bcas2_in.vivo_noRT | Gm12905 | 3.502 | 7.33E-06 | glio |
| Atr_in.vivo_noRT | Mkrn3 | 3.276 | 7.33E-07 | glio |
| Bora_in.vivo_noRT | Gm16283 | 3.412 | 7.33E-07 | glio |
| Chek1_ex.vivo_noRT | Lyl1 | 3.657 | 7.33E-07 | glio |
| Bcas2_in.vivo_noRT | Medag | 3.639 | 7.33E-08 | glio |
| Bora_in.vivo_noRT | Palm3 | 3.868 | 7.33E-08 | glio |
| Hsd17b10_in.vivo_noRT | Tcstv3 | 3.616 | 7.33E-08 | glio |
| culture_effect | Tmem140 | -2.314 | 7.33E-11 | glio |
| Bcas2_in.vivo_noRT | Cys1 | 4.541 | 7.33E-15 | glio |
| Aars2_in.vivo_noRT | Cyp2c23 | 4.804 | 7.33E-16 | glio |

| | | | | |
|---|---|---|---|---|
| Bora_in.vivo_noRT | Gm20711 | 4.936 | 7.33E-17 | glio |
| culture_effect | Pcmtd2 | -2.585 | 7.33E-22 | glio |
| Atr_in.vivo_noRT | B230206H07R | 5.008 | 7.33E-25 | glio |
| culture_effect | Cox5b | 0.303 | 7.34E-03 | glio |
| Interaction_genotype_Aars2 | Olfm2 | -3.072 | 7.34E-03 | glio |
| Interaction_genotype_Aars2 | Gpm6b | 1.625 | 7.34E-03 | glio |
| Interaction_genotype_Bora | Pmp22 | -2.853 | 7.34E-03 | glio |
| Interaction_genotype_Bora | Ppif | 2.231 | 7.34E-03 | glio |
| Interaction_genotype_Bora | Josd2 | -1.103 | 7.34E-03 | glio |
| Interaction_genotype_Rad51 | Irf3 | 1.845 | 7.34E-03 | glio |
| Interaction_genotype_Rpl3 | Elobl | -4.042 | 7.34E-03 | glio |
| Bora_in.vivo_noRT | mt-Nd4 | -0.503 | 7.34E-03 | glio |
| Cenpj_in.vivo_noRT | Bco2 | 2.022 | 7.34E-03 | glio |
| Chek1_in.vivo_noRT | Nphp3 | 1.961 | 7.34E-03 | glio |
| Chek1_ex.vivo_noRT | Ypel4 | 2.444 | 7.34E-03 | glio |
| Fanca_ex.vivo_noRT | Rimklb | 2.124 | 7.34E-03 | glio |
| Interaction_genotype_Aars2 | Gm48772 | -3.929 | 7.34E-04 | glio |
| Interaction_genotype_Bora | Klhl30 | -4.1 | 7.34E-04 | glio |
| Interaction_genotype_Bora | Cdk19 | 2.136 | 7.34E-04 | glio |
| Interaction_genotype_Mtor | Phkg1 | -5.531 | 7.34E-04 | glio |
| Interaction_genotype_Mtor | Rcn1 | 1.715 | 7.34E-04 | glio |
| interaction_RT | Mrpl32 | 0.642 | 7.34E-04 | glio |
| Aars2_in.vivo_noRT | Gm43328 | 2.479 | 7.34E-04 | glio |
| Atp5c1_in.vivo_noRT | Gm11769 | 2.408 | 7.34E-04 | glio |
| Chek1_ex.vivo_noRT | E130215H24R | 2.795 | 7.34E-04 | glio |
| Chek1_ex.vivo_noRT | Tmem88 | 3.079 | 7.34E-04 | glio |
| Hsd17b10_in.vivo_noRT | B230216N24R | 2.768 | 7.34E-04 | glio |
| Bcas2_ex.vivo_noRT | Gm266 | 2.742 | 7.34E-05 | glio |
| Interaction_genotype_Chek1 | Dnm3 | -5.8 | 7.34E-07 | glio |
| Bcas2_in.vivo_noRT | Arhgef19 | 3.628 | 7.34E-07 | glio |
| Aars2_in.vivo_noRT | Kcnab3 | 3.793 | 7.34E-08 | glio |
| Chek1_ex.vivo_noRT | Pla2g2e | 5.545 | 7.34E-09 | glio |
| interaction_RT | Gm26532 | 1.807 | 7.34E-10 | glio |
| Bcas2_in.vivo_noRT | Spink6 | 4.807 | 7.34E-18 | glio |
| Atr_in.vivo_noRT | Ablim2 | 5.001 | 7.34E-25 | glio |
| Interaction_genotype_Aars2 | Itga2b | -3.276 | 7.35E-03 | glio |
| Interaction_genotype_Atr | Krt18 | 4.616 | 7.35E-03 | glio |
| Interaction_genotype_Atr | Mrpl50 | 1.213 | 7.35E-03 | glio |
| Interaction_genotype_Atr | Pnrc1 | 1.215 | 7.35E-03 | glio |
| Interaction_genotype_Mtor | Adam4 | -3.053 | 7.35E-03 | glio |
| Interaction_genotype_Myc | Btbd10 | -2.031 | 7.35E-03 | glio |
| Interaction_genotype_Rad51 | Zfp445 | 1.742 | 7.35E-03 | glio |
| Aars2_in.vivo_noRT | Otub2 | 1.717 | 7.35E-03 | glio |
| Bcas2_in.vivo_noRT | Icmt | 1.403 | 7.35E-03 | glio |
| Bcas2_in.vivo_noRT | Kdm6b | -1.246 | 7.35E-03 | glio |
| Bcas2_ex.vivo_noRT | D630024D03R | 2.155 | 7.35E-03 | glio |
| Chek1_in.vivo_noRT | Wbp4 | -0.978 | 7.35E-03 | glio |
| Chek1_in.vivo_noRT | Hscb | 1.103 | 7.35E-03 | glio |
| Chek1_ex.vivo_noRT | Slc16a2 | 2.505 | 7.35E-03 | glio |
| Interaction_genotype_Mtor | Pagr1a | -3.793 | 7.35E-04 | glio |
| Interaction_genotype_Mtor | Gm7467 | -3.15 | 7.35E-04 | glio |
| Interaction_genotype_Mtor | Gpatch8 | 1.775 | 7.35E-04 | glio |

| | | | | |
|---|---|---|---|---|
| Interaction_genotype_Myc | Fgf8 | -3.323 | 7.35E-04 | glio |
| Interaction_genotype_Myc | D630003M21I | -3.062 | 7.35E-04 | glio |
| Interaction_genotype_Myc | Luc7l3 | 1.473 | 7.35E-04 | glio |
| Bcas2_ex.vivo_noRT | Slc35g3 | 2.342 | 7.35E-04 | glio |
| Bcas2_ex.vivo_noRT | 2300002M23I | 2.742 | 7.35E-04 | glio |
| Chek1_in.vivo_noRT | Lurap1 | 2.395 | 7.35E-04 | glio |
| Atr_in.vivo_noRT | Gm17300 | 2.647 | 7.35E-05 | glio |
| Cyp19a1_ex.vivo_noRT | Gm48617 | 2.959 | 7.35E-05 | glio |
| Cyp19a1_ex.vivo_noRT | Rab27a | 3.689 | 7.35E-05 | glio |
| Bcas2_in.vivo_noRT | Rab11fip1 | 3.513 | 7.35E-06 | glio |
| Aars2_ex.vivo_noRT | H1foo | 3.293 | 7.35E-07 | glio |
| Chek1_in.vivo_noRT | Tmem8b | 3.656 | 7.35E-08 | glio |
| Chek1_in.vivo_noRT | Zfp712 | 3.348 | 7.35E-08 | glio |
| Interaction_genotype_Myc | Dvl3 | -2.14 | 7.36E-03 | glio |
| Aars2_in.vivo_noRT | Tomm40 | -0.812 | 7.36E-03 | glio |
| Atr_in.vivo_noRT | Cdk5r1 | 2.008 | 7.36E-03 | glio |
| Chek1_in.vivo_noRT | Rab43 | 1.816 | 7.36E-03 | glio |
| Chek1_in.vivo_noRT | Tet3 | -1.416 | 7.36E-03 | glio |
| Chek1_ex.vivo_noRT | Cmc2 | -1.245 | 7.36E-03 | glio |
| culture_effect | Zfr2 | -1.916 | 7.36E-04 | glio |
| Interaction_genotype_Atr | Lypd6 | -4.269 | 7.36E-04 | glio |
| Interaction_genotype_Mtor | Lamtor2 | 1.595 | 7.36E-04 | glio |
| Aars2_in.vivo_noRT | Lsm8 | -1.004 | 7.36E-04 | glio |
| Aars2_in.vivo_noRT | Metap2 | -0.902 | 7.36E-04 | glio |
| Aars2_in.vivo_noRT | Emg1 | -0.807 | 7.36E-04 | glio |
| Atp5c1_in.vivo_noRT | Gm15445 | 2.022 | 7.36E-04 | glio |
| Bard1_ex.vivo_noRT | Kcnk7 | 2.814 | 7.36E-04 | glio |
| Bard1_ex.vivo_noRT | Myoz1 | 2.814 | 7.36E-04 | glio |
| Chek1_ex.vivo_noRT | Plppr3 | 3.07 | 7.36E-04 | glio |
| Eif2s1_in.vivo_noRT | Prl3d3 | 1.88 | 7.36E-04 | glio |
| culture_effect | Olfr1189 | -2.19 | 7.36E-05 | glio |
| culture_effect | Dennd5a | 0.638 | 7.36E-05 | glio |
| Interaction_genotype_Myc | Fam219a | -3.778 | 7.36E-05 | glio |
| Interaction_genotype_Nat2 | BC080695 | 3.724 | 7.36E-05 | glio |
| Aars2_ex.vivo_noRT | Gm14207 | 2.499 | 7.36E-05 | glio |
| Bcas2_ex.vivo_noRT | Ildr1 | 2.742 | 7.36E-05 | glio |
| Bora_in.vivo_noRT | Gm15486 | 2.564 | 7.36E-05 | glio |
| Chek1_ex.vivo_noRT | Grin2d | 3.69 | 7.36E-06 | glio |
| interaction_RT | Map1lc3a | 1.04 | 7.36E-07 | glio |
| Atr_in.vivo_noRT | Rhov | 3.347 | 7.36E-07 | glio |
| Hsd17b10_in.vivo_noRT | 1110046J04Ri | 3.616 | 7.36E-07 | glio |
| Hsd17b10_in.vivo_noRT | Gm44658 | 3.084 | 7.36E-07 | glio |
| Interaction_genotype_Mtor | Tmem160 | 2.658 | 7.36E-08 | glio |
| Atr_in.vivo_noRT | Fndc10 | 4.039 | 7.36E-08 | glio |
| Chek1_in.vivo_noRT | Nxph3 | 5.044 | 7.36E-15 | glio |
| Interaction_genotype_Chek1 | Ftl1 | 0.855 | 7.37E-03 | glio |
| Interaction_genotype_Myc | Gins1 | -2.408 | 7.37E-03 | glio |
| Interaction_genotype_Myc | Ube4a | -2.095 | 7.37E-03 | glio |
| interaction_RT | Med7 | 0.659 | 7.37E-03 | glio |
| Bcas2_in.vivo_noRT | Prkd1 | 1.924 | 7.37E-03 | glio |
| Bcas2_in.vivo_noRT | BC039966 | 1.59 | 7.37E-03 | glio |
| Bcas2_in.vivo_noRT | Hmg20b | -0.928 | 7.37E-03 | glio |

| | | | | |
|---|---|---|---|---|
| Bora_in.vivo_noRT | Slc36a4 | 1.933 | 7.37E-03 | glio |
| Bora_in.vivo_noRT | Cd68 | 2.122 | 7.37E-03 | glio |
| Bora_in.vivo_noRT | Jakmip2 | 1.828 | 7.37E-03 | glio |
| Bora_in.vivo_noRT | Gm9887 | 1.923 | 7.37E-03 | glio |
| Bora_in.vivo_noRT | Gm34397 | 1.833 | 7.37E-03 | glio |
| Cenpj_in.vivo_noRT | Stra8 | 2.029 | 7.37E-03 | glio |
| Chek1_in.vivo_noRT | Lmbr1 | 1.521 | 7.37E-03 | glio |
| Chek1_in.vivo_noRT | Acbd6 | -0.953 | 7.37E-03 | glio |
| Chek1_ex.vivo_noRT | S1pr1 | 2.799 | 7.37E-03 | glio |
| Chek1_ex.vivo_noRT | Mfsd4b4 | 2.25 | 7.37E-03 | glio |
| Chek1_ex.vivo_noRT | Leng9 | 2.316 | 7.37E-03 | glio |
| Chek1_ex.vivo_noRT | Tcof1 | -1.224 | 7.37E-03 | glio |
| culture_effect | Fam177a | 1.542 | 7.37E-04 | glio |
| Interaction_genotype_Aars2 | Prkaa2 | -3.796 | 7.37E-04 | glio |
| Interaction_genotype_Aars2 | Gm12905 | -4.537 | 7.37E-04 | glio |
| Interaction_genotype_Aars2 | AA386476 | -4.241 | 7.37E-04 | glio |
| Interaction_genotype_Aars2 | C030005K06R | -4.224 | 7.37E-04 | glio |
| Interaction_genotype_Aars2 | Gm16794 | -4.06 | 7.37E-04 | glio |
| Atr_in.vivo_noRT | Zfp438 | 2.001 | 7.37E-04 | glio |
| Atr_ex.vivo_noRT | Gm47428 | 3.067 | 7.37E-04 | glio |
| Interaction_genotype_Myc | Tubgcp6 | -3.69 | 7.37E-05 | glio |
| Bard1_ex.vivo_noRT | Olfr56 | 2.814 | 7.37E-05 | glio |
| Chek1_ex.vivo_noRT | F630040K05Ri | 3.118 | 7.37E-05 | glio |
| Chek1_ex.vivo_noRT | Zfp275 | -2.264 | 7.37E-05 | glio |
| Interaction_genotype_Mtor | Smtnl1 | -4.099 | 7.37E-06 | glio |
| Aars2_ex.vivo_noRT | Gm38163 | 3.293 | 7.37E-06 | glio |
| Atr_in.vivo_noRT | Mirg | 3.191 | 7.37E-06 | glio |
| Atr_in.vivo_noRT | Ercc6 | 2.479 | 7.37E-06 | glio |
| culture_effect | Camk1d | 2.627 | 7.37E-07 | glio |
| Interaction_genotype_Myc | Anp32b | 2.25 | 7.37E-07 | glio |
| Atp5c1_in.vivo_noRT | Olfr810 | 2.668 | 7.37E-08 | glio |
| Chek1_in.vivo_noRT | Gm20478 | 3.471 | 7.37E-08 | glio |
| Bcas2_in.vivo_noRT | Gm32585 | 4.281 | 7.37E-09 | glio |
| Atr_in.vivo_noRT | Cpne9 | 4.496 | 7.37E-10 | glio |
| Aars2_in.vivo_noRT | Nt5e | 4.384 | 7.37E-13 | glio |
| Bora_in.vivo_noRT | Gstt4 | 4.673 | 7.37E-20 | glio |
| Atr_in.vivo_noRT | Gm35082 | 5.533 | 7.37E-24 | glio |
| Aars2_in.vivo_noRT | Slc19a3 | 5.327 | 7.37E-25 | glio |
| culture_effect | Adamts9 | 1.153 | 7.38E-03 | glio |
| Interaction_genotype_Bcas2 | A930012O16R | -3.692 | 7.38E-03 | glio |
| Interaction_genotype_Mtor | Mlxip | -2.986 | 7.38E-03 | glio |
| Interaction_genotype_Mtor | Nhej1 | -2.966 | 7.38E-03 | glio |
| Aars2_in.vivo_noRT | 9430018G01R | 1.776 | 7.38E-03 | glio |
| Bcas2_ex.vivo_noRT | Alpl | 2.742 | 7.38E-03 | glio |
| Bora_in.vivo_noRT | Atxn1 | 1.736 | 7.38E-03 | glio |
| Bora_in.vivo_noRT | C87436 | 1.19 | 7.38E-03 | glio |
| Hsd17b10_in.vivo_noRT | Mydgf | -0.679 | 7.38E-03 | glio |
| culture_effect | Rap1b | 0.372 | 7.38E-04 | glio |
| Interaction_genotype_Myc | C920021L13Ri | -3.634 | 7.38E-04 | glio |
| Interaction_genotype_Myc | Ufsp1 | -2.762 | 7.38E-04 | glio |
| Atp5c1_in.vivo_noRT | Sprr1a | 2.668 | 7.38E-04 | glio |
| Atp5c1_in.vivo_noRT | Gm47851 | 2.408 | 7.38E-04 | glio |

| | | | | |
|---|---|---|---|---|
| Eif2s1_in.vivo_noRT | Hfm1 | 2.143 | 7.38E-04 | glio |
| Eif2s1_in.vivo_noRT | D630024D03F | 2.143 | 7.38E-04 | glio |
| Hsd17b10_in.vivo_noRT | Reep4 | -1.898 | 7.38E-04 | glio |
| interaction_RT | Fas | 3.679 | 7.38E-05 | glio |
| Atr_in.vivo_noRT | BC028528 | 2.835 | 7.38E-05 | glio |
| Hsd17b10_in.vivo_noRT | Dlx4 | 3.09 | 7.38E-06 | glio |
| culture_effect | Txlng | -0.908 | 7.38E-07 | glio |
| Aars2_in.vivo_noRT | A430035B10R | 3.89 | 7.38E-08 | glio |
| Aars2_in.vivo_noRT | Cyth4 | 3.786 | 7.38E-08 | glio |
| Bora_in.vivo_noRT | Sh2d3c | 4.035 | 7.38E-08 | glio |
| Bora_ex.vivo_noRT | Capn13 | 3.201 | 7.38E-08 | glio |
| Bcas2_in.vivo_noRT | Gm47204 | 4.162 | 7.38E-09 | glio |
| Bcas2_in.vivo_noRT | D030051J21Ri | 3.904 | 7.38E-09 | glio |
| Aars2_in.vivo_noRT | A430106G13R | 4.535 | 7.38E-10 | glio |
| Aars2_in.vivo_noRT | Gm11884 | 4.328 | 7.38E-12 | glio |
| Bora_in.vivo_noRT | Hnf1aos1 | 4.936 | 7.38E-16 | glio |
| culture_effect | Platr16 | -1.831 | 7.39E-03 | glio |
| culture_effect | Rnf214 | 0.571 | 7.39E-03 | glio |
| Interaction_genotype_Atr | Magea10 | -2.991 | 7.39E-03 | glio |
| Interaction_genotype_Atr | Svbp | 1.271 | 7.39E-03 | glio |
| Interaction_genotype_Bora | Asf1a | 1.375 | 7.39E-03 | glio |
| Interaction_genotype_Bora | Tra2b | 1.398 | 7.39E-03 | glio |
| Interaction_genotype_Mtor | Hnrnph3 | 1.721 | 7.39E-03 | glio |
| Interaction_genotype_Myc | Gab3 | -3.063 | 7.39E-03 | glio |
| Bcas2_in.vivo_noRT | A630052C17R | 1.936 | 7.39E-03 | glio |
| Eif2s1_ex.vivo_noRT | Gpr22 | 2.234 | 7.39E-03 | glio |
| Hsd17b10_in.vivo_noRT | Zfp455 | 2.063 | 7.39E-03 | glio |
| Interaction_genotype_Rad21 | Cacng4 | 4.438 | 7.39E-04 | glio |
| Aars2_in.vivo_noRT | Zfpm2 | 2.364 | 7.39E-04 | glio |
| Bcas2_in.vivo_noRT | Foxl2os | 2.299 | 7.39E-04 | glio |
| Chek1_in.vivo_noRT | Sh3bgrl2 | 2.435 | 7.39E-04 | glio |
| Chek1_in.vivo_noRT | Rabep1 | -1.199 | 7.39E-04 | glio |
| Chek1_ex.vivo_noRT | Ahrr | 3.024 | 7.39E-04 | glio |
| Interaction_genotype_Atr | Cavin2 | -4.856 | 7.39E-05 | glio |
| Interaction_genotype_Mtor | Eif4g2 | 1.644 | 7.39E-05 | glio |
| Aars2_in.vivo_noRT | Gm15706 | 2.421 | 7.39E-05 | glio |
| Atr_in.vivo_noRT | 1700001G11R | 2.722 | 7.39E-06 | glio |
| Bora_ex.vivo_noRT | Edn2 | 2.806 | 7.39E-06 | glio |
| Chek1_in.vivo_noRT | 5330426L24Ri | 3.334 | 7.39E-06 | glio |
| Chek1_in.vivo_noRT | Arhgap4 | 3.058 | 7.39E-06 | glio |
| culture_effect | Klk8 | -1.06 | 7.39E-07 | glio |
| culture_effect | 5830454E08Ri | 2.683 | 7.39E-08 | glio |
| Bora_in.vivo_noRT | Gm28424 | 3.925 | 7.39E-09 | glio |
| Atr_in.vivo_noRT | Tmem8b | 4.145 | 7.39E-10 | glio |
| Bcas2_in.vivo_noRT | Gm31152 | 4.285 | 7.39E-12 | glio |
| Aars2_in.vivo_noRT | Armc2 | 4.936 | 7.39E-13 | glio |
| Aars2_in.vivo_noRT | AF067063 | 5.327 | 7.39E-24 | glio |
| culture_effect | Sec24a | -0.477 | 7.40E-03 | glio |
| culture_effect | Pttg1ip | -0.479 | 7.40E-03 | glio |
| Interaction_genotype_Aars2 | D630024D03F | -2.62 | 7.40E-03 | glio |
| Interaction_genotype_Mtor | Nppc | -3.51 | 7.40E-03 | glio |
| Interaction_genotype_Mtor | Lyst | -2.608 | 7.40E-03 | glio |

| | | | | |
|---|---|---|---|---|
| Interaction_genotype_Mtor | Slc20a1 | -2.846 | 7.40E-03 | glio |
| Interaction_genotype_Mtor | Mrpl30 | 1.038 | 7.40E-03 | glio |
| Interaction_genotype_Uqcrc2 | Gm48610 | -4.085 | 7.40E-03 | glio |
| Aars2_in.vivo_noRT | Tbc1d30 | 1.753 | 7.40E-03 | glio |
| Aars2_in.vivo_noRT | Pebp1 | -0.546 | 7.40E-03 | glio |
| Atr_in.vivo_noRT | Ctsh | -1.182 | 7.40E-03 | glio |
| Atr_in.vivo_noRT | Srrm2 | -0.524 | 7.40E-03 | glio |
| Bcas2_in.vivo_noRT | Frmd4a | -1.238 | 7.40E-03 | glio |
| Bora_in.vivo_noRT | Fkbp10 | -1.189 | 7.40E-03 | glio |
| Fanca_ex.vivo_noRT | Gm14929 | 2.124 | 7.40E-03 | glio |
| Fanca_ex.vivo_noRT | Meioc | 2.124 | 7.40E-03 | glio |
| Fanca_ex.vivo_noRT | Tcstv3 | 2.913 | 7.40E-03 | glio |
| Fanca_ex.vivo_noRT | Tmem132c | 2.124 | 7.40E-03 | glio |
| Fanca_ex.vivo_noRT | Gm44693 | 2.124 | 7.40E-03 | glio |
| Hsd17b10_in.vivo_noRT | Spry3 | 2.151 | 7.40E-03 | glio |
| Interaction_genotype_Aars2 | Cog8 | -3.375 | 7.40E-04 | glio |
| Interaction_genotype_Mtor | Plin1 | -4.113 | 7.40E-04 | glio |
| Aars2_in.vivo_noRT | Plxna2 | 2.609 | 7.40E-04 | glio |
| Atr_in.vivo_noRT | Tmod1 | 2.085 | 7.40E-04 | glio |
| Bcas2_in.vivo_noRT | Synm | 2.196 | 7.40E-04 | glio |
| Eif2s1_in.vivo_noRT | Crb3 | 2.143 | 7.40E-04 | glio |
| culture_effect | Xrcc6 | -0.713 | 7.40E-05 | glio |
| Aars2_in.vivo_noRT | Tiprl | -1.298 | 7.40E-05 | glio |
| culture_effect | Gm39556 | -2.845 | 7.40E-06 | glio |
| Interaction_genotype_Myc | Adamts10 | -4.572 | 7.40E-06 | glio |
| Aars2_ex.vivo_noRT | Cox7b2 | 3.629 | 7.40E-06 | glio |
| Bora_in.vivo_noRT | Gm43330 | 3.036 | 7.40E-07 | glio |
| Chek1_in.vivo_noRT | 9430037G07R | 4.135 | 7.40E-09 | glio |
| Aars2_in.vivo_noRT | Ceacam19 | 4.294 | 7.40E-10 | glio |
| Bora_in.vivo_noRT | Nat8f2 | 4.676 | 7.40E-16 | glio |
| Aars2_in.vivo_noRT | Gm26555 | 5.063 | 7.40E-23 | glio |
| culture_effect | Itpr2 | -2.445 | 7.40E-30 | glio |
| Interaction_genotype_Atr | Cenpx | 1.516 | 7.41E-03 | glio |
| Interaction_genotype_Bcas2 | Gm24362 | -3.302 | 7.41E-03 | glio |
| Interaction_genotype_Bcas2 | Lyz2 | -3.716 | 7.41E-03 | glio |
| Interaction_genotype_Bcas2 | D330041H03R | -2.935 | 7.41E-03 | glio |
| Interaction_genotype_Mtor | Pxt1 | -3.158 | 7.41E-03 | glio |
| Interaction_genotype_Mtor | Cyb561a3 | -2.938 | 7.41E-03 | glio |
| Interaction_genotype_Mtor | Clk3 | -1.898 | 7.41E-03 | glio |
| Interaction_genotype_Mtor | Zfr | 1.202 | 7.41E-03 | glio |
| Interaction_genotype_Myc | Papln | -2.941 | 7.41E-03 | glio |
| Interaction_genotype_Myc | Trak1 | -2.542 | 7.41E-03 | glio |
| interaction_RT | Eif2b3 | -0.813 | 7.41E-03 | glio |
| interaction_RT | Pcbp4 | -0.586 | 7.41E-03 | glio |
| Atr_in.vivo_noRT | Rgs2 | 2.094 | 7.41E-03 | glio |
| Bcas2_in.vivo_noRT | Gm36371 | 1.898 | 7.41E-03 | glio |
| Bora_in.vivo_noRT | Prkag2 | 1.418 | 7.41E-03 | glio |
| Bora_in.vivo_noRT | Spry4 | -1.716 | 7.41E-03 | glio |
| Chek1_ex.vivo_noRT | Gstt3 | 2.447 | 7.41E-03 | glio |
| Hsd17b10_in.vivo_noRT | Gm10230 | 2.221 | 7.41E-03 | glio |
| Interaction_genotype_Mtor | Btg1 | 2.013 | 7.41E-04 | glio |
| Aars2_in.vivo_noRT | Msh5 | 2.559 | 7.41E-04 | glio |

| | | | | |
|---|---|---|---|---|
| Atr_in.vivo_noRT | Tmc8 | 2.377 | 7.41E-04 | glio |
| Bcas2_in.vivo_noRT | Nat8f1 | 2.342 | 7.41E-04 | glio |
| Bora_ex.vivo_noRT | Amh | 3.021 | 7.41E-04 | glio |
| Hsd17b10_in.vivo_noRT | Gm47223 | 3.35 | 7.41E-04 | glio |
| Interaction_genotype_Myc | Gjb4 | -4.598 | 7.41E-05 | glio |
| Interaction_genotype_Myc | Vmn1r32 | -3.122 | 7.41E-05 | glio |
| Chek1_in.vivo_noRT | Ampd3 | 2.761 | 7.41E-05 | glio |
| culture_effect | H3f3a | -0.455 | 7.41E-06 | glio |
| Interaction_genotype_Myc | Rps2 | 1.918 | 7.41E-06 | glio |
| Aars2_in.vivo_noRT | Pde4d | 3.151 | 7.41E-06 | glio |
| Chek1_in.vivo_noRT | 2810408A11R | 3.769 | 7.41E-08 | glio |
| Atr_in.vivo_noRT | Clca3a2 | 4.347 | 7.41E-13 | glio |
| Atr_in.vivo_noRT | B930094E09Ri | 5.017 | 7.41E-13 | glio |
| Bcas2_in.vivo_noRT | Cidea | 4.421 | 7.41E-13 | glio |
| culture_effect | Rab19 | -4.166 | 7.41E-14 | glio |
| culture_effect | Dock10 | -2.705 | 7.41E-14 | glio |
| Interaction_genotype_Rpl3 | Gm34885 | -3.835 | 7.42E-03 | glio |
| Interaction_genotype_Rpl3 | Rbpms2 | -3.709 | 7.42E-03 | glio |
| Interaction_genotype_Rpl3 | Gm47758 | 3.681 | 7.42E-03 | glio |
| Aars2_in.vivo_noRT | Jrk | 1.826 | 7.42E-03 | glio |
| Aars2_in.vivo_noRT | Coq9 | -1.328 | 7.42E-03 | glio |
| Atr_in.vivo_noRT | F2r | 1.747 | 7.42E-03 | glio |
| Atr_in.vivo_noRT | Smc1a | -0.849 | 7.42E-03 | glio |
| Atr_ex.vivo_noRT | Epb42 | 2.829 | 7.42E-03 | glio |
| Atr_ex.vivo_noRT | Actb | -0.761 | 7.42E-03 | glio |
| Bcas2_ex.vivo_noRT | Scgb1c1 | 2.349 | 7.42E-03 | glio |
| Bcas2_ex.vivo_noRT | Gprin2 | 2.349 | 7.42E-03 | glio |
| Chek1_in.vivo_noRT | Rps6kl1 | 1.993 | 7.42E-03 | glio |
| Chek1_in.vivo_noRT | Rfc3 | -1.141 | 7.42E-03 | glio |
| Eif2s1_ex.vivo_noRT | Hist1h2ag | 2.234 | 7.42E-03 | glio |
| culture_effect | Mgat4b | 0.61 | 7.42E-04 | glio |
| Interaction_genotype_Atr | Psmc3 | 1.569 | 7.42E-04 | glio |
| Interaction_genotype_Mtor | Tmc7 | -4.282 | 7.42E-04 | glio |
| Interaction_genotype_Myc | Det1 | -2.965 | 7.42E-04 | glio |
| Interaction_genotype_Myc | Gpbp1 | 1.56 | 7.42E-04 | glio |
| interaction_RT | Cox17 | 0.805 | 7.42E-04 | glio |
| Bcas2_in.vivo_noRT | Gm29093 | 2.42 | 7.42E-04 | glio |
| Bcas2_ex.vivo_noRT | Esp15 | 2.742 | 7.42E-04 | glio |
| interaction_RT | Sord | 2.419 | 7.42E-05 | glio |
| Atp5c1_in.vivo_noRT | Serpinb2 | 2.668 | 7.42E-05 | glio |
| Bcas2_in.vivo_noRT | Zdhhc14 | 2.34 | 7.42E-05 | glio |
| Dclre1c_ex.vivo_noRT | D930028M14I | 3.543 | 7.42E-05 | glio |
| Dclre1c_ex.vivo_noRT | AU019990 | 3.08 | 7.42E-05 | glio |
| Bora_in.vivo_noRT | Bfsp2 | 3.564 | 7.42E-06 | glio |
| Bora_in.vivo_noRT | Gm15903 | 3.624 | 7.42E-08 | glio |
| Bcas2_in.vivo_noRT | Gm36757 | 3.943 | 7.42E-09 | glio |
| Aars2_in.vivo_noRT | 5930403N24R | 4.801 | 7.42E-12 | glio |
| Bcas2_in.vivo_noRT | Gm15270 | 4.544 | 7.42E-15 | glio |
| Bcas2_in.vivo_noRT | Serpinb2 | 4.807 | 7.42E-15 | glio |
| culture_effect | Cyp2d22 | -1.839 | 7.43E-03 | glio |
| culture_effect | Ly96 | 0.64 | 7.43E-03 | glio |
| Interaction_genotype_Aars2 | Pldi | -2.728 | 7.43E-03 | glio |

| | | | | |
|---|---|---|---|---|
| Interaction_genotype_Aars2 | Syne3 | -3.567 | 7.43E-03 | glio |
| Interaction_genotype_Aars2 | Sbno1 | 1.373 | 7.43E-03 | glio |
| Interaction_genotype_Myc | Gm26782 | -2.154 | 7.43E-03 | glio |
| Interaction_genotype_Myc | Zwint | 1.215 | 7.43E-03 | glio |
| Interaction_genotype_Pars2 | A330032B11R | -3.381 | 7.43E-03 | glio |
| Interaction_genotype_Pars2 | 1700112J16Ri | -2.855 | 7.43E-03 | glio |
| Interaction_genotype_Pars2 | Npr3 | -3.24 | 7.43E-03 | glio |
| Interaction_genotype_Pars2 | Tlr1 | -3.638 | 7.43E-03 | glio |
| Interaction_genotype_Pars2 | Gm12841 | -2.661 | 7.43E-03 | glio |
| Interaction_genotype_Pars2 | Emb | 2.257 | 7.43E-03 | glio |
| Aars2_in.vivo_noRT | B4galt4 | 1.997 | 7.43E-03 | glio |
| Aars2_in.vivo_noRT | Naxd | -1.016 | 7.43E-03 | glio |
| Aars2_in.vivo_noRT | Atp5h | -0.511 | 7.43E-03 | glio |
| Atp5c1_in.vivo_noRT | 2310067P03R | 2.145 | 7.43E-03 | glio |
| Atp5c1_in.vivo_noRT | A430035B10R | 2.294 | 7.43E-03 | glio |
| Atp5c1_in.vivo_noRT | Prob1 | 2.302 | 7.43E-03 | glio |
| Atr_in.vivo_noRT | Opalin | 2.506 | 7.43E-03 | glio |
| Atr_in.vivo_noRT | Cacna1a | 1.56 | 7.43E-03 | glio |
| Atr_in.vivo_noRT | Mcm3 | -1.204 | 7.43E-03 | glio |
| Atr_in.vivo_noRT | Actg1 | -0.663 | 7.43E-03 | glio |
| Bard1_ex.vivo_noRT | E130102H24R | 2.665 | 7.43E-03 | glio |
| Interaction_genotype_Mtor | Ccdc189 | -4.185 | 7.43E-04 | glio |
| Interaction_genotype_Myc | Gm12794 | -2.729 | 7.43E-04 | glio |
| interaction_RT | Acod1 | 3.462 | 7.43E-04 | glio |
| Aars2_ex.vivo_noRT | Pldi | 2.598 | 7.43E-04 | glio |
| Atr_in.vivo_noRT | Gm47644 | 2.32 | 7.43E-04 | glio |
| Bcas2_in.vivo_noRT | Ttc39c | 1.658 | 7.43E-04 | glio |
| Bcas2_ex.vivo_noRT | Gm10801 | 2.742 | 7.43E-04 | glio |
| Bora_in.vivo_noRT | Cass4 | 2.46 | 7.43E-04 | glio |
| Chek1_ex.vivo_noRT | Kcnj10 | 3.182 | 7.43E-04 | glio |
| Atr_in.vivo_noRT | Cdk16 | -1.517 | 7.43E-05 | glio |
| Bcas2_in.vivo_noRT | Adgrd1 | 2.65 | 7.43E-05 | glio |
| Bcas2_in.vivo_noRT | Crybg2 | 2.8 | 7.43E-05 | glio |
| Chek1_ex.vivo_noRT | Ttll13 | 3.072 | 7.43E-05 | glio |
| Atp5c1_in.vivo_noRT | Apoa5 | 2.668 | 7.43E-08 | glio |
| Atp5c1_in.vivo_noRT | Gm26555 | 2.936 | 7.43E-08 | glio |
| Bora_in.vivo_noRT | Ccr5 | 4.137 | 7.43E-11 | glio |
| Hsd17b10_in.vivo_noRT | Gm6803 | 3.616 | 7.43E-11 | glio |
| Chek1_ex.vivo_noRT | Glyat | 4.045 | 7.43E-12 | glio |
| culture_effect | Gtpbp2 | -2 | 7.43E-19 | glio |
| culture_effect | Rbpms | -1.445 | 7.44E-03 | glio |
| culture_effect | Rps19 | -0.356 | 7.44E-03 | glio |
| Interaction_genotype_Aars2 | Zfp119b | -3.336 | 7.44E-03 | glio |
| Interaction_genotype_Mtor | Hist1h3a | -2.792 | 7.44E-03 | glio |
| Interaction_genotype_Mtor | Gtpbp2 | 1.968 | 7.44E-03 | glio |
| Interaction_genotype_Terf1 | Gm10300 | -4.205 | 7.44E-03 | glio |
| Interaction_genotype_Terf1 | 4930478M13F | -3.851 | 7.44E-03 | glio |
| Interaction_genotype_Terf1 | Gm48933 | -3.999 | 7.44E-03 | glio |
| Aars2_in.vivo_noRT | Cyb5r1 | -1.376 | 7.44E-03 | glio |
| Aars2_ex.vivo_noRT | Rtl9 | 2.582 | 7.44E-03 | glio |
| culture_effect | Asap3 | -2.082 | 7.44E-04 | glio |
| Interaction_genotype_Aars2 | Gm8098 | -4.197 | 7.44E-04 | glio |

| | | | | |
|---|---|---|---|---|
| Interaction_genotype_Aars2 | Gm15127 | -4.148 | 7.44E-04 | glio |
| Atr_in.vivo_noRT | Gm13663 | 2.199 | 7.44E-04 | glio |
| Atr_in.vivo_noRT | Sart3 | -1.568 | 7.44E-04 | glio |
| Chek1_in.vivo_noRT | Plcb4 | 2.457 | 7.44E-04 | glio |
| Interaction_genotype_Mtor | 4932441J04Ri | -4.935 | 7.44E-05 | glio |
| Interaction_genotype_Mtor | Gm38246 | -4.746 | 7.44E-05 | glio |
| Interaction_genotype_Myc | 6330408M09F | -4.761 | 7.44E-05 | glio |
| Interaction_genotype_Myc | Mtcp1 | -4.415 | 7.44E-05 | glio |
| Interaction_genotype_Myc | Msantd2 | -3.564 | 7.44E-05 | glio |
| Interaction_genotype_Myc | Nr2c1 | -3.456 | 7.44E-05 | glio |
| Interaction_genotype_Myc | Gng10 | 1.713 | 7.44E-05 | glio |
| Interaction_genotype_Myc | Rnasek | 1.624 | 7.44E-05 | glio |
| Chek1_ex.vivo_noRT | Prickle2 | 2.9 | 7.44E-05 | glio |
| Chek1_ex.vivo_noRT | Selenok-ps1 | 3.279 | 7.44E-05 | glio |
| Chek1_in.vivo_noRT | Arl4d | 3.195 | 7.44E-06 | glio |
| Hsd17b10_in.vivo_noRT | Gm6566 | 2.691 | 7.44E-06 | glio |
| culture_effect | B230217C12R | 1.243 | 7.44E-07 | glio |
| interaction_RT | Cd47 | 1.362 | 7.44E-08 | glio |
| Chek1_in.vivo_noRT | Prox2 | 4.318 | 7.44E-09 | glio |
| Bora_in.vivo_noRT | Clca3a2 | 3.75 | 7.44E-10 | glio |
| Bora_in.vivo_noRT | Hif3a | 4.818 | 7.44E-12 | glio |
| culture_effect | Sec11a | -0.874 | 7.44E-14 | glio |
| Chek1_in.vivo_noRT | Gm38477 | 4.522 | 7.44E-17 | glio |
| Interaction_genotype_Aars2 | Paip2 | 0.876 | 7.45E-03 | glio |
| Interaction_genotype_Bora | Lncbate10 | -2.522 | 7.45E-03 | glio |
| Interaction_genotype_Bora | 1700023H06R | -3.81 | 7.45E-03 | glio |
| Interaction_genotype_Bora | Ndc80 | 2.222 | 7.45E-03 | glio |
| Interaction_genotype_Cdkn2a | Kcnk7 | 3.378 | 7.45E-03 | glio |
| Interaction_genotype_Pars2 | F630040K05Ri | -3.343 | 7.45E-03 | glio |
| Aars2_in.vivo_noRT | Thap3 | 1.032 | 7.45E-03 | glio |
| Aars2_ex.vivo_noRT | B020031M17F | 3.293 | 7.45E-03 | glio |
| Atr_in.vivo_noRT | Gm15398 | 1.759 | 7.45E-03 | glio |
| Atr_in.vivo_noRT | Glod4 | -1.046 | 7.45E-03 | glio |
| Atr_in.vivo_noRT | Sfpq | -0.961 | 7.45E-03 | glio |
| Bora_in.vivo_noRT | Cd59a | 2.017 | 7.45E-03 | glio |
| Aars2_in.vivo_noRT | Pmp22 | 2.215 | 7.45E-04 | glio |
| Aars2_in.vivo_noRT | Rpl13 | -0.705 | 7.45E-04 | glio |
| Atr_in.vivo_noRT | Cnot4 | -1.101 | 7.45E-04 | glio |
| Atr_ex.vivo_noRT | Lrrc4b | 2.769 | 7.45E-04 | glio |
| Atr_ex.vivo_noRT | Anp32b | -0.778 | 7.45E-04 | glio |
| Bora_in.vivo_noRT | Nos3 | 2.339 | 7.45E-04 | glio |
| Bora_in.vivo_noRT | Faim | -1.286 | 7.45E-04 | glio |
| Interaction_genotype_Aars2 | Gm2115 | -4.27 | 7.45E-05 | glio |
| Interaction_genotype_Mtor | Gm43951 | -3.967 | 7.45E-05 | glio |
| Bora_in.vivo_noRT | Reck | 2.843 | 7.45E-05 | glio |
| Bora_in.vivo_noRT | Pip5k1a | -1.636 | 7.45E-05 | glio |
| Interaction_genotype_Mtor | Fhl4 | -4.781 | 7.45E-06 | glio |
| Aars2_in.vivo_noRT | Gm20536 | 3.43 | 7.45E-07 | glio |
| Bcas2_in.vivo_noRT | Pde11a | 4.278 | 7.45E-09 | glio |
| Chek1_in.vivo_noRT | Gm1123 | 4.136 | 7.45E-09 | glio |
| Aars2_in.vivo_noRT | Tgfb1i1 | 3.994 | 7.45E-10 | glio |
| Bora_in.vivo_noRT | Gm50334 | 4.815 | 7.45E-10 | glio |

| | | | | |
|---|---|---|---|---|
| Chek1_in.vivo_noRT | Serpina3g | 3.847 | 7.45E-10 | glio |
| Chek1_ex.vivo_noRT | Gm49808 | 4.449 | 7.45E-15 | glio |
| Chek1_ex.vivo_noRT | Adam5 | 4.842 | 7.45E-15 | glio |
| Bora_in.vivo_noRT | Slc45a1 | 4.676 | 7.45E-20 | glio |
| culture_effect | Gm12703 | -1.201 | 7.46E-03 | glio |
| Interaction_genotype_Aars2 | Card19 | 1.507 | 7.46E-03 | glio |
| Interaction_genotype_Mtor | Ears2 | -2.417 | 7.46E-03 | glio |
| Interaction_genotype_Mtor | Pcolce2 | 1.867 | 7.46E-03 | glio |
| Interaction_genotype_Myc | Bfsp2 | -3.387 | 7.46E-03 | glio |
| Interaction_genotype_Myc | Pcdhgc5 | -2.772 | 7.46E-03 | glio |
| Interaction_genotype_Myc | Mettl7a1 | -2.459 | 7.46E-03 | glio |
| Interaction_genotype_Myc | Sufu | -2.069 | 7.46E-03 | glio |
| Interaction_genotype_Rpl3 | Cpeb1 | -3.943 | 7.46E-03 | glio |
| Atr_ex.vivo_noRT | Cavin2 | -2.585 | 7.46E-03 | glio |
| Atr_ex.vivo_noRT | Selenop | 1.718 | 7.46E-03 | glio |
| Bcas2_in.vivo_noRT | Pcx | 1.948 | 7.46E-03 | glio |
| Bcas2_in.vivo_noRT | Tmem43 | -1.254 | 7.46E-03 | glio |
| Interaction_genotype_Aars2 | Gm10447 | -3.308 | 7.46E-04 | glio |
| Bcas2_in.vivo_noRT | Gm42372 | 2.807 | 7.46E-04 | glio |
| Chek1_ex.vivo_noRT | Tmem88b | 2.803 | 7.46E-04 | glio |
| culture_effect | Cyth1 | -1.786 | 7.46E-06 | glio |
| interaction_RT | Zscan26 | 1.507 | 7.46E-06 | glio |
| Chek1_in.vivo_noRT | Car11 | 2.816 | 7.46E-06 | glio |
| Chek1_ex.vivo_noRT | Gm45591 | 3.26 | 7.46E-06 | glio |
| Chek1_ex.vivo_noRT | Zfp811 | 3.872 | 7.46E-06 | glio |
| Interaction_genotype_Myc | Set | 2.129 | 7.46E-07 | glio |
| Atp5c1_in.vivo_noRT | Gjd4 | 2.668 | 7.46E-08 | glio |
| Aars2_in.vivo_noRT | Bend7 | 4.548 | 7.46E-12 | glio |
| Chek1_ex.vivo_noRT | 4933400F21Ri | 4.842 | 7.46E-15 | glio |
| Aars2_in.vivo_noRT | Gm26978 | 5.577 | 7.46E-22 | glio |
| Atr_in.vivo_noRT | Slc14a2 | 5.533 | 7.46E-30 | glio |
| culture_effect | Abra | -1.46 | 7.47E-03 | glio |
| culture_effect | Zfp655 | -0.498 | 7.47E-03 | glio |
| Interaction_genotype_Atr | Tph2 | -2.932 | 7.47E-03 | glio |
| Interaction_genotype_Chek1 | Gpr17 | 3.512 | 7.47E-03 | glio |
| Interaction_genotype_Mtor | Slc38a1 | 1.28 | 7.47E-03 | glio |
| Interaction_genotype_Myc | Arhgef40 | -2.545 | 7.47E-03 | glio |
| Interaction_genotype_Rad51 | Krt18 | 5.353 | 7.47E-03 | glio |
| Aars2_ex.vivo_noRT | Calml4 | 2.613 | 7.47E-03 | glio |
| Atr_in.vivo_noRT | AW011738 | 1.568 | 7.47E-03 | glio |
| Eif2s1_in.vivo_noRT | Gm30042 | 1.883 | 7.47E-03 | glio |
| Fanca_ex.vivo_noRT | Pla1a | 2.913 | 7.47E-03 | glio |
| Interaction_genotype_Mtor | Gm19325 | -4.071 | 7.47E-04 | glio |
| Interaction_genotype_Mtor | Pus7l | -3.069 | 7.47E-04 | glio |
| Atr_in.vivo_noRT | Virma | -1.286 | 7.47E-04 | glio |
| Chek1_in.vivo_noRT | Ube2e3 | -1.306 | 7.47E-04 | glio |
| Hsd17b10_in.vivo_noRT | Rmdn1 | -1.879 | 7.47E-04 | glio |
| Aars2_in.vivo_noRT | 1700126G02R | 3.194 | 7.47E-06 | glio |
| Aars2_in.vivo_noRT | 4930542D17R | 3.047 | 7.47E-06 | glio |
| Chek1_ex.vivo_noRT | Gvin1 | 4.447 | 7.47E-07 | glio |
| Bcas2_in.vivo_noRT | Gm34256 | 4.007 | 7.47E-08 | glio |
| Chek1_ex.vivo_noRT | Rbfaos | 3.652 | 7.47E-08 | glio |

| | | | | |
|---|---|---|---|---|
| Chek1_ex.vivo_noRT | 4931431C16R | 3.664 | 7.47E-08 | glio |
| Atr_in.vivo_noRT | Tmem86a | 4.166 | 7.47E-09 | glio |
| Atr_in.vivo_noRT | Spx | 3.807 | 7.47E-09 | glio |
| Aars2_in.vivo_noRT | Gm48933 | 4.325 | 7.47E-10 | glio |
| culture_effect | Snrpa | -0.373 | 7.48E-03 | glio |
| Interaction_genotype_Aars2 | Prl2c2 | -3.212 | 7.48E-03 | glio |
| Interaction_genotype_Aars2 | Zfp551 | -3.285 | 7.48E-03 | glio |
| Interaction_genotype_Aars2 | Adcy3 | -2.598 | 7.48E-03 | glio |
| Interaction_genotype_Bora | Etaa1os | -3.341 | 7.48E-03 | glio |
| interaction_RT | Vgf | 2.04 | 7.48E-03 | glio |
| Aars2_in.vivo_noRT | Exoc3 | -1.161 | 7.48E-03 | glio |
| Atr_in.vivo_noRT | Tbca | -0.496 | 7.48E-03 | glio |
| Bard1_ex.vivo_noRT | Lrrc24 | 2.611 | 7.48E-03 | glio |
| Bard1_ex.vivo_noRT | Cnfn | 3.006 | 7.48E-03 | glio |
| Bard1_ex.vivo_noRT | Cdr2l | -1.329 | 7.48E-03 | glio |
| Bcas2_in.vivo_noRT | Gpr146 | 2.024 | 7.48E-03 | glio |
| Bcas2_ex.vivo_noRT | Cyc1 | -0.836 | 7.48E-03 | glio |
| Bora_in.vivo_noRT | Casp3 | -1.317 | 7.48E-03 | glio |
| Chek1_ex.vivo_noRT | Rps15 | -0.621 | 7.48E-03 | glio |
| Interaction_genotype_Myc | B4galnt4 | -3.64 | 7.48E-04 | glio |
| Interaction_genotype_Myc | Styx | -3.368 | 7.48E-04 | glio |
| Aars2_in.vivo_noRT | Col6a1 | 2.362 | 7.48E-04 | glio |
| Cenpj_ex.vivo_noRT | Gm2022 | 2.591 | 7.48E-04 | glio |
| Aars2_in.vivo_noRT | Cox7a1 | 3.008 | 7.48E-05 | glio |
| Bora_in.vivo_noRT | Gm15728 | 2.984 | 7.48E-05 | glio |
| culture_effect | Marveld1 | 1.148 | 7.48E-06 | glio |
| culture_effect | Map2k3 | 0.68 | 7.48E-06 | glio |
| Interaction_genotype_Myc | Smg6 | -3.468 | 7.48E-06 | glio |
| Interaction_genotype_Myc | Slpi | -5.671 | 7.48E-06 | glio |
| Interaction_genotype_Myc | 9330104G04R | -5.057 | 7.48E-06 | glio |
| Aars2_in.vivo_noRT | Gm26728 | 3.297 | 7.48E-06 | glio |
| Atr_in.vivo_noRT | Syt17 | 3.469 | 7.48E-06 | glio |
| Atr_in.vivo_noRT | Gm26621 | 3.131 | 7.48E-06 | glio |
| Aars2_in.vivo_noRT | Rasl2-9 | 3.889 | 7.48E-07 | glio |
| Chek1_ex.vivo_noRT | Pacrg | 4.817 | 7.48E-10 | glio |
| Interaction_genotype_Chek1 | Pde3a | -3.557 | 7.49E-03 | glio |
| Interaction_genotype_Mtor | Ehhadh | -3.556 | 7.49E-03 | glio |
| Interaction_genotype_Mtor | Hacl1 | -3.028 | 7.49E-03 | glio |
| Interaction_genotype_Mtor | Klk10 | -3.013 | 7.49E-03 | glio |
| Interaction_genotype_Mtor | Wbp4 | 1.459 | 7.49E-03 | glio |
| Interaction_genotype_Mtor | Rpl27a | 0.94 | 7.49E-03 | glio |
| Interaction_genotype_Myc | Fam160b2 | -2.514 | 7.49E-03 | glio |
| Interaction_genotype_Myc | Sec14l2 | -2.35 | 7.49E-03 | glio |
| interaction_RT | Lrrc14 | -0.821 | 7.49E-03 | glio |
| interaction_RT | Snhg15 | 0.803 | 7.49E-03 | glio |
| Atr_in.vivo_noRT | Spin2c | 1.894 | 7.49E-03 | glio |
| Bora_in.vivo_noRT | Atp6v1e1 | -0.697 | 7.49E-03 | glio |
| Chek1_in.vivo_noRT | Ccdc141 | 2.049 | 7.49E-03 | glio |
| Chek1_in.vivo_noRT | Ovca2 | 1.853 | 7.49E-03 | glio |
| Chek1_ex.vivo_noRT | Phf5a | -0.669 | 7.49E-03 | glio |
| Hsd17b10_in.vivo_noRT | Enpp3 | 2.049 | 7.49E-03 | glio |
| Interaction_genotype_Mtor | Agpat1 | -1.973 | 7.49E-04 | glio |

| | | | | |
|---|---|---|---|---|
| Interaction_genotype_Myc | Zfat | -3.228 | 7.49E-04 | glio |
| Atr_in.vivo_noRT | Rplp0 | -0.689 | 7.49E-04 | glio |
| Chek1_in.vivo_noRT | Actr5 | 2.009 | 7.49E-04 | glio |
| Interaction_genotype_Mtor | Gm15728 | -4.87 | 7.49E-05 | glio |
| Interaction_genotype_Myc | 1700040D17R | -4.82 | 7.49E-05 | glio |
| Interaction_genotype_Myc | Pbld1 | -4.744 | 7.49E-05 | glio |
| Interaction_genotype_Myc | Anapc13 | 2.008 | 7.49E-05 | glio |
| Aars2_in.vivo_noRT | 1600002K03R | 3.263 | 7.49E-06 | glio |
| Bcas2_in.vivo_noRT | Tgfb1i1 | 2.964 | 7.49E-06 | glio |
| Bora_in.vivo_noRT | Hoga1 | 3.122 | 7.49E-06 | glio |
| culture_effect | Agfg1 | 0.696 | 7.49E-07 | glio |
| Interaction_genotype_Mtor | Gm9958 | -5.903 | 7.49E-07 | glio |
| Chek1_in.vivo_noRT | Etl4 | 3.168 | 7.49E-08 | glio |
| culture_effect | Klf2 | -1.755 | 7.49E-09 | glio |
| culture_effect | Tmem63a | -1.146 | 7.49E-10 | glio |
| Aars2_in.vivo_noRT | Gm16244 | 4.67 | 7.49E-11 | glio |
| Atr_in.vivo_noRT | Ccr5 | 4.734 | 7.49E-14 | glio |
| Atr_in.vivo_noRT | Klhdc8a | 5.533 | 7.49E-16 | glio |
| Bora_in.vivo_noRT | Gm44646 | 4.936 | 7.49E-18 | glio |
| Interaction_genotype_Atr | Hoxa13 | -3.536 | 7.50E-03 | glio |
| Interaction_genotype_Mtor | Xpc | -2.314 | 7.50E-03 | glio |
| Interaction_genotype_Myc | Lrrc73 | -2.314 | 7.50E-03 | glio |
| Interaction_genotype_Myc | Ddi2 | -1.942 | 7.50E-03 | glio |
| Interaction_genotype_Myc | Mtrf1l | -1.993 | 7.50E-03 | glio |
| Interaction_genotype_Pars2 | Ankrd1 | -4.114 | 7.50E-03 | glio |
| interaction_RT | B4galt6 | 1.356 | 7.50E-03 | glio |
| Aars2_in.vivo_noRT | Eps8l1 | 1.953 | 7.50E-03 | glio |
| Aars2_in.vivo_noRT | Oas2 | 1.801 | 7.50E-03 | glio |
| Aars2_in.vivo_noRT | Mttp | 1.517 | 7.50E-03 | glio |
| Atr_in.vivo_noRT | Pigk | 1.028 | 7.50E-03 | glio |
| Bard1_ex.vivo_noRT | Gm37864 | 2.421 | 7.50E-03 | glio |
| Bora_in.vivo_noRT | Dtx1 | 1.968 | 7.50E-03 | glio |
| Bora_ex.vivo_noRT | Lgals4 | 2.54 | 7.50E-03 | glio |
| Cenpj_in.vivo_noRT | Olfr718-ps1 | 1.758 | 7.50E-03 | glio |
| Cenpj_in.vivo_noRT | Npr3 | 2.022 | 7.50E-03 | glio |
| Cenpj_in.vivo_noRT | Adgrb3 | 2.301 | 7.50E-03 | glio |
| Cenpj_in.vivo_noRT | 9530018F02Ri | 2.464 | 7.50E-03 | glio |
| Cenpj_ex.vivo_noRT | Nat8f4 | 2.601 | 7.50E-03 | glio |
| Chek1_ex.vivo_noRT | Xrcc6 | -1.495 | 7.50E-03 | glio |
| interaction_RT | Rab3a | 1.198 | 7.50E-04 | glio |
| Aars2_in.vivo_noRT | Ttc30a1 | 2.429 | 7.50E-04 | glio |
| Aars2_ex.vivo_noRT | Igdcc4 | 2.899 | 7.50E-04 | glio |
| Aars2_ex.vivo_noRT | Adcy2 | 2.9 | 7.50E-04 | glio |
| Aars2_ex.vivo_noRT | Gm39556 | 3.293 | 7.50E-04 | glio |
| Bora_ex.vivo_noRT | Dcdc2a | 3.201 | 7.50E-05 | glio |
| culture_effect | Atf4 | -0.537 | 7.50E-06 | glio |
| culture_effect | Pir | -1.206 | 7.50E-07 | glio |
| Chek1_in.vivo_noRT | Gm47428 | 3.868 | 7.50E-09 | glio |
| Bcas2_in.vivo_noRT | 4930447F24Ri | 4.547 | 7.50E-12 | glio |
| culture_effect | Epb41l2 | 1.173 | 7.50E-15 | glio |
| Bora_in.vivo_noRT | Dcdc2a | 4.676 | 7.50E-15 | glio |
| Atr_in.vivo_noRT | AI182371 | 4.751 | 7.50E-17 | glio |

| | | | | |
|---|---|---|---|---|
| Bcas2_in.vivo_noRT | Esp15 | 4.807 | 7.50E-17 | glio |
| Bcas2_in.vivo_noRT | Sytl4 | 4.541 | 7.50E-19 | glio |
| Bcas2_in.vivo_noRT | 1700112J16Ri | 4.807 | 7.50E-19 | glio |
| Bcas2_in.vivo_noRT | Clec4d | 4.807 | 7.50E-19 | glio |
| culture_effect | Sdf2 | -1.324 | 7.50E-24 | glio |
| culture_effect | Ypel3 | -0.513 | 7.51E-03 | glio |
| Interaction_genotype_Aars2 | Gm12840 | -3.15 | 7.51E-03 | glio |
| Interaction_genotype_Aars2 | Kcnmb4os2 | -3.134 | 7.51E-03 | glio |
| Interaction_genotype_Atr | Anks6 | -3.014 | 7.51E-03 | glio |
| Interaction_genotype_Myc | Gm49801 | -3.096 | 7.51E-03 | glio |
| Interaction_genotype_Myc | Cyp4f16 | -2.52 | 7.51E-03 | glio |
| Interaction_genotype_Myc | Bod1 | 1.314 | 7.51E-03 | glio |
| Interaction_genotype_Myc | Tmem147 | 1.216 | 7.51E-03 | glio |
| Aars2_in.vivo_noRT | Cpa2 | 1.84 | 7.51E-03 | glio |
| Atr_in.vivo_noRT | Nr3c1 | -1.168 | 7.51E-03 | glio |
| Bora_in.vivo_noRT | Clcn2 | 2.081 | 7.51E-03 | glio |
| Bora_in.vivo_noRT | Zfp438 | 1.65 | 7.51E-03 | glio |
| Chek1_in.vivo_noRT | D330050G23R | 1.629 | 7.51E-03 | glio |
| Chek1_in.vivo_noRT | Psmb7 | -0.676 | 7.51E-03 | glio |
| Chek1_in.vivo_noRT | Zfas1 | -0.923 | 7.51E-03 | glio |
| Chek1_in.vivo_noRT | Mrfap1 | -0.534 | 7.51E-03 | glio |
| Chek1_ex.vivo_noRT | Ddx56 | -1.617 | 7.51E-03 | glio |
| Chek1_ex.vivo_noRT | Ostc | -0.677 | 7.51E-03 | glio |
| culture_effect | Sumf2 | -0.702 | 7.51E-04 | glio |
| Interaction_genotype_Rpl3 | Folr1 | -3.918 | 7.51E-04 | glio |
| Aars2_in.vivo_noRT | Zdhhc1 | 2.197 | 7.51E-04 | glio |
| Atp5c1_in.vivo_noRT | 4833419F23Ri | 2.668 | 7.51E-04 | glio |
| Hsd17b10_in.vivo_noRT | C230037L18Ri | 2.447 | 7.51E-04 | glio |
| Interaction_genotype_Aars2 | Rnf165 | -4.714 | 7.51E-05 | glio |
| interaction_RT | Cox10 | -1.326 | 7.51E-05 | glio |
| Aars2_in.vivo_noRT | Cyb5d1 | 2.791 | 7.51E-05 | glio |
| Aars2_in.vivo_noRT | 4732419C18R | 2.738 | 7.51E-05 | glio |
| Bard1_ex.vivo_noRT | Gm48215 | 2.814 | 7.51E-05 | glio |
| Chek1_in.vivo_noRT | Gm48796 | 2.672 | 7.51E-05 | glio |
| Bora_ex.vivo_noRT | B230206H07R | 2.806 | 7.51E-06 | glio |
| Chek1_in.vivo_noRT | 4833419F23Ri | 5.044 | 7.51E-13 | glio |
| Bcas2_in.vivo_noRT | Gm48215 | 4.807 | 7.51E-20 | glio |
| Bcas2_in.vivo_noRT | Slc19a3 | 4.807 | 7.51E-21 | glio |
| Interaction_genotype_Atr | Klhl30 | -2.806 | 7.52E-03 | glio |
| Interaction_genotype_Mtor | Kctd21 | -3.15 | 7.52E-03 | glio |
| Interaction_genotype_Myc | Tmem42 | -2.174 | 7.52E-03 | glio |
| Interaction_genotype_Pars2 | Gm13288 | -2.988 | 7.52E-03 | glio |
| Interaction_genotype_Pars2 | Accsl | -3.982 | 7.52E-03 | glio |
| Interaction_genotype_Pars2 | Pole | 2.152 | 7.52E-03 | glio |
| Aars2_in.vivo_noRT | Pax9 | 1.833 | 7.52E-03 | glio |
| Aars2_in.vivo_noRT | Rps9 | -0.529 | 7.52E-03 | glio |
| Aars2_ex.vivo_noRT | Thsd1 | 2.311 | 7.52E-03 | glio |
| Atp5c1_in.vivo_noRT | Olfr279 | 2.139 | 7.52E-03 | glio |
| Atr_in.vivo_noRT | Zc3h12b | 1.81 | 7.52E-03 | glio |
| Atr_in.vivo_noRT | Hist1h2bb | 1.879 | 7.52E-03 | glio |
| Bcas2_in.vivo_noRT | Rpl12 | -0.491 | 7.52E-03 | glio |
| Chek1_ex.vivo_noRT | Slc46a2 | 2.307 | 7.52E-03 | glio |

| | | | | |
|---|---|---|---|---|
| Eif2s1_in.vivo_noRT | Spn | 2.144 | 7.52E-03 | glio |
| Hsd17b10_in.vivo_noRT | Pagr1a | 1.932 | 7.52E-03 | glio |
| Interaction_genotype_Mtor | Il2rb | -3.935 | 7.52E-04 | glio |
| Interaction_genotype_Myc | Pard6g | -3.534 | 7.52E-04 | glio |
| Interaction_genotype_Myc | 1700008J07Ri | -3.268 | 7.52E-04 | glio |
| Interaction_genotype_Myc | Fbxw4 | -2.607 | 7.52E-04 | glio |
| Bard1_ex.vivo_noRT | Pou5f1 | 3.203 | 7.52E-04 | glio |
| Chek1_in.vivo_noRT | Abhd18 | 2.505 | 7.52E-04 | glio |
| Chek1_in.vivo_noRT | Pced1a | 1.845 | 7.52E-04 | glio |
| Atr_ex.vivo_noRT | Bpifa3 | 3.164 | 7.52E-05 | glio |
| Bcas2_in.vivo_noRT | Oxld1 | 2.206 | 7.52E-05 | glio |
| Interaction_genotype_Myc | Kctd18 | -4.31 | 7.52E-06 | glio |
| Interaction_genotype_Myc | Eps8l2 | -5.122 | 7.52E-06 | glio |
| Interaction_genotype_Myc | Ngfr | -3.443 | 7.52E-06 | glio |
| Atp5c1_in.vivo_noRT | Prl2c2 | 3.199 | 7.52E-06 | glio |
| Chek1_in.vivo_noRT | Gm48357 | 2.728 | 7.52E-06 | glio |
| culture_effect | Tmem176b | -1.226 | 7.52E-09 | glio |
| culture_effect | Yipf3 | -0.845 | 7.52E-09 | glio |
| Hsd17b10_in.vivo_noRT | Mcmdc2 | 3.887 | 7.52E-11 | glio |
| Chek1_in.vivo_noRT | Olfr171 | 5.044 | 7.52E-20 | glio |
| Chek1_in.vivo_noRT | Gm17203 | 4.904 | 7.52E-22 | glio |
| culture_effect | AY036118 | -3.368 | 7.52E-27 | glio |
| culture_effect | Gm48617 | -1.198 | 7.53E-03 | glio |
| culture_effect | Tmem87b | -0.817 | 7.53E-03 | glio |
| Interaction_genotype_Aars2 | Mpp2 | -3.27 | 7.53E-03 | glio |
| Interaction_genotype_Aars2 | Lmo7 | -2.902 | 7.53E-03 | glio |
| Interaction_genotype_Aars2 | Jdp2 | 1.878 | 7.53E-03 | glio |
| Interaction_genotype_Cdkn2a | Adam5 | 2.839 | 7.53E-03 | glio |
| Interaction_genotype_Myc | Cntnap1 | -2.736 | 7.53E-03 | glio |
| Interaction_genotype_Myc | Atr | -2.493 | 7.53E-03 | glio |
| Interaction_genotype_Myc | Exosc2 | -1.856 | 7.53E-03 | glio |
| Interaction_genotype_Myc | Uqcrb | 1.07 | 7.53E-03 | glio |
| Interaction_genotype_Rad51 | Fbxl12os | -3.731 | 7.53E-03 | glio |
| Aars2_in.vivo_noRT | Mycbp | -1.151 | 7.53E-03 | glio |
| Atp5c1_in.vivo_noRT | Gm13822 | 2.142 | 7.53E-03 | glio |
| Atp5c1_in.vivo_noRT | 4930478M13F | 2.289 | 7.53E-03 | glio |
| Atp5c1_in.vivo_noRT | Igsf23 | 2.32 | 7.53E-03 | glio |
| Atr_in.vivo_noRT | Pdpr | 1.689 | 7.53E-03 | glio |
| Atr_in.vivo_noRT | Ptma | -0.554 | 7.53E-03 | glio |
| Chek1_ex.vivo_noRT | Iws1 | -1.438 | 7.53E-03 | glio |
| Chek1_ex.vivo_noRT | S100a13 | 0.938 | 7.53E-03 | glio |
| Hsd17b10_in.vivo_noRT | Olfml2a | 2.113 | 7.53E-03 | glio |
| Hsd17b10_in.vivo_noRT | Pawr | 1.85 | 7.53E-03 | glio |
| Hsd17b10_in.vivo_noRT | Foxa1 | 2.295 | 7.53E-03 | glio |
| Hsd17b10_in.vivo_noRT | Gm16133 | 2.154 | 7.53E-03 | glio |
| Interaction_genotype_Mtor | Ifi205 | -5.27 | 7.53E-04 | glio |
| Interaction_genotype_Myc | Usp44 | -3.95 | 7.53E-04 | glio |
| Interaction_genotype_Myc | Whrn | -3.849 | 7.53E-04 | glio |
| Interaction_genotype_Myc | Myl12a | 1.403 | 7.53E-04 | glio |
| Atr_in.vivo_noRT | Lurap1 | 2.355 | 7.53E-04 | glio |
| Atr_in.vivo_noRT | D630003M21I | 2.091 | 7.53E-04 | glio |
| Bora_in.vivo_noRT | Crybg2 | 2.412 | 7.53E-04 | glio |

| | | | | |
|---|---|---|---|---|
| Chek1_ex.vivo_noRT | Rab37 | 2.92 | 7.53E-04 | glio |
| culture_effect | Gprc5a | -1.922 | 7.53E-05 | glio |
| culture_effect | Klf3 | 0.747 | 7.53E-05 | glio |
| Atr_in.vivo_noRT | Efr3b | 3.102 | 7.53E-05 | glio |
| Atr_in.vivo_noRT | Rpl10 | -0.757 | 7.53E-05 | glio |
| Interaction_genotype_Mtor | 2900042K21R | -5.211 | 7.53E-06 | glio |
| Bcas2_in.vivo_noRT | 2810457G06R | 3.56 | 7.53E-06 | glio |
| Chek1_in.vivo_noRT | Itga2b | 3.119 | 7.53E-06 | glio |
| Cyp19a1_in.vivo_noRT | Vcan | 3.366 | 7.53E-06 | glio |
| interaction_RT | Ly6e | 1.338 | 7.53E-07 | glio |
| Aars2_in.vivo_noRT | Arhgap4 | 3.341 | 7.53E-07 | glio |
| Bcas2_in.vivo_noRT | 5430400D12R | 3.525 | 7.53E-07 | glio |
| Hsd17b10_in.vivo_noRT | AI463229 | 2.971 | 7.53E-07 | glio |
| Atr_ex.vivo_noRT | Apoa5 | 3.164 | 7.53E-08 | glio |
| Atr_in.vivo_noRT | Gm35986 | 4.485 | 7.53E-12 | glio |
| Bcas2_in.vivo_noRT | Rab27a | 4.799 | 7.53E-15 | glio |
| culture_effect | Copg1 | -0.433 | 7.54E-03 | glio |
| culture_effect | Ankrd17 | -0.408 | 7.54E-03 | glio |
| Interaction_genotype_Atr | Ahrr | -3.359 | 7.54E-03 | glio |
| Interaction_genotype_Mtor | Fam120c | -3.151 | 7.54E-03 | glio |
| Interaction_genotype_Mtor | Dpy30 | 1.293 | 7.54E-03 | glio |
| Interaction_genotype_Mtor | Rps15 | 0.861 | 7.54E-03 | glio |
| Interaction_genotype_Myc | Tcf20 | -1.822 | 7.54E-03 | glio |
| Aars2_in.vivo_noRT | Epor | 1.877 | 7.54E-03 | glio |
| Aars2_in.vivo_noRT | Rpl34 | -0.599 | 7.54E-03 | glio |
| Aars2_ex.vivo_noRT | 4921508D12R | 2.499 | 7.54E-03 | glio |
| Atp5c1_in.vivo_noRT | Ttll13 | 1.935 | 7.54E-03 | glio |
| Atr_ex.vivo_noRT | Hsd17b11 | 2.048 | 7.54E-03 | glio |
| Bcas2_ex.vivo_noRT | Tmem229b | 2.714 | 7.54E-03 | glio |
| Bora_in.vivo_noRT | Sapcd2 | -1.39 | 7.54E-03 | glio |
| Bora_in.vivo_noRT | Cxxc1 | -1.467 | 7.54E-03 | glio |
| Brca2_in.vivo_noRT | Gm41307 | 1.763 | 7.54E-03 | glio |
| Hsd17b10_in.vivo_noRT | A430035B10R | 2.18 | 7.54E-03 | glio |
| culture_effect | Emb | -0.868 | 7.54E-04 | glio |
| Interaction_genotype_Aars2 | Cops8 | 1.775 | 7.54E-04 | glio |
| Interaction_genotype_Myc | Gm43597 | -3.433 | 7.54E-04 | glio |
| Interaction_genotype_Myc | Sat2 | -3.121 | 7.54E-04 | glio |
| Interaction_genotype_Myc | Rpusd1 | -2.792 | 7.54E-04 | glio |
| interaction_RT | Srsf5 | -0.672 | 7.54E-04 | glio |
| Atr_in.vivo_noRT | Zbtb11os1 | 2.321 | 7.54E-04 | glio |
| Atr_ex.vivo_noRT | Hes7 | 3.073 | 7.54E-04 | glio |
| Bora_in.vivo_noRT | Nptx1 | 2.673 | 7.54E-04 | glio |
| Chek1_ex.vivo_noRT | Gm41764 | 2.66 | 7.54E-04 | glio |
| culture_effect | Eif6 | -0.485 | 7.54E-05 | glio |
| Interaction_genotype_Mtor | Bco2 | -4.054 | 7.54E-05 | glio |
| Interaction_genotype_Mtor | Rbp7 | -4.792 | 7.54E-05 | glio |
| Aars2_in.vivo_noRT | Zfp982 | 2.684 | 7.54E-05 | glio |
| Atr_ex.vivo_noRT | Gm13822 | 3.566 | 7.54E-05 | glio |
| Atr_ex.vivo_noRT | Gm34885 | 3.446 | 7.54E-05 | glio |
| Aars2_in.vivo_noRT | Pfkfb1 | 3.165 | 7.54E-06 | glio |
| Atr_in.vivo_noRT | Gm10353 | 3.019 | 7.54E-06 | glio |
| Aars2_in.vivo_noRT | 9530053A07R | 3.513 | 7.54E-07 | glio |

| | | | | |
|---|---|---|---|---|
| Aars2_in.vivo_noRT | Arsj | 3.722 | 7.54E-07 | glio |
| Bcas2_in.vivo_noRT | Gm10655 | 3.622 | 7.54E-07 | glio |
| Bcas2_in.vivo_noRT | 4833407H14R | 4.173 | 7.54E-08 | glio |
| Chek1_in.vivo_noRT | Zdhhc14 | 3.1 | 7.54E-08 | glio |
| Atr_in.vivo_noRT | Olfml2a | 4.03 | 7.54E-09 | glio |
| Bcas2_in.vivo_noRT | Grin2d | 4.028 | 7.54E-10 | glio |
| Bora_in.vivo_noRT | Gm20513 | 3.364 | 7.54E-10 | glio |
| culture_effect | Gm14968 | 3.648 | 7.54E-12 | glio |
| culture_effect | Rps15 | -0.815 | 7.54E-12 | glio |
| Bcas2_in.vivo_noRT | Mettl7a3 | 4.684 | 7.54E-13 | glio |
| Bcas2_in.vivo_noRT | Adam2 | 4.281 | 7.54E-13 | glio |
| Bcas2_in.vivo_noRT | Gem | 4.807 | 7.54E-13 | glio |
| culture_effect | Gm15972 | -1.529 | 7.55E-03 | glio |
| Interaction_genotype_Aars2 | Card10 | -3.395 | 7.55E-03 | glio |
| Interaction_genotype_Atr | Slc35d2 | -3.5 | 7.55E-03 | glio |
| Interaction_genotype_Myc | Usp49 | -2.775 | 7.55E-03 | glio |
| Interaction_genotype_Myc | Grik5 | -2.345 | 7.55E-03 | glio |
| Interaction_genotype_Pars2 | Fam92a | 1.094 | 7.55E-03 | glio |
| Interaction_genotype_Rad51 | 2210417A02R | -4.071 | 7.55E-03 | glio |
| Aars2_in.vivo_noRT | Firre | 1.788 | 7.55E-03 | glio |
| Aars2_in.vivo_noRT | Tm7sf3 | -1.015 | 7.55E-03 | glio |
| Atr_in.vivo_noRT | Ncan | 1.733 | 7.55E-03 | glio |
| Bcas2_in.vivo_noRT | Gm14419 | 1.755 | 7.55E-03 | glio |
| Chek1_in.vivo_noRT | Rps27 | -0.749 | 7.55E-03 | glio |
| Interaction_genotype_Mtor | Glis1 | -3.941 | 7.55E-04 | glio |
| Chek1_in.vivo_noRT | Gm29394 | 2.334 | 7.55E-04 | glio |
| culture_effect | Rfx2 | -2.306 | 7.55E-05 | glio |
| culture_effect | Plxnb3 | -1.112 | 7.55E-05 | glio |
| Interaction_genotype_Mtor | Etaa1os | -4.38 | 7.55E-05 | glio |
| Interaction_genotype_Mtor | Emc6 | 1.837 | 7.55E-05 | glio |
| Aars2_in.vivo_noRT | Kdm1b | 2.324 | 7.55E-05 | glio |
| Bora_in.vivo_noRT | BC051077 | 3.232 | 7.55E-05 | glio |
| Interaction_genotype_Atr | Tex19.1 | 2.409 | 7.55E-06 | glio |
| Interaction_genotype_Myc | 1700029J07Ri | -5.202 | 7.55E-06 | glio |
| Interaction_genotype_Myc | Gm14207 | -3.522 | 7.55E-06 | glio |
| Atr_in.vivo_noRT | Plpp2 | -1.43 | 7.55E-06 | glio |
| Bcas2_in.vivo_noRT | 4930515G01R | 2.925 | 7.55E-06 | glio |
| Chek1_in.vivo_noRT | Trappc13 | -1.953 | 7.55E-06 | glio |
| culture_effect | Syngr3 | -2.517 | 7.55E-07 | glio |
| Bcas2_in.vivo_noRT | Lynx1 | 3.503 | 7.55E-07 | glio |
| culture_effect | Rbm25 | -0.533 | 7.55E-08 | glio |
| Bcas2_in.vivo_noRT | Klra6 | 4.215 | 7.55E-09 | glio |
| Bcas2_in.vivo_noRT | 2810410L24Ri | 4.281 | 7.55E-10 | glio |
| culture_effect | Repin1 | -1.392 | 7.55E-12 | glio |
| Aars2_in.vivo_noRT | Gm45591 | 5.067 | 7.55E-18 | glio |
| Bora_in.vivo_noRT | A930019D19R | 4.936 | 7.55E-25 | glio |
| culture_effect | Sae1 | -0.405 | 7.56E-03 | glio |
| Interaction_genotype_Bcas2 | Cdh10 | -2.952 | 7.56E-03 | glio |
| Interaction_genotype_Bora | Cnpy4 | 1.973 | 7.56E-03 | glio |
| Interaction_genotype_Chek1 | Jade1 | -2.584 | 7.56E-03 | glio |
| Interaction_genotype_Mtor | Ganab | 1.304 | 7.56E-03 | glio |
| Aars2_in.vivo_noRT | Hbegf | -1.129 | 7.56E-03 | glio |

| | | | | |
|---|---|---|---|---|
| Atr_in.vivo_noRT | Tcaim | 1.821 | 7.56E-03 | glio |
| Bora_in.vivo_noRT | Rab7b | 1.919 | 7.56E-03 | glio |
| Chek1_ex.vivo_noRT | 4930506C21R | 2.496 | 7.56E-03 | glio |
| Chek1_ex.vivo_noRT | 2900009J06Ri | 2.418 | 7.56E-03 | glio |
| Chek1_ex.vivo_noRT | Ptk2 | -1.313 | 7.56E-03 | glio |
| Cyp19a1_ex.vivo_noRT | BC049987 | 2.159 | 7.56E-03 | glio |
| Bora_in.vivo_noRT | Gpatch3 | 2.376 | 7.56E-04 | glio |
| Cenpj_in.vivo_noRT | Gm12841 | 2.022 | 7.56E-04 | glio |
| Atr_in.vivo_noRT | Hist1h3d | 2.63 | 7.56E-05 | glio |
| Chek1_in.vivo_noRT | Cwc15 | -1.144 | 7.56E-05 | glio |
| culture_effect | Zfhx2os | 2.359 | 7.56E-06 | glio |
| Interaction_genotype_Myc | Gldc | -5.01 | 7.56E-06 | glio |
| Interaction_genotype_Myc | Fes | -5.163 | 7.56E-06 | glio |
| Interaction_genotype_Myc | Slirp | 2.27 | 7.56E-06 | glio |
| Aars2_in.vivo_noRT | Ngef | 3.318 | 7.56E-08 | glio |
| Hsd17b10_in.vivo_noRT | 4930447K03R | 3.356 | 7.56E-10 | glio |
| Atr_in.vivo_noRT | 9430037G07R | 4.625 | 7.56E-11 | glio |
| Atr_in.vivo_noRT | Bend7 | 4.755 | 7.56E-13 | glio |
| Bora_in.vivo_noRT | D730045A05R | 4.936 | 7.56E-13 | glio |
| Hsd17b10_in.vivo_noRT | Gm13191 | 3.616 | 7.56E-15 | glio |
| Hsd17b10_in.vivo_noRT | 4933416E03Ri | 3.616 | 7.56E-15 | glio |
| Atr_in.vivo_noRT | Col8a1 | 6.043 | 7.56E-19 | glio |
| Chek1_in.vivo_noRT | Dpep3 | 4.511 | 7.56E-20 | glio |
| culture_effect | Naxd | -0.418 | 7.57E-03 | glio |
| Interaction_genotype_Chek1 | Zfp143 | -2.867 | 7.57E-03 | glio |
| Interaction_genotype_Chek1 | Prdx5 | 1.337 | 7.57E-03 | glio |
| Interaction_genotype_Myc | Lca5 | -2.954 | 7.57E-03 | glio |
| Aars2_in.vivo_noRT | Zfp647 | 1.641 | 7.57E-03 | glio |
| Atp5c1_in.vivo_noRT | Cfap20 | -0.554 | 7.57E-03 | glio |
| Bcas2_in.vivo_noRT | Smim8 | -1.092 | 7.57E-03 | glio |
| Bora_in.vivo_noRT | Nif3l1 | -1.162 | 7.57E-03 | glio |
| Bora_in.vivo_noRT | Col2a1 | -2.011 | 7.57E-03 | glio |
| Cenpj_in.vivo_noRT | BC065403 | 2.282 | 7.57E-03 | glio |
| Chek1_ex.vivo_noRT | Ccdc96 | 2.244 | 7.57E-03 | glio |
| Interaction_genotype_Atr | Hist1h2ai | -4.369 | 7.57E-04 | glio |
| Interaction_genotype_Atr | Tal2 | -3.088 | 7.57E-04 | glio |
| Aars2_in.vivo_noRT | B230216N24R | 2.653 | 7.57E-04 | glio |
| Bcas2_in.vivo_noRT | Slc4a8 | 1.996 | 7.57E-04 | glio |
| Bora_in.vivo_noRT | Serinc2 | 2.352 | 7.57E-04 | glio |
| Bora_in.vivo_noRT | Eif2b4 | -1.392 | 7.57E-04 | glio |
| Chek1_in.vivo_noRT | Stard10 | 2.377 | 7.57E-04 | glio |
| Hsd17b10_in.vivo_noRT | Medag | 2.448 | 7.57E-04 | glio |
| Interaction_genotype_Mtor | Adam5 | -3.835 | 7.57E-06 | glio |
| Eif2s1_in.vivo_noRT | Gm49860 | 3.192 | 7.57E-06 | glio |
| Eif2s1_in.vivo_noRT | Hhla1 | -3.249 | 7.57E-06 | glio |
| Bcas2_in.vivo_noRT | Nppc | 4.337 | 7.57E-09 | glio |
| Bora_in.vivo_noRT | Lama5 | 4.414 | 7.57E-10 | glio |
| Chek1_ex.vivo_noRT | Gm12794 | 4.842 | 7.57E-15 | glio |
| Interaction_genotype_Bcas2 | 5730507C01R | -2.733 | 7.58E-03 | glio |
| Interaction_genotype_Bcas2 | 9230112J17Ri | -3.046 | 7.58E-03 | glio |
| Interaction_genotype_Bcas2 | Celf5 | -3.819 | 7.58E-03 | glio |
| Interaction_genotype_Bcas2 | B230311B06R | -3.823 | 7.58E-03 | glio |

| | | | | |
|---|---|---|---|---|
| Interaction_genotype_Bcas2 | Gm49801 | -3.745 | 7.58E-03 | glio |
| Interaction_genotype_Bcas2 | Gm11549 | -3.483 | 7.58E-03 | glio |
| Interaction_genotype_Bcas2 | Gm9970 | -3.727 | 7.58E-03 | glio |
| Interaction_genotype_Bcas2 | Phf1 | -3.103 | 7.58E-03 | glio |
| Interaction_genotype_Bcas2 | Cstb | 1.643 | 7.58E-03 | glio |
| Interaction_genotype_Mtor | Map7d1 | -1.743 | 7.58E-03 | glio |
| Interaction_genotype_Mtor | Vmp1 | 1.64 | 7.58E-03 | glio |
| Interaction_genotype_Myc | Hhex | -2.654 | 7.58E-03 | glio |
| Interaction_genotype_Myc | 1700007L15Ri | -2.22 | 7.58E-03 | glio |
| Interaction_genotype_Myc | Tm9sf3 | 1.149 | 7.58E-03 | glio |
| Interaction_genotype_Myc | Cope | 1.156 | 7.58E-03 | glio |
| interaction_RT | Osr2 | 1.616 | 7.58E-03 | glio |
| interaction_RT | mt-Co2 | 0.583 | 7.58E-03 | glio |
| Aars2_in.vivo_noRT | Arhgef40 | 1.614 | 7.58E-03 | glio |
| Atp5c1_in.vivo_noRT | Gm41442 | 2.157 | 7.58E-03 | glio |
| Atp5c1_in.vivo_noRT | Lamc2 | 2.185 | 7.58E-03 | glio |
| Atr_in.vivo_noRT | Deptor | 1.713 | 7.58E-03 | glio |
| Bcas2_in.vivo_noRT | Pcgf1 | 1.371 | 7.58E-03 | glio |
| Bora_in.vivo_noRT | Ubap2 | -0.922 | 7.58E-03 | glio |
| Chek1_in.vivo_noRT | Spin2c | 1.928 | 7.58E-03 | glio |
| Chek1_in.vivo_noRT | Mef2b | 1.897 | 7.58E-03 | glio |
| Eif2s1_in.vivo_noRT | Gm4858 | 2.143 | 7.58E-03 | glio |
| Eif2s1_ex.vivo_noRT | Rnf165 | 2.521 | 7.58E-03 | glio |
| Interaction_genotype_Aars2 | Lama5 | -4.353 | 7.58E-04 | glio |
| Interaction_genotype_Aars2 | Dpy19l1 | 2.494 | 7.58E-04 | glio |
| Atr_in.vivo_noRT | Spef1 | 2.409 | 7.58E-04 | glio |
| Atr_in.vivo_noRT | Mgst3 | 2.084 | 7.58E-04 | glio |
| Bcas2_in.vivo_noRT | Rpl22l1 | 0.65 | 7.58E-04 | glio |
| Bora_in.vivo_noRT | Zfp114 | 2.463 | 7.58E-04 | glio |
| Interaction_genotype_Mtor | Ltbp2 | -3.702 | 7.58E-05 | glio |
| Bora_in.vivo_noRT | Zfp992 | 2.908 | 7.58E-05 | glio |
| Chek1_in.vivo_noRT | Zbtb8a | 2.982 | 7.58E-05 | glio |
| Interaction_genotype_Mtor | Timm13 | 2.123 | 7.58E-06 | glio |
| Interaction_genotype_Myc | Eif3a | 1.554 | 7.58E-06 | glio |
| Aars2_in.vivo_noRT | Gm50322 | 3.431 | 7.58E-06 | glio |
| Atr_in.vivo_noRT | Plpp7 | 2.884 | 7.58E-06 | glio |
| Bcas2_in.vivo_noRT | Gm34590 | 3.234 | 7.58E-06 | glio |
| Chek1_ex.vivo_noRT | Gm11973 | 3.079 | 7.58E-06 | glio |
| culture_effect | Hps4 | 1.19 | 7.58E-09 | glio |
| Chek1_ex.vivo_noRT | AC154200.1 | 3.872 | 7.58E-09 | glio |
| Aars2_in.vivo_noRT | Trpm3 | 4.805 | 7.58E-12 | glio |
| Aars2_in.vivo_noRT | Gm40557 | 4.804 | 7.58E-12 | glio |
| Bcas2_in.vivo_noRT | Gm10801 | 4.807 | 7.58E-17 | glio |
| culture_effect | Bnip2 | 0.333 | 7.59E-03 | glio |
| Interaction_genotype_Myc | Mterf4 | -2.056 | 7.59E-03 | glio |
| Interaction_genotype_Pars2 | Ccdc71l | -3.238 | 7.59E-03 | glio |
| interaction_RT | Sncaip | 1.966 | 7.59E-03 | glio |
| Aars2_ex.vivo_noRT | Fsd2 | 2.706 | 7.59E-03 | glio |
| Atr_in.vivo_noRT | Dcaf12 | -0.98 | 7.59E-03 | glio |
| Atr_in.vivo_noRT | Ccnc | -0.824 | 7.59E-03 | glio |
| Chek1_ex.vivo_noRT | Tnfsf13os | 2.462 | 7.59E-03 | glio |
| Chek1_ex.vivo_noRT | Gm15473 | 2.585 | 7.59E-03 | glio |

| | | | | |
|---|---|---|---|---|
| Cyp19a1_ex.vivo_noRT | Dpp6 | 2.159 | 7.59E-03 | glio |
| Cyp19a1_ex.vivo_noRT | Serpinb9g | 2.159 | 7.59E-03 | glio |
| Hsd17b10_in.vivo_noRT | A430033K04R | 2.028 | 7.59E-03 | glio |
| Interaction_genotype_Mtor | Nat14 | -3.54 | 7.59E-04 | glio |
| Interaction_genotype_Myc | Prdm10 | -3.366 | 7.59E-04 | glio |
| interaction_RT | Dad1 | 0.66 | 7.59E-04 | glio |
| Atr_in.vivo_noRT | Akr1c12 | 2.362 | 7.59E-04 | glio |
| Aars2_ex.vivo_noRT | Gm35439 | 2.893 | 7.59E-05 | glio |
| Aars2_ex.vivo_noRT | 9930111J21Ri | 3.293 | 7.59E-05 | glio |
| Atr_in.vivo_noRT | Gm43138 | 2.897 | 7.59E-05 | glio |
| Bcas2_ex.vivo_noRT | 1700112J16Ri | 2.937 | 7.59E-05 | glio |
| Interaction_genotype_Mtor | Eif2s1 | 1.99 | 7.59E-06 | glio |
| Aars2_in.vivo_noRT | Rab36 | 3.158 | 7.59E-06 | glio |
| Hsd17b10_in.vivo_noRT | Gm43951 | 2.69 | 7.59E-06 | glio |
| Hsd17b10_in.vivo_noRT | S100a14 | 2.816 | 7.59E-06 | glio |
| Atr_ex.vivo_noRT | Cd200r2 | 3.164 | 7.59E-07 | glio |
| Bora_in.vivo_noRT | Ly6c2 | 4.033 | 7.59E-09 | glio |
| Bora_in.vivo_noRT | A930003A15R | 3.784 | 7.59E-09 | glio |
| Chek1_in.vivo_noRT | Tmem190 | 5.044 | 7.59E-11 | glio |
| Bcas2_in.vivo_noRT | Ripply3 | 4.807 | 7.59E-13 | glio |
| culture_effect | Hbq1b | 3.988 | 7.59E-15 | glio |
| Bcas2_in.vivo_noRT | Gm48008 | 4.543 | 7.59E-16 | glio |
| culture_effect | Morc4 | -1.934 | 7.59E-18 | glio |
| culture_effect | Cldn5 | -1.201 | 7.60E-03 | glio |
| culture_effect | Zfp653 | -1.149 | 7.60E-03 | glio |
| culture_effect | Tmf1 | 0.489 | 7.60E-03 | glio |
| Interaction_genotype_Mtor | Cntnap1 | -2.882 | 7.60E-03 | glio |
| Interaction_genotype_Myc | Sap18b | -2.445 | 7.60E-03 | glio |
| Interaction_genotype_Myc | Plcd1 | -2.275 | 7.60E-03 | glio |
| Atr_in.vivo_noRT | Gm43774 | 1.784 | 7.60E-03 | glio |
| Atr_in.vivo_noRT | Dad1 | -0.717 | 7.60E-03 | glio |
| Bora_in.vivo_noRT | H2-DMa | -1.068 | 7.60E-03 | glio |
| Chek1_in.vivo_noRT | Hba-a1 | 2.402 | 7.60E-03 | glio |
| Chek1_ex.vivo_noRT | E530011L22Ri | 2.257 | 7.60E-03 | glio |
| Hsd17b10_in.vivo_noRT | Rab11fip1 | 2.322 | 7.60E-03 | glio |
| Hsd17b10_in.vivo_noRT | Sphk1 | 2.11 | 7.60E-03 | glio |
| Hsd17b10_in.vivo_noRT | Grk5 | -1.719 | 7.60E-03 | glio |
| Hsd17b10_in.vivo_noRT | Capn10 | 1.235 | 7.60E-03 | glio |
| culture_effect | Clca3a1 | -1.986 | 7.60E-04 | glio |
| culture_effect | Gm15941 | -1.959 | 7.60E-04 | glio |
| culture_effect | Nav2 | 0.582 | 7.60E-04 | glio |
| Aars2_in.vivo_noRT | Adap1 | 2.732 | 7.60E-04 | glio |
| Aars2_in.vivo_noRT | Chic1 | -1.669 | 7.60E-04 | glio |
| Aars2_in.vivo_noRT | Pdhb | -1.186 | 7.60E-04 | glio |
| Chek1_in.vivo_noRT | Fam177a | 2.276 | 7.60E-04 | glio |
| culture_effect | Ucn2 | -2.453 | 7.60E-05 | glio |
| Interaction_genotype_Myc | Sec1 | -4.655 | 7.60E-05 | glio |
| Aars2_in.vivo_noRT | Nptx1 | 3.063 | 7.60E-05 | glio |
| Aars2_in.vivo_noRT | Nabp1 | 2.701 | 7.60E-05 | glio |
| Atr_ex.vivo_noRT | Parvg | 3.774 | 7.60E-05 | glio |
| Chek1_ex.vivo_noRT | 1700011M02F | 4.449 | 7.60E-05 | glio |
| Atr_in.vivo_noRT | Mroh2b | 3.205 | 7.60E-06 | glio |

| | | | | |
|---|---|---|---|---|
| Chek1_in.vivo_noRT | Gm39556 | 3.133 | 7.60E-06 | glio |
| Bora_in.vivo_noRT | Gm8098 | 3.433 | 7.60E-07 | glio |
| culture_effect | Cfap36 | 0.917 | 7.60E-09 | glio |
| culture_effect | BC049715 | 2.458 | 7.60E-10 | glio |
| culture_effect | Psmb4 | -1.395 | 7.60E-26 | glio |
| culture_effect | Gm42670 | 1.238 | 7.61E-03 | glio |
| culture_effect | Dph6 | 0.548 | 7.61E-03 | glio |
| culture_effect | Tbca | 0.25 | 7.61E-03 | glio |
| Interaction_genotype_Atr | Amotl2 | 1.705 | 7.61E-03 | glio |
| Interaction_genotype_Mtor | Gm20075 | -3.033 | 7.61E-03 | glio |
| Interaction_genotype_Myc | Gm19426 | -2.392 | 7.61E-03 | glio |
| interaction_RT | Mfsd11 | 1.105 | 7.61E-03 | glio |
| interaction_RT | Arhgef7 | -0.873 | 7.61E-03 | glio |
| interaction_RT | Camta1 | 0.873 | 7.61E-03 | glio |
| interaction_RT | Plp1 | 1.012 | 7.61E-03 | glio |
| Atr_in.vivo_noRT | Ppp1r16a | 1.455 | 7.61E-03 | glio |
| Atr_ex.vivo_noRT | Lamc2 | 2.215 | 7.61E-03 | glio |
| Bcas2_in.vivo_noRT | Gm10419 | 2.154 | 7.61E-03 | glio |
| Bcas2_in.vivo_noRT | Egfl8 | 2.084 | 7.61E-03 | glio |
| Chek1_in.vivo_noRT | Tcaim | 1.855 | 7.61E-03 | glio |
| Interaction_genotype_Mtor | Gm20300 | -4.04 | 7.61E-04 | glio |
| Interaction_genotype_Myc | Zfp628 | -3.079 | 7.61E-04 | glio |
| Aars2_in.vivo_noRT | Homez | 2.42 | 7.61E-04 | glio |
| Atr_in.vivo_noRT | Acsf2 | 2.121 | 7.61E-04 | glio |
| Atr_in.vivo_noRT | Ano10 | 1.523 | 7.61E-04 | glio |
| Bora_in.vivo_noRT | Hrh2 | 1.926 | 7.61E-04 | glio |
| Chek1_in.vivo_noRT | Gm9993 | 2.228 | 7.61E-04 | glio |
| Interaction_genotype_Mtor | Pfkfb2 | -4.868 | 7.61E-05 | glio |
| Interaction_genotype_Myc | Tedc1 | -3.388 | 7.61E-05 | glio |
| Aars2_in.vivo_noRT | Fzd7 | 2.686 | 7.61E-05 | glio |
| Atr_in.vivo_noRT | St18 | 2.171 | 7.61E-05 | glio |
| Interaction_genotype_Chek1 | Atp6v1d | 2.234 | 7.61E-06 | glio |
| Interaction_genotype_Chek1 | H2-Eb1 | 5.205 | 7.61E-06 | glio |
| Aars2_in.vivo_noRT | 1110013H19R | 3.455 | 7.61E-06 | glio |
| Aars2_in.vivo_noRT | Kif5a | 3.397 | 7.61E-06 | glio |
| Bcas2_in.vivo_noRT | Gm15834 | 3.154 | 7.61E-06 | glio |
| Bora_in.vivo_noRT | Plin1 | 3.223 | 7.61E-06 | glio |
| Bcas2_in.vivo_noRT | Gm9484 | 3.649 | 7.61E-07 | glio |
| Bcas2_in.vivo_noRT | Gm9999 | 3.832 | 7.61E-09 | glio |
| Chek1_in.vivo_noRT | Stx19 | 4.132 | 7.61E-13 | glio |
| Bcas2_in.vivo_noRT | Cldn2 | 4.807 | 7.61E-25 | glio |
| culture_effect | Bpgm | -2.461 | 7.61E-34 | glio |
| culture_effect | Timm17a | 0.301 | 7.62E-03 | glio |
| Interaction_genotype_Aars2 | Gabarap | 0.879 | 7.62E-03 | glio |
| Interaction_genotype_Myc | Cyb5r3 | 1.246 | 7.62E-03 | glio |
| Interaction_genotype_Rad51 | Gm45620 | -3.71 | 7.62E-03 | glio |
| interaction_RT | Dnase1l1 | 1.189 | 7.62E-03 | glio |
| Atp5c1_in.vivo_noRT | AC125149.2 | 1.901 | 7.62E-03 | glio |
| Atr_in.vivo_noRT | Hist1h3i | 1.718 | 7.62E-03 | glio |
| Atr_ex.vivo_noRT | Gm49223 | 2.182 | 7.62E-03 | glio |
| Atr_ex.vivo_noRT | Gm11713 | 2.45 | 7.62E-03 | glio |
| Bard1_ex.vivo_noRT | Dll1 | 3.026 | 7.62E-03 | glio |

| | | | | |
|---|---|---|---|---|
| Cenpj_in.vivo_noRT | Zeb2os | 2.183 | 7.62E-03 | glio |
| Chek1_ex.vivo_noRT | Mov10 | -1.575 | 7.62E-03 | glio |
| Chek1_ex.vivo_noRT | Mad2l1 | -1.421 | 7.62E-03 | glio |
| Eif2s1_in.vivo_noRT | Gm20767 | 2.143 | 7.62E-03 | glio |
| Eif2s1_in.vivo_noRT | Gm36198 | 1.883 | 7.62E-03 | glio |
| Fanca_ex.vivo_noRT | Lhx1os | 2.524 | 7.62E-03 | glio |
| Hsd17b10_in.vivo_noRT | 1700063J08Ri | 2.109 | 7.62E-03 | glio |
| Hsd17b10_in.vivo_noRT | Tec | 1.94 | 7.62E-03 | glio |
| Interaction_genotype_Atr | March7 | 2.001 | 7.62E-04 | glio |
| Interaction_genotype_Bcas2 | 1700029J07Ri | -4.538 | 7.62E-04 | glio |
| Interaction_genotype_Myc | Halr1 | -2.463 | 7.62E-04 | glio |
| Interaction_genotype_Myc | Gm15533 | -2.729 | 7.62E-04 | glio |
| Interaction_genotype_Myc | AC159140.1 | -2.863 | 7.62E-04 | glio |
| Interaction_genotype_Myc | Lyl1 | -3.264 | 7.62E-04 | glio |
| Interaction_genotype_Myc | Bbs1 | -3.833 | 7.62E-04 | glio |
| Interaction_genotype_Myc | Plin1 | -3.904 | 7.62E-04 | glio |
| Interaction_genotype_Myc | Ece2 | -3.641 | 7.62E-04 | glio |
| Interaction_genotype_Myc | Meis3 | -3.173 | 7.62E-04 | glio |
| Interaction_genotype_Myc | Alg3 | -2.793 | 7.62E-04 | glio |
| Atp5c1_in.vivo_noRT | C330024C12R | 2.277 | 7.62E-04 | glio |
| Atr_in.vivo_noRT | Cul9 | 2.602 | 7.62E-04 | glio |
| Chek1_in.vivo_noRT | Gm15614 | 2.399 | 7.62E-04 | glio |
| Interaction_genotype_Mtor | C330013E15Ri | -4.312 | 7.62E-05 | glio |
| Interaction_genotype_Myc | Stard10 | -4.386 | 7.62E-05 | glio |
| interaction_RT | 1500011B03R | 1.008 | 7.62E-05 | glio |
| Aars2_in.vivo_noRT | Zfp658 | 2.799 | 7.62E-05 | glio |
| Aars2_ex.vivo_noRT | Olfr552 | 2.893 | 7.62E-06 | glio |
| Atr_ex.vivo_noRT | Edn2 | 2.769 | 7.62E-06 | glio |
| Bcas2_in.vivo_noRT | Gm19409 | 3.227 | 7.62E-06 | glio |
| Bora_in.vivo_noRT | 4930481B07R | 3.372 | 7.62E-06 | glio |
| Bora_in.vivo_noRT | Speg | 3.559 | 7.62E-07 | glio |
| Bora_in.vivo_noRT | Prr7 | 3.545 | 7.62E-07 | glio |
| Chek1_ex.vivo_noRT | Selenbp1 | 5.243 | 7.62E-08 | glio |
| culture_effect | Vwa1 | -2.482 | 7.62E-11 | glio |
| Chek1_in.vivo_noRT | Gm26978 | 4.784 | 7.62E-17 | glio |
| Atr_in.vivo_noRT | D430001F17R | 5.533 | 7.62E-18 | glio |
| Atr_in.vivo_noRT | Gm47510 | 5.273 | 7.62E-26 | glio |
| Atr_in.vivo_noRT | AU019990 | 5.533 | 7.62E-26 | glio |
| culture_effect | Zfp213 | 0.916 | 7.63E-03 | glio |
| Interaction_genotype_Aars2 | Gm32051 | -3.399 | 7.63E-03 | glio |
| Interaction_genotype_Aars2 | Gm45902 | -3.326 | 7.63E-03 | glio |
| Interaction_genotype_Aars2 | Dlgap1 | -3.014 | 7.63E-03 | glio |
| Interaction_genotype_Aars2 | Tmem43 | 1.877 | 7.63E-03 | glio |
| Interaction_genotype_Aars2 | Ubxn6 | 1.325 | 7.63E-03 | glio |
| Interaction_genotype_Aars2 | Anapc16 | 1.092 | 7.63E-03 | glio |
| Interaction_genotype_Bcas2 | Wnt16 | -3.453 | 7.63E-03 | glio |
| Interaction_genotype_Bcas2 | Rarres1 | -3.486 | 7.63E-03 | glio |
| Interaction_genotype_Bcas2 | Gm14636 | -4.034 | 7.63E-03 | glio |
| Interaction_genotype_Bcas2 | Fbxo44 | 2.184 | 7.63E-03 | glio |
| Interaction_genotype_Bcas2 | Pgls | 0.988 | 7.63E-03 | glio |
| Interaction_genotype_Bora | D930028M14I | -3.222 | 7.63E-03 | glio |
| Interaction_genotype_Bora | Gm2415 | -3.709 | 7.63E-03 | glio |

| | | | | |
|---|---|---|---|---|
| Interaction_genotype_Bora | Klf4 | 2.137 | 7.63E-03 | glio |
| Interaction_genotype_Chek1 | Adgrb3 | -3.629 | 7.63E-03 | glio |
| Interaction_genotype_Chek1 | Rcan2 | -3.833 | 7.63E-03 | glio |
| Interaction_genotype_Chek1 | Gm13663 | -3.417 | 7.63E-03 | glio |
| Interaction_genotype_Chek1 | Jam2 | -3.153 | 7.63E-03 | glio |
| Interaction_genotype_Mtor | Ehd2 | -2.075 | 7.63E-03 | glio |
| Interaction_genotype_Myc | Grb7 | -3.082 | 7.63E-03 | glio |
| Interaction_genotype_Myc | Szt2 | -2.408 | 7.63E-03 | glio |
| Interaction_genotype_Terf1 | Nploc4 | 1.888 | 7.63E-03 | glio |
| interaction_RT | Dancr | 0.957 | 7.63E-03 | glio |
| Aars2_in.vivo_noRT | 1700120C14R | 1.839 | 7.63E-03 | glio |
| Aars2_in.vivo_noRT | Snx6 | -0.907 | 7.63E-03 | glio |
| Aars2_ex.vivo_noRT | Dok2 | 2.317 | 7.63E-03 | glio |
| Atp5c1_in.vivo_noRT | A330015K06R | 2.011 | 7.63E-03 | glio |
| Atp5c1_in.vivo_noRT | Hba-a2 | 2.287 | 7.63E-03 | glio |
| Chek1_in.vivo_noRT | Zfp658 | 1.979 | 7.63E-03 | glio |
| Chek1_in.vivo_noRT | Slc39a13 | 1.273 | 7.63E-03 | glio |
| Bcas2_in.vivo_noRT | Zfand4 | 2.531 | 7.63E-04 | glio |
| Bcas2_in.vivo_noRT | Aig1 | -0.665 | 7.63E-04 | glio |
| Cenpj_ex.vivo_noRT | Plk5 | 2.996 | 7.63E-04 | glio |
| Chek1_ex.vivo_noRT | Gm12280 | 3.364 | 7.63E-04 | glio |
| Interaction_genotype_Atr | Gm43137 | -4.568 | 7.63E-05 | glio |
| Aars2_in.vivo_noRT | Gm16845 | 2.656 | 7.63E-05 | glio |
| Bcas2_ex.vivo_noRT | Dpp6 | 2.742 | 7.63E-05 | glio |
| Bcas2_ex.vivo_noRT | Ccdc125 | 3.125 | 7.63E-05 | glio |
| Bcas2_ex.vivo_noRT | Serpinb9g | 2.742 | 7.63E-05 | glio |
| Bcas2_ex.vivo_noRT | BC049987 | 2.742 | 7.63E-05 | glio |
| Bora_in.vivo_noRT | Zfp982 | 2.706 | 7.63E-05 | glio |
| Chek1_ex.vivo_noRT | Taf9 | 3.469 | 7.63E-05 | glio |
| Chek1_ex.vivo_noRT | Hspe1 | -0.939 | 7.63E-05 | glio |
| Eif2s1_in.vivo_noRT | Gm34059 | 2.143 | 7.63E-05 | glio |
| Eif2s1_in.vivo_noRT | Gm48435 | 2.143 | 7.63E-05 | glio |
| Eif2s1_in.vivo_noRT | Helt | 2.143 | 7.63E-05 | glio |
| Eif2s1_in.vivo_noRT | Gm20625 | 2.143 | 7.63E-05 | glio |
| Eif2s1_in.vivo_noRT | Gm5157 | 2.143 | 7.63E-05 | glio |
| Eif2s1_in.vivo_noRT | Gm44736 | 2.143 | 7.63E-05 | glio |
| Eif2s1_in.vivo_noRT | Fam227b | 2.143 | 7.63E-05 | glio |
| Eif2s1_in.vivo_noRT | Gm20658 | 2.143 | 7.63E-05 | glio |
| culture_effect | Lrrc28 | 1.7 | 7.63E-06 | glio |
| Atr_in.vivo_noRT | Wif1 | 3.527 | 7.63E-06 | glio |
| Interaction_genotype_Atr | Hbq1b | -5.873 | 7.63E-07 | glio |
| Chek1_in.vivo_noRT | Gm11767 | 3.645 | 7.63E-07 | glio |
| Chek1_in.vivo_noRT | Gm10125 | 3.842 | 7.63E-08 | glio |
| Bora_in.vivo_noRT | Alox12 | 4.148 | 7.63E-09 | glio |
| Atr_in.vivo_noRT | Gm1123 | 4.626 | 7.63E-11 | glio |
| Interaction_genotype_Bora | 6330408M09F | -3.769 | 7.64E-03 | glio |
| Interaction_genotype_Bora | BC030343 | -3.553 | 7.64E-03 | glio |
| Interaction_genotype_Bora | Mir670hg | -3.75 | 7.64E-03 | glio |
| Interaction_genotype_Bora | Chd8 | 1.663 | 7.64E-03 | glio |
| Interaction_genotype_Chek1 | 4933429H19R | -3.475 | 7.64E-03 | glio |
| Interaction_genotype_Chek1 | Gm37933 | -3.286 | 7.64E-03 | glio |
| Interaction_genotype_Chek1 | Ahi1 | 2.155 | 7.64E-03 | glio |

| | | | | |
|---|---|---|---|---|
| Interaction_genotype_Mtor | Hsf4 | -3.29 | 7.64E-03 | glio |
| Interaction_genotype_Myc | Pacs1 | -3.018 | 7.64E-03 | glio |
| Interaction_genotype_Myc | Tmem218 | -2.377 | 7.64E-03 | glio |
| Aars2_in.vivo_noRT | Faxc | 1.495 | 7.64E-03 | glio |
| Aars2_in.vivo_noRT | Lysmd2 | 1.176 | 7.64E-03 | glio |
| Abtb1_ex.vivo_noRT | Ap1g2 | -3.177 | 7.64E-03 | glio |
| Abtb1_ex.vivo_noRT | Gm50115 | 3.303 | 7.64E-03 | glio |
| Atr_in.vivo_noRT | Lima1 | -1.157 | 7.64E-03 | glio |
| Atr_ex.vivo_noRT | C4a | 2.376 | 7.64E-03 | glio |
| Bora_in.vivo_noRT | Asb11 | 1.802 | 7.64E-03 | glio |
| Bora_in.vivo_noRT | Tmod1 | 1.717 | 7.64E-03 | glio |
| Bora_in.vivo_noRT | Adipor2 | 1.173 | 7.64E-03 | glio |
| Cenpj_in.vivo_noRT | Hrct1 | 2.022 | 7.64E-03 | glio |
| Chek1_in.vivo_noRT | Sod1 | -0.749 | 7.64E-03 | glio |
| Hsd17b10_in.vivo_noRT | 9530018F02Ri | 2.23 | 7.64E-03 | glio |
| culture_effect | 2010106G04R | 1.947 | 7.64E-04 | glio |
| Interaction_genotype_Mtor | Lrrc75b | -4.275 | 7.64E-04 | glio |
| Atp5c1_in.vivo_noRT | Scel | 1.75 | 7.64E-04 | glio |
| Atr_ex.vivo_noRT | Dnajb13 | 3.178 | 7.64E-04 | glio |
| Chek1_in.vivo_noRT | Phldb3 | 2.458 | 7.64E-04 | glio |
| Chek1_ex.vivo_noRT | Tmem202 | 3.006 | 7.64E-04 | glio |
| Aars2_ex.vivo_noRT | Ltbp2 | 2.897 | 7.64E-05 | glio |
| Atr_in.vivo_noRT | Pls1 | 2.747 | 7.64E-05 | glio |
| culture_effect | Gm34047 | 2.568 | 7.64E-06 | glio |
| Chek1_in.vivo_noRT | Gm2415 | 3.254 | 7.64E-06 | glio |
| Atr_in.vivo_noRT | Strn3 | -1.915 | 7.64E-07 | glio |
| Bcas2_in.vivo_noRT | Jmjd8 | 3.441 | 7.64E-07 | glio |
| Bcas2_in.vivo_noRT | Etl4 | 2.931 | 7.64E-07 | glio |
| Chek1_in.vivo_noRT | Dlx2 | 3.661 | 7.64E-07 | glio |
| Chek1_in.vivo_noRT | AV356131 | 3.76 | 7.64E-08 | glio |
| Chek1_ex.vivo_noRT | Tmeff2 | 5.247 | 7.64E-08 | glio |
| Atr_ex.vivo_noRT | Gm18280 | 4.191 | 7.64E-11 | glio |
| Hsd17b10_in.vivo_noRT | 1700019E08Ri | 3.616 | 7.64E-11 | glio |
| Chek1_ex.vivo_noRT | Gm49504 | 4.443 | 7.64E-12 | glio |
| Bora_in.vivo_noRT | Igf2bp1 | 4.936 | 7.64E-13 | glio |
| Atr_in.vivo_noRT | Gm45338 | 5.533 | 7.64E-31 | glio |
| Interaction_genotype_Atr | Gm16794 | -3.128 | 7.65E-03 | glio |
| Interaction_genotype_Bora | Zfr | 1.063 | 7.65E-03 | glio |
| Interaction_genotype_Rad51 | 4930579K19R | -4.017 | 7.65E-03 | glio |
| Interaction_genotype_Rpl3 | Tecrl | -3.851 | 7.65E-03 | glio |
| Aars2_in.vivo_noRT | Ran | -0.608 | 7.65E-03 | glio |
| Atr_in.vivo_noRT | Vegfd | 1.834 | 7.65E-03 | glio |
| Atr_in.vivo_noRT | Fyco1 | 1.538 | 7.65E-03 | glio |
| Atr_in.vivo_noRT | Ddx54 | -1.091 | 7.65E-03 | glio |
| Bcas2_in.vivo_noRT | 4833417C18R | 1.982 | 7.65E-03 | glio |
| Bcas2_in.vivo_noRT | Bdh2 | 1.981 | 7.65E-03 | glio |
| Bora_in.vivo_noRT | Zfp235 | 1.634 | 7.65E-03 | glio |
| Bora_ex.vivo_noRT | Rorc | 2.971 | 7.65E-03 | glio |
| Brca2_in.vivo_noRT | Gm13398 | 1.763 | 7.65E-03 | glio |
| Brca2_in.vivo_noRT | Gm266 | 1.763 | 7.65E-03 | glio |
| Brca2_in.vivo_noRT | Hbq1b | 2.542 | 7.65E-03 | glio |
| Brca2_in.vivo_noRT | Ildr1 | 1.763 | 7.65E-03 | glio |

| | | | | |
|---|---|---|---|---|
| Cdkn2a_in.vivo_noRT | A930019D19R | -1.675 | 7.65E-03 | glio |
| Chek1_ex.vivo_noRT | Dnajc28 | 2.663 | 7.65E-03 | glio |
| culture_effect | Msrb2 | -1.907 | 7.65E-04 | glio |
| culture_effect | Idi1 | -0.764 | 7.65E-04 | glio |
| culture_effect | Ezr | -0.528 | 7.65E-04 | glio |
| Interaction_genotype_Myc | 1810041H14R | -3.962 | 7.65E-04 | glio |
| Atr_in.vivo_noRT | Eny2 | -0.644 | 7.65E-04 | glio |
| Bcas2_in.vivo_noRT | Rxra | 2.569 | 7.65E-04 | glio |
| Bora_in.vivo_noRT | Ston1 | 2.486 | 7.65E-04 | glio |
| Bora_in.vivo_noRT | Abce1 | -1.016 | 7.65E-04 | glio |
| Bora_ex.vivo_noRT | Gm48161 | 2.808 | 7.65E-04 | glio |
| Bcas2_ex.vivo_noRT | Pcdhb20 | 3.729 | 7.65E-05 | glio |
| Atr_in.vivo_noRT | Tlcd2 | 3.487 | 7.65E-06 | glio |
| Atr_in.vivo_noRT | 3110045C21R | 2.923 | 7.65E-06 | glio |
| Bcas2_in.vivo_noRT | Pate2 | 3.327 | 7.65E-06 | glio |
| culture_effect | Sowahc | -1.156 | 7.65E-07 | glio |
| culture_effect | Rnaseh2a | -0.967 | 7.65E-08 | glio |
| culture_effect | Krcc1 | 1.041 | 7.65E-12 | glio |
| Atr_in.vivo_noRT | Zfp786 | 5.269 | 7.65E-15 | glio |
| culture_effect | Masp2 | -1.469 | 7.66E-03 | glio |
| Interaction_genotype_Aars2 | Hfm1 | -2.609 | 7.66E-03 | glio |
| Interaction_genotype_Aars2 | Prelid3a | -3.269 | 7.66E-03 | glio |
| Interaction_genotype_Aars2 | Pde4a | -3.019 | 7.66E-03 | glio |
| Interaction_genotype_Bcas2 | Tmem236 | -3.171 | 7.66E-03 | glio |
| Interaction_genotype_Mtor | Zfp951 | -2.401 | 7.66E-03 | glio |
| Interaction_genotype_Myc | Gm30042 | -2.469 | 7.66E-03 | glio |
| Interaction_genotype_Myc | 3110001I22Ri | -2.936 | 7.66E-03 | glio |
| Interaction_genotype_Myc | Galnt4 | -2.888 | 7.66E-03 | glio |
| interaction_RT | Tmeff2 | 3.036 | 7.66E-03 | glio |
| interaction_RT | Snrpa | 0.603 | 7.66E-03 | glio |
| interaction_RT | Tmem234 | -0.436 | 7.66E-03 | glio |
| Chek1_in.vivo_noRT | Gm48855 | 1.77 | 7.66E-03 | glio |
| Chek1_in.vivo_noRT | Gmppa | 1.198 | 7.66E-03 | glio |
| Chek1_in.vivo_noRT | Egr1 | -1.719 | 7.66E-03 | glio |
| culture_effect | Slc22a1 | -1.714 | 7.66E-04 | glio |
| Interaction_genotype_Aars2 | 4833419F23Ri | -4.29 | 7.66E-04 | glio |
| Interaction_genotype_Aars2 | 4930505N22R | -3.909 | 7.66E-04 | glio |
| Interaction_genotype_Aars2 | Gm11423 | -3.453 | 7.66E-04 | glio |
| Interaction_genotype_Aars2 | Exosc1 | 1.644 | 7.66E-04 | glio |
| Bora_in.vivo_noRT | Meak7 | 2.443 | 7.66E-04 | glio |
| Bora_in.vivo_noRT | Moap1 | 2.225 | 7.66E-04 | glio |
| Chek1_ex.vivo_noRT | Mfap3l | 3.342 | 7.66E-04 | glio |
| culture_effect | Ccdc24 | -2.238 | 7.66E-05 | glio |
| Interaction_genotype_Mtor | Kank2 | -4.212 | 7.66E-05 | glio |
| Bora_in.vivo_noRT | Vcan | 3.067 | 7.66E-05 | glio |
| Eif2s1_in.vivo_noRT | Olfr810 | 2.143 | 7.66E-05 | glio |
| Atr_in.vivo_noRT | 9330020H09R | 2.892 | 7.66E-06 | glio |
| Bora_in.vivo_noRT | Serpinb6a | 3.248 | 7.66E-06 | glio |
| culture_effect | Abhd8 | 0.875 | 7.66E-07 | glio |
| Bcas2_in.vivo_noRT | Gm13404 | 4.069 | 7.66E-09 | glio |
| Aars2_in.vivo_noRT | Mfap3l | 4.936 | 7.66E-11 | glio |
| Bora_in.vivo_noRT | Pecam1 | 4.927 | 7.66E-12 | glio |

| | | | | |
|---|---|---|---|---|
| Bora_in.vivo_noRT | Smtnl1 | 4.936 | 7.66E-20 | glio |
| Atr_in.vivo_noRT | Gm17203 | 4.883 | 7.66E-22 | glio |
| culture_effect | Gm47510 | -1.194 | 7.67E-03 | glio |
| Interaction_genotype_Aars2 | Gm13404 | -3.386 | 7.67E-03 | glio |
| Interaction_genotype_Aars2 | Zic4 | -3.481 | 7.67E-03 | glio |
| Interaction_genotype_Aars2 | Gm15601 | -3.424 | 7.67E-03 | glio |
| Interaction_genotype_Aars2 | Timm8b | 1.039 | 7.67E-03 | glio |
| Interaction_genotype_Atr | Gm15446 | -2.996 | 7.67E-03 | glio |
| Interaction_genotype_Atr | Ccdc28a | 2.176 | 7.67E-03 | glio |
| Interaction_genotype_Atr | 2410006H16R | 1.193 | 7.67E-03 | glio |
| Interaction_genotype_Pars2 | 6430562O15R | -3.336 | 7.67E-03 | glio |
| Interaction_genotype_Pars2 | Slc19a3 | -2.662 | 7.67E-03 | glio |
| interaction_RT | Specc1 | -0.619 | 7.67E-03 | glio |
| Atr_in.vivo_noRT | Ypel1 | 1.923 | 7.67E-03 | glio |
| Atr_ex.vivo_noRT | Gm44899 | 2.059 | 7.67E-03 | glio |
| Bcas2_in.vivo_noRT | Tppp | 1.726 | 7.67E-03 | glio |
| Bcas2_in.vivo_noRT | Pmepa1 | -1.055 | 7.67E-03 | glio |
| Bora_in.vivo_noRT | Prdm9 | 1.997 | 7.67E-03 | glio |
| Bora_in.vivo_noRT | Rnasel | 1.796 | 7.67E-03 | glio |
| Bora_ex.vivo_noRT | Prox2os | 2.408 | 7.67E-03 | glio |
| Chek1_ex.vivo_noRT | Nbr1 | 1.264 | 7.67E-03 | glio |
| Hsd17b10_in.vivo_noRT | Rnf144b | 2.086 | 7.67E-03 | glio |
| Hsd17b10_in.vivo_noRT | Atl2 | -1.406 | 7.67E-03 | glio |
| Interaction_genotype_Mtor | Etohd2 | -4.256 | 7.67E-04 | glio |
| Interaction_genotype_Mtor | A830082N09R | -4.175 | 7.67E-04 | glio |
| Interaction_genotype_Mtor | Slc25a15 | -3.485 | 7.67E-04 | glio |
| Aars2_in.vivo_noRT | Srp19 | -0.927 | 7.67E-04 | glio |
| Atr_ex.vivo_noRT | Cldn9 | 3.15 | 7.67E-04 | glio |
| culture_effect | Nudt19 | 0.666 | 7.67E-05 | glio |
| Bora_in.vivo_noRT | Cd93 | 2.903 | 7.67E-05 | glio |
| Chek1_in.vivo_noRT | Zglp1 | 3.079 | 7.67E-05 | glio |
| Atp5c1_in.vivo_noRT | Gm30794 | 2.668 | 7.67E-07 | glio |
| culture_effect | Rpl15 | -0.712 | 7.67E-08 | glio |
| Atr_ex.vivo_noRT | Gm49172 | 3.164 | 7.67E-08 | glio |
| Bora_in.vivo_noRT | Gm17227 | 3.554 | 7.67E-08 | glio |
| culture_effect | Gnptg | -1.459 | 7.67E-10 | glio |
| Bora_in.vivo_noRT | Notch4 | 4.203 | 7.67E-10 | glio |
| Bcas2_in.vivo_noRT | Dok2 | 4.281 | 7.67E-13 | glio |
| Aars2_in.vivo_noRT | Gm14002 | 4.805 | 7.67E-14 | glio |
| Bora_in.vivo_noRT | Olfr543 | 4.676 | 7.67E-19 | glio |
| Aars2_in.vivo_noRT | Galnt3 | 1.99 | 7.68E-03 | glio |
| Bard1_ex.vivo_noRT | 9130230L23Ri | 2.421 | 7.68E-03 | glio |
| Bard1_ex.vivo_noRT | Gadd45g | 1.688 | 7.68E-03 | glio |
| Bcas2_ex.vivo_noRT | Smarcad1 | 1.09 | 7.68E-03 | glio |
| Bora_in.vivo_noRT | Coprs | -1.176 | 7.68E-03 | glio |
| Hsd17b10_in.vivo_noRT | Kctd21 | 2.056 | 7.68E-03 | glio |
| Interaction_genotype_Myc | Il20rb | -4.052 | 7.68E-04 | glio |
| Interaction_genotype_Myc | Doc2g | -3.919 | 7.68E-04 | glio |
| Atr_in.vivo_noRT | A930001M01F | 2.373 | 7.68E-04 | glio |
| Atr_in.vivo_noRT | Atxn7l3b | -0.893 | 7.68E-04 | glio |
| Atr_ex.vivo_noRT | 9130230L23Ri | 2.771 | 7.68E-04 | glio |
| Atr_ex.vivo_noRT | Serpinb6b | 3.368 | 7.68E-04 | glio |

| | | | | |
|---|---|---|---|---|
| Fanca_in.vivo_noRT | Clcf1 | 2.244 | 7.68E-04 | glio |
| Aars2_ex.vivo_noRT | Ms4a15 | 3.293 | 7.68E-06 | glio |
| Bora_in.vivo_noRT | Gm36445 | 3.171 | 7.68E-06 | glio |
| culture_effect | Cited2 | 1.433 | 7.68E-11 | glio |
| Aars2_in.vivo_noRT | Gm16853 | 4.293 | 7.68E-11 | glio |
| Hsd17b10_in.vivo_noRT | Tal2 | 3.353 | 7.68E-11 | glio |
| Atr_in.vivo_noRT | Slc23a3 | 4.632 | 7.68E-12 | glio |
| Atr_in.vivo_noRT | Sod3 | 5.533 | 7.68E-12 | glio |
| Atr_in.vivo_noRT | Gm34885 | 5.273 | 7.68E-14 | glio |
| Atr_ex.vivo_noRT | Gm14929 | 4.547 | 7.68E-14 | glio |
| Atr_ex.vivo_noRT | Gm12794 | 4.725 | 7.68E-14 | glio |
| Bora_in.vivo_noRT | Adam12 | 4.676 | 7.68E-14 | glio |
| Bora_in.vivo_noRT | Gm15533 | 4.936 | 7.68E-22 | glio |
| Chek1_in.vivo_noRT | Gjd4 | 5.044 | 7.68E-26 | glio |
| culture_effect | Hexim2 | 0.643 | 7.69E-03 | glio |
| culture_effect | Slc30a5 | -0.521 | 7.69E-03 | glio |
| culture_effect | Bag5 | 0.482 | 7.69E-03 | glio |
| culture_effect | Asap2 | 0.549 | 7.69E-03 | glio |
| Interaction_genotype_Aars2 | A930004D18R | -3.353 | 7.69E-03 | glio |
| Interaction_genotype_Aars2 | Tmem219 | 1.737 | 7.69E-03 | glio |
| Interaction_genotype_Atr | Med19 | 1.158 | 7.69E-03 | glio |
| Interaction_genotype_Chek1 | Nme6 | -1.982 | 7.69E-03 | glio |
| Interaction_genotype_Chek1 | Rdm1 | 2.243 | 7.69E-03 | glio |
| Interaction_genotype_Mtor | Galnt3 | -3.369 | 7.69E-03 | glio |
| Interaction_genotype_Mtor | Catsper2 | -2.943 | 7.69E-03 | glio |
| Interaction_genotype_Mtor | Sar1b | 1.368 | 7.69E-03 | glio |
| Interaction_genotype_Myc | Ikbkg | -2.138 | 7.69E-03 | glio |
| Interaction_genotype_Pars2 | 4833419F23Ri | -3.727 | 7.69E-03 | glio |
| Interaction_genotype_Pars2 | Gm49701 | -3.581 | 7.69E-03 | glio |
| Aars2_in.vivo_noRT | Zdhhc15 | 1.663 | 7.69E-03 | glio |
| Aars2_in.vivo_noRT | Dbf4 | -1.256 | 7.69E-03 | glio |
| Aars2_in.vivo_noRT | Polr2m | -0.813 | 7.69E-03 | glio |
| Atr_in.vivo_noRT | Ube2l3 | -0.751 | 7.69E-03 | glio |
| Bora_ex.vivo_noRT | Neurl3 | 2.802 | 7.69E-03 | glio |
| Chek1_in.vivo_noRT | Fam189b | 2.036 | 7.69E-03 | glio |
| Chek1_in.vivo_noRT | Snx17 | -0.852 | 7.69E-03 | glio |
| Chek1_ex.vivo_noRT | Entpd3 | 2.468 | 7.69E-03 | glio |
| Chek1_ex.vivo_noRT | Imp4 | -1.137 | 7.69E-03 | glio |
| Interaction_genotype_Atr | Kctd11 | -4.325 | 7.69E-04 | glio |
| Bcas2_ex.vivo_noRT | Lrguk | 2.742 | 7.69E-04 | glio |
| Bcas2_ex.vivo_noRT | Gm20275 | 2.742 | 7.69E-04 | glio |
| Bora_in.vivo_noRT | Serp2 | 2.199 | 7.69E-04 | glio |
| Bora_in.vivo_noRT | E130102H24R | 2.349 | 7.69E-04 | glio |
| Chek1_in.vivo_noRT | Sbk1 | 2.486 | 7.69E-04 | glio |
| Chek1_in.vivo_noRT | G730013B05R | -1.642 | 7.69E-04 | glio |
| Eif2s1_in.vivo_noRT | Apoa5 | 2.143 | 7.69E-05 | glio |
| culture_effect | Rpl3l | -2.871 | 7.69E-07 | glio |
| Chek1_in.vivo_noRT | Kcnab3 | 3.51 | 7.69E-07 | glio |
| Chek1_ex.vivo_noRT | Zfp990 | 4.124 | 7.69E-07 | glio |
| Chek1_ex.vivo_noRT | Gm26789 | 3.862 | 7.69E-08 | glio |
| Chek1_ex.vivo_noRT | Iqch | 3.877 | 7.69E-08 | glio |
| Atr_in.vivo_noRT | 1700063J08Ri | 4.026 | 7.69E-09 | glio |

| | | | | |
|---|---|---|---|---|
| Bora_in.vivo_noRT | Rilp | 4.411 | 7.69E-09 | glio |
| Bora_in.vivo_noRT | Apobr | 4.936 | 7.69E-11 | glio |
| Aars2_in.vivo_noRT | BC026762 | 4.947 | 7.69E-14 | glio |
| Interaction_genotype_Aars2 | Ppfia3 | -3.253 | 7.70E-03 | glio |
| Interaction_genotype_Myc | Dubr | -2.36 | 7.70E-03 | glio |
| Aars2_in.vivo_noRT | Dusp4 | 1.852 | 7.70E-03 | glio |
| Aars2_in.vivo_noRT | Tmem143 | 1.59 | 7.70E-03 | glio |
| Atp5c1_in.vivo_noRT | Cgn | 2.142 | 7.70E-03 | glio |
| Atp5c1_in.vivo_noRT | Svopl | 2.015 | 7.70E-03 | glio |
| Atr_in.vivo_noRT | Car5b | 2.022 | 7.70E-03 | glio |
| Bcas2_in.vivo_noRT | Ippk | 1.45 | 7.70E-03 | glio |
| Hsd17b10_in.vivo_noRT | Gm12905 | 2.311 | 7.70E-03 | glio |
| Interaction_genotype_Atr | Rps24 | 1.141 | 7.70E-04 | glio |
| Interaction_genotype_Mtor | Dhx29 | -2.609 | 7.70E-04 | glio |
| Aars2_in.vivo_noRT | Rab30 | 2.145 | 7.70E-04 | glio |
| Atp5c1_in.vivo_noRT | Cyp4f40 | 2.537 | 7.70E-04 | glio |
| Bard1_ex.vivo_noRT | Ceacam16 | 2.814 | 7.70E-04 | glio |
| Bora_in.vivo_noRT | Cox7a1 | 2.618 | 7.70E-04 | glio |
| Cenpj_in.vivo_noRT | Gm12794 | 2.022 | 7.70E-04 | glio |
| Chek1_in.vivo_noRT | Sphk1 | 2.469 | 7.70E-04 | glio |
| Chek1_in.vivo_noRT | Lap3 | -0.992 | 7.70E-04 | glio |
| culture_effect | Wnt10a | 1.885 | 7.70E-05 | glio |
| Interaction_genotype_Atr | Dnajc9 | 2.354 | 7.70E-05 | glio |
| Interaction_genotype_Myc | Dda1 | 1.721 | 7.70E-05 | glio |
| Aars2_in.vivo_noRT | Frg1 | -1.442 | 7.70E-05 | glio |
| Bora_in.vivo_noRT | Zfp988 | 2.804 | 7.70E-05 | glio |
| culture_effect | Psd3 | 1.982 | 7.70E-09 | glio |
| Aars2_in.vivo_noRT | Gpr37l1 | 4.213 | 7.70E-10 | glio |
| Chek1_in.vivo_noRT | Nrros | 4.401 | 7.70E-11 | glio |
| Bora_in.vivo_noRT | Capn9 | 4.936 | 7.70E-15 | glio |
| Interaction_genotype_Mtor | Pgm5 | -3.662 | 7.71E-03 | glio |
| Interaction_genotype_Mtor | Gpam | -2.814 | 7.71E-03 | glio |
| Interaction_genotype_Mtor | Soga3 | -2.261 | 7.71E-03 | glio |
| Interaction_genotype_Myc | St14 | -2.717 | 7.71E-03 | glio |
| Interaction_genotype_Myc | H2-Q7 | 2.627 | 7.71E-03 | glio |
| Interaction_genotype_Myc | Map1b | 1.263 | 7.71E-03 | glio |
| interaction_RT | Klf5 | 0.966 | 7.71E-03 | glio |
| Aars2_in.vivo_noRT | Gm20075 | 1.849 | 7.71E-03 | glio |
| Chek1_ex.vivo_noRT | Mical1 | 2.233 | 7.71E-03 | glio |
| culture_effect | Gm14305 | 1.123 | 7.71E-04 | glio |
| Interaction_genotype_Mtor | Tbcb | 1.605 | 7.71E-04 | glio |
| Chek1_ex.vivo_noRT | Cd59b | 3.071 | 7.71E-04 | glio |
| Interaction_genotype_Myc | Sall3 | -3.662 | 7.71E-05 | glio |
| Interaction_genotype_Mtor | 4930524O07R | -5.263 | 7.71E-06 | glio |
| Aars2_in.vivo_noRT | Gm4189 | 3.125 | 7.71E-06 | glio |
| Bora_ex.vivo_noRT | A930002C04R | 2.802 | 7.71E-06 | glio |
| Interaction_genotype_Mtor | Fbxl2 | -5.334 | 7.71E-07 | glio |
| Hsd17b10_in.vivo_noRT | Cyp4x1 | 3.35 | 7.71E-07 | glio |
| Bcas2_in.vivo_noRT | Isl1 | 4.004 | 7.71E-09 | glio |
| Bora_in.vivo_noRT | Grin3b | 4.414 | 7.71E-11 | glio |
| Chek1_ex.vivo_noRT | Npnt | 4.045 | 7.71E-11 | glio |
| Bora_in.vivo_noRT | Gm5788 | 4.936 | 7.71E-23 | glio |

| | | | | |
|---|---|---|---|---|
| culture_effect | Slc30a4 | -0.627 | 7.72E-03 | glio |
| culture_effect | Tbrg4 | 0.54 | 7.72E-03 | glio |
| Interaction_genotype_Aars2 | C920021L13Ri | -3.212 | 7.72E-03 | glio |
| Interaction_genotype_Aars2 | Mnat1 | 1.5 | 7.72E-03 | glio |
| Interaction_genotype_Mtor | Slc9a9 | -3.371 | 7.72E-03 | glio |
| Interaction_genotype_Myc | Gm34939 | -2.466 | 7.72E-03 | glio |
| Interaction_genotype_Myc | Ppm1f | -2.454 | 7.72E-03 | glio |
| Interaction_genotype_Myc | Rps3a1 | 1.01 | 7.72E-03 | glio |
| Interaction_genotype_Rad9a | Carns1 | -4.518 | 7.72E-03 | glio |
| Bard1_ex.vivo_noRT | Stra8 | 2.419 | 7.72E-03 | glio |
| Chek1_ex.vivo_noRT | Apold1 | 2.83 | 7.72E-03 | glio |
| culture_effect | Cmtm5 | -0.731 | 7.72E-04 | glio |
| Interaction_genotype_Mtor | Gm4755 | -3.717 | 7.72E-04 | glio |
| Interaction_genotype_Mtor | Fkbp1a | 1.312 | 7.72E-04 | glio |
| Aars2_ex.vivo_noRT | Gm43625 | 2.907 | 7.72E-04 | glio |
| Aars2_ex.vivo_noRT | Esm1 | 2.317 | 7.72E-04 | glio |
| Atr_ex.vivo_noRT | Stra8 | 2.769 | 7.72E-04 | glio |
| Chek1_in.vivo_noRT | Gm904 | 2.481 | 7.72E-04 | glio |
| Interaction_genotype_Mtor | Mamdc2 | -4.629 | 7.72E-05 | glio |
| Bora_in.vivo_noRT | Foxa1 | 3.084 | 7.72E-05 | glio |
| Bora_in.vivo_noRT | AA386476 | 2.906 | 7.72E-05 | glio |
| Aars2_in.vivo_noRT | Fgfr1 | 3.689 | 7.72E-06 | glio |
| Bard1_ex.vivo_noRT | Gm34059 | 2.814 | 7.72E-06 | glio |
| Bard1_ex.vivo_noRT | Halr1 | 2.814 | 7.72E-06 | glio |
| Bcas2_in.vivo_noRT | Adcy2 | 3.053 | 7.72E-07 | glio |
| Chek1_ex.vivo_noRT | Me2 | -2.511 | 7.72E-07 | glio |
| Bcas2_in.vivo_noRT | Gm45620 | 4.807 | 7.72E-13 | glio |
| culture_effect | Ap5m1 | -0.707 | 7.73E-03 | glio |
| interaction_RT | Batf2 | 1.567 | 7.73E-03 | glio |
| Aars2_in.vivo_noRT | Ptpn3 | 1.844 | 7.73E-03 | glio |
| Atr_in.vivo_noRT | Maff | -1.488 | 7.73E-03 | glio |
| Bora_in.vivo_noRT | Hspa9 | -0.636 | 7.73E-03 | glio |
| Cdkn2a_in.vivo_noRT | Gm49808 | -1.675 | 7.73E-03 | glio |
| Cdkn2a_in.vivo_noRT | Slc14a2 | -1.675 | 7.73E-03 | glio |
| Cdkn2a_in.vivo_noRT | Gpr17 | -1.922 | 7.73E-03 | glio |
| Cdkn2a_in.vivo_noRT | Anxa8 | 1.903 | 7.73E-03 | glio |
| Hsd17b10_in.vivo_noRT | Firre | 1.896 | 7.73E-03 | glio |
| Interaction_genotype_Aars2 | Prrc1 | 2.129 | 7.73E-04 | glio |
| Interaction_genotype_Myc | Tmem265 | -3.281 | 7.73E-04 | glio |
| Aars2_in.vivo_noRT | Rnasel | 2.186 | 7.73E-04 | glio |
| Atp5c1_in.vivo_noRT | Gm40557 | 2.667 | 7.73E-04 | glio |
| Atr_in.vivo_noRT | Bag4 | -1.595 | 7.73E-04 | glio |
| Bcas2_in.vivo_noRT | Fes | 2.487 | 7.73E-04 | glio |
| Bora_in.vivo_noRT | Cbx2 | 2.188 | 7.73E-04 | glio |
| Chek1_ex.vivo_noRT | Gm47223 | 4.045 | 7.73E-04 | glio |
| Atr_ex.vivo_noRT | Gm1123 | 3.732 | 7.73E-05 | glio |
| culture_effect | Rhobtb3 | 0.957 | 7.73E-07 | glio |
| Interaction_genotype_Myc | Chia1 | -4.059 | 7.73E-07 | glio |
| Atr_in.vivo_noRT | Rpl15 | -1.044 | 7.73E-07 | glio |
| Interaction_genotype_Myc | Sec61b | 2.407 | 7.73E-08 | glio |
| Atr_in.vivo_noRT | Dzank1 | 4.358 | 7.73E-10 | glio |
| Bcas2_in.vivo_noRT | Ephx3 | 4.079 | 7.73E-10 | glio |

| | | | | |
|---|---|---|---|---|
| Chek1_in.vivo_noRT | Slxl1 | 4.265 | 7.73E-10 | glio |
| Bora_in.vivo_noRT | Ramp3 | 4.67 | 7.73E-12 | glio |
| Bcas2_in.vivo_noRT | Gm38534 | 4.807 | 7.73E-13 | glio |
| Bcas2_in.vivo_noRT | 9130230L23Ri | 4.547 | 7.73E-14 | glio |
| Aars2_in.vivo_noRT | 5033417F24Ri | 5.327 | 7.73E-15 | glio |
| Atr_in.vivo_noRT | Scg2 | 5.533 | 7.73E-31 | glio |
| Interaction_genotype_Atr | Mtrr | -2.433 | 7.74E-03 | glio |
| Interaction_genotype_Lmo7 | 1700015F17Ri | -4.257 | 7.74E-03 | glio |
| Interaction_genotype_Mtor | Gm11127 | -3.332 | 7.74E-03 | glio |
| Interaction_genotype_Mtor | Faxc | -2.515 | 7.74E-03 | glio |
| Interaction_genotype_Myc | Syngr4 | -2.992 | 7.74E-03 | glio |
| Interaction_genotype_Myc | Fancl | -2.452 | 7.74E-03 | glio |
| Aars2_in.vivo_noRT | Phf11b | 2.148 | 7.74E-03 | glio |
| Bcas2_in.vivo_noRT | 6430590A07R | 1.53 | 7.74E-03 | glio |
| Bora_in.vivo_noRT | 3110083C13R | 1.933 | 7.74E-03 | glio |
| Chek1_in.vivo_noRT | Mrps18a | -0.785 | 7.74E-03 | glio |
| Chek1_ex.vivo_noRT | Ttf2 | -1.819 | 7.74E-03 | glio |
| Cyp19a1_ex.vivo_noRT | Adam5 | 2.159 | 7.74E-03 | glio |
| culture_effect | Pclo | -0.894 | 7.74E-04 | glio |
| culture_effect | Dnm1l | 0.579 | 7.74E-04 | glio |
| Interaction_genotype_Ifnar1 | Cxcl1 | 4.971 | 7.74E-04 | glio |
| Interaction_genotype_Ifnar1 | Zscan4-ps2 | 3.521 | 7.74E-04 | glio |
| interaction_RT | Bckdk | -0.873 | 7.74E-04 | glio |
| interaction_RT | Tmed4 | 0.744 | 7.74E-04 | glio |
| Bcas2_in.vivo_noRT | AI480526 | 2.386 | 7.74E-04 | glio |
| Bcas2_in.vivo_noRT | Mettl9 | -1.367 | 7.74E-04 | glio |
| Bora_in.vivo_noRT | Btbd17 | 2.905 | 7.74E-04 | glio |
| Bora_in.vivo_noRT | Fkbp15 | 1.592 | 7.74E-04 | glio |
| Hsd17b10_in.vivo_noRT | Gpr37l1 | 2.502 | 7.74E-04 | glio |
| Hsd17b10_in.vivo_noRT | Dusp28 | -1.977 | 7.74E-04 | glio |
| Interaction_genotype_Mtor | Prkag2 | -3.494 | 7.74E-05 | glio |
| Atr_in.vivo_noRT | Gm47689 | 3.144 | 7.74E-06 | glio |
| Bcas2_in.vivo_noRT | Serpinb9b | 3.81 | 7.74E-06 | glio |
| Hsd17b10_in.vivo_noRT | Fam131a | 3.09 | 7.74E-06 | glio |
| Aars2_in.vivo_noRT | Gm15952 | 3.56 | 7.74E-07 | glio |
| Aars2_in.vivo_noRT | Dennd2d | 3.06 | 7.74E-07 | glio |
| Aars2_in.vivo_noRT | Bst1 | 3.224 | 7.74E-07 | glio |
| Interaction_genotype_Myc | Klhl30 | -5.894 | 7.74E-08 | glio |
| Interaction_genotype_Myc | Adam12 | -5.412 | 7.74E-08 | glio |
| Aars2_in.vivo_noRT | Pafah2 | 4.088 | 7.74E-08 | glio |
| Hsd17b10_in.vivo_noRT | Gm28940 | 3.1 | 7.74E-08 | glio |
| Aars2_in.vivo_noRT | Neurl3 | 4.336 | 7.74E-10 | glio |
| Aars2_in.vivo_noRT | Tnxb | 4.174 | 7.74E-10 | glio |
| Atr_in.vivo_noRT | Appbp2os | 4.47 | 7.74E-10 | glio |
| Atr_in.vivo_noRT | Serpina3i | 4.212 | 7.74E-11 | glio |
| Atr_ex.vivo_noRT | Akr1c18 | 4.583 | 7.74E-11 | glio |
| Chek1_in.vivo_noRT | Gm40264 | 4.784 | 7.74E-16 | glio |
| Aars2_in.vivo_noRT | Spi1 | 5.327 | 7.74E-17 | glio |
| culture_effect | Brinp3 | -6.011 | 7.74E-31 | glio |
| Interaction_genotype_Atr | 4933433G15R | -3.304 | 7.75E-03 | glio |
| Interaction_genotype_Mtor | Gm38414 | -3.481 | 7.75E-03 | glio |
| Interaction_genotype_Mtor | Aldh4a1 | -2.831 | 7.75E-03 | glio |

| | | | | |
|---|---|---|---|---|
| Interaction_genotype_Myc | Map3k15 | -2.458 | 7.75E-03 | glio |
| Interaction_genotype_Myc | Fkbp1b | -2.78 | 7.75E-03 | glio |
| Interaction_genotype_Myc | Cep170b | -2.441 | 7.75E-03 | glio |
| interaction_RT | Prkca | -1.281 | 7.75E-03 | glio |
| Atr_in.vivo_noRT | Nono | -0.717 | 7.75E-03 | glio |
| Bora_in.vivo_noRT | Snph | 1.893 | 7.75E-03 | glio |
| Bora_in.vivo_noRT | Arhgef7 | -1.258 | 7.75E-03 | glio |
| Cenpj_ex.vivo_noRT | Tent5c | 2.198 | 7.75E-03 | glio |
| Cenpj_ex.vivo_noRT | Auts2 | 2.591 | 7.75E-03 | glio |
| Cenpj_ex.vivo_noRT | Dio1 | 2.591 | 7.75E-03 | glio |
| Chek1_in.vivo_noRT | Smim1 | 1.594 | 7.75E-03 | glio |
| culture_effect | Pdk3 | 0.57 | 7.75E-04 | glio |
| Interaction_genotype_Mtor | 9330151L19Ri | -3.393 | 7.75E-04 | glio |
| Aars2_in.vivo_noRT | Draxin | 2.378 | 7.75E-04 | glio |
| Aars2_in.vivo_noRT | Sdcbp | -1.071 | 7.75E-04 | glio |
| Atr_in.vivo_noRT | Apobec1 | 2.375 | 7.75E-04 | glio |
| Atr_ex.vivo_noRT | Lrrc24 | 2.976 | 7.75E-04 | glio |
| interaction_RT | Fam107b | 2.199 | 7.75E-05 | glio |
| Bard1_ex.vivo_noRT | Gm45338 | 2.421 | 7.75E-05 | glio |
| Chek1_ex.vivo_noRT | H2-Ab1 | 2.985 | 7.75E-05 | glio |
| Atr_ex.vivo_noRT | B230206H07R | 2.769 | 7.75E-06 | glio |
| Hsd17b10_in.vivo_noRT | Ssh3 | 2.816 | 7.75E-06 | glio |
| Aars2_in.vivo_noRT | Gm50218 | 3.733 | 7.75E-07 | glio |
| Chek1_in.vivo_noRT | Cyth4 | 3.503 | 7.75E-07 | glio |
| culture_effect | Marcks | 1.074 | 7.75E-09 | glio |
| culture_effect | Purb | 0.618 | 7.75E-10 | glio |
| Bora_in.vivo_noRT | Gm43137 | 4.42 | 7.75E-10 | glio |
| Hsd17b10_in.vivo_noRT | Gm13348 | 3.616 | 7.75E-10 | glio |
| Chek1_in.vivo_noRT | Sox11 | 4.397 | 7.75E-11 | glio |
| Aars2_in.vivo_noRT | Gm9929 | 4.548 | 7.75E-13 | glio |
| Aars2_in.vivo_noRT | Gm15127 | 5.067 | 7.75E-14 | glio |
| Bora_in.vivo_noRT | Fbxl13 | 4.676 | 7.75E-15 | glio |
| Bora_in.vivo_noRT | Rassf6 | 4.936 | 7.75E-17 | glio |
| culture_effect | 4930554I06Ri | 1.521 | 7.76E-03 | glio |
| culture_effect | Enpp4 | -0.662 | 7.76E-03 | glio |
| Interaction_genotype_Atr | Adam4 | -3.169 | 7.76E-03 | glio |
| Interaction_genotype_Atr | Gm42726 | -3.27 | 7.76E-03 | glio |
| Interaction_genotype_Atr | BC065397 | -3.717 | 7.76E-03 | glio |
| Interaction_genotype_Bcas2 | Ap3m1 | 1.88 | 7.76E-03 | glio |
| Interaction_genotype_Cdkn2a | Gm266 | 2.835 | 7.76E-03 | glio |
| Interaction_genotype_Chek1 | Pom121 | -2.209 | 7.76E-03 | glio |
| Interaction_genotype_Ifnar1 | Gm49698 | 2.734 | 7.76E-03 | glio |
| Interaction_genotype_Ifnar1 | Olfr881 | 2.734 | 7.76E-03 | glio |
| Interaction_genotype_Mtor | Gsr | 1.389 | 7.76E-03 | glio |
| Interaction_genotype_Myc | 4930506C21R | -3.012 | 7.76E-03 | glio |
| Interaction_genotype_Myc | Dleu7 | -2.999 | 7.76E-03 | glio |
| Interaction_genotype_Myc | Pcp4l1 | -2.865 | 7.76E-03 | glio |
| Interaction_genotype_Myc | Ei24 | -1.533 | 7.76E-03 | glio |
| Interaction_genotype_Rpl3 | Rarres1 | -3.689 | 7.76E-03 | glio |
| Interaction_genotype_Rpl3 | Plcl1 | -3.078 | 7.76E-03 | glio |
| Interaction_genotype_Rpl3 | Tmem179 | -3.707 | 7.76E-03 | glio |
| Aars2_in.vivo_noRT | 6230400D17R | 1.857 | 7.76E-03 | glio |

| | | | | |
|---|---|---|---|---|
| Aars2_in.vivo_noRT | Pars2 | 1.828 | 7.76E-03 | glio |
| Aars2_in.vivo_noRT | Fut10 | 1.552 | 7.76E-03 | glio |
| Aars2_in.vivo_noRT | Braf | 1.163 | 7.76E-03 | glio |
| Atr_in.vivo_noRT | Elp4 | 1.209 | 7.76E-03 | glio |
| Atr_in.vivo_noRT | Rps2 | -0.605 | 7.76E-03 | glio |
| Bcas2_in.vivo_noRT | Dyrk4 | 2.239 | 7.76E-03 | glio |
| Bcas2_in.vivo_noRT | A330023F24Ri | 2.028 | 7.76E-03 | glio |
| Bcas2_in.vivo_noRT | Syt14 | 1.588 | 7.76E-03 | glio |
| Bcas2_in.vivo_noRT | Prr14l | -1.334 | 7.76E-03 | glio |
| Bora_in.vivo_noRT | Chd9 | -1.199 | 7.76E-03 | glio |
| Cenpj_ex.vivo_noRT | Tmem52 | 2.591 | 7.76E-03 | glio |
| Cenpj_ex.vivo_noRT | Gm36043 | 2.591 | 7.76E-03 | glio |
| Cenpj_ex.vivo_noRT | Gm49860 | 2.407 | 7.76E-03 | glio |
| Cenpj_ex.vivo_noRT | Gpr17 | 2.591 | 7.76E-03 | glio |
| Interaction_genotype_Mtor | Ss18l1 | -3.64 | 7.76E-04 | glio |
| Interaction_genotype_Mtor | Adamts14 | -3.266 | 7.76E-04 | glio |
| Interaction_genotype_Myc | Dppa2 | -3.564 | 7.76E-04 | glio |
| Aars2_in.vivo_noRT | Hr | 2.159 | 7.76E-04 | glio |
| Atr_in.vivo_noRT | Nemp2 | 2.587 | 7.76E-04 | glio |
| Interaction_genotype_Myc | Iqgap2 | -3.777 | 7.76E-05 | glio |
| Bora_in.vivo_noRT | Slc25a53 | 3.032 | 7.76E-05 | glio |
| Atr_in.vivo_noRT | Gm14273 | 3.043 | 7.76E-06 | glio |
| Atr_in.vivo_noRT | Gpr135 | 3.092 | 7.76E-06 | glio |
| Chek1_ex.vivo_noRT | Gm26590 | 3.664 | 7.76E-06 | glio |
| Bcas2_in.vivo_noRT | Chordc1 | -1.937 | 7.76E-07 | glio |
| Bcas2_in.vivo_noRT | Ntn1 | 3.905 | 7.76E-11 | glio |
| Atr_in.vivo_noRT | 3930402G23R | 5.146 | 7.76E-13 | glio |
| Bcas2_in.vivo_noRT | Cers3 | 4.275 | 7.76E-15 | glio |
| Chek1_ex.vivo_noRT | Rimklb | 4.842 | 7.76E-16 | glio |
| Bcas2_in.vivo_noRT | Ccdc80 | 4.807 | 7.76E-19 | glio |
| culture_effect | Capn5 | -1.007 | 7.77E-03 | glio |
| Interaction_genotype_Aars2 | Gm36037 | -2.429 | 7.77E-03 | glio |
| Interaction_genotype_Mtor | Slc6a11 | -2.667 | 7.77E-03 | glio |
| Interaction_genotype_Mtor | Sars2 | -2.729 | 7.77E-03 | glio |
| Interaction_genotype_Myc | Zfp788 | -2.043 | 7.77E-03 | glio |
| Interaction_genotype_Myc | Lyrm9 | -1.706 | 7.77E-03 | glio |
| Interaction_genotype_Rad51 | Gtf2a1 | 2.143 | 7.77E-03 | glio |
| interaction_RT | Cd59a | 2.054 | 7.77E-03 | glio |
| Aars2_in.vivo_noRT | Pkm | -0.698 | 7.77E-03 | glio |
| Atr_in.vivo_noRT | Hsd17b12 | -1.018 | 7.77E-03 | glio |
| Bora_in.vivo_noRT | Fcor | 1.927 | 7.77E-03 | glio |
| Bora_ex.vivo_noRT | Gm19409 | 2.705 | 7.77E-03 | glio |
| Chek1_in.vivo_noRT | Gdap1l1 | 2.166 | 7.77E-03 | glio |
| culture_effect | Dcpp1 | -2.261 | 7.77E-04 | glio |
| culture_effect | Osm | -1.884 | 7.77E-04 | glio |
| Interaction_genotype_Myc | Dnaja4 | -3.794 | 7.77E-04 | glio |
| interaction_RT | Cenpb | -0.999 | 7.77E-04 | glio |
| Bora_in.vivo_noRT | Tcf4 | -1.001 | 7.77E-04 | glio |
| culture_effect | Dusp6 | -0.686 | 7.77E-05 | glio |
| Atr_in.vivo_noRT | Fkbp3 | -1.064 | 7.77E-05 | glio |
| Atr_ex.vivo_noRT | Gm12589 | 2.764 | 7.77E-05 | glio |
| Bora_in.vivo_noRT | Cdrt4 | 2.961 | 7.77E-05 | glio |

| | | | | |
|---|---|---|---|---|
| Eif2s1_in.vivo_noRT | Gjd4 | 2.143 | 7.77E-05 | glio |
| Interaction_genotype_Bora | H2-Ab1 | 4.739 | 7.77E-06 | glio |
| Interaction_genotype_Bora | Rnf165 | -5.591 | 7.77E-06 | glio |
| Interaction_genotype_Mtor | Srsf3 | 2.313 | 7.77E-06 | glio |
| Aars2_ex.vivo_noRT | Gm10482 | 3.293 | 7.77E-06 | glio |
| Bcas2_in.vivo_noRT | Gm5861 | 3.759 | 7.77E-06 | glio |
| culture_effect | Icosl | -2.825 | 7.77E-07 | glio |
| culture_effect | 1700012D14R | 2.197 | 7.77E-07 | glio |
| Atr_in.vivo_noRT | Fam71e1 | 3.111 | 7.77E-07 | glio |
| Atr_ex.vivo_noRT | Gjd4 | 3.164 | 7.77E-08 | glio |
| Chek1_ex.vivo_noRT | BB557941 | 4.443 | 7.77E-10 | glio |
| Aars2_in.vivo_noRT | Fam228a | 4.539 | 7.77E-11 | glio |
| culture_effect | Ncam2 | -3.169 | 7.77E-12 | glio |
| Atr_in.vivo_noRT | Rdh12 | 4.888 | 7.77E-14 | glio |
| Bora_in.vivo_noRT | Gm48170 | 4.936 | 7.77E-19 | glio |
| Interaction_genotype_Aars2 | C87436 | -1.826 | 7.78E-03 | glio |
| Interaction_genotype_Atr | Olfr543 | -2.505 | 7.78E-03 | glio |
| Interaction_genotype_Atr | 1700024I08Ri | -2.7 | 7.78E-03 | glio |
| Interaction_genotype_Atr | Nkx2-2os | -3.405 | 7.78E-03 | glio |
| Interaction_genotype_Atr | 4933404O12R | -3.207 | 7.78E-03 | glio |
| Interaction_genotype_Mtor | A630089N07R | -2.809 | 7.78E-03 | glio |
| Interaction_genotype_Mtor | Dmd | -2.36 | 7.78E-03 | glio |
| Interaction_genotype_Myc | Pacrg | -2.732 | 7.78E-03 | glio |
| Interaction_genotype_Myc | Gm20628 | -3.169 | 7.78E-03 | glio |
| Interaction_genotype_Myc | Gm16973 | -3.09 | 7.78E-03 | glio |
| Interaction_genotype_Myc | Rbak | -2.513 | 7.78E-03 | glio |
| Interaction_genotype_Myc | A130014A01R | -2.562 | 7.78E-03 | glio |
| Interaction_genotype_Myc | Pard3b | -2.396 | 7.78E-03 | glio |
| Interaction_genotype_Myc | Ttbk2 | -2.284 | 7.78E-03 | glio |
| Interaction_genotype_Myc | Shkbp1 | -2.343 | 7.78E-03 | glio |
| Bcas2_in.vivo_noRT | Pigv | 2.047 | 7.78E-03 | glio |
| Bora_in.vivo_noRT | B3gat2 | 2.076 | 7.78E-03 | glio |
| Bora_in.vivo_noRT | L3mbtl2 | -1.285 | 7.78E-03 | glio |
| Eif2s1_in.vivo_noRT | Tmem74b | 1.879 | 7.78E-03 | glio |
| Aars2_ex.vivo_noRT | Kcnk13 | 2.505 | 7.78E-04 | glio |
| Bora_in.vivo_noRT | D330050I16Ri | 1.966 | 7.78E-04 | glio |
| Bora_ex.vivo_noRT | Olfr90 | 2.808 | 7.78E-04 | glio |
| Hsd17b10_in.vivo_noRT | Gpr162 | 2.568 | 7.78E-04 | glio |
| Interaction_genotype_Mtor | Ppnr | -4.798 | 7.78E-05 | glio |
| Interaction_genotype_Myc | Xkr8 | -4.327 | 7.78E-05 | glio |
| Atp5c1_in.vivo_noRT | Gm15816 | 2.668 | 7.78E-05 | glio |
| Bcas2_in.vivo_noRT | Insrr | 2.56 | 7.78E-05 | glio |
| Bora_in.vivo_noRT | Etaa1os | 2.72 | 7.78E-05 | glio |
| Chek1_in.vivo_noRT | B230208H11R | 2.887 | 7.78E-05 | glio |
| Chek1_ex.vivo_noRT | Gnptg | 2.123 | 7.78E-05 | glio |
| Interaction_genotype_Myc | Rpl22l1 | 2.002 | 7.78E-06 | glio |
| Aars2_in.vivo_noRT | C1qtnf12 | 3.479 | 7.78E-06 | glio |
| Bcas2_in.vivo_noRT | A530083I20Ri | 3.799 | 7.78E-07 | glio |
| Bcas2_in.vivo_noRT | Ccl8 | 4.281 | 7.78E-08 | glio |
| Bcas2_in.vivo_noRT | Gm30124 | 3.537 | 7.78E-08 | glio |
| Hsd17b10_in.vivo_noRT | Gm26978 | 3.356 | 7.78E-09 | glio |
| Chek1_ex.vivo_noRT | 2810049E08Ri | 4.842 | 7.78E-10 | glio |

| | | | | |
|---|---|---|---|---|
| Aars2_in.vivo_noRT | Gm48772 | 4.419 | 7.78E-11 | glio |
| culture_effect | Clk1 | -1.418 | 7.78E-16 | glio |
| Bcas2_in.vivo_noRT | Strip2 | 4.543 | 7.78E-17 | glio |
| Interaction_genotype_Mtor | Zfp521 | -2.339 | 7.79E-03 | glio |
| Aars2_in.vivo_noRT | Prkce | 1.401 | 7.79E-03 | glio |
| Atp5c1_in.vivo_noRT | Ttc16 | 2.145 | 7.79E-03 | glio |
| Atp5c1_in.vivo_noRT | 4933433G15R | 2.289 | 7.79E-03 | glio |
| Chek1_in.vivo_noRT | Dph6 | -1.289 | 7.79E-03 | glio |
| Chek1_in.vivo_noRT | Aff4 | -0.937 | 7.79E-03 | glio |
| Eif2s1_ex.vivo_noRT | Cxcl5 | 2.234 | 7.79E-03 | glio |
| Aars2_ex.vivo_noRT | Btn1a1 | 2.509 | 7.79E-04 | glio |
| Bora_ex.vivo_noRT | S1pr1 | 3.537 | 7.79E-04 | glio |
| Chek1_in.vivo_noRT | Pou5f1 | 2.259 | 7.79E-04 | glio |
| Interaction_genotype_Atr | Serping1 | 4.124 | 7.79E-05 | glio |
| Aars2_ex.vivo_noRT | Stat4 | 4.16 | 7.79E-06 | glio |
| culture_effect | 1700010B08R | -2.621 | 7.79E-07 | glio |
| Bora_in.vivo_noRT | Dcpp1 | 3.609 | 7.79E-07 | glio |
| Chek1_in.vivo_noRT | Dhrs9 | 3.457 | 7.79E-08 | glio |
| Hsd17b10_in.vivo_noRT | Aqp9 | 3.616 | 7.79E-10 | glio |
| Aars2_in.vivo_noRT | 2810049E08Ri | 4.29 | 7.79E-12 | glio |
| Aars2_in.vivo_noRT | Halr1 | 5.06 | 7.79E-27 | glio |
| Aars2_in.vivo_noRT | Idi2 | 5.327 | 7.79E-27 | glio |
| Interaction_genotype_Mtor | Dnajc27 | -2.707 | 7.80E-03 | glio |
| Interaction_genotype_Mtor | Snapc4 | -2.544 | 7.80E-03 | glio |
| Interaction_genotype_Mtor | Rps11 | 0.844 | 7.80E-03 | glio |
| Interaction_genotype_Rpl3 | 2210406O10R | -3.75 | 7.80E-03 | glio |
| Aars2_ex.vivo_noRT | Gm37986 | 2.495 | 7.80E-03 | glio |
| Atr_in.vivo_noRT | Wdr24 | 1.362 | 7.80E-03 | glio |
| Bcas2_in.vivo_noRT | Ccnb1 | -1.57 | 7.80E-03 | glio |
| Bora_in.vivo_noRT | Sp1 | -1.085 | 7.80E-03 | glio |
| Cenpj_ex.vivo_noRT | Dtx1 | 2.982 | 7.80E-03 | glio |
| Cyp19a1_ex.vivo_noRT | Arhgef16 | 2.559 | 7.80E-03 | glio |
| Cyp19a1_ex.vivo_noRT | 4933400F21Ri | 2.159 | 7.80E-03 | glio |
| Cyp19a1_ex.vivo_noRT | Gm12794 | 2.159 | 7.80E-03 | glio |
| Interaction_genotype_Aars2 | Synpr | -4.597 | 7.80E-04 | glio |
| Interaction_genotype_Aars2 | Gm49864 | -4.04 | 7.80E-04 | glio |
| Interaction_genotype_Rad51 | Rab3ip | 2.154 | 7.80E-04 | glio |
| Aars2_in.vivo_noRT | Elob | -0.691 | 7.80E-04 | glio |
| Atr_in.vivo_noRT | Efna5 | 2.339 | 7.80E-04 | glio |
| Atr_in.vivo_noRT | Sdhaf4 | -1.348 | 7.80E-04 | glio |
| Interaction_genotype_Rad51 | Psme4 | 2.417 | 7.80E-05 | glio |
| Aars2_in.vivo_noRT | Rapgefl1 | 2.729 | 7.80E-05 | glio |
| Cenpj_in.vivo_noRT | C330002G04R | 2.548 | 7.80E-05 | glio |
| Cenpj_in.vivo_noRT | Slc17a6 | 2.022 | 7.80E-05 | glio |
| culture_effect | Acsl5 | -0.684 | 7.80E-06 | glio |
| Atp5c1_in.vivo_noRT | 9430041J12Ri | 2.668 | 7.80E-06 | glio |
| Atr_in.vivo_noRT | Nbeal2 | 3.244 | 7.80E-06 | glio |
| Atr_in.vivo_noRT | Dtx1 | 3.103 | 7.80E-06 | glio |
| Bora_in.vivo_noRT | Ankrd34a | 3.288 | 7.80E-06 | glio |
| Interaction_genotype_Mtor | Gm14929 | -4.099 | 7.80E-07 | glio |
| Chek1_in.vivo_noRT | Reck | 3.48 | 7.80E-07 | glio |
| Chek1_ex.vivo_noRT | Etl4 | 4.842 | 7.80E-11 | glio |

| | | | | |
|---|---|---|---|---|
| culture_effect | Gaa | -1.11 | 7.80E-12 | glio |
| Chek1_in.vivo_noRT | Necab2 | 4.78 | 7.80E-14 | glio |
| Atr_in.vivo_noRT | A930002C04R | 5.273 | 7.80E-27 | glio |
| culture_effect | Cyb5rl | 0.968 | 7.81E-03 | glio |
| Interaction_genotype_Mtor | Fbxo17 | -2.664 | 7.81E-03 | glio |
| Interaction_genotype_Mtor | Adnp2 | -2.265 | 7.81E-03 | glio |
| Interaction_genotype_Mtor | Fam8a1 | -1.916 | 7.81E-03 | glio |
| Interaction_genotype_Myc | Kctd14 | -2.761 | 7.81E-03 | glio |
| Interaction_genotype_Myc | Dhtkd1 | -3.319 | 7.81E-03 | glio |
| Interaction_genotype_Myc | Gtf2e1 | -2.408 | 7.81E-03 | glio |
| Interaction_genotype_Myc | Tpra1 | -2.253 | 7.81E-03 | glio |
| Interaction_genotype_Myc | C130036L24Ri | -2.43 | 7.81E-03 | glio |
| interaction_RT | Ccl19 | 2.3 | 7.81E-03 | glio |
| interaction_RT | Tmem259 | -0.637 | 7.81E-03 | glio |
| interaction_RT | Lima1 | -0.751 | 7.81E-03 | glio |
| Atp5c1_in.vivo_noRT | 3930402G23R | 2.281 | 7.81E-03 | glio |
| Atr_in.vivo_noRT | Pgs1 | -1.237 | 7.81E-03 | glio |
| Atr_in.vivo_noRT | Cdk12 | -0.927 | 7.81E-03 | glio |
| Atr_in.vivo_noRT | Pfdn4 | -0.772 | 7.81E-03 | glio |
| Atr_in.vivo_noRT | Nedd8 | -0.611 | 7.81E-03 | glio |
| Bcas2_in.vivo_noRT | 2810455O05R | 2.031 | 7.81E-03 | glio |
| Bcas2_in.vivo_noRT | Zfp87 | 1.673 | 7.81E-03 | glio |
| Bcas2_in.vivo_noRT | Txndc17 | -0.623 | 7.81E-03 | glio |
| Bora_in.vivo_noRT | A130014A01R | 1.767 | 7.81E-03 | glio |
| Brca2_ex.vivo_noRT | Myoz1 | 2.825 | 7.81E-03 | glio |
| Cenpj_in.vivo_noRT | Gm2115 | 2.022 | 7.81E-03 | glio |
| Chek1_in.vivo_noRT | Ahi1 | -1.248 | 7.81E-03 | glio |
| Fanca_in.vivo_noRT | Klhl32 | -2.703 | 7.81E-03 | glio |
| Interaction_genotype_Mtor | Nsl1 | -3.333 | 7.81E-04 | glio |
| Interaction_genotype_Mtor | Rbm19 | -2.908 | 7.81E-04 | glio |
| Aars2_ex.vivo_noRT | Gm43701 | 2.31 | 7.81E-04 | glio |
| Aars2_ex.vivo_noRT | Ctss | 2.899 | 7.81E-04 | glio |
| Bora_in.vivo_noRT | Bms1 | -1.44 | 7.81E-04 | glio |
| culture_effect | Gm36839 | 2.125 | 7.81E-05 | glio |
| culture_effect | Celf4 | 0.897 | 7.81E-05 | glio |
| Interaction_genotype_Mtor | Grin2d | -4.473 | 7.81E-05 | glio |
| Interaction_genotype_Mtor | Tns4 | -4.246 | 7.81E-05 | glio |
| Interaction_genotype_Mtor | Padi2 | -4.856 | 7.81E-05 | glio |
| Interaction_genotype_Mtor | Mrps24 | 1.691 | 7.81E-05 | glio |
| Bcas2_in.vivo_noRT | Rab40b | 3.07 | 7.81E-06 | glio |
| Chek1_ex.vivo_noRT | Fabp4 | 4.055 | 7.81E-06 | glio |
| culture_effect | 2610035D17R | 1.198 | 7.81E-08 | glio |
| Bcas2_in.vivo_noRT | Pcdhga2 | 3.537 | 7.81E-08 | glio |
| Chek1_in.vivo_noRT | Tmem44 | 3.785 | 7.81E-08 | glio |
| Chek1_ex.vivo_noRT | Ggt1 | 4.842 | 7.81E-09 | glio |
| culture_effect | Ctnnal1 | 1.927 | 7.81E-12 | glio |
| Bcas2_in.vivo_noRT | Plk5 | 4.151 | 7.81E-12 | glio |
| Bcas2_in.vivo_noRT | Upb1 | 4.544 | 7.81E-12 | glio |
| culture_effect | Tmem80 | 1.788 | 7.81E-18 | glio |
| Interaction_genotype_Mtor | Egln2 | -1.618 | 7.82E-03 | glio |
| Interaction_genotype_Mtor | Aurkaip1 | 1.217 | 7.82E-03 | glio |
| Interaction_genotype_Pars2 | Gm10767 | -3.718 | 7.82E-03 | glio |

| | | | | |
|---|---|---|---|---|
| interaction_RT | Sept10 | 0.943 | 7.82E-03 | glio |
| Atr_in.vivo_noRT | Cbx2 | 1.74 | 7.82E-03 | glio |
| Cdkn2a_in.vivo_noRT | Gm4340 | -1.675 | 7.82E-03 | glio |
| Cdkn2a_in.vivo_noRT | Gm16239 | -1.675 | 7.82E-03 | glio |
| Cdkn2a_in.vivo_noRT | Gm13032 | -2.061 | 7.82E-03 | glio |
| Cdkn2a_in.vivo_noRT | Hoxc8 | -1.675 | 7.82E-03 | glio |
| Cdkn2a_in.vivo_noRT | Gm10324 | -1.941 | 7.82E-03 | glio |
| Cdkn2a_in.vivo_noRT | Igsf3 | 1.048 | 7.82E-03 | glio |
| Chek1_in.vivo_noRT | Zfp946 | 1.854 | 7.82E-03 | glio |
| Chek1_ex.vivo_noRT | Mmp23 | 2.469 | 7.82E-03 | glio |
| Chek1_ex.vivo_noRT | Aatk | 2.681 | 7.82E-03 | glio |
| culture_effect | Prex2 | -2.371 | 7.82E-04 | glio |
| culture_effect | Sobp | -1.728 | 7.82E-04 | glio |
| Interaction_genotype_Chek1 | Krt18 | 6.261 | 7.82E-04 | glio |
| Interaction_genotype_Myc | Ccr5 | -3.492 | 7.82E-04 | glio |
| interaction_RT | Bcl3 | 2.486 | 7.82E-04 | glio |
| Chek1_in.vivo_noRT | Rarres2 | 2.366 | 7.82E-04 | glio |
| Chek1_in.vivo_noRT | 1700010I14Ri | 2.374 | 7.82E-04 | glio |
| Chek1_ex.vivo_noRT | Rbp2 | 3.469 | 7.82E-04 | glio |
| Eif2s1_in.vivo_noRT | Gm21319 | 2.143 | 7.82E-04 | glio |
| Hsd17b10_in.vivo_noRT | 1700029J07Ri | 2.59 | 7.82E-04 | glio |
| Aars2_ex.vivo_noRT | Cidea | 3.293 | 7.82E-05 | glio |
| Atr_in.vivo_noRT | Tmem60 | -1.553 | 7.82E-05 | glio |
| Chek1_in.vivo_noRT | Ddx27 | -1.449 | 7.82E-05 | glio |
| Interaction_genotype_Mtor | Gm45669 | -4.624 | 7.82E-06 | glio |
| Hsd17b10_in.vivo_noRT | Tmem190 | 3.616 | 7.82E-06 | glio |
| Hsd17b10_in.vivo_noRT | A330015K06R | 2.959 | 7.82E-06 | glio |
| Aars2_ex.vivo_noRT | Gm128 | 3.676 | 7.82E-07 | glio |
| Aars2_in.vivo_noRT | 2410022M11F | 3.8 | 7.82E-08 | glio |
| culture_effect | Ripk2 | -1.503 | 7.82E-12 | glio |
| Aars2_in.vivo_noRT | Oxt | 4.685 | 7.82E-12 | glio |
| Atr_in.vivo_noRT | Gm50295 | 4.741 | 7.82E-12 | glio |
| Aars2_in.vivo_noRT | 1700017M07F | 5.327 | 7.82E-19 | glio |
| Aars2_in.vivo_noRT | Slc16a4 | 5.063 | 7.82E-19 | glio |
| Atr_in.vivo_noRT | Folr1 | 5.533 | 7.82E-20 | glio |
| Interaction_genotype_Atr | C8g | -3.319 | 7.83E-03 | glio |
| Interaction_genotype_Bcas2 | Trim17 | -2.464 | 7.83E-03 | glio |
| Interaction_genotype_Bcas2 | Gm49860 | -3.047 | 7.83E-03 | glio |
| Interaction_genotype_Bcas2 | Ccl9 | -2.458 | 7.83E-03 | glio |
| Interaction_genotype_Bcas2 | E130215H24R | -3.321 | 7.83E-03 | glio |
| Interaction_genotype_Bcas2 | Ankrd34a | -3.607 | 7.83E-03 | glio |
| Interaction_genotype_Mtor | 2610318N02R | -1.911 | 7.83E-03 | glio |
| Interaction_genotype_Myc | Wars2 | -2.611 | 7.83E-03 | glio |
| Aars2_in.vivo_noRT | Cdk16 | -1.011 | 7.83E-03 | glio |
| Aars2_in.vivo_noRT | Gbp2 | -1.071 | 7.83E-03 | glio |
| Aars2_ex.vivo_noRT | Ppm1d | -2.033 | 7.83E-03 | glio |
| Atr_in.vivo_noRT | Acta2 | 2.198 | 7.83E-03 | glio |
| Atr_in.vivo_noRT | Gm45397 | 1.679 | 7.83E-03 | glio |
| Bard1_in.vivo_noRT | Chia1 | 1.98 | 7.83E-03 | glio |
| Bcas2_in.vivo_noRT | Cysltr2 | 1.803 | 7.83E-03 | glio |
| Bcas2_in.vivo_noRT | Atp5a1 | -0.54 | 7.83E-03 | glio |
| Bora_in.vivo_noRT | Slc8a3 | 1.957 | 7.83E-03 | glio |

| | | | | |
|---|---|---|---|---|
| Bora_in.vivo_noRT | Polh | 1.446 | 7.83E-03 | glio |
| Chek1_in.vivo_noRT | Phgdh | -0.868 | 7.83E-03 | glio |
| culture_effect | Serpinb8 | 1.256 | 7.83E-04 | glio |
| Interaction_genotype_Atr | Nlrx1 | -4.411 | 7.83E-04 | glio |
| Interaction_genotype_Atr | C130040N14R | -4.232 | 7.83E-04 | glio |
| Atr_in.vivo_noRT | Hnrnpa0 | -1.199 | 7.83E-04 | glio |
| Bcas2_in.vivo_noRT | Gm14023 | 2.529 | 7.83E-04 | glio |
| Bcas2_in.vivo_noRT | Pfdn2 | -0.676 | 7.83E-04 | glio |
| Bora_in.vivo_noRT | Gm50203 | 2.304 | 7.83E-04 | glio |
| Chek1_in.vivo_noRT | Hist1h3g | 2.581 | 7.83E-04 | glio |
| Eif2s1_in.vivo_noRT | Gm10447 | 2.143 | 7.83E-04 | glio |
| Interaction_genotype_Myc | Gm30692 | -3.122 | 7.83E-05 | glio |
| Interaction_genotype_Myc | Adprm | -3.53 | 7.83E-05 | glio |
| Aars2_in.vivo_noRT | Gm17501 | 2.904 | 7.83E-05 | glio |
| Aars2_in.vivo_noRT | Yipf4 | -1.103 | 7.83E-05 | glio |
| Bora_in.vivo_noRT | Adam4 | 2.595 | 7.83E-05 | glio |
| Hsd17b10_in.vivo_noRT | St8sia1 | 3.356 | 7.83E-06 | glio |
| Atp5c1_in.vivo_noRT | Ank1 | 2.668 | 7.83E-07 | glio |
| Chek1_in.vivo_noRT | Gm35551 | 3.228 | 7.83E-07 | glio |
| Atr_in.vivo_noRT | Ift27 | -2.449 | 7.83E-08 | glio |
| Bora_in.vivo_noRT | Gm28578 | 4.676 | 7.83E-11 | glio |
| culture_effect | Arpp19 | 0.877 | 7.83E-13 | glio |
| Aars2_in.vivo_noRT | 5430402O13R | 5.327 | 7.83E-14 | glio |
| Chek1_in.vivo_noRT | Klk13 | 4.511 | 7.83E-19 | glio |
| culture_effect | Memo1 | 0.42 | 7.84E-03 | glio |
| Interaction_genotype_Atr | Sipa1l2 | -3.241 | 7.84E-03 | glio |
| Interaction_genotype_Atr | Sorl1 | -3.577 | 7.84E-03 | glio |
| Interaction_genotype_Atr | Neu1 | 1.757 | 7.84E-03 | glio |
| Interaction_genotype_Bcas2 | Gm32250 | -2.656 | 7.84E-03 | glio |
| Interaction_genotype_Bcas2 | 9430041J12Ri | -2.859 | 7.84E-03 | glio |
| Interaction_genotype_Ifnar1 | BC061195 | 2.734 | 7.84E-03 | glio |
| Interaction_genotype_Mtor | Tcim | 2.242 | 7.84E-03 | glio |
| Interaction_genotype_Uqcrb | B3gat3 | 1.634 | 7.84E-03 | glio |
| Interaction_genotype_Uqcrb | Sprr1a | -4.395 | 7.84E-03 | glio |
| Interaction_genotype_Uqcrb | 1700015F17Ri | -4.335 | 7.84E-03 | glio |
| Aars2_in.vivo_noRT | Frg2f1 | 1.671 | 7.84E-03 | glio |
| Bcas2_in.vivo_noRT | Ang | 2.032 | 7.84E-03 | glio |
| Interaction_genotype_Aars2 | Gm15728 | -4.39 | 7.84E-04 | glio |
| Interaction_genotype_Bora | A330084C13R | -4.635 | 7.84E-04 | glio |
| Interaction_genotype_Bora | Esrp2 | -4.191 | 7.84E-04 | glio |
| Interaction_genotype_Myc | Gm33466 | -2.729 | 7.84E-04 | glio |
| Interaction_genotype_Myc | Gm10638 | -3.482 | 7.84E-04 | glio |
| Interaction_genotype_Myc | Cdk18 | -3.142 | 7.84E-04 | glio |
| Interaction_genotype_Myc | Mtmr4 | -2.965 | 7.84E-04 | glio |
| Interaction_genotype_Myc | Map2k3os | -3.017 | 7.84E-04 | glio |
| Aars2_in.vivo_noRT | Psmc3 | -1.119 | 7.84E-04 | glio |
| Atr_in.vivo_noRT | P4ha3 | 2.298 | 7.84E-04 | glio |
| Atr_in.vivo_noRT | Calhm2 | 1.968 | 7.84E-04 | glio |
| Bora_in.vivo_noRT | Lsm6 | -1.025 | 7.84E-04 | glio |
| Chek1_in.vivo_noRT | Gm13203 | 2.544 | 7.84E-04 | glio |
| Chek1_ex.vivo_noRT | Mogat1 | 3.258 | 7.84E-04 | glio |
| Hsd17b10_in.vivo_noRT | Cd59b | 2.571 | 7.84E-04 | glio |

| | | | | |
|---|---|---|---|---|
| culture_effect | Hsd17b11 | -1.496 | 7.84E-05 | glio |
| Atr_in.vivo_noRT | Cysltr2 | 2.529 | 7.84E-05 | glio |
| Bcas2_ex.vivo_noRT | Gm15941 | 3.52 | 7.84E-05 | glio |
| Bora_in.vivo_noRT | Tnp2 | 2.874 | 7.84E-05 | glio |
| Bora_in.vivo_noRT | Saysd1 | -1.901 | 7.84E-05 | glio |
| Interaction_genotype_Mtor | Ccser1 | -4.492 | 7.84E-06 | glio |
| Bcas2_in.vivo_noRT | Cyc1 | -1.138 | 7.84E-06 | glio |
| culture_effect | Efhb | -2.961 | 7.84E-07 | glio |
| Atr_in.vivo_noRT | Asap3 | 3.328 | 7.84E-07 | glio |
| culture_effect | Sgcb | -1.344 | 7.84E-08 | glio |
| Chek1_ex.vivo_noRT | Fam131a | 4.443 | 7.84E-08 | glio |
| Chek1_ex.vivo_noRT | Gm10447 | 3.568 | 7.84E-08 | glio |
| Atr_in.vivo_noRT | Gm49494 | 4.188 | 7.84E-09 | glio |
| Bora_in.vivo_noRT | Amhr2 | 4.584 | 7.84E-09 | glio |
| Bora_in.vivo_noRT | Gm7102 | 4.676 | 7.84E-11 | glio |
| Bora_in.vivo_noRT | Cgn | 4.41 | 7.84E-11 | glio |
| Chek1_ex.vivo_noRT | Boc | 4.449 | 7.84E-13 | glio |
| Atr_in.vivo_noRT | Kpna7 | 5.269 | 7.84E-17 | glio |
| culture_effect | Gm43566 | 0.723 | 7.85E-03 | glio |
| culture_effect | Trip4 | 0.531 | 7.85E-03 | glio |
| culture_effect | Tmem263 | -0.407 | 7.85E-03 | glio |
| Interaction_genotype_Atr | Pxt1 | -3.405 | 7.85E-03 | glio |
| Interaction_genotype_Atr | Abhd15 | -3.568 | 7.85E-03 | glio |
| Interaction_genotype_Atr | Gm26519 | -3.167 | 7.85E-03 | glio |
| Interaction_genotype_Mtor | Hist1h2bn | -3.092 | 7.85E-03 | glio |
| Interaction_genotype_Mtor | Zfp821 | -2.336 | 7.85E-03 | glio |
| Interaction_genotype_Myc | Zfp352 | -2.729 | 7.85E-03 | glio |
| Interaction_genotype_Myc | Zfp141 | -2.568 | 7.85E-03 | glio |
| Interaction_genotype_Myc | Fkbp3 | 1.198 | 7.85E-03 | glio |
| interaction_RT | Irf2bpl | 1.021 | 7.85E-03 | glio |
| interaction_RT | Eif2ak2 | -0.689 | 7.85E-03 | glio |
| Aars2_in.vivo_noRT | 4932422M17F | 1.764 | 7.85E-03 | glio |
| Bora_in.vivo_noRT | Zfp983 | 1.705 | 7.85E-03 | glio |
| Cenpj_in.vivo_noRT | Gm44658 | 2.017 | 7.85E-03 | glio |
| Chek1_in.vivo_noRT | Rmst | 2.064 | 7.85E-03 | glio |
| Chek1_in.vivo_noRT | Nabp1 | 1.903 | 7.85E-03 | glio |
| Interaction_genotype_Myc | BC026762 | -3.637 | 7.85E-04 | glio |
| interaction_RT | Itm2a | 1.786 | 7.85E-04 | glio |
| interaction_RT | Srsf6 | -0.688 | 7.85E-04 | glio |
| Atr_ex.vivo_noRT | Peli3 | 3.077 | 7.85E-04 | glio |
| Atr_ex.vivo_noRT | Chit1 | 3.633 | 7.85E-04 | glio |
| Bcas2_in.vivo_noRT | Hr | 2.188 | 7.85E-04 | glio |
| Bcas2_in.vivo_noRT | Hoxaas3 | 2.03 | 7.85E-04 | glio |
| Bcas2_in.vivo_noRT | Eif4e2 | -0.987 | 7.85E-04 | glio |
| culture_effect | Bcl6 | -2.227 | 7.85E-05 | glio |
| Aars2_in.vivo_noRT | Neto1 | -1.773 | 7.85E-05 | glio |
| Bcas2_ex.vivo_noRT | Gm26542 | 3.715 | 7.85E-05 | glio |
| Chek1_ex.vivo_noRT | Abtb1 | 2.861 | 7.85E-05 | glio |
| Atr_in.vivo_noRT | Camkv | 3.299 | 7.85E-06 | glio |
| culture_effect | Psmc5 | -0.574 | 7.85E-07 | glio |
| Atr_in.vivo_noRT | Map1b | -1.948 | 7.85E-08 | glio |
| Atr_ex.vivo_noRT | 2310003N18R | 4.521 | 7.85E-10 | glio |

| | | | | |
|---|---|---|---|---|
| Bora_in.vivo_noRT | D030068K23R | 4.545 | 7.85E-11 | glio |
| Aars2_in.vivo_noRT | Grm1 | 4.223 | 7.85E-12 | glio |
| Atr_in.vivo_noRT | Gm15080 | 5.041 | 7.85E-12 | glio |
| Bora_in.vivo_noRT | Fam25c | 4.404 | 7.85E-16 | glio |
| Bora_in.vivo_noRT | Fam166a | 4.672 | 7.85E-18 | glio |
| Bora_in.vivo_noRT | Ccdc125 | 4.936 | 7.85E-18 | glio |
| culture_effect | BC037039 | 1.008 | 7.86E-03 | glio |
| Interaction_genotype_Aars2 | Myo1h | -3.396 | 7.86E-03 | glio |
| Interaction_genotype_Aars2 | 5033430I15Ri | -3.042 | 7.86E-03 | glio |
| Interaction_genotype_Aars2 | Atp2a2 | 1.509 | 7.86E-03 | glio |
| Interaction_genotype_Aars2 | Mcrip1 | 1.028 | 7.86E-03 | glio |
| Interaction_genotype_Aars2 | Fkbp3 | 0.987 | 7.86E-03 | glio |
| Interaction_genotype_Mtor | Btbd17 | -3.937 | 7.86E-03 | glio |
| Interaction_genotype_Mtor | Wdr74 | 1.388 | 7.86E-03 | glio |
| Interaction_genotype_Myc | Dtwd2 | -2.615 | 7.86E-03 | glio |
| Aars2_in.vivo_noRT | Ppp1ca | -0.633 | 7.86E-03 | glio |
| Aars2_ex.vivo_noRT | Gm47428 | 2.613 | 7.86E-03 | glio |
| Atr_in.vivo_noRT | Gpsm1 | -1.411 | 7.86E-03 | glio |
| Bcas2_in.vivo_noRT | Rdm1 | -1.233 | 7.86E-03 | glio |
| Chek1_in.vivo_noRT | Fzd7 | 1.893 | 7.86E-03 | glio |
| Chek1_ex.vivo_noRT | A930001C03R | 2.347 | 7.86E-03 | glio |
| Hsd17b10_in.vivo_noRT | Gm36495 | 2.322 | 7.86E-03 | glio |
| Hsd17b10_in.vivo_noRT | Gm27010 | 2.015 | 7.86E-03 | glio |
| culture_effect | Ephx1 | 1.864 | 7.86E-04 | glio |
| Aars2_in.vivo_noRT | Ndufa7 | -0.678 | 7.86E-04 | glio |
| Bora_ex.vivo_noRT | C330024C12R | 2.808 | 7.86E-04 | glio |
| Eif2s1_in.vivo_noRT | Grin3b | 2.682 | 7.86E-04 | glio |
| Aars2_in.vivo_noRT | Catsper2 | 2.736 | 7.86E-05 | glio |
| Hsd17b10_in.vivo_noRT | Chd4 | -0.872 | 7.86E-05 | glio |
| Interaction_genotype_Mtor | 4930417H01R | -5.594 | 7.86E-06 | glio |
| Aars2_in.vivo_noRT | Gm26982 | 3.106 | 7.86E-06 | glio |
| Atr_in.vivo_noRT | Ucn2 | 3.04 | 7.86E-06 | glio |
| Bora_in.vivo_noRT | Gm10602 | 3.529 | 7.86E-06 | glio |
| Aars2_ex.vivo_noRT | Slc14a2 | 3.293 | 7.86E-08 | glio |
| Aars2_in.vivo_noRT | Stab1 | 4.193 | 7.86E-09 | glio |
| Chek1_in.vivo_noRT | Sptbn4 | 4.025 | 7.86E-09 | glio |
| Chek1_in.vivo_noRT | Kcnk7 | 3.995 | 7.86E-12 | glio |
| Bcas2_in.vivo_noRT | 4930555A03R | 4.599 | 7.86E-13 | glio |
| Chek1_in.vivo_noRT | Gm11627 | 5.044 | 7.86E-15 | glio |
| Chek1_in.vivo_noRT | Obscn | 5.044 | 7.86E-16 | glio |
| culture_effect | Gm3448 | 2.838 | 7.86E-18 | glio |
| Atr_in.vivo_noRT | Gm5934 | 5.27 | 7.86E-23 | glio |
| culture_effect | Gm13594 | 1.657 | 7.87E-03 | glio |
| culture_effect | Pcgf6 | -0.426 | 7.87E-03 | glio |
| Interaction_genotype_Myc | Rnf208 | -3.231 | 7.87E-03 | glio |
| Chek1_in.vivo_noRT | St3gal6 | 1.903 | 7.87E-03 | glio |
| Chek1_in.vivo_noRT | Rps26 | -0.548 | 7.87E-03 | glio |
| Chek1_in.vivo_noRT | Rplp2 | -0.497 | 7.87E-03 | glio |
| Cyp19a1_ex.vivo_noRT | Krtcap3 | 2.351 | 7.87E-03 | glio |
| Interaction_genotype_Mtor | Tma7 | 1.467 | 7.87E-04 | glio |
| interaction_RT | Wdr18 | -0.797 | 7.87E-04 | glio |
| Atr_in.vivo_noRT | Fam160b2 | 1.975 | 7.87E-04 | glio |

| | | | | |
|---|---|---|---|---|
| Bora_in.vivo_noRT | Gm42701 | 2.42 | 7.87E-04 | glio |
| Chek1_in.vivo_noRT | BC065397 | 2.698 | 7.87E-04 | glio |
| Eif2s1_in.vivo_noRT | Speer4e | 2.143 | 7.87E-04 | glio |
| Eif2s1_in.vivo_noRT | C430002N11R | 2.143 | 7.87E-04 | glio |
| Eif2s1_in.vivo_noRT | A330049N07R | 1.877 | 7.87E-04 | glio |
| culture_effect | Vill | -1.616 | 7.87E-05 | glio |
| Interaction_genotype_Myc | Slc45a3 | -4.519 | 7.87E-05 | glio |
| Interaction_genotype_Myc | Timm10 | 1.902 | 7.87E-05 | glio |
| Interaction_genotype_Myc | Actb | 1.684 | 7.87E-05 | glio |
| Aars2_in.vivo_noRT | Oxsm | 2.895 | 7.87E-05 | glio |
| Bcas2_in.vivo_noRT | Mettl5os | 2.642 | 7.87E-05 | glio |
| Bcas2_ex.vivo_noRT | Adam5 | 2.742 | 7.87E-05 | glio |
| Bcas2_ex.vivo_noRT | 4933400F21Ri | 2.742 | 7.87E-05 | glio |
| Chek1_ex.vivo_noRT | Fetub | 2.817 | 7.87E-05 | glio |
| Atr_in.vivo_noRT | March11 | 3.26 | 7.87E-06 | glio |
| Bcas2_in.vivo_noRT | Hspa12a | 3.507 | 7.87E-07 | glio |
| Bora_in.vivo_noRT | Cnksr1 | 3.367 | 7.87E-07 | glio |
| Chek1_ex.vivo_noRT | Gm10401 | 3.86 | 7.87E-07 | glio |
| Bora_in.vivo_noRT | Cyp26b1 | 4.011 | 7.87E-08 | glio |
| Bora_in.vivo_noRT | Slc6a11 | 3.504 | 7.87E-10 | glio |
| Bora_in.vivo_noRT | Ankrd33b | 4.42 | 7.87E-10 | glio |
| Bora_in.vivo_noRT | 4933406C10R | 3.9 | 7.87E-10 | glio |
| Aars2_in.vivo_noRT | Elobl | 5.327 | 7.87E-13 | glio |
| Interaction_genotype_Atr | Lbhd2 | -3.606 | 7.88E-03 | glio |
| Interaction_genotype_Bcas2 | Ube4bos1 | -2.987 | 7.88E-03 | glio |
| Chek1_ex.vivo_noRT | Gm11696 | 2.47 | 7.88E-03 | glio |
| Chek1_ex.vivo_noRT | Rbm3 | -0.635 | 7.88E-03 | glio |
| Interaction_genotype_Mtor | Sh2d3c | -4.225 | 7.88E-04 | glio |
| Interaction_genotype_Mtor | Rhbdl1 | -3.948 | 7.88E-04 | glio |
| Interaction_genotype_Pars2 | Prl7c1 | -3.364 | 7.88E-04 | glio |
| Interaction_genotype_Pars2 | Rapgef3os2 | -3.454 | 7.88E-04 | glio |
| Abtb1_ex.vivo_noRT | Trmt61b | 2.95 | 7.88E-04 | glio |
| Atr_ex.vivo_noRT | Serpinb9 | 3.153 | 7.88E-04 | glio |
| Bcas2_in.vivo_noRT | Phkg1 | 3.377 | 7.88E-04 | glio |
| Bcas2_ex.vivo_noRT | Gm34939 | 2.742 | 7.88E-04 | glio |
| Bcas2_ex.vivo_noRT | Nrn1l | 3.125 | 7.88E-04 | glio |
| Bora_in.vivo_noRT | Gm43066 | 2.284 | 7.88E-04 | glio |
| Interaction_genotype_Aars2 | Gm11549 | -4.873 | 7.88E-05 | glio |
| Interaction_genotype_Myc | Etfdh | -3.103 | 7.88E-05 | glio |
| Bcas2_ex.vivo_noRT | Gm12794 | 2.742 | 7.88E-05 | glio |
| Chek1_in.vivo_noRT | Bpifc | 3.024 | 7.88E-05 | glio |
| Chek1_ex.vivo_noRT | Rftn2 | 3.072 | 7.88E-05 | glio |
| Atr_in.vivo_noRT | Gng8 | 3.028 | 7.88E-06 | glio |
| Bcas2_in.vivo_noRT | 0610009L18Ri | 3.296 | 7.88E-06 | glio |
| Atr_in.vivo_noRT | Ppp1r26 | 3.625 | 7.88E-07 | glio |
| Bora_in.vivo_noRT | Tbkbp1 | 3.643 | 7.88E-07 | glio |
| Bora_in.vivo_noRT | BC037704 | 3.412 | 7.88E-07 | glio |
| culture_effect | 1700086O06R | 1.518 | 7.88E-08 | glio |
| culture_effect | Sil1 | -1.258 | 7.88E-08 | glio |
| Atr_in.vivo_noRT | Gm48357 | 3.218 | 7.88E-08 | glio |
| culture_effect | A430106G13R | 3.292 | 7.88E-10 | glio |
| Bora_in.vivo_noRT | Cfb | 3.632 | 7.88E-11 | glio |

| | | | | |
|---|---|---|---|---|
| Bora_in.vivo_noRT | Gm45669 | 4.673 | 7.88E-15 | glio |
| Aars2_in.vivo_noRT | Iqch | 5.063 | 7.88E-18 | glio |
| Aars2_in.vivo_noRT | Serpinb2 | 5.327 | 7.88E-18 | glio |
| culture_effect | Cnmd | -3.986 | 7.88E-25 | glio |
| culture_effect | Nrcam | -0.642 | 7.89E-03 | glio |
| Interaction_genotype_Bora | Gtsf1 | -2.898 | 7.89E-03 | glio |
| Interaction_genotype_Bora | D030068K23R | -3.601 | 7.89E-03 | glio |
| Interaction_genotype_Bora | Eif3a | 0.735 | 7.89E-03 | glio |
| Interaction_genotype_Mtor | Gnaz | -3.387 | 7.89E-03 | glio |
| Interaction_genotype_Mtor | Lamp2 | 1.22 | 7.89E-03 | glio |
| Interaction_genotype_Myc | Plcd4 | -2.794 | 7.89E-03 | glio |
| Interaction_genotype_Myc | Pcdhgb6 | -2.649 | 7.89E-03 | glio |
| Interaction_genotype_Myc | Mkrn3 | -2.899 | 7.89E-03 | glio |
| Interaction_genotype_Myc | Aldh16a1 | -2.298 | 7.89E-03 | glio |
| Interaction_genotype_Myc | Copz1 | 1.253 | 7.89E-03 | glio |
| Atr_in.vivo_noRT | Ercc8 | 1.441 | 7.89E-03 | glio |
| Atr_in.vivo_noRT | Thap12 | -1.172 | 7.89E-03 | glio |
| Bcas2_in.vivo_noRT | Zfp981 | 1.926 | 7.89E-03 | glio |
| Bora_in.vivo_noRT | Ap1g1 | -1.195 | 7.89E-03 | glio |
| Bora_ex.vivo_noRT | Col6a3 | 2.408 | 7.89E-03 | glio |
| Aars2_in.vivo_noRT | 9530068E07Ri | -1.22 | 7.89E-04 | glio |
| Interaction_genotype_Nat2 | Zfp352 | 4.564 | 7.89E-05 | glio |
| culture_effect | Tpsab1 | -2.884 | 7.89E-07 | glio |
| Atr_in.vivo_noRT | Gm12089 | 3.833 | 7.89E-08 | glio |
| Chek1_ex.vivo_noRT | Acoxl | 4.209 | 7.89E-08 | glio |
| Chek1_in.vivo_noRT | 1700047K16R | 4.386 | 7.89E-10 | glio |
| Hsd17b10_in.vivo_noRT | Gm44899 | 3.616 | 7.89E-11 | glio |
| Bcas2_in.vivo_noRT | B3gnt4 | 4.544 | 7.89E-14 | glio |
| Chek1_in.vivo_noRT | Gm33051 | 4.391 | 7.89E-16 | glio |
| culture_effect | Whamm | -0.636 | 7.90E-03 | glio |
| Interaction_genotype_Atr | Gm26930 | -3.583 | 7.90E-03 | glio |
| Interaction_genotype_Atr | Micall2 | -3.305 | 7.90E-03 | glio |
| Interaction_genotype_Bcas2 | Krt18 | 4.921 | 7.90E-03 | glio |
| Interaction_genotype_Bora | Eng | -3.816 | 7.90E-03 | glio |
| Interaction_genotype_Chek1 | Gtf2f1 | 1.992 | 7.90E-03 | glio |
| Interaction_genotype_Ifnar1 | Idi2 | 2.734 | 7.90E-03 | glio |
| Interaction_genotype_Mtor | Tff2 | -3.046 | 7.90E-03 | glio |
| Interaction_genotype_Mtor | Vamp8 | -2.675 | 7.90E-03 | glio |
| Interaction_genotype_Mtor | Gm12059 | -2.988 | 7.90E-03 | glio |
| Interaction_genotype_Mtor | Dtymk | 1.398 | 7.90E-03 | glio |
| Interaction_genotype_Mtor | Ndfip1 | 1.201 | 7.90E-03 | glio |
| Interaction_genotype_Myc | 1110038B12R | 1.167 | 7.90E-03 | glio |
| Bcas2_in.vivo_noRT | Gh | 2.222 | 7.90E-03 | glio |
| Bcas2_ex.vivo_noRT | Tcp11x2 | 2.349 | 7.90E-03 | glio |
| Bora_in.vivo_noRT | Inca1 | 1.797 | 7.90E-03 | glio |
| Chek1_ex.vivo_noRT | Ablim3 | 2.53 | 7.90E-03 | glio |
| Eif2s1_in.vivo_noRT | Gm5129 | 2.397 | 7.90E-03 | glio |
| Hsd17b10_in.vivo_noRT | Slc14a1 | 1.731 | 7.90E-03 | glio |
| Interaction_genotype_Atr | Gatad2a | 1.715 | 7.90E-04 | glio |
| Interaction_genotype_Myc | Ppp1r13l | -3.852 | 7.90E-04 | glio |
| Interaction_genotype_Myc | Slc18b1 | -3.046 | 7.90E-04 | glio |
| Interaction_genotype_Myc | Arpc1b | 1.372 | 7.90E-04 | glio |

| | | | | |
|---|---|---|---|---|
| Atp5c1_in.vivo_noRT | Pla2g2d | 2.405 | 7.90E-04 | glio |
| Atp5c1_in.vivo_noRT | Pls1 | 2.488 | 7.90E-04 | glio |
| Atp5c1_in.vivo_noRT | Fbn2 | -1.635 | 7.90E-04 | glio |
| Chek1_in.vivo_noRT | Tcim | -1.569 | 7.90E-04 | glio |
| Interaction_genotype_Myc | Gm10602 | -4.989 | 7.90E-05 | glio |
| Aars2_in.vivo_noRT | Acad10 | 2.821 | 7.90E-05 | glio |
| Aars2_in.vivo_noRT | Gm9844 | 2.675 | 7.90E-05 | glio |
| Bora_ex.vivo_noRT | Etl4 | 3.201 | 7.90E-05 | glio |
| culture_effect | Gm19705 | 1.201 | 7.90E-06 | glio |
| Aars2_in.vivo_noRT | Psma3 | -0.929 | 7.90E-06 | glio |
| culture_effect | Ripor2 | -3.107 | 7.90E-07 | glio |
| Bcas2_in.vivo_noRT | A930001C03R | 4.154 | 7.90E-10 | glio |
| Bcas2_in.vivo_noRT | Zg16 | 4.154 | 7.90E-10 | glio |
| culture_effect | Igtp | -3.15 | 7.90E-59 | glio |
| Interaction_genotype_Aars2 | Klrk1 | 2.71 | 7.91E-03 | glio |
| Interaction_genotype_Aars2 | Nanos3 | -2.804 | 7.91E-03 | glio |
| Interaction_genotype_Aars2 | 5033406O09R | 1.714 | 7.91E-03 | glio |
| Interaction_genotype_Bora | Elp2 | 2.035 | 7.91E-03 | glio |
| Interaction_genotype_Mtor | Grb7 | -3.291 | 7.91E-03 | glio |
| Interaction_genotype_Mtor | Gprc5b | -3.209 | 7.91E-03 | glio |
| Interaction_genotype_Mtor | Psmd12 | 1.211 | 7.91E-03 | glio |
| Interaction_genotype_Myc | Lrrc8e | -2.724 | 7.91E-03 | glio |
| Interaction_genotype_Myc | Tmem50b | -2.243 | 7.91E-03 | glio |
| Interaction_genotype_Pars2 | Ifi207 | -3.974 | 7.91E-03 | glio |
| Interaction_genotype_Rad51 | Derl2 | 1.533 | 7.91E-03 | glio |
| Interaction_genotype_Rpl3 | Cog8 | -2.926 | 7.91E-03 | glio |
| Interaction_genotype_Terf1 | Vps33b | -3.276 | 7.91E-03 | glio |
| Interaction_genotype_Terf1 | Apoh | -3.967 | 7.91E-03 | glio |
| Interaction_genotype_Terf1 | Gm41318 | -3.94 | 7.91E-03 | glio |
| Interaction_genotype_Terf1 | Ube2g2 | 1.655 | 7.91E-03 | glio |
| Aars2_in.vivo_noRT | Frs3 | 1.899 | 7.91E-03 | glio |
| Aars2_in.vivo_noRT | Nat14 | 1.701 | 7.91E-03 | glio |
| Bcas2_ex.vivo_noRT | Has1 | 2.558 | 7.91E-03 | glio |
| Hsd17b10_in.vivo_noRT | Gm16754 | 2.112 | 7.91E-03 | glio |
| Hsd17b10_in.vivo_noRT | Tbc1d30 | 1.815 | 7.91E-03 | glio |
| Hsd17b10_in.vivo_noRT | Kdm2b | -1.525 | 7.91E-03 | glio |
| Interaction_genotype_Myc | Nipal3 | -3.717 | 7.91E-04 | glio |
| Interaction_genotype_Myc | Ndor1 | -2.995 | 7.91E-04 | glio |
| Interaction_genotype_Myc | Abhd17c | -2.561 | 7.91E-04 | glio |
| Interaction_genotype_Myc | Rps15a | 1.352 | 7.91E-04 | glio |
| Aars2_in.vivo_noRT | Mgarp | 2.295 | 7.91E-04 | glio |
| Bora_in.vivo_noRT | Gm28221 | 2.267 | 7.91E-04 | glio |
| Chek1_ex.vivo_noRT | Ptgds | 3.525 | 7.91E-04 | glio |
| Eif2s1_ex.vivo_noRT | Gm15972 | 3.415 | 7.91E-04 | glio |
| Atr_in.vivo_noRT | Scamp3 | -1.487 | 7.91E-05 | glio |
| Bcas2_in.vivo_noRT | Fbxl22 | 2.691 | 7.91E-05 | glio |
| Bcas2_in.vivo_noRT | Ctss | 2.428 | 7.91E-05 | glio |
| Bora_in.vivo_noRT | Mkrn3 | 2.679 | 7.91E-05 | glio |
| Bora_ex.vivo_noRT | Stra8 | 3.599 | 7.91E-06 | glio |
| Chek1_in.vivo_noRT | Gm49077 | 3.279 | 7.91E-06 | glio |
| Bora_ex.vivo_noRT | Cd7 | 3.201 | 7.91E-07 | glio |
| Bora_ex.vivo_noRT | Gm48764 | 3.201 | 7.91E-07 | glio |

| | | | | |
|---|---|---|---|---|
| Chek1_in.vivo_noRT | Asgr1 | 4.385 | 7.91E-10 | glio |
| culture_effect | Enkd1 | 0.607 | 7.92E-03 | glio |
| culture_effect | Mplkip | 0.5 | 7.92E-03 | glio |
| Interaction_genotype_Aars2 | Ddost | 0.965 | 7.92E-03 | glio |
| Interaction_genotype_Pars2 | Sucnr1 | -2.268 | 7.92E-03 | glio |
| Interaction_genotype_Pars2 | Pik3ap1 | -3.158 | 7.92E-03 | glio |
| Interaction_genotype_Pars2 | Nudt22 | 2.19 | 7.92E-03 | glio |
| Interaction_genotype_Rad51 | A430106G13R | -3.917 | 7.92E-03 | glio |
| interaction_RT | Serping1 | 1.994 | 7.92E-03 | glio |
| interaction_RT | Fa2h | 2.237 | 7.92E-03 | glio |
| Aars2_in.vivo_noRT | 9530059O14R | 1.896 | 7.92E-03 | glio |
| Aars2_in.vivo_noRT | Lgals3 | -0.842 | 7.92E-03 | glio |
| Atr_in.vivo_noRT | Sf3b1 | -0.816 | 7.92E-03 | glio |
| Bcas2_in.vivo_noRT | Cpt1b | 1.926 | 7.92E-03 | glio |
| Bcas2_in.vivo_noRT | Ccdc173 | 1.814 | 7.92E-03 | glio |
| Bcas2_in.vivo_noRT | Ppa2 | -1.202 | 7.92E-03 | glio |
| Bcas2_in.vivo_noRT | Rfc1 | -0.817 | 7.92E-03 | glio |
| Bora_in.vivo_noRT | Tmem223 | 2.007 | 7.92E-03 | glio |
| Bora_in.vivo_noRT | Pimreg | -1.571 | 7.92E-03 | glio |
| Brca2_in.vivo_noRT | Serpinb9g | 1.763 | 7.92E-03 | glio |
| Brca2_in.vivo_noRT | Adh7 | 2.161 | 7.92E-03 | glio |
| Cenpj_ex.vivo_noRT | Gm34776 | 2.198 | 7.92E-03 | glio |
| Hsd17b10_in.vivo_noRT | Rnf208 | 2.224 | 7.92E-03 | glio |
| Interaction_genotype_Atr | B930059L03Ri | -4.344 | 7.92E-04 | glio |
| interaction_RT | 1600020E01Ri | 1.394 | 7.92E-04 | glio |
| Aars2_in.vivo_noRT | Ipp | 1.941 | 7.92E-04 | glio |
| Interaction_genotype_Mtor | Gm45406 | -3.577 | 7.92E-05 | glio |
| culture_effect | Lyrm1 | 2.26 | 7.92E-06 | glio |
| culture_effect | Pomc | 1.609 | 7.92E-06 | glio |
| culture_effect | Asb3 | -0.991 | 7.92E-06 | glio |
| Interaction_genotype_Mtor | Noxred1 | -5.142 | 7.92E-06 | glio |
| Interaction_genotype_Mtor | Gm43653 | -5.566 | 7.92E-07 | glio |
| Interaction_genotype_Mtor | Gm49504 | -4.498 | 7.92E-07 | glio |
| Aars2_ex.vivo_noRT | Zscan4-ps2 | 3.293 | 7.92E-07 | glio |
| Bora_in.vivo_noRT | Mroh7 | 3.412 | 7.92E-07 | glio |
| Atr_in.vivo_noRT | 4930481B07R | 3.969 | 7.92E-08 | glio |
| Atr_in.vivo_noRT | Gm10138 | 3.449 | 7.92E-08 | glio |
| Bora_in.vivo_noRT | Gm42664 | 4.286 | 7.92E-13 | glio |
| Atr_in.vivo_noRT | Gm50138 | 5.011 | 7.92E-18 | glio |
| culture_effect | Wdsub1 | -0.76 | 7.93E-03 | glio |
| Interaction_genotype_Atr | Mfhas1 | -2.9 | 7.93E-03 | glio |
| Interaction_genotype_Atr | Limk1 | -2.009 | 7.93E-03 | glio |
| Interaction_genotype_Atr | Pigk | -1.75 | 7.93E-03 | glio |
| Interaction_genotype_Bcas2 | Gm15326 | -2.629 | 7.93E-03 | glio |
| Interaction_genotype_Bcas2 | C330002G04R | -2.652 | 7.93E-03 | glio |
| Interaction_genotype_Bcas2 | Gm14327 | -3.642 | 7.93E-03 | glio |
| Interaction_genotype_Bcas2 | Cbr3 | -3.748 | 7.93E-03 | glio |
| Interaction_genotype_Bcas2 | 6330415G19R | -3.212 | 7.93E-03 | glio |
| Interaction_genotype_Myc | Zfp229 | -2.836 | 7.93E-03 | glio |
| Interaction_genotype_Pars2 | Mcmdc2 | -3.118 | 7.93E-03 | glio |
| Atr_in.vivo_noRT | B4galt2 | 1.313 | 7.93E-03 | glio |
| Bcas2_ex.vivo_noRT | C430042M11F | 2.793 | 7.93E-03 | glio |

| | | | | |
|---|---|---|---|---|
| Bora_in.vivo_noRT | Ska2 | -1.321 | 7.93E-03 | glio |
| Chek1_in.vivo_noRT | Gm49041 | 1.746 | 7.93E-03 | glio |
| Chek1_in.vivo_noRT | Ssbp4 | -1.425 | 7.93E-03 | glio |
| Fanca_ex.vivo_noRT | Gm13078 | 2.124 | 7.93E-03 | glio |
| Hsd17b10_in.vivo_noRT | Vcan | 2.268 | 7.93E-03 | glio |
| Hsd17b10_in.vivo_noRT | Cd38 | 2.05 | 7.93E-03 | glio |
| Interaction_genotype_Mtor | Nrbp1 | 1.662 | 7.93E-04 | glio |
| Interaction_genotype_Myc | Rbfaos | -3.002 | 7.93E-04 | glio |
| Atp5c1_in.vivo_noRT | Ccr5 | 2.396 | 7.93E-04 | glio |
| Atp5c1_in.vivo_noRT | Ppp1r18os | 2.837 | 7.93E-04 | glio |
| Atr_in.vivo_noRT | Eif4ebp2 | -1.414 | 7.93E-04 | glio |
| Bcas2_in.vivo_noRT | Plek2 | 2.356 | 7.93E-04 | glio |
| Chek1_in.vivo_noRT | Kctd1 | 2.339 | 7.93E-04 | glio |
| Chek1_in.vivo_noRT | Prelid3b | -1.263 | 7.93E-04 | glio |
| Chek1_in.vivo_noRT | Sulf2 | -1.332 | 7.93E-04 | glio |
| Chek1_ex.vivo_noRT | Arhgef15 | 2.953 | 7.93E-04 | glio |
| Chek1_ex.vivo_noRT | Atad3a | -1.446 | 7.93E-04 | glio |
| Interaction_genotype_Mtor | 9630014M24F | -4.487 | 7.93E-05 | glio |
| Interaction_genotype_Mtor | Gm9908 | -5.042 | 7.93E-05 | glio |
| Interaction_genotype_Mtor | Syp | -5.145 | 7.93E-05 | glio |
| Interaction_genotype_Mtor | Gm49479 | -4.586 | 7.93E-05 | glio |
| Interaction_genotype_Mtor | Gm44190 | -3.925 | 7.93E-05 | glio |
| Atr_in.vivo_noRT | Nop58 | -1.042 | 7.93E-05 | glio |
| Interaction_genotype_Myc | Pmp22 | -4.194 | 7.93E-06 | glio |
| Atp5c1_in.vivo_noRT | Oit3 | 3.19 | 7.93E-06 | glio |
| Atp5c1_in.vivo_noRT | Vsig10l | 3.308 | 7.93E-06 | glio |
| Atp5c1_in.vivo_noRT | Armc2 | 3.324 | 7.93E-06 | glio |
| Interaction_genotype_Myc | Baalc | -5.499 | 7.93E-08 | glio |
| Interaction_genotype_Myc | Ldlrad3 | -5.733 | 7.93E-08 | glio |
| Chek1_in.vivo_noRT | Gm48682 | 3.732 | 7.93E-08 | glio |
| Bora_in.vivo_noRT | Npy4r | 3.569 | 7.93E-09 | glio |
| Chek1_ex.vivo_noRT | Gm36757 | 5.243 | 7.93E-10 | glio |
| culture_effect | Vwa5a | -1.557 | 7.93E-11 | glio |
| Chek1_ex.vivo_noRT | Npy1r | 4.842 | 7.93E-13 | glio |
| Interaction_genotype_Atr | D930030I03Ri | -3.409 | 7.94E-03 | glio |
| Interaction_genotype_Mtor | Sbk1 | -3.322 | 7.94E-03 | glio |
| Interaction_genotype_Mtor | Vmn1r139 | -2.886 | 7.94E-03 | glio |
| Interaction_genotype_Mtor | Kmt2e | 1.241 | 7.94E-03 | glio |
| Interaction_genotype_Myc | Cyb5r1 | 1.792 | 7.94E-03 | glio |
| Interaction_genotype_Rpl3 | Wdr74 | 1.286 | 7.94E-03 | glio |
| Interaction_genotype_Terf1 | Gm10399 | -3.597 | 7.94E-03 | glio |
| Interaction_genotype_Terf1 | 4933429H19R | -3.713 | 7.94E-03 | glio |
| Interaction_genotype_Terf1 | 4930426I24Ri | -3.908 | 7.94E-03 | glio |
| Bcas2_in.vivo_noRT | Kdelr2 | -0.743 | 7.94E-03 | glio |
| Chek1_in.vivo_noRT | Mpped2 | 1.958 | 7.94E-03 | glio |
| Hsd17b10_in.vivo_noRT | Hoxa13 | 2.169 | 7.94E-03 | glio |
| Hsd17b10_in.vivo_noRT | Ldlrad3 | 1.901 | 7.94E-03 | glio |
| culture_effect | Rpf1 | -0.526 | 7.94E-04 | glio |
| Interaction_genotype_Mtor | Ccdc142os | -3.852 | 7.94E-04 | glio |
| Interaction_genotype_Myc | Myom1 | -3.951 | 7.94E-04 | glio |
| Interaction_genotype_Rad21 | Zan | 4.84 | 7.94E-04 | glio |
| Aars2_in.vivo_noRT | Arpc2 | -0.86 | 7.94E-04 | glio |

| | | | | |
|---|---|---|---|---|
| Atr_ex.vivo_noRT | Fas | 3.549 | 7.94E-04 | glio |
| Dclre1c_ex.vivo_noRT | Gm30042 | 3.086 | 7.94E-04 | glio |
| Interaction_genotype_Atr | Crabp1 | -4.355 | 7.94E-05 | glio |
| Aars2_in.vivo_noRT | Rhpn2 | 2.483 | 7.94E-05 | glio |
| Atr_in.vivo_noRT | Ccdc142os | 2.745 | 7.94E-05 | glio |
| Bora_in.vivo_noRT | Micall2 | 3.277 | 7.94E-06 | glio |
| Bora_in.vivo_noRT | Flnb | -1.981 | 7.94E-06 | glio |
| culture_effect | Ifit3 | 1.123 | 7.94E-07 | glio |
| Bcas2_in.vivo_noRT | Pgls | -1.196 | 7.94E-07 | glio |
| Atr_in.vivo_noRT | Gm28875 | 3.839 | 7.94E-08 | glio |
| Bcas2_in.vivo_noRT | Fhl4 | 3.556 | 7.94E-08 | glio |
| Bcas2_in.vivo_noRT | Gm27253 | 4.541 | 7.94E-11 | glio |
| culture_effect | Ier3 | 0.596 | 7.95E-03 | glio |
| Interaction_genotype_Aars2 | Gm41409 | -3.046 | 7.95E-03 | glio |
| Interaction_genotype_Aars2 | Ccdc66 | -2.673 | 7.95E-03 | glio |
| Interaction_genotype_Aars2 | Tbc1d20 | 1.759 | 7.95E-03 | glio |
| Interaction_genotype_Atr | Hsf2bp | -3.139 | 7.95E-03 | glio |
| Interaction_genotype_Atr | Foxc2 | -3.491 | 7.95E-03 | glio |
| Interaction_genotype_Atr | Hipk2 | 1.707 | 7.95E-03 | glio |
| Interaction_genotype_Bcas2 | Csta2 | 2.406 | 7.95E-03 | glio |
| Interaction_genotype_Bcas2 | Litaf | 1.593 | 7.95E-03 | glio |
| Interaction_genotype_Chek1 | Tfb2m | 2.124 | 7.95E-03 | glio |
| Interaction_genotype_Mtor | Rad9b | -2.913 | 7.95E-03 | glio |
| Interaction_genotype_Mtor | AV099323 | -2.425 | 7.95E-03 | glio |
| Interaction_genotype_Mtor | 4933434E20Ri | 1.594 | 7.95E-03 | glio |
| Interaction_genotype_Myc | Pde1a | -3.108 | 7.95E-03 | glio |
| Interaction_genotype_Myc | Nphp3 | -2.968 | 7.95E-03 | glio |
| Interaction_genotype_Myc | Rab8b | -2.186 | 7.95E-03 | glio |
| Interaction_genotype_Myc | Wasf3 | -2.073 | 7.95E-03 | glio |
| Interaction_genotype_Myc | Dclre1a | -2.199 | 7.95E-03 | glio |
| Aars2_in.vivo_noRT | 5430431A17R | 1.801 | 7.95E-03 | glio |
| Aars2_in.vivo_noRT | Ppm1h | 1.373 | 7.95E-03 | glio |
| Aars2_in.vivo_noRT | Sem1 | -0.609 | 7.95E-03 | glio |
| Atr_ex.vivo_noRT | Nwd1 | 2.182 | 7.95E-03 | glio |
| Bcas2_in.vivo_noRT | Ncapg | -1.401 | 7.95E-03 | glio |
| Chek1_in.vivo_noRT | Ccdc173 | 1.802 | 7.95E-03 | glio |
| Chek1_ex.vivo_noRT | Tmsb15b2 | 2.761 | 7.95E-03 | glio |
| Hsd17b10_in.vivo_noRT | Gm19325 | 2.173 | 7.95E-03 | glio |
| culture_effect | Tcea2 | 1.761 | 7.95E-04 | glio |
| Interaction_genotype_Mtor | Frmd8os | -4.452 | 7.95E-04 | glio |
| Interaction_genotype_Mtor | Sgsh | -4.196 | 7.95E-04 | glio |
| Interaction_genotype_Mtor | Edn1 | -4.214 | 7.95E-04 | glio |
| Interaction_genotype_Myc | Abca16 | -3.669 | 7.95E-04 | glio |
| Interaction_genotype_Myc | Tubd1 | -3.304 | 7.95E-04 | glio |
| Interaction_genotype_Myc | Ighmbp2 | -3.059 | 7.95E-04 | glio |
| Interaction_genotype_Myc | Axl | 1.505 | 7.95E-04 | glio |
| Aars2_in.vivo_noRT | 6430548M08F | 2.421 | 7.95E-04 | glio |
| Atr_ex.vivo_noRT | Phf11a | 3.159 | 7.95E-04 | glio |
| Bora_in.vivo_noRT | 1700047M11F | 2.209 | 7.95E-04 | glio |
| Interaction_genotype_Mtor | C330024C12R | -4.101 | 7.95E-05 | glio |
| Aars2_in.vivo_noRT | Efna4 | 2.845 | 7.95E-05 | glio |
| Aars2_in.vivo_noRT | Tmem107 | -1.7 | 7.95E-05 | glio |

| | | | | |
|---|---|---|---|---|
| Bard1_ex.vivo_noRT | Gm11583 | 2.814 | 7.95E-05 | glio |
| Bcas2_in.vivo_noRT | Sbk2 | 2.841 | 7.95E-05 | glio |
| culture_effect | Prex1 | -1.413 | 7.95E-06 | glio |
| culture_effect | Pdlim1 | -1.166 | 7.95E-06 | glio |
| Interaction_genotype_Myc | Phtf1os | -5.077 | 7.95E-06 | glio |
| Atp5c1_in.vivo_noRT | Akr1c18 | 2.668 | 7.95E-06 | glio |
| Atp5c1_in.vivo_noRT | 4921504E06Ri | 2.668 | 7.95E-06 | glio |
| Atp5c1_in.vivo_noRT | 4931431C16R | 2.668 | 7.95E-06 | glio |
| Chek1_in.vivo_noRT | Cyp3a13 | 3.381 | 7.95E-06 | glio |
| culture_effect | Adgrg6 | 0.964 | 7.95E-07 | glio |
| Interaction_genotype_Mtor | Gm20515 | -5.755 | 7.95E-07 | glio |
| Aars2_in.vivo_noRT | Rex2 | 3.653 | 7.95E-07 | glio |
| Chek1_ex.vivo_noRT | Myo5c | 4.118 | 7.95E-07 | glio |
| Hsd17b10_in.vivo_noRT | Gm47872 | 3.352 | 7.95E-07 | glio |
| Chek1_ex.vivo_noRT | Ssh3 | 4.045 | 7.95E-08 | glio |
| culture_effect | Cyp2j6 | -1.307 | 7.95E-11 | glio |
| Aars2_in.vivo_noRT | Thnsl2 | 4.159 | 7.95E-15 | glio |
| Atr_in.vivo_noRT | Gm15520 | 4.754 | 7.95E-15 | glio |
| Atr_in.vivo_noRT | Gm37035 | 4.616 | 7.95E-16 | glio |
| Atr_in.vivo_noRT | Gm12610 | 5.004 | 7.95E-19 | glio |
| Cdkn2a_in.vivo_noRT | Cdkn2a | -1.266 | 7.95E-23 | glio |
| culture_effect | Jph4 | 1.601 | 7.96E-03 | glio |
| Interaction_genotype_Chek1 | Fhl4 | -3.449 | 7.96E-03 | glio |
| Interaction_genotype_Pars2 | Gm37800 | -3.338 | 7.96E-03 | glio |
| Interaction_genotype_Pars2 | B4galnt1 | -3.101 | 7.96E-03 | glio |
| Interaction_genotype_Pars2 | Abcf2 | 1.418 | 7.96E-03 | glio |
| Interaction_genotype_Rad51 | Gm35066 | -3.792 | 7.96E-03 | glio |
| Interaction_genotype_Rad51 | D630036H23F | -3.691 | 7.96E-03 | glio |
| Interaction_genotype_Rad51 | Gm10399 | -3.532 | 7.96E-03 | glio |
| Interaction_genotype_Rad51 | Cers4 | 2.637 | 7.96E-03 | glio |
| Interaction_genotype_Rad51 | Srp54a | 2.414 | 7.96E-03 | glio |
| interaction_RT | Serpinh1 | -0.634 | 7.96E-03 | glio |
| Aars2_in.vivo_noRT | Magoh | -0.694 | 7.96E-03 | glio |
| Bcas2_in.vivo_noRT | Smim12 | -1.208 | 7.96E-03 | glio |
| Chek1_in.vivo_noRT | Cntd1 | 1.969 | 7.96E-03 | glio |
| Chek1_in.vivo_noRT | Clec16a | 1.799 | 7.96E-03 | glio |
| Hsd17b10_in.vivo_noRT | Pkhd1 | 2.099 | 7.96E-03 | glio |
| Interaction_genotype_Bora | Gm10010 | -4.412 | 7.96E-04 | glio |
| Interaction_genotype_Bora | Gm9922 | -4.203 | 7.96E-04 | glio |
| Interaction_genotype_Mtor | Dock7 | 1.842 | 7.96E-04 | glio |
| Interaction_genotype_Myc | Ccdc146 | -3.911 | 7.96E-04 | glio |
| Bcas2_in.vivo_noRT | Zfp28 | 2.465 | 7.96E-04 | glio |
| Bcas2_in.vivo_noRT | Abcf2 | -1.39 | 7.96E-04 | glio |
| Chek1_in.vivo_noRT | Gm13663 | 2.221 | 7.96E-04 | glio |
| Interaction_genotype_Myc | Dnase1l2 | -4.371 | 7.96E-05 | glio |
| Chek1_in.vivo_noRT | Stac3 | 2.901 | 7.96E-05 | glio |
| Chek1_in.vivo_noRT | AC125149.3 | 2.639 | 7.96E-05 | glio |
| Aars2_in.vivo_noRT | Abcd2 | 3.488 | 7.96E-06 | glio |
| Bcas2_ex.vivo_noRT | Gm41307 | 3.131 | 7.96E-06 | glio |
| Chek1_in.vivo_noRT | Gm26559 | 3.233 | 7.96E-06 | glio |
| Chek1_in.vivo_noRT | Neurl1a | 3.736 | 7.96E-07 | glio |
| Bora_in.vivo_noRT | Gm9908 | 4.019 | 7.96E-08 | glio |

| | | | | |
|---|---|---|---|---|
| Aars2_in.vivo_noRT | Frmd4b | 4.259 | 7.96E-10 | glio |
| Interaction_genotype_Hsd17b | Acod1 | 5.427 | 7.97E-03 | glio |
| Atr_in.vivo_noRT | Sorbs2 | 1.854 | 7.97E-03 | glio |
| Atr_in.vivo_noRT | Prkaa2 | 1.679 | 7.97E-03 | glio |
| Atr_in.vivo_noRT | Zc3h11a | 1.753 | 7.97E-03 | glio |
| Atr_in.vivo_noRT | Nbeal1 | -1.29 | 7.97E-03 | glio |
| Eif2s1_in.vivo_noRT | Nat8f2 | 1.883 | 7.97E-03 | glio |
| Hsd17b10_in.vivo_noRT | 3110082J24Ri | 2.174 | 7.97E-03 | glio |
| Interaction_genotype_Myc | Haus8 | -2.337 | 7.97E-04 | glio |
| Atr_in.vivo_noRT | Pih1d2 | 2.075 | 7.97E-04 | glio |
| Atr_in.vivo_noRT | Rnd1 | 2.171 | 7.97E-04 | glio |
| Interaction_genotype_Myc | Ablim3 | -4.661 | 7.97E-05 | glio |
| Hsd17b10_in.vivo_noRT | Syp | 3.225 | 7.97E-05 | glio |
| culture_effect | Plgrkt | -0.634 | 7.97E-06 | glio |
| Atr_ex.vivo_noRT | A930002C04R | 2.764 | 7.97E-06 | glio |
| Chek1_in.vivo_noRT | Rpl15 | -0.903 | 7.97E-06 | glio |
| Hsd17b10_in.vivo_noRT | Camk2b | 3.094 | 7.97E-06 | glio |
| Interaction_genotype_Chek1 | Zfpl1 | 2.872 | 7.97E-07 | glio |
| Interaction_genotype_Myc | Ift20 | 2.326 | 7.97E-07 | glio |
| Aars2_in.vivo_noRT | Rasal1 | 3.851 | 7.97E-07 | glio |
| Bcas2_in.vivo_noRT | Gm43258 | 3.411 | 7.97E-07 | glio |
| Chek1_in.vivo_noRT | Ggnbp1 | 3.361 | 7.97E-07 | glio |
| Chek1_ex.vivo_noRT | Gm10432 | 4.045 | 7.97E-07 | glio |
| Chek1_ex.vivo_noRT | Gm4841 | 4.842 | 7.97E-07 | glio |
| Interaction_genotype_Myc | 4930558J18Ri | -5.796 | 7.97E-08 | glio |
| Bora_in.vivo_noRT | Sh3gl3 | 4.024 | 7.97E-09 | glio |
| Chek1_ex.vivo_noRT | Ushbp1 | 4.901 | 7.97E-09 | glio |
| culture_effect | Mlec | 0.955 | 7.97E-12 | glio |
| Aars2_in.vivo_noRT | Cstdc4 | 4.67 | 7.97E-13 | glio |
| Atr_in.vivo_noRT | Prm3 | 5.584 | 7.97E-20 | glio |
| Atr_in.vivo_noRT | Ccser1 | 5.533 | 7.97E-21 | glio |
| Aars2_in.vivo_noRT | Gpc5 | 5.327 | 7.97E-26 | glio |
| Interaction_genotype_Atr | Ubn1 | 1.502 | 7.98E-03 | glio |
| Interaction_genotype_Mtor | Hoxa1 | -3 | 7.98E-03 | glio |
| Interaction_genotype_Mtor | Sprtn | -2.473 | 7.98E-03 | glio |
| Interaction_genotype_Rpl3 | Elavl1 | 1.448 | 7.98E-03 | glio |
| Aars2_in.vivo_noRT | Dnaja4 | 1.929 | 7.98E-03 | glio |
| Aars2_in.vivo_noRT | Arhgap26 | 1.839 | 7.98E-03 | glio |
| Atr_ex.vivo_noRT | Ksr2 | 2.188 | 7.98E-03 | glio |
| Bora_in.vivo_noRT | Cgas | -1.441 | 7.98E-03 | glio |
| Bora_in.vivo_noRT | Ddx3x | -0.757 | 7.98E-03 | glio |
| Bora_in.vivo_noRT | Phf5a | -0.587 | 7.98E-03 | glio |
| Cenpj_in.vivo_noRT | Gm14019 | 2.022 | 7.98E-03 | glio |
| Chek1_in.vivo_noRT | Gm44867 | 1.986 | 7.98E-03 | glio |
| Chek1_in.vivo_noRT | 6430590A07R | 1.515 | 7.98E-03 | glio |
| Cyp19a1_ex.vivo_noRT | Gm15533 | 2.159 | 7.98E-03 | glio |
| Fanca_ex.vivo_noRT | Gm13964 | 2.124 | 7.98E-03 | glio |
| Interaction_genotype_Mtor | Psmb4 | 1.604 | 7.98E-04 | glio |
| Interaction_genotype_Myc | Gm28875 | -3.936 | 7.98E-04 | glio |
| Aars2_in.vivo_noRT | Rpl18a | -0.609 | 7.98E-04 | glio |
| Chek1_in.vivo_noRT | Slc25a53 | 2.607 | 7.98E-04 | glio |
| Eif2s1_in.vivo_noRT | Ifi207 | 2.941 | 7.98E-04 | glio |

| | | | | |
|---|---|---|---|---|
| Interaction_genotype_Myc | Gm12758 | -4.42 | 7.98E-05 | glio |
| Hsd17b10_in.vivo_noRT | Nrn1l | 2.691 | 7.98E-05 | glio |
| culture_effect | Fxyd7 | 1.79 | 7.98E-06 | glio |
| culture_effect | Tmed9 | -0.54 | 7.98E-06 | glio |
| Interaction_genotype_Mtor | Gm17203 | -3.848 | 7.98E-06 | glio |
| Aars2_in.vivo_noRT | Astn2 | 3.239 | 7.98E-07 | glio |
| Chek1_in.vivo_noRT | Gm2762 | 3.471 | 7.98E-08 | glio |
| Chek1_in.vivo_noRT | Kmo | 3.402 | 7.98E-09 | glio |
| Aars2_in.vivo_noRT | Hba-a2 | 4.415 | 7.98E-10 | glio |
| Chek1_in.vivo_noRT | Tlr1 | 4.784 | 7.98E-12 | glio |
| Chek1_ex.vivo_noRT | Gm6803 | 4.842 | 7.98E-13 | glio |
| Bcas2_in.vivo_noRT | Gm11899 | 4.285 | 7.98E-16 | glio |
| Interaction_genotype_Atr | Car5a | -3.02 | 7.99E-03 | glio |
| Interaction_genotype_Bcas2 | Zfpm1 | -3.724 | 7.99E-03 | glio |
| Interaction_genotype_Bcas2 | Stard10 | -3.453 | 7.99E-03 | glio |
| Interaction_genotype_Bcas2 | Atp6v1f | 0.9 | 7.99E-03 | glio |
| Interaction_genotype_Mtor | Mrnip | -2.402 | 7.99E-03 | glio |
| Interaction_genotype_Myc | Tef | -2.512 | 7.99E-03 | glio |
| Interaction_genotype_Myc | Psma5 | 1.437 | 7.99E-03 | glio |
| Bora_in.vivo_noRT | Gm31645 | 1.982 | 7.99E-03 | glio |
| Bora_in.vivo_noRT | Snx3 | -0.513 | 7.99E-03 | glio |
| Dclre1c_ex.vivo_noRT | Pdgfb | 2.528 | 7.99E-03 | glio |
| Atr_in.vivo_noRT | 4930509H03R | 2.386 | 7.99E-04 | glio |
| Bcas2_ex.vivo_noRT | Olfr552 | 2.342 | 7.99E-04 | glio |
| Cenpj_in.vivo_noRT | Gm15533 | 2.022 | 7.99E-04 | glio |
| interaction_RT | D10Wsu102e | -1.231 | 7.99E-06 | glio |
| Aars2_in.vivo_noRT | Gm12971 | 3.219 | 7.99E-06 | glio |
| Bard1_ex.vivo_noRT | Itgbl1 | 3.592 | 7.99E-06 | glio |
| Bard1_ex.vivo_noRT | Helt | 2.814 | 7.99E-06 | glio |
| Bard1_ex.vivo_noRT | Gm20625 | 2.814 | 7.99E-06 | glio |
| Aars2_in.vivo_noRT | 4930455G09R | 4.021 | 7.99E-07 | glio |
| interaction_RT | Atp6v0b | 0.835 | 7.99E-08 | glio |
| Atr_in.vivo_noRT | Ccl20 | 5.533 | 7.99E-08 | glio |
| Hsd17b10_in.vivo_noRT | Cfh | 2.837 | 7.99E-08 | glio |
| Chek1_in.vivo_noRT | Il17b | 4.393 | 7.99E-09 | glio |
| Chek1_in.vivo_noRT | C230037L18Ri | 3.875 | 7.99E-09 | glio |
| Chek1_ex.vivo_noRT | Ngb | 4.842 | 7.99E-09 | glio |
| Atr_in.vivo_noRT | 4921536K21R | 5.273 | 7.99E-15 | glio |
| Atr_in.vivo_noRT | 5430402O13R | 5.533 | 7.99E-15 | glio |
| culture_effect | Hdgf | 0.93 | 7.99E-17 | glio |
| Chek1_in.vivo_noRT | Gm12841 | 5.044 | 7.99E-23 | glio |
| Interaction_genotype_Atr | 4930432K21R | -2.888 | 8.00E-03 | glio |
| Interaction_genotype_Atr | Nbeal1 | 1.794 | 8.00E-03 | glio |
| Interaction_genotype_Bcas2 | Gm47232 | -3.534 | 8.00E-03 | glio |
| Interaction_genotype_Bcas2 | Npnt | -2.596 | 8.00E-03 | glio |
| Interaction_genotype_Bcas2 | Clec12b | -2.46 | 8.00E-03 | glio |
| Interaction_genotype_Bcas2 | Ache | -3.691 | 8.00E-03 | glio |
| Interaction_genotype_Bcas2 | Gm20682 | -3.018 | 8.00E-03 | glio |
| Interaction_genotype_Bcas2 | Gm10655 | -3.626 | 8.00E-03 | glio |
| Interaction_genotype_Bcas2 | D930030I03Ri | -3.694 | 8.00E-03 | glio |
| Interaction_genotype_Bcas2 | 2600006K01R | -3.595 | 8.00E-03 | glio |
| Interaction_genotype_Bcas2 | Cnksr3 | -2.643 | 8.00E-03 | glio |

| | | | | |
|---|---|---|---|---|
| Interaction_genotype_Myc | Stra8 | -2.602 | 8.00E-03 | glio |
| Interaction_genotype_Myc | Kctd21 | -2.938 | 8.00E-03 | glio |
| Interaction_genotype_Pars2 | Psmd2 | 1.063 | 8.00E-03 | glio |
| Aars2_in.vivo_noRT | Tomm7 | -0.785 | 8.00E-03 | glio |
| Aars2_ex.vivo_noRT | Pdgfb | 2.13 | 8.00E-03 | glio |
| Aars2_ex.vivo_noRT | Myd88 | -2.056 | 8.00E-03 | glio |
| Atr_in.vivo_noRT | Smurf1 | 1.76 | 8.00E-03 | glio |
| Atr_in.vivo_noRT | Fdps | -1.242 | 8.00E-03 | glio |
| Chek1_ex.vivo_noRT | Gla | 1.633 | 8.00E-03 | glio |
| Hsd17b10_in.vivo_noRT | Ubxn10 | 1.761 | 8.00E-03 | glio |
| Interaction_genotype_Mtor | Mertk | -3.843 | 8.00E-04 | glio |
| Interaction_genotype_Mtor | Exd2 | -3.614 | 8.00E-04 | glio |
| Interaction_genotype_Mtor | Lgals3bp | 1.885 | 8.00E-04 | glio |
| Atr_in.vivo_noRT | 2310015A10R | 2.488 | 8.00E-04 | glio |
| Atr_ex.vivo_noRT | Dapk2 | 2.367 | 8.00E-04 | glio |
| Chek1_in.vivo_noRT | Hpdl | 2.22 | 8.00E-04 | glio |
| culture_effect | Pcsk1n | -1.093 | 8.00E-05 | glio |
| culture_effect | Cavin1 | 0.933 | 8.00E-06 | glio |
| Interaction_genotype_Mtor | Grcc10 | 2.014 | 8.00E-06 | glio |
| Interaction_genotype_Myc | Rpl35a | 1.515 | 8.00E-06 | glio |
| Atr_in.vivo_noRT | Palb2 | 2.522 | 8.00E-06 | glio |
| Aars2_in.vivo_noRT | Atp5a1 | -1.169 | 8.00E-07 | glio |
| Atr_in.vivo_noRT | AW046200 | 3.475 | 8.00E-07 | glio |
| Hsd17b10_in.vivo_noRT | Tdrd5 | 3.23 | 8.00E-07 | glio |
| Interaction_genotype_Mtor | Zfp786 | -6.367 | 8.00E-08 | glio |
| Atr_in.vivo_noRT | Upk3bl | 3.457 | 8.00E-08 | glio |
| Hsd17b10_in.vivo_noRT | 5730507C01R | 3.093 | 8.00E-09 | glio |
| Aars2_in.vivo_noRT | Gm2016 | 5.327 | 8.00E-24 | glio |
| culture_effect | Arhgap26 | -1.53 | 8.01E-03 | glio |
| culture_effect | Ddx55 | -0.71 | 8.01E-03 | glio |
| culture_effect | Usp48 | -0.489 | 8.01E-03 | glio |
| culture_effect | Abhd17b | -0.362 | 8.01E-03 | glio |
| culture_effect | Wdr74 | -0.361 | 8.01E-03 | glio |
| Interaction_genotype_Atr | Zscan26 | 1.772 | 8.01E-03 | glio |
| Interaction_genotype_Atr | Trmt6 | 1.446 | 8.01E-03 | glio |
| Interaction_genotype_Atr | Skp1a | 1.083 | 8.01E-03 | glio |
| Atp5c1_in.vivo_noRT | Pax5 | 2.351 | 8.01E-03 | glio |
| Bard1_ex.vivo_noRT | AC125149.2 | 2.419 | 8.01E-03 | glio |
| Bard1_ex.vivo_noRT | Plk5 | 2.421 | 8.01E-03 | glio |
| Bora_in.vivo_noRT | Zfp930 | 1.35 | 8.01E-03 | glio |
| Interaction_genotype_Atr | Camkk2 | -3.068 | 8.01E-04 | glio |
| Aars2_in.vivo_noRT | Baiap2 | -1.603 | 8.01E-04 | glio |
| Aars2_in.vivo_noRT | Actl6a | -1.196 | 8.01E-04 | glio |
| Bcas2_in.vivo_noRT | Slc4a11 | 2.547 | 8.01E-04 | glio |
| Bcas2_in.vivo_noRT | Arf2 | -1.694 | 8.01E-04 | glio |
| Interaction_genotype_Myc | Gm11767 | -4.614 | 8.01E-05 | glio |
| Interaction_genotype_Myc | Gm26671 | -3.122 | 8.01E-05 | glio |
| Bcas2_in.vivo_noRT | Gm42701 | 2.809 | 8.01E-05 | glio |
| Bora_in.vivo_noRT | Gm2415 | 2.911 | 8.01E-05 | glio |
| Bora_in.vivo_noRT | Cdc25a | -1.84 | 8.01E-05 | glio |
| Interaction_genotype_Mtor | 1700101I11Ri | -5.758 | 8.01E-06 | glio |
| Interaction_genotype_Mtor | Rab21 | 2.081 | 8.01E-06 | glio |

| | | | | |
|---|---|---|---|---|
| Aars2_in.vivo_noRT | Dlx2 | 3.944 | 8.01E-08 | glio |
| Aars2_in.vivo_noRT | Rnase4 | 3.764 | 8.01E-08 | glio |
| Chek1_in.vivo_noRT | Gm50107 | 3.529 | 8.01E-08 | glio |
| Chek1_in.vivo_noRT | Gm42699 | 3.528 | 8.01E-08 | glio |
| culture_effect | Mtdh | 0.621 | 8.01E-11 | glio |
| Bora_in.vivo_noRT | Hcn3 | 4.294 | 8.01E-12 | glio |
| Aars2_in.vivo_noRT | Vmn2r18 | 5.327 | 8.01E-27 | glio |
| culture_effect | Ndufa3 | 0.467 | 8.02E-03 | glio |
| Interaction_genotype_Aars2 | Fsbp | -3.442 | 8.02E-03 | glio |
| Interaction_genotype_Aars2 | AA388235 | -3.449 | 8.02E-03 | glio |
| Interaction_genotype_Aars2 | Cyb5r1 | 1.91 | 8.02E-03 | glio |
| interaction_RT | Otos | 1.839 | 8.02E-03 | glio |
| Atr_ex.vivo_noRT | Ccr5 | 2.4 | 8.02E-03 | glio |
| Chek1_in.vivo_noRT | Cic | -1.19 | 8.02E-03 | glio |
| Cyp19a1_ex.vivo_noRT | Nrros | 2.943 | 8.02E-03 | glio |
| Hsd17b10_in.vivo_noRT | Gm26581 | 2.221 | 8.02E-03 | glio |
| Interaction_genotype_Atr | Rftn2 | -3.749 | 8.02E-04 | glio |
| Interaction_genotype_Atr | Cdk7 | 1.841 | 8.02E-04 | glio |
| Interaction_genotype_Atr | Map1lc3a | 1.424 | 8.02E-04 | glio |
| Chek1_in.vivo_noRT | Ints8 | -1.42 | 8.02E-04 | glio |
| Interaction_genotype_Bora | Wdr26 | 2.064 | 8.02E-05 | glio |
| Interaction_genotype_Bora | Baalc | -4.49 | 8.02E-05 | glio |
| Interaction_genotype_Bora | Nxph4 | -4.957 | 8.02E-05 | glio |
| Interaction_genotype_Myc | 9430060I03Ri | -4.068 | 8.02E-05 | glio |
| interaction_RT | Ipo4 | -1.099 | 8.02E-05 | glio |
| Atr_in.vivo_noRT | March7 | -1.846 | 8.02E-06 | glio |
| Bard1_ex.vivo_noRT | Gm5157 | 2.814 | 8.02E-06 | glio |
| culture_effect | Cd72 | -3.101 | 8.02E-07 | glio |
| Bora_in.vivo_noRT | Itga7 | 3.506 | 8.02E-07 | glio |
| Bcas2_in.vivo_noRT | Hba-a2 | 3.895 | 8.02E-08 | glio |
| Bora_in.vivo_noRT | Adam33 | 3.779 | 8.02E-08 | glio |
| culture_effect | Rpl13a | -1.442 | 8.02E-27 | glio |
| culture_effect | Zic4 | 1.689 | 8.03E-03 | glio |
| culture_effect | 9330121J05Ri | 1.364 | 8.03E-03 | glio |
| Interaction_genotype_Aars2 | Lst1 | -3.369 | 8.03E-03 | glio |
| Interaction_genotype_Aars2 | 1700081N11R | -3.653 | 8.03E-03 | glio |
| Interaction_genotype_Aars2 | Gm49479 | -3.265 | 8.03E-03 | glio |
| Interaction_genotype_Aars2 | Rbm19 | -2.359 | 8.03E-03 | glio |
| Interaction_genotype_Aars2 | Cd63 | 0.783 | 8.03E-03 | glio |
| Interaction_genotype_Aars2 | Chchd2 | 0.785 | 8.03E-03 | glio |
| Interaction_genotype_Atr | Nr1h3 | -2.79 | 8.03E-03 | glio |
| Interaction_genotype_Bora | Hyal1 | -3.39 | 8.03E-03 | glio |
| Interaction_genotype_Bora | Mis12 | 1.966 | 8.03E-03 | glio |
| Interaction_genotype_Myc | Prss36 | -2.343 | 8.03E-03 | glio |
| Interaction_genotype_Myc | Gys1 | -2.3 | 8.03E-03 | glio |
| Interaction_genotype_Rad51 | Lipt2 | -3.096 | 8.03E-03 | glio |
| Atr_in.vivo_noRT | Mpped2 | 1.918 | 8.03E-03 | glio |
| Bcas2_in.vivo_noRT | 2810001G20R | 1.299 | 8.03E-03 | glio |
| Cenpj_in.vivo_noRT | Hnf1aos1 | 2.022 | 8.03E-03 | glio |
| Hsd17b10_in.vivo_noRT | Sv2a | 2.179 | 8.03E-03 | glio |
| culture_effect | March5 | 0.469 | 8.03E-04 | glio |
| Interaction_genotype_Myc | Ptpn22 | -4.238 | 8.03E-04 | glio |

| | | | | |
|---|---|---|---|---|
| Interaction_genotype_Myc | Dhodh | -2.973 | 8.03E-04 | glio |
| Atr_in.vivo_noRT | Desi2 | -1.316 | 8.03E-04 | glio |
| Atr_ex.vivo_noRT | Plk5 | 2.771 | 8.03E-04 | glio |
| Atr_ex.vivo_noRT | H2-DMb2 | 3.164 | 8.03E-04 | glio |
| Atr_ex.vivo_noRT | 9030612E09Ri | 2.968 | 8.03E-04 | glio |
| Atr_ex.vivo_noRT | Nr4a3 | 3.031 | 8.03E-04 | glio |
| Bcas2_in.vivo_noRT | Bend6 | 2.394 | 8.03E-04 | glio |
| culture_effect | D830025C05R | 2.031 | 8.03E-05 | glio |
| Interaction_genotype_Mtor | A530083I20Ri | -5.725 | 8.03E-06 | glio |
| Interaction_genotype_Mtor | Mettl7a3 | -5.043 | 8.03E-06 | glio |
| Atr_in.vivo_noRT | Gm15879 | 4.04 | 8.03E-06 | glio |
| Atr_ex.vivo_noRT | Crb3 | 3.164 | 8.03E-06 | glio |
| Bora_in.vivo_noRT | Gm26881 | 2.756 | 8.03E-06 | glio |
| Chek1_in.vivo_noRT | Setd4 | 3.098 | 8.03E-06 | glio |
| Aars2_in.vivo_noRT | Gm6225 | 3.651 | 8.03E-07 | glio |
| Aars2_in.vivo_noRT | Gm37170 | 3.484 | 8.03E-07 | glio |
| Aars2_in.vivo_noRT | Itga2b | 3.402 | 8.03E-07 | glio |
| culture_effect | Gm30074 | 2.483 | 8.03E-08 | glio |
| culture_effect | Qdpr | 0.891 | 8.03E-08 | glio |
| Aars2_ex.vivo_noRT | Gm4340 | 3.293 | 8.03E-08 | glio |
| Aars2_ex.vivo_noRT | Gm16239 | 3.293 | 8.03E-08 | glio |
| interaction_RT | Cad | -1.968 | 8.03E-10 | glio |
| culture_effect | Hmgn1 | 0.821 | 8.03E-14 | glio |
| Chek1_in.vivo_noRT | Pkd1l2 | 4.78 | 8.03E-14 | glio |
| Chek1_in.vivo_noRT | Htr1b | 5.044 | 8.03E-14 | glio |
| Aars2_in.vivo_noRT | Krt19 | 5.327 | 8.03E-29 | glio |
| culture_effect | Bbx | -0.392 | 8.04E-03 | glio |
| Interaction_genotype_Atr | Rbfaos | -2.643 | 8.04E-03 | glio |
| Interaction_genotype_Mtor | Mxd4 | 1.772 | 8.04E-03 | glio |
| Interaction_genotype_Mtor | Csrp2 | 2.024 | 8.04E-03 | glio |
| Interaction_genotype_Myc | Msh5 | -3.12 | 8.04E-03 | glio |
| Interaction_genotype_Myc | Farp2 | -2.678 | 8.04E-03 | glio |
| Interaction_genotype_Myc | Hmgb3 | -1.677 | 8.04E-03 | glio |
| Interaction_genotype_Myc | Fdps | 1.863 | 8.04E-03 | glio |
| interaction_RT | Ide | -0.723 | 8.04E-03 | glio |
| Aars2_in.vivo_noRT | Chd6 | -1.128 | 8.04E-03 | glio |
| Atr_in.vivo_noRT | Chp1 | -1.19 | 8.04E-03 | glio |
| Bcas2_in.vivo_noRT | Sgsh | 2.064 | 8.04E-03 | glio |
| Interaction_genotype_Aars2 | Syp | -4.665 | 8.04E-04 | glio |
| Interaction_genotype_Bora | AC154200.1 | -3.505 | 8.04E-04 | glio |
| Interaction_genotype_Mtor | Decr1 | 1.923 | 8.04E-04 | glio |
| Interaction_genotype_Terf1 | Rassf6 | -4.261 | 8.04E-04 | glio |
| Aars2_in.vivo_noRT | Aldh4a1 | 2.133 | 8.04E-04 | glio |
| Atp5c1_in.vivo_noRT | Gm31518 | 2.668 | 8.04E-04 | glio |
| Cenpj_in.vivo_noRT | Slc19a3 | 2.022 | 8.04E-04 | glio |
| Interaction_genotype_Mtor | Hnf1aos1 | -4.103 | 8.04E-05 | glio |
| Aars2_in.vivo_noRT | Ogdhl | 2.772 | 8.04E-05 | glio |
| Interaction_genotype_Myc | Gm48610 | -5.16 | 8.04E-06 | glio |
| Interaction_genotype_Mtor | Gm50394 | -5.671 | 8.04E-07 | glio |
| Chek1_in.vivo_noRT | 1810012K16R | 3.527 | 8.04E-08 | glio |
| Atr_in.vivo_noRT | Tnfrsf4 | 3.947 | 8.04E-09 | glio |
| Chek1_in.vivo_noRT | Kcnk4 | 4.784 | 8.04E-12 | glio |

| | | | | |
|---|---|---|---|---|
| culture_effect | Gm43728 | 1.525 | 8.05E-03 | glio |
| Interaction_genotype_Myc | 1500002F19Ri | -2.841 | 8.05E-03 | glio |
| Interaction_genotype_Myc | Nagpa | -2.608 | 8.05E-03 | glio |
| Interaction_genotype_Myc | U2af1 | 1.088 | 8.05E-03 | glio |
| Aars2_in.vivo_noRT | Sh3yl1 | 1.702 | 8.05E-03 | glio |
| Aars2_in.vivo_noRT | Kdm5a | -0.973 | 8.05E-03 | glio |
| Aars2_ex.vivo_noRT | Krtcap2 | 0.736 | 8.05E-03 | glio |
| Atp5c1_in.vivo_noRT | Dcbld1 | -1.666 | 8.05E-03 | glio |
| Atp5c1_in.vivo_noRT | Cyb5b | -0.827 | 8.05E-03 | glio |
| Atr_in.vivo_noRT | Cpt1c | 1.725 | 8.05E-03 | glio |
| Bcas2_in.vivo_noRT | 1700019L13Ri | 1.978 | 8.05E-03 | glio |
| Bcas2_in.vivo_noRT | Gm38037 | 1.919 | 8.05E-03 | glio |
| Bora_ex.vivo_noRT | Chdh | 2.716 | 8.05E-03 | glio |
| Chek1_in.vivo_noRT | Phactr1 | 1.102 | 8.05E-03 | glio |
| Hsd17b10_in.vivo_noRT | Txnip | -2.52 | 8.05E-03 | glio |
| Interaction_genotype_Aars2 | Ntf5 | -4.442 | 8.05E-04 | glio |
| Aars2_ex.vivo_noRT | Platr31 | 3.693 | 8.05E-04 | glio |
| Atp5c1_in.vivo_noRT | Lhx1os | 2.142 | 8.05E-04 | glio |
| Bora_in.vivo_noRT | Gm29107 | 2.499 | 8.05E-04 | glio |
| Cenpj_in.vivo_noRT | Slc16a4 | 2.287 | 8.05E-04 | glio |
| Cenpj_in.vivo_noRT | Gpr4 | 2.022 | 8.05E-04 | glio |
| Cenpj_in.vivo_noRT | Prl7c1 | 2.022 | 8.05E-04 | glio |
| Cenpj_in.vivo_noRT | Gm33466 | 2.022 | 8.05E-04 | glio |
| Cenpj_ex.vivo_noRT | Gm45406 | 2.591 | 8.05E-04 | glio |
| Chek1_in.vivo_noRT | Tmem40 | 2.699 | 8.05E-04 | glio |
| Chek1_in.vivo_noRT | 2410022M11F | 2.472 | 8.05E-04 | glio |
| Chek1_in.vivo_noRT | Tmem260 | 1.989 | 8.05E-04 | glio |
| Chek1_ex.vivo_noRT | Gm16845 | 2.856 | 8.05E-04 | glio |
| Eif2s1_in.vivo_noRT | Bcl2l15 | 2.414 | 8.05E-04 | glio |
| Eif2s1_in.vivo_noRT | Rimklb | 1.88 | 8.05E-04 | glio |
| Eif2s1_in.vivo_noRT | Myo5c | 2.67 | 8.05E-04 | glio |
| Eif2s1_in.vivo_noRT | Htr1b | 2.666 | 8.05E-04 | glio |
| Bcas2_in.vivo_noRT | Zfp781 | 2.705 | 8.05E-05 | glio |
| Bora_in.vivo_noRT | Hsd17b14 | 2.979 | 8.05E-05 | glio |
| Bora_in.vivo_noRT | Ebf4 | 2.877 | 8.05E-05 | glio |
| Atp5c1_in.vivo_noRT | Gast | 2.668 | 8.05E-06 | glio |
| Atr_in.vivo_noRT | Xlr4c | 3.499 | 8.05E-07 | glio |
| culture_effect | Igfbp6 | 1.722 | 8.05E-08 | glio |
| Interaction_genotype_Mtor | Gm49086 | -5.662 | 8.05E-08 | glio |
| Atr_in.vivo_noRT | Slc41a3 | 3.81 | 8.05E-09 | glio |
| Chek1_ex.vivo_noRT | Spink4 | 4.842 | 8.05E-09 | glio |
| Bcas2_in.vivo_noRT | Akap3 | 4.547 | 8.05E-11 | glio |
| Chek1_in.vivo_noRT | Calhm5 | 4.316 | 8.05E-11 | glio |
| culture_effect | Ifit3b | 1.666 | 8.05E-12 | glio |
| Bcas2_in.vivo_noRT | Gm16685 | 4.149 | 8.05E-13 | glio |
| Bcas2_in.vivo_noRT | Gm35439 | 4.428 | 8.05E-16 | glio |
| Interaction_genotype_Atr | Dnase1l2 | -3.213 | 8.06E-03 | glio |
| Interaction_genotype_Atr | Gm15743 | -3.02 | 8.06E-03 | glio |
| Interaction_genotype_Atr | Abcc10 | -3.349 | 8.06E-03 | glio |
| Interaction_genotype_Atr | Stard8 | -3.011 | 8.06E-03 | glio |
| Aars2_in.vivo_noRT | A930041C12R | 1.908 | 8.06E-03 | glio |
| Aars2_in.vivo_noRT | 4732440D04R | 1.757 | 8.06E-03 | glio |

| | | | | |
|---|---|---|---|---|
| Chek1_in.vivo_noRT | Pltp | 1.791 | 8.06E-03 | glio |
| Interaction_genotype_Aars2 | Gm19531 | -3.811 | 8.06E-04 | glio |
| Interaction_genotype_Mtor | C430014B12R | -3.777 | 8.06E-04 | glio |
| Interaction_genotype_Myc | Bcl9 | -2.525 | 8.06E-04 | glio |
| Bcas2_in.vivo_noRT | Fbxw4 | 1.627 | 8.06E-04 | glio |
| Bcas2_in.vivo_noRT | Hecw2 | 2.922 | 8.06E-05 | glio |
| Bora_in.vivo_noRT | Ctrl | 2.912 | 8.06E-05 | glio |
| Aars2_in.vivo_noRT | Gm34552 | 3.133 | 8.06E-06 | glio |
| Bcas2_in.vivo_noRT | Gm48529 | 3.077 | 8.06E-06 | glio |
| Chek1_in.vivo_noRT | Eif4ebp2 | -1.891 | 8.06E-06 | glio |
| Bora_in.vivo_noRT | Ctss | 2.976 | 8.06E-07 | glio |
| Aars2_in.vivo_noRT | Hoxa6 | 4.294 | 8.06E-11 | glio |
| Aars2_in.vivo_noRT | Milr1 | 4.592 | 8.06E-11 | glio |
| Chek1_ex.vivo_noRT | Lrguk | 4.842 | 8.06E-12 | glio |
| Chek1_ex.vivo_noRT | Gm20275 | 4.842 | 8.06E-12 | glio |
| Chek1_in.vivo_noRT | Stra8 | 4.521 | 8.06E-14 | glio |
| Chek1_ex.vivo_noRT | Kcnk7 | 6.676 | 8.06E-19 | glio |
| culture_effect | Gad1 | -1.201 | 8.07E-03 | glio |
| culture_effect | Aldh6a1 | -1.241 | 8.07E-03 | glio |
| culture_effect | Amph | 0.953 | 8.07E-03 | glio |
| culture_effect | 9530034E10Ri | 0.773 | 8.07E-03 | glio |
| interaction_RT | Renbp | 1.218 | 8.07E-03 | glio |
| interaction_RT | E030030I06Ril | 0.853 | 8.07E-03 | glio |
| interaction_RT | Iqgap1 | -0.538 | 8.07E-03 | glio |
| Aars2_ex.vivo_noRT | Crybb1 | 2.324 | 8.07E-03 | glio |
| Bora_in.vivo_noRT | C3 | 1.934 | 8.07E-03 | glio |
| Bora_in.vivo_noRT | Cby1 | -1.152 | 8.07E-03 | glio |
| Chek1_ex.vivo_noRT | Pde11a | 2.545 | 8.07E-03 | glio |
| Chek1_ex.vivo_noRT | H60b | 2.429 | 8.07E-03 | glio |
| Interaction_genotype_Aars2 | Loxl4 | -3.856 | 8.07E-04 | glio |
| Interaction_genotype_Aars2 | Ephx1 | -4.056 | 8.07E-04 | glio |
| Interaction_genotype_Aars2 | Bcat1 | 2.367 | 8.07E-04 | glio |
| Interaction_genotype_Aars2 | Uchl3 | 1.727 | 8.07E-04 | glio |
| Chek1_in.vivo_noRT | Plekhg5 | 2.323 | 8.07E-04 | glio |
| Chek1_in.vivo_noRT | Trim35 | -1.506 | 8.07E-04 | glio |
| Eif2s1_in.vivo_noRT | Olfr171 | 2.143 | 8.07E-04 | glio |
| Hsd17b10_in.vivo_noRT | 0610009L18Ri | 2.636 | 8.07E-04 | glio |
| Atr_ex.vivo_noRT | Hnf1aos1 | 3.164 | 8.07E-05 | glio |
| Bard1_ex.vivo_noRT | Gm29670 | 2.814 | 8.07E-05 | glio |
| Bcas2_in.vivo_noRT | 2610507I01Ri | 2.539 | 8.07E-05 | glio |
| Chek1_ex.vivo_noRT | Plac1 | 3.48 | 8.07E-05 | glio |
| Chek1_in.vivo_noRT | Gm49479 | 3.206 | 8.07E-06 | glio |
| Aars2_in.vivo_noRT | Ctss | 3.209 | 8.07E-08 | glio |
| Bcas2_in.vivo_noRT | Selp | 3.763 | 8.07E-08 | glio |
| Aars2_in.vivo_noRT | Slc16a6 | 4.68 | 8.07E-11 | glio |
| Bora_in.vivo_noRT | Ifitm5 | 4.414 | 8.07E-11 | glio |
| Atr_in.vivo_noRT | Mettl7b | 5.533 | 8.07E-16 | glio |
| Interaction_genotype_Atr | Wdr70 | 1.275 | 8.08E-03 | glio |
| Interaction_genotype_Bcas2 | Necab2 | -3.295 | 8.08E-03 | glio |
| Interaction_genotype_Bcas2 | Gid4 | 1.571 | 8.08E-03 | glio |
| Interaction_genotype_Mtor | Them4 | -2.349 | 8.08E-03 | glio |
| Interaction_genotype_Mtor | Tnks2 | 1.493 | 8.08E-03 | glio |

| | | | | |
|---|---|---|---|---|
| Interaction_genotype_Mtor | Elof1 | 1.417 | 8.08E-03 | glio |
| interaction_RT | Slc35f1 | -1.267 | 8.08E-03 | glio |
| Bcas2_in.vivo_noRT | Zfp873 | 1.74 | 8.08E-03 | glio |
| Eif2s1_in.vivo_noRT | Pdrg1 | -0.707 | 8.08E-03 | glio |
| Bard1_ex.vivo_noRT | Gm45336 | 2.814 | 8.08E-04 | glio |
| Bora_in.vivo_noRT | Asf1a | -1.172 | 8.08E-04 | glio |
| Chek1_in.vivo_noRT | Eif3e | -0.65 | 8.08E-04 | glio |
| Bcas2_in.vivo_noRT | Tmsb15b2 | 3.087 | 8.08E-05 | glio |
| Bora_in.vivo_noRT | Gm29361 | 2.911 | 8.08E-05 | glio |
| Bora_in.vivo_noRT | Jun | -1.401 | 8.08E-05 | glio |
| Bora_in.vivo_noRT | D230022J07Ri | 3.248 | 8.08E-06 | glio |
| Atr_in.vivo_noRT | Gask1b | 3.738 | 8.08E-07 | glio |
| culture_effect | Dnajb13 | -3.23 | 8.08E-08 | glio |
| Bcas2_in.vivo_noRT | Hist1h4b | 4.102 | 8.08E-08 | glio |
| Bora_in.vivo_noRT | A430057M04F | 3.751 | 8.08E-08 | glio |
| interaction_RT | Atpif1 | 1.031 | 8.08E-09 | glio |
| interaction_RT | Tcn2 | 2.194 | 8.08E-09 | glio |
| Aars2_in.vivo_noRT | H2-Q10 | 4.217 | 8.08E-10 | glio |
| Atr_in.vivo_noRT | Rab14 | -1.83 | 8.08E-11 | glio |
| Bora_in.vivo_noRT | Gm38393 | 4.676 | 8.08E-15 | glio |
| Aars2_in.vivo_noRT | Gm10354 | 5.327 | 8.08E-21 | glio |
| Bora_in.vivo_noRT | Slc19a3 | 4.936 | 8.08E-22 | glio |
| Interaction_genotype_Atr | Ebf4 | -3.446 | 8.09E-03 | glio |
| Interaction_genotype_Mtor | A930001M01F | -3.201 | 8.09E-03 | glio |
| Interaction_genotype_Mtor | Hlcs | -2.909 | 8.09E-03 | glio |
| Interaction_genotype_Mtor | Fbl | 1.169 | 8.09E-03 | glio |
| Cenpj_ex.vivo_noRT | Gbp10 | 2.591 | 8.09E-03 | glio |
| Chek1_in.vivo_noRT | Npepps | -1.189 | 8.09E-03 | glio |
| Interaction_genotype_Myc | 2810021J22Ri | -3.822 | 8.09E-04 | glio |
| Aars2_in.vivo_noRT | Inca1 | 2.187 | 8.09E-04 | glio |
| Atp5c1_in.vivo_noRT | 4930502E18Ri | 2.133 | 8.09E-04 | glio |
| Interaction_genotype_Atr | Zfp983 | -4.213 | 8.09E-05 | glio |
| Interaction_genotype_Atr | Nat8l | -3.458 | 8.09E-05 | glio |
| Interaction_genotype_Atr | Cnksr1 | -4.658 | 8.09E-05 | glio |
| Interaction_genotype_Mtor | Hspb2 | -2.806 | 8.09E-05 | glio |
| Atr_ex.vivo_noRT | Crhbp | 3.069 | 8.09E-05 | glio |
| Bcas2_in.vivo_noRT | 2310008N11R | 3.021 | 8.09E-05 | glio |
| Bcas2_in.vivo_noRT | Tma16 | -1.341 | 8.09E-05 | glio |
| Bora_in.vivo_noRT | C030005K06R | 2.837 | 8.09E-05 | glio |
| culture_effect | C920021L13Ri | 2.174 | 8.09E-06 | glio |
| Aars2_in.vivo_noRT | Nupr1l | 3.546 | 8.09E-06 | glio |
| Bora_ex.vivo_noRT | Crb3 | 3.201 | 8.09E-06 | glio |
| Hsd17b10_in.vivo_noRT | Gm10300 | 3.356 | 8.09E-06 | glio |
| culture_effect | Pfn2 | 0.938 | 8.09E-07 | glio |
| Interaction_genotype_Mtor | Adamts4 | -5.241 | 8.09E-07 | glio |
| Aars2_ex.vivo_noRT | Gm48780 | 3.293 | 8.09E-07 | glio |
| Chek1_ex.vivo_noRT | Acox2 | 3.861 | 8.09E-07 | glio |
| Aars2_in.vivo_noRT | Tbxa2r | 3.487 | 8.09E-08 | glio |
| Aars2_in.vivo_noRT | Pax3 | 3.112 | 8.09E-08 | glio |
| Aars2_in.vivo_noRT | Dcpp3 | 3.809 | 8.09E-09 | glio |
| Bcas2_in.vivo_noRT | Gm32090 | 4.015 | 8.09E-10 | glio |
| Atr_in.vivo_noRT | Ankef1 | 4.794 | 8.09E-12 | glio |

| | | | | |
|---|---|---|---|---|
| Bcas2_in.vivo_noRT | Mpzl2 | 4.547 | 8.09E-16 | glio |
| culture_effect | Metap2 | 0.961 | 8.09E-17 | glio |
| culture_effect | BC005561 | -0.51 | 8.10E-03 | glio |
| Interaction_genotype_Atr | Gm266 | -2.369 | 8.10E-03 | glio |
| Interaction_genotype_Bora | Gm13398 | -2.532 | 8.10E-03 | glio |
| Interaction_genotype_Bora | Sh2d3c | -3.814 | 8.10E-03 | glio |
| Interaction_genotype_Bora | Gfpt1 | 1.615 | 8.10E-03 | glio |
| Interaction_genotype_Mtor | Armc9 | -2.792 | 8.10E-03 | glio |
| Interaction_genotype_Terf1 | BC065403 | -3.644 | 8.10E-03 | glio |
| interaction_RT | Cpq | 0.714 | 8.10E-03 | glio |
| interaction_RT | Rangrf | -0.701 | 8.10E-03 | glio |
| interaction_RT | Ppp4c | 0.65 | 8.10E-03 | glio |
| Aars2_in.vivo_noRT | Gm42869 | 1.867 | 8.10E-03 | glio |
| Aars2_in.vivo_noRT | Nop14 | -0.959 | 8.10E-03 | glio |
| Aars2_in.vivo_noRT | Eif4g2 | -0.609 | 8.10E-03 | glio |
| Aars2_ex.vivo_noRT | Nek4 | -2.257 | 8.10E-03 | glio |
| Atp5c1_in.vivo_noRT | Gm43409 | 2.405 | 8.10E-03 | glio |
| Bard1_ex.vivo_noRT | Fhl1 | 2.104 | 8.10E-03 | glio |
| Bcas2_in.vivo_noRT | Gm47509 | 2.009 | 8.10E-03 | glio |
| Bcas2_in.vivo_noRT | 2410080I02Ri | 1.894 | 8.10E-03 | glio |
| Bora_in.vivo_noRT | D130040H23R | 1.841 | 8.10E-03 | glio |
| Interaction_genotype_Myc | Zfp712 | -3.416 | 8.10E-04 | glio |
| Interaction_genotype_Myc | Srl | -3.812 | 8.10E-04 | glio |
| Interaction_genotype_Myc | Ywhaz | 1.359 | 8.10E-04 | glio |
| Atr_in.vivo_noRT | Dennd3 | 2.489 | 8.10E-04 | glio |
| Bora_in.vivo_noRT | Scml2 | 2.27 | 8.10E-04 | glio |
| culture_effect | Gpd2 | -0.906 | 8.10E-05 | glio |
| culture_effect | Oxr1 | -0.661 | 8.10E-05 | glio |
| Interaction_genotype_Mtor | Hk1os | -4.229 | 8.10E-05 | glio |
| Atr_in.vivo_noRT | Prpf38b | -1.326 | 8.10E-05 | glio |
| Aars2_in.vivo_noRT | Glt8d2 | 3.275 | 8.10E-06 | glio |
| Chek1_ex.vivo_noRT | Gm49579 | 4.055 | 8.10E-08 | glio |
| Bora_in.vivo_noRT | Gm12860 | 3.928 | 8.10E-13 | glio |
| culture_effect | Rcn1 | -1.173 | 8.10E-17 | glio |
| Chek1_in.vivo_noRT | A930033M14R | 5.044 | 8.10E-18 | glio |
| Bcas2_in.vivo_noRT | Cgnl1 | 4.807 | 8.10E-23 | glio |
| Interaction_genotype_Aars2 | Cox18 | 1.483 | 8.11E-03 | glio |
| Interaction_genotype_Aars2 | Rpl36al | 0.773 | 8.11E-03 | glio |
| Interaction_genotype_Mtor | Slc22a5 | -2.976 | 8.11E-03 | glio |
| Interaction_genotype_Mtor | Lratd2 | -2.976 | 8.11E-03 | glio |
| Interaction_genotype_Mtor | E230016M11R | -2.784 | 8.11E-03 | glio |
| Interaction_genotype_Mtor | 4632404H12R | -2.887 | 8.11E-03 | glio |
| Aars2_in.vivo_noRT | Mcm8 | 1.303 | 8.11E-03 | glio |
| Aars2_in.vivo_noRT | Uba5 | -1.146 | 8.11E-03 | glio |
| Atp5c1_in.vivo_noRT | Gm48161 | 1.88 | 8.11E-03 | glio |
| Cenpj_ex.vivo_noRT | Ciita | 2.192 | 8.11E-03 | glio |
| Chek1_in.vivo_noRT | Fancf | 1.605 | 8.11E-03 | glio |
| Chek1_ex.vivo_noRT | EU599041 | 2.549 | 8.11E-03 | glio |
| Chek1_ex.vivo_noRT | Plaat3 | 1.298 | 8.11E-03 | glio |
| Cyp19a1_ex.vivo_noRT | Gpr4 | 2.159 | 8.11E-03 | glio |
| Cyp19a1_ex.vivo_noRT | Gm33466 | 2.159 | 8.11E-03 | glio |
| culture_effect | Pcdhb17 | -2.114 | 8.11E-04 | glio |

| | | | | |
|---|---|---|---|---|
| Aars2_in.vivo_noRT | Hcn1 | 2.584 | 8.11E-04 | glio |
| Aars2_in.vivo_noRT | 1700112D23R | 2.363 | 8.11E-04 | glio |
| Aars2_in.vivo_noRT | Sumo2 | -0.714 | 8.11E-04 | glio |
| Bora_in.vivo_noRT | Uhrf1 | -1.539 | 8.11E-04 | glio |
| Chek1_in.vivo_noRT | Reep6 | 1.867 | 8.11E-04 | glio |
| culture_effect | Cdk9 | -0.698 | 8.11E-05 | glio |
| Interaction_genotype_Myc | Oit3 | -4.291 | 8.11E-05 | glio |
| Interaction_genotype_Myc | Afmid | -4.252 | 8.11E-06 | glio |
| Interaction_genotype_Myc | Mettl15 | -4.004 | 8.11E-06 | glio |
| Interaction_genotype_Myc | Gm14419 | -4.339 | 8.11E-06 | glio |
| Atr_in.vivo_noRT | Tmem88 | 3.051 | 8.11E-06 | glio |
| Bard1_ex.vivo_noRT | Gm44736 | 2.814 | 8.11E-06 | glio |
| Bcas2_in.vivo_noRT | A930033H14R | 3.312 | 8.11E-06 | glio |
| Aars2_in.vivo_noRT | Sdcbp2 | 3.632 | 8.11E-07 | glio |
| Chek1_in.vivo_noRT | Hoxa13 | 3.597 | 8.11E-07 | glio |
| culture_effect | Chit1 | -3.653 | 8.11E-09 | glio |
| Interaction_genotype_Myc | Rpl10a | 2.291 | 8.11E-09 | glio |
| Interaction_genotype_Myc | Nup93 | -4.338 | 8.11E-09 | glio |
| Bcas2_in.vivo_noRT | Ifi214 | 4.016 | 8.11E-09 | glio |
| Atr_in.vivo_noRT | Alpl | 4.349 | 8.11E-11 | glio |
| Bcas2_in.vivo_noRT | Npnt | 4.541 | 8.11E-19 | glio |
| culture_effect | Cd7 | -1.198 | 8.12E-03 | glio |
| culture_effect | Gm9774 | -1.625 | 8.12E-03 | glio |
| culture_effect | Naaa | -0.774 | 8.12E-03 | glio |
| Interaction_genotype_Bora | Saysd1 | 2.116 | 8.12E-03 | glio |
| Interaction_genotype_Mtor | Ebi3 | -3.841 | 8.12E-03 | glio |
| Interaction_genotype_Myc | Pigw | -2.57 | 8.12E-03 | glio |
| Interaction_genotype_Myc | Tarsl2 | -2.318 | 8.12E-03 | glio |
| Interaction_genotype_Myc | Efcab7 | -2.188 | 8.12E-03 | glio |
| Aars2_in.vivo_noRT | Gng3 | 1.829 | 8.12E-03 | glio |
| Aars2_in.vivo_noRT | Hs6st1 | 1.452 | 8.12E-03 | glio |
| Aars2_in.vivo_noRT | Gfm1 | -1.327 | 8.12E-03 | glio |
| Aars2_in.vivo_noRT | Gramd1a | 1.203 | 8.12E-03 | glio |
| Aars2_in.vivo_noRT | Rbmxl1 | -1.016 | 8.12E-03 | glio |
| Atr_in.vivo_noRT | Pnmal2 | 1.876 | 8.12E-03 | glio |
| Atr_in.vivo_noRT | Zfp536 | 1.429 | 8.12E-03 | glio |
| Bcas2_in.vivo_noRT | Zfp963 | 2.2 | 8.12E-03 | glio |
| Bcas2_in.vivo_noRT | 2210011C24R | 1.917 | 8.12E-03 | glio |
| Bora_in.vivo_noRT | Phf13 | -1.357 | 8.12E-03 | glio |
| Chek1_in.vivo_noRT | Pnpo | -1.356 | 8.12E-03 | glio |
| Chek1_in.vivo_noRT | Ostf1 | -0.952 | 8.12E-03 | glio |
| culture_effect | AC241534.1 | 1.803 | 8.12E-04 | glio |
| culture_effect | Gtf2h2 | -0.706 | 8.12E-04 | glio |
| culture_effect | Tsta3 | -0.509 | 8.12E-04 | glio |
| culture_effect | Mrpl12 | 0.373 | 8.12E-04 | glio |
| Interaction_genotype_Rpl3 | Gm49708 | -4.306 | 8.12E-04 | glio |
| Interaction_genotype_Rpl3 | Rcan2 | -4.727 | 8.12E-04 | glio |
| Interaction_genotype_Rpl3 | Olfm2 | -4.664 | 8.12E-04 | glio |
| Interaction_genotype_Rpl3 | Alox12 | -4.646 | 8.12E-04 | glio |
| Interaction_genotype_Rpl3 | 5430402O13R | -4.67 | 8.12E-04 | glio |
| Chek1_ex.vivo_noRT | Gm47972 | 2.985 | 8.12E-04 | glio |
| Interaction_genotype_Myc | Farp1 | -3.435 | 8.12E-05 | glio |

| | | | | |
|---|---|---|---|---|
| Atr_in.vivo_noRT | Carf | 2.76 | 8.12E-05 | glio |
| Aars2_in.vivo_noRT | Poldip2 | -1.587 | 8.12E-06 | glio |
| culture_effect | 9330182L06Ri | -2.51 | 8.12E-07 | glio |
| Atr_in.vivo_noRT | Rpl29 | -0.877 | 8.12E-07 | glio |
| Bcas2_in.vivo_noRT | Ifit1bl1 | 4.275 | 8.12E-07 | glio |
| Chek1_in.vivo_noRT | 1700010B08R | 3.357 | 8.12E-09 | glio |
| Aars2_in.vivo_noRT | Il2rb | 4.321 | 8.12E-11 | glio |
| Chek1_in.vivo_noRT | Gm41409 | 4.512 | 8.12E-13 | glio |
| culture_effect | Wipf1 | -1.32 | 8.12E-14 | glio |
| Chek1_in.vivo_noRT | Ric3 | 4.784 | 8.12E-15 | glio |
| Bora_in.vivo_noRT | Pakap | 4.404 | 8.12E-16 | glio |
| culture_effect | AW209491 | 0.605 | 8.13E-03 | glio |
| Interaction_genotype_Atr | Ildr1 | -2.369 | 8.13E-03 | glio |
| Interaction_genotype_Atr | Firre | -3.151 | 8.13E-03 | glio |
| Interaction_genotype_Atr | Cpe | 1.579 | 8.13E-03 | glio |
| Interaction_genotype_Bcas2 | Vmn1r32 | -2.458 | 8.13E-03 | glio |
| Interaction_genotype_Pars2 | Prdm5 | 1.919 | 8.13E-03 | glio |
| Atr_ex.vivo_noRT | Zfp119a | 1.815 | 8.13E-03 | glio |
| Bcas2_in.vivo_noRT | Phex | 1.857 | 8.13E-03 | glio |
| Bora_ex.vivo_noRT | 4933408J17Ri | 2.417 | 8.13E-03 | glio |
| Chek1_in.vivo_noRT | Angpt2 | 1.796 | 8.13E-03 | glio |
| Cyp19a1_ex.vivo_noRT | Ngb | 2.933 | 8.13E-03 | glio |
| Fanca_ex.vivo_noRT | Gm16933 | 2.124 | 8.13E-03 | glio |
| Fanca_ex.vivo_noRT | Itgbl1 | 2.524 | 8.13E-03 | glio |
| Fanca_ex.vivo_noRT | Lncbate10 | 2.126 | 8.13E-03 | glio |
| Fanca_ex.vivo_noRT | Nat8f2 | 2.524 | 8.13E-03 | glio |
| Fanca_ex.vivo_noRT | Gm47510 | 2.124 | 8.13E-03 | glio |
| Fanca_ex.vivo_noRT | B230311B06R | 3.134 | 8.13E-03 | glio |
| Fanca_ex.vivo_noRT | Gm5788 | 2.124 | 8.13E-03 | glio |
| Fanca_ex.vivo_noRT | Mmp13 | 2.124 | 8.13E-03 | glio |
| Fanca_ex.vivo_noRT | Gm48617 | 2.124 | 8.13E-03 | glio |
| Bcas2_ex.vivo_noRT | Gm30042 | 2.742 | 8.13E-04 | glio |
| Chek1_in.vivo_noRT | P4hb | -0.696 | 8.13E-04 | glio |
| Atp5c1_in.vivo_noRT | Flacc1 | 3.196 | 8.13E-07 | glio |
| Bora_in.vivo_noRT | Elfn2 | 3.162 | 8.13E-07 | glio |
| Atr_in.vivo_noRT | Ahrr | 4.733 | 8.13E-12 | glio |
| Atr_in.vivo_noRT | Gm24362 | 4.49 | 8.13E-13 | glio |
| Atr_in.vivo_noRT | 2010320O07R | 4.883 | 8.13E-13 | glio |
| Atr_in.vivo_noRT | Shisal2a | 5.27 | 8.13E-13 | glio |
| Atr_in.vivo_noRT | Nrn1l | 4.608 | 8.13E-13 | glio |
| Bora_in.vivo_noRT | 8030423F21Ri | 4.407 | 8.13E-14 | glio |
| Chek1_in.vivo_noRT | Gm12703 | 4.777 | 8.13E-22 | glio |
| Interaction_genotype_Atr | Zfp14 | -3.403 | 8.14E-03 | glio |
| Interaction_genotype_Myc | Adap1 | -3.352 | 8.14E-03 | glio |
| Aars2_in.vivo_noRT | Poldip3 | -1.01 | 8.14E-03 | glio |
| Bora_in.vivo_noRT | Tmem255a | 2.028 | 8.14E-03 | glio |
| Brca2_in.vivo_noRT | Gm12841 | 1.763 | 8.14E-03 | glio |
| Cdkn2a_in.vivo_noRT | A330015K06R | -2.332 | 8.14E-03 | glio |
| Cdkn2a_in.vivo_noRT | Myl7 | 1.902 | 8.14E-03 | glio |
| culture_effect | Il11 | 2.033 | 8.14E-04 | glio |
| Interaction_genotype_Myc | 2810410L24Ri | -3.805 | 8.14E-04 | glio |
| Interaction_genotype_Myc | 3010001F23Ri | -3.357 | 8.14E-04 | glio |

| | | | | |
|---|---|---|---|---|
| Interaction_genotype_Myc | Tnfaip6 | -2.79 | 8.14E-04 | glio |
| Interaction_genotype_Myc | Rpl23 | 1.239 | 8.14E-04 | glio |
| interaction_RT | Usp1 | -0.806 | 8.14E-04 | glio |
| Atp5c1_in.vivo_noRT | Necab2 | 2.404 | 8.14E-04 | glio |
| Atp5c1_in.vivo_noRT | 1110046J04Ri | 2.668 | 8.14E-04 | glio |
| Atr_in.vivo_noRT | Cdc42ep2 | 2.456 | 8.14E-04 | glio |
| Atr_ex.vivo_noRT | AC125149.2 | 2.769 | 8.14E-04 | glio |
| Aars2_in.vivo_noRT | Gm43445 | 2.684 | 8.14E-05 | glio |
| Bora_in.vivo_noRT | Il23a | 2.815 | 8.14E-05 | glio |
| Interaction_genotype_Mtor | Icam5 | -5.225 | 8.14E-06 | glio |
| Interaction_genotype_Myc | Gm50203 | -4.621 | 8.14E-06 | glio |
| Interaction_genotype_Myc | Gm13663 | -4.638 | 8.14E-06 | glio |
| Interaction_genotype_Myc | Pebp1 | 1.816 | 8.14E-06 | glio |
| Atr_in.vivo_noRT | Gm11767 | 3.611 | 8.14E-07 | glio |
| Aars2_in.vivo_noRT | Mterf1b | 3.559 | 8.14E-08 | glio |
| Aars2_in.vivo_noRT | 1700011D18R | 4.282 | 8.14E-10 | glio |
| culture_effect | Hgh1 | -0.548 | 8.15E-03 | glio |
| Interaction_genotype_Aars2 | Gm48742 | -2.759 | 8.15E-03 | glio |
| Interaction_genotype_Aars2 | Gm27010 | -3.173 | 8.15E-03 | glio |
| Interaction_genotype_Aars2 | Gm47438 | -3.535 | 8.15E-03 | glio |
| Interaction_genotype_Bcas2 | Gm47622 | -2.64 | 8.15E-03 | glio |
| Interaction_genotype_Bcas2 | Gm20633 | -3.31 | 8.15E-03 | glio |
| Interaction_genotype_Bcas2 | Gm50399 | -3.382 | 8.15E-03 | glio |
| Interaction_genotype_Bcas2 | Cpne8 | -3.612 | 8.15E-03 | glio |
| Interaction_genotype_Bcas2 | Gm47484 | -3.68 | 8.15E-03 | glio |
| Interaction_genotype_Cdkn2a | Serpinb9g | 2.835 | 8.15E-03 | glio |
| Interaction_genotype_Mtor | Ttc39c | -2.302 | 8.15E-03 | glio |
| Interaction_genotype_Myc | Bcor | -2.493 | 8.15E-03 | glio |
| interaction_RT | Ntm | -1.151 | 8.15E-03 | glio |
| Aars2_in.vivo_noRT | Ptov1 | -0.699 | 8.15E-03 | glio |
| Atp5c1_in.vivo_noRT | Slc30a6 | -1.428 | 8.15E-03 | glio |
| Chek1_in.vivo_noRT | Fermt3 | 1.854 | 8.15E-03 | glio |
| Chek1_in.vivo_noRT | Pif1 | 1.84 | 8.15E-03 | glio |
| Chek1_in.vivo_noRT | Mfap3 | -1.133 | 8.15E-03 | glio |
| Chek1_ex.vivo_noRT | Aoc1 | 3.163 | 8.15E-04 | glio |
| interaction_RT | Med28 | 0.647 | 8.15E-05 | glio |
| Atr_in.vivo_noRT | Slc22a5 | 2.553 | 8.15E-05 | glio |
| Chek1_ex.vivo_noRT | Prc1 | -2.202 | 8.15E-05 | glio |
| Chek1_in.vivo_noRT | Gm48832 | 3.028 | 8.15E-06 | glio |
| Atp5c1_in.vivo_noRT | Cd36 | 2.668 | 8.15E-07 | glio |
| Aars2_in.vivo_noRT | Ptgis | 5.327 | 8.15E-15 | glio |
| Bcas2_in.vivo_noRT | Gm5788 | 4.807 | 8.15E-22 | glio |
| culture_effect | Gm9484 | 1.594 | 8.16E-03 | glio |
| culture_effect | 6230400D17R | 1.195 | 8.16E-03 | glio |
| Interaction_genotype_Myc | Gm49135 | -3.063 | 8.16E-03 | glio |
| Interaction_genotype_Myc | Kctd12 | -2.93 | 8.16E-03 | glio |
| Interaction_genotype_Myc | Alg8 | -2.003 | 8.16E-03 | glio |
| interaction_RT | Stpg2 | 2.555 | 8.16E-03 | glio |
| Aars2_in.vivo_noRT | Slc22a21 | 1.953 | 8.16E-03 | glio |
| Aars2_in.vivo_noRT | 2610021A01R | 1.512 | 8.16E-03 | glio |
| Atr_in.vivo_noRT | Irf5 | 2.226 | 8.16E-03 | glio |
| Bard1_ex.vivo_noRT | S100a14 | 2.415 | 8.16E-03 | glio |

| | | | | |
|---|---|---|---|---|
| Bard1_ex.vivo_noRT | Gm39556 | 2.814 | 8.16E-03 | glio |
| Bora_ex.vivo_noRT | Fhad1 | 2.615 | 8.16E-03 | glio |
| Chek1_in.vivo_noRT | Lamp1 | -0.681 | 8.16E-03 | glio |
| Eif2s1_in.vivo_noRT | Tcstv3 | 2.143 | 8.16E-03 | glio |
| Interaction_genotype_Aars2 | Gm10762 | -3.444 | 8.16E-04 | glio |
| Interaction_genotype_Mtor | Tert | -3.896 | 8.16E-04 | glio |
| Interaction_genotype_Myc | Atp6v1f | 1.245 | 8.16E-04 | glio |
| Atr_in.vivo_noRT | Mthfs | 2.089 | 8.16E-04 | glio |
| Bcas2_in.vivo_noRT | Camkv | 2.572 | 8.16E-04 | glio |
| Bcas2_in.vivo_noRT | Ttc38 | 2.407 | 8.16E-04 | glio |
| Bcas2_in.vivo_noRT | Tuba1c | -1.43 | 8.16E-04 | glio |
| Bora_in.vivo_noRT | 1700020L24Ri | 2.276 | 8.16E-04 | glio |
| Hsd17b10_in.vivo_noRT | Nhsl2 | 2.498 | 8.16E-04 | glio |
| culture_effect | Gm20275 | -2.289 | 8.16E-06 | glio |
| culture_effect | C1d | 0.696 | 8.16E-07 | glio |
| Interaction_genotype_Mtor | Oit3 | -5.66 | 8.16E-07 | glio |
| Interaction_genotype_Mtor | Tmem132c | -4.099 | 8.16E-07 | glio |
| Atp5c1_in.vivo_noRT | Gm43701 | 2.668 | 8.16E-07 | glio |
| Hsd17b10_in.vivo_noRT | Skida1 | 3.616 | 8.16E-07 | glio |
| Aars2_in.vivo_noRT | Traf1 | 3.722 | 8.16E-08 | glio |
| Aars2_in.vivo_noRT | Traf3ip3 | 4.263 | 8.16E-10 | glio |
| Aars2_in.vivo_noRT | Gm13054 | 4.801 | 8.16E-15 | glio |
| culture_effect | Arhgap18 | -0.471 | 8.17E-03 | glio |
| Interaction_genotype_Cdkn2a | Ank1 | 2.85 | 8.17E-03 | glio |
| Interaction_genotype_Myc | Rala | 1.157 | 8.17E-03 | glio |
| interaction_RT | Stpg4 | 2.239 | 8.17E-03 | glio |
| Aars2_ex.vivo_noRT | Tmem39b | -1.887 | 8.17E-03 | glio |
| Atr_in.vivo_noRT | Prkaa1 | -1.109 | 8.17E-03 | glio |
| Bora_in.vivo_noRT | Chd8 | -1.145 | 8.17E-03 | glio |
| Chek1_in.vivo_noRT | Trdmt1 | 1.708 | 8.17E-03 | glio |
| Chek1_in.vivo_noRT | Rbms1 | -0.884 | 8.17E-03 | glio |
| culture_effect | Wdr76 | 0.771 | 8.17E-04 | glio |
| interaction_RT | Atic | -0.853 | 8.17E-04 | glio |
| interaction_RT | Bnip3l | 0.822 | 8.17E-04 | glio |
| Bard1_ex.vivo_noRT | 1700010B08R | 2.814 | 8.17E-04 | glio |
| Bcas2_in.vivo_noRT | Slc24a5 | -1.574 | 8.17E-04 | glio |
| Bcas2_in.vivo_noRT | Mkrn1 | -1.1 | 8.17E-04 | glio |
| Bcas2_in.vivo_noRT | Clint1 | -0.751 | 8.17E-04 | glio |
| Chek1_ex.vivo_noRT | Ccl7 | 3.489 | 8.17E-04 | glio |
| culture_effect | Coq8b | -1.045 | 8.17E-05 | glio |
| Interaction_genotype_Myc | Gm30074 | -4.192 | 8.17E-05 | glio |
| Aars2_in.vivo_noRT | Ndufb8 | -0.752 | 8.17E-05 | glio |
| Chek1_in.vivo_noRT | Gm43689 | 2.578 | 8.17E-05 | glio |
| Interaction_genotype_Mtor | Phldb2 | -5.222 | 8.17E-06 | glio |
| Interaction_genotype_Mtor | Hmgb1 | 2.128 | 8.17E-06 | glio |
| Atr_in.vivo_noRT | Gsto2 | 3.159 | 8.17E-06 | glio |
| Atr_in.vivo_noRT | Dhrs11 | 2.909 | 8.17E-06 | glio |
| Chek1_in.vivo_noRT | Hhipl2 | 3.484 | 8.17E-06 | glio |
| Chek1_in.vivo_noRT | Gm16279 | 3.312 | 8.17E-06 | glio |
| culture_effect | Lsm5 | 0.595 | 8.17E-07 | glio |
| Interaction_genotype_Nat2 | Gm5788 | 4.205 | 8.17E-07 | glio |
| Bora_ex.vivo_noRT | Gad1 | 3.201 | 8.17E-07 | glio |

| | | | | |
|---|---|---|---|---|
| Bora_in.vivo_noRT | Gm10484 | 3.935 | 8.17E-08 | glio |
| Bora_in.vivo_noRT | Mamdc2 | 3.626 | 8.17E-08 | glio |
| Bora_in.vivo_noRT | Cyp27b1 | 3.375 | 8.17E-08 | glio |
| Chek1_in.vivo_noRT | Medag | 3.876 | 8.17E-09 | glio |
| Chek1_in.vivo_noRT | Gm10734 | 4.057 | 8.17E-09 | glio |
| Aars2_in.vivo_noRT | Ube2e1 | -2.414 | 8.17E-11 | glio |
| culture_effect | Smim4 | 1.434 | 8.17E-13 | glio |
| Bora_in.vivo_noRT | Gm37800 | 4.936 | 8.17E-15 | glio |
| Atr_in.vivo_noRT | Snrpn | 4.875 | 8.17E-19 | glio |
| Interaction_genotype_Mtor | Mterf1b | -3.094 | 8.18E-03 | glio |
| Interaction_genotype_Myc | Crocc | -2.981 | 8.18E-03 | glio |
| Interaction_genotype_Myc | Pcmt1 | 1.157 | 8.18E-03 | glio |
| Aars2_in.vivo_noRT | Dnhd1 | 1.627 | 8.18E-03 | glio |
| Bard1_ex.vivo_noRT | Rhpn2 | 2.531 | 8.18E-03 | glio |
| Bora_in.vivo_noRT | Ifi208 | 1.991 | 8.18E-03 | glio |
| Interaction_genotype_Mtor | Vim | 1.411 | 8.18E-04 | glio |
| Interaction_genotype_Myc | Slc15a4 | -2.955 | 8.18E-04 | glio |
| Atr_in.vivo_noRT | Adamts9 | 2.261 | 8.18E-04 | glio |
| Bcas2_in.vivo_noRT | Egf | 2.615 | 8.18E-04 | glio |
| Interaction_genotype_Myc | Pdss1 | -3.728 | 8.18E-05 | glio |
| Bora_in.vivo_noRT | Actn4 | -1.227 | 8.18E-05 | glio |
| Interaction_genotype_Mtor | Fbxo45 | -4.254 | 8.18E-06 | glio |
| Bcas2_in.vivo_noRT | Ngef | 2.799 | 8.18E-06 | glio |
| Atr_in.vivo_noRT | Spata45 | 3.888 | 8.18E-09 | glio |
| Chek1_in.vivo_noRT | Prox2os | 3.74 | 8.18E-10 | glio |
| Atr_in.vivo_noRT | K230015D01R | 5.267 | 8.18E-13 | glio |
| Atr_in.vivo_noRT | Efhd1 | 5.017 | 8.18E-13 | glio |
| Bora_in.vivo_noRT | Tdrd5 | 4.55 | 8.18E-13 | glio |
| Bora_in.vivo_noRT | Pygo1 | 4.677 | 8.18E-13 | glio |
| Atr_in.vivo_noRT | Gm5286 | 5.533 | 8.18E-24 | glio |
| Atr_in.vivo_noRT | Gm34776 | 5.533 | 8.18E-24 | glio |
| Atr_in.vivo_noRT | Gm36793 | 5.533 | 8.18E-26 | glio |
| Interaction_genotype_Bora | Fsbp | -3.667 | 8.19E-03 | glio |
| Interaction_genotype_Bora | Cpne9 | -3.793 | 8.19E-03 | glio |
| Interaction_genotype_Bora | Jph4 | -3.479 | 8.19E-03 | glio |
| Interaction_genotype_Bora | Srl | -3.606 | 8.19E-03 | glio |
| Interaction_genotype_Bora | Gm49874 | -3.524 | 8.19E-03 | glio |
| Interaction_genotype_Bora | Cdk1 | 2.147 | 8.19E-03 | glio |
| Interaction_genotype_Mtor | Hist2h2bb | -2.728 | 8.19E-03 | glio |
| Interaction_genotype_Pars2 | Gm36037 | -2.664 | 8.19E-03 | glio |
| Interaction_genotype_Pars2 | D630024D03R | -2.855 | 8.19E-03 | glio |
| Interaction_genotype_Pars2 | Gm50394 | -3.47 | 8.19E-03 | glio |
| Interaction_genotype_Rad21 | Comp | 4.58 | 8.19E-03 | glio |
| Interaction_genotype_Terf1 | D030040B21R | -3.753 | 8.19E-03 | glio |
| Aars2_in.vivo_noRT | H2afv | -0.568 | 8.19E-03 | glio |
| Atr_in.vivo_noRT | 1700066M21R | 1.741 | 8.19E-03 | glio |
| Atr_in.vivo_noRT | Nars | -0.814 | 8.19E-03 | glio |
| Bcas2_in.vivo_noRT | Hcfc1r1 | 2.216 | 8.19E-03 | glio |
| Bcas2_in.vivo_noRT | 2010109A12R | 1.927 | 8.19E-03 | glio |
| Brca2_in.vivo_noRT | Tmem74b | 2.032 | 8.19E-03 | glio |
| Brca2_in.vivo_noRT | Gm12794 | 1.763 | 8.19E-03 | glio |
| Chek1_in.vivo_noRT | Rab27b | 2.102 | 8.19E-03 | glio |

| | | | | |
|---|---|---|---|---|
| Chek1_in.vivo_noRT | Nudt16 | 1.718 | 8.19E-03 | glio |
| Chek1_in.vivo_noRT | Zfp651 | 1.553 | 8.19E-03 | glio |
| Fanca_ex.vivo_noRT | Gm12703 | 2.124 | 8.19E-03 | glio |
| Atp5c1_in.vivo_noRT | Itga5 | -2.132 | 8.19E-04 | glio |
| Bard1_ex.vivo_noRT | Gm15929 | 2.62 | 8.19E-04 | glio |
| Bcas2_in.vivo_noRT | Gzmm | 2.478 | 8.19E-04 | glio |
| Bcas2_in.vivo_noRT | Dock11 | 2.264 | 8.19E-04 | glio |
| Bcas2_in.vivo_noRT | Kif7 | 2.273 | 8.19E-04 | glio |
| Bcas2_in.vivo_noRT | Mcm5 | -1.576 | 8.19E-04 | glio |
| Chek1_in.vivo_noRT | Gm10762 | 2.021 | 8.19E-04 | glio |
| Interaction_genotype_Mtor | Gm30790 | -4.841 | 8.19E-05 | glio |
| Interaction_genotype_Mtor | Arhgef15 | -4.608 | 8.19E-05 | glio |
| Aars2_in.vivo_noRT | Atp8a2 | 2.648 | 8.19E-06 | glio |
| Chek1_in.vivo_noRT | 1700066B19R | 3.067 | 8.19E-06 | glio |
| culture_effect | Sgsm3 | 1.215 | 8.19E-07 | glio |
| Interaction_genotype_Mtor | Gm44693 | -4.099 | 8.19E-07 | glio |
| Interaction_genotype_Myc | D430040D24F | -5.679 | 8.19E-07 | glio |
| Atp5c1_in.vivo_noRT | 9130019O22R | 2.668 | 8.19E-07 | glio |
| Interaction_genotype_Mtor | Capn13 | -4.099 | 8.19E-08 | glio |
| Aars2_in.vivo_noRT | Gm48610 | 4.544 | 8.19E-11 | glio |
| Hsd17b10_in.vivo_noRT | Gm15138 | 3.35 | 8.19E-11 | glio |
| culture_effect | Fam219a | 0.903 | 8.20E-03 | glio |
| culture_effect | Fam162a | 0.402 | 8.20E-03 | glio |
| Interaction_genotype_Aars2 | Gm40040 | -3.419 | 8.20E-03 | glio |
| Interaction_genotype_Aars2 | Gm41760 | -3.22 | 8.20E-03 | glio |
| Interaction_genotype_Aars2 | Gm16046 | -3.187 | 8.20E-03 | glio |
| Interaction_genotype_Aars2 | D930019O06F | -3.317 | 8.20E-03 | glio |
| Interaction_genotype_Aars2 | Asphd2 | -3.015 | 8.20E-03 | glio |
| Interaction_genotype_Aars2 | Fbln5 | 2.04 | 8.20E-03 | glio |
| Interaction_genotype_Aars2 | Ptch1 | 1.931 | 8.20E-03 | glio |
| Interaction_genotype_Aars2 | Zfpl1 | 1.429 | 8.20E-03 | glio |
| Interaction_genotype_Aars2 | Cltb | 1.319 | 8.20E-03 | glio |
| Interaction_genotype_Aars2 | Ndufa12 | 0.801 | 8.20E-03 | glio |
| Interaction_genotype_Atr | Fgd3 | -3.595 | 8.20E-03 | glio |
| Interaction_genotype_Bcas2 | Gm14023 | -3.676 | 8.20E-03 | glio |
| Interaction_genotype_Bcas2 | Xpr1 | 2.33 | 8.20E-03 | glio |
| Interaction_genotype_Myc | Gm11583 | -2.204 | 8.20E-03 | glio |
| Interaction_genotype_Myc | Rps21 | 1.272 | 8.20E-03 | glio |
| interaction_RT | Clint1 | -0.485 | 8.20E-03 | glio |
| Atp5c1_in.vivo_noRT | Gm47232 | 2.145 | 8.20E-03 | glio |
| Atr_in.vivo_noRT | Dlk2 | 1.651 | 8.20E-03 | glio |
| Atr_ex.vivo_noRT | Zbed5 | 2.399 | 8.20E-03 | glio |
| Bora_ex.vivo_noRT | Mgll | 2.808 | 8.20E-03 | glio |
| Chek1_in.vivo_noRT | 4732419C18R | 1.918 | 8.20E-03 | glio |
| Fanca_ex.vivo_noRT | Gm8016 | 2.124 | 8.20E-03 | glio |
| Interaction_genotype_Mtor | Zfp994 | -3.036 | 8.20E-04 | glio |
| Aars2_ex.vivo_noRT | Gm21818 | 3.293 | 8.20E-04 | glio |
| Aars2_ex.vivo_noRT | Dcn | 2.9 | 8.20E-04 | glio |
| Aars2_ex.vivo_noRT | 8030423F21Ri | 2.717 | 8.20E-04 | glio |
| Bard1_ex.vivo_noRT | Fam189a1 | 2.814 | 8.20E-04 | glio |
| Bard1_ex.vivo_noRT | Gm16344 | 2.814 | 8.20E-04 | glio |
| Chek1_in.vivo_noRT | Fhl2 | 2.275 | 8.20E-04 | glio |

| | | | | |
|---|---|---|---|---|
| Chek1_in.vivo_noRT | Rdm1 | -1.558 | 8.20E-04 | glio |
| culture_effect | Lmo7 | 1.511 | 8.20E-05 | glio |
| Interaction_genotype_Myc | Zhx2 | -4.718 | 8.20E-05 | glio |
| Atr_in.vivo_noRT | Mtx2 | -1.164 | 8.20E-05 | glio |
| Atr_ex.vivo_noRT | Ntn1 | 3.164 | 8.20E-05 | glio |
| Aars2_in.vivo_noRT | Dhrs13os | 3.491 | 8.20E-06 | glio |
| culture_effect | N4bp2 | -1.239 | 8.20E-07 | glio |
| Aars2_in.vivo_noRT | Gm39556 | 3.416 | 8.20E-07 | glio |
| Bora_ex.vivo_noRT | 1700019E08Ri | 3.994 | 8.20E-08 | glio |
| Bora_ex.vivo_noRT | Gm13191 | 3.201 | 8.20E-08 | glio |
| Bora_ex.vivo_noRT | 4933416E03Ri | 3.201 | 8.20E-08 | glio |
| Chek1_in.vivo_noRT | 4933421A08R | 4.264 | 8.20E-09 | glio |
| Chek1_in.vivo_noRT | Gm30124 | 3.773 | 8.20E-09 | glio |
| Chek1_in.vivo_noRT | 1700015F17Ri | 4.784 | 8.20E-12 | glio |
| Hsd17b10_in.vivo_noRT | Ankrd22 | 3.616 | 8.20E-14 | glio |
| culture_effect | Slc50a1 | -1.076 | 8.20E-15 | glio |
| Atr_in.vivo_noRT | Gm44646 | 5.533 | 8.20E-22 | glio |
| Bcas2_in.vivo_noRT | Gm26543 | 4.807 | 8.20E-23 | glio |
| culture_effect | Hspb9 | 1.628 | 8.21E-03 | glio |
| culture_effect | St7l | -0.576 | 8.21E-03 | glio |
| Interaction_genotype_Aars2 | Gm31415 | -2.778 | 8.21E-03 | glio |
| Interaction_genotype_Aars2 | Gemin4 | -3.355 | 8.21E-03 | glio |
| Interaction_genotype_Aars2 | Mob3b | -2.758 | 8.21E-03 | glio |
| Interaction_genotype_Aars2 | Mcph1 | -1.917 | 8.21E-03 | glio |
| Interaction_genotype_Aars2 | Zranb1 | 1.667 | 8.21E-03 | glio |
| Interaction_genotype_Aars2 | Prrc2b | 1.458 | 8.21E-03 | glio |
| Interaction_genotype_Mtor | Oas1g | -2.761 | 8.21E-03 | glio |
| Interaction_genotype_Rad51 | Dph5 | 1.721 | 8.21E-03 | glio |
| interaction_RT | Gclm | -0.79 | 8.21E-03 | glio |
| Atr_in.vivo_noRT | Vars | -0.99 | 8.21E-03 | glio |
| Bcas2_ex.vivo_noRT | Arhgef16 | 2.342 | 8.21E-03 | glio |
| Chek1_ex.vivo_noRT | Tyw1 | -1.686 | 8.21E-03 | glio |
| Hsd17b10_in.vivo_noRT | Pmel | -1.872 | 8.21E-03 | glio |
| Interaction_genotype_Mtor | Ccnd1 | 2.284 | 8.21E-04 | glio |
| Interaction_genotype_Pars2 | 2010001K21R | -4.22 | 8.21E-04 | glio |
| Bcas2_in.vivo_noRT | Cyp3a13 | 2.615 | 8.21E-04 | glio |
| Bcas2_in.vivo_noRT | Brip1os | -1.555 | 8.21E-04 | glio |
| Bora_ex.vivo_noRT | Lrrc4b | 2.806 | 8.21E-04 | glio |
| Bora_ex.vivo_noRT | Luzp2 | 2.404 | 8.21E-04 | glio |
| Bora_ex.vivo_noRT | Gm37864 | 2.808 | 8.21E-04 | glio |
| Cenpj_ex.vivo_noRT | Dmrtc1a | 2.591 | 8.21E-04 | glio |
| culture_effect | Fam221a | -1.526 | 8.21E-05 | glio |
| Interaction_genotype_Mtor | Ccl7 | -5.452 | 8.21E-05 | glio |
| Atp5c1_in.vivo_noRT | AC163347.1 | 2.405 | 8.21E-05 | glio |
| Chek1_ex.vivo_noRT | Gm41492 | 3.072 | 8.21E-05 | glio |
| Atr_in.vivo_noRT | Mogat2 | 3.016 | 8.21E-06 | glio |
| Atr_ex.vivo_noRT | Gm8013 | 4.159 | 8.21E-06 | glio |
| Atr_in.vivo_noRT | C330013E15Ri | 3.418 | 8.21E-07 | glio |
| Bcas2_in.vivo_noRT | Sec1 | 3.65 | 8.21E-07 | glio |
| Bcas2_in.vivo_noRT | 1810034E14Ri | 3.452 | 8.21E-07 | glio |
| Chek1_in.vivo_noRT | Cnnm1 | 4.132 | 8.21E-08 | glio |
| Atr_in.vivo_noRT | Fsbp | 4.09 | 8.21E-09 | glio |

| | | | | |
|---|---|---|---|---|
| Interaction_genotype_Bcas2 | Tardbp | 1.045 | 8.22E-03 | glio |
| Interaction_genotype_Myc | Cyp27b1 | -2.713 | 8.22E-03 | glio |
| Interaction_genotype_Myc | Psrc1 | -2.609 | 8.22E-03 | glio |
| Eif2s1_in.vivo_noRT | Fbxl2 | 2.024 | 8.22E-03 | glio |
| Eif2s1_in.vivo_noRT | Gm14379 | 2.15 | 8.22E-03 | glio |
| Eif2s1_in.vivo_noRT | Kif5a | 2.417 | 8.22E-03 | glio |
| Aars2_ex.vivo_noRT | Cxcr6 | 3.293 | 8.22E-04 | glio |
| Chek1_in.vivo_noRT | Wdr5b | 2.357 | 8.22E-04 | glio |
| Chek1_ex.vivo_noRT | Gm31518 | 3.018 | 8.22E-04 | glio |
| Interaction_genotype_Aars2 | Gm13594 | -5.135 | 8.22E-05 | glio |
| Chek1_in.vivo_noRT | Zyx | -1.742 | 8.22E-05 | glio |
| Bcas2_in.vivo_noRT | Pcdhb18 | 3.057 | 8.22E-06 | glio |
| Atr_in.vivo_noRT | Usp44 | 3.544 | 8.22E-07 | glio |
| Bcas2_in.vivo_noRT | Gm15246 | 3.456 | 8.22E-07 | glio |
| Chek1_in.vivo_noRT | Lsp1 | 3.857 | 8.22E-07 | glio |
| culture_effect | 9430060I03Ri | 2.724 | 8.22E-10 | glio |
| interaction_RT | Selenok | 1.177 | 8.22E-10 | glio |
| culture_effect | Gm38037 | 1.218 | 8.23E-03 | glio |
| Interaction_genotype_Bora | Gm32591 | -3.472 | 8.23E-03 | glio |
| Interaction_genotype_Myc | Tmbim6 | 1.078 | 8.23E-03 | glio |
| Aars2_ex.vivo_noRT | Tcf19 | -2.236 | 8.23E-03 | glio |
| Atr_in.vivo_noRT | Gm48960 | 1.782 | 8.23E-03 | glio |
| Atr_in.vivo_noRT | Cabp1 | 1.701 | 8.23E-03 | glio |
| Atr_in.vivo_noRT | Hist3h2a | 1.866 | 8.23E-03 | glio |
| Chek1_in.vivo_noRT | Pfkfb3 | 1.72 | 8.23E-03 | glio |
| Chek1_in.vivo_noRT | Ikbkg | 1.392 | 8.23E-03 | glio |
| Chek1_ex.vivo_noRT | Rab36 | 2.418 | 8.23E-03 | glio |
| Interaction_genotype_Myc | Gm26877 | -3.784 | 8.23E-04 | glio |
| Interaction_genotype_Myc | Gm36371 | -3.451 | 8.23E-04 | glio |
| Interaction_genotype_Myc | S100a10 | 1.844 | 8.23E-04 | glio |
| Hsd17b10_in.vivo_noRT | Slc25a35 | 2.506 | 8.23E-04 | glio |
| Hsd17b10_in.vivo_noRT | Gm15510 | 2.548 | 8.23E-04 | glio |
| Interaction_genotype_Aars2 | Gm49312 | -4.217 | 8.23E-05 | glio |
| Interaction_genotype_Aars2 | Hadha | 2.29 | 8.23E-05 | glio |
| Bcas2_in.vivo_noRT | Gm17655 | 3.143 | 8.23E-05 | glio |
| Bcas2_in.vivo_noRT | Hmgb1 | -0.782 | 8.23E-05 | glio |
| Bora_in.vivo_noRT | Asap3 | 2.727 | 8.23E-05 | glio |
| culture_effect | Lmbr1l | -1.288 | 8.23E-06 | glio |
| culture_effect | Prkar2a | 0.833 | 8.23E-06 | glio |
| Interaction_genotype_Myc | Gm50138 | -4.307 | 8.23E-06 | glio |
| interaction_RT | Phactr1 | -1.279 | 8.23E-06 | glio |
| Chek1_ex.vivo_noRT | Pla2g2f | 4.842 | 8.23E-10 | glio |
| Aars2_in.vivo_noRT | Nrip3 | 4.282 | 8.23E-11 | glio |
| Bcas2_in.vivo_noRT | Cpxm1 | 4.541 | 8.23E-14 | glio |
| Bcas2_in.vivo_noRT | Htr2b | 4.284 | 8.23E-15 | glio |
| Bcas2_in.vivo_noRT | Gm48742 | 5.47 | 8.23E-21 | glio |
| Bcas2_in.vivo_noRT | Gpr4 | 4.807 | 8.23E-21 | glio |
| Bcas2_in.vivo_noRT | Prl7c1 | 4.807 | 8.23E-21 | glio |
| Bcas2_in.vivo_noRT | Gm33466 | 4.807 | 8.23E-21 | glio |
| culture_effect | Clptm1 | -0.415 | 8.24E-03 | glio |
| culture_effect | Dynlt3 | 0.359 | 8.24E-03 | glio |
| Interaction_genotype_Aars2 | Slc25a47 | -3.321 | 8.24E-03 | glio |

| | | | | |
|---|---|---|---|---|
| Interaction_genotype_Aars2 | Rpgr | -3.135 | 8.24E-03 | glio |
| Interaction_genotype_Mtor | Cks1b | 1.564 | 8.24E-03 | glio |
| Interaction_genotype_Myc | Pdcd7 | -1.712 | 8.24E-03 | glio |
| Atp5c1_in.vivo_noRT | Thoc3 | -0.856 | 8.24E-03 | glio |
| Bcas2_in.vivo_noRT | A430018G15R | 1.63 | 8.24E-03 | glio |
| Chek1_in.vivo_noRT | Tubb4a | 2.033 | 8.24E-03 | glio |
| Chek1_in.vivo_noRT | Naa16 | -1.359 | 8.24E-03 | glio |
| Chek1_in.vivo_noRT | Ndufa3 | -1.137 | 8.24E-03 | glio |
| Chek1_in.vivo_noRT | Atp5o | -0.624 | 8.24E-03 | glio |
| Eif2s1_ex.vivo_noRT | Gm2022 | 2.234 | 8.24E-03 | glio |
| Hsd17b10_in.vivo_noRT | Cnfn | 2.221 | 8.24E-03 | glio |
| culture_effect | Taf5 | -0.642 | 8.24E-04 | glio |
| Interaction_genotype_Myc | Lipe | -3.784 | 8.24E-04 | glio |
| Bcas2_ex.vivo_noRT | A330076H08R | 2.742 | 8.24E-04 | glio |
| Bcas2_ex.vivo_noRT | Wdr93 | 3.116 | 8.24E-04 | glio |
| Hsd17b10_in.vivo_noRT | Gm47061 | 2.832 | 8.24E-04 | glio |
| Interaction_genotype_Aars2 | Cdkn2b | 2.629 | 8.24E-05 | glio |
| Interaction_genotype_Mtor | Flnc | -4.739 | 8.24E-05 | glio |
| Interaction_genotype_Mtor | Pard3bos1 | -4.28 | 8.24E-05 | glio |
| Aars2_in.vivo_noRT | Ift27 | -1.715 | 8.24E-05 | glio |
| Chek1_in.vivo_noRT | Grpel2 | -1.908 | 8.24E-06 | glio |
| Chek1_ex.vivo_noRT | Celsr3 | 3.932 | 8.24E-07 | glio |
| Chek1_ex.vivo_noRT | Pnrc1 | 1.546 | 8.24E-07 | glio |
| Bora_in.vivo_noRT | Cxcr2 | 3.624 | 8.24E-08 | glio |
| Atr_in.vivo_noRT | Gm16794 | 3.962 | 8.24E-09 | glio |
| Chek1_in.vivo_noRT | Pcdhga2 | 3.773 | 8.24E-09 | glio |
| culture_effect | Cct8 | -0.713 | 8.24E-10 | glio |
| Chek1_in.vivo_noRT | Has1 | 4.146 | 8.24E-11 | glio |
| Aars2_in.vivo_noRT | Gm45890 | 4.924 | 8.24E-12 | glio |
| Bora_in.vivo_noRT | Gm50394 | 4.67 | 8.24E-12 | glio |
| Atr_in.vivo_noRT | Gm43331 | 4.374 | 8.24E-14 | glio |
| Chek1_ex.vivo_noRT | Gm14929 | 4.842 | 8.24E-16 | glio |
| Interaction_genotype_Aars2 | Gm42984 | -3.328 | 8.25E-03 | glio |
| Interaction_genotype_Bora | Gm31063 | -2.659 | 8.25E-03 | glio |
| Interaction_genotype_Rad51 | Rab10 | 1.127 | 8.25E-03 | glio |
| Aars2_ex.vivo_noRT | Kcnj13 | 2.417 | 8.25E-03 | glio |
| Atr_in.vivo_noRT | 2900009J06Ri | 1.868 | 8.25E-03 | glio |
| Atr_in.vivo_noRT | Cnst | 1.536 | 8.25E-03 | glio |
| Atr_ex.vivo_noRT | Iqch | 2.199 | 8.25E-03 | glio |
| Atr_ex.vivo_noRT | Lrrc46 | 2.583 | 8.25E-03 | glio |
| Bora_in.vivo_noRT | Lrp4 | 1.979 | 8.25E-03 | glio |
| culture_effect | Col5a2 | -1.66 | 8.25E-04 | glio |
| Interaction_genotype_Mtor | Rps6kl1 | -4.083 | 8.25E-04 | glio |
| Atp5c1_ex.vivo_noRT | Pla1a | 3.758 | 8.25E-04 | glio |
| Atp5c1_ex.vivo_noRT | Gjd4 | 2.61 | 8.25E-04 | glio |
| Atr_in.vivo_noRT | Hba-a1 | 2.892 | 8.25E-04 | glio |
| Chek1_in.vivo_noRT | Gtf2ird2 | 2.169 | 8.25E-04 | glio |
| Interaction_genotype_Mtor | Gm11899 | -3.577 | 8.25E-05 | glio |
| Aars2_in.vivo_noRT | Myorg | 2.728 | 8.25E-05 | glio |
| Aars2_in.vivo_noRT | Foxc2 | 2.845 | 8.25E-05 | glio |
| Interaction_genotype_Ifnar1 | Igtp | -1.772 | 8.25E-06 | glio |
| Interaction_genotype_Ifnar1 | Gbp7 | -1.487 | 8.25E-06 | glio |

| | | | | |
|---|---|---|---|---|
| Interaction_genotype_Ifnar1 | Ifi47 | -3.319 | 8.25E-06 | glio |
| Aars2_in.vivo_noRT | Gm48832 | 3.311 | 8.25E-07 | glio |
| Aars2_in.vivo_noRT | Hoxa13 | 3.88 | 8.25E-08 | glio |
| Bcas2_in.vivo_noRT | Gm45441 | 3.748 | 8.25E-08 | glio |
| Aars2_in.vivo_noRT | Gm34256 | 4.526 | 8.25E-10 | glio |
| Aars2_in.vivo_noRT | Gipc2 | 5.063 | 8.25E-12 | glio |
| culture_effect | Mkrn1 | -1.613 | 8.25E-24 | glio |
| Interaction_genotype_Atr | Accsl | -3.567 | 8.26E-03 | glio |
| Interaction_genotype_Bcas2 | Qars | 1.669 | 8.26E-03 | glio |
| Interaction_genotype_Bcas2 | Mcm3 | 1.746 | 8.26E-03 | glio |
| Interaction_genotype_Myc | Kptn | -2.755 | 8.26E-03 | glio |
| Interaction_genotype_Myc | Tbc1d9b | -2.332 | 8.26E-03 | glio |
| Interaction_genotype_Myc | 2700046G09R | -2.263 | 8.26E-03 | glio |
| Interaction_genotype_Myc | Gm9828 | -2.581 | 8.26E-03 | glio |
| Interaction_genotype_Pars2 | Psca | -3.316 | 8.26E-03 | glio |
| Atr_in.vivo_noRT | Ifi213 | -1.482 | 8.26E-03 | glio |
| Bora_in.vivo_noRT | Gm45051 | 1.788 | 8.26E-03 | glio |
| Bora_in.vivo_noRT | Ephx4 | 1.116 | 8.26E-03 | glio |
| Cenpj_in.vivo_noRT | Zfp575 | 2.477 | 8.26E-03 | glio |
| Chek1_in.vivo_noRT | Irf5 | 2.266 | 8.26E-03 | glio |
| Chek1_ex.vivo_noRT | Gm13404 | 2.49 | 8.26E-03 | glio |
| Chek1_ex.vivo_noRT | 2310015A10R | 2.484 | 8.26E-03 | glio |
| Hsd17b10_in.vivo_noRT | Gm10734 | 2.1 | 8.26E-03 | glio |
| Interaction_genotype_Myc | Rbm48 | -2.633 | 8.26E-04 | glio |
| Chek1_in.vivo_noRT | Cdc42ep2 | 2.49 | 8.26E-04 | glio |
| Chek1_in.vivo_noRT | Rbm39 | -0.697 | 8.26E-04 | glio |
| Hsd17b10_in.vivo_noRT | Gm8212 | 2.514 | 8.26E-04 | glio |
| culture_effect | Ppp1ca | 0.422 | 8.26E-05 | glio |
| Interaction_genotype_Mtor | Wdr90 | -4.425 | 8.26E-05 | glio |
| Aars2_in.vivo_noRT | Shroom1 | 2.709 | 8.26E-05 | glio |
| Bcas2_ex.vivo_noRT | Gm15533 | 2.742 | 8.26E-05 | glio |
| Interaction_genotype_Mtor | Cope | 2.132 | 8.26E-06 | glio |
| Chek1_in.vivo_noRT | Astn2 | 2.956 | 8.26E-06 | glio |
| Bcas2_in.vivo_noRT | 2600006K01R | 3.604 | 8.26E-07 | glio |
| Bora_in.vivo_noRT | Phf11a | 3.368 | 8.26E-07 | glio |
| Hsd17b10_in.vivo_noRT | Gm34047 | 3.616 | 8.26E-07 | glio |
| Bcas2_in.vivo_noRT | Gm48933 | 3.805 | 8.26E-08 | glio |
| Chek1_in.vivo_noRT | Gm5624 | 4.054 | 8.26E-10 | glio |
| Atr_in.vivo_noRT | Ms4a6d | 4.745 | 8.26E-16 | glio |
| Atr_in.vivo_noRT | Gm40264 | 5.273 | 8.26E-19 | glio |
| culture_effect | Adra1b | -2.557 | 8.26E-20 | glio |
| Aars2_in.vivo_noRT | C330002G04R | 5.327 | 8.26E-24 | glio |
| Interaction_genotype_Atr | AC159140.1 | -2.503 | 8.27E-03 | glio |
| Interaction_genotype_Atr | Atp6v0b | 0.779 | 8.27E-03 | glio |
| Interaction_genotype_Myc | Slc9a5 | -2.882 | 8.27E-03 | glio |
| Atr_in.vivo_noRT | Gm20274 | 1.582 | 8.27E-03 | glio |
| Atr_ex.vivo_noRT | Akr1b7 | 2.764 | 8.27E-03 | glio |
| Bard1_ex.vivo_noRT | Tdrp | 2.57 | 8.27E-03 | glio |
| Bcas2_in.vivo_noRT | Gm43445 | 1.901 | 8.27E-03 | glio |
| Bcas2_in.vivo_noRT | Tbx2 | 1.63 | 8.27E-03 | glio |
| Chek1_in.vivo_noRT | Gdf11 | 1.507 | 8.27E-03 | glio |
| Cyp19a1_ex.vivo_noRT | Npnt | 2.161 | 8.27E-03 | glio |

| | | | | |
|---|---|---|---|---|
| Cyp19a1_ex.vivo_noRT | Gm15816 | 2.604 | 8.27E-03 | glio |
| Fanca_ex.vivo_noRT | Angptl7 | 2.124 | 8.27E-03 | glio |
| Fanca_ex.vivo_noRT | Cldn5 | 2.124 | 8.27E-03 | glio |
| Interaction_genotype_Myc | Gm49172 | -2.469 | 8.27E-04 | glio |
| Interaction_genotype_Myc | Zkscan4 | -3.871 | 8.27E-04 | glio |
| Interaction_genotype_Myc | Zfp40 | -3.565 | 8.27E-04 | glio |
| Interaction_genotype_Myc | Trmt112 | 1.273 | 8.27E-04 | glio |
| Interaction_genotype_Myc | Rps25 | 1.31 | 8.27E-04 | glio |
| Cdkn2a_in.vivo_noRT | Rbfaos | -2.592 | 8.27E-04 | glio |
| Cdkn2a_in.vivo_noRT | Ablim2 | -2.207 | 8.27E-04 | glio |
| Cdkn2a_in.vivo_noRT | Cntn1 | -2.064 | 8.27E-04 | glio |
| Cdkn2a_in.vivo_noRT | Ivns1abp | 0.956 | 8.27E-04 | glio |
| Cdkn2a_in.vivo_noRT | Trf | 1.411 | 8.27E-04 | glio |
| Chek1_ex.vivo_noRT | 1810012K16R | 2.877 | 8.27E-04 | glio |
| Chek1_ex.vivo_noRT | Rnf144b | 3.079 | 8.27E-04 | glio |
| Hsd17b10_in.vivo_noRT | Lncenc1 | 2.31 | 8.27E-04 | glio |
| culture_effect | Smap1 | 0.515 | 8.27E-05 | glio |
| Interaction_genotype_Mtor | B230206H07R | -3.176 | 8.27E-05 | glio |
| Interaction_genotype_Mtor | Ndufb9 | 1.553 | 8.27E-05 | glio |
| interaction_RT | Samm50 | -0.854 | 8.27E-05 | glio |
| Atr_ex.vivo_noRT | Thnsl2 | 3.164 | 8.27E-06 | glio |
| Bard1_ex.vivo_noRT | Gm20658 | 2.814 | 8.27E-06 | glio |
| Bora_in.vivo_noRT | Cib2 | 3.578 | 8.27E-06 | glio |
| Chek1_ex.vivo_noRT | Rax | 3.689 | 8.27E-06 | glio |
| Aars2_in.vivo_noRT | Itga3 | 3.697 | 8.27E-07 | glio |
| Aars2_in.vivo_noRT | March11 | 3.59 | 8.27E-07 | glio |
| Aars2_in.vivo_noRT | Matk | 3.175 | 8.27E-07 | glio |
| Chek1_in.vivo_noRT | E230025N22R | 3.22 | 8.27E-07 | glio |
| culture_effect | Cotl1 | 1.502 | 8.27E-10 | glio |
| Bora_in.vivo_noRT | BC049987 | 4.153 | 8.27E-17 | glio |
| Interaction_genotype_Atr | Lcat | -3.45 | 8.28E-03 | glio |
| Interaction_genotype_Atr | Zbtb8os | 1.388 | 8.28E-03 | glio |
| Interaction_genotype_Myc | Dnajc5 | -1.519 | 8.28E-03 | glio |
| Aars2_in.vivo_noRT | B230217C12R | 1.355 | 8.28E-03 | glio |
| Aars2_in.vivo_noRT | Rwdd1 | -0.524 | 8.28E-03 | glio |
| Aars2_ex.vivo_noRT | Il12rb1 | 2.505 | 8.28E-03 | glio |
| Bcas2_in.vivo_noRT | H60b | 1.909 | 8.28E-03 | glio |
| Bcas2_in.vivo_noRT | Snta1 | 1.567 | 8.28E-03 | glio |
| Bora_in.vivo_noRT | Haus2 | -1.431 | 8.28E-03 | glio |
| Bora_in.vivo_noRT | Dnm2 | -1.206 | 8.28E-03 | glio |
| Bora_in.vivo_noRT | Thoc7 | -0.994 | 8.28E-03 | glio |
| culture_effect | Tmem101 | -0.749 | 8.28E-04 | glio |
| Interaction_genotype_Mtor | Grip2 | -4.033 | 8.28E-04 | glio |
| Interaction_genotype_Mtor | Serf2 | 1.414 | 8.28E-04 | glio |
| Bora_in.vivo_noRT | Med28 | -0.734 | 8.28E-04 | glio |
| Atr_ex.vivo_noRT | Ier5 | 2.471 | 8.28E-05 | glio |
| Bora_ex.vivo_noRT | Gm12589 | 2.802 | 8.28E-05 | glio |
| Aars2_in.vivo_noRT | Tsn | -1.024 | 8.28E-06 | glio |
| Atr_in.vivo_noRT | Gm49077 | 3.245 | 8.28E-06 | glio |
| Hsd17b10_in.vivo_noRT | Gm21818 | 3.616 | 8.28E-07 | glio |
| Interaction_genotype_Mtor | D430001F17R | -6.044 | 8.28E-08 | glio |
| Chek1_in.vivo_noRT | Tmem179 | 3.885 | 8.28E-08 | glio |

| | | | | |
|---|---|---|---|---|
| Chek1_in.vivo_noRT | Adcy2 | 3.29 | 8.28E-08 | glio |
| Bcas2_in.vivo_noRT | Prl2c3 | 4.807 | 8.28E-14 | glio |
| Atr_in.vivo_noRT | Gm50024 | 5.27 | 8.28E-26 | glio |
| culture_effect | D930016D06F | -0.815 | 8.29E-03 | glio |
| culture_effect | Arl6ip5 | -0.563 | 8.29E-03 | glio |
| Interaction_genotype_Atr | Gm20682 | -2.805 | 8.29E-03 | glio |
| Interaction_genotype_Atr | Ift88 | -2.683 | 8.29E-03 | glio |
| Interaction_genotype_Mtor | Clba1 | -2.416 | 8.29E-03 | glio |
| Interaction_genotype_Myc | D130062J10Ri | -2.797 | 8.29E-03 | glio |
| Interaction_genotype_Myc | Zfp747 | -2.741 | 8.29E-03 | glio |
| Bora_in.vivo_noRT | Ndufa10 | -0.729 | 8.29E-03 | glio |
| Cenpj_ex.vivo_noRT | Gm20732 | 2.591 | 8.29E-03 | glio |
| Cenpj_ex.vivo_noRT | Vgll2 | 2.591 | 8.29E-03 | glio |
| Chek1_in.vivo_noRT | Frk | 1.962 | 8.29E-03 | glio |
| Chek1_in.vivo_noRT | Cbx2 | 1.762 | 8.29E-03 | glio |
| Chek1_ex.vivo_noRT | C230037L18Ri | 2.276 | 8.29E-03 | glio |
| Chek1_ex.vivo_noRT | Ccdc78 | 2.409 | 8.29E-03 | glio |
| Eif2s1_in.vivo_noRT | Pla1a | 2.142 | 8.29E-03 | glio |
| Eif2s1_in.vivo_noRT | Gm49312 | 2.118 | 8.29E-03 | glio |
| culture_effect | Itga8 | 1.031 | 8.29E-04 | glio |
| culture_effect | Tmem245 | 0.611 | 8.29E-04 | glio |
| Interaction_genotype_Myc | Cd36 | -2.729 | 8.29E-04 | glio |
| Interaction_genotype_Myc | 4833417C18R | -3.791 | 8.29E-04 | glio |
| Interaction_genotype_Myc | Hist1h2bn | -3.676 | 8.29E-04 | glio |
| Interaction_genotype_Myc | Rbpms2 | -3.625 | 8.29E-04 | glio |
| Interaction_genotype_Myc | Igsf11 | -3.58 | 8.29E-04 | glio |
| Interaction_genotype_Myc | Gng7 | -3.478 | 8.29E-04 | glio |
| Interaction_genotype_Myc | Ube2n | 1.196 | 8.29E-04 | glio |
| Interaction_genotype_Rad51 | Rassf6 | -3.885 | 8.29E-04 | glio |
| Atr_in.vivo_noRT | Rhobtb3 | -1.591 | 8.29E-04 | glio |
| Atr_in.vivo_noRT | Nhp2 | -0.884 | 8.29E-04 | glio |
| Bcas2_in.vivo_noRT | Dlg5 | 2.294 | 8.29E-04 | glio |
| Atr_in.vivo_noRT | Dennd2c | 2.747 | 8.29E-05 | glio |
| Bcas2_in.vivo_noRT | Pde4d | 3.161 | 8.29E-06 | glio |
| Chek1_ex.vivo_noRT | Sh3gl3 | 3.929 | 8.29E-06 | glio |
| culture_effect | Eps8l2 | 2.624 | 8.29E-07 | glio |
| Bora_in.vivo_noRT | Retn | 4.469 | 8.29E-07 | glio |
| Aars2_in.vivo_noRT | 4930515G01R | 3.444 | 8.29E-08 | glio |
| Aars2_in.vivo_noRT | Rab3b | 4.17 | 8.29E-10 | glio |
| Bcas2_in.vivo_noRT | Cxcl13 | 4.807 | 8.29E-13 | glio |
| Chek1_ex.vivo_noRT | 1700019E08Ri | 4.842 | 8.29E-13 | glio |
| culture_effect | Uqcrc2 | -1.115 | 8.29E-15 | glio |
| Bora_in.vivo_noRT | Angptl7 | 4.07 | 8.29E-17 | glio |
| culture_effect | Sptbn4 | 1.484 | 8.30E-03 | glio |
| culture_effect | Rad51d | 0.569 | 8.30E-03 | glio |
| culture_effect | Smap2 | 0.507 | 8.30E-03 | glio |
| Interaction_genotype_Mtor | Tlcd2 | -3.523 | 8.30E-03 | glio |
| Aars2_in.vivo_noRT | Aldh3b1 | 1.922 | 8.30E-03 | glio |
| Atr_ex.vivo_noRT | 5730507C01R | 1.973 | 8.30E-03 | glio |
| Atr_ex.vivo_noRT | Igfbp7 | 2.367 | 8.30E-03 | glio |
| Bcas2_in.vivo_noRT | Spats1 | 1.976 | 8.30E-03 | glio |
| Chek1_ex.vivo_noRT | Cdo1 | 2.591 | 8.30E-03 | glio |

| | | | | |
|---|---|---|---|---|
| Chek1_ex.vivo_noRT | C2 | 2.059 | 8.30E-03 | glio |
| Chek1_ex.vivo_noRT | Fxn | -1.49 | 8.30E-03 | glio |
| Interaction_genotype_Atr | Foxred2 | -3.248 | 8.30E-04 | glio |
| Bora_ex.vivo_noRT | Gm44646 | 2.615 | 8.30E-04 | glio |
| Aars2_in.vivo_noRT | Fam161b | 2.779 | 8.30E-05 | glio |
| Aars2_in.vivo_noRT | Eif4b | -1.148 | 8.30E-05 | glio |
| Aars2_in.vivo_noRT | Arpc1a | -1.193 | 8.30E-05 | glio |
| Atr_in.vivo_noRT | Anxa11 | 3.115 | 8.30E-05 | glio |
| Atr_in.vivo_noRT | Gm26901 | 2.947 | 8.30E-05 | glio |
| Bcas2_in.vivo_noRT | Iqgap1 | -1.047 | 8.30E-05 | glio |
| Bora_in.vivo_noRT | Tspan3 | -1.421 | 8.30E-06 | glio |
| culture_effect | Gm42997 | 2.397 | 8.30E-07 | glio |
| Bcas2_in.vivo_noRT | Ccl19 | 2.856 | 8.30E-07 | glio |
| Bcas2_in.vivo_noRT | Ocm | 3.029 | 8.30E-07 | glio |
| Atp5c1_in.vivo_noRT | A930019D19R | 2.668 | 8.30E-08 | glio |
| Bcas2_in.vivo_noRT | Col6a3 | 3.771 | 8.30E-10 | glio |
| Aars2_in.vivo_noRT | Spdye4a | 4.669 | 8.30E-11 | glio |
| Aars2_in.vivo_noRT | Gm16685 | 4.668 | 8.30E-16 | glio |
| Interaction_genotype_Aars2 | 4930447K03R | -2.562 | 8.31E-03 | glio |
| Interaction_genotype_Aars2 | Rrs1 | 1.359 | 8.31E-03 | glio |
| Interaction_genotype_Bcas2 | Gm45833 | -4.58 | 8.31E-03 | glio |
| Interaction_genotype_Myc | Tctn3 | -3.169 | 8.31E-03 | glio |
| Interaction_genotype_Myc | Tfap4 | -2.308 | 8.31E-03 | glio |
| interaction_RT | Iah1 | 0.667 | 8.31E-03 | glio |
| Aars2_in.vivo_noRT | Ulk1 | 1.522 | 8.31E-03 | glio |
| Atp5c1_in.vivo_noRT | Gm48265 | 1.885 | 8.31E-03 | glio |
| Atr_in.vivo_noRT | Uap1l1 | -1.31 | 8.31E-03 | glio |
| Hsd17b10_in.vivo_noRT | Gm16174 | 2.226 | 8.31E-03 | glio |
| Hsd17b10_in.vivo_noRT | Pcsk4 | 2.067 | 8.31E-03 | glio |
| culture_effect | Cth | -1.075 | 8.31E-04 | glio |
| Interaction_genotype_Aars2 | Gm10425 | -4.327 | 8.31E-04 | glio |
| Interaction_genotype_Aars2 | Prl3d3 | -2.866 | 8.31E-04 | glio |
| Interaction_genotype_Rad9a | Klrk1 | 4.305 | 8.31E-04 | glio |
| Interaction_genotype_Rad9a | Gm29340 | -4.76 | 8.31E-04 | glio |
| Interaction_genotype_Rad9a | Nxph4 | -5.014 | 8.31E-04 | glio |
| Atr_in.vivo_noRT | Hnrnpa3 | -0.996 | 8.31E-04 | glio |
| Bora_in.vivo_noRT | Azin1 | -1.075 | 8.31E-04 | glio |
| Chek1_ex.vivo_noRT | Gm15743 | 2.754 | 8.31E-04 | glio |
| Aars2_in.vivo_noRT | Atp5g1 | -0.818 | 8.31E-05 | glio |
| Chek1_ex.vivo_noRT | Scarb1 | -2.508 | 8.31E-05 | glio |
| Eif2s1_in.vivo_noRT | A930019D19R | 2.143 | 8.31E-05 | glio |
| interaction_RT | Yaf2 | 0.859 | 8.31E-06 | glio |
| Bcas2_in.vivo_noRT | Krt17 | 5.485 | 8.31E-06 | glio |
| Bcas2_in.vivo_noRT | Gm20513 | 2.71 | 8.31E-07 | glio |
| Bcas2_in.vivo_noRT | Noxred1 | 4.079 | 8.31E-10 | glio |
| Atr_in.vivo_noRT | Gm9922 | 4.398 | 8.31E-11 | glio |
| Aars2_in.vivo_noRT | Ksr2 | 4.289 | 8.31E-14 | glio |
| Aars2_in.vivo_noRT | Zfp786 | 5.063 | 8.31E-14 | glio |
| Aars2_in.vivo_noRT | Rapgef3os2 | 5.327 | 8.31E-24 | glio |
| Aars2_in.vivo_noRT | Gpr4 | 5.327 | 8.31E-25 | glio |
| Aars2_in.vivo_noRT | Prl7c1 | 5.327 | 8.31E-25 | glio |
| Aars2_in.vivo_noRT | Gm33466 | 5.327 | 8.31E-25 | glio |

| | | | | |
|---|---|---|---|---|
| culture_effect | Kcna4 | -1.694 | 8.32E-03 | glio |
| Interaction_genotype_Aars2 | Tedc2 | -2.789 | 8.32E-03 | glio |
| Interaction_genotype_Mtor | Sik1 | -3.311 | 8.32E-03 | glio |
| Interaction_genotype_Mtor | Kctd12 | -3.138 | 8.32E-03 | glio |
| Interaction_genotype_Mtor | Reps1 | -2.046 | 8.32E-03 | glio |
| Aars2_in.vivo_noRT | Gm10130 | 1.525 | 8.32E-03 | glio |
| Aars2_in.vivo_noRT | Samd4b | -1.066 | 8.32E-03 | glio |
| Aars2_in.vivo_noRT | Ndufb6 | -0.648 | 8.32E-03 | glio |
| Atr_in.vivo_noRT | Vat1 | -1.098 | 8.32E-03 | glio |
| Atr_in.vivo_noRT | Uck2 | -0.935 | 8.32E-03 | glio |
| Chek1_ex.vivo_noRT | Sprr1a | 2.47 | 8.32E-03 | glio |
| Chek1_ex.vivo_noRT | Cr1l | -1.415 | 8.32E-03 | glio |
| Chek1_ex.vivo_noRT | Lyar | -0.813 | 8.32E-03 | glio |
| Fanca_ex.vivo_noRT | Dpep3 | 2.124 | 8.32E-03 | glio |
| Fanca_ex.vivo_noRT | Erich6b | 2.124 | 8.32E-03 | glio |
| Fanca_ex.vivo_noRT | Tac2 | 2.124 | 8.32E-03 | glio |
| Fanca_ex.vivo_noRT | Prdm1 | 2.886 | 8.32E-03 | glio |
| Interaction_genotype_Aars2 | Mir670hg | -4.424 | 8.32E-04 | glio |
| Interaction_genotype_Aars2 | Gm14636 | -4.793 | 8.32E-04 | glio |
| Interaction_genotype_Aars2 | Copz1 | 1.59 | 8.32E-04 | glio |
| Interaction_genotype_Aars2 | Guk1 | 1.301 | 8.32E-04 | glio |
| Interaction_genotype_Aars2 | Ndufs6 | 1.213 | 8.32E-04 | glio |
| Interaction_genotype_Myc | Olfr718-ps1 | -2.859 | 8.32E-04 | glio |
| Interaction_genotype_Myc | Pygb | -3.07 | 8.32E-04 | glio |
| Interaction_genotype_Myc | Gemin2 | -2.381 | 8.32E-04 | glio |
| Interaction_genotype_Myc | Bnip2 | 1.42 | 8.32E-04 | glio |
| Bard1_ex.vivo_noRT | Pax5 | 3.464 | 8.32E-04 | glio |
| Bora_in.vivo_noRT | B230209E15Ri | 2.656 | 8.32E-04 | glio |
| Bora_in.vivo_noRT | Alyref2 | 2.022 | 8.32E-04 | glio |
| Chek1_ex.vivo_noRT | Clec1a | 3.079 | 8.32E-04 | glio |
| Atr_in.vivo_noRT | B3gntl1 | 2.683 | 8.32E-05 | glio |
| Bard1_ex.vivo_noRT | Fabp6 | 2.814 | 8.32E-05 | glio |
| Bard1_ex.vivo_noRT | Tcstv1 | 3.213 | 8.32E-05 | glio |
| Chek1_ex.vivo_noRT | Gm28578 | 3.598 | 8.32E-05 | glio |
| Aars2_in.vivo_noRT | Arl4d | 3.478 | 8.32E-07 | glio |
| Aars2_in.vivo_noRT | Ifi207 | 5.063 | 8.32E-12 | glio |
| Bcas2_in.vivo_noRT | Gm39929 | 4.349 | 8.32E-15 | glio |
| Chek1_in.vivo_noRT | Luzp2 | 4.518 | 8.32E-18 | glio |
| culture_effect | Card19 | 1.629 | 8.32E-24 | glio |
| culture_effect | Tslrn1 | 1.067 | 8.33E-03 | glio |
| Interaction_genotype_Lmo7 | Gm49494 | -4.047 | 8.33E-03 | glio |
| Interaction_genotype_Mtor | Vav2 | -2.949 | 8.33E-03 | glio |
| interaction_RT | Bcl6 | 2.339 | 8.33E-03 | glio |
| interaction_RT | Reps2 | -0.765 | 8.33E-03 | glio |
| Bora_in.vivo_noRT | Srsf4 | -1.014 | 8.33E-03 | glio |
| Bora_in.vivo_noRT | Actl6a | -0.94 | 8.33E-03 | glio |
| Chek1_in.vivo_noRT | Gm5141 | 1.931 | 8.33E-03 | glio |
| Chek1_in.vivo_noRT | Fzd5 | 1.888 | 8.33E-03 | glio |
| Chek1_in.vivo_noRT | Gm20756 | 1.8 | 8.33E-03 | glio |
| Chek1_ex.vivo_noRT | Sult2b1 | 2.103 | 8.33E-03 | glio |
| culture_effect | Il6st | 0.761 | 8.33E-04 | glio |
| culture_effect | Rspry1 | -0.578 | 8.33E-04 | glio |

| | | | | |
|---|---|---|---|---|
| Interaction_genotype_Pars2 | Zfp786 | -4.529 | 8.33E-04 | glio |
| Aars2_ex.vivo_noRT | Dreh | 2.899 | 8.33E-04 | glio |
| Eif2s1_in.vivo_noRT | Meioc | 1.88 | 8.33E-04 | glio |
| Eif2s1_in.vivo_noRT | Gm6803 | 2.143 | 8.33E-04 | glio |
| Eif2s1_in.vivo_noRT | Ifitm5 | 2.669 | 8.33E-04 | glio |
| Hsd17b10_in.vivo_noRT | Gm9908 | 2.699 | 8.33E-04 | glio |
| Chek1_in.vivo_noRT | Rspo3 | 3.218 | 8.33E-05 | glio |
| Chek1_in.vivo_noRT | Cenpq | -1.454 | 8.33E-05 | glio |
| Atr_ex.vivo_noRT | Akap14 | 3.164 | 8.33E-06 | glio |
| Chek1_ex.vivo_noRT | Cyp7a1 | 3.866 | 8.33E-06 | glio |
| Aars2_in.vivo_noRT | Usp44 | 3.852 | 8.33E-08 | glio |
| Chek1_in.vivo_noRT | Gm36757 | 3.65 | 8.33E-08 | glio |
| Hsd17b10_in.vivo_noRT | Ccl3 | 3.356 | 8.33E-11 | glio |
| Bora_in.vivo_noRT | S100a14 | 4.136 | 8.33E-12 | glio |
| Chek1_in.vivo_noRT | D930028M14I | 5.044 | 8.33E-16 | glio |
| culture_effect | H2-Eb1 | -12.217 | 8.33E-69 | glio |
| Interaction_genotype_Aars2 | Flicr | -3.271 | 8.34E-03 | glio |
| Interaction_genotype_Mtor | Serpina3g | -2.903 | 8.34E-03 | glio |
| Interaction_genotype_Rad51 | Otos | -3.031 | 8.34E-03 | glio |
| Bard1_ex.vivo_noRT | Gm9999 | 2.636 | 8.34E-03 | glio |
| Eif2s1_in.vivo_noRT | Syp | 2.531 | 8.34E-03 | glio |
| Hsd17b10_in.vivo_noRT | D830035M03I | 2.03 | 8.34E-03 | glio |
| culture_effect | Gm45620 | 1.924 | 8.34E-04 | glio |
| Interaction_genotype_Atr | Stx1b | -3.828 | 8.34E-04 | glio |
| Interaction_genotype_Atr | Pitx3 | -3.35 | 8.34E-04 | glio |
| Interaction_genotype_Myc | Gm45441 | -3.805 | 8.34E-04 | glio |
| Atr_ex.vivo_noRT | S100a14 | 2.764 | 8.34E-04 | glio |
| Bcas2_in.vivo_noRT | F3 | -1.395 | 8.34E-04 | glio |
| Bora_in.vivo_noRT | Trip13 | -1.742 | 8.34E-04 | glio |
| Interaction_genotype_Mtor | Xylb | -4.741 | 8.34E-05 | glio |
| Interaction_genotype_Mtor | Psd3 | -3.881 | 8.34E-05 | glio |
| Aars2_in.vivo_noRT | Slc11a1 | 2.749 | 8.34E-05 | glio |
| Aars2_ex.vivo_noRT | Nmrk2 | 3.293 | 8.34E-05 | glio |
| Aars2_ex.vivo_noRT | C330002G04R | 2.706 | 8.34E-05 | glio |
| Atr_in.vivo_noRT | Klf15 | 2.776 | 8.34E-05 | glio |
| Chek1_in.vivo_noRT | Gm5431 | 2.6 | 8.34E-05 | glio |
| Interaction_genotype_Mtor | St13 | 2.108 | 8.34E-06 | glio |
| Interaction_genotype_Myc | Grk6 | -3.503 | 8.34E-06 | glio |
| Chek1_ex.vivo_noRT | Nell2 | 3.664 | 8.34E-06 | glio |
| Atp5c1_in.vivo_noRT | Gm49172 | 2.408 | 8.34E-07 | glio |
| Atr_in.vivo_noRT | Slco2a1 | 3.535 | 8.34E-08 | glio |
| Interaction_genotype_Aars2 | Rpl28 | 0.765 | 8.35E-03 | glio |
| Interaction_genotype_Atr | Gm15890 | -2.76 | 8.35E-03 | glio |
| Interaction_genotype_Bcas2 | Gm30692 | -2.458 | 8.35E-03 | glio |
| Interaction_genotype_Bcas2 | Osr2 | 3.061 | 8.35E-03 | glio |
| Interaction_genotype_Bcas2 | Ube2cbp | -3.107 | 8.35E-03 | glio |
| Interaction_genotype_Ifnar1 | Glyat | 2.733 | 8.35E-03 | glio |
| Interaction_genotype_Ifnar1 | Tpo | 2.734 | 8.35E-03 | glio |
| Interaction_genotype_Ifnar1 | B2m | -0.883 | 8.35E-03 | glio |
| Interaction_genotype_Mtor | Cadm4 | -2.01 | 8.35E-03 | glio |
| Interaction_genotype_Myc | Dusp12 | -2.108 | 8.35E-03 | glio |
| Interaction_genotype_Myc | Arhgap28 | -2.084 | 8.35E-03 | glio |

| | | | | |
|---|---|---|---|---|
| Interaction_genotype_Myc | Uap1l1 | -1.85 | 8.35E-03 | glio |
| Interaction_genotype_Pars2 | Gm30938 | -3.202 | 8.35E-03 | glio |
| Bard1_ex.vivo_noRT | Bco1 | 2.419 | 8.35E-03 | glio |
| Bcas2_ex.vivo_noRT | Serpinb9 | 2.73 | 8.35E-03 | glio |
| Bora_in.vivo_noRT | Dnajb13 | 1.857 | 8.35E-03 | glio |
| Bora_ex.vivo_noRT | Bbof1 | 2.626 | 8.35E-03 | glio |
| Bora_ex.vivo_noRT | Cyp2d22 | 2.93 | 8.35E-03 | glio |
| Cenpj_in.vivo_noRT | Aftph | 0.693 | 8.35E-03 | glio |
| Chek1_ex.vivo_noRT | Gm47644 | 2.501 | 8.35E-03 | glio |
| Chek1_ex.vivo_noRT | Haglr | 2.667 | 8.35E-03 | glio |
| Chek1_ex.vivo_noRT | Nol3 | 2.227 | 8.35E-03 | glio |
| Chek1_ex.vivo_noRT | Cdk20 | 2.093 | 8.35E-03 | glio |
| culture_effect | 2900089D17R | 1.533 | 8.35E-04 | glio |
| Interaction_genotype_Mtor | Sec61g | 1.899 | 8.35E-04 | glio |
| Interaction_genotype_Myc | Mks1 | -3.22 | 8.35E-04 | glio |
| Interaction_genotype_Myc | Polg2 | -3.06 | 8.35E-04 | glio |
| Atp5c1_in.vivo_noRT | Gm43256 | 2.405 | 8.35E-04 | glio |
| Bora_in.vivo_noRT | Bnip3l | -1.264 | 8.35E-04 | glio |
| culture_effect | 2210417A02R | 2.306 | 8.35E-05 | glio |
| Interaction_genotype_Mtor | Gm27010 | -4.453 | 8.35E-05 | glio |
| Interaction_genotype_Mtor | Gm36839 | -4.595 | 8.35E-05 | glio |
| Atr_ex.vivo_noRT | Rab17 | 3.642 | 8.35E-05 | glio |
| Bora_in.vivo_noRT | Gatad2a | -1.451 | 8.35E-05 | glio |
| Hsd17b10_in.vivo_noRT | Mogat1 | 3.094 | 8.35E-05 | glio |
| Interaction_genotype_Myc | Uqcc2 | 1.902 | 8.35E-06 | glio |
| Bora_in.vivo_noRT | Hnrnpa2b1 | -0.787 | 8.35E-06 | glio |
| Chek1_ex.vivo_noRT | Iqcf1 | 3.754 | 8.35E-06 | glio |
| Aars2_in.vivo_noRT | Cyp2u1 | 3.223 | 8.35E-07 | glio |
| Bora_in.vivo_noRT | B930059L03Ri | 3.751 | 8.35E-07 | glio |
| culture_effect | Csrnp1 | -1.855 | 8.35E-11 | glio |
| Chek1_in.vivo_noRT | Gm42918 | 4.925 | 8.35E-16 | glio |
| Chek1_in.vivo_noRT | Esp24 | 5.044 | 8.35E-22 | glio |
| Interaction_genotype_Aars2 | 1700124L16Ri | -3.448 | 8.36E-03 | glio |
| Interaction_genotype_Aars2 | Nr1h2 | 1.408 | 8.36E-03 | glio |
| Interaction_genotype_Atr | Serpinb9g | -2.369 | 8.36E-03 | glio |
| Interaction_genotype_Bora | Dcaf12 | 1.433 | 8.36E-03 | glio |
| Interaction_genotype_Myc | Med9os | -2.728 | 8.36E-03 | glio |
| interaction_RT | Nop9 | -0.877 | 8.36E-03 | glio |
| interaction_RT | Siah2 | 0.821 | 8.36E-03 | glio |
| Aars2_in.vivo_noRT | Crocc | 1.878 | 8.36E-03 | glio |
| Aars2_in.vivo_noRT | Mtx3 | 1.741 | 8.36E-03 | glio |
| Aars2_in.vivo_noRT | Alkbh7 | -1.225 | 8.36E-03 | glio |
| Atp5c1_in.vivo_noRT | Gm45890 | 2.273 | 8.36E-03 | glio |
| Chek1_in.vivo_noRT | A230056P14R | 1.949 | 8.36E-03 | glio |
| Chek1_in.vivo_noRT | Lrpprc | -1.36 | 8.36E-03 | glio |
| Chek1_ex.vivo_noRT | Bambi | 2.229 | 8.36E-03 | glio |
| culture_effect | Cisd2 | 0.42 | 8.36E-04 | glio |
| Interaction_genotype_Atr | Slc26a6 | -4.05 | 8.36E-04 | glio |
| Interaction_genotype_Atr | Ak1 | -3.819 | 8.36E-04 | glio |
| Interaction_genotype_Mtor | 3110001I22Ri | -3.94 | 8.36E-04 | glio |
| Aars2_in.vivo_noRT | Ubl5 | -0.766 | 8.36E-04 | glio |
| Bcas2_ex.vivo_noRT | Gpr146 | 3.522 | 8.36E-04 | glio |

| | | | | |
|---|---|---|---|---|
| Bcas2_ex.vivo_noRT | Aoc1 | 3.392 | 8.36E-04 | glio |
| Cenpj_ex.vivo_noRT | Gm11899 | 2.591 | 8.36E-04 | glio |
| Chek1_ex.vivo_noRT | Ccr2 | 3.163 | 8.36E-04 | glio |
| Interaction_genotype_Myc | Ahrr | -4.517 | 8.36E-05 | glio |
| Atr_in.vivo_noRT | Mettl5os | 2.594 | 8.36E-05 | glio |
| Atp5c1_in.vivo_noRT | Gm36172 | 2.663 | 8.36E-06 | glio |
| Bcas2_in.vivo_noRT | Olfr920 | 3.057 | 8.36E-06 | glio |
| Bcas2_in.vivo_noRT | Brsk2 | 2.884 | 8.36E-06 | glio |
| Bora_ex.vivo_noRT | Thnsl2 | 3.201 | 8.36E-06 | glio |
| Interaction_genotype_Aars2 | Gm11627 | -5.532 | 8.36E-07 | glio |
| Interaction_genotype_Aars2 | Gm27239 | -5.931 | 8.36E-07 | glio |
| Aars2_ex.vivo_noRT | Ccl3 | 3.293 | 8.36E-07 | glio |
| Bcas2_in.vivo_noRT | Fam13a | 4.007 | 8.36E-09 | glio |
| Chek1_in.vivo_noRT | A330040F15Ri | 4.264 | 8.36E-09 | glio |
| Aars2_in.vivo_noRT | Gm20667 | 4.326 | 8.36E-10 | glio |
| Atr_in.vivo_noRT | Sema4a | 4.49 | 8.36E-10 | glio |
| Chek1_in.vivo_noRT | Gm47204 | 4.398 | 8.36E-10 | glio |
| Chek1_in.vivo_noRT | Gm13404 | 4.306 | 8.36E-10 | glio |
| Atr_in.vivo_noRT | A330076H08R | 6.071 | 8.36E-25 | glio |
| Atr_in.vivo_noRT | Adam5 | 5.269 | 8.36E-25 | glio |
| Interaction_genotype_Bcas2 | Sh3d21 | -3.04 | 8.37E-03 | glio |
| Interaction_genotype_Myc | Fads1 | -1.729 | 8.37E-03 | glio |
| Interaction_genotype_Rpl3 | Gm13986 | -3.889 | 8.37E-03 | glio |
| Aars2_ex.vivo_noRT | Lncenc1 | 2.495 | 8.37E-03 | glio |
| Aars2_ex.vivo_noRT | Ncf1 | 2.312 | 8.37E-03 | glio |
| Atr_in.vivo_noRT | Nudt16 | 1.685 | 8.37E-03 | glio |
| Bora_in.vivo_noRT | Rsph3a | 1.826 | 8.37E-03 | glio |
| Bora_in.vivo_noRT | Snrpe | -0.48 | 8.37E-03 | glio |
| Hsd17b10_in.vivo_noRT | Slc47a2 | 2.031 | 8.37E-03 | glio |
| culture_effect | Rasip1 | -2.185 | 8.37E-04 | glio |
| culture_effect | 2810403D21R | 0.961 | 8.37E-04 | glio |
| culture_effect | Aurkb | -0.765 | 8.37E-04 | glio |
| Aars2_in.vivo_noRT | Mok | 2.436 | 8.37E-04 | glio |
| Bora_in.vivo_noRT | Atp6v0e | -0.688 | 8.37E-04 | glio |
| Chek1_in.vivo_noRT | Cox16 | -1.116 | 8.37E-04 | glio |
| Aars2_in.vivo_noRT | Gm49768 | 2.574 | 8.37E-05 | glio |
| Bard1_ex.vivo_noRT | Olfr810 | 2.814 | 8.37E-06 | glio |
| Bora_ex.vivo_noRT | Akap14 | 3.201 | 8.37E-06 | glio |
| Bora_ex.vivo_noRT | Ccl9 | 2.808 | 8.37E-06 | glio |
| Aars2_ex.vivo_noRT | AU019990 | 3.293 | 8.37E-07 | glio |
| Atr_in.vivo_noRT | 5930403N24R | 5.007 | 8.37E-13 | glio |
| Bcas2_in.vivo_noRT | Rftn2 | 4.416 | 8.37E-13 | glio |
| Aars2_in.vivo_noRT | Olfr171 | 5.327 | 8.37E-22 | glio |
| Aars2_in.vivo_noRT | Gstt4 | 5.063 | 8.37E-23 | glio |
| Interaction_genotype_Mtor | Kptn | -2.96 | 8.38E-03 | glio |
| Interaction_genotype_Mtor | Ndufa1 | 1.414 | 8.38E-03 | glio |
| Interaction_genotype_Myc | Tmem54 | -2.469 | 8.38E-03 | glio |
| Interaction_genotype_Myc | Ttyh3 | -2.713 | 8.38E-03 | glio |
| Interaction_genotype_Myc | Fh1 | -1.636 | 8.38E-03 | glio |
| Aars2_in.vivo_noRT | Haghl | 1.545 | 8.38E-03 | glio |
| Aars2_in.vivo_noRT | Decr1 | -1.039 | 8.38E-03 | glio |
| Atr_in.vivo_noRT | Gpc1 | 1.485 | 8.38E-03 | glio |

| | | | | |
|---|---|---|---|---|
| Bcas2_in.vivo_noRT | Hist2h4 | 1.877 | 8.38E-03 | glio |
| Bcas2_ex.vivo_noRT | Gm48140 | 1.949 | 8.38E-03 | glio |
| Cenpj_in.vivo_noRT | 1700124L16Ri | 2.372 | 8.38E-03 | glio |
| Cenpj_in.vivo_noRT | Cct6b | 2.141 | 8.38E-03 | glio |
| Cenpj_in.vivo_noRT | Ccnj | 2.318 | 8.38E-03 | glio |
| Chek1_ex.vivo_noRT | Gm49164 | 2.577 | 8.38E-03 | glio |
| Cyp19a1_ex.vivo_noRT | Gm26555 | 2.159 | 8.38E-03 | glio |
| Interaction_genotype_Mtor | Ube2n | 1.406 | 8.38E-04 | glio |
| Interaction_genotype_Mtor | Rps25 | 1.146 | 8.38E-04 | glio |
| Interaction_genotype_Myc | Tomm5 | 1.399 | 8.38E-04 | glio |
| Interaction_genotype_Mtor | Prr7 | -4.796 | 8.38E-05 | glio |
| Atr_ex.vivo_noRT | Hspd1 | -1.018 | 8.38E-05 | glio |
| Chek1_ex.vivo_noRT | Mtmr11 | 3.507 | 8.38E-05 | glio |
| Atr_in.vivo_noRT | Gm43328 | 3.195 | 8.38E-06 | glio |
| interaction_RT | Tspan13 | 1.372 | 8.38E-07 | glio |
| Hsd17b10_in.vivo_noRT | Tff2 | 3.356 | 8.38E-07 | glio |
| interaction_RT | Rsrp1 | 1.385 | 8.38E-08 | glio |
| Bora_in.vivo_noRT | Gm30082 | 4.286 | 8.38E-10 | glio |
| Atr_in.vivo_noRT | Capza2 | -1.676 | 8.38E-11 | glio |
| interaction_RT | mt-Nd1 | -1.314 | 8.38E-12 | glio |
| Bcas2_in.vivo_noRT | Gm15816 | 4.807 | 8.38E-15 | glio |
| Bcas2_in.vivo_noRT | Cfh | 4.027 | 8.38E-15 | glio |
| culture_effect | Batf2 | -4.307 | 8.38E-25 | glio |
| Atr_in.vivo_noRT | 1700003E24Ri | 5.533 | 8.38E-31 | glio |
| culture_effect | Slfn3 | 1.144 | 8.39E-03 | glio |
| Interaction_genotype_Mtor | Nkrf | -2.373 | 8.39E-03 | glio |
| Interaction_genotype_Myc | Gm7324 | -2.817 | 8.39E-03 | glio |
| Interaction_genotype_Myc | Dhrs11 | -2.8 | 8.39E-03 | glio |
| Interaction_genotype_Rpl3 | Ccdc85c | -3.658 | 8.39E-03 | glio |
| Atp5c1_in.vivo_noRT | A930003O13R | 2.401 | 8.39E-03 | glio |
| Atr_in.vivo_noRT | Nhej1 | 1.747 | 8.39E-03 | glio |
| Bora_ex.vivo_noRT | Gvin1 | 2.806 | 8.39E-03 | glio |
| Chek1_in.vivo_noRT | Zfp747 | 1.766 | 8.39E-03 | glio |
| Chek1_ex.vivo_noRT | Sult6b1 | 2.424 | 8.39E-03 | glio |
| Interaction_genotype_Aars2 | Mzf1 | -4.019 | 8.39E-04 | glio |
| Interaction_genotype_Aars2 | Tead3 | -4.148 | 8.39E-04 | glio |
| Interaction_genotype_Aars2 | Gm20324 | -3.645 | 8.39E-04 | glio |
| Interaction_genotype_Aars2 | Clec4d | -3.219 | 8.39E-04 | glio |
| Interaction_genotype_Mtor | Nr1h3 | -3.438 | 8.39E-04 | glio |
| Interaction_genotype_Mtor | Gnb2 | 1.6 | 8.39E-04 | glio |
| Bora_in.vivo_noRT | Zfp62 | 1.63 | 8.39E-04 | glio |
| Chek1_ex.vivo_noRT | Gm50394 | 2.85 | 8.39E-04 | glio |
| Eif2s1_in.vivo_noRT | 1700019E08Ri | 2.143 | 8.39E-04 | glio |
| culture_effect | Zc3h14 | 0.564 | 8.39E-05 | glio |
| Atr_in.vivo_noRT | AC132307.1 | 2.505 | 8.39E-05 | glio |
| culture_effect | Eml2 | -1.611 | 8.39E-06 | glio |
| Atr_in.vivo_noRT | Zfp963 | 3.463 | 8.39E-06 | glio |
| Atr_in.vivo_noRT | Rgcc | -2.32 | 8.39E-06 | glio |
| Aars2_in.vivo_noRT | Jmjd8 | 3.961 | 8.39E-09 | glio |
| Atr_in.vivo_noRT | Pkhd1 | 4.016 | 8.39E-09 | glio |
| Bora_in.vivo_noRT | Pcdhgb6 | 3.751 | 8.39E-10 | glio |
| Aars2_in.vivo_noRT | Gm13348 | 5.327 | 8.39E-20 | glio |

| | | | | |
|---|---|---|---|---|
| Atr_in.vivo_noRT | Fam25c | 5.001 | 8.39E-20 | glio |
| culture_effect | Il15 | -0.86 | 8.40E-03 | glio |
| culture_effect | Phf20 | -0.472 | 8.40E-03 | glio |
| Interaction_genotype_Bora | Slc26a6 | -3.415 | 8.40E-03 | glio |
| Interaction_genotype_Bora | Mrpl51 | 0.828 | 8.40E-03 | glio |
| Interaction_genotype_Mtor | Fzd4 | -3.051 | 8.40E-03 | glio |
| Interaction_genotype_Mtor | Nagk | -2.188 | 8.40E-03 | glio |
| Atp5c1_in.vivo_noRT | Ankrd1 | 2.668 | 8.40E-03 | glio |
| Atp5c1_in.vivo_noRT | Rgs9 | 1.868 | 8.40E-03 | glio |
| Atp5c1_in.vivo_noRT | Tfrc | -1.106 | 8.40E-03 | glio |
| Atr_in.vivo_noRT | Uhmk1 | -1.238 | 8.40E-03 | glio |
| Chek1_in.vivo_noRT | Catsper2 | 1.917 | 8.40E-03 | glio |
| Chek1_ex.vivo_noRT | Ncam1 | -1.781 | 8.40E-03 | glio |
| Hsd17b10_in.vivo_noRT | Dnttip1 | -0.818 | 8.40E-03 | glio |
| culture_effect | Cldn10 | -2.156 | 8.40E-04 | glio |
| culture_effect | Gm12925 | 1.988 | 8.40E-04 | glio |
| Interaction_genotype_Atr | Vgll4 | 1.992 | 8.40E-04 | glio |
| Bora_in.vivo_noRT | Arhgap33 | 2.473 | 8.40E-04 | glio |
| Chek1_in.vivo_noRT | Tnnc1 | 2.654 | 8.40E-04 | glio |
| Interaction_genotype_Mtor | Arhgap4 | -4.552 | 8.40E-05 | glio |
| Atr_in.vivo_noRT | Gm45494 | 2.655 | 8.40E-05 | glio |
| Bcas2_in.vivo_noRT | Nr1h4 | 2.758 | 8.40E-05 | glio |
| Chek1_in.vivo_noRT | Zfp637 | -1.547 | 8.40E-05 | glio |
| Chek1_ex.vivo_noRT | Clcc1 | -2.306 | 8.40E-05 | glio |
| Atp5c1_in.vivo_noRT | 4930423D22R | 2.408 | 8.40E-06 | glio |
| Atp5c1_in.vivo_noRT | Boc | 2.404 | 8.40E-06 | glio |
| Bora_in.vivo_noRT | Myzap | 2.615 | 8.40E-06 | glio |
| Bora_in.vivo_noRT | Fkbp9 | -1.508 | 8.40E-06 | glio |
| Interaction_genotype_Myc | Coa3 | 2.342 | 8.40E-07 | glio |
| Interaction_genotype_Myc | Gm15743 | -5.323 | 8.40E-07 | glio |
| Chek1_in.vivo_noRT | Tmem116 | 3.866 | 8.40E-07 | glio |
| Chek1_in.vivo_noRT | Rnase4 | 3.481 | 8.40E-07 | glio |
| Chek1_ex.vivo_noRT | Fsd2 | 4.256 | 8.40E-07 | glio |
| Hsd17b10_in.vivo_noRT | Tent5c | 2.69 | 8.40E-07 | glio |
| culture_effect | Gm43137 | 3.688 | 8.40E-13 | glio |
| Chek1_in.vivo_noRT | Cgnl1 | 5.044 | 8.40E-25 | glio |
| culture_effect | 2810414N06R | 1.437 | 8.41E-03 | glio |
| culture_effect | Ccnd1 | 0.425 | 8.41E-03 | glio |
| Interaction_genotype_Aars2 | 4930524O07R | -3.221 | 8.41E-03 | glio |
| interaction_RT | Bank1 | 1.942 | 8.41E-03 | glio |
| interaction_RT | Gm19684 | 2.765 | 8.41E-03 | glio |
| interaction_RT | S100a1 | 0.733 | 8.41E-03 | glio |
| interaction_RT | Prpf8 | -0.626 | 8.41E-03 | glio |
| interaction_RT | Hspa4l | 0.719 | 8.41E-03 | glio |
| interaction_RT | Mrpl33 | 0.471 | 8.41E-03 | glio |
| interaction_RT | Rps12 | -0.726 | 8.41E-03 | glio |
| Aars2_in.vivo_noRT | Ube2v1 | -0.867 | 8.41E-03 | glio |
| Bcas2_in.vivo_noRT | Vegfd | 1.861 | 8.41E-03 | glio |
| Bcas2_in.vivo_noRT | Kctd6 | 1.822 | 8.41E-03 | glio |
| Chek1_ex.vivo_noRT | Tlr1 | 2.414 | 8.41E-03 | glio |
| Interaction_genotype_Mtor | Ak3 | -2.492 | 8.41E-04 | glio |
| Atp5c1_in.vivo_noRT | 4930505N22R | 2.408 | 8.41E-04 | glio |

| | | | | |
|---|---|---|---|---|
| Cenpj_in.vivo_noRT | Gm30794 | 2.022 | 8.41E-04 | glio |
| Hsd17b10_in.vivo_noRT | Cyp26b1 | 2.691 | 8.41E-04 | glio |
| Interaction_genotype_Chek1 | Stat3 | 2.255 | 8.41E-05 | glio |
| Interaction_genotype_Myc | Itga2 | -3.953 | 8.41E-05 | glio |
| Interaction_genotype_Myc | Rab2a | 1.761 | 8.41E-05 | glio |
| Bcas2_in.vivo_noRT | Ran | -0.859 | 8.41E-05 | glio |
| culture_effect | Eva1b | 1.036 | 8.41E-06 | glio |
| Aars2_ex.vivo_noRT | Gm41307 | 2.9 | 8.41E-06 | glio |
| Abtb1_ex.vivo_noRT | Lce1g | 4.587 | 8.41E-06 | glio |
| culture_effect | Dpf1 | 1.882 | 8.41E-07 | glio |
| Interaction_genotype_Atr | Selenok | 2.016 | 8.41E-07 | glio |
| Atr_in.vivo_noRT | Gm17477 | 3.087 | 8.41E-07 | glio |
| Atr_in.vivo_noRT | Slc35e3 | 2.008 | 8.41E-07 | glio |
| Chek1_in.vivo_noRT | Ccdc184 | 3.71 | 8.41E-07 | glio |
| Chek1_in.vivo_noRT | 3110082J24Ri | 3.602 | 8.41E-07 | glio |
| culture_effect | Pdrg1 | -0.796 | 8.41E-08 | glio |
| Interaction_genotype_Mtor | A430010J10Ri | -7.855 | 8.41E-08 | glio |
| Interaction_genotype_Mtor | Ccl9 | -4.492 | 8.41E-08 | glio |
| Chek1_ex.vivo_noRT | Dusp15 | 5.243 | 8.41E-11 | glio |
| Chek1_in.vivo_noRT | Ccdc162 | 4.515 | 8.41E-16 | glio |
| Bora_in.vivo_noRT | Agap2 | 4.936 | 8.41E-18 | glio |
| Bcas2_in.vivo_noRT | Impg2 | 5.325 | 8.41E-20 | glio |
| Interaction_genotype_Mtor | Trim71 | -3.114 | 8.42E-03 | glio |
| Interaction_genotype_Mtor | Dnajb5 | -2.885 | 8.42E-03 | glio |
| Interaction_genotype_Rad51 | Rnf144a | 1.799 | 8.42E-03 | glio |
| interaction_RT | Cacng5 | 2.56 | 8.42E-03 | glio |
| interaction_RT | Tbrg4 | -0.886 | 8.42E-03 | glio |
| Aars2_in.vivo_noRT | Bak1 | -1.026 | 8.42E-03 | glio |
| Aars2_ex.vivo_noRT | Npr3 | 2.321 | 8.42E-03 | glio |
| Bcas2_in.vivo_noRT | Ly6g6d | 1.977 | 8.42E-03 | glio |
| Bcas2_in.vivo_noRT | Fbxo46 | 1.707 | 8.42E-03 | glio |
| Bcas2_in.vivo_noRT | Exosc3 | -0.792 | 8.42E-03 | glio |
| Bora_in.vivo_noRT | Nebl | -1.553 | 8.42E-03 | glio |
| Bora_ex.vivo_noRT | Il2rb | 2.615 | 8.42E-03 | glio |
| Chek1_in.vivo_noRT | Smim10l1 | -0.996 | 8.42E-03 | glio |
| Chek1_ex.vivo_noRT | Ccdc136 | 2.129 | 8.42E-03 | glio |
| Interaction_genotype_Myc | Usp4 | -2.323 | 8.42E-04 | glio |
| Aars2_in.vivo_noRT | Rab26 | 2.226 | 8.42E-04 | glio |
| Aars2_ex.vivo_noRT | Gm33051 | 2.495 | 8.42E-04 | glio |
| Bora_in.vivo_noRT | 2900052L18Ri | 1.954 | 8.42E-04 | glio |
| Atr_in.vivo_noRT | Snn | 2.798 | 8.42E-05 | glio |
| Interaction_genotype_Mtor | Vax2 | -5.216 | 8.42E-06 | glio |
| Interaction_genotype_Mtor | Gm21761 | -4.099 | 8.42E-06 | glio |
| Atr_in.vivo_noRT | Gm32051 | 3.244 | 8.42E-06 | glio |
| Atr_in.vivo_noRT | Catsperz | 3.059 | 8.42E-06 | glio |
| Aars2_ex.vivo_noRT | mt-Nd1 | 1.038 | 8.42E-07 | glio |
| Atr_in.vivo_noRT | Snph | 3.282 | 8.42E-07 | glio |
| Atr_ex.vivo_noRT | Gm266 | 3.164 | 8.42E-07 | glio |
| Bcas2_in.vivo_noRT | Neurl3 | 3.817 | 8.42E-08 | glio |
| culture_effect | Tpt1 | -0.933 | 8.42E-09 | glio |
| Aars2_in.vivo_noRT | Camkk1 | 4.556 | 8.42E-11 | glio |
| Atr_in.vivo_noRT | Gm41349 | 5.27 | 8.42E-17 | glio |

| | | | | |
|---|---|---|---|---|
| Interaction_genotype_Bcas2 | Cpm | 2.481 | 8.43E-03 | glio |
| Interaction_genotype_Mtor | Gm904 | -3.112 | 8.43E-03 | glio |
| Interaction_genotype_Mtor | Prepl | -2.223 | 8.43E-03 | glio |
| Aars2_in.vivo_noRT | Hnrnph1 | -0.737 | 8.43E-03 | glio |
| Aars2_in.vivo_noRT | Ppib | -0.579 | 8.43E-03 | glio |
| Atr_in.vivo_noRT | Vill | 1.578 | 8.43E-03 | glio |
| Atr_ex.vivo_noRT | Gadd45g | 2.058 | 8.43E-03 | glio |
| Bora_in.vivo_noRT | Pycard | 1.799 | 8.43E-03 | glio |
| Chek1_in.vivo_noRT | Nfatc2 | 2.012 | 8.43E-03 | glio |
| Chek1_in.vivo_noRT | Rpl27a | -0.54 | 8.43E-03 | glio |
| Interaction_genotype_Myc | Zfp865 | -3.021 | 8.43E-04 | glio |
| Interaction_genotype_Myc | 9330020H09R | -2.988 | 8.43E-04 | glio |
| Chek1_in.vivo_noRT | Hmgn5 | -1.37 | 8.43E-04 | glio |
| culture_effect | Atp5f1 | -0.422 | 8.43E-05 | glio |
| Bard1_ex.vivo_noRT | Nkg7 | 3.217 | 8.43E-05 | glio |
| Interaction_genotype_Ifnar1 | Stat1 | -1.605 | 8.43E-06 | glio |
| Interaction_genotype_Mtor | Rspo1 | -5.073 | 8.43E-06 | glio |
| Atr_in.vivo_noRT | Gm19325 | 3.569 | 8.43E-07 | glio |
| Bcas2_in.vivo_noRT | 1700124L16Ri | 3.573 | 8.43E-07 | glio |
| Hsd17b10_in.vivo_noRT | Gm10642 | 3.352 | 8.43E-07 | glio |
| Atr_in.vivo_noRT | Gm34121 | 3.408 | 8.43E-08 | glio |
| Bcas2_in.vivo_noRT | Mir670hg | 4.157 | 8.43E-08 | glio |
| Bora_in.vivo_noRT | Ltb | 3.877 | 8.43E-09 | glio |
| Chek1_ex.vivo_noRT | Adcy2 | 4.449 | 8.43E-09 | glio |
| Chek1_ex.vivo_noRT | Igdcc4 | 4.449 | 8.43E-09 | glio |
| Aars2_in.vivo_noRT | Gpr21 | 4.215 | 8.43E-11 | glio |
| Aars2_in.vivo_noRT | Gm37214 | 5.067 | 8.43E-13 | glio |
| Bora_in.vivo_noRT | Ms4a15 | 4.299 | 8.43E-15 | glio |
| Bora_in.vivo_noRT | Gm19276 | 4.936 | 8.43E-23 | glio |
| Interaction_genotype_Atr | Stac3 | -3.448 | 8.44E-03 | glio |
| Interaction_genotype_Mtor | Gm10401 | -2.879 | 8.44E-03 | glio |
| Interaction_genotype_Terf1 | 4930412M03F | -3.83 | 8.44E-03 | glio |
| interaction_RT | Rasip1 | 2.707 | 8.44E-03 | glio |
| Bcas2_in.vivo_noRT | Zfp882 | 2.119 | 8.44E-03 | glio |
| Bcas2_in.vivo_noRT | Mrpl27 | -0.759 | 8.44E-03 | glio |
| Bora_in.vivo_noRT | 1700063D05R | 2.024 | 8.44E-03 | glio |
| Bora_in.vivo_noRT | Atp11a | 1.892 | 8.44E-03 | glio |
| Bora_in.vivo_noRT | Akr7a5 | -1.187 | 8.44E-03 | glio |
| Bora_in.vivo_noRT | Usp25 | -1.159 | 8.44E-03 | glio |
| Bora_in.vivo_noRT | Nup85 | -1.179 | 8.44E-03 | glio |
| Bora_in.vivo_noRT | Ddx6 | -0.949 | 8.44E-03 | glio |
| Chek1_ex.vivo_noRT | A230087F16Ri | 2.483 | 8.44E-03 | glio |
| Eif2s1_in.vivo_noRT | Tmem54 | 1.883 | 8.44E-03 | glio |
| Eif2s1_in.vivo_noRT | Cebpd | 1.195 | 8.44E-03 | glio |
| Interaction_genotype_Aars2 | Zfp105 | -3.153 | 8.44E-04 | glio |
| Bard1_ex.vivo_noRT | Spi1 | 2.985 | 8.44E-04 | glio |
| Bard1_ex.vivo_noRT | Nrros | 3.219 | 8.44E-04 | glio |
| Bcas2_in.vivo_noRT | Spata45 | 2.37 | 8.44E-04 | glio |
| Bora_in.vivo_noRT | 4930556M19F | 2.289 | 8.44E-04 | glio |
| Aars2_in.vivo_noRT | Stx4a | -1.773 | 8.44E-05 | glio |
| Interaction_genotype_Mtor | 1700086P04R | -5.043 | 8.44E-06 | glio |
| Chek1_in.vivo_noRT | Bpifa3 | 2.63 | 8.44E-06 | glio |

| | | | | |
|---|---|---|---|---|
| Bcas2_in.vivo_noRT | 1700081N11R | 3.954 | 8.44E-07 | glio |
| culture_effect | Chka | -1.057 | 8.44E-08 | glio |
| Bcas2_in.vivo_noRT | Ceacam19 | 3.775 | 8.44E-08 | glio |
| Hsd17b10_in.vivo_noRT | Prap1 | 3.616 | 8.44E-12 | glio |
| Atr_in.vivo_noRT | Oxt | 4.892 | 8.44E-13 | glio |
| Atr_in.vivo_noRT | Tcap | 5.154 | 8.44E-16 | glio |
| Bcas2_in.vivo_noRT | Akap14 | 4.544 | 8.44E-17 | glio |
| Atr_ex.vivo_noRT | Gm34059 | 4.985 | 8.44E-18 | glio |
| Atr_ex.vivo_noRT | Gm13078 | 5.288 | 8.44E-18 | glio |
| Atr_in.vivo_noRT | Gm11899 | 5.011 | 8.44E-21 | glio |
| Interaction_genotype_Atr | Pacc1 | -2.842 | 8.45E-03 | glio |
| Interaction_genotype_Atr | Nfyc | 1.635 | 8.45E-03 | glio |
| Interaction_genotype_Bcas2 | Gm26671 | -2.458 | 8.45E-03 | glio |
| Interaction_genotype_Bora | 4921504E06Ri | -2.823 | 8.45E-03 | glio |
| Interaction_genotype_Bora | Gm31518 | -3.567 | 8.45E-03 | glio |
| Interaction_genotype_Bora | Gm14379 | -3.36 | 8.45E-03 | glio |
| Interaction_genotype_Bora | Efcab9 | -3.386 | 8.45E-03 | glio |
| Interaction_genotype_Bora | Cyth4 | -3.459 | 8.45E-03 | glio |
| Interaction_genotype_Bora | Kctd2 | 1.937 | 8.45E-03 | glio |
| Interaction_genotype_Cdkn2a | Gm12794 | 2.835 | 8.45E-03 | glio |
| interaction_RT | Dcp2 | 0.907 | 8.45E-03 | glio |
| Aars2_in.vivo_noRT | Rpl32 | -0.537 | 8.45E-03 | glio |
| Atp5c1_in.vivo_noRT | A230020J21Ri | 2.268 | 8.45E-03 | glio |
| Bcas2_in.vivo_noRT | Zscan22 | 1.649 | 8.45E-03 | glio |
| Bora_in.vivo_noRT | H2-Eb1 | -0.956 | 8.45E-03 | glio |
| Bora_ex.vivo_noRT | A930033M14F | 2.236 | 8.45E-03 | glio |
| Chek1_in.vivo_noRT | Rapgefl1 | 1.91 | 8.45E-03 | glio |
| Chek1_ex.vivo_noRT | Cenpc1 | -1.487 | 8.45E-03 | glio |
| Hsd17b10_in.vivo_noRT | Gm30790 | 2.1 | 8.45E-03 | glio |
| Hsd17b10_in.vivo_noRT | Elmod3 | 1.883 | 8.45E-03 | glio |
| Interaction_genotype_Mtor | Ptn | -4.757 | 8.45E-04 | glio |
| Interaction_genotype_Myc | Ubxn1 | 1.009 | 8.45E-04 | glio |
| Bora_in.vivo_noRT | Trim66 | 2.224 | 8.45E-04 | glio |
| Chek1_ex.vivo_noRT | Tmem140 | 2.618 | 8.45E-04 | glio |
| Interaction_genotype_Mtor | Fbxo24 | -4.668 | 8.45E-05 | glio |
| Interaction_genotype_Myc | Zfp606 | -3.718 | 8.45E-05 | glio |
| Aars2_in.vivo_noRT | Trim6 | 2.881 | 8.45E-05 | glio |
| Aars2_in.vivo_noRT | Ppfia2 | 2.212 | 8.45E-05 | glio |
| Aars2_in.vivo_noRT | Rpe | -1.756 | 8.45E-05 | glio |
| Bcas2_in.vivo_noRT | Vgll2 | 2.471 | 8.45E-05 | glio |
| Interaction_genotype_Mtor | Bclaf1 | 2.143 | 8.45E-06 | glio |
| Aars2_in.vivo_noRT | Gap43 | 3.881 | 8.45E-07 | glio |
| Aars2_ex.vivo_noRT | Gm36793 | 3.293 | 8.45E-07 | glio |
| Chek1_in.vivo_noRT | Gm10230 | 3.649 | 8.45E-07 | glio |
| Aars2_in.vivo_noRT | Gm50399 | 3.774 | 8.45E-09 | glio |
| culture_effect | Olfm2 | 3.247 | 8.45E-10 | glio |
| Bcas2_in.vivo_noRT | Nrros | 4.165 | 8.45E-10 | glio |
| Atr_in.vivo_noRT | Gm16853 | 4.5 | 8.45E-12 | glio |
| Atr_in.vivo_noRT | Gtsf1l | 3.904 | 8.45E-12 | glio |
| Atr_in.vivo_noRT | Gm41764 | 4.617 | 8.45E-14 | glio |
| Atr_in.vivo_noRT | D730045A05R | 5.533 | 8.45E-16 | glio |
| Atr_in.vivo_noRT | Rftn2 | 5.142 | 8.45E-17 | glio |

| | | | | |
|---|---|---|---|---|
| culture_effect | Fxyd5 | 0.747 | 8.46E-03 | glio |
| Interaction_genotype_Aars2 | Gm33782 | -2.992 | 8.46E-03 | glio |
| Interaction_genotype_Atr | Hyal1 | -3.062 | 8.46E-03 | glio |
| Interaction_genotype_Atr | Pfdn2 | 0.789 | 8.46E-03 | glio |
| Atr_in.vivo_noRT | Syt14 | 1.539 | 8.46E-03 | glio |
| Cenpj_ex.vivo_noRT | Gm41349 | 2.591 | 8.46E-03 | glio |
| Cenpj_ex.vivo_noRT | Serpina3g | 2.591 | 8.46E-03 | glio |
| Chek1_ex.vivo_noRT | Sdc3 | 2.382 | 8.46E-03 | glio |
| Chek1_ex.vivo_noRT | Cntn1 | 2.646 | 8.46E-03 | glio |
| Chek1_ex.vivo_noRT | Zdhhc15 | 2.257 | 8.46E-03 | glio |
| Fanca_ex.vivo_noRT | Agrp | 2.518 | 8.46E-03 | glio |
| Interaction_genotype_Uqcrc2 | Gm27239 | -4.754 | 8.46E-04 | glio |
| Aars2_in.vivo_noRT | 1500002F19Ri | 2.282 | 8.46E-04 | glio |
| Atr_ex.vivo_noRT | Ms4a4d | 3.164 | 8.46E-04 | glio |
| Bora_in.vivo_noRT | Pcdhb22 | 2.28 | 8.46E-04 | glio |
| Eif2s1_in.vivo_noRT | Accsl | 2.937 | 8.46E-04 | glio |
| Eif2s1_in.vivo_noRT | Col8a1 | 2.666 | 8.46E-04 | glio |
| Chek1_in.vivo_noRT | Ldlrad3 | 2.568 | 8.46E-05 | glio |
| Chek1_in.vivo_noRT | Dhx40 | -1.618 | 8.46E-05 | glio |
| Aars2_in.vivo_noRT | 8030456M14F | 3.174 | 8.46E-06 | glio |
| Interaction_genotype_Mtor | Srcin1 | -5.498 | 8.46E-07 | glio |
| Interaction_genotype_Mtor | Gm49864 | -5.683 | 8.46E-07 | glio |
| Interaction_genotype_Mtor | 4930483J18Ri | -5.516 | 8.46E-07 | glio |
| Interaction_genotype_Mtor | Gm27241 | -4.424 | 8.46E-07 | glio |
| Interaction_genotype_Mtor | Slc1a2 | -5.021 | 8.46E-07 | glio |
| Hsd17b10_in.vivo_noRT | Cxcl16 | 3.352 | 8.46E-07 | glio |
| Aars2_in.vivo_noRT | Dnaaf3 | 3.374 | 8.46E-08 | glio |
| Bcas2_in.vivo_noRT | A430035B10R | 3.9 | 8.46E-08 | glio |
| Aars2_in.vivo_noRT | Tcam1 | 4.149 | 8.46E-09 | glio |
| Aars2_in.vivo_noRT | Gm26682 | 4.409 | 8.46E-11 | glio |
| Aars2_in.vivo_noRT | Gm47622 | 5.327 | 8.46E-24 | glio |
| Interaction_genotype_Aars2 | Ubtd2 | -2.355 | 8.47E-03 | glio |
| Interaction_genotype_Chek1 | Gm9958 | -3.734 | 8.47E-03 | glio |
| Interaction_genotype_Mtor | Zfp974 | -3.292 | 8.47E-03 | glio |
| Interaction_genotype_Mtor | Ncoa2 | -2.324 | 8.47E-03 | glio |
| Interaction_genotype_Mtor | Shisa5 | 1.564 | 8.47E-03 | glio |
| Interaction_genotype_Mtor | Actg1 | 1.189 | 8.47E-03 | glio |
| Interaction_genotype_Myc | Dagla | -3.216 | 8.47E-03 | glio |
| Interaction_genotype_Myc | Gpsm1 | -2.174 | 8.47E-03 | glio |
| Interaction_genotype_Pars2 | Dnah10 | -3.227 | 8.47E-03 | glio |
| Interaction_genotype_Rpl3 | Ifit1bl1 | -4.532 | 8.47E-03 | glio |
| Interaction_genotype_Rpl3 | BC030343 | -3.736 | 8.47E-03 | glio |
| Interaction_genotype_Rpl3 | 1700023H06R | -3.993 | 8.47E-03 | glio |
| Interaction_genotype_Rpl3 | Gm47204 | -3.815 | 8.47E-03 | glio |
| Bcas2_in.vivo_noRT | Pqlc2 | 1.651 | 8.47E-03 | glio |
| Bora_in.vivo_noRT | Il1rapl1 | 1.969 | 8.47E-03 | glio |
| Bora_in.vivo_noRT | Maml3 | 1.42 | 8.47E-03 | glio |
| Bora_in.vivo_noRT | Gltp | -1.095 | 8.47E-03 | glio |
| Bora_in.vivo_noRT | Kif5b | -0.668 | 8.47E-03 | glio |
| Bora_in.vivo_noRT | Mrps24 | -0.58 | 8.47E-03 | glio |
| Chek1_in.vivo_noRT | Rasip1 | 2.014 | 8.47E-03 | glio |
| Chek1_in.vivo_noRT | D830050J10Ri | 1.83 | 8.47E-03 | glio |

| | | | | |
|---|---|---|---|---|
| Chek1_in.vivo_noRT | Fan1 | 1.801 | 8.47E-03 | glio |
| Chek1_in.vivo_noRT | Hax1 | -0.815 | 8.47E-03 | glio |
| Chek1_in.vivo_noRT | Arpc2 | -0.64 | 8.47E-03 | glio |
| Cyp19a1_ex.vivo_noRT | Cd36 | 2.159 | 8.47E-03 | glio |
| Cyp19a1_ex.vivo_noRT | Gstt4 | 2.159 | 8.47E-03 | glio |
| Eif2s1_in.vivo_noRT | Nwd1 | 1.879 | 8.47E-03 | glio |
| Hsd17b10_in.vivo_noRT | Lmod1 | 2.185 | 8.47E-03 | glio |
| Hsd17b10_in.vivo_noRT | Pim3 | -0.9 | 8.47E-03 | glio |
| Interaction_genotype_Myc | Nedd8 | 1.248 | 8.47E-04 | glio |
| Bora_in.vivo_noRT | Kcna4 | 2.41 | 8.47E-04 | glio |
| Interaction_genotype_Mtor | Ccdc62 | -4.304 | 8.47E-05 | glio |
| Interaction_genotype_Myc | Sh2d3c | -4.808 | 8.47E-05 | glio |
| Atr_in.vivo_noRT | Rspo3 | 3.178 | 8.47E-05 | glio |
| Atr_in.vivo_noRT | 4921531C22R | 2.692 | 8.47E-05 | glio |
| Bora_in.vivo_noRT | Gm26981 | 2.762 | 8.47E-05 | glio |
| Atr_ex.vivo_noRT | Ildr1 | 3.164 | 8.47E-07 | glio |
| Atr_ex.vivo_noRT | Zfp788 | -3.111 | 8.47E-07 | glio |
| Chek1_in.vivo_noRT | 6330415G19R | 3.737 | 8.47E-08 | glio |
| Chek1_in.vivo_noRT | Gm3604 | 3.761 | 8.47E-08 | glio |
| Aars2_in.vivo_noRT | Gm17210 | 4.267 | 8.47E-10 | glio |
| Chek1_ex.vivo_noRT | Gm35082 | 4.045 | 8.47E-10 | glio |
| Hsd17b10_in.vivo_noRT | Fam166a | 3.352 | 8.47E-10 | glio |
| culture_effect | Rbp2 | -4.229 | 8.47E-11 | glio |
| Bora_in.vivo_noRT | 4930434B07R | 4.673 | 8.47E-15 | glio |
| culture_effect | Cxcr2 | -1.66 | 8.48E-03 | glio |
| culture_effect | Dhx32 | 0.57 | 8.48E-03 | glio |
| Interaction_genotype_Aars2 | Vdac1 | 1.092 | 8.48E-03 | glio |
| Interaction_genotype_Myc | Gm10644 | -3.048 | 8.48E-03 | glio |
| Atr_in.vivo_noRT | Mthfd2 | -1.045 | 8.48E-03 | glio |
| Atr_ex.vivo_noRT | Gm28578 | 2.703 | 8.48E-03 | glio |
| Atr_ex.vivo_noRT | Gm26789 | 2.183 | 8.48E-03 | glio |
| Atr_ex.vivo_noRT | Gm14379 | 2.441 | 8.48E-03 | glio |
| Atr_ex.vivo_noRT | Gm11491 | 2.594 | 8.48E-03 | glio |
| Atr_ex.vivo_noRT | Apold1 | 2.893 | 8.48E-03 | glio |
| Atr_ex.vivo_noRT | Jund | 1.005 | 8.48E-03 | glio |
| Bcas2_in.vivo_noRT | Ldlrad3 | 1.801 | 8.48E-03 | glio |
| Bora_in.vivo_noRT | Tent5d | 1.874 | 8.48E-03 | glio |
| Eif2s1_in.vivo_noRT | 2510046G10R | 1.989 | 8.48E-03 | glio |
| Interaction_genotype_Aars2 | Psmc6 | 1.464 | 8.48E-04 | glio |
| Interaction_genotype_Bora | Gm28874 | -4.409 | 8.48E-04 | glio |
| Chek1_in.vivo_noRT | Mt4 | 3.048 | 8.48E-04 | glio |
| Interaction_genotype_Mtor | Cbs | -4.86 | 8.48E-05 | glio |
| Hsd17b10_in.vivo_noRT | Sult5a1 | 3.084 | 8.48E-05 | glio |
| Hsd17b10_in.vivo_noRT | Cfb | 2.312 | 8.48E-05 | glio |
| Chek1_ex.vivo_noRT | Pitx3 | 3.079 | 8.48E-06 | glio |
| Interaction_genotype_Mtor | Npl | -5.416 | 8.48E-07 | glio |
| Bcas2_in.vivo_noRT | Gm9922 | 3.671 | 8.48E-08 | glio |
| culture_effect | Spata2l | -1.733 | 8.49E-03 | glio |
| culture_effect | Asah2 | -1.031 | 8.49E-03 | glio |
| culture_effect | Ahr | 0.62 | 8.49E-03 | glio |
| Interaction_genotype_Atr | Csgalnact1 | 2.622 | 8.49E-03 | glio |
| Interaction_genotype_Bora | Cdca7 | 1.816 | 8.49E-03 | glio |

| | | | | |
|---|---|---|---|---|
| Interaction_genotype_Bora | Car6 | 1.949 | 8.49E-03 | glio |
| Interaction_genotype_Cdkn2a | Fam131a | 3.769 | 8.49E-03 | glio |
| Interaction_genotype_Mtor | Strap | 1.248 | 8.49E-03 | glio |
| Interaction_genotype_Myc | Gm16685 | -2.47 | 8.49E-03 | glio |
| Interaction_genotype_Myc | Efr3a | 1.142 | 8.49E-03 | glio |
| Interaction_genotype_Myc | Hdlbp | 1.132 | 8.49E-03 | glio |
| Interaction_genotype_Pars2 | Lrp2bp | -3.3 | 8.49E-03 | glio |
| Interaction_genotype_Pars2 | Pldi | -2.963 | 8.49E-03 | glio |
| Interaction_genotype_Pars2 | Gm47540 | -2.661 | 8.49E-03 | glio |
| Interaction_genotype_Rpl3 | Ntn3 | -3.47 | 8.49E-03 | glio |
| Interaction_genotype_Rpl3 | Gm10655 | -3.829 | 8.49E-03 | glio |
| Interaction_genotype_Rpl3 | Trp53bp1 | -2.801 | 8.49E-03 | glio |
| Interaction_genotype_Rpl3 | Nedd9 | 1.907 | 8.49E-03 | glio |
| Interaction_genotype_Rpl3 | Ube2j2 | 1.244 | 8.49E-03 | glio |
| Aars2_in.vivo_noRT | Nob1 | -1.13 | 8.49E-03 | glio |
| Aars2_in.vivo_noRT | Gps1 | -0.905 | 8.49E-03 | glio |
| Bcas2_in.vivo_noRT | N4bp2 | -1.4 | 8.49E-03 | glio |
| Bora_in.vivo_noRT | Mlip | 1.932 | 8.49E-03 | glio |
| Bora_in.vivo_noRT | Tgfb2 | 1.802 | 8.49E-03 | glio |
| Cenpj_in.vivo_noRT | Gm39326 | 2.285 | 8.49E-03 | glio |
| Chek1_in.vivo_noRT | Osbp | -1.088 | 8.49E-03 | glio |
| Chek1_in.vivo_noRT | Elavl1 | -0.921 | 8.49E-03 | glio |
| Chek1_ex.vivo_noRT | A930024E05Ri | 2.525 | 8.49E-03 | glio |
| culture_effect | Ints13 | -0.547 | 8.49E-04 | glio |
| Interaction_genotype_Myc | Gm45669 | -3.254 | 8.49E-04 | glio |
| Interaction_genotype_Myc | Snn | -3.761 | 8.49E-04 | glio |
| Atr_in.vivo_noRT | Lig1 | -1.581 | 8.49E-04 | glio |
| Bcas2_in.vivo_noRT | 1700086P04R | 2.398 | 8.49E-04 | glio |
| Bora_in.vivo_noRT | Gan | 1.645 | 8.49E-04 | glio |
| Chek1_in.vivo_noRT | Irak3 | 2.528 | 8.49E-04 | glio |
| Chek1_in.vivo_noRT | Peli1 | -1.507 | 8.49E-04 | glio |
| culture_effect | Snrpc | -0.436 | 8.49E-05 | glio |
| Interaction_genotype_Mtor | Gm37035 | -3.869 | 8.49E-05 | glio |
| Interaction_genotype_Myc | Npnt | -3.26 | 8.49E-05 | glio |
| Interaction_genotype_Aars2 | Sod3 | -6.035 | 8.49E-06 | glio |
| Aars2_in.vivo_noRT | Sept1 | 3.391 | 8.49E-06 | glio |
| Bard1_ex.vivo_noRT | Apoa5 | 2.814 | 8.49E-06 | glio |
| Chek1_in.vivo_noRT | Stpg2 | 3.095 | 8.49E-06 | glio |
| Atr_in.vivo_noRT | Gm35551 | 3.717 | 8.49E-09 | glio |
| Bcas2_in.vivo_noRT | 4933400F21Ri | 4.016 | 8.49E-16 | glio |
| culture_effect | Cspp1 | -0.569 | 8.50E-03 | glio |
| culture_effect | Micos13 | 0.3 | 8.50E-03 | glio |
| Interaction_genotype_Aars2 | 5930403N24R | -3.377 | 8.50E-03 | glio |
| Interaction_genotype_Atr | Eml6 | -3.415 | 8.50E-03 | glio |
| Interaction_genotype_Atr | 2610016A17R | -3.259 | 8.50E-03 | glio |
| Interaction_genotype_Cdkn2a | A330049N07R | 2.702 | 8.50E-03 | glio |
| Interaction_genotype_Cdkn2a | Gm15533 | 2.835 | 8.50E-03 | glio |
| Interaction_genotype_Cdkn2a | Olfr718-ps1 | 2.978 | 8.50E-03 | glio |
| Interaction_genotype_Chek1 | Irf1 | 2.682 | 8.50E-03 | glio |
| Interaction_genotype_Mtor | Yeats4 | 1.386 | 8.50E-03 | glio |
| Interaction_genotype_Mtor | Hnrnpf | 1.034 | 8.50E-03 | glio |
| Interaction_genotype_Pars2 | Hfm1 | -2.843 | 8.50E-03 | glio |

| | | | | |
|---|---|---|---|---|
| Aars2_in.vivo_noRT | Trnau1ap | -0.853 | 8.50E-03 | glio |
| Atp5c1_ex.vivo_noRT | Gm42670 | -3.031 | 8.50E-03 | glio |
| Atp5c1_ex.vivo_noRT | 4930447K03R | 2.608 | 8.50E-03 | glio |
| Atr_in.vivo_noRT | Fzd5 | 1.848 | 8.50E-03 | glio |
| Atr_in.vivo_noRT | Crtc1 | 1.676 | 8.50E-03 | glio |
| Atr_in.vivo_noRT | Lmbrd2 | -1.134 | 8.50E-03 | glio |
| Bcas2_ex.vivo_noRT | Snhg4.1 | 2.349 | 8.50E-03 | glio |
| Bcas2_ex.vivo_noRT | Gm13398 | 1.945 | 8.50E-03 | glio |
| Bcas2_ex.vivo_noRT | Tmeff2 | 3.129 | 8.50E-03 | glio |
| Bcas2_ex.vivo_noRT | Rasl10a | 2.349 | 8.50E-03 | glio |
| Bcas2_ex.vivo_noRT | H2-DMb2 | 2.742 | 8.50E-03 | glio |
| Bora_in.vivo_noRT | Eml1 | 1.77 | 8.50E-03 | glio |
| Chek1_ex.vivo_noRT | Sqstm1 | 0.959 | 8.50E-03 | glio |
| Cyp19a1_ex.vivo_noRT | 9130019O22R | 2.159 | 8.50E-03 | glio |
| Bora_ex.vivo_noRT | 9130230L23Ri | 2.808 | 8.50E-04 | glio |
| Cenpj_in.vivo_noRT | Ank1 | 2.022 | 8.50E-04 | glio |
| Chek1_in.vivo_noRT | Mtcp1 | 2.354 | 8.50E-04 | glio |
| Chek1_in.vivo_noRT | Cep83os | 2.26 | 8.50E-04 | glio |
| Hsd17b10_in.vivo_noRT | Tnxb | 2.463 | 8.50E-04 | glio |
| Interaction_genotype_Myc | E030037K01Ri | -4.064 | 8.50E-05 | glio |
| Aars2_in.vivo_noRT | 9130015A21R | 4.942 | 8.50E-05 | glio |
| Bora_in.vivo_noRT | Fgfr1 | 3.299 | 8.50E-05 | glio |
| Bora_in.vivo_noRT | Gm9993 | 2.559 | 8.50E-05 | glio |
| culture_effect | Gm49774 | 1.671 | 8.50E-06 | glio |
| Bard1_ex.vivo_noRT | Gm49172 | 2.814 | 8.50E-06 | glio |
| Bard1_ex.vivo_noRT | Gjd4 | 2.814 | 8.50E-06 | glio |
| Bcas2_in.vivo_noRT | Gm26728 | 3.305 | 8.50E-06 | glio |
| Bora_in.vivo_noRT | Gm43727 | 3.109 | 8.50E-06 | glio |
| culture_effect | Cxcr6 | -3.124 | 8.50E-07 | glio |
| Bora_in.vivo_noRT | Dct | 4.133 | 8.50E-07 | glio |
| culture_effect | Ppp1r18os | 2.88 | 8.50E-08 | glio |
| Atr_in.vivo_noRT | D930030I03Ri | 4.294 | 8.50E-09 | glio |
| Chek1_in.vivo_noRT | Fhl4 | 3.793 | 8.50E-09 | glio |
| Aars2_in.vivo_noRT | Gm11491 | 4.285 | 8.50E-10 | glio |
| culture_effect | 2010001A14R | 1.995 | 8.50E-13 | glio |
| culture_effect | Agpat4 | -1.71 | 8.50E-17 | glio |
| Bora_in.vivo_noRT | Olfr56 | 4.936 | 8.50E-21 | glio |
| Atr_in.vivo_noRT | Spink6 | 5.533 | 8.50E-23 | glio |
| Interaction_genotype_Atr | Ltb4r1 | -3.542 | 8.51E-03 | glio |
| Interaction_genotype_Myc | CAAA0114733 | -2.667 | 8.51E-03 | glio |
| Interaction_genotype_Rpl3 | St8sia1 | -3.843 | 8.51E-03 | glio |
| Atr_in.vivo_noRT | Cog3 | 1.605 | 8.51E-03 | glio |
| Bcas2_in.vivo_noRT | Cd38 | 1.941 | 8.51E-03 | glio |
| Bcas2_in.vivo_noRT | Dnlz | -1.099 | 8.51E-03 | glio |
| Chek1_ex.vivo_noRT | Plpp7 | 2.211 | 8.51E-03 | glio |
| Cyp19a1_ex.vivo_noRT | Klk13 | 2.159 | 8.51E-03 | glio |
| Interaction_genotype_Aars2 | Gm5152 | -4.289 | 8.51E-04 | glio |
| Interaction_genotype_Mtor | Akna | -3.955 | 8.51E-04 | glio |
| Interaction_genotype_Myc | AW046200 | -3.76 | 8.51E-04 | glio |
| Atr_in.vivo_noRT | Egln3 | 2.324 | 8.51E-04 | glio |
| Atr_in.vivo_noRT | Rab7 | -1.154 | 8.51E-04 | glio |
| Bard1_ex.vivo_noRT | Cemip | 2.814 | 8.51E-04 | glio |

| | | | | |
|---|---|---|---|---|
| Bora_in.vivo_noRT | 9030407P20R | 2.39 | 8.51E-04 | glio |
| Bora_in.vivo_noRT | Gemin6 | -1.468 | 8.51E-04 | glio |
| Hsd17b10_in.vivo_noRT | 1700025N23R | 2.245 | 8.51E-04 | glio |
| culture_effect | Prune2 | -1.645 | 8.51E-05 | glio |
| culture_effect | Ogt | -0.586 | 8.51E-05 | glio |
| Interaction_genotype_Mtor | Ttc33 | 2.002 | 8.51E-05 | glio |
| Interaction_genotype_Mtor | Prkar1a | 1.718 | 8.51E-05 | glio |
| Interaction_genotype_Myc | Adamts12 | -3.313 | 8.51E-05 | glio |
| Aars2_in.vivo_noRT | Nudcd2 | -0.959 | 8.51E-05 | glio |
| Chek1_in.vivo_noRT | Abi3bp | 2.815 | 8.51E-05 | glio |
| Bora_in.vivo_noRT | Gm15097 | 2.605 | 8.51E-06 | glio |
| Atp5c1_in.vivo_noRT | Gm12800 | 2.668 | 8.51E-07 | glio |
| Aars2_in.vivo_noRT | 4921507P07R | 3.562 | 8.51E-08 | glio |
| Aars2_in.vivo_noRT | Hist2h3c2 | 5.06 | 8.51E-12 | glio |
| Aars2_in.vivo_noRT | Nrn1l | 4.401 | 8.51E-12 | glio |
| Aars2_in.vivo_noRT | Aqp9 | 5.327 | 8.51E-20 | glio |
| Atr_in.vivo_noRT | Strip2 | 5.269 | 8.51E-22 | glio |
| Chek1_in.vivo_noRT | A930002C04R | 4.784 | 8.51E-23 | glio |
| culture_effect | Adam2 | -1.46 | 8.52E-03 | glio |
| Interaction_genotype_Aars2 | Ucma | -3.833 | 8.52E-03 | glio |
| Interaction_genotype_Atr | Gm43581 | -3.012 | 8.52E-03 | glio |
| Interaction_genotype_Atr | 5033430I15Ri | -2.947 | 8.52E-03 | glio |
| Interaction_genotype_Atr | Ing1 | 1.421 | 8.52E-03 | glio |
| Interaction_genotype_Bcas2 | Smim13 | 1.833 | 8.52E-03 | glio |
| Aars2_in.vivo_noRT | Timm8b | -0.77 | 8.52E-03 | glio |
| Atp5c1_in.vivo_noRT | Sema4f | 2.139 | 8.52E-03 | glio |
| Atr_in.vivo_noRT | 9330159F19Ri | 1.796 | 8.52E-03 | glio |
| Atr_in.vivo_noRT | Cenpi | 1.539 | 8.52E-03 | glio |
| Bcas2_ex.vivo_noRT | Nox1 | 2.445 | 8.52E-03 | glio |
| Brca2_in.vivo_noRT | Slc19a3 | 1.763 | 8.52E-03 | glio |
| Chek1_in.vivo_noRT | Ppp1r3f | 2.063 | 8.52E-03 | glio |
| Cyp19a1_ex.vivo_noRT | H1foo | 2.159 | 8.52E-03 | glio |
| Cyp19a1_ex.vivo_noRT | Sytl4 | 2.159 | 8.52E-03 | glio |
| Cyp19a1_ex.vivo_noRT | Gm12800 | 2.159 | 8.52E-03 | glio |
| Cyp19a1_ex.vivo_noRT | 1600014C23R | 3.018 | 8.52E-03 | glio |
| Cyp19a1_ex.vivo_noRT | Gm5890 | 2.159 | 8.52E-03 | glio |
| Cyp19a1_ex.vivo_noRT | Zscan4c | 2.159 | 8.52E-03 | glio |
| Cyp19a1_ex.vivo_noRT | Speer4a | 2.159 | 8.52E-03 | glio |
| Cyp19a1_ex.vivo_noRT | BC080695 | 2.159 | 8.52E-03 | glio |
| culture_effect | Sipa1l2 | 1.687 | 8.52E-04 | glio |
| culture_effect | Hmox2 | -0.691 | 8.52E-04 | glio |
| Interaction_genotype_Mtor | 1500004A13R | -3.964 | 8.52E-04 | glio |
| Interaction_genotype_Mtor | Zbtb7a | 2.013 | 8.52E-04 | glio |
| Interaction_genotype_Myc | Gm12800 | -2.729 | 8.52E-04 | glio |
| Interaction_genotype_Myc | Hoga1 | -3.748 | 8.52E-04 | glio |
| Interaction_genotype_Myc | Pigb | -3.036 | 8.52E-04 | glio |
| Aars2_in.vivo_noRT | Bmerb1 | 2.554 | 8.52E-04 | glio |
| Aars2_in.vivo_noRT | Gm10941 | 2.15 | 8.52E-04 | glio |
| Aars2_in.vivo_noRT | Kctd18 | 1.944 | 8.52E-04 | glio |
| Atr_ex.vivo_noRT | Gm26978 | 2.577 | 8.52E-04 | glio |
| Aars2_in.vivo_noRT | Gm10658 | 2.661 | 8.52E-05 | glio |
| Bard1_ex.vivo_noRT | 4930512B01R | 2.814 | 8.52E-05 | glio |

| | | | | |
|---|---|---|---|---|
| Eif2s1_ex.vivo_noRT | Eif2s1 | -1.297 | 8.52E-08 | glio |
| culture_effect | Mdh1 | -0.726 | 8.52E-09 | glio |
| Bcas2_in.vivo_noRT | Sox11 | 4.161 | 8.52E-10 | glio |
| Bora_in.vivo_noRT | Gm42849 | 4.67 | 8.52E-14 | glio |
| Atr_in.vivo_noRT | Vmn1r91 | 4.88 | 8.52E-15 | glio |
| Chek1_in.vivo_noRT | Crhbp | 5.044 | 8.52E-17 | glio |
| culture_effect | Gm17092 | 2.622 | 8.52E-19 | glio |
| Aars2_in.vivo_noRT | Slc45a1 | 5.067 | 8.52E-23 | glio |
| culture_effect | Arel1 | -0.78 | 8.53E-03 | glio |
| culture_effect | Vps4a | 0.523 | 8.53E-03 | glio |
| Interaction_genotype_Mtor | Nsun7 | -3.546 | 8.53E-03 | glio |
| Interaction_genotype_Mtor | Ly6g6d | -3.184 | 8.53E-03 | glio |
| Interaction_genotype_Myc | Tes3-ps | -2.986 | 8.53E-03 | glio |
| Interaction_genotype_Myc | Ctif | -2.676 | 8.53E-03 | glio |
| Interaction_genotype_Rpl3 | Ankef1 | -3.803 | 8.53E-03 | glio |
| Interaction_genotype_Rpl3 | Gm4419 | -3.142 | 8.53E-03 | glio |
| Aars2_in.vivo_noRT | BC052040 | 1.533 | 8.53E-03 | glio |
| Aars2_in.vivo_noRT | Atp6v1d | -0.872 | 8.53E-03 | glio |
| Aars2_ex.vivo_noRT | Adgrd1 | 2.582 | 8.53E-03 | glio |
| Atr_in.vivo_noRT | Mrpl34 | -0.743 | 8.53E-03 | glio |
| Bora_in.vivo_noRT | Mrps36 | -0.719 | 8.53E-03 | glio |
| Chek1_in.vivo_noRT | Nr4a1 | -1.238 | 8.53E-03 | glio |
| Interaction_genotype_Mtor | Gm21818 | -4.099 | 8.53E-04 | glio |
| Interaction_genotype_Mtor | Dzank1 | -4.123 | 8.53E-04 | glio |
| Interaction_genotype_Mtor | Gm46447 | -3.631 | 8.53E-04 | glio |
| Interaction_genotype_Mtor | 2600006K01R | -4.066 | 8.53E-04 | glio |
| Interaction_genotype_Mtor | Ube2j2 | 1.519 | 8.53E-04 | glio |
| Aars2_in.vivo_noRT | Mmp14 | 2.453 | 8.53E-04 | glio |
| Aars2_ex.vivo_noRT | Slc14a1 | 2.718 | 8.53E-04 | glio |
| Atr_in.vivo_noRT | Lrrc20 | 2.164 | 8.53E-04 | glio |
| Bora_in.vivo_noRT | Gm19605 | 2.389 | 8.53E-04 | glio |
| Interaction_genotype_Mtor | Acss1 | -4.455 | 8.53E-05 | glio |
| Interaction_genotype_Myc | Gm42695 | -4.446 | 8.53E-05 | glio |
| Interaction_genotype_Myc | Tas1r1 | -4.054 | 8.53E-05 | glio |
| Interaction_genotype_Myc | Tmem158 | -3.845 | 8.53E-05 | glio |
| Atr_ex.vivo_noRT | 4921504E06Ri | 2.859 | 8.53E-05 | glio |
| Bora_ex.vivo_noRT | Bpifa3 | 3.201 | 8.53E-05 | glio |
| Chek1_in.vivo_noRT | Nts | 3.001 | 8.53E-05 | glio |
| Eif2s1_in.vivo_noRT | Gm49808 | 2.143 | 8.53E-05 | glio |
| Interaction_genotype_Mtor | Sgtb | -5.29 | 8.53E-06 | glio |
| Bora_in.vivo_noRT | Zfp493 | 3.112 | 8.53E-06 | glio |
| Chek1_ex.vivo_noRT | Maff | 2.516 | 8.53E-06 | glio |
| Chek1_in.vivo_noRT | Gm26519 | 3.857 | 8.53E-07 | glio |
| Hsd17b10_in.vivo_noRT | Bcl2l15 | 3.356 | 8.53E-08 | glio |
| Aars2_in.vivo_noRT | Nhsl2 | 4.208 | 8.53E-10 | glio |
| Chek1_ex.vivo_noRT | Gpr17 | 4.842 | 8.53E-10 | glio |
| Atr_ex.vivo_noRT | Olfr881 | 4.309 | 8.53E-13 | glio |
| Aars2_in.vivo_noRT | Gm16178 | 5.327 | 8.53E-27 | glio |
| Aars2_in.vivo_noRT | Gm364 | 5.327 | 8.53E-27 | glio |
| Interaction_genotype_Bcas2 | Apobr | -3.84 | 8.54E-03 | glio |
| Interaction_genotype_Bcas2 | Prickle4 | -3.332 | 8.54E-03 | glio |
| Interaction_genotype_Bcas2 | Car12 | -3.386 | 8.54E-03 | glio |

| | | | | |
|---|---|---|---|---|
| Interaction_genotype_Bcas2 | Syp | -3.898 | 8.54E-03 | glio |
| Interaction_genotype_Bcas2 | Gm36266 | -3.96 | 8.54E-03 | glio |
| Interaction_genotype_Bcas2 | Smim12 | 1.647 | 8.54E-03 | glio |
| Interaction_genotype_Mtor | 1700066M21F | -2.872 | 8.54E-03 | glio |
| Interaction_genotype_Mtor | Gnai3 | -1.41 | 8.54E-03 | glio |
| Interaction_genotype_Myc | Gm48086 | -3.075 | 8.54E-03 | glio |
| Aars2_in.vivo_noRT | Atg14 | 1.529 | 8.54E-03 | glio |
| Aars2_in.vivo_noRT | Lars | -0.77 | 8.54E-03 | glio |
| Atr_in.vivo_noRT | Setdb2 | 1.584 | 8.54E-03 | glio |
| Bora_in.vivo_noRT | Gm4356 | 2.135 | 8.54E-03 | glio |
| Bora_in.vivo_noRT | Ppfia2 | 1.554 | 8.54E-03 | glio |
| Cyp19a1_ex.vivo_noRT | BB287469 | 2.159 | 8.54E-03 | glio |
| Interaction_genotype_Aars2 | Car14 | -4.027 | 8.54E-04 | glio |
| Interaction_genotype_Myc | Echdc3 | -3.146 | 8.54E-04 | glio |
| Atr_ex.vivo_noRT | Gm48742 | 2.586 | 8.54E-04 | glio |
| Bcas2_in.vivo_noRT | March11 | 2.534 | 8.54E-04 | glio |
| culture_effect | Taco1os | 1.695 | 8.54E-05 | glio |
| Atr_in.vivo_noRT | Gm26670 | 2.583 | 8.54E-05 | glio |
| Atr_ex.vivo_noRT | Gm48265 | 3.164 | 8.54E-05 | glio |
| Chek1_ex.vivo_noRT | Arvcf | 3.747 | 8.54E-05 | glio |
| Chek1_ex.vivo_noRT | Col28a1 | 3.464 | 8.54E-05 | glio |
| Eif2s1_in.vivo_noRT | Slc14a2 | 2.143 | 8.54E-05 | glio |
| Eif2s1_in.vivo_noRT | C330024C12R | 2.806 | 8.54E-05 | glio |
| Eif2s1_in.vivo_noRT | Gm4340 | 2.143 | 8.54E-05 | glio |
| Eif2s1_in.vivo_noRT | Gm16239 | 2.143 | 8.54E-05 | glio |
| Atr_in.vivo_noRT | Egf | 3.342 | 8.54E-06 | glio |
| Chek1_ex.vivo_noRT | Gm45606 | 3.747 | 8.54E-06 | glio |
| Interaction_genotype_Mtor | Lyz2 | -6.131 | 8.54E-07 | glio |
| Aars2_in.vivo_noRT | 1700066B19R | 3.35 | 8.54E-07 | glio |
| Bora_in.vivo_noRT | Gm15927 | 3.935 | 8.54E-09 | glio |
| Chek1_ex.vivo_noRT | Olfr718-ps1 | 4.449 | 8.54E-12 | glio |
| interaction_RT | Nes | -2.148 | 8.54E-16 | glio |
| Chek1_in.vivo_noRT | Gm26543 | 5.044 | 8.54E-25 | glio |
| culture_effect | Zfp958 | -0.676 | 8.55E-03 | glio |
| culture_effect | Nbn | -0.479 | 8.55E-03 | glio |
| Interaction_genotype_Mtor | Casp9 | -2.703 | 8.55E-03 | glio |
| Interaction_genotype_Mtor | Ccdc136 | -2.726 | 8.55E-03 | glio |
| Interaction_genotype_Mtor | Gm50240 | -2.273 | 8.55E-03 | glio |
| Interaction_genotype_Myc | Trim14 | -3.221 | 8.55E-03 | glio |
| Interaction_genotype_Myc | Spata2 | -2.195 | 8.55E-03 | glio |
| Atr_in.vivo_noRT | Prr11 | 1.48 | 8.55E-03 | glio |
| Bard1_ex.vivo_noRT | Olfr1189 | 2.415 | 8.55E-03 | glio |
| Bard1_ex.vivo_noRT | 2010001K21R | 2.443 | 8.55E-03 | glio |
| Bcas2_ex.vivo_noRT | Tcstv1 | 2.342 | 8.55E-03 | glio |
| Bora_in.vivo_noRT | Mturn | 1.811 | 8.55E-03 | glio |
| Bora_in.vivo_noRT | Cdc123 | -0.758 | 8.55E-03 | glio |
| Bora_ex.vivo_noRT | Gm44899 | 2.097 | 8.55E-03 | glio |
| Chek1_in.vivo_noRT | Napsa | 2.002 | 8.55E-03 | glio |
| Chek1_in.vivo_noRT | Ctif | 1.753 | 8.55E-03 | glio |
| Interaction_genotype_Aars2 | C130060C02R | -3.733 | 8.55E-04 | glio |
| Interaction_genotype_Chek1 | Itih5 | 4.751 | 8.55E-04 | glio |
| Interaction_genotype_Chek1 | Cwc15 | 1.501 | 8.55E-04 | glio |

| | | | | |
|---|---|---|---|---|
| Interaction_genotype_Myc | Cox5b | 1.295 | 8.55E-04 | glio |
| Aars2_in.vivo_noRT | Hsdl1 | 1.722 | 8.55E-04 | glio |
| Bora_in.vivo_noRT | Decr1 | -1.324 | 8.55E-04 | glio |
| Bora_in.vivo_noRT | Tmem251 | -1.553 | 8.55E-05 | glio |
| Atr_in.vivo_noRT | Gm16279 | 3.278 | 8.55E-06 | glio |
| Interaction_genotype_Mtor | AV039307 | -6.558 | 8.55E-08 | glio |
| Interaction_genotype_Mtor | Trim17 | -4.498 | 8.55E-08 | glio |
| Aars2_in.vivo_noRT | Pde5a | 4.04 | 8.55E-09 | glio |
| Hsd17b10_in.vivo_noRT | Gm45406 | 3.094 | 8.55E-09 | glio |
| Hsd17b10_in.vivo_noRT | Serpinb2 | 3.616 | 8.55E-09 | glio |
| Bcas2_in.vivo_noRT | Sprr1a | 4.807 | 8.55E-12 | glio |
| Aars2_in.vivo_noRT | Gm5941 | 5.327 | 8.55E-14 | glio |
| Bora_in.vivo_noRT | Lrp2bp | 4.672 | 8.55E-14 | glio |
| Atr_in.vivo_noRT | Hdac9 | 5.267 | 8.55E-15 | glio |
| Bcas2_in.vivo_noRT | Speer4a | 4.541 | 8.55E-19 | glio |
| Bcas2_in.vivo_noRT | 2410017I17Ri | 4.807 | 8.55E-23 | glio |
| culture_effect | Otos | 4.421 | 8.55E-24 | glio |
| culture_effect | Phka2 | 0.908 | 8.56E-03 | glio |
| Interaction_genotype_Aars2 | Ube2l3 | 1.103 | 8.56E-03 | glio |
| Interaction_genotype_Bcas2 | Gm32834 | -2.934 | 8.56E-03 | glio |
| Interaction_genotype_Bcas2 | Ciao1 | 1.523 | 8.56E-03 | glio |
| Interaction_genotype_Bcas2 | Hexa | 1.282 | 8.56E-03 | glio |
| Interaction_genotype_Mtor | Kirrel3 | -3.043 | 8.56E-03 | glio |
| Interaction_genotype_Mtor | Bmi1 | -1.582 | 8.56E-03 | glio |
| Interaction_genotype_Myc | Zc3h6 | -2.902 | 8.56E-03 | glio |
| Interaction_genotype_Myc | Kdm3b | -2.178 | 8.56E-03 | glio |
| Interaction_genotype_Pars2 | Zfp473 | -3.481 | 8.56E-03 | glio |
| Interaction_genotype_Terf1 | Cbr3 | -4.227 | 8.56E-03 | glio |
| interaction_RT | Pip4p1 | 0.855 | 8.56E-03 | glio |
| Aars2_in.vivo_noRT | Gm43303 | 1.721 | 8.56E-03 | glio |
| Aars2_in.vivo_noRT | Lncpint | 1.284 | 8.56E-03 | glio |
| Aars2_in.vivo_noRT | Tjp1 | 1.004 | 8.56E-03 | glio |
| Aars2_in.vivo_noRT | Dut | -0.902 | 8.56E-03 | glio |
| Bora_in.vivo_noRT | Prpf6 | -0.996 | 8.56E-03 | glio |
| Bora_ex.vivo_noRT | Gm49223 | 2.22 | 8.56E-03 | glio |
| Cenpj_ex.vivo_noRT | Ocm | 2.591 | 8.56E-03 | glio |
| Dclre1c_ex.vivo_noRT | Dpp6 | 2.296 | 8.56E-03 | glio |
| Bora_in.vivo_noRT | Gm17501 | 2.513 | 8.56E-04 | glio |
| Bora_in.vivo_noRT | Gm43768 | 2.298 | 8.56E-04 | glio |
| Bora_in.vivo_noRT | Arid3c | 2.181 | 8.56E-04 | glio |
| interaction_RT | Mthfd1 | -1.334 | 8.56E-05 | glio |
| Bcas2_in.vivo_noRT | Arhgap33 | 2.873 | 8.56E-05 | glio |
| Chek1_ex.vivo_noRT | Gm13403 | 3.753 | 8.56E-06 | glio |
| Interaction_genotype_Mtor | Sh3d21 | -5.073 | 8.56E-07 | glio |
| Chek1_in.vivo_noRT | Aldh1a1 | 3.538 | 8.56E-07 | glio |
| Chek1_in.vivo_noRT | Spink10 | 3.248 | 8.56E-07 | glio |
| Aars2_ex.vivo_noRT | Hoxc8 | 3.293 | 8.56E-08 | glio |
| Atr_in.vivo_noRT | Zscan2 | 3.819 | 8.56E-09 | glio |
| Atr_in.vivo_noRT | Igf2bp1 | 5.533 | 8.56E-16 | glio |
| culture_effect | Kcnn4 | -5.002 | 8.56E-19 | glio |
| Atr_in.vivo_noRT | Gm34939 | 5.269 | 8.56E-20 | glio |
| culture_effect | Cd200r2 | -1.198 | 8.57E-03 | glio |

| | | | | |
|---|---|---|---|---|
| culture_effect | Dnph1 | -0.439 | 8.57E-03 | glio |
| Interaction_genotype_Atr | Rab2b | 1.715 | 8.57E-03 | glio |
| Interaction_genotype_Atr | Pmf1 | 1.558 | 8.57E-03 | glio |
| Interaction_genotype_Mtor | Tbpl1 | 1.579 | 8.57E-03 | glio |
| Interaction_genotype_Myc | Gm17655 | -3.2 | 8.57E-03 | glio |
| Interaction_genotype_Myc | Zfp607b | -2.438 | 8.57E-03 | glio |
| Interaction_genotype_Myc | Psph | 1.409 | 8.57E-03 | glio |
| interaction_RT | Dyrk3 | 1.251 | 8.57E-03 | glio |
| Bcas2_in.vivo_noRT | Gm26532 | -1.265 | 8.57E-03 | glio |
| Bcas2_ex.vivo_noRT | Nkg7 | 2.347 | 8.57E-03 | glio |
| Chek1_in.vivo_noRT | Kif2a | -1.105 | 8.57E-03 | glio |
| Eif2s1_in.vivo_noRT | Ppp1r1b | -2.063 | 8.57E-03 | glio |
| Fanca_ex.vivo_noRT | Cd7 | 2.124 | 8.57E-03 | glio |
| Fanca_ex.vivo_noRT | Gm48764 | 2.124 | 8.57E-03 | glio |
| Fanca_ex.vivo_noRT | Gad1 | 2.124 | 8.57E-03 | glio |
| culture_effect | Klhl15 | -0.958 | 8.57E-04 | glio |
| Interaction_genotype_Myc | Trmt61b | -2.729 | 8.57E-04 | glio |
| Aars2_in.vivo_noRT | Tmem82 | 2.565 | 8.57E-04 | glio |
| Atp5c1_in.vivo_noRT | Clca3a1 | 2.396 | 8.57E-04 | glio |
| Atr_in.vivo_noRT | Slc9a5 | 2.322 | 8.57E-04 | glio |
| Chek1_ex.vivo_noRT | Gm34885 | 2.994 | 8.57E-04 | glio |
| Interaction_genotype_Myc | Gm32250 | -3.32 | 8.57E-05 | glio |
| interaction_RT | Sdf2 | 0.788 | 8.57E-05 | glio |
| Bora_in.vivo_noRT | Gm19684 | 2.978 | 8.57E-05 | glio |
| Chek1_ex.vivo_noRT | Gm15903 | 3.38 | 8.57E-05 | glio |
| Interaction_genotype_Mtor | Gm15601 | -5.473 | 8.57E-06 | glio |
| Interaction_genotype_Mtor | Rps6 | 1.425 | 8.57E-06 | glio |
| culture_effect | Dpep2 | -2.889 | 8.57E-07 | glio |
| Interaction_genotype_Mtor | Rasgrp2 | -5.762 | 8.57E-07 | glio |
| Atp5c1_in.vivo_noRT | Trmt61b | 2.668 | 8.57E-07 | glio |
| culture_effect | Flna | 0.854 | 8.57E-09 | glio |
| Bora_in.vivo_noRT | Tslp | 4.291 | 8.57E-09 | glio |
| Hsd17b10_in.vivo_noRT | Gm14540 | 3.093 | 8.57E-09 | glio |
| Chek1_in.vivo_noRT | Brsk2 | 3.899 | 8.57E-10 | glio |
| culture_effect | Slc39a13 | 1.432 | 8.57E-11 | glio |
| Chek1_in.vivo_noRT | Pldi | 5.044 | 8.57E-19 | glio |
| Interaction_genotype_Atr | Car12 | -3.161 | 8.58E-03 | glio |
| Interaction_genotype_Atr | Asb16 | -3.079 | 8.58E-03 | glio |
| Interaction_genotype_Myc | Klf15 | -2.799 | 8.58E-03 | glio |
| Interaction_genotype_Pars2 | 1700016P04R | -3.398 | 8.58E-03 | glio |
| Interaction_genotype_Pars2 | Serpinb6a | -3.763 | 8.58E-03 | glio |
| Atr_ex.vivo_noRT | Plekhf2 | 1.335 | 8.58E-03 | glio |
| Bcas2_in.vivo_noRT | Psma3 | -0.49 | 8.58E-03 | glio |
| Bora_in.vivo_noRT | Trmt44 | 1.547 | 8.58E-03 | glio |
| Bora_in.vivo_noRT | Csnk2a1 | -0.776 | 8.58E-03 | glio |
| Cenpj_in.vivo_noRT | 1110020A21R | 2.37 | 8.58E-03 | glio |
| Chek1_ex.vivo_noRT | Pacs1 | 2.461 | 8.58E-03 | glio |
| culture_effect | Tet3 | -0.792 | 8.58E-04 | glio |
| Interaction_genotype_Myc | Myl7 | -3.887 | 8.58E-04 | glio |
| Interaction_genotype_Myc | Mrpl34 | 1.33 | 8.58E-04 | glio |
| Bcas2_ex.vivo_noRT | Vmn2r54 | 2.742 | 8.58E-04 | glio |
| Bcas2_ex.vivo_noRT | Ipcef1 | 2.742 | 8.58E-04 | glio |

| | | | | |
|---|---|---|---|---|
| Bcas2_ex.vivo_noRT | Gm31152 | 2.949 | 8.58E-04 | glio |
| Bora_in.vivo_noRT | D730045B01R | 2.15 | 8.58E-04 | glio |
| Interaction_genotype_Myc | Fhl1 | -3.44 | 8.58E-05 | glio |
| Chek1_ex.vivo_noRT | Fgf21 | 3.851 | 8.58E-05 | glio |
| Bora_in.vivo_noRT | Gm6598 | 2.942 | 8.58E-06 | glio |
| Bora_in.vivo_noRT | Lmnb1 | -2.361 | 8.58E-07 | glio |
| Atr_in.vivo_noRT | Emid1 | 4.141 | 8.58E-08 | glio |
| culture_effect | Hmgcl | -2.106 | 8.58E-09 | glio |
| Chek1_in.vivo_noRT | 1700030C12R | 4.118 | 8.58E-09 | glio |
| Aars2_in.vivo_noRT | Ephx1 | 4.428 | 8.58E-11 | glio |
| Atr_in.vivo_noRT | Trpm3 | 5.011 | 8.58E-13 | glio |
| Atr_in.vivo_noRT | Gm40557 | 5.011 | 8.58E-13 | glio |
| Bora_in.vivo_noRT | Col19a1 | 4.019 | 8.58E-14 | glio |
| Chek1_in.vivo_noRT | Gm43256 | 4.78 | 8.58E-14 | glio |
| Atr_in.vivo_noRT | 2300002M23F | 4.365 | 8.58E-15 | glio |
| Bcas2_in.vivo_noRT | 1700108F19Ri | 4.543 | 8.58E-16 | glio |
| Bora_in.vivo_noRT | Gm32250 | 4.154 | 8.58E-16 | glio |
| Bora_in.vivo_noRT | Scgb1c1 | 4.936 | 8.58E-17 | glio |
| Interaction_genotype_Aars2 | Sucnr1 | -2.034 | 8.59E-03 | glio |
| Interaction_genotype_Bcas2 | 4930554I06Ri | -3.67 | 8.59E-03 | glio |
| Interaction_genotype_Mtor | Gm28592 | -3.048 | 8.59E-03 | glio |
| Interaction_genotype_Mtor | Hdac10 | -2.756 | 8.59E-03 | glio |
| Interaction_genotype_Mtor | Chchd6 | -2.671 | 8.59E-03 | glio |
| Interaction_genotype_Mtor | Aunip | -2.585 | 8.59E-03 | glio |
| Interaction_genotype_Mtor | Snupn | -2.401 | 8.59E-03 | glio |
| Interaction_genotype_Mtor | Gm48226 | -2.843 | 8.59E-03 | glio |
| Interaction_genotype_Mtor | Ptrh1 | -2.228 | 8.59E-03 | glio |
| Interaction_genotype_Myc | Pms1 | -2.517 | 8.59E-03 | glio |
| Interaction_genotype_Myc | Zfp775 | -2.442 | 8.59E-03 | glio |
| Interaction_genotype_Myc | Ptms | 1.175 | 8.59E-03 | glio |
| Aars2_in.vivo_noRT | Adm | 1.73 | 8.59E-03 | glio |
| Aars2_in.vivo_noRT | Atp5l | -0.59 | 8.59E-03 | glio |
| Atp5c1_ex.vivo_noRT | C1ra | 2.592 | 8.59E-03 | glio |
| Atp5c1_ex.vivo_noRT | Gm11131 | 3.373 | 8.59E-03 | glio |
| Atp5c1_ex.vivo_noRT | Fam166a | 2.61 | 8.59E-03 | glio |
| culture_effect | Cox8a | 0.372 | 8.59E-04 | glio |
| Interaction_genotype_Myc | Gm19541 | -3.651 | 8.59E-04 | glio |
| interaction_RT | Idnk | 1.102 | 8.59E-05 | glio |
| interaction_RT | Larp1 | -0.76 | 8.59E-05 | glio |
| interaction_RT | Tmsb10 | 1.056 | 8.59E-05 | glio |
| Bcas2_in.vivo_noRT | Lbx2 | 3.047 | 8.59E-05 | glio |
| Atp5c1_in.vivo_noRT | Gm38163 | 2.668 | 8.59E-06 | glio |
| Bora_in.vivo_noRT | Gm44091 | 3.017 | 8.59E-06 | glio |
| Chek1_in.vivo_noRT | Gemin4 | 3.298 | 8.59E-06 | glio |
| Chek1_in.vivo_noRT | Gm13283 | 3.152 | 8.59E-06 | glio |
| Atr_in.vivo_noRT | Npl | 3.237 | 8.59E-07 | glio |
| Aars2_in.vivo_noRT | Rdh12 | 4.681 | 8.59E-13 | glio |
| Aars2_in.vivo_noRT | 5330439K02R | 5.06 | 8.59E-13 | glio |
| Aars2_in.vivo_noRT | Oit3 | 5.327 | 8.59E-16 | glio |
| Chek1_in.vivo_noRT | Abcb5 | 5.044 | 8.59E-24 | glio |
| Interaction_genotype_Cdkn2a | Gm35439 | 3.088 | 8.60E-03 | glio |
| Interaction_genotype_Cdkn2a | Gm33466 | 2.835 | 8.60E-03 | glio |

| | | | | |
|---|---|---|---|---|
| Interaction_genotype_Mtor | Calhm2 | -2.294 | 8.60E-03 | glio |
| Interaction_genotype_Rad51 | Nxph4 | -3.649 | 8.60E-03 | glio |
| Interaction_genotype_Rad51 | Gm50295 | -3.769 | 8.60E-03 | glio |
| Interaction_genotype_Rad51 | Klhdc10 | 2.109 | 8.60E-03 | glio |
| Interaction_genotype_Uqcrc2 | 5430402O13R | -4.028 | 8.60E-03 | glio |
| Aars2_in.vivo_noRT | Rps19 | -0.513 | 8.60E-03 | glio |
| Atr_in.vivo_noRT | Psmd8 | -0.808 | 8.60E-03 | glio |
| Atr_ex.vivo_noRT | Acy3 | 2.614 | 8.60E-03 | glio |
| Chek1_in.vivo_noRT | Gm16196 | 1.813 | 8.60E-03 | glio |
| Chek1_in.vivo_noRT | Cd47 | -0.907 | 8.60E-03 | glio |
| Chek1_in.vivo_noRT | Rbm25 | -0.521 | 8.60E-03 | glio |
| Chek1_in.vivo_noRT | Ybx1 | -0.502 | 8.60E-03 | glio |
| Chek1_ex.vivo_noRT | Tmem169 | 2.111 | 8.60E-03 | glio |
| Hsd17b10_in.vivo_noRT | Abcd2 | 2.304 | 8.60E-03 | glio |
| Hsd17b10_in.vivo_noRT | 1700001L19Ri | 1.848 | 8.60E-03 | glio |
| culture_effect | Psme2b | -1.885 | 8.60E-04 | glio |
| culture_effect | Sec16b | -1.302 | 8.60E-04 | glio |
| Interaction_genotype_Aars2 | Gm43581 | -3.849 | 8.60E-04 | glio |
| Interaction_genotype_Bcas2 | Hist1h4b | -4.856 | 8.60E-04 | glio |
| Interaction_genotype_Mtor | BC024978 | -3.344 | 8.60E-04 | glio |
| Interaction_genotype_Mtor | Spag9 | 1.622 | 8.60E-04 | glio |
| Interaction_genotype_Rad21 | Gm5602 | 3.796 | 8.60E-04 | glio |
| Aars2_in.vivo_noRT | Mea1 | -1.284 | 8.60E-04 | glio |
| Atr_in.vivo_noRT | Maats1 | 2.444 | 8.60E-04 | glio |
| Atr_in.vivo_noRT | Nt5c3 | -1.302 | 8.60E-04 | glio |
| Atr_in.vivo_noRT | Hspa4 | -1.192 | 8.60E-04 | glio |
| Atr_in.vivo_noRT | Rps9 | -0.666 | 8.60E-04 | glio |
| Bora_ex.vivo_noRT | Dapk2 | 2.404 | 8.60E-04 | glio |
| interaction_RT | Gm26917 | -1.77 | 8.60E-05 | glio |
| culture_effect | Ubap2 | 0.601 | 8.60E-06 | glio |
| Atr_in.vivo_noRT | Cyp3a13 | 3.341 | 8.60E-06 | glio |
| Hsd17b10_in.vivo_noRT | Krtdap | 3.225 | 8.60E-06 | glio |
| Bcas2_in.vivo_noRT | Gm5624 | 3.818 | 8.60E-09 | glio |
| culture_effect | Alox15 | -3.5 | 8.60E-10 | glio |
| Interaction_genotype_Mtor | Gm15708 | -3.245 | 8.61E-03 | glio |
| Interaction_genotype_Mtor | Pknox2 | -3.347 | 8.61E-03 | glio |
| Interaction_genotype_Mtor | Tnfrsf9 | -3.051 | 8.61E-03 | glio |
| Interaction_genotype_Mtor | 3010003L21Ri | -2.741 | 8.61E-03 | glio |
| Interaction_genotype_Rad51 | Hgs | 1.865 | 8.61E-03 | glio |
| Interaction_genotype_Terf1 | 4930554I06Ri | -4.218 | 8.61E-03 | glio |
| Interaction_genotype_Terf1 | Hba-a2 | -4.088 | 8.61E-03 | glio |
| Atr_in.vivo_noRT | Coro1c | -1.077 | 8.61E-03 | glio |
| Bcas2_in.vivo_noRT | Syk | 1.745 | 8.61E-03 | glio |
| Brca2_in.vivo_noRT | Prl7c1 | 1.763 | 8.61E-03 | glio |
| Brca2_in.vivo_noRT | Gm33466 | 1.763 | 8.61E-03 | glio |
| Hsd17b10_in.vivo_noRT | Gm44190 | 1.891 | 8.61E-03 | glio |
| culture_effect | Sema3a | -1.979 | 8.61E-04 | glio |
| culture_effect | Enpp3 | -2.102 | 8.61E-04 | glio |
| culture_effect | 2010204K13R | 0.539 | 8.61E-04 | glio |
| Interaction_genotype_Atr | Ube2r2 | 1.28 | 8.61E-04 | glio |
| Hsd17b10_in.vivo_noRT | Alpl | 2.431 | 8.61E-04 | glio |
| Interaction_genotype_Mtor | Spi1 | -4.278 | 8.61E-05 | glio |

| | | | | |
|---|---|---|---|---|
| Aars2_in.vivo_noRT | Gm10553 | 2.78 | 8.61E-05 | glio |
| Aars2_in.vivo_noRT | Cox6c | -0.872 | 8.61E-05 | glio |
| Aars2_ex.vivo_noRT | Obscn | 3.293 | 8.61E-05 | glio |
| Bcas2_in.vivo_noRT | Gadl1 | 2.632 | 8.61E-05 | glio |
| Bora_ex.vivo_noRT | Gm21319 | 3.201 | 8.61E-06 | glio |
| Chek1_ex.vivo_noRT | Gm43258 | 3.871 | 8.61E-06 | glio |
| culture_effect | A330084C13R | 2.757 | 8.61E-07 | glio |
| Bora_in.vivo_noRT | Srl | 3.517 | 8.61E-07 | glio |
| Chek1_ex.vivo_noRT | Atp8a2 | 4.055 | 8.61E-08 | glio |
| Aars2_in.vivo_noRT | Camk1d | 4.526 | 8.61E-11 | glio |
| Atr_in.vivo_noRT | Gm32585 | 5.008 | 8.61E-12 | glio |
| Aars2_in.vivo_noRT | Gm10482 | 4.805 | 8.61E-18 | glio |
| Interaction_genotype_Bcas2 | Gm18280 | -2.595 | 8.62E-03 | glio |
| Interaction_genotype_Bcas2 | 5031434O11R | -3.946 | 8.62E-03 | glio |
| Interaction_genotype_Bcas2 | Ptms | 1.104 | 8.62E-03 | glio |
| Interaction_genotype_Myc | Wfdc18 | -3.666 | 8.62E-03 | glio |
| Interaction_genotype_Myc | Golph3l | -2.193 | 8.62E-03 | glio |
| interaction_RT | Ipo13 | -0.963 | 8.62E-03 | glio |
| interaction_RT | Ppp1r14c | 0.739 | 8.62E-03 | glio |
| Atr_in.vivo_noRT | Ciao2b | -0.729 | 8.62E-03 | glio |
| Bard1_ex.vivo_noRT | Gm44646 | 2.227 | 8.62E-03 | glio |
| Bard1_ex.vivo_noRT | Sap30bpos | 2.421 | 8.62E-03 | glio |
| Bora_in.vivo_noRT | Fem1c | -1.291 | 8.62E-03 | glio |
| Bora_in.vivo_noRT | Avpi1 | -1.292 | 8.62E-03 | glio |
| Chek1_ex.vivo_noRT | Rps6 | -0.606 | 8.62E-03 | glio |
| culture_effect | Dus3l | -0.947 | 8.62E-04 | glio |
| Interaction_genotype_Chek1 | Ppp1r2 | 2.111 | 8.62E-04 | glio |
| Interaction_genotype_Chek1 | Lamp1 | 1.326 | 8.62E-04 | glio |
| Interaction_genotype_Myc | Sag | -3.783 | 8.62E-04 | glio |
| Aars2_in.vivo_noRT | Slc36a4 | 2.323 | 8.62E-04 | glio |
| Atr_in.vivo_noRT | Plpp6 | 2.235 | 8.62E-04 | glio |
| Atr_in.vivo_noRT | Smagp | 2.308 | 8.62E-04 | glio |
| Bcas2_in.vivo_noRT | Rit2 | 2.315 | 8.62E-04 | glio |
| Bcas2_in.vivo_noRT | Axin1 | -1.742 | 8.62E-04 | glio |
| Chek1_in.vivo_noRT | Tnfrsf22 | 2.318 | 8.62E-04 | glio |
| Interaction_genotype_Rpl3 | Rhob | 3.18 | 8.62E-05 | glio |
| Chek1_in.vivo_noRT | Rpl10a | -0.654 | 8.62E-05 | glio |
| Interaction_genotype_Mtor | Gm2415 | -5.54 | 8.62E-06 | glio |
| Aars2_in.vivo_noRT | Mill2 | 3.135 | 8.62E-06 | glio |
| Chek1_ex.vivo_noRT | Hist1h2bc | 4.015 | 8.62E-07 | glio |
| Chek1_ex.vivo_noRT | Gpr146 | 5.249 | 8.62E-09 | glio |
| Chek1_ex.vivo_noRT | Slco2a1 | 4.842 | 8.62E-09 | glio |
| Hsd17b10_in.vivo_noRT | Gm47404 | 3.616 | 8.62E-14 | glio |
| culture_effect | Adgrv1 | -0.931 | 8.63E-03 | glio |
| culture_effect | Man2b1 | -0.887 | 8.63E-03 | glio |
| culture_effect | Erbb3 | -0.501 | 8.63E-03 | glio |
| Interaction_genotype_Aars2 | Rtl8b | -2.623 | 8.63E-03 | glio |
| Interaction_genotype_Atr | Zfp600 | -3.223 | 8.63E-03 | glio |
| Interaction_genotype_Myc | Kcnd1 | -2.607 | 8.63E-03 | glio |
| interaction_RT | Gm49890 | 1.861 | 8.63E-03 | glio |
| Atr_in.vivo_noRT | Limk1 | 1.33 | 8.63E-03 | glio |
| Bcas2_in.vivo_noRT | AC168977.2 | 2.053 | 8.63E-03 | glio |

| | | | | |
|---|---|---|---|---|
| Bora_in.vivo_noRT | Nudt3 | -0.859 | 8.63E-03 | glio |
| Cdkn2a_in.vivo_noRT | Gm13856 | -1.675 | 8.63E-03 | glio |
| Cdkn2a_in.vivo_noRT | Gm21936 | -1.675 | 8.63E-03 | glio |
| Cdkn2a_in.vivo_noRT | Gm17203 | -1.798 | 8.63E-03 | glio |
| Cenpj_in.vivo_noRT | Pdgfra | -2.295 | 8.63E-03 | glio |
| Chek1_in.vivo_noRT | Stat4 | 1.879 | 8.63E-03 | glio |
| Chek1_in.vivo_noRT | Col28a1 | 1.844 | 8.63E-03 | glio |
| Chek1_in.vivo_noRT | Vill | 1.596 | 8.63E-03 | glio |
| Hsd17b10_in.vivo_noRT | C430042M11F | 2.087 | 8.63E-03 | glio |
| Hsd17b10_in.vivo_noRT | Maml1 | 1.407 | 8.63E-03 | glio |
| Interaction_genotype_Mtor | Snrpa | 1.731 | 8.63E-04 | glio |
| Atp5c1_in.vivo_noRT | Skida1 | 2.668 | 8.63E-04 | glio |
| Bcas2_in.vivo_noRT | Gm15952 | 2.504 | 8.63E-04 | glio |
| Bcas2_ex.vivo_noRT | Gm36026 | 2.742 | 8.63E-04 | glio |
| Bora_in.vivo_noRT | Susd1 | 2.624 | 8.63E-04 | glio |
| Chek1_in.vivo_noRT | Mgll | 2.339 | 8.63E-04 | glio |
| Chek1_ex.vivo_noRT | Lyz2 | 3.063 | 8.63E-04 | glio |
| Interaction_genotype_Mtor | Fgf1 | -4.65 | 8.63E-05 | glio |
| Interaction_genotype_Myc | Gm15867 | -4.418 | 8.63E-05 | glio |
| Interaction_genotype_Myc | Bace1 | -3.605 | 8.63E-05 | glio |
| Interaction_genotype_Myc | Rps7 | 1.575 | 8.63E-05 | glio |
| Aars2_in.vivo_noRT | Rpl21 | -0.714 | 8.63E-05 | glio |
| Atr_ex.vivo_noRT | Rgs2 | 3.248 | 8.63E-05 | glio |
| Bcas2_in.vivo_noRT | 4930589L23Ri | 2.929 | 8.63E-05 | glio |
| Bora_in.vivo_noRT | Metrn | -1.24 | 8.63E-05 | glio |
| Interaction_genotype_Myc | Daam2 | -4.997 | 8.63E-06 | glio |
| Interaction_genotype_Myc | Rab11fip4os1 | -5.177 | 8.63E-06 | glio |
| Bcas2_in.vivo_noRT | 4932441J04Ri | 3.556 | 8.63E-07 | glio |
| Chek1_in.vivo_noRT | Gm9833 | 3.263 | 8.63E-07 | glio |
| Aars2_in.vivo_noRT | 3110082J24Ri | 3.885 | 8.63E-08 | glio |
| Atr_ex.vivo_noRT | Gm10801 | 3.947 | 8.63E-08 | glio |
| Chek1_ex.vivo_noRT | Pcdhb19 | 4.447 | 8.63E-08 | glio |
| Hsd17b10_in.vivo_noRT | Tmem236 | 3.35 | 8.63E-08 | glio |
| Aars2_in.vivo_noRT | Gm28874 | 4.305 | 8.63E-10 | glio |
| Aars2_in.vivo_noRT | Sult5a1 | 4.794 | 8.63E-11 | glio |
| Bora_in.vivo_noRT | Ncf1 | 4.283 | 8.63E-13 | glio |
| Bcas2_in.vivo_noRT | Ssh3 | 4.536 | 8.63E-14 | glio |
| Chek1_in.vivo_noRT | Myo5c | 5.044 | 8.63E-14 | glio |
| Chek1_ex.vivo_noRT | Gm15533 | 4.842 | 8.63E-15 | glio |
| Atr_in.vivo_noRT | Prickle2 | 5.533 | 8.63E-21 | glio |
| culture_effect | Vipr2 | -1.713 | 8.64E-03 | glio |
| culture_effect | Unc5cl | -1.692 | 8.64E-03 | glio |
| culture_effect | Tmem88 | -1.654 | 8.64E-03 | glio |
| culture_effect | Rusc1 | -0.737 | 8.64E-03 | glio |
| culture_effect | Prkab1 | -0.628 | 8.64E-03 | glio |
| culture_effect | Zfp639 | 0.53 | 8.64E-03 | glio |
| culture_effect | Mepce | -0.46 | 8.64E-03 | glio |
| Interaction_genotype_Mtor | Gm9993 | -2.562 | 8.64E-03 | glio |
| Aars2_in.vivo_noRT | Mtus2 | -1.149 | 8.64E-03 | glio |
| Atp5c1_in.vivo_noRT | Gm12280 | 2.404 | 8.64E-03 | glio |
| Atp5c1_in.vivo_noRT | Ushbp1 | 2.142 | 8.64E-03 | glio |
| Atr_in.vivo_noRT | Mavs | 1.441 | 8.64E-03 | glio |

| | | | | |
|---|---|---|---|---|
| Brca2_ex.vivo_noRT | Sucnr1 | 2.03 | 8.64E-03 | glio |
| Cenpj_in.vivo_noRT | Gm10324 | 1.756 | 8.64E-03 | glio |
| Chek1_ex.vivo_noRT | Rad51ap1 | -1.759 | 8.64E-03 | glio |
| Eif2s1_in.vivo_noRT | Brsk2 | 2.052 | 8.64E-03 | glio |
| Eif2s1_in.vivo_noRT | Fabp5 | 0.848 | 8.64E-03 | glio |
| interaction_RT | Cript | 0.717 | 8.64E-04 | glio |
| Aars2_in.vivo_noRT | Eaf2 | 2.451 | 8.64E-04 | glio |
| Aars2_in.vivo_noRT | Gm10687 | 2.506 | 8.64E-04 | glio |
| Aars2_in.vivo_noRT | Tecpr1 | 1.953 | 8.64E-04 | glio |
| Bcas2_in.vivo_noRT | Grasp | -1.854 | 8.64E-04 | glio |
| Chek1_in.vivo_noRT | Axin2 | 2.474 | 8.64E-04 | glio |
| Chek1_ex.vivo_noRT | Hsd17b11 | 2.483 | 8.64E-04 | glio |
| Aars2_in.vivo_noRT | Gm9776 | 2.702 | 8.64E-05 | glio |
| Hsd17b10_in.vivo_noRT | Tas1r1 | 2.589 | 8.64E-05 | glio |
| Interaction_genotype_Myc | Exoc6 | -4.462 | 8.64E-06 | glio |
| Interaction_genotype_Myc | Gm44702 | -5.218 | 8.64E-06 | glio |
| Aars2_in.vivo_noRT | Slc46a1 | 3.293 | 8.64E-06 | glio |
| Aars2_in.vivo_noRT | Katna1 | -1.737 | 8.64E-06 | glio |
| Bora_in.vivo_noRT | Hmgcs2 | 3.898 | 8.64E-07 | glio |
| culture_effect | Ndufaf4 | 0.772 | 8.64E-10 | glio |
| Atr_in.vivo_noRT | Pcdhgb6 | 4.349 | 8.64E-13 | glio |
| Aars2_in.vivo_noRT | 4921536K21R | 5.067 | 8.64E-14 | glio |
| culture_effect | Cox14 | 1.168 | 8.64E-19 | glio |
| Bora_in.vivo_noRT | Esm1 | 4.672 | 8.64E-20 | glio |
| Interaction_genotype_Bora | Tgs1 | 1.311 | 8.65E-03 | glio |
| Interaction_genotype_Mtor | Puf60 | 1.323 | 8.65E-03 | glio |
| Interaction_genotype_Myc | 0610009E02Ri | -3.157 | 8.65E-03 | glio |
| Atr_in.vivo_noRT | Hbegf | -1.151 | 8.65E-03 | glio |
| Atr_in.vivo_noRT | Serf2 | -0.549 | 8.65E-03 | glio |
| Chek1_in.vivo_noRT | Zkscan5 | 1.566 | 8.65E-03 | glio |
| Eif2s1_in.vivo_noRT | Kcnb1 | 2.345 | 8.65E-03 | glio |
| Interaction_genotype_Myc | Gm29170 | -3.468 | 8.65E-04 | glio |
| Interaction_genotype_Myc | Unc5c | -2.841 | 8.65E-04 | glio |
| Atr_in.vivo_noRT | Gab3 | 2.359 | 8.65E-04 | glio |
| Atr_in.vivo_noRT | Fendrr | 2.362 | 8.65E-04 | glio |
| Chek1_ex.vivo_noRT | Stom | 2.447 | 8.65E-04 | glio |
| Bora_in.vivo_noRT | Ccdc8 | 2.745 | 8.65E-05 | glio |
| Interaction_genotype_Mtor | Col6a1 | -4.049 | 8.65E-06 | glio |
| Interaction_genotype_Mtor | Ube2k | 2.036 | 8.65E-06 | glio |
| Aars2_in.vivo_noRT | Zfp59 | 3.17 | 8.65E-06 | glio |
| Atr_ex.vivo_noRT | Ccl9 | 2.771 | 8.65E-06 | glio |
| Atr_ex.vivo_noRT | Mag | 3.962 | 8.65E-06 | glio |
| culture_effect | Nos1 | 1.157 | 8.65E-07 | glio |
| culture_effect | Ints4 | -1.148 | 8.65E-07 | glio |
| Aars2_ex.vivo_noRT | Spink8 | 3.293 | 8.65E-07 | glio |
| Aars2_ex.vivo_noRT | Psmb11 | 3.293 | 8.65E-07 | glio |
| Bcas2_in.vivo_noRT | Hist1h4f | 4.015 | 8.65E-08 | glio |
| Hsd17b10_in.vivo_noRT | Lrguk | 3.352 | 8.65E-09 | glio |
| Chek1_in.vivo_noRT | Ifi214 | 4.253 | 8.65E-10 | glio |
| culture_effect | Crls1 | 0.409 | 8.66E-03 | glio |
| culture_effect | Mrps5 | 0.427 | 8.66E-03 | glio |
| Aars2_in.vivo_noRT | Ywhaz | -0.619 | 8.66E-03 | glio |

| | | | | |
|---|---|---|---|---|
| Bcas2_in.vivo_noRT | Sgf29 | -1.23 | 8.66E-03 | glio |
| Bcas2_in.vivo_noRT | Riok3 | 0.945 | 8.66E-03 | glio |
| Bora_in.vivo_noRT | Rgs5 | 2.366 | 8.66E-03 | glio |
| Bora_in.vivo_noRT | Nme7 | -1.247 | 8.66E-03 | glio |
| Bora_ex.vivo_noRT | Rslcan18 | 2.414 | 8.66E-03 | glio |
| Interaction_genotype_Mtor | Park7 | 1.61 | 8.66E-04 | glio |
| Interaction_genotype_Myc | Gm15726 | -3.531 | 8.66E-04 | glio |
| Interaction_genotype_Myc | Zfp65 | -3.481 | 8.66E-04 | glio |
| Interaction_genotype_Pars2 | B230319C09R | -4.985 | 8.66E-04 | glio |
| interaction_RT | Gart | -1.043 | 8.66E-04 | glio |
| Atr_in.vivo_noRT | Cdkn1b | -1.49 | 8.66E-04 | glio |
| Bard1_ex.vivo_noRT | 1700017M07F | 2.745 | 8.66E-04 | glio |
| culture_effect | P3h4 | 0.682 | 8.66E-05 | glio |
| culture_effect | Eif4h | -0.54 | 8.66E-05 | glio |
| Interaction_genotype_Mtor | Gm29361 | -4.838 | 8.66E-05 | glio |
| Interaction_genotype_Mtor | Ulk3 | -4.783 | 8.66E-05 | glio |
| Interaction_genotype_Myc | D830025C05R | -4.529 | 8.66E-05 | glio |
| Interaction_genotype_Myc | 5730507C01R | -3.397 | 8.66E-05 | glio |
| Interaction_genotype_Myc | Fam185a | -3.373 | 8.66E-05 | glio |
| Bcas2_ex.vivo_noRT | Gm33466 | 2.742 | 8.66E-05 | glio |
| Interaction_genotype_Aars2 | Wfdc1 | -4.844 | 8.66E-06 | glio |
| Interaction_genotype_Atr | Gm13288 | -4.505 | 8.66E-06 | glio |
| Bcas2_in.vivo_noRT | Slc22a5 | 2.89 | 8.66E-06 | glio |
| Aars2_in.vivo_noRT | Gm26879 | 3.32 | 8.66E-08 | glio |
| Chek1_in.vivo_noRT | Zfp652os | 3.738 | 8.66E-08 | glio |
| Bora_in.vivo_noRT | Gm11696 | 4.07 | 8.66E-09 | glio |
| Bora_in.vivo_noRT | Klk14 | 4.405 | 8.66E-10 | glio |
| Chek1_ex.vivo_noRT | Scgb1c1 | 4.449 | 8.66E-10 | glio |
| Atr_ex.vivo_noRT | Spink6 | 4.547 | 8.66E-11 | glio |
| Interaction_genotype_Mtor | Emc7 | 1.111 | 8.67E-03 | glio |
| Interaction_genotype_Myc | Gm12971 | -3.094 | 8.67E-03 | glio |
| Interaction_genotype_Myc | Chrnb1 | -2.901 | 8.67E-03 | glio |
| Interaction_genotype_Myc | Lrrk2 | -2.21 | 8.67E-03 | glio |
| Interaction_genotype_Myc | Chd1 | 1.216 | 8.67E-03 | glio |
| Aars2_in.vivo_noRT | Tmem176b | -1.386 | 8.67E-03 | glio |
| Aars2_in.vivo_noRT | Stip1 | -0.678 | 8.67E-03 | glio |
| Chek1_in.vivo_noRT | Plin2 | -0.909 | 8.67E-03 | glio |
| Interaction_genotype_Myc | Gm5890 | -2.729 | 8.67E-04 | glio |
| Aars2_in.vivo_noRT | Nr1h2 | -1.247 | 8.67E-04 | glio |
| Aars2_ex.vivo_noRT | Tal2 | 2.311 | 8.67E-04 | glio |
| Atp5c1_in.vivo_noRT | Gm34047 | 2.668 | 8.67E-04 | glio |
| Atp5c1_in.vivo_noRT | Zfp14 | 2.676 | 8.67E-04 | glio |
| Atr_in.vivo_noRT | Smarcd3 | 2.078 | 8.67E-04 | glio |
| Bcas2_in.vivo_noRT | Rad23a | -1.401 | 8.67E-04 | glio |
| Chek1_in.vivo_noRT | Cyb561 | 2.569 | 8.67E-04 | glio |
| Chek1_in.vivo_noRT | Tceal9 | -1.38 | 8.67E-04 | glio |
| Interaction_genotype_Myc | Ndufa6 | 1.664 | 8.67E-05 | glio |
| Chek1_in.vivo_noRT | Cmas | -1.42 | 8.67E-05 | glio |
| Interaction_genotype_Myc | Arhgef19 | -5.179 | 8.67E-06 | glio |
| Atr_in.vivo_noRT | Gm4181 | 3.121 | 8.67E-06 | glio |
| Atr_ex.vivo_noRT | A930019D19R | 3.164 | 8.67E-08 | glio |
| culture_effect | Rasgrp2 | 3.293 | 8.67E-11 | glio |

| | | | | |
|---|---|---|---|---|
| culture_effect | Dnajb14 | 0.518 | 8.68E-03 | glio |
| Interaction_genotype_Aars2 | Zfp575 | -3.509 | 8.68E-03 | glio |
| Interaction_genotype_Aars2 | Elf4 | -2.938 | 8.68E-03 | glio |
| Interaction_genotype_Rpl3 | Gm27253 | -3.642 | 8.68E-03 | glio |
| Interaction_genotype_Rpl3 | Gm26728 | -3.827 | 8.68E-03 | glio |
| Bora_in.vivo_noRT | Naa10 | -0.71 | 8.68E-03 | glio |
| Cenpj_ex.vivo_noRT | Zfp352 | 2.591 | 8.68E-03 | glio |
| Cenpj_ex.vivo_noRT | Pla2g2f | 2.591 | 8.68E-03 | glio |
| Chek1_ex.vivo_noRT | Ankef1 | 2.487 | 8.68E-03 | glio |
| Hsd17b10_in.vivo_noRT | Etl4 | 1.74 | 8.68E-03 | glio |
| Interaction_genotype_Myc | Psmb11 | -2.749 | 8.68E-04 | glio |
| Atp5c1_in.vivo_noRT | Gm21818 | 2.668 | 8.68E-04 | glio |
| Atr_in.vivo_noRT | Gm14488 | 2.561 | 8.68E-04 | glio |
| Bcas2_in.vivo_noRT | Wapl | -1.546 | 8.68E-04 | glio |
| Bora_in.vivo_noRT | Gm10800 | 2.626 | 8.68E-04 | glio |
| Chek1_in.vivo_noRT | Zfp458 | 2.522 | 8.68E-04 | glio |
| culture_effect | Dlg1 | 0.934 | 8.68E-05 | glio |
| Interaction_genotype_Hsd17b | Klrk1 | 4.734 | 8.68E-05 | glio |
| Interaction_genotype_Hsd17b | Klhdc8a | -4.722 | 8.68E-05 | glio |
| Atr_ex.vivo_noRT | Car10 | 3.164 | 8.68E-05 | glio |
| Bora_in.vivo_noRT | Elf3 | 3.419 | 8.68E-06 | glio |
| Bcas2_in.vivo_noRT | Dcpp3 | 3.289 | 8.68E-07 | glio |
| culture_effect | Pcf11 | -1.09 | 8.68E-08 | glio |
| culture_effect | Dpm3 | 0.884 | 8.68E-08 | glio |
| Bcas2_in.vivo_noRT | Aqp5 | 4.544 | 8.68E-09 | glio |
| Hsd17b10_in.vivo_noRT | Ltbp2 | 3.616 | 8.68E-11 | glio |
| Hsd17b10_in.vivo_noRT | Lmx1a | 3.616 | 8.68E-11 | glio |
| Atr_in.vivo_noRT | 9930111J21Ri | 4.353 | 8.68E-13 | glio |
| Atr_in.vivo_noRT | Plk5 | 4.877 | 8.68E-16 | glio |
| culture_effect | Rpn2 | -1.213 | 8.68E-22 | glio |
| Atr_in.vivo_noRT | Ccdc125 | 5.533 | 8.68E-22 | glio |
| culture_effect | Rfc2 | -0.466 | 8.69E-03 | glio |
| culture_effect | Med21 | 0.283 | 8.69E-03 | glio |
| Interaction_genotype_Mtor | Trim14 | -3.427 | 8.69E-03 | glio |
| Interaction_genotype_Mtor | A530041M06F | -3.273 | 8.69E-03 | glio |
| Aars2_ex.vivo_noRT | Bbs2 | 2.355 | 8.69E-03 | glio |
| Bcas2_ex.vivo_noRT | Col5a2 | 2.155 | 8.69E-03 | glio |
| Bora_in.vivo_noRT | 9230112E08Ri | 1.941 | 8.69E-03 | glio |
| Bora_in.vivo_noRT | C530044C16R | 1.861 | 8.69E-03 | glio |
| Bora_in.vivo_noRT | Nop56 | -0.777 | 8.69E-03 | glio |
| Cenpj_in.vivo_noRT | Dnm3 | 2.272 | 8.69E-03 | glio |
| Cenpj_in.vivo_noRT | Hbegf | -0.816 | 8.69E-03 | glio |
| Chek1_in.vivo_noRT | Arid3c | 1.756 | 8.69E-03 | glio |
| Chek1_in.vivo_noRT | Ero1l | -1.206 | 8.69E-03 | glio |
| Chek1_ex.vivo_noRT | Pomc | 1.878 | 8.69E-03 | glio |
| Chek1_ex.vivo_noRT | Klhl28 | 1.583 | 8.69E-03 | glio |
| Chek1_ex.vivo_noRT | Spty2d1 | 1.197 | 8.69E-03 | glio |
| Interaction_genotype_Myc | Zscan4c | -2.729 | 8.69E-04 | glio |
| Aars2_in.vivo_noRT | Cbarp | 2.092 | 8.69E-04 | glio |
| Interaction_genotype_Atr | H2-K1 | 1.808 | 8.69E-05 | glio |
| Chek1_in.vivo_noRT | Ubb | -0.728 | 8.69E-05 | glio |
| Atr_ex.vivo_noRT | Gm39526 | 3.164 | 8.69E-06 | glio |

| | | | | |
|---|---|---|---|---|
| Interaction_genotype_Mtor | Gm37800 | -5.966 | 8.69E-08 | glio |
| Chek1_in.vivo_noRT | Stab1 | 3.91 | 8.69E-08 | glio |
| Chek1_ex.vivo_noRT | Cxcl13 | 4.449 | 8.69E-08 | glio |
| Chek1_in.vivo_noRT | Gm15941 | 3.624 | 8.69E-09 | glio |
| Chek1_ex.vivo_noRT | Myt1 | 4.443 | 8.69E-11 | glio |
| Atr_in.vivo_noRT | Acat3 | 4.741 | 8.69E-18 | glio |
| Interaction_genotype_Aars2 | Rab25 | -2.936 | 8.70E-03 | glio |
| Interaction_genotype_Aars2 | Ptp4a1.1 | -3.214 | 8.70E-03 | glio |
| Interaction_genotype_Aars2 | Ccdc184 | -3.569 | 8.70E-03 | glio |
| Interaction_genotype_Aars2 | Uckl1os | -3.471 | 8.70E-03 | glio |
| Interaction_genotype_Aars2 | Frmd4a | 1.751 | 8.70E-03 | glio |
| Interaction_genotype_Aars2 | Lars | 1.053 | 8.70E-03 | glio |
| Interaction_genotype_Atr | Gm26692 | -3.117 | 8.70E-03 | glio |
| Interaction_genotype_Bcas2 | 2610016A17R | -3.557 | 8.70E-03 | glio |
| Interaction_genotype_Bcas2 | D5Ertd579e | 1.457 | 8.70E-03 | glio |
| Interaction_genotype_Bora | Alyref | 1.094 | 8.70E-03 | glio |
| Interaction_genotype_Mtor | Gm5141 | -3.131 | 8.70E-03 | glio |
| Interaction_genotype_Mtor | Ttll5 | -2.176 | 8.70E-03 | glio |
| Interaction_genotype_Mtor | Ddb1 | 1.308 | 8.70E-03 | glio |
| Interaction_genotype_Rad21 | Gja4 | 4.024 | 8.70E-03 | glio |
| Interaction_genotype_Rad21 | Ablim2 | 2.753 | 8.70E-03 | glio |
| Interaction_genotype_Rad21 | Gm15326 | 2.959 | 8.70E-03 | glio |
| Interaction_genotype_Rad21 | Col5a3 | -2.223 | 8.70E-03 | glio |
| Interaction_genotype_Rad21 | Bpgm | 1.556 | 8.70E-03 | glio |
| interaction_RT | Hspe1-rs1 | -0.935 | 8.70E-03 | glio |
| interaction_RT | Eif4h | -0.615 | 8.70E-03 | glio |
| Aars2_in.vivo_noRT | Wipi2 | -1.196 | 8.70E-03 | glio |
| Aars2_in.vivo_noRT | Kdm4a | -1.224 | 8.70E-03 | glio |
| Aars2_in.vivo_noRT | Rcc2 | -0.788 | 8.70E-03 | glio |
| Bcas2_in.vivo_noRT | Spata2l | 2.015 | 8.70E-03 | glio |
| Bora_in.vivo_noRT | 3300002I08Ri | 1.91 | 8.70E-03 | glio |
| Chek1_in.vivo_noRT | Lpin2 | 1.512 | 8.70E-03 | glio |
| culture_effect | Hmx2 | 0.846 | 8.70E-04 | glio |
| culture_effect | Plpp2 | 0.417 | 8.70E-04 | glio |
| Interaction_genotype_Aars2 | Rbm3os | -4.018 | 8.70E-04 | glio |
| Bcas2_in.vivo_noRT | Smim41 | 2.613 | 8.70E-04 | glio |
| Bcas2_in.vivo_noRT | Ppp1r8 | -1.589 | 8.70E-04 | glio |
| Cenpj_ex.vivo_noRT | Smtnl1 | 2.591 | 8.70E-04 | glio |
| culture_effect | Spop | 0.494 | 8.70E-05 | glio |
| Atp5c1_in.vivo_noRT | Gm13054 | 2.664 | 8.70E-05 | glio |
| Atr_in.vivo_noRT | Pcna | -1.504 | 8.70E-05 | glio |
| Chek1_in.vivo_noRT | Rps6ka4 | 2.615 | 8.70E-05 | glio |
| Chek1_in.vivo_noRT | Phlda1 | -1.27 | 8.70E-05 | glio |
| Interaction_genotype_Atr | Txnip | 5.474 | 8.70E-06 | glio |
| Atr_in.vivo_noRT | Gm36757 | 4.14 | 8.70E-10 | glio |
| culture_effect | Has2 | -1.32 | 8.70E-11 | glio |
| Bora_in.vivo_noRT | Ssh3 | 4.136 | 8.70E-12 | glio |
| Aars2_in.vivo_noRT | Pmch | 4.669 | 8.70E-15 | glio |
| Aars2_in.vivo_noRT | Gm44983 | 4.539 | 8.70E-15 | glio |
| Bcas2_in.vivo_noRT | Adgre5 | 4.547 | 8.70E-15 | glio |
| culture_effect | Tgfbr1 | -0.669 | 8.71E-03 | glio |
| Interaction_genotype_Atr | Pafah2 | -3.526 | 8.71E-03 | glio |

| | | | | |
|---|---|---|---|---|
| Interaction_genotype_Myc | BC147527 | -2.729 | 8.71E-03 | glio |
| interaction_RT | 1700120G11R | 2.789 | 8.71E-03 | glio |
| interaction_RT | Ppl | 1.172 | 8.71E-03 | glio |
| Aars2_in.vivo_noRT | Tmem104 | 1.694 | 8.71E-03 | glio |
| Atr_in.vivo_noRT | Fan1 | 1.761 | 8.71E-03 | glio |
| Chek1_ex.vivo_noRT | Atp5f1 | -0.643 | 8.71E-03 | glio |
| culture_effect | Rhox2g | -0.697 | 8.71E-04 | glio |
| Interaction_genotype_Myc | BC080695 | -2.729 | 8.71E-04 | glio |
| Interaction_genotype_Myc | Tmeff1 | -3.315 | 8.71E-04 | glio |
| Interaction_genotype_Myc | Vars2 | -3.389 | 8.71E-04 | glio |
| Interaction_genotype_Myc | Dlx1 | -3.173 | 8.71E-04 | glio |
| Bora_in.vivo_noRT | Gm32913 | 3.665 | 8.71E-04 | glio |
| Interaction_genotype_Atr | Gm14002 | -4.519 | 8.71E-05 | glio |
| Interaction_genotype_Atr | Gm29083 | -4.493 | 8.71E-05 | glio |
| Interaction_genotype_Mtor | Zfp85os | -4.714 | 8.71E-05 | glio |
| Interaction_genotype_Mtor | Gstm6 | -4.93 | 8.71E-05 | glio |
| Atr_in.vivo_noRT | Dnajc21 | -1.44 | 8.71E-06 | glio |
| Atr_ex.vivo_noRT | Rtl5 | 3.929 | 8.71E-06 | glio |
| Chek1_in.vivo_noRT | Gm12454 | 3.139 | 8.71E-06 | glio |
| Chek1_in.vivo_noRT | Cyp2u1 | 2.94 | 8.71E-06 | glio |
| interaction_RT | Fbln5 | 1.985 | 8.71E-08 | glio |
| Bora_in.vivo_noRT | Txndc2 | 4.157 | 8.71E-14 | glio |
| Chek1_ex.vivo_noRT | Hist1h2ag | 4.842 | 8.71E-14 | glio |
| Bcas2_in.vivo_noRT | Gm48899 | 4.547 | 8.71E-15 | glio |
| Atr_in.vivo_noRT | Pakap | 5.001 | 8.71E-20 | glio |
| culture_effect | Oas2 | -1.442 | 8.72E-03 | glio |
| culture_effect | Zxdb | -0.723 | 8.72E-03 | glio |
| culture_effect | Lamtor5 | 0.349 | 8.72E-03 | glio |
| Interaction_genotype_Myc | Gm36913 | -3.023 | 8.72E-03 | glio |
| Interaction_genotype_Myc | Fam161b | -2.971 | 8.72E-03 | glio |
| Atr_in.vivo_noRT | Smim7 | -0.902 | 8.72E-03 | glio |
| Chek1_ex.vivo_noRT | Gm15708 | 2.493 | 8.72E-03 | glio |
| Chek1_ex.vivo_noRT | Rflnb | 1.836 | 8.72E-03 | glio |
| Hsd17b10_in.vivo_noRT | Ncam2 | -1.907 | 8.72E-03 | glio |
| Hsd17b10_in.vivo_noRT | Nelfb | -1.308 | 8.72E-03 | glio |
| Interaction_genotype_Cdkn2a | Cd36 | 3.634 | 8.72E-04 | glio |
| Aars2_in.vivo_noRT | Wnk2 | 2.411 | 8.72E-04 | glio |
| Aars2_in.vivo_noRT | Pdk2 | 2.426 | 8.72E-04 | glio |
| Aars2_in.vivo_noRT | 2810006K23R | 1.561 | 8.72E-04 | glio |
| Bora_in.vivo_noRT | Wdr27 | 2.108 | 8.72E-04 | glio |
| Aars2_in.vivo_noRT | Susd1 | 3.015 | 8.72E-05 | glio |
| Bora_in.vivo_noRT | Senp6 | -1.524 | 8.72E-05 | glio |
| Bora_in.vivo_noRT | Gm28875 | 3.242 | 8.72E-06 | glio |
| Bora_in.vivo_noRT | Dock11 | 2.927 | 8.72E-06 | glio |
| Bcas2_in.vivo_noRT | Ap4s1 | -2.296 | 8.72E-07 | glio |
| Hsd17b10_in.vivo_noRT | Dntt | 3.616 | 8.72E-07 | glio |
| Bcas2_in.vivo_noRT | Gm42918 | 4.165 | 8.72E-12 | glio |
| Chek1_in.vivo_noRT | Gm16244 | 4.91 | 8.72E-12 | glio |
| Interaction_genotype_Ifnar1 | Gm14929 | 2.734 | 8.73E-03 | glio |
| Interaction_genotype_Mtor | Pglyrp1 | -3.213 | 8.73E-03 | glio |
| Aars2_ex.vivo_noRT | 4931431C16R | 2.114 | 8.73E-03 | glio |
| Aars2_ex.vivo_noRT | Pid1 | 2.505 | 8.73E-03 | glio |

| | | | | |
|---|---|---|---|---|
| Atp5c1_in.vivo_noRT | Lyz2 | 2.282 | 8.73E-03 | glio |
| Atr_in.vivo_noRT | BC048403 | 1.884 | 8.73E-03 | glio |
| Bcas2_in.vivo_noRT | Pcdhga6 | 1.832 | 8.73E-03 | glio |
| Bora_in.vivo_noRT | Tmem68 | -1.095 | 8.73E-03 | glio |
| Bora_ex.vivo_noRT | C4a | 2.414 | 8.73E-03 | glio |
| Chek1_in.vivo_noRT | Commd3 | -0.948 | 8.73E-03 | glio |
| Eif2s1_in.vivo_noRT | F630040K05Ri | 2.02 | 8.73E-03 | glio |
| Interaction_genotype_Mtor | Nipbl | 1.512 | 8.73E-04 | glio |
| Interaction_genotype_Myc | Gm39469 | -3.072 | 8.73E-04 | glio |
| Aars2_in.vivo_noRT | Eml1 | 2.161 | 8.73E-04 | glio |
| Atp5c1_in.vivo_noRT | Flnc | 2.673 | 8.73E-04 | glio |
| Bcas2_in.vivo_noRT | Isoc2a | 2.192 | 8.73E-04 | glio |
| Bora_in.vivo_noRT | Hist1h4j | 2.817 | 8.73E-04 | glio |
| Cenpj_in.vivo_noRT | Cd36 | 2.022 | 8.73E-04 | glio |
| Cenpj_in.vivo_noRT | Gm43701 | 2.022 | 8.73E-04 | glio |
| Cenpj_in.vivo_noRT | 9130019O22R | 2.022 | 8.73E-04 | glio |
| Chek1_in.vivo_noRT | Fendrr | 2.396 | 8.73E-04 | glio |
| Interaction_genotype_Aars2 | B930094E09Ri | -4.655 | 8.73E-05 | glio |
| Interaction_genotype_Mtor | Trim68 | -4.555 | 8.73E-05 | glio |
| Aars2_in.vivo_noRT | Acy3 | 2.833 | 8.73E-05 | glio |
| Bora_in.vivo_noRT | Trim6 | 3.241 | 8.73E-06 | glio |
| Chek1_in.vivo_noRT | Spon1 | 3.19 | 8.73E-06 | glio |
| Interaction_genotype_Mtor | Gm16794 | -5.673 | 8.73E-07 | glio |
| Chek1_ex.vivo_noRT | Gm31415 | 3.547 | 8.73E-07 | glio |
| Bora_in.vivo_noRT | Prtn3 | 4.154 | 8.73E-09 | glio |
| Atr_in.vivo_noRT | Pde11a | 5.004 | 8.73E-12 | glio |
| Aars2_in.vivo_noRT | A330049N07R | 5.06 | 8.73E-25 | glio |
| Interaction_genotype_Aars2 | Gm35867 | -3.188 | 8.74E-03 | glio |
| Interaction_genotype_Myc | 1110013H19R | -3.236 | 8.74E-03 | glio |
| Atr_in.vivo_noRT | Smarca5 | -0.714 | 8.74E-03 | glio |
| Bard1_ex.vivo_noRT | Gm813 | 2.814 | 8.74E-03 | glio |
| Bora_in.vivo_noRT | Hspa12b | 2.129 | 8.74E-03 | glio |
| Bora_in.vivo_noRT | Gm16230 | 1.919 | 8.74E-03 | glio |
| culture_effect | Urod | -0.662 | 8.74E-04 | glio |
| Interaction_genotype_Myc | Rab40b | -3.685 | 8.74E-04 | glio |
| Aars2_in.vivo_noRT | Mrps16 | -0.931 | 8.74E-04 | glio |
| Atr_in.vivo_noRT | Ifitm3 | -0.783 | 8.74E-04 | glio |
| Bora_in.vivo_noRT | Angpt2 | 2.204 | 8.74E-04 | glio |
| Chek1_in.vivo_noRT | Cldn22 | 2.105 | 8.74E-04 | glio |
| Chek1_in.vivo_noRT | Micos10 | -0.68 | 8.74E-04 | glio |
| Aars2_in.vivo_noRT | 6330403L08Ri | 2.666 | 8.74E-05 | glio |
| Atr_in.vivo_noRT | Gm10282 | 2.757 | 8.74E-05 | glio |
| Chek1_in.vivo_noRT | 1810059H22R | 3.085 | 8.74E-05 | glio |
| Chek1_in.vivo_noRT | Glmp | 3.041 | 8.74E-05 | glio |
| Chek1_in.vivo_noRT | Atp8a2 | 2.365 | 8.74E-05 | glio |
| Interaction_genotype_Mtor | Ttc12 | -5.221 | 8.74E-06 | glio |
| Bora_in.vivo_noRT | Cd3eap | -1.586 | 8.74E-06 | glio |
| culture_effect | Gm10638 | 2.386 | 8.74E-07 | glio |
| Hsd17b10_in.vivo_noRT | Gm43653 | 3.353 | 8.74E-07 | glio |
| Bora_in.vivo_noRT | Fabp4 | 4.407 | 8.74E-10 | glio |
| Atr_in.vivo_noRT | Slc7a3 | 4.804 | 8.74E-12 | glio |
| Bcas2_in.vivo_noRT | Tcstv1 | 4.807 | 8.74E-16 | glio |

| | | | | |
|---|---|---|---|---|
| Atr_in.vivo_noRT | Gm5602 | 4.742 | 8.74E-18 | glio |
| Interaction_genotype_Mtor | Usp35 | -3.205 | 8.75E-03 | glio |
| Interaction_genotype_Mtor | Zfp143 | -2.238 | 8.75E-03 | glio |
| Interaction_genotype_Myc | Pxt1 | -2.976 | 8.75E-03 | glio |
| Interaction_genotype_Rpl3 | Hnrnpdl | 1.18 | 8.75E-03 | glio |
| Atr_in.vivo_noRT | Arntl | 1.433 | 8.75E-03 | glio |
| Atr_ex.vivo_noRT | Ggt5 | 2.463 | 8.75E-03 | glio |
| Bard1_ex.vivo_noRT | Gm39459 | 2.727 | 8.75E-03 | glio |
| Bora_in.vivo_noRT | Vash2 | 1.971 | 8.75E-03 | glio |
| Eif2s1_in.vivo_noRT | 2410141K09R | 1.879 | 8.75E-03 | glio |
| Interaction_genotype_Atr | Scpep1 | 1.846 | 8.75E-04 | glio |
| Interaction_genotype_Myc | BB287469 | -2.729 | 8.75E-04 | glio |
| Interaction_genotype_Myc | Polr2g | 1.497 | 8.75E-04 | glio |
| Interaction_genotype_Uqcrc2 | Gm39326 | -4.869 | 8.75E-04 | glio |
| Atr_in.vivo_noRT | Zmiz1 | -1.623 | 8.75E-04 | glio |
| Bcas2_in.vivo_noRT | Rab11fip5 | 2.499 | 8.75E-04 | glio |
| Bora_in.vivo_noRT | Etv1 | -1.549 | 8.75E-04 | glio |
| Atr_ex.vivo_noRT | Prdm1 | 3.554 | 8.75E-05 | glio |
| Hsd17b10_in.vivo_noRT | Rell2 | 2.964 | 8.75E-05 | glio |
| Interaction_genotype_Myc | Ubr1 | -3.771 | 8.75E-06 | glio |
| Interaction_genotype_Myc | Gm10447 | -4.003 | 8.75E-06 | glio |
| Interaction_genotype_Myc | Hspe1 | 2.046 | 8.75E-06 | glio |
| Aars2_in.vivo_noRT | Slpi | 3.501 | 8.75E-06 | glio |
| Bora_ex.vivo_noRT | Gm39526 | 3.201 | 8.75E-06 | glio |
| Interaction_genotype_Mtor | Camk1d | -5.707 | 8.75E-07 | glio |
| Bora_in.vivo_noRT | Gm27010 | 3.335 | 8.75E-07 | glio |
| Chek1_in.vivo_noRT | Traf1 | 3.44 | 8.75E-07 | glio |
| Interaction_genotype_Mtor | Hp1bp3 | 2.69 | 8.75E-08 | glio |
| Interaction_genotype_Mtor | Dnaja1 | 2.638 | 8.75E-08 | glio |
| Interaction_genotype_Mtor | Eef1a1 | 1.762 | 8.75E-08 | glio |
| Aars2_in.vivo_noRT | 4930426I24Ri | 3.671 | 8.75E-08 | glio |
| Interaction_genotype_Rad21 | H2-Eb2 | 5.785 | 8.75E-09 | glio |
| Bcas2_in.vivo_noRT | A430106G13R | 4.545 | 8.75E-10 | glio |
| Bora_in.vivo_noRT | Rhbg | 3.341 | 8.75E-11 | glio |
| Chek1_in.vivo_noRT | Rspo1 | 4.252 | 8.75E-11 | glio |
| Aars2_in.vivo_noRT | Gm50375 | 5.06 | 8.75E-12 | glio |
| Bcas2_in.vivo_noRT | BC147527 | 4.807 | 8.75E-14 | glio |
| Aars2_in.vivo_noRT | Prl2c2 | 5.327 | 8.75E-16 | glio |
| Bcas2_in.vivo_noRT | Pspn | 4.807 | 8.75E-23 | glio |
| culture_effect | Eml5 | -0.884 | 8.76E-03 | glio |
| culture_effect | Chordc1 | 0.398 | 8.76E-03 | glio |
| Interaction_genotype_Lmo7 | Gm48960 | -3.973 | 8.76E-03 | glio |
| Interaction_genotype_Mtor | Slc25a45 | -2.259 | 8.76E-03 | glio |
| Interaction_genotype_Myc | Fgf15 | -3.236 | 8.76E-03 | glio |
| Interaction_genotype_Myc | 1810009A15R | -2.552 | 8.76E-03 | glio |
| Aars2_in.vivo_noRT | Synm | 1.728 | 8.76E-03 | glio |
| Atr_in.vivo_noRT | Arpc2 | -0.661 | 8.76E-03 | glio |
| Atr_ex.vivo_noRT | Atpaf1 | -1.896 | 8.76E-03 | glio |
| Cenpj_ex.vivo_noRT | Strip2 | 2.192 | 8.76E-03 | glio |
| Cenpj_ex.vivo_noRT | Col19a1 | 2.198 | 8.76E-03 | glio |
| Eif2s1_ex.vivo_noRT | Gm45406 | 2.234 | 8.76E-03 | glio |
| Hsd17b10_in.vivo_noRT | Sptbn4 | 2.068 | 8.76E-03 | glio |

| | | | | |
|---|---|---|---|---|
| culture_effect | Shbg | -2.209 | 8.76E-04 | glio |
| culture_effect | Aste1 | -1.129 | 8.76E-04 | glio |
| culture_effect | P2rx7 | 0.806 | 8.76E-04 | glio |
| culture_effect | Mrps2 | 0.717 | 8.76E-04 | glio |
| Interaction_genotype_Atr | Gm15535 | -4.443 | 8.76E-04 | glio |
| Interaction_genotype_Mtor | Nckap1l | -4.022 | 8.76E-04 | glio |
| Atr_in.vivo_noRT | Msrb2 | 2.245 | 8.76E-04 | glio |
| Bard1_ex.vivo_noRT | AF067061 | 2.814 | 8.76E-04 | glio |
| Bcas2_in.vivo_noRT | Dll1 | 2.572 | 8.76E-04 | glio |
| Interaction_genotype_Aars2 | Gm43653 | -4.663 | 8.76E-05 | glio |
| Interaction_genotype_Mtor | Cfap46 | -4.225 | 8.76E-05 | glio |
| Interaction_genotype_Mtor | Rab7 | 2.047 | 8.76E-05 | glio |
| Interaction_genotype_Mtor | Hnrnpa2b1 | 1.622 | 8.76E-05 | glio |
| Atr_in.vivo_noRT | Cacng4 | 2.394 | 8.76E-05 | glio |
| Chek1_ex.vivo_noRT | Jade1 | -2.338 | 8.76E-05 | glio |
| Hsd17b10_in.vivo_noRT | Flicr | 2.816 | 8.76E-05 | glio |
| Aars2_ex.vivo_noRT | Vtn | 3.293 | 8.76E-07 | glio |
| Chek1_in.vivo_noRT | Gchfr | 3.931 | 8.76E-07 | glio |
| Interaction_genotype_Mtor | Imp3 | 2.946 | 8.76E-08 | glio |
| Interaction_genotype_Mtor | Cox6c | 3.008 | 8.76E-08 | glio |
| Interaction_genotype_Mtor | Ccnb1ip1 | -7.09 | 8.76E-08 | glio |
| Interaction_genotype_Mtor | AY074887 | -7.053 | 8.76E-08 | glio |
| Interaction_genotype_Mtor | Ankrd1 | -7.64 | 8.76E-08 | glio |
| Bora_ex.vivo_noRT | Pla1a | 4.786 | 8.76E-08 | glio |
| Bora_ex.vivo_noRT | Cldn2 | 3.201 | 8.76E-08 | glio |
| Aars2_in.vivo_noRT | Mogat1 | 4.805 | 8.76E-11 | glio |
| Atr_in.vivo_noRT | Gpr22 | 5.533 | 8.76E-25 | glio |
| Interaction_genotype_Atr | Zfp335 | -2.525 | 8.77E-03 | glio |
| Interaction_genotype_Bcas2 | Gm15943 | -2.783 | 8.77E-03 | glio |
| Interaction_genotype_Myc | Spast | -1.867 | 8.77E-03 | glio |
| Interaction_genotype_Myc | Cox5a | 1.051 | 8.77E-03 | glio |
| Atr_in.vivo_noRT | Dctpp1 | -0.795 | 8.77E-03 | glio |
| Bcas2_in.vivo_noRT | Gm41609 | 2.121 | 8.77E-03 | glio |
| Cenpj_in.vivo_noRT | 4930570E03Ri | 1.762 | 8.77E-03 | glio |
| Cenpj_in.vivo_noRT | 4930447K03R | 1.762 | 8.77E-03 | glio |
| Cenpj_ex.vivo_noRT | 4732463B04R | 2.198 | 8.77E-03 | glio |
| Cenpj_ex.vivo_noRT | Gm38477 | 2.192 | 8.77E-03 | glio |
| Chek1_ex.vivo_noRT | Ppnr | 2.481 | 8.77E-03 | glio |
| Fanca_ex.vivo_noRT | Gm34885 | 2.795 | 8.77E-03 | glio |
| Interaction_genotype_Atr | Gm49808 | -2.762 | 8.77E-04 | glio |
| Interaction_genotype_Mtor | Eef1b2 | 1.146 | 8.77E-04 | glio |
| Interaction_genotype_Myc | Nudcd2 | 1.277 | 8.77E-04 | glio |
| Aars2_in.vivo_noRT | Gsto2 | 2.415 | 8.77E-04 | glio |
| Aars2_in.vivo_noRT | Acot1 | -1.509 | 8.77E-04 | glio |
| Atr_in.vivo_noRT | Cnot9 | -1.524 | 8.77E-04 | glio |
| culture_effect | Bik | -1.531 | 8.77E-05 | glio |
| Interaction_genotype_Rad51 | Ifi47 | 3.172 | 8.77E-05 | glio |
| Interaction_genotype_Rad51 | Klhdc8a | -4.211 | 8.77E-05 | glio |
| Interaction_genotype_Rad51 | Gm11627 | -4.734 | 8.77E-05 | glio |
| Chek1_ex.vivo_noRT | Gm47428 | 3.363 | 8.77E-05 | glio |
| Atr_in.vivo_noRT | Rplp2 | -0.848 | 8.77E-06 | glio |
| culture_effect | Eif3a | -0.467 | 8.77E-07 | glio |

| | | | | |
|---|---|---|---|---|
| interaction_RT | Tmem140 | 2.447 | 8.77E-07 | glio |
| Atp5c1_in.vivo_noRT | Gm5890 | 2.668 | 8.77E-07 | glio |
| Atp5c1_in.vivo_noRT | Zscan4c | 2.668 | 8.77E-07 | glio |
| Atp5c1_in.vivo_noRT | BC080695 | 2.668 | 8.77E-07 | glio |
| Bora_in.vivo_noRT | BC065403 | 4.42 | 8.77E-11 | glio |
| Bora_in.vivo_noRT | Gm16183 | 4.818 | 8.77E-11 | glio |
| Bora_in.vivo_noRT | Abcg4 | 4.145 | 8.77E-11 | glio |
| Chek1_in.vivo_noRT | Dlx4 | 4.518 | 8.77E-12 | glio |
| Chek1_in.vivo_noRT | Gm26682 | 4.649 | 8.77E-12 | glio |
| culture_effect | Ctsd | -0.956 | 8.77E-13 | glio |
| Chek1_ex.vivo_noRT | H2-Eb2 | 4.842 | 8.77E-13 | glio |
| Hsd17b10_in.vivo_noRT | Gm31717 | 3.616 | 8.77E-14 | glio |
| Atr_in.vivo_noRT | Gm5941 | 5.533 | 8.77E-15 | glio |
| Chek1_in.vivo_noRT | Gm10324 | 4.777 | 8.77E-19 | glio |
| culture_effect | Impdh2 | -1.118 | 8.77E-20 | glio |
| Chek1_in.vivo_noRT | Dscaml1 | 5.044 | 8.77E-25 | glio |
| Chek1_in.vivo_noRT | 2410017I17Ri | 5.044 | 8.77E-25 | glio |
| Interaction_genotype_Aars2 | Nwd1 | -2.752 | 8.78E-03 | glio |
| Interaction_genotype_Aars2 | Spdye4a | -3.455 | 8.78E-03 | glio |
| Interaction_genotype_Aars2 | Cgn | -3.24 | 8.78E-03 | glio |
| Interaction_genotype_Aars2 | Impdh2 | 0.97 | 8.78E-03 | glio |
| Interaction_genotype_Bcas2 | Slc1a2 | -2.987 | 8.78E-03 | glio |
| Interaction_genotype_Bcas2 | 1700012B09R | -4.185 | 8.78E-03 | glio |
| Interaction_genotype_Bcas2 | Gm44672 | -3.432 | 8.78E-03 | glio |
| Interaction_genotype_Myc | Smpd5 | -2.659 | 8.78E-03 | glio |
| Aars2_in.vivo_noRT | Pld3 | -1.098 | 8.78E-03 | glio |
| Atp5c1_in.vivo_noRT | Cfap58 | 2.668 | 8.78E-03 | glio |
| Atr_in.vivo_noRT | Zfp324 | 1.63 | 8.78E-03 | glio |
| Bcas2_in.vivo_noRT | Plpp2 | -0.747 | 8.78E-03 | glio |
| Bora_in.vivo_noRT | Prdx4 | -0.71 | 8.78E-03 | glio |
| Cenpj_in.vivo_noRT | Afp | -2.553 | 8.78E-03 | glio |
| Chek1_in.vivo_noRT | Steap3 | 1.529 | 8.78E-03 | glio |
| Chek1_in.vivo_noRT | Rplp0 | -0.533 | 8.78E-03 | glio |
| Chek1_ex.vivo_noRT | Zbed4 | -1.779 | 8.78E-03 | glio |
| Hsd17b10_in.vivo_noRT | Hsd17b14 | 2.185 | 8.78E-03 | glio |
| Hsd17b10_in.vivo_noRT | Kif16b | -1.39 | 8.78E-03 | glio |
| Aars2_in.vivo_noRT | Slc2a13 | 2.368 | 8.78E-04 | glio |
| Aars2_ex.vivo_noRT | Pstpip2 | 2.499 | 8.78E-04 | glio |
| Bcas2_ex.vivo_noRT | Aqp9 | 2.742 | 8.78E-04 | glio |
| Chek1_in.vivo_noRT | Gm15998 | 2.41 | 8.78E-04 | glio |
| Chek1_ex.vivo_noRT | Rpl6 | -0.808 | 8.78E-04 | glio |
| culture_effect | Tcf25 | 0.446 | 8.78E-05 | glio |
| culture_effect | Ssr2 | -0.466 | 8.78E-05 | glio |
| Interaction_genotype_Aars2 | Slc25a35 | -4.865 | 8.78E-05 | glio |
| Interaction_genotype_Mtor | Cep295nl | -4.61 | 8.78E-05 | glio |
| Aars2_ex.vivo_noRT | Gstp3 | 3.697 | 8.78E-05 | glio |
| Atr_in.vivo_noRT | Ccdc85c | 2.647 | 8.78E-05 | glio |
| Atr_in.vivo_noRT | Prkar1b | 3.194 | 8.78E-06 | glio |
| Bora_ex.vivo_noRT | Speer4e | 3.201 | 8.78E-06 | glio |
| Interaction_genotype_Mtor | Scube3 | -4.428 | 8.78E-07 | glio |
| Interaction_genotype_Mtor | Gm37170 | -5.876 | 8.78E-07 | glio |
| Atp5c1_in.vivo_noRT | Gm47540 | 2.668 | 8.78E-07 | glio |

| | | | | |
|---|---|---|---|---|
| Atp5c1_in.vivo_noRT | BB287469 | 2.668 | 8.78E-07 | glio |
| culture_effect | Sag | 1.273 | 8.79E-03 | glio |
| Interaction_genotype_Aars2 | Gm47061 | -3.507 | 8.79E-03 | glio |
| Interaction_genotype_Atr | Gm15533 | -2.369 | 8.79E-03 | glio |
| Interaction_genotype_Atr | Rpain | 1.439 | 8.79E-03 | glio |
| Interaction_genotype_Bora | Rex2 | -3.781 | 8.79E-03 | glio |
| Interaction_genotype_Bora | Zfyve16 | -2.99 | 8.79E-03 | glio |
| interaction_RT | Gm15938 | 2.579 | 8.79E-03 | glio |
| interaction_RT | Zfp324 | -1.716 | 8.79E-03 | glio |
| Aars2_in.vivo_noRT | Rnpepl1 | 1.079 | 8.79E-03 | glio |
| Aars2_in.vivo_noRT | Isg15 | -1.241 | 8.79E-03 | glio |
| Atr_in.vivo_noRT | Rad51ap1 | -1.474 | 8.79E-03 | glio |
| Bard1_ex.vivo_noRT | Gm21818 | 2.814 | 8.79E-03 | glio |
| Bcas2_in.vivo_noRT | Tmem183a | -1.139 | 8.79E-03 | glio |
| Bora_in.vivo_noRT | Vdr | 2.128 | 8.79E-03 | glio |
| Interaction_genotype_Aars2 | 3110083C13R | -4.064 | 8.79E-04 | glio |
| Interaction_genotype_Bora | Celf1 | 1.68 | 8.79E-04 | glio |
| Interaction_genotype_Mtor | Cul1 | 1.822 | 8.79E-04 | glio |
| Interaction_genotype_Myc | Cpne9 | -4.004 | 8.79E-04 | glio |
| Atr_in.vivo_noRT | Pam16 | -1.096 | 8.79E-04 | glio |
| Bcas2_in.vivo_noRT | Gm16279 | 2.552 | 8.79E-04 | glio |
| Bora_ex.vivo_noRT | AC125149.2 | 2.806 | 8.79E-04 | glio |
| Bora_ex.vivo_noRT | Gm2762 | 2.979 | 8.79E-04 | glio |
| Chek1_ex.vivo_noRT | Junos | 3.012 | 8.79E-04 | glio |
| Chek1_ex.vivo_noRT | Mylip | 2.386 | 8.79E-04 | glio |
| Eif2s1_in.vivo_noRT | Ltbp2 | 2.143 | 8.79E-04 | glio |
| Eif2s1_in.vivo_noRT | Lmx1a | 2.143 | 8.79E-04 | glio |
| Interaction_genotype_Mtor | Ifi207 | -5.012 | 8.79E-05 | glio |
| Interaction_genotype_Myc | Gm12602 | -4.191 | 8.79E-05 | glio |
| Interaction_genotype_Myc | C330002G04R | -3.316 | 8.79E-05 | glio |
| Aars2_in.vivo_noRT | Atp6v0c | -0.795 | 8.79E-05 | glio |
| Bcas2_in.vivo_noRT | Mmp28 | 2.914 | 8.79E-05 | glio |
| Chek1_in.vivo_noRT | Mettl5os | 2.615 | 8.79E-05 | glio |
| Interaction_genotype_Myc | 1700034I23Ri | -3.526 | 8.79E-06 | glio |
| Aars2_in.vivo_noRT | Asphd2 | 3.123 | 8.79E-06 | glio |
| Aars2_in.vivo_noRT | Dock6 | 3.069 | 8.79E-06 | glio |
| Aars2_in.vivo_noRT | Dnpep | -2.072 | 8.79E-06 | glio |
| interaction_RT | Nucks1 | -1.017 | 8.79E-07 | glio |
| Atr_in.vivo_noRT | Rapgef3 | 3.802 | 8.79E-07 | glio |
| Bcas2_in.vivo_noRT | Serpina3g | 3.611 | 8.79E-09 | glio |
| culture_effect | Rpl21 | -0.767 | 8.79E-11 | glio |
| Atr_in.vivo_noRT | Sdr16c5 | 5.269 | 8.79E-16 | glio |
| Chek1_in.vivo_noRT | Gm12794 | 5.044 | 8.79E-23 | glio |
| Atr_in.vivo_noRT | Gm10324 | 5.771 | 8.79E-26 | glio |
| Atr_in.vivo_noRT | Gm4340 | 5.533 | 8.79E-30 | glio |
| Atr_in.vivo_noRT | Gm16239 | 5.533 | 8.79E-30 | glio |
| Interaction_genotype_Atr | Vsig10l | -3.258 | 8.80E-03 | glio |
| Interaction_genotype_Atr | Zfand4 | -3.416 | 8.80E-03 | glio |
| Interaction_genotype_Atr | Polm | -2.908 | 8.80E-03 | glio |
| Interaction_genotype_Atr | Zfp568 | 2.209 | 8.80E-03 | glio |
| Interaction_genotype_Mtor | 1700010I14Ri | -3.11 | 8.80E-03 | glio |
| Interaction_genotype_Myc | Armh3 | -2.612 | 8.80E-03 | glio |

| | | | | |
|---|---|---|---|---|
| Interaction_genotype_Myc | Grina | 0.973 | 8.80E-03 | glio |
| Atr_in.vivo_noRT | Man2a1 | 1.285 | 8.80E-03 | glio |
| Atr_ex.vivo_noRT | Dpep2 | 2.574 | 8.80E-03 | glio |
| Bcas2_in.vivo_noRT | Rpl8 | -0.521 | 8.80E-03 | glio |
| Bcas2_ex.vivo_noRT | Npr2 | 3.2 | 8.80E-03 | glio |
| culture_effect | Azin1 | 0.431 | 8.80E-04 | glio |
| Interaction_genotype_Mtor | 4933408J17Ri | -3.566 | 8.80E-04 | glio |
| Interaction_genotype_Myc | Nfatc1 | -3.521 | 8.80E-04 | glio |
| Bora_in.vivo_noRT | Nckipsd | 2.331 | 8.80E-04 | glio |
| Bora_in.vivo_noRT | Adamts7 | 2.236 | 8.80E-04 | glio |
| Aars2_in.vivo_noRT | Gm10800 | 3.017 | 8.80E-05 | glio |
| Bora_in.vivo_noRT | Vgll3 | 2.815 | 8.80E-05 | glio |
| Chek1_in.vivo_noRT | Ccdc148 | 3.293 | 8.80E-06 | glio |
| culture_effect | Mrpl21 | -0.61 | 8.80E-07 | glio |
| Bora_in.vivo_noRT | Gm16344 | 4.025 | 8.80E-12 | glio |
| Atr_in.vivo_noRT | Postn | 4.899 | 8.80E-13 | glio |
| culture_effect | Piezo2 | -1.775 | 8.80E-20 | glio |
| Aars2_in.vivo_noRT | Luzp2 | 4.801 | 8.80E-20 | glio |
| culture_effect | Usp16 | 0.432 | 8.81E-03 | glio |
| culture_effect | Bud31 | -0.369 | 8.81E-03 | glio |
| Interaction_genotype_Atr | Gm43653 | -3.042 | 8.81E-03 | glio |
| Interaction_genotype_Bora | Gm16222 | -3.979 | 8.81E-03 | glio |
| Interaction_genotype_Mtor | Cep72 | -3.029 | 8.81E-03 | glio |
| Interaction_genotype_Mtor | Trib2 | -2.967 | 8.81E-03 | glio |
| Interaction_genotype_Mtor | Eif5a2 | -2.508 | 8.81E-03 | glio |
| Interaction_genotype_Mtor | 2810454H06R | -2.678 | 8.81E-03 | glio |
| Interaction_genotype_Myc | Btbd8 | -2.278 | 8.81E-03 | glio |
| Aars2_in.vivo_noRT | Tmem219 | -1.226 | 8.81E-03 | glio |
| Aars2_in.vivo_noRT | Ccnd1 | -0.883 | 8.81E-03 | glio |
| Atr_in.vivo_noRT | Ctif | 1.718 | 8.81E-03 | glio |
| Bora_in.vivo_noRT | Purb | -0.596 | 8.81E-03 | glio |
| Cenpj_in.vivo_noRT | Rab27a | 2.025 | 8.81E-03 | glio |
| Chek1_in.vivo_noRT | Rtraf | -0.679 | 8.81E-03 | glio |
| Chek1_ex.vivo_noRT | Gm13625 | 2.269 | 8.81E-03 | glio |
| Chek1_ex.vivo_noRT | Gm35021 | 1.966 | 8.81E-03 | glio |
| Interaction_genotype_Mtor | Qrsl1 | -2.835 | 8.81E-04 | glio |
| Aars2_in.vivo_noRT | Upp2 | 2.127 | 8.81E-04 | glio |
| Bora_in.vivo_noRT | Mfsd4b4 | 2.211 | 8.81E-04 | glio |
| Bora_in.vivo_noRT | Uqcrc2 | -1.035 | 8.81E-04 | glio |
| Interaction_genotype_Mtor | BC051077 | -5.405 | 8.81E-05 | glio |
| Interaction_genotype_Mtor | Ndrg4 | -4.84 | 8.81E-05 | glio |
| interaction_RT | Gm31717 | 2.555 | 8.81E-05 | glio |
| Atr_in.vivo_noRT | Rab42 | 2.929 | 8.81E-05 | glio |
| Atr_in.vivo_noRT | AI429214 | 2.776 | 8.81E-05 | glio |
| Atr_in.vivo_noRT | Txndc17 | -1.081 | 8.81E-05 | glio |
| Interaction_genotype_Myc | Gm30794 | -3.706 | 8.81E-06 | glio |
| Bcas2_in.vivo_noRT | Gm26982 | 3.116 | 8.81E-06 | glio |
| Interaction_genotype_Mtor | Pldi | -4.793 | 8.81E-07 | glio |
| interaction_RT | Ndufv3 | 1.055 | 8.81E-07 | glio |
| culture_effect | Ipcef1 | -2.748 | 8.81E-08 | glio |
| Interaction_genotype_Myc | Ssr4 | 3.284 | 8.81E-10 | glio |
| Interaction_genotype_Myc | Rbm25 | 2.415 | 8.81E-10 | glio |

| | | | | |
|---|---|---|---|---|
| Interaction_genotype_Myc | Nxph4 | -7.112 | 8.81E-10 | glio |
| Bcas2_in.vivo_noRT | Gm31415 | 4.541 | 8.81E-15 | glio |
| Bcas2_in.vivo_noRT | B4galnt1 | 4.547 | 8.81E-15 | glio |
| Chek1_in.vivo_noRT | Gm6803 | 5.044 | 8.81E-20 | glio |
| culture_effect | Flt3l | -1.097 | 8.82E-03 | glio |
| Interaction_genotype_Aars2 | Psmd12 | 0.993 | 8.82E-03 | glio |
| Interaction_genotype_Mtor | Ap1s3 | -3.201 | 8.82E-03 | glio |
| Interaction_genotype_Mtor | Erbin | 1.405 | 8.82E-03 | glio |
| Interaction_genotype_Myc | Spint4 | -3.028 | 8.82E-03 | glio |
| Interaction_genotype_Rpl3 | Jph4 | -3.662 | 8.82E-03 | glio |
| Atr_in.vivo_noRT | Natd1 | 1.527 | 8.82E-03 | glio |
| Bard1_ex.vivo_noRT | Cxcr6 | 2.814 | 8.82E-03 | glio |
| Bard1_ex.vivo_noRT | Opalin | 3.585 | 8.82E-03 | glio |
| Bcas2_in.vivo_noRT | Exosc1 | -0.89 | 8.82E-03 | glio |
| Bcas2_in.vivo_noRT | Grina | -0.686 | 8.82E-03 | glio |
| Bora_in.vivo_noRT | Lfng | 1.946 | 8.82E-03 | glio |
| Bora_ex.vivo_noRT | Dtx1 | 2.808 | 8.82E-03 | glio |
| Chek1_in.vivo_noRT | Hist1h3b | 2.055 | 8.82E-03 | glio |
| Aars2_in.vivo_noRT | Gnl3 | -1.138 | 8.82E-04 | glio |
| Atr_in.vivo_noRT | Cep83os | 2.22 | 8.82E-04 | glio |
| Atr_in.vivo_noRT | Dnaaf2 | -1.419 | 8.82E-04 | glio |
| Bcas2_in.vivo_noRT | Rad51ap1 | -1.884 | 8.82E-04 | glio |
| Bora_in.vivo_noRT | Ifi205 | 3.116 | 8.82E-04 | glio |
| Chek1_in.vivo_noRT | Hcn2 | 2.59 | 8.82E-04 | glio |
| Chek1_in.vivo_noRT | D11Wsu47e | 1.798 | 8.82E-04 | glio |
| Hsd17b10_in.vivo_noRT | Rab3b | 2.46 | 8.82E-04 | glio |
| Interaction_genotype_Myc | 1810019D21R | -4.235 | 8.82E-05 | glio |
| Interaction_genotype_Myc | Vwce | -4.025 | 8.82E-05 | glio |
| Aars2_in.vivo_noRT | Gm15910 | 2.677 | 8.82E-05 | glio |
| Bora_in.vivo_noRT | 5033430I15Ri | 2.757 | 8.82E-05 | glio |
| culture_effect | Pigp | 0.77 | 8.82E-06 | glio |
| culture_effect | Hs2st1 | 0.683 | 8.82E-06 | glio |
| Atr_in.vivo_noRT | Dll1 | 3.299 | 8.82E-06 | glio |
| Atr_in.vivo_noRT | Reck | 3.44 | 8.82E-07 | glio |
| Bcas2_in.vivo_noRT | Amhr2 | 3.931 | 8.82E-07 | glio |
| Bcas2_in.vivo_noRT | Ccdc107 | -1.978 | 8.82E-07 | glio |
| Aars2_in.vivo_noRT | BC028471 | 4.794 | 8.82E-12 | glio |
| Atr_in.vivo_noRT | Gm43625 | 4.228 | 8.82E-12 | glio |
| Interaction_genotype_Bcas2 | Ier2 | 1.743 | 8.83E-03 | glio |
| Interaction_genotype_Chek1 | 5430402O13R | -3.746 | 8.83E-03 | glio |
| Interaction_genotype_Chek1 | Otos | -3.049 | 8.83E-03 | glio |
| Interaction_genotype_Mtor | Prox2os | -2.795 | 8.83E-03 | glio |
| Atr_in.vivo_noRT | Gm16196 | 1.773 | 8.83E-03 | glio |
| Atr_in.vivo_noRT | Gpr137b | 1.779 | 8.83E-03 | glio |
| Atr_in.vivo_noRT | Tmco6 | 1.683 | 8.83E-03 | glio |
| Bcas2_in.vivo_noRT | Col20a1 | 1.97 | 8.83E-03 | glio |
| Bcas2_in.vivo_noRT | C1ql1 | -1.399 | 8.83E-03 | glio |
| Bcas2_in.vivo_noRT | Josd1 | -1.267 | 8.83E-03 | glio |
| Chek1_in.vivo_noRT | Tmem132a | -1.246 | 8.83E-03 | glio |
| Eif2s1_in.vivo_noRT | Tns4 | 2.016 | 8.83E-03 | glio |
| Interaction_genotype_Aars2 | Trpm3 | -4.144 | 8.83E-04 | glio |
| Interaction_genotype_Aars2 | Camk1d | -4.018 | 8.83E-04 | glio |

| | | | | |
|---|---|---|---|---|
| Interaction_genotype_Aars2 | AC163347.1 | -3.353 | 8.83E-04 | glio |
| Interaction_genotype_Aars2 | Ergic3 | 1.476 | 8.83E-04 | glio |
| Interaction_genotype_Chek1 | Gm15706 | -3.865 | 8.83E-04 | glio |
| Aars2_in.vivo_noRT | Gm20712 | 2.358 | 8.83E-04 | glio |
| Aars2_in.vivo_noRT | Snph | 2.283 | 8.83E-04 | glio |
| Aars2_in.vivo_noRT | Cct6a | -0.824 | 8.83E-04 | glio |
| Atr_ex.vivo_noRT | Qpct | 3.333 | 8.83E-04 | glio |
| Atr_ex.vivo_noRT | Rab2b | 1.666 | 8.83E-04 | glio |
| Atr_in.vivo_noRT | Ric8a | -1.814 | 8.83E-05 | glio |
| Atr_in.vivo_noRT | Vdac3 | -1.127 | 8.83E-05 | glio |
| culture_effect | Eif4a3 | -0.749 | 8.83E-09 | glio |
| Atr_in.vivo_noRT | Kctd14 | 3.609 | 8.83E-09 | glio |
| Chek1_in.vivo_noRT | Gm32585 | 4.518 | 8.83E-10 | glio |
| Bora_in.vivo_noRT | E130215H24R | 4.145 | 8.83E-11 | glio |
| Bora_in.vivo_noRT | 1600014C23R | 5.34 | 8.83E-14 | glio |
| Chek1_ex.vivo_noRT | Slc14a2 | 4.842 | 8.83E-17 | glio |
| Interaction_genotype_Chek1 | Batf2 | 3.217 | 8.84E-03 | glio |
| Interaction_genotype_Mtor | Gstp3 | -3.194 | 8.84E-03 | glio |
| Aars2_in.vivo_noRT | Psmb9 | -0.809 | 8.84E-03 | glio |
| Bcas2_in.vivo_noRT | Pcsk4 | 1.958 | 8.84E-03 | glio |
| Bcas2_in.vivo_noRT | Gigyf1 | 1.166 | 8.84E-03 | glio |
| Bcas2_ex.vivo_noRT | Masp2 | 2.349 | 8.84E-03 | glio |
| Bora_ex.vivo_noRT | Gm50295 | 2.481 | 8.84E-03 | glio |
| Cdkn2a_in.vivo_noRT | Ankrd22 | -1.675 | 8.84E-03 | glio |
| Cenpj_ex.vivo_noRT | 2810049E08Ri | 2.591 | 8.84E-03 | glio |
| Chek1_in.vivo_noRT | Jam2 | 1.557 | 8.84E-03 | glio |
| Chek1_in.vivo_noRT | Psme2 | -0.65 | 8.84E-03 | glio |
| Eif2s1_in.vivo_noRT | Slc25a18 | 1.877 | 8.84E-03 | glio |
| Eif2s1_in.vivo_noRT | Prl2c2 | 2.143 | 8.84E-03 | glio |
| Eif2s1_in.vivo_noRT | Gm35439 | 1.763 | 8.84E-03 | glio |
| Hsd17b10_in.vivo_noRT | 4930590J08Ri | 2.111 | 8.84E-03 | glio |
| Hsd17b10_in.vivo_noRT | Zdhhc3 | -0.871 | 8.84E-03 | glio |
| culture_effect | Slc31a1 | -0.531 | 8.84E-04 | glio |
| Atr_in.vivo_noRT | Fzd4 | 2.266 | 8.84E-04 | glio |
| Cenpj_in.vivo_noRT | Gm36037 | 2.022 | 8.84E-04 | glio |
| Cenpj_in.vivo_noRT | Gm12800 | 2.022 | 8.84E-04 | glio |
| Cenpj_in.vivo_noRT | Trmt61b | 2.022 | 8.84E-04 | glio |
| Cenpj_in.vivo_noRT | Gm5890 | 2.022 | 8.84E-04 | glio |
| Cenpj_in.vivo_noRT | Zscan4c | 2.022 | 8.84E-04 | glio |
| Cenpj_in.vivo_noRT | BC080695 | 2.022 | 8.84E-04 | glio |
| Cenpj_in.vivo_noRT | Gpr37l1 | 2.733 | 8.84E-04 | glio |
| Cenpj_in.vivo_noRT | Gm47540 | 2.022 | 8.84E-04 | glio |
| Cenpj_in.vivo_noRT | BB287469 | 2.022 | 8.84E-04 | glio |
| Cenpj_in.vivo_noRT | Tmeff2 | 3.058 | 8.84E-04 | glio |
| Interaction_genotype_Mtor | Nat8l | -3.222 | 8.84E-05 | glio |
| Chek1_ex.vivo_noRT | Gm48855 | 3.279 | 8.84E-05 | glio |
| Interaction_genotype_Myc | Plcl1 | -4.682 | 8.84E-07 | glio |
| culture_effect | Arid3c | -3.044 | 8.84E-08 | glio |
| culture_effect | Lpcat2 | 1.164 | 8.84E-08 | glio |
| Hsd17b10_in.vivo_noRT | Npy1r | 2.971 | 8.84E-08 | glio |
| Aars2_in.vivo_noRT | Sdr16c5 | 4.01 | 8.84E-10 | glio |
| culture_effect | Slc1a3 | -1.502 | 8.84E-13 | glio |

| | | | | |
|---|---|---|---|---|
| Bora_in.vivo_noRT | Gm49808 | 4.936 | 8.84E-25 | glio |
| culture_effect | Adamts7 | 1.241 | 8.85E-03 | glio |
| Interaction_genotype_Lmo7 | Dnm3 | -3.563 | 8.85E-03 | glio |
| Interaction_genotype_Myc | Rbl2 | -2.548 | 8.85E-03 | glio |
| Interaction_genotype_Myc | Dna2 | -2.167 | 8.85E-03 | glio |
| Interaction_genotype_Rad51 | Hbegf | 1.775 | 8.85E-03 | glio |
| Aars2_ex.vivo_noRT | Clec1a | 2.705 | 8.85E-03 | glio |
| Atp5c1_in.vivo_noRT | Dnm3 | 2.142 | 8.85E-03 | glio |
| Atr_in.vivo_noRT | Mrps35 | -0.89 | 8.85E-03 | glio |
| Atr_in.vivo_noRT | Sarnp | -0.648 | 8.85E-03 | glio |
| Bora_in.vivo_noRT | 4930404N11R | 1.799 | 8.85E-03 | glio |
| Chek1_in.vivo_noRT | Prmt3 | -1.273 | 8.85E-03 | glio |
| Atr_in.vivo_noRT | Chmp1b | 1.387 | 8.85E-04 | glio |
| Chek1_in.vivo_noRT | Cd200 | 2.587 | 8.85E-04 | glio |
| Hsd17b10_in.vivo_noRT | Nrap | 2.637 | 8.85E-04 | glio |
| Atp5c1_in.vivo_noRT | Gm48008 | 2.404 | 8.85E-05 | glio |
| Bora_in.vivo_noRT | Tspan33 | 2.842 | 8.85E-05 | glio |
| Bora_ex.vivo_noRT | Ntn1 | 3.201 | 8.85E-05 | glio |
| Chek1_ex.vivo_noRT | Mettl7a3 | 3.248 | 8.85E-05 | glio |
| interaction_RT | Fth1 | 1.281 | 8.85E-06 | glio |
| Aars2_in.vivo_noRT | Sec61b | -0.966 | 8.85E-06 | glio |
| Atr_in.vivo_noRT | Gm15938 | 3.604 | 8.85E-07 | glio |
| Chek1_ex.vivo_noRT | AI182371 | 3.469 | 8.85E-07 | glio |
| Atr_in.vivo_noRT | A730071L15Ri | 4.473 | 8.85E-10 | glio |
| Aars2_in.vivo_noRT | Clca3a2 | 4.14 | 8.85E-12 | glio |
| culture_effect | Gsdmd | -2.351 | 8.85E-18 | glio |
| Atr_in.vivo_noRT | Gm48617 | 5.269 | 8.85E-26 | glio |
| Interaction_genotype_Aars2 | Gm38979 | -3.562 | 8.86E-03 | glio |
| Interaction_genotype_Rpl3 | Apoa1 | -3.826 | 8.86E-03 | glio |
| Interaction_genotype_Rpl3 | Srcin1 | -3.349 | 8.86E-03 | glio |
| Interaction_genotype_Rpl3 | B630019K06R | -2.962 | 8.86E-03 | glio |
| Aars2_in.vivo_noRT | Aig1 | -0.551 | 8.86E-03 | glio |
| Atr_in.vivo_noRT | BC002059 | 1.509 | 8.86E-03 | glio |
| Bcas2_in.vivo_noRT | Lysmd4 | 1.794 | 8.86E-03 | glio |
| Bcas2_ex.vivo_noRT | Acat3 | 2.166 | 8.86E-03 | glio |
| Bcas2_ex.vivo_noRT | Gm16751 | 2.342 | 8.86E-03 | glio |
| Bcas2_ex.vivo_noRT | Zdhhc14 | 2.347 | 8.86E-03 | glio |
| Cdkn2a_in.vivo_noRT | Gm47404 | -1.675 | 8.86E-03 | glio |
| Cdkn2a_in.vivo_noRT | Flicr | -2.475 | 8.86E-03 | glio |
| Cdkn2a_in.vivo_noRT | Gm31717 | -1.675 | 8.86E-03 | glio |
| culture_effect | Gm11549 | 1.904 | 8.86E-04 | glio |
| culture_effect | Ap4m1 | -0.782 | 8.86E-04 | glio |
| Interaction_genotype_Bora | 5930430L01Ri | -4.002 | 8.86E-04 | glio |
| Interaction_genotype_Bora | 4930558J18Ri | -4.01 | 8.86E-04 | glio |
| Interaction_genotype_Bora | Zfp968 | -4.465 | 8.86E-04 | glio |
| Interaction_genotype_Bora | Tssk6 | -3.674 | 8.86E-04 | glio |
| Interaction_genotype_Bora | Edc4 | 2.382 | 8.86E-04 | glio |
| Interaction_genotype_Bora | BC028471 | -4.412 | 8.86E-04 | glio |
| Aars2_in.vivo_noRT | Surf4 | -0.951 | 8.86E-04 | glio |
| Bora_in.vivo_noRT | Trib1 | -1.377 | 8.86E-04 | glio |
| Chek1_in.vivo_noRT | Gm42941 | 2.251 | 8.86E-04 | glio |
| Chek1_in.vivo_noRT | Aig1 | -0.67 | 8.86E-04 | glio |

| | | | | |
|---|---|---|---|---|
| Interaction_genotype_Aars2 | D030040B21R | -4.595 | 8.86E-05 | glio |
| Interaction_genotype_Mtor | Eif1ad | 1.969 | 8.86E-05 | glio |
| Interaction_genotype_Myc | Calr | 2.208 | 8.86E-05 | glio |
| Atr_in.vivo_noRT | Eif3f | -0.956 | 8.86E-05 | glio |
| Bcas2_in.vivo_noRT | Ccdc13 | 2.856 | 8.86E-05 | glio |
| Hsd17b10_in.vivo_noRT | 2310058D17R | 2.968 | 8.86E-05 | glio |
| Atp5c1_in.vivo_noRT | Slc45a1 | 2.408 | 8.86E-06 | glio |
| Chek1_in.vivo_noRT | Bst1 | 2.928 | 8.86E-06 | glio |
| Chek1_in.vivo_noRT | Eng | 3.847 | 8.86E-07 | glio |
| Hsd17b10_in.vivo_noRT | Carns1 | 3.616 | 8.86E-07 | glio |
| Aars2_in.vivo_noRT | Krt16 | 4.617 | 8.86E-08 | glio |
| Chek1_in.vivo_noRT | Efna2 | 4.301 | 8.86E-08 | glio |
| Aars2_ex.vivo_noRT | mt-Cytb | 1.147 | 8.86E-10 | glio |
| Bcas2_in.vivo_noRT | Tlr1 | 4.547 | 8.86E-11 | glio |
| culture_effect | Anp32e | -0.358 | 8.87E-03 | glio |
| Interaction_genotype_Atr | B230206H07R | -2.239 | 8.87E-03 | glio |
| Interaction_genotype_Atr | Gm26728 | -3.32 | 8.87E-03 | glio |
| Interaction_genotype_Ifnar1 | Tmem132c | 2.734 | 8.87E-03 | glio |
| Interaction_genotype_Ifnar1 | Gm44693 | 2.734 | 8.87E-03 | glio |
| Interaction_genotype_Mtor | 1600014C23R | -3.228 | 8.87E-03 | glio |
| Interaction_genotype_Mtor | Vps8 | -2.848 | 8.87E-03 | glio |
| Interaction_genotype_Mtor | Med18 | -2.485 | 8.87E-03 | glio |
| Interaction_genotype_Mtor | Zfp764 | -2.724 | 8.87E-03 | glio |
| Interaction_genotype_Mtor | Nfatc2ip | -2.515 | 8.87E-03 | glio |
| Interaction_genotype_Mtor | Csnk2b | 1.335 | 8.87E-03 | glio |
| Interaction_genotype_Myc | Gm46378 | -3.143 | 8.87E-03 | glio |
| Interaction_genotype_Myc | Cep128 | -2.603 | 8.87E-03 | glio |
| Interaction_genotype_Myc | Pimreg | -2.146 | 8.87E-03 | glio |
| Aars2_ex.vivo_noRT | Hes5 | 3.293 | 8.87E-03 | glio |
| Atr_in.vivo_noRT | Msln | 1.866 | 8.87E-03 | glio |
| Bora_in.vivo_noRT | Nisch | -0.933 | 8.87E-03 | glio |
| Bora_ex.vivo_noRT | Tns4 | 2.404 | 8.87E-03 | glio |
| Cdkn2a_in.vivo_noRT | Gm14742 | -1.675 | 8.87E-03 | glio |
| Cdkn2a_in.vivo_noRT | Zscan4-ps3 | -1.675 | 8.87E-03 | glio |
| Cdkn2a_in.vivo_noRT | Gm4166 | -1.829 | 8.87E-03 | glio |
| Cdkn2a_in.vivo_noRT | A030014E15Ri | -1.675 | 8.87E-03 | glio |
| Cdkn2a_in.vivo_noRT | Gm21731 | -1.675 | 8.87E-03 | glio |
| Interaction_genotype_Atr | 1700023H06R | -4.439 | 8.87E-04 | glio |
| Interaction_genotype_Atr | Gm42849 | -3.767 | 8.87E-04 | glio |
| Interaction_genotype_Atr | Peli1 | 2.071 | 8.87E-04 | glio |
| Interaction_genotype_Myc | Gm26692 | -3.57 | 8.87E-04 | glio |
| Interaction_genotype_Myc | A730063M14F | -3.295 | 8.87E-04 | glio |
| Interaction_genotype_Rpl3 | Matn4 | -4.322 | 8.87E-04 | glio |
| Chek1_ex.vivo_noRT | Rab3d | 2.713 | 8.87E-04 | glio |
| Aars2_in.vivo_noRT | Hspa12b | 3.029 | 8.87E-05 | glio |
| Bora_ex.vivo_noRT | Pmch | 3.201 | 8.87E-05 | glio |
| Bora_in.vivo_noRT | Gm15056 | 3.502 | 8.87E-06 | glio |
| interaction_RT | Iscu | 1.358 | 8.87E-07 | glio |
| interaction_RT | Tcp11l2 | 2.478 | 8.87E-07 | glio |
| Atr_in.vivo_noRT | Zdhhc14 | 3.066 | 8.87E-08 | glio |
| Atr_in.vivo_noRT | AV356131 | 4.25 | 8.87E-10 | glio |
| Bora_in.vivo_noRT | Grin2d | 4.682 | 8.87E-13 | glio |

| | | | | |
|---|---|---|---|---|
| Interaction_genotype_Mtor | Tbc1d2b | -3.047 | 8.88E-03 | glio |
| Interaction_genotype_Mtor | Mydgf | 1.377 | 8.88E-03 | glio |
| Interaction_genotype_Myc | Gm29358 | -3.026 | 8.88E-03 | glio |
| Interaction_genotype_Myc | Stard4 | -2.209 | 8.88E-03 | glio |
| Bora_in.vivo_noRT | Dhcr24 | -1.33 | 8.88E-03 | glio |
| Bora_in.vivo_noRT | Zswim6 | -1.262 | 8.88E-03 | glio |
| culture_effect | Ntn1 | -1.836 | 8.88E-04 | glio |
| Interaction_genotype_Mtor | Cct6b | -3.889 | 8.88E-04 | glio |
| Interaction_genotype_Mtor | Igf2r | -3.856 | 8.88E-04 | glio |
| Interaction_genotype_Mtor | Coa7 | -2.317 | 8.88E-04 | glio |
| Interaction_genotype_Mtor | Plscr1 | 2.316 | 8.88E-04 | glio |
| Bcas2_in.vivo_noRT | Mirg | 2.465 | 8.88E-04 | glio |
| Hsd17b10_in.vivo_noRT | Nadsyn1 | 2.49 | 8.88E-04 | glio |
| Hsd17b10_in.vivo_noRT | Mta1 | -1.362 | 8.88E-04 | glio |
| culture_effect | Cox19 | 0.571 | 8.88E-05 | glio |
| culture_effect | Ubl4a | 0.556 | 8.88E-05 | glio |
| culture_effect | Yaf2 | 0.482 | 8.88E-05 | glio |
| Aars2_ex.vivo_noRT | Vmn1r88 | 3.293 | 8.88E-05 | glio |
| Atp5c1_in.vivo_noRT | Mpzl2 | 2.408 | 8.88E-05 | glio |
| Atp5c1_in.vivo_noRT | Hdac9 | 2.933 | 8.88E-05 | glio |
| Bcas2_in.vivo_noRT | BC037704 | 2.758 | 8.88E-05 | glio |
| Chek1_ex.vivo_noRT | Dll1 | 4.256 | 8.88E-06 | glio |
| culture_effect | Slc48a1 | 1.132 | 8.88E-07 | glio |
| Atr_ex.vivo_noRT | Mogat1 | 4.565 | 8.88E-07 | glio |
| Bcas2_in.vivo_noRT | Gm15941 | 3.387 | 8.88E-08 | glio |
| Bcas2_in.vivo_noRT | Shisal2a | 4.544 | 8.88E-10 | glio |
| Chek1_ex.vivo_noRT | Slc10a1 | 4.449 | 8.88E-11 | glio |
| Chek1_in.vivo_noRT | Gm20732 | 4.273 | 8.88E-12 | glio |
| Aars2_in.vivo_noRT | C130060C02R | 5.063 | 8.88E-16 | glio |
| Bcas2_in.vivo_noRT | Gm19276 | 4.807 | 8.88E-22 | glio |
| Interaction_genotype_Aars2 | Rnasel | -2.906 | 8.89E-03 | glio |
| Interaction_genotype_Bora | Ccdc80 | -2.704 | 8.89E-03 | glio |
| Interaction_genotype_Myc | Fancf | -2.33 | 8.89E-03 | glio |
| Interaction_genotype_Myc | Nfu1 | 1.308 | 8.89E-03 | glio |
| Aars2_in.vivo_noRT | 1700001L19Ri | 1.728 | 8.89E-03 | glio |
| Atr_in.vivo_noRT | Nptxr | 1.415 | 8.89E-03 | glio |
| Atr_in.vivo_noRT | Ggps1 | -1.091 | 8.89E-03 | glio |
| Atr_in.vivo_noRT | Akt1s1 | -1.135 | 8.89E-03 | glio |
| Atr_in.vivo_noRT | Mrpl2 | -0.813 | 8.89E-03 | glio |
| Bard1_ex.vivo_noRT | Kcnd1 | 2.415 | 8.89E-03 | glio |
| Bora_in.vivo_noRT | Gm45902 | 1.925 | 8.89E-03 | glio |
| Bora_in.vivo_noRT | Wbp2 | -0.993 | 8.89E-03 | glio |
| Chek1_in.vivo_noRT | Kcnip3 | 1.778 | 8.89E-03 | glio |
| Chek1_in.vivo_noRT | Cdkn2aipnl | -1.135 | 8.89E-03 | glio |
| Eif2s1_ex.vivo_noRT | Dmrtc1a | 2.234 | 8.89E-03 | glio |
| Interaction_genotype_Myc | Fads6 | -2.998 | 8.89E-04 | glio |
| Interaction_genotype_Myc | Ccdc137 | -2.521 | 8.89E-04 | glio |
| Bora_in.vivo_noRT | Tfam | -1.072 | 8.89E-04 | glio |
| Chek1_in.vivo_noRT | Gm14488 | 2.595 | 8.89E-04 | glio |
| Interaction_genotype_Myc | Gm16046 | -5.029 | 8.89E-06 | glio |
| Interaction_genotype_Myc | 1700055D18R | -5.446 | 8.89E-06 | glio |
| culture_effect | 1700003G18R | 3.024 | 8.89E-07 | glio |

| | | | | |
|---|---|---|---|---|
| Chek1_in.vivo_noRT | Fam13a | 4.243 | 8.89E-10 | glio |
| Hsd17b10_in.vivo_noRT | 2310003N18R | 3.616 | 8.89E-10 | glio |
| Aars2_in.vivo_noRT | Gpr179 | 4.325 | 8.89E-11 | glio |
| Atr_in.vivo_noRT | Hoxa6 | 4.501 | 8.89E-12 | glio |
| Hsd17b10_in.vivo_noRT | Gm14742 | 3.616 | 8.89E-14 | glio |
| Atr_in.vivo_noRT | Gm43137 | 5.555 | 8.89E-15 | glio |
| Bora_in.vivo_noRT | 5730507C01R | 4.413 | 8.89E-17 | glio |
| Bora_in.vivo_noRT | Gpr4 | 4.936 | 8.89E-22 | glio |
| Bora_in.vivo_noRT | Prl7c1 | 4.936 | 8.89E-22 | glio |
| Bora_in.vivo_noRT | Gast | 5.46 | 8.89E-22 | glio |
| Bora_in.vivo_noRT | Gm33466 | 4.936 | 8.89E-22 | glio |
| Bora_in.vivo_noRT | Prl3d3 | 4.673 | 8.89E-22 | glio |
| culture_effect | Seh1l | -0.37 | 8.90E-03 | glio |
| Interaction_genotype_Aars2 | Gm34121 | -2.877 | 8.90E-03 | glio |
| Interaction_genotype_Atr | Sem1 | 0.921 | 8.90E-03 | glio |
| Interaction_genotype_Bcas2 | Slc17a6 | -2.065 | 8.90E-03 | glio |
| Interaction_genotype_Bcas2 | 5033417F24Ri | -3.344 | 8.90E-03 | glio |
| Interaction_genotype_Bcas2 | Gm10131 | -3.432 | 8.90E-03 | glio |
| Interaction_genotype_Bcas2 | Ppp1r18os | -3.569 | 8.90E-03 | glio |
| Interaction_genotype_Mtor | Rpl26 | 0.824 | 8.90E-03 | glio |
| interaction_RT | Cxxc5 | 1.472 | 8.90E-03 | glio |
| Aars2_in.vivo_noRT | Ppp4r2 | -0.898 | 8.90E-03 | glio |
| Aars2_in.vivo_noRT | Anxa1 | -0.83 | 8.90E-03 | glio |
| Bard1_ex.vivo_noRT | Ric3 | 2.421 | 8.90E-03 | glio |
| Bcas2_ex.vivo_noRT | 4930579C12R | 2.347 | 8.90E-03 | glio |
| Brca2_in.vivo_noRT | Gm30794 | 1.763 | 8.90E-03 | glio |
| Chek1_ex.vivo_noRT | Sltm | -0.903 | 8.90E-03 | glio |
| Interaction_genotype_Atr | Baalc | -3.243 | 8.90E-04 | glio |
| Interaction_genotype_Bora | Rarres1 | -4.298 | 8.90E-04 | glio |
| Interaction_genotype_Bora | Fhl1 | -3.37 | 8.90E-04 | glio |
| Interaction_genotype_Myc | Elavl2 | -3.5 | 8.90E-04 | glio |
| Aars2_in.vivo_noRT | Pycard | 2.189 | 8.90E-04 | glio |
| Aars2_ex.vivo_noRT | Cfap58 | 3.863 | 8.90E-04 | glio |
| Atr_in.vivo_noRT | Mfhas1 | 2.124 | 8.90E-04 | glio |
| Bcas2_in.vivo_noRT | Mroh2b | 2.479 | 8.90E-04 | glio |
| Bcas2_in.vivo_noRT | Shmt2 | -1.348 | 8.90E-04 | glio |
| Bora_in.vivo_noRT | Gm20033 | 1.979 | 8.90E-04 | glio |
| Eif2s1_in.vivo_noRT | Ric3 | 2.407 | 8.90E-04 | glio |
| Eif2s1_in.vivo_noRT | Platr20 | 2.928 | 8.90E-04 | glio |
| Interaction_genotype_Mtor | Prdm15 | -4.078 | 8.90E-05 | glio |
| Bcas2_in.vivo_noRT | Mrpl38 | -1.504 | 8.90E-05 | glio |
| Eif2s1_in.vivo_noRT | Hoxc8 | 2.143 | 8.90E-05 | glio |
| Aars2_in.vivo_noRT | Cysltr2 | 2.837 | 8.90E-06 | glio |
| Atr_in.vivo_noRT | Setd4 | 3.058 | 8.90E-06 | glio |
| Bora_in.vivo_noRT | 9330104G04R | 3.456 | 8.90E-07 | glio |
| Chek1_ex.vivo_noRT | 9230112J17Ri | 3.862 | 8.90E-08 | glio |
| Chek1_ex.vivo_noRT | Gm39929 | 3.679 | 8.90E-08 | glio |
| Chek1_in.vivo_noRT | Gm10644 | 4.326 | 8.90E-10 | glio |
| Aars2_in.vivo_noRT | Gm36172 | 4.794 | 8.90E-18 | glio |
| Interaction_genotype_Bcas2 | Kcnk4 | -3.569 | 8.91E-03 | glio |
| Interaction_genotype_Bcas2 | Pnpla2 | 1.719 | 8.91E-03 | glio |
| Interaction_genotype_Myc | Axin2 | -2.936 | 8.91E-03 | glio |

| | | | | |
|---|---|---|---|---|
| Interaction_genotype_Myc | Bbs5 | -2.451 | 8.91E-03 | glio |
| Interaction_genotype_Myc | Prcd | -2.234 | 8.91E-03 | glio |
| Interaction_genotype_Rpl3 | 1700081N11R | -4.162 | 8.91E-03 | glio |
| Interaction_genotype_Rpl3 | Elovl1 | 1.299 | 8.91E-03 | glio |
| interaction_RT | Aebp2 | -0.654 | 8.91E-03 | glio |
| Atr_ex.vivo_noRT | C430042M11F | 2.642 | 8.91E-03 | glio |
| Atr_ex.vivo_noRT | Crebl2 | 1.815 | 8.91E-03 | glio |
| Bora_ex.vivo_noRT | Nwd1 | 2.22 | 8.91E-03 | glio |
| Cdkn2a_in.vivo_noRT | Gm11232 | -1.675 | 8.91E-03 | glio |
| Hsd17b10_in.vivo_noRT | Fam83e | 2.127 | 8.91E-03 | glio |
| Hsd17b10_in.vivo_noRT | Dcakd | -1.27 | 8.91E-03 | glio |
| culture_effect | Gm28592 | -1.986 | 8.91E-04 | glio |
| Interaction_genotype_Myc | Mboat2 | -2.948 | 8.91E-04 | glio |
| Atr_in.vivo_noRT | Sft2d3 | 1.891 | 8.91E-04 | glio |
| Atr_in.vivo_noRT | Tgs1 | -1.148 | 8.91E-04 | glio |
| Atr_in.vivo_noRT | Olig1 | -1.668 | 8.91E-04 | glio |
| Bcas2_in.vivo_noRT | Ifnb1 | 4.807 | 8.91E-04 | glio |
| Bcas2_in.vivo_noRT | Mars | -1.407 | 8.91E-04 | glio |
| Bcas2_in.vivo_noRT | G3bp1 | -0.81 | 8.91E-04 | glio |
| Atr_ex.vivo_noRT | Pcdh15 | 3.164 | 8.91E-05 | glio |
| Bcas2_in.vivo_noRT | Nptx1 | 3.073 | 8.91E-05 | glio |
| Bora_in.vivo_noRT | Haus1 | -1.868 | 8.91E-05 | glio |
| Bora_in.vivo_noRT | Prune2 | 2.337 | 8.91E-05 | glio |
| Aars2_in.vivo_noRT | Gm48796 | 2.955 | 8.91E-06 | glio |
| Aars2_in.vivo_noRT | Rbpms2 | 3.128 | 8.91E-06 | glio |
| Interaction_genotype_Mtor | Wincr1 | -7.371 | 8.91E-07 | glio |
| Interaction_genotype_Mtor | Celsr3 | -5.519 | 8.91E-07 | glio |
| Aars2_in.vivo_noRT | Comp | 4.207 | 8.91E-09 | glio |
| Chek1_ex.vivo_noRT | H2-M10.1 | 4.256 | 8.91E-09 | glio |
| Aars2_in.vivo_noRT | Ccr5 | 4.527 | 8.91E-13 | glio |
| Bcas2_in.vivo_noRT | Slc2a9 | 4.807 | 8.91E-17 | glio |
| Bcas2_in.vivo_noRT | Gm21761 | 4.807 | 8.91E-19 | glio |
| Atr_in.vivo_noRT | Cfh | 4.754 | 8.91E-20 | glio |
| Atr_in.vivo_noRT | Gm15890 | 5.533 | 8.91E-21 | glio |
| Interaction_genotype_Atr | Gm15326 | -2.416 | 8.92E-03 | glio |
| Interaction_genotype_Atr | D030051J21Ri | -3.221 | 8.92E-03 | glio |
| Interaction_genotype_Rpl3 | Efcab9 | -3.569 | 8.92E-03 | glio |
| Interaction_genotype_Rpl3 | Uqcrc1 | 1.576 | 8.92E-03 | glio |
| Atp5c1_in.vivo_noRT | Gm14968 | 2.289 | 8.92E-03 | glio |
| Atp5c1_in.vivo_noRT | Gm15234 | 2.152 | 8.92E-03 | glio |
| Atr_ex.vivo_noRT | 9230112J17Ri | 2.183 | 8.92E-03 | glio |
| Atr_ex.vivo_noRT | CAAA0106612 | 2.37 | 8.92E-03 | glio |
| Atr_ex.vivo_noRT | Olfr1191-ps1 | 2.376 | 8.92E-03 | glio |
| Interaction_genotype_Mtor | Ncaph2 | 1.686 | 8.92E-04 | glio |
| Aars2_in.vivo_noRT | Hspe1 | -0.708 | 8.92E-04 | glio |
| Aars2_in.vivo_noRT | Actb | -0.746 | 8.92E-04 | glio |
| Bcas2_in.vivo_noRT | Gm43813 | 2.266 | 8.92E-04 | glio |
| Chek1_in.vivo_noRT | Ovgp1 | 2.362 | 8.92E-04 | glio |
| Interaction_genotype_Mtor | Aplp2 | 2.09 | 8.92E-05 | glio |
| Aars2_ex.vivo_noRT | Gm20513 | 3.293 | 8.92E-06 | glio |
| Bora_in.vivo_noRT | Cnbd2 | 3.314 | 8.92E-06 | glio |
| Chek1_in.vivo_noRT | Abhd15 | 3.724 | 8.92E-07 | glio |

| | | | | |
|---|---|---|---|---|
| Chek1_ex.vivo_noRT | Dtx1 | 4.449 | 8.92E-07 | glio |
| culture_effect | Sox8 | -2.71 | 8.92E-08 | glio |
| culture_effect | D130020L05R | 1.915 | 8.92E-08 | glio |
| Interaction_genotype_Mtor | Gm47545 | -5.888 | 8.92E-08 | glio |
| Bard1_ex.vivo_noRT | Gm10801 | 4.234 | 8.92E-08 | glio |
| Bcas2_in.vivo_noRT | Alpl | 3.622 | 8.92E-08 | glio |
| Bcas2_in.vivo_noRT | Kcnk4 | 4.547 | 8.92E-11 | glio |
| Hsd17b10_in.vivo_noRT | Fabp6 | 3.616 | 8.92E-12 | glio |
| Aars2_in.vivo_noRT | Lmx1b | 5.054 | 8.92E-13 | glio |
| Aars2_in.vivo_noRT | Gm13684 | 4.225 | 8.92E-13 | glio |
| Chek1_in.vivo_noRT | Gjb5 | 4.577 | 8.92E-14 | glio |
| culture_effect | Apol9a | 0.403 | 8.93E-03 | glio |
| Interaction_genotype_Atr | Gpr4 | -2.369 | 8.93E-03 | glio |
| Interaction_genotype_Atr | Gm41318 | -3.126 | 8.93E-03 | glio |
| Interaction_genotype_Atr | Kyat1 | -3.61 | 8.93E-03 | glio |
| Interaction_genotype_Atr | 2810455O05R | -3.402 | 8.93E-03 | glio |
| Interaction_genotype_Atr | Ppp1cb | 1.281 | 8.93E-03 | glio |
| Interaction_genotype_Mtor | Pcdhga2 | -3.001 | 8.93E-03 | glio |
| Interaction_genotype_Mtor | Dcun1d2 | -2.234 | 8.93E-03 | glio |
| Interaction_genotype_Myc | Gm8104 | -2.854 | 8.93E-03 | glio |
| Interaction_genotype_Myc | Irf2 | 1.431 | 8.93E-03 | glio |
| Aars2_in.vivo_noRT | Elavl2 | 1.768 | 8.93E-03 | glio |
| Aars2_ex.vivo_noRT | Rbfaos | 2.103 | 8.93E-03 | glio |
| Aars2_ex.vivo_noRT | Gm19345 | 2.505 | 8.93E-03 | glio |
| Atr_in.vivo_noRT | Cyb5b | -1.01 | 8.93E-03 | glio |
| Bcas2_in.vivo_noRT | Hist4h4 | 2.01 | 8.93E-03 | glio |
| Chek1_in.vivo_noRT | Tpi1 | -0.671 | 8.93E-03 | glio |
| Chek1_ex.vivo_noRT | Ap3m2 | 1.865 | 8.93E-03 | glio |
| Eif2s1_in.vivo_noRT | Mecom | 2.143 | 8.93E-03 | glio |
| Eif2s1_in.vivo_noRT | Cys1 | 1.877 | 8.93E-03 | glio |
| Hsd17b10_in.vivo_noRT | Gm16794 | 2.045 | 8.93E-03 | glio |
| Interaction_genotype_Aars2 | Gm49086 | -3.597 | 8.93E-04 | glio |
| Interaction_genotype_Bora | Lyz2 | -4.559 | 8.93E-04 | glio |
| Interaction_genotype_Bora | Gm34885 | -4.444 | 8.93E-04 | glio |
| Interaction_genotype_Myc | Gm46404 | -3.969 | 8.93E-04 | glio |
| Atp5c1_in.vivo_noRT | 4930483J18Ri | 2.408 | 8.93E-04 | glio |
| Atp5c1_in.vivo_noRT | Gm49223 | 2.145 | 8.93E-04 | glio |
| Atp5c1_in.vivo_noRT | Dntt | 2.668 | 8.93E-04 | glio |
| Hsd17b10_in.vivo_noRT | Gm16048 | 2.711 | 8.93E-04 | glio |
| Interaction_genotype_Myc | Fhl4 | -4.022 | 8.93E-05 | glio |
| Hsd17b10_in.vivo_noRT | 1700047K16R | 2.958 | 8.93E-05 | glio |
| Aars2_ex.vivo_noRT | Ly6d | 3.293 | 8.93E-06 | glio |
| culture_effect | Dio1 | -2.77 | 8.93E-07 | glio |
| culture_effect | Gm17259 | 2.205 | 8.93E-07 | glio |
| culture_effect | Smc6 | 0.7 | 8.93E-07 | glio |
| culture_effect | Capg | -0.69 | 8.93E-07 | glio |
| Interaction_genotype_Mtor | Selenos | 2.642 | 8.93E-07 | glio |
| Bora_in.vivo_noRT | Ntf5 | 3.641 | 8.93E-07 | glio |
| Bora_in.vivo_noRT | Wdr93 | 4.28 | 8.93E-11 | glio |
| Atr_in.vivo_noRT | Tmem236 | 5.267 | 8.93E-18 | glio |
| Interaction_genotype_Atr | Zfp516 | 1.665 | 8.94E-03 | glio |
| Interaction_genotype_Hsd17k | Gm14968 | -4.239 | 8.94E-03 | glio |

| | | | | |
|---|---|---|---|---|
| Interaction_genotype_Rad51 | Afg1l | -3.83 | 8.94E-03 | glio |
| Bora_in.vivo_noRT | Wdpcp | 1.843 | 8.94E-03 | glio |
| Bora_ex.vivo_noRT | Ksr2 | 2.226 | 8.94E-03 | glio |
| Interaction_genotype_Myc | Gm20682 | -3.165 | 8.94E-04 | glio |
| Interaction_genotype_Myc | Gm44096 | -3.528 | 8.94E-04 | glio |
| Interaction_genotype_Myc | Ccdc13 | -3.783 | 8.94E-04 | glio |
| Aars2_in.vivo_noRT | Apbb3 | 2.094 | 8.94E-04 | glio |
| Aars2_in.vivo_noRT | Efcab7 | 1.662 | 8.94E-04 | glio |
| Interaction_genotype_Mtor | Adamts10 | -4.049 | 8.94E-05 | glio |
| Chek1_in.vivo_noRT | Zc3h18 | -1.45 | 8.94E-05 | glio |
| Atp5c1_in.vivo_noRT | Gstt4 | 2.405 | 8.94E-06 | glio |
| Aars2_in.vivo_noRT | Lynx1 | 4.023 | 8.94E-09 | glio |
| Chek1_in.vivo_noRT | Pde11a | 4.515 | 8.94E-10 | glio |
| Chek1_ex.vivo_noRT | Ocm | 4.842 | 8.94E-10 | glio |
| Atr_in.vivo_noRT | Abcb5 | 5.533 | 8.94E-28 | glio |
| Interaction_genotype_Bcas2 | Prl2c3 | -3.26 | 8.95E-03 | glio |
| Interaction_genotype_Mtor | Acad10 | -3.21 | 8.95E-03 | glio |
| Interaction_genotype_Myc | Dnaaf5 | -2.76 | 8.95E-03 | glio |
| Interaction_genotype_Myc | Akip1 | -2.082 | 8.95E-03 | glio |
| Interaction_genotype_Myc | Trim34a | -1.849 | 8.95E-03 | glio |
| Aars2_in.vivo_noRT | Sap30bp | -1.008 | 8.95E-03 | glio |
| culture_effect | Sf3b1 | -0.431 | 8.95E-04 | glio |
| Interaction_genotype_Mtor | Fndc7 | -4.176 | 8.95E-04 | glio |
| Interaction_genotype_Myc | Gm15133 | -3.684 | 8.95E-04 | glio |
| Interaction_genotype_Myc | Mastl | -3.003 | 8.95E-04 | glio |
| Atp5c1_in.vivo_noRT | Angptl7 | 1.802 | 8.95E-04 | glio |
| Atr_in.vivo_noRT | Sec22c | 2.025 | 8.95E-04 | glio |
| Bard1_ex.vivo_noRT | Sapcd1 | 3.4 | 8.95E-04 | glio |
| Bora_in.vivo_noRT | Lmtk3 | 2.326 | 8.95E-04 | glio |
| Eif2s1_in.vivo_noRT | Slc16a5 | 1.879 | 8.95E-04 | glio |
| Eif2s1_in.vivo_noRT | Gm48170 | 2.143 | 8.95E-04 | glio |
| Aars2_in.vivo_noRT | Gm14326 | 2.818 | 8.95E-05 | glio |
| Atr_in.vivo_noRT | B230206L02Ri | 2.731 | 8.95E-05 | glio |
| Bora_in.vivo_noRT | Rragd | 2.938 | 8.95E-05 | glio |
| Hsd17b10_in.vivo_noRT | Asgr1 | 2.958 | 8.95E-05 | glio |
| Bora_in.vivo_noRT | Tes | -2.225 | 8.95E-06 | glio |
| culture_effect | Map3k5 | -2.7 | 8.95E-07 | glio |
| Interaction_genotype_Mtor | D030068K23R | -5.965 | 8.95E-07 | glio |
| Interaction_genotype_Mtor | Impg2 | -4.887 | 8.95E-07 | glio |
| Bora_in.vivo_noRT | 2610016A17R | 3.883 | 8.95E-08 | glio |
| Bora_in.vivo_noRT | Gm43463 | 4.279 | 8.95E-13 | glio |
| culture_effect | Trim2 | 0.655 | 8.96E-03 | glio |
| Interaction_genotype_Atr | Psmd11 | 0.975 | 8.96E-03 | glio |
| Interaction_genotype_Myc | Efcab2 | -2.61 | 8.96E-03 | glio |
| Interaction_genotype_Myc | Ppat | 1.492 | 8.96E-03 | glio |
| Interaction_genotype_Myc | Cdv3 | 0.97 | 8.96E-03 | glio |
| Interaction_genotype_Rpl3 | Snn | -3.727 | 8.96E-03 | glio |
| Aars2_in.vivo_noRT | Snx2 | -1.018 | 8.96E-03 | glio |
| Aars2_in.vivo_noRT | Dync1li1 | -1.019 | 8.96E-03 | glio |
| Aars2_in.vivo_noRT | Eif4ebp1 | -0.83 | 8.96E-03 | glio |
| Atr_ex.vivo_noRT | Clec4d | 1.979 | 8.96E-03 | glio |
| Atr_ex.vivo_noRT | Aak1 | -1.105 | 8.96E-03 | glio |

| | | | | |
|---|---|---|---|---|
| Bcas2_in.vivo_noRT | Slc29a1 | -1.266 | 8.96E-03 | glio |
| Bcas2_in.vivo_noRT | Tardbp | -0.72 | 8.96E-03 | glio |
| Bora_in.vivo_noRT | Erbb3 | -1.174 | 8.96E-03 | glio |
| Brca2_in.vivo_noRT | Ank1 | 1.763 | 8.96E-03 | glio |
| Chek1_in.vivo_noRT | Entpd5 | 1.846 | 8.96E-03 | glio |
| Chek1_in.vivo_noRT | Ing5 | -1.172 | 8.96E-03 | glio |
| Chek1_ex.vivo_noRT | Gm45767 | 2.424 | 8.96E-03 | glio |
| Cyp19a1_ex.vivo_noRT | Zscan4-ps2 | 2.159 | 8.96E-03 | glio |
| Interaction_genotype_Mtor | Tbkbp1 | -4.181 | 8.96E-04 | glio |
| Interaction_genotype_Myc | Gm49463 | -2.955 | 8.96E-04 | glio |
| Bora_in.vivo_noRT | Pdlim2 | 2.169 | 8.96E-04 | glio |
| culture_effect | Luzp1 | 0.933 | 8.96E-06 | glio |
| Aars2_ex.vivo_noRT | Ccdc162 | 3.293 | 8.96E-06 | glio |
| Atr_in.vivo_noRT | Aldh1a2 | 3.176 | 8.96E-07 | glio |
| Bcas2_in.vivo_noRT | Gm45890 | 3.89 | 8.96E-08 | glio |
| Bcas2_in.vivo_noRT | K230015D01R | 4.541 | 8.96E-10 | glio |
| Atr_in.vivo_noRT | 1700030C12R | 4.608 | 8.96E-11 | glio |
| Hsd17b10_in.vivo_noRT | Gm39929 | 3.679 | 8.96E-11 | glio |
| Aars2_in.vivo_noRT | Gm24362 | 4.284 | 8.96E-12 | glio |
| culture_effect | Ifi206 | -3.843 | 8.96E-13 | glio |
| Chek1_in.vivo_noRT | Nxnl2 | 4.522 | 8.96E-13 | glio |
| Aars2_in.vivo_noRT | Mettl7b | 5.327 | 8.96E-15 | glio |
| Aars2_in.vivo_noRT | Gm48155 | 5.327 | 8.96E-29 | glio |
| culture_effect | Tbcb | -0.329 | 8.97E-03 | glio |
| Interaction_genotype_Atr | Amt | -3.533 | 8.97E-03 | glio |
| Interaction_genotype_Mtor | 4933432I03Ri | -3.231 | 8.97E-03 | glio |
| Interaction_genotype_Myc | Msl3l2 | -2.989 | 8.97E-03 | glio |
| Interaction_genotype_Myc | Nutf2 | -2.91 | 8.97E-03 | glio |
| Interaction_genotype_Myc | Mtss1 | -3.146 | 8.97E-03 | glio |
| Interaction_genotype_Myc | Arv1 | -2.293 | 8.97E-03 | glio |
| Chek1_in.vivo_noRT | Gm15726 | 1.786 | 8.97E-03 | glio |
| Chek1_in.vivo_noRT | Msl1 | -1.002 | 8.97E-03 | glio |
| Interaction_genotype_Mtor | Gm5914 | -3.032 | 8.97E-04 | glio |
| Interaction_genotype_Myc | Thg1l | -2.795 | 8.97E-04 | glio |
| Bora_ex.vivo_noRT | Plk5 | 2.808 | 8.97E-04 | glio |
| Chek1_ex.vivo_noRT | Cyp4v3 | 2.775 | 8.97E-04 | glio |
| Interaction_genotype_Aars2 | Cox4i1 | 1.098 | 8.97E-05 | glio |
| culture_effect | Gpx4 | 0.712 | 8.97E-10 | glio |
| Bora_in.vivo_noRT | Plcl1 | 4.936 | 8.97E-18 | glio |
| culture_effect | Cald1 | 1.767 | 8.97E-23 | glio |
| Interaction_genotype_Aars2 | B430010I23Ri | -3.145 | 8.98E-03 | glio |
| Interaction_genotype_Atr | Lrrc8c | -1.86 | 8.98E-03 | glio |
| Interaction_genotype_Chek1 | Gm11457 | -3.502 | 8.98E-03 | glio |
| Interaction_genotype_Mtor | Hist1h1d | -2.677 | 8.98E-03 | glio |
| Interaction_genotype_Myc | Nkx2-2 | -2.235 | 8.98E-03 | glio |
| Interaction_genotype_Myc | Gm9993 | -2.471 | 8.98E-03 | glio |
| Aars2_ex.vivo_noRT | Gm39929 | 2.13 | 8.98E-03 | glio |
| Bcas2_ex.vivo_noRT | Card9 | 2.342 | 8.98E-03 | glio |
| Hsd17b10_in.vivo_noRT | Aldh1a1 | 2.114 | 8.98E-03 | glio |
| Interaction_genotype_Atr | Glis1 | -4.166 | 8.98E-04 | glio |
| Interaction_genotype_Mtor | 1700012D14R | -3.456 | 8.98E-04 | glio |
| Aars2_ex.vivo_noRT | Gm40264 | 2.706 | 8.98E-04 | glio |

| | | | | |
|---|---|---|---|---|
| Atr_ex.vivo_noRT | A930024E05Ri | 3.062 | 8.98E-04 | glio |
| Bora_in.vivo_noRT | Cyb5d1 | 2.401 | 8.98E-04 | glio |
| Chek1_in.vivo_noRT | Npff | 2.244 | 8.98E-04 | glio |
| Chek1_in.vivo_noRT | Dock5 | 2.025 | 8.98E-04 | glio |
| Chek1_in.vivo_noRT | Myo6 | 2.012 | 8.98E-04 | glio |
| culture_effect | Nqo1 | 1.933 | 8.98E-05 | glio |
| Interaction_genotype_Mtor | Serpinb6a | -4.8 | 8.98E-05 | glio |
| Chek1_in.vivo_noRT | Rab2a | -1.025 | 8.98E-05 | glio |
| Hsd17b10_in.vivo_noRT | D6Ertd527e | 2.834 | 8.98E-05 | glio |
| Bcas2_in.vivo_noRT | Tnc | 3.758 | 8.98E-06 | glio |
| Bcas2_in.vivo_noRT | A230072E10Ri | 3.104 | 8.98E-06 | glio |
| Bora_ex.vivo_noRT | A330049N07R | 2.802 | 8.98E-06 | glio |
| Atr_in.vivo_noRT | Prrg1 | 3.509 | 8.98E-07 | glio |
| interaction_RT | Pdcd4 | 1.714 | 8.98E-09 | glio |
| Aars2_in.vivo_noRT | Slc7a3 | 4.06 | 8.98E-09 | glio |
| Bcas2_in.vivo_noRT | 4833419F23Ri | 4.807 | 8.98E-12 | glio |
| Aars2_in.vivo_noRT | Erich6b | 5.327 | 8.98E-26 | glio |
| Aars2_in.vivo_noRT | Lncbate10 | 5.327 | 8.98E-26 | glio |
| Interaction_genotype_Atr | Gm28731 | -3.427 | 8.99E-03 | glio |
| Aars2_ex.vivo_noRT | Plac8 | 2.718 | 8.99E-03 | glio |
| Atr_in.vivo_noRT | Klf12 | 1.821 | 8.99E-03 | glio |
| Atr_in.vivo_noRT | Lrrc75a | 1.709 | 8.99E-03 | glio |
| Atr_in.vivo_noRT | Zfp532 | 1.38 | 8.99E-03 | glio |
| Atr_in.vivo_noRT | Gtf2a2 | -0.732 | 8.99E-03 | glio |
| Atr_in.vivo_noRT | Cct7 | -0.65 | 8.99E-03 | glio |
| Bcas2_ex.vivo_noRT | Gm34047 | 2.529 | 8.99E-03 | glio |
| Chek1_in.vivo_noRT | Nemp2 | 2.097 | 8.99E-03 | glio |
| Chek1_in.vivo_noRT | Anxa9 | 1.78 | 8.99E-03 | glio |
| Chek1_in.vivo_noRT | Ube2l6 | -0.969 | 8.99E-03 | glio |
| Cyp19a1_ex.vivo_noRT | Card9 | 2.533 | 8.99E-03 | glio |
| Cyp19a1_ex.vivo_noRT | Ccl3 | 2.159 | 8.99E-03 | glio |
| Hsd17b10_in.vivo_noRT | Sgk3 | 2.062 | 8.99E-03 | glio |
| Hsd17b10_in.vivo_noRT | Gm15614 | 2.022 | 8.99E-03 | glio |
| Hsd17b10_in.vivo_noRT | Uggt1 | -1.063 | 8.99E-03 | glio |
| Interaction_genotype_Lmo7 | Txnip | 4.366 | 8.99E-04 | glio |
| Interaction_genotype_Lmo7 | Loxl4 | -4.567 | 8.99E-04 | glio |
| Interaction_genotype_Lmo7 | Gm43581 | -4.526 | 8.99E-04 | glio |
| Interaction_genotype_Lmo7 | 5430402O13R | -4.961 | 8.99E-04 | glio |
| Atr_in.vivo_noRT | Epb42 | 2.412 | 8.99E-04 | glio |
| Bcas2_ex.vivo_noRT | St18 | 2.742 | 8.99E-04 | glio |
| Bora_in.vivo_noRT | Rpgr | 2.276 | 8.99E-04 | glio |
| Bora_in.vivo_noRT | Ypel1 | 2.384 | 8.99E-04 | glio |
| Bora_in.vivo_noRT | Commd8 | -1.351 | 8.99E-04 | glio |
| Bora_ex.vivo_noRT | Ckm | 3.197 | 8.99E-04 | glio |
| Bora_in.vivo_noRT | 9330179D12R | 3.006 | 8.99E-05 | glio |
| Bora_in.vivo_noRT | 1700037C18R | 2.634 | 8.99E-05 | glio |
| Bcas2_in.vivo_noRT | Gm20536 | 3.428 | 8.99E-07 | glio |
| Hsd17b10_in.vivo_noRT | Pnp2 | -2.621 | 8.99E-07 | glio |
| Atr_ex.vivo_noRT | A330076H08R | 3.962 | 8.99E-08 | glio |
| Atr_in.vivo_noRT | Gm48682 | 4.222 | 8.99E-10 | glio |
| Bcas2_in.vivo_noRT | Lrrc4b | 3.896 | 8.99E-11 | glio |
| Atr_in.vivo_noRT | Crhbp | 5.533 | 8.99E-20 | glio |

| | | | | |
|---|---|---|---|---|
| Interaction_genotype_Bcas2 | Vmn2r44 | -2.46 | 9.00E-03 | glio |
| Interaction_genotype_Bcas2 | Rab3b | -3.403 | 9.00E-03 | glio |
| Interaction_genotype_Bcas2 | Gm50107 | -3.269 | 9.00E-03 | glio |
| Interaction_genotype_Myc | 9030612E09Ri | -2.776 | 9.00E-03 | glio |
| Interaction_genotype_Myc | Hist2h4 | -2.596 | 9.00E-03 | glio |
| Atp5c1_in.vivo_noRT | Serpinb9b | 2.615 | 9.00E-03 | glio |
| Bcas2_ex.vivo_noRT | Pde10a | 1.778 | 9.00E-03 | glio |
| Bora_in.vivo_noRT | F8a | 1.328 | 9.00E-03 | glio |
| Chek1_in.vivo_noRT | Tal1 | 1.925 | 9.00E-03 | glio |
| Chek1_in.vivo_noRT | Adcy9 | 1.744 | 9.00E-03 | glio |
| Chek1_ex.vivo_noRT | Fignl2 | 2.639 | 9.00E-03 | glio |
| Hsd17b10_in.vivo_noRT | Stmn4 | 3.1 | 9.00E-03 | glio |
| Interaction_genotype_Bora | Pyy | -3.615 | 9.00E-04 | glio |
| Aars2_ex.vivo_noRT | Gm31063 | 2.369 | 9.00E-04 | glio |
| Aars2_ex.vivo_noRT | Gm43256 | 2.921 | 9.00E-04 | glio |
| Atp5c1_in.vivo_noRT | Carns1 | 2.668 | 9.00E-04 | glio |
| Atr_in.vivo_noRT | Lcat | 3.2 | 9.00E-06 | glio |
| Interaction_genotype_Atr | Dnm3 | -5.412 | 9.00E-07 | glio |
| Interaction_genotype_Atr | Cela1 | 3.141 | 9.00E-07 | glio |
| Interaction_genotype_Atr | Ntn4 | -4.35 | 9.00E-07 | glio |
| Interaction_genotype_Atr | Adgrb3 | -5.797 | 9.00E-07 | glio |
| Interaction_genotype_Atr | Col8a1 | -5.809 | 9.00E-07 | glio |
| Interaction_genotype_Atr | Gm11457 | -5.67 | 9.00E-07 | glio |
| Aars2_in.vivo_noRT | 1700101I11Ri | 3.686 | 9.00E-07 | glio |
| Chek1_in.vivo_noRT | B230319C09R | 3.734 | 9.00E-07 | glio |
| Interaction_genotype_Mtor | Rapgef3os2 | -5.284 | 9.00E-09 | glio |
| interaction_RT | Zfand5 | 1.12 | 9.00E-09 | glio |
| Atr_in.vivo_noRT | Syne3 | 4.277 | 9.00E-09 | glio |
| Atr_in.vivo_noRT | Gm16754 | 4.029 | 9.00E-09 | glio |
| Aars2_in.vivo_noRT | Vsig10l | 4.674 | 9.00E-12 | glio |
| Chek1_ex.vivo_noRT | Vmn2r56 | 4.842 | 9.00E-13 | glio |
| Chek1_in.vivo_noRT | Flacc1 | 5.044 | 9.00E-17 | glio |
| Bora_in.vivo_noRT | Sytl4 | 4.67 | 9.00E-20 | glio |
| Interaction_genotype_Mtor | Gm36445 | -3.214 | 9.01E-03 | glio |
| interaction_RT | Dap3 | -0.675 | 9.01E-03 | glio |
| Aars2_in.vivo_noRT | 1700109H08R | 1.956 | 9.01E-03 | glio |
| Atr_in.vivo_noRT | Zkscan5 | 1.532 | 9.01E-03 | glio |
| Atr_in.vivo_noRT | Nop10 | -0.811 | 9.01E-03 | glio |
| Bard1_ex.vivo_noRT | Serpinb2 | 2.415 | 9.01E-03 | glio |
| Bcas2_in.vivo_noRT | Atmin | 1.368 | 9.01E-03 | glio |
| Bora_in.vivo_noRT | Tspan32 | 1.958 | 9.01E-03 | glio |
| Bora_in.vivo_noRT | Tuba4a | 1.636 | 9.01E-03 | glio |
| Chek1_in.vivo_noRT | Pes1 | -1.046 | 9.01E-03 | glio |
| Chek1_ex.vivo_noRT | Fnbp1l | -1.77 | 9.01E-03 | glio |
| Eif2s1_ex.vivo_noRT | Gm11899 | 2.234 | 9.01E-03 | glio |
| Hsd17b10_in.vivo_noRT | Cdrt4 | 2.162 | 9.01E-03 | glio |
| Hsd17b10_in.vivo_noRT | Tbc1d10a | -1.084 | 9.01E-03 | glio |
| Interaction_genotype_Mtor | Itgbl1 | -3.308 | 9.01E-04 | glio |
| Atr_in.vivo_noRT | Myo19 | 1.866 | 9.01E-04 | glio |
| Chek1_in.vivo_noRT | Sik1 | 2.412 | 9.01E-04 | glio |
| Chek1_in.vivo_noRT | Ptrh1 | 1.874 | 9.01E-04 | glio |
| Interaction_genotype_Mtor | Mtpn | 1.668 | 9.01E-05 | glio |

| | | | | |
|---|---|---|---|---|
| Interaction_genotype_Mtor | Lmx1b | -5.462 | 9.01E-06 | glio |
| Chek1_ex.vivo_noRT | Vmn1r91 | 3.862 | 9.01E-07 | glio |
| culture_effect | Igf2 | -1.436 | 9.01E-09 | glio |
| Bora_in.vivo_noRT | St8sia1 | 4.676 | 9.01E-11 | glio |
| Bcas2_in.vivo_noRT | Gm11769 | 4.547 | 9.01E-13 | glio |
| Atr_in.vivo_noRT | Cd83 | 4.533 | 9.01E-14 | glio |
| Chek1_ex.vivo_noRT | Gpr22 | 4.842 | 9.01E-14 | glio |
| Chek1_in.vivo_noRT | Boc | 4.78 | 9.01E-21 | glio |
| Interaction_genotype_Bora | Gm13270 | -3.593 | 9.02E-03 | glio |
| Interaction_genotype_Mtor | Zbtb34 | -3.14 | 9.02E-03 | glio |
| Interaction_genotype_Myc | Hist1h2ag | -2.197 | 9.02E-03 | glio |
| Interaction_genotype_Myc | Ushbp1 | -2.914 | 9.02E-03 | glio |
| Interaction_genotype_Myc | Upf2 | -1.783 | 9.02E-03 | glio |
| Interaction_genotype_Pars2 | Capn13 | -2.268 | 9.02E-03 | glio |
| Interaction_genotype_Pars2 | Tmem170b | 2.237 | 9.02E-03 | glio |
| Interaction_genotype_Pars2 | Maged2 | 1.371 | 9.02E-03 | glio |
| interaction_RT | Zfhx2os | -2.218 | 9.02E-03 | glio |
| interaction_RT | Crabp2 | 2.018 | 9.02E-03 | glio |
| interaction_RT | Ano6 | 0.762 | 9.02E-03 | glio |
| interaction_RT | Snx6 | 0.612 | 9.02E-03 | glio |
| interaction_RT | G3bp1 | -0.516 | 9.02E-03 | glio |
| Bcas2_in.vivo_noRT | Tnks | -1.206 | 9.02E-03 | glio |
| Bcas2_in.vivo_noRT | Chchd4 | -1.048 | 9.02E-03 | glio |
| Bora_in.vivo_noRT | Pfkm | 1.199 | 9.02E-03 | glio |
| Eif2s1_in.vivo_noRT | Adgre5 | 1.883 | 9.02E-03 | glio |
| Eif2s1_in.vivo_noRT | Gm48899 | 1.883 | 9.02E-03 | glio |
| Eif2s1_in.vivo_noRT | B4galnt1 | 1.883 | 9.02E-03 | glio |
| Eif2s1_in.vivo_noRT | Tmem30c | 2.265 | 9.02E-03 | glio |
| Eif2s1_in.vivo_noRT | Tm9sf2 | -0.719 | 9.02E-03 | glio |
| Eif2s1_in.vivo_noRT | Col9a1 | -0.834 | 9.02E-03 | glio |
| Interaction_genotype_Mtor | Fut8 | -2.604 | 9.02E-04 | glio |
| Aars2_ex.vivo_noRT | Slc15a3 | 3.098 | 9.02E-04 | glio |
| Bcas2_in.vivo_noRT | Slc2a13 | 2.397 | 9.02E-04 | glio |
| Bora_in.vivo_noRT | Zfp772 | 2.45 | 9.02E-04 | glio |
| Chek1_ex.vivo_noRT | AI854703 | 3.001 | 9.02E-04 | glio |
| Chek1_ex.vivo_noRT | Mrpl51 | -0.734 | 9.02E-04 | glio |
| Interaction_genotype_Mtor | Iqcf1 | -4.534 | 9.02E-05 | glio |
| Interaction_genotype_Myc | Gm31415 | -3.758 | 9.02E-05 | glio |
| Aars2_in.vivo_noRT | Spink4 | 2.582 | 9.02E-05 | glio |
| Aars2_in.vivo_noRT | Psmd4 | -0.955 | 9.02E-05 | glio |
| Atr_in.vivo_noRT | Pon3 | 2.783 | 9.02E-05 | glio |
| Chek1_in.vivo_noRT | Zfp418 | 2.988 | 9.02E-05 | glio |
| Bora_in.vivo_noRT | Gpaa1 | -1.873 | 9.02E-06 | glio |
| Hsd17b10_in.vivo_noRT | D830039M14I | 3.35 | 9.02E-06 | glio |
| Aars2_in.vivo_noRT | 5330426L24Ri | 3.617 | 9.02E-07 | glio |
| Interaction_genotype_Mtor | Hist2h3c2 | -6.926 | 9.02E-08 | glio |
| Aars2_in.vivo_noRT | 1110020A21R | 4.376 | 9.02E-10 | glio |
| Atr_in.vivo_noRT | Klra6 | 4.423 | 9.02E-10 | glio |
| Aars2_in.vivo_noRT | Gm43625 | 4.022 | 9.02E-11 | glio |
| Atr_in.vivo_noRT | Mterf1b | 4.275 | 9.02E-11 | glio |
| Bcas2_in.vivo_noRT | Sap30bpos | 3.931 | 9.02E-11 | glio |
| Hsd17b10_in.vivo_noRT | Spink8 | 3.356 | 9.02E-11 | glio |

| | | | | |
|---|---|---|---|---|
| culture_effect | Sncaip | -3.173 | 9.02E-13 | glio |
| Bcas2_in.vivo_noRT | 4732463B04R | 4.543 | 9.02E-17 | glio |
| Chek1_in.vivo_noRT | Pspn | 5.044 | 9.02E-25 | glio |
| Interaction_genotype_Aars2 | 6330408M09F | -3.436 | 9.03E-03 | glio |
| Aars2_in.vivo_noRT | Ccdc124 | -0.78 | 9.03E-03 | glio |
| Atr_in.vivo_noRT | Relt | 1.919 | 9.03E-03 | glio |
| Bard1_ex.vivo_noRT | Rab27a | 2.419 | 9.03E-03 | glio |
| Bora_in.vivo_noRT | Bik | 1.665 | 9.03E-03 | glio |
| Interaction_genotype_Aars2 | Flnc | -4.259 | 9.03E-04 | glio |
| Interaction_genotype_Aars2 | Gm30794 | -3.01 | 9.03E-04 | glio |
| Interaction_genotype_Atr | Glt1d1 | -3.943 | 9.03E-04 | glio |
| Interaction_genotype_Atr | Ube2d3 | 0.955 | 9.03E-04 | glio |
| Bcas2_in.vivo_noRT | Mbd3 | -0.952 | 9.03E-04 | glio |
| Interaction_genotype_Bcas2 | Spink2 | -5.796 | 9.03E-05 | glio |
| Interaction_genotype_Bcas2 | Atpif1 | 1.337 | 9.03E-05 | glio |
| Chek1_in.vivo_noRT | Vma21 | -1.093 | 9.03E-05 | glio |
| Chek1_ex.vivo_noRT | Cyp4x1 | 3.26 | 9.03E-05 | glio |
| Chek1_ex.vivo_noRT | Ly6g6d | 3.377 | 9.03E-05 | glio |
| Interaction_genotype_Mtor | Gm26555 | -3.836 | 9.03E-06 | glio |
| Interaction_genotype_Mtor | Gm5286 | -4.099 | 9.03E-06 | glio |
| Atr_in.vivo_noRT | 2810021J22Ri | 3.214 | 9.03E-06 | glio |
| Chek1_in.vivo_noRT | Myl9 | 3.481 | 9.03E-06 | glio |
| Chek1_in.vivo_noRT | Tead3 | 3.105 | 9.03E-06 | glio |
| Dclre1c_ex.vivo_noRT | Gm2022 | 3.446 | 9.03E-06 | glio |
| Bcas2_in.vivo_noRT | Gm20667 | 3.807 | 9.03E-08 | glio |
| Bcas2_in.vivo_noRT | 4930478M13F | 4.428 | 9.03E-10 | glio |
| culture_effect | Mras | 1.09 | 9.03E-12 | glio |
| Atr_in.vivo_noRT | Gm37214 | 5.273 | 9.03E-14 | glio |
| Chek1_in.vivo_noRT | Gm44658 | 4.512 | 9.03E-14 | glio |
| culture_effect | Mfsd4a | -0.956 | 9.04E-03 | glio |
| Interaction_genotype_Atr | Cisd2 | 1.131 | 9.04E-03 | glio |
| Interaction_genotype_Hsd17b | Pnp2 | 2.726 | 9.04E-03 | glio |
| Interaction_genotype_Hsd17b | BC065403 | -3.87 | 9.04E-03 | glio |
| Interaction_genotype_Hsd17b | Pola2 | 2.184 | 9.04E-03 | glio |
| Interaction_genotype_Hsd17b | Crppa | 3.216 | 9.04E-03 | glio |
| Interaction_genotype_Hsd17b | Rassf6 | -3.472 | 9.04E-03 | glio |
| Interaction_genotype_Hsd17b | Cyp1a1 | -5.307 | 9.04E-03 | glio |
| Interaction_genotype_Mtor | 4930404N11R | -3.025 | 9.04E-03 | glio |
| Interaction_genotype_Myc | Sh3bp5 | -2.966 | 9.04E-03 | glio |
| Interaction_genotype_Myc | C87436 | -1.792 | 9.04E-03 | glio |
| Interaction_genotype_Terf1 | Gm48242 | -3.379 | 9.04E-03 | glio |
| Aars2_in.vivo_noRT | Mis12 | -1.29 | 9.04E-03 | glio |
| Atr_in.vivo_noRT | Sars | -0.758 | 9.04E-03 | glio |
| Bard1_ex.vivo_noRT | Gm21762 | 2.421 | 9.04E-03 | glio |
| Chek1_in.vivo_noRT | Cbx1 | -0.653 | 9.04E-03 | glio |
| Chek1_ex.vivo_noRT | Ucma | 2.792 | 9.04E-03 | glio |
| Interaction_genotype_Aars2 | Olfr552 | -2.947 | 9.04E-04 | glio |
| Interaction_genotype_Atr | Efhd1 | -4.127 | 9.04E-04 | glio |
| Interaction_genotype_Myc | Tbkbp1 | -3.971 | 9.04E-04 | glio |
| Aars2_ex.vivo_noRT | Dll1 | 3.483 | 9.04E-04 | glio |
| Bcas2_in.vivo_noRT | Alpk2 | 3.091 | 9.04E-04 | glio |
| Interaction_genotype_Aars2 | Gm48933 | -4.817 | 9.04E-05 | glio |

| | | | | |
|---|---|---|---|---|
| Bcas2_in.vivo_noRT | Phtf1os | 2.883 | 9.04E-05 | glio |
| Bcas2_in.vivo_noRT | Gm26879 | 2.8 | 9.04E-06 | glio |
| Chek1_in.vivo_noRT | Poln | 3.541 | 9.04E-07 | glio |
| Chek1_in.vivo_noRT | Nppc | 4.574 | 9.04E-10 | glio |
| Chek1_in.vivo_noRT | Rslcan18 | 4.658 | 9.04E-14 | glio |
| Bora_in.vivo_noRT | Vmn2r44 | 4.936 | 9.04E-23 | glio |
| Bora_in.vivo_noRT | Mmp13 | 4.936 | 9.04E-23 | glio |
| culture_effect | B3gnt5 | -1.41 | 9.05E-03 | glio |
| Interaction_genotype_Bora | Nnt | -3.225 | 9.05E-03 | glio |
| Interaction_genotype_Mtor | St6gal1 | -3.32 | 9.05E-03 | glio |
| Interaction_genotype_Mtor | Rab42 | -3.091 | 9.05E-03 | glio |
| Interaction_genotype_Mtor | Jrkl | -2.479 | 9.05E-03 | glio |
| Interaction_genotype_Mtor | Brd4 | 1.3 | 9.05E-03 | glio |
| Interaction_genotype_Myc | Hist1h4k | -3.017 | 9.05E-03 | glio |
| Atp5c1_in.vivo_noRT | Zfp986 | 2.154 | 9.05E-03 | glio |
| Atr_in.vivo_noRT | Ttl | 1.498 | 9.05E-03 | glio |
| Bora_in.vivo_noRT | Ip6k3 | 1.964 | 9.05E-03 | glio |
| culture_effect | Tmem147 | 0.431 | 9.05E-04 | glio |
| Interaction_genotype_Abtb1 | 4933408J17Ri | 4.532 | 9.05E-04 | glio |
| Interaction_genotype_Mtor | Lce1g | -4.097 | 9.05E-04 | glio |
| Interaction_genotype_Mtor | Gm41492 | -3.507 | 9.05E-04 | glio |
| Interaction_genotype_Myc | 4732419C18R | -3.603 | 9.05E-04 | glio |
| Interaction_genotype_Myc | Tcirg1 | -2.754 | 9.05E-04 | glio |
| Interaction_genotype_Myc | Txn2 | 1.4 | 9.05E-04 | glio |
| interaction_RT | Atad3a | -0.823 | 9.05E-04 | glio |
| Aars2_ex.vivo_noRT | Fabp4 | 3.304 | 9.05E-04 | glio |
| Bora_ex.vivo_noRT | Slc5a5 | 3.454 | 9.05E-04 | glio |
| Atr_in.vivo_noRT | Ttbk1 | 2.736 | 9.05E-05 | glio |
| Bcas2_in.vivo_noRT | Tes3-ps | 2.773 | 9.05E-05 | glio |
| Bcas2_in.vivo_noRT | Shd | 2.719 | 9.05E-05 | glio |
| Bcas2_in.vivo_noRT | Icosl | 2.517 | 9.05E-05 | glio |
| Bcas2_in.vivo_noRT | H2-M3 | -2.035 | 9.05E-05 | glio |
| Bora_in.vivo_noRT | Pcdhgb1 | 2.402 | 9.05E-05 | glio |
| Bora_ex.vivo_noRT | Nrros | 3.601 | 9.05E-05 | glio |
| Atr_in.vivo_noRT | 4732491K20R | 2.827 | 9.05E-06 | glio |
| Bcas2_in.vivo_noRT | Ppnr | 3.145 | 9.05E-06 | glio |
| interaction_RT | Lmbrd1 | 1.42 | 9.05E-07 | glio |
| Chek1_in.vivo_noRT | Kctd14 | 3.12 | 9.05E-07 | glio |
| Atr_in.vivo_noRT | Ptpa | -1.893 | 9.05E-08 | glio |
| culture_effect | Dynll2 | 0.727 | 9.05E-09 | glio |
| Bcas2_in.vivo_noRT | 1700015F17Ri | 4.547 | 9.05E-11 | glio |
| culture_effect | Ppp1r18 | 1.414 | 9.05E-14 | glio |
| Aars2_in.vivo_noRT | Gm49223 | 4.804 | 9.05E-17 | glio |
| Aars2_in.vivo_noRT | Gm16741 | 5.327 | 9.05E-17 | glio |
| Aars2_in.vivo_noRT | 9130230L23Ri | 5.067 | 9.05E-17 | glio |
| Interaction_genotype_Atr | Olfr1191-ps1 | -3.016 | 9.06E-03 | glio |
| Interaction_genotype_Mtor | Krtcap2 | 1.187 | 9.06E-03 | glio |
| Interaction_genotype_Myc | Tradd | -2.066 | 9.06E-03 | glio |
| Aars2_in.vivo_noRT | Inpp4a | 1.448 | 9.06E-03 | glio |
| Aars2_in.vivo_noRT | Dnajc19 | -0.819 | 9.06E-03 | glio |
| Cenpj_ex.vivo_noRT | C430002N11R | 2.196 | 9.06E-03 | glio |
| Chek1_ex.vivo_noRT | Gm47509 | 2.46 | 9.06E-03 | glio |

| | | | | |
|---|---|---|---|---|
| culture_effect | Daam1 | 0.711 | 9.06E-04 | glio |
| Interaction_genotype_Atr | Gm46218 | -4.088 | 9.06E-04 | glio |
| Interaction_genotype_Myc | P4ha3 | -3.08 | 9.06E-04 | glio |
| Bora_in.vivo_noRT | Dpp9 | -1.612 | 9.06E-04 | glio |
| Interaction_genotype_Mtor | Tcp11x2 | -3.97 | 9.06E-05 | glio |
| Interaction_genotype_Mtor | Klhdc1 | -4.468 | 9.06E-05 | glio |
| Interaction_genotype_Mtor | 1110006O24R | -4.374 | 9.06E-05 | glio |
| Interaction_genotype_Mtor | Olfm1 | -4.025 | 9.06E-05 | glio |
| Bora_in.vivo_noRT | C1d | -1.477 | 9.06E-05 | glio |
| Bcas2_in.vivo_noRT | Scd3 | 3.419 | 9.06E-06 | glio |
| Hsd17b10_in.vivo_noRT | Kcnd1 | 3.094 | 9.06E-07 | glio |
| Aars2_in.vivo_noRT | Poln | 3.824 | 9.06E-08 | glio |
| Chek1_in.vivo_noRT | A430106G13R | 4.252 | 9.06E-09 | glio |
| Atr_in.vivo_noRT | Bhlhe41 | 5.011 | 9.06E-10 | glio |
| culture_effect | Hspe1-rs1 | 0.548 | 9.07E-03 | glio |
| Interaction_genotype_Mtor | Cdca8 | 1.97 | 9.07E-03 | glio |
| Interaction_genotype_Rad51 | Gm19276 | -2.656 | 9.07E-03 | glio |
| Aars2_in.vivo_noRT | Trim62 | 1.61 | 9.07E-03 | glio |
| Cenpj_in.vivo_noRT | Serpinb2 | 2.022 | 9.07E-03 | glio |
| Cenpj_in.vivo_noRT | Id2 | -1.447 | 9.07E-03 | glio |
| Chek1_ex.vivo_noRT | Map2k3os | 1.733 | 9.07E-03 | glio |
| culture_effect | Asb11 | -1.823 | 9.07E-04 | glio |
| Interaction_genotype_Aars2 | Gm43701 | -3.016 | 9.07E-04 | glio |
| Interaction_genotype_Aars2 | Gm4221 | -4.168 | 9.07E-04 | glio |
| Interaction_genotype_Aars2 | Gm32090 | -3.538 | 9.07E-04 | glio |
| Interaction_genotype_Aars2 | Clcn1 | -3.341 | 9.07E-04 | glio |
| Interaction_genotype_Aars2 | Derl2 | 1.627 | 9.07E-04 | glio |
| Interaction_genotype_Aars2 | Gtf3c6 | 1.615 | 9.07E-04 | glio |
| Interaction_genotype_Aars2 | Nr2c2ap | 1.505 | 9.07E-04 | glio |
| Interaction_genotype_Bcas2 | Gm11973 | -3.562 | 9.07E-04 | glio |
| Interaction_genotype_Myc | 2310003N18R | -3.122 | 9.07E-04 | glio |
| Interaction_genotype_Myc | Ankrd23 | -3.986 | 9.07E-04 | glio |
| Interaction_genotype_Myc | Glt1d1 | -3.507 | 9.07E-04 | glio |
| Interaction_genotype_Myc | Calhm2 | -2.951 | 9.07E-04 | glio |
| Interaction_genotype_Myc | Ilf3 | -1.978 | 9.07E-04 | glio |
| Interaction_genotype_Myc | Mrps23 | 1.468 | 9.07E-04 | glio |
| Interaction_genotype_Myc | Mrpl20 | 1.407 | 9.07E-04 | glio |
| Aars2_in.vivo_noRT | Gm44190 | 2.539 | 9.07E-05 | glio |
| Atr_in.vivo_noRT | Gm36371 | 2.624 | 9.07E-05 | glio |
| Atr_in.vivo_noRT | Firre | 2.532 | 9.07E-05 | glio |
| Atr_ex.vivo_noRT | A930033M14F | 2.997 | 9.07E-05 | glio |
| Atr_ex.vivo_noRT | Gm33973 | 3.164 | 9.07E-05 | glio |
| Bcas2_in.vivo_noRT | Kcnb1 | 2.943 | 9.07E-05 | glio |
| Chek1_in.vivo_noRT | Gm49164 | 2.853 | 9.07E-05 | glio |
| Interaction_genotype_Ifnar1 | Nlrc5 | -3.704 | 9.07E-06 | glio |
| Interaction_genotype_Myc | Gm43701 | -3.711 | 9.07E-06 | glio |
| Hsd17b10_in.vivo_noRT | 7530414M10F | 3.09 | 9.07E-06 | glio |
| Atr_ex.vivo_noRT | Dpp6 | 3.164 | 9.07E-07 | glio |
| Bora_in.vivo_noRT | Enpp3 | 3.37 | 9.07E-07 | glio |
| Chek1_in.vivo_noRT | Apold1 | 3.934 | 9.07E-07 | glio |
| Aars2_in.vivo_noRT | Sv2a | 3.889 | 9.07E-08 | glio |
| Aars2_in.vivo_noRT | Ggt5 | 3.694 | 9.07E-09 | glio |

| | | | | |
|---|---|---|---|---|
| Bora_in.vivo_noRT | Insc | 3.479 | 9.07E-09 | glio |
| Atr_in.vivo_noRT | Nrip3 | 4.489 | 9.07E-12 | glio |
| Atr_in.vivo_noRT | Gm48772 | 4.626 | 9.07E-12 | glio |
| Chek1_in.vivo_noRT | Milr1 | 4.832 | 9.07E-12 | glio |
| Aars2_in.vivo_noRT | H1foo | 4.794 | 9.07E-21 | glio |
| Interaction_genotype_Mtor | Ly6c2 | -3.196 | 9.08E-03 | glio |
| Interaction_genotype_Mtor | Cdc42bpg | -2.884 | 9.08E-03 | glio |
| Interaction_genotype_Mtor | Hdhd3 | -2.712 | 9.08E-03 | glio |
| Interaction_genotype_Myc | Fam53b | -2.482 | 9.08E-03 | glio |
| Interaction_genotype_Myc | Mmab | -2.233 | 9.08E-03 | glio |
| Interaction_genotype_Myc | Prkd3 | -1.877 | 9.08E-03 | glio |
| Aars2_in.vivo_noRT | Gm13470 | 1.726 | 9.08E-03 | glio |
| Aars2_in.vivo_noRT | Slc25a28 | -0.843 | 9.08E-03 | glio |
| Atr_in.vivo_noRT | Ccdc189 | 1.889 | 9.08E-03 | glio |
| Bcas2_in.vivo_noRT | Fam53b | 1.655 | 9.08E-03 | glio |
| Bora_in.vivo_noRT | Phf23 | -0.967 | 9.08E-03 | glio |
| Bora_in.vivo_noRT | Fbxo22 | 0.958 | 9.08E-03 | glio |
| Chek1_in.vivo_noRT | Xrn1 | -1.273 | 9.08E-03 | glio |
| Chek1_in.vivo_noRT | Ctbs | 1.171 | 9.08E-03 | glio |
| Interaction_genotype_Mtor | Prdx4 | 1.857 | 9.08E-04 | glio |
| Interaction_genotype_Aars2 | Gm9732 | 2.273 | 9.08E-05 | glio |
| Interaction_genotype_Atr | Mettl18 | -4.155 | 9.08E-05 | glio |
| Interaction_genotype_Atr | Gm48882 | -4.946 | 9.08E-05 | glio |
| Interaction_genotype_Myc | A530083I20Ri | -5.517 | 9.08E-06 | glio |
| Interaction_genotype_Myc | Eif3l | 2.129 | 9.08E-06 | glio |
| culture_effect | Ppat | -0.759 | 9.08E-07 | glio |
| culture_effect | Col11a2 | -1.485 | 9.08E-21 | glio |
| culture_effect | Ccdc134 | -0.882 | 9.09E-03 | glio |
| Interaction_genotype_Aars2 | Mrpl10 | 1.762 | 9.09E-03 | glio |
| Interaction_genotype_Myc | Slc22a21 | -2.984 | 9.09E-03 | glio |
| Aars2_in.vivo_noRT | Sag | 1.948 | 9.09E-03 | glio |
| Aars2_in.vivo_noRT | Atp9b | 1.336 | 9.09E-03 | glio |
| Atr_in.vivo_noRT | Txnrd2 | 1.594 | 9.09E-03 | glio |
| Bcas2_in.vivo_noRT | Selenok | -0.628 | 9.09E-03 | glio |
| Eif2s1_in.vivo_noRT | BB557941 | 1.883 | 9.09E-03 | glio |
| Eif2s1_in.vivo_noRT | Zmynd10 | 1.883 | 9.09E-03 | glio |
| Interaction_genotype_Aars2 | Gm10501 | -4.328 | 9.09E-04 | glio |
| Interaction_genotype_Aars2 | Gm34047 | -4.111 | 9.09E-04 | glio |
| Interaction_genotype_Aars2 | Gm15952 | -4.153 | 9.09E-04 | glio |
| Interaction_genotype_Aars2 | Dnm3 | -3.375 | 9.09E-04 | glio |
| Interaction_genotype_Aars2 | Parp1 | 1.628 | 9.09E-04 | glio |
| Interaction_genotype_Myc | Zscan4-ps2 | -2.729 | 9.09E-04 | glio |
| Interaction_genotype_Myc | Osgin1 | -3.504 | 9.09E-04 | glio |
| Interaction_genotype_Myc | Gm19391 | -3.091 | 9.09E-04 | glio |
| Atr_in.vivo_noRT | Zfpm2 | 2.318 | 9.09E-04 | glio |
| Atr_ex.vivo_noRT | Sap30bpos | 2.77 | 9.09E-04 | glio |
| Bora_in.vivo_noRT | Tra2b | -1.179 | 9.09E-04 | glio |
| Brca2_ex.vivo_noRT | Gm9758 | 2.82 | 9.09E-04 | glio |
| Brca2_ex.vivo_noRT | Gm48215 | 2.825 | 9.09E-04 | glio |
| Brca2_ex.vivo_noRT | Olfr56 | 2.82 | 9.09E-04 | glio |
| Interaction_genotype_Myc | Col5a3 | -3.377 | 9.09E-05 | glio |
| Bcas2_in.vivo_noRT | Msmo1 | -1.804 | 9.09E-05 | glio |

| | | | | |
|---|---|---|---|---|
| Chek1_in.vivo_noRT | Gm17056 | 2.84 | 9.09E-05 | glio |
| Chek1_in.vivo_noRT | Nsun7 | 3.034 | 9.09E-05 | glio |
| Chek1_in.vivo_noRT | 1700123O20R | -1.644 | 9.09E-05 | glio |
| Atp5c1_in.vivo_noRT | Pitx3 | 2.668 | 9.09E-06 | glio |
| Bora_ex.vivo_noRT | Ptprc | 3.201 | 9.09E-06 | glio |
| Bora_ex.vivo_noRT | Ggt1 | 3.994 | 9.09E-06 | glio |
| Aars2_in.vivo_noRT | Arhgdib | 3.442 | 9.09E-07 | glio |
| Aars2_in.vivo_noRT | Gm30541 | 3.631 | 9.09E-08 | glio |
| Aars2_in.vivo_noRT | Rpl3l | 3.39 | 9.09E-08 | glio |
| Bora_in.vivo_noRT | Gm36026 | 4.285 | 9.09E-14 | glio |
| Bora_in.vivo_noRT | Ube4bos1 | 4.672 | 9.09E-16 | glio |
| Atr_in.vivo_noRT | Bcl2l15 | 5.273 | 9.09E-18 | glio |
| Bora_in.vivo_noRT | Gm4027 | 4.936 | 9.09E-19 | glio |
| Interaction_genotype_Myc | Slc7a3 | -3.027 | 9.10E-03 | glio |
| Interaction_genotype_Myc | Uhrf1bp1 | -2.233 | 9.10E-03 | glio |
| Interaction_genotype_Myc | Churc1 | 1.28 | 9.10E-03 | glio |
| Interaction_genotype_Rad21 | Map7 | 3.86 | 9.10E-03 | glio |
| interaction_RT | Tmem218 | 1.414 | 9.10E-03 | glio |
| Aars2_in.vivo_noRT | BC049715 | 1.79 | 9.10E-03 | glio |
| Chek1_in.vivo_noRT | Zfp689 | 1.556 | 9.10E-03 | glio |
| Chek1_in.vivo_noRT | Sirt6 | -1.341 | 9.10E-03 | glio |
| Cyp19a1_ex.vivo_noRT | AU019990 | 2.159 | 9.10E-03 | glio |
| Cyp19a1_ex.vivo_noRT | Gm48780 | 2.159 | 9.10E-03 | glio |
| interaction_RT | Pak3 | 1.079 | 9.10E-04 | glio |
| Atr_in.vivo_noRT | Snu13 | -0.651 | 9.10E-04 | glio |
| Chek1_in.vivo_noRT | Enho | 2.385 | 9.10E-04 | glio |
| Chek1_in.vivo_noRT | Osgin1 | 2.208 | 9.10E-04 | glio |
| Chek1_in.vivo_noRT | Srp68 | -1.281 | 9.10E-04 | glio |
| culture_effect | Kdm5a | -0.589 | 9.10E-05 | glio |
| Interaction_genotype_Chek1 | Txnip | 6.096 | 9.10E-05 | glio |
| Bcas2_in.vivo_noRT | Gm12689 | 3.098 | 9.10E-05 | glio |
| culture_effect | Rpgrip1 | 0.9 | 9.10E-06 | glio |
| Aars2_in.vivo_noRT | Kcnip3 | 2.852 | 9.10E-06 | glio |
| Aars2_in.vivo_noRT | Ube2l3 | -1.333 | 9.10E-06 | glio |
| Chek1_ex.vivo_noRT | Gm8300 | 4.842 | 9.10E-06 | glio |
| Bcas2_in.vivo_noRT | Gm47223 | 4.541 | 9.10E-07 | glio |
| Bcas2_in.vivo_noRT | Ankef1 | 4.068 | 9.10E-09 | glio |
| Aars2_in.vivo_noRT | Mag | 4.018 | 9.10E-10 | glio |
| Bora_in.vivo_noRT | Alpl | 4.811 | 9.10E-13 | glio |
| Chek1_in.vivo_noRT | 4930505N22R | 4.784 | 9.10E-14 | glio |
| Atr_in.vivo_noRT | 6430562O15R | 5.011 | 9.10E-16 | glio |
| culture_effect | Pdcd11 | 0.424 | 9.11E-03 | glio |
| Interaction_genotype_Hsd17b | Thap8 | -4.24 | 9.11E-03 | glio |
| Interaction_genotype_Mtor | Tmem8b | -3.071 | 9.11E-03 | glio |
| Interaction_genotype_Myc | Ift172 | -2.485 | 9.11E-03 | glio |
| Interaction_genotype_Myc | Itga5 | -2.345 | 9.11E-03 | glio |
| Interaction_genotype_Myc | Ttc5 | -2.05 | 9.11E-03 | glio |
| Interaction_genotype_Myc | Cdk13 | -1.643 | 9.11E-03 | glio |
| Interaction_genotype_Myc | Caprin1 | 1.096 | 9.11E-03 | glio |
| Aars2_in.vivo_noRT | Hdac2 | -0.858 | 9.11E-03 | glio |
| Chek1_ex.vivo_noRT | Cisd1 | -0.739 | 9.11E-03 | glio |
| Hsd17b10_in.vivo_noRT | 1700055D18R | 2.226 | 9.11E-03 | glio |

| | | | | |
|---|---|---|---|---|
| Hsd17b10_in.vivo_noRT | Kcnab3 | 2.082 | 9.11E-03 | glio |
| culture_effect | Gm32585 | 2.013 | 9.11E-04 | glio |
| Interaction_genotype_Atr | Plcb4 | -4.177 | 9.11E-04 | glio |
| Interaction_genotype_Mtor | Vmn1r181 | -3.838 | 9.11E-04 | glio |
| Interaction_genotype_Mtor | Ramp1 | -4.658 | 9.11E-04 | glio |
| Interaction_genotype_Mtor | Lclat1 | -2.292 | 9.11E-04 | glio |
| Interaction_genotype_Myc | Mrpl57 | 1.547 | 9.11E-04 | glio |
| Bora_in.vivo_noRT | Tnni1 | 1.747 | 9.11E-04 | glio |
| Bcas2_in.vivo_noRT | Gm48086 | 2.859 | 9.11E-05 | glio |
| Aars2_in.vivo_noRT | Gm43689 | 2.861 | 9.11E-06 | glio |
| Aars2_in.vivo_noRT | Itgae | 3.21 | 9.11E-06 | glio |
| Chek1_in.vivo_noRT | Dennd2d | 2.777 | 9.11E-06 | glio |
| Aars2_in.vivo_noRT | Stpg2 | 3.378 | 9.11E-07 | glio |
| Chek1_in.vivo_noRT | C430042M11F | 4.038 | 9.11E-09 | glio |
| Atr_in.vivo_noRT | Pcdhga1 | 4.482 | 9.11E-14 | glio |
| Hsd17b10_in.vivo_noRT | Cldn2 | 3.616 | 9.11E-15 | glio |
| Bcas2_in.vivo_noRT | Gm13178 | 4.291 | 9.11E-17 | glio |
| Chek1_in.vivo_noRT | Lrguk | 4.78 | 9.11E-17 | glio |
| Chek1_ex.vivo_noRT | Cldn2 | 4.842 | 9.11E-18 | glio |
| Interaction_genotype_Bora | Gm44096 | -3.33 | 9.12E-03 | glio |
| Interaction_genotype_Chek1 | Gm32090 | -3.299 | 9.12E-03 | glio |
| Interaction_genotype_Chek1 | Dhrs13 | -2.496 | 9.12E-03 | glio |
| Atr_in.vivo_noRT | Epor | 1.831 | 9.12E-03 | glio |
| Atr_ex.vivo_noRT | Pcdhga11 | 2.193 | 9.12E-03 | glio |
| Bard1_ex.vivo_noRT | Pcdhgb5 | 2.421 | 9.12E-03 | glio |
| Bcas2_in.vivo_noRT | Slc7a1 | -1.191 | 9.12E-03 | glio |
| Brca2_in.vivo_noRT | Cd36 | 1.763 | 9.12E-03 | glio |
| Brca2_in.vivo_noRT | Gm43701 | 1.763 | 9.12E-03 | glio |
| Brca2_in.vivo_noRT | 9130019O22R | 1.763 | 9.12E-03 | glio |
| Chek1_in.vivo_noRT | Abhd16a | -1.147 | 9.12E-03 | glio |
| Interaction_genotype_Atr | Gjb4 | -4.238 | 9.12E-04 | glio |
| Atr_in.vivo_noRT | Iars | -1.241 | 9.12E-04 | glio |
| Cenpj_in.vivo_noRT | Tex52 | 2.67 | 9.12E-04 | glio |
| culture_effect | Gde1 | 0.631 | 9.12E-05 | glio |
| interaction_RT | Snhg8 | 1.021 | 9.12E-05 | glio |
| interaction_RT | Tnc | 3.983 | 9.12E-05 | glio |
| Atr_in.vivo_noRT | Ldlrad3 | 2.528 | 9.12E-05 | glio |
| Aars2_in.vivo_noRT | Gm38414 | 3.456 | 9.12E-06 | glio |
| Chek1_ex.vivo_noRT | Egr2 | 3.758 | 9.12E-06 | glio |
| Interaction_genotype_Mtor | Prox2 | -6.226 | 9.12E-07 | glio |
| Chek1_in.vivo_noRT | Cacng5 | 3.193 | 9.12E-07 | glio |
| Chek1_ex.vivo_noRT | Bpifa3 | 4.842 | 9.12E-11 | glio |
| Bora_in.vivo_noRT | Gm48742 | 4.286 | 9.12E-14 | glio |
| culture_effect | Hpse | -1.68 | 9.13E-03 | glio |
| culture_effect | Pax3 | 1.013 | 9.13E-03 | glio |
| Interaction_genotype_Atr | Nectin3 | -2.372 | 9.13E-03 | glio |
| Interaction_genotype_Mtor | Pltp | -2.886 | 9.13E-03 | glio |
| Interaction_genotype_Mtor | Adgra3 | 1.623 | 9.13E-03 | glio |
| Interaction_genotype_Myc | Rab27a | -2.599 | 9.13E-03 | glio |
| Interaction_genotype_Myc | Dennd3 | -3.125 | 9.13E-03 | glio |
| Interaction_genotype_Myc | Tbca | 0.784 | 9.13E-03 | glio |
| Interaction_genotype_Rad51 | Chpt1 | 1.615 | 9.13E-03 | glio |

| | | | | |
|---|---|---|---|---|
| Interaction_genotype_Rpl3 | Gm31518 | -3.75 | 9.13E-03 | glio |
| Interaction_genotype_Rpl3 | BC037704 | -3.465 | 9.13E-03 | glio |
| interaction_RT | Pyroxd2 | 1.937 | 9.13E-03 | glio |
| interaction_RT | Apod | 2.029 | 9.13E-03 | glio |
| Aars2_in.vivo_noRT | Dnajb11 | -1.009 | 9.13E-03 | glio |
| Bcas2_in.vivo_noRT | Gm12940 | 1.665 | 9.13E-03 | glio |
| Bora_in.vivo_noRT | Rnf113a1 | 1.534 | 9.13E-03 | glio |
| Chek1_in.vivo_noRT | Bag1 | -0.662 | 9.13E-03 | glio |
| culture_effect | Utp14b | -1.128 | 9.13E-04 | glio |
| Atr_in.vivo_noRT | Zfp112 | 2.313 | 9.13E-04 | glio |
| Bcas2_in.vivo_noRT | Carmil1 | 2.277 | 9.13E-04 | glio |
| Chek1_in.vivo_noRT | Galk2 | 1.659 | 9.13E-04 | glio |
| Hsd17b10_in.vivo_noRT | Lrrc20 | 2.245 | 9.13E-04 | glio |
| Aars2_in.vivo_noRT | Kctd11 | 2.855 | 9.13E-05 | glio |
| Bora_in.vivo_noRT | Mcph1 | -1.757 | 9.13E-05 | glio |
| Chek1_in.vivo_noRT | Daam2 | 2.871 | 9.13E-05 | glio |
| Hsd17b10_in.vivo_noRT | 1700040D17R | 3.083 | 9.13E-05 | glio |
| culture_effect | Anxa1 | -0.69 | 9.13E-06 | glio |
| Atr_ex.vivo_noRT | Ptprc | 3.164 | 9.13E-06 | glio |
| culture_effect | Fbxo9 | 0.982 | 9.13E-07 | glio |
| Atr_in.vivo_noRT | E230025N22R | 3.186 | 9.13E-07 | glio |
| Atr_ex.vivo_noRT | Serpinb9g | 3.164 | 9.13E-07 | glio |
| Atr_ex.vivo_noRT | BC049987 | 3.164 | 9.13E-07 | glio |
| Bcas2_in.vivo_noRT | Limch1 | 3.769 | 9.13E-07 | glio |
| Aars2_in.vivo_noRT | Gm26766 | 3.631 | 9.13E-08 | glio |
| Chek1_in.vivo_noRT | Fsd2 | 3.661 | 9.13E-08 | glio |
| Aars2_in.vivo_noRT | 5430400D12R | 4.045 | 9.13E-09 | glio |
| culture_effect | Dnajb9 | -0.569 | 9.14E-03 | glio |
| Interaction_genotype_Bora | 1700024I08Ri | -2.849 | 9.14E-03 | glio |
| Bcas2_in.vivo_noRT | Nudt7 | 1.833 | 9.14E-03 | glio |
| Bcas2_ex.vivo_noRT | Tmem35b | 2.238 | 9.14E-03 | glio |
| Chek1_ex.vivo_noRT | Tctn1 | 2.516 | 9.14E-03 | glio |
| Chek1_ex.vivo_noRT | Gm28535 | 2.467 | 9.14E-03 | glio |
| Chek1_ex.vivo_noRT | Lsm5 | -0.792 | 9.14E-03 | glio |
| Eif2s1_in.vivo_noRT | Dok3 | 2.31 | 9.14E-03 | glio |
| Hsd17b10_in.vivo_noRT | Poln | 2.113 | 9.14E-03 | glio |
| culture_effect | Hfe | -1.505 | 9.14E-04 | glio |
| Chek1_in.vivo_noRT | Ece2 | 2.275 | 9.14E-04 | glio |
| Hsd17b10_in.vivo_noRT | Rnft2 | 2.813 | 9.14E-05 | glio |
| Interaction_genotype_Myc | Tmem74b | -4.229 | 9.14E-06 | glio |
| Atr_in.vivo_noRT | Mfsd7a | 3.927 | 9.14E-07 | glio |
| Bcas2_in.vivo_noRT | Spata5 | -2.161 | 9.14E-07 | glio |
| Bora_ex.vivo_noRT | Slc17a6 | 3.201 | 9.14E-09 | glio |
| Bora_in.vivo_noRT | Thap8 | 4.676 | 9.14E-11 | glio |
| Hsd17b10_in.vivo_noRT | Slc16a5 | 3.352 | 9.14E-12 | glio |
| Bora_in.vivo_noRT | Pdlim3 | 4.157 | 9.14E-13 | glio |
| culture_effect | Polr1b | -0.792 | 9.15E-03 | glio |
| Interaction_genotype_Chek1 | 1810013L24Ri | 2.01 | 9.15E-03 | glio |
| Interaction_genotype_Myc | Gm16279 | -3.147 | 9.15E-03 | glio |
| Interaction_genotype_Myc | Wfdc13 | -1.778 | 9.15E-03 | glio |
| Interaction_genotype_Terf1 | 1110046J04Ri | -4.039 | 9.15E-03 | glio |
| Atr_in.vivo_noRT | Entpd5 | 1.806 | 9.15E-03 | glio |

| | | | | |
|---|---|---|---|---|
| Bcas2_in.vivo_noRT | Dcun1d5 | -0.719 | 9.15E-03 | glio |
| Bcas2_in.vivo_noRT | AI662270 | -0.706 | 9.15E-03 | glio |
| Bcas2_ex.vivo_noRT | Dock8 | -2.136 | 9.15E-03 | glio |
| Bcas2_ex.vivo_noRT | Grcc10 | -0.722 | 9.15E-03 | glio |
| Bora_in.vivo_noRT | Rab18 | -0.816 | 9.15E-03 | glio |
| Chek1_in.vivo_noRT | Dnajb5 | 1.713 | 9.15E-03 | glio |
| Chek1_ex.vivo_noRT | Ywhae | -0.564 | 9.15E-03 | glio |
| Hsd17b10_in.vivo_noRT | Tnfrsf4 | 2.03 | 9.15E-03 | glio |
| Hsd17b10_in.vivo_noRT | Gm4755 | 1.859 | 9.15E-03 | glio |
| Interaction_genotype_Aars2 | Gm15743 | -3.828 | 9.15E-04 | glio |
| Interaction_genotype_Myc | Cdkn1c | -3.761 | 9.15E-04 | glio |
| culture_effect | C230037L18Ri | 2.034 | 9.15E-05 | glio |
| Aars2_in.vivo_noRT | Snf8 | -1.203 | 9.15E-05 | glio |
| Bora_in.vivo_noRT | Spry2 | -2.37 | 9.15E-05 | glio |
| culture_effect | Chmp2a | -0.556 | 9.15E-06 | glio |
| Aars2_in.vivo_noRT | Gm13830 | 3.396 | 9.15E-06 | glio |
| Atr_in.vivo_noRT | 2310058D17R | 4.354 | 9.15E-10 | glio |
| Atr_in.vivo_noRT | Cdkal1 | 1.223 | 9.16E-03 | glio |
| Atr_ex.vivo_noRT | Tmem190 | 2.81 | 9.16E-03 | glio |
| Bcas2_in.vivo_noRT | Tpcn1 | 1.6 | 9.16E-03 | glio |
| Bora_in.vivo_noRT | Nf1 | 1.046 | 9.16E-03 | glio |
| Bora_ex.vivo_noRT | 5730507C01R | 2.01 | 9.16E-03 | glio |
| Cenpj_in.vivo_noRT | Fam166a | 1.758 | 9.16E-03 | glio |
| Chek1_ex.vivo_noRT | Gm16244 | 2.471 | 9.16E-03 | glio |
| Cyp19a1_ex.vivo_noRT | Gm36793 | 2.159 | 9.16E-03 | glio |
| Interaction_genotype_Aars2 | 4833418N02R | -4.268 | 9.16E-04 | glio |
| Interaction_genotype_Aars2 | Gm14207 | -2.827 | 9.16E-04 | glio |
| Interaction_genotype_Aars2 | Unc50 | 1.714 | 9.16E-04 | glio |
| Interaction_genotype_Myc | A930029G22R | -3.238 | 9.16E-04 | glio |
| Chek1_in.vivo_noRT | 9030025P20R | 2.571 | 9.16E-04 | glio |
| Chek1_in.vivo_noRT | Srek1 | -1.232 | 9.16E-04 | glio |
| Chek1_ex.vivo_noRT | Wdr46 | -1.74 | 9.16E-04 | glio |
| Hsd17b10_in.vivo_noRT | Gm26766 | 2.447 | 9.16E-04 | glio |
| culture_effect | Cyp51 | -0.767 | 9.16E-05 | glio |
| Interaction_genotype_Nat2 | Tcstv1 | 4.218 | 9.16E-05 | glio |
| Aars2_ex.vivo_noRT | Rslcan18 | 3.282 | 9.16E-05 | glio |
| Bora_ex.vivo_noRT | 4930527F14Ri | 2.806 | 9.16E-05 | glio |
| Bcas2_in.vivo_noRT | Gpr21 | 2.942 | 9.16E-06 | glio |
| Atr_in.vivo_noRT | Proscos | 3.638 | 9.16E-07 | glio |
| culture_effect | Zfp932 | 1.315 | 9.16E-08 | glio |
| Bcas2_in.vivo_noRT | Calhm5 | 4.079 | 9.16E-10 | glio |
| Chek1_ex.vivo_noRT | Gm31063 | 3.918 | 9.16E-10 | glio |
| Atr_in.vivo_noRT | Selp | 4.489 | 9.16E-11 | glio |
| Aars2_in.vivo_noRT | Gm15816 | 5.327 | 9.16E-18 | glio |
| Interaction_genotype_Mtor | Nr2c2ap | 1.35 | 9.17E-03 | glio |
| Interaction_genotype_Myc | Pde6h | -2.501 | 9.17E-03 | glio |
| Interaction_genotype_Myc | Vkorc1l1 | -1.725 | 9.17E-03 | glio |
| Interaction_genotype_Rad51 | B3galt6 | -2.838 | 9.17E-03 | glio |
| Interaction_genotype_Uqcrc2 | D830039M14I | -4.091 | 9.17E-03 | glio |
| Interaction_genotype_Uqcrc2 | Akap3 | -4.031 | 9.17E-03 | glio |
| interaction_RT | Serpinb2 | 2.337 | 9.17E-03 | glio |
| interaction_RT | Cox6c | 0.525 | 9.17E-03 | glio |

| | | | | |
|---|---|---|---|---|
| Aars2_ex.vivo_noRT | Pcdhga1 | 2.324 | 9.17E-03 | glio |
| Bcas2_in.vivo_noRT | Marf1 | 1.231 | 9.17E-03 | glio |
| Bora_in.vivo_noRT | Senp2 | -1.311 | 9.17E-03 | glio |
| Chek1_in.vivo_noRT | Tmem246 | 1.81 | 9.17E-03 | glio |
| Chek1_ex.vivo_noRT | Rpl27 | -0.692 | 9.17E-03 | glio |
| Fanca_ex.vivo_noRT | Zfp143 | -1.919 | 9.17E-03 | glio |
| Hsd17b10_in.vivo_noRT | Gm15492 | 2.08 | 9.17E-03 | glio |
| culture_effect | Calr | -0.488 | 9.17E-04 | glio |
| Interaction_genotype_Aars2 | 1700026N04R | -3.696 | 9.17E-04 | glio |
| interaction_RT | Ftl1 | 0.568 | 9.17E-04 | glio |
| Aars2_in.vivo_noRT | Rbm27 | -1.343 | 9.17E-04 | glio |
| Aars2_ex.vivo_noRT | Gm48170 | 2.495 | 9.17E-04 | glio |
| Aars2_ex.vivo_noRT | Bco2 | 2.732 | 9.17E-04 | glio |
| Bcas2_ex.vivo_noRT | Slc6a11 | 2.742 | 9.17E-04 | glio |
| Bcas2_ex.vivo_noRT | 1700047M11F | 3.131 | 9.17E-04 | glio |
| Bora_ex.vivo_noRT | Lrrc24 | 3.008 | 9.17E-04 | glio |
| Bora_ex.vivo_noRT | H2-DMb2 | 3.201 | 9.17E-04 | glio |
| Chek1_in.vivo_noRT | Zfp112 | 2.347 | 9.17E-04 | glio |
| culture_effect | 1700052K11R | 1.291 | 9.17E-05 | glio |
| culture_effect | Scamp5 | -0.666 | 9.17E-05 | glio |
| Aars2_in.vivo_noRT | A230087F16Ri | 2.703 | 9.17E-05 | glio |
| Bora_in.vivo_noRT | Afg1l | 2.856 | 9.17E-05 | glio |
| Aars2_in.vivo_noRT | Tomm20 | -0.912 | 9.17E-06 | glio |
| Atr_in.vivo_noRT | Mlip | 3.066 | 9.17E-06 | glio |
| Bora_in.vivo_noRT | Farsa | -1.786 | 9.17E-06 | glio |
| Chek1_in.vivo_noRT | Klhdc8b | 3.23 | 9.17E-06 | glio |
| Eif2s1_ex.vivo_noRT | Gm15138 | 3.419 | 9.17E-06 | glio |
| Interaction_genotype_Mtor | Halr1 | -3.833 | 9.17E-07 | glio |
| Hsd17b10_in.vivo_noRT | H3f3aos | 3.498 | 9.17E-07 | glio |
| Atr_in.vivo_noRT | Gm45441 | 4.474 | 9.17E-11 | glio |
| Atr_in.vivo_noRT | Prox2 | 4.807 | 9.17E-11 | glio |
| Bora_in.vivo_noRT | AC154200.1 | 4.936 | 9.17E-19 | glio |
| Interaction_genotype_Atr | Ifi214 | -3.314 | 9.18E-03 | glio |
| Interaction_genotype_Bora | 2310058D17R | -3.395 | 9.18E-03 | glio |
| Interaction_genotype_Mtor | Anxa8 | -3.167 | 9.18E-03 | glio |
| Interaction_genotype_Mtor | Clic4 | 1.463 | 9.18E-03 | glio |
| Interaction_genotype_Myc | Mroh2b | -3.058 | 9.18E-03 | glio |
| Interaction_genotype_Myc | Khnyn | -2.604 | 9.18E-03 | glio |
| interaction_RT | Tnfaip6 | 1.114 | 9.18E-03 | glio |
| Atr_in.vivo_noRT | Sipa1 | 1.524 | 9.18E-03 | glio |
| Bcas2_in.vivo_noRT | B4galt4 | 1.995 | 9.18E-03 | glio |
| Bcas2_in.vivo_noRT | Pars2 | 1.836 | 9.18E-03 | glio |
| Bcas2_ex.vivo_noRT | 1700006H21R | 2.737 | 9.18E-03 | glio |
| Bora_in.vivo_noRT | Cog6 | 1.346 | 9.18E-03 | glio |
| Chek1_in.vivo_noRT | Cish | 1.866 | 9.18E-03 | glio |
| Cyp19a1_ex.vivo_noRT | Vtn | 2.159 | 9.18E-03 | glio |
| Cyp19a1_ex.vivo_noRT | Spink8 | 2.159 | 9.18E-03 | glio |
| Cyp19a1_ex.vivo_noRT | Psmb11 | 2.159 | 9.18E-03 | glio |
| Interaction_genotype_Bcas2 | Ist1 | 2.272 | 9.18E-04 | glio |
| Interaction_genotype_Mtor | Lrrc28 | -3.154 | 9.18E-04 | glio |
| Interaction_genotype_Myc | Exoc4 | -2.181 | 9.18E-04 | glio |
| Aars2_ex.vivo_noRT | Gm813 | 3.293 | 9.18E-04 | glio |

| | | | | |
|---|---|---|---|---|
| Atr_in.vivo_noRT | Wfs1 | 2.31 | 9.18E-04 | glio |
| Atr_in.vivo_noRT | Gm30025 | 1.788 | 9.18E-04 | glio |
| Atr_ex.vivo_noRT | Gm14540 | 2.37 | 9.18E-04 | glio |
| Atr_ex.vivo_noRT | Acod1 | 3.947 | 9.18E-04 | glio |
| Chek1_in.vivo_noRT | 1700123M08F | 2.391 | 9.18E-04 | glio |
| Chek1_in.vivo_noRT | Gm29994 | 2.166 | 9.18E-04 | glio |
| Chek1_in.vivo_noRT | Platr16 | 2.567 | 9.18E-04 | glio |
| Chek1_in.vivo_noRT | Dmpk | 2.31 | 9.18E-04 | glio |
| Chek1_in.vivo_noRT | Sf3b6 | -0.704 | 9.18E-04 | glio |
| culture_effect | Crnde | 0.975 | 9.18E-05 | glio |
| culture_effect | Gjc1 | 0.945 | 9.18E-05 | glio |
| Interaction_genotype_Aars2 | Eif4ebp1 | 1.727 | 9.18E-05 | glio |
| Aars2_ex.vivo_noRT | 1700034I23Ri | 2.495 | 9.18E-05 | glio |
| culture_effect | 4430402I18Ri | -2.371 | 9.18E-06 | glio |
| culture_effect | App | -0.573 | 9.18E-06 | glio |
| Atr_in.vivo_noRT | Fxyd4 | 3.418 | 9.18E-06 | glio |
| culture_effect | Prkci | 1.464 | 9.18E-08 | glio |
| Atr_in.vivo_noRT | Ubc | -1.869 | 9.18E-08 | glio |
| Bora_in.vivo_noRT | Pdzk1ip1 | 3.946 | 9.18E-08 | glio |
| Aars2_in.vivo_noRT | Cbs | 4.137 | 9.18E-09 | glio |
| culture_effect | AC168977.1 | -4.128 | 9.18E-11 | glio |
| Bora_in.vivo_noRT | Oit3 | 4.936 | 9.18E-14 | glio |
| culture_effect | Cbr3 | 1.53 | 9.19E-03 | glio |
| Interaction_genotype_Atr | Gm26978 | -2.696 | 9.19E-03 | glio |
| Interaction_genotype_Bora | Gm42670 | -3.115 | 9.19E-03 | glio |
| Interaction_genotype_Mtor | Gm16196 | -2.545 | 9.19E-03 | glio |
| Interaction_genotype_Mtor | Sbno1 | 1.391 | 9.19E-03 | glio |
| Interaction_genotype_Myc | Homez | -3.011 | 9.19E-03 | glio |
| Aars2_in.vivo_noRT | Ifit3 | -1.687 | 9.19E-03 | glio |
| Aars2_in.vivo_noRT | Brk1 | -0.606 | 9.19E-03 | glio |
| Atr_in.vivo_noRT | Crlf1 | 1.406 | 9.19E-03 | glio |
| Bcas2_ex.vivo_noRT | Gm13684 | 2.348 | 9.19E-03 | glio |
| Bora_in.vivo_noRT | Rtn4ip1 | 1.618 | 9.19E-03 | glio |
| Chek1_in.vivo_noRT | Znhit6 | -1.055 | 9.19E-03 | glio |
| Eif2s1_in.vivo_noRT | Gm31415 | 1.877 | 9.19E-03 | glio |
| Interaction_genotype_Atr | Gm48682 | -3.783 | 9.19E-04 | glio |
| Interaction_genotype_Bcas2 | Smim8 | 1.869 | 9.19E-04 | glio |
| Aars2_in.vivo_noRT | Ripor2 | 2.351 | 9.19E-04 | glio |
| Bcas2_ex.vivo_noRT | Gm12841 | 2.349 | 9.19E-04 | glio |
| Hsd17b10_in.vivo_noRT | BC037032 | 2.698 | 9.19E-04 | glio |
| Interaction_genotype_Mtor | Wdr5b | -4.467 | 9.19E-05 | glio |
| Bcas2_ex.vivo_noRT | Capn13 | 2.742 | 9.19E-06 | glio |
| Chek1_ex.vivo_noRT | Grm1 | 3.467 | 9.19E-06 | glio |
| Chek1_ex.vivo_noRT | Gm31152 | 3.468 | 9.19E-06 | glio |
| Chek1_ex.vivo_noRT | Ascl4 | 4.066 | 9.19E-06 | glio |
| Atr_in.vivo_noRT | Rhbdf2 | 3.213 | 9.19E-07 | glio |
| Atr_in.vivo_noRT | Amd2 | 3.355 | 9.19E-07 | glio |
| Bora_ex.vivo_noRT | Cd200r2 | 3.201 | 9.19E-07 | glio |
| culture_effect | Anxa5 | 0.782 | 9.19E-08 | glio |
| Aars2_in.vivo_noRT | Ccdc184 | 3.993 | 9.19E-08 | glio |
| Bora_in.vivo_noRT | Cyp1a1 | 4.936 | 9.19E-08 | glio |
| culture_effect | Ppid | -0.742 | 9.19E-09 | glio |

| | | | | |
|---|---|---|---|---|
| culture_effect | Fam53a | 1.089 | 9.19E-10 | glio |
| Bora_in.vivo_noRT | Gm48215 | 4.936 | 9.19E-21 | glio |
| Aars2_in.vivo_noRT | Dmrtc1a | 5.327 | 9.19E-23 | glio |
| Aars2_in.vivo_noRT | Gm12589 | 5.327 | 9.19E-23 | glio |
| culture_effect | Il12rb1 | -6.005 | 9.19E-28 | glio |
| culture_effect | Taf1 | -0.384 | 9.20E-03 | glio |
| Interaction_genotype_Aars2 | Gm19426 | -2.687 | 9.20E-03 | glio |
| Interaction_genotype_Bcas2 | Ttbk1 | -3.372 | 9.20E-03 | glio |
| Interaction_genotype_Bora | Gm11273 | -2.95 | 9.20E-03 | glio |
| Interaction_genotype_Bora | Triobp | 1.893 | 9.20E-03 | glio |
| Interaction_genotype_Lmo7 | Gm20635 | -3.974 | 9.20E-03 | glio |
| Interaction_genotype_Lmo7 | A530083I20Ri | -4.479 | 9.20E-03 | glio |
| Interaction_genotype_Lmo7 | Tcea2 | -3.996 | 9.20E-03 | glio |
| Interaction_genotype_Mtor | Tesc | 2.094 | 9.20E-03 | glio |
| Interaction_genotype_Mtor | Arf4 | 1.211 | 9.20E-03 | glio |
| Interaction_genotype_Myc | Timm8a1 | 1.418 | 9.20E-03 | glio |
| Interaction_genotype_Pars2 | Tmem74b | -2.982 | 9.20E-03 | glio |
| Aars2_in.vivo_noRT | Atg3 | -1.176 | 9.20E-03 | glio |
| Aars2_in.vivo_noRT | Suclg1 | -0.858 | 9.20E-03 | glio |
| Hsd17b10_in.vivo_noRT | AW046200 | 2.079 | 9.20E-03 | glio |
| Hsd17b10_in.vivo_noRT | Tmem91 | 2.1 | 9.20E-03 | glio |
| interaction_RT | Poldip3 | -0.836 | 9.20E-04 | glio |
| culture_effect | Rab30 | 1.534 | 9.20E-05 | glio |
| culture_effect | Fkbp1a | 0.375 | 9.20E-05 | glio |
| Interaction_genotype_Ifnar1 | Gjd4 | 3.531 | 9.20E-05 | glio |
| Bcas2_in.vivo_noRT | Zbtb46 | 2.968 | 9.20E-05 | glio |
| Bcas2_in.vivo_noRT | Timm50 | -1.499 | 9.20E-05 | glio |
| Bora_in.vivo_noRT | Gm4221 | 2.832 | 9.20E-05 | glio |
| Bora_in.vivo_noRT | Npl | 2.64 | 9.20E-05 | glio |
| Hsd17b10_in.vivo_noRT | Tnfsf13os | 2.845 | 9.20E-05 | glio |
| Hsd17b10_in.vivo_noRT | Gm11884 | 2.617 | 9.20E-05 | glio |
| Bcas2_in.vivo_noRT | Gm50322 | 3.429 | 9.20E-06 | glio |
| Hsd17b10_in.vivo_noRT | Gprin2 | 2.704 | 9.20E-06 | glio |
| Atr_in.vivo_noRT | Adcy7 | 3.237 | 9.20E-07 | glio |
| Atp5c1_in.vivo_noRT | Slc14a2 | 2.668 | 9.20E-08 | glio |
| culture_effect | Fcf1 | -0.771 | 9.20E-09 | glio |
| Atr_in.vivo_noRT | 5330439K02R | 5.267 | 9.20E-14 | glio |
| Bcas2_in.vivo_noRT | BB557941 | 4.547 | 9.20E-15 | glio |
| culture_effect | Zbtb17 | 0.724 | 9.21E-03 | glio |
| Interaction_genotype_Mtor | B4galt4 | -3.198 | 9.21E-03 | glio |
| Interaction_genotype_Mtor | Trp53inp2 | -2.91 | 9.21E-03 | glio |
| Interaction_genotype_Mtor | Appl1 | -1.965 | 9.21E-03 | glio |
| Interaction_genotype_Mtor | Lactb | -1.88 | 9.21E-03 | glio |
| Interaction_genotype_Myc | Hist1h2bb | -2.892 | 9.21E-03 | glio |
| Interaction_genotype_Pars2 | Pnp | 1.241 | 9.21E-03 | glio |
| Atr_ex.vivo_noRT | Ccnl1 | 1.247 | 9.21E-03 | glio |
| Bard1_ex.vivo_noRT | Ctss | 2.421 | 9.21E-03 | glio |
| Bcas2_in.vivo_noRT | Ubash3a | 2.855 | 9.21E-03 | glio |
| Bcas2_ex.vivo_noRT | Ank1 | 1.958 | 9.21E-03 | glio |
| Bcas2_ex.vivo_noRT | Myzap | 2.342 | 9.21E-03 | glio |
| Bora_ex.vivo_noRT | 4921504E06Ri | 2.113 | 9.21E-03 | glio |
| Cenpj_in.vivo_noRT | Tas1r1 | 2.05 | 9.21E-03 | glio |

| | | | | |
|---|---|---|---|---|
| Interaction_genotype_Atr | Gm15929 | -3.346 | 9.21E-04 | glio |
| Interaction_genotype_Myc | Gm10260 | -4.072 | 9.21E-04 | glio |
| Interaction_genotype_Myc | Kiz | -2.883 | 9.21E-04 | glio |
| Atp5c1_in.vivo_noRT | 1700016P04R | 2.402 | 9.21E-04 | glio |
| Atr_in.vivo_noRT | Dpep2 | 2.205 | 9.21E-04 | glio |
| Atr_in.vivo_noRT | Npff | 2.211 | 9.21E-04 | glio |
| Bcas2_in.vivo_noRT | 9430060I03Ri | 2.325 | 9.21E-04 | glio |
| Cenpj_ex.vivo_noRT | Gm35082 | 2.592 | 9.21E-04 | glio |
| Atp5c1_in.vivo_noRT | 1700108F19Ri | 2.404 | 9.21E-05 | glio |
| culture_effect | Ackr3 | -1.271 | 9.21E-06 | glio |
| Interaction_genotype_Myc | Rasgrp2 | -5.006 | 9.21E-06 | glio |
| Atr_in.vivo_noRT | Ccdc148 | 3.259 | 9.21E-06 | glio |
| Bora_in.vivo_noRT | Wnt6 | 3.761 | 9.21E-06 | glio |
| Bora_in.vivo_noRT | 1700084E18Ri | 3.695 | 9.21E-07 | glio |
| Interaction_genotype_Mtor | Ddost | 2.561 | 9.21E-08 | glio |
| Bcas2_in.vivo_noRT | 1700011D18R | 3.762 | 9.21E-08 | glio |
| Chek1_in.vivo_noRT | Gm47101 | 4.252 | 9.21E-11 | glio |
| Aars2_in.vivo_noRT | Tac2 | 5.327 | 9.21E-26 | glio |
| Interaction_genotype_Atr | Slc47a2 | -3.182 | 9.22E-03 | glio |
| Interaction_genotype_Atr | Trim3 | -2.647 | 9.22E-03 | glio |
| Interaction_genotype_Bcas2 | Bc1 | -3.633 | 9.22E-03 | glio |
| Interaction_genotype_Bora | Stx1b | -3.194 | 9.22E-03 | glio |
| Interaction_genotype_Mtor | Gm47972 | -3.077 | 9.22E-03 | glio |
| Interaction_genotype_Mtor | Gprc5a | -3.111 | 9.22E-03 | glio |
| Interaction_genotype_Myc | Arhgap4 | -2.809 | 9.22E-03 | glio |
| Interaction_genotype_Myc | Lcat | -3.014 | 9.22E-03 | glio |
| Interaction_genotype_Myc | Prelid2 | -2.979 | 9.22E-03 | glio |
| Interaction_genotype_Myc | A730017L22Ri | -2.515 | 9.22E-03 | glio |
| Interaction_genotype_Myc | Ift122 | -2.552 | 9.22E-03 | glio |
| Interaction_genotype_Myc | Tgfbrap1 | -1.899 | 9.22E-03 | glio |
| interaction_RT | Pot1b | 1.29 | 9.22E-03 | glio |
| Aars2_in.vivo_noRT | Nfyb | -1.097 | 9.22E-03 | glio |
| Aars2_in.vivo_noRT | Mtch1 | -0.882 | 9.22E-03 | glio |
| Atr_in.vivo_noRT | Pcdhgc5 | 1.724 | 9.22E-03 | glio |
| Bcas2_in.vivo_noRT | Gm49602 | 1.868 | 9.22E-03 | glio |
| Bcas2_in.vivo_noRT | Set | -0.503 | 9.22E-03 | glio |
| Interaction_genotype_Atr | Gm42984 | -4.173 | 9.22E-04 | glio |
| Bard1_ex.vivo_noRT | Adam2 | 2.814 | 9.22E-04 | glio |
| Interaction_genotype_Bora | Gm27239 | -4.84 | 9.22E-05 | glio |
| culture_effect | Ctsf | -2.21 | 9.22E-06 | glio |
| Interaction_genotype_Myc | Mpp2 | -5.063 | 9.22E-06 | glio |
| Aars2_in.vivo_noRT | Bc1 | 3.299 | 9.22E-06 | glio |
| Atr_in.vivo_noRT | Gja4 | 2.945 | 9.22E-06 | glio |
| Chek1_ex.vivo_noRT | Lrrn4cl | 4.045 | 9.22E-06 | glio |
| culture_effect | Sdhd | -0.893 | 9.22E-11 | glio |
| Bcas2_in.vivo_noRT | Clcn1 | 4.161 | 9.22E-14 | glio |
| Aars2_in.vivo_noRT | Gm40538 | 5.063 | 9.22E-20 | glio |
| culture_effect | Ube2r2 | 0.298 | 9.23E-03 | glio |
| Interaction_genotype_Bcas2 | Gpc5 | -2.46 | 9.23E-03 | glio |
| Interaction_genotype_Myc | Spin4 | -2.278 | 9.23E-03 | glio |
| Interaction_genotype_Myc | Plod2 | 1.848 | 9.23E-03 | glio |
| Interaction_genotype_Myc | Iscu | 1.479 | 9.23E-03 | glio |

| | | | | |
|---|---|---|---|---|
| Interaction_genotype_Nat2 | Gm48780 | 2.937 | 9.23E-03 | glio |
| Atr_in.vivo_noRT | Gm15726 | 1.746 | 9.23E-03 | glio |
| Atr_in.vivo_noRT | Mrtfb | 1.712 | 9.23E-03 | glio |
| Bcas2_in.vivo_noRT | Erfe | 1.793 | 9.23E-03 | glio |
| Bcas2_in.vivo_noRT | Kctd7 | 1.805 | 9.23E-03 | glio |
| Bcas2_in.vivo_noRT | Col9a1 | -1.059 | 9.23E-03 | glio |
| Brca2_in.vivo_noRT | Gm36037 | 1.763 | 9.23E-03 | glio |
| Chek1_in.vivo_noRT | Gng7 | 1.855 | 9.23E-03 | glio |
| Chek1_in.vivo_noRT | Snrpb | -0.504 | 9.23E-03 | glio |
| Hsd17b10_in.vivo_noRT | Cyth4 | 2.076 | 9.23E-03 | glio |
| Hsd17b10_in.vivo_noRT | Slc6a19 | 1.978 | 9.23E-03 | glio |
| Hsd17b10_in.vivo_noRT | Ugt8a | -1.878 | 9.23E-03 | glio |
| culture_effect | Pmp2 | 1.781 | 9.23E-04 | glio |
| Interaction_genotype_Aars2 | A630089N07R | -3.657 | 9.23E-04 | glio |
| Interaction_genotype_Mtor | Gm31645 | -4.138 | 9.23E-04 | glio |
| Interaction_genotype_Myc | Dlk2 | -2.914 | 9.23E-04 | glio |
| Atr_in.vivo_noRT | Ino80dos | 2.444 | 9.23E-04 | glio |
| Interaction_genotype_Mtor | Klhdc2 | 2.092 | 9.23E-05 | glio |
| Interaction_genotype_Mtor | Rer1 | 1.708 | 9.23E-05 | glio |
| Aars2_in.vivo_noRT | Gm14443 | 2.594 | 9.23E-05 | glio |
| Atp5c1_in.vivo_noRT | E030042O20R | 3.196 | 9.23E-05 | glio |
| Interaction_genotype_Myc | Lpin2 | -4.16 | 9.23E-06 | glio |
| Interaction_genotype_Mtor | Gm36198 | -5.394 | 9.23E-08 | glio |
| Interaction_genotype_Mtor | Clec12b | -4.494 | 9.23E-08 | glio |
| Atr_in.vivo_noRT | Sptbn4 | 3.985 | 9.23E-09 | glio |
| interaction_RT | Ly6a | 3.594 | 9.23E-10 | glio |
| Aars2_in.vivo_noRT | BC030343 | 4.29 | 9.23E-10 | glio |
| Bcas2_in.vivo_noRT | BC048671 | 3.506 | 9.23E-10 | glio |
| Chek1_in.vivo_noRT | Zfp808 | 3.927 | 9.23E-11 | glio |
| Aars2_in.vivo_noRT | B3gnt4 | 5.063 | 9.23E-17 | glio |
| culture_effect | 0610009B22R | 0.399 | 9.24E-03 | glio |
| Interaction_genotype_Atr | Creb5 | 1.879 | 9.24E-03 | glio |
| Interaction_genotype_Mtor | Wfdc18 | -3.875 | 9.24E-03 | glio |
| Aars2_in.vivo_noRT | Pfdn5 | -0.507 | 9.24E-03 | glio |
| Bcas2_ex.vivo_noRT | Bmp15 | 2.348 | 9.24E-03 | glio |
| Bcas2_ex.vivo_noRT | Tubb4a | 3.124 | 9.24E-03 | glio |
| Bora_in.vivo_noRT | Zfp597 | 1.776 | 9.24E-03 | glio |
| Bora_in.vivo_noRT | Sgf29 | -1.215 | 9.24E-03 | glio |
| Brca2_in.vivo_noRT | Gm12800 | 1.763 | 9.24E-03 | glio |
| Brca2_in.vivo_noRT | Trmt61b | 1.763 | 9.24E-03 | glio |
| Brca2_in.vivo_noRT | Gm5890 | 1.763 | 9.24E-03 | glio |
| Brca2_in.vivo_noRT | Zscan4c | 1.763 | 9.24E-03 | glio |
| Brca2_in.vivo_noRT | BC080695 | 1.763 | 9.24E-03 | glio |
| Brca2_in.vivo_noRT | Gm47540 | 1.763 | 9.24E-03 | glio |
| Brca2_in.vivo_noRT | BB287469 | 1.763 | 9.24E-03 | glio |
| Chek1_ex.vivo_noRT | Smpdl3a | 2.205 | 9.24E-03 | glio |
| Hsd17b10_in.vivo_noRT | 9330179D12R | 2.207 | 9.24E-03 | glio |
| Interaction_genotype_Myc | Wdr44 | -2.598 | 9.24E-04 | glio |
| Aars2_in.vivo_noRT | 2500004C02R | 1.954 | 9.24E-04 | glio |
| Atr_in.vivo_noRT | Papolg | 1.859 | 9.24E-04 | glio |
| Atr_in.vivo_noRT | Serf1 | -1.045 | 9.24E-04 | glio |
| Bora_in.vivo_noRT | Anxa9 | 2.201 | 9.24E-04 | glio |

| | | | | |
|---|---|---|---|---|
| Chek1_in.vivo_noRT | AC168977.2 | 2.504 | 9.24E-04 | glio |
| Atr_in.vivo_noRT | H6pd | 2.202 | 9.24E-05 | glio |
| culture_effect | Ptms | 0.59 | 9.24E-06 | glio |
| Atr_in.vivo_noRT | A430105J06Ri | 3.025 | 9.24E-06 | glio |
| Aars2_in.vivo_noRT | Gm49479 | 3.489 | 9.24E-07 | glio |
| Bcas2_in.vivo_noRT | Myoz1 | 3.112 | 9.24E-08 | glio |
| Hsd17b10_in.vivo_noRT | Gm15816 | 3.616 | 9.24E-09 | glio |
| Atr_in.vivo_noRT | Drd4 | 4.349 | 9.24E-10 | glio |
| Interaction_genotype_Atr | Gm12602 | -3.038 | 9.25E-03 | glio |
| Interaction_genotype_Atr | Tssk6 | -2.483 | 9.25E-03 | glio |
| Interaction_genotype_Myc | Lrmp | -2.548 | 9.25E-03 | glio |
| interaction_RT | Urah | 1.537 | 9.25E-03 | glio |
| Aars2_ex.vivo_noRT | Gm17210 | 2.706 | 9.25E-03 | glio |
| Atr_ex.vivo_noRT | 4921536K21R | 2.494 | 9.25E-03 | glio |
| Atr_ex.vivo_noRT | Nol11 | -0.792 | 9.25E-03 | glio |
| Bora_ex.vivo_noRT | Apold1 | 2.954 | 9.25E-03 | glio |
| Chek1_ex.vivo_noRT | Gm10516 | 2.409 | 9.25E-03 | glio |
| interaction_RT | Naa10 | -0.692 | 9.25E-04 | glio |
| Aars2_in.vivo_noRT | 3110083C13R | 2.323 | 9.25E-04 | glio |
| Atr_in.vivo_noRT | Rpl7 | -0.651 | 9.25E-04 | glio |
| Chek1_ex.vivo_noRT | Htr1b | 2.887 | 9.25E-04 | glio |
| Hsd17b10_in.vivo_noRT | Gm49767 | 2.578 | 9.25E-04 | glio |
| culture_effect | Rtl1 | -2.231 | 9.25E-05 | glio |
| Interaction_genotype_Bora | Chpf | 2.836 | 9.25E-06 | glio |
| Interaction_genotype_Mtor | 4930481B07R | -5.59 | 9.25E-06 | glio |
| Bcas2_in.vivo_noRT | Kif5a | 3.395 | 9.25E-06 | glio |
| Interaction_genotype_Myc | Gm13272 | -5.714 | 9.25E-08 | glio |
| Atp5c1_in.vivo_noRT | AC154200.1 | 3.19 | 9.25E-08 | glio |
| Atp5c1_in.vivo_noRT | Gm4340 | 2.668 | 9.25E-08 | glio |
| Atp5c1_in.vivo_noRT | Gm16239 | 2.668 | 9.25E-08 | glio |
| Chek1_ex.vivo_noRT | Tmem132c | 4.842 | 9.25E-16 | glio |
| Interaction_genotype_Mtor | Slc9a5 | -3.027 | 9.26E-03 | glio |
| Interaction_genotype_Mtor | Slc29a2 | -3.11 | 9.26E-03 | glio |
| Interaction_genotype_Mtor | Gm16286 | 1.265 | 9.26E-03 | glio |
| Interaction_genotype_Rad51 | Snai1 | -3.969 | 9.26E-03 | glio |
| Interaction_genotype_Rad51 | Srgap1 | -3.627 | 9.26E-03 | glio |
| Interaction_genotype_Rad51 | Gm9958 | -3.74 | 9.26E-03 | glio |
| Interaction_genotype_Rad51 | Wincr1 | -4.736 | 9.26E-03 | glio |
| Atr_in.vivo_noRT | Anxa9 | 1.74 | 9.26E-03 | glio |
| Bora_in.vivo_noRT | Msantd1 | 1.949 | 9.26E-03 | glio |
| Bora_in.vivo_noRT | Zfp607b | 1.616 | 9.26E-03 | glio |
| Bora_in.vivo_noRT | Lyrm9 | 1.226 | 9.26E-03 | glio |
| Bora_ex.vivo_noRT | Iqch | 2.236 | 9.26E-03 | glio |
| Bora_ex.vivo_noRT | Hopx | 2.619 | 9.26E-03 | glio |
| Chek1_in.vivo_noRT | Bop1 | -1.08 | 9.26E-03 | glio |
| Chek1_ex.vivo_noRT | Nupr1 | 1.916 | 9.26E-03 | glio |
| Interaction_genotype_Mtor | Gm15894 | -3.054 | 9.26E-04 | glio |
| Bora_in.vivo_noRT | Dad1 | -0.822 | 9.26E-04 | glio |
| Chek1_in.vivo_noRT | Gm43378 | 2.245 | 9.26E-04 | glio |
| Bard1_ex.vivo_noRT | Kcnip3 | 3.611 | 9.26E-05 | glio |
| culture_effect | Serpinb9 | -2.971 | 9.26E-07 | glio |
| Atr_ex.vivo_noRT | 1700108F19Ri | 3.568 | 9.26E-07 | glio |

| | | | | |
|---|---|---|---|---|
| Bcas2_in.vivo_noRT | Gm44096 | 3.537 | 9.26E-08 | glio |
| Bcas2_in.vivo_noRT | Ldhd | 3.89 | 9.26E-11 | glio |
| Aars2_in.vivo_noRT | Hdac9 | 5.06 | 9.26E-14 | glio |
| Chek1_in.vivo_noRT | Olfr1191-ps1 | 4.917 | 9.26E-15 | glio |
| Chek1_in.vivo_noRT | AF067063 | 5.044 | 9.26E-22 | glio |
| Aars2_in.vivo_noRT | Cd7 | 5.063 | 9.26E-24 | glio |
| Interaction_genotype_Aars2 | Cpxm1 | -2.872 | 9.27E-03 | glio |
| Interaction_genotype_Aars2 | 4933421A08R | -3.46 | 9.27E-03 | glio |
| Interaction_genotype_Bcas2 | Gm37933 | -2.889 | 9.27E-03 | glio |
| Interaction_genotype_Mtor | Myl6 | 1.106 | 9.27E-03 | glio |
| Interaction_genotype_Myc | Tctn1 | -3.017 | 9.27E-03 | glio |
| Interaction_genotype_Myc | Stmn4 | -4.286 | 9.27E-03 | glio |
| Interaction_genotype_Myc | Col20a1 | -2.913 | 9.27E-03 | glio |
| Interaction_genotype_Myc | Spdl1 | -1.902 | 9.27E-03 | glio |
| Interaction_genotype_Myc | Armcx3 | -1.772 | 9.27E-03 | glio |
| Interaction_genotype_Myc | mt-Nd3 | 1.46 | 9.27E-03 | glio |
| interaction_RT | Bex2 | 0.777 | 9.27E-03 | glio |
| Bcas2_in.vivo_noRT | 4930590J08Ri | 2.002 | 9.27E-03 | glio |
| Bcas2_in.vivo_noRT | Cotl1 | 1.38 | 9.27E-03 | glio |
| Bora_in.vivo_noRT | Tmem121 | -1.395 | 9.27E-03 | glio |
| Dclre1c_ex.vivo_noRT | Gm12794 | 2.292 | 9.27E-03 | glio |
| Fanca_ex.vivo_noRT | Cd200r2 | 2.124 | 9.27E-03 | glio |
| Interaction_genotype_Myc | Pdlim4 | -4.016 | 9.27E-04 | glio |
| Interaction_genotype_Myc | Pqbp1 | -1.853 | 9.27E-04 | glio |
| Interaction_genotype_Myc | Cfap36 | 1.73 | 9.27E-04 | glio |
| Bora_ex.vivo_noRT | S100a14 | 2.802 | 9.27E-04 | glio |
| culture_effect | Snai1 | 2.183 | 9.27E-05 | glio |
| Aars2_in.vivo_noRT | Oasl2 | -1.617 | 9.27E-05 | glio |
| culture_effect | Nub1 | -0.69 | 9.27E-06 | glio |
| Bora_in.vivo_noRT | Gm43409 | 4.673 | 9.27E-10 | glio |
| Chek1_ex.vivo_noRT | Gm34939 | 4.842 | 9.27E-12 | glio |
| Atr_in.vivo_noRT | Elobl | 5.533 | 9.27E-14 | glio |
| Bcas2_in.vivo_noRT | AC159140.1 | 4.547 | 9.27E-18 | glio |
| culture_effect | Cct5 | -1.057 | 9.27E-20 | glio |
| culture_effect | Hltf | -0.546 | 9.28E-03 | glio |
| Interaction_genotype_Bcas2 | Slc35g3 | -2.464 | 9.28E-03 | glio |
| Interaction_genotype_Mtor | Ubiad1 | -2.574 | 9.28E-03 | glio |
| Atr_in.vivo_noRT | Zer1 | 1.516 | 9.28E-03 | glio |
| Atr_ex.vivo_noRT | Ncaph2 | -0.957 | 9.28E-03 | glio |
| Bcas2_in.vivo_noRT | E130114P18Ri | 2.055 | 9.28E-03 | glio |
| Bcas2_in.vivo_noRT | Gm15492 | 1.98 | 9.28E-03 | glio |
| Bcas2_in.vivo_noRT | Tert | 1.821 | 9.28E-03 | glio |
| Bora_in.vivo_noRT | Hist1h1b | -2.255 | 9.28E-03 | glio |
| Chek1_ex.vivo_noRT | Zfp14 | 2.488 | 9.28E-03 | glio |
| Chek1_ex.vivo_noRT | Gab2 | 2.278 | 9.28E-03 | glio |
| Chek1_ex.vivo_noRT | Fam111a | -1.876 | 9.28E-03 | glio |
| Chek1_ex.vivo_noRT | Nt5dc2 | -1.783 | 9.28E-03 | glio |
| culture_effect | Immp2l | -1.206 | 9.28E-04 | glio |
| Interaction_genotype_Aars2 | Gprasp1 | -3.678 | 9.28E-04 | glio |
| interaction_RT | C1ra | 2.273 | 9.28E-04 | glio |
| Aars2_in.vivo_noRT | Mrpl20 | -0.918 | 9.28E-04 | glio |
| Aars2_ex.vivo_noRT | Fbxl13 | 2.717 | 9.28E-04 | glio |

| | | | | |
|---|---|---|---|---|
| Eif2s1_in.vivo_noRT | Gm4027 | 2.143 | 9.28E-04 | glio |
| Eif2s1_in.vivo_noRT | AC154200.1 | 2.143 | 9.28E-04 | glio |
| Atr_in.vivo_noRT | Cnn1 | 3.863 | 9.28E-06 | glio |
| Atr_in.vivo_noRT | Cep112it | 3.089 | 9.28E-06 | glio |
| Chek1_in.vivo_noRT | Moap1 | 2.856 | 9.28E-06 | glio |
| Chek1_ex.vivo_noRT | Prl2c2 | 3.664 | 9.28E-06 | glio |
| Interaction_genotype_Mtor | Gm13078 | -4.099 | 9.28E-07 | glio |
| Atr_in.vivo_noRT | Tmem91 | 3.491 | 9.28E-07 | glio |
| Bora_in.vivo_noRT | Zfp968 | 3.559 | 9.28E-07 | glio |
| Chek1_ex.vivo_noRT | Spi1 | 3.871 | 9.28E-07 | glio |
| Chek1_ex.vivo_noRT | Impg2 | 4.055 | 9.28E-09 | glio |
| culture_effect | 4930589L23Ri | 3.032 | 9.28E-10 | glio |
| Chek1_in.vivo_noRT | Vmn2r54 | 3.882 | 9.28E-12 | glio |
| Aars2_in.vivo_noRT | Mecom | 5.327 | 9.28E-16 | glio |
| Bora_in.vivo_noRT | Gm8016 | 4.936 | 9.28E-23 | glio |
| Atr_in.vivo_noRT | Serpinb9g | 5.533 | 9.28E-27 | glio |
| culture_effect | Gm48357 | -1.444 | 9.29E-03 | glio |
| culture_effect | Ssr1 | -0.361 | 9.29E-03 | glio |
| culture_effect | Ctsz | -0.32 | 9.29E-03 | glio |
| Interaction_genotype_Atr | Trappc3 | 1.337 | 9.29E-03 | glio |
| Interaction_genotype_Myc | AW554918 | -2.558 | 9.29E-03 | glio |
| Interaction_genotype_Myc | Zfp770 | -1.484 | 9.29E-03 | glio |
| interaction_RT | Dhrs3 | 2.181 | 9.29E-03 | glio |
| interaction_RT | Il18 | 1.232 | 9.29E-03 | glio |
| interaction_RT | Plaur | 1.016 | 9.29E-03 | glio |
| Bora_ex.vivo_noRT | Gm48242 | 2.206 | 9.29E-03 | glio |
| Cenpj_in.vivo_noRT | Gm15816 | 2.022 | 9.29E-03 | glio |
| Chek1_ex.vivo_noRT | Rnd1 | 2.138 | 9.29E-03 | glio |
| Eif2s1_ex.vivo_noRT | Smtnl1 | 2.234 | 9.29E-03 | glio |
| culture_effect | H2-M10.1 | -1.803 | 9.29E-04 | glio |
| Interaction_genotype_Myc | B230311B06R | -3.958 | 9.29E-04 | glio |
| Atr_ex.vivo_noRT | Gm13449 | 2.376 | 9.29E-04 | glio |
| Atr_ex.vivo_noRT | Kcnd1 | 2.764 | 9.29E-04 | glio |
| Bcas2_in.vivo_noRT | Smim26 | -1.446 | 9.29E-04 | glio |
| Chek1_in.vivo_noRT | Gpr137b | 2.233 | 9.29E-04 | glio |
| culture_effect | 5330439K02R | 2.27 | 9.29E-05 | glio |
| culture_effect | E130317F20Ri | 1.154 | 9.29E-05 | glio |
| Interaction_genotype_Mtor | Gm42664 | -4.035 | 9.29E-05 | glio |
| Interaction_genotype_Mtor | 2610016A17R | -5.359 | 9.29E-06 | glio |
| Aars2_in.vivo_noRT | AC125149.3 | 2.922 | 9.29E-06 | glio |
| Aars2_ex.vivo_noRT | Mpzl2 | 3.293 | 9.29E-06 | glio |
| Atr_in.vivo_noRT | Gm41760 | 3.079 | 9.29E-06 | glio |
| Chek1_in.vivo_noRT | Scn2a | 3.16 | 9.29E-06 | glio |
| culture_effect | Gm36356 | 2.363 | 9.29E-07 | glio |
| Interaction_genotype_Mtor | Gm30262 | -6.152 | 9.29E-07 | glio |
| Atr_in.vivo_noRT | Gm43329 | 3.197 | 9.29E-07 | glio |
| Aars2_in.vivo_noRT | Gm44672 | 4.562 | 9.29E-12 | glio |
| Chek1_ex.vivo_noRT | Gm44693 | 4.842 | 9.29E-16 | glio |
| Bcas2_in.vivo_noRT | Prl3d3 | 4.544 | 9.29E-21 | glio |
| Interaction_genotype_Aars2 | Swsap1 | -2.892 | 9.30E-03 | glio |
| Interaction_genotype_Atr | Gm26916 | -3.175 | 9.30E-03 | glio |
| Interaction_genotype_Atr | Mphosph8 | 1.405 | 9.30E-03 | glio |

| | | | | |
|---|---|---|---|---|
| Aars2_in.vivo_noRT | Mfap5 | 1.971 | 9.30E-03 | glio |
| Bcas2_in.vivo_noRT | Atp2c1 | 1.12 | 9.30E-03 | glio |
| Cdkn2a_in.vivo_noRT | Gm6566 | -2.073 | 9.30E-03 | glio |
| Cdkn2a_in.vivo_noRT | Pspn | -1.675 | 9.30E-03 | glio |
| Eif2s1_in.vivo_noRT | Gm20646 | 2.143 | 9.30E-03 | glio |
| culture_effect | 1700063J08Ri | 1.802 | 9.30E-04 | glio |
| culture_effect | Elp6 | -0.913 | 9.30E-04 | glio |
| culture_effect | St3gal5 | 0.617 | 9.30E-04 | glio |
| Aars2_in.vivo_noRT | Zbtb3 | 2.386 | 9.30E-04 | glio |
| Aars2_in.vivo_noRT | Map3k14 | 2.136 | 9.30E-04 | glio |
| Bcas2_in.vivo_noRT | Frmd4b | 2.423 | 9.30E-04 | glio |
| Interaction_genotype_Myc | Gm48170 | -3.526 | 9.30E-05 | glio |
| Interaction_genotype_Myc | Tollip | -3.193 | 9.30E-05 | glio |
| Interaction_genotype_Myc | H3f3a | 1.367 | 9.30E-05 | glio |
| interaction_RT | 1110008P14R | 0.962 | 9.30E-05 | glio |
| Aars2_in.vivo_noRT | Pcdhb17 | 2.73 | 9.30E-05 | glio |
| Aars2_in.vivo_noRT | Dusp15 | 2.478 | 9.30E-05 | glio |
| Interaction_genotype_Mtor | S100a7a | -5.351 | 9.30E-06 | glio |
| Interaction_genotype_Mtor | Scgb1c1 | -4.492 | 9.30E-06 | glio |
| Interaction_genotype_Mtor | Gstt4 | -3.836 | 9.30E-06 | glio |
| Interaction_genotype_Mtor | Magt1 | 2.502 | 9.30E-06 | glio |
| Interaction_genotype_Mtor | Rdh12 | -5.6 | 9.30E-07 | glio |
| Interaction_genotype_Mtor | Kcnk13 | -4.624 | 9.30E-07 | glio |
| Bcas2_in.vivo_noRT | Gm50399 | 3.255 | 9.30E-07 | glio |
| Hsd17b10_in.vivo_noRT | 5430402O13R | 3.616 | 9.30E-07 | glio |
| Hsd17b10_in.vivo_noRT | 4930447F24Ri | 3.356 | 9.30E-07 | glio |
| Aars2_in.vivo_noRT | 1700040D17R | 4.263 | 9.30E-09 | glio |
| Chek1_in.vivo_noRT | 4930481B07R | 4.258 | 9.30E-09 | glio |
| Bora_in.vivo_noRT | Gm42695 | 4.676 | 9.30E-12 | glio |
| Aars2_in.vivo_noRT | Adam2 | 4.801 | 9.30E-16 | glio |
| Atr_in.vivo_noRT | Ceacam16 | 5.413 | 9.30E-20 | glio |
| culture_effect | Adam5 | -1.198 | 9.31E-03 | glio |
| Interaction_genotype_Atr | Gm50334 | -3.313 | 9.31E-03 | glio |
| Interaction_genotype_Atr | Dusp14 | 1.599 | 9.31E-03 | glio |
| Interaction_genotype_Bora | 1500002C15R | -3.592 | 9.31E-03 | glio |
| Interaction_genotype_Rpl3 | 2310067P03R | -3.567 | 9.31E-03 | glio |
| Interaction_genotype_Rpl3 | Kpna7 | -3.246 | 9.31E-03 | glio |
| interaction_RT | Fem1a | -1.13 | 9.31E-03 | glio |
| interaction_RT | Lbp | 0.995 | 9.31E-03 | glio |
| Atr_ex.vivo_noRT | Col28a1 | 2.569 | 9.31E-03 | glio |
| Atr_ex.vivo_noRT | Zfp985 | -2.026 | 9.31E-03 | glio |
| Bcas2_in.vivo_noRT | Syn3 | 1.954 | 9.31E-03 | glio |
| Bcas2_in.vivo_noRT | Odc1 | -0.907 | 9.31E-03 | glio |
| Chek1_in.vivo_noRT | Atg5 | -0.951 | 9.31E-03 | glio |
| Hsd17b10_in.vivo_noRT | Zfp458 | 2.14 | 9.31E-03 | glio |
| Bcas2_in.vivo_noRT | 2310030G06R | 2.437 | 9.31E-04 | glio |
| Chek1_ex.vivo_noRT | Cd74 | 3.414 | 9.31E-04 | glio |
| Interaction_genotype_Mtor | Ssh3 | -4.096 | 9.31E-05 | glio |
| Interaction_genotype_Mtor | Lrguk | -3.835 | 9.31E-05 | glio |
| Interaction_genotype_Mtor | Gm45589 | -4.642 | 9.31E-05 | glio |
| Interaction_genotype_Mtor | Gm26910 | -4.168 | 9.31E-05 | glio |
| Interaction_genotype_Mtor | Ccdc9b | -3.457 | 9.31E-05 | glio |

| | | | | |
|---|---|---|---|---|
| Interaction_genotype_Mtor | Shisa4 | 2.188 | 9.31E-05 | glio |
| interaction_RT | Hist1h2ak | -2.445 | 9.31E-05 | glio |
| Atr_in.vivo_noRT | Myo1b | 3.149 | 9.31E-05 | glio |
| culture_effect | Trp53 | -0.487 | 9.31E-06 | glio |
| Atr_in.vivo_noRT | Fam71f2 | 3.189 | 9.31E-06 | glio |
| culture_effect | Rsbn1 | -1.054 | 9.31E-08 | glio |
| Atr_in.vivo_noRT | Gm42997 | 3.651 | 9.31E-08 | glio |
| Bora_in.vivo_noRT | Gtsf1l | 3.307 | 9.31E-09 | glio |
| Hsd17b10_in.vivo_noRT | Gm11899 | 3.094 | 9.31E-09 | glio |
| Atr_in.vivo_noRT | Celf5 | 4.47 | 9.31E-10 | glio |
| Hsd17b10_in.vivo_noRT | Vtn | 3.356 | 9.31E-11 | glio |
| Hsd17b10_in.vivo_noRT | Gm48170 | 3.616 | 9.31E-11 | glio |
| Bcas2_in.vivo_noRT | Gm47851 | 4.547 | 9.31E-13 | glio |
| culture_effect | Synrg | 0.477 | 9.32E-03 | glio |
| Interaction_genotype_Atr | Cd36 | -2.369 | 9.32E-03 | glio |
| Interaction_genotype_Atr | Kpna4 | 1.449 | 9.32E-03 | glio |
| Interaction_genotype_Bcas2 | Mrpl38 | 1.442 | 9.32E-03 | glio |
| Interaction_genotype_Chek1 | Sqstm1 | 1.441 | 9.32E-03 | glio |
| Interaction_genotype_Myc | Neurl1b | -3.001 | 9.32E-03 | glio |
| Interaction_genotype_Myc | Myo16 | -2.997 | 9.32E-03 | glio |
| Interaction_genotype_Myc | Tpmt | -2.102 | 9.32E-03 | glio |
| Interaction_genotype_Myc | Zc3h15 | 1.069 | 9.32E-03 | glio |
| Interaction_genotype_Myc | Dynll1 | 0.942 | 9.32E-03 | glio |
| Interaction_genotype_Rpl3 | 9130604C24R | -3.941 | 9.32E-03 | glio |
| Aars2_in.vivo_noRT | 9130221H12R | 1.747 | 9.32E-03 | glio |
| Aars2_in.vivo_noRT | Dpp7 | 1.577 | 9.32E-03 | glio |
| Aars2_in.vivo_noRT | Trmt1l | -1.224 | 9.32E-03 | glio |
| Atp5c1_in.vivo_noRT | Ptpn22 | 2.405 | 9.32E-03 | glio |
| Atp5c1_in.vivo_noRT | Gm16794 | 2.144 | 9.32E-03 | glio |
| Bcas2_in.vivo_noRT | Atg9b | -1.564 | 9.32E-03 | glio |
| Bora_in.vivo_noRT | Capn1 | 1.358 | 9.32E-03 | glio |
| Interaction_genotype_Myc | Rnf217 | -2.59 | 9.32E-04 | glio |
| interaction_RT | Asb3 | 1.197 | 9.32E-04 | glio |
| culture_effect | Pcdhgb5 | -2.445 | 9.32E-06 | glio |
| culture_effect | Ssbp1 | -0.49 | 9.32E-06 | glio |
| Aars2_in.vivo_noRT | Myl7 | 3.199 | 9.32E-06 | glio |
| Atr_in.vivo_noRT | Arhgdib | 3.112 | 9.32E-06 | glio |
| Atr_ex.vivo_noRT | A330049N07R | 2.764 | 9.32E-06 | glio |
| Chek1_in.vivo_noRT | 1700120G11R | 3.329 | 9.32E-06 | glio |
| Interaction_genotype_Mtor | Abcg4 | -5.455 | 9.32E-07 | glio |
| Atr_in.vivo_noRT | Gm26981 | 3.359 | 9.32E-07 | glio |
| Chek1_in.vivo_noRT | Ucma | 4.045 | 9.32E-07 | glio |
| culture_effect | Zfp945 | -1.078 | 9.32E-08 | glio |
| culture_effect | Decr1 | -0.865 | 9.32E-08 | glio |
| Aars2_in.vivo_noRT | Gm5544 | 4.536 | 9.32E-11 | glio |
| Hsd17b10_in.vivo_noRT | Ctla4 | 3.616 | 9.32E-14 | glio |
| Hsd17b10_in.vivo_noRT | Zscan4-ps3 | 3.616 | 9.32E-14 | glio |
| Hsd17b10_in.vivo_noRT | A030014E15Ri | 3.616 | 9.32E-14 | glio |
| Hsd17b10_in.vivo_noRT | Gm21731 | 3.616 | 9.32E-14 | glio |
| culture_effect | Psmb10 | -1.825 | 9.32E-34 | glio |
| Interaction_genotype_Aars2 | Slc1a3 | 1.843 | 9.33E-03 | glio |
| Interaction_genotype_Bora | Siah1b | 2.059 | 9.33E-03 | glio |

| | | | | |
|---|---|---|---|---|
| Interaction_genotype_Mtor | Trim3 | -2.455 | 9.33E-03 | glio |
| Interaction_genotype_Myc | Mdga2 | -2.556 | 9.33E-03 | glio |
| Aars2_in.vivo_noRT | Gpr17 | 1.613 | 9.33E-03 | glio |
| Atp5c1_in.vivo_noRT | Prox2os | 1.892 | 9.33E-03 | glio |
| Atr_in.vivo_noRT | Pkn3 | 1.66 | 9.33E-03 | glio |
| Bard1_ex.vivo_noRT | Osm | 2.415 | 9.33E-03 | glio |
| Bard1_ex.vivo_noRT | Gm26978 | 2.227 | 9.33E-03 | glio |
| Bora_in.vivo_noRT | Syde2 | 1.86 | 9.33E-03 | glio |
| Bora_in.vivo_noRT | Rnd1 | 1.754 | 9.33E-03 | glio |
| Bora_in.vivo_noRT | Nr4a1 | -1.197 | 9.33E-03 | glio |
| Chek1_in.vivo_noRT | Gm12059 | 1.863 | 9.33E-03 | glio |
| Atr_in.vivo_noRT | Rnf8 | -1.56 | 9.33E-04 | glio |
| Bora_ex.vivo_noRT | Slc7a14 | 2.942 | 9.33E-04 | glio |
| Interaction_genotype_Myc | Zfp345 | -3.198 | 9.33E-05 | glio |
| Interaction_genotype_Myc | Snap29 | -3.548 | 9.33E-05 | glio |
| Atp5c1_in.vivo_noRT | Pmp2 | 2.668 | 9.33E-05 | glio |
| culture_effect | Gm30025 | 1.413 | 9.33E-07 | glio |
| culture_effect | Afap1l2 | 1.946 | 9.33E-10 | glio |
| Chek1_in.vivo_noRT | 1700019E08Ri | 5.044 | 9.33E-20 | glio |
| Interaction_genotype_Aars2 | Gm12940 | -2.84 | 9.34E-03 | glio |
| Interaction_genotype_Atr | Scn2a | -3.217 | 9.34E-03 | glio |
| Interaction_genotype_Chek1 | Tnfrsf23 | -4.091 | 9.34E-03 | glio |
| Interaction_genotype_Mtor | Marcks | 1.754 | 9.34E-03 | glio |
| Interaction_genotype_Myc | Rab40c | -2.294 | 9.34E-03 | glio |
| Interaction_genotype_Myc | Caprin2 | -2.323 | 9.34E-03 | glio |
| Atr_in.vivo_noRT | Zrsr2 | -0.775 | 9.34E-03 | glio |
| Bora_in.vivo_noRT | Mapkbp1 | 1.682 | 9.34E-03 | glio |
| Eif2s1_in.vivo_noRT | Iqch | 1.879 | 9.34E-03 | glio |
| Eif2s1_in.vivo_noRT | Klk13 | 1.61 | 9.34E-03 | glio |
| culture_effect | 1700081N11R | 2.101 | 9.34E-04 | glio |
| Interaction_genotype_Atr | Hint2 | 1.824 | 9.34E-04 | glio |
| Interaction_genotype_Mtor | Nhsl2 | -3.952 | 9.34E-04 | glio |
| Aars2_in.vivo_noRT | Fkbp3 | -0.862 | 9.34E-04 | glio |
| culture_effect | Nppb | 2.085 | 9.34E-05 | glio |
| Atr_in.vivo_noRT | Gm12981 | 2.746 | 9.34E-05 | glio |
| Bora_ex.vivo_noRT | Clec12b | 2.806 | 9.34E-06 | glio |
| Bora_ex.vivo_noRT | Olfr171 | 3.201 | 9.34E-06 | glio |
| culture_effect | Fgl2 | -2.852 | 9.34E-07 | glio |
| Interaction_genotype_Mtor | Gm12589 | -4.498 | 9.34E-07 | glio |
| Atr_in.vivo_noRT | 4933433G15R | 3.867 | 9.34E-08 | glio |
| Hsd17b10_in.vivo_noRT | Gm41307 | 3.616 | 9.34E-13 | glio |
| Aars2_in.vivo_noRT | Masp2 | 4.399 | 9.34E-14 | glio |
| Bora_in.vivo_noRT | Prl2c2 | 4.936 | 9.34E-14 | glio |
| Interaction_genotype_Atr | Mtcp1 | -3.146 | 9.35E-03 | glio |
| Interaction_genotype_Bcas2 | Fosb | 2.962 | 9.35E-03 | glio |
| Interaction_genotype_Myc | Atrnl1 | -2.656 | 9.35E-03 | glio |
| Aars2_in.vivo_noRT | Ripk3 | 2.133 | 9.35E-03 | glio |
| Aars2_in.vivo_noRT | Hectd2 | 1.685 | 9.35E-03 | glio |
| Aars2_in.vivo_noRT | Bmp1 | 1.648 | 9.35E-03 | glio |
| Aars2_in.vivo_noRT | Hebp1 | -1.587 | 9.35E-03 | glio |
| Aars2_in.vivo_noRT | Vti1b | -0.924 | 9.35E-03 | glio |
| Aars2_in.vivo_noRT | Rnf130 | -0.951 | 9.35E-03 | glio |

| | | | | |
|---|---|---|---|---|
| Aars2_in.vivo_noRT | Pcbp1 | -0.672 | 9.35E-03 | glio |
| Atr_ex.vivo_noRT | Dennd2d | 2.37 | 9.35E-03 | glio |
| Bcas2_in.vivo_noRT | Etaa1 | -1.356 | 9.35E-03 | glio |
| Interaction_genotype_Mtor | Xist | -3.934 | 9.35E-04 | glio |
| Aars2_ex.vivo_noRT | Ubxn10 | 2.893 | 9.35E-04 | glio |
| Atr_in.vivo_noRT | 1110019D14R | 2.274 | 9.35E-04 | glio |
| Atr_ex.vivo_noRT | Mapk15 | 3.145 | 9.35E-04 | glio |
| Cenpj_in.vivo_noRT | Zscan4-ps2 | 2.022 | 9.35E-04 | glio |
| Chek1_ex.vivo_noRT | Gm14264 | 2.821 | 9.35E-04 | glio |
| Interaction_genotype_Mtor | Crabp1 | -4.586 | 9.35E-06 | glio |
| Interaction_genotype_Myc | Gm14167 | -4.53 | 9.35E-06 | glio |
| Interaction_genotype_Myc | Glyat | -3.526 | 9.35E-06 | glio |
| Interaction_genotype_Myc | Nupl2 | -4.549 | 9.35E-07 | glio |
| Interaction_genotype_Bora | Rftn2 | -3.115 | 9.36E-03 | glio |
| Interaction_genotype_Mtor | Snrnp35 | -2.29 | 9.36E-03 | glio |
| Interaction_genotype_Myc | Hist1h4c | -2.925 | 9.36E-03 | glio |
| Interaction_genotype_Myc | Siae | -2.643 | 9.36E-03 | glio |
| Interaction_genotype_Myc | Nhlrc3 | -2.391 | 9.36E-03 | glio |
| Interaction_genotype_Rpl3 | Mex3c | 2.214 | 9.36E-03 | glio |
| Aars2_in.vivo_noRT | Mgll | 1.843 | 9.36E-03 | glio |
| Bora_in.vivo_noRT | Nlrc5 | -1.391 | 9.36E-03 | glio |
| Bora_in.vivo_noRT | Rrm2b | -1.087 | 9.36E-03 | glio |
| Bora_in.vivo_noRT | Cnot4 | -0.859 | 9.36E-03 | glio |
| Chek1_ex.vivo_noRT | Fam107b | 1.867 | 9.36E-03 | glio |
| culture_effect | Gm43330 | -1.854 | 9.36E-04 | glio |
| culture_effect | Hba-a2 | 1.926 | 9.36E-04 | glio |
| culture_effect | 4930426I24Ri | 1.746 | 9.36E-04 | glio |
| Interaction_genotype_Bcas2 | Gm26910 | -3.999 | 9.36E-04 | glio |
| Interaction_genotype_Mtor | Gm29170 | -3.476 | 9.36E-04 | glio |
| Aars2_in.vivo_noRT | Mapkbp1 | 2.062 | 9.36E-04 | glio |
| Atr_in.vivo_noRT | Gm26797 | 1.971 | 9.36E-04 | glio |
| Chek1_ex.vivo_noRT | Slc15a2 | 3.028 | 9.36E-05 | glio |
| culture_effect | Ephb4 | -1.246 | 9.36E-06 | glio |
| Bora_in.vivo_noRT | Gm32816 | 3.41 | 9.36E-06 | glio |
| Bora_in.vivo_noRT | Gm13986 | 3.678 | 9.36E-07 | glio |
| Atr_in.vivo_noRT | Zfp273 | 3.767 | 9.36E-08 | glio |
| culture_effect | Nova1 | -2.267 | 9.36E-09 | glio |
| culture_effect | Dbf4 | 1.006 | 9.36E-09 | glio |
| Atr_in.vivo_noRT | Zfp575 | 4.281 | 9.36E-09 | glio |
| Atr_in.vivo_noRT | Tmem44 | 4.275 | 9.36E-10 | glio |
| culture_effect | 2010320O07R | 3.232 | 9.36E-11 | glio |
| Aars2_ex.vivo_noRT | Gm13856 | 4.076 | 9.36E-11 | glio |
| Atr_in.vivo_noRT | 1700012B09R | 5.273 | 9.36E-11 | glio |
| Aars2_in.vivo_noRT | Lrfn1 | 4.684 | 9.36E-12 | glio |
| Chek1_in.vivo_noRT | Gm21762 | 4.78 | 9.36E-15 | glio |
| Bora_in.vivo_noRT | Psmb11 | 4.413 | 9.36E-18 | glio |
| culture_effect | Med27 | -2.281 | 9.36E-34 | glio |
| Interaction_genotype_Aars2 | E130208F15Ri | -3.538 | 9.37E-03 | glio |
| Interaction_genotype_Mtor | Washc3 | 1.242 | 9.37E-03 | glio |
| Interaction_genotype_Myc | Tgfb1i1 | -2.783 | 9.37E-03 | glio |
| Aars2_in.vivo_noRT | Ppm1g | -0.856 | 9.37E-03 | glio |
| Atr_in.vivo_noRT | Zswim3 | 1.736 | 9.37E-03 | glio |

| | | | | |
|---|---|---|---|---|
| Atr_in.vivo_noRT | Gramd3 | 1.731 | 9.37E-03 | glio |
| Bcas2_in.vivo_noRT | Tbc1d30 | 1.741 | 9.37E-03 | glio |
| Bcas2_in.vivo_noRT | Atp1b3 | -0.828 | 9.37E-03 | glio |
| Chek1_in.vivo_noRT | Nr2c1 | 1.441 | 9.37E-03 | glio |
| Chek1_in.vivo_noRT | Pigm | 1.269 | 9.37E-03 | glio |
| culture_effect | Apool | -0.672 | 9.37E-04 | glio |
| Interaction_genotype_Mtor | Snph | -3.938 | 9.37E-04 | glio |
| Interaction_genotype_Mtor | Lrrn4cl | -4.316 | 9.37E-04 | glio |
| Interaction_genotype_Mtor | Pde6h | -3.37 | 9.37E-04 | glio |
| Interaction_genotype_Myc | AU019990 | -2.729 | 9.37E-04 | glio |
| Interaction_genotype_Myc | Gm5861 | -4.489 | 9.37E-04 | glio |
| Interaction_genotype_Myc | Hist1h3b | -3.89 | 9.37E-04 | glio |
| Interaction_genotype_Myc | Ppef2 | -3.037 | 9.37E-04 | glio |
| Bcas2_in.vivo_noRT | Plxna2 | 2.607 | 9.37E-04 | glio |
| Bcas2_in.vivo_noRT | Msh5 | 2.557 | 9.37E-04 | glio |
| Bora_in.vivo_noRT | Neurl1a | 2.601 | 9.37E-04 | glio |
| Interaction_genotype_Chek1 | Scube3 | -4.395 | 9.37E-05 | glio |
| Interaction_genotype_Myc | Rdh12 | -4.231 | 9.37E-05 | glio |
| Aars2_in.vivo_noRT | Fbxl12os | 2.686 | 9.37E-05 | glio |
| Atr_in.vivo_noRT | Slc25a47 | 2.771 | 9.37E-05 | glio |
| Aars2_in.vivo_noRT | Ampd3 | 3.044 | 9.37E-06 | glio |
| culture_effect | Brsk2 | -2.857 | 9.37E-07 | glio |
| Atr_in.vivo_noRT | Epsti1 | -2.357 | 9.37E-07 | glio |
| Atr_ex.vivo_noRT | Sucnr1 | 3.164 | 9.37E-09 | glio |
| Chek1_in.vivo_noRT | 4921508D12R | 4.116 | 9.37E-11 | glio |
| Bora_in.vivo_noRT | Gm49860 | 4.936 | 9.37E-17 | glio |
| Interaction_genotype_Aars2 | 5830444B04R | -2.371 | 9.38E-03 | glio |
| Interaction_genotype_Aars2 | Rgs16 | -3.191 | 9.38E-03 | glio |
| Interaction_genotype_Aars2 | Sf3b6 | 0.823 | 9.38E-03 | glio |
| Interaction_genotype_Atr | 9130019O22R | -2.369 | 9.38E-03 | glio |
| Interaction_genotype_Mtor | B230369F24Ri | -2.992 | 9.38E-03 | glio |
| Interaction_genotype_Mtor | Gpn1 | -1.724 | 9.38E-03 | glio |
| Interaction_genotype_Myc | Atp5d | 1.066 | 9.38E-03 | glio |
| Interaction_genotype_Terf1 | Alox12 | -3.96 | 9.38E-03 | glio |
| Interaction_genotype_Terf1 | Pygo1 | -3.69 | 9.38E-03 | glio |
| Aars2_ex.vivo_noRT | Iqcn | 2.495 | 9.38E-03 | glio |
| Hsd17b10_in.vivo_noRT | Prss22 | 2.539 | 9.38E-03 | glio |
| Hsd17b10_in.vivo_noRT | Mboat7 | -1.267 | 9.38E-03 | glio |
| Interaction_genotype_Bora | Tecrl | -4.46 | 9.38E-04 | glio |
| Interaction_genotype_Myc | B130034C11R | -3.07 | 9.38E-04 | glio |
| Interaction_genotype_Myc | S100a11 | 1.831 | 9.38E-04 | glio |
| Aars2_ex.vivo_noRT | Rtl1 | 2.897 | 9.38E-04 | glio |
| Atp5c1_in.vivo_noRT | Celsr3 | 2.408 | 9.38E-04 | glio |
| Chek1_in.vivo_noRT | Fhod3 | 2.527 | 9.38E-04 | glio |
| Chek1_in.vivo_noRT | Amd2 | 2.356 | 9.38E-04 | glio |
| culture_effect | Aldh2 | -0.704 | 9.38E-05 | glio |
| Interaction_genotype_Myc | 1700081N11R | -4.97 | 9.38E-05 | glio |
| Aars2_ex.vivo_noRT | Gm13713 | 2.893 | 9.38E-05 | glio |
| Bcas2_in.vivo_noRT | 1010001B22R | 2.807 | 9.38E-05 | glio |
| Chek1_ex.vivo_noRT | Gm16741 | 3.103 | 9.38E-05 | glio |
| Atr_in.vivo_noRT | Tmem79 | 3.249 | 9.38E-07 | glio |
| Bcas2_in.vivo_noRT | Cyp2u1 | 3.231 | 9.38E-07 | glio |

| | | | | |
|---|---|---|---|---|
| Bora_in.vivo_noRT | Car5a | 3.139 | 9.38E-07 | glio |
| Atr_in.vivo_noRT | Zfp661 | 3.783 | 9.38E-08 | glio |
| culture_effect | Cisd1 | 0.726 | 9.38E-10 | glio |
| Atr_in.vivo_noRT | Prl2c3 | 5.533 | 9.38E-18 | glio |
| Atr_in.vivo_noRT | Akap14 | 5.27 | 9.38E-22 | glio |
| Atr_in.vivo_noRT | Agap2 | 5.533 | 9.38E-22 | glio |
| Bora_in.vivo_noRT | Clec12b | 5.46 | 9.38E-27 | glio |
| Interaction_genotype_Mtor | Bglap2 | -3.17 | 9.39E-03 | glio |
| Interaction_genotype_Mtor | 1810012K16R | -2.963 | 9.39E-03 | glio |
| Interaction_genotype_Mtor | Gm14325 | -3.12 | 9.39E-03 | glio |
| Interaction_genotype_Mtor | Sh2b1 | -1.926 | 9.39E-03 | glio |
| Interaction_genotype_Myc | Zfp985 | -2.273 | 9.39E-03 | glio |
| interaction_RT | Gm4258 | -1.421 | 9.39E-03 | glio |
| Aars2_in.vivo_noRT | Pkp3 | 1.99 | 9.39E-03 | glio |
| Atr_in.vivo_noRT | Smarca4 | -1.008 | 9.39E-03 | glio |
| Bcas2_in.vivo_noRT | Fbxo44 | -1.451 | 9.39E-03 | glio |
| Cdkn2a_in.vivo_noRT | Kcnk13 | -1.938 | 9.39E-03 | glio |
| Interaction_genotype_Aars2 | Col9a1 | 2.391 | 9.39E-04 | glio |
| Interaction_genotype_Myc | Gm43727 | -3.641 | 9.39E-04 | glio |
| culture_effect | Gm9970 | 2.308 | 9.39E-05 | glio |
| Interaction_genotype_Myc | Gm47622 | -3.304 | 9.39E-05 | glio |
| Aars2_in.vivo_noRT | Btbd17 | 3.295 | 9.39E-05 | glio |
| Atr_in.vivo_noRT | Coro1b | 2.903 | 9.39E-06 | glio |
| Aars2_in.vivo_noRT | 5031434O11R | 3.918 | 9.39E-07 | glio |
| Atr_in.vivo_noRT | Cct6b | 3.244 | 9.39E-07 | glio |
| Chek1_ex.vivo_noRT | Flacc1 | 3.652 | 9.39E-07 | glio |
| culture_effect | Gas2l1 | 0.973 | 9.39E-08 | glio |
| Bcas2_in.vivo_noRT | Traf3ip3 | 3.743 | 9.39E-08 | glio |
| Bcas2_in.vivo_noRT | Kmo | 3.165 | 9.39E-08 | glio |
| Atr_in.vivo_noRT | Abcg4 | 4.742 | 9.39E-14 | glio |
| Aars2_in.vivo_noRT | D730045A05R | 5.327 | 9.39E-15 | glio |
| Aars2_in.vivo_noRT | Gm10324 | 5.06 | 9.39E-21 | glio |
| culture_effect | Gm26812 | 1.215 | 9.40E-03 | glio |
| Aars2_in.vivo_noRT | Derl1 | -0.784 | 9.40E-03 | glio |
| Atr_in.vivo_noRT | Ranbp17 | 1.837 | 9.40E-03 | glio |
| Atr_in.vivo_noRT | Sar1a | -0.858 | 9.40E-03 | glio |
| Atr_in.vivo_noRT | Rpp30 | 0.891 | 9.40E-03 | glio |
| Chek1_in.vivo_noRT | Atg3 | -1.189 | 9.40E-03 | glio |
| Chek1_ex.vivo_noRT | Rims3 | 2.18 | 9.40E-03 | glio |
| Chek1_ex.vivo_noRT | Ctu2 | -1.766 | 9.40E-03 | glio |
| Chek1_ex.vivo_noRT | Gbp6 | 1.634 | 9.40E-03 | glio |
| Dclre1c_ex.vivo_noRT | Pdlim3 | 2.659 | 9.40E-03 | glio |
| Eif2s1_ex.vivo_noRT | Gm12280 | 3.172 | 9.40E-03 | glio |
| Aars2_in.vivo_noRT | Fam228b | 2.534 | 9.40E-04 | glio |
| Aars2_in.vivo_noRT | Zfp91 | -0.973 | 9.40E-04 | glio |
| Bcas2_in.vivo_noRT | Ccdc102a | 2.378 | 9.40E-04 | glio |
| culture_effect | Wars | -0.78 | 9.40E-05 | glio |
| Interaction_genotype_Mtor | Spats1 | -4.805 | 9.40E-05 | glio |
| Chek1_in.vivo_noRT | Zfp619 | 2.841 | 9.40E-05 | glio |
| Chek1_in.vivo_noRT | Gm45589 | 2.755 | 9.40E-05 | glio |
| Chek1_in.vivo_noRT | Tlr6 | 3.209 | 9.40E-06 | glio |
| Interaction_genotype_Mtor | Cpeb1 | -6.124 | 9.40E-07 | glio |

| | | | | |
|---|---|---|---|---|
| Interaction_genotype_Mtor | Gm13964 | -4.099 | 9.40E-07 | glio |
| Interaction_genotype_Myc | Sept1 | -6.012 | 9.40E-07 | glio |
| Interaction_genotype_Myc | Gm38534 | -6.406 | 9.40E-09 | glio |
| culture_effect | Lgr6 | 1.558 | 9.41E-03 | glio |
| culture_effect | Prss12 | -0.969 | 9.41E-03 | glio |
| culture_effect | Tmem68 | -0.469 | 9.41E-03 | glio |
| Interaction_genotype_Bcas2 | Ifi207 | -3.771 | 9.41E-03 | glio |
| Interaction_genotype_Myc | 1700061G19R | -2.872 | 9.41E-03 | glio |
| Interaction_genotype_Myc | Polr3f | -1.773 | 9.41E-03 | glio |
| interaction_RT | Nat10 | -1.109 | 9.41E-03 | glio |
| interaction_RT | Lmf2 | -0.736 | 9.41E-03 | glio |
| Atp5c1_in.vivo_noRT | Gm11549 | 2.142 | 9.41E-03 | glio |
| Atp5c1_in.vivo_noRT | Fam13c | 2.151 | 9.41E-03 | glio |
| Atr_in.vivo_noRT | Cyld | 1.1 | 9.41E-03 | glio |
| Bcas2_in.vivo_noRT | Gm50223 | 1.42 | 9.41E-03 | glio |
| Bcas2_in.vivo_noRT | Clcn4 | -1.023 | 9.41E-03 | glio |
| Bcas2_ex.vivo_noRT | Gm38393 | 2.348 | 9.41E-03 | glio |
| culture_effect | Ccdc82 | -1.004 | 9.41E-04 | glio |
| Bcas2_ex.vivo_noRT | BC048671 | 2.742 | 9.41E-04 | glio |
| Chek1_in.vivo_noRT | Ccdc103 | 2.415 | 9.41E-04 | glio |
| Bora_in.vivo_noRT | H2-Q10 | 2.768 | 9.41E-05 | glio |
| Chek1_in.vivo_noRT | Glb1l | 2.792 | 9.41E-05 | glio |
| Aars2_in.vivo_noRT | Gm17300 | 2.95 | 9.41E-06 | glio |
| Atr_ex.vivo_noRT | Olfr171 | 3.164 | 9.41E-06 | glio |
| Bcas2_in.vivo_noRT | Gpr37l1 | 3.693 | 9.41E-08 | glio |
| Chek1_ex.vivo_noRT | Gm42664 | 4.256 | 9.41E-09 | glio |
| Bcas2_in.vivo_noRT | Ms4a6d | 4.019 | 9.41E-12 | glio |
| Chek1_in.vivo_noRT | Gm26789 | 4.126 | 9.41E-13 | glio |
| culture_effect | Gphn | -0.642 | 9.42E-03 | glio |
| Interaction_genotype_Myc | Prkcz | -2.702 | 9.42E-03 | glio |
| Interaction_genotype_Myc | Prkdc | -2.321 | 9.42E-03 | glio |
| Bcas2_in.vivo_noRT | Arhgap26 | 1.847 | 9.42E-03 | glio |
| Bora_in.vivo_noRT | Chp1 | -1.183 | 9.42E-03 | glio |
| Bora_in.vivo_noRT | Tacc3 | -1.35 | 9.42E-03 | glio |
| Interaction_genotype_Mtor | Sema4f | -3.718 | 9.42E-04 | glio |
| Interaction_genotype_Myc | Ggnbp1 | -3.595 | 9.42E-04 | glio |
| Atp5c1_in.vivo_noRT | Gm44702 | 2.658 | 9.42E-04 | glio |
| Atr_ex.vivo_noRT | Efna1 | 2.613 | 9.42E-04 | glio |
| Bora_in.vivo_noRT | Zfp937 | 2.53 | 9.42E-04 | glio |
| Hsd17b10_in.vivo_noRT | Dut | -0.882 | 9.42E-04 | glio |
| Bora_in.vivo_noRT | Chchd1 | -0.957 | 9.42E-05 | glio |
| culture_effect | Igfbp4 | -1.475 | 9.42E-06 | glio |
| Bora_in.vivo_noRT | Fbxl12os | 3.34 | 9.42E-07 | glio |
| Atr_in.vivo_noRT | Gm49785 | 3.836 | 9.42E-08 | glio |
| Bcas2_in.vivo_noRT | 9030612E09Ri | 3.486 | 9.42E-08 | glio |
| Aars2_in.vivo_noRT | Gm15246 | 3.976 | 9.42E-09 | glio |
| culture_effect | Mgll | -4.025 | 9.42E-11 | glio |
| culture_effect | Bola2 | 0.902 | 9.42E-11 | glio |
| Bcas2_in.vivo_noRT | 9430041J12Ri | 4.807 | 9.42E-18 | glio |
| Interaction_genotype_Cdkn2a | Cys1 | 3.231 | 9.43E-03 | glio |
| Interaction_genotype_Cdkn2a | Gm12800 | 2.835 | 9.43E-03 | glio |
| Interaction_genotype_Cdkn2a | Trmt61b | 2.835 | 9.43E-03 | glio |

| | | | | |
|---|---|---|---|---|
| Interaction_genotype_Cdkn2a | Slc16a5 | 2.706 | 9.43E-03 | glio |
| Interaction_genotype_Cdkn2a | Gm5890 | 2.835 | 9.43E-03 | glio |
| Interaction_genotype_Cdkn2a | Zscan4c | 2.835 | 9.43E-03 | glio |
| Interaction_genotype_Cdkn2a | BC080695 | 2.835 | 9.43E-03 | glio |
| Interaction_genotype_Cdkn2a | BB287469 | 2.835 | 9.43E-03 | glio |
| Interaction_genotype_Mtor | Gm38979 | -3.305 | 9.43E-03 | glio |
| Aars2_in.vivo_noRT | Lca5 | 1.842 | 9.43E-03 | glio |
| Aars2_in.vivo_noRT | Pgam1 | -0.708 | 9.43E-03 | glio |
| Atp5c1_in.vivo_noRT | Gm26674 | 2.023 | 9.43E-03 | glio |
| Atr_ex.vivo_noRT | Tdrp | 2.477 | 9.43E-03 | glio |
| Atr_ex.vivo_noRT | Ttyh1 | -2.268 | 9.43E-03 | glio |
| Atr_ex.vivo_noRT | Morn4 | 1.714 | 9.43E-03 | glio |
| Atr_ex.vivo_noRT | Neto1 | -1.444 | 9.43E-03 | glio |
| Atr_ex.vivo_noRT | Ppat | -1.137 | 9.43E-03 | glio |
| Bcas2_in.vivo_noRT | Yars | -1.119 | 9.43E-03 | glio |
| Bora_in.vivo_noRT | Lypd6 | 1.949 | 9.43E-03 | glio |
| Bora_ex.vivo_noRT | Insm1 | 2.614 | 9.43E-03 | glio |
| culture_effect | Hspe1 | 0.395 | 9.43E-04 | glio |
| Interaction_genotype_Aars2 | Gm40557 | -4.194 | 9.43E-04 | glio |
| Interaction_genotype_Aars2 | Atp5g2 | 0.992 | 9.43E-04 | glio |
| Interaction_genotype_Myc | Cebpz | 1.485 | 9.43E-04 | glio |
| Bora_in.vivo_noRT | Zfp287 | 1.894 | 9.43E-04 | glio |
| Interaction_genotype_Mtor | 2610507I01Ri | -3.829 | 9.43E-05 | glio |
| Interaction_genotype_Mtor | Tmem14c | 1.858 | 9.43E-05 | glio |
| Chek1_in.vivo_noRT | 4930554I06Ri | 2.984 | 9.43E-05 | glio |
| culture_effect | Taf1d | -0.633 | 9.43E-06 | glio |
| Atr_in.vivo_noRT | Flrt3 | 2.823 | 9.43E-06 | glio |
| Bora_in.vivo_noRT | Gm26688 | 3.498 | 9.43E-07 | glio |
| Bora_in.vivo_noRT | Sema6b | 3.46 | 9.43E-07 | glio |
| Bcas2_in.vivo_noRT | Gm28874 | 3.785 | 9.43E-08 | glio |
| Interaction_genotype_Aars2 | Vps13a | -1.315 | 9.44E-03 | glio |
| Interaction_genotype_Aars2 | Sfxn1 | 1.319 | 9.44E-03 | glio |
| Interaction_genotype_Aars2 | Slc25a3 | 0.71 | 9.44E-03 | glio |
| Interaction_genotype_Atr | Dok2 | -2.819 | 9.44E-03 | glio |
| Interaction_genotype_Atr | A330015K06R | -2.991 | 9.44E-03 | glio |
| Interaction_genotype_Atr | Tpd52l1 | 1.71 | 9.44E-03 | glio |
| Interaction_genotype_Mtor | 1700063D05R | -3.337 | 9.44E-03 | glio |
| Interaction_genotype_Pars2 | 4930447K03R | -2.796 | 9.44E-03 | glio |
| Aars2_in.vivo_noRT | Eif2ak4 | 1.2 | 9.44E-03 | glio |
| Aars2_in.vivo_noRT | Psph | -0.845 | 9.44E-03 | glio |
| Atp5c1_ex.vivo_noRT | Rslcan18 | 2.981 | 9.44E-03 | glio |
| Atp5c1_ex.vivo_noRT | Crb3 | 2.597 | 9.44E-03 | glio |
| Atr_in.vivo_noRT | Klf2 | -1.838 | 9.44E-03 | glio |
| Atr_in.vivo_noRT | Sec13 | -0.962 | 9.44E-03 | glio |
| Atr_in.vivo_noRT | Rpl31 | -0.516 | 9.44E-03 | glio |
| Bcas2_ex.vivo_noRT | Ms4a4d | 2.742 | 9.44E-03 | glio |
| Chek1_in.vivo_noRT | Ube2d2a | -0.683 | 9.44E-03 | glio |
| Hsd17b10_in.vivo_noRT | Ampd2 | -1.349 | 9.44E-03 | glio |
| Interaction_genotype_Mtor | Qrich2 | -4.056 | 9.44E-04 | glio |
| Interaction_genotype_Mtor | Glb1l | -3.929 | 9.44E-04 | glio |
| Interaction_genotype_Myc | Gm50375 | -4.023 | 9.44E-04 | glio |
| Aars2_in.vivo_noRT | Dnajb13 | 2.247 | 9.44E-04 | glio |

| | | | | |
|---|---|---|---|---|
| Aars2_in.vivo_noRT | Polr2k | -0.868 | 9.44E-04 | glio |
| Aars2_ex.vivo_noRT | Podxl | 2.899 | 9.44E-04 | glio |
| Atr_in.vivo_noRT | Ebf4 | 2.416 | 9.44E-04 | glio |
| culture_effect | Cops3 | -0.596 | 9.44E-05 | glio |
| culture_effect | Tomm5 | 0.517 | 9.44E-06 | glio |
| Atp5c1_in.vivo_noRT | Esm1 | 2.404 | 9.44E-06 | glio |
| Chek1_in.vivo_noRT | Frmd8os | 3.526 | 9.44E-06 | glio |
| Atr_in.vivo_noRT | Ly6d | 4.806 | 9.44E-18 | glio |
| Chek1_in.vivo_noRT | 4930423D22R | 4.784 | 9.44E-21 | glio |
| Interaction_genotype_Aars2 | A330074K22R | -2.924 | 9.45E-03 | glio |
| Interaction_genotype_Atr | Gm26901 | -3.502 | 9.45E-03 | glio |
| Interaction_genotype_Bora | Ttc19 | 1.631 | 9.45E-03 | glio |
| Interaction_genotype_Mtor | 1700056N10R | -2.843 | 9.45E-03 | glio |
| Interaction_genotype_Mtor | Ftl1-ps1 | -1.947 | 9.45E-03 | glio |
| Interaction_genotype_Myc | Chn1 | -2.376 | 9.45E-03 | glio |
| Aars2_in.vivo_noRT | Lmbrd1 | -1.109 | 9.45E-03 | glio |
| Atr_ex.vivo_noRT | Flicr | 2.51 | 9.45E-03 | glio |
| Atr_ex.vivo_noRT | Pfkl | -1.466 | 9.45E-03 | glio |
| Bard1_ex.vivo_noRT | Map3k5 | 2.415 | 9.45E-03 | glio |
| Bcas2_in.vivo_noRT | Snx10 | -0.887 | 9.45E-03 | glio |
| Bora_ex.vivo_noRT | Igfbp7 | 2.404 | 9.45E-03 | glio |
| Interaction_genotype_Chek1 | Ier5 | 3.294 | 9.45E-04 | glio |
| Interaction_genotype_Myc | Gm36793 | -2.729 | 9.45E-04 | glio |
| Atr_ex.vivo_noRT | Serpinb2 | 2.764 | 9.45E-04 | glio |
| Atr_ex.vivo_noRT | Bglap2 | 3.164 | 9.45E-04 | glio |
| Atr_ex.vivo_noRT | Olfr1189 | 2.764 | 9.45E-04 | glio |
| Bora_in.vivo_noRT | Gm14399 | 2.419 | 9.45E-04 | glio |
| Aars2_in.vivo_noRT | Ccdc159 | 2.904 | 9.45E-05 | glio |
| Aars2_ex.vivo_noRT | Gm20705 | 2.897 | 9.45E-05 | glio |
| Bard1_ex.vivo_noRT | Hist1h2ag | 2.814 | 9.45E-05 | glio |
| Bard1_ex.vivo_noRT | Dapk2 | 2.799 | 9.45E-05 | glio |
| Bcas2_in.vivo_noRT | Sept5 | 3.068 | 9.45E-06 | glio |
| Bora_in.vivo_noRT | Rtl9 | 3.495 | 9.45E-08 | glio |
| Bcas2_in.vivo_noRT | Gm16244 | 4.15 | 9.45E-09 | glio |
| Chek1_ex.vivo_noRT | Pla2g2d | 4.842 | 9.45E-10 | glio |
| Bora_in.vivo_noRT | Zfp352 | 4.936 | 9.45E-15 | glio |
| culture_effect | Tapbpl | -3.472 | 9.45E-48 | glio |
| Interaction_genotype_Mtor | Klhl2 | -2.233 | 9.46E-03 | glio |
| Interaction_genotype_Myc | Mmp17 | -2.29 | 9.46E-03 | glio |
| interaction_RT | Adamts1 | 1.028 | 9.46E-03 | glio |
| Bcas2_in.vivo_noRT | Idnk | -1.1 | 9.46E-03 | glio |
| Bora_in.vivo_noRT | Zfp626 | 1.502 | 9.46E-03 | glio |
| Eif2s1_in.vivo_noRT | Mcmdc2 | 1.883 | 9.46E-03 | glio |
| Hsd17b10_in.vivo_noRT | Zbtb11os1 | 1.964 | 9.46E-03 | glio |
| culture_effect | Pcp4l1 | 1.769 | 9.46E-04 | glio |
| Chek1_in.vivo_noRT | Dixdc1 | 2.41 | 9.46E-04 | glio |
| Chek1_ex.vivo_noRT | Car14 | 2.849 | 9.46E-04 | glio |
| culture_effect | Vamp3 | 0.623 | 9.46E-05 | glio |
| Aars2_in.vivo_noRT | Gm15133 | 2.72 | 9.46E-05 | glio |
| interaction_RT | Napsa | 4.092 | 9.46E-06 | glio |
| Chek1_in.vivo_noRT | Slc24a1 | 3.182 | 9.46E-06 | glio |
| Interaction_genotype_Mtor | Zfp385b | 2.898 | 9.46E-07 | glio |

| | | | | |
|---|---|---|---|---|
| Interaction_genotype_Mtor | Rtraf | 2.434 | 9.46E-07 | glio |
| Interaction_genotype_Mtor | Gchfr | -6.467 | 9.46E-07 | glio |
| Interaction_genotype_Mtor | Prdx1 | 3.58 | 9.46E-07 | glio |
| Interaction_genotype_Mtor | Gm15927 | -5.705 | 9.46E-07 | glio |
| Interaction_genotype_Mtor | mt-Co1 | 2.143 | 9.46E-07 | glio |
| Bcas2_in.vivo_noRT | Serpinb9 | 3.268 | 9.46E-07 | glio |
| Hsd17b10_in.vivo_noRT | Gm30042 | 3.356 | 9.46E-09 | glio |
| Bcas2_in.vivo_noRT | E030037K01Ri | 4.275 | 9.46E-12 | glio |
| Chek1_in.vivo_noRT | Podxl | 4.512 | 9.46E-13 | glio |
| Bcas2_in.vivo_noRT | Vmn2r44 | 4.807 | 9.46E-22 | glio |
| Bcas2_in.vivo_noRT | Mmp13 | 4.807 | 9.46E-22 | glio |
| culture_effect | Mrps27 | -0.687 | 9.47E-03 | glio |
| culture_effect | Ttc13 | -0.61 | 9.47E-03 | glio |
| Interaction_genotype_Mtor | Zfp382 | -2.917 | 9.47E-03 | glio |
| Interaction_genotype_Mtor | Brf2 | -2.218 | 9.47E-03 | glio |
| Interaction_genotype_Myc | Cracr2b | -3.132 | 9.47E-03 | glio |
| Interaction_genotype_Myc | Tex15 | -2.788 | 9.47E-03 | glio |
| Interaction_genotype_Myc | Desi2 | 1.338 | 9.47E-03 | glio |
| Interaction_genotype_Myc | Eif5 | 0.995 | 9.47E-03 | glio |
| interaction_RT | Cabp1 | 1.818 | 9.47E-03 | glio |
| interaction_RT | Trp53inp2 | 1.775 | 9.47E-03 | glio |
| interaction_RT | Hus1 | -1.034 | 9.47E-03 | glio |
| Aars2_in.vivo_noRT | Erp27 | 1.927 | 9.47E-03 | glio |
| Aars2_in.vivo_noRT | Paics | -0.589 | 9.47E-03 | glio |
| Atr_in.vivo_noRT | Tmem136 | 1.624 | 9.47E-03 | glio |
| Bcas2_in.vivo_noRT | Galnt3 | 1.988 | 9.47E-03 | glio |
| Bcas2_ex.vivo_noRT | Zfp980 | 2.893 | 9.47E-03 | glio |
| Bora_in.vivo_noRT | Upp2 | 1.724 | 9.47E-03 | glio |
| Chek1_in.vivo_noRT | Gm28535 | 1.861 | 9.47E-03 | glio |
| Chek1_in.vivo_noRT | Cep290 | -1.239 | 9.47E-03 | glio |
| Chek1_ex.vivo_noRT | Slc6a13 | 2.578 | 9.47E-03 | glio |
| culture_effect | Bclaf1 | -0.409 | 9.47E-04 | glio |
| Bcas2_in.vivo_noRT | Cfap298 | -1.27 | 9.47E-04 | glio |
| Eif2s1_in.vivo_noRT | Msantd2 | -2.088 | 9.47E-04 | glio |
| Interaction_genotype_Mtor | Gm15063 | -5.243 | 9.47E-05 | glio |
| Interaction_genotype_Myc | 4930483J18Ri | -4.146 | 9.47E-05 | glio |
| Bora_in.vivo_noRT | Gfpt1 | -1.543 | 9.47E-05 | glio |
| Bora_ex.vivo_noRT | Gm48265 | 3.201 | 9.47E-05 | glio |
| Atr_in.vivo_noRT | Dst | -1.567 | 9.47E-06 | glio |
| Atr_in.vivo_noRT | Rpl10a | -0.764 | 9.47E-06 | glio |
| Bcas2_in.vivo_noRT | 1700066B19R | 3.36 | 9.47E-07 | glio |
| Hsd17b10_in.vivo_noRT | Tcap | 3.237 | 9.47E-07 | glio |
| Bcas2_in.vivo_noRT | A230020J21Ri | 3.88 | 9.47E-08 | glio |
| Aars2_in.vivo_noRT | Hspa12a | 4.026 | 9.47E-09 | glio |
| Bcas2_in.vivo_noRT | Rab37 | 3.889 | 9.47E-09 | glio |
| Bora_in.vivo_noRT | Gngt2 | 4.936 | 9.47E-13 | glio |
| Bora_in.vivo_noRT | Trim16 | 4.676 | 9.47E-14 | glio |
| culture_effect | Shb | 0.659 | 9.48E-03 | glio |
| Interaction_genotype_Mtor | Icam4 | -2.848 | 9.48E-03 | glio |
| Interaction_genotype_Myc | Gm904 | -2.904 | 9.48E-03 | glio |
| Interaction_genotype_Myc | Wdr47 | -2.145 | 9.48E-03 | glio |
| Interaction_genotype_Myc | Deaf1 | -2.172 | 9.48E-03 | glio |

| | | | | |
|---|---|---|---|---|
| interaction_RT | Rptor | -0.99 | 9.48E-03 | glio |
| Atr_in.vivo_noRT | Gtf2e2 | -0.998 | 9.48E-03 | glio |
| Bora_in.vivo_noRT | Slfn9 | -1.305 | 9.48E-03 | glio |
| Bora_ex.vivo_noRT | Nat8f4 | 2.414 | 9.48E-03 | glio |
| Chek1_in.vivo_noRT | Eif3j2 | 1.907 | 9.48E-03 | glio |
| Chek1_in.vivo_noRT | Gm20324 | 1.7 | 9.48E-03 | glio |
| Chek1_in.vivo_noRT | Ranbp17 | 1.871 | 9.48E-03 | glio |
| Chek1_in.vivo_noRT | Cfap97 | -1.296 | 9.48E-03 | glio |
| Chek1_in.vivo_noRT | Syvn1 | -1.26 | 9.48E-03 | glio |
| Chek1_in.vivo_noRT | Etnk1 | -1.056 | 9.48E-03 | glio |
| Hsd17b10_in.vivo_noRT | Scd3 | 2.228 | 9.48E-03 | glio |
| Hsd17b10_in.vivo_noRT | Klhl20 | 1.204 | 9.48E-03 | glio |
| culture_effect | Xlr3a | -0.819 | 9.48E-04 | glio |
| Interaction_genotype_Myc | Adgrv1 | -3.227 | 9.48E-04 | glio |
| Aars2_in.vivo_noRT | Ppig | -1.024 | 9.48E-04 | glio |
| Atr_in.vivo_noRT | Bend3 | 2.327 | 9.48E-04 | glio |
| Atr_in.vivo_noRT | Rexo1 | -1.416 | 9.48E-04 | glio |
| Bora_in.vivo_noRT | Fbxo48 | 2.439 | 9.48E-04 | glio |
| Bora_in.vivo_noRT | 5930430L01Ri | 2.254 | 9.48E-04 | glio |
| Bora_in.vivo_noRT | Il12rb1 | 1.829 | 9.48E-04 | glio |
| Chek1_in.vivo_noRT | Sgms2 | 2.539 | 9.48E-04 | glio |
| Bora_ex.vivo_noRT | Prdm1 | 3.6 | 9.48E-05 | glio |
| Chek1_in.vivo_noRT | Cib2 | 3.162 | 9.48E-05 | glio |
| culture_effect | Zmynd8 | 0.668 | 9.48E-06 | glio |
| Interaction_genotype_Mtor | Nqo1 | -5.382 | 9.48E-06 | glio |
| Interaction_genotype_Mtor | Nckipsd | -5.103 | 9.48E-06 | glio |
| Interaction_genotype_Mtor | 2210417A02R | -6.179 | 9.48E-07 | glio |
| Chek1_ex.vivo_noRT | Dner | 4.115 | 9.48E-07 | glio |
| Bcas2_in.vivo_noRT | Gm11491 | 3.765 | 9.48E-08 | glio |
| Bora_in.vivo_noRT | Gm41764 | 4.019 | 9.48E-11 | glio |
| Bora_in.vivo_noRT | D630036H23R | 4.676 | 9.48E-12 | glio |
| Chek1_in.vivo_noRT | Gm10399 | 4.642 | 9.48E-12 | glio |
| Atr_in.vivo_noRT | E130215H24R | 4.742 | 9.48E-14 | glio |
| Bcas2_in.vivo_noRT | Sucnr1 | 4.807 | 9.48E-26 | glio |
| Interaction_genotype_Aars2 | Adk | 1.477 | 9.49E-03 | glio |
| Interaction_genotype_Aars2 | Tomm6 | 1.008 | 9.49E-03 | glio |
| Interaction_genotype_Aars2 | Psmb6 | 0.98 | 9.49E-03 | glio |
| Interaction_genotype_Atr | B3galt4 | -3.015 | 9.49E-03 | glio |
| Interaction_genotype_Atr | Ctsl | 1.456 | 9.49E-03 | glio |
| Interaction_genotype_Myc | Ntf5 | -3.017 | 9.49E-03 | glio |
| Interaction_genotype_Myc | Spef1 | -3.001 | 9.49E-03 | glio |
| interaction_RT | Ifnar2 | 0.807 | 9.49E-03 | glio |
| Atr_in.vivo_noRT | Dnajb5 | 1.673 | 9.49E-03 | glio |
| Bard1_ex.vivo_noRT | Shbg | 2.883 | 9.49E-03 | glio |
| Bora_ex.vivo_noRT | Akr1b7 | 2.802 | 9.49E-03 | glio |
| Ercc4_ex.vivo_noRT | Crb3 | 2.68 | 9.49E-03 | glio |
| culture_effect | Ttc32 | 0.603 | 9.49E-04 | glio |
| Interaction_genotype_Ifnar1 | Cd83 | 4.183 | 9.49E-04 | glio |
| Aars2_in.vivo_noRT | Gm47438 | 2.583 | 9.49E-04 | glio |
| interaction_RT | Qpct | 3.852 | 9.49E-05 | glio |
| Bard1_ex.vivo_noRT | Gpr22 | 2.814 | 9.49E-05 | glio |
| Bcas2_in.vivo_noRT | Gm43221 | 2.635 | 9.49E-05 | glio |

| | | | | |
|---|---|---|---|---|
| Chek1_ex.vivo_noRT | Papln | 3.363 | 9.49E-05 | glio |
| Interaction_genotype_Myc | Adamts4 | -4.478 | 9.49E-06 | glio |
| Atr_ex.vivo_noRT | Trim17 | 2.764 | 9.49E-06 | glio |
| Bora_in.vivo_noRT | H60c | 3.343 | 9.49E-06 | glio |
| Bora_in.vivo_noRT | Taf9 | 3.046 | 9.49E-06 | glio |
| Atr_in.vivo_noRT | Zfand4 | 3.52 | 9.49E-07 | glio |
| Bcas2_in.vivo_noRT | Gm50295 | 4.015 | 9.49E-09 | glio |
| Interaction_genotype_Myc | mt-Cytb | 2.369 | 9.49E-13 | glio |
| Chek1_ex.vivo_noRT | Gm26671 | 4.449 | 9.49E-14 | glio |
| Chek1_in.vivo_noRT | Hist1h2ag | 4.511 | 9.49E-18 | glio |
| Chek1_in.vivo_noRT | Cldn5 | 4.777 | 9.49E-22 | glio |
| culture_effect | D3Ertd751e | 0.926 | 9.50E-03 | glio |
| Interaction_genotype_Atr | Gm41764 | -2.853 | 9.50E-03 | glio |
| Interaction_genotype_Atr | Gm20455 | -3.321 | 9.50E-03 | glio |
| Interaction_genotype_Atr | Puf60 | 1.179 | 9.50E-03 | glio |
| Interaction_genotype_Mtor | Cacng7 | -2.949 | 9.50E-03 | glio |
| Interaction_genotype_Mtor | Rhot2 | -1.884 | 9.50E-03 | glio |
| Interaction_genotype_Myc | Hdhd3 | -2.5 | 9.50E-03 | glio |
| Interaction_genotype_Myc | Gm26910 | -2.515 | 9.50E-03 | glio |
| interaction_RT | Sema3e | 1.462 | 9.50E-03 | glio |
| Atr_in.vivo_noRT | Sh3bp2 | 1.73 | 9.50E-03 | glio |
| Atr_in.vivo_noRT | Ddias | 1.647 | 9.50E-03 | glio |
| Bcas2_in.vivo_noRT | Gm10825 | 1.978 | 9.50E-03 | glio |
| Bora_in.vivo_noRT | Tnfaip6 | 1.365 | 9.50E-03 | glio |
| Bora_ex.vivo_noRT | Cxcr2 | 2.615 | 9.50E-03 | glio |
| Hsd17b10_in.vivo_noRT | Ctr9 | -1.097 | 9.50E-03 | glio |
| culture_effect | Cdk1 | -0.712 | 9.50E-04 | glio |
| Atp5c1_in.vivo_noRT | Gm5941 | 2.668 | 9.50E-04 | glio |
| Bcas2_in.vivo_noRT | Bloc1s6os | 2.287 | 9.50E-04 | glio |
| Cyp19a1_in.vivo_noRT | Anxa8 | 2.709 | 9.50E-04 | glio |
| culture_effect | Aldh18a1 | -0.832 | 9.50E-05 | glio |
| Interaction_genotype_Myc | Mtpap | -3.143 | 9.50E-05 | glio |
| Interaction_genotype_Mtor | Prodh | -4.713 | 9.50E-06 | glio |
| Aars2_in.vivo_noRT | Col17a1 | 3.276 | 9.50E-06 | glio |
| Aars2_in.vivo_noRT | Shpk | 3.339 | 9.50E-06 | glio |
| Aars2_in.vivo_noRT | Gm5431 | 2.883 | 9.50E-06 | glio |
| Bcas2_in.vivo_noRT | 8030456M14F | 3.183 | 9.50E-06 | glio |
| Hsd17b10_in.vivo_noRT | D030068K23R | 3.225 | 9.50E-06 | glio |
| Bora_in.vivo_noRT | Zfp980 | 3.953 | 9.50E-08 | glio |
| Atp5c1_in.vivo_noRT | Gm36037 | 3.196 | 9.50E-09 | glio |
| Chek1_in.vivo_noRT | Gm49312 | 3.97 | 9.50E-10 | glio |
| culture_effect | Sparc | 1.176 | 9.50E-13 | glio |
| Chek1_in.vivo_noRT | Magea10 | 4.576 | 9.50E-14 | glio |
| Aars2_in.vivo_noRT | Igf2bp1 | 5.327 | 9.50E-15 | glio |
| culture_effect | Zbp1 | -1.511 | 9.50E-17 | glio |
| Interaction_genotype_Myc | Gm45406 | -2.207 | 9.51E-03 | glio |
| Interaction_genotype_Myc | Med22 | -1.725 | 9.51E-03 | glio |
| Interaction_genotype_Myc | Rhbdf1 | -1.75 | 9.51E-03 | glio |
| Aars2_in.vivo_noRT | Pnpla7 | 1.744 | 9.51E-03 | glio |
| Atr_in.vivo_noRT | Ate1 | 1.032 | 9.51E-03 | glio |
| Bora_ex.vivo_noRT | Gm26789 | 2.22 | 9.51E-03 | glio |
| Hsd17b10_in.vivo_noRT | Inafm1 | 1.749 | 9.51E-03 | glio |

| | | | | |
|---|---|---|---|---|
| Hsd17b10_in.vivo_noRT | Slc22a4 | -1.451 | 9.51E-03 | glio |
| culture_effect | Dnah2os | -1.793 | 9.51E-04 | glio |
| Interaction_genotype_Mtor | P4htm | -4.007 | 9.51E-04 | glio |
| Interaction_genotype_Myc | AW146154 | -3.77 | 9.51E-04 | glio |
| Atr_in.vivo_noRT | Rps13 | -0.608 | 9.51E-04 | glio |
| Interaction_genotype_Bcas2 | Rassf6 | -4.301 | 9.51E-05 | glio |
| Interaction_genotype_Myc | Rpl21 | 1.332 | 9.51E-05 | glio |
| Chek1_in.vivo_noRT | Itgb7 | 2.627 | 9.51E-05 | glio |
| Atr_in.vivo_noRT | Bst1 | 2.894 | 9.51E-06 | glio |
| Aars2_in.vivo_noRT | Gm13270 | 5.327 | 9.51E-14 | glio |
| Chek1_in.vivo_noRT | Cox7b2 | 4.576 | 9.51E-14 | glio |
| Interaction_genotype_Mtor | Zg16 | -3.053 | 9.52E-03 | glio |
| Interaction_genotype_Mtor | Zfp882 | -3.383 | 9.52E-03 | glio |
| Interaction_genotype_Mtor | Zdhhc18 | -2.733 | 9.52E-03 | glio |
| Interaction_genotype_Mtor | Srebf1 | -2.09 | 9.52E-03 | glio |
| Interaction_genotype_Mtor | Arid4b | 1.27 | 9.52E-03 | glio |
| Interaction_genotype_Myc | Golga1 | -2.073 | 9.52E-03 | glio |
| interaction_RT | Pttg1ip | 0.751 | 9.52E-03 | glio |
| Aars2_in.vivo_noRT | Eif3m | -0.632 | 9.52E-03 | glio |
| Bcas2_in.vivo_noRT | A930041C12R | 1.916 | 9.52E-03 | glio |
| Bcas2_in.vivo_noRT | Rnf207 | 1.795 | 9.52E-03 | glio |
| Chek1_ex.vivo_noRT | Apod | 2.812 | 9.52E-03 | glio |
| Hsd17b10_in.vivo_noRT | Szt2 | 1.547 | 9.52E-03 | glio |
| culture_effect | Gm16933 | -1.457 | 9.52E-04 | glio |
| Chek1_in.vivo_noRT | Mt1 | 1.04 | 9.52E-04 | glio |
| Chek1_ex.vivo_noRT | Rapgefl1 | 3.072 | 9.52E-04 | glio |
| culture_effect | Zfp931 | 1.289 | 9.52E-06 | glio |
| Interaction_genotype_Myc | 4930524O07R | -5.052 | 9.52E-06 | glio |
| Aars2_in.vivo_noRT | Cldn10 | 3.119 | 9.52E-06 | glio |
| Interaction_genotype_Myc | mt-Nd4 | 2.088 | 9.52E-08 | glio |
| Chek1_in.vivo_noRT | 4930478M13F | 4.664 | 9.52E-11 | glio |
| Interaction_genotype_Atr | Ubald2 | 1.415 | 9.53E-03 | glio |
| Interaction_genotype_Mtor | Naip6 | -3.062 | 9.53E-03 | glio |
| Interaction_genotype_Mtor | Lgalsl | -1.834 | 9.53E-03 | glio |
| Interaction_genotype_Mtor | Chchd2 | 1.021 | 9.53E-03 | glio |
| Interaction_genotype_Myc | Lst1 | -2.891 | 9.53E-03 | glio |
| interaction_RT | Erf | -0.772 | 9.53E-03 | glio |
| Bcas2_in.vivo_noRT | Gldc | 1.899 | 9.53E-03 | glio |
| Interaction_genotype_Mtor | Pmch | -3.441 | 9.53E-04 | glio |
| Interaction_genotype_Mtor | Stip1 | 1.48 | 9.53E-04 | glio |
| Interaction_genotype_Myc | Gm2012 | -3.829 | 9.53E-04 | glio |
| Interaction_genotype_Myc | Stxbp4 | -3.035 | 9.53E-04 | glio |
| Interaction_genotype_Myc | Psma4 | 1.472 | 9.53E-04 | glio |
| Atp5c1_in.vivo_noRT | Ankrd33b | 2.683 | 9.53E-04 | glio |
| Bcas2_in.vivo_noRT | Tsr1 | -1.409 | 9.53E-04 | glio |
| Bcas2_ex.vivo_noRT | Tmem54 | 2.742 | 9.53E-04 | glio |
| Hsd17b10_in.vivo_noRT | Zfp85os | 2.557 | 9.53E-04 | glio |
| Interaction_genotype_Mtor | Aqp3 | -5.798 | 9.53E-05 | glio |
| Interaction_genotype_Mtor | Med9os | -4.436 | 9.53E-05 | glio |
| Interaction_genotype_Myc | Gm36172 | -3.564 | 9.53E-05 | glio |
| Aars2_in.vivo_noRT | Gm45605 | 2.774 | 9.53E-05 | glio |
| Bora_in.vivo_noRT | Pank2 | -1.636 | 9.53E-05 | glio |

| | | | | |
|---|---|---|---|---|
| culture_effect | Prm1 | -2.546 | 9.53E-06 | glio |
| Aars2_in.vivo_noRT | Sgk2 | 3.11 | 9.53E-06 | glio |
| Atr_ex.vivo_noRT | Mpeg1 | 3.761 | 9.53E-06 | glio |
| Atr_ex.vivo_noRT | 1700047K16R | 4.054 | 9.53E-06 | glio |
| culture_effect | Syngr1 | 0.918 | 9.53E-07 | glio |
| Atr_in.vivo_noRT | Kyat1 | 3.68 | 9.53E-07 | glio |
| Bcas2_in.vivo_noRT | Dnajc28 | 3.661 | 9.53E-07 | glio |
| Bcas2_in.vivo_noRT | Dnaaf3 | 3.384 | 9.53E-08 | glio |
| Bora_in.vivo_noRT | Hist2h3c1 | 4.025 | 9.53E-09 | glio |
| Bcas2_in.vivo_noRT | Bmx | 4.019 | 9.53E-11 | glio |
| culture_effect | Apobec3 | -2.142 | 9.53E-32 | glio |
| culture_effect | Rps27l | 1.427 | 9.53E-40 | glio |
| Interaction_genotype_Atr | Trmt61b | -2.369 | 9.54E-03 | glio |
| Interaction_genotype_Mtor | H2-Aa | 2.653 | 9.54E-03 | glio |
| interaction_RT | Dedd2 | 1.216 | 9.54E-03 | glio |
| interaction_RT | Pitrm1 | -0.828 | 9.54E-03 | glio |
| interaction_RT | Ddx18 | -0.63 | 9.54E-03 | glio |
| Aars2_in.vivo_noRT | mt-Nd3 | 0.832 | 9.54E-03 | glio |
| Atr_in.vivo_noRT | Zfp668 | 1.573 | 9.54E-03 | glio |
| Atr_in.vivo_noRT | Abcf2 | -1.072 | 9.54E-03 | glio |
| Bcas2_ex.vivo_noRT | Gm43463 | 2.349 | 9.54E-03 | glio |
| Brca2_in.vivo_noRT | Gm45591 | 2.036 | 9.54E-03 | glio |
| Brca2_in.vivo_noRT | Cys1 | 2.022 | 9.54E-03 | glio |
| culture_effect | Pde11a | 2.013 | 9.54E-04 | glio |
| Interaction_genotype_Bora | Cpeb1 | -4.552 | 9.54E-04 | glio |
| interaction_RT | Gm32122 | 3.006 | 9.54E-04 | glio |
| Bard1_ex.vivo_noRT | Dcdc2a | 2.814 | 9.54E-04 | glio |
| Eif2s1_in.vivo_noRT | Gm47510 | 1.883 | 9.54E-04 | glio |
| culture_effect | Nsdhl | -1.118 | 9.54E-05 | glio |
| Atr_in.vivo_noRT | Higd1a | -1.497 | 9.54E-05 | glio |
| Chek1_in.vivo_noRT | 1700041G16R | 2.789 | 9.54E-05 | glio |
| Interaction_genotype_Myc | Gm15890 | -4.292 | 9.54E-06 | glio |
| Bcas2_in.vivo_noRT | Emilin2 | 3.2 | 9.54E-06 | glio |
| culture_effect | Adgrb3 | 2.493 | 9.54E-07 | glio |
| Atr_ex.vivo_noRT | Nmrk2 | 3.962 | 9.54E-07 | glio |
| culture_effect | Msrb3 | 1.958 | 9.54E-08 | glio |
| culture_effect | Mogs | -1.043 | 9.54E-08 | glio |
| Atr_in.vivo_noRT | Rpl12 | -1.111 | 9.54E-09 | glio |
| Bora_in.vivo_noRT | Gm9970 | 4.291 | 9.54E-09 | glio |
| Chek1_in.vivo_noRT | Rab37 | 4.126 | 9.54E-10 | glio |
| culture_effect | Zbtb26 | 0.699 | 9.55E-03 | glio |
| Interaction_genotype_Bcas2 | A330032B11R | -3.178 | 9.55E-03 | glio |
| Interaction_genotype_Bcas2 | Atg9b | 3.152 | 9.55E-03 | glio |
| Interaction_genotype_Bcas2 | Serf1 | 1.124 | 9.55E-03 | glio |
| Atr_in.vivo_noRT | BC025920 | 1.693 | 9.55E-03 | glio |
| Atr_in.vivo_noRT | Ube2z | -1.153 | 9.55E-03 | glio |
| Bcas2_in.vivo_noRT | Phf11b | 2.146 | 9.55E-03 | glio |
| Interaction_genotype_Terf1 | Prelid3b | 2.037 | 9.55E-04 | glio |
| Interaction_genotype_Terf1 | Klhdc8a | -3.721 | 9.55E-04 | glio |
| Interaction_genotype_Terf1 | Gm2115 | -4.263 | 9.55E-04 | glio |
| Interaction_genotype_Terf1 | Ntn4 | -3.658 | 9.55E-04 | glio |
| Interaction_genotype_Terf1 | 4930524O07R | -4.88 | 9.55E-04 | glio |

| | | | | |
|---|---|---|---|---|
| Interaction_genotype_Terf1 | Snip1 | 2.888 | 9.55E-04 | glio |
| Interaction_genotype_Terf1 | Itga2 | -4.246 | 9.55E-04 | glio |
| Interaction_genotype_Terf1 | Col6a2 | -3.568 | 9.55E-04 | glio |
| Interaction_genotype_Terf1 | Adam12 | -4.152 | 9.55E-04 | glio |
| Interaction_genotype_Terf1 | 6330415G19R | -4.62 | 9.55E-04 | glio |
| Aars2_in.vivo_noRT | Tent5d | 2.268 | 9.55E-04 | glio |
| Aars2_in.vivo_noRT | Ano6 | -1.434 | 9.55E-04 | glio |
| Interaction_genotype_Mtor | 4930540M05F | -4.411 | 9.55E-05 | glio |
| Cenpj_ex.vivo_noRT | Sucnr1 | 2.591 | 9.55E-05 | glio |
| Interaction_genotype_Myc | Calhm5 | -4.77 | 9.55E-06 | glio |
| Atr_in.vivo_noRT | Serp2 | 2.796 | 9.55E-06 | glio |
| Bora_in.vivo_noRT | Tacr2 | 3.504 | 9.55E-07 | glio |
| Bora_in.vivo_noRT | Gm49864 | 3.504 | 9.55E-07 | glio |
| Hsd17b10_in.vivo_noRT | Muc4 | 3.094 | 9.55E-08 | glio |
| Interaction_genotype_Mtor | Gm50321 | -5.995 | 9.55E-09 | glio |
| Interaction_genotype_Mtor | 1700034I23Ri | -4.896 | 9.55E-09 | glio |
| Bcas2_in.vivo_noRT | Sult5a1 | 4.275 | 9.55E-09 | glio |
| Bcas2_in.vivo_noRT | Gm20275 | 3.453 | 9.55E-10 | glio |
| Bora_in.vivo_noRT | Gm10300 | 4.676 | 9.55E-11 | glio |
| culture_effect | Tmod2 | 1.877 | 9.55E-19 | glio |
| Atr_in.vivo_noRT | Flacc1 | 5.533 | 9.55E-20 | glio |
| Interaction_genotype_Ifnar1 | Gm13078 | 2.734 | 9.56E-03 | glio |
| Interaction_genotype_Ifnar1 | Gbp2 | -1.832 | 9.56E-03 | glio |
| Interaction_genotype_Mtor | Lrp6 | 1.387 | 9.56E-03 | glio |
| interaction_RT | Pgp | -0.554 | 9.56E-03 | glio |
| Chek1_ex.vivo_noRT | Kiss1r | 2.53 | 9.56E-03 | glio |
| Chek1_ex.vivo_noRT | H2-Q4 | 1.644 | 9.56E-03 | glio |
| Eif2s1_in.vivo_noRT | Gm15559 | 2.27 | 9.56E-03 | glio |
| culture_effect | C230013L11Ri | 1.998 | 9.56E-04 | glio |
| Interaction_genotype_Atr | Pecam1 | -4.222 | 9.56E-04 | glio |
| Interaction_genotype_Myc | Mmp28 | -3.815 | 9.56E-04 | glio |
| Interaction_genotype_Myc | Lrrc61 | -3.573 | 9.56E-04 | glio |
| Aars2_in.vivo_noRT | Cd59a | 2.407 | 9.56E-04 | glio |
| Aars2_in.vivo_noRT | Smn1 | -1.313 | 9.56E-04 | glio |
| Atr_in.vivo_noRT | Gpi1 | -0.95 | 9.56E-04 | glio |
| Bcas2_in.vivo_noRT | Hmgcs2 | 2.723 | 9.56E-04 | glio |
| Atr_in.vivo_noRT | Gm17056 | 2.8 | 9.56E-05 | glio |
| Atr_in.vivo_noRT | Gm45589 | 2.721 | 9.56E-05 | glio |
| culture_effect | Gm19541 | 1.937 | 9.56E-06 | glio |
| Aars2_in.vivo_noRT | Palm3 | 3.2 | 9.56E-06 | glio |
| Atr_ex.vivo_noRT | Esr1 | 3.943 | 9.56E-06 | glio |
| culture_effect | Gtsf1l | -2.564 | 9.56E-07 | glio |
| Bora_in.vivo_noRT | Nyap2 | 4.024 | 9.56E-09 | glio |
| Hsd17b10_in.vivo_noRT | 4930512B01R | 3.616 | 9.56E-12 | glio |
| Interaction_genotype_Aars2 | Id3 | 1.728 | 9.57E-03 | glio |
| Interaction_genotype_Atr | Kcnab3 | -3.347 | 9.57E-03 | glio |
| Interaction_genotype_Bcas2 | Gm15127 | -3.397 | 9.57E-03 | glio |
| Interaction_genotype_Myc | Mr1 | -2.997 | 9.57E-03 | glio |
| Atr_in.vivo_noRT | P2ry2 | 1.828 | 9.57E-03 | glio |
| Bcas2_ex.vivo_noRT | Gm28535 | 2.677 | 9.57E-03 | glio |
| Bora_in.vivo_noRT | Fbxo4 | -1.232 | 9.57E-03 | glio |
| Brca2_in.vivo_noRT | Zscan4-ps2 | 1.763 | 9.57E-03 | glio |

| | | | | |
|---|---|---|---|---|
| Chek1_in.vivo_noRT | Csnk1d | -1.034 | 9.57E-03 | glio |
| Chek1_ex.vivo_noRT | Hist1h2af | 2.294 | 9.57E-03 | glio |
| Chek1_ex.vivo_noRT | Shoc2 | 1.282 | 9.57E-03 | glio |
| Interaction_genotype_Aars2 | Gm15579 | -4.474 | 9.57E-04 | glio |
| Interaction_genotype_Mtor | Pten | 1.688 | 9.57E-04 | glio |
| Chek1_ex.vivo_noRT | Rfx6 | 2.961 | 9.57E-04 | glio |
| Interaction_genotype_Mtor | Cnksr1 | -4.34 | 9.57E-05 | glio |
| Interaction_genotype_Mtor | Zfp330 | 2.067 | 9.57E-05 | glio |
| Bcas2_in.vivo_noRT | Pcdhb14 | 2.836 | 9.57E-05 | glio |
| Bora_in.vivo_noRT | Fchsd1 | 2.692 | 9.57E-05 | glio |
| Interaction_genotype_Mtor | Btn1a1 | -4.623 | 9.57E-07 | glio |
| Chek1_in.vivo_noRT | 4930426I24Ri | 3.388 | 9.57E-07 | glio |
| Chek1_ex.vivo_noRT | Ccdc63 | 4.132 | 9.57E-07 | glio |
| Atr_in.vivo_noRT | Ngef | 3.525 | 9.57E-09 | glio |
| Bcas2_in.vivo_noRT | Mogat1 | 4.285 | 9.57E-09 | glio |
| Hsd17b10_in.vivo_noRT | Gm11232 | 3.616 | 9.57E-14 | glio |
| Bcas2_in.vivo_noRT | Cd7 | 4.543 | 9.57E-20 | glio |
| culture_effect | Kif3c | 0.871 | 9.58E-03 | glio |
| Atp5c1_in.vivo_noRT | Krt18 | -2.496 | 9.58E-03 | glio |
| Bora_in.vivo_noRT | Vps13c | 1.373 | 9.58E-03 | glio |
| Cenpj_ex.vivo_noRT | 4930553M12F | 2.198 | 9.58E-03 | glio |
| Chek1_in.vivo_noRT | Gm813 | 1.873 | 9.58E-03 | glio |
| Chek1_in.vivo_noRT | Runx2 | 1.732 | 9.58E-03 | glio |
| Chek1_in.vivo_noRT | Nipsnap3b | -1.04 | 9.58E-03 | glio |
| culture_effect | Ypel4 | -1.9 | 9.58E-04 | glio |
| Atr_in.vivo_noRT | Klhl36 | 2.291 | 9.58E-04 | glio |
| Bcas2_ex.vivo_noRT | St14 | 2.95 | 9.58E-04 | glio |
| Hsd17b10_in.vivo_noRT | Gm30124 | 2.346 | 9.58E-04 | glio |
| culture_effect | Foxk1 | -1.345 | 9.58E-06 | glio |
| Atr_in.vivo_noRT | Med9os | 3.107 | 9.58E-06 | glio |
| Bora_in.vivo_noRT | Apoa1 | 3.903 | 9.58E-08 | glio |
| Atr_in.vivo_noRT | Gm10734 | 4.017 | 9.58E-09 | glio |
| Bora_in.vivo_noRT | Gabrd | 4.02 | 9.58E-09 | glio |
| Bora_in.vivo_noRT | Ninj2 | 4.553 | 9.58E-11 | glio |
| culture_effect | Ap1s1 | 1.172 | 9.58E-21 | glio |
| culture_effect | Rtl5 | -1.612 | 9.59E-03 | glio |
| culture_effect | Tsr1 | -0.457 | 9.59E-03 | glio |
| Interaction_genotype_Myc | Gm50431 | -2.654 | 9.59E-03 | glio |
| Interaction_genotype_Terf1 | Gm29340 | -3.532 | 9.59E-03 | glio |
| interaction_RT | Ncaph2 | -0.592 | 9.59E-03 | glio |
| Atr_in.vivo_noRT | Primpol | 1.521 | 9.59E-03 | glio |
| Chek1_in.vivo_noRT | Fut11 | 1.87 | 9.59E-03 | glio |
| Chek1_in.vivo_noRT | Tomm6 | -0.68 | 9.59E-03 | glio |
| Hsd17b10_in.vivo_noRT | Abcc4 | -1.66 | 9.59E-03 | glio |
| Hsd17b10_in.vivo_noRT | Grsf1 | -0.717 | 9.59E-03 | glio |
| culture_effect | Htt | -0.548 | 9.59E-04 | glio |
| Interaction_genotype_Myc | Six5 | -3.307 | 9.59E-04 | glio |
| Cyp19a1_in.vivo_noRT | Apol6 | -1.979 | 9.59E-04 | glio |
| Interaction_genotype_Mtor | 1700122E12Ri | -4.702 | 9.59E-05 | glio |
| Interaction_genotype_Myc | Uqcr11 | 1.918 | 9.59E-05 | glio |
| Aars2_in.vivo_noRT | Bcl7a | 2.792 | 9.59E-05 | glio |
| Bora_in.vivo_noRT | Nupr1l | 3.156 | 9.59E-05 | glio |

| | | | | |
|---|---|---|---|---|
| Bora_in.vivo_noRT | Gm48684 | 3.632 | 9.59E-06 | glio |
| Interaction_genotype_Bora | Cdk11b | 2.409 | 9.59E-07 | glio |
| Bora_in.vivo_noRT | Rax | 3.757 | 9.59E-09 | glio |
| Hsd17b10_in.vivo_noRT | Gm34939 | 3.352 | 9.59E-09 | glio |
| culture_effect | Rps25 | -0.758 | 9.59E-10 | glio |
| Bora_in.vivo_noRT | Gm39459 | 4.286 | 9.59E-10 | glio |
| Chek1_ex.vivo_noRT | Gm30042 | 4.842 | 9.59E-12 | glio |
| Interaction_genotype_Aars2 | Kcnk13 | -2.559 | 9.60E-03 | glio |
| Interaction_genotype_Aars2 | Ptges | -3.449 | 9.60E-03 | glio |
| Interaction_genotype_Aars2 | Acacb | -3.223 | 9.60E-03 | glio |
| Interaction_genotype_Aars2 | Smurf2 | -1.941 | 9.60E-03 | glio |
| Interaction_genotype_Atr | Gm37933 | -2.697 | 9.60E-03 | glio |
| Interaction_genotype_Atr | Ndufaf5 | -2.141 | 9.60E-03 | glio |
| Interaction_genotype_Atr | Higd2a | 1.244 | 9.60E-03 | glio |
| Interaction_genotype_Bora | 4921523L03Ri | -3.63 | 9.60E-03 | glio |
| Interaction_genotype_Bora | 0610040B10R | -2.708 | 9.60E-03 | glio |
| Interaction_genotype_Ifnar1 | Gm13964 | 2.734 | 9.60E-03 | glio |
| Interaction_genotype_Mtor | Mtss1 | -3.355 | 9.60E-03 | glio |
| Interaction_genotype_Mtor | Hikeshi | 1.244 | 9.60E-03 | glio |
| Interaction_genotype_Myc | Gm13054 | -2.599 | 9.60E-03 | glio |
| Interaction_genotype_Myc | Bbs10 | -2.951 | 9.60E-03 | glio |
| Interaction_genotype_Myc | Tpk1 | -2.697 | 9.60E-03 | glio |
| Interaction_genotype_Pars2 | D230022J07Ri | -3.599 | 9.60E-03 | glio |
| Aars2_in.vivo_noRT | Rpl24 | -0.508 | 9.60E-03 | glio |
| Atp5c1_in.vivo_noRT | Gm43660 | 2.028 | 9.60E-03 | glio |
| Atp5c1_in.vivo_noRT | Mgat2 | -0.954 | 9.60E-03 | glio |
| Atr_in.vivo_noRT | Runx2 | 1.699 | 9.60E-03 | glio |
| Atr_in.vivo_noRT | Nob1 | -1.107 | 9.60E-03 | glio |
| Bard1_ex.vivo_noRT | Ubd | 2.421 | 9.60E-03 | glio |
| Chek1_in.vivo_noRT | Spred3 | 1.6 | 9.60E-03 | glio |
| Hsd17b10_in.vivo_noRT | Gm12500 | 1.975 | 9.60E-03 | glio |
| Interaction_genotype_Mtor | Habp4 | -2.909 | 9.60E-04 | glio |
| Aars2_ex.vivo_noRT | Prrt3 | 2.9 | 9.60E-04 | glio |
| Aars2_ex.vivo_noRT | C3 | 3.293 | 9.60E-04 | glio |
| Chek1_in.vivo_noRT | Fam129c | 2.376 | 9.60E-04 | glio |
| Hsd17b10_in.vivo_noRT | Pcdhga2 | 2.346 | 9.60E-04 | glio |
| Aars2_in.vivo_noRT | Mapt | 2.136 | 9.60E-05 | glio |
| Bora_in.vivo_noRT | Mmp19 | 2.742 | 9.60E-05 | glio |
| Bora_in.vivo_noRT | Nmnat3 | 3.296 | 9.60E-06 | glio |
| Aars2_in.vivo_noRT | E430018J23Ri | 3.272 | 9.60E-07 | glio |
| Hsd17b10_in.vivo_noRT | Upb1 | 3.353 | 9.60E-07 | glio |
| Bora_in.vivo_noRT | Gm128 | 4.018 | 9.60E-13 | glio |
| Chek1_in.vivo_noRT | Gm39526 | 4.521 | 9.60E-17 | glio |
| culture_effect | Tmprss5 | -1.645 | 9.61E-03 | glio |
| Interaction_genotype_Atr | Gm5890 | -2.369 | 9.61E-03 | glio |
| Aars2_in.vivo_noRT | Dus1l | -1.166 | 9.61E-03 | glio |
| Atr_in.vivo_noRT | Vasn | 1.767 | 9.61E-03 | glio |
| Atr_in.vivo_noRT | Hnrnpa1 | -0.574 | 9.61E-03 | glio |
| Bora_in.vivo_noRT | Triobp | -1.312 | 9.61E-03 | glio |
| Chek1_in.vivo_noRT | Nampt | -0.872 | 9.61E-03 | glio |
| Chek1_in.vivo_noRT | Sbds | -0.872 | 9.61E-03 | glio |
| Hsd17b10_in.vivo_noRT | Slc27a3 | 2.033 | 9.61E-03 | glio |

| | | | | |
|---|---|---|---|---|
| Interaction_genotype_Atr | Taf1d | 1.407 | 9.61E-04 | glio |
| Atr_in.vivo_noRT | Gpr161 | 2.406 | 9.61E-04 | glio |
| Atr_in.vivo_noRT | Zfp607b | 1.967 | 9.61E-04 | glio |
| Bard1_ex.vivo_noRT | Prap1 | 2.421 | 9.61E-04 | glio |
| Bora_in.vivo_noRT | Ogdhl | 2.381 | 9.61E-04 | glio |
| Bora_in.vivo_noRT | Tinagl1 | -1.532 | 9.61E-04 | glio |
| Eif2s1_in.vivo_noRT | Spink6 | 2.143 | 9.61E-04 | glio |
| Eif2s1_in.vivo_noRT | Dnm3 | 2.671 | 9.61E-04 | glio |
| Eif2s1_in.vivo_noRT | Gm19265 | 2.154 | 9.61E-04 | glio |
| interaction_RT | Acot13 | 1 | 9.61E-05 | glio |
| Bcas2_in.vivo_noRT | Gdf15 | -2.859 | 9.61E-05 | glio |
| Bora_ex.vivo_noRT | Car10 | 3.201 | 9.61E-05 | glio |
| Hsd17b10_in.vivo_noRT | Zfp808 | 2.499 | 9.61E-05 | glio |
| Aars2_in.vivo_noRT | Gm13283 | 3.435 | 9.61E-07 | glio |
| Atr_in.vivo_noRT | Il23a | 3.412 | 9.61E-07 | glio |
| Chek1_in.vivo_noRT | 9530018F02Ri | 3.658 | 9.61E-07 | glio |
| Chek1_in.vivo_noRT | Mterf1b | 3.276 | 9.61E-07 | glio |
| Interaction_genotype_Mtor | Vmn1r32 | -4.492 | 9.61E-08 | glio |
| Interaction_genotype_Mtor | Gm13191 | -4.099 | 9.61E-08 | glio |
| Interaction_genotype_Mtor | 4933416E03Ri | -4.099 | 9.61E-08 | glio |
| Bcas2_in.vivo_noRT | Gm17210 | 3.747 | 9.61E-08 | glio |
| Hsd17b10_in.vivo_noRT | Oit3 | 3.616 | 9.61E-08 | glio |
| Bcas2_in.vivo_noRT | Vpreb1 | 3.904 | 9.61E-09 | glio |
| Chek1_in.vivo_noRT | Gm31518 | 5.044 | 9.61E-13 | glio |
| culture_effect | Rhou | 1.837 | 9.61E-14 | glio |
| Interaction_genotype_Atr | Slc13a5 | -3.135 | 9.62E-03 | glio |
| Interaction_genotype_Bora | Platr4 | -2.75 | 9.62E-03 | glio |
| Interaction_genotype_Mtor | Gucd1 | -2.412 | 9.62E-03 | glio |
| Interaction_genotype_Mtor | Aatf | 1.432 | 9.62E-03 | glio |
| Aars2_in.vivo_noRT | Zfp420 | 1.99 | 9.62E-03 | glio |
| Atr_in.vivo_noRT | Aimp1 | -0.669 | 9.62E-03 | glio |
| Cdkn2a_in.vivo_noRT | Cldn10 | 1.816 | 9.62E-03 | glio |
| Interaction_genotype_Mtor | Gatad1 | 1.59 | 9.62E-04 | glio |
| interaction_RT | Tmcc3 | -1.018 | 9.62E-04 | glio |
| Aars2_ex.vivo_noRT | Cacng4 | 2.9 | 9.62E-04 | glio |
| Atp5c1_in.vivo_noRT | Dnah10 | 2.289 | 9.62E-04 | glio |
| Atp5c1_in.vivo_noRT | Loxl4 | 2.402 | 9.62E-04 | glio |
| Atr_in.vivo_noRT | Ogfod2 | 1.647 | 9.62E-04 | glio |
| Atr_in.vivo_noRT | Aldoa | -0.973 | 9.62E-04 | glio |
| Bora_in.vivo_noRT | Syk | 2.13 | 9.62E-04 | glio |
| culture_effect | 4930412C18R | 2.003 | 9.62E-05 | glio |
| culture_effect | Rassf7 | 1.348 | 9.62E-05 | glio |
| culture_effect | Brms1 | -0.643 | 9.62E-05 | glio |
| Interaction_genotype_Atr | Gramd2 | -4.6 | 9.62E-05 | glio |
| Interaction_genotype_Atr | Eng | -5.22 | 9.62E-05 | glio |
| Interaction_genotype_Mtor | BC049987 | -3.316 | 9.62E-05 | glio |
| Interaction_genotype_Mtor | Wnt4 | -5.219 | 9.62E-05 | glio |
| Interaction_genotype_Mtor | Gm13986 | -4.739 | 9.62E-05 | glio |
| Interaction_genotype_Mtor | Gng7 | -4.369 | 9.62E-05 | glio |
| Atr_ex.vivo_noRT | Gm34776 | 2.77 | 9.62E-05 | glio |
| Bora_in.vivo_noRT | Rbakdn | 2.759 | 9.62E-05 | glio |
| Bora_ex.vivo_noRT | Vmn1r32 | 2.808 | 9.62E-06 | glio |

| | | | | |
|---|---|---|---|---|
| Chek1_ex.vivo_noRT | Tspyl3 | 3.463 | 9.62E-06 | glio |
| Atr_in.vivo_noRT | Ifit1bl2 | 3.84 | 9.62E-08 | glio |
| culture_effect | Rasl12 | -3.982 | 9.62E-12 | glio |
| culture_effect | Dusp8 | 0.691 | 9.63E-03 | glio |
| Interaction_genotype_Mtor | Zfp169 | -3.027 | 9.63E-03 | glio |
| Interaction_genotype_Myc | Btbd3 | -1.722 | 9.63E-03 | glio |
| Aars2_in.vivo_noRT | Gm32031 | 1.665 | 9.63E-03 | glio |
| Atr_in.vivo_noRT | Rps11 | -0.476 | 9.63E-03 | glio |
| Bora_in.vivo_noRT | Adm | 1.728 | 9.63E-03 | glio |
| Cenpj_ex.vivo_noRT | Myt1 | 2.192 | 9.63E-03 | glio |
| Interaction_genotype_Abtb1 | Myoz1 | 4.323 | 9.63E-04 | glio |
| Interaction_genotype_Bora | Zmynd15 | -4.284 | 9.63E-04 | glio |
| Interaction_genotype_Bora | Pitx3 | -3.499 | 9.63E-04 | glio |
| Interaction_genotype_Bora | Cdk2 | 2.686 | 9.63E-04 | glio |
| Interaction_genotype_Bora | Cops8 | 1.789 | 9.63E-04 | glio |
| Interaction_genotype_Mtor | Ttc21b | -2.859 | 9.63E-04 | glio |
| Aars2_in.vivo_noRT | Cdc42bpg | 2.17 | 9.63E-04 | glio |
| Chek1_in.vivo_noRT | Kcnt2 | 2.321 | 9.63E-04 | glio |
| culture_effect | Ckm | -2.412 | 9.63E-05 | glio |
| culture_effect | Thop1 | -0.65 | 9.63E-05 | glio |
| Atp5c1_in.vivo_noRT | 1700024I08Ri | 2.408 | 9.63E-05 | glio |
| Bcas2_in.vivo_noRT | Phf11a | 2.715 | 9.63E-05 | glio |
| Bcas2_in.vivo_noRT | Knop1 | -1.453 | 9.63E-05 | glio |
| Bora_in.vivo_noRT | Gm10516 | 3.373 | 9.63E-07 | glio |
| Atr_in.vivo_noRT | Sgk3 | 3.979 | 9.63E-09 | glio |
| Atr_in.vivo_noRT | 4930432K21R | 3.681 | 9.63E-09 | glio |
| Aars2_in.vivo_noRT | Cfap46 | 4.859 | 9.63E-15 | glio |
| Bcas2_in.vivo_noRT | Gm45591 | 4.547 | 9.63E-15 | glio |
| Interaction_genotype_Aars2 | Ndufb4 | 1.051 | 9.64E-03 | glio |
| Interaction_genotype_Cdkn2a | Lama5 | -3.704 | 9.64E-03 | glio |
| Interaction_genotype_Myc | Gm4858 | -2.729 | 9.64E-03 | glio |
| Interaction_genotype_Myc | D430020J02Ri | -2.493 | 9.64E-03 | glio |
| Interaction_genotype_Myc | Spidr | -2.376 | 9.64E-03 | glio |
| Interaction_genotype_Myc | Mthfsd | -1.982 | 9.64E-03 | glio |
| Atr_in.vivo_noRT | Caskin2 | 1.543 | 9.64E-03 | glio |
| Bcas2_in.vivo_noRT | Pcdh9 | 1.946 | 9.64E-03 | glio |
| Bcas2_ex.vivo_noRT | B230307C23R | -2.136 | 9.64E-03 | glio |
| Bora_in.vivo_noRT | Hnrnpa0 | -0.845 | 9.64E-03 | glio |
| Chek1_in.vivo_noRT | Hdc | 2.002 | 9.64E-03 | glio |
| Chek1_ex.vivo_noRT | Rnf113a1 | 1.949 | 9.64E-03 | glio |
| Eif2s1_ex.vivo_noRT | Gm11769 | 2.643 | 9.64E-03 | glio |
| Interaction_genotype_Myc | Prickle3 | -3.391 | 9.64E-04 | glio |
| Bora_in.vivo_noRT | Ccdc13 | 2.446 | 9.64E-04 | glio |
| Bora_in.vivo_noRT | Cnot6 | -1.304 | 9.64E-04 | glio |
| culture_effect | 4930578M01F | 1.383 | 9.64E-05 | glio |
| culture_effect | Lgmn | -0.991 | 9.64E-05 | glio |
| culture_effect | Skp1a | 0.524 | 9.64E-06 | glio |
| Chek1_ex.vivo_noRT | Il7 | 4.045 | 9.64E-07 | glio |
| Aars2_in.vivo_noRT | Tnfrsf13b | 4.198 | 9.64E-09 | glio |
| Chek1_ex.vivo_noRT | 1700112J16Ri | 4.256 | 9.64E-11 | glio |
| Chek1_in.vivo_noRT | Fam131a | 4.518 | 9.64E-12 | glio |
| Aars2_in.vivo_noRT | Cpxm1 | 5.06 | 9.64E-17 | glio |

| | | | | |
|---|---|---|---|---|
| Aars2_in.vivo_noRT | Gm43915 | 4.794 | 9.64E-17 | glio |
| Aars2_in.vivo_noRT | Ccl9 | 5.327 | 9.64E-27 | glio |
| Interaction_genotype_Atr | Scgb1c1 | -2.762 | 9.65E-03 | glio |
| Interaction_genotype_Bcas2 | Tlnrd1 | 1.84 | 9.65E-03 | glio |
| Interaction_genotype_Lmo7 | Gm35019 | 3.387 | 9.65E-03 | glio |
| Interaction_genotype_Lmo7 | Tlr1 | -4.115 | 9.65E-03 | glio |
| Interaction_genotype_Mtor | Mbd3 | 1.228 | 9.65E-03 | glio |
| Interaction_genotype_Mtor | Pfdn6 | 1.209 | 9.65E-03 | glio |
| Interaction_genotype_Rpl3 | Slc3a2 | 1.157 | 9.65E-03 | glio |
| Aars2_in.vivo_noRT | Chmp4b | -0.912 | 9.65E-03 | glio |
| Aars2_in.vivo_noRT | Csnk1e | 0.753 | 9.65E-03 | glio |
| Aars2_in.vivo_noRT | Rps20 | -0.564 | 9.65E-03 | glio |
| Bcas2_in.vivo_noRT | Faim2 | 1.544 | 9.65E-03 | glio |
| Chek1_in.vivo_noRT | Gm45716 | 1.829 | 9.65E-03 | glio |
| Interaction_genotype_Myc | Hsf2bp | -3.499 | 9.65E-04 | glio |
| Interaction_genotype_Myc | Plekhg5 | -3.556 | 9.65E-04 | glio |
| Atr_in.vivo_noRT | Lias | -1.375 | 9.65E-04 | glio |
| Bora_in.vivo_noRT | Ifitm2 | -0.699 | 9.65E-04 | glio |
| Chek1_in.vivo_noRT | Rps6 | -0.576 | 9.65E-04 | glio |
| culture_effect | Nrbp2 | -1.839 | 9.65E-05 | glio |
| Aars2_in.vivo_noRT | Rbx1 | -0.731 | 9.65E-05 | glio |
| Bora_in.vivo_noRT | 1110013H19R | 3.064 | 9.65E-05 | glio |
| Bora_in.vivo_noRT | Adcy7 | 2.64 | 9.65E-05 | glio |
| Bora_in.vivo_noRT | Tmem79 | 2.652 | 9.65E-05 | glio |
| culture_effect | Oas1g | 1.884 | 9.65E-06 | glio |
| culture_effect | Pigu | -1.046 | 9.65E-06 | glio |
| culture_effect | Acaca | 1.026 | 9.65E-06 | glio |
| Interaction_genotype_Myc | Slc25a35 | -5.024 | 9.65E-06 | glio |
| interaction_RT | Aqp5 | 4.164 | 9.65E-06 | glio |
| Aars2_in.vivo_noRT | B430219N15R | 3.203 | 9.65E-06 | glio |
| Aars2_in.vivo_noRT | Nyap1 | 3.111 | 9.65E-06 | glio |
| Bcas2_in.vivo_noRT | Gm36839 | 3.245 | 9.65E-06 | glio |
| Aars2_in.vivo_noRT | Gm11127 | 3.766 | 9.65E-08 | glio |
| culture_effect | Tma7 | 0.641 | 9.65E-10 | glio |
| culture_effect | Fchsd2 | -1.681 | 9.65E-16 | glio |
| Aars2_in.vivo_noRT | Slc35g3 | 5.327 | 9.65E-26 | glio |
| Interaction_genotype_Bcas2 | Axin1 | 2.132 | 9.66E-03 | glio |
| Interaction_genotype_Bcas2 | Acot8 | 1.978 | 9.66E-03 | glio |
| Atr_ex.vivo_noRT | Whamm | 1.503 | 9.66E-03 | glio |
| Bcas2_in.vivo_noRT | B230216N24R | 2.133 | 9.66E-03 | glio |
| Bcas2_in.vivo_noRT | 4933404O12R | 1.902 | 9.66E-03 | glio |
| Bcas2_in.vivo_noRT | Nrp2 | -1.115 | 9.66E-03 | glio |
| Cenpj_in.vivo_noRT | 9530062K07R | 2.413 | 9.66E-03 | glio |
| Chek1_in.vivo_noRT | Tbc1d22a | 1.428 | 9.66E-03 | glio |
| Chek1_ex.vivo_noRT | Btbd17 | 2.973 | 9.66E-03 | glio |
| Chek1_ex.vivo_noRT | Tlr6 | 2.344 | 9.66E-03 | glio |
| culture_effect | Dpep3 | -1.467 | 9.66E-04 | glio |
| Interaction_genotype_Myc | 4930555A03R | -3.457 | 9.66E-04 | glio |
| Bcas2_in.vivo_noRT | Tex19.2 | 2.343 | 9.66E-04 | glio |
| Bcas2_in.vivo_noRT | Gm26621 | 2.405 | 9.66E-04 | glio |
| Chek1_in.vivo_noRT | Map3k8 | 2.568 | 9.66E-04 | glio |
| Interaction_genotype_Myc | Gpc5 | -3.124 | 9.66E-05 | glio |

| | | | | |
|---|---|---|---|---|
| Atr_in.vivo_noRT | Wdr27 | 2.973 | 9.66E-07 | glio |
| Interaction_genotype_Bcas2 | Col6a2 | -5.406 | 9.66E-08 | glio |
| Chek1_ex.vivo_noRT | Astn2 | 4.447 | 9.66E-08 | glio |
| Aars2_in.vivo_noRT | Gm10399 | 4.395 | 9.66E-11 | glio |
| Bora_in.vivo_noRT | 1700030L20Ri | 4.28 | 9.66E-11 | glio |
| Interaction_genotype_Atr | Krtcap3 | -2.563 | 9.67E-03 | glio |
| Interaction_genotype_Atr | Ccdc62 | -2.967 | 9.67E-03 | glio |
| Interaction_genotype_Bcas2 | Mucl3 | -3.697 | 9.67E-03 | glio |
| Interaction_genotype_Bora | Gm14968 | -3.309 | 9.67E-03 | glio |
| Interaction_genotype_Rad51 | Lbhd2 | -4.337 | 9.67E-03 | glio |
| Interaction_genotype_Rad51 | Asgr1 | -3.663 | 9.67E-03 | glio |
| Interaction_genotype_Rad51 | Brwd3 | 2.483 | 9.67E-03 | glio |
| Atr_in.vivo_noRT | Gm13261 | 1.759 | 9.67E-03 | glio |
| Atr_ex.vivo_noRT | Rdm1 | 1.309 | 9.67E-03 | glio |
| Bora_in.vivo_noRT | Rad51ap1 | -1.476 | 9.67E-03 | glio |
| Chek1_in.vivo_noRT | Set | -0.515 | 9.67E-03 | glio |
| Interaction_genotype_Bora | Crhbp | -3.772 | 9.67E-04 | glio |
| Interaction_genotype_Myc | Ak4 | -2.977 | 9.67E-04 | glio |
| Interaction_genotype_Myc | Golph3 | 1.34 | 9.67E-04 | glio |
| Interaction_genotype_Rpl3 | A330084C13R | -4.818 | 9.67E-04 | glio |
| Aars2_in.vivo_noRT | Naca | -0.569 | 9.67E-04 | glio |
| Atr_in.vivo_noRT | Gm26881 | 2.066 | 9.67E-04 | glio |
| Aars2_in.vivo_noRT | A230072E10Ri | 3.074 | 9.67E-06 | glio |
| culture_effect | Cenpw | 0.841 | 9.67E-08 | glio |
| Interaction_genotype_Mtor | Gm48610 | -6.529 | 9.67E-08 | glio |
| culture_effect | Spsb2 | -0.967 | 9.67E-09 | glio |
| Aars2_in.vivo_noRT | Cnnm1 | 4.952 | 9.67E-11 | glio |
| Interaction_genotype_Atr | Rdh12 | -3.073 | 9.68E-03 | glio |
| Interaction_genotype_Atr | Gm6598 | -2.94 | 9.68E-03 | glio |
| Interaction_genotype_Bora | Lhx9 | -3.637 | 9.68E-03 | glio |
| Interaction_genotype_Myc | Atp5c1 | 1.174 | 9.68E-03 | glio |
| Interaction_genotype_Pars2 | Krt80 | -3.091 | 9.68E-03 | glio |
| interaction_RT | Arhgef28 | 1.959 | 9.68E-03 | glio |
| Aars2_in.vivo_noRT | Ccdc71l | 1.754 | 9.68E-03 | glio |
| Aars2_in.vivo_noRT | Hoxa1 | 1.829 | 9.68E-03 | glio |
| Atr_in.vivo_noRT | Grk5 | 1.486 | 9.68E-03 | glio |
| Atr_ex.vivo_noRT | Sgk1 | 1.526 | 9.68E-03 | glio |
| Bora_in.vivo_noRT | Rrnad1 | 1.453 | 9.68E-03 | glio |
| Chek1_in.vivo_noRT | Swsap1 | 1.675 | 9.68E-03 | glio |
| Interaction_genotype_Myc | Slc36a2 | -3.656 | 9.68E-04 | glio |
| Interaction_genotype_Myc | Zbtb4 | -3.489 | 9.68E-04 | glio |
| Interaction_genotype_Myc | Pqlc2 | -3.188 | 9.68E-04 | glio |
| Aars2_in.vivo_noRT | C230035I16Ri | 2.449 | 9.68E-04 | glio |
| Bora_in.vivo_noRT | Mrps28 | -1.218 | 9.68E-04 | glio |
| Chek1_in.vivo_noRT | Jmjd1c | -1.181 | 9.68E-04 | glio |
| Interaction_genotype_Mtor | Gm46218 | -4.284 | 9.68E-05 | glio |
| Atr_in.vivo_noRT | Tceal1 | 2.617 | 9.68E-05 | glio |
| Atr_in.vivo_noRT | Dapp1 | 2.343 | 9.68E-05 | glio |
| Bcas2_ex.vivo_noRT | Cd36 | 2.742 | 9.68E-05 | glio |
| Bcas2_ex.vivo_noRT | Gstt4 | 2.742 | 9.68E-05 | glio |
| Interaction_genotype_Mtor | Slc10a1 | -4.233 | 9.68E-06 | glio |
| Aars2_in.vivo_noRT | Ndufs8 | -1.474 | 9.68E-06 | glio |

| | | | | |
|---|---|---|---|---|
| Atp5c1_in.vivo_noRT | Sytl4 | 2.402 | 9.68E-06 | glio |
| Hsd17b10_in.vivo_noRT | Aoc1 | 3.356 | 9.68E-06 | glio |
| Aars2_in.vivo_noRT | Gm12454 | 3.422 | 9.68E-07 | glio |
| Chek1_in.vivo_noRT | Hist1h4k | 3.474 | 9.68E-07 | glio |
| Aars2_in.vivo_noRT | H3f3aos | 4.682 | 9.68E-12 | glio |
| Chek1_in.vivo_noRT | Tmem52 | 4.664 | 9.68E-14 | glio |
| Chek1_ex.vivo_noRT | Gm4340 | 4.842 | 9.68E-17 | glio |
| Chek1_ex.vivo_noRT | Gm16239 | 4.842 | 9.68E-17 | glio |
| Interaction_genotype_Bcas2 | F630040K05Ri | -3.139 | 9.69E-03 | glio |
| Interaction_genotype_Mtor | Rsad1 | -2.775 | 9.69E-03 | glio |
| Interaction_genotype_Myc | Gm20767 | -2.729 | 9.69E-03 | glio |
| Interaction_genotype_Myc | Gm43696 | -2.997 | 9.69E-03 | glio |
| Interaction_genotype_Myc | Stub1 | 1.029 | 9.69E-03 | glio |
| Interaction_genotype_Uqcrc2 | Gm15535 | -4.248 | 9.69E-03 | glio |
| Atp5c1_in.vivo_noRT | Slc6a13 | 2.289 | 9.69E-03 | glio |
| Atp5c1_in.vivo_noRT | S100a14 | 1.868 | 9.69E-03 | glio |
| Atr_in.vivo_noRT | Jun | -0.963 | 9.69E-03 | glio |
| Atr_ex.vivo_noRT | Aarsd1 | -1.17 | 9.69E-03 | glio |
| Bcas2_in.vivo_noRT | Prag1 | 1.759 | 9.69E-03 | glio |
| Bcas2_in.vivo_noRT | Slc25a44 | 1.353 | 9.69E-03 | glio |
| Bcas2_in.vivo_noRT | Map2k3 | -1.028 | 9.69E-03 | glio |
| Bora_in.vivo_noRT | Slc30a7 | -1.263 | 9.69E-03 | glio |
| Chek1_ex.vivo_noRT | Gm11963 | 2.386 | 9.69E-03 | glio |
| Chek1_ex.vivo_noRT | Prr13 | 0.995 | 9.69E-03 | glio |
| culture_effect | Rilp | 2.063 | 9.69E-04 | glio |
| Interaction_genotype_Atr | Snhg15 | 2.1 | 9.69E-04 | glio |
| Interaction_genotype_Mtor | Siva1 | 1.671 | 9.69E-04 | glio |
| interaction_RT | Hsdl2 | 1.223 | 9.69E-04 | glio |
| Aars2_in.vivo_noRT | Uqcc2 | -0.79 | 9.69E-04 | glio |
| Atr_ex.vivo_noRT | Rab27a | 2.769 | 9.69E-04 | glio |
| Bard1_ex.vivo_noRT | Etl4 | 2.814 | 9.69E-04 | glio |
| Bcas2_ex.vivo_noRT | Sdr16c5 | 3.145 | 9.69E-04 | glio |
| Chek1_ex.vivo_noRT | Cinp | -1.67 | 9.69E-04 | glio |
| Chek1_ex.vivo_noRT | Cnih1 | -1.276 | 9.69E-04 | glio |
| Interaction_genotype_Myc | Dnajc19 | 1.698 | 9.69E-05 | glio |
| Atr_in.vivo_noRT | Pkdcc | 2.821 | 9.69E-05 | glio |
| Chek1_in.vivo_noRT | Pnmal2 | 2.696 | 9.69E-05 | glio |
| Chek1_in.vivo_noRT | Cldn34c1 | 2.588 | 9.69E-05 | glio |
| Chek1_in.vivo_noRT | Adi1 | -1.886 | 9.69E-05 | glio |
| Chek1_in.vivo_noRT | Plagl1 | -2.141 | 9.69E-05 | glio |
| Chek1_in.vivo_noRT | Rer1 | -0.969 | 9.69E-05 | glio |
| Aars2_ex.vivo_noRT | Rgs4 | 3.293 | 9.69E-07 | glio |
| Aars2_in.vivo_noRT | Lsp1 | 4.14 | 9.69E-08 | glio |
| Hsd17b10_in.vivo_noRT | Prl2c2 | 3.616 | 9.69E-08 | glio |
| culture_effect | Cxcl9 | -6.517 | 9.69E-12 | glio |
| Aars2_in.vivo_noRT | Accsl | 5.067 | 9.69E-12 | glio |
| Aars2_in.vivo_noRT | Gm15903 | 4.551 | 9.69E-12 | glio |
| Interaction_genotype_Mtor | Trdmt1 | -2.652 | 9.70E-03 | glio |
| Interaction_genotype_Myc | Gm11899 | -2.207 | 9.70E-03 | glio |
| Interaction_genotype_Rad51 | Baiap2l2 | -3.506 | 9.70E-03 | glio |
| interaction_RT | Gm12602 | 2.237 | 9.70E-03 | glio |
| interaction_RT | Gnpat | -0.893 | 9.70E-03 | glio |

| | | | | |
|---|---|---|---|---|
| interaction_RT | Cbx3 | 0.561 | 9.70E-03 | glio |
| Aars2_in.vivo_noRT | Gm38115 | 1.886 | 9.70E-03 | glio |
| Aars2_in.vivo_noRT | Gzf1 | 1.246 | 9.70E-03 | glio |
| Aars2_ex.vivo_noRT | Ccdc63 | 2.582 | 9.70E-03 | glio |
| Bard1_ex.vivo_noRT | Cacng4 | 2.421 | 9.70E-03 | glio |
| Bcas2_in.vivo_noRT | Usp6nl | -1.312 | 9.70E-03 | glio |
| Bora_ex.vivo_noRT | Ntrk1 | 2.755 | 9.70E-03 | glio |
| Cyp19a1_ex.vivo_noRT | Rgs4 | 2.159 | 9.70E-03 | glio |
| Cyp19a1_ex.vivo_noRT | Gm13178 | 2.159 | 9.70E-03 | glio |
| Interaction_genotype_Chek1 | Gnptg | 2.757 | 9.70E-04 | glio |
| Interaction_genotype_Mtor | Tmem151a | -3.967 | 9.70E-04 | glio |
| Interaction_genotype_Myc | Zfpm1 | -3.87 | 9.70E-04 | glio |
| Interaction_genotype_Myc | Gm15128 | -2.971 | 9.70E-04 | glio |
| Interaction_genotype_Myc | Cep63 | -2.421 | 9.70E-04 | glio |
| Aars2_in.vivo_noRT | 1500011B03R | -1.367 | 9.70E-04 | glio |
| Bcas2_in.vivo_noRT | Gm47689 | 2.418 | 9.70E-04 | glio |
| Bcas2_in.vivo_noRT | Zfp184 | 2.172 | 9.70E-04 | glio |
| Bcas2_in.vivo_noRT | Ywhah | -0.727 | 9.70E-04 | glio |
| Chek1_in.vivo_noRT | Ankrd23 | 2.536 | 9.70E-04 | glio |
| Hsd17b10_in.vivo_noRT | St14 | 2.23 | 9.70E-04 | glio |
| Bora_in.vivo_noRT | A430010J10Ri | 3.43 | 9.70E-05 | glio |
| Chek1_ex.vivo_noRT | Cct5 | -0.996 | 9.70E-05 | glio |
| Atr_in.vivo_noRT | Tmem240 | 3.805 | 9.70E-07 | glio |
| Hsd17b10_in.vivo_noRT | Gm5941 | 3.616 | 9.70E-07 | glio |
| Bora_in.vivo_noRT | 4930545E07Ri | 3.903 | 9.70E-08 | glio |
| Bcas2_in.vivo_noRT | Gm8016 | 4.807 | 9.70E-22 | glio |
| culture_effect | Ccdc92 | 0.712 | 9.71E-03 | glio |
| culture_effect | Jdp2 | -0.563 | 9.71E-03 | glio |
| Interaction_genotype_Myc | Fam229b | -1.958 | 9.71E-03 | glio |
| interaction_RT | Sbsn | 1.8 | 9.71E-03 | glio |
| interaction_RT | Larp7 | -0.792 | 9.71E-03 | glio |
| interaction_RT | Mtx2 | 0.515 | 9.71E-03 | glio |
| Bora_in.vivo_noRT | Sdhaf1 | -1.072 | 9.71E-03 | glio |
| Chek1_in.vivo_noRT | Wdr19 | 1.854 | 9.71E-03 | glio |
| Fanca_ex.vivo_noRT | Gm266 | 2.124 | 9.71E-03 | glio |
| Fanca_ex.vivo_noRT | Ildr1 | 2.124 | 9.71E-03 | glio |
| Interaction_genotype_Myc | Igsf23 | -3.876 | 9.71E-04 | glio |
| Bard1_ex.vivo_noRT | Slc38a3 | 2.815 | 9.71E-04 | glio |
| Bcas2_in.vivo_noRT | Ints4 | -1.671 | 9.71E-04 | glio |
| Bcas2_ex.vivo_noRT | Slc25a18 | 2.742 | 9.71E-04 | glio |
| Chek1_ex.vivo_noRT | Dok3 | 3.105 | 9.71E-04 | glio |
| Hsd17b10_in.vivo_noRT | Mterf1b | 2.38 | 9.71E-04 | glio |
| culture_effect | Isoc1 | -0.592 | 9.71E-05 | glio |
| Interaction_genotype_Mtor | Gm30042 | -3.839 | 9.71E-05 | glio |
| Interaction_genotype_Mtor | Krtdap | -4.69 | 9.71E-05 | glio |
| Interaction_genotype_Mtor | Sema4a | -4.826 | 9.71E-05 | glio |
| Aars2_in.vivo_noRT | 9330121J05Ri | 2.754 | 9.71E-05 | glio |
| Chek1_in.vivo_noRT | Zfp516 | -1.734 | 9.71E-05 | glio |
| culture_effect | Zfand6 | -0.588 | 9.71E-06 | glio |
| Aars2_in.vivo_noRT | Dynlt1c | 3.523 | 9.71E-06 | glio |
| Bcas2_in.vivo_noRT | Spata33 | 3.283 | 9.71E-06 | glio |
| Bcas2_in.vivo_noRT | Gm9958 | 3.237 | 9.71E-06 | glio |

| | | | | |
|---|---|---|---|---|
| Bcas2_in.vivo_noRT | Fam189a1 | 3.137 | 9.71E-08 | glio |
| Bcas2_in.vivo_noRT | Acoxl | 4.807 | 9.71E-14 | glio |
| culture_effect | Ddrgk1 | -0.345 | 9.72E-03 | glio |
| Interaction_genotype_Atr | Rab18 | 1.123 | 9.72E-03 | glio |
| Interaction_genotype_Bora | H2afv | 0.875 | 9.72E-03 | glio |
| Interaction_genotype_Myc | Ccdc191 | -2.676 | 9.72E-03 | glio |
| Atr_ex.vivo_noRT | Gm12840 | 2.372 | 9.72E-03 | glio |
| Atr_ex.vivo_noRT | Slc38a3 | 2.367 | 9.72E-03 | glio |
| Bard1_ex.vivo_noRT | Gm13054 | 2.419 | 9.72E-03 | glio |
| Cdkn2a_in.vivo_noRT | Edn2 | -1.675 | 9.72E-03 | glio |
| Cdkn2a_in.vivo_noRT | E230025N22R | -2.307 | 9.72E-03 | glio |
| Cdkn2a_in.vivo_noRT | Gm49698 | -1.675 | 9.72E-03 | glio |
| Cdkn2a_in.vivo_noRT | Olfr881 | -1.675 | 9.72E-03 | glio |
| Chek1_ex.vivo_noRT | Fgfr1 | 2.831 | 9.72E-03 | glio |
| culture_effect | Gtf3c4 | -0.758 | 9.72E-04 | glio |
| Bora_in.vivo_noRT | Sept4 | 2.444 | 9.72E-04 | glio |
| Chek1_in.vivo_noRT | Prx | 2.515 | 9.72E-04 | glio |
| Chek1_in.vivo_noRT | Gm35339 | 2.447 | 9.72E-04 | glio |
| Bora_in.vivo_noRT | Gm28050 | 2.898 | 9.72E-05 | glio |
| Bora_in.vivo_noRT | Hyou1 | -1.853 | 9.72E-05 | glio |
| Bora_in.vivo_noRT | BC037032 | 4.018 | 9.72E-08 | glio |
| Hsd17b10_in.vivo_noRT | Gm33051 | 2.963 | 9.72E-08 | glio |
| Atr_in.vivo_noRT | Gm16244 | 4.876 | 9.72E-12 | glio |
| Chek1_ex.vivo_noRT | C4b | 5.051 | 9.72E-13 | glio |
| culture_effect | Tcim | -4.53 | 9.72E-52 | glio |
| culture_effect | Lrrc4b | -1.45 | 9.73E-03 | glio |
| Interaction_genotype_Aars2 | Capn13 | -2.034 | 9.73E-03 | glio |
| Interaction_genotype_Aars2 | Btn1a1 | -2.558 | 9.73E-03 | glio |
| Interaction_genotype_Aars2 | Fzd7 | -3.076 | 9.73E-03 | glio |
| Interaction_genotype_Aars2 | Paxx | -2.303 | 9.73E-03 | glio |
| Interaction_genotype_Aars2 | Nlk | 1.843 | 9.73E-03 | glio |
| Interaction_genotype_Aars2 | Psmd4 | 0.903 | 9.73E-03 | glio |
| Interaction_genotype_Aars2 | Elob | 0.783 | 9.73E-03 | glio |
| Interaction_genotype_Atr | 4732471J01Ri | -3.302 | 9.73E-03 | glio |
| Interaction_genotype_Bora | Pdik1l | -2.444 | 9.73E-03 | glio |
| Interaction_genotype_Myc | Rrp12 | -2.404 | 9.73E-03 | glio |
| Atr_ex.vivo_noRT | Elobl | 2.648 | 9.73E-03 | glio |
| Atr_ex.vivo_noRT | Hyls1 | 1.859 | 9.73E-03 | glio |
| Hsd17b10_in.vivo_noRT | Gm45833 | 2.769 | 9.73E-03 | glio |
| Interaction_genotype_Aars2 | Nat8l | -2.869 | 9.73E-04 | glio |
| Interaction_genotype_Mtor | 9130008F23Ri | -4.403 | 9.73E-04 | glio |
| Chek1_ex.vivo_noRT | Gm50012 | 3.065 | 9.73E-04 | glio |
| Chek1_ex.vivo_noRT | 1700012B07R | 3.286 | 9.73E-04 | glio |
| Interaction_genotype_Mtor | Ogdhl | -4.646 | 9.73E-05 | glio |
| Aars2_in.vivo_noRT | Inafm1 | 2.387 | 9.73E-05 | glio |
| Atr_in.vivo_noRT | Radil | 2.712 | 9.73E-05 | glio |
| Atr_ex.vivo_noRT | Npy1r | 3.164 | 9.73E-06 | glio |
| Bcas2_in.vivo_noRT | Zfp637 | -1.738 | 9.73E-06 | glio |
| Atr_in.vivo_noRT | 4930522L14Ri | 3.515 | 9.73E-07 | glio |
| Chek1_in.vivo_noRT | Il23a | 3.433 | 9.73E-07 | glio |
| Aars2_in.vivo_noRT | A230020J21Ri | 4.399 | 9.73E-10 | glio |
| Atr_in.vivo_noRT | Gm8300 | 5.533 | 9.73E-11 | glio |

| | | | | |
|---|---|---|---|---|
| Chek1_in.vivo_noRT | Sucnr1 | 5.044 | 9.73E-28 | glio |
| culture_effect | Gm10827 | 1.474 | 9.74E-03 | glio |
| culture_effect | Kdm5b | -0.694 | 9.74E-03 | glio |
| Interaction_genotype_Atr | Pus7l | -2.558 | 9.74E-03 | glio |
| Interaction_genotype_Atr | Trir | 0.905 | 9.74E-03 | glio |
| Atp5c1_in.vivo_noRT | Gm28731 | 2.282 | 9.74E-03 | glio |
| Atr_ex.vivo_noRT | Vps13a | -1.183 | 9.74E-03 | glio |
| Chek1_in.vivo_noRT | Veph1 | 1.909 | 9.74E-03 | glio |
| Chek1_in.vivo_noRT | Nfat5 | -1.002 | 9.74E-03 | glio |
| Cyp19a1_ex.vivo_noRT | Olfr90 | 2.54 | 9.74E-03 | glio |
| Eif2s1_in.vivo_noRT | Car14 | 2.143 | 9.74E-03 | glio |
| Interaction_genotype_Aars2 | Stxbp5 | -3.171 | 9.74E-04 | glio |
| Interaction_genotype_Mtor | Tsfm | 1.732 | 9.74E-04 | glio |
| Interaction_genotype_Myc | Qrich2 | -3.847 | 9.74E-04 | glio |
| Interaction_genotype_Myc | Zkscan7 | -3.706 | 9.74E-04 | glio |
| Interaction_genotype_Myc | Atrx | 1.595 | 9.74E-04 | glio |
| Cenpj_ex.vivo_noRT | Gm21761 | 2.591 | 9.74E-04 | glio |
| Chek1_in.vivo_noRT | Gm20939 | 2.245 | 9.74E-04 | glio |
| Eif2s1_in.vivo_noRT | Gm45151 | 2.142 | 9.74E-04 | glio |
| Eif2s1_in.vivo_noRT | Gm50024 | 1.88 | 9.74E-04 | glio |
| Interaction_genotype_Mtor | Gm34939 | -3.835 | 9.74E-05 | glio |
| Interaction_genotype_Mtor | Srp72 | 1.927 | 9.74E-05 | glio |
| Interaction_genotype_Mtor | Ppia | 1.454 | 9.74E-05 | glio |
| Aars2_in.vivo_noRT | Hist1h4c | 2.894 | 9.74E-05 | glio |
| Bora_ex.vivo_noRT | Tent5c | 2.808 | 9.74E-05 | glio |
| Interaction_genotype_Myc | Gng8 | -4.882 | 9.74E-06 | glio |
| Bcas2_in.vivo_noRT | Ache | 3.234 | 9.74E-06 | glio |
| Bcas2_in.vivo_noRT | Anxa8 | 3.081 | 9.74E-06 | glio |
| Chek1_ex.vivo_noRT | Gata3 | 4.31 | 9.74E-06 | glio |
| Atp5c1_in.vivo_noRT | Zscan4-ps2 | 2.668 | 9.74E-07 | glio |
| Interaction_genotype_Mtor | Baalc | -6.021 | 9.74E-09 | glio |
| Chek1_ex.vivo_noRT | Lmx1a | 4.842 | 9.74E-13 | glio |
| culture_effect | Khdrbs2 | -4.964 | 9.74E-15 | glio |
| culture_effect | Plac8 | -4.634 | 9.74E-15 | glio |
| Chek1_ex.vivo_noRT | Gpr4 | 4.842 | 9.74E-15 | glio |
| Chek1_ex.vivo_noRT | Gm33466 | 4.842 | 9.74E-15 | glio |
| Bora_in.vivo_noRT | Slc2a9 | 5.46 | 9.74E-21 | glio |
| culture_effect | Slc25a16 | 0.627 | 9.75E-03 | glio |
| culture_effect | Itsn2 | 0.385 | 9.75E-03 | glio |
| culture_effect | Cdc42se2 | -0.491 | 9.75E-03 | glio |
| Interaction_genotype_Bcas2 | Olfr552 | -2.464 | 9.75E-03 | glio |
| Interaction_genotype_Bcas2 | Lamc2 | -3.198 | 9.75E-03 | glio |
| Interaction_genotype_Bcas2 | Etfrf1 | 1.961 | 9.75E-03 | glio |
| Interaction_genotype_Myc | Lgr4 | -2.429 | 9.75E-03 | glio |
| Interaction_genotype_Mtor | Cyp4f40 | -3.839 | 9.75E-04 | glio |
| Interaction_genotype_Mtor | Kcnb1 | -4.185 | 9.75E-04 | glio |
| Atr_ex.vivo_noRT | Glrp1 | 2.772 | 9.75E-04 | glio |
| Bcas2_in.vivo_noRT | Ccdc9b | 2.167 | 9.75E-04 | glio |
| Bcas2_in.vivo_noRT | Pdk2 | 2.44 | 9.75E-04 | glio |
| Bora_in.vivo_noRT | Lgr6 | 2.728 | 9.75E-04 | glio |
| Cenpj_in.vivo_noRT | AU019990 | 2.022 | 9.75E-04 | glio |
| Interaction_genotype_Myc | Gm45620 | -4.193 | 9.75E-05 | glio |

| | | | | |
|---|---|---|---|---|
| Interaction_genotype_Myc | Gm10131 | -4.36 | 9.75E-05 | glio |
| Interaction_genotype_Myc | C2cd5 | -3.61 | 9.75E-05 | glio |
| Interaction_genotype_Myc | Abhd6 | -3.123 | 9.75E-05 | glio |
| Interaction_genotype_Myc | Saal1 | -3.126 | 9.75E-05 | glio |
| Atr_in.vivo_noRT | Lysmd4 | 2.52 | 9.75E-05 | glio |
| Bcas2_in.vivo_noRT | AC166172.1 | 2.607 | 9.75E-05 | glio |
| Bcas2_ex.vivo_noRT | 9130019O22R | 2.742 | 9.75E-05 | glio |
| Atr_in.vivo_noRT | Acacb | 3.031 | 9.75E-06 | glio |
| Bora_ex.vivo_noRT | Npy1r | 3.201 | 9.75E-06 | glio |
| Bora_ex.vivo_noRT | Gm6803 | 3.201 | 9.75E-06 | glio |
| Bora_ex.vivo_noRT | Ngb | 3.994 | 9.75E-06 | glio |
| culture_effect | Agpat3 | 0.885 | 9.75E-09 | glio |
| Atr_in.vivo_noRT | Rab11fip4os1 | 4.497 | 9.75E-10 | glio |
| Bcas2_in.vivo_noRT | E030042O20R | 4.807 | 9.75E-11 | glio |
| Aars2_in.vivo_noRT | Gm12925 | 4.869 | 9.75E-12 | glio |
| Chek1_in.vivo_noRT | Ush1g | 5.555 | 9.75E-25 | glio |
| Interaction_genotype_Mtor | Rasa4 | -2.666 | 9.76E-03 | glio |
| Interaction_genotype_Myc | Arsk | -2.436 | 9.76E-03 | glio |
| Interaction_genotype_Myc | Elmsan1 | -2.128 | 9.76E-03 | glio |
| Bcas2_in.vivo_noRT | Ganab | -0.728 | 9.76E-03 | glio |
| Chek1_in.vivo_noRT | Zfp945 | -1.218 | 9.76E-03 | glio |
| Chek1_in.vivo_noRT | Rpl24 | -0.505 | 9.76E-03 | glio |
| Interaction_genotype_Mtor | Zfp933 | -3.32 | 9.76E-04 | glio |
| interaction_RT | Trmt112 | 0.58 | 9.76E-04 | glio |
| Atr_in.vivo_noRT | 4930532G15R | 2.137 | 9.76E-04 | glio |
| Atr_in.vivo_noRT | Trmt1 | -1.312 | 9.76E-04 | glio |
| Bcas2_ex.vivo_noRT | Slc19a3 | 2.348 | 9.76E-04 | glio |
| Chek1_in.vivo_noRT | Clcn5 | -1.446 | 9.76E-04 | glio |
| Chek1_in.vivo_noRT | Map2k2 | -0.914 | 9.76E-04 | glio |
| Chek1_ex.vivo_noRT | Kcna4 | 3.075 | 9.76E-04 | glio |
| Chek1_ex.vivo_noRT | Lsm3 | -1.289 | 9.76E-04 | glio |
| Bora_in.vivo_noRT | Gm50322 | 3.041 | 9.76E-05 | glio |
| Bora_in.vivo_noRT | Ptpn2 | -1.196 | 9.76E-05 | glio |
| Atr_in.vivo_noRT | Aldh1l1 | 4.111 | 9.76E-08 | glio |
| Chek1_in.vivo_noRT | Mfap3l | 4.653 | 9.76E-10 | glio |
| Interaction_genotype_Aars2 | Gm14698 | -3.32 | 9.77E-03 | glio |
| Interaction_genotype_Aars2 | Foxc2 | -3.409 | 9.77E-03 | glio |
| Interaction_genotype_Aars2 | Pate2 | -3.316 | 9.77E-03 | glio |
| Interaction_genotype_Aars2 | Ttbk2 | -2.511 | 9.77E-03 | glio |
| Interaction_genotype_Aars2 | Map3k12 | -2.351 | 9.77E-03 | glio |
| Interaction_genotype_Aars2 | Itsn1 | 1.74 | 9.77E-03 | glio |
| Interaction_genotype_Bcas2 | Npr3 | -3.037 | 9.77E-03 | glio |
| Interaction_genotype_Bcas2 | Cd81 | 2.178 | 9.77E-03 | glio |
| Interaction_genotype_Mtor | Gm36757 | -3.105 | 9.77E-03 | glio |
| Interaction_genotype_Mtor | Gm46430 | -2.5 | 9.77E-03 | glio |
| Interaction_genotype_Mtor | Eral1 | -2.504 | 9.77E-03 | glio |
| Interaction_genotype_Mtor | Klhl13 | -2.485 | 9.77E-03 | glio |
| Interaction_genotype_Mtor | Mki67 | 2.281 | 9.77E-03 | glio |
| Interaction_genotype_Myc | Slxl1 | -2.927 | 9.77E-03 | glio |
| Interaction_genotype_Myc | Zfp169 | -2.815 | 9.77E-03 | glio |
| Interaction_genotype_Myc | Castor2 | -2.552 | 9.77E-03 | glio |
| Interaction_genotype_Myc | Mrps33 | 0.916 | 9.77E-03 | glio |

| | | | | |
|---|---|---|---|---|
| Interaction_genotype_Rpl3 | 5430400D12R | -3.675 | 9.77E-03 | glio |
| Interaction_genotype_Rpl3 | Gm43672 | -3.708 | 9.77E-03 | glio |
| Interaction_genotype_Rpl3 | K230015D01R | -3.672 | 9.77E-03 | glio |
| Interaction_genotype_Rpl3 | Fastkd1 | -3.244 | 9.77E-03 | glio |
| Atp5c1_in.vivo_noRT | Slc45a3 | 2.146 | 9.77E-03 | glio |
| Atp5c1_in.vivo_noRT | Ssh3 | 1.868 | 9.77E-03 | glio |
| Atr_in.vivo_noRT | Pde9a | 1.788 | 9.77E-03 | glio |
| Atr_in.vivo_noRT | Rnf170 | 1.257 | 9.77E-03 | glio |
| Bard1_ex.vivo_noRT | Gm15972 | 2.415 | 9.77E-03 | glio |
| Bora_in.vivo_noRT | Map3k6 | 1.971 | 9.77E-03 | glio |
| Brca2_in.vivo_noRT | Gm48899 | 2.026 | 9.77E-03 | glio |
| Brca2_in.vivo_noRT | AU019990 | 1.763 | 9.77E-03 | glio |
| Brca2_in.vivo_noRT | Smpd5 | 2.163 | 9.77E-03 | glio |
| Chek1_ex.vivo_noRT | Dnase2a | 1.696 | 9.77E-03 | glio |
| culture_effect | Dhx29 | 0.71 | 9.77E-04 | glio |
| Interaction_genotype_Myc | Krtcap3 | -2.923 | 9.77E-04 | glio |
| Bora_in.vivo_noRT | Acy3 | 2.443 | 9.77E-04 | glio |
| Hsd17b10_in.vivo_noRT | A330084C13R | 2.698 | 9.77E-04 | glio |
| Interaction_genotype_Myc | Gm15056 | -4.919 | 9.77E-05 | glio |
| Interaction_genotype_Myc | Slc35g3 | -3.128 | 9.77E-05 | glio |
| Interaction_genotype_Myc | Fbxl13 | -3.844 | 9.77E-05 | glio |
| Interaction_genotype_Myc | Oma1 | -3.977 | 9.77E-05 | glio |
| Interaction_genotype_Myc | Rap1b | 1.522 | 9.77E-05 | glio |
| Atr_in.vivo_noRT | 9330160F10Ri | 2.526 | 9.77E-05 | glio |
| Atr_in.vivo_noRT | B130034C11R | 2.311 | 9.77E-05 | glio |
| Atr_in.vivo_noRT | Fbxl6 | -1.539 | 9.77E-05 | glio |
| Chek1_in.vivo_noRT | Serpine2 | 3.313 | 9.77E-05 | glio |
| Bora_in.vivo_noRT | Cdca5 | -2.276 | 9.77E-06 | glio |
| Chek1_in.vivo_noRT | Aga | -2.166 | 9.77E-07 | glio |
| Atr_in.vivo_noRT | Bfsp2 | 4.161 | 9.77E-08 | glio |
| Atr_in.vivo_noRT | Ifnz | 3.699 | 9.77E-08 | glio |
| Chek1_in.vivo_noRT | Selp | 3.999 | 9.77E-09 | glio |
| Chek1_ex.vivo_noRT | Gm15943 | 3.861 | 9.77E-09 | glio |
| Chek1_ex.vivo_noRT | Krtcap3 | 3.857 | 9.77E-09 | glio |
| culture_effect | Hspb6 | 1.934 | 9.77E-12 | glio |
| culture_effect | Gm11884 | -1.534 | 9.78E-03 | glio |
| Interaction_genotype_Aars2 | Iqcc | -2.462 | 9.78E-03 | glio |
| Interaction_genotype_Mtor | Gja1 | -2.436 | 9.78E-03 | glio |
| Interaction_genotype_Mtor | Mtdh | 0.979 | 9.78E-03 | glio |
| Interaction_genotype_Myc | Ldhd | -2.523 | 9.78E-03 | glio |
| Interaction_genotype_Myc | Slc22a23 | -2.187 | 9.78E-03 | glio |
| interaction_RT | Platr16 | 2.67 | 9.78E-03 | glio |
| Aars2_in.vivo_noRT | Dynll1 | -0.533 | 9.78E-03 | glio |
| Interaction_genotype_Myc | Ccser1 | -3.122 | 9.78E-04 | glio |
| Aars2_in.vivo_noRT | Prdm9 | 2.387 | 9.78E-04 | glio |
| Atr_in.vivo_noRT | Mapkapk5 | 2.325 | 9.78E-04 | glio |
| Bora_in.vivo_noRT | Fam161b | 2.389 | 9.78E-04 | glio |
| Bora_in.vivo_noRT | Aebp1 | -1.692 | 9.78E-04 | glio |
| Chek1_in.vivo_noRT | Afg1l | 2.44 | 9.78E-04 | glio |
| Chek1_in.vivo_noRT | Crabp1 | 2.412 | 9.78E-04 | glio |
| Chek1_in.vivo_noRT | Slc3a2 | -0.948 | 9.78E-04 | glio |
| culture_effect | Bdh2 | 1.838 | 9.78E-05 | glio |

| | | | | |
|---|---|---|---|---|
| culture_effect | Vps16 | -1.034 | 9.78E-05 | glio |
| Interaction_genotype_Myc | Nkx2-2os | -4.548 | 9.78E-05 | glio |
| Aars2_in.vivo_noRT | Psmb3 | -1.09 | 9.78E-05 | glio |
| Atp5c1_in.vivo_noRT | Obscn | 2.668 | 9.78E-05 | glio |
| Atr_in.vivo_noRT | Irs4 | 2.933 | 9.78E-05 | glio |
| Bcas2_ex.vivo_noRT | Klk13 | 2.742 | 9.78E-05 | glio |
| Bora_in.vivo_noRT | C130040N14R | 3.289 | 9.78E-06 | glio |
| culture_effect | Gchfr | 2.838 | 9.78E-07 | glio |
| Bora_ex.vivo_noRT | Stfa3 | 3.201 | 9.78E-08 | glio |
| Bora_ex.vivo_noRT | Olfr787 | 3.201 | 9.78E-08 | glio |
| Interaction_genotype_Atr | Gchfr | -3.395 | 9.79E-03 | glio |
| Interaction_genotype_Bcas2 | Wdr13 | 2.105 | 9.79E-03 | glio |
| Interaction_genotype_Mtor | Ppef1 | -3.108 | 9.79E-03 | glio |
| Interaction_genotype_Myc | Mcmdc2 | -2.425 | 9.79E-03 | glio |
| Interaction_genotype_Myc | Usp11 | -3.138 | 9.79E-03 | glio |
| Interaction_genotype_Rad51 | Ddr1 | 1.536 | 9.79E-03 | glio |
| Interaction_genotype_Terf1 | Tslp | -3.975 | 9.79E-03 | glio |
| Interaction_genotype_Terf1 | Casp1 | 3.525 | 9.79E-03 | glio |
| Interaction_genotype_Terf1 | Sod3 | -3.983 | 9.79E-03 | glio |
| Interaction_genotype_Terf1 | Gm47545 | -3.551 | 9.79E-03 | glio |
| Interaction_genotype_Terf1 | Fscn1 | 1.66 | 9.79E-03 | glio |
| Interaction_genotype_Uqcrb | Hba-a2 | -4.355 | 9.79E-03 | glio |
| Interaction_genotype_Uqcrb | Gm29340 | -3.717 | 9.79E-03 | glio |
| Interaction_genotype_Uqcrb | Rcan2 | -4.224 | 9.79E-03 | glio |
| Interaction_genotype_Uqcrb | 5430402O13R | -4.248 | 9.79E-03 | glio |
| Interaction_genotype_Uqcrb | Pck2 | 1.838 | 9.79E-03 | glio |
| Aars2_in.vivo_noRT | Cyp27a1 | 1.731 | 9.79E-03 | glio |
| Aars2_in.vivo_noRT | Zc3h3 | 1.469 | 9.79E-03 | glio |
| Atp5c1_in.vivo_noRT | Creld2 | -0.779 | 9.79E-03 | glio |
| Atr_ex.vivo_noRT | Rtl9 | 2.453 | 9.79E-03 | glio |
| Bcas2_in.vivo_noRT | Srm | -0.822 | 9.79E-03 | glio |
| Atr_in.vivo_noRT | Gm49539 | 2.379 | 9.79E-04 | glio |
| Atr_in.vivo_noRT | Gm15708 | 2.322 | 9.79E-04 | glio |
| Atr_in.vivo_noRT | Adgra3 | -1.377 | 9.79E-04 | glio |
| Atr_in.vivo_noRT | Sox4 | -1.475 | 9.79E-04 | glio |
| Bcas2_ex.vivo_noRT | Stx19 | 2.742 | 9.79E-04 | glio |
| culture_effect | Slc25a24 | 0.784 | 9.79E-05 | glio |
| Eif2s1_in.vivo_noRT | Igf1 | 2.178 | 9.79E-05 | glio |
| culture_effect | Eng | 2.551 | 9.79E-06 | glio |
| culture_effect | Ap5z1 | -0.986 | 9.79E-06 | glio |
| Interaction_genotype_Mtor | Mir155hg | -5.78 | 9.79E-06 | glio |
| Interaction_genotype_Mtor | Psenen | 1.911 | 9.79E-06 | glio |
| Atr_ex.vivo_noRT | Ngb | 3.947 | 9.79E-06 | glio |
| Atr_ex.vivo_noRT | Clec12b | 2.769 | 9.79E-06 | glio |
| Atr_ex.vivo_noRT | Vmn1r32 | 2.771 | 9.79E-06 | glio |
| Atr_ex.vivo_noRT | 1700019E08Ri | 3.164 | 9.79E-06 | glio |
| Atr_ex.vivo_noRT | Sox11 | 3.962 | 9.79E-06 | glio |
| Atr_ex.vivo_noRT | Cct2 | -0.986 | 9.79E-06 | glio |
| Chek1_in.vivo_noRT | Gm37305 | 3.086 | 9.79E-06 | glio |
| Bcas2_in.vivo_noRT | Tnxb | 3.654 | 9.79E-08 | glio |
| Bcas2_in.vivo_noRT | 1700010B08R | 3.12 | 9.79E-08 | glio |
| Chek1_in.vivo_noRT | Dmrtc1a | 5.044 | 9.79E-21 | glio |

| | | | | |
|---|---|---|---|---|
| Chek1_in.vivo_noRT | Gm12589 | 5.044 | 9.79E-21 | glio |
| culture_effect | Rpain | -0.441 | 9.80E-03 | glio |
| Interaction_genotype_Aars2 | 4930423D22R | -2.353 | 9.80E-03 | glio |
| Interaction_genotype_Bora | Ank1 | -2.519 | 9.80E-03 | glio |
| Interaction_genotype_Bora | Dtl | 2.054 | 9.80E-03 | glio |
| Interaction_genotype_Cdkn2a | Gm10482 | 3.097 | 9.80E-03 | glio |
| Interaction_genotype_Lmo7 | Gm32834 | -3.79 | 9.80E-03 | glio |
| Interaction_genotype_Lmo7 | Krt80 | -3.618 | 9.80E-03 | glio |
| Interaction_genotype_Myc | Selenok-ps1 | -2.728 | 9.80E-03 | glio |
| Interaction_genotype_Myc | Kank4os | -2.151 | 9.80E-03 | glio |
| Interaction_genotype_Pars2 | Prl2c2 | -3.447 | 9.80E-03 | glio |
| Interaction_genotype_Pars2 | Gm2449 | -3.102 | 9.80E-03 | glio |
| Interaction_genotype_Terf1 | Gm15127 | -3.859 | 9.80E-03 | glio |
| Atr_in.vivo_noRT | Pphln1 | -1.056 | 9.80E-03 | glio |
| Bcas2_in.vivo_noRT | R3hcc1l | -1.365 | 9.80E-03 | glio |
| Bcas2_in.vivo_noRT | Kdm3b | 1.283 | 9.80E-03 | glio |
| Bora_in.vivo_noRT | Acad11 | 1.592 | 9.80E-03 | glio |
| Eif2s1_in.vivo_noRT | Slc1a2 | 1.879 | 9.80E-03 | glio |
| Hsd17b10_in.vivo_noRT | Insc | 2.159 | 9.80E-04 | glio |
| Interaction_genotype_Mtor | Gem | -4.492 | 9.80E-05 | glio |
| Interaction_genotype_Mtor | Gm16751 | -3.982 | 9.80E-05 | glio |
| Bora_in.vivo_noRT | Cmtm5 | -2.022 | 9.80E-05 | glio |
| Interaction_genotype_Mtor | Pnpla6 | -4.832 | 9.80E-06 | glio |
| Atr_in.vivo_noRT | 5033430I15Ri | 3.354 | 9.80E-07 | glio |
| Atr_ex.vivo_noRT | Adam5 | 3.164 | 9.80E-07 | glio |
| Atr_ex.vivo_noRT | 4933400F21Ri | 3.164 | 9.80E-07 | glio |
| Aars2_in.vivo_noRT | Gpr162 | 5.019 | 9.80E-13 | glio |
| Aars2_in.vivo_noRT | Slc16a7 | 4.215 | 9.80E-13 | glio |
| Bcas2_in.vivo_noRT | Gm15445 | 4.161 | 9.80E-15 | glio |
| Atr_in.vivo_noRT | Gm28940 | 5.017 | 9.80E-19 | glio |
| Interaction_genotype_Aars2 | Gm35439 | -2.564 | 9.81E-03 | glio |
| Interaction_genotype_Mtor | Tro | -3.161 | 9.81E-03 | glio |
| Interaction_genotype_Pars2 | Hist2h3c1 | -3.533 | 9.81E-03 | glio |
| interaction_RT | Cebpd | 1.194 | 9.81E-03 | glio |
| Bcas2_in.vivo_noRT | Aldh3b1 | 1.93 | 9.81E-03 | glio |
| Chek1_in.vivo_noRT | Cct4 | -0.611 | 9.81E-03 | glio |
| Hsd17b10_in.vivo_noRT | Syvn1 | -1.259 | 9.81E-03 | glio |
| Hsd17b10_in.vivo_noRT | Bcas1 | -1.237 | 9.81E-03 | glio |
| Interaction_genotype_Mtor | Cdpf1 | -2.245 | 9.81E-04 | glio |
| Interaction_genotype_Myc | 4930563E22Ri | -3.646 | 9.81E-04 | glio |
| Interaction_genotype_Myc | Rpl37 | 1.529 | 9.81E-04 | glio |
| Interaction_genotype_Myc | Celsr3 | -4.149 | 9.81E-05 | glio |
| Atr_ex.vivo_noRT | Pcdhb21 | 3.567 | 9.81E-05 | glio |
| Interaction_genotype_Myc | Clec4d | -3.914 | 9.81E-06 | glio |
| Interaction_genotype_Myc | Gng5 | 1.73 | 9.81E-06 | glio |
| Aars2_in.vivo_noRT | Gm36371 | 2.954 | 9.81E-06 | glio |
| Bora_in.vivo_noRT | Gm43608 | 3.762 | 9.81E-08 | glio |
| Aars2_in.vivo_noRT | Gm8212 | 4.225 | 9.81E-10 | glio |
| Chek1_in.vivo_noRT | Vpreb1 | 4.141 | 9.81E-10 | glio |
| Chek1_in.vivo_noRT | Gm45606 | 4.784 | 9.81E-13 | glio |
| Interaction_genotype_Mtor | Gm10069 | -2.796 | 9.82E-03 | glio |
| Interaction_genotype_Mtor | Chd4 | 1.24 | 9.82E-03 | glio |

| | | | | |
|---|---|---|---|---|
| Interaction_genotype_Pars2 | Gm13191 | -2.268 | 9.82E-03 | glio |
| Interaction_genotype_Pars2 | 4933416E03Ri | -2.268 | 9.82E-03 | glio |
| Interaction_genotype_Pars2 | Rbm18 | 1.791 | 9.82E-03 | glio |
| Interaction_genotype_Rad51 | Ndel1 | 1.755 | 9.82E-03 | glio |
| Interaction_genotype_Rpl3 | Gm20667 | -3.724 | 9.82E-03 | glio |
| Interaction_genotype_Rpl3 | Carns1 | -3.68 | 9.82E-03 | glio |
| Interaction_genotype_Rpl3 | Layn | -2.59 | 9.82E-03 | glio |
| Interaction_genotype_Rpl3 | Prkaca | 1.741 | 9.82E-03 | glio |
| interaction_RT | Gm20732 | 2.409 | 9.82E-03 | glio |
| Aars2_in.vivo_noRT | Utp18 | -0.828 | 9.82E-03 | glio |
| Atr_in.vivo_noRT | Pou4f1 | 1.386 | 9.82E-03 | glio |
| Atr_in.vivo_noRT | Stub1 | -0.74 | 9.82E-03 | glio |
| Bard1_ex.vivo_noRT | Cnnm1 | 2.973 | 9.82E-03 | glio |
| Bora_in.vivo_noRT | Pym1 | -1.226 | 9.82E-03 | glio |
| Brca2_in.vivo_noRT | Gm36793 | 1.763 | 9.82E-03 | glio |
| Cdkn2a_in.vivo_noRT | BC061195 | -1.675 | 9.82E-03 | glio |
| Chek1_in.vivo_noRT | Mtcl1 | 1.861 | 9.82E-03 | glio |
| Eif2s1_in.vivo_noRT | H1foo | 1.61 | 9.82E-03 | glio |
| culture_effect | Xkr6 | -0.846 | 9.82E-04 | glio |
| Interaction_genotype_Myc | Sema4c | -3.096 | 9.82E-04 | glio |
| Interaction_genotype_Myc | Calm2 | 1.388 | 9.82E-04 | glio |
| Atr_in.vivo_noRT | Paox | 2.174 | 9.82E-04 | glio |
| Bora_in.vivo_noRT | Ocm | 2.095 | 9.82E-04 | glio |
| Chek1_ex.vivo_noRT | Gipc2 | 3.195 | 9.82E-04 | glio |
| Eif2s1_ex.vivo_noRT | Slc17a6 | 2.234 | 9.82E-04 | glio |
| Hsd17b10_in.vivo_noRT | Gm10484 | 2.615 | 9.82E-04 | glio |
| interaction_RT | A530032D15R | 2.848 | 9.82E-05 | glio |
| Bcas2_in.vivo_noRT | 4930513N10R | 3.199 | 9.82E-05 | glio |
| Interaction_genotype_Mtor | Msh5 | -5.657 | 9.82E-06 | glio |
| Interaction_genotype_Mtor | Rnft2 | -5.113 | 9.82E-06 | glio |
| Bcas2_in.vivo_noRT | Nupr1l | 3.544 | 9.82E-06 | glio |
| Atr_in.vivo_noRT | Gm28417 | 3.213 | 9.82E-07 | glio |
| Chek1_ex.vivo_noRT | Sox11 | 4.842 | 9.82E-09 | glio |
| Bora_in.vivo_noRT | Gm13684 | 3.834 | 9.82E-11 | glio |
| Chek1_in.vivo_noRT | Bco2 | 5.044 | 9.82E-17 | glio |
| Atr_in.vivo_noRT | Gm36198 | 5.273 | 9.82E-20 | glio |
| Interaction_genotype_Mtor | Dguok | 1.468 | 9.83E-03 | glio |
| Interaction_genotype_Myc | Cyfip2 | -2.527 | 9.83E-03 | glio |
| Interaction_genotype_Myc | Tbl1x | 1.084 | 9.83E-03 | glio |
| Interaction_genotype_Rpl3 | AC154200.1 | -2.896 | 9.83E-03 | glio |
| Interaction_genotype_Rpl3 | Gm16222 | -4.162 | 9.83E-03 | glio |
| Interaction_genotype_Rpl3 | Gm13272 | -3.29 | 9.83E-03 | glio |
| Interaction_genotype_Rpl3 | A630052C17R | -3.536 | 9.83E-03 | glio |
| Interaction_genotype_Terf1 | Gm16046 | -3.884 | 9.83E-03 | glio |
| Interaction_genotype_Terf1 | Hdac9 | -3.754 | 9.83E-03 | glio |
| Aars2_ex.vivo_noRT | Gtsf1 | 2.13 | 9.83E-03 | glio |
| Aars2_ex.vivo_noRT | Tbc1d22a | -2.059 | 9.83E-03 | glio |
| Bora_in.vivo_noRT | 4930432K21R | 1.795 | 9.83E-03 | glio |
| Cenpj_ex.vivo_noRT | 4930470G03R | 2.198 | 9.83E-03 | glio |
| Chek1_in.vivo_noRT | Armc7 | 1.677 | 9.83E-03 | glio |
| Chek1_in.vivo_noRT | Rpa1 | -1.251 | 9.83E-03 | glio |
| Hsd17b10_in.vivo_noRT | Zfpm1 | 2.1 | 9.83E-03 | glio |

| | | | | |
|---|---|---|---|---|
| culture_effect | GFP | -1.064 | 9.83E-04 | glio |
| Interaction_genotype_Mtor | 6230400D17R | -3.664 | 9.83E-04 | glio |
| Interaction_genotype_Mtor | Smdt1 | 1.608 | 9.83E-04 | glio |
| Aars2_ex.vivo_noRT | Cxcl1 | 3.293 | 9.83E-04 | glio |
| Aars2_ex.vivo_noRT | Egr2 | 3.007 | 9.83E-04 | glio |
| Atr_in.vivo_noRT | Creb3l1 | 2.326 | 9.83E-04 | glio |
| Atr_in.vivo_noRT | Cfap77 | 2.154 | 9.83E-04 | glio |
| Bora_in.vivo_noRT | Gins4 | -1.341 | 9.83E-04 | glio |
| Chek1_in.vivo_noRT | Mapkapk5 | 2.359 | 9.83E-04 | glio |
| Chek1_ex.vivo_noRT | Six4 | 2.263 | 9.83E-04 | glio |
| culture_effect | Gm16283 | 2.012 | 9.83E-05 | glio |
| Aars2_in.vivo_noRT | 2810405F17Ri | 2.64 | 9.83E-05 | glio |
| Atr_in.vivo_noRT | 1700028J19Ri | 2.736 | 9.83E-05 | glio |
| Bora_in.vivo_noRT | Mecom | 4.936 | 9.83E-14 | glio |
| Interaction_genotype_Aars2 | Rgs9 | -2.816 | 9.84E-03 | glio |
| Interaction_genotype_Ifnar1 | Lrrc72 | 3.318 | 9.84E-03 | glio |
| Interaction_genotype_Myc | Ndufaf7 | -1.954 | 9.84E-03 | glio |
| Interaction_genotype_Myc | Yrdc | 1.115 | 9.84E-03 | glio |
| Interaction_genotype_Myc | Mtpn | 0.933 | 9.84E-03 | glio |
| Aars2_in.vivo_noRT | Suox | 1.697 | 9.84E-03 | glio |
| Aars2_in.vivo_noRT | Wdr83os | -0.716 | 9.84E-03 | glio |
| Atr_in.vivo_noRT | Gm44386 | 1.853 | 9.84E-03 | glio |
| Atr_in.vivo_noRT | Eif4g2 | -0.589 | 9.84E-03 | glio |
| Atr_ex.vivo_noRT | Kpna7 | 2.184 | 9.84E-03 | glio |
| Bora_in.vivo_noRT | Fam98b | -1.19 | 9.84E-03 | glio |
| Eif2s1_in.vivo_noRT | Gm50147 | 2.143 | 9.84E-03 | glio |
| culture_effect | Shld2 | 0.789 | 9.84E-04 | glio |
| Interaction_genotype_Aars2 | 1700015F17Ri | -4.098 | 9.84E-04 | glio |
| Interaction_genotype_Mtor | Fam171a2 | -3.873 | 9.84E-04 | glio |
| Aars2_in.vivo_noRT | C3 | 2.325 | 9.84E-04 | glio |
| Atr_ex.vivo_noRT | Pcdhgb5 | 2.77 | 9.84E-04 | glio |
| Cenpj_in.vivo_noRT | Gm36793 | 2.022 | 9.84E-04 | glio |
| Interaction_genotype_Mtor | Gm9929 | -4.289 | 9.84E-05 | glio |
| Interaction_genotype_Mtor | Gdap1l1 | -5.843 | 9.84E-06 | glio |
| Interaction_genotype_Mtor | Gm44752 | -4.69 | 9.84E-06 | glio |
| Chek1_ex.vivo_noRT | Gm14019 | 3.346 | 9.84E-06 | glio |
| Atr_in.vivo_noRT | Abhd15 | 3.684 | 9.84E-07 | glio |
| Atr_in.vivo_noRT | C030005K06R | 3.434 | 9.84E-07 | glio |
| Aars2_in.vivo_noRT | Slc7a14 | 3.556 | 9.84E-08 | glio |
| Atr_in.vivo_noRT | 6330415G19R | 4.226 | 9.84E-10 | glio |
| Atr_in.vivo_noRT | Fsd2 | 4.15 | 9.84E-10 | glio |
| Atr_in.vivo_noRT | 4933421A08R | 4.754 | 9.84E-11 | glio |
| Atr_in.vivo_noRT | Vsig10l | 4.88 | 9.84E-13 | glio |
| Interaction_genotype_Bcas2 | Rad51ap1 | 2.06 | 9.85E-03 | glio |
| Interaction_genotype_Bora | Soga3 | -2.749 | 9.85E-03 | glio |
| Interaction_genotype_Mtor | Caskin2 | -2.581 | 9.85E-03 | glio |
| Interaction_genotype_Mtor | Ech1 | 1.433 | 9.85E-03 | glio |
| Interaction_genotype_Rpl3 | Ppil6 | -3.327 | 9.85E-03 | glio |
| Interaction_genotype_Rpl3 | Gm39326 | -3.541 | 9.85E-03 | glio |
| Aars2_in.vivo_noRT | Ptprn | 1.77 | 9.85E-03 | glio |
| Aars2_in.vivo_noRT | Arhgap32 | 1.214 | 9.85E-03 | glio |
| Atp5c1_in.vivo_noRT | Gm20667 | 2.189 | 9.85E-03 | glio |

| | | | | |
|---|---|---|---|---|
| Atr_in.vivo_noRT | Etfa | -0.924 | 9.85E-03 | glio |
| Atr_in.vivo_noRT | Pdha1 | -0.727 | 9.85E-03 | glio |
| Bcas2_in.vivo_noRT | Dnaja4 | 1.928 | 9.85E-03 | glio |
| Bcas2_in.vivo_noRT | Unc93b1 | 1.243 | 9.85E-03 | glio |
| Bcas2_in.vivo_noRT | Calu | -0.623 | 9.85E-03 | glio |
| Bora_in.vivo_noRT | Zfp97 | 1.912 | 9.85E-03 | glio |
| Bora_in.vivo_noRT | Mrps25 | -0.831 | 9.85E-03 | glio |
| Bora_in.vivo_noRT | Tspan31 | -0.846 | 9.85E-03 | glio |
| Chek1_ex.vivo_noRT | 0610009E02Ri | 2.562 | 9.85E-03 | glio |
| Chek1_ex.vivo_noRT | Stoml1 | 2.027 | 9.85E-03 | glio |
| Cyp19a1_ex.vivo_noRT | Gm9758 | 2.159 | 9.85E-03 | glio |
| Hsd17b10_in.vivo_noRT | Gm42658 | 1.863 | 9.85E-03 | glio |
| Chek1_ex.vivo_noRT | Rbm11 | 2.889 | 9.85E-04 | glio |
| Eif2s1_ex.vivo_noRT | Cyp2d22 | 3.937 | 9.85E-04 | glio |
| Aars2_ex.vivo_noRT | Glyat | 2.495 | 9.85E-05 | glio |
| Bora_ex.vivo_noRT | Pcdh15 | 3.201 | 9.85E-05 | glio |
| Chek1_in.vivo_noRT | Gm11127 | 3.483 | 9.85E-07 | glio |
| Chek1_in.vivo_noRT | Gm9922 | 3.908 | 9.85E-09 | glio |
| Bora_ex.vivo_noRT | Serpinb9g | 4.363 | 9.85E-11 | glio |
| Atr_in.vivo_noRT | Gm13270 | 5.533 | 9.85E-15 | glio |
| Bcas2_in.vivo_noRT | Pyy | 4.807 | 9.85E-17 | glio |
| Chek1_in.vivo_noRT | Stfa3 | 5.044 | 9.85E-27 | glio |
| Chek1_in.vivo_noRT | Olfr787 | 5.044 | 9.85E-27 | glio |
| Interaction_genotype_Aars2 | Mtmr11 | -3.323 | 9.86E-03 | glio |
| Interaction_genotype_Aars2 | Upk3bl | -3.045 | 9.86E-03 | glio |
| Interaction_genotype_Aars2 | Ptpn3 | -3.175 | 9.86E-03 | glio |
| Interaction_genotype_Aars2 | 4921531C22R | -3.064 | 9.86E-03 | glio |
| Interaction_genotype_Aars2 | Cdc123 | 1.057 | 9.86E-03 | glio |
| Interaction_genotype_Chek1 | Snai1 | -3.848 | 9.86E-03 | glio |
| Interaction_genotype_Cyp19a | Vcan | -4.511 | 9.86E-03 | glio |
| Interaction_genotype_Cyp19a | Scel | 3.069 | 9.86E-03 | glio |
| Interaction_genotype_Mtor | Zfp846 | -2.097 | 9.86E-03 | glio |
| Interaction_genotype_Myc | Ccdc112 | -1.948 | 9.86E-03 | glio |
| Bcas2_ex.vivo_noRT | Bglap2 | 2.742 | 9.86E-03 | glio |
| Cyp19a1_ex.vivo_noRT | Esp24 | 2.159 | 9.86E-03 | glio |
| Dclre1c_ex.vivo_noRT | Zfp352 | 2.9 | 9.86E-03 | glio |
| Hsd17b10_in.vivo_noRT | Cntf | 2.02 | 9.86E-03 | glio |
| culture_effect | Acot5 | -2.025 | 9.86E-04 | glio |
| Bora_in.vivo_noRT | Ccdc159 | 2.513 | 9.86E-04 | glio |
| Chek1_ex.vivo_noRT | Hipk4 | 2.763 | 9.86E-04 | glio |
| Chek1_ex.vivo_noRT | Tha1 | 2.752 | 9.86E-04 | glio |
| Interaction_genotype_Rad21 | Pcdh15 | 4.734 | 9.86E-05 | glio |
| Aars2_in.vivo_noRT | 6330408M09F | 4.285 | 9.86E-09 | glio |
| Bcas2_in.vivo_noRT | Ahrr | 4.007 | 9.86E-09 | glio |
| Aars2_in.vivo_noRT | 4930579C12R | 5.327 | 9.86E-19 | glio |
| Chek1_in.vivo_noRT | Gm26555 | 4.78 | 9.86E-21 | glio |
| culture_effect | AW112010 | -6.823 | 9.86E-26 | glio |
| culture_effect | Atp6v0a2 | 0.533 | 9.87E-03 | glio |
| Interaction_genotype_Atr | Slc1a3 | 1.993 | 9.87E-03 | glio |
| Interaction_genotype_Myc | Bc1 | -2.96 | 9.87E-03 | glio |
| Atp5c1_in.vivo_noRT | Arf4 | -0.515 | 9.87E-03 | glio |
| Atr_in.vivo_noRT | Gm35021 | 1.534 | 9.87E-03 | glio |

| | | | | |
|---|---|---|---|---|
| Atr_in.vivo_noRT | Atp5o | -0.626 | 9.87E-03 | glio |
| Interaction_genotype_Aars2 | Gm10354 | -3.329 | 9.87E-04 | glio |
| Interaction_genotype_Myc | Gm5141 | -3.715 | 9.87E-04 | glio |
| Aars2_in.vivo_noRT | Hist1h2af | 2.532 | 9.87E-05 | glio |
| Aars2_in.vivo_noRT | Frk | 2.759 | 9.87E-05 | glio |
| Atr_in.vivo_noRT | 2600006K01R | 2.875 | 9.87E-05 | glio |
| Interaction_genotype_Mtor | Zfp712 | -4.786 | 9.87E-06 | glio |
| Interaction_genotype_Mtor | Sytl4 | -3.833 | 9.87E-06 | glio |
| Atr_in.vivo_noRT | Gm36696 | 2.931 | 9.87E-06 | glio |
| Chek1_ex.vivo_noRT | Gm17173 | 3.738 | 9.87E-06 | glio |
| Chek1_ex.vivo_noRT | 9630014M24F | 3.652 | 9.87E-06 | glio |
| culture_effect | Cbx7 | -1.753 | 9.87E-07 | glio |
| Interaction_genotype_Mtor | 1700029J07Ri | -6.571 | 9.87E-08 | glio |
| Interaction_genotype_Mtor | 4921504E06Ri | -5.187 | 9.87E-08 | glio |
| Interaction_genotype_Mtor | Gm48140 | -4.629 | 9.87E-08 | glio |
| Chek1_in.vivo_noRT | 4833418N02R | 3.865 | 9.87E-08 | glio |
| Chek1_ex.vivo_noRT | Tcp11x2 | 4.449 | 9.87E-10 | glio |
| culture_effect | Fkbp9 | 0.347 | 9.88E-03 | glio |
| Interaction_genotype_Aars2 | Sdcbp2 | -3.449 | 9.88E-03 | glio |
| Interaction_genotype_Aars2 | Leprotl1 | 1.068 | 9.88E-03 | glio |
| Interaction_genotype_Atr | Vps33b | -2.582 | 9.88E-03 | glio |
| Interaction_genotype_Myc | Wasf1 | -2.218 | 9.88E-03 | glio |
| Interaction_genotype_Nat2 | Gm10482 | 3.208 | 9.88E-03 | glio |
| Interaction_genotype_Nat2 | Gm47404 | 2.678 | 9.88E-03 | glio |
| Interaction_genotype_Pars2 | 3830406C13R | 1.872 | 9.88E-03 | glio |
| Atr_in.vivo_noRT | Mid1ip1 | -1.41 | 9.88E-03 | glio |
| Bora_in.vivo_noRT | Dleu7 | 1.874 | 9.88E-03 | glio |
| Chek1_in.vivo_noRT | Ppil4 | -1.181 | 9.88E-03 | glio |
| Hsd17b10_in.vivo_noRT | Rnase4 | 2.054 | 9.88E-03 | glio |
| culture_effect | AC125149.2 | -1.828 | 9.88E-04 | glio |
| culture_effect | Gm9967 | 1.865 | 9.88E-04 | glio |
| culture_effect | Tmem108 | 1.133 | 9.88E-04 | glio |
| Interaction_genotype_Myc | 0610040B10R | -2.903 | 9.88E-04 | glio |
| Interaction_genotype_Uqcrc2 | Gm19276 | -3.459 | 9.88E-04 | glio |
| Aars2_in.vivo_noRT | Sulf2 | -1.308 | 9.88E-04 | glio |
| Aars2_in.vivo_noRT | Becn1 | -0.902 | 9.88E-04 | glio |
| Atr_in.vivo_noRT | Pir | 1.452 | 9.88E-04 | glio |
| Atr_ex.vivo_noRT | Osm | 2.764 | 9.88E-04 | glio |
| Atr_ex.vivo_noRT | Slc26a11 | 2.775 | 9.88E-04 | glio |
| Atr_ex.vivo_noRT | Junb | 1.883 | 9.88E-04 | glio |
| culture_effect | Plcb1 | 1.59 | 9.88E-05 | glio |
| Interaction_genotype_Myc | Cep295nl | -4.401 | 9.88E-05 | glio |
| Interaction_genotype_Myc | 4930434B07R | -3.845 | 9.88E-05 | glio |
| culture_effect | Irak2 | -1.936 | 9.88E-06 | glio |
| Bora_in.vivo_noRT | Gm49785 | 3.239 | 9.88E-06 | glio |
| culture_effect | Smim27 | 1.244 | 9.88E-08 | glio |
| Bora_in.vivo_noRT | 4922502N22R | 3.95 | 9.88E-09 | glio |
| Chek1_ex.vivo_noRT | Cers3 | 3.871 | 9.88E-09 | glio |
| Aars2_in.vivo_noRT | Lst1 | 4.561 | 9.88E-11 | glio |
| Bora_in.vivo_noRT | Npnt | 4.67 | 9.88E-20 | glio |
| Aars2_in.vivo_noRT | Chia1 | 5.327 | 9.88E-26 | glio |
| Interaction_genotype_Aars2 | Prap1 | -2.427 | 9.89E-03 | glio |

| | | | | |
|---|---|---|---|---|
| Interaction_genotype_Atr | 1010001B22R | -3.121 | 9.89E-03 | glio |
| Interaction_genotype_Bora | Gm50394 | -3.307 | 9.89E-03 | glio |
| Interaction_genotype_Bora | Eftud2 | 1.443 | 9.89E-03 | glio |
| Interaction_genotype_Myc | Hars2 | -2.335 | 9.89E-03 | glio |
| interaction_RT | Gm4013 | 1.044 | 9.89E-03 | glio |
| interaction_RT | G6pdx | -0.651 | 9.89E-03 | glio |
| interaction_RT | Srsf9 | 0.518 | 9.89E-03 | glio |
| Atr_in.vivo_noRT | Wdr19 | 1.814 | 9.89E-03 | glio |
| Bcas2_in.vivo_noRT | Rasl12 | 1.76 | 9.89E-03 | glio |
| Interaction_genotype_Myc | 3110082J24Ri | -3.768 | 9.89E-04 | glio |
| Atr_in.vivo_noRT | Ggcx | 1.701 | 9.89E-04 | glio |
| Eif2s1_in.vivo_noRT | Gm48617 | 1.879 | 9.89E-04 | glio |
| culture_effect | Avl9 | -0.753 | 9.89E-05 | glio |
| Interaction_genotype_Mtor | C030037D09R | -4.439 | 9.89E-05 | glio |
| Interaction_genotype_Mtor | Pigh | -3.456 | 9.89E-05 | glio |
| Interaction_genotype_Mtor | Arhgef12 | 1.868 | 9.89E-05 | glio |
| Interaction_genotype_Myc | Ripply3 | -4.293 | 9.89E-05 | glio |
| Interaction_genotype_Myc | 2310033P09R | -2.811 | 9.89E-05 | glio |
| Atr_in.vivo_noRT | Izumo4 | 2.746 | 9.89E-05 | glio |
| Chek1_in.vivo_noRT | Usp11 | 2.945 | 9.89E-05 | glio |
| Chek1_in.vivo_noRT | Upk1a | 3.252 | 9.89E-06 | glio |
| Aars2_in.vivo_noRT | Gm4181 | 3.428 | 9.89E-07 | glio |
| Bora_in.vivo_noRT | Alox8 | 4.204 | 9.89E-07 | glio |
| interaction_RT | Slc30a4 | 1.879 | 9.89E-08 | glio |
| Aars2_in.vivo_noRT | Lrrc46 | 3.679 | 9.89E-08 | glio |
| Atp5c1_in.vivo_noRT | Hoxc8 | 2.668 | 9.89E-08 | glio |
| culture_effect | Plscr2 | 1.554 | 9.89E-09 | glio |
| Atr_in.vivo_noRT | Pglyrp1 | 4.054 | 9.89E-09 | glio |
| Hsd17b10_in.vivo_noRT | Gm20646 | 4.143 | 9.89E-10 | glio |
| Hsd17b10_in.vivo_noRT | Adam30 | 3.764 | 9.89E-10 | glio |
| culture_effect | B3galt2 | -3.884 | 9.89E-12 | glio |
| culture_effect | Timm8b | 0.796 | 9.89E-12 | glio |
| culture_effect | Gxylt2 | 3.096 | 9.89E-22 | glio |
| Chek1_in.vivo_noRT | Gm2016 | 5.044 | 9.89E-22 | glio |
| culture_effect | Btbd8 | 0.8 | 9.90E-03 | glio |
| culture_effect | Asb13 | -0.68 | 9.90E-03 | glio |
| culture_effect | Polr2d | 0.415 | 9.90E-03 | glio |
| Interaction_genotype_Aars2 | Cfap46 | -2.952 | 9.90E-03 | glio |
| Interaction_genotype_Mtor | Zfp217 | -2.822 | 9.90E-03 | glio |
| Interaction_genotype_Myc | Zglp1 | -3.231 | 9.90E-03 | glio |
| Interaction_genotype_Myc | Sybu | -2.669 | 9.90E-03 | glio |
| interaction_RT | Lrp1 | 2.009 | 9.90E-03 | glio |
| interaction_RT | 1700030K09R | -1.352 | 9.90E-03 | glio |
| interaction_RT | Ppia | 0.551 | 9.90E-03 | glio |
| Aars2_in.vivo_noRT | H2-Ab1 | -1.005 | 9.90E-03 | glio |
| Atr_ex.vivo_noRT | Msl3l2 | 2.592 | 9.90E-03 | glio |
| Bard1_ex.vivo_noRT | C3 | 2.814 | 9.90E-03 | glio |
| Bcas2_in.vivo_noRT | Bms1 | -1.128 | 9.90E-03 | glio |
| Bcas2_in.vivo_noRT | Gdi1 | -0.962 | 9.90E-03 | glio |
| Bcas2_in.vivo_noRT | Nudt4 | -0.839 | 9.90E-03 | glio |
| culture_effect | Gm20455 | 1.972 | 9.90E-04 | glio |
| Interaction_genotype_Myc | Nkiras1 | -2.454 | 9.90E-04 | glio |

| | | | | |
|---|---|---|---|---|
| Bard1_ex.vivo_noRT | Gata3 | 3.563 | 9.90E-04 | glio |
| Bora_in.vivo_noRT | Tigd5 | 2.17 | 9.90E-04 | glio |
| Bora_in.vivo_noRT | Nub1 | -1.278 | 9.90E-04 | glio |
| Chek1_in.vivo_noRT | 5830428M24F | 2.39 | 9.90E-04 | glio |
| Eif2s1_in.vivo_noRT | 5033417F24Ri | 2.666 | 9.90E-04 | glio |
| culture_effect | Gm12860 | -1.942 | 9.90E-05 | glio |
| Atr_in.vivo_noRT | A630052C17R | 2.662 | 9.90E-05 | glio |
| Bcas2_in.vivo_noRT | Gm16835 | 2.88 | 9.90E-05 | glio |
| Aars2_in.vivo_noRT | A430010J10Ri | 3.82 | 9.90E-06 | glio |
| Atr_in.vivo_noRT | Efna2 | 4.261 | 9.90E-08 | glio |
| Bora_in.vivo_noRT | Bhlhe41 | 4.414 | 9.90E-08 | glio |
| Chek1_ex.vivo_noRT | Mau2 | -3.003 | 9.90E-08 | glio |
| Bcas2_in.vivo_noRT | 3930402G23R | 4.42 | 9.90E-10 | glio |
| Aars2_in.vivo_noRT | Rbfaos | 4.41 | 9.90E-16 | glio |
| Bora_in.vivo_noRT | Slc14a2 | 4.936 | 9.90E-25 | glio |
| Interaction_genotype_Atr | Fam3c | -1.424 | 9.91E-03 | glio |
| Interaction_genotype_Bcas2 | Prelid3a | -3.418 | 9.91E-03 | glio |
| Interaction_genotype_Bcas2 | Ubl3 | 1.283 | 9.91E-03 | glio |
| Interaction_genotype_Lmo7 | Syp | -4.572 | 9.91E-03 | glio |
| Interaction_genotype_Lmo7 | Plcl1 | -3.369 | 9.91E-03 | glio |
| Interaction_genotype_Mtor | Tapbpl | 1.867 | 9.91E-03 | glio |
| Interaction_genotype_Myc | Pls1 | -2.865 | 9.91E-03 | glio |
| Interaction_genotype_Myc | Eps8l1 | -2.924 | 9.91E-03 | glio |
| Interaction_genotype_Rpl3 | Crhbp | -3.163 | 9.91E-03 | glio |
| Interaction_genotype_Rpl3 | Tcte2 | -3.145 | 9.91E-03 | glio |
| Aars2_in.vivo_noRT | Rpl3 | -0.511 | 9.91E-03 | glio |
| Atp5c1_in.vivo_noRT | Gm8300 | 2.668 | 9.91E-03 | glio |
| Bcas2_ex.vivo_noRT | Gm48161 | 2.349 | 9.91E-03 | glio |
| Chek1_in.vivo_noRT | 2310015A10R | 1.999 | 9.91E-03 | glio |
| Chek1_in.vivo_noRT | Syne1 | -1.213 | 9.91E-03 | glio |
| Chek1_ex.vivo_noRT | Igf2r | 2.331 | 9.91E-03 | glio |
| Eif2s1_in.vivo_noRT | Gm5662 | 2.143 | 9.91E-03 | glio |
| Eif2s1_in.vivo_noRT | Psmb11 | 1.62 | 9.91E-03 | glio |
| Eif2s1_in.vivo_noRT | Serpina3g | 2.001 | 9.91E-03 | glio |
| Eif2s1_in.vivo_noRT | Stx3 | -1.026 | 9.91E-03 | glio |
| Interaction_genotype_Mtor | Clcn3 | 1.899 | 9.91E-04 | glio |
| Atp5c1_in.vivo_noRT | Lce1g | 2.668 | 9.91E-04 | glio |
| Bcas2_in.vivo_noRT | Ccnh | -1.405 | 9.91E-04 | glio |
| Chek1_in.vivo_noRT | Selenbp1 | 2.605 | 9.91E-04 | glio |
| Chek1_in.vivo_noRT | Hist1h2bp | 2.519 | 9.91E-04 | glio |
| Hsd17b10_in.vivo_noRT | Npb | 2.449 | 9.91E-04 | glio |
| culture_effect | Cd151 | -0.63 | 9.91E-05 | glio |
| Interaction_genotype_Atr | Ift27 | 2.602 | 9.91E-05 | glio |
| Interaction_genotype_Myc | Nrde2 | -3.582 | 9.91E-05 | glio |
| Hsd17b10_in.vivo_noRT | Ntn5 | 2.824 | 9.91E-05 | glio |
| culture_effect | Mpc2 | 0.678 | 9.91E-07 | glio |
| Atr_in.vivo_noRT | Fchsd1 | 3.289 | 9.91E-07 | glio |
| Interaction_genotype_Mtor | Gm48242 | -5.462 | 9.91E-08 | glio |
| Bcas2_in.vivo_noRT | Apol7d | 3.905 | 9.91E-08 | glio |
| Bora_in.vivo_noRT | Lrrn3 | 3.48 | 9.91E-08 | glio |
| culture_effect | Fabp5 | 1.124 | 9.91E-09 | glio |
| Atr_in.vivo_noRT | Cacng5 | 3.683 | 9.91E-09 | glio |

| | | | | |
|---|---|---|---|---|
| culture_effect | Ier2 | -1.928 | 9.91E-19 | glio |
| Interaction_genotype_Atr | Ift46 | 1.737 | 9.92E-03 | glio |
| interaction_RT | Cyp4f18 | 3.547 | 9.92E-03 | glio |
| Aars2_in.vivo_noRT | Sltm | -0.816 | 9.92E-03 | glio |
| Aars2_in.vivo_noRT | Sparc | -0.878 | 9.92E-03 | glio |
| Bora_in.vivo_noRT | Brinp3 | 1.442 | 9.92E-03 | glio |
| Bora_in.vivo_noRT | Emc1 | -1.256 | 9.92E-03 | glio |
| Chek1_ex.vivo_noRT | Gpr141 | 2.867 | 9.92E-03 | glio |
| Chek1_ex.vivo_noRT | Elavl1 | -1.046 | 9.92E-03 | glio |
| Chek1_ex.vivo_noRT | Denr | -0.772 | 9.92E-03 | glio |
| Hsd17b10_in.vivo_noRT | Gm14327 | 2.111 | 9.92E-03 | glio |
| Interaction_genotype_Chek1 | Atpif1 | 1.221 | 9.92E-04 | glio |
| Interaction_genotype_Myc | Nxf1 | -2.07 | 9.92E-04 | glio |
| Atp5c1_in.vivo_noRT | Gm50115 | 2.405 | 9.92E-04 | glio |
| Atr_in.vivo_noRT | Them4 | 1.87 | 9.92E-04 | glio |
| Bcas2_in.vivo_noRT | Bag1 | -0.815 | 9.92E-04 | glio |
| Bora_in.vivo_noRT | Adamts4 | 2.128 | 9.92E-04 | glio |
| Eif2s1_in.vivo_noRT | Nrg2 | 2.669 | 9.92E-04 | glio |
| Atr_in.vivo_noRT | Srcin1 | 2.959 | 9.92E-06 | glio |
| Atr_in.vivo_noRT | Tnp2 | 3.471 | 9.92E-07 | glio |
| Atr_in.vivo_noRT | Chd4 | -1.472 | 9.92E-07 | glio |
| Atr_in.vivo_noRT | Zfp960 | 4.176 | 9.92E-09 | glio |
| Bora_in.vivo_noRT | Cpne8 | 4.202 | 9.92E-09 | glio |
| Chek1_in.vivo_noRT | Gm45441 | 3.984 | 9.92E-09 | glio |
| Aars2_in.vivo_noRT | 4833407H14R | 4.692 | 9.92E-10 | glio |
| Atr_in.vivo_noRT | Tnxb | 4.38 | 9.92E-11 | glio |
| Bora_in.vivo_noRT | Gjb4 | 4.803 | 9.92E-12 | glio |
| Interaction_genotype_Bora | Ptges | -3.673 | 9.93E-03 | glio |
| Interaction_genotype_Bora | Dnase1l2 | -3.377 | 9.93E-03 | glio |
| Interaction_genotype_Myc | C030014I23Ri | -2.772 | 9.93E-03 | glio |
| interaction_RT | Dock5 | -1.587 | 9.93E-03 | glio |
| Aars2_in.vivo_noRT | Gm19541 | 1.89 | 9.93E-03 | glio |
| Aars2_in.vivo_noRT | Tlr3 | 1.197 | 9.93E-03 | glio |
| Aars2_ex.vivo_noRT | Gm42918 | 2.321 | 9.93E-03 | glio |
| Atp5c1_in.vivo_noRT | Rdh12 | 2.023 | 9.93E-03 | glio |
| Atp5c1_in.vivo_noRT | Hnrnph3 | -0.995 | 9.93E-03 | glio |
| Bcas2_in.vivo_noRT | Gm16536 | 1.514 | 9.93E-03 | glio |
| Bora_ex.vivo_noRT | Clec4d | 2.016 | 9.93E-03 | glio |
| Chek1_ex.vivo_noRT | Mafk | 1.684 | 9.93E-03 | glio |
| Chek1_ex.vivo_noRT | Dcun1d5 | -0.749 | 9.93E-03 | glio |
| Interaction_genotype_Myc | Jpx | -2.756 | 9.93E-04 | glio |
| Bora_in.vivo_noRT | Kctd11 | 2.465 | 9.93E-04 | glio |
| Bora_ex.vivo_noRT | Gm14540 | 2.407 | 9.93E-04 | glio |
| Interaction_genotype_Mtor | Tent5c | -3.566 | 9.93E-05 | glio |
| Interaction_genotype_Mtor | Txn2 | 1.915 | 9.93E-05 | glio |
| Chek1_in.vivo_noRT | Gba2 | 2.787 | 9.93E-05 | glio |
| Interaction_genotype_Myc | Trim65 | -4.812 | 9.93E-06 | glio |
| Atr_in.vivo_noRT | Zfp992 | 3.505 | 9.93E-07 | glio |
| Atr_in.vivo_noRT | Chl1 | -1.745 | 9.93E-07 | glio |
| Atr_in.vivo_noRT | Prrc2c | -1.397 | 9.93E-07 | glio |
| Bora_in.vivo_noRT | Synpo2l | 4.291 | 9.93E-13 | glio |
| Bora_in.vivo_noRT | Usp17la | 4.936 | 9.93E-13 | glio |

| | | | | |
|---|---|---|---|---|
| culture_effect | Otof | -4.887 | 9.93E-22 | glio |
| culture_effect | Hras | 1.168 | 9.93E-22 | glio |
| Bora_in.vivo_noRT | Olfr1152 | 5.194 | 9.93E-26 | glio |
| culture_effect | G6pc3 | -0.441 | 9.94E-03 | glio |
| culture_effect | Washc3 | -0.334 | 9.94E-03 | glio |
| Interaction_genotype_Bcas2 | Gm37800 | -3.15 | 9.94E-03 | glio |
| Interaction_genotype_Bcas2 | Gm5152 | -3.538 | 9.94E-03 | glio |
| Interaction_genotype_Bcas2 | Gm49701 | -3.366 | 9.94E-03 | glio |
| Interaction_genotype_Bcas2 | Cep57 | 1.499 | 9.94E-03 | glio |
| Interaction_genotype_Bcas2 | Ppil3 | -1.257 | 9.94E-03 | glio |
| Interaction_genotype_Rad51 | Ephx1 | -3.65 | 9.94E-03 | glio |
| Aars2_in.vivo_noRT | Prelid3b | -0.97 | 9.94E-03 | glio |
| Atr_in.vivo_noRT | Batf | 1.88 | 9.94E-03 | glio |
| Atr_in.vivo_noRT | Wdfy2 | 1.615 | 9.94E-03 | glio |
| Bcas2_in.vivo_noRT | Mboat7 | -1.229 | 9.94E-03 | glio |
| Brca2_in.vivo_noRT | Gm4724 | 2.152 | 9.94E-03 | glio |
| Chek1_ex.vivo_noRT | Gm10800 | 2.645 | 9.94E-03 | glio |
| Aars2_in.vivo_noRT | 2610203C22R | 2.473 | 9.94E-04 | glio |
| Aars2_in.vivo_noRT | Star | 2.298 | 9.94E-04 | glio |
| Bcas2_in.vivo_noRT | Piezo1 | -1.505 | 9.94E-04 | glio |
| Chek1_in.vivo_noRT | Gm5577 | 2.2 | 9.94E-04 | glio |
| Chek1_ex.vivo_noRT | Sept5 | 2.961 | 9.94E-04 | glio |
| Hsd17b10_in.vivo_noRT | Gm9833 | 2.356 | 9.94E-04 | glio |
| Atr_in.vivo_noRT | Puf60 | -1.243 | 9.94E-05 | glio |
| Cyp19a1_ex.vivo_noRT | Gm266 | 2.959 | 9.94E-05 | glio |
| culture_effect | Taf1a | -1.397 | 9.94E-06 | glio |
| Interaction_genotype_Aars2 | Scube3 | -4.59 | 9.94E-06 | glio |
| Bora_in.vivo_noRT | 9130204L05Ri | 3.342 | 9.94E-06 | glio |
| Bora_in.vivo_noRT | Ccdc9b | 2.82 | 9.94E-06 | glio |
| Atr_in.vivo_noRT | Ncbp1 | -1.833 | 9.94E-07 | glio |
| Bora_in.vivo_noRT | Gm7008 | 3.471 | 9.94E-07 | glio |
| Atr_in.vivo_noRT | 9130604C24R | 4.067 | 9.94E-08 | glio |
| Bard1_ex.vivo_noRT | Gm48617 | 3.596 | 9.94E-08 | glio |
| Atr_in.vivo_noRT | Gm10644 | 4.292 | 9.94E-10 | glio |
| culture_effect | Cntn1 | -4.564 | 9.94E-15 | glio |
| culture_effect | C4b | -9.081 | 9.94E-58 | glio |
| Interaction_genotype_Atr | Ephx3 | -3.122 | 9.95E-03 | glio |
| Interaction_genotype_Rad51 | Gm49864 | -3.65 | 9.95E-03 | glio |
| Interaction_genotype_Rad51 | Pip4k2c | 2.136 | 9.95E-03 | glio |
| interaction_RT | Sema4b | 1.169 | 9.95E-03 | glio |
| Aars2_in.vivo_noRT | Gusb | -1.044 | 9.95E-03 | glio |
| Dclre1c_ex.vivo_noRT | Pldi | 2.543 | 9.95E-03 | glio |
| Interaction_genotype_Myc | 2310008N11R | -3.999 | 9.95E-04 | glio |
| Aars2_in.vivo_noRT | 4930581F22Ri | 1.975 | 9.95E-04 | glio |
| Bcas2_in.vivo_noRT | Lcat | 2.474 | 9.95E-04 | glio |
| Hsd17b10_in.vivo_noRT | Gm13610 | 2.46 | 9.95E-04 | glio |
| Atr_in.vivo_noRT | Lrp1b | 2.755 | 9.95E-05 | glio |
| Bora_in.vivo_noRT | 1810021B22R | 2.867 | 9.95E-05 | glio |
| Chek1_in.vivo_noRT | Spata1 | 2.883 | 9.95E-05 | glio |
| Hsd17b10_in.vivo_noRT | Cyp2s1 | 3.088 | 9.95E-05 | glio |
| culture_effect | Nfib | 0.604 | 9.95E-06 | glio |
| Hsd17b10_in.vivo_noRT | Nup93 | -1.974 | 9.95E-06 | glio |

| | | | | |
|---|---|---|---|---|
| Aars2_in.vivo_noRT | Olfr1189 | 3.672 | 9.95E-10 | glio |
| Atr_in.vivo_noRT | Ceacam19 | 4.501 | 9.95E-11 | glio |
| Bora_in.vivo_noRT | Nwd1 | 4.672 | 9.95E-16 | glio |
| Bcas2_in.vivo_noRT | Gm30794 | 4.807 | 9.95E-21 | glio |
| Aars2_in.vivo_noRT | Esm1 | 5.063 | 9.95E-23 | glio |
| Bora_in.vivo_noRT | Gm15908 | 4.936 | 9.95E-26 | glio |
| Bora_in.vivo_noRT | A930014E10Ri | 4.936 | 9.95E-26 | glio |
| culture_effect | Cacna1a | 0.844 | 9.96E-03 | glio |
| Interaction_genotype_Bora | Anks6 | -3.17 | 9.96E-03 | glio |
| Interaction_genotype_Myc | 2600006K01R | -2.867 | 9.96E-03 | glio |
| Interaction_genotype_Myc | Osgepl1 | -1.852 | 9.96E-03 | glio |
| Interaction_genotype_Myc | Acsl3 | -1.614 | 9.96E-03 | glio |
| interaction_RT | Abce1 | -0.516 | 9.96E-03 | glio |
| Aars2_in.vivo_noRT | Dcp1a | 1.439 | 9.96E-03 | glio |
| Aars2_in.vivo_noRT | Chchd3 | -0.714 | 9.96E-03 | glio |
| Atp5c1_in.vivo_noRT | 2810410L24Ri | 2.142 | 9.96E-03 | glio |
| Chek1_in.vivo_noRT | Ufc1 | -0.665 | 9.96E-03 | glio |
| Chek1_ex.vivo_noRT | Maco1 | -1.435 | 9.96E-03 | glio |
| Chek1_ex.vivo_noRT | Hspd1 | -0.695 | 9.96E-03 | glio |
| Interaction_genotype_Atr | Gm13212 | -4.25 | 9.96E-04 | glio |
| Interaction_genotype_Atr | 5430400D12R | -4.121 | 9.96E-04 | glio |
| Interaction_genotype_Mtor | Gm48684 | -4.536 | 9.96E-04 | glio |
| Bcas2_in.vivo_noRT | Saxo2 | 2.346 | 9.96E-04 | glio |
| Interaction_genotype_Aars2 | Tmem50a | 1.842 | 9.96E-05 | glio |
| Interaction_genotype_Myc | Hdgf | 1.59 | 9.96E-05 | glio |
| Chek1_in.vivo_noRT | Ficd | 2.2 | 9.96E-05 | glio |
| Chek1_in.vivo_noRT | Eml1 | 2.821 | 9.96E-06 | glio |
| Atr_in.vivo_noRT | C130040N14R | 3.886 | 9.96E-08 | glio |
| Hsd17b10_in.vivo_noRT | Mecom | 3.616 | 9.96E-08 | glio |
| Hsd17b10_in.vivo_noRT | Dpp6 | 2.689 | 9.96E-08 | glio |
| culture_effect | Tacc1 | 1.289 | 9.96E-09 | glio |
| Hsd17b10_in.vivo_noRT | A930033M14F | 3.616 | 9.96E-10 | glio |
| Aars2_in.vivo_noRT | Car12 | 4.228 | 9.96E-11 | glio |
| Chek1_in.vivo_noRT | K230015D01R | 4.777 | 9.96E-11 | glio |
| Interaction_genotype_Myc | Ntn4 | -6.431 | 9.96E-12 | glio |
| Aars2_in.vivo_noRT | Lce1g | 5.327 | 9.96E-14 | glio |
| Chek1_in.vivo_noRT | Gm44899 | 5.044 | 9.96E-20 | glio |
| Interaction_genotype_Aars2 | Ccno | -3.375 | 9.97E-03 | glio |
| Interaction_genotype_Atr | Tmem250-ps | -2.11 | 9.97E-03 | glio |
| Interaction_genotype_Myc | Zfp768 | -1.879 | 9.97E-03 | glio |
| interaction_RT | Tnnt1 | 1.877 | 9.97E-03 | glio |
| interaction_RT | Rrp12 | -1.342 | 9.97E-03 | glio |
| Aars2_in.vivo_noRT | Gclm | -1.225 | 9.97E-03 | glio |
| Aars2_in.vivo_noRT | Aatf | -0.934 | 9.97E-03 | glio |
| Cenpj_in.vivo_noRT | Akap14 | 1.759 | 9.97E-03 | glio |
| Chek1_ex.vivo_noRT | Kpna2 | -2.06 | 9.97E-03 | glio |
| Chek1_ex.vivo_noRT | Cdc16 | -1.336 | 9.97E-03 | glio |
| culture_effect | Rab15 | -2.072 | 9.97E-04 | glio |
| Interaction_genotype_Myc | Ly6g6d | -3.767 | 9.97E-04 | glio |
| Aars2_ex.vivo_noRT | Ripor2 | 3.289 | 9.97E-04 | glio |
| Bora_in.vivo_noRT | Reep5 | -0.899 | 9.97E-04 | glio |
| Bora_in.vivo_noRT | Txnl4a | -0.755 | 9.97E-04 | glio |

| | | | | |
|---|---|---|---|---|
| Eif2s1_in.vivo_noRT | 9430041J12Ri | 2.143 | 9.97E-04 | glio |
| Interaction_genotype_Atr | Synpr | -5.311 | 9.97E-05 | glio |
| Atp5c1_in.vivo_noRT | D930028M14I | 2.668 | 9.97E-05 | glio |
| Bcas2_in.vivo_noRT | Gpd1 | 3.634 | 9.97E-07 | glio |
| Chek1_in.vivo_noRT | 4921507P07R | 3.279 | 9.97E-07 | glio |
| Chek1_in.vivo_noRT | 9030612E09Ri | 3.723 | 9.97E-09 | glio |
| Bcas2_in.vivo_noRT | Igfbp7 | 4.041 | 9.97E-11 | glio |
| Chek1_in.vivo_noRT | Col8a1 | 5.044 | 9.97E-14 | glio |
| Bora_in.vivo_noRT | AC163347.1 | 4.673 | 9.97E-17 | glio |
| Interaction_genotype_Atr | Zscan4-ps2 | -2.369 | 9.98E-03 | glio |
| Interaction_genotype_Hsd17b | Otof | 3.514 | 9.98E-03 | glio |
| Interaction_genotype_Mtor | Scgb1a1 | -3.179 | 9.98E-03 | glio |
| Interaction_genotype_Mtor | Gm13203 | -3.381 | 9.98E-03 | glio |
| Interaction_genotype_Mtor | Exosc4 | 1.336 | 9.98E-03 | glio |
| Interaction_genotype_Myc | Gm38115 | -2.97 | 9.98E-03 | glio |
| Atp5c1_in.vivo_noRT | 4921523L03Ri | 2.28 | 9.98E-03 | glio |
| Atr_in.vivo_noRT | Zfp146 | 1.439 | 9.98E-03 | glio |
| Atr_in.vivo_noRT | Adamts1 | -1.464 | 9.98E-03 | glio |
| Atr_in.vivo_noRT | Rpl37a | -0.691 | 9.98E-03 | glio |
| Chek1_in.vivo_noRT | Dennd3 | 2 | 9.98E-03 | glio |
| Chek1_in.vivo_noRT | 9330151L19Ri | 1.724 | 9.98E-03 | glio |
| Chek1_in.vivo_noRT | Tigd5 | 1.742 | 9.98E-03 | glio |
| Cyp19a1_ex.vivo_noRT | AF067063 | 2.159 | 9.98E-03 | glio |
| Cyp19a1_ex.vivo_noRT | Gm14820 | 2.159 | 9.98E-03 | glio |
| Cyp19a1_ex.vivo_noRT | Gm2016 | 2.159 | 9.98E-03 | glio |
| Cyp19a1_ex.vivo_noRT | H2-Eb2 | 2.159 | 9.98E-03 | glio |
| Aars2_in.vivo_noRT | Ndufa13 | -0.657 | 9.98E-04 | glio |
| Aars2_ex.vivo_noRT | A030005L19Ri | 2.899 | 9.98E-04 | glio |
| Atr_in.vivo_noRT | Pla2g4c | 2.308 | 9.98E-04 | glio |
| Atr_in.vivo_noRT | Gm10863 | 2.138 | 9.98E-04 | glio |
| Bora_in.vivo_noRT | Stat4 | 2.305 | 9.98E-04 | glio |
| culture_effect | Tnfsf12 | 1.443 | 9.98E-06 | glio |
| Atp5c1_in.vivo_noRT | Npnt | 2.402 | 9.98E-06 | glio |
| Hsd17b10_in.vivo_noRT | Cnksr1 | 3.11 | 9.98E-06 | glio |
| Interaction_genotype_Mtor | C130060C02R | -5.805 | 9.98E-08 | glio |
| Aars2_in.vivo_noRT | Tmem116 | 4.149 | 9.98E-08 | glio |
| Bora_in.vivo_noRT | Pla2g2f | 3.826 | 9.98E-10 | glio |
| Chek1_in.vivo_noRT | Nkg7 | 4.784 | 9.98E-16 | glio |
| Interaction_genotype_Atr | Dhodh | -2.527 | 9.99E-03 | glio |
| Interaction_genotype_Atr | Eya4 | -2.304 | 9.99E-03 | glio |
| Interaction_genotype_Myc | Emilin2 | -3.005 | 9.99E-03 | glio |
| Interaction_genotype_Myc | Snrpd3 | 1.299 | 9.99E-03 | glio |
| interaction_RT | Srsf1 | -0.574 | 9.99E-03 | glio |
| Aars2_in.vivo_noRT | Gm9774 | 1.886 | 9.99E-03 | glio |
| Bora_in.vivo_noRT | Fancl | -1.538 | 9.99E-03 | glio |
| Bora_in.vivo_noRT | Sacs | -1.366 | 9.99E-03 | glio |
| Interaction_genotype_Ifnar1 | Gm48215 | 3.531 | 9.99E-04 | glio |
| Interaction_genotype_Mtor | Rimkla | -3.814 | 9.99E-04 | glio |
| Interaction_genotype_Pars2 | Sprr1a | -4.641 | 9.99E-04 | glio |
| Interaction_genotype_Pars2 | Mecom | -4.155 | 9.99E-04 | glio |
| Interaction_genotype_Pars2 | Col8a1 | -4.401 | 9.99E-04 | glio |
| Interaction_genotype_Uqcrc2 | Gm41396 | -5.069 | 9.99E-04 | glio |

| | | | | |
|---|---|---|---|---|
| Aars2_in.vivo_noRT | Tln1 | -1.198 | 9.99E-04 | glio |
| Atr_in.vivo_noRT | BC024978 | 1.994 | 9.99E-04 | glio |
| Chek1_in.vivo_noRT | Cfl1 | -0.668 | 9.99E-04 | glio |
| Interaction_genotype_Myc | Ccnd1 | 2.266 | 9.99E-05 | glio |
| Atr_ex.vivo_noRT | S1pr1 | 3.878 | 9.99E-05 | glio |
| Bcas2_in.vivo_noRT | Gid4 | -1.58 | 9.99E-05 | glio |
| interaction_RT | Fbxo32 | 1.97 | 9.99E-06 | glio |
| Aars2_in.vivo_noRT | Hist1h2ai | 3.23 | 9.99E-06 | glio |
| Atp5c1_in.vivo_noRT | Gm10354 | 2.668 | 9.99E-06 | glio |
| interaction_RT | Rpl23a | -0.986 | 9.99E-07 | glio |
| Chek1_in.vivo_noRT | Tbxa2r | 3.204 | 9.99E-07 | glio |
| Interaction_genotype_Mtor | B930094E09Ri | -6.197 | 9.99E-08 | glio |
| Bora_in.vivo_noRT | 1110017D15R | 3.936 | 9.99E-09 | glio |
| Atr_in.vivo_noRT | Gm26682 | 4.615 | 9.99E-12 | glio |
| Bora_in.vivo_noRT | Myo5c | 5.46 | 9.99E-16 | glio |
| Aars2_in.vivo_noRT | AF067061 | 5.327 | 9.99E-19 | glio |
| Aars2_in.vivo_noRT | Gm6803 | 5.327 | 9.99E-22 | glio |
